# SARS-CoV-2 Envelope-mediated Golgi pH dysregulation interferes with ERAAP retention in cells

**DOI:** 10.1101/2022.11.29.518257

**Authors:** Valerie Vargas-Zapata, Kristina M. Geiger, Dan Tran, Jessica Ma, Xiaowen Mao, Andreas S. Puschnik, Laurent Coscoy

**Affiliations:** Division of Immunology and Molecular Medicine, Department of Molecular and Cell Biology, University of California, Berkeley, Berkeley, CA, 94720, USA; Division of Infectious Diseases and Vaccinology, School of Public Health, University of California, Berkeley, Berkeley, CA 94720, USA; Division of Microbial Biology, Department of Plant and Microbial Biology, University of California, Berkeley, Berkeley, CA 94720, USA; Chan Zuckerberg Biohub, San Francisco, CA 94158, USA

## Abstract

Endoplasmic reticulum (ER) aminopeptidase associated with antigen processing (ERAAP) trims peptide precursors in the ER for presentation by major histocompatibility (MHC)-I molecules to surveying CD8^+^ T-cells. This function allows ERAAP to regulate the nature and quality of the peptide repertoire and, accordingly, the resulting immune responses. We recently showed that infection with murine cytomegalovirus leads to a dramatic loss of ERAAP levels in infected cells. In mice, this loss is associated with the activation of QFL T-cells, a subset of T-cells that monitor ERAAP integrity and eliminate cells experiencing ERAAP dysfunction. In this study, we aimed to identify host factors that regulate ERAAP expression level and determine whether these could be manipulated during viral infections. We performed a CRISPR knockout screen and identified ERp44 as a factor promoting ERAAP retention in the ER. ERp44’s interaction with ERAAP is dependent on the pH gradient between the ER and Golgi. We hypothesized that viruses that disrupt the pH of the secretory pathway interfere with ERAAP retention. Here, we demonstrate that expression of the Envelope (E) protein from Severe Acute Respiratory Syndrome Coronavirus 2 (SARS-CoV-2) leads to Golgi pH neutralization and consequently decrease of ERAAP intracellular levels. Furthermore, SARS-CoV-2-induced ERAAP loss correlates with its release into the extracellular environment. ERAAP’s reliance on ERp44 and a functioning ER/Golgi pH gradient for proper localization and function led us to propose that ERAAP serves as a sensor of disturbances in the secretory pathway during infection and disease.

## Introduction

The antigen presentation pathway (APP) provides a snapshot of the intracellular environment to surveying immune cells in the form of peptides presented on major histocompatibility (MHC) molecules^1^. This pathway plays a crucial role in the response against viruses and cancers^2,3^. Presentation of foreign peptides (e.g. from viruses) or altered self-peptides (e.g. cancer) presented on MHC-I molecules allows cytotoxic CD8^+^ T-cells to recognize and kill infected or cancerous cells, respectively. In this pathway, peptide precursors are generated in the cytoplasm by the proteasome and transported into the endoplasmic reticulum (ER) by the Transporter associated with Antigen Processing (TAP)^4^. Once in the ER, peptides are edited to properly fit into MHC-I molecules. This function is accomplished by the ER aminopeptidase associated with antigen processing (ERAAP) in mice and ER aminopeptidase (ERAP) 1 and 2 in humans^5–7^. We will refer to these collectively as ERAP. ERAP belongs to the M1 family of metallopeptidases, and it trims peptides on their amino (N-) terminus to an approximate length of 8-10 amino acids. ERAP regulates the quality and nature of the peptide repertoire and therefore shapes the resulting immune responses^8,9^. Loss of ERAAP leads to an altered peptide repertoire characterized by changes in the abundance and length of peptide precursors^10^. The peptidome of ERAAP deficient cells is sufficiently different from normal cells that these MHC-I:peptide complexes can elicit an immune response. Indeed, mice immunized with ERAAP KO cells mount a robust CD8^+^ T-cell response that results in the elimination of these cells^11^. Careful analysis of the responding CD8^+^ T-cells revealed that most of them recognize <u>FL</u>9, a self-peptide derived from the ubiquitously expressed host proteins Fam49A and B, that is presented by the non-classical MHC-I molecule, <u>Q</u>a1b. As a result, these cells were named QFL T-cells. Upon recognition of the FL9:Qa1b complex, QFL T-cells can kill target cells and produce inflammatory cytokines^11,12^. So far, FL9 presentation has only been reported in cells experiencing ERAAP dysfunction and thus, it has been suggested that QFL T-cells serve to monitor ERAAP function and eliminate defective cells.

Due to the central role of the APP in eliciting antiviral CD8^+^ T-cell responses, many viruses have evolved to disrupt different components of this pathway^2,13^, including ERAP. Human cytomegalovirus (HCMV) encodes two micro RNAs, miR-UL112-5p and miR-US4-1, that target ERAP1 for degradation and this reduces the production of HCMV antigenic peptides^14,15^. Recently, our group showed that infection with mouse cytomegalovirus (MCMV) also results in ERAAP downregulation^16^. Interestingly, loss of ERAAP during MCMV infection triggers a QFL T-cell response. In MCMV-infected mice, QFLs proliferate, produce inflammatory cytokines, and contribute to viral restriction^16^. In naïve mice, QFL T-cells have an antigen experienced phenotype^17,18^suggesting that actively surveying ERAAP dysfunction, and by extension the integrity of the APP, is an important aspect of immune surveillance. A human QFL T-cell equivalent has not been identified.

Given the significance of ERAAP’s function and the consequences of its loss, we became interested in the cellular pathways that regulate ERAAP in normal cells and whether those could be disrupted during viral infection. Here we show that ERp44 promotes ERAAP retention in the ER, a process regulated by the pH of the ER and Golgi. We demonstrate that the Envelope (E) protein from Severe Acute Respiratory Syndrome Coronavirus 2 (SARS-CoV-2) increases the Golgi pH, leading to a decrease in ERAAP intracellular protein levels and its release into the extracellular environment. Because E’s ability to modulate ion gradients is part of SARS-CoV-2’s life cycle and pathogenesis, we propose that ERAAP could serve as an indicator of the integrity of the secretory pathway, and its loss and/or secretion could signal an altered cellular state like a viral infection.

## Results

### ERAAP dsRed is expressed and properly localized to the ER

Our lab recently described that cells infected with MCMV express low levels of ERAAP^16^. Through this work, we became interested in identifying the cellular pathways that regulate ERAAP protein levels in cells under normal conditions and determine whether these pathways could be targeted during viral infection. To address this question, we performed a genome-wide CRISPR KO screen to identify host factors that regulate ERAAP levels in cells. To this end, a reporter cell line that expresses an ERAAP-dsRed fusion protein was generated to allow monitoring of ERAAP levels by flow cytometry. Human Burkitt lymphoma B (BJAB) cells were transduced to express ERAAP-dsRed and subsequently two cell clones (B2 and D4) that possessed homogenous, stable, and high expression of ERAAP-dsRed were isolated (Figure 1A).

**Figure 1:**
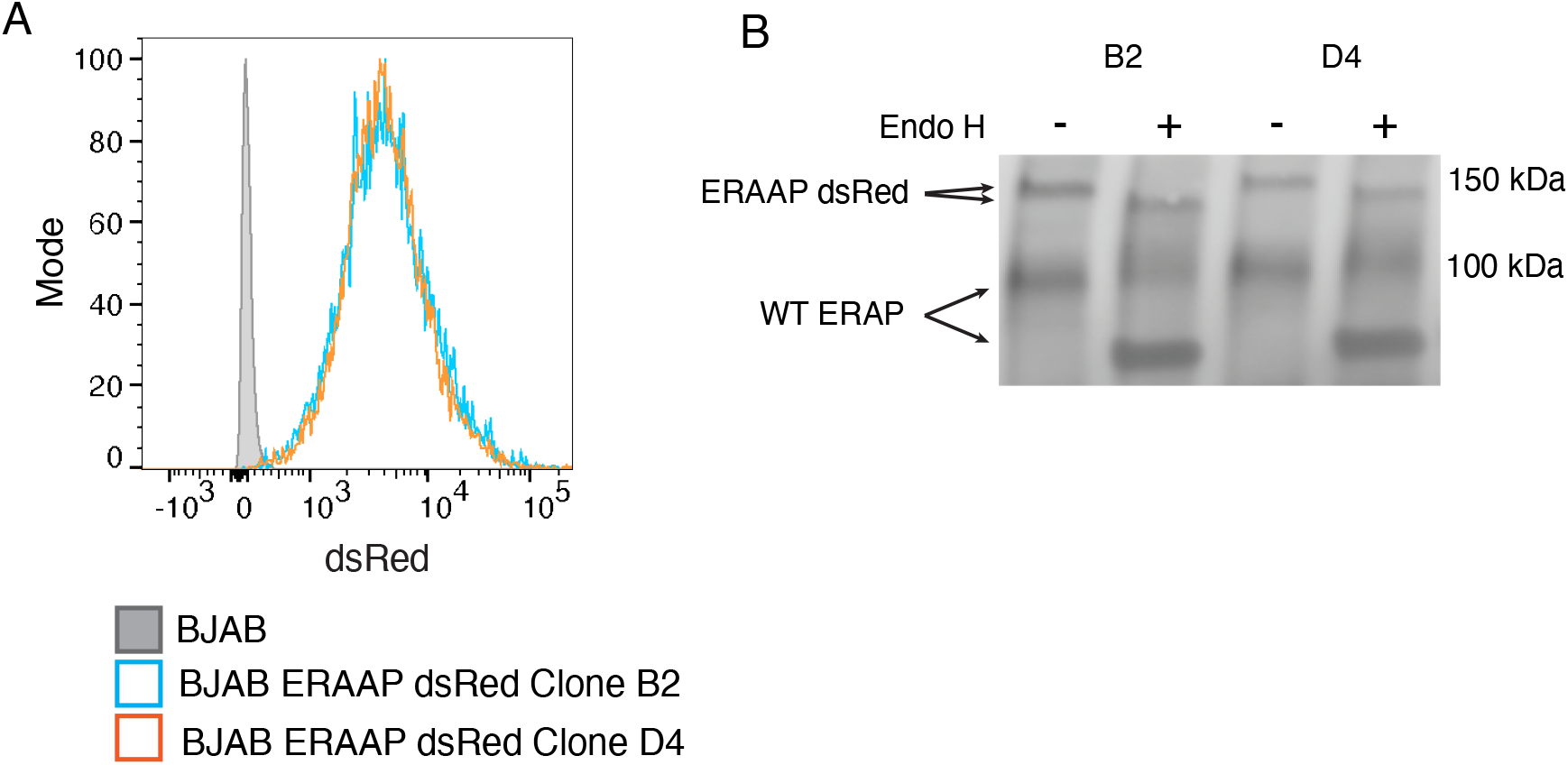
ERAAP dsRed is expressed and properly localized in the ER. A. BJAB cells were transduced to express an ERAAP dsRed fusion. Cells were subcloned and selected for high expression of the construct as measured by flow cytometry. Histogram shows ERAAP dsRed levels in two selected clones, B2 (blue histogram) and D4 (orange histogram), in comparison to non-transduced BJAB cells (grey filled histogram). B. To test for proper ERAAP dsRed localization, Endo H assay was performed. WCL was collected from BJAB ERAAP dsRed clones B2 and D4 and ERAAP was immunoprecipitated using an anti-ERAAP antibody. Purified ERAAP was incubated with Endo H enzyme (Endo H +) or water (Endo H -) as a control. After treatment, ERAAP’s molecular weight was determined by western blot. Shown is a representative ERAAP blot. In non-treated samples, ERAAP dsRed appears as a ~150 kDa band (top set of arrows) while endogenous ERAP (WT ERAP; bottom set of arrows) can be observed as a ~100 kDa band. The appearance of a cleavage product of lower molecular weight indicates Endo H sensitivity.

To confirm the proper ER localization of the ERAAP-dsRed construct, an Endoglycosidase H (Endo H) assay was performed. Glycans added to proteins in the ER are further modified as they transit through the Golgi. These modifications change the sensitivity of the glycan structure to cleavage by enzymes like Endo H. Thus, Endo H sensitivity can be used as an indicator of the general localization within the secretory pathway. ERAAP resides in the ER and should remain sensitive to Endo H cleavage. Indeed, ERAAP-dsRed shows sensitivity to Endo H cleavage as evidenced by the appearance of a lower molecular weight band (shift from ~150 kDa to 130 kDa) in samples treated with the enzyme (Figure 1B) in comparison to untreated samples. Endogenous ERAP was also detected in this gel (~100 kDa) and remained Endo H sensitive suggesting ERAAP-dsRed expression does not interfere with the localization of the endogenous protein. Since both cell clones showed high expression of ERAAP-dsRed and proper ER localization, we selected clone B2 to perform our experiments.

### ERp44 regulates ERAAP levels

To perform the genome-wide CRISPR KO screen, BJAB ERAAP dsRed cells (clone B2) were transduced to express Cas9. These cells were then transduced with the human genome-scale CRISPR KO (GeCKO) v2 library targeting over 19,000 genes (containing 6 guide RNAs (gRNAs) per gene). After selection and recovery, cells with the top and bottom 10-15% dsRed signal were sorted and the gRNAs, present in these cell populations, were identified by sequencing and compared for gene enrichment.

From our hits, ERp44 stood out for its role in modulating the localization and trafficking of different host proteins^19^. Other potential hits obtained from the gRNA library screen mostly corresponded to proteins involved in transcriptional regulation (Supplemental Figure 1). Because of the high likelihood that these factors control ERAAP dsRed transcription from the CMV promoter of our retroviral construct, we decided to only focus on ERp44 in this study.

ERp44’s interaction with ERAP has been reported to regulate ERAP localization by retrieving it from the ER-Golgi intermediate compartment (ERGIC)/cis-Golgi and bringing it back into the ER^20^. To confirm the results of our screen, BJAB ERAAP dsRed cells were transduced with a gRNA targeting ERp44. As expected, cells lacking ERp44 showed a marked decrease in ERAAP protein levels as observed by western blot (Figure 2A) and by flow cytometry (Figure 2B). Knockout of ERAAP did not affect ERp44 levels (Figure 2A).

**Figure 2:**
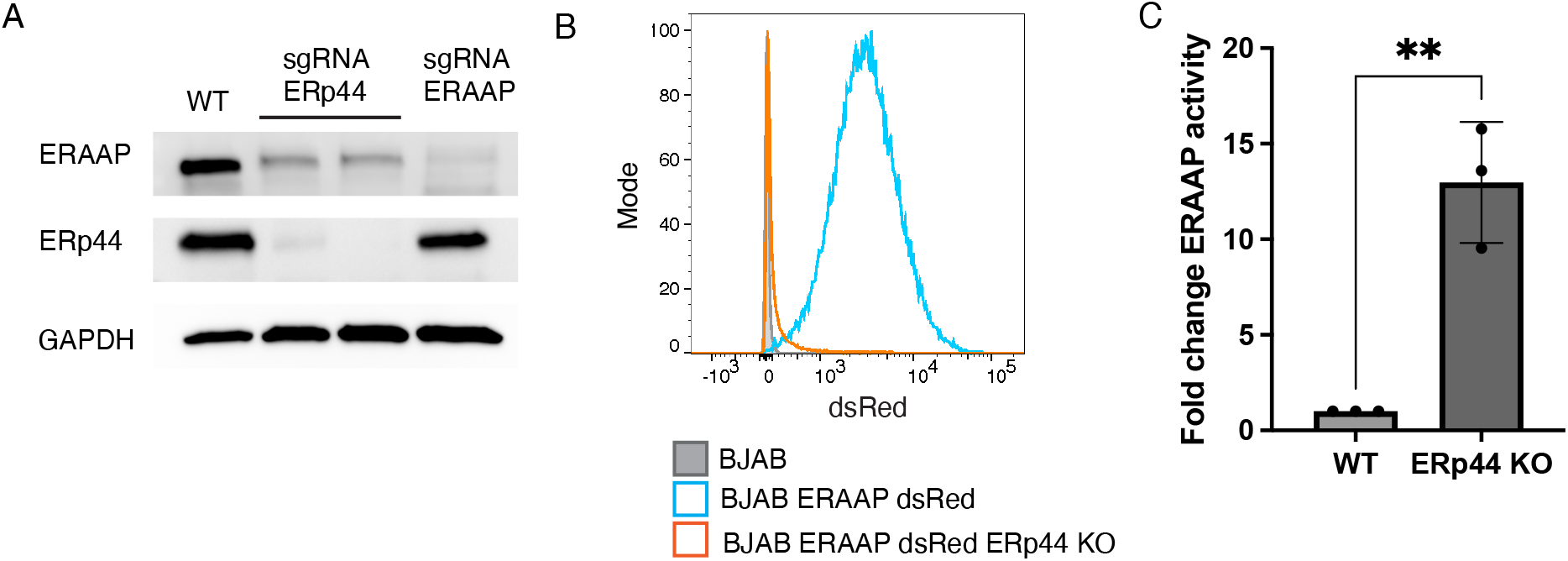
ERp44 regulates ERAAP levels. A. To confirm the results of the CRISPR KO screen, BJAB ERAAP dsRed cells (clone B2) were transfected with two gRNA targeting ERp44 or one gRNA targeting ERAAP as a control. ERAAP levels in ERp44 KO cells were measured by western blot. Representative blot showing ERAAP and ERp44 protein levels in untransfected BJAB ERAAP dsRed (WT) and KO cells. GAPDH is shown as a loading control. B. Histogram showing ERAAP dsRed levels in BJAB ERAAP dsRed (blue histogram) and ERp44 KO cells (orange histogram). Parental BJAB cells are shown as a control (grey filled histogram) C. ERAAP secretion from ERp44 KO cells was measured by LAP assay. ERAAP was immunoprecipitated from the supernatants of BJAB ERAAP dsRed and BJAB ERAAP dsRed ERp44 KO cells. Beads containing ERAAP were incubated with LpNA substrate and LAP activity was measured 8 h after addition of the substrate by measuring optical density (OD) at 410 nm. Background activity coming from media was subtracted from both conditions. Shown is the fold change in LAP activity relative to BJAB ERAAP dsRed (WT) cells. Data represents mean± SD from three independent experiments. Unpaired two-tailed T-test was performed. ** p value < 0.01.

To test if ERAAP was being secreted from BJAB cells lacking ERp44, ERAAP was immunoprecipitated from the supernatant and ERAAP activity was quantified using the leucine aminopeptidase (LAP) activity assay^20^. As predicted, ERp44 KO cells secreted ERAAP as evidenced by increased LAP activity in comparison to parental cells (Figure 2C). ERp44 controls ERAAP retention across a variety of cell lines, both mouse (Supplemental Figure 2) and human^20^ (Figure 2C) suggesting ERp44’s regulation of ERAP is a conserved feature of mammalian cells.

### SARS-CoV-2 Envelope neutralizes the pH of the Golgi

One of the factors regulating ERp44’s function is pH. Dysregulation of the Golgi pH abrogates the ability of ERp44 to interact with its clients and results in their secretion^21^. We reasoned that interfering with the pH homeostasis of the Golgi could have negative consequences for ERAAP function. Interestingly, some viruses encode proteins that modify the pH of intracellular compartments as part of their viral life cycle^22^. In particular, the E proteins from the coronaviruses Infectious Bronchitis virus (IBV) and SARS-CoV-2 have been shown to increase the pH of the Golgi^23,24^. Neutralization of the Golgi space is thought to protect the fusion protein Spike (S) from abnormal cleavage by Golgi resident proteases^23,25^. E oligomerizes into an ion-conductive pore and it is involved in viral assembly, budding, and pathogenesis^26,27^. Viruses lacking E are attenuated^28,29^and E’s ion channel-like activity seems to be particularly important for pathogenesis and SARS-CoV-2-induced pathology^30,31^. Still, much remains to be known about E’s intracellular function and impact on cellular pathways.

We hypothesized that during SARS-CoV-2 infection, E-mediated neutralization of the Golgi and the subsequent loss of ERp44 activity might result in ERAP being secreted rather than being retained into the ER. We decided to first confirm that the expression of SARS-CoV-2 E in cells indeed causes an increase in the pH of the Golgi as previously reported^24^. To that extent, we used the pH reporter pHluorin-TGN38, a Golgi-localized GFP variant that exhibits bimodal excitation at 405 nm and 488 nm^32^. The fluorescence emission from this reporter changes in response to pH. Specifically, the emission ratio of pHluorin (405 nm/488 nm) increases as pH increases. HeLa cells co-transfected with SARS-CoV-2 E showed a higher pHluorin emission ratio in comparison to cells transfected only with the reporter (Figure 3A). Using HeLa cells transfected with the reporter and resuspended in buffers of known pH, we generated a standard curve to convert pHluorin emission ratios to pH values. As expected, cells expressing SARS-CoV-2 E showed neutralization of the Golgi in comparison to control cells. In control cells, the pH of the Golgi was 6.9 and increased to 7.4 in cells expressing SARS-CoV-2 E. As an additional control, we transfected cells with Influenza A virus (IAV) M2, a viral protein known to increase the Golgi pH^23,33–35^, and observed an increased pHluorin emission ratio (Figure 3A) that translated to a Golgi pH of 7.3 (Figure 3B). These data suggest that SARS-CoV-2 E behaves as previously demonstrated by neutralizing the pH of the Golgi.

**Figure 3:**
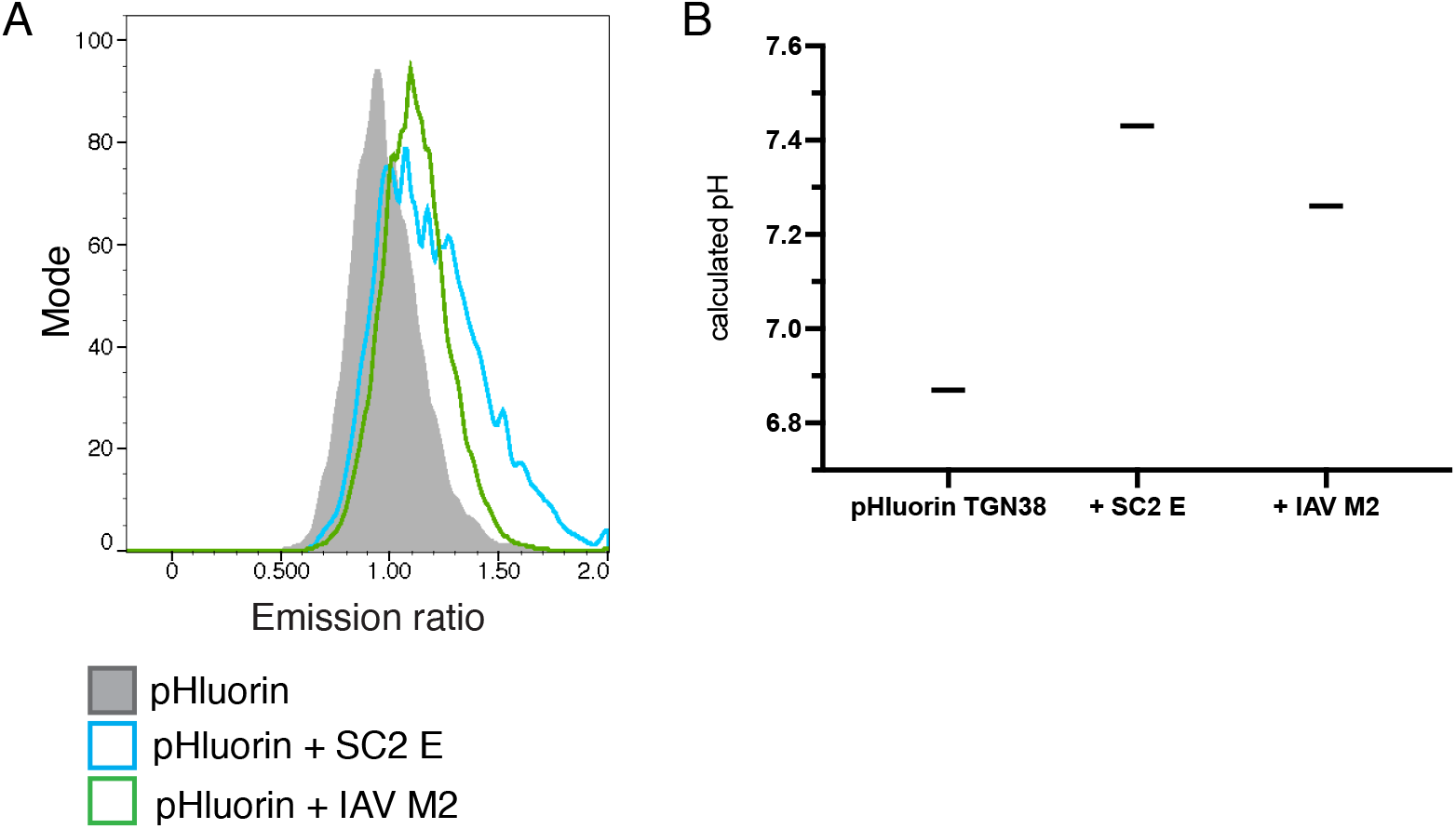
SARS-CoV-2 Envelope neutralizes the pH of the Golgi. A. HeLa cells were transfected with pHluorin-TGN38 alone (grey filled histogram) or co-transfected with pHluorin-TGN38 and SARS-CoV-2 E (blue histogram) or IAV M2 (green histogram). The fluorescence emission of pHluorin-TGN38 was measured by flow cytometry after excitation at 405 nm and 488 nm. Histogram shows the pHIuorin-TGN38 emission ratio (405 nm/488 nm) for each condition. B. Using a pH standard curve, the emission ratios values for each condition were used to calculate the pH of the Golgi. Shown are the calculated pH values for the Golgi space in HeLa cells transfected with pHIuorin-TGN38 alone or co-transfected with SARS-CoV-2 E and IAV M2. Data shown is from one representative experiment. SARS-CoV-2 abbreviated as SC2.

### SARS-CoV-2 Envelope leads to decreased ERAAP levels

Next, we asked if SARS-CoV-2 E expression could result in a reduction in ERAAP’s intracellular levels. To evaluate ERAAP levels specifically in cells expressing SARS-CoV-2 E, we decided to use a co-transfection system. HEK293T cells were chosen as they are easily transfectable, and they express low levels of ERAP endogenously. HEK293T cells were cotransfected with a vector encoding ERAAP-dsRed and another vector encoding GFP and SARS-CoV-2 E (or a vector expressing only GFP as control). 24- and 48-hours post-transfection (hpt), cells were harvested and ERAAP dsRed levels were evaluated by flow cytometry. Cells expressing SARS-CoV-2 E showed decreased ERAAP-dsRed levels in comparison to vector control (Figure 4A). Based on the dsRed geometric mean fluorescence for each population, we found that cells expressing SARS-CoV-2 E have ~40-50% of the levels of ERAAP in control cells (Figure 4B).

**Figure 4:**
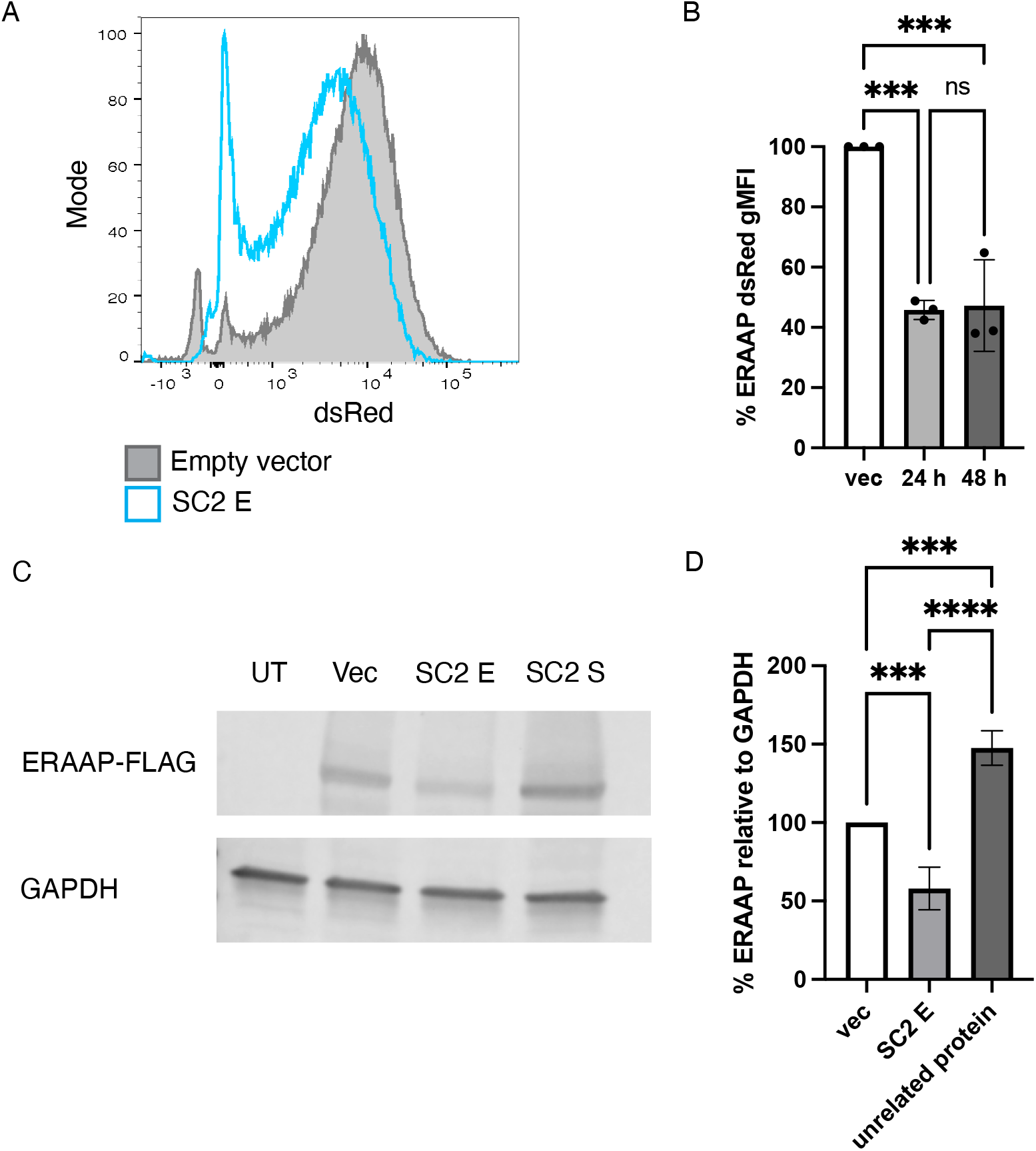
SARS-CoV-2 Envelope leads to decreased ERAAP levels. A. HEK293T cells were co-transfected with ERAAP dsRed and SARS-CoV-2 E (blue histogram), or empty vector control (grey filled histogram) and ERAAP dsRed levels were measured by flow cytometry. Histogram shows ERAAP dsRed levels within transfected cells 24 h post-transfection. B. The geometric mean fluorescence intensity (gMFI) for dsRed was obtained from HEK293T cotransfected with ERAAP dsRed and SARS-CoV-2 E, or empty vector as in A. Graphs shows the percent dsRed gMFI 24 h and 48 h after transfection with SARS-CoV-2 E relative to empty vector across 3 independent experiments. Cells that were co-transfected with empty vector were assumed to represent 100% ERAAP dsRed levels and are shown as a control. Data represents mean± SD. Unpaired One-way ANOVA assuming a Gaussian distribution and Tukey’s multiple comparisons test was performed. *** p value < 0.001, ns=not significant. C. Representative blot showing ERAAP levels in HEK293T co-transfected with ERAAP 3XFlag and SARS-CoV-2 E, SARS-CoV-2 S, or empty vector control. WCL collected 36 h post transfection. GAPDH is shown as a loading control. D. From blots as that shown in C, the band intensity of ERAAP 3XFlag and GAPDH was determined using Image J. The relative ERAAP levels for each condition was obtained after normalizing to respective GAPDH levels. Graph shows the percent ERAAP 3XFlag levels in HEK293T transfected with SARS-CoV-2 E or an unrelated protein (ex. SARS-CoV-2 S). Cells that were co-transfected with ERAAP 3XF and empty vector were assumed to represented 100% ERAAP levels and are shown as a control. Data represents mean ± SD from 5 independent experiments for cells co-transfected with vector or SARS-CoV-2 E and from 3 experiments for unrelated protein. SARS-CoV-2 abbreviated as SC2. Unpaired One-way ANOVA assuming a Gaussian distribution with Tukey’s multiple comparisons test was performed. *** p value < 0.001, **** p value < 0.0001.

The effect of SARS-CoV-2 E on ERAAP protein levels was also confirmed using an alternative method. Cells were co-transfected with a vector encoding a Flag-tagged ERAAP and a vector encoding SARS-CoV-2 E or an empty vector control. 36 hpt, whole cell lysate (WCL) was collected and ERAAP protein levels were evaluated by western blot. As observed before, cells expressing SARS-CoV-2 E have decreased levels of ERAAP. Importantly, this effect is specific to SARS-CoV-2 E as co-transfecting a different viral protein, SARS-CoV-2 S, did not decrease ERAAP levels (Figure 4C). Analysis of ERAAP levels across multiple experiments revealed that cells expressing E have on average ~60% the levels of ERAAP (40% reduction) of cells cotransfected with the empty vector. Cells co-transfected with an unrelated protein did not show any decrease in ERAAP levels (Figure 4D). Taken together, these results indicate that SARS-CoV-2 E causes a loss in ERAAP levels likely as a result of disrupting ERp44-mediated recycling of ERAAP.

### ERAAP is secreted from cells expressing SARS CoV-2 Envelope

We hypothesized that the loss in ERAAP levels observed in cells expressing SARS-CoV-2 E might be the result of ERAAP release into the supernatant as described with cells lacking ERp44^20^ (Figure 2C). To test this, the supernatant of NIH3T3 cells stably expressing ERAAP dsRed and transfected with SARS-CoV-2 E (or empty vector as control) was collected at 48 hpt and a LAP assay was performed.

In cells expressing SARS-CoV-2 E, we observed ~34% ERAAP secretion relative to ERp44 KO cells (Figure 5A). Cells co-transfected with empty vector showed ~13% ERAAP release. This background level of ERAAP secretion is likely due to overexpression of ERAAP in this system which might saturate retention mechanisms^36^. Still, SARS-CoV-2 E induces a significant 2.7-fold increase in ERAAP secretion relative to empty vector control. Importantly the amount of ERAAP being secreted by SARS-CoV-2 E-expressing cells is probably largely underestimated in this assay since only ~20-30% of the cells are transfected with SARS-CoV-2 E (Supplemental Figure 3). Together our results suggest that SARS-CoV-2 E disrupts ERAAP function by preventing the efficient retention of ERAAP inside cells and leading to its secretion.

**Figure 5:**
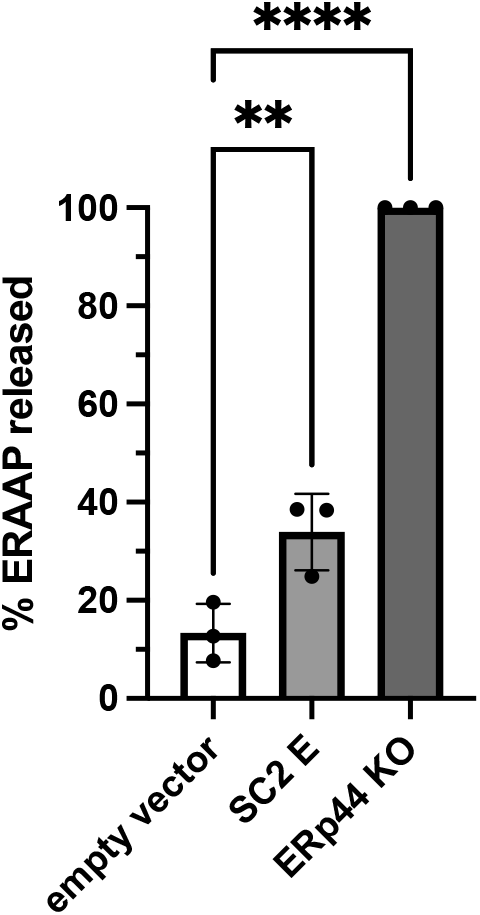
ERAAP is secreted from cells expressing SARS-CoV-2 Envelope. LAP assay was performed on the supernatants of NIH 3T3 ERAAP dsRed transfected with empty vector or SARS-CoV-2 E. Background activity detected in the supernatant of untransfected NIH 3T3 ERAAP dsRed was subtracted from all conditions. LAP activity from the supernatant of NIH 3T3 ERAAP dsRed ERp44 KO cells was measured and assumed to represent 100% ERAAP release. Graph shows the percent of ERAAP released for each condition. Data represents mean ± SD from 3 independent experiments. Unpaired One-way ANOVA assuming a Gaussian distribution with Dunnett’s posttest was performed. ** p value < 0.01, **** p value < 0.0001.

## Discussion

Using a CRISPR KO screen, we identified ERp44 as the main host factor regulating ERAAP retention in cells. ERp44 KO cells showed decreased intracellular ERAAP levels and increased secretion of ERAAP into the extracellular environment. ERp44-mediated retention of ERAAP is in part regulated by pH. Here we report that pH disruption by SARS-CoV-2 E interferes with ERAAP retention in the ER and leads to its secretion. The increase of Golgi pH by E is an important part of the SARS-CoV-2 viral life cycle as it protects the S protein from being inadequately processed by proteases residing in the Golgi^23,25^.

To avoid detection by CD8^+^ T-cells, viruses have evolved a myriad of immune evasion mechanisms aimed at preventing the presentation of antigenic peptides^2,13^. Interfering with ERAAP likely represents another one of these strategies as illustrated by HCMV, and likely MCMV^14–16^. Furthermore, a recent study by Stamatakis, et al., showed that ERAP generates potentially immunogenic SARS-CoV-2 S-derived peptides predicted to bind MHC^37^. Here we show that SARS-CoV-2 E leads to a decrease in ERAP intracellular concentration. This suggests that targeting of the Golgi by SARS-CoV-2 could function to both preserve the integrity of the viral fusion protein S and dampen the generation of virally-derived antigenic peptides through the loss of ERAP. Given the central role of ERAP in modulating T-cell responses, other viruses beyond HCMV, MCMV, and SARS-CoV-2 may have also evolved to disrupt ERAP as part of their immune evasion strategies.

Fortunately, viral immune evasion does not go unchallenged. Cells and the immune system have evolved mechanisms to sense and respond to virally-mediated disturbances. In mice, ERAAP dysfunction is sensed by QFL T-cells. MCMV-induced ERAAP loss results in the presentation of the self-peptide FL9 in the MHC-I molecule Qa1b, the activating ligand of QFL T-cells. In infected mice, QFL T-cells are activated, produce inflammatory cytokines, and can kill cells experiencing ERAAP loss^16^. This exciting observation suggests that ERAAP dysregulation during infection has the potential to be sensed by the immune system. The existence of QFL-like T-cells in humans has yet to be described, but it is tempting to speculate that if a similar immune response exists, it could aid in controlling SARS-CoV-2 infection.

In addition to the loss of intracellular ERAP, we showed that ERAP is secreted from cells expressing SARS-CoV-2 E. Secreted ERAP has been implicated in blood pressure regulation which might have consequences for SARS-CoV-2 infection. ERAP’s potential to modulate the renin-angiotensin-aldosterone system (RAAS), which controls blood pressure, comes from its ability to cleave Angiotensin II (Ang II) into Ang III and Ang IV^38^. Ang II exerts its effects on multiple organs that work in concert to raise blood pressure^39^. To maintain normotension, Ang II levels are carefully regulated. Ang II is cleaved into other angiotensin products that can activate parallel pathways which have the opposite effect of lowering blood pressure^40^. Ang II is normally cleaved by angiotensin-converting enzyme 2 (ACE2). ACE2 has gathered much attention as it is the main receptor SARS-CoV-2 uses to enter cells^41,42^. Upon interaction of SARS-CoV-2 S with ACE2, the virus is internalized causing a temporary depletion of ACE2 at the cell surface, which can result in local increases of Ang II levels^43,44^. Dysregulated RAAS is thought to contribute to the clinical manifestations of COVID-19^45–48^. As recently proposed by D’Amico et al., secreted ERAP could help alleviate some of these effects by lowering the levels of Ang II^47^. Our findings provide the first indication that ERAP can be secreted from SARS-CoV-2 infected cells opening the possibility that ERAP might regulate RAAS during infection. Future studies are needed to address the specific contribution of ERAP as a potential protective factor during SARS-CoV-2 infection. Interestingly, some ERAP polymorphisms predicted to result in lower ERAP activity or loss of function, have been linked to hypertension and blood pressure progression^47^. It is possible that patients with dysfunctional ERAP could be susceptible to more detrimental COVID-19 outcomes because of an inability to dampen Ang II levels. One described ERAP polymorphism, A1533G, results in reduced activity and inability to convert Ang II into its cleavage products^49^. Given these observations, it is certain that the role of ERAP in SARS-CoV-2 infection merits more investigation due to the potential of ERAP to protect or contribute to COVID-19 pathology.

As shown here by us and others, ERAAP localization and retention are regulated by the chaperone ERp44^20^. ERp44 is a member of the protein disulfide isomerase family that cycles between the ER, ERGIC, and cis-Golgi. ERp44 interacts with client proteins, such as ERAP, that lack ER retention motifs (i.e. KDEL) and retrieves them from the Golgi back into the ER. Importantly, ERp44’s function is regulated by pH. In the neutral environment of the ER, ERp44 adopts a close conformation in which its carboxyl (C-) terminal tail occludes the client binding site. In the acidic environment of the ERGIC and cis-Golgi, ERp44 undergoes conformational changes that liberate its C-terminal tail and makes its binding pocket accessible for client binding^19^. The C-terminus of ERp44 contains an RDEL sequence that facilitates the transport of the ERp44-client complex from the Golgi to the ER via KDEL receptors^36^. Similar to what we reported here, increasing the Golgi pH by silencing the Golgi pH regulator (GPHR) results in a decreased ability of ERp44 to bind its clients, and secretion of clients to the extracellular space^21^. In addition to ERAP, ERp44 is involved in regulating other clients including Ero1β, Prx4, and SUMF1^36,50,51^. Secretion of these clients has been reported in cases where interactions with ERp44 are perturbed and so we speculate that these may be also secreted by SARS-CoV-2 Eexpressing cells. Whether these other clients have extracellular functions, and whether their secretion is physiologically relevant remains to be explored.

The observation that pH dysregulation leads to ERAAP loss potentially extends beyond SARS-CoV-2. The E protein belongs to a larger family of virally encoded ion channel-like proteins collectively known as viroporins. Due to their ability to change membrane permeability, viroporins induce a range of physiological changes in the cell and are encoded by viruses across many different families^22,52^ including some that are clinically relevant like IAV, Human Immunodeficiency Virus (HIV), and Hepatitis C Virus (HCV). Of all viroporins, the best studied is IAV M2. M2 is a multifunctional pH-regulated ion channel and similar to SARS-CoV-2 E, M2 localizes to the Golgi membranes where it neutralizes the pH of this organelle^23,33–35^. For M2, the ability to modulate Golgi pH helps protect hemagglutinin from preemptively adopting the low pH conformation needed for fusion^35^. The p7 protein from HCV has also been shown to disrupt H^+^ gradients and increase intracellular pH^53^. It would be interesting to test whether infection with viruses like IAV and HCV could also result in ERAP loss and secretion.

Given that ERAP’s main function is peptide trimming in the ER, it is interesting that this protein lacks an ER retention signal and instead relies on ERp44 for its proper localization and function. This reliance on ERp44 makes ERAP susceptible to different modes of regulation. Indeed, in addition to pH perturbation, ERp44 function is abrogated by disruption of the oxidoreductive state in the ER/Golgi^20,36^, changes in the Zn^2+^ gradient of the secretory pathway^54^, as well as altered trafficking between the Golgi and the ER. This reliance on pH, redox, or Zn^2+^ homeostasis for ERp44 function thus suggests that perturbation of these elements, whether they are mediated by a disease state or an infection, should lead to loss of ERAP. Since the loss of ERAP is associated with the presentation of an alternative immunogenic peptide repertoire on MHC-I molecules, it is expected that perturbation of these pathways would lead to the elimination of the affected cells by the immune system. Notably, because viruses rely on coopting the secretory pathway to produce viral proteins and the assembly of virions, one can reason that it will be difficult for viruses to evade activating pathways that sense perturbations in these compartments. Taken together, we propose that ERAP could serve as a sensor of intracellular disturbances and a link between cellular sensing and activation of the immune system.

## Acknowledgments

We thank Hector Nolla and Alma Valero of the UC Berkeley Cancer Research Lab for their help with flow cytometry. We would like to thank the Shastri lab, Robey lab, and Glaunsinger lab for helpful discussions about this project. The research reported in this manuscript was supported by funding from the National Institute of Allergy and Infectious Diseases of the National Institutes of Health (RO1AI149341).

## Author contributions

VVZ, KG, and LC conceptualized and designed the experiments. VVZ, KG, and AP performed experiments, generated tools, and did data analysis. DT, JM, and XM aided with experiments or generated tools used in this study. VVZ and LC wrote the manuscript.

## Declaration of interests

The authors declare no competing interests.

## Materials and Methods

### Cell lines

HEK293T, HeLa cells, and Phoenix cells were obtained from the UC Berkeley Cell Culture Facility. NIH 3T3, BJAB, and MC57G cells were purchased from ATCC. Except for BJAB, all cells were cultured in DMEM (Gibco, 11995073) containing 10% FBS (VWR, 89510-186) and 1% Pen Strep (Gibco, 15140-122). BJAB cells were cultured in RPMI (Gibco, 11875) containing 10% FBS and 1% Pen Strep. Cells were cultured at 37 °C in 5% CO_2_.

### Generation of ERAAP-dsRed cell lines

To generate stable cell lines expressing ERAAP-dsRed, cells were transduced with retrovirus produced in Phoenix cells. Briefly, Phoenix cells were seeded at 500,000 cells/well in a 6-well plate. 24 hours later, cells were transfected with 2 ug of pQCXIN ERAAP-dsRed using Fugene HD transfection reagent (Promega, E2311) following the manufacturer’s instructions. 48 hours post-transfection, the supernatant was harvested, filtered through a 0.45 μM filter, and added to BJAB or NIH 3T3 target cells.

BJAB transduced to express ERAAP-dsRed were selected with 1 mg/mL neomycin (G418; Corning, 61234RF). After selection, cells were sorted based on high dsRED expression by FACS. Sorted cells were subcloned to identify a clonal population of cells with uniform high expression of ERAAP-dsRed (Clone B2 and D4). HEK293T cells were used to produce lentivirus encoding Cas9 (lentiCas9-Blast, Addgene #52962, gift from Feng Zhang). BJAB ERAAP-dsRed cells were seeded at 1X10^6 cells/well in a 12-well plate and spin-infected with Cas9 expressing lentivirus in presence of 4ug/mL polybrene (Santa Cruz Biotechnology, sc-134220) by centrifugation at 1,000 g for 2 h at 33 °C. 24 h after transduction, the selection was started by culturing cells in media containing 5 ug/mL blasticidin (AG Scientific, B-1247). The selection was continued for 7 days to allow non-transduced cells to die and be removed from the population. BJAB ERAAP-dsRed expressing Cas9 (Clone B2) was selected to perform the genome-wide CRISPR screen.

To generate NIH 3T3 cells expressing ERAAP-dsRed, 200,000 cells/well were seeded in a 6-well plate and supernatant containing retrovirus encoding ERAAP dsRed was added. Cells were spinfected with polybrene at 1,200 rpm for 2 hours. Media was replaced and cells were left to recover overnight. The next day, the selection was started by incubating the transduced cells with media containing 1 mg /mL neomycin (G418). Cells were selected for 10 days and dsRed expression was evaluated by flow cytometry. To isolate a homogenous population of cells expressing ERAAP-dsRed, cells were subcloned. Cell clones were screened by flow cytometry to confirm uniform dsRed expression. Single-cell clone F3 was identified and was used to generate ERp44 KO cells in this background.

### Genome-wide CRISPR KO screen

6.0 × 10^6 BJAB clone B2 cells expressing both Cas9 and ERAAP dsRed were transduced with lentivirus of human GeCKO v2 library (Addgene, #1000000049, gift from Feng Zhang) at an MOI of 0.4 using spin-infection (1,000 g for 2 h at 33 °C). After selection with 3 ug/mL puromycin for 7 days, cells were sorted by flow cytometry based on ERAAP dsRed expression levels. Cells with the top and bottom 10-15% of ERAAP dsRed expression were sorted and collected by FACS for each GeCKO sublibrary (A and B). Genomic DNA was extracted from sorted cells to sequence enriched guide RNA (sgRNA) sequences in each population. Using a two-step nested PCR with KAPA HiFi HotStart ReadyMixPCR Kit (Kapa Biosystems), sgRNA expression cassettes were amplified. PCR products were gel-purified using a QIAquick Gel Purification kit (Qiagen) and sequenced on an Illumina NextSeq 500 using custom sequencing primers. Fastq files were analyzed using the Model-based Analysis of Genome-wide CRISPR/Cas9 Knockout (MaGeCK) method^55^.

### Generation of ERp44 KO cell lines

To generate ERp44 KO cells, cells were first transduced to express Cas9. Generation of BJAB ERAAP-dsRed expressing Cas9 is described above. The same procedure was followed to generate Cas9 expressing NIH 3T3 and MC57G cells.

Guide RNAs targeting ERp44 were cloned into the lentiviral vector pLKO.1 to generate a pLKO.1-ERp44sgRNA lentiviral vector. Sequences for the sgRNAs used are included below.

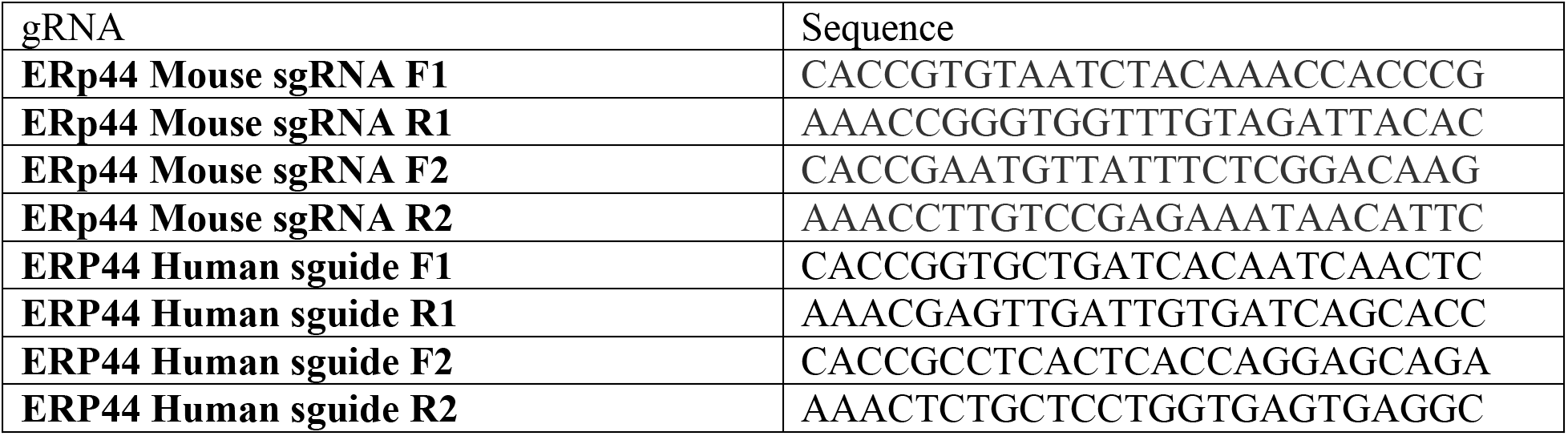

To generate ERp44 KO cell lines for BJAB ERAAP-dsRed (clone B2), NIH 3T3 and MC57G, HEK293T cells were plated at 500,000 cells/well in a 6-well plate. After 24 hours, cells were transfected with 250 ng VSVG, 1,250 ng of psPAX2, and 1,250 ng of pLKO.1-ERp44sgRNA using TransIT-LT1 transfection reagent (Fisher Scientific, MIR2300) following manufacturer’s instructions. After 48 hours, supernatant from transfected HEK293T cells was collected and filtered using a 0.45uM filter. Lentivirus with polybrene was added to target cells plated in a 6-well plate. Cells were spinfected for 2 hours after which fresh media was added to the cells. 24 h later, cells were put into puromycin supplemented media to select for cells expressing ERp44 guide RNA (BJAB ERAAP-dsRed and NIH 3T3-2 ug/mL puromycin, MC57G-4 ug/mL puromycin). After 7 days, KO efficiency was assessed by western blot.

To generate ERP44 KO cells in the NIH 3T3 ERAAP-dsRed background, cells were first transduced to express Cas9. HEK293T cells were transfected with 500 ng VSVG, 500 ng of pRRE, 500 ng of pRev, and 1000 ng of pFUGW-Cas9. 48 h after transfection, lentivirus was harvested and NIH 3T3 ERAAP-dsRed cells (Clone F3) were spinfected as described above. Cells were selected with 5 μg/mL blasticidin for 9 days. The resulting Cas9-expressing NIH 3T3 ERAAP were used to generate ERP44 KO cells. ERp44 mouse sgRNA F1 and R1 were cloned into hLKO to generate hLKO-ERp44sgRNA. NIH 3T3 ERAAP-dsRed cells were transduced with lentivirus encoding ERp44 sgRNA and selected for 9 days in the presence of 200 μg/mL Hygromycin. ERp44 KO was confirmed by western blot.

### Generation of ERAAP KO cells

BJAB ERAAP-dsRed, NIH 3T3, and MC57G expressing Cas9 were transfected with gsRNA targeting ERAAP as described above. gRNAs used:

ERAAP sgRNA F1: CACCGCAGTGGATCAAATTTAACGT
ERAAP sgRNA R1: AAACACGTTAAATTTGATCCACTGC

ERAAP KO splenocytes used for western blot were obtained from ERAAP KO mice that have been previously described (Hammer et al., 2006; Nagarajan et al., 2016).

### Endo H assay

Whole cell lysate (WCL) for BJAB ERAAP-dsRed clones B2 and D4 was made using RIPA lysis buffer (150 mM NaCl, 1% Triton X-100, 5% Na Deoxycholate, 1% SDS, 50 mM Tris pH 8 in water) with Protease inhibitors (Roche, 11836170001). WCL lysate was incubated on ice for 30 min and centrifuged at max speed for 10 min to remove debris. Cleared lysate was transferred to a new tube and anti-ERAAP L1 antibody (1:200 dilution) was added. Samples were incubated rotating overnight at 4 °C. The next day, ERAAP was immunoprecipitated by adding 40 ul of Protein A Dynabeads (Invitrogen, 10001D) to each sample and incubating an additional 2 hr. Using a magnet, beads containing ERAPP were captured. Beads were washed four times with 50 mM Tris, 150 mM NaCl in water. Endo H assay was performed following manufacturer’s instructions (New England Biolabs, P0702S). Following addition of Glycoprotein Denaturing Buffer, each sample was aliquoted into two tubes. 1 ul of EndoH (Endo H +) or 1 ul of water (Endo H -) was then added to each tube. Samples were incubated for 1 h at 37 °C. ERAAP’s glycosylation state was assessed by western blot. Membrane was probed using anti-ERAAP L1 antibody (1:000 dilution). Anti-ERAAP L1 was produced in the Shastri lab.

### ERAAP western blot on cells co-transfected with ERAAP 3XFlag and SARS-CoV-2 E

HEK293T cells were seeded at 1X10^6 cells/well in a 6-well plate. The next day, cells were transfected with 1 ug of plasmid encoding ERAAP 3XFlag (pCDNA3.1 background) and 1 ug of control plasmid (pIRES-GFP), pIRES-GFP SARS-CoV-2 E, or pIRES-GFP SARS-CoV-2 Spike using Fugene HD following manufacturer’s instructions. WCL was collected 36 hours post transfection. Briefly, cells were washed once with PBS (Gibco, 10010023) and detached using trypsin (Gibco, 25300062). Cells were centrifuged at 1,600 rpm for 8 min. Supernatant was carefully removed and cell pellet was resuspended in RIPA lysis buffer with Protease inhibitor. WCL was incubated with 1% Benzonase (Sigma, 1014-25KU) for 15 min at 37 °C. Cleared lysates were collected by centrifuging Benzonase-treated samples for 10 min at 15,000 rpm (4 °C) to pellet remaining genomic DNA.

For western blot, protein concentration was determined using Pierce BCA Protein Assay Kit (Thermo Scientific, 23225) following manufacturer’s instructions. A total of 50 ug of WCL was combined with 4X Laemelli Sample Buffer (BioRad, 1610747) to a final concentration of 1X sample buffer, boiled for 5 min at 100 °C, and loaded into 4-15% Mini-Protean TGX gels (Biorad, 4561084). Gel was run for 1 hr at 120V after which proteins were transferred (1.3 A, 25V, 10 min) into PVDF membranes and blocked for 1 hr at room temperature with 3% bovine serum albumin (BSA) buffer (3% bovine serum albumin, 1 % Triton X-100, 0.5% Tween-20 in PBS). Primary and secondary antibodies were diluted in 3% BSA buffer. Membranes were incubated with the following antibodies: anti-FLAG M2 (Sigma, F1804; 1:1000 dilution), anti-GAPDH 0411 (Santa Cruz Biotechnology, sc-47724; 1:1000 dilution). Primary antibody incubation was done overnight at 4 °C. The next day, the membranes were washed 3 times with PBS for 5 min each and incubated with secondary antibody for 1 hr at room temperature. Secondary antibody used was IRDye 680RD Donkey anti-Mouse IgG (Licor, 926-68072; 1:10000 dilution). Imaging was done using a ChemiDoc MP imager (Biorad). Band intensity analyzed using Image J 1.52q.

### ERAAP western blot on ERp44 KO cell lines

ERp44 KO and WT cell lines (BJAB ERAAP dsRed, NIH 3T3, and MC57G) were lysed in lysis buffer containing 25 mM Tris-HCl (pH 7.6), 150 mM NaCl, 0.1% SDS, 0.5% sodium deoxycholic, and protease inhibitors (Roche) for 30 minutes on ice. Samples were then centrifuged at 20,000xg for 10 min at 4°C to clarify lysate. Lysates were separated by SDS-PAGE and western blot was performed. Membranes were incubated with the following antibodies: anti-ERAAP L1 (1:1000), anti-ERp44 (Invitrogen, #PA5-28484; 1:1000 dilution), anti-GAPDH (Abcam, ab8245; 1:5000 dilution). Rabbit anti-ERAAP L1 was produced in the Shastri lab.

### Golgi pH measurement

pH reporter assay was done as described^23^. Briefly, HeLa cells were seeded at 1X10^6 cells/well in a 6-well plate. The next day, cells were transfected with 2 ug of pHluorin TGN38 alone for generating a standard curve or with 1 ug each of pHluorin TGN38 and pIRES-RFP SARS-CoV-2 E, or pIRES-RFP IAV M2. 18 hours post transfection, media was removed, and cells were washed once with serum free (SF) DMEM. Cells were then incubated for 1 hr at 37 °C with 100 ug/mL cycloheximide (Sigma-Aldrich, C4859) in SF DMEM. Cycloheximide was removed and cells were washed once with PBS, detached using trypsin, and neutralized with ice-cold SF DMEM. Cells were pelleted by spinning 10 min at 1,600 rpm and then washed twice with SF-DMEM. To generate a pH standard curve, cells transfected with pHluorin were resuspended in Na-Ringer buffer (140 mM KCl, 2 mM CaCl_2_, 1 mM MgSO_4_, 1.5 mM K_2_HPO_4_, 10 mM Glucose, 10 mM 2-(N-Morpholino)ethanesulforic acid sodium salt,10 mM HEPES) containing 10 uM monensin (Research Products International, M92100-1.0) and 10 uM nigericin (Cayman Chemical, 11437-5). Cells co-transfected with pHluorin and viral proteins were resuspended in Na-Ringer buffer. Samples were analyzed in the LSR Fortessa Celeste or LSR Fortessa X20 cytometer (BD Biosciences). pH reporter was excited at 405 nm and 408 nm and the emitted fluorescence recorded on FACS Diva software v8.0.1. Data was analyzed in Flow Jo (v10) and the pHluorin emission ratio for each sample was obtained by creating a ratio between the fluorescence emitted after excitation at 405 nm and 488 nm (405 nm/488 nm). The geometric mean of the pHluorin emission ratio was used to obtain a pH value for each sample. The emission ratio of samples resuspended in buffers of known pH was used to generate a standard curve and the formula obtained was used to calculate the luminal Golgi pH of the experimental samples.

### ERAAP dsRed reporter assay

HEK293T cells were seeded at 1X10^6 cells/well in a 6-well plate. The next day, cells were transfected with 1.3 ug of plasmid encoding ERAAP-dsRed (pQCXIP background) and 700 ng of pIRES-GFP, pIRES-GFP SARS-CoV-2 E, or pIRES-GFP SARS-CoV-2 Spike using Fugene HD following manufacturer’s instructions. Cells were harvested for flow cytometry analysis at 24 h and 48 h post-transfection. Briefly, cells were washed with PBS, detached using typsin, and neutralized with DMEM 10% FBS. Cells were centrifuged at 1,600 rpm for 8 min to pellet. Cell pellet was resuspended in flow buffer (3% FBS, 1 mM EDTA in PBS). Cells were washed 2-3 times with flow buffer. Samples were incubated 10 min on ice with viability dye (Invitrogen, 65-0865-14) after which cells were pelleted and resuspended in flow buffer for analysis.

BJAB, BJAB ERAAP-dsRed, and BJAB ERAAP-dsRed ERp44 KO cells were harvested by centrifugation at 1,600 rpm for 8 min. Cell pellets were resuspended in flow buffer (3% FBS, 1 mM EDTA in PBS) and washed 2-3 times before resuspending for analysis.

NIH 3T3, NIH 3T3 ERAAP-dsRed, and NIH 3T3 ERAAP-dsRed ERp44 KO cells were washed with PBS, detached using typsin, and neutralized with DMEM 10% FBS. Cells were centrifuged at 1,600 rpm for 8 min to pellet. Cell pellets were resuspended in flow buffer (3% FBS, 1 mM EDTA in PBS). After 2-3 washes with flow buffer, cells were resuspended in flow buffer for analysis.

Samples were analyzed using the LSR Fortessa Celeste or LSR Fortessa X20 cytometer (BD Biosciences) and data recorded on FACS Diva software v8.0.1. Data was analyzed in Flow Jo (v10).

### ERAAP secretion assay

To evaluate ERAAP secretion from BJAB ERAAP-dsRed ERp44 KO cells, ERAAP was first immunoprecipitated from the supernatant. Cells were seeded at 100,000 cells/well. After 36 h, supernatant was harvested and centrifuged at 1,500 rpm for 5 min to remove cells. Cleared supernatant was incubated with anti-ERAAP L1 antibody (1:50 dilution) for 1 h rotating at 4 °C. ERAAP was immunoprecipitated by adding 50 ul of Protein A Dynabeads (Invitrogen, 10001D) to each sample and incubating for 1 h rotating at 4 °C. Using a magnet, beads containing ERAPP were isolated. Beads were washed four times with PBS and resuspended in 75 uL of PBS. 50 uL of resuspended beads was used for LAP assay.

To measure ERAAP secretion from cells expressing SARS-CoV-2 E, NIH 3T3 ERAAP-dsRed cells were seeded at 200,000 cells/well in a 12-well plate. The next day, cells were transfected with 1 ug of pIRES-GFP (empty vector) or pIRES-GFP SARS-CoV-2 E using Fugene HD following manufacturer’s instructions. Approximately 18 h post transfection, the media was changed to DMEM without phenol red (Gibco, A18967-01). This media was supplemented with 10% FBS, 1% Pen Strep, 1 mM Sodium Pyruvate (Gibco, 11360-070) and 2 mm L-Glutamine (Gibco, 25030-081) to match previous culturing conditions. 48 h after transfection, supernatant was collected. Cell debris was removed by centrifugation at 1,500 rpm for 5 min. Cleared supernatant was moved to a new tube. 50 uL of supernatant was used for LAP assay.

LAP assay was performed by incubating 50 uL of beads containing ERAAP (for BJAB) or 50 uL of supernatant (for NIH 3T3) with 100 ul of 2 mg/mL (in 50 mM Tris, pH 7.6) leucine p-nitroanilide (LpNA; Sigma, L9125). Samples were incubated at 37 °. Cleavage of the peptide substrate results in a colorimetric change that was detected at OD 410 nm. LAP activity was assessed 8 h after addition of the substrate for BJAB cell lines and 19-20 h for NIH 3T3 cell lines.

### Statistical Analysis

Unpaired T-test or Unpaired One-way ANOVA with multiple comparisons analysis was performed in Graph Pad PRISM version 9.4.1 for MacOS. Compiled data is shown as mean±SD.

## Notes

### Competing Interest Statement

The authors have declared no competing interest.

