## Supplemental Figure for "SARS-CoV-2 Envelope-mediated Golgi pH dysregulation interferes with ERAAP retention in cells"

This file includes:

Figure legend for Supplemental Figure 1
Supplemental Figure 2
Supplemental Figure 3

Supplemental Figure 1: **Hits identified in genome-wide CRISPR KO screen**. Hits represent gRNAs enriched in BJAB ERAAP dsRed cells expressing low levels of ERAAP dsRed after transduction with gRNA library. BJAB ERAAP dsRed cells with the top or bottom ~10-15% dsRed signal were sorted, analyzed by sequencing, and compared for gene enrichment. Shown is enrichment analysis for cells showing loss of ERAAP dsRed expression in comparison to high expressors. Enrichment analysis was performed using the MaGeCK method.

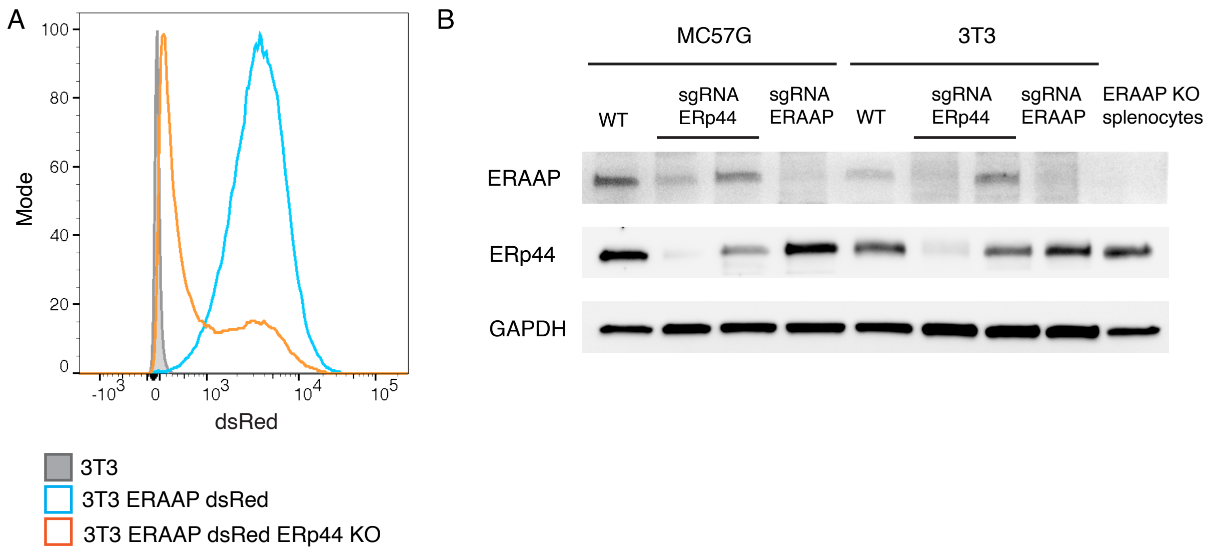


Supplemental Figure 2: **ERp44 regulates ERAAP levels in mouse cells**. A. NIH 3T3 cells were transduced to express ERAAP dsRed and ERp44 was knocked out in this cell line. ERAAP dsRed levels were assessed by flow cytometry. Histogram shows ERAAP dsRed levels in NIH 3T3 ERAAP dsRed (blue histogram) and ERp44 KO cells (orange histogram). Parental NIH 3T3 cells are shown as a control (grey filled histogram). B. MC57G and NIH 3T3 cells were transfected with two gRNA targeting ERp44 or one gRNA targeting ERAAP as a control. ERAAP levels in ERp44 KO cells were measured by western blot. Representative blot showing ERAAP and ERp44 protein levels in untransfected (WT) and KO cells. ERAAP KO splenocytes were included as comparison. GAPDH is shown as a loading control.

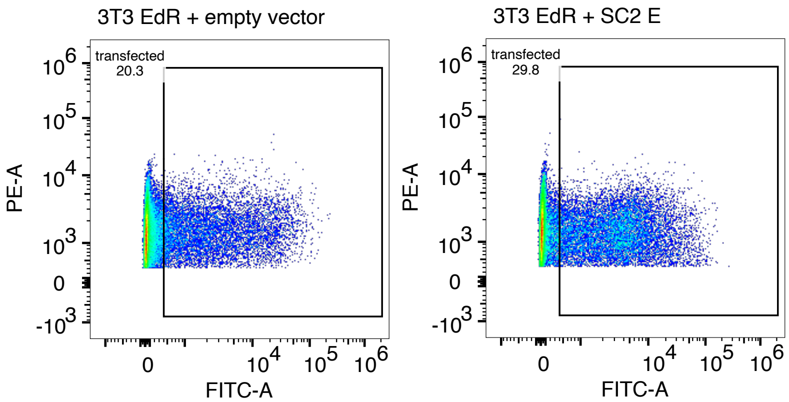


Supplemental Figure 3: **Transfection efficiency of NIH 3T3 ERAAP dsRed cells transfected with SARS-CoV-2 Envelope.** NIH 3T3 ERAAP dsRed cells were transfected with empty vector or SARS-CoV-2 E. The supernatant was collected for LAP assay (Figure 5) and the cells were used to measure transfection efficiency by flow cytometry 48 h post-transfection. Flow plots show the percent of cells that are expressing SARS-CoV-2 E or empty vector (GFP reporter detected with FITC channel) within cells that are expressing the ERAAP dsRed construct (dsRed detected with PE channel).
