## Supplemental Figure 1 for "SARS-CoV-2 Envelope-mediated Golgi pH dysregulation interferes with ERAAP retention in cells"

| id | num | neg\|score | neg\|p-value | neg\|fdr | neg\|rank | neg\|goodsgrna | neg\|lfc | pos\|score | pos\|p-value | pos\|fdr | pos\|rank | pos\|goodsgrna | pos\|lfc |
|---|---|---|---|---|---|---|---|---|---|---|---|---|---|
| TRIM28 | 6 | 1 | 0.99999 | 0.999994 | 21655 | 0 | 12.724 | 9.7158E-17 | 2.2861E-07 | 0.00099 | 1 | 6 | 12.724 |
| ERP44 | 6 | 0.99795 | 0.99793 | 0.999994 | 21580 | 0 | 11.557 | 1.2947E-12 | 2.2861E-07 | 0.00099 | 2 | 6 | 11.557 |
| SPOP | 6 | 0.99903 | 0.99902 | 0.999994 | 21619 | 0 | 8.5488 | 3.4842E-10 | 2.2861E-07 | 0.00099 | 3 | 6 | 8.5488 |
| CCNC | 6 | 0.80567 | 0.85034 | 0.999994 | 17564 | 1 | 10.182 | 1.1113E-08 | 2.2861E-07 | 0.00099 | 4 | 5 | 10.182 |
| MGA | 6 | 0.99995 | 0.99995 | 0.999994 | 21652 | 0 | 8.2785 | 7.4993E-08 | 2.2861E-07 | 0.00099 | 5 | 6 | 8.2785 |
| PAXIP1 | 6 | 0.84935 | 0.87158 | 0.999994 | 18249 | 1 | 6.5322 | 1.6422E-06 | 9.3729E-06 | 0.03041 | 6 | 5 | 6.5322 |
| WDR83 | 6 | 0.96152 | 0.96159 | 0.999994 | 20620 | 0 | 9.1522 | 1.7951E-06 | 9.8301E-06 | 0.03041 | 7 | 5 | 9.1522 |
| ZNF829 | 6 | 0.93692 | 0.93694 | 0.999994 | 20004 | 0 | 6.7114 | 3.1646E-06 | 0.000014859 | 0.040223 | 8 | 4 | 6.7114 |
| UBE2L3 | 6 | 0.86537 | 0.88052 | 0.999994 | 18547 | 1 | 5.5462 | 6.8226E-06 | 0.000040921 | 0.091089 | 9 | 5 | 5.5462 |
| UBE2M | 6 | 0.58245 | 0.727 | 0.999994 | 14712 | 1 | 8.1294 | 9.1327E-06 | 0.000056009 | 0.110261 | 10 | 5 | 8.1294 |
| CHST9 | 4 | 0.99999 | 0.99999 | 0.999994 | 21654 | 0 | 8.9971 | 0.000012035 | 0.000042064 | 0.091089 | 11 | 4 | 8.9971 |
| PPP1R13L | 6 | 0.66471 | 0.78896 | 0.999994 | 15854 | 1 | 7.7787 | 0.000016879 | 0.000099444 | 0.153819 | 12 | 5 | 7.7787 |
| CXorf38 | 5 | 0.99648 | 0.99645 | 0.999994 | 21521 | 0 | 5.7248 | 0.000017008 | 0.000072011 | 0.123762 | 13 | 5 | 5.7248 |
| hsa-mir-6807 | 4 | 0.86194 | 0.86157 | 0.999994 | 18486 | 0 | 9.5479 | 0.000020798 | 0.000074297 | 0.123762 | 14 | 3 | 9.5479 |
| ZNF581 | 5 | 0.48471 | 0.63471 | 0.999994 | 13211 | 1 | 9.3565 | 0.00002794 | 0.00011773 | 0.169967 | 15 | 4 | 9.3565 |
| USP22 | 6 | 0.99997 | 0.99996 | 0.999994 | 21653 | 0 | 6.3995 | 0.000032476 | 0.00018723 | 0.238497 | 16 | 6 | 6.3995 |
| MARCH5 | 5 | 0.98671 | 0.9867 | 0.999994 | 21269 | 0 | 9.3066 | 0.000033716 | 0.00014745 | 0.199567 | 17 | 5 | 9.3066 |
| CELSR1 | 6 | 0.018311 | 0.066506 | 0.93914 | 1377 | 3 | 3.3324 | 0.000039051 | 0.00022381 | 0.251238 | 18 | 3 | 3.3324 |
| SLC27A5 | 5 | 0.33618 | 0.51061 | 0.999994 | 10853 | 2 | 9.1782 | 0.000045813 | 0.00020918 | 0.251238 | 19 | 3 | 9.1782 |
| ELP3 | 5 | 0.98843 | 0.98844 | 0.999994 | 21312 | 0 | 8.3229 | 0.000050605 | 0.00023204 | 0.251238 | 20 | 5 | 8.3229 |
| WDR87 | 6 | 0.99955 | 0.99953 | 0.999994 | 21636 | 0 | 8.0392 | 0.000055014 | 0.0002965 | 0.272483 | 21 | 6 | 8.0392 |
| C5orf51 | 6 | 0.73934 | 0.82476 | 0.999994 | 16728 | 1 | 8.4612 | 0.000055014 | 0.0002965 | 0.272483 | 22 | 5 | 8.4612 |
| DDX39B | 6 | 0.86616 | 0.881 | 0.999994 | 18560 | 1 | 8.7764 | 0.000056264 | 0.00030199 | 0.272483 | 23 | 5 | 8.7764 |
| RING1 | 5 | 0.97609 | 0.97616 | 0.999994 | 20990 | 0 | 5.2669 | 0.000058205 | 0.00026541 | 0.272483 | 24 | 5 | 5.2669 |
| HDLBP | 6 | 0.99529 | 0.99528 | 0.999994 | 21487 | 0 | 8.7471 | 0.00006923 | 0.00037286 | 0.32297 | 25 | 6 | 8.7471 |
| WDR59 | 6 | 0.98335 | 0.98331 | 0.999994 | 21165 | 0 | 8.6916 | 0.000071728 | 0.0003884 | 0.323496 | 26 | 5 | 8.6916 |
| GNAS | 5 | 0.92531 | 0.92555 | 0.999994 | 19752 | 0 | 9.024 | 0.000091005 | 0.00041492 | 0.332783 | 27 | 4 | 9.024 |
| EIF2S2 | 5 | 0.9999 | 0.99991 | 0.999994 | 21651 | 0 | 7.8489 | 0.00010097 | 0.0004611 | 0.350119 | 28 | 5 | 7.8489 |
| NIPBL | 6 | 0.9999 | 0.99989 | 0.999994 | 21650 | 0 | 7.7534 | 0.00010138 | 0.00055803 | 0.389812 | 29 | 6 | 7.7534 |
| ZBTB45 | 5 | 0.24776 | 0.43902 | 0.999994 | 8969 | 1 | 8.9864 | 0.00010276 | 0.00046887 | 0.350119 | 30 | 4 | 8.9864 |
| MAP3K14 | 6 | 0.99988 | 0.99987 | 0.999994 | 21648 | 0 | 7.4638 | 0.00011415 | 0.00062387 | 0.409391 | 31 | 6 | 7.4638 |
| CHUK | 6 | 0.96929 | 0.96929 | 0.999994 | 20809 | 0 | 6.3557 | 0.00011415 | 0.00062387 | 0.409391 | 32 | 5 | 6.3557 |
| HMGB1 | 5 | 0.99988 | 0.99989 | 0.999994 | 21649 | 0 | 7.3593 | 0.00012066 | 0.0005482 | 0.389812 | 33 | 5 | 7.3593 |
| CCND3 | 6 | 0.018663 | 0.067682 | 0.93914 | 1397 | 2 | 2.5756 | 0.00014927 | 0.00081544 | 0.477254 | 34 | 3 | 2.5756 |
| DENND5A | 5 | 0.99985 | 0.99986 | 0.999994 | 21647 | 0 | 6.5204 | 0.00014984 | 0.00069977 | 0.442008 | 35 | 5 | 6.5204 |
| SEC62 | 5 | 0.99772 | 0.9977 | 0.999994 | 21571 | 0 | 7.7672 | 0.00015246 | 0.0007144 | 0.442008 | 36 | 5 | 7.7672 |
| TMED2 | 5 | 0.87892 | 0.87923 | 0.999994 | 18810 | 1 | 2.5853 | 0.00015743 | 0.00073955 | 0.444857 | 37 | 4 | 2.5853 |
| PGP | 6 | 0.98503 | 0.985 | 0.999994 | 21223 | 0 | 4.3368 | 0.00016702 | 0.00091146 | 0.509901 | 38 | 5 | 4.3368 |
| MED13 | 6 | 0.38841 | 0.59493 | 0.999994 | 11773 | 1 | 7.8867 | 0.00017099 | 0.00093798 | 0.509901 | 39 | 5 | 7.8867 |
| EMC6 | 6 | 0.99695 | 0.99694 | 0.999994 | 21537 | 0 | 6.8158 | 0.00017666 | 0.00096678 | 0.509901 | 40 | 6 | 6.8158 |
| SMPDL3A | 6 | 0.99982 | 0.9998 | 0.999994 | 21646 | 0 | 4.6863 | 0.00018148 | 0.0009869 | 0.509901 | 41 | 6 | 4.6863 |
| CSTF2 | 6 | 0.99981 | 0.9998 | 0.999994 | 21645 | 0 | 6.839 | 0.0001873 | 0.0010125 | 0.509901 | 42 | 6 | 6.839 |
| SWT1 | 5 | 0.97386 | 0.9739 | 0.999994 | 20922 | 0 | 3.1506 | 0.00021212 | 0.0010056 | 0.509901 | 43 | 5 | 3.1506 |
| ZNF644 | 6 | 0.99496 | 0.99495 | 0.999994 | 21475 | 0 | 6.1815 | 0.0002128 | 0.0011346 | 0.524099 | 44 | 6 | 6.1815 |
| TOX4 | 6 | 0.88764 | 0.89467 | 0.999994 | 18990 | 1 | 2.9797 | 0.00021491 | 0.0011465 | 0.524099 | 45 | 4 | 2.9797 |
| KDM5C | 6 | 0.99659 | 0.99656 | 0.999994 | 21524 | 0 | 5.8398 | 0.00022239 | 0.001184 | 0.524099 | 46 | 6 | 5.8398 |
| MFN1 | 5 | 0.85603 | 0.8591 | 0.999994 | 18374 | 1 | 8.746 | 0.00022624 | 0.0010715 | 0.524099 | 47 | 4 | 8.746 |
| ZNF345 | 6 | 0.91878 | 0.91927 | 0.999994 | 19603 | 1 | 2.3618 | 0.00023441 | 0.0012461 | 0.524099 | 48 | 4 | 2.3618 |
| ANGEL1 | 5 | 0.95886 | 0.959 | 0.999994 | 20558 | 0 | 8.7306 | 0.00023718 | 0.0011131 | 0.524099 | 49 | 4 | 8.7306 |
| RPN2 | 6 | 0.99976 | 0.99974 | 0.999994 | 21644 | 0 | 6.6867 | 0.00024009 | 0.0012827 | 0.524099 | 50 | 6 | 6.6867 |
| MBOAT7 | 6 | 0.99973 | 0.99971 | 0.999994 | 21642 | 0 | 8.2125 | 0.00025404 | 0.001349 | 0.532462 | 51 | 6 | 8.2125 |

| | | | | | | | | | | | | |
|---|---|---|---|---|---|---|---|---|---|---|---|---|
| PNLIP | 6 | 0.97461 | 0.9745 | 0.999994 | 20945 | 0 | 5.2183 | 0.00026169 | 0.0013979 | 0.532462 | 52 | 5 | 5.2183 |
| CTU1 | 5 | 0.99881 | 0.99883 | 0.999994 | 21612 | 0 | 8.6916 | 0.0002648 | 0.0012493 | 0.524099 | 53 | 5 | 8.6916 |
| NUTM2A | 6 | 0.99973 | 0.99971 | 0.999994 | 21643 | 0 | 6.8457 | 0.00026896 | 0.0014441 | 0.532462 | 54 | 6 | 6.8457 |
| SPEN | 6 | 0.98199 | 0.98192 | 0.999994 | 21136 | 0 | 4.8897 | 0.00028012 | 0.0014999 | 0.532462 | 55 | 5 | 4.8897 |
| XYLT2 | 5 | 0.9997 | 0.9997 | 0.999994 | 21641 | 0 | 6.4099 | 0.00030193 | 0.0014226 | 0.532462 | 56 | 5 | 6.4099 |
| PTGIR | 6 | 0.9997 | 0.99968 | 0.999994 | 21640 | 0 | 4.6327 | 0.00030289 | 0.0016119 | 0.554063 | 57 | 6 | 4.6327 |
| EQTN | 5 | 0.99555 | 0.99552 | 0.999994 | 21491 | 0 | 8.5402 | 0.00031156 | 0.0014702 | 0.532462 | 58 | 5 | 8.5402 |
| TMEM54 | 5 | 0.98831 | 0.9883 | 0.999994 | 21307 | 0 | 7.3324 | 0.00031829 | 0.0014949 | 0.532462 | 59 | 5 | 7.3324 |
| LGALS7 | 6 | 0.99933 | 0.9993 | 0.999994 | 21628 | 0 | 5.9969 | 0.00033116 | 0.0017642 | 0.557992 | 60 | 6 | 5.9969 |
| ZNF541 | 6 | 0.99431 | 0.99428 | 0.999994 | 21455 | 0 | 7.2892 | 0.00033872 | 0.001803 | 0.557992 | 61 | 6 | 7.2892 |
| BCO2 | 4 | 0.99966 | 0.99967 | 0.999994 | 21639 | 0 | 6.5414 | 0.000339 | 0.0012763 | 0.524099 | 62 | 4 | 6.5414 |
| MYO1A | 6 | 0.3985 | 0.60141 | 0.999994 | 11923 | 1 | 5.5703 | 0.00034459 | 0.0018318 | 0.557992 | 63 | 5 | 5.5703 |
| EHMT1 | 6 | 0.97372 | 0.97363 | 0.999994 | 20916 | 0 | 3.4068 | 0.00035249 | 0.0018803 | 0.557992 | 64 | 5 | 3.4068 |
| FLT3LG | 6 | 0.13736 | 0.3097 | 0.999994 | 6118 | 2 | 2.0731 | 0.0003552 | 0.0018917 | 0.557992 | 65 | 3 | 2.0731 |
| FAM187B | 5 | 0.9206 | 0.92079 | 0.999994 | 19641 | 0 | 6.39 | 0.00036592 | 0.0017061 | 0.557992 | 66 | 3 | 6.39 |
| CNTD2 | 6 | 0.32154 | 0.53166 | 0.999994 | 10545 | 1 | 7.8493 | 0.00036615 | 0.0019475 | 0.557992 | 67 | 5 | 7.8493 |
| ADAMTS2 | 6 | 0.27947 | 0.48278 | 0.999994 | 9689 | 3 | 3.3074 | 0.00036753 | 0.0019564 | 0.557992 | 68 | 3 | 3.3074 |
| EXT2 | 6 | 0.022943 | 0.081406 | 0.93914 | 1645 | 2 | 7.9003 | 0.00036977 | 0.0019672 | 0.557992 | 69 | 4 | 7.9003 |
| CCDC61 | 5 | 0.98635 | 0.98637 | 0.999994 | 21256 | 0 | 7.6011 | 0.00037971 | 0.0017623 | 0.557992 | 70 | 5 | 7.6011 |
| RLN3 | 5 | 0.99959 | 0.99959 | 0.999994 | 21638 | 0 | 6.7069 | 0.000411 | 0.0019073 | 0.557992 | 71 | 5 | 6.7069 |
| hsa-mir-604 | 4 | 0.99635 | 0.99636 | 0.999994 | 21514 | 0 | 8.3332 | 0.00041186 | 0.0015488 | 0.540961 | 72 | 4 | 8.3332 |
| FAM129C | 6 | 0.27251 | 0.47466 | 0.999994 | 9528 | 2 | 4.4357 | 0.00041395 | 0.0021875 | 0.599637 | 73 | 3 | 4.4357 |
| HGSNAT | 5 | 0.038361 | 0.11174 | 0.951052 | 2437 | 1 | 8.5362 | 0.00042734 | 0.0019841 | 0.557992 | 74 | 4 | 8.5362 |
| DPAGT1 | 6 | 0.99957 | 0.99955 | 0.999994 | 21637 | 0 | 4.6177 | 0.00043169 | 0.0022744 | 0.608549 | 75 | 6 | 4.6177 |
| PHF6 | 6 | 0.87216 | 0.88466 | 0.999994 | 18678 | 1 | 2.5497 | 0.00043379 | 0.0022831 | 0.608549 | 76 | 4 | 2.5497 |
| CDK8 | 6 | 0.92117 | 0.92147 | 0.999994 | 19656 | 1 | 2.8595 | 0.0004383 | 0.0023044 | 0.608549 | 77 | 5 | 2.8595 |
| IL16 | 6 | 0.46013 | 0.64187 | 0.999994 | 12845 | 2 | 2.7993 | 0.00046274 | 0.0024239 | 0.628045 | 78 | 3 | 2.7993 |
| ALDOA | 4 | 0.99954 | 0.99955 | 0.999994 | 21634 | 0 | 7.9559 | 0.00046438 | 0.0017404 | 0.557992 | 79 | 4 | 7.9559 |
| hsa-mir-3936 | 4 | 0.99954 | 0.99955 | 0.999994 | 21635 | 0 | 7.558 | 0.00046438 | 0.0017404 | 0.557992 | 80 | 4 | 7.558 |
| CREBZF | 5 | 0.99793 | 0.99794 | 0.999994 | 21578 | 0 | 7.6823 | 0.00047034 | 0.0021821 | 0.599637 | 81 | 5 | 7.6823 |
| CPEB4 | 6 | 0.99952 | 0.99951 | 0.999994 | 21633 | 0 | 5.9561 | 0.0004766 | 0.0024843 | 0.628045 | 82 | 6 | 5.9561 |
| CEACAM7 | 6 | 0.945 | 0.94498 | 0.999994 | 20205 | 0 | 4.8941 | 0.00048386 | 0.0025167 | 0.628045 | 83 | 4 | 4.8941 |
| RSPH6A | 6 | 0.99951 | 0.99949 | 0.999994 | 21632 | 0 | 3.7163 | 0.00049135 | 0.0025606 | 0.628045 | 84 | 6 | 3.7163 |
| LZTS1 | 6 | 0.6955 | 0.81065 | 0.999994 | 16262 | 1 | 6.6816 | 0.00049297 | 0.0025689 | 0.628045 | 85 | 5 | 6.6816 |
| ATXN7L2 | 5 | 0.98091 | 0.98101 | 0.999994 | 21116 | 0 | 6.425 | 0.00053023 | 0.0024669 | 0.628045 | 86 | 5 | 6.425 |
| SAMD4B | 6 | 0.74744 | 0.82757 | 0.999994 | 16832 | 1 | 7.1623 | 0.00053216 | 0.0027883 | 0.66353 | 87 | 4 | 7.1623 |
| ZNF503 | 5 | 0.99944 | 0.99944 | 0.999994 | 21631 | 0 | 5.7547 | 0.00055802 | 0.0025812 | 0.628045 | 88 | 5 | 5.7547 |
| RHOJ | 6 | 0.13224 | 0.30318 | 0.999994 | 5980 | 2 | 3.4774 | 0.00057645 | 0.0030128 | 0.665572 | 89 | 4 | 3.4774 |
| KXD1 | 6 | 0.79241 | 0.8447 | 0.999994 | 17381 | 1 | 5.8874 | 0.00058145 | 0.0030453 | 0.665572 | 90 | 5 | 5.8874 |
| CASP12 | 6 | 0.058001 | 0.16146 | 0.967126 | 3337 | 3 | 2.5109 | 0.00058423 | 0.0030581 | 0.665572 | 91 | 3 | 2.5109 |
| OR52W1 | 6 | 0.94529 | 0.94527 | 0.999994 | 20212 | 0 | 7.9027 | 0.00060176 | 0.0031413 | 0.665572 | 92 | 4 | 7.9027 |
| RDH13 | 6 | 0.10884 | 0.26447 | 0.996734 | 5265 | 1 | 6.0673 | 0.00062662 | 0.0032611 | 0.665572 | 93 | 5 | 6.0673 |
| RREB1 | 6 | 0.99937 | 0.99934 | 0.999994 | 21630 | 0 | 4.7079 | 0.0006345 | 0.0032967 | 0.665572 | 94 | 6 | 4.7079 |
| STYK1 | 6 | 0.97853 | 0.97846 | 0.999994 | 21064 | 0 | 4.9829 | 0.00063473 | 0.0032977 | 0.665572 | 95 | 5 | 4.9829 |
| NOVA2 | 5 | 0.99936 | 0.99937 | 0.999994 | 21629 | 0 | 5.1269 | 0.00063706 | 0.0029182 | 0.665572 | 96 | 5 | 5.1269 |
| SLC2A14 | 6 | 0.55053 | 0.70402 | 0.999994 | 14222 | 1 | 0.54192 | 0.00065039 | 0.0033749 | 0.665572 | 97 | 2 | 0.54192 |
| ETV3 | 6 | 0.64685 | 0.77521 | 0.999994 | 15616 | 2 | 4.4554 | 0.00065202 | 0.0033809 | 0.665572 | 98 | 4 | 4.4554 |
| TMEM145 | 6 | 0.45912 | 0.64116 | 0.999994 | 12836 | 1 | 3.0473 | 0.00066213 | 0.0034293 | 0.668582 | 99 | 4 | 3.0473 |
| SH2D7 | 6 | 0.3946 | 0.59889 | 0.999994 | 11867 | 1 | 6.0033 | 0.00066843 | 0.0034579 | 0.668582 | 100 | 4 | 6.0033 |
| FAM71C | 6 | 0.99931 | 0.99928 | 0.999994 | 21627 | 0 | 4.1451 | 0.00068868 | 0.0035542 | 0.675135 | 101 | 6 | 4.1451 |
| ANXA7 | 5 | 0.35932 | 0.52948 | 0.999994 | 11327 | 2 | 0.36392 | 0.00069473 | 0.003187 | 0.665572 | 102 | 2 | 0.36392 |
| HOXC13 | 6 | 0.35976 | 0.57505 | 0.999994 | 11331 | 3 | 3.9749 | 0.0007016 | 0.0036278 | 0.679114 | 103 | 3 | 3.9749 |

| | | | | | | | | | | | | |
|---|---|---|---|---|---|---|---|---|---|---|---|---|
| KCNC4 | 5 | 0.98683 | 0.98681 | 0.999994 | 21273 | 0 | 8.3574 | 0.00071857 | 0.0032881 | 0.665572 | 104 | 5 | 8.3574 |
| SIGLEC12 | 5 | 0.92413 | 0.92436 | 0.999994 | 19719 | 0 | 8.3568 | 0.00072232 | 0.0033004 | 0.665572 | 105 | 3 | 8.3568 |
| FOXE1 | 4 | 0.99927 | 0.99927 | 0.999994 | 21626 | 0 | 7.2863 | 0.00073326 | 0.0027266 | 0.656051 | 106 | 4 | 7.2863 |
| LYPD3 | 6 | 0.99815 | 0.99813 | 0.999994 | 21587 | 0 | 6.5059 | 0.00074877 | 0.0038536 | 0.701265 | 107 | 6 | 6.5059 |
| NonTargetingControlG | 2 | 0.99924 | 0.99922 | 0.999994 | 21625 | 0 | 8.8967 | 0.00076083 | 0.0014267 | 0.532462 | 108 | 2 | 8.8967 |
| MED12 | 5 | 0.97822 | 0.97831 | 0.999994 | 21049 | 0 | 5.7248 | 0.00077761 | 0.0035286 | 0.675135 | 109 | 5 | 5.7248 |
| VMA21 | 4 | 0.99922 | 0.99921 | 0.999994 | 21624 | 0 | 6.6714 | 0.00077935 | 0.002888 | 0.665572 | 110 | 4 | 6.6714 |
| TARS | 4 | 0.96334 | 0.96323 | 0.999994 | 20680 | 0 | 8.568 | 0.00078814 | 0.0029237 | 0.665572 | 111 | 4 | 8.568 |
| BID | 6 | 0.16609 | 0.34582 | 0.999994 | 6885 | 1 | 2.1296 | 0.00082 | 0.0041933 | 0.730812 | 112 | 4 | 2.1296 |
| GJA1 | 5 | 0.99864 | 0.99866 | 0.999994 | 21605 | 0 | 6.8682 | 0.00083965 | 0.003802 | 0.701265 | 113 | 5 | 6.8682 |
| LHB | 6 | 0.85212 | 0.87304 | 0.999994 | 18296 | 1 | 7.4542 | 0.00084354 | 0.0043237 | 0.741562 | 114 | 5 | 7.4542 |
| CTSK | 4 | 0.2466 | 0.38842 | 0.999994 | 8936 | 1 | 4.9399 | 0.00087059 | 0.0032163 | 0.665572 | 115 | 2 | 4.9399 |
| MED29 | 6 | 0.94641 | 0.94638 | 0.999994 | 20241 | 0 | 4.8562 | 0.00087653 | 0.0044942 | 0.741562 | 116 | 4 | 4.8562 |
| RBP1 | 6 | 0.99671 | 0.99668 | 0.999994 | 21528 | 0 | 6.0814 | 0.00087989 | 0.0045125 | 0.741562 | 117 | 6 | 6.0814 |
| TCHH | 6 | 0.9991 | 0.99908 | 0.999994 | 21623 | 0 | 7.166 | 0.00090261 | 0.0046245 | 0.741804 | 118 | 6 | 7.166 |
| FAM105B | 5 | 0.99909 | 0.99911 | 0.999994 | 21622 | 0 | 5.4173 | 0.0009067 | 0.0040576 | 0.720216 | 119 | 5 | 5.4173 |
| FAM208B | 4 | 0.85323 | 0.85278 | 0.999994 | 18316 | 0 | 8.4825 | 0.00090716 | 0.003353 | 0.665572 | 120 | 3 | 8.4825 |
| PSORS1C2 | 5 | 0.0083417 | 0.031293 | 0.862223 | 752 | 2 | 1.0314 | 0.00091894 | 0.0041097 | 0.723537 | 121 | 3 | 1.0314 |
| PELI3 | 6 | 0.99907 | 0.99905 | 0.999994 | 21621 | 0 | 5.7362 | 0.00092803 | 0.0047411 | 0.749404 | 122 | 6 | 5.7362 |
| U2AF1L4 | 6 | 0.99906 | 0.99903 | 0.999994 | 21620 | 0 | 3.3408 | 0.00094488 | 0.0048211 | 0.75546 | 123 | 6 | 3.3408 |
| ZBTB3 | 5 | 0.99762 | 0.99761 | 0.999994 | 21567 | 0 | 7.8329 | 0.00099403 | 0.0044389 | 0.741562 | 124 | 5 | 7.8329 |
| SIGLEC6 | 5 | 0.97877 | 0.97886 | 0.999994 | 21070 | 0 | 1.4887 | 0.0010132 | 0.0045203 | 0.741562 | 125 | 5 | 1.4887 |
| BANF1 | 3 | 0.99898 | 0.99899 | 0.999994 | 21618 | 0 | 7.9312 | 0.0010172 | 0.0028985 | 0.665572 | 126 | 3 | 7.9312 |
| LSR | 6 | 0.99897 | 0.99896 | 0.999994 | 21617 | 0 | 4.5801 | 0.0010277 | 0.0052139 | 0.776285 | 127 | 6 | 4.5801 |
| RGS10 | 5 | 0.99897 | 0.99899 | 0.999994 | 21616 | 0 | 6.4076 | 0.0010293 | 0.0045911 | 0.741804 | 128 | 5 | 6.4076 |
| SLC39A4 | 6 | 0.89051 | 0.8967 | 0.999994 | 19046 | 1 | 6.3311 | 0.0010352 | 0.005255 | 0.776285 | 129 | 5 | 6.3311 |
| SHFM1 | 5 | 0.99896 | 0.99898 | 0.999994 | 21615 | 0 | 4.5924 | 0.0010361 | 0.004624 | 0.741804 | 130 | 5 | 4.5924 |
| LILRB5 | 6 | 0.23649 | 0.43181 | 0.999994 | 8681 | 1 | 7.5719 | 0.0010369 | 0.0052696 | 0.776285 | 131 | 5 | 7.5719 |
| BIN2 | 5 | 0.88462 | 0.88475 | 0.999994 | 18928 | 1 | 1.4006 | 0.0010535 | 0.0047079 | 0.749404 | 132 | 4 | 1.4006 |
| LOC646862 | 4 | 0.98236 | 0.98232 | 0.999994 | 21144 | 0 | 7.8745 | 0.0010659 | 0.0039158 | 0.706642 | 133 | 4 | 7.8745 |
| GNB2L1 | 6 | 0.99892 | 0.99891 | 0.999994 | 21614 | 0 | 5.9076 | 0.0010754 | 0.0054498 | 0.776371 | 134 | 6 | 5.9076 |
| CCDC97 | 6 | 0.50311 | 0.67088 | 0.999994 | 13480 | 1 | 4.1686 | 0.0011016 | 0.0055645 | 0.776371 | 135 | 5 | 4.1686 |
| AMBP | 5 | 0.3666 | 0.53553 | 0.999994 | 11450 | 1 | 7.4584 | 0.0011025 | 0.0049189 | 0.75546 | 136 | 4 | 7.4584 |
| NEUROD1 | 6 | 0.16876 | 0.34919 | 0.999994 | 6952 | 3 | 3.6469 | 0.0011301 | 0.0057099 | 0.776438 | 137 | 3 | 3.6469 |
| RAPGEF2 | 5 | 0.99887 | 0.99888 | 0.999994 | 21613 | 0 | 5.0163 | 0.0011306 | 0.0050401 | 0.768617 | 138 | 5 | 5.0163 |
| SMAP1 | 6 | 0.99821 | 0.99819 | 0.999994 | 21590 | 0 | 6.4115 | 0.0011422 | 0.0057662 | 0.776438 | 139 | 6 | 6.4115 |
| IMPA2 | 6 | 0.28724 | 0.49189 | 0.999994 | 9859 | 3 | 2.9204 | 0.0011458 | 0.0057822 | 0.776438 | 140 | 3 | 2.9204 |
| DCXR | 4 | 0.30255 | 0.44559 | 0.999994 | 10182 | 2 | -0.35656 | 0.0011498 | 0.0042185 | 0.730812 | 141 | 2 | -0.35656 |
| NASP | 6 | 0.081173 | 0.20969 | 0.979587 | 4273 | 3 | 0.86106 | 0.0011543 | 0.0058197 | 0.776438 | 142 | 3 | 0.86106 |
| LONP2 | 6 | 0.57203 | 0.71943 | 0.999994 | 14576 | 1 | 1.4236 | 0.0011591 | 0.0058444 | 0.776438 | 143 | 5 | 1.4236 |
| C14orf119 | 6 | 0.98806 | 0.98802 | 0.999994 | 21301 | 0 | 5.7354 | 0.0011671 | 0.0058855 | 0.777137 | 144 | 6 | 5.7354 |
| FAM216B | 5 | 0.93975 | 0.93993 | 0.999994 | 20070 | 0 | 5.9781 | 0.0011683 | 0.0051983 | 0.776285 | 145 | 4 | 5.9781 |
| EN2 | 5 | 0.90608 | 0.90616 | 0.999994 | 19353 | 0 | 7.1639 | 0.0011813 | 0.0052472 | 0.776285 | 146 | 4 | 7.1639 |
| POTED | 4 | 0.096067 | 0.20803 | 0.979108 | 4818 | 1 | 4.1125 | 0.0011875 | 0.0043621 | 0.741562 | 147 | 3 | 4.1125 |
| ATP10A | 6 | 0.9988 | 0.99879 | 0.999994 | 21611 | 0 | 3.4241 | 0.0011976 | 0.0060295 | 0.780818 | 148 | 6 | 3.4241 |
| SPSB2 | 5 | 0.31732 | 0.49518 | 0.999994 | 10458 | 1 | 8.1534 | 0.0012023 | 0.0053506 | 0.776371 | 149 | 4 | 8.1534 |
| NWD1 | 6 | 0.97896 | 0.97887 | 0.999994 | 21074 | 0 | 5.7592 | 0.0012083 | 0.0060816 | 0.780818 | 150 | 5 | 5.7592 |
| SERPINA10 | 5 | 0.85453 | 0.85791 | 0.999994 | 18340 | 1 | 6.8726 | 0.0012102 | 0.0053903 | 0.776371 | 151 | 4 | 6.8726 |
| AHCYL1 | 6 | 0.9213 | 0.9216 | 0.999994 | 19660 | 1 | 7.5099 | 0.001216 | 0.0061191 | 0.780818 | 152 | 5 | 7.5099 |
| LIM2 | 6 | 0.98684 | 0.98679 | 0.999994 | 21276 | 0 | 4.0027 | 0.0012206 | 0.006142 | 0.780818 | 153 | 6 | 4.0027 |
| SLC25A24 | 4 | 0.67694 | 0.70967 | 0.999994 | 16038 | 1 | 7.8071 | 0.001225 | 0.0044965 | 0.741562 | 154 | 3 | 7.8071 |
| PHLDA3 | 5 | 0.39859 | 0.56238 | 0.999994 | 11925 | 1 | 8.1481 | 0.001227 | 0.0054621 | 0.776371 | 155 | 3 | 8.1481 |

| | | | | | | | | | | | | | |
|---|---|---|---|---|---|---|---|---|---|---|---|---|---|
| B3GNT1 | 5 | 0.13816 | 0.28503 | 0.999632 | 6141 | 2 | 1.8203 | 0.0012319 | 0.0054854 | 0.776371 | 156 | 3 | 1.8203 |
| JMJD4 | 5 | 0.95033 | 0.95051 | 0.999994 | 20343 | 0 | 5.2982 | 0.0012487 | 0.0055673 | 0.776371 | 157 | 4 | 5.2982 |
| FNDC8 | 5 | 0.99875 | 0.99876 | 0.999994 | 21610 | 0 | 7.0586 | 0.0012537 | 0.0055929 | 0.776371 | 158 | 5 | 7.0586 |
| KRTAP10-7 | 6 | 0.87129 | 0.88414 | 0.999994 | 18665 | 1 | 0.52663 | 0.0012722 | 0.006398 | 0.78601 | 159 | 3 | 0.52663 |
| hsa-mir-3185 | 3 | 0.99871 | 0.99872 | 0.999994 | 21609 | 0 | 7.1763 | 0.0012863 | 0.0036378 | 0.679114 | 160 | 3 | 7.1763 |
| NonTargetingControlG | 1 | 0.99871 | 0.99874 | 0.999994 | 21608 | 0 | 10.949 | 0.001287 | 0.0012813 | 0.524099 | 161 | 1 | 10.949 |
| NEK11 | 6 | 0.9987 | 0.99869 | 0.999994 | 21607 | 0 | 5.2951 | 0.001296 | 0.006511 | 0.792107 | 162 | 6 | 5.2951 |
| PHIP | 6 | 0.98383 | 0.98378 | 0.999994 | 21183 | 0 | 7.4068 | 0.0013067 | 0.006559 | 0.79349 | 163 | 5 | 7.4068 |
| BHMT | 6 | 0.40651 | 0.60661 | 0.999994 | 12049 | 2 | 0.70109 | 0.0013287 | 0.006649 | 0.799917 | 164 | 3 | 0.70109 |
| SLC16A11 | 4 | 0.99736 | 0.99736 | 0.999994 | 21558 | 0 | 7.153 | 0.0013382 | 0.0049016 | 0.75546 | 165 | 4 | 7.153 |
| PSKH1 | 6 | 0.31484 | 0.52399 | 0.999994 | 10417 | 1 | 4.7934 | 0.0013443 | 0.0067254 | 0.800124 | 166 | 4 | 4.7934 |
| TOR2A | 5 | 0.59752 | 0.72526 | 0.999994 | 14920 | 1 | 7.9883 | 0.0013496 | 0.0059975 | 0.780818 | 167 | 4 | 7.9883 |
| RNF169 | 6 | 0.92488 | 0.92499 | 0.999994 | 19739 | 1 | 5.6486 | 0.0013514 | 0.0067583 | 0.800124 | 168 | 5 | 5.6486 |
| DACT3 | 3 | 0.99865 | 0.99866 | 0.999994 | 21606 | 0 | 8.3935 | 0.0013536 | 0.0038225 | 0.701265 | 169 | 3 | 8.3935 |
| LAMA2 | 6 | 0.99862 | 0.9986 | 0.999994 | 21604 | 0 | 3.416 | 0.0013848 | 0.0069312 | 0.80013 | 170 | 6 | 3.416 |
| CD3G | 5 | 0.99861 | 0.99863 | 0.999994 | 21603 | 0 | 3.8547 | 0.0013879 | 0.0061639 | 0.780818 | 171 | 5 | 3.8547 |
| PSG8 | 5 | 0.99861 | 0.99863 | 0.999994 | 21602 | 0 | 5.8358 | 0.0013887 | 0.0061658 | 0.780818 | 172 | 5 | 5.8358 |
| TKT | 6 | 0.9986 | 0.99858 | 0.999994 | 21601 | 0 | 3.0695 | 0.0014022 | 0.0070153 | 0.80013 | 173 | 6 | 3.0695 |
| C1orf64 | 5 | 0.30773 | 0.4873 | 0.999994 | 10282 | 2 | 8.0874 | 0.0014181 | 0.0062979 | 0.78601 | 174 | 3 | 8.0874 |
| ATL2 | 6 | 0.0068814 | 0.027994 | 0.855011 | 652 | 4 | -4.7185 | 0.0014183 | 0.007109 | 0.803016 | 175 | 2 | -4.7185 |
| CUX2 | 6 | 0.87062 | 0.88372 | 0.999994 | 18648 | 1 | 5.8933 | 0.001439 | 0.0072055 | 0.803016 | 176 | 5 | 5.8933 |
| DCTPP1 | 6 | 0.25984 | 0.45966 | 0.999994 | 9237 | 1 | 0.89987 | 0.0014417 | 0.0072155 | 0.803016 | 177 | 4 | 0.89987 |
| FKBP2 | 6 | 0.34348 | 0.55663 | 0.999994 | 11020 | 1 | 7.1651 | 0.0014448 | 0.0072302 | 0.803016 | 178 | 4 | 7.1651 |
| ZFP36 | 5 | 0.99855 | 0.99857 | 0.999994 | 21600 | 0 | 5.8779 | 0.0014477 | 0.0064246 | 0.78601 | 179 | 5 | 5.8779 |
| GTF2F1 | 6 | 0.92722 | 0.92728 | 0.999994 | 19795 | 1 | 5.9763 | 0.0014634 | 0.0073065 | 0.803161 | 180 | 5 | 5.9763 |
| SLC1A5 | 6 | 0.9985 | 0.99848 | 0.999994 | 21599 | 0 | 4.0981 | 0.0015018 | 0.0075045 | 0.808507 | 181 | 6 | 4.0981 |
| AHI1 | 4 | 0.90182 | 0.90149 | 0.999994 | 19247 | 0 | 7.6939 | 0.0015265 | 0.0055879 | 0.776371 | 182 | 3 | 7.6939 |
| AHSA1 | 5 | 0.046274 | 0.13143 | 0.95318 | 2817 | 2 | -0.2627 | 0.0015266 | 0.0067647 | 0.800124 | 183 | 2 | -0.2627 |
| CEACAM5 | 6 | 0.9788 | 0.97872 | 0.999994 | 21071 | 0 | 6.2851 | 0.0015302 | 0.0076494 | 0.811952 | 184 | 5 | 6.2851 |
| LRRTM3 | 5 | 0.78134 | 0.80819 | 0.999994 | 17229 | 1 | 6.6635 | 0.0015346 | 0.0067986 | 0.800124 | 185 | 4 | 6.6635 |
| MECP2 | 6 | 0.99845 | 0.99843 | 0.999994 | 21598 | 0 | 5.589 | 0.0015478 | 0.0077404 | 0.811952 | 186 | 6 | 5.589 |
| ZMYM3 | 6 | 0.95004 | 0.95002 | 0.999994 | 20335 | 0 | 7.1427 | 0.0015533 | 0.0077665 | 0.811952 | 187 | 5 | 7.1427 |
| ANKRD50 | 6 | 0.80329 | 0.84927 | 0.999994 | 17529 | 1 | 4.679 | 0.0015604 | 0.0078035 | 0.811952 | 188 | 4 | 4.679 |
| hsa-mir-7855 | 4 | 0.15247 | 0.29171 | 0.999632 | 6520 | 2 | 1.3811 | 0.0015642 | 0.0057227 | 0.776438 | 189 | 2 | 1.3811 |
| hsa-mir-5186 | 4 | 0.82914 | 0.82867 | 0.999994 | 17913 | 0 | 7.8593 | 0.0015668 | 0.0057323 | 0.776438 | 190 | 3 | 7.8593 |
| SYNJ2 | 5 | 0.99843 | 0.99847 | 0.999994 | 21597 | 0 | 3.6393 | 0.0015686 | 0.0069289 | 0.80013 | 191 | 5 | 3.6393 |
| PABPC1L2A | 6 | 0.47613 | 0.65259 | 0.999994 | 13081 | 2 | 4.5728 | 0.0015694 | 0.0078488 | 0.811952 | 192 | 4 | 4.5728 |
| PHC3 | 5 | 0.042679 | 0.12253 | 0.951052 | 2660 | 4 | -0.82836 | 0.0015783 | 0.0069723 | 0.80013 | 193 | 1 | -0.82836 |
| AMN1 | 6 | 0.89334 | 0.89868 | 0.999994 | 19092 | 1 | 5.1441 | 0.0015905 | 0.007953 | 0.812073 | 194 | 5 | 5.1441 |
| LPAR5 | 6 | 0.99841 | 0.99838 | 0.999994 | 21596 | 0 | 5.2083 | 0.0015911 | 0.0079544 | 0.812073 | 195 | 6 | 5.2083 |
| TMED5 | 2 | 0.95677 | 0.95686 | 0.999994 | 20512 | 0 | 8.1003 | 0.0016116 | 0.0030956 | 0.665572 | 196 | 2 | 8.1003 |
| CACNG6 | 6 | 0.99399 | 0.99394 | 0.999994 | 21448 | 0 | 4.5725 | 0.0016119 | 0.0080573 | 0.812073 | 197 | 6 | 4.5725 |
| MED21 | 5 | 0.99836 | 0.99839 | 0.999994 | 21595 | 0 | 5.9833 | 0.0016443 | 0.0072585 | 0.803016 | 198 | 5 | 5.9833 |
| SATB2 | 6 | 0.49901 | 0.66801 | 0.999994 | 13409 | 2 | 2.0407 | 0.0016448 | 0.0082164 | 0.812073 | 199 | 3 | 2.0407 |
| C6orf203 | 5 | 0.98691 | 0.9869 | 0.999994 | 21278 | 0 | 8.0232 | 0.0016458 | 0.0072681 | 0.803016 | 200 | 5 | 8.0232 |
| KMT2A | 6 | 0.066154 | 0.17871 | 0.968171 | 3693 | 1 | 5.9809 | 0.0016656 | 0.008317 | 0.812073 | 201 | 5 | 5.9809 |
| BPNT1 | 6 | 0.050036 | 0.14412 | 0.961983 | 2982 | 2 | -0.060579 | 0.0016677 | 0.0083252 | 0.812073 | 202 | 2 | -0.060579 |
| ZNF382 | 6 | 0.99392 | 0.99388 | 0.999994 | 21446 | 0 | 3.974 | 0.0016679 | 0.008327 | 0.812073 | 203 | 6 | 3.974 |
| UBA3 | 5 | 0.25106 | 0.44161 | 0.999994 | 9035 | 1 | 6.8295 | 0.0016703 | 0.0073733 | 0.806406 | 204 | 4 | 6.8295 |
| SULT2A1 | 6 | 0.98882 | 0.98878 | 0.999994 | 21316 | 0 | 5.376 | 0.0016845 | 0.0084089 | 0.812073 | 205 | 6 | 5.376 |
| IL2RB | 5 | 0.9983 | 0.99832 | 0.999994 | 21594 | 0 | 6.2427 | 0.0017007 | 0.0074999 | 0.808507 | 206 | 5 | 6.2427 |
| SOHLH2 | 6 | 0.99728 | 0.99726 | 0.999994 | 21552 | 0 | 6.7763 | 0.001712 | 0.0085396 | 0.812073 | 207 | 6 | 6.7763 |

| | | | | | | | | | | | | |
|---|---|---|---|---|---|---|---|---|---|---|---|---|
| SPACA4 | 6 | 0.58699 | 0.73037 | 0.999994 | 14776 | 2 | 1.1105 | 0.0017146 | 0.0085511 | 0.812073 | 208 | 3 | 1.1105 |
| SRD5A1 | 3 | 0.99828 | 0.9983 | 0.999994 | 21593 | 0 | 6.9304 | 0.0017199 | 0.0048495 | 0.75546 | 209 | 3 | 6.9304 |
| APLP1 | 6 | 0.99827 | 0.99824 | 0.999994 | 21592 | 0 | 2.9091 | 0.0017333 | 0.0086425 | 0.812073 | 210 | 6 | 2.9091 |
| TMEFF1 | 4 | 0.075997 | 0.17128 | 0.967645 | 4072 | 2 | 2.721 | 0.0017473 | 0.0063509 | 0.78601 | 211 | 2 | 2.721 |
| HSPA1B | 4 | 0.86284 | 0.86246 | 0.999994 | 18504 | 0 | 8.1559 | 0.0017615 | 0.0064017 | 0.78601 | 212 | 3 | 8.1559 |
| PPP2R5D | 5 | 0.82784 | 0.83794 | 0.999994 | 17894 | 1 | 6.9901 | 0.001766 | 0.0077843 | 0.811952 | 213 | 4 | 6.9901 |
| ARG1 | 6 | 0.44639 | 0.63268 | 0.999994 | 12667 | 2 | 1.1999 | 0.0017738 | 0.0088387 | 0.816312 | 214 | 3 | 1.1999 |
| PIGG | 6 | 0.99823 | 0.9982 | 0.999994 | 21591 | 0 | 3.9973 | 0.0017745 | 0.0088428 | 0.816312 | 215 | 6 | 3.9973 |
| STAG2 | 6 | 0.99821 | 0.99818 | 0.999994 | 21589 | 0 | 4.646 | 0.0017907 | 0.0089255 | 0.816312 | 216 | 6 | 4.646 |
| FAM218A | 6 | 0.9892 | 0.98916 | 0.999994 | 21323 | 0 | 3.1919 | 0.0017948 | 0.0089456 | 0.816312 | 217 | 6 | 3.1919 |
| KBTBD3 | 6 | 0.44224 | 0.62992 | 0.999994 | 12614 | 2 | 3.1713 | 0.0018008 | 0.0089717 | 0.816312 | 218 | 4 | 3.1713 |
| APOOL | 5 | 0.91431 | 0.91437 | 0.999994 | 19494 | 0 | 3.5444 | 0.0018137 | 0.0079928 | 0.812073 | 219 | 4 | 3.5444 |
| SFXN4 | 5 | 0.67785 | 0.75663 | 0.999994 | 16057 | 1 | 1.6646 | 0.0018296 | 0.008065 | 0.812073 | 220 | 3 | 1.6646 |
| SEP15 | 5 | 0.99816 | 0.99816 | 0.999994 | 21588 | 0 | 4.7106 | 0.0018445 | 0.0081227 | 0.812073 | 221 | 5 | 4.7106 |
| LOC100131094 | 6 | 0.92407 | 0.92422 | 0.999994 | 19716 | 1 | 7.1001 | 0.0018593 | 0.0092652 | 0.825215 | 222 | 5 | 7.1001 |
| VDAC1 | 6 | 0.96202 | 0.96209 | 0.999994 | 20630 | 0 | 6.1542 | 0.001874 | 0.009343 | 0.825215 | 223 | 5 | 6.1542 |
| LENG8 | 6 | 0.83115 | 0.8621 | 0.999994 | 17949 | 1 | 7.0003 | 0.0018931 | 0.0094426 | 0.825215 | 224 | 4 | 7.0003 |
| TMPRSS12 | 5 | 0.26562 | 0.4533 | 0.999994 | 9362 | 2 | 7.9581 | 0.0019096 | 0.0083942 | 0.812073 | 225 | 3 | 7.9581 |
| OTUD3 | 5 | 0.51702 | 0.66271 | 0.999994 | 13691 | 1 | 8.1207 | 0.0019166 | 0.0084198 | 0.812073 | 226 | 3 | 8.1207 |
| ODF3 | 4 | 0.99807 | 0.99808 | 0.999994 | 21586 | 0 | 4.87 | 0.0019329 | 0.0070203 | 0.80013 | 227 | 4 | 4.87 |
| GDI1 | 6 | 0.78438 | 0.84143 | 0.999994 | 17272 | 1 | 6.9804 | 0.00194 | 0.0096776 | 0.825215 | 228 | 5 | 6.9804 |
| ZNF770 | 6 | 0.99804 | 0.99802 | 0.999994 | 21585 | 0 | 6.7269 | 0.001959 | 0.0097709 | 0.825215 | 229 | 6 | 6.7269 |
| ZC3H18 | 5 | 0.99804 | 0.99805 | 0.999994 | 21584 | 0 | 5.3289 | 0.0019626 | 0.0086046 | 0.812073 | 230 | 5 | 5.3289 |
| PEAK1 | 6 | 0.80075 | 0.84819 | 0.999994 | 17492 | 1 | 7.0448 | 0.0019647 | 0.0098047 | 0.825215 | 231 | 5 | 7.0448 |
| GPS2 | 5 | 0.44862 | 0.6041 | 0.999994 | 12695 | 1 | 7.711 | 0.0019763 | 0.0086613 | 0.812073 | 232 | 4 | 7.711 |
| ARMC4 | 5 | 0.57059 | 0.71005 | 0.999994 | 14550 | 1 | 6.7645 | 0.0019801 | 0.0086786 | 0.812073 | 233 | 4 | 6.7645 |
| CNOT6L | 6 | 0.76219 | 0.83289 | 0.999994 | 17001 | 1 | 7.3669 | 0.0019995 | 0.0099648 | 0.825215 | 234 | 5 | 7.3669 |
| ZNF70 | 6 | 0.20482 | 0.39375 | 0.999994 | 7896 | 1 | 6.0508 | 0.0020102 | 0.010018 | 0.825215 | 235 | 5 | 6.0508 |
| KEAP1 | 6 | 0.99799 | 0.99796 | 0.999994 | 21583 | 0 | 2.1848 | 0.002015 | 0.010035 | 0.825215 | 236 | 6 | 2.1848 |
| ZNF570 | 6 | 0.99798 | 0.99795 | 0.999994 | 21582 | 0 | 3.3674 | 0.0020249 | 0.010086 | 0.825215 | 237 | 6 | 3.3674 |
| MTMR3 | 6 | 0.99797 | 0.99794 | 0.999994 | 21581 | 0 | 6.2245 | 0.0020275 | 0.010095 | 0.825215 | 238 | 6 | 6.2245 |
| NFATC3 | 6 | 0.99794 | 0.99791 | 0.999994 | 21579 | 0 | 3.2186 | 0.0020616 | 0.010258 | 0.825215 | 239 | 6 | 3.2186 |
| NonTargetingControlG | 2 | 0.62186 | 0.62162 | 0.999994 | 15255 | 0 | 5.3777 | 0.0020638 | 0.0039922 | 0.714467 | 240 | 1 | 5.3777 |
| SRP14 | 6 | 0.8824 | 0.89118 | 0.999994 | 18878 | 1 | 5.8235 | 0.0020711 | 0.010302 | 0.825215 | 241 | 5 | 5.8235 |
| CAPN11 | 6 | 0.96661 | 0.96667 | 0.999994 | 20753 | 0 | 4.1583 | 0.0020862 | 0.010364 | 0.825215 | 242 | 4 | 4.1583 |
| CDH23 | 6 | 0.81792 | 0.85584 | 0.999994 | 17744 | 1 | 5.5792 | 0.0020989 | 0.010424 | 0.825215 | 243 | 4 | 5.5792 |
| RAD21 | 4 | 0.99789 | 0.9979 | 0.999994 | 21577 | 0 | 6.1843 | 0.002113 | 0.0076755 | 0.811952 | 244 | 4 | 6.1843 |
| EFHC2 | 6 | 0.5849 | 0.72883 | 0.999994 | 14740 | 2 | 2.5792 | 0.0021195 | 0.010507 | 0.825215 | 245 | 4 | 2.5792 |
| ATP10B | 5 | 0.85698 | 0.8599 | 0.999994 | 18394 | 1 | 6.509 | 0.002134 | 0.0093324 | 0.825215 | 246 | 4 | 6.509 |
| LRRC16A | 6 | 0.99787 | 0.99785 | 0.999994 | 21576 | 0 | 4.5071 | 0.0021349 | 0.010578 | 0.825215 | 247 | 6 | 4.5071 |
| KIAA1919 | 5 | 0.99786 | 0.99787 | 0.999994 | 21575 | 0 | 7.3538 | 0.0021434 | 0.0093745 | 0.825215 | 248 | 5 | 7.3538 |
| C2orf88 | 6 | 0.59164 | 0.73383 | 0.999994 | 14843 | 1 | 0.48699 | 0.002148 | 0.01064 | 0.825215 | 249 | 3 | 0.48699 |
| TM4SF5 | 6 | 0.99784 | 0.99782 | 0.999994 | 21574 | 0 | 3.2367 | 0.0021599 | 0.010698 | 0.825215 | 250 | 6 | 3.2367 |
| PEMT | 6 | 0.78682 | 0.84238 | 0.999994 | 17302 | 1 | 5.097 | 0.0021637 | 0.010716 | 0.825215 | 251 | 4 | 5.097 |
| CPLX1 | 4 | 0.075709 | 0.1707 | 0.967177 | 4062 | 2 | 2.8841 | 0.0021705 | 0.0078739 | 0.811952 | 252 | 2 | 2.8841 |
| MS4A3 | 6 | 0.23187 | 0.42634 | 0.999994 | 8563 | 1 | 5.7698 | 0.0021827 | 0.010805 | 0.825215 | 253 | 5 | 5.7698 |
| CNTN1 | 5 | 0.9015 | 0.90161 | 0.999994 | 19243 | 0 | 1.7166 | 0.0021862 | 0.0095565 | 0.825215 | 254 | 3 | 1.7166 |
| WHAMM | 6 | 0.092477 | 0.23247 | 0.984502 | 4703 | 3 | -0.17031 | 0.0022325 | 0.011039 | 0.829051 | 255 | 3 | -0.17031 |
| GAS6 | 5 | 0.99773 | 0.99772 | 0.999994 | 21573 | 0 | 6.8809 | 0.0022728 | 0.00992 | 0.825215 | 256 | 5 | 6.8809 |
| PRSS21 | 5 | 0.88153 | 0.88173 | 0.999994 | 18861 | 1 | 7.8803 | 0.0022742 | 0.0099259 | 0.825215 | 257 | 4 | 7.8803 |
| ATP1A3 | 6 | 0.54409 | 0.69942 | 0.999994 | 14132 | 2 | 4.5112 | 0.0022817 | 0.011271 | 0.829051 | 258 | 4 | 4.5112 |
| VPS26A | 6 | 0.99772 | 0.9977 | 0.999994 | 21572 | 0 | 5.0989 | 0.0022838 | 0.01128 | 0.829051 | 259 | 6 | 5.0989 |

| | | | | | | | | | | | | | |
|---|---|---|---|---|---|---|---|---|---|---|---|---|---|
| TRAF2 | 5 | 0.97565 | 0.9757 | 0.999994 | 20982 | 0 | 1.399 | 0.0022843 | 0.0099753 | 0.825215 | 260 | 5 | 1.399 |
| GRIK5 | 5 | 0.96718 | 0.96736 | 0.999994 | 20767 | 0 | 7.8782 | 0.0022915 | 0.010005 | 0.825215 | 261 | 4 | 7.8782 |
| JDP2 | 5 | 0.20886 | 0.3878 | 0.999994 | 7999 | 2 | 5.9833 | 0.0023262 | 0.010152 | 0.825215 | 262 | 3 | 5.9833 |
| CLRN3 | 6 | 0.9964 | 0.99638 | 0.999994 | 21516 | 0 | 4.633 | 0.0023311 | 0.011495 | 0.829051 | 263 | 6 | 4.633 |
| VN1R2 | 6 | 0.99766 | 0.99765 | 0.999994 | 21570 | 0 | 4.346 | 0.0023428 | 0.011542 | 0.829051 | 264 | 6 | 4.346 |
| OR6K3 | 6 | 0.6186 | 0.75378 | 0.999994 | 15197 | 1 | 6.911 | 0.0023647 | 0.011642 | 0.829051 | 265 | 4 | 6.911 |
| FAM184B | 4 | 0.99763 | 0.99763 | 0.999994 | 21569 | 0 | 7.0637 | 0.0023692 | 0.0085543 | 0.812073 | 266 | 4 | 7.0637 |
| FBXL18 | 5 | 0.99763 | 0.99762 | 0.999994 | 21568 | 0 | 6.4422 | 0.0023715 | 0.010344 | 0.825215 | 267 | 5 | 6.4422 |
| DNAJC12 | 5 | 0.50657 | 0.65364 | 0.999994 | 13534 | 1 | 7.8647 | 0.0023773 | 0.010366 | 0.825215 | 268 | 4 | 7.8647 |
| SNRNP40 | 6 | 0.8748 | 0.88631 | 0.999994 | 18728 | 1 | 7.0583 | 0.0023896 | 0.011753 | 0.829051 | 269 | 5 | 7.0583 |
| RELB | 6 | 0.9976 | 0.9976 | 0.999994 | 21566 | 0 | 6.063 | 0.0023967 | 0.01179 | 0.829051 | 270 | 6 | 6.063 |
| PNLIPRP2 | 6 | 0.9976 | 0.9976 | 0.999994 | 21565 | 0 | 4.2097 | 0.0023986 | 0.011796 | 0.829051 | 271 | 6 | 4.2097 |
| RFWD2 | 5 | 0.99759 | 0.99758 | 0.999994 | 21564 | 0 | 3.542 | 0.0024076 | 0.01048 | 0.825215 | 272 | 5 | 3.542 |
| RHOB | 4 | 0.71621 | 0.73541 | 0.999994 | 16458 | 1 | 7.6719 | 0.0024087 | 0.0087001 | 0.812073 | 273 | 3 | 7.6719 |
| ZNF584 | 6 | 0.99759 | 0.99759 | 0.999994 | 21563 | 0 | 5.8503 | 0.0024105 | 0.011854 | 0.829051 | 274 | 6 | 5.8503 |
| LGI4 | 6 | 0.88441 | 0.89247 | 0.999994 | 18920 | 1 | 6.2064 | 0.0024126 | 0.011862 | 0.829051 | 275 | 5 | 6.2064 |
| CA12 | 5 | 0.11726 | 0.25316 | 0.994501 | 5548 | 3 | -1.4781 | 0.0024255 | 0.01055 | 0.825215 | 276 | 2 | -1.4781 |
| CST3 | 5 | 0.75886 | 0.79555 | 0.999994 | 16963 | 1 | 0.78271 | 0.0024362 | 0.010589 | 0.825215 | 277 | 3 | 0.78271 |
| CD93 | 6 | 0.99756 | 0.99756 | 0.999994 | 21562 | 0 | 4.3048 | 0.0024421 | 0.012013 | 0.829051 | 278 | 6 | 4.3048 |
| SMURF2 | 6 | 0.38841 | 0.59493 | 0.999994 | 11772 | 2 | 2.3126 | 0.0024439 | 0.012025 | 0.829051 | 279 | 4 | 2.3126 |
| PILRB | 6 | 0.87322 | 0.88531 | 0.999994 | 18700 | 1 | 2.5629 | 0.0024583 | 0.012093 | 0.829051 | 280 | 4 | 2.5629 |
| OR4M1 | 5 | 0.89106 | 0.89118 | 0.999994 | 19055 | 0 | 1.2251 | 0.002459 | 0.010684 | 0.825215 | 281 | 4 | 1.2251 |
| FAU | 3 | 0.98709 | 0.98722 | 0.999994 | 21282 | 0 | 8.7317 | 0.0024913 | 0.0069408 | 0.80013 | 282 | 3 | 8.7317 |
| ATXN3L | 6 | 0.24244 | 0.43891 | 0.999994 | 8845 | 2 | 3.8601 | 0.0025042 | 0.0123 | 0.829051 | 283 | 3 | 3.8601 |
| RIC8A | 6 | 0.99749 | 0.99749 | 0.999994 | 21561 | 0 | 3.4817 | 0.0025063 | 0.012316 | 0.829051 | 284 | 6 | 3.4817 |
| DDI1 | 6 | 0.99749 | 0.99749 | 0.999994 | 21560 | 0 | 3.6618 | 0.0025093 | 0.012329 | 0.829051 | 285 | 6 | 3.6618 |
| PLEKHA1 | 6 | 0.95458 | 0.95464 | 0.999994 | 20444 | 0 | 3.474 | 0.0025159 | 0.012361 | 0.829051 | 286 | 4 | 3.474 |
| TRIM22 | 5 | 0.83374 | 0.84216 | 0.999994 | 17983 | 1 | 6.1806 | 0.0025196 | 0.010912 | 0.825215 | 287 | 4 | 6.1806 |
| RALGPS1 | 5 | 0.97048 | 0.97054 | 0.999994 | 20836 | 0 | 6.1806 | 0.0025196 | 0.010912 | 0.825215 | 288 | 4 | 6.1806 |
| SAP30 | 5 | 0.64969 | 0.745 | 0.999994 | 15654 | 1 | 7.4134 | 0.0025253 | 0.010937 | 0.825215 | 289 | 4 | 7.4134 |
| SMARCAD1 | 6 | 0.97947 | 0.97938 | 0.999994 | 21081 | 0 | 3.1985 | 0.0025318 | 0.012436 | 0.829051 | 290 | 5 | 3.1985 |
| LRFN3 | 6 | 0.9653 | 0.96534 | 0.999994 | 20723 | 0 | 6.5189 | 0.0025449 | 0.012494 | 0.829051 | 291 | 5 | 6.5189 |
| SEMA6A | 6 | 0.65886 | 0.78444 | 0.999994 | 15762 | 2 | 4.3032 | 0.0025496 | 0.012519 | 0.829051 | 292 | 4 | 4.3032 |
| UMODL1 | 5 | 0.083343 | 0.19999 | 0.977164 | 4354 | 2 | 5.5792 | 0.0025666 | 0.011106 | 0.829051 | 293 | 3 | 5.5792 |
| MAP1LC3B2 | 6 | 0.28525 | 0.48956 | 0.999994 | 9817 | 3 | -0.57373 | 0.0025712 | 0.012626 | 0.829051 | 294 | 2 | -0.57373 |
| DDX42 | 5 | 0.33548 | 0.51002 | 0.999994 | 10842 | 1 | 6.1076 | 0.0025754 | 0.011147 | 0.829051 | 295 | 4 | 6.1076 |
| HLA-DMB | 6 | 0.48624 | 0.65941 | 0.999994 | 13231 | 2 | 4.7675 | 0.0025834 | 0.012686 | 0.829051 | 296 | 4 | 4.7675 |
| LDLRAD3 | 6 | 0.99741 | 0.99739 | 0.999994 | 21559 | 0 | 5.2182 | 0.0025897 | 0.01271 | 0.829051 | 297 | 6 | 5.2182 |
| ZNF544 | 5 | 0.60516 | 0.72806 | 0.999994 | 15013 | 1 | 1.2958 | 0.0026137 | 0.011297 | 0.829051 | 298 | 3 | 1.2958 |
| MAGEC2 | 6 | 0.83957 | 0.86639 | 0.999994 | 18079 | 1 | 4.5351 | 0.0026346 | 0.012922 | 0.829051 | 299 | 5 | 4.5351 |
| SNRPA | 5 | 0.9947 | 0.99467 | 0.999994 | 21468 | 0 | 7.1468 | 0.0026372 | 0.011388 | 0.829051 | 300 | 5 | 7.1468 |
| SIPA1L3 | 6 | 0.98644 | 0.98639 | 0.999994 | 21259 | 0 | 5.961 | 0.0026424 | 0.012958 | 0.829051 | 301 | 5 | 5.961 |
| THBS3 | 6 | 0.99736 | 0.99733 | 0.999994 | 21557 | 0 | 6.8275 | 0.0026432 | 0.01296 | 0.829051 | 302 | 6 | 6.8275 |
| USP7 | 6 | 0.3505 | 0.56455 | 0.999994 | 11173 | 1 | 7.0644 | 0.0026465 | 0.012979 | 0.829051 | 303 | 5 | 7.0644 |
| GUCY1A2 | 6 | 0.99734 | 0.99732 | 0.999994 | 21556 | 0 | 3.1839 | 0.0026573 | 0.013024 | 0.829051 | 304 | 6 | 3.1839 |
| SBK2 | 6 | 0.99734 | 0.99732 | 0.999994 | 21555 | 0 | 4.7897 | 0.0026589 | 0.013032 | 0.829051 | 305 | 6 | 4.7897 |
| ZNF507 | 5 | 0.99732 | 0.9973 | 0.999994 | 21554 | 0 | 5.0629 | 0.0026831 | 0.011574 | 0.829051 | 306 | 5 | 5.0629 |
| B3GNT9 | 4 | 0.44014 | 0.58692 | 0.999994 | 12579 | 1 | 7.1302 | 0.0026938 | 0.0096607 | 0.825215 | 307 | 3 | 7.1302 |
| GPR4 | 5 | 0.9899 | 0.98993 | 0.999994 | 21341 | 0 | 7.6925 | 0.0027022 | 0.011645 | 0.829051 | 308 | 5 | 7.6925 |
| CLEC4E | 6 | 0.99729 | 0.99726 | 0.999994 | 21553 | 0 | 4.4694 | 0.0027147 | 0.0133 | 0.829051 | 309 | 6 | 4.4694 |
| IFT81 | 6 | 0.61015 | 0.74744 | 0.999994 | 15074 | 1 | 7.1642 | 0.0027217 | 0.013334 | 0.829051 | 310 | 5 | 7.1642 |
| TBC1D16 | 4 | 0.99727 | 0.99727 | 0.999994 | 21551 | 0 | 5.874 | 0.0027266 | 0.0097791 | 0.825215 | 311 | 4 | 5.874 |

| | | | | | | | | | | | | | |
|---|---|---|---|---|---|---|---|---|---|---|---|---|---|
| LATS2 | 4 | 0.99726 | 0.99725 | 0.999994 | 21550 | 0 | 6.2877 | 0.0027435 | 0.0098445 | 0.825215 | 312 | 4 | 6.2877 |
| PCSK5 | 6 | 0.99725 | 0.99723 | 0.999994 | 21549 | 0 | 5.3345 | 0.0027474 | 0.013443 | 0.829051 | 313 | 6 | 5.3345 |
| ESYT2 | 6 | 0.8389 | 0.86605 | 0.999994 | 18069 | 1 | 0.36647 | 0.0027803 | 0.013612 | 0.829051 | 314 | 3 | 0.36647 |
| TBX18 | 6 | 0.99722 | 0.9972 | 0.999994 | 21548 | 0 | 4.745 | 0.0027835 | 0.013622 | 0.829051 | 315 | 6 | 4.745 |
| DYRK1B | 5 | 0.83541 | 0.84334 | 0.999994 | 18014 | 1 | 7.0133 | 0.0027883 | 0.011989 | 0.829051 | 316 | 4 | 7.0133 |
| SIN3A | 5 | 0.99721 | 0.9972 | 0.999994 | 21547 | 0 | 6.4926 | 0.0027908 | 0.012001 | 0.829051 | 317 | 5 | 6.4926 |
| FARSB | 6 | 0.44403 | 0.63111 | 0.999994 | 12634 | 1 | 6.7279 | 0.0028085 | 0.013742 | 0.829051 | 318 | 5 | 6.7279 |
| IKBKAP | 5 | 0.99718 | 0.99717 | 0.999994 | 21546 | 0 | 5.8608 | 0.0028186 | 0.012111 | 0.829051 | 319 | 5 | 5.8608 |
| PGAM4 | 6 | 0.99718 | 0.99717 | 0.999994 | 21545 | 0 | 5.0543 | 0.0028196 | 0.013792 | 0.829051 | 320 | 6 | 5.0543 |
| SDC2 | 5 | 0.99714 | 0.99712 | 0.999994 | 21544 | 0 | 5.5749 | 0.0028649 | 0.012306 | 0.829051 | 321 | 5 | 5.5749 |
| NR1H4 | 6 | 0.30526 | 0.51286 | 0.999994 | 10238 | 3 | 0.37398 | 0.0028682 | 0.014005 | 0.829051 | 322 | 3 | 0.37398 |
| SAMD12 | 5 | 0.99712 | 0.99711 | 0.999994 | 21543 | 0 | 3.5563 | 0.0028804 | 0.012375 | 0.829051 | 323 | 5 | 3.5563 |
| LINGO1 | 6 | 0.99712 | 0.9971 | 0.999994 | 21542 | 0 | 6.7763 | 0.0028821 | 0.01406 | 0.829051 | 324 | 6 | 6.7763 |
| PLSCR5 | 5 | 0.99246 | 0.99248 | 0.999994 | 21406 | 0 | 6.1768 | 0.0028836 | 0.012389 | 0.829051 | 325 | 5 | 6.1768 |
| CHST6 | 4 | 0.99711 | 0.99711 | 0.999994 | 21541 | 0 | 5.3979 | 0.0028882 | 0.010347 | 0.825215 | 326 | 4 | 5.3979 |
| PKM | 6 | 0.17538 | 0.35734 | 0.999994 | 7120 | 1 | 6.8235 | 0.0028906 | 0.014102 | 0.829051 | 327 | 5 | 6.8235 |
| MOGS | 6 | 0.9971 | 0.99709 | 0.999994 | 21540 | 0 | 2.3672 | 0.0029007 | 0.014148 | 0.829051 | 328 | 6 | 2.3672 |
| KRT74 | 5 | 0.9971 | 0.99709 | 0.999994 | 21539 | 0 | 3.2315 | 0.0029008 | 0.012459 | 0.829051 | 329 | 5 | 3.2315 |
| PRMT6 | 3 | 0.99706 | 0.99708 | 0.999994 | 21538 | 0 | 6.9059 | 0.0029386 | 0.0081524 | 0.812073 | 330 | 3 | 6.9059 |
| VNN2 | 6 | 0.060403 | 0.1666 | 0.967126 | 3442 | 3 | 0.029679 | 0.0029663 | 0.014446 | 0.829051 | 331 | 3 | 0.029679 |
| CALR3 | 5 | 0.92115 | 0.92133 | 0.999994 | 19655 | 0 | 6.4099 | 0.0029706 | 0.012725 | 0.829051 | 332 | 4 | 6.4099 |
| GRK1 | 6 | 0.98439 | 0.98433 | 0.999994 | 21198 | 0 | 1.9718 | 0.0029784 | 0.014504 | 0.829051 | 333 | 5 | 1.9718 |
| PLEKHF1 | 6 | 0.97252 | 0.97245 | 0.999994 | 20883 | 0 | 5.7598 | 0.0029812 | 0.014515 | 0.829051 | 334 | 5 | 5.7598 |
| EPPK1 | 6 | 0.61814 | 0.75343 | 0.999994 | 15191 | 2 | 4.0896 | 0.0030212 | 0.014707 | 0.829051 | 335 | 3 | 4.0896 |
| LEF1 | 6 | 0.94429 | 0.94426 | 0.999994 | 20186 | 0 | 2.9883 | 0.0030227 | 0.014711 | 0.829051 | 336 | 5 | 2.9883 |
| TCEANC | 4 | 0.99695 | 0.99694 | 0.999994 | 21536 | 0 | 6.7713 | 0.0030504 | 0.010925 | 0.825215 | 337 | 4 | 6.7713 |
| AP3D1 | 6 | 0.44032 | 0.62864 | 0.999994 | 12581 | 2 | 2.868 | 0.0030653 | 0.01491 | 0.829051 | 338 | 3 | 2.868 |
| MLL3 | 6 | 0.98314 | 0.98308 | 0.999994 | 21158 | 0 | 2.9249 | 0.0030847 | 0.015001 | 0.829051 | 339 | 5 | 2.9249 |
| ZNF707 | 5 | 0.99691 | 0.99689 | 0.999994 | 21535 | 0 | 7.6074 | 0.0030899 | 0.013186 | 0.829051 | 340 | 5 | 7.6074 |
| DYRK1A | 6 | 0.86466 | 0.8801 | 0.999994 | 18535 | 1 | 6.993 | 0.0030976 | 0.015062 | 0.829051 | 341 | 5 | 6.993 |
| HSF1 | 5 | 0.9969 | 0.99688 | 0.999994 | 21534 | 0 | 3.1527 | 0.003101 | 0.013229 | 0.829051 | 342 | 5 | 3.1527 |
| C15orf61 | 6 | 0.99689 | 0.99688 | 0.999994 | 21533 | 0 | 4.931 | 0.0031074 | 0.015104 | 0.829051 | 343 | 6 | 4.931 |
| ABI3BP | 6 | 0.99688 | 0.99688 | 0.999994 | 21532 | 0 | 2.9318 | 0.0031166 | 0.015143 | 0.829051 | 344 | 6 | 2.9318 |
| KLHDC3 | 6 | 0.86213 | 0.87863 | 0.999994 | 18488 | 1 | 6.8628 | 0.003143 | 0.015255 | 0.829051 | 345 | 4 | 6.8628 |
| ZC3HAV1L | 6 | 0.99203 | 0.99198 | 0.999994 | 21396 | 0 | 6.9713 | 0.0031778 | 0.015407 | 0.829051 | 346 | 6 | 6.9713 |
| MYCBPAP | 6 | 0.99678 | 0.99677 | 0.999994 | 21531 | 0 | 5.1871 | 0.0032157 | 0.015571 | 0.829051 | 347 | 6 | 5.1871 |
| C22orf28 | 6 | 0.82883 | 0.86098 | 0.999994 | 17907 | 1 | 6.7129 | 0.0032256 | 0.015611 | 0.829051 | 348 | 5 | 6.7129 |
| CYFIP1 | 6 | 0.91647 | 0.91717 | 0.999994 | 19549 | 1 | 5.1497 | 0.0032338 | 0.015643 | 0.829051 | 349 | 5 | 5.1497 |
| TRIM34 | 6 | 0.14761 | 0.32264 | 0.999994 | 6393 | 2 | 6.7311 | 0.0032384 | 0.015665 | 0.829051 | 350 | 4 | 6.7311 |
| CSMD3 | 4 | 0.72689 | 0.74291 | 0.999994 | 16567 | 1 | 5.4184 | 0.0032387 | 0.011571 | 0.829051 | 351 | 3 | 5.4184 |
| KRTAP21-2 | 5 | 0.96323 | 0.96338 | 0.999994 | 20675 | 0 | 7.6967 | 0.0032491 | 0.013832 | 0.829051 | 352 | 4 | 7.6967 |
| PRAMEF6 | 2 | 0.97617 | 0.97615 | 0.999994 | 20995 | 0 | 8.444 | 0.0032502 | 0.0062389 | 0.785488 | 353 | 2 | 8.444 |
| PBDC1 | 6 | 0.083255 | 0.21398 | 0.979587 | 4352 | 1 | 6.8881 | 0.0032601 | 0.015757 | 0.829051 | 354 | 5 | 6.8881 |
| RPTOR | 6 | 0.50597 | 0.67287 | 0.999994 | 13523 | 1 | 7.0469 | 0.0032601 | 0.015757 | 0.829051 | 355 | 5 | 7.0469 |
| ATP5I | 6 | 0.99674 | 0.99672 | 0.999994 | 21530 | 0 | 4.7654 | 0.0032624 | 0.015764 | 0.829051 | 356 | 6 | 4.7654 |
| CCDC64B | 5 | 0.31311 | 0.49173 | 0.999994 | 10381 | 1 | 7.7042 | 0.0032628 | 0.013899 | 0.829051 | 357 | 4 | 7.7042 |
| PTPN13 | 4 | 0.96584 | 0.96573 | 0.999994 | 20733 | 0 | 5.0843 | 0.0032706 | 0.011686 | 0.829051 | 358 | 4 | 5.0843 |
| IL15RA | 5 | 0.24165 | 0.43315 | 0.999994 | 8821 | 2 | 0.26808 | 0.0032721 | 0.01393 | 0.829051 | 359 | 2 | 0.26808 |
| TCEB3 | 5 | 0.99672 | 0.9967 | 0.999994 | 21529 | 0 | 7.0807 | 0.0032788 | 0.013954 | 0.829051 | 360 | 5 | 7.0807 |
| ZNF131 | 5 | 0.9967 | 0.99667 | 0.999994 | 21527 | 0 | 4.3741 | 0.0033031 | 0.014042 | 0.829051 | 361 | 5 | 4.3741 |
| PCDHA1 | 6 | 0.21966 | 0.41168 | 0.999994 | 8285 | 3 | -0.67829 | 0.0033049 | 0.015945 | 0.829051 | 362 | 1 | -0.67829 |
| C18orf42 | 6 | 0.99669 | 0.99666 | 0.999994 | 21526 | 0 | 3.3744 | 0.0033062 | 0.015951 | 0.829051 | 363 | 6 | 3.3744 |

| | | | | | | | | | | | | | |
|---|---|---|---|---|---|---|---|---|---|---|---|---|---|
| LRAT | 5 | 0.91827 | 0.91836 | 0.999994 | 19590 | 0 | 1.9792 | 0.0033191 | 0.014114 | 0.829051 | 364 | 3 | 1.9792 |
| MTOR | 5 | 0.87057 | 0.87158 | 0.999994 | 18646 | 1 | 7.6983 | 0.0033222 | 0.014124 | 0.829051 | 365 | 4 | 7.6983 |
| HDC | 6 | 0.99665 | 0.99663 | 0.999994 | 21525 | 0 | 4.4081 | 0.0033466 | 0.016148 | 0.829051 | 366 | 6 | 4.4081 |
| ATG5 | 6 | 0.736 | 0.82363 | 0.999994 | 16676 | 1 | 1.0488 | 0.0033613 | 0.01622 | 0.829051 | 367 | 3 | 1.0488 |
| APOB | 6 | 0.52448 | 0.68564 | 0.999994 | 13809 | 2 | 4.7817 | 0.0033675 | 0.016251 | 0.829051 | 368 | 4 | 4.7817 |
| TTC3 | 6 | 0.43484 | 0.62502 | 0.999994 | 12494 | 1 | 5.5009 | 0.0033737 | 0.016274 | 0.829051 | 369 | 4 | 5.5009 |
| EHMT2 | 6 | 0.97827 | 0.97823 | 0.999994 | 21051 | 0 | 6.0184 | 0.0033875 | 0.016334 | 0.829051 | 370 | 5 | 6.0184 |
| KDM6A | 6 | 0.86178 | 0.87842 | 0.999994 | 18481 | 1 | 5.6165 | 0.0033892 | 0.016342 | 0.829051 | 371 | 5 | 5.6165 |
| GP6 | 6 | 0.99659 | 0.99656 | 0.999994 | 21523 | 0 | 1.8094 | 0.0034143 | 0.016466 | 0.833129 | 372 | 6 | 1.8094 |
| SOS1 | 6 | 0.03613 | 0.11315 | 0.951052 | 2325 | 1 | 5.569 | 0.0034306 | 0.016546 | 0.83326 | 373 | 5 | 5.569 |
| ZNF610 | 6 | 0.62773 | 0.76063 | 0.999994 | 15349 | 1 | 6.7664 | 0.0034428 | 0.016611 | 0.834612 | 374 | 5 | 6.7664 |
| CELA3A | 6 | 0.17487 | 0.3567 | 0.999994 | 7104 | 3 | -0.33227 | 0.0034741 | 0.01675 | 0.83545 | 375 | 3 | -0.33227 |
| HPN | 5 | 0.98626 | 0.98627 | 0.999994 | 21255 | 0 | 6.4524 | 0.0034957 | 0.014827 | 0.829051 | 376 | 5 | 6.4524 |
| PARD3B | 6 | 0.26001 | 0.45985 | 0.999994 | 9243 | 2 | 6.6103 | 0.0035125 | 0.016924 | 0.836675 | 377 | 4 | 6.6103 |
| MED20 | 5 | 0.62008 | 0.73359 | 0.999994 | 15221 | 1 | 5.8608 | 0.0035126 | 0.014882 | 0.829051 | 378 | 4 | 5.8608 |
| NR2C2AP | 6 | 0.99649 | 0.99646 | 0.999994 | 21522 | 0 | 4.5424 | 0.0035143 | 0.016931 | 0.836675 | 379 | 6 | 4.5424 |
| MED13L | 6 | 0.47907 | 0.65455 | 0.999994 | 13130 | 1 | 5.7564 | 0.0035247 | 0.016974 | 0.836675 | 380 | 5 | 5.7564 |
| COPE | 3 | 0.99646 | 0.99652 | 0.999994 | 21520 | 0 | 6.2985 | 0.0035424 | 0.0097983 | 0.825215 | 381 | 3 | 6.2985 |
| PPP1R15A | 6 | 0.99645 | 0.99643 | 0.999994 | 21519 | 0 | 1.9112 | 0.0035462 | 0.017081 | 0.836675 | 382 | 6 | 1.9112 |
| DPH1 | 6 | 0.99629 | 0.99627 | 0.999994 | 21510 | 0 | 7.0138 | 0.0035474 | 0.017089 | 0.836675 | 383 | 6 | 7.0138 |
| NAP1L4 | 5 | 0.79538 | 0.81664 | 0.999994 | 17424 | 1 | -0.39765 | 0.0035542 | 0.015046 | 0.829051 | 384 | 2 | -0.39765 |
| DDB2 | 6 | 0.054958 | 0.15482 | 0.967126 | 3196 | 1 | 5.7637 | 0.0035789 | 0.017247 | 0.836675 | 385 | 5 | 5.7637 |
| FLVCR2 | 5 | 0.30184 | 0.48254 | 0.999994 | 10159 | 2 | 7.6627 | 0.00358 | 0.015156 | 0.829051 | 386 | 3 | 7.6627 |
| MRAS | 6 | 0.99641 | 0.99639 | 0.999994 | 21518 | 0 | 5.8413 | 0.0035865 | 0.017287 | 0.836675 | 387 | 6 | 5.8413 |
| KLK6 | 6 | 0.99641 | 0.99639 | 0.999994 | 21517 | 0 | 2.7737 | 0.0035943 | 0.017323 | 0.836675 | 388 | 6 | 2.7737 |
| CTU2 | 3 | 0.9964 | 0.99646 | 0.999994 | 21515 | 0 | 6.7532 | 0.0036016 | 0.0099588 | 0.825215 | 389 | 3 | 6.7532 |
| CYP2C19 | 6 | 0.99598 | 0.99597 | 0.999994 | 21503 | 0 | 4.8043 | 0.0036092 | 0.017394 | 0.836675 | 390 | 6 | 4.8043 |
| KBTBD6 | 6 | 0.3671 | 0.58141 | 0.999994 | 11460 | 1 | 7.0871 | 0.0036121 | 0.017404 | 0.836675 | 391 | 4 | 7.0871 |
| COL5A1 | 4 | 0.70206 | 0.72577 | 0.999994 | 16327 | 1 | 7.64 | 0.0036255 | 0.01294 | 0.829051 | 392 | 3 | 7.64 |
| SOD3 | 6 | 0.66437 | 0.78869 | 0.999994 | 15846 | 1 | 4.5834 | 0.003627 | 0.017466 | 0.836675 | 393 | 4 | 4.5834 |
| FBXO8 | 6 | 0.42424 | 0.61817 | 0.999994 | 12316 | 2 | 5.1955 | 0.0036284 | 0.017474 | 0.836675 | 394 | 4 | 5.1955 |
| HIST1H1B | 5 | 0.44143 | 0.59798 | 0.999994 | 12602 | 1 | 5.7482 | 0.0036352 | 0.015364 | 0.829051 | 395 | 4 | 5.7482 |
| TEX9 | 6 | 0.99635 | 0.99634 | 0.999994 | 21513 | 0 | 3.4927 | 0.0036518 | 0.017579 | 0.836675 | 396 | 6 | 3.4927 |
| AOC2 | 5 | 0.99635 | 0.99631 | 0.999994 | 21512 | 0 | 4.9089 | 0.0036532 | 0.015435 | 0.829051 | 397 | 5 | 4.9089 |
| CXXC5 | 5 | 0.91185 | 0.91186 | 0.999994 | 19460 | 0 | 7.6506 | 0.0036765 | 0.015528 | 0.829051 | 398 | 3 | 7.6506 |
| OTOF | 5 | 0.96902 | 0.96911 | 0.999994 | 20803 | 0 | 6.1806 | 0.0036767 | 0.015528 | 0.829051 | 399 | 4 | 6.1806 |
| MED19 | 4 | 0.7654 | 0.77207 | 0.999994 | 17034 | 1 | 7.2845 | 0.003677 | 0.013121 | 0.829051 | 400 | 3 | 7.2845 |
| USP12 | 6 | 0.049602 | 0.14317 | 0.960171 | 2960 | 1 | 4.3251 | 0.0036784 | 0.017709 | 0.836675 | 401 | 5 | 4.3251 |
| UNC13C | 6 | 0.2626 | 0.46295 | 0.999994 | 9303 | 2 | 4.7502 | 0.0036784 | 0.017709 | 0.836675 | 402 | 4 | 4.7502 |
| SETBP1 | 6 | 0.9963 | 0.99629 | 0.999994 | 21511 | 0 | 6.0269 | 0.0036963 | 0.017798 | 0.836675 | 403 | 6 | 6.0269 |
| OLIG1 | 6 | 0.070394 | 0.18762 | 0.969543 | 3849 | 3 | 1.4901 | 0.0036997 | 0.017812 | 0.836675 | 404 | 3 | 1.4901 |
| CBX2 | 6 | 0.57832 | 0.72396 | 0.999994 | 14662 | 1 | 7.2296 | 0.003706 | 0.017838 | 0.836675 | 405 | 5 | 7.2296 |
| TPX2 | 5 | 0.97765 | 0.97774 | 0.999994 | 21035 | 0 | 6.7072 | 0.0037198 | 0.015691 | 0.829051 | 406 | 5 | 6.7072 |
| BEST2 | 6 | 0.98951 | 0.98947 | 0.999994 | 21327 | 0 | 5.1274 | 0.0037217 | 0.017909 | 0.836675 | 407 | 6 | 5.1274 |
| DET1 | 6 | 0.99626 | 0.99625 | 0.999994 | 21509 | 0 | 6.68 | 0.0037353 | 0.01797 | 0.836675 | 408 | 6 | 6.68 |
| RIMBP2 | 5 | 0.93397 | 0.93421 | 0.999994 | 19953 | 0 | 2.2413 | 0.0037422 | 0.015776 | 0.829051 | 409 | 4 | 2.2413 |
| LIPF | 6 | 0.96352 | 0.96358 | 0.999994 | 20687 | 0 | 5.4707 | 0.0037683 | 0.018121 | 0.836675 | 410 | 5 | 5.4707 |
| HECTD1 | 5 | 0.73049 | 0.78089 | 0.999994 | 16606 | 1 | 7.6383 | 0.0037815 | 0.015933 | 0.829051 | 411 | 4 | 7.6383 |
| ANKRD33B | 6 | 0.07098 | 0.18884 | 0.969543 | 3873 | 2 | 4.5106 | 0.0037912 | 0.018226 | 0.836675 | 412 | 4 | 4.5106 |
| RSPH9 | 6 | 0.53186 | 0.69087 | 0.999994 | 13933 | 2 | 3.9908 | 0.0037989 | 0.018264 | 0.836675 | 413 | 4 | 3.9908 |
| SPNS2 | 6 | 0.97913 | 0.97903 | 0.999994 | 21076 | 0 | 5.0457 | 0.0038125 | 0.018335 | 0.836675 | 414 | 5 | 5.0457 |
| GSDMA | 6 | 0.9228 | 0.92303 | 0.999994 | 19694 | 1 | 4.5171 | 0.0038226 | 0.018381 | 0.836675 | 415 | 4 | 4.5171 |

| | | | | | | | | | | | | |
|---|---|---|---|---|---|---|---|---|---|---|---|---|
| PAPL | 6 | 0.98366 | 0.98361 | 0.999994 | 21174 | 0 | 4.3044 | 0.0038262 | 0.018397 | 0.836675 | 416 | 5 | 4.3044 |
| NHLRC3 | 5 | 0.56534 | 0.70541 | 0.999994 | 14475 | 2 | 0.58469 | 0.0038362 | 0.01616 | 0.829051 | 417 | 3 | 0.58469 |
| MIF | 6 | 0.89427 | 0.89936 | 0.999994 | 19107 | 1 | 5.3403 | 0.0038456 | 0.018485 | 0.836675 | 418 | 5 | 5.3403 |
| GRAMD1B | 6 | 0.028092 | 0.094295 | 0.945031 | 1909 | 1 | 2.2208 | 0.0038464 | 0.018492 | 0.836675 | 419 | 3 | 2.2208 |
| RECQL5 | 4 | 0.80941 | 0.80975 | 0.999994 | 17623 | 1 | 7.6039 | 0.0038474 | 0.013712 | 0.829051 | 420 | 3 | 7.6039 |
| SPANXE | 2 | 0.99615 | 0.99609 | 0.999994 | 21508 | 0 | 8.2225 | 0.0038513 | 0.0074131 | 0.806682 | 421 | 2 | 8.2225 |
| ANP32A | 6 | 0.99615 | 0.99614 | 0.999994 | 21507 | 0 | 2.2963 | 0.0038521 | 0.018508 | 0.836675 | 422 | 6 | 2.2963 |
| LBX1 | 5 | 0.99614 | 0.99611 | 0.999994 | 21506 | 0 | 2.9792 | 0.0038582 | 0.016257 | 0.829051 | 423 | 5 | 2.9792 |
| ABCE1 | 5 | 0.98659 | 0.98659 | 0.999994 | 21265 | 0 | 6.9665 | 0.0038807 | 0.01634 | 0.829051 | 424 | 5 | 6.9665 |
| PCTP | 5 | 0.90524 | 0.90534 | 0.999994 | 19338 | 0 | 4.5294 | 0.0038832 | 0.016347 | 0.829051 | 425 | 3 | 4.5294 |
| MAMDC4 | 4 | 0.80624 | 0.80686 | 0.999994 | 17574 | 1 | 7.4299 | 0.0039135 | 0.013919 | 0.829051 | 426 | 3 | 7.4299 |
| NXPH4 | 4 | 0.42874 | 0.57525 | 0.999994 | 12391 | 2 | 2.3315 | 0.0039729 | 0.014118 | 0.829051 | 427 | 2 | 2.3315 |
| MED14 | 2 | 0.99603 | 0.99597 | 0.999994 | 21505 | 0 | 8.7867 | 0.0039739 | 0.0076375 | 0.811952 | 428 | 2 | 8.7867 |
| PEX11G | 5 | 0.10195 | 0.22947 | 0.983354 | 5036 | 3 | -1.6055 | 0.0039772 | 0.016724 | 0.83545 | 429 | 2 | -1.6055 |
| ZNF649 | 5 | 0.28355 | 0.46773 | 0.999994 | 9775 | 2 | 7.6074 | 0.0039879 | 0.016769 | 0.83545 | 430 | 3 | 7.6074 |
| SERPINB2 | 5 | 0.37998 | 0.5467 | 0.999994 | 11648 | 2 | 4.1171 | 0.0039921 | 0.016782 | 0.83545 | 431 | 3 | 4.1171 |
| RPS6KB1 | 6 | 0.9865 | 0.98646 | 0.999994 | 21263 | 0 | 6.8352 | 0.0039986 | 0.019151 | 0.849293 | 432 | 5 | 6.8352 |
| C3orf72 | 6 | 0.99599 | 0.99598 | 0.999994 | 21504 | 0 | 2.8496 | 0.0040128 | 0.019217 | 0.849293 | 433 | 6 | 2.8496 |
| MAGEE2 | 4 | 0.99597 | 0.99598 | 0.999994 | 21502 | 0 | 7.2131 | 0.0040307 | 0.014314 | 0.829051 | 434 | 4 | 7.2131 |
| TUB | 6 | 0.51294 | 0.67766 | 0.999994 | 13620 | 2 | 6.8805 | 0.0040435 | 0.019345 | 0.851882 | 435 | 4 | 6.8805 |
| GUCA1B | 4 | 0.93274 | 0.93249 | 0.999994 | 19918 | 0 | 5.9371 | 0.0040481 | 0.014371 | 0.829051 | 436 | 3 | 5.9371 |
| ZBTB14 | 6 | 0.66732 | 0.79104 | 0.999994 | 15882 | 1 | 3.1029 | 0.0040594 | 0.019416 | 0.851882 | 437 | 4 | 3.1029 |
| FOXI3 | 4 | 0.29202 | 0.43485 | 0.999994 | 9953 | 1 | 7.616 | 0.0040827 | 0.014488 | 0.829051 | 438 | 3 | 7.616 |
| GIMAP5 | 6 | 0.97697 | 0.9769 | 0.999994 | 21013 | 0 | 6.1354 | 0.0040942 | 0.019573 | 0.851882 | 439 | 5 | 6.1354 |
| HTATIP2 | 6 | 0.19128 | 0.3771 | 0.999994 | 7520 | 3 | -0.12724 | 0.0040944 | 0.019574 | 0.851882 | 440 | 3 | -0.12724 |
| ERAS | 5 | 0.9959 | 0.99587 | 0.999994 | 21501 | 0 | 3.0938 | 0.0041018 | 0.017237 | 0.836675 | 441 | 5 | 3.0938 |
| NMT2 | 6 | 0.45732 | 0.63992 | 0.999994 | 12812 | 2 | 1.5826 | 0.0041353 | 0.019763 | 0.851882 | 442 | 4 | 1.5826 |
| IFITM3 | 5 | 0.77162 | 0.80263 | 0.999994 | 17109 | 1 | 6.3173 | 0.0041376 | 0.017373 | 0.836675 | 443 | 4 | 6.3173 |
| SP110 | 6 | 0.82539 | 0.85934 | 0.999994 | 17858 | 1 | 1.7442 | 0.0041508 | 0.019838 | 0.851882 | 444 | 5 | 1.7442 |
| NLGN4X | 5 | 0.67262 | 0.7544 | 0.999994 | 15965 | 1 | 0.073499 | 0.0041652 | 0.01749 | 0.836675 | 445 | 1 | 0.073499 |
| KIAA1279 | 6 | 0.92398 | 0.92414 | 0.999994 | 19713 | 1 | 5.6118 | 0.0041714 | 0.019934 | 0.851882 | 446 | 5 | 5.6118 |
| USP9X | 6 | 0.39412 | 0.59858 | 0.999994 | 11863 | 1 | 6.7089 | 0.0042008 | 0.020051 | 0.851882 | 447 | 5 | 6.7089 |
| SPAM1 | 6 | 0.47083 | 0.64901 | 0.999994 | 13016 | 2 | 4.6861 | 0.0042066 | 0.020079 | 0.851882 | 448 | 4 | 4.6861 |
| ZSCAN1 | 6 | 0.94535 | 0.94533 | 0.999994 | 20214 | 0 | 2.9072 | 0.0042119 | 0.020099 | 0.851882 | 449 | 4 | 2.9072 |
| DMKN | 6 | 0.99505 | 0.99504 | 0.999994 | 21479 | 0 | 4.9001 | 0.0042309 | 0.020181 | 0.853037 | 450 | 6 | 4.9001 |
| C19orf12 | 4 | 0.99576 | 0.99577 | 0.999994 | 21500 | 0 | 5.2935 | 0.0042351 | 0.015001 | 0.829051 | 451 | 4 | 5.2935 |
| MAPK14 | 6 | 0.97553 | 0.97545 | 0.999994 | 20977 | 0 | 6.0721 | 0.0042723 | 0.020374 | 0.853037 | 452 | 4 | 6.0721 |
| ZNF497 | 6 | 0.97512 | 0.97503 | 0.999994 | 20956 | 0 | 5.1789 | 0.0042723 | 0.020374 | 0.853037 | 453 | 5 | 5.1789 |
| PLEK2 | 6 | 0.84043 | 0.86684 | 0.999994 | 18093 | 1 | 6.6065 | 0.0042877 | 0.020436 | 0.853037 | 454 | 5 | 6.6065 |
| C14orf183 | 5 | 0.9957 | 0.99568 | 0.999994 | 21499 | 0 | 3.7891 | 0.0042969 | 0.018007 | 0.836675 | 455 | 5 | 3.7891 |
| RUNX2 | 6 | 0.89719 | 0.90149 | 0.999994 | 19164 | 1 | 6.8417 | 0.004298 | 0.020484 | 0.853037 | 456 | 5 | 6.8417 |
| hsa-mir-8085 | 4 | 0.93184 | 0.93158 | 0.999994 | 19896 | 0 | 4.0034 | 0.0043113 | 0.015261 | 0.829051 | 457 | 3 | 4.0034 |
| TPRN | 5 | 0.99569 | 0.99566 | 0.999994 | 21498 | 0 | 5.5158 | 0.0043126 | 0.018064 | 0.836675 | 458 | 5 | 5.5158 |
| RHEBL1 | 4 | 0.90576 | 0.90549 | 0.999994 | 19347 | 0 | 7.2641 | 0.0043147 | 0.015271 | 0.829051 | 459 | 3 | 7.2641 |
| NUMA1 | 6 | 0.2644 | 0.46507 | 0.999994 | 9344 | 3 | 1.0896 | 0.0043199 | 0.020585 | 0.853752 | 460 | 3 | 1.0896 |
| BAIAP2L1 | 6 | 0.27951 | 0.48284 | 0.999994 | 9693 | 1 | 5.3015 | 0.0043348 | 0.020649 | 0.853752 | 461 | 5 | 5.3015 |
| hsa-mir-5690 | 4 | 0.12021 | 0.25066 | 0.992497 | 5636 | 2 | -1.6463 | 0.0043489 | 0.015397 | 0.829051 | 462 | 1 | -1.6463 |
| RAMP2 | 5 | 0.31047 | 0.48951 | 0.999994 | 10336 | 1 | 5.2982 | 0.0043531 | 0.018236 | 0.836675 | 463 | 4 | 5.2982 |
| C1orf131 | 5 | 0.20958 | 0.3888 | 0.999994 | 8019 | 1 | 7.5564 | 0.0043671 | 0.018302 | 0.836675 | 464 | 4 | 7.5564 |
| hsa-mir-4274 | 4 | 0.99563 | 0.99564 | 0.999994 | 21497 | 0 | 5.9953 | 0.0043688 | 0.015466 | 0.829051 | 465 | 4 | 5.9953 |
| C6orf163 | 6 | 0.99563 | 0.99561 | 0.999994 | 21496 | 0 | 3.2016 | 0.0043735 | 0.020825 | 0.853752 | 466 | 6 | 3.2016 |
| RAB20 | 6 | 0.56114 | 0.71162 | 0.999994 | 14391 | 2 | 2.07 | 0.0043763 | 0.020835 | 0.853752 | 467 | 3 | 2.07 |

| | | | | | | | | | | | | | |
|---|---|---|---|---|---|---|---|---|---|---|---|---|---|
| MOAP1 | 5 | 0.99562 | 0.99559 | 0.999994 | 21495 | 0 | 3.9222 | 0.0043826 | 0.018366 | 0.836675 | 468 | 5 | 3.9222 |
| UBAP1 | 6 | 0.63783 | 0.76828 | 0.999994 | 15485 | 2 | 4.2645 | 0.0044017 | 0.020943 | 0.853752 | 469 | 4 | 4.2645 |
| XRCC4 | 6 | 0.98237 | 0.98231 | 0.999994 | 21145 | 0 | 6.7961 | 0.0044033 | 0.020952 | 0.853752 | 470 | 5 | 6.7961 |
| CLEC18B | 4 | 0.99559 | 0.9956 | 0.999994 | 21494 | 0 | 5.9117 | 0.004409 | 0.0156 | 0.829051 | 471 | 4 | 5.9117 |
| TPO | 6 | 0.97066 | 0.97062 | 0.999994 | 20839 | 0 | 3.9314 | 0.0044156 | 0.021001 | 0.853752 | 472 | 5 | 3.9314 |
| MAP4K1 | 6 | 0.99557 | 0.99556 | 0.999994 | 21493 | 0 | 3.1493 | 0.004432 | 0.021074 | 0.853752 | 473 | 6 | 3.1493 |
| PDCL | 6 | 0.90999 | 0.91156 | 0.999994 | 19425 | 1 | 0.97604 | 0.0044326 | 0.021076 | 0.853752 | 474 | 3 | 0.97604 |
| ZMPSTE24 | 6 | 0.44121 | 0.62923 | 0.999994 | 12595 | 2 | 4.1137 | 0.0044364 | 0.021092 | 0.853752 | 475 | 3 | 4.1137 |
| EDDM3A | 5 | 0.99556 | 0.99553 | 0.999994 | 21492 | 0 | 4.5924 | 0.0044392 | 0.018576 | 0.836675 | 476 | 5 | 4.5924 |
| CAMLG | 5 | 0.52372 | 0.66857 | 0.999994 | 13804 | 1 | 1.5218 | 0.0044471 | 0.018613 | 0.836675 | 477 | 4 | 1.5218 |
| PDCD6IP | 6 | 0.92906 | 0.92915 | 0.999994 | 19836 | 0 | 5.2439 | 0.0044967 | 0.021347 | 0.85539 | 478 | 5 | 5.2439 |
| MFAP5 | 6 | 0.99548 | 0.99547 | 0.999994 | 21490 | 0 | 2.8397 | 0.004525 | 0.021485 | 0.85539 | 479 | 6 | 2.8397 |
| NAPSA | 6 | 0.98991 | 0.98986 | 0.999994 | 21342 | 0 | 4.4887 | 0.0045337 | 0.021522 | 0.85539 | 480 | 6 | 4.4887 |
| CROT | 6 | 0.14541 | 0.31988 | 0.999994 | 6335 | 3 | 1.0309 | 0.0045454 | 0.021579 | 0.85539 | 481 | 3 | 1.0309 |
| PPAPDC3 | 6 | 0.73964 | 0.82487 | 0.999994 | 16734 | 1 | 6.4261 | 0.004552 | 0.021606 | 0.85539 | 482 | 5 | 6.4261 |
| C15orf40 | 6 | 0.60403 | 0.74295 | 0.999994 | 14996 | 1 | 6.2561 | 0.0045613 | 0.021646 | 0.85539 | 483 | 5 | 6.2561 |
| EIF2B4 | 4 | 0.99544 | 0.99545 | 0.999994 | 21489 | 0 | 6.752 | 0.0045622 | 0.01614 | 0.829051 | 484 | 4 | 6.752 |
| ZMIZ1 | 5 | 0.33448 | 0.5092 | 0.999994 | 10823 | 2 | 4.8199 | 0.004588 | 0.019167 | 0.849293 | 485 | 3 | 4.8199 |
| CYB5D1 | 5 | 0.9954 | 0.99538 | 0.999994 | 21488 | 0 | 6.4076 | 0.0045966 | 0.019201 | 0.849293 | 486 | 5 | 6.4076 |
| USP35 | 6 | 0.97025 | 0.97022 | 0.999994 | 20829 | 0 | 7.0667 | 0.0046011 | 0.021828 | 0.85539 | 487 | 5 | 7.0667 |
| STK39 | 6 | 0.45948 | 0.64141 | 0.999994 | 12840 | 2 | 6.9359 | 0.0046011 | 0.021828 | 0.85539 | 488 | 4 | 6.9359 |
| SLC22A25 | 6 | 0.29576 | 0.50178 | 0.999994 | 10034 | 2 | 1.1879 | 0.0046073 | 0.021856 | 0.85539 | 489 | 3 | 1.1879 |
| ALPK1 | 6 | 0.50443 | 0.6718 | 0.999994 | 13502 | 1 | 6.942 | 0.004651 | 0.022055 | 0.858991 | 490 | 5 | 6.942 |
| FYCO1 | 5 | 0.85947 | 0.86196 | 0.999994 | 18425 | 1 | 7.517 | 0.0046741 | 0.019525 | 0.851882 | 491 | 3 | 7.517 |
| RHBDL3 | 5 | 0.15064 | 0.30375 | 0.999994 | 6477 | 1 | 1.5746 | 0.0046819 | 0.019551 | 0.851882 | 492 | 3 | 1.5746 |
| PLCB4 | 6 | 0.66153 | 0.78649 | 0.999994 | 15800 | 2 | 4.7181 | 0.0047144 | 0.022345 | 0.862524 | 493 | 4 | 4.7181 |
| CLNK | 5 | 0.25302 | 0.44316 | 0.999994 | 9083 | 2 | 3.5563 | 0.0047247 | 0.019725 | 0.851882 | 494 | 3 | 3.5563 |
| RIF1 | 6 | 0.97649 | 0.97643 | 0.999994 | 21004 | 0 | 4.7744 | 0.0047383 | 0.022452 | 0.863576 | 495 | 5 | 4.7744 |
| PDIA4 | 5 | 0.20024 | 0.37559 | 0.999994 | 7769 | 1 | 6.6037 | 0.0047507 | 0.019815 | 0.851882 | 496 | 4 | 6.6037 |
| FAM96A | 5 | 0.78786 | 0.81203 | 0.999994 | 17320 | 1 | 6.3955 | 0.0047507 | 0.019815 | 0.851882 | 497 | 4 | 6.3955 |
| MDM2 | 5 | 0.99523 | 0.9952 | 0.999994 | 21486 | 0 | 6.1768 | 0.0047748 | 0.019911 | 0.851882 | 498 | 5 | 6.1768 |
| LRRC42 | 6 | 0.99522 | 0.9952 | 0.999994 | 21485 | 0 | 2.3742 | 0.0047807 | 0.022631 | 0.863933 | 499 | 6 | 2.3742 |
| SH3BP1 | 5 | 0.99521 | 0.99519 | 0.999994 | 21484 | 0 | 5.6986 | 0.0047895 | 0.019963 | 0.851882 | 500 | 5 | 5.6986 |
| SLC39A10 | 6 | 0.59156 | 0.73378 | 0.999994 | 14842 | 1 | 4.1635 | 0.0047946 | 0.022685 | 0.863933 | 501 | 5 | 4.1635 |
| NonTargetingControlG | 2 | 0.9952 | 0.99511 | 0.999994 | 21483 | 0 | 8.1601 | 0.0047998 | 0.0092172 | 0.825215 | 502 | 2 | 8.1601 |
| OVCH1 | 6 | 0.99518 | 0.99516 | 0.999994 | 21482 | 0 | 5.8497 | 0.0048217 | 0.022808 | 0.863933 | 503 | 6 | 5.8497 |
| FUT2 | 6 | 0.90324 | 0.9062 | 0.999994 | 19284 | 1 | 5.7724 | 0.0048368 | 0.022878 | 0.863933 | 504 | 5 | 5.7724 |
| SLC5A2 | 4 | 0.47415 | 0.60493 | 0.999994 | 13052 | 1 | 1.314 | 0.0048375 | 0.017092 | 0.836675 | 505 | 3 | 1.314 |
| SLC6A5 | 6 | 0.99515 | 0.99513 | 0.999994 | 21481 | 0 | 3.3997 | 0.0048543 | 0.022958 | 0.863933 | 506 | 6 | 3.3997 |
| NPY1R | 5 | 0.32405 | 0.50063 | 0.999994 | 10600 | 1 | 0.82601 | 0.0048698 | 0.020298 | 0.853037 | 507 | 3 | 0.82601 |
| POGZ | 6 | 0.25279 | 0.45125 | 0.999994 | 9078 | 2 | 3.6338 | 0.0048778 | 0.023048 | 0.863933 | 508 | 4 | 3.6338 |
| GPATCH8 | 6 | 0.30226 | 0.50936 | 0.999994 | 10177 | 3 | 0.099005 | 0.0048834 | 0.023069 | 0.863933 | 509 | 3 | 0.099005 |
| SYT12 | 6 | 0.3148 | 0.52394 | 0.999994 | 10416 | 2 | 6.464 | 0.0048922 | 0.023115 | 0.863933 | 510 | 4 | 6.464 |
| TSFM | 4 | 0.55818 | 0.64371 | 0.999994 | 14337 | 1 | 7.2135 | 0.0049212 | 0.017386 | 0.836675 | 511 | 3 | 7.2135 |
| RABAC1 | 6 | 0.99507 | 0.99506 | 0.999994 | 21480 | 0 | 3.8906 | 0.0049261 | 0.023261 | 0.863933 | 512 | 6 | 3.8906 |
| FAM43A | 6 | 0.091082 | 0.22967 | 0.983354 | 4639 | 2 | 3.9342 | 0.0049393 | 0.02332 | 0.863933 | 513 | 3 | 3.9342 |
| RNH1 | 6 | 0.53772 | 0.69493 | 0.999994 | 14023 | 1 | 0.81582 | 0.0049398 | 0.023323 | 0.863933 | 514 | 3 | 0.81582 |
| C1orf86 | 5 | 0.4654 | 0.61821 | 0.999994 | 12941 | 1 | 5.7477 | 0.0049766 | 0.02073 | 0.853752 | 515 | 4 | 5.7477 |
| CTNND1 | 5 | 0.99499 | 0.99497 | 0.999994 | 21478 | 0 | 7.0303 | 0.0050114 | 0.020867 | 0.853752 | 516 | 5 | 7.0303 |
| P2RY8_X | 6 | 0.99498 | 0.99497 | 0.999994 | 21477 | 0 | 2.5914 | 0.0050173 | 0.023671 | 0.872794 | 517 | 6 | 2.5914 |
| FAT2 | 4 | 0.97331 | 0.9732 | 0.999994 | 20902 | 0 | 7.4088 | 0.0050185 | 0.017691 | 0.836675 | 518 | 4 | 7.4088 |
| CCDC47 | 4 | 0.97789 | 0.97786 | 0.999994 | 21037 | 0 | 7.3338 | 0.0050185 | 0.017691 | 0.836675 | 519 | 4 | 7.3338 |

| | | | | | | | | | | | | |
|---|---|---|---|---|---|---|---|---|---|---|---|---|
| OR6B2 | 6 | 0.99498 | 0.99497 | 0.999994 | 21476 | 0 | 4.4529 | 0.0050235 | 0.023699 | 0.872794 | 520 | 6 | 4.4529 |
| EHBP1L1 | 6 | 0.85136 | 0.87264 | 0.999994 | 18284 | 1 | 3.4934 | 0.0050525 | 0.023817 | 0.874165 | 521 | 4 | 3.4934 |
| DNAJA4 | 5 | 0.64335 | 0.74249 | 0.999994 | 15566 | 1 | 3.1413 | 0.0050654 | 0.021079 | 0.853752 | 522 | 3 | 3.1413 |
| GPR75-ASB3 | 3 | 0.99493 | 0.99504 | 0.999994 | 21474 | 0 | 7.0115 | 0.0050742 | 0.013974 | 0.829051 | 523 | 3 | 7.0115 |
| NTN5 | 6 | 0.18739 | 0.37228 | 0.999994 | 7404 | 3 | -0.19761 | 0.0051088 | 0.024059 | 0.880714 | 524 | 3 | -0.19761 |
| RPS4Y2 | 6 | 0.58303 | 0.72743 | 0.999994 | 14716 | 1 | 6.9857 | 0.0051364 | 0.024187 | 0.880714 | 525 | 5 | 6.9857 |
| hsa-mir-4292 | 4 | 0.98684 | 0.98679 | 0.999994 | 21274 | 0 | 5.8701 | 0.0051599 | 0.018159 | 0.836675 | 526 | 4 | 5.8701 |
| MED16 | 6 | 0.93417 | 0.93424 | 0.999994 | 19957 | 0 | 2.629 | 0.0051606 | 0.024292 | 0.881213 | 527 | 4 | 2.629 |
| P2RX1 | 6 | 0.94388 | 0.94385 | 0.999994 | 20174 | 0 | 0.54293 | 0.0051651 | 0.024311 | 0.881213 | 528 | 3 | 0.54293 |
| C4orf40 | 5 | 0.36529 | 0.53449 | 0.999994 | 11427 | 1 | 7.4641 | 0.0051675 | 0.02148 | 0.85539 | 529 | 4 | 7.4641 |
| FAM229B | 5 | 0.04519 | 0.12878 | 0.951777 | 2772 | 3 | -1.023 | 0.0051794 | 0.021522 | 0.85539 | 530 | 2 | -1.023 |
| UBE2Q2 | 5 | 0.61738 | 0.7326 | 0.999994 | 15176 | 1 | 6.9546 | 0.0051956 | 0.021585 | 0.85539 | 531 | 4 | 6.9546 |
| SPDL1 | 3 | 0.13539 | 0.22962 | 0.983354 | 6062 | 1 | 5.3884 | 0.0052096 | 0.014345 | 0.829051 | 532 | 2 | 5.3884 |
| RBM17 | 5 | 0.44909 | 0.60449 | 0.999994 | 12702 | 1 | 6.0521 | 0.0052312 | 0.021714 | 0.85539 | 533 | 4 | 6.0521 |
| TCEA2 | 5 | 0.99477 | 0.99475 | 0.999994 | 21473 | 0 | 3.3379 | 0.0052346 | 0.021729 | 0.85539 | 534 | 5 | 3.3379 |
| DCAF15 | 6 | 0.99476 | 0.99473 | 0.999994 | 21472 | 0 | 2.2133 | 0.0052352 | 0.024634 | 0.882096 | 535 | 6 | 2.2133 |
| CSTF1 | 3 | 0.99473 | 0.99482 | 0.999994 | 21471 | 0 | 5.6779 | 0.0052726 | 0.01452 | 0.829051 | 536 | 3 | 5.6779 |
| MACF1 | 5 | 0.99473 | 0.99471 | 0.999994 | 21470 | 0 | 2.2475 | 0.005275 | 0.021883 | 0.85539 | 537 | 5 | 2.2475 |
| IFNA8 | 6 | 0.045499 | 0.13416 | 0.95318 | 2791 | 3 | 2.5499 | 0.0052778 | 0.024836 | 0.885141 | 538 | 3 | 2.5499 |
| NFKBIB | 6 | 0.9947 | 0.99467 | 0.999994 | 21469 | 0 | 3.7302 | 0.0053024 | 0.024934 | 0.885141 | 539 | 6 | 3.7302 |
| POLR2L | 4 | 0.99469 | 0.99468 | 0.999994 | 21467 | 0 | 6.1795 | 0.0053052 | 0.018661 | 0.836675 | 540 | 4 | 6.1795 |
| TCF19 | 4 | 0.99469 | 0.99468 | 0.999994 | 21466 | 0 | 5.3601 | 0.0053052 | 0.018661 | 0.836675 | 541 | 4 | 5.3601 |
| ENGASE | 5 | 0.75617 | 0.79411 | 0.999994 | 16932 | 1 | 6.6037 | 0.0053341 | 0.022123 | 0.859479 | 542 | 4 | 6.6037 |
| C3orf62 | 5 | 0.5313 | 0.67522 | 0.999994 | 13917 | 1 | 1.9917 | 0.0053393 | 0.022147 | 0.859479 | 543 | 3 | 1.9917 |
| CXorf64 | 6 | 0.99465 | 0.99462 | 0.999994 | 21465 | 0 | 5.2554 | 0.0053463 | 0.025126 | 0.885141 | 544 | 6 | 5.2554 |
| C2orf49 | 6 | 0.54724 | 0.70167 | 0.999994 | 14170 | 1 | 6.579 | 0.0053532 | 0.025152 | 0.885141 | 545 | 4 | 6.579 |
| SMIM17 | 5 | 0.98725 | 0.98723 | 0.999994 | 21285 | 0 | 5.8558 | 0.0053652 | 0.022251 | 0.861962 | 546 | 5 | 5.8558 |
| FDPS | 3 | 0.99463 | 0.99472 | 0.999994 | 21464 | 0 | 6.7418 | 0.0053688 | 0.014775 | 0.829051 | 547 | 3 | 6.7418 |
| EIF1 | 5 | 0.34038 | 0.51403 | 0.999994 | 10942 | 1 | 6.4592 | 0.0053777 | 0.022296 | 0.862191 | 548 | 4 | 6.4592 |
| HVCN1 | 6 | 0.10178 | 0.25079 | 0.992497 | 5031 | 1 | 1.0682 | 0.0053904 | 0.025316 | 0.885141 | 549 | 3 | 1.0682 |
| DYNLT1 | 5 | 0.98366 | 0.98373 | 0.999994 | 21175 | 0 | 7.0852 | 0.0054027 | 0.022397 | 0.862998 | 550 | 5 | 7.0852 |
| CALML6 | 6 | 0.10461 | 0.25632 | 0.994943 | 5122 | 3 | 2.8696 | 0.0054093 | 0.025393 | 0.885141 | 551 | 3 | 2.8696 |
| NALCN | 6 | 0.26197 | 0.46222 | 0.999994 | 9287 | 1 | 4.0633 | 0.0054101 | 0.025398 | 0.885141 | 552 | 5 | 4.0633 |
| CENPV | 4 | 0.99455 | 0.99453 | 0.999994 | 21463 | 0 | 4.8766 | 0.0054493 | 0.019133 | 0.849293 | 553 | 4 | 4.8766 |
| NDUFAF1 | 6 | 0.84746 | 0.8706 | 0.999994 | 18212 | 1 | 3.8507 | 0.0054663 | 0.025635 | 0.885381 | 554 | 5 | 3.8507 |
| CDC27 | 3 | 0.99453 | 0.99462 | 0.999994 | 21461 | 0 | 6.5071 | 0.0054678 | 0.01502 | 0.829051 | 555 | 3 | 6.5071 |
| RBM8A | 3 | 0.99453 | 0.99462 | 0.999994 | 21462 | 0 | 7.3411 | 0.0054678 | 0.01502 | 0.829051 | 556 | 3 | 7.3411 |
| ONECUT2 | 6 | 0.98385 | 0.98379 | 0.999994 | 21184 | 0 | 4.6954 | 0.0055141 | 0.025859 | 0.890263 | 557 | 5 | 4.6954 |
| CAPN1 | 6 | 0.99447 | 0.99444 | 0.999994 | 21460 | 0 | 4.4641 | 0.0055268 | 0.02591 | 0.890437 | 558 | 6 | 4.4641 |
| C12orf74 | 5 | 0.1605 | 0.31829 | 0.999994 | 6750 | 3 | -1.0789 | 0.005527 | 0.022863 | 0.863933 | 559 | 1 | -1.0789 |
| CDK5R1 | 5 | 0.20298 | 0.37946 | 0.999994 | 7854 | 2 | 3.0801 | 0.0055519 | 0.022968 | 0.863933 | 560 | 3 | 3.0801 |
| FOXJ3 | 6 | 0.9225 | 0.92275 | 0.999994 | 19688 | 1 | 3.1639 | 0.0055593 | 0.026039 | 0.890437 | 561 | 5 | 3.1639 |
| PIK3CB | 6 | 0.19688 | 0.38409 | 0.999994 | 7672 | 2 | 0.39048 | 0.0055821 | 0.026136 | 0.890437 | 562 | 2 | 0.39048 |
| MZF1 | 5 | 0.9944 | 0.99438 | 0.999994 | 21459 | 0 | 2.6655 | 0.0055963 | 0.023133 | 0.863933 | 563 | 5 | 2.6655 |
| SLC17A7 | 6 | 0.99439 | 0.99436 | 0.999994 | 21458 | 0 | 2.2784 | 0.0056143 | 0.02627 | 0.890437 | 564 | 6 | 2.2784 |
| CCL2 | 6 | 0.99438 | 0.99436 | 0.999994 | 21457 | 0 | 2.6875 | 0.0056204 | 0.026292 | 0.890437 | 565 | 6 | 2.6875 |
| USF2 | 5 | 0.95217 | 0.95232 | 0.999994 | 20387 | 0 | 7.4127 | 0.0056336 | 0.023287 | 0.863933 | 566 | 4 | 7.4127 |
| ETHE1 | 5 | 0.46403 | 0.61706 | 0.999994 | 12920 | 2 | 1.0704 | 0.0056475 | 0.023339 | 0.863933 | 567 | 3 | 1.0704 |
| OR2T4 | 5 | 0.8293 | 0.83898 | 0.999994 | 17919 | 1 | 6.5883 | 0.0056475 | 0.023339 | 0.863933 | 568 | 4 | 6.5883 |
| HLA-DOB | 6 | 0.088772 | 0.22506 | 0.983354 | 4545 | 4 | -0.97961 | 0.005672 | 0.026539 | 0.890437 | 569 | 2 | -0.97961 |
| C19orf18 | 6 | 0.73295 | 0.8226 | 0.999994 | 16639 | 1 | 6.2686 | 0.0056806 | 0.026578 | 0.890437 | 570 | 5 | 6.2686 |
| BCL2 | 6 | 0.93699 | 0.93702 | 0.999994 | 20006 | 0 | 6.5522 | 0.0056806 | 0.026578 | 0.890437 | 571 | 4 | 6.5522 |

| | | | | | | | | | | | | |
|---|---|---|---|---|---|---|---|---|---|---|---|---|
| hsa-mir-7153 | 4 | 0.99431 | 0.9943 | 0.999994 | 21456 | 0 | 3.8853 | 0.0056861 | 0.019908 | 0.851882 | 572 | 4 | 3.8853 |
| hsa-mir-4746 | 2 | 0.60586 | 0.60567 | 0.999994 | 15025 | 0 | 4.9993 | 0.0056961 | 0.010913 | 0.825215 | 573 | 1 | 4.9993 |
| CSGALNACT2 | 6 | 0.9943 | 0.99427 | 0.999994 | 21454 | 0 | 1.3566 | 0.0057008 | 0.02667 | 0.890437 | 574 | 6 | 1.3566 |
| AFF2 | 5 | 0.90028 | 0.90039 | 0.999994 | 19225 | 0 | 7.4039 | 0.0057239 | 0.023627 | 0.872794 | 575 | 4 | 7.4039 |
| FGF11 | 6 | 0.61504 | 0.75111 | 0.999994 | 15145 | 1 | 3.1993 | 0.0057283 | 0.026792 | 0.890437 | 576 | 4 | 3.1993 |
| HK2 | 6 | 0.9942 | 0.99417 | 0.999994 | 21453 | 0 | 5.8234 | 0.0057451 | 0.02686 | 0.890437 | 577 | 6 | 5.8234 |
| CLYBL | 4 | 0.95323 | 0.9531 | 0.999994 | 20417 | 0 | 6.9842 | 0.005795 | 0.020268 | 0.853037 | 578 | 4 | 6.9842 |
| LYPD4 | 6 | 0.99419 | 0.99416 | 0.999994 | 21452 | 0 | 1.8208 | 0.0058092 | 0.027147 | 0.890437 | 579 | 6 | 1.8208 |
| TULP2 | 6 | 0.55482 | 0.70709 | 0.999994 | 14280 | 1 | 5.9828 | 0.0058092 | 0.027147 | 0.890437 | 580 | 5 | 5.9828 |
| OR5I1 | 6 | 0.99419 | 0.99416 | 0.999994 | 21451 | 0 | 3.8993 | 0.00581 | 0.027149 | 0.890437 | 581 | 6 | 3.8993 |
| CKM | 6 | 0.99418 | 0.99415 | 0.999994 | 21450 | 0 | 3.1 | 0.0058201 | 0.027187 | 0.890437 | 582 | 6 | 3.1 |
| AGAP11 | 6 | 0.91419 | 0.91519 | 0.999994 | 19492 | 1 | 3.8525 | 0.0058305 | 0.027229 | 0.890437 | 583 | 3 | 3.8525 |
| PTMS | 5 | 0.98923 | 0.98926 | 0.999994 | 21325 | 0 | 5.3289 | 0.0058376 | 0.024091 | 0.880714 | 584 | 5 | 5.3289 |
| hsa-mir-6841 | 4 | 0.78755 | 0.79026 | 0.999994 | 17313 | 1 | 4.0358 | 0.0058513 | 0.020448 | 0.853037 | 585 | 2 | 4.0358 |
| TFAM | 5 | 0.15555 | 0.31104 | 0.999994 | 6609 | 1 | 0.61345 | 0.0058555 | 0.024164 | 0.880714 | 586 | 3 | 0.61345 |
| PACSIN2 | 5 | 0.99414 | 0.99412 | 0.999994 | 21449 | 0 | 3.5149 | 0.0058632 | 0.024199 | 0.880714 | 587 | 5 | 3.5149 |
| hsa-mir-361 | 3 | 0.82779 | 0.82761 | 0.999994 | 17893 | 0 | 8.2926 | 0.0058788 | 0.016102 | 0.829051 | 588 | 2 | 8.2926 |
| NUDCD2 | 6 | 0.021972 | 0.078291 | 0.93914 | 1579 | 3 | -0.57817 | 0.0058971 | 0.027548 | 0.890437 | 589 | 3 | -0.57817 |
| TMED10 | 5 | 0.037264 | 0.10899 | 0.951052 | 2386 | 2 | 0.87996 | 0.0059493 | 0.024538 | 0.882096 | 590 | 3 | 0.87996 |
| ZFP3 | 6 | 0.014814 | 0.055113 | 0.922677 | 1179 | 1 | 2.2243 | 0.0059523 | 0.027786 | 0.890437 | 591 | 4 | 2.2243 |
| C14orf166 | 5 | 0.5493 | 0.69113 | 0.999994 | 14199 | 1 | 6.3358 | 0.0060006 | 0.024733 | 0.883827 | 592 | 4 | 6.3358 |
| AATF | 4 | 0.13337 | 0.27205 | 0.998561 | 6006 | 2 | 3.5409 | 0.0060045 | 0.020943 | 0.853752 | 593 | 2 | 3.5409 |
| SNX22 | 6 | 0.22743 | 0.42099 | 0.999994 | 8464 | 1 | 4.9366 | 0.0060272 | 0.02812 | 0.890437 | 594 | 5 | 4.9366 |
| IPO11 | 6 | 0.99394 | 0.9939 | 0.999994 | 21447 | 0 | 5.3314 | 0.0060577 | 0.028244 | 0.890437 | 595 | 6 | 5.3314 |
| ZNF527 | 6 | 0.99391 | 0.99386 | 0.999994 | 21445 | 0 | 3.3635 | 0.0060923 | 0.028402 | 0.890437 | 596 | 6 | 3.3635 |
| OR5D18 | 6 | 0.99389 | 0.99384 | 0.999994 | 21444 | 0 | 1.6413 | 0.0061052 | 0.028465 | 0.890437 | 597 | 6 | 1.6413 |
| LSM14A | 6 | 0.99389 | 0.99384 | 0.999994 | 21443 | 0 | 2.3118 | 0.0061116 | 0.028494 | 0.890437 | 598 | 6 | 2.3118 |
| SCUBE1 | 4 | 0.72367 | 0.74064 | 0.999994 | 16530 | 1 | 4.4827 | 0.0061145 | 0.021304 | 0.85539 | 599 | 3 | 4.4827 |
| SYT16 | 6 | 0.17026 | 0.35109 | 0.999994 | 6990 | 2 | 3.3198 | 0.0061214 | 0.02854 | 0.890437 | 600 | 3 | 3.3198 |
| DNAJC18 | 6 | 0.80064 | 0.84815 | 0.999994 | 17491 | 1 | 3.8856 | 0.0061214 | 0.02854 | 0.890437 | 601 | 4 | 3.8856 |
| ZNF816 | 6 | 0.66416 | 0.78852 | 0.999994 | 15843 | 1 | 2.07 | 0.0061214 | 0.02854 | 0.890437 | 602 | 3 | 2.07 |
| IGDCC3 | 5 | 0.038589 | 0.11231 | 0.951052 | 2443 | 2 | 2.3587 | 0.0061214 | 0.025197 | 0.885141 | 603 | 3 | 2.3587 |
| OPA3 | 6 | 0.87508 | 0.8865 | 0.999994 | 18739 | 1 | 6.3865 | 0.0061285 | 0.028572 | 0.890437 | 604 | 5 | 6.3865 |
| ACTL7A | 6 | 0.98833 | 0.98829 | 0.999994 | 21308 | 0 | 4.3801 | 0.0061349 | 0.028598 | 0.890437 | 605 | 6 | 4.3801 |
| CHRDL2 | 6 | 0.99385 | 0.9938 | 0.999994 | 21442 | 0 | 2.276 | 0.0061489 | 0.028655 | 0.890437 | 606 | 6 | 2.276 |
| LPO | 5 | 0.99385 | 0.99383 | 0.999994 | 21441 | 0 | 5.6986 | 0.0061519 | 0.025303 | 0.885141 | 607 | 5 | 5.6986 |
| hsa-mir-6729 | 2 | 0.903 | 0.90292 | 0.999994 | 19277 | 0 | 6.4363 | 0.0061662 | 0.011789 | 0.829051 | 608 | 2 | 6.4363 |
| MYADM | 5 | 0.73975 | 0.78555 | 0.999994 | 16736 | 1 | 5.0615 | 0.006168 | 0.025367 | 0.885141 | 609 | 3 | 5.0615 |
| LOC100289187 | 6 | 0.20098 | 0.38906 | 0.999994 | 7790 | 3 | 3.3966 | 0.0061787 | 0.028794 | 0.890437 | 610 | 3 | 3.3966 |
| NUCB1 | 5 | 0.58195 | 0.71969 | 0.999994 | 14704 | 1 | 2.2598 | 0.0061839 | 0.025433 | 0.885141 | 611 | 4 | 2.2598 |
| SLC22A20 | 6 | 0.72616 | 0.82035 | 0.999994 | 16559 | 1 | 6.5919 | 0.006189 | 0.02884 | 0.890437 | 612 | 4 | 6.5919 |
| MKI67 | 6 | 0.90317 | 0.90615 | 0.999994 | 19283 | 1 | 6.2024 | 0.006189 | 0.02884 | 0.890437 | 613 | 4 | 6.2024 |
| GRAMD4 | 5 | 0.8852 | 0.88532 | 0.999994 | 18936 | 1 | 7.3503 | 0.0062131 | 0.025547 | 0.885141 | 614 | 3 | 7.3503 |
| GALR2 | 5 | 0.43371 | 0.59152 | 0.999994 | 12472 | 1 | 7.3503 | 0.0062131 | 0.025547 | 0.885141 | 615 | 4 | 7.3503 |
| ZNF493 | 6 | 0.30605 | 0.51378 | 0.999994 | 10258 | 2 | 6.6036 | 0.0062148 | 0.028946 | 0.890437 | 616 | 4 | 6.6036 |
| NKX2-8 | 5 | 0.98317 | 0.98324 | 0.999994 | 21161 | 0 | 5.8862 | 0.0062308 | 0.025617 | 0.885381 | 617 | 5 | 5.8862 |
| DCAF12 | 6 | 0.99377 | 0.99371 | 0.999994 | 21440 | 0 | 3.657 | 0.0062339 | 0.029027 | 0.890437 | 618 | 6 | 3.657 |
| FAM188B | 6 | 0.00098921 | 0.0050255 | 0.589484 | 163 | 4 | -4.877 | 0.0062349 | 0.029032 | 0.890437 | 619 | 2 | -4.877 |
| GNPDA1 | 6 | 0.94759 | 0.94762 | 0.999994 | 20278 | 0 | 2.6739 | 0.006238 | 0.029044 | 0.890437 | 620 | 5 | 2.6739 |
| NDUFAF4 | 6 | 0.90372 | 0.90657 | 0.999994 | 19298 | 1 | 6.6149 | 0.00625 | 0.02909 | 0.890437 | 621 | 4 | 6.6149 |
| IFLTD1 | 4 | 0.985 | 0.98496 | 0.999994 | 21222 | 0 | 6.9248 | 0.0062615 | 0.021792 | 0.85539 | 622 | 4 | 6.9248 |
| SLC7A1 | 6 | 0.99371 | 0.99366 | 0.999994 | 21439 | 0 | 4.3599 | 0.0062876 | 0.029249 | 0.890437 | 623 | 6 | 4.3599 |

| | | | | | | | | | | | | | |
|---|---|---|---|---|---|---|---|---|---|---|---|---|---|
| SLC35G6 | 6 | 0.9937 | 0.99364 | 0.999994 | 21438 | 0 | 6.2791 | 0.0063041 | 0.029315 | 0.890437 | 624 | 6 | 6.2791 |
| DNAJB12 | 4 | 0.15894 | 0.29839 | 0.999994 | 6699 | 2 | 3.0481 | 0.0063288 | 0.022001 | 0.858434 | 625 | 2 | 3.0481 |
| RMDN3 | 5 | 0.99364 | 0.99362 | 0.999994 | 21437 | 0 | 3.3932 | 0.0063581 | 0.026132 | 0.890437 | 626 | 5 | 3.3932 |
| GCGR | 5 | 0.44005 | 0.59684 | 0.999994 | 12575 | 1 | 7.1451 | 0.0063688 | 0.026173 | 0.890437 | 627 | 4 | 7.1451 |
| TRNT1 | 5 | 0.72802 | 0.7797 | 0.999994 | 16581 | 1 | 5.0615 | 0.0064288 | 0.026386 | 0.890437 | 628 | 4 | 5.0615 |
| VEPH1 | 6 | 0.51142 | 0.67659 | 0.999994 | 13605 | 1 | 6.2487 | 0.0064546 | 0.029972 | 0.890437 | 629 | 4 | 6.2487 |
| MAEL | 6 | 0.99354 | 0.99348 | 0.999994 | 21436 | 0 | 4.0143 | 0.0064643 | 0.030021 | 0.890437 | 630 | 6 | 4.0143 |
| NonTargetingControlG | 2 | 0.99351 | 0.99343 | 0.999994 | 21435 | 0 | 8.2491 | 0.006488 | 0.012396 | 0.829051 | 631 | 2 | 8.2491 |
| NQO1 | 6 | 0.9935 | 0.99345 | 0.999994 | 21434 | 0 | 2.9245 | 0.0064991 | 0.030171 | 0.890437 | 632 | 6 | 2.9245 |
| LIPG | 5 | 0.97592 | 0.97598 | 0.999994 | 20987 | 0 | 5.6402 | 0.0065216 | 0.026734 | 0.890437 | 633 | 5 | 5.6402 |
| IL17RB | 5 | 0.99346 | 0.99344 | 0.999994 | 21433 | 0 | 3.2544 | 0.0065399 | 0.0268 | 0.890437 | 634 | 5 | 3.2544 |
| PAPSS1 | 4 | 0.022921 | 0.066802 | 0.93914 | 1644 | 2 | -2.771 | 0.0065649 | 0.022777 | 0.863933 | 635 | 2 | -2.771 |
| SYNGAP1 | 6 | 0.99343 | 0.99337 | 0.999994 | 21432 | 0 | 4.735 | 0.0065675 | 0.030465 | 0.890437 | 636 | 6 | 4.735 |
| NonTargetingControlG | 2 | 0.99342 | 0.99334 | 0.999994 | 21431 | 0 | 7.867 | 0.0065794 | 0.012571 | 0.829051 | 637 | 2 | 7.867 |
| SELP | 4 | 0.93419 | 0.93398 | 0.999994 | 19958 | 0 | 7.3814 | 0.0066255 | 0.022972 | 0.863933 | 638 | 3 | 7.3814 |
| NUGGC | 6 | 0.99336 | 0.9933 | 0.999994 | 21430 | 0 | 2.7501 | 0.0066369 | 0.030761 | 0.890437 | 639 | 6 | 2.7501 |
| LGSN | 5 | 0.99215 | 0.99217 | 0.999994 | 21399 | 0 | 2.2935 | 0.0066548 | 0.027235 | 0.890437 | 640 | 5 | 2.2935 |
| ARX | 6 | 0.9489 | 0.94889 | 0.999994 | 20307 | 0 | 4.5244 | 0.0066586 | 0.030849 | 0.890437 | 641 | 5 | 4.5244 |
| CCDC130 | 6 | 0.73611 | 0.82366 | 0.999994 | 16678 | 1 | 4.8082 | 0.0066586 | 0.030849 | 0.890437 | 642 | 5 | 4.8082 |
| BCL2L2 | 6 | 0.7774 | 0.83865 | 0.999994 | 17174 | 1 | 3.6886 | 0.0066586 | 0.030849 | 0.890437 | 643 | 5 | 3.6886 |
| SCIN | 6 | 0.64903 | 0.77688 | 0.999994 | 15643 | 1 | 3.4748 | 0.0066586 | 0.030849 | 0.890437 | 644 | 5 | 3.4748 |
| MED1 | 6 | 0.99333 | 0.99328 | 0.999994 | 21429 | 0 | 3.251 | 0.0066668 | 0.03088 | 0.890437 | 645 | 6 | 3.251 |
| NDUFAF3 | 4 | 0.33308 | 0.47687 | 0.999994 | 10791 | 1 | 7.4938 | 0.006682 | 0.023163 | 0.863933 | 646 | 3 | 7.4938 |
| NAGK | 6 | 0.1227 | 0.2908 | 0.999632 | 5730 | 3 | -0.37871 | 0.006685 | 0.03096 | 0.890437 | 647 | 2 | -0.37871 |
| GLYR1 | 6 | 0.98741 | 0.98736 | 0.999994 | 21288 | 0 | 5.3452 | 0.0066855 | 0.030961 | 0.890437 | 648 | 6 | 5.3452 |
| DHX16 | 5 | 0.089244 | 0.20936 | 0.979587 | 4563 | 1 | 7.3095 | 0.0066867 | 0.027351 | 0.890437 | 649 | 4 | 7.3095 |
| CALB1 | 5 | 0.8965 | 0.89662 | 0.999994 | 19152 | 0 | 1.0397 | 0.0067427 | 0.027546 | 0.890437 | 650 | 3 | 1.0397 |
| CLMP | 6 | 0.78332 | 0.84103 | 0.999994 | 17260 | 1 | 3.9298 | 0.006755 | 0.031254 | 0.890437 | 651 | 5 | 3.9298 |
| BSPH1 | 5 | 0.81243 | 0.82754 | 0.999994 | 17658 | 1 | 6.2265 | 0.0067651 | 0.027623 | 0.890437 | 652 | 4 | 6.2265 |
| YY2 | 6 | 0.99281 | 0.99279 | 0.999994 | 21419 | 0 | 3.4335 | 0.0068084 | 0.031474 | 0.890437 | 653 | 6 | 3.4335 |
| IKBKG | 6 | 0.75206 | 0.82921 | 0.999994 | 16879 | 1 | 6.16 | 0.006815 | 0.031501 | 0.890437 | 654 | 5 | 6.16 |
| CC2D1B | 6 | 0.96943 | 0.96942 | 0.999994 | 20817 | 0 | 6.7587 | 0.0068173 | 0.031511 | 0.890437 | 655 | 5 | 6.7587 |
| NR2F6 | 5 | 0.16973 | 0.33174 | 0.999994 | 6979 | 1 | 5.782 | 0.0068199 | 0.02783 | 0.890437 | 656 | 4 | 5.782 |
| KLK14 | 6 | 0.16065 | 0.33906 | 0.999994 | 6757 | 1 | 2.8343 | 0.006834 | 0.031578 | 0.890437 | 657 | 5 | 2.8343 |
| ATP1B3 | 6 | 0.029252 | 0.097024 | 0.945253 | 1970 | 2 | 1.0748 | 0.0068537 | 0.031661 | 0.890437 | 658 | 4 | 1.0748 |
| hsa-mir-8066 | 4 | 0.94582 | 0.94572 | 0.999994 | 20228 | 0 | 7.5205 | 0.0068791 | 0.023793 | 0.874165 | 659 | 4 | 7.5205 |
| ITK | 6 | 0.29051 | 0.49567 | 0.999994 | 9925 | 3 | 0.089515 | 0.00691 | 0.031884 | 0.890437 | 660 | 3 | 0.089515 |
| MAP7D1 | 6 | 0.92882 | 0.92892 | 0.999994 | 19832 | 0 | 3.2223 | 0.0069306 | 0.031968 | 0.890437 | 661 | 5 | 3.2223 |
| NOX5 | 5 | 0.053412 | 0.14861 | 0.965554 | 3136 | 3 | -1.364 | 0.0069342 | 0.028264 | 0.890437 | 662 | 2 | -1.364 |
| FOXD4 | 5 | 0.78158 | 0.80835 | 0.999994 | 17231 | 1 | 5.2982 | 0.0069377 | 0.028278 | 0.890437 | 663 | 4 | 5.2982 |
| KLHL38 | 6 | 0.9621 | 0.96218 | 0.999994 | 20633 | 0 | 4.937 | 0.0069419 | 0.032015 | 0.890437 | 664 | 5 | 4.937 |
| NonTargetingControlG | 2 | 0.99306 | 0.99297 | 0.999994 | 21428 | 0 | 7.498 | 0.006943 | 0.01327 | 0.829051 | 665 | 2 | 7.498 |
| UBE2D1 | 6 | 0.28539 | 0.48974 | 0.999994 | 9820 | 3 | 3.35 | 0.0069458 | 0.032031 | 0.890437 | 666 | 3 | 3.35 |
| BRI3BP | 6 | 0.66229 | 0.78709 | 0.999994 | 15813 | 1 | 4.5528 | 0.0069662 | 0.032118 | 0.890437 | 667 | 5 | 4.5528 |
| ARID1B | 5 | 0.35347 | 0.52475 | 0.999994 | 11223 | 2 | 1.3182 | 0.0069741 | 0.028423 | 0.890437 | 668 | 3 | 1.3182 |
| CHSY1 | 6 | 0.25637 | 0.45556 | 0.999994 | 9158 | 1 | 5.0942 | 0.0069748 | 0.032155 | 0.890437 | 669 | 5 | 5.0942 |
| FAM91A1 | 6 | 0.89148 | 0.89735 | 0.999994 | 19061 | 1 | 6.3805 | 0.0069786 | 0.03217 | 0.890437 | 670 | 5 | 6.3805 |
| MB21D1 | 6 | 0.90289 | 0.90593 | 0.999994 | 19272 | 1 | 5.5184 | 0.0069862 | 0.032201 | 0.890437 | 671 | 5 | 5.5184 |
| RCOR3 | 6 | 0.35162 | 0.56581 | 0.999994 | 11189 | 3 | 3.5289 | 0.0070068 | 0.032296 | 0.890437 | 672 | 3 | 3.5289 |
| H1FX | 6 | 0.20844 | 0.39817 | 0.999994 | 7988 | 1 | 3.5447 | 0.0070072 | 0.032298 | 0.890437 | 673 | 5 | 3.5447 |
| ROPN1L | 6 | 0.99299 | 0.99297 | 0.999994 | 21427 | 0 | 3.7047 | 0.0070095 | 0.032308 | 0.890437 | 674 | 6 | 3.7047 |
| CENPP | 6 | 0.82866 | 0.86092 | 0.999994 | 17905 | 1 | 3.7587 | 0.0070143 | 0.032331 | 0.890437 | 675 | 4 | 3.7587 |

| | | | | | | | | | | | | |
|---|---|---|---|---|---|---|---|---|---|---|---|---|
| hsa-mir-6873 | 2 | 0.99298 | 0.99289 | 0.999994 | 21426 | 0 | 8.5057 | 0.0070202 | 0.013408 | 0.829051 | 676 | 2 | 8.5057 |
| ADIPOR2 | 6 | 0.98165 | 0.98157 | 0.999994 | 21129 | 0 | 5.556 | 0.0070295 | 0.032386 | 0.890437 | 677 | 5 | 5.556 |
| ZC3H13 | 5 | 0.88281 | 0.88298 | 0.999994 | 18888 | 1 | 6.3513 | 0.0070357 | 0.02866 | 0.890437 | 678 | 4 | 6.3513 |
| EIF1AX | 4 | 0.99294 | 0.99292 | 0.999994 | 21425 | 0 | 5.5761 | 0.0070561 | 0.024335 | 0.881213 | 679 | 4 | 5.5761 |
| C12orf45 | 5 | 0.99293 | 0.99293 | 0.999994 | 21424 | 0 | 3.2785 | 0.0070679 | 0.028784 | 0.890437 | 680 | 5 | 3.2785 |
| OR51E2 | 6 | 0.48031 | 0.6554 | 0.999994 | 13146 | 2 | 1.1066 | 0.007072 | 0.032579 | 0.890437 | 681 | 3 | 1.1066 |
| XPO1 | 6 | 0.25312 | 0.45165 | 0.999994 | 9087 | 2 | 4.519 | 0.0070787 | 0.032605 | 0.890437 | 682 | 4 | 4.519 |
| PSIP1 | 6 | 0.98066 | 0.98059 | 0.999994 | 21105 | 0 | 6.1514 | 0.0070799 | 0.032612 | 0.890437 | 683 | 5 | 6.1514 |
| GDAP2 | 6 | 0.64082 | 0.77055 | 0.999994 | 15525 | 1 | 6.3259 | 0.0070879 | 0.032649 | 0.890437 | 684 | 5 | 6.3259 |
| POLR2H | 5 | 0.99291 | 0.99291 | 0.999994 | 21423 | 0 | 5.1479 | 0.0070912 | 0.028877 | 0.890437 | 685 | 5 | 5.1479 |
| SHISA2 | 6 | 0.8901 | 0.89642 | 0.999994 | 19036 | 1 | 3.9609 | 0.0071067 | 0.03273 | 0.890437 | 686 | 5 | 3.9609 |
| NFKBIE | 6 | 0.87602 | 0.88708 | 0.999994 | 18751 | 1 | 6.4238 | 0.007107 | 0.032733 | 0.890437 | 687 | 5 | 6.4238 |
| ATHL1 | 5 | 0.8583 | 0.86096 | 0.999994 | 18410 | 1 | 7.2705 | 0.0071302 | 0.029018 | 0.890437 | 688 | 3 | 7.2705 |
| TFB2M | 5 | 0.99286 | 0.99286 | 0.999994 | 21422 | 0 | 5.6697 | 0.0071445 | 0.029069 | 0.890437 | 689 | 5 | 5.6697 |
| SGK110 | 6 | 0.63362 | 0.76508 | 0.999994 | 15428 | 1 | 5.2561 | 0.0071491 | 0.032917 | 0.890579 | 690 | 5 | 5.2561 |
| POP7 | 4 | 0.99285 | 0.99282 | 0.999994 | 21421 | 0 | 4.2772 | 0.0071493 | 0.024644 | 0.882096 | 691 | 4 | 4.2772 |
| MAPK8 | 5 | 0.99283 | 0.99283 | 0.999994 | 21420 | 0 | 3.404 | 0.0071717 | 0.029157 | 0.890437 | 692 | 5 | 3.404 |
| BAAT | 6 | 0.52801 | 0.68813 | 0.999994 | 13860 | 2 | 6.4639 | 0.0071774 | 0.033027 | 0.890579 | 693 | 4 | 6.4639 |
| ATG16L2 | 6 | 0.40414 | 0.60506 | 0.999994 | 12021 | 2 | 2.0405 | 0.0071912 | 0.033072 | 0.890579 | 694 | 3 | 2.0405 |
| GPR183 | 5 | 0.99278 | 0.99279 | 0.999994 | 21418 | 0 | 4.0801 | 0.0072181 | 0.029316 | 0.890437 | 695 | 5 | 4.0801 |
| ALKBH6 | 5 | 0.99277 | 0.99278 | 0.999994 | 21417 | 0 | 2.3411 | 0.0072327 | 0.029365 | 0.890437 | 696 | 5 | 2.3411 |
| ATIC | 5 | 0.99275 | 0.99276 | 0.999994 | 21416 | 0 | 4.1654 | 0.0072519 | 0.029437 | 0.890437 | 697 | 5 | 4.1654 |
| HPGD | 6 | 0.81801 | 0.85589 | 0.999994 | 17746 | 1 | 1.7165 | 0.0072552 | 0.033324 | 0.893117 | 698 | 4 | 1.7165 |
| NonTargetingControlG | 2 | 0.85246 | 0.85236 | 0.999994 | 18304 | 0 | 5.8633 | 0.0072562 | 0.013839 | 0.829051 | 699 | 2 | 5.8633 |
| DPH6 | 3 | 0.99273 | 0.99282 | 0.999994 | 21415 | 0 | 5.6406 | 0.0072723 | 0.019756 | 0.851882 | 700 | 3 | 5.6406 |
| PLCD1 | 6 | 0.092712 | 0.23292 | 0.984502 | 4710 | 1 | 6.4279 | 0.0072748 | 0.033403 | 0.89322 | 701 | 5 | 6.4279 |
| HSD17B1 | 4 | 0.13241 | 0.27106 | 0.998561 | 5986 | 2 | 2.9609 | 0.0072751 | 0.025052 | 0.885141 | 702 | 2 | 2.9609 |
| C18orf63 | 6 | 0.96914 | 0.96914 | 0.999994 | 20805 | 0 | 4.6784 | 0.007296 | 0.033503 | 0.89322 | 703 | 5 | 4.6784 |
| E4F1 | 5 | 0.99269 | 0.99271 | 0.999994 | 21414 | 0 | 4.9353 | 0.0073107 | 0.029671 | 0.890437 | 704 | 5 | 4.9353 |
| DYDC1 | 4 | 0.99268 | 0.99264 | 0.999994 | 21413 | 0 | 6.0258 | 0.0073186 | 0.025197 | 0.885141 | 705 | 4 | 6.0258 |
| DMRTC2 | 6 | 0.94022 | 0.94018 | 0.999994 | 20090 | 0 | 3.0961 | 0.0073315 | 0.033649 | 0.89322 | 706 | 5 | 3.0961 |
| SH3PXD2A | 6 | 0.97876 | 0.97869 | 0.999994 | 21068 | 0 | 1.5755 | 0.0073599 | 0.033765 | 0.89322 | 707 | 4 | 1.5755 |
| LRRC8A | 6 | 0.76438 | 0.83371 | 0.999994 | 17021 | 1 | 6.7801 | 0.0073649 | 0.033783 | 0.89322 | 708 | 5 | 6.7801 |
| ZCWPW1 | 6 | 0.26637 | 0.4674 | 0.999994 | 9380 | 1 | 4.5572 | 0.0073668 | 0.033788 | 0.89322 | 709 | 4 | 4.5572 |
| HRASLS2 | 6 | 0.99263 | 0.99259 | 0.999994 | 21412 | 0 | 5.2261 | 0.0073734 | 0.033823 | 0.89322 | 710 | 6 | 5.2261 |
| LRRC59 | 4 | 0.99259 | 0.99255 | 0.999994 | 21411 | 0 | 6.6453 | 0.0074115 | 0.025506 | 0.885141 | 711 | 4 | 6.6453 |
| UBE2N | 3 | 0.99259 | 0.99268 | 0.999994 | 21410 | 0 | 6.425 | 0.007415 | 0.020102 | 0.851882 | 712 | 3 | 6.425 |
| hsa-mir-7151 | 2 | 0.94073 | 0.94084 | 0.999994 | 20098 | 0 | 6.9114 | 0.0074253 | 0.01415 | 0.829051 | 713 | 2 | 6.9114 |
| PTOV1 | 6 | 0.99253 | 0.99249 | 0.999994 | 21409 | 0 | 5.0912 | 0.0074656 | 0.034205 | 0.895567 | 714 | 6 | 5.0912 |
| NFE2L1 | 6 | 0.62904 | 0.76162 | 0.999994 | 15364 | 1 | 4.2928 | 0.0074747 | 0.034243 | 0.895567 | 715 | 4 | 4.2928 |
| NPAP1 | 5 | 0.71128 | 0.77167 | 0.999994 | 16410 | 1 | 7.2386 | 0.0075155 | 0.030442 | 0.890437 | 716 | 4 | 7.2386 |
| TPRX1 | 6 | 0.78765 | 0.84272 | 0.999994 | 17315 | 1 | 5.2198 | 0.0075216 | 0.034461 | 0.897383 | 717 | 4 | 5.2198 |
| USP10 | 6 | 0.1384 | 0.31101 | 0.999994 | 6150 | 3 | -0.8409 | 0.0075285 | 0.034492 | 0.897383 | 718 | 2 | -0.8409 |
| L3MBTL1 | 6 | 0.58707 | 0.73043 | 0.999994 | 14777 | 1 | 3.6884 | 0.0075329 | 0.034512 | 0.897383 | 719 | 5 | 3.6884 |
| P2RY1 | 3 | 0.99246 | 0.99256 | 0.999994 | 21408 | 0 | 7.1468 | 0.0075356 | 0.02043 | 0.853037 | 720 | 3 | 7.1468 |
| C9orf152 | 6 | 0.99246 | 0.99242 | 0.999994 | 21407 | 0 | 2.8832 | 0.0075385 | 0.034535 | 0.897383 | 721 | 6 | 2.8832 |
| FAR1 | 6 | 0.99245 | 0.99241 | 0.999994 | 21405 | 0 | 1.8164 | 0.0075462 | 0.034562 | 0.897383 | 722 | 6 | 1.8164 |
| KIAA1009 | 5 | 0.91361 | 0.91368 | 0.999994 | 19484 | 0 | 7.2366 | 0.0075758 | 0.030663 | 0.890437 | 723 | 3 | 7.2366 |
| MAGEC1 | 5 | 0.96109 | 0.96122 | 0.999994 | 20608 | 0 | 7.2366 | 0.0075758 | 0.030663 | 0.890437 | 724 | 4 | 7.2366 |
| FDX1 | 5 | 0.28712 | 0.4706 | 0.999994 | 9854 | 1 | 7.2366 | 0.0075758 | 0.030663 | 0.890437 | 725 | 3 | 7.2366 |
| THG1L | 6 | 0.91791 | 0.91848 | 0.999994 | 19584 | 1 | 5.4535 | 0.0075836 | 0.034733 | 0.897383 | 726 | 5 | 5.4535 |
| ZNF571 | 6 | 0.89262 | 0.89818 | 0.999994 | 19081 | 1 | 5.4495 | 0.0075903 | 0.034759 | 0.897383 | 727 | 5 | 5.4495 |

| | | | | | | | | | | | | |
|---|---|---|---|---|---|---|---|---|---|---|---|---|
| C1R | 6 | 0.99241 | 0.99236 | 0.999994 | 21404 | 0 | 3.1785 | 0.0075935 | 0.034774 | 0.897383 | 728 | 6 | 3.1785 |
| KRT10 | 6 | 0.37253 | 0.5848 | 0.999994 | 11553 | 1 | 6.0625 | 0.0076346 | 0.034945 | 0.897383 | 729 | 5 | 6.0625 |
| OR10H4 | 6 | 0.15784 | 0.33548 | 0.999994 | 6668 | 1 | 4.0428 | 0.007641 | 0.034975 | 0.897383 | 730 | 5 | 4.0428 |
| hsa-mir-3178 | 4 | 0.99235 | 0.99232 | 0.999994 | 21403 | 0 | 4.5467 | 0.0076526 | 0.026292 | 0.890437 | 731 | 4 | 4.5467 |
| LHX2 | 6 | 0.99234 | 0.99231 | 0.999994 | 21402 | 0 | 4.7579 | 0.0076606 | 0.035054 | 0.898105 | 732 | 6 | 4.7579 |
| GPN3 | 5 | 0.73633 | 0.78381 | 0.999994 | 16681 | 1 | 5.9596 | 0.0076928 | 0.031099 | 0.890437 | 733 | 4 | 5.9596 |
| KRTAP3-3 | 5 | 0.955 | 0.95515 | 0.999994 | 20453 | 0 | 6.0535 | 0.007713 | 0.031181 | 0.890437 | 734 | 4 | 6.0535 |
| ZNF563 | 6 | 0.99228 | 0.99225 | 0.999994 | 21401 | 0 | 3.0642 | 0.0077204 | 0.035292 | 0.900122 | 735 | 6 | 3.0642 |
| hsa-mir-192 | 4 | 0.98871 | 0.9887 | 0.999994 | 21315 | 0 | 6.6891 | 0.0077275 | 0.026527 | 0.890437 | 736 | 4 | 6.6891 |
| IL1RN | 6 | 0.48166 | 0.65631 | 0.999994 | 13163 | 1 | 5.9711 | 0.0077276 | 0.035323 | 0.900122 | 737 | 5 | 5.9711 |
| NUFIP2 | 5 | 0.99227 | 0.99228 | 0.999994 | 21400 | 0 | 5.7547 | 0.0077282 | 0.031233 | 0.890437 | 738 | 5 | 5.7547 |
| TSPYL6 | 5 | 0.77261 | 0.80319 | 0.999994 | 17117 | 1 | 1.3478 | 0.007731 | 0.031246 | 0.890437 | 739 | 3 | 1.3478 |
| ELMOD2 | 5 | 0.97467 | 0.97473 | 0.999994 | 20948 | 0 | 7.2223 | 0.0077371 | 0.031269 | 0.890437 | 740 | 5 | 7.2223 |
| MAP2K7 | 6 | 0.28509 | 0.48936 | 0.999994 | 9814 | 3 | 0.81421 | 0.0077534 | 0.035438 | 0.900347 | 741 | 3 | 0.81421 |
| NUTM1 | 6 | 0.89886 | 0.90277 | 0.999994 | 19203 | 1 | 5.1445 | 0.0077682 | 0.035507 | 0.900347 | 742 | 5 | 5.1445 |
| SERPINA5 | 6 | 0.054823 | 0.15455 | 0.967126 | 3191 | 1 | 4.1075 | 0.0077682 | 0.035507 | 0.900347 | 743 | 5 | 4.1075 |
| EPHB4 | 5 | 0.045408 | 0.12932 | 0.952548 | 2784 | 2 | 0.084511 | 0.007772 | 0.031384 | 0.890437 | 744 | 2 | 0.084511 |
| SLC6A4 | 5 | 0.0012487 | 0.0055673 | 0.623123 | 189 | 3 | -4.1233 | 0.0077778 | 0.031399 | 0.890437 | 745 | 2 | -4.1233 |
| NAT6 | 5 | 0.29739 | 0.47895 | 0.999994 | 10077 | 2 | 7.2168 | 0.0077899 | 0.031449 | 0.890437 | 746 | 3 | 7.2168 |
| AOAH | 5 | 0.091127 | 0.21241 | 0.979587 | 4644 | 3 | -1.63 | 0.0078415 | 0.031645 | 0.890437 | 747 | 2 | -1.63 |
| CCDC77 | 4 | 0.71144 | 0.73211 | 0.999994 | 16414 | 1 | 6.8245 | 0.0078674 | 0.02697 | 0.890437 | 748 | 3 | 6.8245 |
| PSMD10 | 6 | 0.93747 | 0.93747 | 0.999994 | 20018 | 0 | 4.5631 | 0.0078711 | 0.035942 | 0.904772 | 749 | 5 | 4.5631 |
| MAGOH | 5 | 0.87804 | 0.8784 | 0.999994 | 18788 | 1 | 7.2068 | 0.0078907 | 0.031835 | 0.890437 | 750 | 3 | 7.2068 |
| GCN1L1 | 6 | 0.99211 | 0.99206 | 0.999994 | 21398 | 0 | 2.1082 | 0.0078949 | 0.036048 | 0.904772 | 751 | 6 | 2.1082 |
| MYBL2 | 5 | 0.97232 | 0.97238 | 0.999994 | 20875 | 0 | 6.0962 | 0.0079121 | 0.031916 | 0.890437 | 752 | 4 | 6.0962 |
| SAPCD1 | 6 | 0.45973 | 0.64159 | 0.999994 | 12844 | 2 | 4.2467 | 0.007922 | 0.036156 | 0.904772 | 753 | 4 | 4.2467 |
| VWA3A | 6 | 0.24955 | 0.44739 | 0.999994 | 9006 | 3 | -0.87324 | 0.007922 | 0.036156 | 0.904772 | 754 | 2 | -0.87324 |
| IGLON5 | 4 | 0.67125 | 0.70613 | 0.999994 | 15947 | 1 | 7.1369 | 0.0079283 | 0.02716 | 0.890437 | 755 | 3 | 7.1369 |
| OCIAD2 | 5 | 0.4929 | 0.64176 | 0.999994 | 13310 | 2 | 0.3508 | 0.0079652 | 0.032115 | 0.890437 | 756 | 2 | 0.3508 |
| hsa-mir-4464 | 4 | 0.99203 | 0.99201 | 0.999994 | 21397 | 0 | 3.4638 | 0.0079666 | 0.027271 | 0.890437 | 757 | 4 | 3.4638 |
| ALDH16A1 | 5 | 0.99199 | 0.99201 | 0.999994 | 21395 | 0 | 2.9671 | 0.0080058 | 0.032273 | 0.890437 | 758 | 5 | 2.9671 |
| PCID2 | 6 | 0.83817 | 0.8657 | 0.999994 | 18058 | 1 | 6.3029 | 0.0080071 | 0.036497 | 0.904772 | 759 | 5 | 6.3029 |
| CLEC2B | 6 | 0.28925 | 0.49419 | 0.999994 | 9897 | 3 | -0.4617 | 0.0080344 | 0.036611 | 0.904772 | 760 | 1 | -0.4617 |
| ENDOU | 6 | 0.93129 | 0.93141 | 0.999994 | 19881 | 0 | 5.8115 | 0.0080356 | 0.036615 | 0.904772 | 761 | 5 | 5.8115 |
| LGALS7B | 4 | 0.99196 | 0.99192 | 0.999994 | 21394 | 0 | 5.3421 | 0.0080433 | 0.027506 | 0.890437 | 762 | 4 | 5.3421 |
| ZNF511 | 5 | 0.5859 | 0.72108 | 0.999994 | 14757 | 1 | 2.7235 | 0.008057 | 0.032481 | 0.890437 | 763 | 3 | 2.7235 |
| ARHGEF37 | 6 | 0.51952 | 0.6822 | 0.999994 | 13739 | 1 | 0.756 | 0.0080589 | 0.036711 | 0.904772 | 764 | 4 | 0.756 |
| MAN1C1 | 4 | 0.14358 | 0.28261 | 0.999632 | 6288 | 1 | 2.8378 | 0.0080652 | 0.027571 | 0.890437 | 765 | 2 | 2.8378 |
| PLCE1 | 5 | 0.99193 | 0.99196 | 0.999994 | 21393 | 0 | 2.0131 | 0.0080664 | 0.032517 | 0.890437 | 766 | 5 | 2.0131 |
| SSX2IP | 6 | 0.2428 | 0.43934 | 0.999994 | 8859 | 2 | -0.36047 | 0.0080906 | 0.03684 | 0.904772 | 767 | 2 | -0.36047 |
| ARHGAP11A | 6 | 0.99191 | 0.99186 | 0.999994 | 21392 | 0 | 3.3058 | 0.0080911 | 0.036843 | 0.904772 | 768 | 6 | 3.3058 |
| FAM211B | 6 | 0.65082 | 0.77824 | 0.999994 | 15670 | 2 | 6.1469 | 0.0080958 | 0.036863 | 0.904772 | 769 | 4 | 6.1469 |
| EFCC1 | 6 | 0.99189 | 0.99184 | 0.999994 | 21391 | 0 | 3.1739 | 0.0081054 | 0.036907 | 0.904772 | 770 | 6 | 3.1739 |
| FRMD3 | 5 | 0.21684 | 0.39885 | 0.999994 | 8187 | 1 | 7.1893 | 0.0081114 | 0.032673 | 0.890437 | 771 | 3 | 7.1893 |
| CFHR3 | 5 | 0.99189 | 0.99192 | 0.999994 | 21390 | 0 | 2.2648 | 0.0081144 | 0.03268 | 0.890437 | 772 | 5 | 2.2648 |
| LRRC57 | 6 | 0.99188 | 0.99183 | 0.999994 | 21389 | 0 | 5.4393 | 0.0081167 | 0.036951 | 0.904772 | 773 | 6 | 5.4393 |
| PHF20L1 | 6 | 0.55424 | 0.70668 | 0.999994 | 14274 | 1 | 3.3473 | 0.0081467 | 0.037069 | 0.904772 | 774 | 3 | 3.3473 |
| ZNF592 | 6 | 0.9908 | 0.99076 | 0.999994 | 21372 | 0 | 5.7413 | 0.0081483 | 0.037078 | 0.904772 | 775 | 6 | 5.7413 |
| HDAC2 | 5 | 0.99184 | 0.99188 | 0.999994 | 21388 | 0 | 5.0005 | 0.0081565 | 0.032845 | 0.890525 | 776 | 5 | 5.0005 |
| DTYMK | 2 | 0.92223 | 0.92229 | 0.999994 | 19682 | 0 | 6.6773 | 0.0081578 | 0.015508 | 0.829051 | 777 | 2 | 6.6773 |
| C1orf63 | 5 | 0.98322 | 0.98328 | 0.999994 | 21162 | 0 | 7.1865 | 0.0081614 | 0.032858 | 0.890525 | 778 | 5 | 7.1865 |
| MATR3 | 6 | 0.98449 | 0.98443 | 0.999994 | 21202 | 0 | 6.3756 | 0.0081618 | 0.037132 | 0.904772 | 779 | 5 | 6.3756 |

| | | | | | | | | | | | | |
|---|---|---|---|---|---|---|---|---|---|---|---|---|
| SEPW1 | 6 | 0.99183 | 0.99179 | 0.999994 | 21387 | 0 | 3.4293 | 0.0081732 | 0.037183 | 0.904772 | 780 | 6 | 3.4293 |
| DUOX1 | 5 | 0.91579 | 0.91589 | 0.999994 | 19533 | 0 | 6.39 | 0.0081861 | 0.032965 | 0.890579 | 781 | 4 | 6.39 |
| LPPR4 | 6 | 0.51858 | 0.68155 | 0.999994 | 13719 | 1 | 0.65422 | 0.0082029 | 0.037305 | 0.904772 | 782 | 3 | 0.65422 |
| CDKN1B | 5 | 0.30826 | 0.48772 | 0.999994 | 10290 | 1 | 2.365 | 0.008209 | 0.033046 | 0.890579 | 783 | 4 | 2.365 |
| MYOT | 6 | 0.14194 | 0.31549 | 0.999994 | 6239 | 3 | -0.65045 | 0.0082194 | 0.037378 | 0.904772 | 784 | 3 | -0.65045 |
| CATSPERB | 6 | 0.67945 | 0.80057 | 0.999994 | 16081 | 2 | -0.24818 | 0.0082194 | 0.037378 | 0.904772 | 785 | 2 | -0.24818 |
| OR6C3 | 5 | 0.69523 | 0.76432 | 0.999994 | 16259 | 1 | 7.1809 | 0.008226 | 0.033106 | 0.890579 | 786 | 3 | 7.1809 |
| FMO5 | 5 | 0.84987 | 0.85418 | 0.999994 | 18260 | 1 | 3.2785 | 0.0082421 | 0.033166 | 0.891071 | 787 | 4 | 3.2785 |
| EIF4A1 | 6 | 0.098213 | 0.24384 | 0.988214 | 4902 | 3 | 0.077423 | 0.0082474 | 0.037486 | 0.904772 | 788 | 3 | 0.077423 |
| ZNF233 | 6 | 0.070581 | 0.18801 | 0.969543 | 3857 | 1 | 3.49 | 0.0082591 | 0.03753 | 0.904772 | 789 | 4 | 3.49 |
| TTC31 | 6 | 0.99174 | 0.9917 | 0.999994 | 21386 | 0 | 1.514 | 0.0082601 | 0.037533 | 0.904772 | 790 | 6 | 1.514 |
| ZBP1 | 6 | 0.99172 | 0.99168 | 0.999994 | 21385 | 0 | 2.9212 | 0.0082788 | 0.037614 | 0.904772 | 791 | 6 | 2.9212 |
| HSPA1A | 5 | 0.064437 | 0.16937 | 0.967126 | 3627 | 2 | 7.1724 | 0.0083438 | 0.033552 | 0.89322 | 792 | 3 | 7.1724 |
| UPK1B | 5 | 0.99165 | 0.99169 | 0.999994 | 21384 | 0 | 2.6283 | 0.0083485 | 0.033566 | 0.89322 | 793 | 5 | 2.6283 |
| SETD1B | 6 | 0.8924 | 0.89802 | 0.999994 | 19077 | 1 | 5.8571 | 0.0083597 | 0.037951 | 0.904772 | 794 | 5 | 5.8571 |
| ANKRD54 | 6 | 0.11991 | 0.28554 | 0.999632 | 5625 | 1 | 3.2689 | 0.0083715 | 0.038002 | 0.904772 | 795 | 4 | 3.2689 |
| VPS4A | 6 | 0.98462 | 0.98455 | 0.999994 | 21212 | 0 | 3.0159 | 0.0083753 | 0.038018 | 0.904772 | 796 | 5 | 3.0159 |
| MSGN1 | 6 | 0.55858 | 0.70981 | 0.999994 | 14342 | 1 | 6.5063 | 0.0083792 | 0.038035 | 0.904772 | 797 | 4 | 6.5063 |
| IL1A | 6 | 0.42942 | 0.62145 | 0.999994 | 12406 | 2 | 0.30346 | 0.0084002 | 0.038127 | 0.904772 | 798 | 3 | 0.30346 |
| MX2 | 5 | 0.58122 | 0.71938 | 0.999994 | 14696 | 1 | 6.2427 | 0.0084068 | 0.03379 | 0.89322 | 799 | 4 | 6.2427 |
| MAPKAPK5 | 6 | 0.30783 | 0.51585 | 0.999994 | 10284 | 2 | 0.71355 | 0.0084276 | 0.038227 | 0.904772 | 800 | 3 | 0.71355 |
| KCNMA1 | 5 | 0.97339 | 0.97344 | 0.999994 | 20906 | 0 | 4.7106 | 0.0084644 | 0.033999 | 0.894584 | 801 | 5 | 4.7106 |
| PGM3 | 6 | 0.77697 | 0.83849 | 0.999994 | 17166 | 1 | 5.9551 | 0.0084654 | 0.038395 | 0.905919 | 802 | 5 | 5.9551 |
| PIH1D2 | 6 | 0.83118 | 0.86211 | 0.999994 | 17950 | 1 | 2.4771 | 0.0084787 | 0.038447 | 0.905919 | 803 | 5 | 2.4771 |
| KMT2D | 6 | 0.9915 | 0.99145 | 0.999994 | 21383 | 0 | 2.1467 | 0.0084956 | 0.038518 | 0.905919 | 804 | 6 | 2.1467 |
| RPL22 | 3 | 0.54198 | 0.57466 | 0.999994 | 14090 | 1 | 8.0729 | 0.0084974 | 0.022898 | 0.863933 | 805 | 2 | 8.0729 |
| GAS7 | 4 | 0.28963 | 0.43242 | 0.999994 | 9904 | 1 | 2.375 | 0.0085149 | 0.029021 | 0.890437 | 806 | 2 | 2.375 |
| MAGEA12 | 6 | 0.70288 | 0.81289 | 0.999994 | 16335 | 1 | 3.6697 | 0.0085165 | 0.038606 | 0.905919 | 807 | 4 | 3.6697 |
| ST3GAL5 | 5 | 0.23297 | 0.42132 | 0.999994 | 8596 | 3 | -0.68674 | 0.0085272 | 0.034215 | 0.895567 | 808 | 2 | -0.68674 |
| NonTargetingControlG | 2 | 0.99147 | 0.99136 | 0.999994 | 21382 | 0 | 8.111 | 0.0085299 | 0.016246 | 0.829051 | 809 | 2 | 8.111 |
| MFSD7 | 6 | 0.000065646 | 0.00035183 | 0.21768 | 34 | 4 | -1.3894 | 0.00854 | 0.038719 | 0.905919 | 810 | 1 | -1.3894 |
| GNA12 | 6 | 0.98963 | 0.98959 | 0.999994 | 21333 | 0 | 6.2228 | 0.0085549 | 0.03878 | 0.905919 | 811 | 6 | 6.2228 |
| DNLZ | 6 | 0.29542 | 0.50136 | 0.999994 | 10025 | 1 | 5.987 | 0.0085549 | 0.03878 | 0.905919 | 812 | 5 | 5.987 |
| SIRT1 | 4 | 0.99138 | 0.99134 | 0.999994 | 21381 | 0 | 4.7334 | 0.0086183 | 0.029351 | 0.890437 | 813 | 4 | 4.7334 |
| DKK4 | 5 | 0.99137 | 0.99142 | 0.999994 | 21380 | 0 | 2.9819 | 0.0086268 | 0.034608 | 0.897383 | 814 | 5 | 2.9819 |
| NonTargetingControlG | 2 | 0.92495 | 0.92507 | 0.999994 | 19741 | 0 | 6.6649 | 0.0086742 | 0.016513 | 0.83326 | 815 | 2 | 6.6649 |
| SPATS1 | 6 | 0.37632 | 0.58724 | 0.999994 | 11598 | 2 | 2.4563 | 0.0086811 | 0.039331 | 0.908838 | 816 | 4 | 2.4563 |
| CT45A1 | 6 | 0.99129 | 0.99125 | 0.999994 | 21379 | 0 | 1.0744 | 0.0087107 | 0.039467 | 0.909688 | 817 | 6 | 1.0744 |
| FXYD5 | 5 | 0.99127 | 0.99133 | 0.999994 | 21378 | 0 | 3.3666 | 0.0087251 | 0.034958 | 0.897383 | 818 | 5 | 3.3666 |
| OSGIN1 | 5 | 0.99127 | 0.99132 | 0.999994 | 21377 | 0 | 3.4529 | 0.0087283 | 0.034968 | 0.897383 | 819 | 5 | 3.4529 |
| EBF1 | 6 | 0.77908 | 0.8393 | 0.999994 | 17198 | 1 | 3.299 | 0.0087646 | 0.039688 | 0.909688 | 820 | 4 | 3.299 |
| MSI2 | 6 | 0.083255 | 0.21398 | 0.979587 | 4351 | 1 | 3.3503 | 0.0087864 | 0.039778 | 0.909688 | 821 | 5 | 3.3503 |
| EXOC3L1 | 6 | 0.9912 | 0.99116 | 0.999994 | 21376 | 0 | 5.5823 | 0.008799 | 0.039834 | 0.909688 | 822 | 6 | 5.5823 |
| TMOD2 | 6 | 0.069817 | 0.18638 | 0.969409 | 3823 | 3 | -0.3512 | 0.0088208 | 0.03993 | 0.909688 | 823 | 3 | -0.3512 |
| BTBD8 | 5 | 0.91418 | 0.91425 | 0.999994 | 19490 | 0 | 2.0749 | 0.0088301 | 0.035332 | 0.900122 | 824 | 4 | 2.0749 |
| LPA | 5 | 0.97179 | 0.97182 | 0.999994 | 20865 | 0 | 6.1426 | 0.0088467 | 0.035393 | 0.900347 | 825 | 4 | 6.1426 |
| HPD | 6 | 0.5907 | 0.73313 | 0.999994 | 14832 | 2 | 6.4213 | 0.0088492 | 0.040047 | 0.909688 | 826 | 4 | 6.4213 |
| PLA2G4F | 6 | 0.99111 | 0.99108 | 0.999994 | 21375 | 0 | 2.3622 | 0.0088914 | 0.040229 | 0.909688 | 827 | 6 | 2.3622 |
| GPR146 | 6 | 0.94933 | 0.94931 | 0.999994 | 20316 | 0 | 6.2132 | 0.0088957 | 0.040247 | 0.909688 | 828 | 4 | 6.2132 |
| NonTargetingControlG | 1 | 0.9911 | 0.99108 | 0.999994 | 21374 | 0 | 9.7087 | 0.0088958 | 0.0088638 | 0.816312 | 829 | 1 | 9.7087 |
| IL25 | 6 | 0.57771 | 0.72351 | 0.999994 | 14654 | 1 | 3.1125 | 0.0089019 | 0.040274 | 0.909688 | 830 | 3 | 3.1125 |
| COLEC10 | 6 | 0.2455 | 0.44257 | 0.999994 | 8910 | 1 | 5.3358 | 0.0089857 | 0.040622 | 0.913215 | 831 | 5 | 5.3358 |

| | | | | | | | | | | | | | |
|---|---|---|---|---|---|---|---|---|---|---|---|---|---|
| ITPRIPL2 | 6 | 0.17229 | 0.35355 | 0.999994 | 7045 | 2 | -0.22901 | 0.0089893 | 0.040638 | 0.913215 | 832 | 2 | -0.22901 |
| PPP1R3E | 6 | 0.96976 | 0.96975 | 0.999994 | 20820 | 0 | 5.8341 | 0.0090042 | 0.040695 | 0.913215 | 833 | 4 | 5.8341 |
| WDR24 | 6 | 0.83576 | 0.86443 | 0.999994 | 18021 | 1 | 5.9269 | 0.0090042 | 0.040695 | 0.913215 | 834 | 4 | 5.9269 |
| NFIB | 5 | 0.95306 | 0.95322 | 0.999994 | 20411 | 0 | 4.5294 | 0.0090582 | 0.036174 | 0.904772 | 835 | 4 | 4.5294 |
| ATAD3A | 5 | 0.87616 | 0.87666 | 0.999994 | 18755 | 1 | 6.1806 | 0.0090582 | 0.036174 | 0.904772 | 836 | 4 | 6.1806 |
| MRPS18C | 4 | 0.015727 | 0.051311 | 0.909275 | 1230 | 1 | 7.0732 | 0.0091029 | 0.030876 | 0.890437 | 837 | 3 | 7.0732 |
| SNPH | 5 | 0.78685 | 0.81141 | 0.999994 | 17303 | 1 | 5.7702 | 0.0091191 | 0.036387 | 0.904772 | 838 | 4 | 5.7702 |
| F8A1 | 6 | 0.97334 | 0.97325 | 0.999994 | 20903 | 0 | 5.8529 | 0.009124 | 0.041208 | 0.917066 | 839 | 5 | 5.8529 |
| COX10 | 3 | 0.73237 | 0.73226 | 0.999994 | 16633 | 0 | 0.66295 | 0.0091366 | 0.024536 | 0.882096 | 840 | 2 | 0.66295 |
| hsa-mir-4673 | 3 | 0.99083 | 0.99093 | 0.999994 | 21373 | 0 | 5.3065 | 0.0091668 | 0.02462 | 0.882096 | 841 | 3 | 5.3065 |
| ZNF215 | 4 | 0.97562 | 0.9755 | 0.999994 | 20981 | 0 | 6.7747 | 0.0091795 | 0.03113 | 0.890437 | 842 | 4 | 6.7747 |
| STARD10 | 6 | 0.53611 | 0.6938 | 0.999994 | 13997 | 1 | 3.5968 | 0.0092138 | 0.041557 | 0.917066 | 843 | 5 | 3.5968 |
| PGS1 | 6 | 0.94892 | 0.9489 | 0.999994 | 20308 | 0 | 4.1918 | 0.0092143 | 0.041557 | 0.917066 | 844 | 5 | 4.1918 |
| KIT | 6 | 0.54557 | 0.7005 | 0.999994 | 14146 | 1 | 6.5511 | 0.0092193 | 0.041579 | 0.917066 | 845 | 4 | 6.5511 |
| C11orf21 | 6 | 0.50191 | 0.67002 | 0.999994 | 13460 | 1 | 5.7547 | 0.0092193 | 0.041579 | 0.917066 | 846 | 5 | 5.7547 |
| OSBPL3 | 6 | 0.72359 | 0.81951 | 0.999994 | 16529 | 1 | 6.6136 | 0.0092193 | 0.041579 | 0.917066 | 847 | 4 | 6.6136 |
| SLC41A3 | 6 | 0.63266 | 0.76434 | 0.999994 | 15408 | 2 | 4.2339 | 0.0092296 | 0.041622 | 0.917066 | 848 | 4 | 4.2339 |
| KCNK17 | 5 | 0.99076 | 0.99081 | 0.999994 | 21371 | 0 | 2.5034 | 0.0092368 | 0.036828 | 0.904772 | 849 | 5 | 2.5034 |
| OVOL2 | 6 | 0.99075 | 0.99071 | 0.999994 | 21370 | 0 | 4.3416 | 0.0092499 | 0.041709 | 0.917066 | 850 | 6 | 4.3416 |
| SMARCE1 | 4 | 0.99075 | 0.9907 | 0.999994 | 21369 | 0 | 4.6417 | 0.0092516 | 0.031364 | 0.890437 | 851 | 4 | 4.6417 |
| TIMM8A | 6 | 0.8119 | 0.85312 | 0.999994 | 17654 | 1 | 6.1739 | 0.0092904 | 0.041884 | 0.917066 | 852 | 5 | 6.1739 |
| ADAM18 | 6 | 0.99071 | 0.99067 | 0.999994 | 21368 | 0 | 1.7232 | 0.0092933 | 0.041896 | 0.917066 | 853 | 6 | 1.7232 |
| PSMC4 | 3 | 0.9907 | 0.99081 | 0.999994 | 21367 | 0 | 5.5479 | 0.0092962 | 0.024967 | 0.885141 | 854 | 3 | 5.5479 |
| HTRA2 | 5 | 0.0086885 | 0.03229 | 0.865899 | 773 | 3 | -2.2003 | 0.0093071 | 0.037102 | 0.904772 | 855 | 2 | -2.2003 |
| TSC22D3 | 6 | 0.71977 | 0.81825 | 0.999994 | 16495 | 1 | 5.8536 | 0.0093077 | 0.041949 | 0.917066 | 856 | 5 | 5.8536 |
| SIAH1 | 6 | 0.068773 | 0.1842 | 0.968526 | 3786 | 2 | 5.7364 | 0.0093077 | 0.041949 | 0.917066 | 857 | 4 | 5.7364 |
| UBXN7 | 6 | 0.88404 | 0.89223 | 0.999994 | 18906 | 1 | 5.6988 | 0.0093172 | 0.041985 | 0.917066 | 858 | 5 | 5.6988 |
| ZNF641 | 6 | 0.99068 | 0.99064 | 0.999994 | 21366 | 0 | 2.6761 | 0.009322 | 0.042004 | 0.917066 | 859 | 6 | 2.6761 |
| ENTPD5 | 5 | 0.098018 | 0.22332 | 0.983354 | 4893 | 3 | -1.6391 | 0.0093229 | 0.037164 | 0.904772 | 860 | 1 | -1.6391 |
| RNF7 | 6 | 0.99067 | 0.99064 | 0.999994 | 21365 | 0 | 3.4538 | 0.0093278 | 0.042029 | 0.917066 | 861 | 6 | 3.4538 |
| ZNF687 | 6 | 0.98638 | 0.98634 | 0.999994 | 21257 | 0 | 3.3874 | 0.0093437 | 0.042099 | 0.917066 | 862 | 5 | 3.3874 |
| RCVRN | 6 | 0.76013 | 0.83213 | 0.999994 | 16983 | 1 | 5.7194 | 0.0093445 | 0.042101 | 0.917066 | 863 | 4 | 5.7194 |
| MEX3B | 4 | 0.89279 | 0.89249 | 0.999994 | 19087 | 0 | 6.4856 | 0.0093448 | 0.031649 | 0.890437 | 864 | 3 | 6.4856 |
| AHDC1 | 4 | 0.31927 | 0.46267 | 0.999994 | 10501 | 1 | 7.4398 | 0.0093478 | 0.031661 | 0.890437 | 865 | 3 | 7.4398 |
| ZNF337 | 6 | 0.99065 | 0.99061 | 0.999994 | 21364 | 0 | 2.8879 | 0.0093531 | 0.042137 | 0.917066 | 866 | 6 | 2.8879 |
| INCA1 | 5 | 0.66622 | 0.75171 | 0.999994 | 15872 | 1 | 7.0942 | 0.0093647 | 0.037303 | 0.904772 | 867 | 4 | 7.0942 |
| RAD18 | 5 | 0.99063 | 0.99067 | 0.999994 | 21363 | 0 | 2.3664 | 0.0093743 | 0.037339 | 0.904772 | 868 | 5 | 2.3664 |
| PANK1 | 4 | 0.99062 | 0.99055 | 0.999994 | 21362 | 0 | 3.9162 | 0.0093828 | 0.031771 | 0.890437 | 869 | 4 | 3.9162 |
| DNAJB1 | 6 | 0.95646 | 0.95653 | 0.999994 | 20502 | 0 | 4.0032 | 0.0093893 | 0.042291 | 0.918424 | 870 | 5 | 4.0032 |
| C20orf201 | 4 | 0.99061 | 0.99054 | 0.999994 | 21361 | 0 | 4.9561 | 0.0093925 | 0.0318 | 0.890437 | 871 | 4 | 4.9561 |
| LILRB2 | 5 | 0.9906 | 0.99064 | 0.999994 | 21360 | 0 | 5.3065 | 0.0093985 | 0.037421 | 0.904772 | 872 | 5 | 5.3065 |
| POGLUT1 | 5 | 0.62114 | 0.73398 | 0.999994 | 15246 | 1 | 4.5878 | 0.0094124 | 0.037473 | 0.904772 | 873 | 4 | 4.5878 |
| TM4SF1 | 5 | 0.073985 | 0.18484 | 0.968526 | 3994 | 2 | 2.5028 | 0.0094164 | 0.037488 | 0.904772 | 874 | 3 | 2.5028 |
| PRTN3 | 5 | 0.58177 | 0.71965 | 0.999994 | 14702 | 1 | 7.0916 | 0.0094194 | 0.037498 | 0.904772 | 875 | 3 | 7.0916 |
| JPH1 | 5 | 0.5421 | 0.68475 | 0.999994 | 14092 | 1 | 7.0916 | 0.0094194 | 0.037498 | 0.904772 | 876 | 3 | 7.0916 |
| HMBS | 6 | 0.88737 | 0.89449 | 0.999994 | 18976 | 1 | 5.881 | 0.0094293 | 0.042455 | 0.918424 | 877 | 5 | 5.881 |
| PCDH19 | 6 | 0.090376 | 0.22824 | 0.983354 | 4608 | 4 | -1.3123 | 0.0094383 | 0.042493 | 0.918424 | 878 | 1 | -1.3123 |
| OR8K3 | 6 | 0.15673 | 0.33409 | 0.999994 | 6645 | 2 | 3.9367 | 0.0094699 | 0.042633 | 0.918424 | 879 | 4 | 3.9367 |
| LUC7L2 | 4 | 0.99051 | 0.99044 | 0.999994 | 21359 | 0 | 3.8765 | 0.0094873 | 0.03208 | 0.890437 | 880 | 4 | 3.8765 |
| NACC2 | 6 | 0.9044 | 0.90711 | 0.999994 | 19309 | 1 | 4.2747 | 0.0094945 | 0.042745 | 0.918424 | 881 | 4 | 4.2747 |
| SSR1 | 6 | 0.9905 | 0.99047 | 0.999994 | 21358 | 0 | 1.3499 | 0.0094952 | 0.042748 | 0.918424 | 882 | 6 | 1.3499 |
| NonTargetingControlG | 1 | 0.99049 | 0.99049 | 0.999994 | 21357 | 0 | 9.6618 | 0.0095087 | 0.0094655 | 0.825215 | 883 | 1 | 9.6618 |

| | | | | | | | | | | | | |
|---|---|---|---|---|---|---|---|---|---|---|---|---|
| COL11A1 | 5 | 0.37048 | 0.53879 | 0.999994 | 11520 | 2 | 1.5753 | 0.00951 | 0.037863 | 0.904772 | 884 | 3 | 1.5753 |
| ZNF862 | 5 | 0.68226 | 0.75856 | 0.999994 | 16121 | 1 | 0.85513 | 0.0095158 | 0.037886 | 0.904772 | 885 | 3 | 0.85513 |
| APBB1 | 4 | 0.30224 | 0.44526 | 0.999994 | 10174 | 1 | 2.3837 | 0.0095263 | 0.032195 | 0.890437 | 886 | 2 | 2.3837 |
| PSME3 | 4 | 0.94442 | 0.94426 | 0.999994 | 20190 | 0 | 7.2663 | 0.0095559 | 0.032298 | 0.890437 | 887 | 4 | 7.2663 |
| CALML5 | 5 | 0.4859 | 0.63572 | 0.999994 | 13228 | 1 | 3.9121 | 0.0095568 | 0.038031 | 0.904772 | 888 | 3 | 3.9121 |
| ZNF8 | 5 | 0.99042 | 0.99045 | 0.999994 | 21356 | 0 | 2.5928 | 0.009583 | 0.03812 | 0.904772 | 889 | 5 | 2.5928 |
| ELSPBP1 | 5 | 0.90469 | 0.90478 | 0.999994 | 19315 | 0 | 6.2746 | 0.0096005 | 0.038182 | 0.904772 | 890 | 4 | 6.2746 |
| FAM120C | 6 | 0.99039 | 0.99036 | 0.999994 | 21355 | 0 | 5.8046 | 0.0096108 | 0.04324 | 0.918424 | 891 | 6 | 5.8046 |
| SVOP | 5 | 0.92803 | 0.92827 | 0.999994 | 19814 | 0 | 7.0762 | 0.0096127 | 0.03823 | 0.904772 | 892 | 3 | 7.0762 |
| GNPDA2 | 6 | 0.069809 | 0.18636 | 0.969409 | 3822 | 1 | 5.0522 | 0.0096153 | 0.04326 | 0.918424 | 893 | 4 | 5.0522 |
| hsa-mir-6755 | 3 | 0.63518 | 0.64485 | 0.999994 | 15447 | 1 | 7.746 | 0.0096285 | 0.025814 | 0.89012 | 894 | 2 | 7.746 |
| PPFIA3 | 6 | 0.99037 | 0.99034 | 0.999994 | 21354 | 0 | 4.0392 | 0.0096332 | 0.043327 | 0.918424 | 895 | 6 | 4.0392 |
| AP2A2 | 6 | 0.77244 | 0.83676 | 0.999994 | 17115 | 1 | 6.1526 | 0.0096397 | 0.043355 | 0.918424 | 896 | 5 | 6.1526 |
| IFITM10 | 6 | 0.99035 | 0.99032 | 0.999994 | 21353 | 0 | 3.6132 | 0.0096474 | 0.043386 | 0.918424 | 897 | 6 | 3.6132 |
| LOC440335 | 6 | 0.98078 | 0.9807 | 0.999994 | 21111 | 0 | 5.0242 | 0.0096678 | 0.043469 | 0.918424 | 898 | 5 | 5.0242 |
| NonTargetingControlG | 2 | 0.99033 | 0.99022 | 0.999994 | 21352 | 0 | 7.2325 | 0.0096698 | 0.018357 | 0.836675 | 899 | 2 | 7.2325 |
| HSPA12B | 5 | 0.1512 | 0.30458 | 0.999994 | 6487 | 2 | 7.0697 | 0.0096942 | 0.038532 | 0.905919 | 900 | 3 | 7.0697 |
| MED23 | 5 | 0.86418 | 0.86594 | 0.999994 | 18526 | 1 | 7.0697 | 0.0096942 | 0.038532 | 0.905919 | 901 | 4 | 7.0697 |
| LRRC49 | 4 | 0.84773 | 0.84727 | 0.999994 | 18214 | 0 | 1.8737 | 0.0096948 | 0.032752 | 0.890437 | 902 | 3 | 1.8737 |
| UNC5A | 5 | 0.63191 | 0.73803 | 0.999994 | 15397 | 1 | 5.6697 | 0.0097111 | 0.038595 | 0.905919 | 903 | 4 | 5.6697 |
| SPARC | 6 | 0.84563 | 0.86958 | 0.999994 | 18180 | 1 | 5.1615 | 0.0097278 | 0.043723 | 0.918424 | 904 | 5 | 5.1615 |
| P2RY2 | 6 | 0.29267 | 0.49816 | 0.999994 | 9968 | 2 | 2.9932 | 0.0097399 | 0.043773 | 0.918424 | 905 | 4 | 2.9932 |
| H3F3A | 3 | 0.98516 | 0.9853 | 0.999994 | 21231 | 0 | 7.9852 | 0.0097419 | 0.026109 | 0.890437 | 906 | 3 | 7.9852 |
| HMGXB3 | 6 | 0.97976 | 0.97968 | 0.999994 | 21085 | 0 | 5.2379 | 0.0097704 | 0.043903 | 0.918424 | 907 | 5 | 5.2379 |
| IGF2BP1 | 5 | 0.36727 | 0.5361 | 0.999994 | 11462 | 2 | 3.1584 | 0.0097907 | 0.038874 | 0.906957 | 908 | 3 | 3.1584 |
| NUTM2G | 6 | 0.60884 | 0.74649 | 0.999994 | 15054 | 2 | 2.3121 | 0.0098071 | 0.044066 | 0.918424 | 909 | 4 | 2.3121 |
| NonTargetingControlG | 2 | 0.51265 | 0.51556 | 0.999994 | 13617 | 1 | 4.4933 | 0.0098098 | 0.018618 | 0.836675 | 910 | 1 | 4.4933 |
| COCH | 5 | 0.99019 | 0.99022 | 0.999994 | 21351 | 0 | 4.1858 | 0.009813 | 0.03895 | 0.906957 | 911 | 5 | 4.1858 |
| DFNA5 | 5 | 0.063649 | 0.1681 | 0.967126 | 3591 | 2 | 1.4654 | 0.0098375 | 0.039043 | 0.908142 | 912 | 3 | 1.4654 |
| KCNIP3 | 6 | 0.99012 | 0.99009 | 0.999994 | 21350 | 0 | 4.5519 | 0.0098767 | 0.044368 | 0.919289 | 913 | 6 | 4.5519 |
| OR10H1 | 6 | 0.55508 | 0.70727 | 0.999994 | 14292 | 2 | 1.3036 | 0.0098873 | 0.044415 | 0.919289 | 914 | 4 | 1.3036 |
| P4HB | 5 | 0.99009 | 0.99013 | 0.999994 | 21349 | 0 | 6.8682 | 0.0099066 | 0.039286 | 0.908838 | 915 | 5 | 6.8682 |
| PDCD1 | 5 | 0.24052 | 0.43163 | 0.999994 | 8784 | 2 | 1.7798 | 0.0099093 | 0.039296 | 0.908838 | 916 | 3 | 1.7798 |
| SLC14A2 | 5 | 0.21953 | 0.40263 | 0.999994 | 8279 | 3 | -0.76357 | 0.009931 | 0.039367 | 0.908838 | 917 | 2 | -0.76357 |
| ZNF335 | 3 | 0.99007 | 0.99019 | 0.999994 | 21348 | 0 | 6.6837 | 0.009931 | 0.026597 | 0.890437 | 918 | 3 | 6.6837 |
| RFPL4A | 6 | 0.99006 | 0.99003 | 0.999994 | 21347 | 0 | 2.611 | 0.0099372 | 0.044593 | 0.919289 | 919 | 6 | 2.611 |
| FUCA2 | 6 | 0.97476 | 0.97465 | 0.999994 | 20950 | 0 | 2.4348 | 0.0099434 | 0.044618 | 0.919289 | 920 | 4 | 2.4348 |
| FAM63B | 6 | 0.99005 | 0.99001 | 0.999994 | 21346 | 0 | 2.5855 | 0.0099518 | 0.044648 | 0.919289 | 921 | 6 | 2.5855 |
| APOE | 6 | 0.49002 | 0.66193 | 0.999994 | 13277 | 1 | 5.872 | 0.009976 | 0.044741 | 0.919289 | 922 | 4 | 5.872 |
| BRI3 | 5 | 0.53317 | 0.67684 | 0.999994 | 13955 | 1 | 4.8677 | 0.0099812 | 0.039571 | 0.909688 | 923 | 4 | 4.8677 |
| TBCD | 5 | 0.80836 | 0.82486 | 0.999994 | 17607 | 1 | 5.1685 | 0.0099812 | 0.039571 | 0.909688 | 924 | 4 | 5.1685 |
| RPL13A | 4 | 0.98623 | 0.98618 | 0.999994 | 21253 | 0 | 6.5069 | 0.009996 | 0.033709 | 0.89322 | 925 | 4 | 6.5069 |
| NonTargetingControlG | 2 | 0.05838 | 0.10321 | 0.946414 | 3358 | 1 | 0.50353 | 0.0099974 | 0.018955 | 0.84631 | 926 | 1 | 0.50353 |
| POLD4 | 5 | 0.81889 | 0.83185 | 0.999994 | 17764 | 1 | 4.0801 | 0.010025 | 0.039729 | 0.909688 | 927 | 4 | 4.0801 |
| CD33 | 6 | 0.21579 | 0.4071 | 0.999994 | 8162 | 1 | 2.9077 | 0.010028 | 0.044948 | 0.919289 | 928 | 5 | 2.9077 |
| ANKRD62 | 6 | 0.025252 | 0.08758 | 0.94395 | 1763 | 4 | -2.1139 | 0.010028 | 0.044948 | 0.919289 | 929 | 1 | -2.1139 |
| FHIT | 6 | 0.98997 | 0.98993 | 0.999994 | 21345 | 0 | 3.3424 | 0.010033 | 0.044972 | 0.919289 | 930 | 6 | 3.3424 |
| NFKBIA | 6 | 0.79129 | 0.84423 | 0.999994 | 17366 | 1 | 5.9011 | 0.010047 | 0.045029 | 0.919289 | 931 | 5 | 5.9011 |
| hsa-mir-6511b-2 | 1 | 0.98993 | 0.98992 | 0.999994 | 21344 | 0 | 9.619 | 0.010074 | 0.010028 | 0.825215 | 932 | 1 | 9.619 |
| MED4 | 4 | 0.93082 | 0.93057 | 0.999994 | 19872 | 0 | 7.2436 | 0.010081 | 0.033978 | 0.894584 | 933 | 3 | 7.2436 |
| hsa-mir-4754 | 4 | 0.98992 | 0.98985 | 0.999994 | 21343 | 0 | 5.6775 | 0.010083 | 0.033983 | 0.894584 | 934 | 4 | 5.6775 |
| ZNF569 | 5 | 0.59643 | 0.72484 | 0.999994 | 14905 | 1 | 1.6104 | 0.010091 | 0.039975 | 0.909688 | 935 | 4 | 1.6104 |

| | | | | | | | | | | | | | |
|---|---|---|---|---|---|---|---|---|---|---|---|---|---|
| SNAPC5 | 5 | 0.12445 | 0.26412 | 0.996734 | 5780 | 2 | 5.8142 | 0.010091 | 0.039975 | 0.909688 | 936 | 3 | 5.8142 |
| RCN3 | 5 | 0.97413 | 0.97417 | 0.999994 | 20932 | 0 | 6.7489 | 0.010091 | 0.039975 | 0.909688 | 937 | 5 | 6.7489 |
| ARHGAP10 | 4 | 0.92746 | 0.92723 | 0.999994 | 19801 | 0 | 4.5075 | 0.010103 | 0.034047 | 0.894772 | 938 | 3 | 4.5075 |
| DHPS | 6 | 0.56029 | 0.71099 | 0.999994 | 14375 | 1 | 5.7444 | 0.010111 | 0.045289 | 0.919289 | 939 | 5 | 5.7444 |
| CCDC53 | 6 | 0.85584 | 0.87509 | 0.999994 | 18364 | 1 | 5.8964 | 0.010111 | 0.045289 | 0.919289 | 940 | 5 | 5.8964 |
| UNC5C | 6 | 0.58873 | 0.73168 | 0.999994 | 14799 | 2 | 6.502 | 0.010111 | 0.045289 | 0.919289 | 941 | 4 | 6.502 |
| CR1L | 6 | 0.12927 | 0.29939 | 0.999994 | 5923 | 3 | 0.6105 | 0.010112 | 0.04529 | 0.919289 | 942 | 3 | 0.6105 |
| ADAM33 | 5 | 0.31953 | 0.49694 | 0.999994 | 10505 | 1 | 0.43831 | 0.010118 | 0.040066 | 0.909688 | 943 | 2 | 0.43831 |
| TCAP | 4 | 0.98987 | 0.9898 | 0.999994 | 21340 | 0 | 4.2068 | 0.010133 | 0.034131 | 0.895567 | 944 | 4 | 4.2068 |
| ATXN7L3 | 6 | 0.47901 | 0.65451 | 0.999994 | 13129 | 1 | 5.0016 | 0.010134 | 0.045391 | 0.919289 | 945 | 5 | 5.0016 |
| KIN | 6 | 0.068773 | 0.1842 | 0.968526 | 3787 | 2 | 4.2018 | 0.010146 | 0.04544 | 0.919289 | 946 | 4 | 4.2018 |
| CASP1 | 5 | 0.98984 | 0.98989 | 0.999994 | 21339 | 0 | 2.7668 | 0.010158 | 0.040215 | 0.909688 | 947 | 5 | 2.7668 |
| C11orf52 | 6 | 0.85049 | 0.87219 | 0.999994 | 18270 | 1 | 4.5382 | 0.010161 | 0.045494 | 0.919289 | 948 | 4 | 4.5382 |
| CGREF1 | 6 | 0.93202 | 0.93214 | 0.999994 | 19900 | 0 | 1.0818 | 0.010168 | 0.045526 | 0.919289 | 949 | 4 | 1.0818 |
| TSEN34 | 6 | 0.98983 | 0.98978 | 0.999994 | 21338 | 0 | 1.2576 | 0.01017 | 0.045535 | 0.919289 | 950 | 6 | 1.2576 |
| GCSAML | 5 | 0.96084 | 0.96098 | 0.999994 | 20598 | 0 | 7.0303 | 0.010235 | 0.040518 | 0.913215 | 951 | 4 | 7.0303 |
| MAP2K1 | 6 | 0.98974 | 0.98969 | 0.999994 | 21337 | 0 | 3.8517 | 0.010255 | 0.045876 | 0.919289 | 952 | 6 | 3.8517 |
| hsa-mir-658 | 4 | 0.98971 | 0.98965 | 0.999994 | 21336 | 0 | 4.6484 | 0.010294 | 0.034653 | 0.897383 | 953 | 4 | 4.6484 |
| CSNK1A1 | 6 | 0.55138 | 0.70462 | 0.999994 | 14232 | 1 | 5.7945 | 0.010302 | 0.046055 | 0.919289 | 954 | 5 | 5.7945 |
| PDCD11 | 4 | 0.98965 | 0.98959 | 0.999994 | 21335 | 0 | 6.1146 | 0.010355 | 0.034848 | 0.897383 | 955 | 4 | 6.1146 |
| IL33 | 6 | 0.98964 | 0.9896 | 0.999994 | 21334 | 0 | 4.154 | 0.010358 | 0.046285 | 0.919289 | 956 | 6 | 4.154 |
| hsa-mir-4430 | 3 | 0.1919 | 0.3025 | 0.999994 | 7544 | 1 | 5.782 | 0.010359 | 0.027702 | 0.890437 | 957 | 2 | 5.782 |
| CDCA4 | 5 | 0.12335 | 0.26242 | 0.995875 | 5748 | 1 | 5.6779 | 0.010367 | 0.040988 | 0.916931 | 958 | 4 | 5.6779 |
| KDELC2 | 5 | 0.25959 | 0.44841 | 0.999994 | 9228 | 1 | 0.98395 | 0.010367 | 0.040988 | 0.916931 | 959 | 3 | 0.98395 |
| RANBP1 | 6 | 0.98963 | 0.98959 | 0.999994 | 21332 | 0 | 5.1122 | 0.010373 | 0.046351 | 0.919289 | 960 | 6 | 5.1122 |
| PRX | 6 | 0.6675 | 0.79119 | 0.999994 | 15884 | 1 | 4.6507 | 0.010373 | 0.046352 | 0.919289 | 961 | 4 | 4.6507 |
| EID2B | 5 | 0.77852 | 0.8066 | 0.999994 | 17189 | 1 | 0.89523 | 0.010399 | 0.041102 | 0.917066 | 962 | 4 | 0.89523 |
| IKBKE | 2 | 0.98959 | 0.9895 | 0.999994 | 21331 | 0 | 7.5479 | 0.010414 | 0.019729 | 0.851882 | 963 | 2 | 7.5479 |
| hsa-mir-3198-1 | 3 | 0.98956 | 0.98969 | 0.999994 | 21329 | 0 | 5.1892 | 0.010439 | 0.027907 | 0.890437 | 964 | 3 | 5.1892 |
| LETM1 | 3 | 0.98956 | 0.98969 | 0.999994 | 21330 | 0 | 5.1053 | 0.010439 | 0.027907 | 0.890437 | 965 | 3 | 5.1053 |
| RNASE6 | 6 | 0.27357 | 0.47594 | 0.999994 | 9558 | 1 | 5.744 | 0.010439 | 0.046614 | 0.919289 | 966 | 4 | 5.744 |
| NMU | 6 | 0.94216 | 0.94213 | 0.999994 | 20129 | 0 | 5.841 | 0.010439 | 0.046614 | 0.919289 | 967 | 5 | 5.841 |
| CXCL14 | 6 | 0.058353 | 0.16222 | 0.967126 | 3356 | 3 | -0.54274 | 0.010448 | 0.046651 | 0.919289 | 968 | 2 | -0.54274 |
| STAP1 | 4 | 0.98955 | 0.98951 | 0.999994 | 21328 | 0 | 5.6775 | 0.01045 | 0.035143 | 0.898494 | 969 | 4 | 5.6775 |
| CABYR | 6 | 0.62559 | 0.75902 | 0.999994 | 15317 | 1 | 4.6132 | 0.010504 | 0.046883 | 0.919289 | 970 | 4 | 4.6132 |
| GABRB3 | 6 | 0.86733 | 0.8817 | 0.999994 | 18584 | 1 | 3.528 | 0.010515 | 0.046921 | 0.919289 | 971 | 4 | 3.528 |
| PZP | 6 | 0.13566 | 0.30751 | 0.999994 | 6072 | 1 | 4.6245 | 0.01056 | 0.047089 | 0.919289 | 972 | 5 | 4.6245 |
| ZNF470 | 5 | 0.043285 | 0.12403 | 0.951052 | 2686 | 2 | -0.21431 | 0.010586 | 0.041799 | 0.917066 | 973 | 1 | -0.21431 |
| CHST3 | 6 | 0.81548 | 0.85473 | 0.999994 | 17706 | 1 | 4.0701 | 0.01063 | 0.047357 | 0.919289 | 974 | 3 | 4.0701 |
| DPP6 | 6 | 0.42616 | 0.6194 | 0.999994 | 12351 | 1 | 3.8583 | 0.01063 | 0.047357 | 0.919289 | 975 | 3 | 3.8583 |
| HLA-A | 6 | 0.68299 | 0.80331 | 0.999994 | 16130 | 2 | 0.6646 | 0.010644 | 0.047419 | 0.919289 | 976 | 3 | 0.6646 |
| FAM156A | 6 | 0.83614 | 0.86461 | 0.999994 | 18023 | 1 | 1.2216 | 0.010644 | 0.047419 | 0.919289 | 977 | 4 | 1.2216 |
| TMEM18 | 6 | 0.98932 | 0.98928 | 0.999994 | 21326 | 0 | 1.7865 | 0.010676 | 0.047546 | 0.919289 | 978 | 6 | 1.7865 |
| CMC1 | 6 | 0.91148 | 0.91281 | 0.999994 | 19455 | 1 | 4.8527 | 0.010709 | 0.047685 | 0.919289 | 979 | 4 | 4.8527 |
| AUNIP | 6 | 0.92567 | 0.92576 | 0.999994 | 19759 | 1 | 5.9981 | 0.010709 | 0.047685 | 0.919289 | 980 | 5 | 5.9981 |
| NMNAT3 | 6 | 0.98923 | 0.98919 | 0.999994 | 21324 | 0 | 2.2013 | 0.010773 | 0.047923 | 0.919289 | 981 | 6 | 2.2013 |
| DND1 | 6 | 0.98141 | 0.98133 | 0.999994 | 21127 | 0 | 6.393 | 0.010779 | 0.047952 | 0.919289 | 982 | 5 | 6.393 |
| NLRP6 | 6 | 0.54922 | 0.70308 | 0.999994 | 14197 | 1 | 5.9001 | 0.010779 | 0.047952 | 0.919289 | 983 | 4 | 5.9001 |
| NT5C2 | 6 | 0.94023 | 0.94019 | 0.999994 | 20091 | 0 | 5.7812 | 0.010779 | 0.047952 | 0.919289 | 984 | 5 | 5.7812 |
| MMP19 | 6 | 0.079414 | 0.20612 | 0.978744 | 4203 | 3 | 0.43735 | 0.010785 | 0.04797 | 0.919289 | 985 | 3 | 0.43735 |
| POU2AF1 | 4 | 0.91436 | 0.91412 | 0.999994 | 19495 | 0 | 5.0883 | 0.010799 | 0.036226 | 0.904772 | 986 | 3 | 5.0883 |
| OR1N1 | 4 | 0.98919 | 0.98916 | 0.999994 | 21322 | 0 | 3.9383 | 0.010812 | 0.036268 | 0.904772 | 987 | 4 | 3.9383 |

| | | | | | | | | | | | | | |
|---|---|---|---|---|---|---|---|---|---|---|---|---|---|
| ZBTB12 | 6 | 0.98915 | 0.98912 | 0.999994 | 21321 | 0 | 4.9656 | 0.010845 | 0.04822 | 0.919864 | 988 | 6 | 4.9656 |
| PATL2 | 6 | 0.33236 | 0.54397 | 0.999994 | 10773 | 1 | 4.7778 | 0.01087 | 0.048326 | 0.920057 | 989 | 5 | 4.7778 |
| PCGF1 | 5 | 0.98913 | 0.98916 | 0.999994 | 21320 | 0 | 4.6527 | 0.010871 | 0.042784 | 0.918424 | 990 | 5 | 4.6527 |
| NonTargetingControlG | 2 | 0.53746 | 0.53887 | 0.999994 | 14020 | 1 | 4.6058 | 0.010879 | 0.020601 | 0.853752 | 991 | 1 | 4.6058 |
| F11 | 6 | 0.29734 | 0.50362 | 0.999994 | 10074 | 1 | 1.509 | 0.010897 | 0.048429 | 0.920057 | 992 | 5 | 1.509 |
| MBNL2 | 6 | 0.98909 | 0.98905 | 0.999994 | 21319 | 0 | 3.024 | 0.010907 | 0.048465 | 0.920057 | 993 | 6 | 3.024 |
| CA11 | 5 | 0.81566 | 0.82973 | 0.999994 | 17708 | 1 | 6.9828 | 0.010929 | 0.043005 | 0.918424 | 994 | 4 | 6.9828 |
| COMMD10 | 5 | 0.93197 | 0.93225 | 0.999994 | 19899 | 0 | 6.9828 | 0.010929 | 0.043005 | 0.918424 | 995 | 4 | 6.9828 |
| ODC1 | 6 | 0.74726 | 0.8275 | 0.999994 | 16830 | 1 | 3.0039 | 0.010939 | 0.048597 | 0.920057 | 996 | 5 | 3.0039 |
| SRPX2 | 6 | 0.28978 | 0.49483 | 0.999994 | 9907 | 1 | 5.0185 | 0.010953 | 0.048648 | 0.920057 | 997 | 5 | 5.0185 |
| IKZF4 | 5 | 0.86277 | 0.86475 | 0.999994 | 18499 | 1 | 6.2408 | 0.010964 | 0.043136 | 0.918424 | 998 | 4 | 6.2408 |
| IFNL1 | 5 | 0.98903 | 0.98908 | 0.999994 | 21318 | 0 | 6.6339 | 0.010966 | 0.043145 | 0.918424 | 999 | 5 | 6.6339 |
| IRF2 | 4 | 0.03206 | 0.08568 | 0.94395 | 2110 | 1 | 6.7316 | 0.010988 | 0.036824 | 0.904772 | 1000 | 3 | 6.7316 |
| IRAK1BP1 | 6 | 0.95348 | 0.95349 | 0.999994 | 20422 | 0 | 5.1343 | 0.011001 | 0.048847 | 0.920057 | 1001 | 4 | 5.1343 |
| CLSPN | 6 | 0.29362 | 0.49927 | 0.999994 | 9989 | 2 | 1.8831 | 0.011009 | 0.048883 | 0.920057 | 1002 | 4 | 1.8831 |
| ESR1 | 6 | 0.36409 | 0.57952 | 0.999994 | 11405 | 2 | 5.9644 | 0.011015 | 0.048903 | 0.920057 | 1003 | 4 | 5.9644 |
| PITHD1 | 5 | 0.49677 | 0.64507 | 0.999994 | 13366 | 2 | 6.9784 | 0.011016 | 0.04332 | 0.918424 | 1004 | 3 | 6.9784 |
| CDK3 | 5 | 0.50203 | 0.6497 | 0.999994 | 13462 | 2 | 6.9784 | 0.011016 | 0.04332 | 0.918424 | 1005 | 3 | 6.9784 |
| TRIM15 | 5 | 0.70195 | 0.76737 | 0.999994 | 16325 | 1 | 6.5252 | 0.01102 | 0.043337 | 0.918424 | 1006 | 3 | 6.5252 |
| TMEM241 | 6 | 0.73758 | 0.82417 | 0.999994 | 16696 | 1 | 3.639 | 0.011042 | 0.049026 | 0.920223 | 1007 | 4 | 3.639 |
| SPANXN4 | 5 | 0.34024 | 0.5139 | 0.999994 | 10938 | 2 | -0.19296 | 0.011053 | 0.043454 | 0.918424 | 1008 | 2 | -0.19296 |
| HIST2H2AB | 6 | 0.77362 | 0.8372 | 0.999994 | 17128 | 1 | -0.080275 | 0.011065 | 0.04911 | 0.920223 | 1009 | 1 | -0.080275 |
| AAR2 | 6 | 0.98401 | 0.98396 | 0.999994 | 21189 | 0 | 6.2986 | 0.011085 | 0.049189 | 0.920223 | 1010 | 5 | 6.2986 |
| ZNF773 | 5 | 0.98888 | 0.98892 | 0.999994 | 21317 | 0 | 2.2756 | 0.011116 | 0.043704 | 0.918424 | 1011 | 5 | 2.2756 |
| CFD | 5 | 0.98358 | 0.98366 | 0.999994 | 21172 | 0 | 4.7106 | 0.011141 | 0.04379 | 0.918424 | 1012 | 5 | 4.7106 |
| GSDMD | 5 | 0.11117 | 0.24377 | 0.988214 | 5337 | 1 | 6.3541 | 0.01117 | 0.043888 | 0.918424 | 1013 | 3 | 6.3541 |
| UBE2G2 | 5 | 0.61749 | 0.73263 | 0.999994 | 15177 | 1 | 6.9634 | 0.011217 | 0.044052 | 0.918424 | 1014 | 4 | 6.9634 |
| TOX2 | 5 | 0.31015 | 0.48925 | 0.999994 | 10327 | 2 | -0.49909 | 0.01124 | 0.044126 | 0.918788 | 1015 | 2 | -0.49909 |
| OMP | 6 | 0.84891 | 0.87135 | 0.999994 | 18238 | 1 | 5.6996 | 0.011253 | 0.04986 | 0.923113 | 1016 | 5 | 5.6996 |
| HPDL | 3 | 0.054351 | 0.12174 | 0.951052 | 3170 | 2 | -3.2541 | 0.011258 | 0.029978 | 0.890437 | 1017 | 1 | -3.2541 |
| LOC200726 | 4 | 0.71987 | 0.73799 | 0.999994 | 16497 | 1 | 6.3108 | 0.011261 | 0.037679 | 0.904772 | 1018 | 3 | 6.3108 |
| ACSS3 | 5 | 0.10413 | 0.23284 | 0.984502 | 5101 | 2 | 0.78416 | 0.011287 | 0.04429 | 0.919289 | 1019 | 3 | 0.78416 |
| CCDC71L | 4 | 0.96601 | 0.96589 | 0.999994 | 20736 | 0 | 5.2408 | 0.011312 | 0.037841 | 0.904772 | 1020 | 4 | 5.2408 |
| ZNF667 | 6 | 0.028099 | 0.094315 | 0.945031 | 1910 | 3 | 1.9228 | 0.011325 | 0.050145 | 0.924139 | 1021 | 3 | 1.9228 |
| PSMC3IP | 5 | 0.86115 | 0.86336 | 0.999994 | 18469 | 1 | 1.702 | 0.011333 | 0.044456 | 0.919289 | 1022 | 4 | 1.702 |
| ZW10 | 6 | 0.85213 | 0.87305 | 0.999994 | 18297 | 1 | 5.4618 | 0.011334 | 0.050173 | 0.924139 | 1023 | 5 | 5.4618 |
| hsa-mir-132 | 4 | 0.066338 | 0.15288 | 0.967126 | 3699 | 2 | -0.41222 | 0.011435 | 0.03821 | 0.904772 | 1024 | 1 | -0.41222 |
| CHMP2A | 5 | 0.98854 | 0.98856 | 0.999994 | 21314 | 0 | 5.0629 | 0.011456 | 0.044875 | 0.919289 | 1025 | 5 | 5.0629 |
| RBX1 | 5 | 0.62194 | 0.73428 | 0.999994 | 15257 | 1 | 4.5924 | 0.011461 | 0.044896 | 0.919289 | 1026 | 4 | 4.5924 |
| CCSAP | 5 | 0.3095 | 0.48874 | 0.999994 | 10316 | 2 | 0.31931 | 0.011473 | 0.044928 | 0.919289 | 1027 | 2 | 0.31931 |
| ZBTB8OS | 6 | 0.81429 | 0.85419 | 0.999994 | 17694 | 1 | 5.3339 | 0.011476 | 0.050719 | 0.924349 | 1028 | 5 | 5.3339 |
| TMEM91 | 6 | 0.90486 | 0.90746 | 0.999994 | 19323 | 1 | 6.0496 | 0.011488 | 0.050773 | 0.924349 | 1029 | 5 | 6.0496 |
| COMMD3-BMI1 | 2 | 0.98851 | 0.98844 | 0.999994 | 21313 | 0 | 7.5126 | 0.011491 | 0.021745 | 0.85539 | 1030 | 2 | 7.5126 |
| IKZF5 | 6 | 0.98751 | 0.98746 | 0.999994 | 21290 | 0 | 4.5674 | 0.011493 | 0.05079 | 0.924349 | 1031 | 6 | 4.5674 |
| ZNF551 | 6 | 0.93171 | 0.93185 | 0.999994 | 19893 | 0 | 4.0432 | 0.011494 | 0.050797 | 0.924349 | 1032 | 4 | 4.0432 |
| POLR2M | 6 | 0.70309 | 0.81295 | 0.999994 | 16340 | 1 | 2.4005 | 0.011513 | 0.050857 | 0.924349 | 1033 | 5 | 2.4005 |
| NonTargetingControlG | 2 | 0.16972 | 0.24046 | 0.987253 | 6978 | 1 | 2.6636 | 0.011545 | 0.021834 | 0.85539 | 1034 | 1 | 2.6636 |
| GPR137 | 6 | 0.63361 | 0.76508 | 0.999994 | 15426 | 2 | 1.1332 | 0.011569 | 0.051086 | 0.924349 | 1035 | 4 | 1.1332 |
| ARFGAP3 | 6 | 0.44745 | 0.63338 | 0.999994 | 12679 | 2 | 0.070369 | 0.011581 | 0.051137 | 0.924349 | 1036 | 3 | 0.070369 |
| OR7G2 | 6 | 0.98842 | 0.98838 | 0.999994 | 21311 | 0 | 3.2341 | 0.011583 | 0.051143 | 0.924349 | 1037 | 6 | 3.2341 |
| KCNG2 | 5 | 0.98841 | 0.98842 | 0.999994 | 21310 | 0 | 2.6389 | 0.01159 | 0.045336 | 0.919289 | 1038 | 5 | 2.6389 |
| SCG3 | 5 | 0.13011 | 0.27275 | 0.998561 | 5936 | 2 | 2.1942 | 0.01159 | 0.045337 | 0.919289 | 1039 | 3 | 2.1942 |

| | | | | | | | | | | | | |
|---|---|---|---|---|---|---|---|---|---|---|---|---|
| FILIP1 | 6 | 0.9884 | 0.98836 | 0.999994 | 21309 | 0 | 3.5964 | 0.011599 | 0.051213 | 0.924349 | 1040 | 6 | 3.5964 |
| hsa-mir-487a | 4 | 0.96698 | 0.96687 | 0.999994 | 20760 | 0 | 7.0847 | 0.0116 | 0.038725 | 0.905919 | 1041 | 4 | 7.0847 |
| FTSJ1 | 6 | 0.90069 | 0.90421 | 0.999994 | 19231 | 1 | 2.8966 | 0.011645 | 0.051393 | 0.924349 | 1042 | 5 | 2.8966 |
| IL17A | 5 | 0.7817 | 0.80842 | 0.999994 | 17235 | 1 | 0.73168 | 0.01166 | 0.045569 | 0.919289 | 1043 | 3 | 0.73168 |
| ST3GAL1 | 6 | 0.59778 | 0.73831 | 0.999994 | 14925 | 2 | -0.17925 | 0.011681 | 0.051529 | 0.925271 | 1044 | 2 | -0.17925 |
| CABP4 | 6 | 0.98831 | 0.98827 | 0.999994 | 21306 | 0 | 3.5561 | 0.01169 | 0.051568 | 0.925271 | 1045 | 6 | 3.5561 |
| ABCD3 | 6 | 0.96262 | 0.96268 | 0.999994 | 20647 | 0 | 3.1026 | 0.011698 | 0.051594 | 0.925271 | 1046 | 4 | 3.1026 |
| APBB1IP | 6 | 0.35334 | 0.56777 | 0.999994 | 11219 | 3 | 3.3413 | 0.011754 | 0.051822 | 0.926769 | 1047 | 3 | 3.3413 |
| OBFC1 | 4 | 0.94368 | 0.94355 | 0.999994 | 20167 | 0 | 7.136 | 0.011766 | 0.039233 | 0.908838 | 1048 | 3 | 7.136 |
| APEX2 | 3 | 0.65951 | 0.66493 | 0.999994 | 15771 | 1 | 2.6281 | 0.011819 | 0.031378 | 0.890437 | 1049 | 2 | 2.6281 |
| IL2 | 6 | 0.1872 | 0.37204 | 0.999994 | 7398 | 3 | 2.9074 | 0.011838 | 0.052145 | 0.926769 | 1050 | 3 | 2.9074 |
| PROB1 | 5 | 0.14905 | 0.30132 | 0.999994 | 6431 | 3 | -1.1687 | 0.011847 | 0.046249 | 0.919289 | 1051 | 2 | -1.1687 |
| IDNK | 6 | 0.28091 | 0.48445 | 0.999994 | 9719 | 2 | 4.0184 | 0.011849 | 0.052193 | 0.926769 | 1052 | 4 | 4.0184 |
| PROS1 | 6 | 0.98814 | 0.9881 | 0.999994 | 21305 | 0 | 1.9444 | 0.011855 | 0.05222 | 0.926769 | 1053 | 6 | 1.9444 |
| IFT57 | 5 | 0.25005 | 0.44082 | 0.999994 | 9020 | 1 | 3.7469 | 0.011857 | 0.046286 | 0.919289 | 1054 | 4 | 3.7469 |
| PLEKHO1 | 5 | 0.98811 | 0.9881 | 0.999994 | 21304 | 0 | 3.6773 | 0.011885 | 0.046387 | 0.919289 | 1055 | 5 | 3.6773 |
| TMBIM4 | 5 | 0.98811 | 0.98809 | 0.999994 | 21303 | 0 | 5.835 | 0.011888 | 0.046395 | 0.919289 | 1056 | 5 | 5.835 |
| DEDD2 | 6 | 0.65618 | 0.78233 | 0.999994 | 15727 | 1 | 6.1557 | 0.011922 | 0.052467 | 0.926769 | 1057 | 5 | 6.1557 |
| ECD | 3 | 0.98807 | 0.98823 | 0.999994 | 21302 | 0 | 6.0743 | 0.011932 | 0.031669 | 0.890437 | 1058 | 3 | 6.0743 |
| NonTargetingControlG | 2 | 0.98805 | 0.98795 | 0.999994 | 21300 | 0 | 7.3756 | 0.011951 | 0.022602 | 0.863933 | 1059 | 2 | 7.3756 |
| MEPE | 5 | 0.92519 | 0.92542 | 0.999994 | 19748 | 0 | 5.835 | 0.011982 | 0.046724 | 0.919289 | 1060 | 4 | 5.835 |
| SNRK | 5 | 0.98498 | 0.98503 | 0.999994 | 21221 | 0 | 4.6527 | 0.011982 | 0.046724 | 0.919289 | 1061 | 5 | 4.6527 |
| AATK | 5 | 0.65678 | 0.74783 | 0.999994 | 15741 | 1 | 6.1084 | 0.011982 | 0.046724 | 0.919289 | 1062 | 4 | 6.1084 |
| BCAS2 | 5 | 0.040884 | 0.11807 | 0.951052 | 2554 | 2 | 6.9136 | 0.011984 | 0.046729 | 0.919289 | 1063 | 3 | 6.9136 |
| HIST1H3J | 6 | 0.92603 | 0.92611 | 0.999994 | 19766 | 1 | 3.5635 | 0.01201 | 0.052814 | 0.926769 | 1064 | 4 | 3.5635 |
| CRTC2 | 6 | 0.62345 | 0.75738 | 0.999994 | 15284 | 2 | 5.5192 | 0.012031 | 0.052909 | 0.926769 | 1065 | 4 | 5.5192 |
| B3GNT4 | 4 | 0.55967 | 0.64446 | 0.999994 | 14365 | 1 | 3.992 | 0.012041 | 0.040079 | 0.909688 | 1066 | 2 | 3.992 |
| hsa-mir-1825 | 3 | 0.98791 | 0.98807 | 0.999994 | 21299 | 0 | 6.1178 | 0.012092 | 0.032076 | 0.890437 | 1067 | 3 | 6.1178 |
| DESI2 | 5 | 0.19598 | 0.36948 | 0.999994 | 7654 | 2 | 0.45924 | 0.012104 | 0.047178 | 0.919289 | 1068 | 2 | 0.45924 |
| hsa-mir-4705 | 4 | 0.82996 | 0.8295 | 0.999994 | 17930 | 0 | 0.42531 | 0.012107 | 0.040286 | 0.909688 | 1069 | 2 | 0.42531 |
| C3orf20 | 5 | 0.46815 | 0.62054 | 0.999994 | 12977 | 2 | 1.1795 | 0.012112 | 0.0472 | 0.919289 | 1070 | 3 | 1.1795 |
| SLC36A2 | 5 | 0.88827 | 0.88838 | 0.999994 | 19002 | 1 | 6.1768 | 0.01218 | 0.047444 | 0.919289 | 1071 | 4 | 6.1768 |
| TCP11 | 6 | 0.34157 | 0.55445 | 0.999994 | 10971 | 2 | 2.1172 | 0.012181 | 0.053498 | 0.926769 | 1072 | 4 | 2.1172 |
| HDAC1 | 6 | 0.98782 | 0.98778 | 0.999994 | 21298 | 0 | 3.3279 | 0.012182 | 0.053505 | 0.926769 | 1073 | 6 | 3.3279 |
| RPL37 | 6 | 0.67358 | 0.79594 | 0.999994 | 15981 | 2 | 5.5996 | 0.012185 | 0.053517 | 0.926769 | 1074 | 4 | 5.5996 |
| RHPN1 | 6 | 0.72669 | 0.82051 | 0.999994 | 16563 | 1 | 6.312 | 0.012185 | 0.053517 | 0.926769 | 1075 | 5 | 6.312 |
| GPR182 | 6 | 0.86818 | 0.88222 | 0.999994 | 18604 | 1 | 5.8673 | 0.012185 | 0.053517 | 0.926769 | 1076 | 5 | 5.8673 |
| CACNA1I | 6 | 0.40168 | 0.60345 | 0.999994 | 11973 | 1 | 1.213 | 0.012185 | 0.05352 | 0.926769 | 1077 | 5 | 1.213 |
| CHMP7 | 5 | 0.98781 | 0.9878 | 0.999994 | 21297 | 0 | 1.884 | 0.012186 | 0.047466 | 0.919289 | 1078 | 5 | 1.884 |
| RAB21 | 5 | 0.12965 | 0.27205 | 0.998561 | 5929 | 1 | 4.3663 | 0.012198 | 0.047505 | 0.919289 | 1079 | 4 | 4.3663 |
| STK32B | 6 | 0.9878 | 0.98776 | 0.999994 | 21296 | 0 | 2.041 | 0.012204 | 0.05359 | 0.926769 | 1080 | 6 | 2.041 |
| CANX | 6 | 0.18388 | 0.36798 | 0.999994 | 7314 | 2 | 4.2402 | 0.012215 | 0.053636 | 0.926769 | 1081 | 4 | 4.2402 |
| hsa-mir-451a | 4 | 0.87769 | 0.87734 | 0.999994 | 18782 | 0 | 0.27178 | 0.01222 | 0.04063 | 0.913215 | 1082 | 2 | 0.27178 |
| CRTC3 | 6 | 0.97462 | 0.97451 | 0.999994 | 20946 | 0 | 3.6207 | 0.012225 | 0.053676 | 0.926769 | 1083 | 4 | 3.6207 |
| CABLES1 | 5 | 0.38868 | 0.55402 | 0.999994 | 11777 | 1 | 6.8934 | 0.012235 | 0.04764 | 0.919289 | 1084 | 4 | 6.8934 |
| S100A5 | 5 | 0.84652 | 0.8516 | 0.999994 | 18201 | 1 | 6.8934 | 0.012235 | 0.04764 | 0.919289 | 1085 | 4 | 6.8934 |
| SOX4 | 6 | 0.98776 | 0.98772 | 0.999994 | 21295 | 0 | 1.4547 | 0.012244 | 0.053744 | 0.926769 | 1086 | 6 | 1.4547 |
| PAGE4 | 5 | 0.46219 | 0.61548 | 0.999994 | 12891 | 2 | 1.5889 | 0.012252 | 0.047693 | 0.919289 | 1087 | 3 | 1.5889 |
| MAP2K5 | 6 | 0.98773 | 0.9877 | 0.999994 | 21294 | 0 | 1.6083 | 0.012267 | 0.05384 | 0.926769 | 1088 | 6 | 1.6083 |
| LMO7 | 6 | 0.58011 | 0.7253 | 0.999994 | 14678 | 2 | 3.2659 | 0.012292 | 0.053943 | 0.926769 | 1089 | 3 | 3.2659 |
| DCP1B | 6 | 0.067528 | 0.18159 | 0.968171 | 3736 | 3 | -0.67734 | 0.012297 | 0.053957 | 0.926769 | 1090 | 2 | -0.67734 |
| ASB2 | 5 | 0.22599 | 0.41165 | 0.999994 | 8425 | 1 | 4.1858 | 0.012304 | 0.047883 | 0.919289 | 1091 | 4 | 4.1858 |

| | | | | | | | | | | | | |
|---|---|---|---|---|---|---|---|---|---|---|---|---|
| NPC2 | 5 | 0.93285 | 0.93314 | 0.999994 | 19923 | 0 | 6.8932 | 0.01232 | 0.047935 | 0.919289 | 1092 | 4 | 6.8932 |
| IGF2BP2 | 5 | 0.48421 | 0.6343 | 0.999994 | 13204 | 2 | 6.8932 | 0.01232 | 0.047935 | 0.919289 | 1093 | 3 | 6.8932 |
| C15orf56 | 6 | 0.98367 | 0.98362 | 0.999994 | 21177 | 0 | 5.4711 | 0.012323 | 0.05407 | 0.926769 | 1094 | 5 | 5.4711 |
| FGF2 | 4 | 0.62697 | 0.6802 | 0.999994 | 15337 | 1 | 3.7238 | 0.012332 | 0.04097 | 0.916931 | 1095 | 3 | 3.7238 |
| PHYHD1 | 5 | 0.98765 | 0.98762 | 0.999994 | 21293 | 0 | 4.686 | 0.012348 | 0.048032 | 0.919667 | 1096 | 5 | 4.686 |
| TSTD2 | 6 | 0.4244 | 0.61828 | 0.999994 | 12323 | 2 | 4.896 | 0.012348 | 0.054156 | 0.926769 | 1097 | 4 | 4.896 |
| MN1 | 6 | 0.064715 | 0.17566 | 0.968171 | 3638 | 4 | -1.637 | 0.012353 | 0.054176 | 0.926769 | 1098 | 2 | -1.637 |
| KRTAP4-7 | 5 | 0.36313 | 0.53266 | 0.999994 | 11390 | 1 | 4.5924 | 0.012366 | 0.048098 | 0.919864 | 1099 | 4 | 4.5924 |
| SEC61A1 | 3 | 0.98763 | 0.98777 | 0.999994 | 21292 | 0 | 5.7269 | 0.012375 | 0.032772 | 0.890437 | 1100 | 3 | 5.7269 |
| PHF16 | 5 | 0.83258 | 0.84133 | 0.999994 | 17967 | 1 | 0.66985 | 0.012403 | 0.048232 | 0.919864 | 1101 | 3 | 0.66985 |
| hsa-mir-1306 | 4 | 0.98759 | 0.98756 | 0.999994 | 21291 | 0 | 3.2816 | 0.012413 | 0.041209 | 0.917066 | 1102 | 4 | 3.2816 |
| HYOU1 | 6 | 0.8782 | 0.88844 | 0.999994 | 18792 | 1 | 6.075 | 0.012437 | 0.054518 | 0.926769 | 1103 | 5 | 6.075 |
| APH1A | 6 | 0.27733 | 0.48032 | 0.999994 | 9641 | 3 | 3.1845 | 0.012463 | 0.054615 | 0.926769 | 1104 | 3 | 3.1845 |
| SPRED3 | 5 | 0.92084 | 0.92102 | 0.999994 | 19650 | 0 | 6.8809 | 0.012499 | 0.048572 | 0.920057 | 1105 | 4 | 6.8809 |
| ACSL5 | 6 | 0.98748 | 0.98744 | 0.999994 | 21289 | 0 | 1.5693 | 0.012515 | 0.054836 | 0.926769 | 1106 | 6 | 1.5693 |
| COX11 | 6 | 0.32748 | 0.53836 | 0.999994 | 10669 | 2 | 4.8497 | 0.012517 | 0.054842 | 0.926769 | 1107 | 4 | 4.8497 |
| KCNA7 | 6 | 0.10722 | 0.26135 | 0.995005 | 5219 | 1 | 3.3579 | 0.012538 | 0.054925 | 0.926769 | 1108 | 5 | 3.3579 |
| TBC1D2 | 5 | 0.61777 | 0.73274 | 0.999994 | 15182 | 1 | 0.38763 | 0.012547 | 0.048735 | 0.920057 | 1109 | 2 | 0.38763 |
| CHPT1 | 6 | 0.55579 | 0.70779 | 0.999994 | 14302 | 1 | 3.739 | 0.012549 | 0.054968 | 0.926769 | 1110 | 3 | 3.739 |
| TCF7 | 6 | 0.39296 | 0.59782 | 0.999994 | 11847 | 2 | 2.1285 | 0.012555 | 0.054992 | 0.926769 | 1111 | 3 | 2.1285 |
| SHROOM1 | 5 | 0.14784 | 0.29954 | 0.999994 | 6404 | 3 | -1.1708 | 0.012566 | 0.048803 | 0.920057 | 1112 | 2 | -1.1708 |
| UBA1 | 3 | 0.7483 | 0.74829 | 0.999994 | 16841 | 0 | 2.8483 | 0.012576 | 0.033266 | 0.892653 | 1113 | 2 | 2.8483 |
| GPR113 | 6 | 0.21596 | 0.4073 | 0.999994 | 8169 | 2 | 3.9702 | 0.012635 | 0.055309 | 0.926769 | 1114 | 4 | 3.9702 |
| DEFB123 | 6 | 0.078377 | 0.20405 | 0.978186 | 4167 | 2 | 3.5674 | 0.012649 | 0.055362 | 0.926769 | 1115 | 4 | 3.5674 |
| APOLD1 | 4 | 0.65389 | 0.69555 | 0.999994 | 15705 | 1 | 6.2443 | 0.012651 | 0.041934 | 0.917066 | 1116 | 3 | 6.2443 |
| ZSCAN30 | 4 | 0.89168 | 0.89142 | 0.999994 | 19066 | 0 | 6.1709 | 0.012651 | 0.041934 | 0.917066 | 1117 | 3 | 6.1709 |
| TMEM129 | 6 | 0.3659 | 0.58064 | 0.999994 | 11440 | 2 | 5.6789 | 0.012662 | 0.055412 | 0.926769 | 1118 | 4 | 5.6789 |
| PGRMC1 | 5 | 0.94002 | 0.9402 | 0.999994 | 20078 | 0 | 6.8682 | 0.012677 | 0.049209 | 0.920223 | 1119 | 4 | 6.8682 |
| KHDC3L | 5 | 0.96783 | 0.96797 | 0.999994 | 20781 | 0 | 1.8915 | 0.012687 | 0.049255 | 0.920284 | 1120 | 3 | 1.8915 |
| TAC4 | 6 | 0.087175 | 0.22182 | 0.983139 | 4484 | 3 | 1.5852 | 0.012706 | 0.055565 | 0.926769 | 1121 | 3 | 1.5852 |
| YIPF4 | 5 | 0.98729 | 0.98725 | 0.999994 | 21287 | 0 | 2.0904 | 0.012711 | 0.049333 | 0.920728 | 1122 | 5 | 2.0904 |
| PMM2 | 6 | 0.10953 | 0.26579 | 0.997535 | 5289 | 1 | 5.9908 | 0.012717 | 0.055608 | 0.926769 | 1123 | 5 | 5.9908 |
| ALG14 | 5 | 0.4473 | 0.60295 | 0.999994 | 12677 | 1 | 4.2513 | 0.012724 | 0.049379 | 0.920728 | 1124 | 4 | 4.2513 |
| SHISA7 | 6 | 0.98727 | 0.98722 | 0.999994 | 21286 | 0 | 4.1376 | 0.012734 | 0.05568 | 0.926769 | 1125 | 6 | 4.1376 |
| TMEM88 | 4 | 0.903 | 0.90269 | 0.999994 | 19278 | 0 | 0.7352 | 0.012743 | 0.042214 | 0.917805 | 1126 | 2 | 0.7352 |
| ZNF175 | 5 | 0.98718 | 0.98715 | 0.999994 | 21284 | 0 | 4.2583 | 0.012819 | 0.049694 | 0.922919 | 1127 | 5 | 4.2583 |
| ZBTB8A | 6 | 0.10069 | 0.24866 | 0.990929 | 4998 | 1 | 1.7323 | 0.012839 | 0.056117 | 0.926769 | 1128 | 4 | 1.7323 |
| TNFRSF10D | 4 | 0.69302 | 0.71991 | 0.999994 | 16235 | 1 | 6.3095 | 0.012845 | 0.042534 | 0.918424 | 1129 | 3 | 6.3095 |
| CD109 | 6 | 0.97528 | 0.97521 | 0.999994 | 20964 | 0 | 4.3673 | 0.012857 | 0.056193 | 0.926769 | 1130 | 5 | 4.3673 |
| FKBP8 | 6 | 0.47221 | 0.64994 | 0.999994 | 13038 | 2 | 5.8097 | 0.012859 | 0.056202 | 0.926769 | 1131 | 4 | 5.8097 |
| CKAP5 | 5 | 0.65892 | 0.7487 | 0.999994 | 15763 | 1 | 4.8199 | 0.012882 | 0.049913 | 0.923113 | 1132 | 4 | 4.8199 |
| MMP2 | 6 | 0.0036779 | 0.016265 | 0.772095 | 419 | 4 | -3.3708 | 0.0129 | 0.05636 | 0.926769 | 1133 | 2 | -3.3708 |
| ZNF805 | 5 | 0.98709 | 0.98707 | 0.999994 | 21283 | 0 | 1.8223 | 0.012911 | 0.050005 | 0.92348 | 1134 | 5 | 1.8223 |
| LOC100127983 | 6 | 0.27576 | 0.47848 | 0.999994 | 9613 | 3 | -0.16809 | 0.012913 | 0.056424 | 0.926769 | 1135 | 2 | -0.16809 |
| OR7D4 | 6 | 0.76142 | 0.8326 | 0.999994 | 16996 | 1 | -0.030839 | 0.012942 | 0.056545 | 0.926769 | 1136 | 2 | -0.030839 |
| TMEM8C | 6 | 0.38113 | 0.59029 | 0.999994 | 11662 | 1 | 5.8474 | 0.012962 | 0.056619 | 0.926769 | 1137 | 5 | 5.8474 |
| TNFRSF13B | 6 | 0.70669 | 0.81406 | 0.999994 | 16381 | 1 | 5.3513 | 0.012962 | 0.056619 | 0.926769 | 1138 | 5 | 5.3513 |
| XPNPEP1 | 4 | 0.63847 | 0.68663 | 0.999994 | 15491 | 1 | 4.3054 | 0.012967 | 0.042898 | 0.918424 | 1139 | 3 | 4.3054 |
| RAB42 | 6 | 0.77675 | 0.8384 | 0.999994 | 17162 | 1 | 1.6524 | 0.012969 | 0.056642 | 0.926769 | 1140 | 4 | 1.6524 |
| WDR38 | 6 | 0.95542 | 0.95548 | 0.999994 | 20464 | 0 | 3.7912 | 0.012972 | 0.056655 | 0.926769 | 1141 | 5 | 3.7912 |
| RASL11A | 5 | 0.95151 | 0.95167 | 0.999994 | 20377 | 0 | 5.8779 | 0.012977 | 0.05023 | 0.924153 | 1142 | 4 | 5.8779 |
| NonTargetingControlG | 2 | 0.987 | 0.98693 | 0.999994 | 21281 | 0 | 6.9818 | 0.013001 | 0.02452 | 0.882096 | 1143 | 2 | 6.9818 |

| | | | | | | | | | | | | | |
|---|---|---|---|---|---|---|---|---|---|---|---|---|---|
| EVL | 6 | 0.021059 | 0.075392 | 0.93914 | 1517 | 4 | -2.3813 | 0.013025 | 0.056851 | 0.926769 | 1144 | 1 | -2.3813 |
| MYCBP | 4 | 0.98697 | 0.98693 | 0.999994 | 21280 | 0 | 3.7499 | 0.013027 | 0.04308 | 0.918424 | 1145 | 4 | 3.7499 |
| CECR5 | 6 | 0.59295 | 0.73476 | 0.999994 | 14861 | 1 | 5.8583 | 0.013051 | 0.056953 | 0.926769 | 1146 | 4 | 5.8583 |
| SLC30A8 | 6 | 0.58101 | 0.72596 | 0.999994 | 14691 | 2 | 5.0428 | 0.013054 | 0.056967 | 0.926769 | 1147 | 4 | 5.0428 |
| ABCA7 | 6 | 0.067569 | 0.18167 | 0.968171 | 3740 | 1 | 3.8139 | 0.013054 | 0.056967 | 0.926769 | 1148 | 3 | 3.8139 |
| OR4C3 | 5 | 0.98695 | 0.98693 | 0.999994 | 21279 | 0 | 4.9089 | 0.013054 | 0.050507 | 0.924349 | 1149 | 5 | 4.9089 |
| DCN | 6 | 0.46625 | 0.64596 | 0.999994 | 12955 | 1 | 2.757 | 0.013058 | 0.056982 | 0.926769 | 1150 | 5 | 2.757 |
| LRRC55 | 6 | 0.048246 | 0.14024 | 0.958716 | 2897 | 1 | 5.2503 | 0.013068 | 0.057023 | 0.926769 | 1151 | 5 | 5.2503 |
| LIN28B | 6 | 0.98643 | 0.98638 | 0.999994 | 21258 | 0 | 2.4422 | 0.013069 | 0.057025 | 0.926769 | 1152 | 5 | 2.4422 |
| ARHGEF17 | 5 | 0.4004 | 0.56385 | 0.999994 | 11956 | 1 | 6.2587 | 0.013091 | 0.050635 | 0.924349 | 1153 | 4 | 6.2587 |
| HMGB4 | 5 | 0.21705 | 0.39915 | 0.999994 | 8196 | 2 | 1.4465 | 0.013107 | 0.050694 | 0.924349 | 1154 | 3 | 1.4465 |
| MEGF9 | 5 | 0.88096 | 0.88119 | 0.999994 | 18852 | 1 | 1.7596 | 0.013114 | 0.05072 | 0.924349 | 1155 | 3 | 1.7596 |
| GKAP1 | 6 | 0.44484 | 0.63166 | 0.999994 | 12645 | 1 | 3.7449 | 0.01312 | 0.057229 | 0.926769 | 1156 | 5 | 3.7449 |
| ALDH1A2 | 6 | 0.88626 | 0.89374 | 0.999994 | 18955 | 1 | 4.3315 | 0.013122 | 0.057242 | 0.926769 | 1157 | 5 | 4.3315 |
| ARHGAP33 | 6 | 0.16441 | 0.34374 | 0.999994 | 6849 | 2 | 2.5027 | 0.013137 | 0.057298 | 0.926769 | 1158 | 4 | 2.5027 |
| BPHL | 6 | 0.052256 | 0.14902 | 0.965566 | 3083 | 1 | 1.246 | 0.013137 | 0.057298 | 0.926769 | 1159 | 5 | 1.246 |
| TMEM194B | 6 | 0.073219 | 0.19345 | 0.975797 | 3965 | 3 | -0.30088 | 0.013137 | 0.057298 | 0.926769 | 1160 | 3 | -0.30088 |
| ORAI1 | 5 | 0.8465 | 0.85159 | 0.999994 | 18200 | 1 | 4.3222 | 0.01314 | 0.050817 | 0.924349 | 1161 | 4 | 4.3222 |
| TCTEX1D2 | 5 | 0.98684 | 0.98682 | 0.999994 | 21277 | 0 | 1.6549 | 0.013157 | 0.050884 | 0.924349 | 1162 | 5 | 1.6549 |
| GPR85 | 6 | 0.0266 | 0.090771 | 0.94395 | 1832 | 3 | 3.1618 | 0.013159 | 0.057383 | 0.926769 | 1163 | 3 | 3.1618 |
| ELP5 | 6 | 0.30769 | 0.51569 | 0.999994 | 10281 | 3 | 3.2019 | 0.013159 | 0.057383 | 0.926769 | 1164 | 3 | 3.2019 |
| JPH4 | 4 | 0.98684 | 0.98679 | 0.999994 | 21275 | 0 | 3.2424 | 0.01316 | 0.04349 | 0.918424 | 1165 | 4 | 3.2424 |
| NOX1 | 6 | 0.77127 | 0.83631 | 0.999994 | 17102 | 1 | 6.0688 | 0.01316 | 0.057387 | 0.926769 | 1166 | 5 | 6.0688 |
| STBD1 | 5 | 0.18952 | 0.36025 | 0.999994 | 7470 | 3 | -0.86903 | 0.013177 | 0.050948 | 0.924349 | 1167 | 2 | -0.86903 |
| TMEM231 | 5 | 0.96233 | 0.96249 | 0.999994 | 20638 | 0 | 5.2669 | 0.01323 | 0.051143 | 0.924349 | 1168 | 4 | 5.2669 |
| GNAO1 | 6 | 0.98677 | 0.98671 | 0.999994 | 21272 | 0 | 3.6892 | 0.013231 | 0.057649 | 0.928172 | 1169 | 6 | 3.6892 |
| VPS18 | 4 | 0.97559 | 0.97547 | 0.999994 | 20980 | 0 | 6.3209 | 0.013241 | 0.043734 | 0.918424 | 1170 | 4 | 6.3209 |
| ZNF446 | 5 | 0.71543 | 0.77366 | 0.999994 | 16453 | 1 | 3.7046 | 0.013254 | 0.051233 | 0.924349 | 1171 | 4 | 3.7046 |
| CASK | 6 | 0.3539 | 0.56841 | 0.999994 | 11233 | 1 | 2.4331 | 0.013269 | 0.057796 | 0.928656 | 1172 | 3 | 2.4331 |
| C10orf67 | 6 | 0.98673 | 0.98668 | 0.999994 | 21271 | 0 | 3.6471 | 0.013274 | 0.057814 | 0.928656 | 1173 | 6 | 3.6471 |
| HOMEZ | 4 | 0.98672 | 0.98667 | 0.999994 | 21270 | 0 | 4.6404 | 0.013276 | 0.043838 | 0.918424 | 1174 | 4 | 4.6404 |
| DONSON | 3 | 0.52103 | 0.56023 | 0.999994 | 13761 | 1 | 7.7728 | 0.013296 | 0.035086 | 0.898105 | 1175 | 2 | 7.7728 |
| LSMD1 | 5 | 0.95554 | 0.95571 | 0.999994 | 20468 | 0 | 3.5324 | 0.013298 | 0.051372 | 0.924349 | 1176 | 4 | 3.5324 |
| TMEM248 | 6 | 0.021581 | 0.077071 | 0.93914 | 1552 | 4 | -2.7901 | 0.013305 | 0.057927 | 0.928945 | 1177 | 2 | -2.7901 |
| ARVCF | 6 | 0.22611 | 0.41942 | 0.999994 | 8427 | 2 | 2.6819 | 0.013312 | 0.057955 | 0.928945 | 1178 | 4 | 2.6819 |
| NonTargetingControlG | 2 | 0.58777 | 0.58779 | 0.999994 | 14788 | 0 | 4.6537 | 0.013325 | 0.025083 | 0.885141 | 1179 | 1 | 4.6537 |
| ZFP37 | 4 | 0.56332 | 0.64634 | 0.999994 | 14437 | 1 | 6.3651 | 0.013346 | 0.044051 | 0.918424 | 1180 | 3 | 6.3651 |
| KIF14 | 4 | 0.98509 | 0.98506 | 0.999994 | 21226 | 0 | 6.3866 | 0.013346 | 0.044051 | 0.918424 | 1181 | 4 | 6.3866 |
| MAP3K7CL | 4 | 0.09067 | 0.19827 | 0.977164 | 4625 | 1 | 6.0528 | 0.013346 | 0.044051 | 0.918424 | 1182 | 3 | 6.0528 |
| ALDH4A1 | 6 | 0.98047 | 0.98038 | 0.999994 | 21103 | 0 | 5.6227 | 0.013359 | 0.058132 | 0.929393 | 1183 | 5 | 5.6227 |
| PSMA6 | 2 | 0.98664 | 0.9866 | 0.999994 | 21267 | 0 | 7.4203 | 0.013361 | 0.025144 | 0.885141 | 1184 | 2 | 7.4203 |
| C9orf72 | 6 | 0.94289 | 0.94287 | 0.999994 | 20149 | 0 | 0.80072 | 0.013389 | 0.058224 | 0.929393 | 1185 | 5 | 0.80072 |
| EMILIN2 | 6 | 0.84908 | 0.87144 | 0.999994 | 18242 | 1 | 2.662 | 0.01339 | 0.058226 | 0.929393 | 1186 | 5 | 2.662 |
| ZDHHC17 | 6 | 0.56557 | 0.71477 | 0.999994 | 14477 | 1 | 0.0022167 | 0.013397 | 0.058251 | 0.929393 | 1187 | 2 | 0.0022167 |
| CAPN2 | 5 | 0.9866 | 0.9866 | 0.999994 | 21266 | 0 | 3.7891 | 0.013402 | 0.051727 | 0.926504 | 1188 | 5 | 3.7891 |
| TRAPPC6A | 6 | 0.38219 | 0.59096 | 0.999994 | 11680 | 2 | 2.6338 | 0.013455 | 0.058434 | 0.929393 | 1189 | 4 | 2.6338 |
| SPHK2 | 6 | 0.78979 | 0.84358 | 0.999994 | 17337 | 1 | 5.2158 | 0.013463 | 0.058462 | 0.929393 | 1190 | 5 | 5.2158 |
| SLC2A10 | 5 | 0.98652 | 0.98653 | 0.999994 | 21264 | 0 | 5.7867 | 0.013479 | 0.051992 | 0.926769 | 1191 | 5 | 5.7867 |
| CTRB1 | 6 | 0.12779 | 0.29747 | 0.999994 | 5867 | 2 | 5.1234 | 0.013485 | 0.058539 | 0.929393 | 1192 | 4 | 5.1234 |
| NAT14 | 5 | 0.58037 | 0.71864 | 0.999994 | 14683 | 1 | 2.8753 | 0.013527 | 0.052156 | 0.926769 | 1193 | 4 | 2.8753 |
| FAM35A | 6 | 0.50598 | 0.67288 | 0.999994 | 13526 | 2 | 5.6169 | 0.013527 | 0.058664 | 0.929393 | 1194 | 4 | 5.6169 |
| ARHGEF9 | 6 | 0.097222 | 0.24191 | 0.987253 | 4867 | 3 | 2.3451 | 0.013529 | 0.058669 | 0.929393 | 1195 | 3 | 2.3451 |

| | | | | | | | | | | | | |
|---|---|---|---|---|---|---|---|---|---|---|---|---|
| SLC12A5 | 6 | 0.49279 | 0.66378 | 0.999994 | 13308 | 2 | 1.0688 | 0.013529 | 0.058669 | 0.929393 | 1196 | 4 | 1.0688 |
| TNIP1 | 3 | 0.98647 | 0.98661 | 0.999994 | 21261 | 0 | 5.7477 | 0.013534 | 0.035666 | 0.903323 | 1197 | 3 | 5.7477 |
| R3HCC1 | 5 | 0.57747 | 0.71609 | 0.999994 | 14649 | 1 | 6.8087 | 0.013569 | 0.052302 | 0.926769 | 1198 | 3 | 6.8087 |
| OR2L3 | 5 | 0.77444 | 0.80423 | 0.999994 | 17136 | 1 | 1.2822 | 0.013574 | 0.052324 | 0.926769 | 1199 | 3 | 1.2822 |
| ZNF827 | 6 | 0.013111 | 0.049486 | 0.906615 | 1072 | 3 | -1.0629 | 0.013576 | 0.05882 | 0.930397 | 1200 | 2 | -1.0629 |
| PHF20 | 6 | 0.14883 | 0.32415 | 0.999994 | 6422 | 2 | 3.6016 | 0.013611 | 0.058927 | 0.930397 | 1201 | 4 | 3.6016 |
| ZNF567 | 6 | 0.96702 | 0.96706 | 0.999994 | 20761 | 0 | 3.1964 | 0.013611 | 0.058927 | 0.930397 | 1202 | 5 | 3.1964 |
| ZNF615 | 6 | 0.77813 | 0.83894 | 0.999994 | 17187 | 1 | 3.5881 | 0.013625 | 0.058971 | 0.930397 | 1203 | 5 | 3.5881 |
| IFNA21 | 5 | 0.89254 | 0.89266 | 0.999994 | 19080 | 0 | 4.6419 | 0.013629 | 0.052498 | 0.926769 | 1204 | 4 | 4.6419 |
| MAP1A | 6 | 0.83243 | 0.86274 | 0.999994 | 17963 | 1 | 4.5303 | 0.013645 | 0.059037 | 0.930397 | 1205 | 4 | 4.5303 |
| GPR1 | 6 | 0.13322 | 0.30443 | 0.999994 | 6001 | 2 | 0.18003 | 0.013668 | 0.059118 | 0.930397 | 1206 | 2 | 0.18003 |
| EYA4 | 6 | 0.92636 | 0.92644 | 0.999994 | 19776 | 1 | 2.0714 | 0.013668 | 0.059118 | 0.930397 | 1207 | 3 | 2.0714 |
| PLEKHH3 | 6 | 0.90733 | 0.90939 | 0.999994 | 19375 | 1 | 4.0512 | 0.013694 | 0.0592 | 0.930397 | 1208 | 5 | 4.0512 |
| SSC5D | 6 | 0.87935 | 0.88918 | 0.999994 | 18815 | 1 | 2.532 | 0.013726 | 0.059294 | 0.930397 | 1209 | 5 | 2.532 |
| ATP5A1 | 5 | 0.98626 | 0.98627 | 0.999994 | 21254 | 0 | 1.4966 | 0.013741 | 0.052898 | 0.926769 | 1210 | 5 | 1.4966 |
| PLCXD3 | 5 | 0.33245 | 0.50754 | 0.999994 | 10779 | 1 | 2.0476 | 0.013783 | 0.053046 | 0.926769 | 1211 | 3 | 2.0476 |
| PPP2R3A | 6 | 0.91608 | 0.91684 | 0.999994 | 19539 | 1 | 3.3032 | 0.013793 | 0.059494 | 0.930397 | 1212 | 5 | 3.3032 |
| SLC16A7 | 5 | 0.68915 | 0.76161 | 0.999994 | 16203 | 1 | 4.0801 | 0.013806 | 0.053131 | 0.926769 | 1213 | 4 | 4.0801 |
| MNT | 6 | 0.87965 | 0.88936 | 0.999994 | 18824 | 1 | 5.0149 | 0.013807 | 0.059545 | 0.930397 | 1214 | 5 | 5.0149 |
| IKZF3 | 6 | 0.13667 | 0.30885 | 0.999994 | 6101 | 3 | -1.0386 | 0.013808 | 0.059547 | 0.930397 | 1215 | 2 | -1.0386 |
| FAM126A | 4 | 0.027581 | 0.076571 | 0.93914 | 1880 | 3 | -2.1096 | 0.013825 | 0.045493 | 0.919289 | 1216 | 1 | -2.1096 |
| ZC3H10 | 6 | 0.6679 | 0.79152 | 0.999994 | 15893 | 1 | 4.9263 | 0.013841 | 0.059651 | 0.930754 | 1217 | 4 | 4.9263 |
| KLF13 | 5 | 0.55942 | 0.7001 | 0.999994 | 14361 | 1 | 6.4099 | 0.013847 | 0.053275 | 0.926769 | 1218 | 4 | 6.4099 |
| INTS4 | 5 | 0.12148 | 0.25959 | 0.994943 | 5683 | 1 | 2.4364 | 0.013847 | 0.053275 | 0.926769 | 1219 | 3 | 2.4364 |
| CDK10 | 5 | 0.17291 | 0.33638 | 0.999994 | 7064 | 1 | 4.7365 | 0.013848 | 0.053277 | 0.926769 | 1220 | 4 | 4.7365 |
| SOHLH1 | 4 | 0.98615 | 0.98611 | 0.999994 | 21251 | 0 | 3.648 | 0.01385 | 0.045578 | 0.919289 | 1221 | 4 | 3.648 |
| ARL4C | 3 | 0.27142 | 0.40142 | 0.999994 | 9510 | 1 | 7.7391 | 0.013859 | 0.036499 | 0.904772 | 1222 | 2 | 7.7391 |
| ZP1 | 6 | 0.13095 | 0.30153 | 0.999994 | 5958 | 2 | 0.40507 | 0.013892 | 0.059813 | 0.931159 | 1223 | 3 | 0.40507 |
| MECR | 4 | 0.92523 | 0.92503 | 0.999994 | 19749 | 0 | 5.896 | 0.013942 | 0.045852 | 0.919289 | 1224 | 3 | 5.896 |
| CXorf65 | 5 | 0.43353 | 0.59138 | 0.999994 | 12471 | 2 | 0.70446 | 0.013946 | 0.053614 | 0.926769 | 1225 | 3 | 0.70446 |
| PRKRIR | 5 | 0.98603 | 0.98606 | 0.999994 | 21249 | 0 | 3.2415 | 0.013969 | 0.053694 | 0.926769 | 1226 | 5 | 3.2415 |
| BANF2 | 6 | 0.75913 | 0.83173 | 0.999994 | 16965 | 1 | 3.1539 | 0.01397 | 0.060061 | 0.931675 | 1227 | 5 | 3.1539 |
| TARS2 | 5 | 0.55096 | 0.69259 | 0.999994 | 14226 | 1 | 5.5158 | 0.014011 | 0.053846 | 0.926769 | 1228 | 4 | 5.5158 |
| C11orf84 | 5 | 0.90953 | 0.90956 | 0.999994 | 19415 | 0 | 1.8999 | 0.014016 | 0.053862 | 0.926769 | 1229 | 3 | 1.8999 |
| IHH | 6 | 0.3675 | 0.58167 | 0.999994 | 11467 | 2 | 0.92918 | 0.014032 | 0.060258 | 0.9326 | 1230 | 3 | 0.92918 |
| ITGA2B | 5 | 0.13623 | 0.28207 | 0.999632 | 6087 | 1 | 6.7763 | 0.014034 | 0.053924 | 0.926769 | 1231 | 4 | 6.7763 |
| MFN2 | 4 | 0.98591 | 0.98588 | 0.999994 | 21248 | 0 | 4.6315 | 0.014093 | 0.046303 | 0.919289 | 1232 | 4 | 4.6315 |
| MAMSTR | 6 | 0.89535 | 0.90016 | 0.999994 | 19134 | 1 | 2.6429 | 0.014104 | 0.060476 | 0.9326 | 1233 | 5 | 2.6429 |
| KCNRG | 4 | 0.34538 | 0.48959 | 0.999994 | 11060 | 1 | 6.0543 | 0.014109 | 0.046344 | 0.919289 | 1234 | 3 | 6.0543 |
| LEKR1 | 5 | 0.98589 | 0.9859 | 0.999994 | 21247 | 0 | 2.7662 | 0.01411 | 0.054191 | 0.926769 | 1235 | 5 | 2.7662 |
| KIAA0247 | 5 | 0.96969 | 0.96975 | 0.999994 | 20819 | 0 | 4.2525 | 0.014122 | 0.054242 | 0.926769 | 1236 | 4 | 4.2525 |
| GATAD2B | 6 | 0.55375 | 0.70635 | 0.999994 | 14265 | 1 | 5.3914 | 0.014127 | 0.060542 | 0.9326 | 1237 | 4 | 5.3914 |
| FAM171A2 | 4 | 0.98586 | 0.98583 | 0.999994 | 21246 | 0 | 3.3772 | 0.014138 | 0.04643 | 0.919289 | 1238 | 4 | 3.3772 |
| WFDC6 | 5 | 0.95374 | 0.95391 | 0.999994 | 20432 | 0 | 4.0088 | 0.014139 | 0.054301 | 0.926769 | 1239 | 4 | 4.0088 |
| MRPL11 | 5 | 0.73672 | 0.78402 | 0.999994 | 16688 | 1 | 6.7756 | 0.014139 | 0.054303 | 0.926769 | 1240 | 4 | 6.7756 |
| SLC6A19 | 6 | 0.069122 | 0.18494 | 0.968526 | 3804 | 3 | -1.0469 | 0.014144 | 0.060589 | 0.9326 | 1241 | 2 | -1.0469 |
| NonTargetingControlG | 2 | 0.9858 | 0.98577 | 0.999994 | 21245 | 0 | 7.6186 | 0.014195 | 0.026692 | 0.890437 | 1242 | 2 | 7.6186 |
| SMOC2 | 5 | 0.13011 | 0.27275 | 0.998561 | 5935 | 1 | 6.221 | 0.014203 | 0.054532 | 0.926769 | 1243 | 4 | 6.221 |
| MEIS3 | 4 | 0.93781 | 0.93767 | 0.999994 | 20025 | 0 | 2.9137 | 0.014255 | 0.046778 | 0.919289 | 1244 | 3 | 2.9137 |
| SUN2 | 6 | 0.50284 | 0.67069 | 0.999994 | 13477 | 1 | 5.2535 | 0.014262 | 0.060958 | 0.9326 | 1245 | 5 | 5.2535 |
| ZNF256 | 5 | 0.98574 | 0.98576 | 0.999994 | 21243 | 0 | 1.3662 | 0.014263 | 0.054733 | 0.926769 | 1246 | 5 | 1.3662 |
| PUM1 | 6 | 0.93217 | 0.93228 | 0.999994 | 19907 | 0 | 1.0581 | 0.014311 | 0.061112 | 0.9326 | 1247 | 3 | 1.0581 |

| | | | | | | | | | | | | |
|---|---|---|---|---|---|---|---|---|---|---|---|---|
| SPANXN5 | 5 | 0.26246 | 0.45071 | 0.999994 | 9294 | 1 | 2.4239 | 0.01438 | 0.055144 | 0.926769 | 1248 | 4 | 2.4239 |
| hsa-mir-3692 | 4 | 0.92615 | 0.92597 | 0.999994 | 19770 | 0 | 6.4922 | 0.014384 | 0.047209 | 0.919289 | 1249 | 3 | 6.4922 |
| PRAMEF8 | 4 | 0.54057 | 0.63514 | 0.999994 | 14068 | 1 | 5.5659 | 0.014385 | 0.047213 | 0.919289 | 1250 | 3 | 5.5659 |
| PTP4A1 | 5 | 0.24108 | 0.43239 | 0.999994 | 8798 | 2 | 2.0758 | 0.014389 | 0.05517 | 0.926769 | 1251 | 3 | 2.0758 |
| RPS7 | 5 | 0.98552 | 0.98556 | 0.999994 | 21240 | 0 | 4.9214 | 0.014479 | 0.055489 | 0.926769 | 1252 | 5 | 4.9214 |
| UBXN2B | 5 | 0.94283 | 0.94302 | 0.999994 | 20146 | 0 | 4.2459 | 0.014493 | 0.055532 | 0.926769 | 1253 | 3 | 4.2459 |
| STX19 | 5 | 0.26839 | 0.45554 | 0.999994 | 9427 | 2 | 1.4959 | 0.014506 | 0.055584 | 0.926769 | 1254 | 3 | 1.4959 |
| CCNE2 | 6 | 0.82592 | 0.85959 | 0.999994 | 17867 | 1 | 6.0735 | 0.014549 | 0.061831 | 0.933948 | 1255 | 5 | 6.0735 |
| C10orf91 | 6 | 0.92307 | 0.92328 | 0.999994 | 19698 | 1 | 4.1991 | 0.014549 | 0.061831 | 0.933948 | 1256 | 5 | 4.1991 |
| ITPA | 5 | 0.74 | 0.78568 | 0.999994 | 16739 | 1 | 3.2396 | 0.014552 | 0.055746 | 0.926769 | 1257 | 4 | 3.2396 |
| VN1R4 | 6 | 0.43094 | 0.62247 | 0.999994 | 12430 | 2 | 2.9337 | 0.014577 | 0.06192 | 0.933948 | 1258 | 4 | 2.9337 |
| SEC31B | 5 | 0.66989 | 0.75325 | 0.999994 | 15929 | 1 | 6.7489 | 0.014599 | 0.055913 | 0.926769 | 1259 | 3 | 6.7489 |
| hsa-mir-4454 | 2 | 0.9854 | 0.98538 | 0.999994 | 21238 | 0 | 7.0404 | 0.0146 | 0.027417 | 0.890437 | 1260 | 2 | 7.0404 |
| F2R | 4 | 0.97194 | 0.97183 | 0.999994 | 20868 | 0 | 5.0937 | 0.014604 | 0.047898 | 0.919289 | 1261 | 4 | 5.0937 |
| USP50 | 6 | 0.2001 | 0.38799 | 0.999994 | 7767 | 3 | 0.10979 | 0.014621 | 0.062049 | 0.933948 | 1262 | 3 | 0.10979 |
| SLC15A1 | 5 | 0.98265 | 0.9827 | 0.999994 | 21151 | 0 | 4.0601 | 0.014644 | 0.05607 | 0.926769 | 1263 | 5 | 4.0601 |
| RYR1 | 6 | 0.9147 | 0.91562 | 0.999994 | 19504 | 1 | 3.6234 | 0.014647 | 0.062135 | 0.933948 | 1264 | 5 | 3.6234 |
| OR2AP1 | 5 | 0.75444 | 0.79319 | 0.999994 | 16903 | 1 | 1.4773 | 0.014669 | 0.056168 | 0.926769 | 1265 | 3 | 1.4773 |
| C1orf177 | 6 | 0.58698 | 0.73036 | 0.999994 | 14775 | 1 | 3.714 | 0.014695 | 0.06229 | 0.933948 | 1266 | 5 | 3.714 |
| HHIP | 6 | 0.0041508 | 0.01803 | 0.795727 | 454 | 2 | -0.39644 | 0.014702 | 0.062315 | 0.933948 | 1267 | 2 | -0.39644 |
| CPN2 | 6 | 0.15265 | 0.32899 | 0.999994 | 6525 | 3 | 3.0871 | 0.014706 | 0.062321 | 0.933948 | 1268 | 3 | 3.0871 |
| PMF1 | 4 | 0.35468 | 0.49903 | 0.999994 | 11244 | 1 | 1.0652 | 0.014721 | 0.048255 | 0.919864 | 1269 | 2 | 1.0652 |
| hsa-mir-3173 | 3 | 0.75111 | 0.75108 | 0.999994 | 16867 | 0 | 0.9665 | 0.014735 | 0.038683 | 0.905919 | 1270 | 2 | 0.9665 |
| IQCH | 6 | 0.46703 | 0.64647 | 0.999994 | 12966 | 2 | 1.5435 | 0.01475 | 0.06246 | 0.933948 | 1271 | 3 | 1.5435 |
| QTRT1 | 5 | 0.98524 | 0.9853 | 0.999994 | 21232 | 0 | 2.3065 | 0.014756 | 0.056484 | 0.926769 | 1272 | 5 | 2.3065 |
| PMPCB | 4 | 0.099453 | 0.21412 | 0.979587 | 4947 | 1 | 6.0581 | 0.014783 | 0.048459 | 0.920057 | 1273 | 3 | 6.0581 |
| DAZ1 | 6 | 0.93067 | 0.9308 | 0.999994 | 19868 | 0 | 0.58518 | 0.014786 | 0.062549 | 0.933948 | 1274 | 4 | 0.58518 |
| hsa-let-7c | 4 | 0.87811 | 0.87776 | 0.999994 | 18789 | 0 | 4.7587 | 0.01481 | 0.048541 | 0.920057 | 1275 | 3 | 4.7587 |
| CSNK1G1 | 5 | 0.36596 | 0.53501 | 0.999994 | 11444 | 2 | 0.77003 | 0.014812 | 0.056685 | 0.926769 | 1276 | 3 | 0.77003 |
| ASPHD2 | 5 | 0.24904 | 0.44002 | 0.999994 | 8993 | 2 | 5.2584 | 0.014832 | 0.056761 | 0.926769 | 1277 | 3 | 5.2584 |
| RNF103-CHMP3 | 3 | 0.98516 | 0.9853 | 0.999994 | 21230 | 0 | 4.0003 | 0.014836 | 0.038925 | 0.906957 | 1278 | 3 | 4.0003 |
| TRPM2 | 5 | 0.3702 | 0.53855 | 0.999994 | 11514 | 2 | 5.1892 | 0.014862 | 0.056859 | 0.926769 | 1279 | 3 | 5.1892 |
| ARIH1 | 6 | 0.27329 | 0.47559 | 0.999994 | 9548 | 2 | 2.2686 | 0.01487 | 0.062809 | 0.933948 | 1280 | 4 | 2.2686 |
| GINS3 | 4 | 0.98513 | 0.9851 | 0.999994 | 21229 | 0 | 6.0319 | 0.014873 | 0.048749 | 0.920057 | 1281 | 4 | 6.0319 |
| NonTargetingControlG | 2 | 0.98512 | 0.98508 | 0.999994 | 21228 | 0 | 8.1016 | 0.014884 | 0.027936 | 0.890437 | 1282 | 2 | 8.1016 |
| DNAAF2 | 5 | 0.23674 | 0.42653 | 0.999994 | 8687 | 1 | 4.3222 | 0.014913 | 0.057034 | 0.926769 | 1283 | 4 | 4.3222 |
| HARBI1 | 6 | 0.56434 | 0.7139 | 0.999994 | 14461 | 2 | 1.651 | 0.014926 | 0.06298 | 0.933948 | 1284 | 4 | 1.651 |
| IMPDH1 | 5 | 0.080351 | 0.19516 | 0.975797 | 4243 | 2 | 6.7304 | 0.014938 | 0.057124 | 0.926769 | 1285 | 3 | 6.7304 |
| BIRC7 | 5 | 0.20111 | 0.37679 | 0.999994 | 7792 | 2 | 6.7304 | 0.014938 | 0.057124 | 0.926769 | 1286 | 3 | 6.7304 |
| ZNF721 | 4 | 0.98505 | 0.98502 | 0.999994 | 21224 | 0 | 6.3046 | 0.014945 | 0.048968 | 0.920223 | 1287 | 4 | 6.3046 |
| NDUFC2 | 6 | 0.21824 | 0.41002 | 0.999994 | 8234 | 3 | 3.0957 | 0.01495 | 0.063045 | 0.933948 | 1288 | 3 | 3.0957 |
| PRR24 | 5 | 0.97671 | 0.97676 | 0.999994 | 21008 | 0 | 4.988 | 0.014972 | 0.057239 | 0.926769 | 1289 | 5 | 4.988 |
| OLR1 | 6 | 0.072054 | 0.19107 | 0.972392 | 3920 | 4 | -1.0271 | 0.014974 | 0.063116 | 0.933948 | 1290 | 2 | -1.0271 |
| SYTL5 | 6 | 0.48073 | 0.65568 | 0.999994 | 13151 | 2 | 0.72379 | 0.014982 | 0.063134 | 0.933948 | 1291 | 3 | 0.72379 |
| DDX26B | 6 | 0.55013 | 0.70373 | 0.999994 | 14213 | 1 | 5.6452 | 0.01502 | 0.063255 | 0.933948 | 1292 | 4 | 5.6452 |
| COL17A1 | 6 | 0.94452 | 0.94449 | 0.999994 | 20192 | 0 | 5.3886 | 0.01502 | 0.063255 | 0.933948 | 1293 | 5 | 5.3886 |
| PAM16 | 5 | 0.97517 | 0.97524 | 0.999994 | 20960 | 0 | 4.5924 | 0.015046 | 0.057503 | 0.927195 | 1294 | 5 | 4.5924 |
| NonTargetingControlG | 2 | 0.063141 | 0.11115 | 0.951052 | 3563 | 1 | 0.43089 | 0.015076 | 0.028298 | 0.890437 | 1295 | 1 | 0.43089 |
| MYH2 | 5 | 0.98485 | 0.98492 | 0.999994 | 21220 | 0 | 1.7189 | 0.015147 | 0.057851 | 0.928656 | 1296 | 5 | 1.7189 |
| C12orf44 | 6 | 0.18914 | 0.37443 | 0.999994 | 7454 | 3 | -0.15128 | 0.015149 | 0.063661 | 0.933948 | 1297 | 3 | -0.15128 |
| TMEM147 | 4 | 0.96287 | 0.96276 | 0.999994 | 20657 | 0 | 5.4757 | 0.015152 | 0.049595 | 0.921876 | 1298 | 4 | 5.4757 |
| CNPPD1 | 3 | 0.98484 | 0.98499 | 0.999994 | 21219 | 0 | 5.3065 | 0.015163 | 0.039712 | 0.909688 | 1299 | 3 | 5.3065 |

| | | | | | | | | | | | | |
|---|---|---|---|---|---|---|---|---|---|---|---|---|
| GGA3 | 6 | 0.87926 | 0.88913 | 0.999994 | 18814 | 1 | 1.0164 | 0.015208 | 0.063836 | 0.933948 | 1300 | 3 | 1.0164 |
| LOX | 5 | 0.98474 | 0.98481 | 0.999994 | 21218 | 0 | 2.147 | 0.015258 | 0.058242 | 0.929393 | 1301 | 5 | 2.147 |
| SLC27A2 | 4 | 0.94516 | 0.94501 | 0.999994 | 20208 | 0 | 2.0184 | 0.015262 | 0.049918 | 0.923113 | 1302 | 4 | 2.0184 |
| KIAA0408 | 5 | 0.6298 | 0.73722 | 0.999994 | 15379 | 1 | 5.6697 | 0.015266 | 0.058273 | 0.929393 | 1303 | 4 | 5.6697 |
| TNFRSF1B | 5 | 0.98471 | 0.98478 | 0.999994 | 21217 | 0 | 1.5712 | 0.015286 | 0.05835 | 0.929393 | 1304 | 5 | 1.5712 |
| GNMT | 5 | 0.10172 | 0.22911 | 0.983354 | 5025 | 2 | 1.3169 | 0.015297 | 0.058385 | 0.929393 | 1305 | 3 | 1.3169 |
| GAMT | 5 | 0.56748 | 0.70732 | 0.999994 | 14505 | 2 | 6.7072 | 0.015308 | 0.058426 | 0.929393 | 1306 | 3 | 6.7072 |
| SMYD3 | 6 | 0.7121 | 0.8158 | 0.999994 | 16421 | 1 | 2.3961 | 0.015317 | 0.064168 | 0.933948 | 1307 | 4 | 2.3961 |
| NADKD1 | 5 | 0.040598 | 0.11738 | 0.951052 | 2542 | 1 | 3.9895 | 0.015345 | 0.058545 | 0.929393 | 1308 | 4 | 3.9895 |
| ZNF526 | 3 | 0.98465 | 0.98481 | 0.999994 | 21215 | 0 | 7.2562 | 0.015349 | 0.040164 | 0.909688 | 1309 | 3 | 7.2562 |
| hsa-mir-4514 | 4 | 0.98465 | 0.9846 | 0.999994 | 21214 | 0 | 6.3491 | 0.015354 | 0.050186 | 0.924139 | 1310 | 4 | 6.3491 |
| NonTargetingControlG | 2 | 0.67746 | 0.67727 | 0.999994 | 16052 | 0 | 4.8923 | 0.015356 | 0.028804 | 0.890437 | 1311 | 1 | 4.8923 |
| UNC5CL | 6 | 0.76389 | 0.83352 | 0.999994 | 17010 | 1 | 5.1253 | 0.015361 | 0.0643 | 0.933948 | 1312 | 5 | 5.1253 |
| ZNF677 | 6 | 0.67302 | 0.79551 | 0.999994 | 15973 | 2 | 2.1837 | 0.015369 | 0.064325 | 0.933948 | 1313 | 3 | 2.1837 |
| KCNMB2 | 6 | 0.070997 | 0.18888 | 0.969543 | 3874 | 4 | -1.4221 | 0.01541 | 0.06447 | 0.933948 | 1314 | 2 | -1.4221 |
| SMG6 | 6 | 0.34771 | 0.56143 | 0.999994 | 11107 | 1 | 5.1537 | 0.015472 | 0.064634 | 0.933948 | 1315 | 5 | 5.1537 |
| POF1B | 4 | 0.98452 | 0.98448 | 0.999994 | 21205 | 0 | 3.1294 | 0.015479 | 0.050567 | 0.924349 | 1316 | 4 | 3.1294 |
| UBL7 | 4 | 0.98452 | 0.98448 | 0.999994 | 21203 | 0 | 5.7767 | 0.015481 | 0.05057 | 0.924349 | 1317 | 4 | 5.7767 |
| BGN | 5 | 0.94245 | 0.94264 | 0.999994 | 20137 | 0 | 2.4611 | 0.015484 | 0.059035 | 0.930397 | 1318 | 4 | 2.4611 |
| CD209 | 6 | 0.29066 | 0.49584 | 0.999994 | 9926 | 2 | 2.4512 | 0.015485 | 0.064669 | 0.933948 | 1319 | 3 | 2.4512 |
| KCTD20 | 6 | 0.9356 | 0.93564 | 0.999994 | 19981 | 0 | 5.0935 | 0.015548 | 0.064894 | 0.933948 | 1320 | 5 | 5.0935 |
| CILP | 6 | 0.98665 | 0.9866 | 0.999994 | 21268 | 0 | 4.5963 | 0.015548 | 0.064894 | 0.933948 | 1321 | 5 | 4.5963 |
| C4BPA | 6 | 0.27688 | 0.47981 | 0.999994 | 9636 | 2 | 3.6855 | 0.015548 | 0.064894 | 0.933948 | 1322 | 3 | 3.6855 |
| MYL1 | 5 | 0.1781 | 0.3439 | 0.999994 | 7179 | 2 | 3.8098 | 0.015554 | 0.059291 | 0.930397 | 1323 | 3 | 3.8098 |
| C19orf47 | 5 | 0.67207 | 0.75418 | 0.999994 | 15956 | 1 | 0.60092 | 0.015554 | 0.059291 | 0.930397 | 1324 | 2 | 0.60092 |
| CREB3L3 | 5 | 0.98443 | 0.98451 | 0.999994 | 21200 | 0 | 2.457 | 0.015571 | 0.059354 | 0.930397 | 1325 | 5 | 2.457 |
| SP6 | 4 | 0.20594 | 0.34682 | 0.999994 | 7924 | 2 | 0.24851 | 0.015578 | 0.050856 | 0.924349 | 1326 | 2 | 0.24851 |
| GPR116 | 5 | 0.98442 | 0.9845 | 0.999994 | 21199 | 0 | 4.8715 | 0.015581 | 0.059391 | 0.930397 | 1327 | 5 | 4.8715 |
| PPRC1 | 6 | 0.85791 | 0.87624 | 0.999994 | 18405 | 1 | 4.6595 | 0.015605 | 0.065068 | 0.933948 | 1328 | 4 | 4.6595 |
| GOLGA8A | 4 | 0.1443 | 0.28334 | 0.999632 | 6310 | 1 | 4.022 | 0.015616 | 0.05097 | 0.924349 | 1329 | 3 | 4.022 |
| FAM83G | 5 | 0.10855 | 0.23973 | 0.987253 | 5260 | 2 | -0.48422 | 0.015624 | 0.059549 | 0.930397 | 1330 | 2 | -0.48422 |
| CYP7A1 | 6 | 0.038843 | 0.11928 | 0.951052 | 2453 | 2 | 3.3212 | 0.015636 | 0.065178 | 0.933948 | 1331 | 3 | 3.3212 |
| NonTargetingControlG | 2 | 0.74009 | 0.73974 | 0.999994 | 16742 | 0 | 5.0872 | 0.015646 | 0.029356 | 0.890437 | 1332 | 1 | 5.0872 |
| ELOVL3 | 6 | 0.88495 | 0.89283 | 0.999994 | 18932 | 1 | 2.5921 | 0.015652 | 0.065228 | 0.933948 | 1333 | 5 | 2.5921 |
| NonTargetingControlG | 2 | 0.98435 | 0.98432 | 0.999994 | 21197 | 0 | 7.7367 | 0.015653 | 0.029368 | 0.890437 | 1334 | 2 | 7.7367 |
| DUSP27 | 5 | 0.86665 | 0.86814 | 0.999994 | 18569 | 1 | 6.1076 | 0.015672 | 0.059715 | 0.930982 | 1335 | 4 | 6.1076 |
| ZNF699 | 5 | 0.98431 | 0.98439 | 0.999994 | 21196 | 0 | 2.7112 | 0.015689 | 0.059775 | 0.931159 | 1336 | 5 | 2.7112 |
| ATL3 | 5 | 0.97363 | 0.97367 | 0.999994 | 20912 | 0 | 4.9089 | 0.01574 | 0.059953 | 0.931675 | 1337 | 5 | 4.9089 |
| hsa-mir-7845 | 4 | 0.90394 | 0.90363 | 0.999994 | 19303 | 0 | 6.808 | 0.01575 | 0.051374 | 0.924349 | 1338 | 3 | 6.808 |
| CEBPB | 4 | 0.98424 | 0.9842 | 0.999994 | 21195 | 0 | 2.51 | 0.015756 | 0.05139 | 0.924349 | 1339 | 4 | 2.51 |
| MLL2 | 2 | 0.98424 | 0.98422 | 0.999994 | 21194 | 0 | 7.1521 | 0.015757 | 0.029556 | 0.890437 | 1340 | 2 | 7.1521 |
| ZNF547 | 5 | 0.083675 | 0.20051 | 0.977164 | 4363 | 1 | 3.6773 | 0.01576 | 0.060021 | 0.931675 | 1341 | 4 | 3.6773 |
| MCRS1 | 5 | 0.90899 | 0.909 | 0.999994 | 19402 | 0 | 5.1053 | 0.01576 | 0.060021 | 0.931675 | 1342 | 4 | 5.1053 |
| RARRES2 | 6 | 0.98616 | 0.98612 | 0.999994 | 21252 | 0 | 4.8224 | 0.015769 | 0.065613 | 0.933948 | 1343 | 5 | 4.8224 |
| CHCHD7 | 6 | 0.093576 | 0.2347 | 0.984971 | 4740 | 3 | -0.42694 | 0.015792 | 0.06568 | 0.933948 | 1344 | 3 | -0.42694 |
| TP53AIP1 | 6 | 0.66865 | 0.79212 | 0.999994 | 15901 | 1 | 5.2729 | 0.015854 | 0.065866 | 0.933948 | 1345 | 4 | 5.2729 |
| LAGE3 | 6 | 0.31311 | 0.52196 | 0.999994 | 10380 | 2 | 5.081 | 0.015854 | 0.065866 | 0.933948 | 1346 | 4 | 5.081 |
| RPA3 | 1 | 0.98414 | 0.98417 | 0.999994 | 21193 | 0 | 9.2695 | 0.015859 | 0.015776 | 0.829051 | 1347 | 1 | 9.2695 |
| GRPEL1 | 5 | 0.98414 | 0.98422 | 0.999994 | 21192 | 0 | 5.8779 | 0.01586 | 0.060348 | 0.9326 | 1348 | 5 | 5.8779 |
| RALGPS2 | 6 | 0.6411 | 0.77076 | 0.999994 | 15530 | 1 | 5.159 | 0.015878 | 0.065935 | 0.933948 | 1349 | 5 | 5.159 |
| MAGEC3 | 5 | 0.24587 | 0.43746 | 0.999994 | 8915 | 1 | 3.5652 | 0.015919 | 0.060549 | 0.9326 | 1350 | 4 | 3.5652 |
| PNPLA4 | 5 | 0.98405 | 0.98412 | 0.999994 | 21191 | 0 | 1.6012 | 0.015949 | 0.060661 | 0.9326 | 1351 | 5 | 1.6012 |

| | | | | | | | | | | | | | |
|---|---|---|---|---|---|---|---|---|---|---|---|---|---|
| RPL26L1 | 5 | 0.54134 | 0.6841 | 0.999994 | 14076 | 2 | 6.5728 | 0.015985 | 0.060777 | 0.9326 | 1352 | 3 | 6.5728 |
| PRICKLE2 | 6 | 0.75871 | 0.83158 | 0.999994 | 16962 | 1 | 4.6612 | 0.015987 | 0.066266 | 0.933948 | 1353 | 4 | 4.6612 |
| SLC25A28 | 6 | 0.44386 | 0.631 | 0.999994 | 12633 | 2 | 5.3454 | 0.015987 | 0.066267 | 0.933948 | 1354 | 4 | 5.3454 |
| FAM217B | 6 | 0.23395 | 0.4288 | 0.999994 | 8619 | 2 | 4.7312 | 0.015992 | 0.066284 | 0.933948 | 1355 | 4 | 4.7312 |
| YEATS4 | 5 | 0.984 | 0.98407 | 0.999994 | 21188 | 0 | 1.2928 | 0.016 | 0.060824 | 0.9326 | 1356 | 5 | 1.2928 |
| ASB3 | 5 | 0.76441 | 0.7986 | 0.999994 | 17022 | 1 | 1.3154 | 0.016042 | 0.060982 | 0.9326 | 1357 | 3 | 1.3154 |
| TMEM140 | 6 | 0.048683 | 0.14122 | 0.959146 | 2918 | 4 | -1.4695 | 0.016043 | 0.066423 | 0.933948 | 1358 | 2 | -1.4695 |
| hsa-mir-485 | 4 | 0.98394 | 0.98391 | 0.999994 | 21187 | 0 | 5.7148 | 0.016057 | 0.052329 | 0.926769 | 1359 | 4 | 5.7148 |
| MAN2C1 | 5 | 0.14046 | 0.28851 | 0.999632 | 6201 | 1 | 6.3363 | 0.016067 | 0.061063 | 0.9326 | 1360 | 4 | 6.3363 |
| PNPLA1 | 5 | 0.61899 | 0.73319 | 0.999994 | 15206 | 1 | 2.1692 | 0.016089 | 0.061137 | 0.9326 | 1361 | 3 | 2.1692 |
| CELF5 | 5 | 0.2829 | 0.46721 | 0.999994 | 9759 | 2 | 6.6635 | 0.016092 | 0.061154 | 0.9326 | 1362 | 3 | 6.6635 |
| TMC4 | 6 | 0.58516 | 0.72902 | 0.999994 | 14744 | 1 | 0.67795 | 0.016099 | 0.066584 | 0.933948 | 1363 | 3 | 0.67795 |
| ENO3 | 5 | 0.90191 | 0.902 | 0.999994 | 19249 | 0 | 3.5387 | 0.016117 | 0.06124 | 0.933254 | 1364 | 4 | 3.5387 |
| hsa-mir-559 | 4 | 0.98388 | 0.98383 | 0.999994 | 21186 | 0 | 2.8668 | 0.016121 | 0.052526 | 0.926769 | 1365 | 4 | 2.8668 |
| SPATC1 | 6 | 0.55743 | 0.709 | 0.999994 | 14321 | 2 | 3.814 | 0.01618 | 0.06683 | 0.933948 | 1366 | 4 | 3.814 |
| MBTPS2 | 5 | 0.98381 | 0.98388 | 0.999994 | 21182 | 0 | 1.817 | 0.016189 | 0.061511 | 0.933948 | 1367 | 5 | 1.817 |
| ARHGEF1 | 5 | 0.48642 | 0.63616 | 0.999994 | 13234 | 2 | 1.3323 | 0.016221 | 0.061622 | 0.933948 | 1368 | 3 | 1.3323 |
| CSRP2 | 6 | 0.90143 | 0.90478 | 0.999994 | 19241 | 1 | 3.7294 | 0.016222 | 0.066953 | 0.933948 | 1369 | 5 | 3.7294 |
| NANP | 6 | 0.3262 | 0.53691 | 0.999994 | 10648 | 3 | 0.62208 | 0.016222 | 0.066953 | 0.933948 | 1370 | 3 | 0.62208 |
| C4orf19 | 6 | 0.83066 | 0.86187 | 0.999994 | 17941 | 1 | 5.5198 | 0.016228 | 0.066971 | 0.933948 | 1371 | 5 | 5.5198 |
| RASD2 | 5 | 0.36423 | 0.53358 | 0.999994 | 11408 | 2 | 6.6597 | 0.016242 | 0.061706 | 0.933948 | 1372 | 3 | 6.6597 |
| PCDHGB4 | 2 | 0.98374 | 0.98372 | 0.999994 | 21181 | 0 | 7.2885 | 0.016259 | 0.03043 | 0.890437 | 1373 | 2 | 7.2885 |
| SSBP1 | 6 | 0.46023 | 0.64194 | 0.999994 | 12848 | 1 | 5.8805 | 0.01626 | 0.067068 | 0.933994 | 1374 | 5 | 5.8805 |
| RPAP3 | 6 | 0.39281 | 0.59772 | 0.999994 | 11845 | 1 | 5.5611 | 0.01626 | 0.067068 | 0.933994 | 1375 | 4 | 5.5611 |
| GPRC5B | 5 | 0.83479 | 0.84291 | 0.999994 | 18001 | 1 | 6.0718 | 0.016269 | 0.061797 | 0.933948 | 1376 | 4 | 6.0718 |
| FAM109A | 5 | 0.050685 | 0.14204 | 0.959216 | 3002 | 2 | 3.3366 | 0.016295 | 0.061883 | 0.933948 | 1377 | 3 | 3.3366 |
| NonTargetingControlG | 2 | 0.98367 | 0.98365 | 0.999994 | 21178 | 0 | 7.2538 | 0.016331 | 0.030565 | 0.890437 | 1378 | 2 | 7.2538 |
| hsa-mir-3619 | 4 | 0.98366 | 0.98362 | 0.999994 | 21176 | 0 | 3.4689 | 0.016336 | 0.053173 | 0.926769 | 1379 | 4 | 3.4689 |
| COL1A1 | 5 | 0.98364 | 0.98372 | 0.999994 | 21173 | 0 | 5.202 | 0.016359 | 0.062112 | 0.933948 | 1380 | 5 | 5.202 |
| MET | 6 | 0.90681 | 0.90897 | 0.999994 | 19367 | 1 | 3.464 | 0.016362 | 0.067381 | 0.934798 | 1381 | 3 | 3.464 |
| ROBO3 | 5 | 0.91556 | 0.91564 | 0.999994 | 19527 | 0 | 1.9225 | 0.016368 | 0.062144 | 0.933948 | 1382 | 3 | 1.9225 |
| LCN10 | 6 | 0.93802 | 0.93799 | 0.999994 | 20028 | 0 | 4.9283 | 0.016376 | 0.067421 | 0.934798 | 1383 | 5 | 4.9283 |
| HLA-G | 6 | 0.94418 | 0.94415 | 0.999994 | 20181 | 0 | 3.5234 | 0.016378 | 0.067427 | 0.934798 | 1384 | 4 | 3.5234 |
| CHRNA2 | 4 | 0.89335 | 0.89306 | 0.999994 | 19093 | 0 | 2.6841 | 0.016398 | 0.053351 | 0.926769 | 1385 | 2 | 2.6841 |
| TUBG1 | 2 | 0.98358 | 0.98355 | 0.999994 | 21171 | 0 | 6.9632 | 0.016422 | 0.030724 | 0.890437 | 1386 | 2 | 6.9632 |
| PABPC1L2B | 2 | 0.69197 | 0.69177 | 0.999994 | 16227 | 0 | 4.9145 | 0.016432 | 0.030738 | 0.890437 | 1387 | 1 | 4.9145 |
| hsa-mir-6748 | 4 | 0.98352 | 0.98348 | 0.999994 | 21170 | 0 | 4.3367 | 0.016483 | 0.053611 | 0.926769 | 1388 | 4 | 4.3367 |
| KCNQ2 | 3 | 0.85388 | 0.85391 | 0.999994 | 18331 | 0 | 7.6157 | 0.016485 | 0.042984 | 0.918424 | 1389 | 2 | 7.6157 |
| GIMAP1 | 6 | 0.8488 | 0.8713 | 0.999994 | 18232 | 1 | -0.68912 | 0.016489 | 0.067762 | 0.935447 | 1390 | 2 | -0.68912 |
| SLC9A3R2 | 5 | 0.43978 | 0.59663 | 0.999994 | 12571 | 1 | 6.6476 | 0.016497 | 0.062613 | 0.933948 | 1391 | 3 | 6.6476 |
| TNFAIP3 | 5 | 0.8407 | 0.84718 | 0.999994 | 18100 | 1 | 3.6179 | 0.016531 | 0.062726 | 0.933948 | 1392 | 4 | 3.6179 |
| HELQ | 5 | 0.98343 | 0.9835 | 0.999994 | 21169 | 0 | 2.8291 | 0.016572 | 0.062881 | 0.933948 | 1393 | 5 | 2.8291 |
| UROD | 3 | 0.64172 | 0.65021 | 0.999994 | 15543 | 1 | 3.1166 | 0.016582 | 0.043216 | 0.918424 | 1394 | 2 | 3.1166 |
| CASP5 | 5 | 0.090468 | 0.21134 | 0.979587 | 4613 | 1 | 4.4873 | 0.016583 | 0.062908 | 0.933948 | 1395 | 4 | 4.4873 |
| hsa-mir-8079 | 4 | 0.98342 | 0.98339 | 0.999994 | 21168 | 0 | 2.5971 | 0.016583 | 0.053921 | 0.926769 | 1396 | 4 | 2.5971 |
| RBMS1 | 6 | 0.30081 | 0.50766 | 0.999994 | 10147 | 3 | -0.34751 | 0.016601 | 0.068113 | 0.935447 | 1397 | 2 | -0.34751 |
| TRAPPC9 | 5 | 0.98338 | 0.98344 | 0.999994 | 21166 | 0 | 6.0521 | 0.016624 | 0.063052 | 0.933948 | 1398 | 5 | 6.0521 |
| CRIPAK | 5 | 0.96289 | 0.96305 | 0.999994 | 20659 | 0 | 6.6354 | 0.016626 | 0.063064 | 0.933948 | 1399 | 4 | 6.6354 |
| C19orf66 | 5 | 0.98334 | 0.9834 | 0.999994 | 21164 | 0 | 2.683 | 0.016661 | 0.063181 | 0.933948 | 1400 | 5 | 2.683 |
| LCLAT1 | 6 | 0.66935 | 0.79266 | 0.999994 | 15920 | 1 | 4.5556 | 0.016706 | 0.068449 | 0.935447 | 1401 | 5 | 4.5556 |
| HIST1H2BB | 6 | 0.65842 | 0.78409 | 0.999994 | 15757 | 1 | 3.7602 | 0.016713 | 0.068466 | 0.935447 | 1402 | 3 | 3.7602 |
| SNCG | 6 | 0.55723 | 0.70885 | 0.999994 | 14318 | 1 | 4.9915 | 0.016726 | 0.068512 | 0.935447 | 1403 | 5 | 4.9915 |

| | | | | | | | | | | | | |
|---|---|---|---|---|---|---|---|---|---|---|---|---|
| RPL36A | 5 | 0.58023 | 0.71852 | 0.999994 | 14681 | 2 | 6.6339 | 0.016748 | 0.063478 | 0.933948 | 1404 | 3 | 6.6339 |
| RGL3 | 5 | 0.16574 | 0.32587 | 0.999994 | 6878 | 2 | 6.6339 | 0.016748 | 0.063478 | 0.933948 | 1405 | 3 | 6.6339 |
| TAS2R7 | 5 | 0.051606 | 0.14427 | 0.961983 | 3050 | 2 | 1.0189 | 0.016787 | 0.063619 | 0.933948 | 1406 | 3 | 1.0189 |
| ISX | 6 | 0.97441 | 0.9743 | 0.999994 | 20938 | 0 | 4.8657 | 0.016794 | 0.068716 | 0.936374 | 1407 | 4 | 4.8657 |
| PHEX | 6 | 0.10372 | 0.2546 | 0.994835 | 5093 | 4 | -0.81764 | 0.016796 | 0.068721 | 0.936374 | 1408 | 1 | -0.81764 |
| UHRF1BP1 | 5 | 0.98316 | 0.98322 | 0.999994 | 21159 | 0 | 4.7663 | 0.016844 | 0.063805 | 0.933948 | 1409 | 5 | 4.7663 |
| RNF224 | 5 | 0.74887 | 0.79026 | 0.999994 | 16849 | 1 | 6.623 | 0.016861 | 0.063863 | 0.933948 | 1410 | 4 | 6.623 |
| NMRK2 | 6 | 0.13661 | 0.30877 | 0.999994 | 6098 | 3 | -0.072541 | 0.01688 | 0.068983 | 0.936996 | 1411 | 3 | -0.072541 |
| hsa-mir-4526 | 4 | 0.86752 | 0.86708 | 0.999994 | 18590 | 0 | 6.538 | 0.016885 | 0.054809 | 0.926769 | 1412 | 3 | 6.538 |
| SLK | 6 | 0.97722 | 0.97716 | 0.999994 | 21022 | 0 | 5.1866 | 0.016915 | 0.069082 | 0.936996 | 1413 | 5 | 5.1866 |
| NDP | 6 | 0.96028 | 0.96034 | 0.999994 | 20590 | 0 | 2.7036 | 0.016935 | 0.069131 | 0.936996 | 1414 | 5 | 2.7036 |
| GABRR1 | 6 | 0.90803 | 0.90996 | 0.999994 | 19384 | 1 | 3.5237 | 0.016939 | 0.069141 | 0.936996 | 1415 | 5 | 3.5237 |
| OR1J4 | 6 | 0.64656 | 0.77499 | 0.999994 | 15610 | 1 | 5.0097 | 0.016961 | 0.0692 | 0.936996 | 1416 | 4 | 5.0097 |
| PREP | 6 | 0.93867 | 0.93863 | 0.999994 | 20044 | 0 | 3.4486 | 0.016964 | 0.06921 | 0.936996 | 1417 | 5 | 3.4486 |
| COX14 | 5 | 0.95803 | 0.95819 | 0.999994 | 20539 | 0 | 6.6189 | 0.016969 | 0.064223 | 0.933948 | 1418 | 4 | 6.6189 |
| EHD1 | 5 | 0.80044 | 0.81978 | 0.999994 | 17486 | 1 | 2.343 | 0.016973 | 0.064235 | 0.933948 | 1419 | 4 | 2.343 |
| DOCK7 | 6 | 0.85668 | 0.87556 | 0.999994 | 18382 | 1 | 3.7409 | 0.016978 | 0.069255 | 0.936996 | 1420 | 5 | 3.7409 |
| SYT3 | 5 | 0.44918 | 0.60456 | 0.999994 | 12704 | 1 | 4.1968 | 0.016978 | 0.064244 | 0.933948 | 1421 | 4 | 4.1968 |
| SNAP91 | 4 | 0.89074 | 0.89045 | 0.999994 | 19049 | 0 | 2.0699 | 0.016994 | 0.055156 | 0.926769 | 1422 | 3 | 2.0699 |
| KRTAP2-2 | 2 | 0.983 | 0.98298 | 0.999994 | 21157 | 0 | 6.9272 | 0.016998 | 0.031744 | 0.890437 | 1423 | 2 | 6.9272 |
| PLCB2 | 6 | 0.98211 | 0.98204 | 0.999994 | 21141 | 0 | 2.8092 | 0.017002 | 0.069327 | 0.936996 | 1424 | 5 | 2.8092 |
| CDC42EP3 | 4 | 0.95145 | 0.95135 | 0.999994 | 20375 | 0 | 5.4995 | 0.017014 | 0.055216 | 0.926769 | 1425 | 4 | 5.4995 |
| CDHR1 | 5 | 0.54194 | 0.6846 | 0.999994 | 14089 | 2 | 3.0656 | 0.017019 | 0.064389 | 0.933948 | 1426 | 3 | 3.0656 |
| MBOAT4 | 4 | 0.98298 | 0.98295 | 0.999994 | 21156 | 0 | 5.3732 | 0.017023 | 0.055239 | 0.926769 | 1427 | 4 | 5.3732 |
| TRIM16L | 5 | 0.85786 | 0.86063 | 0.999994 | 18404 | 1 | 1.2116 | 0.017066 | 0.064561 | 0.933948 | 1428 | 4 | 1.2116 |
| SMAD3 | 6 | 0.96797 | 0.96799 | 0.999994 | 20784 | 0 | 3.2911 | 0.017139 | 0.069748 | 0.937929 | 1429 | 4 | 3.2911 |
| ZSWIM1 | 6 | 0.036624 | 0.11429 | 0.951052 | 2351 | 2 | 1.0292 | 0.017159 | 0.069813 | 0.937929 | 1430 | 3 | 1.0292 |
| hsa-mir-1298 | 4 | 0.98279 | 0.98275 | 0.999994 | 21153 | 0 | 3.2824 | 0.017213 | 0.055804 | 0.926769 | 1431 | 4 | 3.2824 |
| GABRA2 | 6 | 0.97176 | 0.97171 | 0.999994 | 20864 | 0 | 4.7121 | 0.017251 | 0.070078 | 0.937929 | 1432 | 5 | 4.7121 |
| SMARCA1 | 6 | 0.68052 | 0.80143 | 0.999994 | 16100 | 2 | 4.9772 | 0.017252 | 0.070083 | 0.937929 | 1433 | 4 | 4.9772 |
| FOCAD | 6 | 0.85009 | 0.87198 | 0.999994 | 18264 | 1 | 3.5586 | 0.017326 | 0.070306 | 0.937929 | 1434 | 4 | 3.5586 |
| MEF2BNB | 5 | 0.34421 | 0.51717 | 0.999994 | 11033 | 1 | 2.0693 | 0.017339 | 0.065497 | 0.933948 | 1435 | 3 | 2.0693 |
| LCNL1 | 5 | 0.22412 | 0.40901 | 0.999994 | 8382 | 2 | 6.5981 | 0.017339 | 0.065498 | 0.933948 | 1436 | 3 | 6.5981 |
| CNOT2 | 5 | 0.91242 | 0.91244 | 0.999994 | 19470 | 0 | 3.6262 | 0.01738 | 0.065638 | 0.933948 | 1437 | 4 | 3.6262 |
| FBXW10 | 5 | 0.75556 | 0.79378 | 0.999994 | 16926 | 1 | 3.4134 | 0.017382 | 0.065649 | 0.933948 | 1438 | 4 | 3.4134 |
| RAB40A | 5 | 0.97247 | 0.97253 | 0.999994 | 20880 | 0 | 2.0506 | 0.017389 | 0.065668 | 0.933948 | 1439 | 4 | 2.0506 |
| ANP32B | 5 | 0.97376 | 0.9738 | 0.999994 | 20919 | 0 | 3.6421 | 0.017394 | 0.065683 | 0.933948 | 1440 | 5 | 3.6421 |
| BTG1 | 6 | 0.29362 | 0.49927 | 0.999994 | 9988 | 2 | 2.9761 | 0.017427 | 0.0706 | 0.937929 | 1441 | 3 | 2.9761 |
| SIDT2 | 5 | 0.55227 | 0.69375 | 0.999994 | 14248 | 2 | 1.0535 | 0.017443 | 0.065837 | 0.933948 | 1442 | 3 | 1.0535 |
| YIPF2 | 5 | 0.98255 | 0.98262 | 0.999994 | 21150 | 0 | 2.791 | 0.017451 | 0.065869 | 0.933948 | 1443 | 5 | 2.791 |
| DDHD1 | 5 | 0.98255 | 0.98262 | 0.999994 | 21149 | 0 | 1.3514 | 0.017455 | 0.065883 | 0.933948 | 1444 | 5 | 1.3514 |
| CCDC68 | 6 | 0.9673 | 0.96733 | 0.999994 | 20768 | 0 | 5.4201 | 0.017482 | 0.070763 | 0.937929 | 1445 | 5 | 5.4201 |
| ACTL10 | 6 | 0.086093 | 0.21969 | 0.980699 | 4453 | 2 | 5.0102 | 0.017482 | 0.070763 | 0.937929 | 1446 | 4 | 5.0102 |
| CWC27 | 6 | 0.97637 | 0.97631 | 0.999994 | 21002 | 0 | 4.2543 | 0.017497 | 0.070807 | 0.937929 | 1447 | 5 | 4.2543 |
| CCDC147 | 6 | 0.84061 | 0.86693 | 0.999994 | 18099 | 1 | 4.7967 | 0.017512 | 0.070851 | 0.937929 | 1448 | 5 | 4.7967 |
| TRNP1 | 4 | 0.3287 | 0.47236 | 0.999994 | 10694 | 2 | 3.1162 | 0.017515 | 0.056691 | 0.926769 | 1449 | 2 | 3.1162 |
| NonTargetingControlG | 2 | 0.80595 | 0.80557 | 0.999994 | 17572 | 0 | 5.2784 | 0.017526 | 0.032684 | 0.890437 | 1450 | 2 | 5.2784 |
| hsa-mir-22 | 4 | 0.87803 | 0.87768 | 0.999994 | 18787 | 0 | 0.78219 | 0.017552 | 0.056798 | 0.926769 | 1451 | 2 | 0.78219 |
| METAP1D | 6 | 0.27001 | 0.47172 | 0.999994 | 9474 | 3 | 2.7518 | 0.017558 | 0.070988 | 0.937929 | 1452 | 3 | 2.7518 |
| SH3TC1 | 3 | 0.78675 | 0.78667 | 0.999994 | 17300 | 0 | 0.036693 | 0.017561 | 0.045607 | 0.919289 | 1453 | 1 | 0.036693 |
| NTHL1 | 5 | 0.72872 | 0.78003 | 0.999994 | 16588 | 1 | 6.1252 | 0.017566 | 0.066285 | 0.933948 | 1454 | 4 | 6.1252 |
| MAP3K7 | 5 | 0.98243 | 0.98251 | 0.999994 | 21147 | 0 | 2.6144 | 0.017569 | 0.066296 | 0.933948 | 1455 | 5 | 2.6144 |

| | | | | | | | | | | | | |
|---|---|---|---|---|---|---|---|---|---|---|---|---|
| LAMA3 | 5 | 0.73115 | 0.78122 | 0.999994 | 16616 | 1 | 1.136 | 0.017577 | 0.066325 | 0.933948 | 1456 | 3 | 1.136 |
| CAST | 4 | 0.87039 | 0.86997 | 0.999994 | 18642 | 0 | 5.5343 | 0.017594 | 0.056928 | 0.926769 | 1457 | 3 | 5.5343 |
| PIPOX | 5 | 0.9824 | 0.98249 | 0.999994 | 21146 | 0 | 2.1542 | 0.017599 | 0.066388 | 0.933948 | 1458 | 5 | 2.1542 |
| OR52K2 | 5 | 0.75189 | 0.79183 | 0.999994 | 16878 | 1 | 6.5854 | 0.017612 | 0.066429 | 0.933948 | 1459 | 4 | 6.5854 |
| IRGC | 6 | 0.65611 | 0.78229 | 0.999994 | 15726 | 2 | 5.4025 | 0.017618 | 0.071191 | 0.937929 | 1460 | 4 | 5.4025 |
| RNF168 | 6 | 0.58436 | 0.72843 | 0.999994 | 14733 | 2 | 5.6935 | 0.017624 | 0.071213 | 0.937929 | 1461 | 4 | 5.6935 |
| PLA2G4D | 6 | 0.90244 | 0.90557 | 0.999994 | 19259 | 1 | 3.7835 | 0.017624 | 0.071213 | 0.937929 | 1462 | 3 | 3.7835 |
| ALOX12 | 6 | 0.83859 | 0.86591 | 0.999994 | 18063 | 1 | 3.9851 | 0.017647 | 0.07128 | 0.937929 | 1463 | 5 | 3.9851 |
| PALM3 | 6 | 0.19108 | 0.37686 | 0.999994 | 7515 | 3 | 0.43532 | 0.017664 | 0.071335 | 0.937929 | 1464 | 3 | 0.43532 |
| CETN1 | 5 | 0.84342 | 0.84923 | 0.999994 | 18146 | 1 | 2.4812 | 0.01767 | 0.06664 | 0.933948 | 1465 | 4 | 2.4812 |
| ACO2 | 6 | 0.21767 | 0.40932 | 0.999994 | 8215 | 2 | 4.1965 | 0.01774 | 0.071566 | 0.937929 | 1466 | 4 | 4.1965 |
| TRIM41 | 5 | 0.39411 | 0.55863 | 0.999994 | 11859 | 2 | 6.5728 | 0.01777 | 0.066979 | 0.933948 | 1467 | 3 | 6.5728 |
| RADIL | 5 | 0.10995 | 0.24191 | 0.987253 | 5303 | 2 | 6.5728 | 0.01777 | 0.066979 | 0.933948 | 1468 | 3 | 6.5728 |
| CCDC110 | 4 | 0.83651 | 0.83602 | 0.999994 | 18027 | 0 | 0.98214 | 0.017794 | 0.057482 | 0.927195 | 1469 | 3 | 0.98214 |
| HLA-DQA1 | 6 | 0.060741 | 0.16734 | 0.967126 | 3453 | 4 | -1.1732 | 0.017827 | 0.071821 | 0.937929 | 1470 | 2 | -1.1732 |
| EFHC1 | 5 | 0.98216 | 0.98227 | 0.999994 | 21142 | 0 | 1.1992 | 0.017837 | 0.06722 | 0.934798 | 1471 | 5 | 1.1992 |
| CRELD1 | 5 | 0.44622 | 0.60201 | 0.999994 | 12665 | 1 | 3.5501 | 0.017851 | 0.067271 | 0.934798 | 1472 | 4 | 3.5501 |
| SLAMF6 | 6 | 0.34499 | 0.5584 | 0.999994 | 11052 | 2 | 4.0371 | 0.017858 | 0.071914 | 0.937929 | 1473 | 4 | 4.0371 |
| KLHL4 | 6 | 0.96331 | 0.96336 | 0.999994 | 20678 | 0 | 5.2303 | 0.017864 | 0.071933 | 0.937929 | 1474 | 5 | 5.2303 |
| SPINK7 | 6 | 0.93376 | 0.93382 | 0.999994 | 19946 | 0 | 4.5031 | 0.017877 | 0.071977 | 0.937929 | 1475 | 5 | 4.5031 |
| TUBB | 6 | 0.95583 | 0.95591 | 0.999994 | 20477 | 0 | 5.5575 | 0.017883 | 0.071996 | 0.937929 | 1476 | 4 | 5.5575 |
| CYTH2 | 6 | 0.41963 | 0.61516 | 0.999994 | 12247 | 1 | 4.5164 | 0.017886 | 0.072006 | 0.937929 | 1477 | 4 | 4.5164 |
| SUOX | 6 | 0.43123 | 0.62267 | 0.999994 | 12434 | 2 | 3.3624 | 0.017886 | 0.072006 | 0.937929 | 1478 | 3 | 3.3624 |
| HELZ | 6 | 0.67567 | 0.79761 | 0.999994 | 16017 | 2 | 5.4049 | 0.017911 | 0.072067 | 0.937929 | 1479 | 4 | 5.4049 |
| TMED6 | 5 | 0.86104 | 0.86328 | 0.999994 | 18465 | 1 | 6.5726 | 0.017914 | 0.0675 | 0.934798 | 1480 | 4 | 6.5726 |
| PXMP4 | 5 | 0.98208 | 0.9822 | 0.999994 | 21140 | 0 | 2.3664 | 0.017918 | 0.067514 | 0.934798 | 1481 | 5 | 2.3664 |
| B4GALT7 | 6 | 0.95106 | 0.95105 | 0.999994 | 20364 | 0 | 4.3417 | 0.017937 | 0.072146 | 0.937929 | 1482 | 5 | 4.3417 |
| NonTargetingControlG | 2 | 0.39693 | 0.41426 | 0.999994 | 11904 | 1 | 3.8519 | 0.017946 | 0.033451 | 0.89322 | 1483 | 1 | 3.8519 |
| C6orf195 | 5 | 0.0028959 | 0.012439 | 0.733879 | 347 | 1 | 2.3564 | 0.017949 | 0.067609 | 0.935447 | 1484 | 4 | 2.3564 |
| HEATR2 | 6 | 0.91997 | 0.92038 | 0.999994 | 19630 | 1 | 4.9314 | 0.017949 | 0.072186 | 0.937929 | 1485 | 5 | 4.9314 |
| KRTAP10-5 | 6 | 0.62087 | 0.75549 | 0.999994 | 15235 | 2 | 3.8535 | 0.017995 | 0.072322 | 0.937929 | 1486 | 4 | 3.8535 |
| KRT25 | 6 | 0.64983 | 0.77749 | 0.999994 | 15657 | 2 | 3.6748 | 0.018004 | 0.072355 | 0.937929 | 1487 | 3 | 3.6748 |
| POLA2 | 5 | 0.98199 | 0.98211 | 0.999994 | 21137 | 0 | 2.6433 | 0.018009 | 0.067808 | 0.935447 | 1488 | 5 | 2.6433 |
| IRF3 | 5 | 0.96917 | 0.96925 | 0.999994 | 20806 | 0 | 4.7816 | 0.018022 | 0.067852 | 0.935447 | 1489 | 4 | 4.7816 |
| CDV3 | 5 | 0.98196 | 0.98208 | 0.999994 | 21135 | 0 | 1.9326 | 0.018038 | 0.067904 | 0.935447 | 1490 | 5 | 1.9326 |
| HTT | 5 | 0.36382 | 0.53325 | 0.999994 | 11399 | 1 | 6.5598 | 0.018054 | 0.067954 | 0.935447 | 1491 | 3 | 6.5598 |
| ADSSL1 | 6 | 0.94656 | 0.94653 | 0.999994 | 20246 | 0 | 5.6396 | 0.018055 | 0.072499 | 0.937929 | 1492 | 4 | 5.6396 |
| CELA3B | 6 | 0.28593 | 0.49038 | 0.999994 | 9831 | 2 | 0.71723 | 0.018062 | 0.072528 | 0.937929 | 1493 | 3 | 0.71723 |
| PPCS | 3 | 0.70045 | 0.70144 | 0.999994 | 16309 | 1 | 7.5379 | 0.018079 | 0.046895 | 0.919289 | 1494 | 2 | 7.5379 |
| C6orf57 | 5 | 0.92516 | 0.92539 | 0.999994 | 19745 | 0 | 4.1937 | 0.018086 | 0.068073 | 0.935447 | 1495 | 4 | 4.1937 |
| KIAA1644 | 6 | 0.70634 | 0.81397 | 0.999994 | 16376 | 1 | 4.3705 | 0.018087 | 0.072606 | 0.937929 | 1496 | 4 | 4.3705 |
| VSIG4 | 6 | 0.30218 | 0.50926 | 0.999994 | 10168 | 2 | 4.2013 | 0.018087 | 0.072606 | 0.937929 | 1497 | 4 | 4.2013 |
| MRPS33 | 5 | 0.9711 | 0.97114 | 0.999994 | 20850 | 0 | 5.5479 | 0.018088 | 0.068085 | 0.935447 | 1498 | 4 | 5.5479 |
| ZNF865 | 5 | 0.98189 | 0.98201 | 0.999994 | 21134 | 0 | 3.1581 | 0.018108 | 0.06814 | 0.935447 | 1499 | 5 | 3.1581 |
| ZDHHC2 | 6 | 0.31841 | 0.52807 | 0.999994 | 10483 | 1 | 2.3865 | 0.018155 | 0.072808 | 0.937929 | 1500 | 3 | 2.3865 |
| ID3 | 6 | 0.38738 | 0.59425 | 0.999994 | 11757 | 2 | 4.6563 | 0.018155 | 0.072808 | 0.937929 | 1501 | 4 | 4.6563 |
| PRPF18 | 5 | 0.45761 | 0.61166 | 0.999994 | 12815 | 1 | 5.6101 | 0.018163 | 0.068316 | 0.935447 | 1502 | 4 | 5.6101 |
| DNAJB14 | 5 | 0.97712 | 0.9772 | 0.999994 | 21018 | 0 | 1.2973 | 0.018163 | 0.068316 | 0.935447 | 1503 | 5 | 1.2973 |
| CA10 | 5 | 0.93645 | 0.9367 | 0.999994 | 19993 | 0 | 4.1279 | 0.018176 | 0.068367 | 0.935447 | 1504 | 4 | 4.1279 |
| CCSER2 | 5 | 0.45024 | 0.60543 | 0.999994 | 12713 | 2 | -0.12746 | 0.018181 | 0.06838 | 0.935447 | 1505 | 1 | -0.12746 |
| FKBP7 | 5 | 0.64812 | 0.74435 | 0.999994 | 15631 | 1 | 6.5571 | 0.018182 | 0.068381 | 0.935447 | 1506 | 4 | 6.5571 |
| KLRD1 | 6 | 0.091276 | 0.23008 | 0.983704 | 4647 | 1 | 4.3161 | 0.01819 | 0.072897 | 0.938203 | 1507 | 5 | 4.3161 |

| | | | | | | | | | | | | | |
|---|---|---|---|---|---|---|---|---|---|---|---|---|---|
| TPGS2 | 5 | 0.61587 | 0.73202 | 0.999994 | 15157 | 1 | 5.5244 | 0.018201 | 0.068452 | 0.935447 | 1508 | 4 | 5.5244 |
| PLD5 | 6 | 0.55482 | 0.70709 | 0.999994 | 14281 | 1 | 3.9975 | 0.018203 | 0.072942 | 0.938203 | 1509 | 4 | 3.9975 |
| VAT1L | 6 | 0.80285 | 0.84908 | 0.999994 | 17522 | 1 | 3.1759 | 0.018227 | 0.073014 | 0.938203 | 1510 | 5 | 3.1759 |
| WDR1 | 6 | 0.33135 | 0.5428 | 0.999994 | 10752 | 1 | 5.3514 | 0.018255 | 0.07311 | 0.938203 | 1511 | 5 | 5.3514 |
| OXA1L | 5 | 0.96312 | 0.96327 | 0.999994 | 20671 | 0 | 1.8834 | 0.018295 | 0.06876 | 0.936374 | 1512 | 3 | 1.8834 |
| GBP4 | 5 | 0.75638 | 0.7942 | 0.999994 | 16935 | 1 | 6.5468 | 0.018315 | 0.068833 | 0.936374 | 1513 | 4 | 6.5468 |
| LPL | 6 | 0.47978 | 0.65506 | 0.999994 | 13142 | 2 | 0.29918 | 0.018329 | 0.073344 | 0.938203 | 1514 | 3 | 0.29918 |
| OOEP | 6 | 0.97549 | 0.97541 | 0.999994 | 20975 | 0 | 3.1682 | 0.018329 | 0.073345 | 0.938203 | 1515 | 4 | 3.1682 |
| HSPA5 | 3 | 0.98137 | 0.98146 | 0.999994 | 21126 | 0 | 4.8199 | 0.018371 | 0.047567 | 0.919289 | 1516 | 3 | 4.8199 |
| NBAS | 6 | 0.6904 | 0.80917 | 0.999994 | 16214 | 1 | 5.6583 | 0.018409 | 0.073595 | 0.938203 | 1517 | 5 | 5.6583 |
| EID3 | 5 | 0.63406 | 0.7389 | 0.999994 | 15434 | 1 | 6.5412 | 0.01847 | 0.069361 | 0.936996 | 1518 | 3 | 6.5412 |
| C1orf50 | 6 | 0.033576 | 0.10718 | 0.950995 | 2193 | 1 | 1.5179 | 0.018472 | 0.073791 | 0.938203 | 1519 | 4 | 1.5179 |
| BAX | 6 | 0.89672 | 0.90116 | 0.999994 | 19157 | 1 | 4.7909 | 0.018472 | 0.073791 | 0.938203 | 1520 | 5 | 4.7909 |
| hsa-mir-320c-2 | 3 | 0.21899 | 0.33638 | 0.999994 | 8261 | 1 | 5.835 | 0.018483 | 0.047851 | 0.919289 | 1521 | 2 | 5.835 |
| DDX3X | 5 | 0.98152 | 0.98163 | 0.999994 | 21128 | 0 | 1.8557 | 0.018484 | 0.069404 | 0.936998 | 1522 | 5 | 1.8557 |
| L3MBTL2 | 6 | 0.73068 | 0.82184 | 0.999994 | 16608 | 1 | 5.0587 | 0.018496 | 0.073858 | 0.938203 | 1523 | 5 | 5.0587 |
| PRAC1 | 4 | 0.012357 | 0.04105 | 0.891621 | 1019 | 2 | -4.1039 | 0.01852 | 0.059658 | 0.930754 | 1524 | 1 | -4.1039 |
| MRGPRX1 | 6 | 0.13071 | 0.30124 | 0.999994 | 5949 | 1 | 3.4034 | 0.018616 | 0.074223 | 0.938203 | 1525 | 5 | 3.4034 |
| CHD1L | 5 | 0.30206 | 0.4827 | 0.999994 | 10165 | 1 | 4.3741 | 0.01862 | 0.069869 | 0.937929 | 1526 | 4 | 4.3741 |
| hsa-mir-3917 | 4 | 0.80857 | 0.80898 | 0.999994 | 17611 | 1 | 6.3343 | 0.018655 | 0.060055 | 0.931675 | 1527 | 3 | 6.3343 |
| MLPH | 6 | 0.96649 | 0.96654 | 0.999994 | 20749 | 0 | 2.6243 | 0.018724 | 0.074551 | 0.938203 | 1528 | 5 | 2.6243 |
| TMEM52B | 5 | 0.38227 | 0.54861 | 0.999994 | 11681 | 2 | 1.6345 | 0.018739 | 0.070257 | 0.937929 | 1529 | 3 | 1.6345 |
| NonTargetingControlG | 2 | 0.92833 | 0.92848 | 0.999994 | 19821 | 0 | 6.2913 | 0.01874 | 0.034894 | 0.897383 | 1530 | 2 | 6.2913 |
| CETN3 | 4 | 0.98124 | 0.9812 | 0.999994 | 21124 | 0 | 2.4592 | 0.018764 | 0.060364 | 0.9326 | 1531 | 4 | 2.4592 |
| TRIM44 | 6 | 0.19096 | 0.37672 | 0.999994 | 7512 | 3 | 0.025042 | 0.018774 | 0.074691 | 0.938203 | 1532 | 3 | 0.025042 |
| MMRN2 | 6 | 0.97529 | 0.97522 | 0.999994 | 20965 | 0 | 4.8644 | 0.018781 | 0.074713 | 0.938203 | 1533 | 4 | 4.8644 |
| ZNF10 | 5 | 0.98118 | 0.98129 | 0.999994 | 21122 | 0 | 1.5346 | 0.018823 | 0.070527 | 0.937929 | 1534 | 5 | 1.5346 |
| hsa-mir-5010 | 3 | 0.98118 | 0.98125 | 0.999994 | 21121 | 0 | 6.7763 | 0.018825 | 0.048712 | 0.920057 | 1535 | 3 | 6.7763 |
| FAM47A | 6 | 0.25693 | 0.45623 | 0.999994 | 9167 | 2 | 0.98431 | 0.018856 | 0.074939 | 0.938203 | 1536 | 3 | 0.98431 |
| SMIM22 | 6 | 0.93211 | 0.93222 | 0.999994 | 19903 | 0 | 3.0419 | 0.018865 | 0.074967 | 0.938203 | 1537 | 5 | 3.0419 |
| MAT1A | 5 | 0.8039 | 0.82197 | 0.999994 | 17539 | 1 | 1.2512 | 0.018899 | 0.070776 | 0.937929 | 1538 | 3 | 1.2512 |
| HIST3H2A | 5 | 0.018855 | 0.060964 | 0.932932 | 1401 | 2 | 2.0135 | 0.018899 | 0.070776 | 0.937929 | 1539 | 3 | 2.0135 |
| SERPINB6 | 5 | 0.4705 | 0.62259 | 0.999994 | 13010 | 2 | 6.5204 | 0.018919 | 0.070847 | 0.937929 | 1540 | 3 | 6.5204 |
| UGT3A1 | 5 | 0.13091 | 0.27399 | 0.998561 | 5956 | 3 | -1.2588 | 0.018924 | 0.070865 | 0.937929 | 1541 | 1 | -1.2588 |
| MAGT1 | 6 | 0.053283 | 0.15117 | 0.967126 | 3127 | 1 | 3.9105 | 0.018937 | 0.075198 | 0.938203 | 1542 | 4 | 3.9105 |
| EIF2AK1 | 6 | 0.054661 | 0.15418 | 0.967126 | 3181 | 3 | -0.36402 | 0.018942 | 0.075215 | 0.938203 | 1543 | 2 | -0.36402 |
| NRGN | 5 | 0.98106 | 0.98117 | 0.999994 | 21120 | 0 | 1.6361 | 0.018943 | 0.070933 | 0.937929 | 1544 | 5 | 1.6361 |
| NonTargetingControlG | 1 | 0.98105 | 0.98108 | 0.999994 | 21119 | 0 | 9.1152 | 0.018947 | 0.018917 | 0.84631 | 1545 | 1 | 9.1152 |
| hsa-mir-518d | 4 | 0.98105 | 0.981 | 0.999994 | 21118 | 0 | 1.6478 | 0.018954 | 0.060934 | 0.9326 | 1546 | 4 | 1.6478 |
| CBWD3 | 4 | 0.42696 | 0.57338 | 0.999994 | 12364 | 2 | 1.152 | 0.018966 | 0.060972 | 0.9326 | 1547 | 2 | 1.152 |
| hsa-mir-4531 | 4 | 0.82806 | 0.82763 | 0.999994 | 17899 | 0 | 5.7201 | 0.018981 | 0.061013 | 0.9326 | 1548 | 3 | 5.7201 |
| CAPZA1 | 5 | 0.64845 | 0.74448 | 0.999994 | 15634 | 1 | 6.647 | 0.018983 | 0.071071 | 0.937929 | 1549 | 4 | 6.647 |
| HCLS1 | 5 | 0.98096 | 0.98106 | 0.999994 | 21117 | 0 | 3.889 | 0.019037 | 0.071265 | 0.937929 | 1550 | 5 | 3.889 |
| CCDC176 | 5 | 0.97841 | 0.97849 | 0.999994 | 21059 | 0 | 6.509 | 0.019088 | 0.071429 | 0.937929 | 1551 | 5 | 6.509 |
| hsa-mir-3152 | 4 | 0.12015 | 0.25055 | 0.992497 | 5634 | 2 | -0.6289 | 0.019115 | 0.061405 | 0.933948 | 1552 | 2 | -0.6289 |
| PITX1 | 5 | 0.98087 | 0.98096 | 0.999994 | 21114 | 0 | 1.0653 | 0.019132 | 0.071578 | 0.937929 | 1553 | 5 | 1.0653 |
| LPHN2 | 3 | 0.98082 | 0.9809 | 0.999994 | 21113 | 0 | 5.1053 | 0.01918 | 0.049581 | 0.921876 | 1554 | 3 | 5.1053 |
| GAPT | 6 | 0.44988 | 0.63505 | 0.999994 | 12710 | 1 | 4.1969 | 0.019189 | 0.075947 | 0.938203 | 1555 | 4 | 4.1969 |
| MXI1 | 6 | 0.1195 | 0.28479 | 0.999632 | 5613 | 2 | 4.0367 | 0.019189 | 0.075947 | 0.938203 | 1556 | 4 | 4.0367 |
| ADCK3 | 5 | 0.36899 | 0.53751 | 0.999994 | 11490 | 2 | -0.39415 | 0.01921 | 0.07183 | 0.937929 | 1557 | 2 | -0.39415 |
| WDR11 | 5 | 0.40194 | 0.56512 | 0.999994 | 11977 | 1 | 6.5071 | 0.01924 | 0.071927 | 0.937929 | 1558 | 3 | 6.5071 |
| IRS4 | 5 | 0.64902 | 0.74473 | 0.999994 | 15642 | 1 | 6.5071 | 0.01924 | 0.071927 | 0.937929 | 1559 | 4 | 6.5071 |

| | | | | | | | | | | | | | |
|---|---|---|---|---|---|---|---|---|---|---|---|---|---|
| GDAP1L1 | 3 | 0.98071 | 0.9808 | 0.999994 | 21108 | 0 | 3.834 | 0.019286 | 0.049829 | 0.923113 | 1560 | 3 | 3.834 |
| COX19 | 5 | 0.71503 | 0.7735 | 0.999994 | 16450 | 1 | 0.13376 | 0.019296 | 0.072116 | 0.937929 | 1561 | 2 | 0.13376 |
| CEACAM19 | 5 | 0.98068 | 0.98078 | 0.999994 | 21106 | 0 | 1.7962 | 0.019315 | 0.072188 | 0.937929 | 1562 | 5 | 1.7962 |
| CETN2 | 6 | 0.11157 | 0.26973 | 0.998561 | 5351 | 3 | 0.72725 | 0.019315 | 0.076343 | 0.938203 | 1563 | 3 | 0.72725 |
| S100A16 | 6 | 0.081834 | 0.2111 | 0.979587 | 4292 | 4 | -1.0111 | 0.019331 | 0.076395 | 0.938203 | 1564 | 2 | -1.0111 |
| ZCCHC5 | 6 | 0.54841 | 0.70247 | 0.999994 | 14185 | 2 | 5.2792 | 0.019332 | 0.076399 | 0.938203 | 1565 | 4 | 5.2792 |
| KIF1A | 5 | 0.89045 | 0.89056 | 0.999994 | 19043 | 0 | 1.0427 | 0.019342 | 0.072272 | 0.937929 | 1566 | 3 | 1.0427 |
| NCBP2 | 5 | 0.28227 | 0.4667 | 0.999994 | 9744 | 1 | 5.0163 | 0.019344 | 0.072278 | 0.937929 | 1567 | 4 | 5.0163 |
| DRC1 | 5 | 0.39041 | 0.55548 | 0.999994 | 11804 | 1 | 6.4936 | 0.019369 | 0.072369 | 0.937929 | 1568 | 4 | 6.4936 |
| OR51B4 | 6 | 0.045883 | 0.13502 | 0.954328 | 2801 | 3 | -0.28845 | 0.019384 | 0.076558 | 0.938203 | 1569 | 3 | -0.28845 |
| MSANTD4 | 6 | 0.21053 | 0.40074 | 0.999994 | 8035 | 2 | 2.1423 | 0.019384 | 0.076558 | 0.938203 | 1570 | 4 | 2.1423 |
| OR8J3 | 6 | 0.57164 | 0.71914 | 0.999994 | 14569 | 2 | 2.3355 | 0.019384 | 0.076558 | 0.938203 | 1571 | 3 | 2.3355 |
| METAP2 | 6 | 0.97615 | 0.97609 | 0.999994 | 20993 | 0 | 3.063 | 0.01939 | 0.076576 | 0.938203 | 1572 | 5 | 3.063 |
| FABP12 | 6 | 0.92215 | 0.92239 | 0.999994 | 19680 | 1 | 3.7086 | 0.01939 | 0.076576 | 0.938203 | 1573 | 5 | 3.7086 |
| FLAD1 | 6 | 0.2389 | 0.43468 | 0.999994 | 8746 | 1 | 5.5824 | 0.01939 | 0.076576 | 0.938203 | 1574 | 5 | 5.5824 |
| TIE1 | 6 | 0.11549 | 0.27721 | 0.999379 | 5494 | 2 | 1.3993 | 0.019443 | 0.07673 | 0.938203 | 1575 | 4 | 1.3993 |
| LGALS1 | 5 | 0.98053 | 0.98063 | 0.999994 | 21104 | 0 | 1.6299 | 0.01947 | 0.072703 | 0.937929 | 1576 | 5 | 1.6299 |
| CD58 | 6 | 0.35728 | 0.57225 | 0.999994 | 11289 | 3 | 3.1846 | 0.019473 | 0.076828 | 0.938203 | 1577 | 3 | 3.1846 |
| RBM4B | 6 | 0.09206 | 0.23165 | 0.984454 | 4682 | 2 | 3.8324 | 0.019522 | 0.076966 | 0.938203 | 1578 | 4 | 3.8324 |
| LRP10 | 6 | 0.62226 | 0.75649 | 0.999994 | 15263 | 2 | 4.5032 | 0.019533 | 0.076998 | 0.938203 | 1579 | 4 | 4.5032 |
| ARMCX2 | 6 | 0.21658 | 0.40804 | 0.999994 | 8181 | 3 | -0.81705 | 0.019554 | 0.077058 | 0.938203 | 1580 | 2 | -0.81705 |
| TOP3B | 6 | 0.98253 | 0.98247 | 0.999994 | 21148 | 0 | 4.4966 | 0.019594 | 0.077185 | 0.938203 | 1581 | 5 | 4.4966 |
| TMEM163 | 6 | 0.81623 | 0.85508 | 0.999994 | 17717 | 1 | 5.4474 | 0.019594 | 0.077185 | 0.938203 | 1582 | 4 | 5.4474 |
| TAOK2 | 5 | 0.98039 | 0.98047 | 0.999994 | 21101 | 0 | 3.2688 | 0.019614 | 0.073203 | 0.938203 | 1583 | 5 | 3.2688 |
| MAGEB1 | 6 | 0.21053 | 0.40074 | 0.999994 | 8034 | 2 | 1.0574 | 0.019619 | 0.077249 | 0.938203 | 1584 | 3 | 1.0574 |
| CCDC54 | 5 | 0.98038 | 0.98047 | 0.999994 | 21100 | 0 | 1.7038 | 0.01962 | 0.073223 | 0.938203 | 1585 | 5 | 1.7038 |
| SNX11 | 6 | 0.97512 | 0.97504 | 0.999994 | 20957 | 0 | 1.9351 | 0.019665 | 0.077392 | 0.938203 | 1586 | 4 | 1.9351 |
| ZNHIT1 | 4 | 0.98033 | 0.98031 | 0.999994 | 21099 | 0 | 3.6977 | 0.019667 | 0.06299 | 0.933948 | 1587 | 4 | 3.6977 |
| SLC25A26 | 6 | 0.13631 | 0.30837 | 0.999994 | 6092 | 2 | 5.1422 | 0.01967 | 0.077409 | 0.938203 | 1588 | 4 | 5.1422 |
| ZC3H12C | 5 | 0.82805 | 0.83809 | 0.999994 | 17898 | 1 | 1.8316 | 0.019694 | 0.073461 | 0.938203 | 1589 | 3 | 1.8316 |
| DRD2 | 5 | 0.93922 | 0.93941 | 0.999994 | 20054 | 0 | 6.48 | 0.019699 | 0.073479 | 0.938203 | 1590 | 3 | 6.48 |
| FAM124A | 5 | 0.62976 | 0.73721 | 0.999994 | 15378 | 1 | 6.48 | 0.019699 | 0.073479 | 0.938203 | 1591 | 3 | 6.48 |
| EBF4 | 5 | 0.8487 | 0.85326 | 0.999994 | 18227 | 1 | 4.6636 | 0.019703 | 0.073492 | 0.938203 | 1592 | 4 | 4.6636 |
| GPR110 | 6 | 0.96163 | 0.96168 | 0.999994 | 20622 | 0 | 0.70133 | 0.019721 | 0.077577 | 0.938203 | 1593 | 3 | 0.70133 |
| NonTargetingControlG | 2 | 0.46525 | 0.47278 | 0.999994 | 12936 | 1 | 4.1324 | 0.019758 | 0.036733 | 0.904772 | 1594 | 1 | 4.1324 |
| hsa-mir-557 | 4 | 0.98021 | 0.9802 | 0.999994 | 21097 | 0 | 1.8806 | 0.019788 | 0.063334 | 0.933948 | 1595 | 4 | 1.8806 |
| PRICKLE1 | 5 | 0.19498 | 0.36806 | 0.999994 | 7632 | 3 | -0.88577 | 0.019806 | 0.073841 | 0.938203 | 1596 | 2 | -0.88577 |
| IFI44 | 5 | 0.95886 | 0.959 | 0.999994 | 20557 | 0 | 2.829 | 0.019831 | 0.073924 | 0.938203 | 1597 | 3 | 2.829 |
| POGK | 5 | 0.88351 | 0.88365 | 0.999994 | 18897 | 1 | 6.476 | 0.01984 | 0.073954 | 0.938203 | 1598 | 3 | 6.476 |
| WASF3 | 6 | 0.56368 | 0.71343 | 0.999994 | 14445 | 2 | 3.7311 | 0.019853 | 0.077952 | 0.938203 | 1599 | 3 | 3.7311 |
| CEACAM16 | 6 | 0.57303 | 0.72013 | 0.999994 | 14590 | 2 | 0.90286 | 0.019888 | 0.078043 | 0.938203 | 1600 | 4 | 0.90286 |
| PROSC | 6 | 0.54999 | 0.70363 | 0.999994 | 14211 | 2 | 1.8164 | 0.019919 | 0.078126 | 0.938203 | 1601 | 4 | 1.8164 |
| PPP1CC | 6 | 0.51598 | 0.67973 | 0.999994 | 13675 | 1 | 2.9432 | 0.019919 | 0.078126 | 0.938203 | 1602 | 4 | 2.9432 |
| NonTargetingControlG | 2 | 0.58178 | 0.58191 | 0.999994 | 14703 | 1 | 4.5304 | 0.019945 | 0.037074 | 0.904772 | 1603 | 1 | 4.5304 |
| GPR114 | 4 | 0.69465 | 0.72098 | 0.999994 | 16250 | 1 | 1.9334 | 0.019951 | 0.063813 | 0.933948 | 1604 | 2 | 1.9334 |
| AKAP2 | 5 | 0.98005 | 0.98016 | 0.999994 | 21092 | 0 | 2.187 | 0.019951 | 0.074311 | 0.938203 | 1605 | 5 | 2.187 |
| ACTL9 | 5 | 0.10078 | 0.22765 | 0.983354 | 5000 | 2 | -0.0087935 | 0.019968 | 0.074361 | 0.938203 | 1606 | 2 | -0.0087935 |
| PACS2 | 6 | 0.66723 | 0.79096 | 0.999994 | 15881 | 1 | 5.8019 | 0.019981 | 0.078317 | 0.938203 | 1607 | 5 | 5.8019 |
| CYB5R3 | 5 | 0.98 | 0.98011 | 0.999994 | 21091 | 0 | 3.3542 | 0.020001 | 0.074469 | 0.938203 | 1608 | 5 | 3.3542 |
| C9orf24 | 6 | 0.14976 | 0.32533 | 0.999994 | 6454 | 2 | 1.9752 | 0.020012 | 0.078413 | 0.938203 | 1609 | 3 | 1.9752 |
| RNF122 | 4 | 0.97999 | 0.97998 | 0.999994 | 21090 | 0 | 4.0113 | 0.020014 | 0.063982 | 0.933948 | 1610 | 4 | 4.0113 |
| FABP1 | 5 | 0.97999 | 0.98009 | 0.999994 | 21089 | 0 | 2.7149 | 0.020015 | 0.074518 | 0.938203 | 1611 | 5 | 2.7149 |

| | | | | | | | | | | | | | |
|---|---|---|---|---|---|---|---|---|---|---|---|---|---|
| ZIM3 | 6 | 0.90691 | 0.90903 | 0.999994 | 19368 | 1 | 3.3099 | 0.02003 | 0.078485 | 0.938203 | 1612 | 5 | 3.3099 |
| MROH6 | 5 | 0.53919 | 0.68217 | 0.999994 | 14037 | 1 | 0.28732 | 0.020061 | 0.07467 | 0.938203 | 1613 | 2 | 0.28732 |
| ZNF670 | 6 | 0.89045 | 0.89665 | 0.999994 | 19044 | 1 | 4.9011 | 0.020092 | 0.078681 | 0.938203 | 1614 | 4 | 4.9011 |
| VDAC2 | 5 | 0.0069342 | 0.02718 | 0.850392 | 656 | 1 | 6.6235 | 0.020101 | 0.074806 | 0.938203 | 1615 | 4 | 6.6235 |
| UBE2V2 | 4 | 0.81886 | 0.81866 | 0.999994 | 17763 | 1 | 5.9906 | 0.020108 | 0.064282 | 0.933948 | 1616 | 3 | 5.9906 |
| RAB40AL | 4 | 0.97985 | 0.97984 | 0.999994 | 21088 | 0 | 1.7971 | 0.020149 | 0.064404 | 0.933948 | 1617 | 4 | 1.7971 |
| hsa-mir-4311 | 3 | 0.0025294 | 0.0070464 | 0.646564 | 318 | 1 | -0.064351 | 0.020186 | 0.052027 | 0.926769 | 1618 | 1 | -0.064351 |
| CITED2 | 5 | 0.47675 | 0.62782 | 0.999994 | 13096 | 2 | 1.6808 | 0.0202 | 0.075142 | 0.938203 | 1619 | 3 | 1.6808 |
| TSC22D1 | 5 | 0.9798 | 0.9799 | 0.999994 | 21087 | 0 | 1.4235 | 0.020203 | 0.075151 | 0.938203 | 1620 | 5 | 1.4235 |
| TIFA | 6 | 0.82362 | 0.85851 | 0.999994 | 17832 | 1 | 4.826 | 0.020223 | 0.079055 | 0.938203 | 1621 | 5 | 4.826 |
| ZNF180 | 4 | 0.34408 | 0.48826 | 0.999994 | 11030 | 2 | 3.3904 | 0.020226 | 0.064619 | 0.933948 | 1622 | 2 | 3.3904 |
| BHMT2 | 5 | 0.97976 | 0.97987 | 0.999994 | 21086 | 0 | 2.7792 | 0.020237 | 0.07526 | 0.938203 | 1623 | 5 | 2.7792 |
| RNASE2 | 6 | 0.099499 | 0.2464 | 0.98999 | 4954 | 3 | 0.61754 | 0.020241 | 0.079103 | 0.938203 | 1624 | 3 | 0.61754 |
| KRT80 | 6 | 0.83843 | 0.86583 | 0.999994 | 18060 | 1 | 4.4318 | 0.020247 | 0.079125 | 0.938203 | 1625 | 5 | 4.4318 |
| REEP1 | 6 | 0.5206 | 0.68296 | 0.999994 | 13755 | 2 | 3.2322 | 0.020247 | 0.079125 | 0.938203 | 1626 | 3 | 3.2322 |
| TMEM182 | 5 | 0.3698 | 0.5382 | 0.999994 | 11505 | 2 | 2.1277 | 0.02026 | 0.075331 | 0.938203 | 1627 | 3 | 2.1277 |
| NonTargetingControlG | 2 | 0.97973 | 0.97971 | 0.999994 | 21084 | 0 | 7.3279 | 0.020268 | 0.03768 | 0.904772 | 1628 | 2 | 7.3279 |
| GTPBP5 | 5 | 0.62101 | 0.73393 | 0.999994 | 15238 | 1 | 6.4524 | 0.020307 | 0.075479 | 0.938203 | 1629 | 4 | 6.4524 |
| SLC13A4 | 4 | 0.19325 | 0.33373 | 0.999994 | 7581 | 2 | 2.9721 | 0.02032 | 0.064888 | 0.933948 | 1630 | 2 | 2.9721 |
| UNC13A | 6 | 0.51869 | 0.68162 | 0.999994 | 13721 | 2 | 4.7517 | 0.02032 | 0.07934 | 0.938203 | 1631 | 4 | 4.7517 |
| GPSM3 | 5 | 0.80575 | 0.82314 | 0.999994 | 17568 | 1 | 2.7355 | 0.020363 | 0.075675 | 0.938203 | 1632 | 4 | 2.7355 |
| PVALB | 4 | 0.80779 | 0.80825 | 0.999994 | 17593 | 1 | 3.9559 | 0.020408 | 0.065146 | 0.933948 | 1633 | 2 | 3.9559 |
| PRKACG | 6 | 0.98078 | 0.9807 | 0.999994 | 21110 | 0 | 4.4717 | 0.02043 | 0.079682 | 0.938962 | 1634 | 5 | 4.4717 |
| GLUD1 | 5 | 0.52845 | 0.67269 | 0.999994 | 13869 | 2 | 6.4422 | 0.020448 | 0.075954 | 0.938203 | 1635 | 3 | 6.4422 |
| IMPACT | 5 | 0.54015 | 0.68304 | 0.999994 | 14056 | 2 | 6.4422 | 0.020448 | 0.075954 | 0.938203 | 1636 | 3 | 6.4422 |
| CEACAM1 | 6 | 0.95715 | 0.95723 | 0.999994 | 20522 | 0 | 4.5691 | 0.020467 | 0.079785 | 0.938962 | 1637 | 4 | 4.5691 |
| DSCR3 | 4 | 0.97949 | 0.97947 | 0.999994 | 21082 | 0 | 5.4418 | 0.020511 | 0.065452 | 0.933948 | 1638 | 4 | 5.4418 |
| IL32 | 5 | 0.40676 | 0.56917 | 0.999994 | 12058 | 1 | 4.8884 | 0.020532 | 0.076233 | 0.938203 | 1639 | 4 | 4.8884 |
| ARID1A | 5 | 0.21632 | 0.39814 | 0.999994 | 8175 | 1 | 5.1479 | 0.020532 | 0.076233 | 0.938203 | 1640 | 4 | 5.1479 |
| NonTargetingControlG | 2 | 0.85493 | 0.85484 | 0.999994 | 18347 | 0 | 5.5999 | 0.020551 | 0.038172 | 0.904772 | 1641 | 2 | 5.5999 |
| PEX6 | 5 | 0.55746 | 0.69838 | 0.999994 | 14322 | 1 | 0.1897 | 0.020571 | 0.07635 | 0.938203 | 1642 | 2 | 0.1897 |
| ZNF613 | 5 | 0.97942 | 0.97951 | 0.999994 | 21080 | 0 | 1.9354 | 0.020578 | 0.076376 | 0.938203 | 1643 | 5 | 1.9354 |
| PIGA | 6 | 0.29298 | 0.49851 | 0.999994 | 9976 | 2 | 3.9087 | 0.020605 | 0.080194 | 0.938962 | 1644 | 4 | 3.9087 |
| ERI1 | 5 | 0.71236 | 0.77221 | 0.999994 | 16423 | 1 | 4.045 | 0.020633 | 0.076566 | 0.938203 | 1645 | 4 | 4.045 |
| TRERF1 | 4 | 0.53397 | 0.63201 | 0.999994 | 13966 | 1 | 3.9927 | 0.020636 | 0.06582 | 0.933948 | 1646 | 2 | 3.9927 |
| LRRC4 | 5 | 0.2607 | 0.4493 | 0.999994 | 9255 | 1 | 1.8159 | 0.020643 | 0.076604 | 0.938203 | 1647 | 4 | 1.8159 |
| hsa-mir-4487 | 3 | 0.96452 | 0.96463 | 0.999994 | 20713 | 0 | 7.43 | 0.020647 | 0.053166 | 0.926769 | 1648 | 3 | 7.43 |
| TNKS | 6 | 0.87729 | 0.88788 | 0.999994 | 18780 | 1 | 3.901 | 0.02065 | 0.080341 | 0.938962 | 1649 | 5 | 3.901 |
| hsa-mir-4463 | 4 | 0.94596 | 0.94586 | 0.999994 | 20230 | 0 | 6.5016 | 0.020679 | 0.065929 | 0.933948 | 1650 | 4 | 6.5016 |
| ANKS3 | 4 | 0.078618 | 0.17619 | 0.968171 | 4175 | 1 | 4.1228 | 0.020712 | 0.066027 | 0.933948 | 1651 | 2 | 4.1228 |
| ART3 | 5 | 0.97929 | 0.97938 | 0.999994 | 21079 | 0 | 5.4494 | 0.020713 | 0.076837 | 0.938203 | 1652 | 5 | 5.4494 |
| NCR3 | 6 | 0.20795 | 0.39759 | 0.999994 | 7976 | 2 | -0.31248 | 0.020722 | 0.080547 | 0.938962 | 1653 | 2 | -0.31248 |
| UMPS | 6 | 0.96276 | 0.9628 | 0.999994 | 20656 | 0 | 5.4314 | 0.020731 | 0.080577 | 0.938962 | 1654 | 5 | 5.4314 |
| NCOA5 | 5 | 0.8088 | 0.82514 | 0.999994 | 17616 | 1 | -0.35187 | 0.020734 | 0.076916 | 0.938203 | 1655 | 2 | -0.35187 |
| ZNF579 | 5 | 0.97926 | 0.97936 | 0.999994 | 21078 | 0 | 2.1765 | 0.020737 | 0.076929 | 0.938203 | 1656 | 5 | 2.1765 |
| TH | 6 | 0.79924 | 0.84755 | 0.999994 | 17469 | 1 | 4.4605 | 0.020738 | 0.080594 | 0.938962 | 1657 | 4 | 4.4605 |
| CCDC124 | 5 | 0.84878 | 0.85333 | 0.999994 | 18231 | 1 | 6.425 | 0.020751 | 0.076967 | 0.938203 | 1658 | 3 | 6.425 |
| CPNE7 | 5 | 0.24776 | 0.43902 | 0.999994 | 8970 | 1 | 5.3435 | 0.02078 | 0.077057 | 0.938203 | 1659 | 4 | 5.3435 |
| hsa-mir-19b-1 | 4 | 0.91627 | 0.91602 | 0.999994 | 19542 | 0 | 2.5937 | 0.020786 | 0.066221 | 0.933948 | 1660 | 3 | 2.5937 |
| CEP164 | 5 | 0.70488 | 0.76872 | 0.999994 | 16358 | 1 | 3.5171 | 0.020799 | 0.077116 | 0.938203 | 1661 | 4 | 3.5171 |
| EPB42 | 5 | 0.95077 | 0.95094 | 0.999994 | 20351 | 0 | 2.3746 | 0.020826 | 0.077217 | 0.938203 | 1662 | 4 | 2.3746 |
| MLXIP | 6 | 0.41238 | 0.61044 | 0.999994 | 12150 | 2 | 2.3269 | 0.020833 | 0.080887 | 0.939346 | 1663 | 4 | 2.3269 |

| | | | | | | | | | | | | |
|---|---|---|---|---|---|---|---|---|---|---|---|---|
| GRIN2C | 6 | 0.15726 | 0.33478 | 0.999994 | 6654 | 2 | 3.6406 | 0.020852 | 0.080938 | 0.939346 | 1664 | 3 | 3.6406 |
| C20orf173 | 5 | 0.97915 | 0.97924 | 0.999994 | 21077 | 0 | 2.6127 | 0.020854 | 0.077309 | 0.938203 | 1665 | 5 | 2.6127 |
| AWAT2 | 4 | 0.0030026 | 0.010752 | 0.70768 | 358 | 3 | -5.5524 | 0.02086 | 0.06643 | 0.933948 | 1666 | 1 | -5.5524 |
| TM2D3 | 6 | 0.64747 | 0.7757 | 0.999994 | 15623 | 2 | 5.0344 | 0.020878 | 0.081026 | 0.939346 | 1667 | 4 | 5.0344 |
| PEBP1 | 6 | 0.9612 | 0.96125 | 0.999994 | 20609 | 0 | 0.63088 | 0.020972 | 0.081294 | 0.939346 | 1668 | 3 | 0.63088 |
| UPF3B | 6 | 0.77156 | 0.83643 | 0.999994 | 17107 | 1 | 2.4884 | 0.020972 | 0.081294 | 0.939346 | 1669 | 4 | 2.4884 |
| ACOT1 | 5 | 0.41157 | 0.57316 | 0.999994 | 12140 | 2 | 6.4512 | 0.020981 | 0.077736 | 0.938203 | 1670 | 3 | 6.4512 |
| LLGL2 | 5 | 0.97291 | 0.97296 | 0.999994 | 20893 | 0 | 5.6045 | 0.020982 | 0.077736 | 0.938203 | 1671 | 5 | 5.6045 |
| TRAM1 | 6 | 0.61141 | 0.74838 | 0.999994 | 15095 | 2 | 3.1151 | 0.021009 | 0.081407 | 0.939346 | 1672 | 3 | 3.1151 |
| TECR | 4 | 0.94769 | 0.94764 | 0.999994 | 20281 | 0 | 2.4208 | 0.021009 | 0.066867 | 0.933948 | 1673 | 4 | 2.4208 |
| RPL3 | 4 | 0.25725 | 0.39931 | 0.999994 | 9173 | 1 | 5.7234 | 0.021031 | 0.066938 | 0.933948 | 1674 | 3 | 5.7234 |
| PALB2 | 5 | 0.27357 | 0.45966 | 0.999994 | 9557 | 2 | 0.98122 | 0.021035 | 0.077908 | 0.938203 | 1675 | 3 | 0.98122 |
| KCMF1 | 5 | 0.91049 | 0.91052 | 0.999994 | 19440 | 0 | 0.87084 | 0.021035 | 0.077908 | 0.938203 | 1676 | 4 | 0.87084 |
| hsa-mir-4522 | 3 | 0.78787 | 0.78778 | 0.999994 | 17321 | 0 | 7.4127 | 0.021082 | 0.05423 | 0.926769 | 1677 | 2 | 7.4127 |
| RAI14 | 5 | 0.45043 | 0.6056 | 0.999994 | 12717 | 1 | 4.7993 | 0.021083 | 0.078061 | 0.938203 | 1678 | 4 | 4.7993 |
| hsa-mir-6806 | 4 | 0.97891 | 0.97889 | 0.999994 | 21073 | 0 | 4.5491 | 0.021092 | 0.067127 | 0.934215 | 1679 | 4 | 4.5491 |
| LST1 | 5 | 0.43448 | 0.59219 | 0.999994 | 12483 | 2 | 0.75101 | 0.021128 | 0.078219 | 0.938203 | 1680 | 3 | 0.75101 |
| HNRNPH2 | 5 | 0.44697 | 0.60266 | 0.999994 | 12674 | 2 | 4.4204 | 0.021128 | 0.078219 | 0.938203 | 1681 | 3 | 4.4204 |
| TOM1L2 | 6 | 0.7393 | 0.82475 | 0.999994 | 16727 | 1 | 2.7869 | 0.021133 | 0.081791 | 0.939346 | 1682 | 5 | 2.7869 |
| UTP14C | 6 | 0.31598 | 0.5253 | 0.999994 | 10440 | 2 | 2.7037 | 0.02117 | 0.081893 | 0.939346 | 1683 | 4 | 2.7037 |
| NonTargetingControlG | 2 | 0.97883 | 0.97881 | 0.999994 | 21072 | 0 | 6.2932 | 0.021171 | 0.039289 | 0.908838 | 1684 | 2 | 6.2932 |
| NonTargetingControlG | 2 | 0.55915 | 0.55973 | 0.999994 | 14356 | 1 | 4.4037 | 0.021214 | 0.039358 | 0.908838 | 1685 | 1 | 4.4037 |
| NARS | 4 | 0.48184 | 0.60831 | 0.999994 | 13167 | 1 | 6.241 | 0.021217 | 0.067502 | 0.934798 | 1686 | 3 | 6.241 |
| MSANTD3 | 6 | 0.87173 | 0.88439 | 0.999994 | 18672 | 1 | 1.0664 | 0.021222 | 0.082044 | 0.939346 | 1687 | 3 | 1.0664 |
| CCDC85A | 6 | 0.94917 | 0.94915 | 0.999994 | 20312 | 0 | 2.1751 | 0.021239 | 0.082104 | 0.939346 | 1688 | 5 | 2.1751 |
| FDCSP | 5 | 0.36272 | 0.53233 | 0.999994 | 11383 | 1 | 2.6734 | 0.021256 | 0.078643 | 0.938203 | 1689 | 3 | 2.6734 |
| LRRC14 | 5 | 0.82115 | 0.83335 | 0.999994 | 17792 | 1 | 6.4076 | 0.021264 | 0.07867 | 0.938203 | 1690 | 4 | 6.4076 |
| RPS25 | 5 | 0.7565 | 0.79427 | 0.999994 | 16937 | 1 | 6.4076 | 0.021264 | 0.07867 | 0.938203 | 1691 | 4 | 6.4076 |
| SYCN | 6 | 0.68467 | 0.80464 | 0.999994 | 16155 | 1 | 4.2014 | 0.021274 | 0.082202 | 0.939346 | 1692 | 5 | 4.2014 |
| KIR3DL2 | 5 | 0.64821 | 0.74439 | 0.999994 | 15632 | 1 | 4.949 | 0.02128 | 0.078719 | 0.938203 | 1693 | 4 | 4.949 |
| SNTG2 | 5 | 0.94143 | 0.94162 | 0.999994 | 20116 | 0 | 6.3955 | 0.021297 | 0.078779 | 0.938203 | 1694 | 4 | 6.3955 |
| RNF220 | 5 | 0.9787 | 0.97878 | 0.999994 | 21066 | 0 | 3.2369 | 0.021301 | 0.07879 | 0.938203 | 1695 | 5 | 3.2369 |
| LRPAP1 | 6 | 0.044929 | 0.13293 | 0.95318 | 2753 | 2 | 4.1001 | 0.021309 | 0.082309 | 0.939346 | 1696 | 4 | 4.1001 |
| EZR | 6 | 0.15374 | 0.33037 | 0.999994 | 6560 | 1 | 5.1455 | 0.021309 | 0.082309 | 0.939346 | 1697 | 4 | 5.1455 |
| ZNF606 | 6 | 0.93943 | 0.93939 | 0.999994 | 20060 | 0 | 2.7671 | 0.021328 | 0.082368 | 0.939346 | 1698 | 4 | 2.7671 |
| ANKRD13D | 5 | 0.30493 | 0.485 | 0.999994 | 10231 | 2 | -0.10783 | 0.021336 | 0.078899 | 0.938203 | 1699 | 2 | -0.10783 |
| RXRA | 5 | 0.29015 | 0.47303 | 0.999994 | 9911 | 1 | 3.8877 | 0.021341 | 0.078915 | 0.938203 | 1700 | 4 | 3.8877 |
| MED8 | 5 | 0.83612 | 0.84387 | 0.999994 | 18022 | 1 | 4.4318 | 0.021346 | 0.07893 | 0.938203 | 1701 | 4 | 4.4318 |
| CDYL2 | 6 | 0.96055 | 0.96061 | 0.999994 | 20593 | 0 | 2.0993 | 0.021359 | 0.08248 | 0.939346 | 1702 | 5 | 2.0993 |
| RALGAPA1 | 5 | 0.90342 | 0.9035 | 0.999994 | 19290 | 0 | 5.857 | 0.021366 | 0.07899 | 0.938203 | 1703 | 4 | 5.857 |
| ETV2 | 6 | 0.6124 | 0.7491 | 0.999994 | 15115 | 1 | 3.3296 | 0.021396 | 0.082587 | 0.939346 | 1704 | 5 | 3.3296 |
| HUWE1 | 5 | 0.24737 | 0.4387 | 0.999994 | 8958 | 2 | 2.0395 | 0.021398 | 0.079094 | 0.938203 | 1705 | 3 | 2.0395 |
| AKAP17A_X | 6 | 0.96412 | 0.96418 | 0.999994 | 20705 | 0 | 2.5626 | 0.021426 | 0.082685 | 0.939346 | 1706 | 5 | 2.5626 |
| BTD | 6 | 0.15575 | 0.33288 | 0.999994 | 6617 | 1 | 4.968 | 0.021428 | 0.082687 | 0.939346 | 1707 | 5 | 4.968 |
| HBQ1 | 6 | 0.61134 | 0.74833 | 0.999994 | 15093 | 1 | 5.0377 | 0.021428 | 0.082687 | 0.939346 | 1708 | 4 | 5.0377 |
| TMEM74B | 5 | 0.70911 | 0.77067 | 0.999994 | 16396 | 1 | -0.10906 | 0.021475 | 0.079347 | 0.938203 | 1709 | 2 | -0.10906 |
| CFHR5 | 5 | 0.34679 | 0.5193 | 0.999994 | 11089 | 1 | 6.3955 | 0.021485 | 0.079371 | 0.938203 | 1710 | 3 | 6.3955 |
| CORO6 | 5 | 0.13264 | 0.27664 | 0.999379 | 5991 | 1 | 6.3955 | 0.021485 | 0.079371 | 0.938203 | 1711 | 4 | 6.3955 |
| hsa-mir-1287 | 3 | 0.9785 | 0.97858 | 0.999994 | 21063 | 0 | 4.7239 | 0.021497 | 0.055231 | 0.926769 | 1712 | 3 | 4.7239 |
| RAB38 | 6 | 0.96834 | 0.96834 | 0.999994 | 20790 | 0 | 3.3522 | 0.0215 | 0.082903 | 0.939912 | 1713 | 5 | 3.3522 |
| hsa-mir-18a | 4 | 0.97844 | 0.97842 | 0.999994 | 21061 | 0 | 1.9657 | 0.021556 | 0.068478 | 0.935447 | 1714 | 4 | 1.9657 |
| SPIRE1 | 5 | 0.97843 | 0.97851 | 0.999994 | 21060 | 0 | 0.87284 | 0.021567 | 0.079648 | 0.938962 | 1715 | 5 | 0.87284 |

| | | | | | | | | | | | | | |
|---|---|---|---|---|---|---|---|---|---|---|---|---|---|
| RABL6 | 6 | 0.07284 | 0.19266 | 0.974607 | 3954 | 2 | -0.26059 | 0.02163 | 0.083288 | 0.939912 | 1716 | 2 | -0.26059 |
| CTSA | 6 | 0.59508 | 0.73635 | 0.999994 | 14889 | 2 | 1.5731 | 0.02163 | 0.083288 | 0.939912 | 1717 | 4 | 1.5731 |
| PTH | 6 | 0.010294 | 0.040011 | 0.891621 | 889 | 3 | -0.44467 | 0.02163 | 0.083288 | 0.939912 | 1718 | 3 | -0.44467 |
| CHORDC1 | 6 | 0.35357 | 0.56803 | 0.999994 | 11225 | 2 | 5.123 | 0.021646 | 0.083329 | 0.939912 | 1719 | 4 | 5.123 |
| KMT2B | 5 | 0.90495 | 0.90505 | 0.999994 | 19326 | 0 | 2.7054 | 0.021659 | 0.079954 | 0.938962 | 1720 | 4 | 2.7054 |
| MAGEA6 | 5 | 0.94125 | 0.94145 | 0.999994 | 20114 | 0 | 4.5922 | 0.02166 | 0.079959 | 0.938962 | 1721 | 4 | 4.5922 |
| FLT1 | 5 | 0.97833 | 0.97841 | 0.999994 | 21058 | 0 | 1.7565 | 0.021672 | 0.079995 | 0.938962 | 1722 | 5 | 1.7565 |
| GJA10 | 5 | 0.95002 | 0.9502 | 0.999994 | 20334 | 0 | 6.39 | 0.021674 | 0.079998 | 0.938962 | 1723 | 4 | 6.39 |
| ACTRT2 | 4 | 0.19869 | 0.33932 | 0.999994 | 7726 | 2 | 2.8909 | 0.021674 | 0.068839 | 0.936374 | 1724 | 2 | 2.8909 |
| PTRF | 4 | 0.26749 | 0.40971 | 0.999994 | 9408 | 2 | 3.4687 | 0.021674 | 0.068839 | 0.936374 | 1725 | 2 | 3.4687 |
| hsa-mir-638 | 3 | 0.97831 | 0.97838 | 0.999994 | 21056 | 0 | 3.4241 | 0.021688 | 0.05567 | 0.926769 | 1726 | 3 | 3.4241 |
| hsa-mir-1914 | 3 | 0.97831 | 0.97838 | 0.999994 | 21057 | 0 | 5.0163 | 0.021688 | 0.05567 | 0.926769 | 1727 | 3 | 5.0163 |
| PDZD2 | 5 | 0.97831 | 0.97839 | 0.999994 | 21055 | 0 | 2.3409 | 0.02169 | 0.080049 | 0.938962 | 1728 | 5 | 2.3409 |
| WDPCP | 6 | 0.84167 | 0.8675 | 0.999994 | 18117 | 1 | 5.5875 | 0.021693 | 0.083466 | 0.939912 | 1729 | 5 | 5.5875 |
| BHLHE22 | 6 | 0.063144 | 0.1724 | 0.967658 | 3565 | 3 | 2.086 | 0.021693 | 0.083466 | 0.939912 | 1730 | 3 | 2.086 |
| PRMT3 | 6 | 0.74515 | 0.82676 | 0.999994 | 16803 | 1 | 4.8661 | 0.021693 | 0.083466 | 0.939912 | 1731 | 4 | 4.8661 |
| MTRF1 | 4 | 0.91742 | 0.9172 | 0.999994 | 19574 | 0 | 3.4076 | 0.021695 | 0.068903 | 0.936663 | 1732 | 2 | 3.4076 |
| CRADD | 5 | 0.9783 | 0.97838 | 0.999994 | 21054 | 0 | 3.2329 | 0.021699 | 0.080083 | 0.938962 | 1733 | 5 | 3.2329 |
| SUSD3 | 5 | 0.21927 | 0.40225 | 0.999994 | 8270 | 2 | 1.4627 | 0.021707 | 0.080112 | 0.938962 | 1734 | 3 | 1.4627 |
| TNFRSF21 | 6 | 0.64475 | 0.7736 | 0.999994 | 15581 | 1 | 2.489 | 0.021797 | 0.083777 | 0.940296 | 1735 | 5 | 2.489 |
| MGARP | 5 | 0.93252 | 0.93281 | 0.999994 | 19916 | 0 | 6.3809 | 0.021839 | 0.080535 | 0.938962 | 1736 | 3 | 6.3809 |
| ADAT3 | 5 | 0.93617 | 0.9364 | 0.999994 | 19990 | 0 | 6.3809 | 0.021839 | 0.080535 | 0.938962 | 1737 | 4 | 6.3809 |
| NT5M | 6 | 0.1557 | 0.33282 | 0.999994 | 6614 | 3 | 2.6347 | 0.021843 | 0.08391 | 0.940296 | 1738 | 3 | 2.6347 |
| TPRG1L | 5 | 0.001437 | 0.0063797 | 0.636218 | 215 | 1 | 5.5953 | 0.021858 | 0.080606 | 0.938962 | 1739 | 4 | 5.5953 |
| GSX2 | 3 | 0.9086 | 0.90861 | 0.999994 | 19398 | 0 | 7.3773 | 0.021858 | 0.056067 | 0.926769 | 1740 | 3 | 7.3773 |
| OR5C1 | 6 | 0.03698 | 0.11509 | 0.951052 | 2374 | 1 | 3.0161 | 0.021862 | 0.083965 | 0.940296 | 1741 | 5 | 3.0161 |
| SLC7A5 | 6 | 0.62125 | 0.75577 | 0.999994 | 15249 | 1 | 3.8163 | 0.021867 | 0.08398 | 0.940296 | 1742 | 4 | 3.8163 |
| C11orf87 | 4 | 0.9781 | 0.97807 | 0.999994 | 21048 | 0 | 5.7985 | 0.021897 | 0.069449 | 0.937025 | 1743 | 4 | 5.7985 |
| IL31 | 6 | 0.1799 | 0.36302 | 0.999994 | 7219 | 2 | 5.6834 | 0.021918 | 0.084131 | 0.940296 | 1744 | 4 | 5.6834 |
| ZNF444 | 5 | 0.78783 | 0.81201 | 0.999994 | 17318 | 1 | 3.8467 | 0.021938 | 0.080892 | 0.939346 | 1745 | 3 | 3.8467 |
| LBP | 6 | 0.45446 | 0.63802 | 0.999994 | 12763 | 2 | 2.6717 | 0.021939 | 0.084186 | 0.940296 | 1746 | 3 | 2.6717 |
| GDF2 | 5 | 0.97806 | 0.97815 | 0.999994 | 21044 | 0 | 2.4074 | 0.021942 | 0.08091 | 0.939346 | 1747 | 5 | 2.4074 |
| ZNF114 | 6 | 0.83709 | 0.86514 | 0.999994 | 18036 | 1 | 4.8548 | 0.021959 | 0.084246 | 0.940296 | 1748 | 4 | 4.8548 |
| TMEM187 | 6 | 0.9714 | 0.97135 | 0.999994 | 20856 | 0 | 3.733 | 0.021971 | 0.084281 | 0.940296 | 1749 | 5 | 3.733 |
| hsa-mir-130b | 3 | 0.97801 | 0.97807 | 0.999994 | 21040 | 0 | 3.6773 | 0.021993 | 0.056395 | 0.926769 | 1750 | 3 | 3.6773 |
| TUBAL3 | 6 | 0.8339 | 0.86347 | 0.999994 | 17985 | 1 | 4.799 | 0.022012 | 0.084396 | 0.940404 | 1751 | 4 | 4.799 |
| TRPC5 | 5 | 0.46163 | 0.61503 | 0.999994 | 12875 | 2 | 6.3722 | 0.022019 | 0.081155 | 0.939346 | 1752 | 3 | 6.3722 |
| hsa-mir-4707 | 4 | 0.69156 | 0.71897 | 0.999994 | 16222 | 1 | 5.5893 | 0.022067 | 0.069954 | 0.937929 | 1753 | 3 | 5.5893 |
| KDELR1 | 5 | 0.97793 | 0.97801 | 0.999994 | 21038 | 0 | 2.172 | 0.022073 | 0.081343 | 0.939346 | 1754 | 5 | 2.172 |
| PRAC | 5 | 0.58898 | 0.72218 | 0.999994 | 14806 | 1 | 0.26227 | 0.022077 | 0.081352 | 0.939346 | 1755 | 2 | 0.26227 |
| ALMS1 | 6 | 0.77679 | 0.83842 | 0.999994 | 17163 | 1 | 4.6089 | 0.022103 | 0.084667 | 0.940404 | 1756 | 4 | 4.6089 |
| TLE3 | 5 | 0.97789 | 0.97797 | 0.999994 | 21036 | 0 | 2.3738 | 0.022111 | 0.081467 | 0.939346 | 1757 | 5 | 2.3738 |
| DOCK11 | 5 | 0.87945 | 0.87975 | 0.999994 | 18820 | 1 | 3.6773 | 0.022131 | 0.081531 | 0.939346 | 1758 | 4 | 3.6773 |
| HRK | 6 | 0.19663 | 0.38379 | 0.999994 | 7667 | 1 | 3.7435 | 0.022133 | 0.084754 | 0.940435 | 1759 | 4 | 3.7435 |
| hsa-mir-518a-2 | 3 | 0.92703 | 0.92708 | 0.999994 | 19789 | 0 | 7.3684 | 0.022154 | 0.05676 | 0.926769 | 1760 | 3 | 7.3684 |
| CNTNAP1 | 6 | 0.21246 | 0.40308 | 0.999994 | 8072 | 2 | 4.7833 | 0.022162 | 0.084846 | 0.940435 | 1761 | 4 | 4.7833 |
| CAMK2G | 5 | 0.8568 | 0.85974 | 0.999994 | 18384 | 1 | 2.9948 | 0.022193 | 0.081739 | 0.939346 | 1762 | 4 | 2.9948 |
| CALM3 | 6 | 0.95287 | 0.95286 | 0.999994 | 20404 | 0 | 2.5244 | 0.022196 | 0.084941 | 0.940435 | 1763 | 5 | 2.5244 |
| KRTAP10-2 | 6 | 0.23668 | 0.43203 | 0.999994 | 8685 | 2 | 1.2045 | 0.022197 | 0.084945 | 0.940435 | 1764 | 4 | 1.2045 |
| MUC4 | 5 | 0.47052 | 0.6226 | 0.999994 | 13011 | 1 | 1.8116 | 0.022214 | 0.081797 | 0.939346 | 1765 | 4 | 1.8116 |
| CCNL2 | 6 | 0.28327 | 0.4872 | 0.999994 | 9767 | 1 | 4.2213 | 0.022245 | 0.085092 | 0.941573 | 1766 | 4 | 4.2213 |
| RC3H2 | 5 | 0.11547 | 0.25046 | 0.992497 | 5493 | 2 | -0.043568 | 0.02227 | 0.081984 | 0.939346 | 1767 | 2 | -0.043568 |

| | | | | | | | | | | | | | |
|---|---|---|---|---|---|---|---|---|---|---|---|---|---|
| CLN3 | 5 | 0.95936 | 0.9595 | 0.999994 | 20568 | 0 | 6.16 | 0.02229 | 0.082042 | 0.939346 | 1768 | 4 | 6.16 |
| ZFP112 | 6 | 0.87615 | 0.88717 | 0.999994 | 18754 | 1 | 3.4777 | 0.022328 | 0.085351 | 0.942567 | 1769 | 4 | 3.4777 |
| S1PR1 | 6 | 0.22249 | 0.41516 | 0.999994 | 8345 | 3 | 1.7741 | 0.022333 | 0.085364 | 0.942567 | 1770 | 3 | 1.7741 |
| AXDND1 | 5 | 0.90965 | 0.90967 | 0.999994 | 19418 | 0 | 6.3541 | 0.022371 | 0.082315 | 0.939346 | 1771 | 3 | 6.3541 |
| LPIN1 | 5 | 0.91545 | 0.91552 | 0.999994 | 19522 | 0 | 6.3541 | 0.022371 | 0.082315 | 0.939346 | 1772 | 4 | 6.3541 |
| NonTargetingControlG | 2 | 0.97763 | 0.97761 | 0.999994 | 21034 | 0 | 6.5213 | 0.022374 | 0.041464 | 0.917066 | 1773 | 2 | 6.5213 |
| PTPLAD1 | 5 | 0.88714 | 0.88725 | 0.999994 | 18970 | 1 | 0.67584 | 0.022378 | 0.082336 | 0.939346 | 1774 | 3 | 0.67584 |
| FBN1 | 6 | 0.32353 | 0.5339 | 0.999994 | 10584 | 2 | 3.1734 | 0.022414 | 0.085599 | 0.942848 | 1775 | 3 | 3.1734 |
| B9D2 | 5 | 0.3202 | 0.49748 | 0.999994 | 10515 | 2 | 1.8657 | 0.022494 | 0.082737 | 0.939346 | 1776 | 3 | 1.8657 |
| NLRC4 | 6 | 0.97877 | 0.97869 | 0.999994 | 21069 | 0 | 5.114 | 0.022512 | 0.085885 | 0.943068 | 1777 | 5 | 5.114 |
| GABRQ | 6 | 0.53462 | 0.69275 | 0.999994 | 13974 | 2 | 5.4045 | 0.022512 | 0.085885 | 0.943068 | 1778 | 4 | 5.4045 |
| ECM1 | 5 | 0.97748 | 0.97757 | 0.999994 | 21029 | 0 | 4.112 | 0.022516 | 0.082812 | 0.939393 | 1779 | 5 | 4.112 |
| ADCY8 | 6 | 0.34058 | 0.55332 | 0.999994 | 10948 | 2 | 0.59807 | 0.022529 | 0.085935 | 0.943068 | 1780 | 3 | 0.59807 |
| NonTargetingControlG | 2 | 0.4874 | 0.49244 | 0.999994 | 13242 | 1 | 4.2463 | 0.022538 | 0.041763 | 0.917066 | 1781 | 1 | 4.2463 |
| SLC29A2 | 5 | 0.85688 | 0.85982 | 0.999994 | 18389 | 1 | 1.2062 | 0.022587 | 0.083033 | 0.939912 | 1782 | 3 | 1.2062 |
| FBXL15 | 4 | 0.97738 | 0.97733 | 0.999994 | 21027 | 0 | 3.8704 | 0.022621 | 0.071549 | 0.937929 | 1783 | 4 | 3.8704 |
| ASCC2 | 6 | 0.9753 | 0.97523 | 0.999994 | 20966 | 0 | 2.6719 | 0.022647 | 0.086296 | 0.943068 | 1784 | 5 | 2.6719 |
| OPN4 | 5 | 0.97733 | 0.97741 | 0.999994 | 21026 | 0 | 2.4812 | 0.022669 | 0.083307 | 0.939912 | 1785 | 5 | 2.4812 |
| LUC7L3 | 5 | 0.31064 | 0.48966 | 0.999994 | 10340 | 2 | 0.4942 | 0.022669 | 0.083308 | 0.939912 | 1786 | 2 | 0.4942 |
| NonTargetingControlG | 2 | 0.97732 | 0.97729 | 0.999994 | 21025 | 0 | 7.386 | 0.022678 | 0.042014 | 0.917066 | 1787 | 2 | 7.386 |
| OR1J1 | 5 | 0.97729 | 0.97737 | 0.999994 | 21024 | 0 | 1.0515 | 0.022707 | 0.083419 | 0.939912 | 1788 | 5 | 1.0515 |
| TLR4 | 6 | 0.97726 | 0.9772 | 0.999994 | 21023 | 0 | 4.4004 | 0.022708 | 0.086472 | 0.943068 | 1789 | 4 | 4.4004 |
| SNRNP70 | 6 | 0.032854 | 0.10549 | 0.950444 | 2151 | 2 | 3.869 | 0.022715 | 0.086494 | 0.943068 | 1790 | 4 | 3.869 |
| EML5 | 6 | 0.075361 | 0.19791 | 0.977164 | 4047 | 1 | 5.2247 | 0.022715 | 0.086496 | 0.943068 | 1791 | 4 | 5.2247 |
| MAGEL2 | 4 | 0.56199 | 0.64568 | 0.999994 | 14406 | 1 | 3.3858 | 0.022789 | 0.072022 | 0.937929 | 1792 | 3 | 3.3858 |
| VSIG10L | 6 | 0.94599 | 0.94596 | 0.999994 | 20231 | 0 | 1.6217 | 0.022805 | 0.086766 | 0.943068 | 1793 | 4 | 1.6217 |
| hsa-mir-548ai | 4 | 0.97719 | 0.97715 | 0.999994 | 21021 | 0 | 4.082 | 0.022807 | 0.072077 | 0.937929 | 1794 | 4 | 4.082 |
| TATDN2 | 4 | 0.64175 | 0.68857 | 0.999994 | 15545 | 1 | 5.9117 | 0.02281 | 0.072088 | 0.937929 | 1795 | 3 | 5.9117 |
| TUSC1 | 4 | 0.86705 | 0.8666 | 0.999994 | 18574 | 0 | 5.5535 | 0.02281 | 0.072088 | 0.937929 | 1796 | 3 | 5.5535 |
| BCCIP | 5 | 0.099756 | 0.22604 | 0.983354 | 4967 | 3 | -1.4973 | 0.022818 | 0.083772 | 0.940296 | 1797 | 2 | -1.4973 |
| GLO1 | 6 | 0.75137 | 0.82895 | 0.999994 | 16869 | 1 | 4.5339 | 0.022825 | 0.086831 | 0.943068 | 1798 | 5 | 4.5339 |
| RCN2 | 5 | 0.97713 | 0.97721 | 0.999994 | 21019 | 0 | 2.154 | 0.022867 | 0.083948 | 0.940296 | 1799 | 5 | 2.154 |
| OTOG | 6 | 0.78963 | 0.84351 | 0.999994 | 17336 | 1 | 4.4397 | 0.022881 | 0.087 | 0.943068 | 1800 | 4 | 4.4397 |
| GNPNAT1 | 6 | 0.96235 | 0.96244 | 0.999994 | 20639 | 0 | 5.6841 | 0.022881 | 0.087 | 0.943068 | 1801 | 4 | 5.6841 |
| CLIC5 | 5 | 0.97709 | 0.97717 | 0.999994 | 21017 | 0 | 5.8087 | 0.022909 | 0.084093 | 0.940296 | 1802 | 5 | 5.8087 |
| PRKAR1A | 5 | 0.91513 | 0.91521 | 0.999994 | 19516 | 0 | 5.1479 | 0.022924 | 0.084144 | 0.940296 | 1803 | 4 | 5.1479 |
| ZNF468 | 4 | 0.95782 | 0.95779 | 0.999994 | 20533 | 0 | 5.5159 | 0.022932 | 0.072443 | 0.937929 | 1804 | 4 | 5.5159 |
| TM2D1 | 5 | 0.58949 | 0.72235 | 0.999994 | 14813 | 1 | 4.8677 | 0.022934 | 0.084181 | 0.940296 | 1805 | 4 | 4.8677 |
| ZNF140 | 5 | 0.94403 | 0.94424 | 0.999994 | 20178 | 0 | 0.22505 | 0.022937 | 0.084189 | 0.940296 | 1806 | 2 | 0.22505 |
| BAG6 | 5 | 0.95433 | 0.95448 | 0.999994 | 20439 | 0 | 3.6085 | 0.022947 | 0.084237 | 0.940296 | 1807 | 4 | 3.6085 |
| NAA60 | 6 | 0.10117 | 0.2496 | 0.991904 | 5013 | 3 | -0.51239 | 0.022999 | 0.087344 | 0.943068 | 1808 | 3 | -0.51239 |
| ZFPM2 | 6 | 0.62085 | 0.75547 | 0.999994 | 15233 | 2 | 0.15444 | 0.022999 | 0.087344 | 0.943068 | 1809 | 3 | 0.15444 |
| MRPL51 | 5 | 0.38169 | 0.54812 | 0.999994 | 11673 | 2 | 3.9896 | 0.023026 | 0.084517 | 0.940404 | 1810 | 3 | 3.9896 |
| NCOR2 | 6 | 0.95215 | 0.95213 | 0.999994 | 20385 | 0 | 4.3369 | 0.023059 | 0.087515 | 0.943068 | 1811 | 4 | 4.3369 |
| USP43 | 5 | 0.94014 | 0.94032 | 0.999994 | 20084 | 0 | 3.5988 | 0.023059 | 0.084619 | 0.940404 | 1812 | 4 | 3.5988 |
| FAM149B1 | 6 | 0.3514 | 0.56558 | 0.999994 | 11185 | 1 | 2.1626 | 0.0231 | 0.087639 | 0.943068 | 1813 | 4 | 2.1626 |
| TSSK6 | 5 | 0.87575 | 0.87628 | 0.999994 | 18746 | 1 | 4.6924 | 0.023145 | 0.08488 | 0.940435 | 1814 | 4 | 4.6924 |
| PADI1 | 6 | 0.077769 | 0.20285 | 0.977973 | 4140 | 1 | 2.1354 | 0.023164 | 0.087829 | 0.943068 | 1815 | 5 | 2.1354 |
| EOGT | 4 | 0.97683 | 0.97675 | 0.999994 | 21010 | 0 | 5.1569 | 0.023165 | 0.073112 | 0.938203 | 1816 | 4 | 5.1569 |
| NEUROD2 | 6 | 0.64118 | 0.77081 | 0.999994 | 15532 | 1 | 5.2167 | 0.023187 | 0.087892 | 0.943068 | 1817 | 5 | 5.2167 |
| AURKA | 4 | 0.13826 | 0.27712 | 0.999379 | 6146 | 2 | 1.2237 | 0.023253 | 0.07336 | 0.938203 | 1818 | 2 | 1.2237 |
| hsa-mir-5006 | 4 | 0.97667 | 0.97658 | 0.999994 | 21007 | 0 | 2.4505 | 0.023327 | 0.07357 | 0.938203 | 1819 | 4 | 2.4505 |

| | | | | | | | | | | | | | |
|---|---|---|---|---|---|---|---|---|---|---|---|---|---|
| C11orf86 | 6 | 0.92208 | 0.92233 | 0.999994 | 19675 | 1 | 5.5912 | 0.023344 | 0.088373 | 0.943068 | 1820 | 4 | 5.5912 |
| TAF9 | 6 | 0.97046 | 0.97042 | 0.999994 | 20834 | 0 | 4.5528 | 0.023344 | 0.088373 | 0.943068 | 1821 | 5 | 4.5528 |
| GABBR1 | 5 | 0.20082 | 0.37636 | 0.999994 | 7782 | 1 | 1.1771 | 0.023349 | 0.085528 | 0.942567 | 1822 | 3 | 1.1771 |
| OR2L13 | 5 | 0.80518 | 0.82278 | 0.999994 | 17553 | 1 | 2.6723 | 0.023349 | 0.085528 | 0.942567 | 1823 | 4 | 2.6723 |
| GNB5 | 6 | 0.80925 | 0.85192 | 0.999994 | 17620 | 1 | 4.0148 | 0.023378 | 0.088481 | 0.943068 | 1824 | 5 | 4.0148 |
| NonTargetingControlG | 2 | 0.78167 | 0.78117 | 0.999994 | 17233 | 0 | 5.1404 | 0.023386 | 0.043245 | 0.918424 | 1825 | 2 | 5.1404 |
| DST | 6 | 0.81292 | 0.85356 | 0.999994 | 17670 | 1 | 1.3812 | 0.023387 | 0.088511 | 0.943068 | 1826 | 4 | 1.3812 |
| LILRA5 | 6 | 0.58211 | 0.72676 | 0.999994 | 14707 | 1 | 2.5466 | 0.023412 | 0.088579 | 0.943068 | 1827 | 5 | 2.5466 |
| HTR1A | 6 | 0.16753 | 0.34763 | 0.999994 | 6925 | 3 | -0.31298 | 0.023413 | 0.088581 | 0.943068 | 1828 | 3 | -0.31298 |
| SPG7 | 6 | 0.62105 | 0.75561 | 0.999994 | 15240 | 1 | 3.8525 | 0.023413 | 0.088581 | 0.943068 | 1829 | 4 | 3.8525 |
| SYNCRIP | 5 | 0.1108 | 0.24321 | 0.988088 | 5325 | 2 | 0.52113 | 0.023456 | 0.085861 | 0.943068 | 1830 | 2 | 0.52113 |
| FGFRL1 | 5 | 0.97651 | 0.97657 | 0.999994 | 21005 | 0 | 1.9095 | 0.023489 | 0.085971 | 0.943068 | 1831 | 5 | 1.9095 |
| TXLNB | 5 | 0.96057 | 0.9607 | 0.999994 | 20594 | 0 | 3.1166 | 0.023494 | 0.085984 | 0.943068 | 1832 | 4 | 3.1166 |
| RSPO3 | 6 | 0.96098 | 0.96104 | 0.999994 | 20604 | 0 | 3.1048 | 0.023501 | 0.088839 | 0.943068 | 1833 | 5 | 3.1048 |
| GALR1 | 6 | 0.48117 | 0.65599 | 0.999994 | 13154 | 2 | 5.2589 | 0.023517 | 0.088887 | 0.943068 | 1834 | 4 | 5.2589 |
| TFB1M | 6 | 0.18713 | 0.37194 | 0.999994 | 7394 | 2 | 0.048623 | 0.023554 | 0.088986 | 0.943068 | 1835 | 3 | 0.048623 |
| GSPT1 | 4 | 0.96734 | 0.96723 | 0.999994 | 20770 | 0 | 3.8844 | 0.023586 | 0.074324 | 0.938203 | 1836 | 4 | 3.8844 |
| CPNE5 | 6 | 0.98546 | 0.98542 | 0.999994 | 21239 | 0 | 2.7835 | 0.023587 | 0.089084 | 0.943068 | 1837 | 5 | 2.7835 |
| CHI3L2 | 3 | 0.9764 | 0.97649 | 0.999994 | 21003 | 0 | 4.8715 | 0.023598 | 0.060201 | 0.9326 | 1838 | 3 | 4.8715 |
| METTL12 | 6 | 0.13144 | 0.30217 | 0.999994 | 5966 | 3 | -0.13123 | 0.023658 | 0.089261 | 0.943068 | 1839 | 3 | -0.13123 |
| RNPC3 | 5 | 0.57775 | 0.71632 | 0.999994 | 14655 | 1 | 3.235 | 0.023658 | 0.086499 | 0.943068 | 1840 | 4 | 3.235 |
| ADPGK | 6 | 0.3854 | 0.59299 | 0.999994 | 11725 | 1 | 1.9385 | 0.023665 | 0.089279 | 0.943068 | 1841 | 5 | 1.9385 |
| MT1M | 4 | 0.9763 | 0.97619 | 0.999994 | 21001 | 0 | 5.542 | 0.0237 | 0.074666 | 0.938203 | 1842 | 4 | 5.542 |
| TM9SF2 | 6 | 0.28631 | 0.49082 | 0.999994 | 9841 | 2 | 3.4608 | 0.023746 | 0.089527 | 0.943068 | 1843 | 3 | 3.4608 |
| NDUFA12 | 6 | 0.58584 | 0.7295 | 0.999994 | 14756 | 1 | 2.2211 | 0.023749 | 0.089535 | 0.943068 | 1844 | 5 | 2.2211 |
| FMO2 | 5 | 0.82224 | 0.83407 | 0.999994 | 17804 | 1 | 6.453 | 0.023763 | 0.086844 | 0.943068 | 1845 | 3 | 6.453 |
| SMOX | 6 | 0.17818 | 0.36082 | 0.999994 | 7180 | 3 | -0.060764 | 0.023775 | 0.0896 | 0.943068 | 1846 | 3 | -0.060764 |
| ZNF324 | 5 | 0.97621 | 0.97627 | 0.999994 | 20997 | 0 | 1.9832 | 0.023791 | 0.086941 | 0.943068 | 1847 | 5 | 1.9832 |
| PRDM7 | 3 | 0.87937 | 0.87947 | 0.999994 | 18818 | 0 | 7.304 | 0.023792 | 0.06067 | 0.9326 | 1848 | 2 | 7.304 |
| hsa-mir-1193 | 4 | 0.73283 | 0.74726 | 0.999994 | 16638 | 1 | 4.1741 | 0.023805 | 0.074945 | 0.938203 | 1849 | 3 | 4.1741 |
| ABHD12B | 4 | 0.49308 | 0.61325 | 0.999994 | 13312 | 1 | 5.3649 | 0.023808 | 0.074958 | 0.938203 | 1850 | 3 | 5.3649 |
| MEGF8 | 4 | 0.94031 | 0.94019 | 0.999994 | 20094 | 0 | 5.4416 | 0.023808 | 0.074958 | 0.938203 | 1851 | 3 | 5.4416 |
| NonTargetingControlG | 2 | 0.97617 | 0.97615 | 0.999994 | 20994 | 0 | 7.1948 | 0.023827 | 0.044052 | 0.918424 | 1852 | 2 | 7.1948 |
| HIST1H2BK | 2 | 0.97617 | 0.97615 | 0.999994 | 20996 | 0 | 6.8563 | 0.023827 | 0.044052 | 0.918424 | 1853 | 2 | 6.8563 |
| ZCCHC6 | 6 | 0.10564 | 0.25831 | 0.994943 | 5159 | 2 | 1.265 | 0.023831 | 0.089783 | 0.943068 | 1854 | 3 | 1.265 |
| TNNI3 | 5 | 0.97615 | 0.97622 | 0.999994 | 20992 | 0 | 2.9369 | 0.02385 | 0.087128 | 0.943068 | 1855 | 5 | 2.9369 |
| FABP6 | 5 | 0.96854 | 0.96867 | 0.999994 | 20795 | 0 | 3.3399 | 0.023862 | 0.087166 | 0.943068 | 1856 | 4 | 3.3399 |
| CNIH2 | 6 | 0.48756 | 0.6603 | 0.999994 | 13246 | 2 | 1.9052 | 0.023868 | 0.089896 | 0.943068 | 1857 | 3 | 1.9052 |
| NOM1 | 6 | 0.27589 | 0.47864 | 0.999994 | 9614 | 2 | 0.049377 | 0.023868 | 0.089896 | 0.943068 | 1858 | 2 | 0.049377 |
| GRAMD2 | 5 | 0.97612 | 0.97618 | 0.999994 | 20991 | 0 | 2.924 | 0.023883 | 0.087238 | 0.943068 | 1859 | 5 | 2.924 |
| GIPR | 5 | 0.90479 | 0.90488 | 0.999994 | 19319 | 0 | 3.1166 | 0.023907 | 0.08731 | 0.943068 | 1860 | 4 | 3.1166 |
| DUSP28 | 5 | 0.87621 | 0.87671 | 0.999994 | 18756 | 1 | 4.0895 | 0.023913 | 0.087327 | 0.943068 | 1861 | 4 | 4.0895 |
| LIN7B | 6 | 0.28816 | 0.49294 | 0.999994 | 9874 | 2 | 2.6253 | 0.023914 | 0.090017 | 0.943068 | 1862 | 4 | 2.6253 |
| hsa-mir-493 | 4 | 0.97605 | 0.97595 | 0.999994 | 20989 | 0 | 4.0333 | 0.023945 | 0.075354 | 0.938203 | 1863 | 4 | 4.0333 |
| MAST4 | 5 | 0.95804 | 0.9582 | 0.999994 | 20540 | 0 | 4.9089 | 0.024007 | 0.08763 | 0.943068 | 1864 | 4 | 4.9089 |
| DSTN | 5 | 0.45172 | 0.60667 | 0.999994 | 12730 | 2 | 6.2795 | 0.024041 | 0.087745 | 0.943068 | 1865 | 3 | 6.2795 |
| NUSAP1 | 5 | 0.55648 | 0.69751 | 0.999994 | 14309 | 2 | 6.2795 | 0.024041 | 0.087745 | 0.943068 | 1866 | 3 | 6.2795 |
| DKKL1 | 5 | 0.92214 | 0.92231 | 0.999994 | 19679 | 0 | 1.597 | 0.024055 | 0.087784 | 0.943068 | 1867 | 4 | 1.597 |
| KRTAP19-8 | 5 | 0.070855 | 0.17977 | 0.968171 | 3869 | 3 | -1.0818 | 0.024055 | 0.087784 | 0.943068 | 1868 | 2 | -1.0818 |
| XPR1 | 6 | 0.97493 | 0.97484 | 0.999994 | 20953 | 0 | 4.2137 | 0.024108 | 0.090601 | 0.94461 | 1869 | 5 | 4.2137 |
| RELA | 5 | 0.33764 | 0.51181 | 0.999994 | 10890 | 1 | 3.1124 | 0.02411 | 0.087962 | 0.943068 | 1870 | 4 | 3.1124 |
| TMEM9 | 5 | 0.29686 | 0.47851 | 0.999994 | 10063 | 2 | -0.25232 | 0.024134 | 0.088033 | 0.943068 | 1871 | 2 | -0.25232 |

| | | | | | | | | | | | | | |
|---|---|---|---|---|---|---|---|---|---|---|---|---|---|
| ZBTB39 | 5 | 0.21874 | 0.4015 | 0.999994 | 8248 | 3 | -0.76907 | 0.024134 | 0.088033 | 0.943068 | 1872 | 2 | -0.76907 |
| WFDC10B | 5 | 0.81395 | 0.82855 | 0.999994 | 17689 | 1 | 4.8677 | 0.024149 | 0.088086 | 0.943068 | 1873 | 4 | 4.8677 |
| HIST2H2BE | 3 | 0.97581 | 0.9759 | 0.999994 | 20985 | 0 | 3.4241 | 0.02419 | 0.061602 | 0.933948 | 1874 | 3 | 3.4241 |
| CPN1 | 5 | 0.043078 | 0.12352 | 0.951052 | 2680 | 3 | -2.2749 | 0.024206 | 0.088272 | 0.943068 | 1875 | 1 | -2.2749 |
| SLC2A1 | 6 | 0.95259 | 0.9526 | 0.999994 | 20398 | 0 | 3.0068 | 0.024226 | 0.090968 | 0.945479 | 1876 | 5 | 3.0068 |
| SEL1L3 | 5 | 0.52165 | 0.66674 | 0.999994 | 13772 | 1 | 5.6101 | 0.024228 | 0.088334 | 0.943068 | 1877 | 4 | 5.6101 |
| NRSN1 | 5 | 0.96155 | 0.9617 | 0.999994 | 20621 | 0 | 6.2746 | 0.024265 | 0.08845 | 0.943068 | 1878 | 3 | 6.2746 |
| CTNNAL1 | 4 | 0.97126 | 0.97115 | 0.999994 | 20854 | 0 | 5.4797 | 0.024285 | 0.076329 | 0.938203 | 1879 | 4 | 5.4797 |
| MEIOB | 6 | 0.87188 | 0.88449 | 0.999994 | 18675 | 1 | 4.7816 | 0.024293 | 0.091153 | 0.945479 | 1880 | 5 | 4.7816 |
| DDX1 | 5 | 0.9757 | 0.97576 | 0.999994 | 20984 | 0 | 3.889 | 0.024302 | 0.088564 | 0.943068 | 1881 | 5 | 3.889 |
| ZNF432 | 6 | 0.59544 | 0.73661 | 0.999994 | 14897 | 2 | 1.2397 | 0.02433 | 0.091267 | 0.945479 | 1882 | 3 | 1.2397 |
| OR2A2 | 6 | 0.15979 | 0.33797 | 0.999994 | 6725 | 3 | 0.61542 | 0.024338 | 0.091294 | 0.945479 | 1883 | 3 | 0.61542 |
| hsa-mir-521-1 | 4 | 0.97566 | 0.97553 | 0.999994 | 20983 | 0 | 1.8381 | 0.024342 | 0.076501 | 0.938203 | 1884 | 4 | 1.8381 |
| SPATA19 | 5 | 0.25272 | 0.44293 | 0.999994 | 9071 | 2 | 1.3705 | 0.024385 | 0.088823 | 0.943068 | 1885 | 3 | 1.3705 |
| THNSL1 | 6 | 0.97381 | 0.97372 | 0.999994 | 20921 | 0 | 3.1474 | 0.024412 | 0.091504 | 0.945479 | 1886 | 5 | 3.1474 |
| COQ3 | 5 | 0.94091 | 0.94112 | 0.999994 | 20103 | 0 | 3.1082 | 0.024421 | 0.088936 | 0.943068 | 1887 | 4 | 3.1082 |
| OR4A47 | 6 | 0.0057283 | 0.023802 | 0.832699 | 574 | 1 | 3.8434 | 0.024432 | 0.091572 | 0.945479 | 1888 | 5 | 3.8434 |
| TFAP2B | 6 | 0.33158 | 0.54306 | 0.999994 | 10757 | 2 | 0.52817 | 0.024441 | 0.091592 | 0.945479 | 1889 | 3 | 0.52817 |
| ROR2 | 5 | 0.97554 | 0.9756 | 0.999994 | 20978 | 0 | 2.2493 | 0.024462 | 0.089058 | 0.943068 | 1890 | 5 | 2.2493 |
| LEPROTL1 | 6 | 0.72529 | 0.82006 | 0.999994 | 16552 | 1 | 4.1494 | 0.024464 | 0.091661 | 0.945479 | 1891 | 5 | 4.1494 |
| hsa-mir-150 | 4 | 0.97551 | 0.9754 | 0.999994 | 20976 | 0 | 4.445 | 0.024487 | 0.076882 | 0.938203 | 1892 | 4 | 4.445 |
| USP17L2 | 5 | 0.97549 | 0.97554 | 0.999994 | 20974 | 0 | 2.7792 | 0.024515 | 0.089227 | 0.943068 | 1893 | 5 | 2.7792 |
| MRPL33 | 3 | 0.97546 | 0.97554 | 0.999994 | 20973 | 0 | 4.8199 | 0.024543 | 0.062438 | 0.933948 | 1894 | 3 | 4.8199 |
| ACTR2 | 5 | 0.37262 | 0.54056 | 0.999994 | 11555 | 2 | 5.5005 | 0.024544 | 0.08932 | 0.943068 | 1895 | 3 | 5.5005 |
| GPR37 | 5 | 0.93211 | 0.93239 | 0.999994 | 19904 | 0 | 0.85246 | 0.024544 | 0.08932 | 0.943068 | 1896 | 4 | 0.85246 |
| TRIP12 | 6 | 0.10741 | 0.26173 | 0.995257 | 5221 | 4 | -0.83469 | 0.024551 | 0.091925 | 0.946116 | 1897 | 1 | -0.83469 |
| hsa-mir-548aj-1 | 2 | 0.97544 | 0.97543 | 0.999994 | 20972 | 0 | 6.0519 | 0.024557 | 0.045342 | 0.919289 | 1898 | 2 | 6.0519 |
| NonTargetingControlG | 2 | 0.97544 | 0.97543 | 0.999994 | 20971 | 0 | 6.4406 | 0.024557 | 0.045342 | 0.919289 | 1899 | 2 | 6.4406 |
| VASP | 5 | 0.015107 | 0.050507 | 0.909101 | 1196 | 3 | -0.74851 | 0.024575 | 0.089434 | 0.943068 | 1900 | 2 | -0.74851 |
| LGALS9C | 3 | 0.73259 | 0.73247 | 0.999994 | 16636 | 0 | 7.2755 | 0.024591 | 0.062548 | 0.933948 | 1901 | 2 | 7.2755 |
| UGT2A2 | 1 | 0.97537 | 0.9754 | 0.999994 | 20970 | 0 | 8.8893 | 0.024627 | 0.024525 | 0.882096 | 1902 | 1 | 8.8893 |
| CLDND2 | 6 | 0.37253 | 0.5848 | 0.999994 | 11550 | 1 | 3.2216 | 0.024628 | 0.092141 | 0.947138 | 1903 | 5 | 3.2216 |
| FAM122C | 6 | 0.73304 | 0.82263 | 0.999994 | 16641 | 1 | 1.2626 | 0.024635 | 0.092155 | 0.947138 | 1904 | 4 | 1.2626 |
| GGTLC2 | 5 | 0.10667 | 0.2368 | 0.985449 | 5192 | 1 | 1.817 | 0.024645 | 0.08966 | 0.943068 | 1905 | 4 | 1.817 |
| GFRAL | 5 | 0.97535 | 0.97542 | 0.999994 | 20969 | 0 | 2.1129 | 0.024649 | 0.08968 | 0.943068 | 1906 | 5 | 2.1129 |
| PCDHB10 | 5 | 0.97535 | 0.97541 | 0.999994 | 20968 | 0 | 1.7309 | 0.024655 | 0.089702 | 0.943068 | 1907 | 5 | 1.7309 |
| FOXB2 | 5 | 0.97044 | 0.97051 | 0.999994 | 20833 | 0 | 6.1252 | 0.024667 | 0.089749 | 0.943068 | 1908 | 4 | 6.1252 |
| DPH7 | 6 | 0.91532 | 0.91618 | 0.999994 | 19518 | 1 | 5.2409 | 0.024674 | 0.092282 | 0.947514 | 1909 | 4 | 5.2409 |
| TTLL11 | 6 | 0.88225 | 0.89107 | 0.999994 | 18874 | 1 | 1.9205 | 0.024678 | 0.092293 | 0.947514 | 1910 | 5 | 1.9205 |
| UPK2 | 5 | 0.91736 | 0.91745 | 0.999994 | 19572 | 0 | 2.9196 | 0.024701 | 0.089854 | 0.943068 | 1911 | 4 | 2.9196 |
| hsa-mir-517b | 2 | 0.9525 | 0.95251 | 0.999994 | 20395 | 0 | 6.9042 | 0.024709 | 0.045598 | 0.919289 | 1912 | 2 | 6.9042 |
| SLC7A6 | 5 | 0.79971 | 0.81933 | 0.999994 | 17476 | 1 | 0.74364 | 0.024714 | 0.089895 | 0.943068 | 1913 | 3 | 0.74364 |
| B3GNT2 | 5 | 0.97524 | 0.97531 | 0.999994 | 20963 | 0 | 1.5645 | 0.024756 | 0.090029 | 0.943068 | 1914 | 5 | 1.5645 |
| MICU1 | 5 | 0.76374 | 0.79823 | 0.999994 | 17008 | 1 | 4.5833 | 0.024759 | 0.090037 | 0.943068 | 1915 | 4 | 4.5833 |
| ITGB7 | 6 | 0.96209 | 0.96217 | 0.999994 | 20632 | 0 | 2.2228 | 0.024762 | 0.092533 | 0.948217 | 1916 | 5 | 2.2228 |
| TRIM77 | 4 | 0.90656 | 0.90627 | 0.999994 | 19362 | 0 | 0.93133 | 0.024771 | 0.077698 | 0.938203 | 1917 | 2 | 0.93133 |
| FOXQ1 | 6 | 0.43685 | 0.62632 | 0.999994 | 12523 | 1 | 4.3257 | 0.024773 | 0.092567 | 0.948217 | 1918 | 4 | 4.3257 |
| hsa-mir-518c | 4 | 0.97522 | 0.9751 | 0.999994 | 20962 | 0 | 1.6478 | 0.024782 | 0.077734 | 0.938203 | 1919 | 4 | 1.6478 |
| hsa-mir-522 | 3 | 0.91839 | 0.91842 | 0.999994 | 19593 | 0 | 7.1451 | 0.024797 | 0.063024 | 0.933948 | 1920 | 3 | 7.1451 |
| HMG20A | 5 | 0.9752 | 0.97527 | 0.999994 | 20961 | 0 | 1.3795 | 0.0248 | 0.090168 | 0.943068 | 1921 | 5 | 1.3795 |
| APOL1 | 5 | 0.97516 | 0.97523 | 0.999994 | 20959 | 0 | 1.6447 | 0.024839 | 0.090277 | 0.943185 | 1922 | 5 | 1.6447 |
| CSH2 | 5 | 0.95506 | 0.95521 | 0.999994 | 20454 | 0 | 3.3034 | 0.024842 | 0.09029 | 0.943185 | 1923 | 4 | 3.3034 |

| | | | | | | | | | | | | |
|---|---|---|---|---|---|---|---|---|---|---|---|---|
| hsa-mir-4254 | 4 | 0.97516 | 0.97504 | 0.999994 | 20958 | 0 | 4.8422 | 0.024844 | 0.077905 | 0.938203 | 1924 | 4 | 4.8422 |
| PREX1 | 4 | 0.61257 | 0.67218 | 0.999994 | 15120 | 1 | 2.5435 | 0.024864 | 0.077969 | 0.938203 | 1925 | 2 | 2.5435 |
| HOXD10 | 6 | 0.69789 | 0.81137 | 0.999994 | 16276 | 1 | 4.9202 | 0.024877 | 0.092865 | 0.949922 | 1926 | 5 | 4.9202 |
| FMR1NB | 5 | 0.33129 | 0.50661 | 0.999994 | 10751 | 2 | 6.2427 | 0.024883 | 0.090432 | 0.94376 | 1927 | 3 | 6.2427 |
| C1QTNF2 | 4 | 0.9751 | 0.97499 | 0.999994 | 20955 | 0 | 4.4587 | 0.024896 | 0.078075 | 0.938203 | 1928 | 4 | 4.4587 |
| SPRR2A | 6 | 0.45851 | 0.64075 | 0.999994 | 12829 | 1 | 4.5471 | 0.024956 | 0.093102 | 0.950707 | 1929 | 5 | 4.5471 |
| C2CD4D | 4 | 0.36986 | 0.51471 | 0.999994 | 11508 | 1 | 6.1903 | 0.02501 | 0.078391 | 0.938203 | 1930 | 3 | 6.1903 |
| LILRB4 | 5 | 0.97499 | 0.97505 | 0.999994 | 20954 | 0 | 3.8927 | 0.025011 | 0.090853 | 0.945479 | 1931 | 5 | 3.8927 |
| CTNNBIP1 | 4 | 0.26742 | 0.40963 | 0.999994 | 9406 | 2 | 3.1314 | 0.025022 | 0.07843 | 0.938203 | 1932 | 2 | 3.1314 |
| TAS2R1 | 6 | 0.30356 | 0.51086 | 0.999994 | 10206 | 3 | 0.78595 | 0.02505 | 0.093354 | 0.950736 | 1933 | 3 | 0.78595 |
| POC1B-GALNT4 | 2 | 0.83547 | 0.83527 | 0.999994 | 18016 | 0 | 5.3295 | 0.02509 | 0.046278 | 0.919289 | 1934 | 2 | 5.3295 |
| CD6 | 6 | 0.0034124 | 0.015247 | 0.757255 | 397 | 3 | 0.44228 | 0.025105 | 0.09351 | 0.950736 | 1935 | 3 | 0.44228 |
| ZNF681 | 5 | 0.89452 | 0.89464 | 0.999994 | 19114 | 0 | 3.2394 | 0.02511 | 0.091144 | 0.945479 | 1936 | 4 | 3.2394 |
| RAD51AP1 | 5 | 0.42673 | 0.58571 | 0.999994 | 12360 | 2 | 0.92838 | 0.02513 | 0.09121 | 0.945479 | 1937 | 3 | 0.92838 |
| ZBTB43 | 5 | 0.97483 | 0.9749 | 0.999994 | 20952 | 0 | 2.1461 | 0.02517 | 0.091338 | 0.945479 | 1938 | 5 | 2.1461 |
| hsa-mir-4527 | 4 | 0.43711 | 0.5838 | 0.999994 | 12527 | 1 | 6.2158 | 0.025172 | 0.078846 | 0.938203 | 1939 | 3 | 6.2158 |
| ACBD7 | 5 | 0.63171 | 0.73796 | 0.999994 | 15395 | 1 | 2.9444 | 0.02526 | 0.091616 | 0.945479 | 1940 | 4 | 2.9444 |
| GTSF1 | 5 | 0.54129 | 0.68405 | 0.999994 | 14074 | 1 | 3.404 | 0.025261 | 0.091621 | 0.945479 | 1941 | 4 | 3.404 |
| SYNDIG1 | 6 | 0.64067 | 0.77045 | 0.999994 | 15521 | 1 | 3.9754 | 0.025263 | 0.093947 | 0.950736 | 1942 | 4 | 3.9754 |
| COX7C | 6 | 0.73035 | 0.82172 | 0.999994 | 16603 | 1 | 4.2897 | 0.025271 | 0.09397 | 0.950736 | 1943 | 4 | 4.2897 |
| HNRNPUL2 | 5 | 0.70016 | 0.76656 | 0.999994 | 16303 | 1 | 6.2265 | 0.02528 | 0.09168 | 0.945479 | 1944 | 3 | 6.2265 |
| GADD45B | 5 | 0.40586 | 0.56843 | 0.999994 | 12044 | 2 | 2.8884 | 0.025292 | 0.09171 | 0.945479 | 1945 | 3 | 2.8884 |
| PPP1R37 | 5 | 0.75301 | 0.79244 | 0.999994 | 16888 | 1 | 4.0777 | 0.025299 | 0.091732 | 0.945479 | 1946 | 4 | 4.0777 |
| FBLIM1 | 6 | 0.47591 | 0.65244 | 0.999994 | 13079 | 1 | 4.6156 | 0.025301 | 0.094062 | 0.950736 | 1947 | 5 | 4.6156 |
| hsa-mir-1288 | 4 | 0.342 | 0.48608 | 0.999994 | 10981 | 1 | 5.7206 | 0.025322 | 0.079266 | 0.938203 | 1948 | 3 | 5.7206 |
| SMARCB1 | 2 | 0.97462 | 0.97462 | 0.999994 | 20947 | 0 | 6.8441 | 0.025384 | 0.046783 | 0.919289 | 1949 | 2 | 6.8441 |
| CCDC93 | 6 | 0.93649 | 0.93651 | 0.999994 | 19994 | 0 | 0.5756 | 0.025412 | 0.09439 | 0.950736 | 1950 | 3 | 0.5756 |
| RCAN2 | 6 | 0.91107 | 0.91246 | 0.999994 | 19449 | 1 | 1.0215 | 0.025412 | 0.09439 | 0.950736 | 1951 | 3 | 1.0215 |
| BOLL | 4 | 0.044192 | 0.11014 | 0.951052 | 2719 | 3 | -2.0116 | 0.025437 | 0.079595 | 0.938962 | 1952 | 1 | -2.0116 |
| STT3A | 6 | 0.96296 | 0.96302 | 0.999994 | 20661 | 0 | 3.2155 | 0.025438 | 0.094467 | 0.950736 | 1953 | 5 | 3.2155 |
| hsa-mir-7160 | 3 | 0.97453 | 0.97461 | 0.999994 | 20943 | 0 | 6.16 | 0.025472 | 0.064637 | 0.933948 | 1954 | 3 | 6.16 |
| hsa-mir-608 | 3 | 0.97452 | 0.9746 | 0.999994 | 20942 | 0 | 3.8862 | 0.025477 | 0.06465 | 0.933948 | 1955 | 3 | 3.8862 |
| hsa-mir-6790 | 3 | 0.97452 | 0.9746 | 0.999994 | 20941 | 0 | 2.8498 | 0.025485 | 0.064669 | 0.933948 | 1956 | 3 | 2.8498 |
| NPHS2 | 5 | 0.56001 | 0.70065 | 0.999994 | 14370 | 1 | 6.221 | 0.025485 | 0.092367 | 0.947514 | 1957 | 4 | 6.221 |
| LRRC17 | 5 | 0.73779 | 0.78457 | 0.999994 | 16697 | 1 | 6.221 | 0.025485 | 0.092367 | 0.947514 | 1958 | 4 | 6.221 |
| BRINP2 | 6 | 0.7115 | 0.8156 | 0.999994 | 16417 | 1 | 2.1117 | 0.025495 | 0.094609 | 0.950736 | 1959 | 5 | 2.1117 |
| hsa-mir-4737 | 4 | 0.97447 | 0.97436 | 0.999994 | 20940 | 0 | 2.5392 | 0.025525 | 0.079836 | 0.938962 | 1960 | 4 | 2.5392 |
| TTC16 | 6 | 0.88016 | 0.8897 | 0.999994 | 18835 | 1 | 4.4658 | 0.025542 | 0.094768 | 0.950736 | 1961 | 4 | 4.4658 |
| SLC22A18 | 6 | 0.22796 | 0.42163 | 0.999994 | 8477 | 2 | 0.2832 | 0.025548 | 0.094789 | 0.950736 | 1962 | 2 | 0.2832 |
| NUP205 | 6 | 0.28271 | 0.48656 | 0.999994 | 9753 | 1 | 3.7501 | 0.025548 | 0.094789 | 0.950736 | 1963 | 5 | 3.7501 |
| SLC12A1 | 5 | 0.016055 | 0.053187 | 0.916684 | 1248 | 3 | -3.4195 | 0.025592 | 0.092697 | 0.949107 | 1964 | 2 | -3.4195 |
| PRDM11 | 6 | 0.11392 | 0.27418 | 0.998561 | 5444 | 3 | 2.4868 | 0.025593 | 0.094921 | 0.950736 | 1965 | 3 | 2.4868 |
| TUBB3 | 6 | 0.43258 | 0.62355 | 0.999994 | 12453 | 2 | 3.7152 | 0.025593 | 0.094921 | 0.950736 | 1966 | 4 | 3.7152 |
| WNT1 | 3 | 0.091742 | 0.17208 | 0.967658 | 4667 | 1 | 6.139 | 0.025603 | 0.064945 | 0.933948 | 1967 | 2 | 6.139 |
| VAX2 | 5 | 0.1992 | 0.37408 | 0.999994 | 7742 | 3 | -0.88139 | 0.025619 | 0.092789 | 0.949599 | 1968 | 2 | -0.88139 |
| MRPL15 | 4 | 0.97436 | 0.97424 | 0.999994 | 20937 | 0 | 3.9772 | 0.025636 | 0.080157 | 0.938962 | 1969 | 4 | 3.9772 |
| PAGR1 | 6 | 0.16813 | 0.3484 | 0.999994 | 6941 | 1 | 1.6518 | 0.025659 | 0.095122 | 0.950736 | 1970 | 5 | 1.6518 |
| OR2T11 | 5 | 0.93347 | 0.93371 | 0.999994 | 19934 | 0 | 4.7663 | 0.025661 | 0.092919 | 0.95003 | 1971 | 4 | 4.7663 |
| STAR | 4 | 0.26945 | 0.4117 | 0.999994 | 9459 | 1 | 5.8921 | 0.025689 | 0.080319 | 0.938962 | 1972 | 3 | 5.8921 |
| ARL3 | 4 | 0.97426 | 0.97412 | 0.999994 | 20935 | 0 | 4.2674 | 0.025741 | 0.080458 | 0.938962 | 1973 | 4 | 4.2674 |
| OR5AS1 | 6 | 0.91394 | 0.91498 | 0.999994 | 19485 | 1 | 3.5138 | 0.025764 | 0.095416 | 0.950736 | 1974 | 4 | 3.5138 |
| NonTargetingControlG | 2 | 0.85284 | 0.85274 | 0.999994 | 18314 | 0 | 5.4428 | 0.02577 | 0.047473 | 0.919289 | 1975 | 2 | 5.4428 |

| | | | | | | | | | | | | |
|---|---|---|---|---|---|---|---|---|---|---|---|---|
| HSPBP1 | 6 | 0.32766 | 0.53856 | 0.999994 | 10674 | 2 | 4.0222 | 0.025778 | 0.095464 | 0.950736 | 1976 | 4 | 4.0222 |
| HCN2 | 6 | 0.90638 | 0.90866 | 0.999994 | 19358 | 1 | 4.1657 | 0.025778 | 0.095464 | 0.950736 | 1977 | 5 | 4.1657 |
| HTRA4 | 6 | 0.96099 | 0.96105 | 0.999994 | 20605 | 0 | 2.3369 | 0.0258 | 0.095533 | 0.950736 | 1978 | 5 | 2.3369 |
| hsa-mir-4663 | 3 | 0.9742 | 0.97428 | 0.999994 | 20933 | 0 | 5.8862 | 0.025804 | 0.0654 | 0.933948 | 1979 | 3 | 5.8862 |
| FAM19A5 | 4 | 0.92424 | 0.92404 | 0.999994 | 19724 | 0 | 5.7859 | 0.025808 | 0.080656 | 0.939037 | 1980 | 3 | 5.7859 |
| NOD2 | 6 | 0.92246 | 0.92272 | 0.999994 | 19686 | 1 | 1.8611 | 0.025894 | 0.095804 | 0.950736 | 1981 | 5 | 1.8611 |
| FCGBP | 6 | 0.8379 | 0.86556 | 0.999994 | 18050 | 1 | 1.3233 | 0.025894 | 0.095805 | 0.950736 | 1982 | 4 | 1.3233 |
| KIR2DL1 | 6 | 0.89548 | 0.90025 | 0.999994 | 19137 | 1 | 2.2294 | 0.025915 | 0.095865 | 0.950736 | 1983 | 5 | 2.2294 |
| PIP5KL1 | 6 | 0.81096 | 0.85269 | 0.999994 | 17646 | 1 | 2.8177 | 0.025921 | 0.095882 | 0.950736 | 1984 | 5 | 2.8177 |
| ALDH1A1 | 5 | 0.27344 | 0.45954 | 0.999994 | 9551 | 1 | 6.2009 | 0.025927 | 0.093735 | 0.950736 | 1985 | 3 | 6.2009 |
| MED25 | 3 | 0.97406 | 0.97413 | 0.999994 | 20930 | 0 | 3.374 | 0.025935 | 0.065712 | 0.933948 | 1986 | 3 | 3.374 |
| EIF2B1 | 4 | 0.95607 | 0.95598 | 0.999994 | 20485 | 0 | 6.1541 | 0.025977 | 0.081116 | 0.939346 | 1987 | 4 | 6.1541 |
| C17orf102 | 6 | 0.52014 | 0.68263 | 0.999994 | 13748 | 2 | 3.6212 | 0.025985 | 0.096046 | 0.950736 | 1988 | 3 | 3.6212 |
| SLC22A2 | 6 | 0.89066 | 0.89679 | 0.999994 | 19048 | 1 | 4.9168 | 0.025985 | 0.096046 | 0.950736 | 1989 | 5 | 4.9168 |
| TROVE2 | 5 | 0.97401 | 0.97405 | 0.999994 | 20929 | 0 | 2.7397 | 0.025993 | 0.093946 | 0.950736 | 1990 | 5 | 2.7397 |
| TRIM49D1 | 3 | 0.974 | 0.97406 | 0.999994 | 20927 | 0 | 5.1892 | 0.026 | 0.065862 | 0.933948 | 1991 | 3 | 5.1892 |
| STARD13 | 6 | 0.21348 | 0.40432 | 0.999994 | 8101 | 1 | 2.9453 | 0.026002 | 0.0961 | 0.950736 | 1992 | 5 | 2.9453 |
| DYM | 6 | 0.32519 | 0.53577 | 0.999994 | 10625 | 3 | 2.9182 | 0.026009 | 0.09612 | 0.950736 | 1993 | 3 | 2.9182 |
| IFNL3 | 5 | 0.76398 | 0.79836 | 0.999994 | 17014 | 1 | 0.37078 | 0.026031 | 0.094069 | 0.950736 | 1994 | 2 | 0.37078 |
| DIO2 | 5 | 0.97397 | 0.97402 | 0.999994 | 20926 | 0 | 1.8517 | 0.026034 | 0.094077 | 0.950736 | 1995 | 5 | 1.8517 |
| FANCD2OS | 6 | 0.41595 | 0.61272 | 0.999994 | 12191 | 2 | 2.8385 | 0.026047 | 0.096232 | 0.950736 | 1996 | 3 | 2.8385 |
| C2orf50 | 4 | 0.97395 | 0.97382 | 0.999994 | 20925 | 0 | 2.3987 | 0.026047 | 0.081285 | 0.939346 | 1997 | 4 | 2.3987 |
| VSTM2B | 5 | 0.53119 | 0.67513 | 0.999994 | 13915 | 2 | 2.5653 | 0.026053 | 0.094131 | 0.950736 | 1998 | 3 | 2.5653 |
| hsa-mir-7106 | 4 | 0.51439 | 0.62278 | 0.999994 | 13650 | 1 | 5.0028 | 0.026066 | 0.081328 | 0.939346 | 1999 | 3 | 5.0028 |
| MRPS25 | 6 | 0.98613 | 0.98608 | 0.999994 | 21250 | 0 | 4.3894 | 0.026106 | 0.096406 | 0.950736 | 2000 | 5 | 4.3894 |
| KIF1C | 5 | 0.064986 | 0.17022 | 0.967126 | 3651 | 3 | -2.1291 | 0.026121 | 0.094354 | 0.950736 | 2001 | 2 | -2.1291 |
| hsa-mir-637 | 4 | 0.28295 | 0.42557 | 0.999994 | 9762 | 2 | 1.3584 | 0.026121 | 0.081466 | 0.939346 | 2002 | 2 | 1.3584 |
| USHBP1 | 5 | 0.93025 | 0.93053 | 0.999994 | 19860 | 0 | 6.1935 | 0.026154 | 0.094463 | 0.950736 | 2003 | 4 | 6.1935 |
| PNISR | 4 | 0.21299 | 0.35407 | 0.999994 | 8092 | 2 | 2.8192 | 0.026158 | 0.081578 | 0.939346 | 2004 | 2 | 2.8192 |
| ZNF234 | 6 | 0.51042 | 0.6759 | 0.999994 | 13580 | 2 | 4.2898 | 0.026179 | 0.096627 | 0.950736 | 2005 | 4 | 4.2898 |
| hsa-mir-3911 | 4 | 0.84377 | 0.84325 | 0.999994 | 18153 | 0 | 5.5171 | 0.026192 | 0.081657 | 0.939346 | 2006 | 3 | 5.5171 |
| NonTargetingControlG | 2 | 0.97378 | 0.97376 | 0.999994 | 20920 | 0 | 6.3871 | 0.026218 | 0.048252 | 0.919864 | 2007 | 2 | 6.3871 |
| PBX2 | 5 | 0.75252 | 0.79216 | 0.999994 | 16883 | 1 | 5.2042 | 0.026261 | 0.094783 | 0.950736 | 2008 | 3 | 5.2042 |
| CGNL1 | 6 | 0.69801 | 0.81141 | 0.999994 | 16280 | 1 | 2.1727 | 0.026263 | 0.09686 | 0.950736 | 2009 | 5 | 2.1727 |
| hsa-mir-548o-2 | 3 | 0.32892 | 0.44414 | 0.999994 | 10701 | 1 | 7.2168 | 0.026263 | 0.066486 | 0.933948 | 2010 | 2 | 7.2168 |
| hsa-mir-3942 | 3 | 0.97373 | 0.9738 | 0.999994 | 20917 | 0 | 3.338 | 0.026266 | 0.066495 | 0.933948 | 2011 | 3 | 3.338 |
| MAL2 | 5 | 0.86371 | 0.86553 | 0.999994 | 18515 | 1 | 5.9443 | 0.026279 | 0.094845 | 0.950736 | 2012 | 4 | 5.9443 |
| ALDH1A3 | 6 | 0.88614 | 0.89366 | 0.999994 | 18952 | 1 | 4.7403 | 0.026289 | 0.096933 | 0.950736 | 2013 | 5 | 4.7403 |
| OR1L6 | 6 | 0.8884 | 0.89519 | 0.999994 | 19005 | 1 | 4.0897 | 0.026297 | 0.096953 | 0.950736 | 2014 | 5 | 4.0897 |
| ZNF280C | 4 | 0.97368 | 0.97356 | 0.999994 | 20915 | 0 | 4.1864 | 0.026322 | 0.082031 | 0.939346 | 2015 | 4 | 4.1864 |
| ZNF775 | 6 | 0.1059 | 0.25881 | 0.994943 | 5171 | 2 | 1.2864 | 0.026323 | 0.097029 | 0.950736 | 2016 | 4 | 1.2864 |
| C1orf43 | 6 | 0.057922 | 0.16129 | 0.967126 | 3333 | 2 | 4.0736 | 0.026333 | 0.097058 | 0.950736 | 2017 | 4 | 4.0736 |
| WDR41 | 6 | 0.47151 | 0.64945 | 0.999994 | 13022 | 2 | 0.077551 | 0.026333 | 0.097058 | 0.950736 | 2018 | 2 | 0.077551 |
| hsa-mir-6800 | 4 | 0.97366 | 0.97354 | 0.999994 | 20914 | 0 | 2.3201 | 0.026338 | 0.082074 | 0.939346 | 2019 | 4 | 2.3201 |
| M6PR | 6 | 0.27316 | 0.47544 | 0.999994 | 9544 | 1 | 2.7649 | 0.026349 | 0.097107 | 0.950736 | 2020 | 5 | 2.7649 |
| hsa-mir-4763 | 4 | 0.97364 | 0.97352 | 0.999994 | 20913 | 0 | 3.6806 | 0.026364 | 0.082147 | 0.939346 | 2021 | 4 | 3.6806 |
| C14orf166B | 6 | 0.58438 | 0.72844 | 0.999994 | 14735 | 2 | 2.7545 | 0.026379 | 0.097195 | 0.950736 | 2022 | 3 | 2.7545 |
| ACAD8 | 6 | 0.24321 | 0.43982 | 0.999994 | 8869 | 3 | 2.788 | 0.02638 | 0.097198 | 0.950736 | 2023 | 3 | 2.788 |
| PITPNM3 | 6 | 0.79665 | 0.84647 | 0.999994 | 17436 | 1 | 3.6789 | 0.026381 | 0.0972 | 0.950736 | 2024 | 5 | 3.6789 |
| OSTM1 | 5 | 0.86881 | 0.87006 | 0.999994 | 18614 | 1 | 6.1806 | 0.026407 | 0.095256 | 0.950736 | 2025 | 4 | 6.1806 |
| STX7 | 5 | 0.97359 | 0.97363 | 0.999994 | 20910 | 0 | 0.81338 | 0.02641 | 0.095266 | 0.950736 | 2026 | 5 | 0.81338 |
| AP3M1 | 6 | 0.20972 | 0.39977 | 0.999994 | 8021 | 2 | 0.69673 | 0.026443 | 0.09739 | 0.950736 | 2027 | 4 | 0.69673 |

| | | | | | | | | | | | | | |
|---|---|---|---|---|---|---|---|---|---|---|---|---|---|
| hsa-mir-4528 | 1 | 0.97355 | 0.9736 | 0.999994 | 20909 | 0 | 8.8155 | 0.026447 | 0.026383 | 0.890437 | 2028 | 1 | 8.8155 |
| MNAT1 | 6 | 0.85192 | 0.87294 | 0.999994 | 18293 | 1 | 2.8675 | 0.02645 | 0.097412 | 0.950736 | 2029 | 5 | 2.8675 |
| FAM86B1 | 4 | 0.33218 | 0.47598 | 0.999994 | 10769 | 1 | 0.73169 | 0.026546 | 0.082639 | 0.939346 | 2030 | 3 | 0.73169 |
| C16orf11 | 5 | 0.50316 | 0.65068 | 0.999994 | 13482 | 1 | 3.8296 | 0.026553 | 0.095737 | 0.950736 | 2031 | 3 | 3.8296 |
| CMTM7 | 6 | 0.34206 | 0.55501 | 0.999994 | 10986 | 2 | 4.4546 | 0.026586 | 0.097801 | 0.950736 | 2032 | 4 | 4.4546 |
| ARSH | 6 | 0.19706 | 0.38432 | 0.999994 | 7677 | 1 | 5.2288 | 0.026586 | 0.097801 | 0.950736 | 2033 | 5 | 5.2288 |
| ADCK4 | 5 | 0.97337 | 0.97343 | 0.999994 | 20905 | 0 | 0.92307 | 0.026626 | 0.095971 | 0.950736 | 2034 | 5 | 0.92307 |
| ZBBX | 5 | 0.13315 | 0.27743 | 0.999379 | 5998 | 2 | 6.1768 | 0.026648 | 0.096041 | 0.950736 | 2035 | 3 | 6.1768 |
| TNFSF10 | 5 | 0.97335 | 0.9734 | 0.999994 | 20904 | 0 | 3.5579 | 0.026651 | 0.096046 | 0.950736 | 2036 | 5 | 3.5579 |
| MAK16 | 5 | 0.51814 | 0.66369 | 0.999994 | 13713 | 1 | 3.1166 | 0.026671 | 0.096115 | 0.950736 | 2037 | 4 | 3.1166 |
| AJAP1 | 5 | 0.96072 | 0.96086 | 0.999994 | 20597 | 0 | 4.5447 | 0.026671 | 0.096115 | 0.950736 | 2038 | 4 | 4.5447 |
| PRIMA1 | 6 | 0.11665 | 0.27942 | 0.999632 | 5528 | 1 | 3.1704 | 0.026677 | 0.098069 | 0.950736 | 2039 | 5 | 3.1704 |
| KRT17 | 5 | 0.33296 | 0.50795 | 0.999994 | 10787 | 1 | 5.202 | 0.026695 | 0.096191 | 0.950736 | 2040 | 4 | 5.202 |
| NonTargetingControlG | 2 | 0.97329 | 0.97326 | 0.999994 | 20901 | 0 | 6.3741 | 0.026714 | 0.04914 | 0.920223 | 2041 | 2 | 6.3741 |
| PDCL3 | 5 | 0.97324 | 0.97329 | 0.999994 | 20900 | 0 | 1.2095 | 0.02676 | 0.096396 | 0.950736 | 2042 | 5 | 1.2095 |
| SLC22A3 | 6 | 0.617 | 0.75258 | 0.999994 | 15172 | 1 | 4.7027 | 0.026764 | 0.098332 | 0.950736 | 2043 | 5 | 4.7027 |
| PCDHB7 | 6 | 0.17925 | 0.36221 | 0.999994 | 7207 | 2 | 4.1998 | 0.026764 | 0.098332 | 0.950736 | 2044 | 4 | 4.1998 |
| CCDC171 | 6 | 0.8103 | 0.85239 | 0.999994 | 17636 | 1 | 0.81203 | 0.02677 | 0.098344 | 0.950736 | 2045 | 3 | 0.81203 |
| SIGLEC15 | 6 | 0.51687 | 0.68037 | 0.999994 | 13689 | 2 | 4.9213 | 0.026793 | 0.098414 | 0.950983 | 2046 | 4 | 4.9213 |
| KRTAP10-11 | 4 | 0.71279 | 0.73302 | 0.999994 | 16428 | 1 | 5.8407 | 0.026819 | 0.083401 | 0.939912 | 2047 | 3 | 5.8407 |
| PROZ | 6 | 0.92026 | 0.92064 | 0.999994 | 19635 | 1 | 4.2521 | 0.026819 | 0.098492 | 0.95131 | 2048 | 5 | 4.2521 |
| SPATA2 | 5 | 0.44903 | 0.60444 | 0.999994 | 12701 | 2 | 4.0767 | 0.026867 | 0.096729 | 0.950736 | 2049 | 3 | 4.0767 |
| LIN37 | 6 | 0.87488 | 0.88636 | 0.999994 | 18732 | 1 | 1.9205 | 0.026876 | 0.098676 | 0.951383 | 2050 | 5 | 1.9205 |
| LPAR3 | 6 | 0.96071 | 0.96077 | 0.999994 | 20596 | 0 | 3.0762 | 0.026879 | 0.098686 | 0.951383 | 2051 | 5 | 3.0762 |
| PGAM1 | 5 | 0.96934 | 0.96943 | 0.999994 | 20812 | 0 | 2.9342 | 0.026887 | 0.096794 | 0.950736 | 2052 | 4 | 2.9342 |
| SF3A3 | 5 | 0.97248 | 0.97254 | 0.999994 | 20881 | 0 | 4.1539 | 0.026887 | 0.096794 | 0.950736 | 2053 | 4 | 4.1539 |
| NonTargetingControlG | 2 | 0.97311 | 0.97309 | 0.999994 | 20899 | 0 | 6.8491 | 0.026894 | 0.049471 | 0.921154 | 2054 | 2 | 6.8491 |
| HSPB7 | 6 | 0.21874 | 0.4106 | 0.999994 | 8251 | 2 | 3.3627 | 0.026923 | 0.098808 | 0.9514 | 2055 | 4 | 3.3627 |
| FANCE | 4 | 0.27084 | 0.41314 | 0.999994 | 9495 | 1 | 5.7132 | 0.026927 | 0.083699 | 0.940296 | 2056 | 3 | 5.7132 |
| TMEM158 | 6 | 0.27619 | 0.47899 | 0.999994 | 9622 | 1 | 3.4376 | 0.026956 | 0.098915 | 0.951893 | 2057 | 3 | 3.4376 |
| THAP8 | 5 | 0.97304 | 0.9731 | 0.999994 | 20898 | 0 | 2.1684 | 0.026958 | 0.097021 | 0.950736 | 2058 | 5 | 2.1684 |
| SLC25A21 | 6 | 0.17703 | 0.35936 | 0.999994 | 7158 | 3 | -0.07226 | 0.026987 | 0.099005 | 0.951893 | 2059 | 2 | -0.07226 |
| hsa-mir-1269b | 4 | 0.97301 | 0.97289 | 0.999994 | 20897 | 0 | 5.2605 | 0.026994 | 0.083867 | 0.940296 | 2060 | 4 | 5.2605 |
| ZNF746 | 5 | 0.97298 | 0.97303 | 0.999994 | 20896 | 0 | 1.1141 | 0.027016 | 0.097206 | 0.950736 | 2061 | 5 | 1.1141 |
| AQP4 | 5 | 0.97294 | 0.97299 | 0.999994 | 20895 | 0 | 0.80309 | 0.027058 | 0.09734 | 0.950736 | 2062 | 5 | 0.80309 |
| LAMB2 | 5 | 0.90538 | 0.90548 | 0.999994 | 19341 | 0 | 0.51806 | 0.027063 | 0.09736 | 0.950736 | 2063 | 2 | 0.51806 |
| NonTargetingControlG | 2 | 0.97292 | 0.97291 | 0.999994 | 20894 | 0 | 6.0188 | 0.027079 | 0.049777 | 0.923113 | 2064 | 2 | 6.0188 |
| GNAZ | 5 | 0.97289 | 0.97294 | 0.999994 | 20892 | 0 | 3.231 | 0.027108 | 0.097496 | 0.950736 | 2065 | 5 | 3.231 |
| KRTAP12-4 | 5 | 0.97287 | 0.97292 | 0.999994 | 20891 | 0 | 2.2291 | 0.027131 | 0.097565 | 0.950736 | 2066 | 5 | 2.2291 |
| UFSP1 | 5 | 0.33847 | 0.51249 | 0.999994 | 10907 | 2 | 0.7347 | 0.027138 | 0.097588 | 0.950736 | 2067 | 3 | 0.7347 |
| HDDC2 | 6 | 0.51487 | 0.67898 | 0.999994 | 13659 | 2 | 0.47579 | 0.027153 | 0.099482 | 0.951893 | 2068 | 3 | 0.47579 |
| PARG | 4 | 0.68128 | 0.71242 | 0.999994 | 16106 | 1 | 4.4207 | 0.027156 | 0.084325 | 0.940301 | 2069 | 3 | 4.4207 |
| TXNIP | 5 | 0.95682 | 0.95696 | 0.999994 | 20514 | 0 | 3.9895 | 0.027175 | 0.097706 | 0.950736 | 2070 | 4 | 3.9895 |
| NonTargetingControlG | 2 | 0.97278 | 0.97277 | 0.999994 | 20888 | 0 | 7.2183 | 0.027215 | 0.050023 | 0.92348 | 2071 | 2 | 7.2183 |
| C1orf194 | 5 | 0.34897 | 0.52107 | 0.999994 | 11130 | 2 | 0.68666 | 0.02723 | 0.097868 | 0.950736 | 2072 | 3 | 0.68666 |
| B3GNT6 | 5 | 0.97276 | 0.97282 | 0.999994 | 20887 | 0 | 1.6715 | 0.027239 | 0.097897 | 0.950736 | 2073 | 5 | 1.6715 |
| hsa-mir-6766 | 4 | 0.39757 | 0.54321 | 0.999994 | 11911 | 2 | 3.4276 | 0.02728 | 0.084682 | 0.940404 | 2074 | 2 | 3.4276 |
| NKX2-5 | 4 | 0.054113 | 0.12954 | 0.95318 | 3162 | 2 | 0.81902 | 0.02728 | 0.084682 | 0.940404 | 2075 | 2 | 0.81902 |
| INO80D | 5 | 0.2525 | 0.44275 | 0.999994 | 9067 | 2 | -0.24548 | 0.027299 | 0.098074 | 0.950736 | 2076 | 1 | -0.24548 |
| PVRL2 | 6 | 0.93348 | 0.93356 | 0.999994 | 19935 | 0 | 1.4012 | 0.027319 | 0.099955 | 0.951893 | 2077 | 3 | 1.4012 |
| VPS51 | 5 | 0.97268 | 0.97274 | 0.999994 | 20886 | 0 | 2.2618 | 0.027324 | 0.098158 | 0.950736 | 2078 | 5 | 2.2618 |
| RPUSD4 | 4 | 0.049605 | 0.12067 | 0.951052 | 2961 | 1 | 5.6788 | 0.02735 | 0.084867 | 0.940435 | 2079 | 3 | 5.6788 |

| | | | | | | | | | | | | | |
|---|---|---|---|---|---|---|---|---|---|---|---|---|---|
| PLCZ1 | 6 | 0.96967 | 0.96966 | 0.999994 | 20818 | 0 | 4.9658 | 0.027362 | 0.10008 | 0.951893 | 2080 | 5 | 4.9658 |
| SDF2L1 | 6 | 0.80164 | 0.84856 | 0.999994 | 17503 | 1 | 4.3304 | 0.027369 | 0.1001 | 0.951893 | 2081 | 4 | 4.3304 |
| MPP7 | 6 | 0.44787 | 0.63368 | 0.999994 | 12686 | 2 | 3.7752 | 0.027391 | 0.10017 | 0.951893 | 2082 | 4 | 3.7752 |
| COL8A1 | 6 | 0.85325 | 0.87367 | 0.999994 | 18318 | 1 | 2.8462 | 0.027394 | 0.10018 | 0.951893 | 2083 | 5 | 2.8462 |
| DRD4 | 5 | 0.24136 | 0.43276 | 0.999994 | 8810 | 2 | 1.4151 | 0.027481 | 0.098562 | 0.951383 | 2084 | 3 | 1.4151 |
| ALDH8A1 | 6 | 0.26078 | 0.4608 | 0.999994 | 9261 | 1 | 0.85588 | 0.027498 | 0.10048 | 0.951893 | 2085 | 4 | 0.85588 |
| OR51Q1 | 4 | 0.012332 | 0.04097 | 0.891621 | 1014 | 1 | 5.3519 | 0.027499 | 0.085283 | 0.942567 | 2086 | 3 | 5.3519 |
| MRPL37 | 5 | 0.46196 | 0.61531 | 0.999994 | 12884 | 1 | 3.889 | 0.027546 | 0.098719 | 0.951383 | 2087 | 4 | 3.889 |
| PEG3 | 6 | 0.55794 | 0.70938 | 0.999994 | 14333 | 2 | 4.7609 | 0.027549 | 0.10062 | 0.951893 | 2088 | 4 | 4.7609 |
| OTOP3 | 5 | 0.95405 | 0.95422 | 0.999994 | 20436 | 0 | 5.5792 | 0.027579 | 0.098787 | 0.9514 | 2089 | 4 | 5.5792 |
| HLA-DMA | 6 | 0.67237 | 0.795 | 0.999994 | 15959 | 1 | 4.6822 | 0.027609 | 0.10081 | 0.951893 | 2090 | 5 | 4.6822 |
| PDRG1 | 3 | 0.97237 | 0.97244 | 0.999994 | 20878 | 0 | 6.1252 | 0.027629 | 0.06966 | 0.937929 | 2091 | 3 | 6.1252 |
| ZBED6CL | 3 | 0.97237 | 0.97244 | 0.999994 | 20877 | 0 | 4.8253 | 0.027629 | 0.06966 | 0.937929 | 2092 | 3 | 4.8253 |
| hsa-mir-6820 | 3 | 0.97237 | 0.97244 | 0.999994 | 20879 | 0 | 6.476 | 0.027629 | 0.06966 | 0.937929 | 2093 | 3 | 6.476 |
| ZSCAN18 | 4 | 0.95104 | 0.95094 | 0.999994 | 20361 | 0 | 4.8393 | 0.027656 | 0.08572 | 0.943068 | 2094 | 4 | 4.8393 |
| SLC7A10 | 6 | 0.63117 | 0.76323 | 0.999994 | 15388 | 2 | 2.0804 | 0.02768 | 0.101 | 0.951893 | 2095 | 4 | 2.0804 |
| SEZ6L | 6 | 0.96601 | 0.96608 | 0.999994 | 20735 | 0 | 4.8745 | 0.027746 | 0.10119 | 0.951893 | 2096 | 4 | 4.8745 |
| FBXO5 | 5 | 0.90513 | 0.90523 | 0.999994 | 19331 | 0 | 5.2293 | 0.027748 | 0.099181 | 0.951893 | 2097 | 4 | 5.2293 |
| FAM25A | 5 | 0.016531 | 0.054546 | 0.922677 | 1278 | 2 | 0.067608 | 0.02776 | 0.099211 | 0.951893 | 2098 | 2 | 0.067608 |
| STRN | 6 | 0.55108 | 0.70443 | 0.999994 | 14229 | 2 | 2.0709 | 0.027761 | 0.10123 | 0.951893 | 2099 | 3 | 2.0709 |
| RRAS | 6 | 0.92999 | 0.93011 | 0.999994 | 19852 | 0 | 3.0528 | 0.027763 | 0.10124 | 0.951893 | 2100 | 5 | 3.0528 |
| MBD3L1 | 6 | 0.92159 | 0.92188 | 0.999994 | 19668 | 1 | 2.3889 | 0.027798 | 0.10132 | 0.951893 | 2101 | 4 | 2.3889 |
| CRYAA | 6 | 0.56095 | 0.71148 | 0.999994 | 14388 | 2 | 4.4277 | 0.027806 | 0.10134 | 0.951893 | 2102 | 4 | 4.4277 |
| CRYGC | 6 | 0.24923 | 0.447 | 0.999994 | 8999 | 2 | 5.0421 | 0.027806 | 0.10134 | 0.951893 | 2103 | 4 | 5.0421 |
| DCAF13 | 6 | 0.38772 | 0.59449 | 0.999994 | 11762 | 1 | 4.7318 | 0.027806 | 0.10134 | 0.951893 | 2104 | 4 | 4.7318 |
| DDHD2 | 5 | 0.52864 | 0.67286 | 0.999994 | 13874 | 2 | 1.5296 | 0.027806 | 0.099311 | 0.951893 | 2105 | 3 | 1.5296 |
| ARHGEF7 | 5 | 0.79848 | 0.81858 | 0.999994 | 17455 | 1 | 6.1178 | 0.027818 | 0.099348 | 0.951893 | 2106 | 4 | 6.1178 |
| IPCEF1 | 5 | 0.91151 | 0.91154 | 0.999994 | 19456 | 0 | 6.1178 | 0.027818 | 0.099348 | 0.951893 | 2107 | 4 | 6.1178 |
| PDE3A | 5 | 0.47627 | 0.6274 | 0.999994 | 13086 | 2 | 1.6201 | 0.027838 | 0.099396 | 0.951893 | 2108 | 3 | 1.6201 |
| MT1F | 5 | 0.57586 | 0.71469 | 0.999994 | 14630 | 2 | -0.38423 | 0.027838 | 0.099396 | 0.951893 | 2109 | 2 | -0.38423 |
| hsa-mir-34b | 3 | 0.56148 | 0.58848 | 0.999994 | 14395 | 1 | 0.43595 | 0.027839 | 0.070145 | 0.937929 | 2110 | 1 | 0.43595 |
| CYP2C9 | 4 | 0.97214 | 0.97203 | 0.999994 | 20872 | 0 | 1.4 | 0.027863 | 0.086297 | 0.943068 | 2111 | 4 | 1.4 |
| LY96 | 6 | 0.092477 | 0.23247 | 0.984502 | 4701 | 1 | 3.7929 | 0.027868 | 0.10152 | 0.951893 | 2112 | 5 | 3.7929 |
| ENTPD1 | 5 | 0.042512 | 0.12213 | 0.951052 | 2645 | 2 | 1.1106 | 0.027898 | 0.099546 | 0.951893 | 2113 | 3 | 1.1106 |
| LRRC3 | 6 | 0.96532 | 0.96536 | 0.999994 | 20724 | 0 | 3.0724 | 0.027899 | 0.1016 | 0.951893 | 2114 | 5 | 3.0724 |
| NonTargetingControlG | 2 | 0.65755 | 0.65736 | 0.999994 | 15748 | 0 | 4.6389 | 0.027928 | 0.051281 | 0.924349 | 2115 | 1 | 4.6389 |
| SET | 6 | 0.011849 | 0.045223 | 0.891621 | 983 | 1 | 4.2639 | 0.027945 | 0.10173 | 0.951893 | 2116 | 5 | 4.2639 |
| PAG1 | 5 | 0.86777 | 0.8691 | 0.999994 | 18596 | 1 | 0.84492 | 0.027954 | 0.099675 | 0.951893 | 2117 | 3 | 0.84492 |
| SERPINI1 | 4 | 0.97204 | 0.97193 | 0.999994 | 20871 | 0 | 1.9884 | 0.027959 | 0.086552 | 0.943068 | 2118 | 4 | 1.9884 |
| ZNF574 | 5 | 0.56944 | 0.70903 | 0.999994 | 14528 | 2 | 3.0085 | 0.027967 | 0.099708 | 0.951893 | 2119 | 3 | 3.0085 |
| RXFP4 | 6 | 0.1294 | 0.29954 | 0.999994 | 5925 | 1 | 1.2649 | 0.027982 | 0.10184 | 0.951893 | 2120 | 4 | 1.2649 |
| CDR2 | 6 | 0.9484 | 0.94842 | 0.999994 | 20297 | 0 | 4.1089 | 0.027988 | 0.10186 | 0.951893 | 2121 | 5 | 4.1089 |
| IMPG2 | 6 | 0.29714 | 0.50339 | 0.999994 | 10073 | 3 | 2.5891 | 0.028012 | 0.10192 | 0.951893 | 2122 | 3 | 2.5891 |
| ARFIP2 | 5 | 0.91776 | 0.91785 | 0.999994 | 19582 | 0 | 1.9737 | 0.028037 | 0.099882 | 0.951893 | 2123 | 3 | 1.9737 |
| NDUFA7 | 6 | 0.98327 | 0.98321 | 0.999994 | 21163 | 0 | 4.4266 | 0.028081 | 0.10213 | 0.951893 | 2124 | 5 | 4.4266 |
| NME3 | 6 | 0.22784 | 0.42149 | 0.999994 | 8476 | 2 | 4.1036 | 0.028085 | 0.10214 | 0.951893 | 2125 | 4 | 4.1036 |
| PLEKHH1 | 6 | 0.98175 | 0.98167 | 0.999994 | 21131 | 0 | 2.5541 | 0.028104 | 0.10219 | 0.951893 | 2126 | 5 | 2.5541 |
| HS3ST6 | 5 | 0.32693 | 0.50302 | 0.999994 | 10660 | 2 | -0.45016 | 0.028106 | 0.10006 | 0.951893 | 2127 | 2 | -0.45016 |
| PSMB2 | 2 | 0.97189 | 0.97185 | 0.999994 | 20867 | 0 | 6.8226 | 0.028111 | 0.051615 | 0.925271 | 2128 | 2 | 6.8226 |
| DHRSX_X | 6 | 0.74166 | 0.82554 | 0.999994 | 16761 | 1 | 2.1869 | 0.028119 | 0.10224 | 0.951893 | 2129 | 5 | 2.1869 |
| EIF4G2 | 6 | 0.95697 | 0.95705 | 0.999994 | 20519 | 0 | 1.9861 | 0.02812 | 0.10224 | 0.951893 | 2130 | 4 | 1.9861 |
| ANKRD6 | 5 | 0.39322 | 0.5579 | 0.999994 | 11848 | 1 | 2.4936 | 0.02814 | 0.10015 | 0.951893 | 2131 | 3 | 2.4936 |

| | | | | | | | | | | | | | |
|---|---|---|---|---|---|---|---|---|---|---|---|---|---|
| RASSF7 | 6 | 0.83448 | 0.86377 | 0.999994 | 17996 | 1 | 3.7609 | 0.028189 | 0.10245 | 0.952143 | 2132 | 4 | 3.7609 |
| IL8 | 6 | 0.35598 | 0.57076 | 0.999994 | 11268 | 3 | -0.58451 | 0.028231 | 0.10258 | 0.952143 | 2133 | 2 | -0.58451 |
| LMBRD1 | 6 | 0.16716 | 0.34716 | 0.999994 | 6910 | 1 | 1.4454 | 0.028231 | 0.10258 | 0.952143 | 2134 | 4 | 1.4454 |
| hsa-mir-151b | 4 | 0.1792 | 0.31927 | 0.999994 | 7201 | 2 | 3.1379 | 0.028234 | 0.087301 | 0.943068 | 2135 | 2 | 3.1379 |
| GPER1 | 4 | 0.82798 | 0.82755 | 0.999994 | 17897 | 0 | 3.2878 | 0.028264 | 0.087391 | 0.943068 | 2136 | 2 | 3.2878 |
| SYNGR1 | 6 | 0.74461 | 0.82658 | 0.999994 | 16796 | 1 | 3.1572 | 0.028273 | 0.10271 | 0.952143 | 2137 | 5 | 3.1572 |
| hsa-mir-516a-1 | 4 | 0.9717 | 0.9716 | 0.999994 | 20862 | 0 | 2.0518 | 0.028298 | 0.087476 | 0.943068 | 2138 | 4 | 2.0518 |
| CENPC | 3 | 0.63651 | 0.64596 | 0.999994 | 15469 | 1 | 7.1451 | 0.028308 | 0.071236 | 0.937929 | 2139 | 2 | 7.1451 |
| CYB561D1 | 6 | 0.9427 | 0.94269 | 0.999994 | 20144 | 0 | 2.6019 | 0.028313 | 0.10282 | 0.952143 | 2140 | 5 | 2.6019 |
| TTC9B | 6 | 0.94101 | 0.94099 | 0.999994 | 20109 | 0 | 2.2363 | 0.028333 | 0.10289 | 0.952143 | 2141 | 5 | 2.2363 |
| CIDEA | 6 | 0.6438 | 0.77283 | 0.999994 | 15573 | 2 | 0.6574 | 0.028456 | 0.10325 | 0.952231 | 2142 | 3 | 0.6574 |
| NSMAF | 5 | 0.95342 | 0.95358 | 0.999994 | 20420 | 0 | 2.803 | 0.028457 | 0.10088 | 0.951893 | 2143 | 4 | 2.803 |
| EPN2 | 6 | 0.1936 | 0.38002 | 0.999994 | 7589 | 2 | 3.4217 | 0.028479 | 0.10331 | 0.952231 | 2144 | 4 | 3.4217 |
| NonTargetingControlG | 1 | 0.97149 | 0.97151 | 0.999994 | 20858 | 0 | 8.7517 | 0.028512 | 0.028437 | 0.890437 | 2145 | 1 | 8.7517 |
| LAMP2 | 5 | 0.86537 | 0.86698 | 0.999994 | 18546 | 1 | 0.63698 | 0.02852 | 0.10103 | 0.951893 | 2146 | 3 | 0.63698 |
| ZNF621 | 5 | 0.90835 | 0.90839 | 0.999994 | 19392 | 0 | 0.03098 | 0.02852 | 0.10103 | 0.951893 | 2147 | 2 | 0.03098 |
| ULK1 | 6 | 0.03991 | 0.12171 | 0.951052 | 2512 | 1 | 3.4071 | 0.028548 | 0.1035 | 0.952231 | 2148 | 5 | 3.4071 |
| TIMM8B | 5 | 0.79132 | 0.81412 | 0.999994 | 17367 | 1 | 3.3994 | 0.028558 | 0.10111 | 0.951893 | 2149 | 4 | 3.3994 |
| LPAR2 | 6 | 0.27095 | 0.47284 | 0.999994 | 9497 | 1 | 4.2686 | 0.028576 | 0.10359 | 0.952231 | 2150 | 4 | 4.2686 |
| ACPT | 6 | 0.95389 | 0.95392 | 0.999994 | 20434 | 0 | 3.8344 | 0.028594 | 0.10364 | 0.952231 | 2151 | 4 | 3.8344 |
| hsa-mir-6759 | 4 | 0.97133 | 0.97122 | 0.999994 | 20855 | 0 | 3.0044 | 0.028672 | 0.088543 | 0.943068 | 2152 | 4 | 3.0044 |
| CTNNA3 | 6 | 0.68004 | 0.80103 | 0.999994 | 16091 | 2 | 2.2742 | 0.0287 | 0.10396 | 0.952231 | 2153 | 3 | 2.2742 |
| PCMTD1 | 6 | 0.91838 | 0.91891 | 0.999994 | 19592 | 1 | 3.7855 | 0.028711 | 0.10399 | 0.952231 | 2154 | 5 | 3.7855 |
| FAM92B | 6 | 0.93144 | 0.93158 | 0.999994 | 19884 | 0 | 0.38897 | 0.028717 | 0.10401 | 0.952231 | 2155 | 2 | 0.38897 |
| RPS27 | 5 | 0.11988 | 0.25715 | 0.994943 | 5621 | 2 | 0.35554 | 0.028728 | 0.10152 | 0.951893 | 2156 | 2 | 0.35554 |
| FAM46C | 6 | 0.28507 | 0.48934 | 0.999994 | 9813 | 1 | 1.4873 | 0.028755 | 0.10412 | 0.952231 | 2157 | 4 | 1.4873 |
| FHL3 | 5 | 0.9257 | 0.92593 | 0.999994 | 19764 | 0 | 3.4709 | 0.028767 | 0.10162 | 0.951893 | 2158 | 4 | 3.4709 |
| FCHSD1 | 4 | 0.97121 | 0.9711 | 0.999994 | 20851 | 0 | 2.7744 | 0.028788 | 0.088868 | 0.943068 | 2159 | 4 | 2.7744 |
| LTBP4 | 5 | 0.61291 | 0.7309 | 0.999994 | 15121 | 1 | 6.0743 | 0.028789 | 0.10166 | 0.951893 | 2160 | 4 | 6.0743 |
| KLK11 | 5 | 0.84682 | 0.85181 | 0.999994 | 18207 | 1 | 3.5121 | 0.028798 | 0.10169 | 0.951893 | 2161 | 3 | 3.5121 |
| TNRC6B | 6 | 0.7184 | 0.81782 | 0.999994 | 16480 | 1 | 4.7692 | 0.028802 | 0.10426 | 0.952231 | 2162 | 4 | 4.7692 |
| NEK1 | 3 | 0.16611 | 0.26943 | 0.998561 | 6890 | 2 | -1.4496 | 0.028811 | 0.072425 | 0.937929 | 2163 | 1 | -1.4496 |
| ZNF16 | 6 | 0.61407 | 0.7504 | 0.999994 | 15131 | 2 | 4.0197 | 0.028814 | 0.1043 | 0.952231 | 2164 | 4 | 4.0197 |
| PORCN | 6 | 0.31058 | 0.51902 | 0.999994 | 10338 | 2 | 3.5716 | 0.028814 | 0.1043 | 0.952231 | 2165 | 3 | 3.5716 |
| SF3A1 | 6 | 0.071148 | 0.1892 | 0.970085 | 3880 | 3 | 2.1291 | 0.028814 | 0.1043 | 0.952231 | 2166 | 3 | 2.1291 |
| MEIS2 | 6 | 0.29831 | 0.50475 | 0.999994 | 10095 | 3 | 1.1573 | 0.028831 | 0.10434 | 0.952231 | 2167 | 3 | 1.1573 |
| CASQ2 | 6 | 0.16939 | 0.34999 | 0.999994 | 6970 | 3 | 1.1246 | 0.028831 | 0.10434 | 0.952231 | 2168 | 3 | 1.1246 |
| CHST12 | 6 | 0.9759 | 0.97584 | 0.999994 | 20986 | 0 | 3.2901 | 0.028834 | 0.10435 | 0.952231 | 2169 | 5 | 3.2901 |
| GABPB1 | 6 | 0.53769 | 0.6949 | 0.999994 | 14022 | 2 | 4.9002 | 0.028894 | 0.10451 | 0.952231 | 2170 | 4 | 4.9002 |
| RSL24D1 | 3 | 0.85017 | 0.85013 | 0.999994 | 18266 | 0 | 7.124 | 0.028923 | 0.072668 | 0.937929 | 2171 | 2 | 7.124 |
| DBF4B | 4 | 0.97104 | 0.97094 | 0.999994 | 20849 | 0 | 2.4394 | 0.028955 | 0.089293 | 0.943068 | 2172 | 4 | 2.4394 |
| PHYKPL | 1 | 0.97102 | 0.97106 | 0.999994 | 20848 | 0 | 8.7374 | 0.028979 | 0.028895 | 0.890437 | 2173 | 1 | 8.7374 |
| EMC10 | 6 | 0.61386 | 0.75024 | 0.999994 | 15128 | 1 | 3.2301 | 0.028992 | 0.10477 | 0.952231 | 2174 | 5 | 3.2301 |
| KCNK3 | 5 | 0.66157 | 0.74977 | 0.999994 | 15802 | 1 | 5.4733 | 0.028993 | 0.10216 | 0.951893 | 2175 | 4 | 5.4733 |
| ULBP2 | 6 | 0.091772 | 0.23108 | 0.983933 | 4668 | 2 | 3.0471 | 0.028994 | 0.10477 | 0.952231 | 2176 | 3 | 3.0471 |
| IQGAP2 | 6 | 0.90202 | 0.90526 | 0.999994 | 19252 | 1 | 5.2804 | 0.028994 | 0.10477 | 0.952231 | 2177 | 5 | 5.2804 |
| NonTargetingControlG | 2 | 0.97099 | 0.97096 | 0.999994 | 20844 | 0 | 7.4627 | 0.029013 | 0.053222 | 0.926769 | 2178 | 2 | 7.4627 |
| hsa-mir-6786 | 2 | 0.97099 | 0.97096 | 0.999994 | 20845 | 0 | 6.4719 | 0.029013 | 0.053222 | 0.926769 | 2179 | 2 | 6.4719 |
| NonTargetingControlG | 2 | 0.97099 | 0.97096 | 0.999994 | 20846 | 0 | 5.9602 | 0.029013 | 0.053222 | 0.926769 | 2180 | 2 | 5.9602 |
| DDX50 | 6 | 0.5844 | 0.72845 | 0.999994 | 14736 | 1 | 3.0265 | 0.029039 | 0.1049 | 0.952231 | 2181 | 5 | 3.0265 |
| ZFP62 | 6 | 0.97827 | 0.97823 | 0.999994 | 21050 | 0 | 2.6857 | 0.029073 | 0.105 | 0.952231 | 2182 | 5 | 2.6857 |
| NOS1 | 6 | 0.59378 | 0.73538 | 0.999994 | 14875 | 2 | 0.43778 | 0.029086 | 0.10504 | 0.952231 | 2183 | 3 | 0.43778 |

| | | | | | | | | | | | | |
|---|---|---|---|---|---|---|---|---|---|---|---|---|
| HORMAD1 | 4 | 0.43393 | 0.58055 | 0.999994 | 12475 | 2 | 1.1441 | 0.029094 | 0.089677 | 0.943068 | 2184 | 2 | 1.1441 |
| hsa-mir-1827 | 3 | 0.56943 | 0.59422 | 0.999994 | 14527 | 1 | 0.71427 | 0.029116 | 0.073136 | 0.938203 | 2185 | 2 | 0.71427 |
| ABCC10 | 6 | 0.8257 | 0.85948 | 0.999994 | 17864 | 1 | 2.786 | 0.029153 | 0.10523 | 0.952231 | 2186 | 5 | 2.786 |
| TOP3A | 4 | 0.96188 | 0.96177 | 0.999994 | 20626 | 0 | 5.3089 | 0.029175 | 0.089902 | 0.943068 | 2187 | 4 | 5.3089 |
| MICALL2 | 4 | 0.27135 | 0.41367 | 0.999994 | 9506 | 1 | 4.8975 | 0.029175 | 0.089902 | 0.943068 | 2188 | 3 | 4.8975 |
| ALX1 | 6 | 0.34453 | 0.55786 | 0.999994 | 11043 | 3 | 0.91115 | 0.029197 | 0.10534 | 0.952231 | 2189 | 3 | 0.91115 |
| ITM2A | 6 | 0.6572 | 0.78313 | 0.999994 | 15744 | 1 | 4.2433 | 0.029207 | 0.10537 | 0.952231 | 2190 | 4 | 4.2433 |
| OMA1 | 6 | 0.24003 | 0.43601 | 0.999994 | 8772 | 2 | 3.9924 | 0.029207 | 0.10537 | 0.952231 | 2191 | 4 | 3.9924 |
| CSPP1 | 6 | 0.15184 | 0.32797 | 0.999994 | 6502 | 1 | 4.0642 | 0.029207 | 0.10537 | 0.952231 | 2192 | 4 | 4.0642 |
| BTBD17 | 5 | 0.68487 | 0.7597 | 0.999994 | 16157 | 1 | 2.974 | 0.029268 | 0.10279 | 0.952143 | 2193 | 4 | 2.974 |
| ENDOD1 | 4 | 0.575 | 0.65219 | 0.999994 | 14616 | 1 | 4.5039 | 0.029271 | 0.090165 | 0.943068 | 2194 | 3 | 4.5039 |
| SLC22A14 | 4 | 0.84887 | 0.8484 | 0.999994 | 18234 | 0 | 0.10922 | 0.029279 | 0.090191 | 0.943068 | 2195 | 1 | 0.10922 |
| TCF21 | 6 | 0.48183 | 0.65641 | 0.999994 | 13166 | 1 | 4.7348 | 0.02928 | 0.10558 | 0.952231 | 2196 | 4 | 4.7348 |
| BARX1 | 6 | 0.90609 | 0.90843 | 0.999994 | 19354 | 1 | 4.867 | 0.02931 | 0.10567 | 0.952231 | 2197 | 4 | 4.867 |
| TNNT3 | 6 | 0.65585 | 0.78208 | 0.999994 | 15724 | 1 | 4.5482 | 0.02931 | 0.10567 | 0.952231 | 2198 | 5 | 4.5482 |
| FICD | 6 | 0.94833 | 0.94835 | 0.999994 | 20294 | 0 | 4.7045 | 0.02931 | 0.10567 | 0.952231 | 2199 | 4 | 4.7045 |
| DYRK4 | 6 | 0.65796 | 0.78372 | 0.999994 | 15753 | 1 | 3.2353 | 0.029316 | 0.10568 | 0.952231 | 2200 | 5 | 3.2353 |
| OR13C4 | 5 | 0.12943 | 0.27175 | 0.998561 | 5928 | 2 | 1.9181 | 0.029345 | 0.10297 | 0.952231 | 2201 | 3 | 1.9181 |
| PNMA1 | 6 | 0.5326 | 0.69138 | 0.999994 | 13945 | 2 | 2.7398 | 0.029362 | 0.10582 | 0.952231 | 2202 | 3 | 2.7398 |
| CDAN1 | 6 | 0.52957 | 0.68924 | 0.999994 | 13894 | 1 | 3.7243 | 0.029397 | 0.10591 | 0.952231 | 2203 | 5 | 3.7243 |
| GAPDHS | 6 | 0.63283 | 0.76447 | 0.999994 | 15412 | 2 | 3.1691 | 0.029412 | 0.10595 | 0.952231 | 2204 | 4 | 3.1691 |
| IL22RA2 | 6 | 0.89914 | 0.903 | 0.999994 | 19206 | 1 | 2.508 | 0.029417 | 0.10597 | 0.952231 | 2205 | 5 | 2.508 |
| FOSL2 | 4 | 0.60151 | 0.66618 | 0.999994 | 14968 | 1 | 6.2291 | 0.029452 | 0.09066 | 0.94477 | 2206 | 3 | 6.2291 |
| BTBD18 | 6 | 0.33378 | 0.54561 | 0.999994 | 10803 | 1 | 1.5444 | 0.029473 | 0.10613 | 0.952916 | 2207 | 4 | 1.5444 |
| ARPC5L | 6 | 0.4474 | 0.63335 | 0.999994 | 12678 | 1 | 3.9914 | 0.029489 | 0.10617 | 0.952916 | 2208 | 4 | 3.9914 |
| TYRO3 | 5 | 0.84831 | 0.85295 | 0.999994 | 18223 | 1 | 6.0521 | 0.029503 | 0.10332 | 0.952231 | 2209 | 4 | 6.0521 |
| CFL1 | 5 | 0.88024 | 0.8805 | 0.999994 | 18837 | 1 | 6.0521 | 0.029503 | 0.10332 | 0.952231 | 2210 | 3 | 6.0521 |
| TRAPPC11 | 5 | 0.19157 | 0.36317 | 0.999994 | 7530 | 2 | 6.0521 | 0.029503 | 0.10332 | 0.952231 | 2211 | 3 | 6.0521 |
| ZNF154 | 6 | 0.63741 | 0.76795 | 0.999994 | 15479 | 2 | 2.0122 | 0.029588 | 0.10647 | 0.953839 | 2212 | 4 | 2.0122 |
| GALNT15 | 6 | 0.94636 | 0.94632 | 0.999994 | 20239 | 0 | 4.1945 | 0.029621 | 0.10656 | 0.953839 | 2213 | 5 | 4.1945 |
| HLA-DQA2 | 6 | 0.89633 | 0.90088 | 0.999994 | 19149 | 1 | 3.4123 | 0.029645 | 0.10663 | 0.953839 | 2214 | 5 | 3.4123 |
| hsa-mir-4661 | 4 | 0.41767 | 0.56381 | 0.999994 | 12214 | 1 | 3.7036 | 0.029657 | 0.091211 | 0.945479 | 2215 | 2 | 3.7036 |
| MTDH | 6 | 0.29787 | 0.50423 | 0.999994 | 10089 | 3 | 2.8818 | 0.029708 | 0.10679 | 0.953839 | 2216 | 3 | 2.8818 |
| TOR3A | 6 | 0.27446 | 0.47697 | 0.999994 | 9577 | 2 | 2.1713 | 0.029709 | 0.1068 | 0.953839 | 2217 | 3 | 2.1713 |
| CELF6 | 6 | 0.90934 | 0.91102 | 0.999994 | 19410 | 1 | 2.5154 | 0.029727 | 0.10685 | 0.953839 | 2218 | 5 | 2.5154 |
| ZNF486 | 5 | 0.91268 | 0.91271 | 0.999994 | 19472 | 0 | 2.151 | 0.029749 | 0.10391 | 0.952231 | 2219 | 4 | 2.151 |
| C11orf96 | 5 | 0.22036 | 0.40377 | 0.999994 | 8298 | 1 | 4.988 | 0.029763 | 0.10395 | 0.952231 | 2220 | 4 | 4.988 |
| PKD2L1 | 6 | 0.15902 | 0.33698 | 0.999994 | 6704 | 3 | -0.74251 | 0.029776 | 0.10698 | 0.953839 | 2221 | 2 | -0.74251 |
| C17orf62 | 5 | 0.76312 | 0.7979 | 0.999994 | 17005 | 1 | 0.99416 | 0.029832 | 0.10411 | 0.952231 | 2222 | 3 | 0.99416 |
| SNRPG | 4 | 0.96736 | 0.96724 | 0.999994 | 20771 | 0 | 6.2459 | 0.029849 | 0.091706 | 0.945479 | 2223 | 4 | 6.2459 |
| NonTargetingControlG | 2 | 0.39886 | 0.41583 | 0.999994 | 11932 | 1 | 3.8949 | 0.029879 | 0.054746 | 0.926769 | 2224 | 1 | 3.8949 |
| HSPB6 | 3 | 0.83329 | 0.83319 | 0.999994 | 17977 | 0 | 7.0916 | 0.029881 | 0.074885 | 0.938203 | 2225 | 2 | 7.0916 |
| CCDC88B | 6 | 0.10022 | 0.24777 | 0.990682 | 4979 | 2 | 0.169 | 0.029886 | 0.10731 | 0.953839 | 2226 | 3 | 0.169 |
| UBE3D | 6 | 0.43694 | 0.62638 | 0.999994 | 12525 | 2 | 4.2715 | 0.029915 | 0.1074 | 0.953839 | 2227 | 4 | 4.2715 |
| CYP4X1 | 5 | 0.43145 | 0.58963 | 0.999994 | 12438 | 2 | 1.0266 | 0.029947 | 0.10439 | 0.952231 | 2228 | 3 | 1.0266 |
| FGF6 | 6 | 0.067303 | 0.1811 | 0.968171 | 3727 | 3 | -0.69701 | 0.029969 | 0.10755 | 0.953839 | 2229 | 2 | -0.69701 |
| CCL28 | 6 | 0.33796 | 0.55034 | 0.999994 | 10896 | 2 | 1.3901 | 0.029992 | 0.10763 | 0.953839 | 2230 | 4 | 1.3901 |
| TMED8 | 6 | 0.81841 | 0.85607 | 0.999994 | 17754 | 1 | 2.678 | 0.029992 | 0.10763 | 0.953839 | 2231 | 4 | 2.678 |
| UGDH | 6 | 0.3196 | 0.52943 | 0.999994 | 10506 | 1 | 0.46914 | 0.029992 | 0.10763 | 0.953839 | 2232 | 3 | 0.46914 |
| XKR3 | 5 | 0.94301 | 0.9432 | 0.999994 | 20151 | 0 | 1.6052 | 0.030016 | 0.10455 | 0.952231 | 2233 | 4 | 1.6052 |
| YME1L1 | 5 | 0.46941 | 0.62164 | 0.999994 | 12997 | 1 | 4.0956 | 0.030018 | 0.10456 | 0.952231 | 2234 | 4 | 4.0956 |
| HINT3 | 5 | 0.5114 | 0.65789 | 0.999994 | 13604 | 2 | 5.4323 | 0.030023 | 0.10457 | 0.952231 | 2235 | 3 | 5.4323 |

| | | | | | | | | | | | | |
|---|---|---|---|---|---|---|---|---|---|---|---|---|
| CCDC40 | 6 | 0.57138 | 0.71895 | 0.999994 | 14567 | 1 | 1.96 | 0.030024 | 0.10772 | 0.953839 | 2236 | 4 | 1.96 |
| FOXO4 | 6 | 0.64202 | 0.77147 | 0.999994 | 15551 | 1 | 4.2557 | 0.030051 | 0.1078 | 0.953839 | 2237 | 5 | 4.2557 |
| C2CD3 | 6 | 0.9716 | 0.97155 | 0.999994 | 20861 | 0 | 3.1319 | 0.030068 | 0.10785 | 0.953839 | 2238 | 5 | 3.1319 |
| HSF5 | 5 | 0.31166 | 0.4905 | 0.999994 | 10352 | 1 | 2.3522 | 0.030074 | 0.10469 | 0.952231 | 2239 | 4 | 2.3522 |
| UBE3B | 5 | 0.96733 | 0.96751 | 0.999994 | 20769 | 0 | 4.0786 | 0.030112 | 0.10479 | 0.952231 | 2240 | 4 | 4.0786 |
| IFI16 | 6 | 0.82031 | 0.85699 | 0.999994 | 17782 | 1 | 3.3734 | 0.030119 | 0.10799 | 0.953839 | 2241 | 5 | 3.3734 |
| SHOX2 | 4 | 0.56369 | 0.64651 | 0.999994 | 14446 | 1 | 5.3748 | 0.030124 | 0.09242 | 0.94761 | 2242 | 3 | 5.3748 |
| SLC38A7 | 6 | 0.071884 | 0.19069 | 0.972248 | 3914 | 1 | 4.5995 | 0.030141 | 0.10805 | 0.953839 | 2243 | 5 | 4.5995 |
| ZNF823 | 6 | 0.38666 | 0.5938 | 0.999994 | 11745 | 2 | 3.9276 | 0.030141 | 0.10805 | 0.953839 | 2244 | 4 | 3.9276 |
| NonTargetingControlG | 1 | 0.9698 | 0.9699 | 0.999994 | 20821 | 0 | 8.6952 | 0.030204 | 0.03012 | 0.890437 | 2245 | 1 | 8.6952 |
| TMEM126B | 6 | 0.18178 | 0.36539 | 0.999994 | 7265 | 2 | 1.7582 | 0.030225 | 0.10828 | 0.953839 | 2246 | 4 | 1.7582 |
| P2RY11 | 6 | 0.60144 | 0.74101 | 0.999994 | 14967 | 2 | 0.29762 | 0.030273 | 0.10842 | 0.953839 | 2247 | 3 | 0.29762 |
| HSD17B8 | 6 | 0.056574 | 0.15835 | 0.967126 | 3276 | 3 | 2.1273 | 0.030314 | 0.10854 | 0.953839 | 2248 | 3 | 2.1273 |
| KCNH6 | 5 | 0.83415 | 0.84247 | 0.999994 | 17991 | 1 | 6.0163 | 0.030324 | 0.10527 | 0.952231 | 2249 | 3 | 6.0163 |
| hsa-mir-8054 | 4 | 0.40316 | 0.54898 | 0.999994 | 12001 | 1 | 3.6665 | 0.030368 | 0.093085 | 0.950707 | 2250 | 2 | 3.6665 |
| GALP | 6 | 0.88026 | 0.88977 | 0.999994 | 18839 | 1 | 3.849 | 0.030377 | 0.10872 | 0.953839 | 2251 | 5 | 3.849 |
| DLC1 | 5 | 0.20396 | 0.38087 | 0.999994 | 7879 | 3 | -0.72462 | 0.030382 | 0.10541 | 0.952231 | 2252 | 2 | -0.72462 |
| SARNP | 5 | 0.17413 | 0.33813 | 0.999994 | 7086 | 2 | 0.82851 | 0.030382 | 0.10541 | 0.952231 | 2253 | 3 | 0.82851 |
| AGER | 6 | 0.30492 | 0.51243 | 0.999994 | 10230 | 2 | 1.2365 | 0.03041 | 0.10882 | 0.953839 | 2254 | 3 | 1.2365 |
| CUL3 | 6 | 0.92528 | 0.92538 | 0.999994 | 19751 | 1 | 4.4234 | 0.030412 | 0.10882 | 0.953839 | 2255 | 5 | 4.4234 |
| SOX10 | 4 | 0.027304 | 0.076 | 0.93914 | 1873 | 2 | 2.5776 | 0.030421 | 0.093231 | 0.950736 | 2256 | 2 | 2.5776 |
| VWA7 | 6 | 0.64294 | 0.77217 | 0.999994 | 15562 | 2 | 4.6705 | 0.03055 | 0.10921 | 0.953839 | 2257 | 4 | 4.6705 |
| CASC1 | 5 | 0.842 | 0.84818 | 0.999994 | 18123 | 1 | 1.6598 | 0.030568 | 0.10586 | 0.952231 | 2258 | 4 | 1.6598 |
| MC2R | 5 | 0.33346 | 0.50834 | 0.999994 | 10795 | 1 | 6.0066 | 0.03058 | 0.10589 | 0.952231 | 2259 | 3 | 6.0066 |
| TMEM178B | 5 | 0.62925 | 0.73702 | 0.999994 | 15371 | 1 | 2.7317 | 0.030606 | 0.10595 | 0.952231 | 2260 | 4 | 2.7317 |
| COQ2 | 6 | 0.24335 | 0.44001 | 0.999994 | 8872 | 1 | 1.9306 | 0.030611 | 0.10937 | 0.953839 | 2261 | 5 | 1.9306 |
| TNRC6C | 6 | 0.22896 | 0.42282 | 0.999994 | 8502 | 2 | 4.2037 | 0.030676 | 0.10956 | 0.953839 | 2262 | 4 | 4.2037 |
| CLDN8 | 6 | 0.98184 | 0.98176 | 0.999994 | 21132 | 0 | 4.4715 | 0.03068 | 0.10957 | 0.953839 | 2263 | 5 | 4.4715 |
| FAM43B | 5 | 0.34596 | 0.5186 | 0.999994 | 11064 | 2 | 0.042603 | 0.030729 | 0.10624 | 0.952995 | 2264 | 2 | 0.042603 |
| RNF145 | 6 | 0.32561 | 0.53624 | 0.999994 | 10635 | 2 | 3.0318 | 0.030739 | 0.10976 | 0.953839 | 2265 | 4 | 3.0318 |
| ZNF512 | 6 | 0.96045 | 0.96051 | 0.999994 | 20592 | 0 | 4.6332 | 0.030739 | 0.10976 | 0.953839 | 2266 | 5 | 4.6332 |
| ARHGEF6 | 6 | 0.27354 | 0.47589 | 0.999994 | 9555 | 2 | 4.4218 | 0.030763 | 0.10983 | 0.953839 | 2267 | 4 | 4.4218 |
| C4orf22 | 6 | 0.94093 | 0.94091 | 0.999994 | 20104 | 0 | 4.8418 | 0.03078 | 0.10988 | 0.953839 | 2268 | 5 | 4.8418 |
| ZSCAN4 | 6 | 0.64396 | 0.77296 | 0.999994 | 15575 | 1 | 2.029 | 0.030785 | 0.1099 | 0.953839 | 2269 | 5 | 2.029 |
| RAB11FIP5 | 6 | 0.38616 | 0.59347 | 0.999994 | 11734 | 2 | 0.1616 | 0.030824 | 0.10999 | 0.953839 | 2270 | 2 | 0.1616 |
| hsa-mir-8056 | 3 | 0.96917 | 0.96926 | 0.999994 | 20807 | 0 | 4.6527 | 0.030832 | 0.077071 | 0.938203 | 2271 | 3 | 4.6527 |
| PRAMEF22 | 4 | 0.94374 | 0.94361 | 0.999994 | 20172 | 0 | 5.8607 | 0.030837 | 0.094355 | 0.950736 | 2272 | 3 | 5.8607 |
| TMEM123 | 6 | 0.97035 | 0.97032 | 0.999994 | 20831 | 0 | 2.4704 | 0.030859 | 0.1101 | 0.953839 | 2273 | 5 | 2.4704 |
| NonTargetingControlG | 2 | 0.89427 | 0.89415 | 0.999994 | 19108 | 0 | 5.8975 | 0.030891 | 0.056502 | 0.926769 | 2274 | 2 | 5.8975 |
| WTAP | 4 | 0.4133 | 0.55934 | 0.999994 | 12160 | 2 | 3.4581 | 0.030947 | 0.094676 | 0.950736 | 2275 | 2 | 3.4581 |
| IGFL4 | 6 | 0.55669 | 0.70844 | 0.999994 | 14313 | 1 | 3.5392 | 0.030986 | 0.11047 | 0.953839 | 2276 | 5 | 3.5392 |
| LY75-CD302 | 3 | 0.77701 | 0.77696 | 0.999994 | 17167 | 0 | 7.0488 | 0.030992 | 0.077452 | 0.938203 | 2277 | 2 | 7.0488 |
| TRAPPC4 | 3 | 0.4558 | 0.51783 | 0.999994 | 12786 | 1 | 7.0488 | 0.030992 | 0.077452 | 0.938203 | 2278 | 2 | 7.0488 |
| hsa-mir-99a | 4 | 0.3909 | 0.53632 | 0.999994 | 11813 | 1 | 4.4515 | 0.031053 | 0.094977 | 0.950736 | 2279 | 3 | 4.4515 |
| FAM212A | 6 | 0.61768 | 0.75307 | 0.999994 | 15180 | 2 | -0.54903 | 0.03107 | 0.11071 | 0.953839 | 2280 | 2 | -0.54903 |
| ARHGEF3 | 6 | 0.48416 | 0.658 | 0.999994 | 13203 | 2 | 0.0086069 | 0.031072 | 0.11072 | 0.953839 | 2281 | 2 | 0.0086069 |
| hsa-mir-1305 | 2 | 0.22928 | 0.28342 | 0.999632 | 8510 | 1 | 2.9228 | 0.031076 | 0.056822 | 0.926769 | 2282 | 1 | 2.9228 |
| NEUROG3 | 6 | 0.9739 | 0.9738 | 0.999994 | 20924 | 0 | 2.5473 | 0.031095 | 0.11078 | 0.953839 | 2283 | 5 | 2.5473 |
| SUPT20HL1 | 5 | 0.76823 | 0.80071 | 0.999994 | 17064 | 1 | 5.9833 | 0.03111 | 0.10715 | 0.953839 | 2284 | 4 | 5.9833 |
| SLC4A3 | 5 | 0.67475 | 0.75532 | 0.999994 | 16000 | 1 | 5.9833 | 0.03111 | 0.10715 | 0.953839 | 2285 | 4 | 5.9833 |
| hsa-mir-365a | 3 | 0.49498 | 0.54285 | 0.999994 | 13340 | 1 | 0.072937 | 0.031111 | 0.077718 | 0.938203 | 2286 | 1 | 0.072937 |
| SALL3 | 5 | 0.25308 | 0.44321 | 0.999994 | 9086 | 1 | 2.2021 | 0.03112 | 0.10717 | 0.953839 | 2287 | 3 | 2.2021 |

| | | | | | | | | | | | | | |
|---|---|---|---|---|---|---|---|---|---|---|---|---|---|
| PCDHB6 | 5 | 0.43532 | 0.5929 | 0.999994 | 12506 | 2 | 1.0301 | 0.031234 | 0.10743 | 0.953839 | 2288 | 3 | 1.0301 |
| RIT1 | 6 | 0.32766 | 0.53856 | 0.999994 | 10673 | 1 | 2.1437 | 0.031237 | 0.11121 | 0.953839 | 2289 | 4 | 2.1437 |
| RHOT2 | 6 | 0.44707 | 0.63313 | 0.999994 | 12675 | 1 | 4.5692 | 0.031257 | 0.11126 | 0.953839 | 2290 | 4 | 4.5692 |
| BBS10 | 6 | 0.5045 | 0.67185 | 0.999994 | 13503 | 2 | 0.76754 | 0.031292 | 0.11135 | 0.953839 | 2291 | 4 | 0.76754 |
| PLXND1 | 5 | 0.31189 | 0.49069 | 0.999994 | 10356 | 1 | 0.65193 | 0.031304 | 0.10759 | 0.953839 | 2292 | 3 | 0.65193 |
| MR1 | 6 | 0.85693 | 0.87569 | 0.999994 | 18392 | 1 | 1.7638 | 0.031305 | 0.11138 | 0.953839 | 2293 | 5 | 1.7638 |
| SLC12A9 | 4 | 0.91743 | 0.91722 | 0.999994 | 19575 | 0 | 5.7602 | 0.031321 | 0.095738 | 0.950736 | 2294 | 3 | 5.7602 |
| PDPK1 | 4 | 0.96868 | 0.96857 | 0.999994 | 20799 | 0 | 5.5337 | 0.031324 | 0.095746 | 0.950736 | 2295 | 4 | 5.5337 |
| ORMDL2 | 4 | 0.96866 | 0.96856 | 0.999994 | 20798 | 0 | 2.3476 | 0.031335 | 0.09578 | 0.950736 | 2296 | 4 | 2.3476 |
| AURKB | 3 | 0.96862 | 0.96872 | 0.999994 | 20797 | 0 | 6.3809 | 0.03138 | 0.078333 | 0.938203 | 2297 | 3 | 6.3809 |
| DMRTC1 | 6 | 0.47509 | 0.65187 | 0.999994 | 13064 | 1 | 4.0908 | 0.031384 | 0.11161 | 0.953839 | 2298 | 5 | 4.0908 |
| NUDT9 | 5 | 0.28712 | 0.4706 | 0.999994 | 9856 | 2 | 5.9781 | 0.031393 | 0.10781 | 0.953839 | 2299 | 3 | 5.9781 |
| NonTargetingControlG | 2 | 0.9686 | 0.96862 | 0.999994 | 20796 | 0 | 6.9689 | 0.031396 | 0.057391 | 0.926769 | 2300 | 2 | 6.9689 |
| GPR126 | 6 | 0.96823 | 0.96824 | 0.999994 | 20789 | 0 | 3.1902 | 0.031403 | 0.11167 | 0.953839 | 2301 | 5 | 3.1902 |
| ENAH | 6 | 0.96982 | 0.96981 | 0.999994 | 20822 | 0 | 0.45318 | 0.03143 | 0.11174 | 0.953839 | 2302 | 3 | 0.45318 |
| hsa-mir-5008 | 4 | 0.61591 | 0.67405 | 0.999994 | 15159 | 1 | 2.3508 | 0.031434 | 0.096053 | 0.950736 | 2303 | 2 | 2.3508 |
| SPATA17 | 5 | 0.059016 | 0.16043 | 0.967126 | 3381 | 1 | 0.91476 | 0.031442 | 0.10792 | 0.953839 | 2304 | 3 | 0.91476 |
| DOCK3 | 6 | 0.04716 | 0.13786 | 0.957317 | 2861 | 1 | 2.5498 | 0.031447 | 0.11178 | 0.953839 | 2305 | 5 | 2.5498 |
| hsa-mir-7158 | 3 | 0.27064 | 0.40049 | 0.999994 | 9490 | 1 | 7.0361 | 0.031458 | 0.078491 | 0.938203 | 2306 | 2 | 7.0361 |
| hsa-mir-26a-2 | 4 | 0.25579 | 0.3978 | 0.999994 | 9143 | 1 | 4.7286 | 0.031461 | 0.096125 | 0.950736 | 2307 | 3 | 4.7286 |
| CIC | 6 | 0.19706 | 0.38432 | 0.999994 | 7675 | 1 | 2.5046 | 0.031504 | 0.11196 | 0.953839 | 2308 | 5 | 2.5046 |
| BMPR1B | 5 | 0.92229 | 0.92247 | 0.999994 | 19684 | 0 | 6.0759 | 0.031513 | 0.10807 | 0.953839 | 2309 | 3 | 6.0759 |
| hsa-mir-452 | 2 | 0.58109 | 0.58122 | 0.999994 | 14694 | 1 | 4.3319 | 0.031522 | 0.057613 | 0.928172 | 2310 | 1 | 4.3319 |
| FZD5 | 4 | 0.81342 | 0.81352 | 0.999994 | 17682 | 1 | 3.5672 | 0.031549 | 0.096376 | 0.950736 | 2311 | 2 | 3.5672 |
| KIAA1731 | 4 | 0.46048 | 0.5991 | 0.999994 | 12854 | 1 | 3.676 | 0.031549 | 0.096376 | 0.950736 | 2312 | 2 | 3.676 |
| HPS1 | 4 | 0.96844 | 0.96834 | 0.999994 | 20793 | 0 | 2.3086 | 0.031558 | 0.096402 | 0.950736 | 2313 | 4 | 2.3086 |
| CALML4 | 6 | 0.63585 | 0.76679 | 0.999994 | 15456 | 1 | 1.5976 | 0.031568 | 0.11213 | 0.953839 | 2314 | 5 | 1.5976 |
| TMEM86B | 6 | 0.97008 | 0.97005 | 0.999994 | 20825 | 0 | 2.0855 | 0.031569 | 0.11214 | 0.953839 | 2315 | 5 | 2.0855 |
| NonTargetingControlG | 2 | 0.96843 | 0.96845 | 0.999994 | 20791 | 0 | 6.1392 | 0.03157 | 0.057696 | 0.92824 | 2316 | 2 | 6.1392 |
| COQ4 | 6 | 0.89943 | 0.90324 | 0.999994 | 19211 | 1 | 4.4442 | 0.031592 | 0.1122 | 0.953839 | 2317 | 4 | 4.4442 |
| CHRNA10 | 6 | 0.19063 | 0.37632 | 0.999994 | 7503 | 2 | 5.0541 | 0.031592 | 0.1122 | 0.953839 | 2318 | 4 | 5.0541 |
| NPAS1 | 6 | 0.55755 | 0.70908 | 0.999994 | 14326 | 2 | 4.4453 | 0.031592 | 0.1122 | 0.953839 | 2319 | 4 | 4.4453 |
| OVOL3 | 6 | 0.98022 | 0.98012 | 0.999994 | 21098 | 0 | 2.6266 | 0.031633 | 0.11232 | 0.953839 | 2320 | 5 | 2.6266 |
| SHISA3 | 6 | 0.9851 | 0.98507 | 0.999994 | 21227 | 0 | 2.4655 | 0.03165 | 0.11236 | 0.953839 | 2321 | 5 | 2.4655 |
| UBE2C | 6 | 0.035016 | 0.11055 | 0.951052 | 2268 | 3 | -0.71859 | 0.03165 | 0.11236 | 0.953839 | 2322 | 2 | -0.71859 |
| MRGPRX4 | 5 | 0.52647 | 0.67096 | 0.999994 | 13838 | 1 | 3.8042 | 0.031663 | 0.10844 | 0.953839 | 2323 | 4 | 3.8042 |
| FMO4 | 5 | 0.87966 | 0.87994 | 0.999994 | 18825 | 1 | 5.9596 | 0.031688 | 0.1085 | 0.953839 | 2324 | 3 | 5.9596 |
| CHML | 5 | 0.67956 | 0.75736 | 0.999994 | 16083 | 1 | 5.9596 | 0.031688 | 0.1085 | 0.953839 | 2325 | 4 | 5.9596 |
| THBD | 5 | 0.26342 | 0.45151 | 0.999994 | 9324 | 2 | 5.9596 | 0.031688 | 0.1085 | 0.953839 | 2326 | 3 | 5.9596 |
| TMEM121 | 5 | 0.15347 | 0.30799 | 0.999994 | 6548 | 2 | 5.9596 | 0.031688 | 0.1085 | 0.953839 | 2327 | 3 | 5.9596 |
| KLK13 | 6 | 0.94006 | 0.94002 | 0.999994 | 20082 | 0 | 2.9114 | 0.031694 | 0.11249 | 0.954575 | 2328 | 5 | 2.9114 |
| hsa-mir-6504 | 4 | 0.68921 | 0.71744 | 0.999994 | 16204 | 1 | 6.1072 | 0.031815 | 0.097082 | 0.950736 | 2329 | 3 | 6.1072 |
| DGAT2 | 6 | 0.98554 | 0.98552 | 0.999994 | 21241 | 0 | 1.6497 | 0.031823 | 0.11284 | 0.954984 | 2330 | 5 | 1.6497 |
| OASL | 6 | 0.10153 | 0.2503 | 0.992497 | 5020 | 3 | -0.53842 | 0.031843 | 0.11291 | 0.954984 | 2331 | 3 | -0.53842 |
| TRAF7 | 6 | 0.34606 | 0.55959 | 0.999994 | 11069 | 1 | 3.5483 | 0.03186 | 0.11296 | 0.954984 | 2332 | 5 | 3.5483 |
| SOCS2 | 5 | 0.89016 | 0.89029 | 0.999994 | 19041 | 0 | 3.609 | 0.031885 | 0.10897 | 0.953839 | 2333 | 4 | 3.609 |
| CKS1B | 5 | 0.22438 | 0.40938 | 0.999994 | 8390 | 2 | 1.6322 | 0.031893 | 0.10899 | 0.953839 | 2334 | 3 | 1.6322 |
| RNF44 | 5 | 0.92915 | 0.92941 | 0.999994 | 19839 | 0 | 3.2785 | 0.031896 | 0.109 | 0.953839 | 2335 | 4 | 3.2785 |
| TGDS | 3 | 0.57319 | 0.59699 | 0.999994 | 14593 | 1 | 2.1692 | 0.031914 | 0.079536 | 0.938962 | 2336 | 2 | 2.1692 |
| RFXAP | 5 | 0.12824 | 0.26991 | 0.998561 | 5885 | 2 | 2.6671 | 0.031947 | 0.10911 | 0.953839 | 2337 | 3 | 2.6671 |
| ZFAND5 | 6 | 0.40013 | 0.60244 | 0.999994 | 11950 | 2 | 4.5068 | 0.031961 | 0.11324 | 0.954984 | 2338 | 4 | 4.5068 |
| IGSF23 | 5 | 0.49483 | 0.64342 | 0.999994 | 13338 | 1 | 5.9586 | 0.031965 | 0.10915 | 0.953839 | 2339 | 4 | 5.9586 |

| | | | | | | | | | | | | | |
|---|---|---|---|---|---|---|---|---|---|---|---|---|---|
| ANPEP | 5 | 0.41728 | 0.57786 | 0.999994 | 12207 | 1 | 2.8177 | 0.03202 | 0.10928 | 0.953839 | 2340 | 3 | 2.8177 |
| ANTXR1 | 6 | 0.40439 | 0.60523 | 0.999994 | 12026 | 2 | -0.29582 | 0.032022 | 0.11343 | 0.954984 | 2341 | 2 | -0.29582 |
| HOXC12 | 6 | 0.32766 | 0.53856 | 0.999994 | 10675 | 1 | 4.6993 | 0.032036 | 0.11346 | 0.954984 | 2342 | 4 | 4.6993 |
| CCDC34 | 6 | 0.98341 | 0.98336 | 0.999994 | 21167 | 0 | 5.2526 | 0.032041 | 0.11348 | 0.954984 | 2343 | 5 | 5.2526 |
| TXNDC9 | 4 | 0.56734 | 0.64832 | 0.999994 | 14503 | 1 | 4.7309 | 0.032109 | 0.097852 | 0.950736 | 2344 | 3 | 4.7309 |
| TPT1 | 6 | 0.45762 | 0.64012 | 0.999994 | 12816 | 2 | 4.3006 | 0.032135 | 0.11376 | 0.954984 | 2345 | 4 | 4.3006 |
| MSX2 | 6 | 0.97707 | 0.97701 | 0.999994 | 21016 | 0 | 3.5563 | 0.032142 | 0.11377 | 0.954984 | 2346 | 5 | 3.5563 |
| TMEM184B | 5 | 0.88466 | 0.88479 | 0.999994 | 18929 | 1 | 5.6538 | 0.032177 | 0.10963 | 0.953839 | 2347 | 4 | 5.6538 |
| SGCZ | 5 | 0.94783 | 0.94803 | 0.999994 | 20282 | 0 | 2.0805 | 0.032177 | 0.10963 | 0.953839 | 2348 | 4 | 2.0805 |
| RPP25 | 5 | 0.62242 | 0.73446 | 0.999994 | 15265 | 1 | 3.0157 | 0.032181 | 0.10964 | 0.953839 | 2349 | 3 | 3.0157 |
| SNW1 | 5 | 0.42789 | 0.58668 | 0.999994 | 12382 | 1 | 2.8318 | 0.032185 | 0.10966 | 0.953839 | 2350 | 4 | 2.8318 |
| hsa-mir-6783 | 3 | 0.8694 | 0.86946 | 0.999994 | 18627 | 0 | 7.0133 | 0.032195 | 0.08018 | 0.938962 | 2351 | 2 | 7.0133 |
| PPP6R2 | 4 | 0.96778 | 0.96767 | 0.999994 | 20778 | 0 | 2.031 | 0.032224 | 0.098182 | 0.950736 | 2352 | 4 | 2.031 |
| hsa-mir-3617 | 4 | 0.96777 | 0.96767 | 0.999994 | 20777 | 0 | 1.6479 | 0.032229 | 0.09821 | 0.950736 | 2353 | 4 | 1.6479 |
| ADPRHL2 | 5 | 0.94262 | 0.94281 | 0.999994 | 20142 | 0 | 5.9388 | 0.03225 | 0.1098 | 0.953839 | 2354 | 3 | 5.9388 |
| SPIRE2 | 5 | 0.62613 | 0.73588 | 0.999994 | 15326 | 1 | 5.9388 | 0.03225 | 0.1098 | 0.953839 | 2355 | 4 | 5.9388 |
| METRNL | 5 | 0.76098 | 0.79672 | 0.999994 | 16990 | 1 | 4.0801 | 0.032259 | 0.10982 | 0.953839 | 2356 | 4 | 4.0801 |
| C2CD4A | 6 | 0.98527 | 0.98523 | 0.999994 | 21233 | 0 | 2.165 | 0.032291 | 0.11421 | 0.954984 | 2357 | 5 | 2.165 |
| CCDC120 | 5 | 0.89632 | 0.89645 | 0.999994 | 19147 | 0 | 5.359 | 0.032331 | 0.10998 | 0.953839 | 2358 | 4 | 5.359 |
| PEX11A | 6 | 0.31397 | 0.52298 | 0.999994 | 10394 | 2 | 3.7013 | 0.032386 | 0.11447 | 0.954984 | 2359 | 4 | 3.7013 |
| TBC1D12 | 6 | 0.025881 | 0.089093 | 0.94395 | 1795 | 1 | 2.225 | 0.0324 | 0.1145 | 0.954984 | 2360 | 5 | 2.225 |
| CXXC4 | 6 | 0.95939 | 0.95946 | 0.999994 | 20569 | 0 | 2.7914 | 0.032429 | 0.11458 | 0.954984 | 2361 | 5 | 2.7914 |
| hsa-mir-4633 | 2 | 0.96755 | 0.96756 | 0.999994 | 20774 | 0 | 6.1159 | 0.032446 | 0.059206 | 0.930397 | 2362 | 2 | 6.1159 |
| FLOT2 | 6 | 0.89848 | 0.90246 | 0.999994 | 19194 | 1 | 3.7895 | 0.032451 | 0.11465 | 0.954984 | 2363 | 5 | 3.7895 |
| SCN9A | 6 | 0.64212 | 0.77153 | 0.999994 | 15552 | 1 | 0.36588 | 0.032451 | 0.11465 | 0.954984 | 2364 | 3 | 0.36588 |
| LRP3 | 6 | 0.43842 | 0.62733 | 0.999994 | 12546 | 2 | 0.66486 | 0.032476 | 0.11472 | 0.954984 | 2365 | 3 | 0.66486 |
| MSC | 6 | 0.84053 | 0.86689 | 0.999994 | 18097 | 1 | 3.9597 | 0.03258 | 0.11501 | 0.954984 | 2366 | 5 | 3.9597 |
| RPH3AL | 6 | 0.26924 | 0.47083 | 0.999994 | 9450 | 3 | -0.32165 | 0.032601 | 0.11507 | 0.954984 | 2367 | 2 | -0.32165 |
| DPP9 | 6 | 0.57799 | 0.72372 | 0.999994 | 14657 | 2 | 3.1888 | 0.032629 | 0.11514 | 0.954984 | 2368 | 3 | 3.1888 |
| GALNT10 | 6 | 0.61856 | 0.75374 | 0.999994 | 15196 | 2 | 4.7888 | 0.032629 | 0.11514 | 0.954984 | 2369 | 4 | 4.7888 |
| KIAA0430 | 6 | 0.67333 | 0.79575 | 0.999994 | 15976 | 2 | 3.6332 | 0.032629 | 0.11514 | 0.954984 | 2370 | 4 | 3.6332 |
| RFPL2 | 5 | 0.75017 | 0.79095 | 0.999994 | 16857 | 1 | 1.3919 | 0.032659 | 0.11077 | 0.953839 | 2371 | 3 | 1.3919 |
| HEXDC | 6 | 0.73123 | 0.82202 | 0.999994 | 16617 | 1 | 4.2903 | 0.03273 | 0.11542 | 0.954984 | 2372 | 5 | 4.2903 |
| SRGN | 5 | 0.26079 | 0.44939 | 0.999994 | 9263 | 2 | 0.29405 | 0.032751 | 0.111 | 0.953839 | 2373 | 2 | 0.29405 |
| DOM3Z | 6 | 0.535 | 0.69305 | 0.999994 | 13981 | 1 | 1.4035 | 0.032762 | 0.11552 | 0.954984 | 2374 | 4 | 1.4035 |
| CDX2 | 5 | 0.9671 | 0.96728 | 0.999994 | 20764 | 0 | 5.9188 | 0.03281 | 0.11116 | 0.953839 | 2375 | 4 | 5.9188 |
| NCAPH2 | 4 | 0.96718 | 0.96706 | 0.999994 | 20766 | 0 | 3.0299 | 0.03282 | 0.099802 | 0.951893 | 2376 | 4 | 3.0299 |
| RNF152 | 6 | 0.94674 | 0.9467 | 0.999994 | 20256 | 0 | 1.9198 | 0.032844 | 0.11575 | 0.954984 | 2377 | 5 | 1.9198 |
| SSR2 | 6 | 0.63158 | 0.76354 | 0.999994 | 15391 | 2 | 4.6823 | 0.032851 | 0.11577 | 0.954984 | 2378 | 4 | 4.6823 |
| KRT35 | 6 | 0.50045 | 0.669 | 0.999994 | 13428 | 2 | 4.7647 | 0.032851 | 0.11577 | 0.954984 | 2379 | 4 | 4.7647 |
| SPINK8 | 6 | 0.63318 | 0.76474 | 0.999994 | 15418 | 2 | 3.3131 | 0.032851 | 0.11577 | 0.954984 | 2380 | 3 | 3.3131 |
| AGL | 5 | 0.57869 | 0.71717 | 0.999994 | 14666 | 2 | -0.52158 | 0.032868 | 0.11129 | 0.953839 | 2381 | 2 | -0.52158 |
| KPTN | 5 | 0.46359 | 0.61669 | 0.999994 | 12912 | 1 | 3.3864 | 0.032868 | 0.11129 | 0.953839 | 2382 | 4 | 3.3864 |
| FASTKD1 | 6 | 0.86429 | 0.87989 | 0.999994 | 18527 | 1 | 0.91018 | 0.032883 | 0.11586 | 0.954984 | 2383 | 3 | 0.91018 |
| ZER1 | 5 | 0.33817 | 0.51224 | 0.999994 | 10903 | 1 | 5.8358 | 0.032885 | 0.11134 | 0.953839 | 2384 | 4 | 5.8358 |
| ABCD2 | 6 | 0.067503 | 0.18155 | 0.968171 | 3735 | 4 | -1.5373 | 0.032889 | 0.11589 | 0.954984 | 2385 | 2 | -1.5373 |
| TXNDC8 | 5 | 0.54592 | 0.68812 | 0.999994 | 14148 | 1 | 5.8607 | 0.032919 | 0.11143 | 0.953839 | 2386 | 4 | 5.8607 |
| NACA2 | 4 | 0.77291 | 0.77814 | 0.999994 | 17121 | 1 | 3.5403 | 0.032941 | 0.10014 | 0.951893 | 2387 | 2 | 3.5403 |
| PRSS54 | 6 | 0.18475 | 0.36907 | 0.999994 | 7336 | 2 | 2.4418 | 0.032999 | 0.11621 | 0.954984 | 2388 | 3 | 2.4418 |
| CTHRC1 | 5 | 0.90855 | 0.90857 | 0.999994 | 19396 | 0 | 2.2731 | 0.033006 | 0.11165 | 0.953839 | 2389 | 4 | 2.2731 |
| MALT1 | 5 | 0.80773 | 0.82446 | 0.999994 | 17592 | 1 | 2.8646 | 0.033028 | 0.11169 | 0.953839 | 2390 | 4 | 2.8646 |
| ST3GAL2 | 6 | 0.011788 | 0.045018 | 0.891621 | 978 | 3 | -2.2903 | 0.033054 | 0.11638 | 0.954984 | 2391 | 3 | -2.2903 |

| | | | | | | | | | | | | | |
|---|---|---|---|---|---|---|---|---|---|---|---|---|---|
| hsa-mir-6081 | 4 | 0.82653 | 0.82611 | 0.999994 | 17876 | 1 | 3.9992 | 0.033076 | 0.10052 | 0.951893 | 2392 | 2 | 3.9992 |
| KRT28 | 6 | 0.89819 | 0.90225 | 0.999994 | 19186 | 1 | 3.9423 | 0.033083 | 0.11646 | 0.954984 | 2393 | 4 | 3.9423 |
| SYT15 | 6 | 0.41763 | 0.6138 | 0.999994 | 12211 | 2 | 3.4583 | 0.033083 | 0.11646 | 0.954984 | 2394 | 3 | 3.4583 |
| SMC1B | 6 | 0.2459 | 0.44305 | 0.999994 | 8921 | 1 | 4.4213 | 0.033083 | 0.11646 | 0.954984 | 2395 | 5 | 4.4213 |
| TCEB3B | 4 | 0.96686 | 0.96676 | 0.999994 | 20757 | 0 | 1.959 | 0.033143 | 0.1007 | 0.951893 | 2396 | 4 | 1.959 |
| RABEP2 | 4 | 0.17741 | 0.31744 | 0.999994 | 7164 | 1 | 5.8039 | 0.033148 | 0.10072 | 0.951893 | 2397 | 3 | 5.8039 |
| SUSD5 | 6 | 0.45027 | 0.63531 | 0.999994 | 12715 | 2 | 1.5057 | 0.033164 | 0.11668 | 0.954984 | 2398 | 4 | 1.5057 |
| hsa-mir-603 | 4 | 0.96681 | 0.9667 | 0.999994 | 20756 | 0 | 1.9917 | 0.033192 | 0.10084 | 0.951893 | 2399 | 4 | 1.9917 |
| ANGPTL7 | 6 | 0.28456 | 0.48874 | 0.999994 | 9798 | 1 | 2.9514 | 0.033233 | 0.11686 | 0.954984 | 2400 | 5 | 2.9514 |
| TMEM132A | 5 | 0.6351 | 0.73929 | 0.999994 | 15445 | 1 | 1.9422 | 0.033233 | 0.11217 | 0.953839 | 2401 | 3 | 1.9422 |
| MKS1 | 4 | 0.085278 | 0.18841 | 0.969543 | 4422 | 2 | 2.0437 | 0.033234 | 0.10095 | 0.951893 | 2402 | 2 | 2.0437 |
| MYO6 | 6 | 0.97389 | 0.9738 | 0.999994 | 20923 | 0 | 2.0234 | 0.033251 | 0.11692 | 0.954984 | 2403 | 5 | 2.0234 |
| NGB | 6 | 0.30845 | 0.51658 | 0.999994 | 10292 | 2 | 4.9894 | 0.03326 | 0.11694 | 0.954984 | 2404 | 4 | 4.9894 |
| STARD7 | 5 | 0.96753 | 0.9677 | 0.999994 | 20772 | 0 | 4.1171 | 0.033275 | 0.11227 | 0.953839 | 2405 | 4 | 4.1171 |
| PRAMEF5 | 5 | 0.47124 | 0.6232 | 0.999994 | 13018 | 2 | 0.28833 | 0.033279 | 0.11228 | 0.953839 | 2406 | 2 | 0.28833 |
| FPR1 | 6 | 0.8797 | 0.8894 | 0.999994 | 18827 | 1 | 1.7744 | 0.033311 | 0.11708 | 0.955294 | 2407 | 5 | 1.7744 |
| ZNF624 | 4 | 0.96667 | 0.96656 | 0.999994 | 20755 | 0 | 1.8228 | 0.033333 | 0.10123 | 0.951893 | 2408 | 4 | 1.8228 |
| NonTargetingControlG | 2 | 0.96664 | 0.96664 | 0.999994 | 20754 | 0 | 7.1423 | 0.033357 | 0.060785 | 0.9326 | 2409 | 2 | 7.1423 |
| ZNF528 | 6 | 0.33378 | 0.54561 | 0.999994 | 10805 | 1 | 1.7298 | 0.033363 | 0.11723 | 0.956124 | 2410 | 5 | 1.7298 |
| GCNT3 | 6 | 0.96095 | 0.96102 | 0.999994 | 20603 | 0 | 3.9592 | 0.033405 | 0.11735 | 0.956586 | 2411 | 4 | 3.9592 |
| COL21A1 | 6 | 0.97375 | 0.97366 | 0.999994 | 20918 | 0 | 2.4159 | 0.033409 | 0.11737 | 0.956586 | 2412 | 4 | 2.4159 |
| MT1B | 5 | 0.61184 | 0.73048 | 0.999994 | 15106 | 1 | 3.3023 | 0.033422 | 0.1126 | 0.95486 | 2413 | 4 | 3.3023 |
| HM13 | 3 | 0.96651 | 0.96661 | 0.999994 | 20750 | 0 | 3.6666 | 0.033489 | 0.083094 | 0.939912 | 2414 | 3 | 3.6666 |
| MTPN | 5 | 0.023651 | 0.07391 | 0.93914 | 1692 | 2 | -0.37144 | 0.033499 | 0.11279 | 0.954984 | 2415 | 2 | -0.37144 |
| AIRE | 4 | 0.023827 | 0.068724 | 0.93914 | 1701 | 1 | 4.9646 | 0.03352 | 0.10175 | 0.951893 | 2416 | 3 | 4.9646 |
| LENG1 | 4 | 0.96647 | 0.96635 | 0.999994 | 20748 | 0 | 2.5081 | 0.033531 | 0.10178 | 0.951893 | 2417 | 4 | 2.5081 |
| FAM104A | 4 | 0.29258 | 0.43542 | 0.999994 | 9967 | 2 | 0.18357 | 0.033532 | 0.10178 | 0.951893 | 2418 | 2 | 0.18357 |
| PCDHGB2 | 2 | 0.96644 | 0.96644 | 0.999994 | 20747 | 0 | 6.4883 | 0.033564 | 0.061132 | 0.9326 | 2419 | 2 | 6.4883 |
| MTNR1A | 6 | 0.92493 | 0.92504 | 0.999994 | 19740 | 1 | 4.9087 | 0.033589 | 0.11784 | 0.958281 | 2420 | 4 | 4.9087 |
| KCNJ2 | 6 | 0.97797 | 0.97791 | 0.999994 | 21039 | 0 | 3.3589 | 0.033612 | 0.11792 | 0.958281 | 2421 | 5 | 3.3589 |
| KIAA1586 | 6 | 0.94209 | 0.94208 | 0.999994 | 20128 | 0 | 4.09 | 0.033621 | 0.11795 | 0.958281 | 2422 | 4 | 4.09 |
| FGF10 | 6 | 0.86639 | 0.88114 | 0.999994 | 18566 | 1 | 0.47468 | 0.033631 | 0.11798 | 0.958281 | 2423 | 3 | 0.47468 |
| MYH14 | 5 | 0.40878 | 0.57086 | 0.999994 | 12089 | 1 | 0.64665 | 0.033668 | 0.11317 | 0.954984 | 2424 | 3 | 0.64665 |
| ACKR3 | 5 | 0.76072 | 0.79659 | 0.999994 | 16988 | 1 | 1.5424 | 0.033677 | 0.11319 | 0.954984 | 2425 | 3 | 1.5424 |
| MSH3 | 4 | 0.73404 | 0.74809 | 0.999994 | 16653 | 1 | 4.9443 | 0.033678 | 0.10216 | 0.951893 | 2426 | 3 | 4.9443 |
| hsa-mir-4747 | 4 | 0.94149 | 0.94137 | 0.999994 | 20118 | 0 | 5.2336 | 0.033683 | 0.10217 | 0.951893 | 2427 | 3 | 5.2336 |
| RPP40 | 6 | 0.84626 | 0.86994 | 0.999994 | 18192 | 1 | 2.0493 | 0.03373 | 0.11825 | 0.9593 | 2428 | 5 | 2.0493 |
| MAU2 | 4 | 0.96623 | 0.96612 | 0.999994 | 20742 | 0 | 2.585 | 0.033765 | 0.10239 | 0.952143 | 2429 | 4 | 2.585 |
| SPRR2F | 5 | 0.1441 | 0.29397 | 0.999994 | 6301 | 2 | 0.14184 | 0.033783 | 0.11342 | 0.954984 | 2430 | 2 | 0.14184 |
| NonTargetingControlG | 2 | 0.96615 | 0.96616 | 0.999994 | 20740 | 0 | 6.2615 | 0.033852 | 0.061601 | 0.933948 | 2431 | 2 | 6.2615 |
| AMN | 6 | 0.61589 | 0.75177 | 0.999994 | 15158 | 2 | 1.467 | 0.033852 | 0.1186 | 0.9593 | 2432 | 3 | 1.467 |
| TTLL5 | 6 | 0.040727 | 0.12356 | 0.951052 | 2546 | 3 | 1.7008 | 0.033866 | 0.11863 | 0.9593 | 2433 | 3 | 1.7008 |
| C9orf57 | 6 | 0.045976 | 0.13523 | 0.954328 | 2808 | 2 | 3.4118 | 0.033866 | 0.11863 | 0.9593 | 2434 | 3 | 3.4118 |
| OR10S1 | 6 | 0.38897 | 0.59531 | 0.999994 | 11782 | 2 | 3.1815 | 0.033866 | 0.11863 | 0.9593 | 2435 | 3 | 3.1815 |
| SH3GLB1 | 5 | 0.88961 | 0.88973 | 0.999994 | 19029 | 0 | 2.8073 | 0.033893 | 0.11367 | 0.954984 | 2436 | 4 | 2.8073 |
| NonTargetingControlG | 2 | 0.85685 | 0.85679 | 0.999994 | 18387 | 0 | 5.3621 | 0.033896 | 0.061674 | 0.933948 | 2437 | 2 | 5.3621 |
| CTAG2 | 4 | 0.21865 | 0.35991 | 0.999994 | 8246 | 1 | 5.3019 | 0.033933 | 0.10284 | 0.952143 | 2438 | 3 | 5.3019 |
| hsa-mir-4641 | 4 | 0.95871 | 0.95867 | 0.999994 | 20555 | 0 | 4.8386 | 0.033933 | 0.10284 | 0.952143 | 2439 | 4 | 4.8386 |
| KIRREL2 | 6 | 0.96638 | 0.96643 | 0.999994 | 20743 | 0 | 2.0548 | 0.033934 | 0.11884 | 0.959988 | 2440 | 5 | 2.0548 |
| RAPGEF1 | 5 | 0.080461 | 0.19533 | 0.975797 | 4248 | 3 | -1.3442 | 0.033944 | 0.11379 | 0.954984 | 2441 | 1 | -1.3442 |
| CCL27 | 5 | 0.7778 | 0.80617 | 0.999994 | 17180 | 1 | 5.8779 | 0.033968 | 0.11385 | 0.954984 | 2442 | 3 | 5.8779 |
| PMEPA1 | 5 | 0.49568 | 0.64415 | 0.999994 | 13352 | 1 | 3.3115 | 0.033984 | 0.11388 | 0.954984 | 2443 | 4 | 3.3115 |

| | | | | | | | | | | | | | |
|---|---|---|---|---|---|---|---|---|---|---|---|---|---|
| SIDT1 | 6 | 0.7023 | 0.81272 | 0.999994 | 16331 | 1 | 2.8727 | 0.034044 | 0.11914 | 0.960516 | 2444 | 4 | 2.8727 |
| MAST2 | 5 | 0.14054 | 0.28863 | 0.999632 | 6204 | 2 | -0.30316 | 0.034058 | 0.11405 | 0.954984 | 2445 | 2 | -0.30316 |
| PRKCQ | 5 | 0.07811 | 0.19154 | 0.974071 | 4153 | 2 | 5.3435 | 0.034111 | 0.11416 | 0.954984 | 2446 | 3 | 5.3435 |
| EIF4EBP2 | 5 | 0.91917 | 0.91932 | 0.999994 | 19611 | 0 | 1.6905 | 0.034111 | 0.11416 | 0.954984 | 2447 | 3 | 1.6905 |
| HTATSF1 | 6 | 0.24787 | 0.44539 | 0.999994 | 8971 | 2 | 3.4198 | 0.034131 | 0.11936 | 0.961045 | 2448 | 3 | 3.4198 |
| hsa-mir-6809 | 2 | 0.34171 | 0.36935 | 0.999994 | 10975 | 1 | 3.4044 | 0.034192 | 0.062215 | 0.933948 | 2449 | 1 | 3.4044 |
| GAST | 4 | 0.32099 | 0.46441 | 0.999994 | 10533 | 1 | 0.19205 | 0.03423 | 0.10367 | 0.952231 | 2450 | 1 | 0.19205 |
| NonTargetingControlG | 2 | 0.86145 | 0.86145 | 0.999994 | 18474 | 0 | 5.3776 | 0.034276 | 0.062357 | 0.933948 | 2451 | 2 | 5.3776 |
| NOXA1 | 5 | 0.93574 | 0.93597 | 0.999994 | 19986 | 0 | 3.5315 | 0.034279 | 0.11456 | 0.954984 | 2452 | 4 | 3.5315 |
| CNOT6 | 5 | 0.35024 | 0.52213 | 0.999994 | 11163 | 1 | 5.8608 | 0.034285 | 0.11458 | 0.954984 | 2453 | 4 | 5.8608 |
| ECI1 | 5 | 0.9257 | 0.92593 | 0.999994 | 19761 | 0 | 5.8608 | 0.034285 | 0.11458 | 0.954984 | 2454 | 3 | 5.8608 |
| OR9I1 | 6 | 0.45452 | 0.63808 | 0.999994 | 12764 | 1 | 2.5833 | 0.034359 | 0.11997 | 0.961431 | 2455 | 5 | 2.5833 |
| GUK1 | 4 | 0.92604 | 0.92585 | 0.999994 | 19767 | 0 | 4.0431 | 0.034401 | 0.10415 | 0.952231 | 2456 | 2 | 4.0431 |
| GPR97 | 5 | 0.35211 | 0.52363 | 0.999994 | 11196 | 1 | 3.0494 | 0.034406 | 0.11486 | 0.954984 | 2457 | 4 | 3.0494 |
| TMEM106A | 4 | 0.90204 | 0.90171 | 0.999994 | 19253 | 0 | 4.8107 | 0.034417 | 0.10419 | 0.952231 | 2458 | 3 | 4.8107 |
| QSOX1 | 6 | 0.84325 | 0.86835 | 0.999994 | 18144 | 1 | 3.7199 | 0.034424 | 0.12016 | 0.961431 | 2459 | 4 | 3.7199 |
| C10orf114 | 5 | 0.4694 | 0.62163 | 0.999994 | 12996 | 2 | 1.1094 | 0.034425 | 0.1149 | 0.954984 | 2460 | 3 | 1.1094 |
| R3HDML | 6 | 0.62677 | 0.7599 | 0.999994 | 15331 | 2 | -0.5816 | 0.034483 | 0.12034 | 0.961431 | 2461 | 2 | -0.5816 |
| FAM219B | 6 | 0.60117 | 0.74081 | 0.999994 | 14963 | 1 | 2.4857 | 0.034483 | 0.12034 | 0.961431 | 2462 | 4 | 2.4857 |
| SERPINC1 | 6 | 0.1791 | 0.36202 | 0.999994 | 7198 | 3 | -0.52462 | 0.034483 | 0.12034 | 0.961431 | 2463 | 2 | -0.52462 |
| hsa-mir-184 | 3 | 0.86446 | 0.86443 | 0.999994 | 18531 | 0 | 2.4788 | 0.034486 | 0.085343 | 0.942567 | 2464 | 2 | 2.4788 |
| DNAH2 | 4 | 0.69692 | 0.72239 | 0.999994 | 16269 | 1 | 4.4582 | 0.034487 | 0.10438 | 0.952231 | 2465 | 3 | 4.4582 |
| GSC2 | 5 | 0.78434 | 0.80997 | 0.999994 | 17271 | 1 | 3.2075 | 0.034495 | 0.11507 | 0.954984 | 2466 | 4 | 3.2075 |
| LRCH4 | 5 | 0.21189 | 0.39205 | 0.999994 | 8063 | 1 | 5.1519 | 0.034498 | 0.11508 | 0.954984 | 2467 | 4 | 5.1519 |
| NonTargetingControlG | 2 | 0.96549 | 0.96551 | 0.999994 | 20727 | 0 | 6.6668 | 0.034505 | 0.062731 | 0.933948 | 2468 | 2 | 6.6668 |
| TUSC3 | 6 | 0.61806 | 0.75338 | 0.999994 | 15189 | 1 | 3.7773 | 0.034507 | 0.12042 | 0.961504 | 2469 | 5 | 3.7773 |
| FAT1 | 5 | 0.56654 | 0.70649 | 0.999994 | 14490 | 1 | 5.857 | 0.034521 | 0.11514 | 0.954984 | 2470 | 3 | 5.857 |
| SLIT2 | 3 | 0.68667 | 0.68881 | 0.999994 | 16177 | 1 | 2.2923 | 0.034528 | 0.08544 | 0.942567 | 2471 | 2 | 2.2923 |
| GALNTL6 | 5 | 0.25406 | 0.44396 | 0.999994 | 9104 | 2 | 5.8424 | 0.034652 | 0.11543 | 0.954984 | 2472 | 3 | 5.8424 |
| ACTR3C | 6 | 0.081933 | 0.2113 | 0.979587 | 4295 | 3 | -0.74853 | 0.034675 | 0.12089 | 0.962329 | 2473 | 3 | -0.74853 |
| hsa-mir-6071 | 4 | 0.96532 | 0.96521 | 0.999994 | 20725 | 0 | 4.0918 | 0.034677 | 0.10488 | 0.952231 | 2474 | 4 | 4.0918 |
| SLC7A9 | 4 | 0.22551 | 0.36686 | 0.999994 | 8413 | 2 | 3.1045 | 0.034716 | 0.10498 | 0.952231 | 2475 | 2 | 3.1045 |
| ASB6 | 6 | 0.21288 | 0.40359 | 0.999994 | 8087 | 1 | 2.1777 | 0.034721 | 0.12102 | 0.962423 | 2476 | 4 | 2.1777 |
| LOC643355 | 5 | 0.50981 | 0.65646 | 0.999994 | 13574 | 2 | 0.52225 | 0.034746 | 0.11563 | 0.954984 | 2477 | 3 | 0.52225 |
| ARHGAP23 | 5 | 0.86794 | 0.86926 | 0.999994 | 18599 | 1 | 0.6831 | 0.034746 | 0.11563 | 0.954984 | 2478 | 3 | 0.6831 |
| PGBD1 | 6 | 0.12904 | 0.29907 | 0.999994 | 5917 | 2 | 3.8895 | 0.034753 | 0.12112 | 0.96264 | 2479 | 4 | 3.8895 |
| PLEKHA7 | 6 | 0.0050146 | 0.021204 | 0.812607 | 522 | 3 | -3.1914 | 0.034758 | 0.12114 | 0.96264 | 2480 | 2 | -3.1914 |
| RPS27L | 6 | 0.62504 | 0.75862 | 0.999994 | 15307 | 1 | 4.0934 | 0.034777 | 0.12119 | 0.962751 | 2481 | 4 | 4.0934 |
| FRMD6 | 5 | 0.18385 | 0.35216 | 0.999994 | 7313 | 1 | 0.3088 | 0.034838 | 0.11586 | 0.954984 | 2482 | 2 | 0.3088 |
| C21orf91 | 6 | 0.74274 | 0.82592 | 0.999994 | 16775 | 1 | 0.10113 | 0.034868 | 0.12146 | 0.962935 | 2483 | 2 | 0.10113 |
| PHF13 | 5 | 0.030412 | 0.091523 | 0.94395 | 2027 | 4 | -1.6744 | 0.034883 | 0.11596 | 0.954984 | 2484 | 1 | -1.6744 |
| DHX37 | 5 | 0.30726 | 0.48693 | 0.999994 | 10273 | 1 | 2.8731 | 0.03489 | 0.11598 | 0.954984 | 2485 | 4 | 2.8731 |
| TMEM249 | 6 | 0.13458 | 0.30614 | 0.999994 | 6041 | 1 | 4.0383 | 0.03493 | 0.12162 | 0.962935 | 2486 | 4 | 4.0383 |
| FBLN7 | 5 | 0.00022222 | 0.0010482 | 0.352868 | 62 | 4 | -7.0568 | 0.034952 | 0.11613 | 0.954984 | 2487 | 1 | -7.0568 |
| NR2E3 | 5 | 0.82789 | 0.83798 | 0.999994 | 17895 | 1 | -0.085145 | 0.034962 | 0.11615 | 0.954984 | 2488 | 2 | -0.085145 |
| KCNQ4 | 5 | 0.34404 | 0.51703 | 0.999994 | 11029 | 1 | 1.6887 | 0.034962 | 0.11615 | 0.954984 | 2489 | 3 | 1.6887 |
| THAP9 | 5 | 0.93384 | 0.93408 | 0.999994 | 19949 | 0 | 1.0153 | 0.034962 | 0.11615 | 0.954984 | 2490 | 4 | 1.0153 |
| ZNF599 | 6 | 0.40206 | 0.60371 | 0.999994 | 11978 | 1 | 2.5351 | 0.034969 | 0.12173 | 0.962935 | 2491 | 4 | 2.5351 |
| CTXN3 | 6 | 0.41604 | 0.61278 | 0.999994 | 12195 | 2 | 3.7039 | 0.034969 | 0.12173 | 0.962935 | 2492 | 4 | 3.7039 |
| NOL4 | 6 | 0.25205 | 0.45038 | 0.999994 | 9058 | 3 | -0.71662 | 0.035005 | 0.12183 | 0.962935 | 2493 | 2 | -0.71662 |
| FAM81B | 6 | 0.79629 | 0.84632 | 0.999994 | 17432 | 1 | 1.4282 | 0.035005 | 0.12183 | 0.962935 | 2494 | 3 | 1.4282 |
| hsa-mir-3158-1 | 4 | 0.96499 | 0.96488 | 0.999994 | 20719 | 0 | 1.8828 | 0.035006 | 0.10575 | 0.952231 | 2495 | 4 | 1.8828 |

| | | | | | | | | | | | | | |
|---|---|---|---|---|---|---|---|---|---|---|---|---|---|
| CD300A | 5 | 0.80155 | 0.82048 | 0.999994 | 17499 | 1 | 5.835 | 0.035052 | 0.11634 | 0.954984 | 2496 | 3 | 5.835 |
| hsa-mir-1973 | 3 | 0.16022 | 0.2617 | 0.995257 | 6745 | 1 | 6.9182 | 0.03506 | 0.086642 | 0.943068 | 2497 | 2 | 6.9182 |
| DEPDC7 | 5 | 0.81932 | 0.83215 | 0.999994 | 17770 | 1 | 2.8188 | 0.035081 | 0.1164 | 0.954984 | 2498 | 4 | 2.8188 |
| ZNF302 | 6 | 0.93923 | 0.93918 | 0.999994 | 20055 | 0 | 2.5765 | 0.035085 | 0.12206 | 0.962935 | 2499 | 5 | 2.5765 |
| DMWD | 6 | 0.60876 | 0.74643 | 0.999994 | 15053 | 2 | 1.1048 | 0.035087 | 0.12206 | 0.962935 | 2500 | 3 | 1.1048 |
| MMP27 | 6 | 0.19977 | 0.3876 | 0.999994 | 7756 | 3 | -0.52683 | 0.035087 | 0.12206 | 0.962935 | 2501 | 2 | -0.52683 |
| GTF2H2 | 6 | 0.080135 | 0.20757 | 0.978852 | 4235 | 2 | 3.952 | 0.03511 | 0.12214 | 0.962935 | 2502 | 4 | 3.952 |
| HK1 | 5 | 0.59007 | 0.72256 | 0.999994 | 14824 | 1 | 3.9952 | 0.035133 | 0.11652 | 0.954984 | 2503 | 4 | 3.9952 |
| ERMARD | 6 | 0.98462 | 0.98455 | 0.999994 | 21213 | 0 | 2.6093 | 0.035155 | 0.12226 | 0.962935 | 2504 | 5 | 2.6093 |
| TRIM54 | 6 | 0.82691 | 0.86005 | 0.999994 | 17884 | 1 | 3.1041 | 0.035158 | 0.12227 | 0.962935 | 2505 | 5 | 3.1041 |
| hsa-mir-4742 | 4 | 0.71132 | 0.73203 | 0.999994 | 16412 | 1 | 4.4364 | 0.03516 | 0.10618 | 0.952916 | 2506 | 3 | 4.4364 |
| PITRM1 | 6 | 0.52748 | 0.68777 | 0.999994 | 13851 | 2 | 1.5227 | 0.03517 | 0.1223 | 0.962935 | 2507 | 3 | 1.5227 |
| NDUFS6 | 5 | 0.17756 | 0.34312 | 0.999994 | 7168 | 1 | 0.75794 | 0.035204 | 0.11668 | 0.954984 | 2508 | 3 | 0.75794 |
| B4GALT6 | 5 | 0.085165 | 0.20282 | 0.977973 | 4415 | 3 | -1.6207 | 0.035204 | 0.11668 | 0.954984 | 2509 | 1 | -1.6207 |
| C19orf48 | 5 | 0.78492 | 0.81031 | 0.999994 | 17278 | 1 | 2.5607 | 0.035245 | 0.11678 | 0.954984 | 2510 | 4 | 2.5607 |
| C9orf9 | 4 | 0.89499 | 0.89468 | 0.999994 | 19127 | 0 | 3.7246 | 0.035258 | 0.10643 | 0.953839 | 2511 | 3 | 3.7246 |
| AAAS | 5 | 0.8559 | 0.859 | 0.999994 | 18368 | 1 | 3.487 | 0.035296 | 0.11691 | 0.954984 | 2512 | 4 | 3.487 |
| OBP2B | 5 | 0.92455 | 0.92477 | 0.999994 | 19731 | 0 | 3.8261 | 0.035304 | 0.11693 | 0.954984 | 2513 | 4 | 3.8261 |
| KDM5D | 6 | 0.16307 | 0.34211 | 0.999994 | 6820 | 2 | 0.062108 | 0.035307 | 0.12269 | 0.962935 | 2514 | 2 | 0.062108 |
| ANO6 | 6 | 0.20117 | 0.38929 | 0.999994 | 7795 | 2 | 1.5487 | 0.035307 | 0.12269 | 0.962935 | 2515 | 4 | 1.5487 |
| WSCD1 | 5 | 0.32886 | 0.50461 | 0.999994 | 10699 | 1 | 4.7593 | 0.035315 | 0.11695 | 0.954984 | 2516 | 3 | 4.7593 |
| ANXA8L2 | 3 | 0.058598 | 0.12754 | 0.951052 | 3369 | 1 | 6.9059 | 0.035354 | 0.08733 | 0.943068 | 2517 | 2 | 6.9059 |
| GRIN3B | 6 | 0.93311 | 0.9332 | 0.999994 | 19928 | 0 | 4.1906 | 0.035384 | 0.12292 | 0.962935 | 2518 | 4 | 4.1906 |
| NonTargetingControlG | 2 | 0.90571 | 0.9057 | 0.999994 | 19346 | 0 | 5.7547 | 0.035387 | 0.064267 | 0.933948 | 2519 | 2 | 5.7547 |
| OR1D2 | 6 | 0.27446 | 0.47697 | 0.999994 | 9578 | 1 | 3.0857 | 0.035438 | 0.12307 | 0.962935 | 2520 | 3 | 3.0857 |
| ZNF440 | 6 | 0.97598 | 0.97593 | 0.999994 | 20988 | 0 | 4.376 | 0.035438 | 0.12307 | 0.962935 | 2521 | 4 | 4.376 |
| PIGX | 6 | 0.6405 | 0.77034 | 0.999994 | 15517 | 1 | 1.8522 | 0.035472 | 0.12317 | 0.962935 | 2522 | 4 | 1.8522 |
| OR2T12 | 6 | 0.52923 | 0.689 | 0.999994 | 13884 | 2 | 2.1034 | 0.035472 | 0.12317 | 0.962935 | 2523 | 4 | 2.1034 |
| A1CF | 6 | 0.13419 | 0.30565 | 0.999994 | 6027 | 3 | -1.2505 | 0.035472 | 0.12317 | 0.962935 | 2524 | 2 | -1.2505 |
| CREG1 | 6 | 0.90513 | 0.90768 | 0.999994 | 19332 | 1 | 4.0546 | 0.035483 | 0.1232 | 0.962935 | 2525 | 4 | 4.0546 |
| RABGAP1 | 6 | 0.73794 | 0.82429 | 0.999994 | 16701 | 1 | 4.2739 | 0.035483 | 0.1232 | 0.962935 | 2526 | 5 | 4.2739 |
| CMTR2 | 6 | 0.59814 | 0.73856 | 0.999994 | 14928 | 1 | 1.5752 | 0.035551 | 0.12339 | 0.963164 | 2527 | 4 | 1.5752 |
| CHST1 | 5 | 0.88402 | 0.88415 | 0.999994 | 18904 | 1 | 5.8017 | 0.035558 | 0.11749 | 0.957213 | 2528 | 4 | 5.8017 |
| FAIM3 | 5 | 0.072147 | 0.18184 | 0.968322 | 3923 | 3 | -1.579 | 0.035593 | 0.11758 | 0.957238 | 2529 | 2 | -1.579 |
| NLRX1 | 6 | 0.32959 | 0.54075 | 0.999994 | 10712 | 2 | 0.89764 | 0.035609 | 0.12356 | 0.963164 | 2530 | 3 | 0.89764 |
| NAP1L5 | 6 | 0.93876 | 0.93873 | 0.999994 | 20047 | 0 | 2.9511 | 0.035629 | 0.12361 | 0.963212 | 2531 | 3 | 2.9511 |
| MAN1A1 | 5 | 0.85113 | 0.85518 | 0.999994 | 18281 | 1 | 4.5294 | 0.035685 | 0.1178 | 0.958281 | 2532 | 3 | 4.5294 |
| PTPRF | 6 | 0.48769 | 0.66039 | 0.999994 | 13249 | 1 | 3.8335 | 0.035726 | 0.1239 | 0.963705 | 2533 | 4 | 3.8335 |
| CHP1 | 6 | 0.05574 | 0.15652 | 0.967126 | 3235 | 2 | 2.9508 | 0.035726 | 0.1239 | 0.963705 | 2534 | 3 | 2.9508 |
| IFT80 | 5 | 0.96224 | 0.96239 | 0.999994 | 20637 | 0 | 5.8087 | 0.035735 | 0.11792 | 0.958281 | 2535 | 4 | 5.8087 |
| NonTargetingControlG | 2 | 0.96418 | 0.96424 | 0.999994 | 20708 | 0 | 6.1226 | 0.035819 | 0.065003 | 0.933948 | 2536 | 2 | 6.1226 |
| SLC4A1AP | 4 | 0.70798 | 0.7298 | 0.999994 | 16388 | 1 | 5.7003 | 0.035822 | 0.10792 | 0.953839 | 2537 | 3 | 5.7003 |
| NR6A1 | 4 | 0.21173 | 0.35278 | 0.999994 | 8060 | 1 | 4.34 | 0.035823 | 0.10792 | 0.953839 | 2538 | 3 | 4.34 |
| hsa-mir-199a-1 | 3 | 0.96417 | 0.96429 | 0.999994 | 20707 | 0 | 4.5924 | 0.035827 | 0.088377 | 0.943068 | 2539 | 3 | 4.5924 |
| C7orf50 | 4 | 0.80057 | 0.80173 | 0.999994 | 17489 | 1 | 5.2548 | 0.035834 | 0.10795 | 0.953839 | 2540 | 3 | 5.2548 |
| MED30 | 3 | 0.70017 | 0.70119 | 0.999994 | 16304 | 1 | 3.2399 | 0.035853 | 0.088437 | 0.943068 | 2541 | 2 | 3.2399 |
| WEE1 | 3 | 0.14652 | 0.24402 | 0.988309 | 6364 | 2 | -1.6516 | 0.035908 | 0.088568 | 0.943068 | 2542 | 1 | -1.6516 |
| RGP1 | 4 | 0.96407 | 0.96396 | 0.999994 | 20704 | 0 | 1.2299 | 0.035932 | 0.10823 | 0.953839 | 2543 | 4 | 1.2299 |
| TAF11 | 5 | 0.01815 | 0.059043 | 0.931872 | 1370 | 3 | -3.3036 | 0.035937 | 0.11839 | 0.9593 | 2544 | 2 | -3.3036 |
| MRPL23 | 5 | 0.0047979 | 0.020002 | 0.808847 | 507 | 3 | -6.9524 | 0.035975 | 0.11848 | 0.9593 | 2545 | 2 | -6.9524 |
| FBXO39 | 6 | 0.042197 | 0.12684 | 0.951052 | 2624 | 2 | 3.6941 | 0.036007 | 0.12466 | 0.964228 | 2546 | 4 | 3.6941 |
| ZNF501 | 6 | 0.31671 | 0.52613 | 0.999994 | 10446 | 3 | 2.8231 | 0.036007 | 0.12466 | 0.964228 | 2547 | 3 | 2.8231 |

| | | | | | | | | | | | | | |
|---|---|---|---|---|---|---|---|---|---|---|---|---|---|
| RAB34 | 4 | 0.25448 | 0.39652 | 0.999994 | 9115 | 1 | 4.7816 | 0.036017 | 0.10846 | 0.953839 | 2548 | 3 | 4.7816 |
| PSMG2 | 4 | 0.79926 | 0.80057 | 0.999994 | 17470 | 1 | 5.6741 | 0.036017 | 0.10846 | 0.953839 | 2549 | 3 | 5.6741 |
| NDRG4 | 5 | 0.31443 | 0.49282 | 0.999994 | 10407 | 2 | 0.37722 | 0.036028 | 0.1186 | 0.9593 | 2550 | 2 | 0.37722 |
| NonTargetingControlG | 2 | 0.62195 | 0.62171 | 0.999994 | 15258 | 0 | 4.3876 | 0.036082 | 0.065468 | 0.933948 | 2551 | 1 | 4.3876 |
| FGF16 | 5 | 0.73492 | 0.78309 | 0.999994 | 16660 | 1 | 3.0145 | 0.036122 | 0.11883 | 0.959988 | 2552 | 4 | 3.0145 |
| KCTD2 | 6 | 0.35764 | 0.57264 | 0.999994 | 11300 | 3 | -0.38888 | 0.03613 | 0.12499 | 0.964228 | 2553 | 2 | -0.38888 |
| ADCY7 | 5 | 0.22364 | 0.40837 | 0.999994 | 8372 | 1 | 3.9412 | 0.036142 | 0.11888 | 0.959988 | 2554 | 4 | 3.9412 |
| ODF3B | 6 | 0.14046 | 0.3136 | 0.999994 | 6202 | 2 | 0.75949 | 0.036197 | 0.12517 | 0.964228 | 2555 | 4 | 0.75949 |
| EMC7 | 3 | 0.9638 | 0.96393 | 0.999994 | 20701 | 0 | 2.668 | 0.036198 | 0.089208 | 0.943068 | 2556 | 3 | 2.668 |
| MYH1 | 6 | 0.97629 | 0.97624 | 0.999994 | 20999 | 0 | 3.9615 | 0.036273 | 0.12538 | 0.964228 | 2557 | 5 | 3.9615 |
| SCRT1 | 3 | 0.79484 | 0.79471 | 0.999994 | 17411 | 0 | 1.3773 | 0.036295 | 0.089423 | 0.943068 | 2558 | 2 | 1.3773 |
| ZNF18 | 5 | 0.67833 | 0.75684 | 0.999994 | 16065 | 1 | 3.8342 | 0.036301 | 0.11925 | 0.961041 | 2559 | 4 | 3.8342 |
| PDPR | 6 | 0.005672 | 0.023585 | 0.832699 | 568 | 2 | 4.1381 | 0.036305 | 0.12548 | 0.964228 | 2560 | 4 | 4.1381 |
| TPM3 | 6 | 0.43123 | 0.62267 | 0.999994 | 12433 | 1 | 2.1164 | 0.036306 | 0.12548 | 0.964228 | 2561 | 4 | 2.1164 |
| LIN9 | 6 | 0.93978 | 0.93974 | 0.999994 | 20073 | 0 | 3.4497 | 0.03636 | 0.12565 | 0.964228 | 2562 | 5 | 3.4497 |
| ATXN7L1 | 5 | 0.49886 | 0.64692 | 0.999994 | 13408 | 1 | 0.58652 | 0.036392 | 0.11946 | 0.961045 | 2563 | 3 | 0.58652 |
| hsa-mir-6775 | 3 | 0.9636 | 0.96373 | 0.999994 | 20693 | 0 | 3.8927 | 0.036401 | 0.089665 | 0.943068 | 2564 | 3 | 3.8927 |
| OR10AG1 | 6 | 0.87496 | 0.88641 | 0.999994 | 18733 | 1 | 2.5311 | 0.036404 | 0.12576 | 0.964228 | 2565 | 5 | 2.5311 |
| NonTargetingControlG | 2 | 0.9636 | 0.96366 | 0.999994 | 20692 | 0 | 5.4771 | 0.036405 | 0.066042 | 0.933948 | 2566 | 2 | 5.4771 |
| SNRNP200 | 4 | 0.040957 | 0.10369 | 0.946994 | 2560 | 2 | -3.0032 | 0.036414 | 0.10951 | 0.953839 | 2567 | 1 | -3.0032 |
| MRPS23 | 4 | 0.79736 | 0.79885 | 0.999994 | 17442 | 1 | -0.56729 | 0.036414 | 0.10951 | 0.953839 | 2568 | 1 | -0.56729 |
| PDPN | 4 | 0.96356 | 0.96345 | 0.999994 | 20689 | 0 | 3.5004 | 0.036443 | 0.10959 | 0.953839 | 2569 | 4 | 3.5004 |
| hsa-mir-944 | 3 | 0.96355 | 0.96369 | 0.999994 | 20688 | 0 | 2.6821 | 0.036446 | 0.089758 | 0.943068 | 2570 | 3 | 2.6821 |
| LRP8 | 4 | 0.63668 | 0.68557 | 0.999994 | 15470 | 1 | 4.8045 | 0.03646 | 0.10963 | 0.953839 | 2571 | 3 | 4.8045 |
| RIMS3 | 6 | 0.41959 | 0.61513 | 0.999994 | 12246 | 1 | 1.0603 | 0.036504 | 0.12604 | 0.964228 | 2572 | 3 | 1.0603 |
| MYH13 | 6 | 0.66912 | 0.7925 | 0.999994 | 15914 | 1 | 4.1637 | 0.03657 | 0.12622 | 0.964228 | 2573 | 5 | 4.1637 |
| ZSWIM6 | 5 | 0.96144 | 0.96158 | 0.999994 | 20617 | 0 | 2.0862 | 0.036571 | 0.11989 | 0.961431 | 2574 | 4 | 2.0862 |
| MTSS1 | 5 | 0.048669 | 0.13719 | 0.957161 | 2917 | 2 | -0.25195 | 0.036578 | 0.1199 | 0.961431 | 2575 | 2 | -0.25195 |
| MED27 | 3 | 0.96339 | 0.96353 | 0.999994 | 20681 | 0 | 5.6045 | 0.036608 | 0.090108 | 0.943068 | 2576 | 3 | 5.6045 |
| SERPINA7 | 4 | 0.85332 | 0.85287 | 0.999994 | 18321 | 0 | 2.6433 | 0.036652 | 0.11016 | 0.953839 | 2577 | 2 | 2.6433 |
| RPA2 | 2 | 0.96332 | 0.96339 | 0.999994 | 20679 | 0 | 5.9029 | 0.036679 | 0.066514 | 0.933948 | 2578 | 2 | 5.9029 |
| PPID | 4 | 0.96329 | 0.96318 | 0.999994 | 20677 | 0 | 2.0085 | 0.036708 | 0.11031 | 0.953839 | 2579 | 4 | 2.0085 |
| FOXP3 | 5 | 0.9148 | 0.91489 | 0.999994 | 19505 | 0 | 5.5005 | 0.036739 | 0.12029 | 0.961431 | 2580 | 4 | 5.5005 |
| hsa-mir-4449 | 3 | 0.96326 | 0.9634 | 0.999994 | 20676 | 0 | 6.5728 | 0.036745 | 0.090424 | 0.94376 | 2581 | 3 | 6.5728 |
| ZBTB42 | 5 | 0.57564 | 0.71452 | 0.999994 | 14627 | 1 | 5.7702 | 0.036752 | 0.12033 | 0.961431 | 2582 | 4 | 5.7702 |
| KDELC1 | 6 | 0.98019 | 0.98009 | 0.999994 | 21096 | 0 | 2.0133 | 0.036761 | 0.12674 | 0.964228 | 2583 | 5 | 2.0133 |
| SF1 | 6 | 0.0091005 | 0.035878 | 0.875006 | 800 | 4 | -4.7022 | 0.036761 | 0.12674 | 0.964228 | 2584 | 2 | -4.7022 |
| SLAIN2 | 6 | 0.25238 | 0.45077 | 0.999994 | 9066 | 2 | 3.1687 | 0.036761 | 0.12674 | 0.964228 | 2585 | 4 | 3.1687 |
| RAB40C | 5 | 0.95556 | 0.95573 | 0.999994 | 20469 | 0 | 2.4205 | 0.036767 | 0.12036 | 0.961431 | 2586 | 4 | 2.4205 |
| KRTAP19-5 | 6 | 0.43867 | 0.62752 | 0.999994 | 12551 | 1 | 3.6916 | 0.036778 | 0.12679 | 0.964228 | 2587 | 4 | 3.6916 |
| ADCY6 | 4 | 0.1386 | 0.27748 | 0.999379 | 6156 | 1 | 4.9212 | 0.036808 | 0.11057 | 0.953839 | 2588 | 3 | 4.9212 |
| AMIGO1 | 4 | 0.56664 | 0.64798 | 0.999994 | 14491 | 1 | 4.6052 | 0.036842 | 0.11067 | 0.953839 | 2589 | 3 | 4.6052 |
| FAM170B | 5 | 0.16997 | 0.33212 | 0.999994 | 6982 | 2 | -0.11073 | 0.036851 | 0.12056 | 0.961983 | 2590 | 2 | -0.11073 |
| CAPN13 | 6 | 0.79112 | 0.84414 | 0.999994 | 17364 | 1 | 2.0887 | 0.036856 | 0.12702 | 0.964228 | 2591 | 5 | 2.0887 |
| HSP90AB1 | 4 | 0.96313 | 0.96302 | 0.999994 | 20673 | 0 | 2.3824 | 0.036873 | 0.11075 | 0.953839 | 2592 | 4 | 2.3824 |
| NonTargetingControlG | 2 | 0.96308 | 0.96315 | 0.999994 | 20670 | 0 | 6.6293 | 0.036918 | 0.066947 | 0.933948 | 2593 | 2 | 6.6293 |
| RABGGTA | 2 | 0.96308 | 0.96315 | 0.999994 | 20668 | 0 | 5.9244 | 0.036918 | 0.066947 | 0.933948 | 2594 | 2 | 5.9244 |
| BUD31 | 2 | 0.96308 | 0.96315 | 0.999994 | 20669 | 0 | 6.0463 | 0.036918 | 0.066947 | 0.933948 | 2595 | 2 | 6.0463 |
| PINX1 | 6 | 0.90943 | 0.91109 | 0.999994 | 19413 | 1 | 4.3567 | 0.036932 | 0.12722 | 0.964228 | 2596 | 5 | 4.3567 |
| KIAA1199 | 6 | 0.95354 | 0.95355 | 0.999994 | 20424 | 0 | 2.071 | 0.036939 | 0.12724 | 0.964228 | 2597 | 4 | 2.071 |
| NUP50 | 4 | 0.96304 | 0.96292 | 0.999994 | 20664 | 0 | 2.0552 | 0.036959 | 0.11098 | 0.953839 | 2598 | 4 | 2.0552 |
| MCM10 | 6 | 0.96813 | 0.96814 | 0.999994 | 20788 | 0 | 4.5798 | 0.036966 | 0.12731 | 0.964228 | 2599 | 4 | 4.5798 |

| | | | | | | | | | | | | | |
|---|---|---|---|---|---|---|---|---|---|---|---|---|---|
| RGS4 | 6 | 0.34062 | 0.55336 | 0.999994 | 10951 | 3 | 0.33742 | 0.037008 | 0.12742 | 0.964228 | 2600 | 3 | 0.33742 |
| HTR1E | 5 | 0.48115 | 0.63165 | 0.999994 | 13153 | 1 | 1.5929 | 0.037011 | 0.12092 | 0.962329 | 2601 | 3 | 1.5929 |
| RFTN1 | 5 | 0.73055 | 0.78092 | 0.999994 | 16607 | 1 | 1.0156 | 0.037011 | 0.12092 | 0.962329 | 2602 | 3 | 1.0156 |
| AMIGO2 | 4 | 0.3416 | 0.48566 | 0.999994 | 10973 | 1 | 3.568 | 0.037024 | 0.11115 | 0.953839 | 2603 | 2 | 3.568 |
| CLDND1 | 5 | 0.18544 | 0.35446 | 0.999994 | 7352 | 2 | 5.7547 | 0.037034 | 0.12096 | 0.962329 | 2604 | 3 | 5.7547 |
| FAM213B | 5 | 0.95312 | 0.95329 | 0.999994 | 20412 | 0 | 5.7547 | 0.037034 | 0.12096 | 0.962329 | 2605 | 4 | 5.7547 |
| TMED9 | 6 | 0.23585 | 0.43107 | 0.999994 | 8666 | 1 | 3.2246 | 0.037045 | 0.12752 | 0.964228 | 2606 | 5 | 3.2246 |
| C9orf171 | 6 | 0.17102 | 0.35202 | 0.999994 | 7008 | 3 | -0.72621 | 0.037079 | 0.12761 | 0.964228 | 2607 | 2 | -0.72621 |
| AK6 | 5 | 0.94879 | 0.949 | 0.999994 | 20306 | 0 | 4.8199 | 0.037172 | 0.12128 | 0.962897 | 2608 | 4 | 4.8199 |
| hsa-mir-641 | 4 | 0.96195 | 0.96184 | 0.999994 | 20628 | 0 | 4.3584 | 0.037184 | 0.11157 | 0.953839 | 2609 | 4 | 4.3584 |
| SPCS3 | 4 | 0.75922 | 0.76715 | 0.999994 | 16970 | 1 | 4.3482 | 0.037184 | 0.11157 | 0.953839 | 2610 | 3 | 4.3482 |
| MRGBP | 6 | 0.59915 | 0.73931 | 0.999994 | 14943 | 2 | 4.0569 | 0.037188 | 0.1279 | 0.964228 | 2611 | 4 | 4.0569 |
| JKAMP | 4 | 0.74095 | 0.75319 | 0.999994 | 16751 | 1 | -0.29778 | 0.037202 | 0.11162 | 0.953839 | 2612 | 1 | -0.29778 |
| hsa-mir-3658 | 4 | 0.96274 | 0.96264 | 0.999994 | 20654 | 0 | 1.9497 | 0.037262 | 0.11177 | 0.953839 | 2613 | 4 | 1.9497 |
| CSNK2B | 3 | 0.96273 | 0.96287 | 0.999994 | 20653 | 0 | 2.6879 | 0.03727 | 0.091632 | 0.945479 | 2614 | 3 | 2.6879 |
| HAMP | 6 | 0.61894 | 0.75403 | 0.999994 | 15204 | 1 | 4.1125 | 0.037282 | 0.12817 | 0.964228 | 2615 | 5 | 4.1125 |
| APOA1 | 6 | 0.76715 | 0.83476 | 0.999994 | 17053 | 1 | 3.8678 | 0.037302 | 0.12822 | 0.964228 | 2616 | 5 | 3.8678 |
| PXK | 6 | 0.085468 | 0.21845 | 0.980699 | 4428 | 4 | -0.94207 | 0.037337 | 0.12831 | 0.964228 | 2617 | 2 | -0.94207 |
| PPIE | 3 | 0.96252 | 0.96266 | 0.999994 | 20644 | 0 | 6.8425 | 0.037371 | 0.091878 | 0.946116 | 2618 | 3 | 6.8425 |
| TIMM22 | 4 | 0.96262 | 0.96253 | 0.999994 | 20648 | 0 | 2.395 | 0.037377 | 0.11208 | 0.953839 | 2619 | 4 | 2.395 |
| HEMGN | 6 | 0.78504 | 0.8417 | 0.999994 | 17279 | 1 | 2.3987 | 0.037401 | 0.12848 | 0.964228 | 2620 | 5 | 2.3987 |
| PYROXD1 | 4 | 0.83743 | 0.83691 | 0.999994 | 18041 | 0 | 4.2513 | 0.037413 | 0.11217 | 0.953839 | 2621 | 3 | 4.2513 |
| ZNF326 | 4 | 0.96256 | 0.96246 | 0.999994 | 20646 | 0 | 1.108 | 0.037441 | 0.11225 | 0.953839 | 2622 | 4 | 1.108 |
| TGM3 | 6 | 0.67517 | 0.79719 | 0.999994 | 16006 | 1 | 2.975 | 0.037446 | 0.1286 | 0.964228 | 2623 | 5 | 2.975 |
| hsa-mir-371a | 4 | 0.96255 | 0.96245 | 0.999994 | 20645 | 0 | 1.5842 | 0.037451 | 0.11227 | 0.953839 | 2624 | 4 | 1.5842 |
| SNRPA1 | 2 | 0.60299 | 0.60279 | 0.999994 | 14987 | 0 | 4.3193 | 0.03747 | 0.06791 | 0.935447 | 2625 | 1 | 4.3193 |
| BRCA1 | 6 | 0.014479 | 0.054027 | 0.922076 | 1163 | 1 | 4.1053 | 0.037519 | 0.12881 | 0.964228 | 2626 | 5 | 4.1053 |
| ACADS | 6 | 0.89107 | 0.89707 | 0.999994 | 19056 | 1 | 3.608 | 0.037531 | 0.12884 | 0.964228 | 2627 | 4 | 3.608 |
| SEMA3F | 6 | 0.034925 | 0.11034 | 0.951052 | 2265 | 4 | -2.0021 | 0.037556 | 0.12891 | 0.964228 | 2628 | 1 | -2.0021 |
| hsa-mir-563 | 4 | 0.96239 | 0.9623 | 0.999994 | 20641 | 0 | 1.3646 | 0.037605 | 0.11266 | 0.95486 | 2629 | 4 | 1.3646 |
| hsa-mir-4533 | 4 | 0.96238 | 0.96228 | 0.999994 | 20640 | 0 | 2.6193 | 0.037625 | 0.11271 | 0.95487 | 2630 | 4 | 2.6193 |
| CDK11A | 5 | 0.18136 | 0.34861 | 0.999994 | 7261 | 2 | 4.5294 | 0.037634 | 0.12235 | 0.962935 | 2631 | 3 | 4.5294 |
| SCGB3A1 | 5 | 0.11382 | 0.24792 | 0.990682 | 5440 | 1 | 2.9882 | 0.037634 | 0.12235 | 0.962935 | 2632 | 3 | 2.9882 |
| ZFP69 | 5 | 0.34201 | 0.51536 | 0.999994 | 10983 | 2 | -0.41475 | 0.037634 | 0.12235 | 0.962935 | 2633 | 2 | -0.41475 |
| NonTargetingControlG | 2 | 0.81806 | 0.81785 | 0.999994 | 17747 | 0 | 5.0183 | 0.037645 | 0.068201 | 0.935447 | 2634 | 2 | 5.0183 |
| KLK4 | 6 | 0.98385 | 0.9838 | 0.999994 | 21185 | 0 | 1.5121 | 0.037646 | 0.12914 | 0.964228 | 2635 | 5 | 1.5121 |
| SUGP1 | 5 | 0.95688 | 0.95702 | 0.999994 | 20516 | 0 | 5.7269 | 0.037652 | 0.12239 | 0.962935 | 2636 | 4 | 5.7269 |
| LYPD6B | 6 | 0.16317 | 0.34225 | 0.999994 | 6821 | 2 | -0.042174 | 0.037663 | 0.12918 | 0.964228 | 2637 | 2 | -0.042174 |
| IFT20 | 6 | 0.95606 | 0.95615 | 0.999994 | 20484 | 0 | 1.5607 | 0.037665 | 0.12919 | 0.964228 | 2638 | 5 | 1.5607 |
| TCEAL1 | 6 | 0.95583 | 0.95591 | 0.999994 | 20473 | 0 | 4.3923 | 0.037679 | 0.12923 | 0.964228 | 2639 | 5 | 4.3923 |
| SLAMF1 | 6 | 0.18122 | 0.36468 | 0.999994 | 7257 | 1 | -0.078809 | 0.03772 | 0.12934 | 0.964228 | 2640 | 1 | -0.078809 |
| GLT8D2 | 6 | 0.32678 | 0.53757 | 0.999994 | 10658 | 3 | 0.21843 | 0.037775 | 0.12947 | 0.964228 | 2641 | 3 | 0.21843 |
| MORC1 | 5 | 0.65729 | 0.74803 | 0.999994 | 15746 | 1 | 2.448 | 0.037802 | 0.12274 | 0.962935 | 2642 | 4 | 2.448 |
| TUBB4B | 4 | 0.96213 | 0.96202 | 0.999994 | 20634 | 0 | 4.7728 | 0.037867 | 0.11335 | 0.954984 | 2643 | 4 | 4.7728 |
| RNMTL1 | 6 | 0.86593 | 0.88086 | 0.999994 | 18554 | 1 | 3.9957 | 0.037901 | 0.12982 | 0.964228 | 2644 | 4 | 3.9957 |
| FCF1 | 6 | 0.24854 | 0.4462 | 0.999994 | 8981 | 2 | 4.2513 | 0.037901 | 0.12982 | 0.964228 | 2645 | 4 | 4.2513 |
| PYCR2 | 6 | 0.47575 | 0.65232 | 0.999994 | 13073 | 1 | 4.1034 | 0.037901 | 0.12982 | 0.964228 | 2646 | 4 | 4.1034 |
| TLX1NB | 6 | 0.5503 | 0.70386 | 0.999994 | 14217 | 2 | 0.96441 | 0.037912 | 0.12985 | 0.964228 | 2647 | 3 | 0.96441 |
| hsa-mir-548k | 3 | 0.73107 | 0.73098 | 0.999994 | 16615 | 1 | 0.71672 | 0.037922 | 0.093117 | 0.950707 | 2648 | 2 | 0.71672 |
| hsa-mir-518e | 4 | 0.96205 | 0.96194 | 0.999994 | 20631 | 0 | 1.6493 | 0.037948 | 0.11356 | 0.954984 | 2649 | 4 | 1.6493 |
| hsa-mir-4675 | 4 | 0.74862 | 0.75898 | 0.999994 | 16845 | 1 | 5.1476 | 0.037953 | 0.11357 | 0.954984 | 2650 | 3 | 5.1476 |
| SEC61G | 5 | 0.69676 | 0.76501 | 0.999994 | 16268 | 1 | 5.7248 | 0.037955 | 0.1231 | 0.962935 | 2651 | 3 | 5.7248 |

| | | | | | | | | | | | | |
|---|---|---|---|---|---|---|---|---|---|---|---|---|
| LURAP1L | 5 | 0.93863 | 0.93883 | 0.999994 | 20040 | 0 | 0.67587 | 0.037995 | 0.12319 | 0.962935 | 2652 | 3 | 0.67587 |
| KATNA1 | 6 | 0.82857 | 0.86087 | 0.999994 | 17904 | 1 | 1.7693 | 0.038006 | 0.1301 | 0.964228 | 2653 | 5 | 1.7693 |
| AXL | 6 | 0.19283 | 0.37906 | 0.999994 | 7568 | 2 | 0.81179 | 0.038021 | 0.13014 | 0.964228 | 2654 | 4 | 0.81179 |
| CASP4 | 6 | 0.79487 | 0.8457 | 0.999994 | 17412 | 1 | 2.7193 | 0.038021 | 0.13014 | 0.964228 | 2655 | 5 | 2.7193 |
| SEC11A | 5 | 0.9041 | 0.90419 | 0.999994 | 19305 | 0 | 2.4241 | 0.038084 | 0.12339 | 0.963164 | 2656 | 3 | 2.4241 |
| DUPD1 | 6 | 0.19403 | 0.38056 | 0.999994 | 7605 | 2 | 0.95128 | 0.038085 | 0.13032 | 0.964228 | 2657 | 3 | 0.95128 |
| MOB3A | 6 | 0.23187 | 0.42634 | 0.999994 | 8562 | 1 | 2.3029 | 0.038085 | 0.13032 | 0.964228 | 2658 | 5 | 2.3029 |
| hsa-mir-153-2 | 3 | 0.83485 | 0.83474 | 0.999994 | 18002 | 0 | 6.8164 | 0.038126 | 0.093564 | 0.950736 | 2659 | 2 | 6.8164 |
| GPR61 | 6 | 0.23931 | 0.43516 | 0.999994 | 8757 | 2 | 3.2258 | 0.038185 | 0.13061 | 0.964228 | 2660 | 3 | 3.2258 |
| RNLS | 6 | 0.60474 | 0.74349 | 0.999994 | 15006 | 1 | 0.59809 | 0.038251 | 0.13079 | 0.964228 | 2661 | 3 | 0.59809 |
| TRH | 6 | 0.12623 | 0.29553 | 0.999994 | 5830 | 3 | 0.24479 | 0.038251 | 0.13079 | 0.964228 | 2662 | 3 | 0.24479 |
| FAM216A | 6 | 0.078879 | 0.20508 | 0.978186 | 4185 | 1 | 2.3837 | 0.038251 | 0.13079 | 0.964228 | 2663 | 3 | 2.3837 |
| SRP54 | 5 | 0.94144 | 0.94163 | 0.999994 | 20117 | 0 | 5.7015 | 0.038268 | 0.1238 | 0.963705 | 2664 | 4 | 5.7015 |
| NonTargetingControlG | 2 | 0.96173 | 0.96181 | 0.999994 | 20624 | 0 | 6.1927 | 0.038272 | 0.069276 | 0.936996 | 2665 | 2 | 6.1927 |
| IGFLR1 | 6 | 0.98452 | 0.98445 | 0.999994 | 21204 | 0 | 3.537 | 0.038294 | 0.13091 | 0.964228 | 2666 | 5 | 3.537 |
| LIG1 | 5 | 0.12443 | 0.2641 | 0.996734 | 5778 | 1 | 1.2919 | 0.038298 | 0.12386 | 0.963705 | 2667 | 3 | 1.2919 |
| SLC25A10 | 5 | 0.45273 | 0.60752 | 0.999994 | 12744 | 2 | -0.53691 | 0.038298 | 0.12386 | 0.963705 | 2668 | 2 | -0.53691 |
| CDH7 | 5 | 0.080512 | 0.19542 | 0.975797 | 4253 | 3 | -0.70216 | 0.038338 | 0.12396 | 0.963839 | 2669 | 1 | -0.70216 |
| RIMKLA | 6 | 0.91725 | 0.91787 | 0.999994 | 19569 | 1 | 2.6032 | 0.038344 | 0.13104 | 0.964228 | 2670 | 5 | 2.6032 |
| NRIP3 | 6 | 0.12508 | 0.29404 | 0.999994 | 5796 | 1 | 4.745 | 0.038347 | 0.13105 | 0.964228 | 2671 | 5 | 4.745 |
| ZNF587 | 4 | 0.25001 | 0.39193 | 0.999994 | 9017 | 1 | 0.51163 | 0.038357 | 0.11465 | 0.954984 | 2672 | 2 | 0.51163 |
| GRM4 | 6 | 0.65956 | 0.78499 | 0.999994 | 15772 | 2 | 1.4461 | 0.038405 | 0.1312 | 0.964748 | 2673 | 3 | 1.4461 |
| LAIR1 | 5 | 0.30567 | 0.48564 | 0.999994 | 10245 | 2 | 0.17547 | 0.038407 | 0.12412 | 0.964228 | 2674 | 2 | 0.17547 |
| CD14 | 6 | 0.95672 | 0.9568 | 0.999994 | 20509 | 0 | 2.1154 | 0.038427 | 0.13126 | 0.964844 | 2675 | 5 | 2.1154 |
| hsa-mir-3141 | 4 | 0.88731 | 0.88701 | 0.999994 | 18974 | 0 | 4.6821 | 0.038435 | 0.11487 | 0.954984 | 2676 | 3 | 4.6821 |
| OSTN | 6 | 0.65644 | 0.78255 | 0.999994 | 15736 | 2 | 4.1354 | 0.038471 | 0.13137 | 0.964897 | 2677 | 4 | 4.1354 |
| hsa-mir-127 | 4 | 0.96151 | 0.96141 | 0.999994 | 20619 | 0 | 3.5563 | 0.038489 | 0.11501 | 0.954984 | 2678 | 4 | 3.5563 |
| GALNT1 | 6 | 0.80486 | 0.84997 | 0.999994 | 17551 | 1 | 2.4751 | 0.038526 | 0.13153 | 0.965199 | 2679 | 5 | 2.4751 |
| CCKBR | 5 | 0.18459 | 0.3532 | 0.999994 | 7333 | 2 | 5.6986 | 0.038562 | 0.12446 | 0.964228 | 2680 | 3 | 5.6986 |
| ZBTB7A | 6 | 0.96486 | 0.9649 | 0.999994 | 20718 | 0 | 4.7037 | 0.038567 | 0.13163 | 0.965199 | 2681 | 5 | 4.7037 |
| DDAH1 | 6 | 0.34058 | 0.55332 | 0.999994 | 10949 | 1 | 3.0039 | 0.038571 | 0.13164 | 0.965199 | 2682 | 5 | 3.0039 |
| USP28 | 4 | 0.91676 | 0.91653 | 0.999994 | 19557 | 0 | 4.8445 | 0.038592 | 0.11527 | 0.954984 | 2683 | 3 | 4.8445 |
| GOLGA1 | 6 | 0.33568 | 0.54773 | 0.999994 | 10846 | 2 | 1.0226 | 0.038651 | 0.13186 | 0.965685 | 2684 | 4 | 1.0226 |
| hsa-mir-6073 | 3 | 0.96135 | 0.96152 | 0.999994 | 20615 | 0 | 2.0277 | 0.038653 | 0.094764 | 0.950736 | 2685 | 3 | 2.0277 |
| MAD2L1 | 4 | 0.96134 | 0.96124 | 0.999994 | 20614 | 0 | 1.4617 | 0.038656 | 0.11545 | 0.954984 | 2686 | 4 | 1.4617 |
| hsa-mir-4436b-2 | 4 | 0.96134 | 0.96124 | 0.999994 | 20612 | 0 | 2.0886 | 0.038656 | 0.11545 | 0.954984 | 2687 | 4 | 2.0886 |
| CASZ1 | 4 | 0.96134 | 0.96124 | 0.999994 | 20613 | 0 | 2.0868 | 0.038656 | 0.11545 | 0.954984 | 2688 | 4 | 2.0868 |
| ZNF132 | 5 | 0.84449 | 0.85003 | 0.999994 | 18164 | 1 | 3.2372 | 0.038674 | 0.12472 | 0.964228 | 2689 | 4 | 3.2372 |
| ADRA2C | 5 | 0.57231 | 0.71157 | 0.999994 | 14581 | 2 | 5.3289 | 0.038674 | 0.12472 | 0.964228 | 2690 | 3 | 5.3289 |
| KLHL21 | 6 | 0.10216 | 0.25154 | 0.993413 | 5045 | 2 | 0.42406 | 0.038733 | 0.13207 | 0.965913 | 2691 | 2 | 0.42406 |
| TEX261 | 4 | 0.7318 | 0.74649 | 0.999994 | 16624 | 1 | 3.997 | 0.038777 | 0.11577 | 0.954984 | 2692 | 2 | 3.997 |
| C8G | 5 | 0.076841 | 0.18949 | 0.970473 | 4103 | 1 | 4.8715 | 0.038807 | 0.12503 | 0.964228 | 2693 | 3 | 4.8715 |
| NonTargetingControlG | 2 | 0.15586 | 0.23077 | 0.983933 | 6621 | 1 | 1.0594 | 0.03881 | 0.070226 | 0.937929 | 2694 | 1 | 1.0594 |
| ST13 | 5 | 0.85014 | 0.85439 | 0.999994 | 18265 | 1 | 3.1527 | 0.038931 | 0.12534 | 0.964228 | 2695 | 4 | 3.1527 |
| SPANXA2 | 2 | 0.71929 | 0.71899 | 0.999994 | 16488 | 0 | 4.7342 | 0.03894 | 0.070453 | 0.937929 | 2696 | 1 | 4.7342 |
| TTLL3 | 4 | 0.56939 | 0.64933 | 0.999994 | 14526 | 1 | 3.4492 | 0.038959 | 0.11623 | 0.954984 | 2697 | 2 | 3.4492 |
| CBX6 | 5 | 0.57024 | 0.70974 | 0.999994 | 14545 | 1 | 4.9175 | 0.038986 | 0.12546 | 0.964228 | 2698 | 4 | 4.9175 |
| CAMK1 | 5 | 0.64459 | 0.74297 | 0.999994 | 15579 | 1 | 4.1286 | 0.039003 | 0.12549 | 0.964228 | 2699 | 4 | 4.1286 |
| HSPA2 | 6 | 0.50597 | 0.67287 | 0.999994 | 13524 | 1 | 2.5663 | 0.039006 | 0.13282 | 0.965913 | 2700 | 5 | 2.5663 |
| RIPK3 | 5 | 0.93743 | 0.93764 | 0.999994 | 20015 | 0 | 5.3065 | 0.039041 | 0.12558 | 0.964228 | 2701 | 4 | 5.3065 |
| NonTargetingControlG | 2 | 0.96092 | 0.961 | 0.999994 | 20601 | 0 | 6.8835 | 0.039081 | 0.070715 | 0.937929 | 2702 | 2 | 6.8835 |
| NUB1 | 5 | 0.78351 | 0.80946 | 0.999994 | 17261 | 1 | 2.6403 | 0.039092 | 0.12571 | 0.964228 | 2703 | 4 | 2.6403 |

| | | | | | | | | | | | | | |
|---|---|---|---|---|---|---|---|---|---|---|---|---|---|
| OR5W2 | 6 | 0.81487 | 0.85445 | 0.999994 | 17699 | 1 | 2.5444 | 0.039103 | 0.13308 | 0.965913 | 2704 | 5 | 2.5444 |
| GIT1 | 6 | 0.78568 | 0.84194 | 0.999994 | 17288 | 1 | 1.5335 | 0.039128 | 0.13315 | 0.965913 | 2705 | 3 | 1.5335 |
| TANK | 6 | 0.076393 | 0.20006 | 0.977164 | 4087 | 4 | -0.93366 | 0.039171 | 0.13326 | 0.965913 | 2706 | 1 | -0.93366 |
| C10orf107 | 6 | 0.16165 | 0.34035 | 0.999994 | 6781 | 1 | 1.6118 | 0.039208 | 0.13336 | 0.965913 | 2707 | 5 | 1.6118 |
| C1orf115 | 5 | 0.8701 | 0.87116 | 0.999994 | 18637 | 1 | 5.6697 | 0.039244 | 0.12608 | 0.964228 | 2708 | 4 | 5.6697 |
| CAPZB | 5 | 0.88901 | 0.88913 | 0.999994 | 19021 | 0 | 5.6697 | 0.039244 | 0.12608 | 0.964228 | 2709 | 3 | 5.6697 |
| ID2 | 5 | 0.12136 | 0.25944 | 0.994943 | 5678 | 3 | -1.333 | 0.03932 | 0.12625 | 0.964228 | 2710 | 1 | -1.333 |
| SCUBE3 | 6 | 0.51432 | 0.6786 | 0.999994 | 13649 | 2 | -0.39009 | 0.039335 | 0.13371 | 0.965913 | 2711 | 2 | -0.39009 |
| MNDA | 6 | 0.78315 | 0.84095 | 0.999994 | 17253 | 1 | -0.27784 | 0.039335 | 0.13371 | 0.965913 | 2712 | 2 | -0.27784 |
| STRN4 | 5 | 0.75258 | 0.79219 | 0.999994 | 16884 | 1 | 3.1166 | 0.03934 | 0.1263 | 0.964228 | 2713 | 3 | 3.1166 |
| FPGS | 5 | 0.87948 | 0.87977 | 0.999994 | 18821 | 1 | 1.0961 | 0.039388 | 0.12641 | 0.964228 | 2714 | 3 | 1.0961 |
| ZNF460 | 5 | 0.95852 | 0.95867 | 0.999994 | 20546 | 0 | 2.6817 | 0.039405 | 0.12645 | 0.964228 | 2715 | 4 | 2.6817 |
| SDR16C5 | 5 | 0.94134 | 0.94154 | 0.999994 | 20115 | 0 | 4.1084 | 0.039405 | 0.12645 | 0.964228 | 2716 | 4 | 4.1084 |
| TAS2R20 | 6 | 0.96778 | 0.96779 | 0.999994 | 20779 | 0 | 3.5185 | 0.039432 | 0.13397 | 0.965913 | 2717 | 4 | 3.5185 |
| PTH2R | 6 | 0.32232 | 0.53257 | 0.999994 | 10562 | 2 | -0.03648 | 0.039432 | 0.13397 | 0.965913 | 2718 | 2 | -0.03648 |
| NonTargetingControlG | 2 | 0.6922 | 0.69201 | 0.999994 | 16228 | 0 | 4.571 | 0.039467 | 0.071341 | 0.937929 | 2719 | 1 | 4.571 |
| FNBP1L | 6 | 0.80943 | 0.852 | 0.999994 | 17624 | 1 | 1.486 | 0.039477 | 0.13409 | 0.965913 | 2720 | 5 | 1.486 |
| RNF183 | 5 | 0.25478 | 0.44455 | 0.999994 | 9124 | 1 | 1.6738 | 0.03948 | 0.12661 | 0.964228 | 2721 | 3 | 1.6738 |
| SZRD1 | 6 | 0.96272 | 0.96278 | 0.999994 | 20651 | 0 | 3.987 | 0.039485 | 0.13412 | 0.965913 | 2722 | 4 | 3.987 |
| FAM205A | 6 | 0.80137 | 0.84844 | 0.999994 | 17495 | 1 | 3.653 | 0.039485 | 0.13412 | 0.965913 | 2723 | 5 | 3.653 |
| PAK6 | 5 | 0.87717 | 0.87758 | 0.999994 | 18778 | 1 | 2.223 | 0.03952 | 0.1267 | 0.964228 | 2724 | 4 | 2.223 |
| ATP5L2 | 5 | 0.82986 | 0.83939 | 0.999994 | 17926 | 1 | 2.9152 | 0.039539 | 0.12675 | 0.964228 | 2725 | 4 | 2.9152 |
| SPINK2 | 5 | 0.90277 | 0.90286 | 0.999994 | 19266 | 0 | 2.6191 | 0.039543 | 0.12676 | 0.964228 | 2726 | 4 | 2.6191 |
| PGLYRP4 | 5 | 0.19018 | 0.3612 | 0.999994 | 7491 | 3 | -0.87455 | 0.039548 | 0.12677 | 0.964228 | 2727 | 2 | -0.87455 |
| IL17C | 5 | 0.24187 | 0.43345 | 0.999994 | 8831 | 1 | 2.9462 | 0.039577 | 0.12683 | 0.964228 | 2728 | 4 | 2.9462 |
| DARC | 6 | 0.17041 | 0.35127 | 0.999994 | 6992 | 2 | 2.0715 | 0.039588 | 0.1344 | 0.965913 | 2729 | 3 | 2.0715 |
| METTL5 | 6 | 0.29412 | 0.49985 | 0.999994 | 10000 | 3 | 0.35265 | 0.039588 | 0.1344 | 0.965913 | 2730 | 3 | 0.35265 |
| hsa-mir-520b | 3 | 0.96041 | 0.96059 | 0.999994 | 20591 | 0 | 3.4307 | 0.039592 | 0.096915 | 0.950736 | 2731 | 3 | 3.4307 |
| HAUS3 | 5 | 0.55846 | 0.69925 | 0.999994 | 14339 | 2 | 1.4404 | 0.039594 | 0.12688 | 0.964228 | 2732 | 3 | 1.4404 |
| RASL11B | 6 | 0.63267 | 0.76435 | 0.999994 | 15409 | 1 | 3.7381 | 0.039647 | 0.13455 | 0.965913 | 2733 | 5 | 3.7381 |
| CLEC19A | 5 | 0.34229 | 0.5156 | 0.999994 | 10991 | 2 | 4.4873 | 0.039685 | 0.12709 | 0.964228 | 2734 | 3 | 4.4873 |
| DUSP7 | 6 | 0.011235 | 0.043192 | 0.891621 | 942 | 5 | -1.5196 | 0.039691 | 0.13466 | 0.965913 | 2735 | 1 | -1.5196 |
| OR1S2 | 6 | 0.19364 | 0.38007 | 0.999994 | 7590 | 1 | 1.6858 | 0.039719 | 0.13473 | 0.965913 | 2736 | 5 | 1.6858 |
| MPP2 | 4 | 0.13876 | 0.27765 | 0.999519 | 6159 | 1 | 4.7691 | 0.039779 | 0.11842 | 0.9593 | 2737 | 3 | 4.7691 |
| FGFR3 | 6 | 0.88258 | 0.8913 | 0.999994 | 18881 | 1 | 4.4056 | 0.039786 | 0.1349 | 0.965913 | 2738 | 4 | 4.4056 |
| KCNK16 | 6 | 0.010841 | 0.041859 | 0.891621 | 918 | 1 | 3.6878 | 0.039786 | 0.1349 | 0.965913 | 2739 | 4 | 3.6878 |
| FZD4 | 6 | 0.21635 | 0.40776 | 0.999994 | 8177 | 1 | 3.4785 | 0.039786 | 0.1349 | 0.965913 | 2740 | 4 | 3.4785 |
| TTF1 | 4 | 0.3479 | 0.49214 | 0.999994 | 11117 | 1 | 3.3445 | 0.039821 | 0.11852 | 0.9593 | 2741 | 2 | 3.3445 |
| GLTSCR1L | 5 | 0.088928 | 0.20886 | 0.979587 | 4550 | 3 | -1.3882 | 0.039822 | 0.1274 | 0.964228 | 2742 | 1 | -1.3882 |
| NonTargetingControlG | 2 | 0.96016 | 0.96024 | 0.999994 | 20585 | 0 | 5.4338 | 0.039843 | 0.072005 | 0.937929 | 2743 | 2 | 5.4338 |
| TRA2A | 5 | 0.23219 | 0.42024 | 0.999994 | 8576 | 1 | 1.3697 | 0.039845 | 0.12746 | 0.964228 | 2744 | 3 | 1.3697 |
| TCTE3 | 6 | 0.95915 | 0.95923 | 0.999994 | 20560 | 0 | 4.2103 | 0.039869 | 0.13515 | 0.965913 | 2745 | 4 | 4.2103 |
| ABHD8 | 6 | 0.98459 | 0.98452 | 0.999994 | 21209 | 0 | 2.7606 | 0.039902 | 0.13525 | 0.965913 | 2746 | 5 | 2.7606 |
| KRT7 | 6 | 0.063747 | 0.17372 | 0.96793 | 3597 | 1 | 1.5189 | 0.03994 | 0.13535 | 0.965913 | 2747 | 5 | 1.5189 |
| AGO2 | 4 | 0.12379 | 0.25691 | 0.994943 | 5760 | 2 | 2.2875 | 0.039971 | 0.11892 | 0.959988 | 2748 | 2 | 2.2875 |
| THOC7 | 4 | 0.56772 | 0.64849 | 0.999994 | 14510 | 1 | 5.2804 | 0.040043 | 0.11909 | 0.960516 | 2749 | 3 | 5.2804 |
| H3F3C | 5 | 0.15143 | 0.30493 | 0.999994 | 6492 | 3 | -0.70123 | 0.04005 | 0.12793 | 0.964228 | 2750 | 2 | -0.70123 |
| ITGAX | 6 | 0.30891 | 0.51713 | 0.999994 | 10295 | 2 | 2.3473 | 0.04005 | 0.13565 | 0.965913 | 2751 | 3 | 2.3473 |
| TCEB3CL | 4 | 0.77558 | 0.78031 | 0.999994 | 17144 | 1 | 0.19664 | 0.040058 | 0.11914 | 0.960516 | 2752 | 1 | 0.19664 |
| GNGT1 | 6 | 0.18193 | 0.36557 | 0.999994 | 7268 | 2 | 3.8609 | 0.040091 | 0.13576 | 0.965913 | 2753 | 4 | 3.8609 |
| AKT2 | 6 | 0.57187 | 0.7193 | 0.999994 | 14573 | 2 | 0.24909 | 0.040128 | 0.13585 | 0.965913 | 2754 | 2 | 0.24909 |
| CRH | 4 | 0.05836 | 0.13761 | 0.957317 | 3357 | 2 | 1.795 | 0.040172 | 0.11942 | 0.961045 | 2755 | 2 | 1.795 |

| | | | | | | | | | | | | | |
|---|---|---|---|---|---|---|---|---|---|---|---|---|---|
| FBXO18 | 5 | 0.72269 | 0.77715 | 0.999994 | 16523 | 1 | 2.3162 | 0.040174 | 0.1282 | 0.964228 | 2756 | 4 | 2.3162 |
| C2CD2L | 4 | 0.61802 | 0.67522 | 0.999994 | 15186 | 1 | 1.1463 | 0.040186 | 0.11946 | 0.961045 | 2757 | 3 | 1.1463 |
| ZNF384 | 5 | 0.53937 | 0.68234 | 0.999994 | 14041 | 2 | 0.53984 | 0.040187 | 0.12823 | 0.964228 | 2758 | 3 | 0.53984 |
| SDHD | 3 | 0.9598 | 0.95997 | 0.999994 | 20579 | 0 | 2.3753 | 0.040198 | 0.098293 | 0.950736 | 2759 | 3 | 2.3753 |
| XRCC6 | 3 | 0.9598 | 0.95997 | 0.999994 | 20578 | 0 | 4.3741 | 0.040198 | 0.098293 | 0.950736 | 2760 | 3 | 4.3741 |
| PRSS1 | 6 | 0.23187 | 0.42634 | 0.999994 | 8561 | 2 | 0.73395 | 0.040226 | 0.13613 | 0.965913 | 2761 | 3 | 0.73395 |
| C12orf75 | 5 | 0.022294 | 0.070267 | 0.93914 | 1600 | 2 | 2.271 | 0.040278 | 0.12844 | 0.964228 | 2762 | 3 | 2.271 |
| NonTargetingControlG | 2 | 0.95969 | 0.95975 | 0.999994 | 20577 | 0 | 5.182 | 0.040311 | 0.072807 | 0.937929 | 2763 | 2 | 5.182 |
| DUSP9 | 5 | 0.95249 | 0.95264 | 0.999994 | 20394 | 0 | 3.4241 | 0.040337 | 0.12859 | 0.964228 | 2764 | 4 | 3.4241 |
| CUL1 | 4 | 0.12084 | 0.25179 | 0.994 | 5654 | 1 | 4.7517 | 0.040339 | 0.11987 | 0.961431 | 2765 | 3 | 4.7517 |
| C17orf104 | 4 | 0.89813 | 0.89785 | 0.999994 | 19183 | 0 | 5.057 | 0.040352 | 0.11991 | 0.961431 | 2766 | 3 | 5.057 |
| CACNA2D4 | 4 | 0.95963 | 0.95954 | 0.999994 | 20574 | 0 | 2.0934 | 0.040374 | 0.11997 | 0.961431 | 2767 | 4 | 2.0934 |
| LCE1A | 5 | 0.073527 | 0.18415 | 0.968526 | 3971 | 2 | 0.56543 | 0.040392 | 0.1287 | 0.964228 | 2768 | 2 | 0.56543 |
| NT5DC1 | 5 | 0.21032 | 0.38984 | 0.999994 | 8028 | 3 | -0.78236 | 0.040421 | 0.12876 | 0.964228 | 2769 | 2 | -0.78236 |
| SAP18 | 4 | 0.95949 | 0.95941 | 0.999994 | 20572 | 0 | 3.7923 | 0.04051 | 0.12033 | 0.961431 | 2770 | 4 | 3.7923 |
| OR6N2 | 6 | 0.34982 | 0.56379 | 0.999994 | 11153 | 2 | 1.6478 | 0.040511 | 0.13689 | 0.965913 | 2771 | 4 | 1.6478 |
| TMEM170B | 4 | 0.75344 | 0.76274 | 0.999994 | 16894 | 1 | 3.4775 | 0.040512 | 0.12033 | 0.961431 | 2772 | 2 | 3.4775 |
| hsa-mir-718 | 3 | 0.95947 | 0.95963 | 0.999994 | 20570 | 0 | 2.4812 | 0.040532 | 0.099044 | 0.951893 | 2773 | 3 | 2.4812 |
| SCHIP1 | 4 | 0.90475 | 0.90445 | 0.999994 | 19317 | 0 | 4.4601 | 0.04058 | 0.12052 | 0.961971 | 2774 | 3 | 4.4601 |
| NADK2 | 6 | 0.13412 | 0.30555 | 0.999994 | 6024 | 3 | 2.0337 | 0.040593 | 0.13713 | 0.965913 | 2775 | 3 | 2.0337 |
| TAS2R31 | 6 | 0.34965 | 0.5636 | 0.999994 | 11145 | 3 | -0.18922 | 0.040654 | 0.13729 | 0.965913 | 2776 | 2 | -0.18922 |
| LYAR | 6 | 0.29858 | 0.50504 | 0.999994 | 10099 | 1 | 0.69172 | 0.040675 | 0.13736 | 0.965913 | 2777 | 4 | 0.69172 |
| LOC100505549 | 5 | 0.3379 | 0.51202 | 0.999994 | 10893 | 2 | 1.153 | 0.040712 | 0.12942 | 0.964228 | 2778 | 3 | 1.153 |
| CYP26A1 | 5 | 0.9536 | 0.95376 | 0.999994 | 20429 | 0 | 0.64258 | 0.040712 | 0.12942 | 0.964228 | 2779 | 3 | 0.64258 |
| KIF3B | 4 | 0.95927 | 0.95918 | 0.999994 | 20564 | 0 | 1.7665 | 0.040732 | 0.12089 | 0.962329 | 2780 | 4 | 1.7665 |
| hsa-mir-4792 | 4 | 0.95926 | 0.95918 | 0.999994 | 20563 | 0 | 4.5043 | 0.040742 | 0.12092 | 0.962329 | 2781 | 4 | 4.5043 |
| GRIPAP1 | 5 | 0.64572 | 0.74341 | 0.999994 | 15596 | 1 | 2.4492 | 0.040752 | 0.1295 | 0.964228 | 2782 | 4 | 2.4492 |
| IFI30 | 6 | 0.88848 | 0.89525 | 0.999994 | 19009 | 1 | 4.2721 | 0.040774 | 0.13763 | 0.965913 | 2783 | 4 | 4.2721 |
| AKAP5 | 6 | 0.44901 | 0.63447 | 0.999994 | 12700 | 2 | 1.2471 | 0.040784 | 0.13766 | 0.965913 | 2784 | 4 | 1.2471 |
| ACRBP | 6 | 0.63594 | 0.76685 | 0.999994 | 15461 | 1 | 2.2334 | 0.040786 | 0.13766 | 0.965913 | 2785 | 4 | 2.2334 |
| SPPL3 | 5 | 0.2015 | 0.37737 | 0.999994 | 7808 | 3 | -0.77756 | 0.04079 | 0.12959 | 0.964228 | 2786 | 2 | -0.77756 |
| FKBP1A | 6 | 0.8569 | 0.87568 | 0.999994 | 18391 | 1 | 2.3643 | 0.040813 | 0.13773 | 0.965913 | 2787 | 5 | 2.3643 |
| HIATL1 | 5 | 0.80039 | 0.81976 | 0.999994 | 17485 | 1 | 5.2454 | 0.040817 | 0.12966 | 0.964228 | 2788 | 4 | 5.2454 |
| CGB | 6 | 0.89599 | 0.90064 | 0.999994 | 19144 | 1 | 1.6792 | 0.040835 | 0.13779 | 0.965913 | 2789 | 5 | 1.6792 |
| URI1 | 5 | 0.23383 | 0.42251 | 0.999994 | 8614 | 1 | 5.6045 | 0.040899 | 0.12985 | 0.964228 | 2790 | 3 | 5.6045 |
| hsa-mir-3174 | 4 | 0.69235 | 0.71949 | 0.999994 | 16229 | 1 | 4.2534 | 0.040904 | 0.12135 | 0.962897 | 2791 | 3 | 4.2534 |
| DAZL | 6 | 0.59279 | 0.73465 | 0.999994 | 14857 | 2 | 0.11354 | 0.040921 | 0.13802 | 0.965913 | 2792 | 2 | 0.11354 |
| SLC25A51 | 5 | 0.96638 | 0.96656 | 0.999994 | 20745 | 0 | 1.2103 | 0.040931 | 0.12993 | 0.964228 | 2793 | 4 | 1.2103 |
| SPG21 | 6 | 0.85769 | 0.87612 | 0.999994 | 18402 | 1 | 4.0764 | 0.040961 | 0.13813 | 0.965913 | 2794 | 5 | 4.0764 |
| NODAL | 6 | 0.98226 | 0.98219 | 0.999994 | 21143 | 0 | 2.5502 | 0.040978 | 0.13818 | 0.965913 | 2795 | 5 | 2.5502 |
| ZNF595 | 5 | 0.94004 | 0.94022 | 0.999994 | 20080 | 0 | 2.4561 | 0.040982 | 0.13004 | 0.964228 | 2796 | 4 | 2.4561 |
| HAVCR2 | 6 | 0.42925 | 0.62134 | 0.999994 | 12404 | 1 | 2.6643 | 0.040982 | 0.13819 | 0.965913 | 2797 | 4 | 2.6643 |
| ZNF727 | 6 | 0.43165 | 0.62293 | 0.999994 | 12441 | 1 | 2.6075 | 0.040982 | 0.13819 | 0.965913 | 2798 | 4 | 2.6075 |
| TMPRSS15 | 6 | 0.6824 | 0.80288 | 0.999994 | 16124 | 1 | 0.81599 | 0.04103 | 0.13831 | 0.965913 | 2799 | 4 | 0.81599 |
| SH3TC2 | 6 | 0.55788 | 0.70933 | 0.999994 | 14332 | 2 | 1.1638 | 0.04103 | 0.13831 | 0.965913 | 2800 | 4 | 1.1638 |
| STK25 | 6 | 0.63726 | 0.76783 | 0.999994 | 15475 | 1 | 1.4342 | 0.04106 | 0.13838 | 0.965913 | 2801 | 5 | 1.4342 |
| AFG3L2 | 5 | 0.96922 | 0.9693 | 0.999994 | 20808 | 0 | 4.0832 | 0.041077 | 0.13028 | 0.964228 | 2802 | 4 | 4.0832 |
| PRDX2 | 5 | 0.076102 | 0.18831 | 0.969543 | 4077 | 3 | -1.7357 | 0.041099 | 0.13033 | 0.964228 | 2803 | 2 | -1.7357 |
| EBPL | 6 | 0.26868 | 0.47019 | 0.999994 | 9439 | 2 | 0.63255 | 0.041112 | 0.13851 | 0.965913 | 2804 | 3 | 0.63255 |
| hsa-mir-329-2 | 1 | 0.95887 | 0.95906 | 0.999994 | 20559 | 0 | 8.3932 | 0.041127 | 0.041041 | 0.917066 | 2805 | 1 | 8.3932 |
| hsa-mir-2113 | 3 | 0.28417 | 0.4171 | 0.999994 | 9788 | 1 | 0.95407 | 0.041144 | 0.10038 | 0.951893 | 2806 | 2 | 0.95407 |
| LMO2 | 5 | 0.84976 | 0.85409 | 0.999994 | 18256 | 1 | 1.7267 | 0.041168 | 0.13049 | 0.964228 | 2807 | 3 | 1.7267 |

| | | | | | | | | | | | | |
|---|---|---|---|---|---|---|---|---|---|---|---|---|
| LTBR | 5 | 0.82051 | 0.83295 | 0.999994 | 17784 | 1 | 1.6284 | 0.041168 | 0.13049 | 0.964228 | 2808 | 3 | 1.6284 |
| EPX | 4 | 0.95883 | 0.95878 | 0.999994 | 20556 | 0 | 3.4754 | 0.04117 | 0.12204 | 0.962935 | 2809 | 4 | 3.4754 |
| SF3B5 | 4 | 0.80839 | 0.80881 | 0.999994 | 17608 | 1 | 5.1074 | 0.041231 | 0.1222 | 0.962935 | 2810 | 3 | 5.1074 |
| UGT2B7 | 6 | 0.6211 | 0.75565 | 0.999994 | 15242 | 1 | 3.1811 | 0.041246 | 0.13884 | 0.965913 | 2811 | 3 | 3.1811 |
| CSMD1 | 6 | 0.60783 | 0.74575 | 0.999994 | 15045 | 2 | 3.9453 | 0.041246 | 0.13884 | 0.965913 | 2812 | 4 | 3.9453 |
| APPBP2 | 5 | 0.61197 | 0.73053 | 0.999994 | 15109 | 1 | 5.5794 | 0.041247 | 0.13068 | 0.964228 | 2813 | 4 | 5.5794 |
| ZNF706 | 5 | 0.54048 | 0.68333 | 0.999994 | 14063 | 1 | 5.5794 | 0.041247 | 0.13068 | 0.964228 | 2814 | 4 | 5.5794 |
| ARHGEF10L | 6 | 0.95607 | 0.95616 | 0.999994 | 20486 | 0 | 2.4072 | 0.041248 | 0.13884 | 0.965913 | 2815 | 4 | 2.4072 |
| CLEC6A | 6 | 0.71891 | 0.81798 | 0.999994 | 16485 | 1 | 2.333 | 0.041248 | 0.13884 | 0.965913 | 2816 | 4 | 2.333 |
| PPIAL4F | 4 | 0.92213 | 0.92192 | 0.999994 | 19677 | 0 | 4.4464 | 0.041259 | 0.12228 | 0.962935 | 2817 | 3 | 4.4464 |
| hsa-mir-146a | 4 | 0.95869 | 0.95864 | 0.999994 | 20553 | 0 | 2.7109 | 0.041313 | 0.12243 | 0.962935 | 2818 | 4 | 2.7109 |
| TDRD7 | 6 | 0.97806 | 0.978 | 0.999994 | 21045 | 0 | 1.7275 | 0.041318 | 0.13902 | 0.965913 | 2819 | 5 | 1.7275 |
| C1orf189 | 4 | 0.26013 | 0.40224 | 0.999994 | 9246 | 2 | 3.1999 | 0.04133 | 0.12247 | 0.962935 | 2820 | 2 | 3.1999 |
| WAS | 3 | 0.1128 | 0.20001 | 0.977164 | 5398 | 1 | 6.7188 | 0.041361 | 0.10089 | 0.951893 | 2821 | 2 | 6.7188 |
| HRH4 | 6 | 0.36224 | 0.57782 | 0.999994 | 11372 | 2 | -0.3258 | 0.041368 | 0.13916 | 0.965913 | 2822 | 2 | -0.3258 |
| SIMC1 | 6 | 0.9847 | 0.98463 | 0.999994 | 21216 | 0 | 4.2054 | 0.041368 | 0.13916 | 0.965913 | 2823 | 5 | 4.2054 |
| CYP4F8 | 5 | 0.67935 | 0.75727 | 0.999994 | 16080 | 1 | 2.3951 | 0.041374 | 0.13099 | 0.964228 | 2824 | 4 | 2.3951 |
| ATAD3C | 4 | 0.95861 | 0.95856 | 0.999994 | 20552 | 0 | 2.8549 | 0.041386 | 0.12263 | 0.962935 | 2825 | 4 | 2.8549 |
| HSPA8 | 2 | 0.95859 | 0.95866 | 0.999994 | 20551 | 0 | 6.1389 | 0.041407 | 0.074691 | 0.938203 | 2826 | 2 | 6.1389 |
| LOXL4 | 5 | 0.44526 | 0.60123 | 0.999994 | 12652 | 1 | 0.92276 | 0.041408 | 0.13107 | 0.964228 | 2827 | 3 | 0.92276 |
| CCDC106 | 6 | 0.95693 | 0.95701 | 0.999994 | 20517 | 0 | 1.6776 | 0.041432 | 0.13935 | 0.965913 | 2828 | 5 | 1.6776 |
| SHD | 6 | 0.35044 | 0.56448 | 0.999994 | 11169 | 2 | 0.92169 | 0.04147 | 0.13945 | 0.965913 | 2829 | 3 | 0.92169 |
| hsa-mir-579 | 4 | 0.95853 | 0.95847 | 0.999994 | 20548 | 0 | 1.054 | 0.041472 | 0.12286 | 0.962935 | 2830 | 4 | 1.054 |
| CPT1C | 4 | 0.88197 | 0.88159 | 0.999994 | 18868 | 0 | 5.484 | 0.041513 | 0.12296 | 0.962935 | 2831 | 3 | 5.484 |
| CCNO | 6 | 0.94019 | 0.94015 | 0.999994 | 20087 | 0 | 4.1239 | 0.041542 | 0.13965 | 0.965913 | 2832 | 5 | 4.1239 |
| PRKCA | 6 | 0.23173 | 0.42618 | 0.999994 | 8558 | 3 | 2.6396 | 0.041551 | 0.13967 | 0.965913 | 2833 | 3 | 2.6396 |
| GBP3 | 6 | 0.98647 | 0.98642 | 0.999994 | 21262 | 0 | 1.8294 | 0.041581 | 0.13976 | 0.965913 | 2834 | 5 | 1.8294 |
| COMP | 6 | 0.98204 | 0.98197 | 0.999994 | 21139 | 0 | 2.4425 | 0.041585 | 0.13977 | 0.965913 | 2835 | 5 | 2.4425 |
| ABCF3 | 6 | 0.2185 | 0.4103 | 0.999994 | 8240 | 3 | -0.42223 | 0.041585 | 0.13977 | 0.965913 | 2836 | 2 | -0.42223 |
| ARAP1 | 6 | 0.81074 | 0.85259 | 0.999994 | 17643 | 1 | 3.2578 | 0.041585 | 0.13977 | 0.965913 | 2837 | 4 | 3.2578 |
| MANSC1 | 6 | 0.97665 | 0.9766 | 0.999994 | 21006 | 0 | 2.42 | 0.041585 | 0.13977 | 0.965913 | 2838 | 5 | 2.42 |
| UCKL1 | 4 | 0.31193 | 0.45524 | 0.999994 | 10357 | 1 | 1.3315 | 0.041685 | 0.1234 | 0.963164 | 2839 | 3 | 1.3315 |
| SLC25A14 | 5 | 0.12887 | 0.27088 | 0.998561 | 5910 | 2 | 3.714 | 0.041687 | 0.13169 | 0.965199 | 2840 | 3 | 3.714 |
| hsa-mir-5197 | 4 | 0.95828 | 0.95822 | 0.999994 | 20543 | 0 | 4.0052 | 0.041722 | 0.12349 | 0.963164 | 2841 | 4 | 4.0052 |
| DPEP2 | 4 | 0.55535 | 0.6423 | 0.999994 | 14296 | 1 | 4.3664 | 0.041736 | 0.12353 | 0.963164 | 2842 | 3 | 4.3664 |
| hsa-mir-4672 | 4 | 0.19112 | 0.33154 | 0.999994 | 7517 | 1 | 4.5927 | 0.041736 | 0.12353 | 0.963164 | 2843 | 3 | 4.5927 |
| ADAM29 | 6 | 0.66077 | 0.78592 | 0.999994 | 15791 | 1 | 4.1282 | 0.041737 | 0.14017 | 0.965913 | 2844 | 5 | 4.1282 |
| ATR | 5 | 0.79351 | 0.81545 | 0.999994 | 17391 | 1 | 4.5253 | 0.041761 | 0.13186 | 0.965685 | 2845 | 4 | 4.5253 |
| TRPM8 | 6 | 0.4669 | 0.64638 | 0.999994 | 12963 | 2 | 1.1866 | 0.041795 | 0.14032 | 0.965913 | 2846 | 3 | 1.1866 |
| CCDC87 | 6 | 0.92844 | 0.92853 | 0.999994 | 19824 | 0 | 3.5664 | 0.041795 | 0.14032 | 0.965913 | 2847 | 5 | 3.5664 |
| ASS1 | 5 | 0.6028 | 0.7272 | 0.999994 | 14984 | 1 | 3.9895 | 0.041822 | 0.13202 | 0.965913 | 2848 | 4 | 3.9895 |
| MRPL38 | 3 | 0.26402 | 0.39231 | 0.999994 | 9334 | 2 | -0.7547 | 0.041886 | 0.10202 | 0.951893 | 2849 | 1 | -0.7547 |
| CC2D1A | 5 | 0.29031 | 0.47317 | 0.999994 | 9914 | 1 | 5.5534 | 0.041893 | 0.13218 | 0.965913 | 2850 | 3 | 5.5534 |
| CP | 6 | 0.76001 | 0.83207 | 0.999994 | 16982 | 1 | 3.8877 | 0.041915 | 0.14063 | 0.965913 | 2851 | 5 | 3.8877 |
| HDAC8 | 6 | 0.98203 | 0.98196 | 0.999994 | 21138 | 0 | 4.0832 | 0.041915 | 0.14063 | 0.965913 | 2852 | 5 | 4.0832 |
| PARL | 5 | 0.27937 | 0.46436 | 0.999994 | 9687 | 2 | 0.83221 | 0.041919 | 0.13223 | 0.965913 | 2853 | 3 | 0.83221 |
| NonTargetingControlG | 2 | 0.95796 | 0.95803 | 0.999994 | 20538 | 0 | 5.6753 | 0.042039 | 0.075778 | 0.938203 | 2854 | 2 | 5.6753 |
| TDRD9 | 5 | 0.33201 | 0.50717 | 0.999994 | 10767 | 2 | 2.7062 | 0.042046 | 0.13251 | 0.965913 | 2855 | 3 | 2.7062 |
| ADIRF | 5 | 0.0050106 | 0.020865 | 0.811197 | 521 | 1 | 2.3517 | 0.042065 | 0.13255 | 0.965913 | 2856 | 4 | 2.3517 |
| FAM222A | 4 | 0.58643 | 0.65805 | 0.999994 | 14764 | 1 | 4.0816 | 0.042073 | 0.12439 | 0.964228 | 2857 | 3 | 4.0816 |
| TECPR2 | 5 | 0.14198 | 0.29083 | 0.999632 | 6240 | 2 | 0.035535 | 0.042079 | 0.13257 | 0.965913 | 2858 | 2 | 0.035535 |
| FIBCD1 | 5 | 0.81853 | 0.8316 | 0.999994 | 17755 | 1 | 5.022 | 0.042105 | 0.13264 | 0.965913 | 2859 | 4 | 5.022 |

| | | | | | | | | | | | | | |
|---|---|---|---|---|---|---|---|---|---|---|---|---|---|
| RHOBTB3 | 6 | 0.8462 | 0.86991 | 0.999994 | 18190 | 1 | 2.8463 | 0.042146 | 0.14127 | 0.965913 | 2860 | 5 | 2.8463 |
| RHOA | 3 | 0.78982 | 0.78973 | 0.999994 | 17340 | 0 | 1.5797 | 0.042147 | 0.1026 | 0.952143 | 2861 | 2 | 1.5797 |
| KIAA1841 | 6 | 0.93526 | 0.9353 | 0.999994 | 19974 | 0 | 4.6288 | 0.04221 | 0.14144 | 0.965913 | 2862 | 5 | 4.6288 |
| BRD3 | 5 | 0.96524 | 0.96539 | 0.999994 | 20722 | 0 | 4.1731 | 0.042236 | 0.13291 | 0.965913 | 2863 | 4 | 4.1731 |
| HS3ST3B1 | 5 | 0.94245 | 0.94264 | 0.999994 | 20136 | 0 | 1.3396 | 0.042236 | 0.13291 | 0.965913 | 2864 | 4 | 1.3396 |
| MLX | 6 | 0.043731 | 0.1303 | 0.95318 | 2700 | 2 | 0.28398 | 0.042259 | 0.14159 | 0.965913 | 2865 | 3 | 0.28398 |
| hsa-mir-548o | 4 | 0.9577 | 0.95767 | 0.999994 | 20531 | 0 | 1.7103 | 0.042303 | 0.12497 | 0.964228 | 2866 | 4 | 1.7103 |
| ASNA1 | 5 | 0.87326 | 0.87401 | 0.999994 | 18701 | 1 | 2.668 | 0.042307 | 0.13307 | 0.965913 | 2867 | 4 | 2.668 |
| CST8 | 4 | 0.14477 | 0.28384 | 0.999632 | 6318 | 2 | 0.39277 | 0.042324 | 0.12503 | 0.964228 | 2868 | 2 | 0.39277 |
| UCN2 | 6 | 0.112 | 0.27054 | 0.998561 | 5364 | 2 | 3.084 | 0.042368 | 0.14187 | 0.965913 | 2869 | 4 | 3.084 |
| IFT88 | 6 | 0.27572 | 0.47843 | 0.999994 | 9611 | 3 | 0.014046 | 0.042368 | 0.14187 | 0.965913 | 2870 | 3 | 0.014046 |
| KCNMB3 | 6 | 0.061947 | 0.16988 | 0.967126 | 3510 | 3 | -1.3571 | 0.042368 | 0.14187 | 0.965913 | 2871 | 2 | -1.3571 |
| NSUN3 | 6 | 0.91214 | 0.91339 | 0.999994 | 19466 | 1 | 3.1883 | 0.042373 | 0.14189 | 0.965913 | 2872 | 5 | 3.1883 |
| GRAPL | 4 | 0.069284 | 0.15854 | 0.967126 | 3810 | 2 | 0.91149 | 0.042393 | 0.1252 | 0.964228 | 2873 | 2 | 0.91149 |
| hsa-mir-4500 | 4 | 0.5493 | 0.63933 | 0.999994 | 14200 | 1 | 4.871 | 0.042393 | 0.1252 | 0.964228 | 2874 | 3 | 4.871 |
| ZNF804B | 6 | 0.30037 | 0.50714 | 0.999994 | 10139 | 2 | 3.4174 | 0.042409 | 0.14198 | 0.965913 | 2875 | 4 | 3.4174 |
| SPAG5 | 6 | 0.33649 | 0.54866 | 0.999994 | 10860 | 1 | 4.1757 | 0.042414 | 0.14199 | 0.965913 | 2876 | 4 | 4.1757 |
| OR10Q1 | 5 | 0.94505 | 0.94522 | 0.999994 | 20206 | 0 | 5.4444 | 0.042415 | 0.13331 | 0.965913 | 2877 | 4 | 5.4444 |
| GALNS | 4 | 0.95758 | 0.95753 | 0.999994 | 20529 | 0 | 3.4408 | 0.042423 | 0.12528 | 0.964228 | 2878 | 4 | 3.4408 |
| hsa-mir-504 | 3 | 0.9575 | 0.95769 | 0.999994 | 20525 | 0 | 2.6075 | 0.042502 | 0.10337 | 0.952231 | 2879 | 3 | 2.6075 |
| FASTKD2 | 5 | 0.40468 | 0.56744 | 0.999994 | 12031 | 2 | -0.27312 | 0.042512 | 0.13352 | 0.965913 | 2880 | 2 | -0.27312 |
| ADAM22 | 6 | 0.90365 | 0.90651 | 0.999994 | 19296 | 1 | 3.2508 | 0.042518 | 0.14229 | 0.965913 | 2881 | 5 | 3.2508 |
| SLC17A3 | 6 | 0.31129 | 0.51984 | 0.999994 | 10346 | 1 | 3.7019 | 0.042565 | 0.1424 | 0.965913 | 2882 | 5 | 3.7019 |
| TROAP | 4 | 0.74948 | 0.75968 | 0.999994 | 16853 | 1 | 3.9484 | 0.042574 | 0.12568 | 0.964228 | 2883 | 2 | 3.9484 |
| OR52B2 | 6 | 0.97124 | 0.97121 | 0.999994 | 20852 | 0 | 4.0813 | 0.042608 | 0.14252 | 0.965913 | 2884 | 5 | 4.0813 |
| OR10K1 | 6 | 0.085542 | 0.21861 | 0.980699 | 4430 | 2 | 3.2111 | 0.042641 | 0.14262 | 0.965913 | 2885 | 4 | 3.2111 |
| FAM73A | 5 | 0.96366 | 0.96381 | 0.999994 | 20695 | 0 | 2.5146 | 0.042666 | 0.13386 | 0.965913 | 2886 | 4 | 2.5146 |
| hsa-mir-29b-1 | 4 | 0.95733 | 0.95728 | 0.999994 | 20524 | 0 | 3.2194 | 0.042667 | 0.12594 | 0.964228 | 2887 | 4 | 3.2194 |
| KIAA1737 | 6 | 0.21537 | 0.40659 | 0.999994 | 8152 | 2 | 1.4609 | 0.042717 | 0.14285 | 0.965913 | 2888 | 4 | 1.4609 |
| PTH2 | 6 | 0.096227 | 0.23992 | 0.987253 | 4826 | 1 | 1.6977 | 0.042717 | 0.14285 | 0.965913 | 2889 | 3 | 1.6977 |
| TGFBI | 6 | 0.30725 | 0.51519 | 0.999994 | 10272 | 1 | 2.6117 | 0.042717 | 0.14285 | 0.965913 | 2890 | 4 | 2.6117 |
| GPR142 | 4 | 0.35302 | 0.49735 | 0.999994 | 11214 | 1 | 4.6277 | 0.042749 | 0.12617 | 0.964228 | 2891 | 3 | 4.6277 |
| PLOD1 | 6 | 0.1288 | 0.29875 | 0.999994 | 5906 | 2 | 2.4642 | 0.04275 | 0.14294 | 0.965913 | 2892 | 3 | 2.4642 |
| IGHMBP2 | 5 | 0.9307 | 0.93098 | 0.999994 | 19869 | 0 | 3.052 | 0.0428 | 0.13417 | 0.965913 | 2893 | 4 | 3.052 |
| PPP1R7 | 5 | 0.0068827 | 0.027031 | 0.848412 | 653 | 3 | -5.0886 | 0.042815 | 0.13419 | 0.965913 | 2894 | 2 | -5.0886 |
| CTCF | 5 | 0.68305 | 0.7589 | 0.999994 | 16132 | 1 | 4.9089 | 0.042815 | 0.13419 | 0.965913 | 2895 | 4 | 4.9089 |
| PTPRM | 6 | 0.98458 | 0.98451 | 0.999994 | 21208 | 0 | 3.1979 | 0.042816 | 0.1431 | 0.965913 | 2896 | 5 | 3.1979 |
| GPATCH4 | 6 | 0.98008 | 0.97999 | 0.999994 | 21094 | 0 | 3.4488 | 0.042825 | 0.14312 | 0.965913 | 2897 | 4 | 3.4488 |
| ARR3 | 6 | 0.96988 | 0.96985 | 0.999994 | 20823 | 0 | 3.317 | 0.042859 | 0.14321 | 0.965913 | 2898 | 5 | 3.317 |
| NonTargetingControlG | 2 | 0.74854 | 0.74818 | 0.999994 | 16842 | 0 | 4.7048 | 0.04291 | 0.077265 | 0.938203 | 2899 | 1 | 4.7048 |
| BAHD1 | 5 | 0.40843 | 0.57056 | 0.999994 | 12087 | 2 | 5.4149 | 0.042921 | 0.13446 | 0.965913 | 2900 | 3 | 5.4149 |
| PURA | 4 | 0.69806 | 0.72316 | 0.999994 | 16281 | 1 | 3.9725 | 0.042973 | 0.12674 | 0.964228 | 2901 | 3 | 3.9725 |
| CCR7 | 5 | 0.036806 | 0.10789 | 0.951052 | 2365 | 1 | 5.5158 | 0.042975 | 0.13456 | 0.965913 | 2902 | 4 | 5.5158 |
| POLR2J | 5 | 0.79778 | 0.81813 | 0.999994 | 17450 | 1 | 5.5158 | 0.042975 | 0.13456 | 0.965913 | 2903 | 3 | 5.5158 |
| MLL | 5 | 0.54364 | 0.68611 | 0.999994 | 14124 | 1 | 5.5158 | 0.042975 | 0.13456 | 0.965913 | 2904 | 3 | 5.5158 |
| DTD1 | 5 | 0.94549 | 0.94567 | 0.999994 | 20218 | 0 | 5.5158 | 0.042975 | 0.13456 | 0.965913 | 2905 | 4 | 5.5158 |
| TMEM131 | 5 | 0.050996 | 0.14281 | 0.959553 | 3020 | 2 | 0.6722 | 0.043035 | 0.13471 | 0.965913 | 2906 | 3 | 0.6722 |
| hsa-mir-133a-2 | 4 | 0.61478 | 0.6734 | 0.999994 | 15138 | 1 | 5.5346 | 0.043041 | 0.12691 | 0.964228 | 2907 | 3 | 5.5346 |
| NonTargetingControlG | 2 | 0.70175 | 0.70152 | 0.999994 | 16324 | 0 | 4.5659 | 0.043067 | 0.077517 | 0.938203 | 2908 | 1 | 4.5659 |
| KLF7 | 6 | 0.25446 | 0.45327 | 0.999994 | 9111 | 2 | 3.2952 | 0.043075 | 0.14378 | 0.965913 | 2909 | 4 | 3.2952 |
| SYNJ1 | 6 | 0.58945 | 0.73218 | 0.999994 | 14812 | 1 | 1.7216 | 0.043095 | 0.14383 | 0.965913 | 2910 | 5 | 1.7216 |
| OR2B2 | 6 | 0.17507 | 0.35696 | 0.999994 | 7108 | 2 | 0.070304 | 0.043131 | 0.14392 | 0.965913 | 2911 | 2 | 0.070304 |

| | | | | | | | | | | | | | |
|---|---|---|---|---|---|---|---|---|---|---|---|---|---|
| TMEM40 | 5 | 0.36115 | 0.53103 | 0.999994 | 11355 | 2 | -0.16153 | 0.043135 | 0.13495 | 0.965913 | 2912 | 2 | -0.16153 |
| ARHGAP35 | 6 | 0.91203 | 0.91329 | 0.999994 | 19462 | 1 | 1.942 | 0.043136 | 0.14392 | 0.965913 | 2913 | 4 | 1.942 |
| KCNN4 | 6 | 0.67024 | 0.79332 | 0.999994 | 15936 | 1 | 0.99162 | 0.043136 | 0.14392 | 0.965913 | 2914 | 3 | 0.99162 |
| KRTAP5-9 | 6 | 0.50085 | 0.66927 | 0.999994 | 13436 | 2 | 4.5579 | 0.043211 | 0.14413 | 0.965913 | 2915 | 4 | 4.5579 |
| UBL4A | 5 | 0.35374 | 0.52497 | 0.999994 | 11227 | 2 | 0.97231 | 0.04324 | 0.13519 | 0.965913 | 2916 | 3 | 0.97231 |
| ADM5 | 4 | 0.95675 | 0.95666 | 0.999994 | 20510 | 0 | 4.1986 | 0.043251 | 0.12745 | 0.964228 | 2917 | 4 | 4.1986 |
| POLE2 | 3 | 0.15925 | 0.2605 | 0.994943 | 6712 | 1 | 6.6597 | 0.043315 | 0.10522 | 0.952231 | 2918 | 2 | 6.6597 |
| CFTR | 6 | 0.30175 | 0.50877 | 0.999994 | 10156 | 3 | -0.6468 | 0.043322 | 0.14443 | 0.965913 | 2919 | 1 | -0.6468 |
| NDUFB6 | 6 | 0.070288 | 0.18738 | 0.969409 | 3846 | 2 | 3.3664 | 0.043331 | 0.14445 | 0.965913 | 2920 | 4 | 3.3664 |
| AQP12B | 5 | 0.91494 | 0.91503 | 0.999994 | 19510 | 0 | 5.5005 | 0.043336 | 0.1354 | 0.965913 | 2921 | 3 | 5.5005 |
| SPSB1 | 6 | 0.81868 | 0.8562 | 0.999994 | 17760 | 1 | 3.8551 | 0.043337 | 0.14447 | 0.965913 | 2922 | 5 | 3.8551 |
| TAX1BP3 | 6 | 0.19222 | 0.37827 | 0.999994 | 7552 | 1 | 3.763 | 0.043337 | 0.14447 | 0.965913 | 2923 | 4 | 3.763 |
| LRIT3 | 6 | 0.37881 | 0.58882 | 0.999994 | 11637 | 2 | 2.9853 | 0.043337 | 0.14447 | 0.965913 | 2924 | 3 | 2.9853 |
| DPRX | 5 | 0.071522 | 0.18085 | 0.968171 | 3897 | 1 | 2.0693 | 0.043376 | 0.13549 | 0.965913 | 2925 | 4 | 2.0693 |
| TRIM37 | 6 | 0.9305 | 0.93063 | 0.999994 | 19864 | 0 | 3.525 | 0.043388 | 0.14461 | 0.965913 | 2926 | 5 | 3.525 |
| CPNE6 | 4 | 0.95661 | 0.95652 | 0.999994 | 20506 | 0 | 3.2944 | 0.043394 | 0.12781 | 0.964228 | 2927 | 4 | 3.2944 |
| DCUN1D2 | 6 | 0.4097 | 0.60869 | 0.999994 | 12104 | 2 | 2.9668 | 0.043404 | 0.14465 | 0.965913 | 2928 | 4 | 2.9668 |
| PARP10 | 4 | 0.9566 | 0.95651 | 0.999994 | 20504 | 0 | 4.0918 | 0.043405 | 0.12784 | 0.964228 | 2929 | 4 | 4.0918 |
| TFAP2A | 6 | 0.43544 | 0.62542 | 0.999994 | 12507 | 2 | 4.4381 | 0.043418 | 0.1447 | 0.965913 | 2930 | 4 | 4.4381 |
| LIPI | 6 | 0.3299 | 0.5411 | 0.999994 | 10721 | 1 | 1.468 | 0.04342 | 0.1447 | 0.965913 | 2931 | 5 | 1.468 |
| hsa-mir-214 | 4 | 0.95658 | 0.95649 | 0.999994 | 20503 | 0 | 2.0564 | 0.043422 | 0.12788 | 0.964228 | 2932 | 4 | 2.0564 |
| GGN | 6 | 0.8108 | 0.85262 | 0.999994 | 17644 | 1 | 3.5144 | 0.043429 | 0.14473 | 0.965913 | 2933 | 4 | 3.5144 |
| PIK3R2 | 6 | 0.90337 | 0.90631 | 0.999994 | 19288 | 1 | 2.9689 | 0.043461 | 0.14481 | 0.965913 | 2934 | 5 | 2.9689 |
| NPHS1 | 6 | 0.93459 | 0.93468 | 0.999994 | 19959 | 0 | 2.333 | 0.043485 | 0.14487 | 0.965913 | 2935 | 5 | 2.333 |
| C17orf66 | 6 | 0.19739 | 0.38473 | 0.999994 | 7684 | 2 | 0.74458 | 0.043486 | 0.14487 | 0.965913 | 2936 | 3 | 0.74458 |
| SLFN12 | 5 | 0.63516 | 0.73931 | 0.999994 | 15446 | 1 | 2.668 | 0.043535 | 0.13586 | 0.965913 | 2937 | 4 | 2.668 |
| FBN3 | 5 | 0.97235 | 0.97241 | 0.999994 | 20876 | 0 | 3.3734 | 0.043536 | 0.13586 | 0.965913 | 2938 | 4 | 3.3734 |
| KRTAP4-9 | 2 | 0.95645 | 0.95654 | 0.999994 | 20501 | 0 | 5.6862 | 0.04355 | 0.07835 | 0.938203 | 2939 | 2 | 5.6862 |
| DESI1 | 6 | 0.038925 | 0.11947 | 0.951052 | 2456 | 1 | 2.8502 | 0.043552 | 0.14506 | 0.965913 | 2940 | 5 | 2.8502 |
| ABCF2 | 6 | 0.94745 | 0.94747 | 0.999994 | 20276 | 0 | 4.4131 | 0.043561 | 0.14509 | 0.965913 | 2941 | 5 | 4.4131 |
| ARF3 | 5 | 0.042716 | 0.12262 | 0.951052 | 2662 | 1 | 5.5422 | 0.043569 | 0.13593 | 0.965913 | 2942 | 3 | 5.5422 |
| OR10G3 | 6 | 0.89488 | 0.89981 | 0.999994 | 19123 | 1 | 1.8662 | 0.043595 | 0.14517 | 0.965913 | 2943 | 5 | 1.8662 |
| MRPL21 | 4 | 0.12479 | 0.25862 | 0.994943 | 5791 | 1 | 5.416 | 0.043631 | 0.12843 | 0.964228 | 2944 | 3 | 5.416 |
| POTEG | 4 | 0.4051 | 0.55094 | 0.999994 | 12033 | 1 | 4.2908 | 0.043631 | 0.12843 | 0.964228 | 2945 | 3 | 4.2908 |
| HAUS4 | 3 | 0.95636 | 0.95657 | 0.999994 | 20497 | 0 | 2.0691 | 0.043639 | 0.10595 | 0.952231 | 2946 | 3 | 2.0691 |
| KDM4E | 6 | 0.98043 | 0.98033 | 0.999994 | 21102 | 0 | 3.9956 | 0.043647 | 0.14531 | 0.965913 | 2947 | 5 | 3.9956 |
| C10orf12 | 6 | 0.062515 | 0.17109 | 0.96737 | 3537 | 1 | 3.5608 | 0.043647 | 0.14531 | 0.965913 | 2948 | 5 | 3.5608 |
| UGT1A1 | 6 | 0.94871 | 0.94871 | 0.999994 | 20305 | 0 | 1.2798 | 0.043688 | 0.14542 | 0.965913 | 2949 | 5 | 1.2798 |
| ADAM8 | 6 | 0.88592 | 0.8935 | 0.999994 | 18947 | 1 | 3.5805 | 0.043691 | 0.14542 | 0.965913 | 2950 | 4 | 3.5805 |
| hsa-mir-181b-2 | 4 | 0.95629 | 0.9562 | 0.999994 | 20495 | 0 | 2.2358 | 0.043714 | 0.12864 | 0.964228 | 2951 | 4 | 2.2358 |
| FAM65A | 5 | 0.40234 | 0.56547 | 0.999994 | 11984 | 1 | 0.29796 | 0.04374 | 0.13633 | 0.965913 | 2952 | 2 | 0.29796 |
| SAP30L | 4 | 0.95622 | 0.95614 | 0.999994 | 20494 | 0 | 2.3717 | 0.043784 | 0.12882 | 0.964228 | 2953 | 4 | 2.3717 |
| hsa-mir-6877 | 2 | 0.9562 | 0.95629 | 0.999994 | 20492 | 0 | 6.4 | 0.0438 | 0.078781 | 0.938203 | 2954 | 2 | 6.4 |
| NonTargetingControlG | 2 | 0.9562 | 0.95629 | 0.999994 | 20493 | 0 | 6.6424 | 0.0438 | 0.078781 | 0.938203 | 2955 | 2 | 6.6424 |
| HMGA1 | 6 | 0.9637 | 0.96377 | 0.999994 | 20698 | 0 | 1.5743 | 0.043813 | 0.14574 | 0.965913 | 2956 | 5 | 1.5743 |
| ACTR3B | 6 | 0.87692 | 0.88765 | 0.999994 | 18771 | 1 | 1.7432 | 0.043823 | 0.14576 | 0.965913 | 2957 | 5 | 1.7432 |
| NonTargetingControlG | 2 | 0.73922 | 0.7389 | 0.999994 | 16724 | 0 | 4.6751 | 0.043842 | 0.078851 | 0.938203 | 2958 | 1 | 4.6751 |
| HEBP2 | 6 | 0.91546 | 0.91631 | 0.999994 | 19523 | 1 | 2.4498 | 0.043844 | 0.14582 | 0.965913 | 2959 | 5 | 2.4498 |
| CEMP1 | 6 | 0.8771 | 0.88776 | 0.999994 | 18775 | 1 | 2.1504 | 0.043857 | 0.14585 | 0.965913 | 2960 | 5 | 2.1504 |
| hsa-mir-5003 | 4 | 0.76686 | 0.77326 | 0.999994 | 17048 | 1 | 3.6335 | 0.043888 | 0.12912 | 0.964228 | 2961 | 2 | 3.6335 |
| FCGR1A | 3 | 0.95605 | 0.95625 | 0.999994 | 20482 | 0 | 5.5347 | 0.043948 | 0.10664 | 0.953839 | 2962 | 3 | 5.5347 |
| HMGCL | 6 | 0.42427 | 0.61819 | 0.999994 | 12317 | 2 | 0.5437 | 0.043966 | 0.14613 | 0.965913 | 2963 | 3 | 0.5437 |

| | | | | | | | | | | | | | |
|---|---|---|---|---|---|---|---|---|---|---|---|---|---|
| FAM71F2 | 6 | 0.15558 | 0.33268 | 0.999994 | 6610 | 2 | 1.1864 | 0.043966 | 0.14613 | 0.965913 | 2964 | 4 | 1.1864 |
| GDPD1 | 6 | 0.95293 | 0.95292 | 0.999994 | 20408 | 0 | 1.1959 | 0.044058 | 0.14635 | 0.965913 | 2965 | 4 | 1.1959 |
| ATP5B | 4 | 0.62768 | 0.68058 | 0.999994 | 15348 | 1 | 3.8695 | 0.04406 | 0.12956 | 0.964228 | 2966 | 3 | 3.8695 |
| NLRP11 | 4 | 0.95594 | 0.95585 | 0.999994 | 20480 | 0 | 1.2062 | 0.044061 | 0.12956 | 0.964228 | 2967 | 4 | 1.2062 |
| NonTargetingControlG | 2 | 0.27459 | 0.31735 | 0.999994 | 9583 | 1 | 3.182 | 0.044063 | 0.079232 | 0.938203 | 2968 | 1 | 3.182 |
| ARMCX6 | 4 | 0.65322 | 0.69515 | 0.999994 | 15701 | 1 | 2.1145 | 0.044075 | 0.12961 | 0.964228 | 2969 | 3 | 2.1145 |
| PHF19 | 4 | 0.010083 | 0.033985 | 0.867348 | 874 | 2 | -3.4575 | 0.044075 | 0.12961 | 0.964228 | 2970 | 2 | -3.4575 |
| STRC | 5 | 0.56952 | 0.7091 | 0.999994 | 14530 | 2 | 1.4909 | 0.044096 | 0.13714 | 0.965913 | 2971 | 3 | 1.4909 |
| ACADM | 5 | 0.52062 | 0.66585 | 0.999994 | 13756 | 2 | -0.17909 | 0.044096 | 0.13714 | 0.965913 | 2972 | 2 | -0.17909 |
| ZNF559 | 6 | 0.23649 | 0.43181 | 0.999994 | 8682 | 1 | 0.70636 | 0.044112 | 0.1465 | 0.965913 | 2973 | 3 | 0.70636 |
| SLC7A11 | 5 | 0.69179 | 0.76279 | 0.999994 | 16225 | 1 | 2.7298 | 0.04415 | 0.13727 | 0.965913 | 2974 | 4 | 2.7298 |
| YAF2 | 5 | 0.80397 | 0.82201 | 0.999994 | 17541 | 1 | 3.9691 | 0.044163 | 0.1373 | 0.965913 | 2975 | 4 | 3.9691 |
| PTPRK | 5 | 0.83763 | 0.84492 | 0.999994 | 18046 | 1 | 2.0858 | 0.044186 | 0.13734 | 0.965913 | 2976 | 4 | 2.0858 |
| ELF2 | 6 | 0.49682 | 0.66654 | 0.999994 | 13370 | 1 | 0.19272 | 0.044187 | 0.14671 | 0.965913 | 2977 | 2 | 0.19272 |
| PRG3 | 6 | 0.94022 | 0.94017 | 0.999994 | 20089 | 0 | 0.57758 | 0.044187 | 0.14671 | 0.965913 | 2978 | 3 | 0.57758 |
| NDST3 | 6 | 0.44435 | 0.63132 | 0.999994 | 12638 | 2 | 3.0949 | 0.044187 | 0.14671 | 0.965913 | 2979 | 3 | 3.0949 |
| EIF3B | 6 | 0.96019 | 0.96025 | 0.999994 | 20587 | 0 | 3.2648 | 0.044187 | 0.14671 | 0.965913 | 2980 | 5 | 3.2648 |
| SCP2D1 | 6 | 0.16944 | 0.35005 | 0.999994 | 6973 | 3 | 0.63426 | 0.044187 | 0.14671 | 0.965913 | 2981 | 3 | 0.63426 |
| PGBD3 | 6 | 0.59692 | 0.7377 | 0.999994 | 14916 | 2 | 0.46119 | 0.044194 | 0.14672 | 0.965913 | 2982 | 2 | 0.46119 |
| ZSCAN32 | 6 | 0.65048 | 0.77797 | 0.999994 | 15665 | 1 | 1.3544 | 0.044252 | 0.14687 | 0.965913 | 2983 | 5 | 1.3544 |
| PGM2 | 4 | 0.79388 | 0.79579 | 0.999994 | 17394 | 1 | 5.4414 | 0.04427 | 0.13012 | 0.964228 | 2984 | 3 | 5.4414 |
| SNRNP48 | 4 | 0.40449 | 0.55029 | 0.999994 | 12028 | 1 | 4.2047 | 0.04427 | 0.13012 | 0.964228 | 2985 | 3 | 4.2047 |
| WDR25 | 4 | 0.58157 | 0.65557 | 0.999994 | 14701 | 1 | 5.4796 | 0.04427 | 0.13012 | 0.964228 | 2986 | 3 | 5.4796 |
| ADAM21 | 5 | 0.77355 | 0.80371 | 0.999994 | 17126 | 1 | 2.3066 | 0.044301 | 0.13761 | 0.965913 | 2987 | 4 | 2.3066 |
| UTP15 | 3 | 0.12873 | 0.2209 | 0.981851 | 5904 | 1 | 6.6837 | 0.044339 | 0.10752 | 0.953839 | 2988 | 2 | 6.6837 |
| UBXN11 | 6 | 0.37417 | 0.58584 | 0.999994 | 11574 | 2 | 1.538 | 0.044358 | 0.14716 | 0.965913 | 2989 | 4 | 1.538 |
| MRPS28 | 6 | 0.34995 | 0.56394 | 0.999994 | 11157 | 2 | 0.15443 | 0.044358 | 0.14716 | 0.965913 | 2990 | 2 | 0.15443 |
| PSAT1 | 6 | 0.9422 | 0.94218 | 0.999994 | 20131 | 0 | 2.1384 | 0.044362 | 0.14718 | 0.965913 | 2991 | 5 | 2.1384 |
| SMG5 | 5 | 0.60131 | 0.72665 | 0.999994 | 14964 | 1 | 3.9691 | 0.044365 | 0.13776 | 0.965913 | 2992 | 4 | 3.9691 |
| CPEB1 | 4 | 0.45586 | 0.59716 | 0.999994 | 12788 | 1 | 4.3464 | 0.0444 | 0.13045 | 0.964228 | 2993 | 3 | 4.3464 |
| ZFAND4 | 6 | 0.66423 | 0.78858 | 0.999994 | 15844 | 1 | 4.137 | 0.04446 | 0.14742 | 0.965913 | 2994 | 4 | 4.137 |
| PLCB1 | 6 | 0.37736 | 0.58789 | 0.999994 | 11613 | 1 | 3.2341 | 0.044473 | 0.14746 | 0.965913 | 2995 | 5 | 3.2341 |
| PDSS2 | 3 | 0.95551 | 0.95573 | 0.999994 | 20467 | 0 | 6.0163 | 0.044489 | 0.10783 | 0.953839 | 2996 | 3 | 6.0163 |
| OR7D2 | 6 | 0.34618 | 0.55971 | 0.999994 | 11077 | 2 | -0.068504 | 0.044494 | 0.14751 | 0.965913 | 2997 | 2 | -0.068504 |
| RSPH3 | 6 | 0.65624 | 0.78238 | 0.999994 | 15728 | 2 | 4.3684 | 0.044572 | 0.14773 | 0.965913 | 2998 | 4 | 4.3684 |
| MIB1 | 6 | 0.68581 | 0.80555 | 0.999994 | 16169 | 2 | 3.3967 | 0.044586 | 0.14776 | 0.965913 | 2999 | 4 | 3.3967 |
| SLC38A6 | 6 | 0.076499 | 0.20028 | 0.977164 | 4092 | 1 | 3.223 | 0.044591 | 0.14778 | 0.965913 | 3000 | 4 | 3.223 |
| PC | 6 | 0.71875 | 0.81793 | 0.999994 | 16484 | 1 | 4.3718 | 0.044591 | 0.14778 | 0.965913 | 3001 | 4 | 4.3718 |
| PRKAG2 | 6 | 0.97002 | 0.96999 | 0.999994 | 20824 | 0 | 2.8571 | 0.044593 | 0.14778 | 0.965913 | 3002 | 5 | 2.8571 |
| GOLGA6L1 | 5 | 0.56225 | 0.70264 | 0.999994 | 14413 | 1 | 2.3601 | 0.044604 | 0.13831 | 0.965913 | 3003 | 4 | 2.3601 |
| C16orf62 | 5 | 0.33681 | 0.51111 | 0.999994 | 10868 | 2 | 1.7857 | 0.044609 | 0.13833 | 0.965913 | 3004 | 3 | 1.7857 |
| COX6B2 | 5 | 0.94811 | 0.94832 | 0.999994 | 20288 | 0 | 1.9175 | 0.044609 | 0.13833 | 0.965913 | 3005 | 4 | 1.9175 |
| CYP2B6 | 5 | 0.17756 | 0.34312 | 0.999994 | 7169 | 1 | 2.414 | 0.044627 | 0.13837 | 0.965913 | 3006 | 4 | 2.414 |
| ATP6V1E1 | 4 | 0.95537 | 0.95527 | 0.999994 | 20462 | 0 | 1.3822 | 0.044633 | 0.13104 | 0.964228 | 3007 | 4 | 1.3822 |
| CYP3A43 | 5 | 0.35161 | 0.52322 | 0.999994 | 11188 | 1 | 2.162 | 0.044641 | 0.1384 | 0.965913 | 3008 | 4 | 2.162 |
| hsa-mir-6805 | 4 | 0.95535 | 0.95524 | 0.999994 | 20461 | 0 | 1.5508 | 0.044651 | 0.13109 | 0.964228 | 3009 | 4 | 1.5508 |
| LIX1 | 6 | 0.94905 | 0.94904 | 0.999994 | 20310 | 0 | 3.2724 | 0.044661 | 0.14796 | 0.965913 | 3010 | 4 | 3.2724 |
| KRTAP3-1 | 5 | 0.80569 | 0.8231 | 0.999994 | 17565 | 1 | 2.1608 | 0.044673 | 0.13847 | 0.965913 | 3011 | 4 | 2.1608 |
| NonTargetingControlG | 2 | 0.95529 | 0.95538 | 0.999994 | 20458 | 0 | 6.2508 | 0.044707 | 0.08034 | 0.938962 | 3012 | 2 | 6.2508 |
| hsa-mir-5707 | 2 | 0.95529 | 0.95538 | 0.999994 | 20459 | 0 | 6.3655 | 0.044707 | 0.08034 | 0.938962 | 3013 | 2 | 6.3655 |
| OLA1 | 6 | 0.25279 | 0.45125 | 0.999994 | 9074 | 1 | 4.5763 | 0.044711 | 0.1481 | 0.965913 | 3014 | 4 | 4.5763 |
| SUPT6H | 5 | 0.7034 | 0.76804 | 0.999994 | 16343 | 1 | 5.4494 | 0.044731 | 0.13861 | 0.965913 | 3015 | 3 | 5.4494 |

| | | | | | | | | | | | | | |
|---|---|---|---|---|---|---|---|---|---|---|---|---|---|
| SUV39H1 | 6 | 0.2575 | 0.45692 | 0.999994 | 9179 | 1 | 2.797 | 0.044739 | 0.14818 | 0.965913 | 3016 | 5 | 2.797 |
| CEACAM3 | 6 | 0.050709 | 0.14563 | 0.963656 | 3003 | 2 | 3.7999 | 0.04475 | 0.1482 | 0.965913 | 3017 | 4 | 3.7999 |
| RBM33 | 5 | 0.30018 | 0.48122 | 0.999994 | 10136 | 2 | 0.53675 | 0.044763 | 0.13869 | 0.965913 | 3018 | 3 | 0.53675 |
| hsa-mir-124-2 | 4 | 0.95522 | 0.95511 | 0.999994 | 20457 | 0 | 1.7903 | 0.044776 | 0.1314 | 0.964897 | 3019 | 4 | 1.7903 |
| SCGB1D2 | 5 | 0.93192 | 0.93221 | 0.999994 | 19898 | 0 | 0.97549 | 0.044826 | 0.13883 | 0.965913 | 3020 | 3 | 0.97549 |
| CELF4 | 5 | 0.65166 | 0.7458 | 0.999994 | 15682 | 1 | 2.8748 | 0.044826 | 0.13883 | 0.965913 | 3021 | 4 | 2.8748 |
| USP45 | 6 | 0.63274 | 0.7644 | 0.999994 | 15410 | 1 | 3.0711 | 0.044841 | 0.14844 | 0.965913 | 3022 | 5 | 3.0711 |
| KRT77 | 6 | 0.27502 | 0.47761 | 0.999994 | 9593 | 1 | 2.7291 | 0.044841 | 0.14844 | 0.965913 | 3023 | 5 | 2.7291 |
| hsa-mir-633 | 4 | 0.89843 | 0.89814 | 0.999994 | 19192 | 0 | 4.8835 | 0.044898 | 0.13171 | 0.965199 | 3024 | 3 | 4.8835 |
| SH3BP2 | 5 | 0.81238 | 0.82751 | 0.999994 | 17657 | 1 | 2.8679 | 0.044965 | 0.13914 | 0.965913 | 3025 | 4 | 2.8679 |
| RUVBL2 | 6 | 0.88751 | 0.89459 | 0.999994 | 18986 | 1 | 4.2515 | 0.045015 | 0.14888 | 0.965913 | 3026 | 4 | 4.2515 |
| IER3 | 5 | 0.95497 | 0.95513 | 0.999994 | 20452 | 0 | 3.6194 | 0.045016 | 0.13926 | 0.965913 | 3027 | 4 | 3.6194 |
| CNKSR2 | 6 | 0.051137 | 0.14656 | 0.964852 | 3027 | 3 | -0.26055 | 0.045025 | 0.1489 | 0.965913 | 3028 | 2 | -0.26055 |
| APH1B | 6 | 0.44599 | 0.63243 | 0.999994 | 12663 | 1 | 4.3362 | 0.045025 | 0.1489 | 0.965913 | 3029 | 4 | 4.3362 |
| HNF1B | 6 | 0.91623 | 0.91696 | 0.999994 | 19540 | 1 | -0.078203 | 0.045066 | 0.14901 | 0.965913 | 3030 | 2 | -0.078203 |
| IER5L | 6 | 0.45503 | 0.6384 | 0.999994 | 12773 | 2 | 2.078 | 0.045066 | 0.14901 | 0.965913 | 3031 | 3 | 2.078 |
| IL1RL2 | 6 | 0.34836 | 0.56217 | 0.999994 | 11120 | 2 | 2.8943 | 0.045071 | 0.14902 | 0.965913 | 3032 | 3 | 2.8943 |
| OR52I1 | 6 | 0.43891 | 0.62768 | 0.999994 | 12555 | 2 | 4.008 | 0.045071 | 0.14902 | 0.965913 | 3033 | 4 | 4.008 |
| HIST2H2BF | 5 | 0.69505 | 0.76424 | 0.999994 | 16255 | 1 | 4.2459 | 0.045104 | 0.13946 | 0.965913 | 3034 | 3 | 4.2459 |
| hsa-mir-519a-2 | 3 | 0.95489 | 0.95511 | 0.999994 | 20450 | 0 | 3.5803 | 0.04511 | 0.10916 | 0.953839 | 3035 | 3 | 3.5803 |
| RANBP3L | 4 | 0.87706 | 0.87675 | 0.999994 | 18773 | 0 | 1.6215 | 0.045132 | 0.13228 | 0.965913 | 3036 | 3 | 1.6215 |
| ZNF780B | 6 | 0.4132 | 0.61093 | 0.999994 | 12155 | 1 | 3.0084 | 0.045151 | 0.14924 | 0.965913 | 3037 | 5 | 3.0084 |
| FAM131C | 4 | 0.93984 | 0.93973 | 0.999994 | 20076 | 0 | 4.0292 | 0.045189 | 0.13242 | 0.965913 | 3038 | 3 | 4.0292 |
| SPTA1 | 6 | 0.53926 | 0.69602 | 0.999994 | 14040 | 2 | -0.21541 | 0.045249 | 0.1495 | 0.965913 | 3039 | 2 | -0.21541 |
| PLIN3 | 6 | 0.26521 | 0.46604 | 0.999994 | 9354 | 2 | 1.5296 | 0.045249 | 0.1495 | 0.965913 | 3040 | 3 | 1.5296 |
| MIEF2 | 5 | 0.64126 | 0.74167 | 0.999994 | 15536 | 1 | 4.8276 | 0.04525 | 0.1398 | 0.965913 | 3041 | 4 | 4.8276 |
| COPZ2 | 5 | 0.49777 | 0.64594 | 0.999994 | 13393 | 2 | -0.56086 | 0.04525 | 0.1398 | 0.965913 | 3042 | 2 | -0.56086 |
| PAF1 | 5 | 0.72412 | 0.77785 | 0.999994 | 16535 | 1 | 4.5407 | 0.04525 | 0.1398 | 0.965913 | 3043 | 4 | 4.5407 |
| NonTargetingControlG | 2 | 0.87291 | 0.87297 | 0.999994 | 18690 | 0 | 5.3729 | 0.045252 | 0.081214 | 0.939346 | 3044 | 2 | 5.3729 |
| POPDC3 | 5 | 0.59513 | 0.72434 | 0.999994 | 14891 | 1 | 2.506 | 0.045267 | 0.13983 | 0.965913 | 3045 | 4 | 2.506 |
| SULT1E1 | 6 | 0.72014 | 0.81837 | 0.999994 | 16500 | 1 | 1.358 | 0.045286 | 0.1496 | 0.965913 | 3046 | 5 | 1.358 |
| DOK2 | 5 | 0.8292 | 0.83891 | 0.999994 | 17915 | 1 | 5.2908 | 0.045305 | 0.13992 | 0.965913 | 3047 | 4 | 5.2908 |
| DDX17 | 6 | 0.33614 | 0.54824 | 0.999994 | 10850 | 2 | 4.2133 | 0.045321 | 0.1497 | 0.965913 | 3048 | 4 | 4.2133 |
| SLFN13 | 6 | 0.15732 | 0.33485 | 0.999994 | 6655 | 2 | 1.6473 | 0.045374 | 0.14983 | 0.965913 | 3049 | 3 | 1.6473 |
| RBFOX3 | 6 | 0.63029 | 0.76256 | 0.999994 | 15382 | 2 | 3.5029 | 0.045421 | 0.14995 | 0.965913 | 3050 | 4 | 3.5029 |
| PDE7B | 6 | 0.62116 | 0.75571 | 0.999994 | 15247 | 2 | 3.4264 | 0.045421 | 0.14995 | 0.965913 | 3051 | 3 | 3.4264 |
| COX7B2 | 6 | 0.70216 | 0.81268 | 0.999994 | 16329 | 1 | 3.867 | 0.045421 | 0.14995 | 0.965913 | 3052 | 4 | 3.867 |
| CCDC136 | 6 | 0.74213 | 0.82571 | 0.999994 | 16765 | 1 | 3.6468 | 0.045421 | 0.14995 | 0.965913 | 3053 | 5 | 3.6468 |
| RRBP1 | 6 | 0.80924 | 0.85191 | 0.999994 | 17619 | 1 | 4.3576 | 0.045421 | 0.14995 | 0.965913 | 3054 | 5 | 4.3576 |
| PLD1 | 6 | 0.3906 | 0.59635 | 0.999994 | 11809 | 2 | 3.2035 | 0.045421 | 0.14995 | 0.965913 | 3055 | 3 | 3.2035 |
| DDB1 | 5 | 0.63359 | 0.73871 | 0.999994 | 15425 | 1 | 2.913 | 0.045449 | 0.14024 | 0.965913 | 3056 | 4 | 2.913 |
| GJC1 | 5 | 0.32184 | 0.49877 | 0.999994 | 10552 | 1 | 5.4173 | 0.045453 | 0.14025 | 0.965913 | 3057 | 4 | 5.4173 |
| SLC35G1 | 4 | 0.22848 | 0.36994 | 0.999994 | 8489 | 1 | 4.1578 | 0.045462 | 0.13307 | 0.965913 | 3058 | 3 | 4.1578 |
| ECHDC2 | 5 | 0.62895 | 0.7369 | 0.999994 | 15362 | 1 | 0.0033305 | 0.045463 | 0.14027 | 0.965913 | 3059 | 2 | 0.0033305 |
| KBTBD7 | 6 | 0.21662 | 0.40809 | 0.999994 | 8183 | 3 | 0.073596 | 0.045583 | 0.15039 | 0.965913 | 3060 | 3 | 0.073596 |
| AGXT | 6 | 0.50823 | 0.6744 | 0.999994 | 13557 | 1 | 1.5846 | 0.045598 | 0.15043 | 0.965913 | 3061 | 4 | 1.5846 |
| CELF3 | 4 | 0.95438 | 0.95424 | 0.999994 | 20441 | 0 | 2.4281 | 0.045615 | 0.13346 | 0.965913 | 3062 | 4 | 2.4281 |
| KIF4B | 5 | 0.15448 | 0.30947 | 0.999994 | 6578 | 1 | 5.3957 | 0.04562 | 0.14063 | 0.965913 | 3063 | 3 | 5.3957 |
| MED28 | 6 | 0.57641 | 0.72255 | 0.999994 | 14635 | 1 | 2.4148 | 0.045634 | 0.15053 | 0.965913 | 3064 | 5 | 2.4148 |
| RNF146 | 4 | 0.8592 | 0.85879 | 0.999994 | 18417 | 0 | 3.5411 | 0.045669 | 0.13359 | 0.965913 | 3065 | 2 | 3.5411 |
| TM2D2 | 6 | 0.84965 | 0.87174 | 0.999994 | 18253 | 1 | 1.47 | 0.045695 | 0.15068 | 0.965913 | 3066 | 5 | 1.47 |
| hsa-mir-5705 | 4 | 0.3112 | 0.45449 | 0.999994 | 10344 | 1 | 4.2111 | 0.045704 | 0.13367 | 0.965913 | 3067 | 3 | 4.2111 |

| | | | | | | | | | | | | | |
|---|---|---|---|---|---|---|---|---|---|---|---|---|---|
| C4orf36 | 4 | 0.0097884 | 0.033067 | 0.867348 | 858 | 1 | 3.8876 | 0.045704 | 0.13367 | 0.965913 | 3068 | 3 | 3.8876 |
| SLC22A10 | 4 | 0.9543 | 0.95415 | 0.999994 | 20438 | 0 | 3.406 | 0.045705 | 0.13367 | 0.965913 | 3069 | 4 | 3.406 |
| EDA2R | 6 | 0.061603 | 0.16917 | 0.967126 | 3497 | 2 | 3.6274 | 0.045717 | 0.15075 | 0.965913 | 3070 | 4 | 3.6274 |
| RPS8 | 3 | 0.1309 | 0.22374 | 0.983354 | 5955 | 1 | 6.5854 | 0.04578 | 0.11064 | 0.953839 | 3071 | 2 | 6.5854 |
| C15orf32 | 5 | 0.20186 | 0.37791 | 0.999994 | 7820 | 1 | 1.4711 | 0.045785 | 0.141 | 0.965913 | 3072 | 3 | 1.4711 |
| GATA4 | 6 | 0.59316 | 0.73491 | 0.999994 | 14866 | 2 | 4.0395 | 0.0458 | 0.15095 | 0.965913 | 3073 | 4 | 4.0395 |
| PDCD1LG2 | 6 | 0.74705 | 0.82743 | 0.999994 | 16829 | 1 | 3.5048 | 0.0458 | 0.15095 | 0.965913 | 3074 | 4 | 3.5048 |
| ASB1 | 6 | 0.25823 | 0.45779 | 0.999994 | 9202 | 1 | 1.14 | 0.045817 | 0.15099 | 0.965913 | 3075 | 4 | 1.14 |
| PATE3 | 6 | 0.14071 | 0.31394 | 0.999994 | 6209 | 1 | 0.58434 | 0.045827 | 0.15102 | 0.965913 | 3076 | 3 | 0.58434 |
| GDF10 | 5 | 0.34094 | 0.51451 | 0.999994 | 10960 | 1 | 3.4553 | 0.045833 | 0.14112 | 0.965913 | 3077 | 4 | 3.4553 |
| MPP3 | 6 | 0.77607 | 0.83813 | 0.999994 | 17150 | 1 | 3.5881 | 0.045879 | 0.15116 | 0.965913 | 3078 | 5 | 3.5881 |
| C17orf96 | 5 | 0.67638 | 0.75601 | 0.999994 | 16028 | 1 | 0.86173 | 0.045895 | 0.14125 | 0.965913 | 3079 | 3 | 0.86173 |
| CAPN3 | 6 | 0.9338 | 0.93386 | 0.999994 | 19947 | 0 | 4.4956 | 0.045956 | 0.15138 | 0.965913 | 3080 | 5 | 4.4956 |
| RAB39A | 6 | 0.92531 | 0.92541 | 0.999994 | 19753 | 1 | 3.7413 | 0.045956 | 0.15138 | 0.965913 | 3081 | 4 | 3.7413 |
| SIGLECL1 | 6 | 0.88193 | 0.89087 | 0.999994 | 18867 | 1 | 3.7028 | 0.045991 | 0.15146 | 0.965913 | 3082 | 4 | 3.7028 |
| SLCO1B1 | 5 | 0.59093 | 0.72286 | 0.999994 | 14834 | 1 | 2.5781 | 0.046006 | 0.14151 | 0.965913 | 3083 | 4 | 2.5781 |
| ABCB10 | 5 | 0.0073776 | 0.028439 | 0.858757 | 684 | 3 | -0.90693 | 0.046012 | 0.14152 | 0.965913 | 3084 | 2 | -0.90693 |
| SLC10A2 | 5 | 0.84019 | 0.8468 | 0.999994 | 18092 | 1 | 5.479 | 0.046063 | 0.14164 | 0.965913 | 3085 | 4 | 5.479 |
| SLC35A5 | 5 | 0.54253 | 0.68513 | 0.999994 | 14102 | 1 | 2.4619 | 0.046103 | 0.14173 | 0.965913 | 3086 | 4 | 2.4619 |
| MAPK8IP2 | 5 | 0.050013 | 0.14039 | 0.958716 | 2977 | 2 | -0.28464 | 0.046103 | 0.14173 | 0.965913 | 3087 | 1 | -0.28464 |
| SLC7A2 | 5 | 0.77659 | 0.80549 | 0.999994 | 17160 | 1 | 2.3495 | 0.046105 | 0.14173 | 0.965913 | 3088 | 4 | 2.3495 |
| IPO9 | 6 | 0.94564 | 0.94561 | 0.999994 | 20223 | 0 | 2.3031 | 0.046111 | 0.15179 | 0.965913 | 3089 | 5 | 2.3031 |
| hsa-mir-520g | 2 | 0.69667 | 0.69646 | 0.999994 | 16267 | 0 | 4.5122 | 0.046117 | 0.082656 | 0.939346 | 3090 | 1 | 4.5122 |
| SLC17A2 | 6 | 0.71484 | 0.81667 | 0.999994 | 16447 | 1 | 3.6487 | 0.046135 | 0.15185 | 0.965913 | 3091 | 4 | 3.6487 |
| LGR6 | 6 | 0.67961 | 0.80069 | 0.999994 | 16084 | 2 | 0.59071 | 0.046135 | 0.15185 | 0.965913 | 3092 | 3 | 0.59071 |
| UBTD1 | 6 | 0.31203 | 0.52069 | 0.999994 | 10361 | 2 | 2.1764 | 0.046135 | 0.15185 | 0.965913 | 3093 | 4 | 2.1764 |
| GFER | 4 | 0.27461 | 0.41707 | 0.999994 | 9584 | 1 | 4.3022 | 0.046142 | 0.1348 | 0.965913 | 3094 | 3 | 4.3022 |
| TBCC | 2 | 0.95382 | 0.95386 | 0.999994 | 20433 | 0 | 5.3026 | 0.046184 | 0.082765 | 0.939346 | 3095 | 2 | 5.3026 |
| SBSN | 6 | 0.93466 | 0.93474 | 0.999994 | 19960 | 0 | 1.1603 | 0.046208 | 0.15203 | 0.965913 | 3096 | 3 | 1.1603 |
| PIFO | 4 | 0.41493 | 0.56098 | 0.999994 | 12177 | 1 | 0.8082 | 0.046225 | 0.135 | 0.965913 | 3097 | 2 | 0.8082 |
| TBCB | 6 | 0.71354 | 0.81624 | 0.999994 | 16434 | 1 | 4.1953 | 0.046239 | 0.15211 | 0.965913 | 3098 | 5 | 4.1953 |
| RAP2B | 5 | 0.54635 | 0.6885 | 0.999994 | 14158 | 2 | -0.42625 | 0.046285 | 0.14211 | 0.965913 | 3099 | 2 | -0.42625 |
| FFAR2 | 6 | 0.93865 | 0.93861 | 0.999994 | 20041 | 0 | 2.1955 | 0.046324 | 0.15235 | 0.965913 | 3100 | 5 | 2.1955 |
| POLR3E | 5 | 0.70622 | 0.76933 | 0.999994 | 16374 | 1 | 5.3884 | 0.046359 | 0.14226 | 0.965913 | 3101 | 3 | 5.3884 |
| ACE | 6 | 0.0046581 | 0.019909 | 0.808847 | 494 | 2 | 1.178 | 0.046361 | 0.15245 | 0.965913 | 3102 | 3 | 1.178 |
| PODNL1 | 6 | 0.85713 | 0.87581 | 0.999994 | 18395 | 1 | 3.4747 | 0.046421 | 0.1526 | 0.965913 | 3103 | 4 | 3.4747 |
| MUC20 | 3 | 0.95357 | 0.95381 | 0.999994 | 20428 | 0 | 4.4636 | 0.046427 | 0.11207 | 0.953839 | 3104 | 3 | 4.4636 |
| ASTN2 | 6 | 0.94371 | 0.94368 | 0.999994 | 20170 | 0 | 1.8338 | 0.046429 | 0.15262 | 0.965913 | 3105 | 5 | 1.8338 |
| hsa-mir-4466 | 3 | 0.95356 | 0.9538 | 0.999994 | 20426 | 0 | 6.0962 | 0.046442 | 0.1121 | 0.953839 | 3106 | 3 | 6.0962 |
| THBS1 | 4 | 0.95356 | 0.95341 | 0.999994 | 20425 | 0 | 2.1278 | 0.046444 | 0.13555 | 0.965913 | 3107 | 4 | 2.1278 |
| SMIM4 | 5 | 0.10534 | 0.23471 | 0.984971 | 5146 | 3 | -1.623 | 0.046466 | 0.14249 | 0.965913 | 3108 | 2 | -1.623 |
| ALG11 | 3 | 0.9535 | 0.95375 | 0.999994 | 20423 | 0 | 2.8483 | 0.046497 | 0.11221 | 0.953839 | 3109 | 3 | 2.8483 |
| JPH3 | 6 | 0.10242 | 0.25205 | 0.994009 | 5054 | 2 | 2.3504 | 0.046507 | 0.15282 | 0.965913 | 3110 | 3 | 2.3504 |
| GUCY2D | 6 | 0.16295 | 0.34197 | 0.999994 | 6815 | 3 | 1.7854 | 0.046534 | 0.1529 | 0.965913 | 3111 | 3 | 1.7854 |
| PGR | 6 | 0.90482 | 0.90743 | 0.999994 | 19322 | 1 | 0.29337 | 0.046534 | 0.1529 | 0.965913 | 3112 | 3 | 0.29337 |
| EFCAB6 | 6 | 0.67892 | 0.80016 | 0.999994 | 16073 | 1 | 3.9759 | 0.046539 | 0.15292 | 0.965913 | 3113 | 5 | 3.9759 |
| KCNQ3 | 4 | 0.87248 | 0.8721 | 0.999994 | 18684 | 0 | 3.6659 | 0.046552 | 0.13583 | 0.965913 | 3114 | 3 | 3.6659 |
| CYB5B | 4 | 0.38558 | 0.53083 | 0.999994 | 11728 | 1 | 4.4112 | 0.04657 | 0.13587 | 0.965913 | 3115 | 3 | 4.4112 |
| LILRA2 | 6 | 0.43377 | 0.62432 | 0.999994 | 12473 | 2 | 3.2896 | 0.046656 | 0.15322 | 0.965913 | 3116 | 4 | 3.2896 |
| CAPZA3 | 5 | 0.52071 | 0.66592 | 0.999994 | 13758 | 1 | 2.1387 | 0.046688 | 0.14299 | 0.965913 | 3117 | 4 | 2.1387 |
| hsa-mir-6831 | 3 | 0.43854 | 0.5072 | 0.999994 | 12547 | 1 | 1.6868 | 0.046689 | 0.11262 | 0.95486 | 3118 | 2 | 1.6868 |
| GK2 | 5 | 0.50897 | 0.65576 | 0.999994 | 13568 | 2 | 0.24324 | 0.046739 | 0.1431 | 0.965913 | 3119 | 2 | 0.24324 |

| | | | | | | | | | | | | | |
|---|---|---|---|---|---|---|---|---|---|---|---|---|---|
| DCC | 5 | 0.87314 | 0.87389 | 0.999994 | 18695 | 1 | 5.3797 | 0.046752 | 0.14313 | 0.965913 | 3120 | 4 | 5.3797 |
| ZBTB8B | 6 | 0.16165 | 0.34035 | 0.999994 | 6782 | 2 | 4.2206 | 0.04676 | 0.15349 | 0.965913 | 3121 | 4 | 4.2206 |
| VN1R1 | 4 | 0.83693 | 0.83641 | 0.999994 | 18034 | 0 | 1.2717 | 0.046771 | 0.13639 | 0.965913 | 3122 | 3 | 1.2717 |
| GPALPP1 | 5 | 0.88058 | 0.88081 | 0.999994 | 18843 | 1 | 1.0505 | 0.046798 | 0.14323 | 0.965913 | 3123 | 3 | 1.0505 |
| PPP4C | 2 | 0.95319 | 0.95318 | 0.999994 | 20416 | 0 | 5.2822 | 0.046806 | 0.08383 | 0.940296 | 3124 | 2 | 5.2822 |
| EIF4ENIF1 | 5 | 0.62307 | 0.73471 | 0.999994 | 15275 | 1 | 2.2236 | 0.046807 | 0.14325 | 0.965913 | 3125 | 4 | 2.2236 |
| CRYBB3 | 3 | 0.95319 | 0.95344 | 0.999994 | 20415 | 0 | 2.6158 | 0.046808 | 0.11288 | 0.954984 | 3126 | 3 | 2.6158 |
| BCL2L14 | 6 | 0.88767 | 0.89469 | 0.999994 | 18991 | 1 | 3.8198 | 0.04684 | 0.15369 | 0.965913 | 3127 | 4 | 3.8198 |
| FZD6 | 4 | 0.95315 | 0.95301 | 0.999994 | 20413 | 0 | 3.7383 | 0.046852 | 0.13659 | 0.965913 | 3128 | 4 | 3.7383 |
| KLK9 | 6 | 0.91665 | 0.91732 | 0.999994 | 19552 | 1 | 2.7794 | 0.046951 | 0.154 | 0.965913 | 3129 | 5 | 2.7794 |
| OSR1 | 5 | 0.86432 | 0.86607 | 0.999994 | 18529 | 1 | 5.2695 | 0.04697 | 0.14361 | 0.965913 | 3130 | 4 | 5.2695 |
| TYW3 | 6 | 0.2204 | 0.4126 | 0.999994 | 8299 | 2 | 2.9552 | 0.04699 | 0.15411 | 0.965913 | 3131 | 3 | 2.9552 |
| SMAD7 | 6 | 0.45483 | 0.63827 | 0.999994 | 12768 | 2 | 3.9593 | 0.047021 | 0.1542 | 0.965913 | 3132 | 4 | 3.9593 |
| POLD3 | 3 | 0.94724 | 0.94743 | 0.999994 | 20268 | 0 | 6.5468 | 0.047022 | 0.11336 | 0.954984 | 3133 | 3 | 6.5468 |
| ELP2 | 5 | 0.8905 | 0.89061 | 0.999994 | 19045 | 0 | 1.2814 | 0.047024 | 0.14374 | 0.965913 | 3134 | 3 | 1.2814 |
| C1QTNF6 | 4 | 0.76725 | 0.77357 | 0.999994 | 17054 | 1 | 3.9252 | 0.04705 | 0.13708 | 0.965913 | 3135 | 3 | 3.9252 |
| ENPP2 | 6 | 0.9509 | 0.95089 | 0.999994 | 20352 | 0 | 0.77346 | 0.047078 | 0.15434 | 0.965913 | 3136 | 3 | 0.77346 |
| hsa-mir-8074 | 4 | 0.95292 | 0.95277 | 0.999994 | 20407 | 0 | 2.8324 | 0.047081 | 0.13717 | 0.965913 | 3137 | 4 | 2.8324 |
| GOLGA8O | 4 | 0.28148 | 0.42407 | 0.999994 | 9731 | 2 | 2.8597 | 0.047102 | 0.13723 | 0.965913 | 3138 | 2 | 2.8597 |
| CCDC102B | 4 | 0.9529 | 0.95276 | 0.999994 | 20405 | 0 | 2.6992 | 0.047104 | 0.13723 | 0.965913 | 3139 | 4 | 2.6992 |
| C9orf116 | 5 | 0.7937 | 0.81557 | 0.999994 | 17393 | 1 | 5.359 | 0.047118 | 0.14394 | 0.965913 | 3140 | 4 | 5.359 |
| EFEMP2 | 5 | 0.057902 | 0.15861 | 0.967126 | 3332 | 2 | 5.359 | 0.047118 | 0.14394 | 0.965913 | 3141 | 3 | 5.359 |
| PCDHGA8 | 1 | 0.95287 | 0.95303 | 0.999994 | 20403 | 0 | 8.2395 | 0.047133 | 0.047153 | 0.919289 | 3142 | 1 | 8.2395 |
| hsa-mir-1184-3 | 2 | 0.20193 | 0.2635 | 0.996734 | 7824 | 1 | 2.2328 | 0.047167 | 0.084424 | 0.940404 | 3143 | 1 | 2.2328 |
| PRRT2 | 6 | 0.64179 | 0.77129 | 0.999994 | 15546 | 2 | 3.1892 | 0.047168 | 0.15457 | 0.965913 | 3144 | 4 | 3.1892 |
| NIT2 | 4 | 0.75461 | 0.76363 | 0.999994 | 16905 | 1 | 0.041924 | 0.047171 | 0.1374 | 0.965913 | 3145 | 1 | 0.041924 |
| HFM1 | 5 | 0.073586 | 0.18424 | 0.968526 | 3975 | 2 | 5.4661 | 0.047273 | 0.14428 | 0.965913 | 3146 | 3 | 5.4661 |
| NonTargetingControlG | 2 | 0.95272 | 0.95273 | 0.999994 | 20400 | 0 | 5.4732 | 0.047281 | 0.084608 | 0.940404 | 3147 | 2 | 5.4732 |
| DCUN1D1 | 5 | 0.14736 | 0.29882 | 0.999994 | 6386 | 2 | 2.0419 | 0.047283 | 0.14431 | 0.965913 | 3148 | 3 | 2.0419 |
| FAM155B | 5 | 0.87555 | 0.87611 | 0.999994 | 18744 | 1 | 3.4546 | 0.047283 | 0.14431 | 0.965913 | 3149 | 3 | 3.4546 |
| GSKIP | 6 | 0.92469 | 0.92481 | 0.999994 | 19736 | 1 | 2.6179 | 0.047317 | 0.15493 | 0.965913 | 3150 | 5 | 2.6179 |
| ANKS6 | 6 | 0.30475 | 0.51226 | 0.999994 | 10225 | 1 | 2.4926 | 0.047377 | 0.15509 | 0.965913 | 3151 | 4 | 2.4926 |
| FAM178A | 6 | 0.35309 | 0.56749 | 0.999994 | 11217 | 3 | 2.7525 | 0.0474 | 0.15515 | 0.965913 | 3152 | 3 | 2.7525 |
| MYOM3 | 6 | 0.66916 | 0.79252 | 0.999994 | 15916 | 2 | 3.5165 | 0.0474 | 0.15515 | 0.965913 | 3153 | 4 | 3.5165 |
| CALML3 | 6 | 0.093129 | 0.23377 | 0.984971 | 4724 | 2 | 3.35 | 0.0474 | 0.15515 | 0.965913 | 3154 | 3 | 3.35 |
| MBNL3 | 5 | 0.88841 | 0.88852 | 0.999994 | 19006 | 1 | 2.76 | 0.04741 | 0.14459 | 0.965913 | 3155 | 4 | 2.76 |
| NCKAP5L | 6 | 0.69386 | 0.81017 | 0.999994 | 16244 | 1 | 3.7756 | 0.047411 | 0.15517 | 0.965913 | 3156 | 4 | 3.7756 |
| UBA6 | 6 | 0.8107 | 0.85257 | 0.999994 | 17642 | 1 | 2.416 | 0.047433 | 0.15523 | 0.965913 | 3157 | 5 | 2.416 |
| IYD | 6 | 0.21206 | 0.4026 | 0.999994 | 8066 | 2 | -0.39138 | 0.047486 | 0.15536 | 0.965913 | 3158 | 2 | -0.39138 |
| FST | 4 | 0.84406 | 0.84355 | 0.999994 | 18156 | 0 | 4.6916 | 0.047498 | 0.13821 | 0.965913 | 3159 | 3 | 4.6916 |
| hsa-mir-378i | 4 | 0.95247 | 0.95234 | 0.999994 | 20392 | 0 | 1.6137 | 0.047528 | 0.13829 | 0.965913 | 3160 | 4 | 1.6137 |
| COX8C | 5 | 0.036327 | 0.10662 | 0.950995 | 2336 | 2 | 2.0758 | 0.047555 | 0.14492 | 0.965913 | 3161 | 3 | 2.0758 |
| SLC31A1 | 5 | 0.33967 | 0.51344 | 0.999994 | 10923 | 1 | 5.3435 | 0.047564 | 0.14495 | 0.965913 | 3162 | 3 | 5.3435 |
| MUC1 | 5 | 0.39063 | 0.55567 | 0.999994 | 11810 | 1 | 5.3435 | 0.047564 | 0.14495 | 0.965913 | 3163 | 4 | 5.3435 |
| DLK1 | 6 | 0.97446 | 0.97434 | 0.999994 | 20939 | 0 | 3.1229 | 0.047569 | 0.15558 | 0.965913 | 3164 | 5 | 3.1229 |
| FKBPL | 5 | 0.00048686 | 0.0022607 | 0.45324 | 107 | 3 | -8.7382 | 0.047602 | 0.14503 | 0.965913 | 3165 | 2 | -8.7382 |
| CCL4 | 6 | 0.057787 | 0.16099 | 0.967126 | 3327 | 3 | 0.10992 | 0.047648 | 0.15578 | 0.965913 | 3166 | 3 | 0.10992 |
| C3orf84 | 6 | 0.072724 | 0.19241 | 0.974607 | 3950 | 4 | -1.2869 | 0.04773 | 0.15598 | 0.965913 | 3167 | 1 | -1.2869 |
| RBPMS2 | 6 | 0.94488 | 0.94486 | 0.999994 | 20200 | 0 | 1.9767 | 0.047739 | 0.15601 | 0.965913 | 3168 | 4 | 1.9767 |
| TAC1 | 6 | 0.33796 | 0.55034 | 0.999994 | 10900 | 2 | 1.3376 | 0.047739 | 0.15601 | 0.965913 | 3169 | 3 | 1.3376 |
| DHX35 | 4 | 0.64965 | 0.69308 | 0.999994 | 15653 | 1 | 3.6928 | 0.04774 | 0.13883 | 0.965913 | 3170 | 2 | 3.6928 |
| LRRC38 | 6 | 0.27934 | 0.48263 | 0.999994 | 9685 | 1 | 3.2408 | 0.047754 | 0.15604 | 0.965913 | 3171 | 5 | 3.2408 |

| | | | | | | | | | | | | | |
|---|---|---|---|---|---|---|---|---|---|---|---|---|---|
| RPL21 | 3 | 0.077516 | 0.1531 | 0.967126 | 4131 | 1 | 6.5252 | 0.047811 | 0.11509 | 0.954984 | 3172 | 2 | 6.5252 |
| NEUROG1 | 5 | 0.16539 | 0.32536 | 0.999994 | 6869 | 1 | 5.4954 | 0.047823 | 0.14552 | 0.965913 | 3173 | 3 | 5.4954 |
| hsa-mir-4800 | 4 | 0.95216 | 0.95203 | 0.999994 | 20386 | 0 | 2.0317 | 0.047844 | 0.13908 | 0.965913 | 3174 | 4 | 2.0317 |
| PKN1 | 5 | 0.22272 | 0.40708 | 0.999994 | 8351 | 3 | -0.70411 | 0.04785 | 0.14559 | 0.965913 | 3175 | 2 | -0.70411 |
| NonTargetingControlG | 2 | 0.91501 | 0.91506 | 0.999994 | 19512 | 0 | 5.8617 | 0.047857 | 0.08553 | 0.942567 | 3176 | 2 | 5.8617 |
| hsa-mir-134 | 4 | 0.2922 | 0.43503 | 0.999994 | 9956 | 2 | 3.0573 | 0.047885 | 0.13919 | 0.965913 | 3177 | 2 | 3.0573 |
| FAM120AOS | 6 | 0.88589 | 0.89349 | 0.999994 | 18946 | 1 | 4.2904 | 0.04792 | 0.15651 | 0.965913 | 3178 | 4 | 4.2904 |
| HOOK3 | 6 | 0.1705 | 0.3514 | 0.999994 | 6995 | 1 | 2.7324 | 0.047924 | 0.15652 | 0.965913 | 3179 | 5 | 2.7324 |
| EPHB1 | 6 | 0.81316 | 0.85367 | 0.999994 | 17676 | 1 | 4.4667 | 0.047968 | 0.15665 | 0.965913 | 3180 | 5 | 4.4667 |
| ZNF787 | 6 | 0.98401 | 0.98396 | 0.999994 | 21190 | 0 | 3.6332 | 0.048022 | 0.15678 | 0.965913 | 3181 | 5 | 3.6332 |
| RHOH | 5 | 0.59856 | 0.72564 | 0.999994 | 14936 | 1 | 4.8677 | 0.04805 | 0.14603 | 0.965913 | 3182 | 4 | 4.8677 |
| PRKAG3 | 5 | 0.62677 | 0.73611 | 0.999994 | 15332 | 1 | 5.3289 | 0.048096 | 0.14614 | 0.965913 | 3183 | 4 | 5.3289 |
| PSMD11 | 5 | 0.074748 | 0.1861 | 0.969409 | 4027 | 2 | 5.3289 | 0.048096 | 0.14614 | 0.965913 | 3184 | 3 | 5.3289 |
| KIAA0317 | 5 | 0.042352 | 0.12174 | 0.951052 | 2632 | 1 | 5.3289 | 0.048096 | 0.14614 | 0.965913 | 3185 | 4 | 5.3289 |
| KCNH4 | 5 | 0.12794 | 0.26946 | 0.998561 | 5870 | 1 | 5.3289 | 0.048096 | 0.14614 | 0.965913 | 3186 | 3 | 5.3289 |
| RIMBP3 | 6 | 0.95953 | 0.95961 | 0.999994 | 20573 | 0 | 3.6595 | 0.048127 | 0.15705 | 0.965913 | 3187 | 5 | 3.6595 |
| FIGF | 6 | 0.61044 | 0.74766 | 0.999994 | 15078 | 2 | 2.855 | 0.048127 | 0.15705 | 0.965913 | 3188 | 3 | 2.855 |
| MMD | 5 | 0.082443 | 0.19855 | 0.977164 | 4318 | 1 | 2.0931 | 0.04813 | 0.14623 | 0.965913 | 3189 | 4 | 2.0931 |
| AMY2A | 4 | 0.37745 | 0.52252 | 0.999994 | 11616 | 2 | 2.9367 | 0.048189 | 0.13995 | 0.965913 | 3190 | 2 | 2.9367 |
| KCTD10 | 6 | 0.20592 | 0.39508 | 0.999994 | 7922 | 2 | 0.79927 | 0.048191 | 0.15722 | 0.965913 | 3191 | 3 | 0.79927 |
| ISL1 | 6 | 0.032543 | 0.10476 | 0.948754 | 2137 | 2 | 3.4591 | 0.048204 | 0.15726 | 0.965913 | 3192 | 4 | 3.4591 |
| LCAT | 6 | 0.14076 | 0.31399 | 0.999994 | 6211 | 3 | -0.43708 | 0.048218 | 0.15729 | 0.965913 | 3193 | 2 | -0.43708 |
| GABRG2 | 6 | 0.20512 | 0.3941 | 0.999994 | 7900 | 2 | 0.98956 | 0.048218 | 0.15729 | 0.965913 | 3194 | 3 | 0.98956 |
| DEFB127 | 5 | 0.13264 | 0.27664 | 0.999379 | 5990 | 1 | 2.5398 | 0.048235 | 0.14646 | 0.965913 | 3195 | 3 | 2.5398 |
| SEC14L1 | 6 | 0.37243 | 0.58474 | 0.999994 | 11546 | 2 | 4.6105 | 0.048245 | 0.15735 | 0.965913 | 3196 | 4 | 4.6105 |
| SYN1 | 5 | 0.55703 | 0.69801 | 0.999994 | 14317 | 2 | 1.4629 | 0.048249 | 0.1465 | 0.965913 | 3197 | 3 | 1.4629 |
| SLC5A4 | 6 | 0.083562 | 0.21461 | 0.979587 | 4356 | 3 | 0.14419 | 0.048273 | 0.15743 | 0.965913 | 3198 | 3 | 0.14419 |
| UNC119 | 6 | 0.49412 | 0.66466 | 0.999994 | 13327 | 2 | 3.5302 | 0.048339 | 0.15761 | 0.965913 | 3199 | 4 | 3.5302 |
| DYNLRB1 | 5 | 0.6249 | 0.7354 | 0.999994 | 15302 | 1 | 5.3065 | 0.048353 | 0.14672 | 0.965913 | 3200 | 4 | 5.3065 |
| KRTAP9-8 | 5 | 0.93539 | 0.93563 | 0.999994 | 19977 | 0 | 4.2513 | 0.048353 | 0.14672 | 0.965913 | 3201 | 4 | 4.2513 |
| ZNF607 | 6 | 0.19475 | 0.38147 | 0.999994 | 7627 | 1 | 2.5686 | 0.048394 | 0.15776 | 0.965913 | 3202 | 5 | 2.5686 |
| hsa-mir-541 | 4 | 0.91679 | 0.91656 | 0.999994 | 19558 | 0 | 5.0343 | 0.048406 | 0.14052 | 0.965913 | 3203 | 3 | 5.0343 |
| MED6 | 4 | 0.34515 | 0.48936 | 0.999994 | 11054 | 1 | 4.8836 | 0.048406 | 0.14052 | 0.965913 | 3204 | 3 | 4.8836 |
| C11orf48 | 4 | 0.17612 | 0.31614 | 0.999994 | 7136 | 1 | 4.4467 | 0.048407 | 0.14052 | 0.965913 | 3205 | 3 | 4.4467 |
| VIP | 6 | 0.058239 | 0.16197 | 0.967126 | 3349 | 2 | 2.9311 | 0.048411 | 0.1578 | 0.965913 | 3206 | 3 | 2.9311 |
| hsa-mir-548aw | 3 | 0.95154 | 0.95175 | 0.999994 | 20378 | 0 | 5.3435 | 0.048462 | 0.11655 | 0.954984 | 3207 | 3 | 5.3435 |
| ANKRD12 | 5 | 0.049412 | 0.13893 | 0.957514 | 2949 | 1 | 1.1446 | 0.048467 | 0.14694 | 0.965913 | 3208 | 4 | 1.1446 |
| COPB1 | 5 | 0.96102 | 0.96115 | 0.999994 | 20606 | 0 | 5.2293 | 0.048467 | 0.14694 | 0.965913 | 3209 | 3 | 5.2293 |
| GANC | 6 | 0.15327 | 0.32978 | 0.999994 | 6544 | 3 | -0.019782 | 0.048468 | 0.15795 | 0.965913 | 3210 | 3 | -0.019782 |
| RHCE | 6 | 0.66973 | 0.79294 | 0.999994 | 15926 | 2 | 1.3022 | 0.048468 | 0.15795 | 0.965913 | 3211 | 4 | 1.3022 |
| SNX27 | 6 | 0.44541 | 0.63206 | 0.999994 | 12655 | 1 | 3.8279 | 0.048468 | 0.15795 | 0.965913 | 3212 | 5 | 3.8279 |
| ZNF451 | 6 | 0.0084818 | 0.033627 | 0.867348 | 760 | 3 | -0.55702 | 0.04849 | 0.15801 | 0.965913 | 3213 | 2 | -0.55702 |
| NXNL1 | 6 | 0.55103 | 0.70438 | 0.999994 | 14227 | 1 | 4.2761 | 0.048494 | 0.15802 | 0.965913 | 3214 | 4 | 4.2761 |
| PCDHB11 | 6 | 0.48035 | 0.65543 | 0.999994 | 13147 | 2 | 1.3708 | 0.048571 | 0.15823 | 0.965913 | 3215 | 3 | 1.3708 |
| ZSWIM8 | 5 | 0.025961 | 0.079995 | 0.93914 | 1802 | 1 | 2.2449 | 0.048573 | 0.14717 | 0.965913 | 3216 | 4 | 2.2449 |
| SPRY4 | 6 | 0.12978 | 0.3 | 0.999994 | 5930 | 1 | 2.4605 | 0.04866 | 0.15845 | 0.965913 | 3217 | 5 | 2.4605 |
| PPM1N | 5 | 0.0043531 | 0.018236 | 0.795727 | 468 | 1 | 2.0006 | 0.048666 | 0.14736 | 0.965913 | 3218 | 4 | 2.0006 |
| ALYREF | 6 | 0.26436 | 0.46503 | 0.999994 | 9341 | 2 | 1.1452 | 0.048671 | 0.15848 | 0.965913 | 3219 | 3 | 1.1452 |
| FGF9 | 5 | 0.56145 | 0.70191 | 0.999994 | 14394 | 1 | 0.33768 | 0.048675 | 0.14739 | 0.965913 | 3220 | 2 | 0.33768 |
| FAM163A | 5 | 0.46549 | 0.6183 | 0.999994 | 12943 | 2 | 1.9326 | 0.048675 | 0.14739 | 0.965913 | 3221 | 3 | 1.9326 |
| TMEM189 | 6 | 0.92717 | 0.92723 | 0.999994 | 19794 | 1 | 1.7058 | 0.048685 | 0.15851 | 0.965913 | 3222 | 5 | 1.7058 |
| CARD16 | 6 | 0.95583 | 0.95591 | 0.999994 | 20472 | 0 | 3.0261 | 0.048692 | 0.15853 | 0.965913 | 3223 | 5 | 3.0261 |

| | | | | | | | | | | | | |
|---|---|---|---|---|---|---|---|---|---|---|---|---|
| AKAP12 | 6 | 0.47529 | 0.65201 | 0.999994 | 13067 | 2 | 3.8145 | 0.048708 | 0.15857 | 0.965913 | 3224 | 4 | 3.8145 |
| UPK1A | 6 | 0.88114 | 0.89035 | 0.999994 | 18854 | 1 | 2.2674 | 0.048734 | 0.15863 | 0.965913 | 3225 | 4 | 2.2674 |
| WHSC1 | 6 | 0.58744 | 0.7307 | 0.999994 | 14781 | 2 | -0.25055 | 0.048789 | 0.15878 | 0.965913 | 3226 | 2 | -0.25055 |
| PLEC | 6 | 0.066154 | 0.17871 | 0.968171 | 3692 | 1 | 1.4184 | 0.048796 | 0.1588 | 0.965913 | 3227 | 5 | 1.4184 |
| NonTargetingControlG | 2 | 0.95116 | 0.95119 | 0.999994 | 20369 | 0 | 5.7877 | 0.048839 | 0.087182 | 0.943068 | 3228 | 2 | 5.7877 |
| DDX18 | 2 | 0.95116 | 0.95119 | 0.999994 | 20370 | 0 | 5.0883 | 0.048839 | 0.087182 | 0.943068 | 3229 | 2 | 5.0883 |
| C5orf55 | 5 | 0.24934 | 0.44026 | 0.999994 | 9001 | 1 | 4.1855 | 0.048843 | 0.14775 | 0.965913 | 3230 | 4 | 4.1855 |
| MFSD6L | 6 | 0.96811 | 0.96813 | 0.999994 | 20787 | 0 | 2.8532 | 0.048855 | 0.15894 | 0.965913 | 3231 | 4 | 2.8532 |
| SLU7 | 4 | 0.95114 | 0.95105 | 0.999994 | 20367 | 0 | 3.015 | 0.048863 | 0.14169 | 0.965913 | 3232 | 4 | 3.015 |
| SNCB | 5 | 0.15995 | 0.31751 | 0.999994 | 6735 | 3 | -0.63196 | 0.048869 | 0.1478 | 0.965913 | 3233 | 2 | -0.63196 |
| TBC1D31 | 6 | 0.85588 | 0.87511 | 0.999994 | 18366 | 1 | 2.6322 | 0.04887 | 0.15897 | 0.965913 | 3234 | 4 | 2.6322 |
| CERS2 | 6 | 0.94487 | 0.94484 | 0.999994 | 20198 | 0 | 3.647 | 0.048892 | 0.15903 | 0.965913 | 3235 | 5 | 3.647 |
| hsa-mir-512-1 | 4 | 0.95109 | 0.951 | 0.999994 | 20365 | 0 | 2.2302 | 0.048906 | 0.14179 | 0.965913 | 3236 | 4 | 2.2302 |
| DPH2 | 5 | 0.76792 | 0.80056 | 0.999994 | 17059 | 1 | 2.244 | 0.048921 | 0.14791 | 0.965913 | 3237 | 4 | 2.244 |
| hsa-mir-5089 | 3 | 0.95105 | 0.95125 | 0.999994 | 20362 | 0 | 1.9965 | 0.048949 | 0.11758 | 0.957238 | 3238 | 3 | 1.9965 |
| TOMM34 | 4 | 0.67618 | 0.70918 | 0.999994 | 16024 | 1 | 3.6446 | 0.048962 | 0.14193 | 0.965913 | 3239 | 3 | 3.6446 |
| PEX5 | 5 | 0.39335 | 0.558 | 0.999994 | 11852 | 2 | 2.187 | 0.049005 | 0.14808 | 0.965913 | 3240 | 3 | 2.187 |
| OTUD7B | 4 | 0.95098 | 0.95089 | 0.999994 | 20358 | 0 | 3.8868 | 0.049016 | 0.14207 | 0.965913 | 3241 | 4 | 3.8868 |
| YIPF1 | 6 | 0.25115 | 0.4493 | 0.999994 | 9039 | 3 | 0.20908 | 0.04906 | 0.15946 | 0.965913 | 3242 | 3 | 0.20908 |
| RGPD6 | 1 | 0.95094 | 0.95112 | 0.999994 | 20353 | 0 | 8.1913 | 0.049061 | 0.049152 | 0.920223 | 3243 | 1 | 8.1913 |
| CXCR6 | 6 | 0.93008 | 0.93022 | 0.999994 | 19856 | 0 | 3.7437 | 0.049079 | 0.15951 | 0.965913 | 3244 | 5 | 3.7437 |
| POTEE | 6 | 0.034845 | 0.11016 | 0.951052 | 2259 | 4 | -1.4637 | 0.049079 | 0.15951 | 0.965913 | 3245 | 2 | -1.4637 |
| SMAD6 | 6 | 0.55213 | 0.70519 | 0.999994 | 14246 | 2 | 3.7531 | 0.049079 | 0.15951 | 0.965913 | 3246 | 4 | 3.7531 |
| FAN1 | 6 | 0.32139 | 0.53149 | 0.999994 | 10542 | 2 | 3.2229 | 0.049079 | 0.15951 | 0.965913 | 3247 | 4 | 3.2229 |
| SLC24A3 | 5 | 0.46181 | 0.61518 | 0.999994 | 12878 | 2 | 0.92477 | 0.049176 | 0.14845 | 0.965913 | 3248 | 3 | 0.92477 |
| CWC22 | 5 | 0.51096 | 0.65749 | 0.999994 | 13596 | 1 | 5.2348 | 0.049176 | 0.14845 | 0.965913 | 3249 | 4 | 5.2348 |
| hsa-mir-133b | 4 | 0.77873 | 0.78285 | 0.999994 | 17192 | 1 | 4.4842 | 0.049201 | 0.14254 | 0.965913 | 3250 | 3 | 4.4842 |
| ACTN4 | 5 | 0.87136 | 0.87229 | 0.999994 | 18667 | 1 | 2.8564 | 0.049277 | 0.14868 | 0.965913 | 3251 | 4 | 2.8564 |
| NonTargetingControlG | 1 | 0.95069 | 0.95089 | 0.999994 | 20349 | 0 | 8.1844 | 0.049307 | 0.049406 | 0.920728 | 3252 | 1 | 8.1844 |
| EAPP | 6 | 0.77761 | 0.83874 | 0.999994 | 17178 | 1 | 3.1857 | 0.049317 | 0.16011 | 0.965913 | 3253 | 5 | 3.1857 |
| SIGIRR | 6 | 0.54378 | 0.69921 | 0.999994 | 14127 | 1 | 3.7334 | 0.049317 | 0.16011 | 0.965913 | 3254 | 4 | 3.7334 |
| CDPF1 | 6 | 0.68566 | 0.80542 | 0.999994 | 16167 | 2 | 2.1727 | 0.049317 | 0.16011 | 0.965913 | 3255 | 3 | 2.1727 |
| ZNF521 | 5 | 0.93247 | 0.93276 | 0.999994 | 19914 | 0 | 2.9842 | 0.049356 | 0.14885 | 0.965913 | 3256 | 3 | 2.9842 |
| C1RL | 5 | 0.86788 | 0.8692 | 0.999994 | 18598 | 1 | 5.4079 | 0.049376 | 0.14889 | 0.965913 | 3257 | 3 | 5.4079 |
| CCDC148 | 6 | 0.23363 | 0.42843 | 0.999994 | 8609 | 3 | -0.82804 | 0.049385 | 0.16028 | 0.965913 | 3258 | 2 | -0.82804 |
| THADA | 6 | 0.13918 | 0.312 | 0.999994 | 6167 | 1 | 3.3405 | 0.049386 | 0.16028 | 0.965913 | 3259 | 4 | 3.3405 |
| YIF1A | 6 | 0.21798 | 0.4097 | 0.999994 | 8227 | 2 | 4.2775 | 0.049386 | 0.16028 | 0.965913 | 3260 | 4 | 4.2775 |
| IPPK | 5 | 0.030752 | 0.092421 | 0.943968 | 2048 | 3 | -2.4329 | 0.049412 | 0.14898 | 0.965913 | 3261 | 1 | -2.4329 |
| TSPAN33 | 6 | 0.45314 | 0.63716 | 0.999994 | 12747 | 2 | 4.0701 | 0.049421 | 0.16036 | 0.965913 | 3262 | 4 | 4.0701 |
| PNPLA2 | 3 | 0.7854 | 0.78529 | 0.999994 | 17284 | 0 | 6.48 | 0.049426 | 0.11863 | 0.9593 | 3263 | 2 | 6.48 |
| NonTargetingControlG | 2 | 0.95055 | 0.95059 | 0.999994 | 20346 | 0 | 5.1136 | 0.049447 | 0.088229 | 0.943068 | 3264 | 2 | 5.1136 |
| REG1A | 6 | 0.72966 | 0.82148 | 0.999994 | 16594 | 1 | 1.5627 | 0.049494 | 0.16055 | 0.965913 | 3265 | 4 | 1.5627 |
| POU4F3 | 6 | 0.16764 | 0.34778 | 0.999994 | 6929 | 1 | 3.5232 | 0.049498 | 0.16057 | 0.965913 | 3266 | 4 | 3.5232 |
| IFNL2 | 6 | 0.4872 | 0.66006 | 0.999994 | 13239 | 1 | 1.3738 | 0.04957 | 0.16074 | 0.965913 | 3267 | 4 | 1.3738 |
| PRDM6 | 5 | 0.20997 | 0.38932 | 0.999994 | 8024 | 3 | -0.78882 | 0.049571 | 0.1493 | 0.965913 | 3268 | 2 | -0.78882 |
| CD200R1L | 5 | 0.7467 | 0.7891 | 0.999994 | 16825 | 1 | 5.2684 | 0.049574 | 0.14931 | 0.965913 | 3269 | 3 | 5.2684 |
| UBQLN2 | 6 | 0.91045 | 0.91193 | 0.999994 | 19436 | 1 | 3.5539 | 0.049576 | 0.16076 | 0.965913 | 3270 | 5 | 3.5539 |
| hsa-mir-4636 | 3 | 0.50193 | 0.54743 | 0.999994 | 13461 | 1 | -0.28219 | 0.049576 | 0.11894 | 0.959988 | 3271 | 1 | -0.28219 |
| hsa-mir-6749 | 4 | 0.18528 | 0.32552 | 0.999994 | 7351 | 2 | 2.5596 | 0.049585 | 0.14352 | 0.965913 | 3272 | 2 | 2.5596 |
| OLIG3 | 6 | 0.7113 | 0.81553 | 0.999994 | 16411 | 1 | 3.0945 | 0.049594 | 0.1608 | 0.965913 | 3273 | 5 | 3.0945 |
| GPR50 | 6 | 0.2459 | 0.44305 | 0.999994 | 8920 | 3 | 0.17192 | 0.049602 | 0.16082 | 0.965913 | 3274 | 3 | 0.17192 |
| MCUR1 | 6 | 0.82149 | 0.85753 | 0.999994 | 17796 | 1 | 3.2634 | 0.049649 | 0.16096 | 0.965913 | 3275 | 5 | 3.2634 |

| | | | | | | | | | | | | |
|---|---|---|---|---|---|---|---|---|---|---|---|---|
| F5 | 6 | 0.51096 | 0.67629 | 0.999994 | 13597 | 2 | 1.2857 | 0.049656 | 0.16097 | 0.965913 | 3276 | 3 | 1.2857 |
| SLPI | 6 | 0.69176 | 0.80956 | 0.999994 | 16224 | 1 | 3.328 | 0.049809 | 0.16137 | 0.965913 | 3277 | 4 | 3.328 |
| hsa-mir-675 | 4 | 0.95018 | 0.95011 | 0.999994 | 20341 | 0 | 1.9205 | 0.049822 | 0.14414 | 0.965913 | 3278 | 4 | 1.9205 |
| hsa-mir-3660 | 3 | 0.94011 | 0.94022 | 0.999994 | 20083 | 0 | 1.8759 | 0.049823 | 0.11947 | 0.961045 | 3279 | 3 | 1.8759 |
| NonTargetingControlG | 2 | 0.55103 | 0.55184 | 0.999994 | 14228 | 1 | 4.0414 | 0.049853 | 0.088923 | 0.943068 | 3280 | 1 | 4.0414 |
| PSD | 6 | 0.27432 | 0.4768 | 0.999994 | 9571 | 2 | 1.6235 | 0.049873 | 0.16154 | 0.965913 | 3281 | 4 | 1.6235 |
| CUL4B | 6 | 0.51242 | 0.67727 | 0.999994 | 13616 | 2 | 1.0054 | 0.049873 | 0.16154 | 0.965913 | 3282 | 3 | 1.0054 |
| CHDC2 | 6 | 0.90674 | 0.90892 | 0.999994 | 19366 | 1 | 2.8992 | 0.049873 | 0.16154 | 0.965913 | 3283 | 4 | 2.8992 |
| STXBP5 | 4 | 0.95006 | 0.94999 | 0.999994 | 20336 | 0 | 0.99278 | 0.049942 | 0.14443 | 0.965913 | 3284 | 4 | 0.99278 |
| TPD52 | 6 | 0.052553 | 0.14966 | 0.966359 | 3094 | 1 | 1.8761 | 0.049955 | 0.16175 | 0.965913 | 3285 | 5 | 1.8761 |
| BCAM | 5 | 0.6472 | 0.74399 | 0.999994 | 15619 | 1 | 4.3246 | 0.049982 | 0.15024 | 0.965913 | 3286 | 4 | 4.3246 |
| ITGB2 | 5 | 0.64504 | 0.74314 | 0.999994 | 15585 | 1 | 2.5079 | 0.049982 | 0.15024 | 0.965913 | 3287 | 4 | 2.5079 |
| FAM214A | 6 | 0.68301 | 0.80333 | 0.999994 | 16131 | 1 | 3.94 | 0.050001 | 0.16186 | 0.965913 | 3288 | 5 | 3.94 |
| SNAPC3 | 5 | 0.86039 | 0.86272 | 0.999994 | 18452 | 1 | 5.2669 | 0.050003 | 0.1503 | 0.965913 | 3289 | 3 | 5.2669 |
| ARHGEF18 | 6 | 0.42119 | 0.61617 | 0.999994 | 12265 | 1 | 1.692 | 0.050036 | 0.16194 | 0.965913 | 3290 | 5 | 1.692 |
| SSB | 4 | 0.94995 | 0.94988 | 0.999994 | 20331 | 0 | 1.8599 | 0.050045 | 0.14468 | 0.965913 | 3291 | 4 | 1.8599 |
| HNRNPDL | 6 | 0.98009 | 0.97999 | 0.999994 | 21095 | 0 | 2.5768 | 0.050051 | 0.16198 | 0.965913 | 3292 | 5 | 2.5768 |
| FTO | 6 | 0.26315 | 0.46359 | 0.999994 | 9317 | 3 | 2.5345 | 0.050105 | 0.16212 | 0.965913 | 3293 | 3 | 2.5345 |
| RPGRIP1L | 6 | 0.42428 | 0.6182 | 0.999994 | 12318 | 2 | 3.2731 | 0.050105 | 0.16212 | 0.965913 | 3294 | 3 | 3.2731 |
| UNC13B | 6 | 0.076711 | 0.20071 | 0.977164 | 4100 | 2 | 3.0708 | 0.050105 | 0.16212 | 0.965913 | 3295 | 3 | 3.0708 |
| TMEM62 | 6 | 0.95097 | 0.95096 | 0.999994 | 20357 | 0 | 3.906 | 0.050105 | 0.16212 | 0.965913 | 3296 | 4 | 3.906 |
| ATP5J2-PTCD1 | 4 | 0.76592 | 0.77248 | 0.999994 | 17039 | 1 | 4.0466 | 0.050106 | 0.14482 | 0.965913 | 3297 | 3 | 4.0466 |
| ADAMTS6 | 5 | 0.20596 | 0.3837 | 0.999994 | 7925 | 3 | -0.79355 | 0.050144 | 0.15063 | 0.965913 | 3298 | 2 | -0.79355 |
| GATA3 | 6 | 0.96895 | 0.96896 | 0.999994 | 20802 | 0 | 3.2037 | 0.050157 | 0.16226 | 0.965913 | 3299 | 5 | 3.2037 |
| TEDDM1 | 6 | 0.70119 | 0.81238 | 0.999994 | 16318 | 1 | 1.7258 | 0.050167 | 0.16228 | 0.965913 | 3300 | 5 | 1.7258 |
| GSTP1 | 6 | 0.22955 | 0.42352 | 0.999994 | 8521 | 3 | -0.85111 | 0.050198 | 0.16237 | 0.965913 | 3301 | 2 | -0.85111 |
| hsa-mir-8067 | 4 | 0.44977 | 0.59461 | 0.999994 | 12708 | 1 | 1.9488 | 0.050241 | 0.14516 | 0.965913 | 3302 | 3 | 1.9488 |
| AMELX | 6 | 0.97343 | 0.97335 | 0.999994 | 20907 | 0 | 2.0252 | 0.050248 | 0.16251 | 0.965913 | 3303 | 5 | 2.0252 |
| ZNF426 | 6 | 0.56633 | 0.71533 | 0.999994 | 14484 | 1 | 2.8841 | 0.050263 | 0.16254 | 0.965913 | 3304 | 5 | 2.8841 |
| SPARCL1 | 6 | 0.6148 | 0.75094 | 0.999994 | 15139 | 2 | 0.74392 | 0.050276 | 0.16257 | 0.965913 | 3305 | 3 | 0.74392 |
| MYOM1 | 6 | 0.86033 | 0.8776 | 0.999994 | 18448 | 1 | 3.3634 | 0.050276 | 0.16257 | 0.965913 | 3306 | 4 | 3.3634 |
| PCBP3 | 6 | 0.37841 | 0.58856 | 0.999994 | 11628 | 2 | -0.20197 | 0.05028 | 0.16258 | 0.965913 | 3307 | 2 | -0.20197 |
| hsa-mir-3943 | 4 | 0.10211 | 0.21889 | 0.980699 | 5044 | 2 | 2.2125 | 0.050281 | 0.14526 | 0.965913 | 3308 | 2 | 2.2125 |
| ZNF300 | 5 | 0.78852 | 0.81243 | 0.999994 | 17326 | 1 | 2.9948 | 0.050354 | 0.15107 | 0.965913 | 3309 | 4 | 2.9948 |
| CASD1 | 6 | 0.26304 | 0.46347 | 0.999994 | 9315 | 3 | -0.56742 | 0.050361 | 0.16281 | 0.965913 | 3310 | 2 | -0.56742 |
| SORCS1 | 5 | 0.058132 | 0.15897 | 0.967126 | 3343 | 3 | -2.2358 | 0.050388 | 0.15115 | 0.965913 | 3311 | 2 | -2.2358 |
| SGK2 | 6 | 0.54838 | 0.70245 | 0.999994 | 14184 | 2 | -0.079712 | 0.050442 | 0.163 | 0.965913 | 3312 | 1 | -0.079712 |
| CNTN6 | 6 | 0.18573 | 0.37026 | 0.999994 | 7359 | 3 | -0.42947 | 0.050442 | 0.163 | 0.965913 | 3313 | 2 | -0.42947 |
| CENPB | 4 | 0.88199 | 0.88161 | 0.999994 | 18869 | 0 | 4.7088 | 0.050465 | 0.14575 | 0.965913 | 3314 | 3 | 4.7088 |
| FAM151B | 5 | 0.449 | 0.60441 | 0.999994 | 12699 | 1 | 1.8836 | 0.050498 | 0.15139 | 0.965913 | 3315 | 3 | 1.8836 |
| AZIN1 | 6 | 0.61756 | 0.75298 | 0.999994 | 15178 | 1 | 1.8145 | 0.050516 | 0.16318 | 0.965913 | 3316 | 4 | 1.8145 |
| PLA2G12A | 6 | 0.1776 | 0.36008 | 0.999994 | 7171 | 2 | 2.9622 | 0.050518 | 0.16318 | 0.965913 | 3317 | 3 | 2.9622 |
| TBX19 | 6 | 0.42422 | 0.61815 | 0.999994 | 12314 | 2 | 3.9421 | 0.050518 | 0.16318 | 0.965913 | 3318 | 4 | 3.9421 |
| OSBPL5 | 6 | 0.92374 | 0.9239 | 0.999994 | 19712 | 1 | 4.3883 | 0.050518 | 0.16318 | 0.965913 | 3319 | 4 | 4.3883 |
| SGSM2 | 6 | 0.94469 | 0.94466 | 0.999994 | 20195 | 0 | 3.9168 | 0.050518 | 0.16318 | 0.965913 | 3320 | 5 | 3.9168 |
| PATE2 | 6 | 0.14208 | 0.31566 | 0.999994 | 6241 | 1 | 3.5036 | 0.050518 | 0.16318 | 0.965913 | 3321 | 4 | 3.5036 |
| SFXN2 | 6 | 0.12009 | 0.28585 | 0.999632 | 5631 | 4 | -0.73838 | 0.050551 | 0.16328 | 0.965913 | 3322 | 2 | -0.73838 |
| NOX4 | 5 | 0.54144 | 0.68418 | 0.999994 | 14079 | 1 | 0.88406 | 0.050589 | 0.1516 | 0.965913 | 3323 | 4 | 0.88406 |
| BARX2 | 6 | 0.92506 | 0.92516 | 0.999994 | 19744 | 1 | 4.1911 | 0.050645 | 0.16354 | 0.965913 | 3324 | 4 | 4.1911 |
| CAPN5 | 5 | 0.43017 | 0.58856 | 0.999994 | 12415 | 1 | 3.5563 | 0.050651 | 0.15174 | 0.965913 | 3325 | 4 | 3.5563 |
| GBX1 | 5 | 0.15901 | 0.31608 | 0.999994 | 6702 | 2 | 2.4594 | 0.050651 | 0.15174 | 0.965913 | 3326 | 3 | 2.4594 |
| ALG10 | 5 | 0.85591 | 0.85901 | 0.999994 | 18371 | 1 | 2.5627 | 0.05066 | 0.15177 | 0.965913 | 3327 | 4 | 2.5627 |

| | | | | | | | | | | | | |
|---|---|---|---|---|---|---|---|---|---|---|---|---|
| WWC3 | 6 | 0.96932 | 0.96931 | 0.999994 | 20811 | 0 | 1.9112 | 0.050684 | 0.16364 | 0.965913 | 3328 | 5 | 1.9112 |
| hsa-mir-425 | 3 | 0.01812 | 0.04698 | 0.896157 | 1366 | 1 | 6.4422 | 0.050705 | 0.12139 | 0.962897 | 3329 | 2 | 6.4422 |
| NLRP7 | 4 | 0.94929 | 0.94922 | 0.999994 | 20315 | 0 | 1.2557 | 0.050711 | 0.14639 | 0.965913 | 3330 | 4 | 1.2557 |
| PHPT1 | 6 | 0.15813 | 0.33588 | 0.999994 | 6676 | 1 | 2.848 | 0.050713 | 0.16371 | 0.965913 | 3331 | 4 | 2.848 |
| PHACTR3 | 6 | 0.018106 | 0.065863 | 0.93914 | 1365 | 2 | 2.0758 | 0.050713 | 0.16371 | 0.965913 | 3332 | 4 | 2.0758 |
| UHRF2 | 6 | 0.37322 | 0.58524 | 0.999994 | 11562 | 2 | 0.28601 | 0.050771 | 0.16386 | 0.965913 | 3333 | 3 | 0.28601 |
| SWAP70 | 6 | 0.063433 | 0.17303 | 0.96793 | 3574 | 4 | -1.5777 | 0.050771 | 0.16386 | 0.965913 | 3334 | 2 | -1.5777 |
| ASRGL1 | 6 | 0.45367 | 0.63751 | 0.999994 | 12753 | 1 | 1.8961 | 0.050771 | 0.16386 | 0.965913 | 3335 | 4 | 1.8961 |
| SLC26A2 | 6 | 0.5897 | 0.73237 | 0.999994 | 14819 | 2 | 4.192 | 0.050791 | 0.16391 | 0.965913 | 3336 | 4 | 4.192 |
| SRP19 | 4 | 0.31476 | 0.45813 | 0.999994 | 10413 | 1 | 1.2981 | 0.050821 | 0.14666 | 0.965913 | 3337 | 2 | 1.2981 |
| NUP54 | 4 | 0.11426 | 0.24033 | 0.987253 | 5458 | 3 | -1.2832 | 0.050821 | 0.14666 | 0.965913 | 3338 | 1 | -1.2832 |
| NonTargetingControlG | 2 | 0.78992 | 0.78948 | 0.999994 | 17342 | 0 | 4.8183 | 0.050836 | 0.090569 | 0.94461 | 3339 | 2 | 4.8183 |
| JUND | 5 | 0.082582 | 0.19879 | 0.977164 | 4321 | 2 | 0.19297 | 0.050885 | 0.15229 | 0.965913 | 3340 | 2 | 0.19297 |
| TSSK4 | 6 | 0.061952 | 0.16989 | 0.967126 | 3511 | 2 | 3.5284 | 0.0509 | 0.1642 | 0.965913 | 3341 | 4 | 3.5284 |
| CEP76 | 6 | 0.057303 | 0.15993 | 0.967126 | 3305 | 1 | 0.33403 | 0.050903 | 0.16421 | 0.965913 | 3342 | 3 | 0.33403 |
| SREK1 | 6 | 0.84604 | 0.86981 | 0.999994 | 18186 | 1 | 3.4576 | 0.050906 | 0.16422 | 0.965913 | 3343 | 4 | 3.4576 |
| SCAND1 | 4 | 0.11926 | 0.24898 | 0.991255 | 5606 | 2 | 2.1404 | 0.050955 | 0.14701 | 0.965913 | 3344 | 2 | 2.1404 |
| SLC25A6_X | 6 | 0.93591 | 0.93596 | 0.999994 | 19987 | 0 | 1.5847 | 0.051027 | 0.16453 | 0.965913 | 3345 | 4 | 1.5847 |
| FRMD4A | 6 | 0.77953 | 0.83947 | 0.999994 | 17205 | 1 | 1.532 | 0.051027 | 0.16453 | 0.965913 | 3346 | 5 | 1.532 |
| KIAA1549L | 6 | 0.96275 | 0.9628 | 0.999994 | 20655 | 0 | 3.996 | 0.051027 | 0.16453 | 0.965913 | 3347 | 4 | 3.996 |
| OR2Y1 | 6 | 0.1049 | 0.25687 | 0.994943 | 5130 | 2 | 2.8528 | 0.05103 | 0.16454 | 0.965913 | 3348 | 3 | 2.8528 |
| ERN2 | 6 | 0.83893 | 0.86606 | 0.999994 | 18070 | 1 | 3.406 | 0.05103 | 0.16454 | 0.965913 | 3349 | 4 | 3.406 |
| RAB18 | 6 | 0.11503 | 0.27631 | 0.999379 | 5479 | 3 | 1.1102 | 0.051038 | 0.16456 | 0.965913 | 3350 | 3 | 1.1102 |
| PSMC3 | 4 | 0.82538 | 0.82498 | 0.999994 | 17857 | 1 | 3.7412 | 0.051075 | 0.14731 | 0.965913 | 3351 | 3 | 3.7412 |
| FJX1 | 6 | 0.93313 | 0.93322 | 0.999994 | 19929 | 0 | 2.8558 | 0.051086 | 0.16469 | 0.965913 | 3352 | 5 | 2.8558 |
| C14orf28 | 5 | 0.91843 | 0.91853 | 0.999994 | 19594 | 0 | 5.2293 | 0.051098 | 0.15277 | 0.965913 | 3353 | 4 | 5.2293 |
| PET112 | 5 | 0.96638 | 0.96656 | 0.999994 | 20744 | 0 | 5.2293 | 0.051098 | 0.15277 | 0.965913 | 3354 | 4 | 5.2293 |
| GCM1 | 6 | 0.952 | 0.95197 | 0.999994 | 20383 | 0 | 1.6925 | 0.051143 | 0.16485 | 0.965913 | 3355 | 5 | 1.6925 |
| ATF5 | 5 | 0.93125 | 0.93154 | 0.999994 | 19880 | 0 | 2.3626 | 0.051168 | 0.15292 | 0.965913 | 3356 | 4 | 2.3626 |
| IFI35 | 6 | 0.0957 | 0.23891 | 0.987253 | 4804 | 4 | -1.5097 | 0.051174 | 0.16493 | 0.965913 | 3357 | 1 | -1.5097 |
| CCNDBP1 | 5 | 0.91026 | 0.91028 | 0.999994 | 19430 | 0 | 4.1654 | 0.051177 | 0.15294 | 0.965913 | 3358 | 3 | 4.1654 |
| LSM6 | 4 | 0.8874 | 0.8871 | 0.999994 | 18978 | 0 | 4.7794 | 0.051209 | 0.14764 | 0.965913 | 3359 | 3 | 4.7794 |
| KEL | 5 | 0.18918 | 0.35978 | 0.999994 | 7456 | 1 | 5.1053 | 0.051217 | 0.15304 | 0.965913 | 3360 | 3 | 5.1053 |
| NF2 | 6 | 0.97092 | 0.97089 | 0.999994 | 20843 | 0 | 2.6178 | 0.051255 | 0.16514 | 0.965913 | 3361 | 5 | 2.6178 |
| MZB1 | 5 | 0.95785 | 0.95799 | 0.999994 | 20535 | 0 | 5.1341 | 0.051355 | 0.15334 | 0.965913 | 3362 | 3 | 5.1341 |
| hsa-mir-4638 | 2 | 0.94863 | 0.94864 | 0.999994 | 20304 | 0 | 5.4248 | 0.051371 | 0.09146 | 0.945479 | 3363 | 2 | 5.4248 |
| NonTargetingControlG | 2 | 0.94863 | 0.94864 | 0.999994 | 20303 | 0 | 5.7134 | 0.051371 | 0.09146 | 0.945479 | 3364 | 2 | 5.7134 |
| FAM84B | 5 | 0.25974 | 0.44853 | 0.999994 | 9231 | 1 | 2.3724 | 0.05138 | 0.1534 | 0.965913 | 3365 | 4 | 2.3724 |
| DNAH6 | 4 | 0.87155 | 0.87118 | 0.999994 | 18670 | 0 | 1.0163 | 0.051383 | 0.1481 | 0.965913 | 3366 | 3 | 1.0163 |
| MYH6 | 6 | 0.63648 | 0.76725 | 0.999994 | 15468 | 1 | 2.841 | 0.051425 | 0.16558 | 0.965913 | 3367 | 5 | 2.841 |
| C15orf52 | 5 | 0.96194 | 0.96208 | 0.999994 | 20627 | 0 | 3.6876 | 0.051459 | 0.15357 | 0.965913 | 3368 | 4 | 3.6876 |
| OR6B1 | 6 | 0.27446 | 0.47697 | 0.999994 | 9575 | 2 | 0.32742 | 0.051498 | 0.16577 | 0.965913 | 3369 | 3 | 0.32742 |
| CRB3 | 6 | 0.93701 | 0.93704 | 0.999994 | 20007 | 0 | 2.9501 | 0.051543 | 0.16589 | 0.965913 | 3370 | 5 | 2.9501 |
| SMLR1 | 6 | 0.86371 | 0.87954 | 0.999994 | 18514 | 1 | 0.31456 | 0.051553 | 0.16591 | 0.965913 | 3371 | 2 | 0.31456 |
| SLC27A6 | 6 | 0.9444 | 0.94437 | 0.999994 | 20188 | 0 | 1.9401 | 0.051564 | 0.16594 | 0.965913 | 3372 | 5 | 1.9401 |
| KCP | 5 | 0.8025 | 0.8211 | 0.999994 | 17520 | 1 | 2.7883 | 0.051592 | 0.15386 | 0.965913 | 3373 | 4 | 2.7883 |
| UBB | 5 | 0.89824 | 0.89835 | 0.999994 | 19189 | 0 | 5.202 | 0.051612 | 0.15391 | 0.965913 | 3374 | 3 | 5.202 |
| hsa-mir-92a-2 | 4 | 0.32779 | 0.47141 | 0.999994 | 10680 | 2 | 3.027 | 0.051627 | 0.14869 | 0.965913 | 3375 | 2 | 3.027 |
| TMEM254 | 4 | 0.16056 | 0.30005 | 0.999994 | 6752 | 2 | 2.4255 | 0.051627 | 0.14869 | 0.965913 | 3376 | 2 | 2.4255 |
| CPSF3L | 4 | 0.94836 | 0.9483 | 0.999994 | 20295 | 0 | 3.1111 | 0.051644 | 0.14873 | 0.965913 | 3377 | 4 | 3.1111 |
| MMP15 | 6 | 0.57808 | 0.72378 | 0.999994 | 14659 | 2 | 3.3103 | 0.05166 | 0.1662 | 0.965913 | 3378 | 3 | 3.3103 |
| SMCP | 6 | 0.78908 | 0.84329 | 0.999994 | 17332 | 1 | 2.3983 | 0.051677 | 0.16624 | 0.965913 | 3379 | 5 | 2.3983 |

| | | | | | | | | | | | | |
|---|---|---|---|---|---|---|---|---|---|---|---|---|
| NonTargetingControlG | 2 | 0.94832 | 0.94833 | 0.999994 | 20293 | 0 | 6.2943 | 0.051682 | 0.091975 | 0.946184 | 3380 | 2 | 6.2943 |
| TNF | 4 | 0.64762 | 0.69191 | 0.999994 | 15624 | 1 | 3.3035 | 0.051691 | 0.14885 | 0.965913 | 3381 | 3 | 3.3035 |
| CELA2A | 6 | 0.48658 | 0.65964 | 0.999994 | 13235 | 1 | -0.050351 | 0.051699 | 0.16629 | 0.965913 | 3382 | 2 | -0.050351 |
| ZNF536 | 6 | 0.9642 | 0.96425 | 0.999994 | 20709 | 0 | 2.0861 | 0.051708 | 0.16632 | 0.965913 | 3383 | 5 | 2.0861 |
| GPT2 | 4 | 0.94826 | 0.94822 | 0.999994 | 20290 | 0 | 1.6538 | 0.051738 | 0.14897 | 0.965913 | 3384 | 4 | 1.6538 |
| ETS2 | 5 | 0.093231 | 0.21575 | 0.980484 | 4729 | 3 | -1.839 | 0.051764 | 0.15426 | 0.965913 | 3385 | 1 | -1.839 |
| LYG1 | 6 | 0.035219 | 0.11103 | 0.951052 | 2279 | 4 | -2.2877 | 0.051769 | 0.16649 | 0.965913 | 3386 | 2 | -2.2877 |
| ARMC10 | 5 | 0.58979 | 0.72247 | 0.999994 | 14820 | 1 | 2.1543 | 0.051801 | 0.15435 | 0.965913 | 3387 | 4 | 2.1543 |
| SEPSECS | 4 | 0.30598 | 0.44909 | 0.999994 | 10253 | 2 | 2.9054 | 0.051823 | 0.14918 | 0.965913 | 3388 | 2 | 2.9054 |
| KRT73 | 6 | 0.58968 | 0.73236 | 0.999994 | 14818 | 1 | 2.3957 | 0.051831 | 0.16665 | 0.965913 | 3389 | 5 | 2.3957 |
| TIGD2 | 6 | 0.28327 | 0.4872 | 0.999994 | 9768 | 1 | 2.4547 | 0.051863 | 0.16674 | 0.965913 | 3390 | 5 | 2.4547 |
| CCNT2 | 5 | 0.7716 | 0.80262 | 0.999994 | 17108 | 1 | 1.6796 | 0.051884 | 0.15452 | 0.965913 | 3391 | 4 | 1.6796 |
| PRRT3 | 6 | 0.41599 | 0.61275 | 0.999994 | 12193 | 2 | 0.37717 | 0.051904 | 0.16686 | 0.965913 | 3392 | 3 | 0.37717 |
| SULT6B1 | 6 | 0.65417 | 0.7808 | 0.999994 | 15708 | 2 | 3.1008 | 0.051942 | 0.16695 | 0.965913 | 3393 | 4 | 3.1008 |
| TENC1 | 5 | 0.85946 | 0.86195 | 0.999994 | 18424 | 1 | 2.5656 | 0.051944 | 0.15466 | 0.965913 | 3394 | 4 | 2.5656 |
| LY6H | 5 | 0.88535 | 0.88545 | 0.999994 | 18939 | 1 | 2.4122 | 0.051969 | 0.15472 | 0.965913 | 3395 | 4 | 2.4122 |
| KCNA2 | 6 | 0.51949 | 0.68217 | 0.999994 | 13738 | 2 | 2.2773 | 0.051985 | 0.16706 | 0.965913 | 3396 | 3 | 2.2773 |
| DUSP19 | 6 | 0.12834 | 0.29815 | 0.999994 | 5888 | 2 | 3.0305 | 0.052006 | 0.16711 | 0.965913 | 3397 | 4 | 3.0305 |
| SLAMF9 | 5 | 0.92207 | 0.92224 | 0.999994 | 19674 | 0 | 5.1892 | 0.052061 | 0.1549 | 0.965913 | 3398 | 3 | 5.1892 |
| RSF1 | 4 | 0.94794 | 0.94789 | 0.999994 | 20285 | 0 | 1.4865 | 0.052062 | 0.14978 | 0.965913 | 3399 | 4 | 1.4865 |
| CCDC80 | 6 | 0.69897 | 0.8117 | 0.999994 | 16294 | 1 | 1.6212 | 0.052065 | 0.16726 | 0.965913 | 3400 | 5 | 1.6212 |
| STAP2 | 4 | 0.94793 | 0.94788 | 0.999994 | 20284 | 0 | 2.8207 | 0.052075 | 0.14981 | 0.965913 | 3401 | 4 | 2.8207 |
| hsa-mir-199a-2 | 3 | 0.77119 | 0.77122 | 0.999994 | 17100 | 0 | 6.4076 | 0.052082 | 0.12441 | 0.964228 | 3402 | 2 | 6.4076 |
| hsa-mir-4696 | 4 | 0.73937 | 0.752 | 0.999994 | 16729 | 1 | 4.7386 | 0.052093 | 0.14985 | 0.965913 | 3403 | 3 | 4.7386 |
| hsa-mir-3201 | 4 | 0.80594 | 0.8066 | 0.999994 | 17570 | 1 | 3.4867 | 0.052093 | 0.14985 | 0.965913 | 3404 | 2 | 3.4867 |
| ZNF654 | 6 | 0.59074 | 0.73317 | 0.999994 | 14833 | 2 | 1.9185 | 0.052094 | 0.16734 | 0.965913 | 3405 | 4 | 1.9185 |
| MIPOL1 | 3 | 0.9479 | 0.94809 | 0.999994 | 20283 | 0 | 5.1479 | 0.052095 | 0.12444 | 0.964228 | 3406 | 3 | 5.1479 |
| ROPN1B | 5 | 0.87882 | 0.87913 | 0.999994 | 18807 | 1 | 1.1449 | 0.052126 | 0.15504 | 0.965913 | 3407 | 4 | 1.1449 |
| TRPM5 | 6 | 0.92182 | 0.92208 | 0.999994 | 19671 | 1 | 2.9849 | 0.052236 | 0.16767 | 0.965913 | 3408 | 3 | 2.9849 |
| GALNT14 | 6 | 0.20913 | 0.39904 | 0.999994 | 8008 | 2 | 4.1169 | 0.052236 | 0.16767 | 0.965913 | 3409 | 4 | 4.1169 |
| LSM5 | 5 | 0.75727 | 0.79468 | 0.999994 | 16945 | 1 | 2.7673 | 0.052284 | 0.1554 | 0.965913 | 3410 | 3 | 2.7673 |
| SNX31 | 6 | 0.59342 | 0.73511 | 0.999994 | 14871 | 2 | 0.206 | 0.05231 | 0.16785 | 0.965913 | 3411 | 3 | 0.206 |
| YEATS2 | 5 | 0.65 | 0.74512 | 0.999994 | 15659 | 1 | 3.9691 | 0.052318 | 0.15548 | 0.965913 | 3412 | 4 | 3.9691 |
| EPHA2 | 3 | 0.94767 | 0.94786 | 0.999994 | 20280 | 0 | 2.0624 | 0.052332 | 0.12492 | 0.964228 | 3413 | 3 | 2.0624 |
| SLC35E2 | 6 | 0.1761 | 0.35821 | 0.999994 | 7135 | 1 | 3.0001 | 0.052418 | 0.16813 | 0.965913 | 3414 | 4 | 3.0001 |
| PBK | 6 | 0.52705 | 0.68746 | 0.999994 | 13844 | 2 | 3.1333 | 0.052418 | 0.16813 | 0.965913 | 3415 | 4 | 3.1333 |
| TMEM178A | 5 | 0.76689 | 0.79998 | 0.999994 | 17049 | 1 | -0.50222 | 0.052419 | 0.1557 | 0.965913 | 3416 | 2 | -0.50222 |
| HMGCLL1 | 6 | 0.9109 | 0.91231 | 0.999994 | 19445 | 1 | 2.4701 | 0.052439 | 0.16818 | 0.965913 | 3417 | 5 | 2.4701 |
| hsa-mir-3180-1 | 3 | 0.94753 | 0.94774 | 0.999994 | 20277 | 0 | 2.7038 | 0.052467 | 0.12522 | 0.964228 | 3418 | 3 | 2.7038 |
| NPL | 5 | 0.93568 | 0.93592 | 0.999994 | 19982 | 0 | 5.1685 | 0.052505 | 0.1559 | 0.965913 | 3419 | 4 | 5.1685 |
| PRKCH | 5 | 0.017019 | 0.055908 | 0.9235 | 1303 | 2 | 0.0088004 | 0.052509 | 0.15591 | 0.965913 | 3420 | 1 | 0.0088004 |
| FCRL4 | 5 | 0.48442 | 0.63446 | 0.999994 | 13207 | 2 | -0.13717 | 0.052531 | 0.15595 | 0.965913 | 3421 | 2 | -0.13717 |
| RAC3 | 6 | 0.45797 | 0.64038 | 0.999994 | 12821 | 2 | 0.30029 | 0.052553 | 0.16848 | 0.965913 | 3422 | 2 | 0.30029 |
| OR52A1 | 6 | 0.17509 | 0.35699 | 0.999994 | 7112 | 2 | 0.40829 | 0.052553 | 0.16848 | 0.965913 | 3423 | 3 | 0.40829 |
| hsa-mir-4727 | 4 | 0.94742 | 0.94738 | 0.999994 | 20275 | 0 | 4.0875 | 0.052578 | 0.15109 | 0.965913 | 3424 | 4 | 4.0875 |
| SPATC1L | 4 | 0.35928 | 0.50381 | 0.999994 | 11325 | 1 | 3.7239 | 0.052616 | 0.15118 | 0.965913 | 3425 | 2 | 3.7239 |
| LSAMP | 4 | 0.25935 | 0.40146 | 0.999994 | 9223 | 2 | 2.8411 | 0.052616 | 0.15118 | 0.965913 | 3426 | 2 | 2.8411 |
| hsa-mir-1261 | 4 | 0.94737 | 0.94735 | 0.999994 | 20274 | 0 | 3.8385 | 0.052628 | 0.15121 | 0.965913 | 3427 | 4 | 3.8385 |
| NACC1 | 6 | 0.90218 | 0.90538 | 0.999994 | 19256 | 1 | 2.0022 | 0.052638 | 0.16869 | 0.965913 | 3428 | 5 | 2.0022 |
| C1orf162 | 6 | 0.39564 | 0.59957 | 0.999994 | 11883 | 2 | 3.1079 | 0.052654 | 0.16873 | 0.965913 | 3429 | 3 | 3.1079 |
| C14orf169 | 6 | 0.031127 | 0.10147 | 0.946414 | 2066 | 1 | 3.3444 | 0.052654 | 0.16873 | 0.965913 | 3430 | 4 | 3.3444 |
| CHTOP | 6 | 0.95606 | 0.95614 | 0.999994 | 20483 | 0 | 3.8011 | 0.052654 | 0.16873 | 0.965913 | 3431 | 5 | 3.8011 |

| | | | | | | | | | | | | |
|---|---|---|---|---|---|---|---|---|---|---|---|---|
| NCAN | 6 | 0.22576 | 0.419 | 0.999994 | 8419 | 1 | 2.8619 | 0.052664 | 0.16876 | 0.965913 | 3432 | 5 | 2.8619 |
| LRRC37A2 | 4 | 0.74641 | 0.75731 | 0.999994 | 16819 | 1 | 3.8136 | 0.052703 | 0.15137 | 0.965913 | 3433 | 3 | 3.8136 |
| COLCA2 | 5 | 0.40234 | 0.56547 | 0.999994 | 11983 | 1 | 1.0784 | 0.052713 | 0.15633 | 0.965913 | 3434 | 3 | 1.0784 |
| PNMAL2 | 6 | 0.40437 | 0.60521 | 0.999994 | 12025 | 2 | 1.3125 | 0.052752 | 0.16898 | 0.965913 | 3435 | 4 | 1.3125 |
| NonTargetingControlG | 2 | 0.80852 | 0.8082 | 0.999994 | 17610 | 0 | 4.8353 | 0.052865 | 0.093941 | 0.950736 | 3436 | 2 | 4.8353 |
| hsa-mir-548ag-1 | 4 | 0.8016 | 0.80264 | 0.999994 | 17502 | 1 | 3.6639 | 0.052887 | 0.15184 | 0.965913 | 3437 | 3 | 3.6639 |
| KCNJ14 | 5 | 0.91673 | 0.91681 | 0.999994 | 19556 | 0 | 3.1996 | 0.052891 | 0.15673 | 0.965913 | 3438 | 4 | 3.1996 |
| BSCL2 | 5 | 0.71223 | 0.77215 | 0.999994 | 16422 | 1 | 3.5535 | 0.052916 | 0.15679 | 0.965913 | 3439 | 3 | 3.5535 |
| SLC1A7 | 4 | 0.40995 | 0.5559 | 0.999994 | 12107 | 2 | 0.18616 | 0.052923 | 0.15193 | 0.965913 | 3440 | 2 | 0.18616 |
| NonTargetingControlG | 2 | 0.94706 | 0.94708 | 0.999994 | 20266 | 0 | 5.0823 | 0.052937 | 0.094069 | 0.950736 | 3441 | 2 | 5.0823 |
| ZBTB38 | 6 | 0.20075 | 0.38878 | 0.999994 | 7778 | 2 | 3.0604 | 0.052966 | 0.16949 | 0.965913 | 3442 | 4 | 3.0604 |
| hsa-mir-152 | 4 | 0.94703 | 0.947 | 0.999994 | 20265 | 0 | 2.7843 | 0.052968 | 0.15203 | 0.965913 | 3443 | 4 | 2.7843 |
| ATF6 | 5 | 0.96173 | 0.96188 | 0.999994 | 20625 | 0 | 1.666 | 0.052984 | 0.15695 | 0.965913 | 3444 | 3 | 1.666 |
| RPGRIP1 | 5 | 0.95043 | 0.95059 | 0.999994 | 20345 | 0 | 5.1479 | 0.052989 | 0.15696 | 0.965913 | 3445 | 4 | 5.1479 |
| ARHGEF28 | 6 | 0.081069 | 0.20947 | 0.979587 | 4268 | 3 | -0.16542 | 0.053013 | 0.1696 | 0.965913 | 3446 | 2 | -0.16542 |
| TSPAN3 | 6 | 0.76821 | 0.83515 | 0.999994 | 17063 | 1 | 3.9469 | 0.053082 | 0.16977 | 0.965913 | 3447 | 5 | 3.9469 |
| CYB561 | 6 | 0.038855 | 0.11931 | 0.951052 | 2454 | 2 | 3.4556 | 0.053082 | 0.16977 | 0.965913 | 3448 | 4 | 3.4556 |
| GPC6 | 6 | 0.81737 | 0.85559 | 0.999994 | 17736 | 1 | 4.4521 | 0.053082 | 0.16977 | 0.965913 | 3449 | 4 | 4.4521 |
| POLL | 6 | 0.15277 | 0.32915 | 0.999994 | 6528 | 3 | -0.5707 | 0.053094 | 0.16981 | 0.965913 | 3450 | 2 | -0.5707 |
| hsa-mir-7846 | 4 | 0.30262 | 0.44567 | 0.999994 | 10183 | 1 | 3.8665 | 0.053116 | 0.15239 | 0.965913 | 3451 | 3 | 3.8665 |
| hsa-mir-602 | 4 | 0.94684 | 0.9468 | 0.999994 | 20262 | 0 | 0.92465 | 0.053156 | 0.15249 | 0.965913 | 3452 | 4 | 0.92465 |
| OGN | 6 | 0.52712 | 0.68751 | 0.999994 | 13846 | 2 | 0.20111 | 0.053175 | 0.17002 | 0.965913 | 3453 | 3 | 0.20111 |
| CDA | 4 | 0.94681 | 0.94677 | 0.999994 | 20260 | 0 | 4.1526 | 0.053185 | 0.15257 | 0.965913 | 3454 | 4 | 4.1526 |
| RPAP1 | 4 | 0.028633 | 0.07874 | 0.93914 | 1933 | 1 | 3.5125 | 0.05323 | 0.15269 | 0.965913 | 3455 | 3 | 3.5125 |
| SPRED1 | 4 | 0.44528 | 0.59221 | 0.999994 | 12653 | 2 | 2.1965 | 0.053249 | 0.15273 | 0.965913 | 3456 | 2 | 2.1965 |
| PRR5 | 6 | 0.89862 | 0.90258 | 0.999994 | 19196 | 1 | 1.1956 | 0.053257 | 0.1702 | 0.965913 | 3457 | 5 | 1.1956 |
| AGRN | 4 | 0.94672 | 0.94667 | 0.999994 | 20254 | 0 | 1.8208 | 0.053277 | 0.15281 | 0.965913 | 3458 | 4 | 1.8208 |
| MMP17 | 4 | 0.34834 | 0.49259 | 0.999994 | 11119 | 2 | 3.0623 | 0.053284 | 0.15282 | 0.965913 | 3459 | 2 | 3.0623 |
| TIFAB | 4 | 0.88593 | 0.8856 | 0.999994 | 18948 | 0 | 5.3454 | 0.053284 | 0.15282 | 0.965913 | 3460 | 3 | 5.3454 |
| TUBB8 | 6 | 0.86164 | 0.87834 | 0.999994 | 18476 | 1 | 2.057 | 0.05332 | 0.17037 | 0.965913 | 3461 | 5 | 2.057 |
| KLHL13 | 5 | 0.24713 | 0.4385 | 0.999994 | 8953 | 1 | 0.91511 | 0.053367 | 0.1578 | 0.965913 | 3462 | 3 | 0.91511 |
| EXD3 | 6 | 0.78849 | 0.84305 | 0.999994 | 17325 | 1 | 2.4558 | 0.053373 | 0.17051 | 0.965913 | 3463 | 5 | 2.4558 |
| PRR7 | 6 | 0.65033 | 0.77786 | 0.999994 | 15663 | 1 | 2.4048 | 0.053412 | 0.17061 | 0.965913 | 3464 | 4 | 2.4048 |
| CNR2 | 4 | 0.94658 | 0.94653 | 0.999994 | 20248 | 0 | 1.4848 | 0.053415 | 0.15317 | 0.965913 | 3465 | 4 | 1.4848 |
| hsa-mir-3115 | 4 | 0.94656 | 0.9465 | 0.999994 | 20247 | 0 | 1.6996 | 0.053436 | 0.15322 | 0.965913 | 3466 | 4 | 1.6996 |
| C12orf23 | 6 | 0.86496 | 0.88028 | 0.999994 | 18538 | 1 | 2.8784 | 0.053468 | 0.17076 | 0.965913 | 3467 | 5 | 2.8784 |
| ANKRD40 | 6 | 0.67885 | 0.8001 | 0.999994 | 16071 | 2 | 3.7086 | 0.053472 | 0.17076 | 0.965913 | 3468 | 4 | 3.7086 |
| ZNF473 | 6 | 0.93673 | 0.93675 | 0.999994 | 20001 | 0 | 2.0762 | 0.053473 | 0.17076 | 0.965913 | 3469 | 3 | 2.0762 |
| GNRH1 | 6 | 0.25236 | 0.45076 | 0.999994 | 9065 | 1 | 3.7434 | 0.053473 | 0.17077 | 0.965913 | 3470 | 5 | 3.7434 |
| RPS14 | 3 | 0.89871 | 0.89883 | 0.999994 | 19197 | 0 | 1.6094 | 0.053495 | 0.12747 | 0.964228 | 3471 | 3 | 1.6094 |
| SRSF1 | 2 | 0.9465 | 0.94651 | 0.999994 | 20244 | 0 | 5.7736 | 0.0535 | 0.095029 | 0.950736 | 3472 | 2 | 5.7736 |
| NonTargetingControlG | 2 | 0.9465 | 0.94651 | 0.999994 | 20245 | 0 | 5.7517 | 0.0535 | 0.095029 | 0.950736 | 3473 | 2 | 5.7517 |
| hsa-mir-4798 | 4 | 0.94648 | 0.94641 | 0.999994 | 20242 | 0 | 1.0894 | 0.053516 | 0.15342 | 0.965913 | 3474 | 4 | 1.0894 |
| CDK11B | 5 | 0.71465 | 0.7733 | 0.999994 | 16445 | 1 | 5.1342 | 0.053524 | 0.15815 | 0.965913 | 3475 | 4 | 5.1342 |
| GOLGA7B | 5 | 0.16321 | 0.32222 | 0.999994 | 6822 | 1 | 5.1342 | 0.053524 | 0.15815 | 0.965913 | 3476 | 4 | 5.1342 |
| SLC22A12 | 5 | 0.957 | 0.95714 | 0.999994 | 20520 | 0 | 5.1342 | 0.053524 | 0.15815 | 0.965913 | 3477 | 4 | 5.1342 |
| APOBEC3B | 5 | 0.82321 | 0.8347 | 0.999994 | 17828 | 1 | 0.90463 | 0.053531 | 0.15817 | 0.965913 | 3478 | 3 | 0.90463 |
| NonTargetingControlG | 2 | 0.92889 | 0.92903 | 0.999994 | 19834 | 0 | 5.9945 | 0.053544 | 0.095106 | 0.950736 | 3479 | 2 | 5.9945 |
| KYNU | 5 | 0.3434 | 0.51652 | 0.999994 | 11016 | 2 | 1.962 | 0.05357 | 0.15826 | 0.965913 | 3480 | 3 | 1.962 |
| INTS1 | 6 | 0.33771 | 0.55007 | 0.999994 | 10891 | 3 | 2.6384 | 0.053621 | 0.17114 | 0.965913 | 3481 | 3 | 2.6384 |
| SLC25A34 | 6 | 0.64645 | 0.7749 | 0.999994 | 15607 | 2 | 3.1384 | 0.053621 | 0.17114 | 0.965913 | 3482 | 3 | 3.1384 |
| FBXL16 | 6 | 0.30757 | 0.51555 | 0.999994 | 10279 | 2 | 3.5023 | 0.053621 | 0.17114 | 0.965913 | 3483 | 4 | 3.5023 |

| | | | | | | | | | | | | |
|---|---|---|---|---|---|---|---|---|---|---|---|---|
| USP53 | 6 | 0.017716 | 0.064591 | 0.93914 | 1348 | 3 | 0.98771 | 0.053662 | 0.17124 | 0.965913 | 3484 | 3 | 0.98771 |
| MAPK9 | 6 | 0.02597 | 0.08929 | 0.94395 | 1803 | 4 | -2.0389 | 0.053662 | 0.17124 | 0.965913 | 3485 | 2 | -2.0389 |
| PTPRH | 5 | 0.97147 | 0.97152 | 0.999994 | 20857 | 0 | 2.6708 | 0.053724 | 0.1586 | 0.965913 | 3486 | 3 | 2.6708 |
| KIAA0895L | 6 | 0.065326 | 0.17694 | 0.968171 | 3668 | 1 | 2.6194 | 0.053728 | 0.17143 | 0.965913 | 3487 | 5 | 2.6194 |
| SKIL | 6 | 0.49797 | 0.66731 | 0.999994 | 13395 | 1 | 4.2275 | 0.053742 | 0.17147 | 0.965913 | 3488 | 5 | 4.2275 |
| SLC10A5 | 6 | 0.92862 | 0.92872 | 0.999994 | 19828 | 0 | 0.52565 | 0.05377 | 0.17155 | 0.965913 | 3489 | 3 | 0.52565 |
| MKNK2 | 6 | 0.89278 | 0.89829 | 0.999994 | 19086 | 1 | 3.6938 | 0.053774 | 0.17157 | 0.965913 | 3490 | 5 | 3.6938 |
| SNX1 | 4 | 0.50141 | 0.61693 | 0.999994 | 13446 | 1 | 4.4476 | 0.053806 | 0.15414 | 0.965913 | 3491 | 3 | 4.4476 |
| SPSB3 | 6 | 0.053247 | 0.15109 | 0.967126 | 3126 | 4 | -1.3259 | 0.053821 | 0.17168 | 0.965913 | 3492 | 2 | -1.3259 |
| MAGEF1 | 4 | 0.94617 | 0.94608 | 0.999994 | 20237 | 0 | 2.3707 | 0.053831 | 0.15421 | 0.965913 | 3493 | 4 | 2.3707 |
| TP53TG3 | 6 | 0.2628 | 0.46318 | 0.999994 | 9306 | 3 | -0.73616 | 0.053878 | 0.17183 | 0.965913 | 3494 | 2 | -0.73616 |
| KLHL35 | 3 | 0.94612 | 0.94629 | 0.999994 | 20235 | 0 | 4.1654 | 0.053879 | 0.12823 | 0.964228 | 3495 | 3 | 4.1654 |
| CLPTM1 | 4 | 0.94611 | 0.94601 | 0.999994 | 20234 | 0 | 3.6586 | 0.05389 | 0.15436 | 0.965913 | 3496 | 4 | 3.6586 |
| EREG | 4 | 0.66103 | 0.69986 | 0.999994 | 15794 | 1 | 0.63365 | 0.0539 | 0.15438 | 0.965913 | 3497 | 3 | 0.63365 |
| hsa-mir-6743 | 3 | 0.16144 | 0.2633 | 0.996734 | 6773 | 1 | 6.3541 | 0.053926 | 0.12834 | 0.964228 | 3498 | 2 | 6.3541 |
| CLPX | 5 | 0.82135 | 0.83349 | 0.999994 | 17794 | 1 | 3.6179 | 0.053939 | 0.15907 | 0.965913 | 3499 | 4 | 3.6179 |
| EPS15L1 | 5 | 0.47773 | 0.6287 | 0.999994 | 13108 | 2 | -0.48673 | 0.053976 | 0.15916 | 0.965913 | 3500 | 2 | -0.48673 |
| TBC1D10C | 6 | 0.12583 | 0.29503 | 0.999994 | 5813 | 2 | 0.079138 | 0.053986 | 0.17212 | 0.965913 | 3501 | 2 | 0.079138 |
| UBQLN4 | 6 | 0.32522 | 0.5358 | 0.999994 | 10627 | 2 | 3.6021 | 0.054013 | 0.17218 | 0.965913 | 3502 | 4 | 3.6021 |
| NonTargetingControlG | 2 | 0.55349 | 0.55424 | 0.999994 | 14263 | 1 | 4.0068 | 0.054048 | 0.095956 | 0.950736 | 3503 | 1 | 4.0068 |
| TMEM258 | 5 | 0.20052 | 0.37598 | 0.999994 | 7773 | 1 | 3.4241 | 0.054065 | 0.15936 | 0.965913 | 3504 | 4 | 3.4241 |
| SLC22A6 | 6 | 0.9506 | 0.95059 | 0.999994 | 20347 | 0 | 2.1537 | 0.054087 | 0.17237 | 0.965913 | 3505 | 5 | 2.1537 |
| ANKRD46 | 6 | 0.73085 | 0.82189 | 0.999994 | 16611 | 1 | 1.9558 | 0.054087 | 0.17237 | 0.965913 | 3506 | 5 | 1.9558 |
| KCNC2 | 6 | 0.95852 | 0.9586 | 0.999994 | 20547 | 0 | 2.6046 | 0.054087 | 0.17237 | 0.965913 | 3507 | 5 | 2.6046 |
| TMEM51 | 4 | 0.78429 | 0.78748 | 0.999994 | 17270 | 1 | 4.837 | 0.054143 | 0.15499 | 0.965913 | 3508 | 3 | 4.837 |
| AKIRIN2 | 4 | 0.90662 | 0.90634 | 0.999994 | 19363 | 0 | 3.8608 | 0.054143 | 0.15499 | 0.965913 | 3509 | 3 | 3.8608 |
| MFAP1 | 4 | 0.20331 | 0.34405 | 0.999994 | 7861 | 2 | -1.1406 | 0.054153 | 0.155 | 0.965913 | 3510 | 1 | -1.1406 |
| hsa-mir-6753 | 3 | 0.94579 | 0.94595 | 0.999994 | 20227 | 0 | 2.9342 | 0.054206 | 0.12895 | 0.964228 | 3511 | 3 | 2.9342 |
| HSD17B7 | 6 | 0.41416 | 0.61154 | 0.999994 | 12166 | 2 | 3.1457 | 0.054222 | 0.17272 | 0.965913 | 3512 | 4 | 3.1457 |
| C16orf96 | 6 | 0.24442 | 0.44126 | 0.999994 | 8890 | 2 | 1.4208 | 0.054229 | 0.17273 | 0.965913 | 3513 | 3 | 1.4208 |
| TOB1 | 6 | 0.54777 | 0.70204 | 0.999994 | 14174 | 1 | 1.122 | 0.054246 | 0.17277 | 0.965913 | 3514 | 3 | 1.122 |
| RUNDC1 | 3 | 0.94572 | 0.94588 | 0.999994 | 20225 | 0 | 1.9125 | 0.054275 | 0.12911 | 0.964228 | 3515 | 3 | 1.9125 |
| RIPK2 | 5 | 0.43901 | 0.596 | 0.999994 | 12558 | 1 | 0.92803 | 0.054322 | 0.15995 | 0.965913 | 3516 | 3 | 0.92803 |
| hsa-mir-875 | 3 | 0.94568 | 0.94583 | 0.999994 | 20224 | 0 | 5.5464 | 0.054324 | 0.12921 | 0.964228 | 3517 | 3 | 5.5464 |
| ANKDD1B | 6 | 0.88217 | 0.89103 | 0.999994 | 18873 | 1 | 0.42028 | 0.054337 | 0.173 | 0.965913 | 3518 | 3 | 0.42028 |
| GPX3 | 6 | 0.9408 | 0.94078 | 0.999994 | 20100 | 0 | 2.9373 | 0.054351 | 0.17304 | 0.965913 | 3519 | 5 | 2.9373 |
| RBBP8 | 5 | 0.27903 | 0.46407 | 0.999994 | 9673 | 2 | 0.38212 | 0.054382 | 0.16009 | 0.965913 | 3520 | 2 | 0.38212 |
| USP29 | 5 | 0.88927 | 0.88937 | 0.999994 | 19025 | 0 | 3.8551 | 0.054382 | 0.16009 | 0.965913 | 3521 | 3 | 3.8551 |
| PIAS2 | 6 | 0.097027 | 0.24155 | 0.987253 | 4861 | 3 | 1.9825 | 0.054429 | 0.17322 | 0.965913 | 3522 | 3 | 1.9825 |
| RABGAP1L | 6 | 0.4489 | 0.63439 | 0.999994 | 12697 | 1 | 3.5719 | 0.054429 | 0.17322 | 0.965913 | 3523 | 4 | 3.5719 |
| ZFR2 | 6 | 0.22499 | 0.41811 | 0.999994 | 8397 | 2 | 2.4746 | 0.054472 | 0.17333 | 0.965913 | 3524 | 4 | 2.4746 |
| NME5 | 6 | 0.66883 | 0.79227 | 0.999994 | 15905 | 1 | 1.1717 | 0.054472 | 0.17333 | 0.965913 | 3525 | 3 | 1.1717 |
| OPRL1 | 4 | 0.94553 | 0.94541 | 0.999994 | 20219 | 0 | 1.2225 | 0.054475 | 0.15578 | 0.965913 | 3526 | 4 | 1.2225 |
| PRAMEF10 | 5 | 0.38644 | 0.55214 | 0.999994 | 11740 | 1 | 2.1041 | 0.054519 | 0.16039 | 0.965913 | 3527 | 4 | 2.1041 |
| ZKSCAN2 | 5 | 0.97126 | 0.97131 | 0.999994 | 20853 | 0 | 2.6845 | 0.054568 | 0.16051 | 0.965913 | 3528 | 4 | 2.6845 |
| NonTargetingControlG | 2 | 0.94543 | 0.94544 | 0.999994 | 20215 | 0 | 4.6018 | 0.054572 | 0.096867 | 0.950736 | 3529 | 2 | 4.6018 |
| HPS3 | 5 | 0.65124 | 0.74563 | 0.999994 | 15679 | 1 | 5.099 | 0.054621 | 0.16063 | 0.965913 | 3530 | 3 | 5.099 |
| CD59 | 5 | 0.11886 | 0.25563 | 0.994943 | 5597 | 2 | 5.099 | 0.054621 | 0.16063 | 0.965913 | 3531 | 3 | 5.099 |
| BBS4 | 5 | 0.53534 | 0.67874 | 0.999994 | 13985 | 2 | 2.0392 | 0.05463 | 0.16065 | 0.965913 | 3532 | 3 | 2.0392 |
| hsa-mir-107 | 3 | 0.088424 | 0.16765 | 0.967126 | 4532 | 2 | -2.1034 | 0.054653 | 0.12994 | 0.964228 | 3533 | 1 | -2.1034 |
| KIRREL | 6 | 0.42817 | 0.62068 | 0.999994 | 12384 | 2 | 0.4555 | 0.054688 | 0.1739 | 0.966602 | 3534 | 3 | 0.4555 |
| BACE1 | 6 | 0.93673 | 0.93675 | 0.999994 | 20002 | 0 | 1.2823 | 0.054689 | 0.1739 | 0.966602 | 3535 | 5 | 1.2823 |

| | | | | | | | | | | | | |
|---|---|---|---|---|---|---|---|---|---|---|---|---|
| MYO5C | 4 | 0.94531 | 0.94517 | 0.999994 | 20213 | 0 | 1.0522 | 0.054692 | 0.15631 | 0.965913 | 3536 | 4 | 1.0522 |
| UNC119B | 6 | 0.82659 | 0.8599 | 0.999994 | 17879 | 1 | 2.1603 | 0.054705 | 0.17395 | 0.966602 | 3537 | 5 | 2.1603 |
| OR10J1 | 4 | 0.92502 | 0.92482 | 0.999994 | 19743 | 0 | 3.7974 | 0.054721 | 0.15637 | 0.965913 | 3538 | 3 | 3.7974 |
| hsa-mir-342 | 4 | 0.94526 | 0.94512 | 0.999994 | 20211 | 0 | 4.2066 | 0.054743 | 0.15641 | 0.965913 | 3539 | 4 | 4.2066 |
| GPRASP2 | 6 | 0.98646 | 0.98642 | 0.999994 | 21260 | 0 | 1.3745 | 0.054754 | 0.17409 | 0.967011 | 3540 | 5 | 1.3745 |
| DNHD1 | 6 | 0.41429 | 0.61162 | 0.999994 | 12172 | 2 | 1.1036 | 0.054785 | 0.17415 | 0.967011 | 3541 | 3 | 1.1036 |
| MCF2 | 4 | 0.94519 | 0.94504 | 0.999994 | 20209 | 0 | 0.67346 | 0.054812 | 0.15658 | 0.965913 | 3542 | 4 | 0.67346 |
| STARD6 | 6 | 0.95136 | 0.95134 | 0.999994 | 20373 | 0 | 1.009 | 0.054823 | 0.17425 | 0.967011 | 3543 | 4 | 1.009 |
| EIF3K | 6 | 0.34356 | 0.55673 | 0.999994 | 11022 | 3 | -0.54661 | 0.054823 | 0.17425 | 0.967011 | 3544 | 2 | -0.54661 |
| hsa-mir-141 | 4 | 0.70525 | 0.72793 | 0.999994 | 16366 | 1 | 5.1006 | 0.054846 | 0.15667 | 0.965913 | 3545 | 3 | 5.1006 |
| DHRS13 | 4 | 0.94509 | 0.94494 | 0.999994 | 20207 | 0 | 3.5755 | 0.054914 | 0.15684 | 0.965913 | 3546 | 4 | 3.5755 |
| CSRP2BP | 6 | 0.19177 | 0.37771 | 0.999994 | 7537 | 1 | 2.113 | 0.054942 | 0.17454 | 0.967011 | 3547 | 5 | 2.113 |
| OR4K13 | 6 | 0.91293 | 0.91408 | 0.999994 | 19477 | 1 | 3.3798 | 0.054945 | 0.17454 | 0.967011 | 3548 | 4 | 3.3798 |
| IRAK3 | 6 | 0.19283 | 0.37906 | 0.999994 | 7570 | 1 | 3.8455 | 0.054945 | 0.17454 | 0.967011 | 3549 | 5 | 3.8455 |
| ZNF532 | 6 | 0.78577 | 0.84197 | 0.999994 | 17290 | 1 | 3.0189 | 0.054946 | 0.17455 | 0.967011 | 3550 | 4 | 3.0189 |
| ISOC2 | 6 | 0.48258 | 0.65691 | 0.999994 | 13182 | 2 | 1.2542 | 0.054956 | 0.17457 | 0.967011 | 3551 | 3 | 1.2542 |
| PNMA5 | 4 | 0.85827 | 0.85788 | 0.999994 | 18409 | 0 | 4.0335 | 0.054966 | 0.15698 | 0.965913 | 3552 | 3 | 4.0335 |
| MYH7B | 5 | 0.96688 | 0.96706 | 0.999994 | 20758 | 0 | 2.0927 | 0.055001 | 0.16147 | 0.965913 | 3553 | 4 | 2.0927 |
| MYCL1 | 1 | 0.94499 | 0.94521 | 0.999994 | 20204 | 0 | 8.0557 | 0.055006 | 0.054991 | 0.926769 | 3554 | 1 | 8.0557 |
| SYT2 | 4 | 0.4038 | 0.54961 | 0.999994 | 12011 | 2 | 0.85988 | 0.055039 | 0.15716 | 0.965913 | 3555 | 2 | 0.85988 |
| hsa-mir-6793 | 3 | 0.94493 | 0.94506 | 0.999994 | 20202 | 0 | 2.6687 | 0.055066 | 0.13082 | 0.964228 | 3556 | 3 | 2.6687 |
| USP14 | 5 | 0.00080284 | 0.0036397 | 0.555048 | 144 | 4 | -4.0471 | 0.055103 | 0.1617 | 0.965913 | 3557 | 1 | -4.0471 |
| SDE2 | 6 | 0.36429 | 0.57964 | 0.999994 | 11409 | 2 | 1.2497 | 0.055119 | 0.17499 | 0.967616 | 3558 | 3 | 1.2497 |
| NonTargetingControlG | 2 | 0.7515 | 0.75108 | 0.999994 | 16871 | 0 | 4.6186 | 0.055121 | 0.097803 | 0.950736 | 3559 | 1 | 4.6186 |
| SEC22A | 5 | 0.36546 | 0.53462 | 0.999994 | 11430 | 2 | 5.0629 | 0.05513 | 0.16176 | 0.965913 | 3560 | 3 | 5.0629 |
| GTF3C6 | 5 | 0.76438 | 0.79858 | 0.999994 | 17020 | 1 | 5.0629 | 0.05513 | 0.16176 | 0.965913 | 3561 | 4 | 5.0629 |
| SYCP2 | 5 | 0.40192 | 0.56512 | 0.999994 | 11976 | 2 | 5.0629 | 0.05513 | 0.16176 | 0.965913 | 3562 | 3 | 5.0629 |
| UGT2B10 | 6 | 0.59885 | 0.7391 | 0.999994 | 14939 | 1 | 4.0917 | 0.055213 | 0.17524 | 0.968545 | 3563 | 4 | 4.0917 |
| hsa-mir-583 | 4 | 0.37642 | 0.52148 | 0.999994 | 11600 | 1 | 3.8403 | 0.055228 | 0.15761 | 0.965913 | 3564 | 3 | 3.8403 |
| NonTargetingControlG | 2 | 0.94475 | 0.94478 | 0.999994 | 20197 | 0 | 4.8765 | 0.055248 | 0.098018 | 0.950736 | 3565 | 2 | 4.8765 |
| GALNT16 | 6 | 0.14461 | 0.31887 | 0.999994 | 6315 | 2 | 3.2264 | 0.055266 | 0.17538 | 0.968566 | 3566 | 4 | 3.2264 |
| CTNND2 | 4 | 0.94473 | 0.94456 | 0.999994 | 20196 | 0 | 2.4352 | 0.055275 | 0.15773 | 0.965913 | 3567 | 4 | 2.4352 |
| MINPP1 | 6 | 0.6474 | 0.77564 | 0.999994 | 15621 | 1 | 0.37101 | 0.055281 | 0.17542 | 0.968594 | 3568 | 2 | 0.37101 |
| HEY2 | 4 | 0.03437 | 0.090474 | 0.94395 | 2230 | 3 | -3.4344 | 0.055328 | 0.15786 | 0.965913 | 3569 | 1 | -3.4344 |
| BIRC5 | 3 | 0.49679 | 0.54403 | 0.999994 | 13367 | 1 | 4.7933 | 0.055334 | 0.1314 | 0.964897 | 3570 | 2 | 4.7933 |
| ALOX5 | 6 | 0.30218 | 0.50926 | 0.999994 | 10171 | 2 | 4.2871 | 0.055422 | 0.17577 | 0.96929 | 3571 | 4 | 4.2871 |
| TBC1D23 | 6 | 0.16731 | 0.34736 | 0.999994 | 6916 | 2 | 3.0922 | 0.055422 | 0.17577 | 0.96929 | 3572 | 3 | 3.0922 |
| NonTargetingControlG | 2 | 0.50404 | 0.50761 | 0.999994 | 13495 | 1 | 3.7994 | 0.055432 | 0.098337 | 0.950736 | 3573 | 1 | 3.7994 |
| NPBWR1 | 4 | 0.94457 | 0.9444 | 0.999994 | 20194 | 0 | 0.99985 | 0.055434 | 0.1581 | 0.965913 | 3574 | 4 | 0.99985 |
| MCM7 | 5 | 0.44813 | 0.60364 | 0.999994 | 12689 | 2 | 5.0615 | 0.055469 | 0.16248 | 0.965913 | 3575 | 3 | 5.0615 |
| GPR26 | 6 | 0.84113 | 0.86722 | 0.999994 | 18108 | 1 | 3.1524 | 0.055487 | 0.17593 | 0.969457 | 3576 | 4 | 3.1524 |
| TMEM56-RWDD3 | 1 | 0.94443 | 0.94465 | 0.999994 | 20191 | 0 | 8.0418 | 0.055567 | 0.055549 | 0.926769 | 3577 | 1 | 8.0418 |
| RBPJ | 5 | 0.073985 | 0.18484 | 0.968526 | 3995 | 1 | 2.1966 | 0.055573 | 0.16271 | 0.965913 | 3578 | 4 | 2.1966 |
| GMPR | 6 | 0.98371 | 0.98366 | 0.999994 | 21179 | 0 | 3.5515 | 0.055632 | 0.1763 | 0.969889 | 3579 | 5 | 3.5515 |
| PKD1 | 5 | 0.065544 | 0.17112 | 0.96737 | 3674 | 3 | -2.221 | 0.055643 | 0.16288 | 0.965913 | 3580 | 2 | -2.221 |
| NonTargetingControlG | 2 | 0.94077 | 0.94088 | 0.999994 | 20099 | 0 | 6.1937 | 0.055652 | 0.098675 | 0.951383 | 3581 | 2 | 6.1937 |
| TLN2 | 6 | 0.32299 | 0.5333 | 0.999994 | 10575 | 2 | 0.67483 | 0.055669 | 0.17639 | 0.969889 | 3582 | 3 | 0.67483 |
| hsa-mir-6762 | 4 | 0.78179 | 0.78537 | 0.999994 | 17237 | 1 | 4.2269 | 0.055679 | 0.15872 | 0.965913 | 3583 | 3 | 4.2269 |
| C1orf173 | 4 | 0.75841 | 0.7665 | 0.999994 | 16958 | 1 | 3.9426 | 0.055708 | 0.1588 | 0.965913 | 3584 | 3 | 3.9426 |
| PEX19 | 5 | 0.080153 | 0.19482 | 0.975797 | 4236 | 1 | 0.76094 | 0.055711 | 0.16301 | 0.965913 | 3585 | 3 | 0.76094 |
| hsa-mir-6828 | 4 | 0.94428 | 0.94412 | 0.999994 | 20185 | 0 | 3.6704 | 0.055719 | 0.15884 | 0.965913 | 3586 | 4 | 3.6704 |
| DGCR14 | 3 | 0.94427 | 0.94442 | 0.999994 | 20184 | 0 | 2.5001 | 0.055731 | 0.13224 | 0.965913 | 3587 | 3 | 2.5001 |

| | | | | | | | | | | | | | |
|---|---|---|---|---|---|---|---|---|---|---|---|---|---|
| TYR | 3 | 0.45708 | 0.51865 | 0.999994 | 12808 | 1 | 0.42685 | 0.05577 | 0.13233 | 0.965913 | 3588 | 1 | 0.42685 |
| RASEF | 5 | 0.92333 | 0.92352 | 0.999994 | 19704 | 0 | 2.2068 | 0.055771 | 0.16315 | 0.965913 | 3589 | 4 | 2.2068 |
| NR2E1 | 6 | 0.53195 | 0.69092 | 0.999994 | 13936 | 1 | 2.0253 | 0.055772 | 0.17665 | 0.970198 | 3590 | 5 | 2.0253 |
| DNAJC22 | 6 | 0.41947 | 0.61505 | 0.999994 | 12242 | 2 | 4.2836 | 0.055786 | 0.17668 | 0.970198 | 3591 | 4 | 4.2836 |
| hsa-mir-6840 | 4 | 0.94421 | 0.94405 | 0.999994 | 20182 | 0 | 3.2522 | 0.055788 | 0.159 | 0.965913 | 3592 | 4 | 3.2522 |
| FAM131A | 6 | 0.46047 | 0.64209 | 0.999994 | 12853 | 1 | 2.436 | 0.05581 | 0.17675 | 0.970198 | 3593 | 5 | 2.436 |
| CHPF2 | 6 | 0.37099 | 0.58382 | 0.999994 | 11528 | 2 | 1.1957 | 0.055812 | 0.17675 | 0.970198 | 3594 | 4 | 1.1957 |
| SETD7 | 5 | 0.081893 | 0.19767 | 0.977164 | 4294 | 1 | 2.2912 | 0.05582 | 0.16324 | 0.965913 | 3595 | 4 | 2.2912 |
| LCTL | 5 | 0.79743 | 0.8179 | 0.999994 | 17443 | 1 | 4.9428 | 0.055851 | 0.16331 | 0.965913 | 3596 | 4 | 4.9428 |
| NDUFB4 | 3 | 0.94409 | 0.94424 | 0.999994 | 20180 | 0 | 3.0239 | 0.055913 | 0.13263 | 0.965913 | 3597 | 3 | 3.0239 |
| SH3BGRL | 6 | 0.45806 | 0.64044 | 0.999994 | 12822 | 1 | 3.1766 | 0.055918 | 0.17703 | 0.970561 | 3598 | 3 | 3.1766 |
| TTC25 | 5 | 0.011964 | 0.041696 | 0.891621 | 991 | 2 | 5.0814 | 0.055922 | 0.16346 | 0.965913 | 3599 | 3 | 5.0814 |
| KRTAP4-6 | 4 | 0.31883 | 0.46224 | 0.999994 | 10495 | 1 | 3.7204 | 0.055933 | 0.15937 | 0.965913 | 3600 | 2 | 3.7204 |
| hsa-mir-6080 | 3 | 0.94406 | 0.94421 | 0.999994 | 20179 | 0 | 3.5563 | 0.055938 | 0.13267 | 0.965913 | 3601 | 3 | 3.5563 |
| BPIFB2 | 6 | 0.80535 | 0.85018 | 0.999994 | 17557 | 1 | 0.91862 | 0.055941 | 0.17709 | 0.970602 | 3602 | 3 | 0.91862 |
| RBM26 | 5 | 0.0091129 | 0.033475 | 0.867348 | 804 | 4 | -5.5273 | 0.055958 | 0.16355 | 0.965913 | 3603 | 1 | -5.5273 |
| PLCXD1_X | 6 | 0.29276 | 0.49826 | 0.999994 | 9970 | 3 | -0.66673 | 0.05601 | 0.17726 | 0.971262 | 3604 | 1 | -0.66673 |
| hsa-mir-197 | 2 | 0.94397 | 0.94399 | 0.999994 | 20176 | 0 | 4.1177 | 0.056029 | 0.099296 | 0.951893 | 3605 | 2 | 4.1177 |
| NonTargetingControlG | 2 | 0.94397 | 0.94399 | 0.999994 | 20175 | 0 | 6.1427 | 0.056029 | 0.099296 | 0.951893 | 3606 | 2 | 6.1427 |
| ZNF45 | 5 | 0.86076 | 0.86304 | 0.999994 | 18457 | 1 | 1.8773 | 0.056037 | 0.16373 | 0.965913 | 3607 | 4 | 1.8773 |
| HAUS2 | 5 | 0.5089 | 0.65569 | 0.999994 | 13567 | 2 | -0.53379 | 0.056094 | 0.16386 | 0.965913 | 3608 | 2 | -0.53379 |
| ZXDC | 6 | 0.34963 | 0.56359 | 0.999994 | 11144 | 1 | 2.5445 | 0.0561 | 0.17749 | 0.971262 | 3609 | 5 | 2.5445 |
| NOP58 | 5 | 0.324 | 0.50059 | 0.999994 | 10597 | 2 | 5.0259 | 0.05616 | 0.16399 | 0.965913 | 3610 | 3 | 5.0259 |
| HES7 | 6 | 0.0095163 | 0.037337 | 0.881473 | 835 | 3 | 1.5778 | 0.056171 | 0.17769 | 0.971262 | 3611 | 3 | 1.5778 |
| NOVA1 | 6 | 0.22674 | 0.42019 | 0.999994 | 8446 | 2 | -0.071789 | 0.056171 | 0.17769 | 0.971262 | 3612 | 2 | -0.071789 |
| hsa-mir-4498 | 4 | 0.94382 | 0.94367 | 0.999994 | 20173 | 0 | 2.173 | 0.056179 | 0.16 | 0.965913 | 3613 | 4 | 2.173 |
| TAB1 | 5 | 0.29818 | 0.47958 | 0.999994 | 10094 | 2 | 2.3734 | 0.056214 | 0.16409 | 0.965913 | 3614 | 3 | 2.3734 |
| LRRN4CL | 5 | 0.47667 | 0.62775 | 0.999994 | 13094 | 2 | -0.081085 | 0.056251 | 0.16418 | 0.965913 | 3615 | 1 | -0.081085 |
| DACT2 | 6 | 0.9297 | 0.92983 | 0.999994 | 19846 | 0 | 1.1546 | 0.056303 | 0.17802 | 0.971262 | 3616 | 5 | 1.1546 |
| IRGM | 6 | 0.47707 | 0.65322 | 0.999994 | 13101 | 2 | 0.4591 | 0.056306 | 0.17803 | 0.971262 | 3617 | 2 | 0.4591 |
| SYNJ2BP | 6 | 0.89571 | 0.90043 | 0.999994 | 19141 | 1 | 2.1424 | 0.056358 | 0.17816 | 0.971262 | 3618 | 5 | 2.1424 |
| GRM6 | 6 | 0.68699 | 0.80646 | 0.999994 | 16180 | 2 | 0.32991 | 0.05636 | 0.17817 | 0.971262 | 3619 | 3 | 0.32991 |
| PET117 | 6 | 0.57073 | 0.71846 | 0.999994 | 14553 | 2 | 2.7448 | 0.056382 | 0.17823 | 0.971262 | 3620 | 3 | 2.7448 |
| ASPG | 6 | 0.94124 | 0.94122 | 0.999994 | 20113 | 0 | 3.9824 | 0.056382 | 0.17823 | 0.971262 | 3621 | 4 | 3.9824 |
| GDF11 | 6 | 0.027125 | 0.092051 | 0.943968 | 1863 | 3 | -1.3154 | 0.056415 | 0.17832 | 0.971262 | 3622 | 3 | -1.3154 |
| ADRA2A | 5 | 0.35462 | 0.52567 | 0.999994 | 11242 | 1 | 2.6419 | 0.056499 | 0.16472 | 0.965913 | 3623 | 4 | 2.6419 |
| COL14A1 | 5 | 0.27028 | 0.45706 | 0.999994 | 9481 | 1 | 2.7401 | 0.0565 | 0.16472 | 0.965913 | 3624 | 3 | 2.7401 |
| RFWD3 | 5 | 0.13973 | 0.28741 | 0.999632 | 6177 | 2 | 0.70644 | 0.0565 | 0.16472 | 0.965913 | 3625 | 3 | 0.70644 |
| AZI1 | 4 | 0.61879 | 0.67565 | 0.999994 | 15201 | 1 | 3.2588 | 0.056525 | 0.16062 | 0.965913 | 3626 | 3 | 3.2588 |
| DCLK1 | 6 | 0.83172 | 0.86238 | 0.999994 | 17955 | 1 | 1.9293 | 0.056594 | 0.17877 | 0.97162 | 3627 | 5 | 1.9293 |
| SELENBP1 | 6 | 0.1121 | 0.27073 | 0.998561 | 5372 | 3 | -0.32025 | 0.056656 | 0.17894 | 0.97162 | 3628 | 2 | -0.32025 |
| hsa-mir-607 | 3 | 0.40146 | 0.48514 | 0.999994 | 11969 | 1 | 6.2795 | 0.056657 | 0.13417 | 0.965913 | 3629 | 2 | 6.2795 |
| hsa-mir-6864 | 3 | 0.73136 | 0.73127 | 0.999994 | 16618 | 1 | 6.2795 | 0.056657 | 0.13417 | 0.965913 | 3630 | 2 | 6.2795 |
| HSP90B1 | 5 | 0.96289 | 0.96305 | 0.999994 | 20658 | 0 | 2.7439 | 0.056706 | 0.16518 | 0.965913 | 3631 | 4 | 2.7439 |
| TIMM13 | 5 | 0.51677 | 0.66249 | 0.999994 | 13688 | 1 | 5.0163 | 0.056724 | 0.16522 | 0.965913 | 3632 | 3 | 5.0163 |
| MYH11 | 5 | 0.3296 | 0.5052 | 0.999994 | 10715 | 1 | 5.0163 | 0.056724 | 0.16522 | 0.965913 | 3633 | 4 | 5.0163 |
| PRDX1 | 4 | 0.40389 | 0.54971 | 0.999994 | 12016 | 1 | 4.6594 | 0.056729 | 0.16097 | 0.965913 | 3634 | 3 | 4.6594 |
| EFCAB3 | 4 | 0.12385 | 0.25701 | 0.994943 | 5763 | 2 | 2.8473 | 0.056737 | 0.16099 | 0.965913 | 3635 | 2 | 2.8473 |
| MSRA | 5 | 0.81291 | 0.82785 | 0.999994 | 17669 | 1 | 1.9051 | 0.056746 | 0.16527 | 0.965913 | 3636 | 4 | 1.9051 |
| PRKD1 | 6 | 0.13357 | 0.30488 | 0.999994 | 6011 | 3 | -0.68431 | 0.056791 | 0.1793 | 0.97162 | 3637 | 2 | -0.68431 |
| TEX2 | 6 | 0.63568 | 0.76666 | 0.999994 | 15455 | 2 | 4.0986 | 0.056797 | 0.17931 | 0.97162 | 3638 | 4 | 4.0986 |
| ZNF732 | 6 | 0.45912 | 0.64116 | 0.999994 | 12835 | 1 | 3.3811 | 0.056826 | 0.17938 | 0.97162 | 3639 | 5 | 3.3811 |

| | | | | | | | | | | | | | |
|---|---|---|---|---|---|---|---|---|---|---|---|---|---|
| TRIM9 | 5 | 0.062692 | 0.1665 | 0.967126 | 3543 | 1 | 3.49 | 0.056834 | 0.16546 | 0.965913 | 3640 | 4 | 3.49 |
| ZNF33B | 5 | 0.0004927 | 0.0022913 | 0.45324 | 108 | 3 | -8.3993 | 0.056834 | 0.16546 | 0.965913 | 3641 | 2 | -8.3993 |
| TDRD12 | 6 | 0.16842 | 0.34875 | 0.999994 | 6947 | 3 | 0.51327 | 0.056899 | 0.17958 | 0.97162 | 3642 | 3 | 0.51327 |
| ELMO2 | 6 | 0.15558 | 0.33268 | 0.999994 | 6611 | 1 | 1.5127 | 0.056926 | 0.17966 | 0.97162 | 3643 | 5 | 1.5127 |
| ZNF366 | 5 | 0.13629 | 0.28216 | 0.999632 | 6090 | 2 | 0.67003 | 0.056949 | 0.16571 | 0.965913 | 3644 | 3 | 0.67003 |
| SLC22A1 | 4 | 0.25666 | 0.39873 | 0.999994 | 9163 | 1 | 4.0143 | 0.056953 | 0.16133 | 0.965913 | 3645 | 3 | 4.0143 |
| RBM15 | 2 | 0.94302 | 0.94306 | 0.999994 | 20153 | 0 | 4.765 | 0.056977 | 0.10087 | 0.951893 | 3646 | 2 | 4.765 |
| NonTargetingControlG | 2 | 0.94302 | 0.94306 | 0.999994 | 20152 | 0 | 5.9744 | 0.056981 | 0.10087 | 0.951893 | 3647 | 2 | 5.9744 |
| PRPF31 | 5 | 0.17756 | 0.34312 | 0.999994 | 7167 | 1 | 3.876 | 0.057056 | 0.16596 | 0.965913 | 3648 | 3 | 3.876 |
| NICN1 | 5 | 0.24108 | 0.43239 | 0.999994 | 8797 | 1 | 2.2254 | 0.057056 | 0.16596 | 0.965913 | 3649 | 4 | 2.2254 |
| KRT33B | 6 | 0.46223 | 0.64321 | 0.999994 | 12892 | 1 | 2.7841 | 0.057109 | 0.18011 | 0.97162 | 3650 | 5 | 2.7841 |
| FUZ | 6 | 0.43206 | 0.6232 | 0.999994 | 12447 | 2 | 2.8663 | 0.057122 | 0.18014 | 0.97162 | 3651 | 3 | 2.8663 |
| LRRC70 | 6 | 0.88983 | 0.89623 | 0.999994 | 19033 | 1 | 2.5594 | 0.057122 | 0.18014 | 0.97162 | 3652 | 4 | 2.5594 |
| PSG5 | 5 | 0.67508 | 0.75547 | 0.999994 | 16003 | 1 | 0.57997 | 0.057128 | 0.16612 | 0.965913 | 3653 | 3 | 0.57997 |
| ALOX15 | 6 | 0.33537 | 0.54737 | 0.999994 | 10840 | 1 | 0.31797 | 0.057141 | 0.18019 | 0.97162 | 3654 | 3 | 0.31797 |
| PRRG3 | 6 | 0.031513 | 0.10237 | 0.946414 | 2081 | 3 | -0.1959 | 0.057141 | 0.18019 | 0.97162 | 3655 | 2 | -0.1959 |
| H2AFZ | 2 | 0.9428 | 0.94285 | 0.999994 | 20145 | 0 | 5.3692 | 0.057197 | 0.10125 | 0.951893 | 3656 | 2 | 5.3692 |
| WDR81 | 6 | 0.83199 | 0.86251 | 0.999994 | 17958 | 1 | 3.512 | 0.057222 | 0.18038 | 0.97162 | 3657 | 5 | 3.512 |
| ROBO2 | 5 | 0.23777 | 0.4279 | 0.999994 | 8720 | 3 | -0.64001 | 0.057263 | 0.16641 | 0.965913 | 3658 | 2 | -0.64001 |
| FAM107A | 5 | 0.24776 | 0.43902 | 0.999994 | 8967 | 2 | 0.67062 | 0.057263 | 0.16641 | 0.965913 | 3659 | 2 | 0.67062 |
| ZNF830 | 6 | 0.024967 | 0.086938 | 0.94395 | 1756 | 3 | -2.1697 | 0.057303 | 0.18058 | 0.97162 | 3660 | 2 | -2.1697 |
| DNASE1L1 | 5 | 0.96059 | 0.96071 | 0.999994 | 20595 | 0 | 1.8346 | 0.057307 | 0.1665 | 0.965913 | 3661 | 3 | 1.8346 |
| SLCO4A1 | 4 | 0.68996 | 0.71792 | 0.999994 | 16210 | 1 | 1.4277 | 0.057314 | 0.16192 | 0.965913 | 3662 | 3 | 1.4277 |
| CSRNP2 | 6 | 0.10115 | 0.24957 | 0.991904 | 5011 | 3 | 2.5167 | 0.057321 | 0.18062 | 0.97162 | 3663 | 3 | 2.5167 |
| TST | 6 | 0.84221 | 0.8678 | 0.999994 | 18128 | 1 | 3.309 | 0.057321 | 0.18062 | 0.97162 | 3664 | 4 | 3.309 |
| LIF | 6 | 0.23764 | 0.43316 | 0.999994 | 8717 | 2 | 3.1736 | 0.057321 | 0.18062 | 0.97162 | 3665 | 3 | 3.1736 |
| NonTargetingControlG | 2 | 0.61209 | 0.61196 | 0.999994 | 15110 | 0 | 4.1501 | 0.057355 | 0.10153 | 0.951893 | 3666 | 1 | 4.1501 |
| TPPP2 | 5 | 0.88615 | 0.88624 | 0.999994 | 18953 | 1 | 4.988 | 0.057373 | 0.16666 | 0.965913 | 3667 | 3 | 4.988 |
| EIF4EBP1 | 5 | 0.52299 | 0.66792 | 0.999994 | 13794 | 1 | 4.988 | 0.057373 | 0.16666 | 0.965913 | 3668 | 4 | 4.988 |
| PKP4 | 6 | 0.46489 | 0.64502 | 0.999994 | 12932 | 1 | 4.1172 | 0.057386 | 0.1808 | 0.97162 | 3669 | 4 | 4.1172 |
| FHDC1 | 5 | 0.89131 | 0.89144 | 0.999994 | 19058 | 0 | 3.6133 | 0.057387 | 0.16669 | 0.965913 | 3670 | 4 | 3.6133 |
| RPAP2 | 3 | 0.94254 | 0.94266 | 0.999994 | 20139 | 0 | 2.0346 | 0.057462 | 0.13588 | 0.965913 | 3671 | 3 | 2.0346 |
| LARP4B | 6 | 0.64309 | 0.77228 | 0.999994 | 15564 | 2 | 3.4849 | 0.05751 | 0.18111 | 0.97162 | 3672 | 4 | 3.4849 |
| ECHS1 | 5 | 0.95549 | 0.95567 | 0.999994 | 20466 | 0 | 4.9214 | 0.057541 | 0.167 | 0.965913 | 3673 | 4 | 4.9214 |
| RAD51B | 6 | 0.65225 | 0.77936 | 0.999994 | 15691 | 2 | 2.7233 | 0.057542 | 0.18118 | 0.97162 | 3674 | 3 | 2.7233 |
| SRPRB | 6 | 0.56348 | 0.71328 | 0.999994 | 14440 | 1 | 3.361 | 0.057542 | 0.18118 | 0.97162 | 3675 | 4 | 3.361 |
| CALU | 5 | 0.86145 | 0.86362 | 0.999994 | 18475 | 1 | 1.7771 | 0.057553 | 0.16702 | 0.965913 | 3676 | 4 | 1.7771 |
| ARHGEF16 | 6 | 0.79544 | 0.84594 | 0.999994 | 17425 | 1 | 2.2282 | 0.057596 | 0.18133 | 0.97162 | 3677 | 5 | 2.2282 |
| RPL35A | 2 | 0.9424 | 0.94246 | 0.999994 | 20134 | 0 | 5.9345 | 0.057604 | 0.10196 | 0.951893 | 3678 | 2 | 5.9345 |
| hsa-mir-5589 | 4 | 0.76672 | 0.77315 | 0.999994 | 17045 | 1 | 3.4791 | 0.057628 | 0.16246 | 0.965913 | 3679 | 3 | 3.4791 |
| SRRT | 4 | 0.65975 | 0.69911 | 0.999994 | 15775 | 1 | 4.3462 | 0.057628 | 0.16246 | 0.965913 | 3680 | 3 | 4.3462 |
| hsa-mir-1224 | 4 | 0.8223 | 0.82197 | 0.999994 | 17806 | 1 | 4.8422 | 0.057628 | 0.16246 | 0.965913 | 3681 | 3 | 4.8422 |
| hsa-mir-4751 | 4 | 0.79333 | 0.79529 | 0.999994 | 17389 | 1 | 2.7414 | 0.057639 | 0.16247 | 0.965913 | 3682 | 3 | 2.7414 |
| SPAG4 | 6 | 0.41763 | 0.6138 | 0.999994 | 12212 | 1 | 4.1551 | 0.057711 | 0.18162 | 0.97162 | 3683 | 5 | 4.1551 |
| AASDHPPT | 6 | 0.96346 | 0.96352 | 0.999994 | 20684 | 0 | 3.8497 | 0.057723 | 0.18166 | 0.97162 | 3684 | 4 | 3.8497 |
| FANCA | 5 | 0.11739 | 0.25337 | 0.994501 | 5556 | 2 | 4.5196 | 0.057806 | 0.16759 | 0.965913 | 3685 | 3 | 4.5196 |
| hsa-mir-548e | 4 | 0.84864 | 0.84817 | 0.999994 | 18226 | 0 | 3.3018 | 0.057837 | 0.16282 | 0.965913 | 3686 | 2 | 3.3018 |
| NDUFS3 | 5 | 0.025407 | 0.078562 | 0.93914 | 1773 | 2 | 1.0502 | 0.057848 | 0.16768 | 0.965913 | 3687 | 3 | 1.0502 |
| CERCAM | 5 | 0.3864 | 0.55211 | 0.999994 | 11738 | 1 | 4.9696 | 0.057922 | 0.16784 | 0.965913 | 3688 | 4 | 4.9696 |
| OR5A2 | 5 | 0.24154 | 0.433 | 0.999994 | 8815 | 2 | 4.9696 | 0.057922 | 0.16784 | 0.965913 | 3689 | 3 | 4.9696 |
| FAM83C | 5 | 0.084378 | 0.2016 | 0.977164 | 4388 | 1 | 4.9696 | 0.057922 | 0.16784 | 0.965913 | 3690 | 3 | 4.9696 |
| GJB5 | 5 | 0.87899 | 0.87931 | 0.999994 | 18812 | 1 | 4.9696 | 0.057922 | 0.16784 | 0.965913 | 3691 | 4 | 4.9696 |

| | | | | | | | | | | | | |
|---|---|---|---|---|---|---|---|---|---|---|---|---|
| NonTargetingControlG | 2 | 0.94207 | 0.94215 | 0.999994 | 20126 | 0 | 5.6777 | 0.05793 | 0.10248 | 0.952143 | 3692 | 2 | 5.6777 |
| CSTF3 | 2 | 0.94207 | 0.94215 | 0.999994 | 20127 | 0 | 4.1051 | 0.05793 | 0.10248 | 0.952143 | 3693 | 2 | 4.1051 |
| C12orf29 | 4 | 0.93761 | 0.93747 | 0.999994 | 20021 | 0 | 4.9484 | 0.057941 | 0.16301 | 0.965913 | 3694 | 3 | 4.9484 |
| hsa-mir-4306 | 4 | 0.90508 | 0.90478 | 0.999994 | 19330 | 0 | 4.6107 | 0.057941 | 0.16301 | 0.965913 | 3695 | 3 | 4.6107 |
| EPOR | 3 | 0.16659 | 0.27006 | 0.998561 | 6899 | 2 | -1.3913 | 0.057946 | 0.13689 | 0.965913 | 3696 | 1 | -1.3913 |
| SERPINF1 | 6 | 0.96641 | 0.96647 | 0.999994 | 20746 | 0 | 2.0015 | 0.057956 | 0.18228 | 0.97162 | 3697 | 5 | 2.0015 |
| PHB2 | 4 | 0.42979 | 0.57634 | 0.999994 | 12409 | 2 | -0.054983 | 0.057963 | 0.16304 | 0.965913 | 3698 | 1 | -0.054983 |
| ZNF304 | 6 | 0.96443 | 0.96449 | 0.999994 | 20712 | 0 | 3.0912 | 0.057968 | 0.1823 | 0.97162 | 3699 | 5 | 3.0912 |
| MAGI3 | 6 | 0.24776 | 0.44526 | 0.999994 | 8966 | 2 | 0.35071 | 0.05803 | 0.18246 | 0.97162 | 3700 | 3 | 0.35071 |
| ASAP3 | 5 | 0.14694 | 0.29817 | 0.999994 | 6372 | 2 | 2.3794 | 0.058059 | 0.16812 | 0.965913 | 3701 | 3 | 2.3794 |
| KIFC3 | 5 | 0.86084 | 0.8631 | 0.999994 | 18460 | 1 | 0.99632 | 0.058073 | 0.16814 | 0.965913 | 3702 | 3 | 0.99632 |
| C15orf65 | 6 | 0.76832 | 0.83519 | 0.999994 | 17066 | 1 | 1.6409 | 0.058097 | 0.18263 | 0.97162 | 3703 | 5 | 1.6409 |
| SCNN1B | 6 | 0.90117 | 0.90458 | 0.999994 | 19237 | 1 | 0.83161 | 0.05811 | 0.18266 | 0.97162 | 3704 | 3 | 0.83161 |
| TDO2 | 5 | 0.74992 | 0.79083 | 0.999994 | 16856 | 1 | 2.7081 | 0.058124 | 0.16825 | 0.965913 | 3705 | 4 | 2.7081 |
| TTYH1 | 6 | 0.003192 | 0.014435 | 0.753239 | 374 | 1 | 1.4697 | 0.058159 | 0.1828 | 0.97162 | 3706 | 5 | 1.4697 |
| NonTargetingControlG | 2 | 0.5711 | 0.57134 | 0.999994 | 14560 | 1 | 4.0108 | 0.058161 | 0.10287 | 0.952143 | 3707 | 1 | 4.0108 |
| CDKN1A | 6 | 0.13051 | 0.30095 | 0.999994 | 5944 | 2 | 0.077592 | 0.058191 | 0.18288 | 0.97162 | 3708 | 3 | 0.077592 |
| FAM177A1 | 6 | 0.10891 | 0.26461 | 0.996734 | 5268 | 2 | -0.16606 | 0.058239 | 0.183 | 0.97162 | 3709 | 2 | -0.16606 |
| C5orf20 | 5 | 0.36044 | 0.53045 | 0.999994 | 11339 | 1 | 2.1275 | 0.058257 | 0.16854 | 0.965913 | 3710 | 4 | 2.1275 |
| SHOC2 | 6 | 0.94849 | 0.94851 | 0.999994 | 20300 | 0 | 4.286 | 0.058259 | 0.18305 | 0.97162 | 3711 | 4 | 4.286 |
| SKA3 | 6 | 0.14088 | 0.31415 | 0.999994 | 6215 | 2 | 3.1229 | 0.058259 | 0.18305 | 0.97162 | 3712 | 3 | 3.1229 |
| HIST1H4B | 6 | 0.72483 | 0.81992 | 0.999994 | 16547 | 1 | 3.0999 | 0.058259 | 0.18305 | 0.97162 | 3713 | 3 | 3.0999 |
| LRCOL1 | 6 | 0.026932 | 0.091587 | 0.94395 | 1854 | 4 | -1.3018 | 0.058272 | 0.18309 | 0.97162 | 3714 | 2 | -1.3018 |
| RAB2B | 6 | 0.036011 | 0.11288 | 0.951052 | 2314 | 2 | 1.4642 | 0.058272 | 0.18309 | 0.97162 | 3715 | 4 | 1.4642 |
| COL27A1 | 6 | 0.79439 | 0.84549 | 0.999994 | 17400 | 1 | 2.997 | 0.058311 | 0.18319 | 0.97162 | 3716 | 5 | 2.997 |
| EPHX2 | 6 | 0.33236 | 0.54397 | 0.999994 | 10775 | 1 | 3.8263 | 0.058322 | 0.18322 | 0.97162 | 3717 | 4 | 3.8263 |
| C5orf48 | 5 | 0.26475 | 0.45261 | 0.999994 | 9350 | 2 | -0.065469 | 0.058337 | 0.1687 | 0.965913 | 3718 | 2 | -0.065469 |
| PDCD7 | 5 | 0.65371 | 0.74657 | 0.999994 | 15704 | 1 | 3.0031 | 0.058337 | 0.1687 | 0.965913 | 3719 | 3 | 3.0031 |
| ACOX3 | 6 | 0.86741 | 0.88175 | 0.999994 | 18587 | 1 | 2.4115 | 0.058339 | 0.18326 | 0.97162 | 3720 | 5 | 2.4115 |
| IL6R | 5 | 0.73549 | 0.78337 | 0.999994 | 16668 | 1 | 3.5331 | 0.058365 | 0.16875 | 0.965913 | 3721 | 3 | 3.5331 |
| RHPN2 | 6 | 0.97222 | 0.97214 | 0.999994 | 20873 | 0 | 2.296 | 0.058406 | 0.18342 | 0.97162 | 3722 | 4 | 2.296 |
| ZNF568 | 6 | 0.91744 | 0.91804 | 0.999994 | 19576 | 1 | 1.5608 | 0.058443 | 0.18351 | 0.97162 | 3723 | 5 | 1.5608 |
| OTUD7A | 6 | 0.55881 | 0.70997 | 0.999994 | 14347 | 1 | 3.0056 | 0.058448 | 0.18352 | 0.97162 | 3724 | 4 | 3.0056 |
| UBE2E2 | 6 | 0.20483 | 0.39376 | 0.999994 | 7897 | 3 | -0.43915 | 0.058514 | 0.18369 | 0.971846 | 3725 | 2 | -0.43915 |
| ERF | 5 | 0.96659 | 0.96678 | 0.999994 | 20752 | 0 | 4.949 | 0.058518 | 0.16908 | 0.965913 | 3726 | 4 | 4.949 |
| BOD1 | 5 | 0.17554 | 0.34021 | 0.999994 | 7125 | 1 | 4.949 | 0.058518 | 0.16908 | 0.965913 | 3727 | 4 | 4.949 |
| KCNK1 | 5 | 0.82624 | 0.83682 | 0.999994 | 17872 | 1 | 1.8225 | 0.058534 | 0.16911 | 0.965913 | 3728 | 4 | 1.8225 |
| AMDHD2 | 6 | 0.81855 | 0.85614 | 0.999994 | 17756 | 1 | 2.9419 | 0.058559 | 0.18381 | 0.972 | 3729 | 5 | 2.9419 |
| C1orf35 | 6 | 0.32994 | 0.54115 | 0.999994 | 10722 | 2 | 2.3183 | 0.05864 | 0.18401 | 0.972727 | 3730 | 3 | 2.3183 |
| SERINC4 | 6 | 0.27446 | 0.47697 | 0.999994 | 9579 | 2 | 3.843 | 0.058648 | 0.18403 | 0.972727 | 3731 | 4 | 3.843 |
| SNN | 4 | 0.90108 | 0.90076 | 0.999994 | 19236 | 0 | 4.8327 | 0.058756 | 0.16439 | 0.965913 | 3732 | 3 | 4.8327 |
| ZNF354B | 6 | 0.95262 | 0.95263 | 0.999994 | 20399 | 0 | 3.9366 | 0.058764 | 0.18432 | 0.973287 | 3733 | 5 | 3.9366 |
| AIPL1 | 6 | 0.059365 | 0.16441 | 0.967126 | 3398 | 2 | 2.7539 | 0.058764 | 0.18432 | 0.973287 | 3734 | 3 | 2.7539 |
| CATSPERD | 5 | 0.91627 | 0.91636 | 0.999994 | 19544 | 0 | 3.2838 | 0.058773 | 0.16962 | 0.965913 | 3735 | 4 | 3.2838 |
| CCDC73 | 4 | 0.32459 | 0.46816 | 0.999994 | 10609 | 2 | 3.193 | 0.058785 | 0.16443 | 0.965913 | 3736 | 2 | 3.193 |
| hsa-mir-6894 | 4 | 0.70417 | 0.7272 | 0.999994 | 16354 | 1 | 3.8101 | 0.058791 | 0.16444 | 0.965913 | 3737 | 3 | 3.8101 |
| LHX1 | 4 | 0.39243 | 0.5379 | 0.999994 | 11840 | 1 | 4.0616 | 0.058826 | 0.16449 | 0.965913 | 3738 | 3 | 4.0616 |
| HOXB13 | 5 | 0.065375 | 0.17085 | 0.96737 | 3670 | 2 | 2.6365 | 0.058857 | 0.16979 | 0.965913 | 3739 | 3 | 2.6365 |
| SCAMP5 | 5 | 0.83921 | 0.84606 | 0.999994 | 18074 | 1 | 2.4812 | 0.058857 | 0.16979 | 0.965913 | 3740 | 3 | 2.4812 |
| LZIC | 6 | 0.32665 | 0.53742 | 0.999994 | 10656 | 3 | 0.77058 | 0.058859 | 0.18456 | 0.974094 | 3741 | 3 | 0.77058 |
| UQCR10 | 6 | 0.29657 | 0.5027 | 0.999994 | 10055 | 2 | 1.8689 | 0.058859 | 0.18456 | 0.974094 | 3742 | 3 | 1.8689 |
| RAB17 | 5 | 0.68711 | 0.76074 | 0.999994 | 16183 | 1 | 3.8437 | 0.058904 | 0.1699 | 0.965913 | 3743 | 3 | 3.8437 |

| | | | | | | | | | | | | |
|---|---|---|---|---|---|---|---|---|---|---|---|---|
| PANX1 | 5 | 0.18069 | 0.34764 | 0.999994 | 7238 | 3 | -0.94565 | 0.058904 | 0.1699 | 0.965913 | 3744 | 2 | -0.94565 |
| COX5A | 6 | 0.90772 | 0.9097 | 0.999994 | 19379 | 1 | 3.0501 | 0.058908 | 0.18469 | 0.97453 | 3745 | 4 | 3.0501 |
| C6orf120 | 5 | 0.47345 | 0.62504 | 0.999994 | 13048 | 2 | 4.9784 | 0.058912 | 0.16991 | 0.965913 | 3746 | 3 | 4.9784 |
| PDZD11 | 5 | 0.79047 | 0.8136 | 0.999994 | 17355 | 1 | 3.3492 | 0.058921 | 0.16993 | 0.965913 | 3747 | 4 | 3.3492 |
| HIST2H2AC | 3 | 0.42975 | 0.50196 | 0.999994 | 12408 | 1 | 6.221 | 0.058972 | 0.13908 | 0.965913 | 3748 | 2 | 6.221 |
| MAP1S | 6 | 0.7781 | 0.83892 | 0.999994 | 17186 | 1 | 2.3038 | 0.059001 | 0.18491 | 0.97466 | 3749 | 5 | 2.3038 |
| hsa-mir-517a | 3 | 0.94095 | 0.94105 | 0.999994 | 20105 | 0 | 1.6105 | 0.059051 | 0.13924 | 0.965913 | 3750 | 3 | 1.6105 |
| COL19A1 | 6 | 0.4516 | 0.63614 | 0.999994 | 12727 | 2 | 0.5639 | 0.059115 | 0.18519 | 0.97466 | 3751 | 3 | 0.5639 |
| SEC23IP | 4 | 0.8898 | 0.8895 | 0.999994 | 19032 | 0 | 3.9208 | 0.059123 | 0.16503 | 0.965913 | 3752 | 3 | 3.9208 |
| ABCC9 | 4 | 0.045132 | 0.11196 | 0.951052 | 2764 | 2 | 1.7596 | 0.059123 | 0.16503 | 0.965913 | 3753 | 2 | 1.7596 |
| DEFB134 | 4 | 0.26298 | 0.40512 | 0.999994 | 9314 | 2 | 2.7631 | 0.059123 | 0.16503 | 0.965913 | 3754 | 2 | 2.7631 |
| FUT11 | 6 | 0.98534 | 0.9853 | 0.999994 | 21237 | 0 | 3.1249 | 0.059125 | 0.18522 | 0.97466 | 3755 | 5 | 3.1249 |
| ZNF420 | 6 | 0.057707 | 0.16082 | 0.967126 | 3324 | 2 | 1.5275 | 0.059132 | 0.18523 | 0.97466 | 3756 | 4 | 1.5275 |
| SYCE1 | 6 | 0.43215 | 0.62327 | 0.999994 | 12448 | 2 | 1.5289 | 0.059132 | 0.18523 | 0.97466 | 3757 | 4 | 1.5289 |
| HEY1 | 5 | 0.55846 | 0.69925 | 0.999994 | 14340 | 1 | 3.7891 | 0.059161 | 0.17046 | 0.965913 | 3758 | 3 | 3.7891 |
| C11orf16 | 5 | 0.73892 | 0.78512 | 0.999994 | 16720 | 1 | 4.5407 | 0.059161 | 0.17046 | 0.965913 | 3759 | 3 | 4.5407 |
| GABARAPL2 | 6 | 0.59684 | 0.73764 | 0.999994 | 14914 | 1 | 1.5132 | 0.059166 | 0.18531 | 0.97466 | 3760 | 5 | 1.5132 |
| NUDT16 | 6 | 0.62559 | 0.75902 | 0.999994 | 15320 | 1 | 3.837 | 0.059219 | 0.18545 | 0.97466 | 3761 | 5 | 3.837 |
| MLANA | 5 | 0.44114 | 0.59774 | 0.999994 | 12594 | 2 | 4.9214 | 0.059219 | 0.17059 | 0.965913 | 3762 | 3 | 4.9214 |
| FAM47B | 5 | 0.20424 | 0.38126 | 0.999994 | 7884 | 2 | 4.9214 | 0.059219 | 0.17059 | 0.965913 | 3763 | 3 | 4.9214 |
| hsa-mir-6889 | 4 | 0.60073 | 0.66577 | 0.999994 | 14956 | 1 | 4.027 | 0.05922 | 0.1652 | 0.965913 | 3764 | 3 | 4.027 |
| KLHDC8A | 6 | 0.015256 | 0.056555 | 0.9235 | 1205 | 4 | -3.7912 | 0.059294 | 0.18563 | 0.97466 | 3765 | 2 | -3.7912 |
| KCTD12 | 6 | 0.51443 | 0.67868 | 0.999994 | 13652 | 2 | 1.0201 | 0.059294 | 0.18563 | 0.97466 | 3766 | 4 | 1.0201 |
| IFNA6 | 6 | 0.64432 | 0.77325 | 0.999994 | 15577 | 1 | 0.76568 | 0.059401 | 0.18592 | 0.97466 | 3767 | 4 | 0.76568 |
| KIAA0226 | 6 | 0.68246 | 0.80293 | 0.999994 | 16125 | 1 | 3.4375 | 0.05941 | 0.18594 | 0.97466 | 3768 | 4 | 3.4375 |
| GJB4 | 5 | 0.5677 | 0.7075 | 0.999994 | 14509 | 1 | 4.3222 | 0.059442 | 0.17109 | 0.965913 | 3769 | 3 | 4.3222 |
| AGR3 | 6 | 0.26605 | 0.46702 | 0.999994 | 9367 | 3 | 0.24932 | 0.059453 | 0.18605 | 0.97466 | 3770 | 3 | 0.24932 |
| FNIP1 | 5 | 0.42845 | 0.58713 | 0.999994 | 12388 | 1 | 3.5304 | 0.059494 | 0.17121 | 0.965913 | 3771 | 4 | 3.5304 |
| IL11RA | 5 | 0.28282 | 0.46714 | 0.999994 | 9756 | 2 | 0.4001 | 0.05951 | 0.17125 | 0.965913 | 3772 | 2 | 0.4001 |
| OST4 | 4 | 0.11741 | 0.24575 | 0.989866 | 5558 | 1 | 0.53636 | 0.059549 | 0.16574 | 0.965913 | 3773 | 2 | 0.53636 |
| IFNL4 | 4 | 0.51006 | 0.62082 | 0.999994 | 13576 | 1 | 3.2686 | 0.059576 | 0.16579 | 0.965913 | 3774 | 3 | 3.2686 |
| PGPEP1L | 6 | 0.76253 | 0.83301 | 0.999994 | 17003 | 1 | 1.4852 | 0.059634 | 0.18647 | 0.97466 | 3775 | 5 | 1.4852 |
| OR2S2 | 6 | 0.91922 | 0.91969 | 0.999994 | 19615 | 1 | 2.9916 | 0.05967 | 0.18656 | 0.97466 | 3776 | 4 | 2.9916 |
| ADK | 5 | 0.62748 | 0.73636 | 0.999994 | 15343 | 1 | 3.1886 | 0.059674 | 0.1716 | 0.965913 | 3777 | 4 | 3.1886 |
| FSD1 | 6 | 0.9529 | 0.95288 | 0.999994 | 20406 | 0 | 2.9573 | 0.059759 | 0.18679 | 0.97466 | 3778 | 4 | 2.9573 |
| LDLRAD4 | 6 | 0.82378 | 0.85859 | 0.999994 | 17836 | 1 | 3.5377 | 0.059789 | 0.18687 | 0.97466 | 3779 | 5 | 3.5377 |
| CBX8 | 5 | 0.42536 | 0.58455 | 0.999994 | 12342 | 1 | 4.9089 | 0.0598 | 0.17189 | 0.965913 | 3780 | 3 | 4.9089 |
| TUBD1 | 5 | 0.67429 | 0.75512 | 0.999994 | 15993 | 1 | 4.9089 | 0.0598 | 0.17189 | 0.965913 | 3781 | 4 | 4.9089 |
| ATP2A3 | 4 | 0.37507 | 0.52007 | 0.999994 | 11580 | 1 | 3.8288 | 0.059819 | 0.1662 | 0.965913 | 3782 | 3 | 3.8288 |
| C2orf71 | 6 | 0.61501 | 0.75109 | 0.999994 | 15143 | 1 | 3.2213 | 0.059898 | 0.18713 | 0.97466 | 3783 | 5 | 3.2213 |
| ANTXR2 | 4 | 0.0026061 | 0.0093644 | 0.680529 | 324 | 3 | -5.6684 | 0.059909 | 0.16635 | 0.965913 | 3784 | 1 | -5.6684 |
| ZKSCAN4 | 6 | 0.36066 | 0.57605 | 0.999994 | 11346 | 2 | 0.3729 | 0.059912 | 0.18716 | 0.97466 | 3785 | 3 | 0.3729 |
| PARP8 | 5 | 0.80137 | 0.82037 | 0.999994 | 17496 | 1 | 1.6386 | 0.059933 | 0.17219 | 0.965913 | 3786 | 4 | 1.6386 |
| NPY | 6 | 0.95543 | 0.95549 | 0.999994 | 20465 | 0 | 2.3273 | 0.059953 | 0.18727 | 0.97466 | 3787 | 4 | 2.3273 |
| PSMB1 | 3 | 0.93996 | 0.94008 | 0.999994 | 20077 | 0 | 3.714 | 0.060039 | 0.14135 | 0.965913 | 3788 | 3 | 3.714 |
| ITGAM | 6 | 0.56092 | 0.71146 | 0.999994 | 14386 | 2 | 0.44078 | 0.060046 | 0.18751 | 0.97466 | 3789 | 3 | 0.44078 |
| CCT6A | 5 | 0.37155 | 0.53969 | 0.999994 | 11535 | 2 | 0.26655 | 0.060087 | 0.17251 | 0.965913 | 3790 | 2 | 0.26655 |
| TSHZ2 | 5 | 0.86566 | 0.86724 | 0.999994 | 18551 | 1 | 2.2784 | 0.060087 | 0.17251 | 0.965913 | 3791 | 3 | 2.2784 |
| PFKFB1 | 6 | 0.97558 | 0.9755 | 0.999994 | 20979 | 0 | 2.7546 | 0.060122 | 0.18769 | 0.97466 | 3792 | 5 | 2.7546 |
| KRTDAP | 5 | 0.7708 | 0.80215 | 0.999994 | 17092 | 1 | 4.4499 | 0.060138 | 0.17264 | 0.965913 | 3793 | 4 | 4.4499 |
| GOT1 | 5 | 0.68504 | 0.75978 | 0.999994 | 16162 | 1 | 4.0695 | 0.060147 | 0.17266 | 0.965913 | 3794 | 4 | 4.0695 |
| ZNF226 | 6 | 0.47591 | 0.65243 | 0.999994 | 13078 | 2 | 2.0538 | 0.060196 | 0.18788 | 0.97466 | 3795 | 4 | 2.0538 |

| | | | | | | | | | | | | | |
|---|---|---|---|---|---|---|---|---|---|---|---|---|---|
| MSH2 | 6 | 0.91602 | 0.91679 | 0.999994 | 19537 | 1 | 1.2959 | 0.060196 | 0.18788 | 0.97466 | 3796 | 4 | 1.2959 |
| CYP21A2 | 5 | 0.91276 | 0.9128 | 0.999994 | 19474 | 0 | 2.7902 | 0.060209 | 0.1728 | 0.965913 | 3797 | 4 | 2.7902 |
| LAMB4 | 6 | 0.66068 | 0.78584 | 0.999994 | 15789 | 2 | 2.8359 | 0.060268 | 0.18806 | 0.97466 | 3798 | 3 | 2.8359 |
| FAM110B | 4 | 0.59274 | 0.66142 | 0.999994 | 14856 | 1 | 0.48997 | 0.060305 | 0.16701 | 0.965913 | 3799 | 2 | 0.48997 |
| DERL1 | 5 | 0.59579 | 0.72459 | 0.999994 | 14900 | 1 | 0.72117 | 0.060317 | 0.17302 | 0.965913 | 3800 | 3 | 0.72117 |
| NCAPD3 | 6 | 0.88513 | 0.89295 | 0.999994 | 18935 | 1 | 3.9641 | 0.06034 | 0.18825 | 0.97466 | 3801 | 4 | 3.9641 |
| hsa-mir-564 | 2 | 0.93962 | 0.93976 | 0.999994 | 20069 | 0 | 5.6714 | 0.060377 | 0.10654 | 0.953839 | 3802 | 2 | 5.6714 |
| ZNF793 | 6 | 0.90361 | 0.90649 | 0.999994 | 19294 | 1 | 0.92878 | 0.060395 | 0.18838 | 0.97466 | 3803 | 4 | 0.92878 |
| ZC3H3 | 3 | 0.1428 | 0.23921 | 0.987253 | 6264 | 1 | 6.1806 | 0.060428 | 0.14217 | 0.965913 | 3804 | 2 | 6.1806 |
| G6PC | 3 | 0.93938 | 0.93953 | 0.999994 | 20058 | 0 | 6.1806 | 0.060428 | 0.14217 | 0.965913 | 3805 | 3 | 6.1806 |
| KMT2E | 5 | 0.63592 | 0.73961 | 0.999994 | 15459 | 1 | -0.22771 | 0.060448 | 0.1733 | 0.965913 | 3806 | 2 | -0.22771 |
| C6orf7 | 5 | 0.026192 | 0.080615 | 0.93914 | 1814 | 1 | 3.574 | 0.060448 | 0.1733 | 0.965913 | 3807 | 3 | 3.574 |
| THRB | 6 | 0.98069 | 0.98061 | 0.999994 | 21107 | 0 | 2.3741 | 0.060462 | 0.18854 | 0.97466 | 3808 | 4 | 2.3741 |
| LSMEM1 | 6 | 0.47921 | 0.65465 | 0.999994 | 13132 | 2 | 0.85795 | 0.060462 | 0.18854 | 0.97466 | 3809 | 3 | 0.85795 |
| KCNQ1 | 4 | 0.38817 | 0.53349 | 0.999994 | 11768 | 2 | 3.0425 | 0.060576 | 0.16745 | 0.965913 | 3810 | 2 | 3.0425 |
| ME2 | 6 | 0.37253 | 0.5848 | 0.999994 | 11552 | 1 | 3.7092 | 0.060578 | 0.18883 | 0.97466 | 3811 | 4 | 3.7092 |
| PHLPP1 | 6 | 0.94659 | 0.94657 | 0.999994 | 20249 | 0 | 1.7568 | 0.060596 | 0.18888 | 0.97466 | 3812 | 5 | 1.7568 |
| TYK2 | 6 | 0.49116 | 0.66269 | 0.999994 | 13292 | 1 | 3.8025 | 0.060619 | 0.18894 | 0.97466 | 3813 | 4 | 3.8025 |
| FAM46A | 5 | 0.39639 | 0.56055 | 0.999994 | 11894 | 1 | 4.8715 | 0.060633 | 0.1737 | 0.966602 | 3814 | 4 | 4.8715 |
| C6orf223 | 5 | 0.94947 | 0.94966 | 0.999994 | 20321 | 0 | 4.8715 | 0.060633 | 0.1737 | 0.966602 | 3815 | 4 | 4.8715 |
| C9orf47 | 6 | 0.055659 | 0.15634 | 0.967126 | 3228 | 1 | 1.3039 | 0.060637 | 0.18899 | 0.97466 | 3816 | 5 | 1.3039 |
| NEFL | 6 | 0.25125 | 0.44944 | 0.999994 | 9042 | 2 | 3.5445 | 0.060662 | 0.18906 | 0.97466 | 3817 | 4 | 3.5445 |
| TMEM39A | 6 | 0.043458 | 0.1297 | 0.95318 | 2692 | 2 | 3.9166 | 0.060662 | 0.18906 | 0.97466 | 3818 | 4 | 3.9166 |
| IZUMO4 | 6 | 0.2416 | 0.43792 | 0.999994 | 8818 | 1 | 3.2956 | 0.060662 | 0.18906 | 0.97466 | 3819 | 4 | 3.2956 |
| STAG3 | 6 | 0.55892 | 0.71004 | 0.999994 | 14350 | 1 | 2.7572 | 0.060662 | 0.18906 | 0.97466 | 3820 | 5 | 2.7572 |
| ZNF683 | 5 | 0.92073 | 0.92092 | 0.999994 | 19647 | 0 | 2.0344 | 0.06073 | 0.1739 | 0.966602 | 3821 | 4 | 2.0344 |
| TCFL5 | 5 | 0.17552 | 0.34018 | 0.999994 | 7124 | 2 | 0.90325 | 0.060743 | 0.17393 | 0.966602 | 3822 | 3 | 0.90325 |
| FDXACB1 | 6 | 0.94855 | 0.94857 | 0.999994 | 20302 | 0 | 1.9586 | 0.060764 | 0.18932 | 0.97466 | 3823 | 5 | 1.9586 |
| C12orf77 | 6 | 0.80984 | 0.85219 | 0.999994 | 17632 | 1 | 3.24 | 0.060768 | 0.18933 | 0.97466 | 3824 | 4 | 3.24 |
| SLC6A14 | 6 | 0.66059 | 0.78577 | 0.999994 | 15786 | 2 | 2.9914 | 0.060836 | 0.18949 | 0.97466 | 3825 | 4 | 2.9914 |
| hsa-mir-6867 | 3 | 0.93916 | 0.93931 | 0.999994 | 20051 | 0 | 4.2459 | 0.060841 | 0.14305 | 0.965913 | 3826 | 3 | 4.2459 |
| PPARG | 6 | 0.94733 | 0.94734 | 0.999994 | 20271 | 0 | 3.3712 | 0.060864 | 0.18957 | 0.97466 | 3827 | 4 | 3.3712 |
| LTBP2 | 4 | 0.62215 | 0.67756 | 0.999994 | 15262 | 1 | 3.5185 | 0.060869 | 0.16797 | 0.965913 | 3828 | 2 | 3.5185 |
| FZD2 | 4 | 0.63406 | 0.68413 | 0.999994 | 15433 | 1 | 3.5434 | 0.060869 | 0.16797 | 0.965913 | 3829 | 2 | 3.5434 |
| ACAT1 | 6 | 0.45228 | 0.63657 | 0.999994 | 12739 | 2 | 1.4809 | 0.060905 | 0.18968 | 0.97466 | 3830 | 4 | 1.4809 |
| UGT2B11 | 6 | 0.051119 | 0.14652 | 0.964852 | 3024 | 1 | 1.3456 | 0.060938 | 0.18975 | 0.97466 | 3831 | 5 | 1.3456 |
| WASF1 | 6 | 0.97703 | 0.97697 | 0.999994 | 21014 | 0 | 3.1656 | 0.060979 | 0.18986 | 0.97466 | 3832 | 5 | 3.1656 |
| NonTargetingControlG | 1 | 0.93901 | 0.93921 | 0.999994 | 20050 | 0 | 7.9211 | 0.060993 | 0.061023 | 0.9326 | 3833 | 1 | 7.9211 |
| GSTM5 | 5 | 0.28339 | 0.46761 | 0.999994 | 9771 | 1 | -0.017805 | 0.061013 | 0.17453 | 0.967011 | 3834 | 2 | -0.017805 |
| VPS13A | 5 | 0.34679 | 0.5193 | 0.999994 | 11090 | 2 | 4.6527 | 0.061013 | 0.17453 | 0.967011 | 3835 | 3 | 4.6527 |
| G2E3 | 6 | 0.85154 | 0.87272 | 0.999994 | 18287 | 1 | 1.0378 | 0.061017 | 0.18996 | 0.97466 | 3836 | 3 | 1.0378 |
| GFOD2 | 6 | 0.64556 | 0.7742 | 0.999994 | 15592 | 2 | 2.9714 | 0.061093 | 0.19015 | 0.97466 | 3837 | 4 | 2.9714 |
| NLRP12 | 6 | 0.32889 | 0.53995 | 0.999994 | 10700 | 3 | 0.67886 | 0.061093 | 0.19015 | 0.97466 | 3838 | 3 | 0.67886 |
| ECEL1 | 6 | 0.30293 | 0.51014 | 0.999994 | 10192 | 2 | 2.8864 | 0.061098 | 0.19016 | 0.97466 | 3839 | 3 | 2.8864 |
| C8orf4 | 6 | 0.28181 | 0.48549 | 0.999994 | 9737 | 2 | 2.913 | 0.061098 | 0.19016 | 0.97466 | 3840 | 3 | 2.913 |
| GPR132 | 6 | 0.50415 | 0.67161 | 0.999994 | 13497 | 2 | 3.7976 | 0.061098 | 0.19016 | 0.97466 | 3841 | 4 | 3.7976 |
| RPE | 6 | 0.9383 | 0.93825 | 0.999994 | 20033 | 0 | 1.5733 | 0.061113 | 0.1902 | 0.97466 | 3842 | 5 | 1.5733 |
| SAYSD1 | 5 | 0.55297 | 0.69437 | 0.999994 | 14257 | 2 | 4.8677 | 0.061124 | 0.17479 | 0.967058 | 3843 | 3 | 4.8677 |
| hsa-mir-5093 | 4 | 0.41894 | 0.5651 | 0.999994 | 12234 | 2 | 0.52927 | 0.061187 | 0.16852 | 0.965913 | 3844 | 2 | 0.52927 |
| hsa-mir-3186 | 4 | 0.046043 | 0.11371 | 0.951052 | 2810 | 1 | 3.225 | 0.061188 | 0.16852 | 0.965913 | 3845 | 3 | 3.225 |
| hsa-mir-6745 | 3 | 0.53799 | 0.57192 | 0.999994 | 14026 | 1 | 6.16 | 0.061204 | 0.14379 | 0.965913 | 3846 | 2 | 6.16 |
| SLC7A3 | 5 | 0.97063 | 0.97068 | 0.999994 | 20838 | 0 | 4.1171 | 0.061234 | 0.17502 | 0.967616 | 3847 | 4 | 4.1171 |

| | | | | | | | | | | | | | |
|---|---|---|---|---|---|---|---|---|---|---|---|---|---|
| TRIL | 3 | 0.93873 | 0.9389 | 0.999994 | 20046 | 0 | 4.1167 | 0.061266 | 0.14392 | 0.965913 | 3848 | 3 | 4.1167 |
| hsa-mir-4275 | 3 | 0.93873 | 0.93889 | 0.999994 | 20045 | 0 | 2.7254 | 0.06127 | 0.14393 | 0.965913 | 3849 | 3 | 2.7254 |
| CXCL9 | 6 | 0.90827 | 0.91015 | 0.999994 | 19391 | 1 | 3.4305 | 0.061294 | 0.19064 | 0.97466 | 3850 | 4 | 3.4305 |
| CLDN4 | 4 | 0.29702 | 0.43996 | 0.999994 | 10068 | 2 | -0.93305 | 0.061331 | 0.16875 | 0.965913 | 3851 | 1 | -0.93305 |
| ZNF438 | 6 | 0.98576 | 0.98572 | 0.999994 | 21244 | 0 | 3.1285 | 0.061383 | 0.19086 | 0.97466 | 3852 | 5 | 3.1285 |
| LOC283403 | 6 | 0.32634 | 0.53706 | 0.999994 | 10653 | 1 | 2.8125 | 0.061404 | 0.19091 | 0.97466 | 3853 | 4 | 2.8125 |
| CETP | 4 | 0.75813 | 0.76629 | 0.999994 | 16955 | 1 | 3.1079 | 0.061407 | 0.16888 | 0.965913 | 3854 | 3 | 3.1079 |
| MTMR10 | 5 | 0.52613 | 0.67067 | 0.999994 | 13832 | 1 | 4.7322 | 0.061487 | 0.17555 | 0.969019 | 3855 | 3 | 4.7322 |
| NonTargetingControlG | 2 | 0.33504 | 0.36406 | 0.999994 | 10835 | 1 | 3.1956 | 0.061489 | 0.1084 | 0.953839 | 3856 | 1 | 3.1956 |
| hsa-mir-3928 | 3 | 0.93851 | 0.93867 | 0.999994 | 20037 | 0 | 3.3274 | 0.061491 | 0.14437 | 0.965913 | 3857 | 3 | 3.3274 |
| hsa-mir-577 | 4 | 0.3558 | 0.50017 | 0.999994 | 11263 | 1 | 3.1461 | 0.061502 | 0.16905 | 0.965913 | 3858 | 3 | 3.1461 |
| TUBA8 | 4 | 0.60368 | 0.66734 | 0.999994 | 14993 | 1 | 4.1668 | 0.061514 | 0.16907 | 0.965913 | 3859 | 3 | 4.1668 |
| PLEKHF2 | 6 | 0.14947 | 0.32496 | 0.999994 | 6445 | 2 | 3.5141 | 0.061548 | 0.19126 | 0.97466 | 3860 | 4 | 3.5141 |
| ZNF736 | 5 | 0.75158 | 0.79167 | 0.999994 | 16872 | 1 | 0.48692 | 0.06155 | 0.17568 | 0.969272 | 3861 | 2 | 0.48692 |
| DEFB136 | 6 | 0.65833 | 0.78402 | 0.999994 | 15756 | 2 | 3.6321 | 0.061568 | 0.19131 | 0.97466 | 3862 | 4 | 3.6321 |
| HSPA14 | 6 | 0.077571 | 0.20243 | 0.977973 | 4133 | 3 | -1.1357 | 0.061576 | 0.19133 | 0.97466 | 3863 | 3 | -1.1357 |
| CHRNA3 | 5 | 0.75315 | 0.79251 | 0.999994 | 16889 | 1 | 4.813 | 0.061629 | 0.17585 | 0.969446 | 3864 | 3 | 4.813 |
| hsa-mir-4294 | 4 | 0.52363 | 0.62716 | 0.999994 | 13802 | 1 | 3.9612 | 0.061681 | 0.16936 | 0.965913 | 3865 | 3 | 3.9612 |
| STK10 | 6 | 0.23908 | 0.4349 | 0.999994 | 8750 | 2 | 1.5802 | 0.06171 | 0.19166 | 0.97466 | 3866 | 3 | 1.5802 |
| ZNF609 | 6 | 0.88228 | 0.8911 | 0.999994 | 18875 | 1 | 0.24901 | 0.061715 | 0.19168 | 0.97466 | 3867 | 2 | 0.24901 |
| RBM20 | 5 | 0.94696 | 0.94714 | 0.999994 | 20264 | 0 | 2.1708 | 0.061729 | 0.17607 | 0.969889 | 3868 | 3 | 2.1708 |
| SPATA31A4 | 2 | 0.93826 | 0.9384 | 0.999994 | 20031 | 0 | 5.5593 | 0.06174 | 0.10881 | 0.953839 | 3869 | 2 | 5.5593 |
| MEF2B | 6 | 0.076588 | 0.20046 | 0.977164 | 4095 | 3 | 1.6101 | 0.061838 | 0.19196 | 0.97466 | 3870 | 3 | 1.6101 |
| LRIG2 | 5 | 0.018031 | 0.058722 | 0.931872 | 1361 | 2 | 0.35674 | 0.06184 | 0.17631 | 0.969889 | 3871 | 2 | 0.35674 |
| ALG12 | 5 | 0.68185 | 0.75837 | 0.999994 | 16114 | 1 | 1.9597 | 0.061854 | 0.17634 | 0.969889 | 3872 | 4 | 1.9597 |
| DHX38 | 5 | 0.62559 | 0.73568 | 0.999994 | 15322 | 1 | 4.8253 | 0.061858 | 0.17636 | 0.969889 | 3873 | 4 | 4.8253 |
| GPR6 | 6 | 0.029882 | 0.0985 | 0.945253 | 2002 | 4 | -2.9531 | 0.061871 | 0.19204 | 0.97466 | 3874 | 2 | -2.9531 |
| CDC123 | 6 | 0.97827 | 0.97823 | 0.999994 | 21053 | 0 | 1.8185 | 0.061886 | 0.19208 | 0.97466 | 3875 | 5 | 1.8185 |
| LRRC46 | 5 | 0.35668 | 0.52735 | 0.999994 | 11280 | 1 | 0.30399 | 0.061886 | 0.17642 | 0.969889 | 3876 | 2 | 0.30399 |
| NonTargetingControlG | 2 | 0.84271 | 0.84261 | 0.999994 | 18136 | 0 | 4.9349 | 0.061991 | 0.1092 | 0.953839 | 3877 | 2 | 4.9349 |
| GNA13 | 5 | 0.28436 | 0.46838 | 0.999994 | 9791 | 1 | 2.2727 | 0.062021 | 0.17673 | 0.970198 | 3878 | 4 | 2.2727 |
| MARK2 | 6 | 0.24854 | 0.4462 | 0.999994 | 8983 | 2 | 0.0087102 | 0.062059 | 0.19249 | 0.97466 | 3879 | 3 | 0.0087102 |
| C20orf27 | 6 | 0.63821 | 0.76858 | 0.999994 | 15490 | 2 | 3.3587 | 0.062119 | 0.19264 | 0.97466 | 3880 | 4 | 3.3587 |
| PIGW | 6 | 0.16939 | 0.34999 | 0.999994 | 6971 | 2 | 4.3866 | 0.062119 | 0.19264 | 0.97466 | 3881 | 4 | 4.3866 |
| SAT2 | 4 | 0.48257 | 0.60864 | 0.999994 | 13181 | 1 | 4.0514 | 0.062133 | 0.17015 | 0.965913 | 3882 | 3 | 4.0514 |
| GID4 | 4 | 0.084609 | 0.18719 | 0.969409 | 4392 | 2 | 1.6042 | 0.062133 | 0.17015 | 0.965913 | 3883 | 2 | 1.6042 |
| SEPT9 | 5 | 0.084378 | 0.2016 | 0.977164 | 4387 | 1 | 1.0327 | 0.062168 | 0.17704 | 0.970561 | 3884 | 3 | 1.0327 |
| SLC25A31 | 5 | 0.3961 | 0.56032 | 0.999994 | 11891 | 2 | 1.2176 | 0.062168 | 0.17704 | 0.970561 | 3885 | 3 | 1.2176 |
| NonTargetingControlG | 2 | 0.8268 | 0.82663 | 0.999994 | 17883 | 0 | 4.8799 | 0.062192 | 0.10956 | 0.953839 | 3886 | 2 | 4.8799 |
| KDM6B | 4 | 0.295 | 0.43785 | 0.999994 | 10018 | 2 | 0.33707 | 0.06223 | 0.1703 | 0.965913 | 3887 | 2 | 0.33707 |
| HERC1 | 6 | 0.63262 | 0.76431 | 0.999994 | 15407 | 1 | 3.7881 | 0.062233 | 0.19293 | 0.97466 | 3888 | 4 | 3.7881 |
| OR2J2 | 6 | 0.025355 | 0.087828 | 0.94395 | 1770 | 2 | 4.2872 | 0.062233 | 0.19293 | 0.97466 | 3889 | 4 | 4.2872 |
| LHX6 | 4 | 0.44135 | 0.58819 | 0.999994 | 12598 | 1 | 4.8899 | 0.062263 | 0.17036 | 0.965913 | 3890 | 3 | 4.8899 |
| hsa-mir-3686 | 3 | 0.93774 | 0.93787 | 0.999994 | 20024 | 0 | 1.8093 | 0.062265 | 0.14596 | 0.965913 | 3891 | 3 | 1.8093 |
| DPYSL4 | 6 | 0.24398 | 0.44074 | 0.999994 | 8882 | 2 | 3.3443 | 0.062275 | 0.19303 | 0.97466 | 3892 | 4 | 3.3443 |
| LY6G6D | 6 | 0.86184 | 0.87846 | 0.999994 | 18483 | 1 | 3.8019 | 0.06236 | 0.19326 | 0.97466 | 3893 | 5 | 3.8019 |
| DLL4 | 6 | 0.11593 | 0.27805 | 0.999536 | 5512 | 3 | 1.8694 | 0.062383 | 0.19331 | 0.97466 | 3894 | 3 | 1.8694 |
| C10orf120 | 6 | 0.67925 | 0.80043 | 0.999994 | 16079 | 1 | 2.4257 | 0.062399 | 0.19335 | 0.97466 | 3895 | 5 | 2.4257 |
| ZSCAN25 | 5 | 0.75166 | 0.79172 | 0.999994 | 16873 | 1 | 4.8199 | 0.062408 | 0.17753 | 0.971262 | 3896 | 3 | 4.8199 |
| AMER3 | 5 | 0.09877 | 0.22449 | 0.983354 | 4923 | 2 | 4.5188 | 0.062408 | 0.17753 | 0.971262 | 3897 | 3 | 4.5188 |
| EFCAB4A | 4 | 0.33754 | 0.48143 | 0.999994 | 10889 | 1 | 4.9244 | 0.062434 | 0.17065 | 0.965913 | 3898 | 3 | 4.9244 |
| C6orf52 | 5 | 0.36784 | 0.53657 | 0.999994 | 11472 | 1 | 2.0466 | 0.062443 | 0.1776 | 0.971262 | 3899 | 4 | 2.0466 |

| | | | | | | | | | | | | |
|---|---|---|---|---|---|---|---|---|---|---|---|---|
| hsa-mir-4712 | 3 | 0.9375 | 0.93763 | 0.999994 | 20019 | 0 | 2.2526 | 0.062496 | 0.14645 | 0.965913 | 3900 | 3 | 2.2526 |
| LRP1 | 5 | 0.19466 | 0.36758 | 0.999994 | 7623 | 3 | -0.8028 | 0.062503 | 0.17772 | 0.971262 | 3901 | 2 | -0.8028 |
| SYT8 | 5 | 0.2745 | 0.46041 | 0.999994 | 9582 | 1 | 4.7365 | 0.062503 | 0.17772 | 0.971262 | 3902 | 4 | 4.7365 |
| ZFP82 | 6 | 0.98461 | 0.98453 | 0.999994 | 21210 | 0 | 1.8234 | 0.062512 | 0.19364 | 0.97466 | 3903 | 5 | 1.8234 |
| ZCCHC16 | 6 | 0.77972 | 0.83954 | 0.999994 | 17208 | 1 | 0.037084 | 0.062522 | 0.19366 | 0.97466 | 3904 | 2 | 0.037084 |
| SH3D21 | 1 | 0.93745 | 0.93765 | 0.999994 | 20017 | 0 | 7.8908 | 0.062554 | 0.062547 | 0.933948 | 3905 | 1 | 7.8908 |
| GKN2 | 6 | 0.073814 | 0.19469 | 0.975797 | 3987 | 4 | -1.2909 | 0.062568 | 0.19377 | 0.97466 | 3906 | 2 | -1.2909 |
| PACSIN3 | 5 | 0.15932 | 0.31658 | 0.999994 | 6715 | 3 | -0.99443 | 0.06258 | 0.17789 | 0.971262 | 3907 | 2 | -0.99443 |
| TLK1 | 6 | 0.58082 | 0.72582 | 0.999994 | 14689 | 1 | 3.2609 | 0.062632 | 0.19392 | 0.97466 | 3908 | 4 | 3.2609 |
| LYPD6 | 5 | 0.014459 | 0.048657 | 0.902886 | 1161 | 3 | -3.5277 | 0.062647 | 0.17803 | 0.971262 | 3909 | 2 | -3.5277 |
| C10orf137 | 6 | 0.60075 | 0.7405 | 0.999994 | 14958 | 2 | 3.1735 | 0.06268 | 0.19404 | 0.97466 | 3910 | 4 | 3.1735 |
| FCHO1 | 4 | 0.089388 | 0.1959 | 0.975797 | 4573 | 2 | 2.4825 | 0.062685 | 0.17107 | 0.965913 | 3911 | 2 | 2.4825 |
| POLR3A | 4 | 0.8308 | 0.83036 | 0.999994 | 17942 | 0 | 4.49 | 0.062714 | 0.17111 | 0.965913 | 3912 | 3 | 4.49 |
| PROSER1 | 5 | 0.0061839 | 0.024968 | 0.844082 | 603 | 2 | 4.8012 | 0.062745 | 0.17823 | 0.971262 | 3913 | 3 | 4.8012 |
| CHRNE | 5 | 0.38446 | 0.55048 | 0.999994 | 11706 | 1 | 4.4424 | 0.06276 | 0.17827 | 0.971262 | 3914 | 3 | 4.4424 |
| ZSCAN2 | 6 | 0.92835 | 0.92844 | 0.999994 | 19822 | 0 | 0.20954 | 0.062769 | 0.19425 | 0.97466 | 3915 | 2 | 0.20954 |
| GEMIN7 | 4 | 0.23003 | 0.37157 | 0.999994 | 8528 | 2 | -0.50907 | 0.062787 | 0.17123 | 0.965913 | 3916 | 2 | -0.50907 |
| ARHGAP9 | 4 | 0.35741 | 0.50185 | 0.999994 | 11294 | 1 | 0.014897 | 0.062787 | 0.17123 | 0.965913 | 3917 | 1 | 0.014897 |
| hsa-mir-3648 | 2 | 0.93721 | 0.93734 | 0.999994 | 20011 | 0 | 4.3468 | 0.062787 | 0.11055 | 0.953839 | 3918 | 2 | 4.3468 |
| MRPL10 | 5 | 0.51096 | 0.65749 | 0.999994 | 13595 | 1 | 2.187 | 0.062792 | 0.17834 | 0.971262 | 3919 | 4 | 2.187 |
| CRISPLD1 | 6 | 0.21732 | 0.40889 | 0.999994 | 8202 | 2 | -0.15455 | 0.06281 | 0.19435 | 0.97466 | 3920 | 2 | -0.15455 |
| C6orf201 | 5 | 0.25769 | 0.44686 | 0.999994 | 9185 | 2 | 4.8102 | 0.062851 | 0.17848 | 0.971597 | 3921 | 3 | 4.8102 |
| ERAP2 | 6 | 0.98188 | 0.98181 | 0.999994 | 21133 | 0 | 1.1209 | 0.06288 | 0.19452 | 0.97466 | 3922 | 5 | 1.1209 |
| C10orf11 | 4 | 0.74861 | 0.75897 | 0.999994 | 16843 | 1 | 1.1482 | 0.062966 | 0.17155 | 0.965913 | 3923 | 3 | 1.1482 |
| PRDM1 | 6 | 0.83013 | 0.86161 | 0.999994 | 17934 | 1 | 1.3698 | 0.063024 | 0.19487 | 0.97466 | 3924 | 4 | 1.3698 |
| KRTAP2-1 | 6 | 0.6632 | 0.78779 | 0.999994 | 15826 | 1 | 3.2936 | 0.063042 | 0.19491 | 0.97466 | 3925 | 4 | 3.2936 |
| ODF2 | 6 | 0.77706 | 0.83853 | 0.999994 | 17169 | 1 | 0.56375 | 0.063054 | 0.19493 | 0.97466 | 3926 | 3 | 0.56375 |
| BAMBI | 6 | 0.89708 | 0.90142 | 0.999994 | 19161 | 1 | 0.99583 | 0.063054 | 0.19493 | 0.97466 | 3927 | 4 | 0.99583 |
| ESAM | 6 | 0.96167 | 0.96174 | 0.999994 | 20623 | 0 | 1.9584 | 0.063054 | 0.19493 | 0.97466 | 3928 | 3 | 1.9584 |
| DLG4 | 5 | 0.034883 | 0.10297 | 0.946414 | 2262 | 2 | -0.55064 | 0.063137 | 0.1791 | 0.97162 | 3929 | 2 | -0.55064 |
| PKIA | 5 | 0.81934 | 0.83215 | 0.999994 | 17771 | 1 | 3.2302 | 0.063137 | 0.1791 | 0.97162 | 3930 | 4 | 3.2302 |
| CHST14 | 5 | 0.25641 | 0.44584 | 0.999994 | 9160 | 2 | 0.34583 | 0.063137 | 0.1791 | 0.97162 | 3931 | 2 | 0.34583 |
| KIAA1407 | 6 | 0.49533 | 0.66551 | 0.999994 | 13341 | 2 | -0.085914 | 0.063158 | 0.19521 | 0.97466 | 3932 | 2 | -0.085914 |
| FAM206A | 5 | 0.8472 | 0.8521 | 0.999994 | 18210 | 1 | 1.9326 | 0.063165 | 0.17916 | 0.97162 | 3933 | 4 | 1.9326 |
| RAB6C | 5 | 0.15535 | 0.31071 | 0.999994 | 6603 | 1 | 4.7816 | 0.063196 | 0.17923 | 0.97162 | 3934 | 3 | 4.7816 |
| PRLH | 5 | 0.5001 | 0.64797 | 0.999994 | 13424 | 2 | 4.7816 | 0.063196 | 0.17923 | 0.97162 | 3935 | 3 | 4.7816 |
| FYB | 6 | 0.30014 | 0.50687 | 0.999994 | 10133 | 1 | 3.1937 | 0.063237 | 0.19542 | 0.97466 | 3936 | 4 | 3.1937 |
| HS3ST1 | 6 | 0.17391 | 0.35553 | 0.999994 | 7081 | 2 | 3.1895 | 0.063251 | 0.19545 | 0.97466 | 3937 | 3 | 3.1895 |
| C3orf33 | 4 | 0.033752 | 0.089167 | 0.94395 | 2203 | 2 | -0.94888 | 0.063271 | 0.17207 | 0.965913 | 3938 | 1 | -0.94888 |
| MEF2BNB-MEF2B | 3 | 0.29764 | 0.42727 | 0.999994 | 10082 | 1 | 1.9526 | 0.063291 | 0.14807 | 0.965913 | 3939 | 2 | 1.9526 |
| PAX5 | 3 | 0.93669 | 0.93681 | 0.999994 | 19999 | 0 | 4.9696 | 0.063308 | 0.14811 | 0.965913 | 3940 | 3 | 4.9696 |
| C11orf30 | 3 | 0.93667 | 0.93679 | 0.999994 | 19998 | 0 | 5.202 | 0.063326 | 0.14815 | 0.965913 | 3941 | 3 | 5.202 |
| SUDS3 | 4 | 0.93739 | 0.93725 | 0.999994 | 20014 | 0 | 4.7408 | 0.063347 | 0.1722 | 0.965913 | 3942 | 3 | 4.7408 |
| TOMM6 | 5 | 0.75383 | 0.79287 | 0.999994 | 16895 | 1 | 4.7265 | 0.063375 | 0.17963 | 0.97162 | 3943 | 3 | 4.7265 |
| ZBTB49 | 6 | 0.53101 | 0.69025 | 0.999994 | 13911 | 1 | 0.67018 | 0.063385 | 0.1958 | 0.97466 | 3944 | 3 | 0.67018 |
| CLTC | 6 | 0.0743 | 0.19571 | 0.975797 | 4003 | 2 | 2.0353 | 0.063385 | 0.1958 | 0.97466 | 3945 | 3 | 2.0353 |
| PIK3CG | 6 | 0.90494 | 0.90752 | 0.999994 | 19324 | 1 | 1.5148 | 0.063388 | 0.1958 | 0.97466 | 3946 | 5 | 1.5148 |
| LYPLA1 | 4 | 0.91994 | 0.91972 | 0.999994 | 19629 | 0 | 4.085 | 0.063397 | 0.17228 | 0.965913 | 3947 | 3 | 4.085 |
| SBSPON | 5 | 0.012042 | 0.041919 | 0.891621 | 993 | 3 | -3.7721 | 0.063412 | 0.17969 | 0.97162 | 3948 | 2 | -3.7721 |
| PINLYP | 5 | 0.11558 | 0.25062 | 0.992497 | 5499 | 1 | 4.6419 | 0.063431 | 0.17973 | 0.97162 | 3949 | 3 | 4.6419 |
| C17orf74 | 6 | 0.9379 | 0.93788 | 0.999994 | 20027 | 0 | 3.1392 | 0.063437 | 0.19592 | 0.97466 | 3950 | 5 | 3.1392 |
| HBP1 | 6 | 0.56424 | 0.71382 | 0.999994 | 14458 | 2 | 1.4425 | 0.063453 | 0.19595 | 0.97466 | 3951 | 3 | 1.4425 |

| | | | | | | | | | | | | |
|---|---|---|---|---|---|---|---|---|---|---|---|---|
| DMTF1 | 4 | 0.30064 | 0.44368 | 0.999994 | 10144 | 1 | 4.1484 | 0.063467 | 0.1724 | 0.965913 | 3952 | 3 | 4.1484 |
| GSK3B | 6 | 0.94736 | 0.94738 | 0.999994 | 20273 | 0 | 1.593 | 0.06356 | 0.19623 | 0.97466 | 3953 | 5 | 1.593 |
| UHRF1BP1L | 6 | 0.16514 | 0.34463 | 0.999994 | 6866 | 3 | -0.45108 | 0.06356 | 0.19623 | 0.97466 | 3954 | 3 | -0.45108 |
| TUBB2A | 5 | 0.28586 | 0.46956 | 0.999994 | 9828 | 1 | 1.2004 | 0.063565 | 0.18003 | 0.97162 | 3955 | 3 | 1.2004 |
| C6orf1 | 4 | 0.64026 | 0.68767 | 0.999994 | 15514 | 1 | 2.9979 | 0.063577 | 0.17259 | 0.965913 | 3956 | 2 | 2.9979 |
| GZMK | 5 | 0.51938 | 0.66476 | 0.999994 | 13737 | 1 | 0.84082 | 0.063655 | 0.18022 | 0.97162 | 3957 | 3 | 0.84082 |
| DHX58 | 5 | 0.024065 | 0.074993 | 0.93914 | 1712 | 3 | -3.0005 | 0.063655 | 0.18022 | 0.97162 | 3958 | 2 | -3.0005 |
| DTNA | 4 | 0.77282 | 0.77806 | 0.999994 | 17119 | 1 | 0.55174 | 0.063684 | 0.17277 | 0.965913 | 3959 | 2 | 0.55174 |
| DHDH | 6 | 0.42119 | 0.61617 | 0.999994 | 12266 | 1 | 1.1645 | 0.063693 | 0.19658 | 0.97466 | 3960 | 5 | 1.1645 |
| MARCH4 | 6 | 0.41739 | 0.61366 | 0.999994 | 12210 | 2 | 1.9519 | 0.063694 | 0.19658 | 0.97466 | 3961 | 3 | 1.9519 |
| IL3 | 5 | 0.27608 | 0.46168 | 0.999994 | 9619 | 2 | 4.7663 | 0.063742 | 0.1804 | 0.97162 | 3962 | 3 | 4.7663 |
| LRRC30 | 6 | 0.56058 | 0.7112 | 0.999994 | 14379 | 2 | 3.232 | 0.063769 | 0.19674 | 0.97466 | 3963 | 4 | 3.232 |
| SHC3 | 6 | 0.26197 | 0.46222 | 0.999994 | 9288 | 2 | 2.7658 | 0.063788 | 0.19679 | 0.97466 | 3964 | 3 | 2.7658 |
| SERINC2 | 5 | 0.34981 | 0.52175 | 0.999994 | 11152 | 2 | -0.22989 | 0.063811 | 0.18055 | 0.97162 | 3965 | 2 | -0.22989 |
| ARNTL | 6 | 0.93373 | 0.93379 | 0.999994 | 19941 | 0 | 2.8294 | 0.063896 | 0.19705 | 0.97466 | 3966 | 4 | 2.8294 |
| UCP3 | 5 | 0.89295 | 0.89308 | 0.999994 | 19090 | 0 | 0.7367 | 0.063937 | 0.1808 | 0.97162 | 3967 | 3 | 0.7367 |
| FOXR2 | 5 | 0.90958 | 0.90961 | 0.999994 | 19417 | 0 | 3.4923 | 0.063937 | 0.1808 | 0.97162 | 3968 | 3 | 3.4923 |
| SIGLEC5 | 6 | 0.98081 | 0.98073 | 0.999994 | 21112 | 0 | 2.4248 | 0.063988 | 0.19727 | 0.97466 | 3969 | 5 | 2.4248 |
| SLMO2 | 4 | 0.18996 | 0.33033 | 0.999994 | 7482 | 1 | 3.759 | 0.064 | 0.1733 | 0.965913 | 3970 | 3 | 3.759 |
| DDX19B | 5 | 0.91738 | 0.91746 | 0.999994 | 19573 | 0 | 3.4078 | 0.064046 | 0.18103 | 0.97162 | 3971 | 4 | 3.4078 |
| FIZ1 | 4 | 0.92971 | 0.92948 | 0.999994 | 19847 | 0 | 4.0782 | 0.064061 | 0.1734 | 0.966047 | 3972 | 3 | 4.0782 |
| DDX49 | 6 | 0.46949 | 0.64811 | 0.999994 | 12999 | 2 | 2.3147 | 0.064071 | 0.19747 | 0.97466 | 3973 | 4 | 2.3147 |
| HIST1H3G | 6 | 0.28075 | 0.48428 | 0.999994 | 9715 | 2 | 3.1579 | 0.0641 | 0.19754 | 0.97466 | 3974 | 4 | 3.1579 |
| LILRB3 | 6 | 0.92248 | 0.92274 | 0.999994 | 19687 | 1 | 2.3809 | 0.064133 | 0.19761 | 0.97466 | 3975 | 5 | 2.3809 |
| PENK | 6 | 0.60617 | 0.74453 | 0.999994 | 15029 | 1 | 0.83272 | 0.064149 | 0.19764 | 0.97466 | 3976 | 4 | 0.83272 |
| KANK1 | 6 | 0.88129 | 0.89045 | 0.999994 | 18857 | 1 | 0.90013 | 0.064149 | 0.19764 | 0.97466 | 3977 | 3 | 0.90013 |
| SLC44A3 | 6 | 0.098098 | 0.24361 | 0.988088 | 4896 | 1 | 4.2847 | 0.064157 | 0.19766 | 0.97466 | 3978 | 5 | 4.2847 |
| ENOSF1 | 6 | 0.95275 | 0.95275 | 0.999994 | 20401 | 0 | 3.0914 | 0.064164 | 0.19768 | 0.97466 | 3979 | 4 | 3.0914 |
| ZNF710 | 4 | 0.56843 | 0.64884 | 0.999994 | 14516 | 1 | 3.4706 | 0.064224 | 0.1737 | 0.966602 | 3980 | 2 | 3.4706 |
| SEC14L2 | 5 | 0.19384 | 0.36641 | 0.999994 | 7595 | 1 | 1.3182 | 0.064251 | 0.18146 | 0.97162 | 3981 | 4 | 1.3182 |
| hsa-mir-587 | 3 | 0.9357 | 0.93583 | 0.999994 | 19984 | 0 | 1.4488 | 0.064295 | 0.15012 | 0.965913 | 3982 | 3 | 1.4488 |
| OR6X1 | 6 | 0.24619 | 0.44339 | 0.999994 | 8927 | 2 | 2.2379 | 0.064335 | 0.19809 | 0.97466 | 3983 | 3 | 2.2379 |
| TEAD2 | 6 | 0.67638 | 0.79817 | 0.999994 | 16029 | 2 | 2.395 | 0.064335 | 0.19809 | 0.97466 | 3984 | 4 | 2.395 |
| SUCLA2 | 6 | 0.15926 | 0.33727 | 0.999994 | 6713 | 1 | 1.1504 | 0.064335 | 0.19809 | 0.97466 | 3985 | 3 | 1.1504 |
| SERPINB3 | 6 | 0.16848 | 0.34885 | 0.999994 | 6948 | 3 | -0.41409 | 0.064335 | 0.19809 | 0.97466 | 3986 | 2 | -0.41409 |
| CYP46A1 | 6 | 0.96512 | 0.96515 | 0.999994 | 20721 | 0 | 1.5813 | 0.064351 | 0.19812 | 0.97466 | 3987 | 5 | 1.5813 |
| STPG2 | 6 | 0.95769 | 0.95778 | 0.999994 | 20530 | 0 | 3.0637 | 0.064354 | 0.19813 | 0.97466 | 3988 | 4 | 3.0637 |
| SND1 | 6 | 0.93165 | 0.93179 | 0.999994 | 19891 | 0 | 3.2766 | 0.064379 | 0.19819 | 0.97466 | 3989 | 4 | 3.2766 |
| MAP3K2 | 6 | 0.24707 | 0.44443 | 0.999994 | 8948 | 1 | 3.2997 | 0.064379 | 0.19819 | 0.97466 | 3990 | 4 | 3.2997 |
| ZCCHC17 | 6 | 0.039582 | 0.12097 | 0.951052 | 2494 | 2 | 0.06344 | 0.064416 | 0.19827 | 0.97466 | 3991 | 2 | 0.06344 |
| LOC100653515 | 6 | 0.8388 | 0.866 | 0.999994 | 18067 | 1 | 3.5618 | 0.064545 | 0.1986 | 0.97466 | 3992 | 4 | 3.5618 |
| CACNG3 | 6 | 0.15374 | 0.33037 | 0.999994 | 6559 | 3 | 0.27162 | 0.06455 | 0.1986 | 0.97466 | 3993 | 3 | 0.27162 |
| LSM14B | 6 | 0.85458 | 0.87441 | 0.999994 | 18341 | 1 | 1.0772 | 0.06455 | 0.1986 | 0.97466 | 3994 | 3 | 1.0772 |
| OR10A6 | 5 | 0.66364 | 0.75063 | 0.999994 | 15833 | 1 | 4.7365 | 0.064554 | 0.18209 | 0.97162 | 3995 | 3 | 4.7365 |
| CHCHD10 | 5 | 0.14109 | 0.28945 | 0.999632 | 6222 | 2 | 4.7365 | 0.064554 | 0.18209 | 0.97162 | 3996 | 3 | 4.7365 |
| LUZP6 | 5 | 0.41813 | 0.57857 | 0.999994 | 12220 | 2 | -0.39115 | 0.064615 | 0.18222 | 0.97162 | 3997 | 2 | -0.39115 |
| BTN2A2 | 5 | 0.50329 | 0.6508 | 0.999994 | 13485 | 2 | -0.14376 | 0.064615 | 0.18222 | 0.97162 | 3998 | 2 | -0.14376 |
| MED17 | 3 | 0.93536 | 0.93548 | 0.999994 | 19976 | 0 | 4.2589 | 0.064636 | 0.15084 | 0.965913 | 3999 | 3 | 4.2589 |
| ZFY | 6 | 0.94016 | 0.94012 | 0.999994 | 20085 | 0 | 0.94142 | 0.064684 | 0.19892 | 0.97466 | 4000 | 3 | 0.94142 |
| KCNB2 | 6 | 0.25217 | 0.45052 | 0.999994 | 9059 | 3 | -0.3881 | 0.064684 | 0.19892 | 0.97466 | 4001 | 2 | -0.3881 |
| KLHL36 | 4 | 0.33064 | 0.47435 | 0.999994 | 10732 | 1 | 4.2649 | 0.064785 | 0.17465 | 0.967011 | 4002 | 3 | 4.2649 |
| RPS26 | 4 | 0.041649 | 0.10509 | 0.95013 | 2597 | 1 | 4.2721 | 0.064813 | 0.17471 | 0.967011 | 4003 | 3 | 4.2721 |

| | | | | | | | | | | | | | |
|---|---|---|---|---|---|---|---|---|---|---|---|---|---|
| TIMM10B | 4 | 0.69148 | 0.71891 | 0.999994 | 16221 | 1 | 0.97174 | 0.064833 | 0.17474 | 0.967011 | 4004 | 2 | 0.97174 |
| FBXO15 | 6 | 0.68727 | 0.80667 | 0.999994 | 16186 | 2 | 3.5308 | 0.064848 | 0.19932 | 0.97466 | 4005 | 4 | 3.5308 |
| HMGCS2 | 6 | 0.82375 | 0.85857 | 0.999994 | 17834 | 1 | 2.1104 | 0.064867 | 0.19936 | 0.97466 | 4006 | 5 | 2.1104 |
| MED7 | 5 | 0.93531 | 0.93555 | 0.999994 | 19975 | 0 | 1.9134 | 0.064881 | 0.18277 | 0.97162 | 4007 | 4 | 1.9134 |
| CPS1 | 5 | 0.70482 | 0.76869 | 0.999994 | 16357 | 1 | 1.7621 | 0.064893 | 0.1828 | 0.97162 | 4008 | 4 | 1.7621 |
| EGR1 | 5 | 0.63061 | 0.73753 | 0.999994 | 15384 | 1 | 1.8824 | 0.064898 | 0.1828 | 0.97162 | 4009 | 3 | 1.8824 |
| SPC24 | 2 | 0.9351 | 0.9352 | 0.999994 | 19969 | 0 | 3.7624 | 0.064902 | 0.11404 | 0.954984 | 4010 | 2 | 3.7624 |
| C1QL2 | 4 | 0.5649 | 0.64711 | 0.999994 | 14468 | 1 | 3.3929 | 0.064919 | 0.17489 | 0.967344 | 4011 | 2 | 3.3929 |
| HSDL1 | 6 | 0.89685 | 0.90126 | 0.999994 | 19158 | 1 | 0.9077 | 0.064957 | 0.19958 | 0.97466 | 4012 | 3 | 0.9077 |
| KCNK9 | 5 | 0.90887 | 0.90889 | 0.999994 | 19401 | 0 | 3.8087 | 0.064964 | 0.18293 | 0.97162 | 4013 | 4 | 3.8087 |
| ZNF276 | 6 | 0.98461 | 0.98454 | 0.999994 | 21211 | 0 | 1.3502 | 0.064982 | 0.19964 | 0.97466 | 4014 | 5 | 1.3502 |
| OR51E1 | 5 | 0.016601 | 0.054747 | 0.922677 | 1284 | 4 | -1.6368 | 0.064995 | 0.18299 | 0.97162 | 4015 | 1 | -1.6368 |
| KCTD21 | 5 | 0.11175 | 0.2447 | 0.988995 | 5358 | 2 | 2.4513 | 0.065085 | 0.18319 | 0.97162 | 4016 | 3 | 2.4513 |
| HRASLS5 | 6 | 0.92038 | 0.92076 | 0.999994 | 19640 | 1 | 3.3181 | 0.065086 | 0.19988 | 0.97466 | 4017 | 4 | 3.3181 |
| NDST2 | 6 | 0.81285 | 0.85353 | 0.999994 | 17668 | 1 | 2.4457 | 0.065159 | 0.20007 | 0.97466 | 4018 | 5 | 2.4457 |
| NonTargetingControlG | 1 | 0.93483 | 0.93505 | 0.999994 | 19963 | 0 | 7.8398 | 0.065165 | 0.065197 | 0.933948 | 4019 | 1 | 7.8398 |
| NonTargetingControlG | 2 | 0.85653 | 0.85647 | 0.999994 | 18380 | 0 | 5.0394 | 0.065184 | 0.11449 | 0.954984 | 4020 | 2 | 5.0394 |
| NAA40 | 5 | 0.90502 | 0.90512 | 0.999994 | 19329 | 0 | 3.9202 | 0.065185 | 0.1834 | 0.97162 | 4021 | 4 | 3.9202 |
| COMMD6 | 6 | 0.20252 | 0.39095 | 0.999994 | 7844 | 3 | -0.72454 | 0.065192 | 0.20015 | 0.97466 | 4022 | 2 | -0.72454 |
| SERHL2 | 4 | 0.68024 | 0.71173 | 0.999994 | 16094 | 1 | 0.11895 | 0.065209 | 0.17537 | 0.968566 | 4023 | 1 | 0.11895 |
| KPNB1 | 3 | 0.13989 | 0.23548 | 0.985362 | 6181 | 1 | 6.0521 | 0.065214 | 0.15205 | 0.965913 | 4024 | 2 | 6.0521 |
| C3orf52 | 6 | 0.91782 | 0.91838 | 0.999994 | 19583 | 1 | 1.6919 | 0.065214 | 0.2002 | 0.97466 | 4025 | 5 | 1.6919 |
| CCDC153 | 5 | 0.56366 | 0.7039 | 0.999994 | 14444 | 1 | 2.5351 | 0.065219 | 0.18347 | 0.97162 | 4026 | 4 | 2.5351 |
| CLDN9 | 6 | 0.049168 | 0.14228 | 0.959553 | 2939 | 3 | 2.4024 | 0.065281 | 0.20036 | 0.97466 | 4027 | 3 | 2.4024 |
| ADIPOR1 | 6 | 0.16381 | 0.34304 | 0.999994 | 6835 | 2 | 3.7111 | 0.065281 | 0.20036 | 0.97466 | 4028 | 4 | 3.7111 |
| VPS37A | 6 | 0.97905 | 0.97895 | 0.999994 | 21075 | 0 | 3.7488 | 0.065281 | 0.20036 | 0.97466 | 4029 | 5 | 3.7488 |
| NANOS3 | 6 | 0.50886 | 0.67481 | 0.999994 | 13566 | 2 | 3.2536 | 0.065281 | 0.20036 | 0.97466 | 4030 | 4 | 3.2536 |
| HEATR1 | 5 | 0.14021 | 0.28814 | 0.999632 | 6191 | 1 | 4.7106 | 0.0653 | 0.18363 | 0.971789 | 4031 | 4 | 4.7106 |
| CNGA2 | 5 | 0.77781 | 0.80618 | 0.999994 | 17181 | 1 | 2.181 | 0.065353 | 0.18373 | 0.971846 | 4032 | 4 | 2.181 |
| WDR70 | 3 | 0.78158 | 0.78147 | 0.999994 | 17232 | 0 | 3.4241 | 0.065361 | 0.15237 | 0.965913 | 4033 | 2 | 3.4241 |
| GAS2 | 4 | 0.93977 | 0.93967 | 0.999994 | 20072 | 0 | 3.7489 | 0.065373 | 0.17564 | 0.969272 | 4034 | 3 | 3.7489 |
| ZBTB25 | 6 | 0.88988 | 0.89627 | 0.999994 | 19034 | 1 | 1.9338 | 0.065373 | 0.2006 | 0.97466 | 4035 | 5 | 1.9338 |
| SIX3 | 6 | 0.92252 | 0.92277 | 0.999994 | 19690 | 1 | 3.8065 | 0.065481 | 0.20088 | 0.97466 | 4036 | 4 | 3.8065 |
| CHGA | 6 | 0.89012 | 0.89643 | 0.999994 | 19038 | 1 | 3.0553 | 0.065538 | 0.20102 | 0.97466 | 4037 | 3 | 3.0553 |
| hsa-mir-153-1 | 4 | 0.92785 | 0.92762 | 0.999994 | 19811 | 0 | 2.3879 | 0.06555 | 0.17594 | 0.969457 | 4038 | 2 | 2.3879 |
| BAD | 5 | 0.13032 | 0.27309 | 0.998561 | 5940 | 2 | -0.42934 | 0.065554 | 0.18416 | 0.97314 | 4039 | 2 | -0.42934 |
| MYRIP | 6 | 0.87052 | 0.88366 | 0.999994 | 18645 | 1 | 2.9182 | 0.065643 | 0.20129 | 0.97466 | 4040 | 5 | 2.9182 |
| DSP | 6 | 0.28556 | 0.48995 | 0.999994 | 9821 | 3 | 0.41536 | 0.065647 | 0.2013 | 0.97466 | 4041 | 3 | 0.41536 |
| ANKS4B | 6 | 0.60494 | 0.74365 | 0.999994 | 15010 | 1 | 2.0861 | 0.065647 | 0.2013 | 0.97466 | 4042 | 5 | 2.0861 |
| TAF7 | 6 | 0.95934 | 0.95941 | 0.999994 | 20567 | 0 | 3.6937 | 0.065655 | 0.20132 | 0.97466 | 4043 | 4 | 3.6937 |
| SCRT2 | 4 | 0.66029 | 0.69943 | 0.999994 | 15784 | 1 | 4.3287 | 0.065663 | 0.17612 | 0.969889 | 4044 | 3 | 4.3287 |
| PTPRT | 4 | 0.65433 | 0.69581 | 0.999994 | 15709 | 1 | 4.7127 | 0.065685 | 0.17616 | 0.969889 | 4045 | 3 | 4.7127 |
| ACBD4 | 6 | 0.87083 | 0.88385 | 0.999994 | 18652 | 1 | 3.6896 | 0.065727 | 0.2015 | 0.97466 | 4046 | 5 | 3.6896 |
| CYC1 | 6 | 0.47204 | 0.64981 | 0.999994 | 13036 | 1 | 3.4951 | 0.065772 | 0.20159 | 0.97466 | 4047 | 5 | 3.4951 |
| CMTM3 | 6 | 0.026655 | 0.090906 | 0.94395 | 1839 | 3 | -0.093947 | 0.065818 | 0.20171 | 0.97466 | 4048 | 3 | -0.093947 |
| CYTL1 | 6 | 0.08609 | 0.21969 | 0.980699 | 4452 | 3 | -0.76799 | 0.065818 | 0.20171 | 0.97466 | 4049 | 3 | -0.76799 |
| CMKLR1 | 6 | 0.011093 | 0.042716 | 0.891621 | 933 | 1 | 0.99479 | 0.065834 | 0.20176 | 0.97466 | 4050 | 4 | 0.99479 |
| OPRD1 | 6 | 0.68366 | 0.80381 | 0.999994 | 16140 | 2 | 1.1589 | 0.065834 | 0.20176 | 0.97466 | 4051 | 3 | 1.1589 |
| FAM160A2 | 6 | 0.19222 | 0.37827 | 0.999994 | 7553 | 1 | 2.6235 | 0.065842 | 0.20178 | 0.97466 | 4052 | 5 | 2.6235 |
| IL22RA1 | 6 | 0.77604 | 0.83812 | 0.999994 | 17149 | 1 | 3.9873 | 0.065875 | 0.20186 | 0.97466 | 4053 | 4 | 3.9873 |
| GJA9 | 6 | 0.33902 | 0.55155 | 0.999994 | 10916 | 2 | 2.8338 | 0.065875 | 0.20186 | 0.97466 | 4054 | 3 | 2.8338 |
| NonTargetingControlG | 2 | 0.93412 | 0.93421 | 0.999994 | 19956 | 0 | 4.144 | 0.065883 | 0.11564 | 0.954984 | 4055 | 2 | 4.144 |

| | | | | | | | | | | | | | |
|---|---|---|---|---|---|---|---|---|---|---|---|---|---|
| TPH1 | 6 | 0.5372 | 0.69458 | 0.999994 | 14015 | 2 | 1.9175 | 0.065967 | 0.20209 | 0.97466 | 4056 | 3 | 1.9175 |
| WISP1 | 6 | 0.97845 | 0.97839 | 0.999994 | 21062 | 0 | 1.4033 | 0.065988 | 0.20213 | 0.97466 | 4057 | 5 | 1.4033 |
| TRAP1 | 5 | 0.34842 | 0.52062 | 0.999994 | 11121 | 2 | -0.12002 | 0.066022 | 0.18512 | 0.97466 | 4058 | 2 | -0.12002 |
| HNRNPA1L2 | 4 | 0.52083 | 0.62579 | 0.999994 | 13759 | 1 | 3.8297 | 0.06606 | 0.17679 | 0.970198 | 4059 | 3 | 3.8297 |
| PAN3 | 6 | 0.78994 | 0.84364 | 0.999994 | 17343 | 1 | 3.4336 | 0.066078 | 0.20235 | 0.97466 | 4060 | 4 | 3.4336 |
| DPM2 | 5 | 0.96755 | 0.96772 | 0.999994 | 20773 | 0 | 3.4065 | 0.066087 | 0.18525 | 0.97466 | 4061 | 3 | 3.4065 |
| KLK3 | 4 | 0.74565 | 0.75676 | 0.999994 | 16808 | 1 | 2.7964 | 0.06609 | 0.17684 | 0.970231 | 4062 | 3 | 2.7964 |
| ZBTB47 | 5 | 0.92032 | 0.92049 | 0.999994 | 19636 | 0 | 4.69 | 0.066108 | 0.18529 | 0.97466 | 4063 | 3 | 4.69 |
| PIGC | 6 | 0.90375 | 0.9066 | 0.999994 | 19300 | 1 | 2.3853 | 0.066173 | 0.20259 | 0.97466 | 4064 | 5 | 2.3853 |
| DHRS7 | 5 | 0.15244 | 0.3064 | 0.999994 | 6519 | 2 | 0.15076 | 0.066179 | 0.18542 | 0.97466 | 4065 | 2 | 0.15076 |
| SLCO1B7 | 6 | 0.17632 | 0.35847 | 0.999994 | 7141 | 3 | -0.3305 | 0.066181 | 0.20261 | 0.97466 | 4066 | 2 | -0.3305 |
| ZNF44 | 6 | 0.67246 | 0.79508 | 0.999994 | 15962 | 1 | 2.6972 | 0.066226 | 0.20272 | 0.97466 | 4067 | 5 | 2.6972 |
| SFMBT1 | 6 | 0.51053 | 0.67599 | 0.999994 | 13582 | 2 | 2.78 | 0.066226 | 0.20272 | 0.97466 | 4068 | 4 | 2.78 |
| PPP1R35 | 6 | 0.49368 | 0.66438 | 0.999994 | 13320 | 2 | 3.9371 | 0.066226 | 0.20272 | 0.97466 | 4069 | 4 | 3.9371 |
| SYT7 | 6 | 0.52404 | 0.68533 | 0.999994 | 13806 | 2 | 3.4827 | 0.066226 | 0.20272 | 0.97466 | 4070 | 4 | 3.4827 |
| CENPK | 6 | 0.38681 | 0.59389 | 0.999994 | 11749 | 1 | 3.3705 | 0.066226 | 0.20272 | 0.97466 | 4071 | 4 | 3.3705 |
| C12orf49 | 6 | 0.69824 | 0.81147 | 0.999994 | 16285 | 1 | 4.0084 | 0.066226 | 0.20272 | 0.97466 | 4072 | 4 | 4.0084 |
| CALY | 3 | 0.93372 | 0.93383 | 0.999994 | 19940 | 0 | 4.6852 | 0.066276 | 0.15425 | 0.965913 | 4073 | 3 | 4.6852 |
| HIPK1 | 6 | 0.97085 | 0.97082 | 0.999994 | 20841 | 0 | 1.2881 | 0.066284 | 0.20286 | 0.97466 | 4074 | 5 | 1.2881 |
| MEX3C | 6 | 0.95318 | 0.95317 | 0.999994 | 20414 | 0 | 0.57816 | 0.066299 | 0.20289 | 0.97466 | 4075 | 3 | 0.57816 |
| PEPD | 5 | 0.94811 | 0.94832 | 0.999994 | 20287 | 0 | 2.6323 | 0.066303 | 0.18571 | 0.97466 | 4076 | 4 | 2.6323 |
| FAM69B | 5 | 0.057106 | 0.15731 | 0.967126 | 3296 | 2 | 0.088656 | 0.066303 | 0.18571 | 0.97466 | 4077 | 2 | 0.088656 |
| SAMD10 | 5 | 0.4168 | 0.57746 | 0.999994 | 12204 | 2 | 0.08149 | 0.066312 | 0.18573 | 0.97466 | 4078 | 2 | 0.08149 |
| DNAJC24 | 6 | 0.9318 | 0.93193 | 0.999994 | 19895 | 0 | 2.2022 | 0.066389 | 0.20311 | 0.97466 | 4079 | 5 | 2.2022 |
| DNAJC1 | 6 | 0.73187 | 0.82223 | 0.999994 | 16626 | 1 | 1.0994 | 0.066482 | 0.20335 | 0.97466 | 4080 | 5 | 1.0994 |
| CYP4A22 | 5 | 0.86832 | 0.8696 | 0.999994 | 18607 | 1 | 4.5099 | 0.066542 | 0.18621 | 0.97466 | 4081 | 3 | 4.5099 |
| VPS29 | 6 | 0.67476 | 0.79687 | 0.999994 | 16001 | 2 | 3.0566 | 0.066544 | 0.20349 | 0.97466 | 4082 | 4 | 3.0566 |
| ADRBK2 | 6 | 0.082624 | 0.21269 | 0.979587 | 4322 | 2 | 3.5927 | 0.066568 | 0.20353 | 0.97466 | 4083 | 4 | 3.5927 |
| ARAF | 6 | 0.27681 | 0.47972 | 0.999994 | 9634 | 2 | 2.6303 | 0.066597 | 0.20361 | 0.97466 | 4084 | 4 | 2.6303 |
| ANK3 | 6 | 0.13976 | 0.31272 | 0.999994 | 6178 | 3 | -0.9016 | 0.066608 | 0.20363 | 0.97466 | 4085 | 2 | -0.9016 |
| BLVRB | 6 | 0.85462 | 0.87442 | 0.999994 | 18343 | 1 | 3.3956 | 0.066627 | 0.20367 | 0.97466 | 4086 | 4 | 3.3956 |
| TGM5 | 4 | 0.83282 | 0.83235 | 0.999994 | 17969 | 0 | 2.2211 | 0.066642 | 0.17779 | 0.971262 | 4087 | 3 | 2.2211 |
| AKAP6 | 6 | 0.53192 | 0.6909 | 0.999994 | 13935 | 1 | 2.0781 | 0.066643 | 0.20371 | 0.97466 | 4088 | 5 | 2.0781 |
| IBTK | 6 | 0.94977 | 0.94974 | 0.999994 | 20327 | 0 | 1.8529 | 0.066649 | 0.20373 | 0.97466 | 4089 | 5 | 1.8529 |
| CALR | 6 | 0.90825 | 0.91013 | 0.999994 | 19390 | 1 | 1.6518 | 0.066668 | 0.20378 | 0.97466 | 4090 | 5 | 1.6518 |
| KRTAP5-1 | 3 | 0.60911 | 0.62414 | 0.999994 | 15059 | 1 | 3.1166 | 0.066672 | 0.15507 | 0.965913 | 4091 | 2 | 3.1166 |
| hsa-mir-3654 | 3 | 0.92711 | 0.92716 | 0.999994 | 19790 | 0 | 6.173 | 0.066679 | 0.15509 | 0.965913 | 4092 | 3 | 6.173 |
| SIN3B | 5 | 0.74314 | 0.78726 | 0.999994 | 16781 | 1 | 1.6451 | 0.066714 | 0.18661 | 0.97466 | 4093 | 3 | 1.6451 |
| C1QTNF3 | 6 | 0.52854 | 0.68853 | 0.999994 | 13872 | 2 | 2.9091 | 0.066715 | 0.2039 | 0.97466 | 4094 | 4 | 2.9091 |
| NonTargetingControlG | 2 | 0.17676 | 0.24542 | 0.98979 | 7148 | 1 | 1.056 | 0.066733 | 0.11705 | 0.955294 | 4095 | 1 | 1.056 |
| OR52K1 | 5 | 0.33259 | 0.50766 | 0.999994 | 10783 | 2 | 2.5747 | 0.066787 | 0.18678 | 0.97466 | 4096 | 3 | 2.5747 |
| TMEM186 | 6 | 0.84192 | 0.86764 | 0.999994 | 18122 | 1 | 2.9836 | 0.066796 | 0.20411 | 0.97466 | 4097 | 5 | 2.9836 |
| ZC3H6 | 6 | 0.33471 | 0.54662 | 0.999994 | 10826 | 2 | 1.8031 | 0.066822 | 0.20417 | 0.97466 | 4098 | 4 | 1.8031 |
| MDM4 | 6 | 0.56759 | 0.71621 | 0.999994 | 14508 | 2 | 3.858 | 0.066896 | 0.20436 | 0.97466 | 4099 | 4 | 3.858 |
| FAM115C | 6 | 0.81261 | 0.85342 | 0.999994 | 17662 | 1 | 3.6867 | 0.066937 | 0.20445 | 0.97466 | 4100 | 4 | 3.6867 |
| KIAA0391 | 4 | 0.92517 | 0.92497 | 0.999994 | 19746 | 0 | 3.4542 | 0.066938 | 0.17831 | 0.971262 | 4101 | 3 | 3.4542 |
| hsa-mir-4744 | 4 | 0.30015 | 0.44317 | 0.999994 | 10134 | 2 | 2.8616 | 0.066986 | 0.17837 | 0.971262 | 4102 | 2 | 2.8616 |
| TRADD | 5 | 0.42188 | 0.58165 | 0.999994 | 12276 | 1 | 4.6527 | 0.067083 | 0.18739 | 0.97466 | 4103 | 3 | 4.6527 |
| HBM | 6 | 0.34855 | 0.56237 | 0.999994 | 11123 | 3 | 0.16072 | 0.067116 | 0.2049 | 0.97466 | 4104 | 3 | 0.16072 |
| CLIP3 | 6 | 0.62386 | 0.75771 | 0.999994 | 15287 | 2 | 1.7414 | 0.067128 | 0.20493 | 0.97466 | 4105 | 3 | 1.7414 |
| ZFP30 | 6 | 0.3659 | 0.58064 | 0.999994 | 11438 | 2 | 2.8834 | 0.067154 | 0.20499 | 0.97466 | 4106 | 4 | 2.8834 |
| FLYWCH2 | 6 | 0.68353 | 0.80371 | 0.999994 | 16136 | 1 | 3.1009 | 0.06722 | 0.20514 | 0.97466 | 4107 | 5 | 3.1009 |

| | | | | | | | | | | | | |
|---|---|---|---|---|---|---|---|---|---|---|---|---|
| CEACAM21 | 6 | 0.42834 | 0.6208 | 0.999994 | 12387 | 1 | 3.0006 | 0.067263 | 0.20526 | 0.97466 | 4108 | 4 | 3.0006 |
| hsa-mir-4739 | 3 | 0.36891 | 0.46635 | 0.999994 | 11488 | 1 | 4.4636 | 0.067266 | 0.15628 | 0.965913 | 4109 | 2 | 4.4636 |
| EEF1G | 5 | 0.51764 | 0.66325 | 0.999994 | 13704 | 1 | 1.8073 | 0.067309 | 0.18787 | 0.97466 | 4110 | 4 | 1.8073 |
| hsa-let-7e | 4 | 0.87151 | 0.87115 | 0.999994 | 18669 | 0 | 3.3013 | 0.067341 | 0.17898 | 0.97162 | 4111 | 2 | 3.3013 |
| ARHGAP28 | 6 | 0.15575 | 0.33288 | 0.999994 | 6616 | 2 | 2.9504 | 0.067372 | 0.20554 | 0.97466 | 4112 | 4 | 2.9504 |
| WDR20 | 6 | 0.11549 | 0.27721 | 0.999379 | 5496 | 2 | 3.0747 | 0.067383 | 0.20556 | 0.97466 | 4113 | 3 | 3.0747 |
| NFKB2 | 6 | 0.57722 | 0.72316 | 0.999994 | 14646 | 1 | 3.0304 | 0.067383 | 0.20556 | 0.97466 | 4114 | 3 | 3.0304 |
| SLC12A8 | 5 | 0.68111 | 0.75803 | 0.999994 | 16105 | 1 | 2.1233 | 0.067393 | 0.18806 | 0.97466 | 4115 | 3 | 2.1233 |
| CALHM1 | 5 | 0.19829 | 0.37282 | 0.999994 | 7718 | 2 | 1.4594 | 0.067393 | 0.18806 | 0.97466 | 4116 | 3 | 1.4594 |
| OR4X2 | 6 | 0.070641 | 0.18814 | 0.969543 | 3860 | 3 | -1.2624 | 0.067405 | 0.20561 | 0.97466 | 4117 | 2 | -1.2624 |
| VAMP5 | 6 | 0.3505 | 0.56455 | 0.999994 | 11172 | 2 | 1.3916 | 0.067405 | 0.20561 | 0.97466 | 4118 | 3 | 1.3916 |
| RASAL3 | 6 | 0.19004 | 0.37554 | 0.999994 | 7484 | 3 | -0.16 | 0.067436 | 0.20568 | 0.97466 | 4119 | 2 | -0.16 |
| SRBD1 | 5 | 0.70631 | 0.76937 | 0.999994 | 16375 | 1 | 1.3282 | 0.06745 | 0.18819 | 0.97466 | 4120 | 3 | 1.3282 |
| NUDC | 4 | 0.87291 | 0.87256 | 0.999994 | 18691 | 0 | 3.8735 | 0.067466 | 0.17918 | 0.97162 | 4121 | 3 | 3.8735 |
| PLRG1 | 5 | 0.43146 | 0.58964 | 0.999994 | 12439 | 1 | 3.0887 | 0.06748 | 0.18825 | 0.97466 | 4122 | 4 | 3.0887 |
| NonTargetingControlG | 2 | 0.93251 | 0.93265 | 0.999994 | 19915 | 0 | 4.3674 | 0.067486 | 0.11828 | 0.9593 | 4123 | 2 | 4.3674 |
| AVL9 | 6 | 0.048808 | 0.1415 | 0.959146 | 2923 | 4 | -1.9353 | 0.067569 | 0.20599 | 0.97466 | 4124 | 2 | -1.9353 |
| KRTAP9-1 | 5 | 0.52068 | 0.66589 | 0.999994 | 13757 | 2 | -0.36172 | 0.067611 | 0.18852 | 0.97466 | 4125 | 2 | -0.36172 |
| XAF1 | 6 | 0.96406 | 0.96413 | 0.999994 | 20703 | 0 | 2.6955 | 0.067624 | 0.20612 | 0.97466 | 4126 | 5 | 2.6955 |
| hsa-mir-4748 | 3 | 0.27508 | 0.4059 | 0.999994 | 9594 | 1 | 5.9833 | 0.06764 | 0.15707 | 0.965913 | 4127 | 2 | 5.9833 |
| hsa-mir-6088 | 3 | 0.59236 | 0.61128 | 0.999994 | 14853 | 1 | 5.9833 | 0.06764 | 0.15707 | 0.965913 | 4128 | 2 | 5.9833 |
| COX8A | 5 | 0.40109 | 0.56442 | 0.999994 | 11963 | 1 | 4.6419 | 0.067662 | 0.18863 | 0.97466 | 4129 | 3 | 4.6419 |
| ZBTB24 | 5 | 0.78527 | 0.8105 | 0.999994 | 17283 | 1 | 4.6419 | 0.067662 | 0.18863 | 0.97466 | 4130 | 4 | 4.6419 |
| AFTPH | 6 | 0.088336 | 0.22421 | 0.983354 | 4528 | 4 | -1.4003 | 0.067703 | 0.20632 | 0.97466 | 4131 | 2 | -1.4003 |
| SIRT2 | 6 | 0.96379 | 0.96385 | 0.999994 | 20699 | 0 | 1.3878 | 0.067711 | 0.20633 | 0.97466 | 4132 | 5 | 1.3878 |
| IGLL1 | 6 | 0.026102 | 0.089607 | 0.94395 | 1808 | 2 | 2.7025 | 0.067716 | 0.20634 | 0.97466 | 4133 | 3 | 2.7025 |
| OR13C8 | 6 | 0.9037 | 0.90656 | 0.999994 | 19297 | 1 | 2.4969 | 0.067725 | 0.20637 | 0.97466 | 4134 | 3 | 2.4969 |
| OR10J3 | 5 | 0.94951 | 0.9497 | 0.999994 | 20324 | 0 | 0.50071 | 0.067771 | 0.18888 | 0.97466 | 4135 | 2 | 0.50071 |
| CLDN24 | 6 | 0.23599 | 0.43123 | 0.999994 | 8670 | 2 | 3.358 | 0.067839 | 0.20665 | 0.97466 | 4136 | 4 | 3.358 |
| GMPPB | 3 | 0.9321 | 0.93215 | 0.999994 | 19902 | 0 | 4.3741 | 0.067901 | 0.15758 | 0.965913 | 4137 | 3 | 4.3741 |
| PA2G4 | 5 | 0.16198 | 0.32044 | 0.999994 | 6794 | 2 | -0.48864 | 0.067962 | 0.18929 | 0.97466 | 4138 | 1 | -0.48864 |
| DYNC2LI1 | 5 | 0.81011 | 0.82599 | 0.999994 | 17633 | 1 | 4.575 | 0.067991 | 0.18934 | 0.97466 | 4139 | 3 | 4.575 |
| OR1M1 | 5 | 0.82413 | 0.83533 | 0.999994 | 17841 | 1 | 4.7073 | 0.067999 | 0.18936 | 0.97466 | 4140 | 4 | 4.7073 |
| MMP24 | 5 | 0.25149 | 0.44194 | 0.999994 | 9047 | 1 | 0.41327 | 0.068051 | 0.18948 | 0.97466 | 4141 | 2 | 0.41327 |
| FBXO3 | 5 | 0.62825 | 0.73664 | 0.999994 | 15353 | 1 | 1.8704 | 0.068056 | 0.18949 | 0.97466 | 4142 | 4 | 1.8704 |
| SARDH | 5 | 0.82375 | 0.83508 | 0.999994 | 17835 | 1 | 0.50991 | 0.068105 | 0.18958 | 0.97466 | 4143 | 2 | 0.50991 |
| SYNE1 | 5 | 0.69527 | 0.76434 | 0.999994 | 16260 | 1 | 4.279 | 0.068105 | 0.18958 | 0.97466 | 4144 | 4 | 4.279 |
| HCK | 6 | 0.19283 | 0.37906 | 0.999994 | 7571 | 2 | 3.7583 | 0.068154 | 0.2074 | 0.97466 | 4145 | 4 | 3.7583 |
| RDH12 | 6 | 0.87976 | 0.88944 | 0.999994 | 18830 | 1 | 1.7876 | 0.068156 | 0.2074 | 0.97466 | 4146 | 4 | 1.7876 |
| CRLS1 | 6 | 0.96018 | 0.96024 | 0.999994 | 20586 | 0 | 1.9024 | 0.068156 | 0.2074 | 0.97466 | 4147 | 5 | 1.9024 |
| RASL10A | 4 | 0.072375 | 0.16441 | 0.967126 | 3937 | 3 | -1.3149 | 0.068181 | 0.1804 | 0.97162 | 4148 | 1 | -1.3149 |
| MVB12A | 5 | 0.96023 | 0.96037 | 0.999994 | 20589 | 0 | 2.187 | 0.068296 | 0.18999 | 0.97466 | 4149 | 4 | 2.187 |
| NonTargetingControlG | 2 | 0.5716 | 0.57181 | 0.999994 | 14568 | 1 | 3.9445 | 0.068299 | 0.11963 | 0.961431 | 4150 | 1 | 3.9445 |
| NOP14 | 4 | 0.18941 | 0.32978 | 0.999994 | 7467 | 1 | 3.9216 | 0.068346 | 0.18068 | 0.97162 | 4151 | 3 | 3.9216 |
| hsa-mir-3160-1 | 4 | 0.81517 | 0.81519 | 0.999994 | 17701 | 1 | 3.2243 | 0.068346 | 0.18068 | 0.97162 | 4152 | 2 | 3.2243 |
| SPAG11B | 3 | 0.1978 | 0.30987 | 0.999994 | 7702 | 2 | -1.2216 | 0.068372 | 0.15851 | 0.965913 | 4153 | 1 | -1.2216 |
| LTK | 6 | 0.50597 | 0.67287 | 0.999994 | 13525 | 1 | 4.1241 | 0.068425 | 0.20809 | 0.97466 | 4154 | 4 | 4.1241 |
| ANKH | 6 | 0.5209 | 0.68316 | 0.999994 | 13760 | 2 | 0.45508 | 0.068479 | 0.20822 | 0.97466 | 4155 | 3 | 0.45508 |
| ITGB3 | 6 | 0.70346 | 0.81308 | 0.999994 | 16345 | 1 | 1.2186 | 0.068479 | 0.20822 | 0.97466 | 4156 | 3 | 1.2186 |
| NME1 | 6 | 0.9522 | 0.95218 | 0.999994 | 20388 | 0 | 1.8197 | 0.068479 | 0.20822 | 0.97466 | 4157 | 4 | 1.8197 |
| NonTargetingControlG | 2 | 0.93151 | 0.93162 | 0.999994 | 19887 | 0 | 4.4062 | 0.068493 | 0.11996 | 0.961431 | 4158 | 2 | 4.4062 |
| NonTargetingControlG | 2 | 0.93151 | 0.93162 | 0.999994 | 19888 | 0 | 4.4623 | 0.068493 | 0.11996 | 0.961431 | 4159 | 2 | 4.4623 |

| | | | | | | | | | | | | |
|---|---|---|---|---|---|---|---|---|---|---|---|---|
| hsa-mir-6891 | 2 | 0.93151 | 0.93162 | 0.999994 | 19889 | 0 | 4.2647 | 0.068493 | 0.11996 | 0.961431 | 4160 | 2 | 4.2647 |
| PGRMC2 | 5 | 0.013807 | 0.046849 | 0.896157 | 1117 | 1 | 3.1916 | 0.068497 | 0.19042 | 0.97466 | 4161 | 3 | 3.1916 |
| SLC6A12 | 6 | 0.17307 | 0.35451 | 0.999994 | 7067 | 2 | 3.6259 | 0.068513 | 0.20831 | 0.97466 | 4162 | 4 | 3.6259 |
| GSN | 6 | 0.97099 | 0.97096 | 0.999994 | 20847 | 0 | 3.4855 | 0.068513 | 0.20831 | 0.97466 | 4163 | 4 | 3.4855 |
| SPRR1A | 6 | 0.50157 | 0.66977 | 0.999994 | 13450 | 2 | 3.8261 | 0.068513 | 0.20831 | 0.97466 | 4164 | 4 | 3.8261 |
| TRMT10A | 6 | 0.35675 | 0.57162 | 0.999994 | 11283 | 2 | 3.3337 | 0.068513 | 0.20831 | 0.97466 | 4165 | 4 | 3.3337 |
| HILPDA | 6 | 0.21288 | 0.40359 | 0.999994 | 8090 | 1 | 3.5272 | 0.068531 | 0.20835 | 0.97466 | 4166 | 5 | 3.5272 |
| POLR3D | 6 | 0.95236 | 0.95235 | 0.999994 | 20391 | 0 | 4.0725 | 0.06859 | 0.20851 | 0.97466 | 4167 | 4 | 4.0725 |
| TBX21 | 4 | 0.73206 | 0.74667 | 0.999994 | 16629 | 1 | 3.0431 | 0.068607 | 0.18113 | 0.97162 | 4168 | 3 | 3.0431 |
| C21orf58 | 4 | 0.90607 | 0.90578 | 0.999994 | 19352 | 0 | 4.3323 | 0.068607 | 0.18113 | 0.97162 | 4169 | 3 | 4.3323 |
| CUBN | 6 | 0.28001 | 0.48343 | 0.999994 | 9706 | 3 | -0.098453 | 0.068663 | 0.20869 | 0.97466 | 4170 | 3 | -0.098453 |
| C15orf62 | 6 | 0.32793 | 0.53886 | 0.999994 | 10684 | 3 | -0.16533 | 0.068663 | 0.20869 | 0.97466 | 4171 | 2 | -0.16533 |
| DENND4A | 5 | 0.83897 | 0.84588 | 0.999994 | 18071 | 1 | 2.8318 | 0.068675 | 0.19078 | 0.97466 | 4172 | 3 | 2.8318 |
| XK | 4 | 0.23719 | 0.37883 | 0.999994 | 8700 | 1 | 3.1874 | 0.068684 | 0.18127 | 0.97162 | 4173 | 2 | 3.1874 |
| SH3RF3 | 5 | 0.84186 | 0.84808 | 0.999994 | 18120 | 1 | 1.6403 | 0.06874 | 0.19092 | 0.97466 | 4174 | 4 | 1.6403 |
| RTBDN | 6 | 0.12394 | 0.29258 | 0.999632 | 5765 | 1 | 2.5622 | 0.068775 | 0.20894 | 0.97466 | 4175 | 5 | 2.5622 |
| KCNAB1 | 5 | 0.86096 | 0.86321 | 0.999994 | 18462 | 1 | 1.0843 | 0.06878 | 0.191 | 0.97466 | 4176 | 3 | 1.0843 |
| ZNF618 | 6 | 0.019888 | 0.071612 | 0.93914 | 1452 | 1 | 3.8952 | 0.068785 | 0.20896 | 0.97466 | 4177 | 4 | 3.8952 |
| ETV3L | 6 | 0.59424 | 0.73573 | 0.999994 | 14881 | 2 | 3.902 | 0.0688 | 0.20899 | 0.97466 | 4178 | 4 | 3.902 |
| FA2H | 6 | 0.93148 | 0.93163 | 0.999994 | 19886 | 0 | 0.41889 | 0.068828 | 0.20906 | 0.97466 | 4179 | 3 | 0.41889 |
| hsa-mir-891b | 4 | 0.45751 | 0.59785 | 0.999994 | 12814 | 1 | 0.051741 | 0.068843 | 0.18155 | 0.97162 | 4180 | 2 | 0.051741 |
| TTC22 | 5 | 0.93896 | 0.93915 | 0.999994 | 20049 | 0 | 4.3741 | 0.068846 | 0.19115 | 0.97466 | 4181 | 4 | 4.3741 |
| C12orf57 | 6 | 0.88044 | 0.88989 | 0.999994 | 18842 | 1 | 3.678 | 0.068874 | 0.20917 | 0.97466 | 4182 | 4 | 3.678 |
| LHFPL1 | 5 | 0.97149 | 0.97154 | 0.999994 | 20859 | 0 | 1.6583 | 0.068926 | 0.19132 | 0.97466 | 4183 | 4 | 1.6583 |
| RGS11 | 4 | 0.83668 | 0.83618 | 0.999994 | 18029 | 0 | 3.3119 | 0.068931 | 0.18169 | 0.97162 | 4184 | 3 | 3.3119 |
| CYP27B1 | 6 | 0.33718 | 0.54946 | 0.999994 | 10878 | 2 | 3.8228 | 0.06894 | 0.20934 | 0.97466 | 4185 | 4 | 3.8228 |
| IFNE | 6 | 0.5295 | 0.68919 | 0.999994 | 13891 | 2 | 0.73361 | 0.069009 | 0.20952 | 0.97466 | 4186 | 3 | 0.73361 |
| NonTargetingControlG | 2 | 0.14782 | 0.22513 | 0.983354 | 6403 | 1 | 0.70142 | 0.069073 | 0.12087 | 0.962329 | 4187 | 1 | 0.70142 |
| TMED3 | 5 | 0.25773 | 0.44689 | 0.999994 | 9187 | 1 | 0.28502 | 0.069077 | 0.19163 | 0.97466 | 4188 | 2 | 0.28502 |
| CRX | 6 | 0.62344 | 0.75737 | 0.999994 | 15283 | 2 | 0.85702 | 0.069131 | 0.20981 | 0.97466 | 4189 | 3 | 0.85702 |
| C1QL4 | 6 | 0.93544 | 0.93549 | 0.999994 | 19978 | 0 | 2.9175 | 0.069156 | 0.20986 | 0.97466 | 4190 | 3 | 2.9175 |
| C12orf56 | 6 | 0.070803 | 0.18848 | 0.969543 | 3866 | 3 | -1.347 | 0.069169 | 0.20989 | 0.97466 | 4191 | 2 | -1.347 |
| C5AR2 | 6 | 0.95008 | 0.95006 | 0.999994 | 20337 | 0 | 3.1264 | 0.069169 | 0.20989 | 0.97466 | 4192 | 4 | 3.1264 |
| RRNAD1 | 6 | 0.88535 | 0.89311 | 0.999994 | 18940 | 1 | 1.3822 | 0.069179 | 0.20992 | 0.97466 | 4193 | 5 | 1.3822 |
| NUP43 | 4 | 0.23502 | 0.37665 | 0.999994 | 8644 | 2 | -0.10095 | 0.069183 | 0.18213 | 0.97162 | 4194 | 2 | -0.10095 |
| GRIA2 | 6 | 0.62059 | 0.75529 | 0.999994 | 15229 | 1 | 2.9888 | 0.069211 | 0.21 | 0.97466 | 4195 | 5 | 2.9888 |
| ALX4 | 5 | 0.41537 | 0.57625 | 0.999994 | 12183 | 2 | 4.5922 | 0.069233 | 0.19196 | 0.97466 | 4196 | 3 | 4.5922 |
| hsa-mir-190a | 3 | 0.93076 | 0.93081 | 0.999994 | 19870 | 0 | 1.47 | 0.069242 | 0.16031 | 0.965913 | 4197 | 3 | 1.47 |
| CYSLTR1 | 5 | 0.22503 | 0.41028 | 0.999994 | 8398 | 1 | 2.1473 | 0.069252 | 0.192 | 0.97466 | 4198 | 4 | 2.1473 |
| BPI | 5 | 0.16402 | 0.32338 | 0.999994 | 6841 | 1 | 2.187 | 0.069252 | 0.192 | 0.97466 | 4199 | 4 | 2.187 |
| TCTN3 | 5 | 0.83681 | 0.84435 | 0.999994 | 18031 | 1 | 2.7564 | 0.069252 | 0.192 | 0.97466 | 4200 | 4 | 2.7564 |
| WDR36 | 5 | 0.84091 | 0.84735 | 0.999994 | 18104 | 1 | 2.945 | 0.069252 | 0.192 | 0.97466 | 4201 | 4 | 2.945 |
| IFT46 | 6 | 0.74571 | 0.82697 | 0.999994 | 16810 | 1 | 2.5235 | 0.069288 | 0.21018 | 0.97466 | 4202 | 5 | 2.5235 |
| CBY3 | 4 | 0.63903 | 0.68696 | 0.999994 | 15499 | 1 | 4.6516 | 0.069289 | 0.18231 | 0.97162 | 4203 | 3 | 4.6516 |
| ALG1L | 4 | 0.27841 | 0.42095 | 0.999994 | 9662 | 1 | 3.1864 | 0.069314 | 0.18235 | 0.97162 | 4204 | 3 | 3.1864 |
| EEF2 | 3 | 0.21399 | 0.3302 | 0.999994 | 8114 | 1 | 5.9388 | 0.069338 | 0.16051 | 0.965913 | 4205 | 2 | 5.9388 |
| MYD88 | 5 | 0.10028 | 0.22685 | 0.983354 | 4982 | 2 | 0.63649 | 0.069343 | 0.19217 | 0.97466 | 4206 | 3 | 0.63649 |
| FGA | 6 | 0.71557 | 0.8169 | 0.999994 | 16454 | 1 | 3.6576 | 0.069377 | 0.2104 | 0.97466 | 4207 | 5 | 3.6576 |
| TIPIN | 4 | 0.017554 | 0.055439 | 0.922773 | 1344 | 3 | -2.7843 | 0.069379 | 0.18247 | 0.97162 | 4208 | 1 | -2.7843 |
| LACE1 | 6 | 0.063144 | 0.1724 | 0.967658 | 3566 | 3 | 1.397 | 0.069401 | 0.21046 | 0.97466 | 4209 | 3 | 1.397 |
| HAPLN3 | 6 | 0.29641 | 0.50253 | 0.999994 | 10050 | 1 | 2.7126 | 0.069406 | 0.21047 | 0.97466 | 4210 | 4 | 2.7126 |
| ZNF398 | 5 | 0.70134 | 0.7671 | 0.999994 | 16321 | 1 | 1.038 | 0.069407 | 0.1923 | 0.97466 | 4211 | 3 | 1.038 |

| | | | | | | | | | | | | | |
|---|---|---|---|---|---|---|---|---|---|---|---|---|---|
| LHPP | 6 | 0.099638 | 0.24665 | 0.98999 | 4963 | 3 | -0.23311 | 0.069409 | 0.21048 | 0.97466 | 4212 | 3 | -0.23311 |
| SHBG | 6 | 0.38647 | 0.59367 | 0.999994 | 11741 | 2 | 0.35135 | 0.069409 | 0.21048 | 0.97466 | 4213 | 3 | 0.35135 |
| SRGAP2 | 5 | 0.50507 | 0.65236 | 0.999994 | 13513 | 2 | -0.22461 | 0.06941 | 0.1923 | 0.97466 | 4214 | 2 | -0.22461 |
| ACVR1C | 6 | 0.54908 | 0.70298 | 0.999994 | 14195 | 2 | 1.034 | 0.069419 | 0.2105 | 0.97466 | 4215 | 4 | 1.034 |
| NUP210L | 6 | 0.90801 | 0.90994 | 0.999994 | 19383 | 1 | 2.4762 | 0.069502 | 0.21071 | 0.97466 | 4216 | 5 | 2.4762 |
| ZNF516 | 6 | 0.0047708 | 0.020315 | 0.810158 | 504 | 1 | 1.279 | 0.069502 | 0.21071 | 0.97466 | 4217 | 5 | 1.279 |
| SETD8 | 6 | 0.46874 | 0.6476 | 0.999994 | 12988 | 1 | 3.0317 | 0.069502 | 0.21071 | 0.97466 | 4218 | 5 | 3.0317 |
| PROM1 | 5 | 0.96151 | 0.96165 | 0.999994 | 20618 | 0 | 1.3823 | 0.069521 | 0.19254 | 0.97466 | 4219 | 4 | 1.3823 |
| hsa-mir-3666 | 4 | 0.88775 | 0.88744 | 0.999994 | 18993 | 0 | 4.7822 | 0.069524 | 0.18271 | 0.97162 | 4220 | 3 | 4.7822 |
| ZNF467 | 6 | 0.35553 | 0.57027 | 0.999994 | 11259 | 1 | 1.1191 | 0.069622 | 0.211 | 0.97466 | 4221 | 4 | 1.1191 |
| LOC653486 | 6 | 0.26676 | 0.46785 | 0.999994 | 9390 | 2 | 2.3981 | 0.069622 | 0.211 | 0.97466 | 4222 | 4 | 2.3981 |
| TAS2R8 | 6 | 0.32549 | 0.53611 | 0.999994 | 10632 | 2 | 2.6248 | 0.069628 | 0.21102 | 0.97466 | 4223 | 3 | 2.6248 |
| TTLL1 | 6 | 0.54266 | 0.69843 | 0.999994 | 14106 | 1 | 4.0119 | 0.069628 | 0.21102 | 0.97466 | 4224 | 4 | 4.0119 |
| MTG1 | 6 | 0.44057 | 0.62881 | 0.999994 | 12586 | 2 | 3.0519 | 0.069628 | 0.21102 | 0.97466 | 4225 | 3 | 3.0519 |
| hsa-mir-4505 | 4 | 0.17982 | 0.3199 | 0.999994 | 7216 | 2 | -0.28401 | 0.069629 | 0.1829 | 0.97162 | 4226 | 1 | -0.28401 |
| PRRT1 | 5 | 0.748 | 0.78979 | 0.999994 | 16839 | 1 | 1.9721 | 0.069635 | 0.19278 | 0.97466 | 4227 | 4 | 1.9721 |
| GDNF | 6 | 0.81959 | 0.85664 | 0.999994 | 17774 | 1 | 2.2701 | 0.069653 | 0.21108 | 0.97466 | 4228 | 5 | 2.2701 |
| ZNF814 | 6 | 0.98458 | 0.9845 | 0.999994 | 21207 | 0 | 2.5692 | 0.069684 | 0.21116 | 0.97466 | 4229 | 5 | 2.5692 |
| TAAR2 | 5 | 0.93351 | 0.93375 | 0.999994 | 19936 | 0 | 3.1192 | 0.069685 | 0.19287 | 0.97466 | 4230 | 3 | 3.1192 |
| hsa-mir-4439 | 4 | 0.45567 | 0.59708 | 0.999994 | 12785 | 1 | 3.8801 | 0.069763 | 0.18314 | 0.97162 | 4231 | 3 | 3.8801 |
| hsa-mir-3651 | 4 | 0.93374 | 0.93351 | 0.999994 | 19944 | 0 | 3.9139 | 0.069763 | 0.18314 | 0.97162 | 4232 | 3 | 3.9139 |
| C11orf85 | 6 | 0.96413 | 0.96419 | 0.999994 | 20706 | 0 | 2.505 | 0.069845 | 0.21154 | 0.97466 | 4233 | 5 | 2.505 |
| PCDHGA3 | 2 | 0.93013 | 0.93027 | 0.999994 | 19857 | 0 | 5.6978 | 0.069872 | 0.12219 | 0.962935 | 4234 | 2 | 5.6978 |
| ATP12A | 6 | 0.59324 | 0.73496 | 0.999994 | 14867 | 2 | 0.33214 | 0.069887 | 0.21163 | 0.97466 | 4235 | 3 | 0.33214 |
| LIPK | 6 | 0.19543 | 0.38234 | 0.999994 | 7648 | 1 | 3.077 | 0.069887 | 0.21163 | 0.97466 | 4236 | 4 | 3.077 |
| TMEM126A | 6 | 0.45165 | 0.63617 | 0.999994 | 12729 | 2 | 0.63476 | 0.069887 | 0.21163 | 0.97466 | 4237 | 3 | 0.63476 |
| ERH | 3 | 0.3432 | 0.45192 | 0.999994 | 11011 | 1 | 4.6419 | 0.06991 | 0.16167 | 0.965913 | 4238 | 2 | 4.6419 |
| CHMP4A | 6 | 0.032531 | 0.10473 | 0.948754 | 2135 | 2 | 0.096452 | 0.069915 | 0.21169 | 0.97466 | 4239 | 2 | 0.096452 |
| TMEM132E | 6 | 0.9827 | 0.98265 | 0.999994 | 21152 | 0 | 1.1107 | 0.069954 | 0.21179 | 0.97466 | 4240 | 5 | 1.1107 |
| HCN4 | 6 | 0.57175 | 0.71921 | 0.999994 | 14570 | 1 | 3.486 | 0.069973 | 0.21183 | 0.97466 | 4241 | 5 | 3.486 |
| GABBR2 | 6 | 0.78556 | 0.84189 | 0.999994 | 17287 | 1 | 1.4363 | 0.06998 | 0.21185 | 0.97466 | 4242 | 5 | 1.4363 |
| NTAN1 | 5 | 0.80209 | 0.82083 | 0.999994 | 17514 | 1 | 0.1014 | 0.069996 | 0.19353 | 0.97466 | 4243 | 2 | 0.1014 |
| KRT24 | 5 | 0.24776 | 0.43902 | 0.999994 | 8968 | 1 | 2.4812 | 0.069996 | 0.19353 | 0.97466 | 4244 | 3 | 2.4812 |
| SLC25A5 | 5 | 0.51885 | 0.66432 | 0.999994 | 13727 | 1 | 2.4745 | 0.069998 | 0.19353 | 0.97466 | 4245 | 4 | 2.4745 |
| NPY5R | 6 | 0.75231 | 0.8293 | 0.999994 | 16882 | 1 | 2.4772 | 0.069999 | 0.21189 | 0.97466 | 4246 | 5 | 2.4772 |
| HCN3 | 4 | 0.14379 | 0.28283 | 0.999632 | 6292 | 2 | -1.306 | 0.070004 | 0.18356 | 0.97162 | 4247 | 1 | -1.306 |
| MRPL35 | 5 | 0.16156 | 0.31984 | 0.999994 | 6777 | 3 | -1.0305 | 0.070055 | 0.19364 | 0.97466 | 4248 | 1 | -1.0305 |
| MCCC2 | 6 | 0.66638 | 0.79032 | 0.999994 | 15874 | 2 | 2.6598 | 0.070063 | 0.21206 | 0.97466 | 4249 | 3 | 2.6598 |
| OBP2A | 5 | 0.82445 | 0.83555 | 0.999994 | 17844 | 1 | 4.6561 | 0.070084 | 0.1937 | 0.97466 | 4250 | 3 | 4.6561 |
| C1orf172 | 6 | 0.17449 | 0.35622 | 0.999994 | 7098 | 2 | 3.29 | 0.070094 | 0.21213 | 0.97466 | 4251 | 4 | 3.29 |
| B4GALNT3 | 6 | 0.83113 | 0.86209 | 0.999994 | 17947 | 1 | 3.2612 | 0.070113 | 0.21218 | 0.97466 | 4252 | 4 | 3.2612 |
| LRWD1 | 6 | 0.0252 | 0.08747 | 0.94395 | 1762 | 3 | -2.8497 | 0.070155 | 0.21228 | 0.97466 | 4253 | 2 | -2.8497 |
| SQRDL | 5 | 0.80832 | 0.82484 | 0.999994 | 17605 | 1 | 2.4812 | 0.070173 | 0.19388 | 0.97466 | 4254 | 4 | 2.4812 |
| XRCC1 | 6 | 0.97757 | 0.97752 | 0.999994 | 21032 | 0 | 2.8977 | 0.070334 | 0.21274 | 0.97466 | 4255 | 5 | 2.8977 |
| CLU | 6 | 0.1761 | 0.35821 | 0.999994 | 7134 | 1 | 3.4545 | 0.070364 | 0.21281 | 0.97466 | 4256 | 4 | 3.4545 |
| CACNG2 | 6 | 0.22638 | 0.41976 | 0.999994 | 8433 | 2 | 1.1209 | 0.070364 | 0.21281 | 0.97466 | 4257 | 4 | 1.1209 |
| MSH4 | 6 | 0.33749 | 0.54979 | 0.999994 | 10887 | 2 | 2.2112 | 0.070364 | 0.21281 | 0.97466 | 4258 | 4 | 2.2112 |
| LRRC8D | 6 | 0.031947 | 0.10337 | 0.946414 | 2101 | 4 | -2.2668 | 0.070421 | 0.21295 | 0.97466 | 4259 | 2 | -2.2668 |
| VPS13C | 6 | 0.97805 | 0.97799 | 0.999994 | 21043 | 0 | 1.1057 | 0.070425 | 0.21296 | 0.97466 | 4260 | 5 | 1.1057 |
| SHISA8 | 5 | 0.95757 | 0.95771 | 0.999994 | 20528 | 0 | 4.5407 | 0.070435 | 0.19445 | 0.97466 | 4261 | 4 | 4.5407 |
| OR4F6 | 6 | 0.79463 | 0.84561 | 0.999994 | 17405 | 1 | 2.3351 | 0.070483 | 0.2131 | 0.97466 | 4262 | 5 | 2.3351 |
| hsa-mir-6797 | 2 | 0.84425 | 0.84418 | 0.999994 | 18159 | 0 | 5.0288 | 0.070519 | 0.12322 | 0.962935 | 4263 | 2 | 5.0288 |

| | | | | | | | | | | | | | |
|---|---|---|---|---|---|---|---|---|---|---|---|---|---|
| RAX2 | 5 | 0.82656 | 0.83704 | 0.999994 | 17878 | 1 | 0.58844 | 0.070522 | 0.19464 | 0.97466 | 4264 | 3 | 0.58844 |
| FANK1 | 6 | 0.10173 | 0.2507 | 0.992497 | 5028 | 3 | -1.0079 | 0.070527 | 0.21322 | 0.97466 | 4265 | 2 | -1.0079 |
| CSF3 | 3 | 0.16495 | 0.26796 | 0.998561 | 6862 | 2 | -1.5225 | 0.07057 | 0.16301 | 0.965913 | 4266 | 1 | -1.5225 |
| LRRFIP2 | 5 | 0.099774 | 0.22607 | 0.983354 | 4969 | 3 | -0.95813 | 0.070633 | 0.19487 | 0.97466 | 4267 | 1 | -0.95813 |
| HIAT1 | 6 | 0.26276 | 0.46314 | 0.999994 | 9304 | 3 | 2.3153 | 0.070709 | 0.21367 | 0.97466 | 4268 | 3 | 2.3153 |
| RGSL1 | 6 | 0.76448 | 0.83376 | 0.999994 | 17023 | 1 | 3.1626 | 0.070709 | 0.21367 | 0.97466 | 4269 | 5 | 3.1626 |
| KIF26A | 6 | 0.8807 | 0.89007 | 0.999994 | 18845 | 1 | 3.7361 | 0.070709 | 0.21367 | 0.97466 | 4270 | 4 | 3.7361 |
| CSF2RB | 6 | 0.80165 | 0.84857 | 0.999994 | 17504 | 1 | 3.9672 | 0.070709 | 0.21367 | 0.97466 | 4271 | 4 | 3.9672 |
| TAF1A | 4 | 0.094768 | 0.20567 | 0.978603 | 4782 | 3 | -1.3396 | 0.070772 | 0.18485 | 0.97466 | 4272 | 1 | -1.3396 |
| SIM2 | 6 | 0.25094 | 0.44907 | 0.999994 | 9033 | 2 | 3.6486 | 0.070779 | 0.21384 | 0.97466 | 4273 | 4 | 3.6486 |
| TWSG1 | 6 | 0.90297 | 0.90599 | 0.999994 | 19276 | 1 | 1.3172 | 0.070781 | 0.21384 | 0.97466 | 4274 | 5 | 1.3172 |
| CSTF2T | 5 | 0.26839 | 0.45554 | 0.999994 | 9430 | 2 | 1.1504 | 0.070789 | 0.19521 | 0.97466 | 4275 | 3 | 1.1504 |
| RFESD | 5 | 0.45216 | 0.60704 | 0.999994 | 12737 | 2 | -0.0052053 | 0.070789 | 0.19521 | 0.97466 | 4276 | 2 | -0.0052053 |
| RASIP1 | 5 | 0.76804 | 0.80062 | 0.999994 | 17061 | 1 | 1.3136 | 0.070789 | 0.19521 | 0.97466 | 4277 | 4 | 1.3136 |
| MRPS10 | 4 | 0.14292 | 0.28191 | 0.999632 | 6270 | 1 | 2.8156 | 0.070843 | 0.18497 | 0.97466 | 4278 | 3 | 2.8156 |
| SLC2A8 | 5 | 0.1977 | 0.37195 | 0.999994 | 7698 | 2 | 0.50882 | 0.070874 | 0.19538 | 0.97466 | 4279 | 2 | 0.50882 |
| ZNF821 | 6 | 0.45613 | 0.63912 | 0.999994 | 12792 | 1 | 1.9803 | 0.070891 | 0.2141 | 0.97466 | 4280 | 5 | 1.9803 |
| ADCY3 | 6 | 0.86404 | 0.87973 | 0.999994 | 18522 | 1 | 3.8084 | 0.070917 | 0.21417 | 0.97466 | 4281 | 5 | 3.8084 |
| GLTPD2 | 6 | 0.74102 | 0.82533 | 0.999994 | 16752 | 1 | 1.5707 | 0.07092 | 0.21418 | 0.97466 | 4282 | 4 | 1.5707 |
| DERA | 6 | 0.37801 | 0.58831 | 0.999994 | 11624 | 1 | 2.1002 | 0.07093 | 0.2142 | 0.97466 | 4283 | 5 | 2.1002 |
| SHE | 5 | 0.33996 | 0.51368 | 0.999994 | 10930 | 2 | 4.4636 | 0.071011 | 0.19568 | 0.97466 | 4284 | 3 | 4.4636 |
| CLCN5 | 5 | 0.14587 | 0.29659 | 0.999994 | 6348 | 1 | 0.81436 | 0.071011 | 0.19568 | 0.97466 | 4285 | 3 | 0.81436 |
| WNT2B | 3 | 0.66363 | 0.66843 | 0.999994 | 15831 | 1 | 5.8985 | 0.071011 | 0.16391 | 0.965913 | 4286 | 2 | 5.8985 |
| NAA11 | 4 | 0.57996 | 0.65474 | 0.999994 | 14676 | 1 | 3.1541 | 0.071031 | 0.18531 | 0.97466 | 4287 | 2 | 3.1541 |
| OR9Q2 | 6 | 0.12634 | 0.29567 | 0.999994 | 5833 | 2 | 2.7811 | 0.071042 | 0.21447 | 0.97466 | 4288 | 4 | 2.7811 |
| IL4I1 | 6 | 0.97255 | 0.97248 | 0.999994 | 20884 | 0 | 2.2654 | 0.071073 | 0.21454 | 0.97466 | 4289 | 5 | 2.2654 |
| ENTHD1 | 6 | 0.090298 | 0.22808 | 0.983354 | 4605 | 4 | -1.07 | 0.071086 | 0.21458 | 0.97466 | 4290 | 2 | -1.07 |
| RB1 | 6 | 0.047024 | 0.13757 | 0.957317 | 2853 | 3 | -0.025421 | 0.071086 | 0.21458 | 0.97466 | 4291 | 3 | -0.025421 |
| NGFRAP1 | 5 | 0.89824 | 0.89834 | 0.999994 | 19187 | 0 | 3.2785 | 0.071099 | 0.19587 | 0.97466 | 4292 | 4 | 3.2785 |
| BCDIN3D | 6 | 0.98445 | 0.98438 | 0.999994 | 21201 | 0 | 3.9611 | 0.071151 | 0.21474 | 0.97466 | 4293 | 5 | 3.9611 |
| ERGIC3 | 5 | 0.22835 | 0.41493 | 0.999994 | 8486 | 3 | -0.71023 | 0.071161 | 0.19598 | 0.97466 | 4294 | 2 | -0.71023 |
| FAM107B | 6 | 0.94106 | 0.94105 | 0.999994 | 20111 | 0 | 1.2911 | 0.071216 | 0.21489 | 0.97466 | 4295 | 5 | 1.2911 |
| PTCH1 | 5 | 0.66285 | 0.7503 | 0.999994 | 15822 | 1 | 1.1716 | 0.071277 | 0.19624 | 0.97466 | 4296 | 3 | 1.1716 |
| INA | 6 | 0.51863 | 0.68157 | 0.999994 | 13720 | 2 | 3.7896 | 0.071283 | 0.21505 | 0.97466 | 4297 | 4 | 3.7896 |
| ALG9 | 5 | 0.13464 | 0.27969 | 0.999632 | 6042 | 1 | 2.269 | 0.071315 | 0.19632 | 0.97466 | 4298 | 4 | 2.269 |
| CTH | 6 | 0.12635 | 0.29568 | 0.999994 | 5834 | 2 | 3.166 | 0.071336 | 0.21518 | 0.97466 | 4299 | 4 | 3.166 |
| CHST10 | 4 | 0.12598 | 0.26065 | 0.994943 | 5821 | 2 | 1.9631 | 0.071354 | 0.18584 | 0.97466 | 4300 | 2 | 1.9631 |
| MASP1 | 3 | 0.87315 | 0.87326 | 0.999994 | 18696 | 0 | 5.8862 | 0.071409 | 0.16473 | 0.965913 | 4301 | 2 | 5.8862 |
| PDK2 | 5 | 0.82147 | 0.83357 | 0.999994 | 17795 | 1 | 1.6367 | 0.071423 | 0.19654 | 0.97466 | 4302 | 4 | 1.6367 |
| SPATA31A3 | 2 | 0.92857 | 0.92872 | 0.999994 | 19827 | 0 | 4.6414 | 0.071431 | 0.12473 | 0.964228 | 4303 | 2 | 4.6414 |
| SPPL2A | 6 | 0.70639 | 0.81398 | 0.999994 | 16378 | 1 | 1.3016 | 0.071455 | 0.21544 | 0.97466 | 4304 | 3 | 1.3016 |
| RPN1 | 6 | 0.87884 | 0.88884 | 0.999994 | 18808 | 1 | 3.1443 | 0.071464 | 0.21546 | 0.97466 | 4305 | 4 | 3.1443 |
| PBXIP1 | 6 | 0.22875 | 0.42255 | 0.999994 | 8497 | 3 | -0.75163 | 0.071512 | 0.21558 | 0.97466 | 4306 | 2 | -0.75163 |
| BOK | 6 | 0.98294 | 0.9829 | 0.999994 | 21155 | 0 | 1.9126 | 0.071512 | 0.21558 | 0.97466 | 4307 | 4 | 1.9126 |
| CLEC5A | 6 | 0.092947 | 0.23341 | 0.984971 | 4716 | 1 | 3.1126 | 0.071512 | 0.21558 | 0.97466 | 4308 | 4 | 3.1126 |
| SLC30A5 | 4 | 0.28574 | 0.42842 | 0.999994 | 9826 | 2 | -0.97041 | 0.071557 | 0.18617 | 0.97466 | 4309 | 1 | -0.97041 |
| hsa-mir-519a-1 | 3 | 0.92839 | 0.92848 | 0.999994 | 19823 | 0 | 2.8832 | 0.071611 | 0.16515 | 0.965913 | 4310 | 3 | 2.8832 |
| SLC25A42 | 6 | 0.51483 | 0.67895 | 0.999994 | 13658 | 1 | 1.0862 | 0.071614 | 0.21583 | 0.97466 | 4311 | 5 | 1.0862 |
| VAMP1 | 6 | 0.56929 | 0.71744 | 0.999994 | 14524 | 2 | -0.17962 | 0.071618 | 0.21584 | 0.97466 | 4312 | 2 | -0.17962 |
| MSX1 | 6 | 0.8167 | 0.85528 | 0.999994 | 17722 | 1 | 2.7144 | 0.071621 | 0.21584 | 0.97466 | 4313 | 5 | 2.7144 |
| ZNF383 | 6 | 0.25094 | 0.44907 | 0.999994 | 9032 | 2 | 2.3888 | 0.07163 | 0.21587 | 0.97466 | 4314 | 4 | 2.3888 |
| COQ10A | 6 | 0.94923 | 0.94921 | 0.999994 | 20314 | 0 | 1.6501 | 0.071673 | 0.21597 | 0.97466 | 4315 | 5 | 1.6501 |

| | | | | | | | | | | | | | |
|---|---|---|---|---|---|---|---|---|---|---|---|---|---|
| PHAX | 5 | 0.16243 | 0.3211 | 0.999994 | 6806 | 2 | -0.40988 | 0.071677 | 0.19708 | 0.97466 | 4316 | 2 | -0.40988 |
| RPS6KA6 | 6 | 0.62909 | 0.76165 | 0.999994 | 15367 | 1 | 0.85867 | 0.071724 | 0.21609 | 0.97466 | 4317 | 3 | 0.85867 |
| ASAH2 | 6 | 0.19543 | 0.38234 | 0.999994 | 7644 | 2 | 2.6905 | 0.071728 | 0.2161 | 0.97466 | 4318 | 4 | 2.6905 |
| STARD5 | 6 | 0.93357 | 0.93365 | 0.999994 | 19938 | 0 | 3.3597 | 0.071732 | 0.2161 | 0.97466 | 4319 | 5 | 3.3597 |
| hsa-mir-147b | 4 | 0.75665 | 0.76517 | 0.999994 | 16940 | 1 | 4.0718 | 0.071738 | 0.18649 | 0.97466 | 4320 | 3 | 4.0718 |
| CCDC170 | 6 | 0.98533 | 0.98529 | 0.999994 | 21236 | 0 | 1.6814 | 0.071738 | 0.21612 | 0.97466 | 4321 | 5 | 1.6814 |
| hsa-mir-6132 | 3 | 0.92821 | 0.92832 | 0.999994 | 19817 | 0 | 2.451 | 0.071787 | 0.1655 | 0.965913 | 4322 | 3 | 2.451 |
| CELA1 | 6 | 0.39564 | 0.59957 | 0.999994 | 11882 | 1 | 3.9994 | 0.071789 | 0.21624 | 0.97466 | 4323 | 5 | 3.9994 |
| SDCCAG8 | 6 | 0.91638 | 0.91709 | 0.999994 | 19547 | 1 | 2.4353 | 0.071794 | 0.21626 | 0.97466 | 4324 | 3 | 2.4353 |
| LRRC33 | 4 | 0.42684 | 0.57325 | 0.999994 | 12362 | 2 | 0.66712 | 0.071807 | 0.18661 | 0.97466 | 4325 | 2 | 0.66712 |
| RIPPLY2 | 5 | 0.01662 | 0.054807 | 0.922677 | 1286 | 3 | -3.5243 | 0.071816 | 0.19737 | 0.97466 | 4326 | 2 | -3.5243 |
| DTNB | 5 | 0.22649 | 0.41236 | 0.999994 | 8438 | 2 | 0.8226 | 0.071816 | 0.19737 | 0.97466 | 4327 | 3 | 0.8226 |
| TFPI | 6 | 0.59954 | 0.73959 | 0.999994 | 14947 | 2 | 2.5772 | 0.071819 | 0.21632 | 0.97466 | 4328 | 3 | 2.5772 |
| SMIM20 | 6 | 0.95637 | 0.95644 | 0.999994 | 20498 | 0 | 3.7086 | 0.071819 | 0.21632 | 0.97466 | 4329 | 4 | 3.7086 |
| SELRC1 | 6 | 0.25056 | 0.44859 | 0.999994 | 9026 | 3 | 2.2736 | 0.071819 | 0.21632 | 0.97466 | 4330 | 3 | 2.2736 |
| MAP2K2 | 3 | 0.87961 | 0.8797 | 0.999994 | 18823 | 0 | 5.8779 | 0.071862 | 0.16565 | 0.965913 | 4331 | 2 | 5.8779 |
| NonTargetingControlG | 2 | 0.1092 | 0.18424 | 0.968526 | 5274 | 1 | 0.27854 | 0.071936 | 0.12551 | 0.964228 | 4332 | 1 | 0.27854 |
| KIAA0100 | 6 | 0.51718 | 0.68059 | 0.999994 | 13696 | 1 | 1.2717 | 0.071974 | 0.21668 | 0.97466 | 4333 | 5 | 1.2717 |
| hsa-mir-4423 | 4 | 0.87783 | 0.87748 | 0.999994 | 18784 | 0 | 2.2342 | 0.071985 | 0.18688 | 0.97466 | 4334 | 3 | 2.2342 |
| ZNF385A | 6 | 0.64736 | 0.7756 | 0.999994 | 15620 | 2 | 0.15023 | 0.072017 | 0.21678 | 0.97466 | 4335 | 2 | 0.15023 |
| LRRC6 | 6 | 0.96938 | 0.96938 | 0.999994 | 20814 | 0 | 3.4847 | 0.07204 | 0.21684 | 0.97466 | 4336 | 5 | 3.4847 |
| RNF14 | 5 | 0.69425 | 0.76387 | 0.999994 | 16245 | 1 | 2.2846 | 0.072059 | 0.19787 | 0.97466 | 4337 | 4 | 2.2846 |
| KIDINS220 | 6 | 0.89514 | 0.9 | 0.999994 | 19130 | 1 | 1.4509 | 0.072106 | 0.21699 | 0.97466 | 4338 | 5 | 1.4509 |
| ANP32D | 6 | 0.44709 | 0.63314 | 0.999994 | 12676 | 2 | 0.97151 | 0.072157 | 0.21711 | 0.97466 | 4339 | 3 | 0.97151 |
| TMPRSS7 | 6 | 0.053592 | 0.15183 | 0.967126 | 3145 | 4 | -1.8471 | 0.072157 | 0.21711 | 0.97466 | 4340 | 2 | -1.8471 |
| PNN | 5 | 0.35868 | 0.52895 | 0.999994 | 11316 | 1 | 4.4873 | 0.072192 | 0.19813 | 0.97466 | 4341 | 3 | 4.4873 |
| PRELID1 | 5 | 0.023448 | 0.073377 | 0.93914 | 1681 | 2 | 4.4873 | 0.072192 | 0.19813 | 0.97466 | 4342 | 3 | 4.4873 |
| GLIS1 | 4 | 0.0025433 | 0.0091669 | 0.680529 | 319 | 1 | 4.4238 | 0.072246 | 0.18734 | 0.97466 | 4343 | 3 | 4.4238 |
| RPTN | 4 | 0.63922 | 0.68706 | 0.999994 | 15503 | 1 | 4.207 | 0.072246 | 0.18734 | 0.97466 | 4344 | 3 | 4.207 |
| KRTAP10-12 | 4 | 0.80561 | 0.8063 | 0.999994 | 17562 | 1 | 3.8428 | 0.072246 | 0.18734 | 0.97466 | 4345 | 3 | 3.8428 |
| OR2L2 | 5 | 0.41264 | 0.57403 | 0.999994 | 12151 | 2 | 2.4812 | 0.072277 | 0.19833 | 0.97466 | 4346 | 3 | 2.4812 |
| PHC2 | 6 | 0.45706 | 0.63976 | 0.999994 | 12806 | 2 | 2.7468 | 0.072306 | 0.21747 | 0.97466 | 4347 | 3 | 2.7468 |
| MRPS27 | 6 | 0.43723 | 0.62657 | 0.999994 | 12530 | 1 | 2.6018 | 0.07232 | 0.2175 | 0.97466 | 4348 | 4 | 2.6018 |
| SCO2 | 6 | 0.30414 | 0.51154 | 0.999994 | 10212 | 1 | 2.5739 | 0.07232 | 0.2175 | 0.97466 | 4349 | 4 | 2.5739 |
| KCNK5 | 6 | 0.11647 | 0.27908 | 0.999632 | 5524 | 2 | 3.0231 | 0.07232 | 0.2175 | 0.97466 | 4350 | 4 | 3.0231 |
| hsa-mir-6779 | 3 | 0.7166 | 0.71674 | 0.999994 | 16461 | 1 | 5.8608 | 0.072324 | 0.16657 | 0.965913 | 4351 | 2 | 5.8608 |
| NonTargetingControlG | 2 | 0.92767 | 0.92781 | 0.999994 | 19806 | 0 | 4.084 | 0.072334 | 0.12615 | 0.964228 | 4352 | 2 | 4.084 |
| KRT4 | 6 | 0.66462 | 0.78889 | 0.999994 | 15851 | 1 | 1.7366 | 0.072336 | 0.21755 | 0.97466 | 4353 | 5 | 1.7366 |
| FN1 | 6 | 0.16469 | 0.34409 | 0.999994 | 6857 | 3 | -1.5302 | 0.072442 | 0.21778 | 0.97466 | 4354 | 1 | -1.5302 |
| TMEM135 | 5 | 0.27344 | 0.45954 | 0.999994 | 9552 | 1 | 2.9263 | 0.072456 | 0.19873 | 0.97466 | 4355 | 4 | 2.9263 |
| CERS1 | 5 | 0.71146 | 0.77177 | 0.999994 | 16415 | 1 | 1.8588 | 0.072469 | 0.19875 | 0.97466 | 4356 | 4 | 1.8588 |
| CD101 | 5 | 0.31809 | 0.49578 | 0.999994 | 10476 | 2 | -0.17106 | 0.072565 | 0.19894 | 0.97466 | 4357 | 2 | -0.17106 |
| RNF113B | 5 | 0.8525 | 0.85627 | 0.999994 | 18305 | 1 | 4.5424 | 0.072577 | 0.19897 | 0.97466 | 4358 | 3 | 4.5424 |
| POFUT1 | 5 | 0.21776 | 0.40013 | 0.999994 | 8217 | 3 | -0.76684 | 0.072577 | 0.19897 | 0.97466 | 4359 | 2 | -0.76684 |
| PSG1 | 4 | 0.27432 | 0.41675 | 0.999994 | 9572 | 2 | 2.5896 | 0.072596 | 0.18793 | 0.97466 | 4360 | 2 | 2.5896 |
| FXYD7 | 6 | 0.97716 | 0.9771 | 0.999994 | 21020 | 0 | 1.1752 | 0.072653 | 0.21829 | 0.97466 | 4361 | 5 | 1.1752 |
| GGNBP2 | 6 | 0.72198 | 0.81896 | 0.999994 | 16516 | 1 | 2.5382 | 0.072659 | 0.21831 | 0.97466 | 4362 | 5 | 2.5382 |
| AGPAT1 | 6 | 0.85329 | 0.87369 | 0.999994 | 18319 | 1 | 1.696 | 0.072666 | 0.21832 | 0.97466 | 4363 | 5 | 1.696 |
| PDZD9 | 6 | 0.18699 | 0.37177 | 0.999994 | 7387 | 3 | 0.070809 | 0.072679 | 0.21835 | 0.97466 | 4364 | 3 | 0.070809 |
| CNR1 | 6 | 0.86689 | 0.88143 | 0.999994 | 18572 | 1 | 1.7818 | 0.072692 | 0.21839 | 0.97466 | 4365 | 5 | 1.7818 |
| VWA2 | 5 | 0.77579 | 0.80502 | 0.999994 | 17147 | 1 | 0.73329 | 0.072699 | 0.19925 | 0.97466 | 4366 | 3 | 0.73329 |
| NonTargetingControlG | 2 | 0.69716 | 0.69696 | 0.999994 | 16270 | 0 | 4.2911 | 0.072708 | 0.12676 | 0.964228 | 4367 | 1 | 4.2911 |

| | | | | | | | | | | | | | |
|---|---|---|---|---|---|---|---|---|---|---|---|---|---|
| SLC35A3 | 6 | 0.86131 | 0.87815 | 0.999994 | 18473 | 1 | 0.89934 | 0.072732 | 0.21848 | 0.97466 | 4368 | 5 | 0.89934 |
| ATP6AP1 | 2 | 0.92716 | 0.9273 | 0.999994 | 19793 | 0 | 3.2731 | 0.072837 | 0.12696 | 0.964228 | 4369 | 2 | 3.2731 |
| PLD2 | 6 | 0.80965 | 0.8521 | 0.999994 | 17627 | 1 | 3.1302 | 0.072883 | 0.21883 | 0.97466 | 4370 | 4 | 3.1302 |
| KIAA0020 | 6 | 0.77974 | 0.83954 | 0.999994 | 17209 | 1 | 2.6648 | 0.072883 | 0.21883 | 0.97466 | 4371 | 3 | 2.6648 |
| LRP5L | 5 | 0.97173 | 0.97177 | 0.999994 | 20863 | 0 | 2.1324 | 0.072931 | 0.19975 | 0.97466 | 4372 | 4 | 2.1324 |
| DRD3 | 6 | 0.37819 | 0.58843 | 0.999994 | 11627 | 2 | 2.5869 | 0.072984 | 0.21909 | 0.97466 | 4373 | 4 | 2.5869 |
| BPIFC | 5 | 0.75554 | 0.79377 | 0.999994 | 16924 | 1 | 1.6262 | 0.072992 | 0.19988 | 0.97466 | 4374 | 3 | 1.6262 |
| PON3 | 6 | 0.93407 | 0.93414 | 0.999994 | 19955 | 0 | 2.7872 | 0.073088 | 0.21933 | 0.97466 | 4375 | 4 | 2.7872 |
| hsa-mir-4765 | 3 | 0.9269 | 0.92695 | 0.999994 | 19787 | 0 | 1.817 | 0.073104 | 0.16813 | 0.965913 | 4376 | 3 | 1.817 |
| RBPJL | 5 | 0.008788 | 0.032559 | 0.865899 | 779 | 2 | 0.39251 | 0.073142 | 0.20019 | 0.97466 | 4377 | 2 | 0.39251 |
| PI16 | 5 | 0.96943 | 0.96951 | 0.999994 | 20816 | 0 | 4.4636 | 0.073158 | 0.20021 | 0.97466 | 4378 | 4 | 4.4636 |
| GAS2L2 | 5 | 0.7086 | 0.77045 | 0.999994 | 16391 | 1 | 4.4636 | 0.073158 | 0.20021 | 0.97466 | 4379 | 3 | 4.4636 |
| NonTargetingControlG | 2 | 0.5826 | 0.58271 | 0.999994 | 14713 | 1 | 3.9501 | 0.073162 | 0.12748 | 0.964228 | 4380 | 1 | 3.9501 |
| NonTargetingControlG | 2 | 0.92681 | 0.92694 | 0.999994 | 19784 | 0 | 5.3659 | 0.073186 | 0.12753 | 0.964228 | 4381 | 2 | 5.3659 |
| MSRB3 | 6 | 0.26895 | 0.47051 | 0.999994 | 9444 | 3 | -0.051443 | 0.073186 | 0.21956 | 0.97466 | 4382 | 3 | -0.051443 |
| SPATA13 | 6 | 0.39002 | 0.596 | 0.999994 | 11799 | 1 | 2.8689 | 0.073186 | 0.21956 | 0.97466 | 4383 | 4 | 2.8689 |
| RAB30 | 5 | 0.46695 | 0.61953 | 0.999994 | 12964 | 1 | 2.1878 | 0.073266 | 0.20046 | 0.97466 | 4384 | 4 | 2.1878 |
| SLC16A9 | 6 | 0.48763 | 0.66036 | 0.999994 | 13248 | 2 | 0.53507 | 0.073292 | 0.21981 | 0.97466 | 4385 | 3 | 0.53507 |
| CD22 | 5 | 0.84282 | 0.8488 | 0.999994 | 18137 | 1 | 1.0537 | 0.073297 | 0.20052 | 0.97466 | 4386 | 3 | 1.0537 |
| hsa-mir-5693 | 2 | 0.9267 | 0.92683 | 0.999994 | 19782 | 0 | 5.634 | 0.073298 | 0.12771 | 0.964228 | 4387 | 2 | 5.634 |
| hsa-mir-520f | 4 | 0.26459 | 0.40678 | 0.999994 | 9347 | 2 | -0.13724 | 0.073357 | 0.18921 | 0.97466 | 4388 | 2 | -0.13724 |
| KAAG1 | 4 | 0.73341 | 0.74766 | 0.999994 | 16645 | 1 | 1.3744 | 0.073357 | 0.18921 | 0.97466 | 4389 | 2 | 1.3744 |
| PLA2G6 | 5 | 0.92913 | 0.9294 | 0.999994 | 19838 | 0 | 2.9342 | 0.073389 | 0.20073 | 0.97466 | 4390 | 4 | 2.9342 |
| TCEAL6 | 6 | 0.61132 | 0.74831 | 0.999994 | 15092 | 1 | 3.4331 | 0.073399 | 0.22005 | 0.97466 | 4391 | 4 | 3.4331 |
| PCMT1 | 6 | 0.10935 | 0.26544 | 0.997422 | 5282 | 3 | -0.099026 | 0.073478 | 0.22024 | 0.97466 | 4392 | 2 | -0.099026 |
| CASQ1 | 5 | 0.15484 | 0.30998 | 0.999994 | 6592 | 3 | -0.76922 | 0.073497 | 0.20096 | 0.97466 | 4393 | 2 | -0.76922 |
| IDUA | 6 | 0.31857 | 0.52826 | 0.999994 | 10488 | 3 | 0.2768 | 0.073545 | 0.22042 | 0.97466 | 4394 | 3 | 0.2768 |
| FILIP1L | 6 | 0.47966 | 0.65498 | 0.999994 | 13141 | 2 | 0.27532 | 0.073545 | 0.22042 | 0.97466 | 4395 | 2 | 0.27532 |
| GALNT11 | 6 | 0.96787 | 0.96789 | 0.999994 | 20782 | 0 | 1.9008 | 0.07361 | 0.22057 | 0.97466 | 4396 | 4 | 1.9008 |
| COL9A2 | 3 | 0.51885 | 0.55874 | 0.999994 | 13725 | 1 | 4.5199 | 0.07361 | 0.16915 | 0.965913 | 4397 | 2 | 4.5199 |
| TRIO | 3 | 0.92637 | 0.92641 | 0.999994 | 19777 | 0 | 1.329 | 0.073634 | 0.1692 | 0.965913 | 4398 | 3 | 1.329 |
| NLGN4Y | 5 | 0.54785 | 0.6898 | 0.999994 | 14177 | 2 | 1.8181 | 0.07364 | 0.20126 | 0.97466 | 4399 | 3 | 1.8181 |
| GPR108 | 6 | 0.73157 | 0.82213 | 0.999994 | 16622 | 1 | 1.5583 | 0.073743 | 0.2209 | 0.97466 | 4400 | 5 | 1.5583 |
| CYP26C1 | 3 | 0.92622 | 0.92627 | 0.999994 | 19773 | 0 | 3.4636 | 0.073778 | 0.1695 | 0.965913 | 4401 | 3 | 3.4636 |
| TNFAIP2 | 4 | 0.44007 | 0.58685 | 0.999994 | 12576 | 2 | -0.62149 | 0.073802 | 0.18994 | 0.97466 | 4402 | 1 | -0.62149 |
| CLEC3B | 6 | 0.23859 | 0.4343 | 0.999994 | 8739 | 2 | 2.3596 | 0.073863 | 0.22118 | 0.97466 | 4403 | 3 | 2.3596 |
| RNF208 | 6 | 0.49044 | 0.66221 | 0.999994 | 13284 | 2 | 0.96214 | 0.073863 | 0.22118 | 0.97466 | 4404 | 3 | 0.96214 |
| BNIPL | 6 | 0.080435 | 0.20818 | 0.979182 | 4246 | 4 | -1.3622 | 0.073863 | 0.22118 | 0.97466 | 4405 | 2 | -1.3622 |
| S100A14 | 5 | 0.13981 | 0.28754 | 0.999632 | 6180 | 3 | -1.2047 | 0.073874 | 0.20169 | 0.97466 | 4406 | 1 | -1.2047 |
| ASGR2 | 5 | 0.1149 | 0.2496 | 0.991904 | 5473 | 2 | 0.097398 | 0.073874 | 0.20169 | 0.97466 | 4407 | 2 | 0.097398 |
| RASGRF2 | 6 | 0.0085478 | 0.033878 | 0.867348 | 766 | 3 | -3.3589 | 0.073876 | 0.22121 | 0.97466 | 4408 | 2 | -3.3589 |
| PARM1 | 5 | 0.68106 | 0.75801 | 0.999994 | 16104 | 1 | 4.3248 | 0.073894 | 0.20174 | 0.97466 | 4409 | 4 | 4.3248 |
| C14orf1 | 5 | 0.59819 | 0.7255 | 0.999994 | 14929 | 1 | 4.4377 | 0.073902 | 0.20175 | 0.97466 | 4410 | 4 | 4.4377 |
| DEDD | 5 | 0.91713 | 0.91722 | 0.999994 | 19565 | 0 | 1.615 | 0.073911 | 0.20177 | 0.97466 | 4411 | 3 | 1.615 |
| LIMD2 | 5 | 0.047736 | 0.13497 | 0.954328 | 2883 | 2 | 3.0115 | 0.073911 | 0.20177 | 0.97466 | 4412 | 3 | 3.0115 |
| RNF38 | 5 | 0.29809 | 0.47952 | 0.999994 | 10093 | 1 | 2.9466 | 0.073911 | 0.20177 | 0.97466 | 4413 | 3 | 2.9466 |
| PGD | 5 | 0.63316 | 0.73853 | 0.999994 | 15416 | 1 | 1.9522 | 0.073946 | 0.20185 | 0.97466 | 4414 | 4 | 1.9522 |
| SPDYE4 | 4 | 0.79014 | 0.7925 | 0.999994 | 17347 | 1 | 4.3756 | 0.07398 | 0.19025 | 0.97466 | 4415 | 3 | 4.3756 |
| TXNDC12 | 6 | 0.44861 | 0.63419 | 0.999994 | 12694 | 1 | 3.6985 | 0.074061 | 0.22167 | 0.97466 | 4416 | 4 | 3.6985 |
| SRRD | 6 | 0.0019763 | 0.0098582 | 0.680529 | 273 | 5 | -3.7318 | 0.074061 | 0.22167 | 0.97466 | 4417 | 1 | -3.7318 |
| EFNA3 | 4 | 0.41002 | 0.55597 | 0.999994 | 12108 | 1 | 0.9867 | 0.074069 | 0.1904 | 0.97466 | 4418 | 2 | 0.9867 |
| TMEM161B | 6 | 0.92714 | 0.92721 | 0.999994 | 19792 | 1 | 1.7287 | 0.074083 | 0.22173 | 0.97466 | 4419 | 5 | 1.7287 |

| | | | | | | | | | | | | |
|---|---|---|---|---|---|---|---|---|---|---|---|---|
| MAP3K4 | 6 | 0.6467 | 0.77511 | 0.999994 | 15614 | 2 | 2.5997 | 0.074122 | 0.22182 | 0.97466 | 4420 | 4 | 2.5997 |
| hsa-mir-4279 | 4 | 0.83687 | 0.83636 | 0.999994 | 18032 | 0 | 3.7754 | 0.074128 | 0.19049 | 0.97466 | 4421 | 3 | 3.7754 |
| ZNF575 | 3 | 0.8821 | 0.88217 | 0.999994 | 18872 | 0 | 4.8199 | 0.074161 | 0.17026 | 0.965913 | 4422 | 2 | 4.8199 |
| C5orf49 | 6 | 0.92822 | 0.92829 | 0.999994 | 19818 | 0 | 2.3644 | 0.074177 | 0.22195 | 0.97466 | 4423 | 5 | 2.3644 |
| OR10A5 | 6 | 0.082215 | 0.21186 | 0.979587 | 4303 | 2 | 0.93865 | 0.074221 | 0.22206 | 0.97466 | 4424 | 3 | 0.93865 |
| RAB3GAP2 | 6 | 0.13736 | 0.3097 | 0.999994 | 6119 | 3 | 0.23943 | 0.074221 | 0.22206 | 0.97466 | 4425 | 3 | 0.23943 |
| SLC6A7 | 5 | 0.85378 | 0.85729 | 0.999994 | 18328 | 1 | 4.4318 | 0.074297 | 0.2026 | 0.97466 | 4426 | 4 | 4.4318 |
| IDE | 5 | 0.94436 | 0.94455 | 0.999994 | 20187 | 0 | 1.8738 | 0.074298 | 0.2026 | 0.97466 | 4427 | 4 | 1.8738 |
| OR4M2 | 6 | 0.30069 | 0.5075 | 0.999994 | 10145 | 1 | 3.6834 | 0.07432 | 0.2223 | 0.97466 | 4428 | 4 | 3.6834 |
| DEFB115 | 6 | 0.073663 | 0.19437 | 0.975797 | 3978 | 2 | 3.6635 | 0.07435 | 0.22237 | 0.97466 | 4429 | 4 | 3.6635 |
| FKBP10 | 6 | 0.12736 | 0.29693 | 0.999994 | 5859 | 1 | 0.90742 | 0.074422 | 0.22255 | 0.97466 | 4430 | 4 | 0.90742 |
| DAB1 | 6 | 0.91361 | 0.91467 | 0.999994 | 19483 | 1 | 0.11479 | 0.074422 | 0.22255 | 0.97466 | 4431 | 2 | 0.11479 |
| TMEM175 | 6 | 0.21503 | 0.40617 | 0.999994 | 8143 | 2 | 3.9871 | 0.07444 | 0.22259 | 0.97466 | 4432 | 4 | 3.9871 |
| FAH | 5 | 0.43711 | 0.59437 | 0.999994 | 12526 | 1 | 1.7014 | 0.074565 | 0.20316 | 0.97466 | 4433 | 4 | 1.7014 |
| GPI | 6 | 0.80269 | 0.84901 | 0.999994 | 17521 | 1 | 2.2512 | 0.074594 | 0.22295 | 0.97466 | 4434 | 5 | 2.2512 |
| CHD8 | 6 | 0.88701 | 0.89425 | 0.999994 | 18967 | 1 | 1.6311 | 0.074621 | 0.22303 | 0.97466 | 4435 | 5 | 1.6311 |
| SPTY2D1 | 6 | 0.95583 | 0.95591 | 0.999994 | 20471 | 0 | 3.5354 | 0.074657 | 0.22312 | 0.97466 | 4436 | 4 | 3.5354 |
| SUPT16H | 4 | 0.056969 | 0.13498 | 0.954328 | 3291 | 2 | -2.2665 | 0.074728 | 0.19148 | 0.97466 | 4437 | 1 | -2.2665 |
| LRG1 | 6 | 0.023114 | 0.081945 | 0.93914 | 1658 | 4 | -2.1789 | 0.074758 | 0.22336 | 0.97466 | 4438 | 2 | -2.1789 |
| CCL5 | 5 | 0.80521 | 0.8228 | 0.999994 | 17555 | 1 | 0.34341 | 0.074805 | 0.2037 | 0.97466 | 4439 | 2 | 0.34341 |
| CADM4 | 5 | 0.54086 | 0.68368 | 0.999994 | 14071 | 2 | 3.3008 | 0.074805 | 0.2037 | 0.97466 | 4440 | 3 | 3.3008 |
| NonTargetingControlG | 2 | 0.92519 | 0.92532 | 0.999994 | 19747 | 0 | 4.2947 | 0.074812 | 0.13017 | 0.964228 | 4441 | 2 | 4.2947 |
| hsa-mir-320d-1 | 4 | 0.42872 | 0.57523 | 0.999994 | 12390 | 2 | 3.0306 | 0.074821 | 0.19164 | 0.97466 | 4442 | 2 | 3.0306 |
| C1orf233 | 4 | 0.32152 | 0.46497 | 0.999994 | 10544 | 2 | 2.7238 | 0.074821 | 0.19164 | 0.97466 | 4443 | 2 | 2.7238 |
| hsa-mir-6763 | 4 | 0.67647 | 0.70937 | 0.999994 | 16031 | 1 | 4.1318 | 0.074821 | 0.19164 | 0.97466 | 4444 | 3 | 4.1318 |
| TAF5 | 6 | 0.014718 | 0.054826 | 0.922677 | 1172 | 3 | -3.2021 | 0.07483 | 0.22352 | 0.97466 | 4445 | 2 | -3.2021 |
| CXorf40A | 4 | 0.36264 | 0.50723 | 0.999994 | 11381 | 1 | 2.1078 | 0.07487 | 0.19173 | 0.97466 | 4446 | 3 | 2.1078 |
| ZCCHC14 | 6 | 0.66905 | 0.79244 | 0.999994 | 15913 | 1 | 0.93451 | 0.074989 | 0.22391 | 0.97466 | 4447 | 4 | 0.93451 |
| OR2A42 | 2 | 0.92499 | 0.9251 | 0.999994 | 19742 | 0 | 5.1373 | 0.075013 | 0.1305 | 0.964228 | 4448 | 2 | 5.1373 |
| DEFB112 | 6 | 0.96349 | 0.96355 | 0.999994 | 20686 | 0 | 2.6818 | 0.075026 | 0.224 | 0.97466 | 4449 | 5 | 2.6818 |
| FLRT2 | 5 | 0.94425 | 0.94444 | 0.999994 | 20183 | 0 | 0.39928 | 0.075034 | 0.20417 | 0.97466 | 4450 | 2 | 0.39928 |
| ICAM5 | 5 | 0.95854 | 0.95868 | 0.999994 | 20549 | 0 | 0.97017 | 0.075034 | 0.20417 | 0.97466 | 4451 | 3 | 0.97017 |
| ELP6 | 5 | 0.38292 | 0.54915 | 0.999994 | 11692 | 2 | 2.3544 | 0.075034 | 0.20417 | 0.97466 | 4452 | 3 | 2.3544 |
| NUDT4 | 5 | 0.6396 | 0.74102 | 0.999994 | 15510 | 1 | 1.7344 | 0.075063 | 0.20423 | 0.97466 | 4453 | 4 | 1.7344 |
| CLCNKA | 5 | 0.56753 | 0.70736 | 0.999994 | 14507 | 1 | 2.8407 | 0.075102 | 0.20432 | 0.97466 | 4454 | 4 | 2.8407 |
| NKX6-2 | 6 | 0.064058 | 0.17435 | 0.968171 | 3615 | 4 | -1.1739 | 0.075175 | 0.22437 | 0.97466 | 4455 | 2 | -1.1739 |
| STAT6 | 5 | 0.73911 | 0.78523 | 0.999994 | 16723 | 1 | 0.71546 | 0.075248 | 0.20462 | 0.97466 | 4456 | 3 | 0.71546 |
| EVI5 | 6 | 0.8995 | 0.90329 | 0.999994 | 19213 | 1 | 3.2896 | 0.075253 | 0.22454 | 0.97466 | 4457 | 4 | 3.2896 |
| COQ5 | 6 | 0.037337 | 0.1159 | 0.951052 | 2390 | 2 | 2.747 | 0.075253 | 0.22454 | 0.97466 | 4458 | 3 | 2.747 |
| CXCL17 | 6 | 0.80001 | 0.84787 | 0.999994 | 17478 | 1 | 3.8579 | 0.075253 | 0.22454 | 0.97466 | 4459 | 5 | 3.8579 |
| CEP68 | 5 | 0.7453 | 0.78837 | 0.999994 | 16805 | 1 | 4.3946 | 0.075316 | 0.20477 | 0.97466 | 4460 | 3 | 4.3946 |
| ZNF407 | 5 | 0.17457 | 0.3388 | 0.999994 | 7100 | 1 | 2.2565 | 0.075326 | 0.20479 | 0.97466 | 4461 | 4 | 2.2565 |
| AANAT | 6 | 0.95617 | 0.95626 | 0.999994 | 20491 | 0 | 0.90971 | 0.075338 | 0.22476 | 0.97466 | 4462 | 5 | 0.90971 |
| CFC1B | 3 | 0.4002 | 0.48438 | 0.999994 | 11951 | 1 | 5.782 | 0.075369 | 0.1727 | 0.965913 | 4463 | 2 | 5.782 |
| SMU1 | 3 | 0.27928 | 0.41112 | 0.999994 | 9683 | 1 | 5.782 | 0.075369 | 0.1727 | 0.965913 | 4464 | 2 | 5.782 |
| NF1 | 6 | 0.93204 | 0.93216 | 0.999994 | 19901 | 0 | 1.1474 | 0.075384 | 0.22486 | 0.97466 | 4465 | 3 | 1.1474 |
| LMTK2 | 6 | 0.45542 | 0.63866 | 0.999994 | 12781 | 2 | 2.4976 | 0.075389 | 0.22487 | 0.97466 | 4466 | 4 | 2.4976 |
| PSMD8 | 3 | 0.70346 | 0.70428 | 0.999994 | 16346 | 1 | 2.4822 | 0.07541 | 0.17279 | 0.965913 | 4467 | 2 | 2.4822 |
| COL4A3BP | 5 | 0.83159 | 0.84064 | 0.999994 | 17954 | 1 | 1.8071 | 0.075411 | 0.20498 | 0.97466 | 4468 | 4 | 1.8071 |
| RANBP17 | 5 | 0.81037 | 0.82618 | 0.999994 | 17637 | 1 | 1.3273 | 0.075415 | 0.20498 | 0.97466 | 4469 | 3 | 1.3273 |
| ATP2C1 | 5 | 0.75541 | 0.7937 | 0.999994 | 16920 | 1 | 1.1656 | 0.075415 | 0.20498 | 0.97466 | 4470 | 3 | 1.1656 |
| MAG | 6 | 0.9742 | 0.97409 | 0.999994 | 20934 | 0 | 2.2195 | 0.075426 | 0.22497 | 0.97466 | 4471 | 5 | 2.2195 |

| | | | | | | | | | | | | | |
|---|---|---|---|---|---|---|---|---|---|---|---|---|---|
| TACR2 | 5 | 0.83847 | 0.84554 | 0.999994 | 18061 | 1 | 2.1041 | 0.075438 | 0.20503 | 0.97466 | 4472 | 4 | 2.1041 |
| CYP4Z1 | 6 | 0.90356 | 0.90645 | 0.999994 | 19293 | 1 | 1.1039 | 0.07544 | 0.225 | 0.97466 | 4473 | 3 | 1.1039 |
| EPCAM | 6 | 0.76609 | 0.83438 | 0.999994 | 17041 | 1 | 2.091 | 0.075446 | 0.22501 | 0.97466 | 4474 | 5 | 2.091 |
| KALRN | 6 | 0.9693 | 0.9693 | 0.999994 | 20810 | 0 | 1.6138 | 0.075546 | 0.22524 | 0.97466 | 4475 | 5 | 1.6138 |
| UHRF1 | 5 | 0.24622 | 0.43774 | 0.999994 | 8929 | 1 | 4.3222 | 0.07558 | 0.20532 | 0.97466 | 4476 | 4 | 4.3222 |
| EPS8 | 6 | 0.29966 | 0.50631 | 0.999994 | 10121 | 1 | 2.0278 | 0.075589 | 0.22534 | 0.97466 | 4477 | 5 | 2.0278 |
| OR4E2 | 6 | 0.70707 | 0.81419 | 0.999994 | 16383 | 1 | 2.8086 | 0.075601 | 0.22536 | 0.97466 | 4478 | 5 | 2.8086 |
| SMPD4 | 6 | 0.62905 | 0.76162 | 0.999994 | 15366 | 2 | 3.7622 | 0.075626 | 0.22542 | 0.97466 | 4479 | 4 | 3.7622 |
| EMC1 | 4 | 0.73555 | 0.74919 | 0.999994 | 16669 | 1 | 4.1114 | 0.075634 | 0.19303 | 0.97466 | 4480 | 3 | 4.1114 |
| KLHL8 | 6 | 0.93467 | 0.93475 | 0.999994 | 19961 | 0 | 1.6873 | 0.075664 | 0.22552 | 0.97466 | 4481 | 5 | 1.6873 |
| ARHGEF10 | 6 | 0.95332 | 0.95332 | 0.999994 | 20419 | 0 | 2.7294 | 0.075686 | 0.22557 | 0.97466 | 4482 | 4 | 2.7294 |
| SIX1 | 6 | 0.5174 | 0.68072 | 0.999994 | 13699 | 2 | 3.3512 | 0.075686 | 0.22557 | 0.97466 | 4483 | 4 | 3.3512 |
| IL1F10 | 6 | 0.11291 | 0.2723 | 0.998561 | 5400 | 3 | -0.54829 | 0.075705 | 0.22562 | 0.97466 | 4484 | 3 | -0.54829 |
| NonTargetingControlG | 2 | 0.92429 | 0.92438 | 0.999994 | 19726 | 0 | 4.8988 | 0.075712 | 0.13169 | 0.965199 | 4485 | 2 | 4.8988 |
| BIK | 4 | 0.89813 | 0.89785 | 0.999994 | 19182 | 0 | 3.6668 | 0.075724 | 0.19319 | 0.97466 | 4486 | 3 | 3.6668 |
| EIF3I | 4 | 0.73323 | 0.74754 | 0.999994 | 16642 | 1 | 2.8553 | 0.075773 | 0.19328 | 0.97466 | 4487 | 3 | 2.8553 |
| MAGOHB | 3 | 0.78085 | 0.78077 | 0.999994 | 17221 | 0 | 0.96178 | 0.075786 | 0.17355 | 0.966596 | 4488 | 2 | 0.96178 |
| HSPH1 | 6 | 0.8611 | 0.87804 | 0.999994 | 18468 | 1 | 2.6933 | 0.075813 | 0.22589 | 0.97466 | 4489 | 5 | 2.6933 |
| ABCG4 | 5 | 0.38736 | 0.55292 | 0.999994 | 11756 | 1 | 0.88925 | 0.075816 | 0.20581 | 0.97466 | 4490 | 4 | 0.88925 |
| CHMP2B | 4 | 0.075386 | 0.17006 | 0.967126 | 4051 | 3 | -1.1278 | 0.075919 | 0.19353 | 0.97466 | 4491 | 1 | -1.1278 |
| TTC9 | 6 | 0.8246 | 0.85898 | 0.999994 | 17848 | 1 | 3.2806 | 0.076 | 0.22633 | 0.97466 | 4492 | 4 | 3.2806 |
| DNAH9 | 6 | 0.68446 | 0.80447 | 0.999994 | 16151 | 2 | 3.7716 | 0.076 | 0.22633 | 0.97466 | 4493 | 4 | 3.7716 |
| MLLT1 | 6 | 0.92169 | 0.92197 | 0.999994 | 19669 | 1 | 2.8799 | 0.076 | 0.22633 | 0.97466 | 4494 | 3 | 2.8799 |
| NSRP1 | 5 | 0.85949 | 0.86197 | 0.999994 | 18427 | 1 | 2.3959 | 0.076019 | 0.20625 | 0.97466 | 4495 | 4 | 2.3959 |
| CCDC126 | 4 | 0.087963 | 0.19327 | 0.975797 | 4511 | 1 | 3.2495 | 0.076039 | 0.19375 | 0.97466 | 4496 | 2 | 3.2495 |
| C9orf96 | 6 | 0.80821 | 0.85145 | 0.999994 | 17602 | 1 | 2.4219 | 0.076045 | 0.22643 | 0.97466 | 4497 | 5 | 2.4219 |
| SRL | 6 | 0.90927 | 0.91097 | 0.999994 | 19408 | 1 | 2.4499 | 0.076054 | 0.22646 | 0.97466 | 4498 | 4 | 2.4499 |
| POLR1A | 5 | 0.497 | 0.64527 | 0.999994 | 13376 | 2 | 0.54942 | 0.076067 | 0.20635 | 0.97466 | 4499 | 2 | 0.54942 |
| AP1G2 | 6 | 0.016378 | 0.060242 | 0.932932 | 1268 | 2 | 0.22592 | 0.076076 | 0.22651 | 0.97466 | 4500 | 2 | 0.22592 |
| CYB5RL | 6 | 0.096845 | 0.24117 | 0.987253 | 4852 | 4 | -0.9394 | 0.076076 | 0.22651 | 0.97466 | 4501 | 1 | -0.9394 |
| EPHX3 | 6 | 0.02128 | 0.076091 | 0.93914 | 1533 | 5 | -1.0913 | 0.076076 | 0.22651 | 0.97466 | 4502 | 1 | -1.0913 |
| MYOC | 5 | 0.58571 | 0.72101 | 0.999994 | 14754 | 1 | 3.5659 | 0.076091 | 0.2064 | 0.97466 | 4503 | 4 | 3.5659 |
| CHIC1 | 6 | 0.76671 | 0.8346 | 0.999994 | 17044 | 1 | -0.055915 | 0.07614 | 0.22666 | 0.97466 | 4504 | 2 | -0.055915 |
| DNAI2 | 6 | 0.055443 | 0.15589 | 0.967126 | 3217 | 2 | 0.71368 | 0.07614 | 0.22666 | 0.97466 | 4505 | 3 | 0.71368 |
| B9D1 | 6 | 0.82562 | 0.85944 | 0.999994 | 17862 | 1 | 2.4668 | 0.076188 | 0.22677 | 0.97466 | 4506 | 4 | 2.4668 |
| C20orf195 | 6 | 0.95783 | 0.95793 | 0.999994 | 20534 | 0 | 1.644 | 0.076196 | 0.22679 | 0.97466 | 4507 | 5 | 1.644 |
| CHCHD3 | 5 | 0.53985 | 0.68274 | 0.999994 | 14048 | 1 | 0.15916 | 0.076221 | 0.20664 | 0.97466 | 4508 | 2 | 0.15916 |
| HIRA | 5 | 0.22953 | 0.41657 | 0.999994 | 8518 | 2 | -0.16751 | 0.076221 | 0.20664 | 0.97466 | 4509 | 2 | -0.16751 |
| CCDC134 | 5 | 0.90649 | 0.90657 | 0.999994 | 19359 | 0 | 1.8714 | 0.076264 | 0.20674 | 0.97466 | 4510 | 4 | 1.8714 |
| NLRP4 | 6 | 0.9407 | 0.94068 | 0.999994 | 20097 | 0 | 1.1047 | 0.076271 | 0.22697 | 0.97466 | 4511 | 5 | 1.1047 |
| AKR1C1 | 6 | 0.40882 | 0.60814 | 0.999994 | 12090 | 2 | 1.1769 | 0.076277 | 0.22698 | 0.97466 | 4512 | 3 | 1.1769 |
| PUS3 | 6 | 0.54384 | 0.69924 | 0.999994 | 14128 | 2 | 1.0603 | 0.07634 | 0.22713 | 0.97466 | 4513 | 4 | 1.0603 |
| NonTargetingControlG | 1 | 0.92361 | 0.92395 | 0.999994 | 19710 | 0 | 7.6081 | 0.07639 | 0.076397 | 0.938203 | 4514 | 1 | 7.6081 |
| EMR3 | 6 | 0.84364 | 0.86854 | 0.999994 | 18148 | 1 | 1.3025 | 0.076394 | 0.22726 | 0.97466 | 4515 | 5 | 1.3025 |
| ASAH1 | 6 | 0.97806 | 0.978 | 0.999994 | 21046 | 0 | 2.6521 | 0.076406 | 0.22729 | 0.97466 | 4516 | 4 | 2.6521 |
| IPP | 6 | 0.34974 | 0.56371 | 0.999994 | 11150 | 3 | -0.31283 | 0.076406 | 0.22729 | 0.97466 | 4517 | 2 | -0.31283 |
| BCL7C | 5 | 0.05147 | 0.14395 | 0.961589 | 3043 | 2 | 1.3809 | 0.076425 | 0.20707 | 0.97466 | 4518 | 3 | 1.3809 |
| FCRL5 | 6 | 0.98169 | 0.98161 | 0.999994 | 21130 | 0 | 1.5774 | 0.076456 | 0.2274 | 0.97466 | 4519 | 5 | 1.5774 |
| CD44 | 4 | 0.2587 | 0.40081 | 0.999994 | 9208 | 2 | 2.6493 | 0.076456 | 0.19444 | 0.97466 | 4520 | 2 | 2.6493 |
| RAB41 | 5 | 0.75632 | 0.79418 | 0.999994 | 16934 | 1 | 1.9594 | 0.076469 | 0.20716 | 0.97466 | 4521 | 4 | 1.9594 |
| EPHA3 | 5 | 0.11035 | 0.24251 | 0.988088 | 5315 | 1 | 1.3779 | 0.076503 | 0.20723 | 0.97466 | 4522 | 4 | 1.3779 |
| DDIT4 | 4 | 0.44559 | 0.59253 | 0.999994 | 12658 | 1 | 4.077 | 0.076519 | 0.19454 | 0.97466 | 4523 | 3 | 4.077 |

| | | | | | | | | | | | | | |
|---|---|---|---|---|---|---|---|---|---|---|---|---|---|
| LMAN2 | 6 | 0.1469 | 0.32175 | 0.999994 | 6371 | 3 | -0.78814 | 0.076526 | 0.22757 | 0.97466 | 4524 | 2 | -0.78814 |
| B3GNT5 | 6 | 0.052438 | 0.1494 | 0.966183 | 3089 | 4 | -1.3145 | 0.076526 | 0.22757 | 0.97466 | 4525 | 1 | -1.3145 |
| PTPLAD2 | 5 | 0.17271 | 0.33608 | 0.999994 | 7056 | 2 | 2.0807 | 0.076531 | 0.20729 | 0.97466 | 4526 | 3 | 2.0807 |
| MGST1 | 6 | 0.42939 | 0.62144 | 0.999994 | 12405 | 2 | 3.5061 | 0.076574 | 0.22768 | 0.97466 | 4527 | 4 | 3.5061 |
| SCML4 | 6 | 0.90996 | 0.91153 | 0.999994 | 19423 | 1 | 1.5751 | 0.076593 | 0.22773 | 0.97466 | 4528 | 5 | 1.5751 |
| PDAP1 | 5 | 0.33698 | 0.51126 | 0.999994 | 10872 | 1 | 1.6349 | 0.076597 | 0.20742 | 0.97466 | 4529 | 4 | 1.6349 |
| SPTLC2 | 6 | 0.91492 | 0.91582 | 0.999994 | 19509 | 1 | 0.90162 | 0.076617 | 0.22778 | 0.97466 | 4530 | 5 | 0.90162 |
| hsa-mir-649 | 4 | 0.80866 | 0.80907 | 0.999994 | 17613 | 1 | 0.5924 | 0.076647 | 0.19475 | 0.97466 | 4531 | 2 | 0.5924 |
| hsa-mir-591 | 3 | 0.92332 | 0.92337 | 0.999994 | 19702 | 0 | 3.1263 | 0.076677 | 0.17535 | 0.968566 | 4532 | 3 | 3.1263 |
| NonTargetingControlG | 2 | 0.92332 | 0.92341 | 0.999994 | 19701 | 0 | 4.7706 | 0.07668 | 0.13324 | 0.965913 | 4533 | 2 | 4.7706 |
| TMEM45B | 6 | 0.051269 | 0.14686 | 0.964852 | 3037 | 3 | -0.72706 | 0.076685 | 0.22793 | 0.97466 | 4534 | 3 | -0.72706 |
| SNAI2 | 4 | 0.75467 | 0.76368 | 0.999994 | 16906 | 1 | 3.5366 | 0.076708 | 0.19485 | 0.97466 | 4535 | 3 | 3.5366 |
| CAMTA1 | 6 | 0.61591 | 0.75178 | 0.999994 | 15160 | 2 | 2.7298 | 0.076732 | 0.22804 | 0.97466 | 4536 | 4 | 2.7298 |
| UBAC1 | 6 | 0.93398 | 0.93405 | 0.999994 | 19954 | 0 | 3.4581 | 0.07675 | 0.22808 | 0.97466 | 4537 | 4 | 3.4581 |
| DACH1 | 6 | 0.42164 | 0.61648 | 0.999994 | 12272 | 2 | 3.133 | 0.07675 | 0.22808 | 0.97466 | 4538 | 4 | 3.133 |
| SFTPC | 6 | 0.33181 | 0.54334 | 0.999994 | 10762 | 2 | 2.8212 | 0.07675 | 0.22808 | 0.97466 | 4539 | 3 | 2.8212 |
| CEBPD | 6 | 0.92219 | 0.92244 | 0.999994 | 19681 | 1 | 3.0598 | 0.07675 | 0.22808 | 0.97466 | 4540 | 4 | 3.0598 |
| ADAM28 | 6 | 0.095967 | 0.23942 | 0.987253 | 4814 | 2 | 3.3534 | 0.07675 | 0.22808 | 0.97466 | 4541 | 4 | 3.3534 |
| RSRC1 | 6 | 0.12362 | 0.29215 | 0.999632 | 5757 | 2 | 3.0812 | 0.07675 | 0.22808 | 0.97466 | 4542 | 4 | 3.0812 |
| SEMA4F | 5 | 0.19233 | 0.36427 | 0.999994 | 7557 | 1 | 1.7592 | 0.076754 | 0.20773 | 0.97466 | 4543 | 3 | 1.7592 |
| BATF | 6 | 0.81748 | 0.85563 | 0.999994 | 17738 | 1 | 2.6794 | 0.076755 | 0.22809 | 0.97466 | 4544 | 4 | 2.6794 |
| XIAP | 6 | 0.18716 | 0.37199 | 0.999994 | 7396 | 1 | 2.276 | 0.076778 | 0.22814 | 0.97466 | 4545 | 5 | 2.276 |
| TCP11L1 | 6 | 0.080885 | 0.20909 | 0.979587 | 4263 | 3 | -0.60013 | 0.076817 | 0.22824 | 0.97466 | 4546 | 3 | -0.60013 |
| KCNJ15 | 6 | 0.28481 | 0.48904 | 0.999994 | 9805 | 1 | 1.7929 | 0.076827 | 0.22826 | 0.97466 | 4547 | 5 | 1.7929 |
| NonTargetingControlG | 2 | 0.92315 | 0.92323 | 0.999994 | 19699 | 0 | 5.0385 | 0.076847 | 0.1335 | 0.965913 | 4548 | 2 | 5.0385 |
| MYO5B | 5 | 0.90801 | 0.90805 | 0.999994 | 19382 | 0 | 1.637 | 0.076849 | 0.20793 | 0.97466 | 4549 | 4 | 1.637 |
| RECQL | 6 | 0.81927 | 0.85647 | 0.999994 | 17769 | 1 | 2.3751 | 0.076854 | 0.22831 | 0.97466 | 4550 | 5 | 2.3751 |
| LMAN2L | 6 | 0.75511 | 0.83029 | 0.999994 | 16916 | 1 | 1.0838 | 0.07687 | 0.22834 | 0.97466 | 4551 | 3 | 1.0838 |
| NonTargetingControlG | 2 | 0.076119 | 0.13235 | 0.95318 | 4079 | 1 | -0.11404 | 0.07688 | 0.13354 | 0.965913 | 4552 | 1 | -0.11404 |
| TERF2IP | 6 | 0.94948 | 0.94945 | 0.999994 | 20322 | 0 | 1.2029 | 0.076882 | 0.22837 | 0.97466 | 4553 | 5 | 1.2029 |
| NonTargetingControlG | 2 | 0.22482 | 0.28016 | 0.999632 | 8394 | 1 | 2.5027 | 0.077052 | 0.13383 | 0.965913 | 4554 | 1 | 2.5027 |
| CARKD | 6 | 0.59354 | 0.7352 | 0.999994 | 14874 | 1 | 1.4652 | 0.077075 | 0.22882 | 0.97466 | 4555 | 5 | 1.4652 |
| PRDX6 | 6 | 0.28265 | 0.48649 | 0.999994 | 9751 | 3 | 2.3147 | 0.077117 | 0.22892 | 0.97466 | 4556 | 3 | 2.3147 |
| ANKRD66 | 6 | 0.93661 | 0.93662 | 0.999994 | 19996 | 0 | 3.4144 | 0.077178 | 0.22906 | 0.97466 | 4557 | 4 | 3.4144 |
| TANGO2 | 6 | 0.51632 | 0.67997 | 0.999994 | 13685 | 1 | 3.2691 | 0.077188 | 0.22908 | 0.97466 | 4558 | 4 | 3.2691 |
| ZNF324B | 6 | 0.97476 | 0.97466 | 0.999994 | 20951 | 0 | 1.4865 | 0.077213 | 0.22914 | 0.97466 | 4559 | 5 | 1.4865 |
| RNF111 | 6 | 0.53143 | 0.69056 | 0.999994 | 13923 | 2 | 2.1474 | 0.077214 | 0.22914 | 0.97466 | 4560 | 4 | 2.1474 |
| FAM175A | 4 | 0.59813 | 0.66431 | 0.999994 | 14927 | 1 | 0.53344 | 0.077216 | 0.1957 | 0.97466 | 4561 | 2 | 0.53344 |
| GPR21 | 4 | 0.20894 | 0.34989 | 0.999994 | 8000 | 1 | 0.50097 | 0.077216 | 0.1957 | 0.97466 | 4562 | 2 | 0.50097 |
| ACTN1 | 6 | 0.062962 | 0.172 | 0.967658 | 3554 | 4 | -1.5606 | 0.077231 | 0.22917 | 0.97466 | 4563 | 2 | -1.5606 |
| SMKR1 | 6 | 0.51496 | 0.67904 | 0.999994 | 13660 | 2 | 2.0307 | 0.077231 | 0.22917 | 0.97466 | 4564 | 3 | 2.0307 |
| H1FNT | 6 | 0.098261 | 0.24392 | 0.988236 | 4906 | 3 | -0.76594 | 0.077231 | 0.22917 | 0.97466 | 4565 | 3 | -0.76594 |
| WISP2 | 6 | 0.47159 | 0.64951 | 0.999994 | 13023 | 2 | 1.1424 | 0.077231 | 0.22917 | 0.97466 | 4566 | 3 | 1.1424 |
| DEFB105B | 2 | 0.92274 | 0.92282 | 0.999994 | 19693 | 0 | 3.7764 | 0.077255 | 0.13415 | 0.965913 | 4567 | 2 | 3.7764 |
| ITGB4 | 6 | 0.10671 | 0.26036 | 0.994943 | 5198 | 1 | 0.077001 | 0.077293 | 0.22932 | 0.97466 | 4568 | 2 | 0.077001 |
| NEDD9 | 6 | 0.63799 | 0.76842 | 0.999994 | 15486 | 1 | 2.1708 | 0.077295 | 0.22933 | 0.97466 | 4569 | 5 | 2.1708 |
| KCTD15 | 6 | 0.17432 | 0.35603 | 0.999994 | 7091 | 3 | -0.17432 | 0.077373 | 0.22949 | 0.97466 | 4570 | 3 | -0.17432 |
| CD5L | 5 | 0.92445 | 0.92467 | 0.999994 | 19730 | 0 | 2.3808 | 0.077381 | 0.20902 | 0.97466 | 4571 | 3 | 2.3808 |
| MARK4 | 5 | 0.95434 | 0.95449 | 0.999994 | 20440 | 0 | 1.8791 | 0.077417 | 0.2091 | 0.97466 | 4572 | 4 | 1.8791 |
| TMEM125 | 4 | 0.61466 | 0.67334 | 0.999994 | 15137 | 1 | 4.3943 | 0.077443 | 0.19608 | 0.97466 | 4573 | 3 | 4.3943 |
| XPOT | 6 | 0.93744 | 0.93745 | 0.999994 | 20016 | 0 | 1.378 | 0.077454 | 0.22968 | 0.97466 | 4574 | 5 | 1.378 |
| RAB28 | 6 | 0.97436 | 0.97425 | 0.999994 | 20936 | 0 | 1.9645 | 0.077461 | 0.2297 | 0.97466 | 4575 | 5 | 1.9645 |

| | | | | | | | | | | | | | |
|---|---|---|---|---|---|---|---|---|---|---|---|---|---|
| ZNF626 | 5 | 0.86372 | 0.86555 | 0.999994 | 18516 | 1 | 1.8035 | 0.077491 | 0.20925 | 0.97466 | 4576 | 4 | 1.8035 |
| NonTargetingControlG | 2 | 0.6233 | 0.62306 | 0.999994 | 15280 | 0 | 4.0469 | 0.077523 | 0.13458 | 0.965913 | 4577 | 1 | 4.0469 |
| NTNG2 | 6 | 0.58558 | 0.72932 | 0.999994 | 14752 | 2 | 2.918 | 0.07754 | 0.22988 | 0.97466 | 4578 | 3 | 2.918 |
| CNFN | 6 | 0.8426 | 0.86801 | 0.999994 | 18133 | 1 | 3.2753 | 0.07754 | 0.22988 | 0.97466 | 4579 | 4 | 3.2753 |
| SAMD5 | 6 | 0.62466 | 0.75835 | 0.999994 | 15298 | 1 | 2.5925 | 0.07754 | 0.22988 | 0.97466 | 4580 | 3 | 2.5925 |
| RGS13 | 6 | 0.41011 | 0.60898 | 0.999994 | 12111 | 1 | 3.5305 | 0.077548 | 0.2299 | 0.97466 | 4581 | 4 | 3.5305 |
| RANBP10 | 5 | 0.023296 | 0.072978 | 0.93914 | 1670 | 2 | 4.3222 | 0.077585 | 0.20944 | 0.97466 | 4582 | 3 | 4.3222 |
| RABL5 | 5 | 0.9437 | 0.94392 | 0.999994 | 20169 | 0 | 4.3222 | 0.077585 | 0.20944 | 0.97466 | 4583 | 3 | 4.3222 |
| TAPT1 | 5 | 0.57438 | 0.71341 | 0.999994 | 14610 | 1 | 4.3222 | 0.077585 | 0.20944 | 0.97466 | 4584 | 3 | 4.3222 |
| MORF4L2 | 6 | 0.84161 | 0.86747 | 0.999994 | 18115 | 1 | 0.88387 | 0.077621 | 0.23007 | 0.97466 | 4585 | 5 | 0.88387 |
| GNAI3 | 5 | 0.13622 | 0.28206 | 0.999632 | 6086 | 2 | -0.45437 | 0.077636 | 0.20956 | 0.97466 | 4586 | 2 | -0.45437 |
| C8orf58 | 4 | 0.80868 | 0.80908 | 0.999994 | 17614 | 1 | 3.2599 | 0.077694 | 0.19649 | 0.97466 | 4587 | 3 | 3.2599 |
| SFTPA1 | 6 | 0.36694 | 0.5813 | 0.999994 | 11456 | 2 | 0.012991 | 0.077743 | 0.23036 | 0.97466 | 4588 | 2 | 0.012991 |
| AGFG2 | 4 | 0.87689 | 0.87658 | 0.999994 | 18769 | 0 | 3.7077 | 0.077766 | 0.19662 | 0.97466 | 4589 | 3 | 3.7077 |
| NonTargetingControlG | 2 | 0.92223 | 0.92229 | 0.999994 | 19683 | 0 | 4.6716 | 0.07777 | 0.13497 | 0.965913 | 4590 | 2 | 4.6716 |
| CHST5 | 5 | 0.71939 | 0.77556 | 0.999994 | 16490 | 1 | -0.096245 | 0.077857 | 0.21001 | 0.97466 | 4591 | 2 | -0.096245 |
| AR | 6 | 0.35267 | 0.56702 | 0.999994 | 11210 | 3 | 1.2264 | 0.077897 | 0.23072 | 0.97466 | 4592 | 3 | 1.2264 |
| CLLU1OS | 6 | 0.6605 | 0.7857 | 0.999994 | 15785 | 1 | 2.1432 | 0.077897 | 0.23072 | 0.97466 | 4593 | 4 | 2.1432 |
| hsa-mir-4474 | 3 | 0.92202 | 0.92208 | 0.999994 | 19673 | 0 | 2.1801 | 0.077975 | 0.17799 | 0.971262 | 4594 | 3 | 2.1801 |
| NonTargetingControlG | 2 | 0.91445 | 0.91449 | 0.999994 | 19496 | 0 | 5.5946 | 0.077976 | 0.13528 | 0.965913 | 4595 | 2 | 5.5946 |
| MMAA | 6 | 0.087596 | 0.22268 | 0.983354 | 4497 | 4 | -1.1004 | 0.078007 | 0.23098 | 0.97466 | 4596 | 1 | -1.1004 |
| KNTC1 | 6 | 0.86468 | 0.88012 | 0.999994 | 18536 | 1 | 3.5221 | 0.078016 | 0.231 | 0.97466 | 4597 | 5 | 3.5221 |
| REL | 6 | 0.029528 | 0.097676 | 0.945253 | 1986 | 1 | 2.0758 | 0.078065 | 0.2311 | 0.97466 | 4598 | 5 | 2.0758 |
| ZNF257 | 5 | 0.78175 | 0.80845 | 0.999994 | 17236 | 1 | 2.219 | 0.07809 | 0.2105 | 0.97466 | 4599 | 3 | 2.219 |
| SUPT20HL2 | 5 | 0.19924 | 0.37413 | 0.999994 | 7743 | 1 | 0.31946 | 0.07809 | 0.2105 | 0.97466 | 4600 | 2 | 0.31946 |
| CBFA2T2 | 6 | 0.43312 | 0.62391 | 0.999994 | 12465 | 2 | 3.5991 | 0.078135 | 0.23126 | 0.97466 | 4601 | 4 | 3.5991 |
| HOXC8 | 6 | 0.88786 | 0.89481 | 0.999994 | 18996 | 1 | 3.4927 | 0.078135 | 0.23126 | 0.97466 | 4602 | 4 | 3.4927 |
| SOCS6 | 6 | 0.97051 | 0.97046 | 0.999994 | 20837 | 0 | 3.3447 | 0.078135 | 0.23126 | 0.97466 | 4603 | 4 | 3.3447 |
| CEACAM8 | 5 | 0.21659 | 0.3985 | 0.999994 | 8182 | 2 | 1.2213 | 0.078255 | 0.21085 | 0.97466 | 4604 | 3 | 1.2213 |
| RASGRP2 | 6 | 0.92333 | 0.92353 | 0.999994 | 19705 | 1 | 2.4539 | 0.078257 | 0.23155 | 0.97466 | 4605 | 4 | 2.4539 |
| UBE2S | 4 | 0.72675 | 0.74282 | 0.999994 | 16564 | 1 | 3.4619 | 0.078267 | 0.19746 | 0.97466 | 4606 | 2 | 3.4619 |
| hsa-mir-8076 | 2 | 0.59473 | 0.59465 | 0.999994 | 14886 | 0 | 3.938 | 0.078311 | 0.13585 | 0.965913 | 4607 | 1 | 3.938 |
| TTYH3 | 5 | 0.23765 | 0.42774 | 0.999994 | 8718 | 1 | 1.7663 | 0.078318 | 0.21098 | 0.97466 | 4608 | 4 | 1.7663 |
| ZBTB16 | 6 | 0.11057 | 0.26782 | 0.998561 | 5319 | 4 | -0.712 | 0.078324 | 0.23171 | 0.97466 | 4609 | 1 | -0.712 |
| LIPC | 6 | 0.9006 | 0.90415 | 0.999994 | 19230 | 1 | 2.1575 | 0.078354 | 0.23177 | 0.97466 | 4610 | 5 | 2.1575 |
| DEFB135 | 6 | 0.42284 | 0.61727 | 0.999994 | 12295 | 1 | 1.6982 | 0.078368 | 0.2318 | 0.97466 | 4611 | 5 | 1.6982 |
| SEPT11 | 5 | 0.75207 | 0.79193 | 0.999994 | 16880 | 1 | 4.314 | 0.078384 | 0.21111 | 0.97466 | 4612 | 3 | 4.314 |
| ARHGEF40 | 5 | 0.13962 | 0.28726 | 0.999632 | 6175 | 1 | 4.314 | 0.078384 | 0.21111 | 0.97466 | 4613 | 4 | 4.314 |
| GUCY2C | 6 | 0.20431 | 0.39313 | 0.999994 | 7886 | 2 | 3.241 | 0.078407 | 0.23189 | 0.97466 | 4614 | 4 | 3.241 |
| B3GALTL | 5 | 0.21874 | 0.4015 | 0.999994 | 8249 | 1 | 2.8654 | 0.078413 | 0.21116 | 0.97466 | 4615 | 4 | 2.8654 |
| ZBTB22 | 5 | 0.049046 | 0.13806 | 0.957317 | 2933 | 3 | -2.444 | 0.078428 | 0.21119 | 0.97466 | 4616 | 2 | -2.444 |
| TFAP2D | 5 | 0.9599 | 0.96004 | 0.999994 | 20581 | 0 | 3.8211 | 0.078428 | 0.21119 | 0.97466 | 4617 | 4 | 3.8211 |
| SPTBN4 | 5 | 0.907 | 0.90705 | 0.999994 | 19370 | 0 | 1.1413 | 0.078428 | 0.21119 | 0.97466 | 4618 | 3 | 1.1413 |
| hsa-mir-181c | 3 | 0.78507 | 0.78494 | 0.999994 | 17280 | 0 | 5.6986 | 0.078438 | 0.17889 | 0.97162 | 4619 | 2 | 5.6986 |
| MFSD10 | 6 | 0.71432 | 0.8165 | 0.999994 | 16443 | 1 | 2.7262 | 0.078518 | 0.23214 | 0.97466 | 4620 | 5 | 2.7262 |
| HIST1H4A | 5 | 0.45096 | 0.60604 | 0.999994 | 12724 | 1 | 3.9895 | 0.078527 | 0.21137 | 0.97466 | 4621 | 4 | 3.9895 |
| CKLF-CMTM1 | 6 | 0.16081 | 0.33926 | 0.999994 | 6760 | 3 | 0.17078 | 0.078562 | 0.23225 | 0.97466 | 4622 | 3 | 0.17078 |
| PSRC1 | 6 | 0.16392 | 0.34318 | 0.999994 | 6839 | 1 | 1.6775 | 0.078562 | 0.23225 | 0.97466 | 4623 | 4 | 1.6775 |
| hsa-mir-7702 | 4 | 0.77206 | 0.77745 | 0.999994 | 17112 | 1 | 3.4507 | 0.078647 | 0.19811 | 0.97466 | 4624 | 3 | 3.4507 |
| C20orf24 | 6 | 0.92739 | 0.92748 | 0.999994 | 19800 | 1 | 2.6775 | 0.078682 | 0.23253 | 0.97466 | 4625 | 4 | 2.6775 |
| PYHIN1 | 6 | 0.27306 | 0.47532 | 0.999994 | 9542 | 2 | 1.2761 | 0.078694 | 0.23256 | 0.97466 | 4626 | 3 | 1.2761 |
| CCDC117 | 6 | 0.68092 | 0.80174 | 0.999994 | 16102 | 2 | 2.8382 | 0.078721 | 0.23263 | 0.97466 | 4627 | 3 | 2.8382 |

| | | | | | | | | | | | | |
|---|---|---|---|---|---|---|---|---|---|---|---|---|
| TSHR | 6 | 0.95668 | 0.95676 | 0.999994 | 20508 | 0 | 1.6568 | 0.078777 | 0.23276 | 0.97466 | 4628 | 5 | 1.6568 |
| EXOC6 | 5 | 0.84964 | 0.85399 | 0.999994 | 18252 | 1 | 2.605 | 0.078798 | 0.21194 | 0.97466 | 4629 | 4 | 2.605 |
| OR2H1 | 6 | 0.15376 | 0.3304 | 0.999994 | 6561 | 2 | 0.83906 | 0.0788 | 0.23282 | 0.97466 | 4630 | 3 | 0.83906 |
| C8orf33 | 5 | 0.96086 | 0.961 | 0.999994 | 20599 | 0 | 2.1238 | 0.078839 | 0.21202 | 0.97466 | 4631 | 4 | 2.1238 |
| NAA20 | 5 | 0.41051 | 0.57231 | 0.999994 | 12120 | 1 | 1.83 | 0.078846 | 0.21203 | 0.97466 | 4632 | 4 | 1.83 |
| KRT75 | 6 | 0.1057 | 0.25841 | 0.994943 | 5162 | 2 | 0.50376 | 0.07885 | 0.23293 | 0.97466 | 4633 | 3 | 0.50376 |
| CCDC91 | 6 | 0.022655 | 0.080483 | 0.93914 | 1628 | 4 | -1.6389 | 0.07885 | 0.23293 | 0.97466 | 4634 | 2 | -1.6389 |
| NonTargetingControlG | 2 | 0.4435 | 0.45361 | 0.999994 | 12630 | 1 | 3.464 | 0.078854 | 0.13672 | 0.965913 | 4635 | 1 | 3.464 |
| ZNF646 | 5 | 0.41955 | 0.57971 | 0.999994 | 12244 | 1 | 3.6013 | 0.078883 | 0.21211 | 0.97466 | 4636 | 3 | 3.6013 |
| CCL15 | 6 | 0.36066 | 0.57605 | 0.999994 | 11350 | 1 | 0.93242 | 0.078932 | 0.23312 | 0.97466 | 4637 | 4 | 0.93242 |
| TMPRSS11B | 4 | 0.061105 | 0.1429 | 0.959553 | 3472 | 3 | -1.6413 | 0.078937 | 0.19858 | 0.97466 | 4638 | 1 | -1.6413 |
| C17orf107 | 6 | 0.25279 | 0.45125 | 0.999994 | 9076 | 2 | 0.71821 | 0.079038 | 0.23335 | 0.97466 | 4639 | 3 | 0.71821 |
| CCR9 | 5 | 0.21403 | 0.39499 | 0.999994 | 8115 | 2 | -0.024963 | 0.079046 | 0.21245 | 0.97466 | 4640 | 2 | -0.024963 |
| HSPA4L | 6 | 0.2283 | 0.42201 | 0.999994 | 8485 | 1 | 2.1008 | 0.0791 | 0.23348 | 0.97466 | 4641 | 5 | 2.1008 |
| OR2F2 | 5 | 0.091108 | 0.21239 | 0.979587 | 4643 | 3 | -1.4598 | 0.079204 | 0.21277 | 0.97466 | 4642 | 2 | -1.4598 |
| CD164 | 6 | 0.082503 | 0.21246 | 0.979587 | 4319 | 3 | -0.36139 | 0.079209 | 0.23374 | 0.97466 | 4643 | 3 | -0.36139 |
| FGF22 | 6 | 0.93279 | 0.93289 | 0.999994 | 19920 | 0 | 1.5152 | 0.079209 | 0.23374 | 0.97466 | 4644 | 4 | 1.5152 |
| ANKRD16 | 6 | 0.53612 | 0.6938 | 0.999994 | 13999 | 2 | -0.38377 | 0.079209 | 0.23374 | 0.97466 | 4645 | 2 | -0.38377 |
| ARRDC1 | 3 | 0.35616 | 0.45918 | 0.999994 | 11270 | 1 | 5.6294 | 0.07923 | 0.18048 | 0.97162 | 4646 | 2 | 5.6294 |
| RSPO1 | 6 | 0.20136 | 0.38952 | 0.999994 | 7800 | 3 | 0.80433 | 0.079249 | 0.23383 | 0.97466 | 4647 | 3 | 0.80433 |
| ZNF347 | 6 | 0.67611 | 0.79797 | 0.999994 | 16023 | 2 | 3.0069 | 0.079249 | 0.23383 | 0.97466 | 4648 | 4 | 3.0069 |
| ENTPD8 | 6 | 0.344 | 0.55725 | 0.999994 | 11028 | 2 | -0.12314 | 0.079249 | 0.23383 | 0.97466 | 4649 | 2 | -0.12314 |
| ELL2 | 6 | 0.82954 | 0.86131 | 0.999994 | 17920 | 1 | 2.402 | 0.079254 | 0.23385 | 0.97466 | 4650 | 5 | 2.402 |
| OR1F1 | 6 | 0.41591 | 0.61271 | 0.999994 | 12190 | 2 | 3.0823 | 0.079261 | 0.23386 | 0.97466 | 4651 | 4 | 3.0823 |
| DCLRE1A | 6 | 0.96936 | 0.96936 | 0.999994 | 20813 | 0 | 1.6391 | 0.079304 | 0.23395 | 0.97466 | 4652 | 5 | 1.6391 |
| OR8I2 | 5 | 0.72265 | 0.77714 | 0.999994 | 16522 | 1 | 2.0377 | 0.079328 | 0.21302 | 0.97466 | 4653 | 4 | 2.0377 |
| NonTargetingControlG | 1 | 0.92067 | 0.92103 | 0.999994 | 19644 | 0 | 7.5499 | 0.079332 | 0.079367 | 0.938203 | 4654 | 1 | 7.5499 |
| OPRK1 | 6 | 0.5649 | 0.71431 | 0.999994 | 14467 | 2 | 3.159 | 0.07934 | 0.23403 | 0.97466 | 4655 | 4 | 3.159 |
| UTP23 | 6 | 0.65819 | 0.7839 | 0.999994 | 15755 | 1 | 3.002 | 0.07934 | 0.23403 | 0.97466 | 4656 | 4 | 3.002 |
| CA6 | 6 | 0.93239 | 0.93251 | 0.999994 | 19911 | 0 | 3.3313 | 0.07934 | 0.23403 | 0.97466 | 4657 | 4 | 3.3313 |
| CACNA1A | 6 | 0.073919 | 0.19491 | 0.975797 | 3990 | 2 | 2.9341 | 0.07934 | 0.23403 | 0.97466 | 4658 | 3 | 2.9341 |
| FTH1P18 | 5 | 0.87668 | 0.87712 | 0.999994 | 18765 | 1 | 2.9237 | 0.079369 | 0.21312 | 0.97466 | 4659 | 4 | 2.9237 |
| SFI1 | 5 | 0.92643 | 0.92665 | 0.999994 | 19778 | 0 | 2.1999 | 0.079376 | 0.21313 | 0.97466 | 4660 | 4 | 2.1999 |
| SP140 | 4 | 0.0050259 | 0.017719 | 0.795727 | 524 | 3 | -4.8014 | 0.079415 | 0.19938 | 0.97466 | 4661 | 1 | -4.8014 |
| PDHA1 | 6 | 0.63765 | 0.76814 | 0.999994 | 15482 | 1 | 1.7772 | 0.079445 | 0.23427 | 0.97466 | 4662 | 5 | 1.7772 |
| BLOC1S2 | 6 | 0.98529 | 0.98525 | 0.999994 | 21234 | 0 | 2.2853 | 0.079452 | 0.23429 | 0.97466 | 4663 | 5 | 2.2853 |
| ORM1 | 5 | 0.65235 | 0.74606 | 0.999994 | 15692 | 1 | 4.0982 | 0.079455 | 0.21329 | 0.97466 | 4664 | 4 | 4.0982 |
| SSBP4 | 4 | 0.87504 | 0.87469 | 0.999994 | 18737 | 0 | 3.5723 | 0.079463 | 0.19947 | 0.97466 | 4665 | 2 | 3.5723 |
| HP | 6 | 0.68861 | 0.80773 | 0.999994 | 16198 | 2 | 2.6534 | 0.079511 | 0.23443 | 0.97466 | 4666 | 4 | 2.6534 |
| EPS8L1 | 6 | 0.65721 | 0.78314 | 0.999994 | 15745 | 2 | 1.1745 | 0.079513 | 0.23444 | 0.97466 | 4667 | 3 | 1.1745 |
| MYEOV | 6 | 0.78234 | 0.84062 | 0.999994 | 17243 | 1 | 1.5775 | 0.07955 | 0.23452 | 0.97466 | 4668 | 5 | 1.5775 |
| ZNF484 | 6 | 0.50083 | 0.66925 | 0.999994 | 13435 | 1 | 1.5606 | 0.079553 | 0.23453 | 0.97466 | 4669 | 3 | 1.5606 |
| DPPA4 | 6 | 0.12702 | 0.29651 | 0.999994 | 5849 | 2 | 2.5613 | 0.079553 | 0.23453 | 0.97466 | 4670 | 4 | 2.5613 |
| TP53RK | 4 | 0.9322 | 0.93194 | 0.999994 | 19908 | 0 | 3.5498 | 0.079637 | 0.19975 | 0.97466 | 4671 | 3 | 3.5498 |
| TIMP3 | 5 | 0.29226 | 0.47476 | 0.999994 | 9958 | 2 | 0.21219 | 0.079645 | 0.21369 | 0.97466 | 4672 | 2 | 0.21219 |
| FAM71D | 6 | 0.9208 | 0.92115 | 0.999994 | 19648 | 1 | 3.2619 | 0.079687 | 0.23483 | 0.97466 | 4673 | 5 | 3.2619 |
| GPR141 | 6 | 0.073585 | 0.1942 | 0.975797 | 3974 | 3 | -0.51678 | 0.07975 | 0.23497 | 0.97466 | 4674 | 3 | -0.51678 |
| AKIP1 | 5 | 0.90605 | 0.90614 | 0.999994 | 19351 | 0 | 4.2513 | 0.079783 | 0.21398 | 0.97466 | 4675 | 4 | 4.2513 |
| CTNS | 6 | 0.66559 | 0.78968 | 0.999994 | 15865 | 1 | 1.5006 | 0.079812 | 0.23513 | 0.97466 | 4676 | 5 | 1.5006 |
| RAP1GAP2 | 5 | 0.2382 | 0.42848 | 0.999994 | 8731 | 3 | -0.61416 | 0.079822 | 0.21406 | 0.97466 | 4677 | 2 | -0.61416 |
| ICOSLG | 5 | 0.39076 | 0.55576 | 0.999994 | 11812 | 2 | -0.5129 | 0.079822 | 0.21406 | 0.97466 | 4678 | 2 | -0.5129 |
| CXorf57 | 6 | 0.59541 | 0.73659 | 0.999994 | 14896 | 1 | 0.62584 | 0.07984 | 0.23519 | 0.97466 | 4679 | 3 | 0.62584 |

| | | | | | | | | | | | | | |
|---|---|---|---|---|---|---|---|---|---|---|---|---|---|
| NECAB1 | 6 | 0.57687 | 0.7229 | 0.999994 | 14642 | 2 | 2.619 | 0.079864 | 0.23524 | 0.97466 | 4680 | 3 | 2.619 |
| BTBD7 | 6 | 0.96549 | 0.96554 | 0.999994 | 20728 | 0 | 2.9061 | 0.079875 | 0.23526 | 0.97466 | 4681 | 5 | 2.9061 |
| CCDC89 | 6 | 0.35167 | 0.56586 | 0.999994 | 11190 | 2 | -0.027128 | 0.079962 | 0.23546 | 0.97466 | 4682 | 1 | -0.027128 |
| ARID5B | 6 | 0.54742 | 0.70179 | 0.999994 | 14172 | 1 | 0.0098261 | 0.079962 | 0.23546 | 0.97466 | 4683 | 2 | 0.0098261 |
| MCC | 6 | 0.33939 | 0.55195 | 0.999994 | 10920 | 2 | 2.5866 | 0.079964 | 0.23547 | 0.97466 | 4684 | 4 | 2.5866 |
| CCL18 | 5 | 0.96439 | 0.96453 | 0.999994 | 20711 | 0 | 2.8613 | 0.080004 | 0.21443 | 0.97466 | 4685 | 4 | 2.8613 |
| CCDC155 | 6 | 0.95992 | 0.96 | 0.999994 | 20582 | 0 | 1.5529 | 0.080123 | 0.23583 | 0.97466 | 4686 | 5 | 1.5529 |
| UBXN2A | 5 | 0.64323 | 0.74244 | 0.999994 | 15565 | 1 | 2.4812 | 0.080157 | 0.21474 | 0.97466 | 4687 | 3 | 2.4812 |
| hsa-mir-26a-1 | 3 | 0.9198 | 0.91983 | 0.999994 | 19625 | 0 | 1.7113 | 0.080201 | 0.18242 | 0.97162 | 4688 | 3 | 1.7113 |
| UGT1A10 | 3 | 0.91978 | 0.91982 | 0.999994 | 19624 | 0 | 1.1307 | 0.080217 | 0.18245 | 0.97162 | 4689 | 3 | 1.1307 |
| HIVEP3 | 6 | 0.55202 | 0.70511 | 0.999994 | 14243 | 2 | 1.8156 | 0.080234 | 0.2361 | 0.97466 | 4690 | 3 | 1.8156 |
| PRAME | 6 | 0.30475 | 0.51226 | 0.999994 | 10224 | 1 | 2.3749 | 0.080234 | 0.2361 | 0.97466 | 4691 | 3 | 2.3749 |
| CDHR4 | 6 | 0.48955 | 0.66163 | 0.999994 | 13272 | 2 | 0.5654 | 0.080234 | 0.2361 | 0.97466 | 4692 | 3 | 0.5654 |
| LCOR | 6 | 0.92727 | 0.92733 | 0.999994 | 19797 | 1 | 2.5187 | 0.080248 | 0.23614 | 0.97466 | 4693 | 4 | 2.5187 |
| FBXO38 | 6 | 0.44514 | 0.63185 | 0.999994 | 12649 | 1 | 2.4093 | 0.080251 | 0.23614 | 0.97466 | 4694 | 5 | 2.4093 |
| hsa-mir-614 | 3 | 0.91974 | 0.91976 | 0.999994 | 19623 | 0 | 1.5245 | 0.080264 | 0.18254 | 0.97162 | 4695 | 3 | 1.5245 |
| ZNF512B | 4 | 0.89014 | 0.88985 | 0.999994 | 19040 | 0 | 3.5722 | 0.080274 | 0.20072 | 0.97466 | 4696 | 2 | 3.5722 |
| TRPM6 | 5 | 0.23374 | 0.4224 | 0.999994 | 8612 | 3 | -0.61399 | 0.080285 | 0.21501 | 0.97466 | 4697 | 2 | -0.61399 |
| DNAJA1 | 6 | 0.98568 | 0.98565 | 0.999994 | 21242 | 0 | 2.6839 | 0.080289 | 0.23623 | 0.97466 | 4698 | 5 | 2.6839 |
| hsa-mir-4309 | 3 | 0.24611 | 0.37011 | 0.999994 | 8926 | 1 | 5.6402 | 0.080336 | 0.1827 | 0.97162 | 4699 | 2 | 5.6402 |
| PRKG1 | 5 | 0.95928 | 0.95941 | 0.999994 | 20566 | 0 | 0.85502 | 0.080373 | 0.21519 | 0.97466 | 4700 | 3 | 0.85502 |
| POU3F4 | 6 | 0.80466 | 0.84989 | 0.999994 | 17548 | 1 | 2.3211 | 0.080425 | 0.23655 | 0.97466 | 4701 | 5 | 2.3211 |
| RBM42 | 5 | 0.36382 | 0.53325 | 0.999994 | 11398 | 2 | 4.2459 | 0.080438 | 0.21534 | 0.97466 | 4702 | 3 | 4.2459 |
| QRICH2 | 5 | 0.94524 | 0.94542 | 0.999994 | 20210 | 0 | 4.2459 | 0.080438 | 0.21534 | 0.97466 | 4703 | 3 | 4.2459 |
| TSEN2 | 5 | 0.43234 | 0.59037 | 0.999994 | 12451 | 1 | 4.2459 | 0.080438 | 0.21534 | 0.97466 | 4704 | 3 | 4.2459 |
| DNAJC4 | 5 | 0.17588 | 0.34067 | 0.999994 | 7132 | 1 | 4.2459 | 0.080438 | 0.21534 | 0.97466 | 4705 | 4 | 4.2459 |
| KIF26B | 5 | 0.95615 | 0.95632 | 0.999994 | 20490 | 0 | 4.2459 | 0.080438 | 0.21534 | 0.97466 | 4706 | 3 | 4.2459 |
| RBMXL3 | 5 | 0.74222 | 0.78677 | 0.999994 | 16766 | 1 | 4.2459 | 0.080438 | 0.21534 | 0.97466 | 4707 | 3 | 4.2459 |
| MTRR | 6 | 0.05755 | 0.16046 | 0.967126 | 3317 | 4 | -1.4936 | 0.080489 | 0.2367 | 0.97466 | 4708 | 2 | -1.4936 |
| TMEM66 | 5 | 0.95015 | 0.95033 | 0.999994 | 20339 | 0 | 3.9876 | 0.080495 | 0.21545 | 0.97466 | 4709 | 4 | 3.9876 |
| LUM | 5 | 0.92788 | 0.9281 | 0.999994 | 19812 | 0 | 1.4572 | 0.080505 | 0.21547 | 0.97466 | 4710 | 4 | 1.4572 |
| ELP4 | 6 | 0.9736 | 0.97351 | 0.999994 | 20911 | 0 | 3.2714 | 0.080535 | 0.23682 | 0.97466 | 4711 | 5 | 3.2714 |
| BEAN1 | 6 | 0.83387 | 0.86346 | 0.999994 | 17984 | 1 | 1.615 | 0.080585 | 0.23692 | 0.97466 | 4712 | 5 | 1.615 |
| MUC22 | 5 | 0.14143 | 0.28998 | 0.999632 | 6227 | 3 | -1.2615 | 0.080593 | 0.21565 | 0.97466 | 4713 | 1 | -1.2615 |
| FOXJ2 | 6 | 0.2064 | 0.39568 | 0.999994 | 7943 | 3 | 0.81295 | 0.080595 | 0.23693 | 0.97466 | 4714 | 3 | 0.81295 |
| hsa-mir-2110 | 3 | 0.91939 | 0.91942 | 0.999994 | 19617 | 0 | 5.1892 | 0.080612 | 0.18323 | 0.97162 | 4715 | 3 | 5.1892 |
| RTN2 | 4 | 0.82902 | 0.82856 | 0.999994 | 17911 | 0 | 3.7972 | 0.080683 | 0.2014 | 0.97466 | 4716 | 3 | 3.7972 |
| ASB17 | 6 | 0.021666 | 0.07734 | 0.93914 | 1556 | 3 | -0.40672 | 0.080695 | 0.23718 | 0.97466 | 4717 | 3 | -0.40672 |
| hsa-mir-1184-1 | 4 | 0.42585 | 0.57222 | 0.999994 | 12348 | 2 | 0.2207 | 0.080708 | 0.20144 | 0.97466 | 4718 | 2 | 0.2207 |
| CENPBD1 | 6 | 0.51456 | 0.67876 | 0.999994 | 13655 | 2 | 3.4246 | 0.080843 | 0.23751 | 0.97466 | 4719 | 4 | 3.4246 |
| CDH17 | 6 | 0.21018 | 0.40031 | 0.999994 | 8027 | 2 | 3.3721 | 0.080843 | 0.23751 | 0.97466 | 4720 | 4 | 3.3721 |
| GNG13 | 6 | 0.85541 | 0.87486 | 0.999994 | 18356 | 1 | 2.9276 | 0.080843 | 0.23751 | 0.97466 | 4721 | 3 | 2.9276 |
| hsa-mir-3907 | 2 | 0.9191 | 0.91918 | 0.999994 | 19609 | 0 | 4.5808 | 0.080895 | 0.13997 | 0.965913 | 4722 | 2 | 4.5808 |
| ACOT8 | 6 | 0.29863 | 0.5051 | 0.999994 | 10102 | 3 | -0.19256 | 0.08099 | 0.23786 | 0.97466 | 4723 | 2 | -0.19256 |
| DHX34 | 6 | 0.16379 | 0.34301 | 0.999994 | 6834 | 2 | 1.1834 | 0.081 | 0.23789 | 0.97466 | 4724 | 4 | 1.1834 |
| DEFB104A | 6 | 0.70096 | 0.8123 | 0.999994 | 16315 | 1 | 1.4209 | 0.081 | 0.23789 | 0.97466 | 4725 | 3 | 1.4209 |
| MCCD1 | 6 | 0.81336 | 0.85376 | 0.999994 | 17681 | 1 | 1.0169 | 0.081 | 0.23789 | 0.97466 | 4726 | 4 | 1.0169 |
| hsa-mir-122 | 4 | 0.88282 | 0.88245 | 0.999994 | 18889 | 0 | 2.7742 | 0.081018 | 0.20196 | 0.97466 | 4727 | 3 | 2.7742 |
| CDY1 | 6 | 0.76896 | 0.83545 | 0.999994 | 17073 | 1 | 1.5836 | 0.081041 | 0.23799 | 0.97466 | 4728 | 5 | 1.5836 |
| hsa-mir-106a | 4 | 0.68148 | 0.71254 | 0.999994 | 16110 | 1 | 3.6978 | 0.081066 | 0.20204 | 0.97466 | 4729 | 3 | 3.6978 |
| TRAF3IP2 | 5 | 0.044468 | 0.12699 | 0.951052 | 2732 | 2 | 1.3791 | 0.08115 | 0.21682 | 0.97466 | 4730 | 3 | 1.3791 |
| NonTargetingControlG | 2 | 0.91883 | 0.91891 | 0.999994 | 19605 | 0 | 5.4242 | 0.081169 | 0.14044 | 0.965913 | 4731 | 2 | 5.4242 |

| | | | | | | | | | | | | | |
|---|---|---|---|---|---|---|---|---|---|---|---|---|---|
| IGSF3 | 6 | 0.24477 | 0.44168 | 0.999994 | 8896 | 3 | 0.46096 | 0.081227 | 0.23845 | 0.97466 | 4732 | 3 | 0.46096 |
| SCNN1D | 5 | 0.55293 | 0.69435 | 0.999994 | 14256 | 1 | 1.3401 | 0.081243 | 0.21703 | 0.97466 | 4733 | 4 | 1.3401 |
| SRGAP2D | 6 | 0.67725 | 0.79884 | 0.999994 | 16049 | 1 | 1.3027 | 0.081346 | 0.23874 | 0.97466 | 4734 | 3 | 1.3027 |
| DCUN1D5 | 6 | 0.59962 | 0.73965 | 0.999994 | 14948 | 2 | 2.4562 | 0.081415 | 0.23889 | 0.97466 | 4735 | 4 | 2.4562 |
| STIL | 6 | 0.15953 | 0.33763 | 0.999994 | 6720 | 2 | 3.0848 | 0.08144 | 0.23895 | 0.97466 | 4736 | 4 | 3.0848 |
| FGD6 | 6 | 0.92618 | 0.92625 | 0.999994 | 19771 | 1 | 2.9543 | 0.08144 | 0.23895 | 0.97466 | 4737 | 4 | 2.9543 |
| CNEP1R1 | 6 | 0.74 | 0.82499 | 0.999994 | 16740 | 1 | 2.3709 | 0.081604 | 0.23933 | 0.97466 | 4738 | 5 | 2.3709 |
| ZFP1 | 6 | 0.77914 | 0.83933 | 0.999994 | 17200 | 1 | 1.6534 | 0.081636 | 0.23939 | 0.97466 | 4739 | 5 | 1.6534 |
| ZC3H4 | 5 | 0.4377 | 0.59487 | 0.999994 | 12536 | 1 | 4.3412 | 0.081638 | 0.21785 | 0.97466 | 4740 | 3 | 4.3412 |
| IL20RB | 6 | 0.097924 | 0.24326 | 0.988088 | 4891 | 3 | 1.2973 | 0.081681 | 0.2395 | 0.97466 | 4741 | 3 | 1.2973 |
| APOBEC3D | 6 | 0.79881 | 0.84736 | 0.999994 | 17461 | 1 | 2.3993 | 0.081694 | 0.23954 | 0.97466 | 4742 | 4 | 2.3993 |
| GABARAP | 5 | 0.041213 | 0.11893 | 0.951052 | 2572 | 1 | 4.0698 | 0.081734 | 0.21806 | 0.97466 | 4743 | 4 | 4.0698 |
| EPHA7 | 5 | 0.41949 | 0.57966 | 0.999994 | 12243 | 2 | 2.7432 | 0.081739 | 0.21806 | 0.97466 | 4744 | 3 | 2.7432 |
| FGB | 4 | 0.14716 | 0.28624 | 0.999632 | 6380 | 1 | 3.4872 | 0.081762 | 0.2032 | 0.97466 | 4745 | 2 | 3.4872 |
| GLUD2 | 6 | 0.21932 | 0.41127 | 0.999994 | 8272 | 2 | 0.72941 | 0.081833 | 0.23989 | 0.97466 | 4746 | 3 | 0.72941 |
| NonTargetingControlG | 2 | 0.62733 | 0.62714 | 0.999994 | 15341 | 0 | 4.0181 | 0.081837 | 0.1415 | 0.965913 | 4747 | 1 | 4.0181 |
| C11orf80 | 5 | 0.31189 | 0.49069 | 0.999994 | 10355 | 1 | 1.64 | 0.081849 | 0.21829 | 0.97466 | 4748 | 3 | 1.64 |
| DBT | 6 | 0.82003 | 0.85685 | 0.999994 | 17778 | 1 | 1.3212 | 0.081881 | 0.24 | 0.97466 | 4749 | 5 | 1.3212 |
| ZDHHC19 | 5 | 0.91503 | 0.91512 | 0.999994 | 19514 | 0 | 2.0007 | 0.081902 | 0.21841 | 0.97466 | 4750 | 4 | 2.0007 |
| TSSK3 | 4 | 0.47496 | 0.60529 | 0.999994 | 13061 | 1 | 3.0425 | 0.081937 | 0.2035 | 0.97466 | 4751 | 3 | 3.0425 |
| NEBL | 5 | 0.8168 | 0.83046 | 0.999994 | 17725 | 1 | 2.2805 | 0.081951 | 0.21851 | 0.97466 | 4752 | 4 | 2.2805 |
| SLC1A3 | 4 | 0.92333 | 0.92315 | 0.999994 | 19703 | 0 | 3.5775 | 0.081959 | 0.20353 | 0.97466 | 4753 | 3 | 3.5775 |
| EVA1B | 6 | 0.67923 | 0.80041 | 0.999994 | 16078 | 1 | 2.4443 | 0.082033 | 0.24034 | 0.97466 | 4754 | 4 | 2.4443 |
| BCAT1 | 6 | 0.33068 | 0.54204 | 0.999994 | 10734 | 3 | -0.68991 | 0.082033 | 0.24034 | 0.97466 | 4755 | 2 | -0.68991 |
| CLNS1A | 5 | 0.95515 | 0.9553 | 0.999994 | 20455 | 0 | 1.0938 | 0.082091 | 0.21879 | 0.97466 | 4756 | 4 | 1.0938 |
| PHC1 | 5 | 0.42896 | 0.58755 | 0.999994 | 12396 | 2 | -0.57417 | 0.082091 | 0.21879 | 0.97466 | 4757 | 2 | -0.57417 |
| HAO1 | 5 | 0.058387 | 0.15938 | 0.967126 | 3360 | 1 | 1.6785 | 0.082104 | 0.2188 | 0.97466 | 4758 | 4 | 1.6785 |
| RFPL1 | 6 | 0.60263 | 0.7419 | 0.999994 | 14978 | 2 | 3.4121 | 0.082146 | 0.24062 | 0.97466 | 4759 | 4 | 3.4121 |
| LAMTOR5 | 5 | 0.050905 | 0.14258 | 0.959553 | 3013 | 1 | 4.0742 | 0.08216 | 0.21892 | 0.97466 | 4760 | 3 | 4.0742 |
| NKAIN1 | 5 | 0.72186 | 0.77674 | 0.999994 | 16515 | 1 | 4.1858 | 0.08216 | 0.21892 | 0.97466 | 4761 | 3 | 4.1858 |
| EMC9 | 5 | 0.32679 | 0.5029 | 0.999994 | 10659 | 2 | 4.1858 | 0.08216 | 0.21892 | 0.97466 | 4762 | 3 | 4.1858 |
| PPP1R14A | 6 | 0.98005 | 0.97996 | 0.999994 | 21093 | 0 | 2.8521 | 0.082162 | 0.24067 | 0.97466 | 4763 | 5 | 2.8521 |
| NOXRED1 | 6 | 0.013277 | 0.050025 | 0.909101 | 1081 | 1 | 2.9441 | 0.082165 | 0.24068 | 0.97466 | 4764 | 3 | 2.9441 |
| KRT33A | 6 | 0.60554 | 0.74407 | 0.999994 | 15022 | 2 | 2.622 | 0.082165 | 0.24068 | 0.97466 | 4765 | 3 | 2.622 |
| TERF1 | 6 | 0.93303 | 0.93312 | 0.999994 | 19926 | 0 | 2.6807 | 0.082184 | 0.24072 | 0.97466 | 4766 | 5 | 2.6807 |
| TLCD1 | 5 | 0.030545 | 0.091887 | 0.943968 | 2031 | 1 | 2.7351 | 0.082184 | 0.21897 | 0.97466 | 4767 | 4 | 2.7351 |
| KPNA5 | 6 | 0.24334 | 0.44 | 0.999994 | 8871 | 2 | 2.4824 | 0.082221 | 0.2408 | 0.97466 | 4768 | 4 | 2.4824 |
| DISP1 | 6 | 0.89988 | 0.90357 | 0.999994 | 19219 | 1 | 2.5764 | 0.082255 | 0.24089 | 0.97466 | 4769 | 5 | 2.5764 |
| FCAR | 5 | 0.95198 | 0.95213 | 0.999994 | 20382 | 0 | 2.0155 | 0.082271 | 0.21915 | 0.97466 | 4770 | 4 | 2.0155 |
| DAPP1 | 5 | 0.74281 | 0.78708 | 0.999994 | 16776 | 1 | -0.053576 | 0.082289 | 0.21918 | 0.97466 | 4771 | 2 | -0.053576 |
| SPP2 | 5 | 0.21978 | 0.40297 | 0.999994 | 8286 | 2 | 2.4443 | 0.082292 | 0.21919 | 0.97466 | 4772 | 3 | 2.4443 |
| ASZ1 | 5 | 0.40892 | 0.57097 | 0.999994 | 12092 | 2 | -0.060602 | 0.082292 | 0.21919 | 0.97466 | 4773 | 2 | -0.060602 |
| LRP5 | 5 | 0.69864 | 0.76588 | 0.999994 | 16290 | 1 | 2.0879 | 0.082292 | 0.21919 | 0.97466 | 4774 | 3 | 2.0879 |
| ZNF235 | 4 | 0.93223 | 0.93198 | 0.999994 | 19909 | 0 | 2.0081 | 0.082301 | 0.20408 | 0.97466 | 4775 | 3 | 2.0081 |
| ADAL | 4 | 0.043638 | 0.10903 | 0.951052 | 2697 | 1 | 2.1996 | 0.082301 | 0.20408 | 0.97466 | 4776 | 3 | 2.1996 |
| POU3F2 | 6 | 0.48441 | 0.65816 | 0.999994 | 13206 | 1 | 2.6055 | 0.082328 | 0.24106 | 0.97466 | 4777 | 5 | 2.6055 |
| KPNA1 | 6 | 0.098876 | 0.24517 | 0.98979 | 4927 | 1 | 3.0623 | 0.082381 | 0.24118 | 0.97466 | 4778 | 4 | 3.0623 |
| SPATA4 | 6 | 0.29417 | 0.4999 | 0.999994 | 10001 | 3 | 0.45408 | 0.082381 | 0.24118 | 0.97466 | 4779 | 3 | 0.45408 |
| RORB | 5 | 0.92109 | 0.92128 | 0.999994 | 19654 | 0 | 1.2857 | 0.082421 | 0.21946 | 0.97466 | 4780 | 3 | 1.2857 |
| NonTargetingControlG | 2 | 0.91756 | 0.91764 | 0.999994 | 19579 | 0 | 4.7401 | 0.082439 | 0.14249 | 0.965913 | 4781 | 2 | 4.7401 |
| EPN1 | 6 | 0.51709 | 0.68052 | 0.999994 | 13693 | 1 | 1.4919 | 0.082494 | 0.24144 | 0.97466 | 4782 | 5 | 1.4919 |
| METTL1 | 5 | 0.91745 | 0.91753 | 0.999994 | 19577 | 0 | 1.8591 | 0.082564 | 0.21975 | 0.97466 | 4783 | 4 | 1.8591 |

| | | | | | | | | | | | | | |
|---|---|---|---|---|---|---|---|---|---|---|---|---|---|
| PKMYT1 | 3 | 0.26766 | 0.39676 | 0.999994 | 9415 | 1 | 5.5792 | 0.082581 | 0.18718 | 0.97466 | 4784 | 2 | 5.5792 |
| OR8B4 | 6 | 0.85177 | 0.87285 | 0.999994 | 18291 | 1 | 2.8699 | 0.082581 | 0.24165 | 0.97466 | 4785 | 5 | 2.8699 |
| AFF1 | 6 | 0.7429 | 0.82598 | 0.999994 | 16778 | 1 | 2.7625 | 0.082584 | 0.24165 | 0.97466 | 4786 | 4 | 2.7625 |
| RGS16 | 5 | 0.063587 | 0.168 | 0.967126 | 3585 | 2 | 3.9333 | 0.082622 | 0.21987 | 0.97466 | 4787 | 3 | 3.9333 |
| NonTargetingControlG | 2 | 0.88378 | 0.88373 | 0.999994 | 18900 | 0 | 5.1076 | 0.082659 | 0.14283 | 0.965913 | 4788 | 2 | 5.1076 |
| SLFN14 | 4 | 0.50714 | 0.61956 | 0.999994 | 13539 | 1 | 1.4262 | 0.08269 | 0.20474 | 0.97466 | 4789 | 3 | 1.4262 |
| HRC | 5 | 0.95223 | 0.95237 | 0.999994 | 20390 | 0 | 3.8762 | 0.082728 | 0.22009 | 0.97466 | 4790 | 3 | 3.8762 |
| NonTargetingControlG | 1 | 0.91727 | 0.91768 | 0.999994 | 19570 | 0 | 7.4833 | 0.082731 | 0.082695 | 0.939346 | 4791 | 1 | 7.4833 |
| UBXN4 | 6 | 0.01911 | 0.069144 | 0.93914 | 1414 | 3 | 0.54777 | 0.082763 | 0.24205 | 0.97466 | 4792 | 3 | 0.54777 |
| PBOV1 | 6 | 0.66409 | 0.78846 | 0.999994 | 15840 | 2 | -0.15655 | 0.082834 | 0.2422 | 0.97466 | 4793 | 2 | -0.15655 |
| PIK3R6 | 5 | 0.94981 | 0.94999 | 0.999994 | 20328 | 0 | 1.6201 | 0.082837 | 0.22032 | 0.97466 | 4794 | 4 | 1.6201 |
| CHMP4B | 5 | 0.52543 | 0.67006 | 0.999994 | 13824 | 1 | 3.2177 | 0.082922 | 0.22049 | 0.97466 | 4795 | 4 | 3.2177 |
| ASB16 | 5 | 0.18028 | 0.34701 | 0.999994 | 7225 | 2 | 3.9996 | 0.082934 | 0.22051 | 0.97466 | 4796 | 3 | 3.9996 |
| E2F7 | 5 | 0.56006 | 0.7007 | 0.999994 | 14372 | 1 | -0.24308 | 0.082934 | 0.22051 | 0.97466 | 4797 | 2 | -0.24308 |
| KCNK6 | 5 | 0.76111 | 0.79679 | 0.999994 | 16991 | 1 | 0.95238 | 0.082949 | 0.22054 | 0.97466 | 4798 | 4 | 0.95238 |
| hsa-mir-6733 | 4 | 0.66949 | 0.70506 | 0.999994 | 15923 | 1 | 1.89 | 0.083008 | 0.20527 | 0.97466 | 4799 | 2 | 1.89 |
| GCSAM | 6 | 0.87278 | 0.88505 | 0.999994 | 18687 | 1 | 3.0333 | 0.083031 | 0.24266 | 0.97466 | 4800 | 4 | 3.0333 |
| OR6C74 | 6 | 0.091542 | 0.23062 | 0.983933 | 4658 | 3 | 1.6545 | 0.083031 | 0.24266 | 0.97466 | 4801 | 3 | 1.6545 |
| CENPQ | 6 | 0.72057 | 0.8185 | 0.999994 | 16503 | 1 | 1.5929 | 0.083053 | 0.24271 | 0.97466 | 4802 | 5 | 1.5929 |
| KANSL1L | 5 | 0.29937 | 0.48055 | 0.999994 | 10118 | 2 | 1.3498 | 0.083059 | 0.22076 | 0.97466 | 4803 | 3 | 1.3498 |
| MIOX | 6 | 0.084715 | 0.21695 | 0.980699 | 4396 | 3 | 0.7601 | 0.083079 | 0.24277 | 0.97466 | 4804 | 3 | 0.7601 |
| PAQR7 | 5 | 0.48451 | 0.63455 | 0.999994 | 13209 | 2 | 4.1654 | 0.083112 | 0.22086 | 0.97466 | 4805 | 3 | 4.1654 |
| FRMD5 | 5 | 0.27081 | 0.45748 | 0.999994 | 9494 | 1 | 4.1654 | 0.083112 | 0.22086 | 0.97466 | 4806 | 4 | 4.1654 |
| NonTargetingControlG | 2 | 0.91686 | 0.91694 | 0.999994 | 19560 | 0 | 3.9728 | 0.083139 | 0.14358 | 0.965913 | 4807 | 2 | 3.9728 |
| HACE1 | 4 | 0.59748 | 0.66395 | 0.999994 | 14919 | 1 | 0.52855 | 0.083237 | 0.20564 | 0.97466 | 4808 | 2 | 0.52855 |
| SLAMF8 | 6 | 0.97752 | 0.97747 | 0.999994 | 21030 | 0 | 2.7257 | 0.083286 | 0.24327 | 0.97466 | 4809 | 5 | 2.7257 |
| PHKB | 5 | 0.97043 | 0.97049 | 0.999994 | 20832 | 0 | 1.7896 | 0.083294 | 0.22122 | 0.97466 | 4810 | 4 | 1.7896 |
| C14orf159 | 6 | 0.29597 | 0.50202 | 0.999994 | 10037 | 1 | 1.3512 | 0.083315 | 0.24335 | 0.97466 | 4811 | 5 | 1.3512 |
| NOTCH4 | 6 | 0.40565 | 0.60604 | 0.999994 | 12039 | 2 | 2.3252 | 0.083355 | 0.24343 | 0.97466 | 4812 | 4 | 2.3252 |
| HEXA | 6 | 0.33547 | 0.54748 | 0.999994 | 10841 | 2 | 1.4118 | 0.08336 | 0.24344 | 0.97466 | 4813 | 4 | 1.4118 |
| PAK7 | 5 | 0.63097 | 0.73767 | 0.999994 | 15387 | 1 | -0.016762 | 0.083433 | 0.2215 | 0.97466 | 4814 | 2 | -0.016762 |
| GNG4 | 6 | 0.55336 | 0.70608 | 0.999994 | 14261 | 2 | 3.5295 | 0.083463 | 0.24367 | 0.97466 | 4815 | 4 | 3.5295 |
| KRTAP4-3 | 6 | 0.93609 | 0.93613 | 0.999994 | 19989 | 0 | 1.7567 | 0.083469 | 0.24368 | 0.97466 | 4816 | 5 | 1.7567 |
| GALNT9 | 6 | 0.26915 | 0.47073 | 0.999994 | 9449 | 3 | 1.7486 | 0.083478 | 0.24371 | 0.97466 | 4817 | 3 | 1.7486 |
| IL2RA | 6 | 0.85088 | 0.8724 | 0.999994 | 18276 | 1 | 0.048626 | 0.083478 | 0.24371 | 0.97466 | 4818 | 2 | 0.048626 |
| KDM3B | 6 | 3.155E-07 | 1.6003E-06 | 0.00495 | 7 | 4 | -9.0246 | 0.083478 | 0.24371 | 0.97466 | 4819 | 2 | -9.0246 |
| TF | 6 | 0.87453 | 0.88614 | 0.999994 | 18721 | 1 | 1.2881 | 0.083478 | 0.24371 | 0.97466 | 4820 | 4 | 1.2881 |
| hsa-mir-1249 | 3 | 0.18045 | 0.28786 | 0.999632 | 7228 | 1 | 5.5479 | 0.083492 | 0.18899 | 0.97466 | 4821 | 2 | 5.5479 |
| MAGED4 | 6 | 0.23296 | 0.42764 | 0.999994 | 8595 | 1 | 2.2355 | 0.08352 | 0.24381 | 0.97466 | 4822 | 5 | 2.2355 |
| LEPREL1 | 5 | 0.39019 | 0.5553 | 0.999994 | 11801 | 1 | 4.017 | 0.083533 | 0.2217 | 0.97466 | 4823 | 4 | 4.017 |
| E2F8 | 6 | 0.93762 | 0.93762 | 0.999994 | 20022 | 0 | 2.6581 | 0.083571 | 0.24394 | 0.97466 | 4824 | 3 | 2.6581 |
| ABHD15 | 6 | 0.16912 | 0.34964 | 0.999994 | 6962 | 2 | 0.50066 | 0.083571 | 0.24394 | 0.97466 | 4825 | 3 | 0.50066 |
| MYNN | 6 | 0.78216 | 0.84055 | 0.999994 | 17242 | 1 | 1.3506 | 0.083571 | 0.24394 | 0.97466 | 4826 | 5 | 1.3506 |
| TRIM32 | 5 | 0.78127 | 0.80815 | 0.999994 | 17228 | 1 | 2.3438 | 0.083609 | 0.22186 | 0.97466 | 4827 | 3 | 2.3438 |
| hsa-mir-5088 | 3 | 0.91627 | 0.91625 | 0.999994 | 19543 | 0 | 1.9267 | 0.083728 | 0.18944 | 0.97466 | 4828 | 3 | 1.9267 |
| ABHD17B | 5 | 0.61188 | 0.7305 | 0.999994 | 15108 | 1 | 3.9444 | 0.083737 | 0.22212 | 0.97466 | 4829 | 3 | 3.9444 |
| COMMD4 | 5 | 0.62127 | 0.73401 | 0.999994 | 15250 | 1 | 1.3915 | 0.083745 | 0.22213 | 0.97466 | 4830 | 4 | 1.3915 |
| LDOC1L | 6 | 0.94333 | 0.9433 | 0.999994 | 20160 | 0 | 1.44 | 0.083746 | 0.24433 | 0.97466 | 4831 | 5 | 1.44 |
| ZMAT2 | 4 | 0.241 | 0.38267 | 0.999994 | 8793 | 1 | 3.0416 | 0.083765 | 0.2065 | 0.97466 | 4832 | 2 | 3.0416 |
| CACUL1 | 6 | 0.22143 | 0.41383 | 0.999994 | 8314 | 3 | -0.40502 | 0.083781 | 0.24442 | 0.97466 | 4833 | 2 | -0.40502 |
| EYA2 | 6 | 0.030608 | 0.1002 | 0.946394 | 2036 | 3 | -0.54872 | 0.083781 | 0.24442 | 0.97466 | 4834 | 3 | -0.54872 |
| C15orf59 | 5 | 0.51968 | 0.66502 | 0.999994 | 13742 | 2 | 0.38203 | 0.083807 | 0.22225 | 0.97466 | 4835 | 2 | 0.38203 |

| | | | | | | | | | | | | | |
|---|---|---|---|---|---|---|---|---|---|---|---|---|---|
| PTGER1 | 4 | 0.57862 | 0.65404 | 0.999994 | 14665 | 1 | 2.017 | 0.08382 | 0.20659 | 0.97466 | 4836 | 3 | 2.017 |
| AFF3 | 6 | 0.24686 | 0.4442 | 0.999994 | 8942 | 3 | -0.41218 | 0.083886 | 0.24467 | 0.97466 | 4837 | 2 | -0.41218 |
| MAP2K3 | 6 | 0.47433 | 0.65134 | 0.999994 | 13053 | 1 | 3.1624 | 0.083896 | 0.2447 | 0.97466 | 4838 | 4 | 3.1624 |
| MMADHC | 6 | 0.32634 | 0.53706 | 0.999994 | 10652 | 1 | 1.3361 | 0.083896 | 0.2447 | 0.97466 | 4839 | 3 | 1.3361 |
| BAI2 | 6 | 0.046754 | 0.13696 | 0.957161 | 2841 | 2 | 2.7775 | 0.083897 | 0.2447 | 0.97466 | 4840 | 3 | 2.7775 |
| STH | 6 | 0.26762 | 0.46887 | 0.999994 | 9409 | 2 | 2.8728 | 0.083905 | 0.24472 | 0.97466 | 4841 | 4 | 2.8728 |
| MTX3 | 6 | 0.00079503 | 0.0040795 | 0.560963 | 142 | 4 | -5.6784 | 0.083939 | 0.24479 | 0.97466 | 4842 | 1 | -5.6784 |
| CPNE3 | 6 | 0.094119 | 0.23577 | 0.985362 | 4754 | 2 | 1.1493 | 0.084018 | 0.24498 | 0.97466 | 4843 | 3 | 1.1493 |
| MRPS18A | 3 | 0.86594 | 0.86593 | 0.999994 | 18555 | 0 | 5.5272 | 0.084043 | 0.19005 | 0.97466 | 4844 | 2 | 5.5272 |
| CARD8 | 5 | 0.95642 | 0.95657 | 0.999994 | 20500 | 0 | 2.0493 | 0.084077 | 0.2228 | 0.97466 | 4845 | 4 | 2.0493 |
| CWC25 | 5 | 0.67353 | 0.75478 | 0.999994 | 15979 | 1 | 3.1166 | 0.084091 | 0.22283 | 0.97466 | 4846 | 4 | 3.1166 |
| SLC8B1 | 3 | 0.91586 | 0.91584 | 0.999994 | 19534 | 0 | 2.9532 | 0.084135 | 0.19022 | 0.97466 | 4847 | 3 | 2.9532 |
| CEP89 | 6 | 0.96607 | 0.96615 | 0.999994 | 20739 | 0 | 1.1379 | 0.084153 | 0.24529 | 0.97466 | 4848 | 5 | 1.1379 |
| ERVMER34-1 | 6 | 0.59455 | 0.73596 | 0.999994 | 14883 | 1 | 1.0143 | 0.084171 | 0.24532 | 0.97466 | 4849 | 4 | 1.0143 |
| RGCC | 6 | 0.38841 | 0.59493 | 0.999994 | 11771 | 2 | 2.3275 | 0.084173 | 0.24533 | 0.97466 | 4850 | 4 | 2.3275 |
| PRKD2 | 6 | 0.94397 | 0.94395 | 0.999994 | 20177 | 0 | 1.7258 | 0.084179 | 0.24534 | 0.97466 | 4851 | 5 | 1.7258 |
| C8orf86 | 5 | 0.23353 | 0.42211 | 0.999994 | 8605 | 3 | -0.66552 | 0.084246 | 0.22314 | 0.97466 | 4852 | 2 | -0.66552 |
| OR11H1 | 4 | 0.29875 | 0.44172 | 0.999994 | 10108 | 2 | 2.6289 | 0.084253 | 0.20729 | 0.97466 | 4853 | 2 | 2.6289 |
| KCNMB1 | 5 | 0.96358 | 0.96373 | 0.999994 | 20690 | 0 | 3.4241 | 0.084382 | 0.22341 | 0.97466 | 4854 | 4 | 3.4241 |
| DIRAS2 | 5 | 0.8462 | 0.85137 | 0.999994 | 18191 | 1 | 1.3671 | 0.08441 | 0.22348 | 0.97466 | 4855 | 4 | 1.3671 |
| WRN | 6 | 0.63877 | 0.76902 | 0.999994 | 15497 | 2 | 0.73427 | 0.084465 | 0.24599 | 0.97466 | 4856 | 3 | 0.73427 |
| RAB14 | 6 | 0.61011 | 0.74742 | 0.999994 | 15073 | 1 | 2.3036 | 0.08448 | 0.24603 | 0.97466 | 4857 | 5 | 2.3036 |
| OR51B5 | 4 | 0.41825 | 0.5644 | 0.999994 | 12225 | 1 | 3.7728 | 0.08448 | 0.20764 | 0.97466 | 4858 | 3 | 3.7728 |
| SCIMP | 5 | 0.087886 | 0.20719 | 0.978744 | 4506 | 3 | -1.6601 | 0.084488 | 0.22362 | 0.97466 | 4859 | 1 | -1.6601 |
| ZFP2 | 6 | 0.22508 | 0.41821 | 0.999994 | 8400 | 3 | 1.1531 | 0.084506 | 0.2461 | 0.97466 | 4860 | 3 | 1.1531 |
| PGLYRP3 | 6 | 0.031898 | 0.10325 | 0.946414 | 2099 | 1 | 0.96176 | 0.084569 | 0.24624 | 0.97466 | 4861 | 5 | 0.96176 |
| NonTargetingControlG | 2 | 0.52899 | 0.5308 | 0.999994 | 13882 | 1 | 3.7059 | 0.084573 | 0.14585 | 0.965913 | 4862 | 1 | 3.7059 |
| TMEM87A | 6 | 0.53809 | 0.6952 | 0.999994 | 14027 | 2 | 2.6344 | 0.084623 | 0.24637 | 0.97466 | 4863 | 3 | 2.6344 |
| MPND | 6 | 0.23389 | 0.42872 | 0.999994 | 8617 | 3 | 2.2271 | 0.084623 | 0.24637 | 0.97466 | 4864 | 3 | 2.2271 |
| SLC17A8 | 6 | 0.87462 | 0.8862 | 0.999994 | 18724 | 1 | 2.8901 | 0.084623 | 0.24637 | 0.97466 | 4865 | 3 | 2.8901 |
| C4orf26 | 6 | 0.58142 | 0.72624 | 0.999994 | 14699 | 2 | 2.5805 | 0.084623 | 0.24637 | 0.97466 | 4866 | 3 | 2.5805 |
| ALKBH5 | 5 | 0.6816 | 0.75825 | 0.999994 | 16112 | 1 | 1.6179 | 0.084685 | 0.22402 | 0.97466 | 4867 | 3 | 1.6179 |
| CRABP2 | 5 | 0.83954 | 0.84629 | 0.999994 | 18078 | 1 | 1.6458 | 0.084701 | 0.22406 | 0.97466 | 4868 | 4 | 1.6458 |
| MRPL52 | 5 | 0.16229 | 0.3209 | 0.999994 | 6802 | 2 | 4.1171 | 0.084785 | 0.22422 | 0.97466 | 4869 | 3 | 4.1171 |
| CD4 | 5 | 0.14736 | 0.29882 | 0.999994 | 6385 | 2 | 4.1171 | 0.084785 | 0.22422 | 0.97466 | 4870 | 3 | 4.1171 |
| LCE1D | 5 | 0.35027 | 0.52215 | 0.999994 | 11166 | 2 | 4.1171 | 0.084785 | 0.22422 | 0.97466 | 4871 | 3 | 4.1171 |
| MOXD1 | 6 | 0.98456 | 0.98449 | 0.999994 | 21206 | 0 | 1.7117 | 0.08482 | 0.24685 | 0.97466 | 4872 | 5 | 1.7117 |
| MPRIP | 6 | 0.47985 | 0.6551 | 0.999994 | 13143 | 2 | 1.3431 | 0.084858 | 0.24694 | 0.97466 | 4873 | 3 | 1.3431 |
| MADD | 6 | 0.35324 | 0.56764 | 0.999994 | 11218 | 3 | 1.2632 | 0.084858 | 0.24694 | 0.97466 | 4874 | 3 | 1.2632 |
| NonTargetingControlG | 2 | 0.91512 | 0.91517 | 0.999994 | 19515 | 0 | 3.1561 | 0.084883 | 0.14634 | 0.965913 | 4875 | 2 | 3.1561 |
| PRCD | 5 | 0.021852 | 0.069068 | 0.93914 | 1571 | 4 | -1.7118 | 0.084883 | 0.22442 | 0.97466 | 4876 | 1 | -1.7118 |
| ADCY2 | 6 | 0.52301 | 0.68461 | 0.999994 | 13795 | 2 | 1.0612 | 0.084911 | 0.24708 | 0.97466 | 4877 | 3 | 1.0612 |
| SLC22A16 | 6 | 0.14145 | 0.3149 | 0.999994 | 6230 | 2 | 0.90831 | 0.084911 | 0.24708 | 0.97466 | 4878 | 4 | 0.90831 |
| hsa-mir-1302-8 | 4 | 0.078405 | 0.17579 | 0.968171 | 4168 | 1 | 2.1161 | 0.084915 | 0.20836 | 0.97466 | 4879 | 3 | 2.1161 |
| ASB10 | 5 | 0.53688 | 0.68014 | 0.999994 | 14012 | 1 | 2.2802 | 0.084915 | 0.22447 | 0.97466 | 4880 | 4 | 2.2802 |
| OSR2 | 6 | 0.95994 | 0.96001 | 0.999994 | 20583 | 0 | 1.8368 | 0.084945 | 0.24717 | 0.97466 | 4881 | 5 | 1.8368 |
| hsa-mir-93 | 3 | 0.91503 | 0.91502 | 0.999994 | 19513 | 0 | 1.2179 | 0.084969 | 0.19185 | 0.97466 | 4882 | 3 | 1.2179 |
| APOA4 | 6 | 0.13736 | 0.3097 | 0.999994 | 6117 | 2 | 2.888 | 0.084979 | 0.24723 | 0.97466 | 4883 | 4 | 2.888 |
| PIK3C2G | 5 | 0.95661 | 0.95676 | 0.999994 | 20507 | 0 | 2.7637 | 0.084993 | 0.22464 | 0.97466 | 4884 | 4 | 2.7637 |
| PLOD3 | 4 | 0.92904 | 0.92881 | 0.999994 | 19835 | 0 | 2.4913 | 0.085003 | 0.2085 | 0.97466 | 4885 | 3 | 2.4913 |
| PITPNA | 6 | 0.95676 | 0.95683 | 0.999994 | 20511 | 0 | 3.3031 | 0.085004 | 0.24729 | 0.97466 | 4886 | 4 | 3.3031 |
| MYEOV2 | 6 | 0.31484 | 0.52399 | 0.999994 | 10418 | 1 | 1.9776 | 0.085004 | 0.24729 | 0.97466 | 4887 | 5 | 1.9776 |

| | | | | | | | | | | | | | |
|---|---|---|---|---|---|---|---|---|---|---|---|---|---|
| hsa-let-7f-2 | 4 | 0.10654 | 0.22676 | 0.983354 | 5189 | 1 | 4.3114 | 0.085012 | 0.20851 | 0.97466 | 4888 | 3 | 4.3114 |
| NonTargetingControlG | 2 | 0.35444 | 0.37954 | 0.999994 | 11238 | 1 | 3.0435 | 0.085024 | 0.14657 | 0.965913 | 4889 | 1 | 3.0435 |
| NSUN6 | 5 | 0.73493 | 0.7831 | 0.999994 | 16661 | 1 | 3.6894 | 0.085027 | 0.22471 | 0.97466 | 4890 | 3 | 3.6894 |
| CCDC113 | 5 | 0.72983 | 0.78056 | 0.999994 | 16596 | 1 | -0.14294 | 0.085027 | 0.22471 | 0.97466 | 4891 | 2 | -0.14294 |
| ABR | 5 | 0.34961 | 0.52157 | 0.999994 | 11142 | 1 | -0.11577 | 0.085037 | 0.22473 | 0.97466 | 4892 | 2 | -0.11577 |
| WHSC1L1 | 6 | 0.8578 | 0.87618 | 0.999994 | 18403 | 1 | 2.1998 | 0.085046 | 0.24738 | 0.97466 | 4893 | 5 | 2.1998 |
| TJP2 | 6 | 0.41429 | 0.61162 | 0.999994 | 12170 | 1 | 1.5076 | 0.085049 | 0.24739 | 0.97466 | 4894 | 5 | 1.5076 |
| TIMM21 | 6 | 0.56714 | 0.71589 | 0.999994 | 14498 | 1 | 2.7251 | 0.085078 | 0.24746 | 0.97466 | 4895 | 5 | 2.7251 |
| ACTL8 | 4 | 0.33092 | 0.47463 | 0.999994 | 10742 | 1 | 4.0087 | 0.085127 | 0.2087 | 0.97466 | 4896 | 3 | 4.0087 |
| MXRA7 | 4 | 0.39824 | 0.54392 | 0.999994 | 11919 | 2 | 2.8725 | 0.085127 | 0.2087 | 0.97466 | 4897 | 2 | 2.8725 |
| TXLNG | 6 | 0.93962 | 0.93957 | 0.999994 | 20067 | 0 | 1.5955 | 0.085137 | 0.24757 | 0.97466 | 4898 | 5 | 1.5955 |
| DBH | 6 | 0.1205 | 0.28666 | 0.999632 | 5642 | 4 | -1.2249 | 0.0852 | 0.24771 | 0.97466 | 4899 | 1 | -1.2249 |
| KRTAP4-2 | 6 | 0.34228 | 0.55527 | 0.999994 | 10990 | 2 | 2.6856 | 0.0852 | 0.24771 | 0.97466 | 4900 | 3 | 2.6856 |
| STAT3 | 5 | 0.17078 | 0.33326 | 0.999994 | 7001 | 1 | 2.1565 | 0.085229 | 0.22514 | 0.97466 | 4901 | 4 | 2.1565 |
| RRAGC | 5 | 0.90563 | 0.90573 | 0.999994 | 19345 | 0 | 1.5132 | 0.085236 | 0.22516 | 0.97466 | 4902 | 4 | 1.5132 |
| CCDC43 | 4 | 0.86002 | 0.85962 | 0.999994 | 18438 | 0 | 1.5431 | 0.085267 | 0.20894 | 0.97466 | 4903 | 3 | 1.5431 |
| C6orf136 | 6 | 0.83771 | 0.86545 | 0.999994 | 18047 | 1 | 2.6007 | 0.085324 | 0.248 | 0.97466 | 4904 | 4 | 2.6007 |
| EYA3 | 5 | 0.8232 | 0.83469 | 0.999994 | 17827 | 1 | 4.1387 | 0.085353 | 0.22539 | 0.97466 | 4905 | 4 | 4.1387 |
| KRTAP12-2 | 6 | 0.134 | 0.30541 | 0.999994 | 6021 | 3 | -0.43591 | 0.085384 | 0.24813 | 0.97466 | 4906 | 1 | -0.43591 |
| RGS3 | 5 | 0.63254 | 0.73828 | 0.999994 | 15404 | 1 | 1.9311 | 0.085393 | 0.22547 | 0.97466 | 4907 | 4 | 1.9311 |
| ACP5 | 6 | 0.92264 | 0.92287 | 0.999994 | 19692 | 1 | 2.8637 | 0.085433 | 0.24825 | 0.97466 | 4908 | 3 | 2.8637 |
| NXT1 | 6 | 0.22814 | 0.42184 | 0.999994 | 8480 | 2 | 2.8572 | 0.085433 | 0.24825 | 0.97466 | 4909 | 3 | 2.8572 |
| FHL5 | 6 | 0.37458 | 0.5861 | 0.999994 | 11577 | 1 | 3.5989 | 0.085433 | 0.24825 | 0.97466 | 4910 | 4 | 3.5989 |
| DGKG | 6 | 0.49883 | 0.6679 | 0.999994 | 13407 | 2 | 3.5427 | 0.085433 | 0.24825 | 0.97466 | 4911 | 4 | 3.5427 |
| XAGE5 | 5 | 0.47171 | 0.62357 | 0.999994 | 13027 | 1 | 0.71678 | 0.085453 | 0.22559 | 0.97466 | 4912 | 3 | 0.71678 |
| hsa-mir-664b | 4 | 0.77473 | 0.7796 | 0.999994 | 17139 | 1 | 4.0345 | 0.085458 | 0.20924 | 0.97466 | 4913 | 3 | 4.0345 |
| hsa-mir-4709 | 4 | 0.67454 | 0.70817 | 0.999994 | 15997 | 1 | 3.56 | 0.085458 | 0.20924 | 0.97466 | 4914 | 3 | 3.56 |
| ELL | 4 | 0.47564 | 0.60558 | 0.999994 | 13070 | 1 | 4.0225 | 0.085518 | 0.20935 | 0.97466 | 4915 | 3 | 4.0225 |
| AMPD1 | 6 | 0.56234 | 0.71248 | 0.999994 | 14417 | 1 | 2.1278 | 0.085545 | 0.24851 | 0.97466 | 4916 | 5 | 2.1278 |
| TNIP2 | 6 | 0.97257 | 0.9725 | 0.999994 | 20885 | 0 | 1.3218 | 0.085568 | 0.24856 | 0.97466 | 4917 | 4 | 1.3218 |
| LRRC71 | 6 | 0.941 | 0.94098 | 0.999994 | 20108 | 0 | 3.154 | 0.085611 | 0.24866 | 0.97466 | 4918 | 4 | 3.154 |
| OR8U8 | 4 | 0.021721 | 0.064264 | 0.93914 | 1559 | 2 | -3.2135 | 0.085638 | 0.20952 | 0.97466 | 4919 | 2 | -3.2135 |
| DRG1 | 6 | 0.92417 | 0.92432 | 0.999994 | 19722 | 1 | 0.78471 | 0.085642 | 0.24873 | 0.97466 | 4920 | 5 | 0.78471 |
| AOX1 | 6 | 0.12087 | 0.28736 | 0.999632 | 5656 | 3 | 0.26781 | 0.085664 | 0.24877 | 0.97466 | 4921 | 3 | 0.26781 |
| SNX19 | 6 | 0.4221 | 0.6168 | 0.999994 | 12282 | 1 | 2.2121 | 0.085694 | 0.24884 | 0.97466 | 4922 | 5 | 2.2121 |
| VAV3 | 5 | 0.089747 | 0.21018 | 0.979587 | 4584 | 1 | 1.4518 | 0.085701 | 0.22607 | 0.97466 | 4923 | 4 | 1.4518 |
| CNTRL | 5 | 0.83571 | 0.84357 | 0.999994 | 18020 | 1 | 2.8068 | 0.085744 | 0.22617 | 0.97466 | 4924 | 4 | 2.8068 |
| SLC4A8 | 6 | 0.015105 | 0.056057 | 0.9235 | 1194 | 3 | -1.2326 | 0.085804 | 0.24909 | 0.97466 | 4925 | 1 | -1.2326 |
| NonTargetingControlG | 2 | 0.91419 | 0.91424 | 0.999994 | 19491 | 0 | 5.0055 | 0.085814 | 0.1478 | 0.965913 | 4926 | 2 | 5.0055 |
| OR4N4 | 6 | 0.76584 | 0.83427 | 0.999994 | 17037 | 1 | 2.1001 | 0.085935 | 0.2494 | 0.97466 | 4927 | 4 | 2.1001 |
| COX7A1 | 6 | 0.93159 | 0.93174 | 0.999994 | 19890 | 0 | 1.741 | 0.085955 | 0.24944 | 0.97466 | 4928 | 5 | 1.741 |
| SHC2 | 4 | 0.60453 | 0.66779 | 0.999994 | 15000 | 1 | 3.156 | 0.08599 | 0.2101 | 0.97466 | 4929 | 2 | 3.156 |
| TOR1AIP1 | 6 | 0.33103 | 0.54242 | 0.999994 | 10744 | 3 | 0.054271 | 0.086051 | 0.24966 | 0.97466 | 4930 | 3 | 0.054271 |
| ZNF81 | 6 | 0.063959 | 0.17416 | 0.968171 | 3606 | 3 | -0.091748 | 0.086051 | 0.24966 | 0.97466 | 4931 | 3 | -0.091748 |
| TKTL1 | 6 | 0.0036997 | 0.016357 | 0.772095 | 420 | 2 | 2.5506 | 0.086051 | 0.24966 | 0.97466 | 4932 | 4 | 2.5506 |
| TYMP | 6 | 0.9367 | 0.93673 | 0.999994 | 20000 | 0 | 1.9525 | 0.086051 | 0.24966 | 0.97466 | 4933 | 4 | 1.9525 |
| KLHDC10 | 6 | 0.0046665 | 0.01993 | 0.808847 | 495 | 2 | 0.67684 | 0.086051 | 0.24966 | 0.97466 | 4934 | 3 | 0.67684 |
| KIR3DL1 | 6 | 0.34771 | 0.56143 | 0.999994 | 11102 | 2 | 2.2208 | 0.086051 | 0.24966 | 0.97466 | 4935 | 3 | 2.2208 |
| C1orf106 | 6 | 0.35612 | 0.57092 | 0.999994 | 11269 | 3 | -0.3674 | 0.086093 | 0.24976 | 0.97466 | 4936 | 1 | -0.3674 |
| ARFGEF1 | 6 | 0.10419 | 0.25549 | 0.994943 | 5105 | 2 | -0.23871 | 0.086093 | 0.24976 | 0.97466 | 4937 | 2 | -0.23871 |
| PLAC8L1 | 5 | 0.91224 | 0.91228 | 0.999994 | 19468 | 0 | 3.2452 | 0.086103 | 0.22691 | 0.97466 | 4938 | 4 | 3.2452 |
| TNNT1 | 6 | 0.56331 | 0.71317 | 0.999994 | 14436 | 2 | 3.5611 | 0.086128 | 0.24984 | 0.97466 | 4939 | 4 | 3.5611 |

| | | | | | | | | | | | | |
|---|---|---|---|---|---|---|---|---|---|---|---|---|
| GPC5 | 6 | 0.066742 | 0.17992 | 0.968171 | 3714 | 2 | 3.2282 | 0.086128 | 0.24984 | 0.97466 | 4940 | 4 | 3.2282 |
| C4orf29 | 4 | 0.77109 | 0.77666 | 0.999994 | 17098 | 1 | 2.7189 | 0.086184 | 0.21041 | 0.97466 | 4941 | 3 | 2.7189 |
| PPIAL4D | 3 | 0.085894 | 0.16434 | 0.967126 | 4443 | 1 | 0.26463 | 0.086267 | 0.19436 | 0.97466 | 4942 | 1 | 0.26463 |
| OSGEP | 3 | 0.38203 | 0.47395 | 0.999994 | 11678 | 1 | 5.4733 | 0.086268 | 0.19436 | 0.97466 | 4943 | 2 | 5.4733 |
| THAP5 | 4 | 0.040534 | 0.10286 | 0.946414 | 2538 | 1 | 1.5013 | 0.08636 | 0.21068 | 0.97466 | 4944 | 3 | 1.5013 |
| CATSPERG | 5 | 0.8336 | 0.84206 | 0.999994 | 17981 | 1 | 4.0801 | 0.086366 | 0.22745 | 0.97466 | 4945 | 3 | 4.0801 |
| ARMC1 | 5 | 0.56154 | 0.702 | 0.999994 | 14396 | 2 | 4.0801 | 0.086366 | 0.22745 | 0.97466 | 4946 | 3 | 4.0801 |
| C17orf47 | 5 | 0.26958 | 0.45649 | 0.999994 | 9462 | 2 | 4.0801 | 0.086366 | 0.22745 | 0.97466 | 4947 | 3 | 4.0801 |
| OC90 | 5 | 0.37157 | 0.53971 | 0.999994 | 11536 | 2 | 4.0801 | 0.086366 | 0.22745 | 0.97466 | 4948 | 3 | 4.0801 |
| POU5F1B | 4 | 0.13725 | 0.27605 | 0.999379 | 6113 | 2 | 2.9479 | 0.086397 | 0.21074 | 0.97466 | 4949 | 2 | 2.9479 |
| NonTargetingControlG | 2 | 0.9136 | 0.91368 | 0.999994 | 19482 | 0 | 4.3944 | 0.086398 | 0.14876 | 0.965913 | 4950 | 2 | 4.3944 |
| SRPK2 | 6 | 0.61317 | 0.7497 | 0.999994 | 15122 | 1 | 2.7567 | 0.086403 | 0.25047 | 0.97466 | 4951 | 5 | 2.7567 |
| OR51B6 | 6 | 0.49266 | 0.66368 | 0.999994 | 13306 | 2 | 1.6315 | 0.086434 | 0.25054 | 0.97466 | 4952 | 3 | 1.6315 |
| FAM172A | 6 | 0.42593 | 0.61926 | 0.999994 | 12350 | 2 | 0.77683 | 0.086434 | 0.25054 | 0.97466 | 4953 | 3 | 0.77683 |
| SBK1 | 6 | 0.22308 | 0.41585 | 0.999994 | 8359 | 2 | 0.54209 | 0.086439 | 0.25055 | 0.97466 | 4954 | 3 | 0.54209 |
| NonTargetingControlG | 2 | 0.91352 | 0.91359 | 0.999994 | 19480 | 0 | 4.1244 | 0.086483 | 0.1489 | 0.965913 | 4955 | 2 | 4.1244 |
| LAG3 | 5 | 0.0074528 | 0.028669 | 0.858757 | 688 | 2 | -0.2108 | 0.086519 | 0.22777 | 0.97466 | 4956 | 2 | -0.2108 |
| AP4E1 | 5 | 0.9067 | 0.90677 | 0.999994 | 19364 | 0 | 0.6959 | 0.086519 | 0.22777 | 0.97466 | 4957 | 3 | 0.6959 |
| C9orf135 | 6 | 0.98317 | 0.98311 | 0.999994 | 21160 | 0 | 0.8667 | 0.086523 | 0.25073 | 0.97466 | 4958 | 5 | 0.8667 |
| TMX1 | 6 | 0.086077 | 0.21965 | 0.980699 | 4451 | 3 | 1.6976 | 0.086534 | 0.25076 | 0.97466 | 4959 | 3 | 1.6976 |
| ZNF414 | 4 | 0.39561 | 0.54121 | 0.999994 | 11881 | 2 | 1.0001 | 0.086537 | 0.21097 | 0.97466 | 4960 | 2 | 1.0001 |
| ZNF578 | 6 | 0.92414 | 0.9243 | 0.999994 | 19720 | 1 | 1.2704 | 0.086538 | 0.25077 | 0.97466 | 4961 | 5 | 1.2704 |
| HIVEP1 | 5 | 0.83474 | 0.84288 | 0.999994 | 17999 | 1 | 1.5748 | 0.086546 | 0.22783 | 0.97466 | 4962 | 4 | 1.5748 |
| NR1D1 | 5 | 0.13732 | 0.28374 | 0.999632 | 6116 | 1 | 1.5712 | 0.086546 | 0.22783 | 0.97466 | 4963 | 4 | 1.5712 |
| MCTS1 | 5 | 0.35399 | 0.52517 | 0.999994 | 11236 | 1 | 3.024 | 0.086546 | 0.22783 | 0.97466 | 4964 | 4 | 3.024 |
| HSD11B1L | 5 | 0.94455 | 0.94475 | 0.999994 | 20193 | 0 | 2.6831 | 0.086546 | 0.22783 | 0.97466 | 4965 | 4 | 2.6831 |
| SLX4IP | 6 | 0.93495 | 0.93501 | 0.999994 | 19966 | 0 | 2.6154 | 0.086583 | 0.25086 | 0.97466 | 4966 | 4 | 2.6154 |
| DECR1 | 6 | 0.95782 | 0.95792 | 0.999994 | 20532 | 0 | 2.4134 | 0.086583 | 0.25086 | 0.97466 | 4967 | 5 | 2.4134 |
| BNIP3 | 6 | 0.8676 | 0.88187 | 0.999994 | 18592 | 1 | 1.8235 | 0.086591 | 0.25088 | 0.97466 | 4968 | 4 | 1.8235 |
| FBXO10 | 5 | 0.77808 | 0.80634 | 0.999994 | 17185 | 1 | 0.15058 | 0.086594 | 0.22793 | 0.97466 | 4969 | 2 | 0.15058 |
| CREBRF | 5 | 0.91457 | 0.91465 | 0.999994 | 19500 | 0 | 1.8125 | 0.086594 | 0.22793 | 0.97466 | 4970 | 3 | 1.8125 |
| RUVBL1 | 4 | 0.87504 | 0.87469 | 0.999994 | 18738 | 0 | 3.0857 | 0.086617 | 0.21111 | 0.97466 | 4971 | 3 | 3.0857 |
| hsa-mir-548v | 3 | 0.91331 | 0.91331 | 0.999994 | 19479 | 0 | 3.5391 | 0.086691 | 0.19518 | 0.97466 | 4972 | 3 | 3.5391 |
| WDR19 | 6 | 0.41123 | 0.6097 | 0.999994 | 12133 | 2 | 1.2199 | 0.086749 | 0.25124 | 0.97466 | 4973 | 4 | 1.2199 |
| EPHA5 | 6 | 0.83999 | 0.8666 | 0.999994 | 18088 | 1 | 1.6141 | 0.086868 | 0.25152 | 0.97466 | 4974 | 5 | 1.6141 |
| SIKE1 | 5 | 0.038181 | 0.11129 | 0.951052 | 2426 | 2 | 3.9196 | 0.086886 | 0.22855 | 0.97466 | 4975 | 3 | 3.9196 |
| KNDC1 | 5 | 0.74507 | 0.78827 | 0.999994 | 16800 | 1 | -0.27027 | 0.086899 | 0.22857 | 0.97466 | 4976 | 2 | -0.27027 |
| SLC25A35 | 5 | 0.91093 | 0.91095 | 0.999994 | 19446 | 0 | 3.7994 | 0.086904 | 0.22858 | 0.97466 | 4977 | 3 | 3.7994 |
| C10orf131 | 3 | 0.2583 | 0.38528 | 0.999994 | 9204 | 1 | 5.4494 | 0.086919 | 0.19563 | 0.97466 | 4978 | 2 | 5.4494 |
| hsa-mir-4776-1 | 4 | 0.57692 | 0.65317 | 0.999994 | 14644 | 1 | 2.712 | 0.086945 | 0.21162 | 0.97466 | 4979 | 3 | 2.712 |
| COL6A2 | 4 | 0.21865 | 0.35991 | 0.999994 | 8245 | 1 | 2.3288 | 0.086981 | 0.21169 | 0.97466 | 4980 | 3 | 2.3288 |
| RIN3 | 6 | 0.76375 | 0.83348 | 0.999994 | 17009 | 1 | 0.25253 | 0.086985 | 0.2518 | 0.97466 | 4981 | 2 | 0.25253 |
| SNURF | 6 | 0.94996 | 0.94993 | 0.999994 | 20332 | 0 | 1.895 | 0.086985 | 0.2518 | 0.97466 | 4982 | 4 | 1.895 |
| NonTargetingControlG | 2 | 0.030779 | 0.056304 | 0.9235 | 2051 | 1 | -0.78209 | 0.086989 | 0.14972 | 0.965913 | 4983 | 1 | -0.78209 |
| H1F0 | 6 | 0.14324 | 0.31715 | 0.999994 | 6280 | 2 | 0.58211 | 0.087005 | 0.25185 | 0.97466 | 4984 | 3 | 0.58211 |
| KBTBD8 | 6 | 0.94305 | 0.94303 | 0.999994 | 20154 | 0 | 2.1616 | 0.087039 | 0.25192 | 0.97466 | 4985 | 4 | 2.1616 |
| SSX3 | 5 | 0.13483 | 0.27998 | 0.999632 | 6046 | 2 | 3.5863 | 0.087068 | 0.2289 | 0.97466 | 4986 | 3 | 3.5863 |
| TPBGL | 6 | 0.96197 | 0.96205 | 0.999994 | 20629 | 0 | 2.3213 | 0.087116 | 0.25209 | 0.97466 | 4987 | 4 | 2.3213 |
| NPTX2 | 6 | 0.71704 | 0.81736 | 0.999994 | 16469 | 1 | 2.199 | 0.087116 | 0.25209 | 0.97466 | 4988 | 4 | 2.199 |
| TMC7 | 6 | 0.098994 | 0.2454 | 0.98979 | 4931 | 4 | -1.0626 | 0.087124 | 0.25211 | 0.97466 | 4989 | 2 | -1.0626 |
| hsa-mir-4536-1 | 2 | 0.91286 | 0.91292 | 0.999994 | 19475 | 0 | 3.0853 | 0.087145 | 0.14997 | 0.965913 | 4990 | 2 | 3.0853 |
| C15orf54 | 6 | 0.60953 | 0.74699 | 0.999994 | 15066 | 2 | 2.6028 | 0.087159 | 0.25218 | 0.97466 | 4991 | 4 | 2.6028 |

| | | | | | | | | | | | | |
|---|---|---|---|---|---|---|---|---|---|---|---|---|
| NRSN2 | 6 | 0.056683 | 0.15859 | 0.967126 | 3280 | 2 | 2.4158 | 0.087195 | 0.25227 | 0.97466 | 4992 | 3 | 2.4158 |
| SLC2A7 | 6 | 0.29568 | 0.50168 | 0.999994 | 10032 | 2 | 2.4666 | 0.087195 | 0.25227 | 0.97466 | 4993 | 3 | 2.4666 |
| KLF6 | 5 | 0.12358 | 0.2628 | 0.996469 | 5756 | 1 | 3.771 | 0.087231 | 0.22925 | 0.97466 | 4994 | 3 | 3.771 |
| hsa-mir-6810 | 4 | 0.84238 | 0.84183 | 0.999994 | 18131 | 0 | 3.9859 | 0.087251 | 0.21212 | 0.97466 | 4995 | 3 | 3.9859 |
| WNT10B | 6 | 0.0026277 | 0.012342 | 0.733879 | 327 | 2 | 2.3253 | 0.087296 | 0.2525 | 0.97466 | 4996 | 4 | 2.3253 |
| DARS | 4 | 0.86502 | 0.86464 | 0.999994 | 18539 | 0 | 4.52 | 0.087297 | 0.2122 | 0.97466 | 4997 | 3 | 4.52 |
| hsa-mir-433 | 4 | 0.82994 | 0.82949 | 0.999994 | 17929 | 0 | 3.0868 | 0.087297 | 0.2122 | 0.97466 | 4998 | 2 | 3.0868 |
| NOSIP | 6 | 0.9213 | 0.9216 | 0.999994 | 19661 | 1 | 2.0448 | 0.087299 | 0.2525 | 0.97466 | 4999 | 4 | 2.0448 |
| EFTUD1 | 6 | 0.46504 | 0.64512 | 0.999994 | 12934 | 2 | 1.2719 | 0.087302 | 0.25251 | 0.97466 | 5000 | 4 | 1.2719 |
| C10orf55 | 6 | 0.15595 | 0.33313 | 0.999994 | 6626 | 3 | -0.73093 | 0.087302 | 0.25251 | 0.97466 | 5001 | 2 | -0.73093 |
| NME6 | 6 | 0.49984 | 0.66857 | 0.999994 | 13421 | 2 | 2.0079 | 0.087302 | 0.25251 | 0.97466 | 5002 | 3 | 2.0079 |
| ATP6AP2 | 2 | 0.91266 | 0.91272 | 0.999994 | 19471 | 0 | 4.802 | 0.087342 | 0.15028 | 0.965913 | 5003 | 2 | 4.802 |
| hsa-mir-3663 | 4 | 0.1882 | 0.32854 | 0.999994 | 7423 | 2 | -0.97371 | 0.087347 | 0.21229 | 0.97466 | 5004 | 1 | -0.97371 |
| TACC3 | 4 | 0.060341 | 0.14143 | 0.959146 | 3434 | 3 | -2.2057 | 0.087347 | 0.21229 | 0.97466 | 5005 | 1 | -2.2057 |
| ZPLD1 | 3 | 0.57024 | 0.5948 | 0.999994 | 14544 | 1 | 4.5294 | 0.087358 | 0.19649 | 0.97466 | 5006 | 2 | 4.5294 |
| IFT122 | 5 | 0.18919 | 0.35979 | 0.999994 | 7458 | 3 | -0.91038 | 0.087361 | 0.22952 | 0.97466 | 5007 | 2 | -0.91038 |
| ZNF714 | 5 | 0.93353 | 0.93378 | 0.999994 | 19937 | 0 | 0.35554 | 0.087361 | 0.22952 | 0.97466 | 5008 | 2 | 0.35554 |
| C1orf146 | 6 | 0.89846 | 0.90244 | 0.999994 | 19193 | 1 | 2.0649 | 0.087395 | 0.25272 | 0.97466 | 5009 | 5 | 2.0649 |
| CLEC11A | 6 | 0.21964 | 0.41166 | 0.999994 | 8282 | 1 | 2.3707 | 0.087402 | 0.25274 | 0.97466 | 5010 | 5 | 2.3707 |
| SEH1L | 3 | 0.70041 | 0.70141 | 0.999994 | 16308 | 1 | 5.4455 | 0.087407 | 0.19659 | 0.97466 | 5011 | 2 | 5.4455 |
| TAF2 | 4 | 0.63749 | 0.68606 | 0.999994 | 15480 | 1 | 4.1481 | 0.087412 | 0.21239 | 0.97466 | 5012 | 3 | 4.1481 |
| MYLK2 | 5 | 0.95404 | 0.95421 | 0.999994 | 20435 | 0 | 3.8235 | 0.087422 | 0.22965 | 0.97466 | 5013 | 4 | 3.8235 |
| OPALIN | 6 | 0.2178 | 0.40948 | 0.999994 | 8218 | 3 | 0.77733 | 0.087535 | 0.25304 | 0.97466 | 5014 | 3 | 0.77733 |
| ADAM12 | 6 | 0.83967 | 0.86644 | 0.999994 | 18080 | 1 | 2.7213 | 0.087535 | 0.25304 | 0.97466 | 5015 | 4 | 2.7213 |
| PDE2A | 5 | 0.37923 | 0.54607 | 0.999994 | 11638 | 1 | 0.36076 | 0.087559 | 0.22993 | 0.97466 | 5016 | 2 | 0.36076 |
| NFIX | 5 | 0.68416 | 0.75938 | 0.999994 | 16147 | 1 | 1.403 | 0.087563 | 0.22994 | 0.97466 | 5017 | 4 | 1.403 |
| CLDN23 | 6 | 0.10454 | 0.25619 | 0.994943 | 5115 | 2 | 1.686 | 0.087573 | 0.25312 | 0.97466 | 5018 | 4 | 1.686 |
| TDRD5 | 6 | 0.32107 | 0.5311 | 0.999994 | 10536 | 2 | 3.3225 | 0.087579 | 0.25314 | 0.97466 | 5019 | 4 | 3.3225 |
| OXT | 6 | 0.54352 | 0.69902 | 0.999994 | 14120 | 1 | 1.0164 | 0.087606 | 0.2532 | 0.97466 | 5020 | 5 | 1.0164 |
| NonTargetingControlG | 2 | 0.89779 | 0.8977 | 0.999994 | 19174 | 0 | 5.268 | 0.087628 | 0.15076 | 0.965913 | 5021 | 2 | 5.268 |
| RNF182 | 5 | 0.51135 | 0.65783 | 0.999994 | 13602 | 2 | 2.6407 | 0.087632 | 0.23006 | 0.97466 | 5022 | 3 | 2.6407 |
| RPL39 | 2 | 0.91234 | 0.91239 | 0.999994 | 19469 | 0 | 4.8682 | 0.087663 | 0.1508 | 0.965913 | 5023 | 2 | 4.8682 |
| AVEN | 6 | 0.66286 | 0.78754 | 0.999994 | 15823 | 1 | 2.307 | 0.087689 | 0.25339 | 0.97466 | 5024 | 5 | 2.307 |
| CNIH | 6 | 0.65912 | 0.78463 | 0.999994 | 15764 | 2 | 1.1072 | 0.087771 | 0.25358 | 0.97466 | 5025 | 4 | 1.1072 |
| LSM1 | 6 | 0.96344 | 0.96349 | 0.999994 | 20682 | 0 | 1.3154 | 0.087771 | 0.25358 | 0.97466 | 5026 | 4 | 1.3154 |
| UBE2F | 5 | 0.85923 | 0.86174 | 0.999994 | 18418 | 1 | 1.83 | 0.087803 | 0.2304 | 0.97466 | 5027 | 4 | 1.83 |
| ANO8 | 3 | 0.28863 | 0.42245 | 0.999994 | 9888 | 2 | -0.62287 | 0.087824 | 0.19741 | 0.97466 | 5028 | 1 | -0.62287 |
| SCAMP4 | 5 | 0.088106 | 0.20754 | 0.978852 | 4519 | 2 | 3.2312 | 0.087865 | 0.23052 | 0.97466 | 5029 | 3 | 3.2312 |
| SCFD1 | 4 | 0.27906 | 0.4216 | 0.999994 | 9677 | 1 | 0.55724 | 0.087893 | 0.21317 | 0.97466 | 5030 | 2 | 0.55724 |
| PRPF19 | 3 | 0.55269 | 0.58219 | 0.999994 | 14255 | 1 | 5.359 | 0.087904 | 0.19757 | 0.97466 | 5031 | 2 | 5.359 |
| RHNO1 | 5 | 0.46856 | 0.62093 | 0.999994 | 12983 | 1 | -0.13659 | 0.087953 | 0.23069 | 0.97466 | 5032 | 2 | -0.13659 |
| TMEM156 | 6 | 0.54642 | 0.7011 | 0.999994 | 14159 | 1 | 0.93487 | 0.087969 | 0.25404 | 0.97466 | 5033 | 5 | 0.93487 |
| HOXD11 | 5 | 0.28586 | 0.46956 | 0.999994 | 9829 | 1 | 2.0745 | 0.087978 | 0.23074 | 0.97466 | 5034 | 4 | 2.0745 |
| hsa-mir-4452 | 4 | 0.88442 | 0.88406 | 0.999994 | 18922 | 0 | 2.5039 | 0.08799 | 0.21334 | 0.97466 | 5035 | 3 | 2.5039 |
| KIR3DL3 | 6 | 0.85873 | 0.87672 | 0.999994 | 18414 | 1 | 2.2242 | 0.088022 | 0.25418 | 0.97466 | 5036 | 5 | 2.2242 |
| PIK3CA | 6 | 0.66318 | 0.78778 | 0.999994 | 15825 | 1 | 2.2067 | 0.088023 | 0.25419 | 0.97466 | 5037 | 4 | 2.2067 |
| HRG | 6 | 0.20734 | 0.39683 | 0.999994 | 7955 | 3 | -0.013494 | 0.088023 | 0.25419 | 0.97466 | 5038 | 3 | -0.013494 |
| SCN1B | 5 | 0.35934 | 0.52949 | 0.999994 | 11328 | 1 | 1.3293 | 0.088084 | 0.23096 | 0.97466 | 5039 | 3 | 1.3293 |
| GCHFR | 5 | 0.55898 | 0.6997 | 0.999994 | 14353 | 1 | 3.021 | 0.088099 | 0.23098 | 0.97466 | 5040 | 3 | 3.021 |
| TMEM97 | 5 | 0.96306 | 0.96322 | 0.999994 | 20666 | 0 | 3.2677 | 0.088102 | 0.231 | 0.97466 | 5041 | 4 | 3.2677 |
| CMBL | 6 | 0.052236 | 0.14897 | 0.965566 | 3079 | 4 | -1.7959 | 0.088164 | 0.25452 | 0.97466 | 5042 | 2 | -1.7959 |
| TMOD3 | 6 | 0.84093 | 0.86711 | 0.999994 | 18105 | 1 | 1.713 | 0.088164 | 0.25452 | 0.97466 | 5043 | 3 | 1.713 |

| | | | | | | | | | | | | | |
|---|---|---|---|---|---|---|---|---|---|---|---|---|---|
| ANKLE2 | 3 | 0.73613 | 0.73602 | 0.999994 | 16679 | 0 | 0.013486 | 0.088209 | 0.19818 | 0.97466 | 5044 | 1 | 0.013486 |
| RUFY3 | 5 | 0.74693 | 0.78923 | 0.999994 | 16827 | 1 | 0.74132 | 0.088259 | 0.23131 | 0.97466 | 5045 | 3 | 0.74132 |
| hsa-mir-3180-3 | 4 | 0.61027 | 0.67089 | 0.999994 | 15077 | 1 | 3.0879 | 0.088267 | 0.21377 | 0.97466 | 5046 | 2 | 3.0879 |
| VSTM1 | 6 | 0.97704 | 0.97698 | 0.999994 | 21015 | 0 | 3.2628 | 0.088275 | 0.25478 | 0.97466 | 5047 | 4 | 3.2628 |
| ELF5 | 4 | 0.55236 | 0.64085 | 0.999994 | 14249 | 1 | 2.756 | 0.088291 | 0.21382 | 0.97466 | 5048 | 3 | 2.756 |
| KTI12 | 6 | 0.35801 | 0.57307 | 0.999994 | 11304 | 2 | 2.4081 | 0.088305 | 0.25485 | 0.97466 | 5049 | 4 | 2.4081 |
| DTX4 | 6 | 0.89513 | 0.9 | 0.999994 | 19129 | 1 | 0.98796 | 0.088311 | 0.25486 | 0.97466 | 5050 | 5 | 0.98796 |
| RGS12 | 6 | 0.73736 | 0.82409 | 0.999994 | 16691 | 1 | 1.6106 | 0.088318 | 0.25488 | 0.97466 | 5051 | 5 | 1.6106 |
| PTPN9 | 6 | 0.94665 | 0.94662 | 0.999994 | 20252 | 0 | 1.3086 | 0.088321 | 0.25489 | 0.97466 | 5052 | 4 | 1.3086 |
| KLRC2 | 6 | 0.83743 | 0.86532 | 0.999994 | 18040 | 1 | 3.12 | 0.088329 | 0.2549 | 0.97466 | 5053 | 4 | 3.12 |
| hsa-mir-3137 | 3 | 0.41769 | 0.49473 | 0.999994 | 12215 | 1 | 3.1885 | 0.088356 | 0.19844 | 0.97466 | 5054 | 2 | 3.1885 |
| CEACAM4 | 5 | 0.39792 | 0.56182 | 0.999994 | 11915 | 2 | 0.56899 | 0.088369 | 0.23152 | 0.97466 | 5055 | 2 | 0.56899 |
| JAK1 | 6 | 0.95964 | 0.95971 | 0.999994 | 20575 | 0 | 2.2669 | 0.088432 | 0.25515 | 0.97466 | 5056 | 5 | 2.2669 |
| NonTargetingControlG | 2 | 0.59381 | 0.59374 | 0.999994 | 14876 | 0 | 3.8636 | 0.088448 | 0.15204 | 0.965913 | 5057 | 1 | 3.8636 |
| FBXL22 | 6 | 0.59703 | 0.73778 | 0.999994 | 14918 | 1 | 1.4019 | 0.088455 | 0.2552 | 0.97466 | 5058 | 5 | 1.4019 |
| MT4 | 6 | 0.039029 | 0.11971 | 0.951052 | 2463 | 4 | -1.8997 | 0.088504 | 0.25531 | 0.97466 | 5059 | 1 | -1.8997 |
| KIAA0513 | 6 | 0.9149 | 0.9158 | 0.999994 | 19508 | 1 | 1.5101 | 0.088516 | 0.25534 | 0.97466 | 5060 | 5 | 1.5101 |
| GSTO2 | 6 | 0.3675 | 0.58167 | 0.999994 | 11469 | 1 | 1.0803 | 0.088531 | 0.25537 | 0.97466 | 5061 | 5 | 1.0803 |
| SPEM1 | 5 | 0.86106 | 0.86329 | 0.999994 | 18467 | 1 | 1.8219 | 0.088577 | 0.23193 | 0.97466 | 5062 | 4 | 1.8219 |
| NEK9 | 5 | 0.96369 | 0.96384 | 0.999994 | 20696 | 0 | 2.0467 | 0.088607 | 0.23199 | 0.97466 | 5063 | 4 | 2.0467 |
| COL8A2 | 5 | 0.97202 | 0.97206 | 0.999994 | 20869 | 0 | 2.3035 | 0.088709 | 0.23221 | 0.97466 | 5064 | 4 | 2.3035 |
| CDH22 | 4 | 0.36398 | 0.50861 | 0.999994 | 11403 | 2 | 2.785 | 0.088726 | 0.21453 | 0.97466 | 5065 | 2 | 2.785 |
| FOXB1 | 5 | 0.931 | 0.93128 | 0.999994 | 19877 | 0 | 1.7559 | 0.088731 | 0.23226 | 0.97466 | 5066 | 4 | 1.7559 |
| NAA35 | 6 | 0.94684 | 0.9468 | 0.999994 | 20261 | 0 | 2.7653 | 0.088733 | 0.25581 | 0.97466 | 5067 | 4 | 2.7653 |
| ZNF529 | 6 | 0.96294 | 0.96299 | 0.999994 | 20660 | 0 | 3.0554 | 0.088734 | 0.25581 | 0.97466 | 5068 | 4 | 3.0554 |
| BTN3A3 | 6 | 0.14699 | 0.32186 | 0.999994 | 6375 | 2 | 3.26 | 0.088734 | 0.25581 | 0.97466 | 5069 | 4 | 3.26 |
| AKR1C4 | 6 | 0.44607 | 0.63248 | 0.999994 | 12664 | 2 | 2.5178 | 0.088734 | 0.25581 | 0.97466 | 5070 | 3 | 2.5178 |
| SPTLC3 | 6 | 0.37396 | 0.58571 | 0.999994 | 11570 | 2 | 3.013 | 0.088734 | 0.25581 | 0.97466 | 5071 | 4 | 3.013 |
| PTGDS | 5 | 0.36072 | 0.53069 | 0.999994 | 11351 | 1 | 0.7046 | 0.088736 | 0.23227 | 0.97466 | 5072 | 3 | 0.7046 |
| DNMT3B | 5 | 0.17699 | 0.34228 | 0.999994 | 7156 | 2 | 3.6773 | 0.088736 | 0.23227 | 0.97466 | 5073 | 3 | 3.6773 |
| ENOPH1 | 6 | 0.46209 | 0.64312 | 0.999994 | 12889 | 1 | 1.0865 | 0.088745 | 0.25583 | 0.97466 | 5074 | 5 | 1.0865 |
| SCAMP2 | 6 | 0.28123 | 0.48483 | 0.999994 | 9729 | 1 | 2.3738 | 0.088785 | 0.25592 | 0.974731 | 5075 | 4 | 2.3738 |
| BCL3 | 6 | 0.30416 | 0.51156 | 0.999994 | 10214 | 3 | -0.63301 | 0.088792 | 0.25594 | 0.974731 | 5076 | 2 | -0.63301 |
| CABP5 | 5 | 0.75472 | 0.79333 | 0.999994 | 16907 | 1 | 3.9749 | 0.088796 | 0.23238 | 0.97466 | 5077 | 3 | 3.9749 |
| CAPN12 | 6 | 0.32232 | 0.53257 | 0.999994 | 10561 | 1 | 2.2928 | 0.088933 | 0.25626 | 0.974766 | 5078 | 4 | 2.2928 |
| ADORA2A | 6 | 0.10779 | 0.26245 | 0.995875 | 5231 | 2 | 2.2324 | 0.088933 | 0.25626 | 0.974766 | 5079 | 4 | 2.2324 |
| C1QL3 | 6 | 0.058897 | 0.16341 | 0.967126 | 3377 | 3 | -0.4563 | 0.088933 | 0.25626 | 0.974766 | 5080 | 3 | -0.4563 |
| JMY | 5 | 0.3063 | 0.48616 | 0.999994 | 10261 | 2 | -0.098715 | 0.088938 | 0.23267 | 0.97466 | 5081 | 1 | -0.098715 |
| NonTargetingControlG | 2 | 0.38228 | 0.40214 | 0.999994 | 11682 | 1 | 3.0621 | 0.08897 | 0.15287 | 0.965913 | 5082 | 1 | 3.0621 |
| HMGA2 | 5 | 0.57576 | 0.71461 | 0.999994 | 14629 | 1 | 1.9768 | 0.088996 | 0.23278 | 0.97466 | 5083 | 4 | 1.9768 |
| OSBP | 5 | 0.21418 | 0.39521 | 0.999994 | 8122 | 1 | 1.8439 | 0.089003 | 0.2328 | 0.97466 | 5084 | 4 | 1.8439 |
| LAIR2 | 5 | 0.59934 | 0.72592 | 0.999994 | 14945 | 1 | 1.2405 | 0.089032 | 0.23285 | 0.97466 | 5085 | 4 | 1.2405 |
| HTR5A | 6 | 0.53068 | 0.69002 | 0.999994 | 13907 | 2 | 2.0282 | 0.08908 | 0.25657 | 0.974766 | 5086 | 3 | 2.0282 |
| CXorf48 | 6 | 0.26622 | 0.46722 | 0.999994 | 9369 | 2 | 2.3998 | 0.08908 | 0.25657 | 0.974766 | 5087 | 3 | 2.3998 |
| RAB3GAP1 | 5 | 0.3164 | 0.49443 | 0.999994 | 10443 | 1 | 1.1791 | 0.089084 | 0.23295 | 0.97466 | 5088 | 4 | 1.1791 |
| RNF126 | 6 | 0.88698 | 0.89423 | 0.999994 | 18966 | 1 | 3.5395 | 0.089088 | 0.2566 | 0.974766 | 5089 | 4 | 3.5395 |
| COMMD2 | 5 | 0.13125 | 0.27454 | 0.998561 | 5964 | 1 | 1.1301 | 0.089105 | 0.23301 | 0.97466 | 5090 | 4 | 1.1301 |
| GPR37L1 | 5 | 0.25905 | 0.44799 | 0.999994 | 9220 | 1 | 1.4748 | 0.089106 | 0.23301 | 0.97466 | 5091 | 4 | 1.4748 |
| TMEM214 | 6 | 0.73784 | 0.82426 | 0.999994 | 16699 | 1 | 2.5272 | 0.08914 | 0.2567 | 0.974766 | 5092 | 4 | 2.5272 |
| PRRX2 | 5 | 0.78588 | 0.81085 | 0.999994 | 17292 | 1 | 3.7014 | 0.089167 | 0.23313 | 0.97466 | 5093 | 3 | 3.7014 |
| SPATA16 | 6 | 0.026711 | 0.091042 | 0.94395 | 1841 | 1 | 2.7764 | 0.089264 | 0.25699 | 0.974766 | 5094 | 5 | 2.7764 |
| TMEM243 | 6 | 0.83816 | 0.86569 | 0.999994 | 18057 | 1 | 2.1804 | 0.08927 | 0.257 | 0.974766 | 5095 | 4 | 2.1804 |

| | | | | | | | | | | | |
|---|---|---|---|---|---|---|---|---|---|---|---|
| EGLN3 | 5 | 0.42736 | 0.58624 | 0.999994 | 12374 | 2 | 0.32632 | 0.089288 | 0.23335 | 0.97466 | 5096 | 2 | 0.32632 |
| HERC4 | 5 | 0.39254 | 0.55731 | 0.999994 | 11842 | 1 | 2.0261 | 0.08929 | 0.23336 | 0.97466 | 5097 | 4 | 2.0261 |
| MRPL17 | 3 | 0.83336 | 0.83326 | 0.999994 | 17978 | 0 | 2.3076 | 0.089339 | 0.20031 | 0.97466 | 5098 | 2 | 2.3076 |
| ADNP | 6 | 0.36968 | 0.58303 | 0.999994 | 11502 | 2 | 1.4181 | 0.089351 | 0.2572 | 0.974766 | 5099 | 3 | 1.4181 |
| hsa-mir-6794 | 2 | 0.91064 | 0.91068 | 0.999994 | 19443 | 0 | 4.5542 | 0.08936 | 0.15349 | 0.965913 | 5100 | 2 | 4.5542 |
| TTC4 | 6 | 0.98507 | 0.98504 | 0.999994 | 21225 | 0 | 1.3532 | 0.089425 | 0.25736 | 0.974766 | 5101 | 5 | 1.3532 |
| BACH2 | 6 | 0.53189 | 0.69088 | 0.999994 | 13934 | 2 | 2.404 | 0.089443 | 0.25741 | 0.974766 | 5102 | 4 | 2.404 |
| SGPP1 | 6 | 0.66396 | 0.78836 | 0.999994 | 15839 | 1 | 2.0415 | 0.089456 | 0.25743 | 0.974766 | 5103 | 5 | 2.0415 |
| SERAC1 | 6 | 0.20566 | 0.39475 | 0.999994 | 7915 | 2 | 2.3542 | 0.08946 | 0.25744 | 0.974766 | 5104 | 3 | 2.3542 |
| GDPD3 | 6 | 0.35827 | 0.57335 | 0.999994 | 11308 | 2 | 2.4527 | 0.08946 | 0.25744 | 0.974766 | 5105 | 3 | 2.4527 |
| NonTargetingControlG | 2 | 0.27122 | 0.31477 | 0.999994 | 9504 | 1 | 2.4884 | 0.089474 | 0.15368 | 0.965913 | 5106 | 1 | 2.4884 |
| OR1I1 | 5 | 0.010258 | 0.036817 | 0.881473 | 885 | 1 | 1.3401 | 0.089489 | 0.23377 | 0.97466 | 5107 | 3 | 1.3401 |
| CACNG4 | 5 | 0.53137 | 0.67529 | 0.999994 | 13920 | 1 | 3.7666 | 0.089535 | 0.23386 | 0.97466 | 5108 | 4 | 3.7666 |
| OR4C46 | 4 | 0.3235 | 0.46701 | 0.999994 | 10582 | 2 | 2.6167 | 0.08956 | 0.21591 | 0.97466 | 5109 | 2 | 2.6167 |
| SNUPN | 6 | 0.070113 | 0.18701 | 0.969409 | 3834 | 2 | 2.726 | 0.089591 | 0.25775 | 0.974766 | 5110 | 4 | 2.726 |
| PDK3 | 6 | 0.051201 | 0.14671 | 0.964852 | 3033 | 4 | -1.6498 | 0.089604 | 0.25778 | 0.974766 | 5111 | 1 | -1.6498 |
| NOS1AP | 5 | 0.66437 | 0.75092 | 0.999994 | 15847 | 1 | 3.69 | 0.089609 | 0.234 | 0.97466 | 5112 | 4 | 3.69 |
| hsa-let-7d | 3 | 0.80344 | 0.80323 | 0.999994 | 17532 | 0 | 5.3797 | 0.089622 | 0.20087 | 0.97466 | 5113 | 2 | 5.3797 |
| ALDH18A1 | 5 | 0.54174 | 0.68444 | 0.999994 | 14086 | 1 | 3.6643 | 0.089625 | 0.23403 | 0.97466 | 5114 | 4 | 3.6643 |
| TLE6 | 5 | 0.020011 | 0.064111 | 0.93914 | 1461 | 3 | -3.403 | 0.089638 | 0.23405 | 0.97466 | 5115 | 2 | -3.403 |
| DRAP1 | 5 | 0.044968 | 0.12822 | 0.951777 | 2755 | 2 | 0.7669 | 0.089638 | 0.23405 | 0.97466 | 5116 | 3 | 0.7669 |
| GPR135 | 5 | 0.82231 | 0.83412 | 0.999994 | 17807 | 1 | 0.35554 | 0.089638 | 0.23405 | 0.97466 | 5117 | 2 | 0.35554 |
| MGAT5B | 5 | 0.72409 | 0.77783 | 0.999994 | 16534 | 1 | 3.8355 | 0.089644 | 0.23407 | 0.97466 | 5118 | 4 | 3.8355 |
| FBXO33 | 6 | 0.51139 | 0.67657 | 0.999994 | 13603 | 1 | 2.4154 | 0.089656 | 0.2579 | 0.974766 | 5119 | 5 | 2.4154 |
| FKBP15 | 6 | 0.41884 | 0.61461 | 0.999994 | 12232 | 1 | 2.6004 | 0.089678 | 0.25795 | 0.974766 | 5120 | 4 | 2.6004 |
| FAM204A | 6 | 0.023087 | 0.081853 | 0.93914 | 1655 | 3 | -0.33882 | 0.089727 | 0.25807 | 0.974766 | 5121 | 3 | -0.33882 |
| KRTAP20-3 | 6 | 0.029428 | 0.097438 | 0.945253 | 1978 | 2 | 2.4696 | 0.089739 | 0.2581 | 0.974766 | 5122 | 4 | 2.4696 |
| FOXK2 | 4 | 0.72359 | 0.74059 | 0.999994 | 16528 | 1 | 3.914 | 0.089754 | 0.21623 | 0.97466 | 5123 | 3 | 3.914 |
| PTTG1 | 4 | 0.83004 | 0.82957 | 0.999994 | 17932 | 0 | 2.4983 | 0.089761 | 0.21625 | 0.97466 | 5124 | 3 | 2.4983 |
| FRK | 4 | 0.9176 | 0.9174 | 0.999994 | 19580 | 0 | 2.5917 | 0.089769 | 0.21626 | 0.97466 | 5125 | 3 | 2.5917 |
| TRIM45 | 6 | 0.95583 | 0.95591 | 0.999994 | 20475 | 0 | 0.90785 | 0.089778 | 0.25818 | 0.974766 | 5126 | 5 | 0.90785 |
| MAN2A2 | 5 | 0.3474 | 0.51979 | 0.999994 | 11097 | 2 | -0.22582 | 0.089791 | 0.23437 | 0.97466 | 5127 | 1 | -0.22582 |
| MAGEB18 | 6 | 0.54354 | 0.69905 | 0.999994 | 14123 | 2 | 1.4318 | 0.089839 | 0.25832 | 0.974766 | 5128 | 4 | 1.4318 |
| hsa-mir-4316 | 3 | 0.91013 | 0.91016 | 0.999994 | 19426 | 0 | 2.4812 | 0.089865 | 0.20136 | 0.97466 | 5129 | 3 | 2.4812 |
| FGF12 | 4 | 0.010127 | 0.034109 | 0.867348 | 879 | 2 | -4.3068 | 0.08988 | 0.21643 | 0.97466 | 5130 | 1 | -4.3068 |
| FADS1 | 5 | 0.61784 | 0.73277 | 0.999994 | 15183 | 1 | 2.3652 | 0.08988 | 0.23456 | 0.97466 | 5131 | 4 | 2.3652 |
| NABP1 | 5 | 0.051533 | 0.14409 | 0.961983 | 3048 | 3 | -1.6436 | 0.0899 | 0.2346 | 0.97466 | 5132 | 1 | -1.6436 |
| NMUR2 | 6 | 0.64363 | 0.77269 | 0.999994 | 15570 | 2 | 2.5635 | 0.089983 | 0.25864 | 0.974766 | 5133 | 4 | 2.5635 |
| PPM1H | 6 | 0.90191 | 0.90517 | 0.999994 | 19250 | 1 | 2.3851 | 0.090002 | 0.25869 | 0.974766 | 5134 | 5 | 2.3851 |
| TRMT1L | 6 | 0.49587 | 0.66587 | 0.999994 | 13356 | 2 | 1.8623 | 0.090061 | 0.25881 | 0.974766 | 5135 | 4 | 1.8623 |
| PIWIL1 | 4 | 0.023382 | 0.0678 | 0.93914 | 1674 | 1 | 3.3701 | 0.090096 | 0.21678 | 0.97466 | 5136 | 2 | 3.3701 |
| TNS1 | 6 | 0.37253 | 0.5848 | 0.999994 | 11551 | 1 | 2.1277 | 0.0901 | 0.25891 | 0.974766 | 5137 | 4 | 2.1277 |
| CLEC9A | 6 | 0.32929 | 0.54042 | 0.999994 | 10706 | 2 | 2.0208 | 0.0901 | 0.25891 | 0.974766 | 5138 | 4 | 2.0208 |
| REXO1L1 | 6 | 0.1082 | 0.26325 | 0.996734 | 5245 | 2 | 2.6009 | 0.090177 | 0.25908 | 0.974766 | 5139 | 3 | 2.6009 |
| PLCH2 | 6 | 0.057653 | 0.16069 | 0.967126 | 3319 | 2 | 2.3081 | 0.090177 | 0.25908 | 0.974766 | 5140 | 3 | 2.3081 |
| PLD6 | 6 | 0.3027 | 0.50987 | 0.999994 | 10187 | 1 | 3.5397 | 0.090177 | 0.25908 | 0.974766 | 5141 | 4 | 3.5397 |
| SLC35B2 | 6 | 0.18734 | 0.37221 | 0.999994 | 7402 | 3 | 1.9175 | 0.090177 | 0.25908 | 0.974766 | 5142 | 3 | 1.9175 |
| HES3 | 5 | 0.54926 | 0.69109 | 0.999994 | 14198 | 2 | -0.23489 | 0.090228 | 0.23526 | 0.97466 | 5143 | 2 | -0.23489 |
| FAM163B | 5 | 0.93078 | 0.93106 | 0.999994 | 19871 | 0 | 2.7259 | 0.090258 | 0.23531 | 0.97466 | 5144 | 4 | 2.7259 |
| IMPA1 | 4 | 0.23816 | 0.3798 | 0.999994 | 8730 | 2 | -0.48904 | 0.090266 | 0.21707 | 0.97466 | 5145 | 1 | -0.48904 |
| CARHSP1 | 5 | 0.17245 | 0.3357 | 0.999994 | 7047 | 2 | 2.2214 | 0.090345 | 0.2355 | 0.97466 | 5146 | 3 | 2.2214 |
| OR14I1 | 5 | 0.87101 | 0.87197 | 0.999994 | 18654 | 1 | 3.1166 | 0.090345 | 0.2355 | 0.97466 | 5147 | 3 | 3.1166 |

| | | | | | | | | | | | | |
|---|---|---|---|---|---|---|---|---|---|---|---|---|
| ACTR6 | 6 | 0.056063 | 0.15723 | 0.967126 | 3250 | 1 | 3.2821 | 0.090372 | 0.25953 | 0.974766 | 5148 | 5 | 3.2821 |
| RIIAD1 | 6 | 0.24907 | 0.44682 | 0.999994 | 8995 | 1 | 2.9181 | 0.09038 | 0.25954 | 0.974766 | 5149 | 5 | 2.9181 |
| GARNL3 | 5 | 0.72461 | 0.77809 | 0.999994 | 16542 | 1 | 3.4241 | 0.090421 | 0.23565 | 0.97466 | 5150 | 4 | 3.4241 |
| CCR4 | 6 | 0.58768 | 0.73088 | 0.999994 | 14786 | 2 | 1.0593 | 0.090447 | 0.2597 | 0.974766 | 5151 | 3 | 1.0593 |
| NARR | 6 | 0.93925 | 0.93919 | 0.999994 | 20056 | 0 | 1.1136 | 0.090447 | 0.2597 | 0.974766 | 5152 | 4 | 1.1136 |
| GFOD1 | 6 | 0.69453 | 0.81037 | 0.999994 | 16248 | 1 | 3.6016 | 0.09048 | 0.25978 | 0.974766 | 5153 | 5 | 3.6016 |
| DEFB106B | 2 | 0.90943 | 0.90945 | 0.999994 | 19412 | 0 | 4.8112 | 0.090571 | 0.15539 | 0.965913 | 5154 | 2 | 4.8112 |
| PIK3AP1 | 6 | 0.0076912 | 0.03088 | 0.862223 | 706 | 3 | -0.75438 | 0.090571 | 0.25999 | 0.974766 | 5155 | 2 | -0.75438 |
| C9orf40 | 5 | 0.84871 | 0.85327 | 0.999994 | 18228 | 1 | 1.3501 | 0.090731 | 0.23628 | 0.97466 | 5156 | 4 | 1.3501 |
| ZNF785 | 5 | 0.60724 | 0.72879 | 0.999994 | 15039 | 1 | 1.3073 | 0.090731 | 0.23628 | 0.97466 | 5157 | 3 | 1.3073 |
| RSPH10B2 | 5 | 0.42258 | 0.58222 | 0.999994 | 12291 | 1 | 2.3416 | 0.090731 | 0.23628 | 0.97466 | 5158 | 3 | 2.3416 |
| FAM133A | 5 | 0.90879 | 0.90882 | 0.999994 | 19400 | 0 | 1.9326 | 0.090731 | 0.23628 | 0.97466 | 5159 | 4 | 1.9326 |
| DEFB119 | 6 | 0.10262 | 0.25243 | 0.994162 | 5058 | 2 | 2.3157 | 0.090754 | 0.2604 | 0.97477 | 5160 | 4 | 2.3157 |
| LGALS16 | 5 | 0.90519 | 0.9053 | 0.999994 | 19335 | 0 | 3.8453 | 0.090802 | 0.23642 | 0.97466 | 5161 | 4 | 3.8453 |
| DRAM2 | 4 | 0.040278 | 0.10233 | 0.946414 | 2530 | 1 | 3.0054 | 0.090804 | 0.21797 | 0.97466 | 5162 | 3 | 3.0054 |
| ABCC1 | 5 | 0.96584 | 0.96599 | 0.999994 | 20734 | 0 | 1.1173 | 0.09084 | 0.23649 | 0.97466 | 5163 | 3 | 1.1173 |
| TUBG2 | 4 | 0.73989 | 0.7524 | 0.999994 | 16738 | 1 | 3.3837 | 0.090909 | 0.21814 | 0.97466 | 5164 | 3 | 3.3837 |
| PKD2 | 6 | 0.039907 | 0.1217 | 0.951052 | 2510 | 2 | 2.5263 | 0.090928 | 0.26081 | 0.974875 | 5165 | 4 | 2.5263 |
| UPB1 | 6 | 0.9853 | 0.98526 | 0.999994 | 21235 | 0 | 1.8421 | 0.090937 | 0.26083 | 0.974875 | 5166 | 5 | 1.8421 |
| NAT8B | 6 | 0.39453 | 0.59885 | 0.999994 | 11866 | 2 | 2.8125 | 0.090958 | 0.26088 | 0.974875 | 5167 | 3 | 2.8125 |
| OR1C1 | 6 | 0.49688 | 0.66658 | 0.999994 | 13373 | 2 | 3.1932 | 0.090958 | 0.26088 | 0.974875 | 5168 | 4 | 3.1932 |
| MOSPD2 | 6 | 0.06651 | 0.17945 | 0.968171 | 3704 | 4 | -1.6124 | 0.090963 | 0.26089 | 0.974875 | 5169 | 1 | -1.6124 |
| COL5A3 | 4 | 0.92406 | 0.92387 | 0.999994 | 19715 | 0 | 4.0945 | 0.090992 | 0.21828 | 0.97466 | 5170 | 3 | 4.0945 |
| hsa-mir-552 | 3 | 0.18018 | 0.28751 | 0.999632 | 7223 | 2 | -1.3834 | 0.091052 | 0.20367 | 0.97466 | 5171 | 1 | -1.3834 |
| DMPK | 5 | 0.8626 | 0.86459 | 0.999994 | 18496 | 1 | 3.8042 | 0.091123 | 0.23708 | 0.97466 | 5172 | 3 | 3.8042 |
| ZNF668 | 4 | 0.20183 | 0.34253 | 0.999994 | 7819 | 2 | 2.6722 | 0.091127 | 0.2185 | 0.97466 | 5173 | 2 | 2.6722 |
| MANBA | 5 | 0.79878 | 0.81877 | 0.999994 | 17460 | 1 | 0.71106 | 0.091153 | 0.23713 | 0.97466 | 5174 | 3 | 0.71106 |
| IGDCC4 | 6 | 0.86385 | 0.87962 | 0.999994 | 18518 | 1 | 2.492 | 0.091194 | 0.26142 | 0.974875 | 5175 | 5 | 2.492 |
| MYB | 5 | 0.33098 | 0.50635 | 0.999994 | 10743 | 1 | 3.9691 | 0.09121 | 0.23723 | 0.97466 | 5176 | 4 | 3.9691 |
| TUBGCP6 | 5 | 0.81836 | 0.83147 | 0.999994 | 17753 | 1 | 3.9691 | 0.09121 | 0.23723 | 0.97466 | 5177 | 4 | 3.9691 |
| PEX2 | 5 | 0.31732 | 0.49518 | 0.999994 | 10459 | 1 | 0.033401 | 0.091211 | 0.23723 | 0.97466 | 5178 | 1 | 0.033401 |
| RHOV | 6 | 0.89497 | 0.89988 | 0.999994 | 19126 | 1 | 2.4462 | 0.091225 | 0.26149 | 0.974875 | 5179 | 5 | 2.4462 |
| VWA8 | 5 | 0.90033 | 0.90045 | 0.999994 | 19227 | 0 | 2.9044 | 0.091278 | 0.23736 | 0.97466 | 5180 | 4 | 2.9044 |
| CEP120 | 6 | 0.33535 | 0.54735 | 0.999994 | 10839 | 3 | -0.57695 | 0.091302 | 0.26168 | 0.974875 | 5181 | 2 | -0.57695 |
| A2ML1 | 6 | 0.096669 | 0.24081 | 0.987253 | 4845 | 2 | 1.2515 | 0.091302 | 0.26168 | 0.974875 | 5182 | 4 | 1.2515 |
| HCCS | 5 | 0.51289 | 0.65919 | 0.999994 | 13618 | 1 | 1.429 | 0.091338 | 0.23748 | 0.97466 | 5183 | 4 | 1.429 |
| HR | 6 | 0.92648 | 0.92657 | 0.999994 | 19779 | 1 | 0.90379 | 0.091404 | 0.26191 | 0.974875 | 5184 | 5 | 0.90379 |
| CEP85 | 6 | 0.10314 | 0.25345 | 0.994501 | 5072 | 2 | 3.6027 | 0.091453 | 0.26201 | 0.974875 | 5185 | 4 | 3.6027 |
| KLF11 | 6 | 0.075572 | 0.19835 | 0.977164 | 4057 | 1 | 2.1794 | 0.091458 | 0.26203 | 0.974875 | 5186 | 5 | 2.1794 |
| hsa-mir-1206 | 1 | 0.90851 | 0.90883 | 0.999994 | 19395 | 0 | 7.3134 | 0.091485 | 0.091462 | 0.945479 | 5187 | 1 | 7.3134 |
| RGS21 | 4 | 0.20963 | 0.3506 | 0.999994 | 8020 | 1 | 3.1181 | 0.091502 | 0.21908 | 0.97466 | 5188 | 2 | 3.1181 |
| DIMT1 | 4 | 0.30201 | 0.44503 | 0.999994 | 10162 | 1 | 3.1616 | 0.091502 | 0.21908 | 0.97466 | 5189 | 2 | 3.1616 |
| RPL19 | 4 | 0.30547 | 0.44856 | 0.999994 | 10242 | 1 | 4.0004 | 0.091502 | 0.21908 | 0.97466 | 5190 | 3 | 4.0004 |
| APOBEC3A | 5 | 0.8552 | 0.85841 | 0.999994 | 18350 | 1 | 1.307 | 0.091577 | 0.23791 | 0.97466 | 5191 | 4 | 1.307 |
| ELAC2 | 5 | 0.37803 | 0.54511 | 0.999994 | 11625 | 2 | -0.30016 | 0.091582 | 0.23792 | 0.97466 | 5192 | 1 | -0.30016 |
| HIC1 | 5 | 0.13952 | 0.2871 | 0.999632 | 6173 | 3 | -1.1698 | 0.091582 | 0.23792 | 0.97466 | 5193 | 2 | -1.1698 |
| RD3L | 6 | 0.74113 | 0.82537 | 0.999994 | 16754 | 1 | 3.01 | 0.091606 | 0.26237 | 0.974875 | 5194 | 4 | 3.01 |
| SPDYE1 | 6 | 0.00057139 | 0.0029868 | 0.53161 | 115 | 5 | -2.9871 | 0.091616 | 0.26239 | 0.974875 | 5195 | 1 | -2.9871 |
| GFI1B | 5 | 0.15915 | 0.3163 | 0.999994 | 6707 | 1 | 0.42913 | 0.091655 | 0.23807 | 0.97466 | 5196 | 2 | 0.42913 |
| RAB37 | 5 | 0.12443 | 0.2641 | 0.996734 | 5779 | 2 | 3.1813 | 0.091655 | 0.23807 | 0.97466 | 5197 | 3 | 3.1813 |
| SLC52A3 | 5 | 0.08804 | 0.20743 | 0.978852 | 4515 | 2 | 2.6294 | 0.091655 | 0.23807 | 0.97466 | 5198 | 3 | 2.6294 |
| YTHDF1 | 6 | 0.95707 | 0.95715 | 0.999994 | 20521 | 0 | 1.9326 | 0.091661 | 0.2625 | 0.974875 | 5199 | 5 | 1.9326 |

| | | | | | | | | | | | | |
|---|---|---|---|---|---|---|---|---|---|---|---|---|
| DSC1 | 6 | 0.8999 | 0.9036 | 0.999994 | 19220 | 1 | 2.5968 | 0.091669 | 0.26252 | 0.974875 | 5200 | 5 | 2.5968 |
| SIAH3 | 6 | 0.11069 | 0.26804 | 0.998561 | 5324 | 1 | 1.3267 | 0.091762 | 0.26273 | 0.974875 | 5201 | 4 | 1.3267 |
| MRPL3 | 3 | 0.89437 | 0.89442 | 0.999994 | 19110 | 0 | 5.4487 | 0.091765 | 0.20506 | 0.97466 | 5202 | 3 | 5.4487 |
| NCBP1 | 6 | 0.25823 | 0.45779 | 0.999994 | 9201 | 1 | 2.8546 | 0.091821 | 0.26287 | 0.974875 | 5203 | 4 | 2.8546 |
| FRMD7 | 6 | 0.27187 | 0.47392 | 0.999994 | 9518 | 3 | 2.0551 | 0.091821 | 0.26287 | 0.974875 | 5204 | 3 | 2.0551 |
| RPL13 | 3 | 0.10586 | 0.1908 | 0.972248 | 5167 | 1 | 0.35554 | 0.091822 | 0.20518 | 0.97466 | 5205 | 1 | 0.35554 |
| ERCC4 | 5 | 0.053322 | 0.14838 | 0.965554 | 3129 | 1 | 1.2881 | 0.091822 | 0.23841 | 0.97466 | 5206 | 4 | 1.2881 |
| C5AR1 | 6 | 0.97804 | 0.97799 | 0.999994 | 21042 | 0 | 2.6144 | 0.091848 | 0.26293 | 0.974875 | 5207 | 5 | 2.6144 |
| YBEY | 6 | 0.55803 | 0.70944 | 0.999994 | 14334 | 1 | 1.7333 | 0.091877 | 0.26298 | 0.974875 | 5208 | 4 | 1.7333 |
| SLC26A11 | 6 | 0.93212 | 0.93224 | 0.999994 | 19906 | 0 | 1.4271 | 0.091877 | 0.26298 | 0.974875 | 5209 | 4 | 1.4271 |
| ZNF550 | 6 | 0.25984 | 0.45966 | 0.999994 | 9236 | 2 | 1.7778 | 0.091877 | 0.26298 | 0.974875 | 5210 | 3 | 1.7778 |
| CUEDC2 | 4 | 0.056004 | 0.13317 | 0.95318 | 3247 | 2 | 0.99071 | 0.091892 | 0.21973 | 0.97466 | 5211 | 2 | 0.99071 |
| hsa-mir-3198-2 | 1 | 0.90809 | 0.90839 | 0.999994 | 19385 | 0 | 7.304 | 0.091914 | 0.091898 | 0.946116 | 5212 | 1 | 7.304 |
| PTPRG | 5 | 0.76511 | 0.79897 | 0.999994 | 17030 | 1 | 2.4563 | 0.091945 | 0.23863 | 0.97466 | 5213 | 4 | 2.4563 |
| ANXA1 | 5 | 0.10497 | 0.23417 | 0.984971 | 5132 | 2 | -0.4785 | 0.091953 | 0.23866 | 0.97466 | 5214 | 1 | -0.4785 |
| PDE10A | 5 | 0.033256 | 0.098879 | 0.945253 | 2175 | 2 | 0.12511 | 0.091953 | 0.23866 | 0.97466 | 5215 | 2 | 0.12511 |
| HIST1H2BH | 5 | 0.87688 | 0.8773 | 0.999994 | 18768 | 1 | 3.4733 | 0.091975 | 0.2387 | 0.97466 | 5216 | 3 | 3.4733 |
| RGAG1 | 3 | 0.23374 | 0.35477 | 0.999994 | 8613 | 2 | -0.9569 | 0.091994 | 0.20551 | 0.97466 | 5217 | 1 | -0.9569 |
| SCMH1 | 6 | 0.67538 | 0.79738 | 0.999994 | 16010 | 2 | 2.1868 | 0.092056 | 0.26339 | 0.974875 | 5218 | 4 | 2.1868 |
| HSF2 | 5 | 0.17687 | 0.3421 | 0.999994 | 7150 | 2 | 1.8422 | 0.092062 | 0.23889 | 0.97466 | 5219 | 3 | 1.8422 |
| hsa-mir-3187 | 3 | 0.83394 | 0.83383 | 0.999994 | 17987 | 0 | 5.3065 | 0.0921 | 0.2057 | 0.97466 | 5220 | 2 | 5.3065 |
| hsa-mir-4525 | 3 | 0.66339 | 0.66823 | 0.999994 | 15829 | 1 | 5.3065 | 0.0921 | 0.2057 | 0.97466 | 5221 | 2 | 5.3065 |
| RBP5 | 5 | 0.53457 | 0.67809 | 0.999994 | 13973 | 1 | 3.5113 | 0.092134 | 0.23904 | 0.97466 | 5222 | 4 | 3.5113 |
| GTPBP3 | 6 | 0.1195 | 0.28479 | 0.999632 | 5610 | 2 | 0.182 | 0.092141 | 0.26357 | 0.974875 | 5223 | 2 | 0.182 |
| FAM203A | 6 | 0.88738 | 0.8945 | 0.999994 | 18977 | 1 | 0.98811 | 0.092141 | 0.26357 | 0.974875 | 5224 | 4 | 0.98811 |
| C20orf194 | 5 | 0.85951 | 0.86199 | 0.999994 | 18428 | 1 | 1.6557 | 0.0922 | 0.23918 | 0.97466 | 5225 | 4 | 1.6557 |
| GHRH | 5 | 0.88089 | 0.88111 | 0.999994 | 18851 | 1 | 3.4511 | 0.092201 | 0.23918 | 0.97466 | 5226 | 3 | 3.4511 |
| PNKP | 5 | 0.97204 | 0.97208 | 0.999994 | 20870 | 0 | 2.9834 | 0.092201 | 0.23918 | 0.97466 | 5227 | 4 | 2.9834 |
| LYSMD2 | 5 | 0.31783 | 0.49558 | 0.999994 | 10468 | 1 | 3.752 | 0.092204 | 0.23919 | 0.97466 | 5228 | 4 | 3.752 |
| hsa-mir-4643 | 3 | 0.70069 | 0.70165 | 0.999994 | 16311 | 1 | 1.9886 | 0.092206 | 0.20591 | 0.97466 | 5229 | 2 | 1.9886 |
| CLCNKB | 5 | 0.90911 | 0.90911 | 0.999994 | 19404 | 0 | 3.8789 | 0.092236 | 0.23924 | 0.97466 | 5230 | 4 | 3.8789 |
| C10orf53 | 6 | 0.6785 | 0.79983 | 0.999994 | 16068 | 2 | 2.8524 | 0.092242 | 0.26381 | 0.97488 | 5231 | 4 | 2.8524 |
| TARP | 5 | 0.5083 | 0.65516 | 0.999994 | 13558 | 2 | 1.1983 | 0.092346 | 0.23947 | 0.97466 | 5232 | 3 | 1.1983 |
| DBI | 4 | 0.85593 | 0.85552 | 0.999994 | 18372 | 0 | 1.5836 | 0.092355 | 0.22047 | 0.97466 | 5233 | 3 | 1.5836 |
| NonTargetingControlG | 2 | 0.90764 | 0.90762 | 0.999994 | 19378 | 0 | 4.4061 | 0.092364 | 0.15821 | 0.965913 | 5234 | 2 | 4.4061 |
| CNN1 | 3 | 0.25342 | 0.37922 | 0.999994 | 9089 | 2 | -0.84763 | 0.092379 | 0.20622 | 0.97466 | 5235 | 1 | -0.84763 |
| ZNF385D | 6 | 0.15233 | 0.32861 | 0.999994 | 6514 | 3 | 0.56033 | 0.092425 | 0.26422 | 0.975894 | 5236 | 3 | 0.56033 |
| C1orf21 | 5 | 0.84265 | 0.84867 | 0.999994 | 18134 | 1 | 2.7992 | 0.092456 | 0.23969 | 0.97466 | 5237 | 4 | 2.7992 |
| C8orf74 | 5 | 0.66494 | 0.75117 | 0.999994 | 15855 | 1 | 0.1237 | 0.09252 | 0.23981 | 0.97466 | 5238 | 2 | 0.1237 |
| KRTAP17-1 | 6 | 0.31779 | 0.52735 | 0.999994 | 10466 | 2 | 0.91913 | 0.092556 | 0.2645 | 0.976218 | 5239 | 3 | 0.91913 |
| RAP1A | 6 | 0.090258 | 0.228 | 0.983354 | 4601 | 2 | 0.20996 | 0.092556 | 0.2645 | 0.976218 | 5240 | 2 | 0.20996 |
| TMEM107 | 6 | 0.31795 | 0.52754 | 0.999994 | 10471 | 2 | 0.73551 | 0.092581 | 0.26456 | 0.976218 | 5241 | 3 | 0.73551 |
| ALPK2 | 4 | 0.2053 | 0.34611 | 0.999994 | 7903 | 2 | -0.9189 | 0.092583 | 0.22087 | 0.97466 | 5242 | 1 | -0.9189 |
| FOXRED2 | 4 | 0.23962 | 0.38127 | 0.999994 | 8765 | 2 | 0.42797 | 0.092583 | 0.22087 | 0.97466 | 5243 | 2 | 0.42797 |
| HOXD8 | 6 | 0.34859 | 0.56241 | 0.999994 | 11125 | 2 | 2.882 | 0.092592 | 0.26458 | 0.976218 | 5244 | 4 | 2.882 |
| SLC31A2 | 3 | 0.86592 | 0.8659 | 0.999994 | 18553 | 0 | 5.2982 | 0.092687 | 0.20683 | 0.97466 | 5245 | 2 | 5.2982 |
| PRR23A | 5 | 0.19952 | 0.37454 | 0.999994 | 7750 | 2 | 1.9591 | 0.09269 | 0.24015 | 0.97466 | 5246 | 3 | 1.9591 |
| APBA3 | 5 | 0.12613 | 0.26673 | 0.998555 | 5824 | 2 | -0.29235 | 0.09269 | 0.24015 | 0.97466 | 5247 | 2 | -0.29235 |
| C7orf63 | 5 | 0.95124 | 0.95141 | 0.999994 | 20371 | 0 | 1.7496 | 0.092695 | 0.24017 | 0.97466 | 5248 | 4 | 1.7496 |
| PHYH | 6 | 0.26637 | 0.4674 | 0.999994 | 9378 | 2 | 3.4964 | 0.092703 | 0.26483 | 0.976736 | 5249 | 4 | 3.4964 |
| PRPF4 | 4 | 0.27461 | 0.41707 | 0.999994 | 9585 | 1 | 4.4598 | 0.092767 | 0.22117 | 0.97466 | 5250 | 3 | 4.4598 |
| C1orf111 | 5 | 0.65538 | 0.74725 | 0.999994 | 15720 | 1 | 2.4064 | 0.092803 | 0.24037 | 0.97466 | 5251 | 4 | 2.4064 |

| | | | | | | | | | | | | | |
|---|---|---|---|---|---|---|---|---|---|---|---|---|---|
| CYP4F11 | 6 | 0.84301 | 0.86823 | 0.999994 | 18139 | 1 | 2.3087 | 0.092834 | 0.26512 | 0.976736 | 5252 | 3 | 2.3087 |
| NDUFAF2 | 4 | 0.93238 | 0.93212 | 0.999994 | 19910 | 0 | 3.9837 | 0.092925 | 0.22144 | 0.97466 | 5253 | 3 | 3.9837 |
| SYT5 | 5 | 0.90557 | 0.90567 | 0.999994 | 19344 | 0 | 1.2706 | 0.092954 | 0.24067 | 0.97466 | 5254 | 4 | 1.2706 |
| hsa-mir-3915 | 4 | 0.2023 | 0.34305 | 0.999994 | 7838 | 1 | 1.9111 | 0.092968 | 0.22151 | 0.97466 | 5255 | 2 | 1.9111 |
| KCNN2 | 3 | 0.90701 | 0.90706 | 0.999994 | 19371 | 0 | 1.9911 | 0.092988 | 0.20741 | 0.97466 | 5256 | 3 | 1.9911 |
| PROK1 | 6 | 0.41514 | 0.61219 | 0.999994 | 12180 | 2 | 1.6271 | 0.092989 | 0.26549 | 0.977843 | 5257 | 4 | 1.6271 |
| PRG4 | 6 | 0.9241 | 0.92426 | 0.999994 | 19718 | 1 | 1.3055 | 0.093042 | 0.26562 | 0.978056 | 5258 | 5 | 1.3055 |
| EIF5AL1 | 5 | 0.58236 | 0.71983 | 0.999994 | 14711 | 1 | 3.5542 | 0.093043 | 0.24085 | 0.97466 | 5259 | 3 | 3.5542 |
| IST1 | 5 | 0.88175 | 0.88196 | 0.999994 | 18864 | 1 | 1.4472 | 0.093068 | 0.24091 | 0.97466 | 5260 | 4 | 1.4472 |
| PTPRZ1 | 6 | 0.63499 | 0.76611 | 0.999994 | 15444 | 2 | 0.7217 | 0.093077 | 0.26569 | 0.978142 | 5261 | 3 | 0.7217 |
| C4orf48 | 5 | 0.3712 | 0.5394 | 0.999994 | 11531 | 2 | -0.27704 | 0.09308 | 0.24093 | 0.97466 | 5262 | 2 | -0.27704 |
| hsa-mir-1913 | 3 | 0.68453 | 0.68681 | 0.999994 | 16152 | 1 | 5.2684 | 0.093121 | 0.20768 | 0.97466 | 5263 | 2 | 5.2684 |
| RAP2A | 6 | 0.94154 | 0.94152 | 0.999994 | 20121 | 0 | 1.7656 | 0.093136 | 0.26583 | 0.978382 | 5264 | 5 | 1.7656 |
| RNF214 | 5 | 0.010586 | 0.037763 | 0.885019 | 906 | 2 | 1.5901 | 0.093174 | 0.24111 | 0.97466 | 5265 | 3 | 1.5901 |
| TRMT5 | 4 | 0.26107 | 0.40319 | 0.999994 | 9270 | 1 | 4.1585 | 0.093194 | 0.22187 | 0.97466 | 5266 | 3 | 4.1585 |
| SH3BGR | 4 | 0.28228 | 0.42487 | 0.999994 | 9745 | 2 | 2.4976 | 0.093194 | 0.22187 | 0.97466 | 5267 | 2 | 2.4976 |
| FCN2 | 6 | 0.52762 | 0.68788 | 0.999994 | 13854 | 2 | 2.2756 | 0.093219 | 0.26602 | 0.978454 | 5268 | 4 | 2.2756 |
| C16orf89 | 6 | 0.23431 | 0.42923 | 0.999994 | 8624 | 2 | 3.2433 | 0.093289 | 0.26618 | 0.978454 | 5269 | 4 | 3.2433 |
| NonTargetingControlG | 2 | 0.63862 | 0.63845 | 0.999994 | 15495 | 0 | 3.9816 | 0.093293 | 0.15965 | 0.965913 | 5270 | 1 | 3.9816 |
| FTH1 | 5 | 0.046362 | 0.13162 | 0.95318 | 2826 | 3 | -2.1567 | 0.093392 | 0.24154 | 0.97466 | 5271 | 2 | -2.1567 |
| ADH7 | 6 | 0.64026 | 0.77015 | 0.999994 | 15513 | 1 | 1.7906 | 0.093393 | 0.26642 | 0.978454 | 5272 | 5 | 1.7906 |
| PITPNM2 | 5 | 0.10646 | 0.23647 | 0.985449 | 5187 | 1 | 3.1686 | 0.093393 | 0.24154 | 0.97466 | 5273 | 4 | 3.1686 |
| EDDM3B | 6 | 0.84909 | 0.87144 | 0.999994 | 18243 | 1 | 1.5089 | 0.093413 | 0.26647 | 0.978454 | 5274 | 5 | 1.5089 |
| OR2A1 | 5 | 0.88749 | 0.88759 | 0.999994 | 18984 | 1 | 3.9078 | 0.093419 | 0.2416 | 0.97466 | 5275 | 3 | 3.9078 |
| SON | 5 | 0.42536 | 0.58455 | 0.999994 | 12343 | 1 | 1.8603 | 0.093455 | 0.24168 | 0.97466 | 5276 | 4 | 1.8603 |
| KRT27 | 5 | 0.48582 | 0.63566 | 0.999994 | 13226 | 2 | 1.7405 | 0.093455 | 0.24168 | 0.97466 | 5277 | 3 | 1.7405 |
| CAV1 | 5 | 0.61086 | 0.7301 | 0.999994 | 15084 | 1 | 1.2585 | 0.093455 | 0.24168 | 0.97466 | 5278 | 3 | 1.2585 |
| PSD2 | 6 | 0.20795 | 0.39759 | 0.999994 | 7975 | 2 | 0.66054 | 0.093494 | 0.26666 | 0.978454 | 5279 | 3 | 0.66054 |
| SPAG6 | 6 | 0.27069 | 0.47254 | 0.999994 | 9491 | 2 | 0.46666 | 0.093494 | 0.26666 | 0.978454 | 5280 | 3 | 0.46666 |
| NonTargetingControlG | 1 | 0.9065 | 0.90683 | 0.999994 | 19361 | 0 | 7.2755 | 0.093498 | 0.093444 | 0.950736 | 5281 | 1 | 7.2755 |
| hsa-mir-4722 | 1 | 0.9065 | 0.90683 | 0.999994 | 19360 | 0 | 7.2755 | 0.093498 | 0.093444 | 0.950736 | 5282 | 1 | 7.2755 |
| CTSL2 | 6 | 0.9566 | 0.95667 | 0.999994 | 20505 | 0 | 1.2993 | 0.093508 | 0.2667 | 0.978454 | 5283 | 5 | 1.2993 |
| F2 | 5 | 0.8766 | 0.87704 | 0.999994 | 18763 | 1 | 2.2741 | 0.093515 | 0.2418 | 0.97466 | 5284 | 4 | 2.2741 |
| PCGF6 | 6 | 0.28328 | 0.48721 | 0.999994 | 9769 | 2 | 2.1608 | 0.093567 | 0.26685 | 0.978454 | 5285 | 4 | 2.1608 |
| SHISA5 | 4 | 0.33228 | 0.47609 | 0.999994 | 10770 | 2 | 2.6755 | 0.093581 | 0.22249 | 0.97466 | 5286 | 2 | 2.6755 |
| ENTPD7 | 6 | 0.60762 | 0.7456 | 0.999994 | 15042 | 2 | 2.7848 | 0.093587 | 0.2669 | 0.978454 | 5287 | 3 | 2.7848 |
| TMEM232 | 6 | 0.93148 | 0.93162 | 0.999994 | 19885 | 0 | 2.7129 | 0.093587 | 0.2669 | 0.978454 | 5288 | 3 | 2.7129 |
| NonTargetingControlG | 2 | 0.90634 | 0.90634 | 0.999994 | 19357 | 0 | 3.1385 | 0.093664 | 0.16022 | 0.965913 | 5289 | 2 | 3.1385 |
| ZNF548 | 6 | 0.93509 | 0.93514 | 0.999994 | 19968 | 0 | 1.1492 | 0.093714 | 0.26716 | 0.978454 | 5290 | 5 | 1.1492 |
| C11orf71 | 5 | 0.52513 | 0.66978 | 0.999994 | 13819 | 2 | -0.28965 | 0.093817 | 0.24237 | 0.97466 | 5291 | 2 | -0.28965 |
| MED9 | 5 | 0.93277 | 0.93306 | 0.999994 | 19919 | 0 | 2.1347 | 0.093817 | 0.24237 | 0.97466 | 5292 | 3 | 2.1347 |
| KRTAP5-10 | 5 | 0.57533 | 0.71423 | 0.999994 | 14621 | 2 | -0.52341 | 0.093817 | 0.24237 | 0.97466 | 5293 | 2 | -0.52341 |
| OR6C68 | 5 | 0.015788 | 0.052443 | 0.914561 | 1232 | 3 | -3.8117 | 0.093817 | 0.24237 | 0.97466 | 5294 | 2 | -3.8117 |
| NCAM1 | 6 | 0.77589 | 0.83807 | 0.999994 | 17148 | 1 | 2.1997 | 0.093888 | 0.26758 | 0.978454 | 5295 | 5 | 2.1997 |
| ZNF799 | 2 | 0.37975 | 0.4001 | 0.999994 | 11646 | 1 | 3.0802 | 0.093931 | 0.16062 | 0.965913 | 5296 | 1 | 3.0802 |
| CRIP2 | 5 | 0.50566 | 0.65286 | 0.999994 | 13517 | 2 | 3.8927 | 0.093938 | 0.24261 | 0.97466 | 5297 | 3 | 3.8927 |
| MPHOSPH10 | 6 | 0.50879 | 0.67476 | 0.999994 | 13564 | 2 | 0.62798 | 0.093969 | 0.26776 | 0.978454 | 5298 | 3 | 0.62798 |
| EHHADH | 6 | 0.82874 | 0.86095 | 0.999994 | 17906 | 1 | 0.77798 | 0.093969 | 0.26776 | 0.978454 | 5299 | 3 | 0.77798 |
| C8orf47 | 5 | 0.24286 | 0.43482 | 0.999994 | 8860 | 3 | -0.60612 | 0.093981 | 0.24268 | 0.97466 | 5300 | 2 | -0.60612 |
| MCFD2 | 5 | 0.18959 | 0.36035 | 0.999994 | 7472 | 3 | -0.90782 | 0.093981 | 0.24268 | 0.97466 | 5301 | 2 | -0.90782 |
| ADRB3 | 6 | 0.51357 | 0.67809 | 0.999994 | 13632 | 2 | 1.8112 | 0.093989 | 0.26782 | 0.978454 | 5302 | 4 | 1.8112 |
| BMP7 | 6 | 0.40923 | 0.60839 | 0.999994 | 12096 | 1 | 2.8304 | 0.093999 | 0.26784 | 0.978454 | 5303 | 5 | 2.8304 |

| | | | | | | | | | | | | | |
|---|---|---|---|---|---|---|---|---|---|---|---|---|---|
| MCIDAS | 5 | 0.72531 | 0.77842 | 0.999994 | 16553 | 1 | 1.7877 | 0.094124 | 0.24298 | 0.97466 | 5304 | 4 | 1.7877 |
| THAP4 | 4 | 0.24177 | 0.38344 | 0.999994 | 8827 | 1 | 3.329 | 0.094202 | 0.22345 | 0.97466 | 5305 | 3 | 3.329 |
| OXSR1 | 6 | 0.61022 | 0.74749 | 0.999994 | 15076 | 2 | 2.9714 | 0.094203 | 0.26832 | 0.978454 | 5306 | 4 | 2.9714 |
| CORO1A | 5 | 0.54905 | 0.6909 | 0.999994 | 14193 | 2 | 1.7058 | 0.094252 | 0.24324 | 0.97466 | 5307 | 3 | 1.7058 |
| FBXO34 | 5 | 0.2522 | 0.44252 | 0.999994 | 9061 | 2 | 0.31552 | 0.094252 | 0.24324 | 0.97466 | 5308 | 2 | 0.31552 |
| PITX3 | 6 | 0.35514 | 0.56983 | 0.999994 | 11254 | 2 | 0.24096 | 0.094276 | 0.26846 | 0.978454 | 5309 | 3 | 0.24096 |
| COX7A2 | 6 | 0.49717 | 0.66678 | 0.999994 | 13384 | 1 | 0.20116 | 0.094276 | 0.26846 | 0.978454 | 5310 | 3 | 0.20116 |
| NonTargetingControlG | 2 | 0.0079512 | 0.015132 | 0.757255 | 728 | 1 | -1.4375 | 0.09429 | 0.16119 | 0.965913 | 5311 | 1 | -1.4375 |
| GPD1 | 6 | 0.28762 | 0.49234 | 0.999994 | 9867 | 3 | -0.59613 | 0.094343 | 0.2686 | 0.978454 | 5312 | 2 | -0.59613 |
| SECTM1 | 5 | 0.31588 | 0.49401 | 0.999994 | 10439 | 2 | 1.2268 | 0.094351 | 0.24342 | 0.97466 | 5313 | 3 | 1.2268 |
| POSTN | 6 | 0.49066 | 0.66235 | 0.999994 | 13287 | 2 | 3.4109 | 0.094406 | 0.26875 | 0.978454 | 5314 | 4 | 3.4109 |
| TTPAL | 6 | 0.54172 | 0.69776 | 0.999994 | 14084 | 1 | 2.567 | 0.094406 | 0.26875 | 0.978454 | 5315 | 3 | 2.567 |
| VPS26B | 6 | 0.33909 | 0.55163 | 0.999994 | 10918 | 2 | 2.5548 | 0.094406 | 0.26875 | 0.978454 | 5316 | 3 | 2.5548 |
| CHRND | 6 | 0.18297 | 0.36684 | 0.999994 | 7289 | 2 | 2.6534 | 0.094406 | 0.26875 | 0.978454 | 5317 | 3 | 2.6534 |
| SEBOX | 4 | 0.06809 | 0.15622 | 0.967126 | 3751 | 2 | -2.1921 | 0.094422 | 0.2238 | 0.97466 | 5318 | 1 | -2.1921 |
| MFSD8 | 6 | 0.51086 | 0.67622 | 0.999994 | 13591 | 1 | 1.2068 | 0.094429 | 0.26879 | 0.978454 | 5319 | 5 | 1.2068 |
| BCL6B | 6 | 0.049238 | 0.14243 | 0.959553 | 2941 | 4 | -1.8508 | 0.094458 | 0.26886 | 0.978454 | 5320 | 1 | -1.8508 |
| RFT1 | 4 | 0.88638 | 0.88609 | 0.999994 | 18956 | 0 | 2.3921 | 0.0945 | 0.22391 | 0.97466 | 5321 | 3 | 2.3921 |
| BATF2 | 5 | 0.096762 | 0.22133 | 0.983093 | 4849 | 3 | -1.7799 | 0.094525 | 0.24375 | 0.97466 | 5322 | 2 | -1.7799 |
| TRIM62 | 5 | 0.9348 | 0.93504 | 0.999994 | 19962 | 0 | 3.0085 | 0.094525 | 0.24375 | 0.97466 | 5323 | 3 | 3.0085 |
| EGFL8 | 4 | 0.16407 | 0.30368 | 0.999994 | 6843 | 2 | -1.1044 | 0.094545 | 0.22399 | 0.97466 | 5324 | 1 | -1.1044 |
| hsa-mir-520c | 3 | 0.90348 | 0.9035 | 0.999994 | 19292 | 0 | 1.5712 | 0.094551 | 0.21046 | 0.97466 | 5325 | 3 | 1.5712 |
| NonTargetingControlG | 2 | 0.90542 | 0.90537 | 0.999994 | 19342 | 0 | 2.9103 | 0.094584 | 0.16165 | 0.965913 | 5326 | 2 | 2.9103 |
| YPEL3 | 5 | 0.13902 | 0.28637 | 0.999632 | 6163 | 2 | 0.066488 | 0.094585 | 0.24387 | 0.97466 | 5327 | 2 | 0.066488 |
| ZFP14 | 6 | 0.55694 | 0.70863 | 0.999994 | 14316 | 2 | 1.6797 | 0.094588 | 0.26914 | 0.978454 | 5328 | 3 | 1.6797 |
| RIPK1 | 4 | 0.36432 | 0.50895 | 0.999994 | 11412 | 2 | 2.6815 | 0.094682 | 0.22422 | 0.97466 | 5329 | 2 | 2.6815 |
| FTSJ2 | 5 | 0.15237 | 0.30631 | 0.999994 | 6517 | 2 | 3.889 | 0.094691 | 0.24408 | 0.97466 | 5330 | 3 | 3.889 |
| COL4A1 | 5 | 0.021012 | 0.066808 | 0.93914 | 1515 | 1 | 3.889 | 0.094691 | 0.24408 | 0.97466 | 5331 | 4 | 3.889 |
| OR1L1 | 5 | 0.60532 | 0.72811 | 0.999994 | 15017 | 1 | 3.889 | 0.094691 | 0.24408 | 0.97466 | 5332 | 3 | 3.889 |
| ATF4 | 5 | 0.53185 | 0.67569 | 0.999994 | 13932 | 2 | 3.889 | 0.094691 | 0.24408 | 0.97466 | 5333 | 3 | 3.889 |
| PCSK1 | 5 | 0.951 | 0.95117 | 0.999994 | 20360 | 0 | 2.4812 | 0.094719 | 0.24414 | 0.97466 | 5334 | 4 | 2.4812 |
| DVL2 | 6 | 0.46914 | 0.64786 | 0.999994 | 12991 | 2 | 0.75061 | 0.094744 | 0.26951 | 0.978454 | 5335 | 3 | 0.75061 |
| MTUS1 | 6 | 0.037466 | 0.11619 | 0.951052 | 2398 | 3 | -1.8355 | 0.094744 | 0.26951 | 0.978454 | 5336 | 2 | -1.8355 |
| BRWD3 | 5 | 0.66606 | 0.75164 | 0.999994 | 15869 | 1 | 1.4004 | 0.094758 | 0.24421 | 0.97466 | 5337 | 4 | 1.4004 |
| VASH1 | 6 | 0.035801 | 0.1124 | 0.951052 | 2306 | 3 | 0.79333 | 0.094796 | 0.26964 | 0.978454 | 5338 | 3 | 0.79333 |
| SGTA | 6 | 0.031169 | 0.10157 | 0.946414 | 2067 | 4 | -2.3292 | 0.094796 | 0.26964 | 0.978454 | 5339 | 1 | -2.3292 |
| FIS1 | 2 | 0.6689 | 0.66868 | 0.999994 | 15907 | 0 | 4.0606 | 0.094819 | 0.16201 | 0.965913 | 5340 | 1 | 4.0606 |
| hsa-mir-219a-1 | 3 | 0.10685 | 0.19208 | 0.974607 | 5206 | 2 | -1.3671 | 0.094843 | 0.21101 | 0.97466 | 5341 | 1 | -1.3671 |
| ZAR1L | 6 | 0.3169 | 0.52633 | 0.999994 | 10448 | 2 | 3.0941 | 0.09495 | 0.27001 | 0.978454 | 5342 | 4 | 3.0941 |
| ALCAM | 6 | 0.82663 | 0.85992 | 0.999994 | 17880 | 1 | 1.8051 | 0.094972 | 0.27005 | 0.978454 | 5343 | 5 | 1.8051 |
| FOSL1 | 6 | 0.11366 | 0.27373 | 0.998561 | 5430 | 2 | 0.085962 | 0.094979 | 0.27006 | 0.978454 | 5344 | 2 | 0.085962 |
| PSCA | 5 | 0.82072 | 0.83309 | 0.999994 | 17787 | 1 | 1.9219 | 0.09498 | 0.24466 | 0.97466 | 5345 | 4 | 1.9219 |
| CCDC88A | 5 | 0.55482 | 0.69604 | 0.999994 | 14284 | 2 | 0.70391 | 0.094982 | 0.24467 | 0.97466 | 5346 | 3 | 0.70391 |
| HCFC2 | 5 | 0.83108 | 0.84026 | 0.999994 | 17946 | 1 | 3.4354 | 0.094985 | 0.24468 | 0.97466 | 5347 | 3 | 3.4354 |
| HIST1H3E | 6 | 0.80573 | 0.85036 | 0.999994 | 17566 | 1 | 3.1804 | 0.095052 | 0.27023 | 0.978454 | 5348 | 5 | 3.1804 |
| hsa-mir-4532 | 3 | 0.90495 | 0.90498 | 0.999994 | 19325 | 0 | 0.96768 | 0.095052 | 0.21141 | 0.97466 | 5349 | 3 | 0.96768 |
| hsa-mir-3151 | 4 | 0.068539 | 0.15708 | 0.967126 | 3775 | 1 | 2.4938 | 0.095116 | 0.22494 | 0.97466 | 5350 | 3 | 2.4938 |
| TRIM14 | 5 | 0.13996 | 0.28779 | 0.999632 | 6183 | 2 | 3.473 | 0.095123 | 0.24497 | 0.97466 | 5351 | 3 | 3.473 |
| hsa-mir-8065 | 4 | 0.74571 | 0.7568 | 0.999994 | 16809 | 1 | 4.3491 | 0.095128 | 0.22495 | 0.97466 | 5352 | 3 | 4.3491 |
| SART3 | 2 | 0.74508 | 0.74475 | 0.999994 | 16801 | 0 | 4.349 | 0.09516 | 0.16255 | 0.965913 | 5353 | 1 | 4.349 |
| ARMC8 | 6 | 0.14339 | 0.31735 | 0.999994 | 6283 | 2 | 0.81798 | 0.095171 | 0.27049 | 0.978454 | 5354 | 3 | 0.81798 |
| DLX2 | 6 | 0.535 | 0.69305 | 0.999994 | 13980 | 1 | 0.7949 | 0.095171 | 0.27049 | 0.978454 | 5355 | 3 | 0.7949 |

| | | | | | | | | | | | | | |
|---|---|---|---|---|---|---|---|---|---|---|---|---|---|
| TXNL4B | 4 | 0.25001 | 0.39193 | 0.999994 | 9016 | 1 | 2.8933 | 0.095175 | 0.22504 | 0.97466 | 5356 | 2 | 2.8933 |
| ARHGAP5 | 6 | 0.37948 | 0.58926 | 0.999994 | 11643 | 2 | 2.3914 | 0.095175 | 0.2705 | 0.978454 | 5357 | 4 | 2.3914 |
| TIAL1 | 6 | 0.63697 | 0.76763 | 0.999994 | 15472 | 1 | 1.6451 | 0.095188 | 0.27052 | 0.978454 | 5358 | 5 | 1.6451 |
| ZNF583 | 5 | 0.070074 | 0.1785 | 0.968171 | 3829 | 3 | -1.0367 | 0.0952 | 0.24513 | 0.97466 | 5359 | 2 | -1.0367 |
| NDUFS5 | 5 | 0.42789 | 0.58668 | 0.999994 | 12380 | 2 | 3.7749 | 0.095218 | 0.24517 | 0.97466 | 5360 | 3 | 3.7749 |
| ANKRD33 | 4 | 0.26212 | 0.40422 | 0.999994 | 9292 | 2 | -0.35168 | 0.095227 | 0.22511 | 0.97466 | 5361 | 2 | -0.35168 |
| TAPBPL | 6 | 0.71816 | 0.81774 | 0.999994 | 16476 | 1 | 1.1382 | 0.095277 | 0.27074 | 0.978454 | 5362 | 5 | 1.1382 |
| NonTargetingControlG | 2 | 0.90471 | 0.90466 | 0.999994 | 19316 | 0 | 4.7203 | 0.095293 | 0.16276 | 0.965913 | 5363 | 2 | 4.7203 |
| GNG5 | 3 | 0.87628 | 0.87638 | 0.999994 | 18758 | 0 | 5.2293 | 0.095322 | 0.2119 | 0.97466 | 5364 | 2 | 5.2293 |
| PYY | 3 | 0.12029 | 0.20988 | 0.979587 | 5637 | 1 | 5.2293 | 0.095322 | 0.2119 | 0.97466 | 5365 | 2 | 5.2293 |
| CCNE1 | 6 | 0.60225 | 0.7416 | 0.999994 | 14975 | 1 | 3.1354 | 0.095347 | 0.2709 | 0.978454 | 5366 | 4 | 3.1354 |
| NonTargetingControlG | 2 | 0.7756 | 0.77512 | 0.999994 | 17146 | 0 | 4.4114 | 0.095348 | 0.16285 | 0.965913 | 5367 | 2 | 4.4114 |
| SELM | 4 | 0.11446 | 0.24068 | 0.987253 | 5461 | 3 | -0.90266 | 0.095385 | 0.22536 | 0.97466 | 5368 | 1 | -0.90266 |
| FAM126B | 6 | 0.47493 | 0.65176 | 0.999994 | 13060 | 2 | 0.83004 | 0.095421 | 0.27106 | 0.978454 | 5369 | 4 | 0.83004 |
| PAK4 | 6 | 0.35798 | 0.57303 | 0.999994 | 11303 | 2 | 1.789 | 0.095433 | 0.27109 | 0.978454 | 5370 | 3 | 1.789 |
| RAI1 | 6 | 0.88247 | 0.89122 | 0.999994 | 18879 | 1 | 1.4312 | 0.095437 | 0.2711 | 0.978454 | 5371 | 5 | 1.4312 |
| JAG2 | 4 | 0.10041 | 0.21583 | 0.980538 | 4985 | 1 | 2.2076 | 0.09552 | 0.22559 | 0.97466 | 5372 | 3 | 2.2076 |
| hsa-mir-6811 | 4 | 0.93847 | 0.93833 | 0.999994 | 20035 | 0 | 2.9082 | 0.095543 | 0.22563 | 0.97466 | 5373 | 3 | 2.9082 |
| DAP3 | 3 | 0.90442 | 0.90443 | 0.999994 | 19310 | 0 | 1.1927 | 0.095584 | 0.2124 | 0.97466 | 5374 | 3 | 1.1927 |
| LCN12 | 5 | 0.54132 | 0.68407 | 0.999994 | 14075 | 1 | 1.3805 | 0.09561 | 0.2459 | 0.97466 | 5375 | 4 | 1.3805 |
| EIF3F | 5 | 0.95018 | 0.95036 | 0.999994 | 20342 | 0 | 3.7885 | 0.095628 | 0.24593 | 0.97466 | 5376 | 3 | 3.7885 |
| HECTD2 | 3 | 0.039841 | 0.097499 | 0.945253 | 2508 | 2 | -3.124 | 0.09565 | 0.21253 | 0.97466 | 5377 | 1 | -3.124 |
| TECPR1 | 6 | 0.39933 | 0.60194 | 0.999994 | 11937 | 2 | 2.1136 | 0.095687 | 0.27165 | 0.978454 | 5378 | 4 | 2.1136 |
| hsa-mir-5572 | 4 | 0.81814 | 0.81796 | 0.999994 | 17750 | 1 | 2.7395 | 0.095727 | 0.22592 | 0.97466 | 5379 | 3 | 2.7395 |
| CHMP5 | 5 | 0.63644 | 0.73982 | 0.999994 | 15467 | 1 | 0.97218 | 0.0958 | 0.24628 | 0.97466 | 5380 | 4 | 0.97218 |
| ETV4 | 5 | 0.49756 | 0.64574 | 0.999994 | 13391 | 1 | 0.94199 | 0.0958 | 0.24628 | 0.97466 | 5381 | 3 | 0.94199 |
| C19orf81 | 6 | 0.91806 | 0.91861 | 0.999994 | 19587 | 1 | 2.0673 | 0.095813 | 0.27193 | 0.978454 | 5382 | 5 | 2.0673 |
| DIABLO | 6 | 0.13187 | 0.30271 | 0.999994 | 5974 | 2 | 0.27558 | 0.095915 | 0.27214 | 0.978454 | 5383 | 3 | 0.27558 |
| GPR34 | 6 | 0.26197 | 0.46222 | 0.999994 | 9290 | 1 | 0.46953 | 0.095915 | 0.27214 | 0.978454 | 5384 | 3 | 0.46953 |
| HPSE | 6 | 0.97412 | 0.97402 | 0.999994 | 20931 | 0 | 2.196 | 0.09592 | 0.27215 | 0.978454 | 5385 | 5 | 2.196 |
| SLCO3A1 | 6 | 0.85176 | 0.87284 | 0.999994 | 18290 | 1 | 2.3321 | 0.095944 | 0.27221 | 0.978454 | 5386 | 4 | 2.3321 |
| SLC2A2 | 6 | 0.94497 | 0.94496 | 0.999994 | 20203 | 0 | 1.4408 | 0.095944 | 0.27221 | 0.978454 | 5387 | 5 | 1.4408 |
| SP1 | 5 | 0.38593 | 0.5517 | 0.999994 | 11733 | 1 | 3.6317 | 0.095991 | 0.24664 | 0.97466 | 5388 | 4 | 3.6317 |
| GOLGA6L9 | 2 | 0.18523 | 0.25144 | 0.993413 | 7347 | 1 | 0.96205 | 0.096012 | 0.1639 | 0.965913 | 5389 | 1 | 0.96205 |
| ZNF26 | 6 | 0.611 | 0.74807 | 0.999994 | 15087 | 2 | 2.3393 | 0.096026 | 0.27238 | 0.978454 | 5390 | 4 | 2.3393 |
| SNAP25 | 6 | 0.42439 | 0.61828 | 0.999994 | 12321 | 1 | 1.5553 | 0.096049 | 0.27244 | 0.978454 | 5391 | 5 | 1.5553 |
| ERG | 6 | 0.33673 | 0.54893 | 0.999994 | 10867 | 3 | 2.1507 | 0.09606 | 0.27246 | 0.978454 | 5392 | 3 | 2.1507 |
| SUGP2 | 6 | 0.90138 | 0.90474 | 0.999994 | 19239 | 1 | 3.4118 | 0.09606 | 0.27246 | 0.978454 | 5393 | 4 | 3.4118 |
| TOB2 | 6 | 0.17203 | 0.35324 | 0.999994 | 7036 | 2 | 2.6027 | 0.09606 | 0.27246 | 0.978454 | 5394 | 3 | 2.6027 |
| DEFB104B | 1 | 0.90392 | 0.90426 | 0.999994 | 19302 | 0 | 7.2305 | 0.096077 | 0.096042 | 0.950736 | 5395 | 1 | 7.2305 |
| COX6A1 | 5 | 0.15261 | 0.30667 | 0.999994 | 6524 | 2 | 1.2921 | 0.096129 | 0.24692 | 0.97466 | 5396 | 3 | 1.2921 |
| RAB23 | 6 | 0.45189 | 0.63632 | 0.999994 | 12733 | 2 | 2.4266 | 0.096142 | 0.27263 | 0.978454 | 5397 | 4 | 2.4266 |
| MKLN1 | 6 | 0.31494 | 0.5241 | 0.999994 | 10419 | 2 | 2.3554 | 0.096142 | 0.27263 | 0.978454 | 5398 | 3 | 2.3554 |
| ING1 | 6 | 0.83876 | 0.86599 | 0.999994 | 18066 | 1 | 1.3381 | 0.09621 | 0.27281 | 0.978454 | 5399 | 5 | 1.3381 |
| ZNF436 | 4 | 0.023513 | 0.068071 | 0.93914 | 1687 | 3 | -4.9608 | 0.096224 | 0.22671 | 0.97466 | 5400 | 1 | -4.9608 |
| SLC8A2 | 5 | 0.80705 | 0.824 | 0.999994 | 17585 | 1 | 3.3799 | 0.096267 | 0.24718 | 0.97466 | 5401 | 4 | 3.3799 |
| hsa-mir-1587 | 4 | 0.84407 | 0.84355 | 0.999994 | 18157 | 0 | 3.0921 | 0.096267 | 0.22679 | 0.97466 | 5402 | 3 | 3.0921 |
| CDC42BPG | 6 | 0.38168 | 0.59064 | 0.999994 | 11672 | 1 | 2.3193 | 0.096487 | 0.27345 | 0.978454 | 5403 | 3 | 2.3193 |
| EI24 | 6 | 0.052382 | 0.14928 | 0.966106 | 3086 | 2 | 0.79336 | 0.096487 | 0.27345 | 0.978454 | 5404 | 3 | 0.79336 |
| PPT1 | 6 | 0.14394 | 0.31804 | 0.999994 | 6297 | 2 | 1.1322 | 0.096487 | 0.27345 | 0.978454 | 5405 | 3 | 1.1322 |
| ITPRIP | 6 | 0.67013 | 0.79324 | 0.999994 | 15933 | 2 | 2.6068 | 0.096498 | 0.27347 | 0.978454 | 5406 | 4 | 2.6068 |
| SMIM7 | 5 | 0.59254 | 0.72346 | 0.999994 | 14855 | 1 | 0.2604 | 0.096545 | 0.24774 | 0.97466 | 5407 | 2 | 0.2604 |

| | | | | | | | | | | | | |
|---|---|---|---|---|---|---|---|---|---|---|---|---|
| APLN | 5 | 0.24682 | 0.43824 | 0.999994 | 8939 | 2 | 1.71 | 0.096545 | 0.24774 | 0.97466 | 5408 | 3 | 1.71 |
| CD164L2 | 5 | 0.5264 | 0.67091 | 0.999994 | 13833 | 2 | 0.19388 | 0.096548 | 0.24775 | 0.97466 | 5409 | 1 | 0.19388 |
| TRIP10 | 5 | 0.20137 | 0.37718 | 0.999994 | 7801 | 2 | 0.22103 | 0.096548 | 0.24775 | 0.97466 | 5410 | 2 | 0.22103 |
| PEAR1 | 5 | 0.74293 | 0.78715 | 0.999994 | 16779 | 1 | 3.6845 | 0.096651 | 0.24797 | 0.97466 | 5411 | 4 | 3.6845 |
| STXBP4 | 6 | 0.25815 | 0.4577 | 0.999994 | 9198 | 2 | 2.9863 | 0.096677 | 0.27387 | 0.978454 | 5412 | 4 | 2.9863 |
| HSD17B11 | 6 | 0.2203 | 0.41247 | 0.999994 | 8297 | 3 | 0.139 | 0.096677 | 0.27387 | 0.978454 | 5413 | 3 | 0.139 |
| TMPRSS3 | 6 | 0.31354 | 0.52247 | 0.999994 | 10389 | 3 | 0.52749 | 0.096695 | 0.27391 | 0.978454 | 5414 | 3 | 0.52749 |
| MARS | 5 | 0.67427 | 0.75511 | 0.999994 | 15992 | 1 | 2.66 | 0.096724 | 0.24812 | 0.97466 | 5415 | 4 | 2.66 |
| BICC1 | 6 | 0.93815 | 0.93812 | 0.999994 | 20029 | 0 | 2.7648 | 0.096726 | 0.27398 | 0.978454 | 5416 | 4 | 2.7648 |
| hsa-mir-6842 | 4 | 0.73156 | 0.74632 | 0.999994 | 16621 | 1 | 2.968 | 0.096739 | 0.22755 | 0.97466 | 5417 | 3 | 2.968 |
| TSPYL4 | 5 | 0.8116 | 0.82698 | 0.999994 | 17651 | 1 | 0.0038629 | 0.096765 | 0.24821 | 0.97466 | 5418 | 1 | 0.0038629 |
| PLXNB3 | 5 | 0.065751 | 0.17145 | 0.967658 | 3680 | 2 | 1.4833 | 0.096765 | 0.24821 | 0.97466 | 5419 | 3 | 1.4833 |
| hsa-mir-518a-1 | 4 | 0.89143 | 0.89118 | 0.999994 | 19060 | 0 | 1.7512 | 0.096783 | 0.22763 | 0.97466 | 5420 | 3 | 1.7512 |
| ZNF786 | 5 | 0.77372 | 0.8038 | 0.999994 | 17130 | 1 | 1.2274 | 0.096809 | 0.24831 | 0.97466 | 5421 | 4 | 1.2274 |
| hsa-mir-6716 | 4 | 0.58329 | 0.65645 | 0.999994 | 14721 | 1 | 3.339 | 0.096853 | 0.22774 | 0.97466 | 5422 | 2 | 3.339 |
| PRAMEF18 | 4 | 0.38482 | 0.53006 | 0.999994 | 11713 | 1 | 3.0954 | 0.096853 | 0.22774 | 0.97466 | 5423 | 2 | 3.0954 |
| hsa-mir-1539 | 3 | 0.67562 | 0.67881 | 0.999994 | 16015 | 1 | 4.9257 | 0.096874 | 0.21488 | 0.97466 | 5424 | 2 | 4.9257 |
| C6orf48 | 5 | 0.91109 | 0.91112 | 0.999994 | 19451 | 0 | 3.6511 | 0.096891 | 0.24848 | 0.97466 | 5425 | 3 | 3.6511 |
| SLC25A32 | 6 | 0.79612 | 0.84626 | 0.999994 | 17428 | 1 | 2.3738 | 0.096929 | 0.27444 | 0.978454 | 5426 | 4 | 2.3738 |
| CSNK1G2 | 6 | 0.23793 | 0.43351 | 0.999994 | 8723 | 2 | -0.11142 | 0.096929 | 0.27444 | 0.978454 | 5427 | 2 | -0.11142 |
| ATP2B3 | 5 | 0.45735 | 0.61145 | 0.999994 | 12813 | 1 | 0.48425 | 0.096948 | 0.24858 | 0.97466 | 5428 | 2 | 0.48425 |
| HIST1H2BO | 6 | 0.4408 | 0.62896 | 0.999994 | 12587 | 1 | 1.8702 | 0.096955 | 0.2745 | 0.978454 | 5429 | 5 | 1.8702 |
| KCNJ8 | 6 | 0.36139 | 0.57683 | 0.999994 | 11359 | 3 | 2.1967 | 0.096964 | 0.27452 | 0.978454 | 5430 | 3 | 2.1967 |
| LPPR1 | 6 | 0.23105 | 0.42536 | 0.999994 | 8544 | 3 | 1.922 | 0.096964 | 0.27452 | 0.978454 | 5431 | 3 | 1.922 |
| ZNF404 | 6 | 0.020277 | 0.072857 | 0.93914 | 1476 | 2 | 3.052 | 0.096964 | 0.27452 | 0.978454 | 5432 | 4 | 3.052 |
| TCEAL4 | 5 | 0.19147 | 0.36303 | 0.999994 | 7525 | 3 | -0.70485 | 0.097004 | 0.2487 | 0.97466 | 5433 | 2 | -0.70485 |
| KCNJ3 | 6 | 0.64166 | 0.77121 | 0.999994 | 15541 | 2 | -0.018037 | 0.097007 | 0.27463 | 0.978454 | 5434 | 2 | -0.018037 |
| ADAMTSL2 | 5 | 0.94365 | 0.94387 | 0.999994 | 20166 | 0 | 1.518 | 0.097058 | 0.24879 | 0.97466 | 5435 | 4 | 1.518 |
| LYRM1 | 6 | 0.81299 | 0.8536 | 0.999994 | 17672 | 1 | 0.92793 | 0.097085 | 0.2748 | 0.978454 | 5436 | 3 | 0.92793 |
| C12orf50 | 6 | 0.74939 | 0.82823 | 0.999994 | 16852 | 1 | 1.1999 | 0.097093 | 0.27482 | 0.978454 | 5437 | 5 | 1.1999 |
| IFFO2 | 3 | 0.58419 | 0.60509 | 0.999994 | 14730 | 1 | 5.1685 | 0.097138 | 0.21538 | 0.97466 | 5438 | 2 | 5.1685 |
| NRL | 6 | 0.91873 | 0.91923 | 0.999994 | 19601 | 1 | 1.2738 | 0.097167 | 0.27496 | 0.978454 | 5439 | 5 | 1.2738 |
| NonTargetingControlG | 1 | 0.90282 | 0.90317 | 0.999994 | 19270 | 0 | 7.2068 | 0.097175 | 0.097163 | 0.950736 | 5440 | 1 | 7.2068 |
| GRIP1 | 5 | 0.86769 | 0.86901 | 0.999994 | 18594 | 1 | 2.9342 | 0.097206 | 0.24906 | 0.97466 | 5441 | 4 | 2.9342 |
| NonTargetingControlG | 2 | 0.90278 | 0.90269 | 0.999994 | 19268 | 0 | 4.8606 | 0.097223 | 0.16577 | 0.965913 | 5442 | 2 | 4.8606 |
| PCDHGC4 | 2 | 0.68968 | 0.6895 | 0.999994 | 16206 | 0 | 4.1139 | 0.09723 | 0.16578 | 0.965913 | 5443 | 1 | 4.1139 |
| SPAG7 | 6 | 0.36213 | 0.5777 | 0.999994 | 11370 | 1 | 0.75559 | 0.097248 | 0.27512 | 0.978454 | 5444 | 5 | 0.75559 |
| NonTargetingControlG | 2 | 0.90272 | 0.90263 | 0.999994 | 19265 | 0 | 4.7019 | 0.097276 | 0.16585 | 0.965913 | 5445 | 2 | 4.7019 |
| THAP11 | 4 | 0.65193 | 0.69439 | 0.999994 | 15687 | 1 | 2.5746 | 0.097329 | 0.22853 | 0.97466 | 5446 | 3 | 2.5746 |
| RAB5B | 5 | 0.95541 | 0.95557 | 0.999994 | 20463 | 0 | 1.4948 | 0.09733 | 0.24933 | 0.97466 | 5447 | 4 | 1.4948 |
| COX7B | 4 | 0.93656 | 0.93639 | 0.999994 | 19995 | 0 | 4.2766 | 0.097351 | 0.22856 | 0.97466 | 5448 | 3 | 4.2766 |
| RDH10 | 6 | 0.4736 | 0.65084 | 0.999994 | 13050 | 1 | 3.2376 | 0.097395 | 0.27543 | 0.978454 | 5449 | 4 | 3.2376 |
| DSE | 5 | 0.95611 | 0.95627 | 0.999994 | 20489 | 0 | 1.2802 | 0.097401 | 0.24945 | 0.97466 | 5450 | 4 | 1.2802 |
| INTS7 | 5 | 0.52657 | 0.67106 | 0.999994 | 13839 | 1 | 0.14662 | 0.097417 | 0.24949 | 0.97466 | 5451 | 2 | 0.14662 |
| FUT5 | 5 | 0.16898 | 0.33064 | 0.999994 | 6960 | 2 | 0.79605 | 0.097417 | 0.24949 | 0.97466 | 5452 | 3 | 0.79605 |
| INMT | 5 | 0.085767 | 0.20378 | 0.978186 | 4437 | 3 | -1.8331 | 0.097417 | 0.24949 | 0.97466 | 5453 | 1 | -1.8331 |
| C17orf49 | 6 | 0.92876 | 0.92886 | 0.999994 | 19831 | 0 | 2.3806 | 0.097419 | 0.27548 | 0.978454 | 5454 | 5 | 2.3806 |
| NPIPA5 | 1 | 0.90258 | 0.90292 | 0.999994 | 19264 | 0 | 7.2059 | 0.09742 | 0.097392 | 0.950736 | 5455 | 1 | 7.2059 |
| hsa-mir-6505 | 4 | 0.58896 | 0.65937 | 0.999994 | 14804 | 1 | 2.6465 | 0.097446 | 0.22871 | 0.97466 | 5456 | 3 | 2.6465 |
| SREK1IP1 | 6 | 0.11159 | 0.26976 | 0.998561 | 5353 | 2 | 2.07 | 0.097467 | 0.27559 | 0.978454 | 5457 | 4 | 2.07 |
| FAIM | 6 | 0.82257 | 0.85805 | 0.999994 | 17815 | 1 | 2.3752 | 0.097467 | 0.27559 | 0.978454 | 5458 | 4 | 2.3752 |
| HIST1H2AH | 6 | 0.93961 | 0.93956 | 0.999994 | 20066 | 0 | 3.1283 | 0.097485 | 0.27562 | 0.978454 | 5459 | 4 | 3.1283 |

| | | | | | | | | | | | | | |
|---|---|---|---|---|---|---|---|---|---|---|---|---|---|
| SYNE4 | 6 | 0.58898 | 0.73184 | 0.999994 | 14805 | 1 | 2.0377 | 0.097501 | 0.27565 | 0.978454 | 5460 | 5 | 2.0377 |
| KIAA0146 | 6 | 0.6039 | 0.74287 | 0.999994 | 14995 | 1 | 1.0634 | 0.097509 | 0.27567 | 0.978454 | 5461 | 5 | 1.0634 |
| PADI4 | 6 | 0.8442 | 0.86884 | 0.999994 | 18158 | 1 | 1.5549 | 0.097533 | 0.27572 | 0.978454 | 5462 | 5 | 1.5549 |
| SMAD4 | 6 | 0.71668 | 0.81726 | 0.999994 | 16463 | 1 | 2.3573 | 0.097602 | 0.27588 | 0.978454 | 5463 | 3 | 2.3573 |
| EIF4E | 5 | 0.082368 | 0.19843 | 0.977164 | 4310 | 3 | -0.6949 | 0.097644 | 0.24996 | 0.97466 | 5464 | 2 | -0.6949 |
| ATP5O | 5 | 0.74121 | 0.78627 | 0.999994 | 16755 | 1 | 1.5303 | 0.097644 | 0.24996 | 0.97466 | 5465 | 3 | 1.5303 |
| NonTargetingControlG | 1 | 0.90233 | 0.90269 | 0.999994 | 19257 | 0 | 7.1976 | 0.09767 | 0.097638 | 0.950736 | 5466 | 1 | 7.1976 |
| RPS15 | 5 | 0.87115 | 0.8721 | 0.999994 | 18661 | 1 | -0.32472 | 0.097677 | 0.25002 | 0.97466 | 5467 | 2 | -0.32472 |
| LRRC4B | 6 | 0.19543 | 0.38234 | 0.999994 | 7645 | 1 | 1.1531 | 0.097682 | 0.27603 | 0.978454 | 5468 | 3 | 1.1531 |
| SLC11A2 | 6 | 0.53279 | 0.69152 | 0.999994 | 13948 | 1 | 0.7536 | 0.097682 | 0.27603 | 0.978454 | 5469 | 4 | 0.7536 |
| MFSD5 | 6 | 0.57275 | 0.71992 | 0.999994 | 14588 | 2 | 0.15013 | 0.097682 | 0.27603 | 0.978454 | 5470 | 3 | 0.15013 |
| CIZ1 | 5 | 0.96009 | 0.96023 | 0.999994 | 20584 | 0 | 2.1141 | 0.097713 | 0.2501 | 0.97466 | 5471 | 4 | 2.1141 |
| RRAS2 | 5 | 0.61124 | 0.73025 | 0.999994 | 15090 | 1 | 3.8042 | 0.097762 | 0.2502 | 0.97466 | 5472 | 3 | 3.8042 |
| PNRC1 | 6 | 0.22704 | 0.42052 | 0.999994 | 8453 | 1 | 2.1281 | 0.097842 | 0.27629 | 0.978454 | 5473 | 4 | 2.1281 |
| SENP8 | 5 | 0.91752 | 0.9176 | 0.999994 | 19578 | 0 | 2.2806 | 0.097877 | 0.25042 | 0.97466 | 5474 | 4 | 2.2806 |
| TMEM99 | 5 | 0.49855 | 0.64664 | 0.999994 | 13403 | 1 | 1.753 | 0.097894 | 0.25046 | 0.97466 | 5475 | 3 | 1.753 |
| SIM1 | 5 | 0.80826 | 0.8248 | 0.999994 | 17603 | 1 | 2.9133 | 0.097894 | 0.25046 | 0.97466 | 5476 | 3 | 2.9133 |
| DUSP11 | 4 | 0.10253 | 0.21962 | 0.980699 | 5057 | 2 | -1.7107 | 0.097901 | 0.22939 | 0.97466 | 5477 | 1 | -1.7107 |
| PTF1A | 6 | 0.14485 | 0.31919 | 0.999994 | 6319 | 3 | -0.12448 | 0.097917 | 0.27642 | 0.978454 | 5478 | 3 | -0.12448 |
| SCUBE2 | 6 | 0.0022325 | 0.010844 | 0.709416 | 295 | 1 | 1.4279 | 0.097917 | 0.27642 | 0.978454 | 5479 | 3 | 1.4279 |
| OR7G3 | 6 | 0.2295 | 0.42346 | 0.999994 | 8516 | 1 | 3.205 | 0.097962 | 0.27649 | 0.978454 | 5480 | 4 | 3.205 |
| TRAF6 | 5 | 0.90142 | 0.90153 | 0.999994 | 19240 | 0 | 2.0764 | 0.09801 | 0.25067 | 0.97466 | 5481 | 4 | 2.0764 |
| hsa-mir-769 | 4 | 0.83796 | 0.83745 | 0.999994 | 18053 | 0 | 3.0723 | 0.098062 | 0.22964 | 0.97466 | 5482 | 3 | 3.0723 |
| TMEM151B | 6 | 0.14394 | 0.31804 | 0.999994 | 6296 | 1 | 2.6827 | 0.098072 | 0.27668 | 0.978454 | 5483 | 4 | 2.6827 |
| HIST1H2AD | 5 | 0.039012 | 0.1134 | 0.951052 | 2460 | 2 | 0.83177 | 0.09809 | 0.25083 | 0.97466 | 5484 | 3 | 0.83177 |
| PRKCDBP | 3 | 0.9019 | 0.90196 | 0.999994 | 19248 | 0 | 1.9326 | 0.098101 | 0.2172 | 0.97466 | 5485 | 3 | 1.9326 |
| QRSL1 | 5 | 0.9199 | 0.92008 | 0.999994 | 19627 | 0 | 3.5498 | 0.098154 | 0.25095 | 0.97466 | 5486 | 4 | 3.5498 |
| PIK3C3 | 5 | 0.1955 | 0.3688 | 0.999994 | 7651 | 1 | 3.529 | 0.098163 | 0.25097 | 0.97466 | 5487 | 3 | 3.529 |
| ZNF346 | 6 | 0.16689 | 0.34684 | 0.999994 | 6904 | 2 | 2.0988 | 0.098173 | 0.27685 | 0.978454 | 5488 | 4 | 2.0988 |
| GLIPR1L1 | 5 | 0.8673 | 0.86869 | 0.999994 | 18582 | 1 | 3.6368 | 0.098183 | 0.25101 | 0.97466 | 5489 | 3 | 3.6368 |
| NDC1 | 6 | 0.003394 | 0.015165 | 0.757255 | 393 | 3 | -1.7418 | 0.098201 | 0.27689 | 0.978454 | 5490 | 3 | -1.7418 |
| NUF2 | 4 | 0.30201 | 0.44503 | 0.999994 | 10163 | 1 | 4.0813 | 0.098335 | 0.23008 | 0.97466 | 5491 | 3 | 4.0813 |
| SCGB1D1 | 4 | 0.49382 | 0.61357 | 0.999994 | 13322 | 1 | 2.7104 | 0.098359 | 0.23012 | 0.97466 | 5492 | 3 | 2.7104 |
| RPS5 | 4 | 0.038935 | 0.099663 | 0.946394 | 2457 | 2 | 0.77955 | 0.098375 | 0.23015 | 0.97466 | 5493 | 2 | 0.77955 |
| SKA2 | 3 | 0.73755 | 0.73747 | 0.999994 | 16694 | 0 | 0.39048 | 0.098411 | 0.2178 | 0.97466 | 5494 | 1 | 0.39048 |
| PIF1 | 5 | 0.020484 | 0.065375 | 0.93914 | 1483 | 2 | 2.4115 | 0.098459 | 0.25155 | 0.97466 | 5495 | 3 | 2.4115 |
| PPEF2 | 5 | 0.52934 | 0.67346 | 0.999994 | 13889 | 1 | 2.6925 | 0.098528 | 0.25169 | 0.97466 | 5496 | 4 | 2.6925 |
| CXorf66 | 6 | 0.29429 | 0.50004 | 0.999994 | 10002 | 3 | -0.019025 | 0.098616 | 0.27758 | 0.978454 | 5497 | 3 | -0.019025 |
| ARL14EP | 5 | 0.6456 | 0.74336 | 0.999994 | 15593 | 1 | 1.611 | 0.09863 | 0.25188 | 0.97466 | 5498 | 4 | 1.611 |
| MB21D2 | 5 | 0.48123 | 0.63173 | 0.999994 | 13155 | 2 | 0.26209 | 0.098667 | 0.25194 | 0.97466 | 5499 | 2 | 0.26209 |
| CCDC149 | 5 | 0.39632 | 0.5605 | 0.999994 | 11892 | 1 | 3.647 | 0.098667 | 0.25194 | 0.97466 | 5500 | 3 | 3.647 |
| ARHGAP29 | 5 | 0.36596 | 0.53501 | 0.999994 | 11443 | 1 | 1.0646 | 0.098667 | 0.25194 | 0.97466 | 5501 | 3 | 1.0646 |
| DEFB129 | 5 | 0.49174 | 0.64077 | 0.999994 | 13297 | 1 | 1.9367 | 0.098667 | 0.25194 | 0.97466 | 5502 | 3 | 1.9367 |
| GLOD5 | 5 | 0.88081 | 0.88104 | 0.999994 | 18849 | 1 | 1.5925 | 0.098708 | 0.25204 | 0.97466 | 5503 | 4 | 1.5925 |
| SLC6A2 | 6 | 0.27628 | 0.47909 | 0.999994 | 9624 | 3 | 0.21753 | 0.09872 | 0.27774 | 0.978454 | 5504 | 3 | 0.21753 |
| SFXN3 | 4 | 0.84894 | 0.84847 | 0.999994 | 18239 | 0 | 2.549 | 0.098778 | 0.2308 | 0.97466 | 5505 | 3 | 2.549 |
| CCDC59 | 4 | 0.80705 | 0.80759 | 0.999994 | 17586 | 1 | 3.5368 | 0.098778 | 0.2308 | 0.97466 | 5506 | 3 | 3.5368 |
| TNPO2 | 4 | 0.91178 | 0.91157 | 0.999994 | 19458 | 0 | 2.3559 | 0.098778 | 0.2308 | 0.97466 | 5507 | 3 | 2.3559 |
| AQP11 | 6 | 0.032962 | 0.10576 | 0.950444 | 2155 | 3 | -0.96677 | 0.098824 | 0.27793 | 0.978454 | 5508 | 3 | -0.96677 |
| SERINC3 | 6 | 0.097708 | 0.24282 | 0.988088 | 4878 | 2 | 3.1496 | 0.098883 | 0.27803 | 0.978454 | 5509 | 4 | 3.1496 |
| CRYBB1 | 6 | 0.010448 | 0.04055 | 0.891621 | 900 | 3 | 1.8495 | 0.098883 | 0.27803 | 0.978454 | 5510 | 3 | 1.8495 |
| CCDC151 | 6 | 0.5449 | 0.69999 | 0.999994 | 14136 | 1 | 2.817 | 0.098883 | 0.27803 | 0.978454 | 5511 | 4 | 2.817 |

| | | | | | | | | | | | | |
|---|---|---|---|---|---|---|---|---|---|---|---|---|
| AP1G1 | 6 | 0.53456 | 0.69272 | 0.999994 | 13972 | 2 | 3.2412 | 0.098883 | 0.27803 | 0.978454 | 5512 | 4 | 3.2412 |
| CACNG8 | 5 | 0.94974 | 0.94993 | 0.999994 | 20326 | 0 | 1.5779 | 0.098913 | 0.25244 | 0.97466 | 5513 | 4 | 1.5779 |
| MYF5 | 5 | 0.92157 | 0.92174 | 0.999994 | 19667 | 0 | 1.7167 | 0.098929 | 0.25246 | 0.97466 | 5514 | 4 | 1.7167 |
| ETNPPL | 5 | 0.93393 | 0.93417 | 0.999994 | 19951 | 0 | 3.0997 | 0.098992 | 0.25259 | 0.97466 | 5515 | 4 | 3.0997 |
| CCND2 | 5 | 0.95822 | 0.95837 | 0.999994 | 20542 | 0 | 1.2092 | 0.099001 | 0.25261 | 0.97466 | 5516 | 3 | 1.2092 |
| C1orf141 | 3 | 0.87854 | 0.87864 | 0.999994 | 18800 | 0 | 1.128 | 0.099005 | 0.21893 | 0.97466 | 5517 | 2 | 1.128 |
| CENPJ | 5 | 0.93757 | 0.93777 | 0.999994 | 20020 | 0 | 3.7891 | 0.09905 | 0.25271 | 0.97466 | 5518 | 3 | 3.7891 |
| MTMR6 | 5 | 0.95258 | 0.95274 | 0.999994 | 20397 | 0 | 3.7891 | 0.09905 | 0.25271 | 0.97466 | 5519 | 3 | 3.7891 |
| OTOL1 | 5 | 0.03174 | 0.094945 | 0.945031 | 2093 | 1 | 3.7891 | 0.09905 | 0.25271 | 0.97466 | 5520 | 4 | 3.7891 |
| OSER1 | 4 | 0.88746 | 0.88715 | 0.999994 | 18981 | 0 | 0.60647 | 0.099087 | 0.23133 | 0.97466 | 5521 | 2 | 0.60647 |
| CCDC168 | 4 | 0.65751 | 0.69776 | 0.999994 | 15747 | 1 | 2.0558 | 0.099087 | 0.23133 | 0.97466 | 5522 | 3 | 2.0558 |
| FAM135A | 6 | 0.76864 | 0.83532 | 0.999994 | 17070 | 1 | 2.0904 | 0.099109 | 0.27839 | 0.978454 | 5523 | 4 | 2.0904 |
| CYP3A7 | 6 | 0.070092 | 0.18696 | 0.969409 | 3831 | 3 | -0.99344 | 0.099145 | 0.27844 | 0.978454 | 5524 | 2 | -0.99344 |
| CIB2 | 6 | 0.51356 | 0.67809 | 0.999994 | 13631 | 2 | 1.1459 | 0.099145 | 0.27844 | 0.978454 | 5525 | 4 | 1.1459 |
| NonTargetingControlG | 2 | 0.79881 | 0.79837 | 0.999994 | 17462 | 0 | 4.5284 | 0.099164 | 0.16878 | 0.965913 | 5526 | 2 | 4.5284 |
| NYAP2 | 5 | 0.098841 | 0.22461 | 0.983354 | 4925 | 3 | -1.6476 | 0.099174 | 0.25296 | 0.97466 | 5527 | 2 | -1.6476 |
| PRKAR2B | 5 | 0.66317 | 0.75044 | 0.999994 | 15824 | 1 | 1.2828 | 0.099177 | 0.25296 | 0.97466 | 5528 | 4 | 1.2828 |
| OR52N4 | 6 | 0.92422 | 0.92439 | 0.999994 | 19723 | 1 | 3.3262 | 0.099207 | 0.27855 | 0.978454 | 5529 | 4 | 3.3262 |
| GLT6D1 | 6 | 0.85735 | 0.87593 | 0.999994 | 18399 | 1 | 3.2789 | 0.099235 | 0.2786 | 0.978454 | 5530 | 4 | 3.2789 |
| HARS | 3 | 0.16979 | 0.2741 | 0.998561 | 6981 | 1 | 5.1053 | 0.099246 | 0.21939 | 0.97466 | 5531 | 2 | 5.1053 |
| NDUFS7 | 4 | 0.47742 | 0.60636 | 0.999994 | 13105 | 1 | 0.56931 | 0.099261 | 0.23161 | 0.97466 | 5532 | 2 | 0.56931 |
| DEF8 | 4 | 0.40836 | 0.55429 | 0.999994 | 12085 | 1 | 3.6245 | 0.09929 | 0.23165 | 0.97466 | 5533 | 3 | 3.6245 |
| ASCC3 | 5 | 0.89484 | 0.89499 | 0.999994 | 19120 | 0 | 1.5415 | 0.099326 | 0.25323 | 0.97466 | 5534 | 4 | 1.5415 |
| COMMD1 | 6 | 0.0062462 | 0.025717 | 0.845768 | 608 | 3 | -1.5052 | 0.099375 | 0.27883 | 0.978454 | 5535 | 3 | -1.5052 |
| ZMYM1 | 6 | 0.023254 | 0.082391 | 0.940526 | 1667 | 3 | -0.78438 | 0.099394 | 0.27887 | 0.978454 | 5536 | 1 | -0.78438 |
| HSFY2 | 2 | 0.86731 | 0.86732 | 0.999994 | 18583 | 0 | 4.9232 | 0.099397 | 0.16913 | 0.965913 | 5537 | 2 | 4.9232 |
| TMEM64 | 5 | 0.40737 | 0.56968 | 0.999994 | 12064 | 1 | 2.1227 | 0.099402 | 0.25338 | 0.97466 | 5538 | 4 | 2.1227 |
| MPZL1 | 5 | 0.55915 | 0.69986 | 0.999994 | 14355 | 1 | -0.36509 | 0.099413 | 0.2534 | 0.97466 | 5539 | 2 | -0.36509 |
| FETUB | 5 | 0.18199 | 0.3495 | 0.999994 | 7271 | 3 | -0.9052 | 0.099413 | 0.2534 | 0.97466 | 5540 | 2 | -0.9052 |
| hsa-mir-1202 | 4 | 0.16937 | 0.30911 | 0.999994 | 6967 | 2 | 2.5256 | 0.099418 | 0.23187 | 0.97466 | 5541 | 2 | 2.5256 |
| SEPT2 | 5 | 0.91356 | 0.91362 | 0.999994 | 19481 | 0 | 1.2419 | 0.099434 | 0.25344 | 0.97466 | 5542 | 4 | 1.2419 |
| INPP5E | 6 | 0.673 | 0.79549 | 0.999994 | 15971 | 2 | 2.3574 | 0.09944 | 0.27893 | 0.978454 | 5543 | 3 | 2.3574 |
| NonTargetingControlG | 2 | 0.90054 | 0.90046 | 0.999994 | 19229 | 0 | 2.6407 | 0.099458 | 0.16923 | 0.965913 | 5544 | 2 | 2.6407 |
| NXF1 | 5 | 0.92072 | 0.92091 | 0.999994 | 19646 | 0 | 1.3607 | 0.099481 | 0.25354 | 0.97466 | 5545 | 4 | 1.3607 |
| CLCA1 | 6 | 0.65262 | 0.77963 | 0.999994 | 15694 | 2 | 3.4705 | 0.099504 | 0.27904 | 0.978454 | 5546 | 4 | 3.4705 |
| C6orf165 | 5 | 0.61873 | 0.73309 | 0.999994 | 15198 | 1 | 2.121 | 0.099521 | 0.25362 | 0.97466 | 5547 | 4 | 2.121 |
| ICAM1 | 5 | 0.94612 | 0.94631 | 0.999994 | 20236 | 0 | 2.6653 | 0.099529 | 0.25364 | 0.97466 | 5548 | 4 | 2.6653 |
| OR7E24 | 6 | 0.18122 | 0.36468 | 0.999994 | 7255 | 1 | 1.0413 | 0.099552 | 0.27911 | 0.978454 | 5549 | 4 | 1.0413 |
| TRPC4AP | 6 | 0.98373 | 0.98368 | 0.999994 | 21180 | 0 | 1.2489 | 0.099552 | 0.27911 | 0.978454 | 5550 | 4 | 1.2489 |
| DNAJC21 | 6 | 0.79065 | 0.84394 | 0.999994 | 17358 | 1 | 0.3044 | 0.099552 | 0.27911 | 0.978454 | 5551 | 3 | 0.3044 |
| EIF2D | 5 | 0.046127 | 0.13109 | 0.95318 | 2814 | 3 | -2.2006 | 0.09963 | 0.25383 | 0.97466 | 5552 | 2 | -2.2006 |
| SH3YL1 | 5 | 0.21723 | 0.3994 | 0.999994 | 8200 | 3 | -0.74566 | 0.09963 | 0.25383 | 0.97466 | 5553 | 1 | -0.74566 |
| DEFB107B | 2 | 0.57762 | 0.57778 | 0.999994 | 14652 | 1 | 3.7387 | 0.099648 | 0.16951 | 0.965913 | 5554 | 1 | 3.7387 |
| ZNF343 | 5 | 0.9456 | 0.9458 | 0.999994 | 20220 | 0 | 1.6174 | 0.099703 | 0.25399 | 0.97466 | 5555 | 4 | 1.6174 |
| ZNF211 | 6 | 0.34862 | 0.56244 | 0.999994 | 11127 | 2 | 1.9918 | 0.099745 | 0.27941 | 0.978454 | 5556 | 4 | 1.9918 |
| hsa-mir-2355 | 4 | 0.16234 | 0.30187 | 0.999994 | 6804 | 2 | -0.39451 | 0.099749 | 0.23242 | 0.97466 | 5557 | 1 | -0.39451 |
| SPRR2G | 4 | 0.10995 | 0.23278 | 0.984502 | 5306 | 2 | 2.248 | 0.099757 | 0.23244 | 0.97466 | 5558 | 2 | 2.248 |
| RAB6A | 3 | 0.43878 | 0.50735 | 0.999994 | 12553 | 1 | 4.8199 | 0.099809 | 0.22043 | 0.97466 | 5559 | 2 | 4.8199 |
| IMMT | 5 | 0.57815 | 0.71667 | 0.999994 | 14661 | 2 | 0.54658 | 0.099811 | 0.2542 | 0.97466 | 5560 | 2 | 0.54658 |
| PKHD1 | 6 | 0.48328 | 0.65738 | 0.999994 | 13191 | 1 | 1.243 | 0.099835 | 0.27954 | 0.978454 | 5561 | 4 | 1.243 |
| DYNLT3 | 6 | 0.058932 | 0.16348 | 0.967126 | 3378 | 3 | -1.4269 | 0.099835 | 0.27954 | 0.978454 | 5562 | 2 | -1.4269 |
| KIF24 | 6 | 0.13799 | 0.3105 | 0.999994 | 6135 | 2 | 0.3422 | 0.099835 | 0.27954 | 0.978454 | 5563 | 3 | 0.3422 |

| | | | | | | | | | | | | |
|---|---|---|---|---|---|---|---|---|---|---|---|---|
| POLK | 6 | 0.059181 | 0.164 | 0.967126 | 3387 | 2 | -0.13489 | 0.099835 | 0.27954 | 0.978454 | 5564 | 2 | -0.13489 |
| TBX1 | 6 | 0.98134 | 0.98125 | 0.999994 | 21125 | 0 | 2.9163 | 0.099836 | 0.27954 | 0.978454 | 5565 | 4 | 2.9163 |
| UBAP2L | 6 | 0.57078 | 0.71849 | 0.999994 | 14555 | 2 | 2.9435 | 0.099836 | 0.27954 | 0.978454 | 5566 | 4 | 2.9435 |
| hsa-mir-634 | 2 | 0.90016 | 0.90007 | 0.999994 | 19222 | 0 | 3.0718 | 0.099836 | 0.16981 | 0.965913 | 5567 | 2 | 3.0718 |
| PIGM | 5 | 0.36079 | 0.53074 | 0.999994 | 11352 | 1 | 2.7721 | 0.099869 | 0.2543 | 0.97466 | 5568 | 4 | 2.7721 |
| hsa-mir-4456 | 4 | 0.2251 | 0.36646 | 0.999994 | 8402 | 1 | 4.1795 | 0.099891 | 0.23263 | 0.97466 | 5569 | 3 | 4.1795 |
| FOXN4 | 4 | 0.42956 | 0.57611 | 0.999994 | 12407 | 1 | 2.7521 | 0.099906 | 0.23266 | 0.97466 | 5570 | 3 | 2.7521 |
| ADAT2 | 6 | 0.56707 | 0.71584 | 0.999994 | 14497 | 2 | 1.6069 | 0.099916 | 0.27967 | 0.978454 | 5571 | 3 | 1.6069 |
| RRP1B | 5 | 0.2275 | 0.41374 | 0.999994 | 8470 | 1 | 1.8015 | 0.099955 | 0.25447 | 0.97466 | 5572 | 4 | 1.8015 |
| SELT | 5 | 0.96104 | 0.96117 | 0.999994 | 20607 | 0 | 1.3519 | 0.099956 | 0.25447 | 0.97466 | 5573 | 4 | 1.3519 |
| hsa-mir-4291 | 2 | 0.89998 | 0.89989 | 0.999994 | 19221 | 0 | 4.6393 | 0.10002 | 0.17011 | 0.965913 | 5574 | 2 | 4.6393 |
| GJB6 | 6 | 0.055866 | 0.15679 | 0.967126 | 3242 | 4 | -1.6927 | 0.10007 | 0.27992 | 0.978454 | 5575 | 2 | -1.6927 |
| PIWIL3 | 6 | 0.24214 | 0.43855 | 0.999994 | 8836 | 3 | -0.58024 | 0.10007 | 0.27992 | 0.978454 | 5576 | 2 | -0.58024 |
| GPX6 | 6 | 0.55181 | 0.70496 | 0.999994 | 14240 | 1 | 2.2469 | 0.10007 | 0.27993 | 0.978454 | 5577 | 4 | 2.2469 |
| MAS1 | 5 | 0.78454 | 0.81009 | 0.999994 | 17275 | 1 | 1.3794 | 0.10009 | 0.25472 | 0.97466 | 5578 | 4 | 1.3794 |
| CYP17A1 | 5 | 0.84606 | 0.85125 | 0.999994 | 18187 | 1 | 1.8625 | 0.10011 | 0.25476 | 0.97466 | 5579 | 4 | 1.8625 |
| SCGB2A1 | 5 | 0.91717 | 0.91725 | 0.999994 | 19566 | 0 | 3.6718 | 0.10016 | 0.25485 | 0.97466 | 5580 | 3 | 3.6718 |
| ESRRA | 6 | 0.56214 | 0.71234 | 0.999994 | 14410 | 1 | 2.8006 | 0.10018 | 0.2801 | 0.978454 | 5581 | 4 | 2.8006 |
| PLEKHG5 | 6 | 0.34706 | 0.5607 | 0.999994 | 11094 | 2 | 2.6527 | 0.10021 | 0.28013 | 0.978454 | 5582 | 4 | 2.6527 |
| NR3C2 | 5 | 0.18877 | 0.35917 | 0.999994 | 7444 | 2 | 3.6811 | 0.10024 | 0.25502 | 0.97466 | 5583 | 3 | 3.6811 |
| EIF2S1 | 4 | 0.14239 | 0.28136 | 0.999632 | 6249 | 2 | 1.9543 | 0.10026 | 0.23321 | 0.97466 | 5584 | 2 | 1.9543 |
| SPATA32 | 4 | 0.22225 | 0.36356 | 0.999994 | 8337 | 1 | 3.4649 | 0.10026 | 0.23321 | 0.97466 | 5585 | 3 | 3.4649 |
| KRTAP19-6 | 2 | 0.89974 | 0.89965 | 0.999994 | 19217 | 0 | 4.4064 | 0.10026 | 0.17046 | 0.965913 | 5586 | 2 | 4.4064 |
| BLVRA | 6 | 0.010112 | 0.039395 | 0.891621 | 878 | 3 | -1.5438 | 0.1003 | 0.28029 | 0.978454 | 5587 | 1 | -1.5438 |
| OR8D2 | 6 | 0.3691 | 0.58266 | 0.999994 | 11492 | 2 | 0.86945 | 0.10031 | 0.2803 | 0.978454 | 5588 | 4 | 0.86945 |
| LPCAT1 | 6 | 0.59285 | 0.7347 | 0.999994 | 14858 | 2 | 0.040928 | 0.10031 | 0.2803 | 0.978454 | 5589 | 2 | 0.040928 |
| DEFB103B | 1 | 0.89968 | 0.90005 | 0.999994 | 19216 | 0 | 7.1468 | 0.10032 | 0.10034 | 0.951893 | 5590 | 1 | 7.1468 |
| NXNL2 | 4 | 0.12884 | 0.26559 | 0.997535 | 5908 | 2 | 1.3217 | 0.10034 | 0.23335 | 0.97466 | 5591 | 2 | 1.3217 |
| ACD | 4 | 0.73163 | 0.74637 | 0.999994 | 16623 | 1 | 2.9468 | 0.10036 | 0.23338 | 0.97466 | 5592 | 3 | 2.9468 |
| RAPGEFL1 | 4 | 0.80311 | 0.804 | 0.999994 | 17525 | 1 | 4.174 | 0.10036 | 0.23338 | 0.97466 | 5593 | 3 | 4.174 |
| RPL30 | 5 | 0.12165 | 0.25986 | 0.994943 | 5691 | 2 | -0.27568 | 0.10037 | 0.25527 | 0.97466 | 5594 | 2 | -0.27568 |
| hsa-mir-8087 | 3 | 0.89958 | 0.89967 | 0.999994 | 19215 | 0 | 3.3942 | 0.10042 | 0.22162 | 0.97466 | 5595 | 3 | 3.3942 |
| CTTNBP2 | 5 | 0.88846 | 0.88857 | 0.999994 | 19007 | 0 | 1.2451 | 0.10046 | 0.25546 | 0.97466 | 5596 | 4 | 1.2451 |
| CGB1 | 4 | 0.33447 | 0.47827 | 0.999994 | 10821 | 1 | 2.5577 | 0.10048 | 0.23357 | 0.97466 | 5597 | 3 | 2.5577 |
| MRPL27 | 3 | 0.89292 | 0.89295 | 0.999994 | 19088 | 0 | 5.0629 | 0.10049 | 0.22176 | 0.97466 | 5598 | 3 | 5.0629 |
| NonTargetingControlG | 2 | 0.31802 | 0.35067 | 0.999994 | 10473 | 1 | 2.8267 | 0.10049 | 0.17081 | 0.965913 | 5599 | 1 | 2.8267 |
| VWC2 | 6 | 0.45584 | 0.63893 | 0.999994 | 12787 | 2 | 0.099424 | 0.10051 | 0.28063 | 0.978454 | 5600 | 2 | 0.099424 |
| LTV1 | 6 | 0.1279 | 0.29763 | 0.999994 | 5869 | 3 | -0.64749 | 0.10051 | 0.28063 | 0.978454 | 5601 | 2 | -0.64749 |
| IGFL3 | 5 | 0.81858 | 0.83163 | 0.999994 | 17757 | 1 | 1.1966 | 0.10051 | 0.25558 | 0.97466 | 5602 | 4 | 1.1966 |
| ITCH | 6 | 0.28489 | 0.48913 | 0.999994 | 9808 | 3 | -0.23919 | 0.10052 | 0.28065 | 0.978454 | 5603 | 2 | -0.23919 |
| CRP | 5 | 0.95345 | 0.95361 | 0.999994 | 20421 | 0 | 1.8465 | 0.10063 | 0.25579 | 0.97466 | 5604 | 3 | 1.8465 |
| RXFP2 | 6 | 0.22524 | 0.4184 | 0.999994 | 8406 | 3 | 0.75567 | 0.10073 | 0.28096 | 0.978454 | 5605 | 3 | 0.75567 |
| CSNK2A2 | 6 | 0.20005 | 0.38794 | 0.999994 | 7764 | 1 | 0.40839 | 0.10073 | 0.28096 | 0.978454 | 5606 | 3 | 0.40839 |
| AUH | 6 | 0.19673 | 0.38391 | 0.999994 | 7670 | 3 | 0.58826 | 0.10073 | 0.28096 | 0.978454 | 5607 | 3 | 0.58826 |
| NonTargetingControlG | 2 | 0.89917 | 0.89906 | 0.999994 | 19207 | 0 | 2.2878 | 0.10083 | 0.17135 | 0.965913 | 5608 | 2 | 2.2878 |
| CEP250 | 4 | 0.93105 | 0.93081 | 0.999994 | 19879 | 0 | 4.0118 | 0.10085 | 0.23418 | 0.97466 | 5609 | 3 | 4.0118 |
| KLHDC1 | 6 | 0.96894 | 0.96895 | 0.999994 | 20801 | 0 | 1.4728 | 0.10087 | 0.28119 | 0.978454 | 5610 | 4 | 1.4728 |
| RPS6KA1 | 5 | 0.56559 | 0.70565 | 0.999994 | 14478 | 2 | 3.5982 | 0.1009 | 0.25634 | 0.974766 | 5611 | 3 | 3.5982 |
| CEP128 | 5 | 0.092869 | 0.21516 | 0.979587 | 4715 | 1 | 1.4166 | 0.10097 | 0.25648 | 0.974766 | 5612 | 4 | 1.4166 |
| TPSG1 | 5 | 0.32098 | 0.4981 | 0.999994 | 10532 | 1 | 0.8013 | 0.10103 | 0.25658 | 0.974766 | 5613 | 3 | 0.8013 |
| USP24 | 5 | 0.22227 | 0.40646 | 0.999994 | 8339 | 1 | 3.5111 | 0.10103 | 0.25658 | 0.974766 | 5614 | 3 | 3.5111 |
| OR56B1 | 5 | 0.94491 | 0.94508 | 0.999994 | 20201 | 0 | 1.5311 | 0.10104 | 0.2566 | 0.974766 | 5615 | 4 | 1.5311 |

| | | | | | | | | | | | | |
|---|---|---|---|---|---|---|---|---|---|---|---|---|
| OR4L1 | 6 | 0.5791 | 0.72454 | 0.999994 | 14669 | 2 | 1.2872 | 0.10105 | 0.28148 | 0.978454 | 5616 | 4 | 1.2872 |
| HMGCR | 4 | 0.18057 | 0.32067 | 0.999994 | 7235 | 1 | 2.262 | 0.10105 | 0.23452 | 0.97466 | 5617 | 3 | 2.262 |
| MCOLN1 | 6 | 0.38643 | 0.59365 | 0.999994 | 11739 | 2 | 1.7569 | 0.1011 | 0.28158 | 0.978454 | 5618 | 4 | 1.7569 |
| TDRD6 | 5 | 0.23087 | 0.4184 | 0.999994 | 8538 | 2 | 0.42534 | 0.10115 | 0.25682 | 0.974766 | 5619 | 2 | 0.42534 |
| hsa-mir-124-3 | 3 | 0.669 | 0.67301 | 0.999994 | 15911 | 1 | 1.8622 | 0.10115 | 0.22305 | 0.97466 | 5620 | 2 | 1.8622 |
| TRMT61A | 5 | 0.21272 | 0.39321 | 0.999994 | 8081 | 1 | 1.8462 | 0.10115 | 0.25683 | 0.974766 | 5621 | 4 | 1.8462 |
| NonTargetingControlG | 2 | 0.89883 | 0.89872 | 0.999994 | 19200 | 0 | 3.3649 | 0.10117 | 0.17188 | 0.965913 | 5622 | 2 | 3.3649 |
| CMYA5 | 6 | 0.71626 | 0.81712 | 0.999994 | 16459 | 1 | 1.2382 | 0.1012 | 0.28172 | 0.978454 | 5623 | 3 | 1.2382 |
| MAPK1 | 6 | 0.0060594 | 0.025019 | 0.844082 | 597 | 3 | -1.786 | 0.1012 | 0.28172 | 0.978454 | 5624 | 3 | -1.786 |
| NonTargetingControlG | 2 | 0.87603 | 0.87607 | 0.999994 | 18752 | 0 | 4.8832 | 0.10121 | 0.17197 | 0.965913 | 5625 | 2 | 4.8832 |
| ATP11B | 6 | 0.73895 | 0.82463 | 0.999994 | 16721 | 1 | 1.1452 | 0.10126 | 0.28181 | 0.978454 | 5626 | 3 | 1.1452 |
| SLA | 6 | 0.6654 | 0.78953 | 0.999994 | 15859 | 2 | 0.1642 | 0.10126 | 0.28181 | 0.978454 | 5627 | 3 | 0.1642 |
| KLHL42 | 5 | 0.92932 | 0.92958 | 0.999994 | 19841 | 0 | 2.9087 | 0.10126 | 0.25703 | 0.974766 | 5628 | 4 | 2.9087 |
| FANCG | 6 | 0.032297 | 0.1042 | 0.947284 | 2121 | 2 | -0.26415 | 0.10136 | 0.28198 | 0.978454 | 5629 | 2 | -0.26415 |
| C11orf58 | 5 | 0.76279 | 0.79772 | 0.999994 | 17004 | 1 | 0.18827 | 0.10138 | 0.25727 | 0.974766 | 5630 | 2 | 0.18827 |
| ZYG11B | 5 | 0.82157 | 0.83363 | 0.999994 | 17797 | 1 | 2.4384 | 0.10144 | 0.25738 | 0.974766 | 5631 | 3 | 2.4384 |
| LGI3 | 6 | 0.25191 | 0.45021 | 0.999994 | 9055 | 3 | -0.80875 | 0.10147 | 0.28216 | 0.978454 | 5632 | 1 | -0.80875 |
| TMEM130 | 4 | 0.092445 | 0.20148 | 0.977164 | 4699 | 2 | 0.12746 | 0.10149 | 0.23522 | 0.97466 | 5633 | 2 | 0.12746 |
| hsa-mir-4646 | 3 | 0.79751 | 0.79735 | 0.999994 | 17445 | 0 | 5.1866 | 0.10152 | 0.22371 | 0.97466 | 5634 | 2 | 5.1866 |
| LOC643669 | 5 | 0.95828 | 0.95843 | 0.999994 | 20544 | 0 | 1.449 | 0.10153 | 0.25754 | 0.974766 | 5635 | 4 | 1.449 |
| CBLC | 5 | 0.74762 | 0.78959 | 0.999994 | 16834 | 1 | 2.2632 | 0.10156 | 0.2576 | 0.974766 | 5636 | 4 | 2.2632 |
| FOXRED1 | 5 | 0.13308 | 0.27732 | 0.999379 | 5997 | 3 | -2.6337 | 0.10156 | 0.2576 | 0.974766 | 5637 | 2 | -2.6337 |
| GOLGA2 | 5 | 0.47178 | 0.62364 | 0.999994 | 13029 | 2 | -1.1036 | 0.10156 | 0.2576 | 0.974766 | 5638 | 2 | -1.1036 |
| TMEM134 | 5 | 0.1529 | 0.3071 | 0.999994 | 6531 | 2 | 3.4665 | 0.10158 | 0.25764 | 0.974766 | 5639 | 3 | 3.4665 |
| RFFL | 5 | 0.52432 | 0.66909 | 0.999994 | 13807 | 2 | 3.3715 | 0.10161 | 0.25769 | 0.974766 | 5640 | 3 | 3.3715 |
| hsa-mir-548f-5 | 4 | 0.79837 | 0.79978 | 0.999994 | 17453 | 1 | 2.9103 | 0.10164 | 0.23545 | 0.97466 | 5641 | 3 | 2.9103 |
| PAIP2B | 6 | 0.52038 | 0.6828 | 0.999994 | 13753 | 2 | 0.33239 | 0.10165 | 0.28246 | 0.978454 | 5642 | 3 | 0.33239 |
| HMOX1 | 6 | 0.16905 | 0.34955 | 0.999994 | 6961 | 2 | 0.15691 | 0.10165 | 0.28246 | 0.978454 | 5643 | 2 | 0.15691 |
| MMRN1 | 6 | 0.64655 | 0.77499 | 0.999994 | 15609 | 2 | 0.11921 | 0.10165 | 0.28246 | 0.978454 | 5644 | 3 | 0.11921 |
| PKN2 | 6 | 0.18475 | 0.36907 | 0.999994 | 7338 | 2 | 2.0683 | 0.10168 | 0.28251 | 0.978454 | 5645 | 4 | 2.0683 |
| TMEM205 | 6 | 0.37387 | 0.58566 | 0.999994 | 11569 | 1 | 0.79853 | 0.10171 | 0.28254 | 0.978454 | 5646 | 4 | 0.79853 |
| TTC39B | 6 | 0.52897 | 0.68883 | 0.999994 | 13881 | 2 | 2.0671 | 0.10171 | 0.28254 | 0.978454 | 5647 | 4 | 2.0671 |
| RPS11 | 2 | 0.89824 | 0.89814 | 0.999994 | 19188 | 0 | 4.7025 | 0.10176 | 0.17285 | 0.965913 | 5648 | 2 | 4.7025 |
| PPP3R2 | 5 | 0.86942 | 0.87056 | 0.999994 | 18628 | 1 | 3.5443 | 0.10177 | 0.25802 | 0.974766 | 5649 | 3 | 3.5443 |
| PRR13 | 5 | 0.87853 | 0.87886 | 0.999994 | 18799 | 1 | 2.2214 | 0.10178 | 0.25803 | 0.974766 | 5650 | 4 | 2.2214 |
| PEX3 | 6 | 0.19981 | 0.38764 | 0.999994 | 7761 | 2 | 0.90014 | 0.1018 | 0.2827 | 0.978454 | 5651 | 3 | 0.90014 |
| CTSL1 | 4 | 0.050893 | 0.12325 | 0.951052 | 3012 | 1 | 0.5223 | 0.10184 | 0.23578 | 0.97466 | 5652 | 2 | 0.5223 |
| SEC14L3 | 6 | 0.88649 | 0.8939 | 0.999994 | 18959 | 1 | 3.1962 | 0.10188 | 0.28282 | 0.978454 | 5653 | 4 | 3.1962 |
| EIF2AK2 | 6 | 0.1781 | 0.36074 | 0.999994 | 7178 | 3 | 1.9855 | 0.10188 | 0.28282 | 0.978454 | 5654 | 3 | 1.9855 |
| BABAM1 | 6 | 0.078245 | 0.2038 | 0.978186 | 4156 | 2 | 3.1136 | 0.10188 | 0.28282 | 0.978454 | 5655 | 4 | 3.1136 |
| NMB | 5 | 0.068386 | 0.17581 | 0.968171 | 3762 | 1 | 3.4944 | 0.10194 | 0.25834 | 0.974766 | 5656 | 4 | 3.4944 |
| NonTargetingControlG | 2 | 0.85611 | 0.85605 | 0.999994 | 18375 | 0 | 4.8341 | 0.10195 | 0.17313 | 0.965913 | 5657 | 2 | 4.8341 |
| BSN | 5 | 0.84889 | 0.85341 | 0.999994 | 18235 | 1 | 2.02 | 0.10198 | 0.2584 | 0.974766 | 5658 | 4 | 2.02 |
| TRAT1 | 5 | 0.69814 | 0.76565 | 0.999994 | 16283 | 1 | 3.3527 | 0.10201 | 0.25847 | 0.974766 | 5659 | 3 | 3.3527 |
| ZNF582 | 6 | 0.81775 | 0.85576 | 0.999994 | 17742 | 1 | 2.099 | 0.10202 | 0.28305 | 0.978454 | 5660 | 4 | 2.099 |
| PXN | 3 | 0.7956 | 0.79544 | 0.999994 | 17427 | 0 | 0.92449 | 0.10213 | 0.22489 | 0.97466 | 5661 | 2 | 0.92449 |
| hsa-mir-200a | 3 | 0.059875 | 0.12927 | 0.952548 | 3415 | 2 | -2.5007 | 0.10213 | 0.22489 | 0.97466 | 5662 | 1 | -2.5007 |
| ZNF749 | 6 | 0.58104 | 0.72598 | 0.999994 | 14692 | 1 | 2.7136 | 0.10219 | 0.28332 | 0.978454 | 5663 | 4 | 2.7136 |
| KIAA1324 | 6 | 0.29457 | 0.50036 | 0.999994 | 10008 | 2 | 2.3241 | 0.10221 | 0.28335 | 0.978454 | 5664 | 4 | 2.3241 |
| IL36B | 6 | 0.56265 | 0.71269 | 0.999994 | 14424 | 2 | -0.036821 | 0.10223 | 0.28339 | 0.978454 | 5665 | 2 | -0.036821 |
| CNIH3 | 6 | 0.96306 | 0.96312 | 0.999994 | 20667 | 0 | 2.734 | 0.10223 | 0.28339 | 0.978454 | 5666 | 4 | 2.734 |
| KLHL20 | 6 | 0.053067 | 0.1507 | 0.966359 | 3115 | 3 | -0.80021 | 0.10224 | 0.28341 | 0.978454 | 5667 | 2 | -0.80021 |

| | | | | | | | | | | | | |
|---|---|---|---|---|---|---|---|---|---|---|---|---|
| GNAI1 | 6 | 0.34059 | 0.55333 | 0.999994 | 10950 | 3 | 0.97174 | 0.10224 | 0.28341 | 0.978454 | 5668 | 3 | 0.97174 |
| TSSC4 | 6 | 0.44177 | 0.6296 | 0.999994 | 12607 | 2 | -0.46084 | 0.10224 | 0.28341 | 0.978454 | 5669 | 2 | -0.46084 |
| AP5B1 | 5 | 0.80051 | 0.81983 | 0.999994 | 17488 | 1 | 3.714 | 0.10225 | 0.25894 | 0.974766 | 5670 | 3 | 3.714 |
| SMARCA4 | 5 | 0.33346 | 0.50834 | 0.999994 | 10796 | 2 | 3.714 | 0.10225 | 0.25894 | 0.974766 | 5671 | 3 | 3.714 |
| ARL2BP | 5 | 0.28228 | 0.46671 | 0.999994 | 9746 | 2 | 3.714 | 0.10225 | 0.25894 | 0.974766 | 5672 | 3 | 3.714 |
| BIVM | 6 | 0.56157 | 0.71193 | 0.999994 | 14399 | 2 | 1.9591 | 0.10225 | 0.28342 | 0.978454 | 5673 | 4 | 1.9591 |
| OVCH2 | 5 | 0.042561 | 0.12223 | 0.951052 | 2650 | 3 | -2.0692 | 0.10227 | 0.25898 | 0.974766 | 5674 | 2 | -2.0692 |
| FAM192A | 6 | 0.51925 | 0.682 | 0.999994 | 13735 | 1 | 2.6015 | 0.10244 | 0.28374 | 0.978454 | 5675 | 3 | 2.6015 |
| PLA2G2A | 6 | 0.46529 | 0.64531 | 0.999994 | 12937 | 2 | 2.3704 | 0.10245 | 0.28375 | 0.978454 | 5676 | 4 | 2.3704 |
| hsa-mir-4271 | 2 | 0.4794 | 0.48528 | 0.999994 | 13138 | 1 | 3.4584 | 0.10247 | 0.17394 | 0.966602 | 5677 | 1 | 3.4584 |
| PIEZO1 | 5 | 0.46654 | 0.61919 | 0.999994 | 12959 | 1 | 1.9486 | 0.1025 | 0.25942 | 0.974766 | 5678 | 4 | 1.9486 |
| PSMG1 | 4 | 0.27197 | 0.41433 | 0.999994 | 9519 | 1 | 1.8516 | 0.1025 | 0.23683 | 0.97466 | 5679 | 3 | 1.8516 |
| WDR33 | 3 | 0.85259 | 0.85255 | 0.999994 | 18308 | 0 | 5.0163 | 0.1025 | 0.22556 | 0.97466 | 5680 | 2 | 5.0163 |
| FBXL3 | 5 | 0.45871 | 0.61258 | 0.999994 | 12831 | 2 | 1.2942 | 0.10251 | 0.25944 | 0.974766 | 5681 | 3 | 1.2942 |
| FAM173A | 5 | 0.20315 | 0.37972 | 0.999994 | 7858 | 3 | -0.75391 | 0.10251 | 0.25944 | 0.974766 | 5682 | 1 | -0.75391 |
| ATP6V0E1 | 6 | 0.56993 | 0.71789 | 0.999994 | 14539 | 1 | 2.5871 | 0.10256 | 0.28392 | 0.978454 | 5683 | 3 | 2.5871 |
| hsa-mir-8063 | 3 | 0.89739 | 0.89747 | 0.999994 | 19166 | 0 | 2.5626 | 0.10261 | 0.22576 | 0.97466 | 5684 | 3 | 2.5626 |
| ATXN10 | 5 | 0.15347 | 0.30799 | 0.999994 | 6547 | 1 | 3.2729 | 0.10264 | 0.25969 | 0.974766 | 5685 | 3 | 3.2729 |
| C15orf60 | 5 | 0.18527 | 0.3542 | 0.999994 | 7348 | 1 | 2.4812 | 0.10268 | 0.25976 | 0.974766 | 5686 | 4 | 2.4812 |
| SLC5A6 | 6 | 0.59558 | 0.73671 | 0.999994 | 14898 | 2 | 2.1571 | 0.10269 | 0.28414 | 0.978454 | 5687 | 4 | 2.1571 |
| ANKRD45 | 5 | 0.97013 | 0.97018 | 0.999994 | 20827 | 0 | 2.6369 | 0.1027 | 0.25981 | 0.974766 | 5688 | 4 | 2.6369 |
| DCAF10 | 5 | 0.95584 | 0.956 | 0.999994 | 20479 | 0 | 1.3285 | 0.10274 | 0.25988 | 0.974766 | 5689 | 4 | 1.3285 |
| SPP1 | 5 | 0.8224 | 0.83418 | 0.999994 | 17812 | 1 | -0.058619 | 0.10277 | 0.25993 | 0.974766 | 5690 | 1 | -0.058619 |
| SUPT3H | 6 | 0.62889 | 0.76151 | 0.999994 | 15361 | 2 | 0.72482 | 0.10278 | 0.28429 | 0.978454 | 5691 | 4 | 0.72482 |
| COG6 | 6 | 0.88037 | 0.88985 | 0.999994 | 18841 | 1 | 0.94635 | 0.1028 | 0.28431 | 0.978454 | 5692 | 3 | 0.94635 |
| BSDC1 | 6 | 0.3554 | 0.57011 | 0.999994 | 11257 | 2 | -0.044454 | 0.1028 | 0.28431 | 0.978454 | 5693 | 2 | -0.044454 |
| STARD4 | 6 | 0.96939 | 0.96939 | 0.999994 | 20815 | 0 | 1.5525 | 0.1028 | 0.28431 | 0.978454 | 5694 | 3 | 1.5525 |
| SORBS3 | 6 | 0.008877 | 0.035086 | 0.867348 | 784 | 1 | 2.163 | 0.1028 | 0.28431 | 0.978454 | 5695 | 3 | 2.163 |
| ASAH2B | 5 | 0.52281 | 0.66775 | 0.999994 | 13792 | 2 | -0.16316 | 0.10282 | 0.26004 | 0.974766 | 5696 | 2 | -0.16316 |
| EMILIN1 | 5 | 0.10563 | 0.23519 | 0.985362 | 5158 | 3 | -0.87284 | 0.10282 | 0.26004 | 0.974766 | 5697 | 2 | -0.87284 |
| NonTargetingControlG | 2 | 0.7939 | 0.79347 | 0.999994 | 17396 | 0 | 4.4692 | 0.10287 | 0.17453 | 0.967011 | 5698 | 2 | 4.4692 |
| BIRC8 | 6 | 0.28091 | 0.48445 | 0.999994 | 9721 | 2 | 2.0105 | 0.10292 | 0.2845 | 0.978535 | 5699 | 4 | 2.0105 |
| RDH11 | 5 | 0.35675 | 0.52741 | 0.999994 | 11284 | 2 | 0.38702 | 0.10294 | 0.26025 | 0.974766 | 5700 | 2 | 0.38702 |
| NonTargetingControlG | 2 | 0.89703 | 0.89692 | 0.999994 | 19160 | 0 | 2.6981 | 0.10297 | 0.1747 | 0.967011 | 5701 | 2 | 2.6981 |
| AHNAK | 5 | 0.20445 | 0.38156 | 0.999994 | 7889 | 1 | 1.5617 | 0.10299 | 0.26036 | 0.974766 | 5702 | 4 | 1.5617 |
| CASP8AP2 | 5 | 0.43293 | 0.59086 | 0.999994 | 12463 | 1 | 2.7516 | 0.1031 | 0.26057 | 0.974875 | 5703 | 4 | 2.7516 |
| CTC1 | 5 | 0.10697 | 0.23725 | 0.985914 | 5211 | 2 | 3.473 | 0.10313 | 0.26061 | 0.974875 | 5704 | 3 | 3.473 |
| FGF3 | 6 | 0.28569 | 0.49009 | 0.999994 | 9822 | 1 | 3.0469 | 0.10321 | 0.28498 | 0.978871 | 5705 | 4 | 3.0469 |
| HAUS8 | 3 | 0.077382 | 0.15292 | 0.967126 | 4125 | 1 | -0.25259 | 0.10323 | 0.22691 | 0.97466 | 5706 | 1 | -0.25259 |
| ABCA8 | 6 | 0.094484 | 0.23648 | 0.985449 | 4768 | 4 | -1.424 | 0.10325 | 0.28504 | 0.978871 | 5707 | 2 | -1.424 |
| TP53 | 6 | 0.3141 | 0.52315 | 0.999994 | 10399 | 1 | 2.2095 | 0.10328 | 0.2851 | 0.978871 | 5708 | 3 | 2.2095 |
| GPCPD1 | 6 | 0.49739 | 0.66693 | 0.999994 | 13389 | 1 | 2.5437 | 0.10328 | 0.2851 | 0.978871 | 5709 | 4 | 2.5437 |
| CCNF | 4 | 0.80553 | 0.80623 | 0.999994 | 17560 | 1 | 3.1667 | 0.1033 | 0.23812 | 0.97466 | 5710 | 3 | 3.1667 |
| SRP72 | 5 | 0.38554 | 0.5514 | 0.999994 | 11727 | 2 | 3.679 | 0.10331 | 0.26097 | 0.974875 | 5711 | 3 | 3.679 |
| UTP3 | 3 | 0.46248 | 0.52195 | 0.999994 | 12896 | 1 | 4.988 | 0.10332 | 0.22707 | 0.97466 | 5712 | 2 | 4.988 |
| TMEM71 | 5 | 0.13665 | 0.28272 | 0.999632 | 6100 | 2 | 3.2997 | 0.10337 | 0.26109 | 0.974875 | 5713 | 3 | 3.2997 |
| HYLS1 | 5 | 0.74613 | 0.78882 | 0.999994 | 16815 | 1 | 3.5437 | 0.1034 | 0.26113 | 0.974875 | 5714 | 3 | 3.5437 |
| MTERFD1 | 5 | 0.88008 | 0.88033 | 0.999994 | 18833 | 1 | -0.078502 | 0.10342 | 0.26117 | 0.974875 | 5715 | 2 | -0.078502 |
| DRAM1 | 4 | 0.1109 | 0.23446 | 0.984971 | 5329 | 1 | 0.40627 | 0.10342 | 0.23833 | 0.97466 | 5716 | 2 | 0.40627 |
| hsa-mir-6884 | 4 | 0.74781 | 0.75838 | 0.999994 | 16836 | 1 | 2.5601 | 0.10345 | 0.23837 | 0.97466 | 5717 | 3 | 2.5601 |
| STMND1 | 6 | 0.52115 | 0.68333 | 0.999994 | 13763 | 2 | 2.2763 | 0.10346 | 0.28539 | 0.978871 | 5718 | 4 | 2.2763 |
| CD1E | 6 | 0.021661 | 0.07732 | 0.93914 | 1555 | 3 | -0.96753 | 0.10346 | 0.28539 | 0.978871 | 5719 | 3 | -0.96753 |

| | | | | | | | | | | | | | |
|---|---|---|---|---|---|---|---|---|---|---|---|---|---|
| PXDC1 | 6 | 0.049358 | 0.14266 | 0.959553 | 2946 | 2 | 2.2544 | 0.1035 | 0.28547 | 0.978871 | 5720 | 4 | 2.2544 |
| PDHB | 5 | 0.78818 | 0.81223 | 0.999994 | 17322 | 1 | 2.7951 | 0.10353 | 0.26141 | 0.974875 | 5721 | 4 | 2.7951 |
| hsa-mir-876 | 4 | 0.80403 | 0.80481 | 0.999994 | 17543 | 1 | 1.6617 | 0.10358 | 0.23857 | 0.97466 | 5722 | 3 | 1.6617 |
| PDF | 5 | 0.10097 | 0.22795 | 0.983354 | 5006 | 1 | 1.5549 | 0.10365 | 0.26165 | 0.974875 | 5723 | 3 | 1.5549 |
| SIT1 | 6 | 0.054391 | 0.15359 | 0.967126 | 3172 | 3 | 0.95501 | 0.10367 | 0.28572 | 0.978871 | 5724 | 3 | 0.95501 |
| ROGDI | 5 | 0.28818 | 0.47146 | 0.999994 | 9875 | 1 | 1.0828 | 0.10383 | 0.26199 | 0.974875 | 5725 | 3 | 1.0828 |
| EIF5B | 5 | 0.077219 | 0.19009 | 0.970632 | 4118 | 2 | 1.8684 | 0.10383 | 0.26199 | 0.974875 | 5726 | 3 | 1.8684 |
| CLDN22 | 6 | 0.1799 | 0.36302 | 0.999994 | 7217 | 1 | 2.3423 | 0.10399 | 0.28625 | 0.978871 | 5727 | 3 | 2.3423 |
| GLYATL2 | 6 | 0.88879 | 0.89548 | 0.999994 | 19017 | 1 | 2.6507 | 0.10399 | 0.28625 | 0.978871 | 5728 | 3 | 2.6507 |
| HPCAL1 | 5 | 0.21447 | 0.3956 | 0.999994 | 8133 | 2 | 0.89727 | 0.104 | 0.26232 | 0.974875 | 5729 | 3 | 0.89727 |
| hsa-mir-125b-1 | 3 | 0.8014 | 0.80122 | 0.999994 | 17497 | 0 | 4.9696 | 0.104 | 0.22842 | 0.97466 | 5730 | 2 | 4.9696 |
| HABP2 | 6 | 0.64645 | 0.7749 | 0.999994 | 15606 | 2 | 2.2086 | 0.10404 | 0.28632 | 0.978871 | 5731 | 4 | 2.2086 |
| PHKA2 | 6 | 0.21911 | 0.41104 | 0.999994 | 8266 | 3 | -0.19135 | 0.10404 | 0.28632 | 0.978871 | 5732 | 3 | -0.19135 |
| NXT2 | 6 | 0.29641 | 0.50253 | 0.999994 | 10048 | 1 | 2.2266 | 0.10404 | 0.28632 | 0.978871 | 5733 | 4 | 2.2266 |
| CCNB1 | 6 | 0.091727 | 0.23099 | 0.983933 | 4665 | 3 | -0.56658 | 0.10404 | 0.28632 | 0.978871 | 5734 | 2 | -0.56658 |
| PSMD6 | 5 | 0.028038 | 0.085406 | 0.94395 | 1904 | 3 | -3.2402 | 0.10411 | 0.26254 | 0.974875 | 5735 | 1 | -3.2402 |
| TOR1B | 5 | 0.67084 | 0.75366 | 0.999994 | 15942 | 1 | 3.6773 | 0.10412 | 0.26257 | 0.974875 | 5736 | 3 | 3.6773 |
| IER5 | 5 | 0.86014 | 0.86252 | 0.999994 | 18442 | 1 | 3.6773 | 0.10412 | 0.26257 | 0.974875 | 5737 | 3 | 3.6773 |
| ULK2 | 5 | 0.59382 | 0.72388 | 0.999994 | 14877 | 1 | 2.7977 | 0.10413 | 0.26258 | 0.974875 | 5738 | 4 | 2.7977 |
| CBLN4 | 6 | 0.14312 | 0.317 | 0.999994 | 6277 | 2 | 2.968 | 0.10413 | 0.28645 | 0.978871 | 5739 | 4 | 2.968 |
| NonTargetingControlG | 1 | 0.89582 | 0.89618 | 0.999994 | 19143 | 0 | 7.0762 | 0.10418 | 0.1041 | 0.952231 | 5740 | 1 | 7.0762 |
| hsa-mir-4797 | 3 | 0.89542 | 0.8955 | 0.999994 | 19136 | 0 | 1.7093 | 0.10421 | 0.2288 | 0.97466 | 5741 | 3 | 1.7093 |
| PDCD4 | 5 | 0.31155 | 0.49042 | 0.999994 | 10349 | 2 | 0.088738 | 0.10423 | 0.26277 | 0.974875 | 5742 | 2 | 0.088738 |
| ACR | 4 | 0.84058 | 0.84003 | 0.999994 | 18098 | 0 | 4.0125 | 0.10425 | 0.23962 | 0.97466 | 5743 | 3 | 4.0125 |
| SPDYC | 6 | 0.51908 | 0.68188 | 0.999994 | 13731 | 1 | 2.3495 | 0.10426 | 0.28669 | 0.979159 | 5744 | 4 | 2.3495 |
| CRB2 | 5 | 0.9612 | 0.96135 | 0.999994 | 20610 | 0 | 3.2838 | 0.1043 | 0.2629 | 0.974875 | 5745 | 4 | 3.2838 |
| TIPARP | 6 | 0.17705 | 0.35939 | 0.999994 | 7159 | 3 | 0.086022 | 0.10431 | 0.28675 | 0.979159 | 5746 | 3 | 0.086022 |
| CXCL11 | 6 | 0.065005 | 0.17627 | 0.968171 | 3652 | 2 | 2.0518 | 0.10444 | 0.28696 | 0.979159 | 5747 | 4 | 2.0518 |
| CSTA | 4 | 0.28308 | 0.42569 | 0.999994 | 9764 | 2 | 1.0278 | 0.10445 | 0.23996 | 0.97466 | 5748 | 2 | 1.0278 |
| hsa-mir-6769a | 4 | 0.57889 | 0.65419 | 0.999994 | 14668 | 1 | 0.51717 | 0.10445 | 0.23996 | 0.97466 | 5749 | 2 | 0.51717 |
| INADL | 6 | 0.16322 | 0.34231 | 0.999994 | 6823 | 3 | -0.66483 | 0.10449 | 0.28705 | 0.979159 | 5750 | 2 | -0.66483 |
| EPT1 | 6 | 0.1294 | 0.29954 | 0.999994 | 5927 | 1 | 0.77201 | 0.10449 | 0.28705 | 0.979159 | 5751 | 4 | 0.77201 |
| BATF3 | 6 | 0.0488 | 0.14148 | 0.959146 | 2922 | 2 | 0.0045841 | 0.10449 | 0.28705 | 0.979159 | 5752 | 2 | 0.0045841 |
| NEURL | 6 | 0.04264 | 0.12785 | 0.951674 | 2656 | 3 | -0.20527 | 0.10449 | 0.28705 | 0.979159 | 5753 | 2 | -0.20527 |
| CORO1B | 6 | 0.86727 | 0.88166 | 0.999994 | 18580 | 1 | 2.0557 | 0.10451 | 0.2871 | 0.979159 | 5754 | 4 | 2.0557 |
| MRPL55 | 5 | 0.2687 | 0.4558 | 0.999994 | 9440 | 2 | 3.6035 | 0.10461 | 0.26352 | 0.974875 | 5755 | 3 | 3.6035 |
| ODF3L2 | 5 | 0.82116 | 0.83336 | 0.999994 | 17793 | 1 | 1.4732 | 0.10473 | 0.26376 | 0.97488 | 5756 | 4 | 1.4732 |
| BTBD6 | 5 | 0.48409 | 0.6342 | 0.999994 | 13202 | 2 | 1.4755 | 0.10473 | 0.26376 | 0.97488 | 5757 | 3 | 1.4755 |
| PICALM | 6 | 0.27917 | 0.48243 | 0.999994 | 9681 | 2 | 0.552 | 0.10475 | 0.28747 | 0.97947 | 5758 | 3 | 0.552 |
| ITGAL | 5 | 0.54296 | 0.68552 | 0.999994 | 14112 | 2 | 2.9342 | 0.10475 | 0.2638 | 0.97488 | 5759 | 3 | 2.9342 |
| NOTCH2 | 6 | 0.88939 | 0.89591 | 0.999994 | 19027 | 1 | 2.0225 | 0.10487 | 0.28765 | 0.97947 | 5760 | 4 | 2.0225 |
| SMIM6 | 5 | 0.71029 | 0.77121 | 0.999994 | 16401 | 1 | 2.2971 | 0.1049 | 0.26412 | 0.975862 | 5761 | 4 | 2.2971 |
| MAGEA9B | 1 | 0.89509 | 0.89542 | 0.999994 | 19128 | 0 | 7.0586 | 0.10491 | 0.10482 | 0.952231 | 5762 | 1 | 7.0586 |
| hsa-mir-147a | 4 | 0.32438 | 0.46793 | 0.999994 | 10604 | 2 | 1.8235 | 0.10491 | 0.24074 | 0.97466 | 5763 | 2 | 1.8235 |
| MAGEE1 | 6 | 0.32095 | 0.53096 | 0.999994 | 10531 | 1 | 2.5535 | 0.10498 | 0.28782 | 0.979545 | 5764 | 4 | 2.5535 |
| CENPH | 3 | 0.22365 | 0.34221 | 0.999994 | 8374 | 1 | 3.404 | 0.10502 | 0.23034 | 0.97466 | 5765 | 2 | 3.404 |
| LRRC19 | 6 | 0.38959 | 0.59571 | 0.999994 | 11791 | 2 | 2.1104 | 0.10515 | 0.2881 | 0.979689 | 5766 | 4 | 2.1104 |
| NonTargetingControlG | 1 | 0.89484 | 0.89515 | 0.999994 | 19121 | 0 | 7.058 | 0.10516 | 0.10506 | 0.952231 | 5767 | 1 | 7.058 |
| C8orf87 | 6 | 0.20721 | 0.39665 | 0.999994 | 7953 | 2 | 2.7184 | 0.10516 | 0.28811 | 0.979689 | 5768 | 3 | 2.7184 |
| ENPP5 | 6 | 0.42819 | 0.62069 | 0.999994 | 12385 | 2 | 2.3278 | 0.10516 | 0.28811 | 0.979689 | 5769 | 3 | 2.3278 |
| KRTAP5-2 | 6 | 0.83497 | 0.86402 | 0.999994 | 18005 | 1 | 2.6304 | 0.10516 | 0.28811 | 0.979689 | 5770 | 3 | 2.6304 |
| hsa-mir-4644 | 3 | 0.71477 | 0.71493 | 0.999994 | 16446 | 1 | 5.0877 | 0.10522 | 0.23071 | 0.97466 | 5771 | 2 | 5.0877 |

| | | | | | | | | | | | | |
|---|---|---|---|---|---|---|---|---|---|---|---|---|
| SMIM13 | 5 | 0.8949 | 0.89503 | 0.999994 | 19124 | 0 | 0.2521 | 0.10532 | 0.2649 | 0.976736 | 5772 | 2 | 0.2521 |
| ANGEL2 | 4 | 0.90501 | 0.90471 | 0.999994 | 19328 | 0 | 3.3222 | 0.10538 | 0.2415 | 0.97466 | 5773 | 2 | 3.3222 |
| MCL1 | 4 | 0.28527 | 0.42793 | 0.999994 | 9818 | 2 | 2.4596 | 0.10538 | 0.2415 | 0.97466 | 5774 | 2 | 2.4596 |
| hsa-mir-296 | 4 | 0.14793 | 0.28704 | 0.999632 | 6406 | 2 | 2.1527 | 0.10538 | 0.2415 | 0.97466 | 5775 | 2 | 2.1527 |
| ECM2 | 5 | 0.48803 | 0.63752 | 0.999994 | 13256 | 2 | -0.33488 | 0.10542 | 0.26511 | 0.976736 | 5776 | 2 | -0.33488 |
| INTU | 5 | 0.14941 | 0.30189 | 0.999994 | 6442 | 3 | -0.9744 | 0.10542 | 0.26511 | 0.976736 | 5777 | 1 | -0.9744 |
| SLC35A4 | 5 | 0.010726 | 0.038146 | 0.88817 | 913 | 2 | 0.28907 | 0.10542 | 0.26511 | 0.976736 | 5778 | 2 | 0.28907 |
| hsa-mir-3164 | 4 | 0.86228 | 0.86191 | 0.999994 | 18491 | 0 | 0.19788 | 0.10543 | 0.24159 | 0.97466 | 5779 | 1 | 0.19788 |
| ALPP | 6 | 0.93005 | 0.93018 | 0.999994 | 19855 | 0 | 2.4439 | 0.10544 | 0.28856 | 0.979944 | 5780 | 4 | 2.4439 |
| hsa-mir-6895 | 3 | 0.89455 | 0.89462 | 0.999994 | 19115 | 0 | 2.5437 | 0.10545 | 0.23113 | 0.97466 | 5781 | 3 | 2.5437 |
| hsa-mir-6845 | 2 | 0.15453 | 0.22979 | 0.983354 | 6580 | 1 | 0.53776 | 0.10548 | 0.17854 | 0.97162 | 5782 | 1 | 0.53776 |
| hsa-mir-6085 | 4 | 0.89651 | 0.89622 | 0.999994 | 19153 | 0 | 3.3024 | 0.10548 | 0.24166 | 0.97466 | 5783 | 3 | 3.3024 |
| NCOA1 | 6 | 0.03121 | 0.10167 | 0.946414 | 2068 | 2 | 2.699 | 0.10553 | 0.28869 | 0.980082 | 5784 | 4 | 2.699 |
| FAM171B | 6 | 0.2501 | 0.44805 | 0.999994 | 9021 | 3 | 0.60865 | 0.10557 | 0.28876 | 0.980082 | 5785 | 3 | 0.60865 |
| MDP1 | 6 | 0.40373 | 0.60481 | 0.999994 | 12010 | 2 | 2.0004 | 0.10562 | 0.28883 | 0.980082 | 5786 | 4 | 2.0004 |
| SLC18A1 | 5 | 0.29459 | 0.47666 | 0.999994 | 10009 | 2 | 3.396 | 0.10562 | 0.26551 | 0.977843 | 5787 | 3 | 3.396 |
| hsa-mir-6510 | 3 | 0.0030941 | 0.0085803 | 0.680529 | 367 | 1 | 4.9214 | 0.10562 | 0.23143 | 0.97466 | 5788 | 2 | 4.9214 |
| hsa-mir-4314 | 3 | 0.89438 | 0.89443 | 0.999994 | 19111 | 0 | 4.2459 | 0.10562 | 0.23143 | 0.97466 | 5789 | 3 | 4.2459 |
| WWP2 | 6 | 0.10207 | 0.25137 | 0.993413 | 5043 | 3 | -0.90354 | 0.10562 | 0.28884 | 0.980082 | 5790 | 2 | -0.90354 |
| PRKCD | 6 | 0.36066 | 0.57605 | 0.999994 | 11345 | 1 | 2.516 | 0.10565 | 0.28889 | 0.980101 | 5791 | 4 | 2.516 |
| GEMIN2 | 4 | 0.0068588 | 0.023732 | 0.832699 | 649 | 2 | -1.6498 | 0.10566 | 0.24194 | 0.97466 | 5792 | 2 | -1.6498 |
| RBAK-RBAKDN | 1 | 0.89432 | 0.89463 | 0.999994 | 19109 | 0 | 7.0474 | 0.10568 | 0.1056 | 0.952231 | 5793 | 1 | 7.0474 |
| hsa-mir-1283-2 | 4 | 0.30488 | 0.44796 | 0.999994 | 10229 | 1 | 2.2277 | 0.10578 | 0.24214 | 0.97466 | 5794 | 3 | 2.2277 |
| NonTargetingControlG | 2 | 0.89419 | 0.89407 | 0.999994 | 19105 | 0 | 4.7722 | 0.10581 | 0.17904 | 0.97162 | 5795 | 2 | 4.7722 |
| KRT72 | 5 | 0.95255 | 0.9527 | 0.999994 | 20396 | 0 | 0.75052 | 0.10581 | 0.26589 | 0.978382 | 5796 | 4 | 0.75052 |
| ACAP2 | 5 | 0.94026 | 0.94044 | 0.999994 | 20092 | 0 | 0.9758 | 0.10581 | 0.26589 | 0.978382 | 5797 | 4 | 0.9758 |
| VPS9D1 | 5 | 0.2156 | 0.39712 | 0.999994 | 8157 | 1 | -0.096639 | 0.10588 | 0.266 | 0.978454 | 5798 | 2 | -0.096639 |
| NonTargetingControlG | 2 | 0.89411 | 0.89398 | 0.999994 | 19101 | 0 | 3.6634 | 0.10589 | 0.17917 | 0.97162 | 5799 | 2 | 3.6634 |
| TBCCD1 | 5 | 0.45091 | 0.60599 | 0.999994 | 12722 | 2 | 3.5213 | 0.10593 | 0.26612 | 0.978454 | 5800 | 3 | 3.5213 |
| DENR | 6 | 0.070301 | 0.18741 | 0.969409 | 3847 | 3 | 1.2115 | 0.10604 | 0.28952 | 0.980804 | 5801 | 3 | 1.2115 |
| TMEM92 | 6 | 0.1799 | 0.36302 | 0.999994 | 7218 | 2 | 2.2384 | 0.10604 | 0.28952 | 0.980804 | 5802 | 3 | 2.2384 |
| GGT7 | 6 | 0.17867 | 0.36147 | 0.999994 | 7190 | 3 | -0.77972 | 0.10608 | 0.2896 | 0.980804 | 5803 | 2 | -0.77972 |
| ZCRB1 | 6 | 0.11549 | 0.27721 | 0.999379 | 5495 | 3 | 0.21653 | 0.10608 | 0.2896 | 0.980804 | 5804 | 3 | 0.21653 |
| SH2D3C | 6 | 0.55021 | 0.70379 | 0.999994 | 14214 | 2 | 0.62992 | 0.10608 | 0.2896 | 0.980804 | 5805 | 3 | 0.62992 |
| MEX3A | 5 | 0.30437 | 0.48458 | 0.999994 | 10215 | 1 | 2.6564 | 0.1061 | 0.26644 | 0.978454 | 5806 | 4 | 2.6564 |
| PAX4 | 5 | 0.31108 | 0.49002 | 0.999994 | 10341 | 2 | 0.48781 | 0.10613 | 0.26649 | 0.978454 | 5807 | 2 | 0.48781 |
| hsa-mir-4520a | 4 | 0.92825 | 0.92802 | 0.999994 | 19820 | 0 | 4.091 | 0.10616 | 0.24272 | 0.97466 | 5808 | 3 | 4.091 |
| CAPN10 | 6 | 0.47704 | 0.65321 | 0.999994 | 13099 | 2 | 0.96508 | 0.10616 | 0.28973 | 0.980804 | 5809 | 4 | 0.96508 |
| CRISP2 | 6 | 0.31184 | 0.52048 | 0.999994 | 10354 | 1 | -0.43669 | 0.10616 | 0.28973 | 0.980804 | 5810 | 2 | -0.43669 |
| NonTargetingControlG | 2 | 0.8788 | 0.87888 | 0.999994 | 18806 | 0 | 4.9112 | 0.10618 | 0.17959 | 0.97162 | 5811 | 2 | 4.9112 |
| FAM20B | 6 | 0.37576 | 0.58687 | 0.999994 | 11590 | 2 | 2.1481 | 0.10628 | 0.28992 | 0.981141 | 5812 | 4 | 2.1481 |
| NonTargetingControlG | 2 | 0.89371 | 0.89359 | 0.999994 | 19099 | 0 | 4.6888 | 0.10629 | 0.17975 | 0.97162 | 5813 | 2 | 4.6888 |
| ICT1 | 5 | 0.15989 | 0.31741 | 0.999994 | 6732 | 3 | -1.1745 | 0.1063 | 0.26681 | 0.978454 | 5814 | 2 | -1.1745 |
| C4orf27 | 5 | 0.22008 | 0.40336 | 0.999994 | 8291 | 3 | -0.76029 | 0.10631 | 0.26683 | 0.978454 | 5815 | 2 | -0.76029 |
| hsa-mir-363 | 3 | 0.89363 | 0.89368 | 0.999994 | 19097 | 0 | 2.5805 | 0.10637 | 0.23277 | 0.97466 | 5816 | 3 | 2.5805 |
| PRAMEF2 | 5 | 0.35651 | 0.52722 | 0.999994 | 11275 | 2 | 3.3935 | 0.10637 | 0.26696 | 0.978454 | 5817 | 3 | 3.3935 |
| hsa-mir-3945 | 4 | 0.80019 | 0.8014 | 0.999994 | 17480 | 1 | 3.0519 | 0.10649 | 0.24323 | 0.97466 | 5818 | 2 | 3.0519 |
| TRDN | 4 | 0.92868 | 0.92847 | 0.999994 | 19830 | 0 | 3.9961 | 0.10649 | 0.24323 | 0.97466 | 5819 | 3 | 3.9961 |
| GPR63 | 4 | 0.9028 | 0.90249 | 0.999994 | 19269 | 0 | 2.5232 | 0.10651 | 0.24326 | 0.97466 | 5820 | 3 | 2.5232 |
| FAM160B1 | 4 | 0.86077 | 0.86035 | 0.999994 | 18458 | 0 | 4.1179 | 0.10657 | 0.24335 | 0.97466 | 5821 | 3 | 4.1179 |
| GNPAT | 6 | 0.10454 | 0.25619 | 0.994943 | 5116 | 4 | -0.76883 | 0.10658 | 0.29039 | 0.981963 | 5822 | 2 | -0.76883 |
| ARSE | 6 | 0.11425 | 0.27483 | 0.998561 | 5457 | 2 | 1.5364 | 0.10658 | 0.29039 | 0.981963 | 5823 | 3 | 1.5364 |

| | | | | | | | | | | | |
|---|---|---|---|---|---|---|---|---|---|---|---|
| NonTargetingControlG | 2 | 0.38774 | 0.40662 | 0.999994 | 11763 | 1 | 3.0877 | 0.10659 | 0.18019 | 0.97162 | 5824 | 1 | 3.0877 |
| NonTargetingControlG | 2 | 0.4653 | 0.47281 | 0.999994 | 12938 | 1 | 3.3162 | 0.10659 | 0.18019 | 0.97162 | 5825 | 1 | 3.3162 |
| MS4A15 | 6 | 0.26323 | 0.46369 | 0.999994 | 9318 | 2 | 1.8667 | 0.10663 | 0.29048 | 0.982102 | 5826 | 4 | 1.8667 |
| EPB41L1 | 4 | 0.87504 | 0.87469 | 0.999994 | 18736 | 0 | 2.278 | 0.10668 | 0.24352 | 0.97466 | 5827 | 3 | 2.278 |
| ZNF259 | 5 | 0.12148 | 0.25959 | 0.994943 | 5682 | 2 | 3.6179 | 0.10668 | 0.26756 | 0.978454 | 5828 | 3 | 3.6179 |
| NDUFA11 | 6 | 0.067814 | 0.18218 | 0.968526 | 3744 | 2 | 1.8395 | 0.10678 | 0.29074 | 0.982601 | 5829 | 4 | 1.8395 |
| LRRIQ4 | 4 | 0.92546 | 0.92525 | 0.999994 | 19756 | 0 | 1.1656 | 0.10682 | 0.24373 | 0.97466 | 5830 | 3 | 1.1656 |
| UPK3A | 5 | 0.65882 | 0.74866 | 0.999994 | 15761 | 1 | 1.1872 | 0.10697 | 0.26811 | 0.978454 | 5831 | 4 | 1.1872 |
| RAB33A | 6 | 0.39641 | 0.60005 | 0.999994 | 11895 | 2 | 2.5383 | 0.107 | 0.29107 | 0.982859 | 5832 | 3 | 2.5383 |
| ZNF543 | 5 | 0.0068873 | 0.027042 | 0.848412 | 655 | 1 | 0.94975 | 0.10708 | 0.26832 | 0.978454 | 5833 | 3 | 0.94975 |
| UGT8 | 6 | 0.029638 | 0.097938 | 0.945253 | 1989 | 2 | 1.2113 | 0.10709 | 0.29123 | 0.983099 | 5834 | 3 | 1.2113 |
| RHBDF2 | 4 | 0.20299 | 0.34373 | 0.999994 | 7855 | 2 | 2.1066 | 0.10709 | 0.24418 | 0.97466 | 5835 | 2 | 2.1066 |
| DEFB118 | 4 | 0.19398 | 0.33446 | 0.999994 | 7601 | 1 | 2.4138 | 0.10713 | 0.24423 | 0.97466 | 5836 | 3 | 2.4138 |
| IQCK | 6 | 0.92342 | 0.92361 | 0.999994 | 19706 | 1 | 2.0397 | 0.10723 | 0.29144 | 0.983138 | 5837 | 4 | 2.0397 |
| ZNF530 | 6 | 0.9418 | 0.94178 | 0.999994 | 20123 | 0 | 0.020559 | 0.10727 | 0.29151 | 0.983138 | 5838 | 2 | 0.020559 |
| ZFYVE1 | 6 | 0.19544 | 0.38235 | 0.999994 | 7649 | 2 | 0.91182 | 0.10727 | 0.29151 | 0.983138 | 5839 | 4 | 0.91182 |
| CHN2 | 6 | 0.93866 | 0.93862 | 0.999994 | 20043 | 0 | 2.5024 | 0.10728 | 0.29151 | 0.983138 | 5840 | 4 | 2.5024 |
| ELOVL1 | 5 | 0.38593 | 0.5517 | 0.999994 | 11732 | 1 | 1.9826 | 0.1074 | 0.26897 | 0.978454 | 5841 | 3 | 1.9826 |
| ADAM19 | 5 | 0.9484 | 0.9486 | 0.999994 | 20296 | 0 | 0.65765 | 0.1074 | 0.26897 | 0.978454 | 5842 | 3 | 0.65765 |
| CLEC4G | 5 | 0.87866 | 0.87899 | 0.999994 | 18805 | 1 | 2.4704 | 0.1074 | 0.26897 | 0.978454 | 5843 | 3 | 2.4704 |
| VGLL3 | 4 | 0.35824 | 0.5027 | 0.999994 | 11307 | 2 | 0.036369 | 0.10743 | 0.2447 | 0.97466 | 5844 | 2 | 0.036369 |
| ROCK2 | 6 | 0.97029 | 0.97026 | 0.999994 | 20830 | 0 | 2.6152 | 0.10745 | 0.2918 | 0.983225 | 5845 | 4 | 2.6152 |
| FGF14 | 5 | 0.94088 | 0.94109 | 0.999994 | 20102 | 0 | 1.2688 | 0.10747 | 0.26909 | 0.978454 | 5846 | 4 | 1.2688 |
| MFAP2 | 5 | 0.57737 | 0.71602 | 0.999994 | 14648 | 2 | 0.68326 | 0.10749 | 0.26913 | 0.978454 | 5847 | 3 | 0.68326 |
| ZNF280D | 6 | 0.29049 | 0.49565 | 0.999994 | 9921 | 3 | 1.9559 | 0.10755 | 0.29195 | 0.983225 | 5848 | 3 | 1.9559 |
| RWDD1 | 5 | 0.35134 | 0.52301 | 0.999994 | 11184 | 1 | 2.9082 | 0.10757 | 0.26929 | 0.978454 | 5849 | 4 | 2.9082 |
| ORM2 | 4 | 0.87224 | 0.87189 | 0.999994 | 18680 | 0 | 3.6525 | 0.10757 | 0.24492 | 0.97466 | 5850 | 3 | 3.6525 |
| ZNF841 | 5 | 0.69447 | 0.76397 | 0.999994 | 16247 | 1 | 0.98841 | 0.10763 | 0.26941 | 0.978454 | 5851 | 4 | 0.98841 |
| KRT14 | 5 | 0.84638 | 0.8515 | 0.999994 | 18195 | 1 | 1.107 | 0.10768 | 0.26951 | 0.978454 | 5852 | 3 | 1.107 |
| hsa-mir-1246 | 4 | 0.095084 | 0.20627 | 0.978744 | 4787 | 2 | 1.5999 | 0.10768 | 0.2451 | 0.97466 | 5853 | 2 | 1.5999 |
| TEKT3 | 6 | 0.18021 | 0.36343 | 0.999994 | 7224 | 2 | 2.2023 | 0.10772 | 0.29221 | 0.983225 | 5854 | 4 | 2.2023 |
| XCR1 | 6 | 0.91452 | 0.91547 | 0.999994 | 19498 | 1 | 2.2288 | 0.10778 | 0.29233 | 0.983225 | 5855 | 4 | 2.2288 |
| AASDH | 5 | 0.72868 | 0.78002 | 0.999994 | 16587 | 1 | 0.4641 | 0.10779 | 0.26973 | 0.978454 | 5856 | 2 | 0.4641 |
| TAGLN3 | 5 | 0.94096 | 0.94116 | 0.999994 | 20106 | 0 | 2.4786 | 0.10784 | 0.26983 | 0.978454 | 5857 | 4 | 2.4786 |
| ABLIM3 | 5 | 0.95806 | 0.95822 | 0.999994 | 20541 | 0 | 3.2313 | 0.10787 | 0.26989 | 0.978454 | 5858 | 4 | 3.2313 |
| SLC34A3 | 5 | 0.69925 | 0.76613 | 0.999994 | 16299 | 1 | 1.1205 | 0.10787 | 0.26989 | 0.978454 | 5859 | 3 | 1.1205 |
| RHOD | 5 | 0.011613 | 0.040693 | 0.891621 | 967 | 1 | 0.9273 | 0.10787 | 0.26989 | 0.978454 | 5860 | 4 | 0.9273 |
| ATG4D | 5 | 0.053525 | 0.14888 | 0.965566 | 3141 | 2 | 0.65364 | 0.10787 | 0.26989 | 0.978454 | 5861 | 3 | 0.65364 |
| VEGFB | 5 | 0.14338 | 0.29292 | 0.999994 | 6282 | 1 | 0.97041 | 0.10787 | 0.26989 | 0.978454 | 5862 | 3 | 0.97041 |
| FAM168A | 5 | 0.88347 | 0.88362 | 0.999994 | 18896 | 1 | 1.7551 | 0.10787 | 0.26989 | 0.978454 | 5863 | 4 | 1.7551 |
| FFAR3 | 6 | 0.41929 | 0.61491 | 0.999994 | 12240 | 1 | 2.1287 | 0.10788 | 0.29249 | 0.983225 | 5864 | 4 | 2.1287 |
| RASAL2 | 6 | 0.47893 | 0.65445 | 0.999994 | 13127 | 2 | 0.075455 | 0.1079 | 0.29251 | 0.983225 | 5865 | 2 | 0.075455 |
| S100A7 | 6 | 0.10457 | 0.25625 | 0.994943 | 5119 | 4 | -1.1681 | 0.1079 | 0.29251 | 0.983225 | 5866 | 2 | -1.1681 |
| OLAH | 6 | 0.64666 | 0.77507 | 0.999994 | 15612 | 1 | 2.294 | 0.1079 | 0.29251 | 0.983225 | 5867 | 4 | 2.294 |
| hsa-mir-4784 | 4 | 0.43125 | 0.57783 | 0.999994 | 12435 | 1 | 2.3666 | 0.10793 | 0.24549 | 0.97466 | 5868 | 3 | 2.3666 |
| C9orf139 | 5 | 0.094677 | 0.218 | 0.980699 | 4776 | 1 | 3.41 | 0.10794 | 0.27003 | 0.978454 | 5869 | 4 | 3.41 |
| PCCA | 5 | 0.96301 | 0.96316 | 0.999994 | 20662 | 0 | 3.5475 | 0.10801 | 0.27019 | 0.978454 | 5870 | 3 | 3.5475 |
| PDLIM2 | 6 | 0.35849 | 0.57361 | 0.999994 | 11312 | 2 | 2.8504 | 0.10815 | 0.29293 | 0.983225 | 5871 | 4 | 2.8504 |
| KRTAP8-1 | 6 | 0.28695 | 0.49154 | 0.999994 | 9852 | 3 | 1.9166 | 0.10815 | 0.29293 | 0.983225 | 5872 | 3 | 1.9166 |
| TINAGL1 | 6 | 0.15137 | 0.32739 | 0.999994 | 6490 | 2 | 2.2836 | 0.10815 | 0.29293 | 0.983225 | 5873 | 3 | 2.2836 |
| NonTargetingControlG | 2 | 0.89184 | 0.8917 | 0.999994 | 19071 | 0 | 4.9759 | 0.10816 | 0.18266 | 0.97162 | 5874 | 2 | 4.9759 |
| TPD52L3 | 6 | 0.22698 | 0.42045 | 0.999994 | 8452 | 2 | 0.54532 | 0.10817 | 0.29296 | 0.983225 | 5875 | 3 | 0.54532 |

| | | | | | | | | | | | | |
|---|---|---|---|---|---|---|---|---|---|---|---|---|
| ARMC7 | 5 | 0.22649 | 0.41236 | 0.999994 | 8439 | 1 | 2.2973 | 0.10822 | 0.27059 | 0.978454 | 5876 | 4 | 2.2973 |
| PSEN2 | 5 | 0.95752 | 0.95767 | 0.999994 | 20526 | 0 | 2.1362 | 0.10822 | 0.27059 | 0.978454 | 5877 | 4 | 2.1362 |
| PCDHGA5 | 2 | 0.89177 | 0.89163 | 0.999994 | 19069 | 0 | 3.6975 | 0.10823 | 0.18278 | 0.97162 | 5878 | 2 | 3.6975 |
| MATN4 | 6 | 0.32095 | 0.53096 | 0.999994 | 10530 | 2 | 1.9708 | 0.10823 | 0.29306 | 0.983225 | 5879 | 4 | 1.9708 |
| TXNRD3NB | 6 | 0.30964 | 0.51794 | 0.999994 | 10319 | 3 | 0.51067 | 0.10831 | 0.29318 | 0.983225 | 5880 | 3 | 0.51067 |
| SAG | 6 | 0.072017 | 0.19098 | 0.972293 | 3919 | 2 | -0.37578 | 0.10831 | 0.29318 | 0.983225 | 5881 | 2 | -0.37578 |
| HAND1 | 6 | 0.17451 | 0.35625 | 0.999994 | 7099 | 3 | 1.3119 | 0.10831 | 0.29318 | 0.983225 | 5882 | 3 | 1.3119 |
| hsa-mir-513b | 4 | 0.34605 | 0.49027 | 0.999994 | 11066 | 1 | 2.31 | 0.10835 | 0.24616 | 0.97466 | 5883 | 3 | 2.31 |
| hsa-let-7b | 3 | 0.89159 | 0.89163 | 0.999994 | 19063 | 0 | 0.73775 | 0.10841 | 0.23658 | 0.97466 | 5884 | 3 | 0.73775 |
| ITPR2 | 6 | 0.83055 | 0.86182 | 0.999994 | 17940 | 1 | 2.2948 | 0.10846 | 0.29342 | 0.983276 | 5885 | 4 | 2.2948 |
| DHODH | 5 | 0.93962 | 0.93981 | 0.999994 | 20068 | 0 | 3.4732 | 0.10846 | 0.27105 | 0.978454 | 5886 | 4 | 3.4732 |
| NonTargetingControlG | 2 | 0.8915 | 0.89136 | 0.999994 | 19062 | 0 | 3.4416 | 0.1085 | 0.18315 | 0.97162 | 5887 | 2 | 3.4416 |
| GPSM1 | 5 | 0.95099 | 0.95116 | 0.999994 | 20359 | 0 | 1.8343 | 0.10855 | 0.27121 | 0.978454 | 5888 | 4 | 1.8343 |
| PGM1 | 6 | 0.18927 | 0.37461 | 0.999994 | 7461 | 2 | -0.059864 | 0.10856 | 0.29358 | 0.983304 | 5889 | 2 | -0.059864 |
| PRRC2A | 5 | 0.098198 | 0.22359 | 0.983354 | 4900 | 2 | 1.8359 | 0.10863 | 0.27135 | 0.978454 | 5890 | 3 | 1.8359 |
| OBSCN | 5 | 0.21179 | 0.3919 | 0.999994 | 8061 | 2 | 2.465 | 0.10863 | 0.27135 | 0.978454 | 5891 | 3 | 2.465 |
| BRD4 | 6 | 0.33796 | 0.55034 | 0.999994 | 10897 | 2 | 1.9 | 0.10866 | 0.29375 | 0.983309 | 5892 | 4 | 1.9 |
| KRBOX4 | 3 | 0.89133 | 0.89138 | 0.999994 | 19059 | 0 | 2.7864 | 0.10867 | 0.23707 | 0.97466 | 5893 | 3 | 2.7864 |
| hsa-mir-2909 | 4 | 0.42912 | 0.57564 | 0.999994 | 12400 | 2 | 0.097668 | 0.10867 | 0.24668 | 0.97466 | 5894 | 2 | 0.097668 |
| ALDH5A1 | 6 | 0.32788 | 0.53881 | 0.999994 | 10683 | 3 | 0.87598 | 0.10875 | 0.29388 | 0.983309 | 5895 | 3 | 0.87598 |
| ASUN | 6 | 0.34319 | 0.55631 | 0.999994 | 11010 | 2 | 0.71901 | 0.10875 | 0.29388 | 0.983309 | 5896 | 3 | 0.71901 |
| ENO1 | 6 | 0.32522 | 0.5358 | 0.999994 | 10626 | 1 | 1.5115 | 0.10875 | 0.29388 | 0.983309 | 5897 | 4 | 1.5115 |
| MCOLN2 | 5 | 0.28436 | 0.46838 | 0.999994 | 9794 | 1 | 2.8106 | 0.10877 | 0.27164 | 0.978454 | 5898 | 4 | 2.8106 |
| DRAXIN | 5 | 0.036257 | 0.10645 | 0.950995 | 2331 | 1 | 2.2039 | 0.10885 | 0.27178 | 0.978454 | 5899 | 4 | 2.2039 |
| RILPL2 | 3 | 0.87852 | 0.87862 | 0.999994 | 18798 | 0 | 4.8253 | 0.10888 | 0.23746 | 0.97466 | 5900 | 2 | 4.8253 |
| FAM114A2 | 5 | 0.1032 | 0.23137 | 0.984304 | 5073 | 2 | 1.4571 | 0.10889 | 0.27185 | 0.978454 | 5901 | 3 | 1.4571 |
| CEP70 | 5 | 0.26425 | 0.4522 | 0.999994 | 9337 | 2 | -0.11833 | 0.10889 | 0.27185 | 0.978454 | 5902 | 2 | -0.11833 |
| hsa-mir-5585 | 4 | 0.75543 | 0.76427 | 0.999994 | 16922 | 1 | 3.3575 | 0.10897 | 0.24715 | 0.97466 | 5903 | 2 | 3.3575 |
| DNAH12 | 5 | 0.17182 | 0.33478 | 0.999994 | 7033 | 2 | 3.0735 | 0.10904 | 0.27216 | 0.978454 | 5904 | 3 | 3.0735 |
| hsa-mir-1301 | 4 | 0.59663 | 0.66348 | 0.999994 | 14906 | 1 | 2.8031 | 0.10911 | 0.24737 | 0.97466 | 5905 | 3 | 2.8031 |
| HBG2 | 2 | 0.89082 | 0.89066 | 0.999994 | 19051 | 0 | 3.0434 | 0.10918 | 0.18421 | 0.973158 | 5906 | 2 | 3.0434 |
| FEZF2 | 6 | 0.84371 | 0.86858 | 0.999994 | 18151 | 1 | 2.6724 | 0.10922 | 0.29462 | 0.983644 | 5907 | 4 | 2.6724 |
| LRRC36 | 5 | 0.10051 | 0.22722 | 0.983354 | 4991 | 3 | -1.4601 | 0.10927 | 0.27257 | 0.978454 | 5908 | 2 | -1.4601 |
| TMEM119 | 5 | 0.14425 | 0.29419 | 0.999994 | 6308 | 2 | 2.4812 | 0.10927 | 0.27257 | 0.978454 | 5909 | 3 | 2.4812 |
| DEFB103A | 5 | 0.54353 | 0.68603 | 0.999994 | 14121 | 1 | 3.2839 | 0.10932 | 0.27267 | 0.978454 | 5910 | 3 | 3.2839 |
| PRAM1 | 5 | 0.79088 | 0.81386 | 0.999994 | 17362 | 1 | 1.0725 | 0.10932 | 0.27267 | 0.978454 | 5911 | 4 | 1.0725 |
| ANHX | 6 | 0.0048271 | 0.020525 | 0.81027 | 508 | 2 | 3.0545 | 0.10944 | 0.29497 | 0.983787 | 5912 | 4 | 3.0545 |
| SVIP | 6 | 0.39412 | 0.59858 | 0.999994 | 11861 | 1 | 2.9067 | 0.10944 | 0.29497 | 0.983787 | 5913 | 4 | 2.9067 |
| SCGB1D4 | 6 | 0.52332 | 0.68482 | 0.999994 | 13799 | 1 | 2.4275 | 0.1095 | 0.29507 | 0.983787 | 5914 | 4 | 2.4275 |
| PPP1R3F | 6 | 0.35741 | 0.57239 | 0.999994 | 11292 | 1 | 2.4562 | 0.1095 | 0.29507 | 0.983787 | 5915 | 4 | 2.4562 |
| ANKRD30B | 6 | 0.80672 | 0.85081 | 0.999994 | 17579 | 1 | 2.0057 | 0.1095 | 0.29507 | 0.983787 | 5916 | 4 | 2.0057 |
| DCK | 6 | 0.50465 | 0.67195 | 0.999994 | 13507 | 1 | 2.9896 | 0.1095 | 0.29507 | 0.983787 | 5917 | 4 | 2.9896 |
| hsa-mir-3925 | 3 | 0.4832 | 0.5352 | 0.999994 | 13190 | 1 | 4.8199 | 0.10955 | 0.23877 | 0.97466 | 5918 | 2 | 4.8199 |
| RFC3 | 3 | 0.025116 | 0.063796 | 0.937389 | 1760 | 1 | 4.8199 | 0.10955 | 0.23877 | 0.97466 | 5919 | 2 | 4.8199 |
| SPACA3 | 4 | 0.8845 | 0.88415 | 0.999994 | 18924 | 0 | 3.0172 | 0.10962 | 0.24818 | 0.97466 | 5920 | 2 | 3.0172 |
| PPM1J | 4 | 0.39337 | 0.53883 | 0.999994 | 11853 | 1 | 3.2749 | 0.10962 | 0.24818 | 0.97466 | 5921 | 2 | 3.2749 |
| TEX12 | 6 | 0.066001 | 0.17838 | 0.968171 | 3689 | 2 | 2.7298 | 0.10971 | 0.29541 | 0.983965 | 5922 | 4 | 2.7298 |
| NPEPL1 | 5 | 0.92778 | 0.928 | 0.999994 | 19808 | 0 | 3.5563 | 0.10979 | 0.27357 | 0.978454 | 5923 | 3 | 3.5563 |
| SLC45A2 | 5 | 0.33417 | 0.50895 | 0.999994 | 10816 | 2 | 3.5563 | 0.10979 | 0.27357 | 0.978454 | 5924 | 3 | 3.5563 |
| NGF | 4 | 0.8942 | 0.89389 | 0.999994 | 19106 | 0 | 2.3511 | 0.10982 | 0.2485 | 0.97466 | 5925 | 3 | 2.3511 |
| hsa-mir-5196 | 4 | 0.21941 | 0.36067 | 0.999994 | 8273 | 2 | 2.2222 | 0.10993 | 0.24867 | 0.97466 | 5926 | 2 | 2.2222 |
| hsa-mir-6512 | 4 | 0.75643 | 0.76499 | 0.999994 | 16936 | 1 | 3.0384 | 0.10995 | 0.24871 | 0.97466 | 5927 | 2 | 3.0384 |

| | | | | | | | | | | | | |
|---|---|---|---|---|---|---|---|---|---|---|---|---|
| MAGED1 | 3 | 0.061652 | 0.13172 | 0.95318 | 3498 | 1 | 4.8293 | 0.11008 | 0.23975 | 0.97466 | 5928 | 2 | 4.8293 |
| GPR153 | 5 | 0.294 | 0.47616 | 0.999994 | 9997 | 1 | 1.4525 | 0.1101 | 0.27415 | 0.978454 | 5929 | 4 | 1.4525 |
| RAPGEF4 | 6 | 0.63149 | 0.76346 | 0.999994 | 15390 | 1 | 1.7997 | 0.11013 | 0.29608 | 0.984434 | 5930 | 4 | 1.7997 |
| TBXA2R | 5 | 0.11542 | 0.25039 | 0.992497 | 5489 | 3 | -1.3666 | 0.11015 | 0.27423 | 0.978454 | 5931 | 2 | -1.3666 |
| FERMT3 | 5 | 0.80205 | 0.82081 | 0.999994 | 17511 | 1 | 0.95227 | 0.11015 | 0.27423 | 0.978454 | 5932 | 3 | 0.95227 |
| NonTargetingControlG | 2 | 0.86027 | 0.86026 | 0.999994 | 18445 | 0 | 4.7624 | 0.11016 | 0.18571 | 0.97466 | 5933 | 2 | 4.7624 |
| ATF1 | 6 | 0.68037 | 0.8013 | 0.999994 | 16097 | 1 | 0.15388 | 0.11018 | 0.29616 | 0.984541 | 5934 | 2 | 0.15388 |
| SENP6 | 5 | 0.017995 | 0.058632 | 0.931872 | 1360 | 1 | 0.072624 | 0.11022 | 0.27437 | 0.978454 | 5935 | 2 | 0.072624 |
| C1QL1 | 5 | 0.8344 | 0.84267 | 0.999994 | 17993 | 1 | 1.3652 | 0.11024 | 0.27441 | 0.978454 | 5936 | 4 | 1.3652 |
| ZFAND3 | 5 | 0.9602 | 0.96034 | 0.999994 | 20588 | 0 | 1.055 | 0.11025 | 0.27443 | 0.978454 | 5937 | 4 | 1.055 |
| NonTargetingControlG | 2 | 0.88974 | 0.88962 | 0.999994 | 19030 | 0 | 4.0624 | 0.11026 | 0.18585 | 0.97466 | 5938 | 2 | 4.0624 |
| UCK2 | 5 | 0.017763 | 0.057988 | 0.929685 | 1350 | 1 | 2.3735 | 0.1103 | 0.27452 | 0.978454 | 5939 | 4 | 2.3735 |
| RBM18 | 2 | 0.86903 | 0.86903 | 0.999994 | 18618 | 0 | 4.9257 | 0.11031 | 0.18592 | 0.97466 | 5940 | 2 | 4.9257 |
| FAIM2 | 5 | 0.96312 | 0.96328 | 0.999994 | 20672 | 0 | 1.4651 | 0.11033 | 0.27457 | 0.978454 | 5941 | 4 | 1.4651 |
| TMEM47 | 5 | 0.39149 | 0.55639 | 0.999994 | 11822 | 1 | 1.1902 | 0.11036 | 0.27464 | 0.978454 | 5942 | 4 | 1.1902 |
| FSD1L | 6 | 0.096145 | 0.23976 | 0.987253 | 4821 | 3 | -1.3265 | 0.11038 | 0.29649 | 0.984782 | 5943 | 2 | -1.3265 |
| OTUB2 | 5 | 0.23067 | 0.41812 | 0.999994 | 8537 | 3 | -0.6912 | 0.11039 | 0.2747 | 0.978454 | 5944 | 2 | -0.6912 |
| KIAA0586 | 3 | 0.88957 | 0.88961 | 0.999994 | 19028 | 0 | 0.71362 | 0.11043 | 0.24039 | 0.97466 | 5945 | 3 | 0.71362 |
| C9orf117 | 6 | 0.32419 | 0.53467 | 0.999994 | 10602 | 2 | 3.2081 | 0.11052 | 0.29672 | 0.984876 | 5946 | 4 | 3.2081 |
| GTF3C4 | 5 | 0.40401 | 0.56689 | 0.999994 | 12019 | 2 | 3.2279 | 0.11053 | 0.27496 | 0.978454 | 5947 | 3 | 3.2279 |
| WDR4 | 5 | 0.61565 | 0.73195 | 0.999994 | 15154 | 1 | 3.1637 | 0.11054 | 0.27497 | 0.978454 | 5948 | 3 | 3.1637 |
| OR10H2 | 6 | 0.21481 | 0.40594 | 0.999994 | 8139 | 3 | -0.97824 | 0.11056 | 0.29679 | 0.984876 | 5949 | 2 | -0.97824 |
| MYLK3 | 6 | 0.3307 | 0.54205 | 0.999994 | 10737 | 3 | -0.43814 | 0.11059 | 0.29684 | 0.984876 | 5950 | 2 | -0.43814 |
| ZNF177 | 5 | 0.5726 | 0.71182 | 0.999994 | 14585 | 2 | -0.35942 | 0.11067 | 0.27524 | 0.978454 | 5951 | 1 | -0.35942 |
| KIAA1324L | 5 | 0.72383 | 0.77771 | 0.999994 | 16532 | 1 | 3.3058 | 0.11077 | 0.27542 | 0.978454 | 5952 | 3 | 3.3058 |
| ZNF771 | 5 | 0.28304 | 0.46732 | 0.999994 | 9763 | 2 | -0.44493 | 0.11091 | 0.27567 | 0.978454 | 5953 | 1 | -0.44493 |
| NonTargetingControlG | 2 | 0.88908 | 0.88897 | 0.999994 | 19022 | 0 | 4.8207 | 0.11092 | 0.18686 | 0.97466 | 5954 | 2 | 4.8207 |
| UBE2L6 | 6 | 0.2506 | 0.44865 | 0.999994 | 9028 | 2 | 0.48745 | 0.111 | 0.29748 | 0.984876 | 5955 | 3 | 0.48745 |
| BET1L | 6 | 0.041948 | 0.12627 | 0.951052 | 2612 | 3 | -0.25167 | 0.111 | 0.29749 | 0.984876 | 5956 | 2 | -0.25167 |
| ANGPTL5 | 5 | 0.62408 | 0.7351 | 0.999994 | 15291 | 1 | 1.2303 | 0.11105 | 0.27595 | 0.978454 | 5957 | 4 | 1.2303 |
| LGALSL | 5 | 0.21911 | 0.40204 | 0.999994 | 8265 | 1 | 1.3773 | 0.11105 | 0.27595 | 0.978454 | 5958 | 3 | 1.3773 |
| OR8G5 | 5 | 0.17439 | 0.33852 | 0.999994 | 7095 | 1 | 1.4714 | 0.11105 | 0.27595 | 0.978454 | 5959 | 3 | 1.4714 |
| ANGPT1 | 5 | 0.18698 | 0.35665 | 0.999994 | 7386 | 3 | -0.96559 | 0.11105 | 0.27595 | 0.978454 | 5960 | 2 | -0.96559 |
| hsa-mir-4745 | 2 | 0.88894 | 0.88883 | 0.999994 | 19020 | 0 | 3.5943 | 0.11106 | 0.18706 | 0.97466 | 5961 | 2 | 3.5943 |
| CYGB | 5 | 0.97251 | 0.97257 | 0.999994 | 20882 | 0 | 2.1446 | 0.11106 | 0.27596 | 0.978454 | 5962 | 4 | 2.1446 |
| CTDSP1 | 6 | 0.62118 | 0.75572 | 0.999994 | 15248 | 2 | 2.7837 | 0.11108 | 0.29763 | 0.984876 | 5963 | 4 | 2.7837 |
| C10orf35 | 6 | 0.16716 | 0.34716 | 0.999994 | 6911 | 2 | 0.86989 | 0.11115 | 0.29774 | 0.984876 | 5964 | 4 | 0.86989 |
| TRIM13 | 4 | 0.17173 | 0.31158 | 0.999994 | 7030 | 1 | 1.2811 | 0.11118 | 0.25066 | 0.97466 | 5965 | 2 | 1.2811 |
| NonTargetingControlG | 2 | 0.88882 | 0.8887 | 0.999994 | 19018 | 0 | 4.8104 | 0.11118 | 0.18725 | 0.97466 | 5966 | 2 | 4.8104 |
| SIPA1 | 3 | 0.70611 | 0.70676 | 0.999994 | 16373 | 1 | 2.3537 | 0.11119 | 0.2418 | 0.97466 | 5967 | 2 | 2.3537 |
| hsa-mir-186 | 3 | 0.88877 | 0.88883 | 0.999994 | 19016 | 0 | 4.0893 | 0.11123 | 0.24187 | 0.97466 | 5968 | 3 | 4.0893 |
| OCM2 | 3 | 0.88876 | 0.88882 | 0.999994 | 19015 | 0 | 1.587 | 0.11124 | 0.24189 | 0.97466 | 5969 | 3 | 1.587 |
| SLC16A8 | 6 | 0.27565 | 0.47835 | 0.999994 | 9609 | 3 | 0.15389 | 0.11125 | 0.2979 | 0.984876 | 5970 | 3 | 0.15389 |
| BRF2 | 4 | 0.29996 | 0.44296 | 0.999994 | 10129 | 1 | 2.5656 | 0.11126 | 0.2508 | 0.97466 | 5971 | 3 | 2.5656 |
| LEMD3 | 3 | 0.88873 | 0.88879 | 0.999994 | 19013 | 0 | 1.7482 | 0.11127 | 0.24196 | 0.97466 | 5972 | 3 | 1.7482 |
| CCDC179 | 6 | 0.9423 | 0.94227 | 0.999994 | 20133 | 0 | 2.7652 | 0.1113 | 0.29798 | 0.984876 | 5973 | 4 | 2.7652 |
| PAGE2B | 3 | 0.88861 | 0.88867 | 0.999994 | 19010 | 0 | 1.674 | 0.11139 | 0.24219 | 0.97466 | 5974 | 3 | 1.674 |
| MTMR12 | 6 | 0.44297 | 0.63042 | 0.999994 | 12623 | 2 | 0.73911 | 0.11141 | 0.29814 | 0.984876 | 5975 | 4 | 0.73911 |
| TRIM47 | 5 | 0.65634 | 0.74765 | 0.999994 | 15731 | 1 | 3.4701 | 0.11145 | 0.27673 | 0.978454 | 5976 | 3 | 3.4701 |
| C11orf40 | 5 | 0.55832 | 0.69912 | 0.999994 | 14338 | 1 | 3.2785 | 0.11145 | 0.27673 | 0.978454 | 5977 | 4 | 3.2785 |
| GTF2B | 3 | 0.88848 | 0.88854 | 0.999994 | 19008 | 0 | 4.4428 | 0.11152 | 0.24243 | 0.97466 | 5978 | 3 | 4.4428 |
| PIP5K1A | 5 | 0.10814 | 0.23911 | 0.987253 | 5243 | 1 | -0.4299 | 0.11153 | 0.27689 | 0.978454 | 5979 | 2 | -0.4299 |

| | | | | | | | | | | | | | |
|---|---|---|---|---|---|---|---|---|---|---|---|---|---|
| MEI1 | 6 | 0.27562 | 0.47831 | 0.999994 | 9608 | 3 | -0.48273 | 0.11154 | 0.29834 | 0.984876 | 5980 | 2 | -0.48273 |
| GSG1L | 6 | 0.066742 | 0.17992 | 0.968171 | 3713 | 3 | -0.17225 | 0.11154 | 0.29834 | 0.984876 | 5981 | 2 | -0.17225 |
| TMEM150A | 6 | 0.021722 | 0.077504 | 0.93914 | 1560 | 3 | 0.11765 | 0.11154 | 0.29834 | 0.984876 | 5982 | 3 | 0.11765 |
| JAKMIP2 | 6 | 0.18816 | 0.37325 | 0.999994 | 7422 | 2 | 1.8005 | 0.11154 | 0.29834 | 0.984876 | 5983 | 3 | 1.8005 |
| PTGES2 | 6 | 0.28721 | 0.49185 | 0.999994 | 9857 | 2 | 0.95453 | 0.11154 | 0.29834 | 0.984876 | 5984 | 3 | 0.95453 |
| IFNA16 | 4 | 0.93003 | 0.9298 | 0.999994 | 19854 | 0 | 2.3837 | 0.11155 | 0.25124 | 0.97466 | 5985 | 3 | 2.3837 |
| SLC7A13 | 5 | 0.9303 | 0.93058 | 0.999994 | 19861 | 0 | 1.7676 | 0.11157 | 0.27696 | 0.978454 | 5986 | 4 | 1.7676 |
| hsa-mir-520d | 3 | 0.77824 | 0.77819 | 0.999994 | 17188 | 0 | 4.6774 | 0.11161 | 0.24258 | 0.97466 | 5987 | 2 | 4.6774 |
| hsa-mir-8080 | 4 | 0.94217 | 0.94205 | 0.999994 | 20130 | 0 | 2.1018 | 0.11161 | 0.25135 | 0.97466 | 5988 | 3 | 2.1018 |
| hsa-mir-4320 | 4 | 0.43287 | 0.5795 | 0.999994 | 12462 | 2 | 0.23157 | 0.11163 | 0.25137 | 0.97466 | 5989 | 2 | 0.23157 |
| GDF1 | 2 | 0.88835 | 0.88823 | 0.999994 | 19004 | 0 | 3.2104 | 0.11165 | 0.18797 | 0.97466 | 5990 | 2 | 3.2104 |
| GPLD1 | 5 | 0.75913 | 0.7957 | 0.999994 | 16966 | 1 | 1.6976 | 0.11165 | 0.27711 | 0.978454 | 5991 | 4 | 1.6976 |
| CDIPT | 4 | 0.041247 | 0.10426 | 0.947414 | 2575 | 2 | 2.4672 | 0.11166 | 0.25141 | 0.97466 | 5992 | 2 | 2.4672 |
| C11orf73 | 6 | 0.38681 | 0.59389 | 0.999994 | 11748 | 1 | 1.7393 | 0.11169 | 0.29858 | 0.984876 | 5993 | 4 | 1.7393 |
| IQCA1 | 5 | 0.85026 | 0.85448 | 0.999994 | 18267 | 1 | 3.261 | 0.11171 | 0.27724 | 0.978454 | 5994 | 4 | 3.261 |
| BRSK1 | 6 | 0.38325 | 0.59162 | 0.999994 | 11693 | 1 | 1.9098 | 0.11173 | 0.29866 | 0.984876 | 5995 | 4 | 1.9098 |
| NonTargetingControlG | 1 | 0.88826 | 0.88859 | 0.999994 | 19001 | 0 | 6.9304 | 0.11174 | 0.1117 | 0.953839 | 5996 | 1 | 6.9304 |
| NonTargetingControlG | 2 | 0.26432 | 0.30962 | 0.999994 | 9339 | 1 | 2.4504 | 0.11176 | 0.18815 | 0.97466 | 5997 | 1 | 2.4504 |
| LRRC25 | 5 | 0.29476 | 0.47679 | 0.999994 | 10012 | 2 | 3.2264 | 0.11178 | 0.27739 | 0.978454 | 5998 | 3 | 3.2264 |
| ISM1 | 5 | 0.052464 | 0.14633 | 0.964852 | 3092 | 2 | 3.5115 | 0.11178 | 0.27739 | 0.978454 | 5999 | 3 | 3.5115 |
| MLL5 | 5 | 0.61062 | 0.73002 | 0.999994 | 15080 | 1 | 0.33961 | 0.11187 | 0.27758 | 0.978454 | 6000 | 2 | 0.33961 |
| DUXA | 6 | 0.32958 | 0.54074 | 0.999994 | 10711 | 3 | 1.9685 | 0.1119 | 0.29892 | 0.984876 | 6001 | 3 | 1.9685 |
| ATP6V0A2 | 6 | 0.4873 | 0.66012 | 0.999994 | 13241 | 1 | 3.2341 | 0.1119 | 0.29892 | 0.984876 | 6002 | 4 | 3.2341 |
| BACH1 | 6 | 0.67158 | 0.79437 | 0.999994 | 15950 | 2 | 2.5815 | 0.1119 | 0.29892 | 0.984876 | 6003 | 3 | 2.5815 |
| hsa-mir-382 | 3 | 0.45904 | 0.51983 | 0.999994 | 12834 | 1 | 1.4674 | 0.1119 | 0.24312 | 0.97466 | 6004 | 2 | 1.4674 |
| SPATA33 | 3 | 0.71773 | 0.71779 | 0.999994 | 16473 | 1 | 3.404 | 0.1119 | 0.24312 | 0.97466 | 6005 | 2 | 3.404 |
| DPF2 | 4 | 0.035386 | 0.092522 | 0.943968 | 2286 | 3 | -2.0635 | 0.1119 | 0.25183 | 0.97466 | 6006 | 1 | -2.0635 |
| hsa-mir-4730 | 4 | 0.93824 | 0.9381 | 0.999994 | 20030 | 0 | 2.6502 | 0.11194 | 0.25189 | 0.97466 | 6007 | 3 | 2.6502 |
| LPAR4 | 4 | 0.55389 | 0.64163 | 0.999994 | 14268 | 1 | 2.2675 | 0.11195 | 0.2519 | 0.97466 | 6008 | 3 | 2.2675 |
| CCM2L | 6 | 0.30572 | 0.51339 | 0.999994 | 10246 | 3 | -0.40367 | 0.11197 | 0.29905 | 0.984876 | 6009 | 1 | -0.40367 |
| RGPD3 | 4 | 0.45399 | 0.59634 | 0.999994 | 12760 | 1 | 2.4293 | 0.11203 | 0.25201 | 0.97466 | 6010 | 3 | 2.4293 |
| PECAM1 | 6 | 0.18528 | 0.3697 | 0.999994 | 7349 | 3 | -0.16063 | 0.11205 | 0.29916 | 0.984876 | 6011 | 2 | -0.16063 |
| ZHX1 | 5 | 0.095308 | 0.21902 | 0.980699 | 4795 | 2 | 1.464 | 0.11207 | 0.27795 | 0.978454 | 6012 | 3 | 1.464 |
| ZNRF1 | 5 | 0.72414 | 0.77787 | 0.999994 | 16536 | 1 | 1.4575 | 0.1121 | 0.27802 | 0.978454 | 6013 | 4 | 1.4575 |
| TEX37 | 5 | 0.057263 | 0.15756 | 0.967126 | 3302 | 1 | -0.27877 | 0.11215 | 0.2781 | 0.978454 | 6014 | 2 | -0.27877 |
| PRMT8 | 5 | 0.074994 | 0.1865 | 0.969409 | 4035 | 3 | -0.8732 | 0.11215 | 0.2781 | 0.978454 | 6015 | 2 | -0.8732 |
| TMA16 | 5 | 0.21781 | 0.4002 | 0.999994 | 8221 | 1 | 1.817 | 0.11215 | 0.2781 | 0.978454 | 6016 | 4 | 1.817 |
| hsa-mir-8088 | 3 | 0.88785 | 0.88792 | 0.999994 | 18995 | 0 | 1.3951 | 0.11215 | 0.24358 | 0.97466 | 6017 | 3 | 1.3951 |
| NUTM2F | 6 | 0.61096 | 0.74804 | 0.999994 | 15086 | 1 | 1.9318 | 0.11216 | 0.29932 | 0.984876 | 6018 | 4 | 1.9318 |
| WFDC11 | 5 | 0.23279 | 0.42108 | 0.999994 | 8588 | 3 | -0.61783 | 0.11224 | 0.27828 | 0.978454 | 6019 | 2 | -0.61783 |
| MORN3 | 6 | 0.61669 | 0.75236 | 0.999994 | 15167 | 2 | 1.9827 | 0.11227 | 0.29949 | 0.984876 | 6020 | 4 | 1.9827 |
| ANKRD11 | 4 | 0.11711 | 0.24522 | 0.98979 | 5539 | 2 | -1.6539 | 0.11228 | 0.25242 | 0.97466 | 6021 | 1 | -1.6539 |
| NEK6 | 5 | 0.6243 | 0.73517 | 0.999994 | 15294 | 1 | 1.747 | 0.11232 | 0.27845 | 0.978454 | 6022 | 4 | 1.747 |
| EMB | 6 | 0.33497 | 0.54693 | 0.999994 | 10831 | 3 | -0.58895 | 0.11233 | 0.2996 | 0.984876 | 6023 | 2 | -0.58895 |
| CAMK2B | 6 | 0.07881 | 0.20495 | 0.978186 | 4183 | 4 | -1.5071 | 0.11233 | 0.2996 | 0.984876 | 6024 | 1 | -1.5071 |
| MST1 | 6 | 0.46258 | 0.64346 | 0.999994 | 12899 | 2 | 2.5978 | 0.11237 | 0.29966 | 0.984876 | 6025 | 4 | 2.5978 |
| AVPR1B | 6 | 0.016138 | 0.059465 | 0.932932 | 1251 | 4 | -2.7128 | 0.1125 | 0.29988 | 0.984876 | 6026 | 2 | -2.7128 |
| NonTargetingControlG | 2 | 0.88748 | 0.88737 | 0.999994 | 18982 | 0 | 3.7099 | 0.11252 | 0.18934 | 0.97466 | 6027 | 2 | 3.7099 |
| MOB2 | 6 | 0.46267 | 0.64353 | 0.999994 | 12900 | 1 | 3.0083 | 0.11255 | 0.29996 | 0.984876 | 6028 | 4 | 3.0083 |
| MCOLN3 | 5 | 0.2829 | 0.46721 | 0.999994 | 9760 | 1 | 1.1065 | 0.11258 | 0.27892 | 0.978454 | 6029 | 4 | 1.1065 |
| FAM53B | 6 | 0.062875 | 0.17182 | 0.967658 | 3550 | 4 | -1.8997 | 0.11259 | 0.30001 | 0.984876 | 6030 | 2 | -1.8997 |
| hsa-mir-629 | 2 | 0.61252 | 0.61239 | 0.999994 | 15117 | 0 | 3.7734 | 0.11262 | 0.1895 | 0.97466 | 6031 | 1 | 3.7734 |

| | | | | | | | | | | | | |
|---|---|---|---|---|---|---|---|---|---|---|---|---|
| FBXO31 | 6 | 0.25181 | 0.45009 | 0.999994 | 9053 | 3 | -0.76762 | 0.11266 | 0.30013 | 0.984876 | 6032 | 2 | -0.76762 |
| DBX2 | 6 | 0.94048 | 0.94046 | 0.999994 | 20095 | 0 | 3.0053 | 0.11268 | 0.30016 | 0.984876 | 6033 | 4 | 3.0053 |
| KLHL22 | 5 | 0.10068 | 0.22749 | 0.983354 | 4996 | 2 | 1.817 | 0.11269 | 0.2791 | 0.978454 | 6034 | 3 | 1.817 |
| CXorf56 | 5 | 0.59194 | 0.72324 | 0.999994 | 14848 | 1 | 1.2389 | 0.11269 | 0.2791 | 0.978454 | 6035 | 3 | 1.2389 |
| hsa-mir-616 | 4 | 0.17089 | 0.3107 | 0.999994 | 7004 | 1 | 3.0947 | 0.11271 | 0.25311 | 0.97466 | 6036 | 2 | 3.0947 |
| LOC100862671 | 4 | 0.01334 | 0.044028 | 0.891621 | 1088 | 1 | 3.8837 | 0.11279 | 0.25324 | 0.97466 | 6037 | 3 | 3.8837 |
| WBSCR28 | 5 | 0.71057 | 0.77134 | 0.999994 | 16405 | 1 | 1.0623 | 0.11281 | 0.27933 | 0.978454 | 6038 | 4 | 1.0623 |
| LRP1B | 5 | 0.017182 | 0.056363 | 0.9235 | 1310 | 1 | -0.025142 | 0.1129 | 0.27951 | 0.978454 | 6039 | 2 | -0.025142 |
| GPA33 | 6 | 0.21288 | 0.40359 | 0.999994 | 8089 | 1 | 2.5465 | 0.11299 | 0.30067 | 0.984876 | 6040 | 4 | 2.5465 |
| POLR2J2 | 3 | 0.78982 | 0.78973 | 0.999994 | 17339 | 0 | 4.7106 | 0.11307 | 0.2453 | 0.97466 | 6041 | 2 | 4.7106 |
| hsa-mir-33b | 3 | 0.1225 | 0.21278 | 0.979587 | 5720 | 2 | -1.6634 | 0.11307 | 0.24531 | 0.97466 | 6042 | 1 | -1.6634 |
| C20orf197 | 6 | 0.025473 | 0.088106 | 0.94395 | 1775 | 3 | -1.5358 | 0.11312 | 0.30088 | 0.984876 | 6043 | 3 | -1.5358 |
| ZFYVE9 | 6 | 0.0056758 | 0.023601 | 0.832699 | 569 | 4 | -3.1443 | 0.11312 | 0.30088 | 0.984876 | 6044 | 2 | -3.1443 |
| hsa-mir-1290 | 4 | 0.80174 | 0.80275 | 0.999994 | 17508 | 1 | 2.1798 | 0.11317 | 0.25381 | 0.97466 | 6045 | 3 | 2.1798 |
| KIFC2 | 5 | 0.61845 | 0.73299 | 0.999994 | 15194 | 1 | 2.7172 | 0.11323 | 0.28014 | 0.978454 | 6046 | 4 | 2.7172 |
| PHYHIPL | 5 | 0.80365 | 0.82183 | 0.999994 | 17536 | 1 | 1.1784 | 0.11324 | 0.28016 | 0.978454 | 6047 | 4 | 1.1784 |
| DTWD2 | 4 | 0.75734 | 0.76568 | 0.999994 | 16947 | 1 | 3.5867 | 0.11327 | 0.25397 | 0.97466 | 6048 | 3 | 3.5867 |
| CRY2 | 6 | 0.094718 | 0.23695 | 0.985449 | 4780 | 2 | 0.88386 | 0.1133 | 0.30117 | 0.984876 | 6049 | 4 | 0.88386 |
| AMOTL1 | 6 | 0.17575 | 0.35778 | 0.999994 | 7129 | 2 | 0.64221 | 0.1133 | 0.30117 | 0.984876 | 6050 | 3 | 0.64221 |
| ITPR1 | 6 | 0.081021 | 0.20937 | 0.979587 | 4266 | 3 | -0.92986 | 0.1133 | 0.30117 | 0.984876 | 6051 | 3 | -0.92986 |
| NonTargetingControlG | 2 | 0.8867 | 0.88659 | 0.999994 | 18962 | 0 | 3.2337 | 0.1133 | 0.19058 | 0.97466 | 6052 | 2 | 3.2337 |
| NonTargetingControlG | 2 | 0.78783 | 0.78737 | 0.999994 | 17319 | 0 | 4.3864 | 0.11332 | 0.1906 | 0.97466 | 6053 | 2 | 4.3864 |
| FXYD4 | 6 | 0.63555 | 0.76654 | 0.999994 | 15449 | 2 | 0.66915 | 0.11333 | 0.30122 | 0.984876 | 6054 | 3 | 0.66915 |
| GPR82 | 6 | 0.22754 | 0.42112 | 0.999994 | 8471 | 2 | 2.8588 | 0.11333 | 0.30122 | 0.984876 | 6055 | 4 | 2.8588 |
| CDC37L1 | 5 | 0.6952 | 0.7643 | 0.999994 | 16257 | 1 | 2.4578 | 0.11334 | 0.28036 | 0.978454 | 6056 | 4 | 2.4578 |
| PRSS48 | 5 | 0.89559 | 0.89573 | 0.999994 | 19139 | 0 | 3.5791 | 0.11343 | 0.28054 | 0.978454 | 6057 | 4 | 3.5791 |
| ST18 | 6 | 0.093311 | 0.23415 | 0.984971 | 4731 | 3 | 1.3319 | 0.11343 | 0.30139 | 0.985285 | 6058 | 3 | 1.3319 |
| hsa-mir-4491 | 4 | 0.74297 | 0.75474 | 0.999994 | 16780 | 1 | 2.8082 | 0.11347 | 0.25425 | 0.97466 | 6059 | 3 | 2.8082 |
| NonTargetingControlG | 2 | 0.88652 | 0.88642 | 0.999994 | 18960 | 0 | 2.8428 | 0.11348 | 0.19085 | 0.97466 | 6060 | 2 | 2.8428 |
| SRY | 5 | 0.86797 | 0.86928 | 0.999994 | 18600 | 1 | 1.1715 | 0.1135 | 0.28068 | 0.978454 | 6061 | 4 | 1.1715 |
| QKI | 6 | 0.72211 | 0.81901 | 0.999994 | 16518 | 1 | 2.2224 | 0.1136 | 0.30164 | 0.985764 | 6062 | 4 | 2.2224 |
| TMTC2 | 5 | 0.92752 | 0.92775 | 0.999994 | 19803 | 0 | 3.2567 | 0.11361 | 0.28088 | 0.978454 | 6063 | 3 | 3.2567 |
| FGFR1 | 6 | 0.62609 | 0.75938 | 0.999994 | 15325 | 2 | 2.1948 | 0.11362 | 0.30167 | 0.985764 | 6064 | 4 | 2.1948 |
| NOTCH3 | 4 | 0.89752 | 0.89723 | 0.999994 | 19169 | 0 | 4.0806 | 0.11371 | 0.25461 | 0.97466 | 6065 | 3 | 4.0806 |
| LOC100129083 | 5 | 0.87691 | 0.87733 | 0.999994 | 18770 | 1 | 1.132 | 0.11378 | 0.28122 | 0.978454 | 6066 | 3 | 1.132 |
| TPP2 | 5 | 0.63136 | 0.73783 | 0.999994 | 15389 | 1 | 1.6144 | 0.11378 | 0.28122 | 0.978454 | 6067 | 4 | 1.6144 |
| RNF167 | 5 | 0.10613 | 0.23597 | 0.985362 | 5178 | 2 | -0.10738 | 0.11381 | 0.28128 | 0.978454 | 6068 | 2 | -0.10738 |
| FAM60A | 5 | 0.15546 | 0.31089 | 0.999994 | 6606 | 3 | -1.1345 | 0.11381 | 0.28128 | 0.978454 | 6069 | 2 | -1.1345 |
| hsa-mir-4772 | 4 | 0.88025 | 0.87988 | 0.999994 | 18838 | 0 | 3.0373 | 0.11382 | 0.25479 | 0.97466 | 6070 | 2 | 3.0373 |
| hsa-mir-532 | 4 | 0.85397 | 0.85355 | 0.999994 | 18333 | 0 | 3.3333 | 0.11382 | 0.25479 | 0.97466 | 6071 | 2 | 3.3333 |
| ZSCAN10 | 5 | 0.42065 | 0.58061 | 0.999994 | 12260 | 2 | 3.3636 | 0.11385 | 0.28135 | 0.978454 | 6072 | 3 | 3.3636 |
| CCP110 | 4 | 0.32999 | 0.4737 | 0.999994 | 10723 | 2 | -0.85143 | 0.11396 | 0.25499 | 0.97466 | 6073 | 1 | -0.85143 |
| hsa-mir-7515 | 2 | 0.88602 | 0.88591 | 0.999994 | 18950 | 0 | 4.1198 | 0.11398 | 0.19163 | 0.97466 | 6074 | 2 | 4.1198 |
| POT1 | 3 | 0.56993 | 0.59458 | 0.999994 | 14538 | 1 | 0.54779 | 0.114 | 0.24703 | 0.97466 | 6075 | 1 | 0.54779 |
| RPS23 | 3 | 0.8413 | 0.84124 | 0.999994 | 18110 | 0 | 4.69 | 0.11404 | 0.24711 | 0.97466 | 6076 | 2 | 4.69 |
| SYVN1 | 6 | 0.96469 | 0.96473 | 0.999994 | 20715 | 0 | 0.7664 | 0.11404 | 0.30234 | 0.985876 | 6077 | 3 | 0.7664 |
| ZNF25 | 6 | 0.82463 | 0.85899 | 0.999994 | 17849 | 1 | 0.85564 | 0.11404 | 0.30234 | 0.985876 | 6078 | 4 | 0.85564 |
| C9orf131 | 6 | 0.079128 | 0.20556 | 0.978603 | 4194 | 2 | 1.3541 | 0.11404 | 0.30234 | 0.985876 | 6079 | 4 | 1.3541 |
| FASN | 5 | 0.012906 | 0.044324 | 0.891621 | 1055 | 2 | -0.47533 | 0.11408 | 0.28178 | 0.978454 | 6080 | 2 | -0.47533 |
| DNAJC27 | 5 | 0.88427 | 0.8844 | 0.999994 | 18914 | 1 | 1.1458 | 0.11412 | 0.28185 | 0.978454 | 6081 | 4 | 1.1458 |
| SPINT2 | 6 | 0.42173 | 0.61653 | 0.999994 | 12273 | 2 | 2.1641 | 0.11417 | 0.30255 | 0.985876 | 6082 | 4 | 2.1641 |
| hsa-mir-3149 | 2 | 0.88581 | 0.88573 | 0.999994 | 18943 | 0 | 2.809 | 0.11419 | 0.19192 | 0.97466 | 6083 | 2 | 2.809 |

| | | | | | | | | | | | | | |
|---|---|---|---|---|---|---|---|---|---|---|---|---|---|
| ZNF394 | 4 | 0.13688 | 0.27568 | 0.999379 | 6103 | 2 | -0.59678 | 0.11421 | 0.25541 | 0.97466 | 6084 | 2 | -0.59678 |
| ACHE | 6 | 0.60619 | 0.74455 | 0.999994 | 15030 | 1 | 2.2264 | 0.11429 | 0.30273 | 0.985876 | 6085 | 4 | 2.2264 |
| OR1N2 | 6 | 0.9437 | 0.94367 | 0.999994 | 20168 | 0 | 2.8362 | 0.1143 | 0.30275 | 0.985876 | 6086 | 4 | 2.8362 |
| MAPKAPK3 | 6 | 0.89608 | 0.9007 | 0.999994 | 19145 | 1 | 2.4861 | 0.11435 | 0.30282 | 0.985876 | 6087 | 3 | 2.4861 |
| SERTAD2 | 6 | 0.18716 | 0.37199 | 0.999994 | 7397 | 2 | 2.5391 | 0.11435 | 0.30282 | 0.985876 | 6088 | 3 | 2.5391 |
| BAIAP2L2 | 6 | 0.5175 | 0.68079 | 0.999994 | 13701 | 2 | 2.0171 | 0.11435 | 0.30282 | 0.985876 | 6089 | 3 | 2.0171 |
| GUCD1 | 6 | 0.37609 | 0.58708 | 0.999994 | 11595 | 2 | 2.4511 | 0.11435 | 0.30282 | 0.985876 | 6090 | 3 | 2.4511 |
| VGLL2 | 6 | 0.68796 | 0.80722 | 0.999994 | 16195 | 2 | 0.10665 | 0.11435 | 0.30282 | 0.985876 | 6091 | 2 | 0.10665 |
| ZG16B | 6 | 0.33989 | 0.55251 | 0.999994 | 10928 | 3 | -0.36057 | 0.11435 | 0.30282 | 0.985876 | 6092 | 1 | -0.36057 |
| CDC42SE2 | 6 | 0.14065 | 0.31385 | 0.999994 | 6205 | 3 | -0.66559 | 0.11437 | 0.30284 | 0.985876 | 6093 | 3 | -0.66559 |
| DNASE2B | 4 | 0.59683 | 0.66359 | 0.999994 | 14912 | 1 | 3.1525 | 0.11445 | 0.25579 | 0.97466 | 6094 | 2 | 3.1525 |
| TOPBP1 | 4 | 0.38288 | 0.52811 | 0.999994 | 11689 | 2 | 2.5753 | 0.11445 | 0.25579 | 0.97466 | 6095 | 2 | 2.5753 |
| ASPDH | 5 | 0.58098 | 0.71917 | 0.999994 | 14690 | 1 | 1.7564 | 0.1145 | 0.28258 | 0.978454 | 6096 | 4 | 1.7564 |
| ABHD13 | 6 | 0.94441 | 0.94439 | 0.999994 | 20189 | 0 | 0.93057 | 0.11453 | 0.3031 | 0.98595 | 6097 | 3 | 0.93057 |
| PIGB | 5 | 0.016694 | 0.055014 | 0.922677 | 1290 | 3 | -2.1109 | 0.11461 | 0.28279 | 0.978454 | 6098 | 2 | -2.1109 |
| NT5E | 6 | 0.92594 | 0.92603 | 0.999994 | 19765 | 1 | 2.8795 | 0.1147 | 0.30335 | 0.98595 | 6099 | 4 | 2.8795 |
| KRT1 | 6 | 0.11298 | 0.27245 | 0.998561 | 5403 | 2 | 0.92081 | 0.11471 | 0.30337 | 0.98595 | 6100 | 3 | 0.92081 |
| GRSF1 | 5 | 0.96095 | 0.96107 | 0.999994 | 20602 | 0 | 1.9549 | 0.11476 | 0.28305 | 0.978454 | 6101 | 4 | 1.9549 |
| ZFAND1 | 6 | 0.12561 | 0.29476 | 0.999994 | 5809 | 2 | 2.2797 | 0.11482 | 0.30354 | 0.986066 | 6102 | 3 | 2.2797 |
| C22orf42 | 5 | 0.80235 | 0.82099 | 0.999994 | 17518 | 1 | 0.66715 | 0.11483 | 0.28319 | 0.978454 | 6103 | 3 | 0.66715 |
| GALNTL5 | 5 | 0.82454 | 0.83564 | 0.999994 | 17847 | 1 | 0.31858 | 0.11483 | 0.28319 | 0.978454 | 6104 | 2 | 0.31858 |
| hsa-mir-8077 | 2 | 0.88499 | 0.88495 | 0.999994 | 18933 | 0 | 4.9241 | 0.11501 | 0.19319 | 0.97466 | 6105 | 2 | 4.9241 |
| CHRFAM7A | 6 | 0.66029 | 0.78554 | 0.999994 | 15783 | 1 | 2.197 | 0.11506 | 0.30394 | 0.986296 | 6106 | 4 | 2.197 |
| FAM127B | 5 | 0.87404 | 0.87472 | 0.999994 | 18710 | 1 | 1.2896 | 0.11509 | 0.28367 | 0.978454 | 6107 | 4 | 1.2896 |
| ENG | 5 | 0.82038 | 0.83286 | 0.999994 | 17783 | 1 | 1.4623 | 0.1151 | 0.28368 | 0.978454 | 6108 | 4 | 1.4623 |
| MS4A5 | 6 | 0.4768 | 0.65303 | 0.999994 | 13097 | 1 | 3.1611 | 0.11511 | 0.30403 | 0.986296 | 6109 | 4 | 3.1611 |
| KLK1 | 5 | 0.36495 | 0.53418 | 0.999994 | 11420 | 2 | 3.0391 | 0.11513 | 0.28374 | 0.978454 | 6110 | 3 | 3.0391 |
| SLIRP | 4 | 0.64675 | 0.69142 | 0.999994 | 15615 | 1 | 2.9346 | 0.11517 | 0.2569 | 0.974766 | 6111 | 3 | 2.9346 |
| RNASEH2C | 6 | 0.92947 | 0.9296 | 0.999994 | 19842 | 0 | 2.7217 | 0.11517 | 0.30413 | 0.986296 | 6112 | 4 | 2.7217 |
| TCP1 | 4 | 0.17061 | 0.31041 | 0.999994 | 6999 | 2 | 2.2013 | 0.11524 | 0.257 | 0.974766 | 6113 | 2 | 2.2013 |
| SOCS5 | 6 | 0.084751 | 0.21702 | 0.980699 | 4397 | 4 | -1.1593 | 0.11525 | 0.30424 | 0.986296 | 6114 | 1 | -1.1593 |
| PSMD9 | 6 | 0.41164 | 0.60995 | 0.999994 | 12142 | 2 | 2.0857 | 0.11535 | 0.30441 | 0.986296 | 6115 | 3 | 2.0857 |
| VIPAS39 | 5 | 0.11031 | 0.24245 | 0.988088 | 5312 | 2 | -0.51777 | 0.11538 | 0.28423 | 0.978454 | 6116 | 2 | -0.51777 |
| NOP9 | 5 | 0.26684 | 0.4543 | 0.999994 | 9394 | 2 | 2.9886 | 0.11538 | 0.28423 | 0.978454 | 6117 | 3 | 2.9886 |
| LOXL2 | 5 | 0.37246 | 0.54044 | 0.999994 | 11547 | 2 | -0.26942 | 0.11538 | 0.28423 | 0.978454 | 6118 | 2 | -0.26942 |
| SERINC5 | 6 | 0.10516 | 0.25737 | 0.994943 | 5136 | 2 | 2.3329 | 0.11539 | 0.30446 | 0.986296 | 6119 | 3 | 2.3329 |
| EPRS | 5 | 0.48561 | 0.63548 | 0.999994 | 13222 | 1 | 1.9508 | 0.11541 | 0.28428 | 0.978454 | 6120 | 4 | 1.9508 |
| LARP7 | 5 | 0.006433 | 0.025738 | 0.845768 | 621 | 2 | 3.0983 | 0.11543 | 0.28433 | 0.978454 | 6121 | 3 | 3.0983 |
| INSL5 | 6 | 0.86454 | 0.88003 | 0.999994 | 18534 | 1 | 0.40987 | 0.11545 | 0.30457 | 0.986296 | 6122 | 3 | 0.40987 |
| AMBRA1 | 6 | 0.62389 | 0.75773 | 0.999994 | 15288 | 2 | 2.482 | 0.11545 | 0.30457 | 0.986296 | 6123 | 4 | 2.482 |
| NonTargetingControlG | 2 | 0.88452 | 0.88446 | 0.999994 | 18925 | 0 | 4.4596 | 0.11548 | 0.19391 | 0.97466 | 6124 | 2 | 4.4596 |
| LOXL1 | 6 | 0.5103 | 0.67582 | 0.999994 | 13579 | 1 | 2.4063 | 0.1155 | 0.30464 | 0.986316 | 6125 | 3 | 2.4063 |
| ARL5B | 5 | 0.96796 | 0.9681 | 0.999994 | 20783 | 0 | 1.6792 | 0.11551 | 0.28447 | 0.978535 | 6126 | 4 | 1.6792 |
| TEX38 | 5 | 0.5078 | 0.65475 | 0.999994 | 13549 | 1 | 3.1603 | 0.11552 | 0.2845 | 0.978535 | 6127 | 3 | 3.1603 |
| CCDC111 | 5 | 0.063028 | 0.16707 | 0.967126 | 3556 | 2 | 3.3751 | 0.11556 | 0.28456 | 0.978584 | 6128 | 3 | 3.3751 |
| hsa-mir-527 | 1 | 0.88441 | 0.88471 | 0.999994 | 18921 | 0 | 6.8554 | 0.11559 | 0.11553 | 0.954984 | 6129 | 1 | 6.8554 |
| OR1L3 | 5 | 0.040004 | 0.1159 | 0.951052 | 2516 | 3 | -2.6345 | 0.1157 | 0.28484 | 0.978871 | 6130 | 1 | -2.6345 |
| C4orf3 | 5 | 0.26782 | 0.4551 | 0.999994 | 9420 | 1 | 1.4692 | 0.1157 | 0.28484 | 0.978871 | 6131 | 4 | 1.4692 |
| EME2 | 5 | 0.72498 | 0.77826 | 0.999994 | 16548 | 1 | 1.4105 | 0.11571 | 0.28486 | 0.978871 | 6132 | 4 | 1.4105 |
| HES1 | 6 | 0.13612 | 0.30813 | 0.999994 | 6082 | 3 | -0.15359 | 0.11573 | 0.30501 | 0.986316 | 6133 | 3 | -0.15359 |
| KLHDC2 | 6 | 0.53614 | 0.69382 | 0.999994 | 14000 | 2 | 1.0516 | 0.11573 | 0.30501 | 0.986316 | 6134 | 4 | 1.0516 |
| TUBB2B | 6 | 0.43492 | 0.62508 | 0.999994 | 12496 | 2 | 1.178 | 0.11577 | 0.30508 | 0.986316 | 6135 | 4 | 1.178 |

| | | | | | | | | | | | | | |
|---|---|---|---|---|---|---|---|---|---|---|---|---|---|
| G3BP2 | 6 | 0.12268 | 0.29075 | 0.999632 | 5729 | 3 | 0.60839 | 0.11577 | 0.30508 | 0.986316 | 6136 | 3 | 0.60839 |
| SSR4 | 5 | 0.10713 | 0.23749 | 0.986719 | 5215 | 1 | 1.208 | 0.11579 | 0.28501 | 0.978871 | 6137 | 4 | 1.208 |
| PPIP5K1 | 4 | 0.6375 | 0.68608 | 0.999994 | 15481 | 1 | 3.2934 | 0.11585 | 0.25792 | 0.974766 | 6138 | 3 | 3.2934 |
| GPR107 | 6 | 0.46967 | 0.64823 | 0.999994 | 13001 | 1 | 2.0686 | 0.11588 | 0.30526 | 0.986322 | 6139 | 4 | 2.0686 |
| RHOC | 5 | 0.49714 | 0.64541 | 0.999994 | 13381 | 2 | 1.0197 | 0.11598 | 0.28537 | 0.978871 | 6140 | 3 | 1.0197 |
| DCP2 | 6 | 0.68491 | 0.80483 | 0.999994 | 16158 | 1 | 2.2098 | 0.11602 | 0.30548 | 0.986322 | 6141 | 3 | 2.2098 |
| COMTD1 | 6 | 0.40322 | 0.60448 | 0.999994 | 12004 | 2 | 2.7233 | 0.11602 | 0.30548 | 0.986322 | 6142 | 4 | 2.7233 |
| TWIST2 | 5 | 0.20236 | 0.37862 | 0.999994 | 7840 | 3 | -0.73259 | 0.11602 | 0.28544 | 0.978871 | 6143 | 2 | -0.73259 |
| hsa-mir-1185-1 | 4 | 0.49761 | 0.61526 | 0.999994 | 13392 | 1 | 0.19736 | 0.11604 | 0.25822 | 0.974766 | 6144 | 2 | 0.19736 |
| MICALL1 | 5 | 0.91603 | 0.91611 | 0.999994 | 19538 | 0 | 1.9463 | 0.11605 | 0.2855 | 0.978871 | 6145 | 4 | 1.9463 |
| ASGR1 | 4 | 0.90434 | 0.90403 | 0.999994 | 19307 | 0 | 2.6235 | 0.11612 | 0.25834 | 0.974766 | 6146 | 3 | 2.6235 |
| SLC45A1 | 6 | 0.083595 | 0.21467 | 0.979587 | 4358 | 2 | 2.3921 | 0.11623 | 0.30582 | 0.986404 | 6147 | 4 | 2.3921 |
| NAA10 | 5 | 0.90782 | 0.90787 | 0.999994 | 19380 | 0 | 3.4241 | 0.11625 | 0.28587 | 0.978871 | 6148 | 3 | 3.4241 |
| BAK1 | 5 | 0.47496 | 0.62631 | 0.999994 | 13062 | 2 | 3.4241 | 0.11625 | 0.28587 | 0.978871 | 6149 | 3 | 3.4241 |
| ARHGAP11B | 5 | 0.27903 | 0.46407 | 0.999994 | 9674 | 2 | 3.4241 | 0.11625 | 0.28587 | 0.978871 | 6150 | 3 | 3.4241 |
| DERL2 | 6 | 0.30198 | 0.50903 | 0.999994 | 10161 | 2 | 1.7856 | 0.11625 | 0.30585 | 0.986404 | 6151 | 3 | 1.7856 |
| LYZ | 6 | 0.25883 | 0.4585 | 0.999994 | 9212 | 2 | 2.742 | 0.11625 | 0.30585 | 0.986404 | 6152 | 4 | 2.742 |
| hsa-mir-4420 | 4 | 0.58665 | 0.65816 | 0.999994 | 14768 | 1 | 2.2869 | 0.11626 | 0.25857 | 0.974766 | 6153 | 3 | 2.2869 |
| CACNA1C | 5 | 0.93177 | 0.93206 | 0.999994 | 19894 | 0 | 1.2694 | 0.11628 | 0.28591 | 0.978871 | 6154 | 4 | 1.2694 |
| PASD1 | 6 | 0.95096 | 0.95095 | 0.999994 | 20355 | 0 | 1.1556 | 0.11629 | 0.3059 | 0.986404 | 6155 | 4 | 1.1556 |
| HIST1H4I | 6 | 0.078325 | 0.20396 | 0.978186 | 4164 | 3 | -0.14742 | 0.11629 | 0.3059 | 0.986404 | 6156 | 3 | -0.14742 |
| NonTargetingControlG | 2 | 0.88366 | 0.88361 | 0.999994 | 18898 | 0 | 2.3819 | 0.11634 | 0.19519 | 0.97466 | 6157 | 2 | 2.3819 |
| TEX33 | 5 | 0.72123 | 0.77643 | 0.999994 | 16510 | 1 | 1.1881 | 0.11643 | 0.2862 | 0.978871 | 6158 | 3 | 1.1881 |
| IFT43 | 4 | 0.91934 | 0.91913 | 0.999994 | 19616 | 0 | 3.898 | 0.11651 | 0.25897 | 0.974766 | 6159 | 3 | 3.898 |
| CCL25 | 6 | 0.35118 | 0.56533 | 0.999994 | 11179 | 2 | 2.1061 | 0.11659 | 0.30636 | 0.987544 | 6160 | 4 | 2.1061 |
| PITX2 | 5 | 0.28767 | 0.47104 | 0.999994 | 9869 | 2 | 1.3186 | 0.11661 | 0.28654 | 0.978871 | 6161 | 3 | 1.3186 |
| SERPINF2 | 5 | 0.16346 | 0.32259 | 0.999994 | 6826 | 2 | 0.12437 | 0.11661 | 0.28654 | 0.978871 | 6162 | 2 | 0.12437 |
| ZNF808 | 5 | 0.50383 | 0.65125 | 0.999994 | 13491 | 2 | 1.9067 | 0.11661 | 0.28654 | 0.978871 | 6163 | 3 | 1.9067 |
| CDADC1 | 6 | 0.5765 | 0.72261 | 0.999994 | 14637 | 1 | 3.0842 | 0.11682 | 0.30673 | 0.987798 | 6164 | 4 | 3.0842 |
| HSF4 | 5 | 0.24825 | 0.4394 | 0.999994 | 8975 | 1 | 0.98609 | 0.11689 | 0.28702 | 0.979159 | 6165 | 4 | 0.98609 |
| SEPT14 | 6 | 0.050797 | 0.14581 | 0.963846 | 3007 | 4 | -1.3906 | 0.11693 | 0.3069 | 0.987798 | 6166 | 2 | -1.3906 |
| FEM1B | 6 | 0.51712 | 0.68053 | 0.999994 | 13694 | 2 | 1.0986 | 0.11693 | 0.3069 | 0.987798 | 6167 | 4 | 1.0986 |
| OR5T1 | 6 | 0.60227 | 0.74162 | 0.999994 | 14976 | 2 | 2.0329 | 0.11693 | 0.3069 | 0.987798 | 6168 | 4 | 2.0329 |
| TICAM2 | 6 | 0.86521 | 0.88042 | 0.999994 | 18542 | 1 | 2.1171 | 0.11693 | 0.3069 | 0.987798 | 6169 | 3 | 2.1171 |
| TGFB1 | 6 | 0.89271 | 0.89824 | 0.999994 | 19083 | 1 | 1.2204 | 0.11696 | 0.30695 | 0.987798 | 6170 | 4 | 1.2204 |
| CMPK2 | 6 | 0.10482 | 0.25673 | 0.994943 | 5125 | 4 | -0.83025 | 0.11696 | 0.30695 | 0.987798 | 6171 | 2 | -0.83025 |
| IFNA4 | 5 | 0.18501 | 0.35382 | 0.999994 | 7341 | 2 | 0.45353 | 0.11696 | 0.28717 | 0.979159 | 6172 | 2 | 0.45353 |
| PAX6 | 5 | 0.43099 | 0.58925 | 0.999994 | 12431 | 2 | 0.26309 | 0.11696 | 0.28717 | 0.979159 | 6173 | 2 | 0.26309 |
| ASXL2 | 5 | 0.19734 | 0.37145 | 0.999994 | 7682 | 1 | 1.3182 | 0.11701 | 0.28726 | 0.979315 | 6174 | 4 | 1.3182 |
| TOR1A | 5 | 0.72464 | 0.77811 | 0.999994 | 16543 | 1 | 3.1552 | 0.11703 | 0.28731 | 0.979331 | 6175 | 4 | 3.1552 |
| SEC61A2 | 4 | 0.018622 | 0.057715 | 0.929232 | 1395 | 2 | -1.7571 | 0.11707 | 0.25984 | 0.974766 | 6176 | 2 | -1.7571 |
| PTRH2 | 4 | 0.16738 | 0.30707 | 0.999994 | 6919 | 2 | 2.6435 | 0.11708 | 0.25986 | 0.974766 | 6177 | 2 | 2.6435 |
| TGIF2LX | 6 | 0.29284 | 0.49836 | 0.999994 | 9971 | 2 | 1.8545 | 0.11714 | 0.30724 | 0.988343 | 6178 | 4 | 1.8545 |
| MTHFD1L | 5 | 0.086667 | 0.20524 | 0.978342 | 4473 | 2 | 3.1275 | 0.11714 | 0.28753 | 0.97947 | 6179 | 3 | 3.1275 |
| ZNF222 | 5 | 0.91034 | 0.91037 | 0.999994 | 19433 | 0 | 1.0129 | 0.11715 | 0.28754 | 0.97947 | 6180 | 4 | 1.0129 |
| MRPL4 | 3 | 0.14821 | 0.24618 | 0.98999 | 6412 | 1 | 4.5924 | 0.11717 | 0.25166 | 0.97466 | 6181 | 2 | 4.5924 |
| SLC12A3 | 6 | 0.59044 | 0.73293 | 0.999994 | 14829 | 2 | 2.0701 | 0.11718 | 0.3073 | 0.988343 | 6182 | 3 | 2.0701 |
| SERTM1 | 6 | 0.3675 | 0.58167 | 0.999994 | 11468 | 1 | 1.1274 | 0.11718 | 0.30731 | 0.988343 | 6183 | 3 | 1.1274 |
| HLCS | 5 | 0.092057 | 0.21387 | 0.979587 | 4680 | 2 | 0.043024 | 0.11722 | 0.28767 | 0.97947 | 6184 | 2 | 0.043024 |
| MIER2 | 4 | 0.93022 | 0.92999 | 0.999994 | 19859 | 0 | 2.311 | 0.11727 | 0.26014 | 0.974766 | 6185 | 3 | 2.311 |
| C3orf58 | 4 | 0.12109 | 0.25219 | 0.994162 | 5667 | 2 | -1.2288 | 0.11731 | 0.26021 | 0.974766 | 6186 | 1 | -1.2288 |
| PCDHA4 | 2 | 0.88265 | 0.88264 | 0.999994 | 18884 | 0 | 2.4489 | 0.11735 | 0.19672 | 0.97466 | 6187 | 2 | 2.4489 |

| | | | | | | | | | | | | |
|---|---|---|---|---|---|---|---|---|---|---|---|---|
| NonTargetingControlG | 2 | 0.88265 | 0.88264 | 0.999994 | 18883 | 0 | 2.5156 | 0.11735 | 0.19672 | 0.97466 | 6188 | 2 | 2.5156 |
| MBD3L4 | 3 | 0.11566 | 0.20377 | 0.978186 | 5505 | 2 | -2.1744 | 0.11735 | 0.25186 | 0.97466 | 6189 | 1 | -2.1744 |
| SNTB1 | 4 | 0.71976 | 0.73792 | 0.999994 | 16494 | 1 | 2.3591 | 0.11738 | 0.26032 | 0.974766 | 6190 | 3 | 2.3591 |
| GINS2 | 4 | 0.71833 | 0.73689 | 0.999994 | 16479 | 1 | 2.4685 | 0.11749 | 0.26049 | 0.974875 | 6191 | 3 | 2.4685 |
| MYH4 | 5 | 0.96576 | 0.96591 | 0.999994 | 20732 | 0 | 2.0889 | 0.1175 | 0.28817 | 0.979689 | 6192 | 4 | 2.0889 |
| OMG | 6 | 0.97281 | 0.97274 | 0.999994 | 20890 | 0 | 3.0352 | 0.11755 | 0.30789 | 0.988566 | 6193 | 4 | 3.0352 |
| NDEL1 | 6 | 0.005208 | 0.02191 | 0.819701 | 534 | 3 | -0.94574 | 0.11762 | 0.30799 | 0.988566 | 6194 | 2 | -0.94574 |
| C12orf10 | 6 | 0.82008 | 0.85688 | 0.999994 | 17779 | 1 | 1.936 | 0.11762 | 0.30799 | 0.988566 | 6195 | 4 | 1.936 |
| LOC100505841 | 5 | 0.72778 | 0.77958 | 0.999994 | 16578 | 1 | -0.055017 | 0.11762 | 0.2884 | 0.979944 | 6196 | 2 | -0.055017 |
| FAM173B | 5 | 0.5636 | 0.70385 | 0.999994 | 14443 | 2 | 2.2616 | 0.11771 | 0.28857 | 0.979944 | 6197 | 3 | 2.2616 |
| CCR10 | 5 | 0.050013 | 0.14039 | 0.958716 | 2979 | 2 | 0.71374 | 0.11771 | 0.28857 | 0.979944 | 6198 | 3 | 0.71374 |
| GOLGA3 | 5 | 0.42155 | 0.58135 | 0.999994 | 12270 | 1 | 1.6261 | 0.11771 | 0.28858 | 0.979944 | 6199 | 4 | 1.6261 |
| SLC36A1 | 4 | 0.82701 | 0.8266 | 0.999994 | 17887 | 1 | 3.5134 | 0.11772 | 0.26085 | 0.974875 | 6200 | 3 | 3.5134 |
| OR52N1 | 6 | 0.86004 | 0.87745 | 0.999994 | 18439 | 1 | 3.1654 | 0.11778 | 0.30824 | 0.988566 | 6201 | 4 | 3.1654 |
| IL36RN | 6 | 0.12736 | 0.29693 | 0.999994 | 5858 | 2 | 1.7437 | 0.1178 | 0.30827 | 0.988566 | 6202 | 4 | 1.7437 |
| DDX28 | 5 | 0.97017 | 0.97022 | 0.999994 | 20828 | 0 | 3.404 | 0.11796 | 0.28907 | 0.980152 | 6203 | 4 | 3.404 |
| ORAI2 | 2 | 0.48808 | 0.49308 | 0.999994 | 13257 | 1 | 3.369 | 0.11808 | 0.19785 | 0.97466 | 6204 | 1 | 3.369 |
| NonTargetingControlG | 2 | 0.36691 | 0.38963 | 0.999994 | 11455 | 1 | 3.0753 | 0.11808 | 0.19785 | 0.97466 | 6205 | 1 | 3.0753 |
| DLG5 | 5 | 0.1207 | 0.25843 | 0.994943 | 5651 | 3 | -1.3392 | 0.11818 | 0.2895 | 0.980804 | 6206 | 2 | -1.3392 |
| C2orf53 | 4 | 0.0081401 | 0.027828 | 0.855011 | 739 | 1 | 2.3116 | 0.11825 | 0.26167 | 0.974875 | 6207 | 3 | 2.3116 |
| SMYD4 | 6 | 0.84433 | 0.86891 | 0.999994 | 18161 | 1 | 2.1176 | 0.11826 | 0.30899 | 0.988566 | 6208 | 3 | 2.1176 |
| GALE | 6 | 0.18058 | 0.3639 | 0.999994 | 7236 | 3 | 0.62 | 0.11833 | 0.30912 | 0.988566 | 6209 | 3 | 0.62 |
| PSMA3 | 3 | 0.26942 | 0.39897 | 0.999994 | 9457 | 1 | 4.2689 | 0.11835 | 0.25299 | 0.97466 | 6210 | 2 | 4.2689 |
| STXBP3 | 4 | 0.36512 | 0.50977 | 0.999994 | 11424 | 2 | 2.5008 | 0.11837 | 0.26185 | 0.974875 | 6211 | 2 | 2.5008 |
| hsa-mir-525 | 4 | 0.72994 | 0.74515 | 0.999994 | 16597 | 1 | 2.1592 | 0.11837 | 0.26186 | 0.974875 | 6212 | 2 | 2.1592 |
| RPS20 | 4 | 0.051668 | 0.12475 | 0.951052 | 3051 | 2 | -0.40655 | 0.11856 | 0.26215 | 0.974875 | 6213 | 2 | -0.40655 |
| H2AFJ | 6 | 0.56477 | 0.71422 | 0.999994 | 14466 | 2 | 1.2412 | 0.11857 | 0.30948 | 0.988566 | 6214 | 3 | 1.2412 |
| CABLES2 | 5 | 0.28044 | 0.46521 | 0.999994 | 9710 | 2 | 0.048679 | 0.11858 | 0.29024 | 0.981963 | 6215 | 2 | 0.048679 |
| URM1 | 5 | 0.87034 | 0.87137 | 0.999994 | 18641 | 1 | 0.67357 | 0.11866 | 0.29039 | 0.981963 | 6216 | 3 | 0.67357 |
| OSBPL11 | 6 | 0.34332 | 0.55645 | 0.999994 | 11013 | 3 | 1.9457 | 0.11871 | 0.30969 | 0.988566 | 6217 | 3 | 1.9457 |
| FITM2 | 6 | 0.27818 | 0.4813 | 0.999994 | 9658 | 2 | 2.4842 | 0.11871 | 0.30969 | 0.988566 | 6218 | 3 | 2.4842 |
| C9orf62 | 6 | 0.72861 | 0.82114 | 0.999994 | 16585 | 1 | 2.4763 | 0.11871 | 0.30969 | 0.988566 | 6219 | 3 | 2.4763 |
| FAM118A | 6 | 0.68497 | 0.80488 | 0.999994 | 16161 | 2 | 0.019789 | 0.11871 | 0.3097 | 0.988566 | 6220 | 2 | 0.019789 |
| GUF1 | 6 | 0.66862 | 0.7921 | 0.999994 | 15899 | 2 | 2.7603 | 0.11872 | 0.30972 | 0.988566 | 6221 | 4 | 2.7603 |
| hsa-mir-6857 | 4 | 0.38025 | 0.5254 | 0.999994 | 11651 | 2 | 2.6207 | 0.11881 | 0.26256 | 0.974875 | 6222 | 2 | 2.6207 |
| DNAJC7 | 5 | 0.69085 | 0.76237 | 0.999994 | 16217 | 1 | 3.1831 | 0.11887 | 0.29076 | 0.982601 | 6223 | 3 | 3.1831 |
| NonTargetingControlG | 2 | 0.88108 | 0.88111 | 0.999994 | 18853 | 0 | 2.4125 | 0.11892 | 0.19912 | 0.97466 | 6224 | 2 | 2.4125 |
| ZNF608 | 5 | 0.029556 | 0.089316 | 0.94395 | 1987 | 2 | -0.036842 | 0.11896 | 0.29093 | 0.982839 | 6225 | 2 | -0.036842 |
| CLRN1 | 5 | 0.076067 | 0.18826 | 0.969543 | 4075 | 1 | 2.4619 | 0.11903 | 0.29104 | 0.982859 | 6226 | 3 | 2.4619 |
| PIP4K2C | 6 | 0.0019597 | 0.0097759 | 0.680529 | 269 | 5 | -4.9727 | 0.11907 | 0.3103 | 0.988566 | 6227 | 1 | -4.9727 |
| IRAK4 | 6 | 0.60689 | 0.74504 | 0.999994 | 15035 | 2 | 2.6295 | 0.11907 | 0.3103 | 0.988566 | 6228 | 4 | 2.6295 |
| SIAE | 6 | 0.34767 | 0.56139 | 0.999994 | 11100 | 3 | 0.44632 | 0.11907 | 0.3103 | 0.988566 | 6229 | 3 | 0.44632 |
| hsa-mir-567 | 4 | 0.85036 | 0.84988 | 0.999994 | 18269 | 0 | 2.7502 | 0.11908 | 0.26301 | 0.974875 | 6230 | 3 | 2.7502 |
| RINL | 5 | 0.20137 | 0.37718 | 0.999994 | 7802 | 1 | 1.0699 | 0.11908 | 0.29114 | 0.98296 | 6231 | 4 | 1.0699 |
| FBXL17 | 6 | 0.91116 | 0.91252 | 0.999994 | 19453 | 1 | 1.8325 | 0.11909 | 0.31033 | 0.988566 | 6232 | 4 | 1.8325 |
| ADH6 | 6 | 0.24339 | 0.44005 | 0.999994 | 8873 | 2 | 2.1366 | 0.11916 | 0.31045 | 0.988566 | 6233 | 4 | 2.1366 |
| ABCC2 | 4 | 0.87317 | 0.87282 | 0.999994 | 18697 | 0 | 1.221 | 0.1192 | 0.26319 | 0.974875 | 6234 | 2 | 1.221 |
| NUBP1 | 3 | 0.23468 | 0.35595 | 0.999994 | 8636 | 1 | 3.061 | 0.11922 | 0.25397 | 0.97466 | 6235 | 2 | 3.061 |
| LIN54 | 5 | 0.89622 | 0.89634 | 0.999994 | 19146 | 0 | 1.9355 | 0.11928 | 0.29151 | 0.983138 | 6236 | 3 | 1.9355 |
| MLST8 | 5 | 0.22835 | 0.41493 | 0.999994 | 8487 | 2 | 0.073819 | 0.11928 | 0.29151 | 0.983138 | 6237 | 2 | 0.073819 |
| CBR1 | 6 | 0.82062 | 0.85713 | 0.999994 | 17785 | 1 | 2.1972 | 0.11931 | 0.31066 | 0.988566 | 6238 | 4 | 2.1972 |
| ARFGEF2 | 6 | 0.041689 | 0.1257 | 0.951052 | 2600 | 3 | -1.6 | 0.11932 | 0.31069 | 0.988566 | 6239 | 2 | -1.6 |

| | | | | | | | | | | | | | |
|---|---|---|---|---|---|---|---|---|---|---|---|---|---|
| DTX2 | 6 | 0.85591 | 0.87512 | 0.999994 | 18370 | 1 | 2.3486 | 0.11936 | 0.31076 | 0.988566 | 6240 | 3 | 2.3486 |
| TNFSF12 | 5 | 0.79159 | 0.81428 | 0.999994 | 17374 | 1 | 2.9819 | 0.11937 | 0.29168 | 0.983225 | 6241 | 4 | 2.9819 |
| NonTargetingControlG | 2 | 0.88063 | 0.88068 | 0.999994 | 18844 | 0 | 2.1106 | 0.11937 | 0.1998 | 0.97466 | 6242 | 2 | 2.1106 |
| DOCK5 | 5 | 0.94852 | 0.94874 | 0.999994 | 20301 | 0 | 2.5525 | 0.11942 | 0.29177 | 0.983225 | 6243 | 4 | 2.5525 |
| GPR87 | 6 | 0.94246 | 0.94243 | 0.999994 | 20138 | 0 | 2.084 | 0.11943 | 0.31088 | 0.988566 | 6244 | 4 | 2.084 |
| NonTargetingControlG | 2 | 0.82541 | 0.82522 | 0.999994 | 17860 | 0 | 4.4955 | 0.11943 | 0.1999 | 0.97466 | 6245 | 2 | 4.4955 |
| FADS3 | 6 | 0.36084 | 0.57625 | 0.999994 | 11353 | 2 | 1.9179 | 0.11945 | 0.31091 | 0.988566 | 6246 | 4 | 1.9179 |
| ZNF461 | 5 | 0.95105 | 0.95123 | 0.999994 | 20363 | 0 | 1.9538 | 0.11948 | 0.29187 | 0.983225 | 6247 | 4 | 1.9538 |
| DIDO1 | 6 | 0.0068596 | 0.027923 | 0.855011 | 650 | 3 | -0.43821 | 0.11948 | 0.31094 | 0.988566 | 6248 | 3 | -0.43821 |
| STIM2 | 4 | 0.54254 | 0.63609 | 0.999994 | 14104 | 1 | 3.7653 | 0.11948 | 0.26363 | 0.974875 | 6249 | 3 | 3.7653 |
| ATAD1 | 4 | 0.79045 | 0.79276 | 0.999994 | 17351 | 1 | 3.3897 | 0.11948 | 0.26363 | 0.974875 | 6250 | 3 | 3.3897 |
| hsa-mir-6824 | 4 | 0.2424 | 0.38407 | 0.999994 | 8844 | 1 | 2.6754 | 0.11948 | 0.26363 | 0.974875 | 6251 | 3 | 2.6754 |
| SYT14 | 4 | 0.19533 | 0.33589 | 0.999994 | 7643 | 1 | 2.4066 | 0.11948 | 0.26363 | 0.974875 | 6252 | 3 | 2.4066 |
| FAM161B | 4 | 0.075226 | 0.16977 | 0.967126 | 4044 | 1 | 2.0224 | 0.11948 | 0.26363 | 0.974875 | 6253 | 3 | 2.0224 |
| SERF2 | 4 | 0.60083 | 0.66582 | 0.999994 | 14959 | 1 | 3.4385 | 0.11948 | 0.26363 | 0.974875 | 6254 | 3 | 3.4385 |
| LNX1 | 5 | 0.83017 | 0.83961 | 0.999994 | 17935 | 1 | 1.4106 | 0.11961 | 0.29212 | 0.983225 | 6255 | 4 | 1.4106 |
| CRAT | 6 | 0.11565 | 0.27752 | 0.999379 | 5504 | 2 | 2.0001 | 0.11964 | 0.31118 | 0.988566 | 6256 | 4 | 2.0001 |
| CSPG5 | 6 | 0.17975 | 0.36285 | 0.999994 | 7215 | 3 | -0.51123 | 0.11965 | 0.31119 | 0.988566 | 6257 | 2 | -0.51123 |
| ESX1 | 6 | 0.074882 | 0.19692 | 0.977164 | 4031 | 4 | -1.4112 | 0.11968 | 0.31124 | 0.988566 | 6258 | 1 | -1.4112 |
| ZNF816-ZNF321P | 2 | 0.88026 | 0.88032 | 0.999994 | 18840 | 0 | 3.9995 | 0.11974 | 0.20032 | 0.97466 | 6259 | 2 | 3.9995 |
| C6orf141 | 5 | 0.87413 | 0.87481 | 0.999994 | 18712 | 1 | 1.1728 | 0.11976 | 0.29238 | 0.983225 | 6260 | 4 | 1.1728 |
| ERICH1 | 5 | 0.90389 | 0.90398 | 0.999994 | 19301 | 0 | 2.2375 | 0.11978 | 0.29241 | 0.983225 | 6261 | 4 | 2.2375 |
| C20orf26 | 5 | 0.40543 | 0.56807 | 0.999994 | 12037 | 2 | 3.3152 | 0.1198 | 0.29245 | 0.983225 | 6262 | 3 | 3.3152 |
| PKNOX2 | 4 | 0.93374 | 0.93351 | 0.999994 | 19943 | 0 | 3.449 | 0.11983 | 0.26419 | 0.975894 | 6263 | 3 | 3.449 |
| C11orf53 | 5 | 0.7731 | 0.80346 | 0.999994 | 17122 | 1 | 1.3182 | 0.11987 | 0.29259 | 0.983225 | 6264 | 3 | 1.3182 |
| LOC147646 | 6 | 0.41732 | 0.61361 | 0.999994 | 12209 | 2 | 0.97107 | 0.11991 | 0.3116 | 0.988566 | 6265 | 3 | 0.97107 |
| MUC21 | 6 | 0.52468 | 0.68579 | 0.999994 | 13812 | 2 | 0.068934 | 0.11991 | 0.3116 | 0.988566 | 6266 | 2 | 0.068934 |
| ELMO1 | 6 | 0.16885 | 0.3493 | 0.999994 | 6957 | 2 | 0.14394 | 0.11991 | 0.3116 | 0.988566 | 6267 | 2 | 0.14394 |
| PSMA2 | 4 | 0.8079 | 0.80836 | 0.999994 | 17598 | 1 | 3.8877 | 0.11996 | 0.26439 | 0.976183 | 6268 | 3 | 3.8877 |
| CENPT | 6 | 0.84002 | 0.86662 | 0.999994 | 18090 | 1 | 3.1266 | 0.11998 | 0.3117 | 0.988566 | 6269 | 4 | 3.1266 |
| KIAA0556 | 6 | 0.84082 | 0.86705 | 0.999994 | 18102 | 1 | 2.1946 | 0.11998 | 0.3117 | 0.988566 | 6270 | 3 | 2.1946 |
| TRAPPC12 | 6 | 0.38618 | 0.59349 | 0.999994 | 11735 | 2 | 2.1166 | 0.11998 | 0.3117 | 0.988566 | 6271 | 3 | 2.1166 |
| MGAT1 | 6 | 0.54861 | 0.70263 | 0.999994 | 14189 | 2 | 3.0568 | 0.11998 | 0.3117 | 0.988566 | 6272 | 4 | 3.0568 |
| AIF1L | 5 | 0.43475 | 0.59243 | 0.999994 | 12489 | 1 | 1.4239 | 0.12 | 0.29284 | 0.983225 | 6273 | 4 | 1.4239 |
| SIGLEC7 | 5 | 0.94912 | 0.94932 | 0.999994 | 20311 | 0 | 1.1086 | 0.12001 | 0.29286 | 0.983225 | 6274 | 3 | 1.1086 |
| hsa-mir-6130 | 3 | 0.69109 | 0.69272 | 0.999994 | 16219 | 1 | 4.5294 | 0.12017 | 0.25501 | 0.97466 | 6275 | 2 | 4.5294 |
| DDX5 | 6 | 0.62201 | 0.75631 | 0.999994 | 15259 | 2 | 0.89222 | 0.12018 | 0.31198 | 0.988735 | 6276 | 4 | 0.89222 |
| VWC2L | 6 | 0.35122 | 0.56538 | 0.999994 | 11180 | 3 | 1.5064 | 0.12018 | 0.31198 | 0.988735 | 6277 | 3 | 1.5064 |
| UBL4B | 6 | 0.71096 | 0.81543 | 0.999994 | 16409 | 1 | 2.0882 | 0.12018 | 0.31198 | 0.988735 | 6278 | 4 | 2.0882 |
| hsa-mir-3941 | 4 | 0.86613 | 0.86572 | 0.999994 | 18558 | 0 | 2.5537 | 0.12018 | 0.26474 | 0.976633 | 6279 | 3 | 2.5537 |
| SPATS2 | 5 | 0.83254 | 0.84131 | 0.999994 | 17965 | 1 | 1.3182 | 0.12019 | 0.29321 | 0.983225 | 6280 | 4 | 1.3182 |
| NonTargetingControlG | 2 | 0.8798 | 0.87986 | 0.999994 | 18831 | 0 | 4.6445 | 0.1202 | 0.20102 | 0.97466 | 6281 | 2 | 4.6445 |
| FTMT | 5 | 0.95465 | 0.95481 | 0.999994 | 20445 | 0 | 1.0843 | 0.12024 | 0.29331 | 0.983225 | 6282 | 4 | 1.0843 |
| PRAMEF13 | 5 | 0.89715 | 0.89729 | 0.999994 | 19162 | 0 | 1.2366 | 0.12026 | 0.29334 | 0.983225 | 6283 | 4 | 1.2366 |
| PCDH17 | 6 | 0.53923 | 0.696 | 0.999994 | 14039 | 2 | 1.7862 | 0.12034 | 0.31226 | 0.989294 | 6284 | 4 | 1.7862 |
| TNP2 | 5 | 0.58347 | 0.72022 | 0.999994 | 14725 | 1 | 0.3524 | 0.12035 | 0.29351 | 0.983276 | 6285 | 2 | 0.3524 |
| KRT13 | 5 | 0.55894 | 0.69967 | 0.999994 | 14351 | 2 | 0.79663 | 0.12035 | 0.29351 | 0.983276 | 6286 | 3 | 0.79663 |
| IL37 | 4 | 0.19967 | 0.34034 | 0.999994 | 7753 | 1 | 3.2719 | 0.1204 | 0.26511 | 0.976736 | 6287 | 2 | 3.2719 |
| VDAC3 | 4 | 0.78105 | 0.78476 | 0.999994 | 17223 | 1 | 3.1998 | 0.1204 | 0.26511 | 0.976736 | 6288 | 2 | 3.1998 |
| RETN | 5 | 0.9638 | 0.96395 | 0.999994 | 20700 | 0 | 2.8178 | 0.12043 | 0.29365 | 0.983304 | 6289 | 4 | 2.8178 |
| GNG11 | 3 | 0.70638 | 0.70702 | 0.999994 | 16377 | 1 | 2.3387 | 0.12062 | 0.25551 | 0.97466 | 6290 | 2 | 2.3387 |
| ACTR1B | 3 | 0.042862 | 0.1042 | 0.947284 | 2671 | 2 | -1.1509 | 0.12062 | 0.25551 | 0.97466 | 6291 | 1 | -1.1509 |

| | | | | | | | | | | | | | |
|---|---|---|---|---|---|---|---|---|---|---|---|---|---|
| GMPR2 | 6 | 0.22249 | 0.41515 | 0.999994 | 8344 | 2 | 2.4181 | 0.12065 | 0.31276 | 0.989342 | 6292 | 4 | 2.4181 |
| C4A | 6 | 0.34951 | 0.56345 | 0.999994 | 11136 | 2 | -0.2272 | 0.12066 | 0.31277 | 0.989342 | 6293 | 3 | -0.2272 |
| TARM1 | 6 | 0.29377 | 0.49942 | 0.999994 | 9991 | 2 | 0.71555 | 0.12066 | 0.31277 | 0.989342 | 6294 | 3 | 0.71555 |
| CBX3 | 6 | 0.087939 | 0.22338 | 0.983354 | 4509 | 4 | -0.95834 | 0.12066 | 0.31277 | 0.989342 | 6295 | 2 | -0.95834 |
| RTN4RL2 | 5 | 0.8263 | 0.83685 | 0.999994 | 17873 | 1 | 1.5265 | 0.12067 | 0.2941 | 0.983429 | 6296 | 4 | 1.5265 |
| SIX5 | 6 | 0.76757 | 0.83492 | 0.999994 | 17056 | 1 | 1.965 | 0.12079 | 0.313 | 0.989623 | 6297 | 4 | 1.965 |
| KIAA1210 | 5 | 0.94919 | 0.94938 | 0.999994 | 20313 | 0 | 1.7277 | 0.12084 | 0.2944 | 0.983431 | 6298 | 4 | 1.7277 |
| ADH1A | 5 | 0.3889 | 0.55422 | 0.999994 | 11781 | 2 | -0.59488 | 0.12084 | 0.2944 | 0.983431 | 6299 | 1 | -0.59488 |
| AES | 5 | 0.27273 | 0.45898 | 0.999994 | 9531 | 1 | 3.2136 | 0.12085 | 0.29442 | 0.983431 | 6300 | 4 | 3.2136 |
| NonTargetingControlG | 2 | 0.45842 | 0.46672 | 0.999994 | 12828 | 1 | 3.2414 | 0.12087 | 0.20199 | 0.97466 | 6301 | 1 | 3.2414 |
| ELANE | 5 | 0.50364 | 0.6511 | 0.999994 | 13488 | 1 | 2.7684 | 0.12087 | 0.29446 | 0.983431 | 6302 | 3 | 2.7684 |
| ANKMY2 | 6 | 0.68368 | 0.80383 | 0.999994 | 16141 | 2 | 1.8767 | 0.1209 | 0.31316 | 0.989846 | 6303 | 4 | 1.8767 |
| SERTAD1 | 5 | 0.95484 | 0.95501 | 0.999994 | 20448 | 0 | 1.7181 | 0.12106 | 0.29481 | 0.983787 | 6304 | 4 | 1.7181 |
| SURF2 | 6 | 0.72326 | 0.81939 | 0.999994 | 16526 | 1 | -0.20866 | 0.12108 | 0.31345 | 0.989846 | 6305 | 2 | -0.20866 |
| hsa-mir-6769b | 3 | 0.247 | 0.37121 | 0.999994 | 8943 | 2 | -0.881 | 0.12112 | 0.25604 | 0.974766 | 6306 | 1 | -0.881 |
| CD40LG | 6 | 0.41567 | 0.61254 | 0.999994 | 12188 | 2 | 0.77001 | 0.12113 | 0.31353 | 0.989846 | 6307 | 4 | 0.77001 |
| DCX | 5 | 0.21781 | 0.4002 | 0.999994 | 8223 | 1 | 1.2473 | 0.12117 | 0.29504 | 0.983787 | 6308 | 4 | 1.2473 |
| NonTargetingControlG | 2 | 0.7772 | 0.77672 | 0.999994 | 17170 | 0 | 4.3234 | 0.12119 | 0.20252 | 0.97466 | 6309 | 2 | 4.3234 |
| NonTargetingControlG | 2 | 0.032301 | 0.058964 | 0.931872 | 2122 | 1 | -0.96697 | 0.12119 | 0.20252 | 0.97466 | 6310 | 1 | -0.96697 |
| AWAT1 | 5 | 0.61254 | 0.73075 | 0.999994 | 15118 | 1 | 1.0955 | 0.12123 | 0.29513 | 0.983848 | 6311 | 4 | 1.0955 |
| SLC44A2 | 6 | 0.56195 | 0.71221 | 0.999994 | 14405 | 2 | 1.8709 | 0.12132 | 0.31383 | 0.989846 | 6312 | 4 | 1.8709 |
| CREB3 | 4 | 0.78962 | 0.79205 | 0.999994 | 17335 | 1 | 2.2886 | 0.12134 | 0.26659 | 0.978454 | 6313 | 3 | 2.2886 |
| KLK5 | 5 | 0.87385 | 0.87455 | 0.999994 | 18708 | 1 | 1.8405 | 0.12135 | 0.29536 | 0.983965 | 6314 | 4 | 1.8405 |
| TNFRSF10C | 5 | 0.46187 | 0.61522 | 0.999994 | 12882 | 1 | 1.0959 | 0.12143 | 0.2955 | 0.983965 | 6315 | 4 | 1.0959 |
| CASC4 | 6 | 0.18122 | 0.36468 | 0.999994 | 7256 | 1 | 1.7669 | 0.12143 | 0.31402 | 0.989846 | 6316 | 4 | 1.7669 |
| NonTargetingControlG | 2 | 0.87855 | 0.87862 | 0.999994 | 18802 | 0 | 1.7518 | 0.12145 | 0.20289 | 0.97466 | 6317 | 2 | 1.7518 |
| NonTargetingControlG | 1 | 0.87855 | 0.87883 | 0.999994 | 18801 | 0 | 6.735 | 0.12145 | 0.12131 | 0.962897 | 6318 | 1 | 6.735 |
| EDNRA | 6 | 0.36323 | 0.57895 | 0.999994 | 11393 | 2 | 3.0344 | 0.12146 | 0.31407 | 0.989846 | 6319 | 4 | 3.0344 |
| ABI1 | 6 | 0.24266 | 0.43917 | 0.999994 | 8854 | 2 | 3.0464 | 0.12146 | 0.31407 | 0.989846 | 6320 | 4 | 3.0464 |
| TRPM7 | 5 | 0.74499 | 0.78823 | 0.999994 | 16799 | 1 | 1.3484 | 0.12148 | 0.29558 | 0.983965 | 6321 | 4 | 1.3484 |
| SSX2 | 5 | 0.946 | 0.94619 | 0.999994 | 20232 | 0 | 2.4812 | 0.1215 | 0.29562 | 0.983965 | 6322 | 3 | 2.4812 |
| GTPBP2 | 6 | 0.0029765 | 0.013647 | 0.75005 | 353 | 2 | 1.3118 | 0.12159 | 0.31427 | 0.989846 | 6323 | 4 | 1.3118 |
| CGB2 | 2 | 0.87839 | 0.87846 | 0.999994 | 18796 | 0 | 2.9368 | 0.12161 | 0.20314 | 0.97466 | 6324 | 2 | 2.9368 |
| CLDN5 | 4 | 0.42161 | 0.56787 | 0.999994 | 12271 | 1 | 1.811 | 0.12165 | 0.26705 | 0.978454 | 6325 | 3 | 1.811 |
| PRICKLE3 | 5 | 0.53064 | 0.67463 | 0.999994 | 13905 | 1 | 1.7041 | 0.12169 | 0.29598 | 0.984434 | 6326 | 4 | 1.7041 |
| PAEP | 5 | 0.26539 | 0.45312 | 0.999994 | 9357 | 2 | 0.44926 | 0.12171 | 0.29602 | 0.984434 | 6327 | 2 | 0.44926 |
| GABRB2 | 4 | 0.89813 | 0.89785 | 0.999994 | 19180 | 0 | 1.3477 | 0.12183 | 0.26732 | 0.978454 | 6328 | 2 | 1.3477 |
| EFNB1 | 4 | 0.092192 | 0.20102 | 0.977164 | 4687 | 3 | -1.5324 | 0.12183 | 0.26732 | 0.978454 | 6329 | 1 | -1.5324 |
| HIST1H2BD | 4 | 0.80939 | 0.80973 | 0.999994 | 17622 | 1 | 1.9559 | 0.12185 | 0.26735 | 0.978454 | 6330 | 3 | 1.9559 |
| GBP1 | 4 | 0.24262 | 0.38434 | 0.999994 | 8850 | 2 | 2.0902 | 0.12188 | 0.2674 | 0.978454 | 6331 | 2 | 2.0902 |
| MATN1 | 5 | 0.57085 | 0.71028 | 0.999994 | 14557 | 1 | 1.1524 | 0.12195 | 0.29644 | 0.984782 | 6332 | 3 | 1.1524 |
| LHFPL4 | 5 | 0.50575 | 0.65294 | 0.999994 | 13519 | 1 | 1.6063 | 0.12195 | 0.29646 | 0.984782 | 6333 | 4 | 1.6063 |
| NonTargetingControlG | 2 | 0.87793 | 0.87801 | 0.999994 | 18786 | 0 | 3.7942 | 0.12207 | 0.20384 | 0.97466 | 6334 | 2 | 3.7942 |
| SGPL1 | 6 | 0.031344 | 0.10197 | 0.946414 | 2076 | 3 | -1.577 | 0.12208 | 0.31506 | 0.989846 | 6335 | 2 | -1.577 |
| EFNB3 | 6 | 0.86129 | 0.87814 | 0.999994 | 18472 | 1 | 0.62034 | 0.12208 | 0.31506 | 0.989846 | 6336 | 3 | 0.62034 |
| TTLL9 | 3 | 0.13452 | 0.22851 | 0.983354 | 6040 | 1 | -0.097799 | 0.1221 | 0.25716 | 0.974766 | 6337 | 1 | -0.097799 |
| C16orf95 | 5 | 0.20208 | 0.3782 | 0.999994 | 7831 | 1 | 1.393 | 0.12215 | 0.29683 | 0.984876 | 6338 | 4 | 1.393 |
| hsa-mir-4724 | 4 | 0.68141 | 0.7125 | 0.999994 | 16108 | 1 | 2.651 | 0.12219 | 0.26789 | 0.978454 | 6339 | 2 | 2.651 |
| ZNF679 | 6 | 0.06463 | 0.17549 | 0.968171 | 3636 | 4 | -1.6127 | 0.12222 | 0.31529 | 0.989846 | 6340 | 2 | -1.6127 |
| SLC24A4 | 6 | 0.032751 | 0.10524 | 0.95013 | 2147 | 2 | 2.1752 | 0.12222 | 0.31529 | 0.989846 | 6341 | 4 | 2.1752 |
| ZNF586 | 6 | 0.71747 | 0.8175 | 0.999994 | 16472 | 1 | 2.0381 | 0.12222 | 0.31529 | 0.989846 | 6342 | 3 | 2.0381 |
| INTS5 | 5 | 0.074017 | 0.1849 | 0.968526 | 3997 | 3 | -1.1873 | 0.12224 | 0.297 | 0.984876 | 6343 | 2 | -1.1873 |

| | | | | | | | | | | | | |
|---|---|---|---|---|---|---|---|---|---|---|---|---|
| CD81 | 5 | 0.91031 | 0.91033 | 0.999994 | 19432 | 0 | 1.5663 | 0.12224 | 0.297 | 0.984876 | 6344 | 3 | 1.5663 |
| SYT9 | 5 | 0.72429 | 0.77794 | 0.999994 | 16539 | 1 | 1.4348 | 0.12226 | 0.29703 | 0.984876 | 6345 | 4 | 1.4348 |
| MEIG1 | 6 | 0.18716 | 0.37199 | 0.999994 | 7395 | 2 | 2.5477 | 0.12227 | 0.31537 | 0.989846 | 6346 | 4 | 2.5477 |
| FSHB | 6 | 0.57309 | 0.72017 | 0.999994 | 14591 | 2 | 1.9467 | 0.12228 | 0.31538 | 0.989846 | 6347 | 4 | 1.9467 |
| KRT6C | 5 | 0.88917 | 0.88928 | 0.999994 | 19023 | 0 | 1.3426 | 0.12238 | 0.29726 | 0.984876 | 6348 | 4 | 1.3426 |
| SLC39A7 | 3 | 0.20586 | 0.32004 | 0.999994 | 7920 | 1 | 4.4636 | 0.12241 | 0.25753 | 0.974766 | 6349 | 2 | 4.4636 |
| COL6A3 | 6 | 0.53545 | 0.69335 | 0.999994 | 13988 | 2 | 2.8876 | 0.12249 | 0.31568 | 0.989846 | 6350 | 4 | 2.8876 |
| TRIM23 | 5 | 0.1489 | 0.3011 | 0.999994 | 6424 | 1 | 2.9667 | 0.1225 | 0.29748 | 0.984876 | 6351 | 3 | 2.9667 |
| HCFC1R1 | 6 | 0.41429 | 0.61162 | 0.999994 | 12171 | 1 | 1.6259 | 0.12258 | 0.31581 | 0.989846 | 6352 | 4 | 1.6259 |
| CCDC83 | 6 | 0.37058 | 0.58357 | 0.999994 | 11523 | 2 | 3.0303 | 0.12258 | 0.31581 | 0.989846 | 6353 | 4 | 3.0303 |
| ENDOG | 6 | 0.82194 | 0.85775 | 0.999994 | 17801 | 1 | 1.9354 | 0.12262 | 0.31587 | 0.989846 | 6354 | 4 | 1.9354 |
| NonTargetingControlG | 2 | 0.87737 | 0.87743 | 0.999994 | 18781 | 0 | 1.7359 | 0.12263 | 0.20467 | 0.97466 | 6355 | 2 | 1.7359 |
| PEF1 | 5 | 0.77706 | 0.80574 | 0.999994 | 17168 | 1 | 3.0931 | 0.12265 | 0.29776 | 0.984876 | 6356 | 3 | 3.0931 |
| ADCYAP1R1 | 6 | 0.96914 | 0.96913 | 0.999994 | 20804 | 0 | 2.607 | 0.12269 | 0.31598 | 0.989846 | 6357 | 4 | 2.607 |
| CDK19 | 6 | 0.09789 | 0.24318 | 0.988088 | 4889 | 3 | 0.43085 | 0.12278 | 0.3161 | 0.989846 | 6358 | 3 | 0.43085 |
| BTBD1 | 5 | 0.64155 | 0.74178 | 0.999994 | 15539 | 1 | 2.7612 | 0.12279 | 0.298 | 0.984876 | 6359 | 3 | 2.7612 |
| TRIOBP | 5 | 0.14577 | 0.29644 | 0.999994 | 6346 | 2 | 0.040433 | 0.12279 | 0.298 | 0.984876 | 6360 | 2 | 0.040433 |
| NLGN1 | 6 | 0.65863 | 0.78425 | 0.999994 | 15759 | 2 | 2.5591 | 0.12283 | 0.31619 | 0.989846 | 6361 | 4 | 2.5591 |
| USP25 | 4 | 0.77687 | 0.78131 | 0.999994 | 17164 | 1 | 2.1516 | 0.12284 | 0.26892 | 0.978454 | 6362 | 3 | 2.1516 |
| ALAD | 6 | 0.26081 | 0.46084 | 0.999994 | 9264 | 3 | -0.7133 | 0.12287 | 0.31625 | 0.989846 | 6363 | 2 | -0.7133 |
| hsa-mir-1266 | 3 | 0.2226 | 0.34086 | 0.999994 | 8347 | 1 | 2.4093 | 0.12289 | 0.25807 | 0.974766 | 6364 | 2 | 2.4093 |
| ALX3 | 6 | 0.51983 | 0.6824 | 0.999994 | 13745 | 1 | 2.001 | 0.12293 | 0.31636 | 0.989846 | 6365 | 4 | 2.001 |
| KLK12 | 6 | 0.37736 | 0.58789 | 0.999994 | 11615 | 1 | 2.5004 | 0.12297 | 0.31643 | 0.989846 | 6366 | 4 | 2.5004 |
| VPS33A | 6 | 0.79877 | 0.84734 | 0.999994 | 17458 | 1 | 2.8852 | 0.12297 | 0.31643 | 0.989846 | 6367 | 4 | 2.8852 |
| WNT2 | 6 | 0.04861 | 0.14106 | 0.959146 | 2914 | 2 | 2.1636 | 0.12297 | 0.31643 | 0.989846 | 6368 | 4 | 2.1636 |
| PATL1 | 6 | 0.29858 | 0.50504 | 0.999994 | 10100 | 1 | 2.289 | 0.12301 | 0.31648 | 0.989846 | 6369 | 4 | 2.289 |
| SIGLEC11 | 6 | 0.34866 | 0.56249 | 0.999994 | 11128 | 3 | -0.49612 | 0.12306 | 0.31656 | 0.989846 | 6370 | 1 | -0.49612 |
| CPSF2 | 5 | 0.9637 | 0.96384 | 0.999994 | 20697 | 0 | 2.4876 | 0.12317 | 0.2987 | 0.984876 | 6371 | 4 | 2.4876 |
| NonTargetingControlG | 1 | 0.87681 | 0.87713 | 0.999994 | 18767 | 0 | 6.7069 | 0.12319 | 0.12309 | 0.962935 | 6372 | 1 | 6.7069 |
| TRIM60 | 4 | 0.93948 | 0.93939 | 0.999994 | 20062 | 0 | 2.4674 | 0.12327 | 0.26959 | 0.978454 | 6373 | 3 | 2.4674 |
| NonTargetingControlG | 2 | 0.87672 | 0.87675 | 0.999994 | 18766 | 0 | 4.4697 | 0.12328 | 0.20568 | 0.97466 | 6374 | 2 | 4.4697 |
| CXorf23 | 6 | 0.02577 | 0.088818 | 0.94395 | 1789 | 1 | 0.0023606 | 0.12329 | 0.31691 | 0.989846 | 6375 | 2 | 0.0023606 |
| CXCR2 | 6 | 0.26868 | 0.47019 | 0.999994 | 9437 | 2 | 1.8097 | 0.12329 | 0.31691 | 0.989846 | 6376 | 4 | 1.8097 |
| CTSZ | 6 | 0.97855 | 0.97849 | 0.999994 | 21065 | 0 | 2.12 | 0.12329 | 0.31692 | 0.989846 | 6377 | 4 | 2.12 |
| VTA1 | 5 | 0.94018 | 0.94035 | 0.999994 | 20086 | 0 | 0.99137 | 0.12331 | 0.29896 | 0.984876 | 6378 | 4 | 0.99137 |
| FTSJ3 | 4 | 0.33568 | 0.47951 | 0.999994 | 10845 | 1 | 3.749 | 0.12331 | 0.26965 | 0.978454 | 6379 | 3 | 3.749 |
| NonTargetingControlG | 2 | 0.38978 | 0.4083 | 0.999994 | 11793 | 1 | 2.9082 | 0.12336 | 0.2058 | 0.97466 | 6380 | 1 | 2.9082 |
| KIF21A | 5 | 0.49493 | 0.64351 | 0.999994 | 13339 | 1 | 0.76161 | 0.12339 | 0.29912 | 0.984876 | 6381 | 3 | 0.76161 |
| GOLGA5 | 5 | 0.52748 | 0.67186 | 0.999994 | 13850 | 2 | 0.18126 | 0.12339 | 0.29912 | 0.984876 | 6382 | 2 | 0.18126 |
| DHRS2 | 5 | 0.060007 | 0.16211 | 0.967126 | 3421 | 3 | -2.108 | 0.12339 | 0.29912 | 0.984876 | 6383 | 2 | -2.108 |
| hsa-mir-6511a-1 | 4 | 0.9415 | 0.94138 | 0.999994 | 20120 | 0 | 2.8001 | 0.1234 | 0.26979 | 0.978454 | 6384 | 3 | 2.8001 |
| METTL15 | 5 | 0.52141 | 0.66653 | 0.999994 | 13764 | 2 | 3.049 | 0.12343 | 0.2992 | 0.984876 | 6385 | 3 | 3.049 |
| TREX2 | 5 | 0.90199 | 0.90209 | 0.999994 | 19251 | 0 | 2.9585 | 0.12344 | 0.29921 | 0.984876 | 6386 | 3 | 2.9585 |
| METTL16 | 5 | 0.20923 | 0.38832 | 0.999994 | 8011 | 1 | 2.4619 | 0.12351 | 0.29935 | 0.984876 | 6387 | 4 | 2.4619 |
| ACSM3 | 4 | 0.25038 | 0.39229 | 0.999994 | 9023 | 1 | 0.30965 | 0.12351 | 0.26998 | 0.978454 | 6388 | 1 | 0.30965 |
| RPL28 | 5 | 0.81413 | 0.82868 | 0.999994 | 17692 | 1 | 1.2175 | 0.12353 | 0.29939 | 0.984876 | 6389 | 4 | 1.2175 |
| PIANP | 5 | 0.36797 | 0.53667 | 0.999994 | 11474 | 1 | 1.5712 | 0.12354 | 0.29941 | 0.984876 | 6390 | 4 | 1.5712 |
| TAAR1 | 6 | 0.51506 | 0.67912 | 0.999994 | 13664 | 2 | 2.1187 | 0.12358 | 0.31735 | 0.989846 | 6391 | 3 | 2.1187 |
| ZAK | 6 | 0.21288 | 0.40359 | 0.999994 | 8088 | 3 | -0.71141 | 0.12359 | 0.31737 | 0.989846 | 6392 | 2 | -0.71141 |
| HDGF | 5 | 0.29432 | 0.47643 | 0.999994 | 10003 | 1 | 2.854 | 0.12366 | 0.29961 | 0.984876 | 6393 | 3 | 2.854 |
| IL4 | 4 | 0.71827 | 0.73685 | 0.999994 | 16478 | 1 | 3.3865 | 0.12375 | 0.27036 | 0.978454 | 6394 | 3 | 3.3865 |
| ACY3 | 6 | 0.019957 | 0.071819 | 0.93914 | 1458 | 5 | -2.1637 | 0.12384 | 0.31778 | 0.989846 | 6395 | 1 | -2.1637 |

| | | | | | | | | | | | | |
|---|---|---|---|---|---|---|---|---|---|---|---|---|
| FAM26F | 5 | 0.13816 | 0.28503 | 0.999632 | 6142 | 1 | 1.7135 | 0.12385 | 0.29997 | 0.984876 | 6396 | 4 | 1.7135 |
| CNTD1 | 5 | 0.58492 | 0.72073 | 0.999994 | 14741 | 1 | 1.7536 | 0.12398 | 0.30022 | 0.984876 | 6397 | 4 | 1.7536 |
| ZNF331 | 4 | 0.77611 | 0.78072 | 0.999994 | 17152 | 1 | 1.8635 | 0.12399 | 0.27072 | 0.978454 | 6398 | 3 | 1.8635 |
| ARID3A | 5 | 0.7273 | 0.77937 | 0.999994 | 16571 | 1 | 2.8999 | 0.12401 | 0.30026 | 0.984876 | 6399 | 4 | 2.8999 |
| ANKRD13B | 4 | 0.3081 | 0.45129 | 0.999994 | 10286 | 2 | 2.3296 | 0.12402 | 0.27077 | 0.978454 | 6400 | 2 | 2.3296 |
| hsa-mir-643 | 2 | 0.87598 | 0.87602 | 0.999994 | 18750 | 0 | 2.9403 | 0.12402 | 0.2068 | 0.97466 | 6401 | 2 | 2.9403 |
| MYO9A | 6 | 0.06954 | 0.1858 | 0.969409 | 3815 | 4 | -1.2814 | 0.12405 | 0.31809 | 0.989846 | 6402 | 1 | -1.2814 |
| PSMD5 | 6 | 0.62766 | 0.76059 | 0.999994 | 15347 | 2 | -0.32948 | 0.12405 | 0.31809 | 0.989846 | 6403 | 2 | -0.32948 |
| GPT | 6 | 0.11412 | 0.27461 | 0.998561 | 5453 | 3 | -0.6649 | 0.12405 | 0.31809 | 0.989846 | 6404 | 2 | -0.6649 |
| NKAPL | 5 | 0.036462 | 0.10699 | 0.950995 | 2344 | 2 | 2.941 | 0.12405 | 0.30033 | 0.984876 | 6405 | 3 | 2.941 |
| ZNF596 | 6 | 0.20152 | 0.38973 | 0.999994 | 7809 | 2 | 0.97613 | 0.12406 | 0.31812 | 0.989846 | 6406 | 3 | 0.97613 |
| FLG | 5 | 0.85331 | 0.85691 | 0.999994 | 18320 | 1 | 3.4365 | 0.12407 | 0.30037 | 0.984876 | 6407 | 3 | 3.4365 |
| H3F3B | 6 | 0.5325 | 0.69131 | 0.999994 | 13943 | 2 | 3.0637 | 0.12421 | 0.31834 | 0.989846 | 6408 | 4 | 3.0637 |
| OR5M11 | 5 | 0.64937 | 0.74486 | 0.999994 | 15647 | 1 | 3.1198 | 0.12422 | 0.30066 | 0.984876 | 6409 | 4 | 3.1198 |
| hsa-mir-764 | 3 | 0.84828 | 0.84826 | 0.999994 | 18221 | 0 | 0.56457 | 0.12427 | 0.25961 | 0.974766 | 6410 | 1 | 0.56457 |
| TMEM185A | 5 | 0.01306 | 0.044758 | 0.891621 | 1067 | 2 | -0.3618 | 0.1243 | 0.30079 | 0.984876 | 6411 | 2 | -0.3618 |
| CACNB1 | 5 | 0.024298 | 0.075616 | 0.93914 | 1723 | 2 | 0.77703 | 0.1243 | 0.30079 | 0.984876 | 6412 | 3 | 0.77703 |
| WDR91 | 6 | 0.41437 | 0.61166 | 0.999994 | 12173 | 2 | 2.0064 | 0.12432 | 0.31851 | 0.989846 | 6413 | 3 | 2.0064 |
| PRDM5 | 6 | 0.231 | 0.4253 | 0.999994 | 8541 | 3 | -0.26864 | 0.12432 | 0.31851 | 0.989846 | 6414 | 3 | -0.26864 |
| POU1F1 | 6 | 0.84705 | 0.87036 | 0.999994 | 18208 | 1 | 2.0433 | 0.12437 | 0.31858 | 0.989846 | 6415 | 4 | 2.0433 |
| hsa-mir-1179 | 4 | 0.82255 | 0.82222 | 0.999994 | 17814 | 1 | 0.42715 | 0.12441 | 0.27136 | 0.978454 | 6416 | 2 | 0.42715 |
| COX17 | 4 | 0.60291 | 0.66692 | 0.999994 | 14985 | 1 | 2.8956 | 0.12443 | 0.27138 | 0.978454 | 6417 | 3 | 2.8956 |
| ZNF428 | 5 | 0.54321 | 0.68575 | 0.999994 | 14116 | 1 | 0.78868 | 0.12446 | 0.30112 | 0.984876 | 6418 | 3 | 0.78868 |
| DCTN6 | 5 | 0.2126 | 0.39306 | 0.999994 | 8079 | 2 | 3.2838 | 0.1245 | 0.3012 | 0.984876 | 6419 | 3 | 3.2838 |
| DFFB | 5 | 0.28575 | 0.46948 | 0.999994 | 9827 | 2 | 3.2838 | 0.1245 | 0.3012 | 0.984876 | 6420 | 3 | 3.2838 |
| AGPAT6 | 5 | 0.072144 | 0.18183 | 0.968322 | 3922 | 2 | 3.2838 | 0.1245 | 0.3012 | 0.984876 | 6421 | 3 | 3.2838 |
| PRY2 | 2 | 0.87536 | 0.87542 | 0.999994 | 18743 | 0 | 4.4987 | 0.12464 | 0.20777 | 0.97466 | 6422 | 2 | 4.4987 |
| hsa-mir-4540 | 4 | 0.24131 | 0.38297 | 0.999994 | 8808 | 2 | 2.1084 | 0.1247 | 0.27182 | 0.978454 | 6423 | 2 | 2.1084 |
| VGLL1 | 6 | 0.67841 | 0.79976 | 0.999994 | 16067 | 2 | 1.9302 | 0.12475 | 0.31921 | 0.989846 | 6424 | 3 | 1.9302 |
| NonTargetingControlG | 2 | 0.87522 | 0.87529 | 0.999994 | 18740 | 0 | 1.7674 | 0.12478 | 0.20799 | 0.97466 | 6425 | 2 | 1.7674 |
| AKAP3 | 5 | 0.6851 | 0.7598 | 0.999994 | 16164 | 1 | 1.9983 | 0.12479 | 0.30175 | 0.985876 | 6426 | 4 | 1.9983 |
| hsa-mir-8061 | 4 | 0.60865 | 0.67002 | 0.999994 | 15051 | 1 | 2.3931 | 0.12482 | 0.27201 | 0.978454 | 6427 | 3 | 2.3931 |
| EXOC5 | 4 | 0.61547 | 0.67379 | 0.999994 | 15150 | 1 | 2.2585 | 0.12483 | 0.27202 | 0.978454 | 6428 | 3 | 2.2585 |
| KANK2 | 3 | 0.42437 | 0.49872 | 0.999994 | 12319 | 1 | 4.3946 | 0.12493 | 0.26034 | 0.974766 | 6429 | 2 | 4.3946 |
| NUDT3 | 5 | 0.62705 | 0.73621 | 0.999994 | 15338 | 1 | 0.88914 | 0.12496 | 0.30208 | 0.985876 | 6430 | 3 | 0.88914 |
| TMEM201 | 5 | 0.47068 | 0.62273 | 0.999994 | 13013 | 2 | 0.076561 | 0.12496 | 0.30208 | 0.985876 | 6431 | 2 | 0.076561 |
| EFCAB2 | 5 | 0.3666 | 0.53553 | 0.999994 | 11449 | 2 | 0.67892 | 0.12498 | 0.30212 | 0.985876 | 6432 | 3 | 0.67892 |
| RPAIN | 4 | 0.12093 | 0.25194 | 0.994 | 5659 | 2 | -1.4861 | 0.12501 | 0.27228 | 0.978454 | 6433 | 1 | -1.4861 |
| SLITRK2 | 4 | 0.40705 | 0.55295 | 0.999994 | 12062 | 2 | 0.24976 | 0.12501 | 0.27228 | 0.978454 | 6434 | 2 | 0.24976 |
| UNC79 | 6 | 0.95248 | 0.95248 | 0.999994 | 20393 | 0 | 1.4507 | 0.12507 | 0.31971 | 0.989846 | 6435 | 4 | 1.4507 |
| BTBD16 | 6 | 0.33236 | 0.54397 | 0.999994 | 10777 | 1 | 2.5013 | 0.12512 | 0.31979 | 0.989846 | 6436 | 4 | 2.5013 |
| PAQR6 | 6 | 0.45536 | 0.63861 | 0.999994 | 12780 | 1 | -0.071696 | 0.12513 | 0.31981 | 0.989846 | 6437 | 2 | -0.071696 |
| RRAGB | 5 | 0.67187 | 0.75409 | 0.999994 | 15953 | 1 | 1.3171 | 0.12521 | 0.30254 | 0.985876 | 6438 | 4 | 1.3171 |
| TMEM31 | 6 | 0.41563 | 0.61251 | 0.999994 | 12187 | 2 | 2.7593 | 0.12524 | 0.31999 | 0.989846 | 6439 | 4 | 2.7593 |
| CAMK4 | 6 | 0.77688 | 0.83847 | 0.999994 | 17165 | 1 | 2.9407 | 0.12529 | 0.32008 | 0.989846 | 6440 | 4 | 2.9407 |
| hsa-mir-2053 | 4 | 0.65083 | 0.69377 | 0.999994 | 15671 | 1 | 2.5438 | 0.12531 | 0.27278 | 0.978454 | 6441 | 3 | 2.5438 |
| PARPBP | 6 | 0.10591 | 0.25882 | 0.994943 | 5172 | 2 | 1.9933 | 0.12533 | 0.32013 | 0.989846 | 6442 | 3 | 1.9933 |
| SAA2 | 5 | 0.48246 | 0.63281 | 0.999994 | 13177 | 1 | 0.9091 | 0.12535 | 0.3028 | 0.985876 | 6443 | 4 | 0.9091 |
| PFKL | 6 | 0.8467 | 0.87017 | 0.999994 | 18204 | 1 | 0.18467 | 0.12539 | 0.32024 | 0.989846 | 6444 | 2 | 0.18467 |
| ZBTB41 | 6 | 0.39487 | 0.59906 | 0.999994 | 11871 | 2 | 1.9283 | 0.12539 | 0.32024 | 0.989846 | 6445 | 4 | 1.9283 |
| TIMM10 | 4 | 0.43438 | 0.58099 | 0.999994 | 12482 | 1 | 3.7888 | 0.12541 | 0.27295 | 0.978454 | 6446 | 3 | 3.7888 |
| DDX3Y | 6 | 0.88594 | 0.89352 | 0.999994 | 18949 | 1 | 2.7678 | 0.12542 | 0.32028 | 0.989846 | 6447 | 4 | 2.7678 |

| | | | | | | | | | | | | |
|---|---|---|---|---|---|---|---|---|---|---|---|---|
| ZNF878 | 5 | 0.22802 | 0.41448 | 0.999994 | 8478 | 2 | 1.7773 | 0.12544 | 0.30296 | 0.98595 | 6448 | 3 | 1.7773 |
| DBNDD1 | 4 | 0.029171 | 0.07984 | 0.93914 | 1967 | 2 | -3.8455 | 0.12545 | 0.27302 | 0.978454 | 6449 | 1 | -3.8455 |
| hsa-mir-7114 | 2 | 0.87452 | 0.87459 | 0.999994 | 18720 | 0 | 2.283 | 0.12548 | 0.20903 | 0.97466 | 6450 | 2 | 2.283 |
| NonTargetingControlG | 2 | 0.87449 | 0.87456 | 0.999994 | 18719 | 0 | 4.5949 | 0.12551 | 0.20908 | 0.97466 | 6451 | 2 | 4.5949 |
| C10orf10 | 5 | 0.51363 | 0.65977 | 0.999994 | 13635 | 1 | 3.2785 | 0.12559 | 0.30323 | 0.98595 | 6452 | 3 | 3.2785 |
| AMPD2 | 5 | 0.78246 | 0.80885 | 0.999994 | 17245 | 1 | 3.2785 | 0.12559 | 0.30323 | 0.98595 | 6453 | 4 | 3.2785 |
| RABL2B | 5 | 0.82987 | 0.8394 | 0.999994 | 17927 | 1 | 3.1708 | 0.12559 | 0.30323 | 0.98595 | 6454 | 3 | 3.1708 |
| PHF2 | 5 | 0.31588 | 0.49401 | 0.999994 | 10437 | 2 | 3.2785 | 0.12559 | 0.30323 | 0.98595 | 6455 | 3 | 3.2785 |
| LAMC3 | 5 | 0.40177 | 0.56497 | 0.999994 | 11975 | 2 | 3.2785 | 0.12559 | 0.30323 | 0.98595 | 6456 | 3 | 3.2785 |
| TAS2R41 | 6 | 0.0073599 | 0.029678 | 0.862223 | 683 | 2 | 1.9832 | 0.12562 | 0.3206 | 0.989846 | 6457 | 3 | 1.9832 |
| PEX7 | 6 | 0.0018023 | 0.0089781 | 0.680529 | 252 | 4 | -3.5119 | 0.12562 | 0.3206 | 0.989846 | 6458 | 2 | -3.5119 |
| MAFK | 6 | 0.4082 | 0.60771 | 0.999994 | 12082 | 2 | 1.8227 | 0.12562 | 0.3206 | 0.989846 | 6459 | 3 | 1.8227 |
| PFAS | 6 | 0.40839 | 0.60785 | 0.999994 | 12086 | 2 | 2.7962 | 0.12562 | 0.3206 | 0.989846 | 6460 | 4 | 2.7962 |
| C2orf69 | 6 | 0.057545 | 0.16045 | 0.967126 | 3316 | 3 | 0.078521 | 0.12562 | 0.3206 | 0.989846 | 6461 | 3 | 0.078521 |
| TPM1 | 6 | 0.094391 | 0.23631 | 0.985449 | 4764 | 4 | -1.0882 | 0.12567 | 0.32067 | 0.989846 | 6462 | 2 | -1.0882 |
| hsa-mir-3684 | 4 | 0.71786 | 0.73656 | 0.999994 | 16475 | 1 | 0.28841 | 0.12571 | 0.27339 | 0.978454 | 6463 | 1 | 0.28841 |
| NEURL1B | 6 | 0.010394 | 0.040366 | 0.891621 | 895 | 3 | 1.1182 | 0.12572 | 0.32075 | 0.989846 | 6464 | 3 | 1.1182 |
| HSD17B6 | 5 | 0.91954 | 0.91971 | 0.999994 | 19620 | 0 | 1.0768 | 0.12591 | 0.30384 | 0.986296 | 6465 | 4 | 1.0768 |
| OR7G1 | 6 | 0.048509 | 0.14084 | 0.959146 | 2909 | 4 | -1.2733 | 0.12592 | 0.32106 | 0.989846 | 6466 | 1 | -1.2733 |
| CIAO1 | 3 | 0.24956 | 0.37442 | 0.999994 | 9007 | 1 | 4.3741 | 0.12593 | 0.26148 | 0.974875 | 6467 | 2 | 4.3741 |
| NRF1 | 3 | 0.77287 | 0.77286 | 0.999994 | 17120 | 0 | 4.3741 | 0.12593 | 0.26148 | 0.974875 | 6468 | 2 | 4.3741 |
| CYP3A7-CYP3AP1 | 3 | 0.81699 | 0.81681 | 0.999994 | 17727 | 0 | 4.3741 | 0.12593 | 0.26148 | 0.974875 | 6469 | 2 | 4.3741 |
| PEX10 | 6 | 0.85007 | 0.87197 | 0.999994 | 18263 | 1 | 2.0078 | 0.12598 | 0.32115 | 0.989846 | 6470 | 4 | 2.0078 |
| ADAMTS17 | 5 | 0.53259 | 0.67633 | 0.999994 | 13944 | 2 | 1.9766 | 0.12604 | 0.30407 | 0.986296 | 6471 | 3 | 1.9766 |
| FOXP2 | 4 | 0.22744 | 0.36888 | 0.999994 | 8466 | 2 | 2.5855 | 0.12609 | 0.27396 | 0.978454 | 6472 | 2 | 2.5855 |
| RSPO4 | 4 | 0.0013456 | 0.0049276 | 0.589484 | 201 | 2 | -1.9971 | 0.12609 | 0.27396 | 0.978454 | 6473 | 2 | -1.9971 |
| hsa-mir-570 | 4 | 0.36338 | 0.50797 | 0.999994 | 11395 | 1 | 3.2302 | 0.12609 | 0.27396 | 0.978454 | 6474 | 2 | 3.2302 |
| hsa-mir-6773 | 4 | 0.85428 | 0.85386 | 0.999994 | 18337 | 0 | 3.6588 | 0.12609 | 0.27396 | 0.978454 | 6475 | 3 | 3.6588 |
| CCDC159 | 4 | 0.62433 | 0.67878 | 0.999994 | 15295 | 1 | -0.42174 | 0.12612 | 0.27402 | 0.978454 | 6476 | 1 | -0.42174 |
| hsa-mir-409 | 4 | 0.48971 | 0.61175 | 0.999994 | 13274 | 1 | 2.824 | 0.12613 | 0.27403 | 0.978454 | 6477 | 3 | 2.824 |
| LPPR5 | 5 | 0.6992 | 0.76611 | 0.999994 | 16298 | 1 | 1.1286 | 0.12615 | 0.30428 | 0.986296 | 6478 | 4 | 1.1286 |
| TCN1 | 6 | 0.24141 | 0.43767 | 0.999994 | 8811 | 2 | 1.2509 | 0.12617 | 0.32145 | 0.989846 | 6479 | 3 | 1.2509 |
| CXCL6 | 5 | 0.42194 | 0.58171 | 0.999994 | 12279 | 1 | 3.2838 | 0.12617 | 0.30433 | 0.986296 | 6480 | 4 | 3.2838 |
| RGS9BP | 5 | 0.59613 | 0.72472 | 0.999994 | 14903 | 1 | 1.4967 | 0.12618 | 0.30435 | 0.986296 | 6481 | 4 | 1.4967 |
| LARP6 | 5 | 0.33967 | 0.51344 | 0.999994 | 10924 | 2 | 1.1059 | 0.12621 | 0.30439 | 0.986296 | 6482 | 3 | 1.1059 |
| LYVE1 | 6 | 0.10121 | 0.24968 | 0.991904 | 5016 | 2 | 2.9645 | 0.12622 | 0.32152 | 0.989846 | 6483 | 4 | 2.9645 |
| TTK | 6 | 0.075488 | 0.19817 | 0.977164 | 4054 | 2 | 1.2767 | 0.12622 | 0.32152 | 0.989846 | 6484 | 4 | 1.2767 |
| NonTargetingControlG | 1 | 0.87377 | 0.87403 | 0.999994 | 18706 | 0 | 6.6476 | 0.12623 | 0.12613 | 0.964228 | 6485 | 1 | 6.6476 |
| MRPL47 | 5 | 0.33618 | 0.51061 | 0.999994 | 10852 | 1 | 1.0286 | 0.12625 | 0.30447 | 0.986296 | 6486 | 4 | 1.0286 |
| OPLAH | 6 | 0.62857 | 0.76127 | 0.999994 | 15356 | 1 | 2.4763 | 0.12634 | 0.3217 | 0.989846 | 6487 | 3 | 2.4763 |
| DLX6 | 6 | 0.61918 | 0.75421 | 0.999994 | 15209 | 2 | 2.2377 | 0.12634 | 0.3217 | 0.989846 | 6488 | 3 | 2.2377 |
| MITF | 6 | 0.39477 | 0.59899 | 0.999994 | 11869 | 1 | 2.3523 | 0.12634 | 0.3217 | 0.989846 | 6489 | 3 | 2.3523 |
| KLC1 | 6 | 0.0075285 | 0.030301 | 0.862223 | 694 | 2 | 1.9321 | 0.12634 | 0.3217 | 0.989846 | 6490 | 3 | 1.9321 |
| SLC26A10 | 6 | 0.48017 | 0.65532 | 0.999994 | 13145 | 2 | 1.4715 | 0.12634 | 0.3217 | 0.989846 | 6491 | 3 | 1.4715 |
| CREM | 6 | 0.24129 | 0.43753 | 0.999994 | 8807 | 3 | 1.7055 | 0.12634 | 0.3217 | 0.989846 | 6492 | 3 | 1.7055 |
| RNF39 | 6 | 0.21964 | 0.41166 | 0.999994 | 8284 | 2 | 2.585 | 0.12634 | 0.3217 | 0.989846 | 6493 | 4 | 2.585 |
| PPP1R3G | 6 | 0.5439 | 0.6993 | 0.999994 | 14129 | 2 | 2.7507 | 0.12634 | 0.3217 | 0.989846 | 6494 | 4 | 2.7507 |
| COQ7 | 2 | 0.047342 | 0.084706 | 0.94395 | 2871 | 1 | -0.7608 | 0.12634 | 0.21038 | 0.97466 | 6495 | 1 | -0.7608 |
| TOR1AIP2 | 6 | 0.099887 | 0.24712 | 0.99045 | 4972 | 2 | 2.1074 | 0.12642 | 0.32183 | 0.989846 | 6496 | 4 | 2.1074 |
| METTL21B | 5 | 0.43185 | 0.58997 | 0.999994 | 12443 | 1 | 3.1338 | 0.12644 | 0.30481 | 0.986316 | 6497 | 3 | 3.1338 |
| IL17REL | 5 | 0.96706 | 0.96724 | 0.999994 | 20762 | 0 | 1.4568 | 0.12645 | 0.30482 | 0.986316 | 6498 | 4 | 1.4568 |
| HAO2 | 6 | 0.071158 | 0.18922 | 0.970085 | 3881 | 3 | -0.805 | 0.12645 | 0.32188 | 0.989846 | 6499 | 3 | -0.805 |

| | | | | | | | | | | | | |
|---|---|---|---|---|---|---|---|---|---|---|---|---|
| CD320 | 6 | 0.68211 | 0.80267 | 0.999994 | 16117 | 2 | 0.41992 | 0.12645 | 0.32188 | 0.989846 | 6500 | 2 | 0.41992 |
| LUZP2 | 5 | 0.45957 | 0.61329 | 0.999994 | 12841 | 1 | 0.97895 | 0.12649 | 0.30491 | 0.986316 | 6501 | 4 | 0.97895 |
| DHRS12 | 5 | 0.4483 | 0.6038 | 0.999994 | 12692 | 2 | 3.0142 | 0.12649 | 0.30491 | 0.986316 | 6502 | 3 | 3.0142 |
| DOCK10 | 5 | 0.7133 | 0.77267 | 0.999994 | 16433 | 1 | 1.3472 | 0.12657 | 0.30504 | 0.986316 | 6503 | 4 | 1.3472 |
| C5orf42 | 4 | 0.90672 | 0.90645 | 0.999994 | 19365 | 0 | 2.443 | 0.12658 | 0.2747 | 0.978454 | 6504 | 3 | 2.443 |
| NonTargetingControlG | 1 | 0.87342 | 0.87368 | 0.999994 | 18703 | 0 | 6.6354 | 0.12658 | 0.1265 | 0.964228 | 6505 | 1 | 6.6354 |
| TPTE2 | 5 | 0.20524 | 0.38268 | 0.999994 | 7902 | 2 | 0.86317 | 0.12664 | 0.30517 | 0.986316 | 6506 | 3 | 0.86317 |
| POLD1 | 5 | 0.19423 | 0.36698 | 0.999994 | 7611 | 1 | 1.082 | 0.12664 | 0.30518 | 0.986316 | 6507 | 4 | 1.082 |
| E2F6 | 6 | 0.083845 | 0.21518 | 0.979587 | 4371 | 3 | -0.40351 | 0.1267 | 0.32228 | 0.989846 | 6508 | 2 | -0.40351 |
| PFDN5 | 6 | 0.34969 | 0.56364 | 0.999994 | 11147 | 3 | 0.34076 | 0.12674 | 0.32232 | 0.989846 | 6509 | 3 | 0.34076 |
| A3GALT2 | 5 | 0.46807 | 0.62047 | 0.999994 | 12975 | 2 | 3.0559 | 0.1268 | 0.30547 | 0.986322 | 6510 | 3 | 3.0559 |
| SMN2 | 4 | 0.11166 | 0.23581 | 0.985362 | 5356 | 1 | 3.8578 | 0.12683 | 0.27508 | 0.978454 | 6511 | 3 | 3.8578 |
| CCL3 | 6 | 0.054345 | 0.1535 | 0.967126 | 3169 | 2 | -0.14309 | 0.12683 | 0.32246 | 0.989846 | 6512 | 2 | -0.14309 |
| NonTargetingControlG | 2 | 0.13755 | 0.21801 | 0.980699 | 6121 | 1 | 0.2361 | 0.12685 | 0.21113 | 0.97466 | 6513 | 1 | 0.2361 |
| NAA38 | 6 | 0.021635 | 0.077237 | 0.93914 | 1554 | 3 | 0.48978 | 0.12686 | 0.32249 | 0.989846 | 6514 | 3 | 0.48978 |
| SFT2D1 | 6 | 0.17168 | 0.35282 | 0.999994 | 7029 | 3 | 0.73328 | 0.12695 | 0.32265 | 0.989846 | 6515 | 3 | 0.73328 |
| RNGTT | 5 | 0.18369 | 0.35193 | 0.999994 | 7306 | 2 | -0.013224 | 0.12697 | 0.30578 | 0.986404 | 6516 | 2 | -0.013224 |
| DUOXA2 | 6 | 0.753 | 0.82954 | 0.999994 | 16887 | 1 | 1.7892 | 0.12698 | 0.32268 | 0.989846 | 6517 | 4 | 1.7892 |
| ERLIN1 | 5 | 0.44495 | 0.60097 | 0.999994 | 12646 | 1 | 1.3346 | 0.12705 | 0.30592 | 0.986404 | 6518 | 4 | 1.3346 |
| hsa-mir-378c | 4 | 0.91096 | 0.91073 | 0.999994 | 19447 | 0 | 3.107 | 0.12706 | 0.27543 | 0.978454 | 6519 | 3 | 3.107 |
| FAM129B | 4 | 0.022864 | 0.066669 | 0.93914 | 1640 | 1 | 2.3493 | 0.12706 | 0.27543 | 0.978454 | 6520 | 3 | 2.3493 |
| AGAP3 | 3 | 0.030751 | 0.076893 | 0.93914 | 2047 | 2 | -1.5626 | 0.12706 | 0.2627 | 0.974875 | 6521 | 1 | -1.5626 |
| hsa-mir-1257 | 4 | 0.85113 | 0.85065 | 0.999994 | 18282 | 0 | 2.2119 | 0.12709 | 0.27548 | 0.978454 | 6522 | 3 | 2.2119 |
| VCX3A | 2 | 0.79456 | 0.79414 | 0.999994 | 17404 | 0 | 4.3131 | 0.12712 | 0.21156 | 0.97466 | 6523 | 2 | 4.3131 |
| TMEM88B | 4 | 0.78033 | 0.78416 | 0.999994 | 17216 | 1 | 1.9866 | 0.12713 | 0.27555 | 0.978454 | 6524 | 3 | 1.9866 |
| GNB3 | 6 | 0.053082 | 0.15073 | 0.966359 | 3117 | 3 | 0.50673 | 0.1273 | 0.32316 | 0.989881 | 6525 | 3 | 0.50673 |
| VLDLR | 5 | 0.33197 | 0.50714 | 0.999994 | 10766 | 2 | -0.12183 | 0.12741 | 0.3066 | 0.987798 | 6526 | 2 | -0.12183 |
| PDE8B | 6 | 0.022377 | 0.079594 | 0.93914 | 1606 | 3 | 0.14735 | 0.12743 | 0.32336 | 0.989881 | 6527 | 3 | 0.14735 |
| OR13H1 | 6 | 0.37948 | 0.58925 | 0.999994 | 11641 | 2 | 1.0418 | 0.12743 | 0.32336 | 0.989881 | 6528 | 3 | 1.0418 |
| GAK | 4 | 0.34893 | 0.49318 | 0.999994 | 11129 | 1 | 3.4781 | 0.12749 | 0.27613 | 0.978454 | 6529 | 3 | 3.4781 |
| CYTH1 | 4 | 0.49683 | 0.6149 | 0.999994 | 13372 | 1 | 3.2826 | 0.12749 | 0.27613 | 0.978454 | 6530 | 2 | 3.2826 |
| R3HDM4 | 4 | 0.23612 | 0.37776 | 0.999994 | 8675 | 2 | 2.3102 | 0.12749 | 0.27613 | 0.978454 | 6531 | 2 | 2.3102 |
| KIAA0319 | 4 | 0.58216 | 0.65588 | 0.999994 | 14708 | 1 | 3.3189 | 0.12749 | 0.27613 | 0.978454 | 6532 | 3 | 3.3189 |
| hsa-mir-566 | 1 | 0.87249 | 0.87275 | 0.999994 | 18685 | 0 | 6.6189 | 0.12751 | 0.12738 | 0.964228 | 6533 | 1 | 6.6189 |
| hsa-mir-340 | 3 | 0.77977 | 0.77969 | 0.999994 | 17210 | 0 | 4.3222 | 0.12756 | 0.26325 | 0.974875 | 6534 | 2 | 4.3222 |
| NET1 | 5 | 0.93954 | 0.93973 | 0.999994 | 20064 | 0 | 1.4383 | 0.12757 | 0.30688 | 0.987798 | 6535 | 4 | 1.4383 |
| hsa-mir-4640 | 3 | 0.7578 | 0.7578 | 0.999994 | 16950 | 0 | -0.37391 | 0.1276 | 0.26331 | 0.974875 | 6536 | 1 | -0.37391 |
| NonTargetingControlG | 2 | 0.19877 | 0.26119 | 0.994943 | 7728 | 1 | 0.92908 | 0.12763 | 0.21232 | 0.97466 | 6537 | 1 | 0.92908 |
| NPHP4 | 6 | 0.029969 | 0.098712 | 0.945253 | 2006 | 2 | 0.13267 | 0.12769 | 0.32377 | 0.99028 | 6538 | 2 | 0.13267 |
| FBXO45 | 4 | 0.55867 | 0.64396 | 0.999994 | 14344 | 1 | 2.8207 | 0.12777 | 0.27656 | 0.978454 | 6539 | 3 | 2.8207 |
| CRIPT | 5 | 0.34421 | 0.51717 | 0.999994 | 11032 | 1 | 2.5798 | 0.12783 | 0.30734 | 0.988343 | 6540 | 4 | 2.5798 |
| PTPRE | 5 | 0.53519 | 0.67861 | 0.999994 | 13984 | 2 | 1.5651 | 0.12787 | 0.30742 | 0.988455 | 6541 | 3 | 1.5651 |
| TTC23 | 6 | 0.56058 | 0.7112 | 0.999994 | 14378 | 1 | 1.9838 | 0.12798 | 0.32421 | 0.990429 | 6542 | 3 | 1.9838 |
| MMP23B | 6 | 0.12253 | 0.29049 | 0.999632 | 5722 | 3 | -0.42609 | 0.12798 | 0.32421 | 0.990429 | 6543 | 2 | -0.42609 |
| RAB15 | 5 | 0.42789 | 0.58668 | 0.999994 | 12381 | 1 | 1.6842 | 0.12799 | 0.30763 | 0.988566 | 6544 | 4 | 1.6842 |
| ERBB2 | 5 | 0.27023 | 0.45702 | 0.999994 | 9480 | 2 | 0.94079 | 0.128 | 0.30767 | 0.988566 | 6545 | 3 | 0.94079 |
| ZNF737 | 5 | 0.82192 | 0.83386 | 0.999994 | 17800 | 1 | 0.99817 | 0.12801 | 0.30769 | 0.988566 | 6546 | 4 | 0.99817 |
| KLF14 | 5 | 0.96566 | 0.96581 | 0.999994 | 20730 | 0 | 1.8476 | 0.12802 | 0.3077 | 0.988566 | 6547 | 4 | 1.8476 |
| hsa-mir-4524a | 4 | 0.78451 | 0.78766 | 0.999994 | 17274 | 1 | 3.138 | 0.1281 | 0.2771 | 0.978454 | 6548 | 3 | 3.138 |
| SOX30 | 6 | 0.34731 | 0.56099 | 0.999994 | 11096 | 3 | 1.6439 | 0.1281 | 0.3244 | 0.990429 | 6549 | 3 | 1.6439 |
| LCE5A | 6 | 0.92467 | 0.9248 | 0.999994 | 19735 | 1 | 2.2408 | 0.12817 | 0.3245 | 0.990429 | 6550 | 3 | 2.2408 |
| CCL4L2 | 2 | 0.33499 | 0.36402 | 0.999994 | 10832 | 1 | 2.7465 | 0.12821 | 0.2132 | 0.97466 | 6551 | 1 | 2.7465 |

| | | | | | | | | | | | | | |
|---|---|---|---|---|---|---|---|---|---|---|---|---|---|
| ATPIF1 | 6 | 0.30933 | 0.5176 | 0.999994 | 10311 | 2 | 0.27879 | 0.12824 | 0.32461 | 0.990429 | 6552 | 3 | 0.27879 |
| SUPT4H1 | 4 | 0.16644 | 0.30611 | 0.999994 | 6896 | 2 | 1.9122 | 0.12825 | 0.27735 | 0.978454 | 6553 | 2 | 1.9122 |
| BAZ1A | 6 | 0.61812 | 0.75341 | 0.999994 | 15190 | 2 | 2.6036 | 0.12826 | 0.32464 | 0.990429 | 6554 | 4 | 2.6036 |
| KDM4C | 6 | 0.49537 | 0.66553 | 0.999994 | 13345 | 2 | 1.8278 | 0.12829 | 0.32469 | 0.990429 | 6555 | 4 | 1.8278 |
| TPPP3 | 5 | 0.38121 | 0.54771 | 0.999994 | 11664 | 2 | 2.9205 | 0.1283 | 0.30819 | 0.988566 | 6556 | 3 | 2.9205 |
| TMEM185B | 4 | 0.15838 | 0.29784 | 0.999994 | 6682 | 2 | -0.60098 | 0.12832 | 0.27745 | 0.978454 | 6557 | 2 | -0.60098 |
| STX10 | 6 | 0.046294 | 0.13594 | 0.95468 | 2819 | 3 | 0.6255 | 0.12833 | 0.32473 | 0.990429 | 6558 | 3 | 0.6255 |
| ZCCHC4 | 5 | 0.39811 | 0.56199 | 0.999994 | 11917 | 1 | 1.3078 | 0.12843 | 0.30846 | 0.988566 | 6559 | 4 | 1.3078 |
| PRSS42 | 6 | 0.67668 | 0.79839 | 0.999994 | 16036 | 1 | 1.8472 | 0.12844 | 0.3249 | 0.990752 | 6560 | 4 | 1.8472 |
| NonTargetingControlG | 2 | 0.87156 | 0.87159 | 0.999994 | 18671 | 0 | 2.3223 | 0.12844 | 0.21356 | 0.97466 | 6561 | 2 | 2.3223 |
| BRDT | 5 | 0.61773 | 0.73272 | 0.999994 | 15181 | 1 | 1.2159 | 0.12847 | 0.30853 | 0.988566 | 6562 | 4 | 1.2159 |
| NFYC | 3 | 0.67495 | 0.67821 | 0.999994 | 16002 | 1 | 4.314 | 0.12848 | 0.26427 | 0.975922 | 6563 | 2 | 4.314 |
| UBE2O | 5 | 0.86385 | 0.86565 | 0.999994 | 18519 | 1 | 2.7563 | 0.12853 | 0.30862 | 0.988566 | 6564 | 4 | 2.7563 |
| NECAP1 | 6 | 0.24323 | 0.43985 | 0.999994 | 8870 | 2 | 1.5353 | 0.12871 | 0.32532 | 0.991152 | 6565 | 3 | 1.5353 |
| CLIP1 | 6 | 0.64784 | 0.77597 | 0.999994 | 15628 | 1 | 2.9235 | 0.12876 | 0.32539 | 0.991152 | 6566 | 4 | 2.9235 |
| hsa-mir-376a-1 | 4 | 0.005401 | 0.018967 | 0.803792 | 551 | 1 | 2.9947 | 0.12876 | 0.27811 | 0.978454 | 6567 | 3 | 2.9947 |
| CCDC175 | 5 | 0.49826 | 0.64639 | 0.999994 | 13399 | 1 | 1.2697 | 0.12881 | 0.30914 | 0.988566 | 6568 | 4 | 1.2697 |
| NAV1 | 5 | 0.049384 | 0.13886 | 0.957514 | 2947 | 2 | 0.002332 | 0.12881 | 0.30914 | 0.988566 | 6569 | 2 | 0.002332 |
| C1orf228 | 5 | 0.19356 | 0.36603 | 0.999994 | 7587 | 3 | -0.86177 | 0.12881 | 0.30914 | 0.988566 | 6570 | 2 | -0.86177 |
| PRM1 | 5 | 0.95489 | 0.95505 | 0.999994 | 20449 | 0 | 1.049 | 0.12883 | 0.30919 | 0.988566 | 6571 | 4 | 1.049 |
| NonTargetingControlG | 1 | 0.87116 | 0.8714 | 0.999994 | 18662 | 0 | 6.5883 | 0.12884 | 0.12871 | 0.964228 | 6572 | 1 | 6.5883 |
| TGFA | 5 | 0.81246 | 0.82756 | 0.999994 | 17659 | 1 | 1.8633 | 0.12885 | 0.30922 | 0.988566 | 6573 | 4 | 1.8633 |
| MTHFS | 5 | 0.46944 | 0.62167 | 0.999994 | 12998 | 1 | 1.7723 | 0.12888 | 0.30927 | 0.988566 | 6574 | 4 | 1.7723 |
| PRSS37 | 6 | 0.48372 | 0.65769 | 0.999994 | 13196 | 2 | 2.04 | 0.1289 | 0.32563 | 0.991485 | 6575 | 4 | 2.04 |
| POU3F1 | 5 | 0.34961 | 0.52157 | 0.999994 | 11141 | 1 | 3.0437 | 0.129 | 0.30949 | 0.988566 | 6576 | 3 | 3.0437 |
| ISCA2 | 4 | 0.75506 | 0.76397 | 0.999994 | 16914 | 1 | 2.9085 | 0.12902 | 0.2785 | 0.978454 | 6577 | 2 | 2.9085 |
| DCDC1 | 6 | 0.60827 | 0.74606 | 0.999994 | 15048 | 2 | 1.8283 | 0.12905 | 0.32589 | 0.991729 | 6578 | 4 | 1.8283 |
| TECTA | 5 | 0.8423 | 0.84841 | 0.999994 | 18130 | 1 | 3.1559 | 0.12908 | 0.30965 | 0.988566 | 6579 | 3 | 3.1559 |
| TIGD3 | 5 | 0.17442 | 0.33857 | 0.999994 | 7096 | 2 | -0.46677 | 0.12912 | 0.30973 | 0.988566 | 6580 | 2 | -0.46677 |
| ISL2 | 5 | 0.29624 | 0.47801 | 0.999994 | 10042 | 1 | 2.9846 | 0.12913 | 0.30975 | 0.988566 | 6581 | 3 | 2.9846 |
| CLK3 | 6 | 0.96693 | 0.96698 | 0.999994 | 20759 | 0 | 0.97077 | 0.1292 | 0.32612 | 0.991729 | 6582 | 3 | 0.97077 |
| PIM1 | 6 | 0.13829 | 0.31087 | 0.999994 | 6147 | 3 | -1.0421 | 0.1292 | 0.32612 | 0.991729 | 6583 | 2 | -1.0421 |
| STX1A | 5 | 0.38167 | 0.5481 | 0.999994 | 11670 | 1 | 1.0109 | 0.12925 | 0.30998 | 0.988566 | 6584 | 4 | 1.0109 |
| hsa-mir-4698 | 4 | 0.21896 | 0.36021 | 0.999994 | 8259 | 1 | 2.066 | 0.12929 | 0.27889 | 0.978454 | 6585 | 3 | 2.066 |
| SREBF2 | 5 | 0.29624 | 0.47801 | 0.999994 | 10043 | 2 | 2.8524 | 0.1293 | 0.31007 | 0.988566 | 6586 | 3 | 2.8524 |
| ZNF600 | 5 | 0.48907 | 0.6384 | 0.999994 | 13268 | 1 | 0.82963 | 0.1293 | 0.31007 | 0.988566 | 6587 | 4 | 0.82963 |
| TRIM3 | 5 | 0.091233 | 0.21257 | 0.979587 | 4645 | 2 | 2.9128 | 0.12931 | 0.31008 | 0.988566 | 6588 | 3 | 2.9128 |
| DOCK9 | 6 | 0.075686 | 0.19858 | 0.977164 | 4061 | 4 | -1.0912 | 0.1294 | 0.32645 | 0.991729 | 6589 | 2 | -1.0912 |
| MYEF2 | 5 | 0.1955 | 0.3688 | 0.999994 | 7650 | 1 | 2.1027 | 0.12949 | 0.3104 | 0.988566 | 6590 | 4 | 2.1027 |
| MTAP | 5 | 0.54139 | 0.68414 | 0.999994 | 14077 | 1 | 0.20437 | 0.12969 | 0.31078 | 0.988566 | 6591 | 2 | 0.20437 |
| GGACT | 5 | 0.32115 | 0.49824 | 0.999994 | 10539 | 2 | -0.42307 | 0.12969 | 0.31078 | 0.988566 | 6592 | 1 | -0.42307 |
| LRRTM4 | 6 | 0.00854 | 0.033845 | 0.867348 | 765 | 2 | 0.18055 | 0.1297 | 0.32692 | 0.991729 | 6593 | 3 | 0.18055 |
| EHD3 | 5 | 0.56207 | 0.70248 | 0.999994 | 14408 | 2 | 2.7953 | 0.12977 | 0.31091 | 0.988566 | 6594 | 3 | 2.7953 |
| NEDD8-MDP1 | 2 | 0.43454 | 0.44594 | 0.999994 | 12485 | 1 | 3.1179 | 0.1298 | 0.21566 | 0.97466 | 6595 | 1 | 3.1179 |
| NonTargetingControlG | 2 | 0.87017 | 0.87016 | 0.999994 | 18638 | 0 | 4.6516 | 0.12983 | 0.21569 | 0.97466 | 6596 | 2 | 4.6516 |
| MMP11 | 5 | 0.34625 | 0.51884 | 0.999994 | 11079 | 2 | 2.5272 | 0.12984 | 0.31106 | 0.988566 | 6597 | 3 | 2.5272 |
| ARMS2 | 5 | 0.0096504 | 0.035068 | 0.867348 | 851 | 2 | 1.0108 | 0.12984 | 0.31106 | 0.988566 | 6598 | 3 | 1.0108 |
| hsa-mir-593 | 4 | 0.6732 | 0.70734 | 0.999994 | 15975 | 1 | 1.9923 | 0.12987 | 0.27978 | 0.978454 | 6599 | 3 | 1.9923 |
| FAAH2 | 6 | 0.64657 | 0.775 | 0.999994 | 15611 | 2 | 2.7612 | 0.12987 | 0.32716 | 0.991729 | 6600 | 4 | 2.7612 |
| TNFSF13 | 6 | 0.97625 | 0.97619 | 0.999994 | 20998 | 0 | 2.5106 | 0.12987 | 0.32716 | 0.991729 | 6601 | 4 | 2.5106 |
| SEC24C | 6 | 0.2199 | 0.41198 | 0.999994 | 8287 | 3 | 1.6689 | 0.12987 | 0.32716 | 0.991729 | 6602 | 3 | 1.6689 |
| AKT1S1 | 6 | 0.0037303 | 0.016472 | 0.772095 | 424 | 2 | 2.7563 | 0.12987 | 0.32716 | 0.991729 | 6603 | 4 | 2.7563 |

| | | | | | | | | | | | | |
|---|---|---|---|---|---|---|---|---|---|---|---|---|
| C1QB | 6 | 0.35672 | 0.57159 | 0.999994 | 11282 | 3 | 1.8367 | 0.12987 | 0.32716 | 0.991729 | 6604 | 3 | 1.8367 |
| ADAMTS3 | 6 | 0.11982 | 0.28537 | 0.999632 | 5620 | 3 | 1.2188 | 0.12987 | 0.32716 | 0.991729 | 6605 | 3 | 1.2188 |
| XAGE3 | 5 | 0.21228 | 0.39259 | 0.999994 | 8068 | 1 | 2.9309 | 0.12992 | 0.31121 | 0.988566 | 6606 | 3 | 2.9309 |
| C2orf66 | 6 | 0.39327 | 0.59801 | 0.999994 | 11849 | 2 | 0.17393 | 0.13001 | 0.32739 | 0.991729 | 6607 | 2 | 0.17393 |
| OR8K5 | 6 | 0.18295 | 0.36682 | 0.999994 | 7288 | 3 | -0.062193 | 0.13001 | 0.32739 | 0.991729 | 6608 | 3 | -0.062193 |
| BRAT1 | 5 | 0.038705 | 0.1126 | 0.951052 | 2449 | 3 | -1.9993 | 0.13001 | 0.31137 | 0.988566 | 6609 | 2 | -1.9993 |
| CCDC28A | 4 | 0.13785 | 0.27669 | 0.999379 | 6131 | 1 | 2.0367 | 0.13001 | 0.28001 | 0.978454 | 6610 | 3 | 2.0367 |
| BRF1 | 3 | 0.7397 | 0.73962 | 0.999994 | 16735 | 0 | 4.2513 | 0.13008 | 0.26611 | 0.978454 | 6611 | 2 | 4.2513 |
| NonTargetingControlG | 2 | 0.86986 | 0.86986 | 0.999994 | 18632 | 0 | 3.0736 | 0.13014 | 0.21618 | 0.97466 | 6612 | 2 | 3.0736 |
| IDO1 | 6 | 0.91895 | 0.91942 | 0.999994 | 19606 | 1 | 2.7472 | 0.13015 | 0.32762 | 0.991729 | 6613 | 4 | 2.7472 |
| PCDH20 | 6 | 0.37736 | 0.58789 | 0.999994 | 11611 | 2 | 2.7491 | 0.13018 | 0.32766 | 0.991729 | 6614 | 4 | 2.7491 |
| PANX2 | 5 | 0.29246 | 0.47493 | 0.999994 | 9962 | 1 | 0.97498 | 0.13019 | 0.3117 | 0.988566 | 6615 | 4 | 0.97498 |
| MDH1B | 6 | 0.17299 | 0.35443 | 0.999994 | 7065 | 2 | 0.28593 | 0.13026 | 0.32779 | 0.991729 | 6616 | 2 | 0.28593 |
| DNMT3L | 6 | 0.012757 | 0.04829 | 0.90085 | 1050 | 5 | -1.063 | 0.13026 | 0.32779 | 0.991729 | 6617 | 1 | -1.063 |
| TMEM89 | 5 | 0.07201 | 0.18162 | 0.968171 | 3918 | 1 | 0.95409 | 0.13027 | 0.31182 | 0.988735 | 6618 | 4 | 0.95409 |
| SPINT4 | 5 | 0.075181 | 0.18681 | 0.969409 | 4042 | 2 | 2.8561 | 0.13031 | 0.31191 | 0.988735 | 6619 | 3 | 2.8561 |
| hsa-mir-498 | 4 | 0.90296 | 0.90266 | 0.999994 | 19275 | 0 | 0.39252 | 0.13034 | 0.28051 | 0.978454 | 6620 | 1 | 0.39252 |
| TAF6 | 4 | 0.078777 | 0.17648 | 0.968171 | 4180 | 1 | 1.0499 | 0.13034 | 0.28051 | 0.978454 | 6621 | 3 | 1.0499 |
| ATP2B4 | 4 | 0.805 | 0.80573 | 0.999994 | 17552 | 1 | 1.8934 | 0.13036 | 0.28053 | 0.978454 | 6622 | 3 | 1.8934 |
| PLA2G7 | 6 | 0.13406 | 0.30547 | 0.999994 | 6023 | 2 | 0.54928 | 0.13039 | 0.328 | 0.991729 | 6623 | 2 | 0.54928 |
| PCDHGB5 | 2 | 0.14932 | 0.22618 | 0.983354 | 6438 | 1 | 1.2437 | 0.13039 | 0.21656 | 0.97466 | 6624 | 1 | 1.2437 |
| RGS14 | 5 | 0.24261 | 0.43447 | 0.999994 | 8849 | 1 | 1.0598 | 0.13041 | 0.3121 | 0.988962 | 6625 | 3 | 1.0598 |
| DAAM2 | 6 | 0.27792 | 0.48099 | 0.999994 | 9650 | 1 | 2.0052 | 0.13044 | 0.32808 | 0.991729 | 6626 | 4 | 2.0052 |
| CHTF18 | 5 | 0.20271 | 0.37908 | 0.999994 | 7850 | 3 | -0.79027 | 0.13051 | 0.3123 | 0.989294 | 6627 | 1 | -0.79027 |
| FXYD3 | 4 | 0.69536 | 0.72139 | 0.999994 | 16261 | 1 | 1.6467 | 0.13065 | 0.28096 | 0.978454 | 6628 | 3 | 1.6467 |
| SRPR | 5 | 0.052728 | 0.14695 | 0.964901 | 3100 | 2 | 3.0795 | 0.13066 | 0.31259 | 0.989342 | 6629 | 3 | 3.0795 |
| ACOT2 | 5 | 0.75864 | 0.79544 | 0.999994 | 16960 | 1 | 0.6589 | 0.13068 | 0.31261 | 0.989342 | 6630 | 3 | 0.6589 |
| PSMB11 | 6 | 0.14739 | 0.32236 | 0.999994 | 6387 | 2 | 1.8796 | 0.13068 | 0.32848 | 0.991729 | 6631 | 4 | 1.8796 |
| C2CD4C | 5 | 0.87408 | 0.87477 | 0.999994 | 18711 | 1 | 2.0086 | 0.13074 | 0.31273 | 0.989342 | 6632 | 4 | 2.0086 |
| GNG8 | 5 | 0.43483 | 0.59249 | 0.999994 | 12491 | 2 | 1.0945 | 0.13075 | 0.31274 | 0.989342 | 6633 | 3 | 1.0945 |
| NPLOC4 | 4 | 0.14127 | 0.28021 | 0.999632 | 6224 | 1 | 1.8903 | 0.13077 | 0.28116 | 0.978454 | 6634 | 3 | 1.8903 |
| CD24 | 4 | 0.8627 | 0.86233 | 0.999994 | 18498 | 0 | 0.66095 | 0.13081 | 0.28123 | 0.978454 | 6635 | 3 | 0.66095 |
| SLC51A | 5 | 0.55576 | 0.69688 | 0.999994 | 14301 | 1 | 2.9212 | 0.13086 | 0.31296 | 0.989623 | 6636 | 3 | 2.9212 |
| SLC30A2 | 6 | 0.79105 | 0.84411 | 0.999994 | 17363 | 1 | 1.2464 | 0.13087 | 0.32878 | 0.991729 | 6637 | 3 | 1.2464 |
| CRIM1 | 6 | 0.61165 | 0.74854 | 0.999994 | 15100 | 2 | 2.0241 | 0.13087 | 0.32878 | 0.991729 | 6638 | 4 | 2.0241 |
| TMTC1 | 6 | 0.63387 | 0.76527 | 0.999994 | 15430 | 1 | 1.647 | 0.13087 | 0.32878 | 0.991729 | 6639 | 4 | 1.647 |
| FOXO3 | 4 | 0.79754 | 0.79901 | 0.999994 | 17447 | 1 | 2.9668 | 0.13088 | 0.28134 | 0.978454 | 6640 | 2 | 2.9668 |
| MCM3AP | 4 | 0.19827 | 0.3389 | 0.999994 | 7717 | 2 | 2.4287 | 0.13088 | 0.28134 | 0.978454 | 6641 | 2 | 2.4287 |
| COL22A1 | 6 | 0.64045 | 0.77031 | 0.999994 | 15516 | 2 | -0.21419 | 0.13096 | 0.32892 | 0.991729 | 6642 | 2 | -0.21419 |
| PRSS57 | 6 | 0.17379 | 0.35538 | 0.999994 | 7078 | 3 | -0.62401 | 0.13096 | 0.32892 | 0.991729 | 6643 | 2 | -0.62401 |
| PTPRO | 6 | 0.23141 | 0.42578 | 0.999994 | 8554 | 2 | 1.7238 | 0.13096 | 0.32892 | 0.991729 | 6644 | 4 | 1.7238 |
| OLFML1 | 5 | 0.41039 | 0.57221 | 0.999994 | 12117 | 1 | 1.7192 | 0.13102 | 0.31324 | 0.989846 | 6645 | 3 | 1.7192 |
| NonTargetingControlG | 2 | 0.86893 | 0.86891 | 0.999994 | 18616 | 0 | 4.3105 | 0.13107 | 0.21758 | 0.97466 | 6646 | 2 | 4.3105 |
| HECA | 4 | 0.029776 | 0.081052 | 0.93914 | 1996 | 3 | -1.4776 | 0.1311 | 0.28168 | 0.978454 | 6647 | 1 | -1.4776 |
| GPR139 | 6 | 0.017396 | 0.063532 | 0.937389 | 1331 | 3 | -0.77819 | 0.13111 | 0.32914 | 0.991729 | 6648 | 3 | -0.77819 |
| RPL27A | 6 | 0.080331 | 0.20797 | 0.979108 | 4241 | 2 | 2.5096 | 0.13111 | 0.32914 | 0.991729 | 6649 | 4 | 2.5096 |
| MRGPRX3 | 6 | 0.15817 | 0.33593 | 0.999994 | 6678 | 2 | 2.8367 | 0.13111 | 0.32914 | 0.991729 | 6650 | 4 | 2.8367 |
| HS3ST2 | 6 | 0.0019877 | 0.0099117 | 0.680529 | 275 | 4 | -4.2139 | 0.13129 | 0.32944 | 0.992034 | 6651 | 2 | -4.2139 |
| COMMD9 | 6 | 0.47891 | 0.65444 | 0.999994 | 13126 | 1 | 1.7482 | 0.13129 | 0.32944 | 0.992034 | 6652 | 3 | 1.7482 |
| PALLD | 5 | 0.45709 | 0.61124 | 0.999994 | 12809 | 1 | 0.73055 | 0.13133 | 0.31379 | 0.989846 | 6653 | 3 | 0.73055 |
| KHSRP | 5 | 0.060564 | 0.16302 | 0.967126 | 3447 | 3 | -1.5007 | 0.13134 | 0.3138 | 0.989846 | 6654 | 2 | -1.5007 |
| RERGL | 5 | 0.18978 | 0.36062 | 0.999994 | 7477 | 3 | -0.912 | 0.13134 | 0.3138 | 0.989846 | 6655 | 2 | -0.912 |

| | | | | | | | | | | | | | |
|---|---|---|---|---|---|---|---|---|---|---|---|---|---|
| SPHK1 | 5 | 0.019899 | 0.063819 | 0.937389 | 1453 | 1 | 3.035 | 0.13139 | 0.3139 | 0.989846 | 6656 | 3 | 3.035 |
| EGR4 | 6 | 0.055389 | 0.15578 | 0.967126 | 3215 | 1 | 2.2138 | 0.13142 | 0.32962 | 0.992034 | 6657 | 4 | 2.2138 |
| DNMT1 | 6 | 0.0036684 | 0.016228 | 0.772095 | 416 | 3 | -2.8888 | 0.13145 | 0.32969 | 0.992034 | 6658 | 3 | -2.8888 |
| NOL8 | 4 | 0.03704 | 0.095838 | 0.945031 | 2377 | 2 | -1.1715 | 0.13148 | 0.28226 | 0.978454 | 6659 | 2 | -1.1715 |
| NOMO3 | 3 | 0.69359 | 0.69501 | 0.999994 | 16241 | 1 | 0.98695 | 0.13151 | 0.26768 | 0.978454 | 6660 | 2 | 0.98695 |
| BRMS1 | 6 | 0.123 | 0.29136 | 0.999632 | 5739 | 3 | -0.93477 | 0.13157 | 0.32986 | 0.992034 | 6661 | 2 | -0.93477 |
| LRRC31 | 6 | 0.095998 | 0.23947 | 0.987253 | 4816 | 4 | -1.1481 | 0.13157 | 0.32986 | 0.992034 | 6662 | 2 | -1.1481 |
| MRPL50 | 6 | 0.1976 | 0.38498 | 0.999994 | 7691 | 2 | 1.4394 | 0.13157 | 0.32986 | 0.992034 | 6663 | 4 | 1.4394 |
| CENPN | 4 | 0.79136 | 0.79355 | 0.999994 | 17368 | 1 | 1.7778 | 0.13166 | 0.28258 | 0.978454 | 6664 | 3 | 1.7778 |
| BIRC3 | 5 | 0.95125 | 0.95141 | 0.999994 | 20372 | 0 | 1.2322 | 0.13167 | 0.31439 | 0.989846 | 6665 | 4 | 1.2322 |
| NOG | 5 | 0.57574 | 0.71459 | 0.999994 | 14628 | 1 | 1.339 | 0.13167 | 0.31439 | 0.989846 | 6666 | 4 | 1.339 |
| DHX33 | 5 | 0.38508 | 0.551 | 0.999994 | 11719 | 1 | 1.163 | 0.13167 | 0.31439 | 0.989846 | 6667 | 4 | 1.163 |
| DEFB124 | 5 | 0.85583 | 0.85894 | 0.999994 | 18363 | 1 | 0.41485 | 0.13168 | 0.3144 | 0.989846 | 6668 | 2 | 0.41485 |
| TMEM63C | 5 | 0.51623 | 0.66203 | 0.999994 | 13680 | 2 | 2.7254 | 0.13168 | 0.3144 | 0.989846 | 6669 | 3 | 2.7254 |
| KIF23 | 4 | 0.59832 | 0.66442 | 0.999994 | 14931 | 1 | 2.2172 | 0.13177 | 0.28275 | 0.978454 | 6670 | 3 | 2.2172 |
| PLGRKT | 5 | 0.8489 | 0.85341 | 0.999994 | 18237 | 1 | 2.7707 | 0.13185 | 0.31472 | 0.989846 | 6671 | 3 | 2.7707 |
| hsa-mir-548b | 4 | 0.94003 | 0.93993 | 0.999994 | 20079 | 0 | 2.893 | 0.13188 | 0.28292 | 0.978454 | 6672 | 3 | 2.893 |
| ZNF616 | 6 | 0.97808 | 0.97802 | 0.999994 | 21047 | 0 | 2.4164 | 0.13189 | 0.33037 | 0.992034 | 6673 | 4 | 2.4164 |
| TUBB4A | 6 | 0.38841 | 0.59493 | 0.999994 | 11770 | 2 | -0.070985 | 0.1319 | 0.33038 | 0.992034 | 6674 | 2 | -0.070985 |
| OR8D1 | 6 | 0.0029995 | 0.013736 | 0.751291 | 357 | 4 | -1.0402 | 0.1319 | 0.33038 | 0.992034 | 6675 | 2 | -1.0402 |
| CNTLN | 6 | 0.65998 | 0.78531 | 0.999994 | 15779 | 2 | 2.6556 | 0.13195 | 0.33047 | 0.992034 | 6676 | 4 | 2.6556 |
| MAPKAPK2 | 6 | 0.21579 | 0.4071 | 0.999994 | 8164 | 2 | 2.0746 | 0.13201 | 0.33056 | 0.992034 | 6677 | 4 | 2.0746 |
| YWHAH | 5 | 0.054772 | 0.15193 | 0.967126 | 3185 | 3 | -2.1776 | 0.13204 | 0.31508 | 0.989846 | 6678 | 2 | -2.1776 |
| PKDREJ | 6 | 0.16813 | 0.3484 | 0.999994 | 6942 | 2 | 2.0068 | 0.1321 | 0.33071 | 0.992034 | 6679 | 3 | 2.0068 |
| NDN | 6 | 0.62414 | 0.75793 | 0.999994 | 15292 | 1 | 2.093 | 0.13214 | 0.33077 | 0.992034 | 6680 | 4 | 2.093 |
| KBTBD13 | 5 | 0.46141 | 0.61486 | 0.999994 | 12872 | 1 | 0.17743 | 0.13216 | 0.31529 | 0.989846 | 6681 | 2 | 0.17743 |
| SOX3 | 5 | 0.89471 | 0.89484 | 0.999994 | 19118 | 0 | 0.24307 | 0.13216 | 0.31529 | 0.989846 | 6682 | 2 | 0.24307 |
| NonTargetingControlG | 2 | 0.86782 | 0.86783 | 0.999994 | 18597 | 0 | 4.3301 | 0.13218 | 0.21925 | 0.97466 | 6683 | 2 | 4.3301 |
| INSM1 | 6 | 0.81823 | 0.85598 | 0.999994 | 17751 | 1 | 2.4664 | 0.13219 | 0.33086 | 0.992034 | 6684 | 4 | 2.4664 |
| MED31 | 3 | 0.77081 | 0.77082 | 0.999994 | 17094 | 0 | 3.9543 | 0.13221 | 0.26848 | 0.978454 | 6685 | 2 | 3.9543 |
| DSCC1 | 5 | 0.18986 | 0.36074 | 0.999994 | 7480 | 2 | 1.1216 | 0.13228 | 0.31552 | 0.989846 | 6686 | 3 | 1.1216 |
| FOSB | 6 | 0.20382 | 0.39252 | 0.999994 | 7874 | 3 | 0.82947 | 0.1323 | 0.33102 | 0.992034 | 6687 | 3 | 0.82947 |
| MAP3K12 | 6 | 0.23187 | 0.42634 | 0.999994 | 8565 | 3 | -0.47672 | 0.1323 | 0.33102 | 0.992034 | 6688 | 2 | -0.47672 |
| BICD2 | 5 | 0.83815 | 0.84531 | 0.999994 | 18056 | 1 | 2.8508 | 0.13231 | 0.31559 | 0.989846 | 6689 | 3 | 2.8508 |
| DMXL1 | 5 | 0.94547 | 0.94566 | 0.999994 | 20217 | 0 | 1.5707 | 0.13232 | 0.3156 | 0.989846 | 6690 | 4 | 1.5707 |
| C8orf59 | 2 | 0.86767 | 0.86767 | 0.999994 | 18593 | 0 | 4.4968 | 0.13233 | 0.21946 | 0.97466 | 6691 | 2 | 4.4968 |
| KRTAP4-1 | 6 | 0.87114 | 0.88404 | 0.999994 | 18660 | 1 | 2.0173 | 0.13237 | 0.33115 | 0.992034 | 6692 | 4 | 2.0173 |
| FAM154A | 5 | 0.44137 | 0.59793 | 0.999994 | 12599 | 2 | 3.0134 | 0.1324 | 0.31575 | 0.989846 | 6693 | 3 | 3.0134 |
| PGK2 | 5 | 0.96533 | 0.96548 | 0.999994 | 20726 | 0 | 1.8338 | 0.13243 | 0.31579 | 0.989846 | 6694 | 4 | 1.8338 |
| PRKAR2A | 6 | 0.097696 | 0.24281 | 0.988088 | 4877 | 4 | -0.86623 | 0.13243 | 0.33123 | 0.992034 | 6695 | 2 | -0.86623 |
| KRT6B | 6 | 0.50983 | 0.6755 | 0.999994 | 13575 | 1 | 1.63 | 0.13243 | 0.33123 | 0.992034 | 6696 | 4 | 1.63 |
| FBXO44 | 6 | 0.43914 | 0.62783 | 0.999994 | 12561 | 2 | 1.9608 | 0.13243 | 0.33123 | 0.992034 | 6697 | 3 | 1.9608 |
| CDO1 | 6 | 0.34783 | 0.56157 | 0.999994 | 11111 | 2 | 2.0488 | 0.13245 | 0.33126 | 0.992034 | 6698 | 4 | 2.0488 |
| NonTargetingControlG | 2 | 0.86749 | 0.8675 | 0.999994 | 18589 | 0 | 3.8947 | 0.13251 | 0.21974 | 0.97466 | 6699 | 2 | 3.8947 |
| hsa-mir-4422 | 4 | 0.4931 | 0.61326 | 0.999994 | 13314 | 1 | 3.1066 | 0.13252 | 0.28389 | 0.978454 | 6700 | 2 | 3.1066 |
| GNL3 | 4 | 0.1302 | 0.26793 | 0.998561 | 5937 | 2 | 2.3547 | 0.13252 | 0.28389 | 0.978454 | 6701 | 2 | 2.3547 |
| EPN3 | 4 | 0.51367 | 0.62245 | 0.999994 | 13636 | 1 | 2.7675 | 0.13252 | 0.28389 | 0.978454 | 6702 | 2 | 2.7675 |
| HNRNPC | 4 | 0.20334 | 0.34408 | 0.999994 | 7863 | 2 | 2.5593 | 0.13252 | 0.28389 | 0.978454 | 6703 | 2 | 2.5593 |
| hsa-mir-185 | 4 | 0.2977 | 0.44067 | 0.999994 | 10084 | 1 | 3.0024 | 0.13254 | 0.28393 | 0.978454 | 6704 | 3 | 3.0024 |
| HDHD3 | 6 | 0.5296 | 0.68926 | 0.999994 | 13895 | 1 | 1.5217 | 0.13255 | 0.33141 | 0.992091 | 6705 | 4 | 1.5217 |
| MYL10 | 6 | 0.63605 | 0.76693 | 0.999994 | 15464 | 2 | 1.4145 | 0.13255 | 0.33141 | 0.992091 | 6706 | 4 | 1.4145 |
| FAM170A | 5 | 0.25773 | 0.44689 | 0.999994 | 9188 | 2 | 1.7343 | 0.13258 | 0.31607 | 0.989846 | 6707 | 3 | 1.7343 |

| | | | | | | | | | | | | | |
|---|---|---|---|---|---|---|---|---|---|---|---|---|---|
| TAS2R40 | 6 | 0.25427 | 0.45304 | 0.999994 | 9109 | 3 | -0.71322 | 0.1327 | 0.33168 | 0.992121 | 6708 | 2 | -0.71322 |
| SNAI3 | 6 | 0.86746 | 0.88178 | 0.999994 | 18588 | 1 | 0.35595 | 0.1327 | 0.33168 | 0.992121 | 6709 | 3 | 0.35595 |
| TBXAS1 | 5 | 0.77484 | 0.80446 | 0.999994 | 17140 | 1 | 2.1614 | 0.13272 | 0.31633 | 0.989846 | 6710 | 4 | 2.1614 |
| STX8 | 5 | 0.48363 | 0.63378 | 0.999994 | 13193 | 2 | 0.45323 | 0.13287 | 0.31662 | 0.989846 | 6711 | 2 | 0.45323 |
| ZBTB34 | 6 | 0.061594 | 0.16915 | 0.967126 | 3495 | 3 | -0.11038 | 0.13292 | 0.33201 | 0.992196 | 6712 | 3 | -0.11038 |
| ATP6V1G3 | 6 | 0.79149 | 0.84431 | 0.999994 | 17373 | 1 | 0.64342 | 0.13292 | 0.33201 | 0.992196 | 6713 | 3 | 0.64342 |
| CLEC4A | 6 | 0.1345 | 0.30603 | 0.999994 | 6039 | 3 | -0.17975 | 0.13292 | 0.33201 | 0.992196 | 6714 | 3 | -0.17975 |
| HTN3 | 2 | 0.86706 | 0.86707 | 0.999994 | 18575 | 0 | 3.6764 | 0.13294 | 0.22038 | 0.97466 | 6715 | 2 | 3.6764 |
| GORAB | 5 | 0.87485 | 0.87547 | 0.999994 | 18731 | 1 | 0.75622 | 0.13298 | 0.31683 | 0.989846 | 6716 | 3 | 0.75622 |
| NonTargetingControlG | 1 | 0.86699 | 0.86723 | 0.999994 | 18573 | 0 | 6.509 | 0.13301 | 0.13285 | 0.965913 | 6717 | 1 | 6.509 |
| ATOH1 | 6 | 0.42347 | 0.61764 | 0.999994 | 12304 | 2 | 1.504 | 0.13306 | 0.3322 | 0.992251 | 6718 | 3 | 1.504 |
| ZNF213 | 4 | 0.61555 | 0.67384 | 0.999994 | 15152 | 1 | 1.9049 | 0.13309 | 0.28476 | 0.978871 | 6719 | 2 | 1.9049 |
| DCAF5 | 6 | 0.81038 | 0.85242 | 0.999994 | 17638 | 1 | 0.73421 | 0.13313 | 0.33231 | 0.992388 | 6720 | 3 | 0.73421 |
| ST8SIA4 | 5 | 0.18271 | 0.35051 | 0.999994 | 7283 | 1 | 0.70118 | 0.13317 | 0.31719 | 0.989846 | 6721 | 4 | 0.70118 |
| OR2W3 | 5 | 0.43684 | 0.59415 | 0.999994 | 12522 | 1 | -0.015828 | 0.13317 | 0.31719 | 0.989846 | 6722 | 2 | -0.015828 |
| NonTargetingControlG | 2 | 0.86682 | 0.86684 | 0.999994 | 18571 | 0 | 3.0893 | 0.13318 | 0.22073 | 0.97466 | 6723 | 2 | 3.0893 |
| RNF186 | 5 | 0.81606 | 0.82998 | 0.999994 | 17714 | 1 | 3.1527 | 0.13326 | 0.31738 | 0.989846 | 6724 | 3 | 3.1527 |
| GRN | 6 | 0.52685 | 0.68733 | 0.999994 | 13841 | 2 | 2.5493 | 0.13328 | 0.33252 | 0.992526 | 6725 | 4 | 2.5493 |
| ZNF587B | 5 | 0.77736 | 0.80592 | 0.999994 | 17172 | 1 | 1.0677 | 0.13328 | 0.31742 | 0.989846 | 6726 | 4 | 1.0677 |
| GOLGA6L10 | 5 | 0.62374 | 0.73497 | 0.999994 | 15286 | 1 | 2.4023 | 0.13332 | 0.31749 | 0.989846 | 6727 | 4 | 2.4023 |
| POLR2F | 4 | 0.049872 | 0.12123 | 0.951052 | 2968 | 2 | -2.5147 | 0.13337 | 0.28518 | 0.978871 | 6728 | 1 | -2.5147 |
| OR4K1 | 5 | 0.20268 | 0.37905 | 0.999994 | 7849 | 2 | -0.40205 | 0.13338 | 0.31761 | 0.989846 | 6729 | 1 | -0.40205 |
| ZNF490 | 4 | 0.79389 | 0.7958 | 0.999994 | 17395 | 1 | 2.2699 | 0.13352 | 0.28541 | 0.978871 | 6730 | 3 | 2.2699 |
| FSTL3 | 6 | 0.63707 | 0.7677 | 0.999994 | 15473 | 2 | 0.93049 | 0.13352 | 0.3329 | 0.992917 | 6731 | 3 | 0.93049 |
| C5 | 6 | 0.56876 | 0.71706 | 0.999994 | 14521 | 2 | 0.98254 | 0.13352 | 0.3329 | 0.992917 | 6732 | 4 | 0.98254 |
| COL20A1 | 5 | 0.78298 | 0.80916 | 0.999994 | 17250 | 1 | 0.37398 | 0.13355 | 0.31792 | 0.989846 | 6733 | 2 | 0.37398 |
| CABP2 | 6 | 0.39813 | 0.60114 | 0.999994 | 11918 | 2 | 2.3508 | 0.13358 | 0.33299 | 0.992917 | 6734 | 3 | 2.3508 |
| NAT9 | 5 | 0.91803 | 0.91814 | 0.999994 | 19586 | 0 | 0.1289 | 0.13365 | 0.3181 | 0.989846 | 6735 | 2 | 0.1289 |
| ARMCX5-GPRASP2 | 5 | 0.91465 | 0.91472 | 0.999994 | 19503 | 0 | 0.98576 | 0.13366 | 0.31812 | 0.989846 | 6736 | 4 | 0.98576 |
| ARID5A | 6 | 0.027871 | 0.093775 | 0.945031 | 1898 | 2 | 1.9309 | 0.13368 | 0.33314 | 0.992917 | 6737 | 4 | 1.9309 |
| NonTargetingControlG | 2 | 0.51499 | 0.51774 | 0.999994 | 13661 | 1 | 3.3716 | 0.13376 | 0.22164 | 0.97466 | 6738 | 1 | 3.3716 |
| GABRA5 | 4 | 0.90704 | 0.90678 | 0.999994 | 19372 | 0 | 2.4742 | 0.13378 | 0.28581 | 0.978871 | 6739 | 3 | 2.4742 |
| OR5B12 | 5 | 0.58841 | 0.72196 | 0.999994 | 14795 | 1 | 0.82033 | 0.13379 | 0.31836 | 0.989846 | 6740 | 4 | 0.82033 |
| hsa-mir-937 | 3 | 0.66062 | 0.66586 | 0.999994 | 15787 | 1 | 4.1654 | 0.13387 | 0.27037 | 0.978454 | 6741 | 2 | 4.1654 |
| DPPA5 | 5 | 0.56309 | 0.70338 | 0.999994 | 14431 | 2 | 1.4274 | 0.13394 | 0.31863 | 0.989846 | 6742 | 3 | 1.4274 |
| APOBR | 5 | 0.41177 | 0.57332 | 0.999994 | 12143 | 2 | -0.14965 | 0.13394 | 0.31863 | 0.989846 | 6743 | 1 | -0.14965 |
| UBN2 | 6 | 0.83102 | 0.86204 | 0.999994 | 17945 | 1 | 2.0186 | 0.13396 | 0.33358 | 0.992917 | 6744 | 4 | 2.0186 |
| LRRIQ3 | 6 | 0.51523 | 0.67922 | 0.999994 | 13667 | 2 | 1.1682 | 0.13396 | 0.33358 | 0.992917 | 6745 | 3 | 1.1682 |
| USP2 | 6 | 0.60265 | 0.74191 | 0.999994 | 14979 | 2 | 0.7023 | 0.13396 | 0.33358 | 0.992917 | 6746 | 3 | 0.7023 |
| SDR39U1 | 6 | 0.18559 | 0.37008 | 0.999994 | 7357 | 2 | 0.43311 | 0.13396 | 0.33358 | 0.992917 | 6747 | 3 | 0.43311 |
| EMCN | 5 | 0.2106 | 0.39024 | 0.999994 | 8043 | 2 | 2.911 | 0.13398 | 0.31869 | 0.989846 | 6748 | 3 | 2.911 |
| DBP | 6 | 0.44462 | 0.63151 | 0.999994 | 12642 | 1 | 1.5919 | 0.13399 | 0.33362 | 0.992917 | 6749 | 4 | 1.5919 |
| RMDN1 | 6 | 0.85675 | 0.87559 | 0.999994 | 18383 | 1 | 2.285 | 0.13401 | 0.33366 | 0.992917 | 6750 | 4 | 2.285 |
| DEFB4B | 4 | 0.88283 | 0.88246 | 0.999994 | 18890 | 0 | 1.7353 | 0.13409 | 0.28631 | 0.978871 | 6751 | 3 | 1.7353 |
| FAM157A | 4 | 0.059945 | 0.14067 | 0.959146 | 3419 | 1 | 2.9949 | 0.1341 | 0.28633 | 0.978871 | 6752 | 2 | 2.9949 |
| KCNE2 | 5 | 0.16883 | 0.33041 | 0.999994 | 6956 | 3 | -0.97561 | 0.13411 | 0.31892 | 0.989846 | 6753 | 1 | -0.97561 |
| JSRP1 | 4 | 0.28345 | 0.42607 | 0.999994 | 9774 | 2 | -0.0044156 | 0.13412 | 0.28636 | 0.978871 | 6754 | 2 | -0.0044156 |
| BAG4 | 4 | 0.27907 | 0.4216 | 0.999994 | 9678 | 2 | -0.90943 | 0.13412 | 0.28636 | 0.978871 | 6755 | 1 | -0.90943 |
| SUSD2 | 4 | 0.16156 | 0.30109 | 0.999994 | 6776 | 1 | 0.41358 | 0.13412 | 0.28636 | 0.978871 | 6756 | 1 | 0.41358 |
| HSD17B12 | 5 | 0.16349 | 0.32262 | 0.999994 | 6828 | 1 | 1.3103 | 0.13416 | 0.31902 | 0.989846 | 6757 | 4 | 1.3103 |
| RAB5C | 5 | 0.12824 | 0.26991 | 0.998561 | 5884 | 2 | 2.0827 | 0.13426 | 0.3192 | 0.989846 | 6758 | 3 | 2.0827 |
| CDK16 | 5 | 0.3725 | 0.54047 | 0.999994 | 11549 | 1 | 2.899 | 0.13426 | 0.3192 | 0.989846 | 6759 | 3 | 2.899 |

| | | | | | | | | | | | | |
|---|---|---|---|---|---|---|---|---|---|---|---|---|
| SPINK6 | 6 | 0.64853 | 0.7765 | 0.999994 | 15635 | 1 | 1.9024 | 0.13429 | 0.33407 | 0.992933 | 6760 | 3 | 1.9024 |
| RAB26 | 6 | 0.63715 | 0.76776 | 0.999994 | 15474 | 2 | 2.0632 | 0.13429 | 0.33407 | 0.992933 | 6761 | 3 | 2.0632 |
| RBM46 | 6 | 0.00098959 | 0.0050264 | 0.589484 | 164 | 3 | 1.2446 | 0.13429 | 0.33407 | 0.992933 | 6762 | 3 | 1.2446 |
| DNASE2 | 6 | 0.36093 | 0.57634 | 0.999994 | 11354 | 2 | 1.8453 | 0.13432 | 0.33413 | 0.992933 | 6763 | 4 | 1.8453 |
| DDX55 | 6 | 0.35308 | 0.56747 | 0.999994 | 11216 | 1 | 2.7753 | 0.13432 | 0.33413 | 0.992933 | 6764 | 4 | 2.7753 |
| PHGR1 | 6 | 0.50506 | 0.67221 | 0.999994 | 13512 | 2 | 2.3564 | 0.13432 | 0.33413 | 0.992933 | 6765 | 4 | 2.3564 |
| OR2B3 | 6 | 0.34414 | 0.55743 | 0.999994 | 11031 | 2 | 1.9782 | 0.13432 | 0.33413 | 0.992933 | 6766 | 4 | 1.9782 |
| NEU1 | 5 | 0.97085 | 0.97091 | 0.999994 | 20842 | 0 | 2.3967 | 0.13436 | 0.31939 | 0.989846 | 6767 | 4 | 2.3967 |
| MRPS26 | 5 | 0.23219 | 0.42024 | 0.999994 | 8575 | 1 | 0.053003 | 0.13439 | 0.31942 | 0.989846 | 6768 | 2 | 0.053003 |
| PTPRN2 | 5 | 0.17529 | 0.33983 | 0.999994 | 7116 | 2 | -0.37695 | 0.13439 | 0.31942 | 0.989846 | 6769 | 2 | -0.37695 |
| NANOS2 | 5 | 0.18607 | 0.35535 | 0.999994 | 7366 | 2 | 1.9592 | 0.13445 | 0.31954 | 0.989846 | 6770 | 3 | 1.9592 |
| CAV3 | 5 | 0.37845 | 0.54544 | 0.999994 | 11629 | 1 | 1.6377 | 0.13447 | 0.31957 | 0.989846 | 6771 | 4 | 1.6377 |
| OR4F17 | 6 | 0.53917 | 0.69596 | 0.999994 | 14036 | 2 | -0.11545 | 0.13461 | 0.33456 | 0.993364 | 6772 | 1 | -0.11545 |
| ZNF442 | 6 | 0.026711 | 0.091043 | 0.94395 | 1842 | 3 | -0.60199 | 0.13461 | 0.33456 | 0.993364 | 6773 | 3 | -0.60199 |
| LONRF3 | 6 | 0.6407 | 0.77047 | 0.999994 | 15523 | 2 | 2.047 | 0.13462 | 0.33458 | 0.993364 | 6774 | 4 | 2.047 |
| hsa-mir-320c-1 | 4 | 0.91601 | 0.91578 | 0.999994 | 19536 | 0 | 2.192 | 0.13465 | 0.28716 | 0.979159 | 6775 | 3 | 2.192 |
| FAM49B | 5 | 0.96843 | 0.96856 | 0.999994 | 20792 | 0 | 1.308 | 0.13465 | 0.3199 | 0.989846 | 6776 | 4 | 1.308 |
| KLHL23 | 5 | 0.12838 | 0.27014 | 0.998561 | 5891 | 2 | -0.31619 | 0.13466 | 0.31992 | 0.989846 | 6777 | 2 | -0.31619 |
| PRICKLE4 | 5 | 0.34931 | 0.52132 | 0.999994 | 11133 | 2 | 1.1632 | 0.13466 | 0.31992 | 0.989846 | 6778 | 3 | 1.1632 |
| NonTargetingControlG | 2 | 0.86533 | 0.86535 | 0.999994 | 18545 | 0 | 2.6159 | 0.13467 | 0.22296 | 0.97466 | 6779 | 2 | 2.6159 |
| BHLHE40 | 6 | 0.62919 | 0.76172 | 0.999994 | 15370 | 1 | 1.9533 | 0.13472 | 0.33473 | 0.993364 | 6780 | 4 | 1.9533 |
| SCGB1A1 | 5 | 0.68663 | 0.76051 | 0.999994 | 16176 | 1 | 1.3424 | 0.13474 | 0.32006 | 0.989846 | 6781 | 4 | 1.3424 |
| SMAD9 | 6 | 0.43896 | 0.62772 | 0.999994 | 12557 | 2 | 1.5934 | 0.1348 | 0.33486 | 0.993364 | 6782 | 4 | 1.5934 |
| hsa-mir-647 | 4 | 0.60723 | 0.66924 | 0.999994 | 15038 | 1 | 2.3224 | 0.13481 | 0.28742 | 0.97947 | 6783 | 3 | 2.3224 |
| TNNT2 | 5 | 0.017066 | 0.056046 | 0.9235 | 1306 | 2 | 2.7894 | 0.13483 | 0.32022 | 0.989846 | 6784 | 3 | 2.7894 |
| PRPF38A | 3 | 0.55755 | 0.58558 | 0.999994 | 14327 | 1 | 2.668 | 0.13484 | 0.27144 | 0.978454 | 6785 | 2 | 2.668 |
| CYP2R1 | 5 | 0.92621 | 0.92642 | 0.999994 | 19772 | 0 | 3.0504 | 0.13486 | 0.32029 | 0.989846 | 6786 | 3 | 3.0504 |
| SLC17A1 | 5 | 0.070099 | 0.17854 | 0.968171 | 3832 | 2 | 3.0154 | 0.13494 | 0.32044 | 0.989846 | 6787 | 3 | 3.0154 |
| IRX4 | 6 | 0.10237 | 0.25195 | 0.994 | 5053 | 3 | 1.5855 | 0.13494 | 0.33506 | 0.993364 | 6788 | 3 | 1.5855 |
| ZCCHC2 | 4 | 0.92094 | 0.9207 | 0.999994 | 19652 | 0 | 2.7348 | 0.13496 | 0.28763 | 0.97947 | 6789 | 3 | 2.7348 |
| NonTargetingControlG | 2 | 0.86504 | 0.86507 | 0.999994 | 18540 | 0 | 4.1348 | 0.13496 | 0.22339 | 0.97466 | 6790 | 2 | 4.1348 |
| RINT1 | 3 | 0.82291 | 0.82283 | 0.999994 | 17821 | 0 | 4.218 | 0.13497 | 0.27156 | 0.978454 | 6791 | 2 | 4.218 |
| SPTB | 6 | 0.081307 | 0.20996 | 0.979587 | 4277 | 3 | -0.55296 | 0.13508 | 0.33528 | 0.993364 | 6792 | 2 | -0.55296 |
| SNX21 | 5 | 0.056003 | 0.15483 | 0.967126 | 3246 | 1 | 2.968 | 0.13517 | 0.32085 | 0.989846 | 6793 | 4 | 2.968 |
| TTC36 | 5 | 0.43233 | 0.59036 | 0.999994 | 12450 | 2 | 0.83152 | 0.13523 | 0.32096 | 0.989846 | 6794 | 3 | 0.83152 |
| DYSF | 5 | 0.14151 | 0.2901 | 0.999632 | 6231 | 3 | -1.2715 | 0.13523 | 0.32096 | 0.989846 | 6795 | 2 | -1.2715 |
| OR6A2 | 5 | 0.36878 | 0.53734 | 0.999994 | 11485 | 2 | 1.2058 | 0.13525 | 0.32101 | 0.989846 | 6796 | 3 | 1.2058 |
| OR51G2 | 6 | 0.42328 | 0.61754 | 0.999994 | 12298 | 1 | 2.0606 | 0.13528 | 0.3356 | 0.993964 | 6797 | 4 | 2.0606 |
| SEZ6 | 5 | 0.79444 | 0.81605 | 0.999994 | 17402 | 1 | 0.071676 | 0.13532 | 0.32113 | 0.989846 | 6798 | 2 | 0.071676 |
| AGBL2 | 5 | 0.32505 | 0.50145 | 0.999994 | 10620 | 1 | 1.6443 | 0.13535 | 0.32119 | 0.989846 | 6799 | 4 | 1.6443 |
| KRTAP3-2 | 5 | 0.88721 | 0.88732 | 0.999994 | 18972 | 1 | 2.5114 | 0.13536 | 0.32121 | 0.989846 | 6800 | 4 | 2.5114 |
| PRSS22 | 5 | 0.13436 | 0.27929 | 0.999632 | 6034 | 1 | 1.7203 | 0.13548 | 0.32141 | 0.989846 | 6801 | 3 | 1.7203 |
| SCRN3 | 5 | 0.18574 | 0.35491 | 0.999994 | 7360 | 2 | -0.3843 | 0.13548 | 0.32141 | 0.989846 | 6802 | 2 | -0.3843 |
| LCN15 | 6 | 0.19812 | 0.38563 | 0.999994 | 7716 | 3 | -0.23301 | 0.13551 | 0.33595 | 0.994237 | 6803 | 3 | -0.23301 |
| ANKRD30A | 5 | 0.40219 | 0.56534 | 0.999994 | 11981 | 1 | 1.4775 | 0.13561 | 0.32165 | 0.989846 | 6804 | 4 | 1.4775 |
| EN1 | 5 | 0.78511 | 0.8104 | 0.999994 | 17282 | 1 | 1.41 | 0.13566 | 0.32174 | 0.989846 | 6805 | 4 | 1.41 |
| TAS2R14 | 6 | 0.52745 | 0.68774 | 0.999994 | 13849 | 1 | 1.8912 | 0.13567 | 0.33619 | 0.994237 | 6806 | 3 | 1.8912 |
| NonTargetingControlG | 1 | 0.86431 | 0.86458 | 0.999994 | 18528 | 0 | 6.4592 | 0.13569 | 0.13554 | 0.965913 | 6807 | 1 | 6.4592 |
| TNXB | 5 | 0.42917 | 0.58773 | 0.999994 | 12402 | 1 | 2.5583 | 0.13572 | 0.32184 | 0.989846 | 6808 | 4 | 2.5583 |
| WDR77 | 4 | 0.38025 | 0.5254 | 0.999994 | 11650 | 1 | 2.3472 | 0.13583 | 0.28894 | 0.980115 | 6809 | 2 | 2.3472 |
| NFAM1 | 6 | 0.50368 | 0.67127 | 0.999994 | 13490 | 2 | 2.7315 | 0.13587 | 0.33653 | 0.994237 | 6810 | 4 | 2.7315 |
| CHD2 | 6 | 0.047024 | 0.13757 | 0.957317 | 2852 | 2 | 2.0598 | 0.13588 | 0.33653 | 0.994237 | 6811 | 4 | 2.0598 |

| | | | | | | | | | | | | | |
|---|---|---|---|---|---|---|---|---|---|---|---|---|---|
| NAGS | 4 | 0.8903 | 0.89001 | 0.999994 | 19042 | 0 | 2.6979 | 0.13593 | 0.28909 | 0.980152 | 6812 | 3 | 2.6979 |
| CSF3R | 6 | 0.48665 | 0.65969 | 0.999994 | 13236 | 2 | 2.3216 | 0.13596 | 0.33666 | 0.994237 | 6813 | 3 | 2.3216 |
| AXIN2 | 6 | 0.56044 | 0.71109 | 0.999994 | 14377 | 1 | 0.91994 | 0.13598 | 0.33669 | 0.994237 | 6814 | 4 | 0.91994 |
| PALMD | 6 | 0.11544 | 0.27712 | 0.999379 | 5490 | 3 | 0.28066 | 0.13598 | 0.33669 | 0.994237 | 6815 | 3 | 0.28066 |
| CASS4 | 3 | 0.20245 | 0.31577 | 0.999994 | 7842 | 1 | -0.089046 | 0.13601 | 0.27271 | 0.978454 | 6816 | 1 | -0.089046 |
| TMEM235 | 5 | 0.53248 | 0.67623 | 0.999994 | 13942 | 2 | 1.0324 | 0.13606 | 0.32246 | 0.989846 | 6817 | 3 | 1.0324 |
| GPR160 | 5 | 0.88127 | 0.88148 | 0.999994 | 18856 | 1 | 2.1504 | 0.13606 | 0.32246 | 0.989846 | 6818 | 4 | 2.1504 |
| LY6E | 5 | 0.96888 | 0.969 | 0.999994 | 20800 | 0 | 0.94888 | 0.13607 | 0.32247 | 0.989846 | 6819 | 4 | 0.94888 |
| MUTYH | 6 | 0.070311 | 0.18743 | 0.969409 | 3848 | 4 | -1.2982 | 0.13616 | 0.33697 | 0.994237 | 6820 | 2 | -1.2982 |
| RAB22A | 5 | 0.34032 | 0.51398 | 0.999994 | 10940 | 2 | -0.07509 | 0.13619 | 0.3227 | 0.989846 | 6821 | 2 | -0.07509 |
| KCTD3 | 4 | 0.82258 | 0.82225 | 0.999994 | 17816 | 1 | 2.8644 | 0.13623 | 0.28955 | 0.980804 | 6822 | 2 | 2.8644 |
| GNG10 | 5 | 0.69662 | 0.76494 | 0.999994 | 16266 | 1 | 3.073 | 0.13628 | 0.32285 | 0.989846 | 6823 | 4 | 3.073 |
| MED15 | 5 | 0.47762 | 0.6286 | 0.999994 | 13107 | 1 | 3.1166 | 0.13628 | 0.32285 | 0.989846 | 6824 | 3 | 3.1166 |
| CHMP3 | 5 | 0.85658 | 0.85956 | 0.999994 | 18381 | 1 | 3.1166 | 0.13628 | 0.32285 | 0.989846 | 6825 | 3 | 3.1166 |
| NCF2 | 5 | 0.92459 | 0.92481 | 0.999994 | 19733 | 0 | 3.1166 | 0.13628 | 0.32285 | 0.989846 | 6826 | 3 | 3.1166 |
| RPH3A | 5 | 0.9257 | 0.92593 | 0.999994 | 19763 | 0 | 3.1166 | 0.13628 | 0.32285 | 0.989846 | 6827 | 4 | 3.1166 |
| hsa-mir-575 | 3 | 0.07523 | 0.14998 | 0.966359 | 4045 | 2 | -2.6942 | 0.13633 | 0.27306 | 0.978454 | 6828 | 1 | -2.6942 |
| GUCY1A3 | 6 | 0.29406 | 0.49978 | 0.999994 | 9999 | 2 | 2.5079 | 0.13636 | 0.33729 | 0.994237 | 6829 | 4 | 2.5079 |
| CDH20 | 6 | 0.33874 | 0.55123 | 0.999994 | 10912 | 3 | -0.31625 | 0.13638 | 0.33733 | 0.994237 | 6830 | 2 | -0.31625 |
| EIF3E | 6 | 0.37181 | 0.58435 | 0.999994 | 11540 | 2 | 2.0352 | 0.1364 | 0.33734 | 0.994237 | 6831 | 3 | 2.0352 |
| COPS7A | 6 | 0.39056 | 0.59633 | 0.999994 | 11806 | 2 | 2.5627 | 0.13646 | 0.33744 | 0.994237 | 6832 | 4 | 2.5627 |
| TAS1R3 | 5 | 0.28954 | 0.47254 | 0.999994 | 9903 | 1 | 0.73849 | 0.13648 | 0.32324 | 0.989881 | 6833 | 3 | 0.73849 |
| OR10J5 | 5 | 0.22707 | 0.41316 | 0.999994 | 8455 | 3 | -0.67865 | 0.13648 | 0.32324 | 0.989881 | 6834 | 2 | -0.67865 |
| hsa-mir-4794 | 2 | 0.75536 | 0.75492 | 0.999994 | 16919 | 0 | 4.1391 | 0.13651 | 0.22563 | 0.97466 | 6835 | 1 | 4.1391 |
| ZNF17 | 5 | 0.94487 | 0.94505 | 0.999994 | 20199 | 0 | 1.0651 | 0.13659 | 0.32344 | 0.989965 | 6836 | 3 | 1.0651 |
| MED18 | 6 | 0.49938 | 0.66826 | 0.999994 | 13412 | 1 | 0.22682 | 0.1366 | 0.33767 | 0.994237 | 6837 | 2 | 0.22682 |
| FIGNL2 | 5 | 0.72551 | 0.77851 | 0.999994 | 16555 | 1 | 1.9813 | 0.13662 | 0.32349 | 0.989995 | 6838 | 4 | 1.9813 |
| PTK7 | 5 | 0.62303 | 0.73469 | 0.999994 | 15273 | 1 | 0.98936 | 0.13672 | 0.32365 | 0.990175 | 6839 | 4 | 0.98936 |
| CCDC158 | 5 | 0.52535 | 0.66998 | 0.999994 | 13823 | 2 | 0.22954 | 0.13674 | 0.32369 | 0.990175 | 6840 | 2 | 0.22954 |
| UAP1 | 5 | 0.40426 | 0.5671 | 0.999994 | 12024 | 2 | -0.11345 | 0.13674 | 0.32369 | 0.990175 | 6841 | 2 | -0.11345 |
| CRTAC1 | 4 | 0.89242 | 0.89213 | 0.999994 | 19078 | 0 | 2.9795 | 0.13679 | 0.29039 | 0.981963 | 6842 | 3 | 2.9795 |
| NonTargetingControlG | 2 | 0.8632 | 0.86323 | 0.999994 | 18508 | 0 | 4.0853 | 0.1368 | 0.22607 | 0.97466 | 6843 | 2 | 4.0853 |
| MLLT6 | 5 | 0.6201 | 0.7336 | 0.999994 | 15222 | 1 | 3.0322 | 0.13681 | 0.32383 | 0.990342 | 6844 | 3 | 3.0322 |
| LTB4R | 6 | 0.13631 | 0.30837 | 0.999994 | 6091 | 2 | 1.9583 | 0.13684 | 0.33804 | 0.994237 | 6845 | 4 | 1.9583 |
| IFT52 | 6 | 0.5858 | 0.72948 | 0.999994 | 14755 | 2 | 2.6751 | 0.13695 | 0.33822 | 0.994237 | 6846 | 4 | 2.6751 |
| UFC1 | 6 | 0.687 | 0.80647 | 0.999994 | 16181 | 1 | 2.6258 | 0.13695 | 0.33822 | 0.994237 | 6847 | 4 | 2.6258 |
| METTL17 | 6 | 0.75486 | 0.83021 | 0.999994 | 16910 | 1 | 2.0741 | 0.13695 | 0.33822 | 0.994237 | 6848 | 3 | 2.0741 |
| THSD4 | 6 | 0.92739 | 0.92747 | 0.999994 | 19799 | 1 | 2.0929 | 0.13695 | 0.33822 | 0.994237 | 6849 | 3 | 2.0929 |
| INTS2 | 6 | 0.39741 | 0.60069 | 0.999994 | 11910 | 2 | 2.5133 | 0.13697 | 0.33824 | 0.994237 | 6850 | 4 | 2.5133 |
| EFNB2 | 5 | 0.0066529 | 0.026377 | 0.848412 | 638 | 2 | 0.63864 | 0.13699 | 0.32416 | 0.990429 | 6851 | 3 | 0.63864 |
| SPATA18 | 5 | 0.42637 | 0.58543 | 0.999994 | 12354 | 1 | 0.16955 | 0.13703 | 0.32424 | 0.990429 | 6852 | 2 | 0.16955 |
| ODF2L | 5 | 0.56672 | 0.70664 | 0.999994 | 14493 | 2 | -0.30577 | 0.13703 | 0.32424 | 0.990429 | 6853 | 2 | -0.30577 |
| LSM4 | 4 | 0.74512 | 0.75636 | 0.999994 | 16802 | 1 | 3.1382 | 0.13707 | 0.29083 | 0.982652 | 6854 | 2 | 3.1382 |
| OR8B8 | 5 | 0.93515 | 0.93538 | 0.999994 | 19970 | 0 | 1.5469 | 0.13708 | 0.32433 | 0.990429 | 6855 | 4 | 1.5469 |
| HDAC9 | 4 | 0.85389 | 0.85347 | 0.999994 | 18332 | 0 | 0.93207 | 0.13721 | 0.29105 | 0.982859 | 6856 | 2 | 0.93207 |
| KIAA0196 | 5 | 0.58694 | 0.72146 | 0.999994 | 14774 | 1 | 1.2152 | 0.13724 | 0.32463 | 0.990429 | 6857 | 4 | 1.2152 |
| KDM4B | 5 | 0.49984 | 0.64775 | 0.999994 | 13420 | 1 | 1.2499 | 0.13728 | 0.32471 | 0.990429 | 6858 | 3 | 1.2499 |
| ENO4 | 5 | 0.94723 | 0.94741 | 0.999994 | 20267 | 0 | 1.5586 | 0.13728 | 0.32471 | 0.990429 | 6859 | 3 | 1.5586 |
| ACTL7B | 6 | 0.90102 | 0.90446 | 0.999994 | 19235 | 1 | 0.2716 | 0.13731 | 0.33875 | 0.994237 | 6860 | 2 | 0.2716 |
| UBOX5 | 6 | 0.34348 | 0.55664 | 0.999994 | 11021 | 3 | 0.85172 | 0.13731 | 0.33875 | 0.994237 | 6861 | 3 | 0.85172 |
| TTC13 | 6 | 0.66912 | 0.7925 | 0.999994 | 15915 | 2 | 0.72992 | 0.13731 | 0.33875 | 0.994237 | 6862 | 3 | 0.72992 |
| KLKB1 | 6 | 0.87835 | 0.88852 | 0.999994 | 18795 | 1 | 1.6641 | 0.13735 | 0.33881 | 0.994237 | 6863 | 3 | 1.6641 |

| | | | | | | | | | | | |
|---|---|---|---|---|---|---|---|---|---|---|---|
| FXYD1 | 5 | 0.92324 | 0.92343 | 0.999994 | 19700 | 0 | 3.0064 | 0.1374 | 0.32493 | 0.990752 | 6864 | 3 | 3.0064 |
| hsa-mir-4647 | 3 | 0.23141 | 0.35185 | 0.999994 | 8552 | 1 | 4.0801 | 0.13753 | 0.2744 | 0.978454 | 6865 | 2 | 4.0801 |
| FARS2 | 5 | 0.35073 | 0.52252 | 0.999994 | 11177 | 2 | 3.0916 | 0.13753 | 0.32515 | 0.991152 | 6866 | 3 | 3.0916 |
| SURF6 | 5 | 0.96305 | 0.9632 | 0.999994 | 20665 | 0 | 2.3787 | 0.1376 | 0.32529 | 0.991152 | 6867 | 4 | 2.3787 |
| NonTargetingControlG | 2 | 0.86239 | 0.86241 | 0.999994 | 18493 | 0 | 2.4973 | 0.13761 | 0.22731 | 0.97466 | 6868 | 2 | 2.4973 |
| CDH4 | 5 | 0.31414 | 0.49255 | 0.999994 | 10402 | 2 | 1.3253 | 0.13763 | 0.32534 | 0.991152 | 6869 | 3 | 1.3253 |
| DMRTA1 | 6 | 0.4408 | 0.62896 | 0.999994 | 12590 | 1 | 1.9054 | 0.13769 | 0.33932 | 0.994237 | 6870 | 4 | 1.9054 |
| NonTargetingControlG | 2 | 0.26257 | 0.30828 | 0.999994 | 9299 | 1 | 2.4071 | 0.13772 | 0.22749 | 0.97466 | 6871 | 1 | 2.4071 |
| RFX1 | 6 | 0.020985 | 0.075132 | 0.93914 | 1514 | 4 | -1.9143 | 0.13774 | 0.3394 | 0.994237 | 6872 | 2 | -1.9143 |
| JAGN1 | 6 | 0.012773 | 0.048339 | 0.90085 | 1051 | 4 | -2.368 | 0.13774 | 0.3394 | 0.994237 | 6873 | 2 | -2.368 |
| PLK3 | 5 | 0.70111 | 0.76699 | 0.999994 | 16317 | 1 | 1.9536 | 0.13776 | 0.32558 | 0.991477 | 6874 | 4 | 1.9536 |
| KRTAP9-7 | 6 | 0.3141 | 0.52315 | 0.999994 | 10400 | 1 | 2.2062 | 0.1378 | 0.3395 | 0.994237 | 6875 | 4 | 2.2062 |
| HSD3B2 | 6 | 0.19034 | 0.37593 | 0.999994 | 7496 | 2 | 0.80769 | 0.13789 | 0.33964 | 0.994237 | 6876 | 3 | 0.80769 |
| SPATA31A6 | 5 | 0.068386 | 0.17581 | 0.968171 | 3763 | 1 | 1.346 | 0.13791 | 0.32587 | 0.991729 | 6877 | 4 | 1.346 |
| RCOR2 | 5 | 0.92465 | 0.92486 | 0.999994 | 19734 | 0 | 2.11 | 0.13801 | 0.32606 | 0.991729 | 6878 | 3 | 2.11 |
| DENND4B | 5 | 0.022649 | 0.071249 | 0.93914 | 1626 | 2 | -0.2381 | 0.13801 | 0.32606 | 0.991729 | 6879 | 2 | -0.2381 |
| FBXO46 | 6 | 0.2542 | 0.45295 | 0.999994 | 9108 | 1 | 1.1277 | 0.13804 | 0.33986 | 0.994237 | 6880 | 3 | 1.1277 |
| RHEB | 6 | 0.12561 | 0.29476 | 0.999994 | 5810 | 2 | -0.013836 | 0.13809 | 0.33995 | 0.994237 | 6881 | 2 | -0.013836 |
| WFDC12 | 6 | 0.026766 | 0.091176 | 0.94395 | 1847 | 3 | 0.18706 | 0.13809 | 0.33995 | 0.994237 | 6882 | 3 | 0.18706 |
| ZNF34 | 6 | 0.87336 | 0.8854 | 0.999994 | 18702 | 1 | 2.1538 | 0.13809 | 0.33995 | 0.994237 | 6883 | 4 | 2.1538 |
| LARP1B | 6 | 0.65592 | 0.78214 | 0.999994 | 15725 | 1 | 1.8725 | 0.13809 | 0.33995 | 0.994237 | 6884 | 4 | 1.8725 |
| EMP3 | 6 | 0.88714 | 0.89434 | 0.999994 | 18969 | 1 | 0.70129 | 0.13809 | 0.33995 | 0.994237 | 6885 | 3 | 0.70129 |
| MYBPC3 | 4 | 0.90813 | 0.90785 | 0.999994 | 19388 | 0 | 2.9655 | 0.1381 | 0.29241 | 0.983225 | 6886 | 2 | 2.9655 |
| hsa-mir-187 | 4 | 0.087435 | 0.19231 | 0.974607 | 4493 | 1 | 2.1294 | 0.13817 | 0.29252 | 0.983225 | 6887 | 3 | 2.1294 |
| HDAC7 | 5 | 0.73102 | 0.78116 | 0.999994 | 16613 | 1 | 1.9223 | 0.13818 | 0.32637 | 0.991729 | 6888 | 4 | 1.9223 |
| TRAF5 | 5 | 0.7869 | 0.81144 | 0.999994 | 17304 | 1 | 0.31062 | 0.13818 | 0.32637 | 0.991729 | 6889 | 2 | 0.31062 |
| C10orf25 | 5 | 0.95985 | 0.95999 | 0.999994 | 20580 | 0 | 3.0147 | 0.13823 | 0.32648 | 0.991729 | 6890 | 3 | 3.0147 |
| FRG2B | 2 | 0.86175 | 0.86176 | 0.999994 | 18479 | 0 | 2.2784 | 0.13825 | 0.22826 | 0.97466 | 6891 | 2 | 2.2784 |
| MUC12 | 6 | 0.62289 | 0.75697 | 0.999994 | 15271 | 2 | 1.9474 | 0.13828 | 0.34024 | 0.994237 | 6892 | 4 | 1.9474 |
| NonTargetingControlG | 1 | 0.86171 | 0.86198 | 0.999994 | 18478 | 0 | 6.4076 | 0.13829 | 0.13808 | 0.965913 | 6893 | 1 | 6.4076 |
| OR3A3 | 5 | 0.21197 | 0.39217 | 0.999994 | 8064 | 2 | 2.634 | 0.13831 | 0.32663 | 0.991729 | 6894 | 3 | 2.634 |
| DNALI1 | 6 | 0.34008 | 0.55273 | 0.999994 | 10935 | 2 | -0.12302 | 0.13841 | 0.34042 | 0.994237 | 6895 | 2 | -0.12302 |
| ZBTB9 | 6 | 0.048333 | 0.14044 | 0.958716 | 2901 | 3 | 1.0831 | 0.13842 | 0.34043 | 0.994237 | 6896 | 3 | 1.0831 |
| CYP2U1 | 6 | 0.74425 | 0.82645 | 0.999994 | 16791 | 1 | 2.3186 | 0.13842 | 0.34043 | 0.994237 | 6897 | 3 | 2.3186 |
| AZU1 | 6 | 0.67709 | 0.79871 | 0.999994 | 16043 | 2 | 2.2605 | 0.13842 | 0.34043 | 0.994237 | 6898 | 3 | 2.2605 |
| GCNT7 | 6 | 0.7711 | 0.83625 | 0.999994 | 17099 | 1 | 2.2375 | 0.13842 | 0.34043 | 0.994237 | 6899 | 3 | 2.2375 |
| ERI2 | 4 | 0.10432 | 0.22281 | 0.983354 | 5109 | 1 | 3.0394 | 0.13851 | 0.29306 | 0.983225 | 6900 | 2 | 3.0394 |
| NCR1 | 6 | 0.048164 | 0.14007 | 0.958716 | 2895 | 2 | 1.5899 | 0.13866 | 0.34077 | 0.994237 | 6901 | 4 | 1.5899 |
| BRAF | 6 | 0.62311 | 0.75713 | 0.999994 | 15276 | 2 | 1.657 | 0.13866 | 0.34078 | 0.994237 | 6902 | 3 | 1.657 |
| NBPF8 | 4 | 0.80348 | 0.80431 | 0.999994 | 17533 | 1 | 0.94155 | 0.1387 | 0.29336 | 0.983225 | 6903 | 2 | 0.94155 |
| hsa-mir-98 | 4 | 0.15683 | 0.29622 | 0.999994 | 6647 | 2 | -0.33821 | 0.1387 | 0.29336 | 0.983225 | 6904 | 2 | -0.33821 |
| NonTargetingControlG | 1 | 0.86119 | 0.86148 | 0.999994 | 18470 | 0 | 6.3955 | 0.13881 | 0.13859 | 0.965913 | 6905 | 1 | 6.3955 |
| C10orf32 | 6 | 0.1369 | 0.30914 | 0.999994 | 6104 | 3 | -0.28654 | 0.13886 | 0.3411 | 0.994237 | 6906 | 2 | -0.28654 |
| KANK3 | 6 | 0.29393 | 0.49962 | 0.999994 | 9995 | 2 | 1.9685 | 0.13887 | 0.34112 | 0.994237 | 6907 | 4 | 1.9685 |
| hsa-mir-380 | 4 | 0.53499 | 0.63247 | 0.999994 | 13979 | 1 | 1.759 | 0.13887 | 0.29361 | 0.983304 | 6908 | 3 | 1.759 |
| OR2C3 | 6 | 0.33236 | 0.54397 | 0.999994 | 10772 | 2 | 2.0142 | 0.13889 | 0.34115 | 0.994237 | 6909 | 4 | 2.0142 |
| NonTargetingControlG | 2 | 0.33386 | 0.3631 | 0.999994 | 10810 | 1 | 2.6695 | 0.13891 | 0.22925 | 0.97466 | 6910 | 1 | 2.6695 |
| C1orf174 | 6 | 0.51338 | 0.67795 | 0.999994 | 13627 | 1 | 1.7505 | 0.13891 | 0.34118 | 0.994237 | 6911 | 4 | 1.7505 |
| FZD10 | 6 | 0.74076 | 0.82524 | 0.999994 | 16749 | 1 | 1.6822 | 0.13893 | 0.3412 | 0.994237 | 6912 | 4 | 1.6822 |
| FAM21A | 6 | 0.94175 | 0.94173 | 0.999994 | 20122 | 0 | 2.0794 | 0.13899 | 0.34133 | 0.994237 | 6913 | 4 | 2.0794 |
| SRSF5 | 4 | 0.43333 | 0.57996 | 0.999994 | 12468 | 2 | 0.66263 | 0.139 | 0.2938 | 0.983309 | 6914 | 2 | 0.66263 |
| PPP1R27 | 6 | 0.07435 | 0.19581 | 0.975797 | 4005 | 3 | 1.2607 | 0.13903 | 0.34138 | 0.994237 | 6915 | 3 | 1.2607 |

| | | | | | | | | | | | | |
|---|---|---|---|---|---|---|---|---|---|---|---|---|
| ZNF14 | 6 | 0.11729 | 0.28061 | 0.999632 | 5550 | 2 | 2.0198 | 0.13904 | 0.34141 | 0.994237 | 6916 | 3 | 2.0198 |
| DCAF11 | 5 | 0.8426 | 0.84863 | 0.999994 | 18132 | 1 | 1.781 | 0.13905 | 0.32798 | 0.991729 | 6917 | 4 | 1.781 |
| CIDEB | 5 | 0.17881 | 0.3449 | 0.999994 | 7192 | 1 | 0.12737 | 0.13905 | 0.32799 | 0.991729 | 6918 | 2 | 0.12737 |
| GUCA2B | 5 | 0.49428 | 0.64295 | 0.999994 | 13334 | 2 | 0.12788 | 0.13905 | 0.32799 | 0.991729 | 6919 | 2 | 0.12788 |
| C16orf54 | 5 | 0.20652 | 0.38445 | 0.999994 | 7944 | 3 | -0.80947 | 0.13905 | 0.32799 | 0.991729 | 6920 | 2 | -0.80947 |
| S100P | 6 | 0.20429 | 0.3931 | 0.999994 | 7885 | 2 | 1.9344 | 0.1391 | 0.34151 | 0.994392 | 6921 | 3 | 1.9344 |
| PARK7 | 5 | 0.9685 | 0.96864 | 0.999994 | 20794 | 0 | 1.7947 | 0.13911 | 0.32809 | 0.991729 | 6922 | 4 | 1.7947 |
| POLR1D | 5 | 0.55746 | 0.69838 | 0.999994 | 14324 | 2 | -0.25127 | 0.13912 | 0.32811 | 0.991729 | 6923 | 2 | -0.25127 |
| DSTYK | 5 | 0.23674 | 0.42653 | 0.999994 | 8686 | 2 | -0.52809 | 0.13912 | 0.32811 | 0.991729 | 6924 | 2 | -0.52809 |
| IL34 | 5 | 0.89882 | 0.89893 | 0.999994 | 19199 | 0 | 0.88677 | 0.13912 | 0.32811 | 0.991729 | 6925 | 3 | 0.88677 |
| ARL1 | 6 | 0.036306 | 0.11357 | 0.951052 | 2335 | 4 | -2.1817 | 0.13913 | 0.34155 | 0.994392 | 6926 | 2 | -2.1817 |
| THOC5 | 4 | 0.64552 | 0.6907 | 0.999994 | 15591 | 1 | 2.9556 | 0.1392 | 0.29409 | 0.983429 | 6927 | 2 | 2.9556 |
| hsa-mir-4317 | 4 | 0.18701 | 0.32731 | 0.999994 | 7388 | 1 | 3.304 | 0.1392 | 0.29409 | 0.983429 | 6928 | 3 | 3.304 |
| NonTargetingControlG | 2 | 0.44985 | 0.45921 | 0.999994 | 12709 | 1 | 3.2025 | 0.13921 | 0.2297 | 0.97466 | 6929 | 1 | 3.2025 |
| hsa-mir-4469 | 4 | 0.66791 | 0.70411 | 0.999994 | 15894 | 1 | 0.68111 | 0.13924 | 0.29415 | 0.983431 | 6930 | 2 | 0.68111 |
| SRGAP3 | 5 | 0.030668 | 0.092213 | 0.943968 | 2040 | 2 | 2.7992 | 0.13926 | 0.32833 | 0.991729 | 6931 | 3 | 2.7992 |
| PCSK7 | 6 | 0.27171 | 0.47372 | 0.999994 | 9515 | 1 | 1.2934 | 0.13931 | 0.34182 | 0.994392 | 6932 | 3 | 1.2934 |
| MRPL54 | 4 | 0.18332 | 0.32352 | 0.999994 | 7301 | 1 | 2.3962 | 0.1394 | 0.29439 | 0.983431 | 6933 | 3 | 2.3962 |
| SH2D4A | 4 | 0.68096 | 0.71222 | 0.999994 | 16103 | 1 | 1.7935 | 0.13942 | 0.29443 | 0.983431 | 6934 | 3 | 1.7935 |
| OR10H5 | 6 | 0.25997 | 0.4598 | 0.999994 | 9240 | 3 | 0.24994 | 0.13946 | 0.34205 | 0.994392 | 6935 | 3 | 0.24994 |
| PLA2G1B | 6 | 0.68599 | 0.80569 | 0.999994 | 16171 | 1 | 2.8617 | 0.13946 | 0.34205 | 0.994392 | 6936 | 4 | 2.8617 |
| AQP8 | 5 | 0.91498 | 0.91505 | 0.999994 | 19511 | 0 | 0.90166 | 0.13949 | 0.32876 | 0.991729 | 6937 | 4 | 0.90166 |
| TAS2R46 | 5 | 0.74268 | 0.78701 | 0.999994 | 16774 | 1 | 1.2666 | 0.1395 | 0.32878 | 0.991729 | 6938 | 4 | 1.2666 |
| BPIFB3 | 5 | 0.96779 | 0.96794 | 0.999994 | 20780 | 0 | 3.01 | 0.13952 | 0.32881 | 0.991729 | 6939 | 4 | 3.01 |
| ZFPM1 | 4 | 0.01997 | 0.060569 | 0.932932 | 1459 | 2 | -1.1615 | 0.13954 | 0.2946 | 0.983644 | 6940 | 2 | -1.1615 |
| NKAIN3 | 6 | 0.0071852 | 0.029081 | 0.859137 | 670 | 3 | -0.82717 | 0.13961 | 0.34228 | 0.994392 | 6941 | 2 | -0.82717 |
| CSMD2 | 6 | 0.28379 | 0.48781 | 0.999994 | 9779 | 3 | -0.036009 | 0.13961 | 0.34228 | 0.994392 | 6942 | 3 | -0.036009 |
| PHB | 5 | 0.27984 | 0.46474 | 0.999994 | 9699 | 2 | 0.29041 | 0.13962 | 0.32898 | 0.991729 | 6943 | 2 | 0.29041 |
| ERC2 | 5 | 0.33998 | 0.51369 | 0.999994 | 10932 | 2 | 1.4106 | 0.13962 | 0.32898 | 0.991729 | 6944 | 3 | 1.4106 |
| SUMO1 | 5 | 0.27078 | 0.45745 | 0.999994 | 9492 | 2 | 2.7527 | 0.13963 | 0.329 | 0.991729 | 6945 | 3 | 2.7527 |
| ZNF157 | 6 | 0.12761 | 0.29725 | 0.999994 | 5863 | 2 | 2.0519 | 0.13971 | 0.34241 | 0.994392 | 6946 | 4 | 2.0519 |
| hsa-mir-519e | 3 | 0.48367 | 0.53551 | 0.999994 | 13195 | 1 | 0.94051 | 0.13981 | 0.27694 | 0.978454 | 6947 | 2 | 0.94051 |
| TATDN1 | 2 | 0.86005 | 0.86003 | 0.999994 | 18440 | 0 | 2.6284 | 0.13995 | 0.2308 | 0.97466 | 6948 | 2 | 2.6284 |
| WDFY1 | 6 | 0.089984 | 0.22746 | 0.983354 | 4593 | 4 | -1.3135 | 0.13996 | 0.34279 | 0.994392 | 6949 | 2 | -1.3135 |
| KIAA1244 | 6 | 0.67734 | 0.79891 | 0.999994 | 16050 | 1 | 2.1326 | 0.13996 | 0.34279 | 0.994392 | 6950 | 3 | 2.1326 |
| ZBED6 | 5 | 0.040644 | 0.11749 | 0.951052 | 2544 | 2 | 1.7339 | 0.13998 | 0.32965 | 0.992034 | 6951 | 3 | 1.7339 |
| MTHFR | 5 | 0.91043 | 0.91044 | 0.999994 | 19435 | 0 | 1.1963 | 0.13998 | 0.32965 | 0.992034 | 6952 | 3 | 1.1963 |
| RGS19 | 4 | 0.59825 | 0.66437 | 0.999994 | 14930 | 1 | 2.7734 | 0.14 | 0.29533 | 0.983965 | 6953 | 2 | 2.7734 |
| NonTargetingControlG | 1 | 0.85996 | 0.86024 | 0.999994 | 18434 | 0 | 6.3722 | 0.14004 | 0.1398 | 0.965913 | 6954 | 1 | 6.3722 |
| NOP10 | 5 | 0.80087 | 0.82006 | 0.999994 | 17493 | 1 | 2.9607 | 0.14005 | 0.3298 | 0.992034 | 6955 | 3 | 2.9607 |
| NDST1 | 6 | 0.063935 | 0.17411 | 0.968171 | 3604 | 4 | -1.159 | 0.14006 | 0.34294 | 0.994392 | 6956 | 2 | -1.159 |
| TET1 | 6 | 0.63874 | 0.76901 | 0.999994 | 15496 | 2 | 0.45979 | 0.14006 | 0.34294 | 0.994392 | 6957 | 3 | 0.45979 |
| OR8B3 | 6 | 0.014032 | 0.05258 | 0.914561 | 1131 | 4 | -2.1815 | 0.14006 | 0.34294 | 0.994392 | 6958 | 2 | -2.1815 |
| MTMR1 | 5 | 0.35856 | 0.52887 | 0.999994 | 11314 | 2 | 2.7287 | 0.14007 | 0.32982 | 0.992034 | 6959 | 3 | 2.7287 |
| SGK196 | 4 | 0.1596 | 0.29907 | 0.999994 | 6722 | 2 | 1.4932 | 0.14013 | 0.29554 | 0.983965 | 6960 | 2 | 1.4932 |
| NDUFB8 | 4 | 0.41351 | 0.55955 | 0.999994 | 12161 | 2 | 2.5081 | 0.14013 | 0.29554 | 0.983965 | 6961 | 2 | 2.5081 |
| hsa-mir-4503 | 4 | 0.342 | 0.48608 | 0.999994 | 10980 | 1 | 2.1242 | 0.14026 | 0.29574 | 0.984211 | 6962 | 3 | 2.1242 |
| DTX1 | 5 | 0.78404 | 0.80979 | 0.999994 | 17266 | 1 | 1.3944 | 0.14032 | 0.33026 | 0.992034 | 6963 | 4 | 1.3944 |
| TDRKH | 6 | 0.80574 | 0.85037 | 0.999994 | 17567 | 1 | 2.1011 | 0.14037 | 0.34342 | 0.994392 | 6964 | 3 | 2.1011 |
| ZNF85 | 6 | 0.74135 | 0.82544 | 0.999994 | 16757 | 1 | 1.8184 | 0.1404 | 0.34346 | 0.994392 | 6965 | 4 | 1.8184 |
| HDX | 6 | 0.19086 | 0.37658 | 0.999994 | 7511 | 3 | -0.31459 | 0.14041 | 0.34347 | 0.994392 | 6966 | 3 | -0.31459 |
| SOGA2 | 4 | 0.86654 | 0.86612 | 0.999994 | 18567 | 0 | 2.0794 | 0.14043 | 0.296 | 0.984434 | 6967 | 3 | 2.0794 |

| | | | | | | | | | | | | | |
|---|---|---|---|---|---|---|---|---|---|---|---|---|---|
| TGM7 | 6 | 0.26282 | 0.46321 | 0.999994 | 9310 | 1 | 0.25334 | 0.14056 | 0.34371 | 0.994392 | 6968 | 3 | 0.25334 |
| ZFYVE28 | 4 | 0.31037 | 0.45363 | 0.999994 | 10332 | 2 | 0.86847 | 0.14062 | 0.29628 | 0.984782 | 6969 | 2 | 0.86847 |
| NonTargetingControlG | 2 | 0.85933 | 0.8593 | 0.999994 | 18421 | 0 | 3.0956 | 0.14067 | 0.23184 | 0.97466 | 6970 | 2 | 3.0956 |
| THAP10 | 5 | 0.40819 | 0.57036 | 0.999994 | 12081 | 2 | 2.6088 | 0.14073 | 0.33098 | 0.992034 | 6971 | 3 | 2.6088 |
| THAP7 | 5 | 0.90476 | 0.90485 | 0.999994 | 19318 | 0 | 2.4619 | 0.14074 | 0.331 | 0.992034 | 6972 | 4 | 2.4619 |
| PLEKHA2 | 6 | 0.052759 | 0.15008 | 0.966359 | 3103 | 2 | 1.8927 | 0.14074 | 0.344 | 0.994392 | 6973 | 4 | 1.8927 |
| hsa-mir-6789 | 3 | 0.16729 | 0.27092 | 0.998561 | 6915 | 2 | -1.0591 | 0.14075 | 0.27796 | 0.978454 | 6974 | 1 | -1.0591 |
| SEMG1 | 5 | 0.0046895 | 0.019584 | 0.808847 | 497 | 2 | 2.9185 | 0.14075 | 0.33101 | 0.992034 | 6975 | 3 | 2.9185 |
| HOXB1 | 4 | 0.77965 | 0.7836 | 0.999994 | 17207 | 1 | 3.5754 | 0.14078 | 0.2965 | 0.984782 | 6976 | 3 | 3.5754 |
| LBH | 6 | 0.47717 | 0.65328 | 0.999994 | 13103 | 2 | 1.4245 | 0.14081 | 0.34408 | 0.994392 | 6977 | 4 | 1.4245 |
| OR11L1 | 6 | 0.40832 | 0.6078 | 0.999994 | 12084 | 1 | 1.9794 | 0.14087 | 0.34418 | 0.994392 | 6978 | 4 | 1.9794 |
| NUDT19 | 5 | 0.92086 | 0.92103 | 0.999994 | 19651 | 0 | 0.99692 | 0.14087 | 0.33124 | 0.992034 | 6979 | 4 | 0.99692 |
| ZKSCAN8 | 5 | 0.10492 | 0.2341 | 0.984971 | 5131 | 3 | -0.65863 | 0.14088 | 0.33126 | 0.992034 | 6980 | 2 | -0.65863 |
| OR5H6 | 5 | 0.56419 | 0.70436 | 0.999994 | 14457 | 2 | 2.1374 | 0.14088 | 0.33126 | 0.992034 | 6981 | 3 | 2.1374 |
| SERF1A | 5 | 0.32537 | 0.5017 | 0.999994 | 10629 | 2 | -0.22915 | 0.14092 | 0.33133 | 0.992091 | 6982 | 1 | -0.22915 |
| INPP5J | 6 | 0.88588 | 0.89348 | 0.999994 | 18945 | 1 | 1.5808 | 0.14099 | 0.34436 | 0.994392 | 6983 | 4 | 1.5808 |
| GJA4 | 6 | 0.11278 | 0.27206 | 0.998561 | 5397 | 3 | 0.94888 | 0.14099 | 0.34436 | 0.994392 | 6984 | 3 | 0.94888 |
| LGI2 | 6 | 0.75499 | 0.83025 | 0.999994 | 16911 | 1 | 2.3667 | 0.14099 | 0.34436 | 0.994392 | 6985 | 4 | 2.3667 |
| TMEM127 | 5 | 0.39867 | 0.56245 | 0.999994 | 11927 | 2 | 2.5381 | 0.14101 | 0.33149 | 0.992121 | 6986 | 3 | 2.5381 |
| HCAR1 | 6 | 0.43783 | 0.62694 | 0.999994 | 12539 | 2 | 1.6982 | 0.14104 | 0.34446 | 0.994392 | 6987 | 4 | 1.6982 |
| STX18 | 6 | 0.010392 | 0.04036 | 0.891621 | 894 | 2 | 0.073434 | 0.14104 | 0.34446 | 0.994392 | 6988 | 3 | 0.073434 |
| FERMT1 | 6 | 0.96223 | 0.96231 | 0.999994 | 20636 | 0 | 0.16076 | 0.14104 | 0.34446 | 0.994392 | 6989 | 2 | 0.16076 |
| PCYOX1 | 6 | 0.65942 | 0.78487 | 0.999994 | 15769 | 1 | 2.2357 | 0.14104 | 0.34446 | 0.994392 | 6990 | 4 | 2.2357 |
| ZIC5 | 5 | 0.96382 | 0.96397 | 0.999994 | 20702 | 0 | 2.0756 | 0.14113 | 0.3317 | 0.992121 | 6991 | 4 | 2.0756 |
| TNFSF9 | 4 | 0.91311 | 0.91291 | 0.999994 | 19478 | 0 | 2.1643 | 0.1412 | 0.29713 | 0.984876 | 6992 | 3 | 2.1643 |
| ELMOD1 | 6 | 0.10826 | 0.26336 | 0.996734 | 5247 | 2 | 1.6643 | 0.14122 | 0.34474 | 0.994392 | 6993 | 4 | 1.6643 |
| SMN1 | 6 | 0.41111 | 0.60962 | 0.999994 | 12130 | 2 | 1.5956 | 0.14129 | 0.34485 | 0.994392 | 6994 | 4 | 1.5956 |
| MLH3 | 6 | 0.26868 | 0.47019 | 0.999994 | 9438 | 2 | 1.6789 | 0.14131 | 0.34488 | 0.994392 | 6995 | 4 | 1.6789 |
| LARP1 | 5 | 0.19314 | 0.36543 | 0.999994 | 7578 | 2 | 1.6669 | 0.14131 | 0.33205 | 0.992196 | 6996 | 3 | 1.6669 |
| LIPE | 5 | 0.10799 | 0.23887 | 0.987253 | 5239 | 1 | 1.2498 | 0.14132 | 0.33207 | 0.992196 | 6997 | 4 | 1.2498 |
| ZBED1_X | 6 | 0.50805 | 0.67427 | 0.999994 | 13553 | 2 | 0.64468 | 0.14135 | 0.34495 | 0.994392 | 6998 | 3 | 0.64468 |
| FREM1 | 6 | 0.055304 | 0.15561 | 0.967126 | 3212 | 4 | -1.3469 | 0.14135 | 0.34495 | 0.994392 | 6999 | 2 | -1.3469 |
| OR2J3 | 6 | 0.82549 | 0.85938 | 0.999994 | 17861 | 1 | 0.12629 | 0.14135 | 0.34495 | 0.994392 | 7000 | 2 | 0.12629 |
| SLC40A1 | 5 | 0.30304 | 0.48353 | 0.999994 | 10194 | 2 | 3.0394 | 0.14136 | 0.33213 | 0.992196 | 7001 | 3 | 3.0394 |
| CYP2A6 | 6 | 0.78644 | 0.84222 | 0.999994 | 17294 | 1 | 1.5169 | 0.14145 | 0.34509 | 0.994392 | 7002 | 4 | 1.5169 |
| HJURP | 5 | 0.063967 | 0.1686 | 0.967126 | 3608 | 2 | 0.84191 | 0.14149 | 0.33238 | 0.992388 | 7003 | 3 | 0.84191 |
| XPO6 | 5 | 0.60413 | 0.72768 | 0.999994 | 14998 | 1 | 0.57751 | 0.14149 | 0.33238 | 0.992388 | 7004 | 2 | 0.57751 |
| PHF10 | 4 | 0.071396 | 0.16259 | 0.967126 | 3887 | 1 | 2.4941 | 0.14153 | 0.29766 | 0.984876 | 7005 | 3 | 2.4941 |
| DDX51 | 5 | 0.080512 | 0.19542 | 0.975797 | 4252 | 2 | 2.9393 | 0.14154 | 0.33247 | 0.99251 | 7006 | 3 | 2.9393 |
| TMEM252 | 4 | 0.88414 | 0.88379 | 0.999994 | 18911 | 0 | 1.0008 | 0.14154 | 0.29767 | 0.984876 | 7007 | 3 | 1.0008 |
| hsa-mir-4483 | 3 | 0.082733 | 0.16016 | 0.967126 | 4328 | 1 | 3.9895 | 0.14156 | 0.27889 | 0.978454 | 7008 | 2 | 3.9895 |
| PRKACA | 5 | 0.44267 | 0.59906 | 0.999994 | 12619 | 2 | 0.32144 | 0.14161 | 0.33258 | 0.992569 | 7009 | 2 | 0.32144 |
| SAA4 | 4 | 0.50085 | 0.6167 | 0.999994 | 13437 | 1 | 3.4892 | 0.14166 | 0.29784 | 0.984876 | 7010 | 3 | 3.4892 |
| ARGLU1 | 4 | 0.30918 | 0.45237 | 0.999994 | 10306 | 2 | 2.1447 | 0.14168 | 0.29787 | 0.984876 | 7011 | 2 | 2.1447 |
| HNRNPK | 4 | 0.68702 | 0.71605 | 0.999994 | 16182 | 1 | 2.7049 | 0.14168 | 0.29787 | 0.984876 | 7012 | 2 | 2.7049 |
| VASN | 6 | 0.89541 | 0.9002 | 0.999994 | 19135 | 1 | 1.2499 | 0.14171 | 0.34549 | 0.994392 | 7013 | 4 | 1.2499 |
| CDH19 | 6 | 0.005971 | 0.024697 | 0.844082 | 591 | 4 | -3.1174 | 0.14171 | 0.34549 | 0.994392 | 7014 | 2 | -3.1174 |
| COMT | 6 | 0.1888 | 0.37402 | 0.999994 | 7447 | 3 | -0.29191 | 0.14171 | 0.34549 | 0.994392 | 7015 | 3 | -0.29191 |
| NTSR1 | 6 | 0.83545 | 0.86426 | 0.999994 | 18015 | 1 | 2.3339 | 0.14171 | 0.3455 | 0.994392 | 7016 | 4 | 2.3339 |
| KRT71 | 6 | 0.7304 | 0.82173 | 0.999994 | 16605 | 1 | 2.5534 | 0.14172 | 0.34551 | 0.994392 | 7017 | 4 | 2.5534 |
| BRWD1 | 6 | 0.92293 | 0.92315 | 0.999994 | 19695 | 1 | 1.691 | 0.14179 | 0.34561 | 0.994392 | 7018 | 4 | 1.691 |
| BLOC1S6 | 6 | 0.45565 | 0.63881 | 0.999994 | 12784 | 2 | 0.2028 | 0.1418 | 0.34562 | 0.994392 | 7019 | 2 | 0.2028 |

| | | | | | | | | | | | | |
|---|---|---|---|---|---|---|---|---|---|---|---|---|
| PPARGC1B | 6 | 0.89638 | 0.90091 | 0.999994 | 19150 | 1 | 0.79812 | 0.1418 | 0.34562 | 0.994392 | 7020 | 3 | 0.79812 |
| NUDT14 | 6 | 0.15634 | 0.33363 | 0.999994 | 6638 | 2 | 2.0925 | 0.14185 | 0.34569 | 0.994392 | 7021 | 4 | 2.0925 |
| GALNT4 | 6 | 0.71164 | 0.81565 | 0.999994 | 16418 | 1 | 2.4168 | 0.14192 | 0.34582 | 0.994392 | 7022 | 4 | 2.4168 |
| E2F1 | 4 | 0.12072 | 0.25158 | 0.993413 | 5652 | 1 | 2.1621 | 0.14193 | 0.29826 | 0.984876 | 7023 | 2 | 2.1621 |
| MRPS18B | 5 | 0.20955 | 0.38875 | 0.999994 | 8018 | 2 | 0.4673 | 0.14196 | 0.33323 | 0.992917 | 7024 | 2 | 0.4673 |
| LENG9 | 6 | 0.93981 | 0.93977 | 0.999994 | 20075 | 0 | 1.3607 | 0.14198 | 0.3459 | 0.994392 | 7025 | 4 | 1.3607 |
| TSR2 | 3 | 0.72879 | 0.7287 | 0.999994 | 16589 | 1 | 3.9045 | 0.14199 | 0.27933 | 0.978454 | 7026 | 2 | 3.9045 |
| NIPA1 | 5 | 0.012594 | 0.043451 | 0.891621 | 1035 | 1 | 2.7458 | 0.142 | 0.3333 | 0.992917 | 7027 | 3 | 2.7458 |
| PDZD4 | 5 | 0.14526 | 0.29566 | 0.999994 | 6331 | 2 | 1.1623 | 0.14205 | 0.3334 | 0.992917 | 7028 | 3 | 1.1623 |
| C18orf32 | 5 | 0.79004 | 0.81335 | 0.999994 | 17344 | 1 | 1.5427 | 0.14205 | 0.3334 | 0.992917 | 7029 | 4 | 1.5427 |
| TM4SF4 | 5 | 0.81637 | 0.83017 | 0.999994 | 17719 | 1 | 2.8407 | 0.14206 | 0.33341 | 0.992917 | 7030 | 3 | 2.8407 |
| FAM83E | 5 | 0.53624 | 0.67956 | 0.999994 | 14004 | 1 | 2.6265 | 0.14211 | 0.33352 | 0.992917 | 7031 | 3 | 2.6265 |
| MYL2 | 5 | 0.62666 | 0.73607 | 0.999994 | 15328 | 1 | 0.94533 | 0.14213 | 0.33354 | 0.992917 | 7032 | 4 | 0.94533 |
| hsa-mir-6723 | 4 | 0.28279 | 0.42541 | 0.999994 | 9755 | 2 | -0.36022 | 0.14218 | 0.29865 | 0.984876 | 7033 | 2 | -0.36022 |
| NR2F2 | 5 | 0.61838 | 0.73297 | 0.999994 | 15193 | 1 | 0.95991 | 0.14238 | 0.33401 | 0.992933 | 7034 | 4 | 0.95991 |
| MAFB | 6 | 0.57035 | 0.71818 | 0.999994 | 14547 | 1 | 2.0362 | 0.14241 | 0.34658 | 0.994392 | 7035 | 3 | 2.0362 |
| C9orf142 | 6 | 0.4625 | 0.6434 | 0.999994 | 12897 | 1 | 2.0335 | 0.14256 | 0.34683 | 0.994392 | 7036 | 3 | 2.0335 |
| YOD1 | 6 | 0.071494 | 0.18989 | 0.970473 | 3892 | 2 | 1.6551 | 0.14258 | 0.34686 | 0.994392 | 7037 | 4 | 1.6551 |
| OR4S2 | 5 | 0.85401 | 0.85749 | 0.999994 | 18334 | 1 | 1.873 | 0.14261 | 0.33442 | 0.993364 | 7038 | 4 | 1.873 |
| ADCK2 | 6 | 0.33412 | 0.54598 | 0.999994 | 10815 | 3 | -0.44686 | 0.14262 | 0.34694 | 0.994392 | 7039 | 2 | -0.44686 |
| DTHD1 | 6 | 0.88369 | 0.89201 | 0.999994 | 18899 | 1 | 1.1234 | 0.14262 | 0.34694 | 0.994392 | 7040 | 3 | 1.1234 |
| MAZ | 4 | 0.92002 | 0.91981 | 0.999994 | 19632 | 0 | 3.3278 | 0.14269 | 0.29939 | 0.984876 | 7041 | 3 | 3.3278 |
| hsa-mir-6881 | 1 | 0.85724 | 0.8575 | 0.999994 | 18398 | 0 | 6.3173 | 0.14276 | 0.14255 | 0.965913 | 7042 | 1 | 6.3173 |
| ATP2A1 | 5 | 0.11671 | 0.25233 | 0.994162 | 5531 | 1 | 1.1928 | 0.14279 | 0.33475 | 0.993364 | 7043 | 4 | 1.1928 |
| RNF8 | 5 | 0.12054 | 0.25817 | 0.994943 | 5644 | 2 | 0.66563 | 0.14281 | 0.33477 | 0.993364 | 7044 | 3 | 0.66563 |
| NonTargetingControlG | 2 | 0.85718 | 0.85712 | 0.999994 | 18396 | 0 | 1.7848 | 0.14282 | 0.23496 | 0.97466 | 7045 | 2 | 1.7848 |
| SAMD15 | 6 | 0.26637 | 0.4674 | 0.999994 | 9382 | 2 | 1.8494 | 0.14288 | 0.34733 | 0.994392 | 7046 | 3 | 1.8494 |
| ABCF1 | 4 | 0.93485 | 0.93467 | 0.999994 | 19964 | 0 | 2.5951 | 0.1429 | 0.2997 | 0.984876 | 7047 | 3 | 2.5951 |
| DLG1 | 6 | 0.29051 | 0.49567 | 0.999994 | 9924 | 1 | 1.5923 | 0.1429 | 0.34737 | 0.994392 | 7048 | 4 | 1.5923 |
| TRMT112 | 3 | 0.09565 | 0.17729 | 0.968171 | 4803 | 1 | 3.9691 | 0.14291 | 0.28032 | 0.978454 | 7049 | 2 | 3.9691 |
| MPC1 | 5 | 0.79938 | 0.81912 | 0.999994 | 17472 | 1 | 2.9128 | 0.14292 | 0.33496 | 0.993364 | 7050 | 3 | 2.9128 |
| LOC440563 | 4 | 0.2763 | 0.41878 | 0.999994 | 9625 | 2 | 0.45063 | 0.14292 | 0.29975 | 0.984876 | 7051 | 2 | 0.45063 |
| APOA1BP | 6 | 0.49374 | 0.66441 | 0.999994 | 13321 | 2 | -0.23478 | 0.14295 | 0.34743 | 0.994392 | 7052 | 2 | -0.23478 |
| EBF2 | 5 | 0.85341 | 0.857 | 0.999994 | 18322 | 1 | -0.52752 | 0.14307 | 0.33522 | 0.993364 | 7053 | 1 | -0.52752 |
| NonTargetingControlG | 2 | 0.85689 | 0.85683 | 0.999994 | 18390 | 0 | 4.5073 | 0.14311 | 0.23536 | 0.97466 | 7054 | 2 | 4.5073 |
| AIF1 | 6 | 0.40464 | 0.60539 | 0.999994 | 12030 | 2 | 1.8515 | 0.14313 | 0.34769 | 0.994392 | 7055 | 4 | 1.8515 |
| RNMT | 4 | 0.039519 | 0.10084 | 0.946414 | 2492 | 2 | -0.59536 | 0.14316 | 0.30011 | 0.984876 | 7056 | 2 | -0.59536 |
| NRAP | 6 | 0.33478 | 0.54671 | 0.999994 | 10828 | 3 | -0.11895 | 0.1432 | 0.34778 | 0.994392 | 7057 | 2 | -0.11895 |
| THUMPD3 | 6 | 0.047319 | 0.13818 | 0.957317 | 2870 | 4 | -1.5757 | 0.1432 | 0.34778 | 0.994392 | 7058 | 2 | -1.5757 |
| CNDP1 | 6 | 0.12108 | 0.28775 | 0.999632 | 5665 | 4 | -0.88566 | 0.1432 | 0.34778 | 0.994392 | 7059 | 2 | -0.88566 |
| PRSS27 | 6 | 0.26078 | 0.4608 | 0.999994 | 9258 | 2 | -0.48032 | 0.1432 | 0.34778 | 0.994392 | 7060 | 2 | -0.48032 |
| RNF121 | 6 | 0.98122 | 0.98113 | 0.999994 | 21123 | 0 | 1.7795 | 0.14325 | 0.34787 | 0.994392 | 7061 | 4 | 1.7795 |
| LOC100505478 | 5 | 0.82348 | 0.83489 | 0.999994 | 17830 | 1 | 2.2515 | 0.14328 | 0.33562 | 0.993964 | 7062 | 4 | 2.2515 |
| NonTargetingControlG | 2 | 0.34457 | 0.37171 | 0.999994 | 11044 | 1 | 2.7329 | 0.14332 | 0.23568 | 0.97466 | 7063 | 1 | 2.7329 |
| CNTN2 | 5 | 0.34134 | 0.51483 | 0.999994 | 10966 | 2 | 1.4807 | 0.14338 | 0.33579 | 0.994204 | 7064 | 3 | 1.4807 |
| DEFA6 | 5 | 0.364 | 0.5334 | 0.999994 | 11404 | 2 | 0.77423 | 0.14338 | 0.33579 | 0.994204 | 7065 | 3 | 0.77423 |
| SLC12A2 | 5 | 0.040894 | 0.11809 | 0.951052 | 2558 | 1 | 1.0054 | 0.1434 | 0.33584 | 0.994214 | 7066 | 4 | 1.0054 |
| ATP2A2 | 6 | 0.036048 | 0.11297 | 0.951052 | 2322 | 3 | 0.70502 | 0.14342 | 0.34812 | 0.994392 | 7067 | 3 | 0.70502 |
| GPR98 | 6 | 0.16793 | 0.34815 | 0.999994 | 6934 | 2 | 2.017 | 0.14342 | 0.34812 | 0.994392 | 7068 | 3 | 2.017 |
| FRS3 | 6 | 0.03698 | 0.11509 | 0.951052 | 2375 | 3 | 1.0443 | 0.14342 | 0.34812 | 0.994392 | 7069 | 3 | 1.0443 |
| DNTTIP1 | 6 | 0.14264 | 0.31641 | 0.999994 | 6258 | 3 | 1.2439 | 0.14342 | 0.34812 | 0.994392 | 7070 | 3 | 1.2439 |
| GSTA2 | 4 | 0.87665 | 0.87631 | 0.999994 | 18764 | 0 | 1.394 | 0.14343 | 0.30052 | 0.984876 | 7071 | 3 | 1.394 |

| | | | | | | | | | | | | | |
|---|---|---|---|---|---|---|---|---|---|---|---|---|---|
| A4GNT | 6 | 0.25839 | 0.45798 | 0.999994 | 9205 | 3 | -0.74311 | 0.14347 | 0.34819 | 0.994392 | 7072 | 2 | -0.74311 |
| GNE | 6 | 0.58951 | 0.73223 | 0.999994 | 14815 | 1 | 1.0391 | 0.14352 | 0.34827 | 0.994392 | 7073 | 4 | 1.0391 |
| H2AFX | 4 | 0.84268 | 0.84215 | 0.999994 | 18135 | 0 | 1.7667 | 0.14362 | 0.30082 | 0.984876 | 7074 | 3 | 1.7667 |
| OAS2 | 5 | 0.26967 | 0.45657 | 0.999994 | 9467 | 2 | -0.078336 | 0.14363 | 0.33625 | 0.994237 | 7075 | 2 | -0.078336 |
| FRMD8 | 4 | 0.10603 | 0.22588 | 0.983354 | 5177 | 3 | -1.4932 | 0.14366 | 0.30088 | 0.984876 | 7076 | 1 | -1.4932 |
| BRICD5 | 6 | 0.011961 | 0.04558 | 0.891621 | 990 | 3 | -0.44633 | 0.14367 | 0.34849 | 0.994392 | 7077 | 2 | -0.44633 |
| CCDC157 | 5 | 0.0094632 | 0.034524 | 0.867348 | 829 | 2 | 3.0085 | 0.14369 | 0.33635 | 0.994237 | 7078 | 3 | 3.0085 |
| NonTargetingControlG | 2 | 0.63449 | 0.63427 | 0.999994 | 15437 | 0 | 3.733 | 0.14373 | 0.23628 | 0.97466 | 7079 | 1 | 3.733 |
| RAB24 | 4 | 0.86639 | 0.86597 | 0.999994 | 18565 | 0 | 1.2299 | 0.14378 | 0.30105 | 0.984876 | 7080 | 3 | 1.2299 |
| RIMS1 | 5 | 0.24587 | 0.43746 | 0.999994 | 8916 | 1 | 2.2522 | 0.14378 | 0.3365 | 0.994237 | 7081 | 4 | 2.2522 |
| NDUFA10 | 6 | 0.49586 | 0.66586 | 0.999994 | 13354 | 1 | 2.0989 | 0.1438 | 0.3487 | 0.994392 | 7082 | 4 | 2.0989 |
| LRRC66 | 5 | 0.89985 | 0.89997 | 0.999994 | 19218 | 0 | 0.88662 | 0.1438 | 0.33655 | 0.994237 | 7083 | 4 | 0.88662 |
| FCGRT | 6 | 0.86934 | 0.88294 | 0.999994 | 18622 | 1 | 1.7294 | 0.14381 | 0.34871 | 0.994392 | 7084 | 4 | 1.7294 |
| hsa-mir-941-1 | 4 | 0.86283 | 0.86245 | 0.999994 | 18503 | 0 | 3.2434 | 0.14383 | 0.30114 | 0.984876 | 7085 | 3 | 3.2434 |
| SFR1 | 4 | 0.22898 | 0.37046 | 0.999994 | 8503 | 2 | 1.886 | 0.14383 | 0.30114 | 0.984876 | 7086 | 2 | 1.886 |
| MAGIX | 3 | 0.40108 | 0.48491 | 0.999994 | 11961 | 1 | 3.5529 | 0.14399 | 0.28152 | 0.978454 | 7087 | 2 | 3.5529 |
| CHKB | 5 | 0.16556 | 0.3256 | 0.999994 | 6872 | 2 | 2.7851 | 0.14404 | 0.33694 | 0.994237 | 7088 | 3 | 2.7851 |
| TM4SF19 | 6 | 0.46186 | 0.64297 | 0.999994 | 12880 | 1 | 1.8331 | 0.14404 | 0.34905 | 0.994392 | 7089 | 4 | 1.8331 |
| CD79A | 6 | 0.44207 | 0.62979 | 0.999994 | 12611 | 2 | 1.7605 | 0.14407 | 0.34909 | 0.994392 | 7090 | 4 | 1.7605 |
| NonTargetingControlG | 2 | 0.85591 | 0.85584 | 0.999994 | 18369 | 0 | 3.5901 | 0.14409 | 0.23684 | 0.97466 | 7091 | 2 | 3.5901 |
| ANKDD1A | 5 | 0.15399 | 0.30875 | 0.999994 | 6567 | 3 | -1.0425 | 0.14413 | 0.33711 | 0.994237 | 7092 | 2 | -1.0425 |
| CASP8 | 4 | 0.1256 | 0.26004 | 0.994943 | 5808 | 2 | -1.3086 | 0.14416 | 0.30163 | 0.985764 | 7093 | 1 | -1.3086 |
| DCAKD | 5 | 0.8302 | 0.83963 | 0.999994 | 17936 | 1 | 1.8752 | 0.14419 | 0.33722 | 0.994237 | 7094 | 4 | 1.8752 |
| WDR44 | 6 | 0.63601 | 0.7669 | 0.999994 | 15463 | 2 | 1.4667 | 0.14421 | 0.34932 | 0.994392 | 7095 | 4 | 1.4667 |
| BNIP3L | 6 | 0.86617 | 0.88101 | 0.999994 | 18561 | 1 | 2.3753 | 0.14421 | 0.34932 | 0.994392 | 7096 | 4 | 2.3753 |
| EDEM2 | 6 | 0.23702 | 0.43244 | 0.999994 | 8696 | 3 | -0.056695 | 0.14421 | 0.34933 | 0.994392 | 7097 | 3 | -0.056695 |
| CD99_X | 6 | 0.7133 | 0.81617 | 0.999994 | 16432 | 1 | 1.7935 | 0.14434 | 0.34953 | 0.994392 | 7098 | 4 | 1.7935 |
| CCPG1 | 5 | 0.53616 | 0.67949 | 0.999994 | 14001 | 2 | -0.034728 | 0.14442 | 0.33763 | 0.994237 | 7099 | 2 | -0.034728 |
| USP54 | 5 | 0.047964 | 0.13551 | 0.95468 | 2888 | 3 | -1.3942 | 0.14442 | 0.33763 | 0.994237 | 7100 | 1 | -1.3942 |
| FAM177B | 5 | 0.93086 | 0.93114 | 0.999994 | 19874 | 0 | 0.53181 | 0.14442 | 0.33763 | 0.994237 | 7101 | 2 | 0.53181 |
| ARF4 | 6 | 0.93694 | 0.93696 | 0.999994 | 20005 | 0 | 1.5994 | 0.14446 | 0.3497 | 0.994392 | 7102 | 4 | 1.5994 |
| CELSR3 | 4 | 0.9142 | 0.91395 | 0.999994 | 19493 | 0 | 2.8435 | 0.14449 | 0.30212 | 0.985876 | 7103 | 3 | 2.8435 |
| TAL2 | 5 | 0.42024 | 0.58027 | 0.999994 | 12253 | 1 | 0.9611 | 0.14457 | 0.33789 | 0.994237 | 7104 | 4 | 0.9611 |
| ATP4A | 6 | 0.66004 | 0.78535 | 0.999994 | 15780 | 2 | 1.5954 | 0.14458 | 0.34986 | 0.994392 | 7105 | 4 | 1.5954 |
| ZNF556 | 4 | 0.18302 | 0.3232 | 0.999994 | 7291 | 1 | 3.077 | 0.14458 | 0.30225 | 0.985876 | 7106 | 2 | 3.077 |
| KRTAP4-4 | 4 | 0.54618 | 0.6378 | 0.999994 | 14153 | 1 | 3.1087 | 0.14458 | 0.30225 | 0.985876 | 7107 | 2 | 3.1087 |
| ZCCHC18 | 4 | 0.34571 | 0.48993 | 0.999994 | 11063 | 2 | 2.3685 | 0.14458 | 0.30225 | 0.985876 | 7108 | 2 | 2.3685 |
| PHOX2B | 5 | 0.061994 | 0.16539 | 0.967126 | 3512 | 2 | 2.6944 | 0.14464 | 0.33803 | 0.994237 | 7109 | 3 | 2.6944 |
| SPIDR | 3 | 0.11817 | 0.20706 | 0.978744 | 5578 | 2 | -2.0449 | 0.14466 | 0.28225 | 0.978454 | 7110 | 1 | -2.0449 |
| KAT7 | 3 | 0.82959 | 0.82942 | 0.999994 | 17922 | 0 | 0.98172 | 0.14466 | 0.28225 | 0.978454 | 7111 | 2 | 0.98172 |
| GM2A | 6 | 0.52279 | 0.68445 | 0.999994 | 13791 | 2 | 1.5115 | 0.14469 | 0.35004 | 0.994392 | 7112 | 4 | 1.5115 |
| PKHD1L1 | 6 | 0.84952 | 0.87166 | 0.999994 | 18250 | 1 | 1.9309 | 0.14469 | 0.35004 | 0.994392 | 7113 | 4 | 1.9309 |
| NonTargetingControlG | 2 | 0.8553 | 0.8552 | 0.999994 | 18352 | 0 | 2.9111 | 0.1447 | 0.23772 | 0.97466 | 7114 | 2 | 2.9111 |
| DOK6 | 5 | 0.044559 | 0.12723 | 0.951052 | 2740 | 2 | 2.7327 | 0.14472 | 0.33818 | 0.994237 | 7115 | 3 | 2.7327 |
| BFSP1 | 6 | 0.090595 | 0.22869 | 0.983354 | 4620 | 4 | -0.96244 | 0.14474 | 0.35012 | 0.994392 | 7116 | 2 | -0.96244 |
| TMEM190 | 5 | 0.49715 | 0.64542 | 0.999994 | 13383 | 1 | 1.6546 | 0.14476 | 0.33824 | 0.994237 | 7117 | 4 | 1.6546 |
| TUBGCP3 | 4 | 0.55484 | 0.64208 | 0.999994 | 14285 | 1 | 2.3456 | 0.14485 | 0.30265 | 0.985876 | 7118 | 3 | 2.3456 |
| LRRN1 | 5 | 0.051787 | 0.14472 | 0.963012 | 3056 | 1 | 0.45355 | 0.14492 | 0.33853 | 0.994237 | 7119 | 2 | 0.45355 |
| TACC1 | 5 | 0.11958 | 0.25672 | 0.994943 | 5617 | 3 | -1.3786 | 0.14492 | 0.33853 | 0.994237 | 7120 | 2 | -1.3786 |
| SYF2 | 6 | 0.077108 | 0.20151 | 0.977164 | 4110 | 2 | 2.045 | 0.14494 | 0.35043 | 0.994392 | 7121 | 4 | 2.045 |
| SLITRK6 | 6 | 0.22151 | 0.41393 | 0.999994 | 8320 | 3 | 1.4395 | 0.14524 | 0.35088 | 0.994392 | 7122 | 3 | 1.4395 |
| CHST13 | 5 | 0.77948 | 0.80714 | 0.999994 | 17203 | 1 | 1.4036 | 0.14529 | 0.33917 | 0.994237 | 7123 | 4 | 1.4036 |

| | | | | | | | | | | | | | |
|---|---|---|---|---|---|---|---|---|---|---|---|---|---|
| NonTargetingControlG | 2 | 0.85469 | 0.8546 | 0.999994 | 18345 | 0 | 3.811 | 0.14531 | 0.23859 | 0.97466 | 7124 | 2 | 3.811 |
| LAPTM4A | 5 | 0.90815 | 0.90818 | 0.999994 | 19389 | 0 | 0.38247 | 0.14533 | 0.33923 | 0.994237 | 7125 | 2 | 0.38247 |
| SAR1B | 5 | 0.84675 | 0.85176 | 0.999994 | 18205 | 1 | 0.15628 | 0.14533 | 0.33923 | 0.994237 | 7126 | 2 | 0.15628 |
| SIX6 | 5 | 0.32001 | 0.49733 | 0.999994 | 10512 | 1 | 1.8107 | 0.14536 | 0.33928 | 0.994237 | 7127 | 4 | 1.8107 |
| RAB40B | 6 | 0.16093 | 0.33942 | 0.999994 | 6762 | 3 | 0.091753 | 0.14536 | 0.35107 | 0.994392 | 7128 | 3 | 0.091753 |
| KCTD17 | 4 | 0.81371 | 0.81379 | 0.999994 | 17687 | 1 | 3.1951 | 0.14537 | 0.30341 | 0.98595 | 7129 | 3 | 3.1951 |
| VARS | 4 | 0.27882 | 0.42136 | 0.999994 | 9668 | 2 | 2.0573 | 0.14537 | 0.30341 | 0.98595 | 7130 | 2 | 2.0573 |
| BAG5 | 4 | 0.28398 | 0.42661 | 0.999994 | 9783 | 1 | 3.4673 | 0.14537 | 0.30341 | 0.98595 | 7131 | 3 | 3.4673 |
| CHD9 | 6 | 0.29199 | 0.49736 | 0.999994 | 9949 | 2 | 2.1967 | 0.14538 | 0.3511 | 0.994392 | 7132 | 4 | 2.1967 |
| AFAP1L2 | 6 | 0.52466 | 0.68577 | 0.999994 | 13811 | 2 | 1.7647 | 0.14544 | 0.3512 | 0.994392 | 7133 | 3 | 1.7647 |
| TEX15 | 6 | 0.018451 | 0.066995 | 0.93914 | 1385 | 3 | -3.2058 | 0.14544 | 0.3512 | 0.994392 | 7134 | 2 | -3.2058 |
| ABRA | 6 | 0.13691 | 0.30915 | 0.999994 | 6105 | 2 | 0.19154 | 0.14544 | 0.3512 | 0.994392 | 7135 | 3 | 0.19154 |
| ZNF225 | 6 | 0.95921 | 0.9593 | 0.999994 | 20561 | 0 | 1.0999 | 0.14544 | 0.3512 | 0.994392 | 7136 | 4 | 1.0999 |
| hsa-mir-377 | 4 | 0.40633 | 0.55218 | 0.999994 | 12047 | 1 | 2.5925 | 0.14544 | 0.30351 | 0.986066 | 7137 | 3 | 2.5925 |
| OGDHL | 6 | 0.65697 | 0.78296 | 0.999994 | 15742 | 2 | 1.9408 | 0.14546 | 0.35123 | 0.994392 | 7138 | 4 | 1.9408 |
| LRRC18 | 3 | 0.81179 | 0.81156 | 0.999994 | 17652 | 0 | 3.6796 | 0.14546 | 0.28314 | 0.978454 | 7139 | 2 | 3.6796 |
| NonTargetingControlG | 2 | 0.80192 | 0.80153 | 0.999994 | 17510 | 0 | 4.2761 | 0.14552 | 0.23889 | 0.97466 | 7140 | 2 | 4.2761 |
| PDE6D | 5 | 0.28436 | 0.46838 | 0.999994 | 9792 | 1 | 2.5526 | 0.14556 | 0.33966 | 0.994237 | 7141 | 3 | 2.5526 |
| RMND1 | 5 | 0.42343 | 0.58296 | 0.999994 | 12301 | 2 | 2.123 | 0.14556 | 0.33966 | 0.994237 | 7142 | 3 | 2.123 |
| CCAR2 | 5 | 0.28111 | 0.46576 | 0.999994 | 9724 | 2 | 2.6458 | 0.14558 | 0.33969 | 0.994237 | 7143 | 3 | 2.6458 |
| TMTC3 | 6 | 0.25218 | 0.45052 | 0.999994 | 9060 | 3 | 0.69795 | 0.14558 | 0.35141 | 0.994392 | 7144 | 3 | 0.69795 |
| PTGS2 | 6 | 0.94692 | 0.94688 | 0.999994 | 20263 | 0 | 0.84588 | 0.14558 | 0.35141 | 0.994392 | 7145 | 3 | 0.84588 |
| OSBPL7 | 6 | 0.06615 | 0.1787 | 0.968171 | 3691 | 4 | -1.1838 | 0.14558 | 0.35141 | 0.994392 | 7146 | 2 | -1.1838 |
| ZNF705B | 5 | 0.59339 | 0.72375 | 0.999994 | 14870 | 1 | 2.1613 | 0.14562 | 0.33978 | 0.994237 | 7147 | 4 | 2.1613 |
| SLC34A1 | 5 | 0.83689 | 0.8444 | 0.999994 | 18033 | 1 | 1.4829 | 0.14565 | 0.33983 | 0.994237 | 7148 | 4 | 1.4829 |
| CDY1B | 2 | 0.8543 | 0.85423 | 0.999994 | 18338 | 0 | 1.8971 | 0.1457 | 0.23916 | 0.97466 | 7149 | 2 | 1.8971 |
| BCL2L1 | 5 | 0.86312 | 0.86503 | 0.999994 | 18507 | 1 | 2.8818 | 0.14571 | 0.33993 | 0.994237 | 7150 | 3 | 2.8818 |
| COA4 | 5 | 0.58752 | 0.72166 | 0.999994 | 14782 | 1 | 1.064 | 0.14584 | 0.34017 | 0.994237 | 7151 | 4 | 1.064 |
| GART | 6 | 0.33424 | 0.5461 | 0.999994 | 10819 | 3 | 1.647 | 0.14585 | 0.35183 | 0.994392 | 7152 | 3 | 1.647 |
| RBM12B | 6 | 0.61149 | 0.74843 | 0.999994 | 15097 | 1 | 2.1 | 0.14585 | 0.35183 | 0.994392 | 7153 | 3 | 2.1 |
| RAB1B | 6 | 0.3635 | 0.57912 | 0.999994 | 11396 | 2 | 1.9236 | 0.14585 | 0.35183 | 0.994392 | 7154 | 3 | 1.9236 |
| SLX1A | 6 | 0.33856 | 0.55103 | 0.999994 | 10910 | 1 | 1.5993 | 0.14597 | 0.352 | 0.994392 | 7155 | 4 | 1.5993 |
| FLNA | 6 | 0.66073 | 0.78589 | 0.999994 | 15790 | 2 | 1.5904 | 0.14598 | 0.35202 | 0.994392 | 7156 | 4 | 1.5904 |
| TSHZ3 | 5 | 0.41556 | 0.5764 | 0.999994 | 12186 | 1 | 1.5329 | 0.146 | 0.34045 | 0.994237 | 7157 | 4 | 1.5329 |
| ROR1 | 4 | 0.76418 | 0.77114 | 0.999994 | 17018 | 1 | -0.091611 | 0.14602 | 0.30439 | 0.986296 | 7158 | 1 | -0.091611 |
| LRRC2 | 4 | 0.67699 | 0.70969 | 0.999994 | 16040 | 1 | 2.9883 | 0.14604 | 0.30441 | 0.986296 | 7159 | 2 | 2.9883 |
| NonTargetingControlG | 2 | 0.85387 | 0.85379 | 0.999994 | 18330 | 0 | 2.6145 | 0.14613 | 0.23979 | 0.97466 | 7160 | 2 | 2.6145 |
| WNT7B | 6 | 0.34206 | 0.55501 | 0.999994 | 10987 | 2 | 1.6333 | 0.14619 | 0.35235 | 0.994392 | 7161 | 3 | 1.6333 |
| DCAF8L2 | 6 | 0.29349 | 0.4991 | 0.999994 | 9984 | 3 | -0.10714 | 0.14619 | 0.35235 | 0.994392 | 7162 | 3 | -0.10714 |
| CLGN | 5 | 0.43026 | 0.58864 | 0.999994 | 12416 | 2 | 1.0144 | 0.14625 | 0.34086 | 0.994237 | 7163 | 3 | 1.0144 |
| DEF6 | 6 | 0.35741 | 0.57239 | 0.999994 | 11293 | 2 | 0.38444 | 0.1463 | 0.35252 | 0.994392 | 7164 | 3 | 0.38444 |
| GSC | 6 | 0.45706 | 0.63976 | 0.999994 | 12807 | 2 | 0.42487 | 0.1463 | 0.35252 | 0.994392 | 7165 | 3 | 0.42487 |
| ERO1LB | 6 | 0.66234 | 0.78713 | 0.999994 | 15815 | 2 | -0.062747 | 0.1463 | 0.35252 | 0.994392 | 7166 | 2 | -0.062747 |
| TRIM35 | 4 | 0.29866 | 0.44163 | 0.999994 | 10103 | 1 | 3.3379 | 0.14639 | 0.30494 | 0.986316 | 7167 | 3 | 3.3379 |
| PDIK1L | 6 | 0.20788 | 0.3975 | 0.999994 | 7973 | 1 | 2.1076 | 0.14648 | 0.35279 | 0.994392 | 7168 | 3 | 2.1076 |
| SASH3 | 6 | 0.023193 | 0.08219 | 0.940299 | 1663 | 2 | 1.4863 | 0.1465 | 0.35283 | 0.994392 | 7169 | 4 | 1.4863 |
| GABRA1 | 5 | 0.85538 | 0.85857 | 0.999994 | 18355 | 1 | 2.8575 | 0.14651 | 0.34133 | 0.994237 | 7170 | 4 | 2.8575 |
| hsa-mir-6722 | 3 | 0.7702 | 0.7702 | 0.999994 | 17085 | 0 | 3.49 | 0.14652 | 0.28431 | 0.978454 | 7171 | 2 | 3.49 |
| hsa-mir-4251 | 3 | 0.37685 | 0.47096 | 0.999994 | 11606 | 1 | -0.061509 | 0.14652 | 0.28431 | 0.978454 | 7172 | 1 | -0.061509 |
| FAM19A3 | 6 | 0.14115 | 0.3145 | 0.999994 | 6223 | 2 | 2.3195 | 0.14653 | 0.35289 | 0.994392 | 7173 | 4 | 2.3195 |
| RIOK3 | 4 | 0.89471 | 0.89439 | 0.999994 | 19119 | 0 | 1.7224 | 0.14658 | 0.30521 | 0.986316 | 7174 | 3 | 1.7224 |
| NonTargetingControlG | 2 | 0.40675 | 0.42242 | 0.999994 | 12056 | 1 | 2.8957 | 0.14667 | 0.24057 | 0.97466 | 7175 | 1 | 2.8957 |

| | | | | | | | | | | | | | |
|---|---|---|---|---|---|---|---|---|---|---|---|---|---|
| FAM122A | 6 | 0.22293 | 0.41568 | 0.999994 | 8355 | 3 | -0.35389 | 0.14667 | 0.3531 | 0.994392 | 7176 | 3 | -0.35389 |
| SLBP | 5 | 0.48285 | 0.63313 | 0.999994 | 13187 | 2 | 0.018189 | 0.14671 | 0.3417 | 0.994392 | 7177 | 2 | 0.018189 |
| SH3BGRL2 | 5 | 0.9573 | 0.95744 | 0.999994 | 20523 | 0 | 2.211 | 0.14674 | 0.34176 | 0.994392 | 7178 | 3 | 2.211 |
| KLF2 | 5 | 0.90324 | 0.90331 | 0.999994 | 19285 | 0 | -0.40401 | 0.14674 | 0.34176 | 0.994392 | 7179 | 2 | -0.40401 |
| CAMK2D | 5 | 0.92866 | 0.92891 | 0.999994 | 19829 | 0 | 1.3472 | 0.14674 | 0.34176 | 0.994392 | 7180 | 3 | 1.3472 |
| ANKRD49 | 5 | 0.77746 | 0.80598 | 0.999994 | 17176 | 1 | 1.194 | 0.14683 | 0.3419 | 0.994392 | 7181 | 4 | 1.194 |
| CATSPER2 | 6 | 0.43835 | 0.62729 | 0.999994 | 12543 | 2 | 1.977 | 0.14701 | 0.35359 | 0.994392 | 7182 | 3 | 1.977 |
| P2RY12 | 4 | 0.81557 | 0.81556 | 0.999994 | 17707 | 1 | 1.5737 | 0.14706 | 0.30591 | 0.986404 | 7183 | 3 | 1.5737 |
| ST3GAL3 | 6 | 0.44041 | 0.62869 | 0.999994 | 12583 | 2 | 2.6678 | 0.14708 | 0.3537 | 0.994392 | 7184 | 4 | 2.6678 |
| FERMT2 | 5 | 0.64616 | 0.74357 | 0.999994 | 15603 | 1 | 1.7143 | 0.1471 | 0.34237 | 0.994392 | 7185 | 3 | 1.7143 |
| MPI | 6 | 0.1161 | 0.27841 | 0.999632 | 5517 | 3 | -0.58225 | 0.14717 | 0.35384 | 0.994392 | 7186 | 3 | -0.58225 |
| USH1C | 6 | 0.89555 | 0.90031 | 0.999994 | 19138 | 1 | 2.575 | 0.14723 | 0.35392 | 0.994392 | 7187 | 4 | 2.575 |
| LGALS3 | 6 | 0.30979 | 0.51812 | 0.999994 | 10323 | 2 | 1.666 | 0.14724 | 0.35392 | 0.994392 | 7188 | 4 | 1.666 |
| PRKCG | 6 | 0.34195 | 0.55489 | 0.999994 | 10979 | 3 | 0.19848 | 0.14724 | 0.35393 | 0.994392 | 7189 | 3 | 0.19848 |
| KCNJ16 | 6 | 0.068773 | 0.1842 | 0.968526 | 3788 | 3 | -0.56757 | 0.14724 | 0.35393 | 0.994392 | 7190 | 2 | -0.56757 |
| POTEH | 2 | 0.8527 | 0.85259 | 0.999994 | 18311 | 0 | 3.4013 | 0.1473 | 0.2415 | 0.97466 | 7191 | 2 | 3.4013 |
| DPM1 | 5 | 0.53782 | 0.68098 | 0.999994 | 14025 | 2 | -0.56641 | 0.14735 | 0.34283 | 0.994392 | 7192 | 2 | -0.56641 |
| NonTargetingControlG | 2 | 0.85262 | 0.85251 | 0.999994 | 18310 | 0 | 3.5046 | 0.14738 | 0.24162 | 0.97466 | 7193 | 2 | 3.5046 |
| LCE3C | 6 | 0.1518 | 0.32792 | 0.999994 | 6500 | 2 | 1.7645 | 0.14743 | 0.3542 | 0.994392 | 7194 | 4 | 1.7645 |
| ZNF780A | 6 | 0.086819 | 0.22111 | 0.982401 | 4475 | 2 | 1.5964 | 0.14744 | 0.35422 | 0.994392 | 7195 | 4 | 1.5964 |
| DDX11 | 5 | 0.87367 | 0.87439 | 0.999994 | 18705 | 1 | 0.53135 | 0.14746 | 0.34303 | 0.994392 | 7196 | 2 | 0.53135 |
| SNX32 | 5 | 0.56236 | 0.70274 | 0.999994 | 14418 | 2 | 2.2859 | 0.14746 | 0.34303 | 0.994392 | 7197 | 3 | 2.2859 |
| RASL10B | 6 | 0.028369 | 0.094996 | 0.945031 | 1919 | 2 | 1.0325 | 0.14749 | 0.35429 | 0.994392 | 7198 | 3 | 1.0325 |
| ISLR2 | 6 | 0.14509 | 0.31948 | 0.999994 | 6326 | 3 | 0.82555 | 0.14749 | 0.35429 | 0.994392 | 7199 | 3 | 0.82555 |
| MFSD2B | 6 | 0.89754 | 0.90174 | 0.999994 | 19170 | 1 | 0.94317 | 0.14749 | 0.35429 | 0.994392 | 7200 | 3 | 0.94317 |
| CST7 | 4 | 0.40418 | 0.54999 | 0.999994 | 12022 | 1 | 3.2945 | 0.14754 | 0.3066 | 0.987798 | 7201 | 3 | 3.2945 |
| CHURC1 | 6 | 0.8623 | 0.87873 | 0.999994 | 18492 | 1 | 0.38673 | 0.14754 | 0.35436 | 0.994392 | 7202 | 2 | 0.38673 |
| PICK1 | 4 | 0.56178 | 0.64558 | 0.999994 | 14403 | 1 | 1.6719 | 0.14769 | 0.30683 | 0.987798 | 7203 | 3 | 1.6719 |
| CYP2S1 | 6 | 0.1718 | 0.35297 | 0.999994 | 7031 | 3 | 0.83807 | 0.14776 | 0.35471 | 0.994392 | 7204 | 3 | 0.83807 |
| WDR76 | 6 | 0.62684 | 0.75995 | 0.999994 | 15333 | 2 | 1.0393 | 0.14776 | 0.35471 | 0.994392 | 7205 | 4 | 1.0393 |
| ATAD3B | 5 | 0.094024 | 0.21698 | 0.980699 | 4750 | 1 | 1.7266 | 0.14776 | 0.34357 | 0.994392 | 7206 | 4 | 1.7266 |
| LOC100506388 | 6 | 0.36282 | 0.57847 | 0.999994 | 11387 | 2 | 2.0352 | 0.14778 | 0.35474 | 0.994392 | 7207 | 3 | 2.0352 |
| ZNF329 | 5 | 0.91403 | 0.91411 | 0.999994 | 19487 | 0 | 0.74219 | 0.14786 | 0.34375 | 0.994392 | 7208 | 4 | 0.74219 |
| AKAP14 | 5 | 0.7321 | 0.78169 | 0.999994 | 16630 | 1 | 1.5024 | 0.14798 | 0.34395 | 0.994392 | 7209 | 4 | 1.5024 |
| PTRH1 | 6 | 0.34911 | 0.563 | 0.999994 | 11131 | 1 | 1.5594 | 0.14801 | 0.35509 | 0.994392 | 7210 | 4 | 1.5594 |
| CCL21 | 5 | 0.45667 | 0.61088 | 0.999994 | 12801 | 1 | 1.1688 | 0.14803 | 0.34404 | 0.994392 | 7211 | 4 | 1.1688 |
| NonTargetingControlG | 2 | 0.45263 | 0.46163 | 0.999994 | 12742 | 1 | 3.0866 | 0.14803 | 0.24258 | 0.97466 | 7212 | 1 | 3.0866 |
| SMCO3 | 6 | 0.21317 | 0.40394 | 0.999994 | 8093 | 3 | 0.15952 | 0.14803 | 0.35511 | 0.994392 | 7213 | 3 | 0.15952 |
| HIF3A | 6 | 0.056063 | 0.15723 | 0.967126 | 3249 | 1 | 1.8626 | 0.14803 | 0.35511 | 0.994392 | 7214 | 4 | 1.8626 |
| TRIT1 | 5 | 0.88076 | 0.88099 | 0.999994 | 18847 | 1 | 1.0406 | 0.1481 | 0.34416 | 0.994392 | 7215 | 4 | 1.0406 |
| RNF41 | 3 | 0.062374 | 0.13267 | 0.95318 | 3530 | 2 | -3.1851 | 0.14818 | 0.28611 | 0.978871 | 7216 | 1 | -3.1851 |
| ZNF160 | 5 | 0.098593 | 0.22421 | 0.983354 | 4915 | 2 | -0.26847 | 0.14818 | 0.34431 | 0.994392 | 7217 | 2 | -0.26847 |
| SPINK14 | 5 | 0.4612 | 0.6147 | 0.999994 | 12866 | 2 | -0.52149 | 0.14818 | 0.34431 | 0.994392 | 7218 | 2 | -0.52149 |
| ARSI | 6 | 0.40294 | 0.6043 | 0.999994 | 11997 | 2 | 2.2957 | 0.14822 | 0.35538 | 0.994392 | 7219 | 4 | 2.2957 |
| DEFA1 | 6 | 0.66887 | 0.7923 | 0.999994 | 15906 | 1 | 1.8927 | 0.14825 | 0.35542 | 0.994392 | 7220 | 4 | 1.8927 |
| ANAPC2 | 2 | 0.73023 | 0.72997 | 0.999994 | 16600 | 0 | 4.0118 | 0.14827 | 0.24292 | 0.97466 | 7221 | 1 | 4.0118 |
| PPP1R1A | 6 | 0.024055 | 0.084752 | 0.94395 | 1710 | 3 | -1.6857 | 0.14831 | 0.3555 | 0.994392 | 7222 | 3 | -1.6857 |
| NRCAM | 5 | 0.15234 | 0.30624 | 0.999994 | 6515 | 1 | 1.8695 | 0.14836 | 0.34463 | 0.994392 | 7223 | 4 | 1.8695 |
| SLC5A1 | 6 | 0.38784 | 0.59455 | 0.999994 | 11765 | 2 | 1.9754 | 0.14837 | 0.35559 | 0.994392 | 7224 | 4 | 1.9754 |
| LCE2D | 3 | 0.032295 | 0.080408 | 0.93914 | 2120 | 2 | -4.1825 | 0.14842 | 0.28639 | 0.978871 | 7225 | 1 | -4.1825 |
| SKI | 4 | 0.014385 | 0.047213 | 0.896226 | 1159 | 2 | 1.6085 | 0.14844 | 0.30796 | 0.988566 | 7226 | 2 | 1.6085 |
| POU4F2 | 6 | 0.32052 | 0.53049 | 0.999994 | 10523 | 2 | 0.75698 | 0.14846 | 0.35571 | 0.994392 | 7227 | 3 | 0.75698 |

| | | | | | | | | | | | | | |
|---|---|---|---|---|---|---|---|---|---|---|---|---|---|
| STXBP2 | 6 | 0.9735 | 0.97341 | 0.999994 | 20908 | 0 | 1.7774 | 0.14846 | 0.35571 | 0.994392 | 7228 | 4 | 1.7774 |
| APOL3 | 6 | 0.57406 | 0.72086 | 0.999994 | 14604 | 1 | 1.4999 | 0.14849 | 0.35576 | 0.994392 | 7229 | 4 | 1.4999 |
| RPL36A-HNRNPH2 | 2 | 0.8515 | 0.85137 | 0.999994 | 18285 | 0 | 3.7023 | 0.1485 | 0.24326 | 0.97466 | 7230 | 2 | 3.7023 |
| HYDIN | 4 | 0.30126 | 0.44429 | 0.999994 | 10150 | 2 | 0.4205 | 0.1485 | 0.30805 | 0.988566 | 7231 | 2 | 0.4205 |
| IFT74 | 6 | 0.00087723 | 0.0044992 | 0.56977 | 150 | 2 | 0.45716 | 0.14855 | 0.35585 | 0.994392 | 7232 | 3 | 0.45716 |
| MOBP | 5 | 0.059173 | 0.16068 | 0.967126 | 3386 | 1 | 2.187 | 0.14857 | 0.34501 | 0.994392 | 7233 | 3 | 2.187 |
| C15orf26 | 5 | 0.7303 | 0.78079 | 0.999994 | 16601 | 1 | 0.77194 | 0.14867 | 0.34518 | 0.994392 | 7234 | 4 | 0.77194 |
| ZAP70 | 6 | 0.1191 | 0.28403 | 0.999632 | 5604 | 2 | 2.4921 | 0.14874 | 0.35612 | 0.994392 | 7235 | 4 | 2.4921 |
| C2orf62 | 4 | 0.12544 | 0.25977 | 0.994943 | 5803 | 3 | -1.0261 | 0.14875 | 0.30843 | 0.988566 | 7236 | 1 | -1.0261 |
| SYCP3 | 4 | 0.87106 | 0.87068 | 0.999994 | 18655 | 0 | 1.3061 | 0.14875 | 0.30843 | 0.988566 | 7237 | 3 | 1.3061 |
| TMPO | 6 | 0.14763 | 0.32265 | 0.999994 | 6394 | 3 | 0.60146 | 0.1488 | 0.35622 | 0.994392 | 7238 | 3 | 0.60146 |
| OR11A1 | 4 | 0.73879 | 0.75158 | 0.999994 | 16718 | 1 | 2.6059 | 0.14886 | 0.30857 | 0.988566 | 7239 | 3 | 2.6059 |
| NonTargetingControlG | 2 | 0.74059 | 0.74025 | 0.999994 | 16745 | 0 | 3.9954 | 0.14889 | 0.24384 | 0.97466 | 7240 | 1 | 3.9954 |
| KSR2 | 5 | 0.31202 | 0.49082 | 0.999994 | 10360 | 2 | 0.15287 | 0.14892 | 0.34563 | 0.994392 | 7241 | 2 | 0.15287 |
| GCDH | 5 | 0.15114 | 0.30448 | 0.999994 | 6485 | 2 | 0.77794 | 0.14892 | 0.34563 | 0.994392 | 7242 | 3 | 0.77794 |
| CXCR4 | 5 | 0.087361 | 0.20636 | 0.978744 | 4490 | 3 | -1.2362 | 0.14892 | 0.34563 | 0.994392 | 7243 | 2 | -1.2362 |
| SH3D19 | 6 | 0.91488 | 0.91578 | 0.999994 | 19506 | 1 | 1.0137 | 0.14896 | 0.35646 | 0.994392 | 7244 | 4 | 1.0137 |
| GPR27 | 6 | 0.33339 | 0.54516 | 0.999994 | 10792 | 2 | 1.0656 | 0.14896 | 0.35646 | 0.994392 | 7245 | 4 | 1.0656 |
| DOK7 | 6 | 0.23253 | 0.42714 | 0.999994 | 8584 | 2 | 2.053 | 0.14897 | 0.35649 | 0.994392 | 7246 | 3 | 2.053 |
| DCLRE1C | 5 | 0.7617 | 0.7971 | 0.999994 | 16998 | 1 | 1.0213 | 0.14899 | 0.34576 | 0.994392 | 7247 | 4 | 1.0213 |
| PFDN2 | 4 | 0.45903 | 0.59849 | 0.999994 | 12833 | 1 | 1.697 | 0.14902 | 0.30883 | 0.988566 | 7248 | 3 | 1.697 |
| HIST1H2AA | 5 | 0.66874 | 0.75276 | 0.999994 | 15902 | 1 | 2.9342 | 0.1491 | 0.34596 | 0.994392 | 7249 | 3 | 2.9342 |
| RNF157 | 5 | 0.46026 | 0.61388 | 0.999994 | 12849 | 1 | 2.9342 | 0.1491 | 0.34596 | 0.994392 | 7250 | 3 | 2.9342 |
| CCL16 | 6 | 0.19892 | 0.38657 | 0.999994 | 7734 | 3 | -0.55809 | 0.14915 | 0.35674 | 0.994392 | 7251 | 2 | -0.55809 |
| NSUN2 | 6 | 0.31409 | 0.52313 | 0.999994 | 10398 | 2 | -0.39275 | 0.14915 | 0.35674 | 0.994392 | 7252 | 2 | -0.39275 |
| OR4C45 | 6 | 0.80861 | 0.85162 | 0.999994 | 17612 | 1 | 0.91663 | 0.14915 | 0.35674 | 0.994392 | 7253 | 4 | 0.91663 |
| ZNF546 | 6 | 0.36194 | 0.57747 | 0.999994 | 11366 | 3 | -0.47224 | 0.14915 | 0.35674 | 0.994392 | 7254 | 2 | -0.47224 |
| HDGFRP2 | 6 | 0.022992 | 0.081555 | 0.93914 | 1650 | 4 | -2.0803 | 0.14915 | 0.35674 | 0.994392 | 7255 | 2 | -2.0803 |
| ICOS | 5 | 0.29608 | 0.47788 | 0.999994 | 10039 | 2 | -0.54926 | 0.14917 | 0.34608 | 0.994392 | 7256 | 2 | -0.54926 |
| hsa-mir-6892 | 4 | 0.18289 | 0.32306 | 0.999994 | 7287 | 2 | 1.7653 | 0.14926 | 0.30919 | 0.988566 | 7257 | 2 | 1.7653 |
| MED22 | 4 | 0.39007 | 0.53546 | 0.999994 | 11800 | 1 | 3.0263 | 0.14926 | 0.30919 | 0.988566 | 7258 | 2 | 3.0263 |
| ZNF165 | 4 | 0.24598 | 0.38777 | 0.999994 | 8924 | 1 | 3.2693 | 0.14943 | 0.30944 | 0.988566 | 7259 | 3 | 3.2693 |
| LRRCC1 | 5 | 0.10229 | 0.22997 | 0.983417 | 5049 | 2 | -0.47618 | 0.14944 | 0.34659 | 0.994392 | 7260 | 2 | -0.47618 |
| AOC3 | 5 | 0.057481 | 0.15791 | 0.967126 | 3313 | 3 | -2.0398 | 0.14944 | 0.34659 | 0.994392 | 7261 | 1 | -2.0398 |
| DEGS1 | 6 | 0.11097 | 0.26856 | 0.998561 | 5332 | 3 | 0.72264 | 0.14957 | 0.35737 | 0.994392 | 7262 | 3 | 0.72264 |
| DNAJB7 | 5 | 0.80732 | 0.82418 | 0.999994 | 17590 | 1 | 1.2298 | 0.14961 | 0.3469 | 0.994392 | 7263 | 4 | 1.2298 |
| MYF6 | 4 | 0.27249 | 0.41488 | 0.999994 | 9526 | 1 | 3.0323 | 0.14969 | 0.30981 | 0.988566 | 7264 | 3 | 3.0323 |
| RP2 | 3 | 0.8254 | 0.82527 | 0.999994 | 17859 | 0 | 0.30161 | 0.1497 | 0.28778 | 0.979534 | 7265 | 1 | 0.30161 |
| hsa-mir-137 | 3 | 0.018601 | 0.048143 | 0.90085 | 1394 | 2 | -4.843 | 0.1497 | 0.28778 | 0.979534 | 7266 | 1 | -4.843 |
| hsa-mir-548a-3 | 4 | 0.89781 | 0.89753 | 0.999994 | 19175 | 0 | 2.3298 | 0.14972 | 0.30987 | 0.988566 | 7267 | 3 | 2.3298 |
| SLC47A2 | 6 | 0.58422 | 0.72832 | 0.999994 | 14732 | 2 | 0.43801 | 0.14974 | 0.35763 | 0.994392 | 7268 | 3 | 0.43801 |
| GFAP | 6 | 0.87399 | 0.88579 | 0.999994 | 18709 | 1 | 1.9504 | 0.1498 | 0.35771 | 0.994392 | 7269 | 3 | 1.9504 |
| PARP14 | 6 | 0.74929 | 0.8282 | 0.999994 | 16851 | 1 | 1.9533 | 0.14985 | 0.35778 | 0.994392 | 7270 | 4 | 1.9533 |
| OTX1 | 4 | 0.43344 | 0.58007 | 0.999994 | 12469 | 2 | -0.49183 | 0.14987 | 0.3101 | 0.988566 | 7271 | 1 | -0.49183 |
| ZNF229 | 5 | 0.41205 | 0.57355 | 0.999994 | 12146 | 2 | 0.9373 | 0.1499 | 0.3474 | 0.994392 | 7272 | 3 | 0.9373 |
| PIGY | 6 | 0.16015 | 0.33842 | 0.999994 | 6744 | 2 | 0.66309 | 0.15001 | 0.35803 | 0.994392 | 7273 | 4 | 0.66309 |
| ZC2HC1A | 6 | 0.36126 | 0.57671 | 0.999994 | 11358 | 2 | 0.77355 | 0.15001 | 0.35803 | 0.994392 | 7274 | 3 | 0.77355 |
| GCNT2 | 6 | 0.52989 | 0.68946 | 0.999994 | 13899 | 2 | 0.24398 | 0.15001 | 0.35803 | 0.994392 | 7275 | 3 | 0.24398 |
| MIOS | 6 | 0.75914 | 0.83174 | 0.999994 | 16968 | 1 | 2.3617 | 0.15004 | 0.35806 | 0.994392 | 7276 | 4 | 2.3617 |
| OR4C11 | 6 | 0.87583 | 0.88697 | 0.999994 | 18747 | 1 | 1.874 | 0.15005 | 0.35808 | 0.994392 | 7277 | 4 | 1.874 |
| C4orf17 | 4 | 0.055563 | 0.13229 | 0.95318 | 3223 | 2 | 2.2384 | 0.15007 | 0.31039 | 0.988566 | 7278 | 2 | 2.2384 |
| INSL4 | 4 | 0.086667 | 0.19092 | 0.972293 | 4472 | 2 | 0.88229 | 0.15007 | 0.31039 | 0.988566 | 7279 | 2 | 0.88229 |

| | | | | | | | | | | | | | |
|---|---|---|---|---|---|---|---|---|---|---|---|---|---|
| RPL8 | 3 | 0.10987 | 0.19612 | 0.975797 | 5298 | 1 | 3.8042 | 0.15007 | 0.28818 | 0.979689 | 7280 | 2 | 3.8042 |
| SMYD2 | 6 | 0.92236 | 0.92262 | 0.999994 | 19685 | 1 | 2.0585 | 0.15009 | 0.35815 | 0.994392 | 7281 | 4 | 2.0585 |
| ACSS1 | 6 | 0.24411 | 0.4409 | 0.999994 | 8887 | 2 | 1.2355 | 0.15009 | 0.35815 | 0.994392 | 7282 | 4 | 1.2355 |
| KAT2B | 6 | 0.93366 | 0.93373 | 0.999994 | 19939 | 0 | 2.1171 | 0.15009 | 0.35815 | 0.994392 | 7283 | 4 | 2.1171 |
| LRRC1 | 5 | 0.75499 | 0.79348 | 0.999994 | 16912 | 1 | 1.1374 | 0.15023 | 0.348 | 0.994392 | 7284 | 3 | 1.1374 |
| HES5 | 5 | 0.8868 | 0.8869 | 0.999994 | 18964 | 1 | 1.2863 | 0.15024 | 0.34801 | 0.994392 | 7285 | 4 | 1.2863 |
| C2CD4B | 4 | 0.92133 | 0.9211 | 0.999994 | 19663 | 0 | 2.6494 | 0.15025 | 0.31067 | 0.988566 | 7286 | 3 | 2.6494 |
| ROPN1 | 5 | 0.66382 | 0.75071 | 0.999994 | 15837 | 1 | 2.7178 | 0.15027 | 0.34806 | 0.994392 | 7287 | 3 | 2.7178 |
| GBF1 | 6 | 0.81921 | 0.85644 | 0.999994 | 17768 | 1 | 2.5554 | 0.15028 | 0.35843 | 0.994392 | 7288 | 4 | 2.5554 |
| DHDDS | 5 | 0.12233 | 0.26089 | 0.994943 | 5713 | 2 | -0.2598 | 0.15033 | 0.34817 | 0.994392 | 7289 | 1 | -0.2598 |
| NKAIN2 | 6 | 0.58954 | 0.73225 | 0.999994 | 14816 | 2 | 0.60988 | 0.15036 | 0.35855 | 0.994392 | 7290 | 4 | 0.60988 |
| NonTargetingControlG | 2 | 0.84963 | 0.8495 | 0.999994 | 18251 | 0 | 3.5868 | 0.15037 | 0.24598 | 0.97466 | 7291 | 2 | 3.5868 |
| KERA | 6 | 0.87304 | 0.88521 | 0.999994 | 18694 | 1 | 1.7207 | 0.1504 | 0.35863 | 0.994392 | 7292 | 4 | 1.7207 |
| NonTargetingControlG | 2 | 0.82294 | 0.82277 | 0.999994 | 17822 | 0 | 4.3447 | 0.15051 | 0.24618 | 0.97466 | 7293 | 2 | 4.3447 |
| STMN1 | 4 | 0.10561 | 0.22514 | 0.983354 | 5155 | 2 | 1.1258 | 0.15052 | 0.31108 | 0.988566 | 7294 | 2 | 1.1258 |
| G6PC3 | 6 | 0.96659 | 0.96665 | 0.999994 | 20751 | 0 | 1.5939 | 0.15054 | 0.35887 | 0.994392 | 7295 | 4 | 1.5939 |
| CNOT7 | 5 | 0.85853 | 0.86117 | 0.999994 | 18411 | 1 | 0.78062 | 0.15056 | 0.34857 | 0.994392 | 7296 | 3 | 0.78062 |
| EXOG | 5 | 0.9257 | 0.92593 | 0.999994 | 19762 | 0 | 1.4488 | 0.15061 | 0.34865 | 0.994392 | 7297 | 4 | 1.4488 |
| OSGEPL1 | 6 | 0.91844 | 0.91896 | 0.999994 | 19595 | 1 | 0.41167 | 0.15073 | 0.35915 | 0.994392 | 7298 | 3 | 0.41167 |
| CSHL1 | 5 | 0.028428 | 0.086433 | 0.94395 | 1922 | 1 | 1.3558 | 0.15076 | 0.34891 | 0.994392 | 7299 | 4 | 1.3558 |
| hsa-mir-3688-1 | 4 | 0.92908 | 0.92886 | 0.999994 | 19837 | 0 | 1.9484 | 0.15077 | 0.31147 | 0.988566 | 7300 | 3 | 1.9484 |
| CDC37 | 3 | 0.55978 | 0.58723 | 0.999994 | 14367 | 1 | 3.6946 | 0.15085 | 0.28905 | 0.980152 | 7301 | 2 | 3.6946 |
| APRT | 5 | 0.42089 | 0.58081 | 0.999994 | 12263 | 2 | 2.5392 | 0.15086 | 0.34909 | 0.994392 | 7302 | 3 | 2.5392 |
| ZNF429 | 5 | 0.13713 | 0.28345 | 0.999632 | 6111 | 2 | 0.70746 | 0.1509 | 0.34916 | 0.994392 | 7303 | 3 | 0.70746 |
| HAL | 5 | 0.43524 | 0.59283 | 0.999994 | 12504 | 2 | 1.817 | 0.1509 | 0.34916 | 0.994392 | 7304 | 3 | 1.817 |
| RAB8B | 5 | 0.14425 | 0.29419 | 0.999994 | 6309 | 1 | 0.89959 | 0.15096 | 0.34927 | 0.994392 | 7305 | 4 | 0.89959 |
| KIR2DL3 | 5 | 0.93265 | 0.93295 | 0.999994 | 19917 | 0 | 0.83313 | 0.15098 | 0.34932 | 0.994392 | 7306 | 4 | 0.83313 |
| ZACN | 5 | 0.032218 | 0.096155 | 0.945031 | 2112 | 2 | -0.40597 | 0.15104 | 0.34942 | 0.994392 | 7307 | 2 | -0.40597 |
| VPS35 | 6 | 0.67205 | 0.79474 | 0.999994 | 15955 | 2 | 1.6104 | 0.15107 | 0.35965 | 0.994416 | 7308 | 4 | 1.6104 |
| HCAR3 | 6 | 0.89275 | 0.89826 | 0.999994 | 19085 | 1 | 1.7588 | 0.15125 | 0.35994 | 0.994441 | 7309 | 3 | 1.7588 |
| LARS2 | 5 | 0.47807 | 0.62899 | 0.999994 | 13116 | 2 | 2.8668 | 0.15126 | 0.34981 | 0.994392 | 7310 | 3 | 2.8668 |
| PDIA6 | 5 | 0.78586 | 0.81083 | 0.999994 | 17291 | 1 | 1.3595 | 0.15127 | 0.34984 | 0.994392 | 7311 | 4 | 1.3595 |
| WDR74 | 1 | 0.84871 | 0.84899 | 0.999994 | 18229 | 0 | 6.1426 | 0.15129 | 0.15109 | 0.965913 | 7312 | 1 | 6.1426 |
| STK38L | 6 | 0.08798 | 0.22347 | 0.983354 | 4512 | 3 | -1.0299 | 0.15142 | 0.36019 | 0.994562 | 7313 | 2 | -1.0299 |
| IRF2BP2 | 6 | 0.77948 | 0.83946 | 0.999994 | 17204 | 1 | 0.90972 | 0.15142 | 0.36019 | 0.994562 | 7314 | 4 | 0.90972 |
| TMEM151A | 3 | 0.47807 | 0.53191 | 0.999994 | 13115 | 1 | 3.7891 | 0.15146 | 0.28973 | 0.980804 | 7315 | 2 | 3.7891 |
| PTPN23 | 4 | 0.11851 | 0.24763 | 0.990682 | 5590 | 2 | -0.44155 | 0.15149 | 0.31255 | 0.989342 | 7316 | 2 | -0.44155 |
| VNN1 | 6 | 0.67431 | 0.79651 | 0.999994 | 15994 | 1 | 0.94238 | 0.15157 | 0.3604 | 0.994562 | 7317 | 3 | 0.94238 |
| hsa-mir-501 | 3 | 0.29646 | 0.42664 | 0.999994 | 10052 | 1 | 2.5974 | 0.15159 | 0.28987 | 0.981118 | 7318 | 2 | 2.5974 |
| LRTM2 | 5 | 0.67625 | 0.75596 | 0.999994 | 16025 | 1 | -0.18426 | 0.15161 | 0.35045 | 0.994392 | 7319 | 2 | -0.18426 |
| ZNF106 | 5 | 0.91286 | 0.9129 | 0.999994 | 19476 | 0 | 2.8129 | 0.15165 | 0.35052 | 0.994392 | 7320 | 3 | 2.8129 |
| C1orf101 | 6 | 0.63375 | 0.76518 | 0.999994 | 15429 | 1 | 2.3085 | 0.15168 | 0.36058 | 0.994562 | 7321 | 4 | 2.3085 |
| HRCT1 | 5 | 0.94726 | 0.94744 | 0.999994 | 20269 | 0 | 0.7733 | 0.15175 | 0.35069 | 0.994392 | 7322 | 4 | 0.7733 |
| KRT9 | 5 | 0.32765 | 0.50359 | 0.999994 | 10671 | 2 | 2.5764 | 0.15175 | 0.3507 | 0.994392 | 7323 | 3 | 2.5764 |
| PPP1R12C | 6 | 0.59056 | 0.73303 | 0.999994 | 14831 | 1 | 2.0726 | 0.15181 | 0.36076 | 0.994562 | 7324 | 3 | 2.0726 |
| CYBB | 6 | 0.093572 | 0.23469 | 0.984971 | 4739 | 3 | -0.61231 | 0.15181 | 0.36076 | 0.994562 | 7325 | 3 | -0.61231 |
| GPRC6A | 6 | 0.71717 | 0.81741 | 0.999994 | 16470 | 1 | 2.2416 | 0.15191 | 0.36094 | 0.994562 | 7326 | 4 | 2.2416 |
| OR4B1 | 6 | 0.1976 | 0.38498 | 0.999994 | 7692 | 2 | -0.53398 | 0.15199 | 0.36104 | 0.994562 | 7327 | 2 | -0.53398 |
| NonTargetingControlG | 1 | 0.84796 | 0.84826 | 0.999994 | 18217 | 0 | 6.1252 | 0.15204 | 0.15184 | 0.965913 | 7328 | 1 | 6.1252 |
| FSTL5 | 6 | 0.5188 | 0.6817 | 0.999994 | 13723 | 2 | 1.4252 | 0.15205 | 0.36115 | 0.994562 | 7329 | 3 | 1.4252 |
| LIMK1 | 6 | 0.11793 | 0.2818 | 0.999632 | 5571 | 2 | 1.3726 | 0.15205 | 0.36115 | 0.994562 | 7330 | 4 | 1.3726 |
| LCE2B | 6 | 0.53766 | 0.69488 | 0.999994 | 14021 | 2 | 1.1274 | 0.15205 | 0.36115 | 0.994562 | 7331 | 4 | 1.1274 |

| | | | | | | | | | | | | |
|---|---|---|---|---|---|---|---|---|---|---|---|---|
| SSR3 | 5 | 0.44864 | 0.60412 | 0.999994 | 12696 | 2 | 2.5756 | 0.15217 | 0.3514 | 0.994392 | 7332 | 3 | 2.5756 |
| BLMH | 5 | 0.54216 | 0.6848 | 0.999994 | 14095 | 2 | 0.93271 | 0.15217 | 0.3514 | 0.994392 | 7333 | 3 | 0.93271 |
| RABL3 | 5 | 0.90977 | 0.90979 | 0.999994 | 19422 | 0 | 0.51578 | 0.15217 | 0.3514 | 0.994392 | 7334 | 2 | 0.51578 |
| VTN | 5 | 0.22953 | 0.41657 | 0.999994 | 8519 | 2 | 0.47463 | 0.15217 | 0.3514 | 0.994392 | 7335 | 2 | 0.47463 |
| OR6V1 | 6 | 0.91637 | 0.91708 | 0.999994 | 19546 | 1 | 0.55649 | 0.15218 | 0.36133 | 0.994759 | 7336 | 3 | 0.55649 |
| GPR75 | 4 | 0.057098 | 0.13523 | 0.954328 | 3295 | 3 | -1.3995 | 0.15224 | 0.3137 | 0.989846 | 7337 | 1 | -1.3995 |
| GPR150 | 6 | 0.62577 | 0.75916 | 0.999994 | 15323 | 2 | 2.532 | 0.15231 | 0.36153 | 0.994802 | 7338 | 4 | 2.532 |
| DDX46 | 2 | 0.84761 | 0.84751 | 0.999994 | 18213 | 0 | 3.9522 | 0.15239 | 0.24894 | 0.97466 | 7339 | 2 | 3.9522 |
| MAP3K8 | 5 | 0.96708 | 0.96727 | 0.999994 | 20763 | 0 | 0.95973 | 0.1524 | 0.35179 | 0.994392 | 7340 | 4 | 0.95973 |
| UQCRB | 5 | 0.54537 | 0.68762 | 0.999994 | 14143 | 1 | 0.49729 | 0.15246 | 0.35192 | 0.994392 | 7341 | 2 | 0.49729 |
| MLF1 | 6 | 0.14739 | 0.32236 | 0.999994 | 6388 | 2 | 0.89144 | 0.1525 | 0.36182 | 0.994802 | 7342 | 3 | 0.89144 |
| LDHC | 6 | 0.39501 | 0.59915 | 0.999994 | 11873 | 2 | 0.84468 | 0.1525 | 0.36182 | 0.994802 | 7343 | 3 | 0.84468 |
| HSD3B7 | 5 | 0.4997 | 0.64764 | 0.999994 | 13417 | 1 | 1.4264 | 0.15256 | 0.35209 | 0.994392 | 7344 | 4 | 1.4264 |
| EBLN2 | 6 | 0.94317 | 0.94314 | 0.999994 | 20158 | 0 | 0.63259 | 0.15259 | 0.36194 | 0.994908 | 7345 | 4 | 0.63259 |
| TRAF3IP1 | 5 | 0.64122 | 0.74165 | 0.999994 | 15533 | 1 | 1.7807 | 0.15262 | 0.3522 | 0.994392 | 7346 | 4 | 1.7807 |
| TMEM255B | 5 | 0.62602 | 0.73584 | 0.999994 | 15324 | 1 | 2.6085 | 0.15265 | 0.35228 | 0.994392 | 7347 | 3 | 2.6085 |
| hsa-mir-4328 | 4 | 0.77316 | 0.77833 | 0.999994 | 17123 | 1 | 1.0337 | 0.15266 | 0.31431 | 0.989846 | 7348 | 2 | 1.0337 |
| NonTargetingControlG | 2 | 0.054433 | 0.096636 | 0.945031 | 3173 | 1 | -0.81994 | 0.15266 | 0.24932 | 0.97466 | 7349 | 1 | -0.81994 |
| TCERG1L | 5 | 0.93381 | 0.93405 | 0.999994 | 19948 | 0 | 1.0878 | 0.15266 | 0.35228 | 0.994392 | 7350 | 4 | 1.0878 |
| ZCCHC3 | 6 | 0.88922 | 0.89579 | 0.999994 | 19024 | 1 | 1.6271 | 0.1527 | 0.36212 | 0.995271 | 7351 | 4 | 1.6271 |
| C15orf41 | 5 | 0.96303 | 0.96319 | 0.999994 | 20663 | 0 | 1.3156 | 0.15271 | 0.35237 | 0.994392 | 7352 | 4 | 1.3156 |
| RASSF8 | 5 | 0.36385 | 0.53327 | 0.999994 | 11401 | 2 | 2.7576 | 0.15273 | 0.3524 | 0.994392 | 7353 | 3 | 2.7576 |
| SERPINE1 | 4 | 0.33737 | 0.48123 | 0.999994 | 10881 | 1 | 1.6864 | 0.15276 | 0.31446 | 0.989846 | 7354 | 3 | 1.6864 |
| TACSTD2 | 6 | 0.37361 | 0.58548 | 0.999994 | 11564 | 2 | 1.6572 | 0.1528 | 0.36228 | 0.99534 | 7355 | 4 | 1.6572 |
| RTKN2 | 6 | 0.060771 | 0.16741 | 0.967126 | 3458 | 1 | 2.0968 | 0.15281 | 0.36228 | 0.99534 | 7356 | 3 | 2.0968 |
| ROBO1 | 6 | 0.36903 | 0.5826 | 0.999994 | 11491 | 2 | 0.16661 | 0.15287 | 0.36238 | 0.995353 | 7357 | 2 | 0.16661 |
| DEFB131 | 6 | 0.32021 | 0.53012 | 0.999994 | 10516 | 1 | -0.40565 | 0.15287 | 0.36238 | 0.995353 | 7358 | 1 | -0.40565 |
| STK4 | 5 | 0.034746 | 0.10261 | 0.946414 | 2250 | 2 | 2.6846 | 0.15288 | 0.35267 | 0.994392 | 7359 | 3 | 2.6846 |
| CISD3 | 4 | 0.87833 | 0.87797 | 0.999994 | 18794 | 0 | 0.74904 | 0.15296 | 0.31476 | 0.989846 | 7360 | 2 | 0.74904 |
| CNPY2 | 5 | 0.055513 | 0.15368 | 0.967126 | 3221 | 3 | -0.72673 | 0.15302 | 0.3529 | 0.994392 | 7361 | 2 | -0.72673 |
| CNBD1 | 6 | 0.21862 | 0.41044 | 0.999994 | 8242 | 2 | 0.39237 | 0.1531 | 0.3627 | 0.995401 | 7362 | 3 | 0.39237 |
| HYAL4 | 6 | 0.65548 | 0.78183 | 0.999994 | 15721 | 2 | 1.2606 | 0.1531 | 0.3627 | 0.995401 | 7363 | 4 | 1.2606 |
| CDK15 | 6 | 0.52925 | 0.68901 | 0.999994 | 13885 | 2 | 0.65191 | 0.15317 | 0.36281 | 0.995401 | 7364 | 3 | 0.65191 |
| hsa-mir-7975 | 2 | 0.69904 | 0.69885 | 0.999994 | 16296 | 0 | 3.8413 | 0.15319 | 0.25012 | 0.97466 | 7365 | 1 | 3.8413 |
| KCNJ1 | 5 | 0.26962 | 0.45653 | 0.999994 | 9464 | 1 | 0.46218 | 0.15323 | 0.35325 | 0.994392 | 7366 | 2 | 0.46218 |
| PADI2 | 5 | 0.062403 | 0.16604 | 0.967126 | 3531 | 2 | 1.4651 | 0.15329 | 0.35337 | 0.994392 | 7367 | 3 | 1.4651 |
| MMP25 | 5 | 0.84089 | 0.84734 | 0.999994 | 18103 | 1 | 0.16526 | 0.15329 | 0.35337 | 0.994392 | 7368 | 2 | 0.16526 |
| C1QTNF7 | 5 | 0.00068338 | 0.0031349 | 0.537554 | 129 | 2 | -0.33669 | 0.15329 | 0.35337 | 0.994392 | 7369 | 2 | -0.33669 |
| FAM227B | 5 | 0.21689 | 0.39892 | 0.999994 | 8188 | 3 | -0.72728 | 0.15329 | 0.35337 | 0.994392 | 7370 | 2 | -0.72728 |
| MPZL2 | 5 | 0.014133 | 0.047753 | 0.900643 | 1137 | 2 | 0.0080781 | 0.15329 | 0.35337 | 0.994392 | 7371 | 2 | 0.0080781 |
| SFPQ | 4 | 0.0072498 | 0.024978 | 0.844082 | 674 | 3 | -5.2025 | 0.15332 | 0.31531 | 0.989846 | 7372 | 1 | -5.2025 |
| RBP4 | 6 | 0.41363 | 0.61121 | 0.999994 | 12163 | 2 | 1.6256 | 0.15336 | 0.36312 | 0.995401 | 7373 | 4 | 1.6256 |
| AIFM1 | 6 | 0.74252 | 0.82584 | 0.999994 | 16771 | 1 | 2.1387 | 0.1534 | 0.36316 | 0.995401 | 7374 | 4 | 2.1387 |
| MPST | 5 | 0.74179 | 0.78656 | 0.999994 | 16762 | 1 | 0.76173 | 0.15347 | 0.3537 | 0.994392 | 7375 | 4 | 0.76173 |
| NonTargetingControlG | 2 | 0.84653 | 0.84647 | 0.999994 | 18202 | 0 | 4.2579 | 0.15347 | 0.25052 | 0.97466 | 7376 | 2 | 4.2579 |
| TAS2R43 | 6 | 0.58268 | 0.72717 | 0.999994 | 14714 | 2 | 1.813 | 0.1535 | 0.36329 | 0.995401 | 7377 | 4 | 1.813 |
| hsa-mir-889 | 4 | 0.70216 | 0.72584 | 0.999994 | 16328 | 1 | 2.0719 | 0.15353 | 0.31563 | 0.989846 | 7378 | 3 | 2.0719 |
| ZBTB32 | 6 | 0.47509 | 0.65187 | 0.999994 | 13063 | 1 | 1.5511 | 0.15354 | 0.36335 | 0.995401 | 7379 | 4 | 1.5511 |
| LSG1 | 6 | 0.67772 | 0.7992 | 0.999994 | 16055 | 2 | -0.38268 | 0.15354 | 0.36335 | 0.995401 | 7380 | 1 | -0.38268 |
| OR5T3 | 6 | 0.22206 | 0.41465 | 0.999994 | 8334 | 3 | 1.2353 | 0.15354 | 0.36335 | 0.995401 | 7381 | 3 | 1.2353 |
| IL5RA | 6 | 0.35475 | 0.56938 | 0.999994 | 11245 | 1 | 1.6641 | 0.15354 | 0.36335 | 0.995401 | 7382 | 4 | 1.6641 |
| NonTargetingControlG | 2 | 0.80542 | 0.80503 | 0.999994 | 17558 | 0 | 4.2272 | 0.15354 | 0.25063 | 0.97466 | 7383 | 2 | 4.2272 |

| | | | | | | | | | | | | |
|---|---|---|---|---|---|---|---|---|---|---|---|---|
| SOX21 | 2 | 0.84641 | 0.84635 | 0.999994 | 18197 | 0 | 1.5739 | 0.15359 | 0.2507 | 0.97466 | 7384 | 2 | 1.5739 |
| CHRNA7 | 2 | 0.84639 | 0.84634 | 0.999994 | 18196 | 0 | 2.9474 | 0.15361 | 0.25073 | 0.97466 | 7385 | 2 | 2.9474 |
| PCDHB3 | 6 | 0.0077534 | 0.031094 | 0.862223 | 710 | 2 | 2.2841 | 0.15364 | 0.3635 | 0.995401 | 7386 | 4 | 2.2841 |
| ZNF777 | 4 | 0.089913 | 0.19685 | 0.977164 | 4590 | 2 | -0.21716 | 0.15365 | 0.31583 | 0.989846 | 7387 | 2 | -0.21716 |
| ERCC6L | 5 | 0.76015 | 0.79627 | 0.999994 | 16984 | 1 | 2.6362 | 0.15366 | 0.35401 | 0.994392 | 7388 | 3 | 2.6362 |
| ARMC12 | 5 | 0.20675 | 0.38479 | 0.999994 | 7945 | 1 | 1.7086 | 0.15367 | 0.35403 | 0.994392 | 7389 | 4 | 1.7086 |
| NonTargetingControlG | 2 | 0.84629 | 0.84623 | 0.999994 | 18194 | 0 | 3.3325 | 0.15371 | 0.25087 | 0.97466 | 7390 | 2 | 3.3325 |
| ITGAD | 5 | 0.036546 | 0.1072 | 0.950995 | 2347 | 3 | -1.2313 | 0.15374 | 0.35417 | 0.994392 | 7391 | 1 | -1.2313 |
| KIAA0754 | 5 | 0.56738 | 0.70724 | 0.999994 | 14504 | 2 | -0.087341 | 0.15374 | 0.35417 | 0.994392 | 7392 | 1 | -0.087341 |
| hsa-mir-554 | 4 | 0.39243 | 0.5379 | 0.999994 | 11839 | 1 | 2.5056 | 0.15376 | 0.31599 | 0.989846 | 7393 | 3 | 2.5056 |
| CACTIN | 4 | 0.16159 | 0.30112 | 0.999994 | 6778 | 2 | -0.86178 | 0.1538 | 0.31605 | 0.989846 | 7394 | 1 | -0.86178 |
| NonTargetingControlG | 2 | 0.61171 | 0.61158 | 0.999994 | 15102 | 0 | 3.5659 | 0.15391 | 0.25117 | 0.97466 | 7395 | 1 | 3.5659 |
| TRAPPC6B | 6 | 0.084006 | 0.21551 | 0.980236 | 4376 | 4 | -1.2319 | 0.15391 | 0.3639 | 0.995622 | 7396 | 1 | -1.2319 |
| OR14J1 | 6 | 0.94843 | 0.94845 | 0.999994 | 20298 | 0 | 1.4841 | 0.15392 | 0.36391 | 0.995622 | 7397 | 4 | 1.4841 |
| PCSK1N | 3 | 0.64766 | 0.65515 | 0.999994 | 15626 | 1 | 3.2785 | 0.15397 | 0.29248 | 0.983225 | 7398 | 2 | 3.2785 |
| PCCB | 4 | 0.91206 | 0.91186 | 0.999994 | 19463 | 0 | 2.0564 | 0.15399 | 0.31633 | 0.989846 | 7399 | 3 | 2.0564 |
| SNRPN | 6 | 0.23364 | 0.42843 | 0.999994 | 8610 | 2 | 2.2377 | 0.154 | 0.36403 | 0.995622 | 7400 | 4 | 2.2377 |
| MAP1LC3C | 6 | 0.17216 | 0.3534 | 0.999994 | 7041 | 2 | 1.5909 | 0.15403 | 0.36408 | 0.995622 | 7401 | 4 | 1.5909 |
| MYL12A | 5 | 0.71405 | 0.77302 | 0.999994 | 16439 | 1 | 0.98414 | 0.15407 | 0.35477 | 0.994392 | 7402 | 3 | 0.98414 |
| SNX8 | 5 | 0.21947 | 0.40255 | 0.999994 | 8276 | 2 | -0.24189 | 0.15407 | 0.35477 | 0.994392 | 7403 | 2 | -0.24189 |
| CYB561A3 | 6 | 0.024247 | 0.085209 | 0.94395 | 1719 | 2 | 1.7924 | 0.15408 | 0.36414 | 0.995622 | 7404 | 3 | 1.7924 |
| HTR7 | 6 | 0.1947 | 0.38142 | 0.999994 | 7625 | 3 | -0.68637 | 0.15423 | 0.36436 | 0.995622 | 7405 | 2 | -0.68637 |
| MYC | 3 | 0.036791 | 0.090534 | 0.94395 | 2364 | 1 | 0.56221 | 0.15431 | 0.29285 | 0.983225 | 7406 | 1 | 0.56221 |
| NonTargetingControlG | 1 | 0.84564 | 0.84595 | 0.999994 | 18182 | 0 | 6.0743 | 0.15436 | 0.15412 | 0.965913 | 7407 | 1 | 6.0743 |
| MAPK8IP1 | 5 | 0.36578 | 0.53487 | 0.999994 | 11435 | 2 | 2.3546 | 0.15436 | 0.35528 | 0.994392 | 7408 | 3 | 2.3546 |
| LIME1 | 5 | 0.51403 | 0.66011 | 0.999994 | 13641 | 1 | 1.2047 | 0.15439 | 0.35532 | 0.994392 | 7409 | 4 | 1.2047 |
| hsa-mir-3189 | 4 | 0.81706 | 0.81697 | 0.999994 | 17729 | 1 | 1.6709 | 0.15444 | 0.31699 | 0.989846 | 7410 | 3 | 1.6709 |
| DDX19A | 6 | 0.25983 | 0.45964 | 0.999994 | 9235 | 3 | -0.77063 | 0.15446 | 0.36474 | 0.995622 | 7411 | 2 | -0.77063 |
| OR52M1 | 6 | 0.45687 | 0.63962 | 0.999994 | 12803 | 2 | -0.16874 | 0.15446 | 0.36474 | 0.995622 | 7412 | 2 | -0.16874 |
| USP15 | 6 | 0.11894 | 0.28374 | 0.999632 | 5601 | 3 | 0.26247 | 0.15446 | 0.36474 | 0.995622 | 7413 | 3 | 0.26247 |
| GIN1 | 6 | 0.89295 | 0.89841 | 0.999994 | 19089 | 1 | 1.7702 | 0.15446 | 0.36474 | 0.995622 | 7414 | 4 | 1.7702 |
| MMD2 | 6 | 0.10441 | 0.25593 | 0.994943 | 5111 | 3 | 0.0010494 | 0.15446 | 0.36474 | 0.995622 | 7415 | 3 | 0.0010494 |
| OR10G9 | 6 | 0.61169 | 0.74857 | 0.999994 | 15101 | 2 | 1.4415 | 0.15446 | 0.36474 | 0.995622 | 7416 | 4 | 1.4415 |
| APTX | 6 | 0.30127 | 0.50821 | 0.999994 | 10151 | 3 | 1.1227 | 0.15446 | 0.36474 | 0.995622 | 7417 | 3 | 1.1227 |
| PLCD3 | 5 | 0.8432 | 0.84907 | 0.999994 | 18142 | 1 | 1.3629 | 0.15459 | 0.35566 | 0.994392 | 7418 | 3 | 1.3629 |
| hsa-mir-502 | 3 | 0.68029 | 0.683 | 0.999994 | 16095 | 1 | -0.056533 | 0.15463 | 0.29319 | 0.983225 | 7419 | 1 | -0.056533 |
| DDIT4L | 4 | 0.43461 | 0.58122 | 0.999994 | 12486 | 2 | 0.5587 | 0.15464 | 0.3173 | 0.989846 | 7420 | 2 | 0.5587 |
| NMRAL1 | 5 | 0.11046 | 0.24268 | 0.988088 | 5317 | 1 | 1.5712 | 0.15471 | 0.35586 | 0.994392 | 7421 | 4 | 1.5712 |
| SMC5 | 4 | 0.22225 | 0.36356 | 0.999994 | 8338 | 2 | 2.3435 | 0.15474 | 0.31745 | 0.989846 | 7422 | 2 | 2.3435 |
| MTRNR2L2 | 4 | 0.013415 | 0.044272 | 0.891621 | 1093 | 2 | 2.1723 | 0.15474 | 0.31745 | 0.989846 | 7423 | 2 | 2.1723 |
| GTF2I | 6 | 0.66249 | 0.78724 | 0.999994 | 15820 | 2 | 0.27693 | 0.15477 | 0.36521 | 0.996287 | 7424 | 3 | 0.27693 |
| CEACAM6 | 6 | 0.88013 | 0.88968 | 0.999994 | 18834 | 1 | 0.39938 | 0.15477 | 0.36521 | 0.996287 | 7425 | 2 | 0.39938 |
| ZKSCAN5 | 5 | 0.33575 | 0.51023 | 0.999994 | 10847 | 2 | 2.3597 | 0.15478 | 0.35598 | 0.994392 | 7426 | 3 | 2.3597 |
| DEFB113 | 5 | 0.40325 | 0.56625 | 0.999994 | 12006 | 2 | 2.3235 | 0.15487 | 0.35614 | 0.994392 | 7427 | 3 | 2.3235 |
| OR4D10 | 5 | 0.78779 | 0.812 | 0.999994 | 17316 | 1 | 1.4057 | 0.15492 | 0.35623 | 0.994392 | 7428 | 4 | 1.4057 |
| SUMO4 | 6 | 0.93307 | 0.93316 | 0.999994 | 19927 | 0 | 1.5323 | 0.15492 | 0.36545 | 0.996287 | 7429 | 4 | 1.5323 |
| NonTargetingControlG | 2 | 0.24627 | 0.2961 | 0.999994 | 8930 | 1 | 2.1206 | 0.15492 | 0.25267 | 0.97466 | 7430 | 1 | 2.1206 |
| SETMAR | 6 | 0.23486 | 0.42989 | 0.999994 | 8640 | 3 | 1.3473 | 0.15498 | 0.36553 | 0.996287 | 7431 | 3 | 1.3473 |
| DNAJB3 | 6 | 0.31506 | 0.52424 | 0.999994 | 10421 | 2 | 2.6364 | 0.15498 | 0.36553 | 0.996287 | 7432 | 4 | 2.6364 |
| MTMR4 | 5 | 0.14998 | 0.30275 | 0.999994 | 6457 | 1 | 1.2018 | 0.15499 | 0.35634 | 0.994392 | 7433 | 4 | 1.2018 |
| NonTargetingControlG | 2 | 0.84501 | 0.84495 | 0.999994 | 18172 | 0 | 1.5428 | 0.15499 | 0.25279 | 0.97466 | 7434 | 2 | 1.5428 |
| YPEL4 | 5 | 0.41824 | 0.57865 | 0.999994 | 12224 | 2 | 2.5561 | 0.15508 | 0.3565 | 0.994392 | 7435 | 3 | 2.5561 |

| | | | | | | | | | | | | | |
|---|---|---|---|---|---|---|---|---|---|---|---|---|---|
| DCTD | 6 | 0.14046 | 0.3136 | 0.999994 | 6198 | 2 | 1.0396 | 0.15509 | 0.36569 | 0.996287 | 7436 | 3 | 1.0396 |
| hsa-mir-4521 | 4 | 0.92033 | 0.9201 | 0.999994 | 19637 | 0 | 2.8642 | 0.15515 | 0.31805 | 0.989846 | 7437 | 3 | 2.8642 |
| SFMBT2 | 6 | 0.93244 | 0.93255 | 0.999994 | 19912 | 0 | 1.8976 | 0.15518 | 0.36581 | 0.996287 | 7438 | 4 | 1.8976 |
| POLR3G | 5 | 0.94663 | 0.94681 | 0.999994 | 20251 | 0 | 0.93105 | 0.1552 | 0.35672 | 0.994392 | 7439 | 4 | 0.93105 |
| EIF3J | 4 | 0.42137 | 0.56763 | 0.999994 | 12267 | 1 | 2.255 | 0.1552 | 0.31815 | 0.989846 | 7440 | 3 | 2.255 |
| SRA1 | 6 | 0.027285 | 0.09243 | 0.943968 | 1871 | 3 | -0.01732 | 0.15523 | 0.36589 | 0.996287 | 7441 | 3 | -0.01732 |
| KRTAP13-3 | 6 | 0.6677 | 0.79137 | 0.999994 | 15887 | 1 | 0.53753 | 0.15526 | 0.36593 | 0.996287 | 7442 | 3 | 0.53753 |
| hsa-mir-3975 | 3 | 0.53211 | 0.56781 | 0.999994 | 13939 | 1 | -0.51718 | 0.15529 | 0.29394 | 0.983357 | 7443 | 1 | -0.51718 |
| WNT8A | 6 | 0.58673 | 0.73017 | 0.999994 | 14771 | 2 | 1.5343 | 0.1553 | 0.36599 | 0.996287 | 7444 | 4 | 1.5343 |
| hsa-mir-519c | 2 | 0.84457 | 0.84452 | 0.999994 | 18167 | 0 | 3.2701 | 0.15543 | 0.25342 | 0.97466 | 7445 | 2 | 3.2701 |
| ASIC5 | 6 | 0.14357 | 0.31757 | 0.999994 | 6287 | 3 | -1.0963 | 0.15546 | 0.36621 | 0.996708 | 7446 | 2 | -1.0963 |
| AKTIP | 5 | 0.9701 | 0.97016 | 0.999994 | 20826 | 0 | 0.82803 | 0.15546 | 0.35719 | 0.994392 | 7447 | 4 | 0.82803 |
| ZFP92 | 6 | 0.54881 | 0.70278 | 0.999994 | 14191 | 2 | 1.5282 | 0.15547 | 0.36623 | 0.996708 | 7448 | 4 | 1.5282 |
| GSG2 | 4 | 0.43017 | 0.57674 | 0.999994 | 12414 | 1 | 3.1794 | 0.15557 | 0.31866 | 0.989846 | 7449 | 3 | 3.1794 |
| PPWD1 | 4 | 0.74157 | 0.75368 | 0.999994 | 16760 | 1 | 3.5436 | 0.15557 | 0.31866 | 0.989846 | 7450 | 3 | 3.5436 |
| HMGN3 | 5 | 0.90248 | 0.90258 | 0.999994 | 19261 | 0 | 1.6171 | 0.1556 | 0.35742 | 0.994392 | 7451 | 4 | 1.6171 |
| NonTargetingControlG | 2 | 0.84436 | 0.8443 | 0.999994 | 18163 | 0 | 1.5598 | 0.15564 | 0.25375 | 0.97466 | 7452 | 2 | 1.5598 |
| HNRNPD | 6 | 0.49642 | 0.66626 | 0.999994 | 13360 | 2 | 2.4719 | 0.15565 | 0.36649 | 0.997033 | 7453 | 4 | 2.4719 |
| OR2V1 | 5 | 0.36201 | 0.53172 | 0.999994 | 11368 | 1 | 2.3073 | 0.15575 | 0.35769 | 0.994392 | 7454 | 4 | 2.3073 |
| STARD9 | 5 | 0.88471 | 0.88484 | 0.999994 | 18930 | 1 | 2.1237 | 0.15575 | 0.35769 | 0.994392 | 7455 | 3 | 2.1237 |
| ACTA2 | 5 | 0.063125 | 0.16723 | 0.967126 | 3561 | 2 | -0.025273 | 0.15576 | 0.35771 | 0.994392 | 7456 | 2 | -0.025273 |
| FNDC7 | 5 | 0.36423 | 0.53358 | 0.999994 | 11407 | 1 | 1.6208 | 0.15578 | 0.35774 | 0.994392 | 7457 | 4 | 1.6208 |
| GTPBP4 | 4 | 0.012289 | 0.040843 | 0.891621 | 1011 | 2 | -2.6111 | 0.15593 | 0.3192 | 0.989846 | 7458 | 2 | -2.6111 |
| UGT2B15 | 6 | 0.23913 | 0.43496 | 0.999994 | 8751 | 3 | 0.18432 | 0.15594 | 0.36691 | 0.997075 | 7459 | 3 | 0.18432 |
| MIS18A | 5 | 0.069103 | 0.17696 | 0.968171 | 3802 | 2 | 2.8483 | 0.15594 | 0.35806 | 0.994392 | 7460 | 3 | 2.8483 |
| SLC35D1 | 5 | 0.84787 | 0.85259 | 0.999994 | 18216 | 1 | 2.8483 | 0.15594 | 0.35806 | 0.994392 | 7461 | 3 | 2.8483 |
| GRIK2 | 5 | 0.9636 | 0.96375 | 0.999994 | 20694 | 0 | 0.9513 | 0.15595 | 0.35807 | 0.994392 | 7462 | 4 | 0.9513 |
| NonTargetingControlG | 2 | 0.84404 | 0.84397 | 0.999994 | 18155 | 0 | 4.2096 | 0.15596 | 0.25422 | 0.97466 | 7463 | 2 | 4.2096 |
| FAM64A | 6 | 0.83564 | 0.86437 | 0.999994 | 18019 | 1 | 1.1893 | 0.156 | 0.367 | 0.997075 | 7464 | 4 | 1.1893 |
| hsa-mir-101-1 | 4 | 0.68509 | 0.71484 | 0.999994 | 16163 | 1 | 1.9031 | 0.15601 | 0.31932 | 0.989846 | 7465 | 3 | 1.9031 |
| NFKB1 | 6 | 0.48512 | 0.65864 | 0.999994 | 13218 | 2 | 0.53844 | 0.15607 | 0.36712 | 0.997075 | 7466 | 3 | 0.53844 |
| PHF15 | 6 | 0.29051 | 0.49567 | 0.999994 | 9923 | 1 | 1.7635 | 0.15612 | 0.36719 | 0.997075 | 7467 | 3 | 1.7635 |
| PCBD2 | 6 | 0.46625 | 0.64596 | 0.999994 | 12957 | 1 | 2.5774 | 0.15612 | 0.36719 | 0.997075 | 7468 | 4 | 2.5774 |
| GMDS | 6 | 0.89953 | 0.90332 | 0.999994 | 19214 | 1 | 2.2491 | 0.15612 | 0.36719 | 0.997075 | 7469 | 4 | 2.2491 |
| HSPB8 | 6 | 0.24707 | 0.44443 | 0.999994 | 8949 | 1 | 2.3352 | 0.15612 | 0.36719 | 0.997075 | 7470 | 4 | 2.3352 |
| CCDC60 | 5 | 0.1338 | 0.27845 | 0.999632 | 6017 | 2 | 0.77992 | 0.15621 | 0.35853 | 0.994392 | 7471 | 3 | 0.77992 |
| NonTargetingControlG | 2 | 0.84367 | 0.84359 | 0.999994 | 18150 | 0 | 1.8078 | 0.15633 | 0.25475 | 0.97466 | 7472 | 2 | 1.8078 |
| PGGT1B | 5 | 0.19979 | 0.37495 | 0.999994 | 7759 | 1 | 1.0761 | 0.15637 | 0.3588 | 0.994392 | 7473 | 4 | 1.0761 |
| CCDC105 | 5 | 0.95063 | 0.9508 | 0.999994 | 20348 | 0 | 2.6771 | 0.15638 | 0.35881 | 0.994392 | 7474 | 4 | 2.6771 |
| SFRP2 | 6 | 0.97079 | 0.97076 | 0.999994 | 20840 | 0 | 1.5175 | 0.15642 | 0.36762 | 0.997572 | 7475 | 4 | 1.5175 |
| hsa-mir-5739 | 4 | 0.41957 | 0.56577 | 0.999994 | 12245 | 2 | 0.40387 | 0.15643 | 0.31993 | 0.989846 | 7476 | 2 | 0.40387 |
| ACBD3 | 6 | 0.40804 | 0.6076 | 0.999994 | 12076 | 2 | 2.6313 | 0.15644 | 0.36766 | 0.997572 | 7477 | 4 | 2.6313 |
| PPP5C | 6 | 0.85103 | 0.87248 | 0.999994 | 18280 | 1 | 1.942 | 0.15648 | 0.36771 | 0.99759 | 7478 | 4 | 1.942 |
| PCOLCE | 5 | 0.53145 | 0.67535 | 0.999994 | 13924 | 1 | 2.5048 | 0.1565 | 0.35902 | 0.994392 | 7479 | 3 | 2.5048 |
| NRG1 | 5 | 0.90617 | 0.90624 | 0.999994 | 19355 | 0 | 0.93398 | 0.15656 | 0.35913 | 0.994392 | 7480 | 3 | 0.93398 |
| RWDD4 | 5 | 0.51623 | 0.66203 | 0.999994 | 13681 | 2 | 2.4652 | 0.15656 | 0.35913 | 0.994392 | 7481 | 3 | 2.4652 |
| PON1 | 5 | 0.065081 | 0.17039 | 0.967158 | 3656 | 2 | 2.4962 | 0.1566 | 0.35918 | 0.994392 | 7482 | 3 | 2.4962 |
| GYPC | 6 | 0.071565 | 0.19004 | 0.970632 | 3900 | 2 | 1.4895 | 0.15662 | 0.36791 | 0.997833 | 7483 | 4 | 1.4895 |
| CIB4 | 4 | 0.86323 | 0.86287 | 0.999994 | 18510 | 0 | 1.2168 | 0.15666 | 0.3203 | 0.989846 | 7484 | 2 | 1.2168 |
| MDK | 4 | 0.85288 | 0.85243 | 0.999994 | 18315 | 0 | 1.9252 | 0.15666 | 0.3203 | 0.989846 | 7485 | 2 | 1.9252 |
| MTA3 | 6 | 0.96473 | 0.96478 | 0.999994 | 20717 | 0 | 0.88591 | 0.15667 | 0.36798 | 0.997833 | 7486 | 4 | 0.88591 |
| OCA2 | 6 | 0.30519 | 0.51277 | 0.999994 | 10236 | 3 | -0.45621 | 0.15667 | 0.36798 | 0.997833 | 7487 | 2 | -0.45621 |

| | | | | | | | | | | | | | |
|---|---|---|---|---|---|---|---|---|---|---|---|---|---|
| RARS2 | 4 | 0.33648 | 0.48034 | 0.999994 | 10858 | 2 | 2.1787 | 0.15672 | 0.32039 | 0.989846 | 7488 | 2 | 2.1787 |
| HAND2 | 4 | 0.2904 | 0.43322 | 0.999994 | 9919 | 2 | 1.9711 | 0.15672 | 0.32039 | 0.989846 | 7489 | 2 | 1.9711 |
| ERCC8 | 3 | 0.8694 | 0.86946 | 0.999994 | 18624 | 0 | 0.96483 | 0.15673 | 0.29557 | 0.983965 | 7490 | 2 | 0.96483 |
| MAP4 | 5 | 0.88573 | 0.88583 | 0.999994 | 18942 | 1 | 2.3702 | 0.15677 | 0.35947 | 0.994392 | 7491 | 3 | 2.3702 |
| SSX7 | 5 | 0.24274 | 0.43465 | 0.999994 | 8857 | 3 | -0.61649 | 0.15677 | 0.35947 | 0.994392 | 7492 | 1 | -0.61649 |
| TPK1 | 5 | 0.95948 | 0.95962 | 0.999994 | 20571 | 0 | 1.8227 | 0.15678 | 0.3595 | 0.994392 | 7493 | 4 | 1.8227 |
| DEFA4 | 5 | 0.40816 | 0.57034 | 0.999994 | 12079 | 1 | 1.909 | 0.15685 | 0.35962 | 0.994416 | 7494 | 4 | 1.909 |
| TSGA13 | 6 | 0.32021 | 0.53012 | 0.999994 | 10517 | 2 | 2.1558 | 0.15688 | 0.36831 | 0.998341 | 7495 | 3 | 2.1558 |
| ASCL2 | 5 | 0.85567 | 0.85882 | 0.999994 | 18359 | 1 | 1.0285 | 0.15695 | 0.35978 | 0.994441 | 7496 | 4 | 1.0285 |
| LMO1 | 5 | 0.053029 | 0.14769 | 0.965554 | 3112 | 1 | 2.4729 | 0.15695 | 0.35979 | 0.994441 | 7497 | 3 | 2.4729 |
| MCTP1 | 5 | 0.89813 | 0.89824 | 0.999994 | 19184 | 0 | 2.5383 | 0.15698 | 0.35983 | 0.994441 | 7498 | 3 | 2.5383 |
| HLA-DPA1 | 5 | 0.51726 | 0.66292 | 0.999994 | 13697 | 2 | 0.16268 | 0.157 | 0.35987 | 0.994441 | 7499 | 2 | 0.16268 |
| GAL3ST3 | 6 | 0.61679 | 0.75243 | 0.999994 | 15169 | 2 | 1.5992 | 0.15701 | 0.36851 | 0.998341 | 7500 | 3 | 1.5992 |
| PDS5B | 5 | 0.89746 | 0.8976 | 0.999994 | 19167 | 0 | 0.89747 | 0.15711 | 0.36007 | 0.994562 | 7501 | 4 | 0.89747 |
| FLT4 | 5 | 0.81732 | 0.83081 | 0.999994 | 17734 | 1 | 0.91339 | 0.15719 | 0.36022 | 0.994562 | 7502 | 4 | 0.91339 |
| TRIM39 | 5 | 0.023553 | 0.073655 | 0.93914 | 1689 | 4 | -1.6194 | 0.15724 | 0.3603 | 0.994562 | 7503 | 1 | -1.6194 |
| LCT | 6 | 0.42659 | 0.61966 | 0.999994 | 12357 | 2 | 1.7109 | 0.15725 | 0.36888 | 0.998341 | 7504 | 4 | 1.7109 |
| SPACA7 | 6 | 0.88759 | 0.89463 | 0.999994 | 18988 | 1 | 2.668 | 0.15725 | 0.36889 | 0.998341 | 7505 | 4 | 2.668 |
| VDR | 5 | 0.83212 | 0.84102 | 0.999994 | 17961 | 1 | 2.4372 | 0.15732 | 0.36046 | 0.994562 | 7506 | 3 | 2.4372 |
| HADHA | 6 | 0.44055 | 0.62879 | 0.999994 | 12585 | 1 | 1.4946 | 0.15739 | 0.36908 | 0.998341 | 7507 | 4 | 1.4946 |
| HIST1H2AJ | 3 | 0.37003 | 0.46702 | 0.999994 | 11511 | 1 | 3.244 | 0.15742 | 0.29632 | 0.984782 | 7508 | 2 | 3.244 |
| ACTC1 | 6 | 0.49193 | 0.6632 | 0.999994 | 13300 | 2 | 1.7435 | 0.15743 | 0.36914 | 0.998341 | 7509 | 3 | 1.7435 |
| ZNF35 | 6 | 0.7653 | 0.83408 | 0.999994 | 17032 | 1 | 0.74436 | 0.15743 | 0.36914 | 0.998341 | 7510 | 3 | 0.74436 |
| SPAG1 | 6 | 0.19897 | 0.38664 | 0.999994 | 7736 | 2 | 0.063612 | 0.15743 | 0.36914 | 0.998341 | 7511 | 2 | 0.063612 |
| ADAMTS9 | 6 | 0.11991 | 0.28554 | 0.999632 | 5623 | 1 | 2.2333 | 0.15748 | 0.36923 | 0.998341 | 7512 | 4 | 2.2333 |
| KRTAP26-1 | 5 | 0.35593 | 0.52673 | 0.999994 | 11267 | 1 | 0.46141 | 0.15749 | 0.36074 | 0.994562 | 7513 | 2 | 0.46141 |
| ASIC3 | 5 | 0.52199 | 0.66703 | 0.999994 | 13780 | 1 | 1.8315 | 0.15749 | 0.36074 | 0.994562 | 7514 | 3 | 1.8315 |
| SERPINB13 | 5 | 0.56546 | 0.70552 | 0.999994 | 14476 | 2 | 2.7092 | 0.15749 | 0.36074 | 0.994562 | 7515 | 3 | 2.7092 |
| ITIH6 | 6 | 0.92806 | 0.92814 | 0.999994 | 19815 | 0 | 1.665 | 0.15751 | 0.36926 | 0.998341 | 7516 | 4 | 1.665 |
| FCRLB | 6 | 0.9171 | 0.91774 | 0.999994 | 19564 | 1 | 1.5335 | 0.15752 | 0.36928 | 0.998341 | 7517 | 4 | 1.5335 |
| RCC1 | 4 | 0.041313 | 0.1044 | 0.947919 | 2579 | 2 | -0.64374 | 0.15757 | 0.32164 | 0.989846 | 7518 | 2 | -0.64374 |
| CLDN2 | 5 | 0.24278 | 0.43469 | 0.999994 | 8858 | 2 | 2.5924 | 0.15767 | 0.36106 | 0.994562 | 7519 | 3 | 2.5924 |
| LHFPL2 | 5 | 0.92373 | 0.92394 | 0.999994 | 19711 | 0 | 0.30098 | 0.15773 | 0.36117 | 0.994562 | 7520 | 2 | 0.30098 |
| PHF21B | 4 | 0.32452 | 0.46809 | 0.999994 | 10607 | 1 | 1.8698 | 0.15774 | 0.32188 | 0.989846 | 7521 | 3 | 1.8698 |
| CTRL | 6 | 0.16441 | 0.34374 | 0.999994 | 6848 | 2 | 2.4816 | 0.15774 | 0.36961 | 0.9988 | 7522 | 4 | 2.4816 |
| PHLDA2 | 4 | 0.044803 | 0.11133 | 0.951052 | 2749 | 2 | -1.9206 | 0.15779 | 0.32196 | 0.989846 | 7523 | 2 | -1.9206 |
| NonTargetingControlG | 1 | 0.8422 | 0.84246 | 0.999994 | 18127 | 0 | 6.0066 | 0.1578 | 0.15752 | 0.965913 | 7524 | 1 | 6.0066 |
| THNSL2 | 6 | 0.039218 | 0.12013 | 0.951052 | 2479 | 4 | -1.6871 | 0.15784 | 0.36974 | 0.9988 | 7525 | 2 | -1.6871 |
| ANK2 | 6 | 0.11245 | 0.27139 | 0.998561 | 5384 | 4 | -0.91949 | 0.15784 | 0.36974 | 0.9988 | 7526 | 1 | -0.91949 |
| SRRM5 | 6 | 0.90976 | 0.91135 | 0.999994 | 19421 | 1 | 2.4186 | 0.15786 | 0.36979 | 0.9988 | 7527 | 4 | 2.4186 |
| IKBKB | 5 | 0.76548 | 0.79917 | 0.999994 | 17035 | 1 | 1.2376 | 0.15787 | 0.36139 | 0.994786 | 7528 | 4 | 1.2376 |
| NonTargetingControlG | 2 | 0.28358 | 0.32424 | 0.999994 | 9777 | 1 | 2.3137 | 0.1579 | 0.25706 | 0.974766 | 7529 | 1 | 2.3137 |
| C8orf37 | 5 | 0.77447 | 0.80425 | 0.999994 | 17138 | 1 | 1.2867 | 0.1579 | 0.36145 | 0.994802 | 7530 | 4 | 1.2867 |
| SH3GL2 | 6 | 0.54689 | 0.70142 | 0.999994 | 14165 | 1 | 1.8248 | 0.15798 | 0.36998 | 0.9988 | 7531 | 3 | 1.8248 |
| B3GNT8 | 6 | 0.21288 | 0.40359 | 0.999994 | 8086 | 1 | 1.6956 | 0.15802 | 0.37004 | 0.9988 | 7532 | 4 | 1.6956 |
| ADAMTS4 | 6 | 0.22437 | 0.41737 | 0.999994 | 8389 | 2 | 1.2078 | 0.15807 | 0.37011 | 0.9988 | 7533 | 3 | 1.2078 |
| PPP1R3A | 5 | 0.42543 | 0.58462 | 0.999994 | 12344 | 2 | 0.33063 | 0.15808 | 0.36178 | 0.994802 | 7534 | 1 | 0.33063 |
| MAD2L1BP | 5 | 0.10583 | 0.2355 | 0.985362 | 5166 | 1 | 0.2136 | 0.15808 | 0.36178 | 0.994802 | 7535 | 2 | 0.2136 |
| BCL2A1 | 5 | 0.50615 | 0.65329 | 0.999994 | 13529 | 2 | 1.1914 | 0.15808 | 0.36178 | 0.994802 | 7536 | 3 | 1.1914 |
| NPR3 | 6 | 0.04531 | 0.13376 | 0.95318 | 2779 | 2 | 1.6083 | 0.15812 | 0.37019 | 0.9988 | 7537 | 4 | 1.6083 |
| HDGFL1 | 5 | 0.22312 | 0.40765 | 0.999994 | 8361 | 1 | 0.78285 | 0.15812 | 0.36184 | 0.994802 | 7538 | 4 | 0.78285 |
| AKIRIN1 | 5 | 0.32505 | 0.50145 | 0.999994 | 10621 | 1 | 1.0245 | 0.15813 | 0.36186 | 0.994802 | 7539 | 4 | 1.0245 |

| | | | | | | | | | | | | | |
|---|---|---|---|---|---|---|---|---|---|---|---|---|---|
| hsa-mir-5100 | 4 | 0.25618 | 0.39821 | 0.999994 | 9154 | 1 | 1.3653 | 0.15814 | 0.32247 | 0.989846 | 7540 | 3 | 1.3653 |
| WDR5 | 4 | 0.38261 | 0.52784 | 0.999994 | 11685 | 2 | 0.049045 | 0.15814 | 0.32247 | 0.989846 | 7541 | 2 | 0.049045 |
| PRL | 4 | 0.7276 | 0.74343 | 0.999994 | 16576 | 1 | 0.37745 | 0.15814 | 0.32247 | 0.989846 | 7542 | 2 | 0.37745 |
| hsa-mir-202 | 4 | 0.11383 | 0.23958 | 0.987253 | 5441 | 2 | 0.39218 | 0.15814 | 0.32247 | 0.989846 | 7543 | 2 | 0.39218 |
| ARHGAP19 | 6 | 0.87584 | 0.88697 | 0.999994 | 18748 | 1 | 0.27069 | 0.15823 | 0.37036 | 0.9988 | 7544 | 2 | 0.27069 |
| DAK | 6 | 0.26934 | 0.47095 | 0.999994 | 9454 | 1 | 1.9399 | 0.15825 | 0.37039 | 0.9988 | 7545 | 4 | 1.9399 |
| SLC35G3 | 5 | 0.70021 | 0.76658 | 0.999994 | 16305 | 1 | 1.0213 | 0.15835 | 0.36223 | 0.99534 | 7546 | 3 | 1.0213 |
| CCDC69 | 6 | 0.39785 | 0.60097 | 0.999994 | 11914 | 1 | 2.148 | 0.15838 | 0.37058 | 0.998861 | 7547 | 4 | 2.148 |
| GEMIN8 | 5 | 0.94245 | 0.94264 | 0.999994 | 20135 | 0 | 1.7289 | 0.15855 | 0.36255 | 0.995401 | 7548 | 4 | 1.7289 |
| AKAP8 | 6 | 0.41801 | 0.61406 | 0.999994 | 12217 | 2 | 1.6577 | 0.1586 | 0.37091 | 0.998994 | 7549 | 3 | 1.6577 |
| SGOL1 | 4 | 0.31978 | 0.4632 | 0.999994 | 10509 | 1 | 2.2369 | 0.15866 | 0.32323 | 0.989881 | 7550 | 3 | 2.2369 |
| PTGS1 | 4 | 0.49838 | 0.61561 | 0.999994 | 13401 | 1 | 1.8394 | 0.15869 | 0.32329 | 0.989881 | 7551 | 3 | 1.8394 |
| MUL1 | 6 | 0.17566 | 0.35768 | 0.999994 | 7128 | 2 | 0.95538 | 0.15877 | 0.37115 | 0.998994 | 7552 | 4 | 0.95538 |
| NTS | 6 | 0.31871 | 0.52842 | 0.999994 | 10491 | 2 | -0.017542 | 0.15877 | 0.37115 | 0.998994 | 7553 | 2 | -0.017542 |
| FOXR1 | 6 | 0.89947 | 0.90326 | 0.999994 | 19212 | 1 | 1.1936 | 0.15877 | 0.37115 | 0.998994 | 7554 | 4 | 1.1936 |
| NonTargetingControlG | 1 | 0.8412 | 0.84148 | 0.999994 | 18109 | 0 | 5.9833 | 0.1588 | 0.15851 | 0.965913 | 7555 | 1 | 5.9833 |
| BTNL3 | 5 | 0.34612 | 0.51874 | 0.999994 | 11072 | 2 | 1.4246 | 0.15892 | 0.36317 | 0.995401 | 7556 | 3 | 1.4246 |
| CD3E | 5 | 0.29983 | 0.48093 | 0.999994 | 10126 | 2 | -0.41958 | 0.15893 | 0.36318 | 0.995401 | 7557 | 2 | -0.41958 |
| C12orf66 | 5 | 0.78168 | 0.8084 | 0.999994 | 17234 | 1 | 0.71765 | 0.15899 | 0.36329 | 0.995401 | 7558 | 4 | 0.71765 |
| hsa-mir-96 | 3 | 0.85414 | 0.85416 | 0.999994 | 18335 | 0 | 3.2995 | 0.15906 | 0.29815 | 0.984876 | 7559 | 2 | 3.2995 |
| HERPUD2 | 6 | 0.024154 | 0.084996 | 0.94395 | 1715 | 4 | -1.8423 | 0.15909 | 0.37162 | 0.998994 | 7560 | 1 | -1.8423 |
| MTTP | 6 | 0.34974 | 0.56371 | 0.999994 | 11151 | 2 | 1.2468 | 0.15909 | 0.37162 | 0.998994 | 7561 | 4 | 1.2468 |
| PRR15L | 5 | 0.83762 | 0.84491 | 0.999994 | 18045 | 1 | 0.868 | 0.15909 | 0.36346 | 0.995401 | 7562 | 4 | 0.868 |
| SNTA1 | 4 | 0.81864 | 0.81847 | 0.999994 | 17759 | 1 | 1.6349 | 0.15919 | 0.32402 | 0.990429 | 7563 | 3 | 1.6349 |
| hsa-mir-216a | 3 | 0.82231 | 0.8222 | 0.999994 | 17808 | 0 | 3.3053 | 0.15922 | 0.29831 | 0.984876 | 7564 | 2 | 3.3053 |
| hsa-mir-215 | 4 | 0.76538 | 0.77206 | 0.999994 | 17033 | 1 | 3.0669 | 0.15933 | 0.32419 | 0.990429 | 7565 | 3 | 3.0669 |
| KIFC1 | 6 | 0.50459 | 0.67191 | 0.999994 | 13505 | 2 | 1.5334 | 0.15935 | 0.37199 | 0.998994 | 7566 | 4 | 1.5334 |
| TRAPPC5 | 6 | 0.39184 | 0.59713 | 0.999994 | 11829 | 2 | 1.7126 | 0.15937 | 0.37203 | 0.998994 | 7567 | 4 | 1.7126 |
| NonTargetingControlG | 2 | 0.44336 | 0.4535 | 0.999994 | 12627 | 1 | 3.0118 | 0.15938 | 0.25916 | 0.974766 | 7568 | 1 | 3.0118 |
| RUFY1 | 6 | 0.039116 | 0.11991 | 0.951052 | 2470 | 2 | 1.6909 | 0.15939 | 0.37205 | 0.998994 | 7569 | 4 | 1.6909 |
| PRR11 | 5 | 0.90334 | 0.9034 | 0.999994 | 19287 | 0 | 0.93627 | 0.15944 | 0.36413 | 0.995622 | 7570 | 4 | 0.93627 |
| TNFRSF8 | 3 | 0.16549 | 0.26865 | 0.998561 | 6871 | 2 | -1.4144 | 0.15945 | 0.29856 | 0.984876 | 7571 | 1 | -1.4144 |
| ZNF629 | 5 | 0.45373 | 0.60837 | 0.999994 | 12756 | 2 | 2.5181 | 0.15948 | 0.36419 | 0.995622 | 7572 | 3 | 2.5181 |
| ATP5G1 | 6 | 0.28725 | 0.4919 | 0.999994 | 9860 | 1 | 1.373 | 0.15958 | 0.37236 | 0.998994 | 7573 | 4 | 1.373 |
| OR2AG2 | 6 | 0.75105 | 0.82883 | 0.999994 | 16866 | 1 | 1.937 | 0.15958 | 0.37236 | 0.998994 | 7574 | 4 | 1.937 |
| UNC45A | 6 | 0.16117 | 0.33971 | 0.999994 | 6770 | 2 | 1.4752 | 0.1597 | 0.37253 | 0.999063 | 7575 | 4 | 1.4752 |
| SAMD9L | 5 | 0.062354 | 0.16596 | 0.967126 | 3529 | 3 | -0.8548 | 0.1597 | 0.36461 | 0.995622 | 7576 | 2 | -0.8548 |
| ZNF776 | 5 | 0.14094 | 0.28924 | 0.999632 | 6217 | 2 | 2.1684 | 0.1597 | 0.36461 | 0.995622 | 7577 | 3 | 2.1684 |
| KHNYN | 5 | 0.1336 | 0.27814 | 0.999536 | 6012 | 3 | -1.2606 | 0.15981 | 0.36478 | 0.995622 | 7578 | 2 | -1.2606 |
| TMEM200C | 5 | 0.071211 | 0.18035 | 0.968171 | 3883 | 2 | -0.38583 | 0.15981 | 0.36478 | 0.995622 | 7579 | 2 | -0.38583 |
| NonTargetingControlG | 2 | 0.84012 | 0.84005 | 0.999994 | 18091 | 0 | 1.2682 | 0.15988 | 0.25991 | 0.974766 | 7580 | 2 | 1.2682 |
| TSPAN8 | 6 | 0.24266 | 0.43917 | 0.999994 | 8853 | 2 | 2.0588 | 0.15993 | 0.37288 | 0.999346 | 7581 | 3 | 2.0588 |
| NonTargetingControlG | 2 | 0.83998 | 0.8399 | 0.999994 | 18087 | 0 | 2.036 | 0.16002 | 0.26012 | 0.974766 | 7582 | 2 | 2.036 |
| GPN1 | 4 | 0.64775 | 0.69199 | 0.999994 | 15627 | 1 | 1.3737 | 0.16003 | 0.32526 | 0.991152 | 7583 | 2 | 1.3737 |
| MRTO4 | 5 | 0.41865 | 0.57898 | 0.999994 | 12229 | 1 | 2.3227 | 0.16014 | 0.36533 | 0.996287 | 7584 | 4 | 2.3227 |
| SPATA31D4 | 2 | 0.83985 | 0.83977 | 0.999994 | 18084 | 0 | 1.5741 | 0.16015 | 0.26031 | 0.974766 | 7585 | 2 | 1.5741 |
| OR1Q1 | 5 | 0.4902 | 0.63938 | 0.999994 | 13281 | 2 | 0.96316 | 0.16018 | 0.3654 | 0.996287 | 7586 | 3 | 0.96316 |
| AFP | 5 | 0.0631 | 0.16719 | 0.967126 | 3560 | 3 | -2.1449 | 0.16018 | 0.3654 | 0.996287 | 7587 | 1 | -2.1449 |
| ZNF843 | 6 | 0.0038689 | 0.016995 | 0.785648 | 434 | 2 | 0.45169 | 0.16033 | 0.37346 | 0.999517 | 7588 | 3 | 0.45169 |
| HNRNPH1 | 4 | 0.035442 | 0.092625 | 0.943968 | 2290 | 1 | 2.7486 | 0.16047 | 0.32589 | 0.991729 | 7589 | 2 | 2.7486 |
| BLOC1S3 | 5 | 0.51085 | 0.6574 | 0.999994 | 13590 | 2 | -0.032091 | 0.1605 | 0.36594 | 0.996287 | 7590 | 2 | -0.032091 |
| ASCL3 | 5 | 0.89343 | 0.89357 | 0.999994 | 19095 | 0 | 2.3182 | 0.16051 | 0.36597 | 0.996287 | 7591 | 3 | 2.3182 |

| | | | | | | | | | | | | |
|---|---|---|---|---|---|---|---|---|---|---|---|---|
| H2BFM | 6 | 0.3524 | 0.56671 | 0.999994 | 11203 | 2 | 1.6563 | 0.16062 | 0.37391 | 0.999517 | 7592 | 4 | 1.6563 |
| hsa-mir-548aa-2 | 3 | 0.64859 | 0.6559 | 0.999994 | 15637 | 1 | 3.2724 | 0.16062 | 0.29982 | 0.984876 | 7593 | 2 | 3.2724 |
| hsa-mir-4261 | 4 | 0.10995 | 0.23278 | 0.984502 | 5305 | 1 | 2.0673 | 0.16067 | 0.3262 | 0.991729 | 7594 | 3 | 2.0673 |
| hsa-mir-8075 | 2 | 0.8393 | 0.83922 | 0.999994 | 18075 | 0 | 1.1869 | 0.1607 | 0.26108 | 0.974875 | 7595 | 2 | 1.1869 |
| EXOSC7 | 5 | 0.3705 | 0.53881 | 0.999994 | 11521 | 1 | 1.568 | 0.1607 | 0.36629 | 0.996728 | 7596 | 4 | 1.568 |
| SLC30A10 | 6 | 0.64055 | 0.77037 | 0.999994 | 15518 | 2 | 2.054 | 0.16073 | 0.37408 | 0.999517 | 7597 | 4 | 2.054 |
| C1orf56 | 4 | 0.012737 | 0.042197 | 0.891621 | 1049 | 2 | -3.8046 | 0.16074 | 0.32631 | 0.991729 | 7598 | 2 | -3.8046 |
| MPZ | 5 | 0.78554 | 0.81065 | 0.999994 | 17286 | 1 | 2.187 | 0.16075 | 0.36638 | 0.996842 | 7599 | 3 | 2.187 |
| A2M | 6 | 0.26946 | 0.47108 | 0.999994 | 9460 | 2 | 2.5208 | 0.16077 | 0.37414 | 0.999517 | 7600 | 4 | 2.5208 |
| ATPAF1 | 6 | 0.79746 | 0.84681 | 0.999994 | 17444 | 1 | 2.1768 | 0.16077 | 0.37414 | 0.999517 | 7601 | 3 | 2.1768 |
| RAD51D | 6 | 0.33378 | 0.54561 | 0.999994 | 10804 | 2 | 2.0715 | 0.16077 | 0.37414 | 0.999517 | 7602 | 3 | 2.0715 |
| SLC38A1 | 6 | 0.30475 | 0.51226 | 0.999994 | 10223 | 2 | 2.5193 | 0.16077 | 0.37414 | 0.999517 | 7603 | 4 | 2.5193 |
| NRXN3 | 6 | 0.86878 | 0.8826 | 0.999994 | 18613 | 1 | 1.7603 | 0.16077 | 0.37414 | 0.999517 | 7604 | 3 | 1.7603 |
| SHF | 6 | 0.19392 | 0.38042 | 0.999994 | 7599 | 1 | 2.2637 | 0.16077 | 0.37414 | 0.999517 | 7605 | 4 | 2.2637 |
| PPIL6 | 5 | 0.74651 | 0.78901 | 0.999994 | 16821 | 1 | 0.71083 | 0.16089 | 0.36662 | 0.997075 | 7606 | 4 | 0.71083 |
| BET3L | 6 | 0.32048 | 0.53044 | 0.999994 | 10522 | 1 | 1.4061 | 0.16089 | 0.37434 | 0.999517 | 7607 | 4 | 1.4061 |
| ZNF83 | 5 | 0.94942 | 0.9496 | 0.999994 | 20318 | 0 | 2.3549 | 0.16093 | 0.36669 | 0.997075 | 7608 | 4 | 2.3549 |
| TSTD1 | 6 | 0.023387 | 0.082819 | 0.943262 | 1675 | 2 | 0.67077 | 0.16097 | 0.37445 | 0.999517 | 7609 | 3 | 0.67077 |
| IL13RA2 | 6 | 0.94284 | 0.94282 | 0.999994 | 20148 | 0 | 1.626 | 0.16098 | 0.37447 | 0.999517 | 7610 | 4 | 1.626 |
| MCU | 5 | 0.063612 | 0.16804 | 0.967126 | 3587 | 3 | -1.5915 | 0.16099 | 0.36678 | 0.997075 | 7611 | 2 | -1.5915 |
| ATP8B1 | 3 | 0.45193 | 0.51542 | 0.999994 | 12734 | 1 | 1.0119 | 0.16104 | 0.30028 | 0.984876 | 7612 | 2 | 1.0119 |
| KIF4A | 5 | 0.54457 | 0.68693 | 0.999994 | 14134 | 1 | 2.5468 | 0.16108 | 0.36693 | 0.997075 | 7613 | 3 | 2.5468 |
| KIF3C | 6 | 0.095835 | 0.23917 | 0.987253 | 4808 | 3 | -0.32529 | 0.16111 | 0.37468 | 0.999639 | 7614 | 2 | -0.32529 |
| FAM111B | 6 | 0.63222 | 0.76402 | 0.999994 | 15403 | 2 | 1.8874 | 0.16111 | 0.37468 | 0.999639 | 7615 | 4 | 1.8874 |
| RASSF5 | 5 | 0.47279 | 0.62449 | 0.999994 | 13043 | 2 | 2.3622 | 0.16113 | 0.36701 | 0.997075 | 7616 | 3 | 2.3622 |
| NR4A2 | 6 | 0.45045 | 0.63543 | 0.999994 | 12718 | 2 | 1.7938 | 0.16124 | 0.37485 | 0.999639 | 7617 | 4 | 1.7938 |
| KCTD4 | 5 | 0.87298 | 0.87376 | 0.999994 | 18693 | 1 | 1.0128 | 0.16126 | 0.36723 | 0.997075 | 7618 | 4 | 1.0128 |
| STOML3 | 6 | 0.89791 | 0.90203 | 0.999994 | 19177 | 1 | 1.1035 | 0.16126 | 0.37488 | 0.999639 | 7619 | 3 | 1.1035 |
| GMFG | 4 | 0.24353 | 0.38527 | 0.999994 | 8875 | 1 | 2.7948 | 0.16135 | 0.32721 | 0.991729 | 7620 | 2 | 2.7948 |
| HELB | 6 | 0.48106 | 0.65592 | 0.999994 | 13152 | 2 | 2.4926 | 0.16142 | 0.37512 | 0.999639 | 7621 | 4 | 2.4926 |
| KCNK13 | 6 | 0.021252 | 0.076002 | 0.93914 | 1530 | 4 | -2.3893 | 0.16146 | 0.37516 | 0.999639 | 7622 | 2 | -2.3893 |
| RBMS3 | 6 | 0.20239 | 0.39079 | 0.999994 | 7841 | 3 | -0.85529 | 0.16159 | 0.37536 | 0.999639 | 7623 | 2 | -0.85529 |
| C8orf34 | 4 | 0.82655 | 0.82614 | 0.999994 | 17877 | 1 | 3.0146 | 0.16161 | 0.32761 | 0.991729 | 7624 | 3 | 3.0146 |
| TMEM57 | 3 | 0.71036 | 0.71079 | 0.999994 | 16403 | 1 | 3.3298 | 0.16165 | 0.30097 | 0.984876 | 7625 | 2 | 3.3298 |
| GABRB1 | 6 | 0.89877 | 0.9027 | 0.999994 | 19198 | 1 | 1.846 | 0.16177 | 0.37564 | 0.999639 | 7626 | 4 | 1.846 |
| NR5A2 | 5 | 0.015092 | 0.050466 | 0.909101 | 1193 | 3 | -3.5307 | 0.1618 | 0.36818 | 0.99823 | 7627 | 2 | -3.5307 |
| CDYL | 4 | 0.58402 | 0.65682 | 0.999994 | 14729 | 1 | 1.9969 | 0.16182 | 0.32791 | 0.991729 | 7628 | 3 | 1.9969 |
| C2orf76 | 5 | 0.68221 | 0.75854 | 0.999994 | 16120 | 1 | 0.74968 | 0.16188 | 0.36833 | 0.998341 | 7629 | 4 | 0.74968 |
| KATNAL1 | 6 | 0.019693 | 0.071009 | 0.93914 | 1439 | 3 | -0.4545 | 0.16192 | 0.37585 | 0.999639 | 7630 | 2 | -0.4545 |
| NRN1 | 6 | 0.6645 | 0.7888 | 0.999994 | 15849 | 2 | 0.57291 | 0.16192 | 0.37585 | 0.999639 | 7631 | 2 | 0.57291 |
| HBS1L | 6 | 0.53552 | 0.69339 | 0.999994 | 13990 | 2 | 1.7851 | 0.16192 | 0.37585 | 0.999639 | 7632 | 3 | 1.7851 |
| NonTargetingControlG | 1 | 0.83808 | 0.83836 | 0.999994 | 18055 | 0 | 5.9188 | 0.16192 | 0.16164 | 0.965913 | 7633 | 1 | 5.9188 |
| ATP9B | 4 | 0.94264 | 0.94248 | 0.999994 | 20143 | 0 | 1.4604 | 0.16193 | 0.32807 | 0.991729 | 7634 | 3 | 1.4604 |
| ARHGAP44 | 6 | 0.39412 | 0.59858 | 0.999994 | 11860 | 2 | 1.7822 | 0.16198 | 0.37595 | 0.999762 | 7635 | 4 | 1.7822 |
| DOLPP1 | 6 | 0.54908 | 0.70298 | 0.999994 | 14194 | 1 | 1.2318 | 0.16204 | 0.37604 | 0.999762 | 7636 | 4 | 1.2318 |
| HIPK2 | 5 | 0.59982 | 0.72609 | 0.999994 | 14950 | 1 | 2.2415 | 0.16214 | 0.36877 | 0.998341 | 7637 | 3 | 2.2415 |
| SLC19A1 | 5 | 0.20007 | 0.37533 | 0.999994 | 7766 | 3 | -0.82244 | 0.16219 | 0.36884 | 0.998341 | 7638 | 2 | -0.82244 |
| DLST | 5 | 0.75993 | 0.79613 | 0.999994 | 16980 | 1 | 0.58028 | 0.16219 | 0.36884 | 0.998341 | 7639 | 2 | 0.58028 |
| RBM6 | 6 | 0.37553 | 0.58672 | 0.999994 | 11586 | 2 | 1.5822 | 0.16222 | 0.3763 | 0.999964 | 7640 | 4 | 1.5822 |
| FLJ25363 | 5 | 0.92126 | 0.92144 | 0.999994 | 19658 | 0 | 2.4372 | 0.16233 | 0.36912 | 0.998341 | 7641 | 3 | 2.4372 |
| CYSLTR2 | 6 | 0.16609 | 0.34582 | 0.999994 | 6888 | 2 | 1.468 | 0.16234 | 0.37648 | 0.999964 | 7642 | 4 | 1.468 |
| HDAC10 | 5 | 0.96314 | 0.96329 | 0.999994 | 20674 | 0 | 1.6943 | 0.16234 | 0.36913 | 0.998341 | 7643 | 4 | 1.6943 |

| | | | | | | | | | | | | | |
|---|---|---|---|---|---|---|---|---|---|---|---|---|---|
| NonTargetingControlG | 2 | 0.55341 | 0.55416 | 0.999994 | 14262 | 1 | 3.3109 | 0.16237 | 0.26354 | 0.974875 | 7644 | 1 | 3.3109 |
| NonTargetingControlG | 2 | 0.66167 | 0.66145 | 0.999994 | 15805 | 0 | 3.6505 | 0.16237 | 0.26354 | 0.974875 | 7645 | 1 | 3.6505 |
| GSTO1 | 6 | 0.29858 | 0.50504 | 0.999994 | 10098 | 3 | 0.00096563 | 0.16239 | 0.37657 | 1 | 7646 | 3 | 0.00096563 |
| SKA1 | 4 | 0.81828 | 0.81812 | 0.999994 | 17752 | 1 | 2.0715 | 0.1624 | 0.32874 | 0.991729 | 7647 | 3 | 2.0715 |
| RAP1B | 4 | 0.81615 | 0.81615 | 0.999994 | 17715 | 1 | 1.8424 | 0.16254 | 0.32894 | 0.991729 | 7648 | 3 | 1.8424 |
| OR52R1 | 6 | 0.0062027 | 0.025559 | 0.845628 | 605 | 4 | -2.4521 | 0.16255 | 0.37681 | 1 | 7649 | 2 | -2.4521 |
| VAMP4 | 6 | 0.45964 | 0.64152 | 0.999994 | 12842 | 2 | 0.96614 | 0.16255 | 0.37681 | 1 | 7650 | 3 | 0.96614 |
| LRP2BP | 4 | 0.11463 | 0.24097 | 0.987253 | 5467 | 2 | 1.5034 | 0.16256 | 0.32898 | 0.991729 | 7651 | 2 | 1.5034 |
| SLC22A17 | 4 | 0.83737 | 0.83685 | 0.999994 | 18039 | 0 | 3.0034 | 0.16256 | 0.32898 | 0.991729 | 7652 | 3 | 3.0034 |
| HTR1F | 6 | 0.3659 | 0.58064 | 0.999994 | 11437 | 2 | 2.4958 | 0.16272 | 0.37705 | 1 | 7653 | 4 | 2.4958 |
| DEFB4A | 5 | 0.84819 | 0.85284 | 0.999994 | 18220 | 1 | 2.5413 | 0.16275 | 0.36982 | 0.9988 | 7654 | 3 | 2.5413 |
| IFI27L2 | 5 | 0.60782 | 0.72899 | 0.999994 | 15044 | 1 | 2.0496 | 0.16288 | 0.37004 | 0.9988 | 7655 | 4 | 2.0496 |
| ZNF671 | 5 | 0.20311 | 0.37966 | 0.999994 | 7857 | 2 | 2.3508 | 0.16288 | 0.37005 | 0.9988 | 7656 | 3 | 2.3508 |
| CIDEC | 6 | 0.0094383 | 0.037042 | 0.881473 | 823 | 5 | -1.7317 | 0.16292 | 0.37734 | 1 | 7657 | 1 | -1.7317 |
| CEP290 | 5 | 0.6947 | 0.7641 | 0.999994 | 16252 | 1 | 0.88513 | 0.16304 | 0.37031 | 0.9988 | 7658 | 4 | 0.88513 |
| ZNF408 | 5 | 0.131 | 0.27414 | 0.998561 | 5959 | 2 | 0.60677 | 0.16305 | 0.37034 | 0.9988 | 7659 | 3 | 0.60677 |
| RMI1 | 5 | 0.92134 | 0.92152 | 0.999994 | 19664 | 0 | 2.238 | 0.16305 | 0.37034 | 0.9988 | 7660 | 3 | 2.238 |
| CLCA2 | 6 | 0.62559 | 0.75902 | 0.999994 | 15316 | 1 | 1.5741 | 0.16317 | 0.37773 | 1 | 7661 | 4 | 1.5741 |
| CD177 | 6 | 0.018218 | 0.066201 | 0.93914 | 1374 | 1 | 1.6259 | 0.16318 | 0.37773 | 1 | 7662 | 4 | 1.6259 |
| GOLM1 | 4 | 0.92699 | 0.92677 | 0.999994 | 19788 | 0 | 1.7011 | 0.16322 | 0.32996 | 0.992034 | 7663 | 3 | 1.7011 |
| PTGES3 | 5 | 0.018506 | 0.060028 | 0.932932 | 1387 | 1 | 1.1615 | 0.16327 | 0.37071 | 0.998982 | 7664 | 4 | 1.1615 |
| TMEM176A | 6 | 0.070113 | 0.18701 | 0.969409 | 3835 | 2 | 1.8513 | 0.1633 | 0.37792 | 1 | 7665 | 4 | 1.8513 |
| hsa-mir-3677 | 4 | 0.90951 | 0.90923 | 0.999994 | 19414 | 0 | 2.0202 | 0.1633 | 0.33007 | 0.992034 | 7666 | 3 | 2.0202 |
| STRA6 | 6 | 0.53482 | 0.69291 | 0.999994 | 13977 | 2 | 0.46735 | 0.16343 | 0.37811 | 1 | 7667 | 3 | 0.46735 |
| hsa-mir-7-2 | 1 | 0.83656 | 0.83682 | 0.999994 | 18028 | 0 | 5.8862 | 0.16344 | 0.16318 | 0.965913 | 7668 | 1 | 5.8862 |
| ARMC6 | 5 | 0.95578 | 0.95595 | 0.999994 | 20470 | 0 | 2.5266 | 0.16346 | 0.37103 | 0.998994 | 7669 | 4 | 2.5266 |
| KRT38 | 4 | 0.7193 | 0.73757 | 0.999994 | 16489 | 1 | 1.5701 | 0.16347 | 0.3303 | 0.992034 | 7670 | 3 | 1.5701 |
| MEN1 | 5 | 0.031336 | 0.093922 | 0.945031 | 2075 | 2 | 2.3089 | 0.16357 | 0.37121 | 0.998994 | 7671 | 3 | 2.3089 |
| hsa-mir-4788 | 4 | 0.59986 | 0.66527 | 0.999994 | 14951 | 1 | 2.7728 | 0.16362 | 0.33053 | 0.992034 | 7672 | 2 | 2.7728 |
| AGBL3 | 6 | 0.97759 | 0.97753 | 0.999994 | 21033 | 0 | 2.1322 | 0.16369 | 0.37848 | 1 | 7673 | 4 | 2.1322 |
| BCORL1 | 5 | 0.3018 | 0.4825 | 0.999994 | 10158 | 2 | 2.1259 | 0.16378 | 0.37157 | 0.998994 | 7674 | 3 | 2.1259 |
| TMIE | 6 | 0.55034 | 0.70388 | 0.999994 | 14218 | 2 | 1.6336 | 0.16381 | 0.37866 | 1 | 7675 | 4 | 1.6336 |
| HOOK2 | 5 | 0.042287 | 0.12158 | 0.951052 | 2630 | 4 | -1.976 | 0.16384 | 0.37167 | 0.998994 | 7676 | 1 | -1.976 |
| OR52D1 | 6 | 0.1125 | 0.2715 | 0.998561 | 5387 | 2 | 0.72525 | 0.16385 | 0.37872 | 1 | 7677 | 3 | 0.72525 |
| HIGD1B | 5 | 0.79524 | 0.81655 | 0.999994 | 17420 | 1 | 0.65755 | 0.16388 | 0.37173 | 0.998994 | 7678 | 4 | 0.65755 |
| LY6G6C | 6 | 0.0075847 | 0.030503 | 0.862223 | 697 | 3 | 0.24734 | 0.16393 | 0.37883 | 1 | 7679 | 3 | 0.24734 |
| GRAMD1C | 5 | 0.62272 | 0.73457 | 0.999994 | 15270 | 1 | 2.3593 | 0.16394 | 0.37184 | 0.998994 | 7680 | 3 | 2.3593 |
| SPATA8 | 5 | 0.92071 | 0.9209 | 0.999994 | 19645 | 0 | 0.82088 | 0.16397 | 0.37189 | 0.998994 | 7681 | 4 | 0.82088 |
| USE1 | 5 | 0.50072 | 0.64853 | 0.999994 | 13434 | 2 | 1.9074 | 0.16401 | 0.37197 | 0.998994 | 7682 | 3 | 1.9074 |
| NUPR1 | 5 | 0.58536 | 0.72089 | 0.999994 | 14748 | 1 | -0.2591 | 0.16401 | 0.37197 | 0.998994 | 7683 | 2 | -0.2591 |
| C5orf46 | 5 | 0.37845 | 0.54544 | 0.999994 | 11630 | 1 | 0.14624 | 0.16401 | 0.37197 | 0.998994 | 7684 | 2 | 0.14624 |
| KCNS3 | 6 | 0.33186 | 0.54339 | 0.999994 | 10763 | 2 | 2.1465 | 0.1641 | 0.37909 | 1 | 7685 | 4 | 2.1465 |
| FITM1 | 5 | 0.16198 | 0.32044 | 0.999994 | 6792 | 3 | -0.79252 | 0.16411 | 0.37213 | 0.998994 | 7686 | 2 | -0.79252 |
| CA13 | 6 | 0.77439 | 0.83747 | 0.999994 | 17134 | 1 | 0.76076 | 0.16412 | 0.3791 | 1 | 7687 | 4 | 0.76076 |
| TMEM223 | 6 | 0.023065 | 0.081785 | 0.93914 | 1654 | 2 | 1.4555 | 0.16412 | 0.3791 | 1 | 7688 | 3 | 1.4555 |
| RTEL1 | 5 | 0.56432 | 0.70448 | 0.999994 | 14460 | 2 | 2.3486 | 0.16419 | 0.37227 | 0.998994 | 7689 | 3 | 2.3486 |
| MKRN1 | 5 | 0.94374 | 0.94396 | 0.999994 | 20171 | 0 | 1.2862 | 0.16421 | 0.37229 | 0.998994 | 7690 | 4 | 1.2862 |
| HID1 | 5 | 0.49609 | 0.64448 | 0.999994 | 13358 | 1 | 0.75436 | 0.16424 | 0.37235 | 0.998994 | 7691 | 4 | 0.75436 |
| C17orf70 | 5 | 0.52168 | 0.66676 | 0.999994 | 13773 | 2 | 2.7361 | 0.16426 | 0.37238 | 0.998994 | 7692 | 3 | 2.7361 |
| HCST | 5 | 0.88433 | 0.88447 | 0.999994 | 18916 | 1 | 1.8199 | 0.16435 | 0.37254 | 0.999063 | 7693 | 3 | 1.8199 |
| ALLC | 4 | 0.3513 | 0.49565 | 0.999994 | 11183 | 2 | 0.83822 | 0.16435 | 0.33164 | 0.992121 | 7694 | 2 | 0.83822 |
| IFRD2 | 5 | 0.49731 | 0.64556 | 0.999994 | 13386 | 1 | 2.514 | 0.16439 | 0.37261 | 0.999119 | 7695 | 3 | 2.514 |

| | | | | | | | | | | | | | |
|---|---|---|---|---|---|---|---|---|---|---|---|---|---|
| GYS1 | 6 | 0.023775 | 0.08399 | 0.94395 | 1698 | 2 | 2.1065 | 0.16443 | 0.37954 | 1 | 7696 | 4 | 2.1065 |
| MAPKAP1 | 5 | 0.21327 | 0.39398 | 0.999994 | 8094 | 1 | 0.91883 | 0.16444 | 0.37268 | 0.999178 | 7697 | 4 | 0.91883 |
| METTL23 | 6 | 0.50952 | 0.67529 | 0.999994 | 13572 | 2 | 1.3939 | 0.16444 | 0.37955 | 1 | 7698 | 4 | 1.3939 |
| hsa-mir-6079 | 4 | 0.68144 | 0.71251 | 0.999994 | 16109 | 1 | 3.0747 | 0.16449 | 0.33184 | 0.992196 | 7699 | 3 | 3.0747 |
| SSPN | 6 | 0.18835 | 0.37348 | 0.999994 | 7431 | 2 | 0.22289 | 0.16453 | 0.37967 | 1 | 7700 | 3 | 0.22289 |
| ITGA6 | 4 | 0.81808 | 0.81791 | 0.999994 | 17748 | 1 | 2.2332 | 0.16462 | 0.33204 | 0.992196 | 7701 | 2 | 2.2332 |
| ITGA2 | 6 | 0.29598 | 0.50203 | 0.999994 | 10038 | 3 | 0.70272 | 0.16462 | 0.3798 | 1 | 7702 | 3 | 0.70272 |
| ZGPAT | 5 | 0.056454 | 0.15588 | 0.967126 | 3269 | 1 | 1.6077 | 0.16465 | 0.37308 | 0.999517 | 7703 | 4 | 1.6077 |
| ARPC3 | 5 | 0.90501 | 0.9051 | 0.999994 | 19327 | 0 | 1.3269 | 0.16474 | 0.37322 | 0.999517 | 7704 | 3 | 1.3269 |
| hsa-mir-4436b-1 | 3 | 0.53957 | 0.57302 | 0.999994 | 14044 | 1 | 1.779 | 0.16475 | 0.30449 | 0.986296 | 7705 | 2 | 1.779 |
| TCHHL1 | 6 | 0.024884 | 0.086736 | 0.94395 | 1751 | 2 | 0.26175 | 0.16485 | 0.38015 | 1 | 7706 | 2 | 0.26175 |
| SULF1 | 6 | 0.15208 | 0.32829 | 0.999994 | 6506 | 3 | -0.67871 | 0.16485 | 0.38015 | 1 | 7707 | 2 | -0.67871 |
| ZNF75A | 5 | 0.32329 | 0.49999 | 0.999994 | 10580 | 1 | 0.95596 | 0.16493 | 0.37354 | 0.999517 | 7708 | 4 | 0.95596 |
| C12orf73 | 5 | 0.94563 | 0.94582 | 0.999994 | 20221 | 0 | 1.9225 | 0.16493 | 0.37354 | 0.999517 | 7709 | 3 | 1.9225 |
| RNASE12 | 5 | 0.94113 | 0.94133 | 0.999994 | 20112 | 0 | 0.84634 | 0.16496 | 0.37358 | 0.999517 | 7710 | 4 | 0.84634 |
| LOC646498 | 1 | 0.83503 | 0.83527 | 0.999994 | 18007 | 0 | 5.857 | 0.16497 | 0.16477 | 0.965913 | 7711 | 1 | 5.857 |
| NLRP5 | 5 | 0.74751 | 0.78954 | 0.999994 | 16833 | 1 | 2.3281 | 0.16497 | 0.3736 | 0.999517 | 7712 | 3 | 2.3281 |
| OGFRL1 | 6 | 0.66761 | 0.79129 | 0.999994 | 15885 | 1 | 1.8388 | 0.16503 | 0.38041 | 1 | 7713 | 4 | 1.8388 |
| VPS41 | 4 | 0.33731 | 0.48117 | 0.999994 | 10880 | 2 | -0.69106 | 0.16503 | 0.33265 | 0.992621 | 7714 | 1 | -0.69106 |
| ATG10 | 6 | 0.27961 | 0.48296 | 0.999994 | 9695 | 3 | 0.15894 | 0.16505 | 0.38045 | 1 | 7715 | 3 | 0.15894 |
| NDUFAB1 | 5 | 0.2062 | 0.38401 | 0.999994 | 7935 | 1 | 1.9262 | 0.16512 | 0.37387 | 0.999517 | 7716 | 4 | 1.9262 |
| PKNOX1 | 5 | 0.098711 | 0.2244 | 0.983354 | 4921 | 2 | -0.12374 | 0.16513 | 0.37387 | 0.999517 | 7717 | 2 | -0.12374 |
| OR10X1 | 5 | 0.79116 | 0.81403 | 0.999994 | 17365 | 1 | 2.2912 | 0.1652 | 0.37401 | 0.999517 | 7718 | 3 | 2.2912 |
| NonTargetingControlG | 2 | 0.014289 | 0.026864 | 0.848412 | 1148 | 1 | -1.5922 | 0.16525 | 0.26763 | 0.978454 | 7719 | 1 | -1.5922 |
| NonTargetingControlG | 2 | 0.52445 | 0.52655 | 0.999994 | 13808 | 1 | 3.207 | 0.16525 | 0.26763 | 0.978454 | 7720 | 1 | 3.207 |
| NNMT | 5 | 0.14671 | 0.29784 | 0.999994 | 6367 | 2 | 2.2943 | 0.16526 | 0.37412 | 0.999517 | 7721 | 3 | 2.2943 |
| GLIPR1L2 | 6 | 0.4268 | 0.61979 | 0.999994 | 12361 | 2 | 1.2091 | 0.16539 | 0.38094 | 1 | 7722 | 4 | 1.2091 |
| NonTargetingControlG | 2 | 0.8345 | 0.83431 | 0.999994 | 17997 | 0 | 3.6217 | 0.1655 | 0.268 | 0.978454 | 7723 | 2 | 3.6217 |
| hsa-mir-3620 | 3 | 0.62444 | 0.63625 | 0.999994 | 15296 | 1 | 3.4955 | 0.16553 | 0.30533 | 0.986322 | 7724 | 2 | 3.4955 |
| MIXL1 | 6 | 0.059629 | 0.16495 | 0.967126 | 3406 | 3 | 0.48903 | 0.16555 | 0.38116 | 1 | 7725 | 3 | 0.48903 |
| CDKN2A | 6 | 0.15107 | 0.32703 | 0.999994 | 6484 | 2 | 1.7989 | 0.16558 | 0.3812 | 1 | 7726 | 3 | 1.7989 |
| GAREML | 4 | 0.52602 | 0.62827 | 0.999994 | 13831 | 1 | 1.9472 | 0.16562 | 0.3335 | 0.992917 | 7727 | 3 | 1.9472 |
| SEC24B | 5 | 0.19082 | 0.36212 | 0.999994 | 7508 | 2 | 2.3638 | 0.16562 | 0.37473 | 0.999639 | 7728 | 3 | 2.3638 |
| NLRP10 | 6 | 0.61527 | 0.75129 | 0.999994 | 15149 | 2 | 1.9789 | 0.16573 | 0.38143 | 1 | 7729 | 3 | 1.9789 |
| MGRN1 | 6 | 0.95361 | 0.95363 | 0.999994 | 20430 | 0 | 2.0494 | 0.16575 | 0.38146 | 1 | 7730 | 4 | 2.0494 |
| IFIT1 | 6 | 0.45039 | 0.63539 | 0.999994 | 12716 | 2 | 2.0381 | 0.16576 | 0.38147 | 1 | 7731 | 4 | 2.0381 |
| OR11G2 | 6 | 0.0016071 | 0.008028 | 0.680529 | 236 | 4 | -4.9255 | 0.1658 | 0.38152 | 1 | 7732 | 1 | -4.9255 |
| PDZRN4 | 5 | 0.37514 | 0.54269 | 0.999994 | 11584 | 1 | 1.4511 | 0.16588 | 0.37518 | 0.999639 | 7733 | 3 | 1.4511 |
| ANKAR | 5 | 0.4855 | 0.6354 | 0.999994 | 13220 | 2 | -0.028895 | 0.16588 | 0.37518 | 0.999639 | 7734 | 2 | -0.028895 |
| NonTargetingControlG | 2 | 0.83409 | 0.8339 | 0.999994 | 17990 | 0 | 1.6168 | 0.16591 | 0.26857 | 0.978454 | 7735 | 2 | 1.6168 |
| LCE2C | 4 | 0.15582 | 0.29518 | 0.999994 | 6618 | 2 | 1.4453 | 0.16592 | 0.33394 | 0.992933 | 7736 | 2 | 1.4453 |
| ZNF585A | 6 | 0.46625 | 0.64596 | 0.999994 | 12956 | 2 | 1.3993 | 0.16602 | 0.38185 | 1 | 7737 | 3 | 1.3993 |
| CCR8 | 6 | 0.34606 | 0.55959 | 0.999994 | 11068 | 1 | 1.768 | 0.16604 | 0.38187 | 1 | 7738 | 4 | 1.768 |
| WBP1L | 6 | 0.37609 | 0.58708 | 0.999994 | 11593 | 1 | 1.7331 | 0.16606 | 0.38191 | 1 | 7739 | 4 | 1.7331 |
| MTR | 5 | 0.028721 | 0.087183 | 0.94395 | 1939 | 3 | -3.0905 | 0.16606 | 0.37548 | 0.999639 | 7740 | 2 | -3.0905 |
| ABLIM2 | 5 | 0.21519 | 0.39658 | 0.999994 | 8145 | 1 | 0.78456 | 0.16607 | 0.3755 | 0.999639 | 7741 | 4 | 0.78456 |
| OCLM | 2 | 0.81884 | 0.81866 | 0.999994 | 17762 | 0 | 4.361 | 0.16612 | 0.26886 | 0.978454 | 7742 | 2 | 4.361 |
| TRAF3 | 5 | 0.38998 | 0.55511 | 0.999994 | 11798 | 2 | 2.7254 | 0.16615 | 0.37564 | 0.999639 | 7743 | 3 | 2.7254 |
| AP1M2 | 5 | 0.82967 | 0.83924 | 0.999994 | 17924 | 1 | 1.5935 | 0.1662 | 0.37572 | 0.999639 | 7744 | 4 | 1.5935 |
| CTSW | 5 | 0.13209 | 0.27581 | 0.999379 | 5979 | 2 | -0.32486 | 0.16627 | 0.37583 | 0.999639 | 7745 | 1 | -0.32486 |
| ESRP2 | 6 | 0.054823 | 0.15455 | 0.967126 | 3190 | 2 | 0.63304 | 0.16633 | 0.38232 | 1 | 7746 | 3 | 0.63304 |
| C6orf118 | 6 | 0.8844 | 0.89246 | 0.999994 | 18919 | 1 | 1.1279 | 0.16633 | 0.38232 | 1 | 7747 | 4 | 1.1279 |

| | | | | | | | | | | | | |
|---|---|---|---|---|---|---|---|---|---|---|---|---|
| THRSP | 6 | 0.21255 | 0.40319 | 0.999994 | 8077 | 3 | -0.77443 | 0.16633 | 0.38232 | 1 | 7748 | 2 | -0.77443 |
| RYBP | 6 | 0.59191 | 0.73402 | 0.999994 | 14846 | 1 | 1.4174 | 0.16641 | 0.38245 | 1 | 7749 | 4 | 1.4174 |
| RRAGD | 6 | 0.48746 | 0.66023 | 0.999994 | 13243 | 1 | 1.9805 | 0.16655 | 0.38264 | 1 | 7750 | 4 | 1.9805 |
| hsa-let-7i | 4 | 0.91274 | 0.91253 | 0.999994 | 19473 | 0 | 2.5852 | 0.16656 | 0.33486 | 0.993364 | 7751 | 3 | 2.5852 |
| IL1RAP | 5 | 0.78357 | 0.8095 | 0.999994 | 17262 | 1 | 0.92006 | 0.16661 | 0.37643 | 0.999964 | 7752 | 4 | 0.92006 |
| SAMD7 | 4 | 0.88989 | 0.88959 | 0.999994 | 19035 | 0 | 2.2958 | 0.16676 | 0.33516 | 0.993364 | 7753 | 3 | 2.2958 |
| PAIP2 | 6 | 0.94348 | 0.94345 | 0.999994 | 20164 | 0 | 1.0788 | 0.16677 | 0.38296 | 1 | 7754 | 4 | 1.0788 |
| RAB9A | 6 | 0.57814 | 0.72383 | 0.999994 | 14660 | 2 | 1.896 | 0.16677 | 0.38296 | 1 | 7755 | 4 | 1.896 |
| C19orf40 | 4 | 0.74152 | 0.75364 | 0.999994 | 16759 | 1 | 1.7284 | 0.1668 | 0.33522 | 0.993364 | 7756 | 3 | 1.7284 |
| PRRX1 | 4 | 0.62142 | 0.67715 | 0.999994 | 15251 | 1 | 2.4766 | 0.16682 | 0.33525 | 0.993364 | 7757 | 3 | 2.4766 |
| NonTargetingControlG | 2 | 0.83318 | 0.83294 | 0.999994 | 17974 | 0 | 1.2626 | 0.16682 | 0.26984 | 0.978454 | 7758 | 2 | 1.2626 |
| NonTargetingControlG | 2 | 0.83317 | 0.83293 | 0.999994 | 17973 | 0 | 1.4852 | 0.16683 | 0.26986 | 0.978454 | 7759 | 2 | 1.4852 |
| LOC149373 | 4 | 0.59197 | 0.66102 | 0.999994 | 14849 | 1 | 3.1689 | 0.16684 | 0.33527 | 0.993364 | 7760 | 3 | 3.1689 |
| CLVS1 | 6 | 0.58105 | 0.72598 | 0.999994 | 14693 | 2 | 0.12169 | 0.16684 | 0.38306 | 1 | 7761 | 2 | 0.12169 |
| SERPINB8 | 6 | 0.91539 | 0.91625 | 0.999994 | 19520 | 1 | 1.6594 | 0.16685 | 0.38308 | 1 | 7762 | 4 | 1.6594 |
| C5orf64 | 5 | 0.79473 | 0.81623 | 0.999994 | 17409 | 1 | 1.3986 | 0.16686 | 0.37683 | 1 | 7763 | 4 | 1.3986 |
| NPBWR2 | 5 | 0.83173 | 0.84074 | 0.999994 | 17956 | 1 | 1.2408 | 0.16693 | 0.37694 | 1 | 7764 | 4 | 1.2408 |
| TRIM63 | 5 | 0.82614 | 0.83674 | 0.999994 | 17870 | 1 | 2.4416 | 0.16693 | 0.37695 | 1 | 7765 | 3 | 2.4416 |
| ZNF322 | 6 | 0.15 | 0.32563 | 0.999994 | 6458 | 3 | 1.1324 | 0.16694 | 0.38321 | 1 | 7766 | 3 | 1.1324 |
| ATP1A1 | 6 | 0.97468 | 0.97458 | 0.999994 | 20949 | 0 | 2.1969 | 0.16706 | 0.38339 | 1 | 7767 | 4 | 2.1969 |
| NDUFB5 | 6 | 0.29285 | 0.49836 | 0.999994 | 9972 | 2 | 1.1421 | 0.16708 | 0.38343 | 1 | 7768 | 3 | 1.1421 |
| ANKRD37 | 6 | 0.082992 | 0.21343 | 0.979587 | 4338 | 1 | 1.9815 | 0.16711 | 0.38348 | 1 | 7769 | 4 | 1.9815 |
| MSANTD2 | 5 | 0.96571 | 0.96585 | 0.999994 | 20731 | 0 | 0.72103 | 0.16712 | 0.37727 | 1 | 7770 | 4 | 0.72103 |
| HIST1H2AL | 5 | 0.17266 | 0.33601 | 0.999994 | 7055 | 2 | -0.4721 | 0.16716 | 0.37735 | 1 | 7771 | 2 | -0.4721 |
| DLGAP4 | 5 | 0.78665 | 0.81129 | 0.999994 | 17298 | 1 | 0.18216 | 0.16716 | 0.37735 | 1 | 7772 | 2 | 0.18216 |
| NonTargetingControlG | 2 | 0.69116 | 0.69097 | 0.999994 | 16220 | 0 | 3.7278 | 0.16723 | 0.27042 | 0.978454 | 7773 | 1 | 3.7278 |
| FAM155A | 6 | 0.92611 | 0.92618 | 0.999994 | 19768 | 1 | 0.52795 | 0.16727 | 0.38371 | 1 | 7774 | 3 | 0.52795 |
| MRPS21 | 3 | 0.0602 | 0.12974 | 0.95318 | 3430 | 1 | 1.9511 | 0.16736 | 0.30733 | 0.988343 | 7775 | 2 | 1.9511 |
| YPEL1 | 5 | 0.85384 | 0.85735 | 0.999994 | 18329 | 1 | 2.6494 | 0.16739 | 0.37775 | 1 | 7776 | 3 | 2.6494 |
| SEMA7A | 6 | 0.17998 | 0.36313 | 0.999994 | 7222 | 3 | -0.19942 | 0.16742 | 0.38394 | 1 | 7777 | 2 | -0.19942 |
| ABCA5 | 6 | 0.81114 | 0.85276 | 0.999994 | 17647 | 1 | 1.2231 | 0.16742 | 0.38394 | 1 | 7778 | 3 | 1.2231 |
| OR1E2 | 4 | 0.62233 | 0.67767 | 0.999994 | 15264 | 1 | 2.3432 | 0.16743 | 0.33614 | 0.994237 | 7779 | 2 | 2.3432 |
| TUSC5 | 6 | 0.58617 | 0.72975 | 0.999994 | 14759 | 2 | 2.5349 | 0.16753 | 0.3841 | 1 | 7780 | 4 | 2.5349 |
| LSM10 | 6 | 0.039214 | 0.12013 | 0.951052 | 2478 | 3 | 0.12368 | 0.16753 | 0.3841 | 1 | 7781 | 3 | 0.12368 |
| TRIM43B | 4 | 0.11205 | 0.23651 | 0.985449 | 5366 | 3 | -1.0003 | 0.16758 | 0.33636 | 0.994237 | 7782 | 1 | -1.0003 |
| PSAPL1 | 6 | 0.04294 | 0.12854 | 0.951777 | 2675 | 3 | -1.1477 | 0.16767 | 0.3843 | 1 | 7783 | 1 | -1.1477 |
| LOC729020 | 6 | 0.045664 | 0.13452 | 0.953565 | 2795 | 3 | -0.13217 | 0.16768 | 0.38432 | 1 | 7784 | 2 | -0.13217 |
| CHRNA4 | 4 | 0.25951 | 0.40162 | 0.999994 | 9227 | 2 | -0.32268 | 0.16778 | 0.33664 | 0.994237 | 7785 | 2 | -0.32268 |
| ZNF700 | 5 | 0.56426 | 0.70443 | 0.999994 | 14459 | 2 | 2.3389 | 0.1678 | 0.37843 | 1 | 7786 | 3 | 2.3389 |
| LCE4A | 5 | 0.85347 | 0.85704 | 0.999994 | 18323 | 1 | 1.2086 | 0.16782 | 0.37846 | 1 | 7787 | 3 | 1.2086 |
| TOR4A | 6 | 0.2787 | 0.48189 | 0.999994 | 9664 | 3 | -0.36312 | 0.16784 | 0.38454 | 1 | 7788 | 2 | -0.36312 |
| SARS | 3 | 0.5256 | 0.56332 | 0.999994 | 13827 | 1 | 3.2517 | 0.16787 | 0.30788 | 0.988566 | 7789 | 2 | 3.2517 |
| FXR2 | 5 | 0.047041 | 0.13326 | 0.95318 | 2855 | 3 | -1.6403 | 0.1679 | 0.3786 | 1 | 7790 | 2 | -1.6403 |
| PTCHD4 | 4 | 0.79483 | 0.7966 | 0.999994 | 17410 | 1 | 0.29363 | 0.16794 | 0.33688 | 0.994237 | 7791 | 2 | 0.29363 |
| NonTargetingControlG | 2 | 0.83204 | 0.83183 | 0.999994 | 17960 | 0 | 3.7994 | 0.16796 | 0.27148 | 0.978454 | 7792 | 2 | 3.7994 |
| PSG6 | 6 | 0.67773 | 0.79922 | 0.999994 | 16056 | 2 | 1.4395 | 0.16797 | 0.38474 | 1 | 7793 | 4 | 1.4395 |
| COPZ1 | 5 | 0.25539 | 0.44503 | 0.999994 | 9136 | 2 | 2.2388 | 0.16798 | 0.37875 | 1 | 7794 | 3 | 2.2388 |
| PDLIM4 | 4 | 0.4936 | 0.61348 | 0.999994 | 13318 | 1 | 1.7559 | 0.16802 | 0.337 | 0.994237 | 7795 | 2 | 1.7559 |
| HYAL3 | 6 | 0.60864 | 0.74634 | 0.999994 | 15050 | 2 | -0.70728 | 0.16808 | 0.38491 | 1 | 7796 | 1 | -0.70728 |
| PTPRC | 4 | 0.9352 | 0.935 | 0.999994 | 19973 | 0 | 2.9398 | 0.16811 | 0.33712 | 0.994237 | 7797 | 3 | 2.9398 |
| OR11H2 | 5 | 0.34463 | 0.51752 | 0.999994 | 11045 | 2 | 0.076764 | 0.16814 | 0.37903 | 1 | 7798 | 2 | 0.076764 |
| HS3ST5 | 5 | 0.0022482 | 0.009823 | 0.680529 | 298 | 3 | -7.6849 | 0.16825 | 0.37922 | 1 | 7799 | 2 | -7.6849 |

| | | | | | | | | | | | | | |
|---|---|---|---|---|---|---|---|---|---|---|---|---|---|
| SLC25A13 | 5 | 0.78875 | 0.81259 | 0.999994 | 17330 | 1 | 1.1181 | 0.16827 | 0.37927 | 1 | 7800 | 4 | 1.1181 |
| PIGQ | 6 | 0.054792 | 0.15448 | 0.967126 | 3187 | 2 | 1.0262 | 0.1684 | 0.38537 | 1 | 7801 | 3 | 1.0262 |
| NCOA7 | 5 | 0.68512 | 0.75981 | 0.999994 | 16165 | 1 | 2.3328 | 0.1684 | 0.37947 | 1 | 7802 | 3 | 2.3328 |
| LRRC61 | 6 | 0.33389 | 0.54573 | 0.999994 | 10812 | 3 | 1.2201 | 0.16841 | 0.38539 | 1 | 7803 | 3 | 1.2201 |
| hsa-mir-4740 | 4 | 0.85214 | 0.85167 | 0.999994 | 18298 | 0 | 1.4145 | 0.16842 | 0.33759 | 0.994237 | 7804 | 3 | 1.4145 |
| DNAJB9 | 6 | 0.92067 | 0.92102 | 0.999994 | 19643 | 1 | 2.2391 | 0.16842 | 0.38541 | 1 | 7805 | 4 | 2.2391 |
| SPO11 | 4 | 0.90447 | 0.90417 | 0.999994 | 19311 | 0 | 2.3469 | 0.16846 | 0.33764 | 0.994237 | 7806 | 3 | 2.3469 |
| MAGI1 | 6 | 0.02469 | 0.086275 | 0.94395 | 1742 | 1 | 1.5663 | 0.16846 | 0.38547 | 1 | 7807 | 4 | 1.5663 |
| GGT5 | 5 | 0.10986 | 0.24176 | 0.987253 | 5297 | 1 | 0.54942 | 0.16847 | 0.37959 | 1 | 7808 | 2 | 0.54942 |
| OR2D2 | 6 | 0.040183 | 0.12232 | 0.951052 | 2528 | 3 | 0.22471 | 0.16851 | 0.38556 | 1 | 7809 | 3 | 0.22471 |
| OR6P1 | 6 | 0.89568 | 0.90041 | 0.999994 | 19140 | 1 | 1.4904 | 0.16851 | 0.38556 | 1 | 7810 | 4 | 1.4904 |
| hsa-mir-2467 | 3 | 0.20855 | 0.3234 | 0.999994 | 7992 | 1 | 3.2516 | 0.16852 | 0.30857 | 0.988566 | 7811 | 2 | 3.2516 |
| RALYL | 5 | 0.1589 | 0.31592 | 0.999994 | 6696 | 3 | -1.1192 | 0.16856 | 0.37975 | 1 | 7812 | 2 | -1.1192 |
| PDZD3 | 5 | 0.14406 | 0.29392 | 0.999994 | 6300 | 3 | -0.99201 | 0.16856 | 0.37975 | 1 | 7813 | 2 | -0.99201 |
| C17orf100 | 5 | 0.12845 | 0.27024 | 0.998561 | 5896 | 2 | 0.34088 | 0.16856 | 0.37975 | 1 | 7814 | 2 | 0.34088 |
| HYPK | 6 | 0.88513 | 0.89295 | 0.999994 | 18934 | 1 | 1.0371 | 0.16864 | 0.38575 | 1 | 7815 | 4 | 1.0371 |
| MAPK11 | 6 | 0.48144 | 0.65617 | 0.999994 | 13160 | 2 | 2.3073 | 0.16871 | 0.38587 | 1 | 7816 | 4 | 2.3073 |
| APOO | 6 | 0.21253 | 0.40316 | 0.999994 | 8075 | 3 | 1.2796 | 0.16874 | 0.38589 | 1 | 7817 | 3 | 1.2796 |
| GSE1 | 5 | 0.8372 | 0.84462 | 0.999994 | 18037 | 1 | 0.75537 | 0.16877 | 0.38012 | 1 | 7818 | 4 | 0.75537 |
| ALG2 | 4 | 0.018575 | 0.057625 | 0.929232 | 1390 | 2 | -1.3445 | 0.16885 | 0.33824 | 0.994237 | 7819 | 2 | -1.3445 |
| hsa-mir-4726 | 4 | 0.017217 | 0.054719 | 0.922677 | 1315 | 1 | 3.1355 | 0.16885 | 0.33824 | 0.994237 | 7820 | 3 | 3.1355 |
| ZNF500 | 6 | 0.59326 | 0.73498 | 0.999994 | 14869 | 2 | 2.0197 | 0.16889 | 0.38611 | 1 | 7821 | 3 | 2.0197 |
| HSPB11 | 6 | 0.54201 | 0.69797 | 0.999994 | 14091 | 2 | 1.6333 | 0.1689 | 0.38614 | 1 | 7822 | 3 | 1.6333 |
| BCMO1 | 6 | 0.47167 | 0.64957 | 0.999994 | 13024 | 2 | 1.2162 | 0.16912 | 0.38646 | 1 | 7823 | 4 | 1.2162 |
| ZC3H15 | 6 | 0.29051 | 0.49567 | 0.999994 | 9922 | 2 | 1.6448 | 0.1692 | 0.38658 | 1 | 7824 | 4 | 1.6448 |
| NIM1 | 5 | 0.54243 | 0.68504 | 0.999994 | 14101 | 1 | 2.387 | 0.16922 | 0.38092 | 1 | 7825 | 3 | 2.387 |
| C1orf105 | 6 | 0.70036 | 0.81211 | 0.999994 | 16306 | 1 | 1.3077 | 0.16923 | 0.38662 | 1 | 7826 | 4 | 1.3077 |
| EPB41L2 | 6 | 0.94314 | 0.94311 | 0.999994 | 20157 | 0 | 1.3273 | 0.16923 | 0.38662 | 1 | 7827 | 4 | 1.3273 |
| TNKS1BP1 | 6 | 0.11851 | 0.2829 | 0.999632 | 5589 | 3 | -0.095087 | 0.16923 | 0.38662 | 1 | 7828 | 3 | -0.095087 |
| MEAF6 | 6 | 0.15745 | 0.33501 | 0.999994 | 6658 | 2 | 0.48357 | 0.16923 | 0.38662 | 1 | 7829 | 3 | 0.48357 |
| CDCP1 | 6 | 0.013199 | 0.049779 | 0.908143 | 1078 | 4 | -2.1703 | 0.16923 | 0.38662 | 1 | 7830 | 2 | -2.1703 |
| PPAPDC2 | 5 | 0.25106 | 0.44161 | 0.999994 | 9037 | 1 | 1.132 | 0.1693 | 0.38105 | 1 | 7831 | 4 | 1.132 |
| hsa-mir-365b | 4 | 0.8455 | 0.84501 | 0.999994 | 18176 | 0 | 1.929 | 0.1693 | 0.33892 | 0.994237 | 7832 | 3 | 1.929 |
| ATP5J | 5 | 0.20003 | 0.37528 | 0.999994 | 7763 | 2 | 0.9651 | 0.1693 | 0.38105 | 1 | 7833 | 3 | 0.9651 |
| NonTargetingControlG | 2 | 0.43777 | 0.44871 | 0.999994 | 12538 | 1 | 2.8344 | 0.16933 | 0.27343 | 0.978454 | 7834 | 1 | 2.8344 |
| NQO2 | 6 | 0.18528 | 0.3697 | 0.999994 | 7350 | 1 | 1.4986 | 0.16938 | 0.38684 | 1 | 7835 | 3 | 1.4986 |
| ACOT4 | 5 | 0.35504 | 0.526 | 0.999994 | 11252 | 1 | 2.229 | 0.1694 | 0.38123 | 1 | 7836 | 3 | 2.229 |
| FAM26E | 6 | 0.96607 | 0.96614 | 0.999994 | 20738 | 0 | 1.5999 | 0.16941 | 0.38688 | 1 | 7837 | 4 | 1.5999 |
| PHLDB3 | 6 | 0.076446 | 0.20016 | 0.977164 | 4088 | 2 | 1.9711 | 0.1695 | 0.38701 | 1 | 7838 | 4 | 1.9711 |
| hsa-mir-126 | 4 | 0.39576 | 0.54137 | 0.999994 | 11888 | 2 | 0.50631 | 0.16952 | 0.33922 | 0.994237 | 7839 | 2 | 0.50631 |
| hsa-mir-3921 | 4 | 0.068499 | 0.15701 | 0.967126 | 3771 | 2 | -1.9434 | 0.16952 | 0.33922 | 0.994237 | 7840 | 1 | -1.9434 |
| POP1 | 4 | 0.50267 | 0.6175 | 0.999994 | 13474 | 1 | 2.2814 | 0.16953 | 0.33924 | 0.994237 | 7841 | 3 | 2.2814 |
| LIPH | 5 | 0.45486 | 0.60932 | 0.999994 | 12770 | 1 | 0.032022 | 0.16956 | 0.3815 | 1 | 7842 | 2 | 0.032022 |
| NUP160 | 5 | 0.22859 | 0.41526 | 0.999994 | 8494 | 1 | 1.3362 | 0.16961 | 0.38157 | 1 | 7843 | 4 | 1.3362 |
| MAPT | 3 | 0.32462 | 0.44183 | 0.999994 | 10610 | 1 | 3.4241 | 0.16963 | 0.3098 | 0.988566 | 7844 | 2 | 3.4241 |
| DENND2D | 6 | 0.29443 | 0.50022 | 0.999994 | 10005 | 2 | 1.9388 | 0.1697 | 0.3873 | 1 | 7845 | 4 | 1.9388 |
| NGLY1 | 5 | 0.53106 | 0.67501 | 0.999994 | 13912 | 2 | 2.7712 | 0.16976 | 0.38184 | 1 | 7846 | 3 | 2.7712 |
| TICRR | 4 | 0.046229 | 0.11406 | 0.951052 | 2816 | 2 | 0.36371 | 0.16977 | 0.3396 | 0.994237 | 7847 | 2 | 0.36371 |
| ITPKC | 6 | 0.87282 | 0.88507 | 0.999994 | 18688 | 1 | 1.7791 | 0.16982 | 0.3875 | 1 | 7848 | 4 | 1.7791 |
| FAM229A | 5 | 0.80468 | 0.82247 | 0.999994 | 17549 | 1 | 1.3753 | 0.16983 | 0.38196 | 1 | 7849 | 3 | 1.3753 |
| PAQR4 | 6 | 0.12244 | 0.29033 | 0.999632 | 5717 | 4 | -0.84954 | 0.16987 | 0.38758 | 1 | 7850 | 2 | -0.84954 |
| MRFAP1L1 | 6 | 0.81894 | 0.85632 | 0.999994 | 17765 | 1 | 0.5023 | 0.16987 | 0.38758 | 1 | 7851 | 3 | 0.5023 |

| | | | | | | | | | | | | | |
|---|---|---|---|---|---|---|---|---|---|---|---|---|---|
| PRLR | 6 | 0.40956 | 0.6086 | 0.999994 | 12101 | 2 | 1.7889 | 0.16998 | 0.38773 | 1 | 7852 | 4 | 1.7889 |
| ARMC2 | 5 | 0.63518 | 0.73932 | 0.999994 | 15448 | 1 | 2.3774 | 0.17004 | 0.3823 | 1 | 7853 | 3 | 2.3774 |
| CCDC108 | 5 | 0.1147 | 0.24928 | 0.99177 | 5470 | 1 | 2.4619 | 0.1701 | 0.38242 | 1 | 7854 | 3 | 2.4619 |
| ZNF716 | 5 | 0.3455 | 0.51824 | 0.999994 | 11062 | 2 | -0.022886 | 0.1701 | 0.38242 | 1 | 7855 | 2 | -0.022886 |
| EXOC3 | 6 | 0.25173 | 0.45 | 0.999994 | 9051 | 3 | -0.035955 | 0.17014 | 0.38798 | 1 | 7856 | 3 | -0.035955 |
| EPAS1 | 6 | 0.19475 | 0.38147 | 0.999994 | 7628 | 1 | 1.4763 | 0.17014 | 0.38798 | 1 | 7857 | 4 | 1.4763 |
| hsa-mir-4651 | 2 | 0.80829 | 0.80797 | 0.999994 | 17604 | 0 | 4.2624 | 0.17016 | 0.27458 | 0.978454 | 7858 | 2 | 4.2624 |
| MURC | 5 | 0.086485 | 0.20494 | 0.978186 | 4469 | 1 | 2.1552 | 0.17017 | 0.38254 | 1 | 7859 | 4 | 2.1552 |
| HLA-F | 5 | 0.1241 | 0.26359 | 0.996734 | 5772 | 3 | -0.85085 | 0.17025 | 0.38268 | 1 | 7860 | 1 | -0.85085 |
| RIPK4 | 5 | 0.12879 | 0.27077 | 0.998561 | 5905 | 3 | -1.3842 | 0.17025 | 0.38268 | 1 | 7861 | 2 | -1.3842 |
| AVPR2 | 5 | 0.17401 | 0.33796 | 0.999994 | 7085 | 3 | -0.95112 | 0.17025 | 0.38268 | 1 | 7862 | 2 | -0.95112 |
| NPC1 | 6 | 0.033921 | 0.10799 | 0.951052 | 2213 | 3 | 0.022388 | 0.17034 | 0.38827 | 1 | 7863 | 3 | 0.022388 |
| SOX13 | 6 | 0.93714 | 0.93716 | 0.999994 | 20009 | 0 | 2.0742 | 0.17034 | 0.38827 | 1 | 7864 | 3 | 2.0742 |
| ST8SIA6 | 4 | 0.91025 | 0.91001 | 0.999994 | 19429 | 0 | 1.5839 | 0.17041 | 0.34053 | 0.994237 | 7865 | 3 | 1.5839 |
| CAPN6 | 5 | 0.48201 | 0.63242 | 0.999994 | 13171 | 1 | 2.1953 | 0.17044 | 0.38299 | 1 | 7866 | 4 | 2.1953 |
| LOC100287534 | 2 | 0.81595 | 0.81574 | 0.999994 | 17711 | 0 | 4.1572 | 0.17045 | 0.275 | 0.978454 | 7867 | 2 | 4.1572 |
| hsa-mir-3679 | 4 | 0.80635 | 0.80696 | 0.999994 | 17575 | 1 | 1.4957 | 0.17059 | 0.3408 | 0.994237 | 7868 | 3 | 1.4957 |
| RRH | 3 | 0.035839 | 0.088399 | 0.94395 | 2308 | 1 | 3.1606 | 0.17059 | 0.31085 | 0.988566 | 7869 | 2 | 3.1606 |
| WDR7 | 5 | 0.30657 | 0.48637 | 0.999994 | 10266 | 1 | 0.65221 | 0.17063 | 0.38328 | 1 | 7870 | 4 | 0.65221 |
| hsa-mir-4634 | 4 | 0.28621 | 0.42891 | 0.999994 | 9838 | 1 | 2.3682 | 0.17067 | 0.34091 | 0.994237 | 7871 | 3 | 2.3682 |
| COX15 | 5 | 0.68251 | 0.75868 | 0.999994 | 16126 | 1 | 2.3058 | 0.17068 | 0.38336 | 1 | 7872 | 4 | 2.3058 |
| TSPO | 6 | 0.098642 | 0.24469 | 0.988995 | 4919 | 1 | 2.1162 | 0.1707 | 0.3888 | 1 | 7873 | 4 | 2.1162 |
| MTFP1 | 6 | 0.35343 | 0.56787 | 0.999994 | 11220 | 2 | 0.5222 | 0.1707 | 0.3888 | 1 | 7874 | 3 | 0.5222 |
| FGR | 6 | 0.6887 | 0.80779 | 0.999994 | 16199 | 2 | 0.7863 | 0.17074 | 0.38887 | 1 | 7875 | 3 | 0.7863 |
| ELOVL4 | 6 | 0.52035 | 0.68278 | 0.999994 | 13752 | 2 | 1.9931 | 0.17074 | 0.38887 | 1 | 7876 | 4 | 1.9931 |
| hsa-mir-1251 | 4 | 0.39869 | 0.54438 | 0.999994 | 11928 | 2 | 2.2 | 0.17077 | 0.34106 | 0.994237 | 7877 | 2 | 2.2 |
| hsa-mir-330 | 4 | 0.4074 | 0.55329 | 0.999994 | 12065 | 1 | 2.9048 | 0.17077 | 0.34106 | 0.994237 | 7878 | 3 | 2.9048 |
| CCT8 | 4 | 0.15912 | 0.29858 | 0.999994 | 6706 | 2 | 1.2732 | 0.17077 | 0.34106 | 0.994237 | 7879 | 2 | 1.2732 |
| RPP38 | 4 | 0.13495 | 0.27368 | 0.998561 | 6048 | 2 | 1.7305 | 0.17077 | 0.34106 | 0.994237 | 7880 | 2 | 1.7305 |
| NUS1 | 4 | 0.62535 | 0.67933 | 0.999994 | 15312 | 1 | 1.971 | 0.17077 | 0.34106 | 0.994237 | 7881 | 2 | 1.971 |
| hsa-mir-4773-1 | 4 | 0.67471 | 0.70828 | 0.999994 | 15999 | 1 | 2.6388 | 0.17077 | 0.34106 | 0.994237 | 7882 | 2 | 2.6388 |
| hsa-mir-6083 | 4 | 0.93374 | 0.93351 | 0.999994 | 19942 | 0 | 2.9911 | 0.17077 | 0.34106 | 0.994237 | 7883 | 3 | 2.9911 |
| hsa-mir-548q | 2 | 0.82921 | 0.82904 | 0.999994 | 17916 | 0 | 4.1721 | 0.17079 | 0.27548 | 0.978454 | 7884 | 2 | 4.1721 |
| SAMD1 | 6 | 0.95608 | 0.95617 | 0.999994 | 20488 | 0 | 2.1535 | 0.17082 | 0.38898 | 1 | 7885 | 4 | 2.1535 |
| C17orf59 | 5 | 0.035067 | 0.10341 | 0.946414 | 2270 | 3 | -0.74796 | 0.1709 | 0.38376 | 1 | 7886 | 2 | -0.74796 |
| hsa-mir-518b | 4 | 0.92262 | 0.92243 | 0.999994 | 19691 | 0 | 1.6487 | 0.17094 | 0.34131 | 0.994237 | 7887 | 3 | 1.6487 |
| ATP2C2 | 5 | 0.054607 | 0.15154 | 0.967126 | 3180 | 2 | 2.6202 | 0.17101 | 0.38394 | 1 | 7888 | 3 | 2.6202 |
| hsa-mir-4281 | 2 | 0.82895 | 0.82879 | 0.999994 | 17909 | 0 | 2.4865 | 0.17105 | 0.27584 | 0.978454 | 7889 | 2 | 2.4865 |
| CACNA2D1 | 5 | 0.47942 | 0.63013 | 0.999994 | 13139 | 1 | 2.341 | 0.17116 | 0.38419 | 1 | 7890 | 3 | 2.341 |
| NEK10 | 5 | 0.81696 | 0.83056 | 0.999994 | 17726 | 1 | 1.69 | 0.17126 | 0.38436 | 1 | 7891 | 4 | 1.69 |
| LMBR1L | 5 | 0.024121 | 0.075145 | 0.93914 | 1713 | 3 | -3.1571 | 0.17127 | 0.38439 | 1 | 7892 | 1 | -3.1571 |
| ART4 | 5 | 0.46623 | 0.61893 | 0.999994 | 12953 | 2 | -0.45563 | 0.17127 | 0.38439 | 1 | 7893 | 2 | -0.45563 |
| AMBN | 5 | 0.84985 | 0.85416 | 0.999994 | 18259 | 1 | 1.1849 | 0.17127 | 0.38439 | 1 | 7894 | 3 | 1.1849 |
| STT3B | 5 | 0.9081 | 0.90813 | 0.999994 | 19386 | 0 | 1.4853 | 0.17129 | 0.38442 | 1 | 7895 | 4 | 1.4853 |
| LOC100129520 | 6 | 0.47132 | 0.64933 | 0.999994 | 13020 | 2 | 1.8546 | 0.17137 | 0.38979 | 1 | 7896 | 4 | 1.8546 |
| SCN1A | 4 | 0.62546 | 0.6794 | 0.999994 | 15314 | 1 | 0.10066 | 0.17138 | 0.34198 | 0.994392 | 7897 | 2 | 0.10066 |
| FAM3C | 3 | 0.3426 | 0.45161 | 0.999994 | 10996 | 1 | 3.404 | 0.1714 | 0.31169 | 0.988566 | 7898 | 2 | 3.404 |
| TMEM167B | 6 | 0.93347 | 0.93355 | 0.999994 | 19933 | 0 | 1.2389 | 0.17157 | 0.39005 | 1 | 7899 | 4 | 1.2389 |
| AIFM3 | 6 | 0.04861 | 0.14106 | 0.959146 | 2915 | 3 | -0.28855 | 0.17157 | 0.39005 | 1 | 7900 | 2 | -0.28855 |
| GABPB2 | 6 | 0.097239 | 0.24195 | 0.987253 | 4868 | 2 | 0.74128 | 0.17157 | 0.39005 | 1 | 7901 | 3 | 0.74128 |
| VTI1B | 5 | 0.15193 | 0.30568 | 0.999994 | 6504 | 3 | -1.0785 | 0.17159 | 0.38495 | 1 | 7902 | 2 | -1.0785 |
| ZBTB26 | 5 | 0.4244 | 0.58376 | 0.999994 | 12324 | 2 | 0.45369 | 0.17159 | 0.38495 | 1 | 7903 | 2 | 0.45369 |

| | | | | | | | | | | | | |
|---|---|---|---|---|---|---|---|---|---|---|---|---|
| GAREM | 5 | 0.62075 | 0.73383 | 0.999994 | 15231 | 1 | 1.3416 | 0.17164 | 0.38502 | 1 | 7904 | 4 | 1.3416 |
| TANC1 | 5 | 0.55652 | 0.69755 | 0.999994 | 14311 | 1 | 2.6451 | 0.17168 | 0.38507 | 1 | 7905 | 3 | 2.6451 |
| TMPRSS9 | 5 | 0.19082 | 0.36212 | 0.999994 | 7507 | 2 | 2.668 | 0.17168 | 0.38507 | 1 | 7906 | 3 | 2.668 |
| COX4I1 | 5 | 0.24302 | 0.43503 | 0.999994 | 8866 | 2 | 2.6318 | 0.17168 | 0.38507 | 1 | 7907 | 3 | 2.6318 |
| CFP | 5 | 0.11363 | 0.24762 | 0.990682 | 5427 | 1 | 2.668 | 0.17168 | 0.38507 | 1 | 7908 | 3 | 2.668 |
| SLC10A6 | 5 | 0.38987 | 0.55503 | 0.999994 | 11795 | 2 | -0.22399 | 0.17177 | 0.38517 | 1 | 7909 | 2 | -0.22399 |
| TRIM7 | 5 | 0.27429 | 0.46026 | 0.999994 | 9570 | 2 | 0.8917 | 0.17177 | 0.38517 | 1 | 7910 | 3 | 0.8917 |
| CDRT15 | 5 | 0.67739 | 0.75644 | 0.999994 | 16051 | 1 | 0.32521 | 0.17177 | 0.38517 | 1 | 7911 | 2 | 0.32521 |
| ANAPC13 | 6 | 0.96804 | 0.96806 | 0.999994 | 20785 | 0 | 1.9941 | 0.17179 | 0.39038 | 1 | 7912 | 4 | 1.9941 |
| C14orf93 | 6 | 0.86874 | 0.88257 | 0.999994 | 18612 | 1 | 1.0783 | 0.17179 | 0.39038 | 1 | 7913 | 3 | 1.0783 |
| NonTargetingControlG | 2 | 0.82819 | 0.82801 | 0.999994 | 17901 | 0 | 1.8366 | 0.17181 | 0.27691 | 0.978454 | 7914 | 2 | 1.8366 |
| UBE2A | 6 | 0.34169 | 0.55459 | 0.999994 | 10974 | 3 | 1.3757 | 0.1719 | 0.39054 | 1 | 7915 | 3 | 1.3757 |
| NonTargetingControlG | 2 | 0.63275 | 0.63249 | 0.999994 | 15411 | 0 | 3.5174 | 0.17192 | 0.27706 | 0.978454 | 7916 | 1 | 3.5174 |
| DYNC1I1 | 5 | 0.017391 | 0.056946 | 0.925104 | 1330 | 1 | 1.583 | 0.17198 | 0.38541 | 1 | 7917 | 3 | 1.583 |
| MED24 | 5 | 0.22306 | 0.40756 | 0.999994 | 8358 | 3 | -0.67339 | 0.17198 | 0.38541 | 1 | 7918 | 2 | -0.67339 |
| EPM2A | 6 | 0.85929 | 0.87702 | 0.999994 | 18420 | 1 | 1.5092 | 0.17198 | 0.39066 | 1 | 7919 | 4 | 1.5092 |
| DPP4 | 4 | 0.35699 | 0.5014 | 0.999994 | 11288 | 2 | -0.24403 | 0.17206 | 0.34297 | 0.994392 | 7920 | 2 | -0.24403 |
| SCOC | 6 | 0.20058 | 0.38858 | 0.999994 | 7775 | 2 | 0.1567 | 0.17207 | 0.39079 | 1 | 7921 | 3 | 0.1567 |
| ULBP3 | 6 | 0.5967 | 0.73753 | 0.999994 | 14909 | 2 | 1.5411 | 0.17207 | 0.39079 | 1 | 7922 | 4 | 1.5411 |
| RTDR1 | 6 | 0.97226 | 0.97219 | 0.999994 | 20874 | 0 | 2.4082 | 0.1723 | 0.39112 | 1 | 7923 | 4 | 2.4082 |
| CD3EAP | 3 | 0.49016 | 0.53972 | 0.999994 | 13278 | 1 | 1.7768 | 0.17241 | 0.31286 | 0.989484 | 7924 | 2 | 1.7768 |
| NonTargetingControlG | 2 | 0.70969 | 0.70946 | 0.999994 | 16398 | 0 | 3.7494 | 0.17242 | 0.27776 | 0.978454 | 7925 | 1 | 3.7494 |
| HDAC6 | 5 | 0.79912 | 0.81898 | 0.999994 | 17465 | 1 | 2.326 | 0.17254 | 0.38605 | 1 | 7926 | 3 | 2.326 |
| PLEKHB1 | 6 | 0.82695 | 0.86007 | 0.999994 | 17885 | 1 | 1.8288 | 0.17261 | 0.39157 | 1 | 7927 | 4 | 1.8288 |
| MAP3K10 | 6 | 0.95327 | 0.95327 | 0.999994 | 20418 | 0 | 2.3731 | 0.17269 | 0.39168 | 1 | 7928 | 4 | 2.3731 |
| SULT1C3 | 6 | 0.32394 | 0.53438 | 0.999994 | 10595 | 2 | 0.34634 | 0.17269 | 0.39169 | 1 | 7929 | 2 | 0.34634 |
| ERGIC2 | 6 | 0.63951 | 0.76957 | 0.999994 | 15509 | 1 | 0.93994 | 0.17269 | 0.39169 | 1 | 7930 | 4 | 0.93994 |
| RGS17 | 6 | 0.084964 | 0.21746 | 0.980699 | 4407 | 2 | -0.0026848 | 0.1727 | 0.3917 | 1 | 7931 | 2 | -0.0026848 |
| TTLL2 | 4 | 0.86084 | 0.86042 | 0.999994 | 18459 | 0 | 2.095 | 0.17275 | 0.34401 | 0.994392 | 7932 | 3 | 2.095 |
| PRB4 | 3 | 0.3454 | 0.45316 | 0.999994 | 11061 | 1 | 3.0367 | 0.17278 | 0.31326 | 0.989846 | 7933 | 2 | 3.0367 |
| LY6G5B | 6 | 0.21004 | 0.40014 | 0.999994 | 8025 | 3 | 1.0264 | 0.17282 | 0.39188 | 1 | 7934 | 3 | 1.0264 |
| KSR1 | 4 | 0.80428 | 0.80505 | 0.999994 | 17545 | 1 | 2.1017 | 0.17286 | 0.34419 | 0.994392 | 7935 | 3 | 2.1017 |
| NonTargetingControlG | 2 | 0.50847 | 0.51168 | 0.999994 | 13562 | 1 | 3.0939 | 0.17287 | 0.27841 | 0.978454 | 7936 | 1 | 3.0939 |
| LMOD3 | 5 | 0.42619 | 0.58526 | 0.999994 | 12353 | 1 | 0.097287 | 0.17288 | 0.38644 | 1 | 7937 | 2 | 0.097287 |
| SOGA1 | 5 | 0.6408 | 0.7415 | 0.999994 | 15524 | 1 | 2.1996 | 0.17288 | 0.38644 | 1 | 7938 | 3 | 2.1996 |
| TOE1 | 5 | 0.10419 | 0.23293 | 0.984502 | 5104 | 2 | -0.18771 | 0.17288 | 0.38644 | 1 | 7939 | 2 | -0.18771 |
| IGFBP6 | 5 | 0.023234 | 0.072814 | 0.93914 | 1664 | 3 | -0.58114 | 0.17288 | 0.38644 | 1 | 7940 | 2 | -0.58114 |
| SMIM1 | 5 | 0.01531 | 0.051101 | 0.909241 | 1207 | 3 | -0.6763 | 0.17288 | 0.38644 | 1 | 7941 | 2 | -0.6763 |
| ZNF630 | 6 | 0.81518 | 0.85459 | 0.999994 | 17702 | 1 | 1.6811 | 0.17288 | 0.39196 | 1 | 7942 | 3 | 1.6811 |
| OR10AD1 | 6 | 0.41036 | 0.60913 | 0.999994 | 12116 | 1 | 2.2127 | 0.17294 | 0.39206 | 1 | 7943 | 4 | 2.2127 |
| SYT4 | 5 | 0.059538 | 0.16131 | 0.967126 | 3403 | 3 | -0.70029 | 0.17299 | 0.38658 | 1 | 7944 | 2 | -0.70029 |
| ATP2B2 | 5 | 0.13866 | 0.28582 | 0.999632 | 6157 | 2 | 0.56798 | 0.17299 | 0.38658 | 1 | 7945 | 2 | 0.56798 |
| SEMA5B | 6 | 0.056764 | 0.15878 | 0.967126 | 3283 | 3 | 0.15055 | 0.173 | 0.39213 | 1 | 7946 | 3 | 0.15055 |
| hsa-mir-455 | 4 | 0.66111 | 0.6999 | 0.999994 | 15797 | 1 | 2.3739 | 0.17315 | 0.3446 | 0.994392 | 7947 | 2 | 2.3739 |
| SPATA5 | 4 | 0.21102 | 0.35202 | 0.999994 | 8052 | 2 | 1.5462 | 0.17317 | 0.34462 | 0.994392 | 7948 | 2 | 1.5462 |
| SSU72 | 4 | 0.35382 | 0.49818 | 0.999994 | 11230 | 2 | 2.2043 | 0.17317 | 0.34462 | 0.994392 | 7949 | 2 | 2.2043 |
| PRKAA1 | 4 | 0.032023 | 0.085602 | 0.94395 | 2108 | 2 | 0.64994 | 0.17317 | 0.34462 | 0.994392 | 7950 | 2 | 0.64994 |
| TMC5 | 6 | 0.33372 | 0.54555 | 0.999994 | 10800 | 2 | 2.283 | 0.1732 | 0.39241 | 1 | 7951 | 4 | 2.283 |
| NonTargetingControlG | 2 | 0.82673 | 0.82655 | 0.999994 | 17881 | 0 | 3.0718 | 0.17327 | 0.27899 | 0.978454 | 7952 | 2 | 3.0718 |
| DENND6B | 6 | 0.48049 | 0.65552 | 0.999994 | 13149 | 2 | -0.37042 | 0.17336 | 0.39262 | 1 | 7953 | 1 | -0.37042 |
| SYDE1 | 6 | 0.051151 | 0.1466 | 0.964852 | 3029 | 4 | -1.4666 | 0.17336 | 0.39262 | 1 | 7954 | 2 | -1.4666 |
| NonTargetingControlG | 2 | 0.73791 | 0.73761 | 0.999994 | 16700 | 0 | 3.8538 | 0.17339 | 0.27918 | 0.978454 | 7955 | 1 | 3.8538 |

| | | | | | | | | | | | | | |
|---|---|---|---|---|---|---|---|---|---|---|---|---|---|
| PSMD2 | 5 | 0.01011 | 0.03637 | 0.879988 | 877 | 2 | 2.6414 | 0.17343 | 0.38706 | 1 | 7956 | 3 | 2.6414 |
| WASL | 4 | 0.60432 | 0.66768 | 0.999994 | 14999 | 1 | 0.70689 | 0.17348 | 0.34508 | 0.994392 | 7957 | 2 | 0.70689 |
| PIGK | 6 | 0.66336 | 0.78791 | 0.999994 | 15828 | 2 | 0.23461 | 0.17348 | 0.39279 | 1 | 7958 | 3 | 0.23461 |
| AFF4 | 6 | 0.3137 | 0.52267 | 0.999994 | 10391 | 3 | 0.82825 | 0.17348 | 0.39279 | 1 | 7959 | 3 | 0.82825 |
| hsa-mir-372 | 4 | 0.824 | 0.82365 | 0.999994 | 17839 | 1 | 1.6091 | 0.1736 | 0.34526 | 0.994392 | 7960 | 3 | 1.6091 |
| RPLP0 | 5 | 0.31982 | 0.49719 | 0.999994 | 10510 | 2 | 2.3189 | 0.17367 | 0.38733 | 1 | 7961 | 3 | 2.3189 |
| TSPYL2 | 5 | 0.88453 | 0.88466 | 0.999994 | 18926 | 1 | 0.38395 | 0.17368 | 0.38735 | 1 | 7962 | 2 | 0.38395 |
| SLC13A3 | 5 | 0.044558 | 0.12723 | 0.951052 | 2739 | 4 | -1.6136 | 0.17368 | 0.38735 | 1 | 7963 | 1 | -1.6136 |
| ZNF3 | 6 | 0.19753 | 0.3849 | 0.999994 | 7689 | 2 | 1.8635 | 0.17368 | 0.39306 | 1 | 7964 | 3 | 1.8635 |
| OR10P1 | 6 | 0.20909 | 0.39899 | 0.999994 | 8004 | 2 | 2.0076 | 0.1737 | 0.39309 | 1 | 7965 | 4 | 2.0076 |
| FTL | 4 | 0.74354 | 0.75516 | 0.999994 | 16784 | 1 | 1.7972 | 0.1738 | 0.34557 | 0.994392 | 7966 | 3 | 1.7972 |
| NECAB2 | 5 | 0.092302 | 0.21428 | 0.979587 | 4688 | 2 | 0.081605 | 0.17382 | 0.38749 | 1 | 7967 | 2 | 0.081605 |
| C1orf52 | 5 | 0.061405 | 0.16442 | 0.967126 | 3489 | 3 | -2.0359 | 0.17382 | 0.38749 | 1 | 7968 | 2 | -2.0359 |
| GALNT5 | 5 | 0.49019 | 0.63937 | 0.999994 | 13279 | 2 | 2.3636 | 0.17394 | 0.38761 | 1 | 7969 | 3 | 2.3636 |
| WRAP73 | 6 | 0.66411 | 0.78848 | 0.999994 | 15841 | 2 | 1.5998 | 0.17395 | 0.39347 | 1 | 7970 | 4 | 1.5998 |
| POLDIP3 | 6 | 0.49412 | 0.66466 | 0.999994 | 13328 | 1 | 1.7243 | 0.174 | 0.39355 | 1 | 7971 | 4 | 1.7243 |
| NDUFAF6 | 4 | 0.43033 | 0.57688 | 0.999994 | 12418 | 2 | 0.58804 | 0.17407 | 0.34597 | 0.994392 | 7972 | 2 | 0.58804 |
| ACOT6 | 4 | 0.91055 | 0.91031 | 0.999994 | 19441 | 0 | 0.41223 | 0.17407 | 0.34597 | 0.994392 | 7973 | 2 | 0.41223 |
| SLC28A2 | 4 | 0.1886 | 0.32892 | 0.999994 | 7439 | 2 | 0.1902 | 0.17407 | 0.34597 | 0.994392 | 7974 | 2 | 0.1902 |
| PTGES3L | 5 | 0.087799 | 0.20705 | 0.978744 | 4504 | 1 | 2.4834 | 0.1741 | 0.38781 | 1 | 7975 | 3 | 2.4834 |
| RNF222 | 5 | 0.4352 | 0.5928 | 0.999994 | 12503 | 1 | 1.0158 | 0.17415 | 0.38785 | 1 | 7976 | 3 | 1.0158 |
| NonTargetingControlG | 2 | 0.30207 | 0.33846 | 0.999994 | 10166 | 1 | 2.4141 | 0.17416 | 0.28026 | 0.978454 | 7977 | 1 | 2.4141 |
| UBQLN1 | 6 | 0.85648 | 0.87543 | 0.999994 | 18379 | 1 | 2.159 | 0.17417 | 0.39378 | 1 | 7978 | 4 | 2.159 |
| CCNJ | 6 | 0.10618 | 0.25937 | 0.994943 | 5180 | 2 | 1.9738 | 0.17417 | 0.39378 | 1 | 7979 | 3 | 1.9738 |
| MTHFD2 | 6 | 0.16179 | 0.34052 | 0.999994 | 6784 | 3 | 1.0444 | 0.17417 | 0.39378 | 1 | 7980 | 3 | 1.0444 |
| TNFRSF19 | 5 | 0.29171 | 0.47433 | 0.999994 | 9943 | 1 | 2.3934 | 0.17421 | 0.38793 | 1 | 7981 | 3 | 2.3934 |
| hsa-mir-7159 | 4 | 0.53209 | 0.63109 | 0.999994 | 13938 | 1 | 2.0794 | 0.17422 | 0.34619 | 0.994392 | 7982 | 3 | 2.0794 |
| HLA-B | 6 | 0.094176 | 0.23589 | 0.985362 | 4757 | 4 | -1.3257 | 0.17427 | 0.39392 | 1 | 7983 | 2 | -1.3257 |
| KLHDC7B | 6 | 0.05574 | 0.15652 | 0.967126 | 3236 | 2 | 0.4644 | 0.17427 | 0.39392 | 1 | 7984 | 3 | 0.4644 |
| JAKMIP3 | 6 | 0.044226 | 0.13139 | 0.95318 | 2723 | 2 | 0.65726 | 0.17432 | 0.394 | 1 | 7985 | 3 | 0.65726 |
| CUL4A | 6 | 0.12994 | 0.30022 | 0.999994 | 5932 | 2 | 1.5872 | 0.17436 | 0.39406 | 1 | 7986 | 4 | 1.5872 |
| DNAH10 | 5 | 0.87237 | 0.8732 | 0.999994 | 18682 | 1 | 0.53699 | 0.17445 | 0.38822 | 1 | 7987 | 3 | 0.53699 |
| OR8B2 | 6 | 0.50314 | 0.6709 | 0.999994 | 13481 | 2 | 1.5101 | 0.17447 | 0.3942 | 1 | 7988 | 4 | 1.5101 |
| hsa-mir-299 | 4 | 0.33673 | 0.48059 | 0.999994 | 10866 | 2 | -0.76343 | 0.17452 | 0.34664 | 0.994392 | 7989 | 1 | -0.76343 |
| C12orf4 | 5 | 0.3485 | 0.52068 | 0.999994 | 11122 | 2 | 2.0185 | 0.17453 | 0.38831 | 1 | 7990 | 3 | 2.0185 |
| HEPHL1 | 6 | 0.070767 | 0.1884 | 0.969543 | 3864 | 2 | 1.4044 | 0.17457 | 0.39436 | 1 | 7991 | 4 | 1.4044 |
| EIF3G | 4 | 0.7389 | 0.75166 | 0.999994 | 16719 | 1 | 2.7778 | 0.17457 | 0.3467 | 0.994392 | 7992 | 3 | 2.7778 |
| hsa-mir-553 | 2 | 0.78751 | 0.78704 | 0.999994 | 17312 | 0 | 4.0856 | 0.1746 | 0.2809 | 0.978454 | 7993 | 2 | 4.0856 |
| KAT8 | 5 | 0.27748 | 0.46283 | 0.999994 | 9645 | 1 | 1.817 | 0.17466 | 0.38846 | 1 | 7994 | 3 | 1.817 |
| CSNK1D | 5 | 0.8455 | 0.85081 | 0.999994 | 18175 | 1 | 0.7989 | 0.17466 | 0.38846 | 1 | 7995 | 3 | 0.7989 |
| TFF2 | 6 | 0.17266 | 0.35403 | 0.999994 | 7052 | 1 | 0.11371 | 0.17468 | 0.39452 | 1 | 7996 | 2 | 0.11371 |
| MAPK1IP1L | 6 | 0.7182 | 0.81776 | 0.999994 | 16477 | 1 | 1.5375 | 0.17468 | 0.39452 | 1 | 7997 | 3 | 1.5375 |
| F7 | 5 | 0.22057 | 0.40406 | 0.999994 | 8302 | 3 | -0.58423 | 0.17475 | 0.38856 | 1 | 7998 | 2 | -0.58423 |
| TMEM55B | 5 | 0.058016 | 0.15879 | 0.967126 | 3339 | 3 | -2.0466 | 0.17475 | 0.38856 | 1 | 7999 | 2 | -2.0466 |
| NonTargetingControlG | 2 | 0.82523 | 0.82505 | 0.999994 | 17854 | 0 | 1.3089 | 0.17477 | 0.28115 | 0.978454 | 8000 | 2 | 1.3089 |
| CRTC1 | 4 | 0.48771 | 0.61088 | 0.999994 | 13250 | 1 | 2.9809 | 0.17489 | 0.34718 | 0.994392 | 8001 | 3 | 2.9809 |
| hsa-mir-4511 | 3 | 0.84543 | 0.84545 | 0.999994 | 18174 | 0 | 3.0343 | 0.17497 | 0.3157 | 0.989846 | 8002 | 2 | 3.0343 |
| NSMF | 6 | 0.92566 | 0.92575 | 0.999994 | 19758 | 1 | 1.4846 | 0.17499 | 0.39501 | 1 | 8003 | 4 | 1.4846 |
| ANKRD10 | 6 | 0.30238 | 0.50951 | 0.999994 | 10180 | 2 | 0.36608 | 0.175 | 0.39501 | 1 | 8004 | 3 | 0.36608 |
| GDE1 | 6 | 0.16617 | 0.34593 | 0.999994 | 6891 | 3 | -0.47461 | 0.175 | 0.39501 | 1 | 8005 | 3 | -0.47461 |
| SHPRH | 6 | 0.57205 | 0.71944 | 0.999994 | 14577 | 2 | -0.50919 | 0.17501 | 0.39502 | 1 | 8006 | 2 | -0.50919 |
| CA3 | 6 | 0.24656 | 0.44385 | 0.999994 | 8935 | 3 | -0.31042 | 0.17501 | 0.39502 | 1 | 8007 | 2 | -0.31042 |

| | | | | | | | | | | | | |
|---|---|---|---|---|---|---|---|---|---|---|---|---|
| SPATA24 | 6 | 0.16629 | 0.34608 | 0.999994 | 6893 | 2 | 0.54786 | 0.17501 | 0.39502 | 1 | 8008 | 3 | 0.54786 |
| TFCP2 | 6 | 0.52936 | 0.68909 | 0.999994 | 13890 | 2 | 1.682 | 0.17501 | 0.39503 | 1 | 8009 | 4 | 1.682 |
| KISS1R | 4 | 0.039097 | 0.099984 | 0.946394 | 2468 | 2 | 0.095046 | 0.17517 | 0.34759 | 0.994392 | 8010 | 2 | 0.095046 |
| FEN1 | 4 | 0.23114 | 0.37271 | 0.999994 | 8546 | 1 | 3.0334 | 0.17517 | 0.34759 | 0.994392 | 8011 | 3 | 3.0334 |
| PRSS56 | 4 | 0.50297 | 0.61764 | 0.999994 | 13479 | 1 | 2.6443 | 0.17518 | 0.3476 | 0.994392 | 8012 | 3 | 2.6443 |
| NonTargetingControlG | 2 | 0.82482 | 0.82462 | 0.999994 | 17850 | 0 | 3.7785 | 0.17518 | 0.28172 | 0.978454 | 8013 | 2 | 3.7785 |
| hsa-mir-1252 | 3 | 0.8017 | 0.80151 | 0.999994 | 17506 | 0 | 1.4509 | 0.17527 | 0.31602 | 0.989846 | 8014 | 2 | 1.4509 |
| ZNF454 | 6 | 0.01364 | 0.051258 | 0.909241 | 1107 | 2 | 1.4426 | 0.17531 | 0.39547 | 1 | 8015 | 4 | 1.4426 |
| TMEM30A | 6 | 0.48835 | 0.66083 | 0.999994 | 13258 | 1 | 2.1781 | 0.1754 | 0.3956 | 1 | 8016 | 4 | 2.1781 |
| MICU3 | 5 | 0.071522 | 0.18085 | 0.968171 | 3896 | 1 | 2.2073 | 0.17547 | 0.38937 | 1 | 8017 | 3 | 2.2073 |
| NonTargetingControlG | 2 | 0.82452 | 0.82432 | 0.999994 | 17846 | 0 | 1.36 | 0.17548 | 0.28216 | 0.978454 | 8018 | 2 | 1.36 |
| MTRNR2L8 | 2 | 0.82448 | 0.82428 | 0.999994 | 17845 | 0 | 1.5972 | 0.17552 | 0.28222 | 0.978454 | 8019 | 2 | 1.5972 |
| KRT40 | 6 | 0.79702 | 0.84662 | 0.999994 | 17438 | 1 | 0.98297 | 0.17553 | 0.39578 | 1 | 8020 | 3 | 0.98297 |
| GSTT2 | 6 | 0.082439 | 0.21233 | 0.979587 | 4316 | 3 | -0.85345 | 0.17553 | 0.39578 | 1 | 8021 | 3 | -0.85345 |
| hsa-mir-4652 | 4 | 0.30126 | 0.44429 | 0.999994 | 10149 | 2 | 1.7243 | 0.17562 | 0.34827 | 0.994392 | 8022 | 2 | 1.7243 |
| RPS6 | 6 | 0.62559 | 0.75902 | 0.999994 | 15318 | 1 | 1.8681 | 0.1758 | 0.39617 | 1 | 8023 | 4 | 1.8681 |
| FRG2C | 4 | 0.36839 | 0.51317 | 0.999994 | 11478 | 2 | 2.0299 | 0.17586 | 0.34863 | 0.994392 | 8024 | 2 | 2.0299 |
| hsa-mir-4803 | 4 | 0.28287 | 0.42549 | 0.999994 | 9757 | 2 | 1.7667 | 0.17586 | 0.34863 | 0.994392 | 8025 | 2 | 1.7667 |
| PWWP2A | 4 | 0.32372 | 0.46724 | 0.999994 | 10589 | 2 | 1.9313 | 0.17586 | 0.34863 | 0.994392 | 8026 | 2 | 1.9313 |
| PRDM15 | 6 | 0.64429 | 0.77322 | 0.999994 | 15576 | 2 | 0.65343 | 0.17587 | 0.39627 | 1 | 8027 | 3 | 0.65343 |
| CNPY1 | 6 | 0.51383 | 0.67827 | 0.999994 | 13638 | 2 | 0.98162 | 0.17587 | 0.39627 | 1 | 8028 | 3 | 0.98162 |
| FBXW8 | 6 | 0.56653 | 0.71547 | 0.999994 | 14489 | 2 | 0.22434 | 0.17591 | 0.39633 | 1 | 8029 | 3 | 0.22434 |
| NES | 6 | 0.049454 | 0.14286 | 0.959553 | 2953 | 3 | -0.26272 | 0.17591 | 0.39633 | 1 | 8030 | 3 | -0.26272 |
| TNMD | 6 | 0.58519 | 0.72905 | 0.999994 | 14745 | 2 | 0.41328 | 0.17591 | 0.39633 | 1 | 8031 | 3 | 0.41328 |
| YIPF3 | 6 | 0.40009 | 0.60242 | 0.999994 | 11948 | 2 | 1.6347 | 0.17592 | 0.39635 | 1 | 8032 | 4 | 1.6347 |
| CPD | 5 | 0.7844 | 0.81 | 0.999994 | 17273 | 1 | 2.3961 | 0.17597 | 0.38994 | 1 | 8033 | 3 | 2.3961 |
| PCDHGC3 | 2 | 0.82397 | 0.82379 | 0.999994 | 17838 | 0 | 1.7421 | 0.17603 | 0.28294 | 0.978454 | 8034 | 2 | 1.7421 |
| hsa-mir-4308 | 4 | 0.6822 | 0.713 | 0.999994 | 16119 | 1 | 1.5899 | 0.17604 | 0.34889 | 0.994392 | 8035 | 3 | 1.5899 |
| TTLL10 | 6 | 0.23187 | 0.42634 | 0.999994 | 8567 | 2 | 1.8494 | 0.17607 | 0.39656 | 1 | 8036 | 4 | 1.8494 |
| DNASE1 | 4 | 0.54044 | 0.63509 | 0.999994 | 14062 | 1 | 1.3051 | 0.17611 | 0.34898 | 0.994392 | 8037 | 3 | 1.3051 |
| FAXDC2 | 4 | 0.69658 | 0.72216 | 0.999994 | 16265 | 1 | 0.095953 | 0.17616 | 0.34905 | 0.994392 | 8038 | 2 | 0.095953 |
| SUPT7L | 6 | 0.64257 | 0.77188 | 0.999994 | 15556 | 1 | 1.3979 | 0.17619 | 0.39674 | 1 | 8039 | 4 | 1.3979 |
| hsa-mir-32 | 2 | 0.8237 | 0.82352 | 0.999994 | 17833 | 0 | 2.4801 | 0.1763 | 0.28332 | 0.978454 | 8040 | 2 | 2.4801 |
| TSACC | 4 | 0.064808 | 0.14999 | 0.966359 | 3642 | 2 | -3.0155 | 0.17632 | 0.34928 | 0.994392 | 8041 | 1 | -3.0155 |
| TLR6 | 6 | 0.11157 | 0.26973 | 0.998561 | 5350 | 2 | 1.2982 | 0.17635 | 0.39696 | 1 | 8042 | 4 | 1.2982 |
| LOC100288814 | 6 | 0.77777 | 0.83881 | 0.999994 | 17179 | 1 | 1.6177 | 0.17636 | 0.39698 | 1 | 8043 | 4 | 1.6177 |
| TLDC1 | 5 | 0.079711 | 0.19411 | 0.975797 | 4219 | 1 | 1.7524 | 0.17636 | 0.39038 | 1 | 8044 | 3 | 1.7524 |
| SLC39A12 | 5 | 0.78028 | 0.8076 | 0.999994 | 17215 | 1 | 0.56955 | 0.17636 | 0.39038 | 1 | 8045 | 3 | 0.56955 |
| PVR | 5 | 0.076338 | 0.18869 | 0.969543 | 4085 | 3 | -1.715 | 0.17636 | 0.39038 | 1 | 8046 | 2 | -1.715 |
| TG | 4 | 0.50815 | 0.62 | 0.999994 | 13555 | 1 | 2.8735 | 0.17643 | 0.34944 | 0.994392 | 8047 | 3 | 2.8735 |
| CX3CL1 | 6 | 0.021867 | 0.07796 | 0.93914 | 1573 | 2 | 1.5041 | 0.17649 | 0.39716 | 1 | 8048 | 4 | 1.5041 |
| PLEKHG3 | 6 | 0.078809 | 0.20495 | 0.978186 | 4182 | 3 | -0.34752 | 0.17654 | 0.39723 | 1 | 8049 | 3 | -0.34752 |
| TESC | 6 | 0.12491 | 0.29382 | 0.999994 | 5793 | 3 | -1.0344 | 0.1768 | 0.39763 | 1 | 8050 | 2 | -1.0344 |
| VPS4B | 6 | 0.0034777 | 0.015495 | 0.763603 | 402 | 4 | -3.6568 | 0.1768 | 0.39764 | 1 | 8051 | 1 | -3.6568 |
| ITGB1BP2 | 6 | 0.1784 | 0.36111 | 0.999994 | 7184 | 3 | -0.76396 | 0.1768 | 0.39764 | 1 | 8052 | 2 | -0.76396 |
| UBE3A | 5 | 0.33346 | 0.50834 | 0.999994 | 10794 | 2 | 0.98653 | 0.17683 | 0.39091 | 1 | 8053 | 3 | 0.98653 |
| LAMC1 | 5 | 0.89078 | 0.89091 | 0.999994 | 19050 | 0 | 2.3519 | 0.17684 | 0.39092 | 1 | 8054 | 3 | 2.3519 |
| GJD2 | 6 | 0.3226 | 0.53289 | 0.999994 | 10566 | 2 | 1.6913 | 0.17695 | 0.39784 | 1 | 8055 | 3 | 1.6913 |
| NonTargetingControlG | 1 | 0.82301 | 0.8232 | 0.999994 | 17824 | 0 | 5.3819 | 0.17699 | 0.17678 | 0.970198 | 8056 | 1 | 5.3819 |
| NonTargetingControlG | 2 | 0.82299 | 0.82281 | 0.999994 | 17823 | 0 | 3.3791 | 0.17701 | 0.28434 | 0.978454 | 8057 | 2 | 3.3791 |
| ZDHHC14 | 6 | 0.32472 | 0.53525 | 0.999994 | 10613 | 3 | 1.3623 | 0.17715 | 0.39813 | 1 | 8058 | 3 | 1.3623 |
| hsa-mir-599 | 3 | 0.4967 | 0.54398 | 0.999994 | 13365 | 1 | -0.13734 | 0.17717 | 0.31813 | 0.989846 | 8059 | 1 | -0.13734 |

| | | | | | | | | | | | | | |
|---|---|---|---|---|---|---|---|---|---|---|---|---|---|
| MOB4 | 3 | 0.4394 | 0.50773 | 0.999994 | 12566 | 1 | 0.93999 | 0.17717 | 0.31813 | 0.989846 | 8060 | 2 | 0.93999 |
| hsa-mir-600 | 3 | 0.023252 | 0.059389 | 0.932932 | 1665 | 2 | -2.9896 | 0.17717 | 0.31813 | 0.989846 | 8061 | 1 | -2.9896 |
| AEBP2 | 5 | 0.29736 | 0.47892 | 0.999994 | 10076 | 2 | 1.6204 | 0.17727 | 0.39143 | 1 | 8062 | 3 | 1.6204 |
| SCYL3 | 5 | 0.5145 | 0.66053 | 0.999994 | 13653 | 1 | 2.2086 | 0.17727 | 0.39143 | 1 | 8063 | 3 | 2.2086 |
| UROS | 5 | 0.0046349 | 0.019353 | 0.808616 | 488 | 2 | 0.33784 | 0.17727 | 0.39143 | 1 | 8064 | 2 | 0.33784 |
| PIBF1 | 6 | 0.92996 | 0.93008 | 0.999994 | 19850 | 0 | 1.018 | 0.17728 | 0.39833 | 1 | 8065 | 4 | 1.018 |
| NCAPH | 5 | 0.084176 | 0.20128 | 0.977164 | 4381 | 2 | 2.3821 | 0.17734 | 0.39152 | 1 | 8066 | 3 | 2.3821 |
| hsa-mir-4650-2 | 3 | 0.84178 | 0.84171 | 0.999994 | 18118 | 0 | 3.2991 | 0.17736 | 0.31834 | 0.989846 | 8067 | 2 | 3.2991 |
| TBC1D19 | 6 | 0.47633 | 0.65272 | 0.999994 | 13087 | 2 | 1.3865 | 0.17737 | 0.39845 | 1 | 8068 | 4 | 1.3865 |
| STAMBPL1 | 4 | 0.90933 | 0.90905 | 0.999994 | 19409 | 0 | 1.801 | 0.17737 | 0.35078 | 0.994392 | 8069 | 3 | 1.801 |
| FAM71E1 | 6 | 0.86485 | 0.88021 | 0.999994 | 18537 | 1 | 2.2755 | 0.17741 | 0.39852 | 1 | 8070 | 4 | 2.2755 |
| AK1 | 5 | 0.6594 | 0.74889 | 0.999994 | 15767 | 1 | 2.2726 | 0.17741 | 0.3916 | 1 | 8071 | 3 | 2.2726 |
| hsa-mir-211 | 4 | 0.86 | 0.8596 | 0.999994 | 18437 | 0 | 1.0238 | 0.17751 | 0.35098 | 0.994392 | 8072 | 2 | 1.0238 |
| hsa-mir-3190 | 4 | 0.87841 | 0.87805 | 0.999994 | 18797 | 0 | 0.53698 | 0.17751 | 0.35098 | 0.994392 | 8073 | 2 | 0.53698 |
| ST8SIA2 | 6 | 0.17104 | 0.35204 | 0.999994 | 7009 | 3 | -0.74799 | 0.17757 | 0.39876 | 1 | 8074 | 2 | -0.74799 |
| IZUMO3 | 6 | 0.68408 | 0.80416 | 0.999994 | 16145 | 2 | 0.30559 | 0.17757 | 0.39876 | 1 | 8075 | 2 | 0.30559 |
| TTC39A | 6 | 0.19606 | 0.38309 | 0.999994 | 7656 | 3 | 1.0698 | 0.17762 | 0.39884 | 1 | 8076 | 3 | 1.0698 |
| IL23R | 6 | 0.077929 | 0.20317 | 0.97798 | 4146 | 3 | -0.65631 | 0.17764 | 0.39886 | 1 | 8077 | 2 | -0.65631 |
| UBE2R2 | 6 | 0.69866 | 0.8116 | 0.999994 | 16291 | 1 | 0.48033 | 0.17764 | 0.39886 | 1 | 8078 | 3 | 0.48033 |
| CCDC37 | 6 | 0.62843 | 0.76114 | 0.999994 | 15355 | 1 | 1.3383 | 0.17764 | 0.39886 | 1 | 8079 | 4 | 1.3383 |
| GNG3 | 6 | 0.77558 | 0.83795 | 0.999994 | 17145 | 1 | 1.6143 | 0.17764 | 0.39886 | 1 | 8080 | 4 | 1.6143 |
| RLF | 6 | 0.0034787 | 0.015497 | 0.763603 | 403 | 4 | -3.272 | 0.17764 | 0.39886 | 1 | 8081 | 2 | -3.272 |
| IFNGR1 | 6 | 0.47933 | 0.65474 | 0.999994 | 13137 | 2 | 1.3465 | 0.17764 | 0.39887 | 1 | 8082 | 4 | 1.3465 |
| SLC47A1 | 3 | 0.64977 | 0.65691 | 0.999994 | 15656 | 1 | 2.9046 | 0.17768 | 0.31866 | 0.989846 | 8083 | 2 | 2.9046 |
| LMF2 | 5 | 0.14931 | 0.30173 | 0.999994 | 6437 | 3 | -1.1551 | 0.17768 | 0.3919 | 1 | 8084 | 1 | -1.1551 |
| hsa-mir-6771 | 4 | 0.91874 | 0.91854 | 0.999994 | 19602 | 0 | 1.6481 | 0.17768 | 0.35122 | 0.994392 | 8085 | 3 | 1.6481 |
| TGM6 | 6 | 0.51962 | 0.68226 | 0.999994 | 13741 | 2 | 1.4553 | 0.17773 | 0.399 | 1 | 8086 | 3 | 1.4553 |
| CBLL1 | 4 | 0.28506 | 0.42772 | 0.999994 | 9811 | 2 | 1.4199 | 0.17773 | 0.35128 | 0.994392 | 8087 | 2 | 1.4199 |
| ALG10B | 4 | 0.015429 | 0.050406 | 0.909101 | 1210 | 1 | 2.7084 | 0.17777 | 0.35133 | 0.994392 | 8088 | 2 | 2.7084 |
| BCL2L13 | 5 | 0.37514 | 0.54269 | 0.999994 | 11583 | 1 | 2.4756 | 0.17785 | 0.39209 | 1 | 8089 | 3 | 2.4756 |
| hsa-mir-6803 | 4 | 0.79878 | 0.80015 | 0.999994 | 17459 | 1 | 2.6932 | 0.17786 | 0.35145 | 0.994392 | 8090 | 3 | 2.6932 |
| SEMA6C | 6 | 0.88135 | 0.89048 | 0.999994 | 18859 | 1 | 0.64802 | 0.17805 | 0.39951 | 1 | 8091 | 3 | 0.64802 |
| CLPSL2 | 5 | 0.24158 | 0.43306 | 0.999994 | 8817 | 2 | 0.60567 | 0.17808 | 0.39235 | 1 | 8092 | 3 | 0.60567 |
| RPLP1 | 4 | 0.031759 | 0.085086 | 0.94395 | 2095 | 2 | -4.1786 | 0.17819 | 0.35196 | 0.994392 | 8093 | 1 | -4.1786 |
| NonTargetingControlG | 1 | 0.82169 | 0.8219 | 0.999994 | 17799 | 0 | 5.5272 | 0.17831 | 0.17809 | 0.971262 | 8094 | 1 | 5.5272 |
| ZNF84 | 6 | 0.72813 | 0.82098 | 0.999994 | 16583 | 1 | 1.467 | 0.17831 | 0.39988 | 1 | 8095 | 4 | 1.467 |
| C10orf128 | 6 | 0.10925 | 0.26524 | 0.997396 | 5278 | 1 | 2.513 | 0.17831 | 0.39988 | 1 | 8096 | 4 | 2.513 |
| NDUFAF5 | 6 | 0.42085 | 0.61595 | 0.999994 | 12262 | 2 | 1.8395 | 0.17831 | 0.39988 | 1 | 8097 | 4 | 1.8395 |
| IL23A | 6 | 0.5842 | 0.72831 | 0.999994 | 14731 | 2 | 2.1131 | 0.17831 | 0.39988 | 1 | 8098 | 4 | 2.1131 |
| LZTFL1 | 6 | 0.10116 | 0.24959 | 0.991904 | 5012 | 4 | -0.96335 | 0.17839 | 0.4 | 1 | 8099 | 2 | -0.96335 |
| KRTAP9-9 | 4 | 0.099122 | 0.21352 | 0.979587 | 4934 | 2 | 2.1884 | 0.1784 | 0.35226 | 0.994392 | 8100 | 2 | 2.1884 |
| PTPMT1 | 4 | 0.097185 | 0.21006 | 0.979587 | 4865 | 2 | 1.6187 | 0.1784 | 0.35226 | 0.994392 | 8101 | 2 | 1.6187 |
| NCAPG | 5 | 0.064873 | 0.17006 | 0.967126 | 3646 | 3 | -1.0147 | 0.17842 | 0.39271 | 1 | 8102 | 2 | -1.0147 |
| TMEM218 | 5 | 0.084202 | 0.20132 | 0.977164 | 4384 | 1 | 1.9311 | 0.17842 | 0.39271 | 1 | 8103 | 3 | 1.9311 |
| RPUSD1 | 5 | 0.067976 | 0.17513 | 0.968171 | 3749 | 3 | -1.6067 | 0.17842 | 0.39271 | 1 | 8104 | 2 | -1.6067 |
| ARAP3 | 6 | 0.67811 | 0.79952 | 0.999994 | 16062 | 2 | 1.2454 | 0.17845 | 0.40009 | 1 | 8105 | 4 | 1.2454 |
| ATOH7 | 6 | 0.38113 | 0.59029 | 0.999994 | 11663 | 1 | 1.5775 | 0.17845 | 0.40009 | 1 | 8106 | 4 | 1.5775 |
| TCF25 | 6 | 0.36929 | 0.58277 | 0.999994 | 11495 | 2 | 0.4951 | 0.17845 | 0.40009 | 1 | 8107 | 3 | 0.4951 |
| HMCES | 4 | 0.84081 | 0.84026 | 0.999994 | 18101 | 0 | 2.2193 | 0.17846 | 0.35235 | 0.994392 | 8108 | 3 | 2.2193 |
| NAE1 | 4 | 0.42525 | 0.5716 | 0.999994 | 12336 | 1 | 2.2284 | 0.17876 | 0.3528 | 0.994392 | 8109 | 2 | 2.2284 |
| hsa-mir-7843 | 3 | 0.68455 | 0.68682 | 0.999994 | 16153 | 1 | 1.7527 | 0.1788 | 0.31988 | 0.989846 | 8110 | 2 | 1.7527 |
| MEP1A | 6 | 0.97755 | 0.97749 | 0.999994 | 21031 | 0 | 2.2055 | 0.17882 | 0.40062 | 1 | 8111 | 4 | 2.2055 |

| | | | | | | | | | | | | | |
|---|---|---|---|---|---|---|---|---|---|---|---|---|---|
| DNTT | 6 | 0.026103 | 0.089609 | 0.94395 | 1809 | 4 | -1.5146 | 0.17896 | 0.40083 | 1 | 8112 | 1 | -1.5146 |
| SLC25A23 | 6 | 0.070581 | 0.18801 | 0.969543 | 3858 | 4 | -1.3557 | 0.17896 | 0.40083 | 1 | 8113 | 2 | -1.3557 |
| C11orf91 | 5 | 0.4002 | 0.56367 | 0.999994 | 11952 | 2 | 2.187 | 0.17907 | 0.39346 | 1 | 8114 | 3 | 2.187 |
| DGCR6L | 5 | 0.067004 | 0.17353 | 0.96793 | 3721 | 2 | 2.4218 | 0.17907 | 0.39347 | 1 | 8115 | 3 | 2.4218 |
| hsa-mir-6717 | 4 | 0.91533 | 0.91508 | 0.999994 | 19519 | 0 | 1.5762 | 0.1791 | 0.35328 | 0.994392 | 8116 | 3 | 1.5762 |
| C8orf48 | 6 | 0.65787 | 0.78364 | 0.999994 | 15752 | 2 | 1.3662 | 0.17915 | 0.40111 | 1 | 8117 | 4 | 1.3662 |
| OR1G1 | 5 | 0.3296 | 0.5052 | 0.999994 | 10714 | 1 | 2.3322 | 0.17918 | 0.39358 | 1 | 8118 | 3 | 2.3322 |
| SRD5A2 | 6 | 0.26192 | 0.46216 | 0.999994 | 9285 | 3 | -0.06609 | 0.17919 | 0.40117 | 1 | 8119 | 3 | -0.06609 |
| MC5R | 6 | 0.52695 | 0.6874 | 0.999994 | 13842 | 2 | 1.4619 | 0.17919 | 0.40117 | 1 | 8120 | 4 | 1.4619 |
| TLK2 | 6 | 0.007903 | 0.031632 | 0.864893 | 724 | 4 | -1.3099 | 0.17919 | 0.40117 | 1 | 8121 | 2 | -1.3099 |
| hsa-mir-766 | 3 | 0.61698 | 0.63031 | 0.999994 | 15171 | 1 | 3.2785 | 0.17919 | 0.32032 | 0.989846 | 8122 | 2 | 3.2785 |
| hsa-mir-4299 | 4 | 0.72331 | 0.74039 | 0.999994 | 16527 | 1 | 2.009 | 0.17923 | 0.35347 | 0.994392 | 8123 | 3 | 2.009 |
| LYPLA2 | 6 | 0.38932 | 0.59554 | 0.999994 | 11785 | 2 | 1.4517 | 0.17927 | 0.4013 | 1 | 8124 | 4 | 1.4517 |
| ACVR1 | 6 | 0.33695 | 0.5492 | 0.999994 | 10871 | 2 | 1.7604 | 0.17928 | 0.40131 | 1 | 8125 | 4 | 1.7604 |
| TRIM27 | 5 | 0.015065 | 0.050391 | 0.909101 | 1191 | 2 | 2.5535 | 0.17929 | 0.39372 | 1 | 8126 | 3 | 2.5535 |
| hsa-mir-1294 | 4 | 0.91215 | 0.91195 | 0.999994 | 19467 | 0 | 1.2412 | 0.1793 | 0.35357 | 0.994392 | 8127 | 3 | 1.2412 |
| ASCL5 | 6 | 0.5482 | 0.70233 | 0.999994 | 14180 | 2 | 1.3967 | 0.17933 | 0.4014 | 1 | 8128 | 4 | 1.3967 |
| C17orf98 | 5 | 0.02335 | 0.073121 | 0.93914 | 1671 | 3 | -0.96117 | 0.17933 | 0.39377 | 1 | 8129 | 2 | -0.96117 |
| DLEU7 | 5 | 0.14285 | 0.29214 | 0.999632 | 6266 | 3 | -1.1821 | 0.17933 | 0.39377 | 1 | 8130 | 2 | -1.1821 |
| FOXN1 | 5 | 0.057662 | 0.15819 | 0.967126 | 3320 | 2 | -0.50844 | 0.17933 | 0.39377 | 1 | 8131 | 2 | -0.50844 |
| RPL12 | 5 | 0.56243 | 0.70281 | 0.999994 | 14420 | 2 | 0.35975 | 0.17933 | 0.39377 | 1 | 8132 | 2 | 0.35975 |
| ABT1 | 4 | 0.82696 | 0.82656 | 0.999994 | 17886 | 1 | 1.515 | 0.17934 | 0.35363 | 0.994392 | 8133 | 3 | 1.515 |
| SMNDC1 | 4 | 0.036697 | 0.095167 | 0.945031 | 2354 | 2 | -1.8935 | 0.17935 | 0.35363 | 0.994392 | 8134 | 2 | -1.8935 |
| CD69 | 6 | 0.86447 | 0.88 | 0.999994 | 18532 | 1 | 1.7941 | 0.17946 | 0.4016 | 1 | 8135 | 3 | 1.7941 |
| ATF2 | 6 | 0.50397 | 0.67147 | 0.999994 | 13494 | 2 | 1.9264 | 0.17957 | 0.40175 | 1 | 8136 | 4 | 1.9264 |
| hsa-mir-497 | 3 | 0.77011 | 0.77011 | 0.999994 | 17084 | 0 | 2.0013 | 0.17959 | 0.32078 | 0.989846 | 8137 | 2 | 2.0013 |
| C5orf15 | 6 | 0.1832 | 0.36714 | 0.999994 | 7299 | 2 | 2.2589 | 0.17963 | 0.40184 | 1 | 8138 | 4 | 2.2589 |
| CCDC70 | 4 | 0.058739 | 0.13836 | 0.957317 | 3371 | 3 | -1.9398 | 0.17969 | 0.35412 | 0.994392 | 8139 | 1 | -1.9398 |
| SART1 | 4 | 0.45547 | 0.59698 | 0.999994 | 12782 | 1 | 1.9582 | 0.17969 | 0.35412 | 0.994392 | 8140 | 3 | 1.9582 |
| CNTNAP3B | 4 | 0.12693 | 0.26228 | 0.995875 | 5846 | 2 | 1.0711 | 0.1797 | 0.35414 | 0.994392 | 8141 | 2 | 1.0711 |
| ADCY4 | 4 | 0.21746 | 0.35865 | 0.999994 | 8205 | 1 | 2.2506 | 0.1797 | 0.35414 | 0.994392 | 8142 | 2 | 2.2506 |
| ATXN2 | 4 | 0.86998 | 0.86956 | 0.999994 | 18634 | 0 | 2.392 | 0.1797 | 0.35414 | 0.994392 | 8143 | 2 | 2.392 |
| ERN1 | 6 | 0.037446 | 0.11614 | 0.951052 | 2395 | 3 | 0.22219 | 0.17971 | 0.40195 | 1 | 8144 | 3 | 0.22219 |
| CDH8 | 5 | 0.22018 | 0.4035 | 0.999994 | 8294 | 3 | -0.66554 | 0.17973 | 0.39421 | 1 | 8145 | 2 | -0.66554 |
| NonTargetingControlG | 2 | 0.70693 | 0.70672 | 0.999994 | 16382 | 0 | 3.7114 | 0.17976 | 0.28824 | 0.979722 | 8146 | 1 | 3.7114 |
| BYSL | 4 | 0.034891 | 0.09151 | 0.94395 | 2263 | 1 | 3.0521 | 0.1799 | 0.35441 | 0.994392 | 8147 | 3 | 3.0521 |
| SYNC | 6 | 0.34771 | 0.56143 | 0.999994 | 11106 | 1 | 1.3187 | 0.17996 | 0.40232 | 1 | 8148 | 3 | 1.3187 |
| FGG | 6 | 0.71302 | 0.81608 | 0.999994 | 16429 | 1 | 0.46123 | 0.17996 | 0.40232 | 1 | 8149 | 3 | 0.46123 |
| MAATS1 | 5 | 0.39607 | 0.56029 | 0.999994 | 11890 | 2 | 1.5075 | 0.17999 | 0.39452 | 1 | 8150 | 3 | 1.5075 |
| C16orf90 | 5 | 0.24187 | 0.43345 | 0.999994 | 8832 | 1 | 2.4222 | 0.18004 | 0.39457 | 1 | 8151 | 3 | 2.4222 |
| GPR35 | 5 | 0.015528 | 0.051713 | 0.910734 | 1217 | 2 | 1.4509 | 0.18012 | 0.39468 | 1 | 8152 | 3 | 1.4509 |
| hsa-mir-4467 | 2 | 0.73575 | 0.73544 | 0.999994 | 16671 | 0 | 3.8299 | 0.18013 | 0.28876 | 0.980082 | 8153 | 1 | 3.8299 |
| KCNG1 | 6 | 0.85576 | 0.87505 | 0.999994 | 18362 | 1 | 1.611 | 0.18015 | 0.4026 | 1 | 8154 | 4 | 1.611 |
| BCL2L12 | 4 | 0.48232 | 0.60854 | 0.999994 | 13175 | 1 | 2.6309 | 0.18023 | 0.3549 | 0.994392 | 8155 | 2 | 2.6309 |
| EVI2A | 4 | 0.022011 | 0.064863 | 0.93914 | 1584 | 1 | 2.2802 | 0.18027 | 0.35495 | 0.994392 | 8156 | 3 | 2.2802 |
| HIRIP3 | 5 | 0.21276 | 0.39325 | 0.999994 | 8082 | 2 | 0.40442 | 0.18032 | 0.39489 | 1 | 8157 | 2 | 0.40442 |
| hsa-mir-4494 | 4 | 0.87455 | 0.87419 | 0.999994 | 18722 | 0 | 0.64607 | 0.1804 | 0.35515 | 0.994392 | 8158 | 2 | 0.64607 |
| hsa-mir-145 | 4 | 0.27474 | 0.41719 | 0.999994 | 9589 | 2 | 0.2529 | 0.1804 | 0.35515 | 0.994392 | 8159 | 2 | 0.2529 |
| UBR1 | 6 | 0.01727 | 0.063132 | 0.937389 | 1320 | 3 | -0.031787 | 0.18043 | 0.403 | 1 | 8160 | 3 | -0.031787 |
| LILRA1 | 5 | 0.036417 | 0.10687 | 0.950995 | 2341 | 2 | 1.388 | 0.18056 | 0.39516 | 1 | 8161 | 3 | 1.388 |
| OR13D1 | 6 | 0.61129 | 0.74829 | 0.999994 | 15091 | 2 | 1.5334 | 0.18056 | 0.4032 | 1 | 8162 | 4 | 1.5334 |
| MRP63 | 3 | 0.19038 | 0.30056 | 0.999994 | 7498 | 1 | 3.0824 | 0.18071 | 0.32206 | 0.989846 | 8163 | 2 | 3.0824 |

| | | | | | | | | | | | | |
|---|---|---|---|---|---|---|---|---|---|---|---|---|
| PFN4 | 6 | 0.39989 | 0.60229 | 0.999994 | 11947 | 2 | 1.5102 | 0.18073 | 0.40344 | 1 | 8164 | 4 | 1.5102 |
| hsa-mir-8052 | 4 | 0.79072 | 0.79298 | 0.999994 | 17360 | 1 | 2.5446 | 0.1808 | 0.35574 | 0.994392 | 8165 | 2 | 2.5446 |
| ZMYM6NB | 6 | 0.90439 | 0.90709 | 0.999994 | 19308 | 1 | 2.0303 | 0.1809 | 0.40369 | 1 | 8166 | 4 | 2.0303 |
| RGS8 | 5 | 0.023493 | 0.073502 | 0.93914 | 1685 | 2 | 2.417 | 0.18091 | 0.39558 | 1 | 8167 | 3 | 2.417 |
| ST8SIA3 | 5 | 0.51901 | 0.66445 | 0.999994 | 13730 | 1 | 0.35674 | 0.18092 | 0.39558 | 1 | 8168 | 2 | 0.35674 |
| IGFBP1 | 6 | 0.088601 | 0.22474 | 0.983354 | 4539 | 2 | -0.10646 | 0.18093 | 0.40372 | 1 | 8169 | 1 | -0.10646 |
| ZIC2 | 6 | 0.24129 | 0.43753 | 0.999994 | 8805 | 2 | 1.4114 | 0.18112 | 0.404 | 1 | 8170 | 4 | 1.4114 |
| FABP7 | 5 | 0.82401 | 0.83525 | 0.999994 | 17840 | 1 | 2.1619 | 0.18118 | 0.39587 | 1 | 8171 | 3 | 2.1619 |
| PIAS4 | 6 | 0.52642 | 0.68701 | 0.999994 | 13836 | 1 | 1.7131 | 0.1812 | 0.40412 | 1 | 8172 | 4 | 1.7131 |
| FBXL5 | 5 | 0.72533 | 0.77842 | 0.999994 | 16554 | 1 | 1.0017 | 0.18121 | 0.39591 | 1 | 8173 | 3 | 1.0017 |
| TRPV1 | 5 | 0.41731 | 0.57789 | 0.999994 | 12208 | 1 | 1.2959 | 0.18121 | 0.39591 | 1 | 8174 | 3 | 1.2959 |
| NUDT13 | 5 | 0.27415 | 0.46014 | 0.999994 | 9567 | 1 | 0.86788 | 0.18121 | 0.39591 | 1 | 8175 | 3 | 0.86788 |
| SSNA1 | 5 | 0.11063 | 0.24295 | 0.988088 | 5321 | 2 | -0.47914 | 0.18121 | 0.39591 | 1 | 8176 | 2 | -0.47914 |
| CISH | 5 | 0.62047 | 0.73373 | 0.999994 | 15228 | 1 | 0.15327 | 0.18126 | 0.39598 | 1 | 8177 | 2 | 0.15327 |
| GLG1 | 6 | 0.09464 | 0.23678 | 0.985449 | 4773 | 3 | 0.84913 | 0.18129 | 0.40425 | 1 | 8178 | 3 | 0.84913 |
| CEP95 | 6 | 0.3505 | 0.56455 | 0.999994 | 11171 | 2 | 1.6512 | 0.18132 | 0.4043 | 1 | 8179 | 4 | 1.6512 |
| CCNJL | 6 | 0.044194 | 0.13131 | 0.95318 | 2722 | 1 | 1.4774 | 0.18133 | 0.40432 | 1 | 8180 | 4 | 1.4774 |
| KRTAP5-11 | 3 | 0.83234 | 0.83222 | 0.999994 | 17962 | 0 | 3.0857 | 0.18136 | 0.32274 | 0.989846 | 8181 | 2 | 3.0857 |
| TRAM1L1 | 6 | 0.58054 | 0.72563 | 0.999994 | 14686 | 2 | 1.7375 | 0.18149 | 0.40454 | 1 | 8182 | 4 | 1.7375 |
| CYP2F1 | 6 | 0.872 | 0.88457 | 0.999994 | 18676 | 1 | 1.5926 | 0.18158 | 0.40469 | 1 | 8183 | 4 | 1.5926 |
| ATG4C | 6 | 0.045229 | 0.13358 | 0.95318 | 2776 | 2 | 0.17056 | 0.18163 | 0.40476 | 1 | 8184 | 2 | 0.17056 |
| DCAF12L1 | 5 | 0.40971 | 0.57165 | 0.999994 | 12105 | 2 | 1.4281 | 0.18164 | 0.3964 | 1 | 8185 | 3 | 1.4281 |
| RTFDC1 | 6 | 0.74239 | 0.82579 | 0.999994 | 16769 | 1 | 2.027 | 0.18177 | 0.40495 | 1 | 8186 | 3 | 2.027 |
| FAM104B | 6 | 0.63768 | 0.76816 | 0.999994 | 15483 | 2 | 2.2199 | 0.18177 | 0.40495 | 1 | 8187 | 4 | 2.2199 |
| C4orf33 | 6 | 0.96424 | 0.96428 | 0.999994 | 20710 | 0 | 0.83612 | 0.18191 | 0.40515 | 1 | 8188 | 3 | 0.83612 |
| hsa-let-7f-1 | 4 | 0.86597 | 0.86557 | 0.999994 | 18556 | 0 | 1.3413 | 0.18203 | 0.35754 | 0.994392 | 8189 | 3 | 1.3413 |
| DYTN | 5 | 0.034377 | 0.10168 | 0.946414 | 2231 | 2 | 0.046456 | 0.18204 | 0.39683 | 1 | 8190 | 2 | 0.046456 |
| ARHGEF25 | 5 | 0.84835 | 0.85298 | 0.999994 | 18224 | 1 | -0.29354 | 0.18204 | 0.39683 | 1 | 8191 | 2 | -0.29354 |
| HIGD2A | 5 | 0.13905 | 0.28642 | 0.999632 | 6165 | 2 | 0.071034 | 0.18204 | 0.39683 | 1 | 8192 | 2 | 0.071034 |
| GPR89A | 6 | 0.068432 | 0.1835 | 0.968526 | 3766 | 2 | 2.1285 | 0.18209 | 0.40541 | 1 | 8193 | 4 | 2.1285 |
| SYBU | 6 | 0.95147 | 0.95145 | 0.999994 | 20376 | 0 | 1.8725 | 0.18215 | 0.4055 | 1 | 8194 | 4 | 1.8725 |
| RHBDD1 | 6 | 0.42153 | 0.61639 | 0.999994 | 12269 | 2 | 0.53575 | 0.18215 | 0.4055 | 1 | 8195 | 3 | 0.53575 |
| KIAA1984 | 6 | 0.23104 | 0.42534 | 0.999994 | 8542 | 3 | 0.40127 | 0.18215 | 0.4055 | 1 | 8196 | 3 | 0.40127 |
| NUDT18 | 6 | 0.79846 | 0.84722 | 0.999994 | 17454 | 1 | 0.93116 | 0.18215 | 0.4055 | 1 | 8197 | 3 | 0.93116 |
| FASTKD3 | 6 | 0.51353 | 0.67807 | 0.999994 | 13630 | 2 | 0.746 | 0.18215 | 0.4055 | 1 | 8198 | 3 | 0.746 |
| L3MBTL4 | 5 | 0.28436 | 0.46838 | 0.999994 | 9793 | 1 | 2.3348 | 0.18222 | 0.39702 | 1 | 8199 | 3 | 2.3348 |
| CES3 | 6 | 0.27446 | 0.47697 | 0.999994 | 9576 | 2 | 1.8655 | 0.1823 | 0.40572 | 1 | 8200 | 4 | 1.8655 |
| ZNF665 | 5 | 0.68803 | 0.76114 | 0.999994 | 16196 | 1 | 2.2753 | 0.18232 | 0.39712 | 1 | 8201 | 3 | 2.2753 |
| PLAGL2 | 5 | 0.11408 | 0.24831 | 0.990682 | 5451 | 3 | -1.5247 | 0.18234 | 0.39716 | 1 | 8202 | 1 | -1.5247 |
| CST11 | 4 | 0.86713 | 0.86668 | 0.999994 | 18578 | 0 | 1.3542 | 0.18235 | 0.35802 | 0.994392 | 8203 | 3 | 1.3542 |
| FAM50A | 5 | 0.081534 | 0.19708 | 0.977164 | 4283 | 2 | 2.4017 | 0.18237 | 0.39718 | 1 | 8204 | 3 | 2.4017 |
| IFNA13 | 2 | 0.083598 | 0.14429 | 0.961983 | 4359 | 1 | -0.6384 | 0.18241 | 0.29204 | 0.983225 | 8205 | 1 | -0.6384 |
| ANKRD29 | 5 | 0.12534 | 0.2655 | 0.99746 | 5801 | 2 | 2.1815 | 0.18246 | 0.39728 | 1 | 8206 | 3 | 2.1815 |
| ABCA2 | 6 | 0.25304 | 0.45155 | 0.999994 | 9084 | 2 | 1.6727 | 0.18249 | 0.40598 | 1 | 8207 | 3 | 1.6727 |
| SLC45A4 | 2 | 0.8175 | 0.81728 | 0.999994 | 17739 | 0 | 3.5238 | 0.1825 | 0.29216 | 0.983225 | 8208 | 2 | 3.5238 |
| hsa-mir-5697 | 4 | 0.93506 | 0.93488 | 0.999994 | 19967 | 0 | 1.1851 | 0.18253 | 0.35826 | 0.994392 | 8209 | 3 | 1.1851 |
| NDUFB11 | 6 | 0.31716 | 0.52663 | 0.999994 | 10456 | 3 | 0.45442 | 0.18254 | 0.40606 | 1 | 8210 | 3 | 0.45442 |
| NAP1L3 | 6 | 0.88078 | 0.89013 | 0.999994 | 18848 | 1 | 1.8292 | 0.18254 | 0.40606 | 1 | 8211 | 4 | 1.8292 |
| FBXO43 | 6 | 0.29641 | 0.50253 | 0.999994 | 10049 | 1 | 0.55387 | 0.18254 | 0.40606 | 1 | 8212 | 3 | 0.55387 |
| TLR7 | 4 | 0.88433 | 0.88398 | 0.999994 | 18917 | 0 | 3.1821 | 0.18254 | 0.35828 | 0.994392 | 8213 | 3 | 3.1821 |
| OSTF1 | 5 | 0.28586 | 0.46956 | 0.999994 | 9830 | 1 | 2.3896 | 0.18259 | 0.39743 | 1 | 8214 | 3 | 2.3896 |
| hsa-mir-369 | 4 | 0.93058 | 0.93036 | 0.999994 | 19866 | 0 | 2.9655 | 0.1826 | 0.35836 | 0.994392 | 8215 | 3 | 2.9655 |

| | | | | | | | | | | | | |
|---|---|---|---|---|---|---|---|---|---|---|---|---|
| PYCR1 | 6 | 0.51114 | 0.6764 | 0.999994 | 13598 | 2 | 1.291 | 0.18261 | 0.40616 | 1 | 8216 | 4 | 1.291 |
| POLR2C | 1 | 0.81736 | 0.81759 | 0.999994 | 17735 | 0 | 5.4173 | 0.18264 | 0.18244 | 0.97162 | 8217 | 1 | 5.4173 |
| LEPROT | 5 | 0.95631 | 0.95647 | 0.999994 | 20496 | 0 | 2.0656 | 0.18266 | 0.39751 | 1 | 8218 | 3 | 2.0656 |
| ZNF28 | 5 | 0.9476 | 0.94778 | 0.999994 | 20279 | 0 | 1.3504 | 0.18269 | 0.39755 | 1 | 8219 | 3 | 1.3504 |
| DPH3 | 5 | 0.5313 | 0.67523 | 0.999994 | 13918 | 2 | -0.41929 | 0.18269 | 0.39755 | 1 | 8220 | 1 | -0.41929 |
| IL18 | 4 | 0.067054 | 0.15425 | 0.967126 | 3722 | 1 | 2.3646 | 0.1827 | 0.35851 | 0.994392 | 8221 | 2 | 2.3646 |
| NBEAL2 | 4 | 0.3726 | 0.51752 | 0.999994 | 11554 | 2 | -0.10873 | 0.1828 | 0.35865 | 0.994392 | 8222 | 2 | -0.10873 |
| CACNG1 | 4 | 0.82719 | 0.82677 | 0.999994 | 17888 | 0 | 1.3757 | 0.18288 | 0.35877 | 0.994392 | 8223 | 3 | 1.3757 |
| CTAG1B | 2 | 0.81092 | 0.81063 | 0.999994 | 17645 | 0 | 4.1493 | 0.18292 | 0.29275 | 0.983225 | 8224 | 2 | 4.1493 |
| VN1R5 | 5 | 0.85229 | 0.85611 | 0.999994 | 18302 | 1 | 2.257 | 0.18293 | 0.39784 | 1 | 8225 | 3 | 2.257 |
| KNSTRN | 5 | 0.064928 | 0.17014 | 0.967126 | 3649 | 2 | 2.1527 | 0.18296 | 0.39787 | 1 | 8226 | 3 | 2.1527 |
| KIF2C | 5 | 0.7374 | 0.78437 | 0.999994 | 16693 | 1 | 2.1233 | 0.18299 | 0.39789 | 1 | 8227 | 3 | 2.1233 |
| KAZALD1 | 4 | 0.35813 | 0.50259 | 0.999994 | 11306 | 2 | -0.15618 | 0.18311 | 0.35912 | 0.994392 | 8228 | 2 | -0.15618 |
| FABP5 | 6 | 0.11676 | 0.27961 | 0.999632 | 5533 | 3 | 0.71333 | 0.18312 | 0.40688 | 1 | 8229 | 3 | 0.71333 |
| C7orf31 | 6 | 0.95167 | 0.95165 | 0.999994 | 20380 | 0 | 1.1653 | 0.18319 | 0.40698 | 1 | 8230 | 4 | 1.1653 |
| NonTargetingControlG | 1 | 0.81679 | 0.81705 | 0.999994 | 17724 | 0 | 5.4149 | 0.18321 | 0.183 | 0.97162 | 8231 | 1 | 5.4149 |
| TMEM199 | 1 | 0.81679 | 0.81705 | 0.999994 | 17723 | 0 | 5.4149 | 0.18321 | 0.183 | 0.97162 | 8232 | 1 | 5.4149 |
| PACSIN1 | 5 | 0.54596 | 0.68815 | 0.999994 | 14150 | 2 | -0.3542 | 0.18324 | 0.39819 | 1 | 8233 | 2 | -0.3542 |
| NAGPA | 6 | 0.032005 | 0.10351 | 0.946414 | 2105 | 2 | 1.2971 | 0.18329 | 0.40714 | 1 | 8234 | 4 | 1.2971 |
| AP4M1 | 6 | 0.16117 | 0.33971 | 0.999994 | 6769 | 1 | 1.6781 | 0.18332 | 0.40718 | 1 | 8235 | 3 | 1.6781 |
| FBXO48 | 4 | 0.33251 | 0.47631 | 0.999994 | 10781 | 2 | -0.38268 | 0.18337 | 0.35948 | 0.994392 | 8236 | 1 | -0.38268 |
| hsa-mir-3621 | 4 | 0.67604 | 0.70908 | 0.999994 | 16019 | 1 | 2.1931 | 0.18339 | 0.35951 | 0.994392 | 8237 | 2 | 2.1931 |
| POU5F2 | 4 | 0.30327 | 0.44633 | 0.999994 | 10198 | 2 | 1.7835 | 0.18339 | 0.35951 | 0.994392 | 8238 | 2 | 1.7835 |
| ZC3H8 | 4 | 0.028079 | 0.077588 | 0.93914 | 1907 | 1 | 2.2264 | 0.18339 | 0.35951 | 0.994392 | 8239 | 2 | 2.2264 |
| KRTAP10-4 | 4 | 0.1875 | 0.3278 | 0.999994 | 7408 | 1 | 2.813 | 0.18339 | 0.35951 | 0.994392 | 8240 | 3 | 2.813 |
| FATE1 | 6 | 0.38505 | 0.59277 | 0.999994 | 11717 | 2 | 0.71707 | 0.18341 | 0.40732 | 1 | 8241 | 3 | 0.71707 |
| ZNF730 | 6 | 0.73481 | 0.82321 | 0.999994 | 16657 | 1 | 0.60225 | 0.18341 | 0.40732 | 1 | 8242 | 3 | 0.60225 |
| NonTargetingControlG | 1 | 0.81631 | 0.81654 | 0.999994 | 17718 | 0 | 5.5262 | 0.18369 | 0.18349 | 0.97162 | 8243 | 1 | 5.5262 |
| C6orf222 | 6 | 0.64954 | 0.77726 | 0.999994 | 15650 | 2 | 1.364 | 0.18375 | 0.40781 | 1 | 8244 | 3 | 1.364 |
| C20orf112 | 5 | 0.48594 | 0.63576 | 0.999994 | 13229 | 2 | 1.9647 | 0.18379 | 0.3988 | 1 | 8245 | 3 | 1.9647 |
| CNN3 | 6 | 0.39227 | 0.5974 | 0.999994 | 11836 | 2 | 2.3019 | 0.18382 | 0.40792 | 1 | 8246 | 4 | 2.3019 |
| FAM96B | 3 | 0.75514 | 0.75509 | 0.999994 | 16917 | 0 | 2.9272 | 0.18383 | 0.32541 | 0.991152 | 8247 | 2 | 2.9272 |
| CASKIN2 | 3 | 0.67134 | 0.67503 | 0.999994 | 15948 | 1 | 2.9225 | 0.18384 | 0.32543 | 0.991152 | 8248 | 2 | 2.9225 |
| BAHCC1 | 5 | 0.91179 | 0.91179 | 0.999994 | 19459 | 0 | 2.2255 | 0.18386 | 0.39887 | 1 | 8249 | 3 | 2.2255 |
| DPY19L3 | 6 | 0.33649 | 0.54866 | 0.999994 | 10859 | 1 | 0.95501 | 0.18387 | 0.40798 | 1 | 8250 | 3 | 0.95501 |
| RHBDF1 | 6 | 0.064752 | 0.17575 | 0.968171 | 3640 | 2 | -0.040813 | 0.18387 | 0.40798 | 1 | 8251 | 2 | -0.040813 |
| C15orf27 | 5 | 0.65171 | 0.74581 | 0.999994 | 15683 | 1 | 0.15229 | 0.18391 | 0.39893 | 1 | 8252 | 2 | 0.15229 |
| SF3A2 | 4 | 0.20138 | 0.34209 | 0.999994 | 7805 | 2 | 1.2091 | 0.18398 | 0.36037 | 0.994562 | 8253 | 2 | 1.2091 |
| NonTargetingControlG | 2 | 0.81601 | 0.8158 | 0.999994 | 17713 | 0 | 4.1455 | 0.18399 | 0.29429 | 0.983431 | 8254 | 2 | 4.1455 |
| FAM169B | 5 | 0.2062 | 0.38401 | 0.999994 | 7936 | 2 | 0.74272 | 0.18409 | 0.39912 | 1 | 8255 | 3 | 0.74272 |
| BAI3 | 6 | 0.83406 | 0.86356 | 0.999994 | 17988 | 1 | 1.3087 | 0.18411 | 0.40833 | 1 | 8256 | 4 | 1.3087 |
| GAL3ST2 | 5 | 0.49586 | 0.64431 | 0.999994 | 13355 | 2 | 0.016303 | 0.18421 | 0.39925 | 1 | 8257 | 2 | 0.016303 |
| GATAD1 | 5 | 0.071495 | 0.1808 | 0.968171 | 3893 | 3 | -1.9039 | 0.18421 | 0.39925 | 1 | 8258 | 2 | -1.9039 |
| PGAM5 | 5 | 0.22497 | 0.41021 | 0.999994 | 8396 | 3 | -0.66818 | 0.18421 | 0.39925 | 1 | 8259 | 2 | -0.66818 |
| HK3 | 5 | 0.095997 | 0.22009 | 0.980699 | 4815 | 2 | 0.026342 | 0.18421 | 0.39925 | 1 | 8260 | 2 | 0.026342 |
| TMEM191B | 5 | 0.12722 | 0.2684 | 0.998561 | 5852 | 3 | -1.2815 | 0.18421 | 0.39925 | 1 | 8261 | 2 | -1.2815 |
| hsa-mir-146b | 4 | 0.48946 | 0.61164 | 0.999994 | 13271 | 1 | 1.5484 | 0.18421 | 0.36073 | 0.994562 | 8262 | 3 | 1.5484 |
| CAB39L | 6 | 0.030741 | 0.10053 | 0.946414 | 2046 | 1 | 1.3319 | 0.18431 | 0.40863 | 1 | 8263 | 4 | 1.3319 |
| ABCA6 | 6 | 0.56351 | 0.71331 | 0.999994 | 14441 | 2 | 1.5581 | 0.18436 | 0.40871 | 1 | 8264 | 4 | 1.5581 |
| SKIDA1 | 6 | 0.1572 | 0.33468 | 0.999994 | 6653 | 3 | 0.43018 | 0.18437 | 0.40873 | 1 | 8265 | 3 | 0.43018 |
| B3GALT6 | 6 | 0.37948 | 0.58925 | 0.999994 | 11640 | 2 | 0.91787 | 0.18437 | 0.40873 | 1 | 8266 | 3 | 0.91787 |
| GFI1 | 6 | 0.075971 | 0.19918 | 0.977164 | 4070 | 3 | 0.99957 | 0.1844 | 0.40876 | 1 | 8267 | 3 | 0.99957 |

| | | | | | | | | | | | | | |
|---|---|---|---|---|---|---|---|---|---|---|---|---|---|
| SNX33 | 5 | 0.52562 | 0.67021 | 0.999994 | 13828 | 2 | 0.10792 | 0.18443 | 0.3995 | 1 | 8268 | 2 | 0.10792 |
| ZDHHC21 | 6 | 0.39874 | 0.60157 | 0.999994 | 11929 | 2 | 1.324 | 0.18444 | 0.40882 | 1 | 8269 | 3 | 1.324 |
| MEOX1 | 4 | 0.72022 | 0.73825 | 0.999994 | 16501 | 1 | 1.8659 | 0.18449 | 0.36111 | 0.994562 | 8270 | 3 | 1.8659 |
| ABCB7 | 3 | 0.017271 | 0.044885 | 0.891621 | 1323 | 2 | -7.1095 | 0.18449 | 0.32615 | 0.991729 | 8271 | 1 | -7.1095 |
| CA5A | 4 | 0.28872 | 0.43147 | 0.999994 | 9889 | 1 | 2.725 | 0.18451 | 0.36115 | 0.994562 | 8272 | 3 | 2.725 |
| CAMKK2 | 6 | 0.62996 | 0.76229 | 0.999994 | 15380 | 1 | 1.4857 | 0.18452 | 0.40893 | 1 | 8273 | 4 | 1.4857 |
| TAS1R2 | 6 | 0.95039 | 0.95038 | 0.999994 | 20344 | 0 | 2.0195 | 0.18457 | 0.409 | 1 | 8274 | 4 | 2.0195 |
| ZNF513 | 5 | 0.37059 | 0.53889 | 0.999994 | 11524 | 2 | 2.1444 | 0.18458 | 0.39968 | 1 | 8275 | 3 | 2.1444 |
| UTP11L | 6 | 0.057787 | 0.16099 | 0.967126 | 3328 | 2 | 2.1742 | 0.18458 | 0.40902 | 1 | 8276 | 4 | 2.1742 |
| FAM98C | 6 | 0.83677 | 0.86496 | 0.999994 | 18030 | 1 | 0.77808 | 0.18468 | 0.40916 | 1 | 8277 | 4 | 0.77808 |
| BTN3A2 | 5 | 0.13032 | 0.27309 | 0.998561 | 5941 | 2 | 2.2656 | 0.18469 | 0.3998 | 1 | 8278 | 3 | 2.2656 |
| SEMA6D | 5 | 0.24268 | 0.43457 | 0.999994 | 8855 | 3 | -0.58902 | 0.18471 | 0.39983 | 1 | 8279 | 2 | -0.58902 |
| ARL13A | 5 | 0.24116 | 0.43249 | 0.999994 | 8800 | 2 | -0.19682 | 0.18471 | 0.39983 | 1 | 8280 | 1 | -0.19682 |
| TEKT5 | 6 | 0.94826 | 0.94829 | 0.999994 | 20291 | 0 | 1.5659 | 0.18479 | 0.40932 | 1 | 8281 | 4 | 1.5659 |
| TAS2R39 | 6 | 0.26685 | 0.46794 | 0.999994 | 9395 | 3 | -0.43549 | 0.1848 | 0.40934 | 1 | 8282 | 1 | -0.43549 |
| ALPK3 | 3 | 0.6696 | 0.67352 | 0.999994 | 15924 | 1 | 2.6447 | 0.18484 | 0.32654 | 0.991729 | 8283 | 2 | 2.6447 |
| hsa-mir-4530 | 3 | 0.36302 | 0.46299 | 0.999994 | 11389 | 1 | 2.9746 | 0.18489 | 0.3266 | 0.991729 | 8284 | 2 | 2.9746 |
| IARS | 4 | 0.073748 | 0.16697 | 0.967126 | 3982 | 2 | -0.93154 | 0.18501 | 0.36185 | 0.994802 | 8285 | 2 | -0.93154 |
| VEGFC | 6 | 0.11665 | 0.27942 | 0.999632 | 5529 | 1 | -0.016354 | 0.18504 | 0.40967 | 1 | 8286 | 1 | -0.016354 |
| TRIAP1 | 6 | 0.62444 | 0.75818 | 0.999994 | 15297 | 1 | 1.0445 | 0.18504 | 0.40967 | 1 | 8287 | 3 | 1.0445 |
| SEC13 | 3 | 0.098978 | 0.1817 | 0.968171 | 4930 | 1 | 3.0151 | 0.18508 | 0.32679 | 0.991729 | 8288 | 2 | 3.0151 |
| C11orf45 | 6 | 0.84394 | 0.8687 | 0.999994 | 18154 | 1 | 1.6064 | 0.18513 | 0.40978 | 1 | 8289 | 4 | 1.6064 |
| ARMCX4 | 6 | 0.73868 | 0.82454 | 0.999994 | 16716 | 1 | 1.7998 | 0.18514 | 0.4098 | 1 | 8290 | 3 | 1.7998 |
| NDUFC2-KCTD14 | 3 | 0.15998 | 0.26141 | 0.995051 | 6738 | 1 | 2.8447 | 0.18517 | 0.32689 | 0.991729 | 8291 | 2 | 2.8447 |
| LCE2A | 5 | 0.06032 | 0.16263 | 0.967126 | 3433 | 2 | 0.93425 | 0.18521 | 0.40039 | 1 | 8292 | 3 | 0.93425 |
| AKR1B10 | 6 | 0.12465 | 0.29348 | 0.999994 | 5786 | 3 | -0.57177 | 0.18523 | 0.40994 | 1 | 8293 | 2 | -0.57177 |
| CPT2 | 5 | 0.089375 | 0.20958 | 0.979587 | 4571 | 2 | 1.6713 | 0.18525 | 0.40045 | 1 | 8294 | 3 | 1.6713 |
| NonTargetingControlG | 2 | 0.81473 | 0.81453 | 0.999994 | 17698 | 0 | 3.2472 | 0.18527 | 0.29608 | 0.984434 | 8295 | 2 | 3.2472 |
| AUTS2 | 6 | 0.58048 | 0.72558 | 0.999994 | 14685 | 2 | 1.2362 | 0.1853 | 0.41004 | 1 | 8296 | 4 | 1.2362 |
| RALB | 6 | 0.015038 | 0.055841 | 0.9235 | 1190 | 3 | -0.43524 | 0.18545 | 0.41025 | 1 | 8297 | 3 | -0.43524 |
| NETO2 | 4 | 0.039145 | 0.10007 | 0.946394 | 2471 | 2 | -2.1108 | 0.18546 | 0.36249 | 0.995401 | 8298 | 2 | -2.1108 |
| MUC6 | 6 | 0.00065039 | 0.0033749 | 0.541338 | 123 | 5 | -2.4335 | 0.18559 | 0.41047 | 1 | 8299 | 1 | -2.4335 |
| SLC25A41 | 5 | 0.66772 | 0.75235 | 0.999994 | 15888 | 1 | -0.13655 | 0.18563 | 0.40089 | 1 | 8300 | 2 | -0.13655 |
| TAB2 | 5 | 0.734 | 0.78261 | 0.999994 | 16652 | 1 | 0.59203 | 0.18563 | 0.40089 | 1 | 8301 | 2 | 0.59203 |
| CAB39 | 6 | 0.017031 | 0.062376 | 0.937368 | 1305 | 2 | 1.2745 | 0.18576 | 0.4107 | 1 | 8302 | 4 | 1.2745 |
| hsa-mir-6804 | 3 | 0.86818 | 0.8682 | 0.999994 | 18603 | 0 | 3.0692 | 0.18579 | 0.32758 | 0.991729 | 8303 | 2 | 3.0692 |
| hsa-mir-4670 | 4 | 0.91401 | 0.91378 | 0.999994 | 19486 | 0 | 2.6061 | 0.1858 | 0.36298 | 0.995401 | 8304 | 3 | 2.6061 |
| MGLL | 4 | 0.054804 | 0.13083 | 0.95318 | 3189 | 1 | 1.8592 | 0.18581 | 0.363 | 0.995401 | 8305 | 3 | 1.8592 |
| hsa-mir-6758 | 4 | 0.48636 | 0.61031 | 0.999994 | 13233 | 1 | 2.804 | 0.18583 | 0.36303 | 0.995401 | 8306 | 3 | 2.804 |
| INHBB | 6 | 0.64743 | 0.77567 | 0.999994 | 15622 | 2 | -0.052097 | 0.18588 | 0.41087 | 1 | 8307 | 1 | -0.052097 |
| CNNM3 | 6 | 0.28642 | 0.49094 | 0.999994 | 9845 | 2 | 1.3313 | 0.18589 | 0.41089 | 1 | 8308 | 4 | 1.3313 |
| BZW1 | 5 | 0.91811 | 0.9182 | 0.999994 | 19588 | 0 | 0.51776 | 0.18593 | 0.40123 | 1 | 8309 | 2 | 0.51776 |
| TLR5 | 6 | 0.36524 | 0.58022 | 0.999994 | 11426 | 2 | 1.4544 | 0.18595 | 0.41098 | 1 | 8310 | 3 | 1.4544 |
| CLEC12A | 6 | 0.82637 | 0.8598 | 0.999994 | 17874 | 1 | 0.92988 | 0.18595 | 0.41098 | 1 | 8311 | 3 | 0.92988 |
| AQP3 | 4 | 0.76997 | 0.77575 | 0.999994 | 17080 | 1 | 2.9571 | 0.18599 | 0.36325 | 0.995401 | 8312 | 3 | 2.9571 |
| C17orf99 | 6 | 0.59415 | 0.73567 | 0.999994 | 14879 | 1 | 1.594 | 0.18607 | 0.41116 | 1 | 8313 | 4 | 1.594 |
| CYCS | 4 | 0.67113 | 0.70607 | 0.999994 | 15945 | 1 | 1.9563 | 0.1861 | 0.36341 | 0.995401 | 8314 | 3 | 1.9563 |
| NDUFB1 | 5 | 0.37731 | 0.54449 | 0.999994 | 11610 | 2 | 0.67609 | 0.18613 | 0.40144 | 1 | 8315 | 3 | 0.67609 |
| TLR2 | 5 | 0.24919 | 0.44015 | 0.999994 | 8996 | 2 | -0.24638 | 0.18613 | 0.40144 | 1 | 8316 | 2 | -0.24638 |
| CRK | 6 | 0.411 | 0.60955 | 0.999994 | 12129 | 2 | 1.4888 | 0.18616 | 0.41129 | 1 | 8317 | 4 | 1.4888 |
| ZFP41 | 6 | 0.057464 | 0.16028 | 0.967126 | 3312 | 3 | -0.99701 | 0.18616 | 0.41129 | 1 | 8318 | 1 | -0.99701 |
| COL12A1 | 5 | 0.71905 | 0.77541 | 0.999994 | 16486 | 1 | 2.2222 | 0.1862 | 0.40151 | 1 | 8319 | 3 | 2.2222 |

| | | | | | | | | | | | | |
|---|---|---|---|---|---|---|---|---|---|---|---|---|
| KDR | 4 | 0.6374 | 0.68601 | 0.999994 | 15478 | 1 | 2.1961 | 0.18622 | 0.36358 | 0.995496 | 8320 | 3 | 2.1961 |
| AP4S1 | 6 | 0.95754 | 0.95763 | 0.999994 | 20527 | 0 | 1.942 | 0.18629 | 0.41145 | 1 | 8321 | 4 | 1.942 |
| ABHD3 | 5 | 0.30178 | 0.48249 | 0.999994 | 10157 | 2 | 2.2777 | 0.18631 | 0.40165 | 1 | 8322 | 3 | 2.2777 |
| SGK223 | 6 | 0.068432 | 0.1835 | 0.968526 | 3767 | 2 | 1.8152 | 0.18637 | 0.41156 | 1 | 8323 | 4 | 1.8152 |
| hsa-mir-1277 | 4 | 0.27332 | 0.41573 | 0.999994 | 9550 | 1 | 1.677 | 0.18644 | 0.36391 | 0.995622 | 8324 | 3 | 1.677 |
| ZFP69B | 5 | 0.5284 | 0.67264 | 0.999994 | 13867 | 2 | 2.1394 | 0.18653 | 0.40188 | 1 | 8325 | 3 | 2.1394 |
| FAM200A | 5 | 0.79862 | 0.81867 | 0.999994 | 17456 | 1 | 2.2365 | 0.18653 | 0.40189 | 1 | 8326 | 3 | 2.2365 |
| SPHKAP | 6 | 0.061818 | 0.16961 | 0.967126 | 3505 | 1 | 1.0154 | 0.18657 | 0.41183 | 1 | 8327 | 3 | 1.0154 |
| ERCC6 | 2 | 0.33622 | 0.36497 | 0.999994 | 10854 | 1 | 2.3689 | 0.18662 | 0.29794 | 0.984876 | 8328 | 1 | 2.3689 |
| NonTargetingControlG | 2 | 0.20194 | 0.26351 | 0.996734 | 7826 | 1 | 1.505 | 0.18662 | 0.29794 | 0.984876 | 8329 | 1 | 1.505 |
| THSD1 | 6 | 0.65086 | 0.77828 | 0.999994 | 15673 | 2 | 1.5129 | 0.18667 | 0.41198 | 1 | 8330 | 4 | 1.5129 |
| NRDE2 | 5 | 0.65926 | 0.74884 | 0.999994 | 15765 | 1 | 2.4234 | 0.18676 | 0.40215 | 1 | 8331 | 3 | 2.4234 |
| DUSP12 | 6 | 0.47944 | 0.65483 | 0.999994 | 13140 | 2 | 0.2283 | 0.1868 | 0.41215 | 1 | 8332 | 3 | 0.2283 |
| SMIM15 | 6 | 0.20195 | 0.39025 | 0.999994 | 7827 | 2 | 1.0664 | 0.1868 | 0.41215 | 1 | 8333 | 3 | 1.0664 |
| MBP | 6 | 0.60626 | 0.74459 | 0.999994 | 15031 | 2 | 0.49266 | 0.1868 | 0.41215 | 1 | 8334 | 2 | 0.49266 |
| IL12RB1 | 4 | 0.046069 | 0.11376 | 0.951052 | 2813 | 3 | -2.3556 | 0.18687 | 0.36453 | 0.995622 | 8335 | 1 | -2.3556 |
| CCL17 | 4 | 0.9408 | 0.94069 | 0.999994 | 20101 | 0 | 2.5858 | 0.1869 | 0.36458 | 0.995622 | 8336 | 3 | 2.5858 |
| NPS | 5 | 0.0081994 | 0.03088 | 0.862223 | 744 | 1 | 2.1262 | 0.18691 | 0.40232 | 1 | 8337 | 3 | 2.1262 |
| MADCAM1 | 3 | 0.6509 | 0.65782 | 0.999994 | 15675 | 1 | 3.1527 | 0.18693 | 0.32883 | 0.991729 | 8338 | 2 | 3.1527 |
| MSMO1 | 6 | 0.93317 | 0.93325 | 0.999994 | 19930 | 0 | 2.0827 | 0.18695 | 0.41237 | 1 | 8339 | 4 | 2.0827 |
| ALOX15B | 5 | 0.56652 | 0.70647 | 0.999994 | 14488 | 2 | 1.2634 | 0.18703 | 0.40244 | 1 | 8340 | 3 | 1.2634 |
| RNF185 | 6 | 0.90925 | 0.91096 | 0.999994 | 19407 | 1 | 0.1084 | 0.18714 | 0.41262 | 1 | 8341 | 2 | 0.1084 |
| SYNE2 | 6 | 0.14395 | 0.31805 | 0.999994 | 6298 | 3 | -0.34511 | 0.18714 | 0.41262 | 1 | 8342 | 2 | -0.34511 |
| SOX18 | 6 | 0.71721 | 0.81742 | 0.999994 | 16471 | 1 | 0.68782 | 0.18714 | 0.41262 | 1 | 8343 | 4 | 0.68782 |
| TSNARE1 | 5 | 0.77089 | 0.80221 | 0.999994 | 17095 | 1 | -0.13494 | 0.18714 | 0.40256 | 1 | 8344 | 2 | -0.13494 |
| CPE | 6 | 0.87622 | 0.88722 | 0.999994 | 18757 | 1 | 2.085 | 0.18719 | 0.4127 | 1 | 8345 | 4 | 2.085 |
| WTIP | 6 | 0.9097 | 0.91131 | 0.999994 | 19420 | 1 | 1.5043 | 0.18722 | 0.41274 | 1 | 8346 | 4 | 1.5043 |
| hsa-mir-519b | 2 | 0.81277 | 0.81248 | 0.999994 | 17667 | 0 | 1.2864 | 0.18723 | 0.29879 | 0.984876 | 8347 | 2 | 1.2864 |
| MPP4 | 6 | 0.11739 | 0.2808 | 0.999632 | 5555 | 2 | 2.1049 | 0.18725 | 0.41278 | 1 | 8348 | 4 | 2.1049 |
| OR5M10 | 6 | 0.099212 | 0.24583 | 0.989866 | 4938 | 4 | -1.1182 | 0.18731 | 0.41287 | 1 | 8349 | 2 | -1.1182 |
| AACS | 6 | 0.061332 | 0.16861 | 0.967126 | 3487 | 3 | -1.3899 | 0.18735 | 0.41292 | 1 | 8350 | 2 | -1.3899 |
| MARCO | 5 | 0.21942 | 0.40246 | 0.999994 | 8274 | 3 | -0.73295 | 0.18739 | 0.40284 | 1 | 8351 | 2 | -0.73295 |
| PLA2G5 | 6 | 0.42705 | 0.61996 | 0.999994 | 12367 | 2 | 1.7461 | 0.18742 | 0.41303 | 1 | 8352 | 4 | 1.7461 |
| CPZ | 6 | 0.56078 | 0.71134 | 0.999994 | 14382 | 1 | 1.9036 | 0.18745 | 0.41308 | 1 | 8353 | 4 | 1.9036 |
| hsa-mir-6862-2 | 2 | 0.81253 | 0.81225 | 0.999994 | 17661 | 0 | 3.4075 | 0.18747 | 0.29911 | 0.984876 | 8354 | 2 | 3.4075 |
| EPHB3 | 6 | 0.60982 | 0.7472 | 0.999994 | 15069 | 2 | 1.7841 | 0.1875 | 0.41316 | 1 | 8355 | 3 | 1.7841 |
| F12 | 6 | 0.12271 | 0.29083 | 0.999632 | 5731 | 3 | 0.74626 | 0.18754 | 0.41321 | 1 | 8356 | 3 | 0.74626 |
| PNMT | 6 | 0.44236 | 0.62999 | 0.999994 | 12616 | 1 | 0.58985 | 0.18754 | 0.41321 | 1 | 8357 | 3 | 0.58985 |
| CCBP2 | 4 | 0.0039277 | 0.013966 | 0.75278 | 438 | 2 | -4.3256 | 0.1877 | 0.36573 | 0.996287 | 8358 | 1 | -4.3256 |
| hsa-mir-3659 | 4 | 0.81645 | 0.8164 | 0.999994 | 17720 | 1 | 1.775 | 0.1877 | 0.36573 | 0.996287 | 8359 | 3 | 1.775 |
| POMZP3 | 6 | 0.13435 | 0.30585 | 0.999994 | 6033 | 3 | -0.31395 | 0.1878 | 0.41357 | 1 | 8360 | 2 | -0.31395 |
| C12orf42 | 6 | 0.099346 | 0.2461 | 0.98999 | 4943 | 4 | -0.94195 | 0.1878 | 0.41357 | 1 | 8361 | 2 | -0.94195 |
| GPC1 | 6 | 0.22902 | 0.4229 | 0.999994 | 8505 | 3 | 0.3146 | 0.1878 | 0.41357 | 1 | 8362 | 3 | 0.3146 |
| MAGEB17 | 6 | 0.38131 | 0.59041 | 0.999994 | 11667 | 2 | -0.10802 | 0.1878 | 0.41357 | 1 | 8363 | 2 | -0.10802 |
| DEFB128 | 6 | 0.19812 | 0.38562 | 0.999994 | 7715 | 2 | 0.71818 | 0.1878 | 0.41357 | 1 | 8364 | 3 | 0.71818 |
| NRG4 | 5 | 0.21405 | 0.39502 | 0.999994 | 8117 | 2 | 2.2215 | 0.18783 | 0.40332 | 1 | 8365 | 3 | 2.2215 |
| BANK1 | 5 | 0.86999 | 0.87107 | 0.999994 | 18636 | 1 | 0.6761 | 0.18783 | 0.40332 | 1 | 8366 | 3 | 0.6761 |
| CYP2C8 | 5 | 0.3712 | 0.5394 | 0.999994 | 11532 | 1 | 0.4036 | 0.18783 | 0.40332 | 1 | 8367 | 2 | 0.4036 |
| hsa-mir-3202-1 | 4 | 0.54008 | 0.63491 | 0.999994 | 14053 | 1 | 1.9037 | 0.18784 | 0.36592 | 0.996287 | 8368 | 3 | 1.9037 |
| HIPK4 | 6 | 0.95014 | 0.95012 | 0.999994 | 20338 | 0 | 1.1349 | 0.18787 | 0.41368 | 1 | 8369 | 4 | 1.1349 |
| PSD4 | 6 | 0.096123 | 0.23972 | 0.987253 | 4819 | 4 | -0.84114 | 0.1879 | 0.41373 | 1 | 8370 | 1 | -0.84114 |
| RBBP9 | 6 | 0.074814 | 0.19678 | 0.977164 | 4029 | 3 | -0.56217 | 0.1879 | 0.41373 | 1 | 8371 | 2 | -0.56217 |

| | | | | | | | | | | | | | |
|---|---|---|---|---|---|---|---|---|---|---|---|---|---|
| hsa-mir-4507 | 2 | 0.42886 | 0.44111 | 0.999994 | 12392 | 1 | 2.8188 | 0.18797 | 0.29985 | 0.984876 | 8372 | 1 | 2.8188 |
| NonTargetingControlG | 2 | 0.81202 | 0.81174 | 0.999994 | 17655 | 0 | 1.0824 | 0.18798 | 0.29988 | 0.984876 | 8373 | 2 | 1.0824 |
| ADRA1A | 5 | 0.57474 | 0.71371 | 0.999994 | 14613 | 1 | 0.63871 | 0.18809 | 0.40361 | 1 | 8374 | 3 | 0.63871 |
| SCRIB | 5 | 0.070166 | 0.17864 | 0.968171 | 3840 | 2 | -0.55578 | 0.18809 | 0.40361 | 1 | 8375 | 2 | -0.55578 |
| SLC28A1 | 5 | 0.072743 | 0.18285 | 0.968526 | 3951 | 2 | -0.57001 | 0.18809 | 0.40361 | 1 | 8376 | 2 | -0.57001 |
| MYO1H | 6 | 0.6077 | 0.74565 | 0.999994 | 15043 | 1 | 1.6813 | 0.18818 | 0.41415 | 1 | 8377 | 4 | 1.6813 |
| TAF7L | 6 | 0.24875 | 0.44644 | 0.999994 | 8986 | 2 | 1.5271 | 0.18822 | 0.41419 | 1 | 8378 | 4 | 1.5271 |
| CD300C | 6 | 0.87455 | 0.88616 | 0.999994 | 18723 | 1 | 1.6048 | 0.18828 | 0.41427 | 1 | 8379 | 4 | 1.6048 |
| RNASE11 | 6 | 0.25772 | 0.45719 | 0.999994 | 9186 | 2 | 1.643 | 0.18832 | 0.41434 | 1 | 8380 | 4 | 1.643 |
| LGALS8 | 6 | 0.57356 | 0.72051 | 0.999994 | 14601 | 2 | 2.1136 | 0.18839 | 0.41443 | 1 | 8381 | 4 | 2.1136 |
| ROCK1 | 5 | 0.4793 | 0.63004 | 0.999994 | 13134 | 2 | 0.54993 | 0.18841 | 0.40396 | 1 | 8382 | 3 | 0.54993 |
| FAM117B | 4 | 0.0010403 | 0.0038235 | 0.560963 | 173 | 3 | -5.5876 | 0.1885 | 0.36684 | 0.997075 | 8383 | 1 | -5.5876 |
| MS4A8 | 6 | 0.17925 | 0.36221 | 0.999994 | 7206 | 1 | 2.0969 | 0.18856 | 0.41465 | 1 | 8384 | 4 | 2.0969 |
| NonTargetingControlG | 2 | 0.81144 | 0.81115 | 0.999994 | 17650 | 0 | 1.4898 | 0.18856 | 0.30071 | 0.984876 | 8385 | 2 | 1.4898 |
| DPM3 | 6 | 0.39227 | 0.5974 | 0.999994 | 11837 | 1 | 1.2193 | 0.18856 | 0.41466 | 1 | 8386 | 3 | 1.2193 |
| SUV420H2 | 6 | 0.09955 | 0.24649 | 0.98999 | 4956 | 2 | 0.71621 | 0.18856 | 0.41466 | 1 | 8387 | 3 | 0.71621 |
| NonTargetingControlG | 1 | 0.81135 | 0.81152 | 0.999994 | 17649 | 0 | 5.2684 | 0.18865 | 0.18849 | 0.97466 | 8388 | 1 | 5.2684 |
| HNRPLL | 3 | 0.61794 | 0.63107 | 0.999994 | 15184 | 1 | 0.57058 | 0.18867 | 0.33071 | 0.992034 | 8389 | 1 | 0.57058 |
| RTCB | 3 | 0.51822 | 0.5583 | 0.999994 | 13715 | 1 | 1.8508 | 0.18867 | 0.33071 | 0.992034 | 8390 | 2 | 1.8508 |
| SNX16 | 4 | 0.20092 | 0.34162 | 0.999994 | 7788 | 1 | 1.9742 | 0.18879 | 0.36726 | 0.997075 | 8391 | 3 | 1.9742 |
| hsa-mir-506 | 4 | 0.92676 | 0.92656 | 0.999994 | 19783 | 0 | 1.8799 | 0.18887 | 0.36738 | 0.99708 | 8392 | 3 | 1.8799 |
| TRIM5 | 5 | 0.237 | 0.42688 | 0.999994 | 8695 | 3 | -0.62956 | 0.18898 | 0.40461 | 1 | 8393 | 2 | -0.62956 |
| PIK3CD | 5 | 0.0059451 | 0.024262 | 0.837944 | 587 | 4 | -2.4808 | 0.18898 | 0.40461 | 1 | 8394 | 1 | -2.4808 |
| CLCN6 | 6 | 0.16609 | 0.34582 | 0.999994 | 6886 | 1 | 1.1038 | 0.18899 | 0.41529 | 1 | 8395 | 4 | 1.1038 |
| XCL2 | 3 | 0.26034 | 0.38781 | 0.999994 | 9249 | 2 | -0.73954 | 0.18906 | 0.33117 | 0.992034 | 8396 | 1 | -0.73954 |
| AK5 | 5 | 0.36912 | 0.53761 | 0.999994 | 11493 | 1 | 2.4812 | 0.1891 | 0.40476 | 1 | 8397 | 3 | 2.4812 |
| SPIN1 | 5 | 0.055103 | 0.15273 | 0.967126 | 3204 | 2 | 2.4812 | 0.1891 | 0.40476 | 1 | 8398 | 3 | 2.4812 |
| SMYD1 | 5 | 0.29108 | 0.47381 | 0.999994 | 9931 | 2 | 0.45248 | 0.18912 | 0.40479 | 1 | 8399 | 2 | 0.45248 |
| NOL10 | 5 | 0.92127 | 0.92144 | 0.999994 | 19659 | 0 | 1.9672 | 0.18912 | 0.40479 | 1 | 8400 | 3 | 1.9672 |
| AKR1D1 | 6 | 0.86391 | 0.87966 | 0.999994 | 18520 | 1 | -0.048588 | 0.18928 | 0.41571 | 1 | 8401 | 1 | -0.048588 |
| WDR47 | 6 | 0.70235 | 0.81273 | 0.999994 | 16332 | 1 | -0.3345 | 0.18928 | 0.41571 | 1 | 8402 | 2 | -0.3345 |
| ENC1 | 5 | 0.93827 | 0.93846 | 0.999994 | 20032 | 0 | 1.7625 | 0.1894 | 0.40509 | 1 | 8403 | 3 | 1.7625 |
| HIST1H2BM | 6 | 0.58773 | 0.73092 | 0.999994 | 14787 | 2 | 0.93438 | 0.18948 | 0.41599 | 1 | 8404 | 4 | 0.93438 |
| TK1 | 6 | 0.08764 | 0.22276 | 0.983354 | 4499 | 3 | -0.81397 | 0.18948 | 0.41599 | 1 | 8405 | 2 | -0.81397 |
| RALGAPA2 | 6 | 0.45818 | 0.64051 | 0.999994 | 12823 | 2 | 0.03535 | 0.18948 | 0.41599 | 1 | 8406 | 2 | 0.03535 |
| NDUFA8 | 6 | 0.15999 | 0.33822 | 0.999994 | 6740 | 3 | -0.21396 | 0.18948 | 0.41599 | 1 | 8407 | 3 | -0.21396 |
| ABCC11 | 6 | 0.26948 | 0.4711 | 0.999994 | 9461 | 2 | 1.0933 | 0.18948 | 0.41599 | 1 | 8408 | 4 | 1.0933 |
| GC | 6 | 0.72436 | 0.81977 | 0.999994 | 16540 | 1 | 2.1834 | 0.1895 | 0.41602 | 1 | 8409 | 4 | 2.1834 |
| hsa-mir-4289 | 4 | 0.86345 | 0.86309 | 0.999994 | 18513 | 0 | 2.6307 | 0.18965 | 0.3685 | 0.998341 | 8410 | 2 | 2.6307 |
| NPFF | 4 | 0.27834 | 0.42088 | 0.999994 | 9659 | 2 | 1.6845 | 0.18965 | 0.3685 | 0.998341 | 8411 | 2 | 1.6845 |
| SCARA5 | 6 | 0.52511 | 0.6861 | 0.999994 | 13818 | 1 | 1.4975 | 0.18966 | 0.41622 | 1 | 8412 | 3 | 1.4975 |
| MDH1 | 5 | 0.34147 | 0.51494 | 0.999994 | 10969 | 2 | -0.055561 | 0.18968 | 0.40541 | 1 | 8413 | 1 | -0.055561 |
| hsa-mir-7849 | 4 | 0.8137 | 0.81378 | 0.999994 | 17686 | 1 | 0.3362 | 0.1897 | 0.36858 | 0.998341 | 8414 | 1 | 0.3362 |
| HGFAC | 5 | 0.23461 | 0.42359 | 0.999994 | 8630 | 2 | 2.3693 | 0.18971 | 0.40544 | 1 | 8415 | 3 | 2.3693 |
| C11orf63 | 6 | 0.51412 | 0.67846 | 0.999994 | 13645 | 1 | 2.1983 | 0.18975 | 0.41635 | 1 | 8416 | 4 | 2.1983 |
| RABEPK | 5 | 0.72159 | 0.77661 | 0.999994 | 16513 | 1 | 2.0859 | 0.18976 | 0.40549 | 1 | 8417 | 3 | 2.0859 |
| AP2S1 | 5 | 0.43839 | 0.59547 | 0.999994 | 12545 | 2 | 2.1495 | 0.18984 | 0.40558 | 1 | 8418 | 3 | 2.1495 |
| BNIP2 | 6 | 0.40392 | 0.60494 | 0.999994 | 12018 | 1 | 0.45086 | 0.18987 | 0.41655 | 1 | 8419 | 3 | 0.45086 |
| SLC4A9 | 5 | 0.49417 | 0.64286 | 0.999994 | 13333 | 2 | 2.141 | 0.1899 | 0.40565 | 1 | 8420 | 3 | 2.141 |
| HEXIM2 | 3 | 0.8015 | 0.80132 | 0.999994 | 17498 | 0 | 3.1166 | 0.18993 | 0.33214 | 0.992196 | 8421 | 2 | 3.1166 |
| hsa-mir-4711 | 4 | 0.63185 | 0.6829 | 0.999994 | 15396 | 1 | 1.6167 | 0.18995 | 0.36894 | 0.998341 | 8422 | 3 | 1.6167 |
| FBXW9 | 6 | 0.50191 | 0.67002 | 0.999994 | 13458 | 2 | 0.91802 | 0.18998 | 0.41669 | 1 | 8423 | 3 | 0.91802 |

| | | | | | | | | | | | | | |
|---|---|---|---|---|---|---|---|---|---|---|---|---|---|
| NADSYN1 | 5 | 0.099508 | 0.22565 | 0.983354 | 4955 | 2 | 0.36752 | 0.19002 | 0.40579 | 1 | 8424 | 1 | 0.36752 |
| GOLGA6A | 5 | 0.86833 | 0.86961 | 0.999994 | 18608 | 1 | 1.5659 | 0.19002 | 0.40579 | 1 | 8425 | 3 | 1.5659 |
| MAD2L2 | 5 | 0.2863 | 0.46993 | 0.999994 | 9840 | 2 | 0.33752 | 0.19005 | 0.40582 | 1 | 8426 | 2 | 0.33752 |
| SAP30BP | 5 | 0.17957 | 0.34601 | 0.999994 | 7213 | 1 | 1.3455 | 0.19005 | 0.40582 | 1 | 8427 | 3 | 1.3455 |
| TIGD4 | 6 | 0.97532 | 0.97525 | 0.999994 | 20967 | 0 | 2.1819 | 0.19005 | 0.41679 | 1 | 8428 | 4 | 2.1819 |
| NCOA3 | 6 | 0.042988 | 0.12865 | 0.951777 | 2678 | 3 | -1.6558 | 0.19006 | 0.41679 | 1 | 8429 | 2 | -1.6558 |
| CPLX4 | 6 | 0.51185 | 0.67688 | 0.999994 | 13608 | 2 | 1.304 | 0.19009 | 0.41684 | 1 | 8430 | 4 | 1.304 |
| CPM | 6 | 0.9295 | 0.92963 | 0.999994 | 19843 | 0 | 1.7984 | 0.19011 | 0.41688 | 1 | 8431 | 4 | 1.7984 |
| SPATA2L | 4 | 0.53552 | 0.63272 | 0.999994 | 13989 | 1 | 2.5411 | 0.19018 | 0.36926 | 0.998341 | 8432 | 3 | 2.5411 |
| AS3MT | 6 | 0.24524 | 0.44226 | 0.999994 | 8905 | 2 | 1.4713 | 0.19029 | 0.41716 | 1 | 8433 | 4 | 1.4713 |
| SH2D5 | 6 | 0.21267 | 0.40334 | 0.999994 | 8080 | 2 | 0.80588 | 0.1903 | 0.41717 | 1 | 8434 | 3 | 0.80588 |
| SCAMP3 | 4 | 0.51606 | 0.62359 | 0.999994 | 13677 | 1 | 2.363 | 0.19039 | 0.36955 | 0.9988 | 8435 | 2 | 2.363 |
| LGI1 | 6 | 0.7658 | 0.83426 | 0.999994 | 17036 | 1 | 1.4226 | 0.19042 | 0.41735 | 1 | 8436 | 3 | 1.4226 |
| ZBTB21 | 6 | 0.018365 | 0.066697 | 0.93914 | 1379 | 3 | -0.24577 | 0.19043 | 0.41736 | 1 | 8437 | 2 | -0.24577 |
| HSF2BP | 6 | 0.5565 | 0.70831 | 0.999994 | 14310 | 2 | -0.17111 | 0.19043 | 0.41736 | 1 | 8438 | 2 | -0.17111 |
| GLRX5 | 6 | 0.23105 | 0.42535 | 0.999994 | 8543 | 2 | 1.1072 | 0.19044 | 0.41738 | 1 | 8439 | 3 | 1.1072 |
| MARCH8 | 5 | 0.33994 | 0.51366 | 0.999994 | 10929 | 2 | -0.0028704 | 0.19046 | 0.40628 | 1 | 8440 | 2 | -0.0028704 |
| RPL26 | 3 | 0.73032 | 0.7302 | 0.999994 | 16602 | 1 | 1.5208 | 0.19081 | 0.33308 | 0.992917 | 8441 | 2 | 1.5208 |
| PLEKHG7 | 6 | 0.78474 | 0.84159 | 0.999994 | 17276 | 1 | 0.39658 | 0.19087 | 0.41799 | 1 | 8442 | 3 | 0.39658 |
| KCNH7 | 6 | 0.21581 | 0.40712 | 0.999994 | 8167 | 3 | -0.67484 | 0.19087 | 0.41799 | 1 | 8443 | 2 | -0.67484 |
| PIGN | 4 | 0.039876 | 0.10155 | 0.946414 | 2509 | 2 | -0.17981 | 0.19088 | 0.37027 | 0.9988 | 8444 | 2 | -0.17981 |
| NonTargetingControlG | 2 | 0.80911 | 0.80878 | 0.999994 | 17618 | 0 | 2.1592 | 0.19089 | 0.30394 | 0.986296 | 8445 | 2 | 2.1592 |
| ARID3C | 5 | 0.85536 | 0.85855 | 0.999994 | 18354 | 1 | 2.0572 | 0.1909 | 0.40679 | 1 | 8446 | 3 | 2.0572 |
| PHACTR4 | 6 | 0.08785 | 0.22319 | 0.983354 | 4505 | 3 | 0.98761 | 0.19091 | 0.41806 | 1 | 8447 | 3 | 0.98761 |
| AADACL4 | 6 | 0.64562 | 0.77425 | 0.999994 | 15595 | 2 | 2.1211 | 0.19091 | 0.41806 | 1 | 8448 | 4 | 2.1211 |
| OR5K1 | 4 | 0.54053 | 0.63513 | 0.999994 | 14066 | 1 | 1.5255 | 0.19101 | 0.37045 | 0.9988 | 8449 | 3 | 1.5255 |
| RNF135 | 2 | 0.79559 | 0.79515 | 0.999994 | 17426 | 0 | 3.9881 | 0.19102 | 0.30414 | 0.986296 | 8450 | 2 | 3.9881 |
| NonTargetingControlG | 2 | 0.80896 | 0.80863 | 0.999994 | 17617 | 0 | 1.5973 | 0.19104 | 0.30417 | 0.986296 | 8451 | 2 | 1.5973 |
| RAB3C | 6 | 0.76501 | 0.83396 | 0.999994 | 17027 | 1 | 1.1959 | 0.19106 | 0.41828 | 1 | 8452 | 4 | 1.1959 |
| EPHX4 | 5 | 0.36572 | 0.53483 | 0.999994 | 11434 | 2 | 1.2806 | 0.19111 | 0.407 | 1 | 8453 | 3 | 1.2806 |
| SLC17A5 | 5 | 0.9415 | 0.94169 | 0.999994 | 20119 | 0 | 1.5605 | 0.19111 | 0.407 | 1 | 8454 | 3 | 1.5605 |
| C3orf55 | 6 | 0.58665 | 0.73011 | 0.999994 | 14769 | 1 | 1.534 | 0.19111 | 0.41835 | 1 | 8455 | 4 | 1.534 |
| FGF1 | 6 | 0.2925 | 0.49795 | 0.999994 | 9965 | 2 | 1.7892 | 0.19114 | 0.4184 | 1 | 8456 | 4 | 1.7892 |
| UTS2B | 6 | 0.83791 | 0.86557 | 0.999994 | 18052 | 1 | 1.3221 | 0.19115 | 0.41842 | 1 | 8457 | 4 | 1.3221 |
| FNIP2 | 6 | 0.6017 | 0.7412 | 0.999994 | 14970 | 2 | 0.94322 | 0.19116 | 0.41842 | 1 | 8458 | 4 | 0.94322 |
| MLIP | 6 | 0.12902 | 0.29904 | 0.999994 | 5916 | 3 | -0.068567 | 0.19116 | 0.41842 | 1 | 8459 | 3 | -0.068567 |
| GSS | 4 | 0.87133 | 0.87094 | 0.999994 | 18666 | 0 | 2.3487 | 0.19118 | 0.37071 | 0.998982 | 8460 | 2 | 2.3487 |
| SS18 | 6 | 0.12597 | 0.29521 | 0.999994 | 5819 | 3 | -0.069321 | 0.19122 | 0.41851 | 1 | 8461 | 3 | -0.069321 |
| FGFR2 | 5 | 0.51453 | 0.66056 | 0.999994 | 13654 | 2 | 2.2402 | 0.19125 | 0.40715 | 1 | 8462 | 3 | 2.2402 |
| ALOX5AP | 6 | 0.14516 | 0.31957 | 0.999994 | 6327 | 1 | 2.0029 | 0.19126 | 0.41856 | 1 | 8463 | 4 | 2.0029 |
| KDM1B | 5 | 0.82884 | 0.83867 | 0.999994 | 17908 | 1 | 1.1127 | 0.19129 | 0.40721 | 1 | 8464 | 3 | 1.1127 |
| CLEC2L | 5 | 0.083569 | 0.20034 | 0.977164 | 4357 | 3 | -1.6272 | 0.19129 | 0.40721 | 1 | 8465 | 1 | -1.6272 |
| CRABP1 | 6 | 0.64125 | 0.77087 | 0.999994 | 15535 | 1 | 1.542 | 0.19135 | 0.41869 | 1 | 8466 | 4 | 1.542 |
| CTBS | 6 | 0.97046 | 0.97043 | 0.999994 | 20835 | 0 | 1.6573 | 0.19144 | 0.4188 | 1 | 8467 | 4 | 1.6573 |
| KLHL18 | 6 | 0.074901 | 0.19696 | 0.977164 | 4032 | 3 | 0.30459 | 0.19144 | 0.41881 | 1 | 8468 | 3 | 0.30459 |
| YIPF5 | 5 | 0.74424 | 0.78784 | 0.999994 | 16790 | 1 | 2.2549 | 0.19148 | 0.40741 | 1 | 8469 | 3 | 2.2549 |
| NXF5 | 5 | 0.84629 | 0.85143 | 0.999994 | 18193 | 1 | 0.2125 | 0.19155 | 0.40748 | 1 | 8470 | 2 | 0.2125 |
| ANKRD20A1 | 5 | 0.21038 | 0.38991 | 0.999994 | 8030 | 3 | -0.81431 | 0.19155 | 0.40748 | 1 | 8471 | 2 | -0.81431 |
| CLCN4 | 5 | 0.41679 | 0.57745 | 0.999994 | 12203 | 2 | -0.24129 | 0.19155 | 0.40748 | 1 | 8472 | 2 | -0.24129 |
| RTN1 | 5 | 0.23742 | 0.42745 | 0.999994 | 8712 | 2 | 0.18042 | 0.19155 | 0.40748 | 1 | 8473 | 2 | 0.18042 |
| EIF2B3 | 4 | 0.66541 | 0.70258 | 0.999994 | 15860 | 1 | 2.0978 | 0.19157 | 0.37127 | 0.998994 | 8474 | 3 | 2.0978 |
| OR5AP2 | 6 | 0.28611 | 0.49058 | 0.999994 | 9835 | 2 | 1.4549 | 0.19161 | 0.41904 | 1 | 8475 | 4 | 1.4549 |

| | | | | | | | | | | | | | |
|---|---|---|---|---|---|---|---|---|---|---|---|---|---|
| ZBTB48 | 6 | 0.74454 | 0.82655 | 0.999994 | 16795 | 1 | 0.37397 | 0.19165 | 0.4191 | 1 | 8476 | 3 | 0.37397 |
| CITED1 | 6 | 0.24499 | 0.44197 | 0.999994 | 8900 | 3 | 0.30552 | 0.19165 | 0.4191 | 1 | 8477 | 3 | 0.30552 |
| RUNDC3B | 6 | 0.58473 | 0.7287 | 0.999994 | 14739 | 2 | 0.95231 | 0.19165 | 0.4191 | 1 | 8478 | 4 | 0.95231 |
| TFPT | 4 | 0.49026 | 0.612 | 0.999994 | 13282 | 1 | 1.3049 | 0.19171 | 0.37149 | 0.998994 | 8479 | 3 | 1.3049 |
| GAL3ST1 | 5 | 0.58117 | 0.71934 | 0.999994 | 14695 | 1 | 2.4619 | 0.19179 | 0.40777 | 1 | 8480 | 3 | 2.4619 |
| TPGS1 | 6 | 0.25531 | 0.45429 | 0.999994 | 9132 | 1 | 2.0949 | 0.1918 | 0.41932 | 1 | 8481 | 4 | 2.0949 |
| MAP3K3 | 5 | 0.065239 | 0.17064 | 0.967177 | 3661 | 2 | 0.84744 | 0.19183 | 0.40781 | 1 | 8482 | 3 | 0.84744 |
| DAP | 6 | 0.3169 | 0.52633 | 0.999994 | 10451 | 2 | 1.484 | 0.19189 | 0.41945 | 1 | 8483 | 3 | 1.484 |
| ZNF227 | 6 | 0.95583 | 0.95591 | 0.999994 | 20474 | 0 | 0.26363 | 0.19193 | 0.41949 | 1 | 8484 | 2 | 0.26363 |
| EGLN1 | 5 | 0.75657 | 0.7943 | 0.999994 | 16939 | 1 | 0.79192 | 0.19199 | 0.40798 | 1 | 8485 | 3 | 0.79192 |
| hsa-mir-4645 | 4 | 0.87114 | 0.87077 | 0.999994 | 18659 | 0 | 1.2199 | 0.19201 | 0.37192 | 0.998994 | 8486 | 3 | 1.2199 |
| NonTargetingControlG | 2 | 0.80798 | 0.80767 | 0.999994 | 17600 | 0 | 1.0445 | 0.19202 | 0.30548 | 0.986322 | 8487 | 2 | 1.0445 |
| PARP12 | 6 | 0.14968 | 0.32523 | 0.999994 | 6449 | 2 | 1.5111 | 0.19207 | 0.4197 | 1 | 8488 | 4 | 1.5111 |
| TCEA3 | 6 | 0.11718 | 0.28042 | 0.999632 | 5545 | 1 | 1.3321 | 0.19212 | 0.41978 | 1 | 8489 | 4 | 1.3321 |
| NonTargetingControlG | 1 | 0.80786 | 0.80804 | 0.999994 | 17596 | 0 | 5.1892 | 0.19214 | 0.19197 | 0.97466 | 8490 | 1 | 5.1892 |
| USP17L13 | 4 | 0.20756 | 0.3485 | 0.999994 | 7962 | 2 | -0.78606 | 0.19218 | 0.37218 | 0.998994 | 8491 | 2 | -0.78606 |
| OR2A14 | 5 | 0.44095 | 0.59758 | 0.999994 | 12591 | 2 | -0.15618 | 0.19218 | 0.4082 | 1 | 8492 | 2 | -0.15618 |
| NonTargetingControlG | 2 | 0.8078 | 0.8075 | 0.999994 | 17594 | 0 | 1.0742 | 0.1922 | 0.30575 | 0.986404 | 8493 | 2 | 1.0742 |
| hsa-mir-224 | 3 | 0.33656 | 0.44828 | 0.999994 | 10863 | 1 | 2.9838 | 0.1922 | 0.3346 | 0.993364 | 8494 | 2 | 2.9838 |
| MICAL3 | 5 | 0.67018 | 0.75337 | 0.999994 | 15935 | 1 | 1.0224 | 0.19221 | 0.40822 | 1 | 8495 | 3 | 1.0224 |
| SRC | 5 | 0.85522 | 0.85842 | 0.999994 | 18351 | 1 | 2.1272 | 0.19221 | 0.40822 | 1 | 8496 | 3 | 2.1272 |
| THEMIS | 5 | 0.583 | 0.72006 | 0.999994 | 14715 | 1 | -0.12985 | 0.19221 | 0.40822 | 1 | 8497 | 2 | -0.12985 |
| hsa-mir-4777 | 4 | 0.58046 | 0.655 | 0.999994 | 14684 | 1 | 2.1647 | 0.19223 | 0.37225 | 0.998994 | 8498 | 2 | 2.1647 |
| BROX | 6 | 0.65626 | 0.78241 | 0.999994 | 15730 | 2 | 2.0738 | 0.19233 | 0.42009 | 1 | 8499 | 4 | 2.0738 |
| OR6B3 | 6 | 0.86278 | 0.87898 | 0.999994 | 18500 | 1 | 1.9262 | 0.19238 | 0.42016 | 1 | 8500 | 4 | 1.9262 |
| PRODH2 | 6 | 0.45397 | 0.63771 | 0.999994 | 12759 | 2 | 0.92317 | 0.19238 | 0.42016 | 1 | 8501 | 3 | 0.92317 |
| NonTargetingControlG | 2 | 0.32769 | 0.3582 | 0.999994 | 10677 | 1 | 2.3048 | 0.19242 | 0.30607 | 0.98674 | 8502 | 1 | 2.3048 |
| EDN1 | 4 | 0.61148 | 0.67157 | 0.999994 | 15096 | 1 | 2.1104 | 0.19243 | 0.37254 | 0.999063 | 8503 | 3 | 2.1104 |
| WNT8B | 6 | 0.10452 | 0.25616 | 0.994943 | 5114 | 2 | 1.7238 | 0.19251 | 0.42034 | 1 | 8504 | 3 | 1.7238 |
| hsa-mir-642a | 4 | 0.68003 | 0.7116 | 0.999994 | 16090 | 1 | 1.1933 | 0.19274 | 0.37297 | 0.999453 | 8505 | 3 | 1.1933 |
| INSL6 | 6 | 0.17516 | 0.35706 | 0.999994 | 7114 | 2 | 1.6274 | 0.19276 | 0.42069 | 1 | 8506 | 3 | 1.6274 |
| UBN1 | 6 | 0.9461 | 0.94606 | 0.999994 | 20233 | 0 | 0.92273 | 0.19279 | 0.42073 | 1 | 8507 | 3 | 0.92273 |
| hsa-mir-1244-1 | 4 | 0.88297 | 0.88261 | 0.999994 | 18892 | 0 | 1.2269 | 0.19282 | 0.37309 | 0.999517 | 8508 | 3 | 1.2269 |
| AGO3 | 6 | 0.59808 | 0.73852 | 0.999994 | 14926 | 2 | 1.1212 | 0.19283 | 0.42081 | 1 | 8509 | 4 | 1.1212 |
| SLC43A3 | 6 | 0.07607 | 0.19939 | 0.977164 | 4076 | 4 | -1.0968 | 0.19283 | 0.42081 | 1 | 8510 | 2 | -1.0968 |
| TNFSF15 | 5 | 0.74384 | 0.78763 | 0.999994 | 16785 | 1 | 2.0878 | 0.19285 | 0.40896 | 1 | 8511 | 3 | 2.0878 |
| SUV420H1 | 5 | 0.7517 | 0.79174 | 0.999994 | 16874 | 1 | 1.0146 | 0.19298 | 0.4091 | 1 | 8512 | 3 | 1.0146 |
| ZNF101 | 6 | 0.53956 | 0.69623 | 0.999994 | 14043 | 2 | 1.2641 | 0.19306 | 0.42113 | 1 | 8513 | 4 | 1.2641 |
| TCF12 | 5 | 0.058598 | 0.15974 | 0.967126 | 3368 | 3 | -1.8479 | 0.19318 | 0.40933 | 1 | 8514 | 1 | -1.8479 |
| DNAJC15 | 5 | 0.93851 | 0.93871 | 0.999994 | 20038 | 0 | 0.48167 | 0.19318 | 0.40933 | 1 | 8515 | 2 | 0.48167 |
| PRSS12 | 5 | 0.16882 | 0.33039 | 0.999994 | 6955 | 3 | -1.0451 | 0.19318 | 0.40933 | 1 | 8516 | 2 | -1.0451 |
| ADAMTS20 | 5 | 0.40034 | 0.56379 | 0.999994 | 11955 | 1 | -0.21415 | 0.19318 | 0.40933 | 1 | 8517 | 2 | -0.21415 |
| C21orf33 | 6 | 0.8753 | 0.88664 | 0.999994 | 18741 | 1 | 1.7627 | 0.19319 | 0.42133 | 1 | 8518 | 4 | 1.7627 |
| ENOX2 | 6 | 0.22396 | 0.41687 | 0.999994 | 8377 | 1 | 1.3295 | 0.19323 | 0.42139 | 1 | 8519 | 4 | 1.3295 |
| PRDM8 | 6 | 0.14889 | 0.32421 | 0.999994 | 6423 | 3 | 0.63163 | 0.19331 | 0.42149 | 1 | 8520 | 3 | 0.63163 |
| ZNF195 | 4 | 0.67981 | 0.71147 | 0.999994 | 16086 | 1 | 2.05 | 0.19337 | 0.37383 | 0.999517 | 8521 | 2 | 2.05 |
| BDH2 | 6 | 0.97688 | 0.97683 | 0.999994 | 21011 | 0 | 1.5799 | 0.1934 | 0.42162 | 1 | 8522 | 4 | 1.5799 |
| NDOR1 | 5 | 0.05805 | 0.15885 | 0.967126 | 3340 | 2 | 2.3059 | 0.1934 | 0.40958 | 1 | 8523 | 3 | 2.3059 |
| hsa-mir-1237 | 1 | 0.80658 | 0.80677 | 0.999994 | 17578 | 0 | 5.1479 | 0.19342 | 0.19322 | 0.97466 | 8524 | 1 | 5.1479 |
| NonTargetingControlG | 2 | 0.80651 | 0.80615 | 0.999994 | 17577 | 0 | 3.6189 | 0.19349 | 0.30754 | 0.988566 | 8525 | 2 | 3.6189 |
| hsa-mir-676 | 4 | 0.72753 | 0.74339 | 0.999994 | 16575 | 1 | 1.5331 | 0.19351 | 0.37404 | 0.999517 | 8526 | 2 | 1.5331 |
| GATM | 6 | 0.15565 | 0.33276 | 0.999994 | 6613 | 3 | -0.73099 | 0.19354 | 0.42184 | 1 | 8527 | 2 | -0.73099 |

| | | | | | | | | | | | | | |
|---|---|---|---|---|---|---|---|---|---|---|---|---|---|
| TRIP4 | 6 | 0.55458 | 0.70694 | 0.999994 | 14278 | 2 | 0.99224 | 0.19354 | 0.42184 | 1 | 8528 | 4 | 0.99224 |
| MYO5A | 6 | 0.37098 | 0.58382 | 0.999994 | 11526 | 2 | 1.6079 | 0.19354 | 0.42184 | 1 | 8529 | 3 | 1.6079 |
| TSLP | 6 | 0.11177 | 0.27011 | 0.998561 | 5359 | 4 | -0.92647 | 0.19354 | 0.42184 | 1 | 8530 | 2 | -0.92647 |
| C17orf105 | 6 | 0.87319 | 0.8853 | 0.999994 | 18698 | 1 | 1.4284 | 0.19356 | 0.42186 | 1 | 8531 | 4 | 1.4284 |
| COMMD5 | 5 | 0.19401 | 0.36665 | 0.999994 | 7603 | 3 | -0.90269 | 0.1936 | 0.40979 | 1 | 8532 | 2 | -0.90269 |
| LOC100130451 | 4 | 0.53015 | 0.63017 | 0.999994 | 13902 | 1 | 1.1903 | 0.19362 | 0.37417 | 0.999517 | 8533 | 3 | 1.1903 |
| BPIFB1 | 4 | 0.87433 | 0.87398 | 0.999994 | 18715 | 0 | 2.2593 | 0.19366 | 0.37424 | 0.999517 | 8534 | 3 | 2.2593 |
| SMAGP | 4 | 0.27249 | 0.41488 | 0.999994 | 9527 | 1 | 1.2789 | 0.19371 | 0.37429 | 0.999517 | 8535 | 3 | 1.2789 |
| GPR125 | 3 | 0.84982 | 0.8498 | 0.999994 | 18257 | 0 | 1.0229 | 0.19378 | 0.33626 | 0.994237 | 8536 | 2 | 1.0229 |
| OR2L8 | 6 | 0.46159 | 0.64282 | 0.999994 | 12874 | 2 | 1.1962 | 0.19379 | 0.42219 | 1 | 8537 | 4 | 1.1962 |
| TRIM68 | 4 | 0.73908 | 0.75179 | 0.999994 | 16722 | 1 | 2.1975 | 0.1938 | 0.37442 | 0.999517 | 8538 | 2 | 2.1975 |
| ZYG11A | 6 | 0.0062899 | 0.025859 | 0.848212 | 613 | 3 | -0.29417 | 0.19383 | 0.42224 | 1 | 8539 | 2 | -0.29417 |
| TMEM114 | 5 | 0.43723 | 0.59447 | 0.999994 | 12529 | 1 | -0.18145 | 0.19386 | 0.41008 | 1 | 8540 | 2 | -0.18145 |
| IGFBPL1 | 5 | 0.43739 | 0.59461 | 0.999994 | 12533 | 2 | 0.029715 | 0.19386 | 0.41008 | 1 | 8541 | 2 | 0.029715 |
| AIMP2 | 5 | 0.76481 | 0.79881 | 0.999994 | 17025 | 1 | 0.59296 | 0.19386 | 0.41008 | 1 | 8542 | 3 | 0.59296 |
| NPPB | 4 | 0.92822 | 0.92799 | 0.999994 | 19819 | 0 | 1.9811 | 0.19404 | 0.37473 | 0.999639 | 8543 | 3 | 1.9811 |
| hsa-mir-4523 | 2 | 0.8059 | 0.80552 | 0.999994 | 17569 | 0 | 1.358 | 0.1941 | 0.30837 | 0.988566 | 8544 | 2 | 1.358 |
| PAPD4 | 6 | 0.72425 | 0.81973 | 0.999994 | 16538 | 1 | 1.5081 | 0.19415 | 0.4227 | 1 | 8545 | 4 | 1.5081 |
| NEDD4L | 6 | 0.062434 | 0.17093 | 0.96737 | 3532 | 2 | 1.5996 | 0.19415 | 0.4227 | 1 | 8546 | 3 | 1.5996 |
| GPAM | 6 | 0.64173 | 0.77125 | 0.999994 | 15544 | 1 | 1.8947 | 0.1943 | 0.42294 | 1 | 8547 | 4 | 1.8947 |
| PABPN1L | 6 | 0.82095 | 0.85727 | 0.999994 | 17789 | 1 | 1.1009 | 0.1943 | 0.42294 | 1 | 8548 | 3 | 1.1009 |
| AGMO | 6 | 0.82158 | 0.85758 | 0.999994 | 17798 | 1 | 0.60758 | 0.1943 | 0.42294 | 1 | 8549 | 3 | 0.60758 |
| KLF16 | 6 | 0.19265 | 0.37882 | 0.999994 | 7565 | 3 | -0.81875 | 0.1943 | 0.42294 | 1 | 8550 | 2 | -0.81875 |
| BCAS3 | 6 | 0.33473 | 0.54665 | 0.999994 | 10827 | 3 | 1.318 | 0.1943 | 0.42294 | 1 | 8551 | 3 | 1.318 |
| PCDHB16 | 6 | 0.92975 | 0.92988 | 0.999994 | 19848 | 0 | 1.5859 | 0.19439 | 0.42307 | 1 | 8552 | 4 | 1.5859 |
| TAC3 | 4 | 0.83645 | 0.83596 | 0.999994 | 18026 | 0 | 0.20826 | 0.19444 | 0.37533 | 0.999639 | 8553 | 2 | 0.20826 |
| CATSPER3 | 4 | 0.32476 | 0.46833 | 0.999994 | 10614 | 2 | 0.72713 | 0.19444 | 0.37533 | 0.999639 | 8554 | 2 | 0.72713 |
| COL10A1 | 6 | 0.19032 | 0.37591 | 0.999994 | 7495 | 3 | 1.0736 | 0.19448 | 0.4232 | 1 | 8555 | 3 | 1.0736 |
| EPS8L3 | 6 | 0.6784 | 0.79975 | 0.999994 | 16066 | 1 | 1.3611 | 0.19454 | 0.42327 | 1 | 8556 | 4 | 1.3611 |
| hsa-mir-6075 | 4 | 0.66027 | 0.69942 | 0.999994 | 15782 | 1 | 2.2966 | 0.19457 | 0.37552 | 0.999639 | 8557 | 3 | 2.2966 |
| POLR2G | 4 | 0.84164 | 0.84108 | 0.999994 | 18116 | 0 | 2.2318 | 0.19461 | 0.37558 | 0.999639 | 8558 | 2 | 2.2318 |
| GPRIN2 | 5 | 0.17529 | 0.33983 | 0.999994 | 7117 | 2 | 0.36893 | 0.19465 | 0.41096 | 1 | 8559 | 2 | 0.36893 |
| OR14A16 | 5 | 0.012387 | 0.04289 | 0.891621 | 1022 | 4 | -2.6459 | 0.19465 | 0.41096 | 1 | 8560 | 1 | -2.6459 |
| C4orf51 | 5 | 0.86677 | 0.86824 | 0.999994 | 18570 | 1 | 0.28874 | 0.19465 | 0.41096 | 1 | 8561 | 2 | 0.28874 |
| TXNRD2 | 5 | 0.44142 | 0.59797 | 0.999994 | 12600 | 2 | -0.42881 | 0.19465 | 0.41096 | 1 | 8562 | 1 | -0.42881 |
| NUMB | 5 | 0.070766 | 0.17962 | 0.968171 | 3863 | 2 | 0.264 | 0.19465 | 0.41096 | 1 | 8563 | 2 | 0.264 |
| LOC100507462 | 4 | 0.62939 | 0.68154 | 0.999994 | 15375 | 1 | 1.4384 | 0.19467 | 0.37566 | 0.999639 | 8564 | 3 | 1.4384 |
| OR56A5 | 5 | 0.55653 | 0.69756 | 0.999994 | 14312 | 2 | 2.167 | 0.19471 | 0.41104 | 1 | 8565 | 3 | 2.167 |
| GTPBP8 | 6 | 0.044194 | 0.13131 | 0.95318 | 2720 | 3 | -0.28664 | 0.19472 | 0.42355 | 1 | 8566 | 2 | -0.28664 |
| hsa-mir-6720 | 4 | 0.84451 | 0.84401 | 0.999994 | 18166 | 0 | 1.6529 | 0.19473 | 0.37576 | 0.999639 | 8567 | 3 | 1.6529 |
| NFKBID | 6 | 0.55733 | 0.70892 | 0.999994 | 14319 | 2 | 1.9631 | 0.19476 | 0.4236 | 1 | 8568 | 4 | 1.9631 |
| ZNF778 | 5 | 0.10942 | 0.24109 | 0.987253 | 5286 | 3 | -1.6006 | 0.19477 | 0.41109 | 1 | 8569 | 2 | -1.6006 |
| ASH1L | 5 | 0.29161 | 0.47425 | 0.999994 | 9942 | 2 | -0.32827 | 0.19477 | 0.41109 | 1 | 8570 | 2 | -0.32827 |
| VKORC1 | 5 | 0.83494 | 0.84302 | 0.999994 | 18003 | 1 | 0.32284 | 0.19477 | 0.41109 | 1 | 8571 | 2 | 0.32284 |
| CCDC167 | 5 | 0.33833 | 0.51238 | 0.999994 | 10905 | 2 | 0.54942 | 0.19477 | 0.41109 | 1 | 8572 | 2 | 0.54942 |
| AP1AR | 3 | 0.79229 | 0.79218 | 0.999994 | 17378 | 0 | 2.726 | 0.19477 | 0.33732 | 0.994237 | 8573 | 2 | 2.726 |
| TRAPPC3L | 4 | 0.91414 | 0.9139 | 0.999994 | 19489 | 0 | 2.4209 | 0.19477 | 0.37583 | 0.999639 | 8574 | 3 | 2.4209 |
| NonTargetingControlG | 2 | 0.80521 | 0.8048 | 0.999994 | 17556 | 0 | 1.962 | 0.19479 | 0.30929 | 0.988566 | 8575 | 2 | 1.962 |
| PAPPA2 | 6 | 0.19686 | 0.38407 | 0.999994 | 7671 | 2 | 1.5208 | 0.1948 | 0.42366 | 1 | 8576 | 4 | 1.5208 |
| PRKAR1B | 6 | 0.61976 | 0.75465 | 0.999994 | 15216 | 1 | 1.2339 | 0.19495 | 0.42389 | 1 | 8577 | 4 | 1.2339 |
| ATCAY | 4 | 0.80969 | 0.81 | 0.999994 | 17628 | 1 | 1.8857 | 0.195 | 0.37615 | 0.999942 | 8578 | 3 | 1.8857 |
| PARD3 | 3 | 0.28842 | 0.42219 | 0.999994 | 9880 | 1 | -0.43567 | 0.19505 | 0.33762 | 0.994237 | 8579 | 1 | -0.43567 |

| | | | | | | | | | | | | | |
|---|---|---|---|---|---|---|---|---|---|---|---|---|---|
| hsa-mir-7848 | 4 | 0.93212 | 0.93186 | 0.999994 | 19905 | 0 | 1.463 | 0.19506 | 0.37624 | 0.999964 | 8580 | 3 | 1.463 |
| ZNF391 | 6 | 0.87474 | 0.88628 | 0.999994 | 18727 | 1 | 0.51504 | 0.1951 | 0.42412 | 1 | 8581 | 3 | 0.51504 |
| DNAJC5 | 6 | 0.17121 | 0.35225 | 0.999994 | 7013 | 2 | 1.3806 | 0.1951 | 0.42412 | 1 | 8582 | 3 | 1.3806 |
| DUOX2 | 4 | 0.20791 | 0.34884 | 0.999994 | 7974 | 2 | 1.4288 | 0.19522 | 0.37646 | 0.999964 | 8583 | 2 | 1.4288 |
| TMX2 | 4 | 0.10625 | 0.22624 | 0.983354 | 5181 | 2 | 1.7816 | 0.19522 | 0.37646 | 0.999964 | 8584 | 2 | 1.7816 |
| MTERFD3 | 6 | 0.10861 | 0.26404 | 0.996734 | 5261 | 2 | 1.0824 | 0.19524 | 0.4243 | 1 | 8585 | 4 | 1.0824 |
| CCDC42B | 4 | 0.32545 | 0.46905 | 0.999994 | 10630 | 1 | 1.5889 | 0.19524 | 0.37648 | 0.999964 | 8586 | 3 | 1.5889 |
| FOXA3 | 6 | 0.044475 | 0.13194 | 0.95318 | 2733 | 2 | 1.9386 | 0.19529 | 0.42438 | 1 | 8587 | 4 | 1.9386 |
| SORBS1 | 6 | 0.87483 | 0.88633 | 0.999994 | 18730 | 1 | 0.59983 | 0.19529 | 0.42438 | 1 | 8588 | 3 | 0.59983 |
| PAOX | 6 | 0.46609 | 0.64585 | 0.999994 | 12952 | 2 | 0.34273 | 0.19529 | 0.42438 | 1 | 8589 | 2 | 0.34273 |
| EP300 | 6 | 0.073752 | 0.19455 | 0.975797 | 3984 | 2 | 1.2325 | 0.19529 | 0.42438 | 1 | 8590 | 3 | 1.2325 |
| SMG1 | 5 | 0.19782 | 0.37214 | 0.999994 | 7703 | 2 | 1.9083 | 0.19531 | 0.41173 | 1 | 8591 | 3 | 1.9083 |
| SPC25 | 5 | 0.96221 | 0.96236 | 0.999994 | 20635 | 0 | 1.1922 | 0.19542 | 0.41185 | 1 | 8592 | 3 | 1.1922 |
| ANKRD42 | 5 | 0.094469 | 0.21765 | 0.980699 | 4765 | 3 | -1.4734 | 0.19542 | 0.41185 | 1 | 8593 | 2 | -1.4734 |
| OR5P3 | 4 | 0.717 | 0.73595 | 0.999994 | 16468 | 1 | 1.1518 | 0.19573 | 0.37718 | 1 | 8594 | 3 | 1.1518 |
| SSX1 | 6 | 0.88435 | 0.89243 | 0.999994 | 18918 | 1 | 1.4891 | 0.19578 | 0.42507 | 1 | 8595 | 4 | 1.4891 |
| TRPC4 | 6 | 0.4328 | 0.6237 | 0.999994 | 12459 | 2 | 1.8882 | 0.1958 | 0.42509 | 1 | 8596 | 4 | 1.8882 |
| ARHGAP26 | 6 | 0.094709 | 0.23693 | 0.985449 | 4778 | 3 | -0.31174 | 0.19581 | 0.42511 | 1 | 8597 | 2 | -0.31174 |
| GPR68 | 5 | 0.19979 | 0.37495 | 0.999994 | 7760 | 2 | -0.21066 | 0.19586 | 0.41234 | 1 | 8598 | 2 | -0.21066 |
| CORIN | 5 | 0.89656 | 0.89669 | 0.999994 | 19155 | 0 | 1.9288 | 0.19586 | 0.41234 | 1 | 8599 | 3 | 1.9288 |
| MGP | 6 | 0.53363 | 0.6921 | 0.999994 | 13961 | 1 | 1.2958 | 0.19586 | 0.42518 | 1 | 8600 | 4 | 1.2958 |
| C9orf153 | 4 | 0.17287 | 0.3128 | 0.999994 | 7059 | 1 | 1.1663 | 0.19588 | 0.37739 | 1 | 8601 | 3 | 1.1663 |
| SLC25A19 | 5 | 0.070286 | 0.17884 | 0.968171 | 3845 | 3 | -2.0403 | 0.19591 | 0.4124 | 1 | 8602 | 2 | -2.0403 |
| C2orf82 | 6 | 0.8692 | 0.88286 | 0.999994 | 18620 | 1 | 1.9915 | 0.196 | 0.42537 | 1 | 8603 | 4 | 1.9915 |
| NonTargetingControlG | 2 | 0.80399 | 0.80359 | 0.999994 | 17542 | 0 | 3.4202 | 0.19601 | 0.31097 | 0.988566 | 8604 | 2 | 3.4202 |
| NAT1 | 6 | 0.9434 | 0.94336 | 0.999994 | 20161 | 0 | 2.0812 | 0.19606 | 0.42546 | 1 | 8605 | 4 | 2.0812 |
| ZNF674 | 5 | 0.4025 | 0.56561 | 0.999994 | 11989 | 2 | 2.1779 | 0.19608 | 0.4126 | 1 | 8606 | 3 | 2.1779 |
| FBXO27 | 6 | 0.95968 | 0.95976 | 0.999994 | 20576 | 0 | 1.5819 | 0.19608 | 0.42549 | 1 | 8607 | 4 | 1.5819 |
| ACO1 | 6 | 0.36684 | 0.58123 | 0.999994 | 11454 | 2 | 1.1967 | 0.19612 | 0.42556 | 1 | 8608 | 4 | 1.1967 |
| SPACA1 | 4 | 0.21384 | 0.35497 | 0.999994 | 8110 | 1 | 2.8748 | 0.19613 | 0.37778 | 1 | 8609 | 3 | 2.8748 |
| KRTCAP2 | 5 | 0.67954 | 0.75736 | 0.999994 | 16082 | 1 | 2.0998 | 0.19617 | 0.41271 | 1 | 8610 | 3 | 2.0998 |
| ATPBD4 | 5 | 0.94028 | 0.94046 | 0.999994 | 20093 | 0 | 1.8902 | 0.1962 | 0.41274 | 1 | 8611 | 3 | 1.8902 |
| GZMH | 6 | 0.85585 | 0.8751 | 0.999994 | 18365 | 1 | 1.1776 | 0.1962 | 0.42568 | 1 | 8612 | 3 | 1.1776 |
| CBLN1 | 5 | 0.099493 | 0.22562 | 0.983354 | 4951 | 2 | 2.1393 | 0.19621 | 0.41275 | 1 | 8613 | 3 | 2.1393 |
| hsa-mir-4453 | 2 | 0.8037 | 0.80329 | 0.999994 | 17538 | 0 | 2.7233 | 0.1963 | 0.31139 | 0.988566 | 8614 | 2 | 2.7233 |
| TBC1D3F | 1 | 0.80368 | 0.80391 | 0.999994 | 17537 | 0 | 5.0629 | 0.19632 | 0.1961 | 0.97466 | 8615 | 1 | 5.0629 |
| PCSK9 | 6 | 0.64922 | 0.77702 | 0.999994 | 15646 | 1 | 0.82973 | 0.19635 | 0.42588 | 1 | 8616 | 3 | 0.82973 |
| IZUMO1 | 6 | 0.18842 | 0.37355 | 0.999994 | 7432 | 3 | 0.72903 | 0.19636 | 0.42589 | 1 | 8617 | 3 | 0.72903 |
| NGRN | 3 | 0.8489 | 0.84888 | 0.999994 | 18236 | 0 | 3.0621 | 0.19636 | 0.33901 | 0.994237 | 8618 | 2 | 3.0621 |
| NonTargetingControlG | 2 | 0.49837 | 0.50242 | 0.999994 | 13400 | 1 | 2.9903 | 0.19641 | 0.31153 | 0.988566 | 8619 | 1 | 2.9903 |
| PCK2 | 6 | 0.046344 | 0.13605 | 0.95468 | 2823 | 2 | 1.3005 | 0.19641 | 0.42596 | 1 | 8620 | 4 | 1.3005 |
| MED12L | 6 | 0.88395 | 0.89217 | 0.999994 | 18903 | 1 | 1.5133 | 0.19663 | 0.42627 | 1 | 8621 | 4 | 1.5133 |
| RASD1 | 5 | 0.18899 | 0.3595 | 0.999994 | 7450 | 3 | -0.85579 | 0.19671 | 0.41333 | 1 | 8622 | 2 | -0.85579 |
| RHOBTB1 | 5 | 0.13933 | 0.28681 | 0.999632 | 6170 | 3 | -1.1467 | 0.19671 | 0.41333 | 1 | 8623 | 1 | -1.1467 |
| NOV | 4 | 0.8732 | 0.87284 | 0.999994 | 18699 | 0 | 2.7931 | 0.19676 | 0.37871 | 1 | 8624 | 3 | 2.7931 |
| hsa-mir-371b | 1 | 0.80323 | 0.80344 | 0.999994 | 17527 | 0 | 5.0615 | 0.19677 | 0.19655 | 0.97466 | 8625 | 1 | 5.0615 |
| PISD | 4 | 0.63358 | 0.68387 | 0.999994 | 15424 | 1 | 1.6765 | 0.19683 | 0.37879 | 1 | 8626 | 2 | 1.6765 |
| ZSWIM5 | 4 | 0.73924 | 0.7519 | 0.999994 | 16725 | 1 | 2.0426 | 0.19683 | 0.37879 | 1 | 8627 | 3 | 2.0426 |
| GFPT2 | 6 | 0.46034 | 0.64201 | 0.999994 | 12851 | 1 | 1.4398 | 0.19699 | 0.42679 | 1 | 8628 | 4 | 1.4398 |
| KRT76 | 6 | 0.84899 | 0.87139 | 0.999994 | 18240 | 1 | 0.62012 | 0.19699 | 0.42679 | 1 | 8629 | 4 | 0.62012 |
| SLC39A3 | 5 | 0.4133 | 0.57457 | 0.999994 | 12159 | 2 | -0.34983 | 0.19705 | 0.41371 | 1 | 8630 | 2 | -0.34983 |
| ABHD16A | 5 | 0.1622 | 0.32076 | 0.999994 | 6799 | 3 | -1.0975 | 0.19705 | 0.41371 | 1 | 8631 | 2 | -1.0975 |

| | | | | | | | | | | | | |
|---|---|---|---|---|---|---|---|---|---|---|---|---|
| DEGS2 | 6 | 0.85687 | 0.87566 | 0.999994 | 18388 | 1 | 1.8852 | 0.19712 | 0.42698 | 1 | 8632 | 4 | 1.8852 |
| SNAP23 | 4 | 0.91873 | 0.91852 | 0.999994 | 19600 | 0 | 2.8253 | 0.19716 | 0.37926 | 1 | 8633 | 3 | 2.8253 |
| RAD54B | 5 | 0.18084 | 0.34785 | 0.999994 | 7242 | 2 | 2.1453 | 0.19723 | 0.41392 | 1 | 8634 | 3 | 2.1453 |
| CCT3 | 3 | 0.24956 | 0.37442 | 0.999994 | 9009 | 1 | 3.0085 | 0.19726 | 0.33998 | 0.994237 | 8635 | 2 | 3.0085 |
| NUP93 | 3 | 0.65642 | 0.66241 | 0.999994 | 15734 | 1 | 3.0085 | 0.19726 | 0.33998 | 0.994237 | 8636 | 2 | 3.0085 |
| EPB41L4A | 6 | 0.52766 | 0.6879 | 0.999994 | 13856 | 2 | 1.3118 | 0.19727 | 0.4272 | 1 | 8637 | 4 | 1.3118 |
| PARK2 | 6 | 0.070581 | 0.18801 | 0.969543 | 3856 | 2 | 1.8097 | 0.1973 | 0.42722 | 1 | 8638 | 3 | 1.8097 |
| NAALAD2 | 4 | 0.33447 | 0.47827 | 0.999994 | 10822 | 1 | 1.2257 | 0.19733 | 0.37952 | 1 | 8639 | 3 | 1.2257 |
| CDC34 | 6 | 0.11545 | 0.27714 | 0.999379 | 5492 | 1 | 1.939 | 0.19738 | 0.42736 | 1 | 8640 | 4 | 1.939 |
| GRIK3 | 5 | 0.42346 | 0.58298 | 0.999994 | 12303 | 2 | -0.59327 | 0.19744 | 0.41414 | 1 | 8641 | 2 | -0.59327 |
| PUS7 | 5 | 0.0087565 | 0.03248 | 0.865899 | 776 | 3 | -1.1236 | 0.19744 | 0.41414 | 1 | 8642 | 2 | -1.1236 |
| IGSF22 | 5 | 0.10527 | 0.23462 | 0.984971 | 5143 | 2 | 0.73624 | 0.19744 | 0.41414 | 1 | 8643 | 3 | 0.73624 |
| HSPD1 | 4 | 0.42396 | 0.57024 | 0.999994 | 12308 | 1 | 2.0838 | 0.19747 | 0.37972 | 1 | 8644 | 2 | 2.0838 |
| hsa-mir-635 | 4 | 0.57083 | 0.65006 | 0.999994 | 14556 | 1 | 0.13541 | 0.19749 | 0.37973 | 1 | 8645 | 1 | 0.13541 |
| KIF22 | 6 | 0.76203 | 0.83283 | 0.999994 | 17000 | 1 | 0.99764 | 0.19754 | 0.42756 | 1 | 8646 | 4 | 0.99764 |
| PTCH2 | 6 | 0.44935 | 0.63471 | 0.999994 | 12706 | 2 | -0.29126 | 0.19754 | 0.42756 | 1 | 8647 | 2 | -0.29126 |
| CHIC2 | 6 | 0.34345 | 0.5566 | 0.999994 | 11018 | 3 | -0.61169 | 0.19754 | 0.42756 | 1 | 8648 | 2 | -0.61169 |
| SHOX_X | 6 | 0.66386 | 0.78829 | 0.999994 | 15838 | 1 | 0.11049 | 0.19754 | 0.42756 | 1 | 8649 | 2 | 0.11049 |
| PPM1F | 6 | 0.45025 | 0.63529 | 0.999994 | 12714 | 2 | 0.71865 | 0.19754 | 0.42756 | 1 | 8650 | 3 | 0.71865 |
| WNT9B | 6 | 0.43404 | 0.62451 | 0.999994 | 12477 | 2 | 1.1393 | 0.19754 | 0.42756 | 1 | 8651 | 3 | 1.1393 |
| NOMO1 | 6 | 0.052256 | 0.14902 | 0.965566 | 3081 | 2 | 1.6122 | 0.1976 | 0.42766 | 1 | 8652 | 4 | 1.6122 |
| NHLRC1 | 4 | 0.75914 | 0.76708 | 0.999994 | 16967 | 1 | 1.4328 | 0.19765 | 0.37996 | 1 | 8653 | 3 | 1.4328 |
| NonTargetingControlG | 1 | 0.80231 | 0.80254 | 0.999994 | 17517 | 0 | 5.0259 | 0.19769 | 0.19748 | 0.97466 | 8654 | 1 | 5.0259 |
| LRRC8B | 5 | 0.09408 | 0.21706 | 0.980699 | 4753 | 3 | -1.7453 | 0.19773 | 0.41446 | 1 | 8655 | 2 | -1.7453 |
| GBP7 | 4 | 0.17287 | 0.3128 | 0.999994 | 7061 | 1 | 0.59312 | 0.19774 | 0.3801 | 1 | 8656 | 2 | 0.59312 |
| GRAP | 6 | 0.20752 | 0.39704 | 0.999994 | 7961 | 3 | 0.86602 | 0.19776 | 0.42788 | 1 | 8657 | 3 | 0.86602 |
| FZD7 | 5 | 0.4924 | 0.64133 | 0.999994 | 13303 | 2 | -0.49454 | 0.19792 | 0.41469 | 1 | 8658 | 1 | -0.49454 |
| CHRNB2 | 6 | 0.69339 | 0.81003 | 0.999994 | 16239 | 1 | 1.5463 | 0.19793 | 0.42814 | 1 | 8659 | 4 | 1.5463 |
| TPM4 | 6 | 0.50645 | 0.67318 | 0.999994 | 13533 | 2 | 1.3196 | 0.19803 | 0.42828 | 1 | 8660 | 3 | 1.3196 |
| hsa-mir-1469 | 4 | 0.10169 | 0.21814 | 0.980699 | 5024 | 3 | -2.6484 | 0.19803 | 0.38051 | 1 | 8661 | 1 | -2.6484 |
| hsa-mir-526a-1 | 2 | 0.68355 | 0.68339 | 0.999994 | 16138 | 0 | 3.5452 | 0.19806 | 0.31376 | 0.989846 | 8662 | 1 | 3.5452 |
| GSTM3 | 5 | 0.85102 | 0.8551 | 0.999994 | 18279 | 1 | 0.81813 | 0.19807 | 0.41487 | 1 | 8663 | 3 | 0.81813 |
| ME1 | 6 | 0.8827 | 0.89138 | 0.999994 | 18886 | 1 | 0.96491 | 0.19811 | 0.42839 | 1 | 8664 | 4 | 0.96491 |
| WDR90 | 6 | 0.90028 | 0.9039 | 0.999994 | 19226 | 1 | 1.7723 | 0.19813 | 0.42841 | 1 | 8665 | 4 | 1.7723 |
| DNER | 5 | 0.51807 | 0.66363 | 0.999994 | 13711 | 2 | 1.1273 | 0.19817 | 0.41498 | 1 | 8666 | 3 | 1.1273 |
| VMP1 | 5 | 0.22008 | 0.40336 | 0.999994 | 8292 | 3 | -0.76029 | 0.19821 | 0.41503 | 1 | 8667 | 2 | -0.76029 |
| OR56A3 | 6 | 0.97738 | 0.97733 | 0.999994 | 21028 | 0 | 2.0172 | 0.19822 | 0.42856 | 1 | 8668 | 4 | 2.0172 |
| RABGEF1 | 6 | 0.69915 | 0.81175 | 0.999994 | 16297 | 1 | 1.6334 | 0.1983 | 0.42866 | 1 | 8669 | 3 | 1.6334 |
| TFRC | 4 | 0.37677 | 0.52184 | 0.999994 | 11605 | 2 | -0.50335 | 0.19837 | 0.381 | 1 | 8670 | 1 | -0.50335 |
| POLR2J3 | 1 | 0.80157 | 0.8018 | 0.999994 | 17501 | 0 | 5.0163 | 0.19843 | 0.19821 | 0.97466 | 8671 | 1 | 5.0163 |
| POTEM | 1 | 0.80157 | 0.8018 | 0.999994 | 17500 | 0 | 5.0163 | 0.19843 | 0.19821 | 0.97466 | 8672 | 1 | 5.0163 |
| TCF3 | 6 | 0.19447 | 0.38113 | 0.999994 | 7621 | 2 | 0.58218 | 0.19854 | 0.42899 | 1 | 8673 | 3 | 0.58218 |
| CD74 | 6 | 0.018997 | 0.068766 | 0.93914 | 1407 | 4 | -1.3696 | 0.19859 | 0.42905 | 1 | 8674 | 2 | -1.3696 |
| C7 | 6 | 0.64108 | 0.77074 | 0.999994 | 15529 | 2 | 1.4532 | 0.19859 | 0.42905 | 1 | 8675 | 4 | 1.4532 |
| ITPR3 | 6 | 0.26072 | 0.46071 | 0.999994 | 9256 | 3 | -0.60695 | 0.19859 | 0.42905 | 1 | 8676 | 2 | -0.60695 |
| PRKRIP1 | 4 | 0.38542 | 0.53067 | 0.999994 | 11726 | 2 | 1.9269 | 0.19861 | 0.38134 | 1 | 8677 | 2 | 1.9269 |
| BRD9 | 5 | 0.52926 | 0.67339 | 0.999994 | 13886 | 2 | 1.9851 | 0.19869 | 0.41555 | 1 | 8678 | 3 | 1.9851 |
| MRPL2 | 4 | 0.56079 | 0.64506 | 0.999994 | 14383 | 1 | 0.84432 | 0.19873 | 0.3815 | 1 | 8679 | 2 | 0.84432 |
| FBXO16 | 4 | 0.29071 | 0.43353 | 0.999994 | 9927 | 1 | -0.26007 | 0.19873 | 0.3815 | 1 | 8680 | 1 | -0.26007 |
| CKMT2 | 6 | 0.90517 | 0.90771 | 0.999994 | 19334 | 1 | 1.5865 | 0.19877 | 0.42931 | 1 | 8681 | 4 | 1.5865 |
| CD48 | 6 | 0.9464 | 0.94637 | 0.999994 | 20240 | 0 | 0.59624 | 0.19901 | 0.42962 | 1 | 8682 | 3 | 0.59624 |
| NDUFA4L2 | 5 | 0.40764 | 0.56991 | 0.999994 | 12069 | 2 | 1.4767 | 0.19924 | 0.41614 | 1 | 8683 | 3 | 1.4767 |

| | | | | | | | | | | | | |
|---|---|---|---|---|---|---|---|---|---|---|---|---|
| SDAD1 | 5 | 0.18593 | 0.35516 | 0.999994 | 7365 | 2 | -1.2059 | 0.19924 | 0.41614 | 1 | 8684 | 2 | -1.2059 |
| CD163L1 | 5 | 0.4522 | 0.60707 | 0.999994 | 12738 | 1 | 0.213 | 0.19924 | 0.41614 | 1 | 8685 | 2 | 0.213 |
| TXNDC17 | 6 | 0.56079 | 0.71135 | 0.999994 | 14384 | 2 | 2.2434 | 0.19925 | 0.42995 | 1 | 8686 | 4 | 2.2434 |
| NonTargetingControlG | 2 | 0.67281 | 0.67256 | 0.999994 | 15968 | 0 | 3.4976 | 0.19926 | 0.31547 | 0.989846 | 8687 | 1 | 3.4976 |
| C21orf2 | 6 | 0.43617 | 0.6259 | 0.999994 | 12517 | 2 | 1.6629 | 0.19927 | 0.42997 | 1 | 8688 | 4 | 1.6629 |
| ADA | 6 | 0.56946 | 0.71756 | 0.999994 | 14529 | 2 | 1.8034 | 0.19936 | 0.43011 | 1 | 8689 | 4 | 1.8034 |
| C19orf24 | 6 | 0.25871 | 0.45835 | 0.999994 | 9209 | 1 | 1.6045 | 0.19936 | 0.43011 | 1 | 8690 | 4 | 1.6045 |
| LDB2 | 6 | 0.74035 | 0.8251 | 0.999994 | 16743 | 1 | 1.4337 | 0.19939 | 0.43014 | 1 | 8691 | 4 | 1.4337 |
| FCN1 | 5 | 0.27344 | 0.45954 | 0.999994 | 9553 | 2 | 2.1844 | 0.19941 | 0.41634 | 1 | 8692 | 3 | 2.1844 |
| ZNF441 | 6 | 0.942 | 0.94197 | 0.999994 | 20125 | 0 | 0.6689 | 0.19942 | 0.43018 | 1 | 8693 | 4 | 0.6689 |
| MAX | 6 | 0.023145 | 0.082053 | 0.939644 | 1661 | 4 | -1.108 | 0.19942 | 0.43018 | 1 | 8694 | 2 | -1.108 |
| RIC8B | 6 | 0.20267 | 0.39112 | 0.999994 | 7847 | 2 | 1.4677 | 0.19942 | 0.43018 | 1 | 8695 | 4 | 1.4677 |
| NBPF1 | 4 | 0.21255 | 0.35363 | 0.999994 | 8076 | 2 | -0.049843 | 0.19944 | 0.3825 | 1 | 8696 | 2 | -0.049843 |
| hsa-mir-1293 | 4 | 0.2424 | 0.38407 | 0.999994 | 8842 | 1 | 2.7697 | 0.1995 | 0.38258 | 1 | 8697 | 3 | 2.7697 |
| STEAP3 | 6 | 0.14735 | 0.32231 | 0.999994 | 6384 | 3 | 0.47494 | 0.19953 | 0.43033 | 1 | 8698 | 3 | 0.47494 |
| CYB5A | 6 | 0.071033 | 0.18894 | 0.969543 | 3876 | 2 | 1.0133 | 0.19967 | 0.43054 | 1 | 8699 | 3 | 1.0133 |
| MCTP2 | 4 | 0.30359 | 0.44662 | 0.999994 | 10207 | 1 | 2.6554 | 0.19971 | 0.38287 | 1 | 8700 | 3 | 2.6554 |
| LCN8 | 4 | 0.11367 | 0.23929 | 0.987253 | 5431 | 2 | 0.81903 | 0.19971 | 0.38287 | 1 | 8701 | 2 | 0.81903 |
| TMEM242 | 6 | 0.94649 | 0.94646 | 0.999994 | 20243 | 0 | 1.4505 | 0.19971 | 0.43059 | 1 | 8702 | 4 | 1.4505 |
| VPS11 | 3 | 0.31761 | 0.43805 | 0.999994 | 10464 | 1 | 2.9523 | 0.1998 | 0.34275 | 0.994392 | 8703 | 2 | 2.9523 |
| CTR9 | 6 | 0.18309 | 0.367 | 0.999994 | 7293 | 3 | 0.91314 | 0.19981 | 0.43072 | 1 | 8704 | 3 | 0.91314 |
| SLIT1 | 6 | 0.61979 | 0.75467 | 0.999994 | 15217 | 2 | 1.424 | 0.19982 | 0.43073 | 1 | 8705 | 4 | 1.424 |
| PLLP | 6 | 0.94347 | 0.94343 | 0.999994 | 20162 | 0 | 1.2407 | 0.20003 | 0.43101 | 1 | 8706 | 4 | 1.2407 |
| SNED1 | 6 | 0.40219 | 0.60379 | 0.999994 | 11980 | 2 | 1.6051 | 0.2001 | 0.4311 | 1 | 8707 | 4 | 1.6051 |
| FGD2 | 6 | 0.53356 | 0.69205 | 0.999994 | 13959 | 2 | 0.53877 | 0.20016 | 0.43121 | 1 | 8708 | 2 | 0.53877 |
| GINM1 | 6 | 0.73364 | 0.82283 | 0.999994 | 16651 | 1 | 0.63105 | 0.20016 | 0.43121 | 1 | 8709 | 3 | 0.63105 |
| KMT2C | 5 | 0.18271 | 0.35051 | 0.999994 | 7284 | 1 | 1.9143 | 0.20036 | 0.41739 | 1 | 8710 | 3 | 1.9143 |
| TCEB3C | 6 | 0.3933 | 0.59803 | 0.999994 | 11851 | 2 | 1.581 | 0.20037 | 0.4315 | 1 | 8711 | 4 | 1.581 |
| UBXN8 | 6 | 0.46368 | 0.64422 | 0.999994 | 12914 | 1 | 0.57085 | 0.20037 | 0.4315 | 1 | 8712 | 3 | 0.57085 |
| NMUR1 | 6 | 0.059685 | 0.16506 | 0.967126 | 3410 | 4 | -1.5231 | 0.20037 | 0.4315 | 1 | 8713 | 2 | -1.5231 |
| hsa-mir-548ag-2 | 2 | 0.72403 | 0.72376 | 0.999994 | 16533 | 0 | 3.6577 | 0.20048 | 0.31717 | 0.989846 | 8714 | 1 | 3.6577 |
| PIM3 | 6 | 0.19812 | 0.38562 | 0.999994 | 7714 | 2 | 2.176 | 0.20051 | 0.43168 | 1 | 8715 | 4 | 2.176 |
| EMX2 | 6 | 0.62486 | 0.7585 | 0.999994 | 15300 | 2 | 1.5308 | 0.20051 | 0.43168 | 1 | 8716 | 3 | 1.5308 |
| LOC729059 | 2 | 0.61985 | 0.61964 | 0.999994 | 15218 | 0 | 3.3257 | 0.20072 | 0.31751 | 0.989846 | 8717 | 1 | 3.3257 |
| TGS1 | 6 | 0.97279 | 0.97272 | 0.999994 | 20889 | 0 | 1.6937 | 0.20077 | 0.43206 | 1 | 8718 | 4 | 1.6937 |
| NBPF6 | 1 | 0.79922 | 0.79952 | 0.999994 | 17468 | 0 | 4.949 | 0.20078 | 0.20054 | 0.97466 | 8719 | 1 | 4.949 |
| NonTargetingControlG | 2 | 0.79922 | 0.79879 | 0.999994 | 17467 | 0 | 0.99094 | 0.20078 | 0.31759 | 0.989846 | 8720 | 2 | 0.99094 |
| hsa-mir-4288 | 4 | 0.31505 | 0.45842 | 0.999994 | 10420 | 1 | 1.5481 | 0.20085 | 0.38447 | 1 | 8721 | 3 | 1.5481 |
| TLL2 | 6 | 0.96623 | 0.9663 | 0.999994 | 20741 | 0 | 1.4969 | 0.20098 | 0.43236 | 1 | 8722 | 4 | 1.4969 |
| NonTargetingControlG | 2 | 0.79901 | 0.79858 | 0.999994 | 17464 | 0 | 1.8295 | 0.20099 | 0.31788 | 0.989846 | 8723 | 2 | 1.8295 |
| SMG8 | 5 | 0.41697 | 0.5776 | 0.999994 | 12206 | 2 | 2.032 | 0.20103 | 0.41813 | 1 | 8724 | 3 | 2.032 |
| RPS16 | 6 | 0.32634 | 0.53706 | 0.999994 | 10651 | 2 | 1.1305 | 0.20108 | 0.43249 | 1 | 8725 | 3 | 1.1305 |
| CDC6 | 4 | 0.85356 | 0.85312 | 0.999994 | 18325 | 0 | 1.8564 | 0.20121 | 0.38498 | 1 | 8726 | 3 | 1.8564 |
| SETD4 | 6 | 0.91849 | 0.91901 | 0.999994 | 19596 | 1 | 0.86429 | 0.20129 | 0.4328 | 1 | 8727 | 3 | 0.86429 |
| GYS2 | 6 | 0.39068 | 0.59641 | 0.999994 | 11811 | 2 | 0.32732 | 0.20129 | 0.4328 | 1 | 8728 | 3 | 0.32732 |
| LUZP1 | 5 | 0.28519 | 0.46903 | 0.999994 | 9815 | 2 | -0.26996 | 0.20135 | 0.41849 | 1 | 8729 | 2 | -0.26996 |
| ZNF462 | 5 | 0.86262 | 0.86461 | 0.999994 | 18497 | 1 | -0.52564 | 0.20135 | 0.41849 | 1 | 8730 | 2 | -0.52564 |
| PQLC1 | 4 | 0.87971 | 0.87935 | 0.999994 | 18828 | 0 | 1.952 | 0.20157 | 0.38547 | 1 | 8731 | 3 | 1.952 |
| TBC1D8B | 5 | 0.53172 | 0.67558 | 0.999994 | 13929 | 1 | 0.46635 | 0.20158 | 0.41876 | 1 | 8732 | 2 | 0.46635 |
| TENM3 | 5 | 0.15808 | 0.31475 | 0.999994 | 6672 | 1 | -0.31683 | 0.20158 | 0.41876 | 1 | 8733 | 2 | -0.31683 |
| RMND5A | 4 | 0.41599 | 0.56204 | 0.999994 | 12194 | 2 | 1.9266 | 0.2017 | 0.38566 | 1 | 8734 | 2 | 1.9266 |
| POLRMT | 4 | 0.37164 | 0.51655 | 0.999994 | 11539 | 1 | 2.5527 | 0.2017 | 0.38566 | 1 | 8735 | 2 | 2.5527 |

| | | | | | | | | | | | | | |
|---|---|---|---|---|---|---|---|---|---|---|---|---|---|
| SLC27A4 | 5 | 0.53736 | 0.68056 | 0.999994 | 14018 | 1 | -0.18475 | 0.20177 | 0.41898 | 1 | 8736 | 2 | -0.18475 |
| RPS4X | 5 | 0.50053 | 0.64835 | 0.999994 | 13430 | 1 | 0.35554 | 0.20184 | 0.41907 | 1 | 8737 | 1 | 0.35554 |
| C19orf57 | 4 | 0.054967 | 0.13115 | 0.95318 | 3197 | 1 | 2.2153 | 0.20201 | 0.38611 | 1 | 8738 | 3 | 2.2153 |
| AGBL1 | 6 | 0.26528 | 0.46613 | 0.999994 | 9355 | 2 | 1.5441 | 0.20206 | 0.43389 | 1 | 8739 | 4 | 1.5441 |
| SEPT6 | 6 | 0.9763 | 0.97625 | 0.999994 | 21000 | 0 | 1.1557 | 0.20209 | 0.43393 | 1 | 8740 | 4 | 1.1557 |
| TEF | 6 | 0.75542 | 0.8304 | 0.999994 | 16921 | 1 | 1.6637 | 0.20209 | 0.43393 | 1 | 8741 | 3 | 1.6637 |
| PDZK1IP1 | 6 | 0.51885 | 0.68173 | 0.999994 | 13726 | 2 | 1.4641 | 0.20209 | 0.43393 | 1 | 8742 | 3 | 1.4641 |
| MRRF | 6 | 0.3585 | 0.57362 | 0.999994 | 11313 | 2 | 1.6295 | 0.20209 | 0.43393 | 1 | 8743 | 3 | 1.6295 |
| HOXC9 | 5 | 0.21701 | 0.39908 | 0.999994 | 8191 | 3 | -0.72732 | 0.20218 | 0.41944 | 1 | 8744 | 2 | -0.72732 |
| SMURF1 | 6 | 0.13187 | 0.30271 | 0.999994 | 5973 | 2 | -0.18127 | 0.2022 | 0.43409 | 1 | 8745 | 2 | -0.18127 |
| RNF103 | 6 | 0.91407 | 0.91508 | 0.999994 | 19488 | 1 | 1.6065 | 0.2022 | 0.43409 | 1 | 8746 | 4 | 1.6065 |
| ZNF628 | 6 | 0.30933 | 0.5176 | 0.999994 | 10313 | 2 | 1.1831 | 0.20222 | 0.43412 | 1 | 8747 | 4 | 1.1831 |
| ZSCAN31 | 6 | 0.094484 | 0.23648 | 0.985449 | 4770 | 3 | -0.51572 | 0.20222 | 0.43412 | 1 | 8748 | 2 | -0.51572 |
| GOLT1A | 6 | 0.57969 | 0.72498 | 0.999994 | 14673 | 1 | 0.75389 | 0.20222 | 0.43412 | 1 | 8749 | 3 | 0.75389 |
| IER2 | 4 | 0.62881 | 0.68122 | 0.999994 | 15359 | 1 | 1.4372 | 0.20224 | 0.38645 | 1 | 8750 | 3 | 1.4372 |
| TESPA1 | 5 | 0.18729 | 0.35709 | 0.999994 | 7401 | 2 | -0.12222 | 0.20227 | 0.41955 | 1 | 8751 | 2 | -0.12222 |
| BZRAP1 | 4 | 0.36029 | 0.50484 | 0.999994 | 11337 | 1 | 1.2745 | 0.20238 | 0.38665 | 1 | 8752 | 3 | 1.2745 |
| NonTargetingControlG | 2 | 0.79751 | 0.79703 | 0.999994 | 17446 | 0 | 1.9536 | 0.20249 | 0.31996 | 0.989846 | 8753 | 2 | 1.9536 |
| NPAS4 | 4 | 0.82674 | 0.82634 | 0.999994 | 17882 | 1 | 0.57702 | 0.20253 | 0.38687 | 1 | 8754 | 2 | 0.57702 |
| KDELR2 | 4 | 0.43928 | 0.58601 | 0.999994 | 12563 | 2 | 0.6783 | 0.20253 | 0.38687 | 1 | 8755 | 2 | 0.6783 |
| hsa-mir-6499 | 3 | 0.85087 | 0.85082 | 0.999994 | 18275 | 0 | 2.9342 | 0.20255 | 0.34577 | 0.994392 | 8756 | 2 | 2.9342 |
| PCP4 | 6 | 0.1878 | 0.37279 | 0.999994 | 7416 | 1 | 1.6646 | 0.20256 | 0.43461 | 1 | 8757 | 4 | 1.6646 |
| PPP4R1 | 6 | 0.46277 | 0.6436 | 0.999994 | 12902 | 1 | 1.3524 | 0.20265 | 0.43474 | 1 | 8758 | 4 | 1.3524 |
| ZNF134 | 6 | 0.9587 | 0.9588 | 0.999994 | 20554 | 0 | 0.80393 | 0.20265 | 0.43474 | 1 | 8759 | 3 | 0.80393 |
| C10orf99 | 4 | 0.15369 | 0.29299 | 0.999994 | 6557 | 1 | 2.1644 | 0.20266 | 0.38705 | 1 | 8760 | 2 | 2.1644 |
| AHCY | 4 | 0.72115 | 0.73888 | 0.999994 | 16509 | 1 | 2.441 | 0.20266 | 0.38705 | 1 | 8761 | 2 | 2.441 |
| MAPK10 | 4 | 0.16896 | 0.30869 | 0.999994 | 6959 | 1 | 2.1888 | 0.20266 | 0.38705 | 1 | 8762 | 2 | 2.1888 |
| NonTargetingControlG | 2 | 0.7971 | 0.79663 | 0.999994 | 17440 | 0 | 1.3216 | 0.2029 | 0.32055 | 0.989846 | 8763 | 2 | 1.3216 |
| NonTargetingControlG | 2 | 0.7971 | 0.79663 | 0.999994 | 17439 | 0 | 1.7287 | 0.2029 | 0.32055 | 0.989846 | 8764 | 2 | 1.7287 |
| EVA1A | 6 | 0.55056 | 0.70404 | 0.999994 | 14223 | 2 | 1.4002 | 0.20297 | 0.43519 | 1 | 8765 | 3 | 1.4002 |
| WNT16 | 6 | 0.75143 | 0.82897 | 0.999994 | 16870 | 1 | 0.21141 | 0.203 | 0.43523 | 1 | 8766 | 3 | 0.21141 |
| CYP2W1 | 4 | 0.66877 | 0.70463 | 0.999994 | 15904 | 1 | 1.8907 | 0.20306 | 0.3876 | 1 | 8767 | 3 | 1.8907 |
| OR10V1 | 5 | 0.5365 | 0.67981 | 0.999994 | 14009 | 2 | -0.080498 | 0.20306 | 0.42042 | 1 | 8768 | 2 | -0.080498 |
| FAM169A | 5 | 0.056099 | 0.15507 | 0.967126 | 3252 | 3 | -1.9413 | 0.20306 | 0.42042 | 1 | 8769 | 2 | -1.9413 |
| HRASLS | 5 | 0.23093 | 0.41847 | 0.999994 | 8539 | 3 | -0.6674 | 0.20306 | 0.42042 | 1 | 8770 | 2 | -0.6674 |
| EBNA1BP2 | 4 | 0.085055 | 0.18801 | 0.969543 | 4410 | 2 | 0.59401 | 0.20306 | 0.38761 | 1 | 8771 | 2 | 0.59401 |
| GPR33 | 6 | 0.73499 | 0.82328 | 0.999994 | 16662 | 1 | 2.1968 | 0.20313 | 0.4354 | 1 | 8772 | 4 | 2.1968 |
| C22orf39 | 6 | 0.077187 | 0.20166 | 0.977164 | 4115 | 2 | 0.53376 | 0.20318 | 0.43548 | 1 | 8773 | 3 | 0.53376 |
| HRH3 | 6 | 0.28659 | 0.49113 | 0.999994 | 9849 | 3 | 0.22279 | 0.20318 | 0.43548 | 1 | 8774 | 3 | 0.22279 |
| LEO1 | 6 | 0.33851 | 0.55098 | 0.999994 | 10909 | 3 | -0.068725 | 0.20318 | 0.43548 | 1 | 8775 | 2 | -0.068725 |
| NUAK2 | 4 | 0.14213 | 0.28111 | 0.999632 | 6243 | 1 | 1.3704 | 0.20318 | 0.38779 | 1 | 8776 | 3 | 1.3704 |
| hsa-mir-4734 | 3 | 0.048496 | 0.11364 | 0.951052 | 2908 | 1 | 2.8135 | 0.20326 | 0.34656 | 0.994392 | 8777 | 2 | 2.8135 |
| MECOM | 6 | 0.62095 | 0.75554 | 0.999994 | 15236 | 1 | 1.5576 | 0.20329 | 0.43562 | 1 | 8778 | 4 | 1.5576 |
| COL18A1 | 4 | 0.26417 | 0.40632 | 0.999994 | 9336 | 1 | -0.11066 | 0.20331 | 0.38794 | 1 | 8779 | 1 | -0.11066 |
| PDGFRA | 6 | 0.48931 | 0.66147 | 0.999994 | 13270 | 2 | 0.68287 | 0.20333 | 0.43567 | 1 | 8780 | 3 | 0.68287 |
| LANCL3 | 6 | 0.81359 | 0.85388 | 0.999994 | 17685 | 1 | 1.6259 | 0.20334 | 0.4357 | 1 | 8781 | 4 | 1.6259 |
| CYP3A5 | 5 | 0.22993 | 0.41713 | 0.999994 | 8526 | 1 | 1.1279 | 0.20334 | 0.42074 | 1 | 8782 | 3 | 1.1279 |
| NonTargetingControlG | 2 | 0.79653 | 0.79609 | 0.999994 | 17435 | 0 | 2.3364 | 0.20347 | 0.32133 | 0.989846 | 8783 | 2 | 2.3364 |
| NonTargetingControlG | 1 | 0.79651 | 0.79687 | 0.999994 | 17434 | 0 | 4.8715 | 0.20349 | 0.20322 | 0.97466 | 8784 | 1 | 4.8715 |
| TYROBP | 5 | 0.86947 | 0.87062 | 0.999994 | 18630 | 1 | 0.29117 | 0.20349 | 0.4209 | 1 | 8785 | 2 | 0.29117 |
| PSG9 | 6 | 0.1059 | 0.25881 | 0.994943 | 5170 | 2 | 1.4435 | 0.2035 | 0.43592 | 1 | 8786 | 4 | 1.4435 |
| KIAA1456 | 6 | 0.25823 | 0.45779 | 0.999994 | 9200 | 1 | 1.4382 | 0.2036 | 0.43607 | 1 | 8787 | 4 | 1.4382 |

| | | | | | | | | | | | | |
|---|---|---|---|---|---|---|---|---|---|---|---|---|
| hsa-mir-609 | 3 | 0.87416 | 0.8743 | 0.999994 | 18713 | 0 | 2.7792 | 0.2036 | 0.34694 | 0.994392 | 8788 | 2 | 2.7792 |
| MIEF1 | 6 | 0.055821 | 0.15669 | 0.967126 | 3240 | 2 | 1.4935 | 0.20361 | 0.43609 | 1 | 8789 | 3 | 1.4935 |
| AGK | 6 | 0.3854 | 0.59299 | 0.999994 | 11724 | 1 | 1.308 | 0.20365 | 0.43615 | 1 | 8790 | 4 | 1.308 |
| OTOP1 | 6 | 0.44431 | 0.63129 | 0.999994 | 12637 | 2 | 1.9029 | 0.20366 | 0.43617 | 1 | 8791 | 4 | 1.9029 |
| MIER1 | 4 | 0.77861 | 0.78276 | 0.999994 | 17191 | 1 | 1.214 | 0.20368 | 0.38846 | 1 | 8792 | 3 | 1.214 |
| SMC3 | 4 | 0.88306 | 0.88271 | 0.999994 | 18893 | 0 | 2.1383 | 0.20369 | 0.38847 | 1 | 8793 | 2 | 2.1383 |
| GTF2H1 | 6 | 0.90315 | 0.90613 | 0.999994 | 19282 | 1 | 1.3617 | 0.20375 | 0.43629 | 1 | 8794 | 4 | 1.3617 |
| TMCC3 | 5 | 0.62208 | 0.73434 | 0.999994 | 15260 | 1 | 2.1706 | 0.20377 | 0.42122 | 1 | 8795 | 3 | 2.1706 |
| TPRG1 | 6 | 0.022101 | 0.078697 | 0.93914 | 1592 | 4 | -2.3255 | 0.20382 | 0.43639 | 1 | 8796 | 2 | -2.3255 |
| ZBTB6 | 5 | 0.23093 | 0.41847 | 0.999994 | 8540 | 2 | 0.32052 | 0.2039 | 0.42138 | 1 | 8797 | 2 | 0.32052 |
| NUBP2 | 5 | 0.050828 | 0.14239 | 0.959553 | 3008 | 2 | -0.57696 | 0.2039 | 0.42138 | 1 | 8798 | 2 | -0.57696 |
| MESDC2 | 5 | 0.13995 | 0.28776 | 0.999632 | 6182 | 3 | -1.2614 | 0.2039 | 0.42138 | 1 | 8799 | 1 | -1.2614 |
| KIF13B | 6 | 0.6652 | 0.78937 | 0.999994 | 15857 | 2 | 1.548 | 0.20392 | 0.43654 | 1 | 8800 | 3 | 1.548 |
| NonTargetingControlG | 2 | 0.41654 | 0.43057 | 0.999994 | 12200 | 1 | 2.6023 | 0.20393 | 0.32193 | 0.989846 | 8801 | 1 | 2.6023 |
| NonTargetingControlG | 2 | 0.64088 | 0.64067 | 0.999994 | 15528 | 0 | 3.3817 | 0.20393 | 0.32193 | 0.989846 | 8802 | 1 | 3.3817 |
| ZNF658 | 6 | 0.5604 | 0.71106 | 0.999994 | 14376 | 2 | 1.1283 | 0.20393 | 0.43656 | 1 | 8803 | 3 | 1.1283 |
| hsa-mir-125a | 4 | 0.3318 | 0.47557 | 0.999994 | 10760 | 2 | 1.5798 | 0.204 | 0.3889 | 1 | 8804 | 2 | 1.5798 |
| TRMU | 6 | 0.77152 | 0.83641 | 0.999994 | 17106 | 1 | 1.2946 | 0.20406 | 0.43673 | 1 | 8805 | 3 | 1.2946 |
| TXNDC2 | 6 | 0.14976 | 0.32533 | 0.999994 | 6453 | 1 | 1.1766 | 0.20409 | 0.43678 | 1 | 8806 | 3 | 1.1766 |
| LOC730183 | 3 | 0.25639 | 0.3829 | 0.999994 | 9159 | 1 | -1.2059 | 0.20411 | 0.3475 | 0.994392 | 8807 | 1 | -1.2059 |
| hsa-mir-208b | 4 | 0.49705 | 0.615 | 0.999994 | 13378 | 1 | 2.2037 | 0.20422 | 0.38921 | 1 | 8808 | 3 | 2.2037 |
| CHRNB3 | 5 | 0.19295 | 0.36514 | 0.999994 | 7576 | 2 | 0.34891 | 0.20424 | 0.42174 | 1 | 8809 | 2 | 0.34891 |
| SGTB | 5 | 0.36119 | 0.53106 | 0.999994 | 11357 | 2 | 1.6495 | 0.20424 | 0.42174 | 1 | 8810 | 3 | 1.6495 |
| OR4N2 | 5 | 0.56291 | 0.70321 | 0.999994 | 14426 | 2 | 2.1807 | 0.20433 | 0.42183 | 1 | 8811 | 3 | 2.1807 |
| FLJ44313 | 5 | 0.73803 | 0.78468 | 0.999994 | 16705 | 1 | 1.6948 | 0.20433 | 0.42184 | 1 | 8812 | 3 | 1.6948 |
| PYCARD | 5 | 0.098958 | 0.22479 | 0.983354 | 4928 | 3 | -1.5109 | 0.20433 | 0.42184 | 1 | 8813 | 2 | -1.5109 |
| hsa-mir-4802 | 4 | 0.76047 | 0.76816 | 0.999994 | 16986 | 1 | 2.4203 | 0.20433 | 0.38938 | 1 | 8814 | 2 | 2.4203 |
| RPS6KA3 | 6 | 0.84557 | 0.86956 | 0.999994 | 18177 | 1 | 1.8573 | 0.2045 | 0.43736 | 1 | 8815 | 3 | 1.8573 |
| HINT1 | 5 | 0.56504 | 0.70515 | 0.999994 | 14470 | 1 | 2.2268 | 0.20453 | 0.42205 | 1 | 8816 | 3 | 2.2268 |
| CADPS2 | 6 | 0.23793 | 0.43351 | 0.999994 | 8724 | 1 | 1.7375 | 0.20453 | 0.43741 | 1 | 8817 | 4 | 1.7375 |
| HECTD3 | 6 | 0.02311 | 0.081929 | 0.93914 | 1657 | 1 | 1.2692 | 0.20457 | 0.43747 | 1 | 8818 | 4 | 1.2692 |
| MGMT | 5 | 0.13006 | 0.27267 | 0.998561 | 5934 | 3 | -1.4037 | 0.20461 | 0.42215 | 1 | 8819 | 2 | -1.4037 |
| MAP3K5 | 5 | 0.88207 | 0.88225 | 0.999994 | 18871 | 1 | 2.2214 | 0.20461 | 0.42215 | 1 | 8820 | 3 | 2.2214 |
| MAP1LC3B | 6 | 0.13349 | 0.30477 | 0.999994 | 6007 | 3 | -0.58364 | 0.20461 | 0.43754 | 1 | 8821 | 3 | -0.58364 |
| PDE1B | 3 | 0.38185 | 0.47385 | 0.999994 | 11674 | 1 | 2.7297 | 0.20466 | 0.34809 | 0.994392 | 8822 | 2 | 2.7297 |
| ALOX12B | 6 | 0.8761 | 0.88713 | 0.999994 | 18753 | 1 | 1.5605 | 0.20467 | 0.43761 | 1 | 8823 | 3 | 1.5605 |
| ATP5S | 6 | 0.75174 | 0.8291 | 0.999994 | 16875 | 1 | 1.4431 | 0.20471 | 0.43768 | 1 | 8824 | 4 | 1.4431 |
| NonTargetingControlG | 2 | 0.15043 | 0.22694 | 0.983354 | 6467 | 1 | -0.044754 | 0.20479 | 0.32314 | 0.989881 | 8825 | 1 | -0.044754 |
| NonTargetingControlG | 2 | 0.79518 | 0.79473 | 0.999994 | 17419 | 0 | 1.2311 | 0.20482 | 0.32317 | 0.989881 | 8826 | 2 | 1.2311 |
| TM6SF1 | 6 | 0.87936 | 0.88918 | 0.999994 | 18817 | 1 | 1.5641 | 0.20482 | 0.43781 | 1 | 8827 | 3 | 1.5641 |
| OR6F1 | 5 | 0.94227 | 0.94246 | 0.999994 | 20132 | 0 | 1.4378 | 0.20483 | 0.42239 | 1 | 8828 | 3 | 1.4378 |
| SPATA20 | 6 | 0.10351 | 0.25418 | 0.994632 | 5085 | 2 | 1.6735 | 0.20483 | 0.43783 | 1 | 8829 | 3 | 1.6735 |
| CYB5D2 | 6 | 0.13508 | 0.30678 | 0.999994 | 6051 | 2 | 1.983 | 0.20485 | 0.43785 | 1 | 8830 | 4 | 1.983 |
| hsa-mir-2861 | 4 | 0.38743 | 0.53272 | 0.999994 | 11758 | 1 | 2.1347 | 0.20488 | 0.39018 | 1 | 8831 | 3 | 2.1347 |
| RGPD2 | 4 | 0.73225 | 0.74683 | 0.999994 | 16631 | 1 | 0.47708 | 0.20488 | 0.39018 | 1 | 8832 | 2 | 0.47708 |
| PKIB | 6 | 0.54217 | 0.69808 | 0.999994 | 14096 | 2 | 1.1772 | 0.20489 | 0.43792 | 1 | 8833 | 4 | 1.1772 |
| ANAPC5 | 4 | 0.58872 | 0.65925 | 0.999994 | 14798 | 1 | 2.6548 | 0.2049 | 0.3902 | 1 | 8834 | 3 | 2.6548 |
| TXNDC11 | 6 | 0.51129 | 0.67651 | 0.999994 | 13601 | 1 | 1.3911 | 0.20491 | 0.43794 | 1 | 8835 | 4 | 1.3911 |
| PCDHA6 | 2 | 0.79508 | 0.79464 | 0.999994 | 17417 | 0 | 0.85445 | 0.20492 | 0.32332 | 0.989881 | 8836 | 2 | 0.85445 |
| RET | 4 | 0.61005 | 0.67077 | 0.999994 | 15072 | 1 | 1.2284 | 0.20497 | 0.3903 | 1 | 8837 | 3 | 1.2284 |
| UBQLN3 | 6 | 0.22745 | 0.42101 | 0.999994 | 8468 | 2 | 1.1548 | 0.20499 | 0.43806 | 1 | 8838 | 4 | 1.1548 |
| BPTF | 5 | 0.21891 | 0.40175 | 0.999994 | 8257 | 2 | 2.0425 | 0.205 | 0.42259 | 1 | 8839 | 3 | 2.0425 |

| | | | | | | | | | | | | |
|---|---|---|---|---|---|---|---|---|---|---|---|---|
| MRPS12 | 4 | 0.7282 | 0.74385 | 0.999994 | 16584 | 1 | 1.9433 | 0.20501 | 0.39035 | 1 | 8840 | 3 | 1.9433 |
| PTCHD2 | 6 | 0.86065 | 0.87778 | 0.999994 | 18453 | 1 | 1.756 | 0.20505 | 0.43813 | 1 | 8841 | 3 | 1.756 |
| TUFT1 | 6 | 0.46705 | 0.64649 | 0.999994 | 12967 | 2 | 1.5341 | 0.20512 | 0.43824 | 1 | 8842 | 4 | 1.5341 |
| TMEM17 | 4 | 0.102 | 0.21869 | 0.980699 | 5039 | 1 | 2.2594 | 0.20523 | 0.39066 | 1 | 8843 | 3 | 2.2594 |
| PPP1R14D | 6 | 0.077491 | 0.20227 | 0.977973 | 4128 | 3 | 1.0905 | 0.20524 | 0.4384 | 1 | 8844 | 3 | 1.0905 |
| FZD3 | 6 | 0.85003 | 0.87195 | 0.999994 | 18262 | 1 | 1.1814 | 0.20527 | 0.43845 | 1 | 8845 | 4 | 1.1814 |
| FGGY | 4 | 0.86622 | 0.86581 | 0.999994 | 18564 | 0 | 0.87378 | 0.20532 | 0.39079 | 1 | 8846 | 2 | 0.87378 |
| NRROS | 4 | 0.89666 | 0.89637 | 0.999994 | 19156 | 0 | 1.6009 | 0.20532 | 0.39079 | 1 | 8847 | 2 | 1.6009 |
| NonTargetingControlG | 2 | 0.79467 | 0.79425 | 0.999994 | 17407 | 0 | 3.762 | 0.20533 | 0.32389 | 0.990363 | 8848 | 2 | 3.762 |
| SV2C | 5 | 0.26624 | 0.45381 | 0.999994 | 9371 | 2 | 1.1139 | 0.20534 | 0.42296 | 1 | 8849 | 3 | 1.1139 |
| TWIST1 | 5 | 0.035067 | 0.10341 | 0.946414 | 2271 | 2 | -0.14939 | 0.20534 | 0.42296 | 1 | 8850 | 2 | -0.14939 |
| FMNL3 | 5 | 0.38276 | 0.54901 | 0.999994 | 11688 | 2 | 2.1322 | 0.20537 | 0.42301 | 1 | 8851 | 3 | 2.1322 |
| NDUFS1 | 5 | 0.3592 | 0.52938 | 0.999994 | 11323 | 2 | -0.61093 | 0.20542 | 0.42305 | 1 | 8852 | 2 | -0.61093 |
| AKAP1 | 6 | 0.45797 | 0.64038 | 0.999994 | 12820 | 2 | 1.3317 | 0.2055 | 0.43874 | 1 | 8853 | 3 | 1.3317 |
| GDF9 | 6 | 0.35514 | 0.56982 | 0.999994 | 11253 | 1 | 0.86761 | 0.2055 | 0.43875 | 1 | 8854 | 3 | 0.86761 |
| CHEK1 | 3 | 0.013978 | 0.03679 | 0.881473 | 1126 | 1 | 2.7493 | 0.20551 | 0.34898 | 0.994392 | 8855 | 2 | 2.7493 |
| ZBTB1 | 4 | 0.055816 | 0.13279 | 0.95318 | 3239 | 1 | 2.6783 | 0.20552 | 0.39108 | 1 | 8856 | 3 | 2.6783 |
| hsa-mir-601 | 2 | 0.79436 | 0.79394 | 0.999994 | 17399 | 0 | 2.4208 | 0.20564 | 0.3243 | 0.990429 | 8857 | 2 | 2.4208 |
| SVOPL | 5 | 0.1952 | 0.36837 | 0.999994 | 7638 | 3 | -0.82101 | 0.20569 | 0.42336 | 1 | 8858 | 2 | -0.82101 |
| PHF23 | 4 | 0.81983 | 0.81961 | 0.999994 | 17776 | 1 | 2.6839 | 0.2057 | 0.39134 | 1 | 8859 | 3 | 2.6839 |
| hsa-mir-4472-2 | 1 | 0.79428 | 0.7946 | 0.999994 | 17398 | 0 | 4.8199 | 0.20572 | 0.20543 | 0.97466 | 8860 | 1 | 4.8199 |
| ADI1 | 6 | 0.67716 | 0.79876 | 0.999994 | 16046 | 1 | 1.1001 | 0.20573 | 0.43908 | 1 | 8861 | 3 | 1.1001 |
| WIZ | 6 | 0.327 | 0.53782 | 0.999994 | 10661 | 2 | 1.5221 | 0.20573 | 0.43908 | 1 | 8862 | 4 | 1.5221 |
| CEP55 | 6 | 0.0063474 | 0.026065 | 0.848412 | 615 | 2 | 0.81606 | 0.20573 | 0.43908 | 1 | 8863 | 4 | 0.81606 |
| hsa-mir-3182 | 4 | 0.79007 | 0.79244 | 0.999994 | 17345 | 1 | 1.3234 | 0.20574 | 0.3914 | 1 | 8864 | 2 | 1.3234 |
| hsa-mir-6500 | 3 | 0.053837 | 0.12101 | 0.951052 | 3154 | 2 | -3.293 | 0.20582 | 0.34931 | 0.994392 | 8865 | 1 | -3.293 |
| hsa-mir-562 | 3 | 0.62762 | 0.63877 | 0.999994 | 15345 | 1 | 1.4592 | 0.20582 | 0.34931 | 0.994392 | 8866 | 2 | 1.4592 |
| SDPR | 6 | 0.59199 | 0.73408 | 0.999994 | 14851 | 2 | 0.30428 | 0.20585 | 0.43926 | 1 | 8867 | 2 | 0.30428 |
| C19orf10 | 6 | 0.38136 | 0.59044 | 0.999994 | 11668 | 2 | 1.5919 | 0.20586 | 0.43927 | 1 | 8868 | 4 | 1.5919 |
| METTL4 | 6 | 0.69657 | 0.81096 | 0.999994 | 16264 | 1 | 2.0777 | 0.2059 | 0.43933 | 1 | 8869 | 4 | 2.0777 |
| RNASE9 | 6 | 0.17419 | 0.35586 | 0.999994 | 7089 | 2 | 1.4986 | 0.20593 | 0.43938 | 1 | 8870 | 4 | 1.4986 |
| WIPI2 | 6 | 0.19423 | 0.38082 | 0.999994 | 7609 | 2 | 1.3407 | 0.20612 | 0.43966 | 1 | 8871 | 4 | 1.3407 |
| MRS2 | 6 | 0.10868 | 0.26418 | 0.996734 | 5263 | 3 | -0.49873 | 0.20618 | 0.43973 | 1 | 8872 | 2 | -0.49873 |
| ZP4 | 6 | 0.075971 | 0.19918 | 0.977164 | 4071 | 2 | 0.96358 | 0.20618 | 0.43973 | 1 | 8873 | 4 | 0.96358 |
| GRIA4 | 6 | 0.83549 | 0.86428 | 0.999994 | 18017 | 1 | -0.096382 | 0.20618 | 0.43973 | 1 | 8874 | 2 | -0.096382 |
| LTB | 5 | 0.95517 | 0.95532 | 0.999994 | 20456 | 0 | 0.38609 | 0.20628 | 0.424 | 1 | 8875 | 2 | 0.38609 |
| TNFSF12-TNFSF13 | 5 | 0.94946 | 0.94965 | 0.999994 | 20320 | 0 | 1.186 | 0.20628 | 0.424 | 1 | 8876 | 3 | 1.186 |
| CXCL1 | 5 | 0.49584 | 0.64429 | 0.999994 | 13353 | 2 | -0.51266 | 0.20628 | 0.424 | 1 | 8877 | 2 | -0.51266 |
| KIAA1239 | 5 | 0.14973 | 0.30238 | 0.999994 | 6451 | 1 | 1.1922 | 0.20628 | 0.424 | 1 | 8878 | 3 | 1.1922 |
| C22orf46 | 6 | 0.049004 | 0.14191 | 0.959146 | 2929 | 2 | 1.7714 | 0.20631 | 0.4399 | 1 | 8879 | 4 | 1.7714 |
| TRIM46 | 5 | 0.46359 | 0.61669 | 0.999994 | 12911 | 1 | 1.3116 | 0.20634 | 0.42406 | 1 | 8880 | 3 | 1.3116 |
| ATG14 | 5 | 0.79265 | 0.81494 | 0.999994 | 17385 | 1 | 2.2214 | 0.20634 | 0.42406 | 1 | 8881 | 3 | 2.2214 |
| FCAMR | 6 | 0.65277 | 0.77975 | 0.999994 | 15696 | 2 | 1.2051 | 0.20637 | 0.43998 | 1 | 8882 | 4 | 1.2051 |
| HEXIM1 | 4 | 0.12563 | 0.26008 | 0.994943 | 5811 | 2 | 1.3875 | 0.2064 | 0.39236 | 1 | 8883 | 2 | 1.3875 |
| RRAGA | 4 | 0.51874 | 0.62483 | 0.999994 | 13722 | 1 | 2.3727 | 0.2064 | 0.39236 | 1 | 8884 | 2 | 2.3727 |
| PSTK | 4 | 0.34044 | 0.48445 | 0.999994 | 10945 | 1 | 2.7538 | 0.2064 | 0.39236 | 1 | 8885 | 3 | 2.7538 |
| hsa-mir-302d | 4 | 0.33118 | 0.47491 | 0.999994 | 10748 | 2 | 1.7704 | 0.2064 | 0.39236 | 1 | 8886 | 2 | 1.7704 |
| PRY | 6 | 0.57353 | 0.72048 | 0.999994 | 14598 | 1 | 1.6054 | 0.20644 | 0.44008 | 1 | 8887 | 4 | 1.6054 |
| KCNK10 | 6 | 0.028037 | 0.094155 | 0.945031 | 1903 | 3 | 1.0084 | 0.20648 | 0.44013 | 1 | 8888 | 3 | 1.0084 |
| BSG | 6 | 0.29734 | 0.50362 | 0.999994 | 10075 | 3 | 0.0084675 | 0.20653 | 0.44021 | 1 | 8889 | 3 | 0.0084675 |
| HHIPL2 | 6 | 0.50026 | 0.66887 | 0.999994 | 13427 | 2 | 0.13623 | 0.20653 | 0.44021 | 1 | 8890 | 2 | 0.13623 |
| C16orf78 | 6 | 0.038186 | 0.11783 | 0.951052 | 2427 | 2 | 1.7338 | 0.20656 | 0.44026 | 1 | 8891 | 3 | 1.7338 |

| | | | | | | | | | | | | | |
|---|---|---|---|---|---|---|---|---|---|---|---|---|---|
| TIRAP | 6 | 0.12838 | 0.2982 | 0.999994 | 5890 | 2 | 0.37979 | 0.20664 | 0.44039 | 1 | 8892 | 3 | 0.37979 |
| PLEKHA5 | 6 | 0.31809 | 0.52772 | 0.999994 | 10475 | 3 | 0.12384 | 0.20664 | 0.44039 | 1 | 8893 | 3 | 0.12384 |
| NEUROD6 | 6 | 0.10341 | 0.25398 | 0.994632 | 5079 | 3 | -0.17458 | 0.20664 | 0.44039 | 1 | 8894 | 2 | -0.17458 |
| HIGD1C | 6 | 0.90842 | 0.91029 | 0.999994 | 19394 | 1 | 1.0348 | 0.20664 | 0.44039 | 1 | 8895 | 3 | 1.0348 |
| DNMT3A | 6 | 0.8416 | 0.86746 | 0.999994 | 18114 | 1 | 1.4479 | 0.20664 | 0.44039 | 1 | 8896 | 4 | 1.4479 |
| IMMP2L | 6 | 0.49019 | 0.66206 | 0.999994 | 13280 | 2 | 1.1743 | 0.20672 | 0.4405 | 1 | 8897 | 3 | 1.1743 |
| DHFR | 4 | 0.35063 | 0.49496 | 0.999994 | 11175 | 2 | -0.7306 | 0.20674 | 0.39284 | 1 | 8898 | 1 | -0.7306 |
| SLC26A4 | 6 | 0.18437 | 0.36861 | 0.999994 | 7324 | 3 | -0.76066 | 0.20686 | 0.4407 | 1 | 8899 | 2 | -0.76066 |
| BECN1 | 6 | 0.52955 | 0.68923 | 0.999994 | 13893 | 2 | 0.42484 | 0.20686 | 0.4407 | 1 | 8900 | 2 | 0.42484 |
| EYS | 6 | 0.81047 | 0.85247 | 0.999994 | 17639 | 1 | 1.42 | 0.20692 | 0.44078 | 1 | 8901 | 3 | 1.42 |
| FN3KRP | 5 | 0.27175 | 0.45821 | 0.999994 | 9516 | 2 | -0.28875 | 0.2072 | 0.42504 | 1 | 8902 | 1 | -0.28875 |
| PPP1R16B | 5 | 0.51408 | 0.66016 | 0.999994 | 13643 | 2 | 0.6096 | 0.2072 | 0.42504 | 1 | 8903 | 3 | 0.6096 |
| TAT | 5 | 0.047056 | 0.13329 | 0.95318 | 2856 | 3 | -0.91068 | 0.2072 | 0.42504 | 1 | 8904 | 2 | -0.91068 |
| FGF7 | 4 | 0.58863 | 0.65919 | 0.999994 | 14797 | 1 | -0.34131 | 0.20723 | 0.39351 | 1 | 8905 | 1 | -0.34131 |
| CSN2 | 4 | 0.91488 | 0.91464 | 0.999994 | 19507 | 0 | 1.6199 | 0.20723 | 0.39351 | 1 | 8906 | 3 | 1.6199 |
| ITGA8 | 6 | 0.21501 | 0.40616 | 0.999994 | 8142 | 2 | 1.1723 | 0.20723 | 0.4412 | 1 | 8907 | 3 | 1.1723 |
| FAM213A | 6 | 0.2471 | 0.44448 | 0.999994 | 8952 | 2 | 0.72707 | 0.20723 | 0.4412 | 1 | 8908 | 3 | 0.72707 |
| INTS12 | 6 | 0.23522 | 0.43031 | 0.999994 | 8655 | 3 | -0.024022 | 0.20723 | 0.4412 | 1 | 8909 | 3 | -0.024022 |
| CDX1 | 6 | 0.48219 | 0.65665 | 0.999994 | 13173 | 2 | 1.3744 | 0.20728 | 0.44127 | 1 | 8910 | 4 | 1.3744 |
| PHOSPHO1 | 6 | 0.56396 | 0.71362 | 0.999994 | 14451 | 1 | 1.781 | 0.20731 | 0.44132 | 1 | 8911 | 4 | 1.781 |
| ADAM7 | 6 | 0.86029 | 0.87758 | 0.999994 | 18447 | 1 | 1.7457 | 0.20736 | 0.44139 | 1 | 8912 | 3 | 1.7457 |
| CDC42EP5 | 6 | 0.090911 | 0.22934 | 0.983354 | 4634 | 2 | 1.3946 | 0.20736 | 0.4414 | 1 | 8913 | 4 | 1.3946 |
| hsa-mir-1268a | 1 | 0.79262 | 0.79294 | 0.999994 | 17383 | 0 | 4.7663 | 0.20738 | 0.20711 | 0.97466 | 8914 | 1 | 4.7663 |
| MSRB2 | 5 | 0.36238 | 0.53204 | 0.999994 | 11375 | 2 | 1.7139 | 0.20739 | 0.42524 | 1 | 8915 | 3 | 1.7139 |
| CTNNA2 | 5 | 0.41091 | 0.57265 | 0.999994 | 12124 | 2 | -0.55897 | 0.20739 | 0.42524 | 1 | 8916 | 2 | -0.55897 |
| FPR2 | 6 | 0.77029 | 0.83594 | 0.999994 | 17087 | 1 | 1.3333 | 0.20739 | 0.44144 | 1 | 8917 | 4 | 1.3333 |
| LIPJ | 6 | 0.95795 | 0.95804 | 0.999994 | 20537 | 0 | 1.7832 | 0.20746 | 0.44154 | 1 | 8918 | 4 | 1.7832 |
| ATP5SL | 6 | 0.75651 | 0.8308 | 0.999994 | 16938 | 1 | 1.7087 | 0.20747 | 0.44155 | 1 | 8919 | 3 | 1.7087 |
| CWC15 | 3 | 0.031608 | 0.078846 | 0.93914 | 2085 | 2 | -5.9023 | 0.20748 | 0.3511 | 0.994392 | 8920 | 1 | -5.9023 |
| CLIC3 | 6 | 0.6623 | 0.7871 | 0.999994 | 15814 | 2 | 0.54799 | 0.20755 | 0.44167 | 1 | 8921 | 3 | 0.54799 |
| CHST8 | 6 | 0.10788 | 0.26263 | 0.996196 | 5235 | 4 | -0.99395 | 0.20755 | 0.44167 | 1 | 8922 | 2 | -0.99395 |
| hsa-mir-4774 | 4 | 0.67239 | 0.70684 | 0.999994 | 15960 | 1 | 1.1621 | 0.20756 | 0.39399 | 1 | 8923 | 3 | 1.1621 |
| FAM57A | 4 | 0.0082901 | 0.028304 | 0.858439 | 748 | 3 | -4.7742 | 0.20762 | 0.39408 | 1 | 8924 | 1 | -4.7742 |
| GOSR1 | 4 | 0.057332 | 0.13565 | 0.95468 | 3307 | 2 | -0.67568 | 0.20762 | 0.39408 | 1 | 8925 | 2 | -0.67568 |
| EIF2S3 | 4 | 0.0019032 | 0.0069206 | 0.643671 | 261 | 3 | -5.5894 | 0.20762 | 0.39408 | 1 | 8926 | 1 | -5.5894 |
| NonTargetingControlG | 2 | 0.79236 | 0.79192 | 0.999994 | 17379 | 0 | 3.0439 | 0.20764 | 0.32706 | 0.991729 | 8927 | 2 | 3.0439 |
| WDR62 | 6 | 0.88202 | 0.89092 | 0.999994 | 18870 | 1 | 1.1434 | 0.20766 | 0.44183 | 1 | 8928 | 4 | 1.1434 |
| CDKN3 | 6 | 0.17949 | 0.36253 | 0.999994 | 7210 | 1 | 1.2817 | 0.2077 | 0.44189 | 1 | 8929 | 4 | 1.2817 |
| CPSF4L | 6 | 0.57238 | 0.71967 | 0.999994 | 14582 | 2 | 1.0742 | 0.20773 | 0.44193 | 1 | 8930 | 4 | 1.0742 |
| GJB1 | 6 | 0.73355 | 0.8228 | 0.999994 | 16650 | 1 | 1.5314 | 0.20773 | 0.44193 | 1 | 8931 | 4 | 1.5314 |
| EEF1B2 | 6 | 0.037651 | 0.11661 | 0.951052 | 2412 | 4 | -2.0193 | 0.20773 | 0.44193 | 1 | 8932 | 2 | -2.0193 |
| TOPAZ1 | 6 | 0.46737 | 0.64669 | 0.999994 | 12969 | 1 | 1.3539 | 0.20773 | 0.44193 | 1 | 8933 | 4 | 1.3539 |
| ABCC12 | 5 | 0.85975 | 0.86219 | 0.999994 | 18429 | 1 | -0.057527 | 0.20781 | 0.4257 | 1 | 8934 | 2 | -0.057527 |
| RPUSD3 | 5 | 0.14497 | 0.29524 | 0.999994 | 6325 | 3 | -1.2682 | 0.20781 | 0.4257 | 1 | 8935 | 2 | -1.2682 |
| HSD17B13 | 6 | 0.34525 | 0.5587 | 0.999994 | 11055 | 2 | 1.6425 | 0.20781 | 0.44204 | 1 | 8936 | 3 | 1.6425 |
| SLC46A3 | 6 | 0.5769 | 0.72293 | 0.999994 | 14643 | 1 | 1.1589 | 0.20791 | 0.44219 | 1 | 8937 | 4 | 1.1589 |
| TEX35 | 6 | 0.24898 | 0.44671 | 0.999994 | 8991 | 2 | -0.11305 | 0.208 | 0.44232 | 1 | 8938 | 2 | -0.11305 |
| LYNX1 | 6 | 0.38653 | 0.5937 | 0.999994 | 11742 | 2 | 1.0292 | 0.208 | 0.44232 | 1 | 8939 | 3 | 1.0292 |
| RPL10A | 5 | 0.16198 | 0.32044 | 0.999994 | 6795 | 2 | 0.67771 | 0.20818 | 0.42612 | 1 | 8940 | 3 | 0.67771 |
| hsa-mir-1200 | 4 | 0.87829 | 0.87793 | 0.999994 | 18793 | 0 | 1.9038 | 0.20819 | 0.39488 | 1 | 8941 | 3 | 1.9038 |
| FAM26D | 6 | 0.85086 | 0.87238 | 0.999994 | 18274 | 1 | 1.1746 | 0.20825 | 0.44266 | 1 | 8942 | 4 | 1.1746 |
| AGPS | 6 | 0.068432 | 0.1835 | 0.968526 | 3768 | 3 | -0.31738 | 0.20828 | 0.4427 | 1 | 8943 | 3 | -0.31738 |

| | | | | | | | | | | | | | |
|---|---|---|---|---|---|---|---|---|---|---|---|---|---|
| CYP2A13 | 6 | 0.95533 | 0.95538 | 0.999994 | 20460 | 0 | 1.6237 | 0.20833 | 0.44277 | 1 | 8944 | 3 | 1.6237 |
| DNAJB4 | 6 | 0.30879 | 0.51698 | 0.999994 | 10294 | 2 | 1.3572 | 0.20836 | 0.44283 | 1 | 8945 | 4 | 1.3572 |
| SEC24A | 4 | 0.065962 | 0.15215 | 0.967126 | 3687 | 1 | 1.0996 | 0.20851 | 0.39533 | 1 | 8946 | 3 | 1.0996 |
| hsa-mir-6086 | 2 | 0.79145 | 0.79103 | 0.999994 | 17371 | 0 | 2.0508 | 0.20855 | 0.32831 | 0.991729 | 8947 | 2 | 2.0508 |
| RBMXL2 | 5 | 0.00078556 | 0.0035619 | 0.553006 | 139 | 2 | 1.9791 | 0.2086 | 0.4266 | 1 | 8948 | 3 | 1.9791 |
| NR2C1 | 6 | 0.84478 | 0.86914 | 0.999994 | 18170 | 1 | 0.6537 | 0.20865 | 0.44324 | 1 | 8949 | 4 | 0.6537 |
| LRIT1 | 6 | 0.92567 | 0.92576 | 0.999994 | 19760 | 1 | 0.77388 | 0.20865 | 0.44324 | 1 | 8950 | 3 | 0.77388 |
| hsa-mir-940 | 4 | 0.30945 | 0.45265 | 0.999994 | 10315 | 2 | 0.18594 | 0.20873 | 0.39562 | 1 | 8951 | 2 | 0.18594 |
| GPR124 | 6 | 0.98074 | 0.98066 | 0.999994 | 21109 | 0 | 1.1313 | 0.20873 | 0.44336 | 1 | 8952 | 4 | 1.1313 |
| DAO | 6 | 0.40108 | 0.60307 | 0.999994 | 11962 | 2 | 1.9415 | 0.20873 | 0.44337 | 1 | 8953 | 4 | 1.9415 |
| NonTargetingControlG | 2 | 0.54446 | 0.54557 | 0.999994 | 14133 | 1 | 3.0272 | 0.20879 | 0.32863 | 0.991729 | 8954 | 1 | 3.0272 |
| AP1S2 | 6 | 0.16155 | 0.3402 | 0.999994 | 6775 | 3 | -0.57617 | 0.2088 | 0.44347 | 1 | 8955 | 2 | -0.57617 |
| TTC24 | 6 | 0.51331 | 0.67791 | 0.999994 | 13625 | 2 | 0.93018 | 0.2088 | 0.44347 | 1 | 8956 | 4 | 0.93018 |
| C16orf45 | 6 | 0.010141 | 0.039501 | 0.891621 | 881 | 3 | -2.5111 | 0.2088 | 0.44347 | 1 | 8957 | 2 | -2.5111 |
| UHMK1 | 6 | 0.016015 | 0.059084 | 0.931872 | 1246 | 3 | 0.26488 | 0.20883 | 0.44351 | 1 | 8958 | 3 | 0.26488 |
| SLMO1 | 5 | 0.58555 | 0.72094 | 0.999994 | 14751 | 1 | 0.72343 | 0.2089 | 0.42694 | 1 | 8959 | 3 | 0.72343 |
| TAS2R5 | 6 | 0.53964 | 0.69628 | 0.999994 | 14047 | 2 | 1.6274 | 0.2089 | 0.44363 | 1 | 8960 | 4 | 1.6274 |
| WDR5B | 6 | 0.42778 | 0.62043 | 0.999994 | 12378 | 1 | 1.6296 | 0.20896 | 0.4437 | 1 | 8961 | 3 | 1.6296 |
| VAMP7_X | 6 | 0.9313 | 0.93143 | 0.999994 | 19882 | 0 | 1.8635 | 0.20899 | 0.44375 | 1 | 8962 | 4 | 1.8635 |
| TMEM11 | 6 | 0.17745 | 0.35989 | 0.999994 | 7166 | 1 | 1.3607 | 0.20901 | 0.44379 | 1 | 8963 | 4 | 1.3607 |
| NonTargetingControlG | 2 | 0.136 | 0.21691 | 0.980699 | 6077 | 1 | 0.61546 | 0.2092 | 0.32919 | 0.991729 | 8964 | 1 | 0.61546 |
| NonTargetingControlG | 2 | 0.55877 | 0.55935 | 0.999994 | 14346 | 1 | 3.0818 | 0.2092 | 0.32919 | 0.991729 | 8965 | 1 | 3.0818 |
| C14orf79 | 6 | 0.041667 | 0.12565 | 0.951052 | 2599 | 3 | 0.54016 | 0.20923 | 0.44411 | 1 | 8966 | 3 | 0.54016 |
| BEX4 | 6 | 0.15953 | 0.33763 | 0.999994 | 6719 | 2 | 0.19847 | 0.20923 | 0.44411 | 1 | 8967 | 3 | 0.19847 |
| hsa-mir-4720 | 4 | 0.55263 | 0.64098 | 0.999994 | 14253 | 1 | 1.6627 | 0.20924 | 0.39632 | 1 | 8968 | 3 | 1.6627 |
| GNRHR | 6 | 0.46414 | 0.64452 | 0.999994 | 12922 | 2 | 1.1254 | 0.20925 | 0.44414 | 1 | 8969 | 4 | 1.1254 |
| PPP1R9A | 6 | 0.33918 | 0.55172 | 0.999994 | 10919 | 2 | 1.2365 | 0.20929 | 0.44417 | 1 | 8970 | 3 | 1.2365 |
| NFATC1 | 6 | 0.26674 | 0.46782 | 0.999994 | 9388 | 3 | -0.20733 | 0.20933 | 0.44423 | 1 | 8971 | 3 | -0.20733 |
| DYNC1I2 | 5 | 0.093131 | 0.21559 | 0.980236 | 4725 | 2 | 0.21142 | 0.20943 | 0.42754 | 1 | 8972 | 2 | 0.21142 |
| CCL19 | 5 | 0.11757 | 0.25363 | 0.994544 | 5562 | 3 | -1.4555 | 0.20943 | 0.42754 | 1 | 8973 | 2 | -1.4555 |
| PDGFRL | 6 | 0.94951 | 0.94947 | 0.999994 | 20323 | 0 | 1.1339 | 0.20945 | 0.44442 | 1 | 8974 | 4 | 1.1339 |
| IGSF21 | 6 | 0.38368 | 0.59189 | 0.999994 | 11698 | 2 | 1.4801 | 0.20958 | 0.44459 | 1 | 8975 | 4 | 1.4801 |
| ASIC2 | 6 | 0.51728 | 0.68064 | 0.999994 | 13698 | 1 | 1.2436 | 0.20961 | 0.44463 | 1 | 8976 | 4 | 1.2436 |
| CTSS | 6 | 0.87779 | 0.88818 | 0.999994 | 18783 | 1 | 1.3152 | 0.20965 | 0.44469 | 1 | 8977 | 4 | 1.3152 |
| PSMB4 | 2 | 0.017404 | 0.032481 | 0.865899 | 1333 | 1 | -1.7513 | 0.20972 | 0.32991 | 0.992034 | 8978 | 1 | -1.7513 |
| C6orf106 | 6 | 0.46685 | 0.64636 | 0.999994 | 12962 | 2 | 1.9499 | 0.20978 | 0.44487 | 1 | 8979 | 4 | 1.9499 |
| TPRKB | 6 | 0.13525 | 0.30701 | 0.999994 | 6059 | 3 | -1.0729 | 0.2098 | 0.44489 | 1 | 8980 | 2 | -1.0729 |
| CHST2 | 6 | 0.61804 | 0.75336 | 0.999994 | 15187 | 1 | 1.861 | 0.20984 | 0.44494 | 1 | 8981 | 4 | 1.861 |
| CLN5 | 5 | 0.89748 | 0.89762 | 0.999994 | 19168 | 0 | 0.71077 | 0.20986 | 0.42797 | 1 | 8982 | 3 | 0.71077 |
| MS4A10 | 5 | 0.36459 | 0.53388 | 0.999994 | 11414 | 2 | -0.57286 | 0.20986 | 0.42797 | 1 | 8983 | 1 | -0.57286 |
| MVK | 4 | 0.65165 | 0.69424 | 0.999994 | 15681 | 1 | 2.0319 | 0.20992 | 0.39729 | 1 | 8984 | 2 | 2.0319 |
| hsa-mir-3916 | 4 | 0.86221 | 0.86184 | 0.999994 | 18490 | 0 | 2.4319 | 0.20992 | 0.39729 | 1 | 8985 | 2 | 2.4319 |
| MRPS11 | 4 | 0.58622 | 0.65796 | 0.999994 | 14760 | 1 | 2.3454 | 0.20992 | 0.39729 | 1 | 8986 | 2 | 2.3454 |
| HIST1H2AI | 4 | 0.89227 | 0.89199 | 0.999994 | 19074 | 0 | 1.4451 | 0.20998 | 0.39737 | 1 | 8987 | 3 | 1.4451 |
| hsa-mir-8069 | 4 | 0.9031 | 0.90279 | 0.999994 | 19281 | 0 | 2.4059 | 0.20998 | 0.39737 | 1 | 8988 | 3 | 2.4059 |
| PPP1R9B | 6 | 0.25279 | 0.45125 | 0.999994 | 9077 | 1 | 1.7474 | 0.2101 | 0.44531 | 1 | 8989 | 4 | 1.7474 |
| ZNF311 | 5 | 0.19916 | 0.37402 | 0.999994 | 7740 | 3 | -0.79147 | 0.21023 | 0.42839 | 1 | 8990 | 1 | -0.79147 |
| COX16 | 5 | 0.92537 | 0.9256 | 0.999994 | 19755 | 0 | 1.8558 | 0.21023 | 0.42839 | 1 | 8991 | 3 | 1.8558 |
| EMR2 | 5 | 0.23859 | 0.42902 | 0.999994 | 8740 | 2 | 1.917 | 0.21034 | 0.42853 | 1 | 8992 | 3 | 1.917 |
| SAFB2 | 6 | 0.8786 | 0.88868 | 0.999994 | 18803 | 1 | 1.5226 | 0.21035 | 0.44565 | 1 | 8993 | 4 | 1.5226 |
| NonTargetingControlG | 2 | 0.75555 | 0.75511 | 0.999994 | 16925 | 0 | 3.711 | 0.21036 | 0.33079 | 0.992034 | 8994 | 1 | 3.711 |
| C1orf100 | 4 | 0.8118 | 0.81198 | 0.999994 | 17653 | 1 | 1.2512 | 0.21043 | 0.398 | 1 | 8995 | 3 | 1.2512 |

| | | | | | | | | | | | | | |
|---|---|---|---|---|---|---|---|---|---|---|---|---|---|
| ACTL6A | 6 | 0.52589 | 0.68664 | 0.999994 | 13829 | 2 | 1.4269 | 0.21052 | 0.44588 | 1 | 8996 | 4 | 1.4269 |
| ZNHIT6 | 5 | 0.34932 | 0.52133 | 0.999994 | 11134 | 2 | 1.1507 | 0.21061 | 0.42884 | 1 | 8997 | 3 | 1.1507 |
| OR5H14 | 5 | 0.34612 | 0.51874 | 0.999994 | 11071 | 1 | 2.0987 | 0.21061 | 0.42884 | 1 | 8998 | 3 | 2.0987 |
| PDP1 | 5 | 0.85984 | 0.86226 | 0.999994 | 18431 | 1 | 0.31681 | 0.21061 | 0.42884 | 1 | 8999 | 2 | 0.31681 |
| ZNF853 | 5 | 0.045867 | 0.13045 | 0.95318 | 2800 | 3 | -2.1877 | 0.2107 | 0.42894 | 1 | 9000 | 2 | -2.1877 |
| USP11 | 5 | 0.93926 | 0.93945 | 0.999994 | 20057 | 0 | 2.0392 | 0.21077 | 0.42902 | 1 | 9001 | 3 | 2.0392 |
| HAP1 | 5 | 0.36594 | 0.535 | 0.999994 | 11442 | 2 | 1.8809 | 0.21082 | 0.42908 | 1 | 9002 | 3 | 1.8809 |
| PGM2L1 | 6 | 0.096326 | 0.24013 | 0.987253 | 4829 | 2 | 0.9263 | 0.21083 | 0.44631 | 1 | 9003 | 4 | 0.9263 |
| GRID2 | 6 | 0.13684 | 0.30905 | 0.999994 | 6102 | 3 | -0.43611 | 0.21083 | 0.44631 | 1 | 9004 | 3 | -0.43611 |
| MAT2A | 6 | 0.0118 | 0.045055 | 0.891621 | 982 | 3 | -0.36244 | 0.21083 | 0.44631 | 1 | 9005 | 3 | -0.36244 |
| SORBS2 | 3 | 0.11067 | 0.19718 | 0.977164 | 5322 | 1 | -0.027745 | 0.21087 | 0.35471 | 0.994392 | 9006 | 1 | -0.027745 |
| KCNJ6 | 5 | 0.31963 | 0.49702 | 0.999994 | 10508 | 2 | 2.0344 | 0.21087 | 0.42913 | 1 | 9007 | 3 | 2.0344 |
| GRIN2A | 6 | 0.47751 | 0.65351 | 0.999994 | 13106 | 2 | 1.1954 | 0.21095 | 0.44648 | 1 | 9008 | 3 | 1.1954 |
| GRIN2D | 6 | 0.49412 | 0.66466 | 0.999994 | 13329 | 1 | 1.8937 | 0.21098 | 0.44653 | 1 | 9009 | 4 | 1.8937 |
| NonTargetingControlG | 2 | 0.32102 | 0.35299 | 0.999994 | 10534 | 1 | 2.1869 | 0.211 | 0.33164 | 0.992121 | 9010 | 1 | 2.1869 |
| hsa-mir-6885 | 4 | 0.36952 | 0.51436 | 0.999994 | 11498 | 1 | 2.2868 | 0.211 | 0.39886 | 1 | 9011 | 2 | 2.2868 |
| PDE1A | 5 | 0.21327 | 0.39398 | 0.999994 | 8096 | 1 | 1.8488 | 0.21101 | 0.4293 | 1 | 9012 | 3 | 1.8488 |
| HHLA3 | 4 | 0.21546 | 0.3566 | 0.999994 | 8154 | 2 | -0.40908 | 0.21104 | 0.3989 | 1 | 9013 | 2 | -0.40908 |
| PCBD1 | 6 | 0.50596 | 0.67286 | 0.999994 | 13522 | 2 | 1.4698 | 0.21108 | 0.44667 | 1 | 9014 | 3 | 1.4698 |
| CHCHD4 | 6 | 0.010897 | 0.042045 | 0.891621 | 920 | 2 | 1.6276 | 0.21114 | 0.44677 | 1 | 9015 | 4 | 1.6276 |
| EHD2 | 6 | 0.22437 | 0.41737 | 0.999994 | 8388 | 1 | 1.4786 | 0.21122 | 0.44688 | 1 | 9016 | 3 | 1.4786 |
| DNAJC19 | 6 | 0.36257 | 0.57819 | 0.999994 | 11380 | 3 | -0.18984 | 0.21128 | 0.44695 | 1 | 9017 | 2 | -0.18984 |
| MFNG | 6 | 0.18441 | 0.36865 | 0.999994 | 7326 | 3 | -0.74362 | 0.21128 | 0.44695 | 1 | 9018 | 2 | -0.74362 |
| CEP63 | 6 | 0.012622 | 0.047834 | 0.900643 | 1038 | 5 | -1.6785 | 0.21128 | 0.44695 | 1 | 9019 | 1 | -1.6785 |
| TMEM52 | 5 | 0.28854 | 0.47173 | 0.999994 | 9884 | 1 | 0.44961 | 0.21128 | 0.4296 | 1 | 9020 | 2 | 0.44961 |
| DKK1 | 5 | 0.14662 | 0.2977 | 0.999994 | 6365 | 3 | -1.1506 | 0.21128 | 0.4296 | 1 | 9021 | 1 | -1.1506 |
| CARD11 | 5 | 0.063967 | 0.1686 | 0.967126 | 3609 | 2 | 0.80156 | 0.21128 | 0.4296 | 1 | 9022 | 3 | 0.80156 |
| WNT9A | 5 | 0.46394 | 0.61697 | 0.999994 | 12919 | 2 | 0.94333 | 0.21128 | 0.4296 | 1 | 9023 | 3 | 0.94333 |
| METTL3 | 5 | 0.63159 | 0.73792 | 0.999994 | 15392 | 1 | 1.4513 | 0.21128 | 0.4296 | 1 | 9024 | 3 | 1.4513 |
| TSPAN16 | 5 | 0.54652 | 0.68864 | 0.999994 | 14160 | 1 | 0.35278 | 0.21144 | 0.42977 | 1 | 9025 | 2 | 0.35278 |
| SERP2 | 5 | 0.55168 | 0.69324 | 0.999994 | 14236 | 2 | -0.49381 | 0.21144 | 0.42977 | 1 | 9026 | 2 | -0.49381 |
| AKR1C2 | 5 | 0.3278 | 0.50372 | 0.999994 | 10682 | 1 | 0.82577 | 0.21144 | 0.42977 | 1 | 9027 | 3 | 0.82577 |
| TNNI2 | 5 | 0.23728 | 0.42727 | 0.999994 | 8707 | 3 | -0.65476 | 0.21144 | 0.42977 | 1 | 9028 | 2 | -0.65476 |
| VAC14 | 5 | 0.47215 | 0.62395 | 0.999994 | 13037 | 2 | 2.0644 | 0.21147 | 0.42979 | 1 | 9029 | 3 | 2.0644 |
| OR6S1 | 6 | 0.072654 | 0.19228 | 0.974607 | 3946 | 2 | 1.262 | 0.2116 | 0.44737 | 1 | 9030 | 4 | 1.262 |
| hsa-mir-3165 | 4 | 0.89419 | 0.89388 | 0.999994 | 19104 | 0 | 1.221 | 0.2116 | 0.39971 | 1 | 9031 | 3 | 1.221 |
| KBTBD12 | 6 | 0.090055 | 0.22759 | 0.983354 | 4596 | 3 | -1.0851 | 0.2116 | 0.44738 | 1 | 9032 | 2 | -1.0851 |
| DOPEY1 | 6 | 0.42336 | 0.61758 | 0.999994 | 12300 | 1 | 0.73675 | 0.2116 | 0.44738 | 1 | 9033 | 3 | 0.73675 |
| NANS | 6 | 0.10457 | 0.25625 | 0.994943 | 5120 | 3 | -0.90297 | 0.2116 | 0.44738 | 1 | 9034 | 2 | -0.90297 |
| TMEM259 | 6 | 0.36001 | 0.57532 | 0.999994 | 11335 | 3 | 0.1415 | 0.2116 | 0.44738 | 1 | 9035 | 3 | 0.1415 |
| NMBR | 5 | 0.50165 | 0.64935 | 0.999994 | 13453 | 1 | 1.8605 | 0.21162 | 0.42997 | 1 | 9036 | 3 | 1.8605 |
| LRRK1 | 5 | 0.78947 | 0.81301 | 0.999994 | 17334 | 1 | 1.1406 | 0.2118 | 0.43018 | 1 | 9037 | 3 | 1.1406 |
| KANSL2 | 4 | 0.18701 | 0.32731 | 0.999994 | 7389 | 1 | 2.4132 | 0.21189 | 0.40013 | 1 | 9038 | 3 | 2.4132 |
| TMEM14C | 6 | 0.15584 | 0.33298 | 0.999994 | 6620 | 3 | -0.31345 | 0.2119 | 0.44782 | 1 | 9039 | 2 | -0.31345 |
| CACNA1B | 6 | 0.94936 | 0.94933 | 0.999994 | 20317 | 0 | 1.3188 | 0.21194 | 0.44786 | 1 | 9040 | 4 | 1.3188 |
| VBP1 | 6 | 0.028982 | 0.096404 | 0.945031 | 1955 | 3 | -1.6596 | 0.21195 | 0.44789 | 1 | 9041 | 2 | -1.6596 |
| ZNF358 | 4 | 0.23774 | 0.37938 | 0.999994 | 8719 | 2 | 1.4065 | 0.212 | 0.40026 | 1 | 9042 | 2 | 1.4065 |
| ZNF138 | 6 | 0.57119 | 0.7188 | 0.999994 | 14564 | 2 | 1.1009 | 0.21204 | 0.44801 | 1 | 9043 | 4 | 1.1009 |
| FAM211A | 4 | 0.037234 | 0.096248 | 0.945031 | 2384 | 2 | -3.2598 | 0.21205 | 0.40033 | 1 | 9044 | 1 | -3.2598 |
| HIST1H2BI | 4 | 0.64028 | 0.68768 | 0.999994 | 15515 | 1 | 0.5467 | 0.21205 | 0.40033 | 1 | 9045 | 2 | 0.5467 |
| hsa-mir-3664 | 4 | 0.70522 | 0.72791 | 0.999994 | 16365 | 1 | 1.7573 | 0.21208 | 0.40037 | 1 | 9046 | 3 | 1.7573 |
| MTRF1L | 6 | 0.28775 | 0.49248 | 0.999994 | 9871 | 2 | 1.3741 | 0.21209 | 0.44808 | 1 | 9047 | 3 | 1.3741 |

| | | | | | | | | | | | | |
|---|---|---|---|---|---|---|---|---|---|---|---|---|
| PNMA3 | 6 | 0.71138 | 0.81556 | 0.999994 | 16413 | 1 | 1.3675 | 0.21209 | 0.44809 | 1 | 9048 | 4 | 1.3675 |
| RAB11FIP4 | 3 | 0.71376 | 0.71398 | 0.999994 | 16438 | 1 | 1.9168 | 0.21211 | 0.35603 | 0.994392 | 9049 | 2 | 1.9168 |
| EEFSEC | 2 | 0.69877 | 0.69858 | 0.999994 | 16292 | 0 | 3.5216 | 0.21216 | 0.33322 | 0.992917 | 9050 | 1 | 3.5216 |
| GAS1 | 5 | 0.9483 | 0.9485 | 0.999994 | 20292 | 0 | 0.55424 | 0.21222 | 0.43064 | 1 | 9051 | 2 | 0.55424 |
| CKMT1B | 6 | 0.30226 | 0.50936 | 0.999994 | 10178 | 3 | 1.1576 | 0.21225 | 0.44831 | 1 | 9052 | 3 | 1.1576 |
| C2orf42 | 6 | 0.29759 | 0.50391 | 0.999994 | 10080 | 3 | -0.49331 | 0.21227 | 0.44835 | 1 | 9053 | 1 | -0.49331 |
| CHCHD5 | 6 | 0.23027 | 0.4244 | 0.999994 | 8532 | 3 | 0.62483 | 0.21227 | 0.44835 | 1 | 9054 | 3 | 0.62483 |
| COMMD3 | 6 | 0.54957 | 0.70333 | 0.999994 | 14206 | 2 | 0.81363 | 0.21227 | 0.44835 | 1 | 9055 | 3 | 0.81363 |
| ZNF704 | 6 | 0.26388 | 0.46445 | 0.999994 | 9332 | 1 | 1.2263 | 0.21232 | 0.44842 | 1 | 9056 | 4 | 1.2263 |
| MAP3K6 | 6 | 0.029875 | 0.098485 | 0.945253 | 2001 | 3 | 0.61608 | 0.21243 | 0.44857 | 1 | 9057 | 3 | 0.61608 |
| RRN3 | 4 | 0.22062 | 0.36188 | 0.999994 | 8303 | 2 | 1.1151 | 0.21255 | 0.40103 | 1 | 9058 | 2 | 1.1151 |
| MARVELD2 | 4 | 0.92548 | 0.92527 | 0.999994 | 19757 | 0 | 1.3847 | 0.21264 | 0.40116 | 1 | 9059 | 3 | 1.3847 |
| LOC728819 | 6 | 0.12253 | 0.29051 | 0.999632 | 5724 | 3 | -0.66328 | 0.21267 | 0.44892 | 1 | 9060 | 3 | -0.66328 |
| C3orf70 | 6 | 0.022616 | 0.08036 | 0.93914 | 1624 | 4 | -2.6955 | 0.21267 | 0.44892 | 1 | 9061 | 2 | -2.6955 |
| SYDE2 | 5 | 0.82532 | 0.83615 | 0.999994 | 17856 | 1 | 0.82978 | 0.21267 | 0.43113 | 1 | 9062 | 3 | 0.82978 |
| TCL1A | 5 | 0.075181 | 0.18681 | 0.969409 | 4041 | 1 | 0.45811 | 0.21267 | 0.43113 | 1 | 9063 | 2 | 0.45811 |
| OR1L8 | 6 | 0.54043 | 0.69684 | 0.999994 | 14060 | 2 | 1.7735 | 0.21268 | 0.44894 | 1 | 9064 | 3 | 1.7735 |
| SLC25A37 | 4 | 0.62784 | 0.68066 | 0.999994 | 15350 | 1 | 1.8387 | 0.21271 | 0.40124 | 1 | 9065 | 3 | 1.8387 |
| NonTargetingControlG | 2 | 0.78724 | 0.78676 | 0.999994 | 17307 | 0 | 0.97411 | 0.21276 | 0.33405 | 0.992933 | 9066 | 2 | 0.97411 |
| NAALADL1 | 3 | 0.65307 | 0.65964 | 0.999994 | 15699 | 1 | 2.6329 | 0.21277 | 0.35673 | 0.994392 | 9067 | 2 | 2.6329 |
| EIF5A | 3 | 0.21766 | 0.33475 | 0.999994 | 8214 | 1 | 2.7055 | 0.21284 | 0.35681 | 0.994392 | 9068 | 2 | 2.7055 |
| SORD | 6 | 0.33372 | 0.54555 | 0.999994 | 10801 | 2 | 1.7902 | 0.21286 | 0.44917 | 1 | 9069 | 4 | 1.7902 |
| BEND4 | 6 | 0.76425 | 0.83366 | 0.999994 | 17019 | 1 | 0.57594 | 0.21297 | 0.44931 | 1 | 9070 | 3 | 0.57594 |
| SLC2A9 | 6 | 0.076875 | 0.20103 | 0.977164 | 4104 | 2 | 1.5168 | 0.21297 | 0.44931 | 1 | 9071 | 3 | 1.5168 |
| CDR1 | 6 | 0.083755 | 0.21499 | 0.979587 | 4367 | 1 | 0.65355 | 0.21297 | 0.44931 | 1 | 9072 | 3 | 0.65355 |
| KRTAP21-3 | 4 | 0.23824 | 0.37988 | 0.999994 | 8733 | 1 | 1.1173 | 0.21299 | 0.40163 | 1 | 9073 | 3 | 1.1173 |
| RICTOR | 6 | 0.25959 | 0.45938 | 0.999994 | 9230 | 2 | -0.16164 | 0.213 | 0.44935 | 1 | 9074 | 2 | -0.16164 |
| PRAMEF7 | 6 | 0.061775 | 0.16952 | 0.967126 | 3503 | 2 | 1.0828 | 0.213 | 0.44935 | 1 | 9075 | 3 | 1.0828 |
| hsa-mir-1229 | 2 | 0.78196 | 0.78144 | 0.999994 | 17241 | 0 | 3.7656 | 0.21303 | 0.33443 | 0.993364 | 9076 | 2 | 3.7656 |
| hsa-mir-3614 | 3 | 0.74393 | 0.74387 | 0.999994 | 16786 | 0 | 2.4052 | 0.21306 | 0.35708 | 0.994392 | 9077 | 2 | 2.4052 |
| USP21 | 6 | 0.019053 | 0.068951 | 0.93914 | 1411 | 1 | 1.195 | 0.21309 | 0.44948 | 1 | 9078 | 4 | 1.195 |
| SPIN3 | 6 | 0.38989 | 0.59592 | 0.999994 | 11796 | 2 | 1.2068 | 0.21315 | 0.44956 | 1 | 9079 | 4 | 1.2068 |
| OR52E4 | 5 | 0.38503 | 0.55095 | 0.999994 | 11716 | 2 | 1.388 | 0.21317 | 0.43166 | 1 | 9080 | 3 | 1.388 |
| FSIP1 | 5 | 0.26539 | 0.45312 | 0.999994 | 9359 | 2 | -0.55063 | 0.21317 | 0.43166 | 1 | 9081 | 2 | -0.55063 |
| LHX3 | 5 | 0.55169 | 0.69327 | 0.999994 | 14237 | 2 | 0.83043 | 0.21317 | 0.43166 | 1 | 9082 | 3 | 0.83043 |
| FOXO1 | 5 | 0.16296 | 0.32187 | 0.999994 | 6816 | 2 | -0.28207 | 0.21317 | 0.43166 | 1 | 9083 | 2 | -0.28207 |
| hsa-mir-6731 | 3 | 0.23966 | 0.36215 | 0.999994 | 8766 | 1 | 2.6718 | 0.21318 | 0.35722 | 0.994392 | 9084 | 2 | 2.6718 |
| NDUFA9 | 6 | 0.37609 | 0.58708 | 0.999994 | 11594 | 1 | 1.4986 | 0.21322 | 0.44966 | 1 | 9085 | 4 | 1.4986 |
| BIN3 | 6 | 0.44594 | 0.6324 | 0.999994 | 12661 | 2 | 1.8213 | 0.21325 | 0.44971 | 1 | 9086 | 3 | 1.8213 |
| DNM3 | 6 | 0.54335 | 0.69891 | 0.999994 | 14117 | 2 | 1.8049 | 0.21325 | 0.44971 | 1 | 9087 | 3 | 1.8049 |
| FAM184A | 6 | 0.17538 | 0.35734 | 0.999994 | 7119 | 3 | 1.2102 | 0.21325 | 0.44971 | 1 | 9088 | 3 | 1.2102 |
| SPN | 6 | 0.23876 | 0.43451 | 0.999994 | 8741 | 3 | 1.0235 | 0.21325 | 0.44971 | 1 | 9089 | 3 | 1.0235 |
| LYSMD1 | 6 | 0.52695 | 0.6874 | 0.999994 | 13843 | 2 | 1.4424 | 0.21325 | 0.44971 | 1 | 9090 | 3 | 1.4424 |
| SIGLEC9 | 3 | 0.84573 | 0.84577 | 0.999994 | 18183 | 0 | 2.2926 | 0.21335 | 0.3574 | 0.994392 | 9091 | 2 | 2.2926 |
| FBXL4 | 6 | 0.26936 | 0.47099 | 0.999994 | 9456 | 2 | 0.79276 | 0.21336 | 0.44986 | 1 | 9092 | 3 | 0.79276 |
| TLX2 | 4 | 0.90344 | 0.90314 | 0.999994 | 19291 | 0 | 1.3911 | 0.2134 | 0.4022 | 1 | 9093 | 2 | 1.3911 |
| NonTargetingControlG | 2 | 0.78654 | 0.78607 | 0.999994 | 17296 | 0 | 3.5082 | 0.21346 | 0.33499 | 0.993364 | 9094 | 2 | 3.5082 |
| TRAPPC10 | 5 | 0.47448 | 0.62592 | 0.999994 | 13056 | 2 | 1.915 | 0.21349 | 0.43201 | 1 | 9095 | 3 | 1.915 |
| hsa-mir-1284 | 4 | 0.42238 | 0.56865 | 0.999994 | 12290 | 1 | 1.722 | 0.2135 | 0.40236 | 1 | 9096 | 3 | 1.722 |
| CHP2 | 5 | 0.13436 | 0.27929 | 0.999632 | 6035 | 2 | 1.8278 | 0.21351 | 0.43203 | 1 | 9097 | 3 | 1.8278 |
| INVS | 6 | 0.25274 | 0.4512 | 0.999994 | 9072 | 2 | 1.3038 | 0.21352 | 0.45008 | 1 | 9098 | 4 | 1.3038 |
| KDM4D | 5 | 0.47199 | 0.62382 | 0.999994 | 13035 | 2 | 1.8848 | 0.21355 | 0.43206 | 1 | 9099 | 3 | 1.8848 |

| | | | | | | | | | | | | | |
|---|---|---|---|---|---|---|---|---|---|---|---|---|---|
| PCDHGB6 | 2 | 0.78645 | 0.78598 | 0.999994 | 17295 | 0 | 1.6515 | 0.21355 | 0.33512 | 0.993364 | 9100 | 2 | 1.6515 |
| HERC2 | 3 | 0.40279 | 0.48593 | 0.999994 | 11993 | 1 | -0.4454 | 0.21362 | 0.35769 | 0.994392 | 9101 | 1 | -0.4454 |
| INTS8 | 3 | 0.28123 | 0.41349 | 0.999994 | 9728 | 1 | 0.91171 | 0.21362 | 0.35769 | 0.994392 | 9102 | 2 | 0.91171 |
| ZNF597 | 6 | 0.5373 | 0.69463 | 0.999994 | 14016 | 2 | 1.0724 | 0.21363 | 0.45023 | 1 | 9103 | 4 | 1.0724 |
| CTLA4 | 6 | 0.81539 | 0.85469 | 0.999994 | 17705 | 1 | 1.5589 | 0.21369 | 0.45033 | 1 | 9104 | 4 | 1.5589 |
| DES | 6 | 0.56413 | 0.71375 | 0.999994 | 14455 | 1 | 1.6145 | 0.21374 | 0.4504 | 1 | 9105 | 3 | 1.6145 |
| APC2 | 5 | 0.33728 | 0.51152 | 0.999994 | 10879 | 2 | 2.1369 | 0.21377 | 0.43231 | 1 | 9106 | 3 | 2.1369 |
| FAM183A | 4 | 0.81443 | 0.81448 | 0.999994 | 17696 | 1 | 1.7368 | 0.2138 | 0.40276 | 1 | 9107 | 3 | 1.7368 |
| ARHGAP36 | 6 | 0.10503 | 0.25712 | 0.994943 | 5134 | 2 | 1.0865 | 0.2138 | 0.4505 | 1 | 9108 | 3 | 1.0865 |
| CUEDC1 | 6 | 0.34862 | 0.56244 | 0.999994 | 11126 | 1 | 0.44263 | 0.2138 | 0.4505 | 1 | 9109 | 3 | 0.44263 |
| HIST1H4C | 6 | 0.25721 | 0.45656 | 0.999994 | 9170 | 3 | -0.39546 | 0.2138 | 0.4505 | 1 | 9110 | 2 | -0.39546 |
| PELI2 | 6 | 0.49533 | 0.66551 | 0.999994 | 13342 | 1 | 0.90583 | 0.2138 | 0.4505 | 1 | 9111 | 3 | 0.90583 |
| CENPW | 4 | 0.062853 | 0.14622 | 0.964852 | 3549 | 2 | 0.010023 | 0.21384 | 0.40283 | 1 | 9112 | 2 | 0.010023 |
| CIRH1A | 4 | 0.138 | 0.27684 | 0.999379 | 6136 | 2 | 0.91714 | 0.21384 | 0.40283 | 1 | 9113 | 2 | 0.91714 |
| USP3 | 6 | 0.66735 | 0.79107 | 0.999994 | 15883 | 2 | 0.537 | 0.21397 | 0.45073 | 1 | 9114 | 3 | 0.537 |
| HTR6 | 6 | 0.21054 | 0.40075 | 0.999994 | 8038 | 3 | -0.71513 | 0.21397 | 0.45073 | 1 | 9115 | 2 | -0.71513 |
| TDRD1 | 6 | 0.27525 | 0.47788 | 0.999994 | 9597 | 2 | 1.1463 | 0.21397 | 0.45073 | 1 | 9116 | 3 | 1.1463 |
| SNX6 | 6 | 0.66323 | 0.78782 | 0.999994 | 15827 | 2 | 1.0445 | 0.21402 | 0.4508 | 1 | 9117 | 4 | 1.0445 |
| ZNF24 | 5 | 0.75793 | 0.79507 | 0.999994 | 16951 | 1 | 1.3633 | 0.21422 | 0.43277 | 1 | 9118 | 3 | 1.3633 |
| LMOD2 | 5 | 0.18384 | 0.35215 | 0.999994 | 7312 | 3 | -0.90834 | 0.21422 | 0.43277 | 1 | 9119 | 2 | -0.90834 |
| RPS13 | 4 | 0.028738 | 0.078966 | 0.93914 | 1940 | 2 | -1.8345 | 0.21422 | 0.40335 | 1 | 9120 | 2 | -1.8345 |
| ATP1B1 | 6 | 0.6419 | 0.77138 | 0.999994 | 15548 | 2 | 1.1196 | 0.21428 | 0.45114 | 1 | 9121 | 4 | 1.1196 |
| GYPE | 4 | 0.85868 | 0.85829 | 0.999994 | 18412 | 0 | 1.4549 | 0.21437 | 0.40355 | 1 | 9122 | 3 | 1.4549 |
| AGTR1 | 4 | 0.09423 | 0.20471 | 0.978186 | 4761 | 1 | 1.9939 | 0.21442 | 0.40362 | 1 | 9123 | 3 | 1.9939 |
| PLA2G3 | 6 | 0.31333 | 0.52222 | 0.999994 | 10384 | 3 | 1.0844 | 0.21446 | 0.4514 | 1 | 9124 | 3 | 1.0844 |
| ZNF143 | 5 | 0.33618 | 0.51061 | 0.999994 | 10851 | 1 | -0.32634 | 0.21447 | 0.43305 | 1 | 9125 | 2 | -0.32634 |
| MAFF | 5 | 0.39166 | 0.55653 | 0.999994 | 11824 | 2 | 2.3249 | 0.21453 | 0.43313 | 1 | 9126 | 3 | 2.3249 |
| ACTR10 | 3 | 0.17892 | 0.28585 | 0.999632 | 7196 | 1 | 2.4629 | 0.2146 | 0.35871 | 0.994392 | 9127 | 2 | 2.4629 |
| hsa-mir-3714 | 3 | 0.77647 | 0.77643 | 0.999994 | 17158 | 0 | 2.7514 | 0.21469 | 0.35882 | 0.994392 | 9128 | 2 | 2.7514 |
| OFD1 | 5 | 0.12794 | 0.26946 | 0.998561 | 5872 | 2 | 2.1928 | 0.2147 | 0.43331 | 1 | 9129 | 3 | 2.1928 |
| SOST | 4 | 0.59324 | 0.6617 | 0.999994 | 14868 | 1 | 1.674 | 0.21472 | 0.40403 | 1 | 9130 | 3 | 1.674 |
| DMRTB1 | 5 | 0.10305 | 0.23115 | 0.983933 | 5068 | 1 | 0.54942 | 0.21478 | 0.43339 | 1 | 9131 | 2 | 0.54942 |
| GID8 | 5 | 0.34503 | 0.51787 | 0.999994 | 11053 | 2 | -0.50406 | 0.21478 | 0.43339 | 1 | 9132 | 2 | -0.50406 |
| KLHL3 | 5 | 0.89842 | 0.89854 | 0.999994 | 19191 | 0 | 0.35277 | 0.21478 | 0.43339 | 1 | 9133 | 2 | 0.35277 |
| BSND | 3 | 0.50623 | 0.55024 | 0.999994 | 13530 | 1 | 1.6973 | 0.21481 | 0.35895 | 0.994392 | 9134 | 2 | 1.6973 |
| hsa-mir-671 | 3 | 0.068898 | 0.14149 | 0.959146 | 3794 | 1 | 2.4694 | 0.21495 | 0.35909 | 0.994392 | 9135 | 2 | 2.4694 |
| PRSS36 | 6 | 0.18052 | 0.36382 | 0.999994 | 7231 | 3 | -0.73997 | 0.21505 | 0.4522 | 1 | 9136 | 2 | -0.73997 |
| VMO1 | 6 | 0.26591 | 0.46687 | 0.999994 | 9365 | 3 | 0.53309 | 0.21505 | 0.4522 | 1 | 9137 | 3 | 0.53309 |
| SRCIN1 | 6 | 0.022272 | 0.079263 | 0.93914 | 1597 | 3 | -1.9483 | 0.21505 | 0.4522 | 1 | 9138 | 2 | -1.9483 |
| PDCD6 | 6 | 0.3016 | 0.50858 | 0.999994 | 10154 | 3 | 0.38497 | 0.21505 | 0.4522 | 1 | 9139 | 3 | 0.38497 |
| SPRED2 | 6 | 0.6737 | 0.79604 | 0.999994 | 15985 | 2 | 1.1223 | 0.21508 | 0.45224 | 1 | 9140 | 4 | 1.1223 |
| DIO1 | 4 | 0.86819 | 0.86777 | 0.999994 | 18605 | 0 | 1.4071 | 0.21512 | 0.4046 | 1 | 9141 | 3 | 1.4071 |
| LCE1B | 4 | 0.80557 | 0.80626 | 0.999994 | 17561 | 1 | 2.6163 | 0.21516 | 0.40466 | 1 | 9142 | 3 | 2.6163 |
| CLEC16A | 6 | 0.91213 | 0.91338 | 0.999994 | 19465 | 1 | 1.7978 | 0.21516 | 0.45235 | 1 | 9143 | 4 | 1.7978 |
| hsa-mir-661 | 3 | 0.65804 | 0.66371 | 0.999994 | 15754 | 1 | 0.31244 | 0.21523 | 0.35939 | 0.994392 | 9144 | 1 | 0.31244 |
| ADAM23 | 6 | 0.81016 | 0.85233 | 0.999994 | 17634 | 1 | 1.3995 | 0.21524 | 0.45246 | 1 | 9145 | 4 | 1.3995 |
| FAM50B | 6 | 0.18609 | 0.3707 | 0.999994 | 7367 | 2 | 0.57144 | 0.2154 | 0.45267 | 1 | 9146 | 3 | 0.57144 |
| RPS9 | 4 | 0.90167 | 0.90134 | 0.999994 | 19245 | 0 | 2.3648 | 0.2154 | 0.40501 | 1 | 9147 | 2 | 2.3648 |
| FOXA2 | 4 | 0.24532 | 0.38708 | 0.999994 | 8907 | 2 | 1.4122 | 0.2154 | 0.40501 | 1 | 9148 | 2 | 1.4122 |
| NonTargetingControlG | 2 | 0.51091 | 0.51392 | 0.999994 | 13594 | 1 | 2.9063 | 0.21559 | 0.33788 | 0.994237 | 9149 | 1 | 2.9063 |
| hsa-mir-1273a | 2 | 0.11275 | 0.18971 | 0.970473 | 5394 | 1 | -0.49359 | 0.21559 | 0.33788 | 0.994237 | 9150 | 1 | -0.49359 |
| GSTT2B | 4 | 0.09324 | 0.20296 | 0.97798 | 4730 | 2 | -1.8687 | 0.21559 | 0.40528 | 1 | 9151 | 1 | -1.8687 |

| | | | | | | | | | | | | | |
|---|---|---|---|---|---|---|---|---|---|---|---|---|---|
| DDIT3 | 5 | 0.024529 | 0.076214 | 0.93914 | 1731 | 2 | -0.13792 | 0.2157 | 0.43434 | 1 | 9152 | 2 | -0.13792 |
| C6orf99 | 3 | 0.75036 | 0.75031 | 0.999994 | 16858 | 0 | 0.14837 | 0.21572 | 0.35993 | 0.994441 | 9153 | 1 | 0.14837 |
| FAM24A | 4 | 0.039437 | 0.10066 | 0.946414 | 2487 | 1 | 0.99667 | 0.21574 | 0.40548 | 1 | 9154 | 3 | 0.99667 |
| RANBP6 | 6 | 0.54599 | 0.70078 | 0.999994 | 14151 | 2 | 1.148 | 0.21581 | 0.45323 | 1 | 9155 | 4 | 1.148 |
| hsa-mir-639 | 4 | 0.92634 | 0.92614 | 0.999994 | 19775 | 0 | 2.3181 | 0.2159 | 0.4057 | 1 | 9156 | 3 | 2.3181 |
| NonTargetingControlG | 2 | 0.78408 | 0.78355 | 0.999994 | 17268 | 0 | 2.643 | 0.21592 | 0.33833 | 0.994237 | 9157 | 2 | 2.643 |
| DDX43 | 5 | 0.43977 | 0.59661 | 0.999994 | 12570 | 2 | 1.8951 | 0.21595 | 0.43461 | 1 | 9158 | 3 | 1.8951 |
| AQPEP | 5 | 0.38029 | 0.54695 | 0.999994 | 11652 | 2 | 1.8866 | 0.216 | 0.43467 | 1 | 9159 | 3 | 1.8866 |
| TRIM72 | 6 | 0.21244 | 0.40305 | 0.999994 | 8071 | 2 | 1.3299 | 0.21602 | 0.45352 | 1 | 9160 | 4 | 1.3299 |
| GTF2A1L | 6 | 0.02541 | 0.087963 | 0.94395 | 1774 | 3 | -1.3262 | 0.21602 | 0.45352 | 1 | 9161 | 2 | -1.3262 |
| ZFAT | 6 | 0.21353 | 0.40438 | 0.999994 | 8102 | 3 | -0.63943 | 0.21602 | 0.45352 | 1 | 9162 | 2 | -0.63943 |
| CAMK2A | 6 | 0.026422 | 0.09037 | 0.94395 | 1826 | 3 | -0.37743 | 0.21616 | 0.45371 | 1 | 9163 | 3 | -0.37743 |
| SLC36A3 | 6 | 0.15255 | 0.32886 | 0.999994 | 6523 | 2 | 0.73141 | 0.21616 | 0.45371 | 1 | 9164 | 3 | 0.73141 |
| ACP1 | 5 | 0.95097 | 0.95114 | 0.999994 | 20356 | 0 | 0.96805 | 0.21624 | 0.43496 | 1 | 9165 | 3 | 0.96805 |
| CD97 | 5 | 0.76696 | 0.80003 | 0.999994 | 17050 | 1 | 1.124 | 0.21624 | 0.43496 | 1 | 9166 | 3 | 1.124 |
| PPP6R3 | 5 | 0.43511 | 0.59273 | 0.999994 | 12500 | 1 | 0.21588 | 0.21633 | 0.43507 | 1 | 9167 | 2 | 0.21588 |
| ANO10 | 5 | 0.94585 | 0.94605 | 0.999994 | 20229 | 0 | 1.8989 | 0.21634 | 0.43508 | 1 | 9168 | 3 | 1.8989 |
| OGFOD1 | 4 | 0.93169 | 0.93144 | 0.999994 | 19892 | 0 | 2.1124 | 0.21637 | 0.40636 | 1 | 9169 | 3 | 2.1124 |
| PTGDR2 | 4 | 0.54338 | 0.63649 | 0.999994 | 14119 | 1 | 1.4055 | 0.21639 | 0.40639 | 1 | 9170 | 2 | 1.4055 |
| ADORA3 | 5 | 0.38069 | 0.54727 | 0.999994 | 11657 | 2 | 1.3667 | 0.21641 | 0.43515 | 1 | 9171 | 3 | 1.3667 |
| STMN2 | 5 | 0.92688 | 0.92712 | 0.999994 | 19785 | 0 | 1.9546 | 0.21644 | 0.43519 | 1 | 9172 | 3 | 1.9546 |
| NonTargetingControlG | 2 | 0.46816 | 0.47538 | 0.999994 | 12978 | 1 | 2.8007 | 0.21661 | 0.33928 | 0.994237 | 9173 | 1 | 2.8007 |
| FBLN1 | 6 | 0.46185 | 0.64297 | 0.999994 | 12879 | 2 | 0.4737 | 0.21662 | 0.45432 | 1 | 9174 | 3 | 0.4737 |
| EPO | 4 | 0.052166 | 0.12572 | 0.951052 | 3072 | 2 | -1.5008 | 0.21667 | 0.4068 | 1 | 9175 | 2 | -1.5008 |
| AQP2 | 6 | 0.075917 | 0.19908 | 0.977164 | 4068 | 1 | 1.1426 | 0.2167 | 0.45442 | 1 | 9176 | 3 | 1.1426 |
| OPA1 | 4 | 0.29555 | 0.43842 | 0.999994 | 10030 | 1 | 2.1404 | 0.2167 | 0.40683 | 1 | 9177 | 3 | 2.1404 |
| TMSB4Y | 4 | 0.54372 | 0.63665 | 0.999994 | 14126 | 1 | 1.794 | 0.21676 | 0.40691 | 1 | 9178 | 3 | 1.794 |
| NonTargetingControlG | 2 | 0.78324 | 0.78271 | 0.999994 | 17256 | 0 | 3.6819 | 0.21676 | 0.33948 | 0.994237 | 9179 | 2 | 3.6819 |
| CBFA2T3 | 5 | 0.53281 | 0.67653 | 0.999994 | 13949 | 2 | 0.69691 | 0.21679 | 0.43557 | 1 | 9180 | 3 | 0.69691 |
| AHSA2 | 5 | 0.18931 | 0.35996 | 0.999994 | 7463 | 3 | -0.80536 | 0.21679 | 0.43557 | 1 | 9181 | 2 | -0.80536 |
| FIGLA | 5 | 0.40381 | 0.56672 | 0.999994 | 12012 | 2 | 0.66208 | 0.21679 | 0.43557 | 1 | 9182 | 3 | 0.66208 |
| TAS2R30 | 6 | 0.23228 | 0.42682 | 0.999994 | 8579 | 3 | 0.9274 | 0.21682 | 0.45458 | 1 | 9183 | 3 | 0.9274 |
| hsa-mir-3607 | 1 | 0.78318 | 0.78349 | 0.999994 | 17254 | 0 | 4.5081 | 0.21682 | 0.21649 | 0.97466 | 9184 | 1 | 4.5081 |
| YLPM1 | 4 | 0.67435 | 0.70806 | 0.999994 | 15995 | 1 | 1.4913 | 0.21683 | 0.40701 | 1 | 9185 | 3 | 1.4913 |
| PROL1 | 5 | 0.94284 | 0.94303 | 0.999994 | 20147 | 0 | 2.0518 | 0.21688 | 0.43566 | 1 | 9186 | 3 | 2.0518 |
| ZNF703 | 5 | 0.3081 | 0.4876 | 0.999994 | 10287 | 2 | 1.7767 | 0.21691 | 0.43569 | 1 | 9187 | 3 | 1.7767 |
| NonTargetingControlG | 2 | 0.783 | 0.78248 | 0.999994 | 17251 | 0 | 0.71948 | 0.217 | 0.3398 | 0.994237 | 9188 | 2 | 0.71948 |
| LAT2 | 6 | 0.21544 | 0.40668 | 0.999994 | 8153 | 3 | -0.15198 | 0.21701 | 0.45485 | 1 | 9189 | 2 | -0.15198 |
| GTPBP6_X | 6 | 0.15413 | 0.33086 | 0.999994 | 6571 | 3 | -0.92857 | 0.21701 | 0.45485 | 1 | 9190 | 2 | -0.92857 |
| ERP29 | 6 | 0.11289 | 0.27227 | 0.998561 | 5399 | 3 | -0.58 | 0.21701 | 0.45485 | 1 | 9191 | 2 | -0.58 |
| SYPL2 | 6 | 0.2495 | 0.44732 | 0.999994 | 9005 | 2 | 0.68186 | 0.21702 | 0.45486 | 1 | 9192 | 3 | 0.68186 |
| RCBTB1 | 6 | 0.66609 | 0.79009 | 0.999994 | 15870 | 1 | 0.88731 | 0.21702 | 0.45486 | 1 | 9193 | 3 | 0.88731 |
| OPN5 | 6 | 0.29119 | 0.49644 | 0.999994 | 9934 | 1 | 0.22861 | 0.21702 | 0.45486 | 1 | 9194 | 2 | 0.22861 |
| P2RX3 | 6 | 0.36051 | 0.57587 | 0.999994 | 11341 | 3 | 0.94778 | 0.21702 | 0.45486 | 1 | 9195 | 3 | 0.94778 |
| HOXD3 | 3 | 0.74591 | 0.74586 | 0.999994 | 16812 | 0 | 1.1516 | 0.21705 | 0.36134 | 0.994759 | 9196 | 2 | 1.1516 |
| VCX2 | 5 | 0.83757 | 0.84488 | 0.999994 | 18043 | 1 | 1.9653 | 0.21705 | 0.43585 | 1 | 9197 | 3 | 1.9653 |
| REV1 | 6 | 0.58933 | 0.73209 | 0.999994 | 14811 | 2 | 1.4284 | 0.21708 | 0.45493 | 1 | 9198 | 4 | 1.4284 |
| SLC39A5 | 5 | 0.76937 | 0.80134 | 0.999994 | 17076 | 1 | 1.904 | 0.21713 | 0.43595 | 1 | 9199 | 3 | 1.904 |
| EEF2K | 6 | 0.67823 | 0.79961 | 0.999994 | 16064 | 2 | 1.2289 | 0.21714 | 0.455 | 1 | 9200 | 4 | 1.2289 |
| CORO2B | 5 | 0.64356 | 0.74257 | 0.999994 | 15569 | 1 | 0.7121 | 0.21717 | 0.43599 | 1 | 9201 | 3 | 0.7121 |
| NonTargetingControlG | 2 | 0.78256 | 0.78202 | 0.999994 | 17247 | 0 | 1.2615 | 0.21744 | 0.34039 | 0.994237 | 9202 | 2 | 1.2615 |
| RPGR | 6 | 0.96471 | 0.96475 | 0.999994 | 20716 | 0 | 1.6037 | 0.21748 | 0.45547 | 1 | 9203 | 4 | 1.6037 |

| | | | | | | | | | | | | |
|---|---|---|---|---|---|---|---|---|---|---|---|---|
| SLC35G5 | 6 | 0.7614 | 0.8326 | 0.999994 | 16995 | 1 | 1.3594 | 0.21757 | 0.4556 | 1 | 9204 | 4 | 1.3594 |
| HCN1 | 6 | 0.24854 | 0.4462 | 0.999994 | 8982 | 2 | 1.6004 | 0.21761 | 0.45564 | 1 | 9205 | 3 | 1.6004 |
| CCDC107 | 6 | 0.019387 | 0.070024 | 0.93914 | 1425 | 3 | 0.45249 | 0.21761 | 0.45564 | 1 | 9206 | 3 | 0.45249 |
| DSG4 | 4 | 0.29837 | 0.44133 | 0.999994 | 10096 | 1 | 1.582 | 0.21761 | 0.40808 | 1 | 9207 | 3 | 1.582 |
| KRTAP9-3 | 4 | 0.90402 | 0.90371 | 0.999994 | 19304 | 0 | 1.1159 | 0.21765 | 0.40814 | 1 | 9208 | 2 | 1.1159 |
| GSTA1 | 4 | 0.51754 | 0.6243 | 0.999994 | 13703 | 1 | 1.0239 | 0.21765 | 0.40814 | 1 | 9209 | 2 | 1.0239 |
| GDI2 | 5 | 0.27573 | 0.46141 | 0.999994 | 9612 | 2 | 0.35554 | 0.21776 | 0.43664 | 1 | 9210 | 2 | 0.35554 |
| GPR155 | 5 | 0.58034 | 0.71861 | 0.999994 | 14682 | 2 | -0.043431 | 0.21776 | 0.43664 | 1 | 9211 | 2 | -0.043431 |
| TAGLN | 6 | 0.81207 | 0.85319 | 0.999994 | 17656 | 1 | 1.3036 | 0.21792 | 0.45607 | 1 | 9212 | 4 | 1.3036 |
| TMEM132D | 4 | 0.63981 | 0.6874 | 0.999994 | 15512 | 1 | 1.4508 | 0.21792 | 0.4085 | 1 | 9213 | 3 | 1.4508 |
| PTP4A3 | 4 | 0.61109 | 0.67136 | 0.999994 | 15089 | 1 | 2.0303 | 0.21793 | 0.40851 | 1 | 9214 | 2 | 2.0303 |
| PSMD14 | 4 | 0.14941 | 0.28856 | 0.999632 | 6441 | 2 | 1.0036 | 0.21793 | 0.40851 | 1 | 9215 | 2 | 1.0036 |
| C20orf141 | 6 | 0.55265 | 0.70558 | 0.999994 | 14254 | 2 | -0.2663 | 0.21796 | 0.45614 | 1 | 9216 | 2 | -0.2663 |
| SOWAHD | 4 | 0.92612 | 0.92593 | 0.999994 | 19769 | 0 | 2.1758 | 0.218 | 0.40861 | 1 | 9217 | 3 | 2.1758 |
| RAB3B | 6 | 0.92778 | 0.92787 | 0.999994 | 19809 | 1 | 1.1221 | 0.21803 | 0.45624 | 1 | 9218 | 3 | 1.1221 |
| INO80 | 6 | 0.0053153 | 0.022306 | 0.819701 | 541 | 4 | -4.8932 | 0.21803 | 0.45624 | 1 | 9219 | 2 | -4.8932 |
| ZDHHC7 | 6 | 0.50443 | 0.67179 | 0.999994 | 13501 | 2 | 1.5536 | 0.21819 | 0.45644 | 1 | 9220 | 4 | 1.5536 |
| FAM171A1 | 6 | 0.50155 | 0.66975 | 0.999994 | 13449 | 2 | 1.4985 | 0.21828 | 0.45657 | 1 | 9221 | 4 | 1.4985 |
| OR5B2 | 6 | 0.050396 | 0.14491 | 0.963012 | 2991 | 3 | -1.5684 | 0.2184 | 0.45673 | 1 | 9222 | 1 | -1.5684 |
| CRYGA | 5 | 0.29561 | 0.47748 | 0.999994 | 10031 | 2 | -0.31185 | 0.21842 | 0.43736 | 1 | 9223 | 2 | -0.31185 |
| HNRNPR | 5 | 0.31943 | 0.49686 | 0.999994 | 10503 | 2 | -0.23349 | 0.21846 | 0.43741 | 1 | 9224 | 2 | -0.23349 |
| ING2 | 5 | 0.34701 | 0.51947 | 0.999994 | 11093 | 2 | 0.53506 | 0.21855 | 0.4375 | 1 | 9225 | 2 | 0.53506 |
| SLC41A2 | 5 | 0.029777 | 0.089892 | 0.94395 | 1997 | 2 | 1.2087 | 0.21855 | 0.4375 | 1 | 9226 | 3 | 1.2087 |
| PLA2G4B | 6 | 0.26355 | 0.46405 | 0.999994 | 9326 | 2 | 0.71087 | 0.21864 | 0.45706 | 1 | 9227 | 3 | 0.71087 |
| GRIK1 | 6 | 0.67419 | 0.79641 | 0.999994 | 15990 | 2 | 0.054876 | 0.21874 | 0.45721 | 1 | 9228 | 2 | 0.054876 |
| RAB4B | 6 | 0.974 | 0.97391 | 0.999994 | 20928 | 0 | 0.85735 | 0.21874 | 0.45721 | 1 | 9229 | 3 | 0.85735 |
| FAM47E | 6 | 0.7708 | 0.83616 | 0.999994 | 17093 | 1 | 1.0721 | 0.21882 | 0.45733 | 1 | 9230 | 4 | 1.0721 |
| PMF1-BGLAP | 2 | 0.7811 | 0.78062 | 0.999994 | 17224 | 0 | 2.5031 | 0.2189 | 0.34237 | 0.994392 | 9231 | 2 | 2.5031 |
| CLK2 | 5 | 0.50391 | 0.65132 | 0.999994 | 13493 | 1 | 0.3168 | 0.21891 | 0.4379 | 1 | 9232 | 2 | 0.3168 |
| MAST3 | 5 | 0.23642 | 0.4261 | 0.999994 | 8678 | 3 | -0.65147 | 0.21891 | 0.4379 | 1 | 9233 | 2 | -0.65147 |
| TMEM133 | 5 | 0.88647 | 0.88656 | 0.999994 | 18958 | 1 | 0.47285 | 0.21891 | 0.4379 | 1 | 9234 | 2 | 0.47285 |
| ACSM5 | 5 | 0.95221 | 0.95236 | 0.999994 | 20389 | 0 | 0.67062 | 0.21891 | 0.4379 | 1 | 9235 | 2 | 0.67062 |
| RAB6B | 5 | 0.54535 | 0.68761 | 0.999994 | 14142 | 2 | -0.30696 | 0.21891 | 0.4379 | 1 | 9236 | 1 | -0.30696 |
| ZNF729 | 5 | 0.0027078 | 0.011671 | 0.727553 | 331 | 2 | -0.33997 | 0.21891 | 0.4379 | 1 | 9237 | 2 | -0.33997 |
| SIRT7 | 6 | 0.17736 | 0.3598 | 0.999994 | 7162 | 2 | 0.99964 | 0.21897 | 0.45753 | 1 | 9238 | 4 | 0.99964 |
| ACADL | 6 | 0.041249 | 0.12469 | 0.951052 | 2576 | 3 | -0.45054 | 0.21897 | 0.45753 | 1 | 9239 | 3 | -0.45054 |
| RAP1GAP | 5 | 0.57071 | 0.71016 | 0.999994 | 14551 | 1 | 1.5332 | 0.219 | 0.438 | 1 | 9240 | 3 | 1.5332 |
| ARRDC4 | 5 | 0.11949 | 0.25658 | 0.994943 | 5609 | 2 | 0.7836 | 0.219 | 0.438 | 1 | 9241 | 3 | 0.7836 |
| ISG15 | 5 | 0.54049 | 0.68334 | 0.999994 | 14064 | 2 | -0.46175 | 0.219 | 0.438 | 1 | 9242 | 2 | -0.46175 |
| NCF1 | 5 | 0.92154 | 0.92171 | 0.999994 | 19666 | 0 | 2.2214 | 0.21901 | 0.43801 | 1 | 9243 | 3 | 2.2214 |
| KLF1 | 5 | 0.89806 | 0.89818 | 0.999994 | 19179 | 0 | 2.2214 | 0.21901 | 0.43801 | 1 | 9244 | 3 | 2.2214 |
| GPR45 | 6 | 0.013277 | 0.050025 | 0.909101 | 1080 | 1 | 1.1041 | 0.21912 | 0.45773 | 1 | 9245 | 3 | 1.1041 |
| GHRL | 6 | 0.33128 | 0.54271 | 0.999994 | 10750 | 2 | 0.058609 | 0.21912 | 0.45773 | 1 | 9246 | 2 | 0.058609 |
| WNT6 | 5 | 0.47838 | 0.62925 | 0.999994 | 13119 | 1 | 1.6171 | 0.21932 | 0.43836 | 1 | 9247 | 3 | 1.6171 |
| DHCR24 | 4 | 0.21243 | 0.35349 | 0.999994 | 8070 | 2 | 1.2615 | 0.21936 | 0.41052 | 1 | 9248 | 2 | 1.2615 |
| C21orf59 | 4 | 0.010387 | 0.034946 | 0.867348 | 893 | 1 | 2.2413 | 0.21936 | 0.41052 | 1 | 9249 | 2 | 2.2413 |
| THEGL | 6 | 0.23932 | 0.43517 | 0.999994 | 8761 | 3 | -0.66717 | 0.21943 | 0.45816 | 1 | 9250 | 2 | -0.66717 |
| WDR26 | 6 | 0.0099794 | 0.038933 | 0.891621 | 869 | 3 | 0.76359 | 0.21943 | 0.45816 | 1 | 9251 | 3 | 0.76359 |
| CTAGE15 | 6 | 0.20555 | 0.39461 | 0.999994 | 7909 | 3 | -0.059411 | 0.21943 | 0.45816 | 1 | 9252 | 3 | -0.059411 |
| TEC | 6 | 0.93978 | 0.93974 | 0.999994 | 20074 | 0 | 1.3015 | 0.21943 | 0.45817 | 1 | 9253 | 4 | 1.3015 |
| hsa-mir-7704 | 4 | 0.3867 | 0.53197 | 0.999994 | 11746 | 1 | 1.7638 | 0.2195 | 0.41074 | 1 | 9254 | 3 | 1.7638 |
| MARC2 | 6 | 0.95677 | 0.95684 | 0.999994 | 20513 | 0 | 1.5036 | 0.21959 | 0.45839 | 1 | 9255 | 4 | 1.5036 |

| | | | | | | | | | | | | | |
|---|---|---|---|---|---|---|---|---|---|---|---|---|---|
| NonTargetingControlG | 2 | 0.73937 | 0.73903 | 0.999994 | 16730 | 0 | 3.5915 | 0.21971 | 0.34343 | 0.994392 | 9256 | 1 | 3.5915 |
| NonTargetingControlG | 2 | 0.78026 | 0.77976 | 0.999994 | 17214 | 0 | 3.299 | 0.21974 | 0.34347 | 0.994392 | 9257 | 2 | 3.299 |
| LYRM5 | 5 | 0.89885 | 0.89895 | 0.999994 | 19202 | 0 | -0.19104 | 0.21976 | 0.43885 | 1 | 9258 | 2 | -0.19104 |
| EPSTI1 | 5 | 0.27488 | 0.46071 | 0.999994 | 9590 | 2 | 0.21343 | 0.21976 | 0.43885 | 1 | 9259 | 2 | 0.21343 |
| UBD | 5 | 0.47846 | 0.62933 | 0.999994 | 13120 | 1 | 1.2485 | 0.21976 | 0.43885 | 1 | 9260 | 3 | 1.2485 |
| TRIM4 | 5 | 0.198 | 0.37241 | 0.999994 | 7710 | 3 | -0.78686 | 0.21976 | 0.43885 | 1 | 9261 | 2 | -0.78686 |
| FRZB | 6 | 0.087928 | 0.22335 | 0.983354 | 4508 | 1 | 1.5755 | 0.21978 | 0.45867 | 1 | 9262 | 4 | 1.5755 |
| KIAA1683 | 3 | 0.58317 | 0.60432 | 0.999994 | 14719 | 1 | 2.5503 | 0.21978 | 0.36426 | 0.995622 | 9263 | 2 | 2.5503 |
| TRIM64C | 5 | 0.5249 | 0.66959 | 0.999994 | 13815 | 2 | 2.0094 | 0.21979 | 0.43888 | 1 | 9264 | 3 | 2.0094 |
| RELL1 | 5 | 0.84563 | 0.85092 | 0.999994 | 18181 | 1 | 2.048 | 0.21982 | 0.43891 | 1 | 9265 | 3 | 2.048 |
| hsa-mir-4743 | 2 | 0.78015 | 0.77966 | 0.999994 | 17212 | 0 | 0.8372 | 0.21985 | 0.34362 | 0.994392 | 9266 | 2 | 0.8372 |
| RPLP2 | 1 | 0.78015 | 0.7805 | 0.999994 | 17211 | 0 | 4.4318 | 0.21985 | 0.21954 | 0.97466 | 9267 | 1 | 4.4318 |
| SMPX | 6 | 0.76683 | 0.83464 | 0.999994 | 17047 | 1 | 1.6214 | 0.21985 | 0.45877 | 1 | 9268 | 4 | 1.6214 |
| AARS2 | 6 | 0.7008 | 0.81225 | 0.999994 | 16313 | 1 | 1.2564 | 0.21993 | 0.45886 | 1 | 9269 | 4 | 1.2564 |
| WDR54 | 6 | 0.21608 | 0.40745 | 0.999994 | 8173 | 2 | 0.11286 | 0.22001 | 0.45897 | 1 | 9270 | 2 | 0.11286 |
| CCNK | 6 | 0.12353 | 0.29203 | 0.999632 | 5752 | 3 | -1.1042 | 0.22001 | 0.45897 | 1 | 9271 | 1 | -1.1042 |
| BDKRB2 | 5 | 0.026199 | 0.080631 | 0.93914 | 1815 | 2 | 1.9636 | 0.22008 | 0.43917 | 1 | 9272 | 3 | 1.9636 |
| ERVFRD-1 | 5 | 0.24585 | 0.43744 | 0.999994 | 8914 | 2 | 1.7515 | 0.22012 | 0.43921 | 1 | 9273 | 3 | 1.7515 |
| C5orf45 | 5 | 0.074096 | 0.18502 | 0.968626 | 3998 | 2 | 1.9062 | 0.22015 | 0.43925 | 1 | 9274 | 3 | 1.9062 |
| CACNG5 | 6 | 0.11232 | 0.27116 | 0.998561 | 5380 | 3 | 0.50784 | 0.22015 | 0.45917 | 1 | 9275 | 3 | 0.50784 |
| KCTD7 | 5 | 0.48391 | 0.63404 | 0.999994 | 13198 | 2 | 0.32636 | 0.22017 | 0.43928 | 1 | 9276 | 2 | 0.32636 |
| SLC25A48 | 5 | 0.65081 | 0.74546 | 0.999994 | 15669 | 1 | 1.9064 | 0.22018 | 0.43928 | 1 | 9277 | 3 | 1.9064 |
| GSK3A | 6 | 0.44031 | 0.62862 | 0.999994 | 12580 | 2 | 1.5881 | 0.22019 | 0.45923 | 1 | 9278 | 3 | 1.5881 |
| RIPPLY1 | 6 | 0.21053 | 0.40074 | 0.999994 | 8036 | 2 | 1.0534 | 0.22019 | 0.45923 | 1 | 9279 | 4 | 1.0534 |
| ALG8 | 6 | 0.95695 | 0.95702 | 0.999994 | 20518 | 0 | 1.5511 | 0.22019 | 0.45923 | 1 | 9280 | 4 | 1.5511 |
| AMD1 | 6 | 0.066567 | 0.17957 | 0.968171 | 3709 | 3 | -0.18429 | 0.22019 | 0.45923 | 1 | 9281 | 2 | -0.18429 |
| FANCB | 6 | 0.05361 | 0.15188 | 0.967126 | 3147 | 3 | -0.89141 | 0.22019 | 0.45923 | 1 | 9282 | 2 | -0.89141 |
| HOXB7 | 4 | 0.5166 | 0.62383 | 0.999994 | 13686 | 1 | 2.3881 | 0.22021 | 0.41168 | 1 | 9283 | 3 | 2.3881 |
| BCAP31 | 6 | 0.88863 | 0.89536 | 0.999994 | 19011 | 1 | 1.1096 | 0.22022 | 0.45926 | 1 | 9284 | 4 | 1.1096 |
| MYSM1 | 4 | 0.35381 | 0.49817 | 0.999994 | 11228 | 2 | 0.97222 | 0.22028 | 0.41177 | 1 | 9285 | 2 | 0.97222 |
| LETMD1 | 6 | 0.2455 | 0.44257 | 0.999994 | 8911 | 3 | -0.066806 | 0.22031 | 0.45939 | 1 | 9286 | 3 | -0.066806 |
| SNX29 | 6 | 0.8456 | 0.86957 | 0.999994 | 18178 | 1 | 1.2814 | 0.22033 | 0.45942 | 1 | 9287 | 4 | 1.2814 |
| PTCRA | 3 | 0.24364 | 0.36707 | 0.999994 | 8876 | 1 | 1.4822 | 0.22034 | 0.36484 | 0.995675 | 9288 | 2 | 1.4822 |
| RFX5 | 6 | 0.2894 | 0.49439 | 0.999994 | 9898 | 3 | 1.127 | 0.22043 | 0.45955 | 1 | 9289 | 3 | 1.127 |
| ZNF564 | 6 | 0.91863 | 0.91914 | 0.999994 | 19599 | 1 | 1.1658 | 0.22045 | 0.45958 | 1 | 9290 | 4 | 1.1658 |
| RGN | 6 | 0.60493 | 0.74364 | 0.999994 | 15009 | 2 | 1.5956 | 0.22049 | 0.45963 | 1 | 9291 | 4 | 1.5956 |
| METRN | 5 | 0.89267 | 0.8928 | 0.999994 | 19082 | 0 | 1.8599 | 0.22056 | 0.43971 | 1 | 9292 | 3 | 1.8599 |
| CALM1 | 6 | 0.32179 | 0.53197 | 0.999994 | 10550 | 3 | 0.94453 | 0.2206 | 0.45978 | 1 | 9293 | 3 | 0.94453 |
| KRTAP10-10 | 4 | 0.010911 | 0.036571 | 0.881473 | 921 | 2 | 0.82456 | 0.22066 | 0.4123 | 1 | 9294 | 2 | 0.82456 |
| TLL1 | 5 | 0.65008 | 0.74516 | 0.999994 | 15661 | 1 | 1.9089 | 0.22072 | 0.43989 | 1 | 9295 | 3 | 1.9089 |
| TJP1 | 6 | 0.94099 | 0.94098 | 0.999994 | 20107 | 0 | 1.0598 | 0.22077 | 0.45999 | 1 | 9296 | 4 | 1.0598 |
| SEC23B | 6 | 0.3539 | 0.56841 | 0.999994 | 11234 | 2 | 1.0518 | 0.22086 | 0.46011 | 1 | 9297 | 4 | 1.0518 |
| ELN | 4 | 0.055027 | 0.13127 | 0.95318 | 3199 | 2 | -1.5088 | 0.22089 | 0.41262 | 1 | 9298 | 2 | -1.5088 |
| ASCL1 | 4 | 0.53394 | 0.632 | 0.999994 | 13965 | 1 | 0.55498 | 0.22089 | 0.41262 | 1 | 9299 | 2 | 0.55498 |
| MRPL48 | 2 | 0.77911 | 0.7786 | 0.999994 | 17199 | 0 | 2.0143 | 0.22089 | 0.34505 | 0.994392 | 9300 | 2 | 2.0143 |
| TYW1 | 6 | 0.4026 | 0.60408 | 0.999994 | 11991 | 1 | 1.38 | 0.2209 | 0.46017 | 1 | 9301 | 4 | 1.38 |
| TMEM9B | 6 | 0.12987 | 0.30012 | 0.999994 | 5931 | 2 | -0.099327 | 0.22093 | 0.46021 | 1 | 9302 | 2 | -0.099327 |
| CNNM4 | 6 | 0.44634 | 0.63265 | 0.999994 | 12666 | 2 | 0.44396 | 0.22093 | 0.46021 | 1 | 9303 | 3 | 0.44396 |
| SHARPIN | 5 | 0.090252 | 0.21099 | 0.979587 | 4600 | 2 | 2.0497 | 0.22096 | 0.44016 | 1 | 9304 | 3 | 2.0497 |
| NonTargetingControlG | 2 | 0.77903 | 0.77852 | 0.999994 | 17197 | 0 | 0.81428 | 0.22097 | 0.34516 | 0.994392 | 9305 | 2 | 0.81428 |
| NonTargetingControlG | 1 | 0.779 | 0.77933 | 0.999994 | 17195 | 0 | 4.3946 | 0.221 | 0.22067 | 0.97466 | 9306 | 1 | 4.3946 |
| NonTargetingControlG | 1 | 0.779 | 0.77933 | 0.999994 | 17194 | 0 | 4.3946 | 0.221 | 0.22067 | 0.97466 | 9307 | 1 | 4.3946 |

| | | | | | | | | | | | | | |
|---|---|---|---|---|---|---|---|---|---|---|---|---|---|
| TMEM35 | 6 | 0.90593 | 0.9083 | 0.999994 | 19349 | 1 | 1.4929 | 0.221 | 0.46031 | 1 | 9308 | 4 | 1.4929 |
| CCDC112 | 6 | 0.66854 | 0.79203 | 0.999994 | 15897 | 2 | 1.3226 | 0.22102 | 0.46033 | 1 | 9309 | 3 | 1.3226 |
| SF3B4 | 2 | 0.46933 | 0.47641 | 0.999994 | 12994 | 1 | 2.7266 | 0.22106 | 0.34529 | 0.994392 | 9310 | 1 | 2.7266 |
| TMEM245 | 5 | 0.36699 | 0.53587 | 0.999994 | 11458 | 2 | 1.8405 | 0.22119 | 0.44042 | 1 | 9311 | 3 | 1.8405 |
| MRGPRD | 5 | 0.4838 | 0.63393 | 0.999994 | 13197 | 2 | 0.18507 | 0.22119 | 0.44042 | 1 | 9312 | 2 | 0.18507 |
| GPRASP1 | 3 | 0.88135 | 0.88143 | 0.999994 | 18858 | 0 | 2.5922 | 0.22123 | 0.36579 | 0.996287 | 9313 | 2 | 2.5922 |
| QSER1 | 6 | 0.59038 | 0.73288 | 0.999994 | 14828 | 1 | 1.5311 | 0.2213 | 0.46072 | 1 | 9314 | 4 | 1.5311 |
| ARHGAP1 | 6 | 0.54971 | 0.70343 | 0.999994 | 14209 | 2 | 0.1472 | 0.2213 | 0.46072 | 1 | 9315 | 3 | 0.1472 |
| OR56A4 | 6 | 0.35563 | 0.57037 | 0.999994 | 11260 | 2 | 0.60026 | 0.2213 | 0.46072 | 1 | 9316 | 3 | 0.60026 |
| ASB18 | 6 | 0.5454 | 0.70037 | 0.999994 | 14144 | 2 | 1.7271 | 0.22145 | 0.46091 | 1 | 9317 | 3 | 1.7271 |
| BPIFA3 | 6 | 0.54586 | 0.70071 | 0.999994 | 14147 | 2 | 1.6338 | 0.2215 | 0.46099 | 1 | 9318 | 3 | 1.6338 |
| ZNF418 | 6 | 0.23468 | 0.42967 | 0.999994 | 8635 | 1 | 1.8469 | 0.22155 | 0.46105 | 1 | 9319 | 4 | 1.8469 |
| MFSD1 | 6 | 0.61658 | 0.75227 | 0.999994 | 15166 | 2 | 1.7447 | 0.22186 | 0.4615 | 1 | 9320 | 4 | 1.7447 |
| DSCR6 | 6 | 0.094024 | 0.23558 | 0.985362 | 4748 | 4 | -1.1914 | 0.22187 | 0.46152 | 1 | 9321 | 2 | -1.1914 |
| FOXF2 | 4 | 0.89884 | 0.89854 | 0.999994 | 19201 | 0 | 1.1694 | 0.22189 | 0.41401 | 1 | 9322 | 3 | 1.1694 |
| RNF144A | 5 | 0.8782 | 0.87854 | 0.999994 | 18791 | 1 | 1.6717 | 0.22195 | 0.44127 | 1 | 9323 | 3 | 1.6717 |
| SCGN | 5 | 0.29085 | 0.47362 | 0.999994 | 9928 | 2 | 1.8713 | 0.22202 | 0.44134 | 1 | 9324 | 3 | 1.8713 |
| UMOD | 3 | 0.18328 | 0.29143 | 0.999632 | 7300 | 1 | 2.262 | 0.2222 | 0.36681 | 0.997075 | 9325 | 2 | 2.262 |
| GRAP2 | 6 | 0.060403 | 0.1666 | 0.967126 | 3441 | 3 | -0.59689 | 0.22226 | 0.46206 | 1 | 9326 | 3 | -0.59689 |
| NETO1 | 6 | 0.46138 | 0.64268 | 0.999994 | 12868 | 2 | -0.22051 | 0.22226 | 0.46206 | 1 | 9327 | 2 | -0.22051 |
| MUSK | 5 | 0.91921 | 0.91937 | 0.999994 | 19614 | 0 | 1.8548 | 0.22229 | 0.44165 | 1 | 9328 | 3 | 1.8548 |
| OSTC | 4 | 0.53859 | 0.63421 | 0.999994 | 14032 | 1 | 2.2383 | 0.22233 | 0.41466 | 1 | 9329 | 2 | 2.2383 |
| LAMTOR3 | 5 | 0.10855 | 0.23973 | 0.987253 | 5259 | 3 | -0.80014 | 0.22239 | 0.44176 | 1 | 9330 | 2 | -0.80014 |
| ABRACL | 6 | 0.95925 | 0.95934 | 0.999994 | 20562 | 0 | 0.90922 | 0.22253 | 0.46241 | 1 | 9331 | 4 | 0.90922 |
| CHRDL1 | 6 | 0.048707 | 0.14127 | 0.959146 | 2920 | 3 | -0.25116 | 0.22253 | 0.46241 | 1 | 9332 | 3 | -0.25116 |
| NonTargetingControlG | 1 | 0.77741 | 0.77775 | 0.999994 | 17175 | 0 | 4.0015 | 0.22259 | 0.22228 | 0.97466 | 9333 | 1 | 4.0015 |
| KCNIP2 | 5 | 0.77196 | 0.80282 | 0.999994 | 17111 | 1 | 0.4921 | 0.2226 | 0.442 | 1 | 9334 | 2 | 0.4921 |
| DHRS3 | 5 | 0.48139 | 0.63185 | 0.999994 | 13159 | 2 | 0.12261 | 0.2226 | 0.442 | 1 | 9335 | 2 | 0.12261 |
| MEF2C | 3 | 0.058159 | 0.12695 | 0.951052 | 3345 | 2 | -3.2168 | 0.2227 | 0.36735 | 0.99708 | 9336 | 1 | -3.2168 |
| PDCD2L | 6 | 0.039907 | 0.1217 | 0.951052 | 2511 | 3 | 0.85401 | 0.22273 | 0.46269 | 1 | 9337 | 3 | 0.85401 |
| TMEM59 | 6 | 0.27282 | 0.47502 | 0.999994 | 9533 | 2 | 0.41901 | 0.22274 | 0.46271 | 1 | 9338 | 3 | 0.41901 |
| CDKAL1 | 6 | 0.12847 | 0.29832 | 0.999994 | 5897 | 2 | 0.33666 | 0.22274 | 0.46271 | 1 | 9339 | 3 | 0.33666 |
| ARMC9 | 6 | 0.34312 | 0.55624 | 0.999994 | 11008 | 3 | 0.11282 | 0.22274 | 0.46271 | 1 | 9340 | 3 | 0.11282 |
| hsa-mir-4738 | 4 | 0.15031 | 0.2895 | 0.999632 | 6464 | 2 | 0.76561 | 0.223 | 0.41558 | 1 | 9341 | 2 | 0.76561 |
| hsa-mir-3616 | 4 | 0.027602 | 0.076621 | 0.93914 | 1883 | 2 | -1.5631 | 0.223 | 0.41558 | 1 | 9342 | 2 | -1.5631 |
| CCDC86 | 4 | 0.72783 | 0.74359 | 0.999994 | 16579 | 1 | 2.0782 | 0.223 | 0.41558 | 1 | 9343 | 2 | 2.0782 |
| ASMT_X | 6 | 0.95832 | 0.95841 | 0.999994 | 20545 | 0 | 1.8384 | 0.223 | 0.46306 | 1 | 9344 | 4 | 1.8384 |
| KIAA1161 | 5 | 0.2374 | 0.42743 | 0.999994 | 8711 | 3 | -0.62351 | 0.22302 | 0.44248 | 1 | 9345 | 2 | -0.62351 |
| DLAT | 5 | 0.18549 | 0.35453 | 0.999994 | 7353 | 3 | -0.84283 | 0.22302 | 0.44248 | 1 | 9346 | 2 | -0.84283 |
| PIP4K2A | 6 | 0.9121 | 0.91335 | 0.999994 | 19464 | 1 | 1.5789 | 0.22306 | 0.46314 | 1 | 9347 | 3 | 1.5789 |
| BOD1L1 | 6 | 0.039417 | 0.12059 | 0.951052 | 2484 | 2 | 0.056298 | 0.22306 | 0.46314 | 1 | 9348 | 3 | 0.056298 |
| ATP13A2 | 6 | 0.6091 | 0.74668 | 0.999994 | 15058 | 2 | 1.3266 | 0.22309 | 0.46318 | 1 | 9349 | 4 | 1.3266 |
| ZEB2 | 5 | 0.20005 | 0.37531 | 0.999994 | 7765 | 2 | -0.21256 | 0.22328 | 0.44277 | 1 | 9350 | 2 | -0.21256 |
| hsa-mir-1272 | 4 | 0.84903 | 0.84856 | 0.999994 | 18241 | 0 | 1.431 | 0.22334 | 0.41605 | 1 | 9351 | 3 | 1.431 |
| KRTAP13-2 | 5 | 0.4358 | 0.5933 | 0.999994 | 12511 | 1 | 0.35443 | 0.22339 | 0.44288 | 1 | 9352 | 2 | 0.35443 |
| PTPRA | 5 | 0.13765 | 0.28425 | 0.999632 | 6125 | 3 | -1.0559 | 0.22339 | 0.44288 | 1 | 9353 | 2 | -1.0559 |
| TAS2R9 | 6 | 0.24009 | 0.4361 | 0.999994 | 8774 | 3 | -0.15264 | 0.22345 | 0.46368 | 1 | 9354 | 3 | -0.15264 |
| SLC38A9 | 6 | 0.39939 | 0.60198 | 0.999994 | 11939 | 2 | 0.012454 | 0.22345 | 0.46368 | 1 | 9355 | 1 | 0.012454 |
| PRRC2B | 6 | 0.064577 | 0.17538 | 0.968171 | 3634 | 3 | -0.20933 | 0.22345 | 0.46368 | 1 | 9356 | 2 | -0.20933 |
| hsa-mir-6788 | 4 | 0.18866 | 0.32897 | 0.999994 | 7440 | 2 | -0.69387 | 0.2235 | 0.41625 | 1 | 9357 | 1 | -0.69387 |
| NonTargetingControlG | 1 | 0.77647 | 0.77681 | 0.999994 | 17159 | 0 | 4.3222 | 0.22353 | 0.22322 | 0.97466 | 9358 | 1 | 4.3222 |
| TEX13B | 6 | 0.23296 | 0.42764 | 0.999994 | 8593 | 1 | 1.3786 | 0.22353 | 0.46378 | 1 | 9359 | 4 | 1.3786 |

| | | | | | | | | | | | | | |
|---|---|---|---|---|---|---|---|---|---|---|---|---|---|
| NonTargetingControlG | 2 | 0.10634 | 0.17985 | 0.968171 | 5184 | 1 | -0.61351 | 0.22354 | 0.34866 | 0.994392 | 9360 | 1 | -0.61351 |
| PRAMEF21 | 2 | 0.77646 | 0.77597 | 0.999994 | 17157 | 0 | 1.2914 | 0.22354 | 0.34867 | 0.994392 | 9361 | 2 | 1.2914 |
| SLC35D3 | 6 | 0.98089 | 0.98082 | 0.999994 | 21115 | 0 | 1.0802 | 0.22366 | 0.46393 | 1 | 9362 | 4 | 1.0802 |
| GLRX3 | 6 | 0.44821 | 0.63392 | 0.999994 | 12690 | 1 | 1.9057 | 0.22366 | 0.46394 | 1 | 9363 | 4 | 1.9057 |
| SCYL1 | 2 | 0.77633 | 0.77585 | 0.999994 | 17156 | 0 | 2.6737 | 0.22367 | 0.34886 | 0.994392 | 9364 | 2 | 2.6737 |
| DCAF4 | 6 | 0.47853 | 0.6542 | 0.999994 | 13121 | 2 | 1.4256 | 0.22369 | 0.46398 | 1 | 9365 | 3 | 1.4256 |
| CTNNB1 | 6 | 0.62721 | 0.76024 | 0.999994 | 15340 | 2 | 0.61847 | 0.22369 | 0.46398 | 1 | 9366 | 3 | 0.61847 |
| NMNAT2 | 5 | 0.19888 | 0.37365 | 0.999994 | 7731 | 2 | -1.2059 | 0.2237 | 0.44322 | 1 | 9367 | 1 | -1.2059 |
| NonTargetingControlG | 2 | 0.77627 | 0.7758 | 0.999994 | 17155 | 0 | 3.3473 | 0.22373 | 0.34893 | 0.994392 | 9368 | 2 | 3.3473 |
| DNAJB8 | 5 | 0.021465 | 0.068033 | 0.93914 | 1545 | 2 | 2.187 | 0.22373 | 0.44326 | 1 | 9369 | 3 | 2.187 |
| EMC2 | 5 | 0.69087 | 0.76239 | 0.999994 | 16218 | 1 | 2.187 | 0.22373 | 0.44326 | 1 | 9370 | 3 | 2.187 |
| DUOXA1 | 5 | 0.88229 | 0.88248 | 0.999994 | 18876 | 1 | 2.187 | 0.22373 | 0.44326 | 1 | 9371 | 3 | 2.187 |
| hsa-mir-19b-2 | 3 | 0.49345 | 0.54188 | 0.999994 | 13316 | 1 | 2.2194 | 0.22377 | 0.36849 | 0.998341 | 9372 | 2 | 2.2194 |
| EXD1 | 4 | 0.062248 | 0.14509 | 0.963012 | 3521 | 1 | 1.3955 | 0.22378 | 0.41661 | 1 | 9373 | 3 | 1.3955 |
| SGK1 | 6 | 0.16409 | 0.34337 | 0.999994 | 6844 | 3 | 0.2734 | 0.22388 | 0.46423 | 1 | 9374 | 3 | 0.2734 |
| GHRHR | 6 | 0.57412 | 0.7209 | 0.999994 | 14605 | 2 | 0.171 | 0.22396 | 0.46434 | 1 | 9375 | 3 | 0.171 |
| hsa-mir-4313 | 4 | 0.63356 | 0.68386 | 0.999994 | 15423 | 1 | 2.0997 | 0.22398 | 0.41689 | 1 | 9376 | 2 | 2.0997 |
| TMX4 | 5 | 0.24083 | 0.43206 | 0.999994 | 8789 | 3 | -0.59497 | 0.22399 | 0.44353 | 1 | 9377 | 2 | -0.59497 |
| LENEP | 5 | 0.042531 | 0.12217 | 0.951052 | 2648 | 3 | -2.5534 | 0.22399 | 0.44353 | 1 | 9378 | 2 | -2.5534 |
| C10orf105 | 5 | 0.92669 | 0.92691 | 0.999994 | 19781 | 0 | 0.96376 | 0.22399 | 0.44353 | 1 | 9379 | 3 | 0.96376 |
| USP46 | 5 | 0.87175 | 0.87264 | 0.999994 | 18673 | 1 | -0.13609 | 0.22399 | 0.44353 | 1 | 9380 | 2 | -0.13609 |
| CNNM2 | 5 | 0.48867 | 0.63806 | 0.999994 | 13262 | 2 | 0.52243 | 0.22399 | 0.44353 | 1 | 9381 | 2 | 0.52243 |
| GOLGA6L4 | 2 | 0.72094 | 0.72068 | 0.999994 | 16506 | 0 | 3.5659 | 0.22409 | 0.34944 | 0.994392 | 9382 | 1 | 3.5659 |
| PJA2 | 5 | 0.19443 | 0.36724 | 0.999994 | 7620 | 1 | 1.3805 | 0.2241 | 0.44365 | 1 | 9383 | 3 | 1.3805 |
| OR52A5 | 5 | 0.18053 | 0.34739 | 0.999994 | 7232 | 3 | -0.95888 | 0.2241 | 0.44365 | 1 | 9384 | 1 | -0.95888 |
| OR7A5 | 6 | 0.67268 | 0.79524 | 0.999994 | 15966 | 2 | 1.213 | 0.22417 | 0.46463 | 1 | 9385 | 4 | 1.213 |
| SLITRK3 | 4 | 0.8515 | 0.85103 | 0.999994 | 18286 | 0 | 1.1583 | 0.2245 | 0.4176 | 1 | 9386 | 3 | 1.1583 |
| FAM83B | 6 | 0.86127 | 0.87813 | 0.999994 | 18471 | 1 | 1.0581 | 0.22453 | 0.46513 | 1 | 9387 | 4 | 1.0581 |
| GPER | 6 | 0.04264 | 0.12785 | 0.951674 | 2655 | 2 | -0.28095 | 0.22453 | 0.46513 | 1 | 9388 | 2 | -0.28095 |
| PLOD2 | 6 | 0.35679 | 0.57168 | 0.999994 | 11285 | 3 | -0.31549 | 0.22453 | 0.46513 | 1 | 9389 | 2 | -0.31549 |
| TMEM50A | 6 | 0.090551 | 0.22861 | 0.983354 | 4616 | 4 | -1.0698 | 0.22453 | 0.46513 | 1 | 9390 | 2 | -1.0698 |
| GBP2 | 6 | 0.65868 | 0.78429 | 0.999994 | 15760 | 2 | 1.3121 | 0.22454 | 0.46515 | 1 | 9391 | 3 | 1.3121 |
| C16orf70 | 6 | 0.70086 | 0.81227 | 0.999994 | 16314 | 1 | 1.1317 | 0.22455 | 0.46515 | 1 | 9392 | 4 | 1.1317 |
| LOC100287036 | 6 | 0.18657 | 0.37125 | 0.999994 | 7375 | 2 | 1.3035 | 0.22459 | 0.46521 | 1 | 9393 | 4 | 1.3035 |
| BCL7A | 6 | 0.46462 | 0.64483 | 0.999994 | 12927 | 2 | 1.9141 | 0.22461 | 0.46524 | 1 | 9394 | 4 | 1.9141 |
| hsa-mir-7703 | 4 | 0.51813 | 0.62455 | 0.999994 | 13712 | 1 | 1.7827 | 0.22462 | 0.41776 | 1 | 9395 | 2 | 1.7827 |
| CLCN2 | 5 | 0.31062 | 0.48963 | 0.999994 | 10339 | 2 | 0.99941 | 0.22468 | 0.4443 | 1 | 9396 | 3 | 0.99941 |
| hsa-mir-509-1 | 4 | 0.4786 | 0.60689 | 0.999994 | 13122 | 1 | 0.91939 | 0.22479 | 0.41803 | 1 | 9397 | 3 | 0.91939 |
| LLGL1 | 6 | 0.30414 | 0.51154 | 0.999994 | 10213 | 2 | 1.5253 | 0.22483 | 0.46553 | 1 | 9398 | 4 | 1.5253 |
| LEFTY2 | 4 | 0.51767 | 0.62436 | 0.999994 | 13705 | 1 | 1.6577 | 0.22494 | 0.41823 | 1 | 9399 | 3 | 1.6577 |
| hsa-mir-1238 | 3 | 0.19145 | 0.30195 | 0.999994 | 7524 | 2 | -1.2576 | 0.22506 | 0.36983 | 0.9988 | 9400 | 1 | -1.2576 |
| EGR3 | 6 | 0.30218 | 0.50926 | 0.999994 | 10172 | 2 | 1.6851 | 0.22509 | 0.46586 | 1 | 9401 | 3 | 1.6851 |
| KIAA0101 | 5 | 0.49127 | 0.64035 | 0.999994 | 13294 | 2 | 0.94209 | 0.22514 | 0.44482 | 1 | 9402 | 3 | 0.94209 |
| TRPV3 | 4 | 0.34679 | 0.491 | 0.999994 | 11088 | 1 | 1.7901 | 0.22522 | 0.41859 | 1 | 9403 | 2 | 1.7901 |
| hsa-mir-8081 | 4 | 0.93515 | 0.93497 | 0.999994 | 19971 | 0 | 1.1893 | 0.22522 | 0.41859 | 1 | 9404 | 3 | 1.1893 |
| NonTargetingControlG | 2 | 0.48205 | 0.48763 | 0.999994 | 13172 | 1 | 2.8183 | 0.22527 | 0.35105 | 0.994392 | 9405 | 1 | 2.8183 |
| SOWAHB | 6 | 0.19494 | 0.38173 | 0.999994 | 7631 | 2 | 1.6004 | 0.22528 | 0.46613 | 1 | 9406 | 4 | 1.6004 |
| FRMD1 | 6 | 0.90839 | 0.91025 | 0.999994 | 19393 | 1 | 1.025 | 0.22536 | 0.46621 | 1 | 9407 | 4 | 1.025 |
| FRAT2 | 6 | 0.42421 | 0.61815 | 0.999994 | 12313 | 2 | 1.4976 | 0.22537 | 0.46624 | 1 | 9408 | 4 | 1.4976 |
| TLR3 | 6 | 0.15901 | 0.33697 | 0.999994 | 6703 | 2 | 1.8516 | 0.2254 | 0.46628 | 1 | 9409 | 4 | 1.8516 |
| ILDR1 | 6 | 0.060403 | 0.1666 | 0.967126 | 3440 | 2 | 1.5804 | 0.22543 | 0.46632 | 1 | 9410 | 4 | 1.5804 |
| SLC25A17 | 5 | 0.42713 | 0.58605 | 0.999994 | 12369 | 2 | -0.53393 | 0.22544 | 0.44516 | 1 | 9411 | 2 | -0.53393 |

| | | | | | | | | | | | | | |
|---|---|---|---|---|---|---|---|---|---|---|---|---|---|
| RPL23A | 3 | 0.32158 | 0.44019 | 0.999994 | 10546 | 1 | 1.1216 | 0.22563 | 0.37046 | 0.9988 | 9412 | 2 | 1.1216 |
| GPATCH1 | 5 | 0.35075 | 0.52254 | 0.999994 | 11178 | 2 | 1.7429 | 0.22563 | 0.44537 | 1 | 9413 | 3 | 1.7429 |
| LOC100130480 | 5 | 0.3725 | 0.54047 | 0.999994 | 11548 | 1 | 0.69653 | 0.22564 | 0.44538 | 1 | 9414 | 3 | 0.69653 |
| GLDC | 5 | 0.52661 | 0.67109 | 0.999994 | 13840 | 2 | 0.46273 | 0.22564 | 0.44538 | 1 | 9415 | 2 | 0.46273 |
| OR51F2 | 5 | 0.2392 | 0.42985 | 0.999994 | 8753 | 2 | -0.020985 | 0.22564 | 0.44538 | 1 | 9416 | 2 | -0.020985 |
| ARHGEF19 | 4 | 0.22939 | 0.3709 | 0.999994 | 8512 | 2 | 0.73258 | 0.22565 | 0.41917 | 1 | 9417 | 2 | 0.73258 |
| SNAPIN | 4 | 0.17791 | 0.31793 | 0.999994 | 7175 | 2 | -1.1259 | 0.22565 | 0.41917 | 1 | 9418 | 1 | -1.1259 |
| MTRNR2L10 | 3 | 0.37819 | 0.47171 | 0.999994 | 11626 | 1 | 2.336 | 0.22573 | 0.37057 | 0.998861 | 9419 | 2 | 2.336 |
| BPIFB4 | 6 | 0.4064 | 0.60653 | 0.999994 | 12048 | 2 | 1.5326 | 0.22574 | 0.46671 | 1 | 9420 | 4 | 1.5326 |
| CAMTA2 | 6 | 0.90514 | 0.90768 | 0.999994 | 19333 | 1 | 1.3347 | 0.22574 | 0.46671 | 1 | 9421 | 4 | 1.3347 |
| ZBTB46 | 6 | 0.97677 | 0.97673 | 0.999994 | 21009 | 0 | 0.56002 | 0.22574 | 0.46671 | 1 | 9422 | 3 | 0.56002 |
| ZNF813 | 2 | 0.71969 | 0.71937 | 0.999994 | 16493 | 0 | 3.5086 | 0.22585 | 0.35179 | 0.994392 | 9423 | 1 | 3.5086 |
| SMCR7L | 2 | 0.66764 | 0.66744 | 0.999994 | 15886 | 0 | 3.3576 | 0.22585 | 0.35179 | 0.994392 | 9424 | 1 | 3.3576 |
| KCND2 | 6 | 0.078113 | 0.20353 | 0.978186 | 4154 | 3 | -0.94275 | 0.22597 | 0.46704 | 1 | 9425 | 2 | -0.94275 |
| PTPN6 | 6 | 0.23552 | 0.43067 | 0.999994 | 8660 | 2 | 1.0201 | 0.22597 | 0.46704 | 1 | 9426 | 4 | 1.0201 |
| HOXA5 | 4 | 0.092337 | 0.20129 | 0.977164 | 4694 | 2 | 0.65755 | 0.22602 | 0.41969 | 1 | 9427 | 2 | 0.65755 |
| MT3 | 4 | 0.93318 | 0.93294 | 0.999994 | 19931 | 0 | 1.1497 | 0.22604 | 0.4197 | 1 | 9428 | 3 | 1.1497 |
| DISP2 | 4 | 0.78067 | 0.78443 | 0.999994 | 17220 | 1 | 1.4263 | 0.22605 | 0.41971 | 1 | 9429 | 3 | 1.4263 |
| EXOSC8 | 4 | 0.25448 | 0.39652 | 0.999994 | 9113 | 1 | 1.8047 | 0.22611 | 0.4198 | 1 | 9430 | 3 | 1.8047 |
| MAST1 | 5 | 0.51561 | 0.6615 | 0.999994 | 13669 | 2 | -0.21546 | 0.22618 | 0.44597 | 1 | 9431 | 1 | -0.21546 |
| C9orf64 | 5 | 0.38187 | 0.54827 | 0.999994 | 11675 | 2 | 0.074296 | 0.22618 | 0.44597 | 1 | 9432 | 1 | 0.074296 |
| ISY1 | 6 | 0.13144 | 0.30217 | 0.999994 | 5967 | 1 | 1.8046 | 0.22627 | 0.46743 | 1 | 9433 | 4 | 1.8046 |
| CLIC4 | 6 | 0.077779 | 0.20287 | 0.977973 | 4141 | 4 | -1.0022 | 0.22633 | 0.46754 | 1 | 9434 | 2 | -1.0022 |
| PDE6A | 6 | 0.87342 | 0.88544 | 0.999994 | 18704 | 1 | 1.6288 | 0.22637 | 0.46758 | 1 | 9435 | 4 | 1.6288 |
| SLC44A4 | 6 | 0.96768 | 0.96769 | 0.999994 | 20776 | 0 | 1.657 | 0.22646 | 0.46771 | 1 | 9436 | 4 | 1.657 |
| PRELP | 6 | 0.70368 | 0.81315 | 0.999994 | 16348 | 1 | 1.0174 | 0.22647 | 0.46773 | 1 | 9437 | 4 | 1.0174 |
| KLHL29 | 5 | 0.1707 | 0.33313 | 0.999994 | 7000 | 3 | -0.99736 | 0.22653 | 0.44636 | 1 | 9438 | 2 | -0.99736 |
| OR5K3 | 5 | 0.17489 | 0.33925 | 0.999994 | 7105 | 3 | -1.0179 | 0.22653 | 0.44636 | 1 | 9439 | 2 | -1.0179 |
| BLK | 5 | 0.027623 | 0.084316 | 0.94395 | 1885 | 3 | -2.9339 | 0.22658 | 0.44641 | 1 | 9440 | 1 | -2.9339 |
| hsa-mir-6127 | 1 | 0.77335 | 0.77366 | 0.999994 | 17124 | 0 | 4.2459 | 0.22665 | 0.22632 | 0.97466 | 9441 | 1 | 4.2459 |
| NECAB3 | 6 | 0.33246 | 0.54408 | 0.999994 | 10780 | 2 | 1.3996 | 0.22669 | 0.46803 | 1 | 9442 | 4 | 1.3996 |
| CAMKV | 6 | 0.46853 | 0.64744 | 0.999994 | 12982 | 2 | -0.25774 | 0.2267 | 0.46804 | 1 | 9443 | 2 | -0.25774 |
| C16orf71 | 6 | 0.036379 | 0.11374 | 0.951052 | 2339 | 4 | -1.953 | 0.2267 | 0.46804 | 1 | 9444 | 2 | -1.953 |
| KLK8 | 6 | 0.35347 | 0.56791 | 0.999994 | 11221 | 3 | 1.0035 | 0.22675 | 0.46811 | 1 | 9445 | 3 | 1.0035 |
| hsa-mir-151a | 3 | 0.86616 | 0.86616 | 0.999994 | 18559 | 0 | 2.2442 | 0.22679 | 0.37171 | 0.998994 | 9446 | 2 | 2.2442 |
| HGF | 6 | 0.44132 | 0.6293 | 0.999994 | 12597 | 2 | 1.6874 | 0.2268 | 0.46817 | 1 | 9447 | 4 | 1.6874 |
| PAQR5 | 6 | 0.17992 | 0.36305 | 0.999994 | 7221 | 2 | 1.5373 | 0.22689 | 0.46827 | 1 | 9448 | 4 | 1.5373 |
| hsa-mir-412 | 3 | 0.41441 | 0.49278 | 0.999994 | 12174 | 1 | 2.6362 | 0.2269 | 0.37182 | 0.998994 | 9449 | 2 | 2.6362 |
| OR10R2 | 6 | 0.34662 | 0.5602 | 0.999994 | 11086 | 1 | 1.5488 | 0.22696 | 0.46836 | 1 | 9450 | 4 | 1.5488 |
| RPS6KC1 | 5 | 0.54822 | 0.69014 | 0.999994 | 14181 | 2 | -0.5784 | 0.22704 | 0.44689 | 1 | 9451 | 2 | -0.5784 |
| MS4A1 | 5 | 0.11951 | 0.25661 | 0.994943 | 5615 | 2 | 1.1443 | 0.22704 | 0.44689 | 1 | 9452 | 3 | 1.1443 |
| SLC35C2 | 5 | 0.025353 | 0.078409 | 0.93914 | 1769 | 3 | -2.7393 | 0.22704 | 0.44689 | 1 | 9453 | 1 | -2.7393 |
| hsa-mir-6780a | 2 | 0.72752 | 0.72719 | 0.999994 | 16574 | 0 | 3.539 | 0.22706 | 0.35342 | 0.994392 | 9454 | 1 | 3.539 |
| OR2A7 | 2 | 0.7533 | 0.75285 | 0.999994 | 16891 | 0 | 3.6203 | 0.22706 | 0.35342 | 0.994392 | 9455 | 1 | 3.6203 |
| VWA5A | 6 | 0.71583 | 0.81699 | 0.999994 | 16456 | 1 | 0.50127 | 0.22709 | 0.46853 | 1 | 9456 | 3 | 0.50127 |
| PRTG | 6 | 0.53712 | 0.69451 | 0.999994 | 14014 | 1 | 1.0596 | 0.22709 | 0.46853 | 1 | 9457 | 3 | 1.0596 |
| ATXN3 | 6 | 0.35925 | 0.57446 | 0.999994 | 11324 | 3 | 0.98203 | 0.2271 | 0.46856 | 1 | 9458 | 3 | 0.98203 |
| ORC4 | 5 | 0.18691 | 0.35656 | 0.999994 | 7385 | 2 | 1.7708 | 0.2271 | 0.44697 | 1 | 9459 | 3 | 1.7708 |
| MRPL13 | 4 | 0.54934 | 0.63935 | 0.999994 | 14201 | 1 | 1.1626 | 0.22715 | 0.42127 | 1 | 9460 | 3 | 1.1626 |
| LGALS9 | 6 | 0.94679 | 0.94675 | 0.999994 | 20258 | 0 | 1.728 | 0.22725 | 0.46877 | 1 | 9461 | 4 | 1.728 |
| ARHGAP12 | 6 | 0.55537 | 0.70748 | 0.999994 | 14297 | 2 | 1.1168 | 0.22736 | 0.46892 | 1 | 9462 | 4 | 1.1168 |
| CD40 | 6 | 0.67367 | 0.79602 | 0.999994 | 15983 | 2 | 0.99849 | 0.22737 | 0.46894 | 1 | 9463 | 4 | 0.99849 |

| | | | | | | | | | | | | | |
|---|---|---|---|---|---|---|---|---|---|---|---|---|---|
| ATP7A | 6 | 0.23346 | 0.42823 | 0.999994 | 8603 | 3 | -0.63617 | 0.22737 | 0.46894 | 1 | 9464 | 2 | -0.63617 |
| CTAG1A | 6 | 0.6873 | 0.8067 | 0.999994 | 16187 | 1 | 1.2726 | 0.22737 | 0.46894 | 1 | 9465 | 4 | 1.2726 |
| XCL1 | 6 | 0.6719 | 0.79463 | 0.999994 | 15954 | 2 | 0.38118 | 0.22737 | 0.46894 | 1 | 9466 | 3 | 0.38118 |
| ZNF431 | 6 | 0.75333 | 0.82966 | 0.999994 | 16893 | 1 | 1.4945 | 0.2274 | 0.46897 | 1 | 9467 | 3 | 1.4945 |
| OR2T33 | 5 | 0.7351 | 0.78318 | 0.999994 | 16663 | 1 | 1.5388 | 0.22745 | 0.44733 | 1 | 9468 | 3 | 1.5388 |
| RSPO2 | 5 | 0.9236 | 0.9238 | 0.999994 | 19709 | 0 | 1.8137 | 0.22748 | 0.44737 | 1 | 9469 | 3 | 1.8137 |
| ZNF836 | 6 | 0.57763 | 0.72346 | 0.999994 | 14653 | 2 | 1.4741 | 0.22751 | 0.46913 | 1 | 9470 | 3 | 1.4741 |
| STRA13 | 5 | 0.13533 | 0.28073 | 0.999632 | 6060 | 2 | 1.8655 | 0.22751 | 0.4474 | 1 | 9471 | 3 | 1.8655 |
| MAN2A1 | 6 | 0.71204 | 0.81578 | 0.999994 | 16420 | 1 | 1.4223 | 0.22755 | 0.46918 | 1 | 9472 | 4 | 1.4223 |
| C9orf89 | 5 | 0.23224 | 0.42032 | 0.999994 | 8578 | 3 | -0.62868 | 0.2276 | 0.4475 | 1 | 9473 | 2 | -0.62868 |
| SAE1 | 5 | 0.67209 | 0.75419 | 0.999994 | 15957 | 1 | 1.5385 | 0.2276 | 0.4475 | 1 | 9474 | 3 | 1.5385 |
| AREL1 | 5 | 0.84051 | 0.84705 | 0.999994 | 18096 | 1 | 1.8691 | 0.22764 | 0.44753 | 1 | 9475 | 3 | 1.8691 |
| ARHGEF33 | 6 | 0.33649 | 0.54866 | 0.999994 | 10861 | 2 | 1.6537 | 0.22767 | 0.46935 | 1 | 9476 | 4 | 1.6537 |
| hsa-mir-3157 | 4 | 0.93568 | 0.93548 | 0.999994 | 19983 | 0 | 2.395 | 0.22769 | 0.42205 | 1 | 9477 | 3 | 2.395 |
| hsa-mir-3649 | 4 | 0.71678 | 0.73581 | 0.999994 | 16464 | 1 | 2.3402 | 0.22769 | 0.42205 | 1 | 9478 | 2 | 2.3402 |
| hsa-mir-3923 | 4 | 0.71021 | 0.7313 | 0.999994 | 16400 | 1 | 1.9081 | 0.22769 | 0.42205 | 1 | 9479 | 2 | 1.9081 |
| hsa-mir-596 | 4 | 0.005897 | 0.020594 | 0.81027 | 584 | 3 | -5.1525 | 0.2277 | 0.42207 | 1 | 9480 | 1 | -5.1525 |
| hsa-mir-6751 | 4 | 0.36952 | 0.51436 | 0.999994 | 11497 | 1 | 1.2163 | 0.2277 | 0.42207 | 1 | 9481 | 2 | 1.2163 |
| hsa-mir-597 | 4 | 0.92357 | 0.92339 | 0.999994 | 19708 | 0 | 2.0416 | 0.22773 | 0.4221 | 1 | 9482 | 3 | 2.0416 |
| ZNF492 | 2 | 0.77227 | 0.77176 | 0.999994 | 17114 | 0 | 1.3138 | 0.22773 | 0.35432 | 0.994392 | 9483 | 2 | 1.3138 |
| C4orf47 | 6 | 0.79795 | 0.84701 | 0.999994 | 17451 | 1 | 1.1408 | 0.22777 | 0.46949 | 1 | 9484 | 4 | 1.1408 |
| CCDC125 | 5 | 0.51116 | 0.65766 | 0.999994 | 13599 | 2 | -0.29221 | 0.22785 | 0.44777 | 1 | 9485 | 2 | -0.29221 |
| ENOX1 | 5 | 0.53635 | 0.67966 | 0.999994 | 14007 | 2 | 0.53549 | 0.22785 | 0.44777 | 1 | 9486 | 2 | 0.53549 |
| RPS17 | 5 | 0.85222 | 0.85605 | 0.999994 | 18301 | 1 | 1.4568 | 0.22785 | 0.44777 | 1 | 9487 | 3 | 1.4568 |
| CPNE9 | 5 | 0.039229 | 0.11397 | 0.951052 | 2480 | 2 | 0.38761 | 0.22785 | 0.44777 | 1 | 9488 | 1 | 0.38761 |
| hsa-mir-4762 | 3 | 0.83369 | 0.83359 | 0.999994 | 17982 | 0 | 2.5012 | 0.22789 | 0.37287 | 0.999346 | 9489 | 2 | 2.5012 |
| CCL23 | 6 | 0.30923 | 0.51749 | 0.999994 | 10308 | 2 | 1.1043 | 0.2279 | 0.46968 | 1 | 9490 | 4 | 1.1043 |
| MOCS2 | 6 | 0.19707 | 0.38434 | 0.999994 | 7678 | 2 | 0.96813 | 0.22793 | 0.46973 | 1 | 9491 | 4 | 0.96813 |
| SYNPO | 5 | 0.7602 | 0.79629 | 0.999994 | 16985 | 1 | 2.0068 | 0.22801 | 0.44797 | 1 | 9492 | 3 | 2.0068 |
| RIMS4 | 6 | 0.90956 | 0.91119 | 0.999994 | 19416 | 1 | 0.73025 | 0.22805 | 0.46989 | 1 | 9493 | 4 | 0.73025 |
| ACOT9 | 6 | 0.44831 | 0.63399 | 0.999994 | 12693 | 2 | -0.20102 | 0.22805 | 0.46989 | 1 | 9494 | 1 | -0.20102 |
| RPS19BP1 | 3 | 0.16493 | 0.26793 | 0.998561 | 6861 | 1 | -0.19335 | 0.22823 | 0.37322 | 0.999517 | 9495 | 1 | -0.19335 |
| C1D | 5 | 0.09484 | 0.21826 | 0.980699 | 4784 | 2 | 1.803 | 0.22824 | 0.44821 | 1 | 9496 | 3 | 1.803 |
| ALDH6A1 | 6 | 0.44599 | 0.63243 | 0.999994 | 12662 | 1 | 1.8449 | 0.22825 | 0.47015 | 1 | 9497 | 4 | 1.8449 |
| ASTE1 | 6 | 0.96345 | 0.96351 | 0.999994 | 20683 | 0 | 1.7706 | 0.22825 | 0.47015 | 1 | 9498 | 3 | 1.7706 |
| MAP3K13 | 6 | 0.90454 | 0.90721 | 0.999994 | 19312 | 1 | 1.823 | 0.22825 | 0.47015 | 1 | 9499 | 4 | 1.823 |
| hsa-mir-548f-1 | 4 | 0.2236 | 0.36493 | 0.999994 | 8370 | 2 | 0.96125 | 0.22831 | 0.42287 | 1 | 9500 | 2 | 0.96125 |
| LRRC34 | 6 | 0.38142 | 0.59048 | 0.999994 | 11669 | 2 | 0.60043 | 0.22832 | 0.47024 | 1 | 9501 | 3 | 0.60043 |
| KRTAP11-1 | 6 | 0.32825 | 0.53924 | 0.999994 | 10690 | 3 | -0.13886 | 0.22832 | 0.47024 | 1 | 9502 | 2 | -0.13886 |
| RPL35 | 6 | 0.93865 | 0.93862 | 0.999994 | 20042 | 0 | 1.0823 | 0.22832 | 0.47024 | 1 | 9503 | 3 | 1.0823 |
| DVL3 | 5 | 0.16057 | 0.31838 | 0.999994 | 6755 | 2 | 0.73001 | 0.22833 | 0.44829 | 1 | 9504 | 3 | 0.73001 |
| CYP3A4 | 5 | 0.025846 | 0.079697 | 0.93914 | 1794 | 2 | 1.0955 | 0.22833 | 0.44829 | 1 | 9505 | 3 | 1.0955 |
| GTF2F2 | 4 | 0.03821 | 0.098197 | 0.945253 | 2428 | 2 | -1.0128 | 0.22834 | 0.42292 | 1 | 9506 | 2 | -1.0128 |
| ISOC1 | 6 | 0.45091 | 0.63572 | 0.999994 | 12723 | 1 | 1.0394 | 0.22839 | 0.47033 | 1 | 9507 | 4 | 1.0394 |
| ANKRD34B | 6 | 0.20533 | 0.39436 | 0.999994 | 7905 | 3 | -0.42602 | 0.22839 | 0.47033 | 1 | 9508 | 2 | -0.42602 |
| NonTargetingControlG | 1 | 0.77149 | 0.77181 | 0.999994 | 17105 | 0 | 4.1858 | 0.22851 | 0.2282 | 0.97466 | 9509 | 1 | 4.1858 |
| TCTN2 | 5 | 0.15295 | 0.3072 | 0.999994 | 6532 | 2 | 1.8986 | 0.22855 | 0.44853 | 1 | 9510 | 3 | 1.8986 |
| BBX | 6 | 0.35667 | 0.57154 | 0.999994 | 11277 | 1 | 0.97687 | 0.22855 | 0.47057 | 1 | 9511 | 4 | 0.97687 |
| SLC25A38 | 5 | 0.2207 | 0.40422 | 0.999994 | 8306 | 2 | 0.39504 | 0.22858 | 0.44856 | 1 | 9512 | 2 | 0.39504 |
| KIAA1430 | 5 | 0.08779 | 0.20704 | 0.978744 | 4503 | 3 | -1.7628 | 0.22858 | 0.44856 | 1 | 9513 | 2 | -1.7628 |
| NUDT21 | 4 | 0.44195 | 0.58879 | 0.999994 | 12609 | 2 | 1.643 | 0.22871 | 0.42342 | 1 | 9514 | 2 | 1.643 |
| PRSS8 | 5 | 0.7592 | 0.79574 | 0.999994 | 16969 | 1 | 1.8675 | 0.22873 | 0.44872 | 1 | 9515 | 3 | 1.8675 |

| | | | | | | | | | | | | |
|---|---|---|---|---|---|---|---|---|---|---|---|---|
| NKAP | 5 | 0.042071 | 0.12106 | 0.951052 | 2617 | 3 | -2.4809 | 0.22873 | 0.44872 | 1 | 9516 | 2 | -2.4809 |
| LRMP | 5 | 0.15252 | 0.30654 | 0.999994 | 6522 | 3 | -1.0502 | 0.22873 | 0.44872 | 1 | 9517 | 2 | -1.0502 |
| FURIN | 5 | 0.56411 | 0.70429 | 0.999994 | 14454 | 2 | 0.32284 | 0.22873 | 0.44872 | 1 | 9518 | 2 | 0.32284 |
| TEFM | 6 | 0.46206 | 0.64311 | 0.999994 | 12888 | 2 | 1.1367 | 0.22877 | 0.47085 | 1 | 9519 | 3 | 1.1367 |
| NonTargetingControlG | 2 | 0.75176 | 0.75132 | 0.999994 | 16876 | 0 | 3.6164 | 0.22881 | 0.35577 | 0.994392 | 9520 | 1 | 3.6164 |
| CHAT | 6 | 0.37736 | 0.58789 | 0.999994 | 11612 | 2 | 1.6041 | 0.22883 | 0.47095 | 1 | 9521 | 4 | 1.6041 |
| WIBG | 4 | 0.44452 | 0.59144 | 0.999994 | 12641 | 1 | 1.8497 | 0.22884 | 0.4236 | 1 | 9522 | 2 | 1.8497 |
| IQCJ | 6 | 0.67015 | 0.79325 | 0.999994 | 15934 | 2 | 1.0857 | 0.22886 | 0.47098 | 1 | 9523 | 4 | 1.0857 |
| hsa-mir-30a | 4 | 0.82489 | 0.82452 | 0.999994 | 17851 | 1 | 1.359 | 0.22887 | 0.42363 | 1 | 9524 | 3 | 1.359 |
| hsa-mir-7110 | 3 | 0.65497 | 0.66121 | 0.999994 | 15717 | 1 | 2.3184 | 0.22887 | 0.37392 | 0.999517 | 9525 | 2 | 2.3184 |
| TCTEX1D1 | 6 | 0.41128 | 0.60972 | 0.999994 | 12135 | 2 | 1.1332 | 0.229 | 0.47118 | 1 | 9526 | 4 | 1.1332 |
| PCIF1 | 6 | 0.034738 | 0.1099 | 0.951052 | 2249 | 2 | 1.4671 | 0.22912 | 0.47134 | 1 | 9527 | 4 | 1.4671 |
| PSMB10 | 6 | 0.18309 | 0.367 | 0.999994 | 7294 | 2 | 1.065 | 0.22913 | 0.47136 | 1 | 9528 | 3 | 1.065 |
| PJA1 | 5 | 0.0033191 | 0.014114 | 0.75278 | 388 | 2 | 1.916 | 0.22921 | 0.44927 | 1 | 9529 | 3 | 1.916 |
| GGTLC1 | 6 | 0.39227 | 0.5974 | 0.999994 | 11834 | 1 | 0.64755 | 0.22926 | 0.47152 | 1 | 9530 | 3 | 0.64755 |
| ZDHHC15 | 6 | 0.038341 | 0.11818 | 0.951052 | 2433 | 3 | -0.90329 | 0.22926 | 0.47152 | 1 | 9531 | 3 | -0.90329 |
| TSTD3 | 6 | 0.32398 | 0.53442 | 0.999994 | 10596 | 3 | 0.17716 | 0.22926 | 0.47152 | 1 | 9532 | 3 | 0.17716 |
| BLNK | 6 | 0.52268 | 0.68436 | 0.999994 | 13789 | 1 | 0.30177 | 0.22926 | 0.47152 | 1 | 9533 | 3 | 0.30177 |
| DMP1 | 6 | 0.59846 | 0.7388 | 0.999994 | 14935 | 2 | 0.48239 | 0.22926 | 0.47152 | 1 | 9534 | 3 | 0.48239 |
| OTOS | 6 | 0.040893 | 0.12393 | 0.951052 | 2557 | 2 | 1.2581 | 0.22926 | 0.47152 | 1 | 9535 | 4 | 1.2581 |
| ZNF684 | 5 | 0.051477 | 0.14396 | 0.961589 | 3044 | 3 | -2.0796 | 0.22928 | 0.44935 | 1 | 9536 | 2 | -2.0796 |
| MARCH6 | 6 | 0.0063735 | 0.026165 | 0.848412 | 618 | 3 | -2.6226 | 0.2293 | 0.47157 | 1 | 9537 | 3 | -2.6226 |
| TMEM120A | 6 | 0.084319 | 0.21612 | 0.980699 | 4386 | 3 | -0.76336 | 0.2293 | 0.47157 | 1 | 9538 | 3 | -0.76336 |
| SLC9A9 | 6 | 0.046616 | 0.13665 | 0.956777 | 2835 | 4 | -1.629 | 0.2293 | 0.47157 | 1 | 9539 | 2 | -1.629 |
| EFR3A | 6 | 0.19784 | 0.38526 | 0.999994 | 7705 | 2 | 1.7125 | 0.22932 | 0.4716 | 1 | 9540 | 4 | 1.7125 |
| COX5B | 2 | 0.043445 | 0.078156 | 0.93914 | 2691 | 1 | -0.47356 | 0.22941 | 0.35657 | 0.994392 | 9541 | 1 | -0.47356 |
| CDCA8 | 1 | 0.77047 | 0.77081 | 0.999994 | 17089 | 0 | 4.1654 | 0.22953 | 0.22919 | 0.97466 | 9542 | 1 | 4.1654 |
| LYRM2 | 6 | 0.28214 | 0.48588 | 0.999994 | 9741 | 2 | 1.3217 | 0.22962 | 0.47202 | 1 | 9543 | 3 | 1.3217 |
| C12orf76 | 4 | 0.76407 | 0.77106 | 0.999994 | 17015 | 1 | 1.6987 | 0.22966 | 0.42475 | 1 | 9544 | 2 | 1.6987 |
| CHCHD6 | 6 | 0.2438 | 0.44052 | 0.999994 | 8879 | 1 | 1.1547 | 0.22969 | 0.47212 | 1 | 9545 | 4 | 1.1547 |
| NonTargetingControlG | 2 | 0.77026 | 0.76976 | 0.999994 | 17086 | 0 | 2.165 | 0.22974 | 0.35702 | 0.994392 | 9546 | 2 | 2.165 |
| ATN1 | 4 | 0.83997 | 0.83943 | 0.999994 | 18085 | 0 | 1.9848 | 0.22982 | 0.42498 | 1 | 9547 | 3 | 1.9848 |
| NonTargetingControlG | 2 | 0.7701 | 0.76959 | 0.999994 | 17083 | 0 | 2.7756 | 0.2299 | 0.35722 | 0.994392 | 9548 | 2 | 2.7756 |
| SLITRK1 | 6 | 0.73714 | 0.82403 | 0.999994 | 16690 | 1 | 1.4177 | 0.23015 | 0.47275 | 1 | 9549 | 4 | 1.4177 |
| TNFRSF9 | 6 | 0.93134 | 0.93147 | 0.999994 | 19883 | 0 | 0.5282 | 0.23016 | 0.47276 | 1 | 9550 | 3 | 0.5282 |
| SCGB2B2 | 6 | 0.49057 | 0.66229 | 0.999994 | 13286 | 2 | 0.5422 | 0.23016 | 0.47276 | 1 | 9551 | 3 | 0.5422 |
| ESRRG | 6 | 0.077491 | 0.20227 | 0.977973 | 4129 | 2 | 0.92352 | 0.23016 | 0.47276 | 1 | 9552 | 4 | 0.92352 |
| LRRC29 | 6 | 0.010713 | 0.041446 | 0.891621 | 912 | 5 | -1.6516 | 0.23016 | 0.47276 | 1 | 9553 | 1 | -1.6516 |
| FRMPD4 | 6 | 0.00060808 | 0.0031733 | 0.537554 | 120 | 4 | -5.7407 | 0.23016 | 0.47276 | 1 | 9554 | 2 | -5.7407 |
| SALL1 | 6 | 0.31943 | 0.52923 | 0.999994 | 10502 | 3 | -0.4936 | 0.23016 | 0.47276 | 1 | 9555 | 2 | -0.4936 |
| HIVEP2 | 6 | 0.38535 | 0.59296 | 0.999994 | 11722 | 2 | 1.2401 | 0.23023 | 0.47284 | 1 | 9556 | 4 | 1.2401 |
| PDE6C | 6 | 0.11632 | 0.27881 | 0.999632 | 5520 | 2 | 1.3156 | 0.23023 | 0.47285 | 1 | 9557 | 3 | 1.3156 |
| LCE3A | 6 | 0.3663 | 0.5809 | 0.999994 | 11448 | 2 | 1.1835 | 0.23025 | 0.47288 | 1 | 9558 | 3 | 1.1835 |
| CNTNAP5 | 6 | 0.19364 | 0.38007 | 0.999994 | 7591 | 1 | 1.7065 | 0.23026 | 0.47289 | 1 | 9559 | 4 | 1.7065 |
| HYAL1 | 6 | 0.72865 | 0.82115 | 0.999994 | 16586 | 1 | 0.087553 | 0.23027 | 0.47291 | 1 | 9560 | 2 | 0.087553 |
| MTMR11 | 6 | 0.62039 | 0.75514 | 0.999994 | 15226 | 2 | 1.3852 | 0.23027 | 0.47291 | 1 | 9561 | 3 | 1.3852 |
| SLC35B4 | 6 | 0.53321 | 0.69179 | 0.999994 | 13956 | 1 | 1.5102 | 0.23027 | 0.47291 | 1 | 9562 | 3 | 1.5102 |
| SLC46A1 | 5 | 0.43934 | 0.59628 | 0.999994 | 12564 | 1 | 0.82791 | 0.2303 | 0.45046 | 1 | 9563 | 3 | 0.82791 |
| AP3M2 | 5 | 0.52243 | 0.66741 | 0.999994 | 13787 | 2 | 0.81156 | 0.2303 | 0.45046 | 1 | 9564 | 3 | 0.81156 |
| DEXI | 5 | 0.304 | 0.48428 | 0.999994 | 10209 | 2 | 0.37681 | 0.2303 | 0.45046 | 1 | 9565 | 2 | 0.37681 |
| OSBPL2 | 5 | 0.095443 | 0.21924 | 0.980699 | 4800 | 2 | 0.59279 | 0.2303 | 0.45046 | 1 | 9566 | 3 | 0.59279 |
| BLID | 6 | 0.84642 | 0.87003 | 0.999994 | 18198 | 1 | 1.1986 | 0.23033 | 0.47299 | 1 | 9567 | 4 | 1.1986 |

| | | | | | | | | | | | | | |
|---|---|---|---|---|---|---|---|---|---|---|---|---|---|
| hsa-mir-4459 | 1 | 0.76966 | 0.77002 | 0.999994 | 17077 | 0 | 4.2739 | 0.23034 | 0.23002 | 0.97466 | 9568 | 1 | 4.2739 |
| SP2 | 4 | 0.33945 | 0.48342 | 0.999994 | 10921 | 1 | 1.5692 | 0.23042 | 0.42582 | 1 | 9569 | 2 | 1.5692 |
| OPN1SW | 6 | 0.51307 | 0.67775 | 0.999994 | 13622 | 1 | 0.95232 | 0.23045 | 0.47315 | 1 | 9570 | 4 | 0.95232 |
| CGA | 6 | 0.63218 | 0.76399 | 0.999994 | 15402 | 2 | 1.5366 | 0.23058 | 0.47331 | 1 | 9571 | 4 | 1.5366 |
| ACADVL | 4 | 0.2182 | 0.3594 | 0.999994 | 8233 | 2 | 1.04 | 0.23059 | 0.42605 | 1 | 9572 | 2 | 1.04 |
| RFXANK | 6 | 0.91761 | 0.91819 | 0.999994 | 19581 | 1 | 1.2764 | 0.2306 | 0.47334 | 1 | 9573 | 3 | 1.2764 |
| ELOF1 | 5 | 0.81532 | 0.82949 | 0.999994 | 17704 | 1 | 2.1158 | 0.23066 | 0.45084 | 1 | 9574 | 3 | 2.1158 |
| BNC2 | 6 | 0.20339 | 0.39198 | 0.999994 | 7864 | 3 | -0.098316 | 0.23069 | 0.47346 | 1 | 9575 | 3 | -0.098316 |
| SSMEM1 | 6 | 0.096343 | 0.24016 | 0.987253 | 4830 | 2 | 0.96776 | 0.23079 | 0.47361 | 1 | 9576 | 4 | 0.96776 |
| ANKRD18A | 5 | 0.21781 | 0.4002 | 0.999994 | 8222 | 1 | 1.7414 | 0.23081 | 0.45101 | 1 | 9577 | 3 | 1.7414 |
| TMEM43 | 6 | 0.54074 | 0.69706 | 0.999994 | 14069 | 1 | 1.4565 | 0.23085 | 0.4737 | 1 | 9578 | 4 | 1.4565 |
| hsa-mir-3908 | 3 | 0.62488 | 0.63658 | 0.999994 | 15301 | 1 | 1.6381 | 0.23089 | 0.37602 | 0.999762 | 9579 | 2 | 1.6381 |
| NUDT10 | 4 | 0.37771 | 0.5228 | 0.999994 | 11622 | 1 | 1.0877 | 0.23097 | 0.42657 | 1 | 9580 | 3 | 1.0877 |
| hsa-mir-3605 | 4 | 0.021818 | 0.064457 | 0.93914 | 1567 | 2 | -3.0507 | 0.23099 | 0.42659 | 1 | 9581 | 2 | -3.0507 |
| ADCY5 | 6 | 0.19261 | 0.37878 | 0.999994 | 7562 | 2 | 1.1476 | 0.23112 | 0.47407 | 1 | 9582 | 4 | 1.1476 |
| SOLH | 6 | 0.062889 | 0.17185 | 0.967658 | 3551 | 4 | -1.8759 | 0.23114 | 0.4741 | 1 | 9583 | 2 | -1.8759 |
| TENM1 | 6 | 0.043419 | 0.12961 | 0.95318 | 2689 | 3 | -0.81428 | 0.23114 | 0.4741 | 1 | 9584 | 3 | -0.81428 |
| NonTargetingControlG | 2 | 0.14609 | 0.22393 | 0.983354 | 6354 | 1 | -0.23669 | 0.2312 | 0.35896 | 0.994392 | 9585 | 1 | -0.23669 |
| NonTargetingControlG | 2 | 0.69365 | 0.69348 | 0.999994 | 16242 | 0 | 3.3875 | 0.2312 | 0.35896 | 0.994392 | 9586 | 1 | 3.3875 |
| NonTargetingControlG | 2 | 0.093141 | 0.15943 | 0.967126 | 4726 | 1 | -0.79241 | 0.2312 | 0.35896 | 0.994392 | 9587 | 1 | -0.79241 |
| FGF4 | 6 | 0.036048 | 0.11297 | 0.951052 | 2321 | 2 | 1.0925 | 0.23125 | 0.47425 | 1 | 9588 | 4 | 1.0925 |
| MYPOP | 6 | 0.77853 | 0.83909 | 0.999994 | 17190 | 1 | 0.66309 | 0.23126 | 0.47425 | 1 | 9589 | 3 | 0.66309 |
| SUMO3 | 5 | 0.35156 | 0.52318 | 0.999994 | 11187 | 2 | 1.9602 | 0.23141 | 0.45167 | 1 | 9590 | 3 | 1.9602 |
| C19orf71 | 4 | 0.32883 | 0.4725 | 0.999994 | 10697 | 1 | 2.4929 | 0.23144 | 0.42721 | 1 | 9591 | 3 | 2.4929 |
| LRRC23 | 5 | 0.21445 | 0.39558 | 0.999994 | 8130 | 3 | -0.67803 | 0.23146 | 0.45172 | 1 | 9592 | 2 | -0.67803 |
| CCNB2 | 5 | 0.049299 | 0.13867 | 0.957514 | 2942 | 2 | -0.43072 | 0.23146 | 0.45172 | 1 | 9593 | 2 | -0.43072 |
| FAM72D | 2 | 0.76853 | 0.76801 | 0.999994 | 17067 | 0 | 0.74007 | 0.23147 | 0.35929 | 0.994392 | 9594 | 2 | 0.74007 |
| DHRS4 | 6 | 0.27637 | 0.47919 | 0.999994 | 9627 | 2 | 1.855 | 0.23149 | 0.47457 | 1 | 9595 | 4 | 1.855 |
| TTLL6 | 5 | 0.033087 | 0.098455 | 0.945253 | 2166 | 2 | 1.862 | 0.23149 | 0.45177 | 1 | 9596 | 3 | 1.862 |
| KL | 6 | 0.3906 | 0.59635 | 0.999994 | 11807 | 2 | 1.1024 | 0.2315 | 0.47459 | 1 | 9597 | 3 | 1.1024 |
| hsa-mir-8083 | 3 | 0.88409 | 0.88418 | 0.999994 | 18909 | 0 | 2.2635 | 0.23151 | 0.37676 | 1 | 9598 | 2 | 2.2635 |
| PFDN6 | 4 | 0.56436 | 0.64685 | 0.999994 | 14462 | 1 | 1.7452 | 0.23153 | 0.42732 | 1 | 9599 | 3 | 1.7452 |
| C16orf97 | 5 | 0.048122 | 0.13592 | 0.95468 | 2893 | 1 | 1.8809 | 0.23155 | 0.45183 | 1 | 9600 | 3 | 1.8809 |
| PRRC2C | 4 | 0.94363 | 0.94349 | 0.999994 | 20165 | 0 | 1.0386 | 0.23161 | 0.42744 | 1 | 9601 | 3 | 1.0386 |
| ILDR2 | 5 | 0.43481 | 0.59248 | 0.999994 | 12490 | 2 | -0.51258 | 0.23162 | 0.45191 | 1 | 9602 | 1 | -0.51258 |
| CAMSAP2 | 5 | 0.40803 | 0.57023 | 0.999994 | 12075 | 2 | 1.229 | 0.23162 | 0.45191 | 1 | 9603 | 3 | 1.229 |
| KCNB1 | 5 | 0.29243 | 0.4749 | 0.999994 | 9960 | 2 | 1.893 | 0.23165 | 0.45195 | 1 | 9604 | 3 | 1.893 |
| NonTargetingControlG | 2 | 0.42736 | 0.43981 | 0.999994 | 12373 | 1 | 2.6602 | 0.23171 | 0.3596 | 0.994416 | 9605 | 1 | 2.6602 |
| BPGM | 6 | 0.6434 | 0.77251 | 0.999994 | 15568 | 1 | 1.3207 | 0.23171 | 0.47487 | 1 | 9606 | 3 | 1.3207 |
| HAGH | 6 | 0.1121 | 0.27073 | 0.998561 | 5370 | 1 | 1.2549 | 0.23179 | 0.47498 | 1 | 9607 | 4 | 1.2549 |
| KDM8 | 6 | 0.65321 | 0.78007 | 0.999994 | 15700 | 2 | 1.2627 | 0.2318 | 0.47499 | 1 | 9608 | 3 | 1.2627 |
| COLGALT2 | 4 | 0.87286 | 0.87251 | 0.999994 | 18689 | 0 | 1.5662 | 0.2318 | 0.42768 | 1 | 9609 | 3 | 1.5662 |
| CD34 | 6 | 0.68715 | 0.80659 | 0.999994 | 16185 | 1 | 1.1655 | 0.23197 | 0.47521 | 1 | 9610 | 4 | 1.1655 |
| RBM19 | 4 | 0.090178 | 0.19733 | 0.977164 | 4599 | 3 | -1.4684 | 0.23198 | 0.42793 | 1 | 9611 | 1 | -1.4684 |
| CHAF1A | 4 | 0.44122 | 0.58805 | 0.999994 | 12596 | 2 | 1.4098 | 0.23198 | 0.42793 | 1 | 9612 | 2 | 1.4098 |
| ZNF283 | 4 | 0.91645 | 0.9162 | 0.999994 | 19548 | 0 | 0.90158 | 0.23198 | 0.42793 | 1 | 9613 | 2 | 0.90158 |
| SPATA22 | 4 | 0.79321 | 0.79517 | 0.999994 | 17387 | 1 | 1.1739 | 0.23198 | 0.42793 | 1 | 9614 | 2 | 1.1739 |
| IL10RB | 6 | 0.5882 | 0.73128 | 0.999994 | 14794 | 2 | 0.95933 | 0.23198 | 0.47523 | 1 | 9615 | 4 | 0.95933 |
| CPB2 | 6 | 0.4132 | 0.61093 | 0.999994 | 12156 | 2 | 0.2677 | 0.23198 | 0.47523 | 1 | 9616 | 3 | 0.2677 |
| MICAL1 | 6 | 0.037535 | 0.11635 | 0.951052 | 2405 | 3 | -0.80704 | 0.23198 | 0.47523 | 1 | 9617 | 2 | -0.80704 |
| PNLIPRP1 | 6 | 0.5859 | 0.72955 | 0.999994 | 14758 | 2 | 0.45061 | 0.23198 | 0.47523 | 1 | 9618 | 3 | 0.45061 |
| NonTargetingControlG | 1 | 0.76802 | 0.76841 | 0.999994 | 17060 | 0 | 3.83 | 0.23198 | 0.23169 | 0.97466 | 9619 | 1 | 3.83 |

| | | | | | | | | | | | | | |
|---|---|---|---|---|---|---|---|---|---|---|---|---|---|
| MAS1L | 5 | 0.52034 | 0.66559 | 0.999994 | 13751 | 2 | 0.66504 | 0.23207 | 0.45242 | 1 | 9620 | 3 | 0.66504 |
| IVL | 6 | 0.4192 | 0.61485 | 0.999994 | 12237 | 2 | 1.1806 | 0.23213 | 0.47543 | 1 | 9621 | 4 | 1.1806 |
| ZFP64 | 5 | 0.32903 | 0.50475 | 0.999994 | 10703 | 1 | 1.8105 | 0.23215 | 0.4525 | 1 | 9622 | 3 | 1.8105 |
| NonTargetingControlG | 2 | 0.72094 | 0.72068 | 0.999994 | 16507 | 0 | 3.5241 | 0.23221 | 0.36026 | 0.994562 | 9623 | 1 | 3.5241 |
| FGD3 | 3 | 0.22304 | 0.34142 | 0.999994 | 8357 | 1 | 1.466 | 0.23225 | 0.37752 | 1 | 9624 | 2 | 1.466 |
| PPM1D | 5 | 0.92535 | 0.92558 | 0.999994 | 19754 | 0 | 1.8289 | 0.23235 | 0.45273 | 1 | 9625 | 3 | 1.8289 |
| NMI | 6 | 0.28691 | 0.49149 | 0.999994 | 9850 | 1 | 1.1438 | 0.23238 | 0.47577 | 1 | 9626 | 4 | 1.1438 |
| TMPRSS2 | 6 | 0.080331 | 0.20797 | 0.979108 | 4242 | 3 | -0.7179 | 0.23239 | 0.47579 | 1 | 9627 | 2 | -0.7179 |
| OR11H12 | 4 | 0.063092 | 0.14668 | 0.964852 | 3559 | 2 | 0.83288 | 0.23239 | 0.42848 | 1 | 9628 | 2 | 0.83288 |
| hsa-mir-4310 | 4 | 0.88583 | 0.8855 | 0.999994 | 18944 | 0 | 1.0097 | 0.2325 | 0.42863 | 1 | 9629 | 3 | 1.0097 |
| HOXA9 | 6 | 0.0064534 | 0.026479 | 0.848412 | 622 | 5 | -2.3789 | 0.23253 | 0.47598 | 1 | 9630 | 1 | -2.3789 |
| CD8A | 6 | 0.030291 | 0.099463 | 0.945512 | 2024 | 4 | -1.7343 | 0.23253 | 0.47598 | 1 | 9631 | 2 | -1.7343 |
| CSE1L | 5 | 0.68209 | 0.75848 | 0.999994 | 16116 | 1 | 1.7135 | 0.23253 | 0.45293 | 1 | 9632 | 3 | 1.7135 |
| PATE1 | 5 | 0.089375 | 0.20958 | 0.979587 | 4570 | 1 | 0.25425 | 0.23253 | 0.45294 | 1 | 9633 | 2 | 0.25425 |
| OR52L1 | 6 | 0.49848 | 0.66765 | 0.999994 | 13402 | 1 | 1.2513 | 0.23258 | 0.47604 | 1 | 9634 | 4 | 1.2513 |
| CNTNAP2 | 5 | 0.20263 | 0.37896 | 0.999994 | 7845 | 2 | 1.9544 | 0.23259 | 0.45299 | 1 | 9635 | 3 | 1.9544 |
| POLG2 | 5 | 0.091397 | 0.21282 | 0.979587 | 4652 | 3 | -1.676 | 0.23261 | 0.45301 | 1 | 9636 | 2 | -1.676 |
| TTC12 | 5 | 0.014184 | 0.047901 | 0.900643 | 1142 | 4 | -2.8414 | 0.23261 | 0.45301 | 1 | 9637 | 1 | -2.8414 |
| GJC2 | 5 | 0.35284 | 0.52424 | 0.999994 | 11213 | 2 | 0.93854 | 0.23261 | 0.45301 | 1 | 9638 | 3 | 0.93854 |
| ZNF135 | 6 | 0.65992 | 0.78527 | 0.999994 | 15777 | 1 | 1.6696 | 0.23273 | 0.47625 | 1 | 9639 | 4 | 1.6696 |
| EFCAB14 | 5 | 0.39849 | 0.5623 | 0.999994 | 11922 | 2 | 1.7042 | 0.23275 | 0.45317 | 1 | 9640 | 3 | 1.7042 |
| ANO9 | 6 | 0.80422 | 0.8497 | 0.999994 | 17544 | 1 | 1.206 | 0.23283 | 0.47639 | 1 | 9641 | 4 | 1.206 |
| MZT2B | 2 | 0.49143 | 0.4961 | 0.999994 | 13296 | 1 | 2.8215 | 0.23285 | 0.36113 | 0.994562 | 9642 | 1 | 2.8215 |
| SLC18A3 | 6 | 0.024435 | 0.085668 | 0.94395 | 1728 | 4 | -2.9831 | 0.23287 | 0.47643 | 1 | 9643 | 1 | -2.9831 |
| PNPLA7 | 6 | 0.20676 | 0.39613 | 0.999994 | 7946 | 3 | -0.67647 | 0.23287 | 0.47643 | 1 | 9644 | 2 | -0.67647 |
| hsa-mir-1321 | 3 | 0.87812 | 0.87822 | 0.999994 | 18790 | 0 | 2.4487 | 0.23292 | 0.37825 | 1 | 9645 | 2 | 2.4487 |
| SEPT10 | 6 | 0.89095 | 0.89698 | 0.999994 | 19054 | 1 | 1.4713 | 0.23293 | 0.47653 | 1 | 9646 | 3 | 1.4713 |
| ELAVL2 | 6 | 0.69312 | 0.80994 | 0.999994 | 16236 | 1 | 1.1979 | 0.23296 | 0.47657 | 1 | 9647 | 3 | 1.1979 |
| RARS | 6 | 0.3311 | 0.5425 | 0.999994 | 10745 | 3 | -0.10308 | 0.23299 | 0.47662 | 1 | 9648 | 2 | -0.10308 |
| IFNA7 | 5 | 0.046044 | 0.13089 | 0.95318 | 2811 | 3 | -2.6477 | 0.23305 | 0.45349 | 1 | 9649 | 2 | -2.6477 |
| CTTN | 4 | 0.63964 | 0.68731 | 0.999994 | 15511 | 1 | 1.2811 | 0.23305 | 0.42937 | 1 | 9650 | 3 | 1.2811 |
| S100A6 | 6 | 0.21404 | 0.40501 | 0.999994 | 8116 | 2 | 0.45012 | 0.23309 | 0.47674 | 1 | 9651 | 3 | 0.45012 |
| PHTF2 | 6 | 0.56498 | 0.71436 | 0.999994 | 14469 | 1 | 1.0197 | 0.23322 | 0.47691 | 1 | 9652 | 4 | 1.0197 |
| CCDC92 | 4 | 0.036582 | 0.094936 | 0.945031 | 2349 | 2 | -0.53693 | 0.23332 | 0.42974 | 1 | 9653 | 2 | -0.53693 |
| MTRNR2L7 | 4 | 0.30547 | 0.44856 | 0.999994 | 10243 | 1 | 1.7936 | 0.23332 | 0.42974 | 1 | 9654 | 2 | 1.7936 |
| CRYM | 4 | 0.069557 | 0.15906 | 0.967126 | 3817 | 2 | 0.094884 | 0.23332 | 0.42974 | 1 | 9655 | 2 | 0.094884 |
| UQCRQ | 4 | 0.22353 | 0.36486 | 0.999994 | 8366 | 2 | 1.1151 | 0.23332 | 0.42974 | 1 | 9656 | 2 | 1.1151 |
| THAP6 | 6 | 0.39227 | 0.5974 | 0.999994 | 11835 | 2 | 1.4303 | 0.23335 | 0.47707 | 1 | 9657 | 3 | 1.4303 |
| UCP1 | 6 | 0.16609 | 0.34582 | 0.999994 | 6887 | 3 | -0.10707 | 0.23336 | 0.47709 | 1 | 9658 | 3 | -0.10707 |
| C19orf59 | 6 | 0.15078 | 0.32665 | 0.999994 | 6478 | 2 | 0.57374 | 0.23336 | 0.47709 | 1 | 9659 | 3 | 0.57374 |
| ADAMTS10 | 6 | 0.11725 | 0.28056 | 0.999632 | 5547 | 2 | 1.1327 | 0.23338 | 0.47712 | 1 | 9660 | 4 | 1.1327 |
| NUP155 | 3 | 0.19202 | 0.30265 | 0.999994 | 7548 | 2 | -1.1701 | 0.23339 | 0.37876 | 1 | 9661 | 1 | -1.1701 |
| hsa-mir-888 | 3 | 0.8397 | 0.83963 | 0.999994 | 18083 | 0 | 2.4229 | 0.23346 | 0.37883 | 1 | 9662 | 2 | 2.4229 |
| FGD4 | 6 | 0.36964 | 0.583 | 0.999994 | 11500 | 2 | 1.4041 | 0.23347 | 0.47723 | 1 | 9663 | 4 | 1.4041 |
| DMXL2 | 6 | 0.021055 | 0.075382 | 0.93914 | 1516 | 2 | 1.2048 | 0.23353 | 0.47731 | 1 | 9664 | 4 | 1.2048 |
| RNF19A | 4 | 0.28486 | 0.42751 | 0.999994 | 9806 | 1 | 1.1863 | 0.23364 | 0.43021 | 1 | 9665 | 3 | 1.1863 |
| ALB | 5 | 0.59777 | 0.72535 | 0.999994 | 14924 | 1 | 1.712 | 0.23367 | 0.45419 | 1 | 9666 | 3 | 1.712 |
| ANGPTL6 | 5 | 0.88534 | 0.88545 | 0.999994 | 18938 | 1 | 2.025 | 0.23368 | 0.4542 | 1 | 9667 | 3 | 2.025 |
| ZBTB33 | 6 | 0.91883 | 0.91932 | 0.999994 | 19604 | 1 | 1.1824 | 0.23369 | 0.47751 | 1 | 9668 | 3 | 1.1824 |
| PLA2G15 | 6 | 0.29678 | 0.50295 | 0.999994 | 10061 | 3 | 0.31223 | 0.2337 | 0.47753 | 1 | 9669 | 3 | 0.31223 |
| C5orf34 | 4 | 0.26458 | 0.40677 | 0.999994 | 9346 | 1 | 0.65411 | 0.23379 | 0.43043 | 1 | 9670 | 2 | 0.65411 |
| STAT1 | 4 | 0.259 | 0.40112 | 0.999994 | 9215 | 2 | -0.66821 | 0.23379 | 0.43043 | 1 | 9671 | 1 | -0.66821 |

| | | | | | | | | | | | | | |
|---|---|---|---|---|---|---|---|---|---|---|---|---|---|
| SCEL | 5 | 0.40868 | 0.57077 | 0.999994 | 12088 | 2 | -0.24184 | 0.23379 | 0.45432 | 1 | 9672 | 2 | -0.24184 |
| DEFB110 | 5 | 0.09233 | 0.21432 | 0.979587 | 4691 | 3 | -1.6401 | 0.23379 | 0.45432 | 1 | 9673 | 2 | -1.6401 |
| CLDN11 | 5 | 0.4165 | 0.57722 | 0.999994 | 12199 | 2 | 0.32291 | 0.23379 | 0.45432 | 1 | 9674 | 1 | 0.32291 |
| CCDC152 | 5 | 0.5048 | 0.65212 | 0.999994 | 13511 | 1 | 1.6517 | 0.23379 | 0.45432 | 1 | 9675 | 3 | 1.6517 |
| TRPC6 | 6 | 0.41851 | 0.61439 | 0.999994 | 12227 | 2 | 1.4872 | 0.23379 | 0.47764 | 1 | 9676 | 3 | 1.4872 |
| ATPAF2 | 6 | 0.09388 | 0.2353 | 0.985362 | 4746 | 4 | -0.8622 | 0.2338 | 0.47765 | 1 | 9677 | 2 | -0.8622 |
| AFAP1L1 | 6 | 0.26631 | 0.46733 | 0.999994 | 9372 | 3 | 0.27336 | 0.2338 | 0.47765 | 1 | 9678 | 3 | 0.27336 |
| APBA1 | 6 | 0.47514 | 0.6519 | 0.999994 | 13066 | 2 | 0.56031 | 0.2338 | 0.47765 | 1 | 9679 | 3 | 0.56031 |
| GRM5 | 5 | 0.066536 | 0.17272 | 0.96793 | 3706 | 2 | 1.8208 | 0.23387 | 0.4544 | 1 | 9680 | 3 | 1.8208 |
| IGF1R | 6 | 0.16369 | 0.34289 | 0.999994 | 6832 | 2 | 0.9639 | 0.23388 | 0.47775 | 1 | 9681 | 4 | 0.9639 |
| SEC61B | 6 | 0.27064 | 0.47248 | 0.999994 | 9489 | 3 | 1.0365 | 0.23391 | 0.47781 | 1 | 9682 | 3 | 1.0365 |
| NonTargetingControlG | 1 | 0.76608 | 0.76645 | 0.999994 | 17040 | 0 | 3.9687 | 0.23392 | 0.23364 | 0.97466 | 9683 | 1 | 3.9687 |
| hsa-mir-5192 | 3 | 0.47184 | 0.52793 | 0.999994 | 13031 | 1 | -0.1812 | 0.23393 | 0.37936 | 1 | 9684 | 1 | -0.1812 |
| KAL1 | 6 | 0.21995 | 0.41205 | 0.999994 | 8289 | 3 | 0.65343 | 0.23394 | 0.47785 | 1 | 9685 | 3 | 0.65343 |
| hsa-mir-6887 | 3 | 0.87895 | 0.87902 | 0.999994 | 18811 | 0 | 2.3645 | 0.23395 | 0.37937 | 1 | 9686 | 2 | 2.3645 |
| IER3IP1 | 6 | 0.14018 | 0.31326 | 0.999994 | 6190 | 1 | 1.6653 | 0.23397 | 0.47787 | 1 | 9687 | 4 | 1.6653 |
| POTEF | 2 | 0.76509 | 0.76456 | 0.999994 | 17029 | 0 | 3.6416 | 0.23398 | 0.36265 | 0.995401 | 9688 | 2 | 3.6416 |
| DNAL4 | 5 | 0.2765 | 0.46203 | 0.999994 | 9630 | 2 | 1.7957 | 0.23399 | 0.45453 | 1 | 9689 | 3 | 1.7957 |
| RABEP1 | 6 | 0.519 | 0.68183 | 0.999994 | 13729 | 2 | 0.56557 | 0.234 | 0.47791 | 1 | 9690 | 3 | 0.56557 |
| PSME1 | 6 | 0.91709 | 0.91773 | 0.999994 | 19563 | 1 | 0.97152 | 0.234 | 0.47791 | 1 | 9691 | 4 | 0.97152 |
| PCMTD2 | 5 | 0.79874 | 0.81874 | 0.999994 | 17457 | 1 | 1.9196 | 0.23407 | 0.45463 | 1 | 9692 | 3 | 1.9196 |
| MRPS14 | 5 | 0.049144 | 0.13828 | 0.957317 | 2937 | 3 | -2.4357 | 0.23415 | 0.45472 | 1 | 9693 | 1 | -2.4357 |
| VCX | 6 | 0.28607 | 0.49053 | 0.999994 | 9833 | 2 | 1.5637 | 0.23418 | 0.47815 | 1 | 9694 | 3 | 1.5637 |
| KRT78 | 6 | 0.30443 | 0.51189 | 0.999994 | 10216 | 3 | 1.0228 | 0.23437 | 0.4784 | 1 | 9695 | 3 | 1.0228 |
| C5orf54 | 6 | 0.081307 | 0.20996 | 0.979587 | 4278 | 1 | 0.11996 | 0.23443 | 0.47849 | 1 | 9696 | 2 | 0.11996 |
| FMNL1 | 5 | 0.078807 | 0.19264 | 0.974607 | 4181 | 2 | 1.7037 | 0.23448 | 0.45508 | 1 | 9697 | 3 | 1.7037 |
| LMX1B | 5 | 0.14004 | 0.28791 | 0.999632 | 6187 | 3 | -1.2376 | 0.23455 | 0.45515 | 1 | 9698 | 2 | -1.2376 |
| NonTargetingControlG | 2 | 0.51305 | 0.51593 | 0.999994 | 13621 | 1 | 2.7693 | 0.23457 | 0.36347 | 0.995401 | 9699 | 1 | 2.7693 |
| ZNF789 | 6 | 0.31834 | 0.528 | 0.999994 | 10481 | 2 | 1.2745 | 0.23464 | 0.47877 | 1 | 9700 | 4 | 1.2745 |
| GPRIN3 | 5 | 0.21862 | 0.40135 | 0.999994 | 8243 | 3 | -0.75131 | 0.23469 | 0.4553 | 1 | 9701 | 2 | -0.75131 |
| LPIN2 | 6 | 0.12115 | 0.2879 | 0.999632 | 5669 | 1 | 1.1684 | 0.2347 | 0.47885 | 1 | 9702 | 4 | 1.1684 |
| FPR3 | 5 | 0.85162 | 0.85557 | 0.999994 | 18288 | 1 | 1.7837 | 0.2347 | 0.45532 | 1 | 9703 | 3 | 1.7837 |
| WDFY3 | 6 | 0.060771 | 0.16741 | 0.967126 | 3457 | 3 | 0.83929 | 0.23474 | 0.47891 | 1 | 9704 | 3 | 0.83929 |
| ZNF69 | 5 | 0.89832 | 0.89843 | 0.999994 | 19190 | 0 | 1.6414 | 0.23478 | 0.45539 | 1 | 9705 | 3 | 1.6414 |
| WDR6 | 4 | 0.24829 | 0.39017 | 0.999994 | 8976 | 2 | 0.045368 | 0.23482 | 0.43182 | 1 | 9706 | 2 | 0.045368 |
| COL4A6 | 5 | 0.79142 | 0.81418 | 0.999994 | 17369 | 1 | 1.9226 | 0.23485 | 0.45547 | 1 | 9707 | 3 | 1.9226 |
| ZNF423 | 6 | 0.58148 | 0.72629 | 0.999994 | 14700 | 2 | 0.98632 | 0.23487 | 0.47907 | 1 | 9708 | 4 | 0.98632 |
| MPG | 3 | 0.74912 | 0.7491 | 0.999994 | 16850 | 0 | 2.2415 | 0.2349 | 0.38039 | 1 | 9709 | 2 | 2.2415 |
| SNX9 | 6 | 0.50207 | 0.67014 | 0.999994 | 13463 | 2 | 0.65613 | 0.23493 | 0.47915 | 1 | 9710 | 3 | 0.65613 |
| ZNF519 | 6 | 0.32899 | 0.54007 | 0.999994 | 10702 | 2 | 0.85009 | 0.23499 | 0.47925 | 1 | 9711 | 3 | 0.85009 |
| hsa-mir-569 | 4 | 0.79073 | 0.79299 | 0.999994 | 17361 | 1 | 1.214 | 0.235 | 0.43206 | 1 | 9712 | 3 | 1.214 |
| hsa-mir-4701 | 4 | 0.83561 | 0.83513 | 0.999994 | 18018 | 0 | 1.8374 | 0.23502 | 0.43209 | 1 | 9713 | 2 | 1.8374 |
| ZNF726 | 4 | 0.73137 | 0.74618 | 0.999994 | 16619 | 1 | 1.7492 | 0.23506 | 0.43215 | 1 | 9714 | 2 | 1.7492 |
| AMPD3 | 6 | 0.022377 | 0.079594 | 0.93914 | 1607 | 4 | -1.9746 | 0.23529 | 0.47962 | 1 | 9715 | 2 | -1.9746 |
| LGALS13 | 6 | 0.63489 | 0.76604 | 0.999994 | 15443 | 2 | 1.0039 | 0.23534 | 0.4797 | 1 | 9716 | 4 | 1.0039 |
| NonTargetingControlG | 2 | 0.76459 | 0.7641 | 0.999994 | 17024 | 0 | 3.3069 | 0.23541 | 0.3646 | 0.995622 | 9717 | 2 | 3.3069 |
| TMEM143 | 4 | 0.57836 | 0.65391 | 0.999994 | 14663 | 1 | 1.9345 | 0.2355 | 0.43276 | 1 | 9718 | 2 | 1.9345 |
| BPIFB6 | 5 | 0.90018 | 0.90029 | 0.999994 | 19223 | 0 | 1.8441 | 0.23552 | 0.45621 | 1 | 9719 | 3 | 1.8441 |
| ZNF284 | 6 | 0.72568 | 0.82019 | 0.999994 | 16556 | 1 | 1.6042 | 0.23565 | 0.48011 | 1 | 9720 | 4 | 1.6042 |
| DNAJC25 | 6 | 0.019782 | 0.07129 | 0.93914 | 1445 | 4 | -2.4677 | 0.2357 | 0.48018 | 1 | 9721 | 1 | -2.4677 |
| SMDT1 | 4 | 0.17414 | 0.31408 | 0.999994 | 7088 | 2 | -1.2201 | 0.2357 | 0.43304 | 1 | 9722 | 1 | -1.2201 |
| FOXD4L1 | 3 | 0.53645 | 0.57084 | 0.999994 | 14008 | 1 | 2.4803 | 0.23574 | 0.38131 | 1 | 9723 | 2 | 2.4803 |

| | | | | | | | | | | | | |
|---|---|---|---|---|---|---|---|---|---|---|---|---|
| PPIL1 | 6 | 0.74241 | 0.8258 | 0.999994 | 16770 | 1 | 1.1789 | 0.23583 | 0.48035 | 1 | 9724 | 4 | 1.1789 |
| hsa-mir-4685 | 2 | 0.76409 | 0.76361 | 0.999994 | 17017 | 0 | 1.3555 | 0.23591 | 0.36528 | 0.996287 | 9725 | 2 | 1.3555 |
| EAF1 | 6 | 0.04716 | 0.13786 | 0.957317 | 2859 | 2 | 1.3386 | 0.23596 | 0.48053 | 1 | 9726 | 4 | 1.3386 |
| OGFOD2 | 6 | 0.080676 | 0.20868 | 0.979587 | 4258 | 3 | 0.37247 | 0.23597 | 0.48055 | 1 | 9727 | 3 | 0.37247 |
| HIST1H4L | 6 | 0.23592 | 0.43115 | 0.999994 | 8668 | 2 | 1.5905 | 0.23597 | 0.48055 | 1 | 9728 | 3 | 1.5905 |
| C17orf53 | 6 | 0.36244 | 0.57805 | 0.999994 | 11377 | 2 | 1.743 | 0.23597 | 0.48055 | 1 | 9729 | 3 | 1.743 |
| SMPD3 | 6 | 0.71979 | 0.81826 | 0.999994 | 16496 | 1 | 1.7364 | 0.23597 | 0.48055 | 1 | 9730 | 3 | 1.7364 |
| SUPV3L1 | 4 | 0.14914 | 0.28827 | 0.999632 | 6433 | 2 | 0.97614 | 0.23603 | 0.43345 | 1 | 9731 | 2 | 0.97614 |
| CTAGE6 | 4 | 0.75591 | 0.76461 | 0.999994 | 16929 | 1 | 1.8922 | 0.23603 | 0.43345 | 1 | 9732 | 2 | 1.8922 |
| MUC5B | 6 | 0.51989 | 0.68244 | 0.999994 | 13746 | 1 | 0.93754 | 0.23607 | 0.48067 | 1 | 9733 | 4 | 0.93754 |
| ABCA4 | 6 | 0.098611 | 0.24463 | 0.988995 | 4916 | 3 | -1.4052 | 0.23607 | 0.48067 | 1 | 9734 | 2 | -1.4052 |
| GRB2 | 6 | 0.26914 | 0.47073 | 0.999994 | 9448 | 2 | 0.67577 | 0.23607 | 0.48067 | 1 | 9735 | 3 | 0.67577 |
| LAMA4 | 6 | 0.57109 | 0.71873 | 0.999994 | 14559 | 2 | 1.03 | 0.23607 | 0.48067 | 1 | 9736 | 3 | 1.03 |
| hsa-mir-4721 | 3 | 0.78049 | 0.78039 | 0.999994 | 17219 | 0 | 2.3117 | 0.23608 | 0.38167 | 1 | 9737 | 2 | 2.3117 |
| RAB25 | 5 | 0.11461 | 0.24915 | 0.991441 | 5466 | 3 | -1.0054 | 0.23609 | 0.45685 | 1 | 9738 | 2 | -1.0054 |
| AJUBA | 4 | 0.88141 | 0.88103 | 0.999994 | 18860 | 0 | 2.1107 | 0.23612 | 0.43357 | 1 | 9739 | 3 | 2.1107 |
| hsa-mir-8071-1 | 4 | 0.7248 | 0.74143 | 0.999994 | 16546 | 1 | 1.4463 | 0.23616 | 0.43363 | 1 | 9740 | 3 | 1.4463 |
| SNF8 | 5 | 0.64193 | 0.74194 | 0.999994 | 15550 | 1 | 1.9348 | 0.23631 | 0.45709 | 1 | 9741 | 3 | 1.9348 |
| APCDD1L | 6 | 0.021252 | 0.076002 | 0.93914 | 1531 | 3 | -1.245 | 0.23646 | 0.48119 | 1 | 9742 | 2 | -1.245 |
| NFATC4 | 6 | 0.029373 | 0.097313 | 0.945253 | 1977 | 3 | -0.9286 | 0.23646 | 0.48119 | 1 | 9743 | 3 | -0.9286 |
| ST14 | 6 | 0.2952 | 0.50111 | 0.999994 | 10020 | 3 | -0.44518 | 0.23646 | 0.48119 | 1 | 9744 | 1 | -0.44518 |
| ATG4B | 6 | 0.017396 | 0.063532 | 0.937389 | 1332 | 3 | -0.68273 | 0.23646 | 0.48119 | 1 | 9745 | 3 | -0.68273 |
| CMC4 | 5 | 0.52963 | 0.67374 | 0.999994 | 13896 | 2 | 2.0118 | 0.23667 | 0.4575 | 1 | 9746 | 3 | 2.0118 |
| TCEB3CL2 | 1 | 0.7633 | 0.76364 | 0.999994 | 17007 | 0 | 3.9895 | 0.2367 | 0.23644 | 0.97466 | 9747 | 1 | 3.9895 |
| HIST2H2AA4 | 1 | 0.7633 | 0.76364 | 0.999994 | 17006 | 0 | 3.9895 | 0.2367 | 0.23644 | 0.97466 | 9748 | 1 | 3.9895 |
| ITGB1BP1 | 4 | 0.061496 | 0.14363 | 0.961453 | 3493 | 3 | -1.3391 | 0.23672 | 0.43436 | 1 | 9749 | 1 | -1.3391 |
| TMCO5A | 4 | 0.85623 | 0.85582 | 0.999994 | 18377 | 0 | 0.53913 | 0.23672 | 0.43436 | 1 | 9750 | 2 | 0.53913 |
| KRR1 | 4 | 0.11498 | 0.24154 | 0.987253 | 5477 | 3 | -0.86731 | 0.23672 | 0.43436 | 1 | 9751 | 1 | -0.86731 |
| C16orf13 | 6 | 0.40209 | 0.60372 | 0.999994 | 11979 | 2 | 1.1122 | 0.2368 | 0.48162 | 1 | 9752 | 3 | 1.1122 |
| OR5AN1 | 4 | 0.80974 | 0.81006 | 0.999994 | 17631 | 1 | 2.0469 | 0.2369 | 0.4346 | 1 | 9753 | 2 | 2.0469 |
| KCTD11 | 5 | 0.079326 | 0.19347 | 0.975797 | 4202 | 3 | -1.7912 | 0.23693 | 0.45778 | 1 | 9754 | 2 | -1.7912 |
| TNFSF8 | 5 | 0.53987 | 0.68276 | 0.999994 | 14050 | 2 | 1.0023 | 0.23693 | 0.45778 | 1 | 9755 | 3 | 1.0023 |
| ARF5 | 5 | 0.091248 | 0.2126 | 0.979587 | 4646 | 3 | -1.6447 | 0.23693 | 0.45778 | 1 | 9756 | 2 | -1.6447 |
| STXBP1 | 5 | 0.39649 | 0.56063 | 0.999994 | 11897 | 2 | 1.7047 | 0.23699 | 0.45785 | 1 | 9757 | 3 | 1.7047 |
| RNF43 | 6 | 0.23738 | 0.43287 | 0.999994 | 8710 | 2 | 0.84673 | 0.23703 | 0.48193 | 1 | 9758 | 3 | 0.84673 |
| FAM200B | 6 | 0.96273 | 0.96278 | 0.999994 | 20652 | 0 | 1.5024 | 0.23708 | 0.482 | 1 | 9759 | 4 | 1.5024 |
| MMP14 | 6 | 0.42461 | 0.61842 | 0.999994 | 12330 | 1 | 0.46133 | 0.23708 | 0.482 | 1 | 9760 | 2 | 0.46133 |
| DCTN1 | 6 | 0.021139 | 0.075641 | 0.93914 | 1520 | 3 | 0.27149 | 0.23708 | 0.482 | 1 | 9761 | 3 | 0.27149 |
| hsa-mir-936 | 3 | 0.41421 | 0.49266 | 0.999994 | 12167 | 1 | 2.2793 | 0.23709 | 0.38276 | 1 | 9762 | 2 | 2.2793 |
| SLC25A45 | 5 | 0.054804 | 0.152 | 0.967126 | 3188 | 2 | 1.7741 | 0.23714 | 0.458 | 1 | 9763 | 3 | 1.7741 |
| ANXA13 | 5 | 0.36055 | 0.53054 | 0.999994 | 11343 | 2 | 1.7497 | 0.23715 | 0.45803 | 1 | 9764 | 3 | 1.7497 |
| hsa-mir-4731 | 4 | 0.16746 | 0.30714 | 0.999994 | 6922 | 2 | 0.35437 | 0.23716 | 0.43498 | 1 | 9765 | 2 | 0.35437 |
| hsa-mir-802 | 4 | 0.67174 | 0.70644 | 0.999994 | 15951 | 1 | -0.26167 | 0.23716 | 0.43498 | 1 | 9766 | 1 | -0.26167 |
| hsa-mir-217 | 4 | 0.030213 | 0.081921 | 0.93914 | 2020 | 3 | -2.5649 | 0.23716 | 0.43498 | 1 | 9767 | 1 | -2.5649 |
| CASP7 | 4 | 0.23598 | 0.37762 | 0.999994 | 8669 | 2 | 0.73048 | 0.23716 | 0.43498 | 1 | 9768 | 2 | 0.73048 |
| hsa-mir-105-1 | 3 | 0.43773 | 0.50671 | 0.999994 | 12537 | 1 | 1.4263 | 0.23733 | 0.38301 | 1 | 9769 | 2 | 1.4263 |
| HLTF | 5 | 0.44442 | 0.60052 | 0.999994 | 12639 | 2 | 1.7121 | 0.23746 | 0.45834 | 1 | 9770 | 3 | 1.7121 |
| MYH7 | 6 | 0.42206 | 0.61677 | 0.999994 | 12280 | 2 | 1.3385 | 0.23748 | 0.48253 | 1 | 9771 | 4 | 1.3385 |
| hsa-mir-6856 | 3 | 0.6986 | 0.69973 | 0.999994 | 16289 | 1 | 2.3119 | 0.23751 | 0.38319 | 1 | 9772 | 2 | 2.3119 |
| GCA | 6 | 0.23404 | 0.42891 | 0.999994 | 8620 | 3 | -0.52712 | 0.23753 | 0.48259 | 1 | 9773 | 1 | -0.52712 |
| CSF1R | 6 | 0.20148 | 0.38967 | 0.999994 | 7807 | 3 | -0.48586 | 0.23753 | 0.48259 | 1 | 9774 | 1 | -0.48586 |
| GDA | 6 | 0.16291 | 0.34192 | 0.999994 | 6814 | 3 | -0.66149 | 0.23753 | 0.48259 | 1 | 9775 | 2 | -0.66149 |

| | | | | | | | | | | | | | |
|---|---|---|---|---|---|---|---|---|---|---|---|---|---|
| RPL23 | 2 | 0.045141 | 0.081032 | 0.93914 | 2766 | 1 | -1.3946 | 0.23772 | 0.36729 | 0.997075 | 9776 | 1 | -1.3946 |
| DFFA | 6 | 0.63735 | 0.7679 | 0.999994 | 15477 | 2 | 1.3706 | 0.23775 | 0.4829 | 1 | 9777 | 4 | 1.3706 |
| T | 4 | 0.047207 | 0.11598 | 0.951052 | 2865 | 2 | 0.60253 | 0.23778 | 0.43582 | 1 | 9778 | 2 | 0.60253 |
| hsa-mir-3671 | 3 | 0.86513 | 0.86511 | 0.999994 | 18541 | 0 | 2.414 | 0.23783 | 0.38354 | 1 | 9779 | 2 | 2.414 |
| hsa-mir-384 | 4 | 0.8893 | 0.889 | 0.999994 | 19026 | 0 | 1.063 | 0.23792 | 0.436 | 1 | 9780 | 3 | 1.063 |
| ASF1B | 3 | 0.13698 | 0.23167 | 0.984454 | 6108 | 2 | -1.7727 | 0.23795 | 0.38367 | 1 | 9781 | 1 | -1.7727 |
| ZSCAN5B | 5 | 0.23899 | 0.42956 | 0.999994 | 8748 | 3 | -0.59347 | 0.23805 | 0.45895 | 1 | 9782 | 1 | -0.59347 |
| SOX14 | 5 | 0.33512 | 0.50972 | 0.999994 | 10836 | 2 | 1.6743 | 0.23805 | 0.45895 | 1 | 9783 | 3 | 1.6743 |
| CASP2 | 6 | 0.28091 | 0.48445 | 0.999994 | 9720 | 1 | 1.3312 | 0.23808 | 0.48334 | 1 | 9784 | 4 | 1.3312 |
| PCM1 | 5 | 0.033691 | 0.099961 | 0.946394 | 2200 | 1 | 1.8507 | 0.23812 | 0.45904 | 1 | 9785 | 3 | 1.8507 |
| PPL | 6 | 0.95282 | 0.95282 | 0.999994 | 20402 | 0 | 0.91619 | 0.23821 | 0.48352 | 1 | 9786 | 3 | 0.91619 |
| SRD5A3 | 5 | 0.51329 | 0.65952 | 0.999994 | 13623 | 1 | 1.3798 | 0.23822 | 0.45915 | 1 | 9787 | 3 | 1.3798 |
| FAM150B | 5 | 0.35259 | 0.52404 | 0.999994 | 11208 | 1 | 0.44537 | 0.23822 | 0.45915 | 1 | 9788 | 2 | 0.44537 |
| DSN1 | 5 | 0.071722 | 0.18116 | 0.968171 | 3907 | 1 | 0.10105 | 0.23822 | 0.45915 | 1 | 9789 | 2 | 0.10105 |
| GCM2 | 6 | 0.15403 | 0.33073 | 0.999994 | 6570 | 2 | 1.034 | 0.2384 | 0.48378 | 1 | 9790 | 3 | 1.034 |
| hsa-mir-4635 | 4 | 0.04546 | 0.11258 | 0.951052 | 2786 | 2 | 0.39196 | 0.23842 | 0.43664 | 1 | 9791 | 2 | 0.39196 |
| MTL5 | 6 | 0.28172 | 0.48536 | 0.999994 | 9734 | 3 | -0.51348 | 0.23843 | 0.48381 | 1 | 9792 | 1 | -0.51348 |
| CLMN | 5 | 0.18229 | 0.3499 | 0.999994 | 7277 | 2 | 1.1648 | 0.2385 | 0.45945 | 1 | 9793 | 3 | 1.1648 |
| IFIT1B | 5 | 0.085871 | 0.20395 | 0.978186 | 4441 | 1 | 1.7303 | 0.2385 | 0.45945 | 1 | 9794 | 3 | 1.7303 |
| NEK3 | 6 | 0.91681 | 0.91747 | 0.999994 | 19559 | 1 | 1.4488 | 0.23852 | 0.48393 | 1 | 9795 | 4 | 1.4488 |
| CYP2J2 | 6 | 0.018552 | 0.067316 | 0.93914 | 1388 | 2 | 0.98957 | 0.23875 | 0.48424 | 1 | 9796 | 4 | 0.98957 |
| PCDHGA9 | 2 | 0.72943 | 0.72916 | 0.999994 | 16592 | 0 | 3.4678 | 0.23876 | 0.36793 | 0.997833 | 9797 | 1 | 3.4678 |
| PAX9 | 5 | 0.11956 | 0.25668 | 0.994943 | 5616 | 2 | 1.9396 | 0.23881 | 0.45977 | 1 | 9798 | 3 | 1.9396 |
| CLIC2 | 6 | 0.93246 | 0.93256 | 0.999994 | 19913 | 0 | 1.3024 | 0.23882 | 0.48433 | 1 | 9799 | 4 | 1.3024 |
| PLA2G2F | 6 | 0.14373 | 0.31777 | 0.999994 | 6291 | 2 | 1.0337 | 0.23902 | 0.4846 | 1 | 9800 | 4 | 1.0337 |
| ECH1 | 4 | 0.87573 | 0.87539 | 0.999994 | 18745 | 0 | 1.1297 | 0.23903 | 0.43747 | 1 | 9801 | 3 | 1.1297 |
| SENP5 | 6 | 0.61214 | 0.74891 | 0.999994 | 15111 | 2 | 1.3677 | 0.23906 | 0.48468 | 1 | 9802 | 4 | 1.3677 |
| hsa-mir-542 | 4 | 0.7103 | 0.73137 | 0.999994 | 16402 | 1 | 2.0565 | 0.23909 | 0.43755 | 1 | 9803 | 3 | 2.0565 |
| hsa-mir-208a | 1 | 0.76088 | 0.76122 | 0.999994 | 16989 | 0 | 3.8541 | 0.23912 | 0.2389 | 0.97466 | 9804 | 1 | 3.8541 |
| SETDB1 | 5 | 0.000036036 | 0.0001598 | 0.157291 | 23 | 4 | -6.7415 | 0.23915 | 0.46014 | 1 | 9805 | 1 | -6.7415 |
| RHOXF2 | 5 | 0.52341 | 0.66831 | 0.999994 | 13801 | 2 | 1.1401 | 0.23915 | 0.46014 | 1 | 9806 | 3 | 1.1401 |
| ACCSL | 6 | 0.23176 | 0.42621 | 0.999994 | 8560 | 3 | 0.81811 | 0.23915 | 0.48477 | 1 | 9807 | 3 | 0.81811 |
| GTF3C2 | 6 | 0.07724 | 0.20175 | 0.977164 | 4119 | 2 | 1.2707 | 0.23937 | 0.48506 | 1 | 9808 | 3 | 1.2707 |
| TBC1D25 | 6 | 0.36429 | 0.57964 | 0.999994 | 11410 | 2 | 0.88693 | 0.23937 | 0.48506 | 1 | 9809 | 3 | 0.88693 |
| ZNF80 | 6 | 0.027943 | 0.093933 | 0.945031 | 1900 | 3 | -1.765 | 0.2394 | 0.4851 | 1 | 9810 | 2 | -1.765 |
| LELP1 | 4 | 0.69505 | 0.72121 | 0.999994 | 16256 | 1 | 1.9743 | 0.23949 | 0.43807 | 1 | 9811 | 2 | 1.9743 |
| POLR2B | 3 | 0.20951 | 0.32459 | 0.999994 | 8017 | 1 | 2.1786 | 0.23959 | 0.38543 | 1 | 9812 | 2 | 2.1786 |
| MAK | 5 | 0.38981 | 0.55498 | 0.999994 | 11794 | 1 | 0.47469 | 0.23966 | 0.46067 | 1 | 9813 | 3 | 0.47469 |
| WDR35 | 5 | 0.38455 | 0.55057 | 0.999994 | 11707 | 2 | -0.31667 | 0.23966 | 0.46067 | 1 | 9814 | 2 | -0.31667 |
| MMGT1 | 5 | 0.86411 | 0.86589 | 0.999994 | 18524 | 1 | 0.59631 | 0.23966 | 0.46067 | 1 | 9815 | 3 | 0.59631 |
| hsa-mir-526a-2 | 3 | 0.50157 | 0.5472 | 0.999994 | 13451 | 1 | 1.9163 | 0.23968 | 0.38554 | 1 | 9816 | 2 | 1.9163 |
| GFRA4 | 6 | 0.82961 | 0.86135 | 0.999994 | 17923 | 1 | 0.99692 | 0.23996 | 0.48584 | 1 | 9817 | 4 | 0.99692 |
| SEC14L5 | 4 | 0.20208 | 0.34282 | 0.999994 | 7833 | 2 | -0.86726 | 0.23998 | 0.43869 | 1 | 9818 | 1 | -0.86726 |
| ZNF559-ZNF177 | 4 | 0.10901 | 0.2311 | 0.983933 | 5271 | 3 | -1.0139 | 0.23998 | 0.43869 | 1 | 9819 | 1 | -1.0139 |
| RAVER1 | 5 | 0.89234 | 0.89248 | 0.999994 | 19075 | 0 | 1.7724 | 0.23999 | 0.46104 | 1 | 9820 | 3 | 1.7724 |
| ATP1B2 | 6 | 0.76892 | 0.83544 | 0.999994 | 17072 | 1 | 0.98727 | 0.24006 | 0.48599 | 1 | 9821 | 4 | 0.98727 |
| CCDC140 | 6 | 0.53311 | 0.69174 | 0.999994 | 13954 | 2 | 1.1863 | 0.24008 | 0.48602 | 1 | 9822 | 4 | 1.1863 |
| EFEMP1 | 5 | 0.34242 | 0.51571 | 0.999994 | 10994 | 2 | -0.5708 | 0.24009 | 0.46115 | 1 | 9823 | 2 | -0.5708 |
| PFDN4 | 5 | 0.856 | 0.85909 | 0.999994 | 18373 | 1 | 0.62771 | 0.24009 | 0.46115 | 1 | 9824 | 3 | 0.62771 |
| HNF1A | 6 | 0.41465 | 0.61186 | 0.999994 | 12176 | 2 | 1.818 | 0.24024 | 0.48624 | 1 | 9825 | 4 | 1.818 |
| LOC286238 | 6 | 0.10307 | 0.2533 | 0.994501 | 5069 | 2 | 0.97699 | 0.24029 | 0.48631 | 1 | 9826 | 4 | 0.97699 |
| C1orf53 | 4 | 0.17122 | 0.31104 | 0.999994 | 7015 | 1 | 1.5203 | 0.2403 | 0.43914 | 1 | 9827 | 3 | 1.5203 |

| | | | | | | | | | | | | | |
|---|---|---|---|---|---|---|---|---|---|---|---|---|---|
| KIAA0825 | 5 | 0.4567 | 0.61091 | 0.999994 | 12802 | 2 | 1.877 | 0.2403 | 0.46139 | 1 | 9828 | 3 | 1.877 |
| hsa-mir-1283-1 | 4 | 0.69823 | 0.72326 | 0.999994 | 16284 | 1 | 1.3029 | 0.24036 | 0.43923 | 1 | 9829 | 3 | 1.3029 |
| NUCB2 | 5 | 0.024806 | 0.076954 | 0.93914 | 1746 | 3 | -1.342 | 0.24043 | 0.46153 | 1 | 9830 | 2 | -1.342 |
| hsa-mir-3138 | 4 | 0.58905 | 0.65942 | 0.999994 | 14807 | 1 | 2.1475 | 0.24046 | 0.43936 | 1 | 9831 | 3 | 2.1475 |
| TEX26 | 4 | 0.89182 | 0.89155 | 0.999994 | 19070 | 0 | 1.3679 | 0.24054 | 0.43947 | 1 | 9832 | 3 | 1.3679 |
| SPG11 | 6 | 0.68908 | 0.8081 | 0.999994 | 16201 | 2 | 1.4697 | 0.24058 | 0.4867 | 1 | 9833 | 4 | 1.4697 |
| hsa-mir-3159 | 3 | 0.092484 | 0.17307 | 0.96793 | 4704 | 2 | -2.3326 | 0.24063 | 0.38655 | 1 | 9834 | 1 | -2.3326 |
| VCP | 4 | 0.25535 | 0.39736 | 0.999994 | 9134 | 1 | 0.96576 | 0.24066 | 0.43965 | 1 | 9835 | 3 | 0.96576 |
| PCDHGC5 | 1 | 0.75931 | 0.75964 | 0.999994 | 16973 | 0 | 3.8927 | 0.24069 | 0.24048 | 0.97466 | 9836 | 1 | 3.8927 |
| hsa-mir-548i-3 | 1 | 0.75931 | 0.75964 | 0.999994 | 16974 | 0 | 3.8927 | 0.24069 | 0.24048 | 0.97466 | 9837 | 1 | 3.8927 |
| OR1K1 | 6 | 0.42764 | 0.62035 | 0.999994 | 12376 | 2 | 1.0886 | 0.24071 | 0.48689 | 1 | 9838 | 4 | 1.0886 |
| JUN | 5 | 0.022772 | 0.071539 | 0.93914 | 1633 | 2 | 0.77296 | 0.24074 | 0.46186 | 1 | 9839 | 3 | 0.77296 |
| hsa-mir-34a | 4 | 0.36327 | 0.50785 | 0.999994 | 11394 | 2 | -0.50036 | 0.24076 | 0.4398 | 1 | 9840 | 1 | -0.50036 |
| NDNL2 | 6 | 0.000056165 | 0.00030153 | 0.204053 | 29 | 5 | -7.0703 | 0.24079 | 0.487 | 1 | 9841 | 1 | -7.0703 |
| PCDP1 | 6 | 0.45597 | 0.63903 | 0.999994 | 12791 | 2 | 0.53002 | 0.24079 | 0.487 | 1 | 9842 | 3 | 0.53002 |
| SDF2 | 6 | 0.1195 | 0.28479 | 0.999632 | 5614 | 2 | 0.77312 | 0.24082 | 0.48703 | 1 | 9843 | 3 | 0.77312 |
| SCGB1C1 | 4 | 0.80393 | 0.80471 | 0.999994 | 17540 | 1 | 2.2806 | 0.24083 | 0.4399 | 1 | 9844 | 3 | 2.2806 |
| PPP1R15B | 5 | 0.84741 | 0.85227 | 0.999994 | 18211 | 1 | 0.72912 | 0.24088 | 0.46201 | 1 | 9845 | 3 | 0.72912 |
| TMEM141 | 5 | 0.49805 | 0.64619 | 0.999994 | 13397 | 2 | 0.66306 | 0.24088 | 0.46201 | 1 | 9846 | 3 | 0.66306 |
| hsa-mir-935 | 3 | 0.10173 | 0.18535 | 0.968626 | 5029 | 1 | 2.1198 | 0.24093 | 0.38686 | 1 | 9847 | 2 | 2.1198 |
| TRPV5 | 6 | 0.028976 | 0.096393 | 0.945031 | 1953 | 2 | 1.261 | 0.24099 | 0.48727 | 1 | 9848 | 3 | 1.261 |
| HOXA4 | 5 | 0.20587 | 0.38356 | 0.999994 | 7921 | 2 | 0.40777 | 0.24108 | 0.46222 | 1 | 9849 | 2 | 0.40777 |
| ABCB5 | 5 | 0.13717 | 0.28352 | 0.999632 | 6112 | 3 | -1.2503 | 0.24108 | 0.46222 | 1 | 9850 | 2 | -1.2503 |
| STX3 | 4 | 0.45321 | 0.59602 | 0.999994 | 12749 | 1 | 1.4868 | 0.24119 | 0.44039 | 1 | 9851 | 3 | 1.4868 |
| PDGFRB | 5 | 0.46071 | 0.61426 | 0.999994 | 12858 | 1 | 1.899 | 0.24122 | 0.46236 | 1 | 9852 | 3 | 1.899 |
| GML | 6 | 0.91113 | 0.9125 | 0.999994 | 19452 | 1 | 1.2688 | 0.24134 | 0.48772 | 1 | 9853 | 4 | 1.2688 |
| METTL18 | 6 | 0.06297 | 0.17202 | 0.967658 | 3555 | 1 | 0.31046 | 0.24144 | 0.48787 | 1 | 9854 | 3 | 0.31046 |
| DIS3L2 | 6 | 0.51612 | 0.67984 | 0.999994 | 13679 | 2 | 1.5319 | 0.24152 | 0.48797 | 1 | 9855 | 4 | 1.5319 |
| COPA | 4 | 0.86093 | 0.86051 | 0.999994 | 18461 | 0 | 2.3719 | 0.24155 | 0.44086 | 1 | 9856 | 3 | 2.3719 |
| KRTAP12-3 | 5 | 0.036967 | 0.10827 | 0.951052 | 2373 | 2 | 1.6067 | 0.24169 | 0.46288 | 1 | 9857 | 3 | 1.6067 |
| MS4A4A | 6 | 0.091067 | 0.22965 | 0.983354 | 4637 | 1 | 1.1651 | 0.24174 | 0.48828 | 1 | 9858 | 4 | 1.1651 |
| CENPO | 6 | 0.80461 | 0.84986 | 0.999994 | 17547 | 1 | 1.1923 | 0.24182 | 0.48838 | 1 | 9859 | 3 | 1.1923 |
| SELO | 5 | 0.10667 | 0.2368 | 0.985449 | 5193 | 2 | 1.0529 | 0.24182 | 0.46303 | 1 | 9860 | 3 | 1.0529 |
| MAB21L2 | 6 | 0.011085 | 0.042693 | 0.891621 | 932 | 5 | -1.7376 | 0.24187 | 0.48846 | 1 | 9861 | 1 | -1.7376 |
| SYNDIG1L | 5 | 0.4583 | 0.61222 | 0.999994 | 12826 | 2 | 1.1016 | 0.24191 | 0.46313 | 1 | 9862 | 3 | 1.1016 |
| NLRP2 | 6 | 0.45 | 0.63513 | 0.999994 | 12712 | 2 | 1.0536 | 0.24194 | 0.48857 | 1 | 9863 | 4 | 1.0536 |
| ERBB4 | 6 | 0.23897 | 0.43475 | 0.999994 | 8747 | 1 | 1.6218 | 0.24197 | 0.4886 | 1 | 9864 | 4 | 1.6218 |
| OR5H1 | 5 | 0.37771 | 0.54483 | 0.999994 | 11621 | 2 | 1.5929 | 0.24199 | 0.4632 | 1 | 9865 | 3 | 1.5929 |
| hsa-mir-6816 | 3 | 0.86622 | 0.86622 | 0.999994 | 18563 | 0 | 2.4733 | 0.24203 | 0.38804 | 1 | 9866 | 2 | 2.4733 |
| MTO1 | 5 | 0.85253 | 0.8563 | 0.999994 | 18307 | 1 | 1.7596 | 0.24208 | 0.46329 | 1 | 9867 | 3 | 1.7596 |
| REC8 | 4 | 0.48784 | 0.61094 | 0.999994 | 13254 | 1 | 1.5181 | 0.24208 | 0.44158 | 1 | 9868 | 3 | 1.5181 |
| APEH | 6 | 0.95928 | 0.95936 | 0.999994 | 20565 | 0 | 1.3847 | 0.2421 | 0.48876 | 1 | 9869 | 4 | 1.3847 |
| DMTN | 2 | 0.72476 | 0.72445 | 0.999994 | 16545 | 0 | 3.4552 | 0.24211 | 0.36988 | 0.9988 | 9870 | 1 | 3.4552 |
| USP49 | 6 | 0.14845 | 0.32367 | 0.999994 | 6418 | 3 | -0.023461 | 0.24218 | 0.48888 | 1 | 9871 | 3 | -0.023461 |
| PRPF40B | 4 | 0.31402 | 0.45738 | 0.999994 | 10397 | 2 | 1.3381 | 0.24222 | 0.44176 | 1 | 9872 | 2 | 1.3381 |
| POLE3 | 4 | 0.80032 | 0.80151 | 0.999994 | 17483 | 1 | 0.09596 | 0.24223 | 0.44177 | 1 | 9873 | 1 | 0.09596 |
| INPP5D | 6 | 0.78114 | 0.84014 | 0.999994 | 17226 | 1 | 0.16095 | 0.24236 | 0.48914 | 1 | 9874 | 1 | 0.16095 |
| GNS | 6 | 0.18654 | 0.37122 | 0.999994 | 7373 | 2 | 1.168 | 0.2424 | 0.48918 | 1 | 9875 | 4 | 1.168 |
| STX17 | 4 | 0.88728 | 0.88698 | 0.999994 | 18973 | 0 | 1.278 | 0.24247 | 0.44208 | 1 | 9876 | 3 | 1.278 |
| PCDH10 | 6 | 0.22592 | 0.41921 | 0.999994 | 8422 | 1 | 1.8875 | 0.24252 | 0.48934 | 1 | 9877 | 4 | 1.8875 |
| hsa-mir-362 | 4 | 0.81118 | 0.81141 | 0.999994 | 17648 | 1 | 1.7528 | 0.24253 | 0.44217 | 1 | 9878 | 3 | 1.7528 |
| ACOT7 | 6 | 0.20844 | 0.39817 | 0.999994 | 7987 | 2 | 1.4801 | 0.24257 | 0.48939 | 1 | 9879 | 4 | 1.4801 |

| | | | | | | | | | | | | |
|---|---|---|---|---|---|---|---|---|---|---|---|---|
| ROM1 | 6 | 0.124 | 0.29264 | 0.999632 | 5769 | 2 | 0.95207 | 0.24272 | 0.48961 | 1 | 9880 | 4 | 0.95207 |
| HS6ST3 | 5 | 0.84469 | 0.85019 | 0.999994 | 18169 | 1 | 1.4123 | 0.24274 | 0.46402 | 1 | 9881 | 3 | 1.4123 |
| USP9Y | 5 | 0.22658 | 0.41249 | 0.999994 | 8442 | 2 | -0.54614 | 0.24274 | 0.46402 | 1 | 9882 | 2 | -0.54614 |
| C7orf43 | 4 | 0.045278 | 0.11224 | 0.951052 | 2778 | 2 | -0.88968 | 0.24275 | 0.44248 | 1 | 9883 | 2 | -0.88968 |
| PPP1R12B | 5 | 0.69324 | 0.76344 | 0.999994 | 16238 | 1 | 0.31097 | 0.24276 | 0.46403 | 1 | 9884 | 2 | 0.31097 |
| CX3CR1 | 5 | 0.77669 | 0.80553 | 0.999994 | 17161 | 1 | 1.2989 | 0.24276 | 0.46403 | 1 | 9885 | 3 | 1.2989 |
| C17orf72 | 5 | 0.074301 | 0.18536 | 0.968626 | 4004 | 3 | -2.0611 | 0.24276 | 0.46403 | 1 | 9886 | 2 | -2.0611 |
| RMND5B | 6 | 0.23921 | 0.43505 | 0.999994 | 8755 | 2 | 1.2406 | 0.24289 | 0.48983 | 1 | 9887 | 4 | 1.2406 |
| hsa-mir-605 | 3 | 0.73951 | 0.73944 | 0.999994 | 16731 | 0 | 1.0138 | 0.24292 | 0.38901 | 1 | 9888 | 2 | 1.0138 |
| hsa-mir-4791 | 3 | 0.81968 | 0.81954 | 0.999994 | 17775 | 0 | 1.1184 | 0.24292 | 0.38901 | 1 | 9889 | 2 | 1.1184 |
| TIGD7 | 6 | 0.04517 | 0.13345 | 0.95318 | 2770 | 2 | 1.1536 | 0.24297 | 0.48993 | 1 | 9890 | 4 | 1.1536 |
| STX1B | 6 | 0.0019456 | 0.0097037 | 0.680529 | 264 | 4 | -4.8752 | 0.24297 | 0.48993 | 1 | 9891 | 2 | -4.8752 |
| CCDC51 | 6 | 0.081069 | 0.20947 | 0.979587 | 4270 | 2 | 0.29199 | 0.24297 | 0.48993 | 1 | 9892 | 2 | 0.29199 |
| SLC51B | 6 | 0.055426 | 0.15585 | 0.967126 | 3216 | 3 | -1.1816 | 0.24297 | 0.48993 | 1 | 9893 | 2 | -1.1816 |
| SLC5A12 | 6 | 0.51173 | 0.67678 | 0.999994 | 13606 | 2 | 1.3818 | 0.24302 | 0.49 | 1 | 9894 | 4 | 1.3818 |
| ARSJ | 5 | 0.11315 | 0.24685 | 0.98999 | 5412 | 2 | 1.8542 | 0.2431 | 0.4644 | 1 | 9895 | 3 | 1.8542 |
| FAM3A | 6 | 0.055043 | 0.15501 | 0.967126 | 3201 | 4 | -1.7899 | 0.24317 | 0.4902 | 1 | 9896 | 2 | -1.7899 |
| ZNF71 | 5 | 0.83032 | 0.83971 | 0.999994 | 17938 | 1 | 1.1739 | 0.24321 | 0.46453 | 1 | 9897 | 3 | 1.1739 |
| OR6N1 | 5 | 0.81432 | 0.82881 | 0.999994 | 17695 | 1 | 0.78627 | 0.24321 | 0.46453 | 1 | 9898 | 3 | 0.78627 |
| MRPS6 | 4 | 0.77409 | 0.77907 | 0.999994 | 17132 | 1 | 0.96418 | 0.24329 | 0.44319 | 1 | 9899 | 2 | 0.96418 |
| KIAA1614 | 4 | 0.8203 | 0.82006 | 0.999994 | 17781 | 1 | 1.8782 | 0.24331 | 0.44324 | 1 | 9900 | 2 | 1.8782 |
| OR5K2 | 5 | 0.65636 | 0.74766 | 0.999994 | 15732 | 1 | 1.8014 | 0.24338 | 0.46471 | 1 | 9901 | 3 | 1.8014 |
| SGMS2 | 5 | 0.46078 | 0.61432 | 0.999994 | 12859 | 1 | 1.8657 | 0.2434 | 0.46474 | 1 | 9902 | 3 | 1.8657 |
| NLRP8 | 4 | 0.93032 | 0.93009 | 0.999994 | 19862 | 0 | 1.0284 | 0.24343 | 0.44339 | 1 | 9903 | 3 | 1.0284 |
| TEX101 | 6 | 0.098084 | 0.24359 | 0.988088 | 4894 | 2 | 1.3786 | 0.24343 | 0.49057 | 1 | 9904 | 4 | 1.3786 |
| SFTPD | 5 | 0.04869 | 0.13724 | 0.957161 | 2919 | 4 | -1.3888 | 0.24344 | 0.46478 | 1 | 9905 | 1 | -1.3888 |
| hsa-mir-4433 | 4 | 0.83465 | 0.83421 | 0.999994 | 17998 | 0 | 1.8937 | 0.24344 | 0.44341 | 1 | 9906 | 2 | 1.8937 |
| CFHR2 | 6 | 0.38122 | 0.59035 | 0.999994 | 11665 | 2 | 0.97713 | 0.24348 | 0.49062 | 1 | 9907 | 3 | 0.97713 |
| KCNH3 | 4 | 0.70552 | 0.72811 | 0.999994 | 16369 | 1 | 1.3049 | 0.24355 | 0.44357 | 1 | 9908 | 3 | 1.3049 |
| FLJ44635 | 6 | 0.63344 | 0.76494 | 0.999994 | 15420 | 2 | 1.439 | 0.24362 | 0.49081 | 1 | 9909 | 3 | 1.439 |
| EIF3M | 4 | 0.22594 | 0.36731 | 0.999994 | 8423 | 1 | 1.1651 | 0.24365 | 0.44372 | 1 | 9910 | 3 | 1.1651 |
| F13A1 | 6 | 0.51075 | 0.67615 | 0.999994 | 13589 | 2 | 0.99789 | 0.24374 | 0.49096 | 1 | 9911 | 4 | 0.99789 |
| GAN | 4 | 0.15062 | 0.28982 | 0.999632 | 6475 | 2 | 0.32156 | 0.24378 | 0.4439 | 1 | 9912 | 2 | 0.32156 |
| SH3BP5 | 6 | 0.0085102 | 0.033733 | 0.867348 | 763 | 2 | 0.92379 | 0.24379 | 0.49103 | 1 | 9913 | 3 | 0.92379 |
| DNM1L | 6 | 0.65343 | 0.78024 | 0.999994 | 15702 | 2 | 0.036337 | 0.24387 | 0.49112 | 1 | 9914 | 2 | 0.036337 |
| SLC15A4 | 6 | 0.34648 | 0.56005 | 0.999994 | 11082 | 3 | -0.13104 | 0.24387 | 0.49112 | 1 | 9915 | 2 | -0.13104 |
| BNC1 | 6 | 0.19948 | 0.38725 | 0.999994 | 7748 | 2 | 1.5594 | 0.24388 | 0.49115 | 1 | 9916 | 3 | 1.5594 |
| VPS37B | 5 | 0.20579 | 0.38345 | 0.999994 | 7918 | 2 | 1.8167 | 0.244 | 0.46541 | 1 | 9917 | 3 | 1.8167 |
| FYN | 5 | 0.32299 | 0.49974 | 0.999994 | 10576 | 2 | 1.6415 | 0.24404 | 0.46546 | 1 | 9918 | 3 | 1.6415 |
| DDX24 | 5 | 0.035799 | 0.10527 | 0.95013 | 2305 | 2 | 0.018317 | 0.24419 | 0.46564 | 1 | 9919 | 2 | 0.018317 |
| TBC1D15 | 6 | 0.33718 | 0.54946 | 0.999994 | 10877 | 2 | 1.6149 | 0.24429 | 0.49166 | 1 | 9920 | 4 | 1.6149 |
| SEC23A | 6 | 0.40458 | 0.60535 | 0.999994 | 12029 | 2 | 1.2186 | 0.24431 | 0.49168 | 1 | 9921 | 3 | 1.2186 |
| hsa-mir-3133 | 2 | 0.476 | 0.48222 | 0.999994 | 13080 | 1 | 2.6731 | 0.24433 | 0.37122 | 0.998994 | 9922 | 1 | 2.6731 |
| SLC16A5 | 6 | 0.09052 | 0.22854 | 0.983354 | 4615 | 4 | -1.4623 | 0.24434 | 0.49173 | 1 | 9923 | 2 | -1.4623 |
| B3GAT2 | 6 | 0.52492 | 0.68595 | 0.999994 | 13816 | 2 | 1.5633 | 0.24434 | 0.49173 | 1 | 9924 | 4 | 1.5633 |
| MAML2 | 6 | 0.31029 | 0.51868 | 0.999994 | 10330 | 2 | 1.3445 | 0.24458 | 0.49205 | 1 | 9925 | 4 | 1.3445 |
| CS | 5 | 0.31041 | 0.48946 | 0.999994 | 10333 | 2 | 1.6679 | 0.2446 | 0.46611 | 1 | 9926 | 3 | 1.6679 |
| ITM2B | 6 | 0.90466 | 0.90731 | 0.999994 | 19314 | 1 | 0.51126 | 0.24463 | 0.49212 | 1 | 9927 | 3 | 0.51126 |
| DHRS7C | 5 | 0.36791 | 0.53663 | 0.999994 | 11473 | 1 | 0.71164 | 0.2447 | 0.46622 | 1 | 9928 | 3 | 0.71164 |
| RDH5 | 5 | 0.16577 | 0.32591 | 0.999994 | 6879 | 2 | 1.2305 | 0.2447 | 0.46622 | 1 | 9929 | 3 | 1.2305 |
| PRAP1 | 5 | 0.099716 | 0.22598 | 0.983354 | 4965 | 2 | 1.1852 | 0.2447 | 0.46622 | 1 | 9930 | 3 | 1.1852 |
| GPR176 | 6 | 0.63915 | 0.7693 | 0.999994 | 15501 | 2 | 1.6812 | 0.2447 | 0.49222 | 1 | 9931 | 4 | 1.6812 |

| | | | | | | | | | | | | | |
|---|---|---|---|---|---|---|---|---|---|---|---|---|---|
| FAT4 | 5 | 0.96346 | 0.96361 | 0.999994 | 20685 | 0 | 1.1557 | 0.24476 | 0.46627 | 1 | 9932 | 3 | 1.1557 |
| BCHE | 4 | 0.14527 | 0.28433 | 0.999632 | 6332 | 2 | 0.69036 | 0.24483 | 0.44531 | 1 | 9933 | 2 | 0.69036 |
| WRNIP1 | 6 | 0.27263 | 0.47479 | 0.999994 | 9530 | 2 | 1.3885 | 0.24484 | 0.49239 | 1 | 9934 | 3 | 1.3885 |
| L1TD1 | 6 | 0.42068 | 0.61585 | 0.999994 | 12261 | 2 | 1.1364 | 0.24494 | 0.49255 | 1 | 9935 | 3 | 1.1364 |
| GRID2IP | 6 | 0.23683 | 0.4322 | 0.999994 | 8691 | 1 | 0.91121 | 0.24507 | 0.49272 | 1 | 9936 | 4 | 0.91121 |
| EPB41L5 | 6 | 0.30004 | 0.50677 | 0.999994 | 10132 | 3 | -0.10756 | 0.24512 | 0.49278 | 1 | 9937 | 3 | -0.10756 |
| RBMXL1 | 6 | 0.50109 | 0.66946 | 0.999994 | 13439 | 2 | 0.5617 | 0.24512 | 0.49278 | 1 | 9938 | 3 | 0.5617 |
| SELV | 4 | 0.63802 | 0.68636 | 0.999994 | 15487 | 1 | 0.27518 | 0.24514 | 0.44573 | 1 | 9939 | 2 | 0.27518 |
| RPS6KB2 | 5 | 0.12354 | 0.26272 | 0.996368 | 5753 | 3 | -1.2698 | 0.24517 | 0.46673 | 1 | 9940 | 2 | -1.2698 |
| ZNF534 | 6 | 0.9281 | 0.92817 | 0.999994 | 19816 | 0 | 1.0616 | 0.24518 | 0.49287 | 1 | 9941 | 4 | 1.0616 |
| ZNF155 | 6 | 0.31561 | 0.52486 | 0.999994 | 10432 | 2 | 1.3666 | 0.24521 | 0.49292 | 1 | 9942 | 4 | 1.3666 |
| DAZAP1 | 5 | 0.72652 | 0.779 | 0.999994 | 16560 | 1 | 1.7486 | 0.24522 | 0.46678 | 1 | 9943 | 3 | 1.7486 |
| LAS1L | 4 | 0.60515 | 0.66812 | 0.999994 | 15012 | 1 | 1.1669 | 0.24535 | 0.446 | 1 | 9944 | 3 | 1.1669 |
| S100A7A | 5 | 0.36158 | 0.53138 | 0.999994 | 11362 | 2 | -0.4027 | 0.24545 | 0.46704 | 1 | 9945 | 2 | -0.4027 |
| ATP6V0E2 | 6 | 0.26914 | 0.47073 | 0.999994 | 9447 | 2 | 0.15645 | 0.2456 | 0.49344 | 1 | 9946 | 2 | 0.15645 |
| PIR | 6 | 0.28691 | 0.49149 | 0.999994 | 9851 | 2 | 0.89238 | 0.2456 | 0.49344 | 1 | 9947 | 3 | 0.89238 |
| RBM15B | 6 | 0.05782 | 0.16107 | 0.967126 | 3329 | 4 | -1.245 | 0.2456 | 0.49344 | 1 | 9948 | 2 | -1.245 |
| MTIF2 | 6 | 0.24854 | 0.4462 | 0.999994 | 8980 | 2 | 0.73928 | 0.2456 | 0.49344 | 1 | 9949 | 3 | 0.73928 |
| HSFY1 | 6 | 0.54027 | 0.69673 | 0.999994 | 14057 | 1 | 1.0922 | 0.24561 | 0.49345 | 1 | 9950 | 4 | 1.0922 |
| CFDP1 | 6 | 0.61894 | 0.75403 | 0.999994 | 15205 | 1 | 1.3053 | 0.24572 | 0.4936 | 1 | 9951 | 4 | 1.3053 |
| SNX3 | 6 | 0.90519 | 0.90772 | 0.999994 | 19336 | 1 | 1.8114 | 0.24572 | 0.4936 | 1 | 9952 | 4 | 1.8114 |
| ZBTB40 | 6 | 0.96507 | 0.9651 | 0.999994 | 20720 | 0 | 0.81058 | 0.24572 | 0.4936 | 1 | 9953 | 4 | 0.81058 |
| AQP5 | 6 | 0.16703 | 0.347 | 0.999994 | 6905 | 3 | -0.85324 | 0.24572 | 0.4936 | 1 | 9954 | 1 | -0.85324 |
| TSPAN1 | 6 | 0.21098 | 0.40129 | 0.999994 | 8051 | 3 | -0.59161 | 0.24572 | 0.4936 | 1 | 9955 | 2 | -0.59161 |
| HNRNPUL1 | 6 | 0.39698 | 0.6004 | 0.999994 | 11906 | 1 | 0.7226 | 0.24572 | 0.4936 | 1 | 9956 | 3 | 0.7226 |
| CAMSAP1 | 5 | 0.29914 | 0.48036 | 0.999994 | 10116 | 2 | 1.5181 | 0.24592 | 0.46758 | 1 | 9957 | 3 | 1.5181 |
| TMC1 | 4 | 0.11147 | 0.23547 | 0.985362 | 5348 | 2 | -1.4797 | 0.24595 | 0.44681 | 1 | 9958 | 1 | -1.4797 |
| KIF12 | 4 | 0.09074 | 0.1984 | 0.977164 | 4628 | 1 | 0.76942 | 0.24595 | 0.44681 | 1 | 9959 | 2 | 0.76942 |
| TRPC1 | 5 | 0.7671 | 0.80011 | 0.999994 | 17052 | 1 | 1.6427 | 0.24595 | 0.46762 | 1 | 9960 | 3 | 1.6427 |
| TMEM194A | 5 | 0.43514 | 0.59275 | 0.999994 | 12502 | 1 | 1.6604 | 0.24597 | 0.46763 | 1 | 9961 | 3 | 1.6604 |
| TMEM236 | 6 | 0.91028 | 0.9118 | 0.999994 | 19431 | 1 | 1.4642 | 0.24606 | 0.49404 | 1 | 9962 | 4 | 1.4642 |
| hsa-mir-626 | 4 | 0.12115 | 0.2523 | 0.994162 | 5668 | 1 | 2.2003 | 0.2461 | 0.44702 | 1 | 9963 | 3 | 2.2003 |
| DNAJC11 | 4 | 0.018371 | 0.057169 | 0.92665 | 1381 | 2 | 1.2288 | 0.2461 | 0.44702 | 1 | 9964 | 2 | 1.2288 |
| RPS21 | 4 | 0.17521 | 0.31516 | 0.999994 | 7115 | 1 | 2.1857 | 0.2461 | 0.44702 | 1 | 9965 | 2 | 2.1857 |
| ZNF296 | 6 | 0.24256 | 0.43904 | 0.999994 | 8848 | 2 | 1.4046 | 0.24613 | 0.49412 | 1 | 9966 | 4 | 1.4046 |
| SOX1 | 6 | 0.87449 | 0.88612 | 0.999994 | 18718 | 1 | 1.4363 | 0.24623 | 0.49426 | 1 | 9967 | 4 | 1.4363 |
| ALS2 | 5 | 0.53504 | 0.67849 | 0.999994 | 13982 | 1 | 1.616 | 0.24634 | 0.46804 | 1 | 9968 | 3 | 1.616 |
| BLOC1S4 | 4 | 0.22549 | 0.36684 | 0.999994 | 8411 | 2 | -0.96386 | 0.2466 | 0.4477 | 1 | 9969 | 1 | -0.96386 |
| NR2C2 | 6 | 0.98294 | 0.9829 | 0.999994 | 21154 | 0 | 1.5882 | 0.24663 | 0.49482 | 1 | 9970 | 4 | 1.5882 |
| FAM120A | 6 | 0.129 | 0.299 | 0.999994 | 5915 | 3 | -0.68227 | 0.24664 | 0.49483 | 1 | 9971 | 2 | -0.68227 |
| TNFSF11 | 6 | 0.91561 | 0.91645 | 0.999994 | 19530 | 1 | 1.2551 | 0.24664 | 0.49483 | 1 | 9972 | 3 | 1.2551 |
| NAA15 | 5 | 0.83782 | 0.84507 | 0.999994 | 18049 | 1 | 1.0929 | 0.24671 | 0.46845 | 1 | 9973 | 3 | 1.0929 |
| hsa-mir-4465 | 4 | 0.92726 | 0.92704 | 0.999994 | 19796 | 0 | 1.7488 | 0.2468 | 0.44796 | 1 | 9974 | 3 | 1.7488 |
| B4GALT1 | 6 | 0.27615 | 0.47894 | 0.999994 | 9621 | 1 | 1.3702 | 0.24688 | 0.49515 | 1 | 9975 | 4 | 1.3702 |
| ZNF181 | 6 | 0.33512 | 0.5471 | 0.999994 | 10837 | 2 | 1.0343 | 0.24689 | 0.49517 | 1 | 9976 | 4 | 1.0343 |
| RAB1A | 6 | 0.2716 | 0.47359 | 0.999994 | 9513 | 2 | 0.87155 | 0.24691 | 0.49518 | 1 | 9977 | 4 | 0.87155 |
| KRCC1 | 6 | 0.0415 | 0.12528 | 0.951052 | 2591 | 4 | -1.5838 | 0.24691 | 0.49518 | 1 | 9978 | 2 | -1.5838 |
| hsa-mir-1273h | 2 | 0.7444 | 0.74408 | 0.999994 | 16792 | 0 | 3.5192 | 0.24693 | 0.37274 | 0.999216 | 9979 | 1 | 3.5192 |
| MYOZ3 | 5 | 0.086329 | 0.2047 | 0.978186 | 4463 | 2 | -0.52495 | 0.24694 | 0.46871 | 1 | 9980 | 2 | -0.52495 |
| CYSTM1 | 5 | 0.36272 | 0.53233 | 0.999994 | 11385 | 2 | -0.37575 | 0.24694 | 0.46871 | 1 | 9981 | 2 | -0.37575 |
| MYOD1 | 5 | 0.14595 | 0.2967 | 0.999994 | 6349 | 3 | -1.1918 | 0.24694 | 0.46871 | 1 | 9982 | 1 | -1.1918 |
| FBXL14 | 5 | 0.070233 | 0.17876 | 0.968171 | 3843 | 1 | 1.8309 | 0.24698 | 0.46876 | 1 | 9983 | 3 | 1.8309 |

| | | | | | | | | | | | | |
|---|---|---|---|---|---|---|---|---|---|---|---|---|
| OTUD6B | 6 | 0.90858 | 0.91042 | 0.999994 | 19397 | 1 | 1.6208 | 0.24699 | 0.49529 | 1 | 9984 | 4 | 1.6208 |
| POU2F1 | 3 | 0.0051814 | 0.01426 | 0.753188 | 532 | 1 | 2.1815 | 0.24702 | 0.39339 | 1 | 9985 | 2 | 2.1815 |
| hsa-mir-6072 | 4 | 0.91991 | 0.91969 | 0.999994 | 19628 | 0 | 0.84611 | 0.24703 | 0.44825 | 1 | 9986 | 2 | 0.84611 |
| MMS22L | 5 | 0.53132 | 0.67525 | 0.999994 | 13919 | 2 | 1.5475 | 0.24704 | 0.46883 | 1 | 9987 | 3 | 1.5475 |
| EFNA5 | 6 | 0.84342 | 0.86844 | 0.999994 | 18145 | 1 | 1.48 | 0.24715 | 0.49549 | 1 | 9988 | 4 | 1.48 |
| IQCB1 | 6 | 0.68634 | 0.80595 | 0.999994 | 16173 | 2 | 0.85974 | 0.24718 | 0.49553 | 1 | 9989 | 3 | 0.85974 |
| OAT | 6 | 0.20613 | 0.39536 | 0.999994 | 7931 | 3 | -0.6168 | 0.24718 | 0.49553 | 1 | 9990 | 1 | -0.6168 |
| MMP1 | 6 | 0.90242 | 0.90556 | 0.999994 | 19258 | 1 | 1.7085 | 0.2472 | 0.49556 | 1 | 9991 | 4 | 1.7085 |
| CD46 | 5 | 0.18867 | 0.35903 | 0.999994 | 7441 | 3 | -0.86334 | 0.24731 | 0.46912 | 1 | 9992 | 2 | -0.86334 |
| PODXL2 | 4 | 0.55428 | 0.64181 | 0.999994 | 14275 | 1 | 1.4431 | 0.24746 | 0.44885 | 1 | 9993 | 3 | 1.4431 |
| SPAG9 | 4 | 0.049442 | 0.12036 | 0.951052 | 2952 | 1 | 2.0336 | 0.24772 | 0.4492 | 1 | 9994 | 3 | 2.0336 |
| hsa-mir-622 | 3 | 0.17142 | 0.27614 | 0.999379 | 7022 | 1 | -0.061652 | 0.24775 | 0.39417 | 1 | 9995 | 1 | -0.061652 |
| MYT1 | 5 | 0.31196 | 0.49077 | 0.999994 | 10359 | 2 | -0.58277 | 0.24778 | 0.46964 | 1 | 9996 | 2 | -0.58277 |
| CRHR2 | 5 | 0.051945 | 0.1451 | 0.963012 | 3064 | 3 | -1.6713 | 0.24778 | 0.46964 | 1 | 9997 | 2 | -1.6713 |
| FRMPD2 | 5 | 0.71852 | 0.77513 | 0.999994 | 16481 | 1 | 1.4565 | 0.24778 | 0.46964 | 1 | 9998 | 3 | 1.4565 |
| FAM90A1 | 5 | 0.62104 | 0.73394 | 0.999994 | 15239 | 1 | 0.32089 | 0.24778 | 0.46964 | 1 | 9999 | 2 | 0.32089 |
| TRIB1 | 6 | 0.40991 | 0.60884 | 0.999994 | 12106 | 2 | 0.55527 | 0.24785 | 0.49638 | 1 | 10000 | 3 | 0.55527 |
| SNX25 | 4 | 0.74483 | 0.75612 | 0.999994 | 16798 | 1 | 1.0323 | 0.2479 | 0.44941 | 1 | 10001 | 3 | 1.0323 |
| PRKAA2 | 6 | 0.63487 | 0.76602 | 0.999994 | 15442 | 2 | 1.1546 | 0.24792 | 0.49647 | 1 | 10002 | 4 | 1.1546 |
| CARS | 5 | 0.16349 | 0.32262 | 0.999994 | 6827 | 1 | 0.11655 | 0.24795 | 0.46981 | 1 | 10003 | 2 | 0.11655 |
| PPP2R3B_X | 6 | 0.66243 | 0.7872 | 0.999994 | 15818 | 2 | 1.2124 | 0.24795 | 0.49653 | 1 | 10004 | 4 | 1.2124 |
| GDF7 | 4 | 0.85927 | 0.85887 | 0.999994 | 18419 | 0 | 1.9268 | 0.24808 | 0.44967 | 1 | 10005 | 2 | 1.9268 |
| NHLH1 | 6 | 0.33364 | 0.54546 | 0.999994 | 10798 | 2 | 0.61345 | 0.24813 | 0.49675 | 1 | 10006 | 3 | 0.61345 |
| PDE4D | 6 | 0.066782 | 0.18001 | 0.968171 | 3716 | 2 | 0.87111 | 0.24813 | 0.49675 | 1 | 10007 | 3 | 0.87111 |
| RPL3L | 6 | 0.78275 | 0.84079 | 0.999994 | 17248 | 1 | 0.83465 | 0.24813 | 0.49675 | 1 | 10008 | 4 | 0.83465 |
| CTBP2 | 5 | 0.24964 | 0.44049 | 0.999994 | 9010 | 2 | -0.21965 | 0.24818 | 0.47007 | 1 | 10009 | 2 | -0.21965 |
| SMCR8 | 4 | 0.33372 | 0.47749 | 0.999994 | 10799 | 2 | 1.3592 | 0.24821 | 0.44986 | 1 | 10010 | 2 | 1.3592 |
| YTHDC1 | 6 | 0.26092 | 0.46098 | 0.999994 | 9268 | 2 | 1.4514 | 0.24824 | 0.49691 | 1 | 10011 | 3 | 1.4514 |
| SOX15 | 6 | 0.3408 | 0.55357 | 0.999994 | 10954 | 3 | -0.21133 | 0.24825 | 0.49692 | 1 | 10012 | 1 | -0.21133 |
| EEF1A1 | 3 | 0.74861 | 0.7486 | 0.999994 | 16844 | 0 | 2.1365 | 0.24841 | 0.39489 | 1 | 10013 | 2 | 2.1365 |
| A1BG | 6 | 0.49567 | 0.66573 | 0.999994 | 13351 | 2 | 1.5635 | 0.24842 | 0.49715 | 1 | 10014 | 4 | 1.5635 |
| hsa-mir-6832 | 4 | 0.93956 | 0.93946 | 0.999994 | 20065 | 0 | 1.4029 | 0.24845 | 0.45018 | 1 | 10015 | 3 | 1.4029 |
| ZSCAN23 | 5 | 0.75796 | 0.79509 | 0.999994 | 16952 | 1 | 1.4895 | 0.24846 | 0.47035 | 1 | 10016 | 3 | 1.4895 |
| GPN2 | 3 | 0.85698 | 0.85692 | 0.999994 | 18393 | 0 | 0.98288 | 0.24847 | 0.39494 | 1 | 10017 | 2 | 0.98288 |
| PTK6 | 6 | 0.87955 | 0.8893 | 0.999994 | 18822 | 1 | 1.0147 | 0.24849 | 0.49725 | 1 | 10018 | 4 | 1.0147 |
| HMBOX1 | 6 | 0.19189 | 0.37787 | 0.999994 | 7542 | 2 | 0.98909 | 0.24857 | 0.49735 | 1 | 10019 | 3 | 0.98909 |
| OCIAD1 | 6 | 0.38274 | 0.5913 | 0.999994 | 11687 | 1 | 1.2943 | 0.24858 | 0.49737 | 1 | 10020 | 4 | 1.2943 |
| PLEKHG2 | 4 | 0.26632 | 0.40852 | 0.999994 | 9374 | 1 | 1.5694 | 0.24859 | 0.45036 | 1 | 10021 | 3 | 1.5694 |
| RORA | 6 | 0.51595 | 0.67972 | 0.999994 | 13674 | 1 | 0.87043 | 0.2486 | 0.49739 | 1 | 10022 | 4 | 0.87043 |
| ETV6 | 5 | 0.83302 | 0.84166 | 0.999994 | 17971 | 1 | 0.63601 | 0.24863 | 0.47054 | 1 | 10023 | 3 | 0.63601 |
| PRKCB | 6 | 0.44291 | 0.63037 | 0.999994 | 12622 | 2 | 1.3426 | 0.24877 | 0.49762 | 1 | 10024 | 4 | 1.3426 |
| PLP2 | 4 | 0.259 | 0.40112 | 0.999994 | 9216 | 1 | 0.12176 | 0.24883 | 0.45069 | 1 | 10025 | 1 | 0.12176 |
| UBE2G1 | 6 | 0.877 | 0.8877 | 0.999994 | 18772 | 1 | 1.2489 | 0.24884 | 0.49773 | 1 | 10026 | 3 | 1.2489 |
| SCAF1 | 6 | 0.27296 | 0.47519 | 0.999994 | 9541 | 3 | 0.82054 | 0.24888 | 0.49777 | 1 | 10027 | 3 | 0.82054 |
| GNAT2 | 6 | 0.0072474 | 0.029284 | 0.860452 | 673 | 1 | 0.84571 | 0.24892 | 0.49782 | 1 | 10028 | 4 | 0.84571 |
| SPINT1 | 5 | 0.63398 | 0.73887 | 0.999994 | 15432 | 1 | 0.96681 | 0.24917 | 0.47113 | 1 | 10029 | 3 | 0.96681 |
| FBXW2 | 6 | 0.67401 | 0.79627 | 0.999994 | 15988 | 2 | 1.4088 | 0.24917 | 0.49814 | 1 | 10030 | 4 | 1.4088 |
| SNCAIP | 6 | 0.61021 | 0.74749 | 0.999994 | 15075 | 1 | 1.1914 | 0.24919 | 0.49817 | 1 | 10031 | 4 | 1.1914 |
| WDR72 | 5 | 0.93051 | 0.93078 | 0.999994 | 19865 | 0 | 0.61081 | 0.2493 | 0.47125 | 1 | 10032 | 3 | 0.61081 |
| GPR174 | 5 | 0.87065 | 0.87166 | 0.999994 | 18649 | 1 | 1.0006 | 0.2493 | 0.47125 | 1 | 10033 | 3 | 1.0006 |
| LPAR6 | 5 | 0.55488 | 0.6961 | 0.999994 | 14286 | 2 | 1.3059 | 0.2493 | 0.47125 | 1 | 10034 | 3 | 1.3059 |
| GDPD5 | 5 | 0.24179 | 0.43334 | 0.999994 | 8828 | 2 | 0.76102 | 0.2493 | 0.47125 | 1 | 10035 | 3 | 0.76102 |

| | | | | | | | | | | | | |
|---|---|---|---|---|---|---|---|---|---|---|---|---|
| TEX30 | 5 | 0.19187 | 0.36361 | 0.999994 | 7541 | 3 | -0.70435 | 0.2493 | 0.47125 | 1 | 10036 | 2 | -0.70435 |
| NAIP | 5 | 0.55677 | 0.69779 | 0.999994 | 14314 | 1 | 1.6281 | 0.24935 | 0.47131 | 1 | 10037 | 3 | 1.6281 |
| LILRB1 | 3 | 0.73799 | 0.73792 | 0.999994 | 16703 | 0 | 2.2684 | 0.24935 | 0.39589 | 1 | 10038 | 2 | 2.2684 |
| SLC35E2B | 4 | 0.65436 | 0.69584 | 0.999994 | 15710 | 1 | 1.8419 | 0.24936 | 0.45138 | 1 | 10039 | 2 | 1.8419 |
| THEM4 | 6 | 0.20346 | 0.39208 | 0.999994 | 7867 | 2 | 0.1009 | 0.24938 | 0.49842 | 1 | 10040 | 2 | 0.1009 |
| DCHS1 | 6 | 0.54529 | 0.70029 | 0.999994 | 14140 | 2 | -0.018396 | 0.24938 | 0.49842 | 1 | 10041 | 2 | -0.018396 |
| AGTR2 | 6 | 0.30218 | 0.50926 | 0.999994 | 10169 | 2 | 0.31609 | 0.24938 | 0.49842 | 1 | 10042 | 2 | 0.31609 |
| ILVBL | 6 | 0.20302 | 0.39155 | 0.999994 | 7856 | 2 | 1.2682 | 0.24949 | 0.49859 | 1 | 10043 | 4 | 1.2682 |
| NonTargetingControlG | 2 | 0.74073 | 0.7404 | 0.999994 | 16746 | 0 | 3.442 | 0.24955 | 0.3743 | 0.999517 | 10044 | 1 | 3.442 |
| DOCK6 | 6 | 0.16512 | 0.34461 | 0.999994 | 6865 | 1 | 1.2371 | 0.24956 | 0.49868 | 1 | 10045 | 4 | 1.2371 |
| CDSN | 6 | 0.54854 | 0.70257 | 0.999994 | 14188 | 2 | 1.1776 | 0.24966 | 0.49883 | 1 | 10046 | 4 | 1.1776 |
| SCARF2 | 6 | 0.022805 | 0.080962 | 0.93914 | 1634 | 3 | 0.38183 | 0.24969 | 0.49886 | 1 | 10047 | 3 | 0.38183 |
| TCF7L1 | 5 | 0.081232 | 0.1966 | 0.977164 | 4275 | 3 | -1.091 | 0.2497 | 0.47167 | 1 | 10048 | 2 | -1.091 |
| CCL7 | 6 | 0.54052 | 0.6969 | 0.999994 | 14065 | 2 | 1.2263 | 0.24971 | 0.49889 | 1 | 10049 | 4 | 1.2263 |
| AP5S1 | 6 | 0.27898 | 0.48223 | 0.999994 | 9671 | 1 | 0.96376 | 0.24988 | 0.49912 | 1 | 10050 | 4 | 0.96376 |
| SC5D | 6 | 0.12587 | 0.29508 | 0.999994 | 5816 | 3 | -0.35074 | 0.25001 | 0.49929 | 1 | 10051 | 2 | -0.35074 |
| JTB | 5 | 0.12509 | 0.26511 | 0.997396 | 5797 | 2 | -0.018797 | 0.25005 | 0.47205 | 1 | 10052 | 2 | -0.018797 |
| ALDH2 | 6 | 0.47865 | 0.65428 | 0.999994 | 13124 | 1 | 1.2858 | 0.25014 | 0.49947 | 1 | 10053 | 4 | 1.2858 |
| OR4D6 | 5 | 0.30607 | 0.48597 | 0.999994 | 10259 | 2 | 0.30493 | 0.25018 | 0.47219 | 1 | 10054 | 2 | 0.30493 |
| SMARCA2 | 6 | 0.34678 | 0.56037 | 0.999994 | 11087 | 2 | 1.3103 | 0.25019 | 0.49954 | 1 | 10055 | 3 | 1.3103 |
| KIF5B | 4 | 0.87034 | 0.86992 | 0.999994 | 18640 | 0 | 1.5777 | 0.25029 | 0.45263 | 1 | 10056 | 3 | 1.5777 |
| EEF1E1 | 6 | 0.39135 | 0.59681 | 0.999994 | 11818 | 2 | 0.9893 | 0.25031 | 0.49968 | 1 | 10057 | 4 | 0.9893 |
| FAM111A | 5 | 0.84914 | 0.85359 | 0.999994 | 18244 | 1 | 1.657 | 0.25035 | 0.47235 | 1 | 10058 | 3 | 1.657 |
| UNKL | 6 | 0.25091 | 0.44903 | 0.999994 | 9031 | 3 | -0.32678 | 0.25036 | 0.49976 | 1 | 10059 | 2 | -0.32678 |
| SERPINA6 | 6 | 0.60275 | 0.74199 | 0.999994 | 14983 | 2 | 1.34 | 0.25037 | 0.49977 | 1 | 10060 | 3 | 1.34 |
| FOPNL | 5 | 0.18459 | 0.3532 | 0.999994 | 7332 | 1 | 1.5124 | 0.25046 | 0.47246 | 1 | 10061 | 3 | 1.5124 |
| hsa-mir-4318 | 4 | 0.82101 | 0.82073 | 0.999994 | 17790 | 1 | 0.97841 | 0.25049 | 0.45289 | 1 | 10062 | 3 | 0.97841 |
| TOPORS | 6 | 0.068823 | 0.1843 | 0.968526 | 3791 | 3 | 0.73734 | 0.25054 | 0.49999 | 1 | 10063 | 3 | 0.73734 |
| BLCAP | 5 | 0.50257 | 0.65016 | 0.999994 | 13473 | 2 | 0.32614 | 0.25056 | 0.47257 | 1 | 10064 | 2 | 0.32614 |
| CSDC2 | 5 | 0.19979 | 0.37495 | 0.999994 | 7758 | 1 | 1.6808 | 0.25056 | 0.47257 | 1 | 10065 | 3 | 1.6808 |
| AMTN | 5 | 0.031028 | 0.09314 | 0.944268 | 2062 | 1 | 0.26578 | 0.25056 | 0.47257 | 1 | 10066 | 2 | 0.26578 |
| MRM1 | 6 | 0.029126 | 0.09674 | 0.945031 | 1964 | 3 | -1.993 | 0.25057 | 0.50005 | 1 | 10067 | 2 | -1.993 |
| C15orf57 | 6 | 0.76966 | 0.8357 | 0.999994 | 17078 | 1 | 1.4994 | 0.25059 | 0.50007 | 1 | 10068 | 4 | 1.4994 |
| RASGEF1C | 6 | 0.32933 | 0.54047 | 0.999994 | 10707 | 2 | 0.92738 | 0.25061 | 0.5001 | 1 | 10069 | 4 | 0.92738 |
| CTRC | 6 | 0.55548 | 0.70756 | 0.999994 | 14298 | 2 | 0.60602 | 0.25061 | 0.5001 | 1 | 10070 | 3 | 0.60602 |
| GSTZ1 | 6 | 0.43921 | 0.62787 | 0.999994 | 12562 | 2 | 0.53826 | 0.25061 | 0.5001 | 1 | 10071 | 3 | 0.53826 |
| ITPKA | 4 | 0.71537 | 0.73483 | 0.999994 | 16451 | 1 | 1.6793 | 0.25064 | 0.45307 | 1 | 10072 | 3 | 1.6793 |
| EVPLL | 5 | 0.41812 | 0.57855 | 0.999994 | 12219 | 1 | 0.3048 | 0.25078 | 0.4728 | 1 | 10073 | 2 | 0.3048 |
| hsa-mir-526b | 3 | 0.47172 | 0.52786 | 0.999994 | 13028 | 1 | 1.0417 | 0.25085 | 0.3975 | 1 | 10074 | 2 | 1.0417 |
| DPH5 | 6 | 0.59834 | 0.73871 | 0.999994 | 14932 | 2 | 1.2546 | 0.2509 | 0.50049 | 1 | 10075 | 4 | 1.2546 |
| CCL4L1 | 5 | 0.58018 | 0.71847 | 0.999994 | 14679 | 2 | -0.28433 | 0.25097 | 0.473 | 1 | 10076 | 1 | -0.28433 |
| PML | 4 | 0.72301 | 0.74017 | 0.999994 | 16525 | 1 | 2.2491 | 0.2511 | 0.45369 | 1 | 10077 | 3 | 2.2491 |
| MT1H | 4 | 0.89051 | 0.89023 | 0.999994 | 19047 | 0 | 1.8837 | 0.25117 | 0.45379 | 1 | 10078 | 2 | 1.8837 |
| hsa-mir-606 | 3 | 0.15602 | 0.25629 | 0.994943 | 6628 | 1 | -0.046067 | 0.25117 | 0.39784 | 1 | 10079 | 1 | -0.046067 |
| TIMP1 | 3 | 0.67789 | 0.68083 | 0.999994 | 16059 | 1 | 0.35446 | 0.25117 | 0.39784 | 1 | 10080 | 1 | 0.35446 |
| GLCCI1 | 6 | 0.49361 | 0.66433 | 0.999994 | 13319 | 2 | -0.0038531 | 0.25121 | 0.50089 | 1 | 10081 | 2 | -0.0038531 |
| DDX27 | 3 | 0.27036 | 0.40013 | 0.999994 | 9485 | 1 | 2.2979 | 0.25134 | 0.39802 | 1 | 10082 | 2 | 2.2979 |
| MEOX2 | 5 | 0.51806 | 0.66362 | 0.999994 | 13710 | 1 | 1.8243 | 0.25139 | 0.47343 | 1 | 10083 | 3 | 1.8243 |
| ABCB8 | 6 | 0.67722 | 0.79881 | 0.999994 | 16048 | 2 | 1.1892 | 0.25146 | 0.50122 | 1 | 10084 | 4 | 1.1892 |
| PCDH12 | 6 | 0.21081 | 0.40108 | 0.999994 | 8047 | 3 | 0.87157 | 0.2516 | 0.50142 | 1 | 10085 | 3 | 0.87157 |
| PYGL | 5 | 0.47323 | 0.62487 | 0.999994 | 13047 | 2 | -0.51552 | 0.2516 | 0.47367 | 1 | 10086 | 2 | -0.51552 |
| CTDSPL | 5 | 0.11895 | 0.25575 | 0.994943 | 5602 | 2 | 0.18937 | 0.2516 | 0.47367 | 1 | 10087 | 2 | 0.18937 |

| | | | | | | | | | | | | | |
|---|---|---|---|---|---|---|---|---|---|---|---|---|---|
| GPR20 | 5 | 0.32986 | 0.50542 | 0.999994 | 10718 | 2 | 1.6198 | 0.25163 | 0.4737 | 1 | 10088 | 3 | 1.6198 |
| hsa-mir-3162 | 4 | 0.47791 | 0.60658 | 0.999994 | 13113 | 1 | 1.7238 | 0.25179 | 0.45462 | 1 | 10089 | 2 | 1.7238 |
| ZNF577 | 6 | 0.88411 | 0.89229 | 0.999994 | 18910 | 1 | 1.4325 | 0.2518 | 0.50168 | 1 | 10090 | 4 | 1.4325 |
| MBL2 | 6 | 0.20817 | 0.39784 | 0.999994 | 7983 | 3 | -0.92408 | 0.25182 | 0.50169 | 1 | 10091 | 2 | -0.92408 |
| ALDH3A1 | 6 | 0.057048 | 0.1594 | 0.967126 | 3292 | 4 | -1.6013 | 0.25182 | 0.50169 | 1 | 10092 | 2 | -1.6013 |
| CAPNS1 | 6 | 0.84318 | 0.8683 | 0.999994 | 18141 | 1 | 0.96453 | 0.25182 | 0.50169 | 1 | 10093 | 4 | 0.96453 |
| HKR1 | 6 | 0.022499 | 0.079973 | 0.93914 | 1619 | 2 | 0.88022 | 0.25184 | 0.50174 | 1 | 10094 | 4 | 0.88022 |
| TEX29 | 6 | 0.15049 | 0.32626 | 0.999994 | 6473 | 2 | 1.4428 | 0.25191 | 0.50182 | 1 | 10095 | 3 | 1.4428 |
| ATF7IP2 | 5 | 0.20474 | 0.38196 | 0.999994 | 7894 | 2 | 1.1204 | 0.25194 | 0.47404 | 1 | 10096 | 3 | 1.1204 |
| HPCAL4 | 5 | 0.45197 | 0.60689 | 0.999994 | 12735 | 2 | 1.2661 | 0.25194 | 0.47404 | 1 | 10097 | 3 | 1.2661 |
| LOC388849 | 5 | 0.46863 | 0.62097 | 0.999994 | 12984 | 2 | 0.0092545 | 0.25194 | 0.47404 | 1 | 10098 | 2 | 0.0092545 |
| TMEM37 | 5 | 0.03095 | 0.09294 | 0.943968 | 2059 | 2 | 1.6832 | 0.25194 | 0.47404 | 1 | 10099 | 3 | 1.6832 |
| C9orf16 | 4 | 0.8985 | 0.89821 | 0.999994 | 19195 | 0 | 1.7769 | 0.25194 | 0.45484 | 1 | 10100 | 2 | 1.7769 |
| ISYNA1 | 3 | 0.43878 | 0.50735 | 0.999994 | 12552 | 1 | 2.2229 | 0.25195 | 0.39868 | 1 | 10101 | 2 | 2.2229 |
| SERP1 | 6 | 0.012054 | 0.045899 | 0.892434 | 996 | 3 | 0.17473 | 0.25205 | 0.50203 | 1 | 10102 | 3 | 0.17473 |
| PPIAL4E | 4 | 0.83534 | 0.83486 | 0.999994 | 18012 | 0 | 1.187 | 0.25219 | 0.45517 | 1 | 10103 | 3 | 1.187 |
| TMEM237 | 5 | 0.45364 | 0.60829 | 0.999994 | 12752 | 2 | 1.8395 | 0.25219 | 0.47431 | 1 | 10104 | 3 | 1.8395 |
| JRKL | 5 | 0.4189 | 0.57918 | 0.999994 | 12233 | 1 | 1.7296 | 0.25226 | 0.47438 | 1 | 10105 | 3 | 1.7296 |
| KLHL5 | 6 | 0.28253 | 0.48636 | 0.999994 | 9749 | 2 | 1.7058 | 0.25228 | 0.50233 | 1 | 10106 | 4 | 1.7058 |
| LPHN1 | 3 | 0.6467 | 0.65434 | 0.999994 | 15613 | 1 | 1.9728 | 0.2523 | 0.39905 | 1 | 10107 | 2 | 1.9728 |
| INPP4B | 4 | 0.93572 | 0.93552 | 0.999994 | 19985 | 0 | 1.5776 | 0.2523 | 0.45533 | 1 | 10108 | 3 | 1.5776 |
| IL21R | 4 | 0.020786 | 0.062267 | 0.937033 | 1498 | 1 | 1.0367 | 0.25236 | 0.4554 | 1 | 10109 | 3 | 1.0367 |
| PFKFB2 | 5 | 0.18555 | 0.35462 | 0.999994 | 7355 | 1 | 0.83795 | 0.25241 | 0.47455 | 1 | 10110 | 3 | 0.83795 |
| VCY | 4 | 0.10298 | 0.22041 | 0.980699 | 5067 | 2 | 0.38692 | 0.25241 | 0.45547 | 1 | 10111 | 2 | 0.38692 |
| CYLC1 | 5 | 0.83322 | 0.84179 | 0.999994 | 17976 | 1 | 1.4598 | 0.25252 | 0.47466 | 1 | 10112 | 3 | 1.4598 |
| TRIM26 | 4 | 0.33989 | 0.48386 | 0.999994 | 10926 | 1 | 1.0674 | 0.25267 | 0.45581 | 1 | 10113 | 3 | 1.0674 |
| GTF2H2D | 3 | 0.18795 | 0.29744 | 0.999994 | 7419 | 1 | 2.1236 | 0.25269 | 0.39945 | 1 | 10114 | 2 | 2.1236 |
| PGAP3 | 5 | 0.0011074 | 0.00494 | 0.589484 | 180 | 2 | 1.6446 | 0.25269 | 0.47486 | 1 | 10115 | 3 | 1.6446 |
| DAB2 | 5 | 0.10838 | 0.2395 | 0.987253 | 5251 | 3 | -1.4526 | 0.25272 | 0.47489 | 1 | 10116 | 2 | -1.4526 |
| NINJ2 | 6 | 0.84435 | 0.86892 | 0.999994 | 18162 | 1 | 0.98302 | 0.25273 | 0.50293 | 1 | 10117 | 4 | 0.98302 |
| NR1H2 | 6 | 0.48248 | 0.65685 | 0.999994 | 13178 | 1 | 0.93638 | 0.25281 | 0.50303 | 1 | 10118 | 4 | 0.93638 |
| ZNF117 | 4 | 0.87201 | 0.87166 | 0.999994 | 18677 | 0 | 0.50013 | 0.25286 | 0.45608 | 1 | 10119 | 1 | 0.50013 |
| HGC6.3 | 4 | 0.13544 | 0.2742 | 0.998561 | 6067 | 2 | 0.00013813 | 0.25286 | 0.45608 | 1 | 10120 | 2 | 0.00013813 |
| SIX2 | 6 | 0.031392 | 0.10208 | 0.946414 | 2078 | 3 | -0.35339 | 0.253 | 0.5033 | 1 | 10121 | 3 | -0.35339 |
| GCNT1 | 6 | 0.80834 | 0.85151 | 0.999994 | 17606 | 1 | 1.6515 | 0.25301 | 0.50331 | 1 | 10122 | 4 | 1.6515 |
| C17orf77 | 3 | 0.86557 | 0.86556 | 0.999994 | 18550 | 0 | 2.1346 | 0.25301 | 0.3998 | 1 | 10123 | 2 | 2.1346 |
| PLCL1 | 4 | 0.70665 | 0.72891 | 0.999994 | 16379 | 1 | 0.88255 | 0.2531 | 0.45641 | 1 | 10124 | 3 | 0.88255 |
| PGA3 | 6 | 0.46901 | 0.64777 | 0.999994 | 12990 | 2 | 0.28498 | 0.25352 | 0.50399 | 1 | 10125 | 2 | 0.28498 |
| ZGLP1 | 6 | 0.12019 | 0.28604 | 0.999632 | 5635 | 2 | 0.6266 | 0.25352 | 0.50399 | 1 | 10126 | 3 | 0.6266 |
| MRPL41 | 2 | 0.13242 | 0.21444 | 0.979587 | 5987 | 1 | -0.50936 | 0.25352 | 0.37663 | 1 | 10127 | 1 | -0.50936 |
| ETAA1 | 5 | 0.010426 | 0.037284 | 0.881473 | 898 | 2 | 1.4514 | 0.25355 | 0.47579 | 1 | 10128 | 3 | 1.4514 |
| ERCC6L2 | 6 | 0.21703 | 0.40857 | 0.999994 | 8193 | 2 | 1.4898 | 0.25361 | 0.50412 | 1 | 10129 | 4 | 1.4898 |
| DHCR7 | 6 | 0.67308 | 0.79556 | 0.999994 | 15974 | 2 | 1.5067 | 0.25374 | 0.50428 | 1 | 10130 | 4 | 1.5067 |
| hsa-mir-29b-2 | 4 | 0.82222 | 0.82189 | 0.999994 | 17803 | 1 | 1.1518 | 0.25377 | 0.45733 | 1 | 10131 | 3 | 1.1518 |
| SSSCA1 | 5 | 0.95453 | 0.95468 | 0.999994 | 20443 | 0 | 1.4484 | 0.2538 | 0.47604 | 1 | 10132 | 3 | 1.4484 |
| TARSL2 | 5 | 0.91795 | 0.91804 | 0.999994 | 19585 | 0 | 1.4851 | 0.2538 | 0.47604 | 1 | 10133 | 3 | 1.4851 |
| SERGEF | 6 | 0.33796 | 0.55034 | 0.999994 | 10899 | 1 | 1.2794 | 0.25381 | 0.50439 | 1 | 10134 | 3 | 1.2794 |
| SRXN1 | 5 | 0.13962 | 0.28726 | 0.999632 | 6174 | 2 | 1.8061 | 0.25382 | 0.47607 | 1 | 10135 | 3 | 1.8061 |
| UBE2B | 6 | 0.51065 | 0.67608 | 0.999994 | 13586 | 1 | 1.0157 | 0.25383 | 0.50441 | 1 | 10136 | 4 | 1.0157 |
| OR4A15 | 6 | 0.63804 | 0.76846 | 0.999994 | 15489 | 2 | 1.1117 | 0.25389 | 0.50449 | 1 | 10137 | 4 | 1.1117 |
| FMN1 | 6 | 0.86416 | 0.8798 | 0.999994 | 18525 | 1 | 0.42551 | 0.25407 | 0.50472 | 1 | 10138 | 2 | 0.42551 |
| KLRC4 | 6 | 0.50273 | 0.67061 | 0.999994 | 13475 | 2 | 0.6295 | 0.25407 | 0.50472 | 1 | 10139 | 4 | 0.6295 |

| | | | | | | | | | | | | | |
|---|---|---|---|---|---|---|---|---|---|---|---|---|---|
| CTDNEP1 | 5 | 0.38688 | 0.55252 | 0.999994 | 11750 | 2 | 0.72748 | 0.25416 | 0.47642 | 1 | 10140 | 3 | 0.72748 |
| HAUS6 | 5 | 0.081717 | 0.1974 | 0.977164 | 4288 | 3 | -1.9432 | 0.25416 | 0.47642 | 1 | 10141 | 1 | -1.9432 |
| OR1S1 | 5 | 0.41122 | 0.57288 | 0.999994 | 12132 | 2 | 0.0014746 | 0.25416 | 0.47642 | 1 | 10142 | 1 | 0.0014746 |
| HAS1 | 5 | 0.95111 | 0.95128 | 0.999994 | 20366 | 0 | 0.28412 | 0.25427 | 0.47654 | 1 | 10143 | 2 | 0.28412 |
| ZDHHC1 | 6 | 0.3025 | 0.50964 | 0.999994 | 10181 | 2 | 1.3376 | 0.2543 | 0.50502 | 1 | 10144 | 3 | 1.3376 |
| NKIRAS2 | 6 | 0.62253 | 0.75669 | 0.999994 | 15267 | 1 | 0.98656 | 0.2543 | 0.50502 | 1 | 10145 | 3 | 0.98656 |
| CCDC144A | 4 | 0.0075248 | 0.025891 | 0.848212 | 692 | 2 | -4.0109 | 0.25438 | 0.45812 | 1 | 10146 | 2 | -4.0109 |
| C19orf54 | 6 | 0.37099 | 0.58382 | 0.999994 | 11529 | 1 | 1.0434 | 0.25442 | 0.50517 | 1 | 10147 | 4 | 1.0434 |
| RECQL4 | 4 | 0.77612 | 0.78073 | 0.999994 | 17153 | 1 | 1.9182 | 0.25451 | 0.4583 | 1 | 10148 | 2 | 1.9182 |
| BRMS1L | 4 | 0.42053 | 0.56674 | 0.999994 | 12255 | 2 | 1.5631 | 0.25451 | 0.4583 | 1 | 10149 | 2 | 1.5631 |
| SMC1A | 4 | 0.69442 | 0.72083 | 0.999994 | 16246 | 1 | 2.0335 | 0.25451 | 0.4583 | 1 | 10150 | 2 | 2.0335 |
| ETF1 | 4 | 0.30201 | 0.44504 | 0.999994 | 10164 | 2 | 1.3036 | 0.25451 | 0.4583 | 1 | 10151 | 2 | 1.3036 |
| DPYS | 6 | 0.90253 | 0.90565 | 0.999994 | 19262 | 1 | 0.8737 | 0.25458 | 0.50538 | 1 | 10152 | 4 | 0.8737 |
| TTC29 | 5 | 0.92107 | 0.92126 | 0.999994 | 19653 | 0 | 1.5705 | 0.25461 | 0.4769 | 1 | 10153 | 3 | 1.5705 |
| hsa-mir-4729 | 4 | 0.35042 | 0.49471 | 0.999994 | 11168 | 2 | -0.041911 | 0.25465 | 0.45848 | 1 | 10154 | 2 | -0.041911 |
| MEMO1 | 5 | 0.89238 | 0.89252 | 0.999994 | 19076 | 0 | 1.8326 | 0.25465 | 0.47694 | 1 | 10155 | 3 | 1.8326 |
| GRM8 | 6 | 0.32989 | 0.54109 | 0.999994 | 10720 | 2 | 1.8043 | 0.25473 | 0.50556 | 1 | 10156 | 4 | 1.8043 |
| TP53I13 | 6 | 0.14189 | 0.31542 | 0.999994 | 6235 | 3 | 0.65975 | 0.25474 | 0.50559 | 1 | 10157 | 3 | 0.65975 |
| NCAPD2 | 4 | 0.057136 | 0.13529 | 0.954328 | 3299 | 3 | -3.2548 | 0.25491 | 0.45882 | 1 | 10158 | 1 | -3.2548 |
| DAND5 | 5 | 0.43051 | 0.58885 | 0.999994 | 12424 | 2 | 0.71555 | 0.25496 | 0.47729 | 1 | 10159 | 3 | 0.71555 |
| LMO4 | 5 | 0.25287 | 0.44306 | 0.999994 | 9081 | 2 | 0.22764 | 0.25496 | 0.47729 | 1 | 10160 | 2 | 0.22764 |
| TMEM138 | 6 | 0.29895 | 0.50547 | 0.999994 | 10111 | 3 | -0.62595 | 0.255 | 0.50593 | 1 | 10161 | 2 | -0.62595 |
| HOGA1 | 5 | 0.91549 | 0.91556 | 0.999994 | 19524 | 0 | 1.5936 | 0.255 | 0.47734 | 1 | 10162 | 3 | 1.5936 |
| hsa-mir-6128 | 3 | 0.62097 | 0.63343 | 0.999994 | 15237 | 1 | 1.9957 | 0.25503 | 0.40202 | 1 | 10163 | 2 | 1.9957 |
| HNRNPAB | 6 | 0.51469 | 0.67885 | 0.999994 | 13656 | 2 | 1.6147 | 0.25506 | 0.50601 | 1 | 10164 | 4 | 1.6147 |
| RAX | 5 | 0.58216 | 0.71976 | 0.999994 | 14709 | 1 | 1.6214 | 0.25509 | 0.47744 | 1 | 10165 | 3 | 1.6214 |
| KIAA1467 | 6 | 0.010425 | 0.040464 | 0.891621 | 897 | 3 | 0.39895 | 0.25517 | 0.50615 | 1 | 10166 | 3 | 0.39895 |
| ANKRD7 | 6 | 0.024939 | 0.086872 | 0.94395 | 1753 | 2 | 1.2331 | 0.2552 | 0.50619 | 1 | 10167 | 4 | 1.2331 |
| MTBP | 4 | 0.14614 | 0.28523 | 0.999632 | 6357 | 2 | 0.50622 | 0.25531 | 0.45934 | 1 | 10168 | 2 | 0.50622 |
| CLTCL1 | 6 | 0.57775 | 0.72354 | 0.999994 | 14656 | 2 | 0.98897 | 0.25536 | 0.50639 | 1 | 10169 | 4 | 0.98897 |
| ZNF286A | 6 | 0.50637 | 0.67314 | 0.999994 | 13532 | 2 | 1.0949 | 0.25536 | 0.5064 | 1 | 10170 | 4 | 1.0949 |
| CXorf27 | 4 | 0.38094 | 0.52612 | 0.999994 | 11661 | 2 | -0.28945 | 0.25541 | 0.45947 | 1 | 10171 | 1 | -0.28945 |
| ZDHHC23 | 4 | 0.13655 | 0.27535 | 0.999239 | 6097 | 2 | 0.26954 | 0.25541 | 0.45947 | 1 | 10172 | 2 | 0.26954 |
| hsa-mir-190b | 4 | 0.3288 | 0.47246 | 0.999994 | 10695 | 2 | 0.69617 | 0.25541 | 0.45947 | 1 | 10173 | 2 | 0.69617 |
| MEDAG | 6 | 0.72132 | 0.81874 | 0.999994 | 16511 | 1 | 1.1322 | 0.25545 | 0.50652 | 1 | 10174 | 4 | 1.1322 |
| PER3 | 5 | 0.091866 | 0.21355 | 0.979587 | 4674 | 2 | 1.4396 | 0.25547 | 0.47781 | 1 | 10175 | 3 | 1.4396 |
| LHCGR | 5 | 0.16111 | 0.31916 | 0.999994 | 6766 | 2 | 1.1368 | 0.25547 | 0.47781 | 1 | 10176 | 3 | 1.1368 |
| COPS2 | 5 | 0.23513 | 0.42432 | 0.999994 | 8649 | 3 | -0.67213 | 0.25547 | 0.47781 | 1 | 10177 | 2 | -0.67213 |
| CTPS1 | 6 | 0.29342 | 0.49903 | 0.999994 | 9983 | 2 | 1.1588 | 0.25547 | 0.50654 | 1 | 10178 | 3 | 1.1588 |
| ZNF784 | 6 | 0.49281 | 0.6638 | 0.999994 | 13309 | 2 | 1.1768 | 0.25547 | 0.50654 | 1 | 10179 | 3 | 1.1768 |
| LY6G5C | 6 | 0.9671 | 0.96714 | 0.999994 | 20765 | 0 | 1.7738 | 0.25553 | 0.50662 | 1 | 10180 | 4 | 1.7738 |
| GCLC | 6 | 0.081 | 0.20932 | 0.979587 | 4265 | 3 | 0.7637 | 0.25568 | 0.50682 | 1 | 10181 | 3 | 0.7637 |
| hsa-mir-520a | 4 | 0.87107 | 0.87068 | 0.999994 | 18656 | 0 | 2.0555 | 0.25572 | 0.45991 | 1 | 10182 | 3 | 2.0555 |
| FAM103A1 | 5 | 0.84362 | 0.84938 | 0.999994 | 18147 | 1 | 1.5833 | 0.25573 | 0.47811 | 1 | 10183 | 3 | 1.5833 |
| TNIP3 | 3 | 0.48999 | 0.5396 | 0.999994 | 13276 | 1 | -0.14039 | 0.25577 | 0.40279 | 1 | 10184 | 1 | -0.14039 |
| PIH1D1 | 6 | 0.83114 | 0.8621 | 0.999994 | 17948 | 1 | 0.78263 | 0.25578 | 0.50696 | 1 | 10185 | 4 | 0.78263 |
| OR3A2 | 6 | 0.44653 | 0.63278 | 0.999994 | 12670 | 2 | 1.1862 | 0.25578 | 0.50696 | 1 | 10186 | 4 | 1.1862 |
| SLC19A2 | 6 | 0.15866 | 0.33653 | 0.999994 | 6688 | 3 | -0.47187 | 0.25578 | 0.50696 | 1 | 10187 | 3 | -0.47187 |
| NonTargetingControlG | 1 | 0.74414 | 0.74442 | 0.999994 | 16789 | 0 | 3.5563 | 0.25586 | 0.25572 | 0.97466 | 10188 | 1 | 3.5563 |
| TPI1 | 4 | 0.20563 | 0.34645 | 0.999994 | 7914 | 1 | 0.88647 | 0.25591 | 0.46015 | 1 | 10189 | 3 | 0.88647 |
| C2orf43 | 4 | 0.17287 | 0.3128 | 0.999994 | 7060 | 1 | 1.6044 | 0.25593 | 0.46019 | 1 | 10190 | 3 | 1.6044 |
| RELL2 | 4 | 0.18449 | 0.32473 | 0.999994 | 7328 | 1 | 0.78195 | 0.25596 | 0.46022 | 1 | 10191 | 3 | 0.78195 |

| | | | | | | | | | | | | |
|---|---|---|---|---|---|---|---|---|---|---|---|---|
| HOXC6 | 5 | 0.55964 | 0.70032 | 0.999994 | 14363 | 2 | -0.54668 | 0.25597 | 0.47836 | 1 | 10192 | 2 | -0.54668 |
| RPS27A | 6 | 0.83947 | 0.86633 | 0.999994 | 18076 | 1 | 1.6599 | 0.25601 | 0.50727 | 1 | 10193 | 4 | 1.6599 |
| COL11A2 | 6 | 0.05172 | 0.14785 | 0.965554 | 3055 | 2 | 0.41646 | 0.25607 | 0.50735 | 1 | 10194 | 3 | 0.41646 |
| CXCR7 | 5 | 0.41464 | 0.57564 | 0.999994 | 12175 | 1 | 0.8205 | 0.25612 | 0.47853 | 1 | 10195 | 3 | 0.8205 |
| EXTL1 | 5 | 0.11994 | 0.25724 | 0.994943 | 5628 | 3 | -1.4118 | 0.25612 | 0.47853 | 1 | 10196 | 2 | -1.4118 |
| PYGO1 | 6 | 0.026157 | 0.089737 | 0.94395 | 1812 | 3 | 0.75686 | 0.25618 | 0.5075 | 1 | 10197 | 3 | 0.75686 |
| IRG1 | 5 | 0.50933 | 0.65605 | 0.999994 | 13570 | 2 | 1.6387 | 0.2562 | 0.47862 | 1 | 10198 | 3 | 1.6387 |
| KRTAP10-9 | 4 | 0.54153 | 0.63558 | 0.999994 | 14081 | 1 | 1.1019 | 0.25628 | 0.46067 | 1 | 10199 | 3 | 1.1019 |
| POLR1B | 6 | 0.037967 | 0.11731 | 0.951052 | 2421 | 2 | 0.56722 | 0.25634 | 0.50771 | 1 | 10200 | 3 | 0.56722 |
| ADRM1 | 5 | 0.27632 | 0.46189 | 0.999994 | 9626 | 2 | 1.1164 | 0.25643 | 0.47888 | 1 | 10201 | 3 | 1.1164 |
| C1QTNF9B-AS1 | 5 | 0.32068 | 0.49785 | 0.999994 | 10526 | 2 | 0.96201 | 0.25643 | 0.47888 | 1 | 10202 | 3 | 0.96201 |
| TIMM17B | 6 | 0.93303 | 0.93311 | 0.999994 | 19925 | 0 | 1.3123 | 0.25653 | 0.50798 | 1 | 10203 | 3 | 1.3123 |
| SGSM3 | 6 | 0.57318 | 0.72024 | 0.999994 | 14592 | 2 | 1.4561 | 0.25662 | 0.5081 | 1 | 10204 | 4 | 1.4561 |
| RHCG | 6 | 0.80971 | 0.85213 | 0.999994 | 17629 | 1 | 1.4357 | 0.25668 | 0.50817 | 1 | 10205 | 4 | 1.4357 |
| MARCH10 | 6 | 0.014088 | 0.052767 | 0.916123 | 1135 | 3 | 0.77633 | 0.25676 | 0.50828 | 1 | 10206 | 3 | 0.77633 |
| MAB21L1 | 6 | 0.049819 | 0.14365 | 0.961453 | 2965 | 1 | 0.29047 | 0.25679 | 0.50832 | 1 | 10207 | 2 | 0.29047 |
| UTF1 | 6 | 0.53094 | 0.6902 | 0.999994 | 13910 | 2 | -0.048807 | 0.25679 | 0.50832 | 1 | 10208 | 2 | -0.048807 |
| RNF138 | 6 | 0.33021 | 0.54149 | 0.999994 | 10726 | 2 | 0.48585 | 0.25679 | 0.50832 | 1 | 10209 | 3 | 0.48585 |
| UCMA | 6 | 0.011233 | 0.043185 | 0.891621 | 941 | 3 | -1.2979 | 0.25679 | 0.50832 | 1 | 10210 | 2 | -1.2979 |
| PXDN | 6 | 0.32767 | 0.53857 | 0.999994 | 10676 | 2 | 1.0665 | 0.25679 | 0.50832 | 1 | 10211 | 3 | 1.0665 |
| PLXDC1 | 6 | 0.033274 | 0.10647 | 0.950995 | 2176 | 3 | 0.023918 | 0.25679 | 0.50832 | 1 | 10212 | 3 | 0.023918 |
| GSTCD | 6 | 0.26197 | 0.46222 | 0.999994 | 9289 | 2 | 0.46867 | 0.25679 | 0.50832 | 1 | 10213 | 3 | 0.46867 |
| GLTSCR2 | 4 | 0.58562 | 0.65765 | 0.999994 | 14753 | 1 | 1.7226 | 0.2569 | 0.46148 | 1 | 10214 | 2 | 1.7226 |
| POLI | 5 | 0.34389 | 0.51692 | 0.999994 | 11026 | 2 | 0.58867 | 0.25701 | 0.4795 | 1 | 10215 | 2 | 0.58867 |
| HSFX2 | 3 | 0.14832 | 0.24633 | 0.98999 | 6415 | 2 | -1.7255 | 0.25713 | 0.40421 | 1 | 10216 | 1 | -1.7255 |
| hsa-mir-4639 | 4 | 0.62505 | 0.67917 | 0.999994 | 15308 | 1 | 1.2194 | 0.25718 | 0.46187 | 1 | 10217 | 3 | 1.2194 |
| RAB11B | 6 | 0.29901 | 0.50554 | 0.999994 | 10114 | 2 | 1.3821 | 0.25728 | 0.50896 | 1 | 10218 | 3 | 1.3821 |
| SLFN5 | 6 | 0.57564 | 0.72198 | 0.999994 | 14626 | 2 | 0.748 | 0.25754 | 0.5093 | 1 | 10219 | 4 | 0.748 |
| HNRNPM | 6 | 0.22301 | 0.41578 | 0.999994 | 8356 | 2 | 1.0854 | 0.25759 | 0.50936 | 1 | 10220 | 4 | 1.0854 |
| MANBAL | 6 | 0.42482 | 0.61855 | 0.999994 | 12333 | 2 | 1.2738 | 0.25765 | 0.50945 | 1 | 10221 | 3 | 1.2738 |
| USP17L10 | 6 | 0.32717 | 0.53802 | 0.999994 | 10664 | 2 | 1.0679 | 0.25766 | 0.50945 | 1 | 10222 | 4 | 1.0679 |
| MMP7 | 6 | 0.26851 | 0.46997 | 0.999994 | 9434 | 3 | -0.091631 | 0.25781 | 0.50967 | 1 | 10223 | 3 | -0.091631 |
| TES | 4 | 0.38869 | 0.53402 | 0.999994 | 11780 | 2 | 1.4197 | 0.25783 | 0.46273 | 1 | 10224 | 2 | 1.4197 |
| ELOVL7 | 6 | 0.46368 | 0.64422 | 0.999994 | 12916 | 1 | 1.4591 | 0.25784 | 0.50972 | 1 | 10225 | 3 | 1.4591 |
| USP6 | 6 | 0.19892 | 0.38657 | 0.999994 | 7735 | 2 | 1.4466 | 0.25797 | 0.50988 | 1 | 10226 | 4 | 1.4466 |
| FRRS1 | 5 | 0.83475 | 0.84289 | 0.999994 | 18000 | 1 | 1.1615 | 0.25799 | 0.48059 | 1 | 10227 | 3 | 1.1615 |
| CDK18 | 5 | 0.14104 | 0.28939 | 0.999632 | 6220 | 1 | 0.48737 | 0.25799 | 0.48059 | 1 | 10228 | 2 | 0.48737 |
| FBXO42 | 6 | 0.94661 | 0.94659 | 0.999994 | 20250 | 0 | 1.3893 | 0.25808 | 0.51004 | 1 | 10229 | 4 | 1.3893 |
| RSBN1 | 6 | 0.66618 | 0.79015 | 0.999994 | 15871 | 2 | 1.0032 | 0.25814 | 0.51012 | 1 | 10230 | 3 | 1.0032 |
| CD8B | 6 | 0.63849 | 0.7688 | 0.999994 | 15492 | 2 | 1.1491 | 0.25814 | 0.51012 | 1 | 10231 | 3 | 1.1491 |
| ZBTB4 | 6 | 0.33378 | 0.54561 | 0.999994 | 10806 | 1 | 1.0875 | 0.25814 | 0.51012 | 1 | 10232 | 3 | 1.0875 |
| EML3 | 5 | 0.033235 | 0.098827 | 0.945253 | 2173 | 2 | -0.27395 | 0.25822 | 0.48085 | 1 | 10233 | 2 | -0.27395 |
| OR10H3 | 5 | 0.79269 | 0.81496 | 0.999994 | 17386 | 1 | 0.45436 | 0.25822 | 0.48085 | 1 | 10234 | 2 | 0.45436 |
| hsa-mir-6777 | 3 | 0.82246 | 0.82236 | 0.999994 | 17813 | 0 | 2.1187 | 0.25829 | 0.40548 | 1 | 10235 | 2 | 2.1187 |
| CDH15 | 5 | 0.14486 | 0.29507 | 0.999994 | 6320 | 2 | 1.7137 | 0.2583 | 0.48094 | 1 | 10236 | 3 | 1.7137 |
| hsa-mir-135a-2 | 4 | 0.87126 | 0.87088 | 0.999994 | 18663 | 0 | 0.92023 | 0.25831 | 0.46338 | 1 | 10237 | 3 | 0.92023 |
| BTN1A1 | 6 | 0.4955 | 0.66562 | 0.999994 | 13347 | 2 | 0.0078864 | 0.25834 | 0.51036 | 1 | 10238 | 1 | 0.0078864 |
| hsa-mir-548au | 3 | 0.31809 | 0.4383 | 0.999994 | 10477 | 1 | 2.1702 | 0.25842 | 0.40562 | 1 | 10239 | 2 | 2.1702 |
| hsa-mir-4799 | 4 | 0.78982 | 0.79223 | 0.999994 | 17338 | 1 | 2.051 | 0.25851 | 0.46366 | 1 | 10240 | 3 | 2.051 |
| SLC44A5 | 4 | 0.39879 | 0.54448 | 0.999994 | 11930 | 1 | 0.45403 | 0.25852 | 0.46366 | 1 | 10241 | 2 | 0.45403 |
| hsa-mir-1-2 | 4 | 0.87888 | 0.87853 | 0.999994 | 18809 | 0 | 0.7878 | 0.25857 | 0.46372 | 1 | 10242 | 3 | 0.7878 |
| FOXS1 | 6 | 0.32179 | 0.53196 | 0.999994 | 10549 | 2 | 1.6153 | 0.25869 | 0.51081 | 1 | 10243 | 4 | 1.6153 |

| | | | | | | | | | | | | | |
|---|---|---|---|---|---|---|---|---|---|---|---|---|---|
| FNDC3B | 6 | 0.0033613 | 0.015052 | 0.757255 | 391 | 2 | -0.014563 | 0.25871 | 0.51083 | 1 | 10244 | 2 | -0.014563 |
| PTPLA | 6 | 0.78296 | 0.84088 | 0.999994 | 17249 | 1 | 0.57649 | 0.25871 | 0.51083 | 1 | 10245 | 2 | 0.57649 |
| FBXO2 | 5 | 0.95492 | 0.95509 | 0.999994 | 20451 | 0 | 0.85955 | 0.25874 | 0.48144 | 1 | 10246 | 3 | 0.85955 |
| CIB1 | 5 | 0.81302 | 0.82791 | 0.999994 | 17673 | 1 | 1.817 | 0.25874 | 0.48144 | 1 | 10247 | 3 | 1.817 |
| HHLA2 | 5 | 0.055979 | 0.15477 | 0.967126 | 3245 | 3 | -2.0184 | 0.25874 | 0.48144 | 1 | 10248 | 1 | -2.0184 |
| OR2T8 | 6 | 0.67719 | 0.79878 | 0.999994 | 16047 | 1 | 1.1834 | 0.25877 | 0.51091 | 1 | 10249 | 4 | 1.1834 |
| ZNF627 | 5 | 0.48364 | 0.6338 | 0.999994 | 13194 | 2 | 1.6093 | 0.25881 | 0.48151 | 1 | 10250 | 3 | 1.6093 |
| PTGDR | 3 | 0.32602 | 0.44258 | 0.999994 | 10642 | 1 | 2.0152 | 0.25898 | 0.40624 | 1 | 10251 | 2 | 2.0152 |
| WDR49 | 6 | 0.65108 | 0.77845 | 0.999994 | 15677 | 2 | 1.5287 | 0.25902 | 0.51123 | 1 | 10252 | 4 | 1.5287 |
| FAM83H | 6 | 0.93855 | 0.9385 | 0.999994 | 20039 | 0 | 1.1503 | 0.25904 | 0.51126 | 1 | 10253 | 4 | 1.1503 |
| KRTAP19-2 | 4 | 0.14777 | 0.28686 | 0.999632 | 6400 | 2 | -0.22858 | 0.25904 | 0.46436 | 1 | 10254 | 2 | -0.22858 |
| CD276 | 6 | 0.94995 | 0.94992 | 0.999994 | 20330 | 0 | 0.91957 | 0.25916 | 0.51141 | 1 | 10255 | 4 | 0.91957 |
| BMP4 | 6 | 0.079261 | 0.20583 | 0.978744 | 4201 | 4 | -1.0849 | 0.25917 | 0.51143 | 1 | 10256 | 2 | -1.0849 |
| OTP | 6 | 0.20319 | 0.39174 | 0.999994 | 7860 | 1 | 1.6441 | 0.25922 | 0.51149 | 1 | 10257 | 4 | 1.6441 |
| CDC7 | 3 | 0.73484 | 0.73469 | 0.999994 | 16659 | 0 | 1.9552 | 0.25924 | 0.40654 | 1 | 10258 | 2 | 1.9552 |
| ABHD17A | 3 | 0.099162 | 0.18195 | 0.968344 | 4936 | 2 | -2.1852 | 0.25932 | 0.40661 | 1 | 10259 | 1 | -2.1852 |
| RPL38 | 5 | 0.020833 | 0.066322 | 0.93914 | 1500 | 3 | -1.9853 | 0.25951 | 0.48225 | 1 | 10260 | 2 | -1.9853 |
| hsa-mir-514b | 4 | 0.60871 | 0.67005 | 0.999994 | 15052 | 1 | 0.82848 | 0.25961 | 0.46508 | 1 | 10261 | 3 | 0.82848 |
| EBI3 | 6 | 0.091822 | 0.23117 | 0.983933 | 4672 | 3 | -0.52923 | 0.25964 | 0.51202 | 1 | 10262 | 2 | -0.52923 |
| ENTPD6 | 6 | 0.071412 | 0.18973 | 0.970473 | 3889 | 3 | -0.92176 | 0.25967 | 0.51206 | 1 | 10263 | 2 | -0.92176 |
| DLK2 | 5 | 0.37315 | 0.54102 | 0.999994 | 11560 | 1 | 0.97961 | 0.25968 | 0.48243 | 1 | 10264 | 3 | 0.97961 |
| UPF1 | 4 | 0.050061 | 0.12158 | 0.951052 | 2983 | 2 | -0.31893 | 0.25969 | 0.46517 | 1 | 10265 | 2 | -0.31893 |
| FLJ45513 | 4 | 0.26772 | 0.40994 | 0.999994 | 9417 | 2 | -0.41469 | 0.25969 | 0.46518 | 1 | 10266 | 1 | -0.41469 |
| LIMA1 | 4 | 0.048425 | 0.11836 | 0.951052 | 2903 | 2 | -0.0064694 | 0.25969 | 0.46518 | 1 | 10267 | 2 | -0.0064694 |
| CD200 | 6 | 0.44474 | 0.63159 | 0.999994 | 12644 | 2 | 1.0573 | 0.25979 | 0.51222 | 1 | 10268 | 4 | 1.0573 |
| BTG3 | 4 | 0.67094 | 0.70595 | 0.999994 | 15944 | 1 | 1.8388 | 0.25988 | 0.46544 | 1 | 10269 | 3 | 1.8388 |
| OIT3 | 6 | 0.022333 | 0.079457 | 0.93914 | 1603 | 2 | 1.655 | 0.25994 | 0.51242 | 1 | 10270 | 4 | 1.655 |
| MUM1 | 6 | 0.059401 | 0.16449 | 0.967126 | 3399 | 2 | 1.604 | 0.25998 | 0.51247 | 1 | 10271 | 3 | 1.604 |
| PCDH18 | 5 | 0.057662 | 0.15819 | 0.967126 | 3321 | 2 | 1.6361 | 0.25999 | 0.48275 | 1 | 10272 | 3 | 1.6361 |
| DIXDC1 | 6 | 0.62497 | 0.75858 | 0.999994 | 15305 | 2 | 0.89813 | 0.25999 | 0.51248 | 1 | 10273 | 4 | 0.89813 |
| HEBP1 | 6 | 0.22249 | 0.41515 | 0.999994 | 8342 | 2 | 1.2587 | 0.26008 | 0.51259 | 1 | 10274 | 4 | 1.2587 |
| TRPV4 | 6 | 0.069209 | 0.18511 | 0.968626 | 3806 | 3 | -0.71916 | 0.26008 | 0.51259 | 1 | 10275 | 3 | -0.71916 |
| HIST1H4H | 6 | 0.097411 | 0.24227 | 0.988088 | 4871 | 4 | -0.99378 | 0.26008 | 0.51259 | 1 | 10276 | 1 | -0.99378 |
| RHAG | 6 | 0.80061 | 0.84813 | 0.999994 | 17490 | 1 | -0.044717 | 0.26008 | 0.51259 | 1 | 10277 | 2 | -0.044717 |
| PODN | 5 | 0.2317 | 0.41957 | 0.999994 | 8557 | 2 | 1.1048 | 0.26018 | 0.48297 | 1 | 10278 | 3 | 1.1048 |
| B3GALT2 | 6 | 0.007156 | 0.028988 | 0.859137 | 667 | 4 | -2.6938 | 0.26024 | 0.51282 | 1 | 10279 | 2 | -2.6938 |
| TNK2 | 6 | 0.56527 | 0.71457 | 0.999994 | 14471 | 1 | 0.58657 | 0.26024 | 0.51282 | 1 | 10280 | 3 | 0.58657 |
| RYR3 | 4 | 0.27596 | 0.41844 | 0.999994 | 9615 | 1 | 1.4459 | 0.26028 | 0.46598 | 1 | 10281 | 3 | 1.4459 |
| SCN2A | 4 | 0.50904 | 0.62037 | 0.999994 | 13569 | 1 | 0.90316 | 0.26037 | 0.46612 | 1 | 10282 | 3 | 0.90316 |
| hsa-mir-590 | 3 | 0.70122 | 0.70216 | 0.999994 | 16319 | 1 | 2.0165 | 0.2604 | 0.40773 | 1 | 10283 | 2 | 2.0165 |
| SLCO1C1 | 6 | 0.60458 | 0.74338 | 0.999994 | 15003 | 2 | 0.40905 | 0.26054 | 0.51321 | 1 | 10284 | 3 | 0.40905 |
| DCDC2 | 6 | 0.62321 | 0.75721 | 0.999994 | 15279 | 2 | 0.066236 | 0.26054 | 0.51321 | 1 | 10285 | 2 | 0.066236 |
| LUZP4 | 5 | 0.12589 | 0.26635 | 0.998405 | 5817 | 3 | -1.267 | 0.26059 | 0.48341 | 1 | 10286 | 2 | -1.267 |
| FAF1 | 5 | 0.64585 | 0.74345 | 0.999994 | 15599 | 1 | 0.011517 | 0.26059 | 0.48341 | 1 | 10287 | 2 | 0.011517 |
| CT62 | 6 | 0.15555 | 0.33264 | 0.999994 | 6608 | 2 | 1.5927 | 0.2607 | 0.51342 | 1 | 10288 | 4 | 1.5927 |
| ACTRT1 | 6 | 0.33406 | 0.54591 | 0.999994 | 10814 | 2 | 0.81999 | 0.26072 | 0.51344 | 1 | 10289 | 4 | 0.81999 |
| AMY1C | 3 | 0.59664 | 0.61459 | 0.999994 | 14907 | 1 | -0.56303 | 0.26073 | 0.40809 | 1 | 10290 | 1 | -0.56303 |
| CTXN2 | 6 | 0.79528 | 0.84587 | 0.999994 | 17422 | 1 | 1.1135 | 0.26076 | 0.51349 | 1 | 10291 | 3 | 1.1135 |
| MBD3L2 | 4 | 0.86892 | 0.86849 | 0.999994 | 18615 | 0 | 1.0332 | 0.26077 | 0.46663 | 1 | 10292 | 3 | 1.0332 |
| WFDC5 | 5 | 0.21327 | 0.39398 | 0.999994 | 8095 | 1 | 1.5488 | 0.26078 | 0.48363 | 1 | 10293 | 3 | 1.5488 |
| SPRYD7 | 4 | 0.0040481 | 0.014371 | 0.753239 | 443 | 1 | 1.1658 | 0.26081 | 0.46669 | 1 | 10294 | 3 | 1.1658 |
| METTL21C | 6 | 0.66547 | 0.78958 | 0.999994 | 15861 | 2 | 1.0623 | 0.26081 | 0.51358 | 1 | 10295 | 4 | 1.0623 |

| | | | | | | | | | | | | | |
|---|---|---|---|---|---|---|---|---|---|---|---|---|---|
| hsa-mir-3689d-2 | 4 | 0.37453 | 0.51953 | 0.999994 | 11575 | 2 | -0.45977 | 0.26088 | 0.46678 | 1 | 10296 | 1 | -0.45977 |
| hsa-mir-6781 | 4 | 0.41625 | 0.56231 | 0.999994 | 12196 | 2 | 1.5081 | 0.26093 | 0.46685 | 1 | 10297 | 2 | 1.5081 |
| OPCML | 4 | 0.76158 | 0.76907 | 0.999994 | 16997 | 1 | 1.7469 | 0.26095 | 0.46688 | 1 | 10298 | 2 | 1.7469 |
| HMSD | 6 | 0.46036 | 0.64202 | 0.999994 | 12852 | 2 | 1.0608 | 0.26104 | 0.51388 | 1 | 10299 | 3 | 1.0608 |
| hsa-mir-181a-2 | 4 | 0.19075 | 0.33116 | 0.999994 | 7505 | 1 | 1.5948 | 0.26109 | 0.46706 | 1 | 10300 | 2 | 1.5948 |
| hsa-mir-1911 | 4 | 0.58816 | 0.65896 | 0.999994 | 14792 | 1 | 0.79618 | 0.26115 | 0.46713 | 1 | 10301 | 3 | 0.79618 |
| KCNE3 | 4 | 0.46929 | 0.60282 | 0.999994 | 12993 | 1 | 1.0428 | 0.26121 | 0.46721 | 1 | 10302 | 3 | 1.0428 |
| HOXB3 | 5 | 0.10442 | 0.23328 | 0.984913 | 5112 | 3 | -1.6082 | 0.26122 | 0.48409 | 1 | 10303 | 2 | -1.6082 |
| ABHD5 | 5 | 0.42417 | 0.58357 | 0.999994 | 12312 | 2 | 0.56455 | 0.26122 | 0.48409 | 1 | 10304 | 3 | 0.56455 |
| FOXE3 | 5 | 0.0093587 | 0.034213 | 0.867348 | 817 | 4 | -6.027 | 0.26122 | 0.48409 | 1 | 10305 | 1 | -6.027 |
| BRPF3 | 5 | 0.43047 | 0.58882 | 0.999994 | 12423 | 2 | 0.65504 | 0.26126 | 0.48413 | 1 | 10306 | 3 | 0.65504 |
| APOD | 5 | 0.19847 | 0.37308 | 0.999994 | 7724 | 3 | -0.82023 | 0.26126 | 0.48413 | 1 | 10307 | 2 | -0.82023 |
| TIMM17A | 6 | 0.036806 | 0.1147 | 0.951052 | 2366 | 4 | -1.6717 | 0.26127 | 0.51418 | 1 | 10308 | 1 | -1.6717 |
| MAGEA2 | 6 | 0.29872 | 0.5052 | 0.999994 | 10107 | 3 | -0.32131 | 0.26127 | 0.51418 | 1 | 10309 | 2 | -0.32131 |
| PIK3C2B | 6 | 0.40695 | 0.6069 | 0.999994 | 12060 | 2 | 1.1702 | 0.26135 | 0.51429 | 1 | 10310 | 4 | 1.1702 |
| CCDC36 | 6 | 0.56436 | 0.71392 | 0.999994 | 14463 | 2 | 0.4183 | 0.26138 | 0.51434 | 1 | 10311 | 3 | 0.4183 |
| CKMT1A | 6 | 0.14071 | 0.31394 | 0.999994 | 6208 | 3 | -0.11867 | 0.26138 | 0.51434 | 1 | 10312 | 2 | -0.11867 |
| CAPNS2 | 6 | 0.070008 | 0.18678 | 0.969409 | 3827 | 4 | -1.4691 | 0.26138 | 0.51434 | 1 | 10313 | 2 | -1.4691 |
| TBL3 | 5 | 0.095196 | 0.21885 | 0.980699 | 4791 | 2 | 1.8422 | 0.26139 | 0.48427 | 1 | 10314 | 3 | 1.8422 |
| SCARB1 | 6 | 0.21579 | 0.4071 | 0.999994 | 8161 | 2 | 1.4379 | 0.2614 | 0.51436 | 1 | 10315 | 4 | 1.4379 |
| ANKMY1 | 5 | 0.33501 | 0.50962 | 0.999994 | 10833 | 1 | 1.5379 | 0.26141 | 0.48429 | 1 | 10316 | 3 | 1.5379 |
| TRIM58 | 3 | 0.87139 | 0.87146 | 0.999994 | 18668 | 0 | 0.48812 | 0.26142 | 0.40882 | 1 | 10317 | 1 | 0.48812 |
| hsa-mir-1-1 | 4 | 0.59884 | 0.66469 | 0.999994 | 14938 | 1 | 0.94047 | 0.26148 | 0.46754 | 1 | 10318 | 3 | 0.94047 |
| hsa-mir-3646 | 4 | 0.64573 | 0.69082 | 0.999994 | 15597 | 1 | 1.1137 | 0.26154 | 0.46763 | 1 | 10319 | 3 | 1.1137 |
| SLC25A15 | 6 | 0.27541 | 0.47807 | 0.999994 | 9601 | 2 | 1.0572 | 0.26158 | 0.51461 | 1 | 10320 | 4 | 1.0572 |
| MTRNR2L1 | 4 | 0.85251 | 0.85206 | 0.999994 | 18306 | 0 | 1.5923 | 0.26161 | 0.46772 | 1 | 10321 | 2 | 1.5923 |
| MARK3 | 6 | 0.04517 | 0.13345 | 0.95318 | 2768 | 2 | 1.035 | 0.26162 | 0.51466 | 1 | 10322 | 3 | 1.035 |
| RGS1 | 6 | 0.023997 | 0.084597 | 0.94395 | 1708 | 2 | 1.4994 | 0.26163 | 0.51468 | 1 | 10323 | 4 | 1.4994 |
| OTC | 5 | 0.39172 | 0.55659 | 0.999994 | 11825 | 2 | 1.6522 | 0.26167 | 0.48459 | 1 | 10324 | 3 | 1.6522 |
| SNRPD3 | 1 | 0.73829 | 0.73851 | 0.999994 | 16711 | 0 | 3.4241 | 0.26171 | 0.26166 | 0.974875 | 10325 | 1 | 3.4241 |
| HIST2H3C | 1 | 0.73829 | 0.73851 | 0.999994 | 16710 | 0 | 3.4241 | 0.26171 | 0.26166 | 0.974875 | 10326 | 1 | 3.4241 |
| HAGHL | 4 | 0.50747 | 0.61971 | 0.999994 | 13543 | 1 | -0.090384 | 0.26174 | 0.46788 | 1 | 10327 | 1 | -0.090384 |
| USP17L7 | 6 | 0.3033 | 0.51057 | 0.999994 | 10200 | 2 | 1.1638 | 0.26174 | 0.51482 | 1 | 10328 | 4 | 1.1638 |
| ZNF573 | 5 | 0.20761 | 0.38601 | 0.999994 | 7964 | 2 | 1.4878 | 0.26177 | 0.48471 | 1 | 10329 | 3 | 1.4878 |
| hsa-mir-548as | 4 | 0.14614 | 0.28523 | 0.999632 | 6356 | 1 | 0.78055 | 0.26178 | 0.46794 | 1 | 10330 | 3 | 0.78055 |
| hsa-mir-4492 | 2 | 0.43511 | 0.4464 | 0.999994 | 12501 | 1 | 2.4016 | 0.26186 | 0.3816 | 1 | 10331 | 1 | 2.4016 |
| NonTargetingControlG | 2 | 0.15796 | 0.23224 | 0.984502 | 6670 | 1 | 0.4333 | 0.26186 | 0.3816 | 1 | 10332 | 1 | 0.4333 |
| MOG | 4 | 0.054461 | 0.13019 | 0.95318 | 3174 | 3 | -0.6801 | 0.26198 | 0.46819 | 1 | 10333 | 1 | -0.6801 |
| hsa-mir-3166 | 4 | 0.11463 | 0.24097 | 0.987253 | 5468 | 2 | -0.78793 | 0.26198 | 0.46819 | 1 | 10334 | 2 | -0.78793 |
| hsa-mir-196a-1 | 3 | 0.83408 | 0.83397 | 0.999994 | 17989 | 0 | 0.28084 | 0.26201 | 0.40943 | 1 | 10335 | 1 | 0.28084 |
| hsa-mir-4446 | 3 | 0.39329 | 0.48037 | 0.999994 | 11850 | 1 | 1.817 | 0.26201 | 0.40943 | 1 | 10336 | 2 | 1.817 |
| MAML3 | 5 | 0.20023 | 0.37557 | 0.999994 | 7768 | 2 | 1.7611 | 0.26202 | 0.48494 | 1 | 10337 | 3 | 1.7611 |
| FAM76A | 6 | 0.77446 | 0.8375 | 0.999994 | 17137 | 1 | 1.011 | 0.26203 | 0.51519 | 1 | 10338 | 3 | 1.011 |
| CD82 | 6 | 0.15813 | 0.33588 | 0.999994 | 6677 | 2 | 1.0842 | 0.26203 | 0.51519 | 1 | 10339 | 3 | 1.0842 |
| TTC34 | 6 | 0.10052 | 0.24837 | 0.990689 | 4992 | 3 | 0.3407 | 0.26213 | 0.51533 | 1 | 10340 | 3 | 0.3407 |
| LARGE | 6 | 0.016915 | 0.062004 | 0.937033 | 1297 | 4 | -2.1175 | 0.26217 | 0.51538 | 1 | 10341 | 2 | -2.1175 |
| GPR89B | 1 | 0.7378 | 0.738 | 0.999994 | 16698 | 0 | 3.2991 | 0.2622 | 0.26215 | 0.974875 | 10342 | 1 | 3.2991 |
| CD180 | 6 | 0.79166 | 0.84439 | 0.999994 | 17375 | 1 | 1.1908 | 0.26222 | 0.51544 | 1 | 10343 | 4 | 1.1908 |
| SIK2 | 6 | 0.57073 | 0.71846 | 0.999994 | 14552 | 2 | 1.1782 | 0.26228 | 0.51553 | 1 | 10344 | 4 | 1.1782 |
| UBIAD1 | 6 | 0.20195 | 0.39025 | 0.999994 | 7828 | 2 | 1.0363 | 0.26233 | 0.51559 | 1 | 10345 | 4 | 1.0363 |
| GLRA2 | 6 | 0.66256 | 0.7873 | 0.999994 | 15821 | 1 | 1.3371 | 0.26236 | 0.51562 | 1 | 10346 | 4 | 1.3371 |
| hsa-mir-7111 | 3 | 0.25057 | 0.37568 | 0.999994 | 9027 | 1 | 2.0492 | 0.26237 | 0.40981 | 1 | 10347 | 2 | 2.0492 |

| | | | | | | | | | | | | | |
|---|---|---|---|---|---|---|---|---|---|---|---|---|---|
| MYO1F | 6 | 0.96549 | 0.96554 | 0.999994 | 20729 | 0 | 0.97854 | 0.26237 | 0.51563 | 1 | 10348 | 4 | 0.97854 |
| hsa-mir-6839 | 4 | 0.77357 | 0.77866 | 0.999994 | 17127 | 1 | 1.0447 | 0.2624 | 0.46876 | 1 | 10349 | 3 | 1.0447 |
| PPP1R8 | 4 | 0.11096 | 0.23456 | 0.984971 | 5331 | 2 | -1.5654 | 0.26243 | 0.46879 | 1 | 10350 | 1 | -1.5654 |
| SPZ1 | 6 | 0.14302 | 0.31689 | 0.999994 | 6274 | 1 | 1.0354 | 0.26243 | 0.51571 | 1 | 10351 | 3 | 1.0354 |
| PARP1 | 5 | 0.84928 | 0.8537 | 0.999994 | 18246 | 1 | 1.5784 | 0.26243 | 0.48541 | 1 | 10352 | 3 | 1.5784 |
| SRGAP2B | 6 | 0.084021 | 0.21554 | 0.980236 | 4377 | 3 | -0.51867 | 0.2625 | 0.51579 | 1 | 10353 | 2 | -0.51867 |
| SLFN11 | 6 | 0.24944 | 0.44725 | 0.999994 | 9004 | 2 | 0.020619 | 0.2625 | 0.51579 | 1 | 10354 | 2 | 0.020619 |
| LEFTY1 | 6 | 0.13809 | 0.31061 | 0.999994 | 6139 | 2 | 1.3137 | 0.2625 | 0.5158 | 1 | 10355 | 4 | 1.3137 |
| MISP | 2 | 0.35048 | 0.37639 | 0.999994 | 11170 | 1 | 2.0435 | 0.26257 | 0.38203 | 1 | 10356 | 1 | 2.0435 |
| hsa-mir-659 | 4 | 0.80713 | 0.80767 | 0.999994 | 17588 | 1 | 1.8739 | 0.26269 | 0.46915 | 1 | 10357 | 2 | 1.8739 |
| LOC730159 | 4 | 0.069567 | 0.15908 | 0.967126 | 3818 | 2 | -0.14143 | 0.26269 | 0.46915 | 1 | 10358 | 2 | -0.14143 |
| hsa-mir-3192 | 4 | 0.40603 | 0.55186 | 0.999994 | 12045 | 2 | 1.4287 | 0.26269 | 0.46915 | 1 | 10359 | 2 | 1.4287 |
| hsa-mir-4681 | 4 | 0.22144 | 0.36273 | 0.999994 | 8317 | 1 | 1.8076 | 0.26269 | 0.46915 | 1 | 10360 | 2 | 1.8076 |
| DNTTIP2 | 4 | 0.81346 | 0.81356 | 0.999994 | 17683 | 1 | 1.9288 | 0.26269 | 0.46915 | 1 | 10361 | 2 | 1.9288 |
| TMEM200A | 5 | 0.90463 | 0.90471 | 0.999994 | 19313 | 0 | 0.77547 | 0.26273 | 0.48573 | 1 | 10362 | 3 | 0.77547 |
| DDX54 | 5 | 0.1172 | 0.25307 | 0.994501 | 5546 | 2 | 0.69752 | 0.26273 | 0.48573 | 1 | 10363 | 3 | 0.69752 |
| LYN | 5 | 0.033427 | 0.099303 | 0.945253 | 2184 | 2 | 0.35554 | 0.26273 | 0.48573 | 1 | 10364 | 2 | 0.35554 |
| C7orf49 | 5 | 0.40439 | 0.5672 | 0.999994 | 12027 | 2 | 1.7536 | 0.26273 | 0.48573 | 1 | 10365 | 3 | 1.7536 |
| ABCC6 | 6 | 0.59193 | 0.73404 | 0.999994 | 14847 | 2 | 0.031931 | 0.26281 | 0.51619 | 1 | 10366 | 2 | 0.031931 |
| SPAST | 6 | 0.48835 | 0.66083 | 0.999994 | 13259 | 1 | 1.0461 | 0.26281 | 0.51619 | 1 | 10367 | 3 | 1.0461 |
| SDS | 6 | 0.54829 | 0.70239 | 0.999994 | 14183 | 2 | 1.0497 | 0.26282 | 0.5162 | 1 | 10368 | 4 | 1.0497 |
| UNC13D | 5 | 0.061994 | 0.16539 | 0.967126 | 3513 | 2 | 1.6386 | 0.26283 | 0.48584 | 1 | 10369 | 3 | 1.6386 |
| TXNDC5 | 6 | 0.32233 | 0.53258 | 0.999994 | 10564 | 3 | -0.1586 | 0.26285 | 0.51623 | 1 | 10370 | 2 | -0.1586 |
| hsa-mir-4704 | 3 | 0.42783 | 0.5008 | 0.999994 | 12379 | 1 | 1.7742 | 0.26287 | 0.41036 | 1 | 10371 | 2 | 1.7742 |
| RRP7A | 4 | 0.41986 | 0.56606 | 0.999994 | 12251 | 2 | -0.42826 | 0.26292 | 0.46947 | 1 | 10372 | 1 | -0.42826 |
| KCNT2 | 5 | 0.41217 | 0.57365 | 0.999994 | 12148 | 2 | 1.7309 | 0.26306 | 0.48609 | 1 | 10373 | 3 | 1.7309 |
| POMT2 | 5 | 0.17613 | 0.34104 | 0.999994 | 7137 | 3 | -0.63532 | 0.26307 | 0.48611 | 1 | 10374 | 1 | -0.63532 |
| HIBCH | 6 | 0.17203 | 0.35324 | 0.999994 | 7038 | 2 | 1.0571 | 0.26324 | 0.51675 | 1 | 10375 | 4 | 1.0571 |
| PCDHB9 | 6 | 0.61057 | 0.74775 | 0.999994 | 15079 | 2 | 1.3673 | 0.26329 | 0.51681 | 1 | 10376 | 3 | 1.3673 |
| ZBTB7B | 6 | 0.30937 | 0.51764 | 0.999994 | 10314 | 2 | 1.1133 | 0.26331 | 0.51684 | 1 | 10377 | 4 | 1.1133 |
| ZNF783 | 6 | 0.59591 | 0.73696 | 0.999994 | 14901 | 1 | 1.1082 | 0.26337 | 0.51691 | 1 | 10378 | 4 | 1.1082 |
| CRMP1 | 4 | 0.12863 | 0.26525 | 0.997396 | 5901 | 2 | -0.84058 | 0.2634 | 0.47012 | 1 | 10379 | 2 | -0.84058 |
| PRDM13 | 6 | 0.049439 | 0.14282 | 0.959553 | 2951 | 1 | 1.1872 | 0.2634 | 0.51696 | 1 | 10380 | 4 | 1.1872 |
| MAP4K3 | 6 | 0.20127 | 0.38941 | 0.999994 | 7796 | 2 | 0.76474 | 0.26348 | 0.51706 | 1 | 10381 | 4 | 0.76474 |
| RBL2 | 5 | 0.83733 | 0.8447 | 0.999994 | 18038 | 1 | 1.6204 | 0.26356 | 0.48666 | 1 | 10382 | 3 | 1.6204 |
| SPRYD3 | 5 | 0.80169 | 0.82057 | 0.999994 | 17505 | 1 | 0.022001 | 0.26358 | 0.48668 | 1 | 10383 | 1 | 0.022001 |
| PELO | 5 | 0.86582 | 0.8674 | 0.999994 | 18552 | 1 | 1.7731 | 0.2636 | 0.4867 | 1 | 10384 | 3 | 1.7731 |
| LRCH2 | 5 | 0.81465 | 0.82904 | 0.999994 | 17697 | 1 | 1.4121 | 0.26361 | 0.48672 | 1 | 10385 | 3 | 1.4121 |
| C1orf216 | 6 | 0.24999 | 0.44793 | 0.999994 | 9013 | 2 | 1.0684 | 0.26363 | 0.51725 | 1 | 10386 | 3 | 1.0684 |
| C16orf80 | 5 | 0.044587 | 0.12729 | 0.951052 | 2741 | 2 | -0.3951 | 0.26363 | 0.48674 | 1 | 10387 | 2 | -0.3951 |
| ARL10 | 6 | 0.16862 | 0.34902 | 0.999994 | 6950 | 3 | 0.42457 | 0.26366 | 0.51729 | 1 | 10388 | 3 | 0.42457 |
| RS1 | 6 | 0.12214 | 0.28981 | 0.999632 | 5705 | 4 | -1.0308 | 0.26368 | 0.51732 | 1 | 10389 | 1 | -1.0308 |
| CTNNA1 | 4 | 0.51329 | 0.62227 | 0.999994 | 13624 | 1 | 1.2956 | 0.26374 | 0.47055 | 1 | 10390 | 3 | 1.2956 |
| UBR4 | 6 | 0.71956 | 0.81819 | 0.999994 | 16492 | 1 | 1.3012 | 0.26379 | 0.51747 | 1 | 10391 | 3 | 1.3012 |
| IL1R1 | 4 | 0.056429 | 0.13399 | 0.95318 | 3267 | 2 | -0.33209 | 0.26379 | 0.4706 | 1 | 10392 | 2 | -0.33209 |
| ATG4A | 4 | 0.88892 | 0.8886 | 0.999994 | 19019 | 0 | 1.2773 | 0.26393 | 0.47078 | 1 | 10393 | 3 | 1.2773 |
| ZMAT4 | 6 | 0.94005 | 0.94 | 0.999994 | 20081 | 0 | 1.1678 | 0.26405 | 0.5178 | 1 | 10394 | 4 | 1.1678 |
| COLEC11 | 3 | 0.7639 | 0.76389 | 0.999994 | 17011 | 0 | 0.75406 | 0.26411 | 0.41164 | 1 | 10395 | 2 | 0.75406 |
| LYPD8 | 5 | 0.17413 | 0.33813 | 0.999994 | 7087 | 2 | 1.4772 | 0.26421 | 0.48738 | 1 | 10396 | 3 | 1.4772 |
| CCDC160 | 6 | 0.15472 | 0.33164 | 0.999994 | 6587 | 3 | -0.41377 | 0.26423 | 0.51804 | 1 | 10397 | 3 | -0.41377 |
| GK | 6 | 0.49682 | 0.66654 | 0.999994 | 13371 | 1 | 0.54494 | 0.26423 | 0.51804 | 1 | 10398 | 3 | 0.54494 |
| C6orf10 | 6 | 0.28569 | 0.49009 | 0.999994 | 9823 | 2 | 0.9752 | 0.26425 | 0.51807 | 1 | 10399 | 4 | 0.9752 |

| | | | | | | | | | | | | | |
|---|---|---|---|---|---|---|---|---|---|---|---|---|---|
| PARS2 | 5 | 0.53443 | 0.67796 | 0.999994 | 13971 | 2 | -0.34004 | 0.26431 | 0.48749 | 1 | 10400 | 2 | -0.34004 |
| OTUD1 | 5 | 0.51896 | 0.66441 | 0.999994 | 13728 | 2 | -0.020772 | 0.26431 | 0.48749 | 1 | 10401 | 2 | -0.020772 |
| PHRF1 | 5 | 0.75502 | 0.7935 | 0.999994 | 16913 | 1 | 1.5648 | 0.26431 | 0.48749 | 1 | 10402 | 3 | 1.5648 |
| C11orf31 | 6 | 0.95684 | 0.95691 | 0.999994 | 20515 | 0 | 1.4411 | 0.26431 | 0.51816 | 1 | 10403 | 3 | 1.4411 |
| CSH1 | 6 | 0.049087 | 0.14209 | 0.959216 | 2936 | 2 | 0.96268 | 0.26439 | 0.51825 | 1 | 10404 | 3 | 0.96268 |
| LACC1 | 6 | 0.94256 | 0.94254 | 0.999994 | 20140 | 0 | 1.1152 | 0.2644 | 0.51827 | 1 | 10405 | 4 | 1.1152 |
| NonTargetingControlG | 2 | 0.37934 | 0.39976 | 0.999994 | 11639 | 1 | 2.0589 | 0.26446 | 0.38316 | 1 | 10406 | 1 | 2.0589 |
| GCFC2 | 6 | 0.6393 | 0.76941 | 0.999994 | 15504 | 2 | 1.3916 | 0.26467 | 0.51863 | 1 | 10407 | 4 | 1.3916 |
| CLDN10 | 6 | 0.46419 | 0.64456 | 0.999994 | 12923 | 2 | 0.83628 | 0.26478 | 0.51877 | 1 | 10408 | 4 | 0.83628 |
| DLG3 | 6 | 0.96453 | 0.96458 | 0.999994 | 20714 | 0 | 1.2309 | 0.26486 | 0.51887 | 1 | 10409 | 4 | 1.2309 |
| CUZD1 | 6 | 0.29022 | 0.49534 | 0.999994 | 9912 | 3 | -0.59968 | 0.26486 | 0.51887 | 1 | 10410 | 1 | -0.59968 |
| ANXA8L1 | 4 | 0.81863 | 0.81846 | 0.999994 | 17758 | 1 | 0.80615 | 0.26489 | 0.47206 | 1 | 10411 | 3 | 0.80615 |
| ALG6 | 6 | 0.32489 | 0.53544 | 0.999994 | 10615 | 2 | 0.93538 | 0.26499 | 0.51902 | 1 | 10412 | 4 | 0.93538 |
| IGFBP3 | 6 | 0.19522 | 0.38208 | 0.999994 | 7639 | 2 | 0.95564 | 0.265 | 0.51904 | 1 | 10413 | 4 | 0.95564 |
| OR11H4 | 6 | 0.065513 | 0.17736 | 0.968171 | 3673 | 3 | 0.7239 | 0.26509 | 0.51917 | 1 | 10414 | 3 | 0.7239 |
| OR52B6 | 4 | 0.88259 | 0.88222 | 0.999994 | 18882 | 0 | 1.573 | 0.26511 | 0.47234 | 1 | 10415 | 3 | 1.573 |
| OR10A2 | 5 | 0.78874 | 0.81258 | 0.999994 | 17329 | 1 | 1.7905 | 0.26515 | 0.4884 | 1 | 10416 | 3 | 1.7905 |
| CCNYL1 | 5 | 0.81987 | 0.83252 | 0.999994 | 17777 | 1 | 1.6474 | 0.26518 | 0.48843 | 1 | 10417 | 3 | 1.6474 |
| KIF9 | 6 | 0.30457 | 0.51206 | 0.999994 | 10219 | 2 | 0.80357 | 0.26521 | 0.51932 | 1 | 10418 | 4 | 0.80357 |
| EYA1 | 5 | 0.090796 | 0.21187 | 0.979587 | 4630 | 2 | 1.6889 | 0.26528 | 0.48853 | 1 | 10419 | 3 | 1.6889 |
| KIAA1257 | 5 | 0.91107 | 0.91109 | 0.999994 | 19450 | 0 | 1.0234 | 0.26528 | 0.48853 | 1 | 10420 | 3 | 1.0234 |
| FGF19 | 6 | 0.27329 | 0.47559 | 0.999994 | 9549 | 3 | -0.66365 | 0.26532 | 0.51948 | 1 | 10421 | 2 | -0.66365 |
| RXRG | 6 | 0.51626 | 0.67993 | 0.999994 | 13683 | 1 | 1.3149 | 0.26543 | 0.51963 | 1 | 10422 | 4 | 1.3149 |
| KCNJ5 | 6 | 0.22928 | 0.42319 | 0.999994 | 8509 | 1 | 0.82161 | 0.2655 | 0.51971 | 1 | 10423 | 3 | 0.82161 |
| ANXA9 | 5 | 0.56298 | 0.70328 | 0.999994 | 14429 | 1 | 0.61142 | 0.26552 | 0.48879 | 1 | 10424 | 2 | 0.61142 |
| HEATR3 | 5 | 0.54852 | 0.69042 | 0.999994 | 14186 | 2 | -0.14558 | 0.26552 | 0.48879 | 1 | 10425 | 2 | -0.14558 |
| hsa-mir-4323 | 4 | 0.82262 | 0.8223 | 0.999994 | 17817 | 1 | 0.2587 | 0.26556 | 0.47295 | 1 | 10426 | 1 | 0.2587 |
| HIC2 | 4 | 0.87112 | 0.87074 | 0.999994 | 18657 | 0 | 1.8384 | 0.26563 | 0.47304 | 1 | 10427 | 2 | 1.8384 |
| PDE9A | 6 | 0.62254 | 0.7567 | 0.999994 | 15268 | 2 | 0.93899 | 0.26563 | 0.51988 | 1 | 10428 | 3 | 0.93899 |
| ZWINT | 6 | 0.1793 | 0.36229 | 0.999994 | 7208 | 2 | -0.013787 | 0.26563 | 0.51988 | 1 | 10429 | 2 | -0.013787 |
| POU2F2 | 6 | 0.31346 | 0.52238 | 0.999994 | 10385 | 1 | 0.39354 | 0.26563 | 0.51988 | 1 | 10430 | 3 | 0.39354 |
| CERS6 | 5 | 0.3686 | 0.53718 | 0.999994 | 11482 | 1 | 1.6129 | 0.26572 | 0.48902 | 1 | 10431 | 3 | 1.6129 |
| RAB3A | 6 | 0.15688 | 0.33428 | 0.999994 | 6648 | 3 | 0.47359 | 0.26573 | 0.52 | 1 | 10432 | 3 | 0.47359 |
| ARRDC2 | 5 | 0.35013 | 0.52204 | 0.999994 | 11160 | 2 | 1.7611 | 0.26575 | 0.48906 | 1 | 10433 | 3 | 1.7611 |
| GSTA4 | 5 | 0.2757 | 0.46139 | 0.999994 | 9610 | 1 | 1.5915 | 0.2658 | 0.4891 | 1 | 10434 | 3 | 1.5915 |
| BRCC3 | 6 | 0.06058 | 0.16698 | 0.967126 | 3450 | 2 | 0.98446 | 0.26584 | 0.52012 | 1 | 10435 | 4 | 0.98446 |
| ATRX | 6 | 0.00025937 | 0.0013828 | 0.384267 | 72 | 4 | -4.4652 | 0.26585 | 0.52013 | 1 | 10436 | 2 | -4.4652 |
| FCGR3B | 6 | 0.061158 | 0.16824 | 0.967126 | 3482 | 3 | -0.81497 | 0.26585 | 0.52013 | 1 | 10437 | 2 | -0.81497 |
| SLC1A6 | 6 | 0.12235 | 0.29019 | 0.999632 | 5715 | 3 | 0.18863 | 0.26592 | 0.52023 | 1 | 10438 | 3 | 0.18863 |
| FAM102B | 6 | 0.20815 | 0.39782 | 0.999994 | 7982 | 2 | 1.2903 | 0.26598 | 0.52031 | 1 | 10439 | 3 | 1.2903 |
| LAMB3 | 4 | 0.11311 | 0.23838 | 0.987253 | 5410 | 1 | 1.4423 | 0.26602 | 0.47354 | 1 | 10440 | 3 | 1.4423 |
| DNAH11 | 5 | 0.16924 | 0.33102 | 0.999994 | 6965 | 3 | -0.97775 | 0.26617 | 0.48952 | 1 | 10441 | 2 | -0.97775 |
| KIAA0355 | 5 | 0.00035352 | 0.0016544 | 0.402548 | 85 | 2 | -0.46269 | 0.26617 | 0.48952 | 1 | 10442 | 2 | -0.46269 |
| CDKN2AIPNL | 5 | 0.79012 | 0.81339 | 0.999994 | 17346 | 1 | 0.38373 | 0.26617 | 0.48952 | 1 | 10443 | 2 | 0.38373 |
| OPN3 | 5 | 0.14671 | 0.29784 | 0.999994 | 6368 | 2 | 1.1976 | 0.26618 | 0.48954 | 1 | 10444 | 3 | 1.1976 |
| MBOAT2 | 6 | 0.77074 | 0.83612 | 0.999994 | 17091 | 1 | 1.4974 | 0.26626 | 0.52066 | 1 | 10445 | 4 | 1.4974 |
| hsa-mir-1267 | 3 | 0.67459 | 0.67788 | 0.999994 | 15998 | 1 | 1.9646 | 0.26627 | 0.41396 | 1 | 10446 | 2 | 1.9646 |
| OR4K2 | 6 | 0.11067 | 0.268 | 0.998561 | 5323 | 3 | -0.31343 | 0.26631 | 0.52073 | 1 | 10447 | 3 | -0.31343 |
| PHACTR2 | 6 | 0.65023 | 0.7778 | 0.999994 | 15662 | 1 | 0.54491 | 0.26631 | 0.52073 | 1 | 10448 | 3 | 0.54491 |
| PLK4 | 4 | 0.6454 | 0.69063 | 0.999994 | 15590 | 1 | 1.3064 | 0.26635 | 0.47399 | 1 | 10449 | 3 | 1.3064 |
| RGPD1 | 6 | 0.58637 | 0.72989 | 0.999994 | 14763 | 2 | 1.4943 | 0.26637 | 0.52081 | 1 | 10450 | 4 | 1.4943 |
| ARHGEF12 | 5 | 0.40365 | 0.56659 | 0.999994 | 12009 | 2 | 1.493 | 0.26638 | 0.48976 | 1 | 10451 | 3 | 1.493 |

| | | | | | | | | | | | | | |
|---|---|---|---|---|---|---|---|---|---|---|---|---|---|
| TGM1 | 6 | 0.072846 | 0.19267 | 0.974607 | 3955 | 3 | 0.55813 | 0.26638 | 0.52082 | 1 | 10452 | 3 | 0.55813 |
| NLRP9 | 6 | 0.6063 | 0.74462 | 0.999994 | 15032 | 1 | 1.213 | 0.26638 | 0.52082 | 1 | 10453 | 4 | 1.213 |
| PFKFB3 | 6 | 0.81583 | 0.8549 | 0.999994 | 17709 | 1 | 1.2907 | 0.26648 | 0.52096 | 1 | 10454 | 4 | 1.2907 |
| GNB4 | 5 | 0.94197 | 0.94216 | 0.999994 | 20124 | 0 | 1.5795 | 0.26653 | 0.48991 | 1 | 10455 | 3 | 1.5795 |
| ACTN2 | 6 | 0.085935 | 0.21938 | 0.980699 | 4444 | 2 | 1.2643 | 0.26654 | 0.52103 | 1 | 10456 | 3 | 1.2643 |
| SYNPO2 | 5 | 0.39576 | 0.56002 | 0.999994 | 11887 | 1 | 1.4819 | 0.26655 | 0.48993 | 1 | 10457 | 3 | 1.4819 |
| PPP1R16A | 5 | 0.88188 | 0.88208 | 0.999994 | 18866 | 1 | 1.5906 | 0.26664 | 0.49002 | 1 | 10458 | 3 | 1.5906 |
| CASC5 | 5 | 0.080683 | 0.19569 | 0.975797 | 4259 | 2 | 0.51781 | 0.26668 | 0.49006 | 1 | 10459 | 2 | 0.51781 |
| DPCR1 | 5 | 0.21563 | 0.39717 | 0.999994 | 8158 | 3 | -0.76876 | 0.26668 | 0.49006 | 1 | 10460 | 1 | -0.76876 |
| NonTargetingControlG | 2 | 0.44341 | 0.45354 | 0.999994 | 12629 | 1 | 2.3525 | 0.26674 | 0.38452 | 1 | 10461 | 1 | 2.3525 |
| TBL1Y | 6 | 0.1845 | 0.36875 | 0.999994 | 7329 | 2 | 1.5756 | 0.26675 | 0.52128 | 1 | 10462 | 3 | 1.5756 |
| SUZ12 | 4 | 0.19112 | 0.33154 | 0.999994 | 7518 | 1 | 1.7693 | 0.26675 | 0.47452 | 1 | 10463 | 2 | 1.7693 |
| MRPL36 | 4 | 0.36629 | 0.51099 | 0.999994 | 11447 | 2 | 1.2946 | 0.26675 | 0.47452 | 1 | 10464 | 2 | 1.2946 |
| ZNF580 | 6 | 0.84212 | 0.86776 | 0.999994 | 18126 | 1 | 1.2274 | 0.26676 | 0.52131 | 1 | 10465 | 3 | 1.2274 |
| ADAMTS12 | 6 | 0.073239 | 0.19349 | 0.975797 | 3967 | 2 | 1.2811 | 0.26677 | 0.52132 | 1 | 10466 | 3 | 1.2811 |
| TRAF1 | 6 | 0.18349 | 0.36749 | 0.999994 | 7304 | 1 | 1.2513 | 0.26679 | 0.52134 | 1 | 10467 | 3 | 1.2513 |
| hsa-mir-548aj-2 | 4 | 0.91635 | 0.91611 | 0.999994 | 19545 | 0 | 0.87474 | 0.26686 | 0.47466 | 1 | 10468 | 3 | 0.87474 |
| DYRK2 | 6 | 0.31661 | 0.526 | 0.999994 | 10444 | 2 | 1.6657 | 0.26693 | 0.52152 | 1 | 10469 | 4 | 1.6657 |
| SLMAP | 3 | 0.87935 | 0.87945 | 0.999994 | 18816 | 0 | 0.90865 | 0.26695 | 0.41466 | 1 | 10470 | 2 | 0.90865 |
| MOB3C | 3 | 0.25165 | 0.37702 | 0.999994 | 9050 | 1 | -0.27929 | 0.26695 | 0.41466 | 1 | 10471 | 1 | -0.27929 |
| HEPACAM2 | 6 | 0.69018 | 0.80899 | 0.999994 | 16213 | 2 | 0.76954 | 0.26696 | 0.52158 | 1 | 10472 | 3 | 0.76954 |
| CLECL1 | 6 | 0.94574 | 0.94571 | 0.999994 | 20226 | 0 | 0.38368 | 0.26696 | 0.52158 | 1 | 10473 | 2 | 0.38368 |
| MMP10 | 6 | 0.85943 | 0.87711 | 0.999994 | 18423 | 1 | 0.25058 | 0.26696 | 0.52158 | 1 | 10474 | 2 | 0.25058 |
| ICK | 5 | 0.85414 | 0.8576 | 0.999994 | 18336 | 1 | 1.6484 | 0.26715 | 0.4906 | 1 | 10475 | 3 | 1.6484 |
| FUT8 | 4 | 0.84205 | 0.8415 | 0.999994 | 18124 | 0 | 0.94404 | 0.26715 | 0.47504 | 1 | 10476 | 3 | 0.94404 |
| ETFA | 6 | 0.064336 | 0.17492 | 0.968171 | 3624 | 2 | 1.0136 | 0.2672 | 0.52188 | 1 | 10477 | 4 | 1.0136 |
| CYB5R4 | 5 | 0.63199 | 0.73806 | 0.999994 | 15399 | 1 | 0.37042 | 0.2673 | 0.49077 | 1 | 10478 | 2 | 0.37042 |
| FBXL6 | 6 | 0.060009 | 0.16576 | 0.967126 | 3422 | 4 | -1.4152 | 0.26731 | 0.52202 | 1 | 10479 | 2 | -1.4152 |
| WDR75 | 4 | 0.13571 | 0.27448 | 0.998561 | 6074 | 2 | 0.64207 | 0.26735 | 0.4753 | 1 | 10480 | 2 | 0.64207 |
| ZNF267 | 5 | 0.41091 | 0.57265 | 0.999994 | 12126 | 1 | 1.419 | 0.26743 | 0.4909 | 1 | 10481 | 3 | 1.419 |
| STOM | 6 | 0.32653 | 0.53727 | 0.999994 | 10654 | 2 | 0.89577 | 0.26759 | 0.52239 | 1 | 10482 | 4 | 0.89577 |
| CCDC181 | 5 | 0.54679 | 0.68888 | 0.999994 | 14162 | 2 | 1.5169 | 0.26764 | 0.49112 | 1 | 10483 | 3 | 1.5169 |
| LAMA5 | 6 | 0.59901 | 0.73921 | 0.999994 | 14942 | 2 | 1.2176 | 0.26768 | 0.52249 | 1 | 10484 | 3 | 1.2176 |
| HIST1H2AK | 6 | 0.50121 | 0.66953 | 0.999994 | 13440 | 1 | 1.3656 | 0.26768 | 0.52249 | 1 | 10485 | 3 | 1.3656 |
| ATP6V1G1 | 5 | 0.16973 | 0.33174 | 0.999994 | 6980 | 2 | 0.92872 | 0.2677 | 0.49118 | 1 | 10486 | 3 | 0.92872 |
| MTMR2 | 5 | 0.83759 | 0.8449 | 0.999994 | 18044 | 1 | 0.13578 | 0.2677 | 0.49118 | 1 | 10487 | 2 | 0.13578 |
| MAGEA8 | 5 | 0.15564 | 0.31116 | 0.999994 | 6612 | 3 | -1.134 | 0.2677 | 0.49118 | 1 | 10488 | 2 | -1.134 |
| NPVF | 5 | 0.32088 | 0.49801 | 0.999994 | 10528 | 2 | 0.61916 | 0.2677 | 0.49118 | 1 | 10489 | 2 | 0.61916 |
| AKR1A1 | 5 | 0.076338 | 0.18869 | 0.969543 | 4083 | 2 | 1.4781 | 0.2677 | 0.49118 | 1 | 10490 | 3 | 1.4781 |
| AK3 | 5 | 0.30657 | 0.48637 | 0.999994 | 10265 | 2 | -0.054386 | 0.2677 | 0.49118 | 1 | 10491 | 2 | -0.054386 |
| GUCA2A | 5 | 0.79468 | 0.8162 | 0.999994 | 17408 | 1 | 0.85653 | 0.26771 | 0.49119 | 1 | 10492 | 3 | 0.85653 |
| C4orf32 | 4 | 0.87434 | 0.87399 | 0.999994 | 18716 | 0 | 1.8148 | 0.26777 | 0.47585 | 1 | 10493 | 2 | 1.8148 |
| GTF2E1 | 4 | 0.055863 | 0.13288 | 0.95318 | 3241 | 3 | -2.7199 | 0.26783 | 0.47593 | 1 | 10494 | 1 | -2.7199 |
| CCDC41 | 4 | 0.18572 | 0.32598 | 0.999994 | 7358 | 2 | 0.082359 | 0.26783 | 0.47593 | 1 | 10495 | 2 | 0.082359 |
| SPATA31D1 | 6 | 0.93783 | 0.93783 | 0.999994 | 20026 | 0 | 1.0533 | 0.26783 | 0.52273 | 1 | 10496 | 4 | 1.0533 |
| KREMEN1 | 5 | 0.52371 | 0.66857 | 0.999994 | 13803 | 2 | 1.6327 | 0.26789 | 0.49139 | 1 | 10497 | 3 | 1.6327 |
| GNLY | 6 | 0.0015604 | 0.0078035 | 0.677842 | 229 | 3 | -3.8163 | 0.2679 | 0.52281 | 1 | 10498 | 3 | -3.8163 |
| HTRA3 | 6 | 0.0043763 | 0.018864 | 0.803792 | 471 | 3 | -1.2538 | 0.2679 | 0.52281 | 1 | 10499 | 1 | -1.2538 |
| TBX10 | 5 | 0.065241 | 0.17064 | 0.967177 | 3664 | 3 | -1.8824 | 0.26802 | 0.49153 | 1 | 10500 | 2 | -1.8824 |
| hsa-mir-548a-2 | 4 | 0.92478 | 0.92458 | 0.999994 | 19738 | 0 | 1.8903 | 0.26808 | 0.47625 | 1 | 10501 | 3 | 1.8903 |
| ISCU | 2 | 0.33058 | 0.36052 | 0.999994 | 10731 | 1 | 1.9696 | 0.26824 | 0.3854 | 1 | 10502 | 1 | 1.9696 |
| hsa-mir-4632 | 2 | 0.021242 | 0.039411 | 0.891621 | 1529 | 1 | -1.9875 | 0.26824 | 0.3854 | 1 | 10503 | 1 | -1.9875 |

| | | | | | | | | | | | | |
|---|---|---|---|---|---|---|---|---|---|---|---|---|
| CCRL1 | 5 | 0.5632 | 0.70348 | 0.999994 | 14434 | 2 | 0.88086 | 0.26833 | 0.49187 | 1 | 10504 | 3 | 0.88086 |
| C14orf105 | 6 | 0.055254 | 0.15549 | 0.967126 | 3210 | 1 | 0.99429 | 0.26838 | 0.52342 | 1 | 10505 | 4 | 0.99429 |
| hsa-mir-1265 | 4 | 0.26642 | 0.40863 | 0.999994 | 9384 | 2 | 0.85913 | 0.26841 | 0.47667 | 1 | 10506 | 2 | 0.85913 |
| ZNF517 | 4 | 0.1409 | 0.27984 | 0.999632 | 6216 | 2 | 0.13537 | 0.26841 | 0.47668 | 1 | 10507 | 2 | 0.13537 |
| SRFBP1 | 6 | 0.17121 | 0.35225 | 0.999994 | 7014 | 3 | -0.38284 | 0.26842 | 0.52348 | 1 | 10508 | 2 | -0.38284 |
| CCR5 | 6 | 0.45181 | 0.63627 | 0.999994 | 12732 | 2 | 0.48938 | 0.26842 | 0.52348 | 1 | 10509 | 3 | 0.48938 |
| KPRP | 6 | 0.65054 | 0.77802 | 0.999994 | 15666 | 2 | 0.26684 | 0.26842 | 0.52348 | 1 | 10510 | 3 | 0.26684 |
| TENM2 | 6 | 0.27294 | 0.47516 | 0.999994 | 9540 | 3 | -0.11604 | 0.26842 | 0.52348 | 1 | 10511 | 3 | -0.11604 |
| CKS2 | 6 | 0.41014 | 0.609 | 0.999994 | 12112 | 2 | 1.1759 | 0.26842 | 0.52348 | 1 | 10512 | 3 | 1.1759 |
| CWF19L1 | 6 | 0.88797 | 0.89489 | 0.999994 | 18997 | 1 | -0.024328 | 0.26842 | 0.52348 | 1 | 10513 | 2 | -0.024328 |
| EVI2B | 4 | 0.15525 | 0.29462 | 0.999994 | 6601 | 1 | 0.7922 | 0.26845 | 0.47672 | 1 | 10514 | 3 | 0.7922 |
| OR9Q1 | 5 | 0.16714 | 0.3279 | 0.999994 | 6909 | 1 | 0.23851 | 0.26859 | 0.49214 | 1 | 10515 | 2 | 0.23851 |
| ZNF845 | 5 | 0.16198 | 0.32044 | 0.999994 | 6791 | 1 | 0.85113 | 0.26859 | 0.49214 | 1 | 10516 | 3 | 0.85113 |
| PRCC | 6 | 0.011957 | 0.045567 | 0.891621 | 989 | 4 | -2.7708 | 0.26868 | 0.52385 | 1 | 10517 | 2 | -2.7708 |
| ARID3B | 6 | 0.7563 | 0.83072 | 0.999994 | 16933 | 1 | 1.3327 | 0.26884 | 0.52405 | 1 | 10518 | 4 | 1.3327 |
| DPY19L2 | 4 | 0.5048 | 0.61846 | 0.999994 | 13510 | 1 | 1.7407 | 0.26894 | 0.47738 | 1 | 10519 | 3 | 1.7407 |
| OPHN1 | 6 | 0.11032 | 0.26731 | 0.998561 | 5313 | 4 | -0.95798 | 0.26897 | 0.52422 | 1 | 10520 | 2 | -0.95798 |
| OTOR | 6 | 0.16484 | 0.34426 | 0.999994 | 6859 | 3 | -0.76073 | 0.26897 | 0.52422 | 1 | 10521 | 2 | -0.76073 |
| PRKG2 | 6 | 0.66456 | 0.78884 | 0.999994 | 15850 | 2 | 0.79543 | 0.26897 | 0.52422 | 1 | 10522 | 3 | 0.79543 |
| DYNC2H1 | 6 | 0.67706 | 0.79868 | 0.999994 | 16042 | 2 | 0.54176 | 0.26897 | 0.52422 | 1 | 10523 | 3 | 0.54176 |
| PDGFD | 6 | 0.27292 | 0.47514 | 0.999994 | 9538 | 3 | 0.77838 | 0.26924 | 0.52456 | 1 | 10524 | 3 | 0.77838 |
| NonTargetingControlG | 1 | 0.73072 | 0.7309 | 0.999994 | 16609 | 0 | 3.1962 | 0.26928 | 0.26927 | 0.978454 | 10525 | 1 | 3.1962 |
| SCN4A | 4 | 0.16784 | 0.30753 | 0.999994 | 6933 | 2 | 0.70214 | 0.26929 | 0.47785 | 1 | 10526 | 2 | 0.70214 |
| hsa-mir-3188 | 4 | 0.059659 | 0.14013 | 0.958716 | 3409 | 2 | -3.1025 | 0.26941 | 0.47802 | 1 | 10527 | 1 | -3.1025 |
| HACL1 | 4 | 0.29648 | 0.4394 | 0.999994 | 10053 | 2 | 0.42366 | 0.26941 | 0.47802 | 1 | 10528 | 2 | 0.42366 |
| ZNF443 | 6 | 0.48134 | 0.6561 | 0.999994 | 13157 | 2 | 0.72863 | 0.26945 | 0.52483 | 1 | 10529 | 4 | 0.72863 |
| NIPAL2 | 6 | 0.83442 | 0.86374 | 0.999994 | 17994 | 1 | 1.6546 | 0.26946 | 0.52484 | 1 | 10530 | 4 | 1.6546 |
| GTPBP1 | 6 | 0.33236 | 0.54397 | 0.999994 | 10776 | 2 | -0.54643 | 0.26953 | 0.52494 | 1 | 10531 | 2 | -0.54643 |
| ACAD10 | 6 | 0.9328 | 0.9329 | 0.999994 | 19921 | 0 | 0.94767 | 0.26953 | 0.52494 | 1 | 10532 | 4 | 0.94767 |
| GPHB5 | 4 | 0.68132 | 0.71244 | 0.999994 | 16107 | 1 | 1.4739 | 0.26956 | 0.47822 | 1 | 10533 | 3 | 1.4739 |
| HLA-DRA | 6 | 0.1112 | 0.26901 | 0.998561 | 5339 | 2 | 0.83444 | 0.26957 | 0.52498 | 1 | 10534 | 4 | 0.83444 |
| NUMBL | 6 | 0.045038 | 0.13315 | 0.95318 | 2758 | 3 | 0.078257 | 0.26957 | 0.52498 | 1 | 10535 | 3 | 0.078257 |
| LOC100506688 | 6 | 0.31782 | 0.5274 | 0.999994 | 10467 | 2 | 1.3446 | 0.26957 | 0.52498 | 1 | 10536 | 4 | 1.3446 |
| MAGED2 | 6 | 0.37512 | 0.58645 | 0.999994 | 11582 | 2 | 1.3592 | 0.26957 | 0.52498 | 1 | 10537 | 4 | 1.3592 |
| TPSD1 | 5 | 0.15617 | 0.31192 | 0.999994 | 6635 | 3 | -1.1418 | 0.26958 | 0.49318 | 1 | 10538 | 2 | -1.1418 |
| CCNH | 6 | 0.28569 | 0.49009 | 0.999994 | 9825 | 1 | 1.3752 | 0.26964 | 0.52508 | 1 | 10539 | 4 | 1.3752 |
| USP4 | 5 | 0.17756 | 0.34312 | 0.999994 | 7170 | 1 | 1.6204 | 0.26964 | 0.49325 | 1 | 10540 | 3 | 1.6204 |
| PTCD3 | 4 | 0.57009 | 0.64968 | 0.999994 | 14542 | 1 | 1.1135 | 0.26965 | 0.47835 | 1 | 10541 | 3 | 1.1135 |
| HMX3 | 5 | 0.16685 | 0.32747 | 0.999994 | 6903 | 2 | 1.5443 | 0.26969 | 0.49331 | 1 | 10542 | 3 | 1.5443 |
| C15orf43 | 6 | 0.82019 | 0.85694 | 0.999994 | 17780 | 1 | 0.76629 | 0.26981 | 0.52531 | 1 | 10543 | 4 | 0.76629 |
| YAP1 | 6 | 0.37337 | 0.58533 | 0.999994 | 11563 | 2 | 1.4568 | 0.26988 | 0.52538 | 1 | 10544 | 3 | 1.4568 |
| ST3GAL6 | 5 | 0.43463 | 0.59232 | 0.999994 | 12487 | 1 | 0.051996 | 0.26988 | 0.4935 | 1 | 10545 | 2 | 0.051996 |
| QDPR | 5 | 0.032223 | 0.096166 | 0.945031 | 2114 | 2 | 1.2574 | 0.26988 | 0.4935 | 1 | 10546 | 3 | 1.2574 |
| CCDC85B | 5 | 0.15508 | 0.31033 | 0.999994 | 6597 | 2 | 0.77494 | 0.26988 | 0.4935 | 1 | 10547 | 3 | 0.77494 |
| FECH | 5 | 0.45941 | 0.61315 | 0.999994 | 12837 | 2 | 1.6648 | 0.26988 | 0.4935 | 1 | 10548 | 3 | 1.6648 |
| SLC52A2 | 5 | 0.37376 | 0.54154 | 0.999994 | 11568 | 2 | -0.0040952 | 0.26988 | 0.4935 | 1 | 10549 | 2 | -0.0040952 |
| LDHA | 5 | 0.95472 | 0.95487 | 0.999994 | 20447 | 0 | 0.3268 | 0.26997 | 0.4936 | 1 | 10550 | 2 | 0.3268 |
| hsa-mir-645 | 3 | 0.7548 | 0.75474 | 0.999994 | 16909 | 0 | 1.9207 | 0.26997 | 0.41791 | 1 | 10551 | 2 | 1.9207 |
| DTWD1 | 4 | 0.36193 | 0.50652 | 0.999994 | 11365 | 2 | 1.1732 | 0.27001 | 0.47882 | 1 | 10552 | 2 | 1.1732 |
| FAM120B | 4 | 0.81899 | 0.81878 | 0.999994 | 17766 | 1 | 1.7244 | 0.27005 | 0.47889 | 1 | 10553 | 3 | 1.7244 |
| ARFIP1 | 6 | 0.59527 | 0.73649 | 0.999994 | 14893 | 1 | 0.96419 | 0.27005 | 0.52562 | 1 | 10554 | 3 | 0.96419 |
| OR8G1 | 6 | 0.59434 | 0.7358 | 0.999994 | 14882 | 2 | 0.92581 | 0.27007 | 0.52565 | 1 | 10555 | 3 | 0.92581 |

| | | | | | | | | | | | | | |
|---|---|---|---|---|---|---|---|---|---|---|---|---|---|
| RBM23 | 4 | 0.84206 | 0.84151 | 0.999994 | 18125 | 0 | 1.1098 | 0.27008 | 0.47892 | 1 | 10556 | 3 | 1.1098 |
| ADAM9 | 6 | 0.085272 | 0.21805 | 0.980699 | 4421 | 3 | -1.2054 | 0.27014 | 0.52575 | 1 | 10557 | 2 | -1.2054 |
| TMEM116 | 6 | 0.020777 | 0.074478 | 0.93914 | 1494 | 3 | -1.5014 | 0.27014 | 0.52575 | 1 | 10558 | 2 | -1.5014 |
| TEPP | 6 | 0.55071 | 0.70415 | 0.999994 | 14224 | 1 | 1.5015 | 0.2702 | 0.52582 | 1 | 10559 | 4 | 1.5015 |
| hsa-mir-518f | 3 | 0.46375 | 0.52279 | 0.999994 | 12917 | 1 | 1.9514 | 0.27021 | 0.41816 | 1 | 10560 | 2 | 1.9514 |
| AAED1 | 3 | 0.78673 | 0.78664 | 0.999994 | 17299 | 0 | 2.1285 | 0.27022 | 0.41817 | 1 | 10561 | 2 | 2.1285 |
| CIB3 | 6 | 0.20876 | 0.39856 | 0.999994 | 7994 | 2 | 1.292 | 0.27025 | 0.52588 | 1 | 10562 | 3 | 1.292 |
| SHANK1 | 5 | 0.53731 | 0.68051 | 0.999994 | 14017 | 1 | 1.5207 | 0.27028 | 0.49395 | 1 | 10563 | 3 | 1.5207 |
| IGSF8 | 6 | 0.44114 | 0.62919 | 0.999994 | 12593 | 2 | 1.3469 | 0.27032 | 0.52598 | 1 | 10564 | 3 | 1.3469 |
| TAF8 | 3 | 0.16547 | 0.26861 | 0.998561 | 6870 | 1 | -0.37322 | 0.27048 | 0.41843 | 1 | 10565 | 1 | -0.37322 |
| RTN3 | 5 | 0.48479 | 0.63479 | 0.999994 | 13216 | 1 | 0.1474 | 0.2705 | 0.49417 | 1 | 10566 | 2 | 0.1474 |
| TOMM20L | 5 | 0.14521 | 0.29559 | 0.999994 | 6329 | 2 | 0.024562 | 0.2705 | 0.49417 | 1 | 10567 | 2 | 0.024562 |
| LPCAT3 | 5 | 0.56471 | 0.70483 | 0.999994 | 14465 | 2 | 1.4345 | 0.2705 | 0.49417 | 1 | 10568 | 3 | 1.4345 |
| AGAP1 | 6 | 0.012913 | 0.048829 | 0.903125 | 1058 | 3 | -0.26023 | 0.2705 | 0.5262 | 1 | 10569 | 3 | -0.26023 |
| NonTargetingControlG | 2 | 0.47281 | 0.47945 | 0.999994 | 13044 | 1 | 2.4357 | 0.27055 | 0.38676 | 1 | 10570 | 1 | 2.4357 |
| CRLF2_X | 6 | 0.48908 | 0.66131 | 0.999994 | 13269 | 1 | 1.3773 | 0.27057 | 0.52629 | 1 | 10571 | 4 | 1.3773 |
| PLD4 | 6 | 0.089266 | 0.22603 | 0.983354 | 4565 | 2 | 0.60056 | 0.27063 | 0.52637 | 1 | 10572 | 3 | 0.60056 |
| FSCB | 6 | 0.15374 | 0.33037 | 0.999994 | 6558 | 3 | -0.90891 | 0.27063 | 0.52637 | 1 | 10573 | 2 | -0.90891 |
| ZNF197 | 5 | 0.5385 | 0.68157 | 0.999994 | 14030 | 1 | 0.63042 | 0.27066 | 0.49435 | 1 | 10574 | 3 | 0.63042 |
| PREB | 6 | 0.43685 | 0.62632 | 0.999994 | 12524 | 2 | 1.1774 | 0.27068 | 0.52644 | 1 | 10575 | 4 | 1.1774 |
| GIMAP1-GIMAP5 | 4 | 0.55973 | 0.64451 | 0.999994 | 14366 | 1 | 0.46547 | 0.27076 | 0.47985 | 1 | 10576 | 2 | 0.46547 |
| SYCE2 | 5 | 0.79491 | 0.81634 | 0.999994 | 17414 | 1 | 1.7297 | 0.27082 | 0.49452 | 1 | 10577 | 3 | 1.7297 |
| PEX5L | 6 | 0.35441 | 0.56901 | 0.999994 | 11237 | 2 | 1.3871 | 0.27082 | 0.52661 | 1 | 10578 | 3 | 1.3871 |
| hsa-mir-3675 | 4 | 0.76126 | 0.76881 | 0.999994 | 16994 | 1 | 0.79224 | 0.27092 | 0.48006 | 1 | 10579 | 3 | 0.79224 |
| ABHD12 | 6 | 0.18437 | 0.36861 | 0.999994 | 7325 | 2 | -0.52444 | 0.27106 | 0.52691 | 1 | 10580 | 2 | -0.52444 |
| HPSE2 | 6 | 0.91626 | 0.91699 | 0.999994 | 19541 | 1 | 0.61447 | 0.27106 | 0.52691 | 1 | 10581 | 3 | 0.61447 |
| ATAT1 | 5 | 0.3148 | 0.49312 | 0.999994 | 10415 | 2 | -0.45584 | 0.27108 | 0.49481 | 1 | 10582 | 1 | -0.45584 |
| ZNF333 | 5 | 0.0076841 | 0.029338 | 0.860856 | 705 | 3 | -1.3363 | 0.27108 | 0.49481 | 1 | 10583 | 2 | -1.3363 |
| GRP | 5 | 0.007028 | 0.027464 | 0.854935 | 661 | 2 | 0.2287 | 0.27116 | 0.49489 | 1 | 10584 | 2 | 0.2287 |
| C20orf196 | 4 | 0.89495 | 0.89465 | 0.999994 | 19125 | 0 | 1.0953 | 0.27129 | 0.48058 | 1 | 10585 | 3 | 1.0953 |
| ZNF558 | 6 | 0.94544 | 0.94542 | 0.999994 | 20216 | 0 | 1.2081 | 0.27133 | 0.52725 | 1 | 10586 | 4 | 1.2081 |
| hsa-mir-4767 | 4 | 0.30597 | 0.44908 | 0.999994 | 10252 | 2 | 0.85948 | 0.27133 | 0.48063 | 1 | 10587 | 2 | 0.85948 |
| STX4 | 2 | 0.24883 | 0.29802 | 0.999994 | 8989 | 1 | 0.61318 | 0.27134 | 0.38721 | 1 | 10588 | 1 | 0.61318 |
| NonTargetingControlG | 2 | 0.7154 | 0.71512 | 0.999994 | 16452 | 0 | 3.2459 | 0.27134 | 0.38721 | 1 | 10589 | 1 | 3.2459 |
| hsa-mir-3972 | 4 | 0.35698 | 0.5014 | 0.999994 | 11287 | 1 | 1.8172 | 0.27139 | 0.48072 | 1 | 10590 | 3 | 1.8172 |
| GPR15 | 6 | 0.33997 | 0.5526 | 0.999994 | 10931 | 2 | 0.95784 | 0.27142 | 0.52735 | 1 | 10591 | 4 | 0.95784 |
| MAFG | 6 | 0.30675 | 0.5146 | 0.999994 | 10269 | 2 | 1.4647 | 0.27142 | 0.52736 | 1 | 10592 | 3 | 1.4647 |
| hsa-mir-3196 | 3 | 0.7855 | 0.7854 | 0.999994 | 17285 | 0 | 2.187 | 0.27164 | 0.41964 | 1 | 10593 | 2 | 2.187 |
| KANSL3 | 4 | 0.08733 | 0.19213 | 0.974607 | 4488 | 2 | 0.13198 | 0.27165 | 0.48104 | 1 | 10594 | 2 | 0.13198 |
| hsa-mir-1976 | 4 | 0.11132 | 0.2352 | 0.985362 | 5342 | 2 | 0.26737 | 0.27165 | 0.48104 | 1 | 10595 | 2 | 0.26737 |
| ISCA1 | 4 | 0.22827 | 0.36972 | 0.999994 | 8484 | 2 | 1.0086 | 0.27165 | 0.48104 | 1 | 10596 | 2 | 1.0086 |
| FAM198B | 4 | 0.41818 | 0.56434 | 0.999994 | 12222 | 2 | 1.4259 | 0.27165 | 0.48104 | 1 | 10597 | 2 | 1.4259 |
| RP1L1 | 4 | 0.92213 | 0.92192 | 0.999994 | 19678 | 0 | 2.0387 | 0.27165 | 0.48104 | 1 | 10598 | 2 | 2.0387 |
| CD86 | 5 | 0.7606 | 0.79652 | 0.999994 | 16987 | 1 | 0.41291 | 0.2717 | 0.49546 | 1 | 10599 | 2 | 0.41291 |
| APCDD1 | 5 | 0.48273 | 0.63304 | 0.999994 | 13185 | 2 | 1.5626 | 0.27171 | 0.49547 | 1 | 10600 | 3 | 1.5626 |
| PCDHB13 | 5 | 0.037378 | 0.10928 | 0.951052 | 2391 | 3 | -1.5048 | 0.27177 | 0.49553 | 1 | 10601 | 2 | -1.5048 |
| PI15 | 6 | 0.60954 | 0.74699 | 0.999994 | 15067 | 2 | 0.52071 | 0.27208 | 0.52822 | 1 | 10602 | 3 | 0.52071 |
| CLCF1 | 6 | 0.0852 | 0.21791 | 0.980699 | 4418 | 3 | -0.19665 | 0.27208 | 0.52822 | 1 | 10603 | 2 | -0.19665 |
| ZNRF3 | 6 | 0.082781 | 0.21301 | 0.979587 | 4331 | 3 | 0.026787 | 0.27208 | 0.52822 | 1 | 10604 | 3 | 0.026787 |
| GPR83 | 6 | 0.31411 | 0.52316 | 0.999994 | 10401 | 3 | 0.24105 | 0.27208 | 0.52822 | 1 | 10605 | 3 | 0.24105 |
| MAN2B2 | 6 | 0.30933 | 0.5176 | 0.999994 | 10312 | 2 | 0.006269 | 0.27219 | 0.52836 | 1 | 10606 | 2 | 0.006269 |
| DIRAS1 | 6 | 0.018889 | 0.068404 | 0.93914 | 1403 | 4 | -2.0721 | 0.27219 | 0.52836 | 1 | 10607 | 1 | -2.0721 |

| | | | | | | | | | | | | | |
|---|---|---|---|---|---|---|---|---|---|---|---|---|---|
| GYLTL1B | 4 | 0.062158 | 0.14492 | 0.963012 | 3519 | 2 | -1.241 | 0.2723 | 0.4819 | 1 | 10608 | 1 | -1.241 |
| hsa-mir-4725 | 4 | 0.27448 | 0.41693 | 0.999994 | 9580 | 2 | -0.47242 | 0.2723 | 0.4819 | 1 | 10609 | 1 | -0.47242 |
| SIK1 | 6 | 0.097007 | 0.24151 | 0.987253 | 4860 | 2 | 0.79268 | 0.2723 | 0.52848 | 1 | 10610 | 4 | 0.79268 |
| SPIB | 6 | 0.97454 | 0.97443 | 0.999994 | 20944 | 0 | 1.1215 | 0.27238 | 0.52859 | 1 | 10611 | 4 | 1.1215 |
| HMMR | 5 | 0.68698 | 0.76067 | 0.999994 | 16179 | 1 | 0.25786 | 0.27238 | 0.49618 | 1 | 10612 | 2 | 0.25786 |
| OR2A4 | 6 | 0.90759 | 0.90961 | 0.999994 | 19377 | 1 | 1.44 | 0.27247 | 0.5287 | 1 | 10613 | 4 | 1.44 |
| ZSWIM4 | 6 | 0.5421 | 0.69803 | 0.999994 | 14093 | 2 | 0.7829 | 0.27251 | 0.52875 | 1 | 10614 | 4 | 0.7829 |
| hsa-mir-1295a | 4 | 0.10687 | 0.22733 | 0.983354 | 5209 | 1 | 0.65662 | 0.27251 | 0.4822 | 1 | 10615 | 3 | 0.65662 |
| hsa-mir-4659a | 3 | 0.71687 | 0.71698 | 0.999994 | 16465 | 1 | 1.8241 | 0.27255 | 0.42061 | 1 | 10616 | 2 | 1.8241 |
| NBEAL1 | 6 | 0.38775 | 0.59451 | 0.999994 | 11764 | 2 | 1.3205 | 0.27256 | 0.5288 | 1 | 10617 | 3 | 1.3205 |
| ALDOC | 6 | 0.062314 | 0.17068 | 0.967177 | 3525 | 3 | -1.3973 | 0.27269 | 0.52896 | 1 | 10618 | 2 | -1.3973 |
| NonTargetingControlG | 2 | 0.58669 | 0.58673 | 0.999994 | 14770 | 0 | 2.8122 | 0.27278 | 0.38809 | 1 | 10619 | 1 | 2.8122 |
| S1PR4 | 4 | 0.21279 | 0.35388 | 0.999994 | 8083 | 1 | 0.64109 | 0.27283 | 0.48261 | 1 | 10620 | 2 | 0.64109 |
| C3orf79 | 6 | 0.92211 | 0.92236 | 0.999994 | 19676 | 1 | 1.1272 | 0.27292 | 0.52927 | 1 | 10621 | 3 | 1.1272 |
| SMOC1 | 4 | 0.93094 | 0.93071 | 0.999994 | 19876 | 0 | 1.1932 | 0.27302 | 0.48286 | 1 | 10622 | 3 | 1.1932 |
| SPESP1 | 4 | 0.60457 | 0.66781 | 0.999994 | 15002 | 1 | 0.99175 | 0.27303 | 0.48287 | 1 | 10623 | 3 | 0.99175 |
| CHDH | 6 | 0.97827 | 0.97823 | 0.999994 | 21052 | 0 | 1.2738 | 0.27305 | 0.52945 | 1 | 10624 | 4 | 1.2738 |
| AIM2 | 5 | 0.32505 | 0.50145 | 0.999994 | 10623 | 1 | 0.98124 | 0.27306 | 0.49689 | 1 | 10625 | 3 | 0.98124 |
| RAD51 | 5 | 0.12866 | 0.27056 | 0.998561 | 5902 | 2 | 0.25617 | 0.27306 | 0.4969 | 1 | 10626 | 2 | 0.25617 |
| ZMYM6 | 5 | 0.54172 | 0.68442 | 0.999994 | 14085 | 1 | 0.84812 | 0.27306 | 0.4969 | 1 | 10627 | 3 | 0.84812 |
| TMEM38A | 5 | 0.72945 | 0.78039 | 0.999994 | 16593 | 1 | 0.59633 | 0.27306 | 0.4969 | 1 | 10628 | 2 | 0.59633 |
| FDFT1 | 6 | 0.074512 | 0.19615 | 0.975797 | 4016 | 2 | 1.4574 | 0.27306 | 0.52947 | 1 | 10629 | 4 | 1.4574 |
| INS-IGF2 | 3 | 0.45825 | 0.51936 | 0.999994 | 12825 | 1 | 1.8908 | 0.27309 | 0.42117 | 1 | 10630 | 2 | 1.8908 |
| OAF | 6 | 0.005183 | 0.021815 | 0.819701 | 533 | 4 | -2.935 | 0.2731 | 0.52951 | 1 | 10631 | 2 | -2.935 |
| hsa-mir-3175 | 3 | 0.13353 | 0.22722 | 0.983354 | 6009 | 1 | 1.8514 | 0.27315 | 0.42124 | 1 | 10632 | 2 | 1.8514 |
| NXPE4 | 6 | 0.54168 | 0.69774 | 0.999994 | 14083 | 2 | 1.5712 | 0.27326 | 0.52973 | 1 | 10633 | 4 | 1.5712 |
| NT5C3B | 6 | 0.13294 | 0.30409 | 0.999994 | 5994 | 3 | 0.18981 | 0.27329 | 0.52977 | 1 | 10634 | 3 | 0.18981 |
| DNAJC16 | 6 | 0.4132 | 0.61093 | 0.999994 | 12157 | 2 | 1.1008 | 0.2733 | 0.52978 | 1 | 10635 | 3 | 1.1008 |
| C7orf25 | 6 | 0.033625 | 0.1073 | 0.950995 | 2197 | 3 | -1.3558 | 0.2733 | 0.52978 | 1 | 10636 | 2 | -1.3558 |
| IGSF6 | 6 | 0.12035 | 0.28635 | 0.999632 | 5639 | 4 | -0.76467 | 0.2733 | 0.52978 | 1 | 10637 | 2 | -0.76467 |
| PPIF | 6 | 0.53901 | 0.69584 | 0.999994 | 14035 | 2 | 0.85592 | 0.2733 | 0.52978 | 1 | 10638 | 3 | 0.85592 |
| RASSF6 | 6 | 0.068834 | 0.18432 | 0.968526 | 3792 | 3 | -1.3037 | 0.2733 | 0.52978 | 1 | 10639 | 2 | -1.3037 |
| NFATC2 | 6 | 0.22087 | 0.41316 | 0.999994 | 8307 | 2 | 1.1999 | 0.27333 | 0.52982 | 1 | 10640 | 4 | 1.1999 |
| KRTAP21-1 | 6 | 0.58314 | 0.7275 | 0.999994 | 14717 | 2 | 0.89413 | 0.27336 | 0.52986 | 1 | 10641 | 4 | 0.89413 |
| C1QC | 4 | 0.63032 | 0.68203 | 0.999994 | 15383 | 1 | 1.2537 | 0.27341 | 0.48337 | 1 | 10642 | 3 | 1.2537 |
| hsa-mir-4786 | 3 | 0.86621 | 0.86621 | 0.999994 | 18562 | 0 | 0.78924 | 0.27345 | 0.42155 | 1 | 10643 | 2 | 0.78924 |
| hsa-mir-199b | 3 | 0.17642 | 0.28263 | 0.999632 | 7142 | 1 | 1.9141 | 0.27345 | 0.42155 | 1 | 10644 | 2 | 1.9141 |
| hsa-mir-8082 | 3 | 0.16487 | 0.26784 | 0.998561 | 6860 | 2 | -1.4836 | 0.27345 | 0.42155 | 1 | 10645 | 1 | -1.4836 |
| SYNGR2 | 4 | 0.55752 | 0.64339 | 0.999994 | 14325 | 1 | 1.7891 | 0.27346 | 0.48344 | 1 | 10646 | 3 | 1.7891 |
| GRHL1 | 6 | 0.19649 | 0.38362 | 0.999994 | 7661 | 2 | 0.32784 | 0.27353 | 0.53009 | 1 | 10647 | 3 | 0.32784 |
| OR51G1 | 5 | 0.060564 | 0.16302 | 0.967126 | 3446 | 3 | -0.9374 | 0.27369 | 0.49759 | 1 | 10648 | 2 | -0.9374 |
| MAP3K11 | 6 | 0.10349 | 0.25415 | 0.994632 | 5083 | 3 | 0.19395 | 0.27375 | 0.53038 | 1 | 10649 | 3 | 0.19395 |
| PDLIM5 | 6 | 0.4564 | 0.63931 | 0.999994 | 12797 | 2 | 1.0303 | 0.27379 | 0.53043 | 1 | 10650 | 4 | 1.0303 |
| NR1I2 | 6 | 0.25489 | 0.45379 | 0.999994 | 9128 | 3 | -0.28933 | 0.27394 | 0.53062 | 1 | 10651 | 3 | -0.28933 |
| RRP15 | 6 | 0.15232 | 0.32859 | 0.999994 | 6513 | 2 | 0.53208 | 0.27402 | 0.53072 | 1 | 10652 | 3 | 0.53208 |
| KIF21B | 6 | 0.80789 | 0.85131 | 0.999994 | 17597 | 1 | 0.8491 | 0.27407 | 0.53078 | 1 | 10653 | 4 | 0.8491 |
| BTG2 | 6 | 0.67855 | 0.79987 | 0.999994 | 16069 | 2 | 0.47521 | 0.27409 | 0.5308 | 1 | 10654 | 3 | 0.47521 |
| ZNF680 | 5 | 0.12794 | 0.26946 | 0.998561 | 5873 | 3 | -1.0962 | 0.2741 | 0.49802 | 1 | 10655 | 1 | -1.0962 |
| SPTSSA | 5 | 0.88803 | 0.88812 | 0.999994 | 19000 | 1 | 0.33872 | 0.2741 | 0.49802 | 1 | 10656 | 2 | 0.33872 |
| AGPHD1 | 6 | 0.42228 | 0.61693 | 0.999994 | 12287 | 2 | 0.89074 | 0.27413 | 0.53086 | 1 | 10657 | 3 | 0.89074 |
| SPDYE3 | 3 | 0.76998 | 0.76999 | 0.999994 | 17081 | 0 | 0.84667 | 0.27427 | 0.42241 | 1 | 10658 | 2 | 0.84667 |
| IFNA1 | 6 | 0.32042 | 0.53038 | 0.999994 | 10520 | 3 | 0.052832 | 0.2744 | 0.5312 | 1 | 10659 | 3 | 0.052832 |

| | | | | | | | | | | | | | |
|---|---|---|---|---|---|---|---|---|---|---|---|---|---|
| WDR93 | 6 | 0.15596 | 0.33314 | 0.999994 | 6627 | 3 | -0.39312 | 0.2744 | 0.5312 | 1 | 10660 | 3 | -0.39312 |
| ZNF555 | 6 | 0.039814 | 0.12148 | 0.951052 | 2505 | 4 | -1.4602 | 0.2744 | 0.5312 | 1 | 10661 | 2 | -1.4602 |
| MOB3B | 6 | 0.93921 | 0.93915 | 0.999994 | 20053 | 0 | 0.7288 | 0.2744 | 0.5312 | 1 | 10662 | 3 | 0.7288 |
| REEP2 | 4 | 0.054587 | 0.13043 | 0.95318 | 3179 | 1 | 1.1269 | 0.27441 | 0.48466 | 1 | 10663 | 3 | 1.1269 |
| OR4F15 | 5 | 0.51292 | 0.65922 | 0.999994 | 13619 | 2 | -0.50498 | 0.27448 | 0.49843 | 1 | 10664 | 2 | -0.50498 |
| C2orf27A | 5 | 0.12769 | 0.26909 | 0.998561 | 5866 | 2 | 0.97347 | 0.27448 | 0.49843 | 1 | 10665 | 3 | 0.97347 |
| MYO10 | 6 | 0.10676 | 0.26049 | 0.994943 | 5202 | 3 | -0.8232 | 0.27456 | 0.5314 | 1 | 10666 | 2 | -0.8232 |
| CAMKK1 | 6 | 0.62501 | 0.75861 | 0.999994 | 15306 | 1 | 1.5576 | 0.27456 | 0.5314 | 1 | 10667 | 4 | 1.5576 |
| C1orf74 | 6 | 0.62111 | 0.75566 | 0.999994 | 15243 | 2 | -0.43042 | 0.27456 | 0.5314 | 1 | 10668 | 2 | -0.43042 |
| DDX60L | 6 | 0.37099 | 0.58382 | 0.999994 | 11527 | 2 | -0.36019 | 0.27456 | 0.5314 | 1 | 10669 | 2 | -0.36019 |
| CD207 | 6 | 0.90291 | 0.90595 | 0.999994 | 19273 | 1 | 0.91279 | 0.2747 | 0.53158 | 1 | 10670 | 4 | 0.91279 |
| EME1 | 5 | 0.10518 | 0.23448 | 0.984971 | 5138 | 3 | -1.5592 | 0.2747 | 0.49866 | 1 | 10671 | 2 | -1.5592 |
| CT47B1 | 4 | 0.90922 | 0.90894 | 0.999994 | 19406 | 0 | 1.9592 | 0.27475 | 0.4851 | 1 | 10672 | 2 | 1.9592 |
| TRPM4 | 6 | 0.62696 | 0.76004 | 0.999994 | 15335 | 1 | 1.2627 | 0.27478 | 0.53169 | 1 | 10673 | 4 | 1.2627 |
| ARAP2 | 4 | 0.89092 | 0.89062 | 0.999994 | 19053 | 0 | 1.2045 | 0.27478 | 0.48514 | 1 | 10674 | 3 | 1.2045 |
| DHRS1 | 6 | 0.024965 | 0.086934 | 0.94395 | 1755 | 2 | 1.276 | 0.27489 | 0.53183 | 1 | 10675 | 4 | 1.276 |
| ZNF133 | 6 | 0.45664 | 0.63946 | 0.999994 | 12800 | 1 | 1.1246 | 0.27499 | 0.53196 | 1 | 10676 | 3 | 1.1246 |
| hsa-mir-4728 | 2 | 0.39939 | 0.41627 | 0.999994 | 11938 | 1 | 2.2547 | 0.27509 | 0.38954 | 1 | 10677 | 1 | 2.2547 |
| ODF3L1 | 6 | 0.15327 | 0.32978 | 0.999994 | 6543 | 2 | 1.409 | 0.27511 | 0.53211 | 1 | 10678 | 3 | 1.409 |
| PIAS1 | 4 | 0.10114 | 0.21718 | 0.980699 | 5009 | 2 | 0.54421 | 0.27513 | 0.4856 | 1 | 10679 | 2 | 0.54421 |
| SLCO6A1 | 4 | 0.15716 | 0.29658 | 0.999994 | 6652 | 2 | -0.72349 | 0.27513 | 0.4856 | 1 | 10680 | 2 | -0.72349 |
| SPA17 | 6 | 0.78405 | 0.84131 | 0.999994 | 17267 | 1 | 0.98635 | 0.27516 | 0.53217 | 1 | 10681 | 4 | 0.98635 |
| VPRBP | 6 | 0.10839 | 0.26361 | 0.996734 | 5253 | 4 | -1.2403 | 0.27522 | 0.53223 | 1 | 10682 | 2 | -1.2403 |
| ASXL3 | 5 | 0.42482 | 0.58409 | 0.999994 | 12332 | 2 | 1.7403 | 0.2753 | 0.49933 | 1 | 10683 | 3 | 1.7403 |
| RLIM | 5 | 0.37368 | 0.54148 | 0.999994 | 11566 | 2 | 1.7895 | 0.2753 | 0.49933 | 1 | 10684 | 3 | 1.7895 |
| TTC30B | 5 | 0.075799 | 0.18783 | 0.969543 | 4065 | 3 | -1.7563 | 0.27534 | 0.49937 | 1 | 10685 | 2 | -1.7563 |
| hsa-mir-624 | 4 | 0.48627 | 0.61027 | 0.999994 | 13232 | 1 | 0.85961 | 0.27534 | 0.48586 | 1 | 10686 | 3 | 0.85961 |
| hsa-mir-548f-4 | 1 | 0.7246 | 0.72482 | 0.999994 | 16541 | 0 | 3.0188 | 0.2754 | 0.27538 | 0.978454 | 10687 | 1 | 3.0188 |
| AKAP10 | 6 | 0.065593 | 0.17753 | 0.968171 | 3676 | 2 | 1.0869 | 0.27541 | 0.53247 | 1 | 10688 | 4 | 1.0869 |
| ZNF367 | 5 | 0.31047 | 0.48951 | 0.999994 | 10337 | 1 | -0.35786 | 0.27552 | 0.49956 | 1 | 10689 | 2 | -0.35786 |
| NDUFA5 | 4 | 0.11567 | 0.24272 | 0.988088 | 5507 | 2 | 0.18793 | 0.27553 | 0.48611 | 1 | 10690 | 2 | 0.18793 |
| PLAUR | 6 | 0.6508 | 0.77822 | 0.999994 | 15668 | 1 | 0.99456 | 0.27555 | 0.53265 | 1 | 10691 | 4 | 0.99456 |
| CWH43 | 4 | 0.3334 | 0.47718 | 0.999994 | 10793 | 2 | 1.0533 | 0.27569 | 0.48631 | 1 | 10692 | 2 | 1.0533 |
| hsa-mir-6068 | 4 | 0.02675 | 0.074847 | 0.93914 | 1846 | 1 | 1.4606 | 0.27572 | 0.48637 | 1 | 10693 | 3 | 1.4606 |
| MRPS16 | 6 | 0.15997 | 0.33819 | 0.999994 | 6737 | 3 | -0.51106 | 0.27573 | 0.53289 | 1 | 10694 | 2 | -0.51106 |
| TCF7L2 | 6 | 0.31271 | 0.52148 | 0.999994 | 10374 | 2 | 0.83733 | 0.27573 | 0.53289 | 1 | 10695 | 3 | 0.83733 |
| IGSF1 | 6 | 0.22264 | 0.41535 | 0.999994 | 8348 | 3 | -0.073551 | 0.27574 | 0.53289 | 1 | 10696 | 3 | -0.073551 |
| C20orf78 | 6 | 0.41033 | 0.60911 | 0.999994 | 12114 | 2 | 0.19322 | 0.27574 | 0.53289 | 1 | 10697 | 2 | 0.19322 |
| TMPRSS13 | 5 | 0.19004 | 0.36101 | 0.999994 | 7485 | 3 | -0.8587 | 0.27579 | 0.49986 | 1 | 10698 | 2 | -0.8587 |
| DCAF12L2 | 6 | 0.94327 | 0.94325 | 0.999994 | 20159 | 0 | 0.87617 | 0.27583 | 0.533 | 1 | 10699 | 4 | 0.87617 |
| ADAMTS13 | 5 | 0.49068 | 0.63979 | 0.999994 | 13288 | 2 | -0.10983 | 0.27583 | 0.4999 | 1 | 10700 | 2 | -0.10983 |
| IRF1 | 6 | 0.29358 | 0.4992 | 0.999994 | 9987 | 3 | 0.71468 | 0.27585 | 0.53304 | 1 | 10701 | 3 | 0.71468 |
| ATP6V1F | 4 | 0.78626 | 0.78916 | 0.999994 | 17293 | 1 | 1.858 | 0.27595 | 0.48665 | 1 | 10702 | 3 | 1.858 |
| VPS52 | 5 | 0.10753 | 0.23812 | 0.987253 | 5227 | 2 | -0.13297 | 0.27601 | 0.50009 | 1 | 10703 | 2 | -0.13297 |
| TAS2R16 | 6 | 0.12326 | 0.29171 | 0.999632 | 5745 | 2 | 1.425 | 0.27612 | 0.53339 | 1 | 10704 | 4 | 1.425 |
| hsa-mir-154 | 4 | 0.88743 | 0.88713 | 0.999994 | 18980 | 0 | 1.6971 | 0.27612 | 0.48688 | 1 | 10705 | 3 | 1.6971 |
| hsa-mir-3177 | 4 | 0.38329 | 0.52853 | 0.999994 | 11694 | 2 | 1.048 | 0.27614 | 0.4869 | 1 | 10706 | 2 | 1.048 |
| hsa-mir-4752 | 4 | 0.93548 | 0.93528 | 0.999994 | 19979 | 0 | 0.85344 | 0.27622 | 0.48701 | 1 | 10707 | 3 | 0.85344 |
| POU4F1 | 3 | 0.12251 | 0.21278 | 0.979587 | 5721 | 1 | -0.34641 | 0.27627 | 0.4246 | 1 | 10708 | 1 | -0.34641 |
| PPP2R5A | 6 | 0.033723 | 0.10752 | 0.951052 | 2202 | 4 | -1.6688 | 0.2763 | 0.53362 | 1 | 10709 | 1 | -1.6688 |
| CNGA1 | 6 | 0.32021 | 0.53012 | 0.999994 | 10518 | 2 | 0.56134 | 0.2763 | 0.53362 | 1 | 10710 | 3 | 0.56134 |
| PBX1 | 5 | 0.1882 | 0.35839 | 0.999994 | 7424 | 2 | -0.41933 | 0.2765 | 0.50063 | 1 | 10711 | 2 | -0.41933 |

| | | | | | | | | | | | | | |
|---|---|---|---|---|---|---|---|---|---|---|---|---|---|
| TMEM117 | 5 | 0.40145 | 0.56472 | 0.999994 | 11968 | 1 | 0.94742 | 0.2765 | 0.50063 | 1 | 10712 | 3 | 0.94742 |
| ZNF205 | 6 | 0.56156 | 0.71192 | 0.999994 | 14398 | 2 | 1.2463 | 0.27652 | 0.5339 | 1 | 10713 | 4 | 1.2463 |
| ZNF91 | 5 | 0.25106 | 0.44161 | 0.999994 | 9036 | 1 | 1.3737 | 0.27652 | 0.50065 | 1 | 10714 | 3 | 1.3737 |
| ARRDC3 | 6 | 0.57179 | 0.71924 | 0.999994 | 14571 | 2 | 0.75 | 0.27655 | 0.53394 | 1 | 10715 | 4 | 0.75 |
| C1orf227 | 6 | 0.5922 | 0.73422 | 0.999994 | 14852 | 2 | 0.88479 | 0.27665 | 0.53407 | 1 | 10716 | 3 | 0.88479 |
| hsa-mir-6780b | 4 | 0.80961 | 0.80993 | 0.999994 | 17626 | 1 | 1.4051 | 0.27673 | 0.48771 | 1 | 10717 | 3 | 1.4051 |
| ST20 | 2 | 0.74259 | 0.74226 | 0.999994 | 16773 | 0 | 3.365 | 0.27678 | 0.3906 | 1 | 10718 | 1 | 3.365 |
| WFDC10A | 6 | 0.15869 | 0.33656 | 0.999994 | 6689 | 3 | -0.84511 | 0.27699 | 0.5345 | 1 | 10719 | 2 | -0.84511 |
| MTHFSD | 6 | 0.58521 | 0.72907 | 0.999994 | 14746 | 2 | -0.21882 | 0.27699 | 0.5345 | 1 | 10720 | 1 | -0.21882 |
| LMNB1 | 4 | 0.82917 | 0.8287 | 0.999994 | 17914 | 0 | 1.3706 | 0.27705 | 0.48812 | 1 | 10721 | 3 | 1.3706 |
| TMEM253 | 6 | 0.069238 | 0.18517 | 0.968626 | 3807 | 3 | -0.41148 | 0.27711 | 0.53466 | 1 | 10722 | 3 | -0.41148 |
| TMSB10 | 6 | 0.079955 | 0.2072 | 0.978744 | 4232 | 3 | -0.70921 | 0.27711 | 0.53466 | 1 | 10723 | 3 | -0.70921 |
| SYT11 | 6 | 0.83021 | 0.86165 | 0.999994 | 17937 | 1 | 0.4708 | 0.27711 | 0.53466 | 1 | 10724 | 3 | 0.4708 |
| OLFM1 | 6 | 0.12545 | 0.29454 | 0.999994 | 5804 | 3 | 0.24695 | 0.27712 | 0.53467 | 1 | 10725 | 3 | 0.24695 |
| TMSB15A | 5 | 0.51427 | 0.66032 | 0.999994 | 13647 | 2 | 0.87887 | 0.27716 | 0.50135 | 1 | 10726 | 3 | 0.87887 |
| RPL10L | 5 | 0.034749 | 0.10262 | 0.946414 | 2252 | 3 | -2.5878 | 0.27716 | 0.50135 | 1 | 10727 | 1 | -2.5878 |
| COL9A3 | 5 | 0.46113 | 0.61463 | 0.999994 | 12865 | 1 | -0.37027 | 0.27716 | 0.50135 | 1 | 10728 | 2 | -0.37027 |
| hsa-mir-573 | 4 | 0.67243 | 0.70686 | 0.999994 | 15961 | 1 | 0.72469 | 0.27719 | 0.48831 | 1 | 10729 | 3 | 0.72469 |
| NonTargetingControlG | 2 | 0.34498 | 0.37202 | 0.999994 | 11051 | 1 | 1.9818 | 0.27722 | 0.39085 | 1 | 10730 | 1 | 1.9818 |
| TUBGCP5 | 4 | 0.099236 | 0.21372 | 0.979587 | 4940 | 2 | -1.3168 | 0.27733 | 0.48849 | 1 | 10731 | 1 | -1.3168 |
| CKAP2 | 5 | 0.30276 | 0.4833 | 0.999994 | 10189 | 2 | 1.5951 | 0.27736 | 0.50156 | 1 | 10732 | 3 | 1.5951 |
| CCDC24 | 6 | 0.14803 | 0.32317 | 0.999994 | 6409 | 2 | 1.4964 | 0.27741 | 0.53504 | 1 | 10733 | 4 | 1.4964 |
| KIAA0895 | 6 | 0.7141 | 0.81643 | 0.999994 | 16440 | 1 | 1.0974 | 0.27743 | 0.53508 | 1 | 10734 | 4 | 1.0974 |
| MTERFD2 | 6 | 0.23715 | 0.43259 | 0.999994 | 8699 | 3 | -0.34617 | 0.27755 | 0.53523 | 1 | 10735 | 2 | -0.34617 |
| CACNA1E | 6 | 0.63081 | 0.76295 | 0.999994 | 15386 | 2 | 0.72299 | 0.27755 | 0.53523 | 1 | 10736 | 3 | 0.72299 |
| APOL5 | 5 | 0.32369 | 0.50033 | 0.999994 | 10586 | 1 | 1.4643 | 0.27763 | 0.50183 | 1 | 10737 | 3 | 1.4643 |
| HRH1 | 6 | 0.20631 | 0.39557 | 0.999994 | 7941 | 2 | 0.85603 | 0.27778 | 0.53551 | 1 | 10738 | 4 | 0.85603 |
| MAGEB10 | 6 | 0.29858 | 0.50504 | 0.999994 | 10101 | 2 | 0.85748 | 0.27782 | 0.53556 | 1 | 10739 | 4 | 0.85748 |
| C12orf79 | 4 | 0.20786 | 0.3488 | 0.999994 | 7970 | 2 | -0.32417 | 0.27784 | 0.48916 | 1 | 10740 | 1 | -0.32417 |
| KLHL14 | 4 | 0.079132 | 0.17715 | 0.968171 | 4195 | 2 | -1.2031 | 0.27784 | 0.48916 | 1 | 10741 | 1 | -1.2031 |
| MC1R | 4 | 0.93292 | 0.93268 | 0.999994 | 19924 | 0 | 1.246 | 0.27787 | 0.48921 | 1 | 10742 | 3 | 1.246 |
| TBC1D8 | 5 | 0.10588 | 0.23556 | 0.985362 | 5168 | 2 | 0.55251 | 0.27789 | 0.50211 | 1 | 10743 | 3 | 0.55251 |
| CLTB | 5 | 0.36956 | 0.53799 | 0.999994 | 11499 | 2 | 0.54942 | 0.27789 | 0.50211 | 1 | 10744 | 2 | 0.54942 |
| OR5AU1 | 5 | 0.91649 | 0.91658 | 0.999994 | 19550 | 0 | 0.52369 | 0.27789 | 0.50211 | 1 | 10745 | 2 | 0.52369 |
| GPR78 | 6 | 0.29296 | 0.49849 | 0.999994 | 9975 | 2 | -0.48753 | 0.27794 | 0.53572 | 1 | 10746 | 2 | -0.48753 |
| PARP6 | 5 | 0.51236 | 0.65871 | 0.999994 | 13615 | 1 | 1.4322 | 0.27797 | 0.5022 | 1 | 10747 | 3 | 1.4322 |
| DIO3 | 5 | 0.7544 | 0.79316 | 0.999994 | 16901 | 1 | 1.5316 | 0.27809 | 0.50232 | 1 | 10748 | 3 | 1.5316 |
| hsa-mir-4516 | 2 | 0.10838 | 0.183 | 0.968526 | 5252 | 1 | -0.92923 | 0.27818 | 0.39142 | 1 | 10749 | 1 | -0.92923 |
| hsa-mir-4509-1 | 4 | 0.89412 | 0.89382 | 0.999994 | 19103 | 0 | 0.91042 | 0.2782 | 0.48965 | 1 | 10750 | 3 | 0.91042 |
| hsa-mir-4689 | 4 | 0.5301 | 0.63015 | 0.999994 | 13901 | 1 | 1.3205 | 0.27834 | 0.48984 | 1 | 10751 | 3 | 1.3205 |
| hsa-mir-5007 | 4 | 0.16709 | 0.30677 | 0.999994 | 6907 | 2 | -1.0945 | 0.27837 | 0.48988 | 1 | 10752 | 1 | -1.0945 |
| CD47 | 4 | 0.69887 | 0.72367 | 0.999994 | 16293 | 1 | -0.14996 | 0.27837 | 0.48988 | 1 | 10753 | 1 | -0.14996 |
| ANKRD55 | 6 | 0.0084883 | 0.033655 | 0.867348 | 762 | 5 | -2.2763 | 0.27837 | 0.53627 | 1 | 10754 | 1 | -2.2763 |
| GPR171 | 6 | 0.62151 | 0.75594 | 0.999994 | 15253 | 2 | -0.4452 | 0.27837 | 0.53627 | 1 | 10755 | 1 | -0.4452 |
| FGD1 | 6 | 0.52706 | 0.68746 | 0.999994 | 13845 | 1 | 0.29224 | 0.27837 | 0.53627 | 1 | 10756 | 2 | 0.29224 |
| TOM1 | 6 | 0.033996 | 0.10817 | 0.951052 | 2215 | 4 | -2.001 | 0.27845 | 0.53637 | 1 | 10757 | 2 | -2.001 |
| HSDL2 | 6 | 0.32856 | 0.53958 | 0.999994 | 10692 | 3 | 0.24961 | 0.27845 | 0.53637 | 1 | 10758 | 3 | 0.24961 |
| LTB4R2 | 6 | 0.80207 | 0.84874 | 0.999994 | 17513 | 1 | 0.94873 | 0.27845 | 0.53637 | 1 | 10759 | 3 | 0.94873 |
| FBP2 | 6 | 0.4205 | 0.61573 | 0.999994 | 12254 | 2 | -0.040801 | 0.27845 | 0.53637 | 1 | 10760 | 2 | -0.040801 |
| NonTargetingControlG | 1 | 0.72152 | 0.72165 | 0.999994 | 16512 | 0 | 2.7481 | 0.27848 | 0.27839 | 0.978454 | 10761 | 1 | 2.7481 |
| IDH3G | 6 | 0.91958 | 0.92001 | 0.999994 | 19622 | 1 | 1.0646 | 0.27848 | 0.53642 | 1 | 10762 | 4 | 1.0646 |
| G6PD | 2 | 0.21626 | 0.27388 | 0.998561 | 8174 | 1 | 0.28679 | 0.27851 | 0.39162 | 1 | 10763 | 1 | 0.28679 |

| | | | | | | | | | | | | |
|---|---|---|---|---|---|---|---|---|---|---|---|---|
| BAALC | 5 | 0.42099 | 0.58089 | 0.999994 | 12264 | 2 | -0.0065405 | 0.27852 | 0.50281 | 1 | 10764 | 1 | -0.0065405 |
| ADM2 | 3 | 0.29246 | 0.4245 | 0.999994 | 9961 | 1 | 1.7623 | 0.27854 | 0.427 | 1 | 10765 | 2 | 1.7623 |
| SCNN1G | 4 | 0.68215 | 0.71297 | 0.999994 | 16118 | 1 | 0.88459 | 0.27854 | 0.4901 | 1 | 10766 | 3 | 0.88459 |
| NEGR1 | 4 | 0.16764 | 0.30733 | 0.999994 | 6930 | 1 | 0.79411 | 0.2786 | 0.49019 | 1 | 10767 | 3 | 0.79411 |
| C4B_2 | 6 | 0.86868 | 0.88253 | 0.999994 | 18611 | 1 | 0.95969 | 0.27863 | 0.53659 | 1 | 10768 | 4 | 0.95969 |
| KRT5 | 4 | 0.15645 | 0.29582 | 0.999994 | 6640 | 1 | 1.5708 | 0.27876 | 0.49039 | 1 | 10769 | 3 | 1.5708 |
| RPL24 | 3 | 0.45521 | 0.51746 | 0.999994 | 12775 | 1 | 1.9025 | 0.27879 | 0.42729 | 1 | 10770 | 2 | 1.9025 |
| LDHB | 5 | 0.92433 | 0.92455 | 0.999994 | 19727 | 0 | 1.5045 | 0.27883 | 0.50315 | 1 | 10771 | 3 | 1.5045 |
| hsa-mir-23c | 4 | 0.33081 | 0.47452 | 0.999994 | 10739 | 1 | 0.90302 | 0.27884 | 0.4905 | 1 | 10772 | 3 | 0.90302 |
| PLS3 | 6 | 0.46138 | 0.64268 | 0.999994 | 12870 | 2 | 0.91968 | 0.27889 | 0.53692 | 1 | 10773 | 4 | 0.91968 |
| IGFN1 | 6 | 0.22146 | 0.41387 | 0.999994 | 8318 | 3 | -0.66752 | 0.27889 | 0.53693 | 1 | 10774 | 2 | -0.66752 |
| SRM | 6 | 0.78145 | 0.84026 | 0.999994 | 17230 | 1 | 1.4504 | 0.27893 | 0.53698 | 1 | 10775 | 4 | 1.4504 |
| MKL2 | 4 | 0.78508 | 0.78815 | 0.999994 | 17281 | 1 | 0.99876 | 0.27893 | 0.4906 | 1 | 10776 | 3 | 0.99876 |
| ARNT2 | 6 | 0.39685 | 0.60033 | 0.999994 | 11903 | 2 | 1.2181 | 0.27895 | 0.53701 | 1 | 10777 | 4 | 1.2181 |
| TMCO2 | 5 | 0.33088 | 0.50627 | 0.999994 | 10740 | 2 | 1.3182 | 0.27896 | 0.5033 | 1 | 10778 | 3 | 1.3182 |
| CTSF | 4 | 0.93722 | 0.93707 | 0.999994 | 20012 | 0 | 1.4236 | 0.27905 | 0.49077 | 1 | 10779 | 3 | 1.4236 |
| PCDHAC2 | 2 | 0.748 | 0.74763 | 0.999994 | 16838 | 0 | 3.2788 | 0.27911 | 0.39196 | 1 | 10780 | 1 | 3.2788 |
| CCDC65 | 5 | 0.23713 | 0.42706 | 0.999994 | 8698 | 2 | 1.4977 | 0.27913 | 0.50346 | 1 | 10781 | 3 | 1.4977 |
| hsa-mir-5571 | 3 | 0.86894 | 0.86898 | 0.999994 | 18617 | 0 | 1.9951 | 0.27914 | 0.42765 | 1 | 10782 | 2 | 1.9951 |
| PPP1R14C | 6 | 0.72794 | 0.82092 | 0.999994 | 16580 | 1 | 0.76462 | 0.27916 | 0.53728 | 1 | 10783 | 4 | 0.76462 |
| WDR73 | 6 | 0.10462 | 0.25634 | 0.994943 | 5123 | 2 | 1.4216 | 0.27918 | 0.5373 | 1 | 10784 | 3 | 1.4216 |
| IGLL5 | 4 | 0.80792 | 0.80838 | 0.999994 | 17599 | 1 | 0.89327 | 0.2792 | 0.49096 | 1 | 10785 | 3 | 0.89327 |
| ATP6V1B2 | 3 | 0.6055 | 0.62139 | 0.999994 | 15019 | 1 | 1.8436 | 0.27922 | 0.42773 | 1 | 10786 | 2 | 1.8436 |
| SEC16A | 5 | 0.57588 | 0.71471 | 0.999994 | 14631 | 1 | 1.6101 | 0.27928 | 0.50363 | 1 | 10787 | 3 | 1.6101 |
| ACSM2B | 6 | 0.49903 | 0.66803 | 0.999994 | 13410 | 2 | 1.1774 | 0.27937 | 0.53753 | 1 | 10788 | 4 | 1.1774 |
| TPTE | 4 | 0.54524 | 0.63737 | 0.999994 | 14138 | 1 | 1.5003 | 0.27942 | 0.49125 | 1 | 10789 | 3 | 1.5003 |
| COQ9 | 5 | 0.25699 | 0.44629 | 0.999994 | 9168 | 2 | 0.21916 | 0.27954 | 0.50391 | 1 | 10790 | 2 | 0.21916 |
| CDKL4 | 6 | 0.16512 | 0.34461 | 0.999994 | 6864 | 2 | 0.78917 | 0.27962 | 0.53788 | 1 | 10791 | 4 | 0.78917 |
| RASGRF1 | 6 | 0.62014 | 0.75494 | 0.999994 | 15224 | 2 | 0.8853 | 0.27962 | 0.53788 | 1 | 10792 | 3 | 0.8853 |
| OR5M8 | 5 | 0.19829 | 0.37282 | 0.999994 | 7721 | 2 | 1.6953 | 0.27974 | 0.50414 | 1 | 10793 | 3 | 1.6953 |
| KLK2 | 6 | 0.1598 | 0.33799 | 0.999994 | 6727 | 3 | 0.16235 | 0.27976 | 0.53806 | 1 | 10794 | 3 | 0.16235 |
| FTCD | 6 | 0.66982 | 0.79301 | 0.999994 | 15928 | 2 | 0.96752 | 0.27985 | 0.53818 | 1 | 10795 | 4 | 0.96752 |
| ZNF22 | 5 | 0.086178 | 0.20445 | 0.978186 | 4457 | 1 | 1.0647 | 0.27986 | 0.50428 | 1 | 10796 | 3 | 1.0647 |
| NOL11 | 5 | 0.34473 | 0.51761 | 0.999994 | 11047 | 1 | 0.55108 | 0.27986 | 0.50428 | 1 | 10797 | 3 | 0.55108 |
| GNB1L | 4 | 0.55633 | 0.64278 | 0.999994 | 14308 | 1 | 0.71827 | 0.27993 | 0.49189 | 1 | 10798 | 3 | 0.71827 |
| CFHR1 | 5 | 0.77608 | 0.80519 | 0.999994 | 17151 | 1 | 0.66295 | 0.27996 | 0.50438 | 1 | 10799 | 3 | 0.66295 |
| STK19 | 5 | 0.10293 | 0.23096 | 0.983933 | 5064 | 2 | 1.0637 | 0.27996 | 0.50438 | 1 | 10800 | 3 | 1.0637 |
| CD68 | 6 | 0.91559 | 0.91641 | 0.999994 | 19529 | 1 | -0.042345 | 0.27998 | 0.53835 | 1 | 10801 | 2 | -0.042345 |
| NELL2 | 6 | 0.011156 | 0.042928 | 0.891621 | 937 | 4 | -2.9347 | 0.27998 | 0.53835 | 1 | 10802 | 2 | -2.9347 |
| ZHX3 | 4 | 0.30603 | 0.44913 | 0.999994 | 10256 | 1 | 1.7138 | 0.27999 | 0.49197 | 1 | 10803 | 3 | 1.7138 |
| CSPG4 | 4 | 0.73797 | 0.75099 | 0.999994 | 16702 | 1 | 1.3275 | 0.28004 | 0.49203 | 1 | 10804 | 3 | 1.3275 |
| KCTD5 | 5 | 0.76245 | 0.79754 | 0.999994 | 17002 | 1 | 1.4106 | 0.28005 | 0.50448 | 1 | 10805 | 3 | 1.4106 |
| CCDC33 | 6 | 0.2352 | 0.43028 | 0.999994 | 8654 | 2 | 1.423 | 0.28016 | 0.53858 | 1 | 10806 | 3 | 1.423 |
| SGSM1 | 6 | 0.0094832 | 0.03721 | 0.881473 | 831 | 2 | -0.46734 | 0.2802 | 0.53863 | 1 | 10807 | 2 | -0.46734 |
| FUBP1 | 4 | 0.081062 | 0.18065 | 0.968171 | 4267 | 1 | 0.7896 | 0.28024 | 0.4923 | 1 | 10808 | 3 | 0.7896 |
| NonTargetingControlG | 2 | 0.33188 | 0.36154 | 0.999994 | 10764 | 1 | 1.8264 | 0.28028 | 0.39268 | 1 | 10809 | 1 | 1.8264 |
| HTR3D | 6 | 0.47873 | 0.65434 | 0.999994 | 13125 | 2 | 0.90031 | 0.28033 | 0.53878 | 1 | 10810 | 3 | 0.90031 |
| VMAC | 5 | 0.45527 | 0.60967 | 0.999994 | 12778 | 2 | -0.57513 | 0.28046 | 0.5049 | 1 | 10811 | 2 | -0.57513 |
| TOLLIP | 5 | 0.87481 | 0.87544 | 0.999994 | 18729 | 1 | 0.62647 | 0.28046 | 0.5049 | 1 | 10812 | 3 | 0.62647 |
| USP32 | 5 | 0.58664 | 0.72135 | 0.999994 | 14767 | 1 | 0.067309 | 0.28046 | 0.5049 | 1 | 10813 | 2 | 0.067309 |
| SOGA3 | 4 | 0.67579 | 0.70894 | 0.999994 | 16018 | 1 | 0.68601 | 0.28047 | 0.49261 | 1 | 10814 | 3 | 0.68601 |
| POLQ | 6 | 0.045365 | 0.13387 | 0.95318 | 2782 | 2 | 1.1192 | 0.2805 | 0.53899 | 1 | 10815 | 4 | 1.1192 |

| | | | | | | | | | | | | | |
|---|---|---|---|---|---|---|---|---|---|---|---|---|---|
| hsa-mir-6802 | 4 | 0.69476 | 0.72103 | 0.999994 | 16253 | 1 | 1.0949 | 0.28062 | 0.49278 | 1 | 10816 | 3 | 1.0949 |
| CDKL5 | 6 | 0.29542 | 0.50136 | 0.999994 | 10027 | 2 | 1.0156 | 0.2807 | 0.53924 | 1 | 10817 | 3 | 1.0156 |
| ZNF688 | 6 | 0.6494 | 0.77717 | 0.999994 | 15649 | 2 | -0.12224 | 0.2807 | 0.53924 | 1 | 10818 | 1 | -0.12224 |
| hsa-mir-3163 | 3 | 0.71666 | 0.71679 | 0.999994 | 16462 | 1 | 1.571 | 0.28072 | 0.42931 | 1 | 10819 | 2 | 1.571 |
| AAK1 | 4 | 0.089388 | 0.1959 | 0.975797 | 4572 | 2 | 0.74124 | 0.28084 | 0.49308 | 1 | 10820 | 2 | 0.74124 |
| NonTargetingControlG | 2 | 0.37676 | 0.3977 | 0.999994 | 11604 | 1 | 2.0182 | 0.28096 | 0.39312 | 1 | 10821 | 1 | 2.0182 |
| POLA1 | 3 | 0.073569 | 0.14776 | 0.965554 | 3973 | 2 | -2.6619 | 0.2811 | 0.4297 | 1 | 10822 | 1 | -2.6619 |
| MYO7A | 6 | 0.041467 | 0.1252 | 0.951052 | 2589 | 1 | 0.4032 | 0.2811 | 0.53972 | 1 | 10823 | 3 | 0.4032 |
| SLC3A2 | 6 | 0.025118 | 0.087272 | 0.94395 | 1761 | 2 | 0.032159 | 0.2811 | 0.53972 | 1 | 10824 | 2 | 0.032159 |
| MMP20 | 6 | 0.81712 | 0.85546 | 0.999994 | 17730 | 1 | 1.1644 | 0.2811 | 0.53972 | 1 | 10825 | 3 | 1.1644 |
| UQCRH | 4 | 0.78096 | 0.78468 | 0.999994 | 17222 | 1 | 0.66392 | 0.28145 | 0.49388 | 1 | 10826 | 3 | 0.66392 |
| CPA6 | 6 | 0.7556 | 0.83046 | 0.999994 | 16927 | 1 | 1.0927 | 0.28152 | 0.54023 | 1 | 10827 | 3 | 1.0927 |
| SULT1A1 | 4 | 0.79774 | 0.7992 | 0.999994 | 17449 | 1 | 0.25552 | 0.28152 | 0.49396 | 1 | 10828 | 1 | 0.25552 |
| HN1 | 4 | 0.12889 | 0.26567 | 0.997535 | 5912 | 2 | -0.61027 | 0.28152 | 0.49396 | 1 | 10829 | 2 | -0.61027 |
| TSGA10IP | 6 | 0.66169 | 0.7866 | 0.999994 | 15806 | 1 | 0.97788 | 0.28156 | 0.54027 | 1 | 10830 | 4 | 0.97788 |
| CSDE1 | 5 | 0.50221 | 0.64983 | 0.999994 | 13465 | 2 | 0.71498 | 0.28162 | 0.50618 | 1 | 10831 | 3 | 0.71498 |
| SLC6A3 | 5 | 0.3091 | 0.48842 | 0.999994 | 10303 | 2 | 0.0044916 | 0.28162 | 0.50618 | 1 | 10832 | 2 | 0.0044916 |
| CDHR2 | 4 | 0.78324 | 0.78661 | 0.999994 | 17257 | 1 | 1.7144 | 0.28163 | 0.4941 | 1 | 10833 | 2 | 1.7144 |
| UQCRC1 | 4 | 0.10142 | 0.21769 | 0.980699 | 5018 | 1 | 1.978 | 0.28163 | 0.4941 | 1 | 10834 | 2 | 1.978 |
| hsa-mir-1909 | 3 | 0.67809 | 0.68101 | 0.999994 | 16061 | 1 | 1.8787 | 0.28167 | 0.43031 | 1 | 10835 | 2 | 1.8787 |
| AADACL2 | 6 | 0.83392 | 0.86349 | 0.999994 | 17986 | 1 | 1.4887 | 0.28177 | 0.54055 | 1 | 10836 | 4 | 1.4887 |
| VPS36 | 6 | 0.22729 | 0.42083 | 0.999994 | 8462 | 2 | -0.89425 | 0.28181 | 0.54061 | 1 | 10837 | 2 | -0.89425 |
| TMEM69 | 5 | 0.2829 | 0.46721 | 0.999994 | 9761 | 1 | 1.3182 | 0.28184 | 0.5064 | 1 | 10838 | 3 | 1.3182 |
| H2AFY | 5 | 0.83318 | 0.84176 | 0.999994 | 17975 | 1 | 0.67952 | 0.28184 | 0.5064 | 1 | 10839 | 3 | 0.67952 |
| POU6F1 | 5 | 0.43047 | 0.58882 | 0.999994 | 12422 | 2 | -0.16701 | 0.28184 | 0.5064 | 1 | 10840 | 2 | -0.16701 |
| ZNF692 | 6 | 0.52775 | 0.68797 | 0.999994 | 13858 | 2 | 0.69488 | 0.2819 | 0.54073 | 1 | 10841 | 4 | 0.69488 |
| hsa-mir-9-3 | 3 | 0.5882 | 0.60814 | 0.999994 | 14793 | 1 | 1.6045 | 0.28193 | 0.43059 | 1 | 10842 | 2 | 1.6045 |
| CSNK2A3 | 6 | 0.96764 | 0.96765 | 0.999994 | 20775 | 0 | 1.186 | 0.28195 | 0.54079 | 1 | 10843 | 4 | 1.186 |
| WFDC9 | 4 | 0.24621 | 0.38801 | 0.999994 | 8928 | 1 | 0.71515 | 0.28197 | 0.49453 | 1 | 10844 | 3 | 0.71515 |
| hsa-mir-6760 | 3 | 0.40679 | 0.48829 | 0.999994 | 12059 | 1 | 1.8672 | 0.28198 | 0.43065 | 1 | 10845 | 2 | 1.8672 |
| PNP | 6 | 0.32469 | 0.53521 | 0.999994 | 10612 | 1 | 1.1241 | 0.28208 | 0.54097 | 1 | 10846 | 4 | 1.1241 |
| TBL1X | 6 | 0.69939 | 0.81183 | 0.999994 | 16301 | 1 | 0.79587 | 0.28213 | 0.54103 | 1 | 10847 | 4 | 0.79587 |
| UBLCP1 | 6 | 0.95094 | 0.95093 | 0.999994 | 20354 | 0 | 0.99552 | 0.28213 | 0.54103 | 1 | 10848 | 3 | 0.99552 |
| SIRT3 | 5 | 0.14275 | 0.292 | 0.999632 | 6262 | 1 | 0.93167 | 0.28225 | 0.50685 | 1 | 10849 | 3 | 0.93167 |
| TAX1BP1 | 6 | 0.15457 | 0.33144 | 0.999994 | 6583 | 2 | 0.63586 | 0.28226 | 0.54121 | 1 | 10850 | 3 | 0.63586 |
| TGFB2 | 6 | 0.078626 | 0.20457 | 0.978186 | 4176 | 4 | -1.3056 | 0.28226 | 0.54121 | 1 | 10851 | 2 | -1.3056 |
| FAM174B | 6 | 0.50051 | 0.66904 | 0.999994 | 13429 | 1 | 0.36185 | 0.28226 | 0.54121 | 1 | 10852 | 2 | 0.36185 |
| RPL34 | 3 | 0.093929 | 0.17501 | 0.968171 | 4747 | 1 | 1.7977 | 0.28235 | 0.43103 | 1 | 10853 | 2 | 1.7977 |
| SLC16A10 | 6 | 0.62114 | 0.75569 | 0.999994 | 15245 | 2 | 0.95937 | 0.28245 | 0.54145 | 1 | 10854 | 4 | 0.95937 |
| ATP8B4 | 5 | 0.80044 | 0.81978 | 0.999994 | 17487 | 1 | 1.4319 | 0.28251 | 0.50712 | 1 | 10855 | 3 | 1.4319 |
| PTPRD | 5 | 0.9102 | 0.91022 | 0.999994 | 19428 | 0 | 1.4233 | 0.28252 | 0.50713 | 1 | 10856 | 3 | 1.4233 |
| MRPL14 | 5 | 0.18084 | 0.34785 | 0.999994 | 7243 | 1 | 1.5464 | 0.28255 | 0.50717 | 1 | 10857 | 3 | 1.5464 |
| WFIKKN2 | 6 | 0.44207 | 0.62979 | 0.999994 | 12610 | 1 | 1.4597 | 0.28256 | 0.54159 | 1 | 10858 | 4 | 1.4597 |
| OR2D3 | 5 | 0.60583 | 0.72829 | 0.999994 | 15024 | 1 | 0.83882 | 0.28263 | 0.50725 | 1 | 10859 | 3 | 0.83882 |
| CCDC78 | 6 | 0.72698 | 0.82061 | 0.999994 | 16568 | 1 | 0.89679 | 0.28271 | 0.54178 | 1 | 10860 | 4 | 0.89679 |
| FUNDC2 | 6 | 0.34284 | 0.55591 | 0.999994 | 11003 | 3 | 0.77371 | 0.28276 | 0.54183 | 1 | 10861 | 3 | 0.77371 |
| hsa-mir-1269a | 4 | 0.62863 | 0.68113 | 0.999994 | 15357 | 1 | 0.90514 | 0.28277 | 0.49557 | 1 | 10862 | 3 | 0.90514 |
| IFITM2 | 5 | 0.47487 | 0.62623 | 0.999994 | 13059 | 1 | 0.65761 | 0.28282 | 0.50747 | 1 | 10863 | 3 | 0.65761 |
| DIS3 | 4 | 0.82851 | 0.82805 | 0.999994 | 17903 | 0 | 1.571 | 0.28291 | 0.49576 | 1 | 10864 | 2 | 1.571 |
| TLR10 | 4 | 0.07083 | 0.1615 | 0.967126 | 3868 | 3 | -1.986 | 0.28292 | 0.49578 | 1 | 10865 | 1 | -1.986 |
| GTF2IRD2B | 4 | 0.70037 | 0.72462 | 0.999994 | 16307 | 1 | 0.44895 | 0.28292 | 0.49578 | 1 | 10866 | 2 | 0.44895 |
| hsa-mir-1243 | 4 | 0.11495 | 0.24149 | 0.987253 | 5475 | 1 | 0.71063 | 0.28292 | 0.49578 | 1 | 10867 | 2 | 0.71063 |

| | | | | | | | | | | | | | |
|---|---|---|---|---|---|---|---|---|---|---|---|---|---|
| SPATA21 | 6 | 0.3539 | 0.56841 | 0.999994 | 11232 | 2 | 1.0785 | 0.28304 | 0.54218 | 1 | 10868 | 3 | 1.0785 |
| NDUFS2 | 6 | 0.47455 | 0.65151 | 0.999994 | 13057 | 2 | 1.2024 | 0.28304 | 0.54219 | 1 | 10869 | 4 | 1.2024 |
| hsa-mir-5706 | 2 | 0.43268 | 0.44433 | 0.999994 | 12455 | 1 | 2.2879 | 0.28307 | 0.3944 | 1 | 10870 | 1 | 2.2879 |
| C14orf132 | 5 | 0.23199 | 0.41997 | 0.999994 | 8570 | 2 | 1.6603 | 0.28309 | 0.50773 | 1 | 10871 | 3 | 1.6603 |
| GSTK1 | 5 | 0.83914 | 0.846 | 0.999994 | 18073 | 1 | 0.69156 | 0.28314 | 0.5078 | 1 | 10872 | 3 | 0.69156 |
| PLEKHM1 | 5 | 0.70066 | 0.76677 | 0.999994 | 16310 | 1 | 0.53989 | 0.28314 | 0.5078 | 1 | 10873 | 3 | 0.53989 |
| CPSF4 | 6 | 0.06058 | 0.16698 | 0.967126 | 3449 | 3 | -0.25716 | 0.28316 | 0.54232 | 1 | 10874 | 3 | -0.25716 |
| MSL2 | 6 | 0.20625 | 0.3955 | 0.999994 | 7937 | 2 | 0.093418 | 0.28321 | 0.54239 | 1 | 10875 | 2 | 0.093418 |
| EGLN2 | 6 | 0.32012 | 0.53002 | 0.999994 | 10513 | 2 | -0.42575 | 0.28321 | 0.54239 | 1 | 10876 | 2 | -0.42575 |
| FAM194B | 6 | 0.32588 | 0.53655 | 0.999994 | 10640 | 2 | -0.65143 | 0.28321 | 0.54239 | 1 | 10877 | 1 | -0.65143 |
| MLC1 | 6 | 0.32677 | 0.53756 | 0.999994 | 10657 | 2 | 1.0059 | 0.28327 | 0.54247 | 1 | 10878 | 4 | 1.0059 |
| SLC43A2 | 4 | 0.66009 | 0.69932 | 0.999994 | 15781 | 1 | 1.2375 | 0.28328 | 0.49624 | 1 | 10879 | 3 | 1.2375 |
| C22orf29 | 6 | 0.90546 | 0.90792 | 0.999994 | 19343 | 1 | 0.81491 | 0.28332 | 0.54253 | 1 | 10880 | 4 | 0.81491 |
| NPW | 6 | 0.65972 | 0.7851 | 0.999994 | 15774 | 1 | 1.2371 | 0.28334 | 0.54255 | 1 | 10881 | 3 | 1.2371 |
| PYCRL | 5 | 0.53987 | 0.68276 | 0.999994 | 14049 | 1 | 1.3796 | 0.28335 | 0.50804 | 1 | 10882 | 3 | 1.3796 |
| SNIP1 | 3 | 0.13423 | 0.22815 | 0.983354 | 6029 | 2 | -1.6809 | 0.28346 | 0.43219 | 1 | 10883 | 1 | -1.6809 |
| COG1 | 4 | 0.30862 | 0.4518 | 0.999994 | 10293 | 2 | -1.2542 | 0.28351 | 0.49654 | 1 | 10884 | 1 | -1.2542 |
| TMBIM6 | 5 | 0.45525 | 0.60966 | 0.999994 | 12776 | 2 | 1.5236 | 0.28351 | 0.5082 | 1 | 10885 | 3 | 1.5236 |
| DUSP18 | 6 | 0.37736 | 0.58789 | 0.999994 | 11614 | 2 | 1.4478 | 0.28359 | 0.54291 | 1 | 10886 | 4 | 1.4478 |
| ECHDC1 | 6 | 0.89534 | 0.90015 | 0.999994 | 19133 | 1 | 0.9002 | 0.28366 | 0.54299 | 1 | 10887 | 3 | 0.9002 |
| KIAA0226L | 6 | 0.46462 | 0.64483 | 0.999994 | 12928 | 2 | 1.0439 | 0.28366 | 0.54299 | 1 | 10888 | 4 | 1.0439 |
| ASPH | 5 | 0.1067 | 0.23684 | 0.985449 | 5196 | 3 | -0.92565 | 0.28379 | 0.50848 | 1 | 10889 | 2 | -0.92565 |
| C10orf62 | 5 | 0.20787 | 0.38637 | 0.999994 | 7971 | 2 | -0.16852 | 0.28387 | 0.50857 | 1 | 10890 | 2 | -0.16852 |
| APLP2 | 5 | 0.083103 | 0.1996 | 0.977164 | 4342 | 2 | 0.4075 | 0.28387 | 0.50857 | 1 | 10891 | 2 | 0.4075 |
| COL6A5 | 6 | 0.55456 | 0.70692 | 0.999994 | 14277 | 2 | 0.89545 | 0.28407 | 0.54353 | 1 | 10892 | 4 | 0.89545 |
| NDRG1 | 4 | 0.65278 | 0.69489 | 0.999994 | 15697 | 1 | 0.67786 | 0.28411 | 0.4973 | 1 | 10893 | 3 | 0.67786 |
| ZNF766 | 6 | 0.67626 | 0.79808 | 0.999994 | 16026 | 2 | 1.2384 | 0.28414 | 0.54362 | 1 | 10894 | 4 | 1.2384 |
| TALDO1 | 6 | 0.10627 | 0.25953 | 0.994943 | 5182 | 2 | 1.115 | 0.28417 | 0.54367 | 1 | 10895 | 4 | 1.115 |
| SMPDL3B | 5 | 0.62187 | 0.73426 | 0.999994 | 15256 | 1 | 1.3973 | 0.28417 | 0.5089 | 1 | 10896 | 3 | 1.3973 |
| DYNC1H1 | 4 | 0.35382 | 0.49818 | 0.999994 | 11229 | 1 | 1.454 | 0.28424 | 0.49746 | 1 | 10897 | 2 | 1.454 |
| IL20RA | 5 | 0.059322 | 0.16092 | 0.967126 | 3394 | 3 | -1.8138 | 0.28435 | 0.50908 | 1 | 10898 | 2 | -1.8138 |
| LRRC3C | 4 | 0.901 | 0.90068 | 0.999994 | 19234 | 0 | 1.4178 | 0.2845 | 0.49782 | 1 | 10899 | 3 | 1.4178 |
| TAP2 | 5 | 0.17439 | 0.33852 | 0.999994 | 7093 | 1 | 1.2854 | 0.28468 | 0.50943 | 1 | 10900 | 3 | 1.2854 |
| ADAMTS15 | 4 | 0.033552 | 0.088779 | 0.94395 | 2192 | 2 | -3.0264 | 0.28471 | 0.49808 | 1 | 10901 | 1 | -3.0264 |
| hsa-mir-1203 | 3 | 0.68363 | 0.68601 | 0.999994 | 16139 | 1 | 1.703 | 0.28471 | 0.43354 | 1 | 10902 | 2 | 1.703 |
| CXorf49 | 6 | 0.10705 | 0.26102 | 0.994943 | 5213 | 3 | 0.14441 | 0.28476 | 0.54443 | 1 | 10903 | 3 | 0.14441 |
| RPL7A | 4 | 0.72572 | 0.74208 | 0.999994 | 16557 | 1 | 1.0414 | 0.28478 | 0.49818 | 1 | 10904 | 3 | 1.0414 |
| PDCD5 | 5 | 0.40162 | 0.56486 | 0.999994 | 11972 | 1 | 1.6319 | 0.2848 | 0.50955 | 1 | 10905 | 3 | 1.6319 |
| UNG | 5 | 0.014172 | 0.047863 | 0.900643 | 1140 | 4 | -1.7282 | 0.2848 | 0.50955 | 1 | 10906 | 1 | -1.7282 |
| MYBPC1 | 6 | 0.016396 | 0.060295 | 0.932932 | 1270 | 4 | -2.3801 | 0.28481 | 0.54451 | 1 | 10907 | 2 | -2.3801 |
| ALKBH1 | 6 | 0.12597 | 0.29521 | 0.999994 | 5820 | 2 | 0.3472 | 0.28481 | 0.54451 | 1 | 10908 | 3 | 0.3472 |
| ULK4 | 5 | 0.18918 | 0.35978 | 0.999994 | 7457 | 1 | 1.1093 | 0.28482 | 0.50957 | 1 | 10909 | 3 | 1.1093 |
| hsa-mir-1302-5 | 2 | 0.51086 | 0.51388 | 0.999994 | 13592 | 1 | 2.5289 | 0.28489 | 0.39553 | 1 | 10910 | 1 | 2.5289 |
| BCOR | 5 | 0.064548 | 0.16954 | 0.967126 | 3632 | 2 | 0.16289 | 0.2849 | 0.50966 | 1 | 10911 | 2 | 0.16289 |
| ADHFE1 | 3 | 0.21466 | 0.33102 | 0.999994 | 8137 | 1 | 2.0788 | 0.28491 | 0.43372 | 1 | 10912 | 2 | 2.0788 |
| VPS37D | 6 | 0.78715 | 0.84252 | 0.999994 | 17306 | 1 | 1.2256 | 0.28496 | 0.54469 | 1 | 10913 | 4 | 1.2256 |
| SOX2 | 4 | 0.56016 | 0.64474 | 0.999994 | 14373 | 1 | 1.9602 | 0.2851 | 0.49859 | 1 | 10914 | 3 | 1.9602 |
| TMEM196 | 4 | 0.77136 | 0.77688 | 0.999994 | 17103 | 1 | 1.6351 | 0.28522 | 0.49873 | 1 | 10915 | 2 | 1.6351 |
| SLC2A4RG | 4 | 0.81909 | 0.81886 | 0.999994 | 17767 | 1 | 1.5717 | 0.28535 | 0.4989 | 1 | 10916 | 2 | 1.5717 |
| NPDC1 | 6 | 0.73862 | 0.82452 | 0.999994 | 16714 | 1 | 0.77202 | 0.28537 | 0.54522 | 1 | 10917 | 4 | 0.77202 |
| PLEKHA6 | 5 | 0.13854 | 0.28563 | 0.999632 | 6154 | 2 | 0.56311 | 0.28544 | 0.51022 | 1 | 10918 | 3 | 0.56311 |
| CCR1 | 6 | 0.07419 | 0.19548 | 0.975797 | 4000 | 2 | 0.87377 | 0.28545 | 0.54533 | 1 | 10919 | 4 | 0.87377 |

| | | | | | | | | | | | | | |
|---|---|---|---|---|---|---|---|---|---|---|---|---|---|
| TARBP1 | 6 | 0.53895 | 0.69581 | 0.999994 | 14034 | 2 | 0.99159 | 0.28546 | 0.54535 | 1 | 10920 | 4 | 0.99159 |
| KRTAP9-4 | 5 | 0.030645 | 0.092156 | 0.943968 | 2038 | 3 | -3.0689 | 0.28548 | 0.51025 | 1 | 10921 | 2 | -3.0689 |
| TMEM159 | 6 | 0.67666 | 0.79837 | 0.999994 | 16034 | 1 | 0.40285 | 0.28549 | 0.54539 | 1 | 10922 | 3 | 0.40285 |
| JMJD7 | 6 | 0.97187 | 0.97182 | 0.999994 | 20866 | 0 | 1.1628 | 0.28551 | 0.54541 | 1 | 10923 | 4 | 1.1628 |
| NonTargetingControlG | 1 | 0.71445 | 0.71457 | 0.999994 | 16444 | 0 | 2.9091 | 0.28555 | 0.28543 | 0.978871 | 10924 | 1 | 2.9091 |
| hsa-mir-373 | 4 | 0.77208 | 0.77748 | 0.999994 | 17113 | 1 | 1.4223 | 0.28562 | 0.49926 | 1 | 10925 | 2 | 1.4223 |
| EPYC | 4 | 0.74058 | 0.75293 | 0.999994 | 16744 | 1 | 1.2112 | 0.28564 | 0.49929 | 1 | 10926 | 3 | 1.2112 |
| NCKAP5 | 6 | 0.57088 | 0.71857 | 0.999994 | 14558 | 2 | 1.2734 | 0.28568 | 0.54562 | 1 | 10927 | 3 | 1.2734 |
| hsa-mir-494 | 4 | 0.35944 | 0.50396 | 0.999994 | 11329 | 1 | 1.0674 | 0.28584 | 0.49954 | 1 | 10928 | 2 | 1.0674 |
| IL31RA | 6 | 0.31416 | 0.5232 | 0.999994 | 10403 | 2 | 1.4552 | 0.2859 | 0.5459 | 1 | 10929 | 3 | 1.4552 |
| USMG5 | 6 | 0.15767 | 0.33528 | 0.999994 | 6664 | 3 | 0.83178 | 0.28592 | 0.54593 | 1 | 10930 | 3 | 0.83178 |
| CPNE1 | 6 | 0.042162 | 0.12676 | 0.951052 | 2621 | 4 | -1.682 | 0.28593 | 0.54594 | 1 | 10931 | 2 | -1.682 |
| RRP8 | 6 | 0.2164 | 0.40783 | 0.999994 | 8178 | 3 | -0.43038 | 0.28593 | 0.54594 | 1 | 10932 | 2 | -0.43038 |
| SLFN12L | 6 | 0.2683 | 0.46969 | 0.999994 | 9424 | 3 | -0.2972 | 0.28593 | 0.54594 | 1 | 10933 | 2 | -0.2972 |
| GTF3A | 3 | 0.66718 | 0.67148 | 0.999994 | 15880 | 1 | 1.679 | 0.28594 | 0.4348 | 1 | 10934 | 2 | 1.679 |
| hsa-mir-33a | 4 | 0.7773 | 0.78168 | 0.999994 | 17171 | 1 | 0.90559 | 0.28601 | 0.49975 | 1 | 10935 | 3 | 0.90559 |
| RNASEK | 3 | 0.1173 | 0.20592 | 0.978744 | 5551 | 2 | -0.84827 | 0.28604 | 0.4349 | 1 | 10936 | 1 | -0.84827 |
| SPRTN | 5 | 0.25594 | 0.44547 | 0.999994 | 9149 | 2 | -0.17959 | 0.28604 | 0.51084 | 1 | 10937 | 2 | -0.17959 |
| PTPN21 | 6 | 0.53773 | 0.69493 | 0.999994 | 14024 | 2 | 1.0395 | 0.28607 | 0.54613 | 1 | 10938 | 4 | 1.0395 |
| SLC39A11 | 6 | 0.9613 | 0.96135 | 0.999994 | 20611 | 0 | 1.2122 | 0.28609 | 0.54614 | 1 | 10939 | 4 | 1.2122 |
| KLHL26 | 6 | 0.34606 | 0.55959 | 0.999994 | 11070 | 2 | 1.1976 | 0.28617 | 0.54625 | 1 | 10940 | 3 | 1.1976 |
| RLTPR | 4 | 0.54097 | 0.63532 | 0.999994 | 14072 | 1 | 0.70019 | 0.28624 | 0.50004 | 1 | 10941 | 3 | 0.70019 |
| SNX7 | 6 | 0.22632 | 0.41968 | 0.999994 | 8432 | 3 | 0.019081 | 0.28626 | 0.54637 | 1 | 10942 | 3 | 0.019081 |
| OGT | 5 | 0.42985 | 0.58829 | 0.999994 | 12411 | 2 | 0.68636 | 0.2863 | 0.51113 | 1 | 10943 | 3 | 0.68636 |
| ABO | 5 | 0.85351 | 0.85707 | 0.999994 | 18324 | 1 | 0.76075 | 0.2863 | 0.51113 | 1 | 10944 | 3 | 0.76075 |
| CNTF | 5 | 0.79026 | 0.81348 | 0.999994 | 17348 | 1 | 0.57704 | 0.2863 | 0.51113 | 1 | 10945 | 2 | 0.57704 |
| CRYBA2 | 6 | 0.59199 | 0.73408 | 0.999994 | 14850 | 2 | 1.3177 | 0.28635 | 0.54649 | 1 | 10946 | 4 | 1.3177 |
| TCP10L2 | 4 | 0.17543 | 0.3154 | 0.999994 | 7122 | 2 | -0.55821 | 0.28643 | 0.50028 | 1 | 10947 | 2 | -0.55821 |
| SLC24A5 | 6 | 0.1082 | 0.26326 | 0.996734 | 5246 | 2 | 0.4044 | 0.28646 | 0.54663 | 1 | 10948 | 3 | 0.4044 |
| PMP2 | 6 | 0.041948 | 0.12627 | 0.951052 | 2611 | 3 | -0.45355 | 0.28646 | 0.54663 | 1 | 10949 | 2 | -0.45355 |
| CNOT4 | 5 | 0.20137 | 0.37718 | 0.999994 | 7804 | 2 | 1.4408 | 0.2865 | 0.51134 | 1 | 10950 | 3 | 1.4408 |
| VPS37C | 5 | 0.15808 | 0.31475 | 0.999994 | 6673 | 2 | -0.36601 | 0.28656 | 0.5114 | 1 | 10951 | 2 | -0.36601 |
| DAOA | 6 | 0.52929 | 0.68905 | 0.999994 | 13888 | 2 | 1.2235 | 0.2866 | 0.5468 | 1 | 10952 | 4 | 1.2235 |
| SLC5A10 | 6 | 0.096185 | 0.23984 | 0.987253 | 4823 | 2 | 1.2855 | 0.28662 | 0.54684 | 1 | 10953 | 3 | 1.2855 |
| CMTM5 | 6 | 0.016195 | 0.059649 | 0.932932 | 1255 | 5 | -1.0985 | 0.28689 | 0.54717 | 1 | 10954 | 1 | -1.0985 |
| PRPF6 | 6 | 0.45367 | 0.63751 | 0.999994 | 12754 | 2 | 0.79229 | 0.28695 | 0.54724 | 1 | 10955 | 3 | 0.79229 |
| NUDT22 | 5 | 0.039012 | 0.1134 | 0.951052 | 2461 | 2 | 1.4942 | 0.28697 | 0.51186 | 1 | 10956 | 3 | 1.4942 |
| WNT5A | 4 | 0.66687 | 0.70347 | 0.999994 | 15877 | 1 | 0.25304 | 0.28707 | 0.5011 | 1 | 10957 | 1 | 0.25304 |
| OVCA2 | 5 | 0.55389 | 0.69522 | 0.999994 | 14267 | 2 | -0.28995 | 0.28708 | 0.51198 | 1 | 10958 | 2 | -0.28995 |
| TRPC7 | 5 | 0.081769 | 0.19747 | 0.977164 | 4290 | 3 | -1.7381 | 0.28708 | 0.51198 | 1 | 10959 | 1 | -1.7381 |
| CABP7 | 6 | 0.30628 | 0.51405 | 0.999994 | 10260 | 2 | 0.095835 | 0.28731 | 0.54771 | 1 | 10960 | 3 | 0.095835 |
| HIST2H3D | 6 | 0.021963 | 0.078261 | 0.93914 | 1578 | 3 | -0.50222 | 0.28731 | 0.54771 | 1 | 10961 | 3 | -0.50222 |
| HLX | 6 | 0.13836 | 0.31095 | 0.999994 | 6149 | 3 | 0.38416 | 0.28731 | 0.54771 | 1 | 10962 | 3 | 0.38416 |
| C1orf186 | 6 | 0.00099687 | 0.0050666 | 0.589881 | 169 | 5 | -5.466 | 0.28731 | 0.54771 | 1 | 10963 | 1 | -5.466 |
| MED10 | 5 | 0.43914 | 0.59611 | 0.999994 | 12560 | 2 | 1.4785 | 0.28738 | 0.51227 | 1 | 10964 | 3 | 1.4785 |
| PAPOLB | 6 | 0.058302 | 0.16213 | 0.967126 | 3353 | 4 | -1.8161 | 0.2874 | 0.54781 | 1 | 10965 | 2 | -1.8161 |
| TMED7 | 6 | 0.91456 | 0.91551 | 0.999994 | 19499 | 1 | 1.1237 | 0.2874 | 0.54782 | 1 | 10966 | 4 | 1.1237 |
| MYH10 | 5 | 0.34058 | 0.5142 | 0.999994 | 10947 | 2 | 1.4327 | 0.28741 | 0.5123 | 1 | 10967 | 3 | 1.4327 |
| PRR12 | 6 | 0.45379 | 0.63759 | 0.999994 | 12758 | 2 | 0.89673 | 0.28744 | 0.54786 | 1 | 10968 | 4 | 0.89673 |
| hsa-mir-133a-1 | 4 | 0.053049 | 0.12744 | 0.951052 | 3114 | 2 | 0.38819 | 0.28755 | 0.50173 | 1 | 10969 | 2 | 0.38819 |
| HTR2A | 5 | 0.86028 | 0.86264 | 0.999994 | 18446 | 1 | 0.92184 | 0.2876 | 0.51251 | 1 | 10970 | 3 | 0.92184 |
| UBTD2 | 6 | 0.61603 | 0.75186 | 0.999994 | 15162 | 2 | 1.103 | 0.28764 | 0.54814 | 1 | 10971 | 3 | 1.103 |

| | | | | | | | | | | | | | |
|---|---|---|---|---|---|---|---|---|---|---|---|---|---|
| NonTargetingControlG | 2 | 0.65977 | 0.65957 | 0.999994 | 15776 | 0 | 2.8263 | 0.28765 | 0.39723 | 1 | 10972 | 1 | 2.8263 |
| hsa-mir-411 | 4 | 0.91735 | 0.91714 | 0.999994 | 19571 | 0 | 0.92006 | 0.28772 | 0.50195 | 1 | 10973 | 3 | 0.92006 |
| MCM3 | 5 | 0.84718 | 0.85208 | 0.999994 | 18209 | 1 | 0.35136 | 0.28774 | 0.51264 | 1 | 10974 | 2 | 0.35136 |
| N4BP2L2 | 5 | 0.032475 | 0.096814 | 0.945031 | 2134 | 2 | 0.90647 | 0.28774 | 0.51264 | 1 | 10975 | 3 | 0.90647 |
| COPRS | 6 | 0.076689 | 0.20067 | 0.977164 | 4099 | 2 | 1.3494 | 0.28775 | 0.54828 | 1 | 10976 | 3 | 1.3494 |
| CHI3L1 | 6 | 0.036716 | 0.11449 | 0.951052 | 2358 | 4 | -1.5215 | 0.28785 | 0.5484 | 1 | 10977 | 1 | -1.5215 |
| PTBP3 | 6 | 0.20349 | 0.39211 | 0.999994 | 7869 | 3 | 0.078995 | 0.28785 | 0.5484 | 1 | 10978 | 3 | 0.078995 |
| C7orf55 | 5 | 0.56634 | 0.70631 | 0.999994 | 14485 | 2 | 1.6466 | 0.28795 | 0.51287 | 1 | 10979 | 3 | 1.6466 |
| RTN4IP1 | 6 | 0.91818 | 0.91873 | 0.999994 | 19589 | 1 | 0.89623 | 0.28798 | 0.54857 | 1 | 10980 | 4 | 0.89623 |
| ATG2B | 6 | 0.095239 | 0.23799 | 0.987253 | 4792 | 1 | 0.63632 | 0.28811 | 0.54872 | 1 | 10981 | 4 | 0.63632 |
| hsa-mir-203b | 1 | 0.71181 | 0.71192 | 0.999994 | 16419 | 0 | 2.721 | 0.28819 | 0.28809 | 0.979689 | 10982 | 1 | 2.721 |
| PDCD10 | 4 | 0.029064 | 0.079609 | 0.93914 | 1958 | 3 | -2.0805 | 0.2882 | 0.50258 | 1 | 10983 | 1 | -2.0805 |
| FUT6 | 6 | 0.051269 | 0.14686 | 0.964852 | 3038 | 4 | -0.92455 | 0.28821 | 0.54885 | 1 | 10984 | 2 | -0.92455 |
| INIP | 6 | 0.86827 | 0.88228 | 0.999994 | 18606 | 1 | 1.2808 | 0.28821 | 0.54885 | 1 | 10985 | 4 | 1.2808 |
| ZNF540 | 6 | 0.412 | 0.6102 | 0.999994 | 12145 | 1 | 0.8691 | 0.28823 | 0.54886 | 1 | 10986 | 4 | 0.8691 |
| hsa-mir-9-2 | 4 | 0.59523 | 0.66275 | 0.999994 | 14892 | 1 | 0.67678 | 0.28828 | 0.50268 | 1 | 10987 | 3 | 0.67678 |
| ZFYVE21 | 6 | 0.12551 | 0.29464 | 0.999994 | 5805 | 2 | 0.662 | 0.28839 | 0.54908 | 1 | 10988 | 4 | 0.662 |
| SCN3A | 6 | 0.033274 | 0.10647 | 0.950995 | 2177 | 3 | -0.96403 | 0.28846 | 0.54917 | 1 | 10989 | 2 | -0.96403 |
| PNMAL1 | 6 | 0.51118 | 0.67643 | 0.999994 | 13600 | 2 | -0.000097222 | 0.28846 | 0.54917 | 1 | 10990 | 2 | -0.000097222 |
| UCN3 | 6 | 0.25414 | 0.45287 | 0.999994 | 9106 | 3 | 0.44094 | 0.28846 | 0.54917 | 1 | 10991 | 3 | 0.44094 |
| CDC25C | 5 | 0.59132 | 0.723 | 0.999994 | 14838 | 1 | 1.382 | 0.28849 | 0.51341 | 1 | 10992 | 3 | 1.382 |
| NonTargetingControlG | 1 | 0.71146 | 0.71158 | 0.999994 | 16416 | 0 | 2.8769 | 0.28854 | 0.28845 | 0.979944 | 10993 | 1 | 2.8769 |
| RSPH4A | 6 | 0.21815 | 0.40991 | 0.999994 | 8231 | 3 | 0.58052 | 0.28865 | 0.5494 | 1 | 10994 | 3 | 0.58052 |
| MAP4K2 | 5 | 0.46167 | 0.61506 | 0.999994 | 12876 | 2 | -0.0074411 | 0.28874 | 0.51367 | 1 | 10995 | 2 | -0.0074411 |
| CCDC174 | 6 | 0.50742 | 0.67387 | 0.999994 | 13542 | 2 | 0.76256 | 0.28874 | 0.54951 | 1 | 10996 | 3 | 0.76256 |
| TRIM71 | 6 | 0.058975 | 0.16356 | 0.967126 | 3380 | 4 | -1.6684 | 0.28874 | 0.54951 | 1 | 10997 | 2 | -1.6684 |
| ANKRD1 | 6 | 0.11265 | 0.27181 | 0.998561 | 5389 | 3 | -0.57804 | 0.28874 | 0.54951 | 1 | 10998 | 3 | -0.57804 |
| hsa-mir-6129 | 4 | 0.3305 | 0.47421 | 0.999994 | 10730 | 1 | 1.7542 | 0.28884 | 0.5034 | 1 | 10999 | 2 | 1.7542 |
| RNF4 | 2 | 0.52378 | 0.52591 | 0.999994 | 13805 | 1 | 2.4945 | 0.28886 | 0.39794 | 1 | 11000 | 1 | 2.4945 |
| PCDHA2 | 2 | 0.13707 | 0.21767 | 0.980699 | 6109 | 1 | -0.67213 | 0.28886 | 0.39794 | 1 | 11001 | 1 | -0.67213 |
| RHOU | 6 | 0.066957 | 0.18038 | 0.968171 | 3719 | 3 | -1.0296 | 0.28889 | 0.54969 | 1 | 11002 | 2 | -1.0296 |
| KRTAP10-6 | 5 | 0.24066 | 0.43183 | 0.999994 | 8787 | 2 | 1.5493 | 0.28903 | 0.51399 | 1 | 11003 | 3 | 1.5493 |
| PCYT1A | 5 | 0.24682 | 0.43825 | 0.999994 | 8940 | 1 | 0.46563 | 0.28906 | 0.51403 | 1 | 11004 | 2 | 0.46563 |
| TAS2R19 | 6 | 0.94674 | 0.94671 | 0.999994 | 20257 | 0 | 0.8742 | 0.28908 | 0.54993 | 1 | 11005 | 4 | 0.8742 |
| WDR65 | 6 | 0.32904 | 0.54014 | 0.999994 | 10705 | 1 | 0.78883 | 0.28912 | 0.54998 | 1 | 11006 | 4 | 0.78883 |
| TLE2 | 5 | 0.32737 | 0.50337 | 0.999994 | 10665 | 2 | -1.2059 | 0.2892 | 0.51418 | 1 | 11007 | 1 | -1.2059 |
| YTHDF2 | 5 | 0.42186 | 0.58163 | 0.999994 | 12274 | 2 | -0.28902 | 0.28924 | 0.51422 | 1 | 11008 | 2 | -0.28902 |
| RNF130 | 6 | 0.60192 | 0.74135 | 0.999994 | 14971 | 1 | 1.1077 | 0.28927 | 0.55016 | 1 | 11009 | 3 | 1.1077 |
| hsa-mir-323b | 4 | 0.1792 | 0.31927 | 0.999994 | 7203 | 1 | 1.4408 | 0.28927 | 0.50395 | 1 | 11010 | 3 | 1.4408 |
| NCS1 | 6 | 0.33361 | 0.54542 | 0.999994 | 10797 | 3 | 0.66001 | 0.28932 | 0.55024 | 1 | 11011 | 3 | 0.66001 |
| C16orf52 | 5 | 0.63585 | 0.73958 | 0.999994 | 15457 | 1 | 1.3021 | 0.28937 | 0.51436 | 1 | 11012 | 3 | 1.3021 |
| CCNY | 6 | 0.13543 | 0.30722 | 0.999994 | 6064 | 3 | -0.089307 | 0.28942 | 0.55037 | 1 | 11013 | 3 | -0.089307 |
| KRTAP5-7 | 5 | 0.089259 | 0.20938 | 0.979587 | 4564 | 3 | -1.7235 | 0.28943 | 0.51441 | 1 | 11014 | 2 | -1.7235 |
| OR7C1 | 5 | 0.054742 | 0.15185 | 0.967126 | 3184 | 3 | -1.3522 | 0.28947 | 0.51445 | 1 | 11015 | 2 | -1.3522 |
| RSAD2 | 6 | 0.52169 | 0.68368 | 0.999994 | 13774 | 2 | 1.3851 | 0.28949 | 0.55047 | 1 | 11016 | 4 | 1.3851 |
| CSRP3 | 3 | 0.52162 | 0.56061 | 0.999994 | 13771 | 1 | 1.7508 | 0.28953 | 0.43857 | 1 | 11017 | 2 | 1.7508 |
| AP2A1 | 4 | 0.45066 | 0.59497 | 0.999994 | 12721 | 1 | 0.82786 | 0.28963 | 0.50442 | 1 | 11018 | 3 | 0.82786 |
| LAP3 | 4 | 0.67256 | 0.70695 | 0.999994 | 15963 | 1 | 1.068 | 0.28971 | 0.50453 | 1 | 11019 | 2 | 1.068 |
| PRR16 | 4 | 0.89918 | 0.89887 | 0.999994 | 19208 | 0 | 1.2706 | 0.28973 | 0.50455 | 1 | 11020 | 3 | 1.2706 |
| ORC3 | 5 | 0.12735 | 0.26858 | 0.998561 | 5856 | 1 | 1.5526 | 0.28974 | 0.51472 | 1 | 11021 | 3 | 1.5526 |
| ASIC1 | 4 | 0.31302 | 0.45635 | 0.999994 | 10378 | 2 | 0.77369 | 0.28979 | 0.50462 | 1 | 11022 | 2 | 0.77369 |
| USP31 | 6 | 0.54005 | 0.69657 | 0.999994 | 14052 | 2 | 1.3063 | 0.28986 | 0.55093 | 1 | 11023 | 4 | 1.3063 |

| | | | | | | | | | | | | | |
|---|---|---|---|---|---|---|---|---|---|---|---|---|---|
| ZNF835 | 6 | 0.27021 | 0.47198 | 0.999994 | 9478 | 1 | 0.54961 | 0.28987 | 0.55094 | 1 | 11024 | 3 | 0.54961 |
| NonTargetingControlG | 2 | 0.18753 | 0.25303 | 0.994501 | 7409 | 1 | -0.098365 | 0.28988 | 0.39855 | 1 | 11025 | 1 | -0.098365 |
| NonTargetingControlG | 2 | 0.50807 | 0.51132 | 0.999994 | 13554 | 1 | 2.425 | 0.28988 | 0.39855 | 1 | 11026 | 1 | 2.425 |
| KCNK18 | 5 | 0.56668 | 0.70661 | 0.999994 | 14492 | 2 | 1.5374 | 0.28991 | 0.51492 | 1 | 11027 | 3 | 1.5374 |
| TP53TG5 | 6 | 0.45177 | 0.63624 | 0.999994 | 12731 | 1 | 0.37017 | 0.28994 | 0.55102 | 1 | 11028 | 3 | 0.37017 |
| PCGF3 | 6 | 0.3424 | 0.55541 | 0.999994 | 10993 | 2 | 1.1508 | 0.29003 | 0.55113 | 1 | 11029 | 3 | 1.1508 |
| TNFRSF14 | 4 | 0.37406 | 0.51904 | 0.999994 | 11572 | 1 | 0.70358 | 0.29007 | 0.50499 | 1 | 11030 | 3 | 0.70358 |
| PCBP1 | 4 | 0.70324 | 0.72657 | 0.999994 | 16342 | 1 | 0.88411 | 0.29008 | 0.505 | 1 | 11031 | 3 | 0.88411 |
| DOCK1 | 4 | 0.52995 | 0.63009 | 0.999994 | 13900 | 1 | 0.60503 | 0.29015 | 0.50508 | 1 | 11032 | 3 | 0.60503 |
| hsa-mir-383 | 4 | 0.63854 | 0.68667 | 0.999994 | 15493 | 1 | 1.3655 | 0.29027 | 0.50524 | 1 | 11033 | 3 | 1.3655 |
| KLF17 | 5 | 0.52546 | 0.67009 | 0.999994 | 13825 | 2 | -0.59153 | 0.29035 | 0.51541 | 1 | 11034 | 2 | -0.59153 |
| ALG13 | 5 | 0.54149 | 0.68422 | 0.999994 | 14080 | 2 | 0.74314 | 0.29035 | 0.51541 | 1 | 11035 | 3 | 0.74314 |
| ENDOV | 5 | 0.60978 | 0.72972 | 0.999994 | 15068 | 1 | 0.49757 | 0.29035 | 0.51541 | 1 | 11036 | 2 | 0.49757 |
| DPCD | 6 | 0.40392 | 0.60494 | 0.999994 | 12017 | 2 | 0.27123 | 0.29038 | 0.5516 | 1 | 11037 | 2 | 0.27123 |
| TMEM246 | 6 | 0.61796 | 0.75329 | 0.999994 | 15185 | 2 | -0.096957 | 0.29038 | 0.5516 | 1 | 11038 | 1 | -0.096957 |
| KCNIP1 | 4 | 0.79263 | 0.79467 | 0.999994 | 17384 | 1 | 0.93772 | 0.29041 | 0.50543 | 1 | 11039 | 3 | 0.93772 |
| MAGEB5 | 6 | 0.61496 | 0.75105 | 0.999994 | 15142 | 2 | 0.82823 | 0.29045 | 0.5517 | 1 | 11040 | 3 | 0.82823 |
| TOMM70A | 6 | 0.41766 | 0.61383 | 0.999994 | 12213 | 2 | 0.93346 | 0.29062 | 0.55189 | 1 | 11041 | 4 | 0.93346 |
| CIT | 1 | 0.70936 | 0.70945 | 0.999994 | 16397 | 0 | 2.9589 | 0.29064 | 0.29056 | 0.9822 | 11042 | 1 | 2.9589 |
| NOL3 | 6 | 0.026323 | 0.090118 | 0.94395 | 1818 | 2 | 0.872 | 0.29074 | 0.55204 | 1 | 11043 | 3 | 0.872 |
| VPS53 | 3 | 0.0045894 | 0.012688 | 0.733879 | 486 | 1 | 1.9446 | 0.29076 | 0.43985 | 1 | 11044 | 2 | 1.9446 |
| RBM11 | 5 | 0.67998 | 0.75754 | 0.999994 | 16089 | 1 | 0.035307 | 0.29078 | 0.51587 | 1 | 11045 | 2 | 0.035307 |
| LSM7 | 6 | 0.43484 | 0.62502 | 0.999994 | 12493 | 1 | 1.1868 | 0.29082 | 0.55214 | 1 | 11046 | 4 | 1.1868 |
| SAMD3 | 6 | 0.41056 | 0.60926 | 0.999994 | 12121 | 2 | 1.2556 | 0.29086 | 0.55218 | 1 | 11047 | 3 | 1.2556 |
| DPH3P1 | 6 | 0.61632 | 0.75208 | 0.999994 | 15164 | 2 | 0.83149 | 0.29109 | 0.55247 | 1 | 11048 | 4 | 0.83149 |
| DUSP10 | 6 | 0.94904 | 0.94902 | 0.999994 | 20309 | 0 | 1.014 | 0.29111 | 0.5525 | 1 | 11049 | 4 | 1.014 |
| FOLR1 | 4 | 0.93632 | 0.93615 | 0.999994 | 19991 | 0 | 0.71716 | 0.29112 | 0.50634 | 1 | 11050 | 3 | 0.71716 |
| C3orf18 | 5 | 0.73092 | 0.78111 | 0.999994 | 16612 | 1 | 1.6511 | 0.29117 | 0.51629 | 1 | 11051 | 3 | 1.6511 |
| IRS2 | 6 | 0.18944 | 0.37482 | 0.999994 | 7468 | 2 | 1.2448 | 0.29122 | 0.55264 | 1 | 11052 | 4 | 1.2448 |
| OR10K2 | 5 | 0.22976 | 0.41689 | 0.999994 | 8524 | 1 | 0.63549 | 0.29128 | 0.51641 | 1 | 11053 | 3 | 0.63549 |
| TPCN1 | 6 | 0.88071 | 0.89008 | 0.999994 | 18846 | 1 | 1.049 | 0.29135 | 0.55279 | 1 | 11054 | 4 | 1.049 |
| GUCY1B3 | 6 | 0.40759 | 0.60732 | 0.999994 | 12067 | 1 | 0.72112 | 0.29136 | 0.55281 | 1 | 11055 | 4 | 0.72112 |
| ZFR | 6 | 0.33068 | 0.54204 | 0.999994 | 10736 | 1 | 1.3048 | 0.29143 | 0.55289 | 1 | 11056 | 4 | 1.3048 |
| PNO1 | 4 | 0.0026186 | 0.0094134 | 0.680529 | 326 | 3 | -4.266 | 0.29146 | 0.50677 | 1 | 11057 | 1 | -4.266 |
| NPTN | 6 | 0.63201 | 0.76386 | 0.999994 | 15400 | 2 | 0.15916 | 0.29147 | 0.55293 | 1 | 11058 | 3 | 0.15916 |
| KRBA1 | 6 | 0.2452 | 0.44222 | 0.999994 | 8904 | 3 | -0.35756 | 0.29147 | 0.55293 | 1 | 11059 | 2 | -0.35756 |
| SMUG1 | 6 | 0.87224 | 0.88471 | 0.999994 | 18681 | 1 | 0.32063 | 0.29147 | 0.55293 | 1 | 11060 | 2 | 0.32063 |
| PKP2 | 5 | 0.55027 | 0.69197 | 0.999994 | 14215 | 1 | 1.3245 | 0.29154 | 0.5167 | 1 | 11061 | 3 | 1.3245 |
| NAA30 | 6 | 0.035291 | 0.11121 | 0.951052 | 2280 | 4 | -2.26 | 0.29159 | 0.55307 | 1 | 11062 | 1 | -2.26 |
| C1orf127 | 6 | 0.24088 | 0.43707 | 0.999994 | 8791 | 2 | 0.7616 | 0.29159 | 0.55308 | 1 | 11063 | 4 | 0.7616 |
| hsa-mir-3167 | 4 | 0.82361 | 0.82327 | 0.999994 | 17831 | 1 | 0.84746 | 0.29161 | 0.50694 | 1 | 11064 | 3 | 0.84746 |
| F2RL3 | 5 | 0.2171 | 0.39921 | 0.999994 | 8198 | 2 | 0.65857 | 0.29161 | 0.51676 | 1 | 11065 | 3 | 0.65857 |
| TOP1 | 5 | 0.14097 | 0.28929 | 0.999632 | 6219 | 3 | -1.213 | 0.29161 | 0.51676 | 1 | 11066 | 2 | -1.213 |
| FAM9B | 5 | 0.34084 | 0.51442 | 0.999994 | 10955 | 2 | 0.15202 | 0.29161 | 0.51676 | 1 | 11067 | 2 | 0.15202 |
| C3orf43 | 5 | 0.24059 | 0.43173 | 0.999994 | 8785 | 2 | 0.031077 | 0.29161 | 0.51676 | 1 | 11068 | 2 | 0.031077 |
| NRG2 | 5 | 0.30511 | 0.48516 | 0.999994 | 10234 | 1 | 0.45059 | 0.29161 | 0.51676 | 1 | 11069 | 2 | 0.45059 |
| DNAJC5B | 6 | 0.24118 | 0.43741 | 0.999994 | 8802 | 3 | 0.6537 | 0.29162 | 0.55311 | 1 | 11070 | 3 | 0.6537 |
| HABP4 | 5 | 0.93848 | 0.93867 | 0.999994 | 20036 | 0 | 1.6907 | 0.29175 | 0.5169 | 1 | 11071 | 3 | 1.6907 |
| TEAD3 | 5 | 0.0219 | 0.069218 | 0.93914 | 1575 | 4 | -4.6304 | 0.29175 | 0.5169 | 1 | 11072 | 1 | -4.6304 |
| C13orf45 | 5 | 0.082905 | 0.19928 | 0.977164 | 4333 | 2 | 0.59667 | 0.29175 | 0.5169 | 1 | 11073 | 2 | 0.59667 |
| C18orf21 | 4 | 0.41537 | 0.56142 | 0.999994 | 12182 | 2 | 1.3039 | 0.2918 | 0.50719 | 1 | 11074 | 2 | 1.3039 |
| hsa-mir-4327 | 4 | 0.84106 | 0.84051 | 0.999994 | 18107 | 0 | 0.61154 | 0.292 | 0.50745 | 1 | 11075 | 3 | 0.61154 |

| | | | | | | | | | | | | |
|---|---|---|---|---|---|---|---|---|---|---|---|---|
| SETD9 | 5 | 0.39694 | 0.56101 | 0.999994 | 11905 | 2 | 1.3991 | 0.2921 | 0.51728 | 1 | 11076 | 3 | 1.3991 |
| FGF5 | 4 | 0.68753 | 0.71637 | 0.999994 | 16190 | 1 | 1.1622 | 0.29211 | 0.50758 | 1 | 11077 | 3 | 1.1622 |
| UGP2 | 5 | 0.48232 | 0.63269 | 0.999994 | 13174 | 1 | 1.3694 | 0.29235 | 0.51754 | 1 | 11078 | 3 | 1.3694 |
| NonTargetingControlG | 2 | 0.25582 | 0.30324 | 0.999994 | 9144 | 1 | 1.5834 | 0.29237 | 0.4001 | 1 | 11079 | 1 | 1.5834 |
| FLNC | 6 | 0.4624 | 0.64333 | 0.999994 | 12895 | 2 | -0.088342 | 0.29256 | 0.55431 | 1 | 11080 | 2 | -0.088342 |
| EIF3C | 6 | 0.38572 | 0.59319 | 0.999994 | 11729 | 2 | 0.69362 | 0.29256 | 0.55431 | 1 | 11081 | 3 | 0.69362 |
| COPG2 | 6 | 0.51524 | 0.67922 | 0.999994 | 13668 | 1 | 0.89353 | 0.29257 | 0.55433 | 1 | 11082 | 4 | 0.89353 |
| POLR2K | 3 | 0.53993 | 0.57324 | 0.999994 | 14051 | 1 | -1.4517 | 0.29268 | 0.44185 | 1 | 11083 | 1 | -1.4517 |
| ERVV-2 | 5 | 0.11097 | 0.24347 | 0.988088 | 5333 | 2 | 1.3598 | 0.29273 | 0.51796 | 1 | 11084 | 3 | 1.3598 |
| GPR143 | 6 | 0.37547 | 0.58668 | 0.999994 | 11585 | 2 | 1.2978 | 0.29285 | 0.55468 | 1 | 11085 | 4 | 1.2978 |
| HOXD1 | 6 | 0.25636 | 0.45554 | 0.999994 | 9157 | 3 | -0.040366 | 0.29285 | 0.55468 | 1 | 11086 | 3 | -0.040366 |
| SLC35A1 | 6 | 0.2815 | 0.48511 | 0.999994 | 9732 | 3 | 0.21299 | 0.29285 | 0.55468 | 1 | 11087 | 3 | 0.21299 |
| SHPK | 6 | 0.59874 | 0.73902 | 0.999994 | 14937 | 2 | 1.1335 | 0.29285 | 0.55468 | 1 | 11088 | 4 | 1.1335 |
| CRNKL1 | 6 | 0.44152 | 0.62944 | 0.999994 | 12603 | 2 | 0.62437 | 0.29285 | 0.55468 | 1 | 11089 | 3 | 0.62437 |
| KIF5A | 5 | 0.85989 | 0.86231 | 0.999994 | 18432 | 1 | 1.0448 | 0.293 | 0.51824 | 1 | 11090 | 3 | 1.0448 |
| CHID1 | 5 | 0.20567 | 0.38328 | 0.999994 | 7916 | 3 | -0.81824 | 0.293 | 0.51824 | 1 | 11091 | 1 | -0.81824 |
| KCNJ9 | 6 | 0.0053722 | 0.022498 | 0.819701 | 546 | 5 | -5.3036 | 0.29303 | 0.5549 | 1 | 11092 | 1 | -5.3036 |
| AQR | 4 | 0.61176 | 0.67172 | 0.999994 | 15103 | 1 | 1.2146 | 0.29317 | 0.50891 | 1 | 11093 | 3 | 1.2146 |
| MCMDC2 | 6 | 0.49682 | 0.66654 | 0.999994 | 13368 | 2 | 0.80113 | 0.29335 | 0.55531 | 1 | 11094 | 4 | 0.80113 |
| hsa-mir-6726 | 3 | 0.26587 | 0.3946 | 0.999994 | 9363 | 2 | -0.77506 | 0.29341 | 0.44262 | 1 | 11095 | 1 | -0.77506 |
| RPSA | 3 | 0.73342 | 0.73329 | 0.999994 | 16646 | 0 | 0.83941 | 0.29341 | 0.44262 | 1 | 11096 | 2 | 0.83941 |
| NLGN2 | 6 | 0.11687 | 0.27983 | 0.999632 | 5537 | 3 | 0.11627 | 0.29341 | 0.55541 | 1 | 11097 | 3 | 0.11627 |
| OR51T1 | 6 | 0.88493 | 0.89281 | 0.999994 | 18931 | 1 | 0.74598 | 0.29342 | 0.55541 | 1 | 11098 | 4 | 0.74598 |
| PRH2 | 6 | 0.66135 | 0.78635 | 0.999994 | 15799 | 2 | -0.34531 | 0.29348 | 0.55548 | 1 | 11099 | 2 | -0.34531 |
| MTX2 | 6 | 0.14433 | 0.31854 | 0.999994 | 6311 | 3 | -0.10074 | 0.29348 | 0.55548 | 1 | 11100 | 3 | -0.10074 |
| STYX | 5 | 0.075153 | 0.18676 | 0.969409 | 4040 | 3 | -1.7829 | 0.2935 | 0.51875 | 1 | 11101 | 2 | -1.7829 |
| MOCS3 | 5 | 0.7705 | 0.80198 | 0.999994 | 17090 | 1 | 0.8731 | 0.29357 | 0.51882 | 1 | 11102 | 3 | 0.8731 |
| hsa-mir-129-1 | 4 | 0.7371 | 0.75034 | 0.999994 | 16689 | 1 | 1.5622 | 0.2936 | 0.50924 | 1 | 11103 | 2 | 1.5622 |
| FNDC9 | 5 | 0.89466 | 0.89479 | 0.999994 | 19116 | 0 | -0.039373 | 0.2936 | 0.51887 | 1 | 11104 | 2 | -0.039373 |
| NTN3 | 6 | 0.46531 | 0.64533 | 0.999994 | 12939 | 2 | 1.4721 | 0.29374 | 0.55582 | 1 | 11105 | 4 | 1.4721 |
| SPATA31E1 | 6 | 0.89763 | 0.9018 | 0.999994 | 19172 | 1 | 1.3221 | 0.29389 | 0.55601 | 1 | 11106 | 4 | 1.3221 |
| hsa-mir-1270-1 | 4 | 0.42446 | 0.57076 | 0.999994 | 12326 | 2 | 0.47013 | 0.29391 | 0.50945 | 1 | 11107 | 2 | 0.47013 |
| KLHL6 | 6 | 0.2794 | 0.48269 | 0.999994 | 9688 | 3 | 0.69061 | 0.29396 | 0.55608 | 1 | 11108 | 3 | 0.69061 |
| OR4C12 | 6 | 0.19184 | 0.37781 | 0.999994 | 7540 | 3 | 0.40477 | 0.29412 | 0.5563 | 1 | 11109 | 3 | 0.40477 |
| GABRG1 | 6 | 0.2251 | 0.41823 | 0.999994 | 8401 | 2 | 0.70279 | 0.29413 | 0.55631 | 1 | 11110 | 3 | 0.70279 |
| TRIM48 | 5 | 0.21436 | 0.39546 | 0.999994 | 8127 | 2 | 1.2277 | 0.29415 | 0.51946 | 1 | 11111 | 3 | 1.2277 |
| KRT8 | 6 | 0.12293 | 0.29124 | 0.999632 | 5737 | 3 | -0.41483 | 0.2942 | 0.5564 | 1 | 11112 | 2 | -0.41483 |
| HOXB6 | 6 | 0.32296 | 0.53326 | 0.999994 | 10574 | 2 | 0.76817 | 0.2942 | 0.5564 | 1 | 11113 | 4 | 0.76817 |
| TMEM200B | 6 | 0.20191 | 0.3902 | 0.999994 | 7823 | 3 | -0.77276 | 0.2942 | 0.5564 | 1 | 11114 | 2 | -0.77276 |
| NFE2 | 6 | 0.48434 | 0.65812 | 0.999994 | 13205 | 2 | 0.42956 | 0.2942 | 0.5564 | 1 | 11115 | 3 | 0.42956 |
| EMP2 | 5 | 0.90135 | 0.90146 | 0.999994 | 19238 | 0 | 0.84282 | 0.29423 | 0.51954 | 1 | 11116 | 3 | 0.84282 |
| REP15 | 5 | 0.25494 | 0.44468 | 0.999994 | 9130 | 2 | 1.6479 | 0.29425 | 0.51957 | 1 | 11117 | 3 | 1.6479 |
| SULT1A3 | 6 | 0.40061 | 0.60276 | 0.999994 | 11957 | 2 | 0.81041 | 0.29428 | 0.55651 | 1 | 11118 | 4 | 0.81041 |
| SIRPB1 | 6 | 0.1005 | 0.24832 | 0.990682 | 4990 | 3 | 0.073775 | 0.29432 | 0.55656 | 1 | 11119 | 3 | 0.073775 |
| SP8 | 6 | 0.7325 | 0.82245 | 0.999994 | 16635 | 1 | 1.0097 | 0.29438 | 0.55663 | 1 | 11120 | 4 | 1.0097 |
| C17orf85 | 5 | 0.90342 | 0.90349 | 0.999994 | 19289 | 0 | 0.77946 | 0.29454 | 0.51988 | 1 | 11121 | 3 | 0.77946 |
| DTNBP1 | 6 | 0.28398 | 0.48803 | 0.999994 | 9784 | 2 | 0.85713 | 0.29458 | 0.55687 | 1 | 11122 | 3 | 0.85713 |
| CACNA1F | 4 | 0.12861 | 0.26522 | 0.997396 | 5899 | 2 | -0.80313 | 0.2946 | 0.50998 | 1 | 11123 | 2 | -0.80313 |
| DGKZ | 4 | 0.6984 | 0.72336 | 0.999994 | 16288 | 1 | -0.29372 | 0.2946 | 0.50998 | 1 | 11124 | 1 | -0.29372 |
| UCP2 | 4 | 0.092351 | 0.2013 | 0.977164 | 4695 | 3 | -1.2231 | 0.2946 | 0.50998 | 1 | 11125 | 1 | -1.2231 |
| RNF219 | 5 | 0.12842 | 0.27021 | 0.998561 | 5892 | 3 | -1.4263 | 0.2948 | 0.52015 | 1 | 11126 | 1 | -1.4263 |
| FRA10AC1 | 5 | 0.40234 | 0.56547 | 0.999994 | 11985 | 2 | 0.28224 | 0.2948 | 0.52015 | 1 | 11127 | 1 | 0.28224 |

| | | | | | | | | | | | | |
|---|---|---|---|---|---|---|---|---|---|---|---|---|
| RNF17 | 5 | 0.26594 | 0.45356 | 0.999994 | 9366 | 1 | 0.41335 | 0.2948 | 0.52015 | 1 | 11128 | 2 | 0.41335 |
| RTN4 | 6 | 0.22645 | 0.41984 | 0.999994 | 8435 | 2 | 0.89953 | 0.29481 | 0.55717 | 1 | 11129 | 4 | 0.89953 |
| C14orf80 | 6 | 0.061228 | 0.16839 | 0.967126 | 3484 | 2 | 1.1886 | 0.29482 | 0.55717 | 1 | 11130 | 3 | 1.1886 |
| NonTargetingControlG | 1 | 0.70508 | 0.7052 | 0.999994 | 16362 | 0 | 2.9872 | 0.29492 | 0.29487 | 0.983787 | 11131 | 1 | 2.9872 |
| TIMP2 | 5 | 0.31431 | 0.49271 | 0.999994 | 10405 | 2 | 1.4589 | 0.29497 | 0.52035 | 1 | 11132 | 3 | 1.4589 |
| BEND3 | 5 | 0.85631 | 0.85934 | 0.999994 | 18378 | 1 | 1.5061 | 0.295 | 0.52037 | 1 | 11133 | 3 | 1.5061 |
| ZNF837 | 6 | 0.61488 | 0.75099 | 0.999994 | 15140 | 2 | -0.51111 | 0.29502 | 0.55742 | 1 | 11134 | 2 | -0.51111 |
| IFIH1 | 6 | 0.20208 | 0.39042 | 0.999994 | 7834 | 2 | -0.10824 | 0.29502 | 0.55742 | 1 | 11135 | 1 | -0.10824 |
| NOS2 | 6 | 0.066315 | 0.17904 | 0.968171 | 3697 | 2 | -0.0076115 | 0.29502 | 0.55742 | 1 | 11136 | 1 | -0.0076115 |
| PRR5-ARHGAP8 | 3 | 0.85532 | 0.85534 | 0.999994 | 18353 | 0 | 1.7953 | 0.29503 | 0.44441 | 1 | 11137 | 2 | 1.7953 |
| NonTargetingControlG | 2 | 0.55626 | 0.55684 | 0.999994 | 14307 | 1 | 2.5746 | 0.29504 | 0.40173 | 1 | 11138 | 1 | 2.5746 |
| NonTargetingControlG | 2 | 0.47194 | 0.47869 | 0.999994 | 13033 | 1 | 2.3225 | 0.29504 | 0.40173 | 1 | 11139 | 1 | 2.3225 |
| NonTargetingControlG | 2 | 0.39176 | 0.40997 | 0.999994 | 11826 | 1 | 1.9788 | 0.29504 | 0.40173 | 1 | 11140 | 1 | 1.9788 |
| AP1M1 | 6 | 0.85978 | 0.8773 | 0.999994 | 18430 | 1 | 1.1881 | 0.29508 | 0.5575 | 1 | 11141 | 4 | 1.1881 |
| C19orf60 | 3 | 0.10602 | 0.191 | 0.972293 | 5176 | 2 | -2.2985 | 0.29508 | 0.44448 | 1 | 11142 | 1 | -2.2985 |
| KIRREL3 | 6 | 0.48157 | 0.65624 | 0.999994 | 13162 | 2 | 1.4062 | 0.29528 | 0.55775 | 1 | 11143 | 3 | 1.4062 |
| FAM24B | 6 | 0.0096161 | 0.037685 | 0.884154 | 847 | 2 | 1.4771 | 0.29528 | 0.55775 | 1 | 11144 | 3 | 1.4771 |
| FAM159A | 5 | 0.54177 | 0.68447 | 0.999994 | 14087 | 1 | 1.3848 | 0.29531 | 0.52068 | 1 | 11145 | 3 | 1.3848 |
| KLHL28 | 4 | 0.13183 | 0.27046 | 0.998561 | 5972 | 1 | 1.2753 | 0.29533 | 0.5105 | 1 | 11146 | 2 | 1.2753 |
| MLLT4 | 5 | 0.14539 | 0.29585 | 0.999994 | 6334 | 3 | -1.1095 | 0.29535 | 0.52074 | 1 | 11147 | 2 | -1.1095 |
| STUB1 | 6 | 0.13435 | 0.30585 | 0.999994 | 6031 | 2 | 0.078519 | 0.29542 | 0.55792 | 1 | 11148 | 1 | 0.078519 |
| PHTF1 | 6 | 0.15305 | 0.32949 | 0.999994 | 6534 | 2 | 0.70385 | 0.29542 | 0.55792 | 1 | 11149 | 3 | 0.70385 |
| IFRD1 | 6 | 0.34617 | 0.55971 | 0.999994 | 11076 | 3 | 0.23611 | 0.29542 | 0.55792 | 1 | 11150 | 3 | 0.23611 |
| CEACAM20 | 5 | 0.85271 | 0.85645 | 0.999994 | 18312 | 1 | -0.075245 | 0.29551 | 0.5209 | 1 | 11151 | 2 | -0.075245 |
| NonTargetingControlG | 1 | 0.70443 | 0.70453 | 0.999994 | 16356 | 0 | 2.7331 | 0.29557 | 0.29554 | 0.983965 | 11152 | 1 | 2.7331 |
| hsa-mir-1471 | 3 | 0.8572 | 0.85714 | 0.999994 | 18397 | 0 | 1.8612 | 0.29565 | 0.44507 | 1 | 11153 | 2 | 1.8612 |
| hsa-mir-4461 | 4 | 0.14285 | 0.28183 | 0.999632 | 6268 | 1 | 1.6153 | 0.29568 | 0.51076 | 1 | 11154 | 2 | 1.6153 |
| hsa-mir-6801 | 4 | 0.84044 | 0.8399 | 0.999994 | 18094 | 0 | 1.5337 | 0.29568 | 0.51076 | 1 | 11155 | 2 | 1.5337 |
| MMP26 | 4 | 0.62904 | 0.68136 | 0.999994 | 15365 | 1 | 1.3581 | 0.29568 | 0.51076 | 1 | 11156 | 2 | 1.3581 |
| hsa-mir-302a | 3 | 0.42378 | 0.49835 | 0.999994 | 12306 | 1 | -0.074012 | 0.2957 | 0.44512 | 1 | 11157 | 1 | -0.074012 |
| POLM | 6 | 0.0030792 | 0.014032 | 0.75278 | 363 | 4 | -1.6157 | 0.29576 | 0.55835 | 1 | 11158 | 2 | -1.6157 |
| TMLHE | 6 | 0.68794 | 0.8072 | 0.999994 | 16194 | 2 | 1.0211 | 0.2958 | 0.55839 | 1 | 11159 | 4 | 1.0211 |
| OR5M9 | 4 | 0.8397 | 0.83916 | 0.999994 | 18082 | 0 | 0.54552 | 0.29581 | 0.51086 | 1 | 11160 | 2 | 0.54552 |
| CPO | 5 | 0.56717 | 0.70707 | 0.999994 | 14499 | 2 | 0.39901 | 0.2959 | 0.52135 | 1 | 11161 | 2 | 0.39901 |
| IMPG1 | 5 | 0.64377 | 0.74264 | 0.999994 | 15572 | 1 | 1.2768 | 0.2959 | 0.52135 | 1 | 11162 | 3 | 1.2768 |
| CLASP1 | 5 | 0.032949 | 0.098091 | 0.945253 | 2154 | 3 | -1.2663 | 0.2959 | 0.52135 | 1 | 11163 | 1 | -1.2663 |
| GBP5 | 6 | 0.013109 | 0.049478 | 0.906615 | 1071 | 2 | 0.87704 | 0.29591 | 0.55853 | 1 | 11164 | 3 | 0.87704 |
| OVGP1 | 6 | 0.72685 | 0.82056 | 0.999994 | 16565 | 1 | 0.24125 | 0.29591 | 0.55853 | 1 | 11165 | 2 | 0.24125 |
| SERPING1 | 6 | 0.96137 | 0.96142 | 0.999994 | 20616 | 0 | 0.94322 | 0.29591 | 0.55853 | 1 | 11166 | 3 | 0.94322 |
| NonTargetingControlG | 1 | 0.70409 | 0.70418 | 0.999994 | 16353 | 0 | 2.4727 | 0.29591 | 0.29587 | 0.984434 | 11167 | 1 | 2.4727 |
| FAM49A | 6 | 0.90528 | 0.90779 | 0.999994 | 19339 | 1 | 0.67247 | 0.29608 | 0.55875 | 1 | 11168 | 4 | 0.67247 |
| RBBP6 | 5 | 0.48039 | 0.631 | 0.999994 | 13148 | 1 | 1.3293 | 0.2962 | 0.52166 | 1 | 11169 | 3 | 1.3293 |
| TCEAL3 | 5 | 0.17616 | 0.34109 | 0.999994 | 7138 | 2 | -1.4517 | 0.29638 | 0.52184 | 1 | 11170 | 2 | -1.4517 |
| CSN3 | 5 | 0.49309 | 0.64193 | 0.999994 | 13313 | 2 | 0.12939 | 0.2965 | 0.52197 | 1 | 11171 | 2 | 0.12939 |
| NEFM | 5 | 0.26539 | 0.45312 | 0.999994 | 9358 | 2 | 0.4521 | 0.2965 | 0.52197 | 1 | 11172 | 2 | 0.4521 |
| POLR3C | 5 | 0.18116 | 0.34831 | 0.999994 | 7251 | 2 | -0.53516 | 0.2965 | 0.52197 | 1 | 11173 | 2 | -0.53516 |
| PHF3 | 5 | 0.14561 | 0.29621 | 0.999994 | 6340 | 2 | 0.71433 | 0.2965 | 0.52197 | 1 | 11174 | 3 | 0.71433 |
| AIM1 | 5 | 0.013023 | 0.044642 | 0.891621 | 1063 | 2 | 1.518 | 0.29674 | 0.52224 | 1 | 11175 | 3 | 1.518 |
| DYNC1LI2 | 4 | 0.0016019 | 0.0058526 | 0.623123 | 235 | 1 | 1.3626 | 0.297 | 0.51176 | 1 | 11176 | 2 | 1.3626 |
| MAP1LC3A | 5 | 0.11363 | 0.24762 | 0.990682 | 5426 | 1 | 1.295 | 0.29717 | 0.5227 | 1 | 11177 | 3 | 1.295 |
| PAK3 | 6 | 0.91915 | 0.91962 | 0.999994 | 19610 | 1 | 1.1784 | 0.2972 | 0.56013 | 1 | 11178 | 4 | 1.1784 |
| GRAMD1A | 5 | 0.94064 | 0.94084 | 0.999994 | 20096 | 0 | 0.82849 | 0.29735 | 0.52289 | 1 | 11179 | 3 | 0.82849 |

| | | | | | | | | | | | | | |
|---|---|---|---|---|---|---|---|---|---|---|---|---|---|
| KCTD1 | 6 | 0.17632 | 0.35847 | 0.999994 | 7140 | 2 | 1.0505 | 0.29748 | 0.5605 | 1 | 11180 | 4 | 1.0505 |
| GPR148 | 6 | 0.079996 | 0.20727 | 0.978744 | 4233 | 4 | -1.2582 | 0.29755 | 0.56061 | 1 | 11181 | 1 | -1.2582 |
| ATP4B | 6 | 0.20773 | 0.39731 | 0.999994 | 7967 | 2 | 0.0046364 | 0.29755 | 0.56061 | 1 | 11182 | 2 | 0.0046364 |
| GET4 | 5 | 0.045377 | 0.12924 | 0.952548 | 2783 | 2 | 1.5259 | 0.29759 | 0.52315 | 1 | 11183 | 3 | 1.5259 |
| RDH16 | 6 | 0.28218 | 0.48594 | 0.999994 | 9742 | 3 | -0.40555 | 0.29761 | 0.56067 | 1 | 11184 | 2 | -0.40555 |
| METAP1 | 6 | 0.69476 | 0.81044 | 0.999994 | 16254 | 1 | -0.27565 | 0.29761 | 0.56067 | 1 | 11185 | 2 | -0.27565 |
| FAM83D | 6 | 0.58541 | 0.7292 | 0.999994 | 14749 | 2 | 1.0855 | 0.29761 | 0.56067 | 1 | 11186 | 3 | 1.0855 |
| C7orf73 | 5 | 0.10516 | 0.23445 | 0.984971 | 5137 | 2 | -0.54636 | 0.29771 | 0.52327 | 1 | 11187 | 2 | -0.54636 |
| PDE4C | 6 | 0.59608 | 0.7371 | 0.999994 | 14902 | 2 | 1.2627 | 0.29783 | 0.56094 | 1 | 11188 | 3 | 1.2627 |
| PPP1R21 | 4 | 0.43567 | 0.58229 | 0.999994 | 12510 | 2 | 0.61624 | 0.29791 | 0.51243 | 1 | 11189 | 2 | 0.61624 |
| LCE3B | 6 | 0.26521 | 0.46604 | 0.999994 | 9352 | 2 | 0.78136 | 0.29798 | 0.56114 | 1 | 11190 | 4 | 0.78136 |
| KLHL9 | 5 | 0.15428 | 0.30918 | 0.999994 | 6573 | 2 | 1.3488 | 0.298 | 0.52358 | 1 | 11191 | 3 | 1.3488 |
| LOC389895 | 5 | 0.86037 | 0.86271 | 0.999994 | 18451 | 1 | 0.43682 | 0.29809 | 0.52367 | 1 | 11192 | 2 | 0.43682 |
| TMEM220 | 5 | 0.13085 | 0.27391 | 0.998561 | 5953 | 2 | 0.044547 | 0.29809 | 0.52367 | 1 | 11193 | 1 | 0.044547 |
| HS3ST3A1 | 5 | 0.27882 | 0.4639 | 0.999994 | 9667 | 2 | 1.2916 | 0.2981 | 0.52367 | 1 | 11194 | 3 | 1.2916 |
| TMEM104 | 6 | 0.2485 | 0.44615 | 0.999994 | 8979 | 2 | 1.1154 | 0.29812 | 0.56131 | 1 | 11195 | 3 | 1.1154 |
| MORF4L1 | 6 | 0.2949 | 0.50076 | 0.999994 | 10016 | 2 | 0.77755 | 0.29823 | 0.56143 | 1 | 11196 | 4 | 0.77755 |
| NAPA | 3 | 0.1962 | 0.30787 | 0.999994 | 7658 | 1 | -0.43588 | 0.29831 | 0.44791 | 1 | 11197 | 1 | -0.43588 |
| SPRY2 | 3 | 0.41125 | 0.49088 | 0.999994 | 12134 | 1 | 0.97812 | 0.29831 | 0.44791 | 1 | 11198 | 2 | 0.97812 |
| LMCD1 | 6 | 0.8625 | 0.87884 | 0.999994 | 18495 | 1 | 1.0714 | 0.29838 | 0.56161 | 1 | 11199 | 4 | 1.0714 |
| PCDHA11 | 2 | 0.016198 | 0.03033 | 0.862223 | 1256 | 1 | -2.2928 | 0.29845 | 0.40379 | 1 | 11200 | 1 | -2.2928 |
| KIAA1109 | 5 | 0.85215 | 0.85599 | 0.999994 | 18299 | 1 | 1.4625 | 0.29848 | 0.52408 | 1 | 11201 | 3 | 1.4625 |
| KIF5C | 5 | 0.077194 | 0.19005 | 0.970632 | 4116 | 2 | 1.5033 | 0.29869 | 0.52429 | 1 | 11202 | 3 | 1.5033 |
| SRPX | 6 | 0.052443 | 0.14941 | 0.966183 | 3090 | 3 | -1.2503 | 0.29869 | 0.56197 | 1 | 11203 | 3 | -1.2503 |
| TINAG | 5 | 0.29809 | 0.47952 | 0.999994 | 10092 | 2 | 0.33134 | 0.29878 | 0.52439 | 1 | 11204 | 2 | 0.33134 |
| DAZAP2 | 5 | 0.49976 | 0.64769 | 0.999994 | 13419 | 1 | 0.87481 | 0.29878 | 0.52439 | 1 | 11205 | 3 | 0.87481 |
| RHD | 6 | 0.48753 | 0.66028 | 0.999994 | 13244 | 1 | 0.7041 | 0.29889 | 0.56225 | 1 | 11206 | 4 | 0.7041 |
| CPNE2 | 4 | 0.29796 | 0.44093 | 0.999994 | 10090 | 2 | -0.036231 | 0.29905 | 0.51326 | 1 | 11207 | 2 | -0.036231 |
| RFX2 | 5 | 0.026976 | 0.082667 | 0.942677 | 1855 | 2 | -0.446 | 0.29907 | 0.52471 | 1 | 11208 | 2 | -0.446 |
| PIGS | 5 | 0.47807 | 0.62899 | 0.999994 | 13117 | 2 | -0.074443 | 0.29907 | 0.52471 | 1 | 11209 | 1 | -0.074443 |
| HEPH | 5 | 0.21435 | 0.39544 | 0.999994 | 8126 | 3 | -0.79532 | 0.29907 | 0.52471 | 1 | 11210 | 2 | -0.79532 |
| hsa-mir-4686 | 3 | 0.75078 | 0.75074 | 0.999994 | 16862 | 0 | 1.1903 | 0.29916 | 0.44883 | 1 | 11211 | 2 | 1.1903 |
| MCEE | 4 | 0.45054 | 0.59492 | 0.999994 | 12720 | 1 | 1.3379 | 0.29917 | 0.51335 | 1 | 11212 | 2 | 1.3379 |
| LNP1 | 5 | 0.19888 | 0.37365 | 0.999994 | 7732 | 1 | 1.2521 | 0.29918 | 0.52484 | 1 | 11213 | 3 | 1.2521 |
| BUD13 | 4 | 0.19973 | 0.34038 | 0.999994 | 7754 | 2 | 0.64111 | 0.29919 | 0.51336 | 1 | 11214 | 2 | 0.64111 |
| CXorf51A | 6 | 0.19345 | 0.37983 | 0.999994 | 7584 | 1 | 0.6688 | 0.29922 | 0.56266 | 1 | 11215 | 3 | 0.6688 |
| SIPA1L1 | 6 | 0.031347 | 0.10198 | 0.946414 | 2077 | 2 | 0.74802 | 0.29922 | 0.56266 | 1 | 11216 | 3 | 0.74802 |
| BCKDHA | 5 | 0.90523 | 0.90533 | 0.999994 | 19337 | 0 | 1.2363 | 0.29928 | 0.52493 | 1 | 11217 | 3 | 1.2363 |
| RAB11A | 4 | 0.17842 | 0.31847 | 0.999994 | 7187 | 1 | 0.10118 | 0.29938 | 0.5135 | 1 | 11218 | 1 | 0.10118 |
| TUBE1 | 6 | 0.20579 | 0.39491 | 0.999994 | 7919 | 3 | 0.56952 | 0.29939 | 0.56289 | 1 | 11219 | 3 | 0.56952 |
| DAB2IP | 6 | 0.44524 | 0.63193 | 0.999994 | 12651 | 2 | 1.2581 | 0.29955 | 0.56308 | 1 | 11220 | 3 | 1.2581 |
| HNRNPH3 | 5 | 0.033798 | 0.10022 | 0.946394 | 2204 | 3 | -2.3172 | 0.2996 | 0.52527 | 1 | 11221 | 2 | -2.3172 |
| ATP6V1G2 | 5 | 0.50251 | 0.65012 | 0.999994 | 13471 | 2 | 0.35211 | 0.2996 | 0.52527 | 1 | 11222 | 2 | 0.35211 |
| CMTM1 | 5 | 0.50366 | 0.65111 | 0.999994 | 13489 | 2 | 0.45854 | 0.2996 | 0.52527 | 1 | 11223 | 2 | 0.45854 |
| HPX | 5 | 0.41516 | 0.57607 | 0.999994 | 12181 | 2 | 1.3353 | 0.29963 | 0.5253 | 1 | 11224 | 3 | 1.3353 |
| TMSB4X | 3 | 0.2328 | 0.35358 | 0.999994 | 8589 | 2 | -0.91901 | 0.29963 | 0.44933 | 1 | 11225 | 1 | -0.91901 |
| SH3BP5L | 4 | 0.31046 | 0.45372 | 0.999994 | 10335 | 2 | 1.0602 | 0.29986 | 0.51387 | 1 | 11226 | 2 | 1.0602 |
| LMBR1 | 6 | 0.9496 | 0.94955 | 0.999994 | 20325 | 0 | 0.70104 | 0.29987 | 0.56348 | 1 | 11227 | 3 | 0.70104 |
| PTPRS | 5 | 0.016066 | 0.05321 | 0.916684 | 1249 | 1 | 1.3185 | 0.29987 | 0.52556 | 1 | 11228 | 3 | 1.3185 |
| PDZD7 | 5 | 0.87462 | 0.87526 | 0.999994 | 18725 | 1 | 1.2545 | 0.29996 | 0.52565 | 1 | 11229 | 3 | 1.2545 |
| SERPINA9 | 6 | 0.23187 | 0.42634 | 0.999994 | 8568 | 2 | 1.2518 | 0.30001 | 0.56367 | 1 | 11230 | 3 | 1.2518 |
| SLC16A4 | 6 | 0.073967 | 0.19501 | 0.975797 | 3993 | 2 | 0.89273 | 0.30003 | 0.56368 | 1 | 11231 | 3 | 0.89273 |

| | | | | | | | | | | | | | |
|---|---|---|---|---|---|---|---|---|---|---|---|---|---|
| MTSS1L | 6 | 0.28153 | 0.48514 | 0.999994 | 9733 | 2 | 0.90742 | 0.30009 | 0.56377 | 1 | 11232 | 4 | 0.90742 |
| NFKBIZ | 5 | 0.48251 | 0.63286 | 0.999994 | 13180 | 2 | 0.43067 | 0.30011 | 0.5258 | 1 | 11233 | 2 | 0.43067 |
| NonTargetingControlG | 2 | 0.2445 | 0.29481 | 0.999994 | 8892 | 1 | 1.2769 | 0.30017 | 0.40486 | 1 | 11234 | 1 | 1.2769 |
| PPP2R5E | 5 | 0.062312 | 0.1659 | 0.967126 | 3524 | 2 | 0.82079 | 0.30022 | 0.52593 | 1 | 11235 | 3 | 0.82079 |
| ITLN2 | 3 | 0.51702 | 0.55748 | 0.999994 | 13690 | 1 | 1.6644 | 0.30048 | 0.45026 | 1 | 11236 | 2 | 1.6644 |
| WDR60 | 6 | 0.75685 | 0.83092 | 0.999994 | 16941 | 1 | 0.6631 | 0.30056 | 0.56437 | 1 | 11237 | 3 | 0.6631 |
| N4BP1 | 6 | 0.2492 | 0.44696 | 0.999994 | 8997 | 3 | -0.26153 | 0.30056 | 0.56437 | 1 | 11238 | 2 | -0.26153 |
| LOC100287177 | 4 | 0.28854 | 0.43128 | 0.999994 | 9882 | 2 | 0.86121 | 0.30057 | 0.51441 | 1 | 11239 | 2 | 0.86121 |
| CARF | 6 | 0.34175 | 0.55466 | 0.999994 | 10976 | 2 | 0.77191 | 0.30057 | 0.56438 | 1 | 11240 | 4 | 0.77191 |
| hsa-mir-4699 | 4 | 0.092337 | 0.20129 | 0.977164 | 4692 | 1 | 1.434 | 0.30061 | 0.51443 | 1 | 11241 | 2 | 1.434 |
| POLR2I | 3 | 0.85259 | 0.85255 | 0.999994 | 18309 | 0 | 0.94562 | 0.30071 | 0.45051 | 1 | 11242 | 2 | 0.94562 |
| ZNF208 | 6 | 0.95305 | 0.95304 | 0.999994 | 20410 | 0 | 1.1546 | 0.30089 | 0.56477 | 1 | 11243 | 4 | 1.1546 |
| MATN3 | 5 | 0.26294 | 0.4511 | 0.999994 | 9313 | 2 | 1.3838 | 0.30096 | 0.5267 | 1 | 11244 | 3 | 1.3838 |
| PRPSAP1 | 5 | 0.14486 | 0.29507 | 0.999994 | 6321 | 1 | 1.259 | 0.30101 | 0.52674 | 1 | 11245 | 3 | 1.259 |
| NKX2-3 | 5 | 0.88678 | 0.88689 | 0.999994 | 18963 | 1 | 0.092178 | 0.30103 | 0.52677 | 1 | 11246 | 2 | 0.092178 |
| PCYOX1L | 5 | 0.13145 | 0.27482 | 0.998561 | 5968 | 3 | -1.2321 | 0.30103 | 0.52677 | 1 | 11247 | 1 | -1.2321 |
| EEF1D | 5 | 0.35618 | 0.52693 | 0.999994 | 11271 | 2 | -0.091994 | 0.30109 | 0.52682 | 1 | 11248 | 2 | -0.091994 |
| VSX2 | 5 | 0.89011 | 0.89024 | 0.999994 | 19037 | 0 | 0.047232 | 0.30109 | 0.52682 | 1 | 11249 | 2 | 0.047232 |
| ZNF264 | 5 | 0.38971 | 0.5549 | 0.999994 | 11792 | 1 | 1.0851 | 0.30109 | 0.52682 | 1 | 11250 | 3 | 1.0851 |
| GPRC5C | 5 | 0.58732 | 0.72159 | 0.999994 | 14778 | 1 | 1.4268 | 0.30109 | 0.52682 | 1 | 11251 | 3 | 1.4268 |
| ZNRD1 | 2 | 0.37878 | 0.3993 | 0.999994 | 11635 | 1 | 1.8753 | 0.3011 | 0.40541 | 1 | 11252 | 1 | 1.8753 |
| SNAPC2 | 4 | 0.27538 | 0.41784 | 0.999994 | 9599 | 2 | 0.77217 | 0.30118 | 0.51483 | 1 | 11253 | 2 | 0.77217 |
| IQCE | 6 | 0.19424 | 0.38083 | 0.999994 | 7612 | 3 | 0.33309 | 0.3012 | 0.56516 | 1 | 11254 | 3 | 0.33309 |
| KIAA1462 | 6 | 0.50191 | 0.67002 | 0.999994 | 13459 | 2 | 0.80993 | 0.30125 | 0.56521 | 1 | 11255 | 4 | 0.80993 |
| EDEM1 | 6 | 0.41008 | 0.60896 | 0.999994 | 12110 | 2 | 1.0656 | 0.30143 | 0.56544 | 1 | 11256 | 4 | 1.0656 |
| SAMHD1 | 6 | 0.83187 | 0.86246 | 0.999994 | 17957 | 1 | 1.12 | 0.30146 | 0.56548 | 1 | 11257 | 3 | 1.12 |
| hsa-mir-556 | 3 | 0.83269 | 0.83257 | 0.999994 | 17968 | 0 | 1.702 | 0.3016 | 0.45146 | 1 | 11258 | 2 | 1.702 |
| AKR1B1 | 5 | 0.00299 | 0.012795 | 0.733879 | 356 | 1 | -0.25608 | 0.30162 | 0.52741 | 1 | 11259 | 2 | -0.25608 |
| C11orf57 | 5 | 0.8149 | 0.82922 | 0.999994 | 17700 | 1 | -0.32667 | 0.30162 | 0.52741 | 1 | 11260 | 2 | -0.32667 |
| SLC25A22 | 5 | 0.17104 | 0.33366 | 0.999994 | 7010 | 3 | -1.0231 | 0.30162 | 0.52741 | 1 | 11261 | 1 | -1.0231 |
| FAM20C | 5 | 0.087909 | 0.20722 | 0.978744 | 4507 | 3 | -0.66374 | 0.30162 | 0.52741 | 1 | 11262 | 1 | -0.66374 |
| IL12A | 4 | 0.21063 | 0.35162 | 0.999994 | 8044 | 2 | 0.3868 | 0.30165 | 0.51518 | 1 | 11263 | 2 | 0.3868 |
| PAX3 | 5 | 0.52173 | 0.6668 | 0.999994 | 13775 | 1 | 1.4518 | 0.30167 | 0.52746 | 1 | 11264 | 3 | 1.4518 |
| LOC100130370 | 6 | 0.014498 | 0.054081 | 0.922137 | 1165 | 5 | -1.2255 | 0.30182 | 0.56594 | 1 | 11265 | 1 | -1.2255 |
| FCER1A | 6 | 0.82959 | 0.86134 | 0.999994 | 17921 | 1 | 1.0419 | 0.30182 | 0.56594 | 1 | 11266 | 4 | 1.0419 |
| hsa-mir-149 | 3 | 0.15273 | 0.25201 | 0.994009 | 6527 | 1 | 1.6113 | 0.30187 | 0.45174 | 1 | 11267 | 2 | 1.6113 |
| hsa-mir-657 | 3 | 0.28405 | 0.41694 | 0.999994 | 9786 | 2 | -0.61992 | 0.30187 | 0.45174 | 1 | 11268 | 1 | -0.61992 |
| KRTAP1-5 | 5 | 0.014304 | 0.048226 | 0.90085 | 1149 | 2 | 1.2913 | 0.30197 | 0.52779 | 1 | 11269 | 3 | 1.2913 |
| OR52H1 | 5 | 0.16198 | 0.32044 | 0.999994 | 6793 | 2 | 1.4127 | 0.30207 | 0.52788 | 1 | 11270 | 3 | 1.4127 |
| CDK2AP1 | 6 | 0.47664 | 0.65292 | 0.999994 | 13093 | 2 | 1.4076 | 0.30212 | 0.5663 | 1 | 11271 | 4 | 1.4076 |
| RTCA | 6 | 0.23313 | 0.42785 | 0.999994 | 8599 | 2 | 0.85669 | 0.30214 | 0.56633 | 1 | 11272 | 4 | 0.85669 |
| SGOL2 | 5 | 0.14712 | 0.29844 | 0.999994 | 6378 | 3 | -0.91167 | 0.30224 | 0.52806 | 1 | 11273 | 2 | -0.91167 |
| NAGA | 5 | 0.10938 | 0.24103 | 0.987253 | 5285 | 3 | -0.97896 | 0.30224 | 0.52806 | 1 | 11274 | 2 | -0.97896 |
| CDC20B | 5 | 0.050545 | 0.14167 | 0.959146 | 2999 | 4 | -1.262 | 0.30224 | 0.52806 | 1 | 11275 | 1 | -1.262 |
| CEP170B | 6 | 0.87045 | 0.8836 | 0.999994 | 18643 | 1 | 1.0514 | 0.30224 | 0.56644 | 1 | 11276 | 4 | 1.0514 |
| PCDH11X | 6 | 0.34044 | 0.55316 | 0.999994 | 10944 | 3 | 0.69489 | 0.30226 | 0.56647 | 1 | 11277 | 3 | 0.69489 |
| OR5P2 | 4 | 0.79238 | 0.79444 | 0.999994 | 17380 | 1 | 1.3059 | 0.30227 | 0.51563 | 1 | 11278 | 2 | 1.3059 |
| NonTargetingControlG | 1 | 0.69769 | 0.69773 | 0.999994 | 16274 | 0 | 2.7254 | 0.30231 | 0.30231 | 0.985876 | 11279 | 1 | 2.7254 |
| PLA2G12B | 5 | 0.8314 | 0.84051 | 0.999994 | 17952 | 1 | 1.4281 | 0.3024 | 0.52823 | 1 | 11280 | 3 | 1.4281 |
| HSPG2 | 4 | 0.27543 | 0.41788 | 0.999994 | 9602 | 2 | -0.032103 | 0.30241 | 0.51573 | 1 | 11281 | 2 | -0.032103 |
| CSN1S1 | 5 | 0.94563 | 0.94583 | 0.999994 | 20222 | 0 | 0.64697 | 0.30252 | 0.52836 | 1 | 11282 | 3 | 0.64697 |
| SMAD1 | 5 | 0.16057 | 0.31838 | 0.999994 | 6754 | 1 | 1.3008 | 0.30263 | 0.52847 | 1 | 11283 | 3 | 1.3008 |

| | | | | | | | | | | | | |
|---|---|---|---|---|---|---|---|---|---|---|---|---|
| TAMM41 | 6 | 0.27888 | 0.48211 | 0.999994 | 9669 | 3 | 0.64694 | 0.30271 | 0.56701 | 1 | 11284 | 3 | 0.64694 |
| MSI1 | 6 | 0.034405 | 0.1091 | 0.951052 | 2233 | 3 | -0.7389 | 0.30272 | 0.56703 | 1 | 11285 | 3 | -0.7389 |
| NonTargetingControlG | 2 | 0.73523 | 0.73492 | 0.999994 | 16664 | 0 | 3.1343 | 0.30273 | 0.4064 | 1 | 11286 | 1 | 3.1343 |
| FAM101B | 5 | 0.25617 | 0.44565 | 0.999994 | 9153 | 1 | 1.2943 | 0.30274 | 0.52859 | 1 | 11287 | 3 | 1.2943 |
| PVRIG | 5 | 0.14317 | 0.29261 | 0.999632 | 6278 | 2 | 1.5025 | 0.3029 | 0.52877 | 1 | 11288 | 3 | 1.5025 |
| PKIG | 4 | 0.19131 | 0.33173 | 0.999994 | 7522 | 2 | 0.47335 | 0.30301 | 0.51618 | 1 | 11289 | 2 | 0.47335 |
| CRB1 | 6 | 0.58002 | 0.72524 | 0.999994 | 14677 | 1 | 1.1704 | 0.30305 | 0.56745 | 1 | 11290 | 3 | 1.1704 |
| GIMAP2 | 4 | 0.81274 | 0.81289 | 0.999994 | 17665 | 1 | 1.7879 | 0.30312 | 0.51626 | 1 | 11291 | 2 | 1.7879 |
| AMZ2 | 4 | 0.36697 | 0.5117 | 0.999994 | 11457 | 1 | 1.157 | 0.30317 | 0.5163 | 1 | 11292 | 2 | 1.157 |
| MAP2 | 4 | 0.027581 | 0.076571 | 0.93914 | 1881 | 2 | -1.0959 | 0.30319 | 0.51631 | 1 | 11293 | 1 | -1.0959 |
| DBC1 | 3 | 0.38731 | 0.47698 | 0.999994 | 11755 | 1 | 1.6734 | 0.30322 | 0.45315 | 1 | 11294 | 2 | 1.6734 |
| hsa-mir-5188 | 3 | 0.79928 | 0.79913 | 0.999994 | 17471 | 0 | 1.3946 | 0.30324 | 0.45317 | 1 | 11295 | 2 | 1.3946 |
| CYP1A1 | 6 | 0.4994 | 0.66828 | 0.999994 | 13413 | 2 | 0.93119 | 0.3033 | 0.56775 | 1 | 11296 | 4 | 0.93119 |
| ZNF790 | 6 | 0.64499 | 0.77378 | 0.999994 | 15584 | 1 | 0.63307 | 0.30338 | 0.56785 | 1 | 11297 | 4 | 0.63307 |
| OR6C76 | 6 | 0.56072 | 0.71131 | 0.999994 | 14381 | 2 | 0.75869 | 0.3034 | 0.56788 | 1 | 11298 | 4 | 0.75869 |
| PANX3 | 6 | 0.28569 | 0.49009 | 0.999994 | 9824 | 2 | 1.0311 | 0.30342 | 0.56791 | 1 | 11299 | 3 | 1.0311 |
| RAP1GDS1 | 6 | 0.08764 | 0.22276 | 0.983354 | 4500 | 2 | 0.56018 | 0.30343 | 0.56792 | 1 | 11300 | 3 | 0.56018 |
| ZZZ3 | 6 | 0.49951 | 0.66835 | 0.999994 | 13416 | 2 | 0.30644 | 0.30343 | 0.56792 | 1 | 11301 | 3 | 0.30644 |
| TFDP3 | 6 | 0.88415 | 0.89231 | 0.999994 | 18912 | 1 | 0.90328 | 0.30343 | 0.56792 | 1 | 11302 | 4 | 0.90328 |
| MTNR1B | 6 | 0.34084 | 0.55361 | 0.999994 | 10956 | 2 | 0.35245 | 0.30345 | 0.56794 | 1 | 11303 | 3 | 0.35245 |
| MAP4K4 | 6 | 0.022444 | 0.079803 | 0.93914 | 1613 | 5 | -1.0198 | 0.30345 | 0.56794 | 1 | 11304 | 1 | -1.0198 |
| CTDSP2 | 5 | 0.018446 | 0.059846 | 0.932932 | 1383 | 4 | -3.4826 | 0.30346 | 0.52937 | 1 | 11305 | 1 | -3.4826 |
| OXCT2 | 5 | 0.8353 | 0.84327 | 0.999994 | 18011 | 1 | 0.84238 | 0.30346 | 0.52937 | 1 | 11306 | 3 | 0.84238 |
| DENND1A | 6 | 0.26247 | 0.4628 | 0.999994 | 9296 | 2 | 1.2934 | 0.30346 | 0.56795 | 1 | 11307 | 3 | 1.2934 |
| MYH9 | 5 | 0.66218 | 0.75003 | 0.999994 | 15811 | 1 | 1.3184 | 0.30361 | 0.52953 | 1 | 11308 | 3 | 1.3184 |
| ASTN1 | 6 | 0.34483 | 0.55822 | 0.999994 | 11050 | 2 | 0.89857 | 0.30377 | 0.5683 | 1 | 11309 | 4 | 0.89857 |
| SMARCA5 | 5 | 0.00050285 | 0.0023457 | 0.457631 | 110 | 2 | -0.19983 | 0.30379 | 0.52972 | 1 | 11310 | 2 | -0.19983 |
| PWP2 | 5 | 0.059252 | 0.16081 | 0.967126 | 3391 | 2 | 0.1108 | 0.30379 | 0.52972 | 1 | 11311 | 2 | 0.1108 |
| FBXO24 | 5 | 0.86378 | 0.86559 | 0.999994 | 18517 | 1 | 0.45358 | 0.30379 | 0.52972 | 1 | 11312 | 2 | 0.45358 |
| REG4 | 6 | 0.054558 | 0.15395 | 0.967126 | 3178 | 4 | -1.7207 | 0.30391 | 0.56848 | 1 | 11313 | 1 | -1.7207 |
| C18orf25 | 6 | 0.86337 | 0.87934 | 0.999994 | 18512 | 1 | 0.78707 | 0.30395 | 0.56853 | 1 | 11314 | 4 | 0.78707 |
| HEXB | 6 | 0.95466 | 0.95472 | 0.999994 | 20446 | 0 | 0.73081 | 0.304 | 0.5686 | 1 | 11315 | 4 | 0.73081 |
| LBX2 | 5 | 0.60361 | 0.72749 | 0.999994 | 14989 | 1 | 1.3782 | 0.30403 | 0.52994 | 1 | 11316 | 3 | 1.3782 |
| EHD4 | 6 | 0.69295 | 0.80989 | 0.999994 | 16234 | 1 | 1.3011 | 0.30407 | 0.56869 | 1 | 11317 | 4 | 1.3011 |
| RASGEF1A | 6 | 0.61181 | 0.74865 | 0.999994 | 15105 | 1 | 0.93994 | 0.30416 | 0.5688 | 1 | 11318 | 4 | 0.93994 |
| ANK1 | 5 | 0.22887 | 0.41564 | 0.999994 | 8499 | 3 | -0.66203 | 0.30418 | 0.5301 | 1 | 11319 | 2 | -0.66203 |
| TMEM211 | 5 | 0.29484 | 0.47686 | 0.999994 | 10013 | 2 | 0.50491 | 0.30418 | 0.5301 | 1 | 11320 | 2 | 0.50491 |
| SLC22A7 | 6 | 0.01727 | 0.063132 | 0.937389 | 1322 | 3 | -0.77806 | 0.30441 | 0.56912 | 1 | 11321 | 3 | -0.77806 |
| RAD1 | 6 | 0.6326 | 0.7643 | 0.999994 | 15406 | 2 | 1.1646 | 0.30442 | 0.56915 | 1 | 11322 | 4 | 1.1646 |
| UBE2I | 3 | 0.49666 | 0.54395 | 0.999994 | 13364 | 1 | 1.2351 | 0.30455 | 0.45457 | 1 | 11323 | 2 | 1.2351 |
| SLC9A3 | 6 | 0.61994 | 0.75478 | 0.999994 | 15220 | 2 | -0.13075 | 0.30463 | 0.5694 | 1 | 11324 | 1 | -0.13075 |
| CHAC2 | 6 | 0.036706 | 0.11447 | 0.951052 | 2356 | 3 | -0.27478 | 0.30463 | 0.5694 | 1 | 11325 | 3 | -0.27478 |
| hsa-mir-30e | 4 | 0.71239 | 0.73276 | 0.999994 | 16424 | 1 | 0.66861 | 0.30465 | 0.51738 | 1 | 11326 | 2 | 0.66861 |
| UTP18 | 4 | 0.018975 | 0.058484 | 0.931872 | 1406 | 2 | -4.1095 | 0.30465 | 0.51738 | 1 | 11327 | 1 | -4.1095 |
| PROP1 | 6 | 0.092712 | 0.23292 | 0.984502 | 4709 | 2 | 0.71025 | 0.30469 | 0.56949 | 1 | 11328 | 4 | 0.71025 |
| NFKBIL1 | 5 | 0.36382 | 0.53325 | 0.999994 | 11400 | 1 | -0.1341 | 0.30478 | 0.53073 | 1 | 11329 | 2 | -0.1341 |
| ZNF772 | 6 | 0.93192 | 0.93203 | 0.999994 | 19897 | 0 | 0.62216 | 0.3048 | 0.56961 | 1 | 11330 | 4 | 0.62216 |
| CLPSL1 | 3 | 0.73327 | 0.73313 | 0.999994 | 16643 | 0 | 1.6902 | 0.30484 | 0.4549 | 1 | 11331 | 2 | 1.6902 |
| RGS5 | 6 | 0.59307 | 0.73485 | 0.999994 | 14864 | 2 | 0.7313 | 0.30512 | 0.56999 | 1 | 11332 | 4 | 0.7313 |
| LOC81691 | 5 | 0.77363 | 0.80375 | 0.999994 | 17129 | 1 | 1.4589 | 0.30512 | 0.53106 | 1 | 11333 | 3 | 1.4589 |
| SCG5 | 5 | 0.53963 | 0.68255 | 0.999994 | 14046 | 2 | 1.4035 | 0.30517 | 0.53112 | 1 | 11334 | 3 | 1.4035 |
| CLDN14 | 6 | 0.26591 | 0.46687 | 0.999994 | 9364 | 1 | 1.0904 | 0.3052 | 0.57009 | 1 | 11335 | 4 | 1.0904 |

| | | | | | | | | | | | | | |
|---|---|---|---|---|---|---|---|---|---|---|---|---|---|
| TSPAN18 | 4 | 0.17461 | 0.31456 | 0.999994 | 7101 | 1 | 1.388 | 0.30521 | 0.5178 | 1 | 11336 | 2 | 1.388 |
| TCTE1 | 6 | 0.92781 | 0.92789 | 0.999994 | 19810 | 1 | 0.28491 | 0.30527 | 0.57017 | 1 | 11337 | 2 | 0.28491 |
| TMEM8B | 5 | 0.33887 | 0.5128 | 0.999994 | 10914 | 2 | 1.4081 | 0.30528 | 0.53124 | 1 | 11338 | 3 | 1.4081 |
| FBXO11 | 6 | 0.024166 | 0.085031 | 0.94395 | 1717 | 3 | 0.56453 | 0.30532 | 0.57023 | 1 | 11339 | 3 | 0.56453 |
| hsa-mir-124-1 | 3 | 0.30404 | 0.43067 | 0.999994 | 10210 | 1 | 1.4993 | 0.30533 | 0.45542 | 1 | 11340 | 2 | 1.4993 |
| BEND6 | 6 | 0.86931 | 0.88292 | 0.999994 | 18621 | 1 | 1.1711 | 0.30533 | 0.57025 | 1 | 11341 | 3 | 1.1711 |
| C11orf54 | 6 | 0.924 | 0.92416 | 0.999994 | 19714 | 1 | 0.55064 | 0.30534 | 0.57027 | 1 | 11342 | 3 | 0.55064 |
| SREBF1 | 6 | 0.26874 | 0.47026 | 0.999994 | 9441 | 3 | -0.32877 | 0.30534 | 0.57027 | 1 | 11343 | 2 | -0.32877 |
| GPRC5A | 5 | 0.024556 | 0.076291 | 0.93914 | 1733 | 3 | -2.4842 | 0.30538 | 0.53133 | 1 | 11344 | 2 | -2.4842 |
| TAGAP | 5 | 0.33742 | 0.51163 | 0.999994 | 10884 | 1 | 1.1067 | 0.30538 | 0.53133 | 1 | 11345 | 3 | 1.1067 |
| TYRP1 | 5 | 0.48407 | 0.63417 | 0.999994 | 13201 | 1 | 0.24379 | 0.30538 | 0.53133 | 1 | 11346 | 2 | 0.24379 |
| KIF2A | 5 | 0.021935 | 0.069317 | 0.93914 | 1577 | 3 | -3.1051 | 0.30538 | 0.53133 | 1 | 11347 | 1 | -3.1051 |
| STS | 6 | 0.48894 | 0.66121 | 0.999994 | 13266 | 2 | 0.91283 | 0.30542 | 0.57037 | 1 | 11348 | 4 | 0.91283 |
| YBX3 | 5 | 0.8195 | 0.83227 | 0.999994 | 17773 | 1 | 1.3873 | 0.30549 | 0.53145 | 1 | 11349 | 3 | 1.3873 |
| RSU1 | 6 | 0.86169 | 0.87838 | 0.999994 | 18477 | 1 | 0.64507 | 0.3055 | 0.57047 | 1 | 11350 | 3 | 0.64507 |
| MOK | 6 | 0.52223 | 0.68405 | 0.999994 | 13781 | 2 | 0.70015 | 0.3055 | 0.57047 | 1 | 11351 | 4 | 0.70015 |
| HIST1H2BC | 6 | 0.91597 | 0.91676 | 0.999994 | 19535 | 1 | 1.2344 | 0.30566 | 0.57064 | 1 | 11352 | 4 | 1.2344 |
| PPP2R4 | 3 | 0.46662 | 0.52457 | 0.999994 | 12960 | 1 | 1.7471 | 0.30577 | 0.45589 | 1 | 11353 | 2 | 1.7471 |
| ATP6V0A4 | 4 | 0.41072 | 0.55668 | 0.999994 | 12123 | 2 | 0.80364 | 0.30584 | 0.51827 | 1 | 11354 | 2 | 0.80364 |
| DSEL | 6 | 0.10406 | 0.25525 | 0.994943 | 5098 | 4 | -1.2519 | 0.3059 | 0.57097 | 1 | 11355 | 2 | -1.2519 |
| CDH10 | 6 | 0.25113 | 0.44928 | 0.999994 | 9038 | 3 | -0.80686 | 0.3059 | 0.57097 | 1 | 11356 | 1 | -0.80686 |
| TSC22D4 | 6 | 0.00049933 | 0.0026082 | 0.49544 | 109 | 5 | -2.9365 | 0.3059 | 0.57097 | 1 | 11357 | 1 | -2.9365 |
| ESCO2 | 6 | 0.0046404 | 0.019844 | 0.808847 | 489 | 3 | -2.1411 | 0.3059 | 0.57097 | 1 | 11358 | 3 | -2.1411 |
| KIAA1033 | 6 | 0.33744 | 0.54974 | 0.999994 | 10886 | 3 | 0.72075 | 0.30607 | 0.57118 | 1 | 11359 | 3 | 0.72075 |
| PNMA2 | 5 | 0.083015 | 0.19947 | 0.977164 | 4339 | 2 | -0.0055901 | 0.3062 | 0.53221 | 1 | 11360 | 2 | -0.0055901 |
| KRTAP29-1 | 5 | 0.46741 | 0.61993 | 0.999994 | 12970 | 2 | 1.3596 | 0.3062 | 0.53221 | 1 | 11361 | 3 | 1.3596 |
| C8A | 6 | 0.53405 | 0.69237 | 0.999994 | 13967 | 1 | 0.84247 | 0.3062 | 0.57133 | 1 | 11362 | 4 | 0.84247 |
| PNKD | 5 | 0.35762 | 0.52813 | 0.999994 | 11299 | 1 | 1.4023 | 0.3062 | 0.53221 | 1 | 11363 | 3 | 1.4023 |
| KCNN3 | 6 | 0.9392 | 0.93915 | 0.999994 | 20052 | 0 | 1.3884 | 0.30622 | 0.57135 | 1 | 11364 | 4 | 1.3884 |
| LACTB2 | 6 | 0.21751 | 0.40912 | 0.999994 | 8210 | 3 | 0.43739 | 0.30635 | 0.57153 | 1 | 11365 | 3 | 0.43739 |
| CLDN25 | 5 | 0.017298 | 0.056677 | 0.9235 | 1327 | 1 | 0.081211 | 0.30643 | 0.53246 | 1 | 11366 | 2 | 0.081211 |
| NME4 | 5 | 0.79528 | 0.81657 | 0.999994 | 17421 | 1 | 1.2189 | 0.30643 | 0.53246 | 1 | 11367 | 3 | 1.2189 |
| ATXN7L3B | 5 | 0.16736 | 0.32824 | 0.999994 | 6918 | 3 | -1.1116 | 0.30643 | 0.53246 | 1 | 11368 | 2 | -1.1116 |
| WIPF1 | 4 | 0.31399 | 0.45735 | 0.999994 | 10395 | 2 | -0.47937 | 0.30649 | 0.51875 | 1 | 11369 | 1 | -0.47937 |
| GRIN1 | 6 | 0.12054 | 0.28674 | 0.999632 | 5646 | 3 | 0.29352 | 0.3065 | 0.57171 | 1 | 11370 | 3 | 0.29352 |
| PPP4R4 | 5 | 0.35351 | 0.52478 | 0.999994 | 11224 | 1 | 1.3263 | 0.30654 | 0.53256 | 1 | 11371 | 3 | 1.3263 |
| RPS28 | 3 | 0.17262 | 0.27775 | 0.999519 | 7051 | 1 | 1.7294 | 0.30657 | 0.45676 | 1 | 11372 | 2 | 1.7294 |
| AHR | 6 | 0.66704 | 0.79083 | 0.999994 | 15879 | 2 | 1.0129 | 0.30659 | 0.57182 | 1 | 11373 | 3 | 1.0129 |
| hsa-mir-4266 | 3 | 0.044552 | 0.10796 | 0.951052 | 2737 | 2 | -3.7871 | 0.30662 | 0.45681 | 1 | 11374 | 1 | -3.7871 |
| NDUFB2 | 6 | 0.27699 | 0.47994 | 0.999994 | 9638 | 3 | 0.39165 | 0.30664 | 0.57188 | 1 | 11375 | 3 | 0.39165 |
| ZNF79 | 6 | 0.10371 | 0.25457 | 0.994835 | 5091 | 3 | -0.57601 | 0.30671 | 0.57197 | 1 | 11376 | 2 | -0.57601 |
| FKBP9 | 6 | 0.0075268 | 0.030295 | 0.862223 | 693 | 5 | -1.9824 | 0.30671 | 0.57197 | 1 | 11377 | 1 | -1.9824 |
| WDR48 | 6 | 0.42532 | 0.61887 | 0.999994 | 12340 | 2 | 0.29197 | 0.30676 | 0.57204 | 1 | 11378 | 2 | 0.29197 |
| CLCN3 | 6 | 0.84968 | 0.87176 | 0.999994 | 18254 | 1 | 0.49785 | 0.30676 | 0.57204 | 1 | 11379 | 3 | 0.49785 |
| KHDRBS2 | 6 | 0.21802 | 0.40976 | 0.999994 | 8228 | 3 | -0.70194 | 0.30676 | 0.57204 | 1 | 11380 | 2 | -0.70194 |
| WFDC2 | 6 | 0.80952 | 0.85205 | 0.999994 | 17625 | 1 | 0.76424 | 0.30677 | 0.57205 | 1 | 11381 | 4 | 0.76424 |
| NSF | 4 | 0.8249 | 0.82453 | 0.999994 | 17852 | 1 | 1.7169 | 0.30692 | 0.51908 | 1 | 11382 | 2 | 1.7169 |
| PTPRJ | 6 | 0.5013 | 0.66959 | 0.999994 | 13442 | 1 | 1.2248 | 0.30707 | 0.5724 | 1 | 11383 | 3 | 1.2248 |
| SLC16A2 | 5 | 0.011493 | 0.040362 | 0.891621 | 954 | 4 | -2.9861 | 0.30714 | 0.53318 | 1 | 11384 | 1 | -2.9861 |
| CYP19A1 | 5 | 0.82514 | 0.83603 | 0.999994 | 17853 | 1 | 0.56209 | 0.30714 | 0.53318 | 1 | 11385 | 3 | 0.56209 |
| PLCXD2 | 5 | 0.52892 | 0.6731 | 0.999994 | 13880 | 2 | -0.50415 | 0.30714 | 0.53318 | 1 | 11386 | 1 | -0.50415 |
| ANKRD27 | 5 | 0.31732 | 0.49518 | 0.999994 | 10461 | 1 | 1.4672 | 0.3072 | 0.53324 | 1 | 11387 | 3 | 1.4672 |

| | | | | | | | | | | | | | |
|---|---|---|---|---|---|---|---|---|---|---|---|---|---|
| SPOCK2 | 4 | 0.37 | 0.51486 | 0.999994 | 11509 | 2 | 0.70142 | 0.3072 | 0.5193 | 1 | 11388 | 2 | 0.70142 |
| NonTargetingControlG | 2 | 0.22414 | 0.27965 | 0.999632 | 8383 | 1 | 0.89135 | 0.30722 | 0.40913 | 1 | 11389 | 1 | 0.89135 |
| ACSM4 | 6 | 0.18413 | 0.3683 | 0.999994 | 7319 | 3 | -0.61619 | 0.3073 | 0.5727 | 1 | 11390 | 2 | -0.61619 |
| UIMC1 | 5 | 0.18671 | 0.35626 | 0.999994 | 7378 | 1 | 1.3199 | 0.30731 | 0.53336 | 1 | 11391 | 3 | 1.3199 |
| GAGE1 | 6 | 0.97804 | 0.97798 | 0.999994 | 21041 | 0 | 0.58722 | 0.30736 | 0.57279 | 1 | 11392 | 4 | 0.58722 |
| MARVELD3 | 6 | 0.95583 | 0.95591 | 0.999994 | 20478 | 0 | 0.78052 | 0.30738 | 0.5728 | 1 | 11393 | 4 | 0.78052 |
| PHYHIP | 6 | 0.53345 | 0.69197 | 0.999994 | 13958 | 1 | 1.354 | 0.30741 | 0.57284 | 1 | 11394 | 4 | 1.354 |
| RECK | 5 | 0.20923 | 0.38832 | 0.999994 | 8012 | 1 | 1.2768 | 0.30747 | 0.53353 | 1 | 11395 | 3 | 1.2768 |
| TSSK1B | 6 | 0.012633 | 0.047867 | 0.900643 | 1039 | 2 | 1.0095 | 0.3076 | 0.57306 | 1 | 11396 | 4 | 1.0095 |
| AMHR2 | 6 | 0.23187 | 0.42634 | 0.999994 | 8566 | 2 | 0.76553 | 0.30774 | 0.57323 | 1 | 11397 | 4 | 0.76553 |
| METTL20 | 6 | 0.47784 | 0.65374 | 0.999994 | 13110 | 1 | 0.8775 | 0.30775 | 0.57324 | 1 | 11398 | 3 | 0.8775 |
| PPARA | 5 | 0.094677 | 0.218 | 0.980699 | 4774 | 3 | -0.76625 | 0.30775 | 0.53382 | 1 | 11399 | 2 | -0.76625 |
| RBM25 | 4 | 0.56953 | 0.6494 | 0.999994 | 14531 | 1 | 1.4988 | 0.30778 | 0.51973 | 1 | 11400 | 2 | 1.4988 |
| CLEC4C | 6 | 0.037651 | 0.11661 | 0.951052 | 2410 | 2 | 0.7491 | 0.30784 | 0.57335 | 1 | 11401 | 4 | 0.7491 |
| PLAG1 | 6 | 0.008955 | 0.035364 | 0.870224 | 788 | 4 | -4.3474 | 0.308 | 0.57357 | 1 | 11402 | 2 | -4.3474 |
| TBPL2 | 6 | 0.15584 | 0.33298 | 0.999994 | 6619 | 2 | -0.52143 | 0.308 | 0.57357 | 1 | 11403 | 2 | -0.52143 |
| ACAN | 6 | 0.60167 | 0.74118 | 0.999994 | 14969 | 2 | -0.28177 | 0.308 | 0.57357 | 1 | 11404 | 1 | -0.28177 |
| TBC1D20 | 6 | 0.5725 | 0.71975 | 0.999994 | 14584 | 2 | 0.66257 | 0.30802 | 0.57359 | 1 | 11405 | 4 | 0.66257 |
| RALGDS | 6 | 0.15117 | 0.32715 | 0.999994 | 6486 | 3 | -0.4153 | 0.30808 | 0.57366 | 1 | 11406 | 3 | -0.4153 |
| DNAJC13 | 6 | 0.86524 | 0.88044 | 0.999994 | 18543 | 1 | 0.9319 | 0.30809 | 0.57368 | 1 | 11407 | 4 | 0.9319 |
| NonTargetingControlG | 2 | 0.58082 | 0.58097 | 0.999994 | 14688 | 1 | 2.5722 | 0.30813 | 0.40967 | 1 | 11408 | 1 | 2.5722 |
| hsa-mir-3148 | 4 | 0.15034 | 0.28953 | 0.999632 | 6465 | 1 | 0.36709 | 0.30813 | 0.52001 | 1 | 11409 | 2 | 0.36709 |
| CCDC23 | 6 | 0.15767 | 0.33528 | 0.999994 | 6663 | 3 | 0.58321 | 0.30814 | 0.57373 | 1 | 11410 | 3 | 0.58321 |
| ACSL6 | 5 | 0.24236 | 0.4341 | 0.999994 | 8840 | 1 | 0.12529 | 0.30822 | 0.53429 | 1 | 11411 | 2 | 0.12529 |
| TNFSF18 | 6 | 0.94799 | 0.94803 | 0.999994 | 20286 | 0 | 0.97701 | 0.30833 | 0.57397 | 1 | 11412 | 4 | 0.97701 |
| LRRN2 | 5 | 0.23727 | 0.42724 | 0.999994 | 8705 | 2 | 0.20602 | 0.30833 | 0.53442 | 1 | 11413 | 2 | 0.20602 |
| HSD17B14 | 6 | 0.23121 | 0.42556 | 0.999994 | 8547 | 2 | 1.1048 | 0.30843 | 0.5741 | 1 | 11414 | 3 | 1.1048 |
| OR2G2 | 5 | 0.26839 | 0.45554 | 0.999994 | 9429 | 1 | 1.3316 | 0.30857 | 0.53467 | 1 | 11415 | 3 | 1.3316 |
| TRIM11 | 6 | 0.15735 | 0.33488 | 0.999994 | 6657 | 2 | 0.02114 | 0.30869 | 0.57442 | 1 | 11416 | 2 | 0.02114 |
| ZNF669 | 6 | 0.046534 | 0.13647 | 0.956051 | 2832 | 3 | 0.016296 | 0.30884 | 0.57461 | 1 | 11417 | 3 | 0.016296 |
| FXYD6 | 5 | 0.3382 | 0.51226 | 0.999994 | 10904 | 2 | 1.2368 | 0.30891 | 0.53503 | 1 | 11418 | 3 | 1.2368 |
| PAGE1 | 5 | 0.25974 | 0.44853 | 0.999994 | 9232 | 1 | -0.16575 | 0.30891 | 0.53503 | 1 | 11419 | 2 | -0.16575 |
| CXCR1 | 5 | 0.38868 | 0.55402 | 0.999994 | 11779 | 1 | 0.11655 | 0.30891 | 0.53503 | 1 | 11420 | 2 | 0.11655 |
| SLC9B2 | 5 | 0.013317 | 0.04547 | 0.891621 | 1084 | 2 | -0.15199 | 0.30891 | 0.53503 | 1 | 11421 | 1 | -0.15199 |
| CRLF3 | 5 | 0.06875 | 0.17639 | 0.968171 | 3784 | 3 | -2.0533 | 0.30904 | 0.53518 | 1 | 11422 | 2 | -2.0533 |
| hsa-mir-612 | 4 | 0.28723 | 0.42996 | 0.999994 | 9858 | 1 | 1.7293 | 0.30905 | 0.52068 | 1 | 11423 | 2 | 1.7293 |
| PID1 | 6 | 0.020333 | 0.073047 | 0.93914 | 1478 | 2 | -0.011297 | 0.30911 | 0.57492 | 1 | 11424 | 2 | -0.011297 |
| PGAM2 | 6 | 0.013872 | 0.052031 | 0.911383 | 1120 | 5 | -2.203 | 0.30911 | 0.57492 | 1 | 11425 | 1 | -2.203 |
| OR7C2 | 6 | 0.079856 | 0.207 | 0.978744 | 4229 | 2 | 0.74918 | 0.30917 | 0.57499 | 1 | 11426 | 4 | 0.74918 |
| SEMA4G | 5 | 0.49192 | 0.64093 | 0.999994 | 13299 | 2 | 1.2163 | 0.30928 | 0.53543 | 1 | 11427 | 3 | 1.2163 |
| FAM199X | 6 | 0.1781 | 0.36073 | 0.999994 | 7177 | 2 | 0.95012 | 0.30935 | 0.57522 | 1 | 11428 | 4 | 0.95012 |
| ATP8B2 | 5 | 0.19508 | 0.3682 | 0.999994 | 7636 | 3 | -0.89578 | 0.3094 | 0.53556 | 1 | 11429 | 2 | -0.89578 |
| FOXF1 | 6 | 0.069862 | 0.18647 | 0.969409 | 3825 | 2 | 1.3566 | 0.30941 | 0.57529 | 1 | 11430 | 4 | 1.3566 |
| STK32C | 6 | 0.57286 | 0.71999 | 0.999994 | 14589 | 2 | 0.93974 | 0.30943 | 0.57531 | 1 | 11431 | 4 | 0.93974 |
| OR12D2 | 6 | 0.066955 | 0.18038 | 0.968171 | 3718 | 4 | -1.3998 | 0.30948 | 0.57537 | 1 | 11432 | 2 | -1.3998 |
| hsa-mir-302b | 3 | 0.85178 | 0.85176 | 0.999994 | 18292 | 0 | 0.81053 | 0.30967 | 0.46007 | 1 | 11433 | 2 | 0.81053 |
| hsa-mir-4508 | 3 | 0.71074 | 0.71112 | 0.999994 | 16406 | 1 | 0.53652 | 0.30967 | 0.46007 | 1 | 11434 | 1 | 0.53652 |
| ABCB11 | 6 | 0.6567 | 0.78274 | 0.999994 | 15739 | 2 | 1.0385 | 0.30967 | 0.5756 | 1 | 11435 | 3 | 1.0385 |
| CCAR1 | 6 | 0.015596 | 0.05769 | 0.929232 | 1222 | 3 | 0.26812 | 0.30967 | 0.5756 | 1 | 11436 | 3 | 0.26812 |
| STRA8 | 6 | 0.36227 | 0.57785 | 0.999994 | 11373 | 3 | -0.2351 | 0.30967 | 0.5756 | 1 | 11437 | 2 | -0.2351 |
| DFNB59 | 6 | 0.60382 | 0.74281 | 0.999994 | 14994 | 1 | 1.3255 | 0.30972 | 0.57566 | 1 | 11438 | 4 | 1.3255 |
| IDH2 | 5 | 0.35024 | 0.52213 | 0.999994 | 11164 | 1 | 1.2802 | 0.30973 | 0.5359 | 1 | 11439 | 3 | 1.2802 |

| | | | | | | | | | | | | | |
|---|---|---|---|---|---|---|---|---|---|---|---|---|---|
| ACE2 | 6 | 0.67083 | 0.79379 | 0.999994 | 15941 | 2 | 0.29406 | 0.30979 | 0.57575 | 1 | 11440 | 3 | 0.29406 |
| GCSH | 4 | 0.072267 | 0.16422 | 0.967126 | 3928 | 2 | -1.9078 | 0.30999 | 0.5214 | 1 | 11441 | 1 | -1.9078 |
| hsa-mir-3689d-1 | 4 | 0.73738 | 0.75056 | 0.999994 | 16692 | 1 | 0.67857 | 0.30999 | 0.5214 | 1 | 11442 | 2 | 0.67857 |
| OGDH | 4 | 0.32386 | 0.4674 | 0.999994 | 10591 | 1 | 0.95705 | 0.30999 | 0.5214 | 1 | 11443 | 2 | 0.95705 |
| ZNF124 | 6 | 0.011457 | 0.043935 | 0.891621 | 953 | 2 | 1.3468 | 0.31004 | 0.57605 | 1 | 11444 | 4 | 1.3468 |
| ZMYND8 | 6 | 0.72689 | 0.82058 | 0.999994 | 16566 | 1 | 0.58052 | 0.31014 | 0.57619 | 1 | 11445 | 4 | 0.58052 |
| PDHX | 6 | 0.0078817 | 0.031561 | 0.864134 | 720 | 4 | -1.4443 | 0.31023 | 0.57628 | 1 | 11446 | 2 | -1.4443 |
| DNAJC14 | 6 | 0.63298 | 0.76459 | 0.999994 | 15413 | 2 | -0.074748 | 0.31023 | 0.57628 | 1 | 11447 | 2 | -0.074748 |
| ZDHHC18 | 6 | 0.10263 | 0.25245 | 0.994162 | 5059 | 2 | 0.64757 | 0.31033 | 0.57642 | 1 | 11448 | 4 | 0.64757 |
| TDRD10 | 6 | 0.66936 | 0.79267 | 0.999994 | 15921 | 2 | 1.1444 | 0.31033 | 0.57643 | 1 | 11449 | 3 | 1.1444 |
| SACM1L | 5 | 0.070568 | 0.17931 | 0.968171 | 3855 | 3 | -0.85349 | 0.31035 | 0.53655 | 1 | 11450 | 2 | -0.85349 |
| WAPAL | 5 | 0.33151 | 0.50679 | 0.999994 | 10755 | 2 | -0.023506 | 0.31035 | 0.53655 | 1 | 11451 | 2 | -0.023506 |
| SATB1 | 5 | 0.43051 | 0.58885 | 0.999994 | 12425 | 2 | -0.40536 | 0.31035 | 0.53655 | 1 | 11452 | 2 | -0.40536 |
| DSG3 | 6 | 0.17687 | 0.35917 | 0.999994 | 7151 | 1 | 0.72197 | 0.31038 | 0.57649 | 1 | 11453 | 4 | 0.72197 |
| PBX3 | 5 | 0.15462 | 0.30967 | 0.999994 | 6584 | 3 | -0.87453 | 0.31039 | 0.53658 | 1 | 11454 | 2 | -0.87453 |
| CLDN3 | 5 | 0.54605 | 0.68824 | 0.999994 | 14152 | 1 | 1.2724 | 0.31053 | 0.53674 | 1 | 11455 | 3 | 1.2724 |
| CA4 | 6 | 0.093153 | 0.23383 | 0.984971 | 4727 | 2 | 0.64334 | 0.31057 | 0.57672 | 1 | 11456 | 4 | 0.64334 |
| ANKZF1 | 6 | 0.14143 | 0.31486 | 0.999994 | 6228 | 2 | 1.1396 | 0.31059 | 0.57674 | 1 | 11457 | 4 | 1.1396 |
| NonTargetingControlG | 2 | 0.70817 | 0.70792 | 0.999994 | 16390 | 0 | 2.9607 | 0.31066 | 0.41122 | 1 | 11458 | 1 | 2.9607 |
| KRAS | 6 | 0.20592 | 0.39508 | 0.999994 | 7923 | 2 | 0.92904 | 0.31069 | 0.57686 | 1 | 11459 | 4 | 0.92904 |
| ALPI | 6 | 0.76408 | 0.83359 | 0.999994 | 17016 | 1 | 1.3788 | 0.31072 | 0.57692 | 1 | 11460 | 3 | 1.3788 |
| PIP4K2B | 6 | 0.95138 | 0.95136 | 0.999994 | 20374 | 0 | 0.34011 | 0.31079 | 0.57699 | 1 | 11461 | 3 | 0.34011 |
| PLXNA4 | 6 | 0.36856 | 0.58233 | 0.999994 | 11480 | 2 | 1.2737 | 0.31113 | 0.57742 | 1 | 11462 | 3 | 1.2737 |
| MOGAT1 | 5 | 0.036405 | 0.10683 | 0.950995 | 2340 | 4 | -1.9155 | 0.31124 | 0.5375 | 1 | 11463 | 1 | -1.9155 |
| C9orf170 | 5 | 0.04519 | 0.12878 | 0.951777 | 2771 | 2 | 0.25449 | 0.31124 | 0.5375 | 1 | 11464 | 1 | 0.25449 |
| RBM10 | 4 | 0.78329 | 0.78666 | 0.999994 | 17259 | 1 | 1.4757 | 0.31126 | 0.52235 | 1 | 11465 | 2 | 1.4757 |
| CAP1 | 6 | 0.34148 | 0.55436 | 0.999994 | 10970 | 2 | 1.1732 | 0.31133 | 0.57768 | 1 | 11466 | 4 | 1.1732 |
| C9orf169 | 4 | 0.84983 | 0.84936 | 0.999994 | 18258 | 0 | 0.68884 | 0.31136 | 0.52243 | 1 | 11467 | 2 | 0.68884 |
| GBGT1 | 5 | 0.34266 | 0.51591 | 0.999994 | 10998 | 2 | -0.26925 | 0.31149 | 0.53776 | 1 | 11468 | 2 | -0.26925 |
| UBR7 | 5 | 0.050013 | 0.14039 | 0.958716 | 2978 | 2 | -0.1165 | 0.31149 | 0.53776 | 1 | 11469 | 2 | -0.1165 |
| HSPA9 | 5 | 0.19232 | 0.36425 | 0.999994 | 7556 | 2 | -0.014904 | 0.31149 | 0.53776 | 1 | 11470 | 2 | -0.014904 |
| NDC80 | 5 | 0.012087 | 0.042066 | 0.891621 | 998 | 3 | -4.1848 | 0.31149 | 0.53776 | 1 | 11471 | 2 | -4.1848 |
| COL15A1 | 6 | 0.11659 | 0.2793 | 0.999632 | 5525 | 2 | 0.59213 | 0.31149 | 0.57788 | 1 | 11472 | 3 | 0.59213 |
| MAGEB3 | 6 | 0.82235 | 0.85795 | 0.999994 | 17810 | 1 | 0.45033 | 0.31149 | 0.57788 | 1 | 11473 | 3 | 0.45033 |
| COX6A2 | 6 | 0.14697 | 0.32184 | 0.999994 | 6373 | 3 | -0.58714 | 0.31149 | 0.57788 | 1 | 11474 | 2 | -0.58714 |
| CD53 | 6 | 0.25721 | 0.45656 | 0.999994 | 9171 | 3 | -0.6316 | 0.31157 | 0.57797 | 1 | 11475 | 2 | -0.6316 |
| SEMG2 | 6 | 0.32224 | 0.53249 | 0.999994 | 10559 | 3 | -0.60411 | 0.31157 | 0.57797 | 1 | 11476 | 2 | -0.60411 |
| PKN3 | 6 | 0.74224 | 0.82574 | 0.999994 | 16767 | 1 | 0.7245 | 0.31157 | 0.57797 | 1 | 11477 | 3 | 0.7245 |
| CRYZL1 | 6 | 0.36977 | 0.58307 | 0.999994 | 11504 | 2 | 0.53259 | 0.31157 | 0.57797 | 1 | 11478 | 3 | 0.53259 |
| OR4N5 | 5 | 0.14096 | 0.28927 | 0.999632 | 6218 | 2 | 1.2299 | 0.31161 | 0.53787 | 1 | 11479 | 3 | 1.2299 |
| NonTargetingControlG | 2 | 0.14782 | 0.22513 | 0.983354 | 6402 | 1 | -0.70918 | 0.31165 | 0.41185 | 1 | 11480 | 1 | -0.70918 |
| PGK1 | 6 | 0.48268 | 0.65698 | 0.999994 | 13184 | 1 | 1.0324 | 0.31167 | 0.57807 | 1 | 11481 | 3 | 1.0324 |
| PDIA5 | 5 | 0.19019 | 0.36121 | 0.999994 | 7493 | 2 | 1.4037 | 0.31184 | 0.53812 | 1 | 11482 | 3 | 1.4037 |
| MRGPRG | 4 | 0.026602 | 0.074543 | 0.93914 | 1834 | 2 | -0.15201 | 0.31193 | 0.52286 | 1 | 11483 | 2 | -0.15201 |
| FOLR2 | 5 | 0.23583 | 0.42528 | 0.999994 | 8665 | 1 | -0.15964 | 0.31194 | 0.53824 | 1 | 11484 | 2 | -0.15964 |
| CLSTN1 | 6 | 0.3402 | 0.55287 | 0.999994 | 10936 | 2 | 0.58832 | 0.31205 | 0.57854 | 1 | 11485 | 4 | 0.58832 |
| FBXO6 | 6 | 0.1508 | 0.32668 | 0.999994 | 6479 | 1 | 0.71664 | 0.31207 | 0.57856 | 1 | 11486 | 4 | 0.71664 |
| COQ6 | 6 | 0.16957 | 0.35022 | 0.999994 | 6975 | 2 | 0.57871 | 0.3123 | 0.57882 | 1 | 11487 | 3 | 0.57871 |
| POLR3F | 4 | 0.85067 | 0.85019 | 0.999994 | 18272 | 0 | 1.179 | 0.31232 | 0.52314 | 1 | 11488 | 2 | 1.179 |
| FMN2 | 5 | 0.87028 | 0.87132 | 0.999994 | 18639 | 1 | 1.5529 | 0.31248 | 0.53882 | 1 | 11489 | 3 | 1.5529 |
| NonTargetingControlG | 1 | 0.68751 | 0.68749 | 0.999994 | 16189 | 0 | 2.4878 | 0.31249 | 0.31261 | 0.989342 | 11490 | 1 | 2.4878 |
| SKP1 | 5 | 0.28099 | 0.46567 | 0.999994 | 9723 | 2 | -0.37976 | 0.31255 | 0.53889 | 1 | 11491 | 2 | -0.37976 |

| | | | | | | | | | | | | | |
|---|---|---|---|---|---|---|---|---|---|---|---|---|---|
| SOX6 | 5 | 0.5622 | 0.7026 | 0.999994 | 14412 | 1 | 0.45146 | 0.31267 | 0.53901 | 1 | 11492 | 2 | 0.45146 |
| GZMB | 5 | 0.64624 | 0.74361 | 0.999994 | 15604 | 1 | 1.1977 | 0.31294 | 0.5393 | 1 | 11493 | 3 | 1.1977 |
| RNF133 | 6 | 0.30574 | 0.51341 | 0.999994 | 10247 | 3 | -0.26751 | 0.31294 | 0.57964 | 1 | 11494 | 2 | -0.26751 |
| NT5C1A | 6 | 0.80354 | 0.8494 | 0.999994 | 17534 | 1 | 1.1328 | 0.31311 | 0.57984 | 1 | 11495 | 4 | 1.1328 |
| EGR2 | 3 | 0.45617 | 0.51807 | 0.999994 | 12793 | 1 | -0.27156 | 0.31322 | 0.46376 | 1 | 11496 | 1 | -0.27156 |
| SLC25A3 | 4 | 0.79416 | 0.79604 | 0.999994 | 17397 | 1 | 1.8119 | 0.31328 | 0.52386 | 1 | 11497 | 2 | 1.8119 |
| ZFYVE20 | 5 | 0.25162 | 0.44206 | 0.999994 | 9049 | 2 | 0.61852 | 0.3133 | 0.53968 | 1 | 11498 | 3 | 0.61852 |
| DGKK | 5 | 0.18581 | 0.35501 | 0.999994 | 7362 | 3 | -0.8405 | 0.3133 | 0.53968 | 1 | 11499 | 1 | -0.8405 |
| LCN9 | 5 | 0.69798 | 0.76557 | 0.999994 | 16279 | 1 | 0.88362 | 0.31334 | 0.53972 | 1 | 11500 | 3 | 0.88362 |
| GOSR2 | 5 | 0.36001 | 0.53008 | 0.999994 | 11336 | 2 | 1.078 | 0.31334 | 0.53972 | 1 | 11501 | 3 | 1.078 |
| FEZ2 | 5 | 0.1531 | 0.30742 | 0.999994 | 6538 | 3 | -1.0231 | 0.31334 | 0.53972 | 1 | 11502 | 2 | -1.0231 |
| TMEM180 | 5 | 0.54292 | 0.68548 | 0.999994 | 14110 | 2 | 1.1482 | 0.31334 | 0.53972 | 1 | 11503 | 3 | 1.1482 |
| NonTargetingControlG | 2 | 0.25277 | 0.30097 | 0.999994 | 9073 | 1 | 1.0255 | 0.31335 | 0.41288 | 1 | 11504 | 1 | 1.0255 |
| STC1 | 5 | 0.32743 | 0.50341 | 0.999994 | 10666 | 2 | 1.3596 | 0.31339 | 0.53977 | 1 | 11505 | 3 | 1.3596 |
| RPL32 | 3 | 0.66785 | 0.67203 | 0.999994 | 15890 | 1 | 1.7484 | 0.31341 | 0.46396 | 1 | 11506 | 2 | 1.7484 |
| CCDC11 | 6 | 0.30225 | 0.50934 | 0.999994 | 10176 | 3 | 0.33207 | 0.31345 | 0.58027 | 1 | 11507 | 3 | 0.33207 |
| CLPS | 6 | 0.58328 | 0.72761 | 0.999994 | 14720 | 2 | 0.3286 | 0.31345 | 0.58027 | 1 | 11508 | 2 | 0.3286 |
| FAM5B | 3 | 0.49941 | 0.54577 | 0.999994 | 13414 | 1 | 1.569 | 0.31347 | 0.46403 | 1 | 11509 | 2 | 1.569 |
| SLIT3 | 6 | 0.0057776 | 0.023982 | 0.834563 | 576 | 2 | 0.90565 | 0.3135 | 0.58032 | 1 | 11510 | 4 | 0.90565 |
| ZNF506 | 6 | 0.050981 | 0.14621 | 0.964852 | 3017 | 3 | -1.4609 | 0.31354 | 0.58038 | 1 | 11511 | 2 | -1.4609 |
| SCG2 | 6 | 0.64305 | 0.77226 | 0.999994 | 15563 | 2 | 0.17423 | 0.31354 | 0.58038 | 1 | 11512 | 3 | 0.17423 |
| CDH11 | 6 | 0.33065 | 0.542 | 0.999994 | 10733 | 3 | -0.5168 | 0.31354 | 0.58038 | 1 | 11513 | 1 | -0.5168 |
| CDCA3 | 6 | 0.46937 | 0.64803 | 0.999994 | 12995 | 1 | 0.41372 | 0.31354 | 0.58038 | 1 | 11514 | 3 | 0.41372 |
| hsa-mir-378d-1 | 4 | 0.69775 | 0.72294 | 0.999994 | 16275 | 1 | 1.0856 | 0.31358 | 0.52409 | 1 | 11515 | 2 | 1.0856 |
| JMJD8 | 4 | 0.020656 | 0.06202 | 0.937033 | 1490 | 2 | 0.27929 | 0.31358 | 0.52409 | 1 | 11516 | 2 | 0.27929 |
| NLGN3 | 4 | 0.62013 | 0.67642 | 0.999994 | 15223 | 1 | 0.95347 | 0.31358 | 0.52409 | 1 | 11517 | 2 | 0.95347 |
| hsa-mir-375 | 4 | 0.68269 | 0.71332 | 0.999994 | 16127 | 1 | 0.17661 | 0.31358 | 0.52409 | 1 | 11518 | 1 | 0.17661 |
| MLNR | 5 | 0.76192 | 0.79724 | 0.999994 | 16999 | 1 | 1.2424 | 0.31373 | 0.54011 | 1 | 11519 | 3 | 1.2424 |
| TBX15 | 6 | 0.41591 | 0.61271 | 0.999994 | 12189 | 2 | 0.94258 | 0.31381 | 0.58071 | 1 | 11520 | 4 | 0.94258 |
| FBXO9 | 6 | 0.86999 | 0.88332 | 0.999994 | 18635 | 1 | 0.68214 | 0.31382 | 0.58074 | 1 | 11521 | 4 | 0.68214 |
| CD247 | 6 | 0.046294 | 0.13594 | 0.95468 | 2820 | 2 | 0.86263 | 0.31387 | 0.5808 | 1 | 11522 | 4 | 0.86263 |
| SLC22A9 | 4 | 0.058574 | 0.13802 | 0.957317 | 3366 | 2 | -0.74849 | 0.31394 | 0.52434 | 1 | 11523 | 2 | -0.74849 |
| FCN3 | 5 | 0.17164 | 0.33453 | 0.999994 | 7027 | 3 | -1.0259 | 0.31398 | 0.54037 | 1 | 11524 | 2 | -1.0259 |
| MMP8 | 5 | 0.055594 | 0.15387 | 0.967126 | 3225 | 4 | -1.0923 | 0.31398 | 0.54037 | 1 | 11525 | 1 | -1.0923 |
| NonTargetingControlG | 2 | 0.54012 | 0.54139 | 0.999994 | 14055 | 1 | 1.7691 | 0.31415 | 0.41337 | 1 | 11526 | 1 | 1.7691 |
| C16orf55 | 5 | 0.14417 | 0.29407 | 0.999994 | 6304 | 2 | 1.5605 | 0.31424 | 0.54064 | 1 | 11527 | 3 | 1.5605 |
| PDXDC1 | 4 | 0.68574 | 0.71525 | 0.999994 | 16168 | 1 | 0.22739 | 0.31428 | 0.52459 | 1 | 11528 | 1 | 0.22739 |
| NRBP2 | 4 | 0.13621 | 0.27501 | 0.998701 | 6085 | 2 | -0.40314 | 0.31428 | 0.52459 | 1 | 11529 | 2 | -0.40314 |
| NOS3 | 5 | 0.089888 | 0.21043 | 0.979587 | 4588 | 2 | 1.2987 | 0.31432 | 0.54074 | 1 | 11530 | 3 | 1.2987 |
| hsa-mir-198 | 3 | 0.26999 | 0.39971 | 0.999994 | 9473 | 2 | -0.66978 | 0.31434 | 0.46498 | 1 | 11531 | 1 | -0.66978 |
| IRF5 | 4 | 0.072562 | 0.16477 | 0.967126 | 3943 | 3 | -1.5901 | 0.31468 | 0.52488 | 1 | 11532 | 1 | -1.5901 |
| MIEN1 | 5 | 0.0017438 | 0.0076878 | 0.672529 | 243 | 4 | -7.892 | 0.31468 | 0.54112 | 1 | 11533 | 1 | -7.892 |
| TMEM95 | 5 | 0.54225 | 0.68489 | 0.999994 | 14099 | 2 | 0.095162 | 0.31468 | 0.54112 | 1 | 11534 | 2 | 0.095162 |
| CDK20 | 5 | 0.13283 | 0.27694 | 0.999379 | 5992 | 3 | -1.2056 | 0.31468 | 0.54112 | 1 | 11535 | 2 | -1.2056 |
| C9 | 5 | 0.3984 | 0.56222 | 0.999994 | 11921 | 2 | 0.03601 | 0.31468 | 0.54112 | 1 | 11536 | 1 | 0.03601 |
| hsa-mir-3136 | 4 | 0.40305 | 0.54888 | 0.999994 | 12000 | 1 | 1.3373 | 0.31471 | 0.5249 | 1 | 11537 | 2 | 1.3373 |
| FAM194A | 5 | 0.53226 | 0.67603 | 0.999994 | 13940 | 2 | 1.4042 | 0.31494 | 0.54141 | 1 | 11538 | 3 | 1.4042 |
| C12orf43 | 6 | 0.02807 | 0.094237 | 0.945031 | 1906 | 3 | -2.0906 | 0.31502 | 0.58219 | 1 | 11539 | 2 | -2.0906 |
| MSANTD1 | 6 | 0.36251 | 0.57812 | 0.999994 | 11379 | 2 | -0.22262 | 0.31502 | 0.58219 | 1 | 11540 | 2 | -0.22262 |
| RPUSD2 | 6 | 0.86952 | 0.88304 | 0.999994 | 18631 | 1 | 1.1938 | 0.31502 | 0.58219 | 1 | 11541 | 3 | 1.1938 |
| C2orf70 | 6 | 0.13805 | 0.31057 | 0.999994 | 6138 | 2 | -0.024482 | 0.31502 | 0.58219 | 1 | 11542 | 2 | -0.024482 |
| BMI1 | 6 | 0.11994 | 0.2856 | 0.999632 | 5629 | 4 | -0.71218 | 0.31502 | 0.58219 | 1 | 11543 | 2 | -0.71218 |

| | | | | | | | | | | | | | |
|---|---|---|---|---|---|---|---|---|---|---|---|---|---|
| NonTargetingControlG | 2 | 0.69357 | 0.6934 | 0.999994 | 16240 | 0 | 2.8766 | 0.31509 | 0.41393 | 1 | 11544 | 1 | 2.8766 |
| NonTargetingControlG | 2 | 0.37031 | 0.39243 | 0.999994 | 11517 | 1 | 1.9071 | 0.31509 | 0.41393 | 1 | 11545 | 1 | 1.9071 |
| LYSMD3 | 6 | 0.12843 | 0.29826 | 0.999994 | 5894 | 3 | -1.0442 | 0.31509 | 0.58229 | 1 | 11546 | 1 | -1.0442 |
| PPP2R2C | 6 | 0.48706 | 0.65997 | 0.999994 | 13238 | 2 | 0.62526 | 0.31509 | 0.58229 | 1 | 11547 | 3 | 0.62526 |
| TNFRSF13C | 4 | 0.10897 | 0.23104 | 0.983933 | 5270 | 2 | -0.11146 | 0.31512 | 0.5252 | 1 | 11548 | 2 | -0.11146 |
| hsa-mir-4448 | 3 | 0.061215 | 0.13114 | 0.95318 | 3483 | 2 | -3.0714 | 0.31524 | 0.46591 | 1 | 11549 | 1 | -3.0714 |
| FAM127A | 3 | 0.19352 | 0.30451 | 0.999994 | 7585 | 1 | 1.5048 | 0.31531 | 0.46599 | 1 | 11550 | 2 | 1.5048 |
| ZNF354C | 5 | 0.92458 | 0.92479 | 0.999994 | 19732 | 0 | 0.99658 | 0.31536 | 0.54184 | 1 | 11551 | 3 | 0.99658 |
| OR6C65 | 6 | 0.67608 | 0.79794 | 0.999994 | 16020 | 2 | 0.96919 | 0.31538 | 0.58264 | 1 | 11552 | 4 | 0.96919 |
| NonTargetingControlG | 1 | 0.68456 | 0.6846 | 0.999994 | 16154 | 0 | 2.1632 | 0.31544 | 0.31556 | 0.989846 | 11553 | 1 | 2.1632 |
| NR5A1 | 6 | 0.21348 | 0.40432 | 0.999994 | 8100 | 2 | 0.9089 | 0.31554 | 0.58284 | 1 | 11554 | 3 | 0.9089 |
| CCL22 | 6 | 0.7323 | 0.82238 | 0.999994 | 16632 | 1 | 0.81263 | 0.3156 | 0.58291 | 1 | 11555 | 4 | 0.81263 |
| CPEB3 | 6 | 0.097861 | 0.24313 | 0.988088 | 4885 | 4 | -0.94847 | 0.31575 | 0.5831 | 1 | 11556 | 1 | -0.94847 |
| C6orf132 | 6 | 0.0019597 | 0.0097759 | 0.680529 | 270 | 4 | -4.9756 | 0.31575 | 0.5831 | 1 | 11557 | 2 | -4.9756 |
| CTSH | 5 | 0.11217 | 0.24537 | 0.98979 | 5375 | 2 | 1.3172 | 0.31576 | 0.54229 | 1 | 11558 | 3 | 1.3172 |
| PVRL3 | 6 | 0.77426 | 0.83743 | 0.999994 | 17133 | 1 | 0.91664 | 0.31577 | 0.58313 | 1 | 11559 | 4 | 0.91664 |
| IQGAP1 | 5 | 0.87496 | 0.87557 | 0.999994 | 18734 | 1 | 1.1929 | 0.31578 | 0.54232 | 1 | 11560 | 3 | 1.1929 |
| hsa-mir-3689f | 4 | 0.25381 | 0.39585 | 0.999994 | 9099 | 2 | -0.23251 | 0.31586 | 0.52572 | 1 | 11561 | 2 | -0.23251 |
| OR5L2 | 6 | 0.48312 | 0.65728 | 0.999994 | 13189 | 2 | 0.78763 | 0.31592 | 0.58332 | 1 | 11562 | 4 | 0.78763 |
| RASSF1 | 5 | 0.05124 | 0.14339 | 0.960973 | 3034 | 3 | -2.084 | 0.31595 | 0.54248 | 1 | 11563 | 2 | -2.084 |
| C9orf123 | 5 | 0.42906 | 0.58764 | 0.999994 | 12398 | 2 | -0.49694 | 0.31595 | 0.54248 | 1 | 11564 | 2 | -0.49694 |
| PTGER3 | 5 | 0.33115 | 0.50649 | 0.999994 | 10747 | 2 | -0.2179 | 0.31595 | 0.54248 | 1 | 11565 | 2 | -0.2179 |
| ASCL4 | 5 | 0.0070964 | 0.027664 | 0.855011 | 665 | 4 | -4.3773 | 0.31595 | 0.54248 | 1 | 11566 | 1 | -4.3773 |
| hsa-mir-1225 | 2 | 0.19291 | 0.25697 | 0.994943 | 7573 | 1 | 0.54401 | 0.316 | 0.41447 | 1 | 11567 | 1 | 0.54401 |
| NonTargetingControlG | 2 | 0.042569 | 0.076683 | 0.93914 | 2651 | 1 | -1.9157 | 0.316 | 0.41447 | 1 | 11568 | 1 | -1.9157 |
| KIAA0922 | 6 | 0.44159 | 0.62948 | 0.999994 | 12604 | 2 | 0.72049 | 0.3161 | 0.58355 | 1 | 11569 | 4 | 0.72049 |
| SYNPR | 6 | 0.75715 | 0.83102 | 0.999994 | 16943 | 1 | 0.99265 | 0.31624 | 0.58371 | 1 | 11570 | 4 | 0.99265 |
| IFNW1 | 6 | 0.061442 | 0.16884 | 0.967126 | 3490 | 2 | 1.3257 | 0.31628 | 0.58377 | 1 | 11571 | 4 | 1.3257 |
| hsa-mir-4715 | 4 | 0.10765 | 0.22869 | 0.983354 | 5230 | 2 | -0.051691 | 0.31629 | 0.52605 | 1 | 11572 | 2 | -0.051691 |
| TMEM238 | 5 | 0.8276 | 0.83778 | 0.999994 | 17891 | 1 | 1.5712 | 0.31633 | 0.54285 | 1 | 11573 | 3 | 1.5712 |
| NHSL1 | 5 | 0.55246 | 0.69393 | 0.999994 | 14250 | 2 | 1.5712 | 0.31633 | 0.54285 | 1 | 11574 | 3 | 1.5712 |
| HIF1A | 6 | 0.2626 | 0.46293 | 0.999994 | 9302 | 3 | -0.30889 | 0.31645 | 0.58397 | 1 | 11575 | 2 | -0.30889 |
| PPP1R36 | 5 | 0.22186 | 0.40586 | 0.999994 | 8328 | 3 | -0.70118 | 0.31647 | 0.54301 | 1 | 11576 | 2 | -0.70118 |
| RAD54L2 | 5 | 0.79221 | 0.81466 | 0.999994 | 17377 | 1 | 1.109 | 0.31647 | 0.54301 | 1 | 11577 | 3 | 1.109 |
| hsa-mir-3920 | 3 | 0.81408 | 0.81384 | 0.999994 | 17691 | 0 | 1.817 | 0.31649 | 0.4672 | 1 | 11578 | 2 | 1.817 |
| SNRNP35 | 3 | 0.37571 | 0.47029 | 0.999994 | 11589 | 1 | 1.817 | 0.31649 | 0.4672 | 1 | 11579 | 2 | 1.817 |
| CRCP | 3 | 0.70392 | 0.7047 | 0.999994 | 16350 | 1 | 1.817 | 0.31649 | 0.4672 | 1 | 11580 | 2 | 1.817 |
| ATP6V0C | 3 | 0.86015 | 0.86013 | 0.999994 | 18443 | 0 | 1.817 | 0.31649 | 0.4672 | 1 | 11581 | 2 | 1.817 |
| LINS | 6 | 0.57035 | 0.71818 | 0.999994 | 14546 | 2 | 0.80114 | 0.31654 | 0.5841 | 1 | 11582 | 3 | 0.80114 |
| ZBTB10 | 5 | 0.92747 | 0.9277 | 0.999994 | 19802 | 0 | 0.75799 | 0.31655 | 0.54309 | 1 | 11583 | 3 | 0.75799 |
| hsa-mir-5588 | 2 | 0.23201 | 0.28543 | 0.999632 | 8571 | 1 | 0.96317 | 0.31657 | 0.41481 | 1 | 11584 | 1 | 0.96317 |
| SLC25A30 | 6 | 0.68353 | 0.80372 | 0.999994 | 16137 | 2 | 0.54491 | 0.31658 | 0.58413 | 1 | 11585 | 4 | 0.54491 |
| VSNL1 | 6 | 0.085542 | 0.21861 | 0.980699 | 4431 | 1 | 0.59295 | 0.31663 | 0.58419 | 1 | 11586 | 4 | 0.59295 |
| ZNF416 | 6 | 0.32583 | 0.5365 | 0.999994 | 10638 | 2 | 1.269 | 0.31667 | 0.58425 | 1 | 11587 | 3 | 1.269 |
| DCD | 6 | 0.22784 | 0.42149 | 0.999994 | 8475 | 2 | 0.98703 | 0.31668 | 0.58425 | 1 | 11588 | 4 | 0.98703 |
| E2F2 | 4 | 0.44169 | 0.58851 | 0.999994 | 12605 | 2 | 0.21069 | 0.31672 | 0.52635 | 1 | 11589 | 2 | 0.21069 |
| GSTM1 | 6 | 0.25799 | 0.45751 | 0.999994 | 9194 | 2 | 0.38195 | 0.31677 | 0.58435 | 1 | 11590 | 2 | 0.38195 |
| CPA1 | 6 | 0.061562 | 0.16908 | 0.967126 | 3494 | 4 | -1.2425 | 0.31677 | 0.58435 | 1 | 11591 | 2 | -1.2425 |
| PI4K2A | 6 | 0.23793 | 0.43351 | 0.999994 | 8722 | 2 | 1.1486 | 0.31677 | 0.58435 | 1 | 11592 | 3 | 1.1486 |
| OR2G3 | 6 | 0.32454 | 0.53505 | 0.999994 | 10608 | 2 | 0.0025788 | 0.31677 | 0.58435 | 1 | 11593 | 2 | 0.0025788 |
| OR51D1 | 6 | 0.12381 | 0.29241 | 0.999632 | 5761 | 3 | -0.52015 | 0.31677 | 0.58435 | 1 | 11594 | 3 | -0.52015 |
| hsa-mir-181a-1 | 4 | 0.81591 | 0.81591 | 0.999994 | 17710 | 1 | 1.2998 | 0.31691 | 0.5265 | 1 | 11595 | 2 | 1.2998 |

| | | | | | | | | | | | | |
|---|---|---|---|---|---|---|---|---|---|---|---|---|
| CACFD1 | 6 | 0.146 | 0.32062 | 0.999994 | 6351 | 1 | 1.1035 | 0.31702 | 0.58466 | 1 | 11596 | 4 | 1.1035 |
| UBE2J2 | 5 | 0.067547 | 0.17442 | 0.968171 | 3739 | 3 | -1.5833 | 0.31704 | 0.54358 | 1 | 11597 | 2 | -1.5833 |
| NUP107 | 5 | 0.24166 | 0.43316 | 0.999994 | 8822 | 3 | -0.61156 | 0.31704 | 0.54358 | 1 | 11598 | 2 | -0.61156 |
| CEL | 6 | 0.29966 | 0.50631 | 0.999994 | 10122 | 2 | 0.58104 | 0.31707 | 0.58472 | 1 | 11599 | 4 | 0.58104 |
| NARF | 6 | 0.057303 | 0.15993 | 0.967126 | 3304 | 2 | 0.70288 | 0.31709 | 0.58475 | 1 | 11600 | 4 | 0.70288 |
| NPTXR | 4 | 0.44523 | 0.59215 | 0.999994 | 12650 | 1 | 1.2597 | 0.31713 | 0.52666 | 1 | 11601 | 2 | 1.2597 |
| TMEM2 | 6 | 0.58366 | 0.7279 | 0.999994 | 14727 | 2 | 1.0089 | 0.31716 | 0.58484 | 1 | 11602 | 4 | 1.0089 |
| SLC29A1 | 6 | 0.041603 | 0.12551 | 0.951052 | 2595 | 2 | 0.99768 | 0.31718 | 0.58486 | 1 | 11603 | 4 | 0.99768 |
| LRRC14B | 6 | 0.015896 | 0.058706 | 0.931872 | 1239 | 4 | -2.6143 | 0.31721 | 0.5849 | 1 | 11604 | 2 | -2.6143 |
| WDR88 | 6 | 0.43641 | 0.62604 | 0.999994 | 12519 | 2 | 0.2928 | 0.31721 | 0.5849 | 1 | 11605 | 2 | 0.2928 |
| CSRNP3 | 6 | 0.62693 | 0.76003 | 0.999994 | 15334 | 2 | 0.79023 | 0.31721 | 0.5849 | 1 | 11606 | 3 | 0.79023 |
| ZNF713 | 6 | 0.0077535 | 0.031095 | 0.862223 | 711 | 4 | -4.1722 | 0.31721 | 0.5849 | 1 | 11607 | 1 | -4.1722 |
| SELE | 6 | 0.94666 | 0.94662 | 0.999994 | 20253 | 0 | 0.86766 | 0.31721 | 0.5849 | 1 | 11608 | 3 | 0.86766 |
| hsa-mir-3194 | 4 | 0.82531 | 0.82491 | 0.999994 | 17855 | 1 | 1.5528 | 0.31722 | 0.52673 | 1 | 11609 | 2 | 1.5528 |
| TEX14 | 6 | 0.064068 | 0.17437 | 0.968171 | 3616 | 2 | 1.2635 | 0.31727 | 0.58497 | 1 | 11610 | 3 | 1.2635 |
| MPPE1 | 4 | 0.077143 | 0.17342 | 0.96793 | 4111 | 3 | -1.9139 | 0.31734 | 0.52683 | 1 | 11611 | 1 | -1.9139 |
| API5 | 6 | 0.80021 | 0.84796 | 0.999994 | 17481 | 1 | 0.71363 | 0.31747 | 0.5852 | 1 | 11612 | 4 | 0.71363 |
| MYH15 | 5 | 0.817 | 0.83058 | 0.999994 | 17728 | 1 | 1.2237 | 0.31754 | 0.54412 | 1 | 11613 | 3 | 1.2237 |
| TMEM128 | 4 | 0.60132 | 0.66607 | 0.999994 | 14965 | 1 | 1.3539 | 0.31759 | 0.52701 | 1 | 11614 | 2 | 1.3539 |
| CCDC104 | 5 | 0.53198 | 0.67579 | 0.999994 | 13937 | 2 | 0.049158 | 0.3176 | 0.54417 | 1 | 11615 | 1 | 0.049158 |
| SULT2B1 | 6 | 0.20447 | 0.39333 | 0.999994 | 7890 | 3 | -0.49615 | 0.3176 | 0.58537 | 1 | 11616 | 2 | -0.49615 |
| MAGEA11 | 6 | 0.28944 | 0.49443 | 0.999994 | 9899 | 2 | 0.8998 | 0.31762 | 0.58539 | 1 | 11617 | 4 | 0.8998 |
| TCN2 | 6 | 0.17884 | 0.36168 | 0.999994 | 7194 | 2 | 0.20026 | 0.31797 | 0.58585 | 1 | 11618 | 2 | 0.20026 |
| SPIC | 6 | 0.1456 | 0.32013 | 0.999994 | 6339 | 3 | -0.91303 | 0.31797 | 0.58585 | 1 | 11619 | 3 | -0.91303 |
| PUS10 | 5 | 0.24186 | 0.43343 | 0.999994 | 8830 | 3 | -0.59507 | 0.31804 | 0.54461 | 1 | 11620 | 2 | -0.59507 |
| RNASE1 | 5 | 0.2745 | 0.46041 | 0.999994 | 9581 | 1 | 0.16055 | 0.31804 | 0.54461 | 1 | 11621 | 2 | 0.16055 |
| CACYBP | 3 | 0.24752 | 0.37184 | 0.999994 | 8962 | 1 | 1.4881 | 0.31809 | 0.46882 | 1 | 11622 | 2 | 1.4881 |
| GADD45G | 5 | 0.70507 | 0.76881 | 0.999994 | 16361 | 1 | 1.4842 | 0.31809 | 0.54467 | 1 | 11623 | 3 | 1.4842 |
| EIF4A3 | 3 | 0.25544 | 0.38177 | 0.999994 | 9137 | 2 | -0.82244 | 0.31809 | 0.46883 | 1 | 11624 | 1 | -0.82244 |
| TMEM161A | 5 | 0.088325 | 0.20789 | 0.979108 | 4526 | 2 | 1.0798 | 0.31813 | 0.5447 | 1 | 11625 | 3 | 1.0798 |
| HECW2 | 5 | 0.048612 | 0.13707 | 0.957161 | 2916 | 3 | -2.3093 | 0.31821 | 0.54478 | 1 | 11626 | 2 | -2.3093 |
| DUSP13 | 6 | 0.30142 | 0.50837 | 0.999994 | 10152 | 3 | -0.52688 | 0.31826 | 0.5862 | 1 | 11627 | 2 | -0.52688 |
| IBA57 | 6 | 0.19701 | 0.38426 | 0.999994 | 7673 | 3 | -0.95846 | 0.31826 | 0.5862 | 1 | 11628 | 2 | -0.95846 |
| CENPF | 6 | 0.081623 | 0.21064 | 0.979587 | 4286 | 3 | -0.59339 | 0.31826 | 0.5862 | 1 | 11629 | 2 | -0.59339 |
| NIPAL4 | 6 | 0.061158 | 0.16824 | 0.967126 | 3476 | 2 | 0.87485 | 0.31838 | 0.58633 | 1 | 11630 | 4 | 0.87485 |
| ZNF430 | 6 | 0.43597 | 0.62577 | 0.999994 | 12516 | 2 | 0.83053 | 0.31848 | 0.58645 | 1 | 11631 | 3 | 0.83053 |
| FOXP4 | 6 | 0.1866 | 0.37129 | 0.999994 | 7376 | 3 | 0.36171 | 0.31858 | 0.58657 | 1 | 11632 | 3 | 0.36171 |
| RAB36 | 6 | 0.029031 | 0.09651 | 0.945031 | 1957 | 3 | -0.37028 | 0.31859 | 0.58659 | 1 | 11633 | 3 | -0.37028 |
| SALL4 | 6 | 0.15006 | 0.32569 | 0.999994 | 6460 | 2 | -0.22444 | 0.31859 | 0.58659 | 1 | 11634 | 2 | -0.22444 |
| CD27 | 6 | 0.74237 | 0.82579 | 0.999994 | 16768 | 1 | 1.0745 | 0.3187 | 0.58671 | 1 | 11635 | 3 | 1.0745 |
| hsa-mir-6502 | 3 | 0.71314 | 0.71339 | 0.999994 | 16431 | 1 | 1.381 | 0.31886 | 0.46967 | 1 | 11636 | 2 | 1.381 |
| PECR | 6 | 0.25408 | 0.45279 | 0.999994 | 9105 | 2 | 0.83495 | 0.3189 | 0.58695 | 1 | 11637 | 3 | 0.83495 |
| SLC23A2 | 5 | 0.52181 | 0.66687 | 0.999994 | 13777 | 2 | 0.33517 | 0.31892 | 0.54548 | 1 | 11638 | 1 | 0.33517 |
| hsa-mir-15b | 4 | 0.12845 | 0.26496 | 0.997344 | 5895 | 2 | 0.24983 | 0.31892 | 0.52798 | 1 | 11639 | 2 | 0.24983 |
| CIITA | 6 | 0.54354 | 0.69904 | 0.999994 | 14122 | 2 | 0.90254 | 0.31893 | 0.58698 | 1 | 11640 | 4 | 0.90254 |
| hsa-mir-6133 | 4 | 0.27678 | 0.41929 | 0.999994 | 9633 | 2 | 0.65123 | 0.31908 | 0.5281 | 1 | 11641 | 2 | 0.65123 |
| RNF212 | 6 | 0.15575 | 0.33288 | 0.999994 | 6615 | 3 | 0.53575 | 0.31909 | 0.58718 | 1 | 11642 | 3 | 0.53575 |
| SLC17A4 | 6 | 0.047486 | 0.13855 | 0.957336 | 2879 | 3 | 0.092428 | 0.31913 | 0.58723 | 1 | 11643 | 3 | 0.092428 |
| MATN2 | 6 | 0.25898 | 0.45867 | 0.999994 | 9214 | 1 | 0.71406 | 0.31913 | 0.58723 | 1 | 11644 | 4 | 0.71406 |
| EPHX1 | 6 | 0.27472 | 0.47726 | 0.999994 | 9588 | 3 | -0.64452 | 0.31915 | 0.58726 | 1 | 11645 | 2 | -0.64452 |
| DCAF4L1 | 6 | 0.23063 | 0.42481 | 0.999994 | 8536 | 3 | -0.72875 | 0.31915 | 0.58726 | 1 | 11646 | 2 | -0.72875 |
| TMEM179 | 4 | 0.34962 | 0.49387 | 0.999994 | 11143 | 2 | -0.069968 | 0.31932 | 0.52827 | 1 | 11647 | 2 | -0.069968 |

| | | | | | | | | | | | | | |
|---|---|---|---|---|---|---|---|---|---|---|---|---|---|
| CXorf61 | 4 | 0.3385 | 0.48245 | 0.999994 | 10908 | 2 | -0.75351 | 0.31932 | 0.52827 | 1 | 11648 | 1 | -0.75351 |
| hsa-mir-550a-3 | 3 | 0.62479 | 0.63651 | 0.999994 | 15299 | 1 | 1.4838 | 0.31935 | 0.47018 | 1 | 11649 | 2 | 1.4838 |
| ZFYVE26 | 5 | 0.41679 | 0.57745 | 0.999994 | 12202 | 1 | 1.4272 | 0.31941 | 0.54599 | 1 | 11650 | 3 | 1.4272 |
| hsa-mir-3656 | 4 | 0.055139 | 0.1315 | 0.95318 | 3205 | 2 | -0.85673 | 0.31943 | 0.52835 | 1 | 11651 | 2 | -0.85673 |
| SCXA | 6 | 0.72519 | 0.82003 | 0.999994 | 16551 | 1 | 0.88787 | 0.31946 | 0.58765 | 1 | 11652 | 4 | 0.88787 |
| MOSPD1 | 5 | 0.16938 | 0.33122 | 0.999994 | 6968 | 3 | -0.95428 | 0.31954 | 0.54613 | 1 | 11653 | 1 | -0.95428 |
| TEKT4 | 5 | 0.46833 | 0.6207 | 0.999994 | 12979 | 1 | 1.2126 | 0.31954 | 0.54613 | 1 | 11654 | 3 | 1.2126 |
| PSMB3 | 2 | 0.23149 | 0.28505 | 0.999632 | 8556 | 1 | 0.90204 | 0.31954 | 0.41668 | 1 | 11655 | 1 | 0.90204 |
| TRIM56 | 5 | 0.26367 | 0.45171 | 0.999994 | 9329 | 2 | 1.1802 | 0.31956 | 0.54615 | 1 | 11656 | 3 | 1.1802 |
| ELAVL1 | 6 | 0.001876 | 0.0093521 | 0.680529 | 257 | 3 | -3.886 | 0.31957 | 0.58778 | 1 | 11657 | 2 | -3.886 |
| KIAA1432 | 6 | 0.031623 | 0.10262 | 0.946414 | 2087 | 3 | 0.5516 | 0.31958 | 0.58779 | 1 | 11658 | 3 | 0.5516 |
| PRMT2 | 5 | 0.56311 | 0.7034 | 0.999994 | 14433 | 2 | 1.4439 | 0.31959 | 0.54619 | 1 | 11659 | 3 | 1.4439 |
| LGALS4 | 6 | 0.9146 | 0.91553 | 0.999994 | 19502 | 1 | 1.0356 | 0.31963 | 0.58786 | 1 | 11660 | 4 | 1.0356 |
| TBC1D1 | 6 | 0.034457 | 0.10922 | 0.951052 | 2237 | 3 | -1.1485 | 0.31982 | 0.58801 | 1 | 11661 | 3 | -1.1485 |
| PRELID2 | 6 | 0.16031 | 0.33862 | 0.999994 | 6748 | 3 | -0.4386 | 0.31982 | 0.58801 | 1 | 11662 | 1 | -0.4386 |
| VTI1A | 6 | 0.64764 | 0.77582 | 0.999994 | 15625 | 2 | -0.19614 | 0.31982 | 0.58801 | 1 | 11663 | 2 | -0.19614 |
| NonTargetingControlG | 1 | 0.6801 | 0.68008 | 0.999994 | 16092 | 0 | 2.2348 | 0.3199 | 0.32001 | 0.989846 | 11664 | 1 | 2.2348 |
| AGXT2L2 | 6 | 0.20713 | 0.39657 | 0.999994 | 7949 | 3 | 0.30461 | 0.31991 | 0.58809 | 1 | 11665 | 3 | 0.30461 |
| TSPAN7 | 4 | 0.68743 | 0.71631 | 0.999994 | 16188 | 1 | 0.27984 | 0.32 | 0.52877 | 1 | 11666 | 2 | 0.27984 |
| MAP10 | 5 | 0.0094865 | 0.034598 | 0.867348 | 832 | 4 | -2.0016 | 0.3201 | 0.54669 | 1 | 11667 | 1 | -2.0016 |
| ADORA1 | 5 | 0.89448 | 0.89461 | 0.999994 | 19113 | 0 | 0.39271 | 0.3203 | 0.5469 | 1 | 11668 | 2 | 0.39271 |
| ZNF260 | 5 | 0.19082 | 0.36213 | 0.999994 | 7510 | 2 | 0.29142 | 0.32044 | 0.54705 | 1 | 11669 | 2 | 0.29142 |
| TMPRSS5 | 5 | 0.88666 | 0.88675 | 0.999994 | 18961 | 1 | 1.2249 | 0.32046 | 0.54707 | 1 | 11670 | 3 | 1.2249 |
| YIPF7 | 6 | 0.014718 | 0.054826 | 0.922677 | 1173 | 4 | -3.2395 | 0.32054 | 0.58861 | 1 | 11671 | 2 | -3.2395 |
| MAP6D1 | 6 | 0.35199 | 0.56621 | 0.999994 | 11194 | 3 | 0.26931 | 0.32054 | 0.58861 | 1 | 11672 | 3 | 0.26931 |
| CMC2 | 5 | 0.75479 | 0.79338 | 0.999994 | 16908 | 1 | 0.23307 | 0.32056 | 0.54716 | 1 | 11673 | 2 | 0.23307 |
| LGALS9B | 5 | 0.094078 | 0.21706 | 0.980699 | 4752 | 2 | 0.87759 | 0.32066 | 0.54727 | 1 | 11674 | 3 | 0.87759 |
| AMOTL2 | 4 | 0.7413 | 0.75347 | 0.999994 | 16756 | 1 | 0.39368 | 0.32068 | 0.52926 | 1 | 11675 | 2 | 0.39368 |
| DKK2 | 5 | 0.042876 | 0.12301 | 0.951052 | 2672 | 4 | -2.0917 | 0.32072 | 0.54732 | 1 | 11676 | 1 | -2.0917 |
| LOC100288524 | 5 | 0.30331 | 0.48374 | 0.999994 | 10202 | 2 | 0.8438 | 0.32072 | 0.54732 | 1 | 11677 | 3 | 0.8438 |
| PTGES3L-AARSD1 | 2 | 0.66089 | 0.66069 | 0.999994 | 15793 | 0 | 2.7284 | 0.32088 | 0.41747 | 1 | 11678 | 1 | 2.7284 |
| TCF24 | 5 | 0.131 | 0.27414 | 0.998561 | 5960 | 2 | 1.4524 | 0.32106 | 0.54767 | 1 | 11679 | 3 | 1.4524 |
| SRCAP | 5 | 0.52841 | 0.67265 | 0.999994 | 13868 | 1 | 1.365 | 0.32113 | 0.54774 | 1 | 11680 | 3 | 1.365 |
| KLC3 | 5 | 0.85091 | 0.85501 | 0.999994 | 18277 | 1 | 1.4359 | 0.32126 | 0.54789 | 1 | 11681 | 3 | 1.4359 |
| MRPS15 | 6 | 0.051479 | 0.1473 | 0.965554 | 3045 | 4 | -1.7104 | 0.32128 | 0.5892 | 1 | 11682 | 2 | -1.7104 |
| HHIPL1 | 5 | 0.42458 | 0.5839 | 0.999994 | 12328 | 2 | -0.52809 | 0.32129 | 0.54791 | 1 | 11683 | 2 | -0.52809 |
| NPAT | 5 | 0.01753 | 0.057333 | 0.927806 | 1342 | 3 | -3.6804 | 0.32129 | 0.54791 | 1 | 11684 | 2 | -3.6804 |
| WIPF2 | 5 | 0.85164 | 0.85559 | 0.999994 | 18289 | 1 | 0.54942 | 0.32129 | 0.54791 | 1 | 11685 | 2 | 0.54942 |
| SLC25A39 | 6 | 0.78318 | 0.84097 | 0.999994 | 17255 | 1 | 0.19889 | 0.3213 | 0.58921 | 1 | 11686 | 3 | 0.19889 |
| hsa-mir-6756 | 4 | 0.029963 | 0.081411 | 0.93914 | 2004 | 3 | -2.5775 | 0.32133 | 0.52974 | 1 | 11687 | 1 | -2.5775 |
| C1orf229 | 6 | 0.30533 | 0.51294 | 0.999994 | 10241 | 3 | 0.63296 | 0.32142 | 0.58931 | 1 | 11688 | 3 | 0.63296 |
| ATP1A4 | 3 | 0.54366 | 0.57585 | 0.999994 | 14125 | 1 | 1.4606 | 0.32146 | 0.47238 | 1 | 11689 | 2 | 1.4606 |
| CCNG2 | 6 | 0.22357 | 0.41643 | 0.999994 | 8367 | 3 | 0.6454 | 0.32155 | 0.58941 | 1 | 11690 | 3 | 0.6454 |
| ZNF263 | 5 | 0.24735 | 0.43868 | 0.999994 | 8957 | 1 | 1.4288 | 0.32164 | 0.54826 | 1 | 11691 | 3 | 1.4288 |
| ALG1 | 5 | 0.52859 | 0.67281 | 0.999994 | 13873 | 2 | 0.32908 | 0.32193 | 0.54859 | 1 | 11692 | 2 | 0.32908 |
| CCDC38 | 5 | 0.15428 | 0.30918 | 0.999994 | 6574 | 2 | -0.06568 | 0.32193 | 0.54859 | 1 | 11693 | 2 | -0.06568 |
| MPC2 | 5 | 0.17673 | 0.34189 | 0.999994 | 7147 | 1 | -0.18513 | 0.32193 | 0.54859 | 1 | 11694 | 2 | -0.18513 |
| CBFB | 5 | 0.29624 | 0.47801 | 0.999994 | 10044 | 2 | 0.085808 | 0.32195 | 0.54861 | 1 | 11695 | 2 | 0.085808 |
| CHRM2 | 6 | 0.097054 | 0.2416 | 0.987253 | 4862 | 2 | -0.17636 | 0.32202 | 0.58979 | 1 | 11696 | 2 | -0.17636 |
| AGPAT2 | 6 | 0.42725 | 0.6201 | 0.999994 | 12371 | 2 | 0.91212 | 0.32202 | 0.58979 | 1 | 11697 | 3 | 0.91212 |
| MLXIPL | 6 | 0.066315 | 0.17904 | 0.968171 | 3696 | 2 | -0.24812 | 0.32204 | 0.58981 | 1 | 11698 | 2 | -0.24812 |
| TBK1 | 6 | 0.091368 | 0.23028 | 0.983933 | 4651 | 4 | -1.5017 | 0.32204 | 0.58981 | 1 | 11699 | 2 | -1.5017 |

| | | | | | | | | | | | | | |
|---|---|---|---|---|---|---|---|---|---|---|---|---|---|
| OSCP1 | 6 | 0.064577 | 0.17538 | 0.968171 | 3633 | 3 | -0.43354 | 0.32204 | 0.58981 | 1 | 11700 | 3 | -0.43354 |
| AKT3 | 6 | 0.23644 | 0.43175 | 0.999994 | 8680 | 3 | 0.52708 | 0.32204 | 0.58981 | 1 | 11701 | 3 | 0.52708 |
| PALM | 6 | 0.66192 | 0.78678 | 0.999994 | 15809 | 2 | 0.65867 | 0.32204 | 0.58981 | 1 | 11702 | 3 | 0.65867 |
| C3orf22 | 6 | 0.93837 | 0.93833 | 0.999994 | 20034 | 0 | 1.0375 | 0.32211 | 0.58987 | 1 | 11703 | 3 | 1.0375 |
| UNCX | 6 | 0.087168 | 0.22181 | 0.983139 | 4483 | 3 | 0.74056 | 0.32217 | 0.58992 | 1 | 11704 | 3 | 0.74056 |
| ITSN2 | 5 | 0.069476 | 0.17752 | 0.968171 | 3813 | 1 | 1.2175 | 0.32219 | 0.54885 | 1 | 11705 | 3 | 1.2175 |
| DUSP2 | 6 | 0.096383 | 0.24024 | 0.987253 | 4831 | 2 | 0.76619 | 0.32237 | 0.59008 | 1 | 11706 | 3 | 0.76619 |
| PDZD8 | 6 | 0.2969 | 0.50309 | 0.999994 | 10064 | 3 | 0.58853 | 0.32252 | 0.59019 | 1 | 11707 | 3 | 0.58853 |
| SRPK1 | 6 | 0.070999 | 0.18888 | 0.969543 | 3875 | 3 | 0.60158 | 0.32254 | 0.59021 | 1 | 11708 | 3 | 0.60158 |
| IRGQ | 4 | 0.68712 | 0.7161 | 0.999994 | 16184 | 1 | 1.6455 | 0.32258 | 0.53068 | 1 | 11709 | 2 | 1.6455 |
| AKT1 | 6 | 0.21598 | 0.40732 | 0.999994 | 8170 | 3 | -0.83959 | 0.32259 | 0.59026 | 1 | 11710 | 1 | -0.83959 |
| CRLF1 | 6 | 0.0010461 | 0.0053154 | 0.60466 | 174 | 2 | 0.74412 | 0.32259 | 0.59026 | 1 | 11711 | 3 | 0.74412 |
| MAGI2 | 6 | 0.59351 | 0.73518 | 0.999994 | 14873 | 2 | 1.2711 | 0.32267 | 0.59032 | 1 | 11712 | 3 | 1.2711 |
| MRPL22 | 5 | 0.43893 | 0.59594 | 0.999994 | 12556 | 2 | -0.37697 | 0.32272 | 0.54941 | 1 | 11713 | 2 | -0.37697 |
| C3orf35 | 5 | 0.44211 | 0.59858 | 0.999994 | 12613 | 2 | -0.33473 | 0.32272 | 0.54941 | 1 | 11714 | 2 | -0.33473 |
| SLC46A2 | 5 | 0.12794 | 0.26946 | 0.998561 | 5874 | 2 | -0.24671 | 0.32272 | 0.54941 | 1 | 11715 | 2 | -0.24671 |
| KIAA0195 | 5 | 0.065948 | 0.17176 | 0.967658 | 3686 | 3 | -2.1293 | 0.32272 | 0.54941 | 1 | 11716 | 2 | -2.1293 |
| TANGO6 | 3 | 0.17181 | 0.27668 | 0.999379 | 7032 | 1 | 1.4753 | 0.32277 | 0.47377 | 1 | 11717 | 2 | 1.4753 |
| SEPP1 | 6 | 0.012356 | 0.046908 | 0.896157 | 1017 | 2 | 0.70598 | 0.32288 | 0.59046 | 1 | 11718 | 3 | 0.70598 |
| TONSL | 2 | 0.48176 | 0.48736 | 0.999994 | 13165 | 1 | 2.107 | 0.32298 | 0.41882 | 1 | 11719 | 1 | 2.107 |
| C1orf198 | 5 | 0.24877 | 0.43982 | 0.999994 | 8988 | 2 | 1.4099 | 0.32299 | 0.54971 | 1 | 11720 | 3 | 1.4099 |
| hsa-mir-103b-1 | 1 | 0.67698 | 0.67699 | 0.999994 | 16039 | 0 | 2.1352 | 0.32302 | 0.32313 | 0.989881 | 11721 | 1 | 2.1352 |
| ZSCAN29 | 6 | 0.2916 | 0.4969 | 0.999994 | 9941 | 2 | 0.72943 | 0.32306 | 0.5906 | 1 | 11722 | 3 | 0.72943 |
| CCDC116 | 4 | 0.041028 | 0.10382 | 0.947284 | 2562 | 2 | -1.8437 | 0.3232 | 0.53112 | 1 | 11723 | 1 | -1.8437 |
| hsa-mir-6868 | 4 | 0.25864 | 0.40074 | 0.999994 | 9207 | 1 | 1.0077 | 0.3232 | 0.53112 | 1 | 11724 | 2 | 1.0077 |
| hsa-mir-4496 | 4 | 0.77529 | 0.78007 | 0.999994 | 17142 | 1 | 0.68226 | 0.3232 | 0.53112 | 1 | 11725 | 2 | 0.68226 |
| SLC36A4 | 5 | 0.10473 | 0.23376 | 0.984971 | 5124 | 3 | -1.4816 | 0.3234 | 0.55012 | 1 | 11726 | 2 | -1.4816 |
| CRNN | 5 | 0.79891 | 0.81886 | 0.999994 | 17463 | 1 | -0.14293 | 0.3234 | 0.55012 | 1 | 11727 | 2 | -0.14293 |
| SH2D1B | 5 | 0.051687 | 0.14448 | 0.962688 | 3054 | 3 | -2.1007 | 0.3234 | 0.55012 | 1 | 11728 | 1 | -2.1007 |
| METTL22 | 6 | 0.7779 | 0.83885 | 0.999994 | 17183 | 1 | 0.57823 | 0.32341 | 0.59088 | 1 | 11729 | 3 | 0.57823 |
| CAPRIN2 | 6 | 0.18592 | 0.3705 | 0.999994 | 7363 | 3 | -0.85449 | 0.32341 | 0.59088 | 1 | 11730 | 1 | -0.85449 |
| CLEC17A | 6 | 0.54905 | 0.70296 | 0.999994 | 14192 | 1 | 1.0343 | 0.32341 | 0.59088 | 1 | 11731 | 3 | 1.0343 |
| CEND1 | 4 | 0.42441 | 0.57071 | 0.999994 | 12325 | 2 | 1.0017 | 0.32355 | 0.5314 | 1 | 11732 | 2 | 1.0017 |
| hsa-mir-4510 | 3 | 0.34311 | 0.45188 | 0.999994 | 11007 | 1 | 0.79689 | 0.32356 | 0.47465 | 1 | 11733 | 2 | 0.79689 |
| MAGED4B | 3 | 0.67608 | 0.67922 | 0.999994 | 16021 | 1 | 1.5679 | 0.32357 | 0.47466 | 1 | 11734 | 2 | 1.5679 |
| SULT1C4 | 5 | 0.67068 | 0.75359 | 0.999994 | 15940 | 1 | 1.3546 | 0.32365 | 0.55039 | 1 | 11735 | 3 | 1.3546 |
| LRRC8E | 6 | 0.30456 | 0.51205 | 0.999994 | 10218 | 3 | -0.48139 | 0.3238 | 0.59117 | 1 | 11736 | 2 | -0.48139 |
| BCAR3 | 5 | 0.18666 | 0.3562 | 0.999994 | 7377 | 2 | 1.1538 | 0.3239 | 0.55065 | 1 | 11737 | 3 | 1.1538 |
| KRTAP1-3 | 5 | 0.96251 | 0.96266 | 0.999994 | 20643 | 0 | 1.2092 | 0.32395 | 0.5507 | 1 | 11738 | 3 | 1.2092 |
| RNF181 | 5 | 0.5631 | 0.7034 | 0.999994 | 14432 | 2 | 0.16059 | 0.32403 | 0.5508 | 1 | 11739 | 2 | 0.16059 |
| CCL24 | 5 | 0.38717 | 0.55275 | 0.999994 | 11754 | 1 | 0.64535 | 0.32407 | 0.55084 | 1 | 11740 | 3 | 0.64535 |
| KARS | 5 | 0.44754 | 0.60316 | 0.999994 | 12681 | 2 | -0.25914 | 0.32407 | 0.55084 | 1 | 11741 | 2 | -0.25914 |
| IP6K1 | 5 | 0.29192 | 0.47449 | 0.999994 | 9946 | 2 | -0.52953 | 0.32407 | 0.55084 | 1 | 11742 | 2 | -0.52953 |
| RPS6KL1 | 6 | 0.24742 | 0.44487 | 0.999994 | 8959 | 1 | 1.0408 | 0.32412 | 0.59143 | 1 | 11743 | 3 | 1.0408 |
| CADM1 | 6 | 0.8916 | 0.89744 | 0.999994 | 19064 | 1 | 0.89679 | 0.32412 | 0.59143 | 1 | 11744 | 3 | 0.89679 |
| PPFIA2 | 6 | 0.28893 | 0.49383 | 0.999994 | 9893 | 2 | -0.30184 | 0.32412 | 0.59143 | 1 | 11745 | 2 | -0.30184 |
| APOL6 | 6 | 0.55047 | 0.70397 | 0.999994 | 14220 | 2 | 0.83947 | 0.32415 | 0.59145 | 1 | 11746 | 3 | 0.83947 |
| SLC34A2 | 5 | 0.90292 | 0.90302 | 0.999994 | 19274 | 0 | 1.2488 | 0.32418 | 0.55094 | 1 | 11747 | 3 | 1.2488 |
| hsa-mir-99b | 4 | 0.87973 | 0.87937 | 0.999994 | 18829 | 0 | 1.4897 | 0.32418 | 0.53188 | 1 | 11748 | 2 | 1.4897 |
| DNAJC2 | 6 | 0.068823 | 0.1843 | 0.968526 | 3790 | 2 | 0.98735 | 0.32433 | 0.59159 | 1 | 11749 | 3 | 0.98735 |
| GPNMB | 5 | 0.50386 | 0.65128 | 0.999994 | 13492 | 1 | 1.259 | 0.32445 | 0.55123 | 1 | 11750 | 3 | 1.259 |
| MRPL42 | 5 | 0.71951 | 0.77561 | 0.999994 | 16491 | 1 | 1.11 | 0.3245 | 0.55128 | 1 | 11751 | 3 | 1.11 |

| | | | | | | | | | | | | | |
|---|---|---|---|---|---|---|---|---|---|---|---|---|---|
| TRAF3IP3 | 5 | 0.14138 | 0.28989 | 0.999632 | 6226 | 2 | 1.2059 | 0.32455 | 0.55132 | 1 | 11752 | 3 | 1.2059 |
| RAVER2 | 6 | 0.30356 | 0.51086 | 0.999994 | 10204 | 2 | 0.91997 | 0.32465 | 0.59184 | 1 | 11753 | 3 | 0.91997 |
| NonTargetingControlG | 2 | 0.63933 | 0.63916 | 0.999994 | 15506 | 0 | 2.6322 | 0.32482 | 0.41996 | 1 | 11754 | 1 | 2.6322 |
| SLC25A44 | 6 | 0.12815 | 0.29792 | 0.999994 | 5883 | 2 | 0.089076 | 0.32487 | 0.59202 | 1 | 11755 | 2 | 0.089076 |
| MYO9B | 6 | 0.31744 | 0.52694 | 0.999994 | 10462 | 3 | 0.1836 | 0.32487 | 0.59202 | 1 | 11756 | 3 | 0.1836 |
| MANEA | 6 | 0.67179 | 0.79453 | 0.999994 | 15952 | 1 | 0.6645 | 0.32487 | 0.59202 | 1 | 11757 | 3 | 0.6645 |
| SPOCD1 | 6 | 0.58019 | 0.72535 | 0.999994 | 14680 | 2 | 0.077829 | 0.32487 | 0.59202 | 1 | 11758 | 2 | 0.077829 |
| TXNL1 | 5 | 0.16103 | 0.31905 | 0.999994 | 6764 | 3 | -1.0346 | 0.32489 | 0.55167 | 1 | 11759 | 2 | -1.0346 |
| ZNF682 | 6 | 0.6769 | 0.79855 | 0.999994 | 16037 | 2 | 0.76666 | 0.32501 | 0.59212 | 1 | 11760 | 3 | 0.76666 |
| ACSM2A | 5 | 0.18159 | 0.34894 | 0.999994 | 7264 | 3 | -0.95788 | 0.32507 | 0.55185 | 1 | 11761 | 1 | -0.95788 |
| NNT | 5 | 0.20828 | 0.38698 | 0.999994 | 7985 | 3 | -0.80654 | 0.32507 | 0.55185 | 1 | 11762 | 2 | -0.80654 |
| MRPL24 | 5 | 0.33817 | 0.51224 | 0.999994 | 10902 | 2 | 1.1917 | 0.32507 | 0.55185 | 1 | 11763 | 3 | 1.1917 |
| GPAT2 | 5 | 0.59189 | 0.72322 | 0.999994 | 14845 | 1 | 1.2496 | 0.32518 | 0.55196 | 1 | 11764 | 3 | 1.2496 |
| INPP5A | 6 | 0.22524 | 0.4184 | 0.999994 | 8404 | 2 | 0.59204 | 0.32533 | 0.59237 | 1 | 11765 | 3 | 0.59204 |
| CARD10 | 6 | 0.12198 | 0.28949 | 0.999632 | 5701 | 3 | -0.52658 | 0.32533 | 0.59237 | 1 | 11766 | 2 | -0.52658 |
| ATP6V1H | 6 | 0.11749 | 0.281 | 0.999632 | 5560 | 3 | -0.1604 | 0.32533 | 0.59237 | 1 | 11767 | 2 | -0.1604 |
| NonTargetingControlG | 1 | 0.67449 | 0.67453 | 0.999994 | 15996 | 0 | 2.1381 | 0.32551 | 0.32569 | 0.991543 | 11768 | 1 | 2.1381 |
| GYG2 | 4 | 0.76665 | 0.77308 | 0.999994 | 17043 | 1 | 1.3787 | 0.32556 | 0.5329 | 1 | 11769 | 2 | 1.3787 |
| TMEFF2 | 6 | 0.36313 | 0.57882 | 0.999994 | 11391 | 3 | -0.51209 | 0.32563 | 0.59261 | 1 | 11770 | 2 | -0.51209 |
| hsa-mir-6813 | 2 | 0.12122 | 0.20256 | 0.977973 | 5671 | 1 | -1.0905 | 0.32575 | 0.42052 | 1 | 11771 | 1 | -1.0905 |
| FGF21 | 5 | 0.33211 | 0.50725 | 0.999994 | 10768 | 1 | 0.59515 | 0.3259 | 0.55273 | 1 | 11772 | 3 | 0.59515 |
| hsa-mir-5579 | 3 | 0.78491 | 0.78478 | 0.999994 | 17277 | 0 | 0.1583 | 0.32591 | 0.47708 | 1 | 11773 | 1 | 0.1583 |
| ACSL3 | 5 | 0.033439 | 0.099325 | 0.945253 | 2185 | 1 | 1.2243 | 0.32601 | 0.55285 | 1 | 11774 | 3 | 1.2243 |
| TMEM81 | 5 | 0.10529 | 0.23463 | 0.984971 | 5144 | 3 | -1.4374 | 0.32612 | 0.55296 | 1 | 11775 | 2 | -1.4374 |
| SPEF1 | 6 | 0.50693 | 0.67355 | 0.999994 | 13536 | 1 | 1.1437 | 0.32618 | 0.59304 | 1 | 11776 | 3 | 1.1437 |
| PMEL | 5 | 0.43733 | 0.59456 | 0.999994 | 12531 | 2 | 1.324 | 0.32628 | 0.55312 | 1 | 11777 | 3 | 1.324 |
| ATP13A4 | 6 | 0.52718 | 0.68756 | 0.999994 | 13847 | 2 | 0.70509 | 0.3263 | 0.59313 | 1 | 11778 | 3 | 0.70509 |
| MTPAP | 4 | 0.062347 | 0.14529 | 0.963012 | 3528 | 3 | -1.7905 | 0.32645 | 0.53355 | 1 | 11779 | 1 | -1.7905 |
| hsa-mir-200c | 4 | 0.30589 | 0.44899 | 0.999994 | 10250 | 1 | 0.31105 | 0.32645 | 0.53355 | 1 | 11780 | 2 | 0.31105 |
| PSG3 | 4 | 0.64918 | 0.69281 | 0.999994 | 15645 | 1 | 0.76198 | 0.32645 | 0.53355 | 1 | 11781 | 2 | 0.76198 |
| PRSS50 | 4 | 0.42214 | 0.5684 | 0.999994 | 12283 | 2 | -0.41585 | 0.32645 | 0.53355 | 1 | 11782 | 1 | -0.41585 |
| RNF10 | 6 | 0.099499 | 0.2464 | 0.98999 | 4952 | 3 | 0.28482 | 0.3265 | 0.59328 | 1 | 11783 | 3 | 0.28482 |
| PTPRCAP | 3 | 0.080405 | 0.15702 | 0.967126 | 4244 | 2 | -2.424 | 0.32657 | 0.47778 | 1 | 11784 | 1 | -2.424 |
| LSS | 6 | 0.94307 | 0.94305 | 0.999994 | 20156 | 0 | 1.1144 | 0.32676 | 0.59351 | 1 | 11785 | 3 | 1.1144 |
| MYO15A | 5 | 0.90096 | 0.90108 | 0.999994 | 19232 | 0 | 0.91613 | 0.32677 | 0.55365 | 1 | 11786 | 3 | 0.91613 |
| PSORS1C1 | 5 | 0.049299 | 0.13867 | 0.957514 | 2943 | 2 | -0.29773 | 0.32677 | 0.55365 | 1 | 11787 | 2 | -0.29773 |
| DCAF7 | 6 | 0.0017574 | 0.008765 | 0.680529 | 246 | 4 | -4.2209 | 0.32683 | 0.59356 | 1 | 11788 | 2 | -4.2209 |
| GADL1 | 6 | 0.16299 | 0.34201 | 0.999994 | 6819 | 3 | -0.2177 | 0.32683 | 0.59356 | 1 | 11789 | 3 | -0.2177 |
| KIAA1524 | 6 | 0.60059 | 0.74038 | 0.999994 | 14953 | 2 | 0.066918 | 0.32683 | 0.59356 | 1 | 11790 | 2 | 0.066918 |
| CTSE | 5 | 0.93065 | 0.93093 | 0.999994 | 19867 | 0 | 1.17 | 0.32684 | 0.55371 | 1 | 11791 | 3 | 1.17 |
| UBE2V1 | 6 | 0.3505 | 0.56455 | 0.999994 | 11174 | 2 | 0.1867 | 0.32693 | 0.59364 | 1 | 11792 | 3 | 0.1867 |
| ATP11A | 6 | 0.50131 | 0.66959 | 0.999994 | 13443 | 2 | 0.66791 | 0.32693 | 0.59364 | 1 | 11793 | 3 | 0.66791 |
| IBSP | 5 | 0.3651 | 0.53433 | 0.999994 | 11422 | 2 | -0.17234 | 0.32693 | 0.55382 | 1 | 11794 | 2 | -0.17234 |
| BAG3 | 5 | 0.29333 | 0.47562 | 0.999994 | 9982 | 2 | 1.2732 | 0.32695 | 0.55384 | 1 | 11795 | 3 | 1.2732 |
| RNF223 | 5 | 0.59286 | 0.72356 | 0.999994 | 14859 | 1 | 1.2745 | 0.32697 | 0.55385 | 1 | 11796 | 3 | 1.2745 |
| ARRDC5 | 4 | 0.88393 | 0.88359 | 0.999994 | 18902 | 0 | 1.6372 | 0.327 | 0.53398 | 1 | 11797 | 2 | 1.6372 |
| TEX10 | 4 | 0.039378 | 0.10055 | 0.946414 | 2483 | 3 | -2.7204 | 0.3272 | 0.53411 | 1 | 11798 | 1 | -2.7204 |
| DCTN4 | 4 | 0.32816 | 0.47179 | 0.999994 | 10689 | 2 | -0.080481 | 0.3272 | 0.53411 | 1 | 11799 | 2 | -0.080481 |
| ANXA2R | 5 | 0.37755 | 0.5447 | 0.999994 | 11620 | 2 | 0.15189 | 0.32734 | 0.55422 | 1 | 11800 | 1 | 0.15189 |
| SIAH2 | 5 | 0.031645 | 0.094705 | 0.945031 | 2089 | 3 | -2.6131 | 0.32734 | 0.55422 | 1 | 11801 | 2 | -2.6131 |
| POMK | 5 | 0.72605 | 0.77877 | 0.999994 | 16558 | 1 | 0.31653 | 0.32734 | 0.55422 | 1 | 11802 | 1 | 0.31653 |
| ZNF800 | 6 | 0.3051 | 0.51266 | 0.999994 | 10233 | 3 | 0.53113 | 0.32748 | 0.59405 | 1 | 11803 | 3 | 0.53113 |

| | | | | | | | | | | | | | |
|---|---|---|---|---|---|---|---|---|---|---|---|---|---|
| ZNF23 | 6 | 0.42692 | 0.61988 | 0.999994 | 12363 | 1 | 0.89007 | 0.32757 | 0.59413 | 1 | 11804 | 3 | 0.89007 |
| EDEM3 | 6 | 0.80783 | 0.85129 | 0.999994 | 17595 | 1 | 0.54504 | 0.3276 | 0.59415 | 1 | 11805 | 3 | 0.54504 |
| SYN2 | 6 | 0.26711 | 0.46826 | 0.999994 | 9399 | 3 | -0.53001 | 0.3276 | 0.59415 | 1 | 11806 | 2 | -0.53001 |
| THAP1 | 6 | 0.0022824 | 0.011027 | 0.71022 | 303 | 4 | -5.0163 | 0.3276 | 0.59415 | 1 | 11807 | 2 | -5.0163 |
| GDF6 | 6 | 0.088767 | 0.22505 | 0.983354 | 4544 | 3 | -1.3179 | 0.3276 | 0.59415 | 1 | 11808 | 2 | -1.3179 |
| INO80C | 4 | 0.10828 | 0.2298 | 0.983354 | 5248 | 2 | 0.17783 | 0.32763 | 0.53442 | 1 | 11809 | 2 | 0.17783 |
| NonTargetingControlG | 1 | 0.67221 | 0.67226 | 0.999994 | 15958 | 0 | 2.2432 | 0.32779 | 0.32796 | 0.991729 | 11810 | 1 | 2.2432 |
| HHEX | 5 | 0.12843 | 0.27022 | 0.998561 | 5893 | 3 | -1.2914 | 0.32779 | 0.55467 | 1 | 11811 | 2 | -1.2914 |
| DRGX | 5 | 0.47773 | 0.62871 | 0.999994 | 13109 | 1 | 0.65053 | 0.32779 | 0.55467 | 1 | 11812 | 3 | 0.65053 |
| PKLR | 5 | 0.062261 | 0.16582 | 0.967126 | 3523 | 2 | -0.044274 | 0.32779 | 0.55467 | 1 | 11813 | 2 | -0.044274 |
| ATOX1 | 5 | 0.41921 | 0.57943 | 0.999994 | 12238 | 2 | 0.09764 | 0.32779 | 0.55467 | 1 | 11814 | 2 | 0.09764 |
| BCS1L | 5 | 0.19934 | 0.37427 | 0.999994 | 7745 | 2 | 0.35554 | 0.32779 | 0.55467 | 1 | 11815 | 2 | 0.35554 |
| MARCKSL1 | 5 | 0.13779 | 0.28447 | 0.999632 | 6128 | 3 | -1.2754 | 0.32779 | 0.55467 | 1 | 11816 | 2 | -1.2754 |
| BRD2 | 5 | 0.020954 | 0.066648 | 0.93914 | 1511 | 2 | -0.32278 | 0.32779 | 0.55467 | 1 | 11817 | 2 | -0.32278 |
| hsa-mir-6850 | 2 | 0.36562 | 0.38857 | 0.999994 | 11433 | 1 | 1.5773 | 0.32781 | 0.4218 | 1 | 11818 | 1 | 1.5773 |
| hsa-mir-8053 | 4 | 0.047436 | 0.11641 | 0.951052 | 2877 | 2 | -1.424 | 0.32782 | 0.53455 | 1 | 11819 | 2 | -1.424 |
| TBL2 | 5 | 0.21912 | 0.40206 | 0.999994 | 8267 | 3 | -0.75761 | 0.32789 | 0.55477 | 1 | 11820 | 1 | -0.75761 |
| hsa-mir-2278 | 3 | 0.34433 | 0.45257 | 0.999994 | 11035 | 1 | 1.5521 | 0.3281 | 0.47937 | 1 | 11821 | 2 | 1.5521 |
| BCAR1 | 5 | 0.54754 | 0.68952 | 0.999994 | 14173 | 1 | 1.3565 | 0.32812 | 0.55502 | 1 | 11822 | 3 | 1.3565 |
| DPF3 | 5 | 0.35622 | 0.52697 | 0.999994 | 11272 | 2 | 1.3377 | 0.32815 | 0.55506 | 1 | 11823 | 3 | 1.3377 |
| GFRA1 | 6 | 0.34034 | 0.55304 | 0.999994 | 10941 | 3 | -0.51482 | 0.32831 | 0.5947 | 1 | 11824 | 2 | -0.51482 |
| SIRT5 | 6 | 0.0209 | 0.074867 | 0.93914 | 1505 | 3 | -0.28781 | 0.32831 | 0.5947 | 1 | 11825 | 3 | -0.28781 |
| CACHD1 | 6 | 0.64519 | 0.77393 | 0.999994 | 15586 | 2 | 0.27103 | 0.32831 | 0.5947 | 1 | 11826 | 3 | 0.27103 |
| BTBD3 | 6 | 0.03595 | 0.11275 | 0.951052 | 2309 | 3 | -1.5329 | 0.32831 | 0.5947 | 1 | 11827 | 2 | -1.5329 |
| DDO | 5 | 0.35214 | 0.52365 | 0.999994 | 11197 | 2 | 1.2597 | 0.32838 | 0.55531 | 1 | 11828 | 3 | 1.2597 |
| DZIP1 | 4 | 0.18579 | 0.32605 | 0.999994 | 7361 | 2 | -0.0062246 | 0.32839 | 0.53499 | 1 | 11829 | 2 | -0.0062246 |
| EPHA1 | 5 | 0.58068 | 0.71891 | 0.999994 | 14687 | 1 | 0.40886 | 0.32846 | 0.55539 | 1 | 11830 | 2 | 0.40886 |
| OTOP2 | 5 | 0.84876 | 0.85332 | 0.999994 | 18230 | 1 | 0.8051 | 0.3285 | 0.55544 | 1 | 11831 | 3 | 0.8051 |
| GALNT12 | 5 | 0.89411 | 0.89423 | 0.999994 | 19102 | 0 | 0.086695 | 0.3285 | 0.55544 | 1 | 11832 | 2 | 0.086695 |
| SLC43A1 | 5 | 0.96271 | 0.96287 | 0.999994 | 20650 | 0 | 1.0766 | 0.3285 | 0.55544 | 1 | 11833 | 3 | 1.0766 |
| hsa-mir-6827 | 3 | 0.84598 | 0.846 | 0.999994 | 18185 | 0 | 1.4095 | 0.32856 | 0.47985 | 1 | 11834 | 2 | 1.4095 |
| THAP3 | 5 | 0.025753 | 0.079472 | 0.93914 | 1787 | 1 | 1.2812 | 0.32862 | 0.55556 | 1 | 11835 | 3 | 1.2812 |
| UBE2D2 | 5 | 0.022342 | 0.070403 | 0.93914 | 1604 | 2 | -0.30128 | 0.3287 | 0.55564 | 1 | 11836 | 2 | -0.30128 |
| C4orf46 | 6 | 0.21817 | 0.40993 | 0.999994 | 8232 | 2 | 0.15883 | 0.32876 | 0.59503 | 1 | 11837 | 2 | 0.15883 |
| WDR45B | 6 | 0.58693 | 0.73033 | 0.999994 | 14772 | 2 | 0.15854 | 0.32876 | 0.59503 | 1 | 11838 | 2 | 0.15854 |
| TOM1L1 | 5 | 0.16806 | 0.32927 | 0.999994 | 6939 | 2 | 1.3289 | 0.32877 | 0.55572 | 1 | 11839 | 3 | 1.3289 |
| hsa-mir-5584 | 3 | 0.70668 | 0.70729 | 0.999994 | 16380 | 1 | -0.2302 | 0.32889 | 0.48018 | 1 | 11840 | 1 | -0.2302 |
| PIN1 | 6 | 0.1228 | 0.29099 | 0.999632 | 5734 | 1 | 1.1054 | 0.3289 | 0.59513 | 1 | 11841 | 3 | 1.1054 |
| NonTargetingControlG | 2 | 0.22115 | 0.27746 | 0.999379 | 8311 | 1 | 1.0487 | 0.32895 | 0.4225 | 1 | 11842 | 1 | 1.0487 |
| NonTargetingControlG | 2 | 0.69927 | 0.69909 | 0.999994 | 16300 | 0 | 2.789 | 0.32895 | 0.4225 | 1 | 11843 | 1 | 2.789 |
| GLB1 | 5 | 0.24302 | 0.43503 | 0.999994 | 8864 | 1 | -0.095962 | 0.32897 | 0.55593 | 1 | 11844 | 2 | -0.095962 |
| OR8S1 | 6 | 0.16392 | 0.34318 | 0.999994 | 6838 | 3 | 0.43145 | 0.32899 | 0.59521 | 1 | 11845 | 3 | 0.43145 |
| USP30 | 6 | 0.82284 | 0.85817 | 0.999994 | 17820 | 1 | 1.0026 | 0.32922 | 0.59538 | 1 | 11846 | 3 | 1.0026 |
| HAPLN4 | 5 | 0.49725 | 0.6455 | 0.999994 | 13385 | 2 | 1.0993 | 0.32924 | 0.5562 | 1 | 11847 | 3 | 1.0993 |
| CDH3 | 5 | 0.34273 | 0.51597 | 0.999994 | 11000 | 1 | 0.15966 | 0.32924 | 0.5562 | 1 | 11848 | 2 | 0.15966 |
| TDG | 5 | 0.10114 | 0.2282 | 0.983354 | 5008 | 3 | -1.5735 | 0.32924 | 0.5562 | 1 | 11849 | 2 | -1.5735 |
| RGS9 | 5 | 0.061922 | 0.16527 | 0.967126 | 3509 | 3 | -2.1314 | 0.32924 | 0.5562 | 1 | 11850 | 2 | -2.1314 |
| RFX4 | 5 | 0.57329 | 0.71243 | 0.999994 | 14596 | 2 | -0.47108 | 0.32924 | 0.5562 | 1 | 11851 | 2 | -0.47108 |
| FAM110C | 5 | 0.027262 | 0.083405 | 0.94383 | 1869 | 3 | -3.1034 | 0.32925 | 0.55621 | 1 | 11852 | 2 | -3.1034 |
| C17orf75 | 6 | 0.37028 | 0.58338 | 0.999994 | 11516 | 1 | 0.77046 | 0.32931 | 0.59545 | 1 | 11853 | 3 | 0.77046 |
| COQ10B | 6 | 0.2192 | 0.41114 | 0.999994 | 8268 | 2 | 0.16917 | 0.32931 | 0.59545 | 1 | 11854 | 3 | 0.16917 |
| TMEM72 | 6 | 0.41029 | 0.60909 | 0.999994 | 12113 | 2 | 0.67534 | 0.32931 | 0.59545 | 1 | 11855 | 3 | 0.67534 |

| | | | | | | | | | | | | | |
|---|---|---|---|---|---|---|---|---|---|---|---|---|---|
| ZNF781 | 6 | 0.20267 | 0.39112 | 0.999994 | 7848 | 2 | 0.037715 | 0.32931 | 0.59545 | 1 | 11856 | 2 | 0.037715 |
| C11orf49 | 6 | 0.061158 | 0.16824 | 0.967126 | 3478 | 2 | -0.025341 | 0.32931 | 0.59545 | 1 | 11857 | 2 | -0.025341 |
| hsa-mir-3927 | 4 | 0.84188 | 0.84134 | 0.999994 | 18121 | 0 | 1.7176 | 0.32932 | 0.53571 | 1 | 11858 | 2 | 1.7176 |
| RAET1E | 6 | 0.62087 | 0.75549 | 0.999994 | 15234 | 1 | 0.77393 | 0.32933 | 0.59547 | 1 | 11859 | 3 | 0.77393 |
| KCNA4 | 5 | 0.21827 | 0.40086 | 0.999994 | 8236 | 2 | 1.137 | 0.32957 | 0.55653 | 1 | 11860 | 3 | 1.137 |
| WDR13 | 6 | 0.64612 | 0.77466 | 0.999994 | 15602 | 2 | 0.98298 | 0.32973 | 0.59576 | 1 | 11861 | 3 | 0.98298 |
| C1QTNF9B | 3 | 0.30973 | 0.4338 | 0.999994 | 10321 | 1 | 1.439 | 0.32983 | 0.48114 | 1 | 11862 | 2 | 1.439 |
| ARHGAP20 | 4 | 0.18829 | 0.32861 | 0.999994 | 7426 | 1 | 1.1346 | 0.32985 | 0.53609 | 1 | 11863 | 2 | 1.1346 |
| OCSTAMP | 4 | 0.068489 | 0.15699 | 0.967126 | 3770 | 2 | -1.137 | 0.32985 | 0.53609 | 1 | 11864 | 2 | -1.137 |
| GSTT1 | 5 | 0.11232 | 0.24562 | 0.98979 | 5381 | 2 | 1.3124 | 0.32988 | 0.55688 | 1 | 11865 | 3 | 1.3124 |
| RHO | 5 | 0.86336 | 0.86525 | 0.999994 | 18511 | 1 | 1.2239 | 0.32996 | 0.55697 | 1 | 11866 | 3 | 1.2239 |
| UVSSA | 5 | 0.43086 | 0.58913 | 0.999994 | 12427 | 2 | 0.68783 | 0.33003 | 0.55702 | 1 | 11867 | 3 | 0.68783 |
| ANKRD13A | 5 | 0.057172 | 0.15741 | 0.967126 | 3300 | 3 | -1.7798 | 0.33003 | 0.55702 | 1 | 11868 | 1 | -1.7798 |
| DMRT1 | 5 | 0.025478 | 0.078765 | 0.93914 | 1777 | 3 | -2.2961 | 0.33003 | 0.55702 | 1 | 11869 | 1 | -2.2961 |
| DNAJC5G | 5 | 0.38936 | 0.55459 | 0.999994 | 11786 | 2 | 0.89478 | 0.33003 | 0.55702 | 1 | 11870 | 3 | 0.89478 |
| OPN1LW | 5 | 0.26491 | 0.45273 | 0.999994 | 9351 | 2 | 0.45185 | 0.33003 | 0.55702 | 1 | 11871 | 2 | 0.45185 |
| TBC1D17 | 5 | 0.74668 | 0.7891 | 0.999994 | 16824 | 1 | 0.49773 | 0.33003 | 0.55702 | 1 | 11872 | 2 | 0.49773 |
| CXCL16 | 6 | 0.10537 | 0.25778 | 0.994943 | 5147 | 3 | -0.14226 | 0.33006 | 0.59603 | 1 | 11873 | 3 | -0.14226 |
| FXYD6-FXYD2 | 1 | 0.66992 | 0.66995 | 0.999994 | 15930 | 0 | 2.0195 | 0.33008 | 0.33028 | 0.992034 | 11874 | 1 | 2.0195 |
| FOXH1 | 5 | 0.72732 | 0.77937 | 0.999994 | 16572 | 1 | 1.2897 | 0.33009 | 0.5571 | 1 | 11875 | 3 | 1.2897 |
| LYL1 | 6 | 0.23935 | 0.43521 | 0.999994 | 8762 | 3 | 0.43918 | 0.33014 | 0.5961 | 1 | 11876 | 3 | 0.43918 |
| ZNF552 | 2 | 0.13919 | 0.21915 | 0.980699 | 6168 | 1 | -0.93922 | 0.33015 | 0.42329 | 1 | 11877 | 1 | -0.93922 |
| PDGFC | 6 | 0.14258 | 0.31634 | 0.999994 | 6256 | 3 | 0.13481 | 0.33016 | 0.59611 | 1 | 11878 | 3 | 0.13481 |
| IQSEC3 | 6 | 0.56339 | 0.71322 | 0.999994 | 14438 | 2 | 0.7198 | 0.33026 | 0.59621 | 1 | 11879 | 3 | 0.7198 |
| FFAR4 | 6 | 0.13224 | 0.30318 | 0.999994 | 5981 | 2 | 0.68455 | 0.33026 | 0.59621 | 1 | 11880 | 3 | 0.68455 |
| MYOM2 | 6 | 0.31897 | 0.52871 | 0.999994 | 10497 | 3 | -0.53012 | 0.33026 | 0.59621 | 1 | 11881 | 2 | -0.53012 |
| PELI1 | 6 | 0.28091 | 0.48445 | 0.999994 | 9718 | 3 | -0.12653 | 0.33026 | 0.59621 | 1 | 11882 | 3 | -0.12653 |
| PPP1R18 | 6 | 0.022661 | 0.0805 | 0.93914 | 1630 | 3 | -0.81333 | 0.33026 | 0.59621 | 1 | 11883 | 2 | -0.81333 |
| PSMA1 | 5 | 0.35586 | 0.52668 | 0.999994 | 11265 | 2 | 1.1765 | 0.33034 | 0.55734 | 1 | 11884 | 3 | 1.1765 |
| hsa-mir-5687 | 4 | 0.046607 | 0.1148 | 0.951052 | 2834 | 2 | -1.4918 | 0.33036 | 0.53647 | 1 | 11885 | 1 | -1.4918 |
| FAM196B | 5 | 0.43091 | 0.58918 | 0.999994 | 12428 | 2 | 1.0159 | 0.33047 | 0.55749 | 1 | 11886 | 3 | 1.0159 |
| hsa-mir-6503 | 4 | 0.88407 | 0.88372 | 0.999994 | 18908 | 0 | 1.3315 | 0.33058 | 0.53662 | 1 | 11887 | 2 | 1.3315 |
| INPP1 | 6 | 0.22143 | 0.41383 | 0.999994 | 8316 | 2 | 0.32897 | 0.33066 | 0.59653 | 1 | 11888 | 3 | 0.32897 |
| FAM71A | 6 | 0.21556 | 0.40682 | 0.999994 | 8156 | 3 | 0.43026 | 0.33074 | 0.59659 | 1 | 11889 | 3 | 0.43026 |
| RPL5 | 4 | 0.051668 | 0.12475 | 0.951052 | 3052 | 3 | -4.413 | 0.33075 | 0.53675 | 1 | 11890 | 1 | -4.413 |
| CCNI | 6 | 0.36066 | 0.57605 | 0.999994 | 11349 | 2 | 0.95486 | 0.33076 | 0.5966 | 1 | 11891 | 3 | 0.95486 |
| OR13C2 | 6 | 0.27395 | 0.47636 | 0.999994 | 9564 | 2 | 1.0147 | 0.33081 | 0.59665 | 1 | 11892 | 3 | 1.0147 |
| ZMIZ2 | 5 | 0.043189 | 0.12379 | 0.951052 | 2682 | 3 | -1.8232 | 0.33128 | 0.55833 | 1 | 11893 | 2 | -1.8232 |
| hsa-mir-3154 | 3 | 0.24875 | 0.37339 | 0.999994 | 8987 | 1 | 1.4613 | 0.33141 | 0.4828 | 1 | 11894 | 2 | 1.4613 |
| KLF3 | 5 | 0.090329 | 0.21111 | 0.979587 | 4606 | 3 | -1.6491 | 0.33172 | 0.55878 | 1 | 11895 | 1 | -1.6491 |
| LAMTOR4 | 6 | 0.049912 | 0.14385 | 0.961589 | 2972 | 2 | 1.0305 | 0.33173 | 0.59738 | 1 | 11896 | 3 | 1.0305 |
| SETD5 | 6 | 0.15919 | 0.33719 | 0.999994 | 6710 | 3 | -0.60962 | 0.33182 | 0.59745 | 1 | 11897 | 2 | -0.60962 |
| ADAMTS19 | 6 | 0.93553 | 0.93557 | 0.999994 | 19980 | 0 | 0.29713 | 0.33182 | 0.59745 | 1 | 11898 | 2 | 0.29713 |
| HIST1H2AM | 4 | 0.86104 | 0.86063 | 0.999994 | 18466 | 0 | 1.1012 | 0.33191 | 0.53765 | 1 | 11899 | 2 | 1.1012 |
| C2orf54 | 4 | 0.085994 | 0.18971 | 0.970473 | 4446 | 2 | -0.13382 | 0.33199 | 0.53772 | 1 | 11900 | 2 | -0.13382 |
| PTRHD1 | 5 | 0.47673 | 0.6278 | 0.999994 | 13095 | 2 | 0.3103 | 0.33232 | 0.55939 | 1 | 11901 | 2 | 0.3103 |
| SNAP47 | 4 | 0.043693 | 0.10914 | 0.951052 | 2698 | 2 | -0.091011 | 0.33234 | 0.53798 | 1 | 11902 | 2 | -0.091011 |
| RPS3A | 6 | 0.62973 | 0.76213 | 0.999994 | 15377 | 2 | 0.49611 | 0.33236 | 0.59788 | 1 | 11903 | 3 | 0.49611 |
| TEAD4 | 6 | 0.51411 | 0.67846 | 0.999994 | 13644 | 2 | 0.44746 | 0.33236 | 0.59788 | 1 | 11904 | 3 | 0.44746 |
| COL1A2 | 6 | 0.039325 | 0.12037 | 0.951052 | 2481 | 4 | -1.5447 | 0.33236 | 0.59788 | 1 | 11905 | 2 | -1.5447 |
| ZNF417 | 6 | 0.5855 | 0.72926 | 0.999994 | 14750 | 2 | 0.27253 | 0.33236 | 0.59788 | 1 | 11906 | 3 | 0.27253 |
| CNGA4 | 6 | 0.089513 | 0.22653 | 0.983354 | 4578 | 4 | -1.5242 | 0.33236 | 0.59788 | 1 | 11907 | 2 | -1.5242 |

| | | | | | | | | | | | | | |
|---|---|---|---|---|---|---|---|---|---|---|---|---|---|
| TTF2 | 6 | 0.76762 | 0.83494 | 0.999994 | 17058 | 1 | 0.68897 | 0.33236 | 0.59788 | 1 | 11908 | 3 | 0.68897 |
| ZNF594 | 4 | 0.74805 | 0.75855 | 0.999994 | 16840 | 1 | 1.5143 | 0.33238 | 0.53801 | 1 | 11909 | 2 | 1.5143 |
| SLC37A4 | 4 | 0.89654 | 0.89625 | 0.999994 | 19154 | 0 | 1.3545 | 0.33238 | 0.53801 | 1 | 11910 | 2 | 1.3545 |
| EWSR1 | 4 | 0.0059576 | 0.020788 | 0.81027 | 588 | 3 | -5.0289 | 0.33238 | 0.53801 | 1 | 11911 | 1 | -5.0289 |
| MEIS1 | 4 | 0.2108 | 0.3518 | 0.999994 | 8046 | 2 | 0.33366 | 0.33249 | 0.53808 | 1 | 11912 | 2 | 0.33366 |
| ZNF566 | 5 | 0.55044 | 0.69213 | 0.999994 | 14219 | 1 | 1.1829 | 0.3325 | 0.55958 | 1 | 11913 | 3 | 1.1829 |
| CA1 | 6 | 0.081282 | 0.20991 | 0.979587 | 4276 | 3 | -0.89997 | 0.33257 | 0.59804 | 1 | 11914 | 3 | -0.89997 |
| IFNA17 | 6 | 0.94629 | 0.94625 | 0.999994 | 20238 | 0 | 0.52438 | 0.33257 | 0.59804 | 1 | 11915 | 3 | 0.52438 |
| SLC22A11 | 6 | 0.05431 | 0.15342 | 0.967126 | 3167 | 1 | 0.90127 | 0.3326 | 0.59807 | 1 | 11916 | 3 | 0.90127 |
| OLFM3 | 6 | 0.35911 | 0.57431 | 0.999994 | 11322 | 1 | 1.17 | 0.33274 | 0.59817 | 1 | 11917 | 3 | 1.17 |
| OR2AE1 | 4 | 0.17138 | 0.3112 | 0.999994 | 7021 | 2 | 0.33634 | 0.33276 | 0.53828 | 1 | 11918 | 2 | 0.33634 |
| SKAP1 | 5 | 0.41937 | 0.57957 | 0.999994 | 12241 | 2 | 1.3238 | 0.33278 | 0.55985 | 1 | 11919 | 3 | 1.3238 |
| hsa-mir-619 | 2 | 0.41839 | 0.43214 | 0.999994 | 12226 | 1 | 1.8217 | 0.33285 | 0.42497 | 1 | 11920 | 1 | 1.8217 |
| PUSL1 | 5 | 0.021493 | 0.068102 | 0.93914 | 1549 | 4 | -1.3748 | 0.33302 | 0.56009 | 1 | 11921 | 1 | -1.3748 |
| ZNF549 | 5 | 0.14645 | 0.29745 | 0.999994 | 6363 | 3 | -1.0675 | 0.33302 | 0.56009 | 1 | 11922 | 2 | -1.0675 |
| PRB3 | 6 | 0.13728 | 0.30961 | 0.999994 | 6114 | 3 | -0.46781 | 0.33315 | 0.59849 | 1 | 11923 | 3 | -0.46781 |
| DPYSL5 | 4 | 0.10531 | 0.22463 | 0.983354 | 5145 | 2 | 0.97953 | 0.33328 | 0.53865 | 1 | 11924 | 2 | 0.97953 |
| GLYAT | 5 | 0.13616 | 0.28197 | 0.999632 | 6083 | 2 | 1.2802 | 0.3333 | 0.56039 | 1 | 11925 | 3 | 1.2802 |
| SETD3 | 6 | 0.40676 | 0.60678 | 0.999994 | 12057 | 2 | 1.0671 | 0.33339 | 0.5987 | 1 | 11926 | 3 | 1.0671 |
| NonTargetingControlG | 2 | 0.75456 | 0.75411 | 0.999994 | 16904 | 0 | 2.9186 | 0.33347 | 0.42536 | 1 | 11927 | 1 | 2.9186 |
| ZNF764 | 5 | 0.16251 | 0.32122 | 0.999994 | 6810 | 2 | 1.1462 | 0.33351 | 0.56061 | 1 | 11928 | 3 | 1.1462 |
| GALK2 | 6 | 0.73929 | 0.82474 | 0.999994 | 16726 | 1 | 1.0787 | 0.33352 | 0.5988 | 1 | 11929 | 3 | 1.0787 |
| MAP7 | 5 | 0.86711 | 0.86851 | 0.999994 | 18577 | 1 | 1.1191 | 0.33356 | 0.56067 | 1 | 11930 | 3 | 1.1191 |
| NonTargetingControlG | 2 | 0.46966 | 0.47671 | 0.999994 | 13000 | 1 | 1.9199 | 0.33361 | 0.42545 | 1 | 11931 | 1 | 1.9199 |
| CDRT1 | 5 | 0.20487 | 0.38214 | 0.999994 | 7899 | 3 | -0.77481 | 0.3337 | 0.56079 | 1 | 11932 | 2 | -0.77481 |
| YDJC | 5 | 0.32811 | 0.50399 | 0.999994 | 10687 | 2 | 0.084571 | 0.3337 | 0.56079 | 1 | 11933 | 2 | 0.084571 |
| IRAK2 | 3 | 0.28345 | 0.4162 | 0.999994 | 9773 | 1 | 1.0266 | 0.33371 | 0.48519 | 1 | 11934 | 2 | 1.0266 |
| CPLX3 | 4 | 0.046207 | 0.11401 | 0.951052 | 2815 | 2 | 0.68359 | 0.33373 | 0.53897 | 1 | 11935 | 2 | 0.68359 |
| TRMT2B | 5 | 0.5144 | 0.66044 | 0.999994 | 13651 | 2 | 0.37209 | 0.33375 | 0.56084 | 1 | 11936 | 2 | 0.37209 |
| CALCR | 5 | 0.43255 | 0.59054 | 0.999994 | 12452 | 2 | 1.0753 | 0.33388 | 0.56098 | 1 | 11937 | 3 | 1.0753 |
| SLC25A43 | 6 | 0.082381 | 0.21222 | 0.979587 | 4311 | 2 | 0.78288 | 0.3339 | 0.5991 | 1 | 11938 | 3 | 0.78288 |
| RAB33B | 6 | 0.18287 | 0.36672 | 0.999994 | 7285 | 2 | 0.21305 | 0.33391 | 0.59911 | 1 | 11939 | 3 | 0.21305 |
| NTF3 | 6 | 0.34451 | 0.55785 | 0.999994 | 11042 | 2 | 0.27622 | 0.33391 | 0.59911 | 1 | 11940 | 3 | 0.27622 |
| GIP | 5 | 0.3147 | 0.49304 | 0.999994 | 10412 | 1 | 1.1002 | 0.33391 | 0.56103 | 1 | 11941 | 3 | 1.1002 |
| hsa-mir-6876 | 4 | 0.1231 | 0.25569 | 0.994943 | 5740 | 1 | 1.3884 | 0.33393 | 0.53911 | 1 | 11942 | 2 | 1.3884 |
| NonTargetingControlG | 2 | 0.66416 | 0.66391 | 0.999994 | 15842 | 0 | 2.6302 | 0.33413 | 0.42578 | 1 | 11943 | 1 | 2.6302 |
| ZDHHC6 | 5 | 0.95016 | 0.95034 | 0.999994 | 20340 | 0 | 0.54942 | 0.33428 | 0.56142 | 1 | 11944 | 3 | 0.54942 |
| APOH | 5 | 0.29037 | 0.47322 | 0.999994 | 9916 | 2 | 1.0441 | 0.33428 | 0.56142 | 1 | 11945 | 3 | 1.0441 |
| PLAC9 | 6 | 0.31544 | 0.52466 | 0.999994 | 10430 | 2 | 0.47821 | 0.33432 | 0.59942 | 1 | 11946 | 3 | 0.47821 |
| EXTL2 | 6 | 0.13056 | 0.30103 | 0.999994 | 5947 | 3 | -0.091721 | 0.33432 | 0.59942 | 1 | 11947 | 3 | -0.091721 |
| GDF5 | 4 | 0.2214 | 0.36267 | 0.999994 | 8313 | 2 | -0.054791 | 0.33433 | 0.53941 | 1 | 11948 | 2 | -0.054791 |
| RMI2 | 4 | 0.015166 | 0.049636 | 0.907155 | 1200 | 3 | -4.3734 | 0.33433 | 0.53941 | 1 | 11949 | 1 | -4.3734 |
| hsa-mir-4471 | 3 | 0.36976 | 0.46686 | 0.999994 | 11503 | 1 | -0.25854 | 0.33446 | 0.48596 | 1 | 11950 | 1 | -0.25854 |
| OAS3 | 3 | 0.093002 | 0.17377 | 0.96793 | 4720 | 2 | -1.6173 | 0.33446 | 0.48596 | 1 | 11951 | 1 | -1.6173 |
| GOLGA6B | 5 | 0.033235 | 0.098827 | 0.945253 | 2174 | 2 | 1.2967 | 0.33447 | 0.56161 | 1 | 11952 | 3 | 1.2967 |
| hsa-mir-548ah | 1 | 0.66551 | 0.66556 | 0.999994 | 15863 | 0 | 1.9962 | 0.33449 | 0.33461 | 0.993364 | 11953 | 1 | 1.9962 |
| CDH18 | 6 | 0.16881 | 0.34926 | 0.999994 | 6954 | 3 | 0.12285 | 0.33469 | 0.59969 | 1 | 11954 | 3 | 0.12285 |
| PROC | 5 | 0.063677 | 0.16815 | 0.967126 | 3593 | 3 | -1.8627 | 0.33474 | 0.56189 | 1 | 11955 | 2 | -1.8627 |
| DDX60 | 6 | 0.44641 | 0.6327 | 0.999994 | 12669 | 2 | 1.012 | 0.33482 | 0.5998 | 1 | 11956 | 3 | 1.012 |
| FRYL | 5 | 0.0084832 | 0.031698 | 0.865612 | 761 | 4 | -4.6418 | 0.33489 | 0.56204 | 1 | 11957 | 1 | -4.6418 |
| GMPS | 5 | 0.51974 | 0.66507 | 0.999994 | 13743 | 2 | -0.3554 | 0.33489 | 0.56204 | 1 | 11958 | 1 | -0.3554 |
| C6orf70 | 6 | 0.91981 | 0.92023 | 0.999994 | 19626 | 1 | 0.79654 | 0.33489 | 0.59987 | 1 | 11959 | 3 | 0.79654 |

| | | | | | | | | | | | | |
|---|---|---|---|---|---|---|---|---|---|---|---|---|
| S100A3 | 5 | 0.91945 | 0.9196 | 0.999994 | 19618 | 0 | 1.2527 | 0.33491 | 0.56206 | 1 | 11960 | 3 | 1.2527 |
| CNTN4 | 6 | 0.59032 | 0.73284 | 0.999994 | 14827 | 1 | 0.16623 | 0.33494 | 0.5999 | 1 | 11961 | 2 | 0.16623 |
| AP3S2 | 6 | 0.6835 | 0.80369 | 0.999994 | 16135 | 2 | 0.38649 | 0.33494 | 0.5999 | 1 | 11962 | 3 | 0.38649 |
| UTRN | 6 | 0.55812 | 0.7095 | 0.999994 | 14335 | 1 | 0.47767 | 0.33494 | 0.5999 | 1 | 11963 | 3 | 0.47767 |
| TRIM6-TRIM34 | 4 | 0.40149 | 0.54728 | 0.999994 | 11971 | 2 | -0.73362 | 0.33496 | 0.53988 | 1 | 11964 | 1 | -0.73362 |
| C9orf85 | 4 | 0.55181 | 0.64058 | 0.999994 | 14239 | 1 | 1.028 | 0.33496 | 0.53988 | 1 | 11965 | 2 | 1.028 |
| NMRK1 | 6 | 0.041889 | 0.12614 | 0.951052 | 2607 | 4 | -1.6474 | 0.33501 | 0.59995 | 1 | 11966 | 2 | -1.6474 |
| SHROOM4 | 5 | 0.13757 | 0.28413 | 0.999632 | 6123 | 1 | 1.3902 | 0.33502 | 0.56216 | 1 | 11967 | 3 | 1.3902 |
| OAS1 | 5 | 0.19035 | 0.36145 | 0.999994 | 7497 | 2 | 1.2375 | 0.33513 | 0.56228 | 1 | 11968 | 3 | 1.2375 |
| REEP4 | 3 | 0.52483 | 0.56277 | 0.999994 | 13813 | 1 | 1.4103 | 0.33517 | 0.48671 | 1 | 11969 | 2 | 1.4103 |
| hsa-mir-4655 | 3 | 0.2689 | 0.39831 | 0.999994 | 9443 | 1 | 1.7381 | 0.33522 | 0.48675 | 1 | 11970 | 2 | 1.7381 |
| NonTargetingControlG | 1 | 0.6647 | 0.66474 | 0.999994 | 15853 | 0 | 1.9397 | 0.3353 | 0.33542 | 0.993633 | 11971 | 1 | 1.9397 |
| RFPL3 | 6 | 0.5692 | 0.71737 | 0.999994 | 14523 | 1 | 0.83016 | 0.3354 | 0.60023 | 1 | 11972 | 3 | 0.83016 |
| ZNF230 | 6 | 0.63165 | 0.76358 | 0.999994 | 15394 | 2 | 0.66466 | 0.33544 | 0.60026 | 1 | 11973 | 3 | 0.66466 |
| ERVV-1 | 6 | 0.48563 | 0.65898 | 0.999994 | 13225 | 2 | 0.1845 | 0.33559 | 0.60038 | 1 | 11974 | 2 | 0.1845 |
| CTAGE9 | 5 | 0.51212 | 0.65851 | 0.999994 | 13613 | 2 | 0.48012 | 0.3356 | 0.56276 | 1 | 11975 | 2 | 0.48012 |
| BDKRB1 | 5 | 0.23197 | 0.41996 | 0.999994 | 8569 | 3 | -0.66217 | 0.3356 | 0.56276 | 1 | 11976 | 2 | -0.66217 |
| STX2 | 4 | 0.44408 | 0.59099 | 0.999994 | 12635 | 2 | -0.0033175 | 0.33562 | 0.54036 | 1 | 11977 | 2 | -0.0033175 |
| hsa-mir-627 | 4 | 0.42346 | 0.56973 | 0.999994 | 12302 | 2 | -0.48729 | 0.33562 | 0.54036 | 1 | 11978 | 1 | -0.48729 |
| NT5C1B-RDH14 | 3 | 0.53695 | 0.57119 | 0.999994 | 14013 | 1 | 1.4551 | 0.33563 | 0.48719 | 1 | 11979 | 2 | 1.4551 |
| hsa-mir-183 | 3 | 0.71605 | 0.71619 | 0.999994 | 16457 | 1 | 1.3229 | 0.33583 | 0.48741 | 1 | 11980 | 2 | 1.3229 |
| hsa-mir-6784 | 3 | 0.68036 | 0.68307 | 0.999994 | 16096 | 1 | 0.54942 | 0.33583 | 0.48741 | 1 | 11981 | 1 | 0.54942 |
| BIRC2 | 5 | 0.58438 | 0.72054 | 0.999994 | 14734 | 1 | 1.1974 | 0.33586 | 0.56304 | 1 | 11982 | 3 | 1.1974 |
| PLAC1L | 6 | 0.18592 | 0.3705 | 0.999994 | 7364 | 2 | 1.2406 | 0.33593 | 0.60064 | 1 | 11983 | 3 | 1.2406 |
| hsa-mir-585 | 3 | 0.07915 | 0.15531 | 0.967126 | 4196 | 1 | 1.2771 | 0.33607 | 0.48767 | 1 | 11984 | 2 | 1.2771 |
| TPRA1 | 6 | 0.22709 | 0.42058 | 0.999994 | 8457 | 2 | 0.22866 | 0.33611 | 0.60078 | 1 | 11985 | 2 | 0.22866 |
| ZNF251 | 5 | 0.70349 | 0.76808 | 0.999994 | 16347 | 1 | 1.1392 | 0.33619 | 0.56339 | 1 | 11986 | 3 | 1.1392 |
| NonTargetingControlG | 1 | 0.66375 | 0.66377 | 0.999994 | 15834 | 0 | 1.8772 | 0.33625 | 0.33638 | 0.994237 | 11987 | 1 | 1.8772 |
| KLF12 | 5 | 0.095101 | 0.2187 | 0.980699 | 4788 | 2 | 1.1823 | 0.33634 | 0.56355 | 1 | 11988 | 3 | 1.1823 |
| hsa-mir-3118-4 | 2 | 0.68015 | 0.68 | 0.999994 | 16093 | 0 | 2.5061 | 0.33638 | 0.42724 | 1 | 11989 | 1 | 2.5061 |
| DDX52 | 5 | 0.93391 | 0.93415 | 0.999994 | 19950 | 0 | 0.94772 | 0.3364 | 0.56361 | 1 | 11990 | 3 | 0.94772 |
| CCDC144NL | 5 | 0.15271 | 0.30683 | 0.999994 | 6526 | 1 | 0.50162 | 0.33645 | 0.56365 | 1 | 11991 | 3 | 0.50162 |
| ATF6B | 6 | 0.34955 | 0.5635 | 0.999994 | 11138 | 3 | -0.54048 | 0.33647 | 0.60108 | 1 | 11992 | 2 | -0.54048 |
| ABCG1 | 5 | 0.34448 | 0.51738 | 0.999994 | 11039 | 2 | 1.1466 | 0.33652 | 0.56373 | 1 | 11993 | 3 | 1.1466 |
| KCNJ4 | 6 | 0.053342 | 0.15129 | 0.967126 | 3132 | 3 | 0.24961 | 0.33667 | 0.60123 | 1 | 11994 | 3 | 0.24961 |
| RGL2 | 6 | 0.033659 | 0.10737 | 0.950995 | 2199 | 2 | 0.93632 | 0.3367 | 0.60126 | 1 | 11995 | 3 | 0.93632 |
| TAL1 | 6 | 0.050766 | 0.14575 | 0.963702 | 3005 | 4 | -1.6781 | 0.3367 | 0.60126 | 1 | 11996 | 2 | -1.6781 |
| SERPINA1 | 6 | 0.22398 | 0.4169 | 0.999994 | 8378 | 3 | -0.65083 | 0.3367 | 0.60126 | 1 | 11997 | 2 | -0.65083 |
| UBALD2 | 6 | 0.83084 | 0.86195 | 0.999994 | 17944 | 1 | 0.67223 | 0.3367 | 0.60126 | 1 | 11998 | 3 | 0.67223 |
| PEBP4 | 5 | 0.70878 | 0.77052 | 0.999994 | 16394 | 1 | 1.2457 | 0.33677 | 0.564 | 1 | 11999 | 3 | 1.2457 |
| GXYLT1 | 6 | 0.65347 | 0.78026 | 0.999994 | 15703 | 2 | 1.1749 | 0.33683 | 0.60138 | 1 | 12000 | 3 | 1.1749 |
| CDCA5 | 5 | 0.0080418 | 0.03041 | 0.862223 | 734 | 3 | -6.2407 | 0.33693 | 0.56416 | 1 | 12001 | 2 | -6.2407 |
| DEFB132 | 6 | 0.048789 | 0.14145 | 0.959146 | 2921 | 2 | -0.34894 | 0.33696 | 0.60147 | 1 | 12002 | 2 | -0.34894 |
| DUS2L | 2 | 0.5663 | 0.5666 | 0.999994 | 14483 | 1 | 2.3517 | 0.33701 | 0.42763 | 1 | 12003 | 1 | 2.3517 |
| PAXBP1 | 4 | 0.65041 | 0.69353 | 0.999994 | 15664 | 1 | 0.49778 | 0.33708 | 0.54145 | 1 | 12004 | 2 | 0.49778 |
| NHP2 | 4 | 0.080962 | 0.18046 | 0.968171 | 4264 | 3 | -2.0451 | 0.33708 | 0.54145 | 1 | 12005 | 1 | -2.0451 |
| ITGBL1 | 5 | 0.19902 | 0.37384 | 0.999994 | 7738 | 3 | -0.83573 | 0.3372 | 0.56444 | 1 | 12006 | 2 | -0.83573 |
| PSMC5 | 3 | 0.62114 | 0.63358 | 0.999994 | 15244 | 1 | 0.33023 | 0.33728 | 0.48893 | 1 | 12007 | 1 | 0.33023 |
| hsa-mir-6767 | 3 | 0.20417 | 0.31796 | 0.999994 | 7880 | 1 | 1.4036 | 0.33738 | 0.48903 | 1 | 12008 | 2 | 1.4036 |
| FHOD3 | 5 | 0.73447 | 0.78285 | 0.999994 | 16655 | 1 | 0.57914 | 0.33739 | 0.56465 | 1 | 12009 | 3 | 0.57914 |
| TMEM63B | 4 | 0.0072808 | 0.025069 | 0.844082 | 678 | 2 | -2.2974 | 0.33741 | 0.54169 | 1 | 12010 | 2 | -2.2974 |
| C19orf73 | 6 | 0.1231 | 0.29151 | 0.999632 | 5741 | 3 | -0.4745 | 0.33742 | 0.60183 | 1 | 12011 | 2 | -0.4745 |

| | | | | | | | | | | | | |
|---|---|---|---|---|---|---|---|---|---|---|---|---|
| TRPM3 | 6 | 0.30602 | 0.51376 | 0.999994 | 10255 | 2 | 0.182 | 0.33742 | 0.60183 | 1 | 12012 | 2 | 0.182 |
| CYBA | 5 | 0.26839 | 0.45554 | 0.999994 | 9428 | 2 | 1.4389 | 0.33752 | 0.5648 | 1 | 12013 | 3 | 1.4389 |
| HAT1 | 6 | 0.6447 | 0.77356 | 0.999994 | 15580 | 2 | 0.32097 | 0.3376 | 0.60198 | 1 | 12014 | 3 | 0.32097 |
| hsa-mir-1254-1 | 3 | 0.38743 | 0.47705 | 0.999994 | 11759 | 1 | 1.44 | 0.33771 | 0.48937 | 1 | 12015 | 2 | 1.44 |
| FAM124B | 4 | 0.22777 | 0.36923 | 0.999994 | 8474 | 2 | -0.91237 | 0.33777 | 0.54198 | 1 | 12016 | 1 | -0.91237 |
| hsa-mir-4273 | 4 | 0.3404 | 0.4844 | 0.999994 | 10943 | 2 | -0.34025 | 0.33777 | 0.54198 | 1 | 12017 | 1 | -0.34025 |
| TSTA3 | 6 | 0.12345 | 0.29193 | 0.999632 | 5750 | 3 | -0.56789 | 0.33788 | 0.60219 | 1 | 12018 | 2 | -0.56789 |
| OR5D14 | 6 | 0.8106 | 0.85252 | 0.999994 | 17640 | 1 | -0.2974 | 0.33788 | 0.60219 | 1 | 12019 | 2 | -0.2974 |
| FAM186A | 6 | 0.17507 | 0.35696 | 0.999994 | 7111 | 2 | 0.24634 | 0.33788 | 0.60219 | 1 | 12020 | 3 | 0.24634 |
| CAMK1D | 6 | 0.3076 | 0.51559 | 0.999994 | 10280 | 3 | -0.013471 | 0.33788 | 0.60219 | 1 | 12021 | 3 | -0.013471 |
| KRTAP6-3 | 5 | 0.0061193 | 0.024769 | 0.844082 | 600 | 4 | -2.2544 | 0.33795 | 0.56521 | 1 | 12022 | 1 | -2.2544 |
| CCDC178 | 6 | 0.26006 | 0.4599 | 0.999994 | 9244 | 1 | 1.0028 | 0.33796 | 0.60226 | 1 | 12023 | 3 | 1.0028 |
| RHOG | 6 | 0.23468 | 0.42967 | 0.999994 | 8633 | 1 | 1.3283 | 0.33796 | 0.60226 | 1 | 12024 | 3 | 1.3283 |
| MRPS30 | 3 | 0.80213 | 0.80193 | 0.999994 | 17515 | 0 | 1.3188 | 0.3381 | 0.48979 | 1 | 12025 | 2 | 1.3188 |
| MRAP2 | 6 | 0.65134 | 0.77865 | 0.999994 | 15680 | 2 | 0.23003 | 0.33821 | 0.60245 | 1 | 12026 | 3 | 0.23003 |
| NonTargetingControlG | 2 | 0.52534 | 0.52742 | 0.999994 | 13822 | 1 | 2.1788 | 0.33827 | 0.42842 | 1 | 12027 | 1 | 2.1788 |
| NonTargetingControlG | 2 | 0.62807 | 0.62786 | 0.999994 | 15351 | 0 | 2.4995 | 0.33827 | 0.42842 | 1 | 12028 | 1 | 2.4995 |
| KIAA0947 | 6 | 0.063144 | 0.1724 | 0.967658 | 3564 | 2 | 1.0594 | 0.33844 | 0.60263 | 1 | 12029 | 3 | 1.0594 |
| OPTC | 5 | 0.19157 | 0.36317 | 0.999994 | 7529 | 2 | -0.20313 | 0.33852 | 0.56584 | 1 | 12030 | 1 | -0.20313 |
| OSCAR | 5 | 0.79056 | 0.81366 | 0.999994 | 17357 | 1 | 0.90189 | 0.33852 | 0.56584 | 1 | 12031 | 3 | 0.90189 |
| MPEG1 | 4 | 0.70516 | 0.72788 | 0.999994 | 16363 | 1 | 1.7064 | 0.33869 | 0.54268 | 1 | 12032 | 2 | 1.7064 |
| hsa-mir-4801 | 3 | 0.36803 | 0.46587 | 0.999994 | 11476 | 1 | -0.13516 | 0.33885 | 0.49057 | 1 | 12033 | 1 | -0.13516 |
| SNRPF | 3 | 0.73649 | 0.73639 | 0.999994 | 16683 | 0 | 1.3974 | 0.33895 | 0.49066 | 1 | 12034 | 2 | 1.3974 |
| AP5Z1 | 6 | 0.33127 | 0.54271 | 0.999994 | 10749 | 1 | 0.34681 | 0.33896 | 0.60305 | 1 | 12035 | 3 | 0.34681 |
| PUS7L | 6 | 0.10722 | 0.26135 | 0.995005 | 5218 | 4 | -0.76795 | 0.33896 | 0.60305 | 1 | 12036 | 2 | -0.76795 |
| KIAA1377 | 6 | 0.041931 | 0.12623 | 0.951052 | 2610 | 3 | -0.19413 | 0.33896 | 0.60305 | 1 | 12037 | 3 | -0.19413 |
| WNT4 | 6 | 0.40805 | 0.6076 | 0.999994 | 12077 | 2 | 0.098036 | 0.33896 | 0.60305 | 1 | 12038 | 2 | 0.098036 |
| ACER3 | 6 | 0.57665 | 0.72272 | 0.999994 | 14640 | 2 | 0.94769 | 0.33897 | 0.60306 | 1 | 12039 | 3 | 0.94769 |
| SYNJ2BP-COX16 | 1 | 0.66103 | 0.66103 | 0.999994 | 15795 | 0 | 2.1214 | 0.33897 | 0.33914 | 0.994237 | 12040 | 1 | 2.1214 |
| HIGD1A | 4 | 0.049739 | 0.12095 | 0.951052 | 2964 | 3 | -2.1806 | 0.33903 | 0.54292 | 1 | 12041 | 1 | -2.1806 |
| TTC28 | 5 | 0.12927 | 0.27151 | 0.998561 | 5922 | 2 | 0.89821 | 0.33916 | 0.56649 | 1 | 12042 | 3 | 0.89821 |
| LGALS12 | 5 | 0.14946 | 0.30197 | 0.999994 | 6443 | 3 | -1.1522 | 0.33916 | 0.56649 | 1 | 12043 | 2 | -1.1522 |
| CD151 | 5 | 0.026515 | 0.081459 | 0.93914 | 1828 | 2 | -0.29732 | 0.33916 | 0.56649 | 1 | 12044 | 2 | -0.29732 |
| KRT3 | 6 | 0.5121 | 0.67705 | 0.999994 | 13612 | 2 | 0.68113 | 0.3392 | 0.60323 | 1 | 12045 | 3 | 0.68113 |
| USB1 | 6 | 0.30638 | 0.51416 | 0.999994 | 10263 | 3 | -0.30542 | 0.33936 | 0.60335 | 1 | 12046 | 1 | -0.30542 |
| CXCL10 | 3 | 0.26158 | 0.38934 | 0.999994 | 9274 | 1 | 1.3381 | 0.33937 | 0.49108 | 1 | 12047 | 2 | 1.3381 |
| ITGA7 | 5 | 0.91037 | 0.9104 | 0.999994 | 19434 | 0 | 1.0043 | 0.33941 | 0.56674 | 1 | 12048 | 3 | 1.0043 |
| RPS15A | 2 | 0.057484 | 0.10175 | 0.946414 | 3314 | 1 | -1.8999 | 0.33952 | 0.42922 | 1 | 12049 | 1 | -1.8999 |
| TMEM38B | 5 | 0.56375 | 0.70398 | 0.999994 | 14447 | 2 | 0.3503 | 0.33955 | 0.56687 | 1 | 12050 | 2 | 0.3503 |
| KRTAP7-1 | 6 | 0.80803 | 0.85137 | 0.999994 | 17601 | 1 | 1.0767 | 0.33967 | 0.6036 | 1 | 12051 | 3 | 1.0767 |
| PTCHD1 | 3 | 0.72382 | 0.72376 | 0.999994 | 16531 | 1 | 1.4041 | 0.33968 | 0.49141 | 1 | 12052 | 2 | 1.4041 |
| PRNP | 6 | 0.037544 | 0.11638 | 0.951052 | 2406 | 4 | -1.6137 | 0.33972 | 0.60364 | 1 | 12053 | 1 | -1.6137 |
| C11orf65 | 6 | 0.056574 | 0.15835 | 0.967126 | 3275 | 3 | -0.12542 | 0.33976 | 0.60368 | 1 | 12054 | 2 | -0.12542 |
| RFX7 | 5 | 0.64492 | 0.7431 | 0.999994 | 15583 | 1 | 1.0636 | 0.33978 | 0.56709 | 1 | 12055 | 3 | 1.0636 |
| SH3BP4 | 6 | 0.079636 | 0.20658 | 0.978744 | 4214 | 3 | -0.72808 | 0.33982 | 0.60371 | 1 | 12056 | 3 | -0.72808 |
| HSPB2 | 5 | 0.87589 | 0.8764 | 0.999994 | 18749 | 1 | 1.097 | 0.34003 | 0.56735 | 1 | 12057 | 3 | 1.097 |
| TBC1D29 | 5 | 0.18152 | 0.34883 | 0.999994 | 7263 | 2 | 0.25583 | 0.34013 | 0.56746 | 1 | 12058 | 1 | 0.25583 |
| CCNT1 | 4 | 0.88619 | 0.88588 | 0.999994 | 18954 | 0 | 1.189 | 0.34016 | 0.54377 | 1 | 12059 | 2 | 1.189 |
| KCNJ10 | 5 | 0.47621 | 0.62735 | 0.999994 | 13083 | 1 | 0.4304 | 0.34018 | 0.56752 | 1 | 12060 | 2 | 0.4304 |
| UPP1 | 5 | 0.15935 | 0.31662 | 0.999994 | 6716 | 2 | 0.78756 | 0.3404 | 0.56775 | 1 | 12061 | 3 | 0.78756 |
| GBA2 | 5 | 0.2351 | 0.42428 | 0.999994 | 8648 | 1 | 1.2579 | 0.34053 | 0.56787 | 1 | 12062 | 3 | 1.2579 |
| NonTargetingControlG | 2 | 0.14909 | 0.22601 | 0.983354 | 6432 | 1 | -0.91309 | 0.34056 | 0.42988 | 1 | 12063 | 1 | -0.91309 |

| | | | | | | | | | | | | |
|---|---|---|---|---|---|---|---|---|---|---|---|---|
| TRAPPC13 | 5 | 0.07949 | 0.19375 | 0.975797 | 4207 | 3 | -1.6331 | 0.34058 | 0.56793 | 1 | 12064 | 2 | -1.6331 |
| CCT7 | 4 | 0.004842 | 0.017111 | 0.785648 | 510 | 3 | -4.9272 | 0.34061 | 0.54409 | 1 | 12065 | 1 | -4.9272 |
| ZBED2 | 5 | 0.81779 | 0.83111 | 0.999994 | 17743 | 1 | -0.081442 | 0.34062 | 0.56796 | 1 | 12066 | 2 | -0.081442 |
| hsa-mir-4442 | 3 | 0.26635 | 0.3952 | 0.999994 | 9376 | 1 | 1.466 | 0.34067 | 0.49244 | 1 | 12067 | 2 | 1.466 |
| hsa-mir-4486 | 3 | 0.034483 | 0.085334 | 0.94395 | 2240 | 2 | -4.3829 | 0.34067 | 0.49244 | 1 | 12068 | 1 | -4.3829 |
| CDK1 | 3 | 0.22737 | 0.34681 | 0.999994 | 8463 | 2 | -0.9356 | 0.34067 | 0.49244 | 1 | 12069 | 1 | -0.9356 |
| MRPL49 | 3 | 0.47615 | 0.53066 | 0.999994 | 13082 | 1 | 1.2008 | 0.34067 | 0.49244 | 1 | 12070 | 2 | 1.2008 |
| WBSCR17 | 4 | 0.038576 | 0.098954 | 0.945253 | 2442 | 2 | 0.82415 | 0.34073 | 0.54418 | 1 | 12071 | 2 | 0.82415 |
| C20orf144 | 5 | 0.74875 | 0.79019 | 0.999994 | 16847 | 1 | 1.2748 | 0.34077 | 0.56813 | 1 | 12072 | 3 | 1.2748 |
| PDX1 | 6 | 0.090447 | 0.22839 | 0.983354 | 4612 | 3 | -0.79136 | 0.34079 | 0.60449 | 1 | 12073 | 2 | -0.79136 |
| LAMP1 | 6 | 0.91554 | 0.91638 | 0.999994 | 19526 | 1 | 0.4266 | 0.34079 | 0.60449 | 1 | 12074 | 3 | 0.4266 |
| INSRR | 5 | 0.18108 | 0.3482 | 0.999994 | 7249 | 3 | -0.95557 | 0.3408 | 0.56816 | 1 | 12075 | 2 | -0.95557 |
| C17orf51 | 5 | 0.043421 | 0.12435 | 0.951052 | 2690 | 3 | -2.3843 | 0.3408 | 0.56816 | 1 | 12076 | 2 | -2.3843 |
| DUSP15 | 5 | 0.61257 | 0.73076 | 0.999994 | 15119 | 1 | 0.62289 | 0.3408 | 0.56816 | 1 | 12077 | 3 | 0.62289 |
| CRYGB | 6 | 0.19246 | 0.37857 | 0.999994 | 7559 | 3 | 0.50729 | 0.34086 | 0.60454 | 1 | 12078 | 3 | 0.50729 |
| CCDC39 | 6 | 0.16799 | 0.34822 | 0.999994 | 6938 | 3 | 0.47902 | 0.34087 | 0.60456 | 1 | 12079 | 3 | 0.47902 |
| PTEN | 6 | 0.3675 | 0.58167 | 0.999994 | 11466 | 1 | 1.0483 | 0.34091 | 0.60459 | 1 | 12080 | 3 | 1.0483 |
| PIK3R3 | 5 | 0.93281 | 0.93309 | 0.999994 | 19922 | 0 | 0.66982 | 0.34097 | 0.56832 | 1 | 12081 | 3 | 0.66982 |
| GPR157 | 5 | 0.52232 | 0.66732 | 0.999994 | 13783 | 2 | 1.1268 | 0.34101 | 0.56836 | 1 | 12082 | 3 | 1.1268 |
| REN | 6 | 0.059879 | 0.1655 | 0.967126 | 3417 | 3 | -0.88029 | 0.34104 | 0.60469 | 1 | 12083 | 2 | -0.88029 |
| NEIL1 | 5 | 0.69716 | 0.76519 | 0.999994 | 16271 | 1 | 1.2479 | 0.34131 | 0.56867 | 1 | 12084 | 3 | 1.2479 |
| hsa-mir-4723 | 3 | 0.66447 | 0.66914 | 0.999994 | 15848 | 1 | 1.3477 | 0.34159 | 0.4934 | 1 | 12085 | 2 | 1.3477 |
| hsa-mir-877 | 3 | 0.25568 | 0.38206 | 0.999994 | 9140 | 1 | 1.739 | 0.34161 | 0.49341 | 1 | 12086 | 2 | 1.739 |
| PPP1R1B | 6 | 0.1121 | 0.27073 | 0.998561 | 5371 | 3 | -0.66857 | 0.34163 | 0.60516 | 1 | 12087 | 2 | -0.66857 |
| UQCRFS1 | 4 | 0.90364 | 0.90336 | 0.999994 | 19295 | 0 | 1.4178 | 0.34171 | 0.54495 | 1 | 12088 | 2 | 1.4178 |
| GREM1 | 6 | 0.024603 | 0.086076 | 0.94395 | 1736 | 4 | -2.8473 | 0.34174 | 0.60525 | 1 | 12089 | 2 | -2.8473 |
| SMEK1 | 5 | 0.54043 | 0.68329 | 0.999994 | 14061 | 2 | -0.49971 | 0.34177 | 0.56913 | 1 | 12090 | 2 | -0.49971 |
| SOD2 | 5 | 0.083607 | 0.2004 | 0.977164 | 4360 | 3 | -0.81017 | 0.34177 | 0.56913 | 1 | 12091 | 2 | -0.81017 |
| NACA | 5 | 0.19203 | 0.36382 | 0.999994 | 7549 | 2 | 0.53882 | 0.34177 | 0.56913 | 1 | 12092 | 3 | 0.53882 |
| hsa-mir-4778 | 4 | 0.88019 | 0.87982 | 0.999994 | 18836 | 0 | 1.3934 | 0.34187 | 0.54507 | 1 | 12093 | 2 | 1.3934 |
| FCHO2 | 4 | 0.14433 | 0.28338 | 0.999632 | 6312 | 2 | -1.4855 | 0.34187 | 0.54507 | 1 | 12094 | 2 | -1.4855 |
| GAS2L3 | 6 | 0.60699 | 0.74513 | 0.999994 | 15036 | 2 | 0.78604 | 0.34191 | 0.60538 | 1 | 12095 | 3 | 0.78604 |
| PRR15 | 5 | 0.15307 | 0.30737 | 0.999994 | 6536 | 2 | 1.1743 | 0.34197 | 0.56935 | 1 | 12096 | 3 | 1.1743 |
| ATP6V1B1 | 5 | 0.30266 | 0.48321 | 0.999994 | 10186 | 2 | 1.1335 | 0.34214 | 0.56952 | 1 | 12097 | 3 | 1.1335 |
| LHX5 | 5 | 0.52926 | 0.67339 | 0.999994 | 13887 | 1 | -0.11129 | 0.34219 | 0.56957 | 1 | 12098 | 1 | -0.11129 |
| OXLD1 | 6 | 0.044439 | 0.13187 | 0.95318 | 2731 | 2 | -0.1393 | 0.34238 | 0.60575 | 1 | 12099 | 2 | -0.1393 |
| CHIT1 | 3 | 0.8452 | 0.84522 | 0.999994 | 18173 | 0 | 1.4005 | 0.34243 | 0.49427 | 1 | 12100 | 2 | 1.4005 |
| AURKC | 5 | 0.73955 | 0.78545 | 0.999994 | 16732 | 1 | 1.0717 | 0.34257 | 0.56996 | 1 | 12101 | 3 | 1.0717 |
| CDT1 | 2 | 0.49414 | 0.49855 | 0.999994 | 13331 | 1 | 2.0342 | 0.3427 | 0.43128 | 1 | 12102 | 1 | 2.0342 |
| ZNF223 | 5 | 0.32503 | 0.50143 | 0.999994 | 10619 | 2 | 1.1654 | 0.34271 | 0.5701 | 1 | 12103 | 3 | 1.1654 |
| DCUN1D3 | 6 | 0.11425 | 0.27483 | 0.998561 | 5455 | 2 | 0.76447 | 0.3428 | 0.60608 | 1 | 12104 | 3 | 0.76447 |
| MSANTD3-TMEFF1 | 5 | 0.4911 | 0.64018 | 0.999994 | 13290 | 2 | 0.29724 | 0.34288 | 0.57028 | 1 | 12105 | 2 | 0.29724 |
| KRT37 | 5 | 0.058452 | 0.15949 | 0.967126 | 3364 | 3 | -2.1015 | 0.34288 | 0.57028 | 1 | 12106 | 2 | -2.1015 |
| SMARCC2 | 5 | 0.26331 | 0.45142 | 0.999994 | 9322 | 2 | 0.92425 | 0.34288 | 0.57028 | 1 | 12107 | 3 | 0.92425 |
| VSIG10 | 6 | 0.086945 | 0.22137 | 0.983093 | 4476 | 4 | -1.1849 | 0.34289 | 0.60615 | 1 | 12108 | 2 | -1.1849 |
| RAB12 | 6 | 0.063765 | 0.17374 | 0.96793 | 3599 | 3 | -0.9535 | 0.34289 | 0.60615 | 1 | 12109 | 3 | -0.9535 |
| HSPB1 | 6 | 0.50187 | 0.66998 | 0.999994 | 13456 | 2 | 0.78657 | 0.34289 | 0.60615 | 1 | 12110 | 3 | 0.78657 |
| TNFRSF1A | 6 | 0.036011 | 0.11288 | 0.951052 | 2316 | 2 | 0.70652 | 0.3429 | 0.60616 | 1 | 12111 | 3 | 0.70652 |
| ZNF121 | 6 | 0.28834 | 0.49316 | 0.999994 | 9878 | 3 | -0.31558 | 0.343 | 0.60625 | 1 | 12112 | 2 | -0.31558 |
| ALKBH7 | 6 | 0.032016 | 0.10354 | 0.946414 | 2107 | 3 | -1.706 | 0.343 | 0.60625 | 1 | 12113 | 1 | -1.706 |
| IDH1 | 6 | 0.047353 | 0.13826 | 0.957317 | 2873 | 4 | -1.3683 | 0.343 | 0.60625 | 1 | 12114 | 1 | -1.3683 |
| SLC2A13 | 3 | 0.20161 | 0.3147 | 0.999994 | 7811 | 2 | -1.0813 | 0.3431 | 0.49495 | 1 | 12115 | 1 | -1.0813 |

| | | | | | | | | | | | | | |
|---|---|---|---|---|---|---|---|---|---|---|---|---|---|
| QPCT | 5 | 0.09156 | 0.21307 | 0.979587 | 4661 | 2 | -0.54202 | 0.34316 | 0.57056 | 1 | 12116 | 2 | -0.54202 |
| TBC1D21 | 4 | 0.3413 | 0.48534 | 0.999994 | 10964 | 1 | 0.55694 | 0.34328 | 0.54613 | 1 | 12117 | 2 | 0.55694 |
| PTPN2 | 5 | 0.48476 | 0.63476 | 0.999994 | 13213 | 1 | 1.1419 | 0.34329 | 0.57068 | 1 | 12118 | 3 | 1.1419 |
| CELA2B | 6 | 0.41225 | 0.61036 | 0.999994 | 12149 | 2 | 0.93032 | 0.34329 | 0.60647 | 1 | 12119 | 3 | 0.93032 |
| ISLR | 5 | 0.45306 | 0.60779 | 0.999994 | 12746 | 1 | 0.025169 | 0.34349 | 0.57089 | 1 | 12120 | 2 | 0.025169 |
| LIMCH1 | 5 | 0.13081 | 0.27384 | 0.998561 | 5951 | 3 | -1.2052 | 0.34349 | 0.57089 | 1 | 12121 | 2 | -1.2052 |
| NPY4R | 5 | 0.39546 | 0.55978 | 0.999994 | 11877 | 2 | 0.5306 | 0.34349 | 0.57089 | 1 | 12122 | 2 | 0.5306 |
| FCHSD2 | 5 | 0.52015 | 0.66544 | 0.999994 | 13749 | 2 | 1.2169 | 0.34361 | 0.57102 | 1 | 12123 | 3 | 1.2169 |
| OR2T6 | 6 | 0.35771 | 0.57273 | 0.999994 | 11301 | 2 | -0.19377 | 0.34362 | 0.60673 | 1 | 12124 | 2 | -0.19377 |
| IL1RAPL2 | 6 | 0.070897 | 0.18867 | 0.969543 | 3871 | 2 | 0.37912 | 0.34362 | 0.60673 | 1 | 12125 | 3 | 0.37912 |
| PROX1 | 6 | 0.057458 | 0.16026 | 0.967126 | 3311 | 4 | -1.5176 | 0.34362 | 0.60673 | 1 | 12126 | 1 | -1.5176 |
| VANGL1 | 6 | 0.022011 | 0.078411 | 0.93914 | 1585 | 4 | -2.0203 | 0.34362 | 0.60673 | 1 | 12127 | 2 | -2.0203 |
| ARV1 | 6 | 0.17255 | 0.3539 | 0.999994 | 7048 | 3 | 0.066689 | 0.34362 | 0.60673 | 1 | 12128 | 3 | 0.066689 |
| hsa-mir-508 | 4 | 0.22688 | 0.36827 | 0.999994 | 8448 | 2 | 0.78566 | 0.34375 | 0.54648 | 1 | 12129 | 2 | 0.78566 |
| hsa-mir-335 | 4 | 0.67515 | 0.70855 | 0.999994 | 16004 | 1 | 1.4464 | 0.34392 | 0.54661 | 1 | 12130 | 2 | 1.4464 |
| hsa-mir-548ap | 3 | 0.79452 | 0.79438 | 0.999994 | 17403 | 0 | 1.45 | 0.34396 | 0.49586 | 1 | 12131 | 2 | 1.45 |
| DDX21 | 3 | 0.26384 | 0.39208 | 0.999994 | 9331 | 1 | 1.1804 | 0.344 | 0.49591 | 1 | 12132 | 2 | 1.1804 |
| HESX1 | 4 | 0.4077 | 0.55361 | 0.999994 | 12072 | 1 | 0.50766 | 0.34405 | 0.54672 | 1 | 12133 | 2 | 0.50766 |
| PRSS2 | 4 | 0.045806 | 0.11324 | 0.951052 | 2796 | 2 | -0.53005 | 0.34405 | 0.54672 | 1 | 12134 | 2 | -0.53005 |
| NTMT1 | 3 | 0.71257 | 0.71288 | 0.999994 | 16426 | 1 | 1.4691 | 0.34406 | 0.49597 | 1 | 12135 | 2 | 1.4691 |
| POU2F3 | 6 | 0.3918 | 0.5971 | 0.999994 | 11827 | 1 | 0.18828 | 0.34415 | 0.60717 | 1 | 12136 | 3 | 0.18828 |
| P2RX5 | 5 | 0.22953 | 0.41657 | 0.999994 | 8517 | 1 | 0.57444 | 0.34416 | 0.57155 | 1 | 12137 | 3 | 0.57444 |
| ZMYM4 | 5 | 0.14718 | 0.29854 | 0.999994 | 6382 | 2 | 0.76568 | 0.34416 | 0.57155 | 1 | 12138 | 3 | 0.76568 |
| BTBD9 | 5 | 0.94844 | 0.94865 | 0.999994 | 20299 | 0 | 1.3151 | 0.34427 | 0.57167 | 1 | 12139 | 3 | 1.3151 |
| ZNF425 | 6 | 0.92426 | 0.92442 | 0.999994 | 19725 | 1 | 1.0037 | 0.34441 | 0.60739 | 1 | 12140 | 3 | 1.0037 |
| MGAT4B | 5 | 0.29386 | 0.47604 | 0.999994 | 9994 | 2 | 1.218 | 0.34443 | 0.57182 | 1 | 12141 | 3 | 1.218 |
| FAR2 | 3 | 0.61451 | 0.62838 | 0.999994 | 15135 | 1 | 1.1778 | 0.34452 | 0.49646 | 1 | 12142 | 2 | 1.1778 |
| hsa-mir-4506 | 3 | 0.55922 | 0.58682 | 0.999994 | 14358 | 1 | 1.5934 | 0.34458 | 0.49651 | 1 | 12143 | 2 | 1.5934 |
| PSEN1 | 6 | 0.40127 | 0.6032 | 0.999994 | 11966 | 2 | 0.12322 | 0.34465 | 0.60757 | 1 | 12144 | 2 | 0.12322 |
| UGGT2 | 6 | 0.091776 | 0.23108 | 0.983933 | 4669 | 3 | -1.4699 | 0.34465 | 0.60757 | 1 | 12145 | 2 | -1.4699 |
| ERV3-1 | 6 | 0.49398 | 0.66456 | 0.999994 | 13326 | 2 | 0.53574 | 0.34465 | 0.60757 | 1 | 12146 | 3 | 0.53574 |
| RCC2 | 6 | 0.30155 | 0.50853 | 0.999994 | 10153 | 1 | 0.29953 | 0.34465 | 0.60757 | 1 | 12147 | 2 | 0.29953 |
| GPS1 | 6 | 0.012145 | 0.046202 | 0.892434 | 1002 | 3 | -1.7645 | 0.34465 | 0.60757 | 1 | 12148 | 3 | -1.7645 |
| TRIM10 | 6 | 0.30903 | 0.51725 | 0.999994 | 10298 | 3 | -0.65413 | 0.34465 | 0.60757 | 1 | 12149 | 2 | -0.65413 |
| MTRNR2L4 | 5 | 0.23492 | 0.42401 | 0.999994 | 8641 | 2 | 1.1367 | 0.34469 | 0.5721 | 1 | 12150 | 3 | 1.1367 |
| NT5C | 6 | 0.24863 | 0.44631 | 0.999994 | 8985 | 2 | 0.85736 | 0.3447 | 0.60761 | 1 | 12151 | 3 | 0.85736 |
| MXD3 | 5 | 0.23463 | 0.42362 | 0.999994 | 8631 | 3 | -0.67607 | 0.34473 | 0.57214 | 1 | 12152 | 2 | -0.67607 |
| HS6ST2 | 3 | 0.21043 | 0.32576 | 0.999994 | 8032 | 2 | -1.0228 | 0.34478 | 0.49673 | 1 | 12153 | 1 | -1.0228 |
| EML4 | 5 | 0.33112 | 0.50646 | 0.999994 | 10746 | 1 | -0.092997 | 0.3449 | 0.57232 | 1 | 12154 | 2 | -0.092997 |
| ARHGEF4 | 5 | 0.66565 | 0.75147 | 0.999994 | 15866 | 1 | 0.2348 | 0.3449 | 0.57232 | 1 | 12155 | 2 | 0.2348 |
| CYP4A11 | 5 | 0.91831 | 0.9184 | 0.999994 | 19591 | 0 | 0.97573 | 0.3449 | 0.57232 | 1 | 12156 | 3 | 0.97573 |
| ABCD4 | 5 | 0.16461 | 0.32421 | 0.999994 | 6853 | 3 | -1.0493 | 0.34491 | 0.57233 | 1 | 12157 | 2 | -1.0493 |
| hsa-mir-1297 | 2 | 0.29871 | 0.33588 | 0.999994 | 10106 | 1 | 1.248 | 0.3452 | 0.43291 | 1 | 12158 | 1 | 1.248 |
| GALNT3 | 6 | 0.15634 | 0.33363 | 0.999994 | 6637 | 2 | 0.40856 | 0.34521 | 0.60803 | 1 | 12159 | 3 | 0.40856 |
| CLASRP | 4 | 0.49536 | 0.61425 | 0.999994 | 13344 | 1 | 1.2099 | 0.34524 | 0.5476 | 1 | 12160 | 2 | 1.2099 |
| IL9 | 4 | 0.75326 | 0.7626 | 0.999994 | 16890 | 1 | 0.77861 | 0.34556 | 0.54783 | 1 | 12161 | 2 | 0.77861 |
| hsa-mir-4664 | 4 | 0.76997 | 0.77575 | 0.999994 | 17079 | 1 | -0.14136 | 0.34556 | 0.54783 | 1 | 12162 | 1 | -0.14136 |
| hsa-mir-4481 | 3 | 0.85612 | 0.85607 | 0.999994 | 18376 | 0 | 1.2914 | 0.34559 | 0.49755 | 1 | 12163 | 2 | 1.2914 |
| MUC7 | 5 | 0.17439 | 0.33852 | 0.999994 | 7092 | 3 | -1.009 | 0.3456 | 0.57303 | 1 | 12164 | 2 | -1.009 |
| ATRIP | 5 | 0.096949 | 0.22163 | 0.983093 | 4857 | 3 | -1.568 | 0.3456 | 0.57303 | 1 | 12165 | 2 | -1.568 |
| RAD50 | 5 | 0.064069 | 0.16876 | 0.967126 | 3617 | 2 | -0.28609 | 0.3456 | 0.57303 | 1 | 12166 | 1 | -0.28609 |
| SLC44A1 | 5 | 0.078454 | 0.19207 | 0.974607 | 4170 | 1 | -0.12632 | 0.3456 | 0.57303 | 1 | 12167 | 2 | -0.12632 |

| | | | | | | | | | | | | |
|---|---|---|---|---|---|---|---|---|---|---|---|---|
| SYT1 | 5 | 0.26888 | 0.45594 | 0.999994 | 9442 | 2 | 0.20086 | 0.3456 | 0.57303 | 1 | 12168 | 2 | 0.20086 |
| NonTargetingControlG | 1 | 0.65438 | 0.65443 | 0.999994 | 15711 | 0 | 1.774 | 0.34562 | 0.34584 | 0.994392 | 12169 | 1 | 1.774 |
| MS4A6E | 6 | 0.1888 | 0.37402 | 0.999994 | 7446 | 3 | -0.47073 | 0.34562 | 0.60835 | 1 | 12170 | 2 | -0.47073 |
| ZNF791 | 6 | 0.46202 | 0.64308 | 0.999994 | 12887 | 2 | 1.1732 | 0.34562 | 0.60836 | 1 | 12171 | 3 | 1.1732 |
| FADS6 | 5 | 0.53493 | 0.67839 | 0.999994 | 13978 | 2 | 1.2422 | 0.34564 | 0.57306 | 1 | 12172 | 3 | 1.2422 |
| CREB3L4 | 6 | 0.25786 | 0.45735 | 0.999994 | 9192 | 3 | 0.48832 | 0.34573 | 0.60843 | 1 | 12173 | 3 | 0.48832 |
| FRMD4B | 5 | 0.68325 | 0.75898 | 0.999994 | 16133 | 1 | 1.1225 | 0.34579 | 0.5732 | 1 | 12174 | 3 | 1.1225 |
| EFCAB12 | 6 | 0.20677 | 0.39614 | 0.999994 | 7947 | 3 | 0.097445 | 0.34585 | 0.60851 | 1 | 12175 | 3 | 0.097445 |
| TTR | 6 | 0.54852 | 0.70257 | 0.999994 | 14187 | 1 | 0.70805 | 0.34598 | 0.60862 | 1 | 12176 | 3 | 0.70805 |
| CGB7 | 1 | 0.65395 | 0.65401 | 0.999994 | 15706 | 0 | 2.0052 | 0.34605 | 0.34628 | 0.994392 | 12177 | 1 | 2.0052 |
| HSD17B10 | 5 | 0.1182 | 0.25463 | 0.994835 | 5580 | 3 | -1.2958 | 0.34606 | 0.57347 | 1 | 12178 | 2 | -1.2958 |
| PDLIM1 | 6 | 0.6564 | 0.78252 | 0.999994 | 15733 | 2 | 0.57159 | 0.34606 | 0.60868 | 1 | 12179 | 3 | 0.57159 |
| AMH | 6 | 0.27184 | 0.47389 | 0.999994 | 9517 | 2 | 0.069919 | 0.34606 | 0.60868 | 1 | 12180 | 2 | 0.069919 |
| AP3S1 | 6 | 0.91048 | 0.91196 | 0.999994 | 19438 | 1 | -0.073168 | 0.34606 | 0.60868 | 1 | 12181 | 2 | -0.073168 |
| ZDHHC16 | 6 | 0.15322 | 0.32973 | 0.999994 | 6541 | 3 | -0.80018 | 0.34606 | 0.60868 | 1 | 12182 | 2 | -0.80018 |
| hsa-mir-3179-1 | 4 | 0.39494 | 0.54049 | 0.999994 | 11872 | 2 | 0.79311 | 0.34618 | 0.54831 | 1 | 12183 | 2 | 0.79311 |
| SH2D2A | 3 | 0.43483 | 0.50496 | 0.999994 | 12492 | 1 | -0.045195 | 0.34648 | 0.49847 | 1 | 12184 | 1 | -0.045195 |
| hsa-mir-3147 | 3 | 0.015001 | 0.039309 | 0.891621 | 1189 | 2 | -1.4488 | 0.34648 | 0.49847 | 1 | 12185 | 1 | -1.4488 |
| FGF17 | 5 | 0.4939 | 0.64264 | 0.999994 | 13324 | 2 | 0.50944 | 0.34654 | 0.57394 | 1 | 12186 | 2 | 0.50944 |
| SAC3D1 | 6 | 0.41139 | 0.60979 | 0.999994 | 12137 | 2 | 0.33213 | 0.34655 | 0.60906 | 1 | 12187 | 3 | 0.33213 |
| ANKRD9 | 6 | 0.53243 | 0.69126 | 0.999994 | 13941 | 2 | 0.43296 | 0.34655 | 0.60906 | 1 | 12188 | 3 | 0.43296 |
| PNOC | 6 | 0.31855 | 0.52824 | 0.999994 | 10487 | 3 | -0.51714 | 0.34655 | 0.60906 | 1 | 12189 | 2 | -0.51714 |
| SUCNR1 | 6 | 0.24031 | 0.43637 | 0.999994 | 8778 | 3 | 0.34734 | 0.34655 | 0.60906 | 1 | 12190 | 3 | 0.34734 |
| TMEM120B | 6 | 0.060026 | 0.16579 | 0.967126 | 3424 | 3 | -0.062281 | 0.34655 | 0.60906 | 1 | 12191 | 3 | -0.062281 |
| TGFBR1 | 6 | 0.089734 | 0.22699 | 0.983354 | 4582 | 2 | 0.045003 | 0.34655 | 0.60906 | 1 | 12192 | 2 | 0.045003 |
| AZGP1 | 6 | 0.082781 | 0.21301 | 0.979587 | 4329 | 3 | 0.13544 | 0.34655 | 0.60907 | 1 | 12193 | 3 | 0.13544 |
| AMDHD1 | 6 | 0.016575 | 0.06089 | 0.932932 | 1281 | 4 | -1.5672 | 0.34662 | 0.60911 | 1 | 12194 | 2 | -1.5672 |
| ALS2CR11 | 6 | 0.25977 | 0.45958 | 0.999994 | 9233 | 3 | -0.66959 | 0.34662 | 0.60911 | 1 | 12195 | 1 | -0.66959 |
| FCGR3A | 5 | 0.8582 | 0.86089 | 0.999994 | 18408 | 1 | -0.23131 | 0.34667 | 0.57407 | 1 | 12196 | 2 | -0.23131 |
| FAM180B | 6 | 0.057057 | 0.15942 | 0.967126 | 3294 | 2 | 1.0262 | 0.34674 | 0.60921 | 1 | 12197 | 3 | 1.0262 |
| hsa-mir-656 | 4 | 0.35548 | 0.49985 | 0.999994 | 11258 | 2 | -0.13121 | 0.34683 | 0.54878 | 1 | 12198 | 2 | -0.13121 |
| LARP4 | 5 | 0.66596 | 0.7516 | 0.999994 | 15868 | 1 | 0.31995 | 0.347 | 0.57441 | 1 | 12199 | 2 | 0.31995 |
| KIAA2013 | 5 | 0.51851 | 0.66402 | 0.999994 | 13716 | 2 | 0.30771 | 0.347 | 0.57441 | 1 | 12200 | 2 | 0.30771 |
| SYMPK | 5 | 0.43327 | 0.59116 | 0.999994 | 12467 | 2 | 0.53374 | 0.347 | 0.57441 | 1 | 12201 | 2 | 0.53374 |
| hsa-mir-449c | 3 | 0.42527 | 0.49926 | 0.999994 | 12338 | 1 | 1.2751 | 0.34701 | 0.49901 | 1 | 12202 | 2 | 1.2751 |
| hsa-mir-125b-2 | 4 | 0.85504 | 0.85464 | 0.999994 | 18348 | 0 | 1.3376 | 0.34716 | 0.54901 | 1 | 12203 | 2 | 1.3376 |
| ASNSD1 | 6 | 0.46368 | 0.64422 | 0.999994 | 12915 | 1 | 1.0756 | 0.34727 | 0.60964 | 1 | 12204 | 3 | 1.0756 |
| C1orf95 | 4 | 0.15181 | 0.29105 | 0.999632 | 6501 | 2 | -1.2337 | 0.34729 | 0.5491 | 1 | 12205 | 1 | -1.2337 |
| NonTargetingControlG | 2 | 0.44827 | 0.45777 | 0.999994 | 12691 | 1 | 1.8896 | 0.34736 | 0.43428 | 1 | 12206 | 1 | 1.8896 |
| USPL1 | 5 | 0.089888 | 0.21043 | 0.979587 | 4587 | 2 | 1.3465 | 0.3474 | 0.57481 | 1 | 12207 | 3 | 1.3465 |
| hsa-mir-130a | 4 | 0.39564 | 0.54125 | 0.999994 | 11885 | 1 | 1.2498 | 0.34752 | 0.54928 | 1 | 12208 | 2 | 1.2498 |
| MYO18A | 5 | 0.47171 | 0.62357 | 0.999994 | 13025 | 2 | 0.35554 | 0.34755 | 0.57495 | 1 | 12209 | 2 | 0.35554 |
| PF4V1 | 5 | 0.11916 | 0.25608 | 0.994943 | 5605 | 2 | -0.3381 | 0.34755 | 0.57495 | 1 | 12210 | 2 | -0.3381 |
| MBTD1 | 6 | 0.0011529 | 0.0058128 | 0.623123 | 184 | 5 | -6.1748 | 0.34775 | 0.61 | 1 | 12211 | 1 | -6.1748 |
| hsa-mir-203a | 3 | 0.83257 | 0.83244 | 0.999994 | 17966 | 0 | 1.2532 | 0.34778 | 0.49981 | 1 | 12212 | 2 | 1.2532 |
| MTX1 | 4 | 0.012369 | 0.041086 | 0.891621 | 1020 | 2 | -1.6861 | 0.34782 | 0.5495 | 1 | 12213 | 1 | -1.6861 |
| FANCC | 6 | 0.068663 | 0.18399 | 0.968526 | 3779 | 2 | 1.0839 | 0.34788 | 0.6101 | 1 | 12214 | 3 | 1.0839 |
| GPR19 | 5 | 0.69944 | 0.76621 | 0.999994 | 16302 | 1 | -0.15744 | 0.34793 | 0.57534 | 1 | 12215 | 2 | -0.15744 |
| JOSD2 | 5 | 0.47647 | 0.62759 | 0.999994 | 13089 | 1 | 1.1723 | 0.348 | 0.5754 | 1 | 12216 | 3 | 1.1723 |
| RAB11FIP2 | 6 | 0.057114 | 0.15954 | 0.967126 | 3297 | 2 | 0.6242 | 0.34801 | 0.6102 | 1 | 12217 | 3 | 0.6242 |
| TNFSF13B | 6 | 0.92956 | 0.92969 | 0.999994 | 19844 | 0 | 0.99455 | 0.34805 | 0.61024 | 1 | 12218 | 3 | 0.99455 |
| GHSR | 6 | 0.67291 | 0.79542 | 0.999994 | 15970 | 1 | 0.83315 | 0.34807 | 0.61025 | 1 | 12219 | 3 | 0.83315 |

| | | | | | | | | | | | | | |
|---|---|---|---|---|---|---|---|---|---|---|---|---|---|
| NonTargetingControlG | 1 | 0.65171 | 0.65175 | 0.999994 | 15684 | 0 | 1.9344 | 0.34829 | 0.34848 | 0.994392 | 12220 | 1 | 1.9344 |
| hsa-mir-1275 | 3 | 0.47305 | 0.52869 | 0.999994 | 13045 | 1 | 1.2538 | 0.34832 | 0.50034 | 1 | 12221 | 2 | 1.2538 |
| LRR1 | 4 | 0.17467 | 0.31462 | 0.999994 | 7103 | 2 | 0.066278 | 0.34835 | 0.54989 | 1 | 12222 | 2 | 0.066278 |
| hsa-mir-454 | 3 | 0.88286 | 0.88296 | 0.999994 | 18891 | 0 | 1.2787 | 0.34839 | 0.50041 | 1 | 12223 | 2 | 1.2787 |
| MFAP4 | 6 | 0.66172 | 0.78662 | 0.999994 | 15807 | 2 | -0.19502 | 0.34844 | 0.61054 | 1 | 12224 | 2 | -0.19502 |
| CECR2 | 6 | 0.16464 | 0.34402 | 0.999994 | 6856 | 2 | -0.13664 | 0.34844 | 0.61054 | 1 | 12225 | 2 | -0.13664 |
| GHR | 6 | 0.35025 | 0.56426 | 0.999994 | 11165 | 3 | -0.61001 | 0.34844 | 0.61054 | 1 | 12226 | 2 | -0.61001 |
| LASP1 | 6 | 0.061158 | 0.16824 | 0.967126 | 3479 | 2 | 0.90581 | 0.34855 | 0.61062 | 1 | 12227 | 3 | 0.90581 |
| MUC17 | 5 | 0.88404 | 0.88416 | 0.999994 | 18907 | 1 | -0.40617 | 0.34855 | 0.57597 | 1 | 12228 | 2 | -0.40617 |
| FCER1G | 5 | 0.74393 | 0.78768 | 0.999994 | 16787 | 1 | 0.2264 | 0.34855 | 0.57597 | 1 | 12229 | 2 | 0.2264 |
| VSX1 | 5 | 0.42271 | 0.58233 | 0.999994 | 12292 | 2 | 1.2209 | 0.34865 | 0.57606 | 1 | 12230 | 3 | 1.2209 |
| SLC10A3 | 5 | 0.093369 | 0.21596 | 0.980699 | 4734 | 2 | 1.3356 | 0.34872 | 0.57612 | 1 | 12231 | 3 | 1.3356 |
| HIST3H3 | 6 | 0.12684 | 0.2963 | 0.999994 | 5843 | 3 | -0.056376 | 0.34877 | 0.61079 | 1 | 12232 | 3 | -0.056376 |
| GPBP1 | 4 | 0.57543 | 0.65241 | 0.999994 | 14622 | 1 | 0.98874 | 0.34883 | 0.55024 | 1 | 12233 | 2 | 0.98874 |
| hsa-mir-5694 | 4 | 0.74576 | 0.75684 | 0.999994 | 16811 | 1 | -0.31653 | 0.34883 | 0.55024 | 1 | 12234 | 1 | -0.31653 |
| EDAR | 5 | 0.1559 | 0.31155 | 0.999994 | 6623 | 2 | 1.1236 | 0.3489 | 0.57632 | 1 | 12235 | 3 | 1.1236 |
| KIF19 | 3 | 0.84373 | 0.84373 | 0.999994 | 18152 | 0 | 1.187 | 0.34899 | 0.50103 | 1 | 12236 | 2 | 1.187 |
| BHLHB9 | 5 | 0.58209 | 0.71974 | 0.999994 | 14706 | 1 | 1.0026 | 0.34905 | 0.57647 | 1 | 12237 | 3 | 1.0026 |
| CCNB1IP1 | 5 | 0.3125 | 0.49121 | 0.999994 | 10371 | 1 | 0.93623 | 0.34913 | 0.57655 | 1 | 12238 | 3 | 0.93623 |
| MBD2 | 5 | 0.87788 | 0.87825 | 0.999994 | 18785 | 1 | -0.39221 | 0.34913 | 0.57655 | 1 | 12239 | 2 | -0.39221 |
| NREP | 5 | 0.011567 | 0.040572 | 0.891621 | 961 | 2 | -0.29338 | 0.34913 | 0.57655 | 1 | 12240 | 2 | -0.29338 |
| RFC4 | 3 | 0.42861 | 0.50125 | 0.999994 | 12389 | 1 | 0.085086 | 0.34918 | 0.50122 | 1 | 12241 | 1 | 0.085086 |
| GLRA4 | 5 | 0.86549 | 0.86708 | 0.999994 | 18549 | 1 | 1.132 | 0.3492 | 0.57662 | 1 | 12242 | 3 | 1.132 |
| BST1 | 5 | 0.013962 | 0.047263 | 0.896226 | 1125 | 4 | -2.6054 | 0.34926 | 0.57667 | 1 | 12243 | 1 | -2.6054 |
| AGFG1 | 5 | 0.056363 | 0.15566 | 0.967126 | 3262 | 3 | -1.9419 | 0.34926 | 0.57667 | 1 | 12244 | 2 | -1.9419 |
| CLDN19 | 5 | 0.10523 | 0.23455 | 0.984971 | 5141 | 2 | 0.15613 | 0.34926 | 0.57667 | 1 | 12245 | 1 | 0.15613 |
| ZMAT3 | 4 | 0.82277 | 0.82244 | 0.999994 | 17819 | 1 | 1.3677 | 0.34927 | 0.55055 | 1 | 12246 | 2 | 1.3677 |
| IFI44L | 6 | 0.27222 | 0.47435 | 0.999994 | 9523 | 3 | -0.46273 | 0.34935 | 0.61124 | 1 | 12247 | 1 | -0.46273 |
| NRG3 | 6 | 0.14081 | 0.31406 | 0.999994 | 6212 | 3 | -0.99037 | 0.34935 | 0.61124 | 1 | 12248 | 2 | -0.99037 |
| COL4A5 | 6 | 0.62559 | 0.75902 | 0.999994 | 15319 | 1 | 0.99985 | 0.34935 | 0.61124 | 1 | 12249 | 3 | 0.99985 |
| ZC3H12A | 6 | 0.63437 | 0.76566 | 0.999994 | 15436 | 2 | -0.23562 | 0.34935 | 0.61124 | 1 | 12250 | 1 | -0.23562 |
| TRPC5OS | 6 | 0.33737 | 0.54968 | 0.999994 | 10882 | 3 | 0.54627 | 0.34936 | 0.61126 | 1 | 12251 | 3 | 0.54627 |
| KRTAP4-8 | 4 | 0.053122 | 0.12756 | 0.951052 | 3121 | 3 | -2.0711 | 0.34954 | 0.55076 | 1 | 12252 | 1 | -2.0711 |
| GPR88 | 4 | 0.80031 | 0.80151 | 0.999994 | 17482 | 1 | 1.1438 | 0.3496 | 0.55081 | 1 | 12253 | 2 | 1.1438 |
| PRDX5 | 5 | 0.0083399 | 0.031287 | 0.862223 | 751 | 1 | 1.0323 | 0.34964 | 0.57707 | 1 | 12254 | 3 | 1.0323 |
| CLDN17 | 5 | 0.045829 | 0.13036 | 0.95318 | 2798 | 2 | 1.1196 | 0.3497 | 0.57714 | 1 | 12255 | 3 | 1.1196 |
| NARFL | 5 | 0.26001 | 0.44871 | 0.999994 | 9242 | 2 | 1.0525 | 0.34972 | 0.57716 | 1 | 12256 | 3 | 1.0525 |
| CHD7 | 6 | 0.28086 | 0.48441 | 0.999994 | 9716 | 2 | 0.36294 | 0.34977 | 0.61157 | 1 | 12257 | 3 | 0.36294 |
| PPFIA4 | 6 | 0.64111 | 0.77077 | 0.999994 | 15531 | 2 | 1.1932 | 0.34982 | 0.61161 | 1 | 12258 | 3 | 1.1932 |
| SLC38A5 | 5 | 0.56293 | 0.70323 | 0.999994 | 14427 | 2 | -0.14539 | 0.34994 | 0.57739 | 1 | 12259 | 2 | -0.14539 |
| CREBBP | 5 | 0.019753 | 0.063429 | 0.937389 | 1443 | 4 | -2.7569 | 0.34994 | 0.57739 | 1 | 12260 | 1 | -2.7569 |
| AICDA | 5 | 0.69554 | 0.76446 | 0.999994 | 16263 | 1 | -0.30419 | 0.34994 | 0.57739 | 1 | 12261 | 1 | -0.30419 |
| PLEKHA4 | 6 | 0.38485 | 0.59264 | 0.999994 | 11714 | 2 | 0.74843 | 0.34998 | 0.61174 | 1 | 12262 | 3 | 0.74843 |
| PLEKHG1 | 6 | 0.14494 | 0.31929 | 0.999994 | 6324 | 3 | -0.32714 | 0.35 | 0.61175 | 1 | 12263 | 2 | -0.32714 |
| hsa-mir-6765 | 4 | 0.63942 | 0.68718 | 0.999994 | 15507 | 1 | 0.62965 | 0.35021 | 0.55127 | 1 | 12264 | 2 | 0.62965 |
| SMCR9 | 5 | 0.19829 | 0.37282 | 0.999994 | 7722 | 1 | -0.20476 | 0.35033 | 0.5778 | 1 | 12265 | 2 | -0.20476 |
| WDR27 | 5 | 0.56582 | 0.70585 | 0.999994 | 14480 | 1 | 1.1377 | 0.35034 | 0.57781 | 1 | 12266 | 3 | 1.1377 |
| SBF2 | 4 | 0.73481 | 0.74866 | 0.999994 | 16658 | 1 | 1.3927 | 0.35048 | 0.55148 | 1 | 12267 | 2 | 1.3927 |
| GATSL3 | 6 | 0.17728 | 0.35969 | 0.999994 | 7161 | 3 | 0.48444 | 0.3505 | 0.61213 | 1 | 12268 | 3 | 0.48444 |
| ATG12 | 6 | 0.14669 | 0.32151 | 0.999994 | 6366 | 2 | 0.99366 | 0.3506 | 0.6122 | 1 | 12269 | 3 | 0.99366 |
| IL12RB2 | 6 | 0.088231 | 0.224 | 0.983354 | 4522 | 3 | -0.18399 | 0.3506 | 0.6122 | 1 | 12270 | 3 | -0.18399 |
| OBSL1 | 6 | 0.22524 | 0.4184 | 0.999994 | 8405 | 2 | 1.2104 | 0.35062 | 0.61221 | 1 | 12271 | 3 | 1.2104 |

| | | | | | | | | | | | | |
|---|---|---|---|---|---|---|---|---|---|---|---|---|
| FAAH | 5 | 0.91103 | 0.91105 | 0.999994 | 19448 | 0 | 0.60578 | 0.35064 | 0.57815 | 1 | 12272 | 2 | 0.60578 |
| CCDC137 | 6 | 0.2995 | 0.50612 | 0.999994 | 10120 | 3 | -0.02755 | 0.35069 | 0.61227 | 1 | 12273 | 3 | -0.02755 |
| hsa-mir-3143 | 3 | 0.20743 | 0.32202 | 0.999994 | 7959 | 2 | -1.1208 | 0.35074 | 0.50284 | 1 | 12274 | 1 | -1.1208 |
| hsa-mir-4538 | 3 | 0.10097 | 0.18436 | 0.968526 | 5005 | 2 | -2.4069 | 0.35074 | 0.50284 | 1 | 12275 | 1 | -2.4069 |
| HIST1H2BG | 5 | 0.21705 | 0.39915 | 0.999994 | 8195 | 2 | 0.25417 | 0.35079 | 0.57828 | 1 | 12276 | 2 | 0.25417 |
| NARS2 | 4 | 0.32841 | 0.47206 | 0.999994 | 10691 | 2 | 0.25685 | 0.35095 | 0.55184 | 1 | 12277 | 2 | 0.25685 |
| CABIN1 | 4 | 0.29785 | 0.44081 | 0.999994 | 10088 | 1 | 1.1111 | 0.35098 | 0.55187 | 1 | 12278 | 2 | 1.1111 |
| LRRN4 | 5 | 0.40586 | 0.56843 | 0.999994 | 12043 | 1 | 1.329 | 0.35101 | 0.57851 | 1 | 12279 | 3 | 1.329 |
| RBP7 | 5 | 0.45588 | 0.61019 | 0.999994 | 12789 | 2 | -0.48079 | 0.35123 | 0.57873 | 1 | 12280 | 2 | -0.48079 |
| ZC2HC1B | 6 | 0.18378 | 0.36785 | 0.999994 | 7311 | 2 | 1.1586 | 0.35128 | 0.61275 | 1 | 12281 | 3 | 1.1586 |
| DRD1 | 5 | 0.079827 | 0.19431 | 0.975797 | 4227 | 3 | -1.8363 | 0.35134 | 0.57884 | 1 | 12282 | 2 | -1.8363 |
| COL16A1 | 6 | 0.14607 | 0.32072 | 0.999994 | 6353 | 3 | -0.4331 | 0.35134 | 0.61279 | 1 | 12283 | 2 | -0.4331 |
| INHBE | 6 | 0.0021798 | 0.01064 | 0.706752 | 288 | 4 | -5.1126 | 0.35134 | 0.61279 | 1 | 12284 | 1 | -5.1126 |
| OCLN | 4 | 0.84151 | 0.84094 | 0.999994 | 18113 | 0 | 1.5265 | 0.35163 | 0.55234 | 1 | 12285 | 2 | 1.5265 |
| FXN | 3 | 0.44556 | 0.51147 | 0.999994 | 12657 | 1 | 1.3703 | 0.35168 | 0.50387 | 1 | 12286 | 2 | 1.3703 |
| KBTBD2 | 6 | 0.090289 | 0.22806 | 0.983354 | 4604 | 4 | -1.1742 | 0.35178 | 0.61311 | 1 | 12287 | 2 | -1.1742 |
| ZCCHC9 | 6 | 0.29714 | 0.50339 | 0.999994 | 10072 | 3 | -0.54623 | 0.35185 | 0.61316 | 1 | 12288 | 2 | -0.54623 |
| MOGAT2 | 5 | 0.81935 | 0.83216 | 0.999994 | 17772 | 1 | 0.8611 | 0.35192 | 0.57943 | 1 | 12289 | 3 | 0.8611 |
| MON2 | 6 | 0.21408 | 0.40506 | 0.999994 | 8118 | 2 | 0.97497 | 0.35193 | 0.61322 | 1 | 12290 | 3 | 0.97497 |
| TRIM2 | 5 | 0.044695 | 0.12755 | 0.951052 | 2746 | 2 | 1.0283 | 0.35193 | 0.57944 | 1 | 12291 | 3 | 1.0283 |
| OR5F1 | 6 | 0.32204 | 0.53224 | 0.999994 | 10557 | 3 | -0.30121 | 0.35197 | 0.61324 | 1 | 12292 | 2 | -0.30121 |
| hsa-mir-1303 | 2 | 0.13368 | 0.21533 | 0.980018 | 6015 | 1 | -1.1668 | 0.35198 | 0.43719 | 1 | 12293 | 1 | -1.1668 |
| MTFR1 | 6 | 0.12519 | 0.29419 | 0.999994 | 5799 | 3 | -0.81144 | 0.35199 | 0.61326 | 1 | 12294 | 2 | -0.81144 |
| MRC2 | 6 | 0.61733 | 0.75281 | 0.999994 | 15175 | 2 | -0.35835 | 0.35205 | 0.61332 | 1 | 12295 | 2 | -0.35835 |
| TMEM98 | 6 | 0.6141 | 0.75041 | 0.999994 | 15133 | 2 | 0.3407 | 0.35205 | 0.61332 | 1 | 12296 | 3 | 0.3407 |
| STAG1 | 6 | 0.31144 | 0.52003 | 0.999994 | 10348 | 3 | 0.59279 | 0.35206 | 0.61332 | 1 | 12297 | 3 | 0.59279 |
| PLA2G2E | 6 | 0.13601 | 0.30798 | 0.999994 | 6078 | 2 | 0.57515 | 0.3521 | 0.61335 | 1 | 12298 | 3 | 0.57515 |
| PLAU | 5 | 0.079468 | 0.19371 | 0.975797 | 4206 | 3 | -1.4967 | 0.35211 | 0.57962 | 1 | 12299 | 2 | -1.4967 |
| EIF2AK4 | 5 | 0.7508 | 0.79128 | 0.999994 | 16863 | 1 | 1.0243 | 0.35227 | 0.57979 | 1 | 12300 | 3 | 1.0243 |
| ZXDB | 6 | 0.32353 | 0.5339 | 0.999994 | 10583 | 2 | 0.7058 | 0.35229 | 0.61352 | 1 | 12301 | 3 | 0.7058 |
| hsa-mir-1470 | 4 | 0.14805 | 0.28717 | 0.999632 | 6410 | 2 | 0.030795 | 0.35238 | 0.55287 | 1 | 12302 | 2 | 0.030795 |
| KIF18B | 4 | 0.11519 | 0.24192 | 0.987253 | 5484 | 2 | 0.18711 | 0.35238 | 0.55287 | 1 | 12303 | 2 | 0.18711 |
| CARD14 | 4 | 0.25812 | 0.40019 | 0.999994 | 9197 | 2 | 0.57962 | 0.35238 | 0.55287 | 1 | 12304 | 2 | 0.57962 |
| TNS4 | 6 | 0.11157 | 0.26973 | 0.998561 | 5352 | 2 | 1.1 | 0.35254 | 0.61371 | 1 | 12305 | 3 | 1.1 |
| hsa-mir-6848 | 4 | 0.091101 | 0.19906 | 0.977164 | 4642 | 2 | -1.76 | 0.35254 | 0.55299 | 1 | 12306 | 1 | -1.76 |
| hsa-mir-3183 | 4 | 0.57035 | 0.64982 | 0.999994 | 14548 | 1 | 1.5692 | 0.35254 | 0.55299 | 1 | 12307 | 2 | 1.5692 |
| RASGRP4 | 5 | 0.86802 | 0.86934 | 0.999994 | 18601 | 1 | 1.0799 | 0.35257 | 0.58011 | 1 | 12308 | 3 | 1.0799 |
| DNAH8 | 5 | 0.34284 | 0.51605 | 0.999994 | 11004 | 2 | 1.3182 | 0.35264 | 0.58017 | 1 | 12309 | 3 | 1.3182 |
| UBE2E3 | 6 | 0.29994 | 0.50665 | 0.999994 | 10128 | 3 | 0.53901 | 0.35273 | 0.61387 | 1 | 12310 | 3 | 0.53901 |
| S100Z | 5 | 0.15633 | 0.31216 | 0.999994 | 6636 | 3 | -1.0691 | 0.35274 | 0.58028 | 1 | 12311 | 1 | -1.0691 |
| OR4D1 | 5 | 0.36894 | 0.53747 | 0.999994 | 11489 | 2 | 1.101 | 0.35279 | 0.58034 | 1 | 12312 | 3 | 1.101 |
| PCYT2 | 6 | 0.0015773 | 0.0078931 | 0.680529 | 231 | 3 | -4.218 | 0.35286 | 0.61397 | 1 | 12313 | 1 | -4.218 |
| ATP8B3 | 6 | 0.025659 | 0.088569 | 0.94395 | 1783 | 2 | 0.35587 | 0.35286 | 0.61397 | 1 | 12314 | 3 | 0.35587 |
| ELOVL6 | 6 | 0.2708 | 0.47267 | 0.999994 | 9493 | 3 | -0.19206 | 0.35286 | 0.61397 | 1 | 12315 | 2 | -0.19206 |
| SORT1 | 6 | 0.19244 | 0.37855 | 0.999994 | 7558 | 3 | -0.59493 | 0.35286 | 0.61397 | 1 | 12316 | 2 | -0.59493 |
| NonTargetingControlG | 2 | 0.67538 | 0.6751 | 0.999994 | 16009 | 0 | 2.5281 | 0.35299 | 0.4378 | 1 | 12317 | 1 | 2.5281 |
| PDS5A | 5 | 0.22364 | 0.40837 | 0.999994 | 8373 | 1 | -0.2273 | 0.35307 | 0.5806 | 1 | 12318 | 2 | -0.2273 |
| S1PR2 | 6 | 0.7087 | 0.81468 | 0.999994 | 16392 | 1 | 0.38068 | 0.35321 | 0.61423 | 1 | 12319 | 2 | 0.38068 |
| MYL9 | 6 | 0.0041323 | 0.017953 | 0.795727 | 453 | 4 | -3.4685 | 0.35321 | 0.61423 | 1 | 12320 | 2 | -3.4685 |
| SDCCAG3 | 5 | 0.20627 | 0.3841 | 0.999994 | 7939 | 2 | 1.0348 | 0.35321 | 0.58074 | 1 | 12321 | 3 | 1.0348 |
| TMTC4 | 6 | 0.5521 | 0.70517 | 0.999994 | 14245 | 2 | 0.31315 | 0.35353 | 0.6145 | 1 | 12322 | 1 | 0.31315 |
| FAM189B | 6 | 0.21878 | 0.41065 | 0.999994 | 8253 | 3 | -0.75201 | 0.35353 | 0.6145 | 1 | 12323 | 2 | -0.75201 |

| | | | | | | | | | | | | | |
|---|---|---|---|---|---|---|---|---|---|---|---|---|---|
| NCAM2 | 6 | 0.18745 | 0.37235 | 0.999994 | 7406 | 3 | -0.59651 | 0.35353 | 0.6145 | 1 | 12324 | 2 | -0.59651 |
| ZNF645 | 6 | 0.14257 | 0.31632 | 0.999994 | 6255 | 3 | -0.88076 | 0.35353 | 0.6145 | 1 | 12325 | 2 | -0.88076 |
| SESN3 | 6 | 0.090283 | 0.22805 | 0.983354 | 4602 | 2 | -0.10235 | 0.35353 | 0.6145 | 1 | 12326 | 2 | -0.10235 |
| GOLGB1 | 5 | 0.15652 | 0.31245 | 0.999994 | 6642 | 2 | 0.15205 | 0.35354 | 0.58108 | 1 | 12327 | 2 | 0.15205 |
| ARMCX5 | 5 | 0.89297 | 0.8931 | 0.999994 | 19091 | 0 | 0.22417 | 0.35354 | 0.58108 | 1 | 12328 | 2 | 0.22417 |
| OIP5 | 5 | 0.29353 | 0.47578 | 0.999994 | 9986 | 2 | 0.33466 | 0.35359 | 0.58112 | 1 | 12329 | 2 | 0.33466 |
| NonTargetingControlG | 2 | 0.75617 | 0.75575 | 0.999994 | 16931 | 0 | 2.8183 | 0.35362 | 0.4382 | 1 | 12330 | 1 | 2.8183 |
| hsa-mir-5187 | 4 | 0.11089 | 0.23444 | 0.984971 | 5327 | 3 | -0.86469 | 0.35385 | 0.55396 | 1 | 12331 | 1 | -0.86469 |
| TPBG | 6 | 0.024302 | 0.085342 | 0.94395 | 1724 | 3 | -0.54396 | 0.35391 | 0.61479 | 1 | 12332 | 3 | -0.54396 |
| LRTM1 | 5 | 0.52285 | 0.66778 | 0.999994 | 13793 | 2 | 1.1862 | 0.35392 | 0.58147 | 1 | 12333 | 3 | 1.1862 |
| NonTargetingControlG | 1 | 0.64593 | 0.6459 | 0.999994 | 15600 | 0 | 1.8062 | 0.35407 | 0.3542 | 0.994392 | 12334 | 1 | 1.8062 |
| CYHR1 | 6 | 0.63558 | 0.76657 | 0.999994 | 15450 | 1 | 0.2021 | 0.3542 | 0.61503 | 1 | 12335 | 2 | 0.2021 |
| MACC1 | 6 | 0.45706 | 0.63976 | 0.999994 | 12805 | 1 | -0.33037 | 0.3542 | 0.61503 | 1 | 12336 | 2 | -0.33037 |
| ZNF74 | 6 | 0.30974 | 0.51806 | 0.999994 | 10322 | 3 | -0.33626 | 0.3542 | 0.61503 | 1 | 12337 | 2 | -0.33626 |
| NRK | 6 | 0.23514 | 0.43021 | 0.999994 | 8650 | 2 | 0.18649 | 0.3542 | 0.61503 | 1 | 12338 | 2 | 0.18649 |
| SSX4B | 4 | 0.65199 | 0.69443 | 0.999994 | 15688 | 1 | 1.4246 | 0.35425 | 0.55426 | 1 | 12339 | 2 | 1.4246 |
| hsa-mir-1282 | 3 | 0.49546 | 0.54317 | 0.999994 | 13346 | 1 | -0.13056 | 0.35433 | 0.50663 | 1 | 12340 | 1 | -0.13056 |
| BAI1 | 5 | 0.016737 | 0.055141 | 0.922677 | 1292 | 3 | -3.2872 | 0.35437 | 0.58193 | 1 | 12341 | 2 | -3.2872 |
| RC3H1 | 5 | 0.26281 | 0.451 | 0.999994 | 9307 | 2 | 0.9255 | 0.35439 | 0.58194 | 1 | 12342 | 3 | 0.9255 |
| DROSHA | 4 | 0.038485 | 0.098768 | 0.945253 | 2440 | 1 | -0.85051 | 0.35458 | 0.55451 | 1 | 12343 | 1 | -0.85051 |
| TMEM33 | 6 | 0.20632 | 0.39558 | 0.999994 | 7942 | 3 | -0.67711 | 0.35474 | 0.61544 | 1 | 12344 | 2 | -0.67711 |
| PMP22 | 6 | 0.36984 | 0.5831 | 0.999994 | 11507 | 2 | -0.18447 | 0.35474 | 0.61544 | 1 | 12345 | 2 | -0.18447 |
| AGAP2 | 6 | 0.22067 | 0.41292 | 0.999994 | 8304 | 3 | -0.34464 | 0.35474 | 0.61544 | 1 | 12346 | 2 | -0.34464 |
| KLHDC7A | 6 | 0.63637 | 0.76717 | 0.999994 | 15466 | 2 | 0.35838 | 0.35474 | 0.61544 | 1 | 12347 | 3 | 0.35838 |
| NRP1 | 6 | 0.62424 | 0.75802 | 0.999994 | 15293 | 2 | 0.23365 | 0.35475 | 0.61546 | 1 | 12348 | 3 | 0.23365 |
| C15orf38 | 6 | 0.0993 | 0.24601 | 0.98999 | 4941 | 4 | -0.90605 | 0.35475 | 0.61546 | 1 | 12349 | 1 | -0.90605 |
| SURF4 | 5 | 0.3245 | 0.50099 | 0.999994 | 10606 | 1 | 0.92505 | 0.35476 | 0.58231 | 1 | 12350 | 3 | 0.92505 |
| MLF1IP | 5 | 0.24797 | 0.43917 | 0.999994 | 8973 | 2 | 1.0775 | 0.35477 | 0.58233 | 1 | 12351 | 3 | 1.0775 |
| VOPP1 | 6 | 0.016345 | 0.060138 | 0.932932 | 1263 | 3 | 0.054675 | 0.3549 | 0.61557 | 1 | 12352 | 3 | 0.054675 |
| SF3B1 | 5 | 0.19397 | 0.36659 | 0.999994 | 7600 | 3 | -0.87659 | 0.35492 | 0.58247 | 1 | 12353 | 1 | -0.87659 |
| GJC3 | 5 | 0.028958 | 0.08781 | 0.94395 | 1952 | 3 | -2.7155 | 0.35492 | 0.58247 | 1 | 12354 | 2 | -2.7155 |
| hsa-mir-4736 | 3 | 0.85574 | 0.85573 | 0.999994 | 18360 | 0 | 1.2393 | 0.35509 | 0.50742 | 1 | 12355 | 2 | 1.2393 |
| BMS1 | 5 | 0.25005 | 0.44082 | 0.999994 | 9019 | 1 | 1.3104 | 0.35514 | 0.58269 | 1 | 12356 | 3 | 1.3104 |
| hsa-mir-2681 | 4 | 0.82342 | 0.82308 | 0.999994 | 17829 | 1 | 0.58845 | 0.35527 | 0.55501 | 1 | 12357 | 2 | 0.58845 |
| MGST2 | 6 | 0.039198 | 0.12009 | 0.951052 | 2475 | 3 | -0.54576 | 0.35538 | 0.61594 | 1 | 12358 | 2 | -0.54576 |
| MINK1 | 6 | 0.46998 | 0.64843 | 0.999994 | 13007 | 2 | -0.78558 | 0.35538 | 0.61594 | 1 | 12359 | 2 | -0.78558 |
| SLC5A5 | 6 | 0.24898 | 0.44671 | 0.999994 | 8990 | 3 | -0.66213 | 0.35538 | 0.61594 | 1 | 12360 | 2 | -0.66213 |
| KCNG4 | 6 | 0.89647 | 0.90097 | 0.999994 | 19151 | 1 | 0.38677 | 0.35538 | 0.61594 | 1 | 12361 | 3 | 0.38677 |
| NUDT15 | 5 | 0.86298 | 0.86492 | 0.999994 | 18506 | 1 | 1.0861 | 0.35551 | 0.58306 | 1 | 12362 | 3 | 1.0861 |
| FAM210A | 5 | 0.088631 | 0.20838 | 0.979587 | 4541 | 2 | 0.032369 | 0.35551 | 0.58306 | 1 | 12363 | 2 | 0.032369 |
| SECISBP2 | 5 | 0.14855 | 0.30058 | 0.999994 | 6420 | 2 | -0.098887 | 0.35551 | 0.58306 | 1 | 12364 | 2 | -0.098887 |
| HSPB9 | 5 | 0.3262 | 0.50239 | 0.999994 | 10647 | 2 | -0.41344 | 0.35551 | 0.58306 | 1 | 12365 | 2 | -0.41344 |
| DIRC1 | 6 | 0.032008 | 0.10352 | 0.946414 | 2106 | 3 | -0.32462 | 0.35558 | 0.61611 | 1 | 12366 | 3 | -0.32462 |
| hsa-mir-584 | 4 | 0.82926 | 0.82879 | 0.999994 | 17918 | 0 | 1.2456 | 0.35566 | 0.5553 | 1 | 12367 | 2 | 1.2456 |
| hsa-mir-3145 | 3 | 0.24863 | 0.37325 | 0.999994 | 8984 | 1 | 0.21494 | 0.35581 | 0.5082 | 1 | 12368 | 1 | 0.21494 |
| ARHGEF38 | 6 | 0.73821 | 0.82438 | 0.999994 | 16707 | 1 | 1.2029 | 0.35595 | 0.61638 | 1 | 12369 | 3 | 1.2029 |
| TMEM150B | 6 | 0.34786 | 0.56161 | 0.999994 | 11115 | 3 | 0.48716 | 0.35602 | 0.61644 | 1 | 12370 | 3 | 0.48716 |
| AGA | 6 | 0.16375 | 0.34296 | 0.999994 | 6833 | 3 | -0.22462 | 0.35605 | 0.61646 | 1 | 12371 | 3 | -0.22462 |
| NonTargetingControlG | 1 | 0.64393 | 0.64392 | 0.999994 | 15574 | 0 | 1.8463 | 0.35607 | 0.35622 | 0.994392 | 12372 | 1 | 1.8463 |
| TLE4 | 6 | 0.10542 | 0.25788 | 0.994943 | 5151 | 4 | -1.1157 | 0.35616 | 0.61653 | 1 | 12373 | 1 | -1.1157 |
| LCE6A | 6 | 0.0022077 | 0.010742 | 0.70768 | 293 | 4 | -4.6687 | 0.35616 | 0.61653 | 1 | 12374 | 2 | -4.6687 |
| GDAP1 | 5 | 0.027613 | 0.084296 | 0.94395 | 1884 | 2 | -0.34481 | 0.3562 | 0.58378 | 1 | 12375 | 2 | -0.34481 |

| | | | | | | | | | | | | | |
|---|---|---|---|---|---|---|---|---|---|---|---|---|---|
| HPS4 | 6 | 0.053124 | 0.15083 | 0.966359 | 3122 | 3 | -1.1638 | 0.35621 | 0.61657 | 1 | 12376 | 2 | -1.1638 |
| THOC1 | 4 | 0.42736 | 0.5738 | 0.999994 | 12375 | 2 | 0.87126 | 0.35633 | 0.55581 | 1 | 12377 | 2 | 0.87126 |
| XPNPEP3 | 5 | 0.23698 | 0.42686 | 0.999994 | 8694 | 2 | 1.088 | 0.35634 | 0.58394 | 1 | 12378 | 3 | 1.088 |
| CREB5 | 6 | 0.35753 | 0.57252 | 0.999994 | 11297 | 3 | 0.69024 | 0.35661 | 0.61689 | 1 | 12379 | 3 | 0.69024 |
| hsa-mir-4758 | 4 | 0.36148 | 0.50604 | 0.999994 | 11361 | 2 | 0.086082 | 0.35668 | 0.55608 | 1 | 12380 | 2 | 0.086082 |
| ANXA5 | 6 | 0.029105 | 0.096692 | 0.945031 | 1962 | 3 | 0.50453 | 0.35677 | 0.61702 | 1 | 12381 | 3 | 0.50453 |
| METTL21D | 6 | 0.11013 | 0.26694 | 0.998561 | 5309 | 4 | -0.96607 | 0.35685 | 0.61709 | 1 | 12382 | 1 | -0.96607 |
| ABHD16B | 6 | 0.14072 | 0.31396 | 0.999994 | 6210 | 3 | -0.31512 | 0.35685 | 0.61709 | 1 | 12383 | 3 | -0.31512 |
| MYOZ2 | 6 | 0.2372 | 0.43265 | 0.999994 | 8702 | 3 | -0.31505 | 0.35685 | 0.61709 | 1 | 12384 | 1 | -0.31505 |
| hsa-mir-6878 | 4 | 0.07299 | 0.16557 | 0.967126 | 3960 | 2 | -2.2828 | 0.35689 | 0.55625 | 1 | 12385 | 1 | -2.2828 |
| MYT1L | 5 | 0.15351 | 0.30805 | 0.999994 | 6551 | 2 | -0.40216 | 0.35689 | 0.58451 | 1 | 12386 | 2 | -0.40216 |
| APOBEC3C | 5 | 0.25368 | 0.44369 | 0.999994 | 9097 | 2 | -0.5225 | 0.35689 | 0.58451 | 1 | 12387 | 1 | -0.5225 |
| OR10A3 | 5 | 0.24136 | 0.43276 | 0.999994 | 8809 | 3 | -0.62256 | 0.35689 | 0.58451 | 1 | 12388 | 2 | -0.62256 |
| IQSEC2 | 6 | 0.18904 | 0.3743 | 0.999994 | 7452 | 2 | 1.1519 | 0.3569 | 0.61713 | 1 | 12389 | 3 | 1.1519 |
| USP18 | 5 | 0.55746 | 0.69838 | 0.999994 | 14323 | 2 | 1.1298 | 0.35694 | 0.58456 | 1 | 12390 | 3 | 1.1298 |
| hsa-mir-5047 | 3 | 0.3599 | 0.46127 | 0.999994 | 11333 | 1 | 1.2896 | 0.35695 | 0.50938 | 1 | 12391 | 2 | 1.2896 |
| PCDH11Y | 5 | 0.25343 | 0.44349 | 0.999994 | 9091 | 2 | -0.15088 | 0.35695 | 0.58457 | 1 | 12392 | 2 | -0.15088 |
| hsa-mir-1915 | 3 | 0.49414 | 0.54232 | 0.999994 | 13330 | 1 | 1.3876 | 0.357 | 0.50944 | 1 | 12393 | 2 | 1.3876 |
| RCCD1 | 6 | 0.89632 | 0.90087 | 0.999994 | 19148 | 1 | 0.11588 | 0.35701 | 0.61721 | 1 | 12394 | 2 | 0.11588 |
| APPL1 | 5 | 0.73583 | 0.78355 | 0.999994 | 16672 | 1 | 1.221 | 0.35709 | 0.58471 | 1 | 12395 | 3 | 1.221 |
| LCMT2 | 3 | 0.042862 | 0.1042 | 0.947284 | 2670 | 1 | 1.5676 | 0.35721 | 0.50965 | 1 | 12396 | 2 | 1.5676 |
| CACNA2D3 | 4 | 0.19525 | 0.33579 | 0.999994 | 7641 | 2 | 0.17783 | 0.35723 | 0.5565 | 1 | 12397 | 2 | 0.17783 |
| ZCCHC7 | 5 | 0.91569 | 0.91577 | 0.999994 | 19531 | 0 | 1.0487 | 0.35732 | 0.58494 | 1 | 12398 | 3 | 1.0487 |
| APOBEC3H | 5 | 0.88831 | 0.88841 | 0.999994 | 19003 | 1 | -0.02136 | 0.35732 | 0.58494 | 1 | 12399 | 2 | -0.02136 |
| hsa-mir-3118-5 | 1 | 0.64265 | 0.64263 | 0.999994 | 15557 | 0 | 1.675 | 0.35735 | 0.35749 | 0.994392 | 12400 | 1 | 1.675 |
| RAB27A | 5 | 0.52234 | 0.66734 | 0.999994 | 13784 | 2 | 1.0415 | 0.35745 | 0.58507 | 1 | 12401 | 3 | 1.0415 |
| NonTargetingControlG | 2 | 0.61846 | 0.61823 | 0.999994 | 15195 | 0 | 2.3086 | 0.35749 | 0.44072 | 1 | 12402 | 1 | 2.3086 |
| TCIRG1 | 6 | 0.027051 | 0.091885 | 0.943968 | 1859 | 4 | -2.695 | 0.35749 | 0.61759 | 1 | 12403 | 1 | -2.695 |
| hsa-mir-620 | 3 | 0.83536 | 0.83522 | 0.999994 | 18013 | 0 | 1.2079 | 0.35759 | 0.51003 | 1 | 12404 | 2 | 1.2079 |
| C1S | 6 | 0.92035 | 0.92073 | 0.999994 | 19639 | 1 | 0.45063 | 0.35778 | 0.6178 | 1 | 12405 | 2 | 0.45063 |
| HPS5 | 6 | 0.42735 | 0.62016 | 0.999994 | 12372 | 2 | 0.75216 | 0.35785 | 0.61785 | 1 | 12406 | 3 | 0.75216 |
| PALM2 | 5 | 0.23289 | 0.42122 | 0.999994 | 8591 | 1 | -0.013208 | 0.35787 | 0.58547 | 1 | 12407 | 2 | -0.013208 |
| MYBPC2 | 4 | 0.86603 | 0.86562 | 0.999994 | 18557 | 0 | 1.3318 | 0.35798 | 0.55708 | 1 | 12408 | 2 | 1.3318 |
| BPIFA1 | 5 | 0.92061 | 0.9208 | 0.999994 | 19642 | 0 | 1.2518 | 0.35809 | 0.5857 | 1 | 12409 | 3 | 1.2518 |
| L3MBTL3 | 6 | 0.12802 | 0.29776 | 0.999994 | 5878 | 3 | -0.7402 | 0.35814 | 0.61808 | 1 | 12410 | 2 | -0.7402 |
| MUS81 | 5 | 0.01138 | 0.040046 | 0.891621 | 948 | 2 | 0.042567 | 0.35815 | 0.58577 | 1 | 12411 | 2 | 0.042567 |
| UBXN1 | 4 | 0.020731 | 0.062162 | 0.937033 | 1492 | 2 | 0.49293 | 0.35821 | 0.55725 | 1 | 12412 | 2 | 0.49293 |
| ARL15 | 6 | 0.07446 | 0.19605 | 0.975797 | 4011 | 3 | -0.46068 | 0.35841 | 0.61829 | 1 | 12413 | 3 | -0.46068 |
| NOL6 | 5 | 0.2459 | 0.43749 | 0.999994 | 8919 | 2 | 1.1249 | 0.35849 | 0.58611 | 1 | 12414 | 3 | 1.1249 |
| RPF2 | 4 | 0.28744 | 0.43016 | 0.999994 | 9865 | 2 | 0.70294 | 0.35851 | 0.5575 | 1 | 12415 | 2 | 0.70294 |
| TRIM52 | 5 | 0.084176 | 0.20128 | 0.977164 | 4382 | 2 | 0.73387 | 0.35863 | 0.58625 | 1 | 12416 | 3 | 0.73387 |
| TCF4 | 6 | 0.062492 | 0.17104 | 0.96737 | 3536 | 2 | -0.010529 | 0.3587 | 0.61853 | 1 | 12417 | 2 | -0.010529 |
| STK32A | 6 | 0.27795 | 0.48103 | 0.999994 | 9652 | 2 | 0.37872 | 0.3587 | 0.61853 | 1 | 12418 | 3 | 0.37872 |
| SYTL2 | 6 | 0.025355 | 0.087828 | 0.94395 | 1771 | 3 | -0.65726 | 0.3587 | 0.61853 | 1 | 12419 | 3 | -0.65726 |
| RFK | 5 | 0.27756 | 0.46289 | 0.999994 | 9647 | 2 | 1.2359 | 0.35891 | 0.58653 | 1 | 12420 | 3 | 1.2359 |
| ITGB1 | 5 | 0.91855 | 0.91865 | 0.999994 | 19597 | 0 | 1.0533 | 0.35922 | 0.58685 | 1 | 12421 | 3 | 1.0533 |
| GPR149 | 5 | 0.63394 | 0.73885 | 0.999994 | 15431 | 1 | 1.1782 | 0.35932 | 0.58696 | 1 | 12422 | 3 | 1.1782 |
| NonTargetingControlG | 1 | 0.64067 | 0.64062 | 0.999994 | 15522 | 0 | 1.6678 | 0.35933 | 0.35947 | 0.994392 | 12423 | 1 | 1.6678 |
| NAT8 | 5 | 0.56872 | 0.7084 | 0.999994 | 14519 | 2 | 1.0146 | 0.35947 | 0.58711 | 1 | 12424 | 3 | 1.0146 |
| LAT | 4 | 0.16569 | 0.30535 | 0.999994 | 6877 | 2 | -0.84658 | 0.35954 | 0.55827 | 1 | 12425 | 1 | -0.84658 |
| SBNO2 | 4 | 0.20909 | 0.35004 | 0.999994 | 8003 | 2 | -0.16786 | 0.35954 | 0.55827 | 1 | 12426 | 2 | -0.16786 |
| FSCN2 | 4 | 0.65778 | 0.69793 | 0.999994 | 15751 | 1 | 1.2708 | 0.35965 | 0.55836 | 1 | 12427 | 2 | 1.2708 |

| | | | | | | | | | | | | | |
|---|---|---|---|---|---|---|---|---|---|---|---|---|---|
| C1orf51 | 6 | 0.10539 | 0.25782 | 0.994943 | 5150 | 4 | -1.0883 | 0.35974 | 0.61933 | 1 | 12428 | 2 | -1.0883 |
| BCL2L15 | 6 | 0.29542 | 0.50136 | 0.999994 | 10024 | 3 | -0.10797 | 0.35974 | 0.61933 | 1 | 12429 | 2 | -0.10797 |
| FAM167B | 6 | 0.26111 | 0.4612 | 0.999994 | 9272 | 3 | -0.65762 | 0.35974 | 0.61933 | 1 | 12430 | 1 | -0.65762 |
| GNB2 | 5 | 0.54144 | 0.68417 | 0.999994 | 14078 | 2 | 0.12837 | 0.35977 | 0.58743 | 1 | 12431 | 2 | 0.12837 |
| ABCA10 | 6 | 0.44752 | 0.63343 | 0.999994 | 12680 | 1 | -0.1726 | 0.35982 | 0.61939 | 1 | 12432 | 2 | -0.1726 |
| CYP27A1 | 6 | 0.23709 | 0.43251 | 0.999994 | 8697 | 3 | -0.194 | 0.35984 | 0.6194 | 1 | 12433 | 3 | -0.194 |
| HSBP1 | 5 | 0.30331 | 0.48374 | 0.999994 | 10201 | 2 | 1.0153 | 0.35991 | 0.58757 | 1 | 12434 | 3 | 1.0153 |
| PSTPIP2 | 5 | 0.95419 | 0.95435 | 0.999994 | 20437 | 0 | 1.0464 | 0.36004 | 0.58772 | 1 | 12435 | 3 | 1.0464 |
| AGMAT | 6 | 0.50777 | 0.67408 | 0.999994 | 13547 | 2 | 0.20131 | 0.36008 | 0.61959 | 1 | 12436 | 3 | 0.20131 |
| TK2 | 5 | 0.019835 | 0.063649 | 0.937389 | 1450 | 4 | -1.4726 | 0.36013 | 0.5878 | 1 | 12437 | 1 | -1.4726 |
| CAMK2N2 | 5 | 0.19354 | 0.36599 | 0.999994 | 7586 | 3 | -0.84886 | 0.36013 | 0.5878 | 1 | 12438 | 1 | -0.84886 |
| hsa-mir-4300 | 3 | 0.092303 | 0.17282 | 0.96793 | 4689 | 2 | -2.2201 | 0.36014 | 0.51269 | 1 | 12439 | 1 | -2.2201 |
| DDX23 | 3 | 0.86036 | 0.86033 | 0.999994 | 18450 | 0 | 0.81718 | 0.36014 | 0.51269 | 1 | 12440 | 2 | 0.81718 |
| TBC1D10B | 5 | 0.72807 | 0.77973 | 0.999994 | 16582 | 1 | 0.65509 | 0.36019 | 0.58787 | 1 | 12441 | 3 | 0.65509 |
| CCDC14 | 6 | 0.015854 | 0.05857 | 0.931872 | 1235 | 5 | -2.0286 | 0.36025 | 0.61973 | 1 | 12442 | 1 | -2.0286 |
| LDHAL6B | 6 | 0.19957 | 0.38737 | 0.999994 | 7751 | 3 | -0.51694 | 0.36025 | 0.61973 | 1 | 12443 | 2 | -0.51694 |
| CD72 | 6 | 0.6865 | 0.80606 | 0.999994 | 16174 | 2 | -0.21 | 0.3603 | 0.61978 | 1 | 12444 | 2 | -0.21 |
| ZNF107 | 4 | 0.71045 | 0.73146 | 0.999994 | 16404 | 1 | 0.45915 | 0.36031 | 0.55885 | 1 | 12445 | 2 | 0.45915 |
| NRIP1 | 6 | 0.21701 | 0.40854 | 0.999994 | 8192 | 3 | -0.46388 | 0.36038 | 0.61983 | 1 | 12446 | 2 | -0.46388 |
| ESRRB | 6 | 0.025257 | 0.087593 | 0.94395 | 1764 | 2 | 0.50678 | 0.36042 | 0.61987 | 1 | 12447 | 3 | 0.50678 |
| DPY30 | 4 | 0.29253 | 0.43536 | 0.999994 | 9966 | 2 | 0.63375 | 0.3605 | 0.559 | 1 | 12448 | 2 | 0.63375 |
| NonTargetingControlG | 2 | 0.74546 | 0.74513 | 0.999994 | 16806 | 0 | 2.6596 | 0.36052 | 0.44263 | 1 | 12449 | 1 | 2.6596 |
| HCRT | 6 | 0.51746 | 0.68076 | 0.999994 | 13700 | 2 | 0.64527 | 0.36072 | 0.62009 | 1 | 12450 | 3 | 0.64527 |
| MRPL32 | 5 | 0.0075871 | 0.029059 | 0.859137 | 698 | 4 | -3.8461 | 0.36072 | 0.58844 | 1 | 12451 | 1 | -3.8461 |
| TSC22D2 | 6 | 0.37163 | 0.58424 | 0.999994 | 11538 | 2 | -0.03783 | 0.36073 | 0.6201 | 1 | 12452 | 2 | -0.03783 |
| IAH1 | 6 | 0.88731 | 0.89446 | 0.999994 | 18975 | 1 | 0.12448 | 0.36073 | 0.6201 | 1 | 12453 | 1 | 0.12448 |
| PGBD4 | 6 | 0.48563 | 0.65898 | 0.999994 | 13224 | 2 | 0.062057 | 0.36073 | 0.6201 | 1 | 12454 | 2 | 0.062057 |
| ADH5 | 5 | 0.20877 | 0.38766 | 0.999994 | 7997 | 2 | 1.1727 | 0.36075 | 0.58846 | 1 | 12455 | 3 | 1.1727 |
| LCP1 | 5 | 0.86844 | 0.86971 | 0.999994 | 18609 | 1 | 1.1291 | 0.3608 | 0.58851 | 1 | 12456 | 3 | 1.1291 |
| hsa-mir-3176 | 4 | 0.21412 | 0.35527 | 0.999994 | 8119 | 2 | 0.35135 | 0.36083 | 0.55925 | 1 | 12457 | 2 | 0.35135 |
| CPSF6 | 5 | 0.26711 | 0.45452 | 0.999994 | 9398 | 2 | 1.1379 | 0.36087 | 0.58857 | 1 | 12458 | 3 | 1.1379 |
| SERTAD3 | 6 | 0.15457 | 0.33144 | 0.999994 | 6581 | 1 | 0.81647 | 0.36099 | 0.62029 | 1 | 12459 | 3 | 0.81647 |
| NonTargetingControlG | 2 | 0.63562 | 0.63542 | 0.999994 | 15451 | 0 | 2.3309 | 0.36124 | 0.4431 | 1 | 12460 | 1 | 2.3309 |
| CEBPA | 6 | 0.97951 | 0.97942 | 0.999994 | 21083 | 0 | 1.1777 | 0.36131 | 0.62053 | 1 | 12461 | 3 | 1.1777 |
| NOP16 | 5 | 0.037498 | 0.10959 | 0.951052 | 2402 | 2 | 0.99447 | 0.36133 | 0.58903 | 1 | 12462 | 3 | 0.99447 |
| hsa-mir-3618 | 4 | 0.32589 | 0.46947 | 0.999994 | 10641 | 1 | 1.443 | 0.36146 | 0.5597 | 1 | 12463 | 2 | 1.443 |
| OR4D2 | 5 | 0.19471 | 0.36766 | 0.999994 | 7626 | 3 | -0.89057 | 0.3616 | 0.5893 | 1 | 12464 | 2 | -0.89057 |
| hsa-mir-524 | 3 | 0.56167 | 0.58861 | 0.999994 | 14401 | 1 | 1.2125 | 0.36186 | 0.51445 | 1 | 12465 | 2 | 1.2125 |
| C19orf82 | 6 | 0.3539 | 0.56841 | 0.999994 | 11235 | 2 | 0.71166 | 0.36188 | 0.62098 | 1 | 12466 | 3 | 0.71166 |
| AP5M1 | 6 | 0.16558 | 0.34518 | 0.999994 | 6874 | 3 | 0.12503 | 0.36188 | 0.62098 | 1 | 12467 | 3 | 0.12503 |
| NonTargetingControlG | 2 | 0.1218 | 0.20342 | 0.978186 | 5694 | 1 | -0.32785 | 0.36195 | 0.44357 | 1 | 12468 | 1 | -0.32785 |
| NTRK3 | 6 | 0.26051 | 0.46047 | 0.999994 | 9251 | 2 | -0.16561 | 0.36219 | 0.62124 | 1 | 12469 | 2 | -0.16561 |
| HNF4G | 6 | 0.27465 | 0.47718 | 0.999994 | 9587 | 3 | 0.32067 | 0.36236 | 0.62138 | 1 | 12470 | 3 | 0.32067 |
| POPDC2 | 5 | 0.012486 | 0.043166 | 0.891621 | 1029 | 4 | -1.8642 | 0.36247 | 0.59021 | 1 | 12471 | 1 | -1.8642 |
| EID1 | 5 | 0.59668 | 0.72494 | 0.999994 | 14908 | 1 | 1.3182 | 0.3626 | 0.59034 | 1 | 12472 | 3 | 1.3182 |
| LMOD1 | 5 | 0.65114 | 0.74559 | 0.999994 | 15678 | 1 | 1.3182 | 0.3626 | 0.59034 | 1 | 12473 | 3 | 1.3182 |
| KIAA1217 | 5 | 0.89438 | 0.89452 | 0.999994 | 19112 | 0 | 1.3182 | 0.3626 | 0.59034 | 1 | 12474 | 3 | 1.3182 |
| EID2 | 5 | 0.2221 | 0.40619 | 0.999994 | 8335 | 1 | 0.4777 | 0.36262 | 0.59036 | 1 | 12475 | 2 | 0.4777 |
| SLC6A20 | 4 | 0.90579 | 0.90553 | 0.999994 | 19348 | 0 | 0.62254 | 0.36268 | 0.5606 | 1 | 12476 | 2 | 0.62254 |
| GTF2E2 | 2 | 0.15453 | 0.22979 | 0.983354 | 6579 | 1 | -1.038 | 0.36269 | 0.44402 | 1 | 12477 | 1 | -1.038 |
| LRCH1 | 6 | 0.26865 | 0.47014 | 0.999994 | 9435 | 3 | 0.24171 | 0.36276 | 0.62169 | 1 | 12478 | 3 | 0.24171 |
| DCAF16 | 6 | 0.94982 | 0.94979 | 0.999994 | 20329 | 0 | 0.97819 | 0.36276 | 0.62169 | 1 | 12479 | 3 | 0.97819 |

| | | | | | | | | | | | | | | |
|---|---|---|---|---|---|---|---|---|---|---|---|---|---|---|
| CCT6B | 6 | 0.075381 | 0.19796 | 0.977164 | 4049 | 4 | -1.0522 | 0.36303 | 0.62191 | 1 | 12480 | | 1 | -1.0522 |
| RNASEL | 6 | 0.02721 | 0.092246 | 0.943968 | 1866 | 3 | -0.60713 | 0.36303 | 0.62191 | 1 | 12481 | | 2 | -0.60713 |
| AMICA1 | 6 | 0.092999 | 0.23352 | 0.984971 | 4719 | 3 | -1.1469 | 0.36303 | 0.62191 | 1 | 12482 | | 1 | -1.1469 |
| PRAMEF1 | 3 | 0.72468 | 0.72461 | 0.999994 | 16544 | 1 | 1.1205 | 0.36312 | 0.51571 | 1 | 12483 | | 2 | 1.1205 |
| TRIM6 | 6 | 0.023065 | 0.081785 | 0.93914 | 1653 | 2 | 0.98559 | 0.36325 | 0.62208 | 1 | 12484 | | 3 | 0.98559 |
| TBC1D9B | 6 | 0.50458 | 0.67191 | 0.999994 | 13504 | 2 | 0.62519 | 0.36328 | 0.6221 | 1 | 12485 | | 3 | 0.62519 |
| NME1-NME2 | 4 | 0.04275 | 0.10728 | 0.950995 | 2663 | 2 | -2.0568 | 0.3633 | 0.56108 | 1 | 12486 | | 2 | -2.0568 |
| UACA | 5 | 0.83512 | 0.84315 | 0.999994 | 18008 | 1 | 0.19872 | 0.36336 | 0.5911 | 1 | 12487 | | 2 | 0.19872 |
| NLN | 5 | 0.19421 | 0.36695 | 0.999994 | 7607 | 3 | -0.81026 | 0.36341 | 0.59116 | 1 | 12488 | | 2 | -0.81026 |
| TSPY2 | 4 | 0.61087 | 0.67123 | 0.999994 | 15085 | 1 | 0.8304 | 0.36362 | 0.56131 | 1 | 12489 | | 2 | 0.8304 |
| hsa-mir-655 | 4 | 0.23263 | 0.3742 | 0.999994 | 8586 | 2 | -0.32414 | 0.36362 | 0.56131 | 1 | 12490 | | 2 | -0.32414 |
| GSTM2 | 4 | 0.11292 | 0.23804 | 0.987253 | 5401 | 3 | -1.3732 | 0.36362 | 0.56131 | 1 | 12491 | | 1 | -1.3732 |
| hsa-mir-7156 | 4 | 0.63775 | 0.68622 | 0.999994 | 15484 | 1 | 0.85391 | 0.36362 | 0.56131 | 1 | 12492 | | 2 | 0.85391 |
| GABARAPL1 | 4 | 0.039638 | 0.10108 | 0.946414 | 2497 | 2 | -1.1237 | 0.36372 | 0.56139 | 1 | 12493 | | 2 | -1.1237 |
| GTF2H3 | 6 | 0.25132 | 0.44951 | 0.999994 | 9045 | 2 | -0.90897 | 0.36379 | 0.62251 | 1 | 12494 | | 2 | -0.90897 |
| PPFIA1 | 6 | 0.46625 | 0.64596 | 0.999994 | 12954 | 2 | 0.62928 | 0.36379 | 0.62251 | 1 | 12495 | | 3 | 0.62928 |
| BICD1 | 5 | 0.59836 | 0.72557 | 0.999994 | 14934 | 1 | 1.1333 | 0.36381 | 0.59157 | 1 | 12496 | | 3 | 1.1333 |
| USP26 | 5 | 0.62882 | 0.73685 | 0.999994 | 15360 | 1 | 0.94559 | 0.36385 | 0.59161 | 1 | 12497 | | 3 | 0.94559 |
| CCER1 | 4 | 0.53416 | 0.6321 | 0.999994 | 13968 | 1 | 1.2892 | 0.36385 | 0.56149 | 1 | 12498 | | 2 | 1.2892 |
| NonTargetingControlG | 2 | 0.7433 | 0.74298 | 0.999994 | 16782 | 0 | 2.6774 | 0.36388 | 0.44482 | 1 | 12499 | | 1 | 2.6774 |
| TTC38 | 5 | 0.6981 | 0.76563 | 0.999994 | 16282 | 1 | 1.0209 | 0.3639 | 0.59166 | 1 | 12500 | | 3 | 1.0209 |
| hsa-mir-4262 | 4 | 0.42663 | 0.57302 | 0.999994 | 12359 | 2 | 0.85696 | 0.36397 | 0.56158 | 1 | 12501 | | 2 | 0.85696 |
| FBXO47 | 6 | 0.15349 | 0.33005 | 0.999994 | 6550 | 1 | 1.0204 | 0.36403 | 0.62269 | 1 | 12502 | | 3 | 1.0204 |
| NonTargetingControlG | 1 | 0.63588 | 0.63579 | 0.999994 | 15458 | 0 | 1.8803 | 0.36412 | 0.36422 | 0.995622 | 12503 | | 1 | 1.8803 |
| SLCO2A1 | 5 | 0.01969 | 0.063257 | 0.937389 | 1438 | 2 | 1.02 | 0.36417 | 0.59193 | 1 | 12504 | | 3 | 1.02 |
| hsa-mir-578 | 3 | 0.46464 | 0.52335 | 0.999994 | 12930 | 1 | 1.1582 | 0.36422 | 0.51685 | 1 | 12505 | | 2 | 1.1582 |
| ACSS2 | 5 | 0.23359 | 0.42219 | 0.999994 | 8607 | 3 | -0.6795 | 0.36423 | 0.59198 | 1 | 12506 | | 2 | -0.6795 |
| DGAT2L6 | 6 | 0.24587 | 0.44301 | 0.999994 | 8918 | 3 | -0.73439 | 0.36427 | 0.62289 | 1 | 12507 | | 2 | -0.73439 |
| NDUFC1 | 6 | 0.20387 | 0.39259 | 0.999994 | 7877 | 1 | 1.163 | 0.36449 | 0.62306 | 1 | 12508 | | 3 | 1.163 |
| CDK9 | 4 | 0.61758 | 0.67499 | 0.999994 | 15179 | 1 | 1.3013 | 0.36464 | 0.56206 | 1 | 12509 | | 2 | 1.3013 |
| OR13F1 | 5 | 0.56068 | 0.70123 | 0.999994 | 14380 | 2 | 0.45537 | 0.36465 | 0.59241 | 1 | 12510 | | 2 | 0.45537 |
| PTPN22 | 5 | 0.031113 | 0.093361 | 0.945031 | 2065 | 2 | 1.1512 | 0.36478 | 0.59255 | 1 | 12511 | | 3 | 1.1512 |
| FAM188A | 6 | 0.93707 | 0.93709 | 0.999994 | 20008 | 0 | 0.99613 | 0.36491 | 0.62338 | 1 | 12512 | | 3 | 0.99613 |
| hsa-mir-4297 | 3 | 0.83705 | 0.83695 | 0.999994 | 18035 | 0 | 1.3167 | 0.36508 | 0.51773 | 1 | 12513 | | 2 | 1.3167 |
| PROM2 | 6 | 0.053078 | 0.15073 | 0.966359 | 3116 | 4 | -1.4277 | 0.36522 | 0.62361 | 1 | 12514 | | 2 | -1.4277 |
| ATP5G3 | 6 | 0.21696 | 0.40849 | 0.999994 | 8190 | 3 | -0.060665 | 0.36522 | 0.62361 | 1 | 12515 | | 3 | -0.060665 |
| NEK7 | 6 | 0.27197 | 0.47405 | 0.999994 | 9520 | 3 | 0.37852 | 0.36523 | 0.62363 | 1 | 12516 | | 3 | 0.37852 |
| HNRNPCL1 | 6 | 0.34951 | 0.56345 | 0.999994 | 11137 | 3 | -0.095696 | 0.36532 | 0.62369 | 1 | 12517 | | 2 | -0.095696 |
| LRRC41 | 6 | 0.44377 | 0.63094 | 0.999994 | 12631 | 2 | -0.081896 | 0.36532 | 0.62369 | 1 | 12518 | | 2 | -0.081896 |
| EXTL3 | 6 | 0.67917 | 0.80036 | 0.999994 | 16075 | 2 | -0.26084 | 0.36532 | 0.62369 | 1 | 12519 | | 2 | -0.26084 |
| DDX25 | 6 | 0.014255 | 0.053293 | 0.91738 | 1146 | 3 | -0.46869 | 0.36532 | 0.62369 | 1 | 12520 | | 2 | -0.46869 |
| TMPPE | 5 | 0.14749 | 0.29901 | 0.999994 | 6391 | 2 | 0.4826 | 0.36546 | 0.59325 | 1 | 12521 | | 2 | 0.4826 |
| HMX1 | 4 | 0.16794 | 0.30763 | 0.999994 | 6936 | 2 | -0.97402 | 0.36555 | 0.56275 | 1 | 12522 | | 1 | -0.97402 |
| PPAPDC1B | 5 | 0.022841 | 0.071735 | 0.93914 | 1638 | 2 | 1.0197 | 0.36556 | 0.59334 | 1 | 12523 | | 3 | 1.0197 |
| NonTargetingControlG | 2 | 0.080464 | 0.13929 | 0.958457 | 4249 | 1 | -1.8544 | 0.36557 | 0.44589 | 1 | 12524 | | 1 | -1.8544 |
| MLF2 | 5 | 0.070816 | 0.17971 | 0.968171 | 3867 | 3 | -1.1186 | 0.36558 | 0.59337 | 1 | 12525 | | 2 | -1.1186 |
| MYLK | 5 | 0.035366 | 0.10417 | 0.947284 | 2284 | 4 | -1.0149 | 0.36575 | 0.59352 | 1 | 12526 | | 1 | -1.0149 |
| POM121 | 5 | 0.01329 | 0.045399 | 0.891621 | 1082 | 3 | -3.9563 | 0.36575 | 0.59352 | 1 | 12527 | | 1 | -3.9563 |
| TSPAN17 | 6 | 0.099499 | 0.2464 | 0.98999 | 4953 | 3 | -0.62826 | 0.36584 | 0.62411 | 1 | 12528 | | 2 | -0.62826 |
| MAOA | 5 | 0.78124 | 0.80813 | 0.999994 | 17227 | 1 | 0.99966 | 0.36588 | 0.59365 | 1 | 12529 | | 3 | 0.99966 |
| CRYZ | 6 | 0.54681 | 0.70138 | 0.999994 | 14164 | 2 | 0.94264 | 0.36596 | 0.62421 | 1 | 12530 | | 3 | 0.94264 |
| NCL | 5 | 0.081472 | 0.19699 | 0.977164 | 4280 | 3 | -1.8623 | 0.36601 | 0.59377 | 1 | 12531 | | 2 | -1.8623 |

| | | | | | | | | | | | | |
|---|---|---|---|---|---|---|---|---|---|---|---|---|
| hsa-mir-139 | 4 | 0.37472 | 0.51972 | 0.999994 | 11578 | 1 | 1.2113 | 0.36615 | 0.56323 | 1 | 12532 | 2 | 1.2113 |
| FAM19A1 | 6 | 0.37375 | 0.58558 | 0.999994 | 11567 | 2 | 0.36295 | 0.36616 | 0.62437 | 1 | 12533 | 3 | 0.36295 |
| PRR3 | 5 | 0.015107 | 0.050507 | 0.909101 | 1195 | 2 | 0.20031 | 0.36639 | 0.59415 | 1 | 12534 | 1 | 0.20031 |
| CLPB | 5 | 0.18453 | 0.35311 | 0.999994 | 7331 | 3 | -0.83206 | 0.36639 | 0.59415 | 1 | 12535 | 1 | -0.83206 |
| TMEM61 | 6 | 0.28051 | 0.48401 | 0.999994 | 9712 | 3 | -0.21784 | 0.36642 | 0.62456 | 1 | 12536 | 2 | -0.21784 |
| THSD7A | 5 | 0.89251 | 0.89264 | 0.999994 | 19079 | 0 | 0.12834 | 0.3665 | 0.59426 | 1 | 12537 | 2 | 0.12834 |
| KRTAP13-4 | 6 | 0.27388 | 0.47628 | 0.999994 | 9562 | 2 | 0.29189 | 0.36655 | 0.62466 | 1 | 12538 | 3 | 0.29189 |
| PAAF1 | 6 | 0.01766 | 0.064409 | 0.93914 | 1346 | 3 | 0.19711 | 0.36656 | 0.62467 | 1 | 12539 | 3 | 0.19711 |
| STOX1 | 6 | 0.95639 | 0.95646 | 0.999994 | 20499 | 0 | 0.84175 | 0.36656 | 0.62467 | 1 | 12540 | 3 | 0.84175 |
| BRD1 | 5 | 0.32056 | 0.49776 | 0.999994 | 10524 | 2 | 0.99804 | 0.36672 | 0.59449 | 1 | 12541 | 3 | 0.99804 |
| ADAMTS18 | 4 | 0.092306 | 0.20122 | 0.977164 | 4690 | 2 | -0.61555 | 0.36672 | 0.56366 | 1 | 12542 | 2 | -0.61555 |
| RFX6 | 5 | 0.84608 | 0.85126 | 0.999994 | 18188 | 1 | 1.0524 | 0.36673 | 0.5945 | 1 | 12543 | 3 | 1.0524 |
| LMAN1L | 6 | 0.13053 | 0.30097 | 0.999994 | 5945 | 3 | -0.025506 | 0.36694 | 0.62497 | 1 | 12544 | 3 | -0.025506 |
| PLAGL1 | 5 | 0.11591 | 0.25117 | 0.993242 | 5511 | 3 | -1.4426 | 0.36713 | 0.59488 | 1 | 12545 | 2 | -1.4426 |
| NTRK2 | 6 | 0.16713 | 0.34714 | 0.999994 | 6908 | 3 | -0.80755 | 0.36719 | 0.62519 | 1 | 12546 | 2 | -0.80755 |
| hsa-mir-4312 | 2 | 0.59511 | 0.59503 | 0.999994 | 14890 | 0 | 2.1788 | 0.36721 | 0.44698 | 1 | 12547 | 1 | 2.1788 |
| HP1BP3 | 5 | 0.15448 | 0.30947 | 0.999994 | 6577 | 2 | -0.12492 | 0.36735 | 0.59513 | 1 | 12548 | 2 | -0.12492 |
| CGRRF1 | 6 | 0.072229 | 0.19142 | 0.973951 | 3927 | 2 | 0.27982 | 0.36745 | 0.62538 | 1 | 12549 | 2 | 0.27982 |
| CDNF | 6 | 0.24163 | 0.43797 | 0.999994 | 8819 | 2 | -0.18246 | 0.36745 | 0.62538 | 1 | 12550 | 2 | -0.18246 |
| ZFP90 | 6 | 0.34771 | 0.56143 | 0.999994 | 11105 | 2 | 0.39661 | 0.36745 | 0.62538 | 1 | 12551 | 2 | 0.39661 |
| CTSC | 6 | 0.90207 | 0.9053 | 0.999994 | 19254 | 1 | 1.0373 | 0.36749 | 0.62542 | 1 | 12552 | 3 | 1.0373 |
| FOLR3 | 5 | 0.079468 | 0.19371 | 0.975797 | 4205 | 2 | 1.0536 | 0.36769 | 0.59547 | 1 | 12553 | 3 | 1.0536 |
| MAGEB2 | 6 | 0.36443 | 0.57973 | 0.999994 | 11413 | 2 | -0.34279 | 0.36781 | 0.62567 | 1 | 12554 | 2 | -0.34279 |
| POM121L2 | 5 | 0.18188 | 0.34934 | 0.999994 | 7267 | 2 | -0.06657 | 0.36794 | 0.59573 | 1 | 12555 | 1 | -0.06657 |
| TICAM1 | 5 | 0.020919 | 0.066546 | 0.93914 | 1507 | 3 | -1.8934 | 0.36794 | 0.59573 | 1 | 12556 | 2 | -1.8934 |
| H1FOO | 5 | 0.56874 | 0.70841 | 0.999994 | 14520 | 2 | 0.35554 | 0.36794 | 0.59573 | 1 | 12557 | 2 | 0.35554 |
| NonTargetingControlG | 2 | 0.21762 | 0.27488 | 0.998561 | 8212 | 1 | 0.63471 | 0.36803 | 0.44751 | 1 | 12558 | 1 | 0.63471 |
| TM7SF2 | 5 | 0.94823 | 0.94842 | 0.999994 | 20289 | 0 | 0.94526 | 0.36824 | 0.59603 | 1 | 12559 | 3 | 0.94526 |
| hsa-mir-6514 | 3 | 0.091546 | 0.17181 | 0.967658 | 4659 | 2 | -2.4106 | 0.36836 | 0.52113 | 1 | 12560 | 1 | -2.4106 |
| GBA | 5 | 0.26259 | 0.45083 | 0.999994 | 9301 | 2 | 0.44565 | 0.3684 | 0.59618 | 1 | 12561 | 2 | 0.44565 |
| PRDM10 | 5 | 0.012069 | 0.042004 | 0.891621 | 997 | 3 | -1.9652 | 0.3684 | 0.59618 | 1 | 12562 | 2 | -1.9652 |
| DGCR6 | 4 | 0.13663 | 0.27543 | 0.999239 | 6099 | 2 | -1.3278 | 0.36844 | 0.56497 | 1 | 12563 | 1 | -1.3278 |
| ZNF410 | 6 | 0.10191 | 0.25105 | 0.993091 | 5034 | 4 | -1.0386 | 0.36849 | 0.62619 | 1 | 12564 | 2 | -1.0386 |
| SLC38A10 | 6 | 0.1799 | 0.36302 | 0.999994 | 7220 | 2 | 0.34359 | 0.36849 | 0.62619 | 1 | 12565 | 3 | 0.34359 |
| CST9L | 6 | 0.33503 | 0.54699 | 0.999994 | 10834 | 3 | -0.26913 | 0.36849 | 0.62619 | 1 | 12566 | 2 | -0.26913 |
| LRRC7 | 5 | 0.1766 | 0.3417 | 0.999994 | 7146 | 2 | 0.70907 | 0.36853 | 0.59631 | 1 | 12567 | 3 | 0.70907 |
| C14orf177 | 6 | 0.34642 | 0.55998 | 0.999994 | 11081 | 3 | 0.55417 | 0.36868 | 0.62634 | 1 | 12568 | 3 | 0.55417 |
| TTLL12 | 5 | 0.2112 | 0.3911 | 0.999994 | 8053 | 2 | 1.0597 | 0.36881 | 0.5966 | 1 | 12569 | 3 | 1.0597 |
| CHAD | 5 | 0.12124 | 0.25926 | 0.994943 | 5673 | 2 | -0.32321 | 0.369 | 0.59679 | 1 | 12570 | 2 | -0.32321 |
| NEK8 | 5 | 0.2805 | 0.46525 | 0.999994 | 9711 | 1 | 1.0286 | 0.36902 | 0.59681 | 1 | 12571 | 3 | 1.0286 |
| C2orf48 | 6 | 0.57124 | 0.71885 | 0.999994 | 14565 | 1 | 0.58796 | 0.36927 | 0.62679 | 1 | 12572 | 3 | 0.58796 |
| EXOC3L4 | 6 | 0.68446 | 0.80447 | 0.999994 | 16149 | 1 | 0.69025 | 0.36928 | 0.6268 | 1 | 12573 | 3 | 0.69025 |
| PHF8 | 4 | 0.029463 | 0.080433 | 0.93914 | 1983 | 1 | 1.0905 | 0.36929 | 0.56562 | 1 | 12574 | 2 | 1.0905 |
| ADAM2 | 6 | 0.068949 | 0.18457 | 0.968526 | 3796 | 3 | -0.42323 | 0.36939 | 0.62688 | 1 | 12575 | 3 | -0.42323 |
| TTPA | 6 | 0.14145 | 0.3149 | 0.999994 | 6229 | 2 | 0.74807 | 0.36962 | 0.62707 | 1 | 12576 | 3 | 0.74807 |
| hsa-mir-450a-1 | 2 | 0.56004 | 0.56061 | 0.999994 | 14371 | 1 | 2.0419 | 0.36971 | 0.44862 | 1 | 12577 | 1 | 2.0419 |
| TARBP2 | 4 | 0.24479 | 0.38654 | 0.999994 | 8897 | 2 | -0.84201 | 0.36979 | 0.56599 | 1 | 12578 | 1 | -0.84201 |
| HCAR2 | 5 | 0.80708 | 0.82403 | 0.999994 | 17587 | 1 | 1.1495 | 0.3698 | 0.5976 | 1 | 12579 | 3 | 1.1495 |
| AP4B1 | 6 | 0.79489 | 0.8457 | 0.999994 | 17413 | 1 | 0.47339 | 0.36986 | 0.62722 | 1 | 12580 | 2 | 0.47339 |
| STRIP2 | 5 | 0.14082 | 0.28905 | 0.999632 | 6213 | 3 | -1.1004 | 0.36992 | 0.59773 | 1 | 12581 | 2 | -1.1004 |
| COMMD8 | 6 | 0.46138 | 0.64268 | 0.999994 | 12869 | 1 | 0.87951 | 0.36997 | 0.6273 | 1 | 12582 | 3 | 0.87951 |
| LTBP3 | 6 | 0.33378 | 0.54561 | 0.999994 | 10802 | 1 | 0.54891 | 0.36997 | 0.6273 | 1 | 12583 | 3 | 0.54891 |

| | | | | | | | | | | | | | |
|---|---|---|---|---|---|---|---|---|---|---|---|---|---|
| OCRL | 4 | 0.61947 | 0.67604 | 0.999994 | 15213 | 1 | 0.87175 | 0.36998 | 0.56613 | 1 | 12584 | 2 | 0.87175 |
| LCORL | 5 | 0.5256 | 0.6702 | 0.999994 | 13826 | 2 | -0.36566 | 0.37004 | 0.59785 | 1 | 12585 | 2 | -0.36566 |
| MAB21L3 | 3 | 0.8694 | 0.86946 | 0.999994 | 18625 | 0 | 1.3151 | 0.37005 | 0.52292 | 1 | 12586 | 2 | 1.3151 |
| CUTC | 5 | 0.25483 | 0.44459 | 0.999994 | 9126 | 2 | 0.81958 | 0.37026 | 0.59808 | 1 | 12587 | 3 | 0.81958 |
| TMEM198 | 6 | 0.079204 | 0.20571 | 0.978603 | 4197 | 4 | -1.3639 | 0.3703 | 0.62757 | 1 | 12588 | 1 | -1.3639 |
| GLB1L3 | 5 | 0.041286 | 0.1191 | 0.951052 | 2578 | 3 | -2.359 | 0.37034 | 0.59817 | 1 | 12589 | 2 | -2.359 |
| SSRP1 | 5 | 0.0030349 | 0.012981 | 0.733879 | 360 | 3 | -5.4081 | 0.37034 | 0.59817 | 1 | 12590 | 1 | -5.4081 |
| BTLA | 6 | 0.67522 | 0.79723 | 0.999994 | 16008 | 2 | 0.77864 | 0.37056 | 0.62777 | 1 | 12591 | 3 | 0.77864 |
| hsa-mir-4436a | 4 | 0.82104 | 0.82076 | 0.999994 | 17791 | 1 | 1.3997 | 0.37061 | 0.56662 | 1 | 12592 | 2 | 1.3997 |
| APMAP | 5 | 0.51501 | 0.66097 | 0.999994 | 13662 | 2 | 1.0474 | 0.37063 | 0.59847 | 1 | 12593 | 3 | 1.0474 |
| hsa-mir-486 | 4 | 0.63597 | 0.68518 | 0.999994 | 15462 | 1 | 0.76484 | 0.37066 | 0.56665 | 1 | 12594 | 2 | 0.76484 |
| HSPE1 | 1 | 0.62931 | 0.62922 | 0.999994 | 15374 | 0 | 1.6996 | 0.37069 | 0.37077 | 0.998994 | 12595 | 1 | 1.6996 |
| NonTargetingControlG | 1 | 0.62915 | 0.62906 | 0.999994 | 15368 | 0 | 1.5606 | 0.37085 | 0.37093 | 0.998994 | 12596 | 1 | 1.5606 |
| GP9 | 6 | 0.51065 | 0.67608 | 0.999994 | 13585 | 2 | 0.58446 | 0.37091 | 0.62804 | 1 | 12597 | 3 | 0.58446 |
| ZNF365 | 6 | 0.18196 | 0.36561 | 0.999994 | 7270 | 2 | 1.0861 | 0.37097 | 0.62808 | 1 | 12598 | 3 | 1.0861 |
| UBR2 | 5 | 0.57208 | 0.71138 | 0.999994 | 14579 | 2 | 1.0425 | 0.37102 | 0.59887 | 1 | 12599 | 3 | 1.0425 |
| IL5 | 6 | 0.056257 | 0.15764 | 0.967126 | 3256 | 3 | -0.52594 | 0.37102 | 0.62812 | 1 | 12600 | 3 | -0.52594 |
| BEST1 | 5 | 0.81622 | 0.83008 | 0.999994 | 17716 | 1 | 1.221 | 0.37105 | 0.5989 | 1 | 12601 | 3 | 1.221 |
| SYNGR4 | 5 | 0.84829 | 0.85293 | 0.999994 | 18222 | 1 | 0.94098 | 0.37109 | 0.59893 | 1 | 12602 | 3 | 0.94098 |
| DLGAP3 | 3 | 0.061652 | 0.13172 | 0.95318 | 3499 | 1 | -0.10599 | 0.37109 | 0.52398 | 1 | 12603 | 1 | -0.10599 |
| CDCA7 | 5 | 0.46638 | 0.61905 | 0.999994 | 12958 | 2 | 0.61008 | 0.37111 | 0.59894 | 1 | 12604 | 3 | 0.61008 |
| SEC31A | 5 | 0.45116 | 0.6062 | 0.999994 | 12725 | 2 | 0.59486 | 0.37112 | 0.59896 | 1 | 12605 | 3 | 0.59486 |
| LRRC4C | 5 | 0.79035 | 0.81354 | 0.999994 | 17349 | 1 | 1.045 | 0.37128 | 0.59912 | 1 | 12606 | 3 | 1.045 |
| SIGLEC1 | 6 | 0.056882 | 0.15903 | 0.967126 | 3287 | 4 | -1.3174 | 0.37128 | 0.62833 | 1 | 12607 | 2 | -1.3174 |
| WSB1 | 5 | 0.06872 | 0.17635 | 0.968171 | 3781 | 2 | 0.99025 | 0.3713 | 0.59915 | 1 | 12608 | 3 | 0.99025 |
| ABHD4 | 6 | 0.26973 | 0.47141 | 0.999994 | 9469 | 3 | 0.50721 | 0.37166 | 0.62862 | 1 | 12609 | 3 | 0.50721 |
| hsa-mir-1910 | 2 | 0.19738 | 0.26019 | 0.994943 | 7683 | 1 | -0.61382 | 0.37175 | 0.44994 | 1 | 12610 | 1 | -0.61382 |
| CYLC2 | 5 | 0.88006 | 0.88032 | 0.999994 | 18832 | 1 | 0.7796 | 0.37179 | 0.59967 | 1 | 12611 | 3 | 0.7796 |
| FBXW12 | 6 | 0.033096 | 0.10609 | 0.950452 | 2167 | 3 | -1.901 | 0.37183 | 0.62875 | 1 | 12612 | 2 | -1.901 |
| ERO1L | 4 | 0.27009 | 0.41237 | 0.999994 | 9476 | 2 | -0.74742 | 0.37188 | 0.56755 | 1 | 12613 | 1 | -0.74742 |
| C7orf10 | 4 | 0.015844 | 0.051658 | 0.910734 | 1233 | 3 | -4.3165 | 0.37188 | 0.56755 | 1 | 12614 | 1 | -4.3165 |
| NSFL1C | 4 | 0.3176 | 0.46099 | 0.999994 | 10463 | 2 | -0.12876 | 0.37188 | 0.56755 | 1 | 12615 | 2 | -0.12876 |
| TXNDC16 | 6 | 0.58929 | 0.73207 | 0.999994 | 14810 | 2 | -0.17843 | 0.37191 | 0.62882 | 1 | 12616 | 2 | -0.17843 |
| ZNF605 | 6 | 0.25361 | 0.45224 | 0.999994 | 9095 | 3 | -0.43522 | 0.37191 | 0.62882 | 1 | 12617 | 1 | -0.43522 |
| COPG1 | 5 | 0.88279 | 0.88296 | 0.999994 | 18887 | 1 | 0.054499 | 0.37193 | 0.59982 | 1 | 12618 | 2 | 0.054499 |
| RHOXF2B | 5 | 0.36234 | 0.53201 | 0.999994 | 11374 | 2 | 1.1235 | 0.37193 | 0.59982 | 1 | 12619 | 3 | 1.1235 |
| PCGF5 | 5 | 0.89732 | 0.89746 | 0.999994 | 19165 | 0 | 0.37657 | 0.37193 | 0.59982 | 1 | 12620 | 2 | 0.37657 |
| MLLT10 | 5 | 0.74786 | 0.78971 | 0.999994 | 16837 | 1 | 0.54959 | 0.37193 | 0.59982 | 1 | 12621 | 2 | 0.54959 |
| SEPT12 | 5 | 0.062155 | 0.16565 | 0.967126 | 3517 | 3 | -1.4151 | 0.37193 | 0.59982 | 1 | 12622 | 2 | -1.4151 |
| ZNF675 | 5 | 0.85997 | 0.86237 | 0.999994 | 18435 | 1 | 0.53114 | 0.37193 | 0.59982 | 1 | 12623 | 3 | 0.53114 |
| GOLIM4 | 3 | 0.14421 | 0.24102 | 0.987253 | 6306 | 1 | 1.0967 | 0.37208 | 0.52501 | 1 | 12624 | 2 | 1.0967 |
| RPP25L | 3 | 0.127 | 0.21865 | 0.980699 | 5848 | 2 | -1.86 | 0.37211 | 0.52503 | 1 | 12625 | 1 | -1.86 |
| CHKA | 3 | 0.63421 | 0.64406 | 0.999994 | 15435 | 1 | -0.47684 | 0.37211 | 0.52503 | 1 | 12626 | 1 | -0.47684 |
| UPRT | 5 | 0.79643 | 0.8173 | 0.999994 | 17433 | 1 | 0.99173 | 0.37224 | 0.60013 | 1 | 12627 | 3 | 0.99173 |
| CCDC132 | 6 | 0.38212 | 0.59091 | 0.999994 | 11679 | 2 | 0.71628 | 0.37231 | 0.62912 | 1 | 12628 | 3 | 0.71628 |
| ACCS | 5 | 0.029188 | 0.088389 | 0.94395 | 1968 | 2 | 0.94237 | 0.37235 | 0.60024 | 1 | 12629 | 3 | 0.94237 |
| SPRR2D | 4 | 0.15608 | 0.29548 | 0.999994 | 6631 | 2 | 0.62496 | 0.37242 | 0.56794 | 1 | 12630 | 2 | 0.62496 |
| PI4KA | 5 | 0.56326 | 0.70354 | 0.999994 | 14435 | 2 | 1.0532 | 0.37243 | 0.60032 | 1 | 12631 | 3 | 1.0532 |
| SCLY | 5 | 0.70546 | 0.76899 | 0.999994 | 16368 | 1 | 1.13 | 0.37247 | 0.60036 | 1 | 12632 | 3 | 1.13 |
| BTRC | 6 | 0.081728 | 0.21085 | 0.979587 | 4289 | 3 | -0.57442 | 0.37264 | 0.62938 | 1 | 12633 | 2 | -0.57442 |
| CMIP | 6 | 0.22468 | 0.41773 | 0.999994 | 8392 | 2 | 0.61938 | 0.37264 | 0.62938 | 1 | 12634 | 3 | 0.61938 |
| SH3GLB2 | 6 | 0.23793 | 0.43351 | 0.999994 | 8725 | 1 | 0.024567 | 0.37264 | 0.62938 | 1 | 12635 | 2 | 0.024567 |

| | | | | | | | | | | | | | |
|---|---|---|---|---|---|---|---|---|---|---|---|---|---|
| TRIM17 | 6 | 0.019504 | 0.070386 | 0.93914 | 1429 | 5 | -1.2973 | 0.37264 | 0.62938 | 1 | 12636 | 1 | -1.2973 |
| RASAL1 | 6 | 0.64523 | 0.77396 | 0.999994 | 15587 | 2 | 0.14338 | 0.37264 | 0.62938 | 1 | 12637 | 3 | 0.14338 |
| OR51M1 | 5 | 0.35485 | 0.52585 | 0.999994 | 11247 | 2 | 0.45538 | 0.37268 | 0.60056 | 1 | 12638 | 2 | 0.45538 |
| PNCK | 6 | 0.067151 | 0.1808 | 0.968171 | 3723 | 3 | -0.37799 | 0.37274 | 0.62946 | 1 | 12639 | 3 | -0.37799 |
| RILPL1 | 6 | 0.013628 | 0.051212 | 0.909241 | 1106 | 4 | -2.3803 | 0.37274 | 0.62946 | 1 | 12640 | 2 | -2.3803 |
| FABP3 | 6 | 0.25959 | 0.45938 | 0.999994 | 9229 | 2 | 0.042998 | 0.37274 | 0.62946 | 1 | 12641 | 2 | 0.042998 |
| ARL6IP5 | 5 | 0.45644 | 0.61069 | 0.999994 | 12798 | 2 | 1.0586 | 0.37311 | 0.60102 | 1 | 12642 | 3 | 1.0586 |
| IL15 | 6 | 0.34451 | 0.55785 | 0.999994 | 11040 | 2 | 1.1298 | 0.37323 | 0.62984 | 1 | 12643 | 3 | 1.1298 |
| ROMO1 | 5 | 0.25478 | 0.44455 | 0.999994 | 9123 | 2 | 1.0383 | 0.37327 | 0.60117 | 1 | 12644 | 3 | 1.0383 |
| NonTargetingControlG | 2 | 0.73588 | 0.73558 | 0.999994 | 16673 | 0 | 2.5695 | 0.37333 | 0.45096 | 1 | 12645 | 1 | 2.5695 |
| CANT1 | 4 | 0.16111 | 0.30062 | 0.999994 | 6767 | 2 | -0.5722 | 0.37337 | 0.56863 | 1 | 12646 | 2 | -0.5722 |
| KRT86 | 4 | 0.070559 | 0.161 | 0.967126 | 3854 | 3 | -2.1993 | 0.37337 | 0.56863 | 1 | 12647 | 1 | -2.1993 |
| ITPKB | 4 | 0.29973 | 0.44273 | 0.999994 | 10124 | 2 | -0.23071 | 0.37337 | 0.56863 | 1 | 12648 | 2 | -0.23071 |
| CHST15 | 4 | 0.22972 | 0.37124 | 0.999994 | 8522 | 1 | 1.0046 | 0.37337 | 0.56863 | 1 | 12649 | 2 | 1.0046 |
| CLDN12 | 6 | 0.50476 | 0.67203 | 0.999994 | 13509 | 2 | 0.54925 | 0.37341 | 0.62997 | 1 | 12650 | 3 | 0.54925 |
| CAPN7 | 5 | 0.03236 | 0.096514 | 0.945031 | 2125 | 3 | -2.2519 | 0.37347 | 0.60139 | 1 | 12651 | 2 | -2.2519 |
| TTBK2 | 5 | 0.029123 | 0.088222 | 0.94395 | 1963 | 3 | -1.0589 | 0.37347 | 0.60139 | 1 | 12652 | 2 | -1.0589 |
| ZNF41 | 6 | 0.2732 | 0.47548 | 0.999994 | 9545 | 3 | -0.04858 | 0.3735 | 0.63004 | 1 | 12653 | 3 | -0.04858 |
| SPR | 6 | 0.51519 | 0.6792 | 0.999994 | 13666 | 2 | -0.01721 | 0.3735 | 0.63004 | 1 | 12654 | 2 | -0.01721 |
| ZNF562 | 6 | 0.1846 | 0.36888 | 0.999994 | 7334 | 3 | 0.003225 | 0.3735 | 0.63004 | 1 | 12655 | 3 | 0.003225 |
| TMEM222 | 6 | 0.33716 | 0.54944 | 0.999994 | 10875 | 2 | 0.15179 | 0.37361 | 0.63012 | 1 | 12656 | 2 | 0.15179 |
| RPL31 | 5 | 0.48248 | 0.63284 | 0.999994 | 13179 | 1 | 0.95766 | 0.37364 | 0.60157 | 1 | 12657 | 3 | 0.95766 |
| ZNF207 | 4 | 0.00024512 | 0.00093295 | 0.352558 | 69 | 1 | 0.9616 | 0.37367 | 0.56888 | 1 | 12658 | 2 | 0.9616 |
| DNAJC6 | 5 | 0.85221 | 0.85604 | 0.999994 | 18300 | 1 | -0.19956 | 0.37377 | 0.60169 | 1 | 12659 | 1 | -0.19956 |
| ZNF92 | 5 | 0.34473 | 0.51761 | 0.999994 | 11049 | 1 | 0.67508 | 0.37377 | 0.60169 | 1 | 12660 | 3 | 0.67508 |
| C2orf27B | 3 | 0.69522 | 0.69653 | 0.999994 | 16258 | 1 | 1.0368 | 0.37379 | 0.52679 | 1 | 12661 | 2 | 1.0368 |
| NonTargetingControlG | 2 | 0.6761 | 0.67583 | 0.999994 | 16022 | 0 | 2.3667 | 0.37381 | 0.45127 | 1 | 12662 | 1 | 2.3667 |
| MED26 | 4 | 0.041733 | 0.10525 | 0.95013 | 2603 | 2 | -2.2428 | 0.37394 | 0.56909 | 1 | 12663 | 2 | -2.2428 |
| HIST1H1D | 4 | 0.017664 | 0.055656 | 0.9235 | 1347 | 2 | -3.8817 | 0.37399 | 0.56913 | 1 | 12664 | 1 | -3.8817 |
| USP47 | 5 | 0.81023 | 0.82608 | 0.999994 | 17635 | 1 | 1.0956 | 0.374 | 0.60193 | 1 | 12665 | 3 | 1.0956 |
| hsa-mir-6511b-1 | 3 | 0.43792 | 0.50684 | 0.999994 | 12540 | 1 | -0.012715 | 0.37403 | 0.52702 | 1 | 12666 | 1 | -0.012715 |
| hsa-mir-6782 | 3 | 0.53334 | 0.56864 | 0.999994 | 13957 | 1 | -0.049177 | 0.37403 | 0.52702 | 1 | 12667 | 1 | -0.049177 |
| SLC7A6OS | 3 | 0.0037776 | 0.010445 | 0.702432 | 427 | 2 | -8.7985 | 0.37403 | 0.52702 | 1 | 12668 | 1 | -8.7985 |
| SPINK5 | 5 | 0.18934 | 0.36001 | 0.999994 | 7465 | 3 | -0.93772 | 0.37409 | 0.60203 | 1 | 12669 | 2 | -0.93772 |
| PRSS23 | 6 | 0.20948 | 0.39947 | 0.999994 | 8015 | 2 | 0.54286 | 0.3741 | 0.63051 | 1 | 12670 | 3 | 0.54286 |
| hsa-mir-5696 | 3 | 0.44046 | 0.50839 | 0.999994 | 12584 | 1 | 1.0771 | 0.37414 | 0.52713 | 1 | 12671 | 2 | 1.0771 |
| CDH24 | 4 | 0.92798 | 0.92775 | 0.999994 | 19813 | 0 | 1.3137 | 0.37423 | 0.56929 | 1 | 12672 | 2 | 1.3137 |
| PRPSAP2 | 6 | 0.098098 | 0.24361 | 0.988088 | 4895 | 2 | 0.67298 | 0.37425 | 0.63062 | 1 | 12673 | 3 | 0.67298 |
| UBAC2 | 5 | 0.80361 | 0.8218 | 0.999994 | 17535 | 1 | 1.0458 | 0.37429 | 0.60223 | 1 | 12674 | 3 | 1.0458 |
| P2RY10 | 6 | 0.018451 | 0.066995 | 0.93914 | 1384 | 3 | -1.331 | 0.37429 | 0.63065 | 1 | 12675 | 3 | -1.331 |
| ART1 | 6 | 0.34614 | 0.55967 | 0.999994 | 11075 | 3 | -0.41511 | 0.37443 | 0.63075 | 1 | 12676 | 2 | -0.41511 |
| OLFML2A | 6 | 0.50059 | 0.6691 | 0.999994 | 13432 | 2 | 0.97988 | 0.37443 | 0.63075 | 1 | 12677 | 3 | 0.97988 |
| NYX | 6 | 0.90752 | 0.90955 | 0.999994 | 19376 | 1 | 0.81816 | 0.37443 | 0.63075 | 1 | 12678 | 3 | 0.81816 |
| C5orf22 | 5 | 0.86452 | 0.86626 | 0.999994 | 18533 | 1 | 1.0055 | 0.37446 | 0.6024 | 1 | 12679 | 3 | 1.0055 |
| UBR5 | 5 | 0.13253 | 0.27647 | 0.999379 | 5988 | 3 | -1.2702 | 0.37446 | 0.6024 | 1 | 12680 | 2 | -1.2702 |
| ERBB2IP | 5 | 0.82225 | 0.83408 | 0.999994 | 17805 | 1 | -0.012861 | 0.37446 | 0.6024 | 1 | 12681 | 2 | -0.012861 |
| EXOSC5 | 5 | 0.29118 | 0.47389 | 0.999994 | 9933 | 2 | -0.5595 | 0.37446 | 0.6024 | 1 | 12682 | 2 | -0.5595 |
| CD3D | 5 | 0.5766 | 0.71533 | 0.999994 | 14639 | 2 | -0.1214 | 0.37446 | 0.6024 | 1 | 12683 | 2 | -0.1214 |
| VWA9 | 5 | 0.92768 | 0.92792 | 0.999994 | 19807 | 0 | 0.94886 | 0.37454 | 0.60248 | 1 | 12684 | 3 | 0.94886 |
| ALKBH3 | 5 | 0.36968 | 0.5381 | 0.999994 | 11501 | 2 | 0.98161 | 0.37458 | 0.60252 | 1 | 12685 | 3 | 0.98161 |
| PNPT1 | 4 | 0.93374 | 0.93351 | 0.999994 | 19945 | 0 | 1.0386 | 0.37463 | 0.5696 | 1 | 12686 | 2 | 1.0386 |
| TJAP1 | 5 | 0.50418 | 0.65157 | 0.999994 | 13499 | 1 | 1.0024 | 0.37475 | 0.60268 | 1 | 12687 | 3 | 1.0024 |

| | | | | | | | | | | | | |
|---|---|---|---|---|---|---|---|---|---|---|---|---|
| NonTargetingControlG | 2 | 0.26842 | 0.3127 | 0.999994 | 9432 | 1 | 0.7784 | 0.37477 | 0.45187 | 1 | 12688 | 1 | 0.7784 |
| hsa-mir-6507 | 2 | 0.087295 | 0.15021 | 0.966359 | 4486 | 1 | -1.862 | 0.37477 | 0.45187 | 1 | 12689 | 1 | -1.862 |
| KCTD18 | 6 | 0.17493 | 0.35677 | 0.999994 | 7106 | 3 | 0.36589 | 0.37477 | 0.63104 | 1 | 12690 | 3 | 0.36589 |
| ASB8 | 5 | 0.42461 | 0.58393 | 0.999994 | 12331 | 1 | 1.1903 | 0.37485 | 0.60277 | 1 | 12691 | 3 | 1.1903 |
| CAT | 6 | 0.26013 | 0.46 | 0.999994 | 9247 | 2 | 0.11749 | 0.37496 | 0.63119 | 1 | 12692 | 3 | 0.11749 |
| NonTargetingControlG | 1 | 0.62494 | 0.62472 | 0.999994 | 15303 | 0 | 1.817 | 0.37506 | 0.37514 | 0.999639 | 12693 | 1 | 1.817 |
| NonTargetingControlG | 1 | 0.62494 | 0.62472 | 0.999994 | 15304 | 0 | 1.817 | 0.37506 | 0.37514 | 0.999639 | 12694 | 1 | 1.817 |
| PLCH1 | 6 | 0.11527 | 0.27677 | 0.999379 | 5487 | 3 | 0.4769 | 0.37506 | 0.63127 | 1 | 12695 | 3 | 0.4769 |
| MOGAT3 | 5 | 0.0028489 | 0.01224 | 0.733879 | 345 | 2 | -0.44927 | 0.37519 | 0.6031 | 1 | 12696 | 2 | -0.44927 |
| PRPF4B | 5 | 0.12621 | 0.26685 | 0.998555 | 5828 | 2 | -0.22495 | 0.37521 | 0.60313 | 1 | 12697 | 2 | -0.22495 |
| SPPL2B | 5 | 0.0036577 | 0.015453 | 0.763603 | 415 | 4 | -3.6727 | 0.37521 | 0.60313 | 1 | 12698 | 1 | -3.6727 |
| BRSK2 | 6 | 0.10935 | 0.26544 | 0.997422 | 5283 | 1 | 0.99987 | 0.37527 | 0.63143 | 1 | 12699 | 3 | 0.99987 |
| DMGDH | 6 | 0.41429 | 0.61162 | 0.999994 | 12169 | 2 | 0.56628 | 0.37535 | 0.63149 | 1 | 12700 | 3 | 0.56628 |
| INPP4A | 6 | 0.13435 | 0.30585 | 0.999994 | 6032 | 3 | 0.49363 | 0.37537 | 0.6315 | 1 | 12701 | 3 | 0.49363 |
| NonTargetingControlG | 2 | 0.73352 | 0.73322 | 0.999994 | 16649 | 0 | 2.4631 | 0.37545 | 0.45232 | 1 | 12702 | 1 | 2.4631 |
| RAPGEF3 | 5 | 0.036761 | 0.10776 | 0.951052 | 2361 | 2 | 0.50761 | 0.37546 | 0.60336 | 1 | 12703 | 2 | 0.50761 |
| hsa-mir-16-2 | 4 | 0.4982 | 0.61551 | 0.999994 | 13398 | 1 | 0.44943 | 0.37547 | 0.57022 | 1 | 12704 | 2 | 0.44943 |
| HIST1H4E | 4 | 0.042123 | 0.106 | 0.950444 | 2620 | 2 | -1.3483 | 0.37547 | 0.57022 | 1 | 12705 | 1 | -1.3483 |
| SLC6A1 | 6 | 0.096782 | 0.24104 | 0.987253 | 4850 | 3 | 0.34384 | 0.37548 | 0.6316 | 1 | 12706 | 3 | 0.34384 |
| CHEK2 | 6 | 0.29121 | 0.49647 | 0.999994 | 9936 | 3 | 0.25785 | 0.37553 | 0.63163 | 1 | 12707 | 3 | 0.25785 |
| hsa-mir-4690 | 2 | 0.0076131 | 0.014503 | 0.754766 | 699 | 1 | -3.2397 | 0.37561 | 0.45244 | 1 | 12708 | 1 | -3.2397 |
| KCTD13 | 5 | 0.54083 | 0.68366 | 0.999994 | 14070 | 1 | 0.18214 | 0.37572 | 0.60364 | 1 | 12709 | 2 | 0.18214 |
| CYP8B1 | 5 | 0.1042 | 0.23295 | 0.984502 | 5106 | 2 | 1.1097 | 0.37585 | 0.60375 | 1 | 12710 | 3 | 1.1097 |
| AP2M1 | 3 | 0.59127 | 0.61047 | 0.999994 | 14836 | 1 | 1.2223 | 0.37588 | 0.52892 | 1 | 12711 | 2 | 1.2223 |
| hsa-mir-4302 | 4 | 0.46922 | 0.6028 | 0.999994 | 12992 | 1 | 1.1202 | 0.37591 | 0.57057 | 1 | 12712 | 2 | 1.1202 |
| TMEM102 | 5 | 0.69181 | 0.7628 | 0.999994 | 16226 | 1 | 0.32484 | 0.376 | 0.60389 | 1 | 12713 | 2 | 0.32484 |
| LINGO4 | 5 | 0.037498 | 0.10959 | 0.951052 | 2401 | 4 | -2.5632 | 0.376 | 0.60389 | 1 | 12714 | 1 | -2.5632 |
| KCNG3 | 5 | 0.18557 | 0.35466 | 0.999994 | 7356 | 3 | -0.95183 | 0.376 | 0.60389 | 1 | 12715 | 2 | -0.95183 |
| USF1 | 6 | 0.55482 | 0.70709 | 0.999994 | 14282 | 1 | 0.96926 | 0.376 | 0.63201 | 1 | 12716 | 3 | 0.96926 |
| hsa-mir-7977 | 3 | 0.67921 | 0.68204 | 0.999994 | 16077 | 1 | 1.0146 | 0.37607 | 0.5291 | 1 | 12717 | 2 | 1.0146 |
| FEZ1 | 6 | 0.1508 | 0.32668 | 0.999994 | 6481 | 2 | 0.70973 | 0.37608 | 0.63208 | 1 | 12718 | 3 | 0.70973 |
| TAS2R13 | 6 | 0.27898 | 0.48223 | 0.999994 | 9672 | 2 | 0.085513 | 0.3761 | 0.6321 | 1 | 12719 | 3 | 0.085513 |
| PPOX | 6 | 0.14617 | 0.32085 | 0.999994 | 6358 | 3 | -1.0742 | 0.3761 | 0.6321 | 1 | 12720 | 2 | -1.0742 |
| hsa-mir-1256 | 3 | 0.24631 | 0.37037 | 0.999994 | 8931 | 1 | 1.0997 | 0.37612 | 0.52916 | 1 | 12721 | 2 | 1.0997 |
| CYP51A1 | 5 | 0.56103 | 0.70155 | 0.999994 | 14390 | 2 | 0.50453 | 0.3763 | 0.60416 | 1 | 12722 | 3 | 0.50453 |
| ATP6V1D | 6 | 0.00022096 | 0.001178 | 0.368105 | 61 | 4 | -6.7782 | 0.37632 | 0.63227 | 1 | 12723 | 2 | -6.7782 |
| NonTargetingControlG | 2 | 0.27735 | 0.31947 | 0.999994 | 9642 | 1 | 0.98527 | 0.37632 | 0.45288 | 1 | 12724 | 1 | 0.98527 |
| GIF | 6 | 0.35667 | 0.57154 | 0.999994 | 11278 | 3 | 0.47752 | 0.37643 | 0.63235 | 1 | 12725 | 3 | 0.47752 |
| TTC37 | 4 | 0.48469 | 0.60957 | 0.999994 | 13210 | 1 | 0.87357 | 0.37647 | 0.57098 | 1 | 12726 | 2 | 0.87357 |
| EPHA4 | 5 | 0.67302 | 0.75457 | 0.999994 | 15972 | 1 | 0.46412 | 0.3765 | 0.60438 | 1 | 12727 | 2 | 0.46412 |
| MAGEB4 | 6 | 0.9263 | 0.92637 | 0.999994 | 19774 | 1 | 0.83727 | 0.37655 | 0.63244 | 1 | 12728 | 3 | 0.83727 |
| MAFA | 5 | 0.065777 | 0.17149 | 0.967658 | 3682 | 3 | -1.6369 | 0.37668 | 0.60455 | 1 | 12729 | 2 | -1.6369 |
| VAV1 | 5 | 0.017112 | 0.05616 | 0.9235 | 1307 | 2 | 0.90978 | 0.37725 | 0.60514 | 1 | 12730 | 3 | 0.90978 |
| GRB7 | 6 | 0.14252 | 0.31626 | 0.999994 | 6253 | 2 | -0.13169 | 0.37727 | 0.63301 | 1 | 12731 | 2 | -0.13169 |
| CCDC30 | 6 | 0.28507 | 0.48934 | 0.999994 | 9812 | 3 | -0.52747 | 0.37727 | 0.63301 | 1 | 12732 | 1 | -0.52747 |
| SRGAP1 | 6 | 0.20911 | 0.39901 | 0.999994 | 8007 | 3 | -0.064315 | 0.37727 | 0.63301 | 1 | 12733 | 3 | -0.064315 |
| ARHGEF5 | 6 | 0.14768 | 0.32273 | 0.999994 | 6398 | 3 | -0.54601 | 0.37727 | 0.63301 | 1 | 12734 | 3 | -0.54601 |
| PEX1 | 6 | 0.14383 | 0.3179 | 0.999994 | 6293 | 2 | 0.15072 | 0.37727 | 0.63301 | 1 | 12735 | 2 | 0.15072 |
| CALCA | 6 | 0.26989 | 0.47159 | 0.999994 | 9471 | 3 | 0.32858 | 0.37733 | 0.63306 | 1 | 12736 | 3 | 0.32858 |
| hsa-mir-1255b-1 | 1 | 0.62259 | 0.62232 | 0.999994 | 15269 | 0 | 1.5434 | 0.37741 | 0.37749 | 1 | 12737 | 1 | 1.5434 |
| hsa-mir-582 | 3 | 0.68413 | 0.68643 | 0.999994 | 16146 | 1 | 0.74065 | 0.37742 | 0.53049 | 1 | 12738 | 2 | 0.74065 |
| MRC1 | 5 | 0.29451 | 0.47661 | 0.999994 | 10007 | 1 | 0.89135 | 0.37744 | 0.60533 | 1 | 12739 | 3 | 0.89135 |

| | | | | | | | | | | | | |
|---|---|---|---|---|---|---|---|---|---|---|---|---|
| MZT1 | 4 | 0.4255 | 0.57187 | 0.999994 | 12345 | 2 | -0.67389 | 0.37748 | 0.57171 | 1 | 12740 | 1 | -0.67389 |
| PRAMEF16 | 4 | 0.32745 | 0.47107 | 0.999994 | 10667 | 2 | 0.45784 | 0.37748 | 0.57171 | 1 | 12741 | 2 | 0.45784 |
| DPP3 | 6 | 0.15252 | 0.32884 | 0.999994 | 6521 | 3 | -0.79745 | 0.37752 | 0.63321 | 1 | 12742 | 2 | -0.79745 |
| PDHA2 | 6 | 0.61388 | 0.75026 | 0.999994 | 15129 | 2 | 0.65032 | 0.37752 | 0.63321 | 1 | 12743 | 3 | 0.65032 |
| PTPRN | 6 | 0.029342 | 0.097236 | 0.945253 | 1975 | 4 | -2.1302 | 0.37752 | 0.63321 | 1 | 12744 | 2 | -2.1302 |
| hsa-mir-548m | 4 | 0.86708 | 0.86663 | 0.999994 | 18576 | 0 | 1.0316 | 0.37752 | 0.57175 | 1 | 12745 | 2 | 1.0316 |
| RNF187 | 5 | 0.028843 | 0.087495 | 0.94395 | 1944 | 2 | 1.0912 | 0.37761 | 0.60551 | 1 | 12746 | 3 | 1.0912 |
| hsa-mir-4649 | 3 | 0.70498 | 0.70569 | 0.999994 | 16359 | 1 | 1.2281 | 0.37766 | 0.53074 | 1 | 12747 | 2 | 1.2281 |
| hsa-mir-27b | 4 | 0.32284 | 0.46633 | 0.999994 | 10571 | 2 | 0.51618 | 0.37767 | 0.57187 | 1 | 12748 | 2 | 0.51618 |
| GOT1L1 | 5 | 0.37071 | 0.539 | 0.999994 | 11525 | 1 | 0.1329 | 0.37779 | 0.60569 | 1 | 12749 | 1 | 0.1329 |
| CR1 | 5 | 0.10157 | 0.22886 | 0.983354 | 5021 | 3 | -1.5029 | 0.37779 | 0.60569 | 1 | 12750 | 1 | -1.5029 |
| NUDCD3 | 5 | 0.034058 | 0.10087 | 0.946414 | 2220 | 2 | 0.35554 | 0.37786 | 0.60577 | 1 | 12751 | 2 | 0.35554 |
| WAC | 3 | 0.50069 | 0.54661 | 0.999994 | 13433 | 1 | 1.4583 | 0.3779 | 0.53098 | 1 | 12752 | 2 | 1.4583 |
| EDC3 | 6 | 0.33796 | 0.55034 | 0.999994 | 10898 | 2 | 0.68127 | 0.37798 | 0.63357 | 1 | 12753 | 3 | 0.68127 |
| GLCE | 6 | 0.040107 | 0.12214 | 0.951052 | 2521 | 4 | -1.8725 | 0.37799 | 0.63357 | 1 | 12754 | 1 | -1.8725 |
| KRTAP23-1 | 6 | 0.058239 | 0.16197 | 0.967126 | 3350 | 3 | -0.32932 | 0.37799 | 0.63357 | 1 | 12755 | 3 | -0.32932 |
| CCDC19 | 6 | 0.35489 | 0.56954 | 0.999994 | 11249 | 2 | 0.2035 | 0.37799 | 0.63357 | 1 | 12756 | 3 | 0.2035 |
| ATP6V0A1 | 6 | 0.55568 | 0.70771 | 0.999994 | 14299 | 1 | 1.2329 | 0.37808 | 0.63364 | 1 | 12757 | 3 | 1.2329 |
| MYO3A | 6 | 0.94681 | 0.94678 | 0.999994 | 20259 | 0 | 0.89444 | 0.37812 | 0.63367 | 1 | 12758 | 3 | 0.89444 |
| MAF1 | 5 | 0.040757 | 0.11775 | 0.951052 | 2548 | 2 | 0.96424 | 0.37816 | 0.60606 | 1 | 12759 | 3 | 0.96424 |
| TMEM256 | 5 | 0.20947 | 0.38865 | 0.999994 | 8014 | 1 | 0.95829 | 0.37821 | 0.60611 | 1 | 12760 | 3 | 0.95829 |
| RAB7L1 | 5 | 0.50716 | 0.65418 | 0.999994 | 13540 | 2 | 0.9263 | 0.37828 | 0.60618 | 1 | 12761 | 3 | 0.9263 |
| SAPCD2 | 4 | 0.18133 | 0.32149 | 0.999994 | 7260 | 1 | 1.1114 | 0.37852 | 0.5725 | 1 | 12762 | 2 | 1.1114 |
| hsa-mir-4462 | 4 | 0.84296 | 0.84245 | 0.999994 | 18138 | 0 | 1.104 | 0.37871 | 0.57265 | 1 | 12763 | 2 | 1.104 |
| HOXB9 | 4 | 0.027839 | 0.077112 | 0.93914 | 1897 | 1 | 1.018 | 0.37878 | 0.5727 | 1 | 12764 | 2 | 1.018 |
| SCPEP1 | 5 | 0.035571 | 0.10468 | 0.948754 | 2297 | 4 | -1.9893 | 0.37878 | 0.60671 | 1 | 12765 | 1 | -1.9893 |
| CLEC2D | 5 | 0.11207 | 0.24521 | 0.98979 | 5367 | 3 | -1.466 | 0.37878 | 0.60671 | 1 | 12766 | 1 | -1.466 |
| PATZ1 | 5 | 0.52851 | 0.67275 | 0.999994 | 13871 | 1 | -0.35223 | 0.37878 | 0.60671 | 1 | 12767 | 1 | -0.35223 |
| LYZL2 | 3 | 0.62734 | 0.63856 | 0.999994 | 15342 | 1 | 0.31767 | 0.37891 | 0.53202 | 1 | 12768 | 1 | 0.31767 |
| LIPT1 | 6 | 0.0038707 | 0.017003 | 0.785648 | 435 | 4 | -4.8742 | 0.379 | 0.63434 | 1 | 12769 | 2 | -4.8742 |
| C6orf226 | 6 | 0.8601 | 0.87749 | 0.999994 | 18441 | 1 | 0.82058 | 0.379 | 0.63434 | 1 | 12770 | 3 | 0.82058 |
| RGR | 6 | 0.0017807 | 0.0088761 | 0.680529 | 247 | 2 | 0.039591 | 0.379 | 0.63434 | 1 | 12771 | 2 | 0.039591 |
| PYGO2 | 6 | 0.022143 | 0.078833 | 0.93914 | 1594 | 2 | 0.59015 | 0.37923 | 0.63452 | 1 | 12772 | 3 | 0.59015 |
| KCNT1 | 4 | 0.29569 | 0.43858 | 0.999994 | 10033 | 2 | -0.9603 | 0.37931 | 0.57308 | 1 | 12773 | 1 | -0.9603 |
| ARL4A | 3 | 0.059718 | 0.12905 | 0.952525 | 3411 | 2 | -2.8308 | 0.37945 | 0.53257 | 1 | 12774 | 1 | -2.8308 |
| DMD | 5 | 0.20186 | 0.37791 | 0.999994 | 7822 | 2 | -0.36814 | 0.37955 | 0.60745 | 1 | 12775 | 2 | -0.36814 |
| FBXL8 | 5 | 0.070777 | 0.17964 | 0.968171 | 3865 | 3 | -0.7907 | 0.37955 | 0.60745 | 1 | 12776 | 2 | -0.7907 |
| SGCA | 5 | 0.053409 | 0.14861 | 0.965554 | 3135 | 2 | -0.22594 | 0.37955 | 0.60745 | 1 | 12777 | 2 | -0.22594 |
| SCO1 | 5 | 0.52882 | 0.67302 | 0.999994 | 13879 | 1 | 1.1058 | 0.37963 | 0.60752 | 1 | 12778 | 3 | 1.1058 |
| ARL17A | 5 | 0.84785 | 0.85257 | 0.999994 | 18215 | 1 | 1.1692 | 0.37966 | 0.60755 | 1 | 12779 | 3 | 1.1692 |
| hsa-mir-6739 | 3 | 0.5136 | 0.55514 | 0.999994 | 13634 | 1 | 1.0987 | 0.3797 | 0.53281 | 1 | 12780 | 2 | 1.0987 |
| PRDM9 | 4 | 0.40386 | 0.54967 | 0.999994 | 12013 | 2 | 0.77774 | 0.37971 | 0.57339 | 1 | 12781 | 2 | 0.77774 |
| FAM25G | 3 | 0.67384 | 0.6772 | 0.999994 | 15987 | 1 | 1.1429 | 0.37971 | 0.53283 | 1 | 12782 | 2 | 1.1429 |
| UCHL1 | 6 | 0.18782 | 0.37281 | 0.999994 | 7417 | 2 | 0.63534 | 0.3798 | 0.63496 | 1 | 12783 | 3 | 0.63534 |
| ALDH3B2 | 4 | 0.21781 | 0.35899 | 0.999994 | 8220 | 2 | 0.27574 | 0.37983 | 0.57347 | 1 | 12784 | 2 | 0.27574 |
| ALS2CL | 6 | 0.052658 | 0.14988 | 0.966359 | 3098 | 4 | -1.2093 | 0.37991 | 0.63504 | 1 | 12785 | 1 | -1.2093 |
| CDON | 6 | 0.22662 | 0.42004 | 0.999994 | 8444 | 3 | 0.22673 | 0.38005 | 0.63514 | 1 | 12786 | 3 | 0.22673 |
| TMCO4 | 5 | 0.45879 | 0.61264 | 0.999994 | 12832 | 1 | 0.95248 | 0.38021 | 0.6081 | 1 | 12787 | 3 | 0.95248 |
| NonTargetingControlG | 1 | 0.61967 | 0.61938 | 0.999994 | 15214 | 0 | 1.5133 | 0.38033 | 0.3804 | 1 | 12788 | 1 | 1.5133 |
| TIMM23 | 3 | 0.074 | 0.14836 | 0.965554 | 3996 | 1 | 1.2432 | 0.38039 | 0.53353 | 1 | 12789 | 2 | 1.2432 |
| ST5 | 5 | 0.5077 | 0.65466 | 0.999994 | 13546 | 2 | 0.88417 | 0.38058 | 0.60847 | 1 | 12790 | 3 | 0.88417 |
| hsa-mir-760 | 4 | 0.092948 | 0.20244 | 0.977973 | 4717 | 3 | -1.67 | 0.38068 | 0.57411 | 1 | 12791 | 1 | -1.67 |

| | | | | | | | | | | | | | |
|---|---|---|---|---|---|---|---|---|---|---|---|---|---|
| AAGAB | 4 | 0.26258 | 0.4047 | 0.999994 | 9300 | 2 | -0.31782 | 0.38068 | 0.57411 | 1 | 12792 | 2 | -0.31782 |
| U2SURP | 4 | 0.055725 | 0.13263 | 0.95318 | 3233 | 3 | -2.2655 | 0.38068 | 0.57411 | 1 | 12793 | 1 | -2.2655 |
| NTM | 6 | 0.25459 | 0.45343 | 0.999994 | 9120 | 3 | -0.66245 | 0.38073 | 0.63565 | 1 | 12794 | 2 | -0.66245 |
| AKAP9 | 5 | 0.25905 | 0.44799 | 0.999994 | 9219 | 2 | -0.56907 | 0.38073 | 0.60861 | 1 | 12795 | 2 | -0.56907 |
| FBLN5 | 3 | 0.48886 | 0.53885 | 0.999994 | 13265 | 1 | -0.032027 | 0.38079 | 0.53397 | 1 | 12796 | 1 | -0.032027 |
| UNC93A | 6 | 0.59506 | 0.73633 | 0.999994 | 14888 | 2 | 0.48804 | 0.3808 | 0.63571 | 1 | 12797 | 3 | 0.48804 |
| RAB4A | 6 | 0.022111 | 0.078727 | 0.93914 | 1593 | 3 | -0.37826 | 0.38089 | 0.6358 | 1 | 12798 | 3 | -0.37826 |
| BEGAIN | 6 | 0.50149 | 0.66972 | 0.999994 | 13447 | 1 | 0.75484 | 0.38103 | 0.63592 | 1 | 12799 | 3 | 0.75484 |
| TMEM216 | 5 | 0.92349 | 0.92368 | 0.999994 | 19707 | 0 | -0.037104 | 0.38104 | 0.60891 | 1 | 12800 | 2 | -0.037104 |
| FKBP11 | 5 | 0.13294 | 0.2771 | 0.999379 | 5993 | 3 | -0.74686 | 0.38104 | 0.60891 | 1 | 12801 | 1 | -0.74686 |
| PYROXD2 | 6 | 0.0056964 | 0.023681 | 0.832699 | 571 | 2 | 0.70737 | 0.38113 | 0.63599 | 1 | 12802 | 3 | 0.70737 |
| ZNF98 | 1 | 0.61886 | 0.61858 | 0.999994 | 15202 | 0 | 1.4483 | 0.38114 | 0.38124 | 1 | 12803 | 1 | 1.4483 |
| HINT2 | 5 | 0.49415 | 0.64284 | 0.999994 | 13332 | 1 | 0.89503 | 0.38115 | 0.60902 | 1 | 12804 | 3 | 0.89503 |
| ABHD17C | 4 | 0.15811 | 0.29755 | 0.999994 | 6674 | 1 | 1.1659 | 0.38119 | 0.57449 | 1 | 12805 | 2 | 1.1659 |
| HOXB4 | 4 | 0.071557 | 0.16287 | 0.967126 | 3899 | 2 | 0.32482 | 0.38123 | 0.57452 | 1 | 12806 | 2 | 0.32482 |
| NonTargetingControlG | 2 | 0.24152 | 0.29256 | 0.999632 | 8813 | 1 | 0.54183 | 0.38125 | 0.45605 | 1 | 12807 | 1 | 0.54183 |
| PIGU | 6 | 0.00099123 | 0.005036 | 0.589484 | 167 | 4 | -4.9922 | 0.38136 | 0.63617 | 1 | 12808 | 2 | -4.9922 |
| ADPRHL1 | 5 | 0.63304 | 0.73849 | 0.999994 | 15414 | 1 | -0.30875 | 0.38149 | 0.60937 | 1 | 12809 | 2 | -0.30875 |
| HIST1H3C | 6 | 0.25993 | 0.45977 | 0.999994 | 9239 | 3 | -0.23246 | 0.38149 | 0.63628 | 1 | 12810 | 3 | -0.23246 |
| GRK7 | 6 | 0.01205 | 0.045884 | 0.892434 | 995 | 4 | -3.4306 | 0.38152 | 0.6363 | 1 | 12811 | 2 | -3.4306 |
| WDR3 | 6 | 0.061158 | 0.16824 | 0.967126 | 3481 | 3 | 0.058218 | 0.38165 | 0.6364 | 1 | 12812 | 3 | 0.058218 |
| EMG1 | 6 | 0.26282 | 0.46321 | 0.999994 | 9308 | 2 | -0.5142 | 0.38172 | 0.63646 | 1 | 12813 | 2 | -0.5142 |
| WDR89 | 6 | 0.55738 | 0.70895 | 0.999994 | 14320 | 2 | 0.80606 | 0.38181 | 0.63652 | 1 | 12814 | 3 | 0.80606 |
| PRKD3 | 6 | 0.42319 | 0.61749 | 0.999994 | 12297 | 2 | 0.99157 | 0.38186 | 0.63657 | 1 | 12815 | 3 | 0.99157 |
| CNP | 6 | 0.50415 | 0.67161 | 0.999994 | 13498 | 1 | 0.85763 | 0.3819 | 0.63659 | 1 | 12816 | 3 | 0.85763 |
| hsa-mir-4301 | 4 | 0.44782 | 0.59376 | 0.999994 | 12684 | 1 | 1.2501 | 0.38193 | 0.57507 | 1 | 12817 | 2 | 1.2501 |
| SLC7A4 | 5 | 0.4358 | 0.5933 | 0.999994 | 12512 | 2 | -0.23479 | 0.38209 | 0.60999 | 1 | 12818 | 2 | -0.23479 |
| HMCN1 | 6 | 0.40777 | 0.60744 | 0.999994 | 12073 | 2 | 0.11631 | 0.38211 | 0.63675 | 1 | 12819 | 3 | 0.11631 |
| TMEM215 | 6 | 0.14013 | 0.31318 | 0.999994 | 6188 | 3 | 0.076121 | 0.38223 | 0.63685 | 1 | 12820 | 3 | 0.076121 |
| C11orf94 | 4 | 0.65711 | 0.6975 | 0.999994 | 15743 | 1 | 1.1708 | 0.3823 | 0.57531 | 1 | 12821 | 2 | 1.1708 |
| C6orf58 | 4 | 0.077841 | 0.17472 | 0.968171 | 4143 | 2 | -0.89096 | 0.3823 | 0.57531 | 1 | 12822 | 2 | -0.89096 |
| MAF | 5 | 0.14799 | 0.29975 | 0.999994 | 6407 | 2 | -0.14067 | 0.3823 | 0.61021 | 1 | 12823 | 2 | -0.14067 |
| SEC63 | 6 | 0.25342 | 0.45201 | 0.999994 | 9090 | 2 | -0.080151 | 0.38238 | 0.63697 | 1 | 12824 | 2 | -0.080151 |
| CTSB | 6 | 0.12807 | 0.29783 | 0.999994 | 5880 | 3 | -0.70839 | 0.38238 | 0.63697 | 1 | 12825 | 2 | -0.70839 |
| RPA4 | 6 | 0.75432 | 0.83001 | 0.999994 | 16899 | 1 | 0.51606 | 0.38245 | 0.63702 | 1 | 12826 | 3 | 0.51606 |
| ARFGAP1 | 6 | 0.14046 | 0.3136 | 0.999994 | 6197 | 2 | 0.82661 | 0.38245 | 0.63702 | 1 | 12827 | 3 | 0.82661 |
| GNPTG | 5 | 0.19748 | 0.37164 | 0.999994 | 7686 | 3 | -0.82817 | 0.3825 | 0.61041 | 1 | 12828 | 1 | -0.82817 |
| FREM3 | 5 | 0.018599 | 0.060289 | 0.932932 | 1393 | 3 | -2.6182 | 0.3825 | 0.61041 | 1 | 12829 | 2 | -2.6182 |
| STAT5A | 5 | 0.086565 | 0.20507 | 0.978186 | 4470 | 3 | -1.8599 | 0.3825 | 0.61041 | 1 | 12830 | 2 | -1.8599 |
| GLIS2 | 5 | 0.89937 | 0.89947 | 0.999994 | 19210 | 0 | 0.19801 | 0.38256 | 0.61047 | 1 | 12831 | 2 | 0.19801 |
| BUB1B | 4 | 0.54457 | 0.63703 | 0.999994 | 14135 | 1 | 0.94301 | 0.38274 | 0.57564 | 1 | 12832 | 2 | 0.94301 |
| RSAD1 | 6 | 0.34959 | 0.56354 | 0.999994 | 11139 | 3 | 0.46492 | 0.38278 | 0.63726 | 1 | 12833 | 3 | 0.46492 |
| NonTargetingControlG | 1 | 0.61718 | 0.61689 | 0.999994 | 15173 | 0 | 1.6577 | 0.38282 | 0.38294 | 1 | 12834 | 1 | 1.6577 |
| PER2 | 5 | 0.91048 | 0.9105 | 0.999994 | 19439 | 0 | 0.95795 | 0.38307 | 0.61099 | 1 | 12835 | 3 | 0.95795 |
| HSPBAP1 | 3 | 0.72251 | 0.72249 | 0.999994 | 16521 | 1 | 1.022 | 0.38313 | 0.53635 | 1 | 12836 | 2 | 1.022 |
| TFDP2 | 5 | 0.10753 | 0.23813 | 0.987253 | 5228 | 1 | 0.069181 | 0.38327 | 0.6112 | 1 | 12837 | 2 | 0.069181 |
| TAAR5 | 5 | 0.037447 | 0.10946 | 0.951052 | 2397 | 2 | -0.30517 | 0.38327 | 0.6112 | 1 | 12838 | 2 | -0.30517 |
| RRP12 | 5 | 0.080525 | 0.19544 | 0.975797 | 4254 | 3 | -1.9463 | 0.38327 | 0.6112 | 1 | 12839 | 1 | -1.9463 |
| DECR2 | 6 | 0.63592 | 0.76684 | 0.999994 | 15460 | 2 | 1.1212 | 0.38329 | 0.63765 | 1 | 12840 | 3 | 1.1212 |
| C7orf62 | 6 | 0.19163 | 0.37753 | 0.999994 | 7534 | 2 | 0.93972 | 0.38329 | 0.63765 | 1 | 12841 | 3 | 0.93972 |
| UPK3B | 6 | 0.16202 | 0.3408 | 0.999994 | 6797 | 2 | 1.1084 | 0.38329 | 0.63765 | 1 | 12842 | 3 | 1.1084 |
| MKRN3 | 6 | 0.10065 | 0.2486 | 0.990929 | 4995 | 3 | 0.036727 | 0.38329 | 0.63765 | 1 | 12843 | 3 | 0.036727 |

| | | | | | | | | | | | | | |
|---|---|---|---|---|---|---|---|---|---|---|---|---|---|
| hsa-mir-4782 | 4 | 0.12651 | 0.26156 | 0.995257 | 5836 | 1 | 1.1368 | 0.38331 | 0.57606 | 1 | 12844 | 2 | 1.1368 |
| KRT18 | 5 | 0.1052 | 0.2345 | 0.984971 | 5139 | 2 | -0.53786 | 0.38339 | 0.61132 | 1 | 12845 | 2 | -0.53786 |
| DCUN1D4 | 5 | 0.11021 | 0.2423 | 0.988088 | 5311 | 3 | -1.5419 | 0.38339 | 0.61132 | 1 | 12846 | 2 | -1.5419 |
| OR2T3 | 5 | 0.067606 | 0.17451 | 0.968171 | 3741 | 2 | 0.17428 | 0.38339 | 0.61132 | 1 | 12847 | 2 | 0.17428 |
| FGF8 | 5 | 0.19317 | 0.36545 | 0.999994 | 7579 | 3 | -0.59902 | 0.38339 | 0.61132 | 1 | 12848 | 2 | -0.59902 |
| SCML1 | 5 | 0.91662 | 0.91672 | 0.999994 | 19551 | 0 | 0.81821 | 0.38339 | 0.61132 | 1 | 12849 | 3 | 0.81821 |
| NPR2 | 5 | 0.77894 | 0.80684 | 0.999994 | 17193 | 1 | 0.91257 | 0.38339 | 0.61132 | 1 | 12850 | 3 | 0.91257 |
| HS3ST4 | 6 | 0.07964 | 0.20658 | 0.978744 | 4215 | 4 | -1.3766 | 0.38347 | 0.63779 | 1 | 12851 | 2 | -1.3766 |
| EIF4EBP3 | 6 | 0.084883 | 0.21729 | 0.980699 | 4406 | 2 | 0.32263 | 0.38347 | 0.63779 | 1 | 12852 | 2 | 0.32263 |
| GADD45GIP1 | 3 | 0.33656 | 0.44828 | 0.999994 | 10862 | 1 | 0.13545 | 0.38383 | 0.53707 | 1 | 12853 | 1 | 0.13545 |
| NonTargetingControlG | 2 | 0.56858 | 0.56886 | 0.999994 | 14517 | 1 | 1.8659 | 0.38384 | 0.45776 | 1 | 12854 | 1 | 1.8659 |
| hsa-mir-1271 | 3 | 0.86239 | 0.86236 | 0.999994 | 18494 | 0 | 0.99867 | 0.3839 | 0.53715 | 1 | 12855 | 2 | 0.99867 |
| NEU4 | 5 | 0.09135 | 0.21276 | 0.979587 | 4650 | 3 | -1.666 | 0.38401 | 0.61195 | 1 | 12856 | 1 | -1.666 |
| DLX3 | 6 | 0.13898 | 0.31173 | 0.999994 | 6162 | 3 | 0.062727 | 0.38402 | 0.63821 | 1 | 12857 | 3 | 0.062727 |
| GLUL | 5 | 0.31226 | 0.49102 | 0.999994 | 10366 | 2 | 0.3464 | 0.38407 | 0.61201 | 1 | 12858 | 2 | 0.3464 |
| RBM34 | 6 | 0.23141 | 0.42578 | 0.999994 | 8553 | 2 | -0.16585 | 0.38414 | 0.6383 | 1 | 12859 | 1 | -0.16585 |
| ZDHHC8 | 6 | 0.092477 | 0.23247 | 0.984502 | 4702 | 3 | -0.20322 | 0.38414 | 0.6383 | 1 | 12860 | 3 | -0.20322 |
| hsa-mir-3180-2 | 1 | 0.61582 | 0.6155 | 0.999994 | 15156 | 0 | 1.5789 | 0.38418 | 0.38431 | 1 | 12861 | 1 | 1.5789 |
| hsa-mir-1972-1 | 3 | 0.78569 | 0.78559 | 0.999994 | 17289 | 0 | 1.0481 | 0.38425 | 0.53748 | 1 | 12862 | 2 | 1.0481 |
| hsa-mir-4658 | 4 | 0.44581 | 0.59274 | 0.999994 | 12660 | 1 | 1.1557 | 0.38435 | 0.57683 | 1 | 12863 | 2 | 1.1557 |
| hsa-mir-3140 | 4 | 0.84645 | 0.84598 | 0.999994 | 18199 | 0 | 1.2507 | 0.38437 | 0.57684 | 1 | 12864 | 2 | 1.2507 |
| PMAIP1 | 5 | 0.10229 | 0.22997 | 0.983417 | 5048 | 2 | 0.89636 | 0.3845 | 0.61245 | 1 | 12865 | 3 | 0.89636 |
| AMER1 | 5 | 0.74594 | 0.78871 | 0.999994 | 16813 | 1 | 0.3953 | 0.38466 | 0.61262 | 1 | 12866 | 1 | 0.3953 |
| TMEM167A | 5 | 0.18422 | 0.35265 | 0.999994 | 7322 | 3 | -0.90814 | 0.38466 | 0.61262 | 1 | 12867 | 2 | -0.90814 |
| TMEM41B | 5 | 0.17013 | 0.33234 | 0.999994 | 6987 | 2 | 0.51383 | 0.38466 | 0.61262 | 1 | 12868 | 2 | 0.51383 |
| SLC7A7 | 5 | 0.14956 | 0.30214 | 0.999994 | 6447 | 3 | -1.2101 | 0.38466 | 0.61262 | 1 | 12869 | 2 | -1.2101 |
| hsa-mir-4303 | 4 | 0.23305 | 0.37463 | 0.999994 | 8598 | 2 | -0.073389 | 0.3847 | 0.57709 | 1 | 12870 | 2 | -0.073389 |
| XXYLT1 | 4 | 0.80326 | 0.80412 | 0.999994 | 17528 | 1 | 0.17032 | 0.3847 | 0.57709 | 1 | 12871 | 1 | 0.17032 |
| SNRNP27 | 4 | 0.3131 | 0.45643 | 0.999994 | 10379 | 2 | -0.26799 | 0.3847 | 0.57709 | 1 | 12872 | 2 | -0.26799 |
| CBX1 | 4 | 0.10715 | 0.22782 | 0.983354 | 5217 | 2 | -0.65063 | 0.3847 | 0.57709 | 1 | 12873 | 1 | -0.65063 |
| NOTCH2NL | 6 | 0.017946 | 0.06533 | 0.93914 | 1358 | 3 | -1.4814 | 0.38476 | 0.63879 | 1 | 12874 | 3 | -1.4814 |
| LUC7L | 6 | 0.23931 | 0.43516 | 0.999994 | 8758 | 1 | 0.76968 | 0.38476 | 0.63879 | 1 | 12875 | 3 | 0.76968 |
| MPPED2 | 6 | 0.68235 | 0.80284 | 0.999994 | 16123 | 1 | 0.72594 | 0.38481 | 0.63883 | 1 | 12876 | 3 | 0.72594 |
| KRT79 | 6 | 0.60364 | 0.74268 | 0.999994 | 14990 | 2 | -0.53009 | 0.38491 | 0.63891 | 1 | 12877 | 2 | -0.53009 |
| DACH2 | 6 | 0.029142 | 0.096773 | 0.945031 | 1965 | 3 | -0.63371 | 0.38491 | 0.63891 | 1 | 12878 | 2 | -0.63371 |
| PCDH9 | 6 | 0.4408 | 0.62896 | 0.999994 | 12588 | 2 | 0.43563 | 0.38491 | 0.63891 | 1 | 12879 | 2 | 0.43563 |
| SCML2 | 6 | 0.5478 | 0.70206 | 0.999994 | 14176 | 2 | -0.16475 | 0.38491 | 0.63891 | 1 | 12880 | 2 | -0.16475 |
| hsa-mir-6860 | 3 | 0.75391 | 0.75383 | 0.999994 | 16897 | 0 | 0.13929 | 0.38496 | 0.53822 | 1 | 12881 | 1 | 0.13929 |
| NSMCE4A | 3 | 0.18377 | 0.29205 | 0.999632 | 7310 | 2 | -0.69254 | 0.38496 | 0.53822 | 1 | 12882 | 1 | -0.69254 |
| hsa-mir-212 | 3 | 0.84922 | 0.8492 | 0.999994 | 18245 | 0 | 1.1815 | 0.38517 | 0.53843 | 1 | 12883 | 2 | 1.1815 |
| NXPE2 | 5 | 0.85279 | 0.85651 | 0.999994 | 18313 | 1 | 0.91857 | 0.38519 | 0.61317 | 1 | 12884 | 3 | 0.91857 |
| KCTD19 | 5 | 0.20879 | 0.3877 | 0.999994 | 7998 | 3 | -0.54348 | 0.3852 | 0.61318 | 1 | 12885 | 2 | -0.54348 |
| UBTF | 4 | 0.050973 | 0.1234 | 0.951052 | 3015 | 2 | -1.8756 | 0.3853 | 0.57756 | 1 | 12886 | 2 | -1.8756 |
| ZDHHC5 | 4 | 0.33908 | 0.48304 | 0.999994 | 10917 | 2 | 0.69426 | 0.3853 | 0.57756 | 1 | 12887 | 2 | 0.69426 |
| CLEC1B | 5 | 0.18203 | 0.34954 | 0.999994 | 7272 | 3 | -0.91794 | 0.38537 | 0.61334 | 1 | 12888 | 2 | -0.91794 |
| PKDCC | 5 | 0.53543 | 0.67883 | 0.999994 | 13987 | 1 | 0.88328 | 0.38548 | 0.61344 | 1 | 12889 | 3 | 0.88328 |
| ITGA1 | 5 | 0.028489 | 0.086577 | 0.94395 | 1925 | 3 | -2.9773 | 0.38562 | 0.61357 | 1 | 12890 | 2 | -2.9773 |
| TBC1D26 | 4 | 0.85738 | 0.85699 | 0.999994 | 18400 | 0 | 1.4136 | 0.38571 | 0.57783 | 1 | 12891 | 2 | 1.4136 |
| OR4K15 | 5 | 0.32369 | 0.50033 | 0.999994 | 10587 | 1 | 1.0227 | 0.38572 | 0.61368 | 1 | 12892 | 3 | 1.0227 |
| NonTargetingControlG | 2 | 0.44528 | 0.45517 | 0.999994 | 12654 | 1 | 1.4428 | 0.38589 | 0.45909 | 1 | 12893 | 1 | 1.4428 |
| NCK1 | 5 | 0.45139 | 0.60639 | 0.999994 | 12726 | 2 | 0.8703 | 0.38601 | 0.61396 | 1 | 12894 | 3 | 0.8703 |
| C5orf24 | 5 | 0.24028 | 0.43131 | 0.999994 | 8777 | 1 | 0.12038 | 0.38613 | 0.61408 | 1 | 12895 | 2 | 0.12038 |

| | | | | | | | | | | | | |
|---|---|---|---|---|---|---|---|---|---|---|---|---|
| KCNJ12 | 3 | 0.59342 | 0.61212 | 0.999994 | 14872 | 1 | 1.0007 | 0.38614 | 0.53942 | 1 | 12896 | 2 | 1.0007 |
| PSAP | 6 | 0.62669 | 0.75984 | 0.999994 | 15329 | 2 | 0.80517 | 0.38618 | 0.63991 | 1 | 12897 | 3 | 0.80517 |
| GATS | 3 | 0.86193 | 0.86187 | 0.999994 | 18484 | 0 | 1.1028 | 0.38629 | 0.53956 | 1 | 12898 | 2 | 1.1028 |
| SCLT1 | 6 | 0.13367 | 0.30499 | 0.999994 | 6014 | 3 | -0.64251 | 0.3863 | 0.64 | 1 | 12899 | 2 | -0.64251 |
| ANKRD28 | 6 | 0.8384 | 0.8658 | 0.999994 | 18059 | 1 | 0.91112 | 0.3863 | 0.64 | 1 | 12900 | 3 | 0.91112 |
| SYP | 5 | 0.78422 | 0.8099 | 0.999994 | 17269 | 1 | 0.50406 | 0.38644 | 0.6144 | 1 | 12901 | 2 | 0.50406 |
| TATDN3 | 6 | 0.5358 | 0.69358 | 0.999994 | 13993 | 2 | 1.1079 | 0.38651 | 0.64016 | 1 | 12902 | 3 | 1.1079 |
| STX11 | 6 | 0.06847 | 0.18356 | 0.968526 | 3769 | 3 | -1.2091 | 0.38662 | 0.64024 | 1 | 12903 | 2 | -1.2091 |
| CHRNA5 | 4 | 0.42915 | 0.57568 | 0.999994 | 12401 | 2 | 0.72862 | 0.38672 | 0.57859 | 1 | 12904 | 2 | 0.72862 |
| MAPK6 | 6 | 0.35957 | 0.57482 | 0.999994 | 11330 | 3 | -0.14697 | 0.38678 | 0.64037 | 1 | 12905 | 2 | -0.14697 |
| C17orf64 | 5 | 0.027967 | 0.085207 | 0.94395 | 1901 | 1 | 0.46815 | 0.38696 | 0.61493 | 1 | 12906 | 2 | 0.46815 |
| INSIG1 | 5 | 0.18262 | 0.35038 | 0.999994 | 7279 | 3 | -0.89056 | 0.38696 | 0.61493 | 1 | 12907 | 2 | -0.89056 |
| LOC650293 | 5 | 0.008012 | 0.030311 | 0.862223 | 730 | 3 | -1.7412 | 0.38696 | 0.61493 | 1 | 12908 | 1 | -1.7412 |
| KLHL2 | 5 | 0.050013 | 0.14039 | 0.958716 | 2980 | 3 | -1.5471 | 0.38696 | 0.61493 | 1 | 12909 | 2 | -1.5471 |
| hsa-mir-1296 | 4 | 0.92728 | 0.92706 | 0.999994 | 19798 | 0 | 1.4345 | 0.38697 | 0.57878 | 1 | 12910 | 2 | 1.4345 |
| USP6NL | 5 | 0.34967 | 0.52162 | 0.999994 | 11146 | 2 | -0.040538 | 0.38709 | 0.61506 | 1 | 12911 | 2 | -0.040538 |
| hsa-mir-4756 | 3 | 0.0040507 | 0.011186 | 0.714551 | 444 | 2 | -8.7382 | 0.38716 | 0.54045 | 1 | 12912 | 1 | -8.7382 |
| TRHDE | 4 | 0.2548 | 0.39682 | 0.999994 | 9125 | 2 | -0.082511 | 0.38721 | 0.57897 | 1 | 12913 | 2 | -0.082511 |
| hsa-mir-4753 | 4 | 0.4607 | 0.59918 | 0.999994 | 12857 | 1 | 0.8733 | 0.38723 | 0.57898 | 1 | 12914 | 2 | 0.8733 |
| OR51I2 | 5 | 0.21738 | 0.39961 | 0.999994 | 8203 | 2 | 0.90023 | 0.38725 | 0.61519 | 1 | 12915 | 3 | 0.90023 |
| NOX3 | 5 | 0.4443 | 0.60043 | 0.999994 | 12636 | 2 | -0.57062 | 0.38731 | 0.61526 | 1 | 12916 | 2 | -0.57062 |
| ARL5A | 6 | 0.68775 | 0.80706 | 0.999994 | 16192 | 1 | 0.81596 | 0.3874 | 0.64087 | 1 | 12917 | 3 | 0.81596 |
| BBS2 | 6 | 0.35225 | 0.56654 | 0.999994 | 11201 | 3 | -0.27944 | 0.38744 | 0.6409 | 1 | 12918 | 2 | -0.27944 |
| TSPAN14 | 6 | 0.047365 | 0.13828 | 0.957317 | 2874 | 3 | -1.1034 | 0.38744 | 0.6409 | 1 | 12919 | 2 | -1.1034 |
| OR10T2 | 5 | 0.074468 | 0.18561 | 0.969245 | 4012 | 3 | -1.9903 | 0.38749 | 0.61544 | 1 | 12920 | 2 | -1.9903 |
| PSMD3 | 2 | 0.6465 | 0.64627 | 0.999994 | 15608 | 0 | 1.5178 | 0.3876 | 0.46021 | 1 | 12921 | 1 | 1.5178 |
| CEP104 | 4 | 0.28914 | 0.43192 | 0.999994 | 9896 | 2 | -0.42575 | 0.38767 | 0.57932 | 1 | 12922 | 1 | -0.42575 |
| EIF2B5 | 4 | 0.099556 | 0.21432 | 0.979587 | 4957 | 2 | -0.48385 | 0.38773 | 0.57936 | 1 | 12923 | 2 | -0.48385 |
| SKOR1 | 5 | 0.24489 | 0.43662 | 0.999994 | 8898 | 2 | 0.89189 | 0.38776 | 0.61572 | 1 | 12924 | 3 | 0.89189 |
| C5orf63 | 6 | 0.54965 | 0.70338 | 0.999994 | 14207 | 2 | -0.40969 | 0.38777 | 0.64116 | 1 | 12925 | 1 | -0.40969 |
| KDM3A | 6 | 0.095398 | 0.23832 | 0.987253 | 4797 | 3 | -0.94144 | 0.38777 | 0.64116 | 1 | 12926 | 2 | -0.94144 |
| GRM7 | 6 | 0.24129 | 0.43753 | 0.999994 | 8806 | 2 | -0.51746 | 0.38777 | 0.64116 | 1 | 12927 | 2 | -0.51746 |
| ELMO3 | 5 | 0.2545 | 0.44433 | 0.999994 | 9117 | 2 | -0.54313 | 0.38787 | 0.61582 | 1 | 12928 | 1 | -0.54313 |
| TNFAIP8 | 6 | 0.81265 | 0.85344 | 0.999994 | 17663 | 1 | 1.0144 | 0.38798 | 0.64131 | 1 | 12929 | 3 | 1.0144 |
| CALCB | 6 | 0.074512 | 0.19615 | 0.975797 | 4014 | 2 | 0.22211 | 0.3881 | 0.64139 | 1 | 12930 | 3 | 0.22211 |
| NR1I3 | 5 | 0.016609 | 0.054769 | 0.922677 | 1285 | 3 | -3.4667 | 0.38815 | 0.6161 | 1 | 12931 | 2 | -3.4667 |
| RBM24 | 5 | 0.022864 | 0.071809 | 0.93914 | 1642 | 2 | -0.40293 | 0.38817 | 0.61611 | 1 | 12932 | 2 | -0.40293 |
| FAM81A | 4 | 0.62958 | 0.68165 | 0.999994 | 15376 | 1 | 0.63863 | 0.38819 | 0.5797 | 1 | 12933 | 2 | 0.63863 |
| PDCD2 | 4 | 0.050974 | 0.1234 | 0.951052 | 3016 | 3 | -2.3922 | 0.38819 | 0.5797 | 1 | 12934 | 1 | -2.3922 |
| KCNK12 | 6 | 0.95857 | 0.95866 | 0.999994 | 20550 | 0 | 0.93059 | 0.38819 | 0.64147 | 1 | 12935 | 3 | 0.93059 |
| LOC100288255 | 1 | 0.61178 | 0.61137 | 0.999994 | 15104 | 0 | 1.3492 | 0.38822 | 0.38838 | 1 | 12936 | 1 | 1.3492 |
| GFRA3 | 5 | 0.020795 | 0.06621 | 0.93914 | 1499 | 3 | -3.1619 | 0.38823 | 0.61617 | 1 | 12937 | 2 | -3.1619 |
| ST6GALNAC5 | 6 | 0.015167 | 0.05626 | 0.9235 | 1201 | 2 | 0.048493 | 0.38823 | 0.64151 | 1 | 12938 | 2 | 0.048493 |
| CNIH4 | 5 | 0.66897 | 0.75286 | 0.999994 | 15909 | 1 | 0.91017 | 0.38826 | 0.61621 | 1 | 12939 | 3 | 0.91017 |
| SH2B3 | 4 | 0.032413 | 0.086426 | 0.94395 | 2131 | 2 | -1.4907 | 0.38847 | 0.57992 | 1 | 12940 | 2 | -1.4907 |
| BTNL9 | 4 | 0.64939 | 0.69293 | 0.999994 | 15648 | 1 | 0.88022 | 0.38855 | 0.57998 | 1 | 12941 | 2 | 0.88022 |
| TMCO1 | 6 | 0.22411 | 0.41707 | 0.999994 | 8381 | 3 | -0.62464 | 0.38865 | 0.64185 | 1 | 12942 | 1 | -0.62464 |
| ADAM32 | 6 | 0.34071 | 0.55347 | 0.999994 | 10953 | 2 | 0.4397 | 0.38865 | 0.64185 | 1 | 12943 | 3 | 0.4397 |
| C19orf35 | 6 | 0.14613 | 0.32079 | 0.999994 | 6355 | 3 | -0.41206 | 0.38865 | 0.64185 | 1 | 12944 | 3 | -0.41206 |
| FAM181A | 6 | 0.044377 | 0.13173 | 0.95318 | 2729 | 4 | -1.5143 | 0.38865 | 0.64185 | 1 | 12945 | 2 | -1.5143 |
| RBFOX2 | 5 | 0.78187 | 0.80853 | 0.999994 | 17238 | 1 | 0.87409 | 0.38876 | 0.61669 | 1 | 12946 | 3 | 0.87409 |
| C3orf36 | 5 | 0.12396 | 0.26339 | 0.996734 | 5766 | 3 | -1.3999 | 0.38881 | 0.61674 | 1 | 12947 | 1 | -1.3999 |

| | | | | | | | | | | | | | |
|---|---|---|---|---|---|---|---|---|---|---|---|---|---|
| ASTL | 5 | 0.21242 | 0.39281 | 0.999994 | 8069 | 2 | -0.14597 | 0.38881 | 0.61674 | 1 | 12948 | 2 | -0.14597 |
| SFXN5 | 6 | 0.34961 | 0.56356 | 0.999994 | 11140 | 2 | -0.11228 | 0.38882 | 0.64199 | 1 | 12949 | 2 | -0.11228 |
| HGD | 6 | 0.016452 | 0.060476 | 0.932932 | 1274 | 4 | -2.5076 | 0.38892 | 0.64207 | 1 | 12950 | 2 | -2.5076 |
| NonTargetingControlG | 1 | 0.61078 | 0.61038 | 0.999994 | 15081 | 0 | 1.476 | 0.38922 | 0.38936 | 1 | 12951 | 1 | 1.476 |
| TYSND1 | 6 | 0.14699 | 0.32186 | 0.999994 | 6374 | 3 | -0.81335 | 0.38924 | 0.64232 | 1 | 12952 | 1 | -0.81335 |
| NDE1 | 6 | 0.13224 | 0.30318 | 0.999994 | 5982 | 3 | -0.57705 | 0.38924 | 0.64232 | 1 | 12953 | 1 | -0.57705 |
| PRB2 | 3 | 0.55904 | 0.58671 | 0.999994 | 14354 | 1 | 1.3182 | 0.38927 | 0.54259 | 1 | 12954 | 2 | 1.3182 |
| hsa-mir-8059 | 3 | 0.39804 | 0.48311 | 0.999994 | 11916 | 1 | 1.3182 | 0.38927 | 0.54259 | 1 | 12955 | 2 | 1.3182 |
| AHCYL2 | 5 | 0.52836 | 0.67261 | 0.999994 | 13866 | 1 | -0.015895 | 0.38928 | 0.61724 | 1 | 12956 | 2 | -0.015895 |
| hsa-mir-301a | 2 | 0.47191 | 0.47866 | 0.999994 | 13032 | 1 | 1.5465 | 0.38933 | 0.46134 | 1 | 12957 | 1 | 1.5465 |
| GABPA | 3 | 0.52645 | 0.5639 | 0.999994 | 13837 | 1 | -1.4517 | 0.38942 | 0.54275 | 1 | 12958 | 1 | -1.4517 |
| IZUMO2 | 6 | 0.67515 | 0.79718 | 0.999994 | 16005 | 2 | 0.99397 | 0.38947 | 0.64251 | 1 | 12959 | 3 | 0.99397 |
| ELK1 | 5 | 0.75065 | 0.79118 | 0.999994 | 16860 | 1 | 0.76342 | 0.3895 | 0.61744 | 1 | 12960 | 3 | 0.76342 |
| YIF1B | 6 | 0.92526 | 0.92536 | 0.999994 | 19750 | 1 | 0.98203 | 0.38974 | 0.64272 | 1 | 12961 | 3 | 0.98203 |
| TMEM132B | 6 | 0.068423 | 0.18348 | 0.968526 | 3765 | 2 | 0.51867 | 0.38974 | 0.64272 | 1 | 12962 | 3 | 0.51867 |
| NKRF | 5 | 0.8414 | 0.84773 | 0.999994 | 18112 | 1 | -0.15055 | 0.3898 | 0.61775 | 1 | 12963 | 2 | -0.15055 |
| CPB1 | 5 | 0.26433 | 0.45226 | 0.999994 | 9340 | 2 | -0.45966 | 0.38981 | 0.61776 | 1 | 12964 | 2 | -0.45966 |
| LMX1A | 5 | 0.32288 | 0.49965 | 0.999994 | 10573 | 1 | 1.145 | 0.38984 | 0.61779 | 1 | 12965 | 3 | 1.145 |
| OR4K5 | 4 | 0.18706 | 0.32736 | 0.999994 | 7392 | 2 | -0.40244 | 0.38988 | 0.58099 | 1 | 12966 | 2 | -0.40244 |
| CDC42EP2 | 4 | 0.056827 | 0.13472 | 0.954019 | 3284 | 1 | 0.15517 | 0.38988 | 0.58099 | 1 | 12967 | 1 | 0.15517 |
| SPCS2 | 5 | 0.76855 | 0.80091 | 0.999994 | 17068 | 1 | 0.92402 | 0.39 | 0.61795 | 1 | 12968 | 3 | 0.92402 |
| PIAS3 | 6 | 0.92019 | 0.92057 | 0.999994 | 19633 | 1 | 0.58096 | 0.39001 | 0.64293 | 1 | 12969 | 3 | 0.58096 |
| hsa-mir-6865 | 3 | 0.045483 | 0.10944 | 0.951052 | 2789 | 1 | 0.90294 | 0.39004 | 0.54338 | 1 | 12970 | 2 | 0.90294 |
| PPP1R3B | 4 | 0.20715 | 0.34806 | 0.999994 | 7951 | 1 | 1.2115 | 0.39004 | 0.58111 | 1 | 12971 | 2 | 1.2115 |
| LILRA6 | 5 | 0.56619 | 0.70618 | 0.999994 | 14482 | 2 | 0.34227 | 0.39009 | 0.61805 | 1 | 12972 | 2 | 0.34227 |
| ANKRD22 | 6 | 0.61507 | 0.75114 | 0.999994 | 15146 | 2 | 0.68168 | 0.39015 | 0.64304 | 1 | 12973 | 3 | 0.68168 |
| RIN2 | 6 | 0.13164 | 0.30241 | 0.999994 | 5971 | 3 | 0.38603 | 0.39015 | 0.64304 | 1 | 12974 | 3 | 0.38603 |
| ZNF697 | 6 | 0.53141 | 0.69055 | 0.999994 | 13922 | 2 | -0.10289 | 0.39015 | 0.64304 | 1 | 12975 | 2 | -0.10289 |
| IGIP | 6 | 0.10141 | 0.25008 | 0.992497 | 5017 | 2 | 0.040433 | 0.39015 | 0.64304 | 1 | 12976 | 2 | 0.040433 |
| RASSF2 | 6 | 0.36487 | 0.57999 | 0.999994 | 11419 | 2 | 0.87602 | 0.3902 | 0.64308 | 1 | 12977 | 3 | 0.87602 |
| CCL13 | 3 | 0.22165 | 0.33971 | 0.999994 | 8323 | 2 | -0.9396 | 0.39029 | 0.54365 | 1 | 12978 | 1 | -0.9396 |
| WNT3A | 6 | 0.58989 | 0.73251 | 0.999994 | 14821 | 1 | 0.94286 | 0.39047 | 0.64329 | 1 | 12979 | 3 | 0.94286 |
| LRRC8C | 5 | 0.022841 | 0.071735 | 0.93914 | 1639 | 1 | 0.85157 | 0.39049 | 0.61849 | 1 | 12980 | 3 | 0.85157 |
| TRIM29 | 6 | 0.74092 | 0.8253 | 0.999994 | 16750 | 1 | 0.85624 | 0.39057 | 0.64335 | 1 | 12981 | 3 | 0.85624 |
| ATAD2 | 4 | 0.091605 | 0.19995 | 0.977164 | 4663 | 2 | -0.94363 | 0.39059 | 0.58153 | 1 | 12982 | 2 | -0.94363 |
| ABCC4 | 5 | 0.059535 | 0.1613 | 0.967126 | 3402 | 3 | -2.3183 | 0.39067 | 0.61866 | 1 | 12983 | 1 | -2.3183 |
| NonTargetingControlG | 1 | 0.60932 | 0.60894 | 0.999994 | 15063 | 0 | 1.3533 | 0.39068 | 0.3908 | 1 | 12984 | 1 | 1.3533 |
| QPRT | 5 | 0.82601 | 0.83666 | 0.999994 | 17868 | 1 | 0.80339 | 0.39092 | 0.61892 | 1 | 12985 | 3 | 0.80339 |
| TMUB1 | 5 | 0.23526 | 0.42449 | 0.999994 | 8656 | 2 | 1.0036 | 0.39093 | 0.61892 | 1 | 12986 | 3 | 1.0036 |
| LIPM | 6 | 0.28536 | 0.48969 | 0.999994 | 9819 | 3 | -0.49001 | 0.391 | 0.64366 | 1 | 12987 | 1 | -0.49001 |
| SLC24A1 | 5 | 0.55966 | 0.70034 | 0.999994 | 14364 | 1 | 0.89565 | 0.39108 | 0.61907 | 1 | 12988 | 3 | 0.89565 |
| TNK1 | 5 | 0.42582 | 0.58496 | 0.999994 | 12347 | 1 | 0.88235 | 0.3912 | 0.6192 | 1 | 12989 | 3 | 0.88235 |
| HELT | 6 | 0.030195 | 0.099242 | 0.945253 | 2018 | 4 | -2.589 | 0.39161 | 0.64415 | 1 | 12990 | 1 | -2.589 |
| C7orf69 | 6 | 0.084811 | 0.21715 | 0.980699 | 4400 | 3 | -0.54681 | 0.39161 | 0.64415 | 1 | 12991 | 3 | -0.54681 |
| PRAMEF15 | 6 | 0.50817 | 0.67435 | 0.999994 | 13556 | 1 | 0.27371 | 0.39164 | 0.64417 | 1 | 12992 | 3 | 0.27371 |
| CCDC154 | 4 | 0.40734 | 0.55324 | 0.999994 | 12063 | 2 | 0.71736 | 0.39172 | 0.58238 | 1 | 12993 | 2 | 0.71736 |
| hsa-mir-1208 | 4 | 0.025788 | 0.072888 | 0.93914 | 1790 | 2 | -0.93648 | 0.39178 | 0.58242 | 1 | 12994 | 1 | -0.93648 |
| CHD6 | 5 | 0.55613 | 0.69721 | 0.999994 | 14306 | 2 | -0.15553 | 0.39188 | 0.61991 | 1 | 12995 | 2 | -0.15553 |
| VPS13D | 5 | 0.31588 | 0.49401 | 0.999994 | 10438 | 1 | -0.19505 | 0.39188 | 0.61991 | 1 | 12996 | 2 | -0.19505 |
| PCDHB1 | 5 | 0.39486 | 0.55926 | 0.999994 | 11870 | 2 | 0.27302 | 0.39188 | 0.61991 | 1 | 12997 | 2 | 0.27302 |
| CINP | 5 | 0.12659 | 0.2674 | 0.998561 | 5837 | 3 | -1.4454 | 0.39188 | 0.61991 | 1 | 12998 | 2 | -1.4454 |
| GMPPA | 5 | 0.60681 | 0.72864 | 0.999994 | 15034 | 1 | 0.95597 | 0.39199 | 0.62001 | 1 | 12999 | 3 | 0.95597 |

| | | | | | | | | | | | | |
|---|---|---|---|---|---|---|---|---|---|---|---|---|
| GNA15 | 5 | 0.060766 | 0.16334 | 0.967126 | 3456 | 2 | 0.15814 | 0.39201 | 0.62004 | 1 | 13000 | 1 | 0.15814 |
| TM6SF2 | 5 | 0.46136 | 0.61482 | 0.999994 | 12867 | 2 | -0.21607 | 0.39201 | 0.62004 | 1 | 13001 | 2 | -0.21607 |
| PCDHGA1 | 5 | 0.93594 | 0.93619 | 0.999994 | 19988 | 0 | 0.76093 | 0.39208 | 0.62011 | 1 | 13002 | 3 | 0.76093 |
| NKD2 | 5 | 0.36201 | 0.53172 | 0.999994 | 11367 | 2 | 0.91868 | 0.3922 | 0.62023 | 1 | 13003 | 3 | 0.91868 |
| LINGO3 | 4 | 0.58634 | 0.65801 | 0.999994 | 14762 | 1 | 0.94701 | 0.39232 | 0.58283 | 1 | 13004 | 2 | 0.94701 |
| MRPS34 | 4 | 0.072811 | 0.16525 | 0.967126 | 3953 | 2 | -0.7534 | 0.39237 | 0.58287 | 1 | 13005 | 2 | -0.7534 |
| hsa-mir-7108 | 2 | 0.43032 | 0.44229 | 0.999994 | 12417 | 1 | 1.2747 | 0.39251 | 0.46346 | 1 | 13006 | 1 | 1.2747 |
| VRK2 | 4 | 0.24535 | 0.38711 | 0.999994 | 8908 | 2 | -0.99895 | 0.39257 | 0.58302 | 1 | 13007 | 1 | -0.99895 |
| LDHAL6A | 6 | 0.67558 | 0.79754 | 0.999994 | 16014 | 2 | 0.75288 | 0.39268 | 0.64495 | 1 | 13008 | 3 | 0.75288 |
| SGCD | 5 | 0.2056 | 0.3832 | 0.999994 | 7913 | 2 | 0.86395 | 0.39299 | 0.62106 | 1 | 13009 | 3 | 0.86395 |
| NUDT6 | 5 | 0.11134 | 0.24404 | 0.988309 | 5345 | 3 | -0.59511 | 0.39307 | 0.62115 | 1 | 13010 | 2 | -0.59511 |
| hsa-mir-6785 | 2 | 0.052644 | 0.093603 | 0.945031 | 3097 | 1 | -2.415 | 0.39315 | 0.46386 | 1 | 13011 | 1 | -2.415 |
| hsa-mir-3937 | 2 | 0.50293 | 0.50657 | 0.999994 | 13478 | 1 | 1.6677 | 0.39315 | 0.46386 | 1 | 13012 | 1 | 1.6677 |
| LPIN3 | 6 | 0.20549 | 0.39454 | 0.999994 | 7908 | 2 | 0.28674 | 0.39332 | 0.64544 | 1 | 13013 | 3 | 0.28674 |
| SOCS3 | 6 | 0.057868 | 0.16117 | 0.967126 | 3331 | 3 | -0.58486 | 0.39332 | 0.64544 | 1 | 13014 | 3 | -0.58486 |
| KCNH8 | 6 | 0.33665 | 0.54883 | 0.999994 | 10864 | 3 | -0.22642 | 0.39332 | 0.64544 | 1 | 13015 | 2 | -0.22642 |
| SOX8 | 4 | 0.081955 | 0.18232 | 0.968526 | 4297 | 2 | -0.56835 | 0.39343 | 0.58363 | 1 | 13016 | 2 | -0.56835 |
| FAM73B | 5 | 0.18986 | 0.36074 | 0.999994 | 7481 | 2 | 0.97346 | 0.39344 | 0.62148 | 1 | 13017 | 3 | 0.97346 |
| CES4A | 6 | 0.61523 | 0.75126 | 0.999994 | 15148 | 2 | 0.038405 | 0.39352 | 0.6456 | 1 | 13018 | 2 | 0.038405 |
| CRIP3 | 3 | 0.70199 | 0.70287 | 0.999994 | 16326 | 1 | 1.0094 | 0.39357 | 0.54704 | 1 | 13019 | 2 | 1.0094 |
| STK11IP | 4 | 0.080815 | 0.18019 | 0.968171 | 4261 | 2 | -0.64161 | 0.39359 | 0.58376 | 1 | 13020 | 2 | -0.64161 |
| OPTN | 5 | 0.097887 | 0.22313 | 0.983354 | 4886 | 3 | -1.553 | 0.39366 | 0.62169 | 1 | 13021 | 2 | -1.553 |
| MYOF | 5 | 0.29537 | 0.4773 | 0.999994 | 10022 | 2 | -0.60687 | 0.39366 | 0.62169 | 1 | 13022 | 1 | -0.60687 |
| TAF4B | 5 | 0.40171 | 0.56492 | 0.999994 | 11974 | 2 | -0.38054 | 0.39366 | 0.62169 | 1 | 13023 | 2 | -0.38054 |
| OR9A2 | 3 | 0.68037 | 0.68308 | 0.999994 | 16098 | 1 | -0.10471 | 0.394 | 0.54748 | 1 | 13024 | 1 | -0.10471 |
| MLEC | 5 | 0.12058 | 0.25825 | 0.994943 | 5647 | 2 | 0.95741 | 0.39402 | 0.62205 | 1 | 13025 | 3 | 0.95741 |
| HMGN4 | 5 | 0.1049 | 0.23405 | 0.984971 | 5129 | 3 | -1.5399 | 0.39403 | 0.62206 | 1 | 13026 | 2 | -1.5399 |
| STON1 | 4 | 0.79359 | 0.79553 | 0.999994 | 17392 | 1 | 0.35766 | 0.39407 | 0.58413 | 1 | 13027 | 2 | 0.35766 |
| PVRL1 | 6 | 0.26163 | 0.46183 | 0.999994 | 9277 | 3 | 0.37485 | 0.39413 | 0.6461 | 1 | 13028 | 3 | 0.37485 |
| EZH1 | 5 | 0.38202 | 0.5484 | 0.999994 | 11677 | 2 | 0.79974 | 0.39426 | 0.6223 | 1 | 13029 | 3 | 0.79974 |
| OSGIN2 | 6 | 0.9247 | 0.92483 | 0.999994 | 19737 | 1 | 0.51108 | 0.3943 | 0.64623 | 1 | 13030 | 2 | 0.51108 |
| ABLIM1 | 6 | 0.58877 | 0.73171 | 0.999994 | 14800 | 2 | 0.047972 | 0.3943 | 0.64623 | 1 | 13031 | 2 | 0.047972 |
| ST3GAL4 | 6 | 0.63321 | 0.76477 | 0.999994 | 15419 | 2 | 0.30587 | 0.3943 | 0.64623 | 1 | 13032 | 3 | 0.30587 |
| MFAP3 | 5 | 0.13514 | 0.28045 | 0.999632 | 6055 | 2 | 0.91142 | 0.39431 | 0.62235 | 1 | 13033 | 3 | 0.91142 |
| KCNE1L | 5 | 0.091313 | 0.2127 | 0.979587 | 4648 | 3 | -1.8537 | 0.39434 | 0.62238 | 1 | 13034 | 2 | -1.8537 |
| MGAM | 6 | 0.83949 | 0.86634 | 0.999994 | 18077 | 1 | 0.87257 | 0.3944 | 0.64631 | 1 | 13035 | 3 | 0.87257 |
| ZBTB11 | 5 | 0.0139 | 0.047109 | 0.896226 | 1122 | 3 | -1.5139 | 0.39442 | 0.62246 | 1 | 13036 | 2 | -1.5139 |
| IGFL2 | 5 | 0.38459 | 0.5506 | 0.999994 | 11709 | 2 | 0.79742 | 0.39447 | 0.62251 | 1 | 13037 | 3 | 0.79742 |
| hsa-mir-331 | 4 | 0.28744 | 0.43016 | 0.999994 | 9864 | 1 | 0.4497 | 0.39453 | 0.5845 | 1 | 13038 | 2 | 0.4497 |
| NonTargetingControlG | 2 | 0.41982 | 0.43333 | 0.999994 | 12250 | 1 | 1.2589 | 0.39466 | 0.46483 | 1 | 13039 | 1 | 1.2589 |
| CRCT1 | 5 | 0.057056 | 0.15723 | 0.967126 | 3293 | 2 | 0.56468 | 0.39468 | 0.62271 | 1 | 13040 | 3 | 0.56468 |
| SUV39H2 | 6 | 0.049439 | 0.14282 | 0.959553 | 2950 | 4 | -1.7872 | 0.3948 | 0.64663 | 1 | 13041 | 2 | -1.7872 |
| CLEC2A | 5 | 0.81758 | 0.83097 | 0.999994 | 17741 | 1 | 0.82937 | 0.39485 | 0.62286 | 1 | 13042 | 3 | 0.82937 |
| C10orf68 | 5 | 0.68769 | 0.76098 | 0.999994 | 16191 | 1 | 0.94254 | 0.39488 | 0.62291 | 1 | 13043 | 3 | 0.94254 |
| NUFIP1 | 5 | 0.52765 | 0.67199 | 0.999994 | 13855 | 2 | 0.90589 | 0.3949 | 0.62292 | 1 | 13044 | 3 | 0.90589 |
| DZIP1L | 6 | 0.25998 | 0.45982 | 0.999994 | 9241 | 3 | -0.32629 | 0.39492 | 0.64672 | 1 | 13045 | 2 | -0.32629 |
| TPM2 | 6 | 0.056872 | 0.159 | 0.967126 | 3286 | 3 | -0.019877 | 0.39494 | 0.64674 | 1 | 13046 | 3 | -0.019877 |
| hsa-mir-2115 | 4 | 0.12356 | 0.25651 | 0.994943 | 5754 | 1 | 0.049207 | 0.39496 | 0.58482 | 1 | 13047 | 2 | 0.049207 |
| TP53BP1 | 4 | 0.27023 | 0.41253 | 0.999994 | 9479 | 2 | -0.74768 | 0.39496 | 0.58482 | 1 | 13048 | 1 | -0.74768 |
| VSIG8 | 6 | 0.13083 | 0.3014 | 0.999994 | 5952 | 3 | -0.93282 | 0.39505 | 0.64682 | 1 | 13049 | 1 | -0.93282 |
| ZUFSP | 6 | 0.96359 | 0.96365 | 0.999994 | 20691 | 0 | 0.28735 | 0.39505 | 0.64682 | 1 | 13050 | 2 | 0.28735 |
| MST1R | 6 | 0.042511 | 0.12756 | 0.951052 | 2644 | 4 | -1.6331 | 0.39505 | 0.64682 | 1 | 13051 | 1 | -1.6331 |

| | | | | | | | | | | | | | |
|---|---|---|---|---|---|---|---|---|---|---|---|---|---|
| ZDHHC11 | 6 | 0.62408 | 0.75789 | 0.999994 | 15290 | 2 | 0.70529 | 0.39506 | 0.64683 | 1 | 13052 | 3 | 0.70529 |
| PCDH8 | 5 | 0.61516 | 0.73176 | 0.999994 | 15147 | 1 | 0.42292 | 0.39509 | 0.62312 | 1 | 13053 | 2 | 0.42292 |
| CHTF8 | 5 | 0.28992 | 0.47285 | 0.999994 | 9908 | 1 | 0.15465 | 0.39509 | 0.62312 | 1 | 13054 | 2 | 0.15465 |
| UTP14A | 5 | 0.0196 | 0.063031 | 0.937389 | 1434 | 3 | -3.5626 | 0.39509 | 0.62312 | 1 | 13055 | 2 | -3.5626 |
| SMPD2 | 5 | 0.3257 | 0.50199 | 0.999994 | 10636 | 2 | 0.82748 | 0.39511 | 0.62314 | 1 | 13056 | 3 | 0.82748 |
| RPL17 | 3 | 0.84669 | 0.84668 | 0.999994 | 18203 | 0 | 1.0067 | 0.3952 | 0.54871 | 1 | 13057 | 2 | 1.0067 |
| hsa-mir-320b-1 | 3 | 0.073718 | 0.14798 | 0.965554 | 3980 | 2 | -2.7442 | 0.39528 | 0.54879 | 1 | 13058 | 1 | -2.7442 |
| CLP1 | 3 | 0.085161 | 0.16336 | 0.967126 | 4414 | 1 | -0.013492 | 0.39528 | 0.54879 | 1 | 13059 | 1 | -0.013492 |
| TEKT2 | 5 | 0.24724 | 0.43858 | 0.999994 | 8955 | 1 | 0.94254 | 0.39533 | 0.62336 | 1 | 13060 | 3 | 0.94254 |
| AIMP1 | 6 | 0.8818 | 0.89078 | 0.999994 | 18865 | 1 | 0.95279 | 0.39543 | 0.64712 | 1 | 13061 | 3 | 0.95279 |
| PRR23C | 6 | 0.0068113 | 0.027758 | 0.855011 | 647 | 3 | -0.39494 | 0.39547 | 0.64715 | 1 | 13062 | 3 | -0.39494 |
| HRSP12 | 5 | 0.21857 | 0.40126 | 0.999994 | 8241 | 3 | -0.75423 | 0.39558 | 0.62361 | 1 | 13063 | 1 | -0.75423 |
| SLN | 5 | 0.24108 | 0.43239 | 0.999994 | 8796 | 2 | 0.65165 | 0.39558 | 0.62361 | 1 | 13064 | 3 | 0.65165 |
| PLEKHA3 | 5 | 0.1512 | 0.30458 | 0.999994 | 6488 | 2 | 0.25086 | 0.39558 | 0.62361 | 1 | 13065 | 2 | 0.25086 |
| DLD | 4 | 0.20092 | 0.34162 | 0.999994 | 7787 | 1 | 0.79146 | 0.39571 | 0.5854 | 1 | 13066 | 2 | 0.79146 |
| ABCC3 | 4 | 0.21364 | 0.35478 | 0.999994 | 8105 | 2 | 0.0069903 | 0.39576 | 0.58545 | 1 | 13067 | 2 | 0.0069903 |
| LRFN1 | 5 | 0.1977 | 0.37195 | 0.999994 | 7697 | 2 | -0.34192 | 0.39591 | 0.62394 | 1 | 13068 | 2 | -0.34192 |
| TGFB3 | 6 | 0.038954 | 0.11953 | 0.951052 | 2458 | 2 | 0.97081 | 0.39612 | 0.64763 | 1 | 13069 | 3 | 0.97081 |
| CBR3 | 4 | 0.36828 | 0.51304 | 0.999994 | 11477 | 2 | 0.48669 | 0.39612 | 0.58571 | 1 | 13070 | 2 | 0.48669 |
| ACTA1 | 6 | 0.61101 | 0.74807 | 0.999994 | 15088 | 2 | 0.70027 | 0.39619 | 0.64768 | 1 | 13071 | 3 | 0.70027 |
| WSB2 | 4 | 0.27949 | 0.42203 | 0.999994 | 9690 | 2 | 0.56017 | 0.39621 | 0.58577 | 1 | 13072 | 2 | 0.56017 |
| MAP7D3 | 5 | 0.84562 | 0.85091 | 0.999994 | 18179 | 1 | 1.036 | 0.39621 | 0.62423 | 1 | 13073 | 3 | 1.036 |
| TMEM257 | 6 | 0.090719 | 0.22894 | 0.983354 | 4627 | 3 | -0.26679 | 0.39632 | 0.64777 | 1 | 13074 | 3 | -0.26679 |
| SLC2A4 | 6 | 0.39148 | 0.5969 | 0.999994 | 11820 | 1 | 1.0614 | 0.39642 | 0.64785 | 1 | 13075 | 3 | 1.0614 |
| TLCD2 | 5 | 0.85911 | 0.86164 | 0.999994 | 18415 | 1 | 0.50613 | 0.39643 | 0.62444 | 1 | 13076 | 2 | 0.50613 |
| LRP11 | 6 | 0.124 | 0.29264 | 0.999632 | 5770 | 2 | 0.89837 | 0.39649 | 0.6479 | 1 | 13077 | 3 | 0.89837 |
| C11orf82 | 5 | 0.0089702 | 0.033091 | 0.867348 | 791 | 3 | -3.7777 | 0.39653 | 0.62453 | 1 | 13078 | 2 | -3.7777 |
| TMEM191C | 5 | 0.57877 | 0.71725 | 0.999994 | 14667 | 1 | -0.30617 | 0.39653 | 0.62453 | 1 | 13079 | 2 | -0.30617 |
| ABL1 | 5 | 0.93938 | 0.93957 | 0.999994 | 20059 | 0 | 0.39265 | 0.39653 | 0.62453 | 1 | 13080 | 2 | 0.39265 |
| OR13C5 | 5 | 0.0092527 | 0.033897 | 0.867348 | 812 | 3 | -0.88716 | 0.39653 | 0.62453 | 1 | 13081 | 2 | -0.88716 |
| LRRC39 | 5 | 0.46093 | 0.61443 | 0.999994 | 12862 | 2 | -0.39325 | 0.39665 | 0.62465 | 1 | 13082 | 2 | -0.39325 |
| SYTL1 | 6 | 0.22714 | 0.42064 | 0.999994 | 8459 | 3 | 0.16902 | 0.39667 | 0.64804 | 1 | 13083 | 3 | 0.16902 |
| CCND1 | 5 | 0.5775 | 0.71611 | 0.999994 | 14650 | 2 | 0.12088 | 0.39675 | 0.62475 | 1 | 13084 | 2 | 0.12088 |
| COL24A1 | 6 | 0.15457 | 0.33144 | 0.999994 | 6582 | 2 | 0.80508 | 0.39684 | 0.64817 | 1 | 13085 | 3 | 0.80508 |
| hsa-mir-523 | 1 | 0.60296 | 0.6026 | 0.999994 | 14986 | 0 | 1.3764 | 0.39704 | 0.39725 | 1 | 13086 | 1 | 1.3764 |
| ERCC2 | 2 | 0.74259 | 0.74226 | 0.999994 | 16772 | 0 | 2.4358 | 0.39709 | 0.46648 | 1 | 13087 | 1 | 2.4358 |
| RNASE7 | 4 | 0.26727 | 0.40947 | 0.999994 | 9402 | 2 | -0.037655 | 0.3971 | 0.58645 | 1 | 13088 | 2 | -0.037655 |
| PPP6C | 5 | 0.093672 | 0.21644 | 0.980699 | 4741 | 2 | -0.30507 | 0.39716 | 0.62516 | 1 | 13089 | 2 | -0.30507 |
| LPCAT4 | 6 | 0.78375 | 0.8412 | 0.999994 | 17263 | 1 | 0.63749 | 0.39717 | 0.64843 | 1 | 13090 | 3 | 0.63749 |
| ASB11 | 6 | 0.78912 | 0.84331 | 0.999994 | 17333 | 1 | 0.04503 | 0.39722 | 0.64846 | 1 | 13091 | 3 | 0.04503 |
| POFUT2 | 6 | 0.21655 | 0.40802 | 0.999994 | 8180 | 3 | -0.48825 | 0.39722 | 0.64846 | 1 | 13092 | 2 | -0.48825 |
| IQCF5 | 6 | 0.29186 | 0.49721 | 0.999994 | 9945 | 2 | -0.17365 | 0.39722 | 0.64846 | 1 | 13093 | 2 | -0.17365 |
| ARMCX3 | 6 | 0.081491 | 0.21035 | 0.979587 | 4281 | 3 | -0.46981 | 0.39722 | 0.64846 | 1 | 13094 | 2 | -0.46981 |
| COL5A2 | 6 | 0.013702 | 0.051482 | 0.910467 | 1111 | 4 | -2.1782 | 0.39722 | 0.64846 | 1 | 13095 | 2 | -2.1782 |
| OR2L5 | 6 | 0.13796 | 0.31046 | 0.999994 | 6133 | 3 | 0.32553 | 0.39727 | 0.6485 | 1 | 13096 | 3 | 0.32553 |
| RPL11 | 3 | 0.4394 | 0.50773 | 0.999994 | 12567 | 1 | 0.74782 | 0.39733 | 0.55091 | 1 | 13097 | 2 | 0.74782 |
| PAPD5 | 5 | 0.07655 | 0.18905 | 0.969834 | 4093 | 3 | -1.7823 | 0.39733 | 0.62534 | 1 | 13098 | 2 | -1.7823 |
| C1orf210 | 4 | 0.4771 | 0.60622 | 0.999994 | 13102 | 1 | 1.0872 | 0.39744 | 0.58671 | 1 | 13099 | 2 | 1.0872 |
| RSPRY1 | 6 | 0.23296 | 0.42764 | 0.999994 | 8594 | 2 | 0.94888 | 0.39754 | 0.64871 | 1 | 13100 | 3 | 0.94888 |
| SLC38A11 | 4 | 0.27881 | 0.42134 | 0.999994 | 9666 | 2 | 0.3532 | 0.39765 | 0.58686 | 1 | 13101 | 2 | 0.3532 |
| NKIRAS1 | 6 | 0.11566 | 0.27754 | 0.999379 | 5506 | 3 | -0.95858 | 0.39772 | 0.64886 | 1 | 13102 | 2 | -0.95858 |
| CECR6 | 4 | 0.28226 | 0.42485 | 0.999994 | 9743 | 2 | 0.2982 | 0.39775 | 0.58693 | 1 | 13103 | 2 | 0.2982 |

| | | | | | | | | | | | | | |
|---|---|---|---|---|---|---|---|---|---|---|---|---|---|
| KRTAP19-1 | 4 | 0.432 | 0.5786 | 0.999994 | 12446 | 2 | 0.71745 | 0.39795 | 0.58709 | 1 | 13104 | 2 | 0.71745 |
| hsa-mir-210 | 4 | 0.30033 | 0.44336 | 0.999994 | 10137 | 2 | -0.76203 | 0.39797 | 0.58711 | 1 | 13105 | 1 | -0.76203 |
| EPHA6 | 5 | 0.11807 | 0.25444 | 0.994835 | 5575 | 1 | 0.85667 | 0.39797 | 0.62596 | 1 | 13106 | 3 | 0.85667 |
| ARCN1 | 3 | 0.39975 | 0.48412 | 0.999994 | 11946 | 1 | 1.0155 | 0.39801 | 0.5516 | 1 | 13107 | 2 | 1.0155 |
| PITPNM1 | 6 | 0.27995 | 0.48336 | 0.999994 | 9703 | 3 | -0.78779 | 0.39804 | 0.6491 | 1 | 13108 | 1 | -0.78779 |
| COX7A2L | 6 | 0.21964 | 0.41166 | 0.999994 | 8283 | 1 | -0.21888 | 0.39804 | 0.6491 | 1 | 13109 | 2 | -0.21888 |
| NonTargetingControlG | 2 | 0.703 | 0.70276 | 0.999994 | 16338 | 0 | 2.2147 | 0.39804 | 0.46708 | 1 | 13110 | 1 | 2.2147 |
| hsa-mir-3713 | 4 | 0.58785 | 0.6588 | 0.999994 | 14789 | 1 | 1.2196 | 0.39807 | 0.58718 | 1 | 13111 | 2 | 1.2196 |
| ALDH9A1 | 6 | 0.39633 | 0.59999 | 0.999994 | 11893 | 1 | 0.68998 | 0.39828 | 0.64929 | 1 | 13112 | 3 | 0.68998 |
| HADH | 5 | 0.68482 | 0.75968 | 0.999994 | 16156 | 1 | 0.97467 | 0.39837 | 0.62636 | 1 | 13113 | 3 | 0.97467 |
| IP6K2 | 5 | 0.7725 | 0.80312 | 0.999994 | 17116 | 1 | 0.054857 | 0.39837 | 0.62637 | 1 | 13114 | 2 | 0.054857 |
| PIGR | 6 | 0.91696 | 0.9176 | 0.999994 | 19561 | 1 | 0.94005 | 0.39842 | 0.6494 | 1 | 13115 | 3 | 0.94005 |
| ASAP1 | 5 | 0.78749 | 0.81182 | 0.999994 | 17310 | 1 | 0.8181 | 0.39842 | 0.62642 | 1 | 13116 | 3 | 0.8181 |
| CTSO | 6 | 0.14525 | 0.31968 | 0.999994 | 6330 | 3 | -0.23281 | 0.39848 | 0.64947 | 1 | 13117 | 3 | -0.23281 |
| S1PR3 | 6 | 0.11606 | 0.27833 | 0.999632 | 5516 | 1 | 0.98366 | 0.39848 | 0.64947 | 1 | 13118 | 3 | 0.98366 |
| ZNF662 | 6 | 0.016266 | 0.05988 | 0.932932 | 1260 | 3 | -0.11447 | 0.39848 | 0.64947 | 1 | 13119 | 3 | -0.11447 |
| PROX2 | 6 | 0.0064037 | 0.026282 | 0.848412 | 620 | 3 | -0.69912 | 0.39848 | 0.64947 | 1 | 13120 | 3 | -0.69912 |
| ANO3 | 6 | 0.14902 | 0.32439 | 0.999994 | 6428 | 1 | 0.016918 | 0.39848 | 0.64947 | 1 | 13121 | 2 | 0.016918 |
| GMEB1 | 6 | 0.14046 | 0.3136 | 0.999994 | 6199 | 3 | 0.065794 | 0.39848 | 0.64947 | 1 | 13122 | 3 | 0.065794 |
| LIAS | 6 | 0.15984 | 0.33803 | 0.999994 | 6731 | 3 | -0.58371 | 0.39848 | 0.64947 | 1 | 13123 | 2 | -0.58371 |
| MSL3 | 5 | 0.51882 | 0.6643 | 0.999994 | 13724 | 1 | 0.33336 | 0.39867 | 0.62666 | 1 | 13124 | 2 | 0.33336 |
| ELAVL3 | 5 | 0.2106 | 0.39024 | 0.999994 | 8042 | 2 | -0.28253 | 0.39873 | 0.62672 | 1 | 13125 | 2 | -0.28253 |
| hsa-mir-4438 | 4 | 0.47914 | 0.60712 | 0.999994 | 13131 | 1 | 0.78603 | 0.39877 | 0.5877 | 1 | 13126 | 2 | 0.78603 |
| hsa-mir-1302-1 | 3 | 0.7192 | 0.71921 | 0.999994 | 16487 | 1 | 1.0875 | 0.39881 | 0.55242 | 1 | 13127 | 2 | 1.0875 |
| PTX4 | 6 | 0.028859 | 0.096137 | 0.945031 | 1947 | 2 | 0.4888 | 0.39881 | 0.64973 | 1 | 13128 | 2 | 0.4888 |
| SMIM5 | 6 | 0.11352 | 0.27346 | 0.998561 | 5423 | 3 | -0.9614 | 0.39881 | 0.64973 | 1 | 13129 | 2 | -0.9614 |
| EIF1AY | 6 | 0.34935 | 0.56326 | 0.999994 | 11135 | 2 | -0.193 | 0.39881 | 0.64973 | 1 | 13130 | 2 | -0.193 |
| LEMD1 | 6 | 0.259 | 0.45869 | 0.999994 | 9217 | 3 | -0.081896 | 0.39881 | 0.64973 | 1 | 13131 | 3 | -0.081896 |
| PRDM12 | 4 | 0.82898 | 0.82853 | 0.999994 | 17910 | 0 | 0.29701 | 0.39883 | 0.58773 | 1 | 13132 | 1 | 0.29701 |
| KLHL10 | 6 | 0.62085 | 0.75546 | 0.999994 | 15232 | 1 | 0.88573 | 0.39884 | 0.64974 | 1 | 13133 | 3 | 0.88573 |
| SEMA3C | 6 | 0.64788 | 0.776 | 0.999994 | 15629 | 2 | 0.83037 | 0.39886 | 0.64975 | 1 | 13134 | 3 | 0.83037 |
| SERPINA11 | 6 | 0.31796 | 0.52756 | 0.999994 | 10472 | 3 | 0.37097 | 0.3989 | 0.64978 | 1 | 13135 | 3 | 0.37097 |
| ING4 | 6 | 0.61502 | 0.75109 | 0.999994 | 15144 | 2 | 0.98883 | 0.399 | 0.64985 | 1 | 13136 | 3 | 0.98883 |
| ZDHHC9 | 6 | 0.47237 | 0.65004 | 0.999994 | 13039 | 2 | 0.58141 | 0.39902 | 0.64987 | 1 | 13137 | 3 | 0.58141 |
| TNNI3K | 6 | 0.32402 | 0.53446 | 0.999994 | 10598 | 3 | 0.38561 | 0.39905 | 0.6499 | 1 | 13138 | 3 | 0.38561 |
| PRTFDC1 | 5 | 0.45454 | 0.60906 | 0.999994 | 12765 | 1 | 0.87943 | 0.39906 | 0.62704 | 1 | 13139 | 3 | 0.87943 |
| KRT82 | 5 | 0.18485 | 0.35358 | 0.999994 | 7340 | 3 | -0.98667 | 0.39907 | 0.62705 | 1 | 13140 | 2 | -0.98667 |
| ADAMTSL4 | 4 | 0.18057 | 0.32067 | 0.999994 | 7234 | 1 | 0.80407 | 0.39907 | 0.58793 | 1 | 13141 | 2 | 0.80407 |
| ACPP | 5 | 0.19829 | 0.37282 | 0.999994 | 7719 | 2 | -0.15078 | 0.39925 | 0.62722 | 1 | 13142 | 2 | -0.15078 |
| NonTargetingControlG | 2 | 0.25468 | 0.30235 | 0.999994 | 9122 | 1 | 0.37743 | 0.39931 | 0.46792 | 1 | 13143 | 1 | 0.37743 |
| PPM1E | 6 | 0.062729 | 0.17153 | 0.967658 | 3545 | 2 | 0.72233 | 0.39933 | 0.65012 | 1 | 13144 | 3 | 0.72233 |
| CEP57L1 | 5 | 0.29039 | 0.47324 | 0.999994 | 9918 | 2 | 0.98998 | 0.39942 | 0.62737 | 1 | 13145 | 3 | 0.98998 |
| HOXA3 | 5 | 0.026535 | 0.08152 | 0.93914 | 1830 | 4 | -2.8977 | 0.39943 | 0.62738 | 1 | 13146 | 1 | -2.8977 |
| MTF2 | 5 | 0.23742 | 0.42746 | 0.999994 | 8713 | 2 | 0.51064 | 0.39943 | 0.62738 | 1 | 13147 | 2 | 0.51064 |
| USP8 | 6 | 0.14899 | 0.32435 | 0.999994 | 6427 | 3 | -0.059165 | 0.39946 | 0.65023 | 1 | 13148 | 3 | -0.059165 |
| RNF149 | 5 | 0.73827 | 0.7848 | 0.999994 | 16708 | 1 | 0.52921 | 0.39946 | 0.62741 | 1 | 13149 | 2 | 0.52921 |
| BTBD2 | 6 | 0.0064662 | 0.02653 | 0.848412 | 624 | 3 | -0.85665 | 0.39973 | 0.65043 | 1 | 13150 | 3 | -0.85665 |
| ZNF653 | 5 | 0.41798 | 0.57844 | 0.999994 | 12216 | 2 | -0.33881 | 0.39974 | 0.62768 | 1 | 13151 | 2 | -0.33881 |
| GREB1 | 5 | 0.15915 | 0.3163 | 0.999994 | 6708 | 1 | 0.55981 | 0.39975 | 0.62769 | 1 | 13152 | 2 | 0.55981 |
| TMEM206 | 5 | 0.16296 | 0.32187 | 0.999994 | 6818 | 1 | 0.49151 | 0.4001 | 0.62804 | 1 | 13153 | 2 | 0.49151 |
| hsa-mir-5587 | 3 | 0.747 | 0.74695 | 0.999994 | 16828 | 0 | 1.0968 | 0.4001 | 0.55374 | 1 | 13154 | 2 | 1.0968 |
| BHLHE41 | 5 | 0.143 | 0.29236 | 0.999632 | 6272 | 2 | -0.10401 | 0.40018 | 0.62813 | 1 | 13155 | 2 | -0.10401 |

| | | | | | | | | | | | | | |
|---|---|---|---|---|---|---|---|---|---|---|---|---|---|
| RAMP1 | 5 | 0.20355 | 0.38029 | 0.999994 | 7871 | 2 | -0.37558 | 0.40018 | 0.62813 | 1 | 13156 | 1 | -0.37558 |
| DDX31 | 5 | 0.81304 | 0.82792 | 0.999994 | 17674 | 1 | -0.55605 | 0.40018 | 0.62813 | 1 | 13157 | 2 | -0.55605 |
| SNX18 | 5 | 0.34598 | 0.51862 | 0.999994 | 11065 | 2 | 0.44994 | 0.40018 | 0.62813 | 1 | 13158 | 2 | 0.44994 |
| TLN1 | 5 | 0.090595 | 0.21155 | 0.979587 | 4619 | 3 | -0.66496 | 0.40018 | 0.62813 | 1 | 13159 | 1 | -0.66496 |
| DDR1 | 4 | 0.75125 | 0.76103 | 0.999994 | 16868 | 1 | 0.70292 | 0.4002 | 0.58878 | 1 | 13160 | 2 | 0.70292 |
| HIST1H2BJ | 5 | 0.8575 | 0.86033 | 0.999994 | 18401 | 1 | 0.8581 | 0.4002 | 0.62815 | 1 | 13161 | 3 | 0.8581 |
| LONP1 | 3 | 0.44565 | 0.51154 | 0.999994 | 12659 | 1 | 0.54942 | 0.4003 | 0.55393 | 1 | 13162 | 1 | 0.54942 |
| FCGR1B | 3 | 0.11332 | 0.20069 | 0.977164 | 5418 | 2 | -1.8673 | 0.4003 | 0.55393 | 1 | 13163 | 1 | -1.8673 |
| HCFC1 | 3 | 0.14842 | 0.24646 | 0.98999 | 6416 | 2 | -1.7965 | 0.4003 | 0.55393 | 1 | 13164 | 1 | -1.7965 |
| DIP2A | 6 | 0.88716 | 0.89435 | 0.999994 | 18971 | 1 | 0.18779 | 0.40035 | 0.6509 | 1 | 13165 | 2 | 0.18779 |
| C9orf41 | 6 | 0.00042889 | 0.0022593 | 0.45324 | 99 | 4 | -5.3731 | 0.40035 | 0.6509 | 1 | 13166 | 1 | -5.3731 |
| BCL10 | 6 | 0.2656 | 0.46652 | 0.999994 | 9361 | 3 | -0.77701 | 0.40035 | 0.6509 | 1 | 13167 | 2 | -0.77701 |
| USP44 | 6 | 0.079532 | 0.20635 | 0.978744 | 4210 | 3 | -0.85167 | 0.40035 | 0.6509 | 1 | 13168 | 2 | -0.85167 |
| KHDRBS1 | 5 | 0.15504 | 0.31026 | 0.999994 | 6594 | 1 | 0.88738 | 0.40066 | 0.62862 | 1 | 13169 | 3 | 0.88738 |
| P4HA2 | 4 | 0.75819 | 0.76634 | 0.999994 | 16957 | 1 | 1.2191 | 0.40069 | 0.58914 | 1 | 13170 | 2 | 1.2191 |
| DDR2 | 5 | 0.86209 | 0.86415 | 0.999994 | 18487 | 1 | 0.95146 | 0.40073 | 0.62868 | 1 | 13171 | 3 | 0.95146 |
| BACE2 | 5 | 0.46562 | 0.6184 | 0.999994 | 12948 | 1 | 0.48514 | 0.40085 | 0.62881 | 1 | 13172 | 2 | 0.48514 |
| FANCD2 | 5 | 0.05313 | 0.14793 | 0.965554 | 3123 | 3 | -2.0441 | 0.401 | 0.62895 | 1 | 13173 | 2 | -2.0441 |
| C19orf53 | 5 | 0.84812 | 0.85279 | 0.999994 | 18218 | 1 | 0.1969 | 0.401 | 0.62895 | 1 | 13174 | 1 | 0.1969 |
| DNMBP | 5 | 0.031442 | 0.094193 | 0.945031 | 2079 | 3 | -1.3448 | 0.40107 | 0.62902 | 1 | 13175 | 2 | -1.3448 |
| CLEC7A | 4 | 0.4121 | 0.55809 | 0.999994 | 12147 | 2 | 0.65091 | 0.40115 | 0.5895 | 1 | 13176 | 2 | 0.65091 |
| SP7 | 5 | 0.37125 | 0.53943 | 0.999994 | 11533 | 2 | 0.35518 | 0.40127 | 0.62923 | 1 | 13177 | 2 | 0.35518 |
| hsa-mir-3918 | 4 | 0.062912 | 0.14633 | 0.964852 | 3553 | 1 | 1.1154 | 0.4013 | 0.58961 | 1 | 13178 | 2 | 1.1154 |
| ZNF385B | 5 | 0.020084 | 0.064299 | 0.93914 | 1462 | 1 | 0.80356 | 0.40142 | 0.62939 | 1 | 13179 | 3 | 0.80356 |
| SLCO2B1 | 4 | 0.40905 | 0.55498 | 0.999994 | 12094 | 2 | 0.56741 | 0.4015 | 0.58977 | 1 | 13180 | 2 | 0.56741 |
| GLDN | 4 | 0.81275 | 0.8129 | 0.999994 | 17666 | 1 | -0.6083 | 0.40162 | 0.58986 | 1 | 13181 | 1 | -0.6083 |
| hsa-mir-4315-1 | 4 | 0.85094 | 0.85045 | 0.999994 | 18278 | 0 | 0.90743 | 0.40173 | 0.58993 | 1 | 13182 | 2 | 0.90743 |
| SLC2A5 | 5 | 0.23994 | 0.43084 | 0.999994 | 8768 | 1 | 0.28483 | 0.40183 | 0.62977 | 1 | 13183 | 2 | 0.28483 |
| NonTargetingControlG | 2 | 0.639 | 0.63885 | 0.999994 | 15498 | 0 | 1.9902 | 0.40193 | 0.46962 | 1 | 13184 | 1 | 1.9902 |
| ZNF689 | 2 | 0.43469 | 0.44606 | 0.999994 | 12488 | 1 | 1.3493 | 0.40193 | 0.46962 | 1 | 13185 | 1 | 1.3493 |
| hsa-mir-6846 | 3 | 0.87129 | 0.87135 | 0.999994 | 18664 | 0 | 1.078 | 0.40194 | 0.55562 | 1 | 13186 | 2 | 1.078 |
| CC2D2A | 4 | 0.87384 | 0.87348 | 0.999994 | 18707 | 0 | 1.0171 | 0.40204 | 0.59016 | 1 | 13187 | 2 | 1.0171 |
| TBCK | 6 | 0.21604 | 0.40741 | 0.999994 | 8172 | 3 | 0.11259 | 0.40206 | 0.65223 | 1 | 13188 | 3 | 0.11259 |
| ITFG2 | 6 | 0.28834 | 0.49316 | 0.999994 | 9879 | 2 | -0.31158 | 0.40206 | 0.65223 | 1 | 13189 | 1 | -0.31158 |
| ALDH3B1 | 5 | 0.51011 | 0.65672 | 0.999994 | 13578 | 2 | 0.47153 | 0.40224 | 0.6302 | 1 | 13190 | 2 | 0.47153 |
| TRAK1 | 5 | 0.8379 | 0.84513 | 0.999994 | 18051 | 1 | 1.1043 | 0.40225 | 0.63022 | 1 | 13191 | 3 | 1.1043 |
| TSPO2 | 6 | 0.34332 | 0.55645 | 0.999994 | 11014 | 3 | -0.13769 | 0.4023 | 0.65241 | 1 | 13192 | 2 | -0.13769 |
| ZFHX2 | 5 | 0.13757 | 0.28413 | 0.999632 | 6122 | 2 | 0.92995 | 0.40232 | 0.63028 | 1 | 13193 | 3 | 0.92995 |
| hsa-mir-1285-1 | 4 | 0.32862 | 0.47227 | 0.999994 | 10693 | 2 | 0.65615 | 0.40243 | 0.59044 | 1 | 13194 | 2 | 0.65615 |
| RLN1 | 5 | 0.76673 | 0.79989 | 0.999994 | 17046 | 1 | 0.846 | 0.40246 | 0.63041 | 1 | 13195 | 3 | 0.846 |
| hsa-mir-4999 | 4 | 0.2424 | 0.38407 | 0.999994 | 8843 | 1 | 0.88548 | 0.40257 | 0.59055 | 1 | 13196 | 2 | 0.88548 |
| SLC1A1 | 5 | 0.10253 | 0.23036 | 0.983933 | 5056 | 2 | 0.19436 | 0.40261 | 0.63056 | 1 | 13197 | 2 | 0.19436 |
| RAPGEF5 | 5 | 0.073059 | 0.18336 | 0.968526 | 3961 | 3 | -1.8045 | 0.40261 | 0.63056 | 1 | 13198 | 2 | -1.8045 |
| WDTC1 | 5 | 0.099391 | 0.22545 | 0.983354 | 4945 | 3 | -0.83215 | 0.40264 | 0.63058 | 1 | 13199 | 2 | -0.83215 |
| RPP21 | 4 | 0.40229 | 0.54808 | 0.999994 | 11982 | 2 | 0.63286 | 0.40274 | 0.59069 | 1 | 13200 | 2 | 0.63286 |
| FOXI2 | 5 | 0.0089953 | 0.033153 | 0.867348 | 794 | 1 | 0.94776 | 0.40276 | 0.63071 | 1 | 13201 | 3 | 0.94776 |
| TAAR8 | 6 | 0.064818 | 0.17588 | 0.968171 | 3643 | 2 | 0.53385 | 0.40284 | 0.65283 | 1 | 13202 | 3 | 0.53385 |
| SLC24A2 | 5 | 0.49125 | 0.64032 | 0.999994 | 13293 | 1 | 0.86003 | 0.40322 | 0.63116 | 1 | 13203 | 3 | 0.86003 |
| PRKAB2 | 5 | 0.56972 | 0.70927 | 0.999994 | 14533 | 2 | 0.74297 | 0.40331 | 0.63125 | 1 | 13204 | 3 | 0.74297 |
| PSMD1 | 4 | 0.31879 | 0.4622 | 0.999994 | 10493 | 2 | 0.13207 | 0.40337 | 0.59117 | 1 | 13205 | 2 | 0.13207 |
| PRDM4 | 6 | 0.5468 | 0.70137 | 0.999994 | 14163 | 2 | -0.13605 | 0.40359 | 0.65343 | 1 | 13206 | 1 | -0.13605 |
| VSTM5 | 6 | 0.013417 | 0.050497 | 0.909101 | 1094 | 3 | 0.13107 | 0.40359 | 0.65343 | 1 | 13207 | 3 | 0.13107 |

| | | | | | | | | | | | | |
|---|---|---|---|---|---|---|---|---|---|---|---|---|
| CTBP1 | 6 | 0.021835 | 0.077861 | 0.93914 | 1568 | 4 | -2.3692 | 0.40373 | 0.65353 | 1 | 13208 | 2 | -2.3692 |
| IL1RL1 | 5 | 0.9516 | 0.95176 | 0.999994 | 20379 | 0 | 0.88297 | 0.40391 | 0.63184 | 1 | 13209 | 3 | 0.88297 |
| TMEM53 | 5 | 0.2411 | 0.43242 | 0.999994 | 8799 | 2 | -0.59816 | 0.40397 | 0.63188 | 1 | 13210 | 2 | -0.59816 |
| PQBP1 | 5 | 0.21515 | 0.39653 | 0.999994 | 8144 | 1 | 0.8541 | 0.40398 | 0.6319 | 1 | 13211 | 3 | 0.8541 |
| FBF1 | 5 | 0.063147 | 0.16726 | 0.967126 | 3567 | 3 | -1.9467 | 0.40403 | 0.63195 | 1 | 13212 | 2 | -1.9467 |
| CROCC | 6 | 0.65214 | 0.77927 | 0.999994 | 15689 | 2 | 0.68403 | 0.40407 | 0.65381 | 1 | 13213 | 3 | 0.68403 |
| KRTAP4-12 | 5 | 0.097887 | 0.22313 | 0.983354 | 4887 | 2 | 0.87479 | 0.40421 | 0.63214 | 1 | 13214 | 3 | 0.87479 |
| hsa-mir-548ao | 4 | 0.91858 | 0.91836 | 0.999994 | 19598 | 0 | 0.46859 | 0.40424 | 0.59178 | 1 | 13215 | 1 | 0.46859 |
| hsa-mir-3065 | 4 | 0.75771 | 0.76598 | 0.999994 | 16948 | 1 | 0.82582 | 0.40424 | 0.59178 | 1 | 13216 | 2 | 0.82582 |
| B4GALT5 | 5 | 0.91956 | 0.91972 | 0.999994 | 19621 | 0 | 0.54942 | 0.4043 | 0.63222 | 1 | 13217 | 2 | 0.54942 |
| CLEC18C | 3 | 0.42901 | 0.5015 | 0.999994 | 12397 | 1 | 0.99674 | 0.40433 | 0.55809 | 1 | 13218 | 2 | 0.99674 |
| C19orf70 | 6 | 0.042259 | 0.12699 | 0.951052 | 2628 | 3 | 0.13754 | 0.40436 | 0.65401 | 1 | 13219 | 3 | 0.13754 |
| MVP | 6 | 0.24744 | 0.44489 | 0.999994 | 8960 | 3 | -0.39548 | 0.40436 | 0.65401 | 1 | 13220 | 1 | -0.39548 |
| EGFR | 6 | 0.030107 | 0.099045 | 0.945253 | 2014 | 4 | -1.7901 | 0.40436 | 0.65401 | 1 | 13221 | 2 | -1.7901 |
| MYO1D | 5 | 0.0051045 | 0.021234 | 0.812607 | 527 | 1 | 0.83641 | 0.40438 | 0.63229 | 1 | 13222 | 3 | 0.83641 |
| UTS2R | 5 | 0.18517 | 0.35405 | 0.999994 | 7346 | 2 | 0.93213 | 0.40441 | 0.63232 | 1 | 13223 | 3 | 0.93213 |
| UBE2K | 6 | 0.13235 | 0.30334 | 0.999994 | 5985 | 3 | -1.1156 | 0.40445 | 0.65408 | 1 | 13224 | 2 | -1.1156 |
| ABCG2 | 6 | 0.33876 | 0.55126 | 0.999994 | 10913 | 3 | -0.62961 | 0.40445 | 0.65408 | 1 | 13225 | 2 | -0.62961 |
| CCDC109B | 6 | 0.19508 | 0.38191 | 0.999994 | 7635 | 3 | 0.24782 | 0.40445 | 0.65408 | 1 | 13226 | 3 | 0.24782 |
| ZNF625 | 5 | 0.086068 | 0.20427 | 0.978186 | 4448 | 2 | 0.86977 | 0.4045 | 0.6324 | 1 | 13227 | 3 | 0.86977 |
| hsa-mir-3200 | 3 | 0.74526 | 0.74521 | 0.999994 | 16804 | 0 | 0.98107 | 0.40458 | 0.55834 | 1 | 13228 | 2 | 0.98107 |
| PNMA6A | 5 | 0.77935 | 0.80707 | 0.999994 | 17201 | 1 | 0.79705 | 0.40459 | 0.63249 | 1 | 13229 | 3 | 0.79705 |
| PCP4L1 | 5 | 0.60045 | 0.72633 | 0.999994 | 14952 | 1 | 0.80218 | 0.40461 | 0.63251 | 1 | 13230 | 3 | 0.80218 |
| hsa-mir-4790 | 3 | 0.087518 | 0.16647 | 0.967126 | 4495 | 2 | -0.75138 | 0.40474 | 0.55853 | 1 | 13231 | 1 | -0.75138 |
| PIGL | 2 | 0.73821 | 0.7379 | 0.999994 | 16706 | 0 | 2.2978 | 0.4048 | 0.47156 | 1 | 13232 | 1 | 2.2978 |
| NonTargetingControlG | 2 | 0.65863 | 0.65845 | 0.999994 | 15758 | 0 | 2.0531 | 0.4048 | 0.47156 | 1 | 13233 | 1 | 2.0531 |
| LYRM9 | 5 | 0.0012958 | 0.0057634 | 0.623123 | 192 | 2 | 1.0101 | 0.40492 | 0.63282 | 1 | 13234 | 3 | 1.0101 |
| LACTB | 5 | 0.045931 | 0.13061 | 0.95318 | 2803 | 4 | -1.7772 | 0.40493 | 0.63283 | 1 | 13235 | 1 | -1.7772 |
| C1QBP | 5 | 0.86278 | 0.86475 | 0.999994 | 18501 | 1 | 0.047092 | 0.40493 | 0.63283 | 1 | 13236 | 2 | 0.047092 |
| PTPN4 | 5 | 0.24023 | 0.43124 | 0.999994 | 8775 | 3 | -0.59449 | 0.40493 | 0.63283 | 1 | 13237 | 2 | -0.59449 |
| LEPRE1 | 5 | 0.10743 | 0.23796 | 0.987253 | 5222 | 2 | 0.2217 | 0.40494 | 0.63284 | 1 | 13238 | 2 | 0.2217 |
| MCAM | 5 | 0.036509 | 0.1071 | 0.950995 | 2345 | 1 | 0.78132 | 0.40502 | 0.63292 | 1 | 13239 | 3 | 0.78132 |
| GJB7 | 6 | 0.90536 | 0.90785 | 0.999994 | 19340 | 1 | 0.10981 | 0.40515 | 0.65462 | 1 | 13240 | 2 | 0.10981 |
| SCRG1 | 5 | 0.769 | 0.80114 | 0.999994 | 17074 | 1 | 0.85722 | 0.40526 | 0.63316 | 1 | 13241 | 3 | 0.85722 |
| MEPCE | 4 | 0.2762 | 0.41868 | 0.999994 | 9623 | 2 | -0.62176 | 0.40527 | 0.59255 | 1 | 13242 | 1 | -0.62176 |
| ANO5 | 4 | 0.33272 | 0.47652 | 0.999994 | 10785 | 2 | -0.79248 | 0.40527 | 0.59255 | 1 | 13243 | 1 | -0.79248 |
| RILP | 4 | 0.25808 | 0.40015 | 0.999994 | 9196 | 2 | -0.19408 | 0.40527 | 0.59255 | 1 | 13244 | 2 | -0.19408 |
| CCHCR1 | 5 | 0.7324 | 0.78183 | 0.999994 | 16634 | 1 | 0.8058 | 0.40532 | 0.63321 | 1 | 13245 | 3 | 0.8058 |
| MYBPH | 5 | 0.8869 | 0.887 | 0.999994 | 18965 | 1 | 0.83684 | 0.40538 | 0.63328 | 1 | 13246 | 3 | 0.83684 |
| WBP1 | 5 | 0.0065988 | 0.026224 | 0.848412 | 632 | 3 | -6.5311 | 0.40541 | 0.6333 | 1 | 13247 | 2 | -6.5311 |
| NonTargetingControlG | 2 | 0.6709 | 0.67068 | 0.999994 | 15943 | 0 | 2.1097 | 0.40555 | 0.47204 | 1 | 13248 | 1 | 2.1097 |
| TSPAN5 | 5 | 0.3844 | 0.55044 | 0.999994 | 11705 | 2 | 0.93072 | 0.40559 | 0.63348 | 1 | 13249 | 3 | 0.93072 |
| PRAMEF12 | 6 | 0.60944 | 0.74693 | 0.999994 | 15065 | 1 | 0.42582 | 0.40563 | 0.655 | 1 | 13250 | 3 | 0.42582 |
| SEMA3D | 5 | 0.19082 | 0.36212 | 0.999994 | 7509 | 2 | 0.87596 | 0.4058 | 0.63368 | 1 | 13251 | 3 | 0.87596 |
| PIGH | 5 | 0.89519 | 0.89533 | 0.999994 | 19132 | 0 | -0.02629 | 0.4058 | 0.63368 | 1 | 13252 | 2 | -0.02629 |
| RTF1 | 5 | 0.026022 | 0.080162 | 0.93914 | 1807 | 2 | 0.18735 | 0.4058 | 0.63368 | 1 | 13253 | 2 | 0.18735 |
| ORC2 | 6 | 0.13302 | 0.30419 | 0.999994 | 5996 | 3 | 0.034136 | 0.4059 | 0.6552 | 1 | 13254 | 3 | 0.034136 |
| EXO1 | 6 | 0.30683 | 0.51469 | 0.999994 | 10270 | 2 | 0.12071 | 0.40598 | 0.65526 | 1 | 13255 | 3 | 0.12071 |
| GPR128 | 6 | 0.094709 | 0.23693 | 0.985449 | 4777 | 3 | -0.30663 | 0.40598 | 0.65526 | 1 | 13256 | 1 | -0.30663 |
| MANSC4 | 6 | 0.11717 | 0.2804 | 0.999632 | 5540 | 3 | -0.91381 | 0.40598 | 0.65526 | 1 | 13257 | 2 | -0.91381 |
| ATL1 | 4 | 0.02674 | 0.074833 | 0.93914 | 1845 | 3 | -4.693 | 0.4062 | 0.59324 | 1 | 13258 | 1 | -4.693 |
| NonTargetingControlG | 2 | 0.62527 | 0.625 | 0.999994 | 15311 | 0 | 1.9071 | 0.40637 | 0.47257 | 1 | 13259 | 1 | 1.9071 |

| | | | | | | | | | | | | | |
|---|---|---|---|---|---|---|---|---|---|---|---|---|---|
| TMEM234 | 5 | 0.38463 | 0.55064 | 0.999994 | 11710 | 2 | 0.018812 | 0.40646 | 0.63436 | 1 | 13260 | 2 | 0.018812 |
| TFG | 5 | 0.16533 | 0.32527 | 0.999994 | 6868 | 2 | 0.13565 | 0.40656 | 0.63447 | 1 | 13261 | 1 | 0.13565 |
| CAPRIN1 | 5 | 0.33565 | 0.51017 | 0.999994 | 10844 | 1 | 0.27378 | 0.40656 | 0.63447 | 1 | 13262 | 2 | 0.27378 |
| H2AFB1 | 6 | 0.34912 | 0.563 | 0.999994 | 11132 | 2 | 0.17702 | 0.40667 | 0.65581 | 1 | 13263 | 3 | 0.17702 |
| AKR1B15 | 6 | 0.35178 | 0.56598 | 0.999994 | 11192 | 3 | -0.44996 | 0.40667 | 0.65581 | 1 | 13264 | 2 | -0.44996 |
| MDN1 | 3 | 0.59146 | 0.61061 | 0.999994 | 14839 | 1 | 1.0209 | 0.40683 | 0.56067 | 1 | 13265 | 2 | 1.0209 |
| TRMT11 | 6 | 0.23293 | 0.42762 | 0.999994 | 8592 | 3 | -0.57805 | 0.40685 | 0.65595 | 1 | 13266 | 2 | -0.57805 |
| PLK2 | 3 | 0.030363 | 0.075999 | 0.93914 | 2026 | 1 | 0.20141 | 0.40702 | 0.56086 | 1 | 13267 | 1 | 0.20141 |
| SIGLEC14 | 5 | 0.5578 | 0.69866 | 0.999994 | 14331 | 2 | 0.85667 | 0.4071 | 0.63497 | 1 | 13268 | 3 | 0.85667 |
| LMBRD2 | 5 | 0.044249 | 0.12643 | 0.951052 | 2724 | 3 | -2.3055 | 0.40711 | 0.63497 | 1 | 13269 | 2 | -2.3055 |
| EMX1 | 5 | 0.15882 | 0.31581 | 0.999994 | 6693 | 2 | -0.55444 | 0.40711 | 0.63497 | 1 | 13270 | 2 | -0.55444 |
| FAM102A | 4 | 0.8856 | 0.88527 | 0.999994 | 18941 | 0 | 0.92503 | 0.40715 | 0.59397 | 1 | 13271 | 2 | 0.92503 |
| KIAA1045 | 4 | 0.59147 | 0.66074 | 0.999994 | 14840 | 1 | 0.98701 | 0.40717 | 0.59398 | 1 | 13272 | 2 | 0.98701 |
| FIBIN | 4 | 0.93 | 0.92977 | 0.999994 | 19853 | 0 | 1.329 | 0.4072 | 0.59401 | 1 | 13273 | 2 | 1.329 |
| HS1BP3 | 5 | 0.26664 | 0.45414 | 0.999994 | 9387 | 1 | 0.44085 | 0.40723 | 0.63509 | 1 | 13274 | 2 | 0.44085 |
| hsa-mir-25 | 3 | 0.55814 | 0.58604 | 0.999994 | 14336 | 1 | 0.96214 | 0.40724 | 0.56108 | 1 | 13275 | 2 | 0.96214 |
| CTTNBP2NL | 6 | 0.34685 | 0.56047 | 0.999994 | 11092 | 2 | 0.7392 | 0.40749 | 0.65646 | 1 | 13276 | 3 | 0.7392 |
| ADH4 | 6 | 0.024579 | 0.086017 | 0.94395 | 1734 | 4 | -1.4285 | 0.40755 | 0.6565 | 1 | 13277 | 1 | -1.4285 |
| MS4A14 | 6 | 0.11005 | 0.26679 | 0.998555 | 5307 | 4 | -0.98597 | 0.40755 | 0.6565 | 1 | 13278 | 2 | -0.98597 |
| hsa-mir-3132 | 3 | 0.27549 | 0.40644 | 0.999994 | 9607 | 1 | 1.008 | 0.4077 | 0.56153 | 1 | 13279 | 2 | 1.008 |
| CRKL | 4 | 0.28872 | 0.43147 | 0.999994 | 9890 | 1 | 0.84904 | 0.40782 | 0.59448 | 1 | 13280 | 2 | 0.84904 |
| hsa-mir-6738 | 3 | 0.24956 | 0.37442 | 0.999994 | 9008 | 1 | 0.92352 | 0.40784 | 0.56167 | 1 | 13281 | 2 | 0.92352 |
| EML2 | 5 | 0.26962 | 0.45653 | 0.999994 | 9463 | 2 | -0.5373 | 0.4079 | 0.63574 | 1 | 13282 | 1 | -0.5373 |
| NonTargetingControlG | 2 | 0.67752 | 0.67732 | 0.999994 | 16053 | 0 | 1.8992 | 0.40804 | 0.47368 | 1 | 13283 | 1 | 1.8992 |
| ITGB3BP | 4 | 0.099337 | 0.2139 | 0.979587 | 4942 | 2 | -0.8016 | 0.40809 | 0.59468 | 1 | 13284 | 2 | -0.8016 |
| RABIF | 6 | 0.10351 | 0.25418 | 0.994632 | 5084 | 2 | 0.74542 | 0.40816 | 0.65695 | 1 | 13285 | 3 | 0.74542 |
| LTBP1 | 6 | 0.11479 | 0.27585 | 0.999379 | 5472 | 4 | -0.70518 | 0.40818 | 0.65697 | 1 | 13286 | 2 | -0.70518 |
| TMEM168 | 6 | 0.18055 | 0.36385 | 0.999994 | 7233 | 1 | 0.67852 | 0.40821 | 0.65699 | 1 | 13287 | 3 | 0.67852 |
| CCDC25 | 6 | 0.082907 | 0.21325 | 0.979587 | 4335 | 4 | -1.1041 | 0.40847 | 0.65719 | 1 | 13288 | 2 | -1.1041 |
| ELFN2 | 6 | 0.32045 | 0.53041 | 0.999994 | 10521 | 3 | 0.22137 | 0.4087 | 0.65737 | 1 | 13289 | 3 | 0.22137 |
| SPON1 | 6 | 0.22328 | 0.41609 | 0.999994 | 8365 | 3 | 0.12334 | 0.40875 | 0.65743 | 1 | 13290 | 3 | 0.12334 |
| FLNB | 6 | 0.17771 | 0.36023 | 0.999994 | 7173 | 3 | -0.075047 | 0.40875 | 0.65743 | 1 | 13291 | 2 | -0.075047 |
| MRPL19 | 5 | 0.46551 | 0.61831 | 0.999994 | 12945 | 1 | 0.063373 | 0.40881 | 0.63667 | 1 | 13292 | 2 | 0.063373 |
| LY6D | 5 | 0.84459 | 0.85011 | 0.999994 | 18168 | 1 | 0.71711 | 0.40881 | 0.63667 | 1 | 13293 | 3 | 0.71711 |
| DAPL1 | 6 | 0.1473 | 0.32225 | 0.999994 | 6383 | 3 | -0.42184 | 0.40889 | 0.65753 | 1 | 13294 | 3 | -0.42184 |
| STK38 | 6 | 0.11558 | 0.27738 | 0.999379 | 5500 | 2 | 0.70497 | 0.40907 | 0.65767 | 1 | 13295 | 3 | 0.70497 |
| MINA | 6 | 0.18241 | 0.36616 | 0.999994 | 7278 | 3 | 0.012512 | 0.40907 | 0.65767 | 1 | 13296 | 3 | 0.012512 |
| OSBPL8 | 5 | 0.062894 | 0.16684 | 0.967126 | 3552 | 3 | -0.99201 | 0.40907 | 0.63693 | 1 | 13297 | 2 | -0.99201 |
| IKZF1 | 5 | 0.036966 | 0.10827 | 0.951052 | 2372 | 3 | -2.3382 | 0.40907 | 0.63693 | 1 | 13298 | 2 | -2.3382 |
| CTCFL | 5 | 0.76827 | 0.80073 | 0.999994 | 17065 | 1 | 0.90133 | 0.40927 | 0.63715 | 1 | 13299 | 3 | 0.90133 |
| SLC17A6 | 5 | 0.78234 | 0.80879 | 0.999994 | 17244 | 1 | 0.85504 | 0.40932 | 0.6372 | 1 | 13300 | 3 | 0.85504 |
| OR8H2 | 4 | 0.3116 | 0.4549 | 0.999994 | 10351 | 2 | 0.6084 | 0.40938 | 0.59564 | 1 | 13301 | 2 | 0.6084 |
| STK17B | 6 | 0.8244 | 0.85889 | 0.999994 | 17843 | 1 | 0.99254 | 0.40952 | 0.658 | 1 | 13302 | 3 | 0.99254 |
| NFRKB | 4 | 0.059567 | 0.13996 | 0.958716 | 3404 | 2 | -0.0019938 | 0.40972 | 0.59588 | 1 | 13303 | 2 | -0.0019938 |
| APPL2 | 6 | 0.31159 | 0.5202 | 0.999994 | 10350 | 3 | -0.18004 | 0.4098 | 0.65823 | 1 | 13304 | 1 | -0.18004 |
| OR52N5 | 6 | 0.35992 | 0.57521 | 0.999994 | 11334 | 3 | -0.51512 | 0.4098 | 0.65823 | 1 | 13305 | 1 | -0.51512 |
| PSPN | 6 | 0.023254 | 0.082391 | 0.940526 | 1666 | 2 | -0.12033 | 0.4098 | 0.65823 | 1 | 13306 | 2 | -0.12033 |
| L1CAM | 6 | 0.75818 | 0.83139 | 0.999994 | 16956 | 1 | 0.087978 | 0.40991 | 0.65833 | 1 | 13307 | 2 | 0.087978 |
| CYP11B1 | 6 | 0.31013 | 0.5185 | 0.999994 | 10326 | 3 | -0.25217 | 0.40991 | 0.65833 | 1 | 13308 | 2 | -0.25217 |
| MCF2L | 6 | 0.16716 | 0.34717 | 0.999994 | 6912 | 3 | -0.18911 | 0.40991 | 0.65833 | 1 | 13309 | 3 | -0.18911 |
| hsa-mir-3064 | 4 | 0.17097 | 0.31077 | 0.999994 | 7007 | 2 | 0.5033 | 0.41 | 0.59608 | 1 | 13310 | 2 | 0.5033 |
| NonTargetingControlG | 2 | 0.5902 | 0.59022 | 0.999994 | 14826 | 0 | 1.6602 | 0.41007 | 0.47501 | 1 | 13311 | 1 | 1.6602 |

| | | | | | | | | | | | | |
|---|---|---|---|---|---|---|---|---|---|---|---|---|
| hsa-mir-6774 | 4 | 0.85868 | 0.8583 | 0.999994 | 18413 | 0 | 1.1745 | 0.41009 | 0.59615 | 1 | 13312 | 2 | 1.1745 |
| WDR18 | 3 | 0.02552 | 0.064746 | 0.93914 | 1779 | 1 | 0.99903 | 0.41015 | 0.56399 | 1 | 13313 | 2 | 0.99903 |
| TANC2 | 5 | 0.83518 | 0.84319 | 0.999994 | 18009 | 1 | 0.87996 | 0.41024 | 0.63811 | 1 | 13314 | 3 | 0.87996 |
| APOF | 5 | 0.21439 | 0.3955 | 0.999994 | 8129 | 3 | -0.74121 | 0.41029 | 0.63816 | 1 | 13315 | 2 | -0.74121 |
| CILP2 | 4 | 0.032918 | 0.087456 | 0.94395 | 2153 | 2 | -2.0292 | 0.41032 | 0.59631 | 1 | 13316 | 2 | -2.0292 |
| GEM | 6 | 0.46379 | 0.64429 | 0.999994 | 12918 | 2 | 0.56223 | 0.41038 | 0.65868 | 1 | 13317 | 3 | 0.56223 |
| MLL4 | 4 | 0.5665 | 0.64791 | 0.999994 | 14487 | 1 | 0.96786 | 0.4105 | 0.59644 | 1 | 13318 | 2 | 0.96786 |
| hsa-mir-588 | 4 | 0.14136 | 0.28031 | 0.999632 | 6225 | 2 | -1.6669 | 0.4105 | 0.59644 | 1 | 13319 | 1 | -1.6669 |
| hsa-mir-623 | 4 | 0.86099 | 0.86057 | 0.999994 | 18463 | 0 | 0.51732 | 0.4105 | 0.59644 | 1 | 13320 | 2 | 0.51732 |
| BST2 | 4 | 0.85998 | 0.85958 | 0.999994 | 18436 | 0 | 0.38129 | 0.4105 | 0.59644 | 1 | 13321 | 2 | 0.38129 |
| GSTM4 | 4 | 0.81329 | 0.81339 | 0.999994 | 17680 | 1 | 0.66346 | 0.41052 | 0.59645 | 1 | 13322 | 2 | 0.66346 |
| TMEM106C | 6 | 0.076608 | 0.2005 | 0.977164 | 4097 | 3 | -1.0949 | 0.41056 | 0.65883 | 1 | 13323 | 2 | -1.0949 |
| BMP3 | 5 | 0.20208 | 0.3782 | 0.999994 | 7830 | 1 | 0.55294 | 0.41057 | 0.63844 | 1 | 13324 | 3 | 0.55294 |
| FRMPD1 | 5 | 0.30306 | 0.48354 | 0.999994 | 10195 | 2 | 0.33764 | 0.41057 | 0.63844 | 1 | 13325 | 2 | 0.33764 |
| PRSS3 | 6 | 0.052256 | 0.14902 | 0.965566 | 3082 | 1 | 0.85207 | 0.41061 | 0.65886 | 1 | 13326 | 3 | 0.85207 |
| DOLK | 4 | 0.096671 | 0.20914 | 0.979587 | 4846 | 2 | -0.71584 | 0.41069 | 0.59658 | 1 | 13327 | 2 | -0.71584 |
| hsa-mir-516b-1 | 3 | 0.88081 | 0.88086 | 0.999994 | 18850 | 0 | 0.84824 | 0.41075 | 0.56461 | 1 | 13328 | 2 | 0.84824 |
| RNASE4 | 6 | 0.067836 | 0.18223 | 0.968526 | 3745 | 1 | 0.74149 | 0.41085 | 0.65905 | 1 | 13329 | 3 | 0.74149 |
| EPHB2 | 6 | 0.24085 | 0.43703 | 0.999994 | 8790 | 2 | 0.50988 | 0.41109 | 0.65925 | 1 | 13330 | 3 | 0.50988 |
| TMEM183A | 6 | 0.46098 | 0.64242 | 0.999994 | 12863 | 2 | 0.87874 | 0.41112 | 0.65927 | 1 | 13331 | 3 | 0.87874 |
| NonTargetingControlG | 1 | 0.58887 | 0.58857 | 0.999994 | 14803 | 0 | 1.2586 | 0.41113 | 0.4113 | 1 | 13332 | 1 | 1.2586 |
| PALD1 | 6 | 0.082278 | 0.21202 | 0.979587 | 4307 | 4 | -1.0331 | 0.41124 | 0.65936 | 1 | 13333 | 2 | -1.0331 |
| DOC2B | 6 | 0.62047 | 0.75519 | 0.999994 | 15227 | 2 | -0.23434 | 0.41125 | 0.65937 | 1 | 13334 | 2 | -0.23434 |
| C9orf69 | 5 | 0.66654 | 0.75184 | 0.999994 | 15876 | 1 | 0.57788 | 0.41138 | 0.63922 | 1 | 13335 | 3 | 0.57788 |
| hsa-mir-1291 | 4 | 0.27029 | 0.41258 | 0.999994 | 9482 | 1 | 0.88538 | 0.41144 | 0.59718 | 1 | 13336 | 2 | 0.88538 |
| ARHGAP39 | 6 | 0.17858 | 0.36135 | 0.999994 | 7188 | 3 | 0.33008 | 0.41161 | 0.65963 | 1 | 13337 | 3 | 0.33008 |
| NonTargetingControlG | 2 | 0.30833 | 0.34322 | 0.999994 | 10291 | 1 | 0.71094 | 0.41165 | 0.47604 | 1 | 13338 | 1 | 0.71094 |
| CBWD5 | 2 | 0.16755 | 0.23894 | 0.987253 | 6926 | 1 | -0.38144 | 0.41165 | 0.47604 | 1 | 13339 | 1 | -0.38144 |
| hsa-mir-548am | 2 | 0.19212 | 0.2564 | 0.994943 | 7550 | 1 | -1.0138 | 0.41165 | 0.47604 | 1 | 13340 | 1 | -1.0138 |
| NonTargetingControlG | 2 | 0.087142 | 0.14997 | 0.966359 | 4480 | 1 | -2.178 | 0.41165 | 0.47604 | 1 | 13341 | 1 | -2.178 |
| hsa-mir-505 | 4 | 0.35636 | 0.50075 | 0.999994 | 11273 | 1 | 1.1698 | 0.41176 | 0.59741 | 1 | 13342 | 2 | 1.1698 |
| NANOG | 6 | 0.090011 | 0.22752 | 0.983354 | 4594 | 3 | -0.67184 | 0.41193 | 0.65989 | 1 | 13343 | 3 | -0.67184 |
| ZNF622 | 5 | 0.10953 | 0.24126 | 0.987253 | 5287 | 2 | -0.5564 | 0.41194 | 0.63977 | 1 | 13344 | 2 | -0.5564 |
| TSR3 | 6 | 0.098385 | 0.24417 | 0.988309 | 4910 | 4 | -1.0787 | 0.41196 | 0.65992 | 1 | 13345 | 2 | -1.0787 |
| GTDC2 | 4 | 0.056304 | 0.13375 | 0.95318 | 3258 | 3 | -1.3892 | 0.41199 | 0.59759 | 1 | 13346 | 1 | -1.3892 |
| MXD1 | 5 | 0.81296 | 0.82788 | 0.999994 | 17671 | 1 | -0.15243 | 0.41201 | 0.63983 | 1 | 13347 | 1 | -0.15243 |
| CSRP1 | 6 | 0.50883 | 0.67479 | 0.999994 | 13565 | 2 | 0.52891 | 0.41209 | 0.66002 | 1 | 13348 | 3 | 0.52891 |
| EGFL7 | 6 | 0.95115 | 0.95114 | 0.999994 | 20368 | 0 | 0.090561 | 0.41231 | 0.66018 | 1 | 13349 | 2 | 0.090561 |
| MYO1C | 6 | 0.43142 | 0.62279 | 0.999994 | 12437 | 2 | 0.31475 | 0.41231 | 0.66018 | 1 | 13350 | 3 | 0.31475 |
| DMC1 | 6 | 0.079221 | 0.20575 | 0.978603 | 4199 | 3 | -0.33587 | 0.41236 | 0.66023 | 1 | 13351 | 3 | -0.33587 |
| hsa-mir-4493 | 3 | 0.45371 | 0.51651 | 0.999994 | 12755 | 1 | 0.49207 | 0.4124 | 0.56627 | 1 | 13352 | 1 | 0.49207 |
| hsa-mir-328 | 3 | 0.29639 | 0.42661 | 0.999994 | 10046 | 1 | 0.2743 | 0.4124 | 0.56627 | 1 | 13353 | 1 | 0.2743 |
| SPRR3 | 4 | 0.18067 | 0.3208 | 0.999994 | 7237 | 2 | -0.71207 | 0.41244 | 0.5979 | 1 | 13354 | 2 | -0.71207 |
| hsa-mir-1234 | 4 | 0.13605 | 0.27483 | 0.998561 | 6080 | 2 | -1.3164 | 0.41244 | 0.5979 | 1 | 13355 | 2 | -1.3164 |
| ZNF518A | 6 | 0.16363 | 0.34281 | 0.999994 | 6831 | 3 | -0.64605 | 0.41249 | 0.66033 | 1 | 13356 | 2 | -0.64605 |
| CAPN15 | 5 | 0.41159 | 0.57318 | 0.999994 | 12141 | 2 | 0.21953 | 0.41254 | 0.64035 | 1 | 13357 | 2 | 0.21953 |
| CACNA1D | 5 | 0.53381 | 0.67742 | 0.999994 | 13964 | 2 | 0.71556 | 0.4127 | 0.6405 | 1 | 13358 | 3 | 0.71556 |
| PAPSS2 | 5 | 0.50593 | 0.6531 | 0.999994 | 13521 | 2 | 0.36306 | 0.4128 | 0.6406 | 1 | 13359 | 2 | 0.36306 |
| RNF141 | 5 | 0.50007 | 0.64794 | 0.999994 | 13423 | 2 | -0.40311 | 0.41281 | 0.64061 | 1 | 13360 | 1 | -0.40311 |
| C19orf26 | 5 | 0.024643 | 0.07652 | 0.93914 | 1740 | 4 | -2.0705 | 0.41281 | 0.64061 | 1 | 13361 | 1 | -2.0705 |
| hsa-mir-1343 | 4 | 0.29311 | 0.43596 | 0.999994 | 9979 | 1 | 1.2286 | 0.41282 | 0.5982 | 1 | 13362 | 2 | 1.2286 |
| hsa-mir-4693 | 2 | 0.72288 | 0.7226 | 0.999994 | 16524 | 0 | 2.0527 | 0.41284 | 0.47682 | 1 | 13363 | 1 | 2.0527 |

| | | | | | | | | | | | | | |
|---|---|---|---|---|---|---|---|---|---|---|---|---|---|
| NBPF3 | 3 | 0.041886 | 0.10202 | 0.946414 | 2606 | 1 | 1.0037 | 0.41295 | 0.56686 | 1 | 13364 | 2 | 1.0037 |
| GLI2 | 6 | 0.040845 | 0.12383 | 0.951052 | 2552 | 4 | -1.9875 | 0.41309 | 0.66077 | 1 | 13365 | 2 | -1.9875 |
| hsa-mir-551a | 4 | 0.32545 | 0.46905 | 0.999994 | 10631 | 1 | 1.0888 | 0.41312 | 0.59843 | 1 | 13366 | 2 | 1.0888 |
| AIFM2 | 6 | 0.025621 | 0.088474 | 0.94395 | 1781 | 3 | -2.0488 | 0.41314 | 0.66081 | 1 | 13367 | 1 | -2.0488 |
| HLA-E | 6 | 0.042167 | 0.12677 | 0.951052 | 2622 | 4 | -1.5429 | 0.41314 | 0.66081 | 1 | 13368 | 2 | -1.5429 |
| GALM | 5 | 0.54034 | 0.68321 | 0.999994 | 14059 | 2 | 0.64599 | 0.4132 | 0.641 | 1 | 13369 | 3 | 0.64599 |
| KCNC3 | 6 | 0.0099052 | 0.038712 | 0.890863 | 866 | 3 | -2.7035 | 0.41332 | 0.66094 | 1 | 13370 | 3 | -2.7035 |
| SLC45A3 | 5 | 0.27903 | 0.46407 | 0.999994 | 9675 | 2 | -0.10338 | 0.41351 | 0.6413 | 1 | 13371 | 2 | -0.10338 |
| PXDNL | 3 | 0.027748 | 0.069933 | 0.93914 | 1894 | 2 | -3.9431 | 0.41354 | 0.56749 | 1 | 13372 | 1 | -3.9431 |
| GSTA3 | 5 | 0.079667 | 0.19405 | 0.975797 | 4217 | 2 | 0.76062 | 0.41367 | 0.64146 | 1 | 13373 | 3 | 0.76062 |
| PAFAH1B3 | 5 | 0.83342 | 0.84193 | 0.999994 | 17979 | 1 | 0.88568 | 0.41378 | 0.64158 | 1 | 13374 | 3 | 0.88568 |
| TET3 | 5 | 0.26336 | 0.45146 | 0.999994 | 9323 | 1 | 0.76927 | 0.4138 | 0.64159 | 1 | 13375 | 3 | 0.76927 |
| MYCN | 5 | 0.3916 | 0.55648 | 0.999994 | 11823 | 2 | 0.90106 | 0.41388 | 0.64166 | 1 | 13376 | 3 | 0.90106 |
| FKBP14 | 5 | 0.89188 | 0.89202 | 0.999994 | 19072 | 0 | -0.10835 | 0.41392 | 0.6417 | 1 | 13377 | 2 | -0.10835 |
| CALHM3 | 5 | 0.94999 | 0.95016 | 0.999994 | 20333 | 0 | 0.44626 | 0.41392 | 0.6417 | 1 | 13378 | 2 | 0.44626 |
| KCTD9 | 5 | 0.12762 | 0.26898 | 0.998561 | 5864 | 3 | -0.96358 | 0.41392 | 0.6417 | 1 | 13379 | 1 | -0.96358 |
| MLKL | 5 | 0.88606 | 0.88615 | 0.999994 | 18951 | 1 | 0.79412 | 0.41392 | 0.6417 | 1 | 13380 | 3 | 0.79412 |
| SORCS3 | 5 | 0.73589 | 0.78358 | 0.999994 | 16674 | 1 | 0.30646 | 0.41392 | 0.6417 | 1 | 13381 | 2 | 0.30646 |
| BSPRY | 6 | 0.90714 | 0.90924 | 0.999994 | 19373 | 1 | 0.47089 | 0.41398 | 0.66146 | 1 | 13382 | 3 | 0.47089 |
| PIKFYVE | 6 | 0.15824 | 0.33601 | 0.999994 | 6680 | 3 | 0.0033089 | 0.414 | 0.66148 | 1 | 13383 | 3 | 0.0033089 |
| BRD8 | 4 | 0.0044863 | 0.015866 | 0.766918 | 481 | 2 | -3.6077 | 0.41401 | 0.59908 | 1 | 13384 | 2 | -3.6077 |
| HIST1H4J | 4 | 0.033654 | 0.088971 | 0.94395 | 2198 | 2 | -1.7827 | 0.41406 | 0.59911 | 1 | 13385 | 2 | -1.7827 |
| KLRG2 | 4 | 0.4338 | 0.58042 | 0.999994 | 12474 | 2 | -0.35746 | 0.41412 | 0.59916 | 1 | 13386 | 1 | -0.35746 |
| hsa-mir-4425 | 4 | 0.063976 | 0.14836 | 0.965554 | 3610 | 3 | -1.3113 | 0.41412 | 0.59916 | 1 | 13387 | 1 | -1.3113 |
| PCSK2 | 4 | 0.85683 | 0.85643 | 0.999994 | 18385 | 0 | 1.1031 | 0.41413 | 0.59917 | 1 | 13388 | 2 | 1.1031 |
| MDM1 | 5 | 0.2882 | 0.47148 | 0.999994 | 9876 | 2 | 0.58204 | 0.41419 | 0.64194 | 1 | 13389 | 3 | 0.58204 |
| CHMP1B | 6 | 0.028645 | 0.095636 | 0.945031 | 1935 | 2 | 0.47674 | 0.41421 | 0.66164 | 1 | 13390 | 3 | 0.47674 |
| RIOK1 | 3 | 0.21184 | 0.3275 | 0.999994 | 8062 | 1 | 0.90131 | 0.41422 | 0.56817 | 1 | 13391 | 2 | 0.90131 |
| ZBTB5 | 5 | 0.44208 | 0.59855 | 0.999994 | 12612 | 1 | 0.89236 | 0.41437 | 0.64212 | 1 | 13392 | 3 | 0.89236 |
| CA5B | 6 | 0.27327 | 0.47557 | 0.999994 | 9546 | 1 | 0.29639 | 0.41443 | 0.66179 | 1 | 13393 | 2 | 0.29639 |
| GDPD2 | 6 | 0.098819 | 0.24505 | 0.98979 | 4924 | 4 | -0.82574 | 0.41443 | 0.66179 | 1 | 13394 | 2 | -0.82574 |
| C2orf57 | 6 | 0.099557 | 0.2465 | 0.98999 | 4958 | 3 | -0.98138 | 0.41443 | 0.66179 | 1 | 13395 | 2 | -0.98138 |
| ESYT3 | 6 | 0.63316 | 0.76472 | 0.999994 | 15417 | 2 | 0.70158 | 0.41443 | 0.66179 | 1 | 13396 | 3 | 0.70158 |
| CMTM8 | 5 | 0.90332 | 0.90338 | 0.999994 | 19286 | 0 | 0.79908 | 0.41465 | 0.64238 | 1 | 13397 | 3 | 0.79908 |
| RNASEH2A | 6 | 0.14946 | 0.32495 | 0.999994 | 6444 | 3 | -0.42668 | 0.41479 | 0.66208 | 1 | 13398 | 1 | -0.42668 |
| NRARP | 6 | 0.21743 | 0.40904 | 0.999994 | 8204 | 2 | 1.0579 | 0.41503 | 0.66227 | 1 | 13399 | 3 | 1.0579 |
| OARD1 | 6 | 0.10722 | 0.26135 | 0.995005 | 5220 | 3 | 0.017325 | 0.41517 | 0.66238 | 1 | 13400 | 3 | 0.017325 |
| MESP2 | 6 | 0.64084 | 0.77057 | 0.999994 | 15527 | 1 | -0.27397 | 0.41524 | 0.66244 | 1 | 13401 | 2 | -0.27397 |
| REEP3 | 4 | 0.64599 | 0.69097 | 0.999994 | 15601 | 1 | 0.91401 | 0.41533 | 0.60006 | 1 | 13402 | 2 | 0.91401 |
| EPB49 | 3 | 0.6084 | 0.62357 | 0.999994 | 15049 | 1 | 0.84595 | 0.41534 | 0.5693 | 1 | 13403 | 2 | 0.84595 |
| BCL2L10 | 6 | 0.024618 | 0.086107 | 0.94395 | 1738 | 4 | -1.9336 | 0.41535 | 0.66252 | 1 | 13404 | 2 | -1.9336 |
| HPCA | 4 | 0.84813 | 0.84767 | 0.999994 | 18219 | 0 | 1.0985 | 0.41538 | 0.60009 | 1 | 13405 | 2 | 1.0985 |
| RNASE8 | 6 | 0.15002 | 0.32565 | 0.999994 | 6459 | 2 | 0.097183 | 0.41544 | 0.66259 | 1 | 13406 | 3 | 0.097183 |
| LEMD2 | 6 | 0.22749 | 0.42106 | 0.999994 | 8469 | 3 | -0.80502 | 0.41544 | 0.66259 | 1 | 13407 | 2 | -0.80502 |
| TRIML2 | 5 | 0.32184 | 0.49877 | 0.999994 | 10553 | 1 | 0.84326 | 0.41559 | 0.6433 | 1 | 13408 | 3 | 0.84326 |
| TCEAL7 | 4 | 0.063559 | 0.14757 | 0.965554 | 3582 | 2 | -1.6193 | 0.41562 | 0.60026 | 1 | 13409 | 2 | -1.6193 |
| ESF1 | 6 | 0.18902 | 0.37428 | 0.999994 | 7451 | 2 | 0.76598 | 0.41563 | 0.66274 | 1 | 13410 | 3 | 0.76598 |
| XKR8 | 6 | 0.017772 | 0.064773 | 0.93914 | 1352 | 3 | 0.10072 | 0.41563 | 0.66274 | 1 | 13411 | 3 | 0.10072 |
| ARSB | 6 | 0.82988 | 0.86149 | 0.999994 | 17928 | 1 | -0.36239 | 0.41567 | 0.66278 | 1 | 13412 | 2 | -0.36239 |
| LRRFIP1 | 6 | 0.56791 | 0.71646 | 0.999994 | 14512 | 1 | 0.39875 | 0.41582 | 0.6629 | 1 | 13413 | 3 | 0.39875 |
| hsa-mir-548s | 2 | 0.50133 | 0.50514 | 0.999994 | 13444 | 1 | 1.324 | 0.41583 | 0.4788 | 1 | 13414 | 1 | 1.324 |
| IFT140 | 6 | 0.34101 | 0.55381 | 0.999994 | 10962 | 2 | 0.44779 | 0.41602 | 0.66306 | 1 | 13415 | 3 | 0.44779 |

| | | | | | | | | | | | | | |
|---|---|---|---|---|---|---|---|---|---|---|---|---|---|
| STK11 | 5 | 0.27841 | 0.46356 | 0.999994 | 9661 | 2 | 0.25545 | 0.41607 | 0.64378 | 1 | 13416 | 2 | 0.25545 |
| ZNF763 | 5 | 0.83443 | 0.84268 | 0.999994 | 17995 | 1 | 0.9115 | 0.41609 | 0.64381 | 1 | 13417 | 3 | 0.9115 |
| FAM209A | 6 | 0.23753 | 0.43303 | 0.999994 | 8716 | 2 | 0.50889 | 0.41618 | 0.66319 | 1 | 13418 | 3 | 0.50889 |
| ARL6IP1 | 5 | 0.41116 | 0.57284 | 0.999994 | 12131 | 2 | 0.0072343 | 0.41623 | 0.64394 | 1 | 13419 | 1 | 0.0072343 |
| hsa-mir-3125 | 4 | 0.16162 | 0.30115 | 0.999994 | 6780 | 2 | -1.2174 | 0.41624 | 0.60071 | 1 | 13420 | 1 | -1.2174 |
| PLK1 | 4 | 0.02808 | 0.077589 | 0.93914 | 1908 | 3 | -4.4916 | 0.41624 | 0.60071 | 1 | 13421 | 1 | -4.4916 |
| VAPB | 5 | 0.28519 | 0.46903 | 0.999994 | 9816 | 1 | 0.74087 | 0.41625 | 0.64396 | 1 | 13422 | 3 | 0.74087 |
| BRIP1 | 5 | 0.92434 | 0.92457 | 0.999994 | 19728 | 0 | 0.429 | 0.41635 | 0.64407 | 1 | 13423 | 2 | 0.429 |
| NDUFV2 | 5 | 0.12811 | 0.26971 | 0.998561 | 5881 | 2 | 0.80775 | 0.41635 | 0.64407 | 1 | 13424 | 3 | 0.80775 |
| CAPS2 | 6 | 0.010672 | 0.041302 | 0.891621 | 910 | 3 | -0.26548 | 0.4164 | 0.66336 | 1 | 13425 | 2 | -0.26548 |
| ARHGEF2 | 5 | 0.15882 | 0.3158 | 0.999994 | 6692 | 3 | -1.0326 | 0.4164 | 0.64413 | 1 | 13426 | 2 | -1.0326 |
| MDFI | 6 | 0.13836 | 0.31095 | 0.999994 | 6148 | 2 | 0.69587 | 0.41658 | 0.66351 | 1 | 13427 | 3 | 0.69587 |
| PPAT | 5 | 0.15397 | 0.30872 | 0.999994 | 6566 | 2 | 0.26697 | 0.4166 | 0.64435 | 1 | 13428 | 2 | 0.26697 |
| TFF3 | 5 | 0.018285 | 0.059405 | 0.932932 | 1375 | 4 | -2.8545 | 0.4166 | 0.64435 | 1 | 13429 | 1 | -2.8545 |
| NonTargetingControlG | 1 | 0.5834 | 0.58311 | 0.999994 | 14724 | 0 | 1.0661 | 0.4166 | 0.41677 | 1 | 13430 | 1 | 1.0661 |
| NonTargetingControlG | 1 | 0.58333 | 0.58305 | 0.999994 | 14723 | 0 | 1.059 | 0.41667 | 0.41683 | 1 | 13431 | 1 | 1.059 |
| hsa-mir-3926-2 | 1 | 0.58332 | 0.58304 | 0.999994 | 14722 | 0 | 1.312 | 0.41668 | 0.41684 | 1 | 13432 | 1 | 1.312 |
| POR | 6 | 0.23859 | 0.4343 | 0.999994 | 8737 | 2 | 1.0643 | 0.41669 | 0.66359 | 1 | 13433 | 3 | 1.0643 |
| NUP88 | 5 | 0.36596 | 0.53501 | 0.999994 | 11445 | 1 | -0.26193 | 0.41678 | 0.6445 | 1 | 13434 | 2 | -0.26193 |
| SFRP5 | 5 | 0.61321 | 0.73102 | 0.999994 | 15123 | 1 | 0.8646 | 0.41679 | 0.64451 | 1 | 13435 | 3 | 0.8646 |
| PPHLN1 | 6 | 0.00038854 | 0.0020595 | 0.44006 | 93 | 5 | -4.9416 | 0.41689 | 0.66375 | 1 | 13436 | 1 | -4.9416 |
| IDH3A | 6 | 0.90996 | 0.91154 | 0.999994 | 19424 | 1 | 0.28173 | 0.4169 | 0.66376 | 1 | 13437 | 2 | 0.28173 |
| C15orf53 | 6 | 0.34085 | 0.55362 | 0.999994 | 10957 | 3 | -0.044589 | 0.4169 | 0.66376 | 1 | 13438 | 3 | -0.044589 |
| DBNDD2 | 5 | 0.59755 | 0.72528 | 0.999994 | 14921 | 1 | 0.81016 | 0.41708 | 0.6448 | 1 | 13439 | 3 | 0.81016 |
| KCNAB2 | 4 | 0.0023234 | 0.0083988 | 0.680529 | 307 | 3 | -6.8944 | 0.41711 | 0.6014 | 1 | 13440 | 1 | -6.8944 |
| CPPED1 | 6 | 0.25673 | 0.45598 | 0.999994 | 9166 | 2 | 0.5157 | 0.41714 | 0.66393 | 1 | 13441 | 3 | 0.5157 |
| GNRH2 | 6 | 0.14015 | 0.31321 | 0.999994 | 6189 | 3 | -0.57636 | 0.41736 | 0.66412 | 1 | 13442 | 3 | -0.57636 |
| PIWIL4 | 6 | 0.79623 | 0.8463 | 0.999994 | 17431 | 1 | 0.76895 | 0.41736 | 0.66412 | 1 | 13443 | 3 | 0.76895 |
| ULBP1 | 6 | 0.3146 | 0.52371 | 0.999994 | 10410 | 3 | 0.4614 | 0.41736 | 0.66412 | 1 | 13444 | 3 | 0.4614 |
| ABCA1 | 6 | 0.27604 | 0.4788 | 0.999994 | 9618 | 3 | 0.24454 | 0.41737 | 0.66412 | 1 | 13445 | 3 | 0.24454 |
| hsa-mir-558 | 2 | 0.62926 | 0.62904 | 0.999994 | 15373 | 0 | 1.8318 | 0.41738 | 0.47986 | 1 | 13446 | 1 | 1.8318 |
| NonTargetingControlG | 2 | 0.20533 | 0.26598 | 0.997535 | 7904 | 1 | -0.039791 | 0.41738 | 0.47986 | 1 | 13447 | 1 | -0.039791 |
| JAKMIP1 | 6 | 0.27141 | 0.47336 | 0.999994 | 9508 | 3 | 0.32739 | 0.41743 | 0.66416 | 1 | 13448 | 3 | 0.32739 |
| PRRG2 | 5 | 0.55176 | 0.69331 | 0.999994 | 14238 | 1 | 0.33326 | 0.41743 | 0.64512 | 1 | 13449 | 1 | 0.33326 |
| CDKN2B | 5 | 0.052013 | 0.14526 | 0.963012 | 3069 | 2 | -0.47627 | 0.41743 | 0.64512 | 1 | 13450 | 1 | -0.47627 |
| TRAPPC1 | 5 | 0.18188 | 0.34934 | 0.999994 | 7266 | 2 | 0.091541 | 0.41743 | 0.64512 | 1 | 13451 | 2 | 0.091541 |
| hsa-mir-5688 | 3 | 0.78327 | 0.78316 | 0.999994 | 17258 | 0 | 0.83292 | 0.41747 | 0.57146 | 1 | 13452 | 2 | 0.83292 |
| GRIN2B | 6 | 0.29775 | 0.50409 | 0.999994 | 10085 | 2 | 0.80171 | 0.41749 | 0.6642 | 1 | 13453 | 3 | 0.80171 |
| C17orf82 | 4 | 0.060341 | 0.14143 | 0.959146 | 3435 | 2 | 0.53683 | 0.41761 | 0.60179 | 1 | 13454 | 2 | 0.53683 |
| ENHO | 5 | 0.43452 | 0.59222 | 0.999994 | 12484 | 2 | 0.87888 | 0.41764 | 0.64533 | 1 | 13455 | 3 | 0.87888 |
| C12orf60 | 6 | 0.2594 | 0.45914 | 0.999994 | 9224 | 1 | 0.56323 | 0.41803 | 0.66462 | 1 | 13456 | 3 | 0.56323 |
| REM2 | 4 | 0.76394 | 0.77095 | 0.999994 | 17013 | 1 | 0.46189 | 0.41805 | 0.60214 | 1 | 13457 | 2 | 0.46189 |
| PGM5 | 6 | 0.52145 | 0.68352 | 0.999994 | 13766 | 1 | 0.94276 | 0.41818 | 0.66474 | 1 | 13458 | 3 | 0.94276 |
| P2RX4 | 6 | 0.61137 | 0.74835 | 0.999994 | 15094 | 2 | 0.92829 | 0.41823 | 0.66477 | 1 | 13459 | 3 | 0.92829 |
| hsa-mir-191 | 3 | 0.64239 | 0.65076 | 0.999994 | 15553 | 1 | 0.97633 | 0.4183 | 0.57231 | 1 | 13460 | 2 | 0.97633 |
| C7orf65 | 5 | 0.25558 | 0.44519 | 0.999994 | 9138 | 2 | 0.76975 | 0.4183 | 0.646 | 1 | 13461 | 3 | 0.76975 |
| N4BP2L1 | 6 | 0.21095 | 0.40125 | 0.999994 | 8049 | 3 | -0.34078 | 0.41833 | 0.66486 | 1 | 13462 | 2 | -0.34078 |
| SUCO | 5 | 0.033861 | 0.10039 | 0.946394 | 2209 | 3 | -2.6108 | 0.41836 | 0.64606 | 1 | 13463 | 2 | -2.6108 |
| STC2 | 5 | 0.65004 | 0.74514 | 0.999994 | 15660 | 1 | 0.48047 | 0.41836 | 0.64606 | 1 | 13464 | 1 | 0.48047 |
| PACRGL | 5 | 0.55049 | 0.69217 | 0.999994 | 14221 | 2 | 0.037364 | 0.41836 | 0.64606 | 1 | 13465 | 1 | 0.037364 |
| FAM161A | 6 | 0.051267 | 0.14685 | 0.964852 | 3036 | 3 | -1.4417 | 0.41853 | 0.66502 | 1 | 13466 | 2 | -1.4417 |
| HRH2 | 5 | 0.10922 | 0.24078 | 0.987253 | 5277 | 2 | 0.92397 | 0.41865 | 0.64634 | 1 | 13467 | 3 | 0.92397 |

| | | | | | | | | | | | | | |
|---|---|---|---|---|---|---|---|---|---|---|---|---|---|
| GRPEL2 | 6 | 0.18475 | 0.36907 | 0.999994 | 7337 | 2 | 0.50312 | 0.41871 | 0.66516 | 1 | 13468 | 3 | 0.50312 |
| TDP1 | 5 | 0.13709 | 0.28339 | 0.999632 | 6110 | 2 | 1.0884 | 0.41876 | 0.64645 | 1 | 13469 | 3 | 1.0884 |
| FAHD2B | 5 | 0.085341 | 0.20312 | 0.97798 | 4423 | 3 | -1.8705 | 0.4188 | 0.64647 | 1 | 13470 | 2 | -1.8705 |
| HAUS1 | 6 | 0.12142 | 0.28841 | 0.999632 | 5679 | 3 | -0.18493 | 0.41885 | 0.66528 | 1 | 13471 | 3 | -0.18493 |
| JAM2 | 5 | 0.07292 | 0.18314 | 0.968526 | 3959 | 2 | -0.11169 | 0.41891 | 0.64658 | 1 | 13472 | 2 | -0.11169 |
| CER1 | 4 | 0.67984 | 0.71149 | 0.999994 | 16088 | 1 | -0.45985 | 0.41893 | 0.60281 | 1 | 13473 | 1 | -0.45985 |
| PTPN12 | 4 | 0.22641 | 0.36778 | 0.999994 | 8434 | 1 | -0.39093 | 0.41893 | 0.60281 | 1 | 13474 | 1 | -0.39093 |
| OR5V1 | 5 | 0.46806 | 0.62046 | 0.999994 | 12974 | 2 | 0.39274 | 0.41895 | 0.64662 | 1 | 13475 | 2 | 0.39274 |
| GYPB | 5 | 0.53506 | 0.67851 | 0.999994 | 13983 | 1 | 0.8333 | 0.41896 | 0.64662 | 1 | 13476 | 3 | 0.8333 |
| POLR3B | 5 | 0.18308 | 0.35106 | 0.999994 | 7292 | 2 | -0.25365 | 0.41911 | 0.64676 | 1 | 13477 | 2 | -0.25365 |
| COBLL1 | 6 | 0.25942 | 0.45918 | 0.999994 | 9225 | 3 | -0.66909 | 0.41915 | 0.6655 | 1 | 13478 | 2 | -0.66909 |
| EML6 | 5 | 0.016389 | 0.054142 | 0.922455 | 1269 | 4 | -3.2278 | 0.41922 | 0.64686 | 1 | 13479 | 1 | -3.2278 |
| CCL11 | 5 | 0.44809 | 0.60361 | 0.999994 | 12688 | 2 | -0.042643 | 0.41922 | 0.64686 | 1 | 13480 | 2 | -0.042643 |
| YY1AP1 | 5 | 0.11397 | 0.24815 | 0.990682 | 5447 | 3 | -1.2517 | 0.41922 | 0.64686 | 1 | 13481 | 2 | -1.2517 |
| H2AFB2 | 2 | 0.21875 | 0.27573 | 0.999379 | 8252 | 1 | 0.11493 | 0.41927 | 0.48113 | 1 | 13482 | 1 | 0.11493 |
| TMEM171 | 5 | 0.1144 | 0.24881 | 0.991171 | 5460 | 1 | 0.77611 | 0.41931 | 0.64696 | 1 | 13483 | 3 | 0.77611 |
| CDC14A | 4 | 0.10886 | 0.23085 | 0.983933 | 5266 | 3 | -1.2741 | 0.41933 | 0.6031 | 1 | 13484 | 1 | -1.2741 |
| S100B | 5 | 0.051787 | 0.14472 | 0.963012 | 3057 | 2 | -0.023957 | 0.41935 | 0.647 | 1 | 13485 | 2 | -0.023957 |
| CNGA3 | 6 | 0.060855 | 0.16759 | 0.967126 | 3462 | 4 | -1.8645 | 0.41949 | 0.6658 | 1 | 13486 | 2 | -1.8645 |
| KIAA1522 | 5 | 0.19729 | 0.37137 | 0.999994 | 7680 | 3 | -0.85713 | 0.41957 | 0.64719 | 1 | 13487 | 2 | -0.85713 |
| NAPRT1 | 4 | 0.71414 | 0.73396 | 0.999994 | 16441 | 1 | 0.8467 | 0.41959 | 0.60331 | 1 | 13488 | 2 | 0.8467 |
| BRIX1 | 3 | 0.049891 | 0.11555 | 0.951052 | 2971 | 2 | -2.2 | 0.4197 | 0.57381 | 1 | 13489 | 1 | -2.2 |
| PMS2 | 5 | 0.55163 | 0.6932 | 0.999994 | 14234 | 1 | 0.76454 | 0.41987 | 0.64747 | 1 | 13490 | 3 | 0.76454 |
| PTER | 5 | 0.57111 | 0.7105 | 0.999994 | 14562 | 1 | 0.70167 | 0.41992 | 0.64752 | 1 | 13491 | 3 | 0.70167 |
| CHCHD1 | 5 | 0.15753 | 0.31396 | 0.999994 | 6659 | 2 | 0.16513 | 0.41998 | 0.64758 | 1 | 13492 | 2 | 0.16513 |
| KRT36 | 5 | 0.90619 | 0.90627 | 0.999994 | 19356 | 0 | 0.73184 | 0.41999 | 0.64759 | 1 | 13493 | 3 | 0.73184 |
| WBP4 | 5 | 0.11185 | 0.24486 | 0.989317 | 5362 | 3 | -1.5499 | 0.41999 | 0.6476 | 1 | 13494 | 1 | -1.5499 |
| CENPL | 5 | 0.50237 | 0.64998 | 0.999994 | 13468 | 2 | -0.49801 | 0.41999 | 0.6476 | 1 | 13495 | 2 | -0.49801 |
| SMAD2 | 6 | 0.4888 | 0.66112 | 0.999994 | 13263 | 2 | 0.79642 | 0.42004 | 0.66621 | 1 | 13496 | 3 | 0.79642 |
| hsa-mir-5708 | 1 | 0.57988 | 0.57964 | 0.999994 | 14675 | 0 | 1.2668 | 0.42012 | 0.42028 | 1 | 13497 | 1 | 1.2668 |
| MSLN | 5 | 0.9091 | 0.90911 | 0.999994 | 19403 | 0 | 0.80936 | 0.42017 | 0.64775 | 1 | 13498 | 3 | 0.80936 |
| ARL14EPL | 6 | 0.082873 | 0.21318 | 0.979587 | 4332 | 3 | -1.3704 | 0.42017 | 0.6663 | 1 | 13499 | 2 | -1.3704 |
| TXN | 6 | 0.18122 | 0.36468 | 0.999994 | 7258 | 2 | 0.59811 | 0.42017 | 0.6663 | 1 | 13500 | 3 | 0.59811 |
| CXCL12 | 6 | 0.1106 | 0.26787 | 0.998561 | 5320 | 4 | -0.70844 | 0.42017 | 0.6663 | 1 | 13501 | 1 | -0.70844 |
| FARP1 | 6 | 0.42999 | 0.62184 | 0.999994 | 12413 | 2 | -0.11032 | 0.42017 | 0.6663 | 1 | 13502 | 2 | -0.11032 |
| ACP2 | 6 | 0.069249 | 0.1852 | 0.968626 | 3808 | 1 | 0.82454 | 0.42018 | 0.66632 | 1 | 13503 | 3 | 0.82454 |
| CDX4 | 4 | 0.34269 | 0.48679 | 0.999994 | 10999 | 2 | 0.4302 | 0.42024 | 0.6038 | 1 | 13504 | 2 | 0.4302 |
| C3 | 5 | 0.26246 | 0.45071 | 0.999994 | 9295 | 2 | -0.0042469 | 0.42041 | 0.64799 | 1 | 13505 | 2 | -0.0042469 |
| FSIP2 | 5 | 0.051942 | 0.14509 | 0.963012 | 3062 | 3 | -1.5642 | 0.42041 | 0.64799 | 1 | 13506 | 2 | -1.5642 |
| KLRK1 | 6 | 0.35265 | 0.56699 | 0.999994 | 11209 | 3 | -0.15087 | 0.42043 | 0.66651 | 1 | 13507 | 2 | -0.15087 |
| ZBTB2 | 5 | 0.15048 | 0.3035 | 0.999994 | 6472 | 2 | 0.77078 | 0.42044 | 0.64802 | 1 | 13508 | 3 | 0.77078 |
| TMEM230 | 5 | 0.024529 | 0.076214 | 0.93914 | 1732 | 2 | -1.2059 | 0.42055 | 0.64813 | 1 | 13509 | 2 | -1.2059 |
| GJB3 | 5 | 0.43311 | 0.59102 | 0.999994 | 12464 | 2 | -0.17762 | 0.42068 | 0.64823 | 1 | 13510 | 2 | -0.17762 |
| TMEM27 | 5 | 0.028 | 0.085291 | 0.94395 | 1902 | 3 | -3.0557 | 0.42068 | 0.64823 | 1 | 13511 | 2 | -3.0557 |
| C1orf192 | 4 | 0.33145 | 0.47518 | 0.999994 | 10753 | 2 | 0.53876 | 0.42078 | 0.6042 | 1 | 13512 | 2 | 0.53876 |
| TMEM45A | 6 | 0.37126 | 0.58399 | 0.999994 | 11534 | 2 | 0.039398 | 0.42096 | 0.66691 | 1 | 13513 | 3 | 0.039398 |
| SLC9A1 | 6 | 0.10047 | 0.24825 | 0.990682 | 4987 | 3 | -1.1388 | 0.42096 | 0.66691 | 1 | 13514 | 1 | -1.1388 |
| SPATA31A2 | 4 | 0.2013 | 0.342 | 0.999994 | 7799 | 2 | -0.61202 | 0.42096 | 0.60434 | 1 | 13515 | 2 | -0.61202 |
| FAM92A1 | 5 | 0.91551 | 0.91559 | 0.999994 | 19525 | 0 | 0.75808 | 0.42098 | 0.64854 | 1 | 13516 | 3 | 0.75808 |
| NHLRC4 | 6 | 0.15403 | 0.33073 | 0.999994 | 6569 | 2 | 0.86322 | 0.42099 | 0.66694 | 1 | 13517 | 3 | 0.86322 |
| WWTR1 | 5 | 0.4468 | 0.60252 | 0.999994 | 12673 | 1 | 0.66262 | 0.42107 | 0.64862 | 1 | 13518 | 3 | 0.66262 |
| MMP9 | 6 | 0.75593 | 0.83059 | 0.999994 | 16930 | 1 | 0.58401 | 0.42119 | 0.6671 | 1 | 13519 | 2 | 0.58401 |

| | | | | | | | | | | | | | |
|---|---|---|---|---|---|---|---|---|---|---|---|---|---|
| FAM83A | 6 | 0.023581 | 0.083421 | 0.94383 | 1690 | 3 | 0.31557 | 0.42127 | 0.66716 | 1 | 13520 | 3 | 0.31557 |
| DZIP3 | 5 | 0.04399 | 0.1258 | 0.951052 | 2710 | 2 | -0.56885 | 0.42138 | 0.64895 | 1 | 13521 | 2 | -0.56885 |
| KRTAP22-1 | 5 | 0.21667 | 0.3986 | 0.999994 | 8185 | 2 | 0.73146 | 0.42139 | 0.64895 | 1 | 13522 | 3 | 0.73146 |
| OR2Z1 | 4 | 0.20604 | 0.34691 | 0.999994 | 7928 | 2 | 0.40343 | 0.42147 | 0.60472 | 1 | 13523 | 2 | 0.40343 |
| NonTargetingControlG | 2 | 0.028309 | 0.05196 | 0.911383 | 1915 | 1 | -3.1653 | 0.42148 | 0.48261 | 1 | 13524 | 1 | -3.1653 |
| MAGEA3 | 5 | 0.55503 | 0.69623 | 0.999994 | 14290 | 2 | 0.14524 | 0.4215 | 0.64906 | 1 | 13525 | 2 | 0.14524 |
| CASP9 | 5 | 0.25127 | 0.44176 | 0.999994 | 9043 | 2 | 0.70057 | 0.4215 | 0.64906 | 1 | 13526 | 3 | 0.70057 |
| CNST | 5 | 0.5551 | 0.6963 | 0.999994 | 14293 | 1 | 0.79978 | 0.42151 | 0.64906 | 1 | 13527 | 3 | 0.79978 |
| ARPC4 | 6 | 0.15813 | 0.33588 | 0.999994 | 6675 | 2 | 0.73449 | 0.42158 | 0.6674 | 1 | 13528 | 3 | 0.73449 |
| FAM151A | 6 | 0.38288 | 0.59139 | 0.999994 | 11690 | 2 | 0.038161 | 0.42165 | 0.66746 | 1 | 13529 | 2 | 0.038161 |
| ZSCAN16 | 6 | 0.56382 | 0.71353 | 0.999994 | 14448 | 2 | 0.3903 | 0.42165 | 0.66746 | 1 | 13530 | 3 | 0.3903 |
| FASTK | 6 | 0.23729 | 0.43275 | 0.999994 | 8708 | 3 | -0.54963 | 0.42165 | 0.66746 | 1 | 13531 | 2 | -0.54963 |
| FBRSL1 | 5 | 0.71989 | 0.77578 | 0.999994 | 16498 | 1 | 0.15658 | 0.42184 | 0.64941 | 1 | 13532 | 2 | 0.15658 |
| LATS1 | 5 | 0.73531 | 0.78328 | 0.999994 | 16665 | 1 | 0.76828 | 0.42189 | 0.64946 | 1 | 13533 | 3 | 0.76828 |
| MEGF10 | 2 | 0.47564 | 0.48192 | 0.999994 | 13071 | 1 | 1.2654 | 0.4219 | 0.48291 | 1 | 13534 | 1 | 1.2654 |
| SLC5A8 | 5 | 0.79259 | 0.8149 | 0.999994 | 17382 | 1 | -0.044063 | 0.42193 | 0.6495 | 1 | 13535 | 2 | -0.044063 |
| hsa-mir-4700 | 4 | 0.32189 | 0.46533 | 0.999994 | 10555 | 2 | 0.36216 | 0.42205 | 0.60515 | 1 | 13536 | 2 | 0.36216 |
| NonTargetingControlG | 2 | 0.047434 | 0.084855 | 0.94395 | 2876 | 1 | -2.7445 | 0.42211 | 0.48306 | 1 | 13537 | 1 | -2.7445 |
| SCARA3 | 6 | 0.20152 | 0.38973 | 0.999994 | 7810 | 3 | 0.0052068 | 0.42214 | 0.66785 | 1 | 13538 | 3 | 0.0052068 |
| hsa-mir-223 | 3 | 0.20465 | 0.31857 | 0.999994 | 7893 | 2 | -1.1067 | 0.42216 | 0.57632 | 1 | 13539 | 1 | -1.1067 |
| RFC5 | 3 | 0.050868 | 0.11691 | 0.951052 | 3010 | 2 | -4.9308 | 0.42216 | 0.57632 | 1 | 13540 | 1 | -4.9308 |
| PRSS45 | 4 | 0.26707 | 0.40927 | 0.999994 | 9397 | 2 | -0.2867 | 0.42221 | 0.60526 | 1 | 13541 | 1 | -0.2867 |
| OR52J3 | 4 | 0.73343 | 0.74768 | 0.999994 | 16647 | 1 | -0.36563 | 0.42221 | 0.60526 | 1 | 13542 | 1 | -0.36563 |
| FLRT3 | 6 | 0.073239 | 0.19349 | 0.975797 | 3966 | 3 | -0.60906 | 0.42226 | 0.66795 | 1 | 13543 | 2 | -0.60906 |
| ZNF774 | 6 | 0.64125 | 0.77087 | 0.999994 | 15534 | 2 | 0.6124 | 0.42232 | 0.66799 | 1 | 13544 | 3 | 0.6124 |
| MRPL34 | 3 | 0.34367 | 0.45219 | 0.999994 | 11023 | 1 | 0.91866 | 0.42244 | 0.57659 | 1 | 13545 | 2 | 0.91866 |
| SEC11C | 4 | 0.27932 | 0.42186 | 0.999994 | 9684 | 2 | 0.23806 | 0.42245 | 0.60545 | 1 | 13546 | 2 | 0.23806 |
| MYL7 | 6 | 0.36915 | 0.58268 | 0.999994 | 11494 | 2 | 0.34189 | 0.42246 | 0.6681 | 1 | 13547 | 2 | 0.34189 |
| TSEN15 | 6 | 0.0022684 | 0.010977 | 0.71022 | 301 | 5 | -2.6796 | 0.42246 | 0.6681 | 1 | 13548 | 1 | -2.6796 |
| AP3B2 | 6 | 0.78731 | 0.84258 | 0.999994 | 17309 | 1 | 0.29754 | 0.42246 | 0.6681 | 1 | 13549 | 3 | 0.29754 |
| TNIK | 6 | 0.18373 | 0.3678 | 0.999994 | 7308 | 3 | -0.68923 | 0.42246 | 0.6681 | 1 | 13550 | 1 | -0.68923 |
| PAPD7 | 6 | 0.16391 | 0.34317 | 0.999994 | 6837 | 3 | -0.01543 | 0.42251 | 0.66814 | 1 | 13551 | 3 | -0.01543 |
| KIFAP3 | 4 | 0.17577 | 0.31577 | 0.999994 | 7130 | 2 | -0.023778 | 0.42252 | 0.60549 | 1 | 13552 | 2 | -0.023778 |
| REPS1 | 4 | 0.91067 | 0.91044 | 0.999994 | 19444 | 0 | 1.1723 | 0.42262 | 0.60556 | 1 | 13553 | 2 | 1.1723 |
| EFHD2 | 5 | 0.92756 | 0.92779 | 0.999994 | 19804 | 0 | 0.55589 | 0.42269 | 0.65023 | 1 | 13554 | 2 | 0.55589 |
| TCF15 | 5 | 0.080807 | 0.19589 | 0.975797 | 4260 | 2 | 0.11667 | 0.42269 | 0.65023 | 1 | 13555 | 2 | 0.11667 |
| hsa-mir-1250 | 3 | 0.63255 | 0.64271 | 0.999994 | 15405 | 1 | 0.88002 | 0.4228 | 0.57696 | 1 | 13556 | 2 | 0.88002 |
| KRTAP12-1 | 6 | 0.16462 | 0.344 | 0.999994 | 6854 | 3 | -0.5009 | 0.4228 | 0.66836 | 1 | 13557 | 3 | -0.5009 |
| KRTAP5-4 | 5 | 0.61082 | 0.73009 | 0.999994 | 15083 | 1 | 0.81581 | 0.42297 | 0.65054 | 1 | 13558 | 3 | 0.81581 |
| TDRD3 | 5 | 0.0058555 | 0.023982 | 0.834563 | 580 | 2 | 0.80835 | 0.42299 | 0.65055 | 1 | 13559 | 3 | 0.80835 |
| TEX11 | 6 | 0.12551 | 0.29464 | 0.999994 | 5806 | 2 | 0.67319 | 0.42306 | 0.66855 | 1 | 13560 | 3 | 0.67319 |
| C17orf50 | 5 | 0.041322 | 0.1192 | 0.951052 | 2580 | 3 | -2.4769 | 0.42308 | 0.65064 | 1 | 13561 | 2 | -2.4769 |
| OR2T10 | 5 | 0.57804 | 0.71657 | 0.999994 | 14658 | 2 | -0.40836 | 0.42308 | 0.65064 | 1 | 13562 | 2 | -0.40836 |
| TOP2A | 4 | 0.82772 | 0.82729 | 0.999994 | 17892 | 0 | 0.36498 | 0.42315 | 0.60598 | 1 | 13563 | 1 | 0.36498 |
| IRF9 | 4 | 0.3309 | 0.47461 | 0.999994 | 10741 | 2 | -0.65582 | 0.42315 | 0.60598 | 1 | 13564 | 1 | -0.65582 |
| WFIKKN1 | 5 | 0.19503 | 0.36812 | 0.999994 | 7634 | 2 | 0.80142 | 0.42323 | 0.65079 | 1 | 13565 | 3 | 0.80142 |
| GLB1L | 6 | 0.34972 | 0.56369 | 0.999994 | 11149 | 2 | -0.12339 | 0.42336 | 0.66877 | 1 | 13566 | 2 | -0.12339 |
| HAPLN1 | 3 | 0.35885 | 0.46069 | 0.999994 | 11319 | 1 | -0.18756 | 0.42337 | 0.57751 | 1 | 13567 | 1 | -0.18756 |
| P2RX6 | 6 | 0.16968 | 0.35036 | 0.999994 | 6977 | 2 | 0.072519 | 0.42338 | 0.66878 | 1 | 13568 | 2 | 0.072519 |
| hsa-mir-3673 | 2 | 0.33167 | 0.36138 | 0.999994 | 10759 | 1 | 0.75888 | 0.42344 | 0.48397 | 1 | 13569 | 1 | 0.75888 |
| POU3F3 | 5 | 0.30034 | 0.48135 | 0.999994 | 10138 | 2 | -0.46314 | 0.42352 | 0.65107 | 1 | 13570 | 2 | -0.46314 |
| TRAFD1 | 6 | 0.17203 | 0.35324 | 0.999994 | 7037 | 3 | 0.16233 | 0.42363 | 0.66898 | 1 | 13571 | 3 | 0.16233 |

| | | | | | | | | | | | | | |
|---|---|---|---|---|---|---|---|---|---|---|---|---|---|
| TBC1D22A | 6 | 0.23228 | 0.42682 | 0.999994 | 8580 | 1 | 0.88984 | 0.42366 | 0.66901 | 1 | 13572 | 3 | 0.88984 |
| HIST1H4D | 5 | 0.042853 | 0.12295 | 0.951052 | 2669 | 1 | 0.54338 | 0.42377 | 0.65132 | 1 | 13573 | 3 | 0.54338 |
| hsa-mir-4654 | 4 | 0.32814 | 0.47178 | 0.999994 | 10688 | 2 | 0.52555 | 0.4238 | 0.60645 | 1 | 13574 | 2 | 0.52555 |
| SERPINH1 | 4 | 0.16669 | 0.30638 | 0.999994 | 6901 | 1 | 0.8777 | 0.4238 | 0.60645 | 1 | 13575 | 2 | 0.8777 |
| MYOG | 5 | 0.8843 | 0.88444 | 0.999994 | 18915 | 1 | 0.84128 | 0.42381 | 0.65135 | 1 | 13576 | 3 | 0.84128 |
| ACACB | 6 | 0.11873 | 0.2833 | 0.999632 | 5594 | 3 | -0.24011 | 0.42382 | 0.66914 | 1 | 13577 | 3 | -0.24011 |
| HSPB3 | 6 | 0.58203 | 0.72669 | 0.999994 | 14705 | 2 | 0.66463 | 0.42393 | 0.66922 | 1 | 13578 | 3 | 0.66463 |
| FOXD3 | 6 | 0.078594 | 0.2045 | 0.978186 | 4173 | 4 | -1.212 | 0.424 | 0.66928 | 1 | 13579 | 2 | -1.212 |
| SETD1A | 6 | 0.00035249 | 0.0018803 | 0.435802 | 84 | 5 | -3.4383 | 0.424 | 0.66928 | 1 | 13580 | 1 | -3.4383 |
| HSPE1-MOB4 | 6 | 0.064541 | 0.17532 | 0.968171 | 3630 | 3 | -1.518 | 0.424 | 0.66928 | 1 | 13581 | 1 | -1.518 |
| hsa-mir-3657 | 4 | 0.59836 | 0.66444 | 0.999994 | 14933 | 1 | 0.091192 | 0.42403 | 0.60662 | 1 | 13582 | 1 | 0.091192 |
| hsa-mir-302c | 4 | 0.85912 | 0.85871 | 0.999994 | 18416 | 0 | 0.81062 | 0.42403 | 0.60662 | 1 | 13583 | 2 | 0.81062 |
| SERPINA3 | 5 | 0.20082 | 0.37636 | 0.999994 | 7783 | 2 | 0.37638 | 0.42408 | 0.65162 | 1 | 13584 | 2 | 0.37638 |
| CAMK1G | 5 | 0.65097 | 0.74553 | 0.999994 | 15676 | 1 | 0.83732 | 0.42416 | 0.65169 | 1 | 13585 | 3 | 0.83732 |
| FUT3 | 5 | 0.88238 | 0.88257 | 0.999994 | 18877 | 1 | 0.81433 | 0.42418 | 0.65171 | 1 | 13586 | 3 | 0.81433 |
| CASC3 | 6 | 0.64241 | 0.77176 | 0.999994 | 15554 | 2 | 0.82972 | 0.42423 | 0.66947 | 1 | 13587 | 3 | 0.82972 |
| OTX2 | 5 | 0.1774 | 0.3429 | 0.999994 | 7163 | 2 | 0.76711 | 0.42428 | 0.65181 | 1 | 13588 | 3 | 0.76711 |
| ZNF100 | 6 | 0.31221 | 0.5209 | 0.999994 | 10365 | 2 | 0.61985 | 0.42433 | 0.66954 | 1 | 13589 | 3 | 0.61985 |
| NRXN2 | 6 | 0.45559 | 0.63877 | 0.999994 | 12783 | 2 | 0.66564 | 0.42448 | 0.66967 | 1 | 13590 | 3 | 0.66564 |
| CBR4 | 6 | 0.11577 | 0.27777 | 0.999519 | 5510 | 4 | -0.95146 | 0.42452 | 0.6697 | 1 | 13591 | 1 | -0.95146 |
| DUS4L | 6 | 0.64162 | 0.77117 | 0.999994 | 15540 | 2 | 0.19412 | 0.42452 | 0.6697 | 1 | 13592 | 3 | 0.19412 |
| MAVS | 6 | 0.32286 | 0.53317 | 0.999994 | 10572 | 3 | -0.50898 | 0.42452 | 0.6697 | 1 | 13593 | 1 | -0.50898 |
| PHGDH | 6 | 0.044594 | 0.13221 | 0.95318 | 2742 | 4 | -1.5856 | 0.42452 | 0.6697 | 1 | 13594 | 2 | -1.5856 |
| NonTargetingControlG | 1 | 0.57545 | 0.57514 | 0.999994 | 14623 | 0 | 1.3348 | 0.42455 | 0.42473 | 1 | 13595 | 1 | 1.3348 |
| MSMP | 5 | 0.41302 | 0.57434 | 0.999994 | 12153 | 2 | 0.74037 | 0.42456 | 0.65209 | 1 | 13596 | 3 | 0.74037 |
| RHOBTB2 | 6 | 0.24199 | 0.43838 | 0.999994 | 8834 | 2 | 0.52776 | 0.42487 | 0.66995 | 1 | 13597 | 3 | 0.52776 |
| NKX2-2 | 5 | 0.1868 | 0.3564 | 0.999994 | 7383 | 2 | 0.84002 | 0.42496 | 0.65246 | 1 | 13598 | 3 | 0.84002 |
| HES2 | 4 | 0.86193 | 0.86156 | 0.999994 | 18485 | 0 | 1.0897 | 0.42502 | 0.60739 | 1 | 13599 | 2 | 1.0897 |
| KDSR | 4 | 0.10308 | 0.22062 | 0.981405 | 5070 | 2 | -1.4746 | 0.42505 | 0.60742 | 1 | 13600 | 1 | -1.4746 |
| GFM1 | 5 | 0.44143 | 0.59798 | 0.999994 | 12601 | 2 | 0.77373 | 0.42507 | 0.65257 | 1 | 13601 | 3 | 0.77373 |
| ADRB1 | 4 | 0.75283 | 0.76226 | 0.999994 | 16885 | 1 | 1.095 | 0.4251 | 0.60746 | 1 | 13602 | 2 | 1.095 |
| RBM4 | 2 | 0.27925 | 0.32088 | 0.999994 | 9682 | 1 | -0.42062 | 0.42515 | 0.48514 | 1 | 13603 | 1 | -0.42062 |
| USP17L12 | 1 | 0.57485 | 0.57457 | 0.999994 | 14614 | 0 | 0.95 | 0.42515 | 0.42532 | 1 | 13604 | 1 | 0.95 |
| hsa-mir-6719 | 4 | 0.42361 | 0.56989 | 0.999994 | 12305 | 2 | 0.69107 | 0.42518 | 0.60753 | 1 | 13605 | 2 | 0.69107 |
| UST | 5 | 0.25802 | 0.44714 | 0.999994 | 9195 | 2 | 0.78667 | 0.42519 | 0.65269 | 1 | 13606 | 3 | 0.78667 |
| AGTPBP1 | 6 | 0.25823 | 0.45779 | 0.999994 | 9203 | 3 | -0.46706 | 0.42519 | 0.67021 | 1 | 13607 | 1 | -0.46706 |
| USO1 | 6 | 0.93102 | 0.93115 | 0.999994 | 19878 | 0 | 0.42191 | 0.42521 | 0.67023 | 1 | 13608 | 3 | 0.42191 |
| TRMT6 | 6 | 0.033491 | 0.10699 | 0.950995 | 2191 | 3 | -0.34946 | 0.42521 | 0.67023 | 1 | 13609 | 2 | -0.34946 |
| FAM9A | 6 | 0.79046 | 0.84386 | 0.999994 | 17353 | 1 | 0.303 | 0.42521 | 0.67023 | 1 | 13610 | 2 | 0.303 |
| KIF17 | 6 | 0.22359 | 0.41646 | 0.999994 | 8369 | 3 | -0.41284 | 0.42521 | 0.67023 | 1 | 13611 | 2 | -0.41284 |
| DYNC1LI1 | 6 | 0.022818 | 0.081003 | 0.93914 | 1636 | 4 | -1.8219 | 0.42521 | 0.67023 | 1 | 13612 | 2 | -1.8219 |
| CLCA4 | 5 | 0.33151 | 0.50679 | 0.999994 | 10754 | 2 | 0.35554 | 0.42545 | 0.65296 | 1 | 13613 | 2 | 0.35554 |
| DLX4 | 5 | 0.16945 | 0.33134 | 0.999994 | 6974 | 3 | -1.0898 | 0.42545 | 0.65296 | 1 | 13614 | 2 | -1.0898 |
| SPON2 | 6 | 0.31841 | 0.52807 | 0.999994 | 10482 | 2 | 0.95961 | 0.42548 | 0.67042 | 1 | 13615 | 3 | 0.95961 |
| NonTargetingControlG | 1 | 0.57437 | 0.57411 | 0.999994 | 14608 | 0 | 1.3182 | 0.42563 | 0.42581 | 1 | 13616 | 1 | 1.3182 |
| hsa-mir-663b | 1 | 0.57437 | 0.57411 | 0.999994 | 14609 | 0 | 1.3182 | 0.42563 | 0.42581 | 1 | 13617 | 1 | 1.3182 |
| GRASP | 5 | 0.23604 | 0.42556 | 0.999994 | 8672 | 3 | -0.6214 | 0.42565 | 0.65315 | 1 | 13618 | 2 | -0.6214 |
| BIN1 | 6 | 0.058272 | 0.16206 | 0.967126 | 3352 | 3 | 0.057182 | 0.42573 | 0.67061 | 1 | 13619 | 3 | 0.057182 |
| NSA2 | 4 | 0.31353 | 0.45689 | 0.999994 | 10388 | 1 | 1.2198 | 0.42577 | 0.60797 | 1 | 13620 | 2 | 1.2198 |
| GPR162 | 4 | 0.25666 | 0.39873 | 0.999994 | 9164 | 1 | 0.68224 | 0.42589 | 0.60807 | 1 | 13621 | 2 | 0.68224 |
| MARCH7 | 6 | 0.12377 | 0.29235 | 0.999632 | 5758 | 3 | -1.6723 | 0.42598 | 0.67079 | 1 | 13622 | 2 | -1.6723 |
| TC2N | 6 | 0.24999 | 0.44793 | 0.999994 | 9014 | 3 | -0.82761 | 0.42598 | 0.67079 | 1 | 13623 | 2 | -0.82761 |

| | | | | | | | | | | | | | |
|---|---|---|---|---|---|---|---|---|---|---|---|---|---|
| hsa-mir-4741 | 4 | 0.0084301 | 0.028751 | 0.858757 | 756 | 3 | -5.9737 | 0.42615 | 0.60827 | 1 | 13624 | 1 | -5.9737 |
| ERAL1 | 5 | 0.57504 | 0.71397 | 0.999994 | 14618 | 2 | 0.85598 | 0.42623 | 0.65371 | 1 | 13625 | 3 | 0.85598 |
| STOML2 | 5 | 0.089351 | 0.20954 | 0.979587 | 4569 | 3 | -1.7062 | 0.42623 | 0.65371 | 1 | 13626 | 1 | -1.7062 |
| hsa-mir-6761 | 3 | 0.17156 | 0.27632 | 0.999379 | 7024 | 2 | -1.3192 | 0.42628 | 0.58043 | 1 | 13627 | 1 | -1.3192 |
| PRG2 | 5 | 0.14285 | 0.29214 | 0.999632 | 6267 | 2 | 0.89631 | 0.42643 | 0.65391 | 1 | 13628 | 3 | 0.89631 |
| C9orf78 | 6 | 0.057672 | 0.16074 | 0.967126 | 3322 | 3 | 0.44029 | 0.42653 | 0.6712 | 1 | 13629 | 3 | 0.44029 |
| DLGAP2 | 6 | 0.044625 | 0.13226 | 0.95318 | 2743 | 3 | -1.5755 | 0.42669 | 0.67132 | 1 | 13630 | 3 | -1.5755 |
| SGCB | 3 | 0.085691 | 0.16406 | 0.967126 | 4436 | 2 | -2.6651 | 0.42675 | 0.58091 | 1 | 13631 | 1 | -2.6651 |
| C16orf74 | 6 | 0.063739 | 0.1737 | 0.96793 | 3595 | 3 | -0.74195 | 0.42683 | 0.67141 | 1 | 13632 | 2 | -0.74195 |
| PTDSS2 | 6 | 0.13044 | 0.30086 | 0.999994 | 5943 | 3 | -0.63927 | 0.42683 | 0.67141 | 1 | 13633 | 2 | -0.63927 |
| C19orf77 | 6 | 0.31425 | 0.52332 | 0.999994 | 10404 | 2 | 0.65923 | 0.42694 | 0.6715 | 1 | 13634 | 3 | 0.65923 |
| hsa-mir-544b | 4 | 0.14767 | 0.28675 | 0.999632 | 6395 | 2 | -0.40078 | 0.42698 | 0.60891 | 1 | 13635 | 2 | -0.40078 |
| PSMB8 | 6 | 0.86937 | 0.88296 | 0.999994 | 18623 | 1 | 0.64235 | 0.4271 | 0.67162 | 1 | 13636 | 3 | 0.64235 |
| MAN1B1 | 4 | 0.36857 | 0.51336 | 0.999994 | 11481 | 2 | 0.1535 | 0.42714 | 0.60903 | 1 | 13637 | 2 | 0.1535 |
| EGF | 6 | 0.00087442 | 0.0044832 | 0.56977 | 149 | 4 | -4.6955 | 0.42716 | 0.67167 | 1 | 13638 | 2 | -4.6955 |
| ZNF439 | 6 | 0.23539 | 0.43052 | 0.999994 | 8658 | 3 | -0.050666 | 0.42725 | 0.67174 | 1 | 13639 | 3 | -0.050666 |
| DUSP8 | 2 | 0.46744 | 0.47474 | 0.999994 | 12971 | 1 | 1.1446 | 0.42732 | 0.48664 | 1 | 13640 | 1 | 1.1446 |
| WDR55 | 5 | 0.23628 | 0.4259 | 0.999994 | 8677 | 1 | 0.85782 | 0.42733 | 0.65475 | 1 | 13641 | 3 | 0.85782 |
| OR6K2 | 5 | 0.96601 | 0.96617 | 0.999994 | 20737 | 0 | 0.69036 | 0.4274 | 0.65484 | 1 | 13642 | 3 | 0.69036 |
| CD248 | 6 | 0.14781 | 0.32289 | 0.999994 | 6401 | 2 | 0.65479 | 0.42745 | 0.67189 | 1 | 13643 | 3 | 0.65479 |
| hsa-mir-19a | 3 | 0.61905 | 0.63195 | 0.999994 | 15207 | 1 | 0.83 | 0.42749 | 0.58169 | 1 | 13644 | 2 | 0.83 |
| CPXM2 | 6 | 0.31235 | 0.52105 | 0.999994 | 10368 | 3 | -0.40022 | 0.42772 | 0.67209 | 1 | 13645 | 2 | -0.40022 |
| CELF2 | 6 | 0.30528 | 0.51287 | 0.999994 | 10239 | 3 | -0.46445 | 0.42772 | 0.67209 | 1 | 13646 | 1 | -0.46445 |
| OAZ2 | 6 | 0.38094 | 0.59018 | 0.999994 | 11660 | 2 | 0.41246 | 0.42772 | 0.67209 | 1 | 13647 | 3 | 0.41246 |
| ZMYM2 | 6 | 0.17681 | 0.35911 | 0.999994 | 7149 | 3 | -0.58855 | 0.42772 | 0.67209 | 1 | 13648 | 2 | -0.58855 |
| MPP6 | 6 | 0.09405 | 0.23562 | 0.985362 | 4751 | 4 | -0.98293 | 0.42773 | 0.6721 | 1 | 13649 | 2 | -0.98293 |
| PURG | 5 | 0.79914 | 0.819 | 0.999994 | 17466 | 1 | 0.80326 | 0.42784 | 0.65525 | 1 | 13650 | 3 | 0.80326 |
| SLC25A18 | 5 | 0.2175 | 0.39977 | 0.999994 | 8208 | 2 | -0.27941 | 0.42791 | 0.65532 | 1 | 13651 | 2 | -0.27941 |
| MRPL9 | 5 | 0.012615 | 0.043519 | 0.891621 | 1037 | 4 | -3.7339 | 0.42791 | 0.65532 | 1 | 13652 | 1 | -3.7339 |
| TGFB1I1 | 5 | 0.89719 | 0.89733 | 0.999994 | 19163 | 0 | 0.055273 | 0.42791 | 0.65532 | 1 | 13653 | 1 | 0.055273 |
| TEKT1 | 5 | 0.46551 | 0.61831 | 0.999994 | 12944 | 2 | 0.22656 | 0.42792 | 0.65533 | 1 | 13654 | 2 | 0.22656 |
| DIP2B | 5 | 0.21572 | 0.39729 | 0.999994 | 8160 | 3 | -0.73285 | 0.42792 | 0.65533 | 1 | 13655 | 2 | -0.73285 |
| DENND1C | 5 | 0.24236 | 0.4341 | 0.999994 | 8841 | 1 | 0.72064 | 0.428 | 0.65541 | 1 | 13656 | 3 | 0.72064 |
| ARGFX | 6 | 0.57675 | 0.7228 | 0.999994 | 14641 | 2 | 0.40986 | 0.42803 | 0.67232 | 1 | 13657 | 3 | 0.40986 |
| HCRTR1 | 5 | 0.2418 | 0.43336 | 0.999994 | 8829 | 2 | 0.80652 | 0.42817 | 0.65558 | 1 | 13658 | 3 | 0.80652 |
| ENY2 | 6 | 0.26438 | 0.46506 | 0.999994 | 9343 | 3 | -0.040495 | 0.42826 | 0.67249 | 1 | 13659 | 3 | -0.040495 |
| NonTargetingControlG | 2 | 0.29976 | 0.33666 | 0.999994 | 10125 | 1 | 0.55188 | 0.42826 | 0.48728 | 1 | 13660 | 1 | 0.55188 |
| DIS3L | 5 | 0.6348 | 0.73919 | 0.999994 | 15441 | 1 | 0.69164 | 0.42833 | 0.65572 | 1 | 13661 | 3 | 0.69164 |
| JRK | 6 | 0.50105 | 0.66943 | 0.999994 | 13438 | 1 | 0.84217 | 0.42837 | 0.67258 | 1 | 13662 | 3 | 0.84217 |
| LPPR2 | 6 | 0.124 | 0.29264 | 0.999632 | 5768 | 2 | 0.84804 | 0.42841 | 0.6726 | 1 | 13663 | 3 | 0.84804 |
| OR5D16 | 6 | 0.044494 | 0.13198 | 0.95318 | 2736 | 3 | 0.2194 | 0.42844 | 0.67263 | 1 | 13664 | 3 | 0.2194 |
| hsa-mir-6772 | 4 | 0.030013 | 0.081506 | 0.93914 | 2009 | 3 | -2.0885 | 0.42856 | 0.61011 | 1 | 13665 | 1 | -2.0885 |
| FGFBP1 | 6 | 0.25435 | 0.45313 | 0.999994 | 9110 | 3 | 0.29013 | 0.42872 | 0.67282 | 1 | 13666 | 3 | 0.29013 |
| RXRB | 6 | 0.73654 | 0.82382 | 0.999994 | 16685 | 1 | 0.82211 | 0.42878 | 0.67286 | 1 | 13667 | 3 | 0.82211 |
| hsa-mir-519d | 4 | 0.2467 | 0.38852 | 0.999994 | 8938 | 2 | 0.028322 | 0.42885 | 0.61033 | 1 | 13668 | 2 | 0.028322 |
| ILF3 | 5 | 0.89369 | 0.89382 | 0.999994 | 19098 | 0 | 0.35554 | 0.42886 | 0.65625 | 1 | 13669 | 2 | 0.35554 |
| DCHS2 | 5 | 0.30726 | 0.48693 | 0.999994 | 10274 | 2 | 0.34818 | 0.42896 | 0.65635 | 1 | 13670 | 2 | 0.34818 |
| hsa-mir-15a | 4 | 0.77941 | 0.78339 | 0.999994 | 17202 | 1 | 0.034957 | 0.42896 | 0.61041 | 1 | 13671 | 1 | 0.034957 |
| ELAVL4 | 4 | 0.86071 | 0.8603 | 0.999994 | 18454 | 0 | 1.0691 | 0.4291 | 0.61053 | 1 | 13672 | 2 | 1.0691 |
| CORO7 | 6 | 0.054076 | 0.15292 | 0.967126 | 3161 | 4 | -0.97161 | 0.42914 | 0.67313 | 1 | 13673 | 1 | -0.97161 |
| EBP | 6 | 0.85245 | 0.87323 | 0.999994 | 18303 | 1 | -0.024235 | 0.42914 | 0.67313 | 1 | 13674 | 2 | -0.024235 |
| NonTargetingControlG | 2 | 0.25257 | 0.30083 | 0.999994 | 9068 | 1 | -0.57636 | 0.42916 | 0.48792 | 1 | 13675 | 1 | -0.57636 |

| | | | | | | | | | | | | | |
|---|---|---|---|---|---|---|---|---|---|---|---|---|---|
| ELF3 | 6 | 0.6108 | 0.74794 | 0.999994 | 15082 | 2 | -0.077513 | 0.42919 | 0.67317 | 1 | 13676 | 2 | -0.077513 |
| RARG | 6 | 0.51599 | 0.67974 | 0.999994 | 13676 | 2 | 0.11184 | 0.42919 | 0.67317 | 1 | 13677 | 2 | 0.11184 |
| C10orf118 | 6 | 0.25924 | 0.45896 | 0.999994 | 9221 | 2 | 0.12811 | 0.42919 | 0.67317 | 1 | 13678 | 3 | 0.12811 |
| AFMID | 6 | 0.80869 | 0.85166 | 0.999994 | 17615 | 1 | -0.17411 | 0.42919 | 0.67317 | 1 | 13679 | 2 | -0.17411 |
| hsa-mir-581 | 4 | 0.86989 | 0.86946 | 0.999994 | 18633 | 0 | 0.77803 | 0.42944 | 0.6108 | 1 | 13680 | 2 | 0.77803 |
| TMEM55A | 6 | 0.1878 | 0.37279 | 0.999994 | 7415 | 3 | 0.35738 | 0.42944 | 0.67337 | 1 | 13681 | 3 | 0.35738 |
| GPX5 | 5 | 0.95607 | 0.95623 | 0.999994 | 20487 | 0 | 0.71942 | 0.42962 | 0.657 | 1 | 13682 | 3 | 0.71942 |
| MBD5 | 5 | 0.34987 | 0.52181 | 0.999994 | 11156 | 2 | -0.10863 | 0.42967 | 0.65705 | 1 | 13683 | 2 | -0.10863 |
| DDX53 | 5 | 0.01594 | 0.052857 | 0.916123 | 1242 | 3 | -2.1123 | 0.42967 | 0.65705 | 1 | 13684 | 1 | -2.1123 |
| APIP | 5 | 0.11299 | 0.24661 | 0.98999 | 5405 | 2 | 0.68644 | 0.42967 | 0.65705 | 1 | 13685 | 3 | 0.68644 |
| RGL4 | 6 | 0.36538 | 0.5803 | 0.999994 | 11428 | 2 | 0.69596 | 0.42973 | 0.67358 | 1 | 13686 | 3 | 0.69596 |
| MAPK7 | 5 | 0.35174 | 0.52333 | 0.999994 | 11191 | 2 | 0.78447 | 0.42982 | 0.65719 | 1 | 13687 | 3 | 0.78447 |
| NonTargetingControlG | 1 | 0.57018 | 0.56993 | 0.999994 | 14543 | 0 | 1.0029 | 0.42982 | 0.42998 | 1 | 13688 | 1 | 1.0029 |
| TTYH2 | 3 | 0.81732 | 0.81714 | 0.999994 | 17733 | 0 | 0.80422 | 0.42986 | 0.58409 | 1 | 13689 | 2 | 0.80422 |
| SLC28A3 | 5 | 0.027091 | 0.08297 | 0.94336 | 1862 | 2 | 0.76029 | 0.42997 | 0.65734 | 1 | 13690 | 3 | 0.76029 |
| NEU2 | 6 | 0.66377 | 0.78822 | 0.999994 | 15835 | 2 | 0.53228 | 0.42999 | 0.67379 | 1 | 13691 | 3 | 0.53228 |
| MEST | 5 | 0.46092 | 0.61443 | 0.999994 | 12861 | 2 | 0.67182 | 0.42999 | 0.65736 | 1 | 13692 | 3 | 0.67182 |
| OR4D9 | 5 | 0.93086 | 0.93113 | 0.999994 | 19873 | 0 | 0.6308 | 0.43018 | 0.65753 | 1 | 13693 | 3 | 0.6308 |
| ZNF239 | 6 | 0.49778 | 0.66719 | 0.999994 | 13394 | 2 | 0.87607 | 0.43026 | 0.67399 | 1 | 13694 | 3 | 0.87607 |
| TBC1D30 | 5 | 0.020314 | 0.064924 | 0.93914 | 1477 | 2 | 0.95178 | 0.43027 | 0.65761 | 1 | 13695 | 3 | 0.95178 |
| TIPRL | 5 | 0.050508 | 0.14158 | 0.959146 | 2994 | 2 | 0.71517 | 0.43038 | 0.65771 | 1 | 13696 | 3 | 0.71517 |
| POLR3K | 6 | 0.42983 | 0.62173 | 0.999994 | 12410 | 2 | 0.53453 | 0.4304 | 0.67409 | 1 | 13697 | 3 | 0.53453 |
| GUSB | 6 | 0.13855 | 0.31117 | 0.999994 | 6155 | 3 | 0.004988 | 0.43049 | 0.67416 | 1 | 13698 | 3 | 0.004988 |
| IREB2 | 6 | 0.057307 | 0.15993 | 0.967126 | 3306 | 4 | -1.3273 | 0.43049 | 0.67416 | 1 | 13699 | 2 | -1.3273 |
| TSSK2 | 6 | 0.00036753 | 0.0019564 | 0.439257 | 88 | 3 | -0.88501 | 0.43049 | 0.67416 | 1 | 13700 | 2 | -0.88501 |
| PTTG2 | 5 | 0.034746 | 0.10261 | 0.946414 | 2251 | 4 | -1.752 | 0.43056 | 0.65787 | 1 | 13701 | 1 | -1.752 |
| GSPT2 | 5 | 0.83035 | 0.83973 | 0.999994 | 17939 | 1 | 0.96825 | 0.4306 | 0.65791 | 1 | 13702 | 3 | 0.96825 |
| CAMK2N1 | 5 | 0.93394 | 0.93418 | 0.999994 | 19952 | 0 | 0.73147 | 0.43067 | 0.65799 | 1 | 13703 | 3 | 0.73147 |
| CLTA | 5 | 0.61829 | 0.73293 | 0.999994 | 15192 | 1 | 0.75797 | 0.43085 | 0.65814 | 1 | 13704 | 3 | 0.75797 |
| OSM | 4 | 0.23746 | 0.37909 | 0.999994 | 8714 | 2 | -0.95706 | 0.43091 | 0.61191 | 1 | 13705 | 1 | -0.95706 |
| C19orf45 | 4 | 0.34769 | 0.49191 | 0.999994 | 11101 | 2 | -0.015723 | 0.43091 | 0.61191 | 1 | 13706 | 2 | -0.015723 |
| GDF15 | 4 | 0.17027 | 0.31006 | 0.999994 | 6991 | 2 | -1.1653 | 0.43091 | 0.61191 | 1 | 13707 | 1 | -1.1653 |
| XIRP1 | 6 | 0.22274 | 0.41547 | 0.999994 | 8352 | 3 | 0.19726 | 0.43096 | 0.67452 | 1 | 13708 | 3 | 0.19726 |
| IL18RAP | 5 | 0.40287 | 0.56595 | 0.999994 | 11995 | 2 | 0.19507 | 0.431 | 0.65829 | 1 | 13709 | 2 | 0.19507 |
| SNX20 | 5 | 0.19012 | 0.36112 | 0.999994 | 7490 | 3 | -0.8685 | 0.43105 | 0.65834 | 1 | 13710 | 2 | -0.8685 |
| CCNA2 | 2 | 0.1092 | 0.18424 | 0.968526 | 5273 | 1 | -2.0809 | 0.43106 | 0.48914 | 1 | 13711 | 1 | -2.0809 |
| VPREB1 | 4 | 0.55916 | 0.64422 | 0.999994 | 14357 | 1 | 1.1139 | 0.4312 | 0.61213 | 1 | 13712 | 2 | 1.1139 |
| ADAMTS7 | 3 | 0.033214 | 0.082491 | 0.94117 | 2172 | 1 | 0.95195 | 0.43123 | 0.58546 | 1 | 13713 | 2 | 0.95195 |
| DMBT1 | 6 | 0.23931 | 0.43516 | 0.999994 | 8759 | 2 | 0.37458 | 0.43126 | 0.67476 | 1 | 13714 | 3 | 0.37458 |
| ITGB8 | 6 | 0.14339 | 0.31735 | 0.999994 | 6284 | 3 | -0.57058 | 0.4313 | 0.67479 | 1 | 13715 | 2 | -0.57058 |
| PLA2R1 | 5 | 0.40139 | 0.56467 | 0.999994 | 11967 | 2 | 0.65245 | 0.43142 | 0.65871 | 1 | 13716 | 3 | 0.65245 |
| hsa-mir-3127 | 4 | 0.2062 | 0.34708 | 0.999994 | 7933 | 2 | 0.13201 | 0.43143 | 0.6123 | 1 | 13717 | 2 | 0.13201 |
| SLC25A2 | 4 | 0.19293 | 0.33342 | 0.999994 | 7575 | 2 | -0.78879 | 0.43155 | 0.61239 | 1 | 13718 | 2 | -0.78879 |
| LYZL1 | 6 | 0.35988 | 0.57518 | 0.999994 | 11332 | 2 | -0.054987 | 0.43158 | 0.67498 | 1 | 13719 | 2 | -0.054987 |
| KLRG1 | 5 | 0.042102 | 0.12114 | 0.951052 | 2618 | 2 | 0.21402 | 0.43161 | 0.6589 | 1 | 13720 | 2 | 0.21402 |
| MT1A | 5 | 0.5697 | 0.70926 | 0.999994 | 14532 | 2 | 0.77805 | 0.43174 | 0.65904 | 1 | 13721 | 3 | 0.77805 |
| CORO2A | 6 | 0.90938 | 0.91106 | 0.999994 | 19411 | 1 | 0.9292 | 0.43176 | 0.67513 | 1 | 13722 | 3 | 0.9292 |
| AQP9 | 5 | 0.5751 | 0.71402 | 0.999994 | 14619 | 2 | 0.27551 | 0.4318 | 0.65909 | 1 | 13723 | 2 | 0.27551 |
| ACTG2 | 6 | 0.11209 | 0.27072 | 0.998561 | 5369 | 3 | -0.13162 | 0.43211 | 0.67541 | 1 | 13724 | 3 | -0.13162 |
| ABHD1 | 6 | 0.18792 | 0.37295 | 0.999994 | 7418 | 3 | -0.56924 | 0.43217 | 0.67545 | 1 | 13725 | 2 | -0.56924 |
| GPM6A | 6 | 0.29984 | 0.50653 | 0.999994 | 10127 | 3 | -0.31175 | 0.43217 | 0.67545 | 1 | 13726 | 2 | -0.31175 |
| PUS1 | 6 | 0.10024 | 0.24781 | 0.990682 | 4980 | 3 | -1.2786 | 0.43224 | 0.67551 | 1 | 13727 | 2 | -1.2786 |

| | | | | | | | | | | | | | |
|---|---|---|---|---|---|---|---|---|---|---|---|---|---|
| B3GNTL1 | 5 | 0.91722 | 0.9173 | 0.999994 | 19568 | 0 | 0.12653 | 0.43239 | 0.65963 | 1 | 13728 | 2 | 0.12653 |
| FAM166A | 5 | 0.25396 | 0.44389 | 0.999994 | 9103 | 1 | 0.78616 | 0.43241 | 0.65965 | 1 | 13729 | 3 | 0.78616 |
| FNDC3A | 6 | 0.25279 | 0.45125 | 0.999994 | 9080 | 2 | 0.35876 | 0.43247 | 0.67568 | 1 | 13730 | 3 | 0.35876 |
| ACSF3 | 6 | 0.10677 | 0.26051 | 0.994943 | 5203 | 3 | -0.31072 | 0.43249 | 0.6757 | 1 | 13731 | 3 | -0.31072 |
| ACPL2 | 6 | 0.82845 | 0.86082 | 0.999994 | 17902 | 1 | 0.62124 | 0.43258 | 0.67577 | 1 | 13732 | 3 | 0.62124 |
| ADAP2 | 5 | 0.072327 | 0.18213 | 0.968526 | 3933 | 3 | -1.8981 | 0.43275 | 0.65998 | 1 | 13733 | 2 | -1.8981 |
| LXN | 5 | 0.3994 | 0.56301 | 0.999994 | 11940 | 1 | 0.73142 | 0.43281 | 0.66003 | 1 | 13734 | 3 | 0.73142 |
| hsa-mir-8068 | 4 | 0.43749 | 0.58418 | 0.999994 | 12535 | 2 | -0.097741 | 0.43288 | 0.61338 | 1 | 13735 | 1 | -0.097741 |
| hsa-mir-628 | 4 | 0.78657 | 0.78942 | 0.999994 | 17297 | 1 | -0.27301 | 0.43288 | 0.61338 | 1 | 13736 | 1 | -0.27301 |
| C16orf92 | 4 | 0.065119 | 0.15056 | 0.966359 | 3658 | 1 | 0.6922 | 0.43288 | 0.61338 | 1 | 13737 | 2 | 0.6922 |
| DYRK3 | 4 | 0.058771 | 0.13842 | 0.957336 | 3373 | 2 | -2.9062 | 0.43288 | 0.61338 | 1 | 13738 | 1 | -2.9062 |
| ARHGAP32 | 4 | 0.47988 | 0.60745 | 0.999994 | 13144 | 1 | 0.53254 | 0.43288 | 0.61338 | 1 | 13739 | 2 | 0.53254 |
| C16orf58 | 5 | 0.0052454 | 0.021773 | 0.819701 | 536 | 2 | 0.55063 | 0.4329 | 0.66013 | 1 | 13740 | 3 | 0.55063 |
| ABHD6 | 5 | 0.2864 | 0.47 | 0.999994 | 9843 | 2 | 0.76868 | 0.43307 | 0.6603 | 1 | 13741 | 3 | 0.76868 |
| MKI67IP | 3 | 0.26639 | 0.39524 | 0.999994 | 9383 | 2 | -0.77512 | 0.43317 | 0.58743 | 1 | 13742 | 1 | -0.77512 |
| PPAP2B | 6 | 0.25935 | 0.45909 | 0.999994 | 9222 | 2 | -0.37896 | 0.43321 | 0.67626 | 1 | 13743 | 1 | -0.37896 |
| OR51A4 | 6 | 0.053709 | 0.1521 | 0.967126 | 3151 | 3 | -0.75245 | 0.43321 | 0.67626 | 1 | 13744 | 3 | -0.75245 |
| ECSCR | 6 | 0.077167 | 0.20162 | 0.977164 | 4113 | 4 | -1.1802 | 0.43321 | 0.67626 | 1 | 13745 | 2 | -1.1802 |
| FAM9C | 5 | 0.38501 | 0.55093 | 0.999994 | 11715 | 2 | 0.87876 | 0.43334 | 0.66058 | 1 | 13746 | 3 | 0.87876 |
| hsa-mir-6721 | 4 | 0.65651 | 0.69714 | 0.999994 | 15737 | 1 | 0.89534 | 0.43335 | 0.61374 | 1 | 13747 | 2 | 0.89534 |
| hsa-mir-4278 | 4 | 0.079238 | 0.17735 | 0.968171 | 4200 | 2 | 0.48474 | 0.4335 | 0.61386 | 1 | 13748 | 2 | 0.48474 |
| hsa-mir-6826 | 3 | 0.81308 | 0.81284 | 0.999994 | 17675 | 0 | 0.83512 | 0.43376 | 0.58802 | 1 | 13749 | 2 | 0.83512 |
| PRRG4 | 5 | 0.57932 | 0.71772 | 0.999994 | 14670 | 2 | -0.24553 | 0.43377 | 0.66099 | 1 | 13750 | 2 | -0.24553 |
| DDX39A | 3 | 0.69833 | 0.69947 | 0.999994 | 16287 | 1 | 0.89014 | 0.43383 | 0.58808 | 1 | 13751 | 2 | 0.89014 |
| USP1 | 4 | 0.39189 | 0.53733 | 0.999994 | 11830 | 2 | 0.015428 | 0.43389 | 0.61414 | 1 | 13752 | 2 | 0.015428 |
| PRPH2 | 4 | 0.6386 | 0.68671 | 0.999994 | 15494 | 1 | 0.20887 | 0.43389 | 0.61414 | 1 | 13753 | 1 | 0.20887 |
| hsa-mir-3689c | 2 | 0.70131 | 0.70108 | 0.999994 | 16320 | 0 | 1.9722 | 0.43399 | 0.49118 | 1 | 13754 | 1 | 1.9722 |
| RBM45 | 5 | 0.9402 | 0.94037 | 0.999994 | 20088 | 0 | 0.70409 | 0.4341 | 0.66132 | 1 | 13755 | 3 | 0.70409 |
| hsa-mir-4780 | 4 | 0.3124 | 0.45572 | 0.999994 | 10369 | 1 | 0.66632 | 0.43411 | 0.61428 | 1 | 13756 | 2 | 0.66632 |
| TXK | 6 | 0.27688 | 0.47981 | 0.999994 | 9635 | 3 | -0.086678 | 0.43412 | 0.67698 | 1 | 13757 | 2 | -0.086678 |
| OS9 | 6 | 0.21053 | 0.40074 | 0.999994 | 8037 | 2 | 0.4666 | 0.43425 | 0.67708 | 1 | 13758 | 2 | 0.4666 |
| SLC22A18AS | 6 | 0.6576 | 0.78344 | 0.999994 | 15749 | 2 | 0.48753 | 0.43425 | 0.67708 | 1 | 13759 | 3 | 0.48753 |
| RPS29 | 3 | 0.26185 | 0.38969 | 0.999994 | 9283 | 1 | 0.34197 | 0.4343 | 0.58856 | 1 | 13760 | 1 | 0.34197 |
| CCDC180 | 6 | 0.042653 | 0.12789 | 0.951674 | 2658 | 3 | 0.13849 | 0.43438 | 0.67718 | 1 | 13761 | 3 | 0.13849 |
| CCL3L1 | 6 | 0.28972 | 0.49475 | 0.999994 | 9906 | 3 | 0.24758 | 0.4344 | 0.67719 | 1 | 13762 | 3 | 0.24758 |
| KLRC3 | 5 | 0.24154 | 0.43301 | 0.999994 | 8816 | 3 | -0.611 | 0.43457 | 0.66179 | 1 | 13763 | 2 | -0.611 |
| RWDD3 | 6 | 0.60673 | 0.74492 | 0.999994 | 15033 | 2 | 0.60468 | 0.43457 | 0.67733 | 1 | 13764 | 3 | 0.60468 |
| CCDC103 | 6 | 0.099213 | 0.24583 | 0.989866 | 4939 | 2 | 0.83932 | 0.43459 | 0.67734 | 1 | 13765 | 3 | 0.83932 |
| TPCN2 | 6 | 0.95072 | 0.95071 | 0.999994 | 20350 | 0 | 0.91345 | 0.43461 | 0.67736 | 1 | 13766 | 3 | 0.91345 |
| JHDM1D | 6 | 0.073875 | 0.19481 | 0.975797 | 3989 | 4 | -1.6587 | 0.43463 | 0.67738 | 1 | 13767 | 2 | -1.6587 |
| WBP2NL | 5 | 0.17386 | 0.33774 | 0.999994 | 7080 | 3 | -0.88848 | 0.43471 | 0.66193 | 1 | 13768 | 2 | -0.88848 |
| EIF3L | 6 | 0.15232 | 0.32859 | 0.999994 | 6509 | 3 | 0.15306 | 0.43475 | 0.67747 | 1 | 13769 | 3 | 0.15306 |
| NonTargetingControlG | 2 | 0.4382 | 0.44904 | 0.999994 | 12542 | 1 | 0.90357 | 0.4348 | 0.49172 | 1 | 13770 | 1 | 0.90357 |
| PRCP | 5 | 0.12037 | 0.25792 | 0.994943 | 5640 | 3 | -0.89103 | 0.43499 | 0.6622 | 1 | 13771 | 2 | -0.89103 |
| hsa-mir-6886 | 4 | 0.15193 | 0.29118 | 0.999632 | 6503 | 2 | -1.5585 | 0.435 | 0.61494 | 1 | 13772 | 1 | -1.5585 |
| RNF11 | 4 | 0.068092 | 0.15622 | 0.967126 | 3752 | 2 | -1.3222 | 0.435 | 0.61494 | 1 | 13773 | 1 | -1.3222 |
| NonTargetingControlG | 2 | 0.11867 | 0.19876 | 0.977164 | 5593 | 1 | -2.1024 | 0.43517 | 0.49195 | 1 | 13774 | 1 | -2.1024 |
| CLC | 6 | 0.0094038 | 0.03693 | 0.881473 | 820 | 3 | 0.094505 | 0.4352 | 0.67781 | 1 | 13775 | 3 | 0.094505 |
| NTN1 | 6 | 0.0098873 | 0.038656 | 0.890863 | 865 | 2 | 0.41575 | 0.4352 | 0.67781 | 1 | 13776 | 3 | 0.41575 |
| TAS2R10 | 6 | 0.075365 | 0.19792 | 0.977164 | 4048 | 4 | -1.104 | 0.43524 | 0.67784 | 1 | 13777 | 2 | -1.104 |
| EIF5 | 5 | 0.099196 | 0.22516 | 0.983354 | 4937 | 2 | 0.01444 | 0.43532 | 0.66251 | 1 | 13778 | 2 | 0.01444 |
| PIGO | 5 | 0.51909 | 0.66452 | 0.999994 | 13732 | 2 | 0.63895 | 0.43535 | 0.66254 | 1 | 13779 | 3 | 0.63895 |

| | | | | | | | | | | | | |
|---|---|---|---|---|---|---|---|---|---|---|---|---|
| CRY1 | 5 | 0.10652 | 0.23658 | 0.985449 | 5188 | 3 | -1.459 | 0.43548 | 0.66268 | 1 | 13780 | 2 | -1.459 |
| RNF125 | 5 | 0.079822 | 0.1943 | 0.975797 | 4226 | 2 | 0.79187 | 0.43548 | 0.66268 | 1 | 13781 | 3 | 0.79187 |
| LITAF | 5 | 0.24641 | 0.4379 | 0.999994 | 8932 | 1 | 0.71802 | 0.43554 | 0.66273 | 1 | 13782 | 3 | 0.71802 |
| UQCC | 6 | 0.55204 | 0.70513 | 0.999994 | 14244 | 2 | 0.46641 | 0.43561 | 0.67814 | 1 | 13783 | 3 | 0.46641 |
| NAV3 | 6 | 0.91057 | 0.91204 | 0.999994 | 19442 | 1 | 0.46732 | 0.43574 | 0.67824 | 1 | 13784 | 3 | 0.46732 |
| TNFRSF4 | 5 | 0.19081 | 0.3621 | 0.999994 | 7506 | 3 | -0.87679 | 0.43578 | 0.66296 | 1 | 13785 | 2 | -0.87679 |
| CAMKMT | 6 | 0.025991 | 0.089336 | 0.94395 | 1805 | 3 | -1.2224 | 0.43597 | 0.67842 | 1 | 13786 | 2 | -1.2224 |
| DHX32 | 6 | 0.55869 | 0.70988 | 0.999994 | 14345 | 2 | -0.025352 | 0.43597 | 0.67842 | 1 | 13787 | 2 | -0.025352 |
| CA8 | 5 | 0.90784 | 0.90789 | 0.999994 | 19381 | 0 | 0.66934 | 0.43597 | 0.66314 | 1 | 13788 | 3 | 0.66934 |
| STAC3 | 5 | 0.12093 | 0.25878 | 0.994943 | 5658 | 2 | 0.6505 | 0.43599 | 0.66316 | 1 | 13789 | 3 | 0.6505 |
| CARM1 | 5 | 0.071783 | 0.18127 | 0.968171 | 3911 | 3 | -0.66032 | 0.43609 | 0.66325 | 1 | 13790 | 2 | -0.66032 |
| VGLL4 | 6 | 0.31843 | 0.52811 | 0.999994 | 10484 | 2 | 0.7947 | 0.43623 | 0.67862 | 1 | 13791 | 3 | 0.7947 |
| SLC20A1 | 5 | 0.026838 | 0.082317 | 0.940526 | 1848 | 3 | -1.4418 | 0.43637 | 0.66349 | 1 | 13792 | 2 | -1.4418 |
| KLF8 | 6 | 0.68432 | 0.80434 | 0.999994 | 16148 | 1 | 0.90797 | 0.43641 | 0.67874 | 1 | 13793 | 3 | 0.90797 |
| MRPS2 | 2 | 0.65944 | 0.65923 | 0.999994 | 15770 | 0 | 1.6851 | 0.43651 | 0.49289 | 1 | 13794 | 1 | 1.6851 |
| NKD1 | 3 | 0.77126 | 0.7713 | 0.999994 | 17101 | 0 | 0.54942 | 0.43654 | 0.59082 | 1 | 13795 | 1 | 0.54942 |
| NonTargetingControlG | 2 | 0.071482 | 0.1248 | 0.951052 | 3891 | 1 | -2.631 | 0.43661 | 0.49297 | 1 | 13796 | 1 | -2.631 |
| KIAA1107 | 5 | 0.090556 | 0.21149 | 0.979587 | 4618 | 2 | 0.72855 | 0.43662 | 0.66375 | 1 | 13797 | 3 | 0.72855 |
| GPR12 | 5 | 0.019686 | 0.063244 | 0.937389 | 1437 | 3 | -1.4223 | 0.43669 | 0.66382 | 1 | 13798 | 2 | -1.4223 |
| SMARCD2 | 6 | 0.22155 | 0.41397 | 0.999994 | 8322 | 3 | 0.25201 | 0.43678 | 0.67902 | 1 | 13799 | 3 | 0.25201 |
| CLEC3A | 6 | 0.063531 | 0.17324 | 0.96793 | 3579 | 4 | -1.4188 | 0.43678 | 0.67902 | 1 | 13800 | 2 | -1.4188 |
| SDR9C7 | 6 | 0.20385 | 0.39257 | 0.999994 | 7876 | 3 | -0.65614 | 0.43678 | 0.67902 | 1 | 13801 | 2 | -0.65614 |
| RCSD1 | 4 | 0.24598 | 0.38777 | 0.999994 | 8923 | 2 | -0.27822 | 0.43683 | 0.61633 | 1 | 13802 | 1 | -0.27822 |
| TEX22 | 5 | 0.42192 | 0.58169 | 0.999994 | 12278 | 2 | 0.45423 | 0.43699 | 0.66411 | 1 | 13803 | 2 | 0.45423 |
| TRABD | 6 | 0.1273 | 0.29686 | 0.999994 | 5853 | 3 | -0.17884 | 0.43706 | 0.67923 | 1 | 13804 | 3 | -0.17884 |
| hsa-mir-1538 | 4 | 0.072225 | 0.16414 | 0.967126 | 3926 | 2 | -0.95417 | 0.43707 | 0.61653 | 1 | 13805 | 2 | -0.95417 |
| ZNF705D | 4 | 0.12479 | 0.25862 | 0.994943 | 5792 | 1 | 1.2345 | 0.43712 | 0.61657 | 1 | 13806 | 2 | 1.2345 |
| THUMPD2 | 5 | 0.19974 | 0.37489 | 0.999994 | 7755 | 2 | 0.062453 | 0.43716 | 0.66427 | 1 | 13807 | 2 | 0.062453 |
| CYP4F12 | 5 | 0.23461 | 0.42359 | 0.999994 | 8629 | 1 | 0.73548 | 0.43724 | 0.66435 | 1 | 13808 | 3 | 0.73548 |
| POMGNT1 | 6 | 0.053473 | 0.1516 | 0.967126 | 3139 | 2 | 0.10064 | 0.43724 | 0.67936 | 1 | 13809 | 2 | 0.10064 |
| CUL5 | 4 | 0.33711 | 0.48096 | 0.999994 | 10874 | 2 | 0.26705 | 0.43731 | 0.61669 | 1 | 13810 | 2 | 0.26705 |
| PACRG | 5 | 0.35668 | 0.52735 | 0.999994 | 11281 | 1 | 0.67181 | 0.43738 | 0.6645 | 1 | 13811 | 3 | 0.67181 |
| NonTargetingControlG | 1 | 0.56261 | 0.5625 | 0.999994 | 14423 | 0 | 0.96183 | 0.43739 | 0.43755 | 1 | 13812 | 1 | 0.96183 |
| TRIM64B | 4 | 0.61696 | 0.67464 | 0.999994 | 15170 | 1 | 0.99376 | 0.43741 | 0.61677 | 1 | 13813 | 2 | 0.99376 |
| hsa-mir-517c | 1 | 0.56254 | 0.56242 | 0.999994 | 14422 | 0 | 0.92763 | 0.43746 | 0.43762 | 1 | 13814 | 1 | 0.92763 |
| SPANXC | 1 | 0.56251 | 0.56239 | 0.999994 | 14421 | 0 | 0.8916 | 0.43749 | 0.43765 | 1 | 13815 | 1 | 0.8916 |
| ERVW-1 | 6 | 0.93091 | 0.93105 | 0.999994 | 19875 | 0 | 0.0023308 | 0.43767 | 0.67968 | 1 | 13816 | 2 | 0.0023308 |
| TMPRSS4 | 6 | 0.45854 | 0.64077 | 0.999994 | 12830 | 2 | 0.83769 | 0.43767 | 0.67968 | 1 | 13817 | 3 | 0.83769 |
| CD226 | 5 | 0.12479 | 0.26466 | 0.996734 | 5789 | 2 | -0.16926 | 0.43767 | 0.66477 | 1 | 13818 | 2 | -0.16926 |
| NonTargetingControlG | 1 | 0.56229 | 0.56218 | 0.999994 | 14414 | 0 | 0.95657 | 0.43771 | 0.43787 | 1 | 13819 | 1 | 0.95657 |
| THOC2 | 4 | 0.38828 | 0.53361 | 0.999994 | 11769 | 2 | 0.44095 | 0.43771 | 0.61701 | 1 | 13820 | 2 | 0.44095 |
| SMARCD1 | 4 | 0.31333 | 0.45668 | 0.999994 | 10383 | 2 | -0.74276 | 0.43779 | 0.61706 | 1 | 13821 | 1 | -0.74276 |
| SCP2 | 6 | 0.091067 | 0.22965 | 0.983354 | 4638 | 1 | 0.21284 | 0.43794 | 0.67989 | 1 | 13822 | 2 | 0.21284 |
| DHX40 | 6 | 0.67047 | 0.79351 | 0.999994 | 15939 | 2 | 0.87603 | 0.43798 | 0.67991 | 1 | 13823 | 3 | 0.87603 |
| KREMEN2 | 6 | 0.4132 | 0.61093 | 0.999994 | 12154 | 1 | 0.64715 | 0.43801 | 0.67993 | 1 | 13824 | 3 | 0.64715 |
| HBG1 | 5 | 0.77095 | 0.80225 | 0.999994 | 17097 | 1 | 0.45289 | 0.43812 | 0.66519 | 1 | 13825 | 2 | 0.45289 |
| ENO2 | 5 | 0.90912 | 0.90912 | 0.999994 | 19405 | 0 | 0.78692 | 0.43813 | 0.6652 | 1 | 13826 | 3 | 0.78692 |
| ZNF185 | 6 | 0.091581 | 0.23069 | 0.983933 | 4662 | 3 | -0.34866 | 0.43815 | 0.68004 | 1 | 13827 | 2 | -0.34866 |
| AMY2B | 3 | 0.75772 | 0.7577 | 0.999994 | 16949 | 0 | 0.82959 | 0.43827 | 0.59258 | 1 | 13828 | 2 | 0.82959 |
| hsa-mir-4509-2 | 2 | 0.59463 | 0.59455 | 0.999994 | 14885 | 0 | 1.5669 | 0.43836 | 0.49414 | 1 | 13829 | 1 | 1.5669 |
| MPC1L | 5 | 0.22953 | 0.41657 | 0.999994 | 8520 | 1 | 0.69073 | 0.43841 | 0.66547 | 1 | 13830 | 3 | 0.69073 |
| DQX1 | 6 | 0.27798 | 0.48107 | 0.999994 | 9653 | 3 | -0.56993 | 0.43842 | 0.68026 | 1 | 13831 | 2 | -0.56993 |

| | | | | | | | | | | | | | |
|---|---|---|---|---|---|---|---|---|---|---|---|---|---|
| ESCO1 | 6 | 0.12326 | 0.29171 | 0.999632 | 5744 | 3 | -0.13047 | 0.43842 | 0.68026 | 1 | 13832 | 3 | -0.13047 |
| PSTPIP1 | 6 | 0.091485 | 0.23051 | 0.983933 | 4654 | 1 | 0.4726 | 0.43842 | 0.68026 | 1 | 13833 | 3 | 0.4726 |
| ANAPC1 | 4 | 0.098693 | 0.21272 | 0.979587 | 4920 | 2 | -0.53861 | 0.43846 | 0.61758 | 1 | 13834 | 2 | -0.53861 |
| RPS19 | 3 | 0.071405 | 0.14488 | 0.963012 | 3888 | 2 | -2.6655 | 0.43869 | 0.59301 | 1 | 13835 | 1 | -2.6655 |
| hsa-mir-4657 | 3 | 0.28332 | 0.41605 | 0.999994 | 9770 | 2 | -0.64696 | 0.43869 | 0.59301 | 1 | 13836 | 1 | -0.64696 |
| SAGE1 | 6 | 0.29328 | 0.49884 | 0.999994 | 9981 | 2 | 0.38386 | 0.4387 | 0.68047 | 1 | 13837 | 3 | 0.38386 |
| TMEM203 | 5 | 0.22859 | 0.41526 | 0.999994 | 8493 | 1 | 0.71544 | 0.43873 | 0.6658 | 1 | 13838 | 3 | 0.71544 |
| SPTBN1 | 6 | 0.015871 | 0.058622 | 0.931872 | 1236 | 4 | -2.18 | 0.43875 | 0.68051 | 1 | 13839 | 2 | -2.18 |
| ANKS1B | 6 | 0.32904 | 0.54014 | 0.999994 | 10704 | 1 | 0.73236 | 0.43899 | 0.68069 | 1 | 13840 | 3 | 0.73236 |
| NKX2-1 | 6 | 0.22704 | 0.42052 | 0.999994 | 8454 | 2 | 0.68829 | 0.43903 | 0.68072 | 1 | 13841 | 3 | 0.68829 |
| SMTNL2 | 5 | 0.24682 | 0.43825 | 0.999994 | 8941 | 1 | 0.68888 | 0.43904 | 0.6661 | 1 | 13842 | 3 | 0.68888 |
| SCN11A | 5 | 0.89903 | 0.89913 | 0.999994 | 19204 | 0 | 0.45525 | 0.43914 | 0.66621 | 1 | 13843 | 2 | 0.45525 |
| NonTargetingControlG | 2 | 0.3475 | 0.37405 | 0.999994 | 11099 | 1 | 0.83136 | 0.4392 | 0.49473 | 1 | 13844 | 1 | 0.83136 |
| VRK3 | 6 | 0.9154 | 0.91626 | 0.999994 | 19521 | 1 | 0.68326 | 0.43926 | 0.68089 | 1 | 13845 | 3 | 0.68326 |
| NonTargetingControlG | 2 | 0.34266 | 0.37013 | 0.999994 | 10997 | 1 | 0.58675 | 0.4393 | 0.49479 | 1 | 13846 | 1 | 0.58675 |
| LIPT2 | 5 | 0.55116 | 0.69277 | 0.999994 | 14230 | 2 | -0.025684 | 0.43945 | 0.6665 | 1 | 13847 | 1 | -0.025684 |
| HERC6 | 5 | 0.50135 | 0.6491 | 0.999994 | 13445 | 2 | -0.14994 | 0.43945 | 0.6665 | 1 | 13848 | 1 | -0.14994 |
| MUM1L1 | 5 | 0.72052 | 0.77608 | 0.999994 | 16502 | 1 | 0.74929 | 0.43979 | 0.66679 | 1 | 13849 | 3 | 0.74929 |
| hsa-mir-4539 | 4 | 0.040566 | 0.10292 | 0.946414 | 2539 | 2 | -2.8179 | 0.43984 | 0.61863 | 1 | 13850 | 2 | -2.8179 |
| GSX1 | 4 | 0.050168 | 0.12178 | 0.951052 | 2985 | 2 | 0.20867 | 0.43984 | 0.61863 | 1 | 13851 | 2 | 0.20867 |
| ASB12 | 5 | 0.73645 | 0.78388 | 0.999994 | 16682 | 1 | 0.89106 | 0.43986 | 0.66686 | 1 | 13852 | 3 | 0.89106 |
| hsa-mir-5004 | 4 | 0.57601 | 0.65272 | 0.999994 | 14633 | 1 | 1.0895 | 0.43993 | 0.61871 | 1 | 13853 | 2 | 1.0895 |
| ZBTB44 | 6 | 0.33307 | 0.54478 | 0.999994 | 10790 | 3 | -0.18098 | 0.43994 | 0.68142 | 1 | 13854 | 2 | -0.18098 |
| ELAC1 | 5 | 0.41992 | 0.58 | 0.999994 | 12252 | 2 | 0.57019 | 0.43997 | 0.66696 | 1 | 13855 | 3 | 0.57019 |
| PTX3 | 4 | 0.7419 | 0.75393 | 0.999994 | 16763 | 1 | 1.0119 | 0.44003 | 0.61879 | 1 | 13856 | 2 | 1.0119 |
| SPRYD4 | 6 | 0.39547 | 0.59945 | 0.999994 | 11878 | 2 | 0.5829 | 0.4401 | 0.68156 | 1 | 13857 | 3 | 0.5829 |
| MORN2 | 5 | 0.4244 | 0.58376 | 0.999994 | 12322 | 2 | 0.33626 | 0.44015 | 0.66713 | 1 | 13858 | 2 | 0.33626 |
| TNRC6A | 3 | 0.28638 | 0.41974 | 0.999994 | 9842 | 1 | 0.89436 | 0.44022 | 0.59457 | 1 | 13859 | 2 | 0.89436 |
| ARHGDIG | 4 | 0.79053 | 0.79282 | 0.999994 | 17356 | 1 | 0.8645 | 0.44024 | 0.61895 | 1 | 13860 | 2 | 0.8645 |
| hsa-mir-4285 | 3 | 0.7455 | 0.74545 | 0.999994 | 16807 | 0 | 0.74361 | 0.44048 | 0.59483 | 1 | 13861 | 2 | 0.74361 |
| CNPY3 | 6 | 0.33234 | 0.54395 | 0.999994 | 10771 | 2 | 0.46701 | 0.4405 | 0.68189 | 1 | 13862 | 3 | 0.46701 |
| C3orf37 | 5 | 0.87943 | 0.87973 | 0.999994 | 18819 | 1 | 0.47136 | 0.4405 | 0.66748 | 1 | 13863 | 2 | 0.47136 |
| GPR180 | 5 | 0.52873 | 0.67294 | 0.999994 | 13878 | 2 | -0.48068 | 0.4405 | 0.66748 | 1 | 13864 | 2 | -0.48068 |
| POP4 | 5 | 0.50249 | 0.6501 | 0.999994 | 13470 | 2 | -0.06455 | 0.4405 | 0.66748 | 1 | 13865 | 2 | -0.06455 |
| OR5M1 | 5 | 0.20812 | 0.38672 | 0.999994 | 7980 | 3 | -0.74147 | 0.4405 | 0.66748 | 1 | 13866 | 2 | -0.74147 |
| TECRL | 5 | 0.44509 | 0.60109 | 0.999994 | 12647 | 2 | 0.49532 | 0.4405 | 0.66748 | 1 | 13867 | 2 | 0.49532 |
| TSPAN6 | 5 | 0.24036 | 0.43141 | 0.999994 | 8780 | 2 | -0.53788 | 0.4405 | 0.66748 | 1 | 13868 | 2 | -0.53788 |
| CCDC169 | 5 | 0.17769 | 0.34331 | 0.999994 | 7172 | 3 | -0.93272 | 0.4405 | 0.66748 | 1 | 13869 | 2 | -0.93272 |
| HFE2 | 6 | 0.27787 | 0.48093 | 0.999994 | 9649 | 3 | -0.69092 | 0.4406 | 0.68196 | 1 | 13870 | 2 | -0.69092 |
| STAU1 | 4 | 0.71372 | 0.73369 | 0.999994 | 16437 | 1 | 0.93355 | 0.44067 | 0.61927 | 1 | 13871 | 2 | 0.93355 |
| HNF4A | 5 | 0.38167 | 0.5481 | 0.999994 | 11671 | 1 | 0.89391 | 0.44071 | 0.66766 | 1 | 13872 | 3 | 0.89391 |
| MMP16 | 6 | 0.34021 | 0.55288 | 0.999994 | 10937 | 3 | 0.25655 | 0.44071 | 0.68205 | 1 | 13873 | 3 | 0.25655 |
| DPF1 | 6 | 0.93951 | 0.93946 | 0.999994 | 20063 | 0 | 0.61458 | 0.44071 | 0.68205 | 1 | 13874 | 3 | 0.61458 |
| REG3A | 6 | 0.017462 | 0.063741 | 0.937389 | 1336 | 3 | -1.5368 | 0.44078 | 0.6821 | 1 | 13875 | 3 | -1.5368 |
| TCEAL8 | 5 | 0.25124 | 0.44175 | 0.999994 | 9041 | 2 | 0.73306 | 0.44083 | 0.66778 | 1 | 13876 | 3 | 0.73306 |
| CHRNB1 | 4 | 0.031727 | 0.085019 | 0.94395 | 2091 | 3 | -2.1745 | 0.44086 | 0.61942 | 1 | 13877 | 1 | -2.1745 |
| GLI1 | 4 | 0.20451 | 0.34529 | 0.999994 | 7891 | 1 | 0.81297 | 0.44086 | 0.61942 | 1 | 13878 | 2 | 0.81297 |
| SSTR1 | 6 | 0.13966 | 0.31259 | 0.999994 | 6176 | 3 | -0.48328 | 0.4409 | 0.68218 | 1 | 13879 | 3 | -0.48328 |
| PPP2R1B | 6 | 0.51714 | 0.68056 | 0.999994 | 13695 | 2 | -0.17253 | 0.4409 | 0.68218 | 1 | 13880 | 2 | -0.17253 |
| GATA1 | 6 | 0.056629 | 0.15848 | 0.967126 | 3279 | 2 | 0.37779 | 0.4409 | 0.68218 | 1 | 13881 | 3 | 0.37779 |
| ESPNL | 6 | 0.077066 | 0.20142 | 0.977164 | 4107 | 3 | -0.49639 | 0.4409 | 0.68218 | 1 | 13882 | 3 | -0.49639 |
| B3GAT1 | 4 | 0.0027772 | 0.009962 | 0.680529 | 339 | 2 | -2.9709 | 0.44092 | 0.61946 | 1 | 13883 | 2 | -2.9709 |

| | | | | | | | | | | | | |
|---|---|---|---|---|---|---|---|---|---|---|---|---|
| TP53INP1 | 6 | 0.11329 | 0.27303 | 0.998561 | 5416 | 2 | 0.34789 | 0.441 | 0.68226 | 1 | 13884 | 3 | 0.34789 |
| FOXI1 | 6 | 0.16933 | 0.34992 | 0.999994 | 6966 | 3 | -0.082912 | 0.44114 | 0.68237 | 1 | 13885 | 3 | -0.082912 |
| NBL1 | 2 | 0.40324 | 0.41948 | 0.999994 | 12005 | 1 | 0.92053 | 0.44122 | 0.49608 | 1 | 13886 | 1 | 0.92053 |
| TP53BP2 | 6 | 0.48395 | 0.65785 | 0.999994 | 13199 | 2 | 0.90725 | 0.44128 | 0.68247 | 1 | 13887 | 3 | 0.90725 |
| POLH | 6 | 0.12933 | 0.29945 | 0.999994 | 5924 | 3 | -0.13638 | 0.44129 | 0.68249 | 1 | 13888 | 3 | -0.13638 |
| PNPLA5 | 5 | 0.014273 | 0.048135 | 0.90085 | 1147 | 2 | 0.76357 | 0.44132 | 0.66823 | 1 | 13889 | 3 | 0.76357 |
| LHX8 | 4 | 0.014967 | 0.04904 | 0.904562 | 1186 | 2 | -3.1348 | 0.44153 | 0.61993 | 1 | 13890 | 2 | -3.1348 |
| TNFAIP8L3 | 6 | 0.21675 | 0.40824 | 0.999994 | 8186 | 3 | 0.13326 | 0.44155 | 0.68267 | 1 | 13891 | 3 | 0.13326 |
| ZMYND15 | 5 | 0.81528 | 0.82947 | 0.999994 | 17703 | 1 | -0.46942 | 0.44156 | 0.66846 | 1 | 13892 | 2 | -0.46942 |
| IPO7 | 6 | 0.0034114 | 0.015243 | 0.757255 | 396 | 4 | -5.1063 | 0.44161 | 0.68272 | 1 | 13893 | 1 | -5.1063 |
| HTR3C | 6 | 0.24903 | 0.44678 | 0.999994 | 8992 | 3 | -0.22968 | 0.44161 | 0.68272 | 1 | 13894 | 3 | -0.22968 |
| RBPMS | 5 | 0.21283 | 0.39336 | 0.999994 | 8084 | 3 | -0.73772 | 0.44161 | 0.66852 | 1 | 13895 | 2 | -0.73772 |
| TIMM44 | 4 | 0.18509 | 0.32534 | 0.999994 | 7343 | 2 | -0.044324 | 0.44195 | 0.62023 | 1 | 13896 | 2 | -0.044324 |
| CDC45 | 2 | 0.029684 | 0.054417 | 0.922677 | 1992 | 1 | -3.1502 | 0.44196 | 0.49658 | 1 | 13897 | 1 | -3.1502 |
| NonTargetingControlG | 2 | 0.044312 | 0.079667 | 0.93914 | 2727 | 1 | -2.9141 | 0.44196 | 0.49658 | 1 | 13898 | 1 | -2.9141 |
| PCDHA9 | 2 | 0.46502 | 0.47257 | 0.999994 | 12933 | 1 | 1.1015 | 0.44196 | 0.49658 | 1 | 13899 | 1 | 1.1015 |
| ROS1 | 5 | 0.39446 | 0.55896 | 0.999994 | 11864 | 1 | 0.6835 | 0.44197 | 0.66887 | 1 | 13900 | 3 | 0.6835 |
| ZDHHC24 | 5 | 0.34134 | 0.51483 | 0.999994 | 10968 | 1 | 0.64981 | 0.44202 | 0.66892 | 1 | 13901 | 3 | 0.64981 |
| hsa-mir-4679-1 | 1 | 0.55779 | 0.55763 | 0.999994 | 14330 | 0 | 0.95361 | 0.44221 | 0.44232 | 1 | 13902 | 1 | 0.95361 |
| hsa-mir-4650-1 | 3 | 0.73828 | 0.7382 | 0.999994 | 16709 | 0 | 0.77559 | 0.44233 | 0.59671 | 1 | 13903 | 2 | 0.77559 |
| CDK2 | 5 | 0.20918 | 0.38824 | 0.999994 | 8010 | 3 | -0.75202 | 0.44234 | 0.66923 | 1 | 13904 | 2 | -0.75202 |
| MIPEP | 6 | 0.22249 | 0.41515 | 0.999994 | 8343 | 3 | -0.3115 | 0.44238 | 0.68333 | 1 | 13905 | 2 | -0.3115 |
| PCYT1B | 6 | 0.38941 | 0.59561 | 0.999994 | 11787 | 2 | 0.27526 | 0.44238 | 0.68333 | 1 | 13906 | 2 | 0.27526 |
| ORMDL3 | 6 | 0.92767 | 0.92775 | 0.999994 | 19805 | 1 | 0.4513 | 0.44243 | 0.68338 | 1 | 13907 | 3 | 0.4513 |
| ST6GALNAC2 | 6 | 0.93945 | 0.93941 | 0.999994 | 20061 | 0 | 0.61912 | 0.44245 | 0.68339 | 1 | 13908 | 3 | 0.61912 |
| IQCC | 5 | 0.14768 | 0.29929 | 0.999994 | 6397 | 1 | -0.11046 | 0.44246 | 0.66934 | 1 | 13909 | 2 | -0.11046 |
| GLT8D1 | 5 | 0.0022347 | 0.0097654 | 0.680529 | 296 | 4 | -4.1917 | 0.44247 | 0.66936 | 1 | 13910 | 1 | -4.1917 |
| ANKHD1 | 5 | 0.3237 | 0.50034 | 0.999994 | 10588 | 2 | -0.57557 | 0.44248 | 0.66936 | 1 | 13911 | 2 | -0.57557 |
| MAP6 | 6 | 0.45483 | 0.63827 | 0.999994 | 12769 | 1 | 0.95992 | 0.44251 | 0.68344 | 1 | 13912 | 3 | 0.95992 |
| NonTargetingControlG | 2 | 0.046541 | 0.083363 | 0.94383 | 2833 | 1 | -2.9908 | 0.44275 | 0.49714 | 1 | 13913 | 1 | -2.9908 |
| RMDN2 | 5 | 0.087385 | 0.20641 | 0.978744 | 4491 | 2 | 0.7955 | 0.44276 | 0.66962 | 1 | 13914 | 3 | 0.7955 |
| SULT1A4 | 3 | 0.058226 | 0.12704 | 0.951052 | 3348 | 2 | -3.2596 | 0.44295 | 0.59732 | 1 | 13915 | 1 | -3.2596 |
| CMA1 | 5 | 0.20481 | 0.38205 | 0.999994 | 7895 | 2 | 0.77675 | 0.44295 | 0.66981 | 1 | 13916 | 3 | 0.77675 |
| ANKRA2 | 5 | 0.91199 | 0.912 | 0.999994 | 19461 | 0 | 0.61065 | 0.443 | 0.66986 | 1 | 13917 | 3 | 0.61065 |
| DGKI | 5 | 0.95212 | 0.95227 | 0.999994 | 20384 | 0 | 0.64992 | 0.44309 | 0.66994 | 1 | 13918 | 3 | 0.64992 |
| SDC3 | 6 | 0.61401 | 0.75036 | 0.999994 | 15130 | 1 | 0.92506 | 0.44311 | 0.68392 | 1 | 13919 | 3 | 0.92506 |
| ILKAP | 6 | 0.64962 | 0.77733 | 0.999994 | 15652 | 2 | 0.4206 | 0.44321 | 0.68399 | 1 | 13920 | 3 | 0.4206 |
| SDHAF1 | 5 | 0.55527 | 0.69647 | 0.999994 | 14295 | 2 | 0.5297 | 0.44322 | 0.67007 | 1 | 13921 | 3 | 0.5297 |
| CDKL2 | 6 | 0.67283 | 0.79536 | 0.999994 | 15969 | 2 | 0.44426 | 0.44327 | 0.68404 | 1 | 13922 | 3 | 0.44426 |
| FAXC | 5 | 0.18929 | 0.35993 | 0.999994 | 7462 | 2 | 0.64313 | 0.44336 | 0.6702 | 1 | 13923 | 3 | 0.64313 |
| JAM3 | 3 | 0.66551 | 0.67005 | 0.999994 | 15864 | 1 | 0.60232 | 0.44341 | 0.59778 | 1 | 13924 | 2 | 0.60232 |
| OXR1 | 6 | 0.34469 | 0.55806 | 0.999994 | 11046 | 2 | 0.9312 | 0.44342 | 0.68414 | 1 | 13925 | 3 | 0.9312 |
| GPC2 | 5 | 0.78866 | 0.81252 | 0.999994 | 17327 | 1 | 0.66674 | 0.44346 | 0.6703 | 1 | 13926 | 3 | 0.66674 |
| NonTargetingControlG | 2 | 0.050735 | 0.090393 | 0.94395 | 3004 | 1 | -2.8792 | 0.44353 | 0.49768 | 1 | 13927 | 1 | -2.8792 |
| OLFM4 | 4 | 0.80332 | 0.80418 | 0.999994 | 17530 | 1 | 0.63055 | 0.44365 | 0.62152 | 1 | 13928 | 2 | 0.63055 |
| hsa-mir-4669 | 4 | 0.13766 | 0.2765 | 0.999379 | 6126 | 2 | -0.58897 | 0.44365 | 0.62152 | 1 | 13929 | 2 | -0.58897 |
| hsa-mir-337 | 4 | 0.65186 | 0.69435 | 0.999994 | 15685 | 1 | 0.6881 | 0.44365 | 0.62152 | 1 | 13930 | 2 | 0.6881 |
| MRPS36 | 5 | 0.90372 | 0.90381 | 0.999994 | 19299 | 0 | 0.76735 | 0.44371 | 0.67054 | 1 | 13931 | 3 | 0.76735 |
| LEPREL2 | 5 | 0.82196 | 0.83388 | 0.999994 | 17802 | 1 | 0.13929 | 0.44383 | 0.67066 | 1 | 13932 | 2 | 0.13929 |
| DCT | 6 | 0.079812 | 0.20691 | 0.978744 | 4225 | 4 | -1.6117 | 0.44383 | 0.68446 | 1 | 13933 | 2 | -1.6117 |
| BORA | 6 | 0.10747 | 0.26183 | 0.995265 | 5225 | 4 | -0.8577 | 0.44383 | 0.68446 | 1 | 13934 | 2 | -0.8577 |
| MBD6 | 6 | 0.82926 | 0.86118 | 0.999994 | 17917 | 1 | -0.16407 | 0.44383 | 0.68446 | 1 | 13935 | 2 | -0.16407 |

| | | | | | | | | | | | | | |
|---|---|---|---|---|---|---|---|---|---|---|---|---|---|
| OR3A1 | 6 | 0.14493 | 0.31928 | 0.999994 | 6323 | 2 | 0.79428 | 0.44383 | 0.68446 | 1 | 13936 | 3 | 0.79428 |
| HFE | 6 | 0.12799 | 0.29773 | 0.999994 | 5876 | 3 | -0.49622 | 0.44383 | 0.68446 | 1 | 13937 | 2 | -0.49622 |
| CYP4V2 | 5 | 0.89196 | 0.89211 | 0.999994 | 19073 | 0 | 0.69964 | 0.44388 | 0.6707 | 1 | 13938 | 3 | 0.69964 |
| CHMP6 | 3 | 0.15198 | 0.25101 | 0.993091 | 6505 | 2 | -1.6874 | 0.44394 | 0.59831 | 1 | 13939 | 1 | -1.6874 |
| LOC642778 | 1 | 0.55602 | 0.55586 | 0.999994 | 14304 | 0 | 0.83657 | 0.44398 | 0.44408 | 1 | 13940 | 1 | 0.83657 |
| GCKR | 6 | 0.11454 | 0.27539 | 0.999239 | 5464 | 4 | -1.2326 | 0.44412 | 0.68468 | 1 | 13941 | 2 | -1.2326 |
| OTUD6A | 6 | 0.10335 | 0.25387 | 0.994632 | 5077 | 3 | -0.45679 | 0.44412 | 0.68468 | 1 | 13942 | 2 | -0.45679 |
| CCDC82 | 6 | 0.18404 | 0.36818 | 0.999994 | 7317 | 3 | -0.87281 | 0.44412 | 0.68468 | 1 | 13943 | 1 | -0.87281 |
| KATNBL1 | 5 | 0.3125 | 0.49121 | 0.999994 | 10370 | 1 | -0.086171 | 0.44423 | 0.67104 | 1 | 13944 | 2 | -0.086171 |
| SCN2B | 6 | 0.19543 | 0.38234 | 0.999994 | 7647 | 1 | 0.44928 | 0.44425 | 0.68478 | 1 | 13945 | 3 | 0.44928 |
| LRRC16B | 5 | 0.10333 | 0.23159 | 0.984454 | 5076 | 1 | 0.72423 | 0.44434 | 0.67115 | 1 | 13946 | 3 | 0.72423 |
| NonTargetingControlG | 2 | 0.37991 | 0.40023 | 0.999994 | 11647 | 1 | 0.7283 | 0.44445 | 0.49832 | 1 | 13947 | 1 | 0.7283 |
| WDR52 | 6 | 0.35907 | 0.57426 | 0.999994 | 11321 | 3 | 0.41072 | 0.4445 | 0.68498 | 1 | 13948 | 3 | 0.41072 |
| CDC5L | 4 | 0.24849 | 0.39037 | 0.999994 | 8978 | 2 | -0.7845 | 0.44462 | 0.62227 | 1 | 13949 | 1 | -0.7845 |
| HHATL | 4 | 0.071043 | 0.16193 | 0.967126 | 3877 | 3 | -1.5884 | 0.44462 | 0.62227 | 1 | 13950 | 1 | -1.5884 |
| GPATCH3 | 5 | 0.47932 | 0.63005 | 0.999994 | 13135 | 2 | 0.72979 | 0.44464 | 0.67144 | 1 | 13951 | 3 | 0.72979 |
| FEM1A | 6 | 0.25659 | 0.45582 | 0.999994 | 9162 | 2 | -0.56317 | 0.44465 | 0.68511 | 1 | 13952 | 2 | -0.56317 |
| KLHL40 | 6 | 0.52451 | 0.68566 | 0.999994 | 13810 | 2 | 0.42871 | 0.44469 | 0.68514 | 1 | 13953 | 2 | 0.42871 |
| BBS5 | 5 | 0.034509 | 0.10203 | 0.946414 | 2241 | 2 | -0.39152 | 0.4447 | 0.6715 | 1 | 13954 | 2 | -0.39152 |
| MXRA8 | 4 | 0.20897 | 0.34992 | 0.999994 | 8001 | 2 | -0.031451 | 0.44475 | 0.62237 | 1 | 13955 | 2 | -0.031451 |
| REEP6 | 5 | 0.028589 | 0.086842 | 0.94395 | 1929 | 2 | 0.38894 | 0.44479 | 0.67159 | 1 | 13956 | 2 | 0.38894 |
| NCMAP | 5 | 0.0087612 | 0.032494 | 0.865899 | 777 | 3 | -0.78009 | 0.4448 | 0.6716 | 1 | 13957 | 2 | -0.78009 |
| IL6ST | 5 | 0.008293 | 0.031137 | 0.862223 | 749 | 3 | -1.3317 | 0.4448 | 0.6716 | 1 | 13958 | 2 | -1.3317 |
| SSTR3 | 5 | 0.1928 | 0.36494 | 0.999994 | 7567 | 2 | 0.45714 | 0.4448 | 0.6716 | 1 | 13959 | 2 | 0.45714 |
| KRTAP4-5 | 5 | 0.70711 | 0.76973 | 0.999994 | 16384 | 1 | 0.73795 | 0.44483 | 0.67162 | 1 | 13960 | 3 | 0.73795 |
| STRBP | 6 | 0.22597 | 0.41926 | 0.999994 | 8424 | 2 | 0.94293 | 0.44493 | 0.68532 | 1 | 13961 | 3 | 0.94293 |
| SLC16A1 | 5 | 0.68407 | 0.75935 | 0.999994 | 16144 | 1 | 0.67688 | 0.44499 | 0.67176 | 1 | 13962 | 3 | 0.67688 |
| TMPRSS11A | 6 | 0.47076 | 0.64896 | 0.999994 | 13015 | 2 | 0.3647 | 0.44503 | 0.68539 | 1 | 13963 | 3 | 0.3647 |
| CDY2A | 5 | 0.92987 | 0.93015 | 0.999994 | 19849 | 0 | 0.69271 | 0.44504 | 0.67181 | 1 | 13964 | 3 | 0.69271 |
| SMG9 | 6 | 0.403 | 0.60434 | 0.999994 | 11999 | 2 | 0.15891 | 0.4451 | 0.68545 | 1 | 13965 | 2 | 0.15891 |
| RIPPLY3 | 3 | 0.56358 | 0.58999 | 0.999994 | 14442 | 1 | -0.35364 | 0.44521 | 0.59957 | 1 | 13966 | 1 | -0.35364 |
| SHCBP1 | 4 | 0.10361 | 0.22156 | 0.983093 | 5086 | 2 | -0.62063 | 0.44543 | 0.62288 | 1 | 13967 | 2 | -0.62063 |
| NME2 | 4 | 0.26531 | 0.4075 | 0.999994 | 9356 | 2 | -0.41785 | 0.44552 | 0.62295 | 1 | 13968 | 2 | -0.41785 |
| SYN3 | 5 | 0.1559 | 0.31155 | 0.999994 | 6624 | 2 | 0.061786 | 0.44553 | 0.67228 | 1 | 13969 | 2 | 0.061786 |
| TFR2 | 5 | 0.44311 | 0.59943 | 0.999994 | 12626 | 2 | 0.13173 | 0.44553 | 0.67228 | 1 | 13970 | 2 | 0.13173 |
| PRSS16 | 5 | 0.44973 | 0.60501 | 0.999994 | 12707 | 1 | 0.69031 | 0.44553 | 0.67228 | 1 | 13971 | 3 | 0.69031 |
| CD9 | 6 | 0.23997 | 0.43596 | 0.999994 | 8769 | 2 | 0.21967 | 0.44563 | 0.68586 | 1 | 13972 | 2 | 0.21967 |
| IAPP | 6 | 0.87266 | 0.88498 | 0.999994 | 18686 | 1 | -0.12578 | 0.44563 | 0.68586 | 1 | 13973 | 2 | -0.12578 |
| RIC3 | 6 | 0.086239 | 0.21996 | 0.980699 | 4461 | 4 | -1.1863 | 0.44563 | 0.68586 | 1 | 13974 | 2 | -1.1863 |
| VHL | 6 | 0.39365 | 0.59827 | 0.999994 | 11856 | 2 | 0.31991 | 0.44563 | 0.68586 | 1 | 13975 | 3 | 0.31991 |
| PDZRN3 | 5 | 0.22574 | 0.41131 | 0.999994 | 8417 | 2 | 0.74493 | 0.44588 | 0.67263 | 1 | 13976 | 3 | 0.74493 |
| INSC | 5 | 0.025711 | 0.079373 | 0.93914 | 1786 | 4 | -1.8144 | 0.44592 | 0.67266 | 1 | 13977 | 1 | -1.8144 |
| YWHAQ | 5 | 0.0037892 | 0.015972 | 0.770326 | 428 | 1 | 0.70368 | 0.44609 | 0.67281 | 1 | 13978 | 3 | 0.70368 |
| GAD2 | 6 | 0.20148 | 0.38967 | 0.999994 | 7806 | 3 | -0.43751 | 0.44612 | 0.68623 | 1 | 13979 | 2 | -0.43751 |
| LEP | 6 | 0.012354 | 0.0469 | 0.896157 | 1016 | 5 | -1.2812 | 0.44612 | 0.68623 | 1 | 13980 | 1 | -1.2812 |
| N4BP3 | 6 | 0.65942 | 0.78487 | 0.999994 | 15768 | 2 | 0.14122 | 0.44612 | 0.68623 | 1 | 13981 | 2 | 0.14122 |
| CADM3 | 6 | 0.27963 | 0.48299 | 0.999994 | 9696 | 3 | -0.15758 | 0.44612 | 0.68623 | 1 | 13982 | 3 | -0.15758 |
| TOP1MT | 6 | 0.95445 | 0.95451 | 0.999994 | 20442 | 0 | 0.60235 | 0.44612 | 0.68623 | 1 | 13983 | 3 | 0.60235 |
| BDNF | 6 | 0.069649 | 0.18604 | 0.969409 | 3819 | 1 | 0.88584 | 0.44625 | 0.68633 | 1 | 13984 | 3 | 0.88584 |
| NonTargetingControlG | 2 | 0.39031 | 0.40874 | 0.999994 | 11803 | 1 | 0.90393 | 0.44626 | 0.49957 | 1 | 13985 | 1 | 0.90393 |
| NonTargetingControlG | 2 | 0.045482 | 0.081576 | 0.93914 | 2788 | 1 | -2.9297 | 0.44626 | 0.49957 | 1 | 13986 | 1 | -2.9297 |
| NonTargetingControlG | 2 | 0.28766 | 0.32737 | 0.999994 | 9868 | 1 | 0.74458 | 0.44626 | 0.49957 | 1 | 13987 | 1 | 0.74458 |

| | | | | | | | | | | | | | |
|---|---|---|---|---|---|---|---|---|---|---|---|---|---|
| ZNF479 | 4 | 0.14793 | 0.28704 | 0.999632 | 6405 | 1 | 0.040867 | 0.44633 | 0.62358 | 1 | 13988 | 1 | 0.040867 |
| PLIN1 | 3 | 0.42698 | 0.50028 | 0.999994 | 12365 | 1 | 0.27224 | 0.44636 | 0.60071 | 1 | 13989 | 1 | 0.27224 |
| RPL41 | 3 | 0.13732 | 0.2321 | 0.984502 | 6115 | 2 | -1.5802 | 0.44636 | 0.60071 | 1 | 13990 | 1 | -1.5802 |
| hsa-mir-8060 | 3 | 0.19524 | 0.30667 | 0.999994 | 7640 | 2 | -1.2488 | 0.44636 | 0.60071 | 1 | 13991 | 1 | -1.2488 |
| ZNF705A | 5 | 0.86755 | 0.8689 | 0.999994 | 18591 | 1 | 0.78553 | 0.44641 | 0.67312 | 1 | 13992 | 3 | 0.78553 |
| ZNF224 | 5 | 0.56296 | 0.70327 | 0.999994 | 14428 | 2 | 0.68156 | 0.44659 | 0.67331 | 1 | 13993 | 3 | 0.68156 |
| PURB | 5 | 0.28873 | 0.47188 | 0.999994 | 9891 | 2 | 0.75589 | 0.44664 | 0.67335 | 1 | 13994 | 3 | 0.75589 |
| MYO16 | 5 | 0.096143 | 0.22033 | 0.980699 | 4820 | 2 | 0.72351 | 0.44664 | 0.67335 | 1 | 13995 | 3 | 0.72351 |
| OR14C36 | 6 | 0.53685 | 0.69434 | 0.999994 | 14011 | 1 | 0.92319 | 0.44681 | 0.68676 | 1 | 13996 | 3 | 0.92319 |
| AKR7A3 | 4 | 0.23821 | 0.37985 | 0.999994 | 8732 | 1 | 0.53458 | 0.44685 | 0.62399 | 1 | 13997 | 2 | 0.53458 |
| hsa-mir-3919 | 3 | 0.47797 | 0.53185 | 0.999994 | 13114 | 1 | 0.7703 | 0.44686 | 0.60121 | 1 | 13998 | 2 | 0.7703 |
| hsa-mir-6750 | 2 | 0.072318 | 0.12612 | 0.951052 | 3931 | 1 | -2.7945 | 0.4469 | 0.50002 | 1 | 13999 | 1 | -2.7945 |
| TEX19 | 5 | 0.45693 | 0.61111 | 0.999994 | 12804 | 2 | 0.34056 | 0.44704 | 0.67374 | 1 | 14000 | 1 | 0.34056 |
| ANO4 | 5 | 0.03964 | 0.115 | 0.951052 | 2498 | 2 | -0.057011 | 0.44704 | 0.67374 | 1 | 14001 | 2 | -0.057011 |
| ADAMTS5 | 6 | 0.10669 | 0.26033 | 0.994943 | 5194 | 3 | -0.28329 | 0.44704 | 0.68694 | 1 | 14002 | 3 | -0.28329 |
| OR2H2 | 6 | 0.11298 | 0.27245 | 0.998561 | 5404 | 2 | 0.73933 | 0.44712 | 0.68701 | 1 | 14003 | 3 | 0.73933 |
| B3GALT4 | 5 | 0.19423 | 0.36698 | 0.999994 | 7610 | 1 | 0.28016 | 0.44715 | 0.67384 | 1 | 14004 | 2 | 0.28016 |
| ADAT1 | 5 | 0.34256 | 0.51582 | 0.999994 | 10995 | 2 | 0.71931 | 0.44715 | 0.67384 | 1 | 14005 | 3 | 0.71931 |
| NonTargetingControlG | 2 | 0.75041 | 0.74999 | 0.999994 | 16859 | 0 | 1.7761 | 0.44728 | 0.50028 | 1 | 14006 | 1 | 1.7761 |
| C15orf48 | 6 | 0.84324 | 0.86834 | 0.999994 | 18143 | 1 | 0.068656 | 0.44734 | 0.68717 | 1 | 14007 | 2 | 0.068656 |
| SATL1 | 6 | 0.95789 | 0.958 | 0.999994 | 20536 | 0 | 0.77925 | 0.44741 | 0.68722 | 1 | 14008 | 3 | 0.77925 |
| ACADSB | 5 | 0.5723 | 0.71156 | 0.999994 | 14580 | 2 | 0.71532 | 0.4476 | 0.67429 | 1 | 14009 | 3 | 0.71532 |
| PPP1CB | 3 | 0.58801 | 0.60799 | 0.999994 | 14791 | 1 | 0.77081 | 0.44765 | 0.60201 | 1 | 14010 | 2 | 0.77081 |
| OR4K17 | 5 | 0.74284 | 0.7871 | 0.999994 | 16777 | 1 | 0.67087 | 0.44771 | 0.6744 | 1 | 14011 | 3 | 0.67087 |
| GPR179 | 5 | 0.70521 | 0.76887 | 0.999994 | 16364 | 1 | 0.5856 | 0.44775 | 0.67445 | 1 | 14012 | 3 | 0.5856 |
| hsa-mir-450a-2 | 4 | 0.74351 | 0.75514 | 0.999994 | 16783 | 1 | 0.74755 | 0.44784 | 0.62476 | 1 | 14013 | 2 | 0.74755 |
| hsa-mir-2276 | 3 | 0.16028 | 0.26178 | 0.995257 | 6747 | 1 | 0.71921 | 0.44802 | 0.60238 | 1 | 14014 | 2 | 0.71921 |
| SLC22A5 | 6 | 0.095301 | 0.23812 | 0.987253 | 4794 | 4 | -1.3485 | 0.44814 | 0.68779 | 1 | 14015 | 2 | -1.3485 |
| MYLK4 | 6 | 0.37567 | 0.58681 | 0.999994 | 11588 | 2 | -0.31071 | 0.44814 | 0.68779 | 1 | 14016 | 2 | -0.31071 |
| STK35 | 5 | 0.39962 | 0.56318 | 0.999994 | 11944 | 2 | 0.65585 | 0.44815 | 0.67483 | 1 | 14017 | 3 | 0.65585 |
| MAPK15 | 5 | 0.080461 | 0.19533 | 0.975797 | 4247 | 2 | 0.081678 | 0.4482 | 0.67488 | 1 | 14018 | 2 | 0.081678 |
| TNFRSF6B | 5 | 0.34134 | 0.51483 | 0.999994 | 10967 | 1 | -0.043393 | 0.4482 | 0.67488 | 1 | 14019 | 2 | -0.043393 |
| C20orf111 | 6 | 0.65643 | 0.78254 | 0.999994 | 15735 | 1 | 0.66926 | 0.44859 | 0.68813 | 1 | 14020 | 3 | 0.66926 |
| PANK3 | 4 | 0.16722 | 0.30691 | 0.999994 | 6914 | 1 | 0.91218 | 0.44862 | 0.62536 | 1 | 14021 | 2 | 0.91218 |
| hsa-mir-6852 | 4 | 0.097407 | 0.21047 | 0.979587 | 4870 | 3 | -1.5205 | 0.44862 | 0.62536 | 1 | 14022 | 1 | -1.5205 |
| COX4I2 | 4 | 0.24039 | 0.382 | 0.999994 | 8783 | 1 | 0.69604 | 0.44862 | 0.62536 | 1 | 14023 | 2 | 0.69604 |
| hsa-mir-6855 | 3 | 0.16674 | 0.27023 | 0.998561 | 6902 | 2 | -1.2016 | 0.44889 | 0.60325 | 1 | 14024 | 1 | -1.2016 |
| OPRM1 | 6 | 0.16657 | 0.34644 | 0.999994 | 6898 | 3 | -0.39832 | 0.4489 | 0.68838 | 1 | 14025 | 2 | -0.39832 |
| ADAM10 | 6 | 0.19468 | 0.38141 | 0.999994 | 7624 | 3 | -0.73304 | 0.4489 | 0.68838 | 1 | 14026 | 2 | -0.73304 |
| VSIG2 | 6 | 0.39291 | 0.5978 | 0.999994 | 11846 | 2 | 0.09231 | 0.4489 | 0.68838 | 1 | 14027 | 3 | 0.09231 |
| hsa-mir-138-2 | 4 | 0.11178 | 0.23602 | 0.985362 | 5360 | 2 | -0.32975 | 0.44891 | 0.62558 | 1 | 14028 | 2 | -0.32975 |
| ARL13B | 6 | 0.26282 | 0.46321 | 0.999994 | 9309 | 2 | 0.47213 | 0.44896 | 0.68843 | 1 | 14029 | 3 | 0.47213 |
| TUBA3C | 6 | 0.5623 | 0.71245 | 0.999994 | 14415 | 2 | 0.72654 | 0.44907 | 0.68852 | 1 | 14030 | 3 | 0.72654 |
| NEO1 | 5 | 0.89818 | 0.8983 | 0.999994 | 19185 | 0 | 0.60123 | 0.44908 | 0.67575 | 1 | 14031 | 3 | 0.60123 |
| BAGE2 | 6 | 0.76876 | 0.83537 | 0.999994 | 17071 | 1 | 0.58143 | 0.4491 | 0.68854 | 1 | 14032 | 3 | 0.58143 |
| KLF5 | 5 | 0.34783 | 0.52013 | 0.999994 | 11112 | 2 | 0.78946 | 0.44913 | 0.67581 | 1 | 14033 | 3 | 0.78946 |
| GRXCR1 | 5 | 0.021753 | 0.068807 | 0.93914 | 1562 | 2 | 0.60838 | 0.4492 | 0.67588 | 1 | 14034 | 3 | 0.60838 |
| GCC2 | 6 | 0.20271 | 0.39117 | 0.999994 | 7851 | 3 | -0.7141 | 0.44937 | 0.68875 | 1 | 14035 | 2 | -0.7141 |
| TCEB2 | 5 | 0.26079 | 0.44939 | 0.999994 | 9262 | 2 | 0.48795 | 0.44947 | 0.67613 | 1 | 14036 | 2 | 0.48795 |
| CPLX2 | 5 | 0.47684 | 0.6279 | 0.999994 | 13098 | 2 | 0.76068 | 0.44947 | 0.67613 | 1 | 14037 | 3 | 0.76068 |
| TSPAN15 | 5 | 0.23302 | 0.42139 | 0.999994 | 8597 | 2 | -0.1249 | 0.44947 | 0.67613 | 1 | 14038 | 2 | -0.1249 |
| TMEM184A | 5 | 0.048305 | 0.13637 | 0.956051 | 2898 | 3 | -2.1113 | 0.44947 | 0.67613 | 1 | 14039 | 2 | -2.1113 |

| | | | | | | | | | | | | |
|---|---|---|---|---|---|---|---|---|---|---|---|---|
| IGF2BP3 | 6 | 0.66532 | 0.78947 | 0.999994 | 15858 | 1 | 0.41431 | 0.44956 | 0.6889 | 1 | 14040 | 3 | 0.41431 |
| ZNF491 | 6 | 0.43037 | 0.62209 | 0.999994 | 12420 | 2 | -0.097563 | 0.44961 | 0.68893 | 1 | 14041 | 3 | -0.097563 |
| ZNF860 | 6 | 0.38062 | 0.58995 | 0.999994 | 11655 | 2 | 0.65557 | 0.44966 | 0.68898 | 1 | 14042 | 3 | 0.65557 |
| hsa-mir-5094 | 4 | 0.59676 | 0.66356 | 0.999994 | 14911 | 1 | 0.96651 | 0.44969 | 0.62619 | 1 | 14043 | 2 | 0.96651 |
| MSR1 | 5 | 0.90217 | 0.90227 | 0.999994 | 19255 | 0 | 0.51356 | 0.44981 | 0.67645 | 1 | 14044 | 2 | 0.51356 |
| GTF2H5 | 5 | 0.7344 | 0.78282 | 0.999994 | 16654 | 1 | 0.39094 | 0.44981 | 0.67645 | 1 | 14045 | 2 | 0.39094 |
| CEP135 | 5 | 0.19931 | 0.37423 | 0.999994 | 7744 | 3 | -0.75226 | 0.44981 | 0.67645 | 1 | 14046 | 1 | -0.75226 |
| TPD52L2 | 5 | 0.56238 | 0.70276 | 0.999994 | 14419 | 2 | 0.11162 | 0.44981 | 0.67645 | 1 | 14047 | 2 | 0.11162 |
| BAG2 | 6 | 0.49114 | 0.66267 | 0.999994 | 13291 | 2 | 0.28854 | 0.44992 | 0.68917 | 1 | 14048 | 3 | 0.28854 |
| ZNF253 | 4 | 0.2152 | 0.35633 | 0.999994 | 8147 | 1 | 0.27968 | 0.44995 | 0.62639 | 1 | 14049 | 2 | 0.27968 |
| hsa-mir-3124 | 3 | 0.73347 | 0.73333 | 0.999994 | 16648 | 0 | 0.68371 | 0.44995 | 0.60431 | 1 | 14050 | 2 | 0.68371 |
| TMEM25 | 5 | 0.19263 | 0.36469 | 0.999994 | 7563 | 1 | 0.80833 | 0.44996 | 0.67659 | 1 | 14051 | 3 | 0.80833 |
| GTF2IRD1 | 6 | 0.017465 | 0.063753 | 0.937389 | 1339 | 2 | 0.79953 | 0.45029 | 0.68945 | 1 | 14052 | 3 | 0.79953 |
| PLGLB2 | 1 | 0.54955 | 0.54941 | 0.999994 | 14204 | 0 | 1.0878 | 0.45045 | 0.45052 | 1 | 14053 | 1 | 1.0878 |
| CDY2B | 1 | 0.54955 | 0.54941 | 0.999994 | 14205 | 0 | 1.0878 | 0.45045 | 0.45052 | 1 | 14054 | 1 | 1.0878 |
| PGLYRP1 | 6 | 0.13415 | 0.3056 | 0.999994 | 6026 | 2 | -0.047997 | 0.45045 | 0.68957 | 1 | 14055 | 1 | -0.047997 |
| RSC1A1 | 6 | 0.18113 | 0.36457 | 0.999994 | 7250 | 3 | -0.71368 | 0.45045 | 0.68957 | 1 | 14056 | 1 | -0.71368 |
| SLC9A5 | 5 | 0.19771 | 0.37197 | 0.999994 | 7699 | 2 | -0.26385 | 0.4506 | 0.6772 | 1 | 14057 | 2 | -0.26385 |
| MEP1B | 5 | 0.37105 | 0.53927 | 0.999994 | 11530 | 2 | 0.60536 | 0.45072 | 0.67733 | 1 | 14058 | 3 | 0.60536 |
| hsa-mir-7844 | 3 | 0.79047 | 0.79038 | 0.999994 | 17354 | 0 | 0.81226 | 0.45081 | 0.60517 | 1 | 14059 | 2 | 0.81226 |
| NXN | 5 | 0.94105 | 0.94125 | 0.999994 | 20110 | 0 | 0.73994 | 0.45087 | 0.67748 | 1 | 14060 | 3 | 0.73994 |
| GLYATL3 | 6 | 0.07727 | 0.20181 | 0.977164 | 4120 | 4 | -1.174 | 0.45101 | 0.68997 | 1 | 14061 | 2 | -1.174 |
| EFS | 4 | 0.76707 | 0.77341 | 0.999994 | 17051 | 1 | 0.48797 | 0.45107 | 0.62722 | 1 | 14062 | 2 | 0.48797 |
| TAF1 | 4 | 0.097739 | 0.21104 | 0.979587 | 4879 | 2 | -1.5737 | 0.45107 | 0.62722 | 1 | 14063 | 1 | -1.5737 |
| hsa-mir-548az | 4 | 0.32058 | 0.46401 | 0.999994 | 10525 | 1 | 0.57062 | 0.45107 | 0.62722 | 1 | 14064 | 2 | 0.57062 |
| C18orf54 | 6 | 0.30554 | 0.51319 | 0.999994 | 10244 | 2 | 0.79827 | 0.45109 | 0.69003 | 1 | 14065 | 3 | 0.79827 |
| NPFFR2 | 5 | 0.16426 | 0.32369 | 0.999994 | 6846 | 3 | -1.0755 | 0.45112 | 0.67769 | 1 | 14066 | 2 | -1.0755 |
| OR6C2 | 5 | 0.53271 | 0.67643 | 0.999994 | 13947 | 2 | 0.69221 | 0.45135 | 0.6779 | 1 | 14067 | 3 | 0.69221 |
| PALM2-AKAP2 | 1 | 0.54865 | 0.5485 | 0.999994 | 14190 | 0 | 0.83053 | 0.45135 | 0.45144 | 1 | 14068 | 1 | 0.83053 |
| WT1 | 5 | 0.090431 | 0.21128 | 0.979587 | 4611 | 3 | -1.7071 | 0.45142 | 0.67796 | 1 | 14069 | 2 | -1.7071 |
| BAZ2B | 6 | 0.21769 | 0.40935 | 0.999994 | 8216 | 3 | 0.30608 | 0.45146 | 0.6903 | 1 | 14070 | 3 | 0.30608 |
| hsa-mir-5193 | 3 | 0.78047 | 0.78038 | 0.999994 | 17218 | 0 | 0.81064 | 0.45164 | 0.60602 | 1 | 14071 | 2 | 0.81064 |
| RAB8A | 5 | 0.38902 | 0.55432 | 0.999994 | 11783 | 2 | 0.67692 | 0.45166 | 0.6782 | 1 | 14072 | 3 | 0.67692 |
| LCK | 5 | 0.13483 | 0.27998 | 0.999632 | 6045 | 2 | 0.15495 | 0.45167 | 0.6782 | 1 | 14073 | 2 | 0.15495 |
| TAS2R42 | 6 | 0.20788 | 0.3975 | 0.999994 | 7972 | 3 | -0.011033 | 0.45178 | 0.69054 | 1 | 14074 | 3 | -0.011033 |
| OLIG2 | 5 | 0.1769 | 0.34214 | 0.999994 | 7153 | 3 | -0.95495 | 0.45179 | 0.67832 | 1 | 14075 | 2 | -0.95495 |
| TRMT10C | 5 | 0.20052 | 0.37598 | 0.999994 | 7774 | 1 | 0.70383 | 0.45182 | 0.67835 | 1 | 14076 | 3 | 0.70383 |
| MRPL12 | 2 | 0.49342 | 0.4979 | 0.999994 | 13315 | 1 | 1.118 | 0.45205 | 0.50365 | 1 | 14077 | 1 | 1.118 |
| GZF1 | 6 | 0.075504 | 0.19821 | 0.977164 | 4055 | 4 | -1.4639 | 0.45206 | 0.69075 | 1 | 14078 | 2 | -1.4639 |
| CD63 | 6 | 0.60196 | 0.74138 | 0.999994 | 14972 | 1 | 0.90147 | 0.4521 | 0.69079 | 1 | 14079 | 3 | 0.90147 |
| MPPED1 | 5 | 0.3262 | 0.50239 | 0.999994 | 10646 | 2 | 0.67424 | 0.4521 | 0.67861 | 1 | 14080 | 3 | 0.67424 |
| P2RX7 | 6 | 0.15149 | 0.32754 | 0.999994 | 6495 | 2 | 0.724 | 0.45217 | 0.69084 | 1 | 14081 | 3 | 0.724 |
| GOLPH3 | 5 | 0.70097 | 0.76692 | 0.999994 | 16316 | 1 | 0.59215 | 0.45224 | 0.67874 | 1 | 14082 | 3 | 0.59215 |
| hsa-mir-5700 | 4 | 0.37044 | 0.51533 | 0.999994 | 11519 | 2 | 0.54447 | 0.45228 | 0.62815 | 1 | 14083 | 2 | 0.54447 |
| UBXN6 | 5 | 0.87728 | 0.87769 | 0.999994 | 18779 | 1 | 0.45862 | 0.45239 | 0.67886 | 1 | 14084 | 2 | 0.45862 |
| DEFB116 | 5 | 0.56722 | 0.70711 | 0.999994 | 14500 | 2 | 0.57604 | 0.45239 | 0.67886 | 1 | 14085 | 2 | 0.57604 |
| SLC4A7 | 6 | 0.056144 | 0.1574 | 0.967126 | 3253 | 2 | 0.53883 | 0.4525 | 0.69108 | 1 | 14086 | 3 | 0.53883 |
| KCTD6 | 6 | 0.0014334 | 0.0071799 | 0.656035 | 214 | 5 | -2.9494 | 0.45251 | 0.69109 | 1 | 14087 | 1 | -2.9494 |
| KIAA1024 | 6 | 0.069089 | 0.18487 | 0.968526 | 3801 | 2 | 0.33129 | 0.45251 | 0.69109 | 1 | 14088 | 2 | 0.33129 |
| DENND4C | 6 | 0.322 | 0.5322 | 0.999994 | 10556 | 2 | 0.70769 | 0.45257 | 0.69114 | 1 | 14089 | 3 | 0.70769 |
| TTC5 | 5 | 0.024878 | 0.077144 | 0.93914 | 1750 | 3 | -2.6005 | 0.45269 | 0.67917 | 1 | 14090 | 2 | -2.6005 |
| PLAT | 6 | 0.91921 | 0.91969 | 0.999994 | 19613 | 1 | 0.71947 | 0.45276 | 0.69128 | 1 | 14091 | 3 | 0.71947 |

| | | | | | | | | | | | | | |
|---|---|---|---|---|---|---|---|---|---|---|---|---|---|
| IP6K3 | 6 | 0.031953 | 0.10338 | 0.946414 | 2102 | 3 | -0.070716 | 0.45279 | 0.69131 | 1 | 14092 | 3 | -0.070716 |
| MROH2B | 6 | 0.94736 | 0.94738 | 0.999994 | 20272 | 0 | 0.55832 | 0.45288 | 0.69138 | 1 | 14093 | 3 | 0.55832 |
| CPT1A | 6 | 0.064758 | 0.17576 | 0.968171 | 3641 | 4 | -1.2428 | 0.45288 | 0.69138 | 1 | 14094 | 2 | -1.2428 |
| FASTKD5 | 4 | 0.048607 | 0.1187 | 0.951052 | 2912 | 2 | -0.35043 | 0.45295 | 0.62864 | 1 | 14095 | 2 | -0.35043 |
| EFTUD2 | 3 | 0.020088 | 0.051772 | 0.910734 | 1463 | 1 | 0.78105 | 0.45295 | 0.60732 | 1 | 14096 | 2 | 0.78105 |
| KIAA1549 | 6 | 0.39881 | 0.60161 | 0.999994 | 11931 | 2 | 0.85412 | 0.453 | 0.69148 | 1 | 14097 | 3 | 0.85412 |
| NonTargetingControlG | 1 | 0.54692 | 0.54676 | 0.999994 | 14167 | 0 | 0.8549 | 0.45308 | 0.45314 | 1 | 14098 | 1 | 0.8549 |
| RBM22 | 4 | 0.14191 | 0.28087 | 0.999632 | 6237 | 1 | -0.70838 | 0.4531 | 0.62876 | 1 | 14099 | 1 | -0.70838 |
| FOXJ1 | 5 | 0.039023 | 0.11343 | 0.951052 | 2462 | 2 | 0.67123 | 0.45316 | 0.67961 | 1 | 14100 | 3 | 0.67123 |
| AVPR1A | 5 | 0.73341 | 0.78232 | 0.999994 | 16644 | 1 | 0.028438 | 0.45325 | 0.67969 | 1 | 14101 | 2 | 0.028438 |
| FBXO30 | 6 | 0.60551 | 0.74405 | 0.999994 | 15020 | 2 | 0.42029 | 0.45345 | 0.69181 | 1 | 14102 | 3 | 0.42029 |
| GPR55 | 6 | 0.46891 | 0.64772 | 0.999994 | 12989 | 2 | 0.011485 | 0.45348 | 0.69184 | 1 | 14103 | 2 | 0.011485 |
| CRAMP1L | 6 | 0.029746 | 0.098171 | 0.945253 | 1995 | 4 | -1.7884 | 0.45348 | 0.69184 | 1 | 14104 | 1 | -1.7884 |
| PTBP2 | 6 | 0.25279 | 0.45125 | 0.999994 | 9075 | 2 | 0.40204 | 0.45348 | 0.69184 | 1 | 14105 | 3 | 0.40204 |
| RNASEH2B | 5 | 0.71242 | 0.77223 | 0.999994 | 16425 | 1 | 0.69756 | 0.45349 | 0.67994 | 1 | 14106 | 3 | 0.69756 |
| SNX14 | 5 | 0.10415 | 0.23286 | 0.984502 | 5103 | 3 | -1.2409 | 0.45364 | 0.68008 | 1 | 14107 | 1 | -1.2409 |
| SLC16A6 | 5 | 0.61158 | 0.73038 | 0.999994 | 15098 | 1 | 0.53626 | 0.4539 | 0.68033 | 1 | 14108 | 3 | 0.53626 |
| CNGB1 | 5 | 0.16056 | 0.31836 | 0.999994 | 6753 | 2 | 0.74254 | 0.45397 | 0.68039 | 1 | 14109 | 3 | 0.74254 |
| CCDC177 | 5 | 0.40924 | 0.57126 | 0.999994 | 12097 | 1 | 0.69418 | 0.45399 | 0.6804 | 1 | 14110 | 3 | 0.69418 |
| PTCD2 | 5 | 0.16443 | 0.32394 | 0.999994 | 6850 | 1 | 0.82292 | 0.45411 | 0.68052 | 1 | 14111 | 3 | 0.82292 |
| TEX36 | 4 | 0.75522 | 0.76411 | 0.999994 | 16918 | 1 | 0.14212 | 0.45412 | 0.62952 | 1 | 14112 | 1 | 0.14212 |
| CFB | 6 | 0.16599 | 0.34569 | 0.999994 | 6884 | 3 | -0.39325 | 0.45415 | 0.69236 | 1 | 14113 | 3 | -0.39325 |
| CYB561D2 | 5 | 0.21384 | 0.39475 | 0.999994 | 8109 | 2 | 0.64159 | 0.45418 | 0.6806 | 1 | 14114 | 3 | 0.64159 |
| SOX7 | 5 | 0.75727 | 0.79468 | 0.999994 | 16944 | 1 | 0.61773 | 0.45423 | 0.68066 | 1 | 14115 | 3 | 0.61773 |
| TDP2 | 6 | 0.11183 | 0.27024 | 0.998561 | 5361 | 4 | -1.2822 | 0.45424 | 0.69242 | 1 | 14116 | 2 | -1.2822 |
| C17orf58 | 5 | 0.79466 | 0.81619 | 0.999994 | 17406 | 1 | 0.6227 | 0.45434 | 0.68076 | 1 | 14117 | 3 | 0.6227 |
| SLC26A8 | 6 | 0.43957 | 0.62812 | 0.999994 | 12568 | 2 | 0.57075 | 0.45434 | 0.69249 | 1 | 14118 | 3 | 0.57075 |
| SEC22C | 6 | 0.45498 | 0.63837 | 0.999994 | 12772 | 2 | 0.93118 | 0.45434 | 0.69249 | 1 | 14119 | 3 | 0.93118 |
| ZNF483 | 6 | 0.61187 | 0.7487 | 0.999994 | 15107 | 2 | 0.7647 | 0.45434 | 0.69249 | 1 | 14120 | 3 | 0.7647 |
| VWF | 5 | 0.08804 | 0.20743 | 0.978852 | 4516 | 2 | 0.66268 | 0.45439 | 0.68081 | 1 | 14121 | 3 | 0.66268 |
| NPC1L1 | 6 | 0.061847 | 0.16968 | 0.967126 | 3507 | 3 | -1.357 | 0.45451 | 0.69262 | 1 | 14122 | 1 | -1.357 |
| EMD | 6 | 0.26355 | 0.46405 | 0.999994 | 9327 | 2 | 0.62822 | 0.45451 | 0.69262 | 1 | 14123 | 3 | 0.62822 |
| FIGNL1 | 6 | 0.22213 | 0.41472 | 0.999994 | 8336 | 2 | 0.43782 | 0.45492 | 0.69294 | 1 | 14124 | 3 | 0.43782 |
| hsa-mir-3181 | 3 | 0.16498 | 0.26799 | 0.998561 | 6863 | 2 | -0.84079 | 0.45507 | 0.60953 | 1 | 14125 | 1 | -0.84079 |
| MSTO1 | 3 | 0.66895 | 0.67296 | 0.999994 | 15908 | 1 | 0.79242 | 0.45507 | 0.60953 | 1 | 14126 | 2 | 0.79242 |
| hsa-mir-28 | 3 | 0.070518 | 0.14368 | 0.961453 | 3853 | 2 | -2.9249 | 0.45507 | 0.60953 | 1 | 14127 | 1 | -2.9249 |
| SPTLC1 | 4 | 0.36605 | 0.51073 | 0.999994 | 11446 | 1 | 0.22867 | 0.45515 | 0.63033 | 1 | 14128 | 2 | 0.22867 |
| METTL25 | 5 | 0.70397 | 0.76831 | 0.999994 | 16351 | 1 | 0.066703 | 0.45535 | 0.6817 | 1 | 14129 | 2 | 0.066703 |
| IDS | 6 | 0.078615 | 0.20454 | 0.978186 | 4174 | 2 | 0.25929 | 0.45538 | 0.69327 | 1 | 14130 | 3 | 0.25929 |
| SLC26A9 | 6 | 0.0076492 | 0.030736 | 0.862223 | 702 | 5 | -3.4121 | 0.45538 | 0.69327 | 1 | 14131 | 1 | -3.4121 |
| ZNF280A | 6 | 0.0047403 | 0.020211 | 0.808999 | 502 | 4 | -4.7446 | 0.45538 | 0.69327 | 1 | 14132 | 2 | -4.7446 |
| PLA2G2D | 6 | 0.11267 | 0.27183 | 0.998561 | 5391 | 3 | -0.2836 | 0.45541 | 0.69329 | 1 | 14133 | 3 | -0.2836 |
| IL22 | 6 | 0.92849 | 0.92858 | 0.999994 | 19825 | 0 | 0.8013 | 0.45558 | 0.69344 | 1 | 14134 | 3 | 0.8013 |
| C1orf65 | 5 | 0.80034 | 0.81972 | 0.999994 | 17484 | 1 | 0.63287 | 0.45576 | 0.68209 | 1 | 14135 | 3 | 0.63287 |
| hsa-mir-8070 | 3 | 0.6052 | 0.62114 | 0.999994 | 15014 | 1 | 0.63099 | 0.4558 | 0.61028 | 1 | 14136 | 2 | 0.63099 |
| SLC6A17 | 6 | 0.0088348 | 0.034936 | 0.867348 | 781 | 3 | -3.0931 | 0.45587 | 0.69368 | 1 | 14137 | 3 | -3.0931 |
| FUS | 6 | 0.12277 | 0.29093 | 0.999632 | 5732 | 4 | -0.75484 | 0.45587 | 0.69368 | 1 | 14138 | 2 | -0.75484 |
| CXCL13 | 6 | 0.36272 | 0.57835 | 0.999994 | 11384 | 2 | 0.1825 | 0.45587 | 0.69368 | 1 | 14139 | 3 | 0.1825 |
| CLEC10A | 6 | 0.059232 | 0.16411 | 0.967126 | 3390 | 3 | -1.2625 | 0.45587 | 0.69368 | 1 | 14140 | 2 | -1.2625 |
| TMEM169 | 6 | 0.078265 | 0.20383 | 0.978186 | 4158 | 4 | -1.1928 | 0.45587 | 0.69368 | 1 | 14141 | 2 | -1.1928 |
| NDUFB3 | 6 | 0.031012 | 0.1012 | 0.946414 | 2061 | 3 | -0.77473 | 0.45594 | 0.69373 | 1 | 14142 | 3 | -0.77473 |
| hsa-mir-636 | 2 | 0.051295 | 0.091333 | 0.94395 | 3039 | 1 | -2.9466 | 0.45605 | 0.50645 | 1 | 14143 | 1 | -2.9466 |

| | | | | | | | | | | | | | |
|---|---|---|---|---|---|---|---|---|---|---|---|---|---|
| ZCCHC11 | 6 | 0.1241 | 0.29277 | 0.999823 | 5771 | 1 | 0.44 | 0.45606 | 0.69381 | 1 | 14144 | 3 | 0.44 |
| SGCE | 6 | 0.30581 | 0.51351 | 0.999994 | 10249 | 3 | -0.61913 | 0.45606 | 0.69381 | 1 | 14145 | 1 | -0.61913 |
| DSG2 | 6 | 0.50462 | 0.67194 | 0.999994 | 13506 | 2 | -0.087108 | 0.45606 | 0.69381 | 1 | 14146 | 2 | -0.087108 |
| COL9A1 | 6 | 0.33717 | 0.54946 | 0.999994 | 10876 | 3 | 0.26232 | 0.45606 | 0.69381 | 1 | 14147 | 3 | 0.26232 |
| SLC37A3 | 5 | 0.10968 | 0.24149 | 0.987253 | 5294 | 2 | 0.56408 | 0.45608 | 0.68239 | 1 | 14148 | 3 | 0.56408 |
| RGS7BP | 5 | 0.82753 | 0.83773 | 0.999994 | 17890 | 1 | -0.25505 | 0.45619 | 0.68249 | 1 | 14149 | 2 | -0.25505 |
| PTGIS | 5 | 0.27648 | 0.46202 | 0.999994 | 9629 | 1 | 0.65943 | 0.45636 | 0.68267 | 1 | 14150 | 3 | 0.65943 |
| QRFPR | 5 | 0.12502 | 0.26501 | 0.99736 | 5795 | 2 | 0.54567 | 0.4564 | 0.68272 | 1 | 14151 | 2 | 0.54567 |
| hsa-mir-4667 | 3 | 0.40571 | 0.48765 | 0.999994 | 12040 | 1 | 0.8496 | 0.45655 | 0.61104 | 1 | 14152 | 2 | 0.8496 |
| CRYL1 | 4 | 0.92664 | 0.92644 | 0.999994 | 19780 | 0 | 1.0216 | 0.45658 | 0.63142 | 1 | 14153 | 2 | 1.0216 |
| NonTargetingControlG | 2 | 0.68971 | 0.68952 | 0.999994 | 16207 | 0 | 1.5213 | 0.45661 | 0.50683 | 1 | 14154 | 1 | 1.5213 |
| PMFBP1 | 5 | 0.86664 | 0.86813 | 0.999994 | 18568 | 1 | 0.64174 | 0.45663 | 0.68293 | 1 | 14155 | 3 | 0.64174 |
| NMT1 | 3 | 0.029442 | 0.073883 | 0.93914 | 1980 | 2 | -6.0519 | 0.45664 | 0.61113 | 1 | 14156 | 1 | -6.0519 |
| RGPD5 | 3 | 0.67045 | 0.67424 | 0.999994 | 15937 | 1 | -0.17875 | 0.45664 | 0.61113 | 1 | 14157 | 1 | -0.17875 |
| ANKRD2 | 6 | 0.016187 | 0.059618 | 0.932932 | 1253 | 5 | -1.6836 | 0.45689 | 0.69443 | 1 | 14158 | 1 | -1.6836 |
| TRPV6 | 6 | 0.016408 | 0.060333 | 0.932932 | 1271 | 4 | -2.4682 | 0.45689 | 0.69443 | 1 | 14159 | 1 | -2.4682 |
| LCN2 | 6 | 0.013536 | 0.050907 | 0.909101 | 1103 | 3 | -2.7241 | 0.45689 | 0.69443 | 1 | 14160 | 3 | -2.7241 |
| PRPS1 | 6 | 0.89771 | 0.90186 | 0.999994 | 19173 | 1 | 0.47502 | 0.45689 | 0.69443 | 1 | 14161 | 3 | 0.47502 |
| SLC25A52 | 6 | 0.27495 | 0.47753 | 0.999994 | 9591 | 3 | -0.67322 | 0.45689 | 0.69443 | 1 | 14162 | 2 | -0.67322 |
| FAM63A | 6 | 0.80519 | 0.85011 | 0.999994 | 17554 | 1 | 0.83953 | 0.45691 | 0.69445 | 1 | 14163 | 3 | 0.83953 |
| STEAP4 | 6 | 0.3046 | 0.51209 | 0.999994 | 10220 | 3 | -0.34872 | 0.45703 | 0.69455 | 1 | 14164 | 2 | -0.34872 |
| PLCG1 | 5 | 0.2921 | 0.47463 | 0.999994 | 9955 | 2 | 0.48062 | 0.45718 | 0.68348 | 1 | 14165 | 2 | 0.48062 |
| ALDH7A1 | 6 | 0.34658 | 0.56016 | 0.999994 | 11085 | 2 | 0.51236 | 0.45719 | 0.69466 | 1 | 14166 | 2 | 0.51236 |
| P2RX2 | 4 | 0.87653 | 0.8762 | 0.999994 | 18760 | 0 | 0.29139 | 0.4572 | 0.63191 | 1 | 14167 | 1 | 0.29139 |
| C3orf38 | 5 | 0.50575 | 0.65294 | 0.999994 | 13518 | 1 | 0.80203 | 0.45728 | 0.68357 | 1 | 14168 | 3 | 0.80203 |
| ARSF | 6 | 0.555 | 0.70721 | 0.999994 | 14289 | 2 | -0.013879 | 0.45735 | 0.69478 | 1 | 14169 | 2 | -0.013879 |
| NonTargetingControlG | 1 | 0.5426 | 0.54238 | 0.999994 | 14105 | 0 | 0.80457 | 0.4574 | 0.45755 | 1 | 14170 | 1 | 0.80457 |
| ZNF619 | 6 | 0.030676 | 0.10037 | 0.946394 | 2041 | 4 | -1.8793 | 0.45766 | 0.69502 | 1 | 14171 | 1 | -1.8793 |
| C1orf185 | 6 | 0.012386 | 0.047012 | 0.896157 | 1021 | 4 | -1.2269 | 0.45766 | 0.69502 | 1 | 14172 | 2 | -1.2269 |
| DENND1B | 6 | 0.2153 | 0.40652 | 0.999994 | 8150 | 3 | -0.76406 | 0.45766 | 0.69502 | 1 | 14173 | 1 | -0.76406 |
| CASP10 | 6 | 0.061147 | 0.16822 | 0.967126 | 3474 | 3 | 0.26271 | 0.4577 | 0.69505 | 1 | 14174 | 3 | 0.26271 |
| TEK | 3 | 0.75734 | 0.75733 | 0.999994 | 16946 | 0 | 0.54255 | 0.4578 | 0.6123 | 1 | 14175 | 1 | 0.54255 |
| ZNF480 | 5 | 0.59932 | 0.72591 | 0.999994 | 14944 | 1 | -0.022901 | 0.45783 | 0.68407 | 1 | 14176 | 2 | -0.022901 |
| MAGEA4 | 5 | 0.89012 | 0.89025 | 0.999994 | 19039 | 0 | -0.12389 | 0.45783 | 0.68407 | 1 | 14177 | 2 | -0.12389 |
| ARFRP1 | 6 | 0.23859 | 0.4343 | 0.999994 | 8738 | 2 | 0.82222 | 0.45795 | 0.69525 | 1 | 14178 | 3 | 0.82222 |
| hsa-mir-298 | 4 | 0.80563 | 0.80633 | 0.999994 | 17563 | 1 | 0.68173 | 0.45805 | 0.63256 | 1 | 14179 | 2 | 0.68173 |
| C3orf83 | 5 | 0.39196 | 0.55679 | 0.999994 | 11831 | 2 | 0.64233 | 0.45809 | 0.68433 | 1 | 14180 | 3 | 0.64233 |
| CHN1 | 5 | 0.26839 | 0.45554 | 0.999994 | 9431 | 1 | 0.5484 | 0.45826 | 0.68449 | 1 | 14181 | 3 | 0.5484 |
| RETNLB | 6 | 0.50019 | 0.66883 | 0.999994 | 13426 | 1 | 0.56896 | 0.4584 | 0.69561 | 1 | 14182 | 3 | 0.56896 |
| ABAT | 5 | 0.55249 | 0.69395 | 0.999994 | 14251 | 2 | 0.63645 | 0.45842 | 0.68464 | 1 | 14183 | 3 | 0.63645 |
| DOHH | 5 | 0.069659 | 0.17782 | 0.968171 | 3820 | 2 | 0.65574 | 0.45844 | 0.68466 | 1 | 14184 | 3 | 0.65574 |
| PGF | 6 | 0.47735 | 0.65342 | 0.999994 | 13104 | 2 | 0.47874 | 0.45847 | 0.69566 | 1 | 14185 | 3 | 0.47874 |
| CRYAB | 6 | 0.65675 | 0.78279 | 0.999994 | 15740 | 2 | 0.32697 | 0.45847 | 0.69566 | 1 | 14186 | 3 | 0.32697 |
| hsa-mir-3610 | 4 | 0.094178 | 0.20462 | 0.978186 | 4758 | 2 | -0.63329 | 0.45862 | 0.63301 | 1 | 14187 | 2 | -0.63329 |
| ADAM30 | 5 | 0.79146 | 0.81419 | 0.999994 | 17372 | 1 | 0.63016 | 0.45876 | 0.68495 | 1 | 14188 | 3 | 0.63016 |
| WIPI1 | 6 | 0.88803 | 0.89492 | 0.999994 | 18999 | 1 | -0.24164 | 0.45878 | 0.6959 | 1 | 14189 | 2 | -0.24164 |
| ANXA2 | 5 | 0.8876 | 0.88769 | 0.999994 | 18989 | 1 | 0.68739 | 0.45881 | 0.685 | 1 | 14190 | 3 | 0.68739 |
| SPNS1 | 5 | 0.36555 | 0.53468 | 0.999994 | 11431 | 2 | 0.71274 | 0.4589 | 0.68508 | 1 | 14191 | 3 | 0.71274 |
| CSNK1E | 5 | 0.14192 | 0.29073 | 0.999632 | 6238 | 2 | 0.26406 | 0.45891 | 0.68509 | 1 | 14192 | 2 | 0.26406 |
| VAMP8 | 5 | 0.10041 | 0.22707 | 0.983354 | 4984 | 2 | 0.28636 | 0.45891 | 0.68509 | 1 | 14193 | 2 | 0.28636 |
| MST1L | 4 | 0.48604 | 0.61016 | 0.999994 | 13230 | 1 | 0.31052 | 0.45899 | 0.63331 | 1 | 14194 | 1 | 0.31052 |
| SLC38A8 | 6 | 0.12671 | 0.29613 | 0.999994 | 5838 | 2 | -0.16833 | 0.45908 | 0.69613 | 1 | 14195 | 2 | -0.16833 |

| | | | | | | | | | | | | | |
|---|---|---|---|---|---|---|---|---|---|---|---|---|---|
| RAD17 | 4 | 0.088755 | 0.19473 | 0.975797 | 4543 | 2 | -0.0063667 | 0.45909 | 0.63338 | 1 | 14196 | 2 | -0.0063667 |
| MT1E | 5 | 0.038498 | 0.11209 | 0.951052 | 2441 | 2 | 0.33913 | 0.45923 | 0.68539 | 1 | 14197 | 2 | 0.33913 |
| TRIP13 | 5 | 0.47147 | 0.62338 | 0.999994 | 13021 | 2 | -0.15166 | 0.45931 | 0.68546 | 1 | 14198 | 2 | -0.15166 |
| HOXA6 | 6 | 0.67363 | 0.79598 | 0.999994 | 15982 | 2 | 0.46138 | 0.45934 | 0.69633 | 1 | 14199 | 3 | 0.46138 |
| GRWD1 | 6 | 0.25754 | 0.45696 | 0.999994 | 9180 | 3 | -0.26983 | 0.45935 | 0.69635 | 1 | 14200 | 2 | -0.26983 |
| SCN3B | 6 | 0.034057 | 0.10831 | 0.951052 | 2219 | 2 | 0.43264 | 0.45935 | 0.69635 | 1 | 14201 | 3 | 0.43264 |
| HTR3B | 6 | 0.086324 | 0.22014 | 0.980699 | 4462 | 4 | -1.3011 | 0.45935 | 0.69635 | 1 | 14202 | 2 | -1.3011 |
| AGRP | 6 | 0.053015 | 0.15059 | 0.966359 | 3111 | 3 | -1.221 | 0.45937 | 0.69636 | 1 | 14203 | 2 | -1.221 |
| PTPN20A | 6 | 0.21043 | 0.40061 | 0.999994 | 8031 | 3 | 0.071827 | 0.45942 | 0.69639 | 1 | 14204 | 3 | 0.071827 |
| hsa-mir-4447 | 3 | 0.43675 | 0.50612 | 0.999994 | 12521 | 1 | 0.51851 | 0.45943 | 0.61396 | 1 | 14205 | 1 | 0.51851 |
| RBP2 | 6 | 0.07715 | 0.20159 | 0.977164 | 4112 | 4 | -1.2682 | 0.45944 | 0.69641 | 1 | 14206 | 2 | -1.2682 |
| CLEC4D | 6 | 0.66789 | 0.79151 | 0.999994 | 15892 | 2 | 0.83395 | 0.45967 | 0.69658 | 1 | 14207 | 3 | 0.83395 |
| JUNB | 5 | 0.25005 | 0.44082 | 0.999994 | 9018 | 1 | 0.5959 | 0.45978 | 0.68591 | 1 | 14208 | 3 | 0.5959 |
| CYP2A7 | 6 | 0.66903 | 0.79242 | 0.999994 | 15912 | 1 | 0.12244 | 0.45979 | 0.69667 | 1 | 14209 | 2 | 0.12244 |
| PIDD | 5 | 0.0049167 | 0.020491 | 0.81027 | 513 | 2 | -0.50968 | 0.45986 | 0.68599 | 1 | 14210 | 2 | -0.50968 |
| NCKAP1L | 5 | 0.46989 | 0.62206 | 0.999994 | 13005 | 1 | 0.31893 | 0.45986 | 0.68599 | 1 | 14211 | 2 | 0.31893 |
| hsa-mir-4713 | 4 | 0.12626 | 0.26112 | 0.994943 | 5832 | 1 | 0.40622 | 0.45999 | 0.63408 | 1 | 14212 | 2 | 0.40622 |
| MCMBP | 4 | 0.29469 | 0.43755 | 0.999994 | 10011 | 2 | -0.34564 | 0.45999 | 0.63408 | 1 | 14213 | 1 | -0.34564 |
| GAB4 | 5 | 0.83998 | 0.84664 | 0.999994 | 18086 | 1 | 0.65869 | 0.46006 | 0.68619 | 1 | 14214 | 3 | 0.65869 |
| hsa-mir-761 | 3 | 0.78192 | 0.7818 | 0.999994 | 17239 | 0 | -0.26391 | 0.46016 | 0.61469 | 1 | 14215 | 1 | -0.26391 |
| VWCE | 5 | 0.67981 | 0.75747 | 0.999994 | 16085 | 1 | 0.58825 | 0.46018 | 0.68631 | 1 | 14216 | 3 | 0.58825 |
| SLC50A1 | 6 | 0.027674 | 0.093344 | 0.945031 | 1887 | 4 | -1.7611 | 0.46025 | 0.69703 | 1 | 14217 | 1 | -1.7611 |
| NUP62CL | 6 | 0.3308 | 0.54216 | 0.999994 | 10738 | 3 | -0.56304 | 0.46025 | 0.69703 | 1 | 14218 | 1 | -0.56304 |
| THYN1 | 6 | 0.72421 | 0.81972 | 0.999994 | 16537 | 1 | 0.4064 | 0.46025 | 0.69703 | 1 | 14219 | 2 | 0.4064 |
| EBAG9 | 5 | 0.17921 | 0.34547 | 0.999994 | 7204 | 1 | 0.63083 | 0.46029 | 0.68641 | 1 | 14220 | 3 | 0.63083 |
| WBP5 | 6 | 0.0027406 | 0.012772 | 0.733879 | 334 | 3 | 0.1871 | 0.46047 | 0.69719 | 1 | 14221 | 3 | 0.1871 |
| CREB3L2 | 5 | 0.48073 | 0.6313 | 0.999994 | 13150 | 2 | 0.14747 | 0.46053 | 0.68664 | 1 | 14222 | 2 | 0.14747 |
| BCL9 | 5 | 0.24148 | 0.43293 | 0.999994 | 8812 | 2 | 0.58693 | 0.46053 | 0.68664 | 1 | 14223 | 3 | 0.58693 |
| NUCKS1 | 6 | 0.17057 | 0.35147 | 0.999994 | 6997 | 3 | 0.24554 | 0.46055 | 0.69725 | 1 | 14224 | 3 | 0.24554 |
| VCAN | 4 | 0.43733 | 0.58402 | 0.999994 | 12532 | 2 | -0.31631 | 0.46059 | 0.63453 | 1 | 14225 | 1 | -0.31631 |
| DCAF4L2 | 4 | 0.22326 | 0.3646 | 0.999994 | 8363 | 1 | 0.56937 | 0.46065 | 0.63458 | 1 | 14226 | 2 | 0.56937 |
| YIPF6 | 6 | 0.10844 | 0.26371 | 0.996734 | 5254 | 2 | -0.76156 | 0.46077 | 0.6974 | 1 | 14227 | 2 | -0.76156 |
| TM9SF4 | 5 | 0.10616 | 0.23601 | 0.985362 | 5179 | 3 | -1.5431 | 0.46078 | 0.68688 | 1 | 14228 | 2 | -1.5431 |
| RPE65 | 5 | 0.052184 | 0.14565 | 0.963656 | 3074 | 3 | -2.2265 | 0.46078 | 0.68688 | 1 | 14229 | 1 | -2.2265 |
| RBM12 | 6 | 0.040345 | 0.12269 | 0.951052 | 2532 | 3 | -1.4889 | 0.46085 | 0.69747 | 1 | 14230 | 2 | -1.4889 |
| GPAA1 | 6 | 0.33265 | 0.5443 | 0.999994 | 10784 | 2 | 0.90979 | 0.46086 | 0.69748 | 1 | 14231 | 3 | 0.90979 |
| NonTargetingControlG | 2 | 0.16642 | 0.23816 | 0.987253 | 6895 | 1 | -0.88802 | 0.46089 | 0.50989 | 1 | 14232 | 1 | -0.88802 |
| DUSP21 | 5 | 0.13436 | 0.27929 | 0.999632 | 6036 | 2 | 0.6445 | 0.46108 | 0.68718 | 1 | 14233 | 3 | 0.6445 |
| ITM2C | 6 | 0.47268 | 0.65025 | 0.999994 | 13041 | 2 | 0.45959 | 0.4611 | 0.69766 | 1 | 14234 | 3 | 0.45959 |
| UGGT1 | 6 | 0.11373 | 0.27385 | 0.998561 | 5434 | 3 | -1.1793 | 0.4611 | 0.69766 | 1 | 14235 | 1 | -1.1793 |
| MPP1 | 4 | 0.80206 | 0.80304 | 0.999994 | 17512 | 1 | 0.91378 | 0.46117 | 0.63495 | 1 | 14236 | 2 | 0.91378 |
| GJA5 | 5 | 0.077354 | 0.19031 | 0.971257 | 4123 | 3 | -1.2635 | 0.46118 | 0.68728 | 1 | 14237 | 2 | -1.2635 |
| RNF26 | 6 | 0.82396 | 0.85869 | 0.999994 | 17837 | 1 | 0.8461 | 0.46139 | 0.69787 | 1 | 14238 | 3 | 0.8461 |
| SERPINE3 | 6 | 0.3876 | 0.59441 | 0.999994 | 11761 | 2 | 0.089931 | 0.46143 | 0.69789 | 1 | 14239 | 2 | 0.089931 |
| LDOC1 | 6 | 0.29752 | 0.50383 | 0.999994 | 10078 | 3 | -0.44999 | 0.46148 | 0.69794 | 1 | 14240 | 2 | -0.44999 |
| REXO1 | 5 | 0.0054494 | 0.022571 | 0.819701 | 555 | 4 | -2.1431 | 0.46156 | 0.68763 | 1 | 14241 | 1 | -2.1431 |
| ZYX | 5 | 0.12031 | 0.25784 | 0.994943 | 5638 | 3 | -1.223 | 0.46156 | 0.68763 | 1 | 14242 | 2 | -1.223 |
| SUSD1 | 5 | 0.41328 | 0.57455 | 0.999994 | 12158 | 2 | -0.41117 | 0.46156 | 0.68763 | 1 | 14243 | 2 | -0.41117 |
| CMTR1 | 5 | 0.038665 | 0.11249 | 0.951052 | 2447 | 3 | -2.8044 | 0.46156 | 0.68763 | 1 | 14244 | 1 | -2.8044 |
| hsa-mir-1255a | 4 | 0.027025 | 0.07541 | 0.93914 | 1858 | 2 | -1.8897 | 0.46156 | 0.63526 | 1 | 14245 | 2 | -1.8897 |
| SYCP2L | 6 | 0.22812 | 0.42181 | 0.999994 | 8479 | 2 | 0.80088 | 0.4616 | 0.69803 | 1 | 14246 | 3 | 0.80088 |
| PTK2B | 6 | 0.0026809 | 0.012543 | 0.733879 | 329 | 5 | -2.3679 | 0.46187 | 0.69822 | 1 | 14247 | 1 | -2.3679 |

| | | | | | | | | | | | | | |
|---|---|---|---|---|---|---|---|---|---|---|---|---|---|
| CARD18 | 6 | 0.047272 | 0.1381 | 0.957317 | 2867 | 4 | -1.5522 | 0.46187 | 0.69822 | 1 | 14248 | 2 | -1.5522 |
| KRBA2 | 6 | 0.25054 | 0.44856 | 0.999994 | 9025 | 3 | -0.60073 | 0.46187 | 0.69822 | 1 | 14249 | 2 | -0.60073 |
| OR5K4 | 6 | 0.26067 | 0.46065 | 0.999994 | 9254 | 3 | 0.19335 | 0.46187 | 0.69822 | 1 | 14250 | 3 | 0.19335 |
| TSEN54 | 6 | 0.086077 | 0.21965 | 0.980699 | 4449 | 2 | 0.40799 | 0.46194 | 0.69826 | 1 | 14251 | 3 | 0.40799 |
| HLA-C | 6 | 0.65478 | 0.7813 | 0.999994 | 15713 | 2 | 0.16553 | 0.46199 | 0.69831 | 1 | 14252 | 2 | 0.16553 |
| UBE2J1 | 5 | 0.54237 | 0.68498 | 0.999994 | 14100 | 2 | 0.69713 | 0.46205 | 0.68811 | 1 | 14253 | 3 | 0.69713 |
| THRA | 6 | 0.041358 | 0.12494 | 0.951052 | 2581 | 2 | 0.26069 | 0.46217 | 0.69843 | 1 | 14254 | 3 | 0.26069 |
| ADORA2B | 5 | 0.23331 | 0.42178 | 0.999994 | 8602 | 3 | -0.66331 | 0.46217 | 0.68821 | 1 | 14255 | 2 | -0.66331 |
| SERPINB5 | 5 | 0.25396 | 0.44389 | 0.999994 | 9102 | 2 | -0.33246 | 0.46217 | 0.68821 | 1 | 14256 | 2 | -0.33246 |
| CPNE4 | 5 | 0.094677 | 0.218 | 0.980699 | 4775 | 1 | 0.66059 | 0.46219 | 0.68823 | 1 | 14257 | 3 | 0.66059 |
| hsa-mir-3150a | 4 | 0.61987 | 0.67627 | 0.999994 | 15219 | 1 | 0.83845 | 0.46222 | 0.63573 | 1 | 14258 | 2 | 0.83845 |
| IQUB | 6 | 0.20716 | 0.39661 | 0.999994 | 7952 | 2 | 0.45373 | 0.46233 | 0.69856 | 1 | 14259 | 3 | 0.45373 |
| ADO | 4 | 0.43195 | 0.57854 | 0.999994 | 12445 | 2 | -0.078927 | 0.46233 | 0.63582 | 1 | 14260 | 1 | -0.078927 |
| ZNF281 | 4 | 0.13544 | 0.2742 | 0.998561 | 6066 | 2 | 0.13416 | 0.46233 | 0.63582 | 1 | 14261 | 2 | 0.13416 |
| SCARF1 | 5 | 0.63636 | 0.73979 | 0.999994 | 15465 | 1 | 0.61877 | 0.46253 | 0.68856 | 1 | 14262 | 3 | 0.61877 |
| MYL4 | 5 | 0.11671 | 0.25233 | 0.994162 | 5530 | 2 | 0.66142 | 0.46254 | 0.68858 | 1 | 14263 | 3 | 0.66142 |
| CDS1 | 6 | 0.0024526 | 0.011671 | 0.727553 | 313 | 2 | 0.58001 | 0.46262 | 0.69879 | 1 | 14264 | 3 | 0.58001 |
| ALDH1L2 | 6 | 0.014641 | 0.054555 | 0.922677 | 1169 | 3 | -0.7348 | 0.46263 | 0.6988 | 1 | 14265 | 2 | -0.7348 |
| SDCBP | 4 | 0.91669 | 0.91645 | 0.999994 | 19554 | 0 | 0.89726 | 0.46268 | 0.63607 | 1 | 14266 | 2 | 0.89726 |
| LRRIQ1 | 5 | 0.014506 | 0.048787 | 0.903125 | 1166 | 2 | 0.44108 | 0.46288 | 0.6889 | 1 | 14267 | 2 | 0.44108 |
| P4HTM | 5 | 0.57622 | 0.71499 | 0.999994 | 14634 | 2 | 0.091458 | 0.46288 | 0.6889 | 1 | 14268 | 2 | 0.091458 |
| NR0B1 | 5 | 0.28954 | 0.47254 | 0.999994 | 9902 | 1 | -0.077363 | 0.46288 | 0.6889 | 1 | 14269 | 2 | -0.077363 |
| OR52B4 | 5 | 0.34064 | 0.51425 | 0.999994 | 10952 | 2 | -0.39252 | 0.46288 | 0.6889 | 1 | 14270 | 2 | -0.39252 |
| POLR1C | 4 | 0.8132 | 0.81331 | 0.999994 | 17677 | 1 | 0.82697 | 0.46302 | 0.63632 | 1 | 14271 | 2 | 0.82697 |
| FHL1 | 6 | 0.35588 | 0.57066 | 0.999994 | 11266 | 2 | 0.60751 | 0.46305 | 0.69914 | 1 | 14272 | 3 | 0.60751 |
| KRTAP5-5 | 5 | 0.71994 | 0.77579 | 0.999994 | 16499 | 1 | 0.75046 | 0.46314 | 0.68914 | 1 | 14273 | 3 | 0.75046 |
| RPL18A | 3 | 0.27386 | 0.4044 | 0.999994 | 9560 | 1 | 0.78987 | 0.46318 | 0.61774 | 1 | 14274 | 2 | 0.78987 |
| PI3 | 6 | 0.32956 | 0.54072 | 0.999994 | 10710 | 3 | -0.63509 | 0.4632 | 0.69927 | 1 | 14275 | 2 | -0.63509 |
| USP27X | 6 | 0.62926 | 0.76176 | 0.999994 | 15372 | 2 | 0.31142 | 0.4632 | 0.69927 | 1 | 14276 | 2 | 0.31142 |
| ERAP1 | 6 | 0.52532 | 0.68624 | 0.999994 | 13821 | 2 | -0.084879 | 0.4632 | 0.69927 | 1 | 14277 | 2 | -0.084879 |
| ENKD1 | 4 | 0.0058517 | 0.020449 | 0.81027 | 579 | 2 | -0.19341 | 0.46341 | 0.63663 | 1 | 14278 | 2 | -0.19341 |
| MGAT3 | 4 | 0.78726 | 0.79001 | 0.999994 | 17308 | 1 | 1.0734 | 0.46347 | 0.63668 | 1 | 14279 | 2 | 1.0734 |
| CCDC58 | 4 | 0.30603 | 0.44913 | 0.999994 | 10257 | 2 | 0.46862 | 0.46347 | 0.63668 | 1 | 14280 | 2 | 0.46862 |
| TRAPPC2 | 4 | 0.74138 | 0.75353 | 0.999994 | 16758 | 1 | 0.85526 | 0.46347 | 0.63668 | 1 | 14281 | 2 | 0.85526 |
| PRND | 6 | 0.46871 | 0.64757 | 0.999994 | 12985 | 2 | 0.27272 | 0.46361 | 0.69957 | 1 | 14282 | 1 | 0.27272 |
| PIK3IP1 | 6 | 0.33158 | 0.54307 | 0.999994 | 10758 | 3 | -0.27043 | 0.46361 | 0.69957 | 1 | 14283 | 1 | -0.27043 |
| KDM5B | 5 | 0.92416 | 0.92438 | 0.999994 | 19721 | 0 | 0.58314 | 0.46362 | 0.6896 | 1 | 14284 | 3 | 0.58314 |
| NonTargetingControlG | 2 | 0.21995 | 0.2766 | 0.999379 | 8288 | 1 | -0.12928 | 0.46367 | 0.51188 | 1 | 14285 | 1 | -0.12928 |
| HIPK3 | 5 | 0.93663 | 0.93687 | 0.999994 | 19997 | 0 | 0.40347 | 0.46372 | 0.68969 | 1 | 14286 | 2 | 0.40347 |
| OR1B1 | 6 | 0.43595 | 0.62575 | 0.999994 | 12513 | 2 | 0.84975 | 0.4638 | 0.69971 | 1 | 14287 | 3 | 0.84975 |
| HERPUD1 | 5 | 0.29651 | 0.47823 | 0.999994 | 10054 | 2 | 0.54207 | 0.46385 | 0.68983 | 1 | 14288 | 3 | 0.54207 |
| POM121C | 4 | 0.7653 | 0.772 | 0.999994 | 17031 | 1 | 0.9504 | 0.46387 | 0.63699 | 1 | 14289 | 2 | 0.9504 |
| NonTargetingControlG | 1 | 0.53611 | 0.53588 | 0.999994 | 13998 | 0 | 0.77907 | 0.46389 | 0.46406 | 1 | 14290 | 1 | 0.77907 |
| SFT2D2 | 5 | 0.45318 | 0.60789 | 0.999994 | 12748 | 2 | 0.64255 | 0.46392 | 0.6899 | 1 | 14291 | 3 | 0.64255 |
| NonTargetingControlG | 2 | 0.46436 | 0.472 | 0.999994 | 12925 | 1 | 0.88123 | 0.46397 | 0.5121 | 1 | 14292 | 1 | 0.88123 |
| NANOS1 | 6 | 0.88873 | 0.89543 | 0.999994 | 19014 | 1 | 0.81023 | 0.46428 | 0.70008 | 1 | 14293 | 3 | 0.81023 |
| SGK3 | 6 | 0.34031 | 0.553 | 0.999994 | 10939 | 3 | -0.40613 | 0.46448 | 0.70022 | 1 | 14294 | 2 | -0.40613 |
| S100G | 5 | 0.10799 | 0.23887 | 0.987253 | 5238 | 1 | 0.31363 | 0.4645 | 0.69046 | 1 | 14295 | 2 | 0.31363 |
| CCDC146 | 5 | 0.1754 | 0.34 | 0.999994 | 7121 | 3 | -0.94007 | 0.4645 | 0.69046 | 1 | 14296 | 2 | -0.94007 |
| SASS6 | 5 | 0.12008 | 0.25748 | 0.994943 | 5630 | 3 | -0.94638 | 0.46451 | 0.69048 | 1 | 14297 | 2 | -0.94638 |
| EIF3H | 4 | 0.042543 | 0.10683 | 0.950995 | 2649 | 2 | -3.1895 | 0.46452 | 0.63746 | 1 | 14298 | 1 | -3.1895 |
| hsa-mir-4424 | 4 | 0.29777 | 0.44074 | 0.999994 | 10086 | 2 | -0.37087 | 0.46452 | 0.63746 | 1 | 14299 | 2 | -0.37087 |

| | | | | | | | | | | | | | |
|---|---|---|---|---|---|---|---|---|---|---|---|---|---|
| C1orf158 | 6 | 0.20424 | 0.39303 | 0.999994 | 7882 | 1 | 0.57706 | 0.46455 | 0.70028 | 1 | 14300 | 3 | 0.57706 |
| CDK17 | 6 | 0.55929 | 0.7103 | 0.999994 | 14359 | 2 | 0.61076 | 0.4646 | 0.70031 | 1 | 14301 | 3 | 0.61076 |
| WNK4 | 5 | 0.66637 | 0.75177 | 0.999994 | 15873 | 1 | 0.6405 | 0.46464 | 0.69059 | 1 | 14302 | 3 | 0.6405 |
| ZNF720 | 5 | 0.32505 | 0.50145 | 0.999994 | 10622 | 1 | 0.63216 | 0.46484 | 0.69076 | 1 | 14303 | 3 | 0.63216 |
| NonTargetingControlG | 2 | 0.53961 | 0.54091 | 0.999994 | 14045 | 1 | 1.0094 | 0.4649 | 0.51273 | 1 | 14304 | 1 | 1.0094 |
| ZNF709 | 6 | 0.90144 | 0.9048 | 0.999994 | 19242 | 1 | 0.49687 | 0.46495 | 0.70057 | 1 | 14305 | 3 | 0.49687 |
| CASR | 5 | 0.35463 | 0.52567 | 0.999994 | 11243 | 2 | 0.62247 | 0.46496 | 0.69089 | 1 | 14306 | 3 | 0.62247 |
| C1orf123 | 5 | 0.056341 | 0.15561 | 0.967126 | 3260 | 2 | 0.58147 | 0.46506 | 0.69098 | 1 | 14307 | 3 | 0.58147 |
| FOXG1 | 6 | 0.22649 | 0.41989 | 0.999994 | 8441 | 3 | 0.18226 | 0.46511 | 0.70069 | 1 | 14308 | 3 | 0.18226 |
| hsa-mir-514a-1 | 3 | 0.77521 | 0.77518 | 0.999994 | 17141 | 0 | 0.59191 | 0.46513 | 0.61972 | 1 | 14309 | 2 | 0.59191 |
| CIAPIN1 | 5 | 0.18301 | 0.35095 | 0.999994 | 7290 | 1 | 0.17557 | 0.46531 | 0.69122 | 1 | 14310 | 2 | 0.17557 |
| ORAOV1 | 4 | 0.73986 | 0.75237 | 0.999994 | 16737 | 1 | 1.0005 | 0.46532 | 0.63809 | 1 | 14311 | 2 | 1.0005 |
| WDR43 | 6 | 0.0013623 | 0.0068136 | 0.643671 | 204 | 4 | -6.7637 | 0.46538 | 0.70092 | 1 | 14312 | 2 | -6.7637 |
| MARCH11 | 5 | 0.13907 | 0.28644 | 0.999632 | 6166 | 3 | -1.2144 | 0.4654 | 0.6913 | 1 | 14313 | 2 | -1.2144 |
| ANXA10 | 5 | 0.50938 | 0.65608 | 0.999994 | 13571 | 1 | 0.091523 | 0.4654 | 0.6913 | 1 | 14314 | 1 | 0.091523 |
| ERC1 | 5 | 0.34273 | 0.51597 | 0.999994 | 11001 | 1 | -0.25475 | 0.4654 | 0.6913 | 1 | 14315 | 2 | -0.25475 |
| SSX5 | 5 | 0.11315 | 0.24685 | 0.98999 | 5411 | 3 | -1.3101 | 0.4654 | 0.6913 | 1 | 14316 | 2 | -1.3101 |
| TMEM44 | 5 | 0.026352 | 0.081027 | 0.93914 | 1822 | 3 | -3.0347 | 0.4654 | 0.6913 | 1 | 14317 | 2 | -3.0347 |
| MVD | 3 | 0.79538 | 0.79523 | 0.999994 | 17423 | 0 | 0.61877 | 0.46567 | 0.62026 | 1 | 14318 | 2 | 0.61877 |
| hsa-mir-4519 | 4 | 0.15991 | 0.2994 | 0.999994 | 6733 | 1 | 0.49917 | 0.4659 | 0.63853 | 1 | 14319 | 2 | 0.49917 |
| PPP2CA | 3 | 0.20225 | 0.3155 | 0.999994 | 7837 | 2 | -0.90362 | 0.4659 | 0.62048 | 1 | 14320 | 1 | -0.90362 |
| TCEB1 | 3 | 0.055239 | 0.12297 | 0.951052 | 3209 | 2 | -2.4564 | 0.4659 | 0.62048 | 1 | 14321 | 1 | -2.4564 |
| DYDC2 | 4 | 0.33989 | 0.48386 | 0.999994 | 10927 | 1 | 1.0154 | 0.46591 | 0.63853 | 1 | 14322 | 2 | 1.0154 |
| MIS12 | 3 | 0.55495 | 0.5838 | 0.999994 | 14287 | 1 | 0.68883 | 0.46606 | 0.62066 | 1 | 14323 | 2 | 0.68883 |
| RNF128 | 5 | 0.24921 | 0.44016 | 0.999994 | 8998 | 2 | 0.62968 | 0.46618 | 0.69203 | 1 | 14324 | 3 | 0.62968 |
| NonTargetingControlG | 1 | 0.53366 | 0.53342 | 0.999994 | 13962 | 0 | 0.81521 | 0.46634 | 0.46638 | 1 | 14325 | 1 | 0.81521 |
| RERG | 5 | 0.83967 | 0.84639 | 0.999994 | 18081 | 1 | 0.55781 | 0.46637 | 0.6922 | 1 | 14326 | 3 | 0.55781 |
| APOA2 | 5 | 0.022486 | 0.070813 | 0.93914 | 1617 | 4 | -1.3953 | 0.46638 | 0.69221 | 1 | 14327 | 1 | -1.3953 |
| ITGA3 | 5 | 0.061915 | 0.16525 | 0.967126 | 3508 | 3 | -1.9856 | 0.46638 | 0.69221 | 1 | 14328 | 1 | -1.9856 |
| FAM83F | 5 | 0.066315 | 0.17236 | 0.967658 | 3698 | 3 | -1.793 | 0.46638 | 0.69222 | 1 | 14329 | 2 | -1.793 |
| HOXA13 | 6 | 0.33667 | 0.54886 | 0.999994 | 10865 | 3 | -0.24956 | 0.46682 | 0.70203 | 1 | 14330 | 2 | -0.24956 |
| CENPC1 | 4 | 0.085461 | 0.18874 | 0.969543 | 4427 | 2 | -0.4825 | 0.46685 | 0.63922 | 1 | 14331 | 1 | -0.4825 |
| hsa-mir-4295 | 4 | 0.39199 | 0.53744 | 0.999994 | 11832 | 2 | 0.091389 | 0.46685 | 0.63922 | 1 | 14332 | 2 | 0.091389 |
| NonTargetingControlG | 2 | 0.35574 | 0.38062 | 0.999994 | 11262 | 1 | 0.64336 | 0.46693 | 0.51419 | 1 | 14333 | 1 | 0.64336 |
| hsa-mir-3120 | 2 | 0.11732 | 0.19669 | 0.977164 | 5552 | 1 | -2.2446 | 0.46693 | 0.51419 | 1 | 14334 | 1 | -2.2446 |
| PRKX | 5 | 0.79721 | 0.81777 | 0.999994 | 17441 | 1 | 0.53495 | 0.46693 | 0.69275 | 1 | 14335 | 3 | 0.53495 |
| MDC1 | 6 | 0.20075 | 0.38878 | 0.999994 | 7777 | 2 | 0.60834 | 0.467 | 0.70217 | 1 | 14336 | 3 | 0.60834 |
| NANOGNB | 5 | 0.86542 | 0.86702 | 0.999994 | 18548 | 1 | 0.54429 | 0.46702 | 0.69284 | 1 | 14337 | 3 | 0.54429 |
| RAB27B | 5 | 0.13562 | 0.28119 | 0.999632 | 6071 | 1 | -0.36805 | 0.46726 | 0.69305 | 1 | 14338 | 1 | -0.36805 |
| TPPP | 5 | 0.36542 | 0.53459 | 0.999994 | 11429 | 2 | -1.4517 | 0.46726 | 0.69305 | 1 | 14339 | 1 | -1.4517 |
| GGCX | 5 | 0.67118 | 0.75381 | 0.999994 | 15946 | 1 | 0.57651 | 0.46739 | 0.69317 | 1 | 14340 | 3 | 0.57651 |
| MS4A6A | 6 | 0.25761 | 0.45706 | 0.999994 | 9184 | 3 | 0.24308 | 0.46741 | 0.70246 | 1 | 14341 | 3 | 0.24308 |
| XRCC2 | 6 | 0.073199 | 0.19341 | 0.975797 | 3964 | 4 | -1.0437 | 0.46742 | 0.70247 | 1 | 14342 | 2 | -1.0437 |
| TMEM65 | 6 | 0.10714 | 0.2612 | 0.994943 | 5216 | 3 | -1.1735 | 0.46742 | 0.70247 | 1 | 14343 | 2 | -1.1735 |
| PKP3 | 6 | 0.30218 | 0.50926 | 0.999994 | 10170 | 2 | -0.064951 | 0.46742 | 0.70247 | 1 | 14344 | 2 | -0.064951 |
| C16orf59 | 6 | 0.20267 | 0.39112 | 0.999994 | 7846 | 3 | -0.13169 | 0.46742 | 0.70247 | 1 | 14345 | 3 | -0.13169 |
| TUBA3E | 6 | 0.099083 | 0.24557 | 0.98979 | 4933 | 3 | -0.58916 | 0.46742 | 0.70247 | 1 | 14346 | 2 | -0.58916 |
| ECT2L | 5 | 0.88639 | 0.88649 | 0.999994 | 18957 | 1 | 0.69414 | 0.46747 | 0.69325 | 1 | 14347 | 3 | 0.69414 |
| LMAN1 | 6 | 0.25871 | 0.45835 | 0.999994 | 9210 | 2 | 0.90762 | 0.46749 | 0.70252 | 1 | 14348 | 3 | 0.90762 |
| KIF1B | 6 | 0.85993 | 0.8774 | 0.999994 | 18433 | 1 | 0.74201 | 0.46764 | 0.70264 | 1 | 14349 | 3 | 0.74201 |
| RAB31 | 6 | 0.14564 | 0.32018 | 0.999994 | 6341 | 3 | -0.22986 | 0.46774 | 0.70274 | 1 | 14350 | 3 | -0.22986 |
| ZNF652 | 6 | 0.54288 | 0.69858 | 0.999994 | 14109 | 2 | -0.24267 | 0.4678 | 0.70277 | 1 | 14351 | 2 | -0.24267 |

| | | | | | | | | | | | | | |
|---|---|---|---|---|---|---|---|---|---|---|---|---|---|
| C1QTNF9 | 6 | 0.040343 | 0.12268 | 0.951052 | 2531 | 4 | -1.9534 | 0.4678 | 0.70277 | 1 | 14352 | 1 | -1.9534 |
| TUBA3D | 4 | 0.018179 | 0.056755 | 0.924086 | 1373 | 2 | -3.5679 | 0.4679 | 0.64003 | 1 | 14353 | 2 | -3.5679 |
| RNF150 | 4 | 0.68541 | 0.71505 | 0.999994 | 16166 | 1 | 0.41635 | 0.4679 | 0.64003 | 1 | 14354 | 2 | 0.41635 |
| CARD9 | 4 | 0.44675 | 0.59332 | 0.999994 | 12671 | 1 | 0.40861 | 0.4679 | 0.64003 | 1 | 14355 | 2 | 0.40861 |
| TMEM251 | 5 | 0.57503 | 0.71396 | 0.999994 | 14617 | 2 | 0.60394 | 0.46794 | 0.69369 | 1 | 14356 | 3 | 0.60394 |
| NonTargetingControlG | 2 | 0.63931 | 0.63914 | 0.999994 | 15505 | 0 | 1.3581 | 0.46801 | 0.51498 | 1 | 14357 | 1 | 1.3581 |
| SLC4A10 | 6 | 0.055551 | 0.15614 | 0.967126 | 3222 | 2 | 0.56359 | 0.4681 | 0.703 | 1 | 14358 | 3 | 0.56359 |
| hsa-mir-3909 | 3 | 0.70555 | 0.70622 | 0.999994 | 16370 | 1 | 0.67478 | 0.46823 | 0.62283 | 1 | 14359 | 2 | 0.67478 |
| PPP6R1 | 6 | 0.25361 | 0.45224 | 0.999994 | 9096 | 2 | 0.37928 | 0.46841 | 0.70321 | 1 | 14360 | 3 | 0.37928 |
| ZNF614 | 6 | 0.19948 | 0.38725 | 0.999994 | 7747 | 3 | -0.67673 | 0.46846 | 0.70325 | 1 | 14361 | 2 | -0.67673 |
| ACYP1 | 5 | 0.91918 | 0.91933 | 0.999994 | 19612 | 0 | 0.37485 | 0.46846 | 0.6942 | 1 | 14362 | 2 | 0.37485 |
| KRT32 | 5 | 0.19384 | 0.36641 | 0.999994 | 7597 | 2 | 0.54732 | 0.46861 | 0.69436 | 1 | 14363 | 3 | 0.54732 |
| FAM156B | 2 | 0.3995 | 0.41637 | 0.999994 | 11942 | 1 | 0.5901 | 0.46865 | 0.51543 | 1 | 14364 | 1 | 0.5901 |
| PSME2 | 5 | 0.3619 | 0.53164 | 0.999994 | 11364 | 2 | 0.57799 | 0.46866 | 0.6944 | 1 | 14365 | 3 | 0.57799 |
| KCNV1 | 6 | 0.51623 | 0.67992 | 0.999994 | 13682 | 2 | 0.64453 | 0.46872 | 0.70346 | 1 | 14366 | 3 | 0.64453 |
| GLYATL1 | 6 | 0.58846 | 0.73148 | 0.999994 | 14796 | 2 | -0.17586 | 0.46874 | 0.70347 | 1 | 14367 | 2 | -0.17586 |
| KCNH2 | 5 | 0.40547 | 0.56811 | 0.999994 | 12038 | 2 | 0.11633 | 0.46874 | 0.69448 | 1 | 14368 | 2 | 0.11633 |
| AARD | 3 | 0.65514 | 0.66134 | 0.999994 | 15719 | 1 | 0.6867 | 0.46878 | 0.62341 | 1 | 14369 | 2 | 0.6867 |
| MSH5 | 5 | 0.8445 | 0.85004 | 0.999994 | 18165 | 1 | 0.56855 | 0.46879 | 0.69452 | 1 | 14370 | 3 | 0.56855 |
| HIST1H2AG | 5 | 0.018065 | 0.058811 | 0.931872 | 1363 | 2 | 0.54994 | 0.4688 | 0.69453 | 1 | 14371 | 3 | 0.54994 |
| OR10G8 | 5 | 0.21705 | 0.39915 | 0.999994 | 8197 | 1 | 0.21848 | 0.46883 | 0.69455 | 1 | 14372 | 2 | 0.21848 |
| C1orf112 | 5 | 0.294 | 0.47616 | 0.999994 | 9998 | 2 | 0.57575 | 0.46891 | 0.69462 | 1 | 14373 | 3 | 0.57575 |
| TMEM14B | 3 | 0.7465 | 0.74646 | 0.999994 | 16820 | 0 | 0.52196 | 0.46893 | 0.62356 | 1 | 14374 | 2 | 0.52196 |
| DPP7 | 4 | 0.66818 | 0.70427 | 0.999994 | 15895 | 1 | 0.84003 | 0.46894 | 0.64082 | 1 | 14375 | 2 | 0.84003 |
| GSR | 4 | 0.86845 | 0.86804 | 0.999994 | 18610 | 0 | 0.86293 | 0.46905 | 0.64091 | 1 | 14376 | 2 | 0.86293 |
| FGFBP3 | 5 | 0.074738 | 0.18608 | 0.969409 | 4026 | 2 | 0.64968 | 0.46909 | 0.6948 | 1 | 14377 | 3 | 0.64968 |
| SLC25A29 | 6 | 0.25615 | 0.45529 | 0.999994 | 9152 | 3 | -0.83005 | 0.46909 | 0.70373 | 1 | 14378 | 2 | -0.83005 |
| ARHGAP27 | 5 | 0.39831 | 0.56215 | 0.999994 | 11920 | 2 | 0.59551 | 0.4691 | 0.69482 | 1 | 14379 | 3 | 0.59551 |
| RHOXF1 | 4 | 0.23519 | 0.37685 | 0.999994 | 8653 | 1 | 0.48944 | 0.46913 | 0.64096 | 1 | 14380 | 2 | 0.48944 |
| TMCO6 | 5 | 0.63008 | 0.73733 | 0.999994 | 15381 | 1 | 0.67705 | 0.46937 | 0.69508 | 1 | 14381 | 3 | 0.67705 |
| FAM45A | 6 | 0.68189 | 0.80251 | 0.999994 | 16115 | 2 | -0.022969 | 0.46941 | 0.704 | 1 | 14382 | 2 | -0.022969 |
| hsa-mir-4451 | 4 | 0.51817 | 0.62457 | 0.999994 | 13714 | 1 | 0.84383 | 0.46957 | 0.64131 | 1 | 14383 | 2 | 0.84383 |
| ZNF217 | 6 | 0.012111 | 0.046091 | 0.892434 | 999 | 5 | -4.1481 | 0.46971 | 0.70422 | 1 | 14384 | 1 | -4.1481 |
| UBE2E1 | 6 | 0.020546 | 0.073717 | 0.93914 | 1485 | 4 | -2.195 | 0.46971 | 0.70422 | 1 | 14385 | 1 | -2.195 |
| GP5 | 6 | 0.019547 | 0.070524 | 0.93914 | 1430 | 4 | -2.5586 | 0.46971 | 0.70422 | 1 | 14386 | 2 | -2.5586 |
| PRKRA | 6 | 0.0884 | 0.22435 | 0.983354 | 4531 | 4 | -0.73204 | 0.46971 | 0.70422 | 1 | 14387 | 2 | -0.73204 |
| SRRM4 | 6 | 0.53089 | 0.69016 | 0.999994 | 13908 | 2 | -0.27106 | 0.46971 | 0.70422 | 1 | 14388 | 2 | -0.27106 |
| CATSPER1 | 5 | 0.052171 | 0.14561 | 0.963656 | 3073 | 1 | -0.23064 | 0.46973 | 0.69542 | 1 | 14389 | 2 | -0.23064 |
| RLN2 | 5 | 0.37159 | 0.53973 | 0.999994 | 11537 | 2 | 0.37914 | 0.46994 | 0.69562 | 1 | 14390 | 2 | 0.37914 |
| PHF17 | 4 | 0.29868 | 0.44165 | 0.999994 | 10105 | 2 | 0.08934 | 0.46994 | 0.64157 | 1 | 14391 | 2 | 0.08934 |
| GALC | 4 | 0.28121 | 0.42379 | 0.999994 | 9726 | 2 | 0.42718 | 0.46997 | 0.6416 | 1 | 14392 | 2 | 0.42718 |
| HOXC5 | 6 | 0.39717 | 0.60053 | 0.999994 | 11908 | 2 | 0.025998 | 0.47 | 0.70445 | 1 | 14393 | 2 | 0.025998 |
| hsa-mir-4517 | 4 | 0.83305 | 0.83258 | 0.999994 | 17972 | 0 | 0.76691 | 0.47008 | 0.64168 | 1 | 14394 | 2 | 0.76691 |
| hsa-mir-4429 | 4 | 0.31671 | 0.46009 | 0.999994 | 10447 | 2 | -0.22406 | 0.47008 | 0.64168 | 1 | 14395 | 2 | -0.22406 |
| hsa-mir-4270 | 3 | 0.51566 | 0.55656 | 0.999994 | 13670 | 1 | -0.46439 | 0.47013 | 0.62476 | 1 | 14396 | 1 | -0.46439 |
| hsa-mir-4732 | 3 | 0.0010321 | 0.0029397 | 0.530487 | 172 | 1 | -0.57606 | 0.47013 | 0.62476 | 1 | 14397 | 1 | -0.57606 |
| hsa-mir-4653 | 4 | 0.57595 | 0.65269 | 0.999994 | 14632 | 1 | 0.89201 | 0.47019 | 0.64178 | 1 | 14398 | 2 | 0.89201 |
| SURF1 | 5 | 0.70263 | 0.76768 | 0.999994 | 16334 | 1 | 0.53017 | 0.47021 | 0.69588 | 1 | 14399 | 3 | 0.53017 |
| PRRC1 | 5 | 0.53836 | 0.68145 | 0.999994 | 14028 | 2 | 0.81069 | 0.4703 | 0.69598 | 1 | 14400 | 3 | 0.81069 |
| OR2M4 | 6 | 0.22576 | 0.419 | 0.999994 | 8418 | 2 | 0.26712 | 0.47042 | 0.70476 | 1 | 14401 | 3 | 0.26712 |
| TPP1 | 5 | 0.23561 | 0.42495 | 0.999994 | 8662 | 3 | -0.62955 | 0.47056 | 0.69623 | 1 | 14402 | 2 | -0.62955 |
| WWC1 | 6 | 0.42616 | 0.6194 | 0.999994 | 12352 | 1 | 0.43772 | 0.47058 | 0.70489 | 1 | 14403 | 2 | 0.43772 |

| | | | | | | | | | | | | | |
|---|---|---|---|---|---|---|---|---|---|---|---|---|---|
| ITGA9 | 6 | 0.021972 | 0.078291 | 0.93914 | 1580 | 2 | 0.76825 | 0.47061 | 0.70491 | 1 | 14404 | 3 | 0.76825 |
| SLC6A11 | 6 | 0.86101 | 0.87798 | 0.999994 | 18464 | 1 | 0.76826 | 0.47069 | 0.70499 | 1 | 14405 | 3 | 0.76826 |
| FOXM1 | 6 | 0.19105 | 0.37682 | 0.999994 | 7513 | 3 | -0.083259 | 0.47071 | 0.705 | 1 | 14406 | 3 | -0.083259 |
| EPB41L4B | 4 | 0.10332 | 0.22104 | 0.982265 | 5075 | 2 | -0.61713 | 0.47079 | 0.64223 | 1 | 14407 | 2 | -0.61713 |
| WDR83OS | 5 | 0.57695 | 0.71564 | 0.999994 | 14645 | 1 | 0.5822 | 0.47092 | 0.69655 | 1 | 14408 | 3 | 0.5822 |
| MRGPRX2 | 5 | 0.31544 | 0.49364 | 0.999994 | 10427 | 2 | -0.23774 | 0.47114 | 0.69674 | 1 | 14409 | 2 | -0.23774 |
| AMER2 | 5 | 0.56156 | 0.70201 | 0.999994 | 14397 | 2 | 0.53054 | 0.47126 | 0.69685 | 1 | 14410 | 3 | 0.53054 |
| OR52E6 | 6 | 0.20213 | 0.39049 | 0.999994 | 7835 | 3 | -0.79449 | 0.47126 | 0.70541 | 1 | 14411 | 2 | -0.79449 |
| PLG | 6 | 0.11842 | 0.28274 | 0.999632 | 5587 | 3 | -0.90321 | 0.47136 | 0.70548 | 1 | 14412 | 2 | -0.90321 |
| GSTA5 | 5 | 0.14586 | 0.29658 | 0.999994 | 6347 | 3 | -1.0736 | 0.47142 | 0.69701 | 1 | 14413 | 2 | -1.0736 |
| NAP1L1 | 6 | 0.91571 | 0.91652 | 0.999994 | 19532 | 1 | 0.69609 | 0.47148 | 0.70558 | 1 | 14414 | 3 | 0.69609 |
| XKR9 | 6 | 0.64889 | 0.77677 | 0.999994 | 15640 | 2 | 0.018472 | 0.4715 | 0.70559 | 1 | 14415 | 2 | 0.018472 |
| TP53TG3B | 2 | 0.71687 | 0.71657 | 0.999994 | 16466 | 0 | 1.4356 | 0.47158 | 0.51745 | 1 | 14416 | 1 | 1.4356 |
| hsa-mir-500b | 3 | 0.41903 | 0.49556 | 0.999994 | 12235 | 1 | 0.52781 | 0.47166 | 0.6263 | 1 | 14417 | 2 | 0.52781 |
| KRTAP13-1 | 5 | 0.294 | 0.47616 | 0.999994 | 9996 | 1 | 0.031076 | 0.47169 | 0.69726 | 1 | 14418 | 2 | 0.031076 |
| PTPN3 | 5 | 0.22236 | 0.40658 | 0.999994 | 8340 | 3 | -0.68219 | 0.47171 | 0.69727 | 1 | 14419 | 1 | -0.68219 |
| AHSP | 5 | 0.35566 | 0.5265 | 0.999994 | 11261 | 2 | -0.14129 | 0.47171 | 0.69727 | 1 | 14420 | 1 | -0.14129 |
| NonTargetingControlG | 1 | 0.52825 | 0.52802 | 0.999994 | 13863 | 0 | 0.71931 | 0.47175 | 0.47175 | 1 | 14421 | 1 | 0.71931 |
| HTR3E | 4 | 0.31601 | 0.45938 | 0.999994 | 10441 | 2 | 0.1386 | 0.47187 | 0.64306 | 1 | 14422 | 2 | 0.1386 |
| ITFG1 | 5 | 0.92252 | 0.92272 | 0.999994 | 19689 | 0 | 0.38142 | 0.472 | 0.69754 | 1 | 14423 | 2 | 0.38142 |
| NDST4 | 4 | 0.089095 | 0.19534 | 0.975797 | 4557 | 2 | -0.81911 | 0.47212 | 0.64324 | 1 | 14424 | 2 | -0.81911 |
| ADAMTS8 | 6 | 0.22581 | 0.41906 | 0.999994 | 8421 | 2 | 0.22932 | 0.47212 | 0.70606 | 1 | 14425 | 3 | 0.22932 |
| ARL9 | 6 | 0.00093448 | 0.0047768 | 0.588856 | 156 | 4 | -4.9522 | 0.47212 | 0.70606 | 1 | 14426 | 2 | -4.9522 |
| NXF3 | 6 | 0.007135 | 0.028913 | 0.859137 | 666 | 3 | -1.2092 | 0.47212 | 0.70606 | 1 | 14427 | 2 | -1.2092 |
| PDIA3 | 6 | 0.15403 | 0.33073 | 0.999994 | 6568 | 2 | 0.10596 | 0.47212 | 0.70606 | 1 | 14428 | 3 | 0.10596 |
| NonTargetingControlG | 2 | 0.39667 | 0.41404 | 0.999994 | 11900 | 1 | 0.49166 | 0.47217 | 0.51786 | 1 | 14429 | 1 | 0.49166 |
| NOP56 | 3 | 0.75441 | 0.75434 | 0.999994 | 16902 | 0 | 0.67151 | 0.47218 | 0.62684 | 1 | 14430 | 2 | 0.67151 |
| hsa-mir-548f-2 | 4 | 0.88527 | 0.88494 | 0.999994 | 18937 | 0 | 0.29253 | 0.47222 | 0.64333 | 1 | 14431 | 1 | 0.29253 |
| hsa-mir-617 | 4 | 0.85435 | 0.85392 | 0.999994 | 18339 | 0 | 0.34919 | 0.47222 | 0.64333 | 1 | 14432 | 2 | 0.34919 |
| hsa-mir-3117 | 4 | 0.41093 | 0.55689 | 0.999994 | 12127 | 1 | 0.14952 | 0.47222 | 0.64333 | 1 | 14433 | 2 | 0.14952 |
| ADNP2 | 5 | 0.86776 | 0.86909 | 0.999994 | 18595 | 1 | 0.54808 | 0.47223 | 0.69775 | 1 | 14434 | 3 | 0.54808 |
| HDAC4 | 6 | 0.057196 | 0.1597 | 0.967126 | 3301 | 4 | -1.5307 | 0.47223 | 0.70616 | 1 | 14435 | 2 | -1.5307 |
| TBC1D9 | 5 | 0.039685 | 0.11512 | 0.951052 | 2499 | 1 | 0.57529 | 0.4723 | 0.69784 | 1 | 14436 | 3 | 0.57529 |
| PPIAL4G | 3 | 0.8694 | 0.86946 | 0.999994 | 18626 | 0 | 0.60996 | 0.47235 | 0.62701 | 1 | 14437 | 2 | 0.60996 |
| NBR1 | 6 | 0.91445 | 0.9154 | 0.999994 | 19497 | 1 | 0.93732 | 0.47249 | 0.70637 | 1 | 14438 | 3 | 0.93732 |
| PARVG | 6 | 0.29201 | 0.49739 | 0.999994 | 9952 | 3 | 0.14963 | 0.47251 | 0.70639 | 1 | 14439 | 3 | 0.14963 |
| NonTargetingControlG | 2 | 0.30575 | 0.3413 | 0.999994 | 10248 | 1 | 0.20999 | 0.47254 | 0.5181 | 1 | 14440 | 1 | 0.20999 |
| EFNA2 | 5 | 0.13594 | 0.28166 | 0.999632 | 6076 | 3 | -1.3475 | 0.4727 | 0.69819 | 1 | 14441 | 1 | -1.3475 |
| HKDC1 | 5 | 0.26624 | 0.45381 | 0.999994 | 9370 | 2 | 0.55491 | 0.47309 | 0.69857 | 1 | 14442 | 3 | 0.55491 |
| AURKAIP1 | 6 | 0.057678 | 0.16075 | 0.967126 | 3323 | 4 | -1.7856 | 0.47311 | 0.70683 | 1 | 14443 | 2 | -1.7856 |
| TAF1D | 6 | 0.080329 | 0.20796 | 0.979108 | 4240 | 4 | -0.9446 | 0.47311 | 0.70683 | 1 | 14444 | 2 | -0.9446 |
| TMEM74 | 6 | 0.2311 | 0.42541 | 0.999994 | 8545 | 2 | 0.36912 | 0.47311 | 0.70683 | 1 | 14445 | 3 | 0.36912 |
| CCDC172 | 4 | 0.81063 | 0.81088 | 0.999994 | 17641 | 1 | 0.86798 | 0.47313 | 0.64403 | 1 | 14446 | 2 | 0.86798 |
| TPSAB1 | 6 | 0.028611 | 0.095553 | 0.945031 | 1931 | 4 | -1.8549 | 0.47331 | 0.70698 | 1 | 14447 | 1 | -1.8549 |
| ODF4 | 6 | 0.067916 | 0.18238 | 0.968526 | 3748 | 3 | -0.82088 | 0.47331 | 0.70698 | 1 | 14448 | 3 | -0.82088 |
| CD5 | 6 | 0.013455 | 0.050623 | 0.909101 | 1098 | 4 | -1.8905 | 0.47331 | 0.70698 | 1 | 14449 | 1 | -1.8905 |
| PROSER2 | 6 | 0.91998 | 0.92039 | 0.999994 | 19631 | 1 | 0.73355 | 0.47331 | 0.70698 | 1 | 14450 | 3 | 0.73355 |
| FIP1L1 | 4 | 0.13257 | 0.27122 | 0.998561 | 5989 | 2 | -0.60077 | 0.47338 | 0.64423 | 1 | 14451 | 2 | -0.60077 |
| MAP7D2 | 4 | 0.12583 | 0.26044 | 0.994943 | 5814 | 3 | -1.0347 | 0.47338 | 0.64423 | 1 | 14452 | 1 | -1.0347 |
| MASTL | 4 | 0.072457 | 0.16457 | 0.967126 | 3939 | 3 | -1.3858 | 0.47338 | 0.64423 | 1 | 14453 | 1 | -1.3858 |
| MAOB | 6 | 0.5232 | 0.68474 | 0.999994 | 13796 | 2 | 0.4147 | 0.47339 | 0.70704 | 1 | 14454 | 3 | 0.4147 |
| FTSJD2 | 4 | 0.00098253 | 0.003615 | 0.555048 | 161 | 2 | -4.1541 | 0.4734 | 0.64424 | 1 | 14455 | 2 | -4.1541 |

| | | | | | | | | | | | | |
|---|---|---|---|---|---|---|---|---|---|---|---|---|
| HSD3B1 | 5 | 0.14235 | 0.29142 | 0.999632 | 6247 | 2 | 0.71656 | 0.47362 | 0.69905 | 1 | 14456 | 3 | 0.71656 |
| ATP13A1 | 5 | 0.81598 | 0.82992 | 0.999994 | 17712 | 1 | 0.059614 | 0.47371 | 0.69915 | 1 | 14457 | 2 | 0.059614 |
| WNT3 | 5 | 0.10093 | 0.22788 | 0.983354 | 5004 | 3 | -1.5072 | 0.47371 | 0.69915 | 1 | 14458 | 1 | -1.5072 |
| RIT2 | 5 | 0.51197 | 0.65838 | 0.999994 | 13610 | 2 | -0.27972 | 0.47371 | 0.69915 | 1 | 14459 | 1 | -0.27972 |
| NOB1 | 5 | 0.022966 | 0.072114 | 0.93914 | 1647 | 3 | -3.4904 | 0.47371 | 0.69915 | 1 | 14460 | 1 | -3.4904 |
| WDR96 | 5 | 0.063028 | 0.16707 | 0.967126 | 3557 | 1 | 0.58985 | 0.47371 | 0.69915 | 1 | 14461 | 2 | 0.58985 |
| HIST1H1T | 5 | 0.88741 | 0.88751 | 0.999994 | 18979 | 1 | -0.36482 | 0.47371 | 0.69915 | 1 | 14462 | 2 | -0.36482 |
| TMEM174 | 5 | 0.3686 | 0.53719 | 0.999994 | 11483 | 2 | -1.3892 | 0.47371 | 0.69915 | 1 | 14463 | 2 | -1.3892 |
| ZCCHC10 | 5 | 0.71559 | 0.77373 | 0.999994 | 16455 | 1 | 0.39185 | 0.47371 | 0.69915 | 1 | 14464 | 1 | 0.39185 |
| TCP10 | 5 | 0.14026 | 0.28822 | 0.999632 | 6193 | 2 | -0.21753 | 0.47386 | 0.69928 | 1 | 14465 | 2 | -0.21753 |
| KRTAP10-8 | 6 | 0.33693 | 0.54917 | 0.999994 | 10870 | 3 | 0.2247 | 0.47388 | 0.70744 | 1 | 14466 | 3 | 0.2247 |
| WDR16 | 5 | 0.57945 | 0.71784 | 0.999994 | 14671 | 1 | -0.01198 | 0.47405 | 0.69947 | 1 | 14467 | 2 | -0.01198 |
| GNA14 | 6 | 0.20744 | 0.39694 | 0.999994 | 7960 | 3 | -0.62793 | 0.47408 | 0.7076 | 1 | 14468 | 2 | -0.62793 |
| RNF2 | 6 | 0.46985 | 0.64835 | 0.999994 | 13003 | 2 | 0.7364 | 0.47411 | 0.70763 | 1 | 14469 | 3 | 0.7364 |
| ERMAP | 5 | 0.75332 | 0.79261 | 0.999994 | 16892 | 1 | 0.63827 | 0.47415 | 0.69957 | 1 | 14470 | 3 | 0.63827 |
| DNAI1 | 4 | 0.31537 | 0.45873 | 0.999994 | 10425 | 2 | 0.22586 | 0.47421 | 0.64485 | 1 | 14471 | 2 | 0.22586 |
| WTH3DI | 6 | 0.19008 | 0.37559 | 0.999994 | 7487 | 2 | 0.36307 | 0.47421 | 0.70769 | 1 | 14472 | 3 | 0.36307 |
| ARHGAP4 | 6 | 0.10012 | 0.24758 | 0.990682 | 4977 | 2 | 0.39806 | 0.47435 | 0.70781 | 1 | 14473 | 3 | 0.39806 |
| HTR4 | 5 | 0.30266 | 0.48321 | 0.999994 | 10185 | 1 | 0.60526 | 0.47436 | 0.69978 | 1 | 14474 | 3 | 0.60526 |
| FCGR2B | 6 | 0.043855 | 0.13057 | 0.95318 | 2706 | 2 | -0.28234 | 0.47442 | 0.70786 | 1 | 14475 | 2 | -0.28234 |
| hsa-mir-448 | 3 | 0.24994 | 0.37487 | 0.999994 | 9012 | 1 | 0.71434 | 0.47443 | 0.62912 | 1 | 14476 | 2 | 0.71434 |
| CCBE1 | 5 | 0.10685 | 0.23706 | 0.985687 | 5207 | 2 | 0.58677 | 0.47447 | 0.69988 | 1 | 14477 | 3 | 0.58677 |
| hsa-mir-182 | 4 | 0.30043 | 0.44346 | 0.999994 | 10141 | 2 | 0.076658 | 0.47453 | 0.6451 | 1 | 14478 | 2 | 0.076658 |
| IFITM1 | 6 | 0.10178 | 0.25079 | 0.992497 | 5032 | 3 | -0.057119 | 0.47454 | 0.70797 | 1 | 14479 | 3 | -0.057119 |
| RFX8 | 6 | 0.11038 | 0.26744 | 0.998561 | 5316 | 4 | -1.2082 | 0.47454 | 0.70797 | 1 | 14480 | 2 | -1.2082 |
| SPAG8 | 6 | 0.088537 | 0.22461 | 0.983354 | 4536 | 4 | -1.3477 | 0.47454 | 0.70797 | 1 | 14481 | 1 | -1.3477 |
| TRIM36 | 6 | 0.20763 | 0.39719 | 0.999994 | 7965 | 3 | -0.14363 | 0.47454 | 0.70797 | 1 | 14482 | 3 | -0.14363 |
| RNF25 | 6 | 0.30043 | 0.5072 | 0.999994 | 10142 | 3 | -0.38061 | 0.47454 | 0.70797 | 1 | 14483 | 2 | -0.38061 |
| CD200R1 | 6 | 0.029942 | 0.098656 | 0.945253 | 2003 | 3 | -1.6869 | 0.47454 | 0.70797 | 1 | 14484 | 2 | -1.6869 |
| GNGT2 | 6 | 0.35128 | 0.56545 | 0.999994 | 11181 | 3 | 0.26163 | 0.47458 | 0.70799 | 1 | 14485 | 3 | 0.26163 |
| PRB1 | 6 | 0.14845 | 0.32367 | 0.999994 | 6417 | 1 | 0.60709 | 0.47462 | 0.70802 | 1 | 14486 | 3 | 0.60709 |
| hsa-mir-548j | 3 | 0.82265 | 0.82257 | 0.999994 | 17818 | 0 | 0.57525 | 0.47462 | 0.6293 | 1 | 14487 | 2 | 0.57525 |
| NonTargetingControlG | 2 | 0.68784 | 0.68766 | 0.999994 | 16193 | 0 | 1.3481 | 0.4747 | 0.51961 | 1 | 14488 | 1 | 1.3481 |
| PNLIPRP3 | 5 | 0.80485 | 0.82256 | 0.999994 | 17550 | 1 | 0.55933 | 0.47476 | 0.70013 | 1 | 14489 | 2 | 0.55933 |
| PCDHA5 | 2 | 0.39968 | 0.41651 | 0.999994 | 11945 | 1 | 0.50856 | 0.47519 | 0.51996 | 1 | 14490 | 1 | 0.50856 |
| hsa-mir-4284 | 3 | 0.56723 | 0.59263 | 0.999994 | 14501 | 1 | 0.6336 | 0.4752 | 0.62988 | 1 | 14491 | 2 | 0.6336 |
| TBCE | 5 | 0.34612 | 0.51874 | 0.999994 | 11073 | 2 | 0.55935 | 0.47521 | 0.70056 | 1 | 14492 | 3 | 0.55935 |
| NonTargetingControlG | 2 | 0.016198 | 0.03033 | 0.862223 | 1257 | 1 | -3.722 | 0.47523 | 0.51999 | 1 | 14493 | 1 | -3.722 |
| hsa-mir-615 | 3 | 0.2847 | 0.41776 | 0.999994 | 9802 | 1 | 0.4164 | 0.47526 | 0.62993 | 1 | 14494 | 1 | 0.4164 |
| HIST1H4G | 6 | 0.1976 | 0.38498 | 0.999994 | 7690 | 2 | 0.54258 | 0.47531 | 0.70853 | 1 | 14495 | 3 | 0.54258 |
| RAB39B | 5 | 0.18626 | 0.35561 | 0.999994 | 7368 | 3 | -0.91149 | 0.47531 | 0.70066 | 1 | 14496 | 2 | -0.91149 |
| TMEM30B | 6 | 0.17376 | 0.35533 | 0.999994 | 7076 | 3 | -1.4034 | 0.4754 | 0.7086 | 1 | 14497 | 2 | -1.4034 |
| OR4S1 | 4 | 0.044683 | 0.11109 | 0.951052 | 2745 | 2 | -1.749 | 0.47544 | 0.64582 | 1 | 14498 | 2 | -1.749 |
| ZP3 | 5 | 0.2967 | 0.47838 | 0.999994 | 10058 | 2 | 0.46404 | 0.47551 | 0.70083 | 1 | 14499 | 2 | 0.46404 |
| AMACR | 5 | 0.37292 | 0.54083 | 0.999994 | 11557 | 2 | -0.50927 | 0.47554 | 0.70088 | 1 | 14500 | 2 | -0.50927 |
| PEX12 | 6 | 0.036721 | 0.1145 | 0.951052 | 2360 | 2 | 0.65115 | 0.47556 | 0.70872 | 1 | 14501 | 3 | 0.65115 |
| GSDMB | 5 | 0.24252 | 0.43434 | 0.999994 | 8847 | 3 | -0.59865 | 0.47564 | 0.70094 | 1 | 14502 | 1 | -0.59865 |
| CLVS2 | 5 | 0.30669 | 0.48647 | 0.999994 | 10267 | 2 | 0.11464 | 0.47564 | 0.70094 | 1 | 14503 | 2 | 0.11464 |
| GGT1 | 4 | 0.20318 | 0.34392 | 0.999994 | 7859 | 1 | -0.18125 | 0.47567 | 0.646 | 1 | 14504 | 1 | -0.18125 |
| ZIK1 | 6 | 0.54914 | 0.70301 | 0.999994 | 14196 | 1 | 0.29239 | 0.47585 | 0.70893 | 1 | 14505 | 2 | 0.29239 |
| LMF1 | 6 | 0.085673 | 0.21885 | 0.980699 | 4433 | 3 | -0.88003 | 0.47621 | 0.70921 | 1 | 14506 | 2 | -0.88003 |
| DHX15 | 2 | 0.3302 | 0.36022 | 0.999994 | 10724 | 1 | -0.39492 | 0.47637 | 0.52079 | 1 | 14507 | 1 | -0.39492 |

| | | | | | | | | | | | | |
|---|---|---|---|---|---|---|---|---|---|---|---|---|
| NonTargetingControlG | 2 | 0.094568 | 0.16162 | 0.967126 | 4771 | 1 | -2.5542 | 0.47637 | 0.52079 | 1 | 14508 | 1 | -2.5542 |
| NonTargetingControlG | 2 | 0.54317 | 0.54433 | 0.999994 | 14115 | 1 | 1.1486 | 0.47637 | 0.52079 | 1 | 14509 | 1 | 1.1486 |
| NDRG2 | 5 | 0.043172 | 0.12374 | 0.951052 | 2681 | 2 | 0.026555 | 0.47653 | 0.70139 | 1 | 14510 | 2 | 0.026555 |
| C3orf14 | 5 | 0.16104 | 0.31906 | 0.999994 | 6765 | 2 | -0.41485 | 0.4766 | 0.70142 | 1 | 14511 | 1 | -0.41485 |
| BTG4 | 5 | 0.52828 | 0.67254 | 0.999994 | 13865 | 2 | 0.37956 | 0.4766 | 0.70142 | 1 | 14512 | 2 | 0.37956 |
| PLTP | 5 | 0.22574 | 0.41131 | 0.999994 | 8416 | 2 | 0.32795 | 0.4766 | 0.70142 | 1 | 14513 | 2 | 0.32795 |
| HIST1H2BN | 5 | 0.036791 | 0.10785 | 0.951052 | 2363 | 4 | -1.0442 | 0.4766 | 0.70142 | 1 | 14514 | 1 | -1.0442 |
| NonTargetingControlG | 1 | 0.52333 | 0.52314 | 0.999994 | 13800 | 0 | 0.59262 | 0.47667 | 0.47668 | 1 | 14515 | 1 | 0.59262 |
| ZNF664-FAM101A | 3 | 0.089434 | 0.16897 | 0.967126 | 4575 | 2 | -2.3723 | 0.47668 | 0.63139 | 1 | 14516 | 1 | -2.3723 |
| hsa-mir-543 | 1 | 0.52322 | 0.52304 | 0.999994 | 13798 | 0 | 0.62041 | 0.47678 | 0.47678 | 1 | 14517 | 1 | 0.62041 |
| hsa-mir-548i-2 | 2 | 0.4075 | 0.42304 | 0.999994 | 12066 | 1 | 0.63994 | 0.47679 | 0.52107 | 1 | 14518 | 1 | 0.63994 |
| TMEM255A | 6 | 0.087467 | 0.22242 | 0.983354 | 4494 | 3 | -0.64396 | 0.47679 | 0.70962 | 1 | 14519 | 3 | -0.64396 |
| KPNA7 | 6 | 0.032348 | 0.10431 | 0.947507 | 2123 | 2 | 0.48085 | 0.47681 | 0.70964 | 1 | 14520 | 3 | 0.48085 |
| C8orf82 | 3 | 0.72071 | 0.72069 | 0.999994 | 16504 | 1 | 0.55885 | 0.47691 | 0.63162 | 1 | 14521 | 2 | 0.55885 |
| HADHB | 4 | 0.69013 | 0.71804 | 0.999994 | 16212 | 1 | 0.38975 | 0.47696 | 0.64698 | 1 | 14522 | 1 | 0.38975 |
| BMPR1A | 6 | 0.91558 | 0.91641 | 0.999994 | 19528 | 1 | 0.61085 | 0.47698 | 0.70977 | 1 | 14523 | 3 | 0.61085 |
| FNTB | 6 | 0.065631 | 0.17762 | 0.968171 | 3678 | 3 | -1.2574 | 0.4771 | 0.70986 | 1 | 14524 | 2 | -1.2574 |
| HS6ST1 | 6 | 0.052193 | 0.14889 | 0.965566 | 3075 | 2 | 0.36026 | 0.4771 | 0.70986 | 1 | 14525 | 2 | 0.36026 |
| LAX1 | 6 | 0.064894 | 0.17605 | 0.968171 | 3647 | 3 | -0.99676 | 0.4771 | 0.70986 | 1 | 14526 | 3 | -0.99676 |
| TBPL1 | 6 | 0.26989 | 0.47158 | 0.999994 | 9470 | 3 | -0.16853 | 0.4771 | 0.70986 | 1 | 14527 | 1 | -0.16853 |
| BAZ2A | 6 | 0.25784 | 0.45733 | 0.999994 | 9191 | 3 | 0.095598 | 0.47728 | 0.71001 | 1 | 14528 | 3 | 0.095598 |
| NonTargetingControlG | 1 | 0.5227 | 0.52254 | 0.999994 | 13790 | 0 | 0.70464 | 0.4773 | 0.4773 | 1 | 14529 | 1 | 0.70464 |
| P2RY13 | 6 | 0.00098518 | 0.0050031 | 0.589484 | 162 | 3 | -1.0539 | 0.47736 | 0.71007 | 1 | 14530 | 3 | -1.0539 |
| NonTargetingControlG | 2 | 0.097822 | 0.16672 | 0.967126 | 4882 | 1 | -2.764 | 0.47757 | 0.52161 | 1 | 14531 | 1 | -2.764 |
| CCBL1 | 6 | 0.16393 | 0.34319 | 0.999994 | 6840 | 2 | 0.6914 | 0.47757 | 0.71022 | 1 | 14532 | 3 | 0.6914 |
| CDHR5 | 6 | 0.25597 | 0.45507 | 0.999994 | 9150 | 3 | -0.21493 | 0.47778 | 0.71039 | 1 | 14533 | 3 | -0.21493 |
| TLX3 | 5 | 0.18118 | 0.34835 | 0.999994 | 7252 | 3 | -0.95791 | 0.47779 | 0.70208 | 1 | 14534 | 2 | -0.95791 |
| DBN1 | 6 | 0.35385 | 0.56836 | 0.999994 | 11231 | 3 | 0.06295 | 0.47795 | 0.71052 | 1 | 14535 | 3 | 0.06295 |
| HOXA10 | 5 | 0.32402 | 0.5006 | 0.999994 | 10599 | 2 | 0.22224 | 0.47802 | 0.7022 | 1 | 14536 | 2 | 0.22224 |
| hsa-mir-3944 | 3 | 0.017225 | 0.044776 | 0.891621 | 1316 | 1 | 0.57114 | 0.47803 | 0.63272 | 1 | 14537 | 2 | 0.57114 |
| SLC25A25 | 6 | 0.032047 | 0.1036 | 0.94662 | 2109 | 3 | -1.5594 | 0.47807 | 0.71061 | 1 | 14538 | 3 | -1.5594 |
| KRT31 | 6 | 0.082781 | 0.21301 | 0.979587 | 4330 | 2 | 0.81416 | 0.4781 | 0.71064 | 1 | 14539 | 3 | 0.81416 |
| NonTargetingControlG | 1 | 0.52184 | 0.52167 | 0.999994 | 13778 | 0 | 0.67202 | 0.47816 | 0.47813 | 1 | 14540 | 1 | 0.67202 |
| CUTA | 4 | 0.22551 | 0.36686 | 0.999994 | 8412 | 1 | 0.14497 | 0.47818 | 0.64791 | 1 | 14541 | 1 | 0.14497 |
| TMOD4 | 4 | 0.050511 | 0.12249 | 0.951052 | 2995 | 2 | -1.6359 | 0.47818 | 0.64791 | 1 | 14542 | 2 | -1.6359 |
| SEPT7 | 5 | 0.70885 | 0.77055 | 0.999994 | 16395 | 1 | 0.26863 | 0.47831 | 0.70235 | 1 | 14543 | 2 | 0.26863 |
| RPL10 | 3 | 0.12191 | 0.21198 | 0.979587 | 5700 | 2 | -1.7137 | 0.47834 | 0.63304 | 1 | 14544 | 1 | -1.7137 |
| GPHN | 6 | 0.033717 | 0.10751 | 0.951052 | 2201 | 4 | -1.8626 | 0.47842 | 0.71088 | 1 | 14545 | 2 | -1.8626 |
| TAOK1 | 6 | 0.55949 | 0.71044 | 0.999994 | 14362 | 2 | 0.61297 | 0.47843 | 0.71088 | 1 | 14546 | 3 | 0.61297 |
| PERP | 5 | 0.062552 | 0.16627 | 0.967126 | 3538 | 2 | -0.483 | 0.47851 | 0.70246 | 1 | 14547 | 2 | -0.483 |
| GJD3 | 2 | 0.68822 | 0.68804 | 0.999994 | 16197 | 0 | 1.4637 | 0.4787 | 0.52242 | 1 | 14548 | 1 | 1.4637 |
| TRPT1 | 6 | 0.057826 | 0.16109 | 0.967126 | 3330 | 3 | -0.68821 | 0.47873 | 0.71111 | 1 | 14549 | 3 | -0.68821 |
| TTC21A | 5 | 0.13492 | 0.28013 | 0.999632 | 6047 | 3 | -1.276 | 0.47874 | 0.70258 | 1 | 14550 | 2 | -1.276 |
| GALNT6 | 5 | 0.032366 | 0.096533 | 0.945031 | 2126 | 4 | -1.0472 | 0.47874 | 0.70258 | 1 | 14551 | 1 | -1.0472 |
| XRCC3 | 5 | 0.71269 | 0.77237 | 0.999994 | 16427 | 1 | 0.47984 | 0.47876 | 0.70258 | 1 | 14552 | 2 | 0.47984 |
| PI4K2B | 5 | 0.39387 | 0.55842 | 0.999994 | 11857 | 2 | -0.42872 | 0.47884 | 0.70263 | 1 | 14553 | 2 | -0.42872 |
| hsa-mir-4307 | 4 | 0.84001 | 0.83947 | 0.999994 | 18089 | 0 | 0.77383 | 0.47885 | 0.64839 | 1 | 14554 | 2 | 0.77383 |
| ZNF839 | 6 | 0.24503 | 0.442 | 0.999994 | 8901 | 3 | 0.13714 | 0.47893 | 0.71124 | 1 | 14555 | 3 | 0.13714 |
| ADAD1 | 6 | 0.64171 | 0.77123 | 0.999994 | 15542 | 2 | 0.49235 | 0.479 | 0.7113 | 1 | 14556 | 3 | 0.49235 |
| LMNB2 | 5 | 0.1528 | 0.30696 | 0.999994 | 6529 | 3 | -1.0375 | 0.47901 | 0.70272 | 1 | 14557 | 2 | -1.0375 |
| ADAM11 | 6 | 0.24415 | 0.44094 | 0.999994 | 8888 | 3 | -0.17254 | 0.47902 | 0.71133 | 1 | 14558 | 3 | -0.17254 |
| TBC1D22B | 6 | 0.085673 | 0.21885 | 0.980699 | 4432 | 2 | -0.17788 | 0.47902 | 0.71133 | 1 | 14559 | 2 | -0.17788 |

| | | | | | | | | | | | | | |
|---|---|---|---|---|---|---|---|---|---|---|---|---|---|
| NonTargetingControlG | 2 | 0.59889 | 0.59876 | 0.999994 | 14940 | 0 | 1.1477 | 0.47906 | 0.5227 | 1 | 14560 | 1 | 1.1477 |
| NonTargetingControlG | 2 | 0.61221 | 0.61208 | 0.999994 | 15112 | 0 | 1.2228 | 0.47912 | 0.52274 | 1 | 14561 | 1 | 1.2228 |
| CDK2AP2 | 5 | 0.21951 | 0.40259 | 0.999994 | 8278 | 3 | -0.71862 | 0.47926 | 0.70285 | 1 | 14562 | 2 | -0.71862 |
| AMFR | 4 | 0.2074 | 0.34833 | 0.999994 | 7957 | 2 | 0.11704 | 0.47945 | 0.64886 | 1 | 14563 | 2 | 0.11704 |
| COL4A2 | 5 | 0.3879 | 0.55339 | 0.999994 | 11766 | 1 | 0.60299 | 0.47965 | 0.70305 | 1 | 14564 | 2 | 0.60299 |
| CCR2 | 5 | 0.11988 | 0.25715 | 0.994943 | 5622 | 3 | -0.98412 | 0.47972 | 0.70309 | 1 | 14565 | 2 | -0.98412 |
| AKAP7 | 5 | 0.13646 | 0.28243 | 0.999632 | 6096 | 3 | -1.173 | 0.47981 | 0.70314 | 1 | 14566 | 2 | -1.173 |
| PPA1 | 6 | 0.044194 | 0.13131 | 0.95318 | 2721 | 1 | 0.80703 | 0.47982 | 0.71198 | 1 | 14567 | 3 | 0.80703 |
| LRRC20 | 6 | 0.47007 | 0.6485 | 0.999994 | 13008 | 2 | -0.031407 | 0.47986 | 0.71201 | 1 | 14568 | 2 | -0.031407 |
| LRRC26 | 6 | 0.032998 | 0.10585 | 0.950444 | 2158 | 3 | -0.59736 | 0.47992 | 0.71206 | 1 | 14569 | 2 | -0.59736 |
| APC | 6 | 0.92177 | 0.92204 | 0.999994 | 19670 | 1 | 0.07537 | 0.47992 | 0.71206 | 1 | 14570 | 2 | 0.07537 |
| CUX1 | 4 | 0.70984 | 0.73105 | 0.999994 | 16399 | 1 | 0.56581 | 0.47995 | 0.64924 | 1 | 14571 | 2 | 0.56581 |
| hsa-mir-1289-1 | 4 | 0.0034534 | 0.012344 | 0.733879 | 400 | 2 | -3.919 | 0.48013 | 0.64939 | 1 | 14572 | 2 | -3.919 |
| NonTargetingControlG | 1 | 0.51974 | 0.51956 | 0.999994 | 13744 | 0 | 0.67277 | 0.48026 | 0.48024 | 1 | 14573 | 1 | 0.67277 |
| MCM5 | 4 | 0.0081918 | 0.027986 | 0.855011 | 743 | 3 | -5.069 | 0.48031 | 0.64953 | 1 | 14574 | 1 | -5.069 |
| ZFP36L2 | 6 | 0.0013667 | 0.006836 | 0.643671 | 207 | 5 | -3.1051 | 0.4806 | 0.71257 | 1 | 14575 | 1 | -3.1051 |
| ANP32E | 6 | 0.1975 | 0.38487 | 0.999994 | 7687 | 3 | -0.78335 | 0.4806 | 0.71257 | 1 | 14576 | 2 | -0.78335 |
| PCGF2 | 6 | 0.011689 | 0.044696 | 0.891621 | 971 | 5 | -1.3527 | 0.4806 | 0.71257 | 1 | 14577 | 1 | -1.3527 |
| ZNF268 | 5 | 0.51776 | 0.66336 | 0.999994 | 13707 | 1 | 0.1745 | 0.48065 | 0.70362 | 1 | 14578 | 2 | 0.1745 |
| NPM2 | 6 | 0.25224 | 0.4506 | 0.999994 | 9063 | 3 | 0.09234 | 0.48073 | 0.71268 | 1 | 14579 | 3 | 0.09234 |
| ACBD6 | 5 | 0.34473 | 0.51761 | 0.999994 | 11048 | 2 | 0.097318 | 0.4809 | 0.70374 | 1 | 14580 | 1 | 0.097318 |
| SAFB | 5 | 0.019318 | 0.062241 | 0.937033 | 1421 | 3 | -3.3614 | 0.4809 | 0.70374 | 1 | 14581 | 2 | -3.3614 |
| PF4 | 6 | 0.31544 | 0.52466 | 0.999994 | 10428 | 1 | 0.65517 | 0.48091 | 0.71282 | 1 | 14582 | 3 | 0.65517 |
| ECI2 | 6 | 0.14829 | 0.32349 | 0.999994 | 6414 | 3 | -0.57519 | 0.48131 | 0.71311 | 1 | 14583 | 2 | -0.57519 |
| SHROOM2 | 6 | 0.49947 | 0.66832 | 0.999994 | 13415 | 2 | 0.8355 | 0.48139 | 0.71318 | 1 | 14584 | 3 | 0.8355 |
| ALDOB | 6 | 0.045041 | 0.13315 | 0.95318 | 2759 | 4 | -1.9262 | 0.48148 | 0.71324 | 1 | 14585 | 1 | -1.9262 |
| FAM20A | 6 | 0.075636 | 0.19848 | 0.977164 | 4058 | 3 | -1.5598 | 0.48148 | 0.71324 | 1 | 14586 | 2 | -1.5598 |
| DGKB | 6 | 0.34208 | 0.55504 | 0.999994 | 10988 | 3 | 0.33189 | 0.48149 | 0.71325 | 1 | 14587 | 3 | 0.33189 |
| IRF8 | 5 | 0.6419 | 0.74193 | 0.999994 | 15549 | 1 | 0.0030769 | 0.48176 | 0.7042 | 1 | 14588 | 2 | 0.0030769 |
| ANXA6 | 5 | 0.12245 | 0.26107 | 0.994943 | 5719 | 2 | -0.56978 | 0.48176 | 0.7042 | 1 | 14589 | 1 | -0.56978 |
| hsa-mir-5095 | 3 | 0.48447 | 0.53602 | 0.999994 | 13208 | 1 | 0.68617 | 0.48177 | 0.63572 | 1 | 14590 | 1 | 0.68617 |
| ODAM | 6 | 0.65305 | 0.77995 | 0.999994 | 15698 | 2 | 0.42511 | 0.48179 | 0.71347 | 1 | 14591 | 3 | 0.42511 |
| NonTargetingControlG | 2 | 0.11097 | 0.18694 | 0.969409 | 5334 | 1 | -2.4874 | 0.48197 | 0.52477 | 1 | 14592 | 1 | -2.4874 |
| MDGA1 | 6 | 0.060986 | 0.16786 | 0.967126 | 3467 | 3 | -0.28927 | 0.482 | 0.71361 | 1 | 14593 | 3 | -0.28927 |
| BCAP29 | 6 | 0.031642 | 0.10267 | 0.946414 | 2088 | 3 | -0.88851 | 0.48214 | 0.71372 | 1 | 14594 | 3 | -0.88851 |
| TSPAN19 | 6 | 0.27098 | 0.47288 | 0.999994 | 9498 | 3 | -0.14062 | 0.48232 | 0.71386 | 1 | 14595 | 3 | -0.14062 |
| B3GALNT1 | 6 | 0.037651 | 0.11661 | 0.951052 | 2411 | 3 | -0.49325 | 0.48247 | 0.71398 | 1 | 14596 | 3 | -0.49325 |
| CCDC27 | 6 | 0.11632 | 0.27881 | 0.999632 | 5522 | 3 | -0.1386 | 0.48251 | 0.71401 | 1 | 14597 | 3 | -0.1386 |
| CA7 | 4 | 0.5837 | 0.65664 | 0.999994 | 14728 | 1 | 0.58455 | 0.48255 | 0.65122 | 1 | 14598 | 2 | 0.58455 |
| PRDM2 | 6 | 0.11567 | 0.27755 | 0.999379 | 5508 | 2 | 0.52134 | 0.48257 | 0.71406 | 1 | 14599 | 3 | 0.52134 |
| L2HGDH | 5 | 0.40654 | 0.56901 | 0.999994 | 12051 | 2 | 0.044276 | 0.48261 | 0.70464 | 1 | 14600 | 2 | 0.044276 |
| HIST1H3I | 4 | 0.70292 | 0.72635 | 0.999994 | 16337 | 1 | 0.60047 | 0.48279 | 0.65139 | 1 | 14601 | 2 | 0.60047 |
| EXOSC2 | 4 | 0.088613 | 0.19448 | 0.975797 | 4540 | 2 | -1.7705 | 0.4828 | 0.6514 | 1 | 14602 | 2 | -1.7705 |
| PRKAG1 | 6 | 0.1145 | 0.27531 | 0.999239 | 5462 | 3 | -0.53266 | 0.48282 | 0.71425 | 1 | 14603 | 3 | -0.53266 |
| BLZF1 | 6 | 0.11836 | 0.28263 | 0.999632 | 5584 | 3 | -0.47707 | 0.48288 | 0.71429 | 1 | 14604 | 3 | -0.47707 |
| POMP | 6 | 0.03624 | 0.1134 | 0.951052 | 2330 | 3 | 0.13692 | 0.48296 | 0.71435 | 1 | 14605 | 3 | 0.13692 |
| NUDT2 | 6 | 0.63687 | 0.76755 | 0.999994 | 15471 | 1 | 0.76914 | 0.48304 | 0.71441 | 1 | 14606 | 3 | 0.76914 |
| NonTargetingControlG | 2 | 0.32747 | 0.35804 | 0.999994 | 10668 | 1 | 0.29859 | 0.48311 | 0.52561 | 1 | 14607 | 1 | 0.29859 |
| HOOK1 | 5 | 0.48561 | 0.63548 | 0.999994 | 13223 | 2 | 0.099564 | 0.48334 | 0.70501 | 1 | 14608 | 2 | 0.099564 |
| METTL21A | 5 | 0.080273 | 0.19502 | 0.975797 | 4238 | 3 | -0.7158 | 0.48346 | 0.70507 | 1 | 14609 | 1 | -0.7158 |
| HEATR5B | 5 | 0.80246 | 0.82107 | 0.999994 | 17519 | 1 | 0.043751 | 0.48346 | 0.70507 | 1 | 14610 | 1 | 0.043751 |
| NKX6-1 | 4 | 0.67651 | 0.70939 | 0.999994 | 16032 | 1 | 1.1735 | 0.48348 | 0.65191 | 1 | 14611 | 2 | 1.1735 |

| | | | | | | | | | | | | | |
|---|---|---|---|---|---|---|---|---|---|---|---|---|---|
| DAD1 | 3 | 0.25742 | 0.38416 | 0.999994 | 9177 | 2 | -0.83647 | 0.48355 | 0.63647 | 1 | 14612 | 1 | -0.83647 |
| ZNF576 | 5 | 0.2253 | 0.4107 | 0.999994 | 8408 | 2 | -0.14856 | 0.48388 | 0.70529 | 1 | 14613 | 2 | -0.14856 |
| hsa-mir-874 | 4 | 0.097477 | 0.21061 | 0.979587 | 4872 | 3 | -1.7245 | 0.48396 | 0.65225 | 1 | 14614 | 1 | -1.7245 |
| hsa-mir-6861 | 4 | 0.0056639 | 0.019829 | 0.808847 | 567 | 2 | -1.0033 | 0.48396 | 0.65225 | 1 | 14615 | 1 | -1.0033 |
| VTCN1 | 6 | 0.16888 | 0.34934 | 0.999994 | 6958 | 2 | 0.40648 | 0.4842 | 0.71531 | 1 | 14616 | 3 | 0.40648 |
| KLHL12 | 6 | 0.095134 | 0.2378 | 0.987253 | 4789 | 4 | -0.98769 | 0.4842 | 0.71531 | 1 | 14617 | 2 | -0.98769 |
| NDUFV1 | 6 | 0.01906 | 0.068979 | 0.93914 | 1412 | 3 | -2.359 | 0.4842 | 0.71531 | 1 | 14618 | 2 | -2.359 |
| ODF1 | 6 | 0.2054 | 0.39443 | 0.999994 | 7906 | 3 | -0.040713 | 0.48425 | 0.71535 | 1 | 14619 | 3 | -0.040713 |
| KCNF1 | 6 | 0.019734 | 0.071153 | 0.93914 | 1441 | 3 | -2.3438 | 0.48426 | 0.71537 | 1 | 14620 | 3 | -2.3438 |
| MROH7 | 6 | 0.67705 | 0.79868 | 0.999994 | 16041 | 1 | 0.17983 | 0.48426 | 0.71537 | 1 | 14621 | 2 | 0.17983 |
| SYCE1L | 6 | 0.007715 | 0.030961 | 0.862223 | 709 | 4 | -3.5496 | 0.48426 | 0.71537 | 1 | 14622 | 2 | -3.5496 |
| hsa-mir-7974 | 4 | 0.42294 | 0.56921 | 0.999994 | 12296 | 2 | 0.51226 | 0.48438 | 0.65258 | 1 | 14623 | 2 | 0.51226 |
| UBE4B | 5 | 0.033204 | 0.098746 | 0.945253 | 2171 | 4 | -2.1289 | 0.48454 | 0.70563 | 1 | 14624 | 1 | -2.1289 |
| FAM193B | 5 | 0.84994 | 0.85423 | 0.999994 | 18261 | 1 | -0.45674 | 0.48454 | 0.70563 | 1 | 14625 | 2 | -0.45674 |
| GCK | 6 | 0.2818 | 0.48547 | 0.999994 | 9736 | 3 | 0.089894 | 0.48468 | 0.71567 | 1 | 14626 | 3 | 0.089894 |
| TULP1 | 5 | 0.90285 | 0.90294 | 0.999994 | 19271 | 0 | 0.0035468 | 0.48496 | 0.70585 | 1 | 14627 | 2 | 0.0035468 |
| CDR2L | 5 | 0.46809 | 0.62049 | 0.999994 | 12976 | 2 | -0.49089 | 0.48496 | 0.70585 | 1 | 14628 | 2 | -0.49089 |
| GRAMD3 | 5 | 0.41868 | 0.57901 | 0.999994 | 12230 | 2 | -0.37254 | 0.48496 | 0.70585 | 1 | 14629 | 2 | -0.37254 |
| CHRM5 | 5 | 0.19829 | 0.37282 | 0.999994 | 7720 | 1 | 0.41007 | 0.48496 | 0.70585 | 1 | 14630 | 2 | 0.41007 |
| hsa-mir-744 | 3 | 0.52153 | 0.56056 | 0.999994 | 13770 | 1 | -0.52114 | 0.48499 | 0.63707 | 1 | 14631 | 1 | -0.52114 |
| FAM196A | 6 | 0.35023 | 0.56425 | 0.999994 | 11162 | 2 | 0.6349 | 0.48501 | 0.71592 | 1 | 14632 | 3 | 0.6349 |
| KCNIP4 | 6 | 0.15026 | 0.32594 | 0.999994 | 6463 | 3 | -0.0058794 | 0.48506 | 0.71596 | 1 | 14633 | 3 | -0.0058794 |
| C15orf39 | 6 | 0.000014599 | 0.000088928 | 0.114686 | 15 | 4 | -5.8767 | 0.48506 | 0.71596 | 1 | 14634 | 1 | -5.8767 |
| DLGAP5 | 6 | 0.051503 | 0.14737 | 0.965554 | 3046 | 4 | -1.4859 | 0.48551 | 0.7163 | 1 | 14635 | 2 | -1.4859 |
| PHF12 | 5 | 0.45595 | 0.61026 | 0.999994 | 12790 | 2 | -0.07069 | 0.48558 | 0.70617 | 1 | 14636 | 1 | -0.07069 |
| FAM69A | 5 | 0.45824 | 0.61217 | 0.999994 | 12824 | 2 | -0.23894 | 0.48563 | 0.70619 | 1 | 14637 | 2 | -0.23894 |
| NonTargetingControlG | 1 | 0.51431 | 0.51418 | 0.999994 | 13648 | 0 | 0.6079 | 0.48569 | 0.48569 | 1 | 14638 | 1 | 0.6079 |
| CPEB2 | 6 | 0.12529 | 0.29433 | 0.999994 | 5800 | 3 | -0.01628 | 0.48569 | 0.71644 | 1 | 14639 | 3 | -0.01628 |
| C1GALT1C1 | 6 | 0.031548 | 0.10244 | 0.946414 | 2082 | 3 | -1.3389 | 0.48572 | 0.71647 | 1 | 14640 | 2 | -1.3389 |
| GLIS3 | 6 | 0.15634 | 0.33363 | 0.999994 | 6639 | 2 | 0.55189 | 0.48573 | 0.71648 | 1 | 14641 | 3 | 0.55189 |
| NonTargetingControlG | 2 | 0.10485 | 0.17756 | 0.968171 | 5126 | 1 | -2.5574 | 0.48606 | 0.5277 | 1 | 14642 | 1 | -2.5574 |
| OR51V1 | 5 | 0.29468 | 0.47673 | 0.999994 | 10010 | 2 | -0.27333 | 0.48613 | 0.70644 | 1 | 14643 | 2 | -0.27333 |
| NCCRP1 | 6 | 0.92025 | 0.92063 | 0.999994 | 19634 | 1 | 0.4334 | 0.48614 | 0.71681 | 1 | 14644 | 3 | 0.4334 |
| FAM222B | 6 | 0.89517 | 0.90003 | 0.999994 | 19131 | 1 | 0.36742 | 0.48618 | 0.71684 | 1 | 14645 | 2 | 0.36742 |
| SEMA3G | 5 | 3.2337E-06 | 0.000014859 | 0.026815 | 12 | 3 | -10.625 | 0.48642 | 0.70658 | 1 | 14646 | 1 | -10.625 |
| DDX6 | 6 | 0.034013 | 0.10821 | 0.951052 | 2216 | 4 | -1.7242 | 0.48653 | 0.71712 | 1 | 14647 | 2 | -1.7242 |
| TGFBRAP1 | 6 | 0.071819 | 0.19056 | 0.971868 | 3912 | 4 | -1.1329 | 0.48657 | 0.71715 | 1 | 14648 | 1 | -1.1329 |
| FNDC4 | 6 | 0.33157 | 0.54304 | 0.999994 | 10756 | 3 | 0.30042 | 0.4866 | 0.71717 | 1 | 14649 | 3 | 0.30042 |
| ABCB1 | 4 | 0.2074 | 0.34833 | 0.999994 | 7958 | 2 | -0.089421 | 0.48663 | 0.65435 | 1 | 14650 | 2 | -0.089421 |
| RAB11FIP1 | 6 | 0.61458 | 0.75077 | 0.999994 | 15136 | 2 | 0.47078 | 0.48664 | 0.7172 | 1 | 14651 | 3 | 0.47078 |
| ASPM | 6 | 0.90481 | 0.90741 | 0.999994 | 19320 | 1 | 0.43906 | 0.48665 | 0.7172 | 1 | 14652 | 3 | 0.43906 |
| NonTargetingControlG | 2 | 0.0039467 | 0.0075886 | 0.670742 | 439 | 1 | -3.5999 | 0.48684 | 0.52825 | 1 | 14653 | 1 | -3.5999 |
| GNAT1 | 6 | 0.24167 | 0.43801 | 0.999994 | 8824 | 3 | 0.1216 | 0.4871 | 0.71756 | 1 | 14654 | 3 | 0.1216 |
| NonTargetingControlG | 2 | 0.31446 | 0.34795 | 0.999994 | 10408 | 1 | 0.10641 | 0.48716 | 0.52848 | 1 | 14655 | 1 | 0.10641 |
| FAM47C | 3 | 0.51501 | 0.5561 | 0.999994 | 13663 | 1 | -0.11513 | 0.48721 | 0.63794 | 1 | 14656 | 1 | -0.11513 |
| NIPAL3 | 6 | 0.095609 | 0.23873 | 0.987253 | 4802 | 4 | -0.99995 | 0.48728 | 0.7177 | 1 | 14657 | 2 | -0.99995 |
| hsa-mir-4280 | 4 | 0.77355 | 0.77864 | 0.999994 | 17125 | 1 | 0.46701 | 0.48729 | 0.65484 | 1 | 14658 | 2 | 0.46701 |
| POP5 | 4 | 0.28617 | 0.42887 | 0.999994 | 9836 | 2 | 0.16394 | 0.48734 | 0.65487 | 1 | 14659 | 2 | 0.16394 |
| GP1BB | 5 | 0.046702 | 0.13243 | 0.95318 | 2838 | 4 | -1.7535 | 0.48742 | 0.70712 | 1 | 14660 | 1 | -1.7535 |
| CCDC135 | 5 | 0.42236 | 0.58205 | 0.999994 | 12288 | 2 | 0.03165 | 0.48742 | 0.70712 | 1 | 14661 | 2 | 0.03165 |
| HOXD9 | 5 | 0.11718 | 0.25305 | 0.994501 | 5544 | 3 | -1.2883 | 0.48742 | 0.70712 | 1 | 14662 | 1 | -1.2883 |
| ZNF623 | 6 | 0.069542 | 0.1858 | 0.969409 | 3816 | 3 | 0.0070511 | 0.48744 | 0.71781 | 1 | 14663 | 3 | 0.0070511 |

| | | | | | | | | | | | | | |
|---|---|---|---|---|---|---|---|---|---|---|---|---|---|
| SLC6A6 | 6 | 0.001335 | 0.0066843 | 0.643671 | 199 | 5 | -3.2178 | 0.48744 | 0.71781 | 1 | 14664 | 1 | -3.2178 |
| NSD1 | 6 | 0.33459 | 0.54649 | 0.999994 | 10824 | 3 | 0.2319 | 0.4877 | 0.71802 | 1 | 14665 | 3 | 0.2319 |
| hsa-mir-1245a | 4 | 0.2862 | 0.4289 | 0.999994 | 9837 | 2 | 0.048289 | 0.4877 | 0.65514 | 1 | 14666 | 2 | 0.048289 |
| LRRD1 | 6 | 0.34771 | 0.56143 | 0.999994 | 11103 | 2 | 0.4832 | 0.48774 | 0.71805 | 1 | 14667 | 3 | 0.4832 |
| DCAF6 | 6 | 0.6686 | 0.79208 | 0.999994 | 15898 | 2 | 0.36922 | 0.4878 | 0.71809 | 1 | 14668 | 3 | 0.36922 |
| MAGEH1 | 5 | 0.21391 | 0.39483 | 0.999994 | 8112 | 2 | 0.54942 | 0.48813 | 0.70748 | 1 | 14669 | 2 | 0.54942 |
| NonTargetingControlG | 2 | 0.37563 | 0.39678 | 0.999994 | 11587 | 1 | 0.29171 | 0.48822 | 0.52927 | 1 | 14670 | 1 | 0.29171 |
| ICA1 | 6 | 0.051931 | 0.14835 | 0.965554 | 3061 | 3 | -0.21925 | 0.48837 | 0.71851 | 1 | 14671 | 2 | -0.21925 |
| SLC35F6 | 6 | 0.22697 | 0.42044 | 0.999994 | 8451 | 3 | -0.54898 | 0.48841 | 0.71854 | 1 | 14672 | 2 | -0.54898 |
| MYL6B | 6 | 0.21453 | 0.4056 | 0.999994 | 8134 | 3 | -0.1313 | 0.48846 | 0.71857 | 1 | 14673 | 3 | -0.1313 |
| ASNS | 4 | 0.013172 | 0.043521 | 0.891621 | 1075 | 2 | 0.44127 | 0.48858 | 0.65581 | 1 | 14674 | 2 | 0.44127 |
| hsa-mir-496 | 4 | 0.26681 | 0.40901 | 0.999994 | 9392 | 2 | -0.78473 | 0.48862 | 0.65585 | 1 | 14675 | 1 | -0.78473 |
| hsa-mir-5191 | 4 | 0.23505 | 0.37669 | 0.999994 | 8645 | 2 | -0.64694 | 0.48862 | 0.65585 | 1 | 14676 | 1 | -0.64694 |
| NR2F1 | 5 | 0.046829 | 0.13274 | 0.95318 | 2843 | 3 | -2.4927 | 0.48863 | 0.70773 | 1 | 14677 | 1 | -2.4927 |
| C3orf27 | 5 | 0.07268 | 0.18274 | 0.968526 | 3947 | 3 | -1.6278 | 0.48863 | 0.70773 | 1 | 14678 | 2 | -1.6278 |
| TRO | 5 | 0.21286 | 0.39339 | 0.999994 | 8085 | 2 | -0.30117 | 0.48863 | 0.70773 | 1 | 14679 | 2 | -0.30117 |
| SDHAF2 | 5 | 0.13628 | 0.28215 | 0.999632 | 6088 | 3 | -1.3759 | 0.48863 | 0.70773 | 1 | 14680 | 2 | -1.3759 |
| DIAPH3 | 5 | 0.35234 | 0.52382 | 0.999994 | 11202 | 2 | -0.41947 | 0.48863 | 0.70773 | 1 | 14681 | 1 | -0.41947 |
| FKBP1B | 5 | 0.049384 | 0.13886 | 0.957514 | 2948 | 2 | 0.18423 | 0.48863 | 0.70773 | 1 | 14682 | 2 | 0.18423 |
| NonTargetingControlG | 2 | 0.34202 | 0.36961 | 0.999994 | 10984 | 1 | 0.53524 | 0.48863 | 0.52957 | 1 | 14683 | 1 | 0.53524 |
| UPP2 | 5 | 0.29946 | 0.48063 | 0.999994 | 10119 | 2 | -0.054554 | 0.48883 | 0.70783 | 1 | 14684 | 2 | -0.054554 |
| FRS2 | 6 | 0.9244 | 0.92455 | 0.999994 | 19729 | 1 | 0.15677 | 0.48898 | 0.719 | 1 | 14685 | 2 | 0.15677 |
| NonTargetingControlG | 2 | 0.72908 | 0.7288 | 0.999994 | 16590 | 0 | 1.5513 | 0.48938 | 0.53009 | 1 | 14686 | 1 | 1.5513 |
| HTR1D | 5 | 0.12735 | 0.26858 | 0.998561 | 5855 | 2 | -0.14583 | 0.48939 | 0.70813 | 1 | 14687 | 2 | -0.14583 |
| GAL | 6 | 0.009096 | 0.035864 | 0.875006 | 799 | 5 | -2.0397 | 0.48939 | 0.71929 | 1 | 14688 | 1 | -2.0397 |
| CD300LG | 6 | 0.042441 | 0.12739 | 0.951052 | 2639 | 4 | -1.6151 | 0.48939 | 0.71929 | 1 | 14689 | 2 | -1.6151 |
| ID4 | 6 | 0.084382 | 0.21626 | 0.980699 | 4389 | 4 | -1.2221 | 0.48942 | 0.71931 | 1 | 14690 | 2 | -1.2221 |
| hsa-mir-495 | 4 | 0.79344 | 0.79539 | 0.999994 | 17390 | 1 | 0.62573 | 0.48964 | 0.65661 | 1 | 14691 | 2 | 0.62573 |
| RRAD | 5 | 0.32107 | 0.49817 | 0.999994 | 10537 | 2 | -0.33861 | 0.48971 | 0.70831 | 1 | 14692 | 2 | -0.33861 |
| LYPD5 | 5 | 0.23176 | 0.41964 | 0.999994 | 8559 | 3 | -0.63816 | 0.48974 | 0.70831 | 1 | 14693 | 2 | -0.63816 |
| KIAA0141 | 5 | 0.073386 | 0.1839 | 0.968526 | 3970 | 1 | 0.099564 | 0.48974 | 0.70831 | 1 | 14694 | 2 | 0.099564 |
| PLEKHA8 | 6 | 0.48137 | 0.65612 | 0.999994 | 13158 | 2 | 0.60438 | 0.49 | 0.71974 | 1 | 14695 | 3 | 0.60438 |
| hsa-mir-548an | 4 | 0.15552 | 0.29488 | 0.999994 | 6607 | 1 | 0.43104 | 0.49007 | 0.65693 | 1 | 14696 | 2 | 0.43104 |
| GLP1R | 6 | 0.33383 | 0.54567 | 0.999994 | 10807 | 2 | 0.63143 | 0.4901 | 0.71981 | 1 | 14697 | 3 | 0.63143 |
| NonTargetingControlG | 2 | 0.35258 | 0.37805 | 0.999994 | 11205 | 1 | 0.34476 | 0.4903 | 0.53077 | 1 | 14698 | 1 | 0.34476 |
| BBS7 | 5 | 0.036441 | 0.10693 | 0.950995 | 2343 | 3 | -2.7429 | 0.49033 | 0.70863 | 1 | 14699 | 2 | -2.7429 |
| LYRM4 | 4 | 0.059063 | 0.13899 | 0.957632 | 3383 | 2 | -1.0132 | 0.49043 | 0.65721 | 1 | 14700 | 2 | -1.0132 |
| CADM2 | 6 | 0.47386 | 0.65102 | 0.999994 | 13051 | 2 | 0.32459 | 0.49045 | 0.72007 | 1 | 14701 | 3 | 0.32459 |
| FAM69C | 6 | 0.040107 | 0.12214 | 0.951052 | 2522 | 2 | 0.73249 | 0.49051 | 0.72011 | 1 | 14702 | 3 | 0.73249 |
| FZR1 | 5 | 0.27395 | 0.45997 | 0.999994 | 9563 | 1 | 0.35554 | 0.49053 | 0.70874 | 1 | 14703 | 2 | 0.35554 |
| ASL | 6 | 0.25944 | 0.4592 | 0.999994 | 9226 | 3 | 0.024327 | 0.49059 | 0.72017 | 1 | 14704 | 3 | 0.024327 |
| CLPP | 5 | 0.0095329 | 0.034733 | 0.867348 | 840 | 4 | -4.4385 | 0.49068 | 0.70882 | 1 | 14705 | 1 | -4.4385 |
| KLHDC4 | 5 | 0.019815 | 0.063596 | 0.937389 | 1448 | 3 | -3.0969 | 0.49068 | 0.70882 | 1 | 14706 | 1 | -3.0969 |
| CEP72 | 5 | 0.042452 | 0.12199 | 0.951052 | 2640 | 3 | -2.4363 | 0.49068 | 0.70882 | 1 | 14707 | 1 | -2.4363 |
| GTF2H2C | 2 | 0.7292 | 0.72892 | 0.999994 | 16591 | 0 | 1.5272 | 0.49081 | 0.53112 | 1 | 14708 | 1 | 1.5272 |
| C9orf37 | 6 | 0.4197 | 0.6152 | 0.999994 | 12248 | 2 | -0.20008 | 0.49121 | 0.72063 | 1 | 14709 | 2 | -0.20008 |
| DCLK3 | 6 | 0.090333 | 0.22815 | 0.983354 | 4607 | 4 | -0.91955 | 0.49121 | 0.72063 | 1 | 14710 | 2 | -0.91955 |
| CDKL3 | 6 | 0.11157 | 0.26972 | 0.998561 | 5349 | 3 | -1.0313 | 0.49121 | 0.72063 | 1 | 14711 | 2 | -1.0313 |
| ZFHX3 | 6 | 0.16387 | 0.34311 | 0.999994 | 6836 | 3 | -0.81684 | 0.49121 | 0.72063 | 1 | 14712 | 2 | -0.81684 |
| CH25H | 3 | 0.13797 | 0.23293 | 0.984502 | 6134 | 2 | -1.8841 | 0.49122 | 0.63961 | 1 | 14713 | 1 | -1.8841 |
| EMC4 | 3 | 0.69164 | 0.69321 | 0.999994 | 16223 | 1 | 0.51299 | 0.49122 | 0.63961 | 1 | 14714 | 1 | 0.51299 |
| MTERF | 6 | 0.5911 | 0.73342 | 0.999994 | 14835 | 2 | 0.11593 | 0.49136 | 0.72075 | 1 | 14715 | 2 | 0.11593 |

| | | | | | | | | | | | | | |
|---|---|---|---|---|---|---|---|---|---|---|---|---|---|
| NT5C1B | 5 | 0.41876 | 0.57907 | 0.999994 | 12231 | 2 | 0.082054 | 0.49171 | 0.70941 | 1 | 14716 | 2 | 0.082054 |
| RAPSN | 5 | 0.22431 | 0.40929 | 0.999994 | 8386 | 3 | -0.71639 | 0.49171 | 0.70941 | 1 | 14717 | 1 | -0.71639 |
| RABL2A | 5 | 0.28732 | 0.47076 | 0.999994 | 9861 | 2 | 0.11273 | 0.49171 | 0.70941 | 1 | 14718 | 2 | 0.11273 |
| HEATR4 | 5 | 0.77797 | 0.80626 | 0.999994 | 17184 | 1 | 0.12726 | 0.49171 | 0.70941 | 1 | 14719 | 2 | 0.12726 |
| RNF217 | 6 | 0.0093822 | 0.03685 | 0.881473 | 818 | 2 | 0.68089 | 0.49172 | 0.72102 | 1 | 14720 | 3 | 0.68089 |
| SPACA5 | 6 | 0.57111 | 0.71874 | 0.999994 | 14561 | 1 | 0.42113 | 0.49185 | 0.72112 | 1 | 14721 | 3 | 0.42113 |
| hsa-mir-7157 | 4 | 0.097127 | 0.20995 | 0.979587 | 4864 | 3 | -1.2907 | 0.49196 | 0.65843 | 1 | 14722 | 1 | -1.2907 |
| hsa-mir-568 | 4 | 0.060809 | 0.14232 | 0.959553 | 3460 | 3 | -1.433 | 0.49196 | 0.65843 | 1 | 14723 | 1 | -1.433 |
| hsa-mir-539 | 2 | 0.74444 | 0.74412 | 0.999994 | 16794 | 0 | 1.4399 | 0.49214 | 0.53206 | 1 | 14724 | 1 | 1.4399 |
| hsa-mir-4482 | 4 | 0.86403 | 0.86365 | 0.999994 | 18521 | 0 | 0.72712 | 0.49216 | 0.65858 | 1 | 14725 | 2 | 0.72712 |
| ARF1 | 6 | 0.061377 | 0.16871 | 0.967126 | 3488 | 3 | -1.1693 | 0.49234 | 0.72146 | 1 | 14726 | 2 | -1.1693 |
| MILR1 | 6 | 0.11596 | 0.27813 | 0.999536 | 5514 | 2 | 0.073821 | 0.49244 | 0.72154 | 1 | 14727 | 3 | 0.073821 |
| hsa-mir-1185-2 | 1 | 0.50751 | 0.50738 | 0.999994 | 13544 | 0 | 0.60177 | 0.49249 | 0.4925 | 1 | 14728 | 0 | 0.60177 |
| KCTD14 | 5 | 0.41648 | 0.5772 | 0.999994 | 12198 | 2 | 0.39207 | 0.49267 | 0.7099 | 1 | 14729 | 2 | 0.39207 |
| hsa-mir-8073 | 4 | 0.79952 | 0.80079 | 0.999994 | 17474 | 1 | 0.70213 | 0.49276 | 0.65904 | 1 | 14730 | 2 | 0.70213 |
| ACP6 | 6 | 0.61971 | 0.75461 | 0.999994 | 15215 | 2 | 0.63063 | 0.4929 | 0.7219 | 1 | 14731 | 3 | 0.63063 |
| FKBP3 | 5 | 0.16112 | 0.31918 | 0.999994 | 6768 | 3 | -0.91715 | 0.49317 | 0.71015 | 1 | 14732 | 2 | -0.91715 |
| CLK1 | 6 | 0.13031 | 0.30069 | 0.999994 | 5939 | 3 | -0.28728 | 0.49337 | 0.72226 | 1 | 14733 | 3 | -0.28728 |
| DHRS11 | 6 | 0.90154 | 0.90488 | 0.999994 | 19244 | 1 | 0.41243 | 0.49356 | 0.72239 | 1 | 14734 | 3 | 0.41243 |
| RASGRP3 | 6 | 0.1905 | 0.37614 | 0.999994 | 7499 | 3 | -0.34081 | 0.49356 | 0.72239 | 1 | 14735 | 2 | -0.34081 |
| MRPS24 | 5 | 0.0078007 | 0.029678 | 0.862223 | 714 | 4 | -6.2873 | 0.49356 | 0.71038 | 1 | 14736 | 1 | -6.2873 |
| DLG2 | 5 | 0.42216 | 0.58188 | 0.999994 | 12284 | 1 | 0.15149 | 0.49356 | 0.71038 | 1 | 14737 | 1 | 0.15149 |
| NSUN7 | 6 | 0.098383 | 0.24416 | 0.988309 | 4908 | 1 | 0.57556 | 0.49359 | 0.72241 | 1 | 14738 | 3 | 0.57556 |
| GATC | 6 | 0.22359 | 0.41645 | 0.999994 | 8368 | 2 | -0.5414 | 0.49363 | 0.72244 | 1 | 14739 | 2 | -0.5414 |
| BCR | 6 | 0.005595 | 0.023285 | 0.830713 | 564 | 2 | 0.42708 | 0.49363 | 0.72244 | 1 | 14740 | 3 | 0.42708 |
| RNF6 | 6 | 0.016133 | 0.059447 | 0.932932 | 1250 | 2 | 0.30188 | 0.49363 | 0.72244 | 1 | 14741 | 2 | 0.30188 |
| SOD1 | 3 | 0.10931 | 0.19537 | 0.975797 | 5281 | 2 | -1.9227 | 0.49369 | 0.64065 | 1 | 14742 | 1 | -1.9227 |
| NonTargetingControlG | 1 | 0.50629 | 0.50616 | 0.999994 | 13531 | 0 | 0.53577 | 0.49371 | 0.49372 | 1 | 14743 | 0 | 0.53577 |
| NonTargetingControlG | 2 | 0.29698 | 0.33453 | 0.999994 | 10066 | 1 | -0.13468 | 0.49378 | 0.53326 | 1 | 14744 | 1 | -0.13468 |
| hsa-mir-7-3 | 4 | 0.4291 | 0.57563 | 0.999994 | 12399 | 2 | 0.32318 | 0.494 | 0.65995 | 1 | 14745 | 2 | 0.32318 |
| TSPAN11 | 5 | 0.14046 | 0.28851 | 0.999632 | 6200 | 1 | 0.38008 | 0.49428 | 0.71075 | 1 | 14746 | 2 | 0.38008 |
| PPT2 | 5 | 0.91718 | 0.91726 | 0.999994 | 19567 | 0 | 0.35141 | 0.49434 | 0.71078 | 1 | 14747 | 2 | 0.35141 |
| KCTD16 | 4 | 0.16722 | 0.30691 | 0.999994 | 6913 | 2 | -0.13176 | 0.49436 | 0.66022 | 1 | 14748 | 2 | -0.13176 |
| MKNK1 | 5 | 0.15594 | 0.3116 | 0.999994 | 6625 | 3 | -1.0853 | 0.49457 | 0.7109 | 1 | 14749 | 2 | -1.0853 |
| TCERG1 | 4 | 0.37797 | 0.52306 | 0.999994 | 11623 | 2 | 0.34992 | 0.49462 | 0.66042 | 1 | 14750 | 2 | 0.34992 |
| GLRA3 | 6 | 0.10929 | 0.26532 | 0.997406 | 5279 | 4 | -0.81559 | 0.49465 | 0.7232 | 1 | 14751 | 1 | -0.81559 |
| RHBDL2 | 6 | 0.056934 | 0.15915 | 0.967126 | 3290 | 4 | -1.3627 | 0.49465 | 0.7232 | 1 | 14752 | 1 | -1.3627 |
| PEX26 | 5 | 0.15883 | 0.31582 | 0.999994 | 6694 | 2 | 0.30309 | 0.49469 | 0.71096 | 1 | 14753 | 2 | 0.30309 |
| GINS4 | 2 | 0.53941 | 0.54071 | 0.999994 | 14042 | 1 | 1.0182 | 0.49478 | 0.53399 | 1 | 14754 | 1 | 1.0182 |
| ARHGAP17 | 6 | 0.22872 | 0.42252 | 0.999994 | 8496 | 2 | 0.79678 | 0.4949 | 0.72338 | 1 | 14755 | 3 | 0.79678 |
| ITGA10 | 6 | 0.037878 | 0.11712 | 0.951052 | 2420 | 4 | -1.6894 | 0.49508 | 0.72352 | 1 | 14756 | 2 | -1.6894 |
| GPBAR1 | 5 | 0.83003 | 0.8395 | 0.999994 | 17931 | 1 | 0.50615 | 0.49513 | 0.71119 | 1 | 14757 | 2 | 0.50615 |
| FGL2 | 5 | 0.1702 | 0.33245 | 0.999994 | 6989 | 3 | -0.91796 | 0.49523 | 0.71124 | 1 | 14758 | 2 | -0.91796 |
| TCTN1 | 5 | 0.13506 | 0.28033 | 0.999632 | 6050 | 3 | -1.0329 | 0.49532 | 0.71128 | 1 | 14759 | 2 | -1.0329 |
| TTBK1 | 6 | 0.08418 | 0.21585 | 0.980538 | 4383 | 3 | -0.097937 | 0.49532 | 0.7237 | 1 | 14760 | 3 | -0.097937 |
| AGO4 | 6 | 0.29199 | 0.49736 | 0.999994 | 9951 | 2 | 0.2777 | 0.49533 | 0.72372 | 1 | 14761 | 3 | 0.2777 |
| EVX1 | 5 | 0.1763 | 0.34128 | 0.999994 | 7139 | 2 | 0.38577 | 0.49536 | 0.71131 | 1 | 14762 | 2 | 0.38577 |
| MESDC1 | 6 | 0.049848 | 0.14372 | 0.961453 | 2967 | 3 | -1.2405 | 0.49551 | 0.72385 | 1 | 14763 | 3 | -1.2405 |
| CXCR3 | 6 | 0.060972 | 0.16783 | 0.967126 | 3465 | 3 | -0.7715 | 0.49551 | 0.72385 | 1 | 14764 | 3 | -0.7715 |
| PLXNB2 | 4 | 0.032465 | 0.086529 | 0.94395 | 2132 | 2 | -3.0634 | 0.49573 | 0.66129 | 1 | 14765 | 2 | -3.0634 |
| S1PR5 | 4 | 0.29554 | 0.43841 | 0.999994 | 10029 | 2 | -0.41693 | 0.49573 | 0.66129 | 1 | 14766 | 1 | -0.41693 |
| MRAP | 5 | 0.47069 | 0.62273 | 0.999994 | 13014 | 2 | 0.15709 | 0.49582 | 0.71155 | 1 | 14767 | 2 | 0.15709 |

| | | | | | | | | | | | | | |
|---|---|---|---|---|---|---|---|---|---|---|---|---|---|
| hsa-mir-6858 | 2 | 0.3091 | 0.34382 | 0.999994 | 10302 | 1 | 0.037095 | 0.49582 | 0.53476 | 1 | 14768 | 1 | 0.037095 |
| NonTargetingControlG | 2 | 0.2812 | 0.32242 | 0.999994 | 9725 | 1 | -0.287 | 0.49607 | 0.53494 | 1 | 14769 | 1 | -0.287 |
| hsa-mir-6770-3 | 2 | 0.42437 | 0.43722 | 0.999994 | 12320 | 1 | 0.48789 | 0.4963 | 0.5351 | 1 | 14770 | 1 | 0.48789 |
| LOC100507003 | 6 | 0.11969 | 0.28514 | 0.999632 | 5618 | 3 | -0.99873 | 0.49634 | 0.7245 | 1 | 14771 | 2 | -0.99873 |
| SDHA | 6 | 0.043233 | 0.12919 | 0.952548 | 2685 | 3 | -1.2616 | 0.49634 | 0.7245 | 1 | 14772 | 2 | -1.2616 |
| OR4Q3 | 5 | 0.49643 | 0.64478 | 0.999994 | 13361 | 2 | 0.38542 | 0.49643 | 0.71187 | 1 | 14773 | 2 | 0.38542 |
| OR5L1 | 4 | 0.65762 | 0.69782 | 0.999994 | 15750 | 1 | 0.92075 | 0.4965 | 0.66188 | 1 | 14774 | 2 | 0.92075 |
| NOC2L | 4 | 0.004612 | 0.016305 | 0.772095 | 487 | 2 | -3.7443 | 0.49654 | 0.6619 | 1 | 14775 | 2 | -3.7443 |
| CT45A5 | 3 | 0.082952 | 0.16045 | 0.967126 | 4336 | 2 | -2.6163 | 0.49661 | 0.64185 | 1 | 14776 | 1 | -2.6163 |
| NBPF9 | 4 | 0.27949 | 0.42204 | 0.999994 | 9692 | 2 | 0.061067 | 0.49675 | 0.66206 | 1 | 14777 | 2 | 0.061067 |
| PTCHD3 | 6 | 0.15084 | 0.32674 | 0.999994 | 6482 | 3 | -0.20799 | 0.4968 | 0.72487 | 1 | 14778 | 3 | -0.20799 |
| NonTargetingControlG | 2 | 0.6735 | 0.67325 | 0.999994 | 15978 | 0 | 1.3316 | 0.49686 | 0.53552 | 1 | 14779 | 1 | 1.3316 |
| hsa-mir-942 | 4 | 0.529 | 0.62967 | 0.999994 | 13883 | 1 | 0.31919 | 0.49689 | 0.66216 | 1 | 14780 | 2 | 0.31919 |
| DHX8 | 4 | 0.028611 | 0.078691 | 0.93914 | 1930 | 3 | -4.5616 | 0.49689 | 0.66216 | 1 | 14781 | 1 | -4.5616 |
| LRRC10B | 4 | 0.1832 | 0.32338 | 0.999994 | 7298 | 2 | -0.42484 | 0.49689 | 0.66216 | 1 | 14782 | 2 | -0.42484 |
| ANAPC16 | 5 | 0.073265 | 0.1837 | 0.968526 | 3968 | 3 | -1.8403 | 0.49704 | 0.71219 | 1 | 14783 | 2 | -1.8403 |
| SLC33A1 | 6 | 0.062652 | 0.17137 | 0.967658 | 3541 | 4 | -1.4186 | 0.49718 | 0.72516 | 1 | 14784 | 2 | -1.4186 |
| ARHGAP18 | 6 | 0.23661 | 0.43196 | 0.999994 | 8684 | 3 | -0.82543 | 0.49718 | 0.72516 | 1 | 14785 | 2 | -0.82543 |
| ADCYAP1 | 4 | 0.4104 | 0.55634 | 0.999994 | 12118 | 1 | 0.64318 | 0.49723 | 0.66242 | 1 | 14786 | 2 | 0.64318 |
| TNFRSF11A | 5 | 0.40244 | 0.56556 | 0.999994 | 11988 | 2 | 0.157 | 0.49727 | 0.71232 | 1 | 14787 | 2 | 0.157 |
| CERKL | 6 | 0.13606 | 0.30805 | 0.999994 | 6081 | 3 | -0.39323 | 0.49742 | 0.72535 | 1 | 14788 | 3 | -0.39323 |
| hsa-mir-6829 | 2 | 0.59153 | 0.59152 | 0.999994 | 14841 | 0 | 0.99862 | 0.49754 | 0.53604 | 1 | 14789 | 1 | 0.99862 |
| GSDMC | 6 | 0.92689 | 0.92696 | 0.999994 | 19786 | 1 | 0.72347 | 0.49757 | 0.72546 | 1 | 14790 | 3 | 0.72347 |
| NBN | 6 | 0.096669 | 0.24081 | 0.987253 | 4844 | 3 | -0.15389 | 0.49777 | 0.72561 | 1 | 14791 | 3 | -0.15389 |
| PRMT7 | 6 | 0.18404 | 0.36818 | 0.999994 | 7316 | 2 | 0.68003 | 0.49779 | 0.72562 | 1 | 14792 | 3 | 0.68003 |
| TNFSF4 | 6 | 0.33744 | 0.54974 | 0.999994 | 10885 | 1 | 0.78233 | 0.49783 | 0.72565 | 1 | 14793 | 3 | 0.78233 |
| SLC4A2 | 4 | 0.033144 | 0.087925 | 0.94395 | 2169 | 3 | -4.5009 | 0.4979 | 0.66292 | 1 | 14794 | 1 | -4.5009 |
| ACAD11 | 6 | 0.23404 | 0.42891 | 0.999994 | 8621 | 2 | 0.72509 | 0.49795 | 0.72574 | 1 | 14795 | 3 | 0.72509 |
| CLSTN2 | 5 | 0.33868 | 0.51266 | 0.999994 | 10911 | 2 | -0.037845 | 0.49806 | 0.71274 | 1 | 14796 | 2 | -0.037845 |
| DNPH1 | 4 | 0.93038 | 0.93015 | 0.999994 | 19863 | 0 | 0.89331 | 0.49809 | 0.66306 | 1 | 14797 | 2 | 0.89331 |
| DUSP23 | 6 | 0.21766 | 0.40931 | 0.999994 | 8213 | 3 | -0.83806 | 0.4981 | 0.72585 | 1 | 14798 | 2 | -0.83806 |
| AMPH | 6 | 0.014011 | 0.052506 | 0.914561 | 1129 | 5 | -3.4338 | 0.4981 | 0.72585 | 1 | 14799 | 1 | -3.4338 |
| hsa-mir-4718 | 4 | 0.61383 | 0.6729 | 0.999994 | 15127 | 1 | 0.96973 | 0.49822 | 0.66317 | 1 | 14800 | 2 | 0.96973 |
| SDK1 | 5 | 0.015219 | 0.050837 | 0.909101 | 1203 | 2 | -0.48874 | 0.49824 | 0.71284 | 1 | 14801 | 2 | -0.48874 |
| UBC | 6 | 0.10644 | 0.25986 | 0.994943 | 5186 | 2 | 0.43307 | 0.49849 | 0.72614 | 1 | 14802 | 3 | 0.43307 |
| ART5 | 4 | 0.13201 | 0.27064 | 0.998561 | 5977 | 2 | -1.1051 | 0.49854 | 0.66341 | 1 | 14803 | 1 | -1.1051 |
| AQP6 | 5 | 0.50709 | 0.65411 | 0.999994 | 13538 | 2 | -0.5514 | 0.4988 | 0.71315 | 1 | 14804 | 2 | -0.5514 |
| GIPC2 | 6 | 0.14934 | 0.32479 | 0.999994 | 6439 | 3 | -0.96018 | 0.49883 | 0.7264 | 1 | 14805 | 1 | -0.96018 |
| hsa-mir-6791 | 4 | 0.31387 | 0.45722 | 0.999994 | 10393 | 2 | 0.38976 | 0.49886 | 0.66367 | 1 | 14806 | 2 | 0.38976 |
| CENPE | 3 | 0.65624 | 0.66226 | 0.999994 | 15729 | 1 | -0.15319 | 0.49891 | 0.64281 | 1 | 14807 | 1 | -0.15319 |
| hsa-mir-181b-1 | 3 | 0.46254 | 0.52199 | 0.999994 | 12898 | 1 | 0.012709 | 0.49891 | 0.64281 | 1 | 14808 | 1 | 0.012709 |
| PBX4 | 3 | 0.32576 | 0.44243 | 0.999994 | 10637 | 1 | 0.19543 | 0.49891 | 0.64281 | 1 | 14809 | 1 | 0.19543 |
| KRTAP5-3 | 6 | 0.12245 | 0.29035 | 0.999632 | 5718 | 3 | -0.35557 | 0.49899 | 0.72652 | 1 | 14810 | 3 | -0.35557 |
| ZNF319 | 5 | 0.34612 | 0.51874 | 0.999994 | 11074 | 2 | 0.54942 | 0.49901 | 0.71327 | 1 | 14811 | 2 | 0.54942 |
| C5orf60 | 5 | 0.28472 | 0.46867 | 0.999994 | 9803 | 2 | -0.15605 | 0.49909 | 0.71331 | 1 | 14812 | 2 | -0.15605 |
| SPTBN2 | 6 | 0.26179 | 0.46202 | 0.999994 | 9282 | 3 | -0.19207 | 0.49914 | 0.72664 | 1 | 14813 | 3 | -0.19207 |
| VAMP3 | 6 | 0.12174 | 0.28902 | 0.999632 | 5693 | 1 | 0.040455 | 0.49915 | 0.72665 | 1 | 14814 | 2 | 0.040455 |
| WDR92 | 4 | 0.0021569 | 0.0078255 | 0.677842 | 285 | 2 | -4.7363 | 0.49927 | 0.66398 | 1 | 14815 | 1 | -4.7363 |
| HIST2H2AA3 | 5 | 0.8909 | 0.89103 | 0.999994 | 19052 | 0 | 0.35554 | 0.49935 | 0.71343 | 1 | 14816 | 2 | 0.35554 |
| TSKS | 5 | 0.67375 | 0.75488 | 0.999994 | 15986 | 1 | 0.48904 | 0.49935 | 0.71343 | 1 | 14817 | 2 | 0.48904 |
| ATP6V1C1 | 5 | 0.098545 | 0.22413 | 0.983354 | 4913 | 3 | -1.335 | 0.49935 | 0.71343 | 1 | 14818 | 2 | -1.335 |
| MYADML2 | 5 | 0.48786 | 0.63739 | 0.999994 | 13255 | 2 | 0.021902 | 0.49935 | 0.71343 | 1 | 14819 | 1 | 0.021902 |

| | | | | | | | | | | | | | |
|---|---|---|---|---|---|---|---|---|---|---|---|---|---|
| TCHP | 5 | 0.79959 | 0.81925 | 0.999994 | 17475 | 1 | 0.40647 | 0.4994 | 0.71345 | 1 | 14820 | 2 | 0.40647 |
| RCAN3 | 6 | 0.3628 | 0.57846 | 0.999994 | 11386 | 3 | -0.057417 | 0.49962 | 0.72701 | 1 | 14821 | 2 | -0.057417 |
| ASB5 | 6 | 0.75226 | 0.82928 | 0.999994 | 16881 | 1 | 0.45651 | 0.4997 | 0.72707 | 1 | 14822 | 3 | 0.45651 |
| KIF16B | 6 | 0.042464 | 0.12745 | 0.951052 | 2641 | 4 | -2.0189 | 0.4997 | 0.72707 | 1 | 14823 | 2 | -2.0189 |
| SGPP2 | 6 | 0.25531 | 0.45429 | 0.999994 | 9133 | 2 | -0.41442 | 0.4997 | 0.72707 | 1 | 14824 | 2 | -0.41442 |
| CLDN7 | 6 | 0.019554 | 0.070549 | 0.93914 | 1431 | 2 | 0.683 | 0.49984 | 0.72717 | 1 | 14825 | 3 | 0.683 |
| PIWIL2 | 5 | 0.013489 | 0.045933 | 0.892434 | 1100 | 4 | -2.0182 | 0.49997 | 0.71374 | 1 | 14826 | 1 | -2.0182 |
| ANKS1A | 6 | 0.30326 | 0.51051 | 0.999994 | 10197 | 3 | -0.53275 | 0.5001 | 0.72737 | 1 | 14827 | 2 | -0.53275 |
| AKNA | 6 | 0.22816 | 0.42186 | 0.999994 | 8482 | 3 | -0.2688 | 0.50026 | 0.72749 | 1 | 14828 | 3 | -0.2688 |
| STON2 | 4 | 0.84139 | 0.84082 | 0.999994 | 18111 | 0 | 0.5103 | 0.50026 | 0.66473 | 1 | 14829 | 2 | 0.5103 |
| COA3 | 5 | 0.58315 | 0.72011 | 0.999994 | 14718 | 1 | 0.18323 | 0.50028 | 0.7139 | 1 | 14830 | 2 | 0.18323 |
| NonTargetingControlG | 1 | 0.49971 | 0.4997 | 0.999994 | 13418 | 0 | 0.46507 | 0.50029 | 0.50026 | 1 | 14831 | 0 | 0.46507 |
| PHACTR1 | 4 | 0.60547 | 0.66827 | 0.999994 | 15018 | 1 | 0.46093 | 0.50039 | 0.66484 | 1 | 14832 | 2 | 0.46093 |
| hsa-mir-521-2 | 3 | 0.46267 | 0.52207 | 0.999994 | 12901 | 1 | 0.48966 | 0.50041 | 0.64343 | 1 | 14833 | 1 | 0.48966 |
| TRAPPC3 | 5 | 0.55186 | 0.6934 | 0.999994 | 14242 | 1 | 0.54942 | 0.50054 | 0.71406 | 1 | 14834 | 2 | 0.54942 |
| DDRGK1 | 5 | 0.0449 | 0.12805 | 0.951777 | 2752 | 3 | -1.2901 | 0.50054 | 0.71406 | 1 | 14835 | 2 | -1.2901 |
| DNAJB11 | 6 | 0.075952 | 0.19914 | 0.977164 | 4069 | 4 | -1.2401 | 0.50056 | 0.72772 | 1 | 14836 | 2 | -1.2401 |
| GPR22 | 6 | 0.25591 | 0.45499 | 0.999994 | 9148 | 2 | 0.30397 | 0.50073 | 0.72785 | 1 | 14837 | 3 | 0.30397 |
| GCAT | 6 | 0.95369 | 0.95371 | 0.999994 | 20431 | 0 | 0.24968 | 0.50073 | 0.72785 | 1 | 14838 | 3 | 0.24968 |
| TM4SF18 | 6 | 0.063636 | 0.17347 | 0.96793 | 3588 | 3 | -1.3304 | 0.50073 | 0.72785 | 1 | 14839 | 2 | -1.3304 |
| ARNTL2 | 6 | 0.67259 | 0.79518 | 0.999994 | 15964 | 2 | 0.23097 | 0.50077 | 0.72787 | 1 | 14840 | 3 | 0.23097 |
| C12orf61 | 5 | 0.60887 | 0.7294 | 0.999994 | 15055 | 1 | -0.13416 | 0.50089 | 0.71424 | 1 | 14841 | 2 | -0.13416 |
| NonTargetingControlG | 2 | 0.68679 | 0.68663 | 0.999994 | 16178 | 0 | 1.4001 | 0.50113 | 0.53859 | 1 | 14842 | 1 | 1.4001 |
| PHOSPHO2-KLHL23 | 1 | 0.49882 | 0.49879 | 0.999994 | 13406 | 0 | 0.54004 | 0.50118 | 0.50117 | 1 | 14843 | 0 | 0.54004 |
| CORT | 6 | 0.023443 | 0.083001 | 0.94336 | 1680 | 3 | -0.71284 | 0.50122 | 0.72823 | 1 | 14844 | 2 | -0.71284 |
| LBR | 6 | 0.093396 | 0.23431 | 0.984971 | 4737 | 3 | 0.049353 | 0.50133 | 0.7283 | 1 | 14845 | 3 | 0.049353 |
| NMNAT1 | 1 | 0.49866 | 0.49862 | 0.999994 | 13404 | 0 | 0.49478 | 0.50134 | 0.50132 | 1 | 14846 | 0 | 0.49478 |
| TMUB2 | 6 | 0.83082 | 0.86194 | 0.999994 | 17943 | 1 | 0.56973 | 0.50135 | 0.72831 | 1 | 14847 | 3 | 0.56973 |
| hsa-mir-708 | 2 | 0.70797 | 0.70773 | 0.999994 | 16387 | 0 | 1.484 | 0.50136 | 0.53876 | 1 | 14848 | 1 | 1.484 |
| MARCKS | 6 | 0.26204 | 0.4623 | 0.999994 | 9291 | 3 | -0.014617 | 0.50141 | 0.72836 | 1 | 14849 | 3 | -0.014617 |
| NonTargetingControlG | 2 | 0.42459 | 0.4374 | 0.999994 | 12329 | 1 | 0.49724 | 0.50143 | 0.53881 | 1 | 14850 | 1 | 0.49724 |
| CRHBP | 4 | 0.40273 | 0.54854 | 0.999994 | 11992 | 2 | 0.30434 | 0.5015 | 0.66564 | 1 | 14851 | 2 | 0.30434 |
| SKIV2L | 5 | 0.41051 | 0.5723 | 0.999994 | 12119 | 2 | -0.23811 | 0.50162 | 0.71464 | 1 | 14852 | 1 | -0.23811 |
| COL4A4 | 5 | 0.019913 | 0.063849 | 0.937389 | 1455 | 4 | -1.5706 | 0.50162 | 0.71464 | 1 | 14853 | 1 | -1.5706 |
| hsa-mir-6090 | 2 | 0.19615 | 0.25931 | 0.994943 | 7657 | 1 | -1.5857 | 0.50173 | 0.53904 | 1 | 14854 | 1 | -1.5857 |
| TMPRSS11D | 4 | 0.017464 | 0.05525 | 0.922677 | 1337 | 2 | -2.4853 | 0.50182 | 0.66588 | 1 | 14855 | 2 | -2.4853 |
| EMILIN3 | 4 | 0.031693 | 0.084944 | 0.94395 | 2090 | 3 | -3.2252 | 0.50182 | 0.66588 | 1 | 14856 | 1 | -3.2252 |
| SEMA4D | 4 | 0.43865 | 0.58535 | 0.999994 | 12550 | 1 | -0.077108 | 0.50182 | 0.66588 | 1 | 14857 | 1 | -0.077108 |
| hsa-mir-4283-1 | 4 | 0.32338 | 0.46688 | 0.999994 | 10581 | 1 | 0.49719 | 0.50182 | 0.66588 | 1 | 14858 | 2 | 0.49719 |
| S100A4 | 6 | 0.26606 | 0.46704 | 0.999994 | 9368 | 3 | 0.13107 | 0.50182 | 0.72867 | 1 | 14859 | 3 | 0.13107 |
| WDR61 | 5 | 0.032638 | 0.097248 | 0.945253 | 2141 | 3 | -2.671 | 0.50192 | 0.71481 | 1 | 14860 | 2 | -2.671 |
| hsa-mir-4512 | 3 | 0.27142 | 0.40141 | 0.999994 | 9509 | 2 | -0.73942 | 0.50203 | 0.64409 | 1 | 14861 | 1 | -0.73942 |
| ZNF524 | 6 | 0.81799 | 0.85588 | 0.999994 | 17745 | 1 | 0.25345 | 0.5021 | 0.72891 | 1 | 14862 | 3 | 0.25345 |
| MPV17L | 4 | 0.45519 | 0.59686 | 0.999994 | 12774 | 1 | 0.9321 | 0.5021 | 0.66611 | 1 | 14863 | 2 | 0.9321 |
| USP51 | 6 | 0.00012767 | 0.0006952 | 0.309052 | 42 | 5 | -8.0021 | 0.50219 | 0.72897 | 1 | 14864 | 1 | -8.0021 |
| PNPLA3 | 6 | 0.59421 | 0.7357 | 0.999994 | 14880 | 1 | 0.80907 | 0.50227 | 0.72903 | 1 | 14865 | 3 | 0.80907 |
| hsa-mir-1247 | 4 | 0.38505 | 0.53031 | 0.999994 | 11718 | 2 | 0.22518 | 0.50253 | 0.66643 | 1 | 14866 | 2 | 0.22518 |
| KLHL32 | 5 | 0.11267 | 0.24611 | 0.98999 | 5390 | 2 | -0.37435 | 0.50257 | 0.71516 | 1 | 14867 | 2 | -0.37435 |
| RPS6KA5 | 6 | 0.33496 | 0.54693 | 0.999994 | 10830 | 1 | 0.11827 | 0.50257 | 0.72928 | 1 | 14868 | 2 | 0.11827 |
| SLC29A3 | 4 | 0.037092 | 0.095966 | 0.945031 | 2381 | 2 | 0.2454 | 0.50258 | 0.66646 | 1 | 14869 | 2 | 0.2454 |
| FAM98B | 5 | 0.30826 | 0.48772 | 0.999994 | 10288 | 2 | 0.52791 | 0.50266 | 0.71521 | 1 | 14870 | 2 | 0.52791 |
| ZSCAN9 | 5 | 0.18082 | 0.34782 | 0.999994 | 7240 | 3 | -0.87083 | 0.50266 | 0.71521 | 1 | 14871 | 2 | -0.87083 |

| | | | | | | | | | | | | |
|---|---|---|---|---|---|---|---|---|---|---|---|---|
| TRIM55 | 5 | 0.074416 | 0.18552 | 0.969035 | 4008 | 3 | -0.91213 | 0.50266 | 0.71521 | 1 | 14872 | 1 | -0.91213 |
| APITD1-CORT | 2 | 0.66067 | 0.66047 | 0.999994 | 15788 | 0 | 1.194 | 0.50272 | 0.53974 | 1 | 14873 | 1 | 1.194 |
| hsa-mir-6792 | 3 | 0.22384 | 0.34243 | 0.999994 | 8376 | 2 | -1.0649 | 0.50278 | 0.64439 | 1 | 14874 | 1 | -1.0649 |
| ST20-MTHFS | 1 | 0.49712 | 0.4971 | 0.999994 | 13380 | 0 | 0.40578 | 0.50288 | 0.50285 | 1 | 14875 | 0 | 0.40578 |
| CYBRD1 | 6 | 0.26934 | 0.47096 | 0.999994 | 9455 | 3 | -0.72246 | 0.50288 | 0.72951 | 1 | 14876 | 2 | -0.72246 |
| ALPL | 5 | 0.26712 | 0.45454 | 0.999994 | 9400 | 1 | 0.60711 | 0.50289 | 0.71533 | 1 | 14877 | 2 | 0.60711 |
| C19orf21 | 6 | 0.16274 | 0.3417 | 0.999994 | 6813 | 3 | -0.18979 | 0.50293 | 0.72954 | 1 | 14878 | 3 | -0.18979 |
| RND2 | 5 | 0.85935 | 0.86185 | 0.999994 | 18422 | 1 | 0.42762 | 0.50293 | 0.71535 | 1 | 14879 | 2 | 0.42762 |
| ARL17B | 1 | 0.49705 | 0.49703 | 0.999994 | 13377 | 0 | 0.51707 | 0.50295 | 0.50292 | 1 | 14880 | 0 | 0.51707 |
| OR13C3 | 6 | 0.81397 | 0.85404 | 0.999994 | 17690 | 1 | 0.56398 | 0.50299 | 0.72958 | 1 | 14881 | 3 | 0.56398 |
| PTPN7 | 6 | 0.31776 | 0.52732 | 0.999994 | 10465 | 3 | 0.139 | 0.50307 | 0.72965 | 1 | 14882 | 3 | 0.139 |
| LNX2 | 6 | 0.11633 | 0.27883 | 0.999632 | 5523 | 4 | -0.98527 | 0.50309 | 0.72967 | 1 | 14883 | 1 | -0.98527 |
| TMEM100 | 6 | 0.068366 | 0.18336 | 0.968526 | 3759 | 4 | -1.4152 | 0.50309 | 0.72967 | 1 | 14884 | 1 | -1.4152 |
| FAM86A | 6 | 0.24172 | 0.43806 | 0.999994 | 8826 | 3 | 0.15593 | 0.50309 | 0.72967 | 1 | 14885 | 3 | 0.15593 |
| CD1C | 6 | 0.56187 | 0.71214 | 0.999994 | 14404 | 2 | -0.094846 | 0.50309 | 0.72967 | 1 | 14886 | 2 | -0.094846 |
| LECT2 | 6 | 0.0209 | 0.074867 | 0.93914 | 1504 | 3 | -0.48697 | 0.50325 | 0.7298 | 1 | 14887 | 2 | -0.48697 |
| C8B | 6 | 0.12225 | 0.29002 | 0.999632 | 5711 | 3 | -0.28761 | 0.50326 | 0.72981 | 1 | 14888 | 3 | -0.28761 |
| GRID1 | 6 | 0.31582 | 0.5251 | 0.999994 | 10435 | 3 | 0.026608 | 0.50334 | 0.72987 | 1 | 14889 | 3 | 0.026608 |
| OR5D13 | 5 | 0.23356 | 0.42215 | 0.999994 | 8606 | 2 | 0.59023 | 0.50346 | 0.71564 | 1 | 14890 | 2 | 0.59023 |
| SLC7A8 | 5 | 0.22649 | 0.41236 | 0.999994 | 8440 | 2 | 0.68617 | 0.50356 | 0.7157 | 1 | 14891 | 2 | 0.68617 |
| RRP9 | 1 | 0.49633 | 0.4963 | 0.999994 | 13359 | 0 | 0.45913 | 0.50367 | 0.50361 | 1 | 14892 | 0 | 0.45913 |
| LCN1 | 6 | 0.13363 | 0.30494 | 0.999994 | 6013 | 1 | 0.73289 | 0.50384 | 0.73024 | 1 | 14893 | 3 | 0.73289 |
| GALNT18 | 6 | 0.16734 | 0.34741 | 0.999994 | 6917 | 3 | -0.23663 | 0.50388 | 0.73027 | 1 | 14894 | 3 | -0.23663 |
| NonTargetingControlG | 2 | 0.64825 | 0.64804 | 0.999994 | 15633 | 0 | 1.217 | 0.50394 | 0.54061 | 1 | 14895 | 1 | 1.217 |
| NACAD | 6 | 0.14577 | 0.32035 | 0.999994 | 6345 | 2 | 0.15072 | 0.50396 | 0.73032 | 1 | 14896 | 2 | 0.15072 |
| NUDT12 | 6 | 0.090283 | 0.22805 | 0.983354 | 4603 | 1 | 0.65899 | 0.504 | 0.73035 | 1 | 14897 | 3 | 0.65899 |
| TBC1D10A | 6 | 0.89467 | 0.89966 | 0.999994 | 19117 | 1 | -0.069055 | 0.50412 | 0.73045 | 1 | 14898 | 2 | -0.069055 |
| CXorf51B | 2 | 0.30907 | 0.3438 | 0.999994 | 10300 | 1 | 0.080195 | 0.50423 | 0.54083 | 1 | 14899 | 1 | 0.080195 |
| FMO1 | 4 | 0.23986 | 0.38147 | 0.999994 | 8767 | 2 | -0.64781 | 0.50427 | 0.66776 | 1 | 14900 | 1 | -0.64781 |
| PLEKHH2 | 4 | 0.8674 | 0.86697 | 0.999994 | 18586 | 0 | 0.16563 | 0.50427 | 0.66776 | 1 | 14901 | 1 | 0.16563 |
| NonTargetingControlG | 2 | 0.52872 | 0.53055 | 0.999994 | 13877 | 1 | 0.74478 | 0.5043 | 0.54087 | 1 | 14902 | 1 | 0.74478 |
| PCBP2 | 5 | 0.23289 | 0.42122 | 0.999994 | 8590 | 2 | -0.053618 | 0.50459 | 0.71623 | 1 | 14903 | 1 | -0.053618 |
| NonTargetingControlG | 2 | 0.41555 | 0.42971 | 0.999994 | 12185 | 1 | 0.41504 | 0.50463 | 0.54112 | 1 | 14904 | 1 | 0.41504 |
| WDYHV1 | 6 | 0.10589 | 0.25879 | 0.994943 | 5169 | 4 | -1.0813 | 0.50469 | 0.73088 | 1 | 14905 | 2 | -1.0813 |
| TTC33 | 6 | 0.91046 | 0.91195 | 0.999994 | 19437 | 1 | 0.54959 | 0.5047 | 0.73088 | 1 | 14906 | 3 | 0.54959 |
| COL3A1 | 6 | 0.083139 | 0.21372 | 0.979587 | 4344 | 4 | -1.1918 | 0.5047 | 0.73088 | 1 | 14907 | 2 | -1.1918 |
| RBM48 | 6 | 0.25 | 0.44795 | 0.999994 | 9015 | 3 | -0.6495 | 0.5047 | 0.73088 | 1 | 14908 | 2 | -0.6495 |
| RCBTB2 | 6 | 0.010482 | 0.040669 | 0.891621 | 902 | 3 | -3.0389 | 0.5048 | 0.73095 | 1 | 14909 | 2 | -3.0389 |
| ACAA1 | 6 | 0.069738 | 0.18623 | 0.969409 | 3821 | 2 | 0.50013 | 0.50489 | 0.73102 | 1 | 14910 | 3 | 0.50013 |
| CCDC127 | 5 | 0.50796 | 0.65489 | 0.999994 | 13550 | 1 | -0.2313 | 0.5049 | 0.71639 | 1 | 14911 | 2 | -0.2313 |
| hsa-mir-3529 | 4 | 0.41193 | 0.55791 | 0.999994 | 12144 | 2 | 0.29814 | 0.50493 | 0.66828 | 1 | 14912 | 2 | 0.29814 |
| DOCK8 | 6 | 0.35252 | 0.56685 | 0.999994 | 11204 | 3 | -0.11473 | 0.50504 | 0.73112 | 1 | 14913 | 2 | -0.11473 |
| GRINA | 4 | 0.79497 | 0.79674 | 0.999994 | 17416 | 1 | 0.87986 | 0.50511 | 0.66843 | 1 | 14914 | 2 | 0.87986 |
| hsa-mir-3121 | 4 | 0.39182 | 0.53727 | 0.999994 | 11828 | 2 | 0.20764 | 0.50513 | 0.66844 | 1 | 14915 | 2 | 0.20764 |
| SLC20A2 | 5 | 0.12184 | 0.26013 | 0.994943 | 5697 | 3 | -1.3512 | 0.50522 | 0.71658 | 1 | 14916 | 2 | -1.3512 |
| PLD3 | 6 | 0.10687 | 0.26069 | 0.994943 | 5208 | 4 | -1.3051 | 0.50524 | 0.73128 | 1 | 14917 | 2 | -1.3051 |
| AFAP1 | 4 | 0.31667 | 0.46005 | 0.999994 | 10445 | 2 | -0.84044 | 0.50545 | 0.66868 | 1 | 14918 | 1 | -0.84044 |
| KCNC1 | 6 | 0.10669 | 0.26033 | 0.994943 | 5195 | 2 | 0.14487 | 0.50548 | 0.73147 | 1 | 14919 | 3 | 0.14487 |
| GALNT7 | 5 | 0.23023 | 0.41753 | 0.999994 | 8529 | 2 | -0.60676 | 0.5056 | 0.71678 | 1 | 14920 | 2 | -0.60676 |
| UGT2B4 | 5 | 0.51187 | 0.65829 | 0.999994 | 13609 | 2 | -0.42451 | 0.50571 | 0.71684 | 1 | 14921 | 1 | -0.42451 |
| CLASP2 | 6 | 0.35849 | 0.57361 | 0.999994 | 11311 | 2 | -0.23936 | 0.50576 | 0.73171 | 1 | 14922 | 2 | -0.23936 |
| NonTargetingControlG | 2 | 0.75562 | 0.75519 | 0.999994 | 16928 | 0 | 1.4287 | 0.50588 | 0.54201 | 1 | 14923 | 1 | 1.4287 |

| | | | | | | | | | | | | | |
|---|---|---|---|---|---|---|---|---|---|---|---|---|---|
| SLC9A2 | 6 | 0.6335 | 0.76499 | 0.999994 | 15421 | 2 | 0.56177 | 0.50589 | 0.7318 | 1 | 14924 | 3 | 0.56177 |
| hsa-mir-544a | 4 | 0.87862 | 0.87826 | 0.999994 | 18804 | 0 | 0.68797 | 0.50595 | 0.66907 | 1 | 14925 | 2 | 0.68797 |
| GPHA2 | 5 | 0.28712 | 0.4706 | 0.999994 | 9855 | 2 | 0.23146 | 0.50596 | 0.71697 | 1 | 14926 | 2 | 0.23146 |
| TMC3 | 5 | 0.61725 | 0.73255 | 0.999994 | 15174 | 1 | 0.5204 | 0.506 | 0.71699 | 1 | 14927 | 2 | 0.5204 |
| BTN2A1 | 6 | 0.94258 | 0.94256 | 0.999994 | 20141 | 0 | 0.80493 | 0.50613 | 0.73199 | 1 | 14928 | 3 | 0.80493 |
| CCR6 | 5 | 0.49739 | 0.64562 | 0.999994 | 13388 | 1 | -0.16057 | 0.50614 | 0.71706 | 1 | 14929 | 2 | -0.16057 |
| hsa-mir-4766 | 4 | 0.62318 | 0.67816 | 0.999994 | 15277 | 1 | 0.56166 | 0.50623 | 0.6693 | 1 | 14930 | 2 | 0.56166 |
| EDARADD | 6 | 0.48259 | 0.65693 | 0.999994 | 13183 | 2 | 0.35394 | 0.50626 | 0.73208 | 1 | 14931 | 3 | 0.35394 |
| GCOM1 | 6 | 0.83868 | 0.86595 | 0.999994 | 18064 | 1 | 0.47654 | 0.50635 | 0.73215 | 1 | 14932 | 3 | 0.47654 |
| RND3 | 6 | 0.34986 | 0.56382 | 0.999994 | 11155 | 3 | -0.080668 | 0.50639 | 0.73217 | 1 | 14933 | 2 | -0.080668 |
| TECTB | 6 | 0.58357 | 0.72783 | 0.999994 | 14726 | 2 | -0.15954 | 0.50645 | 0.73222 | 1 | 14934 | 1 | -0.15954 |
| PARD6G | 6 | 0.58764 | 0.73085 | 0.999994 | 14785 | 2 | 0.13098 | 0.50645 | 0.73222 | 1 | 14935 | 2 | 0.13098 |
| C22orf15 | 6 | 0.27988 | 0.48327 | 0.999994 | 9702 | 3 | -0.68993 | 0.50645 | 0.73222 | 1 | 14936 | 1 | -0.68993 |
| ITGA11 | 4 | 0.20597 | 0.34685 | 0.999994 | 7926 | 2 | -0.26175 | 0.50648 | 0.66949 | 1 | 14937 | 2 | -0.26175 |
| hsa-mir-1276 | 4 | 0.4257 | 0.57206 | 0.999994 | 12346 | 2 | 0.35534 | 0.50656 | 0.66956 | 1 | 14938 | 2 | 0.35534 |
| COX6B1 | 4 | 0.020905 | 0.062516 | 0.937389 | 1506 | 3 | -4.8414 | 0.50667 | 0.66964 | 1 | 14939 | 1 | -4.8414 |
| RALBP1 | 5 | 0.11343 | 0.24729 | 0.990682 | 5420 | 2 | 0.27054 | 0.5067 | 0.71738 | 1 | 14940 | 2 | 0.27054 |
| GAD1 | 5 | 0.02596 | 0.07999 | 0.93914 | 1801 | 4 | -1.5602 | 0.5067 | 0.71738 | 1 | 14941 | 1 | -1.5602 |
| SPG20 | 5 | 0.21749 | 0.39975 | 0.999994 | 8207 | 2 | -0.15012 | 0.5067 | 0.71738 | 1 | 14942 | 2 | -0.15012 |
| hsa-mir-7162 | 2 | 0.039762 | 0.071869 | 0.93914 | 2503 | 1 | -3.5551 | 0.5068 | 0.54272 | 1 | 14943 | 1 | -3.5551 |
| BTBD11 | 6 | 0.34236 | 0.55536 | 0.999994 | 10992 | 2 | 0.59082 | 0.50697 | 0.73263 | 1 | 14944 | 3 | 0.59082 |
| CXCL3 | 4 | 0.70308 | 0.72646 | 0.999994 | 16339 | 1 | 0.50977 | 0.50709 | 0.66996 | 1 | 14945 | 2 | 0.50977 |
| BEST4 | 4 | 0.42802 | 0.5745 | 0.999994 | 12383 | 2 | 0.37241 | 0.50718 | 0.67004 | 1 | 14946 | 2 | 0.37241 |
| DAW1 | 6 | 0.058462 | 0.16244 | 0.967126 | 3365 | 4 | -1.6512 | 0.50729 | 0.73285 | 1 | 14947 | 2 | -1.6512 |
| hsa-mir-2054 | 1 | 0.49265 | 0.49264 | 0.999994 | 13304 | 0 | 0.54942 | 0.50735 | 0.50723 | 1 | 14948 | 0 | 0.54942 |
| hsa-mir-7978 | 1 | 0.49265 | 0.49264 | 0.999994 | 13305 | 0 | 0.54942 | 0.50735 | 0.50723 | 1 | 14949 | 0 | 0.54942 |
| ANKRD18B | 6 | 0.039007 | 0.11966 | 0.951052 | 2459 | 2 | 0.43381 | 0.50736 | 0.73291 | 1 | 14950 | 3 | 0.43381 |
| XIRP2 | 5 | 0.22029 | 0.40365 | 0.999994 | 8296 | 2 | -0.58963 | 0.50745 | 0.71778 | 1 | 14951 | 2 | -0.58963 |
| SUSD4 | 6 | 0.3182 | 0.52785 | 0.999994 | 10479 | 3 | -0.60866 | 0.50764 | 0.7331 | 1 | 14952 | 2 | -0.60866 |
| KRTAP19-3 | 6 | 0.11809 | 0.2821 | 0.999632 | 5576 | 3 | -0.55467 | 0.50764 | 0.7331 | 1 | 14953 | 2 | -0.55467 |
| TBC1D2B | 5 | 0.16239 | 0.32104 | 0.999994 | 6805 | 2 | 0.31034 | 0.5077 | 0.71792 | 1 | 14954 | 2 | 0.31034 |
| SAR1A | 6 | 0.23468 | 0.42967 | 0.999994 | 8634 | 3 | 0.039916 | 0.50773 | 0.73318 | 1 | 14955 | 3 | 0.039916 |
| CLDN18 | 4 | 0.082395 | 0.18314 | 0.968526 | 4313 | 3 | -2.2026 | 0.50783 | 0.67054 | 1 | 14956 | 1 | -2.2026 |
| SAV1 | 4 | 0.11734 | 0.24563 | 0.98979 | 5553 | 3 | -0.91935 | 0.50783 | 0.67054 | 1 | 14957 | 1 | -0.91935 |
| ELMSAN1 | 4 | 0.04015 | 0.10209 | 0.946414 | 2525 | 3 | -2.5891 | 0.50783 | 0.67054 | 1 | 14958 | 1 | -2.5891 |
| PRAMEF4 | 5 | 0.69269 | 0.76318 | 0.999994 | 16231 | 1 | 0.12919 | 0.50786 | 0.718 | 1 | 14959 | 2 | 0.12919 |
| FN3K | 6 | 0.13825 | 0.31081 | 0.999994 | 6145 | 3 | -0.32304 | 0.50791 | 0.7333 | 1 | 14960 | 3 | -0.32304 |
| GLTP | 6 | 0.25756 | 0.45699 | 0.999994 | 9182 | 3 | -0.020285 | 0.50796 | 0.73334 | 1 | 14961 | 3 | -0.020285 |
| CLINT1 | 6 | 0.094832 | 0.23719 | 0.985914 | 4783 | 4 | -1.0556 | 0.50811 | 0.73345 | 1 | 14962 | 2 | -1.0556 |
| MROH9 | 6 | 0.0022601 | 0.010939 | 0.71022 | 300 | 4 | -4.0654 | 0.50848 | 0.73373 | 1 | 14963 | 2 | -4.0654 |
| FUT7 | 6 | 0.84178 | 0.86756 | 0.999994 | 18119 | 1 | 0.43924 | 0.50871 | 0.73391 | 1 | 14964 | 3 | 0.43924 |
| SRP68 | 6 | 0.26772 | 0.469 | 0.999994 | 9416 | 3 | 0.17318 | 0.50871 | 0.73391 | 1 | 14965 | 3 | 0.17318 |
| GPR112 | 6 | 0.44514 | 0.63185 | 0.999994 | 12648 | 2 | 0.57193 | 0.50871 | 0.73391 | 1 | 14966 | 3 | 0.57193 |
| TCEAL2 | 6 | 0.21897 | 0.41087 | 0.999994 | 8260 | 2 | 0.37046 | 0.50883 | 0.73399 | 1 | 14967 | 3 | 0.37046 |
| CHRM3 | 6 | 0.94306 | 0.94304 | 0.999994 | 20155 | 0 | 0.37819 | 0.50883 | 0.73399 | 1 | 14968 | 1 | 0.37819 |
| KIAA0319L | 6 | 0.16063 | 0.33904 | 0.999994 | 6756 | 3 | -1.0541 | 0.50883 | 0.73399 | 1 | 14969 | 2 | -1.0541 |
| KTN1 | 4 | 0.50221 | 0.61729 | 0.999994 | 13466 | 1 | 0.79399 | 0.50904 | 0.67144 | 1 | 14970 | 2 | 0.79399 |
| SH3KBP1 | 4 | 0.40293 | 0.54876 | 0.999994 | 11996 | 2 | -0.11189 | 0.5091 | 0.67149 | 1 | 14971 | 2 | -0.11189 |
| NME7 | 6 | 0.086198 | 0.21989 | 0.980699 | 4459 | 4 | -0.72821 | 0.5094 | 0.73442 | 1 | 14972 | 2 | -0.72821 |
| MSRB1 | 4 | 0.83749 | 0.83697 | 0.999994 | 18042 | 0 | 0.58195 | 0.50943 | 0.67175 | 1 | 14973 | 2 | 0.58195 |
| CA9 | 6 | 0.11767 | 0.28134 | 0.999632 | 5564 | 4 | -0.83863 | 0.50962 | 0.73457 | 1 | 14974 | 2 | -0.83863 |
| hsa-mir-6883 | 4 | 0.12798 | 0.2641 | 0.996734 | 5875 | 2 | -0.57812 | 0.50974 | 0.67198 | 1 | 14975 | 2 | -0.57812 |

| | | | | | | | | | | | | | |
|---|---|---|---|---|---|---|---|---|---|---|---|---|---|
| FANCI | 6 | 0.10089 | 0.24905 | 0.991255 | 5003 | | 4 | -1.2784 | 0.50977 | 0.73469 | 1 | 14976 | 2 | -1.2784 |
| ACTR3 | 6 | 0.087344 | 0.22218 | 0.983354 | 4489 | | 3 | -0.27718 | 0.51006 | 0.73491 | 1 | 14977 | 3 | -0.27718 |
| PPP1R32 | 6 | 0.17832 | 0.36101 | 0.999994 | 7182 | | 3 | -0.55637 | 0.51006 | 0.73491 | 1 | 14978 | 1 | -0.55637 |
| SIL1 | 6 | 0.044084 | 0.13106 | 0.95318 | 2715 | | 4 | -1.9713 | 0.51006 | 0.73491 | 1 | 14979 | 2 | -1.9713 |
| GPR115 | 5 | 0.013946 | 0.047224 | 0.896226 | 1124 | | 2 | 0.16834 | 0.51015 | 0.71922 | 1 | 14980 | 2 | 0.16834 |
| TSC1 | 5 | 0.37954 | 0.54634 | 0.999994 | 11644 | | 2 | -0.14389 | 0.51015 | 0.71922 | 1 | 14981 | 2 | -0.14389 |
| INF2 | 5 | 0.0043595 | 0.018266 | 0.795727 | 469 | | 3 | -1.5561 | 0.51015 | 0.71922 | 1 | 14982 | 2 | -1.5561 |
| NLE1 | 5 | 0.10658 | 0.23667 | 0.985449 | 5190 | | 3 | -1.6069 | 0.51015 | 0.71922 | 1 | 14983 | 1 | -1.6069 |
| NonTargetingControlG | 2 | 0.66822 | 0.668 | 0.999994 | 15896 | | 0 | 1.2004 | 0.51019 | 0.54519 | 1 | 14984 | 1 | 1.2004 |
| MCM6 | 4 | 0.29025 | 0.43307 | 0.999994 | 9913 | | 1 | -0.66332 | 0.51024 | 0.67237 | 1 | 14985 | 1 | -0.66332 |
| KRT12 | 6 | 0.22689 | 0.42036 | 0.999994 | 8449 | | 3 | -0.092476 | 0.51031 | 0.7351 | 1 | 14986 | 3 | -0.092476 |
| OR7A10 | 6 | 0.27379 | 0.47617 | 0.999994 | 9559 | | 3 | -0.23281 | 0.51041 | 0.73517 | 1 | 14987 | 2 | -0.23281 |
| INTS6 | 5 | 0.24165 | 0.43315 | 0.999994 | 8820 | | 3 | -0.56224 | 0.51048 | 0.7194 | 1 | 14988 | 2 | -0.56224 |
| ASB9 | 5 | 0.18887 | 0.35933 | 0.999994 | 7449 | | 3 | -0.90099 | 0.51048 | 0.7194 | 1 | 14989 | 2 | -0.90099 |
| BTF3L4 | 4 | 0.8353 | 0.83482 | 0.999994 | 18010 | | 0 | 0.7309 | 0.51079 | 0.67279 | 1 | 14990 | 2 | 0.7309 |
| hsa-mir-4437 | 4 | 0.40121 | 0.54699 | 0.999994 | 11964 | | 1 | 0.58745 | 0.51089 | 0.67286 | 1 | 14991 | 2 | 0.58745 |
| SLC16A12 | 6 | 0.27233 | 0.47446 | 0.999994 | 9524 | | 3 | -0.62412 | 0.511 | 0.73562 | 1 | 14992 | 2 | -0.62412 |
| MAGEA9 | 5 | 0.059807 | 0.16177 | 0.967126 | 3414 | | 3 | -1.9857 | 0.51123 | 0.71981 | 1 | 14993 | 1 | -1.9857 |
| MREG | 6 | 0.039992 | 0.12189 | 0.951052 | 2515 | | 3 | -0.34988 | 0.51124 | 0.7358 | 1 | 14994 | 2 | -0.34988 |
| TRIM65 | 3 | 0.20802 | 0.32275 | 0.999994 | 7977 | | 2 | -1.0843 | 0.51151 | 0.64804 | 1 | 14995 | 1 | -1.0843 |
| RCOR1 | 4 | 0.29447 | 0.43734 | 0.999994 | 10006 | | 1 | 0.31152 | 0.51153 | 0.67337 | 1 | 14996 | 2 | 0.31152 |
| MLLT11 | 4 | 0.85589 | 0.85548 | 0.999994 | 18367 | | 0 | 0.078124 | 0.51153 | 0.67337 | 1 | 14997 | 1 | 0.078124 |
| FAM212B | 6 | 0.8291 | 0.86111 | 0.999994 | 17912 | | 1 | 0.62196 | 0.51173 | 0.73615 | 1 | 14998 | 3 | 0.62196 |
| GLS | 6 | 0.094484 | 0.23648 | 0.985449 | 4769 | | 3 | -0.43897 | 0.51175 | 0.73617 | 1 | 14999 | 3 | -0.43897 |
| MFHAS1 | 6 | 0.67645 | 0.79823 | 0.999994 | 16030 | | 2 | 0.35869 | 0.51179 | 0.7362 | 1 | 15000 | 3 | 0.35869 |
| PSMG3 | 4 | 0.75989 | 0.7677 | 0.999994 | 16979 | | 1 | 1.0153 | 0.51179 | 0.67354 | 1 | 15001 | 2 | 1.0153 |
| TMED1 | 5 | 0.3953 | 0.55965 | 0.999994 | 11876 | | 2 | -0.33662 | 0.51188 | 0.72017 | 1 | 15002 | 2 | -0.33662 |
| TRIM21 | 5 | 0.060491 | 0.16291 | 0.967126 | 3444 | | 2 | 0.42499 | 0.51213 | 0.72029 | 1 | 15003 | 1 | 0.42499 |
| TBCEL | 5 | 0.077327 | 0.19026 | 0.971257 | 4122 | | 2 | -0.43185 | 0.51213 | 0.72029 | 1 | 15004 | 2 | -0.43185 |
| RAC2 | 5 | 0.021336 | 0.067692 | 0.93914 | 1536 | | 4 | -0.93701 | 0.51213 | 0.72029 | 1 | 15005 | 1 | -0.93701 |
| HIST2H3A | 5 | 0.010062 | 0.036232 | 0.879595 | 873 | | 4 | -2.076 | 0.51213 | 0.72029 | 1 | 15006 | 1 | -2.076 |
| TMED7-TICAM2 | 2 | 0.29351 | 0.33189 | 0.999994 | 9985 | | 1 | -0.18991 | 0.51214 | 0.54666 | 1 | 15007 | 1 | -0.18991 |
| CPNE8 | 1 | 0.48784 | 0.48789 | 0.999994 | 13253 | | 0 | 0.39953 | 0.51216 | 0.51203 | 1 | 15008 | 0 | 0.39953 |
| OR7A17 | 6 | 0.11092 | 0.26847 | 0.998561 | 5330 | | 2 | 0.49254 | 0.51241 | 0.73669 | 1 | 15009 | 3 | 0.49254 |
| SLC38A2 | 6 | 0.15006 | 0.32569 | 0.999994 | 6461 | | 1 | 0.69456 | 0.51243 | 0.7367 | 1 | 15010 | 3 | 0.69456 |
| TCF23 | 4 | 0.80299 | 0.80388 | 0.999994 | 17524 | | 1 | 0.97798 | 0.51248 | 0.67409 | 1 | 15011 | 2 | 0.97798 |
| PLBD2 | 6 | 0.18829 | 0.37341 | 0.999994 | 7427 | | 3 | -0.5191 | 0.51256 | 0.7368 | 1 | 15012 | 2 | -0.5191 |
| RRP1 | 6 | 0.0059832 | 0.024748 | 0.844082 | 592 | | 3 | -3.814 | 0.51256 | 0.7368 | 1 | 15013 | 2 | -3.814 |
| NUDT17 | 6 | 0.34771 | 0.56143 | 0.999994 | 11104 | | 2 | 0.74982 | 0.51256 | 0.73681 | 1 | 15014 | 3 | 0.74982 |
| PCNP | 6 | 0.15614 | 0.33338 | 0.999994 | 6634 | | 3 | -0.23623 | 0.51262 | 0.73686 | 1 | 15015 | 2 | -0.23623 |
| APOBEC4 | 6 | 0.016601 | 0.060976 | 0.932932 | 1283 | | 3 | -1.7144 | 0.5127 | 0.73692 | 1 | 15016 | 3 | -1.7144 |
| CXorf58 | 6 | 0.79071 | 0.84397 | 0.999994 | 17359 | | 1 | 0.79493 | 0.51276 | 0.73697 | 1 | 15017 | 3 | 0.79493 |
| DSC3 | 6 | 0.68978 | 0.80867 | 0.999994 | 16209 | | 2 | 0.6053 | 0.51282 | 0.73701 | 1 | 15018 | 3 | 0.6053 |
| FAM132A | 6 | 0.24062 | 0.43674 | 0.999994 | 8786 | | 3 | 0.051474 | 0.51284 | 0.73703 | 1 | 15019 | 3 | 0.051474 |
| hsa-mir-30c-1 | 4 | 0.80113 | 0.80223 | 0.999994 | 17494 | | 1 | -0.068041 | 0.51284 | 0.67436 | 1 | 15020 | 1 | -0.068041 |
| hsa-mir-4445 | 3 | 0.01962 | 0.050619 | 0.909101 | 1435 | | 2 | -5.0491 | 0.51328 | 0.64881 | 1 | 15021 | 1 | -5.0491 |
| IFNB1 | 6 | 0.21073 | 0.40098 | 0.999994 | 8045 | | 3 | -0.49346 | 0.51349 | 0.7375 | 1 | 15022 | 1 | -0.49346 |
| SELPLG | 4 | 0.033806 | 0.089276 | 0.94395 | 2205 | | 2 | -2.8021 | 0.51361 | 0.67497 | 1 | 15023 | 2 | -2.8021 |
| ADRB2 | 6 | 0.24707 | 0.44443 | 0.999994 | 8946 | | 3 | -0.096041 | 0.51376 | 0.73771 | 1 | 15024 | 2 | -0.096041 |
| BPY2C | 2 | 0.66975 | 0.6695 | 0.999994 | 15927 | | 0 | 1.1196 | 0.51392 | 0.54794 | 1 | 15025 | 1 | 1.1196 |
| ABP1 | 6 | 0.66939 | 0.79269 | 0.999994 | 15922 | | 2 | 0.59011 | 0.51404 | 0.73793 | 1 | 15026 | 3 | 0.59011 |
| hsa-mir-300 | 4 | 0.56416 | 0.64675 | 0.999994 | 14456 | | 1 | 0.83265 | 0.51405 | 0.67531 | 1 | 15027 | 2 | 0.83265 |

| | | | | | | | | | | | | |
|---|---|---|---|---|---|---|---|---|---|---|---|---|
| hsa-mir-6501 | 2 | 0.63196 | 0.63171 | 0.999994 | 15398 | 0 | 1.159 | 0.51416 | 0.54814 | 1 | 15028 | 1 | 1.159 |
| ALG5 | 5 | 0.077381 | 0.19035 | 0.971257 | 4124 | 3 | -1.1724 | 0.5143 | 0.72143 | 1 | 15029 | 2 | -1.1724 |
| TAGLN2 | 5 | 0.03346 | 0.09938 | 0.945253 | 2187 | 3 | -2.7495 | 0.5143 | 0.72143 | 1 | 15030 | 2 | -2.7495 |
| ZNF12 | 6 | 0.07728 | 0.20184 | 0.977164 | 4121 | 3 | -0.68553 | 0.51441 | 0.7382 | 1 | 15031 | 2 | -0.68553 |
| GABRD | 5 | 0.47531 | 0.62659 | 0.999994 | 13068 | 2 | -0.37922 | 0.51442 | 0.7215 | 1 | 15032 | 2 | -0.37922 |
| PSMA7 | 5 | 0.012704 | 0.043778 | 0.891621 | 1047 | 3 | -5.5806 | 0.51442 | 0.7215 | 1 | 15033 | 1 | -5.5806 |
| NSG1 | 5 | 0.1979 | 0.37224 | 0.999994 | 7708 | 1 | -0.10247 | 0.51442 | 0.7215 | 1 | 15034 | 2 | -0.10247 |
| MAT2B | 6 | 0.31231 | 0.521 | 0.999994 | 10367 | 3 | -0.16793 | 0.51452 | 0.73829 | 1 | 15035 | 2 | -0.16793 |
| PCDH7 | 6 | 0.22986 | 0.42389 | 0.999994 | 8525 | 3 | -0.014854 | 0.5146 | 0.73835 | 1 | 15036 | 3 | -0.014854 |
| IPO4 | 5 | 0.53286 | 0.67658 | 0.999994 | 13950 | 2 | -0.31676 | 0.5148 | 0.72169 | 1 | 15037 | 2 | -0.31676 |
| CAGE1 | 4 | 0.83348 | 0.833 | 0.999994 | 17980 | 0 | -0.19039 | 0.51487 | 0.67595 | 1 | 15038 | 1 | -0.19039 |
| AKAP8L | 6 | 0.25776 | 0.45722 | 0.999994 | 9190 | 3 | -0.53168 | 0.51493 | 0.7386 | 1 | 15039 | 2 | -0.53168 |
| SAMM50 | 4 | 0.83625 | 0.83575 | 0.999994 | 18025 | 0 | 0.44243 | 0.51509 | 0.67611 | 1 | 15040 | 2 | 0.44243 |
| ISG20 | 6 | 0.57053 | 0.71832 | 0.999994 | 14549 | 2 | 0.66563 | 0.5152 | 0.7388 | 1 | 15041 | 3 | 0.66563 |
| DYNLRB2 | 6 | 0.075747 | 0.19871 | 0.977164 | 4063 | 3 | -1.1532 | 0.5153 | 0.73887 | 1 | 15042 | 2 | -1.1532 |
| KANK4 | 6 | 0.045474 | 0.13411 | 0.95318 | 2787 | 2 | 0.29216 | 0.51539 | 0.73894 | 1 | 15043 | 3 | 0.29216 |
| ABCA12 | 6 | 0.0056196 | 0.023371 | 0.832392 | 565 | 5 | -3.1315 | 0.51544 | 0.73898 | 1 | 15044 | 1 | -3.1315 |
| RASA1 | 6 | 0.014199 | 0.053119 | 0.916684 | 1143 | 3 | -0.35487 | 0.51544 | 0.73898 | 1 | 15045 | 2 | -0.35487 |
| ZNF277 | 6 | 0.01665 | 0.061158 | 0.933313 | 1287 | 4 | -2.7836 | 0.51544 | 0.73898 | 1 | 15046 | 2 | -2.7836 |
| GABRA3 | 6 | 0.3465 | 0.56007 | 0.999994 | 11083 | 3 | 0.14635 | 0.51557 | 0.73909 | 1 | 15047 | 3 | 0.14635 |
| GLI3 | 6 | 0.020838 | 0.07467 | 0.93914 | 1501 | 4 | -1.9296 | 0.51588 | 0.73933 | 1 | 15048 | 2 | -1.9296 |
| OR12D3 | 5 | 0.41091 | 0.57265 | 0.999994 | 12125 | 2 | 0.37888 | 0.5159 | 0.72227 | 1 | 15049 | 2 | 0.37888 |
| hsa-mir-1180 | 4 | 0.61678 | 0.67454 | 0.999994 | 15168 | 1 | 0.6927 | 0.51597 | 0.67681 | 1 | 15050 | 2 | 0.6927 |
| hsa-mir-205 | 4 | 0.41971 | 0.5659 | 0.999994 | 12249 | 2 | -0.2899 | 0.51597 | 0.67681 | 1 | 15051 | 1 | -0.2899 |
| GLB1L2 | 6 | 0.22504 | 0.41817 | 0.999994 | 8399 | 3 | -0.29427 | 0.5162 | 0.73957 | 1 | 15052 | 3 | -0.29427 |
| NonTargetingControlG | 2 | 0.12554 | 0.20913 | 0.979587 | 5807 | 1 | -1.8469 | 0.51623 | 0.54966 | 1 | 15053 | 1 | -1.8469 |
| UGT3A2 | 6 | 0.32228 | 0.53252 | 0.999994 | 10560 | 3 | -0.074225 | 0.51635 | 0.73967 | 1 | 15054 | 3 | -0.074225 |
| C2orf44 | 6 | 0.048321 | 0.14041 | 0.958716 | 2899 | 4 | -1.7094 | 0.51635 | 0.73967 | 1 | 15055 | 2 | -1.7094 |
| ARL8A | 6 | 0.3251 | 0.53567 | 0.999994 | 10624 | 3 | -0.36259 | 0.51635 | 0.73967 | 1 | 15056 | 1 | -0.36259 |
| ZNFX1 | 6 | 0.13113 | 0.30177 | 0.999994 | 5961 | 3 | -0.69949 | 0.51635 | 0.73967 | 1 | 15057 | 3 | -0.69949 |
| USP17L15 | 6 | 0.054364 | 0.15353 | 0.967126 | 3171 | 4 | -0.96679 | 0.51635 | 0.73967 | 1 | 15058 | 1 | -0.96679 |
| TSNAXIP1 | 5 | 0.48773 | 0.63727 | 0.999994 | 13251 | 2 | -0.026182 | 0.5165 | 0.72258 | 1 | 15059 | 2 | -0.026182 |
| AADAC | 5 | 0.17648 | 0.34153 | 0.999994 | 7143 | 3 | -0.98872 | 0.5165 | 0.72258 | 1 | 15060 | 2 | -0.98872 |
| EHF | 5 | 0.072188 | 0.1819 | 0.968322 | 3925 | 3 | -1.644 | 0.5165 | 0.72258 | 1 | 15061 | 2 | -1.644 |
| BRCA2 | 4 | 0.80316 | 0.80403 | 0.999994 | 17526 | 1 | 0.74741 | 0.51656 | 0.67725 | 1 | 15062 | 2 | 0.74741 |
| SLC15A3 | 6 | 0.68594 | 0.80565 | 0.999994 | 16170 | 2 | 0.072264 | 0.51677 | 0.73999 | 1 | 15063 | 2 | 0.072264 |
| GEMIN5 | 3 | 0.053595 | 0.12071 | 0.951052 | 3146 | 2 | -4.8301 | 0.51701 | 0.65044 | 1 | 15064 | 1 | -4.8301 |
| hsa-mir-489 | 4 | 0.57844 | 0.65395 | 0.999994 | 14664 | 1 | 0.75998 | 0.51705 | 0.67764 | 1 | 15065 | 2 | 0.75998 |
| hsa-mir-4497 | 4 | 0.42391 | 0.57018 | 0.999994 | 12307 | 1 | 0.56573 | 0.51734 | 0.67787 | 1 | 15066 | 2 | 0.56573 |
| VIM | 6 | 0.0014417 | 0.0072155 | 0.656523 | 216 | 3 | -0.44145 | 0.51735 | 0.74044 | 1 | 15067 | 2 | -0.44145 |
| BLM | 5 | 0.020954 | 0.066648 | 0.93914 | 1512 | 2 | 0.40503 | 0.51735 | 0.72304 | 1 | 15068 | 2 | 0.40503 |
| NonTargetingControlG | 2 | 0.39769 | 0.41488 | 0.999994 | 11912 | 1 | 0.43609 | 0.51741 | 0.55053 | 1 | 15069 | 1 | 0.43609 |
| SRRM2 | 6 | 0.20164 | 0.38986 | 0.999994 | 7813 | 2 | 0.44506 | 0.51765 | 0.74067 | 1 | 15070 | 3 | 0.44506 |
| SLC22A8 | 5 | 0.34335 | 0.51647 | 0.999994 | 11015 | 1 | 0.21161 | 0.51767 | 0.72321 | 1 | 15071 | 2 | 0.21161 |
| hsa-mir-548y | 3 | 0.80729 | 0.80712 | 0.999994 | 17589 | 0 | 0.36233 | 0.51777 | 0.65076 | 1 | 15072 | 1 | 0.36233 |
| RFPL4AL1 | 4 | 0.021157 | 0.063073 | 0.937389 | 1524 | 1 | 0.69424 | 0.51791 | 0.67835 | 1 | 15073 | 2 | 0.69424 |
| C2orf73 | 6 | 0.19948 | 0.38725 | 0.999994 | 7749 | 2 | 0.38031 | 0.51798 | 0.74094 | 1 | 15074 | 3 | 0.38031 |
| hsa-mir-4460 | 4 | 0.05345 | 0.12823 | 0.951777 | 3138 | 2 | -2.3473 | 0.51841 | 0.67872 | 1 | 15075 | 2 | -2.3473 |
| TBC1D5 | 6 | 0.22823 | 0.42194 | 0.999994 | 8483 | 3 | -0.55645 | 0.51841 | 0.74127 | 1 | 15076 | 2 | -0.55645 |
| INPP5K | 6 | 0.26197 | 0.46222 | 0.999994 | 9286 | 2 | -0.19369 | 0.51841 | 0.74127 | 1 | 15077 | 2 | -0.19369 |
| CPAMD8 | 5 | 0.57447 | 0.71348 | 0.999994 | 14611 | 2 | -0.30499 | 0.51855 | 0.72368 | 1 | 15078 | 2 | -0.30499 |
| COG4 | 5 | 0.0089665 | 0.033077 | 0.867348 | 790 | 4 | -2.3015 | 0.51856 | 0.72369 | 1 | 15079 | 1 | -2.3015 |

| | | | | | | | | | | | | | |
|---|---|---|---|---|---|---|---|---|---|---|---|---|---|
| ZNF189 | 5 | 0.20058 | 0.37605 | 0.999994 | 7776 | 2 | -0.18841 | 0.51856 | 0.72369 | 1 | 15080 | 2 | -0.18841 |
| COPS3 | 4 | 0.04995 | 0.12137 | 0.951052 | 2973 | 3 | -3.5095 | 0.51871 | 0.67896 | 1 | 15081 | 1 | -3.5095 |
| DBNL | 4 | 0.45656 | 0.59743 | 0.999994 | 12799 | 1 | 0.2643 | 0.51871 | 0.67896 | 1 | 15082 | 2 | 0.2643 |
| hsa-mir-6082 | 4 | 0.39148 | 0.53691 | 0.999994 | 11821 | 2 | -0.049754 | 0.51871 | 0.67896 | 1 | 15083 | 2 | -0.049754 |
| KLHL33 | 4 | 0.013993 | 0.046008 | 0.892434 | 1127 | 3 | -2.5279 | 0.51871 | 0.67896 | 1 | 15084 | 1 | -2.5279 |
| GARS | 3 | 0.095926 | 0.17765 | 0.968171 | 4811 | 2 | -2.113 | 0.51875 | 0.65119 | 1 | 15085 | 1 | -2.113 |
| LCN6 | 6 | 0.021094 | 0.075501 | 0.93914 | 1518 | 3 | 0.019635 | 0.51877 | 0.74154 | 1 | 15086 | 3 | 0.019635 |
| FAM13A | 6 | 0.79493 | 0.84572 | 0.999994 | 17415 | 1 | -0.047971 | 0.51885 | 0.74162 | 1 | 15087 | 2 | -0.047971 |
| NonTargetingControlG | 2 | 0.62302 | 0.62281 | 0.999994 | 15272 | 0 | 1.1492 | 0.51895 | 0.55169 | 1 | 15088 | 1 | 1.1492 |
| ABTB2 | 6 | 0.39254 | 0.59756 | 0.999994 | 11843 | 2 | 0.38436 | 0.51915 | 0.74183 | 1 | 15089 | 3 | 0.38436 |
| LCA5 | 6 | 0.24152 | 0.43782 | 0.999994 | 8814 | 3 | 0.14091 | 0.51919 | 0.74186 | 1 | 15090 | 3 | 0.14091 |
| SH3RF2 | 6 | 0.19058 | 0.37624 | 0.999994 | 7501 | 3 | -0.69844 | 0.51923 | 0.74189 | 1 | 15091 | 2 | -0.69844 |
| MITD1 | 6 | 0.036297 | 0.11354 | 0.951052 | 2334 | 4 | -1.5601 | 0.51923 | 0.74189 | 1 | 15092 | 2 | -1.5601 |
| SLC25A47 | 5 | 0.23947 | 0.43021 | 0.999994 | 8764 | 2 | -0.54561 | 0.51951 | 0.7242 | 1 | 15093 | 1 | -0.54561 |
| NonTargetingControlG | 2 | 0.60568 | 0.60551 | 0.999994 | 15023 | 0 | 0.9268 | 0.51971 | 0.55225 | 1 | 15094 | 1 | 0.9268 |
| GPANK1 | 6 | 0.34775 | 0.56148 | 0.999994 | 11109 | 3 | 0.17673 | 0.51989 | 0.74241 | 1 | 15095 | 3 | 0.17673 |
| GBX2 | 4 | 0.019029 | 0.058603 | 0.931872 | 1409 | 3 | -2.7137 | 0.52005 | 0.67998 | 1 | 15096 | 1 | -2.7137 |
| SERPINA4 | 4 | 0.1554 | 0.29477 | 0.999994 | 6604 | 2 | -0.6107 | 0.52005 | 0.67998 | 1 | 15097 | 2 | -0.6107 |
| NEU3 | 6 | 0.16915 | 0.34969 | 0.999994 | 6964 | 3 | -0.29137 | 0.52008 | 0.74255 | 1 | 15098 | 3 | -0.29137 |
| PLXNA3 | 5 | 0.15428 | 0.30918 | 0.999994 | 6575 | 2 | -0.43766 | 0.52009 | 0.72451 | 1 | 15099 | 2 | -0.43766 |
| ENPP7 | 6 | 0.060362 | 0.16651 | 0.967126 | 3436 | 3 | -1.2347 | 0.52018 | 0.74264 | 1 | 15100 | 2 | -1.2347 |
| OR9K2 | 6 | 0.26467 | 0.46541 | 0.999994 | 9349 | 2 | 0.54763 | 0.5203 | 0.74273 | 1 | 15101 | 3 | 0.54763 |
| CST2 | 6 | 0.31787 | 0.52746 | 0.999994 | 10469 | 2 | -0.045314 | 0.5203 | 0.74273 | 1 | 15102 | 2 | -0.045314 |
| PSME4 | 6 | 0.21906 | 0.41098 | 0.999994 | 8263 | 3 | -0.32296 | 0.5203 | 0.74273 | 1 | 15103 | 3 | -0.32296 |
| NKX2-4 | 6 | 0.043015 | 0.12871 | 0.951777 | 2679 | 4 | -1.6372 | 0.5203 | 0.74273 | 1 | 15104 | 2 | -1.6372 |
| GUCY2F | 6 | 0.62524 | 0.75879 | 0.999994 | 15310 | 2 | 0.30915 | 0.52035 | 0.74277 | 1 | 15105 | 3 | 0.30915 |
| MPP5 | 6 | 0.070181 | 0.18715 | 0.969409 | 3841 | 3 | 0.13676 | 0.52037 | 0.74279 | 1 | 15106 | 3 | 0.13676 |
| ZNF711 | 6 | 0.55979 | 0.71064 | 0.999994 | 14368 | 2 | 0.42259 | 0.52039 | 0.7428 | 1 | 15107 | 3 | 0.42259 |
| PSMG4 | 4 | 0.38749 | 0.53278 | 0.999994 | 11760 | 1 | 0.94397 | 0.52051 | 0.68035 | 1 | 15108 | 2 | 0.94397 |
| MAPK3 | 5 | 0.010021 | 0.036125 | 0.877982 | 871 | 4 | -1.9272 | 0.52056 | 0.72477 | 1 | 15109 | 1 | -1.9272 |
| SERINC1 | 5 | 0.92083 | 0.92101 | 0.999994 | 19649 | 0 | 0.24689 | 0.52056 | 0.72477 | 1 | 15110 | 2 | 0.24689 |
| ZBTB17 | 5 | 0.083184 | 0.19973 | 0.977164 | 4346 | 2 | -0.26077 | 0.52058 | 0.72478 | 1 | 15111 | 2 | -0.26077 |
| hsa-mir-613 | 4 | 0.70155 | 0.72541 | 0.999994 | 16322 | 1 | 0.74012 | 0.52066 | 0.68046 | 1 | 15112 | 2 | 0.74012 |
| MAP3K1 | 6 | 0.0014779 | 0.0073815 | 0.665213 | 221 | 3 | -0.43128 | 0.52068 | 0.74302 | 1 | 15113 | 3 | -0.43128 |
| NonTargetingControlG | 2 | 0.43286 | 0.44447 | 0.999994 | 12461 | 1 | 0.4495 | 0.52091 | 0.55311 | 1 | 15114 | 1 | 0.4495 |
| BOLA2 | 6 | 0.61163 | 0.74853 | 0.999994 | 15099 | 2 | 0.36159 | 0.52092 | 0.74317 | 1 | 15115 | 3 | 0.36159 |
| MGAT5 | 6 | 0.18979 | 0.37523 | 0.999994 | 7478 | 2 | 0.7705 | 0.52111 | 0.74331 | 1 | 15116 | 3 | 0.7705 |
| STARD8 | 6 | 0.26355 | 0.46405 | 0.999994 | 9328 | 1 | 0.56883 | 0.52113 | 0.74333 | 1 | 15117 | 3 | 0.56883 |
| hsa-mir-1279 | 2 | 0.23127 | 0.2849 | 0.999632 | 8549 | 1 | -0.61347 | 0.5212 | 0.55332 | 1 | 15118 | 1 | -0.61347 |
| hsa-mir-6854 | 4 | 0.31808 | 0.46149 | 0.999994 | 10474 | 2 | -0.47799 | 0.52132 | 0.68099 | 1 | 15119 | 1 | -0.47799 |
| hsa-mir-155 | 4 | 0.075441 | 0.17017 | 0.967126 | 4052 | 3 | -1.7761 | 0.52132 | 0.68099 | 1 | 15120 | 1 | -1.7761 |
| hsa-mir-6768 | 4 | 0.88975 | 0.88945 | 0.999994 | 19031 | 0 | 0.46286 | 0.52132 | 0.68099 | 1 | 15121 | 1 | 0.46286 |
| NonTargetingControlG | 2 | 0.31883 | 0.35129 | 0.999994 | 10494 | 1 | 0.030276 | 0.52133 | 0.55342 | 1 | 15122 | 1 | 0.030276 |
| JAZF1 | 6 | 0.036898 | 0.11491 | 0.951052 | 2369 | 3 | 0.11091 | 0.52146 | 0.74357 | 1 | 15123 | 3 | 0.11091 |
| IGBP1 | 4 | 0.015277 | 0.049963 | 0.909101 | 1206 | 2 | -3.3385 | 0.52152 | 0.68114 | 1 | 15124 | 2 | -3.3385 |
| OCEL1 | 2 | 0.23908 | 0.29076 | 0.999632 | 8749 | 1 | -0.56233 | 0.52176 | 0.55375 | 1 | 15125 | 1 | -0.56233 |
| NFIL3 | 6 | 0.15057 | 0.32637 | 0.999994 | 6474 | 3 | 0.21844 | 0.52183 | 0.74386 | 1 | 15126 | 3 | 0.21844 |
| CDCP2 | 6 | 0.19784 | 0.38526 | 0.999994 | 7704 | 1 | 0.20913 | 0.5221 | 0.74407 | 1 | 15127 | 3 | 0.20913 |
| hsa-mir-511 | 4 | 0.62509 | 0.67919 | 0.999994 | 15309 | 1 | 0.50603 | 0.52224 | 0.68169 | 1 | 15128 | 2 | 0.50603 |
| KCNU1 | 6 | 0.26282 | 0.46321 | 0.999994 | 9311 | 2 | -0.17116 | 0.52231 | 0.74422 | 1 | 15129 | 2 | -0.17116 |
| VKORC1L1 | 5 | 0.19641 | 0.37011 | 0.999994 | 7660 | 2 | 0.25761 | 0.5224 | 0.72575 | 1 | 15130 | 1 | 0.25761 |
| HDAC3 | 5 | 0.00043202 | 0.0020037 | 0.439257 | 100 | 3 | -8.8041 | 0.5224 | 0.72575 | 1 | 15131 | 1 | -8.8041 |

| | | | | | | | | | | | | | |
|---|---|---|---|---|---|---|---|---|---|---|---|---|---|
| PSMC6 | 6 | 0.008618 | 0.03414 | 0.867348 | 768 | 4 | -4.1623 | 0.5224 | 0.74428 | 1 | 15132 | 2 | -4.1623 |
| TRIP6 | 6 | 0.29754 | 0.50385 | 0.999994 | 10079 | 2 | 0.28855 | 0.52246 | 0.74433 | 1 | 15133 | 2 | 0.28855 |
| SMEK2 | 5 | 0.25068 | 0.44131 | 0.999994 | 9029 | 2 | -0.20775 | 0.52251 | 0.7258 | 1 | 15134 | 2 | -0.20775 |
| NAAA | 6 | 0.46179 | 0.64293 | 0.999994 | 12877 | 2 | -0.13521 | 0.52262 | 0.74444 | 1 | 15135 | 2 | -0.13521 |
| SIGLEC10 | 4 | 0.43182 | 0.57841 | 0.999994 | 12442 | 1 | 0.92664 | 0.52267 | 0.68203 | 1 | 15136 | 2 | 0.92664 |
| C7orf26 | 4 | 0.1055 | 0.22496 | 0.983354 | 5153 | 3 | -1.466 | 0.52271 | 0.68207 | 1 | 15137 | 1 | -1.466 |
| RFC1 | 4 | 0.068525 | 0.15706 | 0.967126 | 3773 | 3 | -1.4929 | 0.52271 | 0.68207 | 1 | 15138 | 1 | -1.4929 |
| DRD5 | 6 | 0.22975 | 0.42377 | 0.999994 | 8523 | 3 | 0.0025451 | 0.52275 | 0.74455 | 1 | 15139 | 3 | 0.0025451 |
| TRDMT1 | 6 | 0.33068 | 0.54204 | 0.999994 | 10735 | 2 | 0.48123 | 0.52277 | 0.74456 | 1 | 15140 | 3 | 0.48123 |
| SRSF2 | 3 | 0.055783 | 0.12371 | 0.951052 | 3237 | 1 | 0.39399 | 0.52279 | 0.65292 | 1 | 15141 | 1 | 0.39399 |
| hsa-mir-6074 | 3 | 0.74074 | 0.74066 | 0.999994 | 16748 | 0 | 0.15537 | 0.52279 | 0.65292 | 1 | 15142 | 1 | 0.15537 |
| hsa-mir-4504 | 4 | 0.29996 | 0.44296 | 0.999994 | 10131 | 1 | 0.78626 | 0.52311 | 0.68238 | 1 | 15143 | 2 | 0.78626 |
| OR2M3 | 5 | 0.71868 | 0.77522 | 0.999994 | 16482 | 1 | 0.37282 | 0.52311 | 0.72612 | 1 | 15144 | 2 | 0.37282 |
| C5orf28 | 5 | 0.29246 | 0.47493 | 0.999994 | 9963 | 2 | 0.27893 | 0.5233 | 0.72623 | 1 | 15145 | 2 | 0.27893 |
| PTPRQ | 6 | 0.061228 | 0.16839 | 0.967126 | 3485 | 3 | -1.2255 | 0.52344 | 0.74506 | 1 | 15146 | 2 | -1.2255 |
| TBX22 | 6 | 0.36248 | 0.57809 | 0.999994 | 11378 | 3 | -0.51802 | 0.52344 | 0.74506 | 1 | 15147 | 2 | -0.51802 |
| SLCO4C1 | 6 | 0.52509 | 0.68608 | 0.999994 | 13817 | 2 | 0.48942 | 0.52344 | 0.74506 | 1 | 15148 | 3 | 0.48942 |
| OR2AT4 | 6 | 0.067528 | 0.18159 | 0.968171 | 3737 | 3 | -0.61098 | 0.52344 | 0.74506 | 1 | 15149 | 1 | -0.61098 |
| CARD17 | 6 | 0.83138 | 0.86221 | 0.999994 | 17951 | 1 | 0.553 | 0.52349 | 0.7451 | 1 | 15150 | 3 | 0.553 |
| hsa-mir-3118-2 | 1 | 0.47648 | 0.47647 | 0.999994 | 13090 | 0 | 0.2989 | 0.52352 | 0.52334 | 1 | 15151 | 0 | 0.2989 |
| HSH2D | 5 | 0.2087 | 0.38754 | 0.999994 | 7993 | 3 | -0.81309 | 0.5236 | 0.7264 | 1 | 15152 | 1 | -0.81309 |
| CDC14B | 5 | 0.56566 | 0.70571 | 0.999994 | 14479 | 2 | -0.099735 | 0.5236 | 0.7264 | 1 | 15153 | 1 | -0.099735 |
| PCDHA10 | 2 | 0.50778 | 0.51106 | 0.999994 | 13548 | 1 | 0.63123 | 0.52361 | 0.55511 | 1 | 15154 | 1 | 0.63123 |
| LINGO2 | 6 | 0.016228 | 0.059753 | 0.932932 | 1259 | 4 | -2.0804 | 0.52371 | 0.74527 | 1 | 15155 | 2 | -2.0804 |
| hsa-mir-4255 | 4 | 0.4848 | 0.60962 | 0.999994 | 13217 | 1 | 0.23632 | 0.5238 | 0.68291 | 1 | 15156 | 2 | 0.23632 |
| hsa-mir-4674 | 4 | 0.18423 | 0.32447 | 0.999994 | 7323 | 2 | -0.66166 | 0.5238 | 0.68291 | 1 | 15157 | 1 | -0.66166 |
| NonTargetingControlG | 2 | 0.75546 | 0.75502 | 0.999994 | 16923 | 0 | 1.4547 | 0.52382 | 0.55527 | 1 | 15158 | 1 | 1.4547 |
| FAM150A | 5 | 0.20377 | 0.38059 | 0.999994 | 7873 | 2 | -0.34716 | 0.52385 | 0.72653 | 1 | 15159 | 2 | -0.34716 |
| GANAB | 6 | 0.0090454 | 0.035675 | 0.873925 | 797 | 3 | -0.094628 | 0.52395 | 0.74546 | 1 | 15160 | 3 | -0.094628 |
| ZNF485 | 6 | 0.65489 | 0.78138 | 0.999994 | 15715 | 2 | -0.52672 | 0.52397 | 0.74548 | 1 | 15161 | 2 | -0.52672 |
| AGBL4 | 6 | 0.19118 | 0.37698 | 0.999994 | 7519 | 3 | -0.20109 | 0.52403 | 0.74553 | 1 | 15162 | 3 | -0.20109 |
| MRPL1 | 5 | 0.36747 | 0.53627 | 0.999994 | 11465 | 1 | 0.052301 | 0.52403 | 0.72664 | 1 | 15163 | 2 | 0.052301 |
| KLF15 | 6 | 0.010491 | 0.040702 | 0.891621 | 903 | 3 | -0.6155 | 0.52405 | 0.74555 | 1 | 15164 | 3 | -0.6155 |
| ITFG3 | 5 | 0.51209 | 0.65847 | 0.999994 | 13611 | 2 | 0.068414 | 0.52411 | 0.72668 | 1 | 15165 | 2 | 0.068414 |
| SULT1C2 | 6 | 0.50191 | 0.67002 | 0.999994 | 13457 | 1 | 0.77101 | 0.52415 | 0.74563 | 1 | 15166 | 3 | 0.77101 |
| TXNRD3 | 6 | 0.28832 | 0.49314 | 0.999994 | 9877 | 2 | 0.49171 | 0.52426 | 0.7457 | 1 | 15167 | 3 | 0.49171 |
| GPKOW | 3 | 0.020897 | 0.053771 | 0.921203 | 1503 | 2 | -3.5467 | 0.52438 | 0.65358 | 1 | 15168 | 1 | -3.5467 |
| RASA4 | 6 | 0.14306 | 0.31693 | 0.999994 | 6275 | 2 | 0.26599 | 0.52471 | 0.74605 | 1 | 15169 | 3 | 0.26599 |
| COL13A1 | 4 | 0.087434 | 0.19231 | 0.974607 | 4492 | 3 | -1.0135 | 0.5248 | 0.6837 | 1 | 15170 | 1 | -1.0135 |
| LCE1E | 4 | 0.59895 | 0.66474 | 0.999994 | 14941 | 1 | 0.19113 | 0.5248 | 0.6837 | 1 | 15171 | 2 | 0.19113 |
| OR8H3 | 5 | 0.1586 | 0.31549 | 0.999994 | 6687 | 3 | -0.82113 | 0.52483 | 0.72705 | 1 | 15172 | 2 | -0.82113 |
| BMP15 | 5 | 0.071722 | 0.18116 | 0.968171 | 3906 | 1 | 0.23383 | 0.52484 | 0.72706 | 1 | 15173 | 2 | 0.23383 |
| TAF15 | 6 | 0.47564 | 0.65225 | 0.999994 | 13072 | 2 | 0.17188 | 0.52492 | 0.74621 | 1 | 15174 | 1 | 0.17188 |
| KIF15 | 6 | 0.5385 | 0.69547 | 0.999994 | 14029 | 2 | 0.13352 | 0.52492 | 0.74621 | 1 | 15175 | 3 | 0.13352 |
| USP19 | 6 | 0.19787 | 0.38531 | 0.999994 | 7707 | 3 | -0.12785 | 0.52492 | 0.74621 | 1 | 15176 | 3 | -0.12785 |
| ARF6 | 5 | 0.0013428 | 0.0059692 | 0.623123 | 200 | 3 | -4.7919 | 0.52501 | 0.72714 | 1 | 15177 | 2 | -4.7919 |
| SLC35D2 | 6 | 0.15851 | 0.33634 | 0.999994 | 6685 | 3 | -0.22345 | 0.52507 | 0.74633 | 1 | 15178 | 3 | -0.22345 |
| GOLGA8B | 3 | 0.16342 | 0.26597 | 0.997535 | 6825 | 2 | -1.5725 | 0.52517 | 0.65394 | 1 | 15179 | 1 | -1.5725 |
| RBFOX1 | 6 | 0.18924 | 0.37456 | 0.999994 | 7459 | 3 | -0.4701 | 0.52522 | 0.74644 | 1 | 15180 | 2 | -0.4701 |
| OXCT1 | 4 | 0.17825 | 0.3183 | 0.999994 | 7181 | 2 | -0.35293 | 0.52526 | 0.68408 | 1 | 15181 | 2 | -0.35293 |
| SPRR2E | 2 | 0.15015 | 0.22673 | 0.983354 | 6462 | 1 | -2.3757 | 0.5254 | 0.55648 | 1 | 15182 | 1 | -2.3757 |
| EXOSC1 | 6 | 0.06072 | 0.1673 | 0.967126 | 3452 | 3 | -0.80479 | 0.52573 | 0.74684 | 1 | 15183 | 3 | -0.80479 |

| | | | | | | | | | | | | | |
|---|---|---|---|---|---|---|---|---|---|---|---|---|---|
| UBE2Q1 | 6 | 0.056306 | 0.15775 | 0.967126 | 3259 | 3 | 0.1747 | 0.52577 | 0.74687 | 1 | 15184 | 3 | 0.1747 |
| PCDHB8 | 5 | 0.24302 | 0.43503 | 0.999994 | 8868 | 2 | -0.064855 | 0.52578 | 0.72755 | 1 | 15185 | 2 | -0.064855 |
| ZNF317 | 6 | 0.49279 | 0.66378 | 0.999994 | 13307 | 1 | 0.67164 | 0.52581 | 0.7469 | 1 | 15186 | 3 | 0.67164 |
| PAN2 | 5 | 0.20734 | 0.38562 | 0.999994 | 7954 | 3 | -0.76763 | 0.52592 | 0.72763 | 1 | 15187 | 2 | -0.76763 |
| GBE1 | 6 | 0.58995 | 0.73255 | 0.999994 | 14823 | 1 | 0.4811 | 0.52604 | 0.74707 | 1 | 15188 | 3 | 0.4811 |
| ARHGAP24 | 6 | 0.10525 | 0.25754 | 0.994943 | 5142 | 4 | -0.94063 | 0.52605 | 0.74708 | 1 | 15189 | 2 | -0.94063 |
| hsa-mir-1226 | 4 | 0.17426 | 0.31421 | 0.999994 | 7090 | 2 | 0.063514 | 0.52606 | 0.6847 | 1 | 15190 | 2 | 0.063514 |
| NonTargetingControlG | 2 | 0.34527 | 0.37225 | 0.999994 | 11056 | 1 | 0.047339 | 0.52609 | 0.55698 | 1 | 15191 | 1 | 0.047339 |
| CRYGD | 5 | 0.4577 | 0.61173 | 0.999994 | 12817 | 1 | 0.20779 | 0.52616 | 0.72777 | 1 | 15192 | 2 | 0.20779 |
| RDX | 4 | 0.83498 | 0.83453 | 0.999994 | 18006 | 0 | 0.1839 | 0.52617 | 0.68479 | 1 | 15193 | 1 | 0.1839 |
| hsa-mir-548ae-1 | 4 | 0.89813 | 0.89785 | 0.999994 | 19181 | 0 | 0.76217 | 0.52617 | 0.68479 | 1 | 15194 | 2 | 0.76217 |
| NonTargetingControlG | 2 | 0.59689 | 0.59677 | 0.999994 | 14915 | 0 | 0.93035 | 0.52623 | 0.55708 | 1 | 15195 | 1 | 0.93035 |
| GNB1 | 6 | 0.35486 | 0.56951 | 0.999994 | 11248 | 2 | 0.65299 | 0.52627 | 0.74723 | 1 | 15196 | 3 | 0.65299 |
| NPEPPS | 6 | 0.03597 | 0.1128 | 0.951052 | 2311 | 3 | -1.1805 | 0.52635 | 0.74729 | 1 | 15197 | 3 | -1.1805 |
| CSAG1 | 5 | 0.19384 | 0.36641 | 0.999994 | 7596 | 2 | -0.48771 | 0.52656 | 0.72799 | 1 | 15198 | 2 | -0.48771 |
| NonTargetingControlG | 2 | 0.75803 | 0.75758 | 0.999994 | 16953 | 0 | 1.3376 | 0.52656 | 0.55733 | 1 | 15199 | 1 | 1.3376 |
| TMEM165 | 5 | 0.17351 | 0.33725 | 0.999994 | 7074 | 2 | 0.52127 | 0.52663 | 0.72802 | 1 | 15200 | 2 | 0.52127 |
| ENTHD2 | 6 | 0.65993 | 0.78527 | 0.999994 | 15778 | 2 | 0.55628 | 0.52663 | 0.74753 | 1 | 15201 | 3 | 0.55628 |
| SNRPE | 5 | 0.0079321 | 0.030077 | 0.862223 | 726 | 3 | -6.263 | 0.52676 | 0.72811 | 1 | 15202 | 2 | -6.263 |
| IFITM5 | 6 | 0.1112 | 0.26901 | 0.998561 | 5340 | 3 | -0.31361 | 0.52678 | 0.74765 | 1 | 15203 | 2 | -0.31361 |
| hsa-mir-3678 | 2 | 0.50602 | 0.50942 | 0.999994 | 13527 | 1 | 0.64415 | 0.52684 | 0.55755 | 1 | 15204 | 1 | 0.64415 |
| AARSD1 | 6 | 0.87715 | 0.88779 | 0.999994 | 18777 | 1 | 0.46899 | 0.52687 | 0.74772 | 1 | 15205 | 3 | 0.46899 |
| NonTargetingControlG | 2 | 0.41821 | 0.43199 | 0.999994 | 12223 | 1 | 0.36266 | 0.52692 | 0.5576 | 1 | 15206 | 1 | 0.36266 |
| OR8U1 | 6 | 0.42189 | 0.61664 | 0.999994 | 12277 | 2 | 0.32678 | 0.52724 | 0.74801 | 1 | 15207 | 3 | 0.32678 |
| NonTargetingControlG | 2 | 0.68228 | 0.68215 | 0.999994 | 16122 | 0 | 1.2438 | 0.52726 | 0.55786 | 1 | 15208 | 1 | 1.2438 |
| GDPGP1 | 4 | 0.009314 | 0.031565 | 0.864134 | 815 | 3 | -5.5335 | 0.52733 | 0.68565 | 1 | 15209 | 1 | -5.5335 |
| RFPL4B | 6 | 0.011748 | 0.044896 | 0.891621 | 975 | 5 | -1.7345 | 0.52738 | 0.74811 | 1 | 15210 | 1 | -1.7345 |
| ZNF285 | 6 | 0.32317 | 0.53352 | 0.999994 | 10578 | 3 | -0.43431 | 0.52738 | 0.74811 | 1 | 15211 | 1 | -0.43431 |
| ARHGAP21 | 5 | 0.067405 | 0.17418 | 0.968171 | 3732 | 2 | -0.20851 | 0.5274 | 0.72844 | 1 | 15212 | 2 | -0.20851 |
| LSM11 | 6 | 0.090398 | 0.22828 | 0.983354 | 4609 | 4 | -1.4635 | 0.52741 | 0.74813 | 1 | 15213 | 2 | -1.4635 |
| NIPSNAP3B | 6 | 0.085689 | 0.21889 | 0.980699 | 4435 | 2 | -0.25365 | 0.52741 | 0.74813 | 1 | 15214 | 2 | -0.25365 |
| LCP2 | 5 | 0.0015049 | 0.0066687 | 0.643671 | 223 | 4 | -4.9332 | 0.52751 | 0.72851 | 1 | 15215 | 1 | -4.9332 |
| NonTargetingControlG | 2 | 0.27121 | 0.31476 | 0.999994 | 9503 | 1 | -0.31627 | 0.52761 | 0.55812 | 1 | 15216 | 1 | -0.31627 |
| TARDBP | 5 | 0.025483 | 0.078785 | 0.93914 | 1778 | 3 | -2.6709 | 0.52762 | 0.72856 | 1 | 15217 | 2 | -2.6709 |
| LOC388813 | 2 | 0.60988 | 0.60972 | 0.999994 | 15071 | 0 | 0.90366 | 0.52766 | 0.55815 | 1 | 15218 | 1 | 0.90366 |
| ATP11C | 6 | 0.26696 | 0.46809 | 0.999994 | 9396 | 1 | 0.6459 | 0.52793 | 0.74853 | 1 | 15219 | 3 | 0.6459 |
| SLC9A3R1 | 4 | 0.7883 | 0.79091 | 0.999994 | 17323 | 1 | 0.39732 | 0.52794 | 0.68612 | 1 | 15220 | 2 | 0.39732 |
| hsa-mir-7161 | 4 | 0.36464 | 0.50927 | 0.999994 | 11415 | 2 | 0.23613 | 0.52803 | 0.68618 | 1 | 15221 | 2 | 0.23613 |
| SENP7 | 6 | 0.30919 | 0.51743 | 0.999994 | 10307 | 2 | 0.68921 | 0.52803 | 0.7486 | 1 | 15222 | 3 | 0.68921 |
| hsa-mir-424 | 4 | 0.23325 | 0.37483 | 0.999994 | 8601 | 2 | -0.65508 | 0.52804 | 0.68619 | 1 | 15223 | 1 | -0.65508 |
| ADRBK1 | 4 | 0.40093 | 0.54669 | 0.999994 | 11958 | 1 | 0.8875 | 0.52807 | 0.68622 | 1 | 15224 | 2 | 0.8875 |
| NonTargetingControlG | 2 | 0.63907 | 0.63892 | 0.999994 | 15500 | 0 | 0.88223 | 0.52814 | 0.55851 | 1 | 15225 | 1 | 0.88223 |
| FLII | 4 | 0.27349 | 0.4159 | 0.999994 | 9554 | 2 | -0.1578 | 0.5282 | 0.68633 | 1 | 15226 | 2 | -0.1578 |
| ADAP1 | 6 | 0.90246 | 0.90558 | 0.999994 | 19260 | 1 | 0.48172 | 0.52822 | 0.74874 | 1 | 15227 | 3 | 0.48172 |
| AGR2 | 6 | 0.029023 | 0.096497 | 0.945031 | 1956 | 4 | -1.9581 | 0.52833 | 0.74883 | 1 | 15228 | 2 | -1.9581 |
| CSF2 | 6 | 0.014144 | 0.052934 | 0.916293 | 1138 | 3 | -0.55625 | 0.52833 | 0.74883 | 1 | 15229 | 2 | -0.55625 |
| TSPAN9 | 6 | 0.017612 | 0.064244 | 0.93914 | 1345 | 5 | -2.1266 | 0.52833 | 0.74883 | 1 | 15230 | 1 | -2.1266 |
| NUAK1 | 6 | 0.26408 | 0.4647 | 0.999994 | 9335 | 2 | 0.2813 | 0.52833 | 0.74883 | 1 | 15231 | 2 | 0.2813 |
| CBLN3 | 6 | 0.049958 | 0.14396 | 0.961589 | 2974 | 4 | -1.4159 | 0.52833 | 0.74883 | 1 | 15232 | 2 | -1.4159 |
| ACOT13 | 6 | 0.13383 | 0.30521 | 0.999994 | 6018 | 2 | 0.66465 | 0.52834 | 0.74884 | 1 | 15233 | 3 | 0.66465 |
| GRK6 | 6 | 0.34332 | 0.55645 | 0.999994 | 11012 | 1 | 0.713 | 0.5284 | 0.74887 | 1 | 15234 | 3 | 0.713 |
| C20orf166 | 6 | 0.31543 | 0.52465 | 0.999994 | 10426 | 3 | 0.11419 | 0.52845 | 0.74892 | 1 | 15235 | 3 | 0.11419 |

| | | | | | | | | | | | | | |
|---|---|---|---|---|---|---|---|---|---|---|---|---|---|
| ANGPTL2 | 4 | 0.56387 | 0.6466 | 0.999994 | 14449 | 1 | 0.19384 | 0.52851 | 0.68655 | 1 | 15236 | 2 | 0.19384 |
| NAT10 | 4 | 0.036186 | 0.09411 | 0.945031 | 2328 | 3 | -3.8041 | 0.52851 | 0.68655 | 1 | 15237 | 1 | -3.8041 |
| TBX5 | 5 | 0.11225 | 0.2455 | 0.98979 | 5379 | 3 | -1.3751 | 0.52855 | 0.72907 | 1 | 15238 | 1 | -1.3751 |
| SRSF8 | 5 | 0.00027929 | 0.0013152 | 0.384267 | 75 | 3 | -0.69003 | 0.52855 | 0.72907 | 1 | 15239 | 2 | -0.69003 |
| RPL36AL | 5 | 0.11159 | 0.24445 | 0.988898 | 5354 | 3 | -1.0033 | 0.52855 | 0.72907 | 1 | 15240 | 2 | -1.0033 |
| ANKRD39 | 5 | 0.87631 | 0.87679 | 0.999994 | 18759 | 1 | 0.21067 | 0.5286 | 0.7291 | 1 | 15241 | 2 | 0.21067 |
| IFI27 | 6 | 0.29867 | 0.50513 | 0.999994 | 10104 | 3 | 0.12788 | 0.52873 | 0.74913 | 1 | 15242 | 3 | 0.12788 |
| IGFBP7 | 6 | 0.46874 | 0.6476 | 0.999994 | 12987 | 2 | 0.22383 | 0.52878 | 0.74916 | 1 | 15243 | 2 | 0.22383 |
| MXRA5 | 6 | 0.35007 | 0.56407 | 0.999994 | 11159 | 2 | 0.10541 | 0.52884 | 0.74921 | 1 | 15244 | 3 | 0.10541 |
| PCLO | 6 | 0.1431 | 0.31698 | 0.999994 | 6276 | 2 | 0.35552 | 0.5289 | 0.74925 | 1 | 15245 | 3 | 0.35552 |
| hsa-mir-6513 | 4 | 0.13151 | 0.27011 | 0.998561 | 5970 | 2 | 0.035678 | 0.5289 | 0.68686 | 1 | 15246 | 2 | 0.035678 |
| NTF4 | 6 | 0.044475 | 0.13194 | 0.95318 | 2734 | 2 | 0.60685 | 0.52905 | 0.74936 | 1 | 15247 | 3 | 0.60685 |
| RDM1 | 6 | 0.028755 | 0.095897 | 0.945031 | 1941 | 2 | 0.73463 | 0.52909 | 0.74939 | 1 | 15248 | 3 | 0.73463 |
| VIPR1 | 6 | 0.75994 | 0.83205 | 0.999994 | 16981 | 1 | 0.04547 | 0.52927 | 0.74953 | 1 | 15249 | 2 | 0.04547 |
| PYGB | 6 | 0.01374 | 0.051589 | 0.910734 | 1113 | 4 | -2.6637 | 0.52937 | 0.7496 | 1 | 15250 | 2 | -2.6637 |
| SLC32A1 | 6 | 0.10084 | 0.24896 | 0.991255 | 5001 | 3 | -0.90446 | 0.52937 | 0.7496 | 1 | 15251 | 2 | -0.90446 |
| EXOSC10 | 6 | 0.059658 | 0.16501 | 0.967126 | 3408 | 4 | -1.4744 | 0.52937 | 0.7496 | 1 | 15252 | 1 | -1.4744 |
| MAPRE2 | 6 | 0.32206 | 0.53228 | 0.999994 | 10558 | 2 | -0.48597 | 0.52937 | 0.7496 | 1 | 15253 | 1 | -0.48597 |
| COX6C | 4 | 0.27544 | 0.4179 | 0.999994 | 9603 | 2 | 0.094144 | 0.52946 | 0.68727 | 1 | 15254 | 2 | 0.094144 |
| LRRTM2 | 6 | 0.59311 | 0.73487 | 0.999994 | 14865 | 2 | 0.15381 | 0.5295 | 0.7497 | 1 | 15255 | 3 | 0.15381 |
| hsa-mir-10b | 4 | 0.64182 | 0.68861 | 0.999994 | 15547 | 1 | 0.75921 | 0.52957 | 0.68736 | 1 | 15256 | 2 | 0.75921 |
| NR1H3 | 4 | 0.10195 | 0.21859 | 0.980699 | 5035 | 2 | -1.1017 | 0.52961 | 0.68739 | 1 | 15257 | 2 | -1.1017 |
| MFRP | 4 | 0.11886 | 0.24826 | 0.990682 | 5596 | 3 | -0.79764 | 0.52961 | 0.68739 | 1 | 15258 | 1 | -0.79764 |
| NDUFA2 | 4 | 0.054317 | 0.12991 | 0.95318 | 3168 | 3 | -2.0153 | 0.52961 | 0.68739 | 1 | 15259 | 1 | -2.0153 |
| BRINP1 | 5 | 0.93518 | 0.93542 | 0.999994 | 19972 | 0 | 0.47188 | 0.52969 | 0.72968 | 1 | 15260 | 2 | 0.47188 |
| TBC1D24 | 5 | 0.0071619 | 0.027847 | 0.855011 | 669 | 3 | -4.9129 | 0.52969 | 0.72968 | 1 | 15261 | 2 | -4.9129 |
| LYPD1 | 5 | 0.18806 | 0.35818 | 0.999994 | 7420 | 3 | -0.72212 | 0.52969 | 0.72968 | 1 | 15262 | 2 | -0.72212 |
| NHS | 5 | 0.7703 | 0.80187 | 0.999994 | 17088 | 1 | 0.24719 | 0.52969 | 0.72968 | 1 | 15263 | 2 | 0.24719 |
| GPR137B | 5 | 0.038858 | 0.11301 | 0.951052 | 2455 | 3 | -2.8418 | 0.52969 | 0.72968 | 1 | 15264 | 1 | -2.8418 |
| ACBD5 | 6 | 0.073941 | 0.19495 | 0.975797 | 3992 | 3 | -1.5428 | 0.52974 | 0.74987 | 1 | 15265 | 2 | -1.5428 |
| POLN | 6 | 0.12054 | 0.28674 | 0.999632 | 5645 | 3 | -0.67757 | 0.52979 | 0.7499 | 1 | 15266 | 2 | -0.67757 |
| hsa-mir-3670-1 | 3 | 0.31364 | 0.4359 | 0.999994 | 10390 | 1 | 0.54988 | 0.52993 | 0.65605 | 1 | 15267 | 1 | 0.54988 |
| RAD23A | 5 | 0.06669 | 0.173 | 0.96793 | 3712 | 3 | -1.691 | 0.53006 | 0.72986 | 1 | 15268 | 2 | -1.691 |
| ZNF880 | 6 | 0.14422 | 0.31839 | 0.999994 | 6307 | 3 | -0.40205 | 0.53015 | 0.75018 | 1 | 15269 | 3 | -0.40205 |
| VCPIP1 | 6 | 0.15299 | 0.32942 | 0.999994 | 6533 | 3 | -0.29084 | 0.53017 | 0.75018 | 1 | 15270 | 3 | -0.29084 |
| KLHL24 | 6 | 0.030466 | 0.099869 | 0.946394 | 2029 | 2 | 0.57459 | 0.53019 | 0.7502 | 1 | 15271 | 3 | 0.57459 |
| TXNDC15 | 6 | 0.89487 | 0.8998 | 0.999994 | 19122 | 1 | 0.21003 | 0.53025 | 0.75025 | 1 | 15272 | 3 | 0.21003 |
| KBTBD11 | 5 | 0.11382 | 0.24792 | 0.990682 | 5439 | 3 | -0.66796 | 0.53029 | 0.73 | 1 | 15273 | 1 | -0.66796 |
| SLC22A24 | 5 | 0.048575 | 0.13698 | 0.957161 | 2911 | 2 | 0.31064 | 0.53035 | 0.73003 | 1 | 15274 | 2 | 0.31064 |
| hsa-mir-370 | 4 | 0.15137 | 0.29061 | 0.999632 | 6489 | 2 | -0.37897 | 0.53037 | 0.68796 | 1 | 15275 | 2 | -0.37897 |
| PTCD1 | 3 | 0.03308 | 0.082197 | 0.940299 | 2164 | 2 | -4.1957 | 0.53037 | 0.65624 | 1 | 15276 | 1 | -4.1957 |
| GAP43 | 4 | 0.0012454 | 0.0045774 | 0.576301 | 188 | 2 | -4.3184 | 0.53041 | 0.68799 | 1 | 15277 | 2 | -4.3184 |
| MGME1 | 6 | 0.074161 | 0.19542 | 0.975797 | 3999 | 4 | -1.0465 | 0.53068 | 0.75057 | 1 | 15278 | 2 | -1.0465 |
| MRPS7 | 4 | 0.10004 | 0.21518 | 0.979587 | 4974 | 2 | -0.70958 | 0.53069 | 0.68818 | 1 | 15279 | 2 | -0.70958 |
| ZNF48 | 5 | 0.12998 | 0.27255 | 0.998561 | 5933 | 3 | -1.2406 | 0.53078 | 0.73028 | 1 | 15280 | 2 | -1.2406 |
| ENTPD3 | 5 | 0.90428 | 0.90437 | 0.999994 | 19306 | 0 | -0.2805 | 0.53078 | 0.73028 | 1 | 15281 | 2 | -0.2805 |
| ANKRD34A | 5 | 0.53064 | 0.67463 | 0.999994 | 13906 | 1 | 0.39727 | 0.53078 | 0.73028 | 1 | 15282 | 2 | 0.39727 |
| NIN | 4 | 0.18104 | 0.32117 | 0.999994 | 7248 | 2 | -0.66769 | 0.53089 | 0.68832 | 1 | 15283 | 1 | -0.66769 |
| APOC4 | 6 | 0.38623 | 0.59352 | 0.999994 | 11737 | 2 | 0.2346 | 0.53104 | 0.75085 | 1 | 15284 | 3 | 0.2346 |
| TCTEX1D4 | 6 | 0.0027234 | 0.012715 | 0.733879 | 332 | 5 | -4.2644 | 0.53111 | 0.7509 | 1 | 15285 | 1 | -4.2644 |
| RAB10 | 6 | 0.52153 | 0.68357 | 0.999994 | 13769 | 2 | 0.3239 | 0.53121 | 0.75099 | 1 | 15286 | 3 | 0.3239 |
| MROH8 | 6 | 0.0061322 | 0.025297 | 0.84412 | 601 | 4 | -2.6702 | 0.53128 | 0.75103 | 1 | 15287 | 2 | -2.6702 |

| | | | | | | | | | | | | | |
|---|---|---|---|---|---|---|---|---|---|---|---|---|---|
| EEF1A2 | 5 | 0.19161 | 0.36322 | 0.999994 | 7533 | 3 | -0.88619 | 0.53133 | 0.73059 | 1 | 15288 | 2 | -0.88619 |
| hsa-mir-4684 | 4 | 0.34181 | 0.48587 | 0.999994 | 10977 | 2 | 0.12254 | 0.53134 | 0.68869 | 1 | 15289 | 2 | 0.12254 |
| ARHGEF35 | 5 | 0.03263 | 0.097231 | 0.945253 | 2140 | 3 | -2.5698 | 0.53136 | 0.7306 | 1 | 15290 | 2 | -2.5698 |
| CHGB | 6 | 0.86323 | 0.87926 | 0.999994 | 18509 | 1 | 0.40394 | 0.53153 | 0.75122 | 1 | 15291 | 2 | 0.40394 |
| AP1S1 | 6 | 0.19222 | 0.37827 | 0.999994 | 7555 | 2 | -0.36242 | 0.53156 | 0.75124 | 1 | 15292 | 2 | -0.36242 |
| C11orf83 | 6 | 0.4386 | 0.62747 | 0.999994 | 12549 | 2 | 0.83594 | 0.53156 | 0.75124 | 1 | 15293 | 3 | 0.83594 |
| UROC1 | 5 | 0.12329 | 0.26233 | 0.995875 | 5746 | 3 | -1.3356 | 0.53173 | 0.73079 | 1 | 15294 | 2 | -1.3356 |
| CCDC122 | 6 | 0.088819 | 0.22516 | 0.983354 | 4546 | 2 | 0.44288 | 0.53179 | 0.75142 | 1 | 15295 | 3 | 0.44288 |
| ARSA | 5 | 0.23407 | 0.42286 | 0.999994 | 8622 | 3 | -0.66067 | 0.53181 | 0.73083 | 1 | 15296 | 2 | -0.66067 |
| ZNF330 | 6 | 0.90255 | 0.90567 | 0.999994 | 19263 | 1 | 0.031383 | 0.53181 | 0.75143 | 1 | 15297 | 2 | 0.031383 |
| SCN5A | 6 | 0.19978 | 0.38761 | 0.999994 | 7757 | 3 | -0.52976 | 0.53183 | 0.75144 | 1 | 15298 | 2 | -0.52976 |
| PPP1R14B | 5 | 0.1179 | 0.25416 | 0.994632 | 5570 | 1 | 0.48166 | 0.53185 | 0.73085 | 1 | 15299 | 2 | 0.48166 |
| PRIM1 | 3 | 0.049978 | 0.11568 | 0.951052 | 2976 | 2 | -3.6315 | 0.53186 | 0.65688 | 1 | 15300 | 1 | -3.6315 |
| NINL | 3 | 0.72204 | 0.722 | 0.999994 | 16517 | 1 | 0.3834 | 0.53186 | 0.65688 | 1 | 15301 | 1 | 0.3834 |
| hsa-mir-3122 | 3 | 0.78397 | 0.78387 | 0.999994 | 17265 | 0 | -0.1052 | 0.53186 | 0.65688 | 1 | 15302 | 1 | -0.1052 |
| DOK5 | 6 | 0.29199 | 0.49736 | 0.999994 | 9950 | 2 | 0.50604 | 0.53187 | 0.75147 | 1 | 15303 | 3 | 0.50604 |
| FOXC2 | 6 | 0.072898 | 0.19277 | 0.97489 | 3957 | 3 | -1.2139 | 0.53187 | 0.75147 | 1 | 15304 | 2 | -1.2139 |
| KNCN | 5 | 0.063948 | 0.16857 | 0.967126 | 3605 | 3 | -2.1837 | 0.53189 | 0.73087 | 1 | 15305 | 2 | -2.1837 |
| CCZ1B | 4 | 0.1989 | 0.33953 | 0.999994 | 7733 | 2 | -0.21199 | 0.53191 | 0.68915 | 1 | 15306 | 2 | -0.21199 |
| PPTC7 | 6 | 0.26637 | 0.4674 | 0.999994 | 9381 | 2 | 0.47193 | 0.53196 | 0.75153 | 1 | 15307 | 3 | 0.47193 |
| hsa-mir-7976 | 4 | 0.53151 | 0.6308 | 0.999994 | 13925 | 1 | 0.88512 | 0.53214 | 0.68935 | 1 | 15308 | 2 | 0.88512 |
| hsa-mir-6764 | 4 | 0.31326 | 0.45659 | 0.999994 | 10382 | 2 | -0.5787 | 0.53214 | 0.68935 | 1 | 15309 | 1 | -0.5787 |
| hsa-mir-4260 | 4 | 0.39143 | 0.53685 | 0.999994 | 11819 | 2 | -0.21415 | 0.53214 | 0.68935 | 1 | 15310 | 1 | -0.21415 |
| EED | 4 | 0.36562 | 0.51028 | 0.999994 | 11432 | 1 | 0.6189 | 0.53227 | 0.68944 | 1 | 15311 | 2 | 0.6189 |
| PPP1R42 | 6 | 0.15777 | 0.33542 | 0.999994 | 6667 | 3 | -0.23226 | 0.53227 | 0.75176 | 1 | 15312 | 3 | -0.23226 |
| FOXC1 | 6 | 0.056901 | 0.15908 | 0.967126 | 3288 | 4 | -1.3747 | 0.53232 | 0.7518 | 1 | 15313 | 1 | -1.3747 |
| SLC18B1 | 5 | 0.35522 | 0.52614 | 0.999994 | 11255 | 2 | -0.42751 | 0.53246 | 0.73118 | 1 | 15314 | 2 | -0.42751 |
| NonTargetingControlG | 2 | 0.062822 | 0.1106 | 0.951052 | 3548 | 1 | -3.4528 | 0.53248 | 0.5618 | 1 | 15315 | 1 | -3.4528 |
| NonTargetingControlG | 1 | 0.46745 | 0.46751 | 0.999994 | 12972 | 0 | 0.23017 | 0.53255 | 0.53233 | 1 | 15316 | 0 | 0.23017 |
| MXD4 | 5 | 0.37753 | 0.54467 | 0.999994 | 11619 | 2 | -0.26204 | 0.53261 | 0.73127 | 1 | 15317 | 2 | -0.26204 |
| LCE3D | 6 | 0.01029 | 0.039999 | 0.891621 | 888 | 5 | -3.5196 | 0.53288 | 0.75222 | 1 | 15318 | 1 | -3.5196 |
| TACR3 | 6 | 0.094119 | 0.23577 | 0.985362 | 4755 | 2 | 0.14247 | 0.53288 | 0.75222 | 1 | 15319 | 3 | 0.14247 |
| hsa-mir-1233-1 | 4 | 0.75384 | 0.76305 | 0.999994 | 16896 | 1 | 0.17069 | 0.53301 | 0.69005 | 1 | 15320 | 1 | 0.17069 |
| OR2AK2 | 5 | 0.19403 | 0.36668 | 0.999994 | 7604 | 3 | -0.84308 | 0.53302 | 0.73149 | 1 | 15321 | 2 | -0.84308 |
| CXADR | 6 | 0.31787 | 0.52746 | 0.999994 | 10470 | 2 | 0.34065 | 0.53307 | 0.75236 | 1 | 15322 | 3 | 0.34065 |
| MARCH9 | 5 | 0.57269 | 0.7119 | 0.999994 | 14587 | 2 | 0.21848 | 0.53348 | 0.73174 | 1 | 15323 | 2 | 0.21848 |
| HMGN2 | 4 | 0.21056 | 0.35155 | 0.999994 | 8041 | 2 | 0.18272 | 0.53356 | 0.69051 | 1 | 15324 | 2 | 0.18272 |
| WBP11 | 4 | 0.076488 | 0.1722 | 0.967658 | 4091 | 3 | -1.0421 | 0.53357 | 0.69051 | 1 | 15325 | 1 | -1.0421 |
| CDH26 | 6 | 0.046474 | 0.13632 | 0.956051 | 2828 | 2 | 0.73709 | 0.53365 | 0.75281 | 1 | 15326 | 3 | 0.73709 |
| KRTAP16-1 | 6 | 0.92197 | 0.92223 | 0.999994 | 19672 | 1 | 0.69181 | 0.53368 | 0.75283 | 1 | 15327 | 3 | 0.69181 |
| NSL1 | 4 | 0.22566 | 0.36701 | 0.999994 | 8415 | 2 | -0.18182 | 0.53387 | 0.69076 | 1 | 15328 | 2 | -0.18182 |
| UGT2A3 | 6 | 0.67635 | 0.79815 | 0.999994 | 16027 | 2 | -0.11148 | 0.53388 | 0.75297 | 1 | 15329 | 2 | -0.11148 |
| FAM149A | 6 | 0.51359 | 0.67811 | 0.999994 | 13633 | 1 | -0.1068 | 0.53388 | 0.75297 | 1 | 15330 | 2 | -0.1068 |
| RCHY1 | 6 | 0.54531 | 0.7003 | 0.999994 | 14141 | 1 | 0.53325 | 0.53397 | 0.75304 | 1 | 15331 | 3 | 0.53325 |
| OR6T1 | 6 | 0.87433 | 0.88601 | 0.999994 | 18714 | 1 | 0.70596 | 0.53408 | 0.75312 | 1 | 15332 | 3 | 0.70596 |
| PLEKHG4 | 5 | 0.15483 | 0.30997 | 0.999994 | 6589 | 3 | -1.0165 | 0.53409 | 0.73206 | 1 | 15333 | 2 | -1.0165 |
| SYAP1 | 5 | 0.033411 | 0.099267 | 0.945253 | 2183 | 4 | -1.516 | 0.53409 | 0.73206 | 1 | 15334 | 1 | -1.516 |
| PLCD4 | 5 | 0.056554 | 0.15613 | 0.967126 | 3273 | 4 | -1.9246 | 0.53409 | 0.73206 | 1 | 15335 | 1 | -1.9246 |
| INO80E | 6 | 0.099487 | 0.24638 | 0.98999 | 4949 | 3 | -0.63314 | 0.5341 | 0.75314 | 1 | 15336 | 3 | -0.63314 |
| NonTargetingControlG | 2 | 0.73203 | 0.73175 | 0.999994 | 16628 | 0 | 1.4284 | 0.53434 | 0.5632 | 1 | 15337 | 1 | 1.4284 |
| RP9 | 6 | 0.18773 | 0.3727 | 0.999994 | 7414 | 3 | -0.53861 | 0.53438 | 0.75334 | 1 | 15338 | 2 | -0.53861 |
| MZT2A | 5 | 0.30054 | 0.4815 | 0.999994 | 10143 | 2 | -0.33778 | 0.53439 | 0.73222 | 1 | 15339 | 2 | -0.33778 |

| | | | | | | | | | | | | | |
|---|---|---|---|---|---|---|---|---|---|---|---|---|---|
| RAMP3 | 6 | 0.15528 | 0.33232 | 0.999994 | 6602 | 3 | -0.57061 | 0.53453 | 0.75345 | 1 | 15340 | 2 | -0.57061 |
| ZXDA | 4 | 0.4181 | 0.56425 | 0.999994 | 12218 | 2 | 0.2927 | 0.53454 | 0.69129 | 1 | 15341 | 2 | 0.2927 |
| OR52E8 | 6 | 0.28402 | 0.48808 | 0.999994 | 9785 | 1 | 0.020173 | 0.53459 | 0.7535 | 1 | 15342 | 2 | 0.020173 |
| SNRPC | 4 | 0.027711 | 0.076845 | 0.93914 | 1889 | 3 | -4.1179 | 0.53461 | 0.69134 | 1 | 15343 | 1 | -4.1179 |
| DDX47 | 4 | 0.35358 | 0.49796 | 0.999994 | 11226 | 2 | -0.18006 | 0.53461 | 0.69134 | 1 | 15344 | 2 | -0.18006 |
| SPINK13 | 4 | 0.24733 | 0.38916 | 0.999994 | 8956 | 2 | -0.18219 | 0.5348 | 0.69148 | 1 | 15345 | 2 | -0.18219 |
| COL4A3 | 5 | 0.029997 | 0.090472 | 0.94395 | 2008 | 4 | -1.7388 | 0.535 | 0.73253 | 1 | 15346 | 1 | -1.7388 |
| PCED1B | 5 | 0.18918 | 0.35978 | 0.999994 | 7455 | 2 | -0.54011 | 0.535 | 0.73253 | 1 | 15347 | 2 | -0.54011 |
| MOCOS | 5 | 0.12012 | 0.25755 | 0.994943 | 5632 | 3 | -1.3261 | 0.535 | 0.73253 | 1 | 15348 | 1 | -1.3261 |
| CYP1B1 | 5 | 0.22624 | 0.41201 | 0.999994 | 8430 | 3 | -0.72115 | 0.535 | 0.73253 | 1 | 15349 | 1 | -0.72115 |
| CD302 | 6 | 0.23384 | 0.42868 | 0.999994 | 8615 | 3 | -0.69593 | 0.535 | 0.75381 | 1 | 15350 | 1 | -0.69593 |
| ATXN2L | 6 | 0.052 | 0.1485 | 0.965554 | 3067 | 4 | -1.5822 | 0.535 | 0.75381 | 1 | 15351 | 1 | -1.5822 |
| ACYP2 | 6 | 0.25573 | 0.45479 | 0.999994 | 9142 | 2 | 0.40885 | 0.535 | 0.75381 | 1 | 15352 | 2 | 0.40885 |
| NonTargetingControlG | 2 | 0.36675 | 0.38949 | 0.999994 | 11453 | 1 | 0.13709 | 0.53503 | 0.56372 | 1 | 15353 | 1 | 0.13709 |
| hsa-mir-376c | 2 | 0.45052 | 0.4598 | 0.999994 | 12719 | 1 | 0.35611 | 0.53508 | 0.56376 | 1 | 15354 | 1 | 0.35611 |
| hsa-mir-6844 | 1 | 0.46488 | 0.46501 | 0.999994 | 12931 | 0 | 0.19033 | 0.53512 | 0.53489 | 1 | 15355 | 0 | 0.19033 |
| CALCOCO2 | 6 | 0.26436 | 0.46503 | 0.999994 | 9342 | 1 | 0.25094 | 0.53514 | 0.75392 | 1 | 15356 | 3 | 0.25094 |
| CLDN16 | 5 | 0.12288 | 0.26171 | 0.995257 | 5735 | 3 | -1.3099 | 0.53516 | 0.73262 | 1 | 15357 | 2 | -1.3099 |
| SYNPO2L | 4 | 0.33605 | 0.47989 | 0.999994 | 10849 | 2 | 0.055326 | 0.53522 | 0.69179 | 1 | 15358 | 2 | 0.055326 |
| CD70 | 6 | 0.57948 | 0.72482 | 0.999994 | 14672 | 2 | 0.48129 | 0.53526 | 0.754 | 1 | 15359 | 3 | 0.48129 |
| B4GALT4 | 6 | 0.60139 | 0.74097 | 0.999994 | 14966 | 2 | 0.59629 | 0.53526 | 0.754 | 1 | 15360 | 3 | 0.59629 |
| BDP1 | 4 | 0.46022 | 0.59899 | 0.999994 | 12846 | 1 | 0.42356 | 0.53533 | 0.69188 | 1 | 15361 | 2 | 0.42356 |
| NonTargetingControlG | 2 | 0.57192 | 0.57211 | 0.999994 | 14574 | 1 | 0.76362 | 0.53535 | 0.56397 | 1 | 15362 | 1 | 0.76362 |
| MYL5 | 6 | 0.77959 | 0.83949 | 0.999994 | 17206 | 1 | 0.66572 | 0.53559 | 0.75423 | 1 | 15363 | 3 | 0.66572 |
| NonTargetingControlG | 1 | 0.46428 | 0.46443 | 0.999994 | 12924 | 0 | 0.23423 | 0.53572 | 0.5355 | 1 | 15364 | 0 | 0.23423 |
| MNS1 | 5 | 0.1189 | 0.25569 | 0.994943 | 5600 | 2 | -0.11046 | 0.53583 | 0.73297 | 1 | 15365 | 2 | -0.11046 |
| ANKRD34C | 5 | 0.46577 | 0.61852 | 0.999994 | 12949 | 1 | 0.29978 | 0.53592 | 0.73303 | 1 | 15366 | 2 | 0.29978 |
| EBLN1 | 6 | 0.033361 | 0.10668 | 0.950995 | 2179 | 3 | -2.0362 | 0.53593 | 0.75447 | 1 | 15367 | 3 | -2.0362 |
| hsa-mir-640 | 4 | 0.27029 | 0.41258 | 0.999994 | 9483 | 2 | -0.56354 | 0.536 | 0.69241 | 1 | 15368 | 1 | -0.56354 |
| PIRT | 6 | 0.088987 | 0.2255 | 0.983354 | 4552 | 3 | -1.082 | 0.53605 | 0.75456 | 1 | 15369 | 2 | -1.082 |
| RAD23B | 6 | 0.0035446 | 0.015758 | 0.766139 | 409 | 5 | -1.4039 | 0.53605 | 0.75456 | 1 | 15370 | 1 | -1.4039 |
| TBC1D7 | 5 | 0.20555 | 0.38312 | 0.999994 | 7910 | 2 | -0.41374 | 0.53607 | 0.73311 | 1 | 15371 | 2 | -0.41374 |
| RPRD1B | 6 | 0.19475 | 0.38147 | 0.999994 | 7629 | 1 | 0.50464 | 0.53616 | 0.75464 | 1 | 15372 | 2 | 0.50464 |
| NonTargetingControlG | 2 | 0.078989 | 0.13693 | 0.957161 | 4189 | 1 | -3.0658 | 0.53623 | 0.56461 | 1 | 15373 | 1 | -3.0658 |
| ATP13A5 | 6 | 0.88382 | 0.89209 | 0.999994 | 18901 | 1 | 0.65256 | 0.53634 | 0.75479 | 1 | 15374 | 3 | 0.65256 |
| hsa-mir-3922 | 4 | 0.3829 | 0.52813 | 0.999994 | 11691 | 1 | 0.72879 | 0.53658 | 0.69288 | 1 | 15375 | 2 | 0.72879 |
| C11orf93 | 6 | 0.65217 | 0.77929 | 0.999994 | 15690 | 2 | 0.49603 | 0.53674 | 0.7551 | 1 | 15376 | 3 | 0.49603 |
| SFN | 6 | 0.063756 | 0.17373 | 0.96793 | 3598 | 4 | -1.3226 | 0.53675 | 0.7551 | 1 | 15377 | 2 | -1.3226 |
| APBB3 | 4 | 0.19359 | 0.33408 | 0.999994 | 7588 | 1 | 0.33043 | 0.53698 | 0.69321 | 1 | 15378 | 1 | 0.33043 |
| hsa-mir-1197 | 4 | 0.25598 | 0.39801 | 0.999994 | 9151 | 1 | -0.075558 | 0.53698 | 0.69321 | 1 | 15379 | 1 | -0.075558 |
| DDA1 | 4 | 0.20092 | 0.34162 | 0.999994 | 7789 | 1 | 0.060919 | 0.53698 | 0.69321 | 1 | 15380 | 1 | 0.060919 |
| GJD4 | 4 | 0.042343 | 0.10644 | 0.950995 | 2631 | 3 | -2.716 | 0.53698 | 0.69321 | 1 | 15381 | 1 | -2.716 |
| MED11 | 4 | 0.021807 | 0.064438 | 0.93914 | 1566 | 3 | -2.4582 | 0.53698 | 0.69321 | 1 | 15382 | 1 | -2.4582 |
| GPR101 | 5 | 0.87071 | 0.87171 | 0.999994 | 18651 | 1 | 0.080426 | 0.53708 | 0.73367 | 1 | 15383 | 2 | 0.080426 |
| FBXW5 | 6 | 0.0022043 | 0.010732 | 0.70768 | 291 | 5 | -4.0538 | 0.53712 | 0.75536 | 1 | 15384 | 1 | -4.0538 |
| FAM181B | 5 | 0.16071 | 0.31858 | 0.999994 | 6758 | 2 | -0.34207 | 0.53717 | 0.73372 | 1 | 15385 | 2 | -0.34207 |
| LPPR3 | 5 | 0.07466 | 0.18595 | 0.969409 | 4021 | 3 | -1.8846 | 0.53732 | 0.7338 | 1 | 15386 | 2 | -1.8846 |
| SNTN | 6 | 0.66921 | 0.79256 | 0.999994 | 15917 | 2 | 0.5186 | 0.53745 | 0.7556 | 1 | 15387 | 3 | 0.5186 |
| TADA2B | 3 | 0.5023 | 0.54768 | 0.999994 | 13467 | 1 | -0.28959 | 0.53749 | 0.65936 | 1 | 15388 | 1 | -0.28959 |
| NonTargetingControlG | 2 | 0.67045 | 0.67022 | 0.999994 | 15938 | 0 | 1.0606 | 0.53763 | 0.56567 | 1 | 15389 | 1 | 1.0606 |
| SUN1 | 5 | 0.62842 | 0.73671 | 0.999994 | 15354 | 1 | 0.11293 | 0.53764 | 0.73398 | 1 | 15390 | 2 | 0.11293 |
| RPL9 | 2 | 0.098099 | 0.16716 | 0.967126 | 4897 | 1 | -2.9412 | 0.53765 | 0.56568 | 1 | 15391 | 1 | -2.9412 |

| | | | | | | | | | | | | |
|---|---|---|---|---|---|---|---|---|---|---|---|---|
| ARL5C | 5 | 0.1784 | 0.34431 | 0.999994 | 7186 | 2 | 0.066784 | 0.53775 | 0.73404 | 1 | 15392 | 2 | 0.066784 |
| B3GNT7 | 6 | 0.066655 | 0.17975 | 0.968171 | 3710 | 4 | -1.3232 | 0.53784 | 0.75591 | 1 | 15393 | 2 | -1.3232 |
| RNF144B | 6 | 0.08152 | 0.21041 | 0.979587 | 4282 | 2 | 0.37952 | 0.53786 | 0.75592 | 1 | 15394 | 3 | 0.37952 |
| COA1 | 6 | 0.13224 | 0.30318 | 0.999994 | 5983 | 2 | -0.015881 | 0.53795 | 0.75599 | 1 | 15395 | 2 | -0.015881 |
| AK8 | 6 | 0.53267 | 0.69143 | 0.999994 | 13946 | 2 | 0.59128 | 0.53803 | 0.75606 | 1 | 15396 | 3 | 0.59128 |
| DKK3 | 4 | 0.35476 | 0.49913 | 0.999994 | 11246 | 1 | 0.36897 | 0.53814 | 0.69414 | 1 | 15397 | 2 | 0.36897 |
| FRAT1 | 6 | 0.21424 | 0.40524 | 0.999994 | 8124 | 2 | 0.37163 | 0.53817 | 0.75617 | 1 | 15398 | 3 | 0.37163 |
| RBP3 | 6 | 0.045433 | 0.13402 | 0.95318 | 2785 | 3 | -1.2009 | 0.5382 | 0.75619 | 1 | 15399 | 2 | -1.2009 |
| BLOC1S1 | 6 | 0.27288 | 0.4751 | 0.999994 | 9536 | 2 | 0.49417 | 0.53822 | 0.75621 | 1 | 15400 | 3 | 0.49417 |
| SCD5 | 5 | 0.44919 | 0.60456 | 0.999994 | 12705 | 2 | -0.097428 | 0.53823 | 0.73431 | 1 | 15401 | 2 | -0.097428 |
| TNNI1 | 6 | 0.92148 | 0.92177 | 0.999994 | 19665 | 1 | 0.59244 | 0.53824 | 0.75623 | 1 | 15402 | 3 | 0.59244 |
| ZC3H11A | 5 | 0.77903 | 0.80689 | 0.999994 | 17196 | 1 | 0.13253 | 0.53835 | 0.73436 | 1 | 15403 | 1 | 0.13253 |
| ALS2CR8 | 5 | 0.090514 | 0.21142 | 0.979587 | 4614 | 3 | -1.7627 | 0.53835 | 0.73436 | 1 | 15404 | 1 | -1.7627 |
| GTPBP10 | 6 | 0.33523 | 0.54723 | 0.999994 | 10838 | 3 | -0.51168 | 0.53843 | 0.75637 | 1 | 15405 | 2 | -0.51168 |
| CDC73 | 6 | 0.046754 | 0.13696 | 0.957161 | 2840 | 3 | -0.52687 | 0.53857 | 0.75646 | 1 | 15406 | 3 | -0.52687 |
| hsa-mir-642b | 1 | 0.46141 | 0.46156 | 0.999994 | 12871 | 0 | 0.21543 | 0.53859 | 0.53833 | 1 | 15407 | 0 | 0.21543 |
| EIF2AK3 | 6 | 0.69011 | 0.80893 | 0.999994 | 16211 | 2 | 0.16117 | 0.53893 | 0.75671 | 1 | 15408 | 3 | 0.16117 |
| SH2D1A | 6 | 0.54104 | 0.69728 | 0.999994 | 14073 | 2 | 0.19343 | 0.53899 | 0.75675 | 1 | 15409 | 3 | 0.19343 |
| PMM1 | 6 | 0.43033 | 0.62206 | 0.999994 | 12419 | 2 | 0.58052 | 0.53909 | 0.75684 | 1 | 15410 | 3 | 0.58052 |
| CERS3 | 6 | 0.089129 | 0.22575 | 0.983354 | 4560 | 3 | -0.61734 | 0.53916 | 0.75689 | 1 | 15411 | 1 | -0.61734 |
| KATNB1 | 6 | 0.17005 | 0.35082 | 0.999994 | 6984 | 2 | -0.063559 | 0.53916 | 0.75689 | 1 | 15412 | 1 | -0.063559 |
| NUP85 | 4 | 0.0069974 | 0.024155 | 0.836935 | 659 | 3 | -4.7804 | 0.53938 | 0.69508 | 1 | 15413 | 1 | -4.7804 |
| TMEM221 | 6 | 0.22628 | 0.41963 | 0.999994 | 8431 | 3 | -0.065697 | 0.53945 | 0.75711 | 1 | 15414 | 3 | -0.065697 |
| CPQ | 6 | 0.60984 | 0.74722 | 0.999994 | 15070 | 2 | 0.31573 | 0.53953 | 0.75717 | 1 | 15415 | 3 | 0.31573 |
| FAM228A | 6 | 0.53598 | 0.69371 | 0.999994 | 13995 | 2 | 0.22602 | 0.53972 | 0.75731 | 1 | 15416 | 3 | 0.22602 |
| SPTBN5 | 6 | 0.0050713 | 0.021397 | 0.81457 | 525 | 3 | -0.52995 | 0.53982 | 0.7574 | 1 | 15417 | 3 | -0.52995 |
| NonTargetingControlG | 2 | 0.075539 | 0.13138 | 0.95318 | 4056 | 1 | -3.3606 | 0.54002 | 0.56749 | 1 | 15418 | 1 | -3.3606 |
| AIDA | 6 | 0.041368 | 0.12496 | 0.951052 | 2582 | 3 | -0.15852 | 0.54009 | 0.75759 | 1 | 15419 | 3 | -0.15852 |
| TM9SF1 | 6 | 0.92998 | 0.93011 | 0.999994 | 19851 | 0 | 0.52841 | 0.54011 | 0.75761 | 1 | 15420 | 3 | 0.52841 |
| BBS12 | 6 | 0.010626 | 0.04115 | 0.891621 | 907 | 5 | -2.4689 | 0.54017 | 0.75765 | 1 | 15421 | 1 | -2.4689 |
| C17orf112 | 6 | 0.0089851 | 0.035468 | 0.870441 | 792 | 5 | -2.7394 | 0.54017 | 0.75765 | 1 | 15422 | 1 | -2.7394 |
| C1orf151-NBL1 | 6 | 0.10893 | 0.26464 | 0.996734 | 5269 | 4 | -1.016 | 0.54026 | 0.75772 | 1 | 15423 | 2 | -1.016 |
| C13orf35 | 6 | 0.23261 | 0.42724 | 0.999994 | 8585 | 3 | -0.45559 | 0.54037 | 0.75781 | 1 | 15424 | 2 | -0.45559 |
| POLR2D | 4 | 0.0033569 | 0.011991 | 0.729405 | 390 | 3 | -6.5337 | 0.54042 | 0.69589 | 1 | 15425 | 1 | -6.5337 |
| MIA3 | 4 | 0.53092 | 0.63051 | 0.999994 | 13909 | 1 | -0.16473 | 0.54042 | 0.69589 | 1 | 15426 | 1 | -0.16473 |
| THOC6 | 6 | 0.8969 | 0.90129 | 0.999994 | 19159 | 1 | 0.5834 | 0.54083 | 0.75816 | 1 | 15427 | 3 | 0.5834 |
| CA14 | 6 | 0.78679 | 0.84236 | 0.999994 | 17301 | 1 | 0.64945 | 0.54107 | 0.75834 | 1 | 15428 | 3 | 0.64945 |
| IL17B | 6 | 0.086111 | 0.21973 | 0.980699 | 4455 | 4 | -1.2917 | 0.54135 | 0.75853 | 1 | 15429 | 2 | -1.2917 |
| DR1 | 6 | 0.25823 | 0.45779 | 0.999994 | 9199 | 3 | -0.77506 | 0.54135 | 0.75853 | 1 | 15430 | 2 | -0.77506 |
| GNG12 | 6 | 0.24297 | 0.43954 | 0.999994 | 8862 | 2 | 0.41814 | 0.54135 | 0.75853 | 1 | 15431 | 2 | 0.41814 |
| ETFB | 6 | 0.443 | 0.63044 | 0.999994 | 12624 | 2 | -0.26248 | 0.54135 | 0.75853 | 1 | 15432 | 2 | -0.26248 |
| ANAPC10 | 6 | 0.076949 | 0.20118 | 0.977164 | 4105 | 3 | -0.7096 | 0.54135 | 0.75853 | 1 | 15433 | 2 | -0.7096 |
| CLEC4F | 6 | 0.13423 | 0.3057 | 0.999994 | 6028 | 3 | -0.83946 | 0.54135 | 0.75853 | 1 | 15434 | 2 | -0.83946 |
| SULT1B1 | 6 | 0.87916 | 0.88906 | 0.999994 | 18813 | 1 | 0.26616 | 0.54136 | 0.75854 | 1 | 15435 | 3 | 0.26616 |
| RNASE13 | 6 | 0.0086523 | 0.034269 | 0.867348 | 771 | 2 | 0.50283 | 0.54138 | 0.75856 | 1 | 15436 | 3 | 0.50283 |
| NonTargetingControlG | 2 | 0.66235 | 0.66212 | 0.999994 | 15816 | 0 | 0.98731 | 0.54152 | 0.56867 | 1 | 15437 | 1 | 0.98731 |
| A4GALT | 3 | 0.26997 | 0.39968 | 0.999994 | 9472 | 2 | -0.67788 | 0.54154 | 0.66111 | 1 | 15438 | 1 | -0.67788 |
| OXSM | 6 | 0.83156 | 0.86231 | 0.999994 | 17953 | 1 | -0.029292 | 0.54181 | 0.7589 | 1 | 15439 | 2 | -0.029292 |
| KIR2DL4 | 6 | 0.64365 | 0.77271 | 0.999994 | 15571 | 2 | 0.34895 | 0.54184 | 0.75892 | 1 | 15440 | 3 | 0.34895 |
| SMTN | 6 | 0.18088 | 0.36428 | 0.999994 | 7244 | 2 | 0.20177 | 0.54187 | 0.75895 | 1 | 15441 | 3 | 0.20177 |
| hsa-mir-4485 | 3 | 0.09775 | 0.18005 | 0.968171 | 4880 | 2 | -2.4449 | 0.54191 | 0.66127 | 1 | 15442 | 1 | -2.4449 |
| C12orf71 | 6 | 0.43998 | 0.62839 | 0.999994 | 12573 | 2 | -0.04604 | 0.54193 | 0.75899 | 1 | 15443 | 2 | -0.04604 |

| | | | | | | | | | | | | |
|---|---|---|---|---|---|---|---|---|---|---|---|---|
| DMRT2 | 5 | 0.011964 | 0.041696 | 0.891621 | 992 | 2 | -0.52707 | 0.54193 | 0.7363 | 1 | 15444 | 1 | -0.52707 |
| PRPS2 | 5 | 0.0019531 | 0.0085666 | 0.680529 | 267 | 3 | -7.8017 | 0.54193 | 0.7363 | 1 | 15445 | 1 | -7.8017 |
| DNM1 | 4 | 0.79618 | 0.79782 | 0.999994 | 17429 | 1 | 0.86246 | 0.54195 | 0.69711 | 1 | 15446 | 2 | 0.86246 |
| FAM131B | 4 | 0.70225 | 0.7259 | 0.999994 | 16330 | 1 | 0.43868 | 0.54203 | 0.69718 | 1 | 15447 | 2 | 0.43868 |
| GTSF1L | 6 | 0.060019 | 0.16578 | 0.967126 | 3423 | 3 | -1.1391 | 0.54229 | 0.75925 | 1 | 15448 | 2 | -1.1391 |
| UBAP1L | 6 | 0.1953 | 0.38218 | 0.999994 | 7642 | 2 | -0.097794 | 0.54243 | 0.75936 | 1 | 15449 | 2 | -0.097794 |
| SRSF12 | 6 | 0.66898 | 0.79238 | 0.999994 | 15910 | 2 | 0.50889 | 0.54243 | 0.75936 | 1 | 15450 | 2 | 0.50889 |
| CNGB3 | 6 | 0.32256 | 0.53284 | 0.999994 | 10565 | 3 | -0.6915 | 0.54243 | 0.75936 | 1 | 15451 | 2 | -0.6915 |
| SLC35G2 | 6 | 0.26739 | 0.46859 | 0.999994 | 9405 | 3 | -0.44517 | 0.54243 | 0.75936 | 1 | 15452 | 2 | -0.44517 |
| hsa-mir-4421 | 3 | 0.13521 | 0.22939 | 0.983354 | 6057 | 2 | -1.93 | 0.54263 | 0.6616 | 1 | 15453 | 1 | -1.93 |
| hsa-mir-4440 | 4 | 0.37321 | 0.51815 | 0.999994 | 11561 | 1 | 0.35298 | 0.54264 | 0.69765 | 1 | 15454 | 2 | 0.35298 |
| FBXW4 | 4 | 0.22153 | 0.36282 | 0.999994 | 8321 | 2 | 0.19063 | 0.54277 | 0.69775 | 1 | 15455 | 2 | 0.19063 |
| hsa-mir-5691 | 4 | 0.76658 | 0.77303 | 0.999994 | 17042 | 1 | 0.39474 | 0.5428 | 0.69778 | 1 | 15456 | 2 | 0.39474 |
| SLC25A16 | 6 | 0.23472 | 0.42973 | 0.999994 | 8637 | 3 | -0.27696 | 0.54286 | 0.75969 | 1 | 15457 | 2 | -0.27696 |
| AMY1A | 6 | 0.5056 | 0.6726 | 0.999994 | 13516 | 1 | 0.54369 | 0.54293 | 0.75974 | 1 | 15458 | 3 | 0.54369 |
| LDLRAP1 | 5 | 0.042648 | 0.12245 | 0.951052 | 2657 | 3 | -1.5656 | 0.54313 | 0.73696 | 1 | 15459 | 2 | -1.5656 |
| MPL | 6 | 0.36475 | 0.57992 | 0.999994 | 11416 | 2 | 0.43133 | 0.54318 | 0.75994 | 1 | 15460 | 3 | 0.43133 |
| FAM228B | 5 | 0.25149 | 0.44194 | 0.999994 | 9046 | 1 | -0.45364 | 0.54323 | 0.73702 | 1 | 15461 | 1 | -0.45364 |
| COLQ | 5 | 0.041399 | 0.1194 | 0.951052 | 2584 | 3 | -2.1662 | 0.54323 | 0.73702 | 1 | 15462 | 2 | -2.1662 |
| MC3R | 5 | 0.80431 | 0.82222 | 0.999994 | 17546 | 1 | 0.54942 | 0.54323 | 0.73702 | 1 | 15463 | 2 | 0.54942 |
| GUCA1A | 5 | 0.080637 | 0.19561 | 0.975797 | 4256 | 2 | 0.57612 | 0.54323 | 0.73702 | 1 | 15464 | 2 | 0.57612 |
| SF3B14 | 5 | 0.039374 | 0.11433 | 0.951052 | 2482 | 4 | -0.92106 | 0.54323 | 0.73702 | 1 | 15465 | 1 | -0.92106 |
| GIPC1 | 4 | 0.068092 | 0.15622 | 0.967126 | 3753 | 1 | 0.41031 | 0.54324 | 0.69812 | 1 | 15466 | 2 | 0.41031 |
| GLA | 5 | 0.8917 | 0.89183 | 0.999994 | 19068 | 0 | 0.60519 | 0.54337 | 0.73709 | 1 | 15467 | 2 | 0.60519 |
| NonTargetingControlG | 2 | 0.16626 | 0.23806 | 0.987253 | 6892 | 1 | -2.2758 | 0.54345 | 0.57012 | 1 | 15468 | 1 | -2.2758 |
| hsa-mir-6509 | 4 | 0.38375 | 0.52897 | 0.999994 | 11700 | 2 | 0.24592 | 0.54351 | 0.69831 | 1 | 15469 | 2 | 0.24592 |
| ASMTL_X | 6 | 0.10456 | 0.25623 | 0.994943 | 5118 | 4 | -1.1276 | 0.54352 | 0.76019 | 1 | 15470 | 2 | -1.1276 |
| C9orf66 | 6 | 0.035166 | 0.11091 | 0.951052 | 2278 | 3 | -1.5253 | 0.54354 | 0.76021 | 1 | 15471 | 2 | -1.5253 |
| DUSP26 | 6 | 0.042684 | 0.12796 | 0.951704 | 2661 | 4 | -2.0941 | 0.54354 | 0.76021 | 1 | 15472 | 1 | -2.0941 |
| TMEM212 | 4 | 0.089083 | 0.19532 | 0.975797 | 4555 | 2 | -0.98821 | 0.54355 | 0.69833 | 1 | 15473 | 2 | -0.98821 |
| NonTargetingControlG | 2 | 0.13783 | 0.21822 | 0.980699 | 6130 | 1 | -2.6473 | 0.54359 | 0.57024 | 1 | 15474 | 1 | -2.6473 |
| ENPEP | 6 | 0.28456 | 0.48874 | 0.999994 | 9799 | 2 | 0.68302 | 0.54368 | 0.76031 | 1 | 15475 | 3 | 0.68302 |
| NTN4 | 6 | 0.87238 | 0.8848 | 0.999994 | 18683 | 1 | 0.59307 | 0.54372 | 0.76035 | 1 | 15476 | 3 | 0.59307 |
| APOBEC3A_B | 2 | 0.077831 | 0.13505 | 0.954328 | 4142 | 1 | -3.3921 | 0.54405 | 0.57059 | 1 | 15477 | 1 | -3.3921 |
| NCEH1 | 6 | 0.053013 | 0.15059 | 0.966359 | 3110 | 1 | 0.21061 | 0.54412 | 0.76066 | 1 | 15478 | 3 | 0.21061 |
| GPR151 | 4 | 0.060382 | 0.14151 | 0.959146 | 3437 | 2 | -1.5945 | 0.5444 | 0.699 | 1 | 15479 | 2 | -1.5945 |
| C7orf61 | 5 | 0.80852 | 0.82496 | 0.999994 | 17609 | 1 | 0.50676 | 0.54443 | 0.73765 | 1 | 15480 | 1 | 0.50676 |
| PRSS35 | 6 | 0.42888 | 0.62111 | 0.999994 | 12394 | 1 | 0.029166 | 0.54451 | 0.76092 | 1 | 15481 | 2 | 0.029166 |
| SMCR7 | 1 | 0.45534 | 0.45546 | 0.999994 | 12779 | 0 | 0.16691 | 0.54466 | 0.54449 | 1 | 15482 | 0 | 0.16691 |
| RAPH1 | 6 | 0.092455 | 0.23243 | 0.984502 | 4700 | 4 | -1.0648 | 0.54469 | 0.76105 | 1 | 15483 | 2 | -1.0648 |
| ANKEF1 | 6 | 0.55311 | 0.70591 | 0.999994 | 14259 | 2 | 0.70667 | 0.54479 | 0.76113 | 1 | 15484 | 3 | 0.70667 |
| JMJD1C | 6 | 0.10575 | 0.25852 | 0.994943 | 5165 | 2 | 0.45939 | 0.54481 | 0.76114 | 1 | 15485 | 3 | 0.45939 |
| hsa-mir-660 | 4 | 0.29801 | 0.44096 | 0.999994 | 10091 | 1 | 0.59579 | 0.54519 | 0.69963 | 1 | 15486 | 2 | 0.59579 |
| JARID2 | 6 | 0.30265 | 0.50982 | 0.999994 | 10184 | 3 | -0.15474 | 0.54533 | 0.76155 | 1 | 15487 | 2 | -0.15474 |
| RPS2 | 4 | 0.49755 | 0.61523 | 0.999994 | 13390 | 1 | 0.11833 | 0.54539 | 0.69979 | 1 | 15488 | 1 | 0.11833 |
| hsa-mir-1302-4 | 4 | 0.76587 | 0.77245 | 0.999994 | 17038 | 1 | -0.24608 | 0.54539 | 0.69979 | 1 | 15489 | 1 | -0.24608 |
| SCRN2 | 4 | 0.85555 | 0.85514 | 0.999994 | 18358 | 0 | 0.087374 | 0.54539 | 0.69979 | 1 | 15490 | 1 | 0.087374 |
| RPL36 | 4 | 0.0022559 | 0.0081693 | 0.680529 | 299 | 3 | -6.8523 | 0.54539 | 0.69979 | 1 | 15491 | 1 | -6.8523 |
| ABCC8 | 6 | 0.033824 | 0.10775 | 0.951052 | 2207 | 2 | 0.22998 | 0.54542 | 0.76162 | 1 | 15492 | 3 | 0.22998 |
| KRTAP20-1 | 5 | 0.17224 | 0.33539 | 0.999994 | 7042 | 2 | -0.24915 | 0.54555 | 0.73827 | 1 | 15493 | 1 | -0.24915 |
| PDE4B | 6 | 0.00035521 | 0.0018917 | 0.435802 | 86 | 4 | -4.3767 | 0.5456 | 0.76176 | 1 | 15494 | 1 | -4.3767 |
| PDCL2 | 6 | 0.26659 | 0.46765 | 0.999994 | 9386 | 3 | -0.57915 | 0.5456 | 0.76176 | 1 | 15495 | 2 | -0.57915 |

| | | | | | | | | | | | | | |
|---|---|---|---|---|---|---|---|---|---|---|---|---|---|
| DHX30 | 6 | 0.54949 | 0.70326 | 0.999994 | 14203 | 2 | -0.37262 | 0.5456 | 0.76176 | 1 | 15496 | 2 | -0.37262 |
| HIP1 | 6 | 0.56276 | 0.71277 | 0.999994 | 14425 | 2 | 0.21498 | 0.54606 | 0.76209 | 1 | 15497 | 2 | 0.21498 |
| NonTargetingControlG | 2 | 0.68497 | 0.68481 | 0.999994 | 16160 | 0 | 1.0856 | 0.54634 | 0.57232 | 1 | 15498 | 1 | 1.0856 |
| hsa-mir-6815 | 4 | 0.053871 | 0.12906 | 0.952525 | 3155 | 2 | -1.4089 | 0.54638 | 0.70057 | 1 | 15499 | 2 | -1.4089 |
| HEPACAM | 6 | 0.57757 | 0.72342 | 0.999994 | 14651 | 2 | 0.62021 | 0.54646 | 0.7624 | 1 | 15500 | 3 | 0.62021 |
| CUL7 | 6 | 0.12242 | 0.2903 | 0.999632 | 5716 | 4 | -0.68175 | 0.54649 | 0.76242 | 1 | 15501 | 1 | -0.68175 |
| DLX1 | 6 | 0.20127 | 0.38941 | 0.999994 | 7797 | 2 | 0.58509 | 0.54649 | 0.76242 | 1 | 15502 | 3 | 0.58509 |
| PRR9 | 6 | 0.30602 | 0.51376 | 0.999994 | 10254 | 2 | 0.011562 | 0.54649 | 0.76242 | 1 | 15503 | 2 | 0.011562 |
| RSRC2 | 6 | 0.0039489 | 0.017297 | 0.785648 | 440 | 4 | -2.8222 | 0.54649 | 0.76242 | 1 | 15504 | 1 | -2.8222 |
| TMEM67 | 6 | 0.5276 | 0.68786 | 0.999994 | 13853 | 1 | 0.58935 | 0.54652 | 0.76244 | 1 | 15505 | 3 | 0.58935 |
| OR10C1 | 5 | 0.014042 | 0.047496 | 0.899064 | 1132 | 3 | -4.0603 | 0.54652 | 0.7388 | 1 | 15506 | 2 | -4.0603 |
| MICAL2 | 6 | 0.19479 | 0.38154 | 0.999994 | 7630 | 3 | -0.25266 | 0.54661 | 0.76252 | 1 | 15507 | 3 | -0.25266 |
| CHAMP1 | 6 | 0.81667 | 0.85526 | 0.999994 | 17721 | 1 | 0.77175 | 0.54673 | 0.7626 | 1 | 15508 | 3 | 0.77175 |
| SRSF11 | 5 | 0.047561 | 0.13453 | 0.953565 | 2880 | 4 | -1.4179 | 0.54677 | 0.73894 | 1 | 15509 | 1 | -1.4179 |
| ZMYND19 | 5 | 0.14023 | 0.28817 | 0.999632 | 6192 | 3 | -1.0691 | 0.54677 | 0.73894 | 1 | 15510 | 2 | -1.0691 |
| MUCL1 | 5 | 0.91528 | 0.91536 | 0.999994 | 19517 | 0 | 0.30173 | 0.54687 | 0.73899 | 1 | 15511 | 2 | 0.30173 |
| OR5B17 | 6 | 0.30775 | 0.51575 | 0.999994 | 10283 | 2 | 0.63622 | 0.54696 | 0.76277 | 1 | 15512 | 3 | 0.63622 |
| NCAPG2 | 3 | 0.071498 | 0.14501 | 0.963012 | 3894 | 2 | -3.0002 | 0.54697 | 0.66352 | 1 | 15513 | 1 | -3.0002 |
| hsa-mir-6744 | 4 | 0.22688 | 0.36827 | 0.999994 | 8447 | 2 | 0.1662 | 0.54706 | 0.7011 | 1 | 15514 | 2 | 0.1662 |
| ZRANB1 | 4 | 0.012186 | 0.040524 | 0.891621 | 1008 | 2 | -4.3972 | 0.54726 | 0.70126 | 1 | 15515 | 1 | -4.3972 |
| TSPAN4 | 6 | 0.90604 | 0.90839 | 0.999994 | 19350 | 1 | 0.54984 | 0.54727 | 0.76298 | 1 | 15516 | 3 | 0.54984 |
| ACAP1 | 5 | 0.88255 | 0.88273 | 0.999994 | 18880 | 1 | 0.085948 | 0.54735 | 0.73924 | 1 | 15517 | 2 | 0.085948 |
| CLDN6 | 6 | 0.13404 | 0.30545 | 0.999994 | 6022 | 2 | 0.35449 | 0.54739 | 0.76307 | 1 | 15518 | 3 | 0.35449 |
| JAK2 | 6 | 0.052501 | 0.14954 | 0.966317 | 3093 | 4 | -2.5714 | 0.54739 | 0.76307 | 1 | 15519 | 2 | -2.5714 |
| SACS | 6 | 0.24372 | 0.44044 | 0.999994 | 8878 | 3 | -0.072006 | 0.54742 | 0.76309 | 1 | 15520 | 3 | -0.072006 |
| hsa-mir-664a | 4 | 0.47863 | 0.6069 | 0.999994 | 13123 | 1 | 0.35814 | 0.54754 | 0.70149 | 1 | 15521 | 2 | 0.35814 |
| PRMT5 | 5 | 0.47171 | 0.62357 | 0.999994 | 13026 | 1 | 0.54942 | 0.54787 | 0.73953 | 1 | 15522 | 2 | 0.54942 |
| TUBA1A | 5 | 0.16806 | 0.32927 | 0.999994 | 6940 | 3 | -0.58793 | 0.54787 | 0.73953 | 1 | 15523 | 2 | -0.58793 |
| hsa-mir-548x-2 | 4 | 0.34443 | 0.48862 | 0.999994 | 11038 | 1 | 0.77617 | 0.5479 | 0.70177 | 1 | 15524 | 2 | 0.77617 |
| AEN | 4 | 0.21258 | 0.35366 | 0.999994 | 8078 | 2 | -0.16459 | 0.54796 | 0.70182 | 1 | 15525 | 2 | -0.16459 |
| OR52E2 | 5 | 0.39345 | 0.55808 | 0.999994 | 11854 | 2 | -0.16891 | 0.54809 | 0.73964 | 1 | 15526 | 2 | -0.16891 |
| FBXO41 | 4 | 0.62338 | 0.67827 | 0.999994 | 15281 | 1 | 0.41517 | 0.54815 | 0.70199 | 1 | 15527 | 2 | 0.41517 |
| GPR119 | 6 | 0.082176 | 0.21179 | 0.979587 | 4302 | 2 | 0.3737 | 0.54828 | 0.76371 | 1 | 15528 | 3 | 0.3737 |
| PYGM | 6 | 0.49714 | 0.66676 | 0.999994 | 13382 | 1 | 0.34703 | 0.54834 | 0.76376 | 1 | 15529 | 3 | 0.34703 |
| RPS6KA4 | 6 | 0.75804 | 0.83134 | 0.999994 | 16954 | 1 | 0.71391 | 0.54841 | 0.76381 | 1 | 15530 | 3 | 0.71391 |
| OR9A4 | 5 | 0.44008 | 0.59687 | 0.999994 | 12577 | 2 | 0.4284 | 0.54846 | 0.73985 | 1 | 15531 | 2 | 0.4284 |
| WWP1 | 6 | 0.12736 | 0.29693 | 0.999994 | 5857 | 1 | 0.1289 | 0.54852 | 0.7639 | 1 | 15532 | 3 | 0.1289 |
| SECISBP2L | 6 | 0.94728 | 0.94729 | 0.999994 | 20270 | 0 | 0.20834 | 0.54852 | 0.7639 | 1 | 15533 | 2 | 0.20834 |
| MKKS | 6 | 0.21096 | 0.40126 | 0.999994 | 8050 | 3 | -0.56043 | 0.54852 | 0.7639 | 1 | 15534 | 1 | -0.56043 |
| hsa-mir-4770 | 4 | 0.46098 | 0.59929 | 0.999994 | 12864 | 1 | 0.6338 | 0.54867 | 0.70239 | 1 | 15535 | 2 | 0.6338 |
| RNASEH1 | 4 | 0.40812 | 0.55403 | 0.999994 | 12078 | 1 | 0.7564 | 0.54889 | 0.70257 | 1 | 15536 | 2 | 0.7564 |
| MRGPRF | 6 | 0.25584 | 0.45492 | 0.999994 | 9145 | 2 | 0.39512 | 0.54892 | 0.7642 | 1 | 15537 | 3 | 0.39512 |
| POMC | 6 | 0.14343 | 0.31741 | 0.999994 | 6285 | 3 | -0.51611 | 0.549 | 0.76426 | 1 | 15538 | 2 | -0.51611 |
| ADAM15 | 5 | 0.0091825 | 0.033682 | 0.867348 | 808 | 3 | -1.0474 | 0.54901 | 0.74015 | 1 | 15539 | 2 | -1.0474 |
| TRIM16 | 5 | 0.063406 | 0.1677 | 0.967126 | 3573 | 3 | -2.0903 | 0.54901 | 0.74015 | 1 | 15540 | 1 | -2.0903 |
| PDZK1 | 5 | 0.040884 | 0.11807 | 0.951052 | 2553 | 3 | -1.0251 | 0.54901 | 0.74015 | 1 | 15541 | 1 | -1.0251 |
| KHDC1L | 6 | 0.1508 | 0.32668 | 0.999994 | 6480 | 2 | -0.20728 | 0.54939 | 0.76455 | 1 | 15542 | 2 | -0.20728 |
| ZBED4 | 6 | 0.42643 | 0.61956 | 0.999994 | 12355 | 2 | -0.24162 | 0.54945 | 0.76459 | 1 | 15543 | 3 | -0.24162 |
| LMTK3 | 6 | 0.10401 | 0.25515 | 0.994943 | 5096 | 3 | -0.84435 | 0.54959 | 0.76472 | 1 | 15544 | 2 | -0.84435 |
| FAM179A | 6 | 0.053107 | 0.15079 | 0.966359 | 3120 | 3 | -1.045 | 0.54962 | 0.76474 | 1 | 15545 | 3 | -1.045 |
| CCDC8 | 6 | 0.75857 | 0.83152 | 0.999994 | 16959 | 1 | 0.42819 | 0.54962 | 0.76474 | 1 | 15546 | 2 | 0.42819 |
| PEX16 | 4 | 0.87067 | 0.87024 | 0.999994 | 18650 | 0 | 0.40979 | 0.5497 | 0.70322 | 1 | 15547 | 2 | 0.40979 |

| | | | | | | | | | | | | |
|---|---|---|---|---|---|---|---|---|---|---|---|---|
| PGC | 5 | 0.068896 | 0.17664 | 0.968171 | 3793 | 3 | -1.9998 | 0.54992 | 0.74067 | 1 | 15548 | 2 | -1.9998 |
| ADIPOQ | 5 | 0.16245 | 0.32113 | 0.999994 | 6807 | 2 | -0.16369 | 0.55015 | 0.7408 | 1 | 15549 | 1 | -0.16369 |
| CELSR2 | 4 | 0.30166 | 0.44469 | 0.999994 | 10155 | 2 | 0.12427 | 0.55029 | 0.70366 | 1 | 15550 | 2 | 0.12427 |
| hsa-mir-2682 | 4 | 0.41817 | 0.56433 | 0.999994 | 12221 | 2 | 0.35767 | 0.55041 | 0.70375 | 1 | 15551 | 2 | 0.35767 |
| ZNF362 | 6 | 0.2705 | 0.47232 | 0.999994 | 9487 | 2 | 0.19385 | 0.5505 | 0.76541 | 1 | 15552 | 3 | 0.19385 |
| ARHGAP31 | 6 | 0.36317 | 0.57888 | 0.999994 | 11392 | 3 | -0.31415 | 0.5505 | 0.76541 | 1 | 15553 | 1 | -0.31415 |
| KIAA1147 | 6 | 0.009819 | 0.038404 | 0.889591 | 861 | 5 | -2.2147 | 0.5505 | 0.76541 | 1 | 15554 | 1 | -2.2147 |
| NonTargetingControlG | 2 | 0.071264 | 0.12445 | 0.951052 | 3886 | 1 | -3.3004 | 0.55052 | 0.57545 | 1 | 15555 | 1 | -3.3004 |
| C11orf88 | 4 | 0.80288 | 0.80378 | 0.999994 | 17523 | 1 | 0.62022 | 0.55069 | 0.70397 | 1 | 15556 | 2 | 0.62022 |
| CEBPG | 6 | 0.30711 | 0.51501 | 0.999994 | 10271 | 1 | 0.42789 | 0.55071 | 0.76557 | 1 | 15557 | 3 | 0.42789 |
| PRDM16 | 5 | 0.059587 | 0.16138 | 0.967126 | 3405 | 3 | -1.9574 | 0.5511 | 0.74132 | 1 | 15558 | 2 | -1.9574 |
| PHF21A | 5 | 0.42886 | 0.58747 | 0.999994 | 12393 | 2 | -0.60444 | 0.5511 | 0.74132 | 1 | 15559 | 2 | -0.60444 |
| GIMD1 | 4 | 0.23719 | 0.37883 | 0.999994 | 8701 | 2 | -0.58402 | 0.5513 | 0.70444 | 1 | 15560 | 1 | -0.58402 |
| ANKHD1-EIF4EBP3 | 4 | 0.17946 | 0.31953 | 0.999994 | 7209 | 2 | -1.2792 | 0.5513 | 0.70444 | 1 | 15561 | 1 | -1.2792 |
| NSUN5 | 4 | 0.71366 | 0.73365 | 0.999994 | 16436 | 1 | -0.042607 | 0.5513 | 0.70444 | 1 | 15562 | 1 | -0.042607 |
| POLR2E | 4 | 0.29224 | 0.43507 | 0.999994 | 9957 | 2 | -0.78917 | 0.5513 | 0.70444 | 1 | 15563 | 1 | -0.78917 |
| TCOF1 | 4 | 0.063559 | 0.14757 | 0.965554 | 3581 | 3 | -1.2422 | 0.5513 | 0.70444 | 1 | 15564 | 1 | -1.2422 |
| hsa-mir-4783 | 4 | 0.0051737 | 0.018207 | 0.795727 | 530 | 3 | -5.6154 | 0.5513 | 0.70444 | 1 | 15565 | 1 | -5.6154 |
| hsa-mir-3612 | 4 | 0.87656 | 0.87622 | 0.999994 | 18762 | 0 | 0.2383 | 0.5513 | 0.70444 | 1 | 15566 | 1 | 0.2383 |
| OR6C1 | 6 | 0.002571 | 0.012129 | 0.732963 | 321 | 4 | -3.158 | 0.55131 | 0.76602 | 1 | 15567 | 2 | -3.158 |
| FMNL2 | 5 | 0.055728 | 0.15419 | 0.967126 | 3234 | 4 | -0.79245 | 0.55138 | 0.74147 | 1 | 15568 | 1 | -0.79245 |
| CSF2RA_X | 6 | 0.46546 | 0.64543 | 0.999994 | 12942 | 2 | 0.19136 | 0.5515 | 0.76615 | 1 | 15569 | 3 | 0.19136 |
| TMEM213 | 6 | 0.21962 | 0.41164 | 0.999994 | 8281 | 3 | -0.7837 | 0.55155 | 0.76619 | 1 | 15570 | 1 | -0.7837 |
| ATP6V0D2 | 6 | 0.64141 | 0.771 | 0.999994 | 15538 | 2 | -0.14522 | 0.55155 | 0.76619 | 1 | 15571 | 2 | -0.14522 |
| TRUB2 | 6 | 0.10957 | 0.26588 | 0.997535 | 5292 | 4 | -1.0896 | 0.55155 | 0.76619 | 1 | 15572 | 1 | -1.0896 |
| CAPS | 6 | 0.065787 | 0.17793 | 0.968171 | 3683 | 2 | 0.030559 | 0.55155 | 0.76619 | 1 | 15573 | 2 | 0.030559 |
| NKX3-2 | 6 | 0.03772 | 0.11676 | 0.951052 | 2416 | 3 | -0.30457 | 0.55155 | 0.76619 | 1 | 15574 | 2 | -0.30457 |
| TSR1 | 3 | 0.079747 | 0.15617 | 0.967126 | 4222 | 2 | -2.8274 | 0.55161 | 0.66561 | 1 | 15575 | 1 | -2.8274 |
| ITIH2 | 6 | 0.30596 | 0.5137 | 0.999994 | 10251 | 3 | -0.61036 | 0.55162 | 0.76625 | 1 | 15576 | 2 | -0.61036 |
| TMPRSS11F | 6 | 0.20424 | 0.39303 | 0.999994 | 7883 | 2 | 0.34084 | 0.55174 | 0.76635 | 1 | 15577 | 3 | 0.34084 |
| TMEM204 | 5 | 0.32493 | 0.50135 | 0.999994 | 10617 | 2 | 0.16883 | 0.55204 | 0.74182 | 1 | 15578 | 2 | 0.16883 |
| RPS18 | 1 | 0.44794 | 0.44802 | 0.999994 | 12687 | 0 | 0.094124 | 0.55206 | 0.55187 | 1 | 15579 | 0 | 0.094124 |
| ZNF280B | 6 | 0.14676 | 0.3216 | 0.999994 | 6369 | 3 | -0.8542 | 0.55215 | 0.76666 | 1 | 15580 | 2 | -0.8542 |
| TMC8 | 6 | 0.23041 | 0.42457 | 0.999994 | 8535 | 2 | 0.49696 | 0.55227 | 0.76675 | 1 | 15581 | 3 | 0.49696 |
| HENMT1 | 6 | 0.44988 | 0.63505 | 0.999994 | 12711 | 1 | 0.71752 | 0.55236 | 0.76682 | 1 | 15582 | 3 | 0.71752 |
| MOCS1 | 5 | 0.096191 | 0.2204 | 0.980699 | 4825 | 3 | -1.5262 | 0.55243 | 0.74204 | 1 | 15583 | 2 | -1.5262 |
| CCNB3 | 4 | 0.36116 | 0.50571 | 0.999994 | 11356 | 2 | -0.52982 | 0.5525 | 0.70537 | 1 | 15584 | 1 | -0.52982 |
| NonTargetingControlG | 2 | 0.5673 | 0.56762 | 0.999994 | 14502 | 1 | 0.58399 | 0.55257 | 0.57702 | 1 | 15585 | 1 | 0.58399 |
| RP1 | 5 | 0.071665 | 0.18108 | 0.968171 | 3903 | 3 | -1.9539 | 0.55259 | 0.74213 | 1 | 15586 | 1 | -1.9539 |
| CYP11A1 | 5 | 0.25354 | 0.44359 | 0.999994 | 9092 | 2 | 0.06615 | 0.55259 | 0.74213 | 1 | 15587 | 1 | 0.06615 |
| INSM2 | 5 | 0.028857 | 0.087539 | 0.94395 | 1946 | 4 | -1.1788 | 0.55259 | 0.74213 | 1 | 15588 | 1 | -1.1788 |
| TMEM184C | 5 | 0.30826 | 0.48772 | 0.999994 | 10289 | 1 | -0.26234 | 0.55259 | 0.74213 | 1 | 15589 | 2 | -0.26234 |
| FAM174A | 5 | 0.3079 | 0.48743 | 0.999994 | 10285 | 2 | -0.22077 | 0.55259 | 0.74213 | 1 | 15590 | 1 | -0.22077 |
| SNAI1 | 4 | 0.097831 | 0.2112 | 0.979587 | 4884 | 2 | -0.30492 | 0.55267 | 0.70552 | 1 | 15591 | 2 | -0.30492 |
| S100A2 | 5 | 0.035497 | 0.1045 | 0.948066 | 2294 | 3 | -0.69071 | 0.55267 | 0.74218 | 1 | 15592 | 2 | -0.69071 |
| NonTargetingControlG | 2 | 0.58734 | 0.58738 | 0.999994 | 14779 | 0 | 0.70535 | 0.55273 | 0.57715 | 1 | 15593 | 1 | 0.70535 |
| IL24 | 6 | 0.35033 | 0.56437 | 0.999994 | 11167 | 3 | -1.0522 | 0.55282 | 0.76718 | 1 | 15594 | 2 | -1.0522 |
| EXO5 | 6 | 0.099757 | 0.24687 | 0.98999 | 4968 | 2 | -0.040351 | 0.55282 | 0.76718 | 1 | 15595 | 2 | -0.040351 |
| CHST4 | 6 | 0.46535 | 0.64535 | 0.999994 | 12940 | 2 | 0.064481 | 0.55282 | 0.76718 | 1 | 15596 | 2 | 0.064481 |
| hsa-mir-4789 | 4 | 0.73537 | 0.74907 | 0.999994 | 16667 | 1 | 0.31247 | 0.55283 | 0.70564 | 1 | 15597 | 2 | 0.31247 |
| TTC8 | 6 | 0.035307 | 0.11126 | 0.951052 | 2281 | 2 | 0.38783 | 0.55292 | 0.76727 | 1 | 15598 | 3 | 0.38783 |
| CBY1 | 6 | 0.0017284 | 0.0086169 | 0.680529 | 242 | 4 | -5.7082 | 0.55294 | 0.76727 | 1 | 15599 | 2 | -5.7082 |

| | | | | | | | | | | | | | |
|---|---|---|---|---|---|---|---|---|---|---|---|---|---|
| LRRC52 | 6 | 0.10852 | 0.26385 | 0.996734 | 5257 | 4 | -1.0732 | 0.55337 | 0.76758 | 1 | 15600 | 1 | -1.0732 |
| KRT15 | 6 | 0.083786 | 0.21506 | 0.979587 | 4368 | 4 | -1.5846 | 0.5534 | 0.7676 | 1 | 15601 | 2 | -1.5846 |
| NonTargetingControlG | 1 | 0.4464 | 0.4465 | 0.999994 | 12668 | 0 | 0.096105 | 0.5536 | 0.55344 | 1 | 15602 | 0 | 0.096105 |
| CHRNA6 | 4 | 0.011359 | 0.037981 | 0.887249 | 947 | 3 | -4.3712 | 0.55367 | 0.70632 | 1 | 15603 | 1 | -4.3712 |
| hsa-mir-3913-1 | 4 | 0.19391 | 0.3344 | 0.999994 | 7598 | 1 | 0.46718 | 0.55376 | 0.7064 | 1 | 15604 | 2 | 0.46718 |
| PRH1 | 5 | 0.23921 | 0.42987 | 0.999994 | 8754 | 3 | -0.59207 | 0.55382 | 0.74283 | 1 | 15605 | 2 | -0.59207 |
| ABCC5 | 5 | 0.55692 | 0.69791 | 0.999994 | 14315 | 1 | 0.18115 | 0.55382 | 0.74283 | 1 | 15606 | 2 | 0.18115 |
| ATRNL1 | 5 | 0.69826 | 0.76571 | 0.999994 | 16286 | 1 | 0.52093 | 0.55382 | 0.74283 | 1 | 15607 | 2 | 0.52093 |
| TNFAIP1 | 5 | 0.0067651 | 0.02669 | 0.848412 | 644 | 4 | -2.8421 | 0.55382 | 0.74283 | 1 | 15608 | 1 | -2.8421 |
| CAPSL | 5 | 0.085093 | 0.2027 | 0.977973 | 4413 | 3 | -1.8407 | 0.55382 | 0.74283 | 1 | 15609 | 1 | -1.8407 |
| AHSG | 5 | 0.4607 | 0.61425 | 0.999994 | 12856 | 2 | -0.54317 | 0.55382 | 0.74283 | 1 | 15610 | 1 | -0.54317 |
| S100A8 | 5 | 0.047853 | 0.13526 | 0.954328 | 2884 | 4 | -1.1733 | 0.55382 | 0.74283 | 1 | 15611 | 1 | -1.1733 |
| CCNL1 | 6 | 0.041272 | 0.12475 | 0.951052 | 2577 | 4 | -1.498 | 0.55398 | 0.76804 | 1 | 15612 | 2 | -1.498 |
| CPA5 | 6 | 0.20372 | 0.3924 | 0.999994 | 7872 | 3 | -0.14353 | 0.55398 | 0.76804 | 1 | 15613 | 1 | -0.14353 |
| CRTAM | 6 | 0.028571 | 0.095453 | 0.945031 | 1928 | 4 | -2.6794 | 0.5541 | 0.76812 | 1 | 15614 | 2 | -2.6794 |
| NonTargetingControlG | 2 | 0.21198 | 0.27077 | 0.998561 | 8065 | 1 | -1.7229 | 0.55422 | 0.57829 | 1 | 15615 | 1 | -1.7229 |
| ZNF740 | 5 | 0.87292 | 0.8737 | 0.999994 | 18692 | 1 | 0.45185 | 0.55438 | 0.74314 | 1 | 15616 | 2 | 0.45185 |
| MPDU1 | 6 | 0.11277 | 0.27204 | 0.998561 | 5396 | 3 | -0.15613 | 0.55453 | 0.76847 | 1 | 15617 | 3 | -0.15613 |
| CDH1 | 6 | 0.15344 | 0.33 | 0.999994 | 6546 | 3 | -0.48761 | 0.55464 | 0.76855 | 1 | 15618 | 2 | -0.48761 |
| NT5DC3 | 4 | 0.19327 | 0.33375 | 0.999994 | 7583 | 1 | 0.55217 | 0.55466 | 0.70707 | 1 | 15619 | 2 | 0.55217 |
| NonTargetingControlG | 2 | 0.36766 | 0.39026 | 0.999994 | 11470 | 1 | 0.071991 | 0.55481 | 0.57874 | 1 | 15620 | 1 | 0.071991 |
| IL9R_X | 6 | 0.56119 | 0.71165 | 0.999994 | 14392 | 2 | 0.3441 | 0.55484 | 0.7687 | 1 | 15621 | 3 | 0.3441 |
| DOT1L | 4 | 0.014054 | 0.046178 | 0.892434 | 1133 | 3 | -4.663 | 0.55491 | 0.70725 | 1 | 15622 | 1 | -4.663 |
| hsa-mir-631 | 4 | 0.42723 | 0.57365 | 0.999994 | 12370 | 2 | -0.091618 | 0.55491 | 0.70725 | 1 | 15623 | 1 | -0.091618 |
| OR2T1 | 4 | 0.37399 | 0.51898 | 0.999994 | 11571 | 2 | -0.66342 | 0.55491 | 0.70725 | 1 | 15624 | 1 | -0.66342 |
| DENND5B | 6 | 0.054469 | 0.15375 | 0.967126 | 3175 | 4 | -1.8823 | 0.55495 | 0.76879 | 1 | 15625 | 2 | -1.8823 |
| EGFLAM | 4 | 0.30327 | 0.44633 | 0.999994 | 10199 | 2 | -0.070811 | 0.55496 | 0.70729 | 1 | 15626 | 2 | -0.070811 |
| TXN2 | 6 | 0.21603 | 0.40739 | 0.999994 | 8171 | 3 | -0.15061 | 0.55501 | 0.76883 | 1 | 15627 | 3 | -0.15061 |
| NonTargetingControlG | 2 | 0.66933 | 0.6691 | 0.999994 | 15919 | 0 | 0.97194 | 0.5551 | 0.57894 | 1 | 15628 | 1 | 0.97194 |
| KCNH5 | 6 | 0.022583 | 0.080239 | 0.93914 | 1623 | 3 | 0.12126 | 0.55518 | 0.76894 | 1 | 15629 | 3 | 0.12126 |
| XRN2 | 5 | 0.0040733 | 0.017128 | 0.785648 | 447 | 3 | -7.1379 | 0.55519 | 0.74362 | 1 | 15630 | 1 | -7.1379 |
| SEMA6B | 5 | 0.49534 | 0.64384 | 0.999994 | 13343 | 2 | 0.27163 | 0.55519 | 0.74362 | 1 | 15631 | 2 | 0.27163 |
| PPIB | 5 | 0.14453 | 0.2946 | 0.999994 | 6314 | 3 | -1.2674 | 0.55519 | 0.74362 | 1 | 15632 | 2 | -1.2674 |
| EIF4G3 | 6 | 0.27839 | 0.48156 | 0.999994 | 9660 | 1 | 0.34232 | 0.55525 | 0.76898 | 1 | 15633 | 2 | 0.34232 |
| ADAR | 5 | 0.37051 | 0.53882 | 0.999994 | 11522 | 2 | -0.56142 | 0.55536 | 0.7437 | 1 | 15634 | 2 | -0.56142 |
| RSG1 | 6 | 0.29972 | 0.50638 | 0.999994 | 10123 | 3 | 0.028936 | 0.55541 | 0.76911 | 1 | 15635 | 3 | 0.028936 |
| TNNC2 | 5 | 0.097887 | 0.22313 | 0.983354 | 4888 | 3 | -1.5189 | 0.55563 | 0.74385 | 1 | 15636 | 2 | -1.5189 |
| OR2T34 | 5 | 0.57354 | 0.71265 | 0.999994 | 14600 | 2 | 0.25646 | 0.55576 | 0.74393 | 1 | 15637 | 2 | 0.25646 |
| NonTargetingControlG | 2 | 0.20714 | 0.26729 | 0.998561 | 7950 | 1 | -0.73923 | 0.55592 | 0.57955 | 1 | 15638 | 1 | -0.73923 |
| CPXCR1 | 6 | 0.43093 | 0.62246 | 0.999994 | 12429 | 1 | 0.071372 | 0.55605 | 0.7696 | 1 | 15639 | 2 | 0.071372 |
| RGS2 | 6 | 0.041052 | 0.12428 | 0.951052 | 2564 | 2 | -0.43019 | 0.55605 | 0.7696 | 1 | 15640 | 2 | -0.43019 |
| RPS4Y1 | 6 | 0.60529 | 0.74389 | 0.999994 | 15016 | 2 | 0.52738 | 0.55605 | 0.7696 | 1 | 15641 | 3 | 0.52738 |
| HPS6 | 6 | 0.062157 | 0.17034 | 0.967126 | 3518 | 4 | -1.7325 | 0.55605 | 0.7696 | 1 | 15642 | 1 | -1.7325 |
| GOLGA7 | 5 | 0.041214 | 0.11893 | 0.951052 | 2573 | 3 | -2.1682 | 0.55607 | 0.7441 | 1 | 15643 | 2 | -2.1682 |
| NCDN | 4 | 0.08269 | 0.18369 | 0.968526 | 4326 | 2 | -1.1806 | 0.5562 | 0.70827 | 1 | 15644 | 2 | -1.1806 |
| ACN9 | 6 | 0.17137 | 0.35244 | 0.999994 | 7018 | 2 | 0.047882 | 0.55622 | 0.76971 | 1 | 15645 | 2 | 0.047882 |
| ZSCAN20 | 6 | 0.067305 | 0.18111 | 0.968171 | 3728 | 4 | -0.87162 | 0.55622 | 0.76971 | 1 | 15646 | 2 | -0.87162 |
| HBEGF | 6 | 0.004455 | 0.019153 | 0.805137 | 478 | 3 | -0.097253 | 0.55622 | 0.76971 | 1 | 15647 | 3 | -0.097253 |
| NonTargetingControlG | 2 | 0.58452 | 0.58458 | 0.999994 | 14737 | 0 | 0.64135 | 0.55646 | 0.57997 | 1 | 15648 | 1 | 0.64135 |
| BBIP1 | 5 | 0.21928 | 0.40227 | 0.999994 | 8271 | 3 | -0.70613 | 0.55649 | 0.74433 | 1 | 15649 | 1 | -0.70613 |
| FAM32A | 4 | 0.17082 | 0.31063 | 0.999994 | 7003 | 2 | -1.1038 | 0.55661 | 0.7086 | 1 | 15650 | 1 | -1.1038 |
| UBALD1 | 3 | 0.030114 | 0.075417 | 0.93914 | 2016 | 2 | -3.2581 | 0.55671 | 0.66786 | 1 | 15651 | 1 | -3.2581 |

| | | | | | | | | | | | | | |
|---|---|---|---|---|---|---|---|---|---|---|---|---|---|
| hsa-mir-20b | 4 | 0.29378 | 0.43664 | 0.999994 | 9992 | 1 | 0.72574 | 0.55679 | 0.70873 | 1 | 15652 | 2 | 0.72574 |
| WARS | 4 | 0.38655 | 0.53183 | 0.999994 | 11743 | 1 | 0.39761 | 0.55697 | 0.70888 | 1 | 15653 | 2 | 0.39761 |
| DAPK2 | 5 | 0.18271 | 0.35051 | 0.999994 | 7282 | 2 | -0.41901 | 0.55719 | 0.74472 | 1 | 15654 | 2 | -0.41901 |
| TBX6 | 5 | 0.04519 | 0.12878 | 0.951777 | 2773 | 2 | 0.036861 | 0.55719 | 0.74472 | 1 | 15655 | 2 | 0.036861 |
| KRT16 | 5 | 0.068935 | 0.1767 | 0.968171 | 3795 | 3 | -1.2599 | 0.55719 | 0.74472 | 1 | 15656 | 2 | -1.2599 |
| SDHC | 5 | 0.19012 | 0.36111 | 0.999994 | 7489 | 2 | 0.31446 | 0.55719 | 0.74472 | 1 | 15657 | 2 | 0.31446 |
| NUP210 | 5 | 0.86297 | 0.8649 | 0.999994 | 18505 | 1 | 0.028946 | 0.55719 | 0.74472 | 1 | 15658 | 2 | 0.028946 |
| RAB7A | 5 | 0.20806 | 0.38664 | 0.999994 | 7979 | 3 | -0.74987 | 0.55719 | 0.74472 | 1 | 15659 | 2 | -0.74987 |
| PNPLA8 | 5 | 0.032269 | 0.096286 | 0.945031 | 2118 | 2 | -0.22954 | 0.55719 | 0.74472 | 1 | 15660 | 2 | -0.22954 |
| KDM5A | 6 | 0.14033 | 0.31344 | 0.999994 | 6194 | 3 | -0.50466 | 0.55749 | 0.77071 | 1 | 15661 | 3 | -0.50466 |
| TSPAN12 | 6 | 0.12253 | 0.2905 | 0.999632 | 5723 | 4 | -0.80195 | 0.55763 | 0.77082 | 1 | 15662 | 2 | -0.80195 |
| PTDSS1 | 6 | 0.15232 | 0.32859 | 0.999994 | 6511 | 3 | -0.005722 | 0.55767 | 0.77085 | 1 | 15663 | 3 | -0.005722 |
| KRT26 | 4 | 0.30309 | 0.44615 | 0.999994 | 10196 | 2 | -0.079842 | 0.55773 | 0.70948 | 1 | 15664 | 2 | -0.079842 |
| ACACA | 6 | 0.32412 | 0.53459 | 0.999994 | 10601 | 1 | 0.49207 | 0.55795 | 0.77108 | 1 | 15665 | 3 | 0.49207 |
| NXPE1 | 6 | 0.14895 | 0.3243 | 0.999994 | 6425 | 3 | -1.0138 | 0.55799 | 0.77111 | 1 | 15666 | 1 | -1.0138 |
| CCDC129 | 6 | 0.01937 | 0.069968 | 0.93914 | 1424 | 5 | -2.3792 | 0.55799 | 0.77111 | 1 | 15667 | 1 | -2.3792 |
| EPHA8 | 4 | 0.17881 | 0.31889 | 0.999994 | 7193 | 1 | 0.47819 | 0.55807 | 0.70975 | 1 | 15668 | 2 | 0.47819 |
| ZNF488 | 5 | 0.26845 | 0.4556 | 0.999994 | 9433 | 2 | -0.26978 | 0.55826 | 0.74529 | 1 | 15669 | 2 | -0.26978 |
| APOC1 | 6 | 0.15143 | 0.32747 | 0.999994 | 6493 | 3 | -0.23763 | 0.55836 | 0.77136 | 1 | 15670 | 2 | -0.23763 |
| SPDEF | 4 | 0.058847 | 0.13855 | 0.957336 | 3376 | 1 | 0.62248 | 0.55854 | 0.71011 | 1 | 15671 | 2 | 0.62248 |
| RGAG4 | 4 | 0.92301 | 0.92284 | 0.999994 | 19697 | 0 | 0.7722 | 0.55871 | 0.71025 | 1 | 15672 | 2 | 0.7722 |
| NLK | 5 | 0.23421 | 0.42304 | 0.999994 | 8623 | 3 | -0.63608 | 0.55876 | 0.74558 | 1 | 15673 | 1 | -0.63608 |
| NEURL2 | 5 | 0.033061 | 0.098381 | 0.945253 | 2163 | 4 | -1.3186 | 0.55876 | 0.74558 | 1 | 15674 | 1 | -1.3186 |
| SETD2 | 5 | 0.023785 | 0.07426 | 0.93914 | 1699 | 3 | -2.8002 | 0.55886 | 0.74563 | 1 | 15675 | 2 | -2.8002 |
| SLCO1B3 | 6 | 0.47322 | 0.65058 | 0.999994 | 13046 | 2 | 0.31827 | 0.55888 | 0.77174 | 1 | 15676 | 3 | 0.31827 |
| hsa-mir-3611 | 1 | 0.44099 | 0.44113 | 0.999994 | 12592 | 0 | 0.046701 | 0.55901 | 0.55886 | 1 | 15677 | 0 | 0.046701 |
| TFIP11 | 4 | 0.058306 | 0.13752 | 0.957317 | 3354 | 3 | -3.313 | 0.55903 | 0.71051 | 1 | 15678 | 1 | -3.313 |
| FUK | 6 | 0.48473 | 0.65838 | 0.999994 | 13212 | 2 | 0.26553 | 0.55908 | 0.77187 | 1 | 15679 | 2 | 0.26553 |
| PDXK | 5 | 0.069337 | 0.17731 | 0.968171 | 3811 | 3 | -2.095 | 0.55918 | 0.74579 | 1 | 15680 | 2 | -2.095 |
| hsa-mir-3938 | 4 | 0.39949 | 0.5452 | 0.999994 | 11941 | 2 | 0.26887 | 0.55924 | 0.71068 | 1 | 15681 | 2 | 0.26887 |
| hsa-mir-4796 | 4 | 0.77737 | 0.78173 | 0.999994 | 17173 | 1 | 0.23422 | 0.55938 | 0.71078 | 1 | 15682 | 2 | 0.23422 |
| ELK3 | 6 | 0.78888 | 0.8432 | 0.999994 | 17331 | 1 | 0.73171 | 0.55939 | 0.77209 | 1 | 15683 | 3 | 0.73171 |
| HOXD12 | 6 | 0.24098 | 0.43717 | 0.999994 | 8792 | 3 | 0.08536 | 0.55939 | 0.77209 | 1 | 15684 | 3 | 0.08536 |
| PPP3CC | 6 | 0.17904 | 0.36193 | 0.999994 | 7197 | 1 | 0.77264 | 0.55939 | 0.77209 | 1 | 15685 | 3 | 0.77264 |
| SGSH | 6 | 0.84365 | 0.86855 | 0.999994 | 18149 | 1 | 0.69411 | 0.55939 | 0.77209 | 1 | 15686 | 3 | 0.69411 |
| LRRC10 | 6 | 0.088557 | 0.22465 | 0.983354 | 4537 | 2 | 0.37224 | 0.55939 | 0.77209 | 1 | 15687 | 3 | 0.37224 |
| GPR52 | 6 | 0.39564 | 0.59957 | 0.999994 | 11884 | 2 | 0.24372 | 0.55942 | 0.77212 | 1 | 15688 | 2 | 0.24372 |
| XKR7 | 2 | 0.23932 | 0.29094 | 0.999632 | 8760 | 1 | -1.6002 | 0.55952 | 0.5824 | 1 | 15689 | 1 | -1.6002 |
| VAX1 | 6 | 0.076196 | 0.19964 | 0.977164 | 4082 | 2 | 0.44263 | 0.55961 | 0.77224 | 1 | 15690 | 3 | 0.44263 |
| KIAA1429 | 4 | 0.7839 | 0.78715 | 0.999994 | 17264 | 1 | 0.7516 | 0.5597 | 0.71102 | 1 | 15691 | 2 | 0.7516 |
| WARS2 | 6 | 0.0028387 | 0.013135 | 0.733879 | 344 | 5 | -2.8855 | 0.55974 | 0.77234 | 1 | 15692 | 1 | -2.8855 |
| ALK | 6 | 0.11565 | 0.27751 | 0.999379 | 5503 | 4 | -0.96518 | 0.55974 | 0.77234 | 1 | 15693 | 2 | -0.96518 |
| ARHGAP15 | 5 | 0.22026 | 0.40362 | 0.999994 | 8295 | 3 | -0.63903 | 0.55991 | 0.74621 | 1 | 15694 | 2 | -0.63903 |
| hsa-mir-4768 | 4 | 0.83251 | 0.83204 | 0.999994 | 17964 | 0 | 0.6089 | 0.55997 | 0.71126 | 1 | 15695 | 2 | 0.6089 |
| NonTargetingControlG | 2 | 0.55462 | 0.55532 | 0.999994 | 14279 | 1 | 0.60635 | 0.56001 | 0.58279 | 1 | 15696 | 1 | 0.60635 |
| SUCLG2 | 6 | 0.66773 | 0.79139 | 0.999994 | 15889 | 1 | 0.20201 | 0.56026 | 0.77272 | 1 | 15697 | 3 | 0.20201 |
| MDGA2 | 6 | 0.17875 | 0.36157 | 0.999994 | 7191 | 3 | -0.2699 | 0.56029 | 0.77275 | 1 | 15698 | 3 | -0.2699 |
| TULP3 | 6 | 0.026268 | 0.089985 | 0.94395 | 1816 | 4 | -1.6286 | 0.56033 | 0.77277 | 1 | 15699 | 2 | -1.6286 |
| PER1 | 4 | 0.58657 | 0.65812 | 0.999994 | 14766 | 1 | 0.74382 | 0.56044 | 0.71161 | 1 | 15700 | 2 | 0.74382 |
| ZNF696 | 4 | 0.049165 | 0.11978 | 0.951052 | 2938 | 3 | -1.7136 | 0.56044 | 0.71162 | 1 | 15701 | 1 | -1.7136 |
| ZDHHC12 | 4 | 0.11137 | 0.23529 | 0.985362 | 5346 | 2 | -1.207 | 0.56044 | 0.71162 | 1 | 15702 | 1 | -1.207 |
| NRP2 | 4 | 0.19838 | 0.33901 | 0.999994 | 7723 | 2 | -1.1466 | 0.56044 | 0.71162 | 1 | 15703 | 1 | -1.1466 |

| | | | | | | | | | | | | | |
|---|---|---|---|---|---|---|---|---|---|---|---|---|---|
| KIF20A | 5 | 0.23516 | 0.42436 | 0.999994 | 8651 | 3 | -0.66852 | 0.56056 | 0.74654 | 1 | 15704 | 2 | -0.66852 |
| MIF4GD | 5 | 0.40121 | 0.56452 | 0.999994 | 11965 | 2 | 0.016024 | 0.56056 | 0.74654 | 1 | 15705 | 2 | 0.016024 |
| CD2BP2 | 5 | 0.51998 | 0.66529 | 0.999994 | 13747 | 2 | -0.46372 | 0.56056 | 0.74654 | 1 | 15706 | 1 | -0.46372 |
| CRYGS | 6 | 0.35867 | 0.57382 | 0.999994 | 11315 | 3 | -0.49204 | 0.56056 | 0.77296 | 1 | 15707 | 2 | -0.49204 |
| C2CD2 | 6 | 0.32766 | 0.53856 | 0.999994 | 10672 | 2 | 0.1984 | 0.56075 | 0.77309 | 1 | 15708 | 2 | 0.1984 |
| ACOX2 | 6 | 0.088327 | 0.2242 | 0.983354 | 4527 | 4 | -1.2851 | 0.56081 | 0.77314 | 1 | 15709 | 2 | -1.2851 |
| TAF1C | 6 | 0.0058938 | 0.024425 | 0.842094 | 583 | 4 | -2.4747 | 0.56081 | 0.77314 | 1 | 15710 | 1 | -2.4747 |
| hsa-mir-329-1 | 4 | 0.88868 | 0.88836 | 0.999994 | 19012 | 0 | 0.51824 | 0.56095 | 0.71202 | 1 | 15711 | 2 | 0.51824 |
| hsa-mir-138-1 | 4 | 0.37861 | 0.52373 | 0.999994 | 11632 | 2 | 0.092458 | 0.56096 | 0.71203 | 1 | 15712 | 2 | 0.092458 |
| IGSF11 | 6 | 0.052905 | 0.15036 | 0.966359 | 3106 | 1 | 0.58705 | 0.56113 | 0.77339 | 1 | 15713 | 3 | 0.58705 |
| FSTL4 | 5 | 0.29895 | 0.48022 | 0.999994 | 10110 | 2 | 0.43121 | 0.56123 | 0.74692 | 1 | 15714 | 2 | 0.43121 |
| CCL20 | 6 | 0.4583 | 0.6406 | 0.999994 | 12827 | 2 | 0.29255 | 0.5613 | 0.77352 | 1 | 15715 | 3 | 0.29255 |
| CCDC96 | 6 | 0.31217 | 0.52085 | 0.999994 | 10363 | 3 | 0.054822 | 0.56159 | 0.77373 | 1 | 15716 | 3 | 0.054822 |
| APLNR | 5 | 0.063726 | 0.16822 | 0.967126 | 3594 | 3 | -2.7019 | 0.56159 | 0.74711 | 1 | 15717 | 2 | -2.7019 |
| NOLC1 | 5 | 0.11664 | 0.25224 | 0.994162 | 5527 | 3 | -1.1384 | 0.56159 | 0.74711 | 1 | 15718 | 1 | -1.1384 |
| BASP1 | 5 | 0.18633 | 0.35573 | 0.999994 | 7370 | 3 | -0.64414 | 0.56161 | 0.74712 | 1 | 15719 | 2 | -0.64414 |
| RTP1 | 6 | 0.25195 | 0.45026 | 0.999994 | 9056 | 3 | -0.18158 | 0.56187 | 0.77396 | 1 | 15720 | 3 | -0.18158 |
| ZMYND12 | 6 | 0.87969 | 0.88939 | 0.999994 | 18826 | 1 | 0.34825 | 0.56187 | 0.77396 | 1 | 15721 | 2 | 0.34825 |
| GPRIN1 | 6 | 0.015569 | 0.057597 | 0.929232 | 1220 | 5 | -1.4237 | 0.56187 | 0.77396 | 1 | 15722 | 1 | -1.4237 |
| hsa-mir-4529 | 4 | 0.85796 | 0.85757 | 0.999994 | 18406 | 0 | 0.39562 | 0.56197 | 0.71281 | 1 | 15723 | 2 | 0.39562 |
| EXOC2 | 6 | 0.020166 | 0.072513 | 0.93914 | 1466 | 3 | -0.04906 | 0.56201 | 0.77406 | 1 | 15724 | 3 | -0.04906 |
| TBC1D13 | 6 | 0.56987 | 0.71785 | 0.999994 | 14536 | 2 | 0.60684 | 0.56206 | 0.77411 | 1 | 15725 | 3 | 0.60684 |
| TRIM43 | 4 | 0.02674 | 0.074833 | 0.93914 | 1844 | 3 | -4.4444 | 0.56211 | 0.71291 | 1 | 15726 | 1 | -4.4444 |
| TUBGCP4 | 4 | 0.01069 | 0.035873 | 0.875006 | 911 | 3 | -5.2104 | 0.56211 | 0.71291 | 1 | 15727 | 1 | -5.2104 |
| OR1E1 | 6 | 0.059363 | 0.1644 | 0.967126 | 3397 | 4 | -1.1392 | 0.56214 | 0.77416 | 1 | 15728 | 2 | -1.1392 |
| FAF2 | 6 | 0.9373 | 0.9373 | 0.999994 | 20013 | 0 | 0.62723 | 0.5622 | 0.77421 | 1 | 15729 | 3 | 0.62723 |
| hsa-mir-6787 | 3 | 0.30224 | 0.42973 | 0.999994 | 10175 | 1 | 0.19844 | 0.56221 | 0.67034 | 1 | 15730 | 1 | 0.19844 |
| RBMY1D | 2 | 0.2989 | 0.33601 | 0.999994 | 10109 | 1 | -0.39685 | 0.56249 | 0.58467 | 1 | 15731 | 1 | -0.39685 |
| LOC643802 | 6 | 0.60474 | 0.74349 | 0.999994 | 15005 | 1 | 0.25173 | 0.56257 | 0.77447 | 1 | 15732 | 3 | 0.25173 |
| DPYSL2 | 5 | 0.19379 | 0.36634 | 0.999994 | 7593 | 3 | -0.88683 | 0.56263 | 0.74769 | 1 | 15733 | 2 | -0.88683 |
| C20orf202 | 5 | 0.089616 | 0.20997 | 0.979587 | 4581 | 3 | -0.96984 | 0.56263 | 0.74769 | 1 | 15734 | 1 | -0.96984 |
| PDYN | 6 | 0.52987 | 0.68944 | 0.999994 | 13898 | 2 | 0.24667 | 0.56288 | 0.7747 | 1 | 15735 | 3 | 0.24667 |
| C5orf58 | 6 | 0.12858 | 0.29845 | 0.999994 | 5898 | 3 | -1.1273 | 0.56288 | 0.7747 | 1 | 15736 | 1 | -1.1273 |
| LRRC56 | 4 | 0.7686 | 0.77463 | 0.999994 | 17069 | 1 | 0.44185 | 0.56292 | 0.71352 | 1 | 15737 | 2 | 0.44185 |
| NonTargetingControlG | 2 | 0.15149 | 0.22768 | 0.983354 | 6496 | 1 | -2.5621 | 0.56293 | 0.58502 | 1 | 15738 | 1 | -2.5621 |
| PAX7 | 4 | 0.087094 | 0.19168 | 0.9742 | 4479 | 2 | -1.3043 | 0.56297 | 0.71356 | 1 | 15739 | 2 | -1.3043 |
| CPVL | 6 | 0.21494 | 0.40607 | 0.999994 | 8141 | 3 | -0.19542 | 0.563 | 0.77479 | 1 | 15740 | 3 | -0.19542 |
| ZNF43 | 6 | 0.096461 | 0.24039 | 0.987253 | 4837 | 2 | 0.70829 | 0.56311 | 0.77487 | 1 | 15741 | 3 | 0.70829 |
| hsa-mir-3130-1 | 4 | 0.66158 | 0.70018 | 0.999994 | 15803 | 1 | 0.57677 | 0.56331 | 0.71384 | 1 | 15742 | 2 | 0.57677 |
| GSG1 | 4 | 0.73629 | 0.74973 | 0.999994 | 16680 | 1 | 0.56064 | 0.56334 | 0.71386 | 1 | 15743 | 2 | 0.56064 |
| hsa-mir-548i-4 | 2 | 0.67886 | 0.67867 | 0.999994 | 16072 | 0 | 0.97362 | 0.56335 | 0.58534 | 1 | 15744 | 1 | 0.97362 |
| SMCO2 | 6 | 0.065884 | 0.17815 | 0.968171 | 3684 | 3 | -1.5761 | 0.56345 | 0.77511 | 1 | 15745 | 1 | -1.5761 |
| FAM162A | 6 | 0.88462 | 0.8926 | 0.999994 | 18927 | 1 | 0.48153 | 0.56353 | 0.77516 | 1 | 15746 | 3 | 0.48153 |
| PRKCZ | 6 | 0.32388 | 0.53431 | 0.999994 | 10593 | 3 | -0.17031 | 0.56357 | 0.77519 | 1 | 15747 | 2 | -0.17031 |
| NPRL3 | 4 | 0.12416 | 0.25753 | 0.994943 | 5773 | 2 | -1.358 | 0.56376 | 0.71418 | 1 | 15748 | 1 | -1.358 |
| hsa-mir-4677 | 4 | 0.58884 | 0.65932 | 0.999994 | 14802 | 1 | -0.095051 | 0.56376 | 0.71418 | 1 | 15749 | 1 | -0.095051 |
| hsa-mir-4324 | 4 | 0.27936 | 0.4219 | 0.999994 | 9686 | 2 | -0.59792 | 0.56376 | 0.71418 | 1 | 15750 | 1 | -0.59792 |
| hsa-mir-665 | 4 | 0.12566 | 0.26012 | 0.994943 | 5812 | 3 | -0.89027 | 0.56376 | 0.71418 | 1 | 15751 | 1 | -0.89027 |
| PET100 | 2 | 0.56833 | 0.56862 | 0.999994 | 14515 | 1 | 0.57319 | 0.56397 | 0.58582 | 1 | 15752 | 1 | 0.57319 |
| GPR64 | 6 | 0.005503 | 0.022984 | 0.827176 | 558 | 3 | -3.117 | 0.564 | 0.77552 | 1 | 15753 | 3 | -3.117 |
| FUT4 | 6 | 0.19222 | 0.37827 | 0.999994 | 7554 | 2 | 0.047458 | 0.56404 | 0.77556 | 1 | 15754 | 2 | 0.047458 |
| KAT5 | 6 | 0.0084791 | 0.033615 | 0.867348 | 759 | 5 | -2.6358 | 0.56404 | 0.77556 | 1 | 15755 | 1 | -2.6358 |

| | | | | | | | | | | | | | |
|---|---|---|---|---|---|---|---|---|---|---|---|---|---|
| NTSR2 | 6 | 0.20109 | 0.38919 | 0.999994 | 7791 | 3 | -0.86331 | 0.56404 | 0.77556 | 1 | 15756 | 2 | -0.86331 |
| IDI2 | 6 | 0.29273 | 0.49822 | 0.999994 | 9969 | 3 | -0.51268 | 0.56404 | 0.77556 | 1 | 15757 | 1 | -0.51268 |
| CEP152 | 6 | 0.19252 | 0.37867 | 0.999994 | 7560 | 3 | -0.93073 | 0.56413 | 0.77562 | 1 | 15758 | 2 | -0.93073 |
| MND1 | 5 | 0.43627 | 0.59369 | 0.999994 | 12518 | 2 | -0.77488 | 0.56426 | 0.74859 | 1 | 15759 | 2 | -0.77488 |
| RNF123 | 4 | 0.1259 | 0.26054 | 0.994943 | 5818 | 1 | 0.77613 | 0.56445 | 0.71473 | 1 | 15760 | 2 | 0.77613 |
| LMLN | 4 | 0.73104 | 0.74593 | 0.999994 | 16614 | 1 | 0.55011 | 0.56463 | 0.71487 | 1 | 15761 | 2 | 0.55011 |
| KLF9 | 6 | 0.066555 | 0.17954 | 0.968171 | 3708 | 3 | -0.91708 | 0.56478 | 0.77611 | 1 | 15762 | 3 | -0.91708 |
| hsa-mir-663a | 1 | 0.43505 | 0.4352 | 0.999994 | 12499 | 0 | -1.2059 | 0.56495 | 0.56483 | 1 | 15763 | 0 | -1.2059 |
| UBASH3B | 4 | 0.34159 | 0.48565 | 0.999994 | 10972 | 2 | -0.031173 | 0.56508 | 0.71519 | 1 | 15764 | 2 | -0.031173 |
| G3BP1 | 4 | 0.84428 | 0.84377 | 0.999994 | 18160 | 0 | 0.50143 | 0.56513 | 0.71524 | 1 | 15765 | 2 | 0.50143 |
| C11orf74 | 4 | 0.16028 | 0.29977 | 0.999994 | 6746 | 2 | -1.0211 | 0.56513 | 0.71524 | 1 | 15766 | 1 | -1.0211 |
| MIP | 6 | 0.57414 | 0.72092 | 0.999994 | 14606 | 2 | 0.38372 | 0.56529 | 0.7765 | 1 | 15767 | 3 | 0.38372 |
| STAU2 | 6 | 0.64578 | 0.77438 | 0.999994 | 15598 | 2 | 0.31111 | 0.56531 | 0.77652 | 1 | 15768 | 2 | 0.31111 |
| NonTargetingControlG | 2 | 0.09182 | 0.15736 | 0.967126 | 4671 | 1 | -3.1579 | 0.56533 | 0.58688 | 1 | 15769 | 1 | -3.1579 |
| RIMS2 | 6 | 0.67519 | 0.7972 | 0.999994 | 16007 | 2 | 0.545 | 0.56535 | 0.77654 | 1 | 15770 | 3 | 0.545 |
| POTEC | 5 | 0.40661 | 0.56905 | 0.999994 | 12053 | 2 | -0.21048 | 0.56537 | 0.74922 | 1 | 15771 | 2 | -0.21048 |
| ZNF620 | 6 | 0.39785 | 0.60097 | 0.999994 | 11913 | 1 | 0.59506 | 0.56547 | 0.77663 | 1 | 15772 | 3 | 0.59506 |
| IL17F | 6 | 0.26349 | 0.46399 | 0.999994 | 9325 | 2 | 0.20784 | 0.56552 | 0.77667 | 1 | 15773 | 3 | 0.20784 |
| NonTargetingControlG | 2 | 0.40484 | 0.42085 | 0.999994 | 12032 | 1 | 0.38465 | 0.56553 | 0.58702 | 1 | 15774 | 1 | 0.38465 |
| EXOC1 | 6 | 0.0012798 | 0.0064342 | 0.636218 | 191 | 3 | -1.1618 | 0.56566 | 0.77676 | 1 | 15775 | 3 | -1.1618 |
| C6orf89 | 6 | 0.90692 | 0.90904 | 0.999994 | 19369 | 1 | 0.36724 | 0.56571 | 0.7768 | 1 | 15776 | 3 | 0.36724 |
| PLAC4 | 5 | 0.80594 | 0.82326 | 0.999994 | 17571 | 1 | -0.46604 | 0.56592 | 0.74952 | 1 | 15777 | 2 | -0.46604 |
| RBMY1B | 1 | 0.43402 | 0.4342 | 0.999994 | 12476 | 0 | -0.018132 | 0.56598 | 0.56584 | 1 | 15778 | 0 | -0.018132 |
| TMEM261 | 5 | 0.029964 | 0.090378 | 0.94395 | 2005 | 3 | -2.1024 | 0.56604 | 0.74958 | 1 | 15779 | 1 | -2.1024 |
| MBD1 | 5 | 0.092368 | 0.21438 | 0.979587 | 4696 | 3 | -1.4541 | 0.56604 | 0.74958 | 1 | 15780 | 2 | -1.4541 |
| ZNF334 | 6 | 0.91014 | 0.91169 | 0.999994 | 19427 | 1 | 0.47868 | 0.56611 | 0.7771 | 1 | 15781 | 3 | 0.47868 |
| ZKSCAN3 | 5 | 0.59574 | 0.72457 | 0.999994 | 14899 | 1 | -0.43929 | 0.56621 | 0.74968 | 1 | 15782 | 2 | -0.43929 |
| PTPRR | 4 | 0.34227 | 0.48637 | 0.999994 | 10989 | 2 | -0.031407 | 0.56622 | 0.7161 | 1 | 15783 | 2 | -0.031407 |
| THTPA | 6 | 0.013788 | 0.051745 | 0.910734 | 1116 | 4 | -2.4718 | 0.56642 | 0.77734 | 1 | 15784 | 2 | -2.4718 |
| MYO19 | 6 | 0.32392 | 0.53436 | 0.999994 | 10594 | 3 | -0.45502 | 0.56642 | 0.77734 | 1 | 15785 | 1 | -0.45502 |
| ALKBH2 | 6 | 0.076132 | 0.19952 | 0.977164 | 4080 | 4 | -1.4793 | 0.56642 | 0.77734 | 1 | 15786 | 2 | -1.4793 |
| UEVLD | 6 | 0.33797 | 0.55035 | 0.999994 | 10901 | 3 | -0.39174 | 0.56642 | 0.77734 | 1 | 15787 | 2 | -0.39174 |
| LOC728392 | 6 | 0.65454 | 0.7811 | 0.999994 | 15712 | 2 | 0.11773 | 0.56642 | 0.77734 | 1 | 15788 | 2 | 0.11773 |
| NUP98 | 6 | 0.000043399 | 0.00024255 | 0.18112 | 24 | 4 | -8.9491 | 0.56642 | 0.77734 | 1 | 15789 | 1 | -8.9491 |
| MRI1 | 4 | 0.16127 | 0.30078 | 0.999994 | 6772 | 1 | 0.67283 | 0.56654 | 0.71636 | 1 | 15790 | 2 | 0.67283 |
| hsa-mir-4755 | 4 | 0.15754 | 0.29696 | 0.999994 | 6661 | 2 | -0.18054 | 0.56666 | 0.71644 | 1 | 15791 | 2 | -0.18054 |
| FUT10 | 4 | 0.1016 | 0.21797 | 0.980699 | 5022 | 1 | 0.37829 | 0.56666 | 0.71644 | 1 | 15792 | 1 | 0.37829 |
| NAT2 | 5 | 0.22709 | 0.41318 | 0.999994 | 8456 | 3 | -0.69336 | 0.56666 | 0.74992 | 1 | 15793 | 2 | -0.69336 |
| hsa-mir-4637 | 3 | 0.13937 | 0.23479 | 0.984971 | 6171 | 2 | -1.6984 | 0.56672 | 0.67241 | 1 | 15794 | 1 | -1.6984 |
| SSTR4 | 5 | 0.44002 | 0.59682 | 0.999994 | 12574 | 2 | -0.55117 | 0.5671 | 0.75017 | 1 | 15795 | 2 | -0.55117 |
| WNK3 | 4 | 0.035515 | 0.092765 | 0.943968 | 2295 | 1 | 0.39614 | 0.56721 | 0.71687 | 1 | 15796 | 2 | 0.39614 |
| IGSF9 | 6 | 0.29096 | 0.49617 | 0.999994 | 9930 | 3 | -0.18936 | 0.56752 | 0.77817 | 1 | 15797 | 2 | -0.18936 |
| PIP | 5 | 0.013487 | 0.045929 | 0.892434 | 1099 | 4 | -2.2531 | 0.56756 | 0.7504 | 1 | 15798 | 1 | -2.2531 |
| EPM2AIP1 | 5 | 0.13413 | 0.27894 | 0.999632 | 6025 | 3 | -1.2972 | 0.56774 | 0.75049 | 1 | 15799 | 2 | -1.2972 |
| SPAG17 | 5 | 0.11248 | 0.24586 | 0.989866 | 5386 | 2 | 0.25958 | 0.56787 | 0.75056 | 1 | 15800 | 2 | 0.25958 |
| SMARCC1 | 6 | 0.019019 | 0.068835 | 0.93914 | 1408 | 3 | -1.083 | 0.56797 | 0.77851 | 1 | 15801 | 2 | -1.083 |
| SNX13 | 6 | 0.021493 | 0.076789 | 0.93914 | 1548 | 5 | -1.3622 | 0.56849 | 0.77887 | 1 | 15802 | 1 | -1.3622 |
| BZW2 | 6 | 0.041167 | 0.12454 | 0.951052 | 2568 | 4 | -1.1679 | 0.56849 | 0.77887 | 1 | 15803 | 1 | -1.1679 |
| SUB1 | 4 | 0.14824 | 0.28735 | 0.999632 | 6413 | 2 | -0.58181 | 0.56857 | 0.71792 | 1 | 15804 | 2 | -0.58181 |
| ONECUT1 | 4 | 0.084632 | 0.18723 | 0.969409 | 4394 | 2 | -0.28672 | 0.56857 | 0.71792 | 1 | 15805 | 2 | -0.28672 |
| PES1 | 4 | 0.10565 | 0.22521 | 0.983354 | 5160 | 2 | -1.6713 | 0.56857 | 0.71792 | 1 | 15806 | 2 | -1.6713 |
| OR2AG1 | 4 | 0.071703 | 0.16316 | 0.967126 | 3905 | 2 | -1.8663 | 0.56867 | 0.718 | 1 | 15807 | 1 | -1.8663 |

| | | | | | | | | | | | | | |
|---|---|---|---|---|---|---|---|---|---|---|---|---|---|
| GCG | 6 | 0.039807 | 0.12147 | 0.951052 | 2504 | 3 | -1.6007 | 0.56872 | 0.77904 | 1 | 15808 | 2 | -1.6007 |
| ERMN | 6 | 0.50833 | 0.67447 | 0.999994 | 13559 | 1 | 0.16856 | 0.56873 | 0.77905 | 1 | 15809 | 3 | 0.16856 |
| SDC1 | 5 | 0.39049 | 0.55555 | 0.999994 | 11805 | 2 | 0.12442 | 0.56884 | 0.75109 | 1 | 15810 | 1 | 0.12442 |
| PVRL4 | 5 | 0.1347 | 0.27979 | 0.999632 | 6043 | 3 | -1.2372 | 0.56884 | 0.75109 | 1 | 15811 | 1 | -1.2372 |
| STEAP1B | 5 | 0.20847 | 0.38723 | 0.999994 | 7989 | 3 | -0.75758 | 0.56884 | 0.75109 | 1 | 15812 | 2 | -0.75758 |
| SRSF10 | 5 | 0.0044414 | 0.01859 | 0.801598 | 476 | 4 | -4.939 | 0.56884 | 0.75109 | 1 | 15813 | 1 | -4.939 |
| CRHR1 | 4 | 0.46323 | 0.60023 | 0.999994 | 12909 | 1 | 0.55523 | 0.56885 | 0.71815 | 1 | 15814 | 1 | 0.55523 |
| PROK2 | 6 | 0.019136 | 0.069208 | 0.93914 | 1416 | 3 | -0.62251 | 0.56909 | 0.77933 | 1 | 15815 | 2 | -0.62251 |
| MYOCD | 4 | 0.8431 | 0.84258 | 0.999994 | 18140 | 0 | 0.39162 | 0.56923 | 0.71844 | 1 | 15816 | 2 | 0.39162 |
| KBTBD4 | 4 | 0.80333 | 0.80419 | 0.999994 | 17531 | 1 | 0.42386 | 0.56929 | 0.71848 | 1 | 15817 | 2 | 0.42386 |
| LRGUK | 5 | 0.1405 | 0.28857 | 0.999632 | 6203 | 3 | -1.0883 | 0.56943 | 0.75141 | 1 | 15818 | 2 | -1.0883 |
| KRTAP4-11 | 6 | 0.041051 | 0.12427 | 0.951052 | 2563 | 3 | -1.4052 | 0.56951 | 0.77966 | 1 | 15819 | 2 | -1.4052 |
| NonTargetingControlG | 2 | 0.035313 | 0.064134 | 0.93914 | 2282 | 1 | -3.8034 | 0.56953 | 0.59017 | 1 | 15820 | 1 | -3.8034 |
| MPZL3 | 6 | 0.55495 | 0.70719 | 0.999994 | 14288 | 2 | 0.51897 | 0.56954 | 0.77969 | 1 | 15821 | 3 | 0.51897 |
| ATP6V1C2 | 6 | 0.17157 | 0.35267 | 0.999994 | 7025 | 2 | 0.36483 | 0.5696 | 0.77972 | 1 | 15822 | 2 | 0.36483 |
| NAT16 | 5 | 0.29246 | 0.47493 | 0.999994 | 9964 | 1 | 0.35554 | 0.56961 | 0.7515 | 1 | 15823 | 2 | 0.35554 |
| NADK | 5 | 0.041914 | 0.12069 | 0.951052 | 2608 | 2 | 0.091718 | 0.56981 | 0.75161 | 1 | 15824 | 2 | 0.091718 |
| APOPT1 | 6 | 0.90099 | 0.90445 | 0.999994 | 19233 | 1 | 0.56128 | 0.56983 | 0.77988 | 1 | 15825 | 3 | 0.56128 |
| SLC2A3 | 6 | 0.44467 | 0.63154 | 0.999994 | 12643 | 2 | 0.73572 | 0.56989 | 0.77992 | 1 | 15826 | 3 | 0.73572 |
| ZNF250 | 6 | 0.29619 | 0.50226 | 0.999994 | 10040 | 3 | -0.3612 | 0.56996 | 0.77996 | 1 | 15827 | 2 | -0.3612 |
| DNAJC3 | 5 | 0.2242 | 0.40912 | 0.999994 | 8384 | 1 | -0.071629 | 0.57 | 0.75171 | 1 | 15828 | 1 | -0.071629 |
| C11orf68 | 5 | 0.21399 | 0.39494 | 0.999994 | 8113 | 1 | -0.16033 | 0.57 | 0.75171 | 1 | 15829 | 2 | -0.16033 |
| ST6GALNAC6 | 6 | 0.44306 | 0.63048 | 0.999994 | 12625 | 2 | 0.30903 | 0.57 | 0.78 | 1 | 15830 | 3 | 0.30903 |
| PLB1 | 6 | 0.2029 | 0.3914 | 0.999994 | 7853 | 2 | 0.29876 | 0.57013 | 0.78011 | 1 | 15831 | 3 | 0.29876 |
| GATA6 | 6 | 0.26779 | 0.46909 | 0.999994 | 9419 | 2 | 0.3421 | 0.57031 | 0.78023 | 1 | 15832 | 3 | 0.3421 |
| NBPF15 | 4 | 0.80695 | 0.80751 | 0.999994 | 17583 | 1 | 0.43949 | 0.57051 | 0.71943 | 1 | 15833 | 2 | 0.43949 |
| FLT3 | 6 | 0.088866 | 0.22527 | 0.983354 | 4547 | 3 | -0.7577 | 0.57069 | 0.78052 | 1 | 15834 | 2 | -0.7577 |
| RAB5A | 6 | 0.2807 | 0.48422 | 0.999994 | 9714 | 3 | -0.14555 | 0.57069 | 0.78052 | 1 | 15835 | 2 | -0.14555 |
| LRBA | 6 | 0.26447 | 0.46516 | 0.999994 | 9345 | 3 | -0.25074 | 0.57069 | 0.78052 | 1 | 15836 | 2 | -0.25074 |
| NonTargetingControlG | 2 | 0.68657 | 0.68641 | 0.999994 | 16175 | 0 | 1.0033 | 0.57089 | 0.59121 | 1 | 15837 | 1 | 1.0033 |
| P4HA1 | 6 | 0.32275 | 0.53305 | 0.999994 | 10570 | 3 | 0.10251 | 0.57096 | 0.78076 | 1 | 15838 | 3 | 0.10251 |
| ARHGEF11 | 5 | 0.068531 | 0.17605 | 0.968171 | 3774 | 3 | -1.732 | 0.57109 | 0.75229 | 1 | 15839 | 1 | -1.732 |
| SLC39A8 | 6 | 0.64284 | 0.77209 | 0.999994 | 15559 | 2 | 0.53029 | 0.5711 | 0.78084 | 1 | 15840 | 3 | 0.53029 |
| NonTargetingControlG | 2 | 0.4636 | 0.47133 | 0.999994 | 12913 | 1 | 0.28757 | 0.57119 | 0.59143 | 1 | 15841 | 1 | 0.28757 |
| CPA2 | 6 | 0.1356 | 0.30743 | 0.999994 | 6070 | 2 | 0.57129 | 0.57121 | 0.78094 | 1 | 15842 | 3 | 0.57129 |
| TFAP2E | 4 | 0.006273 | 0.021832 | 0.819701 | 611 | 3 | -4.1799 | 0.57123 | 0.72 | 1 | 15843 | 1 | -4.1799 |
| NXPH3 | 4 | 0.15369 | 0.29299 | 0.999994 | 6556 | 2 | -0.14919 | 0.57143 | 0.72018 | 1 | 15844 | 2 | -0.14919 |
| MBIP | 2 | 0.69285 | 0.69267 | 0.999994 | 16232 | 0 | 0.91736 | 0.57154 | 0.59169 | 1 | 15845 | 1 | 0.91736 |
| CKAP2L | 4 | 0.056495 | 0.1341 | 0.95318 | 3270 | 2 | -1.5714 | 0.57162 | 0.72031 | 1 | 15846 | 2 | -1.5714 |
| FAM114A1 | 6 | 0.44012 | 0.62849 | 0.999994 | 12578 | 2 | 0.45502 | 0.57163 | 0.78123 | 1 | 15847 | 3 | 0.45502 |
| IL2RG | 6 | 0.17147 | 0.35256 | 0.999994 | 7023 | 2 | 0.44751 | 0.57172 | 0.7813 | 1 | 15848 | 3 | 0.44751 |
| CSNK1G3 | 6 | 0.16234 | 0.34121 | 0.999994 | 6803 | 2 | 0.63348 | 0.57172 | 0.7813 | 1 | 15849 | 3 | 0.63348 |
| EDN3 | 5 | 0.054905 | 0.15223 | 0.967126 | 3195 | 3 | -2.2351 | 0.57177 | 0.75269 | 1 | 15850 | 2 | -2.2351 |
| NHEJ1 | 6 | 0.18757 | 0.37251 | 0.999994 | 7410 | 3 | -0.31478 | 0.57191 | 0.78145 | 1 | 15851 | 3 | -0.31478 |
| GIMAP7 | 6 | 0.37972 | 0.5894 | 0.999994 | 11645 | 2 | -0.15483 | 0.57199 | 0.78151 | 1 | 15852 | 2 | -0.15483 |
| COTL1 | 6 | 0.50726 | 0.67376 | 0.999994 | 13541 | 2 | 0.44357 | 0.57214 | 0.78162 | 1 | 15853 | 3 | 0.44357 |
| DPP8 | 6 | 0.81811 | 0.85593 | 0.999994 | 17749 | 1 | -0.032246 | 0.57226 | 0.78172 | 1 | 15854 | 2 | -0.032246 |
| F13B | 5 | 0.037492 | 0.10958 | 0.951052 | 2400 | 2 | 0.41871 | 0.5723 | 0.75297 | 1 | 15855 | 2 | 0.41871 |
| BMP8B | 5 | 0.90812 | 0.90814 | 0.999994 | 19387 | 0 | 0.05721 | 0.5723 | 0.75297 | 1 | 15856 | 1 | 0.05721 |
| TACR1 | 5 | 0.085341 | 0.20312 | 0.97798 | 4424 | 3 | -0.80009 | 0.5723 | 0.75297 | 1 | 15857 | 2 | -0.80009 |
| MCF2L2 | 5 | 0.16249 | 0.32119 | 0.999994 | 6808 | 2 | 0.049029 | 0.5724 | 0.75303 | 1 | 15858 | 2 | 0.049029 |
| ATOH8 | 4 | 0.4066 | 0.55246 | 0.999994 | 12052 | 2 | -0.064359 | 0.57275 | 0.72119 | 1 | 15859 | 1 | -0.064359 |

| | | | | | | | | | | | | |
|---|---|---|---|---|---|---|---|---|---|---|---|---|
| SCGB2A2 | 6 | 0.84614 | 0.86987 | 0.999994 | 18189 | 1 | 0.35844 | 0.57279 | 0.7821 | 1 | 15860 | 3 | 0.35844 |
| KLHL25 | 6 | 0.56897 | 0.7172 | 0.999994 | 14522 | 2 | 0.14441 | 0.5728 | 0.78211 | 1 | 15861 | 3 | 0.14441 |
| CLIP2 | 6 | 0.095568 | 0.23864 | 0.987253 | 4801 | 4 | -1.4742 | 0.57292 | 0.7822 | 1 | 15862 | 2 | -1.4742 |
| UTS2 | 6 | 0.12215 | 0.28983 | 0.999632 | 5707 | 2 | 0.074566 | 0.57292 | 0.7822 | 1 | 15863 | 2 | 0.074566 |
| ST6GAL2 | 6 | 0.3122 | 0.5209 | 0.999994 | 10364 | 3 | -0.60627 | 0.57292 | 0.7822 | 1 | 15864 | 1 | -0.60627 |
| C14orf142 | 6 | 0.18927 | 0.3746 | 0.999994 | 7460 | 3 | -0.91192 | 0.57294 | 0.78221 | 1 | 15865 | 2 | -0.91192 |
| FAM189A2 | 6 | 0.082381 | 0.21222 | 0.979587 | 4312 | 3 | -0.13055 | 0.57297 | 0.78224 | 1 | 15866 | 3 | -0.13055 |
| ANGPTL1 | 5 | 0.48862 | 0.63801 | 0.999994 | 13261 | 2 | 0.089182 | 0.57299 | 0.75334 | 1 | 15867 | 2 | 0.089182 |
| CDC16 | 4 | 0.078278 | 0.17554 | 0.968171 | 4160 | 2 | -1.3148 | 0.57305 | 0.72143 | 1 | 15868 | 2 | -1.3148 |
| GPR31 | 6 | 0.096316 | 0.24011 | 0.987253 | 4828 | 4 | -1.3035 | 0.57325 | 0.78246 | 1 | 15869 | 2 | -1.3035 |
| DDI2 | 6 | 0.97155 | 0.97151 | 0.999994 | 20860 | 0 | 0.64437 | 0.57345 | 0.78262 | 1 | 15870 | 3 | 0.64437 |
| GHDC | 6 | 0.038745 | 0.11906 | 0.951052 | 2450 | 3 | -1.6587 | 0.57349 | 0.78265 | 1 | 15871 | 2 | -1.6587 |
| CDC42EP4 | 5 | 0.48145 | 0.63191 | 0.999994 | 13161 | 2 | -0.16803 | 0.57353 | 0.75363 | 1 | 15872 | 1 | -0.16803 |
| B4GALNT2 | 5 | 0.28122 | 0.46586 | 0.999994 | 9727 | 2 | -0.46257 | 0.57362 | 0.75368 | 1 | 15873 | 2 | -0.46257 |
| NonTargetingControlG | 2 | 0.43226 | 0.44396 | 0.999994 | 12449 | 1 | 0.17111 | 0.57362 | 0.59335 | 1 | 15874 | 1 | 0.17111 |
| DIRC2 | 4 | 0.23028 | 0.37183 | 0.999994 | 8534 | 2 | -0.070821 | 0.5737 | 0.72195 | 1 | 15875 | 2 | -0.070821 |
| PSG4 | 6 | 0.09339 | 0.2343 | 0.984971 | 4736 | 3 | -0.6646 | 0.57376 | 0.78285 | 1 | 15876 | 2 | -0.6646 |
| POLB | 5 | 0.60891 | 0.72941 | 0.999994 | 15056 | 1 | 0.50833 | 0.57378 | 0.75378 | 1 | 15877 | 2 | 0.50833 |
| ZNF292 | 6 | 0.30933 | 0.5176 | 0.999994 | 10310 | 2 | -0.16335 | 0.57379 | 0.78288 | 1 | 15878 | 3 | -0.16335 |
| hsa-mir-7112-1 | 4 | 0.59758 | 0.664 | 0.999994 | 14922 | 1 | 0.7291 | 0.57386 | 0.72207 | 1 | 15879 | 2 | 0.7291 |
| NUDT11 | 6 | 0.12154 | 0.28863 | 0.999632 | 5688 | 4 | -1.3012 | 0.57392 | 0.78298 | 1 | 15880 | 2 | -1.3012 |
| DYX1C1 | 6 | 0.31544 | 0.52466 | 0.999994 | 10431 | 1 | 0.44456 | 0.57397 | 0.78301 | 1 | 15881 | 3 | 0.44456 |
| KHDC1 | 6 | 0.92711 | 0.92718 | 0.999994 | 19791 | 1 | 0.63016 | 0.57399 | 0.78303 | 1 | 15882 | 3 | 0.63016 |
| SMYD5 | 5 | 0.095833 | 0.21984 | 0.980699 | 4807 | 3 | -1.6614 | 0.574 | 0.75389 | 1 | 15883 | 2 | -1.6614 |
| LRIG1 | 6 | 0.51632 | 0.67997 | 0.999994 | 13684 | 2 | 0.46744 | 0.57401 | 0.78304 | 1 | 15884 | 3 | 0.46744 |
| OR8H1 | 6 | 0.39672 | 0.60024 | 0.999994 | 11901 | 2 | -0.42631 | 0.57406 | 0.78309 | 1 | 15885 | 2 | -0.42631 |
| hsa-mir-6862-1 | 4 | 0.060558 | 0.14184 | 0.959146 | 3445 | 3 | -1.5536 | 0.57406 | 0.72223 | 1 | 15886 | 1 | -1.5536 |
| EIF4G1 | 4 | 0.22308 | 0.36442 | 0.999994 | 8360 | 2 | -0.82369 | 0.57406 | 0.72223 | 1 | 15887 | 1 | -0.82369 |
| KIAA1211L | 5 | 0.17537 | 0.33995 | 0.999994 | 7118 | 3 | -1.0543 | 0.57419 | 0.75401 | 1 | 15888 | 2 | -1.0543 |
| FAM134C | 5 | 0.032178 | 0.096053 | 0.945031 | 2111 | 4 | -2.6975 | 0.57431 | 0.75407 | 1 | 15889 | 1 | -2.6975 |
| FAM71B | 4 | 0.79327 | 0.79523 | 0.999994 | 17388 | 1 | 0.70656 | 0.57436 | 0.72248 | 1 | 15890 | 2 | 0.70656 |
| CEP85L | 6 | 0.11793 | 0.2818 | 0.999632 | 5572 | 2 | 0.060042 | 0.57439 | 0.78334 | 1 | 15891 | 2 | 0.060042 |
| KLRF2 | 6 | 0.79941 | 0.84762 | 0.999994 | 17473 | 1 | 0.36797 | 0.57448 | 0.7834 | 1 | 15892 | 3 | 0.36797 |
| hsa-mir-3662 | 3 | 0.40964 | 0.48996 | 0.999994 | 12102 | 1 | -0.2067 | 0.57455 | 0.67608 | 1 | 15893 | 1 | -0.2067 |
| RAD51C | 6 | 0.0095506 | 0.037449 | 0.881473 | 844 | 2 | 0.49384 | 0.57456 | 0.78346 | 1 | 15894 | 3 | 0.49384 |
| SLC26A5 | 6 | 0.12068 | 0.28701 | 0.999632 | 5650 | 4 | -0.83844 | 0.57459 | 0.78348 | 1 | 15895 | 2 | -0.83844 |
| PNPO | 6 | 0.19725 | 0.38454 | 0.999994 | 7679 | 3 | 0.021915 | 0.5746 | 0.78349 | 1 | 15896 | 3 | 0.021915 |
| AHRR | 4 | 0.28252 | 0.42513 | 0.999994 | 9748 | 1 | 0.29336 | 0.57484 | 0.72285 | 1 | 15897 | 2 | 0.29336 |
| FOLH1 | 6 | 0.53372 | 0.69216 | 0.999994 | 13963 | 2 | 0.18249 | 0.57493 | 0.78373 | 1 | 15898 | 2 | 0.18249 |
| MINOS1 | 5 | 0.013193 | 0.04513 | 0.891621 | 1077 | 4 | -1.8199 | 0.57506 | 0.75447 | 1 | 15899 | 1 | -1.8199 |
| MRPL39 | 5 | 0.030843 | 0.092666 | 0.943968 | 2053 | 3 | -1.1982 | 0.57506 | 0.75447 | 1 | 15900 | 1 | -1.1982 |
| SDCBP2 | 5 | 0.80599 | 0.82329 | 0.999994 | 17573 | 1 | -0.33041 | 0.57506 | 0.75447 | 1 | 15901 | 2 | -0.33041 |
| C22orf23 | 3 | 0.70381 | 0.70461 | 0.999994 | 16349 | 1 | 0.26112 | 0.57511 | 0.67636 | 1 | 15902 | 1 | 0.26112 |
| ALAS2 | 5 | 0.22169 | 0.40561 | 0.999994 | 8325 | 3 | -0.71597 | 0.57519 | 0.75454 | 1 | 15903 | 2 | -0.71597 |
| SST | 4 | 0.27544 | 0.4179 | 0.999994 | 9605 | 2 | -0.0088474 | 0.57525 | 0.72315 | 1 | 15904 | 2 | -0.0088474 |
| FUT9 | 6 | 0.36066 | 0.57605 | 0.999994 | 11348 | 2 | 0.51371 | 0.57526 | 0.78397 | 1 | 15905 | 3 | 0.51371 |
| NonTargetingControlG | 2 | 0.62823 | 0.628 | 0.999994 | 15352 | 0 | 0.75144 | 0.57534 | 0.59472 | 1 | 15906 | 1 | 0.75144 |
| NonTargetingControlG | 2 | 0.021503 | 0.039881 | 0.891621 | 1550 | 1 | -4.1846 | 0.57543 | 0.59479 | 1 | 15907 | 1 | -4.1846 |
| NonTargetingControlG | 1 | 0.42455 | 0.42473 | 0.999994 | 12327 | 0 | -0.097228 | 0.57545 | 0.57514 | 1 | 15908 | 0 | -0.097228 |
| DNAL1 | 2 | 0.59458 | 0.59451 | 0.999994 | 14884 | 0 | 0.69749 | 0.57548 | 0.59483 | 1 | 15909 | 1 | 0.69749 |
| PPAP2A | 6 | 0.067459 | 0.18145 | 0.968171 | 3734 | 3 | -1.1387 | 0.57554 | 0.78417 | 1 | 15910 | 2 | -1.1387 |
| NOTO | 6 | 0.040887 | 0.12392 | 0.951052 | 2555 | 2 | 0.39186 | 0.57554 | 0.78417 | 1 | 15911 | 3 | 0.39186 |

| | | | | | | | | | | | | |
|---|---|---|---|---|---|---|---|---|---|---|---|---|
| SPAG16 | 6 | 0.041464 | 0.12519 | 0.951052 | 2588 | 3 | -1.1445 | 0.57565 | 0.78426 | 1 | 15912 | 3 | -1.1445 |
| SAMD8 | 6 | 0.70075 | 0.81224 | 0.999994 | 16312 | 1 | 0.52847 | 0.57581 | 0.78439 | 1 | 15913 | 3 | 0.52847 |
| SEMA3B | 6 | 0.050503 | 0.14514 | 0.963012 | 2993 | 2 | 0.13694 | 0.57609 | 0.78459 | 1 | 15914 | 2 | 0.13694 |
| HOXB2 | 5 | 0.31858 | 0.49618 | 0.999994 | 10489 | 2 | 0.65682 | 0.57628 | 0.75515 | 1 | 15915 | 2 | 0.65682 |
| SYPL1 | 5 | 0.37604 | 0.54344 | 0.999994 | 11592 | 2 | -0.42262 | 0.57628 | 0.75515 | 1 | 15916 | 1 | -0.42262 |
| IPO5 | 5 | 0.16521 | 0.32509 | 0.999994 | 6867 | 3 | -0.99271 | 0.57628 | 0.75515 | 1 | 15917 | 1 | -0.99271 |
| hsa-mir-3910-1 | 4 | 0.80192 | 0.80292 | 0.999994 | 17509 | 1 | 0.48784 | 0.57634 | 0.72401 | 1 | 15918 | 2 | 0.48784 |
| TMEM79 | 6 | 0.28098 | 0.48453 | 0.999994 | 9722 | 3 | -0.31808 | 0.57643 | 0.78484 | 1 | 15919 | 1 | -0.31808 |
| SNX4 | 6 | 0.21798 | 0.4097 | 0.999994 | 8226 | 3 | -0.026832 | 0.57652 | 0.78491 | 1 | 15920 | 3 | -0.026832 |
| HMOX2 | 6 | 0.66358 | 0.78808 | 0.999994 | 15830 | 2 | 0.095409 | 0.57658 | 0.78495 | 1 | 15921 | 3 | 0.095409 |
| ACAT2 | 6 | 0.81712 | 0.85546 | 0.999994 | 17731 | 1 | 0.50171 | 0.5766 | 0.78497 | 1 | 15922 | 3 | 0.50171 |
| C3orf80 | 4 | 0.81267 | 0.81282 | 0.999994 | 17664 | 1 | 0.028342 | 0.57675 | 0.72432 | 1 | 15923 | 1 | 0.028342 |
| F9 | 6 | 0.78035 | 0.83979 | 0.999994 | 17217 | 1 | 0.61908 | 0.57682 | 0.78514 | 1 | 15924 | 3 | 0.61908 |
| ATG13 | 6 | 0.26057 | 0.46052 | 0.999994 | 9252 | 2 | -0.31859 | 0.57687 | 0.78518 | 1 | 15925 | 2 | -0.31859 |
| FGF13 | 5 | 0.18512 | 0.35397 | 0.999994 | 7344 | 2 | -0.018886 | 0.57694 | 0.75552 | 1 | 15926 | 2 | -0.018886 |
| MCAT | 6 | 0.022661 | 0.0805 | 0.93914 | 1631 | 3 | -0.5037 | 0.57703 | 0.7853 | 1 | 15927 | 3 | -0.5037 |
| EPB41 | 4 | 0.040461 | 0.10271 | 0.946414 | 2534 | 1 | 0.60354 | 0.57707 | 0.72457 | 1 | 15928 | 2 | 0.60354 |
| GNAT3 | 5 | 0.3262 | 0.50239 | 0.999994 | 10645 | 1 | 0.080051 | 0.57708 | 0.7556 | 1 | 15929 | 2 | 0.080051 |
| CNPY4 | 6 | 0.15209 | 0.3283 | 0.999994 | 6507 | 3 | -0.36621 | 0.57711 | 0.78535 | 1 | 15930 | 3 | -0.36621 |
| RNASET2 | 6 | 0.10201 | 0.25126 | 0.993254 | 5041 | 3 | -0.65739 | 0.57718 | 0.7854 | 1 | 15931 | 3 | -0.65739 |
| TBX2 | 4 | 0.56217 | 0.64578 | 0.999994 | 14411 | 1 | 0.43581 | 0.57727 | 0.72471 | 1 | 15932 | 2 | 0.43581 |
| KIAA0930 | 6 | 0.32705 | 0.53789 | 0.999994 | 10663 | 1 | 0.48481 | 0.57728 | 0.78548 | 1 | 15933 | 3 | 0.48481 |
| MTHFD2L | 4 | 0.38907 | 0.53442 | 0.999994 | 11784 | 2 | 0.039142 | 0.57729 | 0.72472 | 1 | 15934 | 2 | 0.039142 |
| FAM157B | 3 | 0.25356 | 0.37938 | 0.999994 | 9093 | 2 | -0.78185 | 0.5773 | 0.67737 | 1 | 15935 | 1 | -0.78185 |
| TRIP11 | 6 | 0.655 | 0.78146 | 0.999994 | 15718 | 2 | 0.15471 | 0.57731 | 0.78551 | 1 | 15936 | 3 | 0.15471 |
| PEX13 | 5 | 0.053409 | 0.14861 | 0.965554 | 3134 | 2 | -0.27161 | 0.57734 | 0.75574 | 1 | 15937 | 1 | -0.27161 |
| CD96 | 5 | 0.93326 | 0.93351 | 0.999994 | 19932 | 0 | 0.12685 | 0.57734 | 0.75574 | 1 | 15938 | 2 | 0.12685 |
| ZNF593 | 6 | 0.08236 | 0.21217 | 0.979587 | 4308 | 3 | -1.1557 | 0.57737 | 0.78555 | 1 | 15939 | 2 | -1.1557 |
| MYLPF | 6 | 0.068097 | 0.18276 | 0.968526 | 3754 | 3 | -1.5992 | 0.57744 | 0.7856 | 1 | 15940 | 2 | -1.5992 |
| hsa-mir-4666b | 4 | 0.20604 | 0.34692 | 0.999994 | 7929 | 2 | -0.35085 | 0.57749 | 0.72488 | 1 | 15941 | 2 | -0.35085 |
| TGFBR2 | 6 | 0.068956 | 0.18458 | 0.968526 | 3797 | 1 | 0.41241 | 0.57771 | 0.78579 | 1 | 15942 | 3 | 0.41241 |
| RPRM | 6 | 0.23686 | 0.43224 | 0.999994 | 8693 | 3 | -0.19715 | 0.57771 | 0.78579 | 1 | 15943 | 3 | -0.19715 |
| hsa-mir-516a-2 | 1 | 0.42222 | 0.4224 | 0.999994 | 12286 | 0 | -0.097701 | 0.57778 | 0.57747 | 1 | 15944 | 0 | -0.097701 |
| hsa-mir-6822 | 4 | 0.19768 | 0.33829 | 0.999994 | 7695 | 2 | -0.25302 | 0.57784 | 0.72516 | 1 | 15945 | 2 | -0.25302 |
| BAGE | 5 | 0.49916 | 0.64718 | 0.999994 | 13411 | 1 | 0.33043 | 0.57784 | 0.75598 | 1 | 15946 | 2 | 0.33043 |
| HDGFRP3 | 6 | 0.66363 | 0.78811 | 0.999994 | 15832 | 2 | 0.50919 | 0.57788 | 0.78591 | 1 | 15947 | 3 | 0.50919 |
| LIG3 | 6 | 0.063784 | 0.17378 | 0.96793 | 3600 | 3 | -0.815 | 0.57792 | 0.78594 | 1 | 15948 | 3 | -0.815 |
| CLK4 | 6 | 0.02881 | 0.096029 | 0.945031 | 1943 | 1 | 0.51579 | 0.57799 | 0.786 | 1 | 15949 | 3 | 0.51579 |
| FCRL3 | 4 | 0.30092 | 0.44396 | 0.999994 | 10148 | 2 | -0.085464 | 0.57827 | 0.72549 | 1 | 15950 | 2 | -0.085464 |
| SPANXF1 | 4 | 0.0059583 | 0.020791 | 0.81027 | 589 | 2 | -4.5236 | 0.57828 | 0.72551 | 1 | 15951 | 1 | -4.5236 |
| MRPL16 | 4 | 0.48723 | 0.61068 | 0.999994 | 13240 | 1 | -0.39063 | 0.57828 | 0.72551 | 1 | 15952 | 1 | -0.39063 |
| ACRV1 | 6 | 0.25051 | 0.44852 | 0.999994 | 9024 | 2 | 0.064338 | 0.57836 | 0.78628 | 1 | 15953 | 3 | 0.064338 |
| FAM154B | 6 | 0.52642 | 0.68701 | 0.999994 | 13834 | 2 | -0.1219 | 0.57838 | 0.7863 | 1 | 15954 | 2 | -0.1219 |
| PARP11 | 6 | 0.030559 | 0.10009 | 0.946394 | 2032 | 4 | -1.8643 | 0.57846 | 0.78635 | 1 | 15955 | 2 | -1.8643 |
| JOSD1 | 6 | 0.10069 | 0.24866 | 0.990929 | 4997 | 2 | 0.010243 | 0.57846 | 0.78635 | 1 | 15956 | 2 | 0.010243 |
| TTC26 | 6 | 0.24789 | 0.44541 | 0.999994 | 8972 | 3 | -0.69632 | 0.57846 | 0.78635 | 1 | 15957 | 1 | -0.69632 |
| hsa-mir-6879 | 1 | 0.42151 | 0.4217 | 0.999994 | 12268 | 0 | -0.10014 | 0.57849 | 0.57821 | 1 | 15958 | 0 | -0.10014 |
| hsa-mir-34c | 4 | 0.81416 | 0.81422 | 0.999994 | 17693 | 1 | 0.26992 | 0.5785 | 0.72567 | 1 | 15959 | 2 | 0.26992 |
| NOA1 | 4 | 0.0014135 | 0.0051663 | 0.595086 | 212 | 1 | 0.65519 | 0.57861 | 0.72577 | 1 | 15960 | 2 | 0.65519 |
| UGCG | 6 | 0.80547 | 0.85024 | 0.999994 | 17559 | 1 | 0.56191 | 0.57867 | 0.78652 | 1 | 15961 | 3 | 0.56191 |
| CCDC81 | 6 | 0.84886 | 0.87133 | 0.999994 | 18233 | 1 | 0.41353 | 0.57877 | 0.7866 | 1 | 15962 | 3 | 0.41353 |
| PIK3R5 | 5 | 0.033373 | 0.099162 | 0.945253 | 2180 | 4 | -1.0096 | 0.57878 | 0.75654 | 1 | 15963 | 1 | -1.0096 |

| | | | | | | | | | | | | |
|---|---|---|---|---|---|---|---|---|---|---|---|---|
| TTC27 | 5 | 0.0021447 | 0.0093791 | 0.680529 | 284 | 4 | -6.8082 | 0.57878 | 0.75654 | 1 | 15964 | 1 | -6.8082 |
| SLTM | 2 | 0.75075 | 0.75033 | 0.999994 | 16861 | 0 | 1.056 | 0.57884 | 0.59748 | 1 | 15965 | 1 | 1.056 |
| ENPP6 | 6 | 0.09423 | 0.236 | 0.985362 | 4762 | 4 | -1.5517 | 0.57885 | 0.78666 | 1 | 15966 | 2 | -1.5517 |
| QARS | 3 | 0.037619 | 0.092423 | 0.943968 | 2409 | 1 | 0.45499 | 0.57909 | 0.6782 | 1 | 15967 | 1 | 0.45499 |
| ANKRD32 | 4 | 0.83623 | 0.83573 | 0.999994 | 18024 | 0 | 0.47796 | 0.57913 | 0.72619 | 1 | 15968 | 2 | 0.47796 |
| hsa-mir-6874 | 4 | 0.14541 | 0.28448 | 0.999632 | 6336 | 2 | -0.62185 | 0.57915 | 0.72622 | 1 | 15969 | 2 | -0.62185 |
| hsa-mir-491 | 4 | 0.069183 | 0.15834 | 0.967126 | 3805 | 2 | -0.33871 | 0.57918 | 0.72624 | 1 | 15970 | 2 | -0.33871 |
| TBKBP1 | 6 | 0.58968 | 0.73236 | 0.999994 | 14817 | 1 | 0.58474 | 0.57922 | 0.78695 | 1 | 15971 | 3 | 0.58474 |
| CCDC94 | 4 | 0.19732 | 0.33794 | 0.999994 | 7681 | 2 | -0.59727 | 0.57948 | 0.72647 | 1 | 15972 | 2 | -0.59727 |
| RNASE3 | 5 | 0.019462 | 0.062642 | 0.937389 | 1427 | 3 | -2.7406 | 0.57955 | 0.75696 | 1 | 15973 | 2 | -2.7406 |
| HTR3A | 6 | 0.26637 | 0.4674 | 0.999994 | 9379 | 2 | -0.32748 | 0.57975 | 0.78735 | 1 | 15974 | 2 | -0.32748 |
| REST | 6 | 0.098178 | 0.24378 | 0.988214 | 4899 | 4 | -1.092 | 0.57975 | 0.78735 | 1 | 15975 | 1 | -1.092 |
| TGM4 | 6 | 0.4605 | 0.64212 | 0.999994 | 12855 | 2 | -0.21483 | 0.57975 | 0.78735 | 1 | 15976 | 2 | -0.21483 |
| TMEM144 | 6 | 0.013682 | 0.051413 | 0.910337 | 1110 | 5 | -1.5564 | 0.57975 | 0.78735 | 1 | 15977 | 1 | -1.5564 |
| FSBP | 6 | 0.014126 | 0.052882 | 0.916123 | 1136 | 4 | -2.7366 | 0.57975 | 0.78735 | 1 | 15978 | 2 | -2.7366 |
| GRM1 | 6 | 0.10371 | 0.25457 | 0.994835 | 5092 | 4 | -1.1533 | 0.57985 | 0.78743 | 1 | 15979 | 2 | -1.1533 |
| hsa-mir-4695 | 4 | 0.036961 | 0.095684 | 0.945031 | 2371 | 3 | -2.6131 | 0.58007 | 0.72693 | 1 | 15980 | 1 | -2.6131 |
| OTUD4 | 4 | 0.064372 | 0.14913 | 0.965739 | 3626 | 3 | -1.3505 | 0.58007 | 0.72693 | 1 | 15981 | 1 | -1.3505 |
| CLSTN3 | 4 | 0.19191 | 0.33236 | 0.999994 | 7545 | 1 | 0.0042019 | 0.58007 | 0.72693 | 1 | 15982 | 1 | 0.0042019 |
| PSKH2 | 5 | 0.2389 | 0.42943 | 0.999994 | 8745 | 2 | -1.2059 | 0.5801 | 0.75726 | 1 | 15983 | 2 | -1.2059 |
| XRCC5 | 5 | 0.036601 | 0.10734 | 0.950995 | 2350 | 3 | -2.0147 | 0.5801 | 0.75726 | 1 | 15984 | 2 | -2.0147 |
| HSPA1L | 5 | 0.21784 | 0.40025 | 0.999994 | 8225 | 3 | -0.7198 | 0.58013 | 0.75727 | 1 | 15985 | 2 | -0.7198 |
| MRPL43 | 3 | 0.13558 | 0.22985 | 0.983354 | 6068 | 1 | 0.30775 | 0.58016 | 0.6787 | 1 | 15986 | 1 | 0.30775 |
| JUP | 6 | 0.22857 | 0.42235 | 0.999994 | 8490 | 2 | 0.4422 | 0.58041 | 0.78783 | 1 | 15987 | 3 | 0.4422 |
| ZC3H7B | 5 | 0.26325 | 0.45137 | 0.999994 | 9319 | 2 | 0.44218 | 0.58056 | 0.7575 | 1 | 15988 | 2 | 0.44218 |
| BEST3 | 6 | 0.048354 | 0.14048 | 0.958716 | 2902 | 4 | -1.834 | 0.58075 | 0.78808 | 1 | 15989 | 2 | -1.834 |
| PSMB9 | 6 | 0.015622 | 0.057774 | 0.929498 | 1225 | 4 | -2.841 | 0.58075 | 0.78808 | 1 | 15990 | 2 | -2.841 |
| ACAP3 | 6 | 0.11991 | 0.28554 | 0.999632 | 5626 | 2 | -0.39768 | 0.58075 | 0.78808 | 1 | 15991 | 2 | -0.39768 |
| LOR | 6 | 0.19442 | 0.38106 | 0.999994 | 7619 | 3 | -0.15948 | 0.58075 | 0.78808 | 1 | 15992 | 3 | -0.15948 |
| STK40 | 6 | 0.96263 | 0.96269 | 0.999994 | 20649 | 0 | 0.36271 | 0.58091 | 0.78822 | 1 | 15993 | 3 | 0.36271 |
| hsa-mir-939 | 2 | 0.31827 | 0.35088 | 0.999994 | 10480 | 1 | -0.41425 | 0.58091 | 0.59915 | 1 | 15994 | 1 | -0.41425 |
| hsa-mir-6843 | 3 | 0.082269 | 0.15954 | 0.967126 | 4306 | 2 | -2.6122 | 0.58105 | 0.67911 | 1 | 15995 | 1 | -2.6122 |
| SSTR2 | 5 | 0.065241 | 0.17064 | 0.967177 | 3663 | 2 | 0.20497 | 0.58119 | 0.75785 | 1 | 15996 | 2 | 0.20497 |
| LYPD2 | 5 | 0.19176 | 0.36345 | 0.999994 | 7536 | 3 | -0.92707 | 0.58133 | 0.75793 | 1 | 15997 | 2 | -0.92707 |
| SMC4 | 4 | 0.81324 | 0.81335 | 0.999994 | 17679 | 1 | 0.3372 | 0.58138 | 0.72798 | 1 | 15998 | 1 | 0.3372 |
| DUSP3 | 4 | 0.20092 | 0.34162 | 0.999994 | 7786 | 2 | -0.70934 | 0.58138 | 0.72798 | 1 | 15999 | 1 | -0.70934 |
| CDH16 | 4 | 0.16463 | 0.30427 | 0.999994 | 6855 | 1 | -0.086366 | 0.58138 | 0.72798 | 1 | 16000 | 1 | -0.086366 |
| IMPDH2 | 4 | 0.019561 | 0.059716 | 0.932932 | 1432 | 1 | 0.32887 | 0.58143 | 0.72801 | 1 | 16001 | 2 | 0.32887 |
| hsa-mir-422a | 2 | 0.75903 | 0.75857 | 0.999994 | 16964 | 0 | 1.0335 | 0.58145 | 0.59955 | 1 | 16002 | 1 | 1.0335 |
| MFSD3 | 5 | 0.065241 | 0.17064 | 0.967177 | 3662 | 2 | 0.13275 | 0.58146 | 0.758 | 1 | 16003 | 1 | 0.13275 |
| ATP5H | 5 | 0.3226 | 0.4994 | 0.999994 | 10568 | 2 | 0.22343 | 0.58146 | 0.758 | 1 | 16004 | 2 | 0.22343 |
| MYLIP | 5 | 0.23727 | 0.42724 | 0.999994 | 8704 | 3 | -0.6237 | 0.58167 | 0.75812 | 1 | 16005 | 2 | -0.6237 |
| TMEM177 | 6 | 0.46455 | 0.64479 | 0.999994 | 12926 | 2 | -0.40592 | 0.5819 | 0.78895 | 1 | 16006 | 2 | -0.40592 |
| AGPAT4 | 6 | 0.0019501 | 0.0097256 | 0.680529 | 266 | 1 | 0.24375 | 0.5819 | 0.78895 | 1 | 16007 | 2 | 0.24375 |
| SLC39A14 | 4 | 0.79681 | 0.79837 | 0.999994 | 17437 | 1 | 0.58896 | 0.582 | 0.72846 | 1 | 16008 | 2 | 0.58896 |
| PCNXL2 | 6 | 0.18657 | 0.37125 | 0.999994 | 7374 | 2 | 0.38304 | 0.58208 | 0.78908 | 1 | 16009 | 3 | 0.38304 |
| BIRC6 | 6 | 0.48275 | 0.65702 | 0.999994 | 13186 | 2 | 0.16175 | 0.58214 | 0.78912 | 1 | 16010 | 3 | 0.16175 |
| QPCTL | 6 | 0.2056 | 0.39467 | 0.999994 | 7912 | 1 | 0.4506 | 0.58258 | 0.78946 | 1 | 16011 | 3 | 0.4506 |
| hsa-mir-3144 | 4 | 0.87654 | 0.8762 | 0.999994 | 18761 | 0 | 0.55712 | 0.5826 | 0.72889 | 1 | 16012 | 2 | 0.55712 |
| HNRNPL | 6 | 0.27999 | 0.4834 | 0.999994 | 9704 | 1 | 0.57003 | 0.58266 | 0.78952 | 1 | 16013 | 3 | 0.57003 |
| TESK2 | 5 | 0.55376 | 0.69511 | 0.999994 | 14266 | 1 | -0.16568 | 0.58273 | 0.75872 | 1 | 16014 | 2 | -0.16568 |
| PION | 2 | 0.31711 | 0.34997 | 0.999994 | 10455 | 1 | -0.39177 | 0.5828 | 0.60062 | 1 | 16015 | 1 | -0.39177 |

| | | | | | | | | | | | | | |
|---|---|---|---|---|---|---|---|---|---|---|---|---|---|
| KRTCAP3 | 6 | 0.15606 | 0.33327 | 0.999994 | 6629 | 3 | -0.4655 | 0.58292 | 0.78972 | 1 | 16016 | 3 | -0.4655 |
| RBM28 | 6 | 0.66123 | 0.78627 | 0.999994 | 15798 | 2 | 0.33657 | 0.58309 | 0.78984 | 1 | 16017 | 3 | 0.33657 |
| SERPIND1 | 6 | 0.85948 | 0.87714 | 0.999994 | 18426 | 1 | 0.6857 | 0.58309 | 0.78984 | 1 | 16018 | 3 | 0.6857 |
| EPB41L3 | 6 | 0.040074 | 0.12208 | 0.951052 | 2520 | 4 | -1.5635 | 0.58312 | 0.78987 | 1 | 16019 | 2 | -1.5635 |
| MYCL | 6 | 0.0042063 | 0.018232 | 0.795727 | 457 | 5 | -1.9897 | 0.58312 | 0.78987 | 1 | 16020 | 1 | -1.9897 |
| PAK1 | 6 | 0.083203 | 0.21386 | 0.979587 | 4348 | 3 | -1.1606 | 0.58312 | 0.78987 | 1 | 16021 | 1 | -1.1606 |
| COL25A1 | 3 | 0.74626 | 0.74621 | 0.999994 | 16816 | 0 | -0.13678 | 0.58313 | 0.68009 | 1 | 16022 | 1 | -0.13678 |
| UBE2W | 6 | 0.26242 | 0.46274 | 0.999994 | 9293 | 2 | 0.30906 | 0.5833 | 0.79002 | 1 | 16023 | 3 | 0.30906 |
| SCCPDH | 6 | 0.54949 | 0.70326 | 0.999994 | 14202 | 2 | 0.13104 | 0.58338 | 0.79007 | 1 | 16024 | 3 | 0.13104 |
| NonTargetingControlG | 2 | 0.45527 | 0.46392 | 0.999994 | 12777 | 1 | 0.19397 | 0.58354 | 0.60119 | 1 | 16025 | 1 | 0.19397 |
| hsa-mir-5682 | 4 | 0.33242 | 0.47623 | 0.999994 | 10778 | 2 | 0.044195 | 0.58359 | 0.72966 | 1 | 16026 | 2 | 0.044195 |
| hsa-mir-3168 | 4 | 0.40409 | 0.54992 | 0.999994 | 12020 | 2 | -0.19162 | 0.58374 | 0.72979 | 1 | 16027 | 1 | -0.19162 |
| NID2 | 4 | 0.67714 | 0.70977 | 0.999994 | 16045 | 1 | 0.17052 | 0.58374 | 0.72979 | 1 | 16028 | 1 | 0.17052 |
| C3orf17 | 4 | 0.025815 | 0.072953 | 0.93914 | 1792 | 2 | -3.6264 | 0.58374 | 0.72979 | 1 | 16029 | 1 | -3.6264 |
| FAM133B | 4 | 0.24394 | 0.38571 | 0.999994 | 8880 | 2 | -1.0791 | 0.58374 | 0.72979 | 1 | 16030 | 1 | -1.0791 |
| OR2M2 | 5 | 0.91666 | 0.91674 | 0.999994 | 19553 | 0 | 0.11388 | 0.58375 | 0.75927 | 1 | 16031 | 2 | 0.11388 |
| RIN1 | 6 | 0.61806 | 0.75337 | 0.999994 | 15188 | 2 | 0.28481 | 0.58398 | 0.79055 | 1 | 16032 | 3 | 0.28481 |
| FLG2 | 5 | 0.1559 | 0.31155 | 0.999994 | 6622 | 2 | -0.30289 | 0.58399 | 0.75941 | 1 | 16033 | 2 | -0.30289 |
| DTX3 | 6 | 0.0056409 | 0.023458 | 0.832699 | 566 | 3 | -2.6294 | 0.584 | 0.79056 | 1 | 16034 | 2 | -2.6294 |
| IRX2 | 5 | 0.86215 | 0.86419 | 0.999994 | 18489 | 1 | -0.42108 | 0.58403 | 0.75944 | 1 | 16035 | 1 | -0.42108 |
| COLEC12 | 6 | 0.027729 | 0.093467 | 0.945031 | 1892 | 4 | -3.2254 | 0.58404 | 0.79059 | 1 | 16036 | 2 | -3.2254 |
| STK36 | 6 | 0.18196 | 0.36561 | 0.999994 | 7269 | 2 | 0.11773 | 0.5841 | 0.79065 | 1 | 16037 | 3 | 0.11773 |
| DPY19L4 | 5 | 0.84677 | 0.85177 | 0.999994 | 18206 | 1 | -0.018083 | 0.58411 | 0.75948 | 1 | 16038 | 2 | -0.018083 |
| C17orf80 | 4 | 0.60211 | 0.6665 | 0.999994 | 14974 | 1 | 0.438 | 0.58412 | 0.73011 | 1 | 16039 | 2 | 0.438 |
| hsa-mir-1263 | 4 | 0.3788 | 0.52394 | 0.999994 | 11636 | 1 | 0.73128 | 0.58428 | 0.73024 | 1 | 16040 | 2 | 0.73128 |
| CCDC12 | 6 | 0.39897 | 0.6017 | 0.999994 | 11934 | 2 | -0.38099 | 0.58481 | 0.79117 | 1 | 16041 | 2 | -0.38099 |
| VEZT | 6 | 0.0074916 | 0.030149 | 0.862223 | 691 | 3 | -3.8684 | 0.58481 | 0.79117 | 1 | 16042 | 2 | -3.8684 |
| SCAF4 | 6 | 0.029485 | 0.097568 | 0.945253 | 1984 | 4 | -1.6353 | 0.58481 | 0.79117 | 1 | 16043 | 2 | -1.6353 |
| STXBP6 | 6 | 0.34436 | 0.55767 | 0.999994 | 11036 | 3 | 0.056238 | 0.58481 | 0.79117 | 1 | 16044 | 3 | 0.056238 |
| SLC5A3 | 6 | 0.0065725 | 0.026922 | 0.848412 | 631 | 2 | -0.12831 | 0.58481 | 0.79117 | 1 | 16045 | 2 | -0.12831 |
| CBLB | 6 | 0.39412 | 0.59858 | 0.999994 | 11862 | 2 | 0.38078 | 0.5849 | 0.79124 | 1 | 16046 | 3 | 0.38078 |
| PARP9 | 6 | 0.036691 | 0.11444 | 0.951052 | 2353 | 4 | -1.717 | 0.58502 | 0.79133 | 1 | 16047 | 1 | -1.717 |
| SBF1 | 6 | 0.13801 | 0.31052 | 0.999994 | 6137 | 2 | 0.62157 | 0.58502 | 0.79133 | 1 | 16048 | 3 | 0.62157 |
| ST6GALNAC1 | 6 | 0.3689 | 0.58252 | 0.999994 | 11487 | 2 | 0.41918 | 0.58526 | 0.79151 | 1 | 16049 | 3 | 0.41918 |
| CCDC62 | 5 | 0.0060267 | 0.024501 | 0.842094 | 594 | 3 | -6.6485 | 0.58527 | 0.76014 | 1 | 16050 | 2 | -6.6485 |
| SLC30A9 | 5 | 0.084176 | 0.20128 | 0.977164 | 4380 | 3 | -1.7148 | 0.5854 | 0.76022 | 1 | 16051 | 2 | -1.7148 |
| ZG16 | 4 | 0.067376 | 0.15487 | 0.967126 | 3731 | 3 | -1.231 | 0.58564 | 0.73134 | 1 | 16052 | 1 | -1.231 |
| IRF7 | 6 | 0.14086 | 0.31412 | 0.999994 | 6214 | 3 | -0.19062 | 0.58569 | 0.79182 | 1 | 16053 | 3 | -0.19062 |
| hsa-mir-586 | 2 | 0.62901 | 0.62879 | 0.999994 | 15363 | 0 | 0.62757 | 0.58578 | 0.60295 | 1 | 16054 | 1 | 0.62757 |
| PIGT | 6 | 0.86914 | 0.88282 | 0.999994 | 18619 | 1 | 0.59404 | 0.58582 | 0.79191 | 1 | 16055 | 3 | 0.59404 |
| FAM21B | 6 | 0.07376 | 0.19458 | 0.975797 | 3985 | 4 | -1.2278 | 0.58584 | 0.79192 | 1 | 16056 | 2 | -1.2278 |
| GABRA6 | 6 | 0.70504 | 0.81357 | 0.999994 | 16360 | 1 | 0.59797 | 0.58589 | 0.79197 | 1 | 16057 | 3 | 0.59797 |
| NDUFB7 | 4 | 0.65252 | 0.69474 | 0.999994 | 15693 | 1 | 0.48623 | 0.58593 | 0.73157 | 1 | 16058 | 2 | 0.48623 |
| UBR3 | 6 | 0.79443 | 0.84552 | 0.999994 | 17401 | 1 | 0.47612 | 0.58593 | 0.79201 | 1 | 16059 | 3 | 0.47612 |
| EFCAB7 | 6 | 0.20814 | 0.3978 | 0.999994 | 7981 | 2 | 0.59885 | 0.58604 | 0.7921 | 1 | 16060 | 3 | 0.59885 |
| TUFM | 2 | 0.3302 | 0.36022 | 0.999994 | 10725 | 1 | -1.1226 | 0.58609 | 0.6032 | 1 | 16061 | 1 | -1.1226 |
| NFATC2IP | 6 | 0.05112 | 0.14652 | 0.964852 | 3025 | 3 | -1.2267 | 0.58611 | 0.79215 | 1 | 16062 | 2 | -1.2267 |
| ZNF471 | 5 | 0.12794 | 0.26946 | 0.998561 | 5871 | 2 | 0.49109 | 0.58615 | 0.76066 | 1 | 16063 | 2 | 0.49109 |
| TACC2 | 4 | 0.033044 | 0.087711 | 0.94395 | 2161 | 3 | -2.166 | 0.58622 | 0.7318 | 1 | 16064 | 1 | -2.166 |
| ACTRT3 | 4 | 0.42186 | 0.56811 | 0.999994 | 12275 | 2 | 0.24428 | 0.58622 | 0.7318 | 1 | 16065 | 2 | 0.24428 |
| IL1B | 4 | 0.31477 | 0.45815 | 0.999994 | 10414 | 2 | -0.63089 | 0.58626 | 0.73182 | 1 | 16066 | 1 | -0.63089 |
| FBXW7 | 4 | 0.049043 | 0.11955 | 0.951052 | 2932 | 1 | 0.59256 | 0.58642 | 0.73194 | 1 | 16067 | 2 | 0.59256 |

| | | | | | | | | | | | | | |
|---|---|---|---|---|---|---|---|---|---|---|---|---|---|
| NonTargetingControlG | 2 | 0.040067 | 0.0724 | 0.93914 | 2518 | 1 | -3.7755 | 0.58652 | 0.60355 | 1 | 16068 | 1 | -3.7755 |
| NonTargetingControlG | 2 | 0.70341 | 0.70319 | 0.999994 | 16344 | 0 | 0.9471 | 0.58655 | 0.60357 | 1 | 16069 | 1 | 0.9471 |
| DHX57 | 5 | 0.045251 | 0.12893 | 0.952273 | 2777 | 3 | -2.3831 | 0.58655 | 0.7609 | 1 | 16070 | 2 | -2.3831 |
| BOC | 4 | 0.27098 | 0.41329 | 0.999994 | 9499 | 2 | -0.22359 | 0.58656 | 0.73205 | 1 | 16071 | 2 | -0.22359 |
| C2orf47 | 6 | 0.60738 | 0.74542 | 0.999994 | 15040 | 2 | 0.072346 | 0.58668 | 0.79257 | 1 | 16072 | 3 | 0.072346 |
| PATE4 | 6 | 0.54633 | 0.70104 | 0.999994 | 14156 | 2 | 0.50622 | 0.5867 | 0.79258 | 1 | 16073 | 3 | 0.50622 |
| ANO1 | 6 | 0.0073004 | 0.029473 | 0.862223 | 679 | 3 | -3.3812 | 0.58671 | 0.79259 | 1 | 16074 | 2 | -3.3812 |
| GCH1 | 6 | 0.06837 | 0.18337 | 0.968526 | 3760 | 2 | 0.65953 | 0.58672 | 0.79259 | 1 | 16075 | 3 | 0.65953 |
| ANAPC15 | 5 | 0.60088 | 0.72649 | 0.999994 | 14960 | 1 | 0.17621 | 0.58676 | 0.76101 | 1 | 16076 | 2 | 0.17621 |
| SCAND3 | 6 | 0.1915 | 0.37738 | 0.999994 | 7526 | 3 | -0.12706 | 0.58677 | 0.79263 | 1 | 16077 | 2 | -0.12706 |
| PCDHB14 | 4 | 0.60599 | 0.66856 | 0.999994 | 15027 | 1 | 0.65497 | 0.58696 | 0.73237 | 1 | 16078 | 2 | 0.65497 |
| ATP5F1 | 5 | 0.24169 | 0.4332 | 0.999994 | 8825 | 2 | -0.59714 | 0.58715 | 0.76124 | 1 | 16079 | 2 | -0.59714 |
| TFPI2 | 6 | 0.29661 | 0.50274 | 0.999994 | 10057 | 3 | -0.14007 | 0.58717 | 0.79295 | 1 | 16080 | 3 | -0.14007 |
| hsa-mir-4252 | 4 | 0.02794 | 0.077307 | 0.93914 | 1899 | 2 | -3.309 | 0.58718 | 0.73255 | 1 | 16081 | 2 | -3.309 |
| hsa-mir-4688 | 4 | 0.02341 | 0.06785 | 0.93914 | 1679 | 3 | -2.8749 | 0.58718 | 0.73255 | 1 | 16082 | 1 | -2.8749 |
| WRB | 4 | 0.73531 | 0.74903 | 0.999994 | 16666 | 1 | -0.13148 | 0.58718 | 0.73255 | 1 | 16083 | 1 | -0.13148 |
| PPP1R11 | 5 | 0.0096032 | 0.03493 | 0.867348 | 845 | 2 | 0.23656 | 0.58726 | 0.7613 | 1 | 16084 | 2 | 0.23656 |
| LRP6 | 6 | 0.096903 | 0.24129 | 0.987253 | 4853 | 3 | -0.13557 | 0.58744 | 0.79315 | 1 | 16085 | 2 | -0.13557 |
| GINS1 | 2 | 0.10539 | 0.17841 | 0.968171 | 5149 | 1 | -3.0205 | 0.58744 | 0.60427 | 1 | 16086 | 1 | -3.0205 |
| TMEM41A | 6 | 0.21492 | 0.40606 | 0.999994 | 8140 | 3 | -0.17101 | 0.58771 | 0.79336 | 1 | 16087 | 3 | -0.17101 |
| OR2M5 | 6 | 0.52741 | 0.68771 | 0.999994 | 13848 | 2 | 0.17075 | 0.58772 | 0.79336 | 1 | 16088 | 2 | 0.17075 |
| TSPAN2 | 5 | 0.19308 | 0.36533 | 0.999994 | 7577 | 3 | -0.67654 | 0.58777 | 0.76156 | 1 | 16089 | 1 | -0.67654 |
| LHX4 | 5 | 0.060147 | 0.16235 | 0.967126 | 3426 | 3 | -2.2053 | 0.58777 | 0.76156 | 1 | 16090 | 2 | -2.2053 |
| ZNF169 | 5 | 0.15969 | 0.31714 | 0.999994 | 6723 | 3 | -1.1083 | 0.58777 | 0.76156 | 1 | 16091 | 2 | -1.1083 |
| KRTAP9-6 | 5 | 0.19785 | 0.37218 | 0.999994 | 7706 | 3 | -0.83472 | 0.58777 | 0.76156 | 1 | 16092 | 2 | -0.83472 |
| LSM2 | 3 | 0.26108 | 0.38872 | 0.999994 | 9271 | 2 | -0.75062 | 0.58794 | 0.68235 | 1 | 16093 | 1 | -0.75062 |
| FARSA | 3 | 0.077757 | 0.15343 | 0.967126 | 4138 | 2 | -3.5125 | 0.58794 | 0.68235 | 1 | 16094 | 1 | -3.5125 |
| KRT222 | 6 | 0.03635 | 0.11368 | 0.951052 | 2338 | 4 | -1.1303 | 0.58802 | 0.7936 | 1 | 16095 | 2 | -1.1303 |
| PFDN1 | 6 | 0.24467 | 0.44157 | 0.999994 | 8893 | 2 | -0.23568 | 0.58802 | 0.7936 | 1 | 16096 | 2 | -0.23568 |
| ANLN | 6 | 0.064313 | 0.17487 | 0.968171 | 3621 | 3 | -0.48323 | 0.58802 | 0.7936 | 1 | 16097 | 1 | -0.48323 |
| NonTargetingControlG | 2 | 0.38465 | 0.40412 | 0.999994 | 11711 | 1 | -0.048029 | 0.58812 | 0.6048 | 1 | 16098 | 1 | -0.048029 |
| HSD17B3 | 5 | 0.572 | 0.7113 | 0.999994 | 14575 | 2 | -0.021698 | 0.58836 | 0.76189 | 1 | 16099 | 2 | -0.021698 |
| KIAA2022 | 5 | 0.23025 | 0.41756 | 0.999994 | 8530 | 2 | 0.47128 | 0.5884 | 0.76191 | 1 | 16100 | 2 | 0.47128 |
| SDK2 | 6 | 0.51608 | 0.67981 | 0.999994 | 13678 | 2 | -0.052291 | 0.58847 | 0.79394 | 1 | 16101 | 2 | -0.052291 |
| MID2 | 6 | 0.49682 | 0.66654 | 0.999994 | 13369 | 2 | 0.43667 | 0.58848 | 0.79395 | 1 | 16102 | 3 | 0.43667 |
| hsa-mir-7107 | 3 | 0.28173 | 0.41409 | 0.999994 | 9735 | 2 | -0.66959 | 0.58854 | 0.68264 | 1 | 16103 | 1 | -0.66959 |
| hsa-mir-6853 | 3 | 0.063469 | 0.13414 | 0.95318 | 3575 | 2 | -3.0072 | 0.58856 | 0.68265 | 1 | 16104 | 1 | -3.0072 |
| TTC9C | 4 | 0.53875 | 0.63429 | 0.999994 | 14033 | 1 | 0.30965 | 0.58858 | 0.73368 | 1 | 16105 | 2 | 0.30965 |
| OR2T27 | 4 | 0.69725 | 0.7226 | 0.999994 | 16272 | 1 | 0.69747 | 0.58859 | 0.73369 | 1 | 16106 | 2 | 0.69747 |
| LAMP3 | 6 | 0.0029099 | 0.013391 | 0.739733 | 348 | 2 | -0.043513 | 0.58865 | 0.79409 | 1 | 16107 | 2 | -0.043513 |
| GAGE2A | 1 | 0.4113 | 0.41148 | 0.999994 | 12136 | 0 | -0.16962 | 0.5887 | 0.58841 | 1 | 16108 | 0 | -0.16962 |
| hsa-mir-4256 | 4 | 0.42416 | 0.57044 | 0.999994 | 12311 | 2 | 0.15313 | 0.58883 | 0.73388 | 1 | 16109 | 2 | 0.15313 |
| hsa-mir-8058 | 4 | 0.31961 | 0.46303 | 0.999994 | 10507 | 1 | 0.32041 | 0.58888 | 0.73392 | 1 | 16110 | 2 | 0.32041 |
| MRPL28 | 4 | 0.16581 | 0.30547 | 0.999994 | 6881 | 2 | -1.1948 | 0.58894 | 0.73397 | 1 | 16111 | 1 | -1.1948 |
| NonTargetingControlG | 1 | 0.41096 | 0.41114 | 0.999994 | 12128 | 0 | -0.15137 | 0.58904 | 0.58873 | 1 | 16112 | 0 | -0.15137 |
| PCDHB15 | 6 | 0.090631 | 0.22875 | 0.983354 | 4623 | 3 | -1.4801 | 0.58909 | 0.79438 | 1 | 16113 | 2 | -1.4801 |
| RAI2 | 5 | 0.058212 | 0.1591 | 0.967126 | 3347 | 3 | -2.2623 | 0.5891 | 0.76232 | 1 | 16114 | 1 | -2.2623 |
| SLC25A12 | 5 | 0.22815 | 0.41467 | 0.999994 | 8481 | 3 | -0.70689 | 0.5891 | 0.76232 | 1 | 16115 | 1 | -0.70689 |
| WDSUB1 | 5 | 0.46597 | 0.61871 | 0.999994 | 12950 | 2 | 0.39153 | 0.5891 | 0.76232 | 1 | 16116 | 2 | 0.39153 |
| BMPR2 | 4 | 0.3575 | 0.50194 | 0.999994 | 11295 | 1 | 0.27795 | 0.58917 | 0.73415 | 1 | 16117 | 2 | 0.27795 |
| EEA1 | 6 | 0.85544 | 0.87488 | 0.999994 | 18357 | 1 | 0.50095 | 0.58921 | 0.79446 | 1 | 16118 | 3 | 0.50095 |
| LOC152586 | 6 | 0.17095 | 0.35193 | 0.999994 | 7006 | 3 | -0.87672 | 0.58925 | 0.7945 | 1 | 16119 | 2 | -0.87672 |

| | | | | | | | | | | | | | |
|---|---|---|---|---|---|---|---|---|---|---|---|---|---|
| TMEM233 | 6 | 0.058336 | 0.16219 | 0.967126 | 3355 | 4 | -1.6487 | 0.58926 | 0.7945 | 1 | 16120 | 1 | -1.6487 |
| SORL1 | 6 | 0.058439 | 0.1624 | 0.967126 | 3363 | 3 | -0.81292 | 0.58927 | 0.79451 | 1 | 16121 | 3 | -0.81292 |
| ADARB1 | 5 | 0.093741 | 0.21654 | 0.980699 | 4744 | 1 | 0.54571 | 0.58942 | 0.76249 | 1 | 16122 | 2 | 0.54571 |
| RAB35 | 6 | 0.044475 | 0.13194 | 0.95318 | 2735 | 2 | 0.18536 | 0.58942 | 0.79462 | 1 | 16123 | 3 | 0.18536 |
| TBC1D28 | 3 | 0.2515 | 0.37684 | 0.999994 | 9048 | 1 | -0.33604 | 0.58961 | 0.68314 | 1 | 16124 | 1 | -0.33604 |
| CHAF1B | 4 | 0.32946 | 0.47317 | 0.999994 | 10709 | 1 | 0.52711 | 0.58971 | 0.73462 | 1 | 16125 | 2 | 0.52711 |
| ABCB4 | 4 | 0.03666 | 0.095093 | 0.945031 | 2352 | 3 | -1.8133 | 0.58975 | 0.73465 | 1 | 16126 | 1 | -1.8133 |
| hsa-mir-16-1 | 3 | 0.65663 | 0.66259 | 0.999994 | 15738 | 1 | -0.54342 | 0.58976 | 0.68322 | 1 | 16127 | 1 | -0.54342 |
| RHOQ | 6 | 0.33181 | 0.54334 | 0.999994 | 10761 | 3 | -0.16549 | 0.59022 | 0.79521 | 1 | 16128 | 2 | -0.16549 |
| FARP2 | 5 | 0.082223 | 0.19818 | 0.977164 | 4304 | 2 | 0.40058 | 0.59024 | 0.76293 | 1 | 16129 | 2 | 0.40058 |
| hsa-mir-3142 | 2 | 0.49195 | 0.49655 | 0.999994 | 13301 | 1 | 0.28472 | 0.59031 | 0.60652 | 1 | 16130 | 1 | 0.28472 |
| hsa-mir-320d-2 | 2 | 0.47591 | 0.48214 | 0.999994 | 13077 | 1 | 0.24666 | 0.59035 | 0.60655 | 1 | 16131 | 1 | 0.24666 |
| HMHB1 | 4 | 0.82982 | 0.82936 | 0.999994 | 17925 | 0 | 0.67118 | 0.59041 | 0.73515 | 1 | 16132 | 2 | 0.67118 |
| NOSTRIN | 4 | 0.22286 | 0.36418 | 0.999994 | 8353 | 2 | -1.2368 | 0.59041 | 0.73515 | 1 | 16133 | 1 | -1.2368 |
| ZNF419 | 5 | 0.85203 | 0.85589 | 0.999994 | 18295 | 1 | 0.073399 | 0.59049 | 0.76309 | 1 | 16134 | 2 | 0.073399 |
| hsa-mir-4441 | 2 | 0.55259 | 0.55335 | 0.999994 | 14252 | 1 | 0.43674 | 0.59054 | 0.6067 | 1 | 16135 | 1 | 0.43674 |
| ASB13 | 6 | 0.11889 | 0.28362 | 0.999632 | 5598 | 4 | -0.85332 | 0.59058 | 0.79551 | 1 | 16136 | 1 | -0.85332 |
| LAYN | 6 | 0.23922 | 0.43506 | 0.999994 | 8756 | 3 | -0.13628 | 0.59061 | 0.79553 | 1 | 16137 | 3 | -0.13628 |
| hsa-mir-6830 | 4 | 0.095935 | 0.20779 | 0.979108 | 4813 | 2 | -1.1152 | 0.59093 | 0.73556 | 1 | 16138 | 2 | -1.1152 |
| ANAPC4 | 3 | 0.41139 | 0.49096 | 0.999994 | 12138 | 1 | 0.37798 | 0.59096 | 0.68378 | 1 | 16139 | 1 | 0.37798 |
| HHAT | 6 | 0.028981 | 0.096403 | 0.945031 | 1954 | 4 | -1.5477 | 0.59105 | 0.79585 | 1 | 16140 | 2 | -1.5477 |
| RWDD2B | 4 | 0.87504 | 0.87469 | 0.999994 | 18735 | 0 | 0.55251 | 0.5911 | 0.73571 | 1 | 16141 | 2 | 0.55251 |
| UBE2H | 6 | 0.28488 | 0.48912 | 0.999994 | 9807 | 2 | 0.2293 | 0.59125 | 0.79598 | 1 | 16142 | 3 | 0.2293 |
| TMCO3 | 6 | 0.60272 | 0.74197 | 0.999994 | 14982 | 2 | 0.28908 | 0.59127 | 0.796 | 1 | 16143 | 3 | 0.28908 |
| LIG4 | 6 | 0.87218 | 0.88467 | 0.999994 | 18679 | 1 | 0.19763 | 0.59132 | 0.79603 | 1 | 16144 | 2 | 0.19763 |
| C1orf27 | 6 | 0.015164 | 0.056252 | 0.9235 | 1199 | 5 | -2.1843 | 0.59134 | 0.79605 | 1 | 16145 | 1 | -2.1843 |
| HLA-DRB1 | 5 | 0.4115 | 0.57311 | 0.999994 | 12139 | 2 | 0.29642 | 0.59145 | 0.76365 | 1 | 16146 | 2 | 0.29642 |
| LRRC37A3 | 6 | 0.20487 | 0.39379 | 0.999994 | 7898 | 3 | -0.71406 | 0.59147 | 0.79615 | 1 | 16147 | 2 | -0.71406 |
| CNTN3 | 6 | 0.42649 | 0.6196 | 0.999994 | 12356 | 2 | 0.23729 | 0.59151 | 0.79619 | 1 | 16148 | 3 | 0.23729 |
| C6orf15 | 5 | 0.071168 | 0.18027 | 0.968171 | 3882 | 3 | -0.70722 | 0.59164 | 0.76377 | 1 | 16149 | 1 | -0.70722 |
| hsa-mir-6125 | 1 | 0.40826 | 0.40845 | 0.999994 | 12083 | 0 | -0.18907 | 0.59174 | 0.59146 | 1 | 16150 | 0 | -0.18907 |
| EFCAB9 | 5 | 0.11767 | 0.25379 | 0.994632 | 5565 | 3 | -1.4582 | 0.59189 | 0.76392 | 1 | 16151 | 2 | -1.4582 |
| MAP4K5 | 4 | 0.40625 | 0.5521 | 0.999994 | 12046 | 2 | 0.09798 | 0.592 | 0.73647 | 1 | 16152 | 2 | 0.09798 |
| TAF13 | 6 | 0.0070316 | 0.028524 | 0.858757 | 662 | 4 | -3.299 | 0.59205 | 0.79656 | 1 | 16153 | 2 | -3.299 |
| SLC2A12 | 6 | 0.088499 | 0.22454 | 0.983354 | 4534 | 2 | -0.43093 | 0.59205 | 0.79656 | 1 | 16154 | 2 | -0.43093 |
| USP17L4 | 2 | 0.60367 | 0.60346 | 0.999994 | 14992 | 0 | 0.63512 | 0.59214 | 0.60796 | 1 | 16155 | 1 | 0.63512 |
| BLOC1S5 | 6 | 0.25589 | 0.45496 | 0.999994 | 9147 | 3 | -0.17958 | 0.59232 | 0.79676 | 1 | 16156 | 3 | -0.17958 |
| ARL8B | 6 | 0.025825 | 0.088968 | 0.94395 | 1793 | 3 | -0.17587 | 0.59239 | 0.79681 | 1 | 16157 | 3 | -0.17587 |
| TGIF2-C20orf24 | 1 | 0.4076 | 0.40781 | 0.999994 | 12068 | 0 | -0.21469 | 0.5924 | 0.59211 | 1 | 16158 | 0 | -0.21469 |
| NonTargetingControlG | 2 | 0.19423 | 0.25794 | 0.994943 | 7608 | 1 | -1.1696 | 0.59258 | 0.6083 | 1 | 16159 | 1 | -1.1696 |
| GPIHBP1 | 6 | 0.1352 | 0.30693 | 0.999994 | 6056 | 3 | -0.98899 | 0.59274 | 0.79707 | 1 | 16160 | 2 | -0.98899 |
| CAMSAP3 | 4 | 0.17461 | 0.31456 | 0.999994 | 7102 | 2 | -0.24781 | 0.59276 | 0.73708 | 1 | 16161 | 1 | -0.24781 |
| ABCA9 | 5 | 0.1915 | 0.36307 | 0.999994 | 7527 | 3 | -0.94138 | 0.59289 | 0.76448 | 1 | 16162 | 2 | -0.94138 |
| FAM118B | 6 | 0.61893 | 0.75402 | 0.999994 | 15203 | 2 | 0.14032 | 0.59295 | 0.79722 | 1 | 16163 | 3 | 0.14032 |
| MORC4 | 5 | 0.013348 | 0.045555 | 0.891621 | 1089 | 4 | -1.8631 | 0.59301 | 0.76455 | 1 | 16164 | 1 | -1.8631 |
| THY1 | 5 | 0.026442 | 0.081269 | 0.93914 | 1827 | 4 | -2.4296 | 0.59301 | 0.76455 | 1 | 16165 | 1 | -2.4296 |
| PPAP2C | 4 | 0.37869 | 0.52384 | 0.999994 | 11634 | 2 | 0.023623 | 0.59314 | 0.73739 | 1 | 16166 | 2 | 0.023623 |
| ANGPT2 | 6 | 0.61416 | 0.75046 | 0.999994 | 15134 | 2 | -0.10256 | 0.5933 | 0.79748 | 1 | 16167 | 2 | -0.10256 |
| PLVAP | 6 | 0.032811 | 0.10539 | 0.95013 | 2150 | 4 | -1.9176 | 0.5933 | 0.79748 | 1 | 16168 | 1 | -1.9176 |
| hsa-mir-4282 | 4 | 0.62697 | 0.6802 | 0.999994 | 15336 | 1 | 0.24742 | 0.59333 | 0.73754 | 1 | 16169 | 2 | 0.24742 |
| CHRM4 | 6 | 0.40013 | 0.60244 | 0.999994 | 11949 | 1 | 0.61022 | 0.59333 | 0.79751 | 1 | 16170 | 3 | 0.61022 |
| FBN2 | 6 | 0.50698 | 0.67358 | 0.999994 | 13537 | 2 | 0.59556 | 0.59338 | 0.79756 | 1 | 16171 | 3 | 0.59556 |

| | | | | | | | | | | | | | |
|---|---|---|---|---|---|---|---|---|---|---|---|---|---|
| OR5AC2 | 4 | 0.15062 | 0.28982 | 0.999632 | 6476 | 1 | 0.3495 | 0.59346 | 0.73765 | 1 | 16172 | 2 | 0.3495 |
| CFLAR | 4 | 0.063559 | 0.14757 | 0.965554 | 3584 | 2 | -0.34226 | 0.5935 | 0.73768 | 1 | 16173 | 2 | -0.34226 |
| CD274 | 5 | 0.86179 | 0.8639 | 0.999994 | 18482 | 1 | 0.25443 | 0.59354 | 0.76485 | 1 | 16174 | 2 | 0.25443 |
| SLC24A6 | 4 | 0.92958 | 0.92936 | 0.999994 | 19845 | 0 | 0.77906 | 0.59355 | 0.73772 | 1 | 16175 | 2 | 0.77906 |
| ZBED3 | 6 | 0.063131 | 0.17236 | 0.967658 | 3562 | 4 | -1.6747 | 0.5936 | 0.79772 | 1 | 16176 | 2 | -1.6747 |
| ICA1L | 6 | 0.19105 | 0.37683 | 0.999994 | 7514 | 2 | 0.04127 | 0.5936 | 0.79772 | 1 | 16177 | 3 | 0.04127 |
| ARHGAP22 | 6 | 0.19654 | 0.38368 | 0.999994 | 7663 | 3 | -0.28225 | 0.5936 | 0.79772 | 1 | 16178 | 2 | -0.28225 |
| C8orf31 | 5 | 0.74864 | 0.79013 | 0.999994 | 16846 | 1 | -0.018997 | 0.59363 | 0.76489 | 1 | 16179 | 2 | -0.018997 |
| KAT6A | 6 | 0.20851 | 0.39827 | 0.999994 | 7990 | 3 | -0.3415 | 0.59385 | 0.7979 | 1 | 16180 | 3 | -0.3415 |
| LY6K | 6 | 0.42774 | 0.62041 | 0.999994 | 12377 | 1 | 0.39665 | 0.59387 | 0.79792 | 1 | 16181 | 3 | 0.39665 |
| TMEM229B | 5 | 0.4292 | 0.58776 | 0.999994 | 12403 | 1 | 0.092117 | 0.59402 | 0.76509 | 1 | 16182 | 2 | 0.092117 |
| ARMCX1 | 5 | 0.28499 | 0.46888 | 0.999994 | 9809 | 1 | 0.53071 | 0.59402 | 0.76509 | 1 | 16183 | 2 | 0.53071 |
| VPS54 | 5 | 0.038308 | 0.11161 | 0.951052 | 2431 | 3 | -2.5996 | 0.59402 | 0.76509 | 1 | 16184 | 1 | -2.5996 |
| NCKIPSD | 5 | 0.015607 | 0.051931 | 0.911383 | 1223 | 4 | -0.94549 | 0.59402 | 0.76509 | 1 | 16185 | 1 | -0.94549 |
| SLC9A4 | 5 | 0.049488 | 0.1391 | 0.957704 | 2955 | 3 | -2.2635 | 0.59413 | 0.76516 | 1 | 16186 | 2 | -2.2635 |
| hsa-mir-1323 | 4 | 0.8497 | 0.84922 | 0.999994 | 18255 | 0 | 0.63598 | 0.59419 | 0.73822 | 1 | 16187 | 2 | 0.63598 |
| LRRC43 | 6 | 0.59298 | 0.73479 | 0.999994 | 14862 | 1 | -0.099712 | 0.59426 | 0.79822 | 1 | 16188 | 2 | -0.099712 |
| IVD | 6 | 0.43595 | 0.62575 | 0.999994 | 12515 | 2 | 0.53916 | 0.59431 | 0.79825 | 1 | 16189 | 3 | 0.53916 |
| SERPINA12 | 6 | 0.56027 | 0.71097 | 0.999994 | 14374 | 2 | 0.08523 | 0.59438 | 0.7983 | 1 | 16190 | 2 | 0.08523 |
| SLC6A13 | 6 | 0.91902 | 0.91949 | 0.999994 | 19607 | 1 | 0.072602 | 0.59438 | 0.7983 | 1 | 16191 | 1 | 0.072602 |
| BMP1 | 6 | 0.55594 | 0.70789 | 0.999994 | 14303 | 2 | -0.27102 | 0.59438 | 0.7983 | 1 | 16192 | 1 | -0.27102 |
| FBXL12 | 6 | 0.46847 | 0.64741 | 0.999994 | 12981 | 1 | 0.62568 | 0.59438 | 0.7983 | 1 | 16193 | 3 | 0.62568 |
| CYP1A2 | 6 | 0.84097 | 0.86713 | 0.999994 | 18106 | 1 | 0.61169 | 0.59471 | 0.79853 | 1 | 16194 | 3 | 0.61169 |
| hsa-mir-5091 | 2 | 0.10455 | 0.17714 | 0.968171 | 5117 | 1 | -2.5017 | 0.59482 | 0.61008 | 1 | 16195 | 1 | -2.5017 |
| COLGALT1 | 6 | 0.11749 | 0.281 | 0.999632 | 5561 | 2 | 0.47751 | 0.59492 | 0.79869 | 1 | 16196 | 3 | 0.47751 |
| OR56A1 | 6 | 0.30903 | 0.51725 | 0.999994 | 10299 | 3 | -0.022222 | 0.59495 | 0.79871 | 1 | 16197 | 3 | -0.022222 |
| LRRC32 | 6 | 0.14802 | 0.32316 | 0.999994 | 6408 | 3 | -0.38014 | 0.59504 | 0.79878 | 1 | 16198 | 2 | -0.38014 |
| PEX14 | 5 | 0.15489 | 0.31006 | 0.999994 | 6593 | 3 | -0.84277 | 0.59508 | 0.7657 | 1 | 16199 | 2 | -0.84277 |
| IFI6 | 5 | 0.055211 | 0.15299 | 0.967126 | 3208 | 3 | -0.60945 | 0.59513 | 0.76573 | 1 | 16200 | 1 | -0.60945 |
| EDF1 | 6 | 0.019348 | 0.069902 | 0.93914 | 1423 | 4 | -3.6068 | 0.59518 | 0.79888 | 1 | 16201 | 2 | -3.6068 |
| MYO3B | 5 | 0.38415 | 0.55021 | 0.999994 | 11703 | 2 | 0.28056 | 0.59523 | 0.76579 | 1 | 16202 | 2 | 0.28056 |
| PPIL2 | 4 | 0.0081764 | 0.027945 | 0.855011 | 740 | 3 | -5.2723 | 0.59535 | 0.73915 | 1 | 16203 | 1 | -5.2723 |
| SLFNL1 | 4 | 0.86026 | 0.85987 | 0.999994 | 18444 | 0 | 0.42639 | 0.59535 | 0.73915 | 1 | 16204 | 1 | 0.42639 |
| ZNF589 | 5 | 0.276 | 0.46161 | 0.999994 | 9617 | 2 | 0.16359 | 0.5954 | 0.76588 | 1 | 16205 | 2 | 0.16359 |
| MAN2B1 | 6 | 0.39359 | 0.59823 | 0.999994 | 11855 | 2 | 0.39005 | 0.59543 | 0.79906 | 1 | 16206 | 3 | 0.39005 |
| OMD | 6 | 0.29045 | 0.49561 | 0.999994 | 9920 | 3 | -0.23932 | 0.5955 | 0.79911 | 1 | 16207 | 2 | -0.23932 |
| CKAP4 | 6 | 0.042966 | 0.1286 | 0.951777 | 2676 | 4 | -2.1904 | 0.59555 | 0.79915 | 1 | 16208 | 2 | -2.1904 |
| NIT1 | 6 | 0.030107 | 0.099046 | 0.945253 | 2015 | 2 | 0.20586 | 0.59555 | 0.79915 | 1 | 16209 | 3 | 0.20586 |
| ACSL1 | 6 | 0.10993 | 0.26656 | 0.998525 | 5301 | 3 | -1.273 | 0.59558 | 0.79916 | 1 | 16210 | 2 | -1.273 |
| SUGT1 | 3 | 0.028719 | 0.072202 | 0.93914 | 1938 | 2 | -6.1018 | 0.59561 | 0.68608 | 1 | 16211 | 1 | -6.1018 |
| ZNHIT2 | 4 | 0.24107 | 0.38273 | 0.999994 | 8795 | 2 | -0.078815 | 0.59568 | 0.73941 | 1 | 16212 | 2 | -0.078815 |
| FAM178B | 6 | 0.12805 | 0.2978 | 0.999994 | 5879 | 2 | 0.35407 | 0.59571 | 0.79927 | 1 | 16213 | 3 | 0.35407 |
| NonTargetingControlG | 2 | 0.39855 | 0.41558 | 0.999994 | 11924 | 1 | -0.053909 | 0.5959 | 0.61096 | 1 | 16214 | 1 | -0.053909 |
| NAP1L2 | 5 | 0.43327 | 0.59116 | 0.999994 | 12466 | 1 | -0.49253 | 0.59599 | 0.76623 | 1 | 16215 | 2 | -0.49253 |
| TNFRSF11B | 6 | 0.44336 | 0.63067 | 0.999994 | 12628 | 2 | -0.095693 | 0.59602 | 0.79951 | 1 | 16216 | 2 | -0.095693 |
| hsa-mir-592 | 3 | 0.13537 | 0.22959 | 0.983354 | 6061 | 2 | -1.8683 | 0.59604 | 0.68626 | 1 | 16217 | 1 | -1.8683 |
| hsa-mir-4499 | 1 | 0.40389 | 0.40407 | 0.999994 | 12015 | 0 | -0.23578 | 0.59611 | 0.5958 | 1 | 16218 | 0 | -0.23578 |
| hsa-mir-378b | 3 | 0.55574 | 0.58432 | 0.999994 | 14300 | 1 | -0.030522 | 0.59612 | 0.6863 | 1 | 16219 | 1 | -0.030522 |
| PXMP2 | 5 | 0.012407 | 0.042947 | 0.891621 | 1023 | 3 | -2.351 | 0.59627 | 0.7664 | 1 | 16220 | 1 | -2.351 |
| RGS18 | 5 | 0.0043141 | 0.018069 | 0.795727 | 465 | 3 | -7.0728 | 0.59627 | 0.7664 | 1 | 16221 | 1 | -7.0728 |
| GRHL2 | 5 | 0.21363 | 0.39447 | 0.999994 | 8104 | 3 | -0.69862 | 0.59627 | 0.7664 | 1 | 16222 | 1 | -0.69862 |
| KRTAP15-1 | 5 | 0.17864 | 0.34466 | 0.999994 | 7189 | 2 | -0.040834 | 0.59634 | 0.76644 | 1 | 16223 | 2 | -0.040834 |

| | | | | | | | | | | | | | |
|---|---|---|---|---|---|---|---|---|---|---|---|---|---|
| hsa-mir-3935 | 4 | 0.17328 | 0.3132 | 0.999994 | 7069 | 2 | -0.54937 | 0.5964 | 0.73996 | 1 | 16224 | 2 | -0.54937 |
| PIGV | 5 | 0.025033 | 0.077563 | 0.93914 | 1757 | 3 | -3.2353 | 0.59664 | 0.76662 | 1 | 16225 | 2 | -3.2353 |
| TERT | 6 | 0.7469 | 0.82738 | 0.999994 | 16826 | 1 | -0.016682 | 0.59668 | 0.79999 | 1 | 16226 | 2 | -0.016682 |
| UPK3BL | 6 | 0.098478 | 0.24437 | 0.988879 | 4912 | 4 | -1.3373 | 0.59674 | 0.80004 | 1 | 16227 | 2 | -1.3373 |
| GRIA3 | 5 | 0.1584 | 0.31519 | 0.999994 | 6683 | 3 | -1.1594 | 0.59701 | 0.76685 | 1 | 16228 | 2 | -1.1594 |
| B4GALT3 | 6 | 0.032256 | 0.1041 | 0.947284 | 2115 | 4 | -1.8645 | 0.59705 | 0.80027 | 1 | 16229 | 2 | -1.8645 |
| THBS2 | 5 | 0.28037 | 0.46514 | 0.999994 | 9709 | 2 | 0.084988 | 0.59708 | 0.76689 | 1 | 16230 | 2 | 0.084988 |
| NonTargetingControlG | 1 | 0.40284 | 0.40302 | 0.999994 | 11994 | 0 | -0.23825 | 0.59716 | 0.59683 | 1 | 16231 | 0 | -0.23825 |
| PUM2 | 6 | 0.15998 | 0.33821 | 0.999994 | 6739 | 3 | -0.7904 | 0.5973 | 0.80044 | 1 | 16232 | 2 | -0.7904 |
| PMCH | 5 | 0.50172 | 0.64941 | 0.999994 | 13454 | 1 | -0.19807 | 0.59749 | 0.76711 | 1 | 16233 | 2 | -0.19807 |
| AARS | 4 | 0.16868 | 0.3084 | 0.999994 | 6951 | 1 | 0.21956 | 0.59749 | 0.74083 | 1 | 16234 | 1 | 0.21956 |
| hsa-mir-548t | 3 | 0.68603 | 0.6882 | 0.999994 | 16172 | 1 | -0.03978 | 0.59758 | 0.68702 | 1 | 16235 | 1 | -0.03978 |
| RPL7L1 | 3 | 0.030694 | 0.076759 | 0.93914 | 2043 | 2 | -4.6327 | 0.59758 | 0.68702 | 1 | 16236 | 1 | -4.6327 |
| FSHR | 6 | 0.33793 | 0.5503 | 0.999994 | 10894 | 3 | -0.329 | 0.5977 | 0.80073 | 1 | 16237 | 2 | -0.329 |
| LYZL6 | 5 | 0.28479 | 0.46872 | 0.999994 | 9804 | 2 | -0.28285 | 0.59771 | 0.76724 | 1 | 16238 | 2 | -0.28285 |
| GAGE10 | 4 | 0.83774 | 0.83723 | 0.999994 | 18048 | 0 | 0.6056 | 0.59774 | 0.74102 | 1 | 16239 | 2 | 0.6056 |
| ZNF850 | 6 | 0.40025 | 0.60252 | 0.999994 | 11953 | 2 | 0.46624 | 0.59782 | 0.80082 | 1 | 16240 | 3 | 0.46624 |
| ASPA | 6 | 0.5814 | 0.72623 | 0.999994 | 14698 | 2 | 0.56384 | 0.59787 | 0.80085 | 1 | 16241 | 3 | 0.56384 |
| KMO | 5 | 0.19877 | 0.37348 | 0.999994 | 7729 | 2 | 0.28796 | 0.59799 | 0.7674 | 1 | 16242 | 2 | 0.28796 |
| MBLAC1 | 6 | 0.59676 | 0.73758 | 0.999994 | 14910 | 2 | 0.59583 | 0.59832 | 0.80118 | 1 | 16243 | 3 | 0.59583 |
| CABP1 | 4 | 0.036331 | 0.0944 | 0.945031 | 2337 | 2 | -1.8696 | 0.59837 | 0.74153 | 1 | 16244 | 2 | -1.8696 |
| VPREB3 | 6 | 0.28641 | 0.49093 | 0.999994 | 9844 | 3 | -0.11265 | 0.59871 | 0.8015 | 1 | 16245 | 3 | -0.11265 |
| NonTargetingControlG | 2 | 0.49027 | 0.49507 | 0.999994 | 13283 | 1 | 0.17496 | 0.59879 | 0.61322 | 1 | 16246 | 1 | 0.17496 |
| NonTargetingControlG | 1 | 0.40106 | 0.4012 | 0.999994 | 11960 | 0 | -0.25444 | 0.59894 | 0.59861 | 1 | 16247 | 0 | -0.25444 |
| SAP25 | 6 | 0.1926 | 0.37876 | 0.999994 | 7561 | 3 | -0.29297 | 0.59898 | 0.8017 | 1 | 16248 | 1 | -0.29297 |
| ZNF350 | 6 | 0.15523 | 0.33225 | 0.999994 | 6600 | 3 | -0.30031 | 0.59898 | 0.8017 | 1 | 16249 | 2 | -0.30031 |
| PGBD2 | 6 | 0.25739 | 0.45678 | 0.999994 | 9176 | 3 | -0.22359 | 0.59898 | 0.8017 | 1 | 16250 | 2 | -0.22359 |
| IL7R | 6 | 0.96089 | 0.96096 | 0.999994 | 20600 | 0 | 0.1718 | 0.59898 | 0.8017 | 1 | 16251 | 2 | 0.1718 |
| FAM185A | 6 | 0.72752 | 0.82078 | 0.999994 | 16573 | 1 | 0.5675 | 0.59904 | 0.80174 | 1 | 16252 | 3 | 0.5675 |
| PCDH1 | 5 | 0.47807 | 0.62899 | 0.999994 | 13118 | 1 | -0.39962 | 0.59913 | 0.76806 | 1 | 16253 | 2 | -0.39962 |
| MAN1A2 | 5 | 0.31564 | 0.4938 | 0.999994 | 10433 | 2 | 0.44169 | 0.59924 | 0.76812 | 1 | 16254 | 2 | 0.44169 |
| hsa-mir-5704 | 4 | 0.90277 | 0.90246 | 0.999994 | 19267 | 0 | 0.52986 | 0.59946 | 0.74241 | 1 | 16255 | 2 | 0.52986 |
| R3HCC1L | 6 | 0.14278 | 0.3166 | 0.999994 | 6263 | 2 | -0.48428 | 0.59947 | 0.80204 | 1 | 16256 | 2 | -0.48428 |
| SERPINE2 | 4 | 0.30892 | 0.45209 | 0.999994 | 10296 | 2 | -0.64245 | 0.59963 | 0.74255 | 1 | 16257 | 1 | -0.64245 |
| PPARGC1A | 6 | 0.46553 | 0.64548 | 0.999994 | 12946 | 1 | 0.63015 | 0.59966 | 0.80219 | 1 | 16258 | 3 | 0.63015 |
| SPRR1B | 6 | 0.02887 | 0.096158 | 0.945031 | 1948 | 3 | -1.4105 | 0.59971 | 0.80223 | 1 | 16259 | 3 | -1.4105 |
| hsa-mir-6077-1 | 4 | 0.10078 | 0.2165 | 0.980699 | 4999 | 2 | -1.2446 | 0.59973 | 0.74264 | 1 | 16260 | 1 | -1.2446 |
| LRIG3 | 6 | 0.3027 | 0.50987 | 0.999994 | 10188 | 2 | 0.58224 | 0.59975 | 0.80226 | 1 | 16261 | 3 | 0.58224 |
| SOCS1 | 5 | 0.27748 | 0.46283 | 0.999994 | 9646 | 1 | -0.065829 | 0.59991 | 0.7685 | 1 | 16262 | 2 | -0.065829 |
| ZADH2 | 4 | 0.77137 | 0.77689 | 0.999994 | 17104 | 1 | 0.43791 | 0.60004 | 0.74287 | 1 | 16263 | 2 | 0.43791 |
| DALRD3 | 6 | 0.14699 | 0.32186 | 0.999994 | 6376 | 3 | -0.42808 | 0.60009 | 0.80251 | 1 | 16264 | 2 | -0.42808 |
| DPT | 4 | 0.21416 | 0.3553 | 0.999994 | 8121 | 2 | -0.40515 | 0.60021 | 0.74302 | 1 | 16265 | 2 | -0.40515 |
| NonTargetingControlG | 2 | 0.54012 | 0.54139 | 0.999994 | 14054 | 1 | 0.53564 | 0.60032 | 0.61442 | 1 | 16266 | 1 | 0.53564 |
| KLHDC9 | 6 | 0.15659 | 0.33391 | 0.999994 | 6643 | 3 | -0.12481 | 0.60033 | 0.80268 | 1 | 16267 | 3 | -0.12481 |
| DGKH | 5 | 0.14317 | 0.29261 | 0.999632 | 6279 | 2 | 0.29792 | 0.60035 | 0.76875 | 1 | 16268 | 2 | 0.29792 |
| C7orf33 | 6 | 0.34426 | 0.55756 | 0.999994 | 11034 | 3 | -0.081253 | 0.60042 | 0.80274 | 1 | 16269 | 2 | -0.081253 |
| KDM2A | 4 | 0.015485 | 0.050584 | 0.909101 | 1215 | 3 | -2.5396 | 0.60061 | 0.74337 | 1 | 16270 | 1 | -2.5396 |
| DEK | 4 | 0.0068276 | 0.023633 | 0.832699 | 648 | 3 | -4.4814 | 0.60061 | 0.74337 | 1 | 16271 | 1 | -4.4814 |
| TGM2 | 4 | 0.40653 | 0.55239 | 0.999994 | 12050 | 1 | -0.0071046 | 0.60061 | 0.74337 | 1 | 16272 | 1 | -0.0071046 |
| NonTargetingControlG | 1 | 0.39929 | 0.39946 | 0.999994 | 11936 | 0 | -0.26572 | 0.60071 | 0.6004 | 1 | 16273 | 0 | -0.26572 |
| NDUFB9 | 4 | 0.34094 | 0.48494 | 0.999994 | 10959 | 1 | 0.4516 | 0.60072 | 0.74346 | 1 | 16274 | 2 | 0.4516 |
| NonTargetingControlG | 1 | 0.39919 | 0.39935 | 0.999994 | 11935 | 0 | -0.26139 | 0.60081 | 0.60051 | 1 | 16275 | 0 | -0.26139 |

| | | | | | | | | | | | | |
|---|---|---|---|---|---|---|---|---|---|---|---|---|
| MYO1B | 6 | 0.29897 | 0.50549 | 0.999994 | 10113 | 2 | 0.39925 | 0.6009 | 0.80309 | 1 | 16276 | 3 | 0.39925 |
| TMX3 | 6 | 0.058164 | 0.16181 | 0.967126 | 3346 | 2 | -0.40267 | 0.60103 | 0.8032 | 1 | 16277 | 2 | -0.40267 |
| PDE4A | 6 | 0.04887 | 0.14162 | 0.959146 | 2924 | 4 | -1.4948 | 0.60103 | 0.8032 | 1 | 16278 | 2 | -1.4948 |
| FAM227A | 6 | 0.011391 | 0.043728 | 0.891621 | 949 | 3 | -2.0862 | 0.60103 | 0.8032 | 1 | 16279 | 2 | -2.0862 |
| hsa-mir-4444-1 | 4 | 0.44897 | 0.59427 | 0.999994 | 12698 | 1 | 0.66719 | 0.60103 | 0.7437 | 1 | 16280 | 2 | 0.66719 |
| ARPC1B | 6 | 0.27388 | 0.47628 | 0.999994 | 9561 | 2 | 0.46144 | 0.60107 | 0.80324 | 1 | 16281 | 3 | 0.46144 |
| hsa-mir-890 | 4 | 0.90877 | 0.90849 | 0.999994 | 19399 | 0 | 0.50295 | 0.60117 | 0.74381 | 1 | 16282 | 2 | 0.50295 |
| NonTargetingControlG | 2 | 0.57647 | 0.57661 | 0.999994 | 14636 | 1 | 0.3855 | 0.60139 | 0.61529 | 1 | 16283 | 1 | 0.3855 |
| HOMER2 | 4 | 0.52179 | 0.62625 | 0.999994 | 13776 | 1 | 0.68081 | 0.6014 | 0.74399 | 1 | 16284 | 2 | 0.68081 |
| hsa-mir-4479 | 3 | 0.35001 | 0.4557 | 0.999994 | 11158 | 1 | 0.12092 | 0.60141 | 0.68886 | 1 | 16285 | 1 | 0.12092 |
| RIBC1 | 5 | 0.89758 | 0.89771 | 0.999994 | 19171 | 0 | 0.12541 | 0.60148 | 0.76941 | 1 | 16286 | 2 | 0.12541 |
| GAB2 | 6 | 0.069263 | 0.18523 | 0.968626 | 3809 | 3 | -0.079503 | 0.60162 | 0.80365 | 1 | 16287 | 3 | -0.079503 |
| NOMO2 | 2 | 0.48758 | 0.49261 | 0.999994 | 13247 | 1 | 0.18899 | 0.60173 | 0.61558 | 1 | 16288 | 1 | 0.18899 |
| SRMS | 5 | 0.14068 | 0.28885 | 0.999632 | 6207 | 3 | -1.3382 | 0.60177 | 0.76959 | 1 | 16289 | 2 | -1.3382 |
| NR1D2 | 6 | 0.47651 | 0.65284 | 0.999994 | 13091 | 2 | -0.42009 | 0.60184 | 0.80381 | 1 | 16290 | 2 | -0.42009 |
| MYOZ1 | 5 | 0.083288 | 0.1999 | 0.977164 | 4353 | 3 | -1.8458 | 0.60184 | 0.76962 | 1 | 16291 | 1 | -1.8458 |
| PPP2R2B | 5 | 0.46317 | 0.61633 | 0.999994 | 12908 | 2 | 0.32966 | 0.60184 | 0.76962 | 1 | 16292 | 1 | 0.32966 |
| COA6 | 5 | 0.0010603 | 0.0047351 | 0.588856 | 177 | 4 | -5.3293 | 0.60184 | 0.76962 | 1 | 16293 | 1 | -5.3293 |
| GPR84 | 5 | 0.24292 | 0.43489 | 0.999994 | 8861 | 3 | -0.56202 | 0.60184 | 0.76962 | 1 | 16294 | 2 | -0.56202 |
| RYK | 5 | 0.20137 | 0.37718 | 0.999994 | 7803 | 3 | -0.70645 | 0.60192 | 0.76966 | 1 | 16295 | 2 | -0.70645 |
| GXYLT2 | 4 | 0.031784 | 0.085137 | 0.94395 | 2096 | 3 | -1.8367 | 0.60199 | 0.74445 | 1 | 16296 | 1 | -1.8367 |
| CXorf36 | 6 | 0.084866 | 0.21726 | 0.980699 | 4404 | 3 | -0.31498 | 0.60204 | 0.80396 | 1 | 16297 | 3 | -0.31498 |
| LILRA3 | 6 | 0.097 | 0.24149 | 0.987253 | 4859 | 4 | -1.0072 | 0.60207 | 0.80398 | 1 | 16298 | 2 | -1.0072 |
| SLC12A6 | 6 | 0.12214 | 0.28981 | 0.999632 | 5706 | 2 | 0.52263 | 0.60217 | 0.80405 | 1 | 16299 | 3 | 0.52263 |
| PPP3CA | 4 | 0.42423 | 0.57052 | 0.999994 | 12315 | 1 | 0.68622 | 0.60229 | 0.74468 | 1 | 16300 | 2 | 0.68622 |
| NonTargetingControlG | 2 | 0.68159 | 0.68149 | 0.999994 | 16111 | 0 | 0.75971 | 0.60231 | 0.61606 | 1 | 16301 | 1 | 0.75971 |
| hsa-mir-3135a | 2 | 0.44549 | 0.45535 | 0.999994 | 12656 | 1 | 0.02956 | 0.60233 | 0.61607 | 1 | 16302 | 1 | 0.02956 |
| ADCK1 | 6 | 0.58786 | 0.73103 | 0.999994 | 14790 | 2 | 0.54486 | 0.60246 | 0.80429 | 1 | 16303 | 3 | 0.54486 |
| CIR1 | 6 | 0.29303 | 0.49856 | 0.999994 | 9977 | 3 | -0.72503 | 0.60249 | 0.80431 | 1 | 16304 | 2 | -0.72503 |
| MAGEA10 | 5 | 0.49564 | 0.64412 | 0.999994 | 13350 | 2 | 0.13706 | 0.60251 | 0.77002 | 1 | 16305 | 2 | 0.13706 |
| NonTargetingControlG | 2 | 0.26931 | 0.31337 | 0.999994 | 9452 | 1 | -0.58182 | 0.60254 | 0.61624 | 1 | 16306 | 1 | -0.58182 |
| ZDBF2 | 6 | 0.77094 | 0.8362 | 0.999994 | 17096 | 1 | 0.64773 | 0.60263 | 0.80442 | 1 | 16307 | 3 | 0.64773 |
| ATP5G2 | 6 | 0.18834 | 0.37347 | 0.999994 | 7430 | 3 | -0.69683 | 0.60263 | 0.80442 | 1 | 16308 | 2 | -0.69683 |
| hsa-mir-6715a | 4 | 0.76501 | 0.77179 | 0.999994 | 17028 | 1 | -0.10879 | 0.6027 | 0.74504 | 1 | 16309 | 1 | -0.10879 |
| MBTPS1 | 5 | 0.13436 | 0.27929 | 0.999632 | 6037 | 3 | -0.93726 | 0.60286 | 0.77022 | 1 | 16310 | 2 | -0.93726 |
| YARS2 | 5 | 0.17094 | 0.3335 | 0.999994 | 7005 | 2 | -0.34428 | 0.60294 | 0.77027 | 1 | 16311 | 2 | -0.34428 |
| ZSWIM7 | 5 | 0.16593 | 0.32613 | 0.999994 | 6882 | 3 | -0.94927 | 0.60295 | 0.77027 | 1 | 16312 | 2 | -0.94927 |
| OR13A1 | 6 | 0.19543 | 0.38234 | 0.999994 | 7646 | 3 | -0.043508 | 0.60313 | 0.80481 | 1 | 16313 | 3 | -0.043508 |
| PRADC1 | 6 | 0.085186 | 0.21789 | 0.980699 | 4416 | 4 | -1.4855 | 0.60317 | 0.80483 | 1 | 16314 | 2 | -1.4855 |
| DUSP6 | 4 | 0.30632 | 0.44944 | 0.999994 | 10262 | 1 | 0.21443 | 0.60325 | 0.74547 | 1 | 16315 | 2 | 0.21443 |
| SFTA3 | 5 | 0.1323 | 0.27613 | 0.999379 | 5984 | 1 | 0.06709 | 0.60328 | 0.77046 | 1 | 16316 | 2 | 0.06709 |
| ANKRD26 | 5 | 0.030468 | 0.091696 | 0.94395 | 2030 | 4 | -1.986 | 0.60328 | 0.77046 | 1 | 16317 | 1 | -1.986 |
| TIMD4 | 5 | 0.091555 | 0.21306 | 0.979587 | 4660 | 3 | -1.5493 | 0.60328 | 0.77046 | 1 | 16318 | 1 | -1.5493 |
| SLC22A4 | 6 | 0.93021 | 0.93034 | 0.999994 | 19858 | 0 | 0.19918 | 0.6036 | 0.80517 | 1 | 16319 | 2 | 0.19918 |
| MTMR9 | 6 | 0.50358 | 0.67122 | 0.999994 | 13487 | 1 | 0.56992 | 0.60378 | 0.80529 | 1 | 16320 | 3 | 0.56992 |
| MBD3L3 | 5 | 0.34048 | 0.51411 | 0.999994 | 10946 | 2 | -0.4929 | 0.60411 | 0.77093 | 1 | 16321 | 2 | -0.4929 |
| APOC2 | 6 | 0.39959 | 0.60209 | 0.999994 | 11943 | 2 | 0.05351 | 0.60411 | 0.80554 | 1 | 16322 | 3 | 0.05351 |
| SPERT | 5 | 0.33462 | 0.50932 | 0.999994 | 10825 | 2 | -0.39992 | 0.60413 | 0.77094 | 1 | 16323 | 2 | -0.39992 |
| CTPS2 | 6 | 0.030576 | 0.10013 | 0.946394 | 2035 | 3 | -1.8325 | 0.60424 | 0.80565 | 1 | 16324 | 2 | -1.8325 |
| NonTargetingControlG | 2 | 0.50558 | 0.50901 | 0.999994 | 13515 | 1 | 0.19401 | 0.60425 | 0.61763 | 1 | 16325 | 1 | 0.19401 |
| XYLB | 4 | 0.39664 | 0.54227 | 0.999994 | 11899 | 2 | -0.038159 | 0.60434 | 0.74635 | 1 | 16326 | 2 | -0.038159 |
| TSPY3 | 1 | 0.39556 | 0.39576 | 0.999994 | 11879 | 0 | -0.28208 | 0.60444 | 0.60405 | 1 | 16327 | 0 | -0.28208 |

| | | | | | | | | | | | | | |
|---|---|---|---|---|---|---|---|---|---|---|---|---|---|
| ZNF57 | 4 | 0.12455 | 0.25821 | 0.994943 | 5783 | 2 | -0.66006 | 0.60447 | 0.74644 | 1 | 16328 | 2 | -0.66006 |
| MAPKBP1 | 6 | 0.44914 | 0.63456 | 0.999994 | 12703 | 2 | -0.22632 | 0.60447 | 0.80581 | 1 | 16329 | 2 | -0.22632 |
| PLGLB1 | 6 | 0.083156 | 0.21375 | 0.979587 | 4345 | 4 | -0.87151 | 0.60447 | 0.80581 | 1 | 16330 | 1 | -0.87151 |
| ATP5C1 | 6 | 0.074416 | 0.19595 | 0.975797 | 4009 | 4 | -1.1598 | 0.60447 | 0.80581 | 1 | 16331 | 1 | -1.1598 |
| NLRC3 | 6 | 0.016358 | 0.060182 | 0.932932 | 1265 | 5 | -0.89484 | 0.60447 | 0.80581 | 1 | 16332 | 1 | -0.89484 |
| LGALS2 | 6 | 0.19623 | 0.38331 | 0.999994 | 7659 | 2 | -0.12883 | 0.6046 | 0.8059 | 1 | 16333 | 2 | -0.12883 |
| GRHL3 | 6 | 0.00050545 | 0.0026342 | 0.49604 | 112 | 3 | -0.091018 | 0.60465 | 0.80594 | 1 | 16334 | 2 | -0.091018 |
| hsa-mir-4269 | 3 | 0.22661 | 0.34586 | 0.999994 | 8443 | 2 | -1.0115 | 0.60489 | 0.69055 | 1 | 16335 | 1 | -1.0115 |
| NonTargetingControlG | 2 | 0.51087 | 0.51389 | 0.999994 | 13593 | 1 | 0.3475 | 0.60503 | 0.61828 | 1 | 16336 | 1 | 0.3475 |
| NonTargetingControlG | 2 | 0.29628 | 0.33398 | 0.999994 | 10045 | 1 | -0.69874 | 0.60515 | 0.61838 | 1 | 16337 | 1 | -0.69874 |
| PNPLA6 | 5 | 0.6193 | 0.7333 | 0.999994 | 15211 | 1 | -0.54508 | 0.60527 | 0.77158 | 1 | 16338 | 2 | -0.54508 |
| DEFB106A | 4 | 0.083184 | 0.18462 | 0.968526 | 4347 | 2 | -0.66874 | 0.6053 | 0.74709 | 1 | 16339 | 1 | -0.66874 |
| C8orf46 | 4 | 0.43858 | 0.58528 | 0.999994 | 12548 | 2 | -0.4589 | 0.6053 | 0.74709 | 1 | 16340 | 1 | -0.4589 |
| SPATA31A5 | 2 | 0.20043 | 0.26241 | 0.995875 | 7771 | 1 | -0.638 | 0.60532 | 0.61851 | 1 | 16341 | 1 | -0.638 |
| RRM2 | 3 | 0.18674 | 0.2959 | 0.999994 | 7380 | 2 | -2.2204 | 0.60546 | 0.69083 | 1 | 16342 | 1 | -2.2204 |
| KLHL11 | 5 | 0.93892 | 0.93912 | 0.999994 | 20048 | 0 | 0.33899 | 0.60546 | 0.77169 | 1 | 16343 | 2 | 0.33899 |
| GORASP1 | 5 | 0.1805 | 0.34734 | 0.999994 | 7230 | 3 | -0.9236 | 0.60546 | 0.77169 | 1 | 16344 | 1 | -0.9236 |
| CDK7 | 4 | 0.86035 | 0.85995 | 0.999994 | 18449 | 0 | 0.50879 | 0.6055 | 0.74725 | 1 | 16345 | 2 | 0.50879 |
| DRP2 | 5 | 0.18404 | 0.35242 | 0.999994 | 7318 | 2 | 0.047803 | 0.60564 | 0.77178 | 1 | 16346 | 2 | 0.047803 |
| NonTargetingControlG | 2 | 0.072436 | 0.12632 | 0.951052 | 3938 | 1 | -3.657 | 0.60564 | 0.61878 | 1 | 16347 | 1 | -3.657 |
| NonTargetingControlG | 2 | 0.25787 | 0.3048 | 0.999994 | 9193 | 1 | -0.7037 | 0.60597 | 0.61904 | 1 | 16348 | 1 | -0.7037 |
| ATG16L1 | 5 | 0.31732 | 0.49518 | 0.999994 | 10460 | 1 | 0.53088 | 0.60597 | 0.77197 | 1 | 16349 | 2 | 0.53088 |
| hsa-mir-106b | 4 | 0.81246 | 0.81262 | 0.999994 | 17660 | 1 | 0.27172 | 0.60611 | 0.74775 | 1 | 16350 | 2 | 0.27172 |
| SNX30 | 4 | 0.12815 | 0.26441 | 0.996734 | 5882 | 2 | -0.44123 | 0.60612 | 0.74775 | 1 | 16351 | 2 | -0.44123 |
| ELTD1 | 6 | 0.62362 | 0.75751 | 0.999994 | 15285 | 2 | 0.57046 | 0.60619 | 0.80706 | 1 | 16352 | 3 | 0.57046 |
| hsa-mir-4735 | 4 | 0.34538 | 0.48959 | 0.999994 | 11059 | 2 | -0.10252 | 0.60619 | 0.74781 | 1 | 16353 | 2 | -0.10252 |
| CORO7-PAM16 | 4 | 0.21473 | 0.35587 | 0.999994 | 8138 | 2 | -0.3338 | 0.60671 | 0.74823 | 1 | 16354 | 2 | -0.3338 |
| RARRES3 | 6 | 0.40513 | 0.60571 | 0.999994 | 12034 | 2 | 0.28223 | 0.60674 | 0.80744 | 1 | 16355 | 3 | 0.28223 |
| UGT2B28 | 4 | 0.29489 | 0.43774 | 0.999994 | 10014 | 2 | -0.36192 | 0.60677 | 0.74829 | 1 | 16356 | 2 | -0.36192 |
| KCND1 | 6 | 0.70876 | 0.8147 | 0.999994 | 16393 | 1 | 0.11777 | 0.6068 | 0.80748 | 1 | 16357 | 3 | 0.11777 |
| IRX3 | 5 | 0.12466 | 0.26446 | 0.996734 | 5787 | 3 | -1.4038 | 0.60688 | 0.77249 | 1 | 16358 | 1 | -1.4038 |
| OR5AK2 | 6 | 0.079632 | 0.20657 | 0.978744 | 4213 | 4 | -1.2753 | 0.60694 | 0.80759 | 1 | 16359 | 2 | -1.2753 |
| FAM89A | 6 | 0.037568 | 0.11643 | 0.951052 | 2407 | 4 | -2.0523 | 0.60694 | 0.80759 | 1 | 16360 | 2 | -2.0523 |
| ADAM20 | 6 | 0.67148 | 0.79429 | 0.999994 | 15949 | 1 | -0.23525 | 0.60694 | 0.80759 | 1 | 16361 | 2 | -0.23525 |
| BCKDHB | 6 | 0.30478 | 0.51229 | 0.999994 | 10227 | 3 | -0.31944 | 0.60695 | 0.80759 | 1 | 16362 | 2 | -0.31944 |
| DHRS4L2 | 2 | 0.55887 | 0.55943 | 0.999994 | 14349 | 1 | 0.38487 | 0.60697 | 0.61986 | 1 | 16363 | 1 | 0.38487 |
| USP20 | 4 | 0.42221 | 0.56847 | 0.999994 | 12285 | 1 | 0.20637 | 0.60712 | 0.74857 | 1 | 16364 | 2 | 0.20637 |
| hsa-mir-302e | 4 | 0.73662 | 0.74997 | 0.999994 | 16686 | 1 | 0.4144 | 0.60722 | 0.74864 | 1 | 16365 | 2 | 0.4144 |
| hsa-mir-8055 | 4 | 0.71355 | 0.73358 | 0.999994 | 16435 | 1 | 0.59537 | 0.60729 | 0.74871 | 1 | 16366 | 2 | 0.59537 |
| TINF2 | 3 | 0.22578 | 0.34484 | 0.999994 | 8420 | 2 | -0.94667 | 0.60748 | 0.6918 | 1 | 16367 | 1 | -0.94667 |
| MSS51 | 6 | 0.2542 | 0.45295 | 0.999994 | 9107 | 2 | 0.41179 | 0.60768 | 0.80813 | 1 | 16368 | 3 | 0.41179 |
| KRT81 | 5 | 0.2062 | 0.38401 | 0.999994 | 7934 | 1 | 0.48904 | 0.60777 | 0.77299 | 1 | 16369 | 2 | 0.48904 |
| DEFB114 | 6 | 0.43276 | 0.62367 | 0.999994 | 12456 | 2 | 0.59587 | 0.60778 | 0.8082 | 1 | 16370 | 3 | 0.59587 |
| ZIC1 | 6 | 0.12731 | 0.29687 | 0.999994 | 5854 | 3 | -0.50051 | 0.608 | 0.80836 | 1 | 16371 | 3 | -0.50051 |
| RAG2 | 3 | 0.1907 | 0.30096 | 0.999994 | 7504 | 2 | -1.2983 | 0.60802 | 0.69207 | 1 | 16372 | 1 | -1.2983 |
| PCDHA8 | 2 | 0.71091 | 0.71065 | 0.999994 | 16408 | 0 | 0.90267 | 0.60806 | 0.62076 | 1 | 16373 | 1 | 0.90267 |
| ZC2HC1C | 5 | 0.12801 | 0.26957 | 0.998561 | 5877 | 2 | 0.22266 | 0.60807 | 0.77316 | 1 | 16374 | 2 | 0.22266 |
| GPM6B | 5 | 0.50333 | 0.65084 | 0.999994 | 13486 | 2 | 0.62898 | 0.60807 | 0.77316 | 1 | 16375 | 2 | 0.62898 |
| RASSF9 | 4 | 0.4091 | 0.55504 | 0.999994 | 12095 | 1 | 0.59436 | 0.60813 | 0.74939 | 1 | 16376 | 2 | 0.59436 |
| GPR152 | 6 | 0.036432 | 0.11386 | 0.951052 | 2342 | 2 | 0.10875 | 0.60815 | 0.80847 | 1 | 16377 | 3 | 0.10875 |
| RALY | 6 | 0.069084 | 0.18486 | 0.968526 | 3800 | 3 | -0.51264 | 0.60816 | 0.80848 | 1 | 16378 | 2 | -0.51264 |
| SPAG11A | 5 | 0.082975 | 0.19941 | 0.977164 | 4337 | 3 | -1.6037 | 0.60824 | 0.77327 | 1 | 16379 | 2 | -1.6037 |

| | | | | | | | | | | | | | |
|---|---|---|---|---|---|---|---|---|---|---|---|---|---|
| PPM1K | 5 | 0.36242 | 0.53208 | 0.999994 | 11376 | 2 | 0.26868 | 0.60828 | 0.77328 | 1 | 16380 | 2 | 0.26868 |
| PRODH | 5 | 0.26377 | 0.4518 | 0.999994 | 9330 | 2 | 0.10121 | 0.6083 | 0.7733 | 1 | 16381 | 2 | 0.10121 |
| HELLS | 4 | 0.34342 | 0.48755 | 0.999994 | 11017 | 1 | 0.77645 | 0.60857 | 0.74974 | 1 | 16382 | 2 | 0.77645 |
| NonTargetingControlG | 2 | 0.61563 | 0.61544 | 0.999994 | 15153 | 0 | 0.5421 | 0.60875 | 0.62132 | 1 | 16383 | 1 | 0.5421 |
| PCDH15 | 6 | 0.27644 | 0.47928 | 0.999994 | 9628 | 3 | -0.37439 | 0.60876 | 0.80892 | 1 | 16384 | 2 | -0.37439 |
| ADD3 | 6 | 0.97694 | 0.97687 | 0.999994 | 21012 | 0 | 0.51098 | 0.60877 | 0.80893 | 1 | 16385 | 3 | 0.51098 |
| PPP2CB | 3 | 0.36522 | 0.46425 | 0.999994 | 11425 | 1 | -0.54352 | 0.60884 | 0.69246 | 1 | 16386 | 1 | -0.54352 |
| NonTargetingControlG | 1 | 0.39116 | 0.39129 | 0.999994 | 11817 | 0 | -0.32582 | 0.60884 | 0.60847 | 1 | 16387 | 0 | -0.32582 |
| ESR2 | 6 | 0.18912 | 0.37441 | 0.999994 | 7453 | 2 | 0.24144 | 0.60887 | 0.809 | 1 | 16388 | 3 | 0.24144 |
| hsa-mir-3939 | 4 | 0.2929 | 0.43576 | 0.999994 | 9973 | 2 | -0.32695 | 0.60909 | 0.75015 | 1 | 16389 | 2 | -0.32695 |
| hsa-mir-4434 | 4 | 0.38474 | 0.52999 | 0.999994 | 11712 | 1 | -0.15714 | 0.60909 | 0.75015 | 1 | 16390 | 1 | -0.15714 |
| TAP1 | 5 | 0.018887 | 0.061047 | 0.932932 | 1402 | 3 | -1.1417 | 0.6093 | 0.77387 | 1 | 16391 | 1 | -1.1417 |
| EVI5L | 5 | 0.050227 | 0.14091 | 0.959146 | 2987 | 3 | -1.307 | 0.6093 | 0.77387 | 1 | 16392 | 1 | -1.307 |
| PABPC3 | 5 | 0.25118 | 0.44169 | 0.999994 | 9040 | 2 | -0.271 | 0.6093 | 0.77387 | 1 | 16393 | 2 | -0.271 |
| TRIM8 | 5 | 0.26164 | 0.45007 | 0.999994 | 9278 | 2 | -0.61415 | 0.6093 | 0.77387 | 1 | 16394 | 1 | -0.61415 |
| NonTargetingControlG | 2 | 0.034757 | 0.063145 | 0.937389 | 2255 | 1 | -3.9583 | 0.60945 | 0.62188 | 1 | 16395 | 1 | -3.9583 |
| MOV10L1 | 6 | 0.019902 | 0.071653 | 0.93914 | 1454 | 4 | -1.9917 | 0.60949 | 0.80947 | 1 | 16396 | 1 | -1.9917 |
| HDHD2 | 6 | 0.64956 | 0.77728 | 0.999994 | 15651 | 2 | 0.14912 | 0.60949 | 0.80947 | 1 | 16397 | 2 | 0.14912 |
| MAPRE1 | 6 | 0.28659 | 0.49113 | 0.999994 | 9848 | 3 | -0.15917 | 0.60949 | 0.80947 | 1 | 16398 | 2 | -0.15917 |
| DGKQ | 6 | 0.11222 | 0.27095 | 0.998561 | 5376 | 4 | -0.86054 | 0.60949 | 0.80947 | 1 | 16399 | 2 | -0.86054 |
| WDR66 | 4 | 0.40671 | 0.55258 | 0.999994 | 12055 | 1 | 0.40654 | 0.60975 | 0.75069 | 1 | 16400 | 2 | 0.40654 |
| VENTX | 4 | 0.7811 | 0.7848 | 0.999994 | 17225 | 1 | 0.32222 | 0.60975 | 0.75069 | 1 | 16401 | 2 | 0.32222 |
| SMIM11 | 5 | 0.11208 | 0.24522 | 0.98979 | 5368 | 3 | -1.4395 | 0.60978 | 0.77414 | 1 | 16402 | 2 | -1.4395 |
| KCND3 | 6 | 0.11404 | 0.27444 | 0.998561 | 5450 | 3 | -0.69576 | 0.60986 | 0.80974 | 1 | 16403 | 3 | -0.69576 |
| hsa-mir-148a | 3 | 0.3522 | 0.45696 | 0.999994 | 11198 | 1 | 0.42601 | 0.60991 | 0.69299 | 1 | 16404 | 1 | 0.42601 |
| hsa-mir-548h-2 | 2 | 0.40584 | 0.42166 | 0.999994 | 12042 | 1 | -0.010256 | 0.61005 | 0.62239 | 1 | 16405 | 1 | -0.010256 |
| RPL6 | 2 | 0.26162 | 0.30757 | 0.999994 | 9276 | 1 | -0.80749 | 0.61011 | 0.62244 | 1 | 16406 | 1 | -0.80749 |
| hsa-mir-219a-2 | 4 | 0.8934 | 0.89311 | 0.999994 | 19094 | 0 | 0.059419 | 0.6102 | 0.75105 | 1 | 16407 | 1 | 0.059419 |
| MIA | 4 | 0.23028 | 0.37183 | 0.999994 | 8533 | 2 | -0.0044697 | 0.6102 | 0.75106 | 1 | 16408 | 2 | -0.0044697 |
| FSCN3 | 6 | 0.25261 | 0.45104 | 0.999994 | 9069 | 3 | -0.085332 | 0.61021 | 0.80999 | 1 | 16409 | 3 | -0.085332 |
| RPL7 | 4 | 0.10971 | 0.23235 | 0.984502 | 5295 | 2 | -1.7774 | 0.61029 | 0.75112 | 1 | 16410 | 1 | -1.7774 |
| SPATA6 | 5 | 0.15477 | 0.30987 | 0.999994 | 6588 | 3 | -1.0328 | 0.61041 | 0.7745 | 1 | 16411 | 2 | -1.0328 |
| FXR1 | 5 | 0.11788 | 0.25412 | 0.994632 | 5569 | 3 | -1.524 | 0.61044 | 0.77452 | 1 | 16412 | 2 | -1.524 |
| DMRTA2 | 5 | 0.041796 | 0.1204 | 0.951052 | 2604 | 4 | -1.1138 | 0.61044 | 0.77452 | 1 | 16413 | 1 | -1.1138 |
| SESTD1 | 5 | 0.67544 | 0.75562 | 0.999994 | 16011 | 1 | 0.41706 | 0.61044 | 0.77452 | 1 | 16414 | 1 | 0.41706 |
| MAPK12 | 5 | 0.23145 | 0.4192 | 0.999994 | 8555 | 3 | -0.69519 | 0.61044 | 0.77452 | 1 | 16415 | 2 | -0.69519 |
| TAS2R38 | 6 | 0.34384 | 0.55708 | 0.999994 | 11025 | 3 | -0.047403 | 0.61071 | 0.81036 | 1 | 16416 | 3 | -0.047403 |
| hsa-mir-6735 | 3 | 0.28861 | 0.42243 | 0.999994 | 9887 | 2 | -0.5876 | 0.6108 | 0.69341 | 1 | 16417 | 1 | -0.5876 |
| hsa-mir-346 | 3 | 0.22712 | 0.34649 | 0.999994 | 8458 | 2 | -0.97463 | 0.6108 | 0.69341 | 1 | 16418 | 1 | -0.97463 |
| HOXB5 | 6 | 0.33236 | 0.54397 | 0.999994 | 10774 | 2 | 0.1066 | 0.6108 | 0.81043 | 1 | 16419 | 3 | 0.1066 |
| C1QTNF8 | 5 | 0.56814 | 0.70789 | 0.999994 | 14514 | 2 | 0.35996 | 0.61096 | 0.77482 | 1 | 16420 | 2 | 0.35996 |
| NonTargetingControlG | 2 | 0.63463 | 0.6344 | 0.999994 | 15438 | 0 | 0.58658 | 0.61102 | 0.62321 | 1 | 16421 | 1 | 0.58658 |
| ZFP36L1 | 4 | 0.028523 | 0.078504 | 0.93914 | 1926 | 3 | -3.5218 | 0.61102 | 0.75172 | 1 | 16422 | 1 | -3.5218 |
| LRRC37A | 6 | 0.1705 | 0.3514 | 0.999994 | 6994 | 2 | 0.28607 | 0.61105 | 0.8106 | 1 | 16423 | 3 | 0.28607 |
| SEPT4 | 4 | 0.10273 | 0.21995 | 0.980699 | 5060 | 1 | 0.45949 | 0.61111 | 0.75179 | 1 | 16424 | 2 | 0.45949 |
| OR10A4 | 5 | 0.16773 | 0.32876 | 0.999994 | 6931 | 3 | -0.99217 | 0.61143 | 0.7751 | 1 | 16425 | 2 | -0.99217 |
| ZNF396 | 6 | 0.88422 | 0.89235 | 0.999994 | 18913 | 1 | 0.50514 | 0.61147 | 0.81089 | 1 | 16426 | 3 | 0.50514 |
| ACSM1 | 6 | 0.466 | 0.64578 | 0.999994 | 12951 | 2 | -0.023513 | 0.61147 | 0.8109 | 1 | 16427 | 3 | -0.023513 |
| NonTargetingControlG | 2 | 0.6316 | 0.63136 | 0.999994 | 15393 | 0 | 0.55183 | 0.61151 | 0.62361 | 1 | 16428 | 1 | 0.55183 |
| SEPN1 | 5 | 0.40234 | 0.56547 | 0.999994 | 11986 | 2 | 0.19532 | 0.61154 | 0.77516 | 1 | 16429 | 2 | 0.19532 |
| AMY1B | 2 | 0.52039 | 0.52277 | 0.999994 | 13754 | 1 | -0.46571 | 0.61158 | 0.62367 | 1 | 16430 | 1 | -0.46571 |
| CCDC22 | 4 | 0.8673 | 0.86687 | 0.999994 | 18581 | 0 | 0.47868 | 0.6117 | 0.75228 | 1 | 16431 | 1 | 0.47868 |

| | | | | | | | | | | | | | |
|---|---|---|---|---|---|---|---|---|---|---|---|---|---|
| MYL3 | 5 | 0.33304 | 0.50801 | 0.999994 | 10789 | 2 | -0.58478 | 0.61177 | 0.77529 | 1 | 16432 | 1 | -0.58478 |
| GPR17 | 4 | 0.15982 | 0.2993 | 0.999994 | 6730 | 2 | -1.3273 | 0.61182 | 0.75238 | 1 | 16433 | 1 | -1.3273 |
| hsa-mir-3912 | 4 | 0.73182 | 0.74651 | 0.999994 | 16625 | 1 | 0.30564 | 0.61183 | 0.75239 | 1 | 16434 | 2 | 0.30564 |
| FAM84A | 6 | 0.05198 | 0.14845 | 0.965554 | 3065 | 3 | -1.5435 | 0.61186 | 0.81119 | 1 | 16435 | 2 | -1.5435 |
| DCST1 | 6 | 0.051945 | 0.14838 | 0.965554 | 3063 | 4 | -1.4 | 0.61186 | 0.81119 | 1 | 16436 | 2 | -1.4 |
| C1orf54 | 6 | 0.056543 | 0.15828 | 0.967126 | 3272 | 4 | -1.4715 | 0.61186 | 0.81119 | 1 | 16437 | 2 | -1.4715 |
| FAM21C | 4 | 0.16997 | 0.30975 | 0.999994 | 6983 | 2 | -1.0584 | 0.61203 | 0.75255 | 1 | 16438 | 1 | -1.0584 |
| HSBP1L1 | 5 | 0.04905 | 0.13807 | 0.957317 | 2934 | 2 | 0.31515 | 0.61204 | 0.77545 | 1 | 16439 | 2 | 0.31515 |
| DHH | 5 | 0.0027813 | 0.01196 | 0.729405 | 340 | 3 | -7.5036 | 0.6122 | 0.77554 | 1 | 16440 | 2 | -7.5036 |
| SPTAN1 | 6 | 0.11254 | 0.2716 | 0.998561 | 5388 | 3 | -0.91206 | 0.61234 | 0.81152 | 1 | 16441 | 2 | -0.91206 |
| KRTAP25-1 | 4 | 0.66214 | 0.70052 | 0.999994 | 15810 | 1 | 0.28664 | 0.61239 | 0.75281 | 1 | 16442 | 2 | 0.28664 |
| PPEF1 | 5 | 0.14355 | 0.29317 | 0.999994 | 6286 | 2 | -0.055909 | 0.61243 | 0.77567 | 1 | 16443 | 2 | -0.055909 |
| SLC52A1 | 6 | 0.54032 | 0.69677 | 0.999994 | 14058 | 2 | 0.011342 | 0.61265 | 0.81178 | 1 | 16444 | 2 | 0.011342 |
| LAMB1 | 5 | 0.20382 | 0.38067 | 0.999994 | 7875 | 3 | -0.86105 | 0.61267 | 0.77581 | 1 | 16445 | 2 | -0.86105 |
| BHLHA15 | 4 | 0.23484 | 0.37647 | 0.999994 | 8639 | 1 | 0.51594 | 0.61268 | 0.75304 | 1 | 16446 | 1 | 0.51594 |
| DEFB125 | 5 | 0.092726 | 0.21494 | 0.979587 | 4711 | 2 | 0.24827 | 0.61279 | 0.77589 | 1 | 16447 | 2 | 0.24827 |
| CCDC63 | 4 | 0.37644 | 0.5215 | 0.999994 | 11601 | 1 | 0.45764 | 0.61282 | 0.75315 | 1 | 16448 | 2 | 0.45764 |
| TMPRSS6 | 6 | 0.32937 | 0.54052 | 0.999994 | 10708 | 3 | -0.34731 | 0.61285 | 0.81193 | 1 | 16449 | 2 | -0.34731 |
| PDE8A | 6 | 0.26326 | 0.46372 | 0.999994 | 9320 | 3 | -0.11423 | 0.61292 | 0.81199 | 1 | 16450 | 3 | -0.11423 |
| EDA | 5 | 0.0003014 | 0.0014217 | 0.384267 | 80 | 2 | -0.4098 | 0.61296 | 0.776 | 1 | 16451 | 2 | -0.4098 |
| ZFYVE19 | 5 | 0.56996 | 0.70947 | 0.999994 | 14540 | 2 | 0.28366 | 0.61306 | 0.77606 | 1 | 16452 | 2 | 0.28366 |
| CD84 | 6 | 0.39109 | 0.59664 | 0.999994 | 11816 | 2 | -0.11914 | 0.61312 | 0.81213 | 1 | 16453 | 2 | -0.11914 |
| TMEM239 | 6 | 0.061158 | 0.16824 | 0.967126 | 3480 | 3 | -1.0313 | 0.61312 | 0.81213 | 1 | 16454 | 2 | -1.0313 |
| KCNS2 | 6 | 0.36066 | 0.57605 | 0.999994 | 11347 | 2 | -0.18503 | 0.61312 | 0.81213 | 1 | 16455 | 1 | -0.18503 |
| NonTargetingControlG | 2 | 0.66994 | 0.6697 | 0.999994 | 15931 | 0 | 0.69884 | 0.61318 | 0.62497 | 1 | 16456 | 1 | 0.69884 |
| STK24 | 6 | 0.53162 | 0.6907 | 0.999994 | 13926 | 2 | 0.24899 | 0.61322 | 0.81222 | 1 | 16457 | 3 | 0.24899 |
| TNC | 6 | 0.24467 | 0.44157 | 0.999994 | 8894 | 2 | 0.039144 | 0.61325 | 0.81224 | 1 | 16458 | 3 | 0.039144 |
| HAX1 | 5 | 0.030101 | 0.090749 | 0.94395 | 2013 | 2 | 0.57876 | 0.61331 | 0.77619 | 1 | 16459 | 2 | 0.57876 |
| DDOST | 4 | 0.049558 | 0.12057 | 0.951052 | 2958 | 2 | -2.0199 | 0.61333 | 0.75358 | 1 | 16460 | 1 | -2.0199 |
| YARS | 4 | 0.043194 | 0.10814 | 0.951052 | 2683 | 3 | -2.9207 | 0.61333 | 0.75358 | 1 | 16461 | 1 | -2.9207 |
| UVRAG | 4 | 0.48167 | 0.60824 | 0.999994 | 13164 | 1 | 0.48066 | 0.61333 | 0.75358 | 1 | 16462 | 2 | 0.48066 |
| SPATA31C1 | 6 | 0.58917 | 0.73197 | 0.999994 | 14809 | 2 | -0.18689 | 0.61347 | 0.81241 | 1 | 16463 | 2 | -0.18689 |
| hsa-mir-6778 | 3 | 0.2598 | 0.38718 | 0.999994 | 9234 | 2 | -0.81607 | 0.61348 | 0.69477 | 1 | 16464 | 1 | -0.81607 |
| GCLM | 6 | 0.036277 | 0.11349 | 0.951052 | 2332 | 4 | -1.6282 | 0.61366 | 0.81254 | 1 | 16465 | 1 | -1.6282 |
| KIF25 | 4 | 0.31691 | 0.46029 | 0.999994 | 10452 | 2 | -0.22852 | 0.6138 | 0.75398 | 1 | 16466 | 2 | -0.22852 |
| hsa-mir-6757 | 4 | 0.87715 | 0.87683 | 0.999994 | 18776 | 0 | 0.57558 | 0.61381 | 0.75399 | 1 | 16467 | 2 | 0.57558 |
| EPS15 | 6 | 0.30502 | 0.51257 | 0.999994 | 10232 | 3 | -0.14881 | 0.61384 | 0.81268 | 1 | 16468 | 3 | -0.14881 |
| PRR4 | 6 | 0.053441 | 0.15153 | 0.967126 | 3137 | 3 | -1.7034 | 0.61384 | 0.81269 | 1 | 16469 | 2 | -1.7034 |
| RPL14 | 4 | 0.84932 | 0.84883 | 0.999994 | 18247 | 0 | 0.63763 | 0.61386 | 0.75402 | 1 | 16470 | 2 | 0.63763 |
| NFIC | 6 | 0.5874 | 0.73068 | 0.999994 | 14780 | 2 | 0.13311 | 0.61388 | 0.81271 | 1 | 16471 | 3 | 0.13311 |
| ARPC4-TTLL3 | 2 | 0.67354 | 0.67328 | 0.999994 | 15980 | 0 | 0.72199 | 0.61397 | 0.62562 | 1 | 16472 | 1 | 0.72199 |
| IQCG | 6 | 0.081834 | 0.2111 | 0.979587 | 4293 | 2 | 0.2442 | 0.61401 | 0.81282 | 1 | 16473 | 3 | 0.2442 |
| NonTargetingControlG | 2 | 0.47244 | 0.47912 | 0.999994 | 13040 | 1 | 0.23697 | 0.61411 | 0.62573 | 1 | 16474 | 1 | 0.23697 |
| KLF4 | 3 | 0.10958 | 0.19574 | 0.975797 | 5293 | 1 | -0.16379 | 0.61411 | 0.6951 | 1 | 16475 | 1 | -0.16379 |
| C14orf182 | 6 | 0.022875 | 0.081201 | 0.93914 | 1643 | 4 | -2.2409 | 0.61424 | 0.81299 | 1 | 16476 | 1 | -2.2409 |
| BFAR | 6 | 0.071094 | 0.18909 | 0.969834 | 3879 | 4 | -1.4333 | 0.61424 | 0.81299 | 1 | 16477 | 1 | -1.4333 |
| RBM44 | 4 | 0.016156 | 0.052399 | 0.914561 | 1252 | 1 | 0.66557 | 0.61431 | 0.75439 | 1 | 16478 | 2 | 0.66557 |
| INPP5B | 5 | 0.12479 | 0.26466 | 0.996734 | 5790 | 2 | -0.52536 | 0.61454 | 0.7769 | 1 | 16479 | 1 | -0.52536 |
| GPR133 | 6 | 0.5953 | 0.73652 | 0.999994 | 14894 | 2 | -0.22546 | 0.61461 | 0.81327 | 1 | 16480 | 2 | -0.22546 |
| MAGEA2B | 1 | 0.38538 | 0.38554 | 0.999994 | 11723 | 0 | -0.34579 | 0.61462 | 0.6143 | 1 | 16481 | 0 | -0.34579 |
| ANKFN1 | 4 | 0.63946 | 0.6872 | 0.999994 | 15508 | 1 | -0.65637 | 0.61469 | 0.75467 | 1 | 16482 | 1 | -0.65637 |
| C6 | 6 | 0.20196 | 0.39027 | 0.999994 | 7829 | 2 | 0.13555 | 0.61489 | 0.81346 | 1 | 16483 | 3 | 0.13555 |

| | | | | | | | | | | | | | |
|---|---|---|---|---|---|---|---|---|---|---|---|---|---|
| CREG2 | 6 | 0.030919 | 0.10097 | 0.946414 | 2057 | 4 | -1.9679 | 0.61492 | 0.81348 | 1 | 16484 | 1 | -1.9679 |
| UNC93B1 | 6 | 0.28453 | 0.48871 | 0.999994 | 9796 | 3 | -0.1382 | 0.61504 | 0.81357 | 1 | 16485 | 3 | -0.1382 |
| NonTargetingControlG | 2 | 0.48961 | 0.49448 | 0.999994 | 13273 | 1 | 0.11524 | 0.61514 | 0.62657 | 1 | 16486 | 1 | 0.11524 |
| C1orf110 | 6 | 0.24654 | 0.44382 | 0.999994 | 8934 | 3 | -0.059895 | 0.61515 | 0.81365 | 1 | 16487 | 3 | -0.059895 |
| OR5H15 | 5 | 0.29584 | 0.4777 | 0.999994 | 10035 | 2 | 0.022234 | 0.61519 | 0.77725 | 1 | 16488 | 2 | 0.022234 |
| hsa-mir-3171 | 2 | 0.062712 | 0.11043 | 0.951052 | 3544 | 1 | -3.7419 | 0.61519 | 0.62662 | 1 | 16489 | 1 | -3.7419 |
| LSMEM2 | 6 | 0.23127 | 0.42562 | 0.999994 | 8550 | 3 | -0.29869 | 0.61522 | 0.8137 | 1 | 16490 | 3 | -0.29869 |
| OLFML3 | 6 | 0.24701 | 0.44437 | 0.999994 | 8944 | 3 | -0.60968 | 0.61523 | 0.81371 | 1 | 16491 | 2 | -0.60968 |
| RTN4RL1 | 5 | 0.15389 | 0.3086 | 0.999994 | 6563 | 2 | -0.50112 | 0.61534 | 0.77734 | 1 | 16492 | 2 | -0.50112 |
| PRPF8 | 4 | 0.054201 | 0.1297 | 0.95318 | 3164 | 3 | -1.8338 | 0.61536 | 0.75521 | 1 | 16493 | 1 | -1.8338 |
| STRADB | 6 | 0.16182 | 0.34057 | 0.999994 | 6785 | 3 | -0.6549 | 0.61541 | 0.81385 | 1 | 16494 | 2 | -0.6549 |
| CEBPE | 6 | 0.015128 | 0.056132 | 0.9235 | 1197 | 4 | -2.5831 | 0.61541 | 0.81385 | 1 | 16495 | 1 | -2.5831 |
| AUP1 | 6 | 0.085079 | 0.2177 | 0.980699 | 4412 | 3 | -1.119 | 0.61541 | 0.81385 | 1 | 16496 | 2 | -1.119 |
| ERBB3 | 5 | 0.50846 | 0.65532 | 0.999994 | 13561 | 2 | 0.14123 | 0.61543 | 0.7774 | 1 | 16497 | 2 | 0.14123 |
| SOCS4 | 5 | 0.18704 | 0.35673 | 0.999994 | 7391 | 3 | -0.92544 | 0.61561 | 0.77751 | 1 | 16498 | 2 | -0.92544 |
| NABP2 | 6 | 0.075031 | 0.19722 | 0.977164 | 4037 | 3 | -0.50947 | 0.61568 | 0.81404 | 1 | 16499 | 2 | -0.50947 |
| hsa-mir-4427 | 4 | 0.39243 | 0.5379 | 0.999994 | 11841 | 1 | 0.18505 | 0.61584 | 0.75561 | 1 | 16500 | 2 | 0.18505 |
| CLEC4M | 5 | 0.88315 | 0.88332 | 0.999994 | 18894 | 1 | 0.27507 | 0.61588 | 0.77766 | 1 | 16501 | 2 | 0.27507 |
| TSNAX | 6 | 0.65084 | 0.77825 | 0.999994 | 15672 | 2 | 0.12151 | 0.61588 | 0.8142 | 1 | 16502 | 3 | 0.12151 |
| LIN52 | 4 | 0.034689 | 0.091082 | 0.94395 | 2248 | 2 | -0.34976 | 0.61594 | 0.75569 | 1 | 16503 | 2 | -0.34976 |
| HES6 | 4 | 0.0067989 | 0.023538 | 0.832699 | 646 | 3 | -4.886 | 0.61599 | 0.75573 | 1 | 16504 | 1 | -4.886 |
| ZNF136 | 6 | 0.3659 | 0.58064 | 0.999994 | 11436 | 1 | 0.48781 | 0.61601 | 0.81429 | 1 | 16505 | 3 | 0.48781 |
| HSPA4 | 6 | 0.19983 | 0.38767 | 0.999994 | 7762 | 3 | -0.49956 | 0.61609 | 0.81434 | 1 | 16506 | 2 | -0.49956 |
| NonTargetingControlG | 1 | 0.38391 | 0.38402 | 0.999994 | 11702 | 0 | -0.38628 | 0.61609 | 0.61578 | 1 | 16507 | 0 | -0.38628 |
| MTIF3 | 5 | 0.13902 | 0.28637 | 0.999632 | 6164 | 1 | 0.4512 | 0.61619 | 0.77786 | 1 | 16508 | 2 | 0.4512 |
| L3HYPDH | 6 | 0.09206 | 0.23165 | 0.984454 | 4681 | 3 | -0.32638 | 0.61627 | 0.81447 | 1 | 16509 | 3 | -0.32638 |
| HIST1H2BF | 4 | 0.072466 | 0.16459 | 0.967126 | 3940 | 3 | -1.8548 | 0.6163 | 0.75599 | 1 | 16510 | 1 | -1.8548 |
| PDE4DIP | 4 | 0.020599 | 0.06191 | 0.937033 | 1487 | 2 | -3.4451 | 0.6165 | 0.75614 | 1 | 16511 | 2 | -3.4451 |
| ATP5D | 6 | 0.1294 | 0.29954 | 0.999994 | 5926 | 3 | -0.75317 | 0.61656 | 0.81467 | 1 | 16512 | 2 | -0.75317 |
| FEM1C | 6 | 0.11928 | 0.28438 | 0.999632 | 5607 | 4 | -1.3303 | 0.61658 | 0.81469 | 1 | 16513 | 1 | -1.3303 |
| ATP6AP1L | 6 | 0.068773 | 0.1842 | 0.968526 | 3789 | 4 | -1.3933 | 0.61658 | 0.81469 | 1 | 16514 | 2 | -1.3933 |
| KRT20 | 5 | 0.18551 | 0.35456 | 0.999994 | 7354 | 3 | -0.7361 | 0.61662 | 0.7781 | 1 | 16515 | 2 | -0.7361 |
| QSOX2 | 6 | 0.21447 | 0.40553 | 0.999994 | 8132 | 2 | 0.39014 | 0.61669 | 0.81477 | 1 | 16516 | 3 | 0.39014 |
| NonTargetingControlG | 2 | 0.43163 | 0.44341 | 0.999994 | 12440 | 1 | 0.065128 | 0.61678 | 0.62793 | 1 | 16517 | 1 | 0.065128 |
| MST4 | 6 | 0.37458 | 0.5861 | 0.999994 | 11576 | 2 | 0.23296 | 0.61687 | 0.8149 | 1 | 16518 | 3 | 0.23296 |
| CCNG1 | 5 | 0.34203 | 0.51537 | 0.999994 | 10985 | 2 | 0.46778 | 0.61699 | 0.77832 | 1 | 16519 | 2 | 0.46778 |
| RIOK2 | 4 | 0.17607 | 0.31609 | 0.999994 | 7133 | 2 | -0.65514 | 0.617 | 0.75656 | 1 | 16520 | 2 | -0.65514 |
| GALNT8 | 5 | 0.73852 | 0.78492 | 0.999994 | 16713 | 1 | -0.52837 | 0.61702 | 0.77834 | 1 | 16521 | 2 | -0.52837 |
| NonTargetingControlG | 2 | 0.46289 | 0.4707 | 0.999994 | 12904 | 1 | 0.058753 | 0.61713 | 0.62823 | 1 | 16522 | 1 | 0.058753 |
| PDE1C | 4 | 0.31266 | 0.45599 | 0.999994 | 10373 | 2 | -0.87152 | 0.61714 | 0.75667 | 1 | 16523 | 1 | -0.87152 |
| ARC | 4 | 0.30969 | 0.45292 | 0.999994 | 10320 | 2 | -0.61474 | 0.61714 | 0.75667 | 1 | 16524 | 1 | -0.61474 |
| ZDHHC20 | 4 | 0.18396 | 0.32418 | 0.999994 | 7315 | 2 | -0.88217 | 0.61714 | 0.75667 | 1 | 16525 | 1 | -0.88217 |
| UXT | 4 | 0.039421 | 0.10064 | 0.946414 | 2485 | 3 | -4.041 | 0.61714 | 0.75667 | 1 | 16526 | 1 | -4.041 |
| RAN | 4 | 0.38848 | 0.53382 | 0.999994 | 11775 | 1 | 0.34902 | 0.61714 | 0.75667 | 1 | 16527 | 1 | 0.34902 |
| CRYBA1 | 5 | 0.39452 | 0.559 | 0.999994 | 11865 | 2 | 0.43789 | 0.61718 | 0.77842 | 1 | 16528 | 2 | 0.43789 |
| NonTargetingControlG | 2 | 0.50844 | 0.51166 | 0.999994 | 13560 | 1 | 0.22511 | 0.61719 | 0.62827 | 1 | 16529 | 1 | 0.22511 |
| WNK1 | 6 | 0.083713 | 0.2149 | 0.979587 | 4366 | 4 | -1.555 | 0.61722 | 0.81515 | 1 | 16530 | 2 | -1.555 |
| SLC9C1 | 4 | 0.91909 | 0.91888 | 0.999994 | 19608 | 0 | 0.58608 | 0.61724 | 0.75676 | 1 | 16531 | 2 | 0.58608 |
| ANAPC7 | 5 | 0.34771 | 0.52003 | 0.999994 | 11108 | 2 | 0.23216 | 0.6174 | 0.77856 | 1 | 16532 | 2 | 0.23216 |
| NonTargetingControlG | 1 | 0.38257 | 0.38268 | 0.999994 | 11684 | 0 | -0.38368 | 0.61743 | 0.61715 | 1 | 16533 | 0 | -0.38368 |
| SV2A | 6 | 0.9429 | 0.94287 | 0.999994 | 20150 | 0 | 0.22025 | 0.6175 | 0.81536 | 1 | 16534 | 3 | 0.22025 |
| C16orf86 | 4 | 0.73959 | 0.75216 | 0.999994 | 16733 | 1 | 0.22984 | 0.61752 | 0.75699 | 1 | 16535 | 2 | 0.22984 |

| | | | | | | | | | | | | | |
|---|---|---|---|---|---|---|---|---|---|---|---|---|---|
| SLC30A6 | 5 | 0.45352 | 0.6082 | 0.999994 | 12750 | 1 | 0.33016 | 0.61754 | 0.77863 | 1 | 16536 | 2 | 0.33016 |
| RTP4 | 6 | 0.43939 | 0.628 | 0.999994 | 12565 | 2 | 0.1444 | 0.61764 | 0.81546 | 1 | 16537 | 3 | 0.1444 |
| ST7 | 5 | 0.49392 | 0.64265 | 0.999994 | 13325 | 2 | 0.53466 | 0.61764 | 0.77869 | 1 | 16538 | 2 | 0.53466 |
| hsa-mir-2277 | 4 | 0.33635 | 0.4802 | 0.999994 | 10855 | 2 | -0.20866 | 0.61766 | 0.7571 | 1 | 16539 | 2 | -0.20866 |
| NonTargetingControlG | 2 | 0.26729 | 0.31185 | 0.999994 | 9403 | 1 | -0.69114 | 0.61767 | 0.62865 | 1 | 16540 | 1 | -0.69114 |
| PADI6 | 4 | 0.3349 | 0.47872 | 0.999994 | 10829 | 2 | -0.17401 | 0.6177 | 0.75714 | 1 | 16541 | 2 | -0.17401 |
| BUB3 | 6 | 0.092868 | 0.23324 | 0.984913 | 4713 | 4 | -1.1656 | 0.61784 | 0.8156 | 1 | 16542 | 2 | -1.1656 |
| STRIP1 | 6 | 0.02634 | 0.090165 | 0.94395 | 1819 | 4 | -2.2316 | 0.61784 | 0.8156 | 1 | 16543 | 1 | -2.2316 |
| NonTargetingControlG | 2 | 0.5015 | 0.5053 | 0.999994 | 13448 | 1 | 0.17449 | 0.61799 | 0.62891 | 1 | 16544 | 1 | 0.17449 |
| LZTS2 | 5 | 0.016307 | 0.053908 | 0.921374 | 1262 | 4 | -3.7335 | 0.61805 | 0.77891 | 1 | 16545 | 1 | -3.7335 |
| CHRNA9 | 5 | 0.089506 | 0.20979 | 0.979587 | 4577 | 3 | -1.6926 | 0.61805 | 0.77891 | 1 | 16546 | 1 | -1.6926 |
| SSH3 | 5 | 0.23548 | 0.42478 | 0.999994 | 8659 | 3 | -0.66716 | 0.61805 | 0.77891 | 1 | 16547 | 2 | -0.66716 |
| FAM167A | 4 | 0.026884 | 0.07513 | 0.93914 | 1851 | 2 | -3.2968 | 0.61836 | 0.75766 | 1 | 16548 | 2 | -3.2968 |
| ERI3 | 5 | 0.0053862 | 0.022336 | 0.819701 | 549 | 2 | 0.10683 | 0.61864 | 0.77925 | 1 | 16549 | 2 | 0.10683 |
| UNC5B | 5 | 0.86527 | 0.86689 | 0.999994 | 18544 | 1 | 0.057298 | 0.61864 | 0.77925 | 1 | 16550 | 2 | 0.057298 |
| RAB43 | 5 | 0.11826 | 0.25474 | 0.994835 | 5581 | 2 | 0.47128 | 0.61864 | 0.77925 | 1 | 16551 | 2 | 0.47128 |
| WFS1 | 5 | 0.80765 | 0.8244 | 0.999994 | 17591 | 1 | 0.14624 | 0.61864 | 0.77925 | 1 | 16552 | 2 | 0.14624 |
| SH2B1 | 2 | 0.38267 | 0.40248 | 0.999994 | 11686 | 1 | -0.31245 | 0.61889 | 0.62967 | 1 | 16553 | 1 | -0.31245 |
| CNTROB | 6 | 0.67797 | 0.79942 | 0.999994 | 16060 | 1 | 0.14053 | 0.61894 | 0.81641 | 1 | 16554 | 3 | 0.14053 |
| BARD1 | 6 | 0.076649 | 0.20058 | 0.977164 | 4098 | 3 | -1.0836 | 0.61907 | 0.8165 | 1 | 16555 | 2 | -1.0836 |
| CCDC66 | 6 | 0.090842 | 0.2292 | 0.983354 | 4632 | 3 | -1.0501 | 0.61907 | 0.8165 | 1 | 16556 | 1 | -1.0501 |
| EFCAB4B | 6 | 0.12497 | 0.29389 | 0.999994 | 5794 | 3 | -0.6697 | 0.61907 | 0.8165 | 1 | 16557 | 1 | -0.6697 |
| NonTargetingControlG | 2 | 0.12266 | 0.20472 | 0.978186 | 5728 | 1 | -3.1001 | 0.61923 | 0.62996 | 1 | 16558 | 1 | -3.1001 |
| SSH1 | 5 | 0.13571 | 0.28129 | 0.999632 | 6073 | 3 | -1.2029 | 0.61925 | 0.77958 | 1 | 16559 | 2 | -1.2029 |
| SCAI | 6 | 0.24036 | 0.43644 | 0.999994 | 8781 | 3 | -0.22482 | 0.61927 | 0.81664 | 1 | 16560 | 3 | -0.22482 |
| MEGF11 | 4 | 0.063559 | 0.14757 | 0.965554 | 3583 | 2 | -0.77509 | 0.61928 | 0.75842 | 1 | 16561 | 2 | -0.77509 |
| FBXO32 | 5 | 0.073763 | 0.1845 | 0.968526 | 3986 | 1 | 0.53773 | 0.61963 | 0.77979 | 1 | 16562 | 2 | 0.53773 |
| CNTFR | 3 | 0.19744 | 0.3094 | 0.999994 | 7685 | 2 | -1.1553 | 0.61969 | 0.69787 | 1 | 16563 | 1 | -1.1553 |
| CCRN4L | 3 | 0.72511 | 0.72505 | 0.999994 | 16550 | 1 | -0.3146 | 0.61973 | 0.69789 | 1 | 16564 | 1 | -0.3146 |
| CMAS | 5 | 0.29368 | 0.47589 | 0.999994 | 9990 | 2 | 0.49948 | 0.61991 | 0.77997 | 1 | 16565 | 2 | 0.49948 |
| PPP5D1 | 6 | 0.071959 | 0.19086 | 0.972248 | 3915 | 4 | -1.527 | 0.61991 | 0.81709 | 1 | 16566 | 2 | -1.527 |
| LPHN3 | 6 | 0.26743 | 0.46864 | 0.999994 | 9407 | 3 | -0.36584 | 0.61991 | 0.81709 | 1 | 16567 | 2 | -0.36584 |
| ZNF891 | 5 | 0.54708 | 0.68912 | 0.999994 | 14169 | 2 | 0.55633 | 0.62013 | 0.78009 | 1 | 16568 | 2 | 0.55633 |
| hsa-mir-548a-1 | 4 | 0.12324 | 0.25593 | 0.994943 | 5743 | 2 | -1.0249 | 0.62018 | 0.75914 | 1 | 16569 | 2 | -1.0249 |
| OR2V2 | 5 | 0.82236 | 0.83416 | 0.999994 | 17811 | 1 | 0.33339 | 0.62032 | 0.7802 | 1 | 16570 | 2 | 0.33339 |
| OR51L1 | 6 | 0.0011909 | 0.0059966 | 0.623123 | 186 | 3 | -0.7048 | 0.62033 | 0.8174 | 1 | 16571 | 2 | -0.7048 |
| F2RL2 | 6 | 0.56204 | 0.71227 | 0.999994 | 14407 | 2 | -0.0078746 | 0.62033 | 0.8174 | 1 | 16572 | 2 | -0.0078746 |
| GLE1 | 2 | 0.50424 | 0.50778 | 0.999994 | 13500 | 1 | 0.1527 | 0.62039 | 0.63092 | 1 | 16573 | 1 | 0.1527 |
| CNKSR1 | 5 | 0.28415 | 0.46823 | 0.999994 | 9787 | 2 | -0.070708 | 0.62053 | 0.78032 | 1 | 16574 | 1 | -0.070708 |
| ZNF93 | 5 | 0.055889 | 0.15456 | 0.967126 | 3243 | 4 | -0.73925 | 0.62053 | 0.78032 | 1 | 16575 | 1 | -0.73925 |
| KRTAP1-1 | 5 | 0.54391 | 0.68635 | 0.999994 | 14130 | 1 | -0.097488 | 0.62053 | 0.78032 | 1 | 16576 | 1 | -0.097488 |
| SYNM | 5 | 0.1482 | 0.30005 | 0.999994 | 6411 | 3 | -0.64801 | 0.62068 | 0.78041 | 1 | 16577 | 2 | -0.64801 |
| STXBP5L | 4 | 0.15174 | 0.29098 | 0.999632 | 6499 | 1 | 0.2143 | 0.62069 | 0.75956 | 1 | 16578 | 2 | 0.2143 |
| ABL2 | 6 | 0.6734 | 0.7958 | 0.999994 | 15977 | 2 | 0.29417 | 0.62092 | 0.81783 | 1 | 16579 | 3 | 0.29417 |
| RIMKLB | 4 | 0.036957 | 0.095678 | 0.945031 | 2370 | 3 | -4.1418 | 0.62105 | 0.75986 | 1 | 16580 | 1 | -4.1418 |
| FAM195A | 4 | 0.017455 | 0.055232 | 0.922677 | 1335 | 3 | -2.5047 | 0.62105 | 0.75986 | 1 | 16581 | 1 | -2.5047 |
| SUFU | 4 | 0.31874 | 0.46215 | 0.999994 | 10492 | 2 | -0.42151 | 0.62105 | 0.75986 | 1 | 16582 | 1 | -0.42151 |
| MYBBP1A | 3 | 0.14619 | 0.2436 | 0.988088 | 6359 | 2 | -1.7522 | 0.62108 | 0.69855 | 1 | 16583 | 1 | -1.7522 |
| SPATA12 | 5 | 0.95181 | 0.95196 | 0.999994 | 20381 | 0 | 0.5115 | 0.62113 | 0.78066 | 1 | 16584 | 2 | 0.5115 |
| LOC283710 | 4 | 0.19768 | 0.33829 | 0.999994 | 7696 | 2 | -0.35513 | 0.62117 | 0.75997 | 1 | 16585 | 2 | -0.35513 |
| C12orf52 | 6 | 0.5742 | 0.72097 | 0.999994 | 14607 | 2 | 0.42578 | 0.62141 | 0.81818 | 1 | 16586 | 3 | 0.42578 |
| hsa-mir-1322 | 1 | 0.37855 | 0.37862 | 0.999994 | 11631 | 0 | -0.43121 | 0.62145 | 0.62117 | 1 | 16587 | 0 | -0.43121 |

| | | | | | | | | | | | | |
|---|---|---|---|---|---|---|---|---|---|---|---|---|
| WDR63 | 6 | 0.95357 | 0.95359 | 0.999994 | 20427 | 0 | 0.32072 | 0.62154 | 0.81827 | 1 | 16588 | 3 | 0.32072 |
| hsa-mir-302f | 2 | 0.12424 | 0.20718 | 0.978744 | 5774 | 1 | -3.0533 | 0.62157 | 0.63191 | 1 | 16589 | 1 | -3.0533 |
| NCOA2 | 6 | 0.88403 | 0.89222 | 0.999994 | 18905 | 1 | 0.34365 | 0.62172 | 0.8184 | 1 | 16590 | 3 | 0.34365 |
| FSTL1 | 6 | 0.14967 | 0.32521 | 0.999994 | 6448 | 3 | -0.78202 | 0.62173 | 0.8184 | 1 | 16591 | 2 | -0.78202 |
| CYP20A1 | 6 | 0.0097189 | 0.038044 | 0.887545 | 854 | 4 | -2.7153 | 0.62173 | 0.8184 | 1 | 16592 | 1 | -2.7153 |
| hsa-mir-7705 | 2 | 0.12871 | 0.21183 | 0.979587 | 5903 | 1 | -3.0559 | 0.6218 | 0.63209 | 1 | 16593 | 1 | -3.0559 |
| CCIN | 6 | 0.027787 | 0.093591 | 0.945031 | 1895 | 4 | -1.7227 | 0.62184 | 0.81849 | 1 | 16594 | 2 | -1.7227 |
| ACOT12 | 4 | 0.25487 | 0.39689 | 0.999994 | 9127 | 2 | -0.52021 | 0.622 | 0.76062 | 1 | 16595 | 2 | -0.52021 |
| ADRA1B | 6 | 0.3169 | 0.52633 | 0.999994 | 10450 | 2 | 0.57635 | 0.62207 | 0.81866 | 1 | 16596 | 3 | 0.57635 |
| LURAP1 | 5 | 0.10995 | 0.24191 | 0.987253 | 5304 | 2 | -0.28709 | 0.62213 | 0.78125 | 1 | 16597 | 1 | -0.28709 |
| CABS1 | 6 | 0.48984 | 0.66182 | 0.999994 | 13275 | 2 | -0.15281 | 0.62225 | 0.81879 | 1 | 16598 | 2 | -0.15281 |
| RASL12 | 6 | 0.011063 | 0.04262 | 0.891621 | 928 | 3 | -2.7878 | 0.62246 | 0.81895 | 1 | 16599 | 2 | -2.7878 |
| MARS2 | 6 | 0.6448 | 0.77363 | 0.999994 | 15582 | 2 | 0.10983 | 0.62247 | 0.81896 | 1 | 16600 | 3 | 0.10983 |
| THEG | 6 | 0.10172 | 0.25068 | 0.992497 | 5026 | 3 | -0.78347 | 0.62254 | 0.819 | 1 | 16601 | 2 | -0.78347 |
| AGGF1 | 6 | 0.337 | 0.54924 | 0.999994 | 10873 | 3 | -0.27189 | 0.6226 | 0.81904 | 1 | 16602 | 2 | -0.27189 |
| hsa-mir-5087 | 2 | 0.4836 | 0.48904 | 0.999994 | 13192 | 1 | 0.13351 | 0.62271 | 0.63288 | 1 | 16603 | 1 | 0.13351 |
| SF3B3 | 5 | 0.34783 | 0.52013 | 0.999994 | 11113 | 1 | 0.37682 | 0.62279 | 0.78162 | 1 | 16604 | 2 | 0.37682 |
| hsa-mir-3128 | 4 | 0.17796 | 0.31799 | 0.999994 | 7176 | 2 | -0.34182 | 0.62283 | 0.7613 | 1 | 16605 | 2 | -0.34182 |
| SLC10A7 | 5 | 0.20851 | 0.38729 | 0.999994 | 7991 | 2 | 0.20136 | 0.62287 | 0.78166 | 1 | 16606 | 2 | 0.20136 |
| VCL | 4 | 0.86739 | 0.86695 | 0.999994 | 18585 | 0 | -0.088773 | 0.62287 | 0.76134 | 1 | 16607 | 1 | -0.088773 |
| NPR1 | 6 | 0.085824 | 0.21915 | 0.980699 | 4439 | 2 | 0.0043762 | 0.62293 | 0.8193 | 1 | 16608 | 3 | 0.0043762 |
| PTAFR | 6 | 0.028132 | 0.094391 | 0.945031 | 1911 | 3 | -1.8154 | 0.62305 | 0.8194 | 1 | 16609 | 1 | -1.8154 |
| KIF7 | 4 | 0.11595 | 0.24321 | 0.988088 | 5513 | 2 | -0.92879 | 0.62305 | 0.7615 | 1 | 16610 | 2 | -0.92879 |
| P2RY4 | 6 | 0.46994 | 0.64841 | 0.999994 | 13006 | 2 | 0.084931 | 0.62309 | 0.81943 | 1 | 16611 | 3 | 0.084931 |
| CCDC142 | 6 | 0.22871 | 0.42252 | 0.999994 | 8495 | 3 | -0.2866 | 0.62318 | 0.81949 | 1 | 16612 | 3 | -0.2866 |
| MELK | 6 | 0.27151 | 0.4735 | 0.999994 | 9512 | 1 | 0.36036 | 0.62324 | 0.81953 | 1 | 16613 | 3 | 0.36036 |
| TRAK2 | 6 | 0.018162 | 0.066028 | 0.93914 | 1372 | 3 | -1.4537 | 0.62337 | 0.81963 | 1 | 16614 | 3 | -1.4537 |
| USP16 | 6 | 0.68903 | 0.80805 | 0.999994 | 16200 | 2 | 0.54224 | 0.6236 | 0.8198 | 1 | 16615 | 3 | 0.54224 |
| SAMD9 | 6 | 0.57361 | 0.72055 | 0.999994 | 14603 | 2 | -0.23552 | 0.62371 | 0.81989 | 1 | 16616 | 3 | -0.23552 |
| C19orf38 | 6 | 0.096799 | 0.24107 | 0.987253 | 4851 | 2 | 0.29082 | 0.6238 | 0.81996 | 1 | 16617 | 2 | 0.29082 |
| CLN6 | 6 | 0.45442 | 0.638 | 0.999994 | 12762 | 2 | 0.5111 | 0.62388 | 0.82001 | 1 | 16618 | 3 | 0.5111 |
| ZFP28 | 6 | 0.1894 | 0.37477 | 0.999994 | 7466 | 3 | -0.35402 | 0.6239 | 0.82002 | 1 | 16619 | 3 | -0.35402 |
| BMF | 5 | 0.7676 | 0.80038 | 0.999994 | 17057 | 1 | 0.26449 | 0.62399 | 0.78233 | 1 | 16620 | 2 | 0.26449 |
| PTH1R | 5 | 0.17157 | 0.33443 | 0.999994 | 7026 | 2 | -0.17388 | 0.62399 | 0.78233 | 1 | 16621 | 1 | -0.17388 |
| PKD1L3 | 6 | 0.83298 | 0.86303 | 0.999994 | 17970 | 1 | 0.45888 | 0.62411 | 0.82018 | 1 | 16622 | 3 | 0.45888 |
| INS | 4 | 0.18878 | 0.3291 | 0.999994 | 7445 | 2 | -0.73228 | 0.62428 | 0.76245 | 1 | 16623 | 2 | -0.73228 |
| ZNF90 | 4 | 0.73757 | 0.75071 | 0.999994 | 16695 | 1 | 0.51194 | 0.62432 | 0.76248 | 1 | 16624 | 2 | 0.51194 |
| hsa-mir-5702 | 4 | 0.037397 | 0.09657 | 0.945031 | 2392 | 2 | -2.6137 | 0.6244 | 0.76255 | 1 | 16625 | 1 | -2.6137 |
| ERCC1 | 6 | 0.60552 | 0.74406 | 0.999994 | 15021 | 2 | 0.52602 | 0.62443 | 0.82041 | 1 | 16626 | 2 | 0.52602 |
| LETM2 | 6 | 0.30218 | 0.50926 | 0.999994 | 10173 | 2 | 0.03601 | 0.62443 | 0.82041 | 1 | 16627 | 1 | 0.03601 |
| TET2 | 6 | 0.34347 | 0.55662 | 0.999994 | 11019 | 3 | -0.096428 | 0.62453 | 0.82049 | 1 | 16628 | 2 | -0.096428 |
| NonTargetingControlG | 2 | 0.71501 | 0.71473 | 0.999994 | 16449 | 0 | 0.84618 | 0.62454 | 0.63438 | 1 | 16629 | 1 | 0.84618 |
| AKAP4 | 5 | 0.45631 | 0.61057 | 0.999994 | 12795 | 2 | 0.40455 | 0.62461 | 0.78269 | 1 | 16630 | 2 | 0.40455 |
| ST6GALNAC4 | 6 | 0.15466 | 0.33156 | 0.999994 | 6585 | 2 | -0.28998 | 0.62474 | 0.82064 | 1 | 16631 | 2 | -0.28998 |
| TIAM1 | 5 | 0.17439 | 0.33852 | 0.999994 | 7094 | 1 | -0.33818 | 0.62482 | 0.7828 | 1 | 16632 | 2 | -0.33818 |
| BCAS1 | 6 | 0.057941 | 0.16133 | 0.967126 | 3335 | 3 | -0.65577 | 0.62484 | 0.82071 | 1 | 16633 | 2 | -0.65577 |
| ATE1 | 6 | 0.67546 | 0.79744 | 0.999994 | 16012 | 1 | 0.46182 | 0.62495 | 0.82081 | 1 | 16634 | 3 | 0.46182 |
| TSKU | 4 | 0.013027 | 0.043079 | 0.891621 | 1064 | 3 | -5.0732 | 0.62497 | 0.763 | 1 | 16635 | 1 | -5.0732 |
| CD7 | 5 | 0.1582 | 0.31491 | 0.999994 | 6679 | 3 | -1.143 | 0.62498 | 0.7829 | 1 | 16636 | 2 | -1.143 |
| hsa-mir-4455 | 3 | 0.61877 | 0.63172 | 0.999994 | 15200 | 1 | -0.53832 | 0.625 | 0.70058 | 1 | 16637 | 1 | -0.53832 |
| NonTargetingControlG | 1 | 0.37474 | 0.37484 | 0.999994 | 11579 | 0 | -0.4447 | 0.62526 | 0.62506 | 1 | 16638 | 0 | -0.4447 |
| NonTargetingControlG | 2 | 0.023172 | 0.042862 | 0.891621 | 1662 | 1 | -4.3938 | 0.62532 | 0.63501 | 1 | 16639 | 1 | -4.3938 |

| | | | | | | | | | | | | | |
|---|---|---|---|---|---|---|---|---|---|---|---|---|---|
| VAPA | 6 | 0.029714 | 0.098103 | 0.945253 | 1994 | 4 | -1.7282 | 0.62536 | 0.82112 | 1 | 16640 | 2 | -1.7282 |
| S100A9 | 6 | 0.54819 | 0.70231 | 0.999994 | 14179 | 2 | -0.39237 | 0.62536 | 0.82112 | 1 | 16641 | 2 | -0.39237 |
| CXXC1 | 5 | 0.000022619 | 0.000095329 | 0.114686 | 18 | 4 | -10.054 | 0.62542 | 0.78316 | 1 | 16642 | 1 | -10.054 |
| DSC2 | 6 | 0.020221 | 0.072682 | 0.93914 | 1472 | 3 | -0.09562 | 0.62546 | 0.82119 | 1 | 16643 | 3 | -0.09562 |
| AGPAT9 | 6 | 0.49449 | 0.66493 | 0.999994 | 13336 | 1 | 0.26339 | 0.62566 | 0.82133 | 1 | 16644 | 3 | 0.26339 |
| LOC100130539 | 6 | 0.10314 | 0.25345 | 0.994501 | 5071 | 3 | -0.40018 | 0.62586 | 0.82146 | 1 | 16645 | 3 | -0.40018 |
| SRSF9 | 6 | 0.37271 | 0.58492 | 0.999994 | 11556 | 2 | 0.069524 | 0.6259 | 0.82148 | 1 | 16646 | 2 | 0.069524 |
| FAM214B | 6 | 0.10061 | 0.24852 | 0.990929 | 4994 | 4 | -1.3532 | 0.62598 | 0.82154 | 1 | 16647 | 2 | -1.3532 |
| BAG1 | 4 | 0.11992 | 0.25014 | 0.992497 | 5627 | 3 | -1.0792 | 0.62605 | 0.76388 | 1 | 16648 | 1 | -1.0792 |
| ZNF655 | 6 | 0.035746 | 0.11228 | 0.951052 | 2303 | 2 | -0.26999 | 0.62615 | 0.82166 | 1 | 16649 | 2 | -0.26999 |
| WNT11 | 5 | 0.090551 | 0.21148 | 0.979587 | 4617 | 3 | -1.4349 | 0.62627 | 0.78365 | 1 | 16650 | 2 | -1.4349 |
| ESM1 | 5 | 0.28198 | 0.46646 | 0.999994 | 9740 | 1 | 0.20725 | 0.62645 | 0.78373 | 1 | 16651 | 2 | 0.20725 |
| LAMP5 | 5 | 0.40934 | 0.57134 | 0.999994 | 12099 | 1 | 0.39981 | 0.62646 | 0.78374 | 1 | 16652 | 2 | 0.39981 |
| ZNF397 | 5 | 0.16739 | 0.32829 | 0.999994 | 6920 | 2 | 0.29338 | 0.62653 | 0.78378 | 1 | 16653 | 2 | 0.29338 |
| MYO7B | 3 | 0.24409 | 0.36763 | 0.999994 | 8883 | 2 | -2.1605 | 0.62664 | 0.70139 | 1 | 16654 | 1 | -2.1605 |
| NFAT5 | 6 | 0.20701 | 0.39644 | 0.999994 | 7948 | 3 | -0.090889 | 0.62668 | 0.82205 | 1 | 16655 | 3 | -0.090889 |
| RORC | 5 | 0.00071181 | 0.0032533 | 0.541338 | 131 | 3 | -6.3574 | 0.62671 | 0.78388 | 1 | 16656 | 2 | -6.3574 |
| ZFP57 | 5 | 0.11718 | 0.25304 | 0.994501 | 5541 | 2 | -0.21957 | 0.62671 | 0.78388 | 1 | 16657 | 2 | -0.21957 |
| TMEM101 | 5 | 0.008799 | 0.032589 | 0.865899 | 780 | 3 | -4.4234 | 0.62679 | 0.78393 | 1 | 16658 | 2 | -4.4234 |
| AKR1C3 | 6 | 0.62918 | 0.76171 | 0.999994 | 15369 | 2 | -0.098788 | 0.62684 | 0.82215 | 1 | 16659 | 2 | -0.098788 |
| ESRP1 | 5 | 0.052126 | 0.14551 | 0.963601 | 3071 | 3 | -1.1234 | 0.62689 | 0.78398 | 1 | 16660 | 2 | -1.1234 |
| HSD11B1 | 6 | 0.10542 | 0.25788 | 0.994943 | 5152 | 3 | -1.2514 | 0.62701 | 0.82227 | 1 | 16661 | 3 | -1.2514 |
| SPEF2 | 6 | 0.31029 | 0.51868 | 0.999994 | 10331 | 2 | 0.20439 | 0.62705 | 0.82231 | 1 | 16662 | 3 | 0.20439 |
| hsa-mir-3667 | 4 | 0.32386 | 0.4674 | 0.999994 | 10590 | 1 | -0.10314 | 0.62709 | 0.76468 | 1 | 16663 | 1 | -0.10314 |
| BVES | 4 | 0.075884 | 0.17105 | 0.96737 | 4067 | 3 | -1.9395 | 0.62709 | 0.76468 | 1 | 16664 | 1 | -1.9395 |
| ZSCAN22 | 6 | 0.25736 | 0.45674 | 0.999994 | 9174 | 2 | -0.034808 | 0.62719 | 0.82242 | 1 | 16665 | 2 | -0.034808 |
| RPS6KA2 | 6 | 0.021757 | 0.077609 | 0.93914 | 1563 | 5 | -1.789 | 0.62719 | 0.82242 | 1 | 16666 | 1 | -1.789 |
| SLC17A9 | 6 | 0.38425 | 0.59225 | 0.999994 | 11704 | 2 | -0.27858 | 0.62719 | 0.82242 | 1 | 16667 | 1 | -0.27858 |
| ZNF148 | 6 | 0.007072 | 0.028681 | 0.858757 | 664 | 4 | -2.2273 | 0.62719 | 0.82242 | 1 | 16668 | 2 | -2.2273 |
| hsa-mir-216b | 4 | 0.20951 | 0.35045 | 0.999994 | 8016 | 2 | -0.54476 | 0.62739 | 0.76493 | 1 | 16669 | 2 | -0.54476 |
| hsa-mir-487b | 4 | 0.6093 | 0.67036 | 0.999994 | 15062 | 1 | 0.30628 | 0.62739 | 0.76493 | 1 | 16670 | 2 | 0.30628 |
| NonTargetingControlG | 2 | 0.51754 | 0.52011 | 0.999994 | 13702 | 1 | 0.17892 | 0.62751 | 0.6369 | 1 | 16671 | 1 | 0.17892 |
| HAUS5 | 3 | 0.28325 | 0.41597 | 0.999994 | 9765 | 2 | -0.64634 | 0.62768 | 0.70191 | 1 | 16672 | 1 | -0.64634 |
| PLSCR1 | 6 | 0.086171 | 0.21984 | 0.980699 | 4456 | 4 | -0.96655 | 0.62786 | 0.82292 | 1 | 16673 | 2 | -0.96655 |
| TREX1 | 6 | 0.06364 | 0.17348 | 0.96793 | 3589 | 3 | -0.52574 | 0.62825 | 0.8232 | 1 | 16674 | 2 | -0.52574 |
| GPR32 | 5 | 0.54689 | 0.68897 | 0.999994 | 14166 | 2 | 0.54942 | 0.62839 | 0.78484 | 1 | 16675 | 2 | 0.54942 |
| SPRN | 6 | 0.35665 | 0.57152 | 0.999994 | 11276 | 3 | -0.034551 | 0.62843 | 0.82333 | 1 | 16676 | 3 | -0.034551 |
| APBA2 | 6 | 0.11236 | 0.27124 | 0.998561 | 5382 | 4 | -1.1249 | 0.62849 | 0.82337 | 1 | 16677 | 2 | -1.1249 |
| hsa-mir-3126 | 4 | 0.29785 | 0.44081 | 0.999994 | 10087 | 1 | 0.081613 | 0.62858 | 0.76592 | 1 | 16678 | 2 | 0.081613 |
| YJEFN3 | 5 | 0.50607 | 0.65321 | 0.999994 | 13528 | 2 | -0.15189 | 0.62859 | 0.78496 | 1 | 16679 | 2 | -0.15189 |
| ZKSCAN7 | 6 | 0.041003 | 0.12418 | 0.951052 | 2561 | 3 | -0.68498 | 0.62861 | 0.82347 | 1 | 16680 | 3 | -0.68498 |
| PRUNE | 6 | 0.011319 | 0.043485 | 0.891621 | 944 | 4 | -4.4393 | 0.62864 | 0.8235 | 1 | 16681 | 2 | -4.4393 |
| MFSD4 | 6 | 0.22947 | 0.42342 | 0.999994 | 8515 | 3 | -0.15112 | 0.62914 | 0.82384 | 1 | 16682 | 3 | -0.15112 |
| ABCG8 | 6 | 0.002498 | 0.011844 | 0.729405 | 316 | 5 | -2.5576 | 0.6293 | 0.82396 | 1 | 16683 | 1 | -2.5576 |
| NHLH2 | 3 | 0.17288 | 0.27808 | 0.999536 | 7062 | 1 | 0.37462 | 0.62938 | 0.70278 | 1 | 16684 | 1 | 0.37462 |
| CCDC13 | 4 | 0.10196 | 0.21861 | 0.980699 | 5037 | 2 | -1.1856 | 0.62942 | 0.76661 | 1 | 16685 | 2 | -1.1856 |
| CLEC12B | 6 | 0.20215 | 0.39051 | 0.999994 | 7836 | 3 | -0.3532 | 0.62944 | 0.82408 | 1 | 16686 | 3 | -0.3532 |
| PPIL3 | 6 | 0.078776 | 0.20488 | 0.978186 | 4179 | 3 | -0.058388 | 0.62952 | 0.82413 | 1 | 16687 | 3 | -0.058388 |
| ID1 | 5 | 0.094223 | 0.21729 | 0.980699 | 4760 | 3 | -1.2838 | 0.62955 | 0.78554 | 1 | 16688 | 2 | -1.2838 |
| ORC5 | 5 | 0.049503 | 0.13913 | 0.957704 | 2957 | 2 | -1.4517 | 0.62955 | 0.78554 | 1 | 16689 | 1 | -1.4517 |
| TGFBR3L | 5 | 0.018159 | 0.059067 | 0.931872 | 1371 | 4 | -1.5639 | 0.62955 | 0.78554 | 1 | 16690 | 1 | -1.5639 |
| RAB9B | 5 | 0.40328 | 0.56628 | 0.999994 | 12007 | 1 | -0.077367 | 0.62955 | 0.78554 | 1 | 16691 | 2 | -0.077367 |

| | | | | | | | | | | | | | |
|---|---|---|---|---|---|---|---|---|---|---|---|---|---|
| ASH2L | 4 | 0.0023234 | 0.0083988 | 0.680529 | 306 | 3 | -6.9133 | 0.62956 | 0.76673 | 1 | 16692 | 1 | -6.9133 |
| SAMD4A | 6 | 0.2136 | 0.40447 | 0.999994 | 8103 | 2 | 0.073529 | 0.62958 | 0.82418 | 1 | 16693 | 2 | 0.073529 |
| FAM153B | 4 | 0.64875 | 0.69256 | 0.999994 | 15638 | 1 | 0.094806 | 0.62971 | 0.76684 | 1 | 16694 | 2 | 0.094806 |
| LMO3 | 6 | 0.20349 | 0.39211 | 0.999994 | 7868 | 3 | -0.12628 | 0.62975 | 0.82431 | 1 | 16695 | 3 | -0.12628 |
| NonTargetingControlG | 2 | 0.66191 | 0.6617 | 0.999994 | 15808 | 0 | 0.63285 | 0.62994 | 0.63892 | 1 | 16696 | 1 | 0.63285 |
| TLE1 | 6 | 0.3102 | 0.51859 | 0.999994 | 10329 | 3 | -0.07269 | 0.62998 | 0.82447 | 1 | 16697 | 3 | -0.07269 |
| KCNK4 | 4 | 0.63565 | 0.685 | 0.999994 | 15453 | 1 | 0.40617 | 0.63001 | 0.76708 | 1 | 16698 | 2 | 0.40617 |
| SLC1A4 | 4 | 0.25707 | 0.39914 | 0.999994 | 9169 | 2 | -0.35473 | 0.63007 | 0.76712 | 1 | 16699 | 2 | -0.35473 |
| PSG2 | 6 | 0.24203 | 0.43842 | 0.999994 | 8835 | 2 | 0.34427 | 0.63012 | 0.82457 | 1 | 16700 | 2 | 0.34427 |
| LOC643037 | 4 | 0.27982 | 0.42236 | 0.999994 | 9698 | 2 | -0.37666 | 0.63013 | 0.76717 | 1 | 16701 | 2 | -0.37666 |
| NonTargetingControlG | 1 | 0.36983 | 0.36994 | 0.999994 | 11506 | 0 | -0.492 | 0.63017 | 0.63006 | 1 | 16702 | 0 | -0.492 |
| NonTargetingControlG | 2 | 0.616 | 0.6158 | 0.999994 | 15161 | 0 | 0.43089 | 0.63029 | 0.63923 | 1 | 16703 | 1 | 0.43089 |
| TIAM2 | 6 | 0.11842 | 0.28274 | 0.999632 | 5586 | 4 | -0.7518 | 0.63043 | 0.82481 | 1 | 16704 | 1 | -0.7518 |
| FAM207A | 6 | 0.015851 | 0.058564 | 0.931872 | 1234 | 4 | -2.629 | 0.63043 | 0.82481 | 1 | 16705 | 2 | -2.629 |
| DENND2A | 4 | 0.013421 | 0.044288 | 0.891621 | 1096 | 3 | -3.1053 | 0.63046 | 0.76743 | 1 | 16706 | 1 | -3.1053 |
| INSR | 6 | 0.62107 | 0.75563 | 0.999994 | 15241 | 1 | 0.52241 | 0.63055 | 0.8249 | 1 | 16707 | 3 | 0.52241 |
| TMEM8A | 6 | 0.17227 | 0.35353 | 0.999994 | 7044 | 2 | -0.48738 | 0.63058 | 0.82493 | 1 | 16708 | 2 | -0.48738 |
| SEL1L2 | 6 | 0.62763 | 0.76055 | 0.999994 | 15346 | 1 | 0.50907 | 0.63061 | 0.82495 | 1 | 16709 | 3 | 0.50907 |
| OR4P4 | 6 | 0.11336 | 0.27317 | 0.998561 | 5419 | 3 | -0.76813 | 0.63062 | 0.82496 | 1 | 16710 | 3 | -0.76813 |
| FBXO25 | 4 | 0.40103 | 0.54679 | 0.999994 | 11959 | 1 | 0.33351 | 0.63066 | 0.76759 | 1 | 16711 | 2 | 0.33351 |
| CREB1 | 6 | 0.053082 | 0.15073 | 0.966359 | 3118 | 4 | -1.2301 | 0.63071 | 0.82503 | 1 | 16712 | 2 | -1.2301 |
| BOD1L2 | 6 | 0.089405 | 0.22632 | 0.983354 | 4574 | 3 | -0.79018 | 0.63073 | 0.82505 | 1 | 16713 | 3 | -0.79018 |
| CDC25B | 5 | 0.35755 | 0.52807 | 0.999994 | 11298 | 2 | -0.40326 | 0.63087 | 0.78629 | 1 | 16714 | 2 | -0.40326 |
| YTHDF3 | 6 | 0.53418 | 0.69246 | 0.999994 | 13969 | 2 | 0.1583 | 0.63088 | 0.82514 | 1 | 16715 | 3 | 0.1583 |
| ZNF202 | 5 | 0.3926 | 0.55737 | 0.999994 | 11844 | 2 | 0.14271 | 0.63112 | 0.78644 | 1 | 16716 | 2 | 0.14271 |
| KITLG | 5 | 0.056606 | 0.15625 | 0.967126 | 3278 | 3 | -1.9097 | 0.63112 | 0.78644 | 1 | 16717 | 2 | -1.9097 |
| ERCC5 | 5 | 0.026345 | 0.081004 | 0.93914 | 1821 | 4 | -1.7485 | 0.63112 | 0.78644 | 1 | 16718 | 1 | -1.7485 |
| WWOX | 6 | 0.37648 | 0.58733 | 0.999994 | 11602 | 2 | 0.070556 | 0.63134 | 0.82547 | 1 | 16719 | 2 | 0.070556 |
| ZNF182 | 6 | 0.1425 | 0.31624 | 0.999994 | 6252 | 2 | 0.31267 | 0.63142 | 0.82552 | 1 | 16720 | 3 | 0.31267 |
| hsa-mir-4472-1 | 4 | 0.78762 | 0.79033 | 0.999994 | 17314 | 1 | 0.57777 | 0.63143 | 0.76821 | 1 | 16721 | 2 | 0.57777 |
| XG | 3 | 0.29673 | 0.42678 | 0.999994 | 10059 | 1 | 0.38621 | 0.63145 | 0.70384 | 1 | 16722 | 1 | 0.38621 |
| LRRC27 | 6 | 0.073737 | 0.19452 | 0.975797 | 3981 | 4 | -1.2466 | 0.63151 | 0.82559 | 1 | 16723 | 2 | -1.2466 |
| SLC19A3 | 5 | 0.013387 | 0.045664 | 0.892434 | 1090 | 3 | -1.0364 | 0.63171 | 0.78677 | 1 | 16724 | 2 | -1.0364 |
| TNR | 4 | 0.41676 | 0.56286 | 0.999994 | 12201 | 2 | -0.000126 | 0.63173 | 0.76845 | 1 | 16725 | 2 | -0.000126 |
| PPARD | 4 | 0.050793 | 0.12305 | 0.951052 | 3006 | 3 | -1.5993 | 0.63174 | 0.76845 | 1 | 16726 | 1 | -1.5993 |
| VPS25 | 4 | 0.31989 | 0.46332 | 0.999994 | 10511 | 2 | -0.6086 | 0.63174 | 0.76845 | 1 | 16727 | 1 | -0.6086 |
| hsa-mir-8089 | 3 | 0.78194 | 0.78182 | 0.999994 | 17240 | 0 | -0.065107 | 0.63175 | 0.704 | 1 | 16728 | 1 | -0.065107 |
| hsa-mir-488 | 4 | 0.83202 | 0.83155 | 0.999994 | 17959 | 0 | 0.11087 | 0.63179 | 0.7685 | 1 | 16729 | 2 | 0.11087 |
| COX20 | 4 | 0.81748 | 0.81736 | 0.999994 | 17737 | 1 | 0.58392 | 0.63249 | 0.76905 | 1 | 16730 | 2 | 0.58392 |
| THAP2 | 4 | 0.19327 | 0.33375 | 0.999994 | 7582 | 1 | 0.17876 | 0.63253 | 0.76908 | 1 | 16731 | 2 | 0.17876 |
| NonTargetingControlG | 2 | 0.35308 | 0.37845 | 0.999994 | 11215 | 1 | -0.28146 | 0.63256 | 0.6411 | 1 | 16732 | 1 | -0.28146 |
| NonTargetingControlG | 2 | 0.31637 | 0.34938 | 0.999994 | 10442 | 1 | -0.51787 | 0.63258 | 0.64112 | 1 | 16733 | 1 | -0.51787 |
| NonTargetingControlG | 2 | 0.51065 | 0.51369 | 0.999994 | 13584 | 1 | 0.18666 | 0.63259 | 0.64113 | 1 | 16734 | 1 | 0.18666 |
| USP34 | 5 | 0.028539 | 0.08671 | 0.94395 | 1927 | 3 | -2.9285 | 0.6326 | 0.7873 | 1 | 16735 | 2 | -2.9285 |
| PRKCSH | 6 | 0.059759 | 0.16524 | 0.967126 | 3412 | 4 | -1.1992 | 0.63262 | 0.82603 | 1 | 16736 | 2 | -1.1992 |
| ASCC1 | 6 | 0.11689 | 0.27987 | 0.999632 | 5538 | 4 | -0.97386 | 0.63266 | 0.82604 | 1 | 16737 | 2 | -0.97386 |
| TPD52L1 | 4 | 0.42706 | 0.57348 | 0.999994 | 12368 | 2 | 0.021144 | 0.63269 | 0.76921 | 1 | 16738 | 2 | 0.021144 |
| OR2C1 | 5 | 0.25621 | 0.44568 | 0.999994 | 9155 | 2 | 0.17837 | 0.6327 | 0.78736 | 1 | 16739 | 2 | 0.17837 |
| KRT39 | 6 | 0.24004 | 0.43603 | 0.999994 | 8773 | 2 | -0.25463 | 0.63272 | 0.82606 | 1 | 16740 | 2 | -0.25463 |
| AK7 | 5 | 0.030308 | 0.091263 | 0.94395 | 2025 | 4 | -1.2461 | 0.63276 | 0.78741 | 1 | 16741 | 1 | -1.2461 |
| HTR1B | 5 | 0.15893 | 0.31595 | 0.999994 | 6697 | 3 | -1.1163 | 0.63276 | 0.78741 | 1 | 16742 | 2 | -1.1163 |
| GP1BA | 5 | 0.049965 | 0.14025 | 0.958716 | 2975 | 4 | -1.8567 | 0.63276 | 0.78741 | 1 | 16743 | 1 | -1.8567 |

| | | | | | | | | | | | | | |
|---|---|---|---|---|---|---|---|---|---|---|---|---|---|
| hsa-mir-1258 | 4 | 0.85363 | 0.85319 | 0.999994 | 18327 | 0 | 0.31679 | 0.63292 | 0.76938 | 1 | 16744 | 2 | 0.31679 |
| FAM221A | 5 | 0.50255 | 0.65014 | 0.999994 | 13472 | 2 | 0.43921 | 0.63305 | 0.78757 | 1 | 16745 | 2 | 0.43921 |
| TNFRSF18 | 5 | 0.26809 | 0.45531 | 0.999994 | 9422 | 2 | 0.11843 | 0.63309 | 0.7876 | 1 | 16746 | 2 | 0.11843 |
| hsa-mir-6870 | 4 | 0.54212 | 0.63587 | 0.999994 | 14094 | 1 | 0.47351 | 0.63315 | 0.76955 | 1 | 16747 | 2 | 0.47351 |
| CCDC74A | 3 | 0.13319 | 0.22677 | 0.983354 | 5999 | 2 | -1.7867 | 0.63327 | 0.70478 | 1 | 16748 | 1 | -1.7867 |
| YTHDC2 | 5 | 0.52866 | 0.67287 | 0.999994 | 13875 | 2 | 0.17411 | 0.63339 | 0.78778 | 1 | 16749 | 2 | 0.17411 |
| REPIN1 | 5 | 0.40425 | 0.5671 | 0.999994 | 12023 | 1 | 0.16253 | 0.63344 | 0.78781 | 1 | 16750 | 2 | 0.16253 |
| ATP8A1 | 5 | 0.64083 | 0.74151 | 0.999994 | 15526 | 1 | 0.54287 | 0.63346 | 0.78782 | 1 | 16751 | 2 | 0.54287 |
| UNC5D | 6 | 0.55886 | 0.71 | 0.999994 | 14348 | 2 | -0.27902 | 0.63349 | 0.82637 | 1 | 16752 | 2 | -0.27902 |
| GRM2 | 6 | 0.20804 | 0.39768 | 0.999994 | 7978 | 3 | -0.69885 | 0.63352 | 0.82639 | 1 | 16753 | 2 | -0.69885 |
| ZNF30 | 6 | 0.40322 | 0.60448 | 0.999994 | 12003 | 2 | -0.035808 | 0.63359 | 0.82641 | 1 | 16754 | 2 | -0.035808 |
| C8orf22 | 5 | 0.45461 | 0.60912 | 0.999994 | 12766 | 2 | 0.083163 | 0.63372 | 0.78797 | 1 | 16755 | 2 | 0.083163 |
| PFKM | 5 | 0.41508 | 0.57601 | 0.999994 | 12178 | 2 | -0.3083 | 0.63399 | 0.78814 | 1 | 16756 | 1 | -0.3083 |
| EVPL | 5 | 0.14531 | 0.29574 | 0.999994 | 6333 | 2 | 0.46608 | 0.63399 | 0.78814 | 1 | 16757 | 2 | 0.46608 |
| GLRB | 5 | 0.60748 | 0.72887 | 0.999994 | 15041 | 1 | 0.30251 | 0.63399 | 0.78814 | 1 | 16758 | 2 | 0.30251 |
| POMT1 | 5 | 0.040268 | 0.11655 | 0.951052 | 2529 | 3 | -2.5639 | 0.63399 | 0.78814 | 1 | 16759 | 1 | -2.5639 |
| FUT1 | 4 | 0.78021 | 0.78406 | 0.999994 | 17213 | 1 | 0.60272 | 0.63414 | 0.77033 | 1 | 16760 | 2 | 0.60272 |
| PTPRU | 4 | 0.28191 | 0.4245 | 0.999994 | 9739 | 1 | 0.40444 | 0.63416 | 0.77034 | 1 | 16761 | 2 | 0.40444 |
| FER | 6 | 0.07793 | 0.20317 | 0.97798 | 4147 | 4 | -1.2646 | 0.63431 | 0.82669 | 1 | 16762 | 1 | -1.2646 |
| COA5 | 3 | 0.27257 | 0.40283 | 0.999994 | 9529 | 2 | -0.68989 | 0.63432 | 0.70533 | 1 | 16763 | 1 | -0.68989 |
| MFSD11 | 6 | 0.42895 | 0.62115 | 0.999994 | 12395 | 2 | 0.038279 | 0.63434 | 0.8267 | 1 | 16764 | 2 | 0.038279 |
| hsa-mir-374a | 4 | 0.080421 | 0.17944 | 0.968171 | 4245 | 2 | -1.7518 | 0.63487 | 0.77092 | 1 | 16765 | 1 | -1.7518 |
| NonTargetingControlG | 2 | 0.059149 | 0.1045 | 0.948066 | 3385 | 1 | -3.8483 | 0.63489 | 0.6431 | 1 | 16766 | 1 | -3.8483 |
| MVB12B | 4 | 0.42331 | 0.56959 | 0.999994 | 12299 | 1 | 0.63294 | 0.63489 | 0.77094 | 1 | 16767 | 2 | 0.63294 |
| hsa-mir-1307 | 4 | 0.77442 | 0.77935 | 0.999994 | 17135 | 1 | 0.51305 | 0.63512 | 0.77111 | 1 | 16768 | 2 | 0.51305 |
| ASXL1 | 6 | 0.014754 | 0.054935 | 0.922677 | 1175 | 4 | -3.5542 | 0.63515 | 0.827 | 1 | 16769 | 2 | -3.5542 |
| MAML1 | 5 | 0.0097986 | 0.035493 | 0.870441 | 859 | 4 | -5.964 | 0.63519 | 0.78882 | 1 | 16770 | 1 | -5.964 |
| AKAP13 | 6 | 0.19183 | 0.3778 | 0.999994 | 7539 | 3 | -0.53843 | 0.63534 | 0.82708 | 1 | 16771 | 2 | -0.53843 |
| KRTAP24-1 | 4 | 0.61407 | 0.67302 | 0.999994 | 15132 | 1 | 0.43257 | 0.63538 | 0.77132 | 1 | 16772 | 2 | 0.43257 |
| IFNAR1 | 6 | 0.052932 | 0.15042 | 0.966359 | 3108 | 4 | -1.8318 | 0.63561 | 0.82718 | 1 | 16773 | 2 | -1.8318 |
| UTY | 6 | 0.018331 | 0.066584 | 0.93914 | 1378 | 4 | -2.9232 | 0.63561 | 0.82718 | 1 | 16774 | 2 | -2.9232 |
| SPATA9 | 6 | 0.96804 | 0.96806 | 0.999994 | 20786 | 0 | 0.33143 | 0.63561 | 0.82718 | 1 | 16775 | 2 | 0.33143 |
| CYP4B1 | 6 | 0.023973 | 0.084532 | 0.94395 | 1706 | 4 | -2.4811 | 0.63561 | 0.82718 | 1 | 16776 | 2 | -2.4811 |
| PSMB7 | 4 | 0.38592 | 0.53116 | 0.999994 | 11731 | 2 | 0.037565 | 0.63562 | 0.7715 | 1 | 16777 | 2 | 0.037565 |
| NonTargetingControlG | 2 | 0.40386 | 0.42 | 0.999994 | 12014 | 1 | -0.14409 | 0.63567 | 0.64379 | 1 | 16778 | 1 | -0.14409 |
| PODXL | 4 | 0.35803 | 0.50248 | 0.999994 | 11305 | 2 | -0.16287 | 0.63581 | 0.77166 | 1 | 16779 | 2 | -0.16287 |
| NonTargetingControlG | 2 | 0.46201 | 0.46989 | 0.999994 | 12886 | 1 | -0.13282 | 0.63581 | 0.64393 | 1 | 16780 | 1 | -0.13282 |
| RRS1 | 3 | 0.18121 | 0.28883 | 0.999632 | 7254 | 2 | -0.93329 | 0.63586 | 0.70612 | 1 | 16781 | 1 | -0.93329 |
| hsa-mir-6506 | 3 | 0.22093 | 0.3388 | 0.999994 | 8308 | 2 | -0.94259 | 0.63618 | 0.7063 | 1 | 16782 | 1 | -0.94259 |
| hsa-mir-6798 | 4 | 0.26631 | 0.40852 | 0.999994 | 9373 | 2 | -1.1006 | 0.6362 | 0.77198 | 1 | 16783 | 2 | -1.1006 |
| PRAMEF17 | 4 | 0.15769 | 0.29712 | 0.999994 | 6666 | 2 | -0.76959 | 0.63632 | 0.77208 | 1 | 16784 | 2 | -0.76959 |
| CSAD | 5 | 0.31381 | 0.49229 | 0.999994 | 10392 | 2 | -0.19824 | 0.63636 | 0.78952 | 1 | 16785 | 2 | -0.19824 |
| RUNX1T1 | 5 | 0.16092 | 0.31889 | 0.999994 | 6761 | 3 | -1.0807 | 0.63645 | 0.78957 | 1 | 16786 | 1 | -1.0807 |
| SAP130 | 5 | 0.11106 | 0.24361 | 0.988088 | 5335 | 2 | 0.070116 | 0.63645 | 0.78957 | 1 | 16787 | 2 | 0.070116 |
| NTNG1 | 5 | 0.32165 | 0.49862 | 0.999994 | 10547 | 2 | 0.57703 | 0.63645 | 0.78957 | 1 | 16788 | 2 | 0.57703 |
| OR4K14 | 5 | 0.41383 | 0.57501 | 0.999994 | 12165 | 2 | -0.56885 | 0.63645 | 0.78957 | 1 | 16789 | 1 | -0.56885 |
| CDC26 | 5 | 0.29706 | 0.47868 | 0.999994 | 10069 | 2 | 0.079231 | 0.63645 | 0.78957 | 1 | 16790 | 2 | 0.079231 |
| SSX4 | 5 | 0.73302 | 0.78213 | 0.999994 | 16640 | 1 | -0.20489 | 0.63646 | 0.78958 | 1 | 16791 | 2 | -0.20489 |
| PACS1 | 4 | 0.42533 | 0.57169 | 0.999994 | 12341 | 1 | -0.016176 | 0.6365 | 0.77221 | 1 | 16792 | 1 | -0.016176 |
| hsa-mir-29a | 4 | 0.23787 | 0.37951 | 0.999994 | 8721 | 2 | -0.47426 | 0.6365 | 0.77221 | 1 | 16793 | 1 | -0.47426 |
| ACOT11 | 6 | 0.035537 | 0.11179 | 0.951052 | 2296 | 4 | -2.0445 | 0.63654 | 0.82755 | 1 | 16794 | 2 | -2.0445 |
| DUS2 | 6 | 0.5976 | 0.73817 | 0.999994 | 14923 | 2 | 0.30827 | 0.63678 | 0.82765 | 1 | 16795 | 1 | 0.30827 |

| | | | | | | | | | | | | | |
|---|---|---|---|---|---|---|---|---|---|---|---|---|---|
| LRIT2 | 6 | 0.64881 | 0.77671 | 0.999994 | 15639 | 2 | 0.060616 | 0.63678 | 0.82765 | 1 | 16796 | 2 | 0.060616 |
| ZCWPW2 | 6 | 0.31706 | 0.52652 | 0.999994 | 10454 | 3 | -0.59248 | 0.63678 | 0.82765 | 1 | 16797 | 1 | -0.59248 |
| PPP1CA | 6 | 0.0096955 | 0.037953 | 0.887249 | 853 | 5 | -1.9918 | 0.63678 | 0.82765 | 1 | 16798 | 1 | -1.9918 |
| C1orf168 | 6 | 0.49597 | 0.66594 | 0.999994 | 13357 | 2 | -0.074142 | 0.63678 | 0.82765 | 1 | 16799 | 2 | -0.074142 |
| INSL3 | 6 | 0.68972 | 0.80863 | 0.999994 | 16208 | 2 | 0.2276 | 0.63678 | 0.82765 | 1 | 16800 | 2 | 0.2276 |
| BCKDK | 6 | 0.39862 | 0.60149 | 0.999994 | 11926 | 2 | -0.08067 | 0.63678 | 0.82765 | 1 | 16801 | 2 | -0.08067 |
| BEX1 | 6 | 0.87049 | 0.88364 | 0.999994 | 18644 | 1 | 0.073257 | 0.63678 | 0.82765 | 1 | 16802 | 2 | 0.073257 |
| FCRL6 | 6 | 0.25984 | 0.45966 | 0.999994 | 9238 | 1 | 0.1163 | 0.63678 | 0.82765 | 1 | 16803 | 2 | 0.1163 |
| CCSER1 | 4 | 0.17656 | 0.31659 | 0.999994 | 7145 | 1 | 0.51579 | 0.63707 | 0.77268 | 1 | 16804 | 2 | 0.51579 |
| DNASE1L2 | 5 | 0.18447 | 0.35302 | 0.999994 | 7327 | 2 | 0.36669 | 0.63712 | 0.78996 | 1 | 16805 | 2 | 0.36669 |
| ARG2 | 5 | 0.19439 | 0.36717 | 0.999994 | 7618 | 3 | -0.89629 | 0.63717 | 0.79 | 1 | 16806 | 1 | -0.89629 |
| HIST1H2AE | 5 | 0.56697 | 0.70687 | 0.999994 | 14496 | 2 | -0.42897 | 0.63732 | 0.79008 | 1 | 16807 | 2 | -0.42897 |
| SMG7 | 3 | 0.078266 | 0.15414 | 0.967126 | 4159 | 2 | -0.80956 | 0.63733 | 0.70689 | 1 | 16808 | 1 | -0.80956 |
| RAG1 | 5 | 0.20186 | 0.37791 | 0.999994 | 7821 | 2 | 0.21177 | 0.63739 | 0.79012 | 1 | 16809 | 2 | 0.21177 |
| TPST2 | 4 | 0.7595 | 0.76737 | 0.999994 | 16977 | 1 | 0.4766 | 0.63743 | 0.77299 | 1 | 16810 | 2 | 0.4766 |
| hsa-mir-548l | 4 | 0.17196 | 0.31182 | 0.999994 | 7035 | 2 | -0.3165 | 0.63746 | 0.77301 | 1 | 16811 | 2 | -0.3165 |
| RRM2B | 5 | 0.132 | 0.27568 | 0.999379 | 5976 | 3 | -1.3675 | 0.63769 | 0.7903 | 1 | 16812 | 2 | -1.3675 |
| NonTargetingControlG | 2 | 0.26288 | 0.3085 | 0.999994 | 9312 | 1 | -0.70651 | 0.63771 | 0.64556 | 1 | 16813 | 1 | -0.70651 |
| FOXD4L2 | 6 | 0.22668 | 0.42011 | 0.999994 | 8445 | 3 | -0.46132 | 0.63789 | 0.8281 | 1 | 16814 | 2 | -0.46132 |
| hsa-mir-4450 | 4 | 0.52826 | 0.62931 | 0.999994 | 13864 | 1 | 0.30635 | 0.63811 | 0.77352 | 1 | 16815 | 2 | 0.30635 |
| AVPI1 | 5 | 0.69467 | 0.76407 | 0.999994 | 16251 | 1 | 0.49306 | 0.63814 | 0.79058 | 1 | 16816 | 2 | 0.49306 |
| DDX20 | 4 | 0.094737 | 0.2056 | 0.978603 | 4781 | 2 | -1.6076 | 0.63815 | 0.77355 | 1 | 16817 | 1 | -1.6076 |
| NonTargetingControlG | 1 | 0.36177 | 0.36194 | 0.999994 | 11363 | 0 | -0.52105 | 0.63823 | 0.63816 | 1 | 16818 | 0 | -0.52105 |
| SIRT6 | 5 | 0.38868 | 0.55402 | 0.999994 | 11778 | 1 | 0.32971 | 0.63833 | 0.79068 | 1 | 16819 | 2 | 0.32971 |
| PHLPP2 | 6 | 0.016719 | 0.061376 | 0.935982 | 1291 | 4 | -2.3223 | 0.63836 | 0.82829 | 1 | 16820 | 2 | -2.3223 |
| CCT5 | 3 | 0.60231 | 0.61892 | 0.999994 | 14977 | 1 | 0.32609 | 0.63851 | 0.70749 | 1 | 16821 | 1 | 0.32609 |
| LRRC37B | 5 | 0.43487 | 0.59253 | 0.999994 | 12495 | 2 | -0.46934 | 0.63857 | 0.79083 | 1 | 16822 | 2 | -0.46934 |
| hsa-mir-3978 | 4 | 0.88163 | 0.88126 | 0.999994 | 18862 | 0 | 0.32308 | 0.63859 | 0.77392 | 1 | 16823 | 2 | 0.32308 |
| FANCF | 4 | 0.21414 | 0.35528 | 0.999994 | 8120 | 2 | -0.46257 | 0.63883 | 0.77411 | 1 | 16824 | 2 | -0.46257 |
| RDH14 | 6 | 0.023087 | 0.081853 | 0.93914 | 1656 | 4 | -3.6529 | 0.63911 | 0.82856 | 1 | 16825 | 1 | -3.6529 |
| SCAF11 | 6 | 0.0089301 | 0.03527 | 0.86993 | 786 | 5 | -2.185 | 0.63911 | 0.82856 | 1 | 16826 | 1 | -2.185 |
| BRK1 | 4 | 0.049315 | 0.12011 | 0.951052 | 2944 | 3 | -1.7924 | 0.63914 | 0.77436 | 1 | 16827 | 1 | -1.7924 |
| SLAIN1 | 6 | 0.18831 | 0.37343 | 0.999994 | 7429 | 2 | -0.44556 | 0.63926 | 0.82861 | 1 | 16828 | 2 | -0.44556 |
| hsa-mir-3973 | 4 | 0.31915 | 0.46255 | 0.999994 | 10500 | 2 | -0.3384 | 0.63942 | 0.77458 | 1 | 16829 | 1 | -0.3384 |
| RPP14 | 5 | 0.038353 | 0.11172 | 0.951052 | 2435 | 3 | -1.838 | 0.63947 | 0.79134 | 1 | 16830 | 1 | -1.838 |
| C6orf47 | 5 | 0.57324 | 0.7124 | 0.999994 | 14594 | 2 | -0.18417 | 0.63947 | 0.79134 | 1 | 16831 | 1 | -0.18417 |
| BTK | 5 | 0.011172 | 0.039457 | 0.891621 | 939 | 3 | -3.5315 | 0.63947 | 0.79134 | 1 | 16832 | 1 | -3.5315 |
| CCDC3 | 4 | 0.82618 | 0.82576 | 0.999994 | 17871 | 1 | -0.57364 | 0.63951 | 0.77465 | 1 | 16833 | 1 | -0.57364 |
| PCDHA13 | 2 | 0.082578 | 0.1427 | 0.959553 | 4320 | 1 | -3.5971 | 0.63955 | 0.64712 | 1 | 16834 | 1 | -3.5971 |
| TVP23C | 4 | 0.28257 | 0.42517 | 0.999994 | 9750 | 2 | -0.48089 | 0.64007 | 0.7751 | 1 | 16835 | 1 | -0.48089 |
| NAIF1 | 4 | 0.0089027 | 0.030245 | 0.862223 | 785 | 3 | -4.7085 | 0.64011 | 0.77513 | 1 | 16836 | 1 | -4.7085 |
| hsa-mir-6718 | 4 | 0.66583 | 0.70284 | 0.999994 | 15867 | 1 | 0.54477 | 0.64017 | 0.77519 | 1 | 16837 | 2 | 0.54477 |
| hsa-mir-26b | 3 | 0.56102 | 0.58814 | 0.999994 | 14389 | 1 | -0.15747 | 0.6402 | 0.70837 | 1 | 16838 | 1 | -0.15747 |
| USP39 | 5 | 0.40259 | 0.56569 | 0.999994 | 11990 | 2 | -0.52483 | 0.64032 | 0.79186 | 1 | 16839 | 2 | -0.52483 |
| MICALCL | 6 | 0.028378 | 0.095015 | 0.945031 | 1920 | 4 | -2.2789 | 0.64036 | 0.82906 | 1 | 16840 | 1 | -2.2789 |
| FMO3 | 6 | 0.1195 | 0.28479 | 0.999632 | 5611 | 3 | -1.1377 | 0.6404 | 0.82907 | 1 | 16841 | 2 | -1.1377 |
| NonTargetingControlG | 2 | 0.027016 | 0.04968 | 0.907155 | 1857 | 1 | -4.3872 | 0.64057 | 0.64798 | 1 | 16842 | 1 | -4.3872 |
| ATP7B | 5 | 0.27692 | 0.46238 | 0.999994 | 9637 | 2 | -0.11369 | 0.64063 | 0.79204 | 1 | 16843 | 2 | -0.11369 |
| RNFT2 | 4 | 0.12133 | 0.25263 | 0.994481 | 5676 | 2 | -1.087 | 0.64065 | 0.77558 | 1 | 16844 | 2 | -1.087 |
| TRPA1 | 5 | 0.18084 | 0.34785 | 0.999994 | 7241 | 1 | 0.064858 | 0.64066 | 0.79205 | 1 | 16845 | 1 | 0.064858 |
| ZNF2 | 5 | 0.15305 | 0.30734 | 0.999994 | 6535 | 3 | -1.1437 | 0.64066 | 0.79205 | 1 | 16846 | 1 | -1.1437 |
| DEFA5 | 5 | 0.24302 | 0.43503 | 0.999994 | 8865 | 2 | -0.23292 | 0.64066 | 0.79205 | 1 | 16847 | 1 | -0.23292 |

| | | | | | | | | | | | | |
|---|---|---|---|---|---|---|---|---|---|---|---|---|
| NonTargetingControlG | 2 | 0.54053 | 0.54183 | 0.999994 | 14067 | 1 | 0.20455 | 0.64069 | 0.64807 | 1 | 16848 | 1 | 0.20455 |
| DERL3 | 5 | 0.46083 | 0.61436 | 0.999994 | 12860 | 2 | -0.39496 | 0.64087 | 0.79217 | 1 | 16849 | 2 | -0.39496 |
| hsa-mir-6863 | 4 | 0.28645 | 0.42914 | 0.999994 | 9846 | 2 | -0.32193 | 0.64089 | 0.77578 | 1 | 16850 | 2 | -0.32193 |
| hsa-mir-891a | 3 | 0.064739 | 0.13584 | 0.95468 | 3639 | 2 | -1.5212 | 0.64089 | 0.70874 | 1 | 16851 | 1 | -1.5212 |
| GTF2H4 | 3 | 0.1268 | 0.21838 | 0.980699 | 5841 | 2 | -0.87182 | 0.64089 | 0.70874 | 1 | 16852 | 1 | -0.87182 |
| hsa-mir-4417 | 3 | 0.25568 | 0.38206 | 0.999994 | 9141 | 1 | 0.025715 | 0.64089 | 0.70874 | 1 | 16853 | 1 | 0.025715 |
| PRC1 | 4 | 0.32106 | 0.46448 | 0.999994 | 10535 | 2 | -0.85289 | 0.641 | 0.77587 | 1 | 16854 | 1 | -0.85289 |
| ZCCHC24 | 4 | 0.070021 | 0.15998 | 0.967126 | 3828 | 2 | -2.4026 | 0.64114 | 0.776 | 1 | 16855 | 2 | -2.4026 |
| SLC8A1 | 4 | 0.85061 | 0.85012 | 0.999994 | 18271 | 0 | 0.55715 | 0.64122 | 0.77607 | 1 | 16856 | 2 | 0.55715 |
| C19orf80 | 4 | 0.54336 | 0.63648 | 0.999994 | 14118 | 1 | 0.40646 | 0.64135 | 0.77617 | 1 | 16857 | 2 | 0.40646 |
| hsa-mir-4524b | 1 | 0.35849 | 0.35861 | 0.999994 | 11310 | 0 | -0.57787 | 0.64151 | 0.6415 | 1 | 16858 | 0 | -0.57787 |
| hsa-mir-4475 | 4 | 0.74631 | 0.75723 | 0.999994 | 16817 | 1 | 0.15986 | 0.64156 | 0.77634 | 1 | 16859 | 2 | 0.15986 |
| DISC1 | 3 | 0.07463 | 0.14918 | 0.965773 | 4020 | 2 | -2.7164 | 0.64157 | 0.70906 | 1 | 16860 | 1 | -2.7164 |
| SLC9A6 | 5 | 0.021838 | 0.069023 | 0.93914 | 1569 | 3 | -2.1433 | 0.64178 | 0.7927 | 1 | 16861 | 2 | -2.1433 |
| KLLN | 6 | 0.53679 | 0.69429 | 0.999994 | 14010 | 2 | -0.00061277 | 0.64191 | 0.82965 | 1 | 16862 | 2 | -0.00061277 |
| ZPBP2 | 6 | 0.13925 | 0.31207 | 0.999994 | 6169 | 3 | -1.0073 | 0.64191 | 0.82965 | 1 | 16863 | 2 | -1.0073 |
| NonTargetingControlG | 2 | 0.47589 | 0.48213 | 0.999994 | 13076 | 1 | 0.12427 | 0.64207 | 0.64923 | 1 | 16864 | 1 | 0.12427 |
| RBM47 | 5 | 0.83868 | 0.84569 | 0.999994 | 18065 | 1 | -0.26285 | 0.64209 | 0.79287 | 1 | 16865 | 1 | -0.26285 |
| NonTargetingControlG | 2 | 0.13852 | 0.21869 | 0.980699 | 6153 | 1 | -3.06 | 0.64217 | 0.64933 | 1 | 16866 | 1 | -3.06 |
| FAM65B | 5 | 0.2272 | 0.41334 | 0.999994 | 8460 | 3 | -0.68353 | 0.64231 | 0.79298 | 1 | 16867 | 2 | -0.68353 |
| hsa-mir-6808 | 4 | 0.11311 | 0.23837 | 0.987253 | 5409 | 3 | -1.333 | 0.64232 | 0.77692 | 1 | 16868 | 1 | -1.333 |
| KCNH1 | 4 | 0.046742 | 0.11506 | 0.951052 | 2839 | 3 | -1.7153 | 0.64232 | 0.77692 | 1 | 16869 | 1 | -1.7153 |
| LZTR1 | 6 | 0.24933 | 0.44713 | 0.999994 | 9000 | 3 | -0.42758 | 0.6425 | 0.82986 | 1 | 16870 | 2 | -0.42758 |
| OCM | 6 | 0.10637 | 0.25971 | 0.994943 | 5185 | 4 | -0.99138 | 0.64257 | 0.82989 | 1 | 16871 | 2 | -0.99138 |
| FBXL20 | 5 | 0.23389 | 0.4226 | 0.999994 | 8618 | 2 | 0.058069 | 0.64278 | 0.79326 | 1 | 16872 | 2 | 0.058069 |
| CORO1C | 5 | 0.56174 | 0.70219 | 0.999994 | 14402 | 2 | -0.2492 | 0.64294 | 0.79336 | 1 | 16873 | 2 | -0.2492 |
| PCNX | 5 | 0.88769 | 0.88778 | 0.999994 | 18992 | 1 | 0.20777 | 0.643 | 0.79339 | 1 | 16874 | 2 | 0.20777 |
| PITPNC1 | 4 | 0.32136 | 0.46479 | 0.999994 | 10541 | 2 | -0.81785 | 0.64306 | 0.77753 | 1 | 16875 | 1 | -0.81785 |
| hsa-mir-572 | 4 | 0.60601 | 0.66856 | 0.999994 | 15028 | 1 | 0.54326 | 0.64313 | 0.77759 | 1 | 16876 | 2 | 0.54326 |
| APOBEC3F | 6 | 0.0027688 | 0.012869 | 0.733879 | 336 | 4 | -4.2272 | 0.64315 | 0.83012 | 1 | 16877 | 1 | -4.2272 |
| TRAPPC8 | 6 | 0.21463 | 0.40572 | 0.999994 | 8136 | 2 | 0.067104 | 0.64315 | 0.83012 | 1 | 16878 | 1 | 0.067104 |
| SPSB4 | 6 | 0.15224 | 0.32848 | 0.999994 | 6508 | 3 | -0.92617 | 0.64321 | 0.83014 | 1 | 16879 | 2 | -0.92617 |
| NonTargetingControlG | 2 | 0.55165 | 0.55245 | 0.999994 | 14235 | 1 | 0.34558 | 0.6436 | 0.65051 | 1 | 16880 | 1 | 0.34558 |
| NonTargetingControlG | 1 | 0.35639 | 0.35654 | 0.999994 | 11274 | 0 | -0.60663 | 0.64361 | 0.64359 | 1 | 16881 | 0 | -0.60663 |
| CYP39A1 | 5 | 0.87114 | 0.87209 | 0.999994 | 18658 | 1 | 0.16082 | 0.64379 | 0.79385 | 1 | 16882 | 1 | 0.16082 |
| TIGD6 | 5 | 0.0041177 | 0.017299 | 0.785648 | 448 | 2 | -0.57841 | 0.64379 | 0.79385 | 1 | 16883 | 2 | -0.57841 |
| hsa-mir-128-1 | 4 | 0.50013 | 0.6164 | 0.999994 | 13425 | 1 | 0.0099988 | 0.64405 | 0.77835 | 1 | 16884 | 2 | 0.0099988 |
| hsa-mir-4330 | 2 | 0.62184 | 0.62161 | 0.999994 | 15254 | 0 | 0.38876 | 0.64412 | 0.65097 | 1 | 16885 | 1 | 0.38876 |
| NUBPL | 6 | 0.061838 | 0.16966 | 0.967126 | 3506 | 3 | -1.0162 | 0.64414 | 0.83052 | 1 | 16886 | 2 | -1.0162 |
| C2orf78 | 6 | 0.51771 | 0.68094 | 0.999994 | 13706 | 1 | 0.38347 | 0.64421 | 0.83054 | 1 | 16887 | 1 | 0.38347 |
| DCDC2B | 6 | 0.075676 | 0.19857 | 0.977164 | 4060 | 3 | -1.494 | 0.64421 | 0.83054 | 1 | 16888 | 2 | -1.494 |
| B2M | 5 | 0.38621 | 0.55196 | 0.999994 | 11736 | 1 | 0.4918 | 0.64427 | 0.79411 | 1 | 16889 | 2 | 0.4918 |
| NonTargetingControlG | 2 | 0.25458 | 0.30228 | 0.999994 | 9119 | 1 | -0.98242 | 0.64435 | 0.65117 | 1 | 16890 | 1 | -0.98242 |
| NonTargetingControlG | 2 | 0.66498 | 0.66468 | 0.999994 | 15856 | 0 | 0.67168 | 0.64455 | 0.65134 | 1 | 16891 | 1 | 0.67168 |
| hsa-mir-3665 | 2 | 0.23643 | 0.28877 | 0.999632 | 8679 | 1 | -1.9548 | 0.64458 | 0.65137 | 1 | 16892 | 1 | -1.9548 |
| ALKBH8 | 5 | 0.54495 | 0.68725 | 0.999994 | 14137 | 2 | 0.088 | 0.64472 | 0.79439 | 1 | 16893 | 2 | 0.088 |
| NonTargetingControlG | 1 | 0.35523 | 0.35537 | 0.999994 | 11256 | 0 | -0.56237 | 0.64477 | 0.64478 | 1 | 16894 | 0 | -0.56237 |
| RBMY1F | 1 | 0.35491 | 0.35504 | 0.999994 | 11250 | 1 | -0.62698 | 0.64509 | 0.64509 | 1 | 16895 | 0 | -0.62698 |
| CMPK1 | 2 | 0.52253 | 0.52474 | 0.999994 | 13788 | 1 | 0.065963 | 0.64522 | 0.65188 | 1 | 16896 | 1 | 0.065963 |
| ADAMDEC1 | 6 | 0.07885 | 0.20502 | 0.978186 | 4184 | 3 | -1.0768 | 0.64541 | 0.83103 | 1 | 16897 | 2 | -1.0768 |
| hsa-mir-3118-1 | 1 | 0.35456 | 0.35471 | 0.999994 | 11240 | 1 | -0.60861 | 0.64544 | 0.64541 | 1 | 16898 | 0 | -0.60861 |
| MME | 5 | 0.40026 | 0.56372 | 0.999994 | 11954 | 2 | -0.28535 | 0.64547 | 0.79483 | 1 | 16899 | 1 | -0.28535 |

| | | | | | | | | | | | | |
|---|---|---|---|---|---|---|---|---|---|---|---|---|
| NonTargetingControlG | 2 | 0.67754 | 0.67734 | 0.999994 | 16054 | 0 | 0.61976 | 0.6455 | 0.65211 | 1 | 16900 | 1 | 0.61976 |
| GOLPH3L | 6 | 0.0034103 | 0.015236 | 0.757255 | 395 | 5 | -2.2274 | 0.64567 | 0.83113 | 1 | 16901 | 1 | -2.2274 |
| RANBP9 | 6 | 0.22201 | 0.41459 | 0.999994 | 8332 | 3 | -0.54004 | 0.64567 | 0.83113 | 1 | 16902 | 2 | -0.54004 |
| APLF | 6 | 0.053554 | 0.15177 | 0.967126 | 3143 | 3 | -0.3181 | 0.64567 | 0.83113 | 1 | 16903 | 1 | -0.3181 |
| SCN7A | 6 | 0.015892 | 0.058697 | 0.931872 | 1238 | 5 | -1.8967 | 0.64567 | 0.83113 | 1 | 16904 | 1 | -1.8967 |
| hsa-mir-7973-1 | 4 | 0.90969 | 0.90942 | 0.999994 | 19419 | 0 | 0.51068 | 0.6457 | 0.77966 | 1 | 16905 | 2 | 0.51068 |
| CACNA1G | 6 | 0.15415 | 0.33087 | 0.999994 | 6572 | 3 | -0.25485 | 0.64573 | 0.83116 | 1 | 16906 | 2 | -0.25485 |
| WLS | 4 | 0.12352 | 0.25643 | 0.994943 | 5751 | 3 | -0.94187 | 0.64574 | 0.77969 | 1 | 16907 | 1 | -0.94187 |
| NonTargetingControlG | 2 | 0.18335 | 0.2501 | 0.992497 | 7302 | 1 | -2.5409 | 0.64583 | 0.65241 | 1 | 16908 | 1 | -2.5409 |
| RGS6 | 6 | 0.097204 | 0.24187 | 0.987253 | 4866 | 3 | -1.0344 | 0.64601 | 0.83127 | 1 | 16909 | 2 | -1.0344 |
| NonTargetingControlG | 2 | 0.48195 | 0.48753 | 0.999994 | 13168 | 1 | -0.010324 | 0.64603 | 0.65261 | 1 | 16910 | 1 | -0.010324 |
| SDF4 | 6 | 0.67983 | 0.80085 | 0.999994 | 16087 | 2 | 0.12324 | 0.64609 | 0.83131 | 1 | 16911 | 2 | 0.12324 |
| ZFAND6 | 5 | 0.039839 | 0.1155 | 0.951052 | 2507 | 4 | -1.8869 | 0.64616 | 0.79524 | 1 | 16912 | 1 | -1.8869 |
| UBE4A | 3 | 0.38386 | 0.47501 | 0.999994 | 11701 | 1 | -0.2537 | 0.64642 | 0.71162 | 1 | 16913 | 1 | -0.2537 |
| hsa-mir-4749 | 4 | 0.87708 | 0.87677 | 0.999994 | 18774 | 0 | 0.40983 | 0.64646 | 0.78028 | 1 | 16914 | 2 | 0.40983 |
| GRB14 | 6 | 0.19869 | 0.38633 | 0.999994 | 7727 | 2 | -0.3566 | 0.6465 | 0.83146 | 1 | 16915 | 2 | -0.3566 |
| OR2A12 | 5 | 0.11047 | 0.2427 | 0.988088 | 5318 | 3 | -0.90799 | 0.64678 | 0.79562 | 1 | 16916 | 2 | -0.90799 |
| ZBTB18 | 5 | 0.2044 | 0.38148 | 0.999994 | 7888 | 3 | -0.81689 | 0.64686 | 0.79567 | 1 | 16917 | 1 | -0.81689 |
| ILK | 6 | 0.06018 | 0.16612 | 0.967126 | 3428 | 4 | -1.6205 | 0.64693 | 0.83163 | 1 | 16918 | 2 | -1.6205 |
| AQP7 | 6 | 0.044083 | 0.13106 | 0.95318 | 2714 | 4 | -1.3281 | 0.64708 | 0.83169 | 1 | 16919 | 2 | -1.3281 |
| METTL7A | 5 | 0.13512 | 0.28043 | 0.999632 | 6053 | 3 | -1.1815 | 0.64743 | 0.796 | 1 | 16920 | 2 | -1.1815 |
| hsa-mir-770 | 3 | 0.11685 | 0.20533 | 0.978342 | 5535 | 2 | -1.3419 | 0.64744 | 0.71216 | 1 | 16921 | 1 | -1.3419 |
| LOC100506422 | 6 | 0.065753 | 0.17787 | 0.968171 | 3681 | 3 | -0.60356 | 0.64758 | 0.83189 | 1 | 16922 | 2 | -0.60356 |
| hsa-mir-5011 | 4 | 0.10297 | 0.2204 | 0.980699 | 5066 | 2 | -0.11873 | 0.64761 | 0.78118 | 1 | 16923 | 2 | -0.11873 |
| hsa-mir-6818 | 4 | 0.45792 | 0.598 | 0.999994 | 12819 | 1 | -0.090196 | 0.64761 | 0.78118 | 1 | 16924 | 1 | -0.090196 |
| CPSF1 | 1 | 0.35207 | 0.35221 | 0.999994 | 11195 | 1 | -0.64826 | 0.64793 | 0.64793 | 1 | 16925 | 0 | -0.64826 |
| hsa-mir-376b | 3 | 0.59966 | 0.6169 | 0.999994 | 14949 | 1 | -0.33136 | 0.64797 | 0.71244 | 1 | 16926 | 1 | -0.33136 |
| USP33 | 5 | 0.26277 | 0.45096 | 0.999994 | 9305 | 1 | 0.018228 | 0.64797 | 0.79632 | 1 | 16927 | 2 | 0.018228 |
| hsa-mir-767 | 4 | 0.75186 | 0.76151 | 0.999994 | 16877 | 1 | 0.2925 | 0.64798 | 0.78149 | 1 | 16928 | 2 | 0.2925 |
| IGF1 | 5 | 0.39401 | 0.55853 | 0.999994 | 11858 | 1 | 0.31054 | 0.64802 | 0.79635 | 1 | 16929 | 2 | 0.31054 |
| PP2D1 | 6 | 0.51048 | 0.67596 | 0.999994 | 13581 | 2 | 0.0040559 | 0.64807 | 0.83209 | 1 | 16930 | 2 | 0.0040559 |
| C1QA | 5 | 0.43746 | 0.59468 | 0.999994 | 12534 | 1 | 0.43449 | 0.64817 | 0.79643 | 1 | 16931 | 2 | 0.43449 |
| ENPP4 | 5 | 0.097893 | 0.22314 | 0.983354 | 4890 | 3 | -1.479 | 0.64827 | 0.7965 | 1 | 16932 | 2 | -1.479 |
| XPA | 6 | 0.29906 | 0.50561 | 0.999994 | 10115 | 3 | -0.5136 | 0.64838 | 0.8322 | 1 | 16933 | 2 | -0.5136 |
| ANXA3 | 6 | 0.27986 | 0.48326 | 0.999994 | 9701 | 3 | -0.24241 | 0.64838 | 0.8322 | 1 | 16934 | 2 | -0.24241 |
| MGEA5 | 6 | 0.00954 | 0.03742 | 0.881473 | 842 | 5 | -1.7546 | 0.64838 | 0.8322 | 1 | 16935 | 1 | -1.7546 |
| RBM5 | 6 | 0.033109 | 0.10612 | 0.950452 | 2168 | 4 | -1.3765 | 0.64838 | 0.8322 | 1 | 16936 | 2 | -1.3765 |
| KLC4 | 5 | 0.2114 | 0.39134 | 0.999994 | 8057 | 2 | -1.4517 | 0.64839 | 0.79657 | 1 | 16937 | 2 | -1.4517 |
| TNFAIP8L1 | 5 | 0.07292 | 0.18314 | 0.968526 | 3958 | 3 | -1.6029 | 0.64841 | 0.79658 | 1 | 16938 | 2 | -1.6029 |
| HPGDS | 5 | 0.3207 | 0.49787 | 0.999994 | 10527 | 2 | 0.16896 | 0.64841 | 0.79658 | 1 | 16939 | 2 | 0.16896 |
| MTG2 | 1 | 0.35068 | 0.35084 | 0.999994 | 11176 | 1 | -0.60574 | 0.64932 | 0.64933 | 1 | 16940 | 0 | -0.60574 |
| SAMD13 | 5 | 0.93488 | 0.93512 | 0.999994 | 19965 | 0 | 0.31419 | 0.64939 | 0.79717 | 1 | 16941 | 2 | 0.31419 |
| NonTargetingControlG | 2 | 0.33443 | 0.36356 | 0.999994 | 10820 | 1 | -0.62629 | 0.64952 | 0.65563 | 1 | 16942 | 1 | -0.62629 |
| hsa-mir-324 | 3 | 0.096513 | 0.17843 | 0.968171 | 4839 | 2 | -2.2036 | 0.64953 | 0.71327 | 1 | 16943 | 1 | -2.2036 |
| NYAP1 | 6 | 0.13497 | 0.30663 | 0.999994 | 6049 | 3 | -0.95543 | 0.64963 | 0.83268 | 1 | 16944 | 2 | -0.95543 |
| CHD3 | 6 | 0.081069 | 0.20947 | 0.979587 | 4269 | 3 | -0.93954 | 0.64965 | 0.83269 | 1 | 16945 | 2 | -0.93954 |
| WWC2 | 3 | 0.27816 | 0.40976 | 0.999994 | 9656 | 1 | 0.52676 | 0.64971 | 0.71336 | 1 | 16946 | 1 | 0.52676 |
| DAPK3 | 5 | 0.22621 | 0.41196 | 0.999994 | 8429 | 3 | -0.66902 | 0.64972 | 0.79738 | 1 | 16947 | 2 | -0.66902 |
| KIAA1328 | 5 | 0.039034 | 0.11345 | 0.951052 | 2464 | 3 | -2.7258 | 0.64972 | 0.79738 | 1 | 16948 | 1 | -2.7258 |
| IRX1 | 5 | 0.29759 | 0.47912 | 0.999994 | 10081 | 2 | -0.44006 | 0.64972 | 0.79738 | 1 | 16949 | 1 | -0.44006 |
| CBLN2 | 5 | 0.17125 | 0.33394 | 0.999994 | 7017 | 3 | -1.0711 | 0.64972 | 0.79738 | 1 | 16950 | 1 | -1.0711 |
| LNPEP | 5 | 0.55079 | 0.69244 | 0.999994 | 14225 | 1 | 0.1485 | 0.64972 | 0.79738 | 1 | 16951 | 2 | 0.1485 |

| | | | | | | | | | | | | | |
|---|---|---|---|---|---|---|---|---|---|---|---|---|---|
| PIN4 | 4 | 0.80684 | 0.80741 | 0.999994 | 17581 | 1 | 0.61355 | 0.64972 | 0.78288 | 1 | 16952 | 2 | 0.61355 |
| TVP23A | 4 | 0.078245 | 0.17547 | 0.968171 | 4157 | 2 | -0.17811 | 0.6499 | 0.78302 | 1 | 16953 | 2 | -0.17811 |
| USP37 | 5 | 0.17258 | 0.33589 | 0.999994 | 7049 | 2 | 0.13949 | 0.65017 | 0.79765 | 1 | 16954 | 2 | 0.13949 |
| hsa-mir-6165 | 1 | 0.34983 | 0.35001 | 0.999994 | 11154 | 1 | -0.65849 | 0.65017 | 0.65018 | 1 | 16955 | 0 | -0.65849 |
| OR4F29 | 2 | 0.18965 | 0.25461 | 0.994835 | 7476 | 1 | -1.6467 | 0.6503 | 0.65632 | 1 | 16956 | 1 | -1.6467 |
| ARHGEF39 | 3 | 0.73863 | 0.73855 | 0.999994 | 16715 | 0 | 0.34547 | 0.65042 | 0.71371 | 1 | 16957 | 1 | 0.34547 |
| FOXL1 | 5 | 0.082443 | 0.19855 | 0.977164 | 4317 | 1 | 0.10633 | 0.65053 | 0.79785 | 1 | 16958 | 1 | 0.10633 |
| hsa-mir-548h-4 | 2 | 0.45947 | 0.46766 | 0.999994 | 12839 | 1 | -0.1086 | 0.65062 | 0.65663 | 1 | 16959 | 1 | -0.1086 |
| CXorf30 | 6 | 0.4922 | 0.66337 | 0.999994 | 13302 | 2 | -0.28586 | 0.65078 | 0.83312 | 1 | 16960 | 2 | -0.28586 |
| hsa-mir-548ax | 3 | 0.45719 | 0.51871 | 0.999994 | 12810 | 1 | -0.31579 | 0.65088 | 0.71395 | 1 | 16961 | 1 | -0.31579 |
| TMEM14A | 6 | 0.059349 | 0.16437 | 0.967126 | 3396 | 4 | -1.3339 | 0.65096 | 0.8332 | 1 | 16962 | 1 | -1.3339 |
| PRAF2 | 6 | 0.16565 | 0.34527 | 0.999994 | 6876 | 3 | -0.87283 | 0.65096 | 0.8332 | 1 | 16963 | 1 | -0.87283 |
| ONECUT3 | 3 | 0.089741 | 0.16939 | 0.967126 | 4583 | 2 | -2.2215 | 0.65097 | 0.71399 | 1 | 16964 | 1 | -2.2215 |
| ZNF214 | 5 | 0.1686 | 0.33006 | 0.999994 | 6949 | 3 | -0.71274 | 0.65113 | 0.79823 | 1 | 16965 | 2 | -0.71274 |
| GPD2 | 6 | 0.10119 | 0.24964 | 0.991904 | 5015 | 3 | -0.90161 | 0.65136 | 0.83335 | 1 | 16966 | 2 | -0.90161 |
| PIH1D3 | 5 | 0.21056 | 0.39018 | 0.999994 | 8040 | 3 | -0.77194 | 0.65136 | 0.79835 | 1 | 16967 | 2 | -0.77194 |
| MSL1 | 4 | 0.039437 | 0.10066 | 0.946414 | 2486 | 2 | -0.033487 | 0.65159 | 0.78436 | 1 | 16968 | 2 | -0.033487 |
| RASA4B | 3 | 0.20089 | 0.31378 | 0.999994 | 7785 | 2 | -1.2093 | 0.65163 | 0.71434 | 1 | 16969 | 1 | -1.2093 |
| TCP11X2 | 5 | 0.16448 | 0.32402 | 0.999994 | 6852 | 2 | 0.34017 | 0.65173 | 0.79858 | 1 | 16970 | 2 | 0.34017 |
| VPS8 | 5 | 0.28146 | 0.46606 | 0.999994 | 9730 | 2 | -0.44138 | 0.65173 | 0.79858 | 1 | 16971 | 1 | -0.44138 |
| RAET1L | 5 | 0.34679 | 0.5193 | 0.999994 | 11091 | 2 | -0.31012 | 0.65173 | 0.79858 | 1 | 16972 | 2 | -0.31012 |
| PPM1L | 4 | 0.075102 | 0.16955 | 0.967126 | 4039 | 3 | -1.3465 | 0.65203 | 0.78471 | 1 | 16973 | 1 | -1.3465 |
| ACIN1 | 4 | 0.37308 | 0.51802 | 0.999994 | 11559 | 2 | -0.015638 | 0.65206 | 0.78473 | 1 | 16974 | 2 | -0.015638 |
| MRFAP1 | 4 | 0.46562 | 0.60127 | 0.999994 | 12947 | 1 | 0.31503 | 0.65209 | 0.78475 | 1 | 16975 | 2 | 0.31503 |
| MRPS17 | 6 | 0.013545 | 0.050941 | 0.909101 | 1105 | 4 | -2.6425 | 0.65221 | 0.83369 | 1 | 16976 | 1 | -2.6425 |
| UPF2 | 4 | 0.8958 | 0.8955 | 0.999994 | 19142 | 0 | 0.35188 | 0.65234 | 0.78497 | 1 | 16977 | 2 | 0.35188 |
| FRY | 4 | 0.72661 | 0.74271 | 0.999994 | 16562 | 1 | 0.56722 | 0.65241 | 0.78503 | 1 | 16978 | 2 | 0.56722 |
| KRIT1 | 4 | 0.099732 | 0.21465 | 0.979587 | 4966 | 1 | 0.3282 | 0.65246 | 0.78506 | 1 | 16979 | 2 | 0.3282 |
| RGPD4 | 3 | 0.18311 | 0.29122 | 0.999632 | 7295 | 2 | -1.1913 | 0.65258 | 0.71485 | 1 | 16980 | 1 | -1.1913 |
| MUC16 | 6 | 0.17745 | 0.35989 | 0.999994 | 7165 | 3 | -0.5138 | 0.65286 | 0.83394 | 1 | 16981 | 2 | -0.5138 |
| YRDC | 5 | 0.2389 | 0.42942 | 0.999994 | 8744 | 2 | 0.16657 | 0.65303 | 0.79931 | 1 | 16982 | 2 | 0.16657 |
| GMFB | 5 | 0.31817 | 0.49586 | 0.999994 | 10478 | 1 | 0.056359 | 0.65304 | 0.79932 | 1 | 16983 | 2 | 0.056359 |
| NonTargetingControlG | 2 | 0.33387 | 0.36311 | 0.999994 | 10811 | 1 | -0.60483 | 0.65306 | 0.65876 | 1 | 16984 | 1 | -0.60483 |
| ACAA2 | 6 | 0.20464 | 0.39353 | 0.999994 | 7892 | 2 | 0.16215 | 0.65314 | 0.83404 | 1 | 16985 | 2 | 0.16215 |
| ERP27 | 5 | 0.7474 | 0.78949 | 0.999994 | 16831 | 1 | 0.4014 | 0.65333 | 0.79949 | 1 | 16986 | 2 | 0.4014 |
| GIMAP6 | 6 | 0.29181 | 0.49715 | 0.999994 | 9944 | 3 | -0.20771 | 0.65338 | 0.83414 | 1 | 16987 | 2 | -0.20771 |
| BAGE3 | 2 | 0.60926 | 0.6091 | 0.999994 | 15061 | 0 | 0.32809 | 0.65339 | 0.65905 | 1 | 16988 | 1 | 0.32809 |
| OR6Q1 | 6 | 0.27288 | 0.4751 | 0.999994 | 9534 | 2 | -0.069886 | 0.65349 | 0.83418 | 1 | 16989 | 1 | -0.069886 |
| CENPA | 6 | 0.043295 | 0.12932 | 0.952548 | 2687 | 3 | -0.60892 | 0.65349 | 0.83418 | 1 | 16990 | 2 | -0.60892 |
| ZBTB7C | 6 | 0.051173 | 0.14665 | 0.964852 | 3031 | 4 | -1.777 | 0.65349 | 0.83418 | 1 | 16991 | 2 | -1.777 |
| CPSF3 | 3 | 0.11685 | 0.20533 | 0.978342 | 5536 | 2 | -1.8653 | 0.65362 | 0.71542 | 1 | 16992 | 1 | -1.8653 |
| TOMM40L | 5 | 0.12149 | 0.25961 | 0.994943 | 5685 | 3 | -1.3791 | 0.65373 | 0.79971 | 1 | 16993 | 2 | -1.3791 |
| HERC5 | 5 | 0.27427 | 0.46024 | 0.999994 | 9569 | 2 | -0.27376 | 0.65376 | 0.79973 | 1 | 16994 | 2 | -0.27376 |
| SYNGR3 | 5 | 0.18741 | 0.35726 | 0.999994 | 7405 | 3 | -0.81903 | 0.65378 | 0.79974 | 1 | 16995 | 2 | -0.81903 |
| hsa-mir-632 | 4 | 0.053225 | 0.12778 | 0.951674 | 3125 | 3 | -2.2125 | 0.65384 | 0.78616 | 1 | 16996 | 1 | -2.2125 |
| hsa-mir-873 | 4 | 0.15094 | 0.29017 | 0.999632 | 6483 | 2 | -0.57164 | 0.65384 | 0.78616 | 1 | 16997 | 1 | -0.57164 |
| RBM27 | 4 | 0.032389 | 0.086377 | 0.94395 | 2127 | 3 | -2.5369 | 0.65384 | 0.78616 | 1 | 16998 | 1 | -2.5369 |
| MALSU1 | 4 | 0.46722 | 0.60192 | 0.999994 | 12968 | 1 | 0.48878 | 0.65402 | 0.78631 | 1 | 16999 | 2 | 0.48878 |
| RASA2 | 6 | 0.50754 | 0.67394 | 0.999994 | 13545 | 1 | 0.095112 | 0.6541 | 0.83443 | 1 | 17000 | 2 | 0.095112 |
| GIMAP8 | 4 | 0.14331 | 0.28233 | 0.999632 | 6281 | 2 | -0.6236 | 0.65411 | 0.78637 | 1 | 17001 | 2 | -0.6236 |
| NonTargetingControlG | 2 | 0.32705 | 0.3577 | 0.999994 | 10662 | 1 | -0.50465 | 0.65441 | 0.65993 | 1 | 17002 | 1 | -0.50465 |
| C1orf159 | 5 | 0.026646 | 0.081804 | 0.93914 | 1838 | 4 | -1.5099 | 0.65457 | 0.80024 | 1 | 17003 | 1 | -1.5099 |

| | | | | | | | | | | | | | |
|---|---|---|---|---|---|---|---|---|---|---|---|---|---|
| hsa-mir-4418 | 4 | 0.59615 | 0.66322 | 0.999994 | 14904 | 1 | 0.065591 | 0.65468 | 0.78683 | 1 | 17004 | 2 | 0.065591 |
| THEMIS2 | 6 | 0.12754 | 0.29715 | 0.999994 | 5862 | 3 | -1.1636 | 0.65472 | 0.83468 | 1 | 17005 | 1 | -1.1636 |
| ZNF879 | 6 | 0.10569 | 0.25841 | 0.994943 | 5161 | 4 | -0.87933 | 0.65472 | 0.83468 | 1 | 17006 | 1 | -0.87933 |
| IL20 | 6 | 0.055956 | 0.15698 | 0.967126 | 3244 | 4 | -0.90605 | 0.65472 | 0.83468 | 1 | 17007 | 2 | -0.90605 |
| CSK | 6 | 0.00076436 | 0.0039199 | 0.560963 | 136 | 4 | -5.3736 | 0.65472 | 0.83468 | 1 | 17008 | 1 | -5.3736 |
| hsa-mir-21 | 4 | 0.029344 | 0.080189 | 0.93914 | 1976 | 3 | -2.0738 | 0.65514 | 0.7872 | 1 | 17009 | 1 | -2.0738 |
| COL28A1 | 4 | 0.31443 | 0.45779 | 0.999994 | 10406 | 2 | -0.26307 | 0.65523 | 0.78727 | 1 | 17010 | 2 | -0.26307 |
| NonTargetingControlG | 2 | 0.5276 | 0.52951 | 0.999994 | 13852 | 1 | 0.21893 | 0.65533 | 0.66075 | 1 | 17011 | 1 | 0.21893 |
| C2orf72 | 6 | 0.36744 | 0.58163 | 0.999994 | 11464 | 2 | 0.0014017 | 0.65543 | 0.83496 | 1 | 17012 | 2 | 0.0014017 |
| OR4F5 | 4 | 0.0027315 | 0.0097983 | 0.680529 | 333 | 2 | -0.278 | 0.65546 | 0.78746 | 1 | 17013 | 2 | -0.278 |
| MMEL1 | 5 | 0.016787 | 0.055285 | 0.922677 | 1294 | 2 | 0.47332 | 0.65552 | 0.80081 | 1 | 17014 | 2 | 0.47332 |
| MCHR2 | 6 | 0.024385 | 0.085545 | 0.94395 | 1727 | 3 | -0.67499 | 0.65556 | 0.835 | 1 | 17015 | 2 | -0.67499 |
| NonTargetingControlG | 1 | 0.34437 | 0.34465 | 0.999994 | 11037 | 1 | -0.69815 | 0.65563 | 0.65566 | 1 | 17016 | 0 | -0.69815 |
| hsa-mir-144 | 4 | 0.70429 | 0.7273 | 0.999994 | 16355 | 1 | 0.087296 | 0.65575 | 0.78771 | 1 | 17017 | 2 | 0.087296 |
| ABHD10 | 5 | 0.16444 | 0.32396 | 0.999994 | 6851 | 3 | -0.92336 | 0.65575 | 0.80095 | 1 | 17018 | 2 | -0.92336 |
| NonTargetingControlG | 2 | 0.024168 | 0.044666 | 0.891621 | 1718 | 1 | -4.4419 | 0.65582 | 0.66114 | 1 | 17019 | 1 | -4.4419 |
| C14orf180 | 5 | 0.11521 | 0.25006 | 0.992497 | 5485 | 3 | -0.88324 | 0.6559 | 0.80105 | 1 | 17020 | 1 | -0.88324 |
| HINFP | 5 | 0.00010855 | 0.00049539 | 0.275324 | 36 | 4 | -8.2976 | 0.6559 | 0.80105 | 1 | 17021 | 1 | -8.2976 |
| C7orf41 | 6 | 0.041052 | 0.12428 | 0.951052 | 2565 | 3 | -0.95482 | 0.65606 | 0.83519 | 1 | 17022 | 2 | -0.95482 |
| GPR156 | 6 | 0.042517 | 0.12757 | 0.951052 | 2646 | 4 | -1.4922 | 0.65606 | 0.83519 | 1 | 17023 | 2 | -1.4922 |
| CDKN1C | 6 | 0.024601 | 0.086067 | 0.94395 | 1735 | 4 | -2.5081 | 0.65606 | 0.83519 | 1 | 17024 | 2 | -2.5081 |
| FAM19A4 | 6 | 0.27399 | 0.4764 | 0.999994 | 9566 | 2 | -0.17581 | 0.65606 | 0.83519 | 1 | 17025 | 2 | -0.17581 |
| NonTargetingControlG | 1 | 0.34393 | 0.3442 | 0.999994 | 11027 | 1 | -0.68827 | 0.65607 | 0.65611 | 1 | 17026 | 0 | -0.68827 |
| FTSJD1 | 3 | 0.76732 | 0.76732 | 0.999994 | 17055 | 0 | 0.2905 | 0.65627 | 0.71681 | 1 | 17027 | 1 | 0.2905 |
| BRPF1 | 5 | 0.28906 | 0.47217 | 0.999994 | 9894 | 2 | 0.23512 | 0.65636 | 0.80133 | 1 | 17028 | 2 | 0.23512 |
| BTN3A1 | 5 | 0.5422 | 0.68484 | 0.999994 | 14098 | 1 | 0.46529 | 0.65643 | 0.80137 | 1 | 17029 | 2 | 0.46529 |
| hsa-mir-4697 | 2 | 0.25897 | 0.30559 | 0.999994 | 9213 | 1 | -1.0809 | 0.65674 | 0.66195 | 1 | 17030 | 1 | -1.0809 |
| PRR5L | 4 | 0.12066 | 0.25146 | 0.993413 | 5649 | 3 | -1.1047 | 0.6568 | 0.7886 | 1 | 17031 | 1 | -1.1047 |
| ARIH2OS | 5 | 0.23611 | 0.42566 | 0.999994 | 8674 | 3 | -0.61296 | 0.65695 | 0.80167 | 1 | 17032 | 2 | -0.61296 |
| NonTargetingControlG | 1 | 0.34292 | 0.34319 | 0.999994 | 11006 | 1 | -0.68528 | 0.65708 | 0.65711 | 1 | 17033 | 0 | -0.68528 |
| hsa-mir-6734 | 1 | 0.3428 | 0.34305 | 0.999994 | 11002 | 1 | -0.68871 | 0.6572 | 0.65723 | 1 | 17034 | 0 | -0.68871 |
| SLC9A7 | 5 | 0.025278 | 0.078204 | 0.93914 | 1765 | 3 | -0.85768 | 0.65721 | 0.80183 | 1 | 17035 | 1 | -0.85768 |
| STMN3 | 5 | 0.54253 | 0.68513 | 0.999994 | 14103 | 2 | 0.57452 | 0.65725 | 0.80186 | 1 | 17036 | 2 | 0.57452 |
| ELFN1 | 6 | 0.43044 | 0.62214 | 0.999994 | 12421 | 2 | -0.32592 | 0.65726 | 0.83566 | 1 | 17037 | 1 | -0.32592 |
| SH3GL3 | 6 | 0.33397 | 0.54582 | 0.999994 | 10813 | 3 | -0.21263 | 0.65726 | 0.83566 | 1 | 17038 | 2 | -0.21263 |
| hsa-mir-548g | 2 | 0.68386 | 0.6837 | 0.999994 | 16142 | 0 | 0.62362 | 0.65735 | 0.66249 | 1 | 17039 | 1 | 0.62362 |
| NonTargetingControlG | 2 | 0.54557 | 0.54662 | 0.999994 | 14145 | 1 | 0.12615 | 0.6575 | 0.66261 | 1 | 17040 | 1 | 0.12615 |
| C6orf211 | 6 | 0.1215 | 0.28858 | 0.999632 | 5687 | 4 | -0.79266 | 0.6576 | 0.83581 | 1 | 17041 | 2 | -0.79266 |
| hsa-mir-3156-3 | 3 | 0.71696 | 0.71707 | 0.999994 | 16467 | 1 | -0.38854 | 0.65768 | 0.71754 | 1 | 17042 | 1 | -0.38854 |
| hsa-mir-670 | 4 | 0.65088 | 0.6938 | 0.999994 | 15674 | 1 | 0.44525 | 0.65774 | 0.78937 | 1 | 17043 | 2 | 0.44525 |
| hsa-mir-2052 | 2 | 0.57239 | 0.57258 | 0.999994 | 14583 | 1 | 0.25425 | 0.65775 | 0.66284 | 1 | 17044 | 1 | 0.25425 |
| FADS2 | 5 | 0.87092 | 0.87188 | 0.999994 | 18653 | 1 | 0.30561 | 0.6578 | 0.80217 | 1 | 17045 | 1 | 0.30561 |
| RB1CC1 | 4 | 0.74442 | 0.75582 | 0.999994 | 16793 | 1 | 0.068652 | 0.65785 | 0.78945 | 1 | 17046 | 1 | 0.068652 |
| FTHL17 | 6 | 0.66245 | 0.78721 | 0.999994 | 15819 | 2 | -0.13934 | 0.65793 | 0.83594 | 1 | 17047 | 2 | -0.13934 |
| C12orf40 | 4 | 0.90173 | 0.9014 | 0.999994 | 19246 | 0 | 0.51264 | 0.65823 | 0.78974 | 1 | 17048 | 2 | 0.51264 |
| SMAP2 | 4 | 0.20917 | 0.35012 | 0.999994 | 8009 | 2 | -0.70553 | 0.65854 | 0.79001 | 1 | 17049 | 2 | -0.70553 |
| TRABD2B | 5 | 0.038658 | 0.11247 | 0.951052 | 2446 | 2 | 0.096521 | 0.65855 | 0.80261 | 1 | 17050 | 2 | 0.096521 |
| MEF2A | 6 | 0.64853 | 0.7765 | 0.999994 | 15636 | 2 | -0.29725 | 0.65859 | 0.83621 | 1 | 17051 | 2 | -0.29725 |
| FBXL2 | 6 | 0.19448 | 0.38115 | 0.999994 | 7622 | 3 | -0.63232 | 0.65861 | 0.83622 | 1 | 17052 | 2 | -0.63232 |
| TRIB3 | 5 | 0.0042408 | 0.017786 | 0.795727 | 460 | 4 | -2.3918 | 0.65861 | 0.80265 | 1 | 17053 | 1 | -2.3918 |
| NCOR1 | 5 | 0.023853 | 0.07444 | 0.93914 | 1704 | 4 | -1.3435 | 0.65861 | 0.80265 | 1 | 17054 | 1 | -1.3435 |
| PANK4 | 5 | 0.009418 | 0.034375 | 0.867348 | 821 | 3 | -0.61541 | 0.65866 | 0.80268 | 1 | 17055 | 2 | -0.61541 |

| | | | | | | | | | | | | | |
|---|---|---|---|---|---|---|---|---|---|---|---|---|---|
| NonTargetingControlG | 2 | 0.36431 | 0.38753 | 0.999994 | 11411 | 1 | -0.45073 | 0.65873 | 0.66368 | 1 | 17056 | 1 | -0.45073 |
| FAHD2A | 6 | 0.9377 | 0.9377 | 0.999994 | 20023 | 0 | -0.05294 | 0.65876 | 0.83628 | 1 | 17057 | 2 | -0.05294 |
| NonTargetingControlG | 2 | 0.52868 | 0.53051 | 0.999994 | 13876 | 1 | 0.10761 | 0.65887 | 0.6638 | 1 | 17058 | 1 | 0.10761 |
| EXOSC3 | 4 | 0.31588 | 0.45925 | 0.999994 | 10436 | 2 | -0.40157 | 0.65891 | 0.79032 | 1 | 17059 | 1 | -0.40157 |
| NARG2 | 6 | 0.029195 | 0.096892 | 0.945132 | 1969 | 4 | -1.8848 | 0.65905 | 0.8364 | 1 | 17060 | 2 | -1.8848 |
| ST6GAL1 | 5 | 0.4151 | 0.57602 | 0.999994 | 12179 | 2 | -0.39471 | 0.65927 | 0.80305 | 1 | 17061 | 2 | -0.39471 |
| FOXD4L5 | 4 | 0.75687 | 0.76532 | 0.999994 | 16942 | 1 | -0.01803 | 0.65935 | 0.7907 | 1 | 17062 | 2 | -0.01803 |
| PGAP2 | 4 | 0.7218 | 0.73932 | 0.999994 | 16514 | 1 | 0.2969 | 0.65939 | 0.79073 | 1 | 17063 | 2 | 0.2969 |
| hsa-mir-3161 | 4 | 0.48397 | 0.60926 | 0.999994 | 13200 | 1 | 0.35817 | 0.65941 | 0.79075 | 1 | 17064 | 2 | 0.35817 |
| hsa-mir-7-1 | 4 | 0.32179 | 0.46524 | 0.999994 | 10548 | 2 | -0.26475 | 0.6595 | 0.79081 | 1 | 17065 | 2 | -0.26475 |
| hsa-mir-1254-2 | 2 | 0.6895 | 0.68932 | 0.999994 | 16205 | 0 | 0.59472 | 0.65957 | 0.66439 | 1 | 17066 | 1 | 0.59472 |
| WBSCR22 | 4 | 0.20085 | 0.34154 | 0.999994 | 7784 | 2 | -0.76999 | 0.65992 | 0.79117 | 1 | 17067 | 2 | -0.76999 |
| SLC41A1 | 6 | 0.13631 | 0.30837 | 0.999994 | 6093 | 2 | 0.11895 | 0.65998 | 0.83679 | 1 | 17068 | 1 | 0.11895 |
| WDR46 | 1 | 0.33999 | 0.34017 | 0.999994 | 10933 | 1 | -0.71712 | 0.66001 | 0.66004 | 1 | 17069 | 0 | -0.71712 |
| C16orf93 | 5 | 0.92035 | 0.92052 | 0.999994 | 19638 | 0 | 0.047527 | 0.66015 | 0.80357 | 1 | 17070 | 2 | 0.047527 |
| MARK1 | 4 | 0.61226 | 0.67201 | 0.999994 | 15113 | 1 | 0.41491 | 0.66017 | 0.79137 | 1 | 17071 | 2 | 0.41491 |
| hsa-mir-6737 | 3 | 0.21882 | 0.33616 | 0.999994 | 8256 | 2 | -1.0546 | 0.66039 | 0.71897 | 1 | 17072 | 1 | -1.0546 |
| SLC35E1 | 4 | 0.21437 | 0.3555 | 0.999994 | 8128 | 2 | -1.0682 | 0.66055 | 0.79169 | 1 | 17073 | 1 | -1.0682 |
| MTRNR2L3 | 4 | 0.2008 | 0.34149 | 0.999994 | 7780 | 2 | -0.80151 | 0.66055 | 0.79169 | 1 | 17074 | 1 | -0.80151 |
| THOP1 | 4 | 0.074434 | 0.16827 | 0.967126 | 4010 | 2 | -1.4861 | 0.66055 | 0.79169 | 1 | 17075 | 1 | -1.4861 |
| F10 | 4 | 0.38575 | 0.53099 | 0.999994 | 11730 | 1 | -0.024114 | 0.66055 | 0.79169 | 1 | 17076 | 1 | -0.024114 |
| LOC100130301 | 5 | 0.56932 | 0.70892 | 0.999994 | 14525 | 2 | 0.042969 | 0.66064 | 0.80388 | 1 | 17077 | 2 | 0.042969 |
| PDLIM7 | 6 | 0.035723 | 0.11223 | 0.951052 | 2302 | 3 | -1.8517 | 0.66079 | 0.83709 | 1 | 17078 | 2 | -1.8517 |
| SLC23A1 | 5 | 0.32969 | 0.50527 | 0.999994 | 10716 | 2 | -0.48536 | 0.66079 | 0.80397 | 1 | 17079 | 1 | -0.48536 |
| CNKSR3 | 5 | 0.66242 | 0.75013 | 0.999994 | 15817 | 1 | 0.54942 | 0.66099 | 0.80409 | 1 | 17080 | 2 | 0.54942 |
| CEP19 | 6 | 0.18225 | 0.36598 | 0.999994 | 7276 | 3 | -0.90916 | 0.66099 | 0.83717 | 1 | 17081 | 2 | -0.90916 |
| CXorf21 | 6 | 0.048001 | 0.1397 | 0.958716 | 2889 | 1 | 0.291 | 0.66099 | 0.83717 | 1 | 17082 | 2 | 0.291 |
| LRRC45 | 6 | 0.3209 | 0.5309 | 0.999994 | 10529 | 3 | -0.50406 | 0.66099 | 0.83717 | 1 | 17083 | 1 | -0.50406 |
| RPL22L1 | 4 | 0.85462 | 0.85421 | 0.999994 | 18344 | 0 | 0.64485 | 0.66127 | 0.79226 | 1 | 17084 | 2 | 0.64485 |
| WSCD2 | 5 | 0.77552 | 0.80486 | 0.999994 | 17143 | 1 | 0.30958 | 0.66128 | 0.80425 | 1 | 17085 | 2 | 0.30958 |
| CLN8 | 6 | 0.044178 | 0.13128 | 0.95318 | 2717 | 4 | -1.4344 | 0.66137 | 0.83733 | 1 | 17086 | 2 | -1.4344 |
| TTC39C | 6 | 0.11545 | 0.27714 | 0.999379 | 5491 | 3 | -0.19697 | 0.66141 | 0.83733 | 1 | 17087 | 2 | -0.19697 |
| IRF4 | 5 | 0.38807 | 0.55352 | 0.999994 | 11767 | 2 | 0.40123 | 0.66148 | 0.80437 | 1 | 17088 | 2 | 0.40123 |
| hsa-mir-2114 | 4 | 0.33897 | 0.48292 | 0.999994 | 10915 | 2 | -0.73225 | 0.66185 | 0.79273 | 1 | 17089 | 1 | -0.73225 |
| SLC35E3 | 5 | 0.082685 | 0.19895 | 0.977164 | 4325 | 2 | -0.41792 | 0.66192 | 0.80464 | 1 | 17090 | 2 | -0.41792 |
| VSTM2L | 5 | 0.43528 | 0.59287 | 0.999994 | 12505 | 2 | -0.4197 | 0.66192 | 0.80464 | 1 | 17091 | 2 | -0.4197 |
| ANGPTL3 | 5 | 0.042352 | 0.12174 | 0.951052 | 2633 | 2 | 0.33083 | 0.66195 | 0.80466 | 1 | 17092 | 2 | 0.33083 |
| STK33 | 5 | 0.03227 | 0.096288 | 0.945031 | 2119 | 4 | -1.941 | 0.66195 | 0.80466 | 1 | 17093 | 1 | -1.941 |
| BOLA1 | 5 | 0.14002 | 0.28788 | 0.999632 | 6185 | 2 | 0.098218 | 0.66195 | 0.80466 | 1 | 17094 | 2 | 0.098218 |
| BCAS4 | 5 | 0.80972 | 0.82573 | 0.999994 | 17630 | 1 | 0.18564 | 0.66195 | 0.80466 | 1 | 17095 | 2 | 0.18564 |
| EIF1AD | 5 | 0.13077 | 0.27378 | 0.998561 | 5950 | 3 | -1.3398 | 0.66195 | 0.80466 | 1 | 17096 | 1 | -1.3398 |
| F3 | 6 | 0.011196 | 0.043059 | 0.891621 | 940 | 5 | -2.5722 | 0.66211 | 0.83761 | 1 | 17097 | 1 | -2.5722 |
| RCL1 | 6 | 0.043972 | 0.13083 | 0.95318 | 2709 | 4 | -2.0083 | 0.66211 | 0.83761 | 1 | 17098 | 1 | -2.0083 |
| hsa-mir-1262 | 4 | 0.1792 | 0.31927 | 0.999994 | 7202 | 1 | 0.21652 | 0.66239 | 0.79318 | 1 | 17099 | 2 | 0.21652 |
| TTC19 | 4 | 0.10634 | 0.2264 | 0.983354 | 5183 | 2 | -1.2153 | 0.66275 | 0.79347 | 1 | 17100 | 1 | -1.2153 |
| ARHGEF15 | 4 | 0.74884 | 0.75916 | 0.999994 | 16848 | 1 | 0.021399 | 0.66279 | 0.7935 | 1 | 17101 | 2 | 0.021399 |
| hsa-mir-421 | 4 | 0.13337 | 0.27205 | 0.998561 | 6005 | 2 | -0.80577 | 0.66282 | 0.79352 | 1 | 17102 | 2 | -0.80577 |
| ZNF254 | 4 | 0.60526 | 0.66817 | 0.999994 | 15015 | 1 | 0.50203 | 0.66299 | 0.79367 | 1 | 17103 | 2 | 0.50203 |
| hsa-mir-449b | 3 | 0.054776 | 0.12233 | 0.951052 | 3186 | 1 | 0.58466 | 0.66302 | 0.7204 | 1 | 17104 | 1 | 0.58466 |
| RRM1 | 3 | 0.52952 | 0.56603 | 0.999994 | 13892 | 1 | -0.37448 | 0.66305 | 0.72042 | 1 | 17105 | 1 | -0.37448 |
| hsa-mir-7850 | 4 | 0.35258 | 0.49692 | 0.999994 | 11206 | 2 | -0.26299 | 0.66311 | 0.79377 | 1 | 17106 | 2 | -0.26299 |
| SHROOM3 | 5 | 0.13616 | 0.28198 | 0.999632 | 6084 | 3 | -1.2056 | 0.66328 | 0.80546 | 1 | 17107 | 1 | -1.2056 |

| | | | | | | | | | | | | | |
|---|---|---|---|---|---|---|---|---|---|---|---|---|---|
| STAC | 5 | 0.37652 | 0.54384 | 0.999994 | 11603 | 2 | -0.063844 | 0.66328 | 0.80546 | 1 | 17108 | 1 | -0.063844 |
| C5orf30 | 5 | 0.39703 | 0.56108 | 0.999994 | 11907 | 2 | -0.43858 | 0.66328 | 0.80546 | 1 | 17109 | 1 | -0.43858 |
| TMEM60 | 5 | 0.17703 | 0.34233 | 0.999994 | 7157 | 2 | 0.021286 | 0.66328 | 0.80546 | 1 | 17110 | 1 | 0.021286 |
| N6AMT1 | 5 | 0.021735 | 0.068753 | 0.93914 | 1561 | 3 | -1.4247 | 0.66328 | 0.80546 | 1 | 17111 | 1 | -1.4247 |
| hsa-mir-4287 | 4 | 0.85123 | 0.85076 | 0.999994 | 18283 | 0 | 0.52745 | 0.66352 | 0.79411 | 1 | 17112 | 2 | 0.52745 |
| hsa-mir-762 | 4 | 0.58628 | 0.65798 | 0.999994 | 14761 | 1 | -0.083612 | 0.66354 | 0.79413 | 1 | 17113 | 1 | -0.083612 |
| ING3 | 4 | 0.006502 | 0.022572 | 0.819701 | 626 | 2 | -3.3222 | 0.66369 | 0.79424 | 1 | 17114 | 2 | -3.3222 |
| hsa-mir-4265 | 4 | 0.35156 | 0.49591 | 0.999994 | 11186 | 1 | 0.49498 | 0.66372 | 0.79426 | 1 | 17115 | 2 | 0.49498 |
| NonTargetingControlG | 2 | 0.29095 | 0.32985 | 0.999994 | 9929 | 1 | -0.79077 | 0.66375 | 0.66804 | 1 | 17116 | 1 | -0.79077 |
| CCDC90B | 4 | 0.18452 | 0.32477 | 0.999994 | 7330 | 2 | -0.74171 | 0.66382 | 0.79434 | 1 | 17117 | 2 | -0.74171 |
| AVP | 5 | 0.20082 | 0.37636 | 0.999994 | 7781 | 1 | 0.3131 | 0.66384 | 0.80579 | 1 | 17118 | 2 | 0.3131 |
| NKG7 | 4 | 0.10673 | 0.22708 | 0.983354 | 5199 | 2 | -1.7729 | 0.66394 | 0.79444 | 1 | 17119 | 1 | -1.7729 |
| RNF207 | 5 | 0.13821 | 0.28513 | 0.999632 | 6144 | 3 | -0.73243 | 0.66397 | 0.80587 | 1 | 17120 | 2 | -0.73243 |
| hsa-mir-7856 | 4 | 0.26737 | 0.40958 | 0.999994 | 9404 | 2 | -0.76187 | 0.66424 | 0.79467 | 1 | 17121 | 1 | -0.76187 |
| RAB11FIP3 | 5 | 0.36664 | 0.53557 | 0.999994 | 11452 | 2 | 0.16901 | 0.66425 | 0.80604 | 1 | 17122 | 2 | 0.16901 |
| hsa-mir-7152 | 4 | 0.92299 | 0.92282 | 0.999994 | 19696 | 0 | 0.43052 | 0.6643 | 0.79474 | 1 | 17123 | 2 | 0.43052 |
| ATP5L | 5 | 0.44763 | 0.60323 | 0.999994 | 12682 | 2 | 0.10975 | 0.66444 | 0.80616 | 1 | 17124 | 2 | 0.10975 |
| hsa-mir-3606 | 4 | 0.38518 | 0.53046 | 0.999994 | 11721 | 1 | 0.54395 | 0.66462 | 0.795 | 1 | 17125 | 2 | 0.54395 |
| ADAMTSL3 | 5 | 0.18955 | 0.36029 | 0.999994 | 7471 | 3 | -0.89799 | 0.66465 | 0.80628 | 1 | 17126 | 2 | -0.89799 |
| SERPINB1 | 6 | 0.046294 | 0.13594 | 0.95468 | 2818 | 4 | -1.3074 | 0.66469 | 0.83865 | 1 | 17127 | 2 | -1.3074 |
| MCM2 | 6 | 0.0001991 | 0.0010678 | 0.352868 | 56 | 5 | -7.4073 | 0.66469 | 0.83865 | 1 | 17128 | 1 | -7.4073 |
| CEP170 | 6 | 0.044274 | 0.13149 | 0.95318 | 2725 | 4 | -1.7135 | 0.66469 | 0.83865 | 1 | 17129 | 2 | -1.7135 |
| SLC4A5 | 5 | 0.035959 | 0.10567 | 0.950444 | 2310 | 4 | -1.4051 | 0.66471 | 0.80631 | 1 | 17130 | 1 | -1.4051 |
| OR56B4 | 5 | 0.033449 | 0.099353 | 0.945253 | 2186 | 3 | -2.9066 | 0.66471 | 0.80631 | 1 | 17131 | 1 | -2.9066 |
| KIF3A | 6 | 0.15545 | 0.33251 | 0.999994 | 6605 | 3 | -0.67215 | 0.66516 | 0.83884 | 1 | 17132 | 2 | -0.67215 |
| hsa-mir-4501 | 2 | 0.2215 | 0.2777 | 0.999519 | 8319 | 1 | -1.1743 | 0.66516 | 0.66923 | 1 | 17133 | 1 | -1.1743 |
| BPIFA2 | 5 | 0.16942 | 0.33129 | 0.999994 | 6972 | 2 | -0.39806 | 0.66517 | 0.80659 | 1 | 17134 | 2 | -0.39806 |
| FAM3D | 5 | 0.28266 | 0.467 | 0.999994 | 9752 | 2 | -0.17416 | 0.66525 | 0.80663 | 1 | 17135 | 2 | -0.17416 |
| DACT1 | 6 | 0.053567 | 0.15179 | 0.967126 | 3144 | 4 | -1.4375 | 0.66532 | 0.83891 | 1 | 17136 | 2 | -1.4375 |
| hsa-mir-580 | 3 | 0.53617 | 0.57066 | 0.999994 | 14002 | 1 | 0.17359 | 0.66542 | 0.72173 | 1 | 17137 | 1 | 0.17359 |
| PAK1IP1 | 5 | 0.064485 | 0.16943 | 0.967126 | 3629 | 3 | -2.2175 | 0.66556 | 0.80683 | 1 | 17138 | 2 | -2.2175 |
| STAB2 | 5 | 0.1174 | 0.25338 | 0.994501 | 5557 | 3 | -1.2148 | 0.66565 | 0.80688 | 1 | 17139 | 1 | -1.2148 |
| CD300LD | 5 | 0.27293 | 0.45915 | 0.999994 | 9539 | 2 | -0.035311 | 0.66565 | 0.80688 | 1 | 17140 | 2 | -0.035311 |
| SUMF1 | 5 | 0.19562 | 0.36895 | 0.999994 | 7653 | 2 | -0.55964 | 0.66571 | 0.80691 | 1 | 17141 | 2 | -0.55964 |
| CCDC102A | 5 | 0.84497 | 0.85039 | 0.999994 | 18171 | 1 | 0.043001 | 0.66576 | 0.80694 | 1 | 17142 | 2 | 0.043001 |
| COPS6 | 6 | 0.0070209 | 0.028486 | 0.858757 | 660 | 5 | -3.8013 | 0.6661 | 0.83922 | 1 | 17143 | 1 | -3.8013 |
| ZDHHC22 | 6 | 0.25179 | 0.45007 | 0.999994 | 9052 | 3 | -0.54955 | 0.6661 | 0.83922 | 1 | 17144 | 2 | -0.54955 |
| EAF2 | 6 | 0.078314 | 0.20393 | 0.978186 | 4163 | 4 | -1.5306 | 0.6661 | 0.83922 | 1 | 17145 | 2 | -1.5306 |
| hsa-mir-3940 | 2 | 0.49665 | 0.50085 | 0.999994 | 13363 | 1 | -0.015615 | 0.66622 | 0.67018 | 1 | 17146 | 1 | -0.015615 |
| RALA | 5 | 0.031615 | 0.094614 | 0.945031 | 2086 | 3 | -2.2467 | 0.66626 | 0.80723 | 1 | 17147 | 1 | -2.2467 |
| NonTargetingControlG | 2 | 0.67006 | 0.66983 | 0.999994 | 15932 | 0 | 0.58717 | 0.66633 | 0.67028 | 1 | 17148 | 1 | 0.58717 |
| MIS18BP1 | 5 | 0.49554 | 0.64404 | 0.999994 | 13348 | 2 | 0.63063 | 0.66634 | 0.80729 | 1 | 17149 | 2 | 0.63063 |
| hsa-mir-507 | 4 | 0.73608 | 0.74958 | 0.999994 | 16677 | 1 | 0.10902 | 0.66639 | 0.79643 | 1 | 17150 | 2 | 0.10902 |
| SEZ6L2 | 5 | 0.64971 | 0.745 | 0.999994 | 15655 | 1 | 0.46947 | 0.66645 | 0.80736 | 1 | 17151 | 2 | 0.46947 |
| hsa-mir-374b | 4 | 0.36861 | 0.51341 | 0.999994 | 11484 | 2 | -0.15908 | 0.66656 | 0.79656 | 1 | 17152 | 2 | -0.15908 |
| SMIM9 | 6 | 0.026643 | 0.090884 | 0.94395 | 1837 | 3 | -1.5409 | 0.6668 | 0.8395 | 1 | 17153 | 2 | -1.5409 |
| ZNF728 | 4 | 0.58452 | 0.65708 | 0.999994 | 14738 | 1 | 0.1296 | 0.66682 | 0.79677 | 1 | 17154 | 2 | 0.1296 |
| C14orf37 | 6 | 0.23003 | 0.4241 | 0.999994 | 8527 | 3 | -0.83878 | 0.66686 | 0.83953 | 1 | 17155 | 1 | -0.83878 |
| NonTargetingControlG | 1 | 0.33303 | 0.33315 | 0.999994 | 10788 | 1 | -0.76681 | 0.66697 | 0.66702 | 1 | 17156 | 0 | -0.76681 |
| PRR25 | 4 | 0.91952 | 0.91931 | 0.999994 | 19619 | 0 | 0.54507 | 0.66736 | 0.79722 | 1 | 17157 | 2 | 0.54507 |
| ATP1B4 | 4 | 0.0096265 | 0.032527 | 0.865899 | 848 | 2 | -3.9532 | 0.66744 | 0.79729 | 1 | 17158 | 2 | -3.9532 |
| ZNF7 | 6 | 0.1335 | 0.30478 | 0.999994 | 6008 | 2 | 0.10618 | 0.6675 | 0.8398 | 1 | 17159 | 2 | 0.10618 |

| | | | | | | | | | | | | | |
|---|---|---|---|---|---|---|---|---|---|---|---|---|---|
| VAV2 | 6 | 0.47114 | 0.64921 | 0.999994 | 13017 | 2 | -0.31464 | 0.6675 | 0.8398 | 1 | 17160 | 1 | -0.31464 |
| ZNF765 | 6 | 0.16706 | 0.34703 | 0.999994 | 6906 | 3 | -0.35049 | 0.6675 | 0.8398 | 1 | 17161 | 2 | -0.35049 |
| KLF10 | 6 | 0.026688 | 0.090985 | 0.94395 | 1840 | 3 | -2.0686 | 0.6675 | 0.8398 | 1 | 17162 | 2 | -2.0686 |
| BCL9L | 6 | 0.21838 | 0.41018 | 0.999994 | 8237 | 3 | -0.81814 | 0.6675 | 0.8398 | 1 | 17163 | 2 | -0.81814 |
| ZNRF2 | 5 | 0.0041182 | 0.017301 | 0.785648 | 449 | 3 | -1.8365 | 0.66756 | 0.80802 | 1 | 17164 | 1 | -1.8365 |
| ZNF320 | 5 | 0.18073 | 0.3477 | 0.999994 | 7239 | 3 | -0.91075 | 0.66756 | 0.80802 | 1 | 17165 | 1 | -0.91075 |
| PRPF38B | 5 | 0.054308 | 0.1508 | 0.966359 | 3166 | 4 | -0.82847 | 0.66756 | 0.80802 | 1 | 17166 | 1 | -0.82847 |
| hsa-mir-6508 | 4 | 0.017742 | 0.055816 | 0.9235 | 1349 | 3 | -2.3448 | 0.66758 | 0.79741 | 1 | 17167 | 1 | -2.3448 |
| ELF1 | 4 | 0.018127 | 0.056649 | 0.9235 | 1367 | 3 | -3.2576 | 0.66758 | 0.79741 | 1 | 17168 | 1 | -3.2576 |
| HUS1 | 4 | 0.032256 | 0.086091 | 0.94395 | 2116 | 3 | -3.1969 | 0.66758 | 0.79741 | 1 | 17169 | 1 | -3.1969 |
| hsa-mir-4515 | 4 | 0.25672 | 0.3988 | 0.999994 | 9165 | 2 | -0.67657 | 0.66758 | 0.79741 | 1 | 17170 | 1 | -0.67657 |
| KCNA1 | 4 | 0.056927 | 0.13491 | 0.954328 | 3289 | 3 | -1.5752 | 0.66758 | 0.79741 | 1 | 17171 | 1 | -1.5752 |
| IFFO1 | 4 | 0.09619 | 0.20826 | 0.97933 | 4824 | 3 | -1.4868 | 0.66758 | 0.79741 | 1 | 17172 | 1 | -1.4868 |
| CAPN8 | 4 | 0.29996 | 0.44296 | 0.999994 | 10130 | 2 | -0.45599 | 0.66758 | 0.79741 | 1 | 17173 | 1 | -0.45599 |
| hsa-mir-3184 | 4 | 0.82591 | 0.82549 | 0.999994 | 17865 | 1 | 0.20371 | 0.66768 | 0.79749 | 1 | 17174 | 2 | 0.20371 |
| S100A10 | 5 | 0.24839 | 0.4395 | 0.999994 | 8977 | 2 | 0.35617 | 0.66772 | 0.80812 | 1 | 17175 | 2 | 0.35617 |
| CDCA2 | 5 | 0.020954 | 0.066648 | 0.93914 | 1510 | 3 | -2.1987 | 0.66775 | 0.80813 | 1 | 17176 | 2 | -2.1987 |
| SMIM12 | 5 | 0.47444 | 0.62589 | 0.999994 | 13055 | 2 | 0.26029 | 0.66843 | 0.80854 | 1 | 17177 | 2 | 0.26029 |
| RAD54L | 5 | 0.19945 | 0.37444 | 0.999994 | 7746 | 2 | -0.034504 | 0.66859 | 0.80864 | 1 | 17178 | 2 | -0.034504 |
| TCTA | 5 | 0.59684 | 0.72501 | 0.999994 | 14913 | 1 | 0.3832 | 0.66859 | 0.80864 | 1 | 17179 | 2 | 0.3832 |
| C2 | 5 | 0.56167 | 0.70211 | 0.999994 | 14400 | 2 | -0.2072 | 0.6687 | 0.80871 | 1 | 17180 | 2 | -0.2072 |
| MYPN | 6 | 0.27999 | 0.4834 | 0.999994 | 9705 | 2 | 0.084904 | 0.66871 | 0.84032 | 1 | 17181 | 2 | 0.084904 |
| VPS45 | 3 | 0.025756 | 0.065293 | 0.93914 | 1788 | 1 | -0.56492 | 0.66872 | 0.7235 | 1 | 17182 | 1 | -0.56492 |
| hsa-mir-6849 | 2 | 0.73083 | 0.73056 | 0.999994 | 16610 | 0 | 0.83783 | 0.66874 | 0.67247 | 1 | 17183 | 1 | 0.83783 |
| CACNB4 | 6 | 0.021364 | 0.07638 | 0.93914 | 1537 | 5 | -2.5182 | 0.66882 | 0.84036 | 1 | 17184 | 1 | -2.5182 |
| CYTH4 | 6 | 0.20345 | 0.39206 | 0.999994 | 7866 | 3 | -0.54774 | 0.66882 | 0.84036 | 1 | 17185 | 2 | -0.54774 |
| PQLC2 | 6 | 0.89123 | 0.89718 | 0.999994 | 19057 | 1 | 0.15644 | 0.66882 | 0.84036 | 1 | 17186 | 2 | 0.15644 |
| NonTargetingControlG | 2 | 0.25749 | 0.30452 | 0.999994 | 9178 | 1 | -1.203 | 0.66898 | 0.67267 | 1 | 17187 | 1 | -1.203 |
| BPY2 | 6 | 0.6665 | 0.7904 | 0.999994 | 15875 | 1 | 0.44547 | 0.66903 | 0.84045 | 1 | 17188 | 2 | 0.44547 |
| UBAP2 | 5 | 0.31117 | 0.49011 | 0.999994 | 10343 | 2 | 0.46651 | 0.6691 | 0.80895 | 1 | 17189 | 2 | 0.46651 |
| EIF2B2 | 5 | 0.050263 | 0.14099 | 0.959146 | 2988 | 3 | -0.88584 | 0.6691 | 0.80895 | 1 | 17190 | 2 | -0.88584 |
| NAB2 | 5 | 0.23122 | 0.41887 | 0.999994 | 8548 | 1 | 0.48156 | 0.6691 | 0.80895 | 1 | 17191 | 2 | 0.48156 |
| ICAM4 | 5 | 0.024631 | 0.076486 | 0.93914 | 1739 | 4 | -1.9615 | 0.6691 | 0.80895 | 1 | 17192 | 1 | -1.9615 |
| NonTargetingControlG | 2 | 0.69065 | 0.69046 | 0.999994 | 16215 | 0 | 0.55171 | 0.66936 | 0.67302 | 1 | 17193 | 1 | 0.55171 |
| hsa-mir-516b-2 | 2 | 0.67367 | 0.67342 | 0.999994 | 15984 | 0 | 0.50898 | 0.66961 | 0.67323 | 1 | 17194 | 1 | 0.50898 |
| NonTargetingControlG | 1 | 0.33029 | 0.33049 | 0.999994 | 10727 | 1 | -0.851 | 0.66971 | 0.66975 | 1 | 17195 | 0 | -0.851 |
| hsa-let-7a-2 | 4 | 0.28744 | 0.43016 | 0.999994 | 9863 | 2 | -0.28307 | 0.66977 | 0.79815 | 1 | 17196 | 1 | -0.28307 |
| MORN5 | 4 | 0.1605 | 0.3 | 0.999994 | 6751 | 2 | -1.1758 | 0.66984 | 0.79818 | 1 | 17197 | 1 | -1.1758 |
| hsa-mir-4775 | 3 | 0.050981 | 0.11708 | 0.951052 | 3018 | 2 | -3.5161 | 0.66991 | 0.72412 | 1 | 17198 | 1 | -3.5161 |
| NonTargetingControlG | 2 | 0.048942 | 0.087371 | 0.94395 | 2926 | 1 | -4.0014 | 0.6701 | 0.67367 | 1 | 17199 | 1 | -4.0014 |
| SYTL4 | 6 | 0.022999 | 0.081581 | 0.93914 | 1651 | 4 | -0.77667 | 0.67044 | 0.841 | 1 | 17200 | 2 | -0.77667 |
| PLEKHG6 | 6 | 0.27674 | 0.47964 | 0.999994 | 9632 | 3 | -0.75836 | 0.67054 | 0.84105 | 1 | 17201 | 2 | -0.75836 |
| TFE3 | 6 | 0.093356 | 0.23423 | 0.984971 | 4733 | 3 | -0.90463 | 0.67056 | 0.84106 | 1 | 17202 | 2 | -0.90463 |
| WFDC13 | 5 | 0.35868 | 0.52895 | 0.999994 | 11317 | 2 | -0.47221 | 0.67062 | 0.80991 | 1 | 17203 | 2 | -0.47221 |
| NDFIP1 | 5 | 0.32115 | 0.49824 | 0.999994 | 10538 | 2 | 0.41985 | 0.67086 | 0.81005 | 1 | 17204 | 2 | 0.41985 |
| IL27 | 5 | 0.067437 | 0.17424 | 0.968171 | 3733 | 3 | -2.0799 | 0.67086 | 0.81005 | 1 | 17205 | 1 | -2.0799 |
| TGIF1 | 5 | 0.39226 | 0.55705 | 0.999994 | 11833 | 2 | 0.068966 | 0.67086 | 0.81005 | 1 | 17206 | 1 | 0.068966 |
| PSD3 | 5 | 0.34119 | 0.51471 | 0.999994 | 10963 | 2 | -0.22814 | 0.67099 | 0.81013 | 1 | 17207 | 2 | -0.22814 |
| LFNG | 6 | 0.26078 | 0.4608 | 0.999994 | 9260 | 2 | -0.22582 | 0.67108 | 0.84126 | 1 | 17208 | 2 | -0.22582 |
| FOXD4L4 | 1 | 0.32882 | 0.329 | 0.999994 | 10696 | 1 | -0.83342 | 0.67118 | 0.67121 | 1 | 17209 | 0 | -0.83342 |
| MAGEB16 | 6 | 0.23224 | 0.42677 | 0.999994 | 8577 | 3 | -0.76527 | 0.67139 | 0.84137 | 1 | 17210 | 2 | -0.76527 |
| MLLT3 | 5 | 0.10143 | 0.22865 | 0.983354 | 5019 | 3 | -1.4955 | 0.67154 | 0.81045 | 1 | 17211 | 2 | -1.4955 |

| | | | | | | | | | | | | |
|---|---|---|---|---|---|---|---|---|---|---|---|---|
| NonTargetingControlG | 2 | 0.26171 | 0.30764 | 0.999994 | 9279 | 1 | -1.2434 | 0.67163 | 0.67504 | 1 | 17212 | 1 | -1.2434 |
| SV2B | 6 | 0.40147 | 0.60333 | 0.999994 | 11970 | 2 | 0.18475 | 0.67173 | 0.84152 | 1 | 17213 | 2 | 0.18475 |
| APCS | 6 | 0.48584 | 0.65913 | 0.999994 | 13227 | 2 | -0.23819 | 0.67186 | 0.84157 | 1 | 17214 | 2 | -0.23819 |
| COMMD7 | 6 | 0.56805 | 0.71656 | 0.999994 | 14513 | 2 | -0.050404 | 0.67214 | 0.84168 | 1 | 17215 | 2 | -0.050404 |
| hsa-mir-1264 | 3 | 0.061313 | 0.13125 | 0.95318 | 3486 | 2 | -1.6121 | 0.67223 | 0.72539 | 1 | 17216 | 1 | -1.6121 |
| NonTargetingControlG | 1 | 0.32776 | 0.32793 | 0.999994 | 10679 | 1 | -0.8175 | 0.67224 | 0.67228 | 1 | 17217 | 0 | -0.8175 |
| TRABD2A | 5 | 0.49653 | 0.64486 | 0.999994 | 13362 | 2 | -0.21361 | 0.67233 | 0.81093 | 1 | 17218 | 1 | -0.21361 |
| OR4C13 | 5 | 0.009978 | 0.035998 | 0.876861 | 868 | 4 | -2.1435 | 0.67233 | 0.81093 | 1 | 17219 | 1 | -2.1435 |
| RAPGEF6 | 5 | 0.042428 | 0.12192 | 0.951052 | 2638 | 3 | -2.189 | 0.67233 | 0.81093 | 1 | 17220 | 1 | -2.189 |
| NonTargetingControlG | 2 | 0.10286 | 0.17451 | 0.968171 | 5063 | 1 | -3.5257 | 0.673 | 0.67624 | 1 | 17221 | 1 | -3.5257 |
| MAGEA10-MAGEA5 | 5 | 0.13321 | 0.27752 | 0.999379 | 6000 | 2 | -0.25939 | 0.67309 | 0.81138 | 1 | 17222 | 2 | -0.25939 |
| OR4C6 | 6 | 0.05653 | 0.15825 | 0.967126 | 3271 | 4 | -1.3228 | 0.67313 | 0.84208 | 1 | 17223 | 2 | -1.3228 |
| HIST1H4K | 1 | 0.32663 | 0.32679 | 0.999994 | 10655 | 1 | -0.81488 | 0.67337 | 0.67345 | 1 | 17224 | 0 | -0.81488 |
| CTSV | 3 | 0.027714 | 0.069858 | 0.93914 | 1890 | 2 | -4.6582 | 0.67346 | 0.72602 | 1 | 17225 | 1 | -4.6582 |
| NonTargetingControlG | 2 | 0.49441 | 0.49881 | 0.999994 | 13335 | 1 | -0.084688 | 0.67354 | 0.67673 | 1 | 17226 | 1 | -0.084688 |
| TMEM14E | 6 | 0.004286 | 0.018528 | 0.801598 | 463 | 5 | -4.361 | 0.67384 | 0.84238 | 1 | 17227 | 1 | -4.361 |
| ATXN7 | 6 | 0.44381 | 0.63097 | 0.999994 | 12632 | 2 | 0.34026 | 0.67384 | 0.84238 | 1 | 17228 | 2 | 0.34026 |
| PTTG1IP | 5 | 0.23999 | 0.43091 | 0.999994 | 8770 | 3 | -0.63204 | 0.67405 | 0.81193 | 1 | 17229 | 2 | -0.63204 |
| MS4A12 | 5 | 0.19157 | 0.36317 | 0.999994 | 7531 | 3 | -0.68053 | 0.67424 | 0.81206 | 1 | 17230 | 2 | -0.68053 |
| NonTargetingControlG | 2 | 0.58907 | 0.58907 | 0.999994 | 14808 | 0 | 0.37433 | 0.67431 | 0.67741 | 1 | 17231 | 1 | 0.37433 |
| OR2F1 | 6 | 0.18418 | 0.36835 | 0.999994 | 7321 | 3 | -0.70198 | 0.67431 | 0.84255 | 1 | 17232 | 2 | -0.70198 |
| hsa-mir-651 | 3 | 0.59015 | 0.60963 | 0.999994 | 14825 | 1 | 0.18115 | 0.6744 | 0.72658 | 1 | 17233 | 1 | 0.18115 |
| ANKRD53 | 5 | 0.50796 | 0.65489 | 0.999994 | 13551 | 2 | 0.021602 | 0.67464 | 0.81231 | 1 | 17234 | 2 | 0.021602 |
| SRSF4 | 5 | 0.040057 | 0.11604 | 0.951052 | 2517 | 3 | -2.7768 | 0.67477 | 0.8124 | 1 | 17235 | 2 | -2.7768 |
| JPH2 | 5 | 0.014319 | 0.048271 | 0.90085 | 1152 | 3 | -1.3675 | 0.67491 | 0.81249 | 1 | 17236 | 2 | -1.3675 |
| SPATA6L | 6 | 0.17776 | 0.3603 | 0.999994 | 7174 | 3 | -0.91035 | 0.67503 | 0.84287 | 1 | 17237 | 1 | -0.91035 |
| SH3BGRL3 | 6 | 0.071246 | 0.18941 | 0.970473 | 3885 | 4 | -0.86684 | 0.67503 | 0.84287 | 1 | 17238 | 1 | -0.86684 |
| GPR137C | 6 | 0.19051 | 0.37616 | 0.999994 | 7500 | 3 | -0.47856 | 0.67517 | 0.84292 | 1 | 17239 | 2 | -0.47856 |
| METTL9 | 4 | 0.26821 | 0.41041 | 0.999994 | 9423 | 2 | -0.66578 | 0.67519 | 0.79984 | 1 | 17240 | 1 | -0.66578 |
| hsa-mir-548ae-2 | 1 | 0.32442 | 0.32458 | 0.999994 | 10605 | 1 | -0.8695 | 0.67558 | 0.67558 | 1 | 17241 | 0 | -0.8695 |
| STOX2 | 5 | 0.90301 | 0.9031 | 0.999994 | 19279 | 0 | 0.1539 | 0.67571 | 0.81296 | 1 | 17242 | 2 | 0.1539 |
| EIF5A2 | 5 | 0.091741 | 0.21335 | 0.979587 | 4666 | 3 | -1.4765 | 0.67575 | 0.81298 | 1 | 17243 | 1 | -1.4765 |
| OR5B21 | 6 | 0.65075 | 0.77818 | 0.999994 | 15667 | 2 | 0.36324 | 0.67579 | 0.84319 | 1 | 17244 | 2 | 0.36324 |
| TRIM49 | 6 | 0.065467 | 0.17725 | 0.968171 | 3672 | 3 | -0.32716 | 0.67597 | 0.84326 | 1 | 17245 | 2 | -0.32716 |
| KIF18A | 6 | 0.54525 | 0.70026 | 0.999994 | 14139 | 2 | -0.16213 | 0.67625 | 0.84338 | 1 | 17246 | 2 | -0.16213 |
| SOCS7 | 6 | 0.1268 | 0.29625 | 0.999994 | 5842 | 3 | -0.23519 | 0.67625 | 0.84338 | 1 | 17247 | 2 | -0.23519 |
| INTS3 | 4 | 0.07904 | 0.17697 | 0.968171 | 4192 | 2 | -1.9139 | 0.6763 | 0.80018 | 1 | 17248 | 1 | -1.9139 |
| hsa-mir-3650 | 4 | 0.1275 | 0.26326 | 0.996734 | 5860 | 3 | -0.66726 | 0.6763 | 0.80018 | 1 | 17249 | 1 | -0.66726 |
| FMR1 | 4 | 0.098476 | 0.21234 | 0.979587 | 4911 | 3 | -0.91962 | 0.6763 | 0.80018 | 1 | 17250 | 1 | -0.91962 |
| FANCM | 5 | 0.0074212 | 0.028578 | 0.858757 | 686 | 2 | -0.51361 | 0.67642 | 0.81337 | 1 | 17251 | 2 | -0.51361 |
| AP2B1 | 5 | 0.010515 | 0.037551 | 0.882924 | 904 | 3 | -5.3755 | 0.67651 | 0.81342 | 1 | 17252 | 2 | -5.3755 |
| NonTargetingControlG | 2 | 0.1847 | 0.25108 | 0.993091 | 7335 | 1 | -2.6507 | 0.67656 | 0.67946 | 1 | 17253 | 1 | -2.6507 |
| OR5T2 | 6 | 0.063544 | 0.17327 | 0.96793 | 3580 | 3 | -1.1769 | 0.67661 | 0.84352 | 1 | 17254 | 2 | -1.1769 |
| PPY | 5 | 0.76392 | 0.79833 | 0.999994 | 17012 | 1 | 0.46098 | 0.67669 | 0.81352 | 1 | 17255 | 2 | 0.46098 |
| FAM122B | 5 | 0.24944 | 0.44034 | 0.999994 | 9003 | 2 | 0.096402 | 0.67678 | 0.81358 | 1 | 17256 | 2 | 0.096402 |
| NonTargetingControlG | 2 | 0.69242 | 0.69223 | 0.999994 | 16230 | 0 | 0.50839 | 0.67698 | 0.67986 | 1 | 17257 | 1 | 0.50839 |
| NonTargetingControlG | 2 | 0.43418 | 0.44563 | 0.999994 | 12480 | 1 | -0.42102 | 0.67723 | 0.68008 | 1 | 17258 | 1 | -0.42102 |
| LEPR | 5 | 0.052893 | 0.14735 | 0.965554 | 3105 | 1 | -0.033303 | 0.67723 | 0.81384 | 1 | 17259 | 2 | -0.033303 |
| ALDH3A2 | 5 | 0.87447 | 0.87512 | 0.999994 | 18717 | 1 | 0.093768 | 0.67726 | 0.81385 | 1 | 17260 | 1 | 0.093768 |
| ZNF565 | 5 | 0.61936 | 0.73331 | 0.999994 | 15212 | 1 | 0.11272 | 0.67726 | 0.81385 | 1 | 17261 | 2 | 0.11272 |
| STAM2 | 5 | 0.054039 | 0.15014 | 0.966359 | 3159 | 4 | -0.90071 | 0.67726 | 0.81385 | 1 | 17262 | 1 | -0.90071 |
| PROCA1 | 5 | 0.046404 | 0.13172 | 0.95318 | 2827 | 3 | -2.3341 | 0.67726 | 0.81385 | 1 | 17263 | 2 | -2.3341 |

| | | | | | | | | | | | | |
|---|---|---|---|---|---|---|---|---|---|---|---|---|
| FKRP | 5 | 0.55416 | 0.69545 | 0.999994 | 14271 | 1 | 0.53639 | 0.67732 | 0.81389 | 1 | 17264 | 2 | 0.53639 |
| SHKBP1 | 5 | 0.16939 | 0.33125 | 0.999994 | 6969 | 3 | -0.95493 | 0.67734 | 0.8139 | 1 | 17265 | 2 | -0.95493 |
| NRD1 | 6 | 0.07708 | 0.20145 | 0.977164 | 4108 | 4 | -1.4415 | 0.67736 | 0.84381 | 1 | 17266 | 2 | -1.4415 |
| ACOX1 | 6 | 0.22935 | 0.42328 | 0.999994 | 8511 | 3 | -0.53099 | 0.67738 | 0.84383 | 1 | 17267 | 1 | -0.53099 |
| GPR56 | 6 | 0.022056 | 0.078545 | 0.93914 | 1588 | 4 | -2.0169 | 0.67738 | 0.84383 | 1 | 17268 | 2 | -2.0169 |
| C9orf91 | 6 | 0.015485 | 0.057293 | 0.927806 | 1214 | 3 | -0.3664 | 0.67738 | 0.84383 | 1 | 17269 | 2 | -0.3664 |
| RANGAP1 | 6 | 0.0080242 | 0.032049 | 0.865899 | 731 | 4 | -3.266 | 0.67738 | 0.84383 | 1 | 17270 | 1 | -3.266 |
| DCTN5 | 4 | 0.22326 | 0.3646 | 0.999994 | 8364 | 2 | -0.56836 | 0.67745 | 0.80054 | 1 | 17271 | 1 | -0.56836 |
| RHBDD2 | 5 | 0.07164 | 0.18103 | 0.968171 | 3902 | 3 | -1.7662 | 0.67766 | 0.81411 | 1 | 17272 | 2 | -1.7662 |
| hsa-mir-1182 | 4 | 0.020782 | 0.062259 | 0.937033 | 1497 | 3 | -4.9366 | 0.67768 | 0.80062 | 1 | 17273 | 1 | -4.9366 |
| SLC39A1 | 5 | 0.76904 | 0.80117 | 0.999994 | 17075 | 1 | 0.22018 | 0.67817 | 0.81441 | 1 | 17274 | 2 | 0.22018 |
| METTL7B | 5 | 0.52026 | 0.66554 | 0.999994 | 13750 | 2 | 0.11977 | 0.67819 | 0.81443 | 1 | 17275 | 2 | 0.11977 |
| NonTargetingControlG | 2 | 0.71777 | 0.71748 | 0.999994 | 16474 | 0 | 0.5698 | 0.67833 | 0.68108 | 1 | 17276 | 1 | 0.5698 |
| HSP90AA1 | 5 | 0.045341 | 0.12915 | 0.952548 | 2780 | 4 | -0.80678 | 0.67852 | 0.81462 | 1 | 17277 | 1 | -0.80678 |
| NonTargetingControlG | 1 | 0.32145 | 0.32155 | 0.999994 | 10543 | 1 | -0.89932 | 0.67855 | 0.67856 | 1 | 17278 | 0 | -0.89932 |
| MID1 | 5 | 0.53609 | 0.67942 | 0.999994 | 13996 | 2 | 0.23616 | 0.67858 | 0.81466 | 1 | 17279 | 2 | 0.23616 |
| MPO | 5 | 0.11779 | 0.25397 | 0.994632 | 5566 | 3 | -1.5074 | 0.67867 | 0.81471 | 1 | 17280 | 1 | -1.5074 |
| hsa-mir-221 | 4 | 0.75923 | 0.76716 | 0.999994 | 16971 | 1 | 0.1388 | 0.6787 | 0.80094 | 1 | 17281 | 1 | 0.1388 |
| TSN | 6 | 0.091826 | 0.23118 | 0.983933 | 4673 | 4 | -1.3995 | 0.6787 | 0.84438 | 1 | 17282 | 1 | -1.3995 |
| LRCH3 | 6 | 0.12162 | 0.28878 | 0.999632 | 5689 | 4 | -0.61419 | 0.6787 | 0.84438 | 1 | 17283 | 1 | -0.61419 |
| TENM4 | 5 | 0.24587 | 0.43746 | 0.999994 | 8917 | 2 | 0.098776 | 0.67877 | 0.81477 | 1 | 17284 | 2 | 0.098776 |
| APP | 6 | 0.40522 | 0.60575 | 0.999994 | 12036 | 2 | -0.023789 | 0.67922 | 0.84458 | 1 | 17285 | 2 | -0.023789 |
| HDDC3 | 6 | 0.065299 | 0.17689 | 0.968171 | 3666 | 2 | -0.116 | 0.67926 | 0.84459 | 1 | 17286 | 2 | -0.116 |
| GPD1L | 5 | 0.18138 | 0.34864 | 0.999994 | 7262 | 2 | -0.61257 | 0.67939 | 0.81515 | 1 | 17287 | 2 | -0.61257 |
| NFX1 | 5 | 0.076487 | 0.18893 | 0.969543 | 4090 | 3 | -0.60057 | 0.6794 | 0.81516 | 1 | 17288 | 1 | -0.60057 |
| HMGN5 | 5 | 0.0076227 | 0.029156 | 0.859467 | 700 | 4 | -3.683 | 0.6794 | 0.81516 | 1 | 17289 | 1 | -3.683 |
| MOB1B | 6 | 0.10048 | 0.24827 | 0.990682 | 4988 | 2 | -0.039773 | 0.67944 | 0.84466 | 1 | 17290 | 2 | -0.039773 |
| ABCD1 | 4 | 0.082255 | 0.18288 | 0.968526 | 4305 | 3 | -1.2119 | 0.67954 | 0.80121 | 1 | 17291 | 1 | -1.2119 |
| HLA-DQB1 | 5 | 0.27857 | 0.4637 | 0.999994 | 9663 | 2 | -0.093462 | 0.67954 | 0.81525 | 1 | 17292 | 2 | -0.093462 |
| KIF6 | 5 | 0.52186 | 0.66691 | 0.999994 | 13779 | 1 | 0.24605 | 0.67955 | 0.81526 | 1 | 17293 | 2 | 0.24605 |
| KRT84 | 5 | 0.12883 | 0.27083 | 0.998561 | 5907 | 2 | -0.063234 | 0.67958 | 0.81528 | 1 | 17294 | 2 | -0.063234 |
| NonTargetingControlG | 2 | 0.28326 | 0.32399 | 0.999994 | 9766 | 1 | -1.1084 | 0.67974 | 0.68239 | 1 | 17295 | 1 | -1.1084 |
| NonTargetingControlG | 2 | 0.73871 | 0.7384 | 0.999994 | 16717 | 0 | 0.6166 | 0.67976 | 0.68241 | 1 | 17296 | 1 | 0.6166 |
| hsa-mir-1228 | 3 | 0.047303 | 0.11194 | 0.951052 | 2869 | 2 | -5.0735 | 0.67982 | 0.72967 | 1 | 17297 | 1 | -5.0735 |
| SH3GL1 | 6 | 0.10216 | 0.25154 | 0.993413 | 5046 | 3 | -0.83325 | 0.67999 | 0.84489 | 1 | 17298 | 2 | -0.83325 |
| ADAM17 | 6 | 0.01215 | 0.046221 | 0.892434 | 1003 | 5 | -2.2356 | 0.67999 | 0.84489 | 1 | 17299 | 1 | -2.2356 |
| CEP44 | 5 | 0.18827 | 0.35846 | 0.999994 | 7425 | 3 | -0.80596 | 0.68001 | 0.81553 | 1 | 17300 | 2 | -0.80596 |
| TTLL7 | 6 | 0.025881 | 0.089094 | 0.94395 | 1796 | 4 | -1.8208 | 0.68006 | 0.84492 | 1 | 17301 | 2 | -1.8208 |
| SIRPD | 6 | 0.29698 | 0.50319 | 0.999994 | 10067 | 2 | 0.0012806 | 0.68007 | 0.84492 | 1 | 17302 | 2 | 0.0012806 |
| MAP2K4 | 5 | 0.42496 | 0.58421 | 0.999994 | 12335 | 2 | 0.28419 | 0.68019 | 0.81564 | 1 | 17303 | 2 | 0.28419 |
| NOL7 | 5 | 0.78871 | 0.81256 | 0.999994 | 17328 | 1 | -0.09782 | 0.68065 | 0.81592 | 1 | 17304 | 1 | -0.09782 |
| TIMM50 | 5 | 0.35129 | 0.52297 | 0.999994 | 11182 | 2 | -0.46176 | 0.68065 | 0.81592 | 1 | 17305 | 2 | -0.46176 |
| THEM6 | 5 | 0.5134 | 0.6596 | 0.999994 | 13628 | 1 | 0.17767 | 0.68065 | 0.81592 | 1 | 17306 | 2 | 0.17767 |
| OR6K6 | 6 | 0.14518 | 0.31959 | 0.999994 | 6328 | 3 | -0.91549 | 0.68079 | 0.84522 | 1 | 17307 | 2 | -0.91549 |
| FBXO40 | 6 | 0.041235 | 0.12466 | 0.951052 | 2574 | 4 | -2.1238 | 0.68092 | 0.84528 | 1 | 17308 | 2 | -2.1238 |
| KCNA3 | 6 | 0.084269 | 0.21603 | 0.980699 | 4385 | 4 | -1.2417 | 0.68145 | 0.84549 | 1 | 17309 | 1 | -1.2417 |
| SERPINB10 | 6 | 0.038678 | 0.11893 | 0.951052 | 2448 | 4 | -1.6585 | 0.68145 | 0.84549 | 1 | 17310 | 1 | -1.6585 |
| CNBD2 | 6 | 0.00057301 | 0.002995 | 0.53161 | 116 | 5 | -6.5052 | 0.68145 | 0.84549 | 1 | 17311 | 1 | -6.5052 |
| ZNF561 | 6 | 0.27703 | 0.47997 | 0.999994 | 9639 | 3 | -1.1807 | 0.68145 | 0.84549 | 1 | 17312 | 2 | -1.1807 |
| PLA2G4A | 6 | 0.83005 | 0.86157 | 0.999994 | 17933 | 1 | -0.076916 | 0.68145 | 0.84549 | 1 | 17313 | 2 | -0.076916 |
| EMP1 | 6 | 0.045977 | 0.13523 | 0.954328 | 2809 | 3 | -1.0572 | 0.68154 | 0.84554 | 1 | 17314 | 2 | -1.0572 |
| EXOSC4 | 4 | 0.036018 | 0.093781 | 0.945031 | 2317 | 3 | -4.0084 | 0.68156 | 0.80185 | 1 | 17315 | 1 | -4.0084 |

| | | | | | | | | | | | | |
|---|---|---|---|---|---|---|---|---|---|---|---|---|
| CRYBB2 | 5 | 0.21011 | 0.38953 | 0.999994 | 8026 | 2 | -0.28923 | 0.6817 | 0.81657 | 1 | 17316 | 1 | -0.28923 |
| FNBP1 | 6 | 0.11525 | 0.27675 | 0.999379 | 5486 | 4 | -0.86831 | 0.68188 | 0.84568 | 1 | 17317 | 2 | -0.86831 |
| FAM19A2 | 5 | 0.093175 | 0.21565 | 0.980236 | 4728 | 3 | -1.4664 | 0.68208 | 0.81682 | 1 | 17318 | 2 | -1.4664 |
| ZCCHC13 | 4 | 0.092736 | 0.20203 | 0.97784 | 4712 | 3 | -1.2562 | 0.6821 | 0.80203 | 1 | 17319 | 1 | -1.2562 |
| ZWILCH | 5 | 0.12617 | 0.26678 | 0.998555 | 5825 | 2 | 0.28095 | 0.68222 | 0.81691 | 1 | 17320 | 2 | 0.28095 |
| SCTR | 6 | 0.055157 | 0.15526 | 0.967126 | 3206 | 4 | -1.6604 | 0.68232 | 0.84585 | 1 | 17321 | 2 | -1.6604 |
| ZZEF1 | 5 | 0.081945 | 0.19775 | 0.977164 | 4296 | 3 | -1.7465 | 0.6825 | 0.81708 | 1 | 17322 | 2 | -1.7465 |
| PRRG1 | 5 | 0.50575 | 0.65294 | 0.999994 | 13520 | 2 | -0.5395 | 0.6825 | 0.81708 | 1 | 17323 | 1 | -0.5395 |
| DNAH3 | 6 | 0.051125 | 0.14653 | 0.964852 | 3026 | 4 | -1.2513 | 0.68252 | 0.84593 | 1 | 17324 | 2 | -1.2513 |
| SRPK3 | 5 | 0.21865 | 0.40139 | 0.999994 | 8247 | 3 | -0.69098 | 0.68263 | 0.81716 | 1 | 17325 | 2 | -0.69098 |
| CXCL2 | 5 | 0.052937 | 0.14745 | 0.965554 | 3109 | 3 | -1.9604 | 0.68271 | 0.81721 | 1 | 17326 | 2 | -1.9604 |
| LRRK2 | 6 | 0.016435 | 0.060421 | 0.932932 | 1272 | 4 | -3.4698 | 0.68303 | 0.84612 | 1 | 17327 | 2 | -3.4698 |
| hsa-mir-549a | 4 | 0.403 | 0.54883 | 0.999994 | 11998 | 2 | -0.50127 | 0.68306 | 0.80234 | 1 | 17328 | 1 | -0.50127 |
| NonTargetingControlG | 2 | 0.53621 | 0.53767 | 0.999994 | 14003 | 1 | 0.077274 | 0.68308 | 0.68546 | 1 | 17329 | 1 | 0.077274 |
| hsa-mir-4253 | 3 | 0.2829 | 0.41554 | 0.999994 | 9758 | 2 | -0.62432 | 0.68317 | 0.73161 | 1 | 17330 | 1 | -0.62432 |
| PLIN5 | 4 | 0.072058 | 0.16383 | 0.967126 | 3921 | 3 | -1.7308 | 0.68333 | 0.80243 | 1 | 17331 | 1 | -1.7308 |
| PDILT | 6 | 0.11831 | 0.28252 | 0.999632 | 5583 | 4 | -0.83191 | 0.68337 | 0.84625 | 1 | 17332 | 2 | -0.83191 |
| GALK1 | 6 | 0.065024 | 0.17631 | 0.968171 | 3655 | 4 | -1.8975 | 0.68349 | 0.8463 | 1 | 17333 | 1 | -1.8975 |
| LOC256021 | 6 | 0.088007 | 0.22353 | 0.983354 | 4514 | 3 | -0.66155 | 0.68349 | 0.8463 | 1 | 17334 | 1 | -0.66155 |
| TSG101 | 6 | 0.20127 | 0.38941 | 0.999994 | 7798 | 3 | -0.44099 | 0.68349 | 0.8463 | 1 | 17335 | 1 | -0.44099 |
| APOL2 | 5 | 0.0054331 | 0.022511 | 0.819701 | 554 | 2 | 0.51209 | 0.68386 | 0.81791 | 1 | 17336 | 2 | 0.51209 |
| PARD6A | 5 | 0.04165 | 0.12002 | 0.951052 | 2598 | 3 | -2.4272 | 0.68387 | 0.81792 | 1 | 17337 | 2 | -2.4272 |
| ITGA5 | 5 | 0.01814 | 0.05902 | 0.931872 | 1369 | 4 | -1.964 | 0.68387 | 0.81792 | 1 | 17338 | 1 | -1.964 |
| NonTargetingControlG | 2 | 0.16741 | 0.23884 | 0.987253 | 6921 | 1 | -2.862 | 0.68388 | 0.68617 | 1 | 17339 | 1 | -2.862 |
| TDGF1 | 5 | 0.71313 | 0.77258 | 0.999994 | 16430 | 1 | 0.28583 | 0.68389 | 0.81793 | 1 | 17340 | 2 | 0.28583 |
| hsa-mir-3668 | 2 | 0.63802 | 0.63785 | 0.999994 | 15488 | 0 | 0.33712 | 0.68403 | 0.68631 | 1 | 17341 | 1 | 0.33712 |
| TMEM48 | 3 | 0.14421 | 0.24102 | 0.987253 | 6305 | 2 | -1.8262 | 0.68417 | 0.73216 | 1 | 17342 | 1 | -1.8262 |
| C10orf2 | 5 | 0.19649 | 0.37021 | 0.999994 | 7662 | 2 | 0.50752 | 0.68424 | 0.81814 | 1 | 17343 | 2 | 0.50752 |
| METTL2B | 4 | 0.58495 | 0.6573 | 0.999994 | 14742 | 1 | 0.086749 | 0.68456 | 0.80282 | 1 | 17344 | 1 | 0.086749 |
| hsa-mir-3153 | 4 | 0.69384 | 0.72045 | 0.999994 | 16243 | 1 | 0.15144 | 0.68456 | 0.80282 | 1 | 17345 | 1 | 0.15144 |
| hsa-mir-4257 | 3 | 0.11853 | 0.20756 | 0.978852 | 5591 | 2 | -0.72747 | 0.68476 | 0.73251 | 1 | 17346 | 1 | -0.72747 |
| C7orf72 | 6 | 0.24231 | 0.43876 | 0.999994 | 8838 | 3 | -0.37426 | 0.68493 | 0.8469 | 1 | 17347 | 2 | -0.37426 |
| CCNA1 | 6 | 0.013045 | 0.049272 | 0.906088 | 1066 | 3 | -3.1653 | 0.68493 | 0.8469 | 1 | 17348 | 1 | -3.1653 |
| HOMER3 | 6 | 0.31907 | 0.52883 | 0.999994 | 10499 | 3 | -0.1191 | 0.68493 | 0.8469 | 1 | 17349 | 2 | -0.1191 |
| PAM | 6 | 0.0085227 | 0.03378 | 0.867348 | 764 | 5 | -3.4999 | 0.68493 | 0.8469 | 1 | 17350 | 1 | -3.4999 |
| LPP | 5 | 0.1515 | 0.30503 | 0.999994 | 6497 | 3 | -1.0777 | 0.68502 | 0.81862 | 1 | 17351 | 2 | -1.0777 |
| PTPN14 | 5 | 0.027738 | 0.084609 | 0.94395 | 1893 | 4 | -2.31 | 0.68515 | 0.81871 | 1 | 17352 | 1 | -2.31 |
| BBC3 | 6 | 0.16477 | 0.34419 | 0.999994 | 6858 | 3 | -0.59764 | 0.68518 | 0.84701 | 1 | 17353 | 2 | -0.59764 |
| ADAMTSL1 | 5 | 0.66079 | 0.74946 | 0.999994 | 15792 | 1 | 0.53303 | 0.68519 | 0.81873 | 1 | 17354 | 2 | 0.53303 |
| TMEM179B | 5 | 0.38074 | 0.5473 | 0.999994 | 11659 | 1 | 0.31446 | 0.68531 | 0.81881 | 1 | 17355 | 2 | 0.31446 |
| TSPEAR | 5 | 0.083851 | 0.20076 | 0.977164 | 4373 | 3 | -0.77078 | 0.68537 | 0.81884 | 1 | 17356 | 2 | -0.77078 |
| RASSF4 | 6 | 0.31859 | 0.52828 | 0.999994 | 10490 | 3 | -0.1764 | 0.6858 | 0.84726 | 1 | 17357 | 2 | -0.1764 |
| CYP2E1 | 5 | 0.19428 | 0.36703 | 0.999994 | 7613 | 3 | -0.76848 | 0.68582 | 0.81911 | 1 | 17358 | 1 | -0.76848 |
| SMARCD3 | 5 | 0.064955 | 0.17018 | 0.967126 | 3650 | 3 | -0.94215 | 0.68582 | 0.81911 | 1 | 17359 | 1 | -0.94215 |
| PPAPDC1A | 5 | 0.55333 | 0.69472 | 0.999994 | 14260 | 2 | -0.7042 | 0.68582 | 0.81911 | 1 | 17360 | 1 | -0.7042 |
| ATP6V0B | 3 | 0.18933 | 0.29921 | 0.999994 | 7464 | 1 | 0.42457 | 0.68587 | 0.73317 | 1 | 17361 | 1 | 0.42457 |
| EIF3CL | 1 | 0.31401 | 0.31415 | 0.999994 | 10396 | 1 | -0.97237 | 0.68599 | 0.686 | 1 | 17362 | 0 | -0.97237 |
| QTRTD1 | 5 | 0.72971 | 0.78052 | 0.999994 | 16595 | 1 | 0.22128 | 0.68599 | 0.81921 | 1 | 17363 | 2 | 0.22128 |
| PBRM1 | 6 | 0.011048 | 0.042573 | 0.891621 | 926 | 4 | -2.4995 | 0.68612 | 0.84738 | 1 | 17364 | 2 | -2.4995 |
| PHOX2A | 6 | 0.1708 | 0.35176 | 0.999994 | 7002 | 3 | -0.28508 | 0.68612 | 0.84738 | 1 | 17365 | 2 | -0.28508 |
| PABPC4 | 5 | 0.16761 | 0.32859 | 0.999994 | 6928 | 3 | -0.66085 | 0.68615 | 0.8193 | 1 | 17366 | 2 | -0.66085 |
| CAPG | 4 | 0.055283 | 0.13176 | 0.95318 | 3211 | 3 | -1.7404 | 0.68618 | 0.80334 | 1 | 17367 | 1 | -1.7404 |

| | | | | | | | | | | | | | |
|---|---|---|---|---|---|---|---|---|---|---|---|---|---|
| PCDHA3 | 2 | 0.58522 | 0.58529 | 0.999994 | 14747 | 0 | 0.20855 | 0.68634 | 0.68842 | 1 | 17368 | 1 | 0.20855 |
| TMEM192 | 5 | 0.12885 | 0.27085 | 0.998561 | 5909 | 3 | -0.82241 | 0.68646 | 0.81949 | 1 | 17369 | 2 | -0.82241 |
| AP1S3 | 5 | 0.090913 | 0.21207 | 0.979587 | 4635 | 3 | -1.5769 | 0.68657 | 0.81956 | 1 | 17370 | 2 | -1.5769 |
| TMEM170A | 6 | 0.029077 | 0.096625 | 0.945031 | 1960 | 3 | -2.0697 | 0.68662 | 0.84757 | 1 | 17371 | 1 | -2.0697 |
| SUMF2 | 5 | 0.012649 | 0.043613 | 0.891621 | 1041 | 3 | -3.6731 | 0.68677 | 0.8197 | 1 | 17372 | 2 | -3.6731 |
| UBA52 | 3 | 0.14462 | 0.24156 | 0.987253 | 6317 | 2 | -1.5082 | 0.6868 | 0.73372 | 1 | 17373 | 1 | -1.5082 |
| hsa-mir-4480 | 4 | 0.076477 | 0.17218 | 0.967658 | 4089 | 2 | -1.8153 | 0.68686 | 0.80355 | 1 | 17374 | 1 | -1.8153 |
| PCDHGA10 | 2 | 0.50212 | 0.50585 | 0.999994 | 13464 | 1 | -0.058843 | 0.68692 | 0.68896 | 1 | 17375 | 1 | -0.058843 |
| HCRTR2 | 5 | 0.15046 | 0.30347 | 0.999994 | 6469 | 1 | -0.097658 | 0.68715 | 0.81995 | 1 | 17376 | 2 | -0.097658 |
| U2AF2 | 5 | 0.16191 | 0.32035 | 0.999994 | 6788 | 3 | -1.1499 | 0.6873 | 0.82006 | 1 | 17377 | 2 | -1.1499 |
| LIN28A | 6 | 0.071751 | 0.19042 | 0.971392 | 3909 | 3 | -0.20945 | 0.68738 | 0.84789 | 1 | 17378 | 2 | -0.20945 |
| RPRML | 4 | 0.35776 | 0.50221 | 0.999994 | 11302 | 2 | -0.38194 | 0.68746 | 0.80374 | 1 | 17379 | 1 | -0.38194 |
| hsa-mir-941-2 | 2 | 0.34535 | 0.37232 | 0.999994 | 11058 | 1 | -0.63054 | 0.68759 | 0.68954 | 1 | 17380 | 1 | -0.63054 |
| LYZL4 | 4 | 0.063935 | 0.14828 | 0.965554 | 3603 | 3 | -2.1655 | 0.68775 | 0.80383 | 1 | 17381 | 1 | -2.1655 |
| GLP2R | 6 | 0.32621 | 0.53692 | 0.999994 | 10649 | 3 | -0.17729 | 0.68784 | 0.84807 | 1 | 17382 | 2 | -0.17729 |
| ABCB6 | 5 | 0.79767 | 0.81805 | 0.999994 | 17448 | 1 | 0.48137 | 0.68808 | 0.82053 | 1 | 17383 | 2 | 0.48137 |
| KIAA1958 | 6 | 0.54595 | 0.70076 | 0.999994 | 14149 | 2 | -0.24607 | 0.68826 | 0.84824 | 1 | 17384 | 2 | -0.24607 |
| SNAPC1 | 6 | 0.12698 | 0.29646 | 0.999994 | 5847 | 2 | -0.40498 | 0.68837 | 0.84828 | 1 | 17385 | 2 | -0.40498 |
| OR10W1 | 5 | 0.17344 | 0.33715 | 0.999994 | 7072 | 3 | -0.82171 | 0.68839 | 0.82072 | 1 | 17386 | 2 | -0.82171 |
| IMP4 | 3 | 0.092493 | 0.17308 | 0.96793 | 4705 | 2 | -2.5792 | 0.6885 | 0.73467 | 1 | 17387 | 1 | -2.5792 |
| NDUFV3 | 4 | 0.12863 | 0.26525 | 0.997396 | 5900 | 3 | -1.5064 | 0.68859 | 0.8041 | 1 | 17388 | 1 | -1.5064 |
| IFNK | 6 | 0.25537 | 0.45436 | 0.999994 | 9135 | 3 | -0.44865 | 0.68859 | 0.84838 | 1 | 17389 | 2 | -0.44865 |
| NonTargetingControlG | 2 | 0.45214 | 0.4612 | 0.999994 | 12736 | 1 | -0.19941 | 0.68866 | 0.69052 | 1 | 17390 | 1 | -0.19941 |
| hsa-mir-1236 | 3 | 0.033606 | 0.083365 | 0.94383 | 2196 | 2 | -5.7703 | 0.68871 | 0.73477 | 1 | 17391 | 1 | -5.7703 |
| HYKK | 3 | 0.087167 | 0.16599 | 0.967126 | 4482 | 2 | -2.3431 | 0.68901 | 0.73494 | 1 | 17392 | 1 | -2.3431 |
| SHISA6 | 5 | 0.6556 | 0.74735 | 0.999994 | 15722 | 1 | -0.21934 | 0.68912 | 0.82113 | 1 | 17393 | 2 | -0.21934 |
| SPANXD | 4 | 0.51235 | 0.62185 | 0.999994 | 13614 | 1 | -0.22434 | 0.68915 | 0.8043 | 1 | 17394 | 1 | -0.22434 |
| MOB1A | 6 | 0.055605 | 0.15623 | 0.967126 | 3226 | 4 | -1.0912 | 0.68919 | 0.84863 | 1 | 17395 | 1 | -1.0912 |
| hsa-mir-410 | 3 | 0.17226 | 0.27728 | 0.999379 | 7043 | 1 | -0.0054062 | 0.68957 | 0.73524 | 1 | 17396 | 1 | -0.0054062 |
| R3HDM2 | 6 | 0.18651 | 0.37118 | 0.999994 | 7372 | 3 | -0.93467 | 0.68969 | 0.84886 | 1 | 17397 | 1 | -0.93467 |
| STK31 | 5 | 0.0096036 | 0.034931 | 0.867348 | 846 | 3 | -0.96084 | 0.68986 | 0.8216 | 1 | 17398 | 2 | -0.96084 |
| hsa-mir-4432 | 2 | 0.64525 | 0.64499 | 0.999994 | 15588 | 0 | 0.27613 | 0.68998 | 0.69172 | 1 | 17399 | 1 | 0.27613 |
| PLXNA1 | 5 | 0.20994 | 0.38928 | 0.999994 | 8022 | 3 | -0.79894 | 0.68999 | 0.82168 | 1 | 17400 | 2 | -0.79894 |
| hsa-mir-7155 | 4 | 0.29768 | 0.44064 | 0.999994 | 10083 | 2 | -0.31393 | 0.68999 | 0.80459 | 1 | 17401 | 1 | -0.31393 |
| ST8SIA5 | 6 | 0.081834 | 0.2111 | 0.979587 | 4291 | 2 | -0.67667 | 0.69023 | 0.84908 | 1 | 17402 | 2 | -0.67667 |
| LDB3 | 6 | 0.041929 | 0.12622 | 0.951052 | 2609 | 3 | -1.4151 | 0.69037 | 0.84913 | 1 | 17403 | 2 | -1.4151 |
| hsa-mir-6812 | 1 | 0.30954 | 0.3096 | 0.999994 | 10318 | 1 | -0.85015 | 0.69046 | 0.69049 | 1 | 17404 | 0 | -0.85015 |
| SLC35E4 | 5 | 0.22859 | 0.41526 | 0.999994 | 8491 | 2 | -0.40244 | 0.6909 | 0.82224 | 1 | 17405 | 1 | -0.40244 |
| PRPF40A | 5 | 0.32491 | 0.50134 | 0.999994 | 10616 | 2 | -0.30104 | 0.6909 | 0.82224 | 1 | 17406 | 1 | -0.30104 |
| ZKSCAN1 | 5 | 0.15753 | 0.31396 | 0.999994 | 6660 | 3 | -0.65354 | 0.6909 | 0.82224 | 1 | 17407 | 1 | -0.65354 |
| PTGR2 | 5 | 0.13028 | 0.27302 | 0.998561 | 5938 | 3 | -1.3129 | 0.6909 | 0.82224 | 1 | 17408 | 1 | -1.3129 |
| NonTargetingControlG | 1 | 0.30907 | 0.30911 | 0.999994 | 10301 | 1 | -1.0027 | 0.69093 | 0.69097 | 1 | 17409 | 0 | -1.0027 |
| OR51S1 | 4 | 0.36512 | 0.50977 | 0.999994 | 11423 | 2 | -0.60111 | 0.69106 | 0.80494 | 1 | 17410 | 1 | -0.60111 |
| LOC154872 | 5 | 0.088471 | 0.20812 | 0.979182 | 4533 | 3 | -1.652 | 0.69111 | 0.82237 | 1 | 17411 | 2 | -1.652 |
| TMEM225 | 5 | 0.59483 | 0.72424 | 0.999994 | 14887 | 1 | 0.0024081 | 0.69127 | 0.82247 | 1 | 17412 | 2 | 0.0024081 |
| LECT1 | 5 | 0.0085739 | 0.031954 | 0.865899 | 767 | 3 | -2.8665 | 0.69149 | 0.82261 | 1 | 17413 | 2 | -2.8665 |
| CD83 | 6 | 0.094221 | 0.23598 | 0.985362 | 4759 | 4 | -1.086 | 0.69167 | 0.84968 | 1 | 17414 | 2 | -1.086 |
| OR8A1 | 5 | 0.045041 | 0.12841 | 0.951777 | 2760 | 3 | -2.4992 | 0.69168 | 0.82274 | 1 | 17415 | 1 | -2.4992 |
| DPYSL3 | 5 | 0.5653 | 0.70537 | 0.999994 | 14474 | 2 | 0.56464 | 0.6917 | 0.82275 | 1 | 17416 | 2 | 0.56464 |
| NonTargetingControlG | 2 | 0.5281 | 0.52998 | 0.999994 | 13862 | 1 | 0.13365 | 0.69182 | 0.69342 | 1 | 17417 | 1 | 0.13365 |
| DOK1 | 5 | 0.2327 | 0.42096 | 0.999994 | 8587 | 3 | -0.66802 | 0.69185 | 0.82285 | 1 | 17418 | 1 | -0.66802 |
| FAM189A1 | 6 | 0.037486 | 0.11624 | 0.951052 | 2399 | 3 | -1.5409 | 0.69198 | 0.84981 | 1 | 17419 | 2 | -1.5409 |

| | | | | | | | | | | | | | |
|---|---|---|---|---|---|---|---|---|---|---|---|---|---|
| DCAF8L1 | 5 | 0.020618 | 0.065733 | 0.93914 | 1489 | 3 | -1.6653 | 0.69209 | 0.823 | 1 | 17420 | 2 | -1.6653 |
| RPS3 | 5 | 0.079888 | 0.1944 | 0.975797 | 4230 | 1 | 0.43335 | 0.69217 | 0.82304 | 1 | 17421 | 2 | 0.43335 |
| KIAA1468 | 5 | 0.25101 | 0.44156 | 0.999994 | 9034 | 2 | 0.54137 | 0.69218 | 0.82305 | 1 | 17422 | 2 | 0.54137 |
| PCBP4 | 6 | 0.003325 | 0.01491 | 0.755841 | 389 | 4 | -3.3841 | 0.69221 | 0.8499 | 1 | 17423 | 2 | -3.3841 |
| CDC25A | 6 | 0.008305 | 0.033013 | 0.867348 | 750 | 5 | -2.4875 | 0.69221 | 0.8499 | 1 | 17424 | 1 | -2.4875 |
| SALL2 | 6 | 0.25012 | 0.44808 | 0.999994 | 9022 | 3 | -0.21113 | 0.69221 | 0.8499 | 1 | 17425 | 2 | -0.21113 |
| NonTargetingControlG | 2 | 0.23388 | 0.28684 | 0.999632 | 8616 | 1 | -1.4835 | 0.69222 | 0.6938 | 1 | 17426 | 1 | -1.4835 |
| CDC42SE1 | 5 | 0.51588 | 0.66173 | 0.999994 | 13673 | 1 | 0.3711 | 0.69223 | 0.82308 | 1 | 17427 | 2 | 0.3711 |
| HS2ST1 | 5 | 0.0079183 | 0.030036 | 0.862223 | 725 | 2 | 0.52052 | 0.69233 | 0.82314 | 1 | 17428 | 2 | 0.52052 |
| TMBIM1 | 6 | 0.10744 | 0.26179 | 0.995257 | 5223 | 4 | -1.1277 | 0.69236 | 0.84996 | 1 | 17429 | 2 | -1.1277 |
| PROKR2 | 5 | 0.50974 | 0.65639 | 0.999994 | 13573 | 1 | 0.078262 | 0.6924 | 0.82319 | 1 | 17430 | 2 | 0.078262 |
| TGOLN2 | 6 | 0.92408 | 0.92424 | 0.999994 | 19717 | 1 | -0.20967 | 0.69245 | 0.84999 | 1 | 17431 | 2 | -0.20967 |
| NonTargetingControlG | 2 | 0.7359 | 0.7356 | 0.999994 | 16675 | 0 | 0.67973 | 0.69249 | 0.69403 | 1 | 17432 | 1 | 0.67973 |
| FAM89B | 5 | 0.052728 | 0.14695 | 0.964901 | 3101 | 3 | -1.0283 | 0.69253 | 0.82327 | 1 | 17433 | 2 | -1.0283 |
| hsa-mir-483 | 1 | 0.30746 | 0.30749 | 0.999994 | 10276 | 1 | -2.2988 | 0.69254 | 0.69255 | 1 | 17434 | 0 | -2.2988 |
| LOC100129924 | 6 | 0.33556 | 0.54759 | 0.999994 | 10843 | 3 | -0.44995 | 0.69264 | 0.85007 | 1 | 17435 | 2 | -0.44995 |
| SSFA2 | 5 | 0.51805 | 0.66361 | 0.999994 | 13709 | 2 | -0.51098 | 0.69279 | 0.82342 | 1 | 17436 | 2 | -0.51098 |
| DENND3 | 5 | 0.012286 | 0.042609 | 0.891621 | 1010 | 3 | -1.2872 | 0.69281 | 0.82343 | 1 | 17437 | 2 | -1.2872 |
| HSD11B2 | 5 | 0.13148 | 0.27487 | 0.998561 | 5969 | 3 | -1.3727 | 0.69281 | 0.82343 | 1 | 17438 | 2 | -1.3727 |
| hsa-mir-345 | 2 | 0.079505 | 0.13772 | 0.957317 | 4209 | 1 | -3.8488 | 0.69293 | 0.69447 | 1 | 17439 | 1 | -3.8488 |
| PRKACB | 5 | 0.053409 | 0.14861 | 0.965554 | 3133 | 2 | -0.22198 | 0.69304 | 0.82358 | 1 | 17440 | 2 | -0.22198 |
| C2orf91 | 5 | 0.065239 | 0.17064 | 0.967177 | 3660 | 2 | -0.048781 | 0.69315 | 0.82364 | 1 | 17441 | 2 | -0.048781 |
| hsa-mir-7853 | 2 | 0.62672 | 0.6265 | 0.999994 | 15330 | 0 | 0.24011 | 0.6933 | 0.69481 | 1 | 17442 | 1 | 0.24011 |
| NonTargetingControlG | 2 | 0.66862 | 0.66841 | 0.999994 | 15900 | 0 | 0.43415 | 0.69331 | 0.69482 | 1 | 17443 | 1 | 0.43415 |
| TMEM154 | 5 | 0.11541 | 0.25038 | 0.992497 | 5488 | 3 | -1.3617 | 0.6935 | 0.82383 | 1 | 17444 | 1 | -1.3617 |
| NonTargetingControlG | 2 | 0.55413 | 0.55485 | 0.999994 | 14270 | 1 | -0.016105 | 0.6935 | 0.69499 | 1 | 17445 | 1 | -0.016105 |
| SMPD1 | 5 | 0.72218 | 0.77691 | 0.999994 | 16519 | 1 | 0.42331 | 0.69368 | 0.82395 | 1 | 17446 | 2 | 0.42331 |
| MTCH1 | 5 | 0.45238 | 0.60721 | 0.999994 | 12741 | 2 | -0.43025 | 0.69396 | 0.82412 | 1 | 17447 | 2 | -0.43025 |
| CDH9 | 6 | 0.54271 | 0.69846 | 0.999994 | 14107 | 1 | 0.43071 | 0.69409 | 0.85071 | 1 | 17448 | 2 | 0.43071 |
| BNIP1 | 5 | 0.27949 | 0.46444 | 0.999994 | 9691 | 2 | -0.11633 | 0.69425 | 0.82431 | 1 | 17449 | 1 | -0.11633 |
| DNAAF3 | 5 | 0.24395 | 0.43585 | 0.999994 | 8881 | 2 | -0.36307 | 0.69425 | 0.82431 | 1 | 17450 | 2 | -0.36307 |
| SLC25A46 | 4 | 0.36881 | 0.51362 | 0.999994 | 11486 | 1 | 0.034316 | 0.69428 | 0.80599 | 1 | 17451 | 1 | 0.034316 |
| hsa-mir-6515 | 3 | 0.19018 | 0.30032 | 0.999994 | 7492 | 2 | -1.2673 | 0.69451 | 0.73813 | 1 | 17452 | 1 | -1.2673 |
| IFT27 | 6 | 0.33036 | 0.54167 | 0.999994 | 10728 | 3 | -0.45536 | 0.69466 | 0.85094 | 1 | 17453 | 2 | -0.45536 |
| hsa-mir-5009 | 4 | 0.39525 | 0.54083 | 0.999994 | 11874 | 2 | -0.61559 | 0.69467 | 0.80612 | 1 | 17454 | 1 | -0.61559 |
| PPP1R1C | 6 | 0.2289 | 0.42275 | 0.999994 | 8500 | 3 | -0.65921 | 0.69472 | 0.85097 | 1 | 17455 | 2 | -0.65921 |
| TMEM86A | 6 | 0.084849 | 0.21723 | 0.980699 | 4402 | 2 | 0.12126 | 0.69472 | 0.85097 | 1 | 17456 | 2 | 0.12126 |
| ANKRD13C | 6 | 0.32388 | 0.53431 | 0.999994 | 10592 | 2 | 0.38395 | 0.69473 | 0.85097 | 1 | 17457 | 2 | 0.38395 |
| PANK2 | 4 | 0.089925 | 0.19688 | 0.977164 | 4591 | 3 | -1.4573 | 0.69477 | 0.80616 | 1 | 17458 | 1 | -1.4573 |
| LOC100130348 | 6 | 0.069498 | 0.18571 | 0.969409 | 3814 | 3 | -0.86537 | 0.69488 | 0.85103 | 1 | 17459 | 2 | -0.86537 |
| FOXP1 | 6 | 0.19293 | 0.37919 | 0.999994 | 7574 | 3 | -0.62 | 0.69495 | 0.85106 | 1 | 17460 | 2 | -0.62 |
| NonTargetingControlG | 2 | 0.40817 | 0.42359 | 0.999994 | 12080 | 1 | -0.43334 | 0.69499 | 0.69634 | 1 | 17461 | 1 | -0.43334 |
| CCDC85C | 5 | 0.15648 | 0.31239 | 0.999994 | 6641 | 3 | -0.73401 | 0.69509 | 0.82484 | 1 | 17462 | 1 | -0.73401 |
| RNF213 | 5 | 0.19213 | 0.36396 | 0.999994 | 7551 | 3 | -0.9145 | 0.69509 | 0.82484 | 1 | 17463 | 1 | -0.9145 |
| SPATA25 | 5 | 0.38123 | 0.54772 | 0.999994 | 11666 | 2 | -0.46181 | 0.69509 | 0.82484 | 1 | 17464 | 1 | -0.46181 |
| ESYT1 | 6 | 0.050579 | 0.14533 | 0.963012 | 3000 | 4 | -1.541 | 0.69512 | 0.85113 | 1 | 17465 | 2 | -1.541 |
| TUSC2 | 5 | 0.7053 | 0.76891 | 0.999994 | 16367 | 1 | 0.48965 | 0.69514 | 0.82486 | 1 | 17466 | 2 | 0.48965 |
| VIPR2 | 5 | 0.11841 | 0.25496 | 0.994943 | 5585 | 2 | 0.37193 | 0.69528 | 0.82495 | 1 | 17467 | 2 | 0.37193 |
| NPY2R | 5 | 0.19431 | 0.36706 | 0.999994 | 7614 | 3 | -0.79968 | 0.69534 | 0.82498 | 1 | 17468 | 2 | -0.79968 |
| NonTargetingControlG | 2 | 0.21127 | 0.27026 | 0.998561 | 8054 | 1 | -1.6152 | 0.69549 | 0.69681 | 1 | 17469 | 1 | -1.6152 |
| NIP7 | 2 | 0.024941 | 0.046002 | 0.892434 | 1754 | 1 | -4.506 | 0.69555 | 0.69687 | 1 | 17470 | 1 | -4.506 |
| PRLHR | 6 | 0.49354 | 0.66429 | 0.999994 | 13317 | 1 | 0.1001 | 0.69569 | 0.85136 | 1 | 17471 | 2 | 0.1001 |

| | | | | | | | | | | | | |
|---|---|---|---|---|---|---|---|---|---|---|---|---|
| IFNA5 | 5 | 0.81757 | 0.83096 | 0.999994 | 17740 | 1 | -0.34697 | 0.69572 | 0.82521 | 1 | 17472 | 2 | -0.34697 |
| IL18R1 | 6 | 0.11373 | 0.27384 | 0.998561 | 5433 | 4 | -1.2357 | 0.69576 | 0.8514 | 1 | 17473 | 2 | -1.2357 |
| PABPC5 | 5 | 0.15894 | 0.31597 | 0.999994 | 6698 | 2 | 0.39208 | 0.69587 | 0.82529 | 1 | 17474 | 2 | 0.39208 |
| ADPRM | 6 | 0.13781 | 0.31027 | 0.999994 | 6129 | 3 | -1.0992 | 0.69628 | 0.85162 | 1 | 17475 | 2 | -1.0992 |
| RGS7 | 6 | 0.021444 | 0.076635 | 0.93914 | 1544 | 4 | -2.0214 | 0.69628 | 0.85162 | 1 | 17476 | 2 | -2.0214 |
| ISG20L2 | 6 | 0.03833 | 0.11814 | 0.951052 | 2432 | 4 | -1.8587 | 0.69628 | 0.85162 | 1 | 17477 | 1 | -1.8587 |
| WFDC8 | 6 | 0.4684 | 0.64735 | 0.999994 | 12980 | 2 | -0.22205 | 0.6963 | 0.85163 | 1 | 17478 | 2 | -0.22205 |
| CYP2C18 | 5 | 0.11758 | 0.25365 | 0.994544 | 5563 | 2 | 0.0080922 | 0.69631 | 0.82556 | 1 | 17479 | 2 | 0.0080922 |
| EXOSC9 | 5 | 0.11519 | 0.25004 | 0.992497 | 5483 | 2 | -0.026166 | 0.69643 | 0.82563 | 1 | 17480 | 1 | -0.026166 |
| XBP1 | 5 | 0.0013501 | 0.0059993 | 0.623123 | 203 | 4 | -3.1276 | 0.69643 | 0.82563 | 1 | 17481 | 1 | -3.1276 |
| CELF1 | 4 | 0.0036328 | 0.012969 | 0.733879 | 413 | 3 | -5.3818 | 0.69684 | 0.80681 | 1 | 17482 | 1 | -5.3818 |
| CTAGE1 | 5 | 0.098221 | 0.22363 | 0.983354 | 4903 | 3 | -1.631 | 0.69687 | 0.82589 | 1 | 17483 | 2 | -1.631 |
| hsa-mir-4694 | 3 | 0.16342 | 0.26597 | 0.997535 | 6824 | 1 | -0.23063 | 0.697 | 0.73962 | 1 | 17484 | 1 | -0.23063 |
| MGAT4C | 6 | 0.33963 | 0.55223 | 0.999994 | 10922 | 3 | -0.49704 | 0.69707 | 0.85195 | 1 | 17485 | 2 | -0.49704 |
| NonTargetingControlG | 2 | 0.181 | 0.24842 | 0.990718 | 7246 | 1 | -1.6441 | 0.69748 | 0.69865 | 1 | 17486 | 1 | -1.6441 |
| NonTargetingControlG | 2 | 0.74657 | 0.74622 | 0.999994 | 16823 | 0 | 0.69757 | 0.69753 | 0.69869 | 1 | 17487 | 1 | 0.69757 |
| PHKG1 | 5 | 0.10413 | 0.23284 | 0.984502 | 5102 | 3 | -1.3245 | 0.69755 | 0.82633 | 1 | 17488 | 2 | -1.3245 |
| MTCP1 | 6 | 0.2146 | 0.40568 | 0.999994 | 8135 | 3 | -0.71008 | 0.69782 | 0.85229 | 1 | 17489 | 2 | -0.71008 |
| ARHGDIA | 6 | 0.12042 | 0.28649 | 0.999632 | 5641 | 3 | -0.59749 | 0.69782 | 0.85229 | 1 | 17490 | 2 | -0.59749 |
| HOXA1 | 6 | 0.010457 | 0.040584 | 0.891621 | 901 | 4 | -2.3106 | 0.69782 | 0.85229 | 1 | 17491 | 1 | -2.3106 |
| LOC100144595 | 6 | 0.038627 | 0.11881 | 0.951052 | 2445 | 4 | -1.4655 | 0.69782 | 0.85229 | 1 | 17492 | 1 | -1.4655 |
| MCM9 | 5 | 0.64802 | 0.74432 | 0.999994 | 15630 | 1 | 0.39068 | 0.69787 | 0.82653 | 1 | 17493 | 1 | 0.39068 |
| N4BP2 | 6 | 0.021707 | 0.077455 | 0.93914 | 1557 | 4 | -1.4363 | 0.69787 | 0.8523 | 1 | 17494 | 2 | -1.4363 |
| hsa-mir-1912 | 2 | 0.36147 | 0.3852 | 0.999994 | 11360 | 1 | -0.79349 | 0.69825 | 0.69936 | 1 | 17495 | 1 | -0.79349 |
| hsa-mir-3131 | 4 | 0.050973 | 0.1234 | 0.951052 | 3014 | 3 | -3.4491 | 0.69826 | 0.80726 | 1 | 17496 | 1 | -3.4491 |
| GALT | 5 | 0.025278 | 0.078204 | 0.93914 | 1766 | 2 | -0.31288 | 0.69829 | 0.82678 | 1 | 17497 | 2 | -0.31288 |
| C9orf114 | 3 | 0.2949 | 0.42582 | 0.999994 | 10015 | 1 | -0.2818 | 0.69833 | 0.74041 | 1 | 17498 | 1 | -0.2818 |
| TEP1 | 6 | 0.35273 | 0.56708 | 0.999994 | 11211 | 3 | -0.068852 | 0.69834 | 0.85249 | 1 | 17499 | 2 | -0.068852 |
| DSCAML1 | 5 | 0.16037 | 0.3181 | 0.999994 | 6749 | 3 | -1.029 | 0.69835 | 0.82681 | 1 | 17500 | 2 | -1.029 |
| PAQR3 | 6 | 0.030677 | 0.10037 | 0.946394 | 2042 | 4 | -1.3837 | 0.69837 | 0.8525 | 1 | 17501 | 2 | -1.3837 |
| RBBP7 | 6 | 0.0348 | 0.11005 | 0.951052 | 2256 | 3 | -2.6494 | 0.6984 | 0.85251 | 1 | 17502 | 2 | -2.6494 |
| NINJ1 | 6 | 0.039527 | 0.12085 | 0.951052 | 2493 | 2 | -0.22133 | 0.69849 | 0.85255 | 1 | 17503 | 2 | -0.22133 |
| RPRD2 | 6 | 0.00060057 | 0.0031372 | 0.537554 | 118 | 5 | -5.0977 | 0.69855 | 0.85257 | 1 | 17504 | 1 | -5.0977 |
| TMEM173 | 6 | 0.072846 | 0.19267 | 0.974607 | 3956 | 3 | -0.54091 | 0.69876 | 0.85267 | 1 | 17505 | 2 | -0.54091 |
| NFXL1 | 5 | 0.17006 | 0.33224 | 0.999994 | 6985 | 3 | -1.0863 | 0.69878 | 0.82707 | 1 | 17506 | 2 | -1.0863 |
| C16orf3 | 5 | 0.041998 | 0.12089 | 0.951052 | 2613 | 3 | -1.5111 | 0.69884 | 0.82709 | 1 | 17507 | 2 | -1.5111 |
| VPS16 | 5 | 0.18813 | 0.35829 | 0.999994 | 7421 | 2 | -0.11124 | 0.69884 | 0.82709 | 1 | 17508 | 2 | -0.11124 |
| FBXL19 | 5 | 0.69295 | 0.7633 | 0.999994 | 16233 | 1 | 0.061989 | 0.69884 | 0.82709 | 1 | 17509 | 1 | 0.061989 |
| OR8D4 | 5 | 0.46302 | 0.6162 | 0.999994 | 12905 | 2 | 0.15799 | 0.69891 | 0.82714 | 1 | 17510 | 2 | 0.15799 |
| NonTargetingControlG | 2 | 0.065941 | 0.11574 | 0.951052 | 3685 | 1 | -3.1836 | 0.699 | 0.70009 | 1 | 17511 | 1 | -3.1836 |
| C4orf6 | 5 | 0.49737 | 0.6456 | 0.999994 | 13387 | 2 | -0.070581 | 0.69921 | 0.82732 | 1 | 17512 | 2 | -0.070581 |
| hsa-mir-1273g | 1 | 0.3007 | 0.3007 | 0.999994 | 10146 | 1 | -0.99445 | 0.6993 | 0.69935 | 1 | 17513 | 0 | -0.99445 |
| INHBC | 6 | 0.23479 | 0.42981 | 0.999994 | 8638 | 3 | -0.76656 | 0.69944 | 0.85294 | 1 | 17514 | 1 | -0.76656 |
| CHL1 | 6 | 0.061158 | 0.16824 | 0.967126 | 3477 | 2 | 0.064344 | 0.69944 | 0.85294 | 1 | 17515 | 2 | 0.064344 |
| IL10RA | 5 | 0.19324 | 0.36557 | 0.999994 | 7580 | 2 | 0.54997 | 0.69977 | 0.82768 | 1 | 17516 | 2 | 0.54997 |
| ZC4H2 | 5 | 0.43966 | 0.59653 | 0.999994 | 12569 | 1 | 0.29137 | 0.6998 | 0.8277 | 1 | 17517 | 2 | 0.29137 |
| GBP6 | 6 | 0.26021 | 0.46011 | 0.999994 | 9248 | 3 | -0.72766 | 0.69981 | 0.8531 | 1 | 17518 | 2 | -0.72766 |
| NUDT7 | 5 | 0.15992 | 0.31746 | 0.999994 | 6734 | 3 | -1.021 | 0.69981 | 0.82771 | 1 | 17519 | 2 | -1.021 |
| hsa-mir-4750 | 1 | 0.30017 | 0.30016 | 0.999994 | 10135 | 1 | -1.1385 | 0.69983 | 0.69987 | 1 | 17520 | 0 | -1.1385 |
| SOAT1 | 5 | 0.18702 | 0.35671 | 0.999994 | 7390 | 3 | -0.95195 | 0.69992 | 0.82778 | 1 | 17521 | 2 | -0.95195 |
| SEMA4B | 5 | 0.73039 | 0.78084 | 0.999994 | 16604 | 1 | 0.26131 | 0.69997 | 0.82781 | 1 | 17522 | 2 | 0.26131 |
| TMEM244 | 5 | 0.03095 | 0.09294 | 0.943968 | 2060 | 3 | -1.7915 | 0.70011 | 0.8279 | 1 | 17523 | 2 | -1.7915 |

| | | | | | | | | | | | | | |
|---|---|---|---|---|---|---|---|---|---|---|---|---|---|
| NCR3LG1 | 5 | 0.10459 | 0.23356 | 0.984971 | 5121 | 3 | -1.0994 | 0.70026 | 0.82799 | 1 | 17524 | 2 | -1.0994 |
| TMEM70 | 6 | 0.18366 | 0.3677 | 0.999994 | 7305 | 3 | -1.0191 | 0.70045 | 0.85337 | 1 | 17525 | 2 | -1.0191 |
| ECSIT | 5 | 0.19663 | 0.37043 | 0.999994 | 7666 | 2 | -0.35502 | 0.70052 | 0.82816 | 1 | 17526 | 2 | -0.35502 |
| hsa-mir-5002 | 4 | 0.25082 | 0.39275 | 0.999994 | 9030 | 2 | -0.94177 | 0.7006 | 0.80801 | 1 | 17527 | 1 | -0.94177 |
| RTTN | 6 | 0.017881 | 0.065127 | 0.93914 | 1355 | 5 | -1.7318 | 0.70064 | 0.85345 | 1 | 17528 | 1 | -1.7318 |
| CD1D | 6 | 0.027251 | 0.092345 | 0.943968 | 1867 | 4 | -1.8155 | 0.70064 | 0.85345 | 1 | 17529 | 2 | -1.8155 |
| LPGAT1 | 6 | 0.031852 | 0.10315 | 0.946414 | 2097 | 2 | 0.22799 | 0.70079 | 0.85353 | 1 | 17530 | 2 | 0.22799 |
| KIAA0753 | 5 | 0.022649 | 0.071249 | 0.93914 | 1627 | 3 | -0.88336 | 0.7008 | 0.82832 | 1 | 17531 | 2 | -0.88336 |
| NonTargetingControlG | 2 | 0.065339 | 0.11474 | 0.951052 | 3669 | 1 | -4.0163 | 0.70083 | 0.70178 | 1 | 17532 | 1 | -4.0163 |
| ITGAE | 5 | 0.11435 | 0.24873 | 0.991042 | 5459 | 3 | -1.2709 | 0.70113 | 0.82853 | 1 | 17533 | 2 | -1.2709 |
| PRSS46 | 6 | 0.032698 | 0.10511 | 0.95013 | 2144 | 4 | -2.6022 | 0.70117 | 0.8537 | 1 | 17534 | 1 | -2.6022 |
| YY1 | 4 | 0.54187 | 0.63576 | 0.999994 | 14088 | 1 | -0.099603 | 0.70123 | 0.80824 | 1 | 17535 | 1 | -0.099603 |
| NonTargetingControlG | 2 | 0.51705 | 0.51966 | 0.999994 | 13692 | 1 | -0.10319 | 0.70126 | 0.70218 | 1 | 17536 | 1 | -0.10319 |
| TAPBP | 5 | 0.21432 | 0.3954 | 0.999994 | 8125 | 3 | -0.75233 | 0.70132 | 0.82866 | 1 | 17537 | 2 | -0.75233 |
| PMVK | 4 | 0.11814 | 0.24698 | 0.990269 | 5577 | 3 | -1.4738 | 0.70141 | 0.80828 | 1 | 17538 | 1 | -1.4738 |
| ENAM | 4 | 0.052905 | 0.12716 | 0.951052 | 3107 | 2 | -3.0643 | 0.70149 | 0.80832 | 1 | 17539 | 1 | -3.0643 |
| LRRC73 | 5 | 0.89167 | 0.8918 | 0.999994 | 19065 | 0 | 0.092465 | 0.70152 | 0.82878 | 1 | 17540 | 2 | 0.092465 |
| hsa-mir-4759 | 3 | 0.55866 | 0.58643 | 0.999994 | 14343 | 1 | 0.46352 | 0.70157 | 0.74231 | 1 | 17541 | 1 | 0.46352 |
| KRT19 | 3 | 0.46409 | 0.52301 | 0.999994 | 12921 | 1 | 0.29367 | 0.70164 | 0.74234 | 1 | 17542 | 1 | 0.29367 |
| C11orf24 | 6 | 0.20766 | 0.39723 | 0.999994 | 7966 | 3 | -1.4043 | 0.70182 | 0.85397 | 1 | 17543 | 2 | -1.4043 |
| DPPA2 | 6 | 0.083137 | 0.21372 | 0.979587 | 4343 | 4 | -1.4722 | 0.70182 | 0.85397 | 1 | 17544 | 2 | -1.4722 |
| BBS1 | 5 | 0.035081 | 0.10345 | 0.946414 | 2272 | 4 | -2.3686 | 0.70184 | 0.82898 | 1 | 17545 | 1 | -2.3686 |
| NonTargetingControlG | 2 | 0.66876 | 0.66854 | 0.999994 | 15903 | 0 | 0.36078 | 0.7022 | 0.70304 | 1 | 17546 | 1 | 0.36078 |
| NELL1 | 5 | 0.095933 | 0.22 | 0.980699 | 4812 | 3 | -1.667 | 0.7023 | 0.82927 | 1 | 17547 | 2 | -1.667 |
| FSD2 | 6 | 0.30755 | 0.51553 | 0.999994 | 10278 | 3 | -0.61003 | 0.70233 | 0.85418 | 1 | 17548 | 2 | -0.61003 |
| hsa-mir-7851 | 2 | 0.76123 | 0.76073 | 0.999994 | 16992 | 0 | 0.67178 | 0.70246 | 0.70327 | 1 | 17549 | 1 | 0.67178 |
| NonTargetingControlG | 2 | 0.50866 | 0.51187 | 0.999994 | 13563 | 1 | -0.12244 | 0.70264 | 0.70343 | 1 | 17550 | 1 | -0.12244 |
| DZANK1 | 6 | 0.3659 | 0.58064 | 0.999994 | 11439 | 2 | -0.38234 | 0.70275 | 0.85437 | 1 | 17551 | 2 | -0.38234 |
| PGLS | 4 | 0.19433 | 0.33482 | 0.999994 | 7616 | 2 | -0.72781 | 0.7029 | 0.80879 | 1 | 17552 | 1 | -0.72781 |
| NonTargetingControlG | 2 | 0.66425 | 0.664 | 0.999994 | 15845 | 0 | 0.33364 | 0.70293 | 0.70371 | 1 | 17553 | 1 | 0.33364 |
| CCBL2 | 6 | 0.034185 | 0.10859 | 0.951052 | 2224 | 3 | -1.7171 | 0.70306 | 0.85448 | 1 | 17554 | 1 | -1.7171 |
| TNPO1 | 6 | 0.086443 | 0.22038 | 0.980699 | 4467 | 4 | -1.1278 | 0.70306 | 0.85448 | 1 | 17555 | 1 | -1.1278 |
| CDH6 | 6 | 0.84225 | 0.86782 | 0.999994 | 18129 | 1 | -0.062397 | 0.70306 | 0.85448 | 1 | 17556 | 2 | -0.062397 |
| AK4 | 5 | 0.54625 | 0.68841 | 0.999994 | 14154 | 2 | 0.27883 | 0.70329 | 0.82988 | 1 | 17557 | 2 | 0.27883 |
| CHADL | 6 | 0.18686 | 0.37163 | 0.999994 | 7384 | 3 | -0.52088 | 0.70339 | 0.85464 | 1 | 17558 | 2 | -0.52088 |
| SEC16B | 5 | 0.071608 | 0.18099 | 0.968171 | 3901 | 3 | -2.0726 | 0.70354 | 0.83004 | 1 | 17559 | 1 | -2.0726 |
| hsa-mir-376a-2 | 3 | 0.70817 | 0.70869 | 0.999994 | 16389 | 1 | 0.41101 | 0.70354 | 0.7435 | 1 | 17560 | 1 | 0.41101 |
| CENPI | 6 | 0.060999 | 0.1679 | 0.967126 | 3469 | 4 | -1.908 | 0.70363 | 0.85475 | 1 | 17561 | 2 | -1.908 |
| CYP26B1 | 6 | 0.61926 | 0.75428 | 0.999994 | 15210 | 2 | -0.48205 | 0.70367 | 0.85477 | 1 | 17562 | 2 | -0.48205 |
| hsa-mir-4473 | 4 | 0.4089 | 0.55484 | 0.999994 | 12091 | 2 | -0.32122 | 0.70379 | 0.80911 | 1 | 17563 | 1 | -0.32122 |
| NonTargetingControlG | 1 | 0.29594 | 0.29589 | 0.999994 | 10036 | 1 | -1.0457 | 0.70406 | 0.70415 | 1 | 17564 | 0 | -1.0457 |
| TGIF2LY | 6 | 0.03604 | 0.11295 | 0.951052 | 2320 | 4 | -1.7645 | 0.70426 | 0.85502 | 1 | 17565 | 2 | -1.7645 |
| ZNF341 | 6 | 0.36735 | 0.58157 | 0.999994 | 11463 | 2 | -0.074415 | 0.70436 | 0.85506 | 1 | 17566 | 2 | -0.074415 |
| SOWAHC | 5 | 0.0047091 | 0.019659 | 0.808847 | 499 | 3 | -3.1386 | 0.70446 | 0.8306 | 1 | 17567 | 1 | -3.1386 |
| CERS5 | 5 | 0.031601 | 0.094589 | 0.945031 | 2083 | 4 | -1.0726 | 0.70446 | 0.8306 | 1 | 17568 | 1 | -1.0726 |
| TCEANC2 | 5 | 0.38688 | 0.55252 | 0.999994 | 11751 | 1 | 0.49449 | 0.70446 | 0.8306 | 1 | 17569 | 2 | 0.49449 |
| COPS7B | 5 | 0.24101 | 0.43229 | 0.999994 | 8794 | 3 | -0.56616 | 0.70446 | 0.8306 | 1 | 17570 | 1 | -0.56616 |
| PPCDC | 5 | 0.45621 | 0.61048 | 0.999994 | 12794 | 2 | -0.034007 | 0.70446 | 0.8306 | 1 | 17571 | 1 | -0.034007 |
| ADAMTS14 | 5 | 0.048882 | 0.13769 | 0.957317 | 2925 | 4 | -1.0276 | 0.70446 | 0.8306 | 1 | 17572 | 1 | -1.0276 |
| hsa-mir-423 | 1 | 0.29546 | 0.29542 | 0.999994 | 10028 | 1 | -1.2 | 0.70454 | 0.70463 | 1 | 17573 | 0 | -1.2 |
| SPHAR | 1 | 0.29536 | 0.2953 | 0.999994 | 10021 | 1 | -1.1068 | 0.70464 | 0.70473 | 1 | 17574 | 0 | -1.1068 |
| S100A12 | 5 | 0.92926 | 0.92953 | 0.999994 | 19840 | 0 | 0.41963 | 0.70467 | 0.83073 | 1 | 17575 | 2 | 0.41963 |

| | | | | | | | | | | | | | |
|---|---|---|---|---|---|---|---|---|---|---|---|---|---|
| BCL2L2-PABPN1 | 6 | 0.020953 | 0.075035 | 0.93914 | 1509 | 5 | -1.9832 | 0.70476 | 0.85523 | 1 | 17576 | 1 | -1.9832 |
| CNN2 | 6 | 0.14902 | 0.32439 | 0.999994 | 6429 | 2 | -0.1094 | 0.70476 | 0.85523 | 1 | 17577 | 2 | -0.1094 |
| FUBP3 | 6 | 0.096544 | 0.24056 | 0.987253 | 4841 | 4 | -0.91229 | 0.70476 | 0.85523 | 1 | 17578 | 1 | -0.91229 |
| ACOXL | 6 | 0.93678 | 0.9368 | 0.999994 | 20003 | 0 | 0.2265 | 0.70476 | 0.85523 | 1 | 17579 | 2 | 0.2265 |
| DDT | 6 | 0.0010501 | 0.0053332 | 0.60466 | 176 | 5 | -1.9952 | 0.70476 | 0.85523 | 1 | 17580 | 1 | -1.9952 |
| TMEM26 | 6 | 0.091485 | 0.23051 | 0.983933 | 4655 | 3 | -0.65838 | 0.70476 | 0.85523 | 1 | 17581 | 1 | -0.65838 |
| NonTargetingControlG | 2 | 0.63921 | 0.63905 | 0.999994 | 15502 | 0 | 0.31723 | 0.70501 | 0.70569 | 1 | 17582 | 1 | 0.31723 |
| KRTAP19-4 | 5 | 0.20162 | 0.37755 | 0.999994 | 7812 | 2 | 0.48068 | 0.70512 | 0.83102 | 1 | 17583 | 2 | 0.48068 |
| E2F5 | 6 | 0.015972 | 0.058953 | 0.931872 | 1244 | 3 | -2.0915 | 0.70529 | 0.85546 | 1 | 17584 | 2 | -2.0915 |
| RNF151 | 6 | 0.55005 | 0.70367 | 0.999994 | 14212 | 2 | -0.2808 | 0.70534 | 0.85548 | 1 | 17585 | 2 | -0.2808 |
| ZNF701 | 3 | 0.1913 | 0.30176 | 0.999994 | 7521 | 2 | -1.1004 | 0.70543 | 0.74461 | 1 | 17586 | 1 | -1.1004 |
| DAPK1 | 6 | 0.26076 | 0.46077 | 0.999994 | 9257 | 3 | -0.70608 | 0.70548 | 0.85554 | 1 | 17587 | 2 | -0.70608 |
| GAA | 5 | 0.089821 | 0.2103 | 0.979587 | 4586 | 3 | -1.7361 | 0.70553 | 0.83129 | 1 | 17588 | 2 | -1.7361 |
| GAGE12H | 2 | 0.05838 | 0.10321 | 0.946414 | 3359 | 1 | -4.0894 | 0.70557 | 0.7062 | 1 | 17589 | 1 | -4.0894 |
| HIBADH | 6 | 0.079717 | 0.20672 | 0.978744 | 4220 | 4 | -1.2738 | 0.70564 | 0.85562 | 1 | 17590 | 2 | -1.2738 |
| SLC18A2 | 5 | 0.86945 | 0.8706 | 0.999994 | 18629 | 1 | 0.00029969 | 0.70588 | 0.83149 | 1 | 17591 | 2 | 0.00029969 |
| CDH12 | 5 | 0.10367 | 0.23211 | 0.984502 | 5088 | 3 | -1.5255 | 0.70588 | 0.83149 | 1 | 17592 | 2 | -1.5255 |
| FAM195B | 5 | 0.48885 | 0.63821 | 0.999994 | 13264 | 2 | -0.58381 | 0.70588 | 0.83149 | 1 | 17593 | 2 | -0.58381 |
| ST8SIA1 | 6 | 0.55221 | 0.70525 | 0.999994 | 14247 | 2 | -0.528 | 0.70601 | 0.85578 | 1 | 17594 | 1 | -0.528 |
| NonTargetingControlG | 2 | 0.222 | 0.27806 | 0.999536 | 8331 | 1 | -1.4407 | 0.70613 | 0.70672 | 1 | 17595 | 1 | -1.4407 |
| SLC6A18 | 4 | 0.14235 | 0.28132 | 0.999632 | 6246 | 2 | -0.83291 | 0.70621 | 0.80992 | 1 | 17596 | 1 | -0.83291 |
| ETV1 | 2 | 0.21152 | 0.27044 | 0.998561 | 8058 | 1 | -1.619 | 0.7063 | 0.7069 | 1 | 17597 | 1 | -1.619 |
| MRO | 4 | 0.079917 | 0.17855 | 0.968171 | 4231 | 3 | -1.3059 | 0.70632 | 0.80996 | 1 | 17598 | 1 | -1.3059 |
| SLC35A2 | 5 | 0.53583 | 0.67918 | 0.999994 | 13994 | 2 | 0.39187 | 0.70652 | 0.83188 | 1 | 17599 | 2 | 0.39187 |
| MEX3D | 5 | 0.24116 | 0.43249 | 0.999994 | 8801 | 3 | -0.63109 | 0.70652 | 0.83188 | 1 | 17600 | 1 | -0.63109 |
| PSMD12 | 5 | 0.0082305 | 0.030965 | 0.862223 | 747 | 3 | -6.213 | 0.70652 | 0.83188 | 1 | 17601 | 1 | -6.213 |
| NonTargetingControlG | 1 | 0.29328 | 0.29319 | 0.999994 | 9980 | 1 | -0.90385 | 0.70672 | 0.70681 | 1 | 17602 | 0 | -0.90385 |
| DCSTAMP | 5 | 0.21036 | 0.38989 | 0.999994 | 8029 | 3 | -0.79244 | 0.70689 | 0.83211 | 1 | 17603 | 2 | -0.79244 |
| MMP3 | 5 | 0.13543 | 0.28087 | 0.999632 | 6065 | 3 | -1.1827 | 0.70708 | 0.83223 | 1 | 17604 | 2 | -1.1827 |
| LTA | 5 | 0.17165 | 0.33455 | 0.999994 | 7028 | 3 | -0.96239 | 0.70711 | 0.83225 | 1 | 17605 | 2 | -0.96239 |
| XKR6 | 5 | 0.14301 | 0.29237 | 0.999632 | 6273 | 3 | -1.1825 | 0.70714 | 0.83226 | 1 | 17606 | 2 | -1.1825 |
| CCL8 | 6 | 0.074962 | 0.19708 | 0.977164 | 4034 | 4 | -1.3494 | 0.70718 | 0.85627 | 1 | 17607 | 2 | -1.3494 |
| MYH8 | 4 | 0.88123 | 0.88084 | 0.999994 | 18855 | 0 | 0.23565 | 0.70731 | 0.81031 | 1 | 17608 | 1 | 0.23565 |
| SLC26A7 | 6 | 0.099059 | 0.24553 | 0.98979 | 4932 | 4 | -0.86321 | 0.70753 | 0.85642 | 1 | 17609 | 2 | -0.86321 |
| NonTargetingControlG | 2 | 0.54973 | 0.55062 | 0.999994 | 14210 | 1 | 0.010334 | 0.70761 | 0.70813 | 1 | 17610 | 1 | 0.010334 |
| MMACHC | 4 | 0.072326 | 0.16433 | 0.967126 | 3932 | 3 | -1.7301 | 0.70776 | 0.81046 | 1 | 17611 | 1 | -1.7301 |
| IFNA2 | 6 | 0.64691 | 0.77525 | 0.999994 | 15617 | 2 | 0.13052 | 0.7078 | 0.85654 | 1 | 17612 | 2 | 0.13052 |
| hsa-mir-4660 | 3 | 0.24645 | 0.37055 | 0.999994 | 8933 | 1 | 0.18124 | 0.70781 | 0.74603 | 1 | 17613 | 1 | 0.18124 |
| GLT1D1 | 5 | 0.0037422 | 0.015776 | 0.766139 | 425 | 2 | 0.10448 | 0.70787 | 0.83274 | 1 | 17614 | 2 | 0.10448 |
| ILF2 | 5 | 0.17957 | 0.34601 | 0.999994 | 7212 | 2 | -0.62137 | 0.70792 | 0.83277 | 1 | 17615 | 2 | -0.62137 |
| AGAP4 | 5 | 0.72653 | 0.77901 | 0.999994 | 16561 | 1 | 0.53773 | 0.70793 | 0.83278 | 1 | 17616 | 2 | 0.53773 |
| CPOX | 2 | 0.075212 | 0.13084 | 0.95318 | 4043 | 1 | -3.908 | 0.70802 | 0.70851 | 1 | 17617 | 1 | -3.908 |
| CD1B | 5 | 0.17272 | 0.33609 | 0.999994 | 7057 | 3 | -0.94486 | 0.70811 | 0.8329 | 1 | 17618 | 2 | -0.94486 |
| RNF175 | 6 | 0.047202 | 0.13795 | 0.957317 | 2864 | 2 | 0.52999 | 0.70822 | 0.85673 | 1 | 17619 | 2 | 0.52999 |
| CDHR3 | 6 | 0.10694 | 0.2608 | 0.994943 | 5210 | 3 | -0.68658 | 0.70822 | 0.85673 | 1 | 17620 | 2 | -0.68658 |
| NonTargetingControlG | 2 | 0.29012 | 0.32922 | 0.999994 | 9910 | 1 | -1.1518 | 0.70822 | 0.7087 | 1 | 17621 | 1 | -1.1518 |
| KDM2B | 4 | 0.04015 | 0.10209 | 0.946414 | 2526 | 2 | -0.92372 | 0.70825 | 0.81063 | 1 | 17622 | 1 | -0.92372 |
| C6orf25 | 3 | 0.2361 | 0.35772 | 0.999994 | 8673 | 2 | -0.89691 | 0.70828 | 0.74632 | 1 | 17623 | 1 | -0.89691 |
| NonTargetingControlG | 2 | 0.74763 | 0.74727 | 0.999994 | 16835 | 0 | 0.65601 | 0.70857 | 0.70903 | 1 | 17624 | 1 | 0.65601 |
| STARD3NL | 6 | 0.29199 | 0.49736 | 0.999994 | 9948 | 2 | 0.054878 | 0.70858 | 0.85687 | 1 | 17625 | 2 | 0.054878 |
| KCNK7 | 6 | 0.084072 | 0.21564 | 0.980236 | 4379 | 4 | -1.1124 | 0.70858 | 0.85687 | 1 | 17626 | 1 | -1.1124 |
| TIGD5 | 6 | 0.063321 | 0.17279 | 0.96793 | 3570 | 4 | -1.1533 | 0.70858 | 0.85687 | 1 | 17627 | 2 | -1.1533 |

| | | | | | | | | | | | | | |
|---|---|---|---|---|---|---|---|---|---|---|---|---|---|
| REPS2 | 5 | 0.0053505 | 0.022192 | 0.819701 | 545 | 4 | -2.7379 | 0.70859 | 0.8332 | 1 | 17628 | 1 | -2.7379 |
| NSUN4 | 5 | 0.06097 | 0.16369 | 0.967126 | 3464 | 2 | -0.23322 | 0.70859 | 0.8332 | 1 | 17629 | 1 | -0.23322 |
| RAB32 | 5 | 0.1979 | 0.37224 | 0.999994 | 7709 | 2 | -0.4759 | 0.70859 | 0.8332 | 1 | 17630 | 1 | -0.4759 |
| CALD1 | 5 | 0.03758 | 0.10981 | 0.951052 | 2408 | 4 | -1.0824 | 0.70859 | 0.8332 | 1 | 17631 | 1 | -1.0824 |
| PIP5K1B | 5 | 0.83899 | 0.8459 | 0.999994 | 18072 | 1 | 0.096856 | 0.70859 | 0.8332 | 1 | 17632 | 1 | 0.096856 |
| NUP62 | 4 | 0.13336 | 0.27204 | 0.998561 | 6004 | 2 | -1.0484 | 0.70879 | 0.81081 | 1 | 17633 | 1 | -1.0484 |
| KIAA1704 | 2 | 0.63361 | 0.63337 | 0.999994 | 15427 | 0 | 0.22693 | 0.7088 | 0.70925 | 1 | 17634 | 1 | 0.22693 |
| SPATA3 | 4 | 0.032694 | 0.086986 | 0.94395 | 2143 | 3 | -1.9681 | 0.70885 | 0.81082 | 1 | 17635 | 1 | -1.9681 |
| CDCA7L | 6 | 0.23683 | 0.4322 | 0.999994 | 8692 | 3 | -0.68488 | 0.70891 | 0.85702 | 1 | 17636 | 2 | -0.68488 |
| NonTargetingControlG | 2 | 0.020094 | 0.037363 | 0.881473 | 1464 | 1 | -4.7061 | 0.70896 | 0.70941 | 1 | 17637 | 1 | -4.7061 |
| ARL11 | 6 | 0.0852 | 0.21791 | 0.980699 | 4417 | 2 | -0.020101 | 0.70913 | 0.85712 | 1 | 17638 | 2 | -0.020101 |
| IPO13 | 3 | 0.19197 | 0.30259 | 0.999994 | 7546 | 2 | -1.3266 | 0.70957 | 0.74712 | 1 | 17639 | 1 | -1.3266 |
| DRG2 | 2 | 0.47276 | 0.47939 | 0.999994 | 13042 | 1 | -0.26043 | 0.70959 | 0.71002 | 1 | 17640 | 1 | -0.26043 |
| FAM160B2 | 6 | 0.097824 | 0.24306 | 0.988088 | 4883 | 4 | -1.2607 | 0.70973 | 0.85739 | 1 | 17641 | 2 | -1.2607 |
| TRIM49B | 2 | 0.69767 | 0.69745 | 0.999994 | 16273 | 0 | 0.47081 | 0.70975 | 0.71016 | 1 | 17642 | 1 | 0.47081 |
| MCM8 | 5 | 0.060393 | 0.16275 | 0.967126 | 3438 | 3 | -0.83138 | 0.71004 | 0.83412 | 1 | 17643 | 2 | -0.83138 |
| SIPA1L2 | 6 | 0.091099 | 0.22971 | 0.983354 | 4641 | 3 | -0.96852 | 0.71023 | 0.8576 | 1 | 17644 | 2 | -0.96852 |
| ZFP106 | 6 | 0.0047619 | 0.020284 | 0.810158 | 503 | 5 | -2.4525 | 0.71023 | 0.8576 | 1 | 17645 | 1 | -2.4525 |
| NDUFAF7 | 4 | 0.059017 | 0.13889 | 0.957514 | 3382 | 2 | -1.8389 | 0.71033 | 0.81132 | 1 | 17646 | 1 | -1.8389 |
| GYPA | 5 | 0.45163 | 0.6066 | 0.999994 | 12728 | 2 | -0.39502 | 0.71034 | 0.83429 | 1 | 17647 | 1 | -0.39502 |
| TSPY8 | 1 | 0.28965 | 0.28957 | 0.999632 | 9905 | 1 | -1.099 | 0.71035 | 0.71047 | 1 | 17648 | 0 | -1.099 |
| ADCY9 | 6 | 0.34828 | 0.56207 | 0.999994 | 11118 | 3 | -0.15388 | 0.71063 | 0.85778 | 1 | 17649 | 2 | -0.15388 |
| hsa-mir-135b | 4 | 0.33385 | 0.47763 | 0.999994 | 10809 | 2 | -0.63913 | 0.71071 | 0.81144 | 1 | 17650 | 1 | -0.63913 |
| TESK1 | 6 | 0.016191 | 0.059635 | 0.932932 | 1254 | 4 | -1.8543 | 0.71077 | 0.85784 | 1 | 17651 | 2 | -1.8543 |
| STARD3 | 4 | 0.031313 | 0.084163 | 0.94395 | 2074 | 3 | -3.9211 | 0.71077 | 0.81146 | 1 | 17652 | 1 | -3.9211 |
| TLR1 | 5 | 0.21445 | 0.39559 | 0.999994 | 8131 | 2 | -0.48717 | 0.711 | 0.83469 | 1 | 17653 | 2 | -0.48717 |
| SLC35F3 | 5 | 0.52321 | 0.66812 | 0.999994 | 13797 | 2 | 0.34899 | 0.711 | 0.83469 | 1 | 17654 | 2 | 0.34899 |
| DPEP3 | 5 | 0.363 | 0.53254 | 0.999994 | 11388 | 2 | 0.1486 | 0.711 | 0.83469 | 1 | 17655 | 2 | 0.1486 |
| PCDHB2 | 5 | 0.31849 | 0.49611 | 0.999994 | 10485 | 2 | -0.12522 | 0.711 | 0.83469 | 1 | 17656 | 1 | -0.12522 |
| SFRP1 | 5 | 0.011146 | 0.039387 | 0.891621 | 935 | 3 | -0.80597 | 0.711 | 0.83469 | 1 | 17657 | 1 | -0.80597 |
| TXLNA | 6 | 0.65578 | 0.78203 | 0.999994 | 15723 | 2 | 0.36829 | 0.71117 | 0.858 | 1 | 17658 | 2 | 0.36829 |
| C10orf113 | 5 | 0.47621 | 0.62735 | 0.999994 | 13084 | 1 | 0.00092159 | 0.7112 | 0.83481 | 1 | 17659 | 2 | 0.00092159 |
| hsa-mir-4488 | 3 | 0.11122 | 0.19789 | 0.977164 | 5341 | 2 | -2.0749 | 0.71138 | 0.74819 | 1 | 17660 | 1 | -2.0749 |
| NonTargetingControlG | 2 | 0.37021 | 0.39235 | 0.999994 | 11515 | 1 | -0.74274 | 0.71145 | 0.71177 | 1 | 17661 | 1 | -0.74274 |
| C12orf39 | 5 | 0.18341 | 0.35153 | 0.999994 | 7303 | 3 | -0.92415 | 0.71153 | 0.83503 | 1 | 17662 | 2 | -0.92415 |
| EFCAB11 | 5 | 0.029282 | 0.088622 | 0.94395 | 1973 | 4 | -1.9707 | 0.7116 | 0.83508 | 1 | 17663 | 1 | -1.9707 |
| hsa-mir-4329 | 2 | 0.74074 | 0.74041 | 0.999994 | 16747 | 0 | 0.66971 | 0.71171 | 0.71203 | 1 | 17664 | 1 | 0.66971 |
| BRAP | 5 | 0.48777 | 0.63731 | 0.999994 | 13252 | 2 | -0.53357 | 0.71177 | 0.83518 | 1 | 17665 | 2 | -0.53357 |
| OR10G7 | 5 | 0.13867 | 0.28583 | 0.999632 | 6158 | 3 | -1.232 | 0.71179 | 0.83519 | 1 | 17666 | 2 | -1.232 |
| SLC11A1 | 6 | 0.024055 | 0.084752 | 0.94395 | 1711 | 4 | -3.463 | 0.71187 | 0.85831 | 1 | 17667 | 1 | -3.463 |
| VANGL2 | 4 | 0.30465 | 0.44772 | 0.999994 | 10222 | 2 | -0.16926 | 0.7119 | 0.81184 | 1 | 17668 | 1 | -0.16926 |
| NEDD4 | 5 | 0.90023 | 0.90034 | 0.999994 | 19224 | 0 | 0.38331 | 0.7119 | 0.83527 | 1 | 17669 | 2 | 0.38331 |
| USP17L24 | 2 | 0.59698 | 0.59686 | 0.999994 | 14917 | 0 | 0.043982 | 0.71196 | 0.71227 | 1 | 17670 | 1 | 0.043982 |
| hsa-mir-193b | 4 | 0.73269 | 0.74716 | 0.999994 | 16637 | 1 | 0.33906 | 0.71198 | 0.81186 | 1 | 17671 | 1 | 0.33906 |
| NonTargetingControlG | 2 | 0.14978 | 0.22649 | 0.983354 | 6455 | 1 | -3.2028 | 0.71227 | 0.71257 | 1 | 17672 | 1 | -3.2028 |
| hsa-mir-4476 | 3 | 0.24709 | 0.37132 | 0.999994 | 8951 | 2 | -0.83079 | 0.71239 | 0.7488 | 1 | 17673 | 1 | -0.83079 |
| hsa-mir-1253 | 4 | 0.82813 | 0.8277 | 0.999994 | 17900 | 0 | -0.15067 | 0.71241 | 0.81202 | 1 | 17674 | 1 | -0.15067 |
| ANKRD20A2 | 1 | 0.28746 | 0.28734 | 0.999632 | 9866 | 1 | -1.009 | 0.71254 | 0.71265 | 1 | 17675 | 0 | -1.009 |
| SLC9B1 | 5 | 0.15365 | 0.30823 | 0.999994 | 6553 | 3 | -1.0549 | 0.71272 | 0.83579 | 1 | 17676 | 2 | -1.0549 |
| hsa-mir-548h-1 | 4 | 0.058745 | 0.13837 | 0.957317 | 3372 | 3 | -2.7459 | 0.71301 | 0.81223 | 1 | 17677 | 1 | -2.7459 |
| GNL2 | 4 | 0.45376 | 0.59626 | 0.999994 | 12757 | 1 | -0.566 | 0.71306 | 0.81224 | 1 | 17678 | 1 | -0.566 |
| NOL9 | 6 | 0.12106 | 0.28774 | 0.999632 | 5664 | 3 | -0.31275 | 0.7131 | 0.8588 | 1 | 17679 | 2 | -0.31275 |

| | | | | | | | | | | | | | |
|---|---|---|---|---|---|---|---|---|---|---|---|---|---|
| LIX1L | 5 | 0.11084 | 0.24327 | 0.988088 | 5326 | 2 | 0.27294 | 0.7131 | 0.83603 | 1 | 17680 | 2 | 0.27294 |
| NAMPT | 5 | 0.11805 | 0.2544 | 0.994835 | 5574 | 3 | -0.82311 | 0.71312 | 0.83605 | 1 | 17681 | 1 | -0.82311 |
| CLEC18A | 5 | 0.053976 | 0.14999 | 0.966359 | 3157 | 4 | -0.8154 | 0.71312 | 0.83605 | 1 | 17682 | 1 | -0.8154 |
| ZNF750 | 6 | 0.56527 | 0.71457 | 0.999994 | 14472 | 1 | 0.13861 | 0.71323 | 0.85886 | 1 | 17683 | 1 | 0.13861 |
| RANBP2 | 6 | 0.0511 | 0.14647 | 0.964852 | 3023 | 4 | -1.3562 | 0.71323 | 0.85886 | 1 | 17684 | 1 | -1.3562 |
| CHCHD2 | 6 | 0.00065894 | 0.0034124 | 0.541338 | 125 | 5 | -3.3497 | 0.71323 | 0.85886 | 1 | 17685 | 1 | -3.3497 |
| PARP16 | 5 | 0.45972 | 0.61339 | 0.999994 | 12843 | 1 | -0.13692 | 0.7133 | 0.83616 | 1 | 17686 | 2 | -0.13692 |
| hsa-mir-5699 | 4 | 0.034658 | 0.091022 | 0.94395 | 2247 | 2 | -2.8941 | 0.71339 | 0.81237 | 1 | 17687 | 1 | -2.8941 |
| RBM3 | 6 | 0.20168 | 0.38991 | 0.999994 | 7814 | 3 | -0.61523 | 0.71351 | 0.85898 | 1 | 17688 | 2 | -0.61523 |
| GNL3L | 4 | 0.052295 | 0.12598 | 0.951052 | 3084 | 3 | -2.7927 | 0.71355 | 0.81243 | 1 | 17689 | 1 | -2.7927 |
| AZI2 | 6 | 0.016953 | 0.062113 | 0.937033 | 1299 | 4 | -1.8326 | 0.71365 | 0.85904 | 1 | 17690 | 2 | -1.8326 |
| ZNF792 | 5 | 0.7142 | 0.77309 | 0.999994 | 16442 | 1 | 0.32217 | 0.71368 | 0.83639 | 1 | 17691 | 2 | 0.32217 |
| ELOVL5 | 6 | 0.21666 | 0.40813 | 0.999994 | 8184 | 3 | -0.32418 | 0.71368 | 0.85906 | 1 | 17692 | 2 | -0.32418 |
| hsa-mir-550a-2 | 1 | 0.28629 | 0.28618 | 0.999632 | 9839 | 1 | -1.2099 | 0.71371 | 0.71381 | 1 | 17693 | 0 | -1.2099 |
| PLA2G4C | 6 | 0.10044 | 0.2482 | 0.990682 | 4986 | 4 | -1.0638 | 0.71373 | 0.85908 | 1 | 17694 | 2 | -1.0638 |
| METTL2A | 5 | 0.10953 | 0.24126 | 0.987253 | 5288 | 2 | -0.37729 | 0.71383 | 0.83648 | 1 | 17695 | 2 | -0.37729 |
| OR10G2 | 5 | 0.2294 | 0.41638 | 0.999994 | 8513 | 3 | -0.6204 | 0.71396 | 0.83657 | 1 | 17696 | 2 | -0.6204 |
| DUS3L | 4 | 0.052414 | 0.12622 | 0.951052 | 3088 | 2 | -2.611 | 0.71438 | 0.81272 | 1 | 17697 | 1 | -2.611 |
| RBBP5 | 4 | 0.090792 | 0.19849 | 0.977164 | 4629 | 2 | -1.6042 | 0.71438 | 0.81272 | 1 | 17698 | 1 | -1.6042 |
| KCNV2 | 4 | 0.084809 | 0.18756 | 0.969543 | 4399 | 1 | 0.17353 | 0.71438 | 0.81272 | 1 | 17699 | 1 | 0.17353 |
| ZNF804A | 6 | 0.85194 | 0.87295 | 0.999994 | 18294 | 1 | -0.20657 | 0.71456 | 0.85944 | 1 | 17700 | 2 | -0.20657 |
| NCK2 | 6 | 0.0011947 | 0.006014 | 0.623123 | 187 | 4 | -4.6383 | 0.71474 | 0.85952 | 1 | 17701 | 1 | -4.6383 |
| SQLE | 5 | 0.024666 | 0.076583 | 0.93914 | 1741 | 3 | -1.9238 | 0.71477 | 0.83703 | 1 | 17702 | 2 | -1.9238 |
| hsa-mir-765 | 3 | 0.36481 | 0.46403 | 0.999994 | 11417 | 1 | -0.032061 | 0.71487 | 0.75027 | 1 | 17703 | 1 | -0.032061 |
| FDX1L | 2 | 0.38046 | 0.40067 | 0.999994 | 11654 | 1 | -0.63857 | 0.71511 | 0.71525 | 1 | 17704 | 1 | -0.63857 |
| MCPH1 | 5 | 0.21379 | 0.39468 | 0.999994 | 8107 | 2 | -0.50036 | 0.71513 | 0.83726 | 1 | 17705 | 2 | -0.50036 |
| hsa-mir-1972-2 | 1 | 0.28453 | 0.28439 | 0.999632 | 9797 | 1 | -1.2303 | 0.71547 | 0.71559 | 1 | 17706 | 0 | -1.2303 |
| TYMS | 5 | 0.24027 | 0.4313 | 0.999994 | 8776 | 3 | -0.60263 | 0.71559 | 0.83755 | 1 | 17707 | 1 | -0.60263 |
| FBXO17 | 5 | 0.74008 | 0.78571 | 0.999994 | 16741 | 1 | -0.20678 | 0.71559 | 0.83755 | 1 | 17708 | 2 | -0.20678 |
| CXCR5 | 6 | 0.11161 | 0.2698 | 0.998561 | 5355 | 4 | -0.80711 | 0.71572 | 0.85996 | 1 | 17709 | 2 | -0.80711 |
| SCN8A | 6 | 0.23252 | 0.42713 | 0.999994 | 8583 | 3 | -0.54364 | 0.71578 | 0.85998 | 1 | 17710 | 2 | -0.54364 |
| CLCN7 | 5 | 0.096428 | 0.22077 | 0.98185 | 4833 | 3 | -0.83921 | 0.71582 | 0.83771 | 1 | 17711 | 2 | -0.83921 |
| hsa-mir-662 | 3 | 0.047183 | 0.11178 | 0.951052 | 2862 | 2 | -3.573 | 0.71583 | 0.75088 | 1 | 17712 | 1 | -3.573 |
| CKLF | 4 | 0.85509 | 0.85468 | 0.999994 | 18349 | 0 | -0.080774 | 0.71583 | 0.81325 | 1 | 17713 | 1 | -0.080774 |
| GMCL1 | 6 | 0.22068 | 0.41293 | 0.999994 | 8305 | 1 | 0.37326 | 0.71588 | 0.86002 | 1 | 17714 | 2 | 0.37326 |
| SETX | 6 | 0.054202 | 0.15319 | 0.967126 | 3165 | 2 | -0.042881 | 0.71602 | 0.86008 | 1 | 17715 | 1 | -0.042881 |
| PHKA1 | 6 | 0.077769 | 0.20285 | 0.977973 | 4139 | 1 | -0.42743 | 0.71602 | 0.86008 | 1 | 17716 | 2 | -0.42743 |
| ZC3H14 | 6 | 0.23726 | 0.43271 | 0.999994 | 8703 | 3 | -0.47671 | 0.71602 | 0.86008 | 1 | 17717 | 2 | -0.47671 |
| NonTargetingControlG | 1 | 0.28394 | 0.28381 | 0.999632 | 9782 | 1 | -1.205 | 0.71606 | 0.71619 | 1 | 17718 | 0 | -1.205 |
| ZFP42 | 5 | 0.53468 | 0.67819 | 0.999994 | 13975 | 1 | -0.023189 | 0.71616 | 0.83791 | 1 | 17719 | 2 | -0.023189 |
| GAGE12B | 2 | 0.18848 | 0.25375 | 0.994632 | 7434 | 1 | -1.7559 | 0.71636 | 0.71645 | 1 | 17720 | 1 | -1.7559 |
| TAS2R60 | 6 | 0.11498 | 0.27622 | 0.999379 | 5478 | 2 | -0.063232 | 0.7164 | 0.86024 | 1 | 17721 | 2 | -0.063232 |
| GGCT | 6 | 0.36386 | 0.57937 | 0.999994 | 11402 | 2 | -0.035923 | 0.71641 | 0.86025 | 1 | 17722 | 2 | -0.035923 |
| hsa-mir-3118-3 | 1 | 0.28358 | 0.28344 | 0.999632 | 9776 | 1 | -1.1749 | 0.71642 | 0.71654 | 1 | 17723 | 0 | -1.1749 |
| SASH1 | 6 | 0.15232 | 0.32859 | 0.999994 | 6510 | 2 | -0.23765 | 0.71644 | 0.86026 | 1 | 17724 | 2 | -0.23765 |
| TCEAL5 | 5 | 0.77277 | 0.80328 | 0.999994 | 17118 | 1 | 0.54942 | 0.71664 | 0.83817 | 1 | 17725 | 2 | 0.54942 |
| SPINK4 | 5 | 0.46187 | 0.61522 | 0.999994 | 12881 | 2 | -0.56689 | 0.71683 | 0.83829 | 1 | 17726 | 2 | -0.56689 |
| NPRL2 | 6 | 0.0018849 | 0.0093992 | 0.680529 | 259 | 5 | -4.4767 | 0.71709 | 0.86054 | 1 | 17727 | 1 | -4.4767 |
| LEPREL4 | 6 | 0.30356 | 0.51086 | 0.999994 | 10205 | 2 | 0.32939 | 0.71714 | 0.86057 | 1 | 17728 | 2 | 0.32939 |
| AADAT | 5 | 0.048434 | 0.13666 | 0.956777 | 2904 | 3 | -1.9317 | 0.71714 | 0.83849 | 1 | 17729 | 2 | -1.9317 |
| XPO4 | 4 | 0.076593 | 0.17239 | 0.967658 | 4096 | 3 | -1.1666 | 0.71741 | 0.81378 | 1 | 17730 | 1 | -1.1666 |
| IKBIP | 3 | 0.54732 | 0.57839 | 0.999994 | 14171 | 1 | 0.0054762 | 0.71743 | 0.75185 | 1 | 17731 | 1 | 0.0054762 |

| | | | | | | | | | | | | | |
|---|---|---|---|---|---|---|---|---|---|---|---|---|---|
| DGKE | 5 | 0.36044 | 0.53045 | 0.999994 | 11340 | 2 | -0.40591 | 0.7175 | 0.83872 | 1 | 17732 | 2 | -0.40591 |
| RNASE10 | 6 | 0.066422 | 0.17928 | 0.968171 | 3702 | 2 | -0.1647 | 0.71752 | 0.86072 | 1 | 17733 | 2 | -0.1647 |
| NonTargetingControlG | 2 | 0.54301 | 0.54418 | 0.999994 | 14113 | 1 | -0.046767 | 0.71753 | 0.71756 | 1 | 17734 | 1 | -0.046767 |
| TAF1B | 2 | 0.7587 | 0.75826 | 0.999994 | 16961 | 0 | 0.69039 | 0.71754 | 0.71757 | 1 | 17735 | 1 | 0.69039 |
| TSPYL5 | 5 | 0.039751 | 0.11528 | 0.951052 | 2501 | 4 | -2.0711 | 0.71775 | 0.83888 | 1 | 17736 | 1 | -2.0711 |
| IQGAP3 | 5 | 0.1037 | 0.23215 | 0.984502 | 5090 | 3 | -1.5301 | 0.71775 | 0.83888 | 1 | 17737 | 1 | -1.5301 |
| TRIM64 | 5 | 0.42397 | 0.58341 | 0.999994 | 12309 | 2 | -0.0091725 | 0.71775 | 0.83888 | 1 | 17738 | 1 | -0.0091725 |
| NonTargetingControlG | 2 | 0.20518 | 0.26586 | 0.997535 | 7901 | 1 | -1.517 | 0.71788 | 0.7179 | 1 | 17739 | 1 | -1.517 |
| VCX3B | 3 | 0.3677 | 0.46568 | 0.999994 | 11471 | 1 | 0.38647 | 0.71796 | 0.75218 | 1 | 17740 | 1 | 0.38647 |
| CLDN1 | 6 | 0.067351 | 0.18122 | 0.968171 | 3730 | 4 | -1.2494 | 0.71796 | 0.86091 | 1 | 17741 | 1 | -1.2494 |
| RAB13 | 6 | 0.29121 | 0.49647 | 0.999994 | 9937 | 2 | -0.22911 | 0.71796 | 0.86091 | 1 | 17742 | 1 | -0.22911 |
| B4GALT2 | 6 | 0.16558 | 0.34518 | 0.999994 | 6873 | 3 | -0.18445 | 0.71796 | 0.86091 | 1 | 17743 | 2 | -0.18445 |
| AQP10 | 6 | 0.058845 | 0.1633 | 0.967126 | 3375 | 3 | -0.44643 | 0.71796 | 0.86091 | 1 | 17744 | 1 | -0.44643 |
| FAM134A | 6 | 0.20045 | 0.38841 | 0.999994 | 7772 | 3 | -0.39678 | 0.71819 | 0.86102 | 1 | 17745 | 2 | -0.39678 |
| RFX3 | 6 | 0.44784 | 0.63366 | 0.999994 | 12685 | 2 | 0.10651 | 0.71829 | 0.86106 | 1 | 17746 | 2 | 0.10651 |
| PXT1 | 5 | 0.20247 | 0.37877 | 0.999994 | 7843 | 3 | -0.81046 | 0.71871 | 0.83947 | 1 | 17747 | 2 | -0.81046 |
| C17orf89 | 5 | 0.41911 | 0.57934 | 0.999994 | 12236 | 2 | 0.31463 | 0.71909 | 0.83972 | 1 | 17748 | 2 | 0.31463 |
| SCT | 6 | 0.066542 | 0.17952 | 0.968171 | 3707 | 4 | -1.9077 | 0.7191 | 0.86142 | 1 | 17749 | 2 | -1.9077 |
| TMEM217 | 6 | 0.032265 | 0.10412 | 0.947284 | 2117 | 4 | -1.9883 | 0.71924 | 0.86148 | 1 | 17750 | 2 | -1.9883 |
| COG2 | 6 | 0.012169 | 0.04628 | 0.892434 | 1005 | 4 | -3.5292 | 0.71924 | 0.86148 | 1 | 17751 | 1 | -3.5292 |
| ACLY | 5 | 0.062577 | 0.16631 | 0.967126 | 3539 | 2 | 0.066594 | 0.71927 | 0.83982 | 1 | 17752 | 2 | 0.066594 |
| MFGE8 | 5 | 0.35668 | 0.52735 | 0.999994 | 11279 | 2 | -0.44393 | 0.7193 | 0.83984 | 1 | 17753 | 2 | -0.44393 |
| GAR1 | 4 | 0.042988 | 0.10775 | 0.951052 | 2677 | 3 | -2.9961 | 0.71944 | 0.81452 | 1 | 17754 | 1 | -2.9961 |
| RAD51AP2 | 5 | 0.0035072 | 0.014864 | 0.755578 | 407 | 2 | 0.43283 | 0.71952 | 0.83998 | 1 | 17755 | 2 | 0.43283 |
| PLAC8 | 5 | 0.069103 | 0.17696 | 0.968171 | 3803 | 2 | -0.28825 | 0.7197 | 0.84008 | 1 | 17756 | 2 | -0.28825 |
| EBF3 | 4 | 0.098245 | 0.21193 | 0.979587 | 4905 | 2 | -1.7808 | 0.7199 | 0.81469 | 1 | 17757 | 1 | -1.7808 |
| ZRANB2 | 5 | 0.0096498 | 0.035067 | 0.867348 | 849 | 3 | -0.84918 | 0.71996 | 0.84024 | 1 | 17758 | 2 | -0.84918 |
| IL10 | 6 | 0.57353 | 0.72049 | 0.999994 | 14599 | 2 | 0.38692 | 0.72006 | 0.86183 | 1 | 17759 | 2 | 0.38692 |
| TSC2 | 5 | 0.073085 | 0.1834 | 0.968526 | 3962 | 2 | -0.30173 | 0.72025 | 0.84042 | 1 | 17760 | 2 | -0.30173 |
| hsa-mir-3119-1 | 4 | 0.12183 | 0.25345 | 0.994501 | 5696 | 3 | -0.98861 | 0.72036 | 0.81486 | 1 | 17761 | 1 | -0.98861 |
| RNF165 | 5 | 0.049592 | 0.13933 | 0.958457 | 2959 | 4 | -1.6878 | 0.72045 | 0.84056 | 1 | 17762 | 1 | -1.6878 |
| NonTargetingControlG | 2 | 0.34622 | 0.37302 | 0.999994 | 11078 | 1 | -0.90728 | 0.72055 | 0.72051 | 1 | 17763 | 1 | -0.90728 |
| C19orf67 | 5 | 0.13847 | 0.28552 | 0.999632 | 6151 | 1 | 0.030634 | 0.72066 | 0.8407 | 1 | 17764 | 2 | 0.030634 |
| ABTB1 | 6 | 0.24774 | 0.44523 | 0.999994 | 8965 | 3 | -0.60002 | 0.72073 | 0.86211 | 1 | 17765 | 2 | -0.60002 |
| GALR3 | 6 | 0.019916 | 0.071688 | 0.93914 | 1456 | 5 | -1.8723 | 0.72073 | 0.86211 | 1 | 17766 | 1 | -1.8723 |
| HIST1H2BL | 6 | 0.017296 | 0.063213 | 0.937389 | 1326 | 4 | -3.7398 | 0.72073 | 0.86211 | 1 | 17767 | 2 | -3.7398 |
| DNAJB5 | 6 | 0.0063564 | 0.0261 | 0.848412 | 617 | 5 | -4.5969 | 0.72073 | 0.86211 | 1 | 17768 | 1 | -4.5969 |
| ASB4 | 6 | 0.015676 | 0.057977 | 0.929685 | 1227 | 4 | -1.2805 | 0.72088 | 0.86218 | 1 | 17769 | 2 | -1.2805 |
| CHM | 3 | 0.25328 | 0.37905 | 0.999994 | 9088 | 2 | -0.85246 | 0.721 | 0.75408 | 1 | 17770 | 1 | -0.85246 |
| MUC15 | 5 | 0.16296 | 0.32187 | 0.999994 | 6817 | 2 | -0.56781 | 0.72106 | 0.84097 | 1 | 17771 | 2 | -0.56781 |
| NonTargetingControlG | 2 | 0.56677 | 0.56708 | 0.999994 | 14494 | 1 | -0.019585 | 0.72137 | 0.7213 | 1 | 17772 | 1 | -0.019585 |
| SLC22A23 | 6 | 0.034751 | 0.10993 | 0.951052 | 2253 | 4 | -2.206 | 0.72139 | 0.86239 | 1 | 17773 | 1 | -2.206 |
| MNF1 | 6 | 0.12219 | 0.28989 | 0.999632 | 5709 | 4 | -1.1044 | 0.72144 | 0.86241 | 1 | 17774 | 2 | -1.1044 |
| C10orf129 | 6 | 0.30476 | 0.51227 | 0.999994 | 10226 | 3 | -0.58831 | 0.72149 | 0.86243 | 1 | 17775 | 2 | -0.58831 |
| ENTPD2 | 5 | 0.15447 | 0.30945 | 0.999994 | 6576 | 3 | -1.1269 | 0.72156 | 0.84127 | 1 | 17776 | 2 | -1.1269 |
| NDUFS8 | 5 | 0.047151 | 0.13351 | 0.95318 | 2857 | 3 | -2.41 | 0.72167 | 0.84133 | 1 | 17777 | 1 | -2.41 |
| PRDX3 | 6 | 0.5865 | 0.72998 | 0.999994 | 14765 | 2 | -0.73699 | 0.72181 | 0.86258 | 1 | 17778 | 2 | -0.73699 |
| NonTargetingControlG | 2 | 0.74977 | 0.74937 | 0.999994 | 16855 | 0 | 0.62427 | 0.72183 | 0.72173 | 1 | 17779 | 1 | 0.62427 |
| DEPTOR | 5 | 0.27816 | 0.46336 | 0.999994 | 9657 | 2 | 0.024254 | 0.72183 | 0.84143 | 1 | 17780 | 2 | 0.024254 |
| hsa-mir-6124 | 3 | 0.27009 | 0.39983 | 0.999994 | 9477 | 1 | 0.23708 | 0.72194 | 0.75466 | 1 | 17781 | 1 | 0.23708 |
| MTFMT | 6 | 0.20181 | 0.39008 | 0.999994 | 7818 | 3 | -0.84919 | 0.72196 | 0.86264 | 1 | 17782 | 2 | -0.84919 |
| SFT2D3 | 4 | 0.052216 | 0.12582 | 0.951052 | 3078 | 3 | -2.3541 | 0.72197 | 0.81538 | 1 | 17783 | 1 | -2.3541 |

| | | | | | | | | | | | | | |
|---|---|---|---|---|---|---|---|---|---|---|---|---|---|
| TPH2 | 6 | 0.088318 | 0.22418 | 0.983354 | 4525 | 4 | -1.5145 | 0.72198 | 0.86265 | 1 | 17784 | 2 | -1.5145 |
| MANF | 6 | 0.045229 | 0.13358 | 0.95318 | 2775 | 4 | -1.3688 | 0.72202 | 0.86266 | 1 | 17785 | 2 | -1.3688 |
| HN1L | 5 | 0.19767 | 0.37192 | 0.999994 | 7694 | 3 | -0.87512 | 0.72202 | 0.84156 | 1 | 17786 | 2 | -0.87512 |
| ARSG | 5 | 0.014459 | 0.048657 | 0.902886 | 1162 | 2 | -0.2289 | 0.72211 | 0.84161 | 1 | 17787 | 2 | -0.2289 |
| PCNXL3 | 5 | 0.022365 | 0.070462 | 0.93914 | 1605 | 3 | -3.3845 | 0.72211 | 0.84161 | 1 | 17788 | 2 | -3.3845 |
| RWDD2A | 6 | 0.44777 | 0.63361 | 0.999994 | 12683 | 2 | 0.047195 | 0.72227 | 0.86278 | 1 | 17789 | 2 | 0.047195 |
| FGF23 | 4 | 0.11504 | 0.24165 | 0.987253 | 5480 | 1 | 0.48879 | 0.72243 | 0.81554 | 1 | 17790 | 1 | 0.48879 |
| LRSAM1 | 3 | 0.70717 | 0.70773 | 0.999994 | 16385 | 1 | -0.26232 | 0.72245 | 0.75499 | 1 | 17791 | 1 | -0.26232 |
| hsa-mir-5194 | 4 | 0.79046 | 0.79277 | 0.999994 | 17352 | 1 | 0.35896 | 0.72251 | 0.81556 | 1 | 17792 | 1 | 0.35896 |
| TRIM50 | 6 | 0.1195 | 0.28479 | 0.999632 | 5612 | 3 | -0.92243 | 0.72274 | 0.863 | 1 | 17793 | 2 | -0.92243 |
| CES1 | 6 | 0.93639 | 0.93641 | 0.999994 | 19992 | 0 | 0.298 | 0.72284 | 0.86304 | 1 | 17794 | 2 | 0.298 |
| CNTNAP3 | 3 | 0.17365 | 0.27908 | 0.999632 | 7075 | 2 | -1.234 | 0.72299 | 0.75532 | 1 | 17795 | 1 | -1.234 |
| EIF4E1B | 6 | 0.026621 | 0.09082 | 0.94395 | 1835 | 4 | -2.1432 | 0.72325 | 0.86321 | 1 | 17796 | 2 | -2.1432 |
| ITIH1 | 6 | 0.032001 | 0.10349 | 0.946414 | 2104 | 4 | -1.5954 | 0.72328 | 0.86323 | 1 | 17797 | 1 | -1.5954 |
| GNAQ | 6 | 0.042895 | 0.12844 | 0.951777 | 2673 | 4 | -1.4677 | 0.72328 | 0.86323 | 1 | 17798 | 2 | -1.4677 |
| FAM110A | 6 | 0.073752 | 0.19455 | 0.975797 | 3983 | 4 | -1.1607 | 0.72328 | 0.86323 | 1 | 17799 | 1 | -1.1607 |
| SPOPL | 6 | 0.18646 | 0.37113 | 0.999994 | 7371 | 3 | -0.84654 | 0.72328 | 0.86323 | 1 | 17800 | 1 | -0.84654 |
| CYP11B2 | 5 | 0.021617 | 0.068445 | 0.93914 | 1553 | 4 | -3.2293 | 0.72339 | 0.84241 | 1 | 17801 | 1 | -3.2293 |
| DIRAS3 | 6 | 0.072267 | 0.1915 | 0.974071 | 3929 | 4 | -1.084 | 0.72356 | 0.86336 | 1 | 17802 | 2 | -1.084 |
| hsa-mir-4477a | 2 | 0.56639 | 0.56668 | 0.999994 | 14486 | 1 | -0.046907 | 0.72373 | 0.72362 | 1 | 17803 | 1 | -0.046907 |
| GPX1 | 5 | 0.25269 | 0.44291 | 0.999994 | 9070 | 2 | -0.55105 | 0.72389 | 0.84272 | 1 | 17804 | 2 | -0.55105 |
| TBATA | 6 | 0.17507 | 0.35696 | 0.999994 | 7110 | 2 | -0.18292 | 0.72404 | 0.86357 | 1 | 17805 | 2 | -0.18292 |
| CDC42 | 6 | 0.085882 | 0.21926 | 0.980699 | 4442 | 4 | -1.0886 | 0.72406 | 0.86358 | 1 | 17806 | 2 | -1.0886 |
| MRPS5 | 5 | 0.31732 | 0.49518 | 0.999994 | 10457 | 2 | 0.12371 | 0.72428 | 0.84295 | 1 | 17807 | 2 | 0.12371 |
| ZNF274 | 6 | 0.10596 | 0.25892 | 0.994943 | 5174 | 4 | -1.3307 | 0.72456 | 0.86379 | 1 | 17808 | 2 | -1.3307 |
| SPANXB1 | 5 | 0.20996 | 0.3893 | 0.999994 | 8023 | 3 | -0.75798 | 0.72462 | 0.84315 | 1 | 17809 | 1 | -0.75798 |
| PDLIM3 | 6 | 0.086393 | 0.22028 | 0.980699 | 4464 | 4 | -1.3885 | 0.72464 | 0.86383 | 1 | 17810 | 2 | -1.3885 |
| REG1B | 5 | 0.12685 | 0.2678 | 0.998561 | 5844 | 3 | -1.3395 | 0.72475 | 0.84322 | 1 | 17811 | 2 | -1.3395 |
| PRPF3 | 5 | 0.57135 | 0.71071 | 0.999994 | 14566 | 1 | -0.24822 | 0.72477 | 0.84324 | 1 | 17812 | 1 | -0.24822 |
| POU5F1 | 5 | 0.14385 | 0.29361 | 0.999994 | 6295 | 3 | -1.3025 | 0.72517 | 0.84352 | 1 | 17813 | 2 | -1.3025 |
| H2AFB3 | 2 | 0.48756 | 0.49259 | 0.999994 | 13245 | 1 | -0.32816 | 0.72521 | 0.72502 | 1 | 17814 | 1 | -0.32816 |
| C1orf226 | 5 | 0.077092 | 0.18989 | 0.970473 | 4109 | 3 | -1.7672 | 0.72526 | 0.84358 | 1 | 17815 | 1 | -1.7672 |
| TNKS2 | 5 | 0.10279 | 0.23077 | 0.983933 | 5062 | 3 | -1.4255 | 0.72574 | 0.84387 | 1 | 17816 | 2 | -1.4255 |
| SP9 | 5 | 0.47351 | 0.62508 | 0.999994 | 13049 | 2 | 0.17896 | 0.72578 | 0.8439 | 1 | 17817 | 1 | 0.17896 |
| BMX | 5 | 0.37692 | 0.54416 | 0.999994 | 11607 | 2 | -0.36677 | 0.72585 | 0.84393 | 1 | 17818 | 1 | -0.36677 |
| hsa-mir-610 | 3 | 0.27516 | 0.40601 | 0.999994 | 9596 | 2 | -0.65425 | 0.72597 | 0.75717 | 1 | 17819 | 1 | -0.65425 |
| PHLDA1 | 6 | 0.2472 | 0.4446 | 0.999994 | 8954 | 2 | 0.10699 | 0.72614 | 0.86448 | 1 | 17820 | 2 | 0.10699 |
| BMP10 | 6 | 0.85034 | 0.87211 | 0.999994 | 18268 | 1 | 0.49215 | 0.72614 | 0.86448 | 1 | 17821 | 2 | 0.49215 |
| NonTargetingControlG | 2 | 0.51514 | 0.51785 | 0.999994 | 13665 | 1 | -0.22784 | 0.72621 | 0.72597 | 1 | 17822 | 1 | -0.22784 |
| WISP3 | 5 | 0.84586 | 0.85109 | 0.999994 | 18184 | 1 | 0.26102 | 0.72668 | 0.84446 | 1 | 17823 | 2 | 0.26102 |
| PASK | 5 | 0.10441 | 0.23326 | 0.984913 | 5110 | 2 | 0.093477 | 0.72669 | 0.84447 | 1 | 17824 | 2 | 0.093477 |
| SNCA | 6 | 0.0087385 | 0.034594 | 0.867348 | 775 | 3 | -1.4122 | 0.72677 | 0.86475 | 1 | 17825 | 2 | -1.4122 |
| FADD | 6 | 0.012555 | 0.047604 | 0.899802 | 1032 | 5 | -1.4083 | 0.72677 | 0.86475 | 1 | 17826 | 1 | -1.4083 |
| SNX12 | 6 | 0.011007 | 0.042426 | 0.891621 | 924 | 4 | -3.2743 | 0.72685 | 0.86479 | 1 | 17827 | 2 | -3.2743 |
| hsa-mir-6882 | 1 | 0.27311 | 0.27313 | 0.998561 | 9543 | 1 | -1.2774 | 0.72689 | 0.72708 | 1 | 17828 | 0 | -1.2774 |
| NonTargetingControlG | 2 | 0.61552 | 0.61533 | 0.999994 | 15151 | 0 | 0.18221 | 0.72694 | 0.72667 | 1 | 17829 | 1 | 0.18221 |
| DCBLD1 | 3 | 0.69789 | 0.69904 | 0.999994 | 16277 | 1 | -0.56822 | 0.72702 | 0.75785 | 1 | 17830 | 1 | -0.56822 |
| NonTargetingControlG | 1 | 0.27289 | 0.2729 | 0.998561 | 9537 | 1 | -1.3649 | 0.72711 | 0.72729 | 1 | 17831 | 0 | -1.3649 |
| hsa-mir-1285-2 | 3 | 0.13041 | 0.22309 | 0.983354 | 5942 | 2 | -1.8979 | 0.72725 | 0.75799 | 1 | 17832 | 1 | -1.8979 |
| WDR53 | 6 | 0.42237 | 0.61698 | 0.999994 | 12289 | 2 | -0.099275 | 0.72727 | 0.86496 | 1 | 17833 | 2 | -0.099275 |
| PLSCR2 | 5 | 0.5193 | 0.66469 | 0.999994 | 13736 | 1 | -0.14109 | 0.7275 | 0.84496 | 1 | 17834 | 2 | -0.14109 |
| SLURP1 | 5 | 0.37191 | 0.53999 | 0.999994 | 11541 | 2 | -0.32365 | 0.72754 | 0.84498 | 1 | 17835 | 2 | -0.32365 |

| | | | | | | | | | | | | | |
|---|---|---|---|---|---|---|---|---|---|---|---|---|---|
| F2RL1 | 5 | 0.096461 | 0.22082 | 0.98185 | 4836 | 1 | -0.49219 | 0.72767 | 0.84506 | 1 | 17836 | 1 | -0.49219 |
| SLC48A1 | 6 | 0.36933 | 0.5828 | 0.999994 | 11496 | 2 | 0.13419 | 0.72787 | 0.86524 | 1 | 17837 | 1 | 0.13419 |
| KCNQ5 | 6 | 0.43719 | 0.62655 | 0.999994 | 12528 | 2 | 0.25987 | 0.72794 | 0.86527 | 1 | 17838 | 2 | 0.25987 |
| FZD1 | 6 | 0.0061926 | 0.025523 | 0.845628 | 604 | 4 | -4.641 | 0.72799 | 0.86529 | 1 | 17839 | 2 | -4.641 |
| CTF1 | 5 | 0.96242 | 0.96259 | 0.999994 | 20642 | 0 | 0.51949 | 0.72811 | 0.84536 | 1 | 17840 | 2 | 0.51949 |
| OR5M3 | 6 | 0.82319 | 0.85832 | 0.999994 | 17826 | 1 | 0.34912 | 0.72817 | 0.86538 | 1 | 17841 | 2 | 0.34912 |
| SLC1A2 | 6 | 0.89168 | 0.8975 | 0.999994 | 19067 | 1 | 0.30789 | 0.7282 | 0.86539 | 1 | 17842 | 2 | 0.30789 |
| HSD17B4 | 4 | 0.005739 | 0.020087 | 0.808847 | 575 | 3 | -1.4095 | 0.72823 | 0.81761 | 1 | 17843 | 1 | -1.4095 |
| COG5 | 6 | 0.0287 | 0.095766 | 0.945031 | 1936 | 2 | -0.085735 | 0.72827 | 0.86542 | 1 | 17844 | 2 | -0.085735 |
| hsa-mir-500a | 4 | 0.74477 | 0.75607 | 0.999994 | 16797 | 1 | -0.44219 | 0.72828 | 0.81763 | 1 | 17845 | 1 | -0.44219 |
| FAM175B | 5 | 0.40238 | 0.5655 | 0.999994 | 11987 | 2 | 0.45425 | 0.72845 | 0.84557 | 1 | 17846 | 2 | 0.45425 |
| NonTargetingControlG | 2 | 0.36726 | 0.38992 | 0.999994 | 11461 | 1 | -0.71434 | 0.72845 | 0.72821 | 1 | 17847 | 1 | -0.71434 |
| PPM1G | 5 | 0.01594 | 0.052857 | 0.916123 | 1243 | 3 | -2.9271 | 0.72849 | 0.84561 | 1 | 17848 | 1 | -2.9271 |
| CYP27C1 | 5 | 0.034883 | 0.10297 | 0.946414 | 2261 | 3 | -2.3047 | 0.72849 | 0.84561 | 1 | 17849 | 2 | -2.3047 |
| TCP11L2 | 6 | 0.57265 | 0.71986 | 0.999994 | 14586 | 2 | -0.42766 | 0.72851 | 0.86553 | 1 | 17850 | 2 | -0.42766 |
| ACER1 | 5 | 0.49075 | 0.63985 | 0.999994 | 13289 | 2 | -0.41484 | 0.72866 | 0.84572 | 1 | 17851 | 2 | -0.41484 |
| NonTargetingControlG | 1 | 0.27133 | 0.27132 | 0.998561 | 9505 | 1 | -1.3125 | 0.72867 | 0.72884 | 1 | 17852 | 0 | -1.3125 |
| NDNF | 5 | 0.21782 | 0.40022 | 0.999994 | 8224 | 2 | 0.25424 | 0.72868 | 0.84573 | 1 | 17853 | 2 | 0.25424 |
| DLEC1 | 5 | 0.087668 | 0.20685 | 0.978744 | 4501 | 2 | -0.077591 | 0.72875 | 0.84578 | 1 | 17854 | 2 | -0.077591 |
| C1orf61 | 5 | 0.4342 | 0.59194 | 0.999994 | 12481 | 1 | 0.028172 | 0.72881 | 0.84581 | 1 | 17855 | 2 | 0.028172 |
| hsa-mir-297 | 1 | 0.27116 | 0.27114 | 0.998561 | 9502 | 1 | -1.3092 | 0.72884 | 0.72901 | 1 | 17856 | 0 | -1.3092 |
| NonTargetingControlG | 1 | 0.27111 | 0.27109 | 0.998561 | 9501 | 1 | -1.427 | 0.72889 | 0.72905 | 1 | 17857 | 0 | -1.427 |
| hsa-mir-20a | 1 | 0.27102 | 0.27099 | 0.998561 | 9500 | 1 | -1.4256 | 0.72898 | 0.72915 | 1 | 17858 | 0 | -1.4256 |
| CTGF | 6 | 0.21781 | 0.40948 | 0.999994 | 8219 | 3 | -0.4073 | 0.72912 | 0.86579 | 1 | 17859 | 2 | -0.4073 |
| FRAS1 | 6 | 0.58992 | 0.73253 | 0.999994 | 14822 | 1 | 0.30139 | 0.72915 | 0.86581 | 1 | 17860 | 2 | 0.30139 |
| GLS2 | 5 | 0.17057 | 0.33296 | 0.999994 | 6996 | 3 | -1.01 | 0.72919 | 0.84607 | 1 | 17861 | 2 | -1.01 |
| MFAP3L | 6 | 0.21747 | 0.40908 | 0.999994 | 8206 | 2 | 0.38131 | 0.72928 | 0.86585 | 1 | 17862 | 2 | 0.38131 |
| UBQLNL | 6 | 0.072306 | 0.19158 | 0.974071 | 3930 | 3 | -1.0593 | 0.72946 | 0.86592 | 1 | 17863 | 2 | -1.0593 |
| ZFP91 | 6 | 0.03772 | 0.11676 | 0.951052 | 2415 | 4 | -0.93108 | 0.72947 | 0.86593 | 1 | 17864 | 2 | -0.93108 |
| RAD9A | 4 | 0.072627 | 0.16489 | 0.967126 | 3945 | 3 | -1.6042 | 0.72957 | 0.81812 | 1 | 17865 | 1 | -1.6042 |
| GFPT1 | 4 | 0.042012 | 0.10579 | 0.950444 | 2614 | 3 | -2.1844 | 0.72957 | 0.81812 | 1 | 17866 | 1 | -2.1844 |
| RESP18 | 5 | 0.56399 | 0.70419 | 0.999994 | 14453 | 2 | 0.20956 | 0.72958 | 0.84629 | 1 | 17867 | 1 | 0.20956 |
| HEG1 | 6 | 0.64337 | 0.77249 | 0.999994 | 15567 | 2 | -0.084804 | 0.72962 | 0.86599 | 1 | 17868 | 2 | -0.084804 |
| EXPH5 | 5 | 0.54969 | 0.69147 | 0.999994 | 14208 | 1 | -0.064356 | 0.7297 | 0.84637 | 1 | 17869 | 2 | -0.064356 |
| FFAR1 | 5 | 0.86072 | 0.86301 | 0.999994 | 18455 | 1 | -0.35047 | 0.72974 | 0.8464 | 1 | 17870 | 1 | -0.35047 |
| hsa-mir-4305 | 2 | 0.55126 | 0.55208 | 0.999994 | 14231 | 1 | -0.095406 | 0.72976 | 0.72954 | 1 | 17871 | 1 | -0.095406 |
| ZNF846 | 6 | 0.24129 | 0.43753 | 0.999994 | 8804 | 3 | -0.45419 | 0.72984 | 0.86609 | 1 | 17872 | 2 | -0.45419 |
| C1QTNF1 | 5 | 0.00071951 | 0.0032913 | 0.541338 | 133 | 4 | -8.5271 | 0.72984 | 0.84645 | 1 | 17873 | 1 | -8.5271 |
| hsa-mir-6847 | 3 | 0.26835 | 0.39762 | 0.999994 | 9426 | 2 | -0.73036 | 0.72986 | 0.75962 | 1 | 17874 | 1 | -0.73036 |
| hsa-mir-378a | 3 | 0.26086 | 0.38845 | 0.999994 | 9265 | 1 | 0.32346 | 0.73002 | 0.75972 | 1 | 17875 | 1 | 0.32346 |
| NonTargetingControlG | 2 | 0.51401 | 0.51681 | 0.999994 | 13640 | 1 | -0.17155 | 0.73028 | 0.73005 | 1 | 17876 | 1 | -0.17155 |
| KIAA2018 | 6 | 0.090621 | 0.22873 | 0.983354 | 4621 | 4 | -0.9218 | 0.73029 | 0.86629 | 1 | 17877 | 2 | -0.9218 |
| hsa-mir-4259 | 3 | 0.23795 | 0.36002 | 0.999994 | 8727 | 1 | -0.52681 | 0.73032 | 0.7599 | 1 | 17878 | 1 | -0.52681 |
| hsa-mir-4761 | 4 | 0.23847 | 0.38011 | 0.999994 | 8735 | 2 | -1.0039 | 0.73034 | 0.8184 | 1 | 17879 | 1 | -1.0039 |
| SLC23A3 | 6 | 0.067533 | 0.1816 | 0.968171 | 3738 | 4 | -1.2576 | 0.73053 | 0.8664 | 1 | 17880 | 1 | -1.2576 |
| hsa-mir-3672 | 1 | 0.26943 | 0.26943 | 0.998561 | 9458 | 1 | -1.2884 | 0.73057 | 0.73074 | 1 | 17881 | 0 | -1.2884 |
| ANKRD65 | 5 | 0.075823 | 0.18786 | 0.969543 | 4066 | 2 | 0.49344 | 0.73058 | 0.84693 | 1 | 17882 | 2 | 0.49344 |
| MAP2K6 | 6 | 0.078298 | 0.20391 | 0.978186 | 4161 | 1 | 0.24157 | 0.73063 | 0.86644 | 1 | 17883 | 2 | 0.24157 |
| SPIN2B | 1 | 0.26933 | 0.26932 | 0.998561 | 9453 | 1 | -1.3081 | 0.73067 | 0.73085 | 1 | 17884 | 0 | -1.3081 |
| TBC1D14 | 6 | 0.24707 | 0.44443 | 0.999994 | 8947 | 1 | 0.3585 | 0.73099 | 0.8666 | 1 | 17885 | 2 | 0.3585 |
| MIIP | 6 | 0.06365 | 0.17351 | 0.96793 | 3592 | 4 | -1.1739 | 0.73111 | 0.86665 | 1 | 17886 | 2 | -1.1739 |
| NKX1-2 | 5 | 0.47933 | 0.63006 | 0.999994 | 13136 | 1 | 0.4186 | 0.73115 | 0.8473 | 1 | 17887 | 2 | 0.4186 |

| | | | | | | | | | | | | |
|---|---|---|---|---|---|---|---|---|---|---|---|---|
| PAH | 6 | 0.044305 | 0.13155 | 0.95318 | 2726 | 4 | -1.6566 | 0.73117 | 0.86669 | 1 | 17888 | 2 | -1.6566 |
| FAM3B | 5 | 0.083631 | 0.20044 | 0.977164 | 4361 | 3 | -1.6318 | 0.7316 | 0.84759 | 1 | 17889 | 1 | -1.6318 |
| DCAF8 | 5 | 0.88446 | 0.88459 | 0.999994 | 18923 | 1 | -0.12878 | 0.73171 | 0.84765 | 1 | 17890 | 2 | -0.12878 |
| SVEP1 | 6 | 0.065155 | 0.17658 | 0.968171 | 3659 | 3 | -1.2627 | 0.73189 | 0.867 | 1 | 17891 | 2 | -1.2627 |
| KIAA1191 | 5 | 0.06467 | 0.16975 | 0.967126 | 3637 | 3 | -1.8614 | 0.73208 | 0.84789 | 1 | 17892 | 2 | -1.8614 |
| hsa-mir-449a | 3 | 0.26306 | 0.39115 | 0.999994 | 9316 | 2 | -0.75561 | 0.73217 | 0.76106 | 1 | 17893 | 1 | -0.75561 |
| OR9G4 | 6 | 0.86176 | 0.87841 | 0.999994 | 18480 | 1 | 0.096251 | 0.73224 | 0.86715 | 1 | 17894 | 2 | 0.096251 |
| hsa-mir-561 | 4 | 0.41697 | 0.56307 | 0.999994 | 12205 | 1 | -0.22368 | 0.73232 | 0.81912 | 1 | 17895 | 1 | -0.22368 |
| VEGFA | 6 | 0.26521 | 0.46604 | 0.999994 | 9353 | 2 | 0.079708 | 0.73245 | 0.86724 | 1 | 17896 | 2 | 0.079708 |
| TAOK3 | 6 | 0.25225 | 0.45061 | 0.999994 | 9064 | 2 | -0.31579 | 0.73257 | 0.86729 | 1 | 17897 | 1 | -0.31579 |
| ACKR2 | 6 | 0.18631 | 0.37095 | 0.999994 | 7369 | 3 | -0.77767 | 0.73266 | 0.86733 | 1 | 17898 | 2 | -0.77767 |
| DNAH14 | 6 | 0.075764 | 0.19875 | 0.977164 | 4064 | 4 | -1.1143 | 0.73291 | 0.86744 | 1 | 17899 | 2 | -1.1143 |
| NonTargetingControlG | 2 | 0.48554 | 0.49076 | 0.999994 | 13221 | 1 | -0.32238 | 0.73322 | 0.73294 | 1 | 17900 | 1 | -0.32238 |
| AEBP1 | 6 | 0.10931 | 0.26535 | 0.997406 | 5280 | 3 | -1.0148 | 0.73328 | 0.86761 | 1 | 17901 | 1 | -1.0148 |
| TFDP1 | 6 | 0.011425 | 0.043835 | 0.891621 | 951 | 5 | -3.4535 | 0.73328 | 0.86761 | 1 | 17902 | 1 | -3.4535 |
| SULT4A1 | 5 | 0.407 | 0.56936 | 0.999994 | 12061 | 2 | 0.48035 | 0.73333 | 0.84871 | 1 | 17903 | 2 | 0.48035 |
| NonTargetingControlG | 1 | 0.26653 | 0.2665 | 0.998525 | 9385 | 1 | -1.4468 | 0.73347 | 0.73364 | 1 | 17904 | 0 | -1.4468 |
| NonTargetingControlG | 2 | 0.68446 | 0.68431 | 0.999994 | 16150 | 0 | 0.30486 | 0.73355 | 0.73326 | 1 | 17905 | 1 | 0.30486 |
| hsa-mir-5684 | 2 | 0.75964 | 0.75915 | 0.999994 | 16978 | 0 | 0.62828 | 0.73375 | 0.73345 | 1 | 17906 | 1 | 0.62828 |
| hsa-mir-3681 | 3 | 0.30911 | 0.43347 | 0.999994 | 10304 | 1 | 0.24585 | 0.734 | 0.76223 | 1 | 17907 | 1 | 0.24585 |
| KRBOX1 | 6 | 0.26835 | 0.46976 | 0.999994 | 9425 | 3 | -0.67632 | 0.73404 | 0.86794 | 1 | 17908 | 2 | -0.67632 |
| CD1A | 5 | 0.24302 | 0.43503 | 0.999994 | 8867 | 2 | 0.29897 | 0.73408 | 0.8492 | 1 | 17909 | 2 | 0.29897 |
| PHLDB1 | 6 | 0.01291 | 0.048814 | 0.903125 | 1057 | 3 | -2.6473 | 0.7342 | 0.86801 | 1 | 17910 | 2 | -2.6473 |
| KY | 4 | 0.14776 | 0.28685 | 0.999632 | 6399 | 2 | -1.053 | 0.73425 | 0.81979 | 1 | 17911 | 1 | -1.053 |
| CRELD2 | 5 | 0.36846 | 0.53706 | 0.999994 | 11479 | 1 | 0.24432 | 0.73444 | 0.84943 | 1 | 17912 | 2 | 0.24432 |
| PDIA2 | 5 | 0.073836 | 0.18461 | 0.968526 | 3988 | 3 | -3.3431 | 0.73448 | 0.84945 | 1 | 17913 | 2 | -3.3431 |
| PEX11B | 5 | 0.31348 | 0.49202 | 0.999994 | 10386 | 2 | -0.44211 | 0.73448 | 0.84945 | 1 | 17914 | 1 | -0.44211 |
| SLC22A13 | 6 | 0.11726 | 0.28056 | 0.999632 | 5549 | 4 | -1.0997 | 0.73476 | 0.86827 | 1 | 17915 | 2 | -1.0997 |
| DSCR4 | 6 | 0.46462 | 0.64483 | 0.999994 | 12929 | 1 | 0.24814 | 0.73476 | 0.86827 | 1 | 17916 | 2 | 0.24814 |
| C11orf42 | 5 | 0.062435 | 0.16609 | 0.967126 | 3533 | 3 | -2.1427 | 0.73524 | 0.84994 | 1 | 17917 | 2 | -2.1427 |
| HELZ2 | 6 | 0.29542 | 0.50136 | 0.999994 | 10026 | 2 | 0.15956 | 0.73525 | 0.86847 | 1 | 17918 | 2 | 0.15956 |
| hsa-mir-6742 | 3 | 0.53029 | 0.56657 | 0.999994 | 13903 | 1 | -0.56597 | 0.73531 | 0.76306 | 1 | 17919 | 1 | -0.56597 |
| ARRB2 | 5 | 0.095443 | 0.21924 | 0.980699 | 4799 | 3 | -1.3209 | 0.73539 | 0.85004 | 1 | 17920 | 2 | -1.3209 |
| hsa-mir-3622b | 1 | 0.2646 | 0.2645 | 0.996734 | 9348 | 1 | -1.4579 | 0.7354 | 0.73556 | 1 | 17921 | 0 | -1.4579 |
| NRBF2 | 6 | 0.11119 | 0.26899 | 0.998561 | 5338 | 3 | -1.131 | 0.73545 | 0.86857 | 1 | 17922 | 2 | -1.131 |
| ZFAND2A | 6 | 0.26679 | 0.46787 | 0.999994 | 9391 | 2 | -0.12719 | 0.73561 | 0.86865 | 1 | 17923 | 2 | -0.12719 |
| RAB3D | 5 | 0.0092681 | 0.033943 | 0.867348 | 814 | 3 | -4.6965 | 0.73564 | 0.85019 | 1 | 17924 | 2 | -4.6965 |
| FAM166B | 6 | 0.016991 | 0.062234 | 0.937033 | 1302 | 2 | -0.034371 | 0.7357 | 0.86869 | 1 | 17925 | 2 | -0.034371 |
| hsa-mir-30d | 3 | 0.12227 | 0.21247 | 0.979587 | 5712 | 2 | -1.9812 | 0.73575 | 0.76337 | 1 | 17926 | 1 | -1.9812 |
| NEIL2 | 6 | 0.31522 | 0.52443 | 0.999994 | 10423 | 3 | -0.2304 | 0.73578 | 0.86873 | 1 | 17927 | 2 | -0.2304 |
| hsa-mir-1304 | 2 | 0.4106 | 0.42559 | 0.999994 | 12122 | 1 | -0.66228 | 0.73579 | 0.73549 | 1 | 17928 | 1 | -0.66228 |
| FLI1 | 4 | 0.62035 | 0.67654 | 0.999994 | 15225 | 1 | -0.24832 | 0.73594 | 0.82042 | 1 | 17929 | 1 | -0.24832 |
| PRR19 | 5 | 0.026623 | 0.081743 | 0.93914 | 1836 | 3 | -2.5558 | 0.73602 | 0.85044 | 1 | 17930 | 2 | -2.5558 |
| PINK1 | 3 | 0.46153 | 0.52137 | 0.999994 | 12873 | 1 | 0.13289 | 0.73618 | 0.76367 | 1 | 17931 | 1 | 0.13289 |
| SIVA1 | 6 | 0.019699 | 0.071029 | 0.93914 | 1440 | 5 | -1.5614 | 0.73645 | 0.86902 | 1 | 17932 | 1 | -1.5614 |
| RBCK1 | 6 | 0.25385 | 0.45253 | 0.999994 | 9100 | 3 | -0.19384 | 0.73645 | 0.86902 | 1 | 17933 | 2 | -0.19384 |
| EML1 | 5 | 0.15393 | 0.30866 | 0.999994 | 6565 | 3 | -1.2027 | 0.73658 | 0.85079 | 1 | 17934 | 1 | -1.2027 |
| SF3B2 | 5 | 0.39643 | 0.56058 | 0.999994 | 11896 | 2 | 0.21677 | 0.73666 | 0.85085 | 1 | 17935 | 2 | 0.21677 |
| SH3RF1 | 5 | 0.37722 | 0.54441 | 0.999994 | 11608 | 2 | -0.52428 | 0.73669 | 0.85087 | 1 | 17936 | 1 | -0.52428 |
| NonTargetingControlG | 2 | 0.38199 | 0.40189 | 0.999994 | 11676 | 1 | -0.75689 | 0.73687 | 0.73655 | 1 | 17937 | 0 | -0.75689 |
| ADCK5 | 5 | 0.38029 | 0.54695 | 0.999994 | 11653 | 1 | 0.44252 | 0.73687 | 0.851 | 1 | 17938 | 2 | 0.44252 |
| NonTargetingControlG | 2 | 0.061489 | 0.1084 | 0.951052 | 3491 | 1 | -4.2009 | 0.73701 | 0.7367 | 1 | 17939 | 0 | -4.2009 |

| | | | | | | | | | | | | | |
|---|---|---|---|---|---|---|---|---|---|---|---|---|---|
| NPAS3 | 6 | 0.22744 | 0.421 | 0.999994 | 8467 | 3 | -0.74618 | 0.73716 | 0.86935 | 1 | 17940 | 1 | -0.74618 |
| CHST11 | 6 | 0.15936 | 0.33739 | 0.999994 | 6717 | 3 | -0.77773 | 0.73721 | 0.86937 | 1 | 17941 | 2 | -0.77773 |
| ATRAID | 5 | 0.10555 | 0.23506 | 0.985362 | 5154 | 2 | -0.42685 | 0.73732 | 0.85129 | 1 | 17942 | 1 | -0.42685 |
| COG8 | 5 | 0.46678 | 0.6194 | 0.999994 | 12961 | 2 | 0.56212 | 0.73732 | 0.85129 | 1 | 17943 | 2 | 0.56212 |
| CCDC88C | 5 | 0.052216 | 0.14573 | 0.963702 | 3077 | 2 | 0.41338 | 0.73732 | 0.85129 | 1 | 17944 | 2 | 0.41338 |
| FAM86C1 | 3 | 0.10218 | 0.18594 | 0.969409 | 5047 | 2 | -0.70487 | 0.73738 | 0.76446 | 1 | 17945 | 1 | -0.70487 |
| RNF170 | 5 | 0.39888 | 0.56262 | 0.999994 | 11933 | 1 | 0.21154 | 0.73747 | 0.85139 | 1 | 17946 | 2 | 0.21154 |
| FNTA | 4 | 0.071764 | 0.16328 | 0.967126 | 3910 | 3 | -1.4456 | 0.73749 | 0.82102 | 1 | 17947 | 1 | -1.4456 |
| TRIM69 | 6 | 0.0042187 | 0.018276 | 0.795727 | 458 | 3 | -0.64466 | 0.73759 | 0.86953 | 1 | 17948 | 2 | -0.64466 |
| TLX1 | 6 | 0.54778 | 0.70204 | 0.999994 | 14175 | 2 | -0.11995 | 0.73776 | 0.86961 | 1 | 17949 | 2 | -0.11995 |
| CCDC114 | 5 | 0.67711 | 0.75632 | 0.999994 | 16044 | 1 | 0.29273 | 0.73782 | 0.85161 | 1 | 17950 | 2 | 0.29273 |
| ZMYND11 | 5 | 0.19658 | 0.37035 | 0.999994 | 7664 | 3 | -0.81115 | 0.73792 | 0.85167 | 1 | 17951 | 2 | -0.81115 |
| RAD9B | 4 | 0.062458 | 0.14548 | 0.963601 | 3534 | 3 | -2.2118 | 0.73803 | 0.8212 | 1 | 17952 | 1 | -2.2118 |
| TTL | 5 | 0.07481 | 0.18621 | 0.969409 | 4028 | 3 | -1.9918 | 0.73817 | 0.85185 | 1 | 17953 | 2 | -1.9918 |
| VWA5B2 | 5 | 0.37004 | 0.5384 | 0.999994 | 11512 | 2 | 0.64367 | 0.73822 | 0.85187 | 1 | 17954 | 2 | 0.64367 |
| GHITM | 5 | 0.37199 | 0.54007 | 0.999994 | 11542 | 2 | -0.10867 | 0.73834 | 0.85194 | 1 | 17955 | 2 | -0.10867 |
| LOC100130880 | 5 | 0.80006 | 0.81954 | 0.999994 | 17479 | 1 | 0.43114 | 0.7384 | 0.85198 | 1 | 17956 | 2 | 0.43114 |
| AHNAK2 | 4 | 0.052616 | 0.12662 | 0.951052 | 3096 | 2 | -2.2607 | 0.73851 | 0.82136 | 1 | 17957 | 1 | -2.2607 |
| PPP1R10 | 4 | 0.14745 | 0.28652 | 0.999632 | 6390 | 2 | -1.3333 | 0.73851 | 0.82136 | 1 | 17958 | 1 | -1.3333 |
| ZNF782 | 6 | 0.08856 | 0.22466 | 0.983354 | 4538 | 4 | -1.1629 | 0.73859 | 0.86997 | 1 | 17959 | 2 | -1.1629 |
| hsa-mir-23a | 3 | 0.26674 | 0.39566 | 0.999994 | 9389 | 2 | -0.73869 | 0.73893 | 0.76547 | 1 | 17960 | 1 | -0.73869 |
| NonTargetingControlG | 2 | 0.75507 | 0.75463 | 0.999994 | 16915 | 0 | 0.48723 | 0.739 | 0.73868 | 1 | 17961 | 0 | 0.48723 |
| ORMDL1 | 6 | 0.11425 | 0.27483 | 0.998561 | 5456 | 3 | -0.40418 | 0.73908 | 0.87019 | 1 | 17962 | 2 | -0.40418 |
| GPSM2 | 5 | 0.1706 | 0.33301 | 0.999994 | 6998 | 3 | -1.0777 | 0.73919 | 0.8525 | 1 | 17963 | 2 | -1.0777 |
| TYW1B | 5 | 0.20626 | 0.38409 | 0.999994 | 7938 | 3 | -0.79102 | 0.73921 | 0.85251 | 1 | 17964 | 1 | -0.79102 |
| CCT2 | 2 | 0.055652 | 0.098675 | 0.945253 | 3227 | 1 | -4.2899 | 0.73926 | 0.73892 | 1 | 17965 | 0 | -4.2899 |
| ANKRD23 | 5 | 0.051578 | 0.1442 | 0.961983 | 3049 | 3 | -2.1714 | 0.73929 | 0.85257 | 1 | 17966 | 2 | -2.1714 |
| IK | 6 | 0.048967 | 0.14183 | 0.959146 | 2927 | 4 | -2.0277 | 0.73929 | 0.87028 | 1 | 17967 | 2 | -2.0277 |
| NonTargetingControlG | 2 | 0.51068 | 0.5137 | 0.999994 | 13587 | 1 | -0.15759 | 0.73932 | 0.73899 | 1 | 17968 | 0 | -0.15759 |
| SEPT8 | 5 | 0.062593 | 0.16634 | 0.967126 | 3540 | 3 | -2.0516 | 0.73937 | 0.85262 | 1 | 17969 | 2 | -2.0516 |
| FHL2 | 6 | 0.031729 | 0.10287 | 0.946414 | 2092 | 3 | -1.7683 | 0.73938 | 0.87031 | 1 | 17970 | 1 | -1.7683 |
| NonTargetingControlG | 1 | 0.26058 | 0.2605 | 0.994943 | 9253 | 1 | -1.5111 | 0.73942 | 0.73968 | 1 | 17971 | 0 | -1.5111 |
| GFRA2 | 2 | 0.087034 | 0.1498 | 0.966359 | 4478 | 1 | -3.1562 | 0.7396 | 0.73926 | 1 | 17972 | 0 | -3.1562 |
| HARS2 | 5 | 0.031249 | 0.093695 | 0.945031 | 2071 | 3 | -2.6387 | 0.73997 | 0.85298 | 1 | 17973 | 2 | -2.6387 |
| NonTargetingControlG | 2 | 0.10561 | 0.17874 | 0.968171 | 5156 | 1 | -3.792 | 0.73997 | 0.73962 | 1 | 17974 | 0 | -3.792 |
| ARPP19 | 5 | 0.55604 | 0.69713 | 0.999994 | 14305 | 2 | -0.4922 | 0.73998 | 0.853 | 1 | 17975 | 2 | -0.4922 |
| NonTargetingControlG | 2 | 0.48289 | 0.4884 | 0.999994 | 13188 | 1 | -0.37407 | 0.73999 | 0.73965 | 1 | 17976 | 0 | -0.37407 |
| LRRC15 | 6 | 0.070501 | 0.18784 | 0.969543 | 3851 | 4 | -1.1893 | 0.7402 | 0.87067 | 1 | 17977 | 2 | -1.1893 |
| hsa-mir-103a-2 | 4 | 0.48478 | 0.60961 | 0.999994 | 13215 | 1 | 0.19672 | 0.74025 | 0.82202 | 1 | 17978 | 1 | 0.19672 |
| YWHAB | 6 | 0.11351 | 0.27344 | 0.998561 | 5422 | 2 | 0.33086 | 0.74026 | 0.8707 | 1 | 17979 | 2 | 0.33086 |
| ARL4D | 6 | 0.046517 | 0.13642 | 0.956051 | 2830 | 3 | -1.6569 | 0.7403 | 0.87072 | 1 | 17980 | 2 | -1.6569 |
| DEPDC1 | 5 | 0.12384 | 0.26319 | 0.996734 | 5762 | 3 | -1.4047 | 0.74037 | 0.85322 | 1 | 17981 | 2 | -1.4047 |
| PRDX4 | 6 | 0.079794 | 0.20686 | 0.978744 | 4224 | 4 | -1.5819 | 0.74043 | 0.87077 | 1 | 17982 | 1 | -1.5819 |
| hsa-mir-30c-2 | 4 | 0.16879 | 0.30852 | 0.999994 | 6953 | 2 | -1.3779 | 0.74047 | 0.8221 | 1 | 17983 | 1 | -1.3779 |
| PTGER2 | 5 | 0.052464 | 0.14633 | 0.964852 | 3091 | 3 | -2.3789 | 0.74048 | 0.8533 | 1 | 17984 | 2 | -2.3789 |
| PHF1 | 5 | 0.44179 | 0.5983 | 0.999994 | 12608 | 1 | 0.072849 | 0.7409 | 0.85356 | 1 | 17985 | 2 | 0.072849 |
| CCK | 5 | 0.0050986 | 0.021208 | 0.812607 | 526 | 4 | -6.8737 | 0.7409 | 0.85356 | 1 | 17986 | 1 | -6.8737 |
| RNF13 | 5 | 0.089276 | 0.2094 | 0.979587 | 4566 | 3 | -0.77762 | 0.74105 | 0.85366 | 1 | 17987 | 2 | -0.77762 |
| KIAA0232 | 6 | 0.018133 | 0.065939 | 0.93914 | 1368 | 3 | -0.71182 | 0.74114 | 0.87109 | 1 | 17988 | 2 | -0.71182 |
| ELMOD3 | 6 | 0.25723 | 0.45658 | 0.999994 | 9172 | 3 | -0.76229 | 0.74114 | 0.87109 | 1 | 17989 | 1 | -0.76229 |
| MRPL46 | 6 | 0.03316 | 0.10624 | 0.950812 | 2170 | 4 | -0.89786 | 0.74114 | 0.87109 | 1 | 17990 | 1 | -0.89786 |
| RPS10 | 3 | 0.8208 | 0.82065 | 0.999994 | 17788 | 0 | 0.15226 | 0.74122 | 0.76694 | 1 | 17991 | 1 | 0.15226 |

| | | | | | | | | | | | | | |
|---|---|---|---|---|---|---|---|---|---|---|---|---|---|
| NonTargetingControlG | 2 | 0.51662 | 0.51925 | 0.999994 | 13687 | 1 | -0.24188 | 0.74145 | 0.74114 | 1 | 17992 | 0 | -0.24188 |
| TWISTNB | 4 | 0.22135 | 0.36263 | 0.999994 | 8312 | 2 | -1.1551 | 0.74163 | 0.82256 | 1 | 17993 | 1 | -1.1551 |
| FSCN1 | 5 | 0.93976 | 0.93994 | 0.999994 | 20071 | 0 | 0.42417 | 0.74164 | 0.85404 | 1 | 17994 | 1 | 0.42417 |
| LHFP | 6 | 0.34451 | 0.55785 | 0.999994 | 11041 | 2 | 0.0049207 | 0.74167 | 0.87132 | 1 | 17995 | 2 | 0.0049207 |
| OR5AR1 | 4 | 0.2152 | 0.35633 | 0.999994 | 8146 | 1 | 0.40679 | 0.74178 | 0.82262 | 1 | 17996 | 1 | 0.40679 |
| NonTargetingControlG | 2 | 0.51426 | 0.51705 | 0.999994 | 13646 | 1 | -0.28263 | 0.7419 | 0.74159 | 1 | 17997 | 0 | -0.28263 |
| NonTargetingControlG | 2 | 0.14751 | 0.2249 | 0.983354 | 6392 | 1 | -2.0165 | 0.74197 | 0.74166 | 1 | 17998 | 0 | -2.0165 |
| MRPS31 | 5 | 0.15321 | 0.30758 | 0.999994 | 6540 | 3 | -1.1189 | 0.74209 | 0.85433 | 1 | 17999 | 2 | -1.1189 |
| CCS | 5 | 0.3296 | 0.5052 | 0.999994 | 10713 | 1 | -0.26976 | 0.7421 | 0.85434 | 1 | 18000 | 2 | -0.26976 |
| hsa-mir-222 | 4 | 0.2625 | 0.40461 | 0.999994 | 9297 | 1 | -0.38816 | 0.74239 | 0.82286 | 1 | 18001 | 1 | -0.38816 |
| PHKG2 | 3 | 0.26635 | 0.3952 | 0.999994 | 9377 | 2 | -0.67225 | 0.74248 | 0.76776 | 1 | 18002 | 1 | -0.67225 |
| hsa-mir-6869 | 3 | 0.1292 | 0.22153 | 0.983093 | 5920 | 1 | -0.26142 | 0.7425 | 0.76778 | 1 | 18003 | 1 | -0.26142 |
| SLC39A2 | 6 | 0.0016145 | 0.0080692 | 0.680529 | 237 | 4 | -5.3057 | 0.74259 | 0.87174 | 1 | 18004 | 2 | -5.3057 |
| EFR3B | 6 | 0.17266 | 0.35403 | 0.999994 | 7054 | 2 | -0.38962 | 0.74263 | 0.87176 | 1 | 18005 | 2 | -0.38962 |
| DMBX1 | 4 | 0.082905 | 0.1841 | 0.968526 | 4334 | 3 | -1.1646 | 0.74269 | 0.82297 | 1 | 18006 | 1 | -1.1646 |
| NonTargetingControlG | 2 | 0.66548 | 0.66518 | 0.999994 | 15862 | 0 | 0.22273 | 0.74278 | 0.74245 | 1 | 18007 | 0 | 0.22273 |
| NonTargetingControlG | 2 | 0.17376 | 0.2433 | 0.988088 | 7077 | 1 | -2.983 | 0.74281 | 0.74248 | 1 | 18008 | 0 | -2.983 |
| SLC16A3 | 4 | 0.78712 | 0.7899 | 0.999994 | 17305 | 1 | 0.35986 | 0.74291 | 0.82306 | 1 | 18009 | 1 | 0.35986 |
| EPG5 | 5 | 0.04144 | 0.1195 | 0.951052 | 2586 | 4 | -1.2609 | 0.743 | 0.85492 | 1 | 18010 | 1 | -1.2609 |
| MKX | 5 | 0.61369 | 0.73119 | 0.999994 | 15126 | 1 | 0.33527 | 0.743 | 0.85492 | 1 | 18011 | 2 | 0.33527 |
| FAM127C | 5 | 0.41276 | 0.57413 | 0.999994 | 12152 | 2 | 0.053855 | 0.74308 | 0.85497 | 1 | 18012 | 2 | 0.053855 |
| hsa-mir-6732 | 3 | 0.05926 | 0.12841 | 0.951777 | 3392 | 2 | -2.86 | 0.74309 | 0.76816 | 1 | 18013 | 1 | -2.86 |
| SYT17 | 5 | 0.23352 | 0.42208 | 0.999994 | 8604 | 3 | -0.68207 | 0.74343 | 0.8552 | 1 | 18014 | 2 | -0.68207 |
| GH2 | 5 | 0.49798 | 0.64611 | 0.999994 | 13396 | 2 | 0.40036 | 0.74349 | 0.85523 | 1 | 18015 | 2 | 0.40036 |
| ZNF248 | 5 | 0.20193 | 0.37802 | 0.999994 | 7825 | 2 | 0.35141 | 0.74352 | 0.85526 | 1 | 18016 | 2 | 0.35141 |
| CDH2 | 4 | 0.23517 | 0.37682 | 0.999994 | 8652 | 2 | -0.89331 | 0.7437 | 0.82337 | 1 | 18017 | 1 | -0.89331 |
| MSH6 | 6 | 0.11225 | 0.27102 | 0.998561 | 5378 | 3 | -0.17416 | 0.74371 | 0.87223 | 1 | 18018 | 2 | -0.17416 |
| hsa-mir-3960 | 1 | 0.25626 | 0.25613 | 0.994943 | 9156 | 1 | -1.5551 | 0.74374 | 0.744 | 1 | 18019 | 0 | -1.5551 |
| PCOLCE2 | 5 | 0.36506 | 0.53429 | 0.999994 | 11421 | 1 | 0.56175 | 0.74391 | 0.85553 | 1 | 18020 | 2 | 0.56175 |
| EOMES | 6 | 0.08902 | 0.22555 | 0.983354 | 4553 | 3 | -1.0507 | 0.74409 | 0.87239 | 1 | 18021 | 2 | -1.0507 |
| CYLD | 5 | 0.50124 | 0.649 | 0.999994 | 13441 | 1 | 0.45849 | 0.74411 | 0.85565 | 1 | 18022 | 2 | 0.45849 |
| hsa-mir-4484 | 3 | 0.67662 | 0.67969 | 0.999994 | 16033 | 1 | 0.44948 | 0.74424 | 0.76891 | 1 | 18023 | 1 | 0.44948 |
| NonTargetingControlG | 2 | 0.22201 | 0.27807 | 0.999536 | 8333 | 1 | -1.6587 | 0.74433 | 0.74401 | 1 | 18024 | 0 | -1.6587 |
| NonTargetingControlG | 2 | 0.31299 | 0.34687 | 0.999994 | 10376 | 1 | -1.1376 | 0.74434 | 0.74402 | 1 | 18025 | 0 | -1.1376 |
| CMTM4 | 6 | 0.13819 | 0.31074 | 0.999994 | 6143 | 3 | -1.0755 | 0.74464 | 0.87263 | 1 | 18026 | 1 | -1.0755 |
| CDC42EP1 | 6 | 0.01028 | 0.039965 | 0.891621 | 887 | 4 | -1.4055 | 0.74464 | 0.87263 | 1 | 18027 | 2 | -1.4055 |
| hsa-mir-6747 | 4 | 0.031111 | 0.083761 | 0.94395 | 2064 | 3 | -3.6316 | 0.74475 | 0.82375 | 1 | 18028 | 1 | -3.6316 |
| PCF11 | 4 | 0.80678 | 0.80735 | 0.999994 | 17580 | 1 | 0.3337 | 0.74475 | 0.82375 | 1 | 18029 | 1 | 0.3337 |
| RPS24 | 4 | 0.079211 | 0.17729 | 0.968171 | 4198 | 3 | -1.4703 | 0.74475 | 0.82375 | 1 | 18030 | 1 | -1.4703 |
| TADA2A | 5 | 0.17399 | 0.33792 | 0.999994 | 7084 | 3 | -0.97647 | 0.74477 | 0.85611 | 1 | 18031 | 1 | -0.97647 |
| HIST1H4F | 5 | 0.18746 | 0.35733 | 0.999994 | 7407 | 3 | -0.87727 | 0.74485 | 0.85616 | 1 | 18032 | 1 | -0.87727 |
| GPRC5D | 6 | 0.32612 | 0.53682 | 0.999994 | 10644 | 2 | 0.13945 | 0.74487 | 0.87272 | 1 | 18033 | 2 | 0.13945 |
| MUT | 6 | 0.19761 | 0.38499 | 0.999994 | 7693 | 2 | 0.45815 | 0.74496 | 0.87276 | 1 | 18034 | 2 | 0.45815 |
| IDH3B | 5 | 0.034398 | 0.10173 | 0.946414 | 2232 | 4 | -1.0304 | 0.74507 | 0.85631 | 1 | 18035 | 1 | -1.0304 |
| ANXA4 | 6 | 0.39584 | 0.5997 | 0.999994 | 11889 | 2 | 0.24135 | 0.74509 | 0.87282 | 1 | 18036 | 2 | 0.24135 |
| SAMSN1 | 6 | 0.076116 | 0.19949 | 0.977164 | 4078 | 4 | -1.0128 | 0.7453 | 0.87292 | 1 | 18037 | 2 | -1.0128 |
| HDAC11 | 6 | 0.09541 | 0.23835 | 0.987253 | 4798 | 4 | -0.87859 | 0.74532 | 0.87293 | 1 | 18038 | 1 | -0.87859 |
| MAP3K9 | 5 | 0.51585 | 0.6617 | 0.999994 | 13671 | 2 | -0.50927 | 0.74536 | 0.85648 | 1 | 18039 | 2 | -0.50927 |
| TNFRSF17 | 5 | 0.028328 | 0.08618 | 0.94395 | 1917 | 4 | -2.4639 | 0.74545 | 0.85654 | 1 | 18040 | 1 | -2.4639 |
| MASP2 | 5 | 0.026527 | 0.081498 | 0.93914 | 1829 | 4 | -1.7775 | 0.7458 | 0.85676 | 1 | 18041 | 1 | -1.7775 |
| VCAM1 | 6 | 0.078706 | 0.20473 | 0.978186 | 4177 | 4 | -1.0182 | 0.74583 | 0.87316 | 1 | 18042 | 1 | -1.0182 |
| LILRA4 | 5 | 0.51587 | 0.66172 | 0.999994 | 13672 | 2 | 0.42821 | 0.7459 | 0.85682 | 1 | 18043 | 2 | 0.42821 |

| | | | | | | | | | | | | | |
|---|---|---|---|---|---|---|---|---|---|---|---|---|---|
| hsa-mir-4426 | 2 | 0.54695 | 0.54797 | 0.999994 | 14168 | 1 | -0.14795 | 0.74594 | 0.7456 | 1 | 18044 | 0 | -0.14795 |
| PARP2 | 5 | 0.0015676 | 0.0069257 | 0.643671 | 230 | 4 | -3.2542 | 0.74599 | 0.85687 | 1 | 18045 | 1 | -3.2542 |
| hsa-mir-6076 | 2 | 0.21705 | 0.27447 | 0.998561 | 8194 | 1 | -1.8315 | 0.7462 | 0.74587 | 1 | 18046 | 0 | -1.8315 |
| ING5 | 5 | 0.1005 | 0.22721 | 0.983354 | 4989 | 3 | -1.4836 | 0.74623 | 0.85703 | 1 | 18047 | 2 | -1.4836 |
| hsa-mir-548p | 1 | 0.2537 | 0.25354 | 0.994544 | 9098 | 1 | -1.5014 | 0.7463 | 0.74659 | 1 | 18048 | 0 | -1.5014 |
| HBZ | 6 | 0.17266 | 0.35403 | 0.999994 | 7053 | 2 | -0.073232 | 0.74631 | 0.87336 | 1 | 18049 | 1 | -0.073232 |
| hsa-mir-3974 | 3 | 0.055372 | 0.12316 | 0.951052 | 3214 | 2 | -3.3658 | 0.74646 | 0.7704 | 1 | 18050 | 1 | -3.3658 |
| TRIM39-RPP21 | 2 | 0.7384 | 0.7381 | 0.999994 | 16712 | 0 | 0.39986 | 0.74646 | 0.74612 | 1 | 18051 | 0 | 0.39986 |
| COL2A1 | 6 | 0.0050258 | 0.021239 | 0.812607 | 523 | 5 | -2.0866 | 0.74647 | 0.87344 | 1 | 18052 | 1 | -2.0866 |
| ITGB5 | 5 | 0.75093 | 0.79133 | 0.999994 | 16864 | 1 | -0.098006 | 0.74649 | 0.85719 | 1 | 18053 | 2 | -0.098006 |
| hsa-mir-367 | 4 | 0.43416 | 0.58076 | 0.999994 | 12479 | 2 | -0.075882 | 0.74684 | 0.82456 | 1 | 18054 | 1 | -0.075882 |
| CD52 | 5 | 0.43554 | 0.59309 | 0.999994 | 12508 | 2 | -0.44128 | 0.74685 | 0.85743 | 1 | 18055 | 2 | -0.44128 |
| NonTargetingControlG | 1 | 0.25305 | 0.25289 | 0.994501 | 9085 | 1 | -1.4333 | 0.74695 | 0.74724 | 1 | 18056 | 0 | -1.4333 |
| MSN | 5 | 0.096157 | 0.22034 | 0.980699 | 4822 | 2 | 0.095108 | 0.74715 | 0.85763 | 1 | 18057 | 2 | 0.095108 |
| NonTargetingControlG | 2 | 0.60266 | 0.60246 | 0.999994 | 14980 | 0 | 0.0024574 | 0.74715 | 0.7468 | 1 | 18058 | 0 | 0.0024574 |
| PLXNC1 | 4 | 0.76125 | 0.7688 | 0.999994 | 16993 | 1 | -0.070593 | 0.74719 | 0.8247 | 1 | 18059 | 1 | -0.070593 |
| STX16 | 5 | 0.82069 | 0.83307 | 0.999994 | 17786 | 1 | 0.32247 | 0.74723 | 0.85769 | 1 | 18060 | 2 | 0.32247 |
| MALL | 6 | 0.32779 | 0.53872 | 0.999994 | 10681 | 3 | -0.43288 | 0.74723 | 0.87379 | 1 | 18061 | 2 | -0.43288 |
| IL13RA1 | 5 | 0.11556 | 0.2506 | 0.992497 | 5498 | 3 | -1.2386 | 0.74727 | 0.85772 | 1 | 18062 | 2 | -1.2386 |
| C9orf84 | 6 | 0.10396 | 0.25506 | 0.994943 | 5094 | 3 | -0.90637 | 0.74751 | 0.87392 | 1 | 18063 | 2 | -0.90637 |
| FGF20 | 5 | 0.462 | 0.61533 | 0.999994 | 12885 | 2 | -0.53029 | 0.74759 | 0.85795 | 1 | 18064 | 1 | -0.53029 |
| RERE | 5 | 0.021437 | 0.067962 | 0.93914 | 1543 | 4 | -1.8244 | 0.74759 | 0.85795 | 1 | 18065 | 1 | -1.8244 |
| UBA7 | 6 | 0.12187 | 0.28927 | 0.999632 | 5698 | 2 | -0.3663 | 0.7476 | 0.87395 | 1 | 18066 | 1 | -0.3663 |
| TUBB6 | 6 | 0.1178 | 0.28158 | 0.999632 | 5567 | 4 | -1.1129 | 0.7476 | 0.87395 | 1 | 18067 | 1 | -1.1129 |
| CATSPER4 | 5 | 0.074229 | 0.18525 | 0.968626 | 4001 | 2 | -0.0040138 | 0.74768 | 0.85801 | 1 | 18068 | 2 | -0.0040138 |
| PWWP2B | 6 | 0.13129 | 0.30198 | 0.999994 | 5965 | 3 | -0.69274 | 0.74774 | 0.87402 | 1 | 18069 | 2 | -0.69274 |
| XKRY | 6 | 0.56208 | 0.7123 | 0.999994 | 14409 | 2 | -0.12795 | 0.74776 | 0.87402 | 1 | 18070 | 2 | -0.12795 |
| UNC50 | 5 | 0.21055 | 0.39016 | 0.999994 | 8039 | 2 | -0.092396 | 0.74782 | 0.85809 | 1 | 18071 | 2 | -0.092396 |
| EVA1C | 6 | 0.26158 | 0.46178 | 0.999994 | 9275 | 3 | -0.21335 | 0.74785 | 0.87407 | 1 | 18072 | 2 | -0.21335 |
| NonTargetingControlG | 2 | 0.4971 | 0.50129 | 0.999994 | 13379 | 1 | -0.30625 | 0.74797 | 0.7476 | 1 | 18073 | 0 | -0.30625 |
| C22orf26 | 2 | 0.62339 | 0.62314 | 0.999994 | 15282 | 0 | 0.057553 | 0.74799 | 0.74762 | 1 | 18074 | 0 | 0.057553 |
| OLFM2 | 5 | 0.15793 | 0.31452 | 0.999994 | 6669 | 3 | -0.94573 | 0.74807 | 0.85828 | 1 | 18075 | 2 | -0.94573 |
| hsa-mir-4760 | 3 | 0.75926 | 0.75926 | 0.999994 | 16972 | 0 | -0.24753 | 0.74812 | 0.77152 | 1 | 18076 | 1 | -0.24753 |
| TRIM38 | 5 | 0.021487 | 0.068084 | 0.93914 | 1547 | 3 | -3.0265 | 0.7482 | 0.85837 | 1 | 18077 | 2 | -3.0265 |
| WDFY2 | 6 | 0.26762 | 0.46887 | 0.999994 | 9413 | 2 | 0.28183 | 0.74825 | 0.87425 | 1 | 18078 | 2 | 0.28183 |
| ARNT | 5 | 0.063967 | 0.1686 | 0.967126 | 3607 | 2 | -0.1632 | 0.74825 | 0.85839 | 1 | 18079 | 2 | -0.1632 |
| hsa-mir-7706 | 3 | 0.27151 | 0.40153 | 0.999994 | 9511 | 2 | -0.73488 | 0.74825 | 0.7716 | 1 | 18080 | 1 | -0.73488 |
| PLAC1 | 6 | 0.24559 | 0.44267 | 0.999994 | 8913 | 3 | -0.37099 | 0.74833 | 0.87429 | 1 | 18081 | 2 | -0.37099 |
| TRPC3 | 5 | 0.044868 | 0.12798 | 0.951704 | 2750 | 4 | -1.6746 | 0.74834 | 0.85845 | 1 | 18082 | 1 | -1.6746 |
| C8orf76 | 6 | 0.60061 | 0.7404 | 0.999994 | 14954 | 2 | -0.088438 | 0.7484 | 0.87432 | 1 | 18083 | 2 | -0.088438 |
| hsa-mir-5000 | 4 | 0.63735 | 0.68598 | 0.999994 | 15476 | 1 | 0.18792 | 0.74853 | 0.8252 | 1 | 18084 | 1 | 0.18792 |
| CHPF | 5 | 0.69456 | 0.76402 | 0.999994 | 16249 | 1 | 0.099004 | 0.74877 | 0.85871 | 1 | 18085 | 2 | 0.099004 |
| HGS | 5 | 0.17924 | 0.34552 | 0.999994 | 7205 | 3 | -0.86671 | 0.74883 | 0.85875 | 1 | 18086 | 1 | -0.86671 |
| NonTargetingControlG | 2 | 0.73664 | 0.73633 | 0.999994 | 16687 | 0 | 0.4521 | 0.74886 | 0.74848 | 1 | 18087 | 0 | 0.4521 |
| C1orf109 | 6 | 0.042755 | 0.12813 | 0.951777 | 2664 | 3 | -1.9673 | 0.74891 | 0.87452 | 1 | 18088 | 2 | -1.9673 |
| FUCA1 | 6 | 0.08669 | 0.22085 | 0.98185 | 4474 | 4 | -1.1378 | 0.74891 | 0.87452 | 1 | 18089 | 1 | -1.1378 |
| ENTPD4 | 6 | 0.003877 | 0.017028 | 0.785648 | 436 | 4 | -4.4405 | 0.74891 | 0.87452 | 1 | 18090 | 2 | -4.4405 |
| KLHL31 | 6 | 0.049023 | 0.14196 | 0.959171 | 2931 | 4 | -1.5921 | 0.74891 | 0.87452 | 1 | 18091 | 1 | -1.5921 |
| BEX5 | 5 | 0.14255 | 0.2917 | 0.999632 | 6254 | 3 | -0.99705 | 0.74933 | 0.85905 | 1 | 18092 | 2 | -0.99705 |
| KIAA1804 | 5 | 0.021383 | 0.067805 | 0.93914 | 1539 | 2 | 0.26551 | 0.74935 | 0.85906 | 1 | 18093 | 2 | 0.26551 |
| COPB2 | 3 | 0.033903 | 0.084035 | 0.94395 | 2212 | 2 | -4.0433 | 0.74936 | 0.77232 | 1 | 18094 | 1 | -4.0433 |
| PCSK4 | 6 | 0.19432 | 0.38094 | 0.999994 | 7615 | 3 | -0.71967 | 0.74939 | 0.87473 | 1 | 18095 | 2 | -0.71967 |

| | | | | | | | | | | | | | |
|---|---|---|---|---|---|---|---|---|---|---|---|---|---|
| DLX5 | 5 | 0.17288 | 0.33634 | 0.999994 | 7063 | 3 | -1.014 | 0.74956 | 0.85919 | 1 | 18096 | 2 | -1.014 |
| CNTN5 | 5 | 0.4621 | 0.61542 | 0.999994 | 12890 | 2 | -0.43878 | 0.74981 | 0.85935 | 1 | 18097 | 2 | -0.43878 |
| hsa-mir-5092 | 2 | 0.24235 | 0.29319 | 0.999994 | 8839 | 1 | -1.6163 | 0.74993 | 0.74952 | 1 | 18098 | 0 | -1.6163 |
| NCALD | 6 | 0.21253 | 0.40316 | 0.999994 | 8074 | 3 | -0.39082 | 0.75006 | 0.87503 | 1 | 18099 | 1 | -0.39082 |
| MRPL18 | 4 | 0.00012154 | 0.0004739 | 0.275324 | 40 | 3 | -9.4506 | 0.75011 | 0.82586 | 1 | 18100 | 1 | -9.4506 |
| NEXN | 5 | 0.04352 | 0.12461 | 0.951052 | 2694 | 3 | -1.1951 | 0.75015 | 0.85956 | 1 | 18101 | 2 | -1.1951 |
| FES | 5 | 0.14221 | 0.29118 | 0.999632 | 6245 | 2 | -0.43048 | 0.75015 | 0.85957 | 1 | 18102 | 1 | -0.43048 |
| RASGRP1 | 4 | 0.1229 | 0.25534 | 0.994943 | 5736 | 2 | -1.4318 | 0.75016 | 0.82588 | 1 | 18103 | 1 | -1.4318 |
| hsa-mir-103a-1 | 4 | 0.070681 | 0.16123 | 0.967126 | 3862 | 3 | -1.5006 | 0.75021 | 0.8259 | 1 | 18104 | 1 | -1.5006 |
| PSENEN | 6 | 0.3906 | 0.59635 | 0.999994 | 11808 | 1 | 0.025849 | 0.75025 | 0.87512 | 1 | 18105 | 1 | 0.025849 |
| C1QTNF4 | 4 | 0.36201 | 0.50662 | 0.999994 | 11369 | 2 | -0.3838 | 0.75029 | 0.82593 | 1 | 18106 | 1 | -0.3838 |
| ENSA | 5 | 0.34855 | 0.52073 | 0.999994 | 11124 | 2 | 0.052505 | 0.75037 | 0.85971 | 1 | 18107 | 2 | 0.052505 |
| CST6 | 2 | 0.58878 | 0.5888 | 0.999994 | 14801 | 0 | 0.001047 | 0.75047 | 0.75005 | 1 | 18108 | 0 | 0.001047 |
| ATXN1L | 4 | 0.6727 | 0.70703 | 0.999994 | 15967 | 1 | 0.13928 | 0.75055 | 0.82603 | 1 | 18109 | 1 | 0.13928 |
| SPDYE2 | 5 | 0.13207 | 0.27577 | 0.999379 | 5978 | 3 | -1.3674 | 0.75068 | 0.8599 | 1 | 18110 | 1 | -1.3674 |
| OR10Z1 | 5 | 0.51855 | 0.66406 | 0.999994 | 13717 | 2 | 0.2867 | 0.75072 | 0.85993 | 1 | 18111 | 2 | 0.2867 |
| SMIM3 | 6 | 0.011859 | 0.045255 | 0.891621 | 984 | 4 | -2.5958 | 0.75076 | 0.87535 | 1 | 18112 | 1 | -2.5958 |
| TBL1XR1 | 6 | 0.82798 | 0.86059 | 0.999994 | 17896 | 1 | 0.18811 | 0.75076 | 0.87535 | 1 | 18113 | 2 | 0.18811 |
| hsa-mir-4716 | 3 | 0.14168 | 0.23783 | 0.987253 | 6233 | 2 | -1.76 | 0.75079 | 0.77328 | 1 | 18114 | 1 | -1.76 |
| VPS13B | 6 | 0.21879 | 0.41066 | 0.999994 | 8254 | 3 | -0.43898 | 0.75108 | 0.8755 | 1 | 18115 | 2 | -0.43898 |
| C9orf43 | 6 | 0.10362 | 0.25438 | 0.994835 | 5087 | 3 | -1.3481 | 0.75127 | 0.87558 | 1 | 18116 | 2 | -1.3481 |
| SLC2A6 | 6 | 0.41034 | 0.60912 | 0.999994 | 12115 | 2 | 0.046305 | 0.75142 | 0.87565 | 1 | 18117 | 2 | 0.046305 |
| SRF | 6 | 0.070255 | 0.18731 | 0.969409 | 3844 | 4 | -1.1663 | 0.75203 | 0.87595 | 1 | 18118 | 1 | -1.1663 |
| SERPINB9 | 6 | 0.67563 | 0.79759 | 0.999994 | 16016 | 2 | 0.014018 | 0.75203 | 0.87595 | 1 | 18119 | 1 | 0.014018 |
| CYP2D6 | 6 | 0.040112 | 0.12216 | 0.951052 | 2523 | 4 | -2.205 | 0.75206 | 0.87597 | 1 | 18120 | 2 | -2.205 |
| CD163 | 5 | 0.032613 | 0.097182 | 0.945253 | 2139 | 3 | -0.6958 | 0.7521 | 0.86082 | 1 | 18121 | 2 | -0.6958 |
| LRRC58 | 5 | 0.14629 | 0.2972 | 0.999994 | 6361 | 2 | -0.09746 | 0.7521 | 0.86082 | 1 | 18122 | 2 | -0.09746 |
| EFNA1 | 6 | 0.06391 | 0.17405 | 0.968171 | 3602 | 4 | -1.085 | 0.75227 | 0.87607 | 1 | 18123 | 2 | -1.085 |
| PCDHGA7 | 2 | 0.6036 | 0.60339 | 0.999994 | 14988 | 0 | -0.048583 | 0.7523 | 0.75185 | 1 | 18124 | 0 | -0.048583 |
| DNAJA2 | 6 | 0.27502 | 0.47761 | 0.999994 | 9592 | 2 | -0.22167 | 0.75247 | 0.87616 | 1 | 18125 | 2 | -0.22167 |
| HNRNPA3 | 5 | 0.031239 | 0.093668 | 0.945031 | 2070 | 4 | -3.1022 | 0.75264 | 0.86119 | 1 | 18126 | 1 | -3.1022 |
| NonTargetingControlG | 2 | 0.65488 | 0.65472 | 0.999994 | 15714 | 0 | 0.11375 | 0.75279 | 0.75233 | 1 | 18127 | 0 | 0.11375 |
| SPATA5L1 | 5 | 0.082368 | 0.19843 | 0.977164 | 4309 | 2 | 0.18567 | 0.75285 | 0.86132 | 1 | 18128 | 2 | 0.18567 |
| ZNF735 | 5 | 0.90482 | 0.9049 | 0.999994 | 19321 | 0 | 0.50465 | 0.75299 | 0.86141 | 1 | 18129 | 2 | 0.50465 |
| NENF | 6 | 0.11323 | 0.2729 | 0.998561 | 5413 | 4 | -0.77603 | 0.75304 | 0.87643 | 1 | 18130 | 2 | -0.77603 |
| NonTargetingControlG | 2 | 0.49556 | 0.49986 | 0.999994 | 13349 | 1 | -0.34689 | 0.75311 | 0.75265 | 1 | 18131 | 0 | -0.34689 |
| AMMECR1 | 5 | 0.78848 | 0.8124 | 0.999994 | 17324 | 1 | 0.26562 | 0.75344 | 0.86169 | 1 | 18132 | 2 | 0.26562 |
| APOL4 | 5 | 0.00021493 | 0.0010184 | 0.352868 | 59 | 3 | -8.2837 | 0.75359 | 0.86179 | 1 | 18133 | 2 | -8.2837 |
| hsa-mir-1281 | 2 | 0.53634 | 0.5378 | 0.999994 | 14006 | 1 | -0.27788 | 0.75388 | 0.75343 | 1 | 18134 | 0 | -0.27788 |
| GABRP | 6 | 0.11379 | 0.27395 | 0.998561 | 5437 | 4 | -0.9637 | 0.7539 | 0.87682 | 1 | 18135 | 1 | -0.9637 |
| FOXD2 | 6 | 0.0053172 | 0.022313 | 0.819701 | 542 | 5 | -3.146 | 0.7539 | 0.87682 | 1 | 18136 | 1 | -3.146 |
| ST7L | 6 | 0.087622 | 0.22272 | 0.983354 | 4498 | 4 | -1.2246 | 0.7539 | 0.87682 | 1 | 18137 | 1 | -1.2246 |
| PAK2 | 6 | 0.4623 | 0.64326 | 0.999994 | 12894 | 2 | -0.16568 | 0.7539 | 0.87682 | 1 | 18138 | 2 | -0.16568 |
| RNF114 | 6 | 0.11305 | 0.27256 | 0.998561 | 5406 | 3 | -1.2498 | 0.7539 | 0.87682 | 1 | 18139 | 1 | -1.2498 |
| FAM179B | 6 | 0.90718 | 0.90927 | 0.999994 | 19374 | 1 | -0.16365 | 0.7539 | 0.87682 | 1 | 18140 | 1 | -0.16365 |
| OSBPL1A | 6 | 0.00038007 | 0.002017 | 0.439257 | 90 | 4 | -2.3928 | 0.75408 | 0.87691 | 1 | 18141 | 2 | -2.3928 |
| STEAP2 | 6 | 0.01621 | 0.059694 | 0.932932 | 1258 | 3 | -0.64884 | 0.75419 | 0.87696 | 1 | 18142 | 2 | -0.64884 |
| DDX59 | 5 | 0.38688 | 0.55252 | 0.999994 | 11752 | 1 | 0.43629 | 0.75437 | 0.8623 | 1 | 18143 | 1 | 0.43629 |
| PCDHA7 | 2 | 0.4718 | 0.47858 | 0.999994 | 13030 | 1 | -0.4765 | 0.75448 | 0.75403 | 1 | 18144 | 0 | -0.4765 |
| WBP2 | 5 | 0.00013357 | 0.00061427 | 0.302318 | 44 | 4 | -6.5605 | 0.7545 | 0.86238 | 1 | 18145 | 1 | -6.5605 |
| ELOVL2 | 4 | 0.07187 | 0.16347 | 0.967126 | 3913 | 2 | -2.0944 | 0.75453 | 0.82758 | 1 | 18146 | 1 | -2.0944 |
| ZNF200 | 6 | 0.025936 | 0.089216 | 0.94395 | 1800 | 4 | -1.706 | 0.75457 | 0.87712 | 1 | 18147 | 2 | -1.706 |

| | | | | | | | | | | | | |
|---|---|---|---|---|---|---|---|---|---|---|---|---|
| SAA2-SAA4 | 4 | 0.12356 | 0.25651 | 0.994943 | 5755 | 2 | -1.3272 | 0.75458 | 0.82759 | 1 | 18148 | 1 | -1.3272 |
| SOX11 | 5 | 0.055809 | 0.15436 | 0.967126 | 3238 | 4 | -0.75732 | 0.75476 | 0.86255 | 1 | 18149 | 1 | -0.75732 |
| RNF115 | 5 | 0.092869 | 0.21516 | 0.979587 | 4714 | 3 | -0.76825 | 0.7548 | 0.86257 | 1 | 18150 | 2 | -0.76825 |
| CSNK1A1L | 6 | 0.60206 | 0.74146 | 0.999994 | 14973 | 2 | 0.24171 | 0.75484 | 0.87726 | 1 | 18151 | 2 | 0.24171 |
| SHMT1 | 5 | 0.17137 | 0.33413 | 0.999994 | 7019 | 2 | -0.6004 | 0.75486 | 0.86261 | 1 | 18152 | 2 | -0.6004 |
| FCGR2A | 5 | 0.18317 | 0.35119 | 0.999994 | 7297 | 3 | -0.93696 | 0.75487 | 0.86262 | 1 | 18153 | 2 | -0.93696 |
| XRN1 | 5 | 0.036866 | 0.10802 | 0.951052 | 2368 | 2 | -0.087225 | 0.75498 | 0.86268 | 1 | 18154 | 2 | -0.087225 |
| LOC441155 | 6 | 0.29714 | 0.50339 | 0.999994 | 10071 | 3 | -0.7549 | 0.75538 | 0.8775 | 1 | 18155 | 2 | -0.7549 |
| RPL18 | 2 | 0.54293 | 0.54411 | 0.999994 | 14111 | 1 | -0.17048 | 0.75549 | 0.75505 | 1 | 18156 | 0 | -0.17048 |
| NonTargetingControlG | 2 | 0.57342 | 0.57359 | 0.999994 | 14597 | 1 | -0.089982 | 0.75553 | 0.75509 | 1 | 18157 | 0 | -0.089982 |
| C2orf68 | 6 | 0.10802 | 0.26289 | 0.99665 | 5241 | 4 | -1.0252 | 0.75559 | 0.87759 | 1 | 18158 | 1 | -1.0252 |
| PMS1 | 6 | 0.23364 | 0.42844 | 0.999994 | 8611 | 3 | -0.59968 | 0.75559 | 0.87759 | 1 | 18159 | 1 | -0.59968 |
| GZMM | 6 | 0.24551 | 0.44258 | 0.999994 | 8912 | 3 | -0.81267 | 0.75559 | 0.87759 | 1 | 18160 | 2 | -0.81267 |
| MRVI1 | 5 | 0.01721 | 0.056443 | 0.9235 | 1312 | 3 | -3.4033 | 0.75584 | 0.86327 | 1 | 18161 | 1 | -3.4033 |
| S100A1 | 5 | 0.88785 | 0.88795 | 0.999994 | 18994 | 1 | -0.085102 | 0.75584 | 0.86327 | 1 | 18162 | 1 | -0.085102 |
| C16orf46 | 3 | 0.28236 | 0.41488 | 0.999994 | 9747 | 2 | -0.61725 | 0.75593 | 0.77669 | 1 | 18163 | 1 | -0.61725 |
| hsa-mir-3615 | 3 | 0.22902 | 0.34885 | 0.999994 | 8506 | 2 | -1.0179 | 0.75593 | 0.77669 | 1 | 18164 | 1 | -1.0179 |
| hsa-mir-1205 | 3 | 0.052404 | 0.11904 | 0.951052 | 3087 | 2 | -0.88911 | 0.75593 | 0.77669 | 1 | 18165 | 1 | -0.88911 |
| hsa-mir-6736 | 3 | 0.16819 | 0.27205 | 0.998561 | 6944 | 2 | -1.4861 | 0.75593 | 0.77669 | 1 | 18166 | 1 | -1.4861 |
| OR5J2 | 5 | 0.44177 | 0.59827 | 0.999994 | 12606 | 2 | -0.13403 | 0.75604 | 0.86339 | 1 | 18167 | 2 | -0.13403 |
| TTN | 5 | 0.14242 | 0.29151 | 0.999632 | 6250 | 3 | -1.1891 | 0.75626 | 0.86354 | 1 | 18168 | 2 | -1.1891 |
| SOS2 | 6 | 0.26773 | 0.469 | 0.999994 | 9418 | 3 | -0.35622 | 0.75638 | 0.87796 | 1 | 18169 | 2 | -0.35622 |
| AADACL3 | 4 | 0.13446 | 0.27319 | 0.998561 | 6038 | 1 | -0.047466 | 0.75648 | 0.8283 | 1 | 18170 | 1 | -0.047466 |
| SIRPB2 | 6 | 0.61491 | 0.75101 | 0.999994 | 15141 | 2 | -0.2202 | 0.7566 | 0.87807 | 1 | 18171 | 2 | -0.2202 |
| RYR2 | 6 | 0.098625 | 0.24465 | 0.988995 | 4917 | 4 | -1.3567 | 0.75668 | 0.87811 | 1 | 18172 | 2 | -1.3567 |
| CSRNP1 | 6 | 0.45237 | 0.63662 | 0.999994 | 12740 | 2 | -0.26953 | 0.75673 | 0.87813 | 1 | 18173 | 1 | -0.26953 |
| SEPT3 | 5 | 0.47582 | 0.62701 | 0.999994 | 13075 | 2 | -0.12271 | 0.75675 | 0.86385 | 1 | 18174 | 1 | -0.12271 |
| hsa-mir-1248 | 4 | 0.67817 | 0.71042 | 0.999994 | 16063 | 1 | 0.13417 | 0.7569 | 0.82847 | 1 | 18175 | 1 | 0.13417 |
| MIER3 | 5 | 0.60815 | 0.72913 | 0.999994 | 15047 | 1 | 0.13282 | 0.75707 | 0.86407 | 1 | 18176 | 1 | 0.13282 |
| UBL5 | 2 | 0.012566 | 0.023723 | 0.832699 | 1033 | 1 | -5.0775 | 0.7573 | 0.75686 | 1 | 18177 | 0 | -5.0775 |
| CDC23 | 2 | 0.23534 | 0.28795 | 0.999632 | 8657 | 1 | -1.6759 | 0.7573 | 0.75686 | 1 | 18178 | 0 | -1.6759 |
| hsa-mir-4290 | 2 | 0.73018 | 0.72993 | 0.999994 | 16599 | 0 | 0.40893 | 0.7573 | 0.75686 | 1 | 18179 | 0 | 0.40893 |
| GLTSCR1 | 4 | 0.27007 | 0.41235 | 0.999994 | 9475 | 2 | -0.92961 | 0.7574 | 0.82866 | 1 | 18180 | 1 | -0.92961 |
| PFKFB4 | 6 | 0.059504 | 0.16471 | 0.967126 | 3401 | 4 | -1.7456 | 0.7574 | 0.87842 | 1 | 18181 | 2 | -1.7456 |
| ZNF184 | 6 | 0.095048 | 0.23763 | 0.987137 | 4786 | 4 | -1.1973 | 0.7574 | 0.87842 | 1 | 18182 | 1 | -1.1973 |
| SNRPB | 6 | 0.00016419 | 0.00089546 | 0.352558 | 50 | 4 | -7.1621 | 0.7574 | 0.87842 | 1 | 18183 | 1 | -7.1621 |
| HSPA13 | 6 | 0.084699 | 0.21692 | 0.980699 | 4395 | 3 | -0.33543 | 0.7574 | 0.87842 | 1 | 18184 | 1 | -0.33543 |
| MYH3 | 4 | 0.31701 | 0.4604 | 0.999994 | 10453 | 2 | -0.7294 | 0.75747 | 0.82869 | 1 | 18185 | 1 | -0.7294 |
| DYNLL2 | 6 | 0.61236 | 0.74907 | 0.999994 | 15114 | 2 | 0.085591 | 0.75749 | 0.87847 | 1 | 18186 | 2 | 0.085591 |
| TBX4 | 5 | 0.02637 | 0.081076 | 0.93914 | 1823 | 4 | -2.4521 | 0.75822 | 0.86486 | 1 | 18187 | 1 | -2.4521 |
| ZNF415 | 5 | 0.05348 | 0.14877 | 0.965566 | 3140 | 3 | -0.88154 | 0.75844 | 0.86502 | 1 | 18188 | 2 | -0.88154 |
| ZNF585B | 6 | 0.11376 | 0.2739 | 0.998561 | 5435 | 3 | -0.82986 | 0.75847 | 0.87892 | 1 | 18189 | 1 | -0.82986 |
| UCHL3 | 6 | 0.66697 | 0.79076 | 0.999994 | 15878 | 2 | -0.01152 | 0.75851 | 0.87894 | 1 | 18190 | 2 | -0.01152 |
| GPR25 | 5 | 0.017256 | 0.056572 | 0.9235 | 1318 | 4 | -2.2766 | 0.75854 | 0.86508 | 1 | 18191 | 1 | -2.2766 |
| NonTargetingControlG | 2 | 0.56093 | 0.56145 | 0.999994 | 14387 | 1 | -0.18547 | 0.75854 | 0.75809 | 1 | 18192 | 0 | -0.18547 |
| SPPL2C | 4 | 0.27135 | 0.41367 | 0.999994 | 9507 | 1 | -0.14511 | 0.75874 | 0.8292 | 1 | 18193 | 1 | -0.14511 |
| C4B | 1 | 0.24125 | 0.24105 | 0.987253 | 8803 | 1 | -1.464 | 0.75875 | 0.75907 | 1 | 18194 | 0 | -1.464 |
| TMEM59L | 5 | 0.064002 | 0.16865 | 0.967126 | 3611 | 3 | -2.0974 | 0.75878 | 0.86524 | 1 | 18195 | 1 | -2.0974 |
| NOXO1 | 6 | 0.075031 | 0.19722 | 0.977164 | 4036 | 4 | -1.3259 | 0.75888 | 0.87912 | 1 | 18196 | 2 | -1.3259 |
| OR4F16 | 5 | 0.3946 | 0.55906 | 0.999994 | 11868 | 2 | -0.084024 | 0.75896 | 0.86536 | 1 | 18197 | 2 | -0.084024 |
| STOML1 | 5 | 0.34783 | 0.52013 | 0.999994 | 11114 | 1 | 0.35554 | 0.75899 | 0.86538 | 1 | 18198 | 2 | 0.35554 |
| LHFPL3 | 5 | 0.85805 | 0.86078 | 0.999994 | 18407 | 1 | 0.13025 | 0.75902 | 0.8654 | 1 | 18199 | 2 | 0.13025 |

| | | | | | | | | | | | | | |
|---|---|---|---|---|---|---|---|---|---|---|---|---|---|
| C17orf78 | 5 | 0.31017 | 0.48927 | 0.999994 | 10328 | 2 | 0.14437 | 0.75905 | 0.86542 | 1 | 18200 | 1 | 0.14437 |
| CRTAP | 6 | 0.024275 | 0.08528 | 0.94395 | 1720 | 4 | -2.6283 | 0.75906 | 0.87921 | 1 | 18201 | 2 | -2.6283 |
| DSPP | 4 | 0.108 | 0.2293 | 0.983354 | 5240 | 3 | -1.0923 | 0.75907 | 0.82934 | 1 | 18202 | 1 | -1.0923 |
| IRX6 | 6 | 0.22484 | 0.41794 | 0.999994 | 8395 | 3 | -0.36809 | 0.75909 | 0.87922 | 1 | 18203 | 2 | -0.36809 |
| hsa-mir-650 | 2 | 0.11132 | 0.18746 | 0.969409 | 5343 | 1 | -3.739 | 0.75933 | 0.75886 | 1 | 18204 | 0 | -3.739 |
| DOC2A | 6 | 0.067883 | 0.18231 | 0.968526 | 3747 | 4 | -1.1671 | 0.75938 | 0.87935 | 1 | 18205 | 2 | -1.1671 |
| RHBDD3 | 4 | 0.092337 | 0.20129 | 0.977164 | 4693 | 1 | 0.43652 | 0.75946 | 0.82951 | 1 | 18206 | 1 | 0.43652 |
| NonTargetingControlG | 1 | 0.24036 | 0.24016 | 0.987253 | 8779 | 1 | -1.7279 | 0.75964 | 0.75996 | 1 | 18207 | 0 | -1.7279 |
| hsa-mir-7852 | 3 | 0.67918 | 0.68202 | 0.999994 | 16076 | 1 | 0.29842 | 0.76019 | 0.77962 | 1 | 18208 | 1 | 0.29842 |
| VILL | 4 | 0.090059 | 0.19712 | 0.977164 | 4597 | 3 | -1.0774 | 0.76021 | 0.82981 | 1 | 18209 | 1 | -1.0774 |
| SPICE1 | 6 | 0.05739 | 0.16013 | 0.967126 | 3310 | 3 | -1.0536 | 0.76026 | 0.87978 | 1 | 18210 | 1 | -1.0536 |
| HNRNPA0 | 6 | 0.10956 | 0.26586 | 0.997535 | 5291 | 4 | -1.1114 | 0.7603 | 0.8798 | 1 | 18211 | 2 | -1.1114 |
| PADI3 | 5 | 0.03719 | 0.10882 | 0.951052 | 2383 | 3 | -2.3564 | 0.76039 | 0.86632 | 1 | 18212 | 2 | -2.3564 |
| LOC402160 | 6 | 0.07269 | 0.19235 | 0.974607 | 3948 | 3 | -1.1482 | 0.76074 | 0.88 | 1 | 18213 | 2 | -1.1482 |
| PDDC1 | 5 | 0.5679 | 0.70767 | 0.999994 | 14511 | 2 | 0.040827 | 0.76081 | 0.86659 | 1 | 18214 | 2 | 0.040827 |
| TREM1 | 6 | 0.23025 | 0.42437 | 0.999994 | 8531 | 3 | -0.28772 | 0.76081 | 0.88004 | 1 | 18215 | 2 | -0.28772 |
| TEAD1 | 6 | 0.090806 | 0.22913 | 0.983354 | 4631 | 3 | -0.31277 | 0.76117 | 0.8802 | 1 | 18216 | 2 | -0.31277 |
| NFIA | 5 | 0.30038 | 0.48138 | 0.999994 | 10140 | 1 | 0.25357 | 0.76122 | 0.86685 | 1 | 18217 | 2 | 0.25357 |
| hsa-mir-378e | 4 | 0.80697 | 0.80752 | 0.999994 | 17584 | 1 | 0.020327 | 0.76137 | 0.83029 | 1 | 18218 | 1 | 0.020327 |
| BGLAP | 5 | 0.89803 | 0.89815 | 0.999994 | 19178 | 0 | -0.33157 | 0.76143 | 0.86699 | 1 | 18219 | 1 | -0.33157 |
| H2AFY2 | 6 | 0.013388 | 0.050399 | 0.909101 | 1091 | 4 | -3.1824 | 0.76148 | 0.88036 | 1 | 18220 | 2 | -3.1824 |
| CST9 | 5 | 0.046522 | 0.13201 | 0.95318 | 2831 | 4 | -1.1147 | 0.76161 | 0.8671 | 1 | 18221 | 1 | -1.1147 |
| APOM | 5 | 0.24188 | 0.43347 | 0.999994 | 8833 | 3 | -0.60958 | 0.76178 | 0.86723 | 1 | 18222 | 2 | -0.60958 |
| OR4C15 | 6 | 0.303 | 0.51022 | 0.999994 | 10193 | 3 | -0.21917 | 0.76186 | 0.88054 | 1 | 18223 | 2 | -0.21917 |
| hsa-mir-379 | 4 | 0.259 | 0.40112 | 0.999994 | 9218 | 2 | -0.72006 | 0.76193 | 0.83052 | 1 | 18224 | 1 | -0.72006 |
| NonTargetingControlG | 2 | 0.61345 | 0.61329 | 0.999994 | 15124 | 0 | -0.0010455 | 0.76196 | 0.76147 | 1 | 18225 | 0 | -0.0010455 |
| hsa-mir-4656 | 4 | 0.19851 | 0.33916 | 0.999994 | 7725 | 2 | -0.93659 | 0.76204 | 0.83058 | 1 | 18226 | 1 | -0.93659 |
| CHMP4C | 5 | 0.10185 | 0.22931 | 0.983354 | 5033 | 2 | -0.46518 | 0.76222 | 0.86753 | 1 | 18227 | 1 | -0.46518 |
| NonTargetingControlG | 2 | 0.23241 | 0.28572 | 0.999632 | 8581 | 1 | -1.395 | 0.76226 | 0.76179 | 1 | 18228 | 0 | -1.395 |
| EPC1 | 6 | 0.01727 | 0.063132 | 0.937389 | 1321 | 3 | -1.0937 | 0.76228 | 0.88075 | 1 | 18229 | 2 | -1.0937 |
| POLR3GL | 5 | 0.065018 | 0.17028 | 0.967126 | 3654 | 3 | -0.83459 | 0.7623 | 0.86758 | 1 | 18230 | 2 | -0.83459 |
| SPINK1 | 6 | 0.67418 | 0.7964 | 0.999994 | 15989 | 1 | 0.0093057 | 0.76233 | 0.88078 | 1 | 18231 | 2 | 0.0093057 |
| OR4X1 | 6 | 0.091941 | 0.2314 | 0.984304 | 4676 | 4 | -1.0043 | 0.76255 | 0.88087 | 1 | 18232 | 2 | -1.0043 |
| NUDT1 | 5 | 0.028705 | 0.087141 | 0.94395 | 1937 | 2 | -0.46926 | 0.76265 | 0.8678 | 1 | 18233 | 2 | -0.46926 |
| NonTargetingControlG | 2 | 0.62244 | 0.62222 | 0.999994 | 15266 | 0 | 0.01251 | 0.76271 | 0.76222 | 1 | 18234 | 0 | 0.01251 |
| SPI1 | 6 | 0.011779 | 0.04498 | 0.891621 | 976 | 5 | -2.1482 | 0.76311 | 0.88114 | 1 | 18235 | 1 | -2.1482 |
| ARL6 | 5 | 0.21728 | 0.39948 | 0.999994 | 8201 | 2 | -0.38049 | 0.76329 | 0.86821 | 1 | 18236 | 2 | -0.38049 |
| SEC14L6 | 5 | 0.53633 | 0.67965 | 0.999994 | 14005 | 2 | 0.56579 | 0.76345 | 0.86831 | 1 | 18237 | 2 | 0.56579 |
| CCDC166 | 5 | 0.89384 | 0.89397 | 0.999994 | 19100 | 0 | 0.35554 | 0.76347 | 0.86833 | 1 | 18238 | 2 | 0.35554 |
| XAGE2 | 5 | 0.078313 | 0.19187 | 0.974607 | 4162 | 2 | 0.3964 | 0.7636 | 0.86842 | 1 | 18239 | 1 | 0.3964 |
| hsa-mir-27a | 3 | 0.22366 | 0.34221 | 0.999994 | 8375 | 2 | -0.87791 | 0.76379 | 0.78204 | 1 | 18240 | 1 | -0.87791 |
| KCNE1 | 6 | 0.0047144 | 0.020114 | 0.808847 | 500 | 2 | 0.035832 | 0.76397 | 0.88153 | 1 | 18241 | 2 | 0.035832 |
| RAD21L1 | 5 | 0.014972 | 0.050122 | 0.909101 | 1187 | 1 | 0.087536 | 0.76397 | 0.86867 | 1 | 18242 | 2 | 0.087536 |
| FBXO4 | 6 | 0.041569 | 0.12544 | 0.951052 | 2594 | 3 | -1.7226 | 0.7642 | 0.88164 | 1 | 18243 | 1 | -1.7226 |
| TDRP | 4 | 0.29117 | 0.434 | 0.999994 | 9932 | 2 | -0.84315 | 0.7644 | 0.83151 | 1 | 18244 | 1 | -0.84315 |
| CHRD | 5 | 0.00023559 | 0.0011081 | 0.352868 | 67 | 3 | -9.1107 | 0.76443 | 0.86896 | 1 | 18245 | 2 | -9.1107 |
| hsa-mir-490 | 4 | 0.26126 | 0.40338 | 0.999994 | 9273 | 2 | -0.8988 | 0.76476 | 0.83164 | 1 | 18246 | 1 | -0.8988 |
| MX1 | 6 | 0.1226 | 0.29061 | 0.999632 | 5727 | 3 | -0.98524 | 0.76484 | 0.88193 | 1 | 18247 | 1 | -0.98524 |
| DGKD | 6 | 0.92856 | 0.92865 | 0.999994 | 19826 | 0 | 0.37765 | 0.76509 | 0.88206 | 1 | 18248 | 2 | 0.37765 |
| KRTAP6-2 | 5 | 0.041509 | 0.11966 | 0.951052 | 2592 | 2 | -0.32284 | 0.76513 | 0.86941 | 1 | 18249 | 2 | -0.32284 |
| TXNRD1 | 6 | 0.42529 | 0.61885 | 0.999994 | 12339 | 2 | 0.16791 | 0.76524 | 0.88213 | 1 | 18250 | 2 | 0.16791 |
| hsa-mir-520h | 1 | 0.23466 | 0.23437 | 0.984971 | 8632 | 1 | -1.7649 | 0.76534 | 0.7657 | 1 | 18251 | 0 | -1.7649 |

| | | | | | | | | | | | | |
|---|---|---|---|---|---|---|---|---|---|---|---|---|
| TMEM181 | 5 | 0.089069 | 0.20907 | 0.979587 | 4554 | 3 | -1.0346 | 0.76548 | 0.86965 | 1 | 18252 | 2 | -1.0346 |
| TMEM139 | 5 | 0.099391 | 0.22545 | 0.983354 | 4946 | 2 | 0.070534 | 0.76571 | 0.8698 | 1 | 18253 | 2 | 0.070534 |
| PARP15 | 6 | 0.27433 | 0.47681 | 0.999994 | 9573 | 3 | -0.66415 | 0.76572 | 0.88236 | 1 | 18254 | 2 | -0.66415 |
| OR2M7 | 6 | 0.047941 | 0.13957 | 0.958716 | 2887 | 3 | -0.73014 | 0.76572 | 0.88236 | 1 | 18255 | 2 | -0.73014 |
| TRIM31 | 6 | 0.068977 | 0.18463 | 0.968526 | 3798 | 4 | -1.2129 | 0.76587 | 0.88243 | 1 | 18256 | 2 | -1.2129 |
| DEPDC1B | 5 | 0.11493 | 0.24965 | 0.991904 | 5474 | 3 | -1.0231 | 0.76598 | 0.86998 | 1 | 18257 | 2 | -1.0231 |
| C7orf71 | 6 | 0.56457 | 0.71407 | 0.999994 | 14464 | 2 | -0.2649 | 0.76611 | 0.88254 | 1 | 18258 | 1 | -0.2649 |
| CD37 | 5 | 0.13601 | 0.28176 | 0.999632 | 6079 | 3 | -1.2598 | 0.76625 | 0.87016 | 1 | 18259 | 2 | -1.2598 |
| hsa-mir-6741 | 3 | 0.27355 | 0.40403 | 0.999994 | 9556 | 1 | -0.30669 | 0.76634 | 0.7838 | 1 | 18260 | 1 | -0.30669 |
| SRR | 6 | 0.046673 | 0.13678 | 0.957161 | 2837 | 4 | -1.6687 | 0.76644 | 0.8827 | 1 | 18261 | 2 | -1.6687 |
| PGPEP1 | 3 | 0.18221 | 0.29009 | 0.999632 | 7275 | 2 | -1.45 | 0.76656 | 0.78395 | 1 | 18262 | 1 | -1.45 |
| LPAR1 | 4 | 0.7594 | 0.76729 | 0.999994 | 16976 | 1 | -0.17891 | 0.76661 | 0.83239 | 1 | 18263 | 1 | -0.17891 |
| HIST1H3D | 6 | 0.34096 | 0.55376 | 0.999994 | 10961 | 2 | -0.050582 | 0.76669 | 0.88281 | 1 | 18264 | 2 | -0.050582 |
| hsa-mir-6746 | 3 | 0.049494 | 0.11501 | 0.951052 | 2956 | 2 | -0.79878 | 0.7667 | 0.78403 | 1 | 18265 | 1 | -0.79878 |
| CDK5RAP3 | 6 | 0.063868 | 0.17398 | 0.968171 | 3601 | 4 | -1.6425 | 0.76671 | 0.88281 | 1 | 18266 | 2 | -1.6425 |
| MLYCD | 6 | 0.08265 | 0.21274 | 0.979587 | 4324 | 4 | -1.1687 | 0.76672 | 0.88282 | 1 | 18267 | 2 | -1.1687 |
| TTC23L | 5 | 0.02379 | 0.074275 | 0.93914 | 1700 | 4 | -1.6641 | 0.76674 | 0.87051 | 1 | 18268 | 1 | -1.6641 |
| ABI3 | 2 | 0.60924 | 0.60908 | 0.999994 | 15060 | 0 | -0.071817 | 0.76707 | 0.76652 | 1 | 18269 | 0 | -0.071817 |
| hsa-mir-1324 | 4 | 0.089458 | 0.19603 | 0.975797 | 4576 | 3 | -1.9884 | 0.76717 | 0.83259 | 1 | 18270 | 1 | -1.9884 |
| HYAL2 | 6 | 0.08929 | 0.22609 | 0.983354 | 4567 | 3 | -0.83929 | 0.76725 | 0.88306 | 1 | 18271 | 2 | -0.83929 |
| MTMR8 | 5 | 0.21881 | 0.4016 | 0.999994 | 8255 | 3 | -0.64628 | 0.76727 | 0.87086 | 1 | 18272 | 2 | -0.64628 |
| MRPL40 | 4 | 0.034582 | 0.090878 | 0.94395 | 2245 | 3 | -3.6027 | 0.76727 | 0.83263 | 1 | 18273 | 1 | -3.6027 |
| HES4 | 5 | 0.22439 | 0.40939 | 0.999994 | 8391 | 1 | 0.27299 | 0.76737 | 0.87093 | 1 | 18274 | 1 | 0.27299 |
| DUSP16 | 6 | 0.18416 | 0.36833 | 0.999994 | 7320 | 2 | -0.46433 | 0.76748 | 0.88318 | 1 | 18275 | 2 | -0.46433 |
| NAPG | 1 | 0.23249 | 0.23221 | 0.984502 | 8582 | 1 | -1.7121 | 0.76751 | 0.76794 | 1 | 18276 | 0 | -1.7121 |
| NonTargetingControlG | 2 | 0.43996 | 0.45058 | 0.999994 | 12572 | 1 | -0.53034 | 0.76778 | 0.76725 | 1 | 18277 | 0 | -0.53034 |
| CA2 | 6 | 0.087991 | 0.2235 | 0.983354 | 4513 | 4 | -1.3974 | 0.76801 | 0.88344 | 1 | 18278 | 2 | -1.3974 |
| HBD | 5 | 0.022018 | 0.069541 | 0.93914 | 1586 | 4 | -2.2854 | 0.76806 | 0.87139 | 1 | 18279 | 1 | -2.2854 |
| hsa-mir-6089-1 | 3 | 0.62545 | 0.63702 | 0.999994 | 15313 | 1 | -0.59413 | 0.7681 | 0.78502 | 1 | 18280 | 1 | -0.59413 |
| PAGE2 | 5 | 0.12143 | 0.25954 | 0.994943 | 5681 | 3 | -1.3299 | 0.76821 | 0.87149 | 1 | 18281 | 2 | -1.3299 |
| ADAMTS16 | 5 | 0.016275 | 0.053818 | 0.921281 | 1261 | 3 | -1.5662 | 0.76834 | 0.87157 | 1 | 18282 | 1 | -1.5662 |
| PIK3R1 | 5 | 0.067714 | 0.17469 | 0.968171 | 3743 | 3 | -1.8191 | 0.76837 | 0.87159 | 1 | 18283 | 1 | -1.8191 |
| FOXN3 | 6 | 0.0049926 | 0.021122 | 0.812607 | 517 | 4 | -4.1422 | 0.7684 | 0.88364 | 1 | 18284 | 1 | -4.1422 |
| IL19 | 6 | 0.31544 | 0.52466 | 0.999994 | 10429 | 2 | -0.023898 | 0.7684 | 0.88364 | 1 | 18285 | 2 | -0.023898 |
| GULP1 | 6 | 0.096922 | 0.24133 | 0.987253 | 4855 | 4 | -0.97719 | 0.7684 | 0.88364 | 1 | 18286 | 1 | -0.97719 |
| SLC25A1 | 4 | 0.0095324 | 0.032217 | 0.865899 | 838 | 3 | -4.6062 | 0.76859 | 0.83316 | 1 | 18287 | 1 | -4.6062 |
| TRANK1 | 5 | 0.0014496 | 0.00643 | 0.636218 | 218 | 3 | -7.3728 | 0.76899 | 0.87202 | 1 | 18288 | 2 | -7.3728 |
| RBM43 | 6 | 0.099634 | 0.24664 | 0.98999 | 4962 | 4 | -0.82213 | 0.76907 | 0.88396 | 1 | 18289 | 2 | -0.82213 |
| NonTargetingControlG | 2 | 0.53166 | 0.5334 | 0.999994 | 13927 | 1 | -0.3072 | 0.76911 | 0.76862 | 1 | 18290 | 0 | -0.3072 |
| SLC4A4 | 6 | 0.59943 | 0.73952 | 0.999994 | 14946 | 2 | -0.28504 | 0.76944 | 0.88414 | 1 | 18291 | 2 | -0.28504 |
| KIAA0368 | 5 | 0.56232 | 0.70271 | 0.999994 | 14416 | 1 | 0.45012 | 0.76948 | 0.87234 | 1 | 18292 | 2 | 0.45012 |
| YWHAG | 6 | 0.21826 | 0.41004 | 0.999994 | 8235 | 3 | -0.49327 | 0.76948 | 0.88416 | 1 | 18293 | 2 | -0.49327 |
| SQSTM1 | 6 | 0.0043199 | 0.018658 | 0.801598 | 466 | 4 | -1.3264 | 0.76951 | 0.88417 | 1 | 18294 | 1 | -1.3264 |
| GJA3 | 6 | 0.020394 | 0.073241 | 0.93914 | 1482 | 4 | -2.2272 | 0.76967 | 0.88425 | 1 | 18295 | 2 | -2.2272 |
| NonTargetingControlG | 2 | 0.003269 | 0.0062828 | 0.636218 | 380 | 1 | -5.6819 | 0.7697 | 0.76921 | 1 | 18296 | 0 | -5.6819 |
| PLEKHO2 | 6 | 0.3169 | 0.52633 | 0.999994 | 10449 | 3 | -0.35851 | 0.77025 | 0.88451 | 1 | 18297 | 2 | -0.35851 |
| NonTargetingControlG | 2 | 0.70261 | 0.70235 | 0.999994 | 16333 | 0 | 0.21582 | 0.77027 | 0.76977 | 1 | 18298 | 0 | 0.21582 |
| ARL14 | 6 | 0.15678 | 0.33416 | 0.999994 | 6646 | 2 | -0.39675 | 0.77033 | 0.88455 | 1 | 18299 | 1 | -0.39675 |
| MAL | 5 | 0.20171 | 0.37769 | 0.999994 | 7815 | 2 | -0.41781 | 0.77041 | 0.87298 | 1 | 18300 | 1 | -0.41781 |
| OR6M1 | 6 | 0.85361 | 0.87388 | 0.999994 | 18326 | 1 | 0.04692 | 0.77041 | 0.88458 | 1 | 18301 | 2 | 0.04692 |
| hsa-mir-4298 | 3 | 0.55932 | 0.58689 | 0.999994 | 14360 | 1 | 0.33796 | 0.77061 | 0.78681 | 1 | 18302 | 1 | 0.33796 |
| CHST7 | 6 | 0.066929 | 0.18032 | 0.968171 | 3717 | 3 | -0.67685 | 0.77075 | 0.88475 | 1 | 18303 | 2 | -0.67685 |

| | | | | | | | | | | | | |
|---|---|---|---|---|---|---|---|---|---|---|---|---|
| RPA1 | 1 | 0.22915 | 0.22883 | 0.983354 | 8508 | 1 | -1.6297 | 0.77085 | 0.77119 | 1 | 18304 | 0 | -1.6297 |
| PRR14L | 5 | 0.06484 | 0.17002 | 0.967126 | 3644 | 3 | -1.9233 | 0.77089 | 0.8733 | 1 | 18305 | 2 | -1.9233 |
| NonTargetingControlG | 1 | 0.22892 | 0.22861 | 0.983354 | 8501 | 1 | -1.8331 | 0.77108 | 0.7714 | 1 | 18306 | 0 | -1.8331 |
| GABRA4 | 6 | 0.061493 | 0.16894 | 0.967126 | 3492 | 4 | -1.2443 | 0.77117 | 0.88493 | 1 | 18307 | 1 | -1.2443 |
| hsa-mir-378d-2 | 4 | 0.32501 | 0.4686 | 0.999994 | 10618 | 2 | -0.59458 | 0.77117 | 0.83422 | 1 | 18308 | 1 | -0.59458 |
| NonTargetingControlG | 1 | 0.22883 | 0.22852 | 0.983354 | 8498 | 1 | -1.6747 | 0.77117 | 0.77149 | 1 | 18309 | 0 | -1.6747 |
| ASB14 | 6 | 0.75437 | 0.83003 | 0.999994 | 16900 | 1 | -0.19963 | 0.77118 | 0.88494 | 1 | 18310 | 2 | -0.19963 |
| DBF4 | 4 | 0.24755 | 0.38941 | 0.999994 | 8963 | 2 | -0.98242 | 0.7712 | 0.83423 | 1 | 18311 | 1 | -0.98242 |
| SMCO4 | 6 | 0.25754 | 0.45696 | 0.999994 | 9181 | 2 | -0.27873 | 0.77133 | 0.885 | 1 | 18312 | 2 | -0.27873 |
| GPR18 | 6 | 0.06099 | 0.16787 | 0.967126 | 3468 | 3 | -1.3787 | 0.77134 | 0.88501 | 1 | 18313 | 2 | -1.3787 |
| NonTargetingControlG | 2 | 0.63305 | 0.63279 | 0.999994 | 15415 | 0 | 0.0080926 | 0.77145 | 0.77097 | 1 | 18314 | 0 | 0.0080926 |
| CTRB2 | 2 | 0.43281 | 0.44443 | 0.999994 | 12460 | 1 | -0.64824 | 0.77174 | 0.77125 | 1 | 18315 | 0 | -0.64824 |
| hsa-mir-3689a | 4 | 0.30651 | 0.44964 | 0.999994 | 10264 | 2 | -0.43996 | 0.77176 | 0.83448 | 1 | 18316 | 1 | -0.43996 |
| GIGYF2 | 5 | 0.049478 | 0.13908 | 0.957704 | 2954 | 4 | -0.918 | 0.77183 | 0.87391 | 1 | 18317 | 1 | -0.918 |
| PIGZ | 5 | 0.088694 | 0.20848 | 0.979587 | 4542 | 3 | -1.1833 | 0.77185 | 0.87392 | 1 | 18318 | 2 | -1.1833 |
| SSX2B | 2 | 0.0032314 | 0.006206 | 0.636218 | 376 | 1 | -5.6992 | 0.77186 | 0.77138 | 1 | 18319 | 0 | -5.6992 |
| UFM1 | 5 | 0.04606 | 0.13092 | 0.95318 | 2812 | 3 | -2.0773 | 0.77189 | 0.87395 | 1 | 18320 | 2 | -2.0773 |
| hsa-mir-3934 | 4 | 0.28004 | 0.42257 | 0.999994 | 9707 | 2 | -0.44273 | 0.77196 | 0.83454 | 1 | 18321 | 1 | -0.44273 |
| POC1B | 6 | 0.15885 | 0.33676 | 0.999994 | 6695 | 3 | -0.91662 | 0.77201 | 0.8853 | 1 | 18322 | 2 | -0.91662 |
| SS18L2 | 3 | 0.0031798 | 0.008819 | 0.680529 | 372 | 2 | -8.9223 | 0.77218 | 0.7879 | 1 | 18323 | 1 | -8.9223 |
| ZNHIT3 | 5 | 0.91132 | 0.91134 | 0.999994 | 19454 | 0 | 0.14129 | 0.77225 | 0.87419 | 1 | 18324 | 2 | 0.14129 |
| AIG1 | 6 | 0.60712 | 0.74523 | 0.999994 | 15037 | 2 | 0.31592 | 0.77227 | 0.88543 | 1 | 18325 | 2 | 0.31592 |
| PLIN2 | 6 | 0.16095 | 0.33944 | 0.999994 | 6763 | 2 | 0.033724 | 0.77227 | 0.88543 | 1 | 18326 | 2 | 0.033724 |
| hsa-mir-218-2 | 4 | 0.0017573 | 0.0063834 | 0.636218 | 245 | 3 | -7.6959 | 0.77278 | 0.83485 | 1 | 18327 | 1 | -7.6959 |
| SPTSSB | 6 | 0.61907 | 0.75413 | 0.999994 | 15208 | 2 | -0.41669 | 0.77281 | 0.88567 | 1 | 18328 | 2 | -0.41669 |
| HIST1H3H | 6 | 0.21757 | 0.4092 | 0.999994 | 8211 | 3 | -0.32932 | 0.77294 | 0.88573 | 1 | 18329 | 2 | -0.32932 |
| YPEL2 | 6 | 0.12135 | 0.28829 | 0.999632 | 5677 | 3 | -0.81218 | 0.77297 | 0.88574 | 1 | 18330 | 2 | -0.81218 |
| HPRT1 | 6 | 0.79835 | 0.84718 | 0.999994 | 17452 | 1 | 0.30727 | 0.77308 | 0.8858 | 1 | 18331 | 2 | 0.30727 |
| SLAMF7 | 5 | 0.46986 | 0.62203 | 0.999994 | 13004 | 2 | -0.19154 | 0.77327 | 0.87487 | 1 | 18332 | 2 | -0.19154 |
| HOPX | 4 | 0.48537 | 0.60989 | 0.999994 | 13219 | 1 | 0.46669 | 0.7733 | 0.83506 | 1 | 18333 | 1 | 0.46669 |
| UBE2U | 5 | 0.81382 | 0.82846 | 0.999994 | 17688 | 1 | -0.14734 | 0.77384 | 0.87525 | 1 | 18334 | 1 | -0.14734 |
| NOL12 | 3 | 0.27199 | 0.40211 | 0.999994 | 9521 | 2 | -0.71269 | 0.77387 | 0.78909 | 1 | 18335 | 1 | -0.71269 |
| NT5C3A | 6 | 0.27984 | 0.48323 | 0.999994 | 9700 | 2 | 0.19403 | 0.77392 | 0.8862 | 1 | 18336 | 2 | 0.19403 |
| IGFALS | 3 | 0.57358 | 0.59726 | 0.999994 | 14602 | 1 | -0.005364 | 0.77397 | 0.78916 | 1 | 18337 | 1 | -0.005364 |
| TAS2R50 | 5 | 0.19263 | 0.36469 | 0.999994 | 7564 | 3 | -0.72966 | 0.77402 | 0.87536 | 1 | 18338 | 2 | -0.72966 |
| LSM3 | 4 | 0.028632 | 0.078739 | 0.93914 | 1932 | 3 | -2.5257 | 0.77405 | 0.83537 | 1 | 18339 | 1 | -2.5257 |
| ADD2 | 5 | 0.55162 | 0.69318 | 0.999994 | 14233 | 2 | -0.44756 | 0.77412 | 0.87543 | 1 | 18340 | 1 | -0.44756 |
| ARSK | 5 | 0.40968 | 0.57161 | 0.999994 | 12103 | 1 | 0.25445 | 0.77424 | 0.87552 | 1 | 18341 | 2 | 0.25445 |
| IL27RA | 3 | 0.20577 | 0.31994 | 0.999994 | 7917 | 2 | -1.1785 | 0.7743 | 0.78941 | 1 | 18342 | 1 | -1.1785 |
| BEX2 | 6 | 0.34131 | 0.55415 | 0.999994 | 10965 | 3 | -0.40382 | 0.77431 | 0.88639 | 1 | 18343 | 2 | -0.40382 |
| NonTargetingControlG | 2 | 0.38068 | 0.40083 | 0.999994 | 11656 | 1 | -0.94857 | 0.7744 | 0.77393 | 1 | 18344 | 0 | -0.94857 |
| KRT85 | 3 | 0.59539 | 0.61364 | 0.999994 | 14895 | 1 | -0.33405 | 0.7745 | 0.78954 | 1 | 18345 | 1 | -0.33405 |
| STK3 | 6 | 0.12202 | 0.28957 | 0.999632 | 5702 | 3 | -1.0113 | 0.77453 | 0.88649 | 1 | 18346 | 2 | -1.0113 |
| TBP | 5 | 0.049697 | 0.13959 | 0.958716 | 2963 | 3 | -2.4018 | 0.7746 | 0.87576 | 1 | 18347 | 1 | -2.4018 |
| YBX2 | 5 | 0.90034 | 0.90046 | 0.999994 | 19228 | 0 | -0.0031142 | 0.7746 | 0.87576 | 1 | 18348 | 1 | -0.0031142 |
| RBM7 | 5 | 0.060822 | 0.16345 | 0.967126 | 3461 | 3 | -2.0429 | 0.77463 | 0.87577 | 1 | 18349 | 2 | -2.0429 |
| ARSD | 4 | 0.14412 | 0.28316 | 0.999632 | 6302 | 2 | -1.1806 | 0.77466 | 0.83562 | 1 | 18350 | 1 | -1.1806 |
| hsa-mir-193a | 3 | 0.22521 | 0.34412 | 0.999994 | 8403 | 2 | -1.072 | 0.77471 | 0.78968 | 1 | 18351 | 1 | -1.072 |
| NID1 | 6 | 0.12838 | 0.2982 | 0.999994 | 5889 | 3 | -0.84154 | 0.7748 | 0.88662 | 1 | 18352 | 2 | -0.84154 |
| TSHB | 4 | 0.37868 | 0.52381 | 0.999994 | 11633 | 2 | -0.75032 | 0.77483 | 0.83568 | 1 | 18353 | 1 | -0.75032 |
| CXCL5 | 5 | 0.10203 | 0.22959 | 0.983354 | 5042 | 2 | -0.18326 | 0.77486 | 0.87592 | 1 | 18354 | 2 | -0.18326 |
| NUDT8 | 5 | 0.0068848 | 0.027036 | 0.848412 | 654 | 3 | -1.2586 | 0.77513 | 0.87611 | 1 | 18355 | 2 | -1.2586 |

| | | | | | | | | | | | | | |
|---|---|---|---|---|---|---|---|---|---|---|---|---|---|
| MDH2 | 4 | 0.01569 | 0.0512 | 0.909241 | 1229 | 2 | -1.8161 | 0.77529 | 0.83589 | 1 | 18356 | 1 | -1.8161 |
| C19orf44 | 6 | 0.044058 | 0.131 | 0.95318 | 2713 | 2 | 0.20287 | 0.77531 | 0.88687 | 1 | 18357 | 1 | 0.20287 |
| BFSP2 | 6 | 0.38681 | 0.59389 | 0.999994 | 11747 | 2 | 0.40083 | 0.77539 | 0.88691 | 1 | 18358 | 2 | 0.40083 |
| hsa-mir-378j | 2 | 0.12691 | 0.21055 | 0.979587 | 5845 | 1 | -3.6621 | 0.77543 | 0.77496 | 1 | 18359 | 0 | -3.6621 |
| IGFL1 | 5 | 0.3723 | 0.54031 | 0.999994 | 11543 | 2 | -0.47511 | 0.77547 | 0.87634 | 1 | 18360 | 2 | -0.47511 |
| PSMA8 | 6 | 0.024355 | 0.08547 | 0.94395 | 1726 | 4 | -2.4329 | 0.7755 | 0.88696 | 1 | 18361 | 2 | -2.4329 |
| MRPS35 | 5 | 0.1057 | 0.23528 | 0.985362 | 5163 | 3 | -1.6491 | 0.77555 | 0.8764 | 1 | 18362 | 1 | -1.6491 |
| KRT23 | 5 | 0.38992 | 0.55506 | 0.999994 | 11797 | 2 | -0.40569 | 0.77555 | 0.8764 | 1 | 18363 | 2 | -0.40569 |
| LOC391322 | 6 | 0.84933 | 0.87157 | 0.999994 | 18248 | 1 | 0.27256 | 0.77562 | 0.88701 | 1 | 18364 | 2 | 0.27256 |
| KCNA6 | 6 | 0.035664 | 0.11209 | 0.951052 | 2300 | 3 | -0.50625 | 0.7758 | 0.8871 | 1 | 18365 | 1 | -0.50625 |
| RUNX3 | 5 | 0.753 | 0.79242 | 0.999994 | 16886 | 1 | 0.29951 | 0.77588 | 0.87662 | 1 | 18366 | 2 | 0.29951 |
| FDXR | 5 | 0.010363 | 0.037099 | 0.881473 | 891 | 4 | -2.9411 | 0.77602 | 0.87671 | 1 | 18367 | 1 | -2.9411 |
| C12orf68 | 5 | 0.22859 | 0.41526 | 0.999994 | 8492 | 3 | -0.60908 | 0.77614 | 0.8768 | 1 | 18368 | 2 | -0.60908 |
| hsa-mir-6875 | 3 | 0.1464 | 0.24386 | 0.988214 | 6362 | 1 | 0.2282 | 0.77639 | 0.79086 | 1 | 18369 | 1 | 0.2282 |
| NonTargetingControlG | 2 | 0.04702 | 0.084168 | 0.94395 | 2851 | 1 | -3.74 | 0.77643 | 0.77595 | 1 | 18370 | 0 | -3.74 |
| RPP30 | 6 | 0.26078 | 0.4608 | 0.999994 | 9259 | 3 | -0.089953 | 0.77651 | 0.88742 | 1 | 18371 | 2 | -0.089953 |
| SERBP1 | 6 | 0.019924 | 0.071718 | 0.93914 | 1457 | 4 | -2.7844 | 0.77652 | 0.88743 | 1 | 18372 | 1 | -2.7844 |
| RAP2C | 6 | 0.089768 | 0.22705 | 0.983354 | 4585 | 3 | -1.2893 | 0.77652 | 0.88743 | 1 | 18373 | 2 | -1.2893 |
| APBB2 | 6 | 0.8628 | 0.87899 | 0.999994 | 18502 | 1 | 0.18963 | 0.77652 | 0.88743 | 1 | 18374 | 2 | 0.18963 |
| NonTargetingControlG | 2 | 0.37636 | 0.39736 | 0.999994 | 11599 | 1 | -0.88157 | 0.77665 | 0.77616 | 1 | 18375 | 0 | -0.88157 |
| HEATR6 | 3 | 0.025901 | 0.065627 | 0.93914 | 1798 | 2 | -4.6514 | 0.77665 | 0.79104 | 1 | 18376 | 1 | -4.6514 |
| NonTargetingControlG | 2 | 0.64062 | 0.64042 | 0.999994 | 15520 | 0 | -0.00148 | 0.77666 | 0.77617 | 1 | 18377 | 0 | -0.00148 |
| OR2T2 | 4 | 0.058144 | 0.1372 | 0.957161 | 3344 | 2 | -1.804 | 0.77669 | 0.83645 | 1 | 18378 | 1 | -1.804 |
| QRFP | 6 | 0.47509 | 0.65187 | 0.999994 | 13065 | 1 | 0.19455 | 0.77677 | 0.88756 | 1 | 18379 | 2 | 0.19455 |
| IKZF2 | 6 | 0.040463 | 0.12294 | 0.951052 | 2535 | 4 | -1.6628 | 0.77697 | 0.88766 | 1 | 18380 | 2 | -1.6628 |
| TNFRSF10B | 6 | 0.13199 | 0.30287 | 0.999994 | 5975 | 2 | -0.00049671 | 0.77714 | 0.88774 | 1 | 18381 | 2 | -0.00049671 |
| NonTargetingControlG | 1 | 0.2227 | 0.2224 | 0.983354 | 8350 | 1 | -1.8475 | 0.7773 | 0.77764 | 1 | 18382 | 0 | -1.8475 |
| SNRPD1 | 2 | 0.096756 | 0.16505 | 0.967126 | 4848 | 1 | -3.931 | 0.77746 | 0.77697 | 1 | 18383 | 0 | -3.931 |
| PRR21 | 5 | 0.023724 | 0.074086 | 0.93914 | 1696 | 2 | -0.46258 | 0.77746 | 0.87769 | 1 | 18384 | 2 | -0.46258 |
| THRAP3 | 5 | 0.0095402 | 0.034754 | 0.867348 | 843 | 3 | -4.6556 | 0.7775 | 0.87772 | 1 | 18385 | 1 | -4.6556 |
| SGMS1 | 5 | 0.049666 | 0.1395 | 0.958716 | 2962 | 4 | -0.75521 | 0.77751 | 0.87773 | 1 | 18386 | 1 | -0.75521 |
| TMA7 | 3 | 0.60072 | 0.6177 | 0.999994 | 14955 | 1 | -0.40862 | 0.77754 | 0.79169 | 1 | 18387 | 1 | -0.40862 |
| CLRN2 | 5 | 0.29659 | 0.4783 | 0.999994 | 10056 | 2 | 0.3471 | 0.77759 | 0.87778 | 1 | 18388 | 2 | 0.3471 |
| ACER2 | 4 | 0.012649 | 0.041927 | 0.891621 | 1042 | 3 | -1.4608 | 0.77764 | 0.83684 | 1 | 18389 | 1 | -1.4608 |
| UTP20 | 5 | 0.0047979 | 0.020002 | 0.808847 | 506 | 3 | -6.9524 | 0.77777 | 0.87792 | 1 | 18390 | 2 | -6.9524 |
| HIST1H3A | 5 | 0.14486 | 0.29507 | 0.999994 | 6322 | 1 | -0.23767 | 0.77783 | 0.87796 | 1 | 18391 | 1 | -0.23767 |
| TMEM202 | 4 | 0.36707 | 0.5118 | 0.999994 | 11459 | 2 | -0.60014 | 0.77789 | 0.83694 | 1 | 18392 | 1 | -0.60014 |
| SOX17 | 4 | 0.76814 | 0.77427 | 0.999994 | 17062 | 1 | 0.16409 | 0.77792 | 0.83696 | 1 | 18393 | 1 | 0.16409 |
| KIAA0907 | 6 | 0.1068 | 0.26057 | 0.994943 | 5204 | 4 | -0.80483 | 0.77794 | 0.88812 | 1 | 18394 | 2 | -0.80483 |
| WDR85 | 4 | 0.49694 | 0.61495 | 0.999994 | 13375 | 1 | 0.047624 | 0.77812 | 0.83704 | 1 | 18395 | 1 | 0.047624 |
| ZNF37A | 6 | 0.011003 | 0.042409 | 0.891621 | 923 | 2 | 0.14067 | 0.77818 | 0.88824 | 1 | 18396 | 2 | 0.14067 |
| NAPEPLD | 6 | 0.030771 | 0.10061 | 0.946414 | 2049 | 4 | -1.6735 | 0.77818 | 0.88824 | 1 | 18397 | 1 | -1.6735 |
| NonTargetingControlG | 2 | 0.68038 | 0.68024 | 0.999994 | 16099 | 0 | 0.1223 | 0.77821 | 0.77771 | 1 | 18398 | 0 | 0.1223 |
| SEMA3E | 5 | 0.078423 | 0.19203 | 0.974607 | 4169 | 3 | -1.9293 | 0.77824 | 0.87824 | 1 | 18399 | 1 | -1.9293 |
| RFC2 | 6 | 0.049004 | 0.14191 | 0.959146 | 2928 | 2 | 0.10918 | 0.77824 | 0.88827 | 1 | 18400 | 2 | 0.10918 |
| CAAP1 | 5 | 0.88172 | 0.88193 | 0.999994 | 18863 | 1 | 0.51133 | 0.77843 | 0.87837 | 1 | 18401 | 2 | 0.51133 |
| ZFPL1 | 5 | 0.046823 | 0.13273 | 0.95318 | 2842 | 3 | -2.6112 | 0.7785 | 0.87841 | 1 | 18402 | 2 | -2.6112 |
| FAM105A | 3 | 0.133 | 0.22654 | 0.983354 | 5995 | 2 | -1.7427 | 0.7785 | 0.79238 | 1 | 18403 | 1 | -1.7427 |
| MAD1L1 | 5 | 0.078747 | 0.19254 | 0.974607 | 4178 | 3 | -1.6927 | 0.77874 | 0.87857 | 1 | 18404 | 2 | -1.6927 |
| OSBP2 | 5 | 0.067201 | 0.17387 | 0.968129 | 3725 | 3 | -1.7802 | 0.77875 | 0.87858 | 1 | 18405 | 2 | -1.7802 |
| CGGBP1 | 6 | 0.88706 | 0.89428 | 0.999994 | 18968 | 1 | -0.019268 | 0.77875 | 0.88852 | 1 | 18406 | 2 | -0.019268 |
| FLOT1 | 6 | 0.32806 | 0.53901 | 0.999994 | 10685 | 3 | -0.45797 | 0.77881 | 0.88855 | 1 | 18407 | 2 | -0.45797 |

| | | | | | | | | | | | | | |
|---|---|---|---|---|---|---|---|---|---|---|---|---|---|
| NPHP3 | 6 | 0.2076 | 0.39716 | 0.999994 | 7963 | 3 | -0.40675 | 0.77885 | 0.88857 | 1 | 18408 | 2 | -0.40675 |
| OR2T29 | 5 | 0.11718 | 0.25304 | 0.994501 | 5542 | 2 | -0.076324 | 0.7789 | 0.87868 | 1 | 18409 | 2 | -0.076324 |
| ATP6V1A | 2 | 0.2078 | 0.26774 | 0.998561 | 7969 | 1 | -2.7701 | 0.77896 | 0.77845 | 1 | 18410 | 0 | -2.7701 |
| LRRC3B | 5 | 0.12105 | 0.25897 | 0.994943 | 5663 | 3 | -1.331 | 0.77901 | 0.87876 | 1 | 18411 | 1 | -1.331 |
| TRIML1 | 5 | 0.29208 | 0.47461 | 0.999994 | 9954 | 2 | -1.4517 | 0.77903 | 0.87876 | 1 | 18412 | 1 | -1.4517 |
| GUCA1C | 5 | 0.88326 | 0.88341 | 0.999994 | 18895 | 1 | 0.22734 | 0.77903 | 0.87876 | 1 | 18413 | 2 | 0.22734 |
| PTHLH | 6 | 0.0066559 | 0.027214 | 0.850392 | 639 | 2 | 0.1291 | 0.77903 | 0.88865 | 1 | 18414 | 2 | 0.1291 |
| DDX56 | 3 | 0.18964 | 0.29959 | 0.999994 | 7474 | 2 | -1.1224 | 0.77908 | 0.79278 | 1 | 18415 | 1 | -1.1224 |
| NonTargetingControlG | 2 | 0.45266 | 0.46165 | 0.999994 | 12743 | 1 | -0.50572 | 0.77944 | 0.77895 | 1 | 18416 | 0 | -0.50572 |
| HOXA7 | 6 | 0.74403 | 0.82638 | 0.999994 | 16788 | 1 | -0.078505 | 0.77947 | 0.88886 | 1 | 18417 | 2 | -0.078505 |
| CISD2 | 5 | 0.80173 | 0.8206 | 0.999994 | 17507 | 1 | 0.5131 | 0.77948 | 0.87908 | 1 | 18418 | 2 | 0.5131 |
| ZNF141 | 6 | 0.14973 | 0.32528 | 0.999994 | 6450 | 3 | -1.0461 | 0.77964 | 0.88892 | 1 | 18419 | 1 | -1.0461 |
| CFL2 | 6 | 0.10849 | 0.26381 | 0.996734 | 5256 | 4 | -0.73149 | 0.77995 | 0.88906 | 1 | 18420 | 2 | -0.73149 |
| PTGER4 | 5 | 0.88802 | 0.88812 | 0.999994 | 18998 | 1 | 0.35944 | 0.77998 | 0.87944 | 1 | 18421 | 2 | 0.35944 |
| CCT8L2 | 5 | 0.31112 | 0.49005 | 0.999994 | 10342 | 2 | 0.271 | 0.78006 | 0.87949 | 1 | 18422 | 2 | 0.271 |
| FLCN | 4 | 0.070507 | 0.16089 | 0.967126 | 3852 | 3 | -2.4987 | 0.78019 | 0.83789 | 1 | 18423 | 1 | -2.4987 |
| SLC16A13 | 6 | 0.065326 | 0.17694 | 0.968171 | 3667 | 3 | -0.44491 | 0.78022 | 0.88919 | 1 | 18424 | 2 | -0.44491 |
| APOC3 | 5 | 0.22199 | 0.40604 | 0.999994 | 8330 | 3 | -0.65584 | 0.78033 | 0.87966 | 1 | 18425 | 2 | -0.65584 |
| LY75 | 5 | 0.27748 | 0.46283 | 0.999994 | 9644 | 2 | 0.14554 | 0.78037 | 0.87969 | 1 | 18426 | 2 | 0.14554 |
| NonTargetingControlG | 2 | 0.16797 | 0.23925 | 0.987253 | 6937 | 1 | -2.4815 | 0.78041 | 0.77991 | 1 | 18427 | 0 | -2.4815 |
| hsa-mir-1289-2 | 3 | 0.52772 | 0.56478 | 0.999994 | 13857 | 1 | 0.12499 | 0.78044 | 0.79375 | 1 | 18428 | 1 | 0.12499 |
| BRS3 | 6 | 0.036814 | 0.11473 | 0.951052 | 2367 | 4 | -1.7595 | 0.78048 | 0.88932 | 1 | 18429 | 2 | -1.7595 |
| PDE3B | 6 | 0.083647 | 0.21477 | 0.979587 | 4362 | 3 | -0.76216 | 0.78049 | 0.88933 | 1 | 18430 | 2 | -0.76216 |
| DPEP1 | 5 | 0.74207 | 0.7867 | 0.999994 | 16764 | 1 | 0.28774 | 0.78058 | 0.87982 | 1 | 18431 | 2 | 0.28774 |
| hsa-mir-3976 | 4 | 0.86404 | 0.86366 | 0.999994 | 18523 | 0 | 0.27285 | 0.78062 | 0.83807 | 1 | 18432 | 1 | 0.27285 |
| USP13 | 5 | 0.012641 | 0.043591 | 0.891621 | 1040 | 3 | -2.5543 | 0.78065 | 0.87987 | 1 | 18433 | 2 | -2.5543 |
| NME8 | 5 | 0.053048 | 0.14773 | 0.965554 | 3113 | 3 | -1.6822 | 0.78066 | 0.87987 | 1 | 18434 | 1 | -1.6822 |
| SPATA31A1 | 5 | 0.3121 | 0.49089 | 0.999994 | 10362 | 2 | -0.25738 | 0.78066 | 0.87987 | 1 | 18435 | 1 | -0.25738 |
| WRAP53 | 5 | 0.056856 | 0.15685 | 0.967126 | 3285 | 3 | -2.3886 | 0.78066 | 0.87987 | 1 | 18436 | 1 | -2.3886 |
| TRAF4 | 6 | 0.009445 | 0.037068 | 0.881473 | 825 | 5 | -3.3133 | 0.78084 | 0.8895 | 1 | 18437 | 1 | -3.3133 |
| GKN1 | 6 | 0.3519 | 0.56612 | 0.999994 | 11193 | 2 | 0.063258 | 0.78084 | 0.8895 | 1 | 18438 | 1 | 0.063258 |
| RTKN | 6 | 0.080867 | 0.20905 | 0.979587 | 4262 | 4 | -1.2799 | 0.78084 | 0.8895 | 1 | 18439 | 2 | -1.2799 |
| RBM38 | 6 | 0.0076592 | 0.030768 | 0.862223 | 703 | 5 | -3.5996 | 0.78084 | 0.8895 | 1 | 18440 | 1 | -3.5996 |
| TGFBR3 | 6 | 0.19898 | 0.38664 | 0.999994 | 7737 | 3 | -0.83359 | 0.78084 | 0.8895 | 1 | 18441 | 2 | -0.83359 |
| hsa-mir-653 | 3 | 0.26036 | 0.38785 | 0.999994 | 9250 | 2 | -0.79283 | 0.78089 | 0.79408 | 1 | 18442 | 1 | -0.79283 |
| hsa-mir-204 | 4 | 0.19514 | 0.33569 | 0.999994 | 7637 | 2 | -0.89188 | 0.78096 | 0.83821 | 1 | 18443 | 1 | -0.89188 |
| PTPN18 | 4 | 0.60458 | 0.66782 | 0.999994 | 15004 | 1 | -0.31339 | 0.781 | 0.83822 | 1 | 18444 | 1 | -0.31339 |
| OR10G4 | 6 | 0.60456 | 0.74336 | 0.999994 | 15001 | 2 | -0.16455 | 0.78105 | 0.88961 | 1 | 18445 | 2 | -0.16455 |
| TMEM82 | 5 | 0.063493 | 0.16783 | 0.967126 | 3578 | 2 | 0.12533 | 0.78113 | 0.8802 | 1 | 18446 | 2 | 0.12533 |
| UQCR11 | 5 | 0.15391 | 0.30863 | 0.999994 | 6564 | 3 | -1.1943 | 0.78117 | 0.88022 | 1 | 18447 | 2 | -1.1943 |
| BTBD10 | 6 | 0.29119 | 0.49645 | 0.999994 | 9935 | 3 | -0.2967 | 0.78127 | 0.88972 | 1 | 18448 | 2 | -0.2967 |
| CCDC64 | 5 | 0.28389 | 0.46802 | 0.999994 | 9780 | 1 | 0.10975 | 0.78135 | 0.88034 | 1 | 18449 | 2 | 0.10975 |
| KIAA1024L | 5 | 0.16421 | 0.32362 | 0.999994 | 6845 | 3 | -1.0637 | 0.78136 | 0.88036 | 1 | 18450 | 2 | -1.0637 |
| ZMAT5 | 5 | 0.11172 | 0.24465 | 0.988995 | 5357 | 3 | -1.1128 | 0.7814 | 0.88038 | 1 | 18451 | 1 | -1.1128 |
| hsa-mir-545 | 4 | 0.31902 | 0.46244 | 0.999994 | 10498 | 2 | -0.61884 | 0.78142 | 0.83842 | 1 | 18452 | 1 | -0.61884 |
| UBA5 | 4 | 0.42061 | 0.56683 | 0.999994 | 12257 | 2 | -0.24191 | 0.78149 | 0.83845 | 1 | 18453 | 1 | -0.24191 |
| ZNF672 | 4 | 0.078331 | 0.17564 | 0.968171 | 4165 | 3 | -1.457 | 0.78164 | 0.83851 | 1 | 18454 | 1 | -1.457 |
| OSBPL6 | 6 | 0.045136 | 0.13337 | 0.95318 | 2765 | 4 | -1.508 | 0.78165 | 0.8899 | 1 | 18455 | 2 | -1.508 |
| KIAA1715 | 6 | 0.067223 | 0.18094 | 0.968171 | 3726 | 2 | -0.18482 | 0.78191 | 0.89002 | 1 | 18456 | 2 | -0.18482 |
| TEX28 | 3 | 0.012604 | 0.03334 | 0.867348 | 1036 | 2 | -1.9624 | 0.78214 | 0.79498 | 1 | 18457 | 1 | -1.9624 |
| FAM193A | 5 | 0.050828 | 0.14239 | 0.959553 | 3009 | 3 | -1.3187 | 0.78227 | 0.88098 | 1 | 18458 | 2 | -1.3187 |
| CLIP4 | 6 | 0.031742 | 0.1029 | 0.946414 | 2094 | 4 | -1.7206 | 0.78228 | 0.89019 | 1 | 18459 | 2 | -1.7206 |

| | | | | | | | | | | | | | |
|---|---|---|---|---|---|---|---|---|---|---|---|---|---|
| RETSAT | 5 | 0.13432 | 0.27922 | 0.999632 | 6030 | 2 | 0.17816 | 0.78232 | 0.88102 | 1 | 18460 | 1 | 0.17816 |
| OR5B3 | 5 | 0.86073 | 0.86302 | 0.999994 | 18456 | 1 | -0.119 | 0.78241 | 0.88108 | 1 | 18461 | 2 | -0.119 |
| SARS2 | 5 | 0.2754 | 0.46116 | 0.999994 | 9600 | 2 | -0.48339 | 0.78249 | 0.88113 | 1 | 18462 | 1 | -0.48339 |
| hsa-mir-6084 | 2 | 0.45479 | 0.46351 | 0.999994 | 12767 | 1 | -0.66002 | 0.78251 | 0.78197 | 1 | 18463 | 0 | -0.66002 |
| CACNA1S | 4 | 0.048607 | 0.1187 | 0.951052 | 2913 | 1 | 0.097663 | 0.78271 | 0.83896 | 1 | 18464 | 1 | 0.097663 |
| hsa-mir-4708 | 3 | 0.18965 | 0.29961 | 0.999994 | 7475 | 2 | -1.3714 | 0.78274 | 0.79541 | 1 | 18465 | 1 | -1.3714 |
| MYO1E | 6 | 0.0076762 | 0.030831 | 0.862223 | 704 | 4 | -2.1057 | 0.78283 | 0.89047 | 1 | 18466 | 2 | -2.1057 |
| PCDHGA12 | 2 | 0.37411 | 0.39552 | 0.999994 | 11573 | 1 | -0.91382 | 0.78287 | 0.78234 | 1 | 18467 | 0 | -0.91382 |
| MMS19 | 5 | 0.019747 | 0.06341 | 0.937389 | 1442 | 3 | -3.5413 | 0.78291 | 0.88141 | 1 | 18468 | 1 | -3.5413 |
| DNAJC8 | 5 | 0.15139 | 0.30488 | 0.999994 | 6491 | 3 | -1.0323 | 0.78291 | 0.88141 | 1 | 18469 | 1 | -1.0323 |
| NLRP13 | 6 | 0.23435 | 0.42927 | 0.999994 | 8625 | 3 | -1.229 | 0.78291 | 0.89051 | 1 | 18470 | 1 | -1.229 |
| GCC1 | 6 | 0.021139 | 0.075641 | 0.93914 | 1521 | 3 | -1.5271 | 0.78299 | 0.89055 | 1 | 18471 | 1 | -1.5271 |
| TMEM108 | 4 | 0.31949 | 0.4629 | 0.999994 | 10504 | 2 | -0.027679 | 0.78303 | 0.83909 | 1 | 18472 | 1 | -0.027679 |
| ACTL6B | 4 | 0.028141 | 0.077709 | 0.93914 | 1913 | 3 | -2.536 | 0.7831 | 0.83913 | 1 | 18473 | 1 | -2.536 |
| SLC8A3 | 6 | 0.16759 | 0.34771 | 0.999994 | 6927 | 3 | -0.75347 | 0.78327 | 0.89068 | 1 | 18474 | 2 | -0.75347 |
| KIAA1143 | 6 | 0.024762 | 0.08644 | 0.94395 | 1744 | 3 | -1.4831 | 0.78332 | 0.89071 | 1 | 18475 | 1 | -1.4831 |
| IRX5 | 5 | 0.3637 | 0.53316 | 0.999994 | 11397 | 2 | 0.5883 | 0.78342 | 0.88176 | 1 | 18476 | 2 | 0.5883 |
| HIST1H2AB | 6 | 0.33587 | 0.54793 | 0.999994 | 10848 | 3 | -0.46904 | 0.78343 | 0.89076 | 1 | 18477 | 2 | -0.46904 |
| NonTargetingControlG | 2 | 0.50162 | 0.50541 | 0.999994 | 13452 | 1 | -0.46545 | 0.78344 | 0.78292 | 1 | 18478 | 0 | -0.46545 |
| IL17RE | 5 | 0.25189 | 0.44227 | 0.999994 | 9054 | 2 | -0.58821 | 0.78346 | 0.88179 | 1 | 18479 | 1 | -0.58821 |
| RASGEF1B | 4 | 0.61634 | 0.67428 | 0.999994 | 15165 | 1 | -0.21005 | 0.78346 | 0.83928 | 1 | 18480 | 1 | -0.21005 |
| TMIGD1 | 4 | 0.40362 | 0.54944 | 0.999994 | 12008 | 1 | 0.22163 | 0.7836 | 0.83934 | 1 | 18481 | 1 | 0.22163 |
| ARHGAP6 | 4 | 0.27045 | 0.41273 | 0.999994 | 9486 | 2 | -0.89947 | 0.78372 | 0.83938 | 1 | 18482 | 1 | -0.89947 |
| DTD2 | 6 | 0.838 | 0.86561 | 0.999994 | 18054 | 1 | -0.26211 | 0.78375 | 0.89092 | 1 | 18483 | 1 | -0.26211 |
| SLC30A7 | 5 | 0.4824 | 0.63277 | 0.999994 | 13176 | 1 | 0.12229 | 0.78392 | 0.8821 | 1 | 18484 | 2 | 0.12229 |
| PCDHGB1 | 2 | 0.48197 | 0.48755 | 0.999994 | 13170 | 1 | -0.52948 | 0.78393 | 0.78341 | 1 | 18485 | 0 | -0.52948 |
| ZSWIM3 | 6 | 0.033034 | 0.10595 | 0.950444 | 2159 | 4 | -1.9085 | 0.78394 | 0.89102 | 1 | 18486 | 2 | -1.9085 |
| C22orf31 | 5 | 0.24528 | 0.43695 | 0.999994 | 8906 | 1 | 0.40617 | 0.78396 | 0.88213 | 1 | 18487 | 2 | 0.40617 |
| ZNF518B | 5 | 0.14237 | 0.29145 | 0.999632 | 6248 | 3 | -0.75985 | 0.78404 | 0.88218 | 1 | 18488 | 2 | -0.75985 |
| SLC25A11 | 4 | 0.009751 | 0.032941 | 0.867348 | 857 | 2 | -1.8872 | 0.78406 | 0.83952 | 1 | 18489 | 1 | -1.8872 |
| ZDHHC13 | 6 | 0.30522 | 0.51281 | 0.999994 | 10237 | 3 | -0.41363 | 0.78406 | 0.89107 | 1 | 18490 | 2 | -0.41363 |
| PPIL4 | 5 | 0.14706 | 0.29834 | 0.999994 | 6377 | 3 | -1.1723 | 0.78413 | 0.88225 | 1 | 18491 | 2 | -1.1723 |
| IL4R | 5 | 0.19774 | 0.372 | 0.999994 | 7701 | 3 | -0.82175 | 0.7844 | 0.88243 | 1 | 18492 | 1 | -0.82175 |
| NFYA | 3 | 0.063645 | 0.13436 | 0.953565 | 3590 | 2 | -2.5563 | 0.7844 | 0.79661 | 1 | 18493 | 1 | -2.5563 |
| PAWR | 6 | 0.5675 | 0.71616 | 0.999994 | 14506 | 2 | -0.36209 | 0.78452 | 0.89129 | 1 | 18494 | 1 | -0.36209 |
| EXOC6B | 5 | 0.0042947 | 0.018002 | 0.795727 | 464 | 4 | -2.495 | 0.78455 | 0.88254 | 1 | 18495 | 1 | -2.495 |
| OXTR | 5 | 0.47196 | 0.62379 | 0.999994 | 13034 | 2 | 0.38599 | 0.78468 | 0.88263 | 1 | 18496 | 2 | 0.38599 |
| LIMD1 | 6 | 0.91152 | 0.91285 | 0.999994 | 19457 | 1 | -0.021155 | 0.78482 | 0.89142 | 1 | 18497 | 2 | -0.021155 |
| PPIA | 5 | 0.45361 | 0.60826 | 0.999994 | 12751 | 2 | -0.26401 | 0.78489 | 0.88276 | 1 | 18498 | 1 | -0.26401 |
| GGA2 | 6 | 0.30208 | 0.50915 | 0.999994 | 10167 | 3 | -0.38946 | 0.7849 | 0.89146 | 1 | 18499 | 1 | -0.38946 |
| hsa-mir-5586 | 4 | 0.042581 | 0.10692 | 0.950995 | 2653 | 3 | -2.3036 | 0.78492 | 0.83987 | 1 | 18500 | 1 | -2.3036 |
| hsa-mir-4428 | 4 | 0.295 | 0.43785 | 0.999994 | 10017 | 2 | -1.448 | 0.78492 | 0.83987 | 1 | 18501 | 1 | -1.448 |
| hsa-mir-1273f | 4 | 0.024281 | 0.069717 | 0.93914 | 1722 | 3 | -3.5012 | 0.78492 | 0.83987 | 1 | 18502 | 1 | -3.5012 |
| hsa-mir-1231 | 4 | 0.82592 | 0.8255 | 0.999994 | 17866 | 1 | 0.064402 | 0.78492 | 0.83987 | 1 | 18503 | 1 | 0.064402 |
| TGIF2 | 6 | 0.53858 | 0.69552 | 0.999994 | 14031 | 2 | 0.38301 | 0.78507 | 0.89154 | 1 | 18504 | 2 | 0.38301 |
| hsa-let-7a-3 | 3 | 0.18372 | 0.29199 | 0.999632 | 7307 | 2 | -1.3278 | 0.7852 | 0.79718 | 1 | 18505 | 1 | -1.3278 |
| hsa-mir-3691 | 3 | 0.35222 | 0.45697 | 0.999994 | 11200 | 1 | -0.066081 | 0.78522 | 0.7972 | 1 | 18506 | 1 | -0.066081 |
| NonTargetingControlG | 2 | 0.28792 | 0.32756 | 0.999994 | 9872 | 1 | -1.3104 | 0.78529 | 0.78478 | 1 | 18507 | 0 | -1.3104 |
| KCNJ11 | 6 | 0.0051311 | 0.021614 | 0.819701 | 529 | 5 | -2.9771 | 0.78532 | 0.89165 | 1 | 18508 | 1 | -2.9771 |
| RSPH10B | 5 | 0.011551 | 0.040529 | 0.891621 | 956 | 3 | -5.7296 | 0.78535 | 0.88306 | 1 | 18509 | 1 | -5.7296 |
| NonTargetingControlG | 2 | 0.16175 | 0.23491 | 0.985287 | 6783 | 1 | -3.2921 | 0.78538 | 0.7849 | 1 | 18510 | 0 | -3.2921 |
| PRKCI | 6 | 0.28845 | 0.4933 | 0.999994 | 9881 | 2 | -0.28323 | 0.78546 | 0.89172 | 1 | 18511 | 2 | -0.28323 |

| | | | | | | | | | | | | | |
|---|---|---|---|---|---|---|---|---|---|---|---|---|---|
| DUSP4 | 5 | 0.47899 | 0.62978 | 0.999994 | 13128 | 2 | -0.50148 | 0.7855 | 0.88316 | 1 | 18512 | 1 | -0.50148 |
| HBE1 | 6 | 0.098976 | 0.24536 | 0.98979 | 4929 | 4 | -0.72645 | 0.78552 | 0.89175 | 1 | 18513 | 2 | -0.72645 |
| PSG11 | 6 | 0.046513 | 0.13641 | 0.956051 | 2829 | 4 | -1.4251 | 0.78556 | 0.89177 | 1 | 18514 | 2 | -1.4251 |
| PLEKHB2 | 5 | 0.050521 | 0.14161 | 0.959146 | 2998 | 2 | -0.55631 | 0.78561 | 0.88323 | 1 | 18515 | 1 | -0.55631 |
| MIA2 | 6 | 0.74654 | 0.82725 | 0.999994 | 16822 | 1 | 0.14267 | 0.7858 | 0.89187 | 1 | 18516 | 2 | 0.14267 |
| G0S2 | 5 | 0.4779 | 0.62885 | 0.999994 | 13111 | 1 | 0.2898 | 0.78582 | 0.88337 | 1 | 18517 | 2 | 0.2898 |
| MRPL20 | 5 | 0.097651 | 0.22276 | 0.983354 | 4875 | 3 | -1.6596 | 0.78586 | 0.88341 | 1 | 18518 | 1 | -1.6596 |
| hsa-mir-548ay | 4 | 0.081579 | 0.18163 | 0.968171 | 4285 | 3 | -1.5942 | 0.78587 | 0.84027 | 1 | 18519 | 1 | -1.5942 |
| TSPAN32 | 5 | 0.89927 | 0.89937 | 0.999994 | 19209 | 0 | 0.24948 | 0.78594 | 0.88347 | 1 | 18520 | 2 | 0.24948 |
| SH2D3A | 5 | 0.16198 | 0.32044 | 0.999994 | 6790 | 2 | 0.10593 | 0.78605 | 0.88354 | 1 | 18521 | 2 | 0.10593 |
| NEDD8 | 2 | 0.50057 | 0.50443 | 0.999994 | 13431 | 1 | -0.41872 | 0.78609 | 0.78563 | 1 | 18522 | 0 | -0.41872 |
| PPP2R2A | 5 | 0.3759 | 0.54333 | 0.999994 | 11591 | 2 | -0.54928 | 0.78627 | 0.8837 | 1 | 18523 | 1 | -0.54928 |
| RFNG | 3 | 0.43914 | 0.50758 | 0.999994 | 12559 | 1 | -0.18073 | 0.78647 | 0.79814 | 1 | 18524 | 1 | -0.18073 |
| ARMC3 | 3 | 0.74635 | 0.74631 | 0.999994 | 16818 | 0 | 0.57501 | 0.78648 | 0.79814 | 1 | 18525 | 1 | 0.57501 |
| C3AR1 | 5 | 0.50281 | 0.65037 | 0.999994 | 13476 | 2 | -0.2941 | 0.78657 | 0.8839 | 1 | 18526 | 2 | -0.2941 |
| NonTargetingControlG | 1 | 0.21342 | 0.21308 | 0.979587 | 8098 | 1 | -2.0533 | 0.78658 | 0.78689 | 1 | 18527 | 0 | -2.0533 |
| OR52I2 | 4 | 0.36801 | 0.51278 | 0.999994 | 11475 | 2 | -0.73638 | 0.78666 | 0.84062 | 1 | 18528 | 1 | -0.73638 |
| TIGD1 | 4 | 0.080214 | 0.17907 | 0.968171 | 4237 | 2 | -2.6301 | 0.7867 | 0.84064 | 1 | 18529 | 1 | -2.6301 |
| DEFB107A | 3 | 0.23945 | 0.36188 | 0.999994 | 8763 | 1 | -0.21663 | 0.78676 | 0.79836 | 1 | 18530 | 1 | -0.21663 |
| FGL1 | 6 | 0.27288 | 0.4751 | 0.999994 | 9535 | 3 | -0.50041 | 0.78677 | 0.89231 | 1 | 18531 | 2 | -0.50041 |
| PCDHB5 | 6 | 0.094157 | 0.23585 | 0.985362 | 4756 | 3 | -1.1861 | 0.78686 | 0.89235 | 1 | 18532 | 1 | -1.1861 |
| NonTargetingControlG | 2 | 0.60593 | 0.60575 | 0.999994 | 15026 | 0 | -0.15672 | 0.78686 | 0.78639 | 1 | 18533 | 0 | -0.15672 |
| ZNF708 | 6 | 0.065006 | 0.17627 | 0.968171 | 3653 | 4 | -1.2479 | 0.78692 | 0.89239 | 1 | 18534 | 2 | -1.2479 |
| GMIP | 5 | 0.4206 | 0.58058 | 0.999994 | 12256 | 2 | 0.45843 | 0.78708 | 0.88425 | 1 | 18535 | 2 | 0.45843 |
| EIF6 | 5 | 0.12452 | 0.26424 | 0.996734 | 5782 | 2 | -1.4517 | 0.78727 | 0.88437 | 1 | 18536 | 2 | -1.4517 |
| KLB | 6 | 0.12014 | 0.28595 | 0.999632 | 5633 | 4 | -0.68195 | 0.78741 | 0.8926 | 1 | 18537 | 2 | -0.68195 |
| HERC3 | 6 | 0.68401 | 0.8041 | 0.999994 | 16143 | 2 | 0.045709 | 0.78762 | 0.89272 | 1 | 18538 | 1 | 0.045709 |
| VEZF1 | 6 | 0.0013784 | 0.0068991 | 0.643671 | 208 | 5 | -5.079 | 0.7877 | 0.89276 | 1 | 18539 | 1 | -5.079 |
| FOXL2 | 5 | 0.2545 | 0.44433 | 0.999994 | 9116 | 2 | 0.028701 | 0.78772 | 0.88468 | 1 | 18540 | 2 | 0.028701 |
| AKR1E2 | 6 | 0.53168 | 0.69076 | 0.999994 | 13928 | 2 | -0.021972 | 0.78778 | 0.89281 | 1 | 18541 | 2 | -0.021972 |
| NonTargetingControlG | 2 | 0.42065 | 0.43403 | 0.999994 | 12258 | 1 | -0.71528 | 0.78788 | 0.78742 | 1 | 18542 | 0 | -0.71528 |
| HIST1H1E | 4 | 0.0025984 | 0.0093393 | 0.680529 | 322 | 3 | -6.655 | 0.78811 | 0.84123 | 1 | 18543 | 1 | -6.655 |
| NR3C1 | 6 | 0.29641 | 0.50253 | 0.999994 | 10047 | 2 | 0.25009 | 0.78826 | 0.89304 | 1 | 18544 | 2 | 0.25009 |
| LOC100129216 | 4 | 0.09297 | 0.20248 | 0.977973 | 4718 | 3 | -1.8741 | 0.78832 | 0.84132 | 1 | 18545 | 1 | -1.8741 |
| PAGE5 | 4 | 0.35018 | 0.49446 | 0.999994 | 11161 | 2 | -0.64415 | 0.78838 | 0.84134 | 1 | 18546 | 1 | -0.64415 |
| hsa-mir-6851 | 3 | 0.7348 | 0.73465 | 0.999994 | 16656 | 0 | -0.44908 | 0.78839 | 0.79956 | 1 | 18547 | 1 | -0.44908 |
| PPIG | 4 | 0.86816 | 0.86774 | 0.999994 | 18602 | 0 | -0.0019081 | 0.78839 | 0.84135 | 1 | 18548 | 1 | -0.0019081 |
| KIAA1671 | 5 | 0.023493 | 0.073502 | 0.93914 | 1684 | 2 | 0.40585 | 0.78842 | 0.88516 | 1 | 18549 | 2 | 0.40585 |
| FAM65C | 6 | 0.14285 | 0.31667 | 0.999994 | 6265 | 3 | -0.54883 | 0.78854 | 0.89316 | 1 | 18550 | 2 | -0.54883 |
| NNAT | 4 | 0.33252 | 0.47633 | 0.999994 | 10782 | 2 | -0.62807 | 0.78864 | 0.84146 | 1 | 18551 | 1 | -0.62807 |
| hsa-mir-6727 | 1 | 0.21135 | 0.21103 | 0.979587 | 8056 | 1 | -2.1132 | 0.78865 | 0.78897 | 1 | 18552 | 0 | -2.1132 |
| CXorf22 | 5 | 0.043889 | 0.12553 | 0.951052 | 2707 | 3 | -2.3659 | 0.78872 | 0.88536 | 1 | 18553 | 2 | -2.3659 |
| NonTargetingControlG | 2 | 0.00098975 | 0.0018652 | 0.435802 | 165 | 1 | -6.1606 | 0.78889 | 0.78845 | 1 | 18554 | 0 | -6.1606 |
| PTPN1 | 6 | 0.60938 | 0.74689 | 0.999994 | 15064 | 1 | 0.21659 | 0.78891 | 0.89333 | 1 | 18555 | 2 | 0.21659 |
| hsa-mir-520e | 4 | 0.22551 | 0.36686 | 0.999994 | 8414 | 2 | -1.1085 | 0.78904 | 0.84164 | 1 | 18556 | 1 | -1.1085 |
| NonTargetingControlG | 2 | 0.68298 | 0.68284 | 0.999994 | 16129 | 0 | 0.10445 | 0.78914 | 0.7887 | 1 | 18557 | 0 | 0.10445 |
| TTC40 | 6 | 0.1898 | 0.37525 | 0.999994 | 7479 | 1 | -0.073988 | 0.78915 | 0.89344 | 1 | 18558 | 2 | -0.073988 |
| RPL29 | 6 | 0.0016897 | 0.0084349 | 0.680529 | 239 | 4 | -3.7084 | 0.78915 | 0.89344 | 1 | 18559 | 1 | -3.7084 |
| NonTargetingControlG | 2 | 0.57112 | 0.57135 | 0.999994 | 14563 | 1 | -0.26042 | 0.78957 | 0.78912 | 1 | 18560 | 0 | -0.26042 |
| USP17L3 | 4 | 0.40321 | 0.54905 | 0.999994 | 12002 | 2 | -0.43405 | 0.78962 | 0.84189 | 1 | 18561 | 1 | -0.43405 |
| ATP5J2 | 5 | 0.088872 | 0.20878 | 0.979587 | 4548 | 3 | -0.98411 | 0.78976 | 0.88605 | 1 | 18562 | 2 | -0.98411 |
| CTAGE4 | 5 | 0.2142 | 0.39523 | 0.999994 | 8123 | 2 | -0.54953 | 0.78981 | 0.88609 | 1 | 18563 | 1 | -0.54953 |

| | | | | | | | | | | | | | |
|---|---|---|---|---|---|---|---|---|---|---|---|---|---|
| hsa-mir-1302-7 | 3 | 0.24606 | 0.37006 | 0.999994 | 8925 | 2 | -0.83332 | 0.78992 | 0.80067 | 1 | 18564 | 1 | -0.83332 |
| NonTargetingControlG | 2 | 0.59307 | 0.59301 | 0.999994 | 14863 | 0 | -0.18255 | 0.7902 | 0.78977 | 1 | 18565 | 0 | -0.18255 |
| hsa-mir-143 | 3 | 0.043384 | 0.10538 | 0.95013 | 2688 | 2 | -2.351 | 0.79025 | 0.80092 | 1 | 18566 | 1 | -2.351 |
| AAMDC | 5 | 0.21092 | 0.39068 | 0.999994 | 8048 | 2 | -0.28129 | 0.7903 | 0.88641 | 1 | 18567 | 2 | -0.28129 |
| DPPA3 | 6 | 0.56599 | 0.71508 | 0.999994 | 14481 | 2 | -0.022834 | 0.79039 | 0.89406 | 1 | 18568 | 2 | -0.022834 |
| DGKA | 5 | 0.0009316 | 0.0041723 | 0.564633 | 155 | 4 | -5.5359 | 0.79039 | 0.88649 | 1 | 18569 | 1 | -5.5359 |
| DEAF1 | 6 | 0.089525 | 0.22656 | 0.983354 | 4579 | 2 | 0.13185 | 0.79043 | 0.89407 | 1 | 18570 | 2 | 0.13185 |
| EVC2 | 5 | 0.2585 | 0.44754 | 0.999994 | 9206 | 1 | -0.38344 | 0.79049 | 0.88655 | 1 | 18571 | 1 | -0.38344 |
| PSPH | 6 | 0.78749 | 0.84265 | 0.999994 | 17311 | 1 | 0.1344 | 0.79057 | 0.89414 | 1 | 18572 | 2 | 0.1344 |
| BIVM-ERCC5 | 3 | 0.37948 | 0.47245 | 0.999994 | 11642 | 1 | -0.4326 | 0.79064 | 0.80121 | 1 | 18573 | 1 | -0.4326 |
| ZNF236 | 5 | 0.024851 | 0.077065 | 0.93914 | 1748 | 4 | -1.7063 | 0.79065 | 0.88667 | 1 | 18574 | 1 | -1.7063 |
| hsa-mir-5698 | 4 | 0.22184 | 0.36313 | 0.999994 | 8327 | 2 | -0.62409 | 0.79076 | 0.84237 | 1 | 18575 | 1 | -0.62409 |
| YWHAZ | 6 | 0.066662 | 0.17976 | 0.968171 | 3711 | 3 | -0.79289 | 0.79081 | 0.89426 | 1 | 18576 | 2 | -0.79289 |
| RAF1 | 6 | 0.11981 | 0.28535 | 0.999632 | 5619 | 4 | -1.2004 | 0.79081 | 0.89426 | 1 | 18577 | 1 | -1.2004 |
| XDH | 6 | 0.010526 | 0.040817 | 0.891621 | 905 | 5 | -1.2827 | 0.79081 | 0.89426 | 1 | 18578 | 1 | -1.2827 |
| MTHFD1 | 6 | 0.0088622 | 0.035033 | 0.867348 | 783 | 4 | -3.7448 | 0.79081 | 0.89426 | 1 | 18579 | 1 | -3.7448 |
| SERPINB12 | 5 | 0.45724 | 0.61136 | 0.999994 | 12811 | 2 | -0.23886 | 0.79093 | 0.88687 | 1 | 18580 | 2 | -0.23886 |
| NLRC5 | 3 | 0.27549 | 0.40644 | 0.999994 | 9606 | 1 | -0.32529 | 0.79094 | 0.80144 | 1 | 18581 | 1 | -0.32529 |
| hsa-mir-3653 | 4 | 0.32423 | 0.46779 | 0.999994 | 10603 | 2 | -0.79612 | 0.79097 | 0.84246 | 1 | 18582 | 1 | -0.79612 |
| TTC32 | 6 | 0.079665 | 0.20662 | 0.978744 | 4216 | 4 | -1.0482 | 0.79104 | 0.89437 | 1 | 18583 | 2 | -1.0482 |
| METTL13 | 5 | 0.088389 | 0.20799 | 0.979108 | 4529 | 2 | 0.5518 | 0.79111 | 0.88699 | 1 | 18584 | 2 | 0.5518 |
| NonTargetingControlG | 2 | 0.56683 | 0.56714 | 0.999994 | 14495 | 1 | -0.27657 | 0.79117 | 0.79074 | 1 | 18585 | 0 | -0.27657 |
| NonTargetingControlG | 2 | 0.36064 | 0.38453 | 0.999994 | 11344 | 1 | -0.95893 | 0.79119 | 0.79075 | 1 | 18586 | 0 | -0.95893 |
| NonTargetingControlG | 1 | 0.20876 | 0.20847 | 0.979587 | 7995 | 1 | -1.9939 | 0.79124 | 0.79156 | 1 | 18587 | 0 | -1.9939 |
| hsa-mir-618 | 4 | 0.13893 | 0.27782 | 0.999534 | 6160 | 2 | -0.93078 | 0.7913 | 0.84262 | 1 | 18588 | 1 | -0.93078 |
| RDH8 | 6 | 0.24974 | 0.44762 | 0.999994 | 9011 | 3 | -0.10392 | 0.79138 | 0.89455 | 1 | 18589 | 2 | -0.10392 |
| PPAN-P2RY11 | 1 | 0.20842 | 0.20814 | 0.979182 | 7986 | 1 | -1.9833 | 0.79158 | 0.7919 | 1 | 18590 | 0 | -1.9833 |
| hsa-mir-4470 | 3 | 0.59294 | 0.61174 | 0.999994 | 14860 | 1 | -0.28713 | 0.79163 | 0.80195 | 1 | 18591 | 1 | -0.28713 |
| ADSS | 6 | 0.055039 | 0.155 | 0.967126 | 3200 | 3 | -1.5227 | 0.79168 | 0.8947 | 1 | 18592 | 1 | -1.5227 |
| RPL37A | 2 | 0.1916 | 0.25601 | 0.994943 | 7532 | 1 | -2.2213 | 0.79184 | 0.79142 | 1 | 18593 | 0 | -2.2213 |
| FAM129A | 6 | 0.0054663 | 0.022843 | 0.824431 | 556 | 4 | -1.272 | 0.79188 | 0.8948 | 1 | 18594 | 2 | -1.272 |
| SEC24D | 5 | 0.57184 | 0.71116 | 0.999994 | 14572 | 2 | -0.18517 | 0.7919 | 0.88753 | 1 | 18595 | 2 | -0.18517 |
| MPHOSPH6 | 6 | 0.0081823 | 0.032583 | 0.865899 | 742 | 4 | -1.8871 | 0.79212 | 0.89492 | 1 | 18596 | 1 | -1.8871 |
| TIMP4 | 6 | 0.313 | 0.52182 | 0.999994 | 10377 | 2 | -0.29477 | 0.79213 | 0.89493 | 1 | 18597 | 2 | -0.29477 |
| LTN1 | 5 | 0.12921 | 0.2714 | 0.998561 | 5921 | 3 | -1.2542 | 0.79232 | 0.88782 | 1 | 18598 | 2 | -1.2542 |
| TRMT1 | 5 | 0.43879 | 0.59581 | 0.999994 | 12554 | 2 | 0.49094 | 0.79244 | 0.8879 | 1 | 18599 | 2 | 0.49094 |
| TTI2 | 5 | 0.020946 | 0.06662 | 0.93914 | 1508 | 4 | -3.2187 | 0.79253 | 0.88797 | 1 | 18600 | 1 | -3.2187 |
| VPS72 | 5 | 0.10563 | 0.23518 | 0.985362 | 5157 | 3 | -1.5074 | 0.79253 | 0.88797 | 1 | 18601 | 1 | -1.5074 |
| FAM46D | 6 | 0.066381 | 0.17918 | 0.968171 | 3700 | 4 | -1.2623 | 0.79255 | 0.89511 | 1 | 18602 | 2 | -1.2623 |
| UBE2T | 6 | 0.28501 | 0.48928 | 0.999994 | 9810 | 3 | -0.58878 | 0.79261 | 0.89514 | 1 | 18603 | 2 | -0.58878 |
| KRT6A | 6 | 0.28793 | 0.49269 | 0.999994 | 9873 | 3 | -0.40149 | 0.79278 | 0.89523 | 1 | 18604 | 2 | -0.40149 |
| HYI | 5 | 0.5918 | 0.72318 | 0.999994 | 14844 | 1 | 0.42763 | 0.79281 | 0.88817 | 1 | 18605 | 2 | 0.42763 |
| IFIT3 | 6 | 0.38357 | 0.59181 | 0.999994 | 11697 | 2 | -0.32191 | 0.79289 | 0.89527 | 1 | 18606 | 2 | -0.32191 |
| TOMM20 | 2 | 0.19112 | 0.25567 | 0.994943 | 7516 | 1 | -2.9949 | 0.79292 | 0.7925 | 1 | 18607 | 0 | -2.9949 |
| hsa-mir-4785 | 3 | 0.055062 | 0.12272 | 0.951052 | 3202 | 2 | -3.0198 | 0.79315 | 0.80306 | 1 | 18608 | 1 | -3.0198 |
| GOLGA6C | 3 | 0.68909 | 0.6909 | 0.999994 | 16202 | 1 | -0.48538 | 0.7933 | 0.80317 | 1 | 18609 | 1 | -0.48538 |
| OR2T5 | 4 | 0.029272 | 0.080043 | 0.93914 | 1971 | 3 | -2.0923 | 0.79331 | 0.8435 | 1 | 18610 | 1 | -2.0923 |
| CYP4F2 | 3 | 0.1735 | 0.27888 | 0.999632 | 7073 | 2 | -0.77941 | 0.79337 | 0.80322 | 1 | 18611 | 1 | -0.77941 |
| B3GAT3 | 4 | 0.090062 | 0.19712 | 0.977164 | 4598 | 3 | -2.1619 | 0.79342 | 0.84356 | 1 | 18612 | 1 | -2.1619 |
| ZNF75D | 6 | 0.087151 | 0.22178 | 0.983139 | 4481 | 4 | -0.99264 | 0.79347 | 0.89555 | 1 | 18613 | 2 | -0.99264 |
| PLA2G16 | 6 | 0.14218 | 0.3158 | 0.999994 | 6244 | 2 | 0.12595 | 0.79364 | 0.89562 | 1 | 18614 | 2 | 0.12595 |
| DDAH2 | 6 | 0.22005 | 0.41217 | 0.999994 | 8290 | 3 | -0.83368 | 0.79368 | 0.89565 | 1 | 18615 | 1 | -0.83368 |

| | | | | | | | | | | | | | |
|---|---|---|---|---|---|---|---|---|---|---|---|---|---|
| SEPHS1 | 6 | 0.0027758 | 0.0129 | 0.733879 | 337 | 5 | -2.5395 | 0.79368 | 0.89565 | 1 | 18616 | 1 | -2.5395 |
| EFCAB13 | 6 | 0.038104 | 0.11762 | 0.951052 | 2425 | 2 | 0.0098964 | 0.79368 | 0.89565 | 1 | 18617 | 1 | 0.0098964 |
| HNRNPA2B1 | 6 | 1.0048E-06 | 5.7152E-06 | 0.012376 | 9 | 5 | -8.2275 | 0.79368 | 0.89565 | 1 | 18618 | 1 | -8.2275 |
| NonTargetingControlG | 2 | 0.30464 | 0.34041 | 0.999994 | 10221 | 1 | -1.2932 | 0.79376 | 0.79334 | 1 | 18619 | 0 | -1.2932 |
| AP3B1 | 5 | 0.64714 | 0.74396 | 0.999994 | 15618 | 1 | 0.35795 | 0.79384 | 0.88887 | 1 | 18620 | 2 | 0.35795 |
| DSCAM | 5 | 0.1827 | 0.3505 | 0.999994 | 7281 | 2 | -0.36618 | 0.79398 | 0.88895 | 1 | 18621 | 1 | -0.36618 |
| CASC10 | 1 | 0.20602 | 0.20572 | 0.978603 | 7927 | 1 | -2.0775 | 0.79398 | 0.79429 | 1 | 18622 | 0 | -2.0775 |
| NonTargetingControlG | 2 | 0.1712 | 0.24147 | 0.987253 | 7012 | 1 | -2.0827 | 0.79417 | 0.79374 | 1 | 18623 | 0 | -2.0827 |
| KRT83 | 5 | 0.45944 | 0.61318 | 0.999994 | 12838 | 2 | 0.54608 | 0.79418 | 0.88911 | 1 | 18624 | 2 | 0.54608 |
| RNF32 | 6 | 0.019326 | 0.069827 | 0.93914 | 1422 | 4 | -2.8698 | 0.79457 | 0.89609 | 1 | 18625 | 2 | -2.8698 |
| ABCG5 | 5 | 0.011497 | 0.040374 | 0.891621 | 955 | 3 | -4.2385 | 0.79468 | 0.88946 | 1 | 18626 | 1 | -4.2385 |
| IL17RA | 6 | 0.13628 | 0.30835 | 0.999994 | 6089 | 3 | -0.42907 | 0.79469 | 0.89614 | 1 | 18627 | 2 | -0.42907 |
| NonTargetingControlG | 2 | 0.35347 | 0.37876 | 0.999994 | 11222 | 1 | -1.093 | 0.79506 | 0.79461 | 1 | 18628 | 0 | -1.093 |
| NonTargetingControlG | 2 | 0.64061 | 0.64041 | 0.999994 | 15519 | 0 | -0.057924 | 0.79507 | 0.79463 | 1 | 18629 | 0 | -0.057924 |
| EMR1 | 4 | 0.3053 | 0.44839 | 0.999994 | 10240 | 2 | -0.46575 | 0.79517 | 0.84435 | 1 | 18630 | 1 | -0.46575 |
| OR1J2 | 5 | 0.14286 | 0.29216 | 0.999632 | 6269 | 3 | -1.1604 | 0.79533 | 0.8899 | 1 | 18631 | 2 | -1.1604 |
| FKBP4 | 6 | 0.00878 | 0.034739 | 0.867348 | 778 | 5 | -2.3174 | 0.79541 | 0.89649 | 1 | 18632 | 1 | -2.3174 |
| XRCC6BP1 | 6 | 0.68494 | 0.80485 | 0.999994 | 16159 | 2 | -0.010366 | 0.79543 | 0.89649 | 1 | 18633 | 2 | -0.010366 |
| NonTargetingControlG | 2 | 0.69318 | 0.69301 | 0.999994 | 16237 | 0 | 0.10191 | 0.79557 | 0.79513 | 1 | 18634 | 0 | 0.10191 |
| RCAN1 | 6 | 0.26683 | 0.46791 | 0.999994 | 9393 | 3 | -0.75855 | 0.79563 | 0.89659 | 1 | 18635 | 2 | -0.75855 |
| SRRM1 | 3 | 0.79044 | 0.79034 | 0.999994 | 17350 | 0 | -0.11662 | 0.79577 | 0.805 | 1 | 18636 | 1 | -0.11662 |
| PLIN4 | 5 | 0.023382 | 0.073209 | 0.93914 | 1673 | 2 | 0.27985 | 0.79582 | 0.89021 | 1 | 18637 | 1 | 0.27985 |
| VWA1 | 2 | 0.54407 | 0.54521 | 0.999994 | 14131 | 1 | -0.39484 | 0.79584 | 0.79542 | 1 | 18638 | 0 | -0.39484 |
| SCRN1 | 5 | 0.15709 | 0.31328 | 0.999994 | 6651 | 3 | -1.125 | 0.79608 | 0.89028 | 1 | 18639 | 1 | -1.125 |
| TBC1D3 | 6 | 0.35738 | 0.57234 | 0.999994 | 11290 | 3 | -0.12874 | 0.79634 | 0.89695 | 1 | 18640 | 2 | -0.12874 |
| MRE11A | 6 | 0.10011 | 0.24756 | 0.990682 | 4976 | 4 | -0.98853 | 0.79634 | 0.89695 | 1 | 18641 | 1 | -0.98853 |
| C2orf16 | 6 | 0.011335 | 0.043527 | 0.891621 | 945 | 5 | -2.9289 | 0.79634 | 0.89695 | 1 | 18642 | 1 | -2.9289 |
| TAAR9 | 6 | 0.34292 | 0.556 | 0.999994 | 11005 | 3 | -0.35702 | 0.79634 | 0.89695 | 1 | 18643 | 1 | -0.35702 |
| SLC3A1 | 6 | 0.057998 | 0.16145 | 0.967126 | 3336 | 4 | -1.5169 | 0.79634 | 0.89695 | 1 | 18644 | 1 | -1.5169 |
| PLSCR3 | 6 | 0.32551 | 0.53613 | 0.999994 | 10633 | 2 | -0.68492 | 0.79634 | 0.89695 | 1 | 18645 | 2 | -0.68492 |
| NonTargetingControlG | 1 | 0.20354 | 0.20326 | 0.97798 | 7870 | 1 | -2.1721 | 0.79646 | 0.79682 | 1 | 18646 | 0 | -2.1721 |
| VIT | 6 | 0.63472 | 0.76591 | 0.999994 | 15440 | 2 | 0.10052 | 0.79665 | 0.8971 | 1 | 18647 | 2 | 0.10052 |
| FAM78A | 3 | 0.0009925 | 0.0028199 | 0.517578 | 168 | 2 | -9.6827 | 0.79676 | 0.80575 | 1 | 18648 | 1 | -9.6827 |
| ZC3HC1 | 6 | 0.0017852 | 0.0089017 | 0.680529 | 249 | 4 | -4.8883 | 0.79688 | 0.89721 | 1 | 18649 | 2 | -4.8883 |
| PROKR1 | 5 | 0.059778 | 0.16171 | 0.967126 | 3413 | 3 | -2.0134 | 0.79691 | 0.89048 | 1 | 18650 | 1 | -2.0134 |
| NonTargetingControlG | 2 | 0.44229 | 0.45256 | 0.999994 | 12615 | 1 | -0.73225 | 0.79715 | 0.79667 | 1 | 18651 | 0 | -0.73225 |
| ACSBG1 | 4 | 0.2846 | 0.42726 | 0.999994 | 9800 | 2 | -0.72631 | 0.79723 | 0.84525 | 1 | 18652 | 1 | -0.72631 |
| NonTargetingControlG | 2 | 0.6079 | 0.60772 | 0.999994 | 15046 | 0 | -0.18138 | 0.79726 | 0.7968 | 1 | 18653 | 0 | -0.18138 |
| NonTargetingControlG | 2 | 0.43261 | 0.44428 | 0.999994 | 12454 | 1 | -0.68164 | 0.79733 | 0.79686 | 1 | 18654 | 0 | -0.68164 |
| WIPF3 | 5 | 0.13813 | 0.28498 | 0.999632 | 6140 | 2 | -0.39907 | 0.79736 | 0.8906 | 1 | 18655 | 1 | -0.39907 |
| NonTargetingControlG | 2 | 0.68276 | 0.68264 | 0.999994 | 16128 | 0 | 0.064948 | 0.79752 | 0.79704 | 1 | 18656 | 0 | 0.064948 |
| FAM159B | 2 | 0.56387 | 0.56426 | 0.999994 | 14450 | 1 | -0.31685 | 0.79754 | 0.79707 | 1 | 18657 | 0 | -0.31685 |
| FAM135B | 5 | 0.22729 | 0.41345 | 0.999994 | 8461 | 3 | -0.68648 | 0.7977 | 0.89068 | 1 | 18658 | 1 | -0.68648 |
| NPHP1 | 6 | 0.035982 | 0.11283 | 0.951052 | 2312 | 4 | -1.8982 | 0.79776 | 0.89765 | 1 | 18659 | 2 | -1.8982 |
| KIAA1211 | 6 | 0.018366 | 0.066701 | 0.93914 | 1380 | 4 | -3.075 | 0.79778 | 0.89766 | 1 | 18660 | 2 | -3.075 |
| CD38 | 6 | 0.28462 | 0.48882 | 0.999994 | 9801 | 2 | -0.053346 | 0.79786 | 0.89769 | 1 | 18661 | 2 | -0.053346 |
| FAP | 3 | 0.16249 | 0.26474 | 0.996878 | 6809 | 1 | 0.076937 | 0.79793 | 0.80662 | 1 | 18662 | 1 | 0.076937 |
| TNPO3 | 5 | 0.05657 | 0.15616 | 0.967126 | 3274 | 4 | -0.89724 | 0.79823 | 0.89082 | 1 | 18663 | 1 | -0.89724 |
| ARID4B | 6 | 0.073929 | 0.19494 | 0.975797 | 3991 | 4 | -1.0376 | 0.79824 | 0.89789 | 1 | 18664 | 2 | -1.0376 |
| hsa-mir-196a-2 | 4 | 0.32269 | 0.46618 | 0.999994 | 10569 | 2 | -0.40496 | 0.79831 | 0.84576 | 1 | 18665 | 1 | -0.40496 |
| ZFC3H1 | 6 | 0.0066045 | 0.027026 | 0.848412 | 633 | 3 | -3.561 | 0.79832 | 0.89792 | 1 | 18666 | 2 | -3.561 |
| NonTargetingControlG | 2 | 0.40802 | 0.42347 | 0.999994 | 12074 | 1 | -0.74591 | 0.79836 | 0.79791 | 1 | 18667 | 0 | -0.74591 |

| | | | | | | | | | | | | | |
|---|---|---|---|---|---|---|---|---|---|---|---|---|---|
| C12orf5 | 5 | 0.46284 | 0.61606 | 0.999994 | 12903 | 2 | -0.32947 | 0.79854 | 0.8909 | 1 | 18668 | 1 | -0.32947 |
| PFKP | 6 | 0.048116 | 0.13995 | 0.958716 | 2892 | 3 | -0.47303 | 0.79864 | 0.89809 | 1 | 18669 | 2 | -0.47303 |
| ALS2CR12 | 6 | 0.059213 | 0.16406 | 0.967126 | 3389 | 4 | -0.82918 | 0.79874 | 0.89813 | 1 | 18670 | 2 | -0.82918 |
| AMMECR1L | 3 | 0.4756 | 0.53033 | 0.999994 | 13069 | 1 | 0.063455 | 0.79874 | 0.80725 | 1 | 18671 | 1 | 0.063455 |
| BANP | 5 | 0.55506 | 0.69627 | 0.999994 | 14291 | 2 | -0.67575 | 0.79879 | 0.89097 | 1 | 18672 | 1 | -0.67575 |
| COG3 | 4 | 0.06433 | 0.14906 | 0.965566 | 3622 | 2 | -1.9214 | 0.79886 | 0.84601 | 1 | 18673 | 1 | -1.9214 |
| ZNF812 | 6 | 0.077558 | 0.20241 | 0.977973 | 4132 | 2 | -0.031461 | 0.79887 | 0.89819 | 1 | 18674 | 2 | -0.031461 |
| TRA2B | 6 | 0.0012784 | 0.0064282 | 0.636218 | 190 | 5 | -4.122 | 0.79895 | 0.89823 | 1 | 18675 | 1 | -4.122 |
| ATMIN | 5 | 0.21535 | 0.3968 | 0.999994 | 8151 | 3 | -0.75193 | 0.79909 | 0.89104 | 1 | 18676 | 1 | -0.75193 |
| TSPYL1 | 5 | 0.20555 | 0.38312 | 0.999994 | 7911 | 2 | 0.06345 | 0.79929 | 0.89109 | 1 | 18677 | 1 | 0.06345 |
| ZNF554 | 5 | 0.12205 | 0.26046 | 0.994943 | 5704 | 2 | -0.25556 | 0.79933 | 0.8911 | 1 | 18678 | 1 | -0.25556 |
| TNN | 4 | 0.032612 | 0.08683 | 0.94395 | 2138 | 3 | -2.9852 | 0.7994 | 0.84624 | 1 | 18679 | 1 | -2.9852 |
| GABRE | 5 | 0.95294 | 0.9531 | 0.999994 | 20409 | 0 | 0.23337 | 0.79985 | 0.89123 | 1 | 18680 | 1 | 0.23337 |
| hsa-mir-378h | 4 | 0.14359 | 0.28263 | 0.999632 | 6289 | 2 | -0.90611 | 0.79985 | 0.84643 | 1 | 18681 | 1 | -0.90611 |
| SMIM14 | 6 | 0.0080902 | 0.032261 | 0.865899 | 737 | 5 | -2.4195 | 0.80013 | 0.89881 | 1 | 18682 | 1 | -2.4195 |
| ZNF768 | 6 | 0.27064 | 0.47248 | 0.999994 | 9488 | 3 | -0.47086 | 0.80015 | 0.89881 | 1 | 18683 | 1 | -0.47086 |
| hsa-mir-4662a | 4 | 0.11246 | 0.23722 | 0.985914 | 5385 | 3 | -0.78118 | 0.80018 | 0.84658 | 1 | 18684 | 1 | -0.78118 |
| FAM117A | 6 | 0.0018089 | 0.0090106 | 0.680529 | 253 | 4 | -5.0077 | 0.80028 | 0.89887 | 1 | 18685 | 2 | -5.0077 |
| SYNRG | 6 | 0.32751 | 0.53839 | 0.999994 | 10670 | 3 | -0.09643 | 0.80031 | 0.89889 | 1 | 18686 | 2 | -0.09643 |
| hsa-mir-4795 | 3 | 0.034458 | 0.085281 | 0.94395 | 2238 | 2 | -0.78 | 0.8004 | 0.80851 | 1 | 18687 | 1 | -0.78 |
| hsa-mir-129-2 | 4 | 0.094622 | 0.2054 | 0.978342 | 4772 | 3 | -1.2612 | 0.80043 | 0.8467 | 1 | 18688 | 1 | -1.2612 |
| INSIG2 | 6 | 0.19706 | 0.38432 | 0.999994 | 7676 | 2 | -0.43176 | 0.80053 | 0.89899 | 1 | 18689 | 2 | -0.43176 |
| CSNK2A1 | 5 | 0.020544 | 0.065535 | 0.93914 | 1484 | 3 | -3.4213 | 0.80062 | 0.89141 | 1 | 18690 | 1 | -3.4213 |
| HTR2B | 5 | 0.22764 | 0.41391 | 0.999994 | 8472 | 3 | -0.7116 | 0.80068 | 0.89143 | 1 | 18691 | 1 | -0.7116 |
| NonTargetingControlG | 1 | 0.19917 | 0.19893 | 0.977164 | 7741 | 1 | -2.1705 | 0.80083 | 0.80107 | 1 | 18692 | 0 | -2.1705 |
| MTRNR2L6 | 6 | 0.02754 | 0.093036 | 0.943968 | 1879 | 4 | -2.0656 | 0.80089 | 0.89916 | 1 | 18693 | 2 | -2.0656 |
| KLHL7 | 6 | 0.23187 | 0.42634 | 0.999994 | 8564 | 3 | -0.9909 | 0.80094 | 0.89919 | 1 | 18694 | 2 | -0.9909 |
| PPP1R3C | 6 | 0.17124 | 0.3523 | 0.999994 | 7016 | 3 | -0.54749 | 0.80104 | 0.89923 | 1 | 18695 | 2 | -0.54749 |
| BAP1 | 6 | 0.00023613 | 0.0012562 | 0.377819 | 68 | 5 | -6.659 | 0.80118 | 0.8993 | 1 | 18696 | 1 | -6.659 |
| TNFRSF12A | 5 | 0.33967 | 0.51344 | 0.999994 | 10925 | 2 | -0.52215 | 0.80119 | 0.89158 | 1 | 18697 | 1 | -0.52215 |
| FCRL2 | 6 | 0.021226 | 0.075915 | 0.93914 | 1528 | 5 | -1.1994 | 0.80119 | 0.89931 | 1 | 18698 | 1 | -1.1994 |
| ZNF598 | 4 | 0.026884 | 0.07513 | 0.93914 | 1852 | 3 | -4.0822 | 0.8012 | 0.84705 | 1 | 18699 | 1 | -4.0822 |
| OR1D5 | 6 | 0.062812 | 0.17169 | 0.967658 | 3547 | 4 | -1.3792 | 0.80121 | 0.89932 | 1 | 18700 | 1 | -1.3792 |
| SOWAHA | 4 | 0.091986 | 0.20064 | 0.977164 | 4679 | 3 | -2.2931 | 0.80158 | 0.84721 | 1 | 18701 | 1 | -2.2931 |
| KPNA4 | 5 | 0.17114 | 0.33379 | 0.999994 | 7011 | 3 | -1.0705 | 0.80159 | 0.89166 | 1 | 18702 | 1 | -1.0705 |
| CRBN | 5 | 0.056397 | 0.15573 | 0.967126 | 3264 | 4 | -0.93371 | 0.80159 | 0.89166 | 1 | 18703 | 1 | -0.93371 |
| ZNF266 | 6 | 0.26104 | 0.46111 | 0.999994 | 9269 | 3 | -0.53938 | 0.80179 | 0.89961 | 1 | 18704 | 2 | -0.53938 |
| hsa-mir-6799 | 3 | 0.28875 | 0.42257 | 0.999994 | 9892 | 1 | -0.061804 | 0.80184 | 0.80959 | 1 | 18705 | 1 | -0.061804 |
| DOK3 | 5 | 0.17649 | 0.34156 | 0.999994 | 7144 | 2 | -0.55483 | 0.80206 | 0.89179 | 1 | 18706 | 1 | -0.55483 |
| LIMK2 | 6 | 0.016489 | 0.060602 | 0.932932 | 1276 | 3 | -0.75877 | 0.80223 | 0.89983 | 1 | 18707 | 1 | -0.75877 |
| FOXK1 | 6 | 0.0044326 | 0.019062 | 0.804634 | 474 | 3 | -1.0121 | 0.80223 | 0.89983 | 1 | 18708 | 1 | -1.0121 |
| CISD1 | 6 | 0.072762 | 0.1925 | 0.974607 | 3952 | 4 | -1.084 | 0.80223 | 0.89983 | 1 | 18709 | 1 | -1.084 |
| SBDS | 6 | 0.021715 | 0.077484 | 0.93914 | 1558 | 4 | -2.8803 | 0.80223 | 0.89983 | 1 | 18710 | 1 | -2.8803 |
| PRKAB1 | 6 | 0.023484 | 0.083125 | 0.94383 | 1683 | 5 | -1.2253 | 0.80223 | 0.89983 | 1 | 18711 | 1 | -1.2253 |
| CD36 | 6 | 0.21904 | 0.41094 | 0.999994 | 8262 | 3 | -0.17548 | 0.80227 | 0.89985 | 1 | 18712 | 2 | -0.17548 |
| HECW1 | 5 | 0.0062936 | 0.025317 | 0.84412 | 614 | 4 | -3.09 | 0.80228 | 0.89184 | 1 | 18713 | 1 | -3.09 |
| RNPS1 | 4 | 0.25448 | 0.39652 | 0.999994 | 9114 | 2 | -0.41245 | 0.80249 | 0.8476 | 1 | 18714 | 1 | -0.41245 |
| NonTargetingControlG | 2 | 0.3093 | 0.34397 | 0.999994 | 10309 | 1 | -1.3247 | 0.80253 | 0.80214 | 1 | 18715 | 0 | -1.3247 |
| hsa-mir-3169 | 3 | 0.63075 | 0.64126 | 0.999994 | 15385 | 1 | 0.25788 | 0.80287 | 0.81039 | 1 | 18716 | 1 | 0.25788 |
| hsa-mir-515-1 | 3 | 0.78251 | 0.78242 | 0.999994 | 17246 | 0 | 0.25325 | 0.80322 | 0.81066 | 1 | 18717 | 1 | 0.25325 |
| FLVCR1 | 6 | 0.08121 | 0.20978 | 0.979587 | 4274 | 4 | -1.2286 | 0.80341 | 0.90045 | 1 | 18718 | 2 | -1.2286 |
| PTPDC1 | 3 | 0.2654 | 0.39403 | 0.999994 | 9360 | 2 | -0.76076 | 0.80343 | 0.81082 | 1 | 18719 | 1 | -0.76076 |

| | | | | | | | | | | | | | |
|---|---|---|---|---|---|---|---|---|---|---|---|---|---|
| NonTargetingControlG | 2 | 0.5634 | 0.5638 | 0.999994 | 14439 | 1 | -0.34935 | 0.80347 | 0.80304 | 1 | 18720 | 0 | -0.34935 |
| MYRF | 4 | 0.32129 | 0.46471 | 0.999994 | 10540 | 2 | -0.52422 | 0.80379 | 0.84818 | 1 | 18721 | 1 | -0.52422 |
| NonTargetingControlG | 2 | 0.65926 | 0.65907 | 0.999994 | 15766 | 0 | -0.001424 | 0.80384 | 0.80342 | 1 | 18722 | 0 | -0.001424 |
| SIGLEC8 | 6 | 0.26973 | 0.47141 | 0.999994 | 9468 | 2 | 0.45558 | 0.80394 | 0.90072 | 1 | 18723 | 2 | 0.45558 |
| C1orf87 | 6 | 0.24036 | 0.43644 | 0.999994 | 8782 | 3 | -0.79452 | 0.80415 | 0.90084 | 1 | 18724 | 1 | -0.79452 |
| hsa-mir-4671 | 3 | 0.045359 | 0.10927 | 0.951052 | 2781 | 2 | -1.1147 | 0.80459 | 0.81171 | 1 | 18725 | 1 | -1.1147 |
| TRIM73 | 4 | 0.04118 | 0.10413 | 0.947284 | 2570 | 3 | -2.6544 | 0.8046 | 0.84858 | 1 | 18726 | 1 | -2.6544 |
| NonTargetingControlG | 2 | 0.58128 | 0.58142 | 0.999994 | 14697 | 1 | -0.28517 | 0.8048 | 0.80437 | 1 | 18727 | 0 | -0.28517 |
| SDSL | 5 | 0.023296 | 0.072978 | 0.93914 | 1669 | 3 | -1.7909 | 0.80499 | 0.89253 | 1 | 18728 | 1 | -1.7909 |
| C12orf65 | 5 | 0.030179 | 0.090945 | 0.94395 | 2017 | 4 | -1.2337 | 0.80499 | 0.89253 | 1 | 18729 | 1 | -1.2337 |
| NUP188 | 5 | 0.50467 | 0.65199 | 0.999994 | 13508 | 2 | -0.31192 | 0.80499 | 0.89253 | 1 | 18730 | 1 | -0.31192 |
| UNC45B | 5 | 0.80695 | 0.82394 | 0.999994 | 17582 | 1 | 0.069993 | 0.80499 | 0.89253 | 1 | 18731 | 1 | 0.069993 |
| hsa-mir-2117 | 3 | 0.16258 | 0.26485 | 0.997104 | 6811 | 2 | -1.3986 | 0.8053 | 0.81227 | 1 | 18732 | 1 | -1.3986 |
| OAZ3 | 5 | 0.48127 | 0.63176 | 0.999994 | 13156 | 2 | -0.55278 | 0.80556 | 0.89269 | 1 | 18733 | 1 | -0.55278 |
| GEMIN4 | 2 | 0.28344 | 0.32413 | 0.999994 | 9772 | 1 | -1.4596 | 0.80562 | 0.80525 | 1 | 18734 | 0 | -1.4596 |
| LOC100129361 | 5 | 0.40516 | 0.56785 | 0.999994 | 12035 | 2 | -0.094493 | 0.8057 | 0.89273 | 1 | 18735 | 1 | -0.094493 |
| TRMT12 | 5 | 0.010092 | 0.036323 | 0.879988 | 876 | 4 | -2.399 | 0.80586 | 0.89277 | 1 | 18736 | 1 | -2.399 |
| DXO | 3 | 0.66154 | 0.66666 | 0.999994 | 15801 | 1 | -0.2976 | 0.80594 | 0.81278 | 1 | 18737 | 1 | -0.2976 |
| hsa-mir-938 | 3 | 0.19806 | 0.31021 | 0.999994 | 7712 | 2 | -1.1911 | 0.80597 | 0.8128 | 1 | 18738 | 1 | -1.1911 |
| hsa-mir-555 | 4 | 0.11475 | 0.24117 | 0.987253 | 5471 | 3 | -0.88423 | 0.80609 | 0.84925 | 1 | 18739 | 1 | -0.88423 |
| TMEM160 | 5 | 0.114 | 0.24819 | 0.990682 | 5448 | 2 | 0.3108 | 0.80611 | 0.89283 | 1 | 18740 | 1 | 0.3108 |
| CEP112 | 6 | 0.09196 | 0.23144 | 0.984304 | 4677 | 4 | -0.74683 | 0.80615 | 0.90185 | 1 | 18741 | 1 | -0.74683 |
| CCDC6 | 6 | 0.4356 | 0.62553 | 0.999994 | 12509 | 2 | -0.28608 | 0.80615 | 0.90185 | 1 | 18742 | 2 | -0.28608 |
| ZRANB3 | 5 | 0.03948 | 0.11459 | 0.951052 | 2489 | 4 | -1.0807 | 0.80631 | 0.89289 | 1 | 18743 | 1 | -1.0807 |
| DHRS4L1 | 6 | 0.63565 | 0.76663 | 0.999994 | 15454 | 1 | -0.29179 | 0.80636 | 0.90196 | 1 | 18744 | 2 | -0.29179 |
| CAPN14 | 5 | 0.039628 | 0.11497 | 0.951052 | 2496 | 4 | -1.639 | 0.80651 | 0.89294 | 1 | 18745 | 1 | -1.639 |
| PYURF | 4 | 0.1212 | 0.25239 | 0.994162 | 5670 | 2 | -0.85173 | 0.80666 | 0.8495 | 1 | 18746 | 1 | -0.85173 |
| NonTargetingControlG | 2 | 0.30233 | 0.33865 | 0.999994 | 10179 | 1 | -1.278 | 0.80668 | 0.80632 | 1 | 18747 | 0 | -1.278 |
| ATAD2B | 5 | 0.11509 | 0.24989 | 0.992497 | 5481 | 3 | -1.4254 | 0.80673 | 0.893 | 1 | 18748 | 1 | -1.4254 |
| CST5 | 5 | 0.15672 | 0.31274 | 0.999994 | 6644 | 3 | -1.0944 | 0.80693 | 0.89306 | 1 | 18749 | 1 | -1.0944 |
| LCMT1 | 6 | 0.059928 | 0.16559 | 0.967126 | 3418 | 4 | -1.2817 | 0.80707 | 0.90231 | 1 | 18750 | 2 | -1.2817 |
| CYTH3 | 6 | 0.00079862 | 0.0040978 | 0.560963 | 143 | 4 | -6.7664 | 0.80713 | 0.90234 | 1 | 18751 | 1 | -6.7664 |
| RSL1D1 | 4 | 0.054891 | 0.13101 | 0.95318 | 3194 | 2 | -3.3003 | 0.80733 | 0.8498 | 1 | 18752 | 1 | -3.3003 |
| ZNF449 | 6 | 0.2451 | 0.44209 | 0.999994 | 8902 | 3 | -0.41546 | 0.80753 | 0.90255 | 1 | 18753 | 2 | -0.41546 |
| PAFAH2 | 6 | 0.58235 | 0.72694 | 0.999994 | 14710 | 2 | 0.27125 | 0.80757 | 0.90256 | 1 | 18754 | 2 | 0.27125 |
| ADIG | 3 | 0.60479 | 0.62084 | 0.999994 | 15007 | 1 | 0.33597 | 0.80787 | 0.8143 | 1 | 18755 | 1 | 0.33597 |
| METTL24 | 5 | 0.09537 | 0.21913 | 0.980699 | 4796 | 3 | -1.0085 | 0.80799 | 0.89335 | 1 | 18756 | 1 | -1.0085 |
| TNP1 | 5 | 0.0051258 | 0.021319 | 0.814212 | 528 | 4 | -3.2701 | 0.80812 | 0.89339 | 1 | 18757 | 1 | -3.2701 |
| IL17RD | 6 | 0.078002 | 0.20332 | 0.97798 | 4151 | 4 | -1.3813 | 0.80834 | 0.90296 | 1 | 18758 | 2 | -1.3813 |
| PAPPA | 6 | 0.047006 | 0.13753 | 0.957317 | 2850 | 4 | -1.9595 | 0.80834 | 0.90296 | 1 | 18759 | 1 | -1.9595 |
| FAM109B | 6 | 0.16667 | 0.34657 | 0.999994 | 6900 | 3 | -0.8865 | 0.80834 | 0.90296 | 1 | 18760 | 2 | -0.8865 |
| C9orf92 | 5 | 0.16914 | 0.3309 | 0.999994 | 6963 | 3 | -1.0593 | 0.80834 | 0.89344 | 1 | 18761 | 1 | -1.0593 |
| THEM5 | 6 | 0.057377 | 0.1601 | 0.967126 | 3309 | 3 | -1.4064 | 0.8084 | 0.90299 | 1 | 18762 | 2 | -1.4064 |
| ARRB1 | 5 | 0.0058745 | 0.024048 | 0.834563 | 581 | 4 | -4.1135 | 0.80872 | 0.89353 | 1 | 18763 | 1 | -4.1135 |
| NonTargetingControlG | 2 | 0.59129 | 0.59127 | 0.999994 | 14837 | 0 | -0.26672 | 0.80881 | 0.8085 | 1 | 18764 | 0 | -0.26672 |
| CLUL1 | 6 | 0.025985 | 0.089317 | 0.94395 | 1804 | 3 | -2.7409 | 0.80882 | 0.9032 | 1 | 18765 | 2 | -2.7409 |
| SELK | 4 | 0.84851 | 0.84805 | 0.999994 | 18225 | 0 | 0.13333 | 0.80887 | 0.85051 | 1 | 18766 | 1 | 0.13333 |
| SHCBP1L | 6 | 0.28856 | 0.49342 | 0.999994 | 9885 | 3 | -0.32826 | 0.80917 | 0.90339 | 1 | 18767 | 2 | -0.32826 |
| hsa-mir-548ar | 2 | 0.021857 | 0.040524 | 0.891621 | 1572 | 1 | -5.1109 | 0.80945 | 0.80913 | 1 | 18768 | 0 | -5.1109 |
| CD28 | 6 | 0.024136 | 0.084949 | 0.94395 | 1714 | 3 | -0.90021 | 0.80948 | 0.90354 | 1 | 18769 | 1 | -0.90021 |
| PRM3 | 6 | 0.086103 | 0.21972 | 0.980699 | 4454 | 4 | -0.93215 | 0.80948 | 0.90354 | 1 | 18770 | 1 | -0.93215 |
| BRE | 6 | 0.42703 | 0.61994 | 0.999994 | 12366 | 2 | -0.3319 | 0.80948 | 0.90354 | 1 | 18771 | 1 | -0.3319 |

| | | | | | | | | | | | | | |
|---|---|---|---|---|---|---|---|---|---|---|---|---|---|
| OR2K2 | 5 | 0.17692 | 0.34216 | 0.999994 | 7154 | 3 | -0.93671 | 0.80952 | 0.89375 | 1 | 18772 | 1 | -0.93671 |
| C1QTNF5 | 4 | 0.37016 | 0.51503 | 0.999994 | 11513 | 2 | -0.39073 | 0.80961 | 0.85086 | 1 | 18773 | 1 | -0.39073 |
| NEIL3 | 4 | 0.009441 | 0.031944 | 0.865899 | 824 | 3 | -5.0581 | 0.80986 | 0.85099 | 1 | 18774 | 1 | -5.0581 |
| PLN | 1 | 0.19006 | 0.18991 | 0.970473 | 7486 | 1 | -2.4264 | 0.80994 | 0.8101 | 1 | 18775 | 0 | -2.4264 |
| SAAL1 | 6 | 0.014803 | 0.055078 | 0.922677 | 1177 | 3 | -2.5803 | 0.80995 | 0.90381 | 1 | 18776 | 1 | -2.5803 |
| hsa-mir-1227 | 2 | 0.095196 | 0.16261 | 0.967126 | 4790 | 1 | -4.1229 | 0.80997 | 0.80966 | 1 | 18777 | 0 | -4.1229 |
| hsa-mir-8062 | 4 | 0.12917 | 0.26616 | 0.998057 | 5919 | 2 | -1.1193 | 0.81002 | 0.85107 | 1 | 18778 | 1 | -1.1193 |
| NFASC | 5 | 0.022292 | 0.070265 | 0.93914 | 1599 | 4 | -1.4523 | 0.81003 | 0.89388 | 1 | 18779 | 1 | -1.4523 |
| NPNT | 6 | 0.10705 | 0.26103 | 0.994943 | 5214 | 4 | -0.8629 | 0.81015 | 0.90391 | 1 | 18780 | 2 | -0.8629 |
| SPANXN2 | 5 | 0.43278 | 0.59074 | 0.999994 | 12457 | 2 | -0.60028 | 0.8102 | 0.89393 | 1 | 18781 | 1 | -0.60028 |
| FNDC1 | 5 | 0.38956 | 0.55477 | 0.999994 | 11790 | 2 | -0.52558 | 0.81038 | 0.89397 | 1 | 18782 | 1 | -0.52558 |
| hsa-mir-6776 | 3 | 0.075666 | 0.15059 | 0.966359 | 4059 | 2 | -2.1931 | 0.81052 | 0.8164 | 1 | 18783 | 1 | -2.1931 |
| hsa-mir-4779 | 3 | 0.48681 | 0.53752 | 0.999994 | 13237 | 1 | -0.49653 | 0.81052 | 0.8164 | 1 | 18784 | 1 | -0.49653 |
| RARA | 6 | 0.55514 | 0.7073 | 0.999994 | 14294 | 2 | 0.25028 | 0.81059 | 0.90412 | 1 | 18785 | 2 | 0.25028 |
| MTCH2 | 6 | 0.23886 | 0.43462 | 0.999994 | 8743 | 3 | -0.69691 | 0.81069 | 0.90417 | 1 | 18786 | 2 | -0.69691 |
| hsa-mir-548ab | 2 | 0.57205 | 0.57224 | 0.999994 | 14578 | 1 | -0.95406 | 0.81073 | 0.81043 | 1 | 18787 | 0 | -0.95406 |
| hsa-mir-4495 | 2 | 0.43104 | 0.4429 | 0.999994 | 12432 | 1 | -1.4705 | 0.81073 | 0.81043 | 1 | 18788 | 0 | -1.4705 |
| NBPF24 | 2 | 0.16221 | 0.23524 | 0.985362 | 6800 | 1 | -3.2294 | 0.81073 | 0.81043 | 1 | 18789 | 0 | -3.2294 |
| RANGRF | 6 | 0.66104 | 0.78613 | 0.999994 | 15796 | 2 | -0.35337 | 0.81091 | 0.9043 | 1 | 18790 | 2 | -0.35337 |
| NonTargetingControlG | 2 | 0.5253 | 0.52738 | 0.999994 | 13820 | 1 | -0.48017 | 0.81107 | 0.81076 | 1 | 18791 | 0 | -0.48017 |
| SUPT20H | 4 | 0.11561 | 0.24263 | 0.988088 | 5501 | 3 | -1.217 | 0.81107 | 0.85154 | 1 | 18792 | 1 | -1.217 |
| TAF9B | 6 | 0.21842 | 0.41022 | 0.999994 | 8238 | 1 | 0.024387 | 0.81107 | 0.90437 | 1 | 18793 | 2 | 0.024387 |
| NonTargetingControlG | 1 | 0.18883 | 0.18866 | 0.969543 | 7448 | 1 | -2.4018 | 0.81117 | 0.81135 | 1 | 18794 | 0 | -2.4018 |
| NonTargetingControlG | 2 | 0.64277 | 0.64256 | 0.999994 | 15558 | 0 | -0.12581 | 0.81118 | 0.81087 | 1 | 18795 | 0 | -0.12581 |
| DUX2 | 1 | 0.18876 | 0.1886 | 0.969543 | 7443 | 1 | -2.1811 | 0.81124 | 0.81141 | 1 | 18796 | 0 | -2.1811 |
| NonTargetingControlG | 1 | 0.1887 | 0.18854 | 0.969543 | 7442 | 1 | -3.8144 | 0.8113 | 0.81146 | 1 | 18797 | 0 | -3.8144 |
| PLSCR4 | 6 | 0.20343 | 0.39204 | 0.999994 | 7865 | 3 | -0.78341 | 0.81133 | 0.9045 | 1 | 18798 | 2 | -0.78341 |
| NonTargetingControlG | 2 | 0.44276 | 0.45297 | 0.999994 | 12621 | 1 | -0.80887 | 0.81142 | 0.81113 | 1 | 18799 | 0 | -0.80887 |
| KLRF1 | 6 | 0.17687 | 0.35917 | 0.999994 | 7152 | 3 | -0.5179 | 0.81152 | 0.90458 | 1 | 18800 | 2 | -0.5179 |
| LEAP2 | 4 | 0.77395 | 0.77898 | 0.999994 | 17131 | 1 | 0.027876 | 0.81156 | 0.85177 | 1 | 18801 | 1 | 0.027876 |
| NonTargetingControlG | 2 | 0.51342 | 0.51628 | 0.999994 | 13629 | 1 | -0.53512 | 0.8116 | 0.81131 | 1 | 18802 | 0 | -0.53512 |
| NonTargetingControlG | 2 | 0.67666 | 0.67642 | 0.999994 | 16035 | 0 | -0.021755 | 0.81177 | 0.81148 | 1 | 18803 | 0 | -0.021755 |
| ARPC2 | 6 | 0.0019794 | 0.0098733 | 0.680529 | 274 | 4 | -4.1668 | 0.81178 | 0.90471 | 1 | 18804 | 2 | -4.1668 |
| C7orf55-LUC7L2 | 5 | 0.039066 | 0.11353 | 0.951052 | 2466 | 4 | -1.026 | 0.81183 | 0.89436 | 1 | 18805 | 1 | -1.026 |
| hsa-mir-1302-6 | 3 | 0.50317 | 0.54824 | 0.999994 | 13483 | 1 | -0.23839 | 0.81213 | 0.81767 | 1 | 18806 | 1 | -0.23839 |
| HOXC4 | 3 | 0.55027 | 0.58047 | 0.999994 | 14216 | 1 | 0.17217 | 0.81235 | 0.81785 | 1 | 18807 | 1 | 0.17217 |
| ZCCHC8 | 6 | 0.087296 | 0.22208 | 0.983354 | 4487 | 4 | -1.2553 | 0.81267 | 0.90514 | 1 | 18808 | 1 | -1.2553 |
| SPATS2L | 6 | 0.22268 | 0.41539 | 0.999994 | 8349 | 3 | -0.65936 | 0.81268 | 0.90515 | 1 | 18809 | 1 | -0.65936 |
| LACRT | 6 | 0.12295 | 0.29128 | 0.999632 | 5738 | 4 | -1.0158 | 0.81287 | 0.90525 | 1 | 18810 | 2 | -1.0158 |
| PEG10 | 5 | 0.12052 | 0.25814 | 0.994943 | 5643 | 3 | -1.4783 | 0.81301 | 0.89466 | 1 | 18811 | 1 | -1.4783 |
| MYO1G | 5 | 0.07471 | 0.18604 | 0.969409 | 4023 | 3 | -1.8258 | 0.81301 | 0.89466 | 1 | 18812 | 1 | -1.8258 |
| TMEM106B | 5 | 0.15981 | 0.31731 | 0.999994 | 6729 | 2 | -0.46129 | 0.81301 | 0.89466 | 1 | 18813 | 1 | -0.46129 |
| OR11H6 | 6 | 0.22743 | 0.42099 | 0.999994 | 8465 | 3 | -0.24685 | 0.81304 | 0.90534 | 1 | 18814 | 2 | -0.24685 |
| NonTargetingControlG | 2 | 0.57977 | 0.57991 | 0.999994 | 14674 | 1 | -0.3355 | 0.81304 | 0.81278 | 1 | 18815 | 0 | -0.3355 |
| CDS2 | 4 | 0.40903 | 0.55496 | 0.999994 | 12093 | 1 | 0.201 | 0.81318 | 0.85254 | 1 | 18816 | 1 | 0.201 |
| C1orf122 | 4 | 0.051834 | 0.12508 | 0.951052 | 3058 | 3 | -1.8313 | 0.81318 | 0.85254 | 1 | 18817 | 1 | -1.8313 |
| TTC1 | 4 | 0.0717 | 0.16316 | 0.967126 | 3904 | 2 | -0.7883 | 0.81318 | 0.85254 | 1 | 18818 | 1 | -0.7883 |
| AAMP | 6 | 0.20117 | 0.38929 | 0.999994 | 7794 | 3 | -0.44959 | 0.81333 | 0.90549 | 1 | 18819 | 2 | -0.44959 |
| DNAJC30 | 6 | 0.62559 | 0.75902 | 0.999994 | 15321 | 2 | 0.29108 | 0.81337 | 0.90551 | 1 | 18820 | 2 | 0.29108 |
| EIF4A2 | 6 | 0.00028925 | 0.0015479 | 0.388244 | 77 | 4 | -4.0864 | 0.81343 | 0.90554 | 1 | 18821 | 2 | -4.0864 |
| PRDM14 | 5 | 0.23618 | 0.42576 | 0.999994 | 8676 | 3 | -0.60786 | 0.8135 | 0.89481 | 1 | 18822 | 1 | -0.60786 |
| NTPCR | 6 | 0.30364 | 0.51095 | 0.999994 | 10208 | 3 | -0.37242 | 0.81352 | 0.90559 | 1 | 18823 | 2 | -0.37242 |

| | | | | | | | | | | | | | |
|---|---|---|---|---|---|---|---|---|---|---|---|---|---|
| RD3 | 6 | 0.028877 | 0.096173 | 0.945031 | 1949 | 4 | -1.7538 | 0.81375 | 0.9057 | 1 | 18824 | 2 | -1.7538 |
| CAND1 | 6 | 0.0025148 | 0.011909 | 0.729405 | 317 | 4 | -2.1832 | 0.81388 | 0.90577 | 1 | 18825 | 2 | -2.1832 |
| DIAPH1 | 5 | 0.075292 | 0.187 | 0.969409 | 4046 | 3 | -1.0294 | 0.81391 | 0.89492 | 1 | 18826 | 1 | -1.0294 |
| SDC4 | 5 | 0.39095 | 0.55592 | 0.999994 | 11815 | 2 | 0.23543 | 0.81397 | 0.89494 | 1 | 18827 | 1 | 0.23543 |
| FIG4 | 3 | 0.15612 | 0.25643 | 0.994943 | 6633 | 1 | 0.42794 | 0.81404 | 0.81918 | 1 | 18828 | 1 | 0.42794 |
| FAM5C | 3 | 0.095288 | 0.17681 | 0.968171 | 4793 | 2 | -2.3824 | 0.81407 | 0.81921 | 1 | 18829 | 1 | -2.3824 |
| ALPPL2 | 6 | 0.045499 | 0.13416 | 0.95318 | 2790 | 4 | -1.6657 | 0.81414 | 0.90589 | 1 | 18830 | 2 | -1.6657 |
| BUB1 | 6 | 0.11874 | 0.28332 | 0.999632 | 5595 | 2 | -0.17992 | 0.8142 | 0.90592 | 1 | 18831 | 2 | -0.17992 |
| FAM25C | 4 | 0.62209 | 0.67753 | 0.999994 | 15261 | 1 | -0.32326 | 0.81426 | 0.85307 | 1 | 18832 | 1 | -0.32326 |
| MICU2 | 5 | 0.024276 | 0.07556 | 0.93914 | 1721 | 4 | -1.7818 | 0.81444 | 0.89507 | 1 | 18833 | 1 | -1.7818 |
| BRINP3 | 6 | 0.21579 | 0.4071 | 0.999994 | 8163 | 2 | -0.069645 | 0.81446 | 0.90608 | 1 | 18834 | 1 | -0.069645 |
| CYR61 | 6 | 0.18502 | 0.36938 | 0.999994 | 7342 | 3 | -0.51145 | 0.81453 | 0.90611 | 1 | 18835 | 2 | -0.51145 |
| NUP35 | 3 | 0.010752 | 0.02872 | 0.858757 | 915 | 2 | -1.4666 | 0.81504 | 0.82003 | 1 | 18836 | 1 | -1.4666 |
| NonTargetingControlG | 2 | 0.1001 | 0.17022 | 0.967126 | 4975 | 1 | -3.3468 | 0.81511 | 0.8149 | 1 | 18837 | 0 | -3.3468 |
| NFE2L2 | 6 | 0.29641 | 0.50253 | 0.999994 | 10051 | 2 | 0.34917 | 0.81527 | 0.90647 | 1 | 18838 | 2 | 0.34917 |
| XAGE1B | 2 | 0.50691 | 0.51026 | 0.999994 | 13535 | 1 | -0.54917 | 0.8153 | 0.81509 | 1 | 18839 | 0 | -0.54917 |
| MTFR2 | 4 | 0.12255 | 0.25472 | 0.994835 | 5726 | 3 | -1.7268 | 0.81549 | 0.85364 | 1 | 18840 | 1 | -1.7268 |
| TTC30A | 6 | 0.0010933 | 0.0055293 | 0.623123 | 178 | 4 | -2.3775 | 0.81556 | 0.90663 | 1 | 18841 | 2 | -2.3775 |
| IGF2R | 3 | 0.55482 | 0.58371 | 0.999994 | 14283 | 1 | 0.48125 | 0.81564 | 0.82051 | 1 | 18842 | 1 | 0.48125 |
| ARHGAP8 | 6 | 0.94944 | 0.94941 | 0.999994 | 20319 | 0 | 0.15742 | 0.81566 | 0.90667 | 1 | 18843 | 2 | 0.15742 |
| MTUS2 | 6 | 0.23727 | 0.43274 | 0.999994 | 8706 | 3 | -0.33856 | 0.81567 | 0.90668 | 1 | 18844 | 2 | -0.33856 |
| NonTargetingControlG | 2 | 0.51922 | 0.52169 | 0.999994 | 13734 | 1 | -0.51934 | 0.81572 | 0.8155 | 1 | 18845 | 0 | -0.51934 |
| NXPE3 | 5 | 0.011159 | 0.039415 | 0.891621 | 938 | 4 | -1.9516 | 0.81577 | 0.89543 | 1 | 18846 | 1 | -1.9516 |
| TRUB1 | 6 | 0.03836 | 0.11821 | 0.951052 | 2436 | 4 | -2.0822 | 0.81584 | 0.90677 | 1 | 18847 | 1 | -2.0822 |
| RNF5 | 4 | 0.023985 | 0.069067 | 0.93914 | 1707 | 3 | -3.0957 | 0.81585 | 0.85382 | 1 | 18848 | 1 | -3.0957 |
| ALAS1 | 6 | 0.00079046 | 0.0040539 | 0.560963 | 141 | 3 | -2.5009 | 0.81605 | 0.90688 | 1 | 18849 | 2 | -2.5009 |
| KLRC1 | 6 | 0.09339 | 0.2343 | 0.984971 | 4735 | 3 | -0.66985 | 0.81617 | 0.90693 | 1 | 18850 | 2 | -0.66985 |
| hsa-mir-100 | 3 | 0.31526 | 0.43677 | 0.999994 | 10424 | 1 | 0.50982 | 0.81621 | 0.82099 | 1 | 18851 | 1 | 0.50982 |
| hsa-mir-551b | 4 | 0.15973 | 0.2992 | 0.999994 | 6724 | 2 | -0.89683 | 0.81623 | 0.85401 | 1 | 18852 | 1 | -0.89683 |
| SLITRK4 | 4 | 0.019077 | 0.058714 | 0.931872 | 1413 | 2 | -3.2954 | 0.81629 | 0.85404 | 1 | 18853 | 1 | -3.2954 |
| DTL | 5 | 0.063237 | 0.16741 | 0.967126 | 3569 | 3 | -1.3898 | 0.81637 | 0.89558 | 1 | 18854 | 1 | -1.3898 |
| NonTargetingControlG | 2 | 0.62635 | 0.62612 | 0.999994 | 15327 | 0 | -0.19192 | 0.81638 | 0.81615 | 1 | 18855 | 0 | -0.19192 |
| YPEL5 | 5 | 0.41852 | 0.57887 | 0.999994 | 12228 | 2 | -0.2292 | 0.81654 | 0.89562 | 1 | 18856 | 1 | -0.2292 |
| C10orf71 | 3 | 0.7762 | 0.77614 | 0.999994 | 17154 | 0 | -0.21843 | 0.81664 | 0.82135 | 1 | 18857 | 1 | -0.21843 |
| UBE2D3 | 6 | 0.035364 | 0.1114 | 0.951052 | 2283 | 4 | -2.2773 | 0.81667 | 0.90718 | 1 | 18858 | 1 | -2.2773 |
| GPR3 | 6 | 0.24223 | 0.43866 | 0.999994 | 8837 | 3 | -0.40417 | 0.81677 | 0.90722 | 1 | 18859 | 2 | -0.40417 |
| NXF2 | 6 | 0.020224 | 0.072689 | 0.93914 | 1473 | 4 | -2.653 | 0.81681 | 0.90725 | 1 | 18860 | 1 | -2.653 |
| TEX264 | 6 | 0.05221 | 0.14892 | 0.965566 | 3076 | 4 | -1.7816 | 0.81685 | 0.90727 | 1 | 18861 | 1 | -1.7816 |
| ABHD2 | 6 | 0.58763 | 0.73084 | 0.999994 | 14784 | 1 | 0.35278 | 0.81686 | 0.90727 | 1 | 18862 | 2 | 0.35278 |
| CTSL | 5 | 0.13368 | 0.27827 | 0.999632 | 6016 | 3 | -1.0872 | 0.81709 | 0.89576 | 1 | 18863 | 1 | -1.0872 |
| WDFY4 | 6 | 0.037521 | 0.11633 | 0.951052 | 2404 | 3 | -1.7282 | 0.81716 | 0.90741 | 1 | 18864 | 1 | -1.7282 |
| ZFAND2B | 6 | 0.014341 | 0.053581 | 0.919407 | 1156 | 5 | -3.5313 | 0.81721 | 0.90743 | 1 | 18865 | 1 | -3.5313 |
| PTP4A2 | 6 | 0.11325 | 0.27295 | 0.998561 | 5414 | 3 | -0.52803 | 0.81726 | 0.90746 | 1 | 18866 | 2 | -0.52803 |
| OR4C16 | 6 | 0.063479 | 0.17312 | 0.96793 | 3577 | 3 | -0.73095 | 0.81745 | 0.90756 | 1 | 18867 | 1 | -0.73095 |
| FAM219A | 6 | 0.5224 | 0.68417 | 0.999994 | 13786 | 2 | 0.17823 | 0.81755 | 0.90761 | 1 | 18868 | 2 | 0.17823 |
| ASPSCR1 | 5 | 0.028176 | 0.085784 | 0.94395 | 1914 | 3 | -2.5808 | 0.81778 | 0.89595 | 1 | 18869 | 1 | -2.5808 |
| KIAA1755 | 6 | 0.094715 | 0.23695 | 0.985449 | 4779 | 4 | -1.4048 | 0.81789 | 0.9078 | 1 | 18870 | 1 | -1.4048 |
| hsa-mir-6859-1 | 1 | 0.18209 | 0.18187 | 0.968322 | 7273 | 1 | -2.3449 | 0.81791 | 0.81814 | 1 | 18871 | 0 | -2.3449 |
| IFNLR1 | 5 | 0.051086 | 0.14301 | 0.95983 | 3021 | 3 | -0.92671 | 0.818 | 0.896 | 1 | 18872 | 1 | -0.92671 |
| SFTPB | 6 | 0.038812 | 0.1192 | 0.951052 | 2452 | 4 | -1.6082 | 0.81802 | 0.90787 | 1 | 18873 | 2 | -1.6082 |
| ZNF174 | 6 | 0.0054259 | 0.022699 | 0.820617 | 553 | 5 | -2.0972 | 0.81819 | 0.90796 | 1 | 18874 | 1 | -2.0972 |
| TNFAIP8L2 | 6 | 0.1569 | 0.3343 | 0.999994 | 6649 | 3 | -0.52377 | 0.81822 | 0.90797 | 1 | 18875 | 1 | -0.52377 |

| | | | | | | | | | | | | |
|---|---|---|---|---|---|---|---|---|---|---|---|---|
| PDGFA | 6 | 0.51917 | 0.68195 | 0.999994 | 13733 | 2 | -0.051567 | 0.81831 | 0.90802 | 1 | 18876 | 1 | -0.051567 |
| NonTargetingControlG | 2 | 0.52109 | 0.52339 | 0.999994 | 13762 | 1 | -0.55684 | 0.8184 | 0.8182 | 1 | 18877 | 0 | -0.55684 |
| EXT1 | 5 | 0.07014 | 0.1786 | 0.968171 | 3837 | 3 | -1.8168 | 0.81857 | 0.89615 | 1 | 18878 | 1 | -1.8168 |
| NSMCE1 | 5 | 0.0097332 | 0.035311 | 0.86993 | 856 | 4 | -2.846 | 0.81857 | 0.89615 | 1 | 18879 | 1 | -2.846 |
| METTL11B | 6 | 0.97872 | 0.97865 | 0.999994 | 21067 | 0 | 0.47417 | 0.81861 | 0.90817 | 1 | 18880 | 2 | 0.47417 |
| UAP1L1 | 4 | 0.3807 | 0.52587 | 0.999994 | 11658 | 2 | -0.68599 | 0.81864 | 0.85514 | 1 | 18881 | 1 | -0.68599 |
| BCAN | 4 | 0.034305 | 0.090334 | 0.94395 | 2229 | 3 | -3.6312 | 0.81868 | 0.85516 | 1 | 18882 | 1 | -3.6312 |
| hsa-mir-503 | 4 | 0.28705 | 0.42976 | 0.999994 | 9853 | 2 | -0.83962 | 0.81879 | 0.85521 | 1 | 18883 | 1 | -0.83962 |
| GPR65 | 6 | 0.22772 | 0.42134 | 0.999994 | 8473 | 3 | -0.60776 | 0.81881 | 0.90826 | 1 | 18884 | 2 | -0.60776 |
| NEB | 6 | 0.16609 | 0.34582 | 0.999994 | 6889 | 3 | -0.7702 | 0.81885 | 0.90828 | 1 | 18885 | 2 | -0.7702 |
| hsa-mir-3180-5 | 3 | 0.725 | 0.72494 | 0.999994 | 16549 | 1 | 0.54942 | 0.81886 | 0.82317 | 1 | 18886 | 1 | 0.54942 |
| ZNF514 | 6 | 0.10232 | 0.25185 | 0.994 | 5052 | 4 | -0.84035 | 0.81887 | 0.90829 | 1 | 18887 | 2 | -0.84035 |
| MOSPD3 | 6 | 0.15232 | 0.32859 | 0.999994 | 6512 | 3 | -0.62407 | 0.8189 | 0.9083 | 1 | 18888 | 2 | -0.62407 |
| LDLRAD1 | 6 | 0.1377 | 0.31013 | 0.999994 | 6127 | 3 | -1.0239 | 0.81898 | 0.90834 | 1 | 18889 | 2 | -1.0239 |
| NonTargetingControlG | 1 | 0.18096 | 0.18075 | 0.968171 | 7245 | 1 | -2.6039 | 0.81904 | 0.81927 | 1 | 18890 | 0 | -2.6039 |
| UFSP2 | 6 | 0.014481 | 0.054034 | 0.922076 | 1164 | 4 | -4.0503 | 0.81909 | 0.9084 | 1 | 18891 | 1 | -4.0503 |
| hsa-mir-5683 | 4 | 0.82232 | 0.82199 | 0.999994 | 17809 | 1 | 0.24679 | 0.81928 | 0.85546 | 1 | 18892 | 1 | 0.24679 |
| OR6C6 | 6 | 0.0016677 | 0.0083252 | 0.680529 | 238 | 4 | -4.2614 | 0.81939 | 0.90857 | 1 | 18893 | 1 | -4.2614 |
| hsa-mir-31 | 2 | 0.08404 | 0.14498 | 0.963012 | 4378 | 1 | -3.5507 | 0.81954 | 0.81937 | 1 | 18894 | 0 | -3.5507 |
| hsa-mir-5001 | 1 | 0.18032 | 0.18009 | 0.968171 | 7226 | 1 | -2.2498 | 0.81968 | 0.81988 | 1 | 18895 | 0 | -2.2498 |
| hsa-mir-218-1 | 4 | 0.12514 | 0.25923 | 0.994943 | 5798 | 3 | -0.92481 | 0.81982 | 0.85573 | 1 | 18896 | 1 | -0.92481 |
| SLC15A2 | 6 | 0.36213 | 0.5777 | 0.999994 | 11371 | 2 | -0.66261 | 0.82005 | 0.90893 | 1 | 18897 | 2 | -0.66261 |
| ZNF844 | 6 | 0.85073 | 0.87232 | 0.999994 | 18273 | 1 | 0.16915 | 0.8203 | 0.90904 | 1 | 18898 | 1 | 0.16915 |
| ATG2A | 6 | 0.011071 | 0.04265 | 0.891621 | 930 | 4 | -3.3064 | 0.8203 | 0.90904 | 1 | 18899 | 1 | -3.3064 |
| UBL3 | 5 | 0.15241 | 0.30635 | 0.999994 | 6518 | 3 | -1.0101 | 0.8206 | 0.8967 | 1 | 18900 | 1 | -1.0101 |
| ATG9A | 6 | 0.08888 | 0.22529 | 0.983354 | 4549 | 4 | -1.1567 | 0.82072 | 0.90926 | 1 | 18901 | 1 | -1.1567 |
| VCY1B | 1 | 0.17918 | 0.17894 | 0.968171 | 7199 | 1 | -2.3675 | 0.82082 | 0.82102 | 1 | 18902 | 0 | -2.3675 |
| hsa-mir-652 | 3 | 0.82313 | 0.82305 | 0.999994 | 17825 | 0 | 0.089877 | 0.82111 | 0.82505 | 1 | 18903 | 1 | 0.089877 |
| RABGGTB | 3 | 0.17445 | 0.28009 | 0.999632 | 7097 | 2 | -1.5021 | 0.82124 | 0.82517 | 1 | 18904 | 1 | -1.5021 |
| ZNF695 | 5 | 0.032711 | 0.097456 | 0.945253 | 2145 | 3 | -2.7668 | 0.82125 | 0.89686 | 1 | 18905 | 1 | -2.7668 |
| LGALS14 | 6 | 0.088396 | 0.22434 | 0.983354 | 4530 | 4 | -0.95238 | 0.82142 | 0.90961 | 1 | 18906 | 2 | -0.95238 |
| MEF2D | 6 | 0.01371 | 0.051504 | 0.910467 | 1112 | 4 | -2.3486 | 0.82159 | 0.90971 | 1 | 18907 | 2 | -2.3486 |
| NonTargetingControlG | 1 | 0.1784 | 0.17819 | 0.968171 | 7185 | 1 | -2.2758 | 0.8216 | 0.8218 | 1 | 18908 | 0 | -2.2758 |
| MBD3L5 | 1 | 0.17839 | 0.17818 | 0.968171 | 7183 | 1 | -2.6295 | 0.82161 | 0.82181 | 1 | 18909 | 0 | -2.6295 |
| CHMP1A | 6 | 0.0051761 | 0.021783 | 0.819701 | 531 | 4 | -3.2443 | 0.82161 | 0.90972 | 1 | 18910 | 2 | -3.2443 |
| ADRA2B | 5 | 0.23459 | 0.42356 | 0.999994 | 8628 | 3 | -0.66409 | 0.82169 | 0.89699 | 1 | 18911 | 1 | -0.66409 |
| MEGF6 | 6 | 0.058574 | 0.16269 | 0.967126 | 3367 | 4 | -1.276 | 0.82188 | 0.90987 | 1 | 18912 | 2 | -1.276 |
| C19orf25 | 6 | 0.06825 | 0.1831 | 0.968526 | 3757 | 4 | -1.5537 | 0.82192 | 0.90988 | 1 | 18913 | 1 | -1.5537 |
| BSX | 6 | 0.0044057 | 0.018966 | 0.803792 | 472 | 5 | -2.4681 | 0.82194 | 0.9099 | 1 | 18914 | 1 | -2.4681 |
| DEFB133 | 2 | 0.66382 | 0.66358 | 0.999994 | 15836 | 0 | -0.10853 | 0.82208 | 0.82191 | 1 | 18915 | 0 | -0.10853 |
| OR4A16 | 6 | 0.037131 | 0.11542 | 0.951052 | 2382 | 4 | -1.6979 | 0.82228 | 0.91009 | 1 | 18916 | 2 | -1.6979 |
| SAMD14 | 6 | 0.065967 | 0.17832 | 0.968171 | 3688 | 3 | -0.15482 | 0.82235 | 0.91013 | 1 | 18917 | 2 | -0.15482 |
| DIP2C | 6 | 0.015369 | 0.05693 | 0.925104 | 1209 | 4 | -3.4495 | 0.82284 | 0.91037 | 1 | 18918 | 1 | -3.4495 |
| CCDC9 | 6 | 0.010262 | 0.039908 | 0.891621 | 886 | 3 | -4.1187 | 0.82284 | 0.91037 | 1 | 18919 | 1 | -4.1187 |
| D2HGDH | 6 | 0.082413 | 0.21228 | 0.979587 | 4314 | 3 | -1.5635 | 0.82284 | 0.91037 | 1 | 18920 | 1 | -1.5635 |
| CDK5RAP2 | 6 | 0.19028 | 0.37586 | 0.999994 | 7494 | 3 | -0.62515 | 0.82284 | 0.91037 | 1 | 18921 | 1 | -0.62515 |
| C6orf62 | 6 | 0.055567 | 0.15616 | 0.967126 | 3224 | 3 | -1.3246 | 0.82315 | 0.91053 | 1 | 18922 | 2 | -1.3246 |
| HMGB3 | 5 | 0.26176 | 0.45017 | 0.999994 | 9280 | 1 | -0.067817 | 0.82316 | 0.89738 | 1 | 18923 | 1 | -0.067817 |
| LRRC28 | 6 | 0.14922 | 0.32464 | 0.999994 | 6435 | 2 | -0.076817 | 0.82347 | 0.91069 | 1 | 18924 | 2 | -0.076817 |
| DUSP5 | 6 | 0.27905 | 0.48229 | 0.999994 | 9676 | 3 | -0.78028 | 0.82372 | 0.91082 | 1 | 18925 | 2 | -0.78028 |
| CTXN1 | 6 | 0.037514 | 0.1163 | 0.951052 | 2403 | 3 | -2.0302 | 0.82384 | 0.91089 | 1 | 18926 | 1 | -2.0302 |
| OXER1 | 3 | 0.17549 | 0.28146 | 0.999632 | 7123 | 2 | -1.4045 | 0.82387 | 0.82739 | 1 | 18927 | 1 | -1.4045 |

| | | | | | | | | | | | | | |
|---|---|---|---|---|---|---|---|---|---|---|---|---|---|
| GPATCH2L | 5 | 0.23217 | 0.42022 | 0.999994 | 8574 | 2 | -0.41567 | 0.82391 | 0.89757 | 1 | 18928 | 1 | -0.41567 |
| ATP1A2 | 5 | 0.015361 | 0.05123 | 0.909241 | 1208 | 3 | -3.662 | 0.82402 | 0.89761 | 1 | 18929 | 1 | -3.662 |
| ERRFI1 | 5 | 0.71874 | 0.77526 | 0.999994 | 16483 | 1 | -0.48152 | 0.82427 | 0.89768 | 1 | 18930 | 1 | -0.48152 |
| NonTargetingControlG | 1 | 0.17562 | 0.17539 | 0.968171 | 7127 | 1 | -2.575 | 0.82438 | 0.82455 | 1 | 18931 | 0 | -2.575 |
| SMIM18 | 6 | 0.2609 | 0.46095 | 0.999994 | 9267 | 3 | -0.68959 | 0.82444 | 0.9112 | 1 | 18932 | 1 | -0.68959 |
| H6PD | 6 | 0.17507 | 0.35696 | 0.999994 | 7109 | 2 | 0.40441 | 0.82452 | 0.91124 | 1 | 18933 | 2 | 0.40441 |
| NonTargetingControlG | 2 | 0.058115 | 0.10279 | 0.946414 | 3342 | 1 | -4.5944 | 0.82461 | 0.82442 | 1 | 18934 | 0 | -4.5944 |
| CREB3L1 | 5 | 0.20736 | 0.38564 | 0.999994 | 7956 | 2 | -0.48355 | 0.82461 | 0.89777 | 1 | 18935 | 1 | -0.48355 |
| NonTargetingControlG | 2 | 0.2165 | 0.27407 | 0.998561 | 8179 | 1 | -2.002 | 0.82471 | 0.82451 | 1 | 18936 | 0 | -2.002 |
| ANKFY1 | 6 | 0.64138 | 0.77097 | 0.999994 | 15537 | 2 | -0.014812 | 0.82476 | 0.91137 | 1 | 18937 | 1 | -0.014812 |
| NonTargetingControlG | 1 | 0.17514 | 0.17493 | 0.968171 | 7113 | 1 | -4.0603 | 0.82486 | 0.82505 | 1 | 18938 | 0 | -4.0603 |
| DHTKD1 | 6 | 0.50319 | 0.67093 | 0.999994 | 13484 | 2 | -0.013947 | 0.8249 | 0.91145 | 1 | 18939 | 2 | -0.013947 |
| IFNG | 6 | 0.26762 | 0.46887 | 0.999994 | 9411 | 2 | 0.17221 | 0.82491 | 0.91146 | 1 | 18940 | 2 | 0.17221 |
| C20orf96 | 5 | 0.004794 | 0.019985 | 0.808847 | 505 | 4 | -2.6812 | 0.82497 | 0.89787 | 1 | 18941 | 1 | -2.6812 |
| CMTM6 | 6 | 0.88749 | 0.89458 | 0.999994 | 18983 | 1 | 0.060309 | 0.8251 | 0.91156 | 1 | 18942 | 2 | 0.060309 |
| hsa-mir-548c | 3 | 0.82564 | 0.82549 | 0.999994 | 17863 | 0 | 0.12532 | 0.82511 | 0.82838 | 1 | 18943 | 1 | 0.12532 |
| ZNF76 | 6 | 0.059079 | 0.16379 | 0.967126 | 3384 | 1 | 0.23629 | 0.82526 | 0.91165 | 1 | 18944 | 2 | 0.23629 |
| EFHB | 5 | 0.0075581 | 0.028976 | 0.859137 | 696 | 4 | -3.0329 | 0.82528 | 0.89794 | 1 | 18945 | 1 | -3.0329 |
| FAM110D | 5 | 0.15849 | 0.31532 | 0.999994 | 6684 | 2 | -0.52515 | 0.82528 | 0.89794 | 1 | 18946 | 1 | -0.52515 |
| HOXB8 | 5 | 0.032971 | 0.098159 | 0.945253 | 2156 | 4 | -1.5244 | 0.82528 | 0.89794 | 1 | 18947 | 1 | -1.5244 |
| SETD6 | 6 | 0.0055756 | 0.023229 | 0.83009 | 563 | 4 | -4.2102 | 0.82535 | 0.91169 | 1 | 18948 | 2 | -4.2102 |
| SLC13A2 | 5 | 0.0029429 | 0.012619 | 0.733879 | 349 | 4 | -3.1748 | 0.82553 | 0.89801 | 1 | 18949 | 1 | -3.1748 |
| AMOT | 6 | 0.30742 | 0.51538 | 0.999994 | 10275 | 3 | -0.41122 | 0.82557 | 0.91181 | 1 | 18950 | 2 | -0.41122 |
| DIAPH2 | 6 | 0.033051 | 0.10599 | 0.950444 | 2162 | 4 | -2.3313 | 0.82559 | 0.91182 | 1 | 18951 | 1 | -2.3313 |
| TAS2R3 | 4 | 0.60902 | 0.67022 | 0.999994 | 15057 | 1 | -0.071514 | 0.82561 | 0.85855 | 1 | 18952 | 1 | -0.071514 |
| TAAR6 | 4 | 0.10796 | 0.22923 | 0.983354 | 5237 | 3 | -1.6178 | 0.82563 | 0.85856 | 1 | 18953 | 1 | -1.6178 |
| KAZN | 5 | 0.21633 | 0.39815 | 0.999994 | 8176 | 3 | -0.69449 | 0.82572 | 0.89807 | 1 | 18954 | 1 | -0.69449 |
| PLXNA2 | 6 | 0.53141 | 0.69055 | 0.999994 | 13921 | 1 | 0.20555 | 0.82576 | 0.91192 | 1 | 18955 | 2 | 0.20555 |
| NUDT16L1 | 3 | 0.28422 | 0.41716 | 0.999994 | 9789 | 1 | -0.34903 | 0.82588 | 0.82902 | 1 | 18956 | 1 | -0.34903 |
| NonTargetingControlG | 2 | 0.32807 | 0.3585 | 0.999994 | 10686 | 1 | -1.4054 | 0.8259 | 0.82572 | 1 | 18957 | 0 | -1.4054 |
| MFSD6 | 6 | 0.51391 | 0.67834 | 0.999994 | 13639 | 2 | -0.008299 | 0.82605 | 0.91208 | 1 | 18958 | 2 | -0.008299 |
| MEFV | 6 | 0.038761 | 0.1191 | 0.951052 | 2451 | 4 | -0.96424 | 0.82608 | 0.91209 | 1 | 18959 | 2 | -0.96424 |
| SLC39A13 | 6 | 0.126 | 0.29525 | 0.999994 | 5822 | 2 | 0.27892 | 0.82622 | 0.91217 | 1 | 18960 | 1 | 0.27892 |
| SCNM1 | 6 | 0.069411 | 0.18553 | 0.969035 | 3812 | 3 | -1.3791 | 0.82622 | 0.91217 | 1 | 18961 | 1 | -1.3791 |
| AGAP5 | 2 | 0.34776 | 0.37425 | 0.999994 | 11110 | 1 | -1.2103 | 0.82631 | 0.82613 | 1 | 18962 | 0 | -1.2103 |
| C19orf52 | 3 | 0.24708 | 0.37132 | 0.999994 | 8950 | 1 | 0.18853 | 0.82661 | 0.82964 | 1 | 18963 | 1 | 0.18853 |
| hsa-mir-3669 | 1 | 0.17333 | 0.17311 | 0.96793 | 7071 | 1 | -2.6138 | 0.82667 | 0.8269 | 1 | 18964 | 0 | -2.6138 |
| KRTAP10-1 | 6 | 0.042068 | 0.12654 | 0.951052 | 2616 | 4 | -2.0027 | 0.82667 | 0.91238 | 1 | 18965 | 2 | -2.0027 |
| CCR3 | 6 | 0.25564 | 0.45469 | 0.999994 | 9139 | 3 | -0.39019 | 0.82668 | 0.91239 | 1 | 18966 | 2 | -0.39019 |
| IL11 | 6 | 0.706 | 0.81386 | 0.999994 | 16372 | 1 | 0.33994 | 0.82673 | 0.91242 | 1 | 18967 | 2 | 0.33994 |
| TMEM229A | 6 | 0.070165 | 0.18711 | 0.969409 | 3839 | 4 | -1.2753 | 0.82688 | 0.91249 | 1 | 18968 | 2 | -1.2753 |
| PDE6G | 6 | 0.058429 | 0.16237 | 0.967126 | 3362 | 4 | -1.3932 | 0.82699 | 0.91255 | 1 | 18969 | 2 | -1.3932 |
| LIPN | 6 | 0.078932 | 0.20519 | 0.978342 | 4187 | 4 | -1.4096 | 0.827 | 0.91256 | 1 | 18970 | 1 | -1.4096 |
| NonTargetingControlG | 2 | 0.64987 | 0.64962 | 0.999994 | 15658 | 0 | -0.13759 | 0.82708 | 0.8269 | 1 | 18971 | 0 | -0.13759 |
| THUMPD1 | 5 | 0.41357 | 0.57479 | 0.999994 | 12162 | 2 | -0.58833 | 0.8272 | 0.89848 | 1 | 18972 | 1 | -0.58833 |
| P2RY14 | 6 | 0.12622 | 0.29552 | 0.999994 | 5829 | 2 | -0.30987 | 0.82724 | 0.9127 | 1 | 18973 | 2 | -0.30987 |
| SLC4A1 | 6 | 0.11401 | 0.27438 | 0.998561 | 5449 | 4 | -0.93044 | 0.82726 | 0.91271 | 1 | 18974 | 1 | -0.93044 |
| hsa-mir-4733 | 4 | 0.54806 | 0.63874 | 0.999994 | 14178 | 1 | -0.31745 | 0.82728 | 0.85939 | 1 | 18975 | 1 | -0.31745 |
| PDXP | 3 | 0.53573 | 0.57034 | 0.999994 | 13991 | 1 | 0.445 | 0.82739 | 0.83028 | 1 | 18976 | 1 | 0.445 |
| NonTargetingControlG | 2 | 0.62068 | 0.62046 | 0.999994 | 15230 | 0 | -0.26234 | 0.8274 | 0.82722 | 1 | 18977 | 0 | -0.26234 |
| NonTargetingControlG | 2 | 0.55365 | 0.5544 | 0.999994 | 14264 | 1 | -0.45442 | 0.82753 | 0.82737 | 1 | 18978 | 0 | -0.45442 |
| CPA3 | 5 | 0.089122 | 0.20917 | 0.979587 | 4559 | 3 | -1.4991 | 0.82758 | 0.8986 | 1 | 18979 | 1 | -1.4991 |

| | | | | | | | | | | | | | |
|---|---|---|---|---|---|---|---|---|---|---|---|---|---|
| SCD | 4 | 0.04788 | 0.1173 | 0.951052 | 2886 | 3 | -3.791 | 0.82758 | 0.85954 | 1 | 18980 | 1 | -3.791 |
| FER1L6 | 6 | 0.1643 | 0.3436 | 0.999994 | 6847 | 3 | -1.0338 | 0.82759 | 0.91288 | 1 | 18981 | 2 | -1.0338 |
| hsa-mir-759 | 3 | 0.27209 | 0.40223 | 0.999994 | 9522 | 2 | -0.64321 | 0.82783 | 0.83065 | 1 | 18982 | 1 | -0.64321 |
| hsa-mir-4703 | 1 | 0.17216 | 0.17192 | 0.967658 | 7040 | 1 | -2.6042 | 0.82784 | 0.82804 | 1 | 18983 | 0 | -2.6042 |
| DOK4 | 6 | 0.10431 | 0.25572 | 0.994943 | 5107 | 4 | -1.0855 | 0.82798 | 0.91308 | 1 | 18984 | 1 | -1.0855 |
| ANKRD44 | 6 | 0.00396 | 0.017334 | 0.785648 | 441 | 3 | -4.1471 | 0.82802 | 0.9131 | 1 | 18985 | 2 | -4.1471 |
| PROCR | 6 | 0.042376 | 0.12724 | 0.951052 | 2634 | 3 | -1.7275 | 0.82807 | 0.91312 | 1 | 18986 | 1 | -1.7275 |
| USP36 | 4 | 0.017334 | 0.054983 | 0.922677 | 1328 | 2 | -4.1434 | 0.82845 | 0.85997 | 1 | 18987 | 1 | -4.1434 |
| RUNDC3A | 6 | 0.59243 | 0.73439 | 0.999994 | 14854 | 2 | 0.25654 | 0.82846 | 0.91333 | 1 | 18988 | 2 | 0.25654 |
| AXIN1 | 3 | 0.52226 | 0.56104 | 0.999994 | 13782 | 1 | 0.048263 | 0.82854 | 0.83125 | 1 | 18989 | 1 | 0.048263 |
| MTA2 | 6 | 0.13894 | 0.31167 | 0.999994 | 6161 | 3 | -0.57178 | 0.82856 | 0.91339 | 1 | 18990 | 2 | -0.57178 |
| CT45A2 | 1 | 0.17138 | 0.17114 | 0.96737 | 7020 | 1 | -2.6358 | 0.82862 | 0.82881 | 1 | 18991 | 0 | -2.6358 |
| hsa-mir-136 | 4 | 0.066535 | 0.15327 | 0.967126 | 3705 | 3 | -1.452 | 0.82871 | 0.86009 | 1 | 18992 | 1 | -1.452 |
| POC5 | 6 | 0.15922 | 0.33723 | 0.999994 | 6711 | 3 | -0.84151 | 0.82879 | 0.91351 | 1 | 18993 | 2 | -0.84151 |
| CNNM1 | 6 | 0.15348 | 0.33005 | 0.999994 | 6549 | 3 | -0.87782 | 0.829 | 0.91361 | 1 | 18994 | 1 | -0.87782 |
| TRHR | 6 | 0.11034 | 0.26735 | 0.998561 | 5314 | 3 | -0.95601 | 0.82924 | 0.91373 | 1 | 18995 | 2 | -0.95601 |
| PRR23B | 4 | 0.66925 | 0.7049 | 0.999994 | 15918 | 1 | 0.21616 | 0.82945 | 0.86047 | 1 | 18996 | 1 | 0.21616 |
| REXO4 | 5 | 0.15881 | 0.31579 | 0.999994 | 6691 | 3 | -1.0963 | 0.82961 | 0.89918 | 1 | 18997 | 1 | -1.0963 |
| ADAD2 | 3 | 0.24937 | 0.37418 | 0.999994 | 9002 | 2 | -0.79109 | 0.82969 | 0.83225 | 1 | 18998 | 1 | -0.79109 |
| STK17A | 5 | 0.043812 | 0.12535 | 0.951052 | 2704 | 4 | -1.8234 | 0.82972 | 0.89922 | 1 | 18999 | 1 | -1.8234 |
| TSPY1 | 6 | 0.61623 | 0.75201 | 0.999994 | 15163 | 2 | 0.24297 | 0.82981 | 0.91405 | 1 | 19000 | 2 | 0.24297 |
| MFI2 | 4 | 0.070156 | 0.16022 | 0.967126 | 3838 | 3 | -2.1388 | 0.82983 | 0.86067 | 1 | 19001 | 1 | -2.1388 |
| FAM217A | 6 | 0.94348 | 0.94345 | 0.999994 | 20163 | 0 | 0.14938 | 0.82991 | 0.91411 | 1 | 19002 | 2 | 0.14938 |
| C11orf1 | 5 | 0.42525 | 0.58446 | 0.999994 | 12337 | 2 | -0.5913 | 0.82991 | 0.89926 | 1 | 19003 | 1 | -0.5913 |
| hsa-mir-4537 | 4 | 0.17196 | 0.31182 | 0.999994 | 7034 | 2 | -1.015 | 0.83001 | 0.86076 | 1 | 19004 | 1 | -1.015 |
| DHRS7B | 6 | 0.19421 | 0.3808 | 0.999994 | 7606 | 3 | -0.64595 | 0.83007 | 0.91419 | 1 | 19005 | 2 | -0.64595 |
| NUDT5 | 6 | 0.11819 | 0.28227 | 0.999632 | 5579 | 4 | -0.88078 | 0.83021 | 0.91425 | 1 | 19006 | 1 | -0.88078 |
| SS18L1 | 6 | 0.13938 | 0.31225 | 0.999994 | 6172 | 3 | -0.58427 | 0.83027 | 0.91429 | 1 | 19007 | 2 | -0.58427 |
| hsa-mir-4719 | 1 | 0.16961 | 0.16939 | 0.967126 | 6976 | 1 | -4.1691 | 0.83039 | 0.83062 | 1 | 19008 | 0 | -4.1691 |
| METTL14 | 4 | 0.28859 | 0.43134 | 0.999994 | 9886 | 2 | -0.95487 | 0.83052 | 0.86102 | 1 | 19009 | 1 | -0.95487 |
| hsa-mir-595 | 4 | 0.060479 | 0.14169 | 0.959146 | 3443 | 3 | -3.3837 | 0.83056 | 0.86104 | 1 | 19010 | 1 | -3.3837 |
| AFM | 6 | 0.15471 | 0.33162 | 0.999994 | 6586 | 3 | -0.57761 | 0.83065 | 0.9145 | 1 | 19011 | 2 | -0.57761 |
| SLC25A4 | 6 | 0.077955 | 0.20321 | 0.97798 | 4149 | 3 | -1.2165 | 0.83068 | 0.91452 | 1 | 19012 | 2 | -1.2165 |
| SCAMP1 | 6 | 0.026886 | 0.091479 | 0.94395 | 1853 | 3 | -0.39629 | 0.83069 | 0.91452 | 1 | 19013 | 1 | -0.39629 |
| TP53TG3D | 3 | 0.14572 | 0.243 | 0.988088 | 6343 | 2 | -1.1212 | 0.83074 | 0.83321 | 1 | 19014 | 1 | -1.1212 |
| C14orf178 | 5 | 0.0028281 | 0.012151 | 0.732963 | 343 | 4 | -2.9859 | 0.83096 | 0.89956 | 1 | 19015 | 1 | -2.9859 |
| BMP2K | 6 | 0.012915 | 0.048837 | 0.903125 | 1059 | 3 | -1.8913 | 0.83099 | 0.91469 | 1 | 19016 | 2 | -1.8913 |
| CDKN2C | 5 | 0.3627 | 0.53232 | 0.999994 | 11382 | 2 | -0.46523 | 0.83108 | 0.89959 | 1 | 19017 | 1 | -0.46523 |
| UBE2NL | 4 | 0.45438 | 0.5965 | 0.999994 | 12761 | 1 | 0.31315 | 0.83121 | 0.86138 | 1 | 19018 | 1 | 0.31315 |
| LOC158434 | 4 | 0.20208 | 0.34282 | 0.999994 | 7832 | 1 | -0.10735 | 0.83133 | 0.86144 | 1 | 19019 | 1 | -0.10735 |
| OR13C9 | 6 | 0.64451 | 0.77341 | 0.999994 | 15578 | 2 | 0.12292 | 0.83144 | 0.91494 | 1 | 19020 | 2 | 0.12292 |
| C2orf40 | 4 | 0.37627 | 0.52134 | 0.999994 | 11597 | 2 | -0.50922 | 0.83149 | 0.86152 | 1 | 19021 | 1 | -0.50922 |
| COIL | 5 | 0.09694 | 0.22162 | 0.983093 | 4856 | 3 | -0.62652 | 0.83164 | 0.89977 | 1 | 19022 | 1 | -0.62652 |
| VSIG1 | 6 | 0.065112 | 0.17649 | 0.968171 | 3657 | 2 | 0.015759 | 0.83187 | 0.91517 | 1 | 19023 | 1 | 0.015759 |
| hsa-mir-6880 | 1 | 0.16794 | 0.16773 | 0.967126 | 6935 | 1 | -2.8239 | 0.83206 | 0.83231 | 1 | 19024 | 0 | -2.8239 |
| hsa-mir-6070 | 3 | 0.39558 | 0.48171 | 0.999994 | 11880 | 1 | 0.46712 | 0.83211 | 0.83436 | 1 | 19025 | 1 | 0.46712 |
| hsa-mir-548ac | 2 | 0.63468 | 0.63445 | 0.999994 | 15439 | 0 | -0.23983 | 0.83222 | 0.832 | 1 | 19026 | 0 | -0.23983 |
| OR1L4 | 6 | 0.9167 | 0.91737 | 0.999994 | 19555 | 1 | 0.20824 | 0.83249 | 0.9155 | 1 | 19027 | 2 | 0.20824 |
| FAM221B | 6 | 0.1135 | 0.27342 | 0.998561 | 5421 | 4 | -0.72427 | 0.83268 | 0.91561 | 1 | 19028 | 2 | -0.72427 |
| IGSF5 | 6 | 0.35221 | 0.5665 | 0.999994 | 11199 | 3 | -0.32106 | 0.8327 | 0.91562 | 1 | 19029 | 2 | -0.32106 |
| CBX4 | 4 | 0.058427 | 0.13775 | 0.957317 | 3361 | 3 | -1.6 | 0.83272 | 0.86214 | 1 | 19030 | 1 | -1.6 |
| hsa-mir-4764 | 4 | 0.43349 | 0.58012 | 0.999994 | 12470 | 2 | -0.032263 | 0.83308 | 0.86231 | 1 | 19031 | 1 | -0.032263 |

| | | | | | | | | | | | | | |
|---|---|---|---|---|---|---|---|---|---|---|---|---|---|
| OSBPL9 | 2 | 0.091488 | 0.15685 | 0.967126 | 4656 | 1 | -4.2707 | 0.83328 | 0.83304 | 1 | 19032 | 0 | -4.2707 |
| SCNN1A | 6 | 0.1032 | 0.25357 | 0.994544 | 5074 | 2 | 0.27022 | 0.83347 | 0.91603 | 1 | 19033 | 2 | 0.27022 |
| BCL11B | 6 | 0.005384 | 0.022543 | 0.819701 | 547 | 5 | -3.5626 | 0.83386 | 0.91624 | 1 | 19034 | 1 | -3.5626 |
| NKX6-3 | 4 | 0.22051 | 0.36176 | 0.999994 | 8301 | 2 | -1.2718 | 0.83387 | 0.86271 | 1 | 19035 | 1 | -1.2718 |
| SSUH2 | 3 | 0.46194 | 0.52163 | 0.999994 | 12883 | 1 | 0.38676 | 0.83403 | 0.83603 | 1 | 19036 | 1 | 0.38676 |
| TMPRSS11E | 5 | 0.056434 | 0.15583 | 0.967126 | 3268 | 4 | -1.3582 | 0.83413 | 0.90048 | 1 | 19037 | 1 | -1.3582 |
| CERK | 6 | 0.08806 | 0.22365 | 0.983354 | 4517 | 4 | -1.1192 | 0.83414 | 0.9164 | 1 | 19038 | 2 | -1.1192 |
| NonTargetingControlG | 1 | 0.16581 | 0.16561 | 0.967126 | 6880 | 1 | -2.1587 | 0.83419 | 0.83446 | 1 | 19039 | 0 | -2.1587 |
| NTRK1 | 5 | 0.21345 | 0.39422 | 0.999994 | 8099 | 3 | -0.72904 | 0.83421 | 0.90051 | 1 | 19040 | 1 | -0.72904 |
| PCDHGA2 | 2 | 0.35741 | 0.38198 | 0.999994 | 11291 | 1 | -1.0894 | 0.83422 | 0.83403 | 1 | 19041 | 0 | -1.0894 |
| DHRS9 | 6 | 0.089948 | 0.22739 | 0.983354 | 4592 | 4 | -1.3267 | 0.83429 | 0.91647 | 1 | 19042 | 1 | -1.3267 |
| NonTargetingControlG | 2 | 0.58694 | 0.58696 | 0.999994 | 14773 | 0 | -0.39951 | 0.83438 | 0.83419 | 1 | 19043 | 0 | -0.39951 |
| RARRES1 | 6 | 0.041713 | 0.12575 | 0.951052 | 2601 | 4 | -1.8884 | 0.8345 | 0.91659 | 1 | 19044 | 2 | -1.8884 |
| MCHR1 | 3 | 0.015921 | 0.041582 | 0.891621 | 1240 | 2 | -7.2311 | 0.83452 | 0.83646 | 1 | 19045 | 1 | -7.2311 |
| TRAPPC2L | 6 | 0.043855 | 0.13057 | 0.95318 | 2705 | 3 | -0.81276 | 0.83454 | 0.91661 | 1 | 19046 | 1 | -0.81276 |
| EMC3 | 6 | 0.57495 | 0.72149 | 0.999994 | 14615 | 2 | -0.10709 | 0.83456 | 0.91662 | 1 | 19047 | 2 | -0.10709 |
| ZNF146 | 6 | 0.24003 | 0.43601 | 0.999994 | 8771 | 3 | -0.081443 | 0.83457 | 0.91662 | 1 | 19048 | 2 | -0.081443 |
| ACKR4 | 4 | 0.89272 | 0.89242 | 0.999994 | 19084 | 0 | 0.39849 | 0.83458 | 0.86309 | 1 | 19049 | 1 | 0.39849 |
| hsa-mir-484 | 3 | 0.0098533 | 0.026391 | 0.848412 | 864 | 2 | -1.1629 | 0.83464 | 0.83657 | 1 | 19050 | 1 | -1.1629 |
| INHBA | 5 | 0.023468 | 0.073428 | 0.93914 | 1682 | 4 | -1.3358 | 0.83469 | 0.90063 | 1 | 19051 | 1 | -1.3358 |
| CCRL2 | 5 | 0.017768 | 0.058001 | 0.929685 | 1351 | 3 | -3.7885 | 0.83469 | 0.90063 | 1 | 19052 | 1 | -3.7885 |
| PHF14 | 5 | 0.40764 | 0.56991 | 0.999994 | 12070 | 2 | 0.25489 | 0.83469 | 0.90063 | 1 | 19053 | 1 | 0.25489 |
| SPECC1L | 6 | 0.034648 | 0.10969 | 0.951052 | 2246 | 3 | -1.3758 | 0.8347 | 0.91669 | 1 | 19054 | 2 | -1.3758 |
| NonTargetingControlG | 2 | 0.2437 | 0.29421 | 0.999994 | 8877 | 1 | -1.8575 | 0.83471 | 0.83451 | 1 | 19055 | 0 | -1.8575 |
| SLC6A15 | 6 | 0.30899 | 0.5172 | 0.999994 | 10297 | 3 | -0.88298 | 0.83482 | 0.91674 | 1 | 19056 | 2 | -0.88298 |
| STK16 | 6 | 0.65273 | 0.77972 | 0.999994 | 15695 | 2 | -0.49124 | 0.83525 | 0.91697 | 1 | 19057 | 1 | -0.49124 |
| C5orf52 | 5 | 0.26762 | 0.45493 | 0.999994 | 9414 | 2 | -0.4783 | 0.83561 | 0.9009 | 1 | 19058 | 1 | -0.4783 |
| DUT | 2 | 0.14462 | 0.2229 | 0.983354 | 6316 | 1 | -4.3339 | 0.83562 | 0.83542 | 1 | 19059 | 0 | -4.3339 |
| ERLEC1 | 6 | 0.15164 | 0.32774 | 0.999994 | 6498 | 3 | -0.65986 | 0.83583 | 0.91728 | 1 | 19060 | 2 | -0.65986 |
| SNX5 | 6 | 0.079545 | 0.20638 | 0.978744 | 4211 | 3 | -0.57596 | 0.83585 | 0.9173 | 1 | 19061 | 1 | -0.57596 |
| TOMM40 | 4 | 0.31006 | 0.45329 | 0.999994 | 10325 | 2 | -0.66779 | 0.83594 | 0.86378 | 1 | 19062 | 1 | -0.66779 |
| ERMP1 | 4 | 0.28434 | 0.42697 | 0.999994 | 9790 | 2 | -0.57856 | 0.83611 | 0.86386 | 1 | 19063 | 1 | -0.57856 |
| TTC14 | 5 | 0.10348 | 0.23182 | 0.984502 | 5082 | 3 | -1.4378 | 0.83612 | 0.90105 | 1 | 19064 | 1 | -1.4378 |
| LONRF2 | 5 | 0.094024 | 0.21698 | 0.980699 | 4749 | 3 | -1.4605 | 0.83625 | 0.90108 | 1 | 19065 | 1 | -1.4605 |
| KIAA1551 | 6 | 0.004678 | 0.01997 | 0.808847 | 496 | 5 | -3.0773 | 0.83627 | 0.91753 | 1 | 19066 | 1 | -3.0773 |
| ZNF219 | 6 | 0.096626 | 0.24073 | 0.987253 | 4842 | 3 | -1.0161 | 0.83631 | 0.91755 | 1 | 19067 | 2 | -1.0161 |
| LAPTM4B | 6 | 0.063739 | 0.1737 | 0.96793 | 3596 | 3 | -1.0904 | 0.83632 | 0.91756 | 1 | 19068 | 2 | -1.0904 |
| GAPDH | 3 | 0.014324 | 0.037651 | 0.884154 | 1153 | 2 | -7.3835 | 0.8364 | 0.83809 | 1 | 19069 | 1 | -7.3835 |
| DEFB130 | 6 | 0.064545 | 0.17533 | 0.968171 | 3631 | 4 | -1.5746 | 0.83674 | 0.9178 | 1 | 19070 | 1 | -1.5746 |
| GRIN3A | 5 | 0.16007 | 0.31768 | 0.999994 | 6743 | 2 | 0.1358 | 0.83698 | 0.90127 | 1 | 19071 | 1 | 0.1358 |
| ZNF660 | 6 | 0.034271 | 0.10878 | 0.951052 | 2227 | 4 | -1.5015 | 0.83698 | 0.91793 | 1 | 19072 | 1 | -1.5015 |
| FRRS1L | 6 | 0.3593 | 0.57453 | 0.999994 | 11326 | 3 | -0.22558 | 0.83728 | 0.91809 | 1 | 19073 | 2 | -0.22558 |
| FKBP5 | 6 | 0.062066 | 0.17014 | 0.967126 | 3515 | 3 | -0.6697 | 0.8373 | 0.91811 | 1 | 19074 | 2 | -0.6697 |
| LPCAT2 | 5 | 0.031936 | 0.09545 | 0.945031 | 2100 | 4 | -1.5319 | 0.83733 | 0.90137 | 1 | 19075 | 1 | -1.5319 |
| OR51A7 | 4 | 0.32302 | 0.46651 | 0.999994 | 10577 | 1 | -0.18536 | 0.83734 | 0.86447 | 1 | 19076 | 1 | -0.18536 |
| ARIH2 | 6 | 0.73006 | 0.82161 | 0.999994 | 16598 | 1 | 0.23539 | 0.83735 | 0.91813 | 1 | 19077 | 2 | 0.23539 |
| UQCRHL | 5 | 0.18858 | 0.3589 | 0.999994 | 7438 | 3 | -0.79635 | 0.83736 | 0.90138 | 1 | 19078 | 1 | -0.79635 |
| PSMC1 | 3 | 0.58509 | 0.60579 | 0.999994 | 14743 | 1 | -0.50418 | 0.83737 | 0.83897 | 1 | 19079 | 1 | -0.50418 |
| BAIAP3 | 5 | 0.50413 | 0.65152 | 0.999994 | 13496 | 2 | -0.56034 | 0.83765 | 0.90146 | 1 | 19080 | 1 | -0.56034 |
| NonTargetingControlG | 1 | 0.16224 | 0.16199 | 0.967126 | 6801 | 1 | -2.4332 | 0.83776 | 0.83804 | 1 | 19081 | 0 | -2.4332 |
| NonTargetingControlG | 2 | 0.49135 | 0.49602 | 0.999994 | 13295 | 1 | -0.70044 | 0.8378 | 0.83775 | 1 | 19082 | 0 | -0.70044 |
| POLG | 6 | 0.35273 | 0.56709 | 0.999994 | 11212 | 2 | -0.091732 | 0.83783 | 0.91839 | 1 | 19083 | 2 | -0.091732 |

| | | | | | | | | | | | | | |
|---|---|---|---|---|---|---|---|---|---|---|---|---|---|
| ME3 | 6 | 0.022809 | 0.080978 | 0.93914 | 1635 | 4 | -1.5062 | 0.83787 | 0.91842 | 1 | 19084 | 2 | -1.5062 |
| PSMB6 | 4 | 0.003281 | 0.011719 | 0.727553 | 383 | 3 | -6.3448 | 0.83794 | 0.86478 | 1 | 19085 | 1 | -6.3448 |
| hsa-mir-101-2 | 4 | 0.08911 | 0.19537 | 0.975797 | 4558 | 3 | -1.5891 | 0.83803 | 0.86483 | 1 | 19086 | 1 | -1.5891 |
| NLRP3 | 5 | 0.8875 | 0.8876 | 0.999994 | 18985 | 1 | -0.066884 | 0.8381 | 0.9016 | 1 | 19087 | 1 | -0.066884 |
| ESPN | 5 | 0.1487 | 0.30079 | 0.999994 | 6421 | 1 | -0.40254 | 0.83813 | 0.90161 | 1 | 19088 | 1 | -0.40254 |
| TMEM209 | 4 | 0.040698 | 0.10317 | 0.946414 | 2545 | 3 | -2.3822 | 0.83814 | 0.86488 | 1 | 19089 | 1 | -2.3822 |
| ABI2 | 6 | 0.15323 | 0.32973 | 0.999994 | 6542 | 2 | -0.06416 | 0.83819 | 0.91858 | 1 | 19090 | 2 | -0.06416 |
| hsa-mir-4435-2 | 3 | 0.11378 | 0.20129 | 0.977164 | 5436 | 2 | -1.3631 | 0.83822 | 0.83972 | 1 | 19091 | 1 | -1.3631 |
| TAF1L | 6 | 0.32987 | 0.54107 | 0.999994 | 10719 | 2 | 0.34167 | 0.83837 | 0.91868 | 1 | 19092 | 2 | 0.34167 |
| NAV2 | 6 | 0.23678 | 0.43215 | 0.999994 | 8689 | 3 | -0.75044 | 0.83842 | 0.91871 | 1 | 19093 | 2 | -0.75044 |
| NonTargetingControlG | 2 | 0.4328 | 0.44442 | 0.999994 | 12458 | 1 | -0.86319 | 0.83862 | 0.83856 | 1 | 19094 | 0 | -0.86319 |
| PAFAH1B1 | 4 | 0.6027 | 0.66682 | 0.999994 | 14981 | 1 | -0.18559 | 0.83874 | 0.86518 | 1 | 19095 | 1 | -0.18559 |
| SLC38A4 | 6 | 0.15997 | 0.33819 | 0.999994 | 6736 | 1 | -0.10839 | 0.83879 | 0.91888 | 1 | 19096 | 2 | -0.10839 |
| ADH1B | 6 | 0.091517 | 0.23057 | 0.983933 | 4657 | 4 | -1.1215 | 0.83887 | 0.91892 | 1 | 19097 | 1 | -1.1215 |
| hsa-mir-301b | 3 | 0.053988 | 0.12122 | 0.951052 | 3158 | 2 | -2.9653 | 0.83908 | 0.8405 | 1 | 19098 | 1 | -2.9653 |
| SLC10A4 | 6 | 0.017144 | 0.062729 | 0.937389 | 1308 | 5 | -2.1898 | 0.83909 | 0.91905 | 1 | 19099 | 1 | -2.1898 |
| LRRTM1 | 4 | 0.29695 | 0.43989 | 0.999994 | 10065 | 2 | -0.61757 | 0.83913 | 0.86539 | 1 | 19100 | 1 | -0.61757 |
| hsa-mir-3674 | 3 | 0.2044 | 0.31823 | 0.999994 | 7887 | 1 | 0.057777 | 0.83914 | 0.84056 | 1 | 19101 | 1 | 0.057777 |
| AMZ1 | 6 | 0.0098486 | 0.038515 | 0.89046 | 863 | 5 | -1.6102 | 0.83916 | 0.91909 | 1 | 19102 | 1 | -1.6102 |
| VIL1 | 6 | 0.62393 | 0.75776 | 0.999994 | 15289 | 2 | -0.14299 | 0.8393 | 0.91916 | 1 | 19103 | 1 | -0.14299 |
| WNT10A | 6 | 0.036697 | 0.11445 | 0.951052 | 2355 | 3 | -1.3844 | 0.83944 | 0.91924 | 1 | 19104 | 2 | -1.3844 |
| NonTargetingControlG | 2 | 0.14744 | 0.22485 | 0.983354 | 6389 | 1 | -3.6995 | 0.83971 | 0.83963 | 1 | 19105 | 0 | -3.6995 |
| GRM3 | 4 | 0.75098 | 0.76083 | 0.999994 | 16865 | 1 | -0.30969 | 0.83983 | 0.86577 | 1 | 19106 | 1 | -0.30969 |
| EFCAB1 | 4 | 0.46872 | 0.60259 | 0.999994 | 12986 | 1 | -0.40692 | 0.8399 | 0.86581 | 1 | 19107 | 1 | -0.40692 |
| hsa-mir-668 | 4 | 0.028467 | 0.078392 | 0.93914 | 1924 | 3 | -1.1271 | 0.83993 | 0.86582 | 1 | 19108 | 1 | -1.1271 |
| FHOD1 | 6 | 0.035581 | 0.1119 | 0.951052 | 2299 | 3 | -0.62899 | 0.83995 | 0.91953 | 1 | 19109 | 2 | -0.62899 |
| CCDC50 | 5 | 0.956 | 0.95616 | 0.999994 | 20481 | 0 | 0.4707 | 0.83997 | 0.90216 | 1 | 19110 | 1 | 0.4707 |
| SFSWAP | 4 | 0.062335 | 0.14526 | 0.963012 | 3527 | 2 | -3.0856 | 0.83997 | 0.86585 | 1 | 19111 | 1 | -3.0856 |
| USP17L5 | 1 | 0.15999 | 0.15975 | 0.967126 | 6741 | 1 | -4.3648 | 0.84001 | 0.84029 | 1 | 19112 | 0 | -4.3648 |
| AHCTF1 | 6 | 0.53308 | 0.69171 | 0.999994 | 13953 | 2 | -0.41717 | 0.84005 | 0.91958 | 1 | 19113 | 1 | -0.41717 |
| PSMD4 | 6 | 0.072362 | 0.19169 | 0.9742 | 3934 | 3 | -1.1005 | 0.84005 | 0.91958 | 1 | 19114 | 1 | -1.1005 |
| KRTAP6-1 | 6 | 0.1351 | 0.30681 | 0.999994 | 6052 | 3 | -0.74397 | 0.84044 | 0.91982 | 1 | 19115 | 2 | -0.74397 |
| ACVR1B | 6 | 0.085043 | 0.21763 | 0.980699 | 4409 | 2 | 0.27506 | 0.84048 | 0.91984 | 1 | 19116 | 2 | 0.27506 |
| ZNF99 | 5 | 0.010739 | 0.038185 | 0.88817 | 914 | 4 | -2.5421 | 0.84061 | 0.90234 | 1 | 19117 | 1 | -2.5421 |
| hsa-mir-4296 | 3 | 0.096742 | 0.17873 | 0.968171 | 4847 | 2 | -2.2303 | 0.84061 | 0.84189 | 1 | 19118 | 1 | -2.2303 |
| hsa-mir-3926-1 | 3 | 0.72703 | 0.72696 | 0.999994 | 16569 | 1 | -0.50493 | 0.8409 | 0.84214 | 1 | 19119 | 1 | -0.50493 |
| SIX4 | 5 | 0.11329 | 0.24707 | 0.990435 | 5415 | 3 | -1.2017 | 0.8409 | 0.90244 | 1 | 19120 | 1 | -1.2017 |
| FCRL1 | 6 | 0.093121 | 0.23376 | 0.984971 | 4722 | 4 | -1.2568 | 0.84093 | 0.9201 | 1 | 19121 | 2 | -1.2568 |
| TIGIT | 6 | 0.45496 | 0.63835 | 0.999994 | 12771 | 2 | -0.38697 | 0.84125 | 0.92029 | 1 | 19122 | 2 | -0.38697 |
| AK2 | 5 | 0.18727 | 0.35706 | 0.999994 | 7400 | 3 | -0.93875 | 0.84145 | 0.90261 | 1 | 19123 | 1 | -0.93875 |
| NonTargetingControlG | 1 | 0.15854 | 0.15826 | 0.967126 | 6686 | 1 | -2.9896 | 0.84146 | 0.84173 | 1 | 19124 | 0 | -2.9896 |
| KLHL1 | 5 | 0.011738 | 0.041059 | 0.891621 | 974 | 4 | -2.7438 | 0.84156 | 0.90264 | 1 | 19125 | 1 | -2.7438 |
| KCNA10 | 6 | 0.02917 | 0.096838 | 0.945031 | 1966 | 4 | -2.1945 | 0.84178 | 0.92058 | 1 | 19126 | 2 | -2.1945 |
| TAZ | 6 | 0.255 | 0.45391 | 0.999994 | 9131 | 3 | -0.25818 | 0.84206 | 0.92074 | 1 | 19127 | 2 | -0.25818 |
| WDR34 | 5 | 0.34652 | 0.51907 | 0.999994 | 11084 | 2 | -0.49098 | 0.84207 | 0.9028 | 1 | 19128 | 1 | -0.49098 |
| MNX1 | 4 | 0.074912 | 0.16918 | 0.967126 | 4033 | 3 | -2.0467 | 0.84209 | 0.86695 | 1 | 19129 | 1 | -2.0467 |
| LRTOMT | 6 | 0.075092 | 0.19735 | 0.977164 | 4038 | 3 | -1.0282 | 0.84211 | 0.92077 | 1 | 19130 | 2 | -1.0282 |
| TBRG1 | 5 | 0.050887 | 0.14253 | 0.959553 | 3011 | 4 | -1.057 | 0.84235 | 0.90289 | 1 | 19131 | 1 | -1.057 |
| NonTargetingControlG | 2 | 0.037303 | 0.067619 | 0.93914 | 2388 | 1 | -4.9701 | 0.84269 | 0.84258 | 1 | 19132 | 0 | -4.9701 |
| LIPA | 6 | 0.26762 | 0.46887 | 0.999994 | 9410 | 1 | 0.28972 | 0.84276 | 0.92113 | 1 | 19133 | 2 | 0.28972 |
| TFEB | 6 | 0.012935 | 0.048902 | 0.903562 | 1060 | 4 | -4.2079 | 0.84277 | 0.92114 | 1 | 19134 | 2 | -4.2079 |
| HOXA11 | 6 | 0.072468 | 0.1919 | 0.974607 | 3941 | 4 | -1.2855 | 0.84285 | 0.92119 | 1 | 19135 | 2 | -1.2855 |

| | | | | | | | | | | | | | |
|---|---|---|---|---|---|---|---|---|---|---|---|---|---|
| CACNB3 | 5 | 0.47128 | 0.62322 | 0.999994 | 13019 | 2 | -0.26301 | 0.84333 | 0.90317 | 1 | 19136 | 1 | -0.26301 |
| hsa-mir-3914-1 | 4 | 0.85576 | 0.85535 | 0.999994 | 18361 | 0 | 0.015148 | 0.84334 | 0.86763 | 1 | 19137 | 1 | 0.015148 |
| EARS2 | 6 | 0.058003 | 0.16146 | 0.967126 | 3338 | 1 | 0.042941 | 0.84337 | 0.92146 | 1 | 19138 | 2 | 0.042941 |
| KRT34 | 5 | 0.041208 | 0.11891 | 0.951052 | 2571 | 4 | -1.5943 | 0.84344 | 0.90321 | 1 | 19139 | 1 | -1.5943 |
| VPS28 | 3 | 0.53121 | 0.5672 | 0.999994 | 13916 | 1 | 0.31173 | 0.84345 | 0.84449 | 1 | 19140 | 1 | 0.31173 |
| RTP3 | 6 | 0.47929 | 0.65471 | 0.999994 | 13133 | 2 | 0.37176 | 0.84353 | 0.92155 | 1 | 19141 | 1 | 0.37176 |
| MSMB | 5 | 0.016682 | 0.054977 | 0.922677 | 1288 | 4 | -2.9862 | 0.84366 | 0.90328 | 1 | 19142 | 1 | -2.9862 |
| PLA1A | 5 | 0.011945 | 0.041644 | 0.891621 | 988 | 3 | -3.5151 | 0.84367 | 0.90328 | 1 | 19143 | 1 | -3.5151 |
| TBC1D4 | 6 | 0.015205 | 0.056393 | 0.9235 | 1202 | 2 | 0.15231 | 0.8438 | 0.92169 | 1 | 19144 | 2 | 0.15231 |
| FAM180A | 5 | 0.027252 | 0.083384 | 0.94383 | 1868 | 3 | -1.9427 | 0.84386 | 0.90334 | 1 | 19145 | 1 | -1.9427 |
| hsa-mir-3914-2 | 1 | 0.15609 | 0.15582 | 0.967126 | 6632 | 1 | -2.5752 | 0.84391 | 0.84419 | 1 | 19146 | 0 | -2.5752 |
| SERPINB4 | 3 | 0.73652 | 0.73642 | 0.999994 | 16684 | 0 | -0.35125 | 0.84408 | 0.84507 | 1 | 19147 | 1 | -0.35125 |
| JAG1 | 6 | 0.46701 | 0.64646 | 0.999994 | 12965 | 2 | 0.054221 | 0.84424 | 0.92193 | 1 | 19148 | 2 | 0.054221 |
| hsa-mir-6890 | 3 | 0.53743 | 0.57153 | 0.999994 | 14019 | 1 | -0.044034 | 0.84428 | 0.84525 | 1 | 19149 | 1 | -0.044034 |
| NonTargetingControlG | 2 | 0.51792 | 0.52048 | 0.999994 | 13708 | 1 | -0.64454 | 0.84431 | 0.84425 | 1 | 19150 | 0 | -0.64454 |
| hsa-mir-5581 | 3 | 0.17045 | 0.27495 | 0.998666 | 6993 | 1 | 0.085805 | 0.84436 | 0.84532 | 1 | 19151 | 1 | 0.085805 |
| NonTargetingControlG | 2 | 0.53431 | 0.53591 | 0.999994 | 13970 | 1 | -0.63615 | 0.84453 | 0.84448 | 1 | 19152 | 0 | -0.63615 |
| hsa-mir-4293 | 4 | 0.61353 | 0.67273 | 0.999994 | 15125 | 1 | -0.37251 | 0.84458 | 0.86833 | 1 | 19153 | 1 | -0.37251 |
| PARVB | 6 | 0.10755 | 0.26199 | 0.995544 | 5229 | 4 | -0.76202 | 0.84469 | 0.92216 | 1 | 19154 | 2 | -0.76202 |
| SH2D4B | 6 | 0.29038 | 0.49551 | 0.999994 | 9917 | 3 | -0.32823 | 0.8447 | 0.92216 | 1 | 19155 | 2 | -0.32823 |
| SPINK9 | 6 | 0.30483 | 0.51234 | 0.999994 | 10228 | 2 | -0.48402 | 0.84484 | 0.92223 | 1 | 19156 | 2 | -0.48402 |
| hsa-mir-7641-2 | 1 | 0.15509 | 0.15482 | 0.967126 | 6598 | 1 | -2.3857 | 0.84491 | 0.8452 | 1 | 19157 | 0 | -2.3857 |
| MIB2 | 5 | 0.31258 | 0.49127 | 0.999994 | 10372 | 2 | -0.60954 | 0.84504 | 0.90369 | 1 | 19158 | 1 | -0.60954 |
| CHSY3 | 6 | 0.84047 | 0.86686 | 0.999994 | 18095 | 1 | 0.30971 | 0.84511 | 0.92238 | 1 | 19159 | 2 | 0.30971 |
| ZNRF4 | 5 | 0.65191 | 0.74588 | 0.999994 | 15686 | 1 | -0.33166 | 0.84511 | 0.90371 | 1 | 19160 | 1 | -0.33166 |
| HIST4H4 | 6 | 0.14106 | 0.31438 | 0.999994 | 6221 | 2 | -0.050282 | 0.84531 | 0.92247 | 1 | 19161 | 2 | -0.050282 |
| RNF113A | 3 | 0.049219 | 0.11462 | 0.951052 | 2940 | 2 | -3.6543 | 0.84556 | 0.84641 | 1 | 19162 | 1 | -3.6543 |
| CLHC1 | 6 | 0.3028 | 0.50998 | 0.999994 | 10190 | 3 | -0.24377 | 0.84557 | 0.92262 | 1 | 19163 | 2 | -0.24377 |
| BBS9 | 5 | 0.52599 | 0.67055 | 0.999994 | 13830 | 2 | -0.49896 | 0.8456 | 0.90387 | 1 | 19164 | 1 | -0.49896 |
| KIF27 | 6 | 0.085206 | 0.21792 | 0.980699 | 4419 | 4 | -0.90568 | 0.84566 | 0.92267 | 1 | 19165 | 2 | -0.90568 |
| NRAS | 6 | 0.026103 | 0.089609 | 0.94395 | 1810 | 3 | -0.35546 | 0.84584 | 0.92278 | 1 | 19166 | 2 | -0.35546 |
| CBL | 5 | 0.04238 | 0.12181 | 0.951052 | 2635 | 4 | -1.1628 | 0.84588 | 0.90396 | 1 | 19167 | 1 | -1.1628 |
| TPMT | 6 | 0.0042663 | 0.018454 | 0.800859 | 461 | 4 | -3.1413 | 0.84593 | 0.92282 | 1 | 19168 | 2 | -3.1413 |
| PPP3R1 | 5 | 0.047041 | 0.13326 | 0.95318 | 2854 | 3 | -1.4663 | 0.84603 | 0.904 | 1 | 19169 | 1 | -1.4663 |
| SMIM19 | 6 | 0.95583 | 0.95591 | 0.999994 | 20476 | 0 | 0.12474 | 0.84604 | 0.92289 | 1 | 19170 | 2 | 0.12474 |
| TREML4 | 6 | 0.078514 | 0.20432 | 0.978186 | 4172 | 4 | -1.1984 | 0.84621 | 0.92298 | 1 | 19171 | 2 | -1.1984 |
| hsa-mir-5695 | 3 | 0.51472 | 0.55593 | 0.999994 | 13657 | 1 | 0.54942 | 0.84625 | 0.84703 | 1 | 19172 | 1 | 0.54942 |
| NonTargetingControlG | 1 | 0.15355 | 0.1533 | 0.967126 | 6552 | 1 | -3.0459 | 0.84645 | 0.84677 | 1 | 19173 | 0 | -3.0459 |
| hsa-mir-3977 | 4 | 0.40583 | 0.55169 | 0.999994 | 12041 | 2 | -0.44096 | 0.84658 | 0.86944 | 1 | 19174 | 1 | -0.44096 |
| RPRD1A | 5 | 0.85324 | 0.85686 | 0.999994 | 18317 | 1 | -0.049112 | 0.84673 | 0.90421 | 1 | 19175 | 1 | -0.049112 |
| ZBTB37 | 5 | 0.87467 | 0.8753 | 0.999994 | 18726 | 1 | -0.3719 | 0.84674 | 0.90421 | 1 | 19176 | 1 | -0.3719 |
| SNX24 | 6 | 0.09776 | 0.24294 | 0.988088 | 4881 | 4 | -0.85597 | 0.84682 | 0.92332 | 1 | 19177 | 1 | -0.85597 |
| DEC1 | 4 | 0.26013 | 0.40224 | 0.999994 | 9245 | 2 | -0.54409 | 0.84697 | 0.86965 | 1 | 19178 | 1 | -0.54409 |
| FAM136A | 6 | 0.079503 | 0.20629 | 0.978744 | 4208 | 4 | -1.0908 | 0.84705 | 0.92345 | 1 | 19179 | 2 | -1.0908 |
| ZDHHC3 | 6 | 0.50801 | 0.67425 | 0.999994 | 13552 | 2 | 0.32846 | 0.84735 | 0.92364 | 1 | 19180 | 1 | 0.32846 |
| KATNAL2 | 6 | 0.03908 | 0.11983 | 0.951052 | 2467 | 4 | -1.6568 | 0.84735 | 0.92364 | 1 | 19181 | 1 | -1.6568 |
| hsa-mir-4276 | 4 | 0.051845 | 0.1251 | 0.951052 | 3059 | 3 | -1.7884 | 0.84747 | 0.86993 | 1 | 19182 | 1 | -1.7884 |
| SLC22A15 | 3 | 0.27712 | 0.40846 | 0.999994 | 9640 | 2 | -0.70142 | 0.84747 | 0.84817 | 1 | 19183 | 1 | -0.70142 |
| DCST2 | 6 | 0.5686 | 0.71694 | 0.999994 | 14518 | 2 | 0.084788 | 0.84749 | 0.92371 | 1 | 19184 | 2 | 0.084788 |
| C22orf24 | 6 | 0.064048 | 0.17433 | 0.968171 | 3614 | 3 | -0.65196 | 0.8475 | 0.92371 | 1 | 19185 | 2 | -0.65196 |
| PRUNE2 | 6 | 0.54823 | 0.70234 | 0.999994 | 14182 | 2 | 0.0094529 | 0.84776 | 0.92384 | 1 | 19186 | 2 | 0.0094529 |
| RBL1 | 5 | 0.64535 | 0.74326 | 0.999994 | 15589 | 1 | 0.014804 | 0.8478 | 0.90454 | 1 | 19187 | 1 | 0.014804 |

| | | | | | | | | | | | | | |
|---|---|---|---|---|---|---|---|---|---|---|---|---|---|
| CYB5R2 | 6 | 0.2172 | 0.40875 | 0.999994 | 8199 | 3 | -0.27486 | 0.84815 | 0.92404 | 1 | 19188 | 2 | -0.27486 |
| IGFBP4 | 6 | 0.16192 | 0.34068 | 0.999994 | 6789 | 3 | -0.44182 | 0.84817 | 0.92405 | 1 | 19189 | 2 | -0.44182 |
| IFT172 | 6 | 0.2801 | 0.48355 | 0.999994 | 9708 | 3 | -0.71964 | 0.8482 | 0.92406 | 1 | 19190 | 2 | -0.71964 |
| hsa-mir-892a | 4 | 0.55897 | 0.64412 | 0.999994 | 14352 | 1 | 0.046382 | 0.84822 | 0.87035 | 1 | 19191 | 1 | 0.046382 |
| RASSF3 | 5 | 0.28364 | 0.4678 | 0.999994 | 9778 | 2 | 0.050665 | 0.84829 | 0.90469 | 1 | 19192 | 1 | 0.050665 |
| NKAIN4 | 4 | 0.88752 | 0.88721 | 0.999994 | 18987 | 0 | 0.36688 | 0.84838 | 0.87044 | 1 | 19193 | 1 | 0.36688 |
| WNT5B | 6 | 0.59386 | 0.73544 | 0.999994 | 14878 | 2 | -0.46325 | 0.8485 | 0.92423 | 1 | 19194 | 2 | -0.46325 |
| DGAT1 | 6 | 0.041834 | 0.12603 | 0.951052 | 2605 | 3 | -2.5196 | 0.84851 | 0.92423 | 1 | 19195 | 2 | -2.5196 |
| PAMR1 | 6 | 0.23734 | 0.43283 | 0.999994 | 8709 | 3 | -0.77035 | 0.84852 | 0.92423 | 1 | 19196 | 1 | -0.77035 |
| NAGLU | 6 | 0.11782 | 0.28161 | 0.999632 | 5568 | 2 | -0.0011919 | 0.84852 | 0.92423 | 1 | 19197 | 1 | -0.0011919 |
| ZPBP | 6 | 0.012117 | 0.046109 | 0.892434 | 1000 | 4 | -2.3885 | 0.84852 | 0.92423 | 1 | 19198 | 1 | -2.3885 |
| TRIM59 | 6 | 0.13397 | 0.30538 | 0.999994 | 6020 | 3 | -0.58617 | 0.84852 | 0.92423 | 1 | 19199 | 1 | -0.58617 |
| GATSL1 | 5 | 0.069979 | 0.17834 | 0.968171 | 3826 | 3 | -2.1144 | 0.84854 | 0.90478 | 1 | 19200 | 1 | -2.1144 |
| NonTargetingControlG | 2 | 0.45638 | 0.46487 | 0.999994 | 12796 | 1 | -0.83461 | 0.84875 | 0.84864 | 1 | 19201 | 0 | -0.83461 |
| C16orf72 | 6 | 0.054844 | 0.15459 | 0.967126 | 3192 | 4 | -1.4131 | 0.84875 | 0.92436 | 1 | 19202 | 2 | -1.4131 |
| SNRPB2 | 6 | 0.49877 | 0.66785 | 0.999994 | 13405 | 2 | -0.12376 | 0.84886 | 0.92442 | 1 | 19203 | 2 | -0.12376 |
| NPIPL3 | 3 | 0.64287 | 0.65117 | 0.999994 | 15560 | 1 | -0.3195 | 0.84901 | 0.84958 | 1 | 19204 | 1 | -0.3195 |
| CCDC74B | 4 | 0.080603 | 0.17979 | 0.968171 | 4255 | 3 | -1.6211 | 0.84906 | 0.87082 | 1 | 19205 | 1 | -1.6211 |
| B4GALNT1 | 5 | 0.11186 | 0.24487 | 0.989317 | 5363 | 3 | -1.3861 | 0.84908 | 0.90496 | 1 | 19206 | 1 | -1.3861 |
| S100A11 | 4 | 0.015559 | 0.050801 | 0.909101 | 1219 | 2 | -3.3692 | 0.84911 | 0.87084 | 1 | 19207 | 1 | -3.3692 |
| LDHD | 6 | 0.11991 | 0.28554 | 0.999632 | 5624 | 3 | -0.27317 | 0.84926 | 0.92466 | 1 | 19208 | 2 | -0.27317 |
| LYRM7 | 5 | 0.17506 | 0.3395 | 0.999994 | 7107 | 3 | -0.98858 | 0.84933 | 0.90503 | 1 | 19209 | 1 | -0.98858 |
| CCDC150 | 6 | 0.012445 | 0.04722 | 0.896226 | 1025 | 4 | -3.0005 | 0.84933 | 0.9247 | 1 | 19210 | 1 | -3.0005 |
| NonTargetingControlG | 2 | 0.62321 | 0.62299 | 0.999994 | 15278 | 0 | -0.35779 | 0.84941 | 0.84928 | 1 | 19211 | 0 | -0.35779 |
| CARTPT | 6 | 0.039151 | 0.11998 | 0.951052 | 2472 | 4 | -1.7148 | 0.84957 | 0.92483 | 1 | 19212 | 2 | -1.7148 |
| CAPN9 | 5 | 0.4424 | 0.59882 | 0.999994 | 12617 | 2 | -0.048448 | 0.84961 | 0.90513 | 1 | 19213 | 1 | -0.048448 |
| IL3RA_X | 6 | 0.5215 | 0.68355 | 0.999994 | 13768 | 2 | -0.33586 | 0.84967 | 0.92489 | 1 | 19214 | 1 | -0.33586 |
| NonTargetingControlG | 2 | 0.3965 | 0.41391 | 0.999994 | 11898 | 1 | -1.1674 | 0.84986 | 0.84973 | 1 | 19215 | 0 | -1.1674 |
| WFDC3 | 5 | 0.54658 | 0.68869 | 0.999994 | 14161 | 2 | 0.14626 | 0.84993 | 0.90522 | 1 | 19216 | 1 | 0.14626 |
| NonTargetingControlG | 2 | 0.37233 | 0.39411 | 0.999994 | 11544 | 1 | -1.1819 | 0.84998 | 0.84984 | 1 | 19217 | 0 | -1.1819 |
| ATM | 6 | 0.89906 | 0.90294 | 0.999994 | 19205 | 1 | 0.15751 | 0.85012 | 0.92515 | 1 | 19218 | 2 | 0.15751 |
| TTC7B | 6 | 0.27395 | 0.47636 | 0.999994 | 9565 | 3 | -0.68051 | 0.85013 | 0.92516 | 1 | 19219 | 2 | -0.68051 |
| NDFIP2 | 6 | 0.10103 | 0.24933 | 0.991784 | 5007 | 4 | -1.0544 | 0.85016 | 0.92517 | 1 | 19220 | 2 | -1.0544 |
| CHERP | 3 | 0.084823 | 0.16291 | 0.967126 | 4401 | 2 | -2.3985 | 0.8502 | 0.85066 | 1 | 19221 | 1 | -2.3985 |
| hsa-mir-1273d | 1 | 0.14974 | 0.14952 | 0.966317 | 6452 | 1 | -2.781 | 0.85026 | 0.85054 | 1 | 19222 | 0 | -2.781 |
| ASIP | 6 | 0.91459 | 0.91552 | 0.999994 | 19501 | 1 | -0.17609 | 0.85035 | 0.92528 | 1 | 19223 | 1 | -0.17609 |
| HNRNPU | 3 | 0.051856 | 0.11829 | 0.951052 | 3060 | 2 | -4.8949 | 0.85051 | 0.85094 | 1 | 19224 | 1 | -4.8949 |
| MSTN | 5 | 0.14396 | 0.29378 | 0.999994 | 6299 | 3 | -1.0793 | 0.85063 | 0.90546 | 1 | 19225 | 1 | -1.0793 |
| SESN2 | 3 | 0.17919 | 0.2862 | 0.999632 | 7200 | 2 | -1.3958 | 0.8508 | 0.85122 | 1 | 19226 | 1 | -1.3958 |
| hsa-mir-513a-1 | 3 | 0.82438 | 0.82426 | 0.999994 | 17842 | 0 | 0.18111 | 0.85087 | 0.85128 | 1 | 19227 | 1 | 0.18111 |
| FCRLA | 6 | 0.054739 | 0.15437 | 0.967126 | 3183 | 4 | -1.3291 | 0.85091 | 0.92558 | 1 | 19228 | 2 | -1.3291 |
| hsa-mir-3156-1 | 3 | 0.28063 | 0.41276 | 0.999994 | 9713 | 2 | -0.66216 | 0.85102 | 0.85142 | 1 | 19229 | 1 | -0.66216 |
| RBFA | 4 | 0.32885 | 0.47252 | 0.999994 | 10698 | 2 | -0.91207 | 0.85104 | 0.87192 | 1 | 19230 | 1 | -0.91207 |
| PCDHGA11 | 3 | 0.72099 | 0.72097 | 0.999994 | 16508 | 1 | 0.41724 | 0.85105 | 0.85146 | 1 | 19231 | 1 | 0.41724 |
| hsa-mir-200b | 4 | 0.056358 | 0.13387 | 0.95318 | 3261 | 3 | -3.6094 | 0.85115 | 0.87198 | 1 | 19232 | 1 | -3.6094 |
| LRRN3 | 6 | 0.033934 | 0.10803 | 0.951052 | 2214 | 2 | 0.2792 | 0.85116 | 0.92572 | 1 | 19233 | 2 | 0.2792 |
| NonTargetingControlG | 2 | 0.15332 | 0.22895 | 0.983354 | 6545 | 1 | -3.6981 | 0.8513 | 0.85118 | 1 | 19234 | 0 | -3.6981 |
| UBXN10 | 6 | 0.10035 | 0.24801 | 0.990682 | 4983 | 4 | -0.80342 | 0.85132 | 0.92581 | 1 | 19235 | 1 | -0.80342 |
| hsa-mir-4319 | 4 | 0.69791 | 0.72305 | 0.999994 | 16278 | 1 | -0.2861 | 0.85133 | 0.87209 | 1 | 19236 | 1 | -0.2861 |
| CSTL1 | 5 | 0.24076 | 0.43196 | 0.999994 | 8788 | 3 | -0.6194 | 0.85141 | 0.9057 | 1 | 19237 | 1 | -0.6194 |
| VRTN | 6 | 0.025327 | 0.087765 | 0.94395 | 1767 | 3 | -0.28191 | 0.85147 | 0.92589 | 1 | 19238 | 2 | -0.28191 |
| DKC1 | 4 | 0.13055 | 0.26852 | 0.998561 | 5946 | 3 | -0.84771 | 0.8515 | 0.87218 | 1 | 19239 | 1 | -0.84771 |

| | | | | | | | | | | | | |
|---|---|---|---|---|---|---|---|---|---|---|---|---|
| RSBN1L | 6 | 0.74604 | 0.82708 | 0.999994 | 16814 | 1 | 0.41091 | 0.8515 | 0.92591 | 1 | 19240 | 2 | 0.41091 |
| RAD52 | 3 | 0.25775 | 0.38461 | 0.999994 | 9189 | 2 | -0.81934 | 0.85151 | 0.85192 | 1 | 19241 | 1 | -0.81934 |
| NonTargetingControlG | 2 | 0.60074 | 0.60056 | 0.999994 | 14957 | 0 | -0.44303 | 0.85158 | 0.85145 | 1 | 19242 | 0 | -0.44303 |
| ACTBL2 | 6 | 0.07977 | 0.20682 | 0.978744 | 4223 | 4 | -1.0137 | 0.8516 | 0.92596 | 1 | 19243 | 2 | -1.0137 |
| PKD2L2 | 4 | 0.06874 | 0.15749 | 0.967126 | 3783 | 3 | -1.8982 | 0.85172 | 0.87231 | 1 | 19244 | 1 | -1.8982 |
| hsa-mir-921 | 3 | 0.52851 | 0.56534 | 0.999994 | 13870 | 1 | -0.20184 | 0.85184 | 0.85223 | 1 | 19245 | 1 | -0.20184 |
| TMSB15B | 3 | 0.53575 | 0.57035 | 0.999994 | 13992 | 1 | -0.31429 | 0.85193 | 0.85232 | 1 | 19246 | 1 | -0.31429 |
| hsa-mir-1302-10 | 4 | 0.060974 | 0.14265 | 0.959553 | 3466 | 3 | -1.4154 | 0.85219 | 0.87259 | 1 | 19247 | 1 | -1.4154 |
| NonTargetingControlG | 2 | 0.23498 | 0.28768 | 0.999632 | 8642 | 1 | -2.0693 | 0.8522 | 0.8521 | 1 | 19248 | 0 | -2.0693 |
| NonTargetingControlG | 2 | 0.22648 | 0.28135 | 0.999632 | 8437 | 1 | -2.0403 | 0.8523 | 0.85219 | 1 | 19249 | 0 | -2.0403 |
| SCGB3A2 | 6 | 0.43649 | 0.62611 | 0.999994 | 12520 | 2 | -0.3166 | 0.85252 | 0.92648 | 1 | 19250 | 1 | -0.3166 |
| NCF4 | 6 | 0.033361 | 0.10668 | 0.950995 | 2178 | 3 | -2.5167 | 0.85259 | 0.92652 | 1 | 19251 | 2 | -2.5167 |
| NonTargetingControlG | 2 | 0.61246 | 0.61233 | 0.999994 | 15116 | 0 | -0.37145 | 0.85264 | 0.85253 | 1 | 19252 | 0 | -0.37145 |
| REEP5 | 6 | 0.04517 | 0.13345 | 0.95318 | 2769 | 4 | -1.7571 | 0.85267 | 0.92657 | 1 | 19253 | 2 | -1.7571 |
| PTGFRN | 6 | 0.65491 | 0.78139 | 0.999994 | 15716 | 2 | -0.20032 | 0.85268 | 0.92657 | 1 | 19254 | 2 | -0.20032 |
| SFTPA2 | 3 | 0.56397 | 0.59027 | 0.999994 | 14452 | 1 | 0.52101 | 0.85269 | 0.85302 | 1 | 19255 | 1 | 0.52101 |
| SPOCK1 | 6 | 0.2435 | 0.44018 | 0.999994 | 8874 | 2 | 0.41443 | 0.85272 | 0.9266 | 1 | 19256 | 2 | 0.41443 |
| TVP23B | 6 | 0.16642 | 0.34625 | 0.999994 | 6894 | 3 | -0.25711 | 0.85278 | 0.92663 | 1 | 19257 | 2 | -0.25711 |
| SPOCK3 | 5 | 0.11364 | 0.24763 | 0.990682 | 5428 | 3 | -1.3705 | 0.85288 | 0.90618 | 1 | 19258 | 1 | -1.3705 |
| C19orf33 | 6 | 0.012415 | 0.047115 | 0.896226 | 1024 | 4 | -2.7569 | 0.85298 | 0.92673 | 1 | 19259 | 2 | -2.7569 |
| POLR1E | 4 | 0.1023 | 0.21922 | 0.980699 | 5051 | 3 | -1.7523 | 0.85306 | 0.87308 | 1 | 19260 | 1 | -1.7523 |
| IL26 | 6 | 0.11672 | 0.27955 | 0.999632 | 5532 | 3 | -0.76136 | 0.85306 | 0.92677 | 1 | 19261 | 2 | -0.76136 |
| TRIM74 | 4 | 0.28191 | 0.4245 | 0.999994 | 9738 | 1 | 0.28598 | 0.8531 | 0.8731 | 1 | 19262 | 1 | 0.28598 |
| NR4A1 | 6 | 0.12535 | 0.29441 | 0.999994 | 5802 | 3 | -0.51817 | 0.85315 | 0.92683 | 1 | 19263 | 2 | -0.51817 |
| ERCC3 | 4 | 0.11748 | 0.24588 | 0.989866 | 5559 | 2 | -2.24 | 0.85351 | 0.87334 | 1 | 19264 | 1 | -2.24 |
| LRPPRC | 5 | 0.055712 | 0.15415 | 0.967126 | 3231 | 4 | -1.057 | 0.85353 | 0.90637 | 1 | 19265 | 1 | -1.057 |
| IL18BP | 6 | 0.037446 | 0.11614 | 0.951052 | 2396 | 4 | -1.6875 | 0.85369 | 0.92711 | 1 | 19266 | 1 | -1.6875 |
| STRADA | 6 | 0.29432 | 0.50009 | 0.999994 | 10004 | 3 | -0.6268 | 0.85375 | 0.92715 | 1 | 19267 | 1 | -0.6268 |
| hsa-mir-378g | 4 | 0.46774 | 0.60216 | 0.999994 | 12973 | 1 | 0.00019986 | 0.85382 | 0.87353 | 1 | 19268 | 1 | 0.00019986 |
| KCNN1 | 6 | 0.058694 | 0.16295 | 0.967126 | 3370 | 4 | -1.6339 | 0.85417 | 0.9274 | 1 | 19269 | 2 | -1.6339 |
| CRISP1 | 5 | 0.16753 | 0.32847 | 0.999994 | 6924 | 3 | -1.0594 | 0.85422 | 0.90659 | 1 | 19270 | 1 | -1.0594 |
| PLEKHD1 | 6 | 0.94673 | 0.94669 | 0.999994 | 20255 | 0 | 0.095686 | 0.85425 | 0.92744 | 1 | 19271 | 2 | 0.095686 |
| TMEM5 | 4 | 0.57553 | 0.65246 | 0.999994 | 14624 | 1 | 0.38984 | 0.8544 | 0.87387 | 1 | 19272 | 1 | 0.38984 |
| hsa-mir-142 | 3 | 0.0029812 | 0.0082589 | 0.680529 | 355 | 2 | -8.9664 | 0.85457 | 0.85486 | 1 | 19273 | 1 | -8.9664 |
| TNFRSF25 | 6 | 0.088499 | 0.22454 | 0.983354 | 4535 | 4 | -0.93355 | 0.85465 | 0.92765 | 1 | 19274 | 1 | -0.93355 |
| UGT2B17 | 4 | 0.46304 | 0.60015 | 0.999994 | 12907 | 1 | -0.33291 | 0.85466 | 0.87401 | 1 | 19275 | 1 | -0.33291 |
| FANCL | 6 | 0.052743 | 0.15004 | 0.966359 | 3102 | 4 | -1.293 | 0.85466 | 0.92766 | 1 | 19276 | 2 | -1.293 |
| XAGE1A | 6 | 0.37364 | 0.5855 | 0.999994 | 11565 | 2 | -0.19544 | 0.8547 | 0.92767 | 1 | 19277 | 1 | -0.19544 |
| CC2D2B | 5 | 0.18945 | 0.36015 | 0.999994 | 7469 | 2 | -0.58247 | 0.85487 | 0.9068 | 1 | 19278 | 1 | -0.58247 |
| IFNA10 | 6 | 0.43504 | 0.62516 | 0.999994 | 12498 | 2 | -0.13405 | 0.85496 | 0.92781 | 1 | 19279 | 2 | -0.13405 |
| LANCL1 | 6 | 0.1883 | 0.37342 | 0.999994 | 7428 | 3 | -0.25606 | 0.85509 | 0.9279 | 1 | 19280 | 1 | -0.25606 |
| LGR4 | 6 | 0.0031386 | 0.014246 | 0.753188 | 371 | 5 | -5.5033 | 0.85522 | 0.92797 | 1 | 19281 | 1 | -5.5033 |
| hsa-mir-6795 | 3 | 0.71658 | 0.71672 | 0.999994 | 16460 | 1 | 0.36257 | 0.85522 | 0.85546 | 1 | 19282 | 1 | 0.36257 |
| MAP1B | 6 | 0.011567 | 0.044286 | 0.891621 | 962 | 4 | -2.6024 | 0.85536 | 0.92805 | 1 | 19283 | 2 | -2.6024 |
| TIAF1 | 3 | 0.67789 | 0.68083 | 0.999994 | 16058 | 1 | -0.60276 | 0.8554 | 0.85565 | 1 | 19284 | 1 | -0.60276 |
| HAS2 | 6 | 0.14001 | 0.31303 | 0.999994 | 6184 | 3 | -0.68204 | 0.85544 | 0.92809 | 1 | 19285 | 1 | -0.68204 |
| WASH1 | 5 | 0.099961 | 0.22635 | 0.983354 | 4973 | 3 | -2.1127 | 0.85556 | 0.90702 | 1 | 19286 | 1 | -2.1127 |
| NELFB | 5 | 2.6102E-06 | 0.000012116 | 0.023852 | 11 | 4 | -10.691 | 0.85556 | 0.90702 | 1 | 19287 | 1 | -10.691 |
| RBM14 | 5 | 0.05619 | 0.15526 | 0.967126 | 3255 | 3 | -1.4517 | 0.85556 | 0.90702 | 1 | 19288 | 1 | -1.4517 |
| GGPS1 | 4 | 0.38944 | 0.5348 | 0.999994 | 11788 | 1 | 0.097013 | 0.85557 | 0.87456 | 1 | 19289 | 1 | 0.097013 |
| hsa-mir-3689b | 3 | 0.75932 | 0.75932 | 0.999994 | 16975 | 0 | 0.26188 | 0.85561 | 0.85584 | 1 | 19290 | 1 | 0.26188 |
| CALB2 | 4 | 0.01728 | 0.054864 | 0.922677 | 1324 | 3 | -2.8173 | 0.85571 | 0.87464 | 1 | 19291 | 1 | -2.8173 |

| | | | | | | | | | | | | | |
|---|---|---|---|---|---|---|---|---|---|---|---|---|---|
| SFTA2 | 6 | 0.92883 | 0.92893 | 0.999994 | 19833 | 0 | -0.011894 | 0.85571 | 0.92825 | 1 | 19292 | 2 | -0.011894 |
| EIF4B | 5 | 0.0041295 | 0.017342 | 0.785648 | 452 | 4 | -2.6146 | 0.85583 | 0.90712 | 1 | 19293 | 1 | -2.6146 |
| SWSAP1 | 5 | 0.0063546 | 0.025512 | 0.845628 | 616 | 4 | -5.2003 | 0.85584 | 0.90712 | 1 | 19294 | 1 | -5.2003 |
| RPIA | 5 | 0.0066518 | 0.026372 | 0.848412 | 637 | 4 | -2.7147 | 0.85595 | 0.90716 | 1 | 19295 | 1 | -2.7147 |
| AREG | 6 | 0.013063 | 0.049338 | 0.906138 | 1068 | 4 | -2.8728 | 0.85601 | 0.92841 | 1 | 19296 | 1 | -2.8728 |
| R3HDM1 | 6 | 0.030212 | 0.099276 | 0.945253 | 2019 | 4 | -1.6996 | 0.85608 | 0.92846 | 1 | 19297 | 2 | -1.6996 |
| CSTB | 5 | 0.43816 | 0.59526 | 0.999994 | 12541 | 2 | -0.60247 | 0.85614 | 0.90722 | 1 | 19298 | 1 | -0.60247 |
| KIAA1598 | 6 | 0.22845 | 0.4222 | 0.999994 | 8488 | 3 | -1.2727 | 0.85625 | 0.92855 | 1 | 19299 | 2 | -1.2727 |
| GLRX | 6 | 0.41631 | 0.61296 | 0.999994 | 12197 | 2 | -0.36293 | 0.8564 | 0.92864 | 1 | 19300 | 2 | -0.36293 |
| UFL1 | 6 | 0.22478 | 0.41786 | 0.999994 | 8393 | 3 | -0.67579 | 0.85642 | 0.92865 | 1 | 19301 | 2 | -0.67579 |
| TMEM87B | 6 | 0.11368 | 0.27376 | 0.998561 | 5432 | 4 | -1.0382 | 0.85642 | 0.92865 | 1 | 19302 | 1 | -1.0382 |
| RIMBP3C | 2 | 0.57075 | 0.57099 | 0.999994 | 14554 | 1 | -0.5641 | 0.85656 | 0.8565 | 1 | 19303 | 0 | -0.5641 |
| hsa-mir-4535 | 4 | 0.4283 | 0.57479 | 0.999994 | 12386 | 2 | -0.14868 | 0.85658 | 0.87514 | 1 | 19304 | 1 | -0.14868 |
| NonTargetingControlG | 2 | 0.53479 | 0.53636 | 0.999994 | 13976 | 1 | -0.68257 | 0.85663 | 0.85658 | 1 | 19305 | 0 | -0.68257 |
| RUSC2 | 6 | 0.063345 | 0.17285 | 0.96793 | 3571 | 3 | -0.60602 | 0.85674 | 0.92883 | 1 | 19306 | 2 | -0.60602 |
| MKRN2 | 6 | 0.38459 | 0.59247 | 0.999994 | 11708 | 2 | -0.21785 | 0.85682 | 0.92887 | 1 | 19307 | 2 | -0.21785 |
| VARS2 | 5 | 0.012961 | 0.044471 | 0.891621 | 1061 | 3 | -3.9899 | 0.857 | 0.9075 | 1 | 19308 | 1 | -3.9899 |
| ZC3H7A | 6 | 0.43133 | 0.62273 | 0.999994 | 12436 | 2 | -0.043472 | 0.857 | 0.92898 | 1 | 19309 | 2 | -0.043472 |
| NonTargetingControlG | 1 | 0.14297 | 0.14276 | 0.959553 | 6271 | 1 | -3.2086 | 0.85703 | 0.85727 | 1 | 19310 | 0 | -3.2086 |
| GGA1 | 3 | 0.31348 | 0.43582 | 0.999994 | 10387 | 1 | 0.40254 | 0.85726 | 0.85734 | 1 | 19311 | 1 | 0.40254 |
| GIMAP4 | 5 | 0.011557 | 0.040544 | 0.891621 | 957 | 4 | -2.2455 | 0.85754 | 0.90768 | 1 | 19312 | 1 | -2.2455 |
| OR2T35 | 5 | 0.2175 | 0.39977 | 0.999994 | 8209 | 2 | -0.0052661 | 0.85762 | 0.90771 | 1 | 19313 | 1 | -0.0052661 |
| SKAP2 | 6 | 0.053342 | 0.15129 | 0.967126 | 3131 | 3 | -0.73873 | 0.85762 | 0.92935 | 1 | 19314 | 2 | -0.73873 |
| HNRNPA1 | 3 | 0.071216 | 0.14463 | 0.963012 | 3884 | 1 | -0.53753 | 0.85767 | 0.85774 | 1 | 19315 | 1 | -0.53753 |
| C14orf2 | 6 | 0.31892 | 0.52865 | 0.999994 | 10496 | 2 | -0.52899 | 0.85768 | 0.92938 | 1 | 19316 | 2 | -0.52899 |
| CRISPLD2 | 6 | 0.0080483 | 0.032128 | 0.865899 | 735 | 4 | -3.8966 | 0.8578 | 0.92945 | 1 | 19317 | 2 | -3.8966 |
| NonTargetingControlG | 2 | 0.21553 | 0.27337 | 0.998561 | 8155 | 1 | -2.0616 | 0.85789 | 0.85783 | 1 | 19318 | 0 | -2.0616 |
| TCAIM | 6 | 0.18288 | 0.36674 | 0.999994 | 7286 | 2 | -0.32279 | 0.85795 | 0.92953 | 1 | 19319 | 2 | -0.32279 |
| NOTUM | 6 | 0.0032772 | 0.014735 | 0.755578 | 382 | 5 | -2.3002 | 0.85798 | 0.92955 | 1 | 19320 | 1 | -2.3002 |
| C7orf29 | 6 | 0.5392 | 0.69598 | 0.999994 | 14038 | 2 | 0.20845 | 0.85821 | 0.92966 | 1 | 19321 | 2 | 0.20845 |
| LY9 | 6 | 0.045007 | 0.13309 | 0.95318 | 2757 | 4 | -1.8837 | 0.85839 | 0.92978 | 1 | 19322 | 1 | -1.8837 |
| HPR | 6 | 0.0118 | 0.045055 | 0.891621 | 981 | 4 | -1.7539 | 0.85855 | 0.92988 | 1 | 19323 | 1 | -1.7539 |
| NonTargetingControlG | 2 | 0.46029 | 0.46838 | 0.999994 | 12850 | 1 | -0.83142 | 0.85865 | 0.85858 | 1 | 19324 | 0 | -0.83142 |
| FAM160A1 | 6 | 0.037857 | 0.11706 | 0.951052 | 2419 | 2 | -0.11787 | 0.85872 | 0.92997 | 1 | 19325 | 2 | -0.11787 |
| TMEM208 | 5 | 0.027718 | 0.084553 | 0.94395 | 1891 | 4 | -2.0681 | 0.85875 | 0.90809 | 1 | 19326 | 1 | -2.0681 |
| PTPN5 | 4 | 0.032539 | 0.086679 | 0.94395 | 2136 | 3 | -3.972 | 0.85882 | 0.87646 | 1 | 19327 | 1 | -3.972 |
| hsa-mir-3129 | 3 | 0.43404 | 0.5045 | 0.999994 | 12478 | 1 | 0.39198 | 0.85885 | 0.85891 | 1 | 19328 | 1 | 0.39198 |
| CST1 | 6 | 0.18849 | 0.37363 | 0.999994 | 7436 | 1 | -0.097686 | 0.8589 | 0.93008 | 1 | 19329 | 2 | -0.097686 |
| VSTM2A | 5 | 0.042261 | 0.12152 | 0.951052 | 2629 | 3 | -1.3009 | 0.85905 | 0.90819 | 1 | 19330 | 1 | -1.3009 |
| KLHL34 | 4 | 0.37036 | 0.51524 | 0.999994 | 11518 | 2 | -0.6226 | 0.8591 | 0.87661 | 1 | 19331 | 1 | -0.6226 |
| PCDHAC1 | 2 | 0.55181 | 0.5526 | 0.999994 | 14241 | 1 | -0.6183 | 0.85911 | 0.85906 | 1 | 19332 | 0 | -0.6183 |
| NAT8L | 5 | 0.063589 | 0.168 | 0.967126 | 3586 | 3 | -2.0206 | 0.85923 | 0.90825 | 1 | 19333 | 1 | -2.0206 |
| hsa-mir-6817 | 4 | 0.32184 | 0.46528 | 0.999994 | 10551 | 1 | -0.087007 | 0.85931 | 0.87675 | 1 | 19334 | 1 | -0.087007 |
| STYXL1 | 5 | 0.15046 | 0.30347 | 0.999994 | 6471 | 3 | -0.97468 | 0.85941 | 0.90832 | 1 | 19335 | 1 | -0.97468 |
| AGXT2L1 | 5 | 0.087941 | 0.20727 | 0.978744 | 4510 | 3 | -1.7755 | 0.85956 | 0.90836 | 1 | 19336 | 1 | -1.7755 |
| CMSS1 | 5 | 0.018908 | 0.06111 | 0.933245 | 1404 | 3 | -3.6262 | 0.8597 | 0.90841 | 1 | 19337 | 1 | -3.6262 |
| MCCC1 | 5 | 0.10906 | 0.24054 | 0.987253 | 5272 | 2 | -0.19492 | 0.85985 | 0.90846 | 1 | 19338 | 1 | -0.19492 |
| hsa-mir-509-3 | 2 | 0.11275 | 0.18971 | 0.970473 | 5395 | 1 | -4.1506 | 0.85989 | 0.85987 | 1 | 19339 | 0 | -4.1506 |
| RPS10-NUDT3 | 1 | 0.14003 | 0.13979 | 0.958716 | 6186 | 1 | -3.501 | 0.85997 | 0.86026 | 1 | 19340 | 0 | -3.501 |
| hsa-let-7a-1 | 4 | 0.14848 | 0.2876 | 0.999632 | 6419 | 2 | -1.0073 | 0.85998 | 0.87713 | 1 | 19341 | 1 | -1.0073 |
| hsa-mir-5689 | 3 | 0.23589 | 0.35746 | 0.999994 | 8667 | 2 | -0.97913 | 0.85998 | 0.86002 | 1 | 19342 | 1 | -0.97913 |
| LIMS3L | 2 | 0.35458 | 0.37966 | 0.999994 | 11241 | 1 | -1.3801 | 0.86002 | 0.86001 | 1 | 19343 | 0 | -1.3801 |

| | | | | | | | | | | | | | |
|---|---|---|---|---|---|---|---|---|---|---|---|---|---|
| CLEC14A | 6 | 0.00076347 | 0.0039172 | 0.560963 | 135 | 4 | -1.7822 | 0.86002 | 0.9307 | 1 | 19344 | 1 | -1.7822 |
| CNTNAP4 | 5 | 0.036288 | 0.10652 | 0.950995 | 2333 | 4 | -2.2337 | 0.86013 | 0.90857 | 1 | 19345 | 1 | -2.2337 |
| CDKL1 | 6 | 0.026395 | 0.090298 | 0.94395 | 1824 | 4 | -2.2526 | 0.86018 | 0.93079 | 1 | 19346 | 1 | -2.2526 |
| SSBP3 | 6 | 0.0076322 | 0.03067 | 0.862223 | 701 | 5 | -2.6699 | 0.86024 | 0.93082 | 1 | 19347 | 1 | -2.6699 |
| DCTN2 | 3 | 0.53291 | 0.56835 | 0.999994 | 13952 | 1 | -0.049653 | 0.86026 | 0.86029 | 1 | 19348 | 1 | -0.049653 |
| TUBA1C | 5 | 0.006029 | 0.024508 | 0.842094 | 595 | 4 | -2.6427 | 0.86035 | 0.90863 | 1 | 19349 | 1 | -2.6427 |
| OR1A2 | 6 | 0.36035 | 0.57572 | 0.999994 | 11338 | 3 | -0.52324 | 0.86045 | 0.93093 | 1 | 19350 | 2 | -0.52324 |
| IGSF10 | 6 | 0.2336 | 0.42839 | 0.999994 | 8608 | 2 | -0.42616 | 0.86052 | 0.93098 | 1 | 19351 | 2 | -0.42616 |
| ATF7 | 6 | 0.86444 | 0.87998 | 0.999994 | 18530 | 1 | 0.18471 | 0.86055 | 0.93099 | 1 | 19352 | 2 | 0.18471 |
| SLC25A36 | 6 | 0.025397 | 0.087933 | 0.94395 | 1772 | 4 | -1.5712 | 0.86079 | 0.93113 | 1 | 19353 | 1 | -1.5712 |
| REXO2 | 3 | 0.2522 | 0.37771 | 0.999994 | 9062 | 1 | -1.4517 | 0.86084 | 0.86086 | 1 | 19354 | 1 | -1.4517 |
| FKBP6 | 4 | 0.83496 | 0.83451 | 0.999994 | 18004 | 0 | 0.16545 | 0.86085 | 0.87763 | 1 | 19355 | 1 | 0.16545 |
| PAIP1 | 5 | 0.15929 | 0.31653 | 0.999994 | 6714 | 3 | -1.174 | 0.86086 | 0.9088 | 1 | 19356 | 1 | -1.174 |
| RNF31 | 6 | 0.27442 | 0.47692 | 0.999994 | 9574 | 3 | -0.52075 | 0.86104 | 0.93127 | 1 | 19357 | 1 | -0.52075 |
| ADD1 | 5 | 0.078955 | 0.19287 | 0.975143 | 4188 | 3 | -1.8062 | 0.86112 | 0.90889 | 1 | 19358 | 1 | -1.8062 |
| PFN1 | 4 | 0.03921 | 0.10019 | 0.946394 | 2476 | 2 | -4.3139 | 0.86113 | 0.87779 | 1 | 19359 | 1 | -4.3139 |
| NAB1 | 5 | 0.04217 | 0.1213 | 0.951052 | 2623 | 3 | -0.77625 | 0.86114 | 0.90889 | 1 | 19360 | 1 | -0.77625 |
| DCLK2 | 6 | 0.26389 | 0.46447 | 0.999994 | 9333 | 2 | -0.09282 | 0.86132 | 0.93143 | 1 | 19361 | 2 | -0.09282 |
| MYO18B | 6 | 0.21874 | 0.4106 | 0.999994 | 8250 | 3 | -0.72383 | 0.86147 | 0.93151 | 1 | 19362 | 2 | -0.72383 |
| HMG20B | 3 | 0.086411 | 0.16503 | 0.967126 | 4465 | 2 | -2.1941 | 0.86152 | 0.86151 | 1 | 19363 | 1 | -2.1941 |
| C11orf70 | 5 | 0.48476 | 0.63477 | 0.999994 | 13214 | 2 | -0.6072 | 0.86159 | 0.90905 | 1 | 19364 | 1 | -0.6072 |
| IRF2BPL | 6 | 0.0057082 | 0.023731 | 0.832699 | 572 | 4 | -3.4281 | 0.86164 | 0.9316 | 1 | 19365 | 2 | -3.4281 |
| hsa-mir-2116 | 4 | 0.011665 | 0.038924 | 0.891621 | 969 | 3 | -3.8844 | 0.86167 | 0.87811 | 1 | 19366 | 1 | -3.8844 |
| CCDC71 | 6 | 0.012345 | 0.046874 | 0.896157 | 1015 | 4 | -2.6001 | 0.86168 | 0.93163 | 1 | 19367 | 2 | -2.6001 |
| IL1RAPL1 | 6 | 0.61567 | 0.7516 | 0.999994 | 15155 | 2 | -0.46302 | 0.86173 | 0.93165 | 1 | 19368 | 2 | -0.46302 |
| PPIC | 6 | 0.60111 | 0.74077 | 0.999994 | 14961 | 2 | 0.36056 | 0.86181 | 0.93169 | 1 | 19369 | 2 | 0.36056 |
| hsa-mir-29c | 4 | 0.15504 | 0.2944 | 0.999994 | 6595 | 2 | -1.3738 | 0.86213 | 0.87839 | 1 | 19370 | 1 | -1.3738 |
| hsa-mir-6893 | 2 | 0.12888 | 0.21195 | 0.979587 | 5911 | 1 | -3.2529 | 0.86223 | 0.86224 | 1 | 19371 | 0 | -3.2529 |
| IFNA14 | 6 | 0.31467 | 0.52379 | 0.999994 | 10411 | 3 | -0.35293 | 0.86237 | 0.93199 | 1 | 19372 | 1 | -0.35293 |
| CNIH1 | 4 | 0.2201 | 0.36135 | 0.999994 | 8293 | 2 | -0.88188 | 0.86241 | 0.87855 | 1 | 19373 | 1 | -0.88188 |
| NonTargetingControlG | 1 | 0.13743 | 0.13722 | 0.957161 | 6120 | 1 | -3.0122 | 0.86257 | 0.86284 | 1 | 19374 | 0 | -3.0122 |
| HOXC10 | 6 | 0.046344 | 0.13605 | 0.95468 | 2824 | 3 | -0.5959 | 0.86258 | 0.93211 | 1 | 19375 | 1 | -0.5959 |
| EFHD1 | 5 | 0.079059 | 0.19303 | 0.975741 | 4193 | 3 | -1.7844 | 0.86276 | 0.90944 | 1 | 19376 | 1 | -1.7844 |
| hsa-mir-1255b-2 | 3 | 0.39679 | 0.48241 | 0.999994 | 11902 | 1 | 0.54942 | 0.86286 | 0.86283 | 1 | 19377 | 1 | 0.54942 |
| RPL27 | 5 | 0.24449 | 0.43631 | 0.999994 | 8891 | 2 | -0.27648 | 0.8629 | 0.90949 | 1 | 19378 | 1 | -0.27648 |
| MTMR7 | 5 | 0.080109 | 0.19475 | 0.975797 | 4234 | 2 | -1.4517 | 0.86291 | 0.90949 | 1 | 19379 | 1 | -1.4517 |
| hsa-mir-92b | 2 | 0.32034 | 0.35246 | 0.999994 | 10519 | 1 | -1.5327 | 0.86315 | 0.86317 | 1 | 19380 | 0 | -1.5327 |
| PON2 | 5 | 0.012043 | 0.041925 | 0.891621 | 994 | 4 | -2.4635 | 0.8634 | 0.90966 | 1 | 19381 | 1 | -2.4635 |
| CASKIN1 | 4 | 0.27879 | 0.42133 | 0.999994 | 9665 | 2 | -0.59054 | 0.86354 | 0.87922 | 1 | 19382 | 1 | -0.59054 |
| GPR39 | 5 | 0.10369 | 0.23214 | 0.984502 | 5089 | 3 | -1.66 | 0.86373 | 0.90977 | 1 | 19383 | 1 | -1.66 |
| CARD6 | 5 | 0.34747 | 0.51985 | 0.999994 | 11098 | 2 | -0.54785 | 0.86376 | 0.90978 | 1 | 19384 | 1 | -0.54785 |
| CSF1 | 3 | 0.54634 | 0.57769 | 0.999994 | 14157 | 1 | 0.47603 | 0.86377 | 0.86372 | 1 | 19385 | 1 | 0.47603 |
| TSHZ1 | 6 | 0.01792 | 0.065245 | 0.93914 | 1357 | 3 | -1.1627 | 0.86377 | 0.93277 | 1 | 19386 | 1 | -1.1627 |
| hsa-mir-6833 | 3 | 0.28646 | 0.41983 | 0.999994 | 9847 | 2 | -0.63342 | 0.86378 | 0.86373 | 1 | 19387 | 1 | -0.63342 |
| NMD3 | 5 | 0.000064111 | 0.0002933 | 0.204053 | 33 | 4 | -8.0639 | 0.8638 | 0.90979 | 1 | 19388 | 1 | -8.0639 |
| COPS5 | 5 | 0.047374 | 0.13407 | 0.95318 | 2875 | 3 | -1.3841 | 0.86387 | 0.90981 | 1 | 19389 | 1 | -1.3841 |
| EIF2A | 4 | 0.04505 | 0.1118 | 0.951052 | 2761 | 3 | -1.5843 | 0.864 | 0.87949 | 1 | 19390 | 1 | -1.5843 |
| TMEM136 | 6 | 0.16264 | 0.34158 | 0.999994 | 6812 | 3 | -0.72973 | 0.86401 | 0.93291 | 1 | 19391 | 2 | -0.72973 |
| FERD3L | 6 | 0.52965 | 0.6893 | 0.999994 | 13897 | 2 | 0.072118 | 0.86419 | 0.93303 | 1 | 19392 | 1 | 0.072118 |
| hsa-mir-548n | 2 | 0.16354 | 0.23617 | 0.985449 | 6829 | 1 | -2.7216 | 0.86419 | 0.86425 | 1 | 19393 | 0 | -2.7216 |
| STAC2 | 6 | 0.65414 | 0.78078 | 0.999994 | 15707 | 2 | -0.1545 | 0.86435 | 0.93312 | 1 | 19394 | 2 | -0.1545 |
| hsa-mir-548f-3 | 3 | 0.47473 | 0.52979 | 0.999994 | 13058 | 1 | 0.30184 | 0.86441 | 0.86439 | 1 | 19395 | 1 | 0.30184 |

| | | | | | | | | | | | | | |
|---|---|---|---|---|---|---|---|---|---|---|---|---|---|
| SYTL3 | 6 | 0.15523 | 0.33225 | 0.999994 | 6599 | 2 | -0.21975 | 0.86468 | 0.93331 | 1 | 19396 | 1 | -0.21975 |
| FRG2 | 6 | 0.2294 | 0.42334 | 0.999994 | 8514 | 3 | -0.55116 | 0.86476 | 0.93334 | 1 | 19397 | 2 | -0.55116 |
| CACNG7 | 6 | 0.32555 | 0.53617 | 0.999994 | 10634 | 2 | -0.43486 | 0.86479 | 0.93336 | 1 | 19398 | 1 | -0.43486 |
| EFCAB5 | 3 | 0.022298 | 0.057096 | 0.92665 | 1601 | 2 | -5.0325 | 0.86482 | 0.86479 | 1 | 19399 | 1 | -5.0325 |
| UNC80 | 6 | 0.56991 | 0.71788 | 0.999994 | 14537 | 2 | -0.19763 | 0.86487 | 0.9334 | 1 | 19400 | 1 | -0.19763 |
| NonTargetingControlG | 1 | 0.13513 | 0.13497 | 0.954328 | 6054 | 1 | -3.6639 | 0.86487 | 0.86512 | 1 | 19401 | 0 | -3.6639 |
| MGAT4A | 5 | 0.13579 | 0.2814 | 0.999632 | 6075 | 3 | -1.239 | 0.86496 | 0.91019 | 1 | 19402 | 1 | -1.239 |
| NonTargetingControlG | 2 | 0.39026 | 0.4087 | 0.999994 | 11802 | 1 | -1.1511 | 0.86496 | 0.865 | 1 | 19403 | 0 | -1.1511 |
| ABHD14A | 5 | 0.033081 | 0.098442 | 0.945253 | 2165 | 3 | -2.8249 | 0.86503 | 0.91021 | 1 | 19404 | 1 | -2.8249 |
| H2AFV | 4 | 0.045934 | 0.1135 | 0.951052 | 2804 | 3 | -2.0059 | 0.86516 | 0.88019 | 1 | 19405 | 1 | -2.0059 |
| ITLN1 | 6 | 0.11898 | 0.28382 | 0.999632 | 5603 | 3 | -0.9689 | 0.86524 | 0.93362 | 1 | 19406 | 1 | -0.9689 |
| IDI1 | 3 | 0.01229 | 0.032556 | 0.865899 | 1012 | 2 | -7.5841 | 0.86535 | 0.86533 | 1 | 19407 | 0 | -7.5841 |
| LOC339862 | 6 | 0.11677 | 0.27964 | 0.999632 | 5534 | 4 | -0.74427 | 0.86539 | 0.9337 | 1 | 19408 | 2 | -0.74427 |
| CNOT8 | 6 | 0.30193 | 0.50899 | 0.999994 | 10160 | 3 | -0.47001 | 0.8654 | 0.93371 | 1 | 19409 | 2 | -0.47001 |
| TREML2 | 6 | 0.013511 | 0.050823 | 0.909101 | 1101 | 4 | -3.7448 | 0.86548 | 0.93375 | 1 | 19410 | 1 | -3.7448 |
| AGPAT3 | 6 | 0.031446 | 0.1022 | 0.946414 | 2080 | 4 | -1.9393 | 0.86554 | 0.9338 | 1 | 19411 | 2 | -1.9393 |
| SOAT2 | 4 | 0.01891 | 0.058345 | 0.931872 | 1405 | 2 | -2.7286 | 0.86554 | 0.88042 | 1 | 19412 | 1 | -2.7286 |
| NonTargetingControlG | 2 | 0.011798 | 0.022314 | 0.819701 | 980 | 1 | -5.6345 | 0.86565 | 0.86568 | 1 | 19413 | 0 | -5.6345 |
| NMS | 6 | 0.050382 | 0.14488 | 0.963012 | 2990 | 4 | -1.531 | 0.86565 | 0.93386 | 1 | 19414 | 2 | -1.531 |
| hsa-mir-4668 | 3 | 0.1356 | 0.22987 | 0.983354 | 6069 | 2 | -1.4751 | 0.86582 | 0.86581 | 1 | 19415 | 0 | -1.4751 |
| ADRA1D | 4 | 0.55762 | 0.64344 | 0.999994 | 14328 | 1 | -0.012753 | 0.86594 | 0.88065 | 1 | 19416 | 1 | -0.012753 |
| SPDYE6 | 1 | 0.13397 | 0.13379 | 0.95318 | 6019 | 1 | -3.4913 | 0.86603 | 0.86629 | 1 | 19417 | 0 | -3.4913 |
| TELO2 | 4 | 0.074673 | 0.16872 | 0.967126 | 4022 | 2 | -2.8656 | 0.86629 | 0.88086 | 1 | 19418 | 1 | -2.8656 |
| NEURL4 | 5 | 0.11365 | 0.24765 | 0.990682 | 5429 | 3 | -1.5037 | 0.86646 | 0.91075 | 1 | 19419 | 1 | -1.5037 |
| PIK3C2A | 5 | 0.11456 | 0.24906 | 0.991255 | 5465 | 3 | -1.257 | 0.8665 | 0.91076 | 1 | 19420 | 1 | -1.257 |
| NonTargetingControlG | 2 | 0.52788 | 0.52978 | 0.999994 | 13859 | 1 | -0.74031 | 0.86665 | 0.86666 | 1 | 19421 | 0 | -0.74031 |
| hsa-mir-5582 | 2 | 0.42208 | 0.43528 | 0.999994 | 12281 | 1 | -1.0663 | 0.86668 | 0.8667 | 1 | 19422 | 0 | -1.0663 |
| GLRA1 | 6 | 0.099708 | 0.24678 | 0.98999 | 4964 | 3 | -1.0083 | 0.86676 | 0.9345 | 1 | 19423 | 1 | -1.0083 |
| PCP2 | 5 | 0.013747 | 0.046677 | 0.896157 | 1114 | 4 | -3.2298 | 0.86681 | 0.91088 | 1 | 19424 | 1 | -3.2298 |
| NEUROD4 | 6 | 0.10277 | 0.25273 | 0.994501 | 5061 | 4 | -1.0006 | 0.86698 | 0.93463 | 1 | 19425 | 2 | -1.0006 |
| BMP6 | 3 | 0.56998 | 0.59462 | 0.999994 | 14541 | 1 | 0.39439 | 0.867 | 0.86698 | 1 | 19426 | 0 | 0.39439 |
| P2RY6 | 6 | 0.017285 | 0.063179 | 0.937389 | 1325 | 3 | -2.7545 | 0.86703 | 0.93465 | 1 | 19427 | 2 | -2.7545 |
| OR5R1 | 5 | 0.32467 | 0.50114 | 0.999994 | 10611 | 1 | -0.29564 | 0.8671 | 0.91097 | 1 | 19428 | 1 | -0.29564 |
| NonTargetingControlG | 2 | 0.54271 | 0.5439 | 0.999994 | 14108 | 1 | -0.67865 | 0.86722 | 0.86723 | 1 | 19429 | 0 | -0.67865 |
| HTN1 | 3 | 0.044704 | 0.1083 | 0.951052 | 2747 | 2 | -3.3641 | 0.86732 | 0.86731 | 1 | 19430 | 0 | -3.3641 |
| NonTargetingControlG | 2 | 0.55768 | 0.55824 | 0.999994 | 14329 | 1 | -0.61511 | 0.86736 | 0.86736 | 1 | 19431 | 0 | -0.61511 |
| hsa-mir-4283-2 | 2 | 0.32366 | 0.35508 | 0.999994 | 10585 | 1 | -1.624 | 0.86739 | 0.8674 | 1 | 19432 | 0 | -1.624 |
| PRHOXNB | 6 | 0.67914 | 0.80034 | 0.999994 | 16074 | 2 | -0.064424 | 0.86755 | 0.93495 | 1 | 19433 | 2 | -0.064424 |
| ZNF639 | 6 | 0.06018 | 0.16612 | 0.967126 | 3427 | 2 | -0.16642 | 0.86756 | 0.93496 | 1 | 19434 | 1 | -0.16642 |
| NIPSNAP1 | 6 | 0.097054 | 0.2416 | 0.987253 | 4863 | 3 | -0.38242 | 0.86757 | 0.93496 | 1 | 19435 | 2 | -0.38242 |
| AMT | 5 | 0.48838 | 0.63781 | 0.999994 | 13260 | 2 | -0.38492 | 0.8676 | 0.91113 | 1 | 19436 | 1 | -0.38492 |
| CLOCK | 3 | 0.21387 | 0.33005 | 0.999994 | 8111 | 2 | -1.1661 | 0.86764 | 0.86765 | 1 | 19437 | 0 | -1.1661 |
| MBNL1 | 3 | 0.12102 | 0.21079 | 0.979587 | 5662 | 2 | -1.8387 | 0.86767 | 0.86767 | 1 | 19438 | 0 | -1.8387 |
| TMEM207 | 6 | 0.042105 | 0.12663 | 0.951052 | 2619 | 3 | -1.6761 | 0.86779 | 0.93509 | 1 | 19439 | 1 | -1.6761 |
| EDN2 | 6 | 0.0091674 | 0.036109 | 0.877982 | 807 | 5 | -4.0885 | 0.86788 | 0.93513 | 1 | 19440 | 1 | -4.0885 |
| SLC5A9 | 5 | 0.013336 | 0.045516 | 0.891621 | 1087 | 4 | -1.5821 | 0.86791 | 0.91125 | 1 | 19441 | 1 | -1.5821 |
| hsa-mir-4325 | 4 | 0.72771 | 0.7435 | 0.999994 | 16577 | 1 | 0.17427 | 0.86791 | 0.88184 | 1 | 19442 | 1 | 0.17427 |
| NonTargetingControlG | 2 | 0.24166 | 0.29267 | 0.999632 | 8823 | 1 | -2.1416 | 0.86792 | 0.86793 | 1 | 19443 | 0 | -2.1416 |
| PPBP | 6 | 0.040924 | 0.124 | 0.951052 | 2559 | 4 | -1.5627 | 0.86804 | 0.93524 | 1 | 19444 | 2 | -1.5627 |
| hsa-mir-611 | 3 | 0.039108 | 0.095817 | 0.945031 | 2469 | 2 | -3.8433 | 0.86815 | 0.86817 | 1 | 19445 | 0 | -3.8433 |
| ZNF611 | 4 | 0.34378 | 0.48794 | 0.999994 | 11024 | 2 | -0.59328 | 0.86815 | 0.882 | 1 | 19446 | 1 | -0.59328 |
| NDUFA4 | 5 | 0.022052 | 0.069639 | 0.93914 | 1587 | 4 | -1.3967 | 0.86821 | 0.91136 | 1 | 19447 | 1 | -1.3967 |

| | | | | | | | | | | | | |
|---|---|---|---|---|---|---|---|---|---|---|---|---|
| FAS | 6 | 0.78986 | 0.84361 | 0.999994 | 17341 | 1 | -0.12391 | 0.86834 | 0.93542 | 1 | 19448 | 2 | -0.12391 |
| OR4D11 | 6 | 0.18377 | 0.36785 | 0.999994 | 7309 | 3 | -0.5318 | 0.86844 | 0.93547 | 1 | 19449 | 2 | -0.5318 |
| NHLRC2 | 6 | 0.0086361 | 0.034212 | 0.867348 | 770 | 5 | -2.8901 | 0.86859 | 0.93556 | 1 | 19450 | 1 | -2.8901 |
| FAM101A | 5 | 0.050612 | 0.14185 | 0.959146 | 3001 | 3 | -0.6658 | 0.86865 | 0.91152 | 1 | 19451 | 1 | -0.6658 |
| PAFAH1B2 | 5 | 0.32184 | 0.49877 | 0.999994 | 10554 | 2 | 0.34177 | 0.86866 | 0.91152 | 1 | 19452 | 1 | 0.34177 |
| HOXD13 | 6 | 0.04716 | 0.13786 | 0.957317 | 2860 | 3 | -0.62054 | 0.86871 | 0.93563 | 1 | 19453 | 2 | -0.62054 |
| ZNF557 | 6 | 0.01943 | 0.070156 | 0.93914 | 1426 | 4 | -1.2387 | 0.86892 | 0.93575 | 1 | 19454 | 2 | -1.2387 |
| CD99L2 | 6 | 0.068631 | 0.18392 | 0.968526 | 3778 | 4 | -1.438 | 0.86897 | 0.93579 | 1 | 19455 | 2 | -1.438 |
| CTSD | 6 | 0.29852 | 0.50499 | 0.999994 | 10097 | 3 | -0.6734 | 0.86904 | 0.93583 | 1 | 19456 | 2 | -0.6734 |
| PCDHA12 | 2 | 0.037682 | 0.068271 | 0.93914 | 2414 | 1 | -5.0879 | 0.86925 | 0.86923 | 1 | 19457 | 0 | -5.0879 |
| DPYD | 6 | 0.31458 | 0.5237 | 0.999994 | 10409 | 3 | -0.50227 | 0.86928 | 0.93597 | 1 | 19458 | 1 | -0.50227 |
| KNOP1 | 5 | 0.24497 | 0.43668 | 0.999994 | 8899 | 2 | 0.18055 | 0.86931 | 0.91174 | 1 | 19459 | 1 | 0.18055 |
| hsa-mir-499b | 3 | 0.44251 | 0.50964 | 0.999994 | 12618 | 1 | 0.18716 | 0.86932 | 0.86938 | 1 | 19460 | 0 | 0.18716 |
| CMTM2 | 5 | 0.2931 | 0.47544 | 0.999994 | 9978 | 2 | 0.30979 | 0.86932 | 0.91175 | 1 | 19461 | 1 | 0.30979 |
| NonTargetingControlG | 1 | 0.13065 | 0.1305 | 0.95318 | 5948 | 1 | -3.1057 | 0.86935 | 0.86956 | 1 | 19462 | 0 | -3.1057 |
| ACVR2A | 6 | 0.92121 | 0.92152 | 0.999994 | 19657 | 1 | 0.13878 | 0.86936 | 0.93601 | 1 | 19463 | 1 | 0.13878 |
| SLC37A2 | 6 | 0.014926 | 0.055481 | 0.922773 | 1184 | 4 | -1.6933 | 0.86967 | 0.93619 | 1 | 19464 | 1 | -1.6933 |
| NAA25 | 5 | 0.042207 | 0.12138 | 0.951052 | 2625 | 4 | -1.2067 | 0.87012 | 0.91202 | 1 | 19465 | 1 | -1.2067 |
| MATK | 6 | 0.047615 | 0.13882 | 0.957514 | 2881 | 3 | -1.8577 | 0.87025 | 0.93654 | 1 | 19466 | 2 | -1.8577 |
| C10orf88 | 4 | 0.49993 | 0.6163 | 0.999994 | 13422 | 1 | -0.09379 | 0.87028 | 0.88332 | 1 | 19467 | 1 | -0.09379 |
| hsa-mir-4431 | 2 | 0.42661 | 0.43919 | 0.999994 | 12358 | 1 | -1.19 | 0.87036 | 0.87036 | 1 | 19468 | 0 | -1.19 |
| IGF2 | 6 | 0.10165 | 0.25055 | 0.992497 | 5023 | 3 | -0.71767 | 0.8704 | 0.93663 | 1 | 19469 | 2 | -0.71767 |
| hsa-mir-24-2 | 3 | 0.070111 | 0.14314 | 0.960171 | 3833 | 2 | -2.7992 | 0.87048 | 0.87055 | 1 | 19470 | 0 | -2.7992 |
| hsa-mir-6069 | 4 | 0.074728 | 0.16882 | 0.967126 | 4024 | 1 | -0.4056 | 0.87051 | 0.88346 | 1 | 19471 | 1 | -0.4056 |
| HDHD1 | 6 | 0.57456 | 0.72122 | 0.999994 | 14612 | 2 | 0.02825 | 0.87063 | 0.93677 | 1 | 19472 | 2 | 0.02825 |
| SEMA4A | 6 | 0.1088 | 0.2644 | 0.996734 | 5264 | 2 | -0.068723 | 0.87067 | 0.93679 | 1 | 19473 | 2 | -0.068723 |
| OVOL1 | 5 | 0.47659 | 0.62768 | 0.999994 | 13092 | 2 | -0.50475 | 0.87072 | 0.91225 | 1 | 19474 | 1 | -0.50475 |
| MAP3K19 | 5 | 0.066276 | 0.17229 | 0.967658 | 3695 | 3 | -1.9829 | 0.87074 | 0.91225 | 1 | 19475 | 1 | -1.9829 |
| SFXN1 | 6 | 0.2169 | 0.40841 | 0.999994 | 8189 | 3 | -0.68364 | 0.87079 | 0.93686 | 1 | 19476 | 2 | -0.68364 |
| NEK5 | 5 | 0.18671 | 0.35626 | 0.999994 | 7379 | 2 | -0.57145 | 0.87093 | 0.91231 | 1 | 19477 | 1 | -0.57145 |
| STMN4 | 6 | 0.011905 | 0.045397 | 0.891621 | 985 | 4 | -0.96004 | 0.87096 | 0.93696 | 1 | 19478 | 2 | -0.96004 |
| ZNF831 | 6 | 0.18475 | 0.36907 | 0.999994 | 7339 | 3 | -0.47954 | 0.87104 | 0.937 | 1 | 19479 | 1 | -0.47954 |
| EIF4H | 6 | 0.0072365 | 0.029246 | 0.860452 | 672 | 4 | -3.4726 | 0.87104 | 0.937 | 1 | 19480 | 1 | -3.4726 |
| DCLRE1B | 6 | 0.060748 | 0.16736 | 0.967126 | 3454 | 4 | -1.3458 | 0.87104 | 0.937 | 1 | 19481 | 1 | -1.3458 |
| UXS1 | 6 | 0.0073061 | 0.029494 | 0.862223 | 680 | 4 | -3.114 | 0.87104 | 0.937 | 1 | 19482 | 1 | -3.114 |
| PSMA4 | 4 | 0.013323 | 0.043979 | 0.891621 | 1085 | 3 | -5.8348 | 0.87108 | 0.88384 | 1 | 19483 | 1 | -5.8348 |
| UPF3A | 3 | 0.30911 | 0.43347 | 0.999994 | 10305 | 1 | 0.46088 | 0.87108 | 0.87114 | 1 | 19484 | 0 | 0.46088 |
| SPRR4 | 6 | 0.14415 | 0.31829 | 0.999994 | 6303 | 2 | 0.2998 | 0.87112 | 0.93704 | 1 | 19485 | 2 | 0.2998 |
| GDF3 | 6 | 0.79145 | 0.84429 | 0.999994 | 17370 | 1 | 0.12887 | 0.87121 | 0.93709 | 1 | 19486 | 2 | 0.12887 |
| C10orf82 | 5 | 0.1981 | 0.37256 | 0.999994 | 7713 | 3 | -0.83522 | 0.87121 | 0.91241 | 1 | 19487 | 1 | -0.83522 |
| LMNA | 5 | 0.18763 | 0.35758 | 0.999994 | 7411 | 2 | -0.14646 | 0.87141 | 0.91248 | 1 | 19488 | 1 | -0.14646 |
| ZNF221 | 6 | 0.35899 | 0.57417 | 0.999994 | 11320 | 2 | 0.44278 | 0.87144 | 0.93722 | 1 | 19489 | 2 | 0.44278 |
| TRMT61B | 5 | 0.004837 | 0.02017 | 0.808847 | 509 | 3 | -1.0333 | 0.87147 | 0.9125 | 1 | 19490 | 1 | -1.0333 |
| KRTAP5-6 | 5 | 0.48195 | 0.63236 | 0.999994 | 13169 | 2 | -0.57967 | 0.8716 | 0.91254 | 1 | 19491 | 1 | -0.57967 |
| ETFDH | 6 | 0.070869 | 0.18861 | 0.969543 | 3870 | 4 | -1.1398 | 0.87165 | 0.93733 | 1 | 19492 | 1 | -1.1398 |
| SHISA9 | 6 | 0.012162 | 0.046255 | 0.892434 | 1004 | 2 | 0.14816 | 0.87183 | 0.93743 | 1 | 19493 | 2 | 0.14816 |
| FAM186B | 6 | 0.097622 | 0.24266 | 0.988088 | 4874 | 4 | -1.1424 | 0.87187 | 0.93746 | 1 | 19494 | 2 | -1.1424 |
| NCR2 | 6 | 0.062492 | 0.17104 | 0.96737 | 3535 | 2 | 0.095834 | 0.87189 | 0.93747 | 1 | 19495 | 2 | 0.095834 |
| GPX2 | 6 | 0.35873 | 0.57389 | 0.999994 | 11318 | 3 | -0.41813 | 0.8719 | 0.93747 | 1 | 19496 | 1 | -0.41813 |
| BTC | 6 | 0.25491 | 0.4538 | 0.999994 | 9129 | 3 | -0.51228 | 0.87196 | 0.93751 | 1 | 19497 | 2 | -0.51228 |
| PIM2 | 4 | 0.19667 | 0.3373 | 0.999994 | 7669 | 1 | 0.23811 | 0.87198 | 0.8844 | 1 | 19498 | 1 | 0.23811 |
| OR9G1 | 6 | 0.074393 | 0.19591 | 0.975797 | 4006 | 4 | -1.2497 | 0.872 | 0.93753 | 1 | 19499 | 2 | -1.2497 |

| | | | | | | | | | | | | |
|---|---|---|---|---|---|---|---|---|---|---|---|---|
| GOPC | 6 | 0.33742 | 0.54972 | 0.999994 | 10883 | 3 | -0.57852 | 0.87207 | 0.93758 | 1 | 19500 | 2 | -0.57852 |
| hsa-mir-3139 | 3 | 0.14368 | 0.24036 | 0.987253 | 6290 | 1 | 0.23671 | 0.87221 | 0.87231 | 1 | 19501 | 0 | 0.23671 |
| PPP1R26 | 6 | 0.067329 | 0.18117 | 0.968171 | 3729 | 4 | -1.1482 | 0.87233 | 0.93774 | 1 | 19502 | 2 | -1.1482 |
| NonTargetingControlG | 1 | 0.12754 | 0.12741 | 0.951052 | 5861 | 1 | -3.5309 | 0.87246 | 0.87272 | 1 | 19503 | 0 | -3.5309 |
| PPP2R5B | 5 | 0.0091124 | 0.033473 | 0.867348 | 803 | 4 | -6.0649 | 0.87256 | 0.91287 | 1 | 19504 | 1 | -6.0649 |
| SRP9 | 4 | 0.0083758 | 0.028583 | 0.858757 | 754 | 3 | -4.6732 | 0.87265 | 0.88482 | 1 | 19505 | 1 | -4.6732 |
| GLI4 | 6 | 0.095735 | 0.23898 | 0.987253 | 4805 | 3 | -1.0761 | 0.87265 | 0.93793 | 1 | 19506 | 1 | -1.0761 |
| S100A13 | 4 | 0.18998 | 0.33036 | 0.999994 | 7483 | 2 | -0.69158 | 0.87272 | 0.88487 | 1 | 19507 | 1 | -0.69158 |
| TCP10L | 6 | 0.70593 | 0.81384 | 0.999994 | 16371 | 1 | 0.16764 | 0.87278 | 0.938 | 1 | 19508 | 2 | 0.16764 |
| RASA3 | 6 | 0.0082029 | 0.032655 | 0.866598 | 745 | 3 | -1.0611 | 0.87291 | 0.93808 | 1 | 19509 | 2 | -1.0611 |
| NGDN | 5 | 0.004995 | 0.020804 | 0.81027 | 518 | 4 | -3.2391 | 0.87298 | 0.91303 | 1 | 19510 | 1 | -3.2391 |
| PGLYRP2 | 5 | 0.04076 | 0.11776 | 0.951052 | 2549 | 4 | -1.6895 | 0.87304 | 0.91305 | 1 | 19511 | 1 | -1.6895 |
| C7orf34 | 5 | 0.493 | 0.64183 | 0.999994 | 13311 | 2 | -0.21029 | 0.87307 | 0.91306 | 1 | 19512 | 1 | -0.21029 |
| KCNAB3 | 6 | 0.011704 | 0.044738 | 0.891621 | 972 | 4 | -2.5683 | 0.87309 | 0.93818 | 1 | 19513 | 2 | -2.5683 |
| NAF1 | 6 | 0.080645 | 0.20861 | 0.979587 | 4257 | 3 | -0.37203 | 0.87315 | 0.93822 | 1 | 19514 | 2 | -0.37203 |
| hsa-mir-6131 | 3 | 0.11132 | 0.19805 | 0.977164 | 5344 | 1 | 0.08452 | 0.87319 | 0.8733 | 1 | 19515 | 0 | 0.08452 |
| PLCL2 | 6 | 0.0023454 | 0.011258 | 0.716138 | 309 | 4 | -1.8688 | 0.8732 | 0.93826 | 1 | 19516 | 2 | -1.8688 |
| WNT7A | 5 | 0.017252 | 0.056562 | 0.9235 | 1317 | 3 | -3.5735 | 0.87334 | 0.91316 | 1 | 19517 | 1 | -3.5735 |
| NOC4L | 4 | 0.0036715 | 0.013103 | 0.733879 | 418 | 3 | -6.8451 | 0.87336 | 0.88526 | 1 | 19518 | 1 | -6.8451 |
| METTL10 | 4 | 0.24598 | 0.38777 | 0.999994 | 8922 | 2 | -0.38185 | 0.87343 | 0.8853 | 1 | 19519 | 1 | -0.38185 |
| RGMB | 3 | 0.63353 | 0.64351 | 0.999994 | 15422 | 1 | -0.11703 | 0.87352 | 0.87364 | 1 | 19520 | 0 | -0.11703 |
| P4HA3 | 6 | 0.43595 | 0.62575 | 0.999994 | 12514 | 2 | 0.31727 | 0.87352 | 0.93845 | 1 | 19521 | 2 | 0.31727 |
| RPL17-C18orf32 | 1 | 0.12638 | 0.1263 | 0.951052 | 5835 | 1 | -5.2266 | 0.87362 | 0.87388 | 1 | 19522 | 0 | -5.2266 |
| hsa-mir-1181 | 3 | 0.030694 | 0.076759 | 0.93914 | 2044 | 2 | -5.7119 | 0.87367 | 0.87379 | 1 | 19523 | 0 | -5.7119 |
| EDIL3 | 4 | 0.035577 | 0.0929 | 0.943968 | 2298 | 3 | -1.9577 | 0.87369 | 0.88547 | 1 | 19524 | 1 | -1.9577 |
| NonTargetingControlG | 2 | 0.26327 | 0.30879 | 0.999994 | 9321 | 1 | -1.8217 | 0.87369 | 0.87376 | 1 | 19525 | 0 | -1.8217 |
| POLR3H | 3 | 0.70317 | 0.704 | 0.999994 | 16341 | 1 | 0.39389 | 0.87375 | 0.87388 | 1 | 19526 | 0 | 0.39389 |
| hsa-mir-8078 | 1 | 0.12618 | 0.12606 | 0.951052 | 5826 | 1 | -5.2324 | 0.87382 | 0.8741 | 1 | 19527 | 0 | -5.2324 |
| NIPSNAP3A | 6 | 0.093129 | 0.23377 | 0.984971 | 4723 | 1 | -0.22109 | 0.87397 | 0.93873 | 1 | 19528 | 2 | -0.22109 |
| TOMM5 | 6 | 0.11843 | 0.28276 | 0.999632 | 5588 | 4 | -0.69796 | 0.87408 | 0.93879 | 1 | 19529 | 1 | -0.69796 |
| hsa-mir-320e | 3 | 0.66785 | 0.67203 | 0.999994 | 15891 | 1 | 0.24325 | 0.87411 | 0.87425 | 1 | 19530 | 0 | 0.24325 |
| ZNF747 | 4 | 0.059337 | 0.13954 | 0.958716 | 3395 | 3 | -2.0124 | 0.87415 | 0.88575 | 1 | 19531 | 1 | -2.0124 |
| CTSG | 5 | 0.0062787 | 0.025267 | 0.84412 | 612 | 4 | -2.396 | 0.87424 | 0.91349 | 1 | 19532 | 1 | -2.396 |
| TRPV2 | 3 | 0.19134 | 0.30182 | 0.999994 | 7523 | 1 | -0.037716 | 0.87428 | 0.87442 | 1 | 19533 | 0 | -0.037716 |
| SRI | 4 | 0.08462 | 0.18721 | 0.969409 | 4393 | 3 | -1.745 | 0.8743 | 0.88585 | 1 | 19534 | 1 | -1.745 |
| hsa-mir-5681a | 4 | 0.23658 | 0.37821 | 0.999994 | 8683 | 2 | -0.61976 | 0.87431 | 0.88585 | 1 | 19535 | 1 | -0.61976 |
| ARHGEF26 | 6 | 0.22143 | 0.41383 | 0.999994 | 8315 | 2 | -0.2773 | 0.87432 | 0.93893 | 1 | 19536 | 2 | -0.2773 |
| DNAJB2 | 5 | 0.40952 | 0.57148 | 0.999994 | 12100 | 1 | -0.43165 | 0.87455 | 0.9136 | 1 | 19537 | 1 | -0.43165 |
| ENPP1 | 5 | 0.23579 | 0.42521 | 0.999994 | 8664 | 2 | -0.46608 | 0.87456 | 0.9136 | 1 | 19538 | 1 | -0.46608 |
| GH1 | 6 | 0.25279 | 0.45125 | 0.999994 | 9079 | 2 | -0.025364 | 0.87456 | 0.93906 | 1 | 19539 | 2 | -0.025364 |
| KANSL1 | 4 | 0.0020653 | 0.0074954 | 0.665213 | 281 | 3 | -8.0506 | 0.87459 | 0.88602 | 1 | 19540 | 1 | -8.0506 |
| ACMSD | 5 | 0.15311 | 0.30742 | 0.999994 | 6539 | 3 | -0.60407 | 0.87461 | 0.91362 | 1 | 19541 | 1 | -0.60407 |
| hsa-mir-1537 | 3 | 0.52146 | 0.56052 | 0.999994 | 13767 | 1 | -0.04147 | 0.87474 | 0.87489 | 1 | 19542 | 0 | -0.04147 |
| PTAR1 | 6 | 0.10087 | 0.24902 | 0.991255 | 5002 | 4 | -1.2284 | 0.87474 | 0.93918 | 1 | 19543 | 1 | -1.2284 |
| ANAPC11 | 4 | 0.20613 | 0.34701 | 0.999994 | 7932 | 2 | -1.1665 | 0.87481 | 0.88616 | 1 | 19544 | 1 | -1.1665 |
| WDR86 | 4 | 0.13696 | 0.27575 | 0.999379 | 6106 | 2 | -1.104 | 0.87485 | 0.88619 | 1 | 19545 | 1 | -1.104 |
| hsa-mir-6126 | 3 | 0.27612 | 0.40723 | 0.999994 | 9620 | 1 | 0.48323 | 0.87487 | 0.875 | 1 | 19546 | 0 | 0.48323 |
| RARB | 6 | 0.15899 | 0.33694 | 0.999994 | 6701 | 2 | -0.0051121 | 0.8749 | 0.93927 | 1 | 19547 | 2 | -0.0051121 |
| MACROD1 | 6 | 0.076771 | 0.20082 | 0.977164 | 4102 | 3 | -1.2061 | 0.87496 | 0.93931 | 1 | 19548 | 2 | -1.2061 |
| HSD17B2 | 5 | 0.53179 | 0.67563 | 0.999994 | 13930 | 2 | -0.48195 | 0.87498 | 0.91376 | 1 | 19549 | 1 | -0.48195 |
| TIA1 | 6 | 0.03542 | 0.11152 | 0.951052 | 2288 | 4 | -1.49 | 0.87511 | 0.93941 | 1 | 19550 | 1 | -1.49 |
| hsa-mir-378f | 3 | 0.25462 | 0.38077 | 0.999994 | 9121 | 2 | -0.75424 | 0.87511 | 0.87525 | 1 | 19551 | 0 | -0.75424 |

| | | | | | | | | | | | | | |
|---|---|---|---|---|---|---|---|---|---|---|---|---|---|
| NonTargetingControlG | 1 | 0.12473 | 0.12463 | 0.951052 | 5788 | 1 | -3.8373 | 0.87527 | 0.87561 | 1 | 19552 | 0 | -3.8373 |
| SPINT3 | 6 | 0.03222 | 0.10402 | 0.947284 | 2113 | 4 | -1.7367 | 0.8753 | 0.93951 | 1 | 19553 | 1 | -1.7367 |
| KRTAP10-3 | 6 | 0.33042 | 0.54173 | 0.999994 | 10729 | 3 | -0.474 | 0.87545 | 0.9396 | 1 | 19554 | 1 | -0.474 |
| CALN1 | 6 | 0.010425 | 0.040464 | 0.891621 | 896 | 4 | -2.2114 | 0.87554 | 0.93965 | 1 | 19555 | 1 | -2.2114 |
| NonTargetingControlG | 2 | 0.12436 | 0.20737 | 0.978852 | 5777 | 2 | -1.4712 | 0.87564 | 0.87569 | 1 | 19556 | 0 | -1.4712 |
| LHFPL5 | 5 | 0.22177 | 0.40575 | 0.999994 | 8326 | 3 | -0.69387 | 0.87565 | 0.91401 | 1 | 19557 | 1 | -0.69387 |
| NonTargetingControlG | 2 | 0.099478 | 0.16926 | 0.967126 | 4948 | 2 | -4.383 | 0.87567 | 0.87572 | 1 | 19558 | 0 | -4.383 |
| ASB7 | 5 | 0.016922 | 0.055649 | 0.9235 | 1298 | 4 | -3.5166 | 0.87568 | 0.91402 | 1 | 19559 | 1 | -3.5166 |
| hsa-mir-3123 | 3 | 0.19151 | 0.30202 | 0.999994 | 7528 | 2 | -1.2566 | 0.87571 | 0.87585 | 1 | 19560 | 0 | -1.2566 |
| HLA-DOA | 6 | 0.085487 | 0.21849 | 0.980699 | 4429 | 3 | -1.2349 | 0.87578 | 0.93979 | 1 | 19561 | 2 | -1.2349 |
| DEFB105A | 4 | 0.43061 | 0.57718 | 0.999994 | 12426 | 2 | -0.26208 | 0.87584 | 0.88686 | 1 | 19562 | 1 | -0.26208 |
| HIP1R | 5 | 0.021838 | 0.069023 | 0.93914 | 1570 | 3 | -2.9693 | 0.87596 | 0.91411 | 1 | 19563 | 1 | -2.9693 |
| ZNF502 | 6 | 0.031997 | 0.10348 | 0.946414 | 2103 | 3 | -1.8197 | 0.87599 | 0.9399 | 1 | 19564 | 2 | -1.8197 |
| GAGE12J | 1 | 0.12379 | 0.1237 | 0.951052 | 5759 | 1 | -3.2529 | 0.87621 | 0.87654 | 1 | 19565 | 0 | -3.2529 |
| MAPK8IP3 | 3 | 0.2789 | 0.41066 | 0.999994 | 9670 | 2 | -0.62105 | 0.87623 | 0.87634 | 1 | 19566 | 0 | -0.62105 |
| ECHDC3 | 5 | 0.21373 | 0.3946 | 0.999994 | 8106 | 2 | 0.28379 | 0.87634 | 0.91424 | 1 | 19567 | 1 | 0.28379 |
| hsa-mir-630 | 4 | 0.14544 | 0.28451 | 0.999632 | 6337 | 2 | -1.172 | 0.87636 | 0.88721 | 1 | 19568 | 1 | -1.172 |
| CCDC15 | 6 | 0.1906 | 0.37628 | 0.999994 | 7502 | 3 | -0.49833 | 0.87638 | 0.94015 | 1 | 19569 | 1 | -0.49833 |
| MRPS9 | 3 | 0.66222 | 0.66722 | 0.999994 | 15812 | 1 | 0.25606 | 0.87641 | 0.8765 | 1 | 19570 | 0 | 0.25606 |
| RGS20 | 5 | 0.47044 | 0.62254 | 0.999994 | 13009 | 2 | -0.22258 | 0.87641 | 0.91426 | 1 | 19571 | 1 | -0.22258 |
| TEX13A | 6 | 0.56527 | 0.71457 | 0.999994 | 14473 | 2 | 0.3172 | 0.87644 | 0.94018 | 1 | 19572 | 2 | 0.3172 |
| KRTAP9-2 | 3 | 0.091001 | 0.17108 | 0.96737 | 4636 | 2 | -2.5577 | 0.87649 | 0.87658 | 1 | 19573 | 0 | -2.5577 |
| NonTargetingControlG | 2 | 0.11465 | 0.19264 | 0.974607 | 5469 | 2 | -4.241 | 0.87651 | 0.87656 | 1 | 19574 | 0 | -4.241 |
| hsa-mir-5692a-1 | 2 | 0.12343 | 0.20593 | 0.978744 | 5749 | 2 | -2.0453 | 0.87657 | 0.87661 | 1 | 19575 | 0 | -2.0453 |
| UFD1L | 2 | 0.074414 | 0.12952 | 0.95318 | 4007 | 2 | -3.9448 | 0.87663 | 0.87667 | 1 | 19576 | 0 | -3.9448 |
| RBBP8NL | 6 | 0.21579 | 0.4071 | 0.999994 | 8165 | 3 | -0.57931 | 0.8767 | 0.94033 | 1 | 19577 | 2 | -0.57931 |
| NBEA | 6 | 0.20079 | 0.38883 | 0.999994 | 7779 | 3 | -0.82732 | 0.87672 | 0.94035 | 1 | 19578 | 2 | -0.82732 |
| TRMT13 | 2 | 0.12313 | 0.20543 | 0.978342 | 5742 | 2 | -0.93993 | 0.87687 | 0.8769 | 1 | 19579 | 0 | -0.93993 |
| ZNF445 | 4 | 0.8546 | 0.85418 | 0.999994 | 18342 | 0 | -0.16216 | 0.87688 | 0.88757 | 1 | 19580 | 1 | -0.16216 |
| OSMR | 6 | 0.31133 | 0.5199 | 0.999994 | 10347 | 3 | -0.33656 | 0.87705 | 0.94054 | 1 | 19581 | 2 | -0.33656 |
| ASPHD1 | 5 | 0.021378 | 0.067797 | 0.93914 | 1538 | 4 | -1.5349 | 0.87717 | 0.91455 | 1 | 19582 | 1 | -1.5349 |
| GON4L | 5 | 0.46339 | 0.61651 | 0.999994 | 12910 | 2 | -0.27384 | 0.87717 | 0.91455 | 1 | 19583 | 1 | -0.27384 |
| FBXO7 | 6 | 0.63565 | 0.76663 | 0.999994 | 15452 | 2 | -0.14239 | 0.87719 | 0.94063 | 1 | 19584 | 2 | -0.14239 |
| ST6GALNAC3 | 6 | 0.01402 | 0.052539 | 0.914561 | 1130 | 5 | -1.3482 | 0.87733 | 0.94071 | 1 | 19585 | 1 | -1.3482 |
| MMAB | 6 | 0.32985 | 0.54106 | 0.999994 | 10717 | 2 | 0.15056 | 0.87739 | 0.94075 | 1 | 19586 | 2 | 0.15056 |
| F8A3 | 1 | 0.12254 | 0.12243 | 0.951052 | 5725 | 1 | -3.3997 | 0.87746 | 0.87777 | 1 | 19587 | 0 | -3.3997 |
| C6orf170 | 6 | 0.9372 | 0.93721 | 0.999994 | 20010 | 0 | -0.23008 | 0.8775 | 0.94081 | 1 | 19588 | 1 | -0.23008 |
| RPL39L | 4 | 0.045642 | 0.11293 | 0.951052 | 2794 | 3 | -1.834 | 0.87751 | 0.88798 | 1 | 19589 | 1 | -1.834 |
| hsa-mir-5189 | 3 | 0.79176 | 0.79166 | 0.999994 | 17376 | 0 | 0.10771 | 0.87754 | 0.87761 | 1 | 19590 | 0 | 0.10771 |
| SOX5 | 4 | 0.012696 | 0.04207 | 0.891621 | 1046 | 3 | -3.1733 | 0.87762 | 0.88804 | 1 | 19591 | 1 | -3.1733 |
| NonTargetingControlG | 2 | 0.12222 | 0.20407 | 0.978186 | 5710 | 2 | -0.81067 | 0.87778 | 0.87785 | 1 | 19592 | 0 | -0.81067 |
| NonTargetingControlG | 2 | 0.12218 | 0.20403 | 0.978186 | 5708 | 2 | -0.91294 | 0.87782 | 0.87789 | 1 | 19593 | 0 | -0.91294 |
| PLEKHM2 | 5 | 0.22106 | 0.40472 | 0.999994 | 8310 | 3 | -0.74579 | 0.87782 | 0.91478 | 1 | 19594 | 1 | -0.74579 |
| C3orf67 | 6 | 0.28947 | 0.49447 | 0.999994 | 9901 | 3 | -0.51198 | 0.87788 | 0.94102 | 1 | 19595 | 2 | -0.51198 |
| ZRSR2 | 3 | 0.26725 | 0.39626 | 0.999994 | 9401 | 1 | -0.016002 | 0.87805 | 0.87814 | 1 | 19596 | 0 | -0.016002 |
| hsa-mir-432 | 4 | 0.34969 | 0.49393 | 0.999994 | 11148 | 1 | 0.2467 | 0.87808 | 0.88834 | 1 | 19597 | 1 | 0.2467 |
| NEK2 | 6 | 0.00043788 | 0.0023023 | 0.45324 | 103 | 5 | -5.4471 | 0.8782 | 0.94111 | 1 | 19598 | 1 | -5.4471 |
| GATA2 | 4 | 0.081346 | 0.1812 | 0.968171 | 4279 | 2 | -0.57079 | 0.8784 | 0.88855 | 1 | 19599 | 1 | -0.57079 |
| PRR22 | 4 | 0.022085 | 0.06502 | 0.93914 | 1591 | 3 | -0.90995 | 0.87847 | 0.88859 | 1 | 19600 | 1 | -0.90995 |
| RXFP1 | 6 | 0.088296 | 0.22414 | 0.983354 | 4523 | 4 | -1.494 | 0.87849 | 0.94116 | 1 | 19601 | 1 | -1.494 |
| IL36A | 4 | 0.12457 | 0.25825 | 0.994943 | 5784 | 2 | -1.3089 | 0.87854 | 0.88864 | 1 | 19602 | 1 | -1.3089 |
| EXOSC6 | 3 | 0.16007 | 0.26152 | 0.995257 | 6742 | 2 | -1.4766 | 0.87861 | 0.8787 | 1 | 19603 | 0 | -1.4766 |

| | | | | | | | | | | | | | |
|---|---|---|---|---|---|---|---|---|---|---|---|---|---|
| SLC13A1 | 6 | 0.06203 | 0.17005 | 0.967126 | 3514 | 4 | -1.7226 | 0.8787 | 0.9412 | 1 | 19604 | 1 | -1.7226 |
| GK5 | 6 | 0.66159 | 0.78653 | 0.999994 | 15804 | 2 | -0.27686 | 0.87872 | 0.94121 | 1 | 19605 | 1 | -0.27686 |
| hsa-mir-323a | 4 | 0.046661 | 0.1149 | 0.951052 | 2836 | 3 | -2.8798 | 0.87875 | 0.88877 | 1 | 19606 | 1 | -2.8798 |
| hsa-mir-920 | 4 | 0.3151 | 0.45848 | 0.999994 | 10422 | 2 | -0.937 | 0.87883 | 0.88883 | 1 | 19607 | 1 | -0.937 |
| NonTargetingControlG | 2 | 0.12094 | 0.2021 | 0.977967 | 5660 | 2 | -0.90304 | 0.87906 | 0.87914 | 1 | 19608 | 0 | -0.90304 |
| SLC30A3 | 4 | 0.82612 | 0.8257 | 0.999994 | 17869 | 1 | -0.20451 | 0.87909 | 0.88898 | 1 | 19609 | 1 | -0.20451 |
| IL12B | 6 | 0.059879 | 0.1655 | 0.967126 | 3416 | 3 | -0.60277 | 0.8794 | 0.94134 | 1 | 19610 | 1 | -0.60277 |
| C21orf62 | 4 | 0.14262 | 0.2816 | 0.999632 | 6257 | 2 | -1.3603 | 0.8795 | 0.88926 | 1 | 19611 | 1 | -1.3603 |
| RRP36 | 3 | 0.067644 | 0.1398 | 0.958716 | 3742 | 1 | -0.019152 | 0.87964 | 0.87972 | 1 | 19612 | 0 | -0.019152 |
| DOCK4 | 5 | 0.33796 | 0.51207 | 0.999994 | 10895 | 2 | -0.55968 | 0.87971 | 0.91549 | 1 | 19613 | 1 | -0.55968 |
| SESN1 | 5 | 0.54218 | 0.68482 | 0.999994 | 14097 | 2 | 0.3656 | 0.88031 | 0.91571 | 1 | 19614 | 1 | 0.3656 |
| hsa-mir-1260a | 3 | 0.76501 | 0.76498 | 0.999994 | 17026 | 0 | -0.25677 | 0.88043 | 0.88048 | 1 | 19615 | 0 | -0.25677 |
| BCAT2 | 3 | 0.20824 | 0.32302 | 0.999994 | 7984 | 2 | -1.0665 | 0.88043 | 0.88048 | 1 | 19616 | 0 | -1.0665 |
| hsa-mir-4419b | 3 | 0.16403 | 0.26674 | 0.998555 | 6842 | 2 | -0.6113 | 0.88043 | 0.88048 | 1 | 19617 | 0 | -0.6113 |
| PSMD13 | 4 | 0.21865 | 0.35991 | 0.999994 | 8244 | 2 | -0.85593 | 0.88065 | 0.89 | 1 | 19618 | 1 | -0.85593 |
| PARP3 | 5 | 0.08601 | 0.20418 | 0.978186 | 4447 | 3 | -1.7396 | 0.88067 | 0.91585 | 1 | 19619 | 1 | -1.7396 |
| TKTL2 | 6 | 0.0058462 | 0.024221 | 0.837859 | 578 | 3 | -1.0098 | 0.88071 | 0.94162 | 1 | 19620 | 1 | -1.0098 |
| AGT | 4 | 0.12092 | 0.25191 | 0.994 | 5657 | 3 | -0.77158 | 0.88072 | 0.89004 | 1 | 19621 | 1 | -0.77158 |
| PFN2 | 6 | 0.19189 | 0.37787 | 0.999994 | 7543 | 1 | -0.050326 | 0.88073 | 0.94162 | 1 | 19622 | 1 | -0.050326 |
| MACROD2 | 4 | 0.7223 | 0.73967 | 0.999994 | 16520 | 1 | 0.16792 | 0.88079 | 0.89008 | 1 | 19623 | 1 | 0.16792 |
| C9orf50 | 5 | 0.57517 | 0.71409 | 0.999994 | 14620 | 2 | 0.11957 | 0.88082 | 0.91591 | 1 | 19624 | 1 | 0.11957 |
| POMGNT2 | 4 | 0.17319 | 0.31311 | 0.999994 | 7068 | 2 | -0.88826 | 0.88083 | 0.8901 | 1 | 19625 | 1 | -0.88826 |
| hsa-mir-6516 | 3 | 0.23574 | 0.35728 | 0.999994 | 8663 | 1 | -0.077701 | 0.88113 | 0.8812 | 1 | 19626 | 0 | -0.077701 |
| PPFIBP2 | 6 | 0.044875 | 0.13281 | 0.95318 | 2751 | 3 | -0.44117 | 0.88147 | 0.94176 | 1 | 19627 | 1 | -0.44117 |
| hsa-mir-148b | 4 | 0.30348 | 0.44653 | 0.999994 | 10203 | 2 | -0.56026 | 0.88152 | 0.89058 | 1 | 19628 | 1 | -0.56026 |
| ARHGAP25 | 6 | 0.003484 | 0.015515 | 0.763603 | 405 | 5 | -2.1678 | 0.882 | 0.94187 | 1 | 19629 | 1 | -2.1678 |
| HMGB2 | 6 | 0.36485 | 0.57998 | 0.999994 | 11418 | 2 | -0.13356 | 0.88205 | 0.94188 | 1 | 19630 | 1 | -0.13356 |
| ZNF648 | 5 | 0.0020763 | 0.0090759 | 0.680529 | 282 | 4 | -6.3019 | 0.88207 | 0.91639 | 1 | 19631 | 1 | -6.3019 |
| CERS4 | 5 | 0.51178 | 0.65821 | 0.999994 | 13607 | 2 | -0.87389 | 0.88208 | 0.91639 | 1 | 19632 | 1 | -0.87389 |
| IFIT5 | 6 | 0.019194 | 0.069383 | 0.93914 | 1418 | 3 | -2.476 | 0.88227 | 0.94193 | 1 | 19633 | 1 | -2.476 |
| GAB3 | 6 | 0.07094 | 0.18875 | 0.969543 | 3872 | 4 | -1.7523 | 0.88228 | 0.94193 | 1 | 19634 | 1 | -1.7523 |
| ZNF33A | 6 | 0.017023 | 0.062351 | 0.937368 | 1304 | 4 | -2.3491 | 0.88229 | 0.94193 | 1 | 19635 | 1 | -2.3491 |
| RND1 | 5 | 0.16815 | 0.32941 | 0.999994 | 6943 | 3 | -0.97082 | 0.88233 | 0.91649 | 1 | 19636 | 1 | -0.97082 |
| FUOM | 6 | 0.0024668 | 0.011726 | 0.727553 | 315 | 4 | -4.8541 | 0.88244 | 0.94197 | 1 | 19637 | 1 | -4.8541 |
| MTRNR2L5 | 4 | 0.10888 | 0.23087 | 0.983933 | 5267 | 3 | -1.662 | 0.88257 | 0.89131 | 1 | 19638 | 1 | -1.662 |
| PRSS58 | 3 | 0.091976 | 0.17239 | 0.967658 | 4678 | 2 | -2.0055 | 0.88269 | 0.88277 | 1 | 19639 | 0 | -2.0055 |
| FH | 6 | 0.0020341 | 0.010099 | 0.68773 | 277 | 5 | -4.3852 | 0.88277 | 0.94203 | 1 | 19640 | 1 | -4.3852 |
| PM20D2 | 6 | 0.053088 | 0.15075 | 0.966359 | 3119 | 4 | -1.2075 | 0.88315 | 0.9421 | 1 | 19641 | 1 | -1.2075 |
| hsa-mir-892b | 4 | 0.81879 | 0.81859 | 0.999994 | 17761 | 1 | 0.02672 | 0.88319 | 0.89174 | 1 | 19642 | 1 | 0.02672 |
| LRFN2 | 5 | 0.028847 | 0.087505 | 0.94395 | 1945 | 3 | -3.1706 | 0.88321 | 0.91684 | 1 | 19643 | 1 | -3.1706 |
| CLCN1 | 6 | 0.024929 | 0.086846 | 0.94395 | 1752 | 3 | -1.9663 | 0.88322 | 0.94211 | 1 | 19644 | 1 | -1.9663 |
| EIF3A | 3 | 0.045114 | 0.10891 | 0.951052 | 2763 | 2 | -3.1879 | 0.88325 | 0.88334 | 1 | 19645 | 0 | -3.1879 |
| hsa-mir-7847 | 3 | 0.7951 | 0.79496 | 0.999994 | 17418 | 0 | 0.31141 | 0.88327 | 0.88338 | 1 | 19646 | 0 | 0.31141 |
| AGTRAP | 6 | 0.017883 | 0.065132 | 0.93914 | 1356 | 4 | -1.5752 | 0.88329 | 0.94213 | 1 | 19647 | 1 | -1.5752 |
| NonTargetingControlG | 2 | 0.11662 | 0.19563 | 0.975797 | 5526 | 2 | -0.87928 | 0.88338 | 0.88334 | 1 | 19648 | 0 | -0.87928 |
| ZNF318 | 5 | 0.37626 | 0.5436 | 0.999994 | 11596 | 2 | 0.18083 | 0.88348 | 0.91693 | 1 | 19649 | 1 | 0.18083 |
| GADD45A | 4 | 0.342 | 0.48608 | 0.999994 | 10982 | 2 | -0.096142 | 0.88351 | 0.89196 | 1 | 19650 | 1 | -0.096142 |
| SPDYA | 5 | 0.041489 | 0.11962 | 0.951052 | 2590 | 3 | -2.7289 | 0.88363 | 0.91698 | 1 | 19651 | 1 | -2.7289 |
| NonTargetingControlG | 2 | 0.11632 | 0.19516 | 0.975797 | 5521 | 2 | -1.2505 | 0.88368 | 0.88363 | 1 | 19652 | 0 | -1.2505 |
| IL6 | 4 | 0.10175 | 0.21825 | 0.980699 | 5030 | 2 | -0.79024 | 0.88379 | 0.89214 | 1 | 19653 | 1 | -0.79024 |
| NonTargetingControlG | 2 | 0.116 | 0.19471 | 0.975797 | 5515 | 2 | -0.74195 | 0.884 | 0.88395 | 1 | 19654 | 0 | -0.74195 |
| ECE2 | 3 | 0.2269 | 0.34623 | 0.999994 | 8450 | 2 | -0.96263 | 0.88402 | 0.88412 | 1 | 19655 | 0 | -0.96263 |

| | | | | | | | | | | | | | |
|---|---|---|---|---|---|---|---|---|---|---|---|---|---|
| HMGN1 | 5 | 0.017267 | 0.056598 | 0.9235 | 1319 | 3 | -3.78 | 0.88416 | 0.91719 | 1 | 19656 | 1 | -3.78 |
| SOSTDC1 | 3 | 0.70166 | 0.70256 | 0.999994 | 16323 | 1 | -0.060383 | 0.88424 | 0.88435 | 1 | 19657 | 0 | -0.060383 |
| LRFN4 | 5 | 0.13118 | 0.27441 | 0.998561 | 5962 | 3 | -1.2898 | 0.88426 | 0.91723 | 1 | 19658 | 1 | -1.2898 |
| RELT | 5 | 0.042487 | 0.12207 | 0.951052 | 2642 | 4 | -2.5763 | 0.88428 | 0.91724 | 1 | 19659 | 1 | -2.5763 |
| CLUAP1 | 5 | 0.028313 | 0.086141 | 0.94395 | 1916 | 3 | -0.73531 | 0.88445 | 0.91732 | 1 | 19660 | 1 | -0.73531 |
| CPA4 | 5 | 0.20173 | 0.37772 | 0.999994 | 7816 | 3 | -0.8684 | 0.88447 | 0.91732 | 1 | 19661 | 1 | -0.8684 |
| PDE11A | 6 | 0.0033026 | 0.014818 | 0.755578 | 385 | 5 | -2.0819 | 0.88466 | 0.9424 | 1 | 19662 | 1 | -2.0819 |
| OR13G1 | 6 | 0.050199 | 0.14448 | 0.962688 | 2986 | 4 | -1.7515 | 0.88466 | 0.9424 | 1 | 19663 | 1 | -1.7515 |
| CFH | 5 | 0.01904 | 0.061478 | 0.936877 | 1410 | 3 | -1.2871 | 0.88468 | 0.9174 | 1 | 19664 | 1 | -1.2871 |
| XYLT1 | 5 | 0.01911 | 0.06167 | 0.937033 | 1415 | 4 | -1.4276 | 0.88474 | 0.91742 | 1 | 19665 | 1 | -1.4276 |
| NUTF2 | 3 | 0.03239 | 0.080622 | 0.93914 | 2128 | 2 | -4.3832 | 0.88481 | 0.88493 | 1 | 19666 | 0 | -4.3832 |
| GMEB2 | 5 | 0.037257 | 0.10898 | 0.951052 | 2385 | 4 | -1.347 | 0.88515 | 0.91758 | 1 | 19667 | 1 | -1.347 |
| ARID2 | 6 | 0.015736 | 0.058177 | 0.931817 | 1231 | 4 | -2.7307 | 0.88528 | 0.94254 | 1 | 19668 | 1 | -2.7307 |
| SGIP1 | 5 | 0.047862 | 0.13527 | 0.954328 | 2885 | 4 | -2.0198 | 0.88546 | 0.91771 | 1 | 19669 | 1 | -2.0198 |
| IGFBP2 | 3 | 0.023844 | 0.060797 | 0.932932 | 1702 | 2 | -4.749 | 0.88552 | 0.88561 | 1 | 19670 | 0 | -4.749 |
| SUN3 | 6 | 0.0049961 | 0.021136 | 0.812607 | 519 | 4 | -1.8991 | 0.88565 | 0.94261 | 1 | 19671 | 1 | -1.8991 |
| hsa-mir-548ba | 1 | 0.1141 | 0.11404 | 0.951052 | 5452 | 1 | -4.2671 | 0.8859 | 0.88622 | 1 | 19672 | 0 | -4.2671 |
| NonTargetingControlG | 1 | 0.11389 | 0.11385 | 0.951052 | 5443 | 1 | -4.1552 | 0.88611 | 0.88644 | 1 | 19673 | 0 | -4.1552 |
| NEDD1 | 5 | 0.19805 | 0.37248 | 0.999994 | 7711 | 3 | -0.85269 | 0.88618 | 0.91798 | 1 | 19674 | 1 | -0.85269 |
| ITGB6 | 6 | 0.083822 | 0.21512 | 0.979587 | 4370 | 4 | -0.99602 | 0.88623 | 0.94274 | 1 | 19675 | 1 | -0.99602 |
| hsa-mir-30b | 3 | 0.27916 | 0.41098 | 0.999994 | 9680 | 2 | -0.65696 | 0.88623 | 0.88629 | 1 | 19676 | 0 | -0.65696 |
| hsa-mir-8084 | 4 | 0.42585 | 0.57222 | 0.999994 | 12349 | 2 | -0.21459 | 0.88636 | 0.89393 | 1 | 19677 | 1 | -0.21459 |
| TSSC1 | 4 | 0.073543 | 0.1666 | 0.967126 | 3972 | 3 | -2.0215 | 0.88655 | 0.89406 | 1 | 19678 | 1 | -2.0215 |
| OR51A2 | 5 | 0.24246 | 0.43425 | 0.999994 | 8846 | 3 | -0.53748 | 0.88712 | 0.91838 | 1 | 19679 | 1 | -0.53748 |
| SHMT2 | 4 | 0.092181 | 0.201 | 0.977164 | 4686 | 3 | -1.2637 | 0.88714 | 0.89445 | 1 | 19680 | 1 | -1.2637 |
| ETS1 | 5 | 0.55849 | 0.69927 | 0.999994 | 14341 | 1 | 0.03185 | 0.88722 | 0.91841 | 1 | 19681 | 1 | 0.03185 |
| GRB10 | 6 | 0.13323 | 0.30444 | 0.999994 | 6002 | 3 | -0.68784 | 0.88724 | 0.94296 | 1 | 19682 | 1 | -0.68784 |
| NonTargetingControlG | 2 | 0.11269 | 0.18962 | 0.970473 | 5392 | 2 | -3.2219 | 0.88731 | 0.8872 | 1 | 19683 | 0 | -3.2219 |
| ACRC | 3 | 0.68061 | 0.68328 | 0.999994 | 16101 | 1 | -0.16062 | 0.88741 | 0.88749 | 1 | 19684 | 0 | -0.16062 |
| TMEM56 | 6 | 0.24514 | 0.44213 | 0.999994 | 8903 | 2 | -0.2077 | 0.88744 | 0.943 | 1 | 19685 | 1 | -0.2077 |
| SCFD2 | 4 | 0.38956 | 0.53493 | 0.999994 | 11789 | 2 | -0.6793 | 0.88747 | 0.89469 | 1 | 19686 | 1 | -0.6793 |
| NonTargetingControlG | 1 | 0.11239 | 0.11234 | 0.951052 | 5383 | 1 | -4.0246 | 0.88761 | 0.88792 | 1 | 19687 | 0 | -4.0246 |
| CPT1B | 5 | 0.0018229 | 0.008039 | 0.680529 | 254 | 4 | -3.9654 | 0.88768 | 0.9186 | 1 | 19688 | 1 | -3.9654 |
| hsa-mir-548aa-1 | 4 | 0.25452 | 0.39656 | 0.999994 | 9118 | 2 | -0.56348 | 0.88784 | 0.89494 | 1 | 19689 | 1 | -0.56348 |
| ZC3HAV1 | 4 | 0.074479 | 0.16836 | 0.967126 | 4013 | 2 | -0.98624 | 0.88786 | 0.89495 | 1 | 19690 | 1 | -0.98624 |
| hsa-mir-4490 | 4 | 0.3479 | 0.49213 | 0.999994 | 11116 | 2 | -0.7086 | 0.88792 | 0.89499 | 1 | 19691 | 1 | -0.7086 |
| WASF2 | 5 | 0.14598 | 0.29675 | 0.999994 | 6350 | 3 | -1.1347 | 0.88805 | 0.91874 | 1 | 19692 | 1 | -1.1347 |
| KCNJ18 | 3 | 0.1079 | 0.19347 | 0.975797 | 5236 | 2 | -1.9618 | 0.88807 | 0.88815 | 1 | 19693 | 0 | -1.9618 |
| AVIL | 6 | 0.31129 | 0.51984 | 0.999994 | 10345 | 2 | 0.18405 | 0.88812 | 0.94314 | 1 | 19694 | 1 | 0.18405 |
| CBS | 6 | 0.064857 | 0.17597 | 0.968171 | 3645 | 4 | -1.2496 | 0.88834 | 0.9432 | 1 | 19695 | 1 | -1.2496 |
| DLL3 | 6 | 0.016559 | 0.060826 | 0.932932 | 1280 | 5 | -2.2535 | 0.88868 | 0.94328 | 1 | 19696 | 1 | -2.2535 |
| hsa-mir-6835 | 2 | 0.11111 | 0.18714 | 0.969409 | 5336 | 2 | -2.9675 | 0.88889 | 0.88878 | 1 | 19697 | 0 | -2.9675 |
| UBE2QL1 | 4 | 0.077003 | 0.17316 | 0.96793 | 4106 | 2 | -1.008 | 0.88929 | 0.89595 | 1 | 19698 | 1 | -1.008 |
| hsa-mir-4666a | 3 | 0.52809 | 0.56505 | 0.999994 | 13861 | 1 | -0.43222 | 0.88933 | 0.88936 | 1 | 19699 | 0 | -0.43222 |
| POLE4 | 5 | 0.088315 | 0.20788 | 0.979108 | 4524 | 3 | -1.6727 | 0.88952 | 0.91932 | 1 | 19700 | 1 | -1.6727 |
| GRIA1 | 4 | 0.49452 | 0.61386 | 0.999994 | 13337 | 1 | 0.1427 | 0.88964 | 0.89619 | 1 | 19701 | 1 | 0.1427 |
| KIAA1967 | 6 | 0.023521 | 0.083234 | 0.94383 | 1688 | 4 | -2.4544 | 0.88979 | 0.94355 | 1 | 19702 | 1 | -2.4544 |
| DCBLD2 | 6 | 0.51011 | 0.67569 | 0.999994 | 13577 | 2 | -0.27393 | 0.89 | 0.94359 | 1 | 19703 | 1 | -0.27393 |
| NonTargetingControlG | 2 | 0.10989 | 0.18528 | 0.968626 | 5299 | 2 | -1.4821 | 0.89011 | 0.88996 | 1 | 19704 | 0 | -1.4821 |
| SYNE3 | 5 | 0.2351 | 0.42428 | 0.999994 | 8647 | 2 | -0.2867 | 0.8902 | 0.91959 | 1 | 19705 | 1 | -0.2867 |
| TMCC1 | 6 | 0.12205 | 0.28964 | 0.999632 | 5703 | 4 | -0.69478 | 0.8904 | 0.94367 | 1 | 19706 | 1 | -0.69478 |
| hsa-mir-10a | 4 | 0.009424 | 0.031901 | 0.865899 | 822 | 3 | -5.5106 | 0.89055 | 0.89685 | 1 | 19707 | 1 | -5.5106 |

| | | | | | | | | | | | | | |
|---|---|---|---|---|---|---|---|---|---|---|---|---|---|
| hsa-mir-194-2 | 4 | 0.29679 | 0.43972 | 0.999994 | 10062 | 2 | -0.48164 | 0.89058 | 0.89687 | 1 | 19708 | 1 | -0.48164 |
| TTLL4 | 5 | 0.28089 | 0.46558 | 0.999994 | 9717 | 2 | -0.17237 | 0.89073 | 0.91981 | 1 | 19709 | 1 | -0.17237 |
| hsa-mir-4264 | 2 | 0.10921 | 0.18425 | 0.968526 | 5276 | 2 | -1.0093 | 0.89079 | 0.89063 | 1 | 19710 | 0 | -1.0093 |
| NonTargetingControlG | 2 | 0.096298 | 0.16436 | 0.967126 | 4827 | 2 | -4.5373 | 0.89087 | 0.89072 | 1 | 19711 | 0 | -4.5373 |
| HAUS7 | 3 | 0.57657 | 0.59948 | 0.999994 | 14638 | 1 | -0.1805 | 0.89107 | 0.89112 | 1 | 19712 | 0 | -0.1805 |
| MRPL44 | 4 | 0.035019 | 0.091758 | 0.94395 | 2269 | 3 | -2.0489 | 0.89117 | 0.8973 | 1 | 19713 | 1 | -2.0489 |
| NAPB | 4 | 0.0053379 | 0.018771 | 0.803321 | 543 | 3 | -6.2201 | 0.89117 | 0.8973 | 1 | 19714 | 1 | -6.2201 |
| COX18 | 4 | 0.43495 | 0.58155 | 0.999994 | 12497 | 2 | -0.24818 | 0.89117 | 0.8973 | 1 | 19715 | 1 | -0.24818 |
| ZNF691 | 4 | 0.55455 | 0.64194 | 0.999994 | 14276 | 1 | -0.1672 | 0.89117 | 0.8973 | 1 | 19716 | 1 | -0.1672 |
| FBL | 4 | 0.10117 | 0.21722 | 0.980699 | 5014 | 3 | -1.3171 | 0.89117 | 0.8973 | 1 | 19717 | 1 | -1.3171 |
| PCDHGB7 | 2 | 0.10848 | 0.18313 | 0.968526 | 5255 | 2 | -1.0688 | 0.89152 | 0.89137 | 1 | 19718 | 0 | -1.0688 |
| DDX10 | 1 | 0.10838 | 0.10828 | 0.951052 | 5250 | 1 | -3.9833 | 0.89162 | 0.89195 | 1 | 19719 | 0 | -3.9833 |
| C9orf156 | 6 | 0.011562 | 0.044271 | 0.891621 | 958 | 5 | -2.762 | 0.89164 | 0.94395 | 1 | 19720 | 1 | -2.762 |
| LY86 | 3 | 0.036033 | 0.088849 | 0.94395 | 2319 | 2 | -2.1596 | 0.89202 | 0.89207 | 1 | 19721 | 0 | -2.1596 |
| TMEM109 | 5 | 0.77753 | 0.80602 | 0.999994 | 17177 | 1 | -0.18055 | 0.89218 | 0.92044 | 1 | 19722 | 1 | -0.18055 |
| ATP10D | 6 | 0.013643 | 0.051267 | 0.909241 | 1108 | 4 | -2.2065 | 0.8924 | 0.94413 | 1 | 19723 | 1 | -2.2065 |
| hsa-mir-3199-1 | 4 | 0.70794 | 0.72977 | 0.999994 | 16386 | 1 | -0.12868 | 0.89249 | 0.89826 | 1 | 19724 | 1 | -0.12868 |
| NonTargetingControlG | 2 | 0.10749 | 0.18164 | 0.968171 | 5226 | 2 | -1.5713 | 0.89251 | 0.89237 | 1 | 19725 | 0 | -1.5713 |
| SGCG | 4 | 0.049057 | 0.11957 | 0.951052 | 2935 | 2 | -2.5919 | 0.89255 | 0.89829 | 1 | 19726 | 1 | -2.5919 |
| RNPEP | 6 | 0.012239 | 0.046515 | 0.895138 | 1009 | 5 | -1.5121 | 0.89269 | 0.94419 | 1 | 19727 | 1 | -1.5121 |
| PHF11 | 6 | 0.068613 | 0.18388 | 0.968526 | 3777 | 4 | -1.6713 | 0.89294 | 0.94425 | 1 | 19728 | 1 | -1.6713 |
| NonTargetingControlG | 2 | 0.10674 | 0.18044 | 0.968171 | 5201 | 2 | -1.3448 | 0.89326 | 0.89314 | 1 | 19729 | 0 | -1.3448 |
| NonTargetingControlG | 2 | 0.1067 | 0.18039 | 0.968171 | 5197 | 2 | -1.2422 | 0.8933 | 0.89317 | 1 | 19730 | 0 | -1.2422 |
| hsa-mir-492 | 4 | 0.05567 | 0.13251 | 0.95318 | 3229 | 2 | -3.2458 | 0.89345 | 0.89893 | 1 | 19731 | 1 | -3.2458 |
| GLMN | 5 | 0.46225 | 0.61553 | 0.999994 | 12893 | 1 | -0.31918 | 0.89348 | 0.92098 | 1 | 19732 | 1 | -0.31918 |
| ARPP21 | 5 | 0.068387 | 0.17581 | 0.968171 | 3764 | 3 | -1.6747 | 0.89352 | 0.921 | 1 | 19733 | 1 | -1.6747 |
| PLS1 | 6 | 0.094472 | 0.23646 | 0.985449 | 4766 | 4 | -1.1902 | 0.89363 | 0.9444 | 1 | 19734 | 1 | -1.1902 |
| hsa-mir-3683 | 3 | 0.091915 | 0.17232 | 0.967658 | 4675 | 1 | 0.21045 | 0.89369 | 0.89374 | 1 | 19735 | 0 | 0.21045 |
| CACNA1H | 6 | 0.23501 | 0.43008 | 0.999994 | 8643 | 3 | -0.41832 | 0.89384 | 0.94445 | 1 | 19736 | 1 | -0.41832 |
| hsa-mir-934 | 3 | 0.095914 | 0.17763 | 0.968171 | 4810 | 1 | -0.20142 | 0.8941 | 0.89414 | 1 | 19737 | 0 | -0.20142 |
| MTF1 | 6 | 0.0041224 | 0.017922 | 0.795727 | 450 | 5 | -2.7331 | 0.89417 | 0.94453 | 1 | 19738 | 1 | -2.7331 |
| NIPAL1 | 6 | 0.0096563 | 0.037821 | 0.885421 | 852 | 5 | -2.945 | 0.89421 | 0.94454 | 1 | 19739 | 1 | -2.945 |
| PHLDB2 | 6 | 0.0018734 | 0.0093393 | 0.680529 | 256 | 5 | -3.1607 | 0.89432 | 0.94456 | 1 | 19740 | 1 | -3.1607 |
| hsa-mir-4691 | 3 | 0.73146 | 0.73137 | 0.999994 | 16620 | 1 | 0.15392 | 0.8944 | 0.89445 | 1 | 19741 | 0 | 0.15392 |
| SBNO1 | 5 | 0.0060397 | 0.024538 | 0.842094 | 596 | 4 | -5.9023 | 0.89444 | 0.92138 | 1 | 19742 | 1 | -5.9023 |
| IQCF1 | 5 | 0.08647 | 0.20492 | 0.978186 | 4468 | 3 | -1.4722 | 0.89455 | 0.92142 | 1 | 19743 | 1 | -1.4722 |
| KRTAP20-2 | 6 | 0.038345 | 0.11818 | 0.951052 | 2434 | 4 | -1.5937 | 0.89496 | 0.94471 | 1 | 19744 | 1 | -1.5937 |
| BMP5 | 5 | 0.55989 | 0.70055 | 0.999994 | 14369 | 2 | 0.16134 | 0.89517 | 0.92168 | 1 | 19745 | 1 | 0.16134 |
| JAK3 | 4 | 0.19177 | 0.33222 | 0.999994 | 7538 | 2 | -0.79636 | 0.89533 | 0.90034 | 1 | 19746 | 1 | -0.79636 |
| HOXC11 | 4 | 0.035434 | 0.092612 | 0.943968 | 2289 | 3 | -4.126 | 0.89534 | 0.90034 | 1 | 19747 | 1 | -4.126 |
| TMEM219 | 4 | 0.029565 | 0.080637 | 0.93914 | 1988 | 3 | -2.3543 | 0.89554 | 0.90047 | 1 | 19748 | 1 | -2.3543 |
| STAT2 | 6 | 0.60487 | 0.7436 | 0.999994 | 15008 | 2 | -0.54309 | 0.89561 | 0.94487 | 1 | 19749 | 1 | -0.54309 |
| hsa-mir-8086 | 1 | 0.10411 | 0.10403 | 0.947284 | 5100 | 1 | -5.9939 | 0.89589 | 0.89623 | 1 | 19750 | 0 | -5.9939 |
| ABHD11 | 5 | 0.01948 | 0.062692 | 0.937389 | 1428 | 4 | -1.7357 | 0.8959 | 0.922 | 1 | 19751 | 1 | -1.7357 |
| STEAP1 | 6 | 0.90303 | 0.90603 | 0.999994 | 19280 | 1 | -0.24167 | 0.89606 | 0.94497 | 1 | 19752 | 1 | -0.24167 |
| hsa-mir-3195 | 2 | 0.054121 | 0.096086 | 0.945031 | 3163 | 2 | -5.027 | 0.89615 | 0.89606 | 1 | 19753 | 0 | -5.027 |
| SSTR5 | 6 | 0.64639 | 0.77487 | 0.999994 | 15605 | 2 | 0.2468 | 0.89622 | 0.945 | 1 | 19754 | 1 | 0.2468 |
| NonTargetingControlG | 2 | 0.013334 | 0.025102 | 0.844082 | 1086 | 2 | -6.4042 | 0.89641 | 0.89633 | 1 | 19755 | 0 | -6.4042 |
| NonTargetingControlG | 1 | 0.10342 | 0.10335 | 0.946414 | 5080 | 1 | -6.0232 | 0.89658 | 0.89691 | 1 | 19756 | 0 | -6.0232 |
| PABPC1L | 5 | 0.19274 | 0.36485 | 0.999994 | 7566 | 3 | -0.84458 | 0.8967 | 0.92232 | 1 | 19757 | 1 | -0.84458 |
| hsa-mir-5680 | 3 | 0.1738 | 0.27926 | 0.999632 | 7079 | 2 | -1.3583 | 0.89673 | 0.89684 | 1 | 19758 | 0 | -1.3583 |
| EXOC4 | 6 | 0.065453 | 0.17722 | 0.968171 | 3671 | 3 | -1.1927 | 0.89673 | 0.94513 | 1 | 19759 | 1 | -1.1927 |

| | | | | | | | | | | | | | |
|---|---|---|---|---|---|---|---|---|---|---|---|---|---|
| SLC14A1 | 6 | 0.0095278 | 0.037378 | 0.881473 | 836 | 4 | -3.7376 | 0.89685 | 0.94515 | 1 | 19760 | 1 | -3.7376 |
| hsa-mir-325 | 4 | 0.74107 | 0.75328 | 0.999994 | 16753 | 1 | 0.1424 | 0.89701 | 0.90156 | 1 | 19761 | 1 | 0.1424 |
| SEPHS2 | 3 | 0.15756 | 0.25828 | 0.994943 | 6662 | 2 | -1.4526 | 0.89702 | 0.89712 | 1 | 19762 | 0 | -1.4526 |
| HLA-DPB1 | 3 | 0.738 | 0.73793 | 0.999994 | 16704 | 0 | 0.077977 | 0.89706 | 0.89716 | 1 | 19763 | 0 | 0.077977 |
| FAM115A | 5 | 0.033402 | 0.099242 | 0.945253 | 2182 | 4 | -2.7447 | 0.89715 | 0.92251 | 1 | 19764 | 1 | -2.7447 |
| hsa-mir-6730 | 3 | 0.14681 | 0.2444 | 0.988879 | 6370 | 2 | -1.7576 | 0.89743 | 0.89751 | 1 | 19765 | 0 | -1.7576 |
| HMGXB4 | 5 | 0.24411 | 0.43599 | 0.999994 | 8886 | 2 | -0.40769 | 0.8975 | 0.92264 | 1 | 19766 | 1 | -0.40769 |
| SDHB | 5 | 0.033835 | 0.10032 | 0.946394 | 2208 | 4 | -3.3431 | 0.89752 | 0.92265 | 1 | 19767 | 1 | -3.3431 |
| GEN1 | 3 | 0.077208 | 0.15268 | 0.967126 | 4117 | 1 | -0.42512 | 0.89753 | 0.8976 | 1 | 19768 | 0 | -0.42512 |
| hsa-mir-4787 | 3 | 0.75409 | 0.75403 | 0.999994 | 16898 | 0 | 0.060671 | 0.89754 | 0.89761 | 1 | 19769 | 0 | 0.060671 |
| IGJ | 5 | 0.17554 | 0.34021 | 0.999994 | 7126 | 2 | 0.18049 | 0.89763 | 0.9227 | 1 | 19770 | 1 | 0.18049 |
| MRPS22 | 5 | 0.1138 | 0.24788 | 0.990682 | 5438 | 2 | -0.22421 | 0.89767 | 0.92272 | 1 | 19771 | 1 | -0.22421 |
| NonTargetingControlG | 2 | 0.10229 | 0.17367 | 0.96793 | 5050 | 2 | -3.3438 | 0.89771 | 0.89761 | 1 | 19772 | 0 | -3.3438 |
| ANG | 5 | 0.049828 | 0.13992 | 0.958716 | 2966 | 4 | -1.0724 | 0.89773 | 0.92275 | 1 | 19773 | 1 | -1.0724 |
| ZHX2 | 5 | 0.044988 | 0.12828 | 0.951777 | 2756 | 4 | -1.2212 | 0.89779 | 0.92277 | 1 | 19774 | 1 | -1.2212 |
| hsa-mir-648 | 3 | 0.2475 | 0.37182 | 0.999994 | 8961 | 2 | -0.83932 | 0.89779 | 0.89788 | 1 | 19775 | 0 | -0.83932 |
| HOXD4 | 5 | 0.10486 | 0.234 | 0.984971 | 5127 | 2 | 0.15229 | 0.89796 | 0.92285 | 1 | 19776 | 1 | 0.15229 |
| DUSP1 | 3 | 0.7962 | 0.79605 | 0.999994 | 17430 | 0 | 0.070564 | 0.89813 | 0.89823 | 1 | 19777 | 0 | 0.070564 |
| SH2B2 | 5 | 0.64292 | 0.74232 | 0.999994 | 15561 | 1 | -0.11855 | 0.89829 | 0.92298 | 1 | 19778 | 1 | -0.11855 |
| hsa-mir-339 | 4 | 0.098586 | 0.21253 | 0.979587 | 4914 | 3 | -1.2879 | 0.89831 | 0.90253 | 1 | 19779 | 1 | -1.2879 |
| LTF | 5 | 0.25587 | 0.44542 | 0.999994 | 9146 | 2 | -0.30342 | 0.8988 | 0.92319 | 1 | 19780 | 1 | -0.30342 |
| PSMC2 | 3 | 0.010307 | 0.027568 | 0.855011 | 890 | 2 | -7.8017 | 0.89885 | 0.89898 | 1 | 19781 | 0 | -7.8017 |
| MDFIC | 5 | 0.054529 | 0.15134 | 0.967126 | 3177 | 4 | -0.93482 | 0.8989 | 0.92323 | 1 | 19782 | 1 | -0.93482 |
| RNF180 | 4 | 0.35694 | 0.50137 | 0.999994 | 11286 | 2 | -0.59694 | 0.8989 | 0.90296 | 1 | 19783 | 1 | -0.59694 |
| hsa-mir-548u | 3 | 0.22255 | 0.34081 | 0.999994 | 8346 | 2 | -1.0948 | 0.89896 | 0.89908 | 1 | 19784 | 0 | -1.0948 |
| GYG1 | 5 | 0.070126 | 0.17858 | 0.968171 | 3836 | 2 | -0.25518 | 0.8991 | 0.92333 | 1 | 19785 | 1 | -0.25518 |
| ANKK1 | 4 | 0.1102 | 0.23321 | 0.984913 | 5310 | 2 | -0.96142 | 0.89944 | 0.90337 | 1 | 19786 | 1 | -0.96142 |
| OR51F1 | 5 | 0.057785 | 0.1584 | 0.967126 | 3326 | 3 | -2.0059 | 0.89964 | 0.92357 | 1 | 19787 | 1 | -2.0059 |
| hsa-mir-24-1 | 4 | 0.089528 | 0.19615 | 0.975797 | 4580 | 3 | -0.83561 | 0.89968 | 0.90355 | 1 | 19788 | 1 | -0.83561 |
| MAEA | 4 | 0.53117 | 0.63064 | 0.999994 | 13914 | 1 | -0.4771 | 0.8997 | 0.90357 | 1 | 19789 | 1 | -0.4771 |
| RBMY1J | 2 | 0.034424 | 0.062591 | 0.937389 | 2234 | 2 | -5.3431 | 0.89984 | 0.89976 | 1 | 19790 | 0 | -5.3431 |
| CEP57 | 5 | 0.039062 | 0.11353 | 0.951052 | 2465 | 4 | -0.89279 | 0.90004 | 0.92374 | 1 | 19791 | 1 | -0.89279 |
| TMEM105 | 5 | 0.013107 | 0.044884 | 0.891621 | 1070 | 3 | -3.6163 | 0.90021 | 0.92381 | 1 | 19792 | 1 | -3.6163 |
| hsa-mir-6825 | 3 | 0.6255 | 0.63706 | 0.999994 | 15315 | 1 | -0.38049 | 0.90024 | 0.90031 | 1 | 19793 | 0 | -0.38049 |
| NHP2L1 | 2 | 0.099589 | 0.16943 | 0.967126 | 4961 | 2 | -4.0101 | 0.90041 | 0.90032 | 1 | 19794 | 0 | -4.0101 |
| hsa-mir-6078 | 4 | 0.80222 | 0.80317 | 0.999994 | 17516 | 1 | -0.2161 | 0.90062 | 0.90423 | 1 | 19795 | 1 | -0.2161 |
| hsa-mir-95 | 4 | 0.16776 | 0.30745 | 0.999994 | 6932 | 2 | -1.367 | 0.90063 | 0.90424 | 1 | 19796 | 1 | -1.367 |
| hsa-mir-3156-2 | 1 | 0.099348 | 0.099343 | 0.945253 | 4944 | 1 | -6.1874 | 0.90065 | 0.90101 | 1 | 19797 | 0 | -6.1874 |
| PPP3CB | 6 | 0.041174 | 0.12454 | 0.951052 | 2569 | 4 | -2.2804 | 0.90068 | 0.94608 | 1 | 19798 | 1 | -2.2804 |
| PLBD1 | 5 | 0.2798 | 0.46471 | 0.999994 | 9697 | 2 | -0.20659 | 0.9008 | 0.92407 | 1 | 19799 | 1 | -0.20659 |
| MOS | 4 | 0.021513 | 0.063811 | 0.937389 | 1551 | 2 | -4.0033 | 0.90084 | 0.90441 | 1 | 19800 | 1 | -4.0033 |
| TAF6L | 4 | 0.027502 | 0.076408 | 0.93914 | 1877 | 3 | -4.7118 | 0.90087 | 0.90444 | 1 | 19801 | 1 | -4.7118 |
| NonTargetingControlG | 2 | 0.099126 | 0.16871 | 0.967126 | 4935 | 2 | -1.6443 | 0.90087 | 0.90082 | 1 | 19802 | 0 | -1.6443 |
| hsa-mir-1268b | 3 | 0.053972 | 0.1212 | 0.951052 | 3156 | 2 | -1.1298 | 0.90092 | 0.90099 | 1 | 19803 | 0 | -1.1298 |
| MRPL30 | 4 | 0.099489 | 0.21418 | 0.979587 | 4950 | 3 | -1.1946 | 0.90105 | 0.90458 | 1 | 19804 | 1 | -1.1946 |
| NonTargetingControlG | 2 | 0.098871 | 0.16833 | 0.967126 | 4926 | 2 | -2.8134 | 0.90113 | 0.90107 | 1 | 19805 | 0 | -2.8134 |
| REG3G | 6 | 0.024165 | 0.085029 | 0.94395 | 1716 | 4 | -2.0343 | 0.90133 | 0.94623 | 1 | 19806 | 1 | -2.0343 |
| KRTAP2-3 | 1 | 0.098627 | 0.098606 | 0.945253 | 4918 | 1 | -6.213 | 0.90137 | 0.90174 | 1 | 19807 | 0 | -6.213 |
| SLA2 | 4 | 0.029641 | 0.080791 | 0.93914 | 1990 | 3 | -2.6422 | 0.90139 | 0.90483 | 1 | 19808 | 1 | -2.6422 |
| hsa-mir-1273c | 4 | 0.4077 | 0.55361 | 0.999994 | 12071 | 2 | -0.92738 | 0.90143 | 0.90487 | 1 | 19809 | 1 | -0.92738 |
| ATAD5 | 5 | 0.020181 | 0.064572 | 0.93914 | 1470 | 4 | -1.6087 | 0.90157 | 0.9244 | 1 | 19810 | 1 | -1.6087 |
| NonTargetingControlG | 2 | 0.098326 | 0.16751 | 0.967126 | 4907 | 2 | -0.94128 | 0.90167 | 0.90159 | 1 | 19811 | 0 | -0.94128 |

| | | | | | | | | | | | | | |
|---|---|---|---|---|---|---|---|---|---|---|---|---|---|
| NonTargetingControlG | 2 | 0.098125 | 0.1672 | 0.967126 | 4898 | 2 | -1.5657 | 0.90188 | 0.90178 | 1 | 19812 | 0 | -1.5657 |
| LOC100996485 | 6 | 0.073621 | 0.19428 | 0.975797 | 3976 | 4 | -1.0432 | 0.90191 | 0.94638 | 1 | 19813 | 1 | -1.0432 |
| CITED4 | 6 | 0.04284 | 0.12832 | 0.951777 | 2667 | 3 | -1.8938 | 0.90191 | 0.94638 | 1 | 19814 | 1 | -1.8938 |
| FGD5 | 5 | 0.061817 | 0.16508 | 0.967126 | 3504 | 3 | -1.8833 | 0.90192 | 0.92456 | 1 | 19815 | 1 | -1.8833 |
| RBKS | 4 | 0.092396 | 0.20139 | 0.977164 | 4697 | 3 | -1.3769 | 0.90202 | 0.9053 | 1 | 19816 | 1 | -1.3769 |
| PREX2 | 5 | 0.21134 | 0.39127 | 0.999994 | 8055 | 3 | -0.71036 | 0.90203 | 0.92461 | 1 | 19817 | 1 | -0.71036 |
| ZNF286B | 5 | 0.0090174 | 0.033214 | 0.867348 | 795 | 4 | -6.08 | 0.90209 | 0.92464 | 1 | 19818 | 1 | -6.08 |
| B3GALT5 | 5 | 0.21524 | 0.39665 | 0.999994 | 8148 | 3 | -0.6849 | 0.90225 | 0.9247 | 1 | 19819 | 1 | -0.6849 |
| ZIC4 | 5 | 0.43188 | 0.58999 | 0.999994 | 12444 | 2 | -0.23869 | 0.90226 | 0.92471 | 1 | 19820 | 1 | -0.23869 |
| MYBPHL | 4 | 0.096993 | 0.20971 | 0.979587 | 4858 | 2 | -0.65975 | 0.90233 | 0.90553 | 1 | 19821 | 1 | -0.65975 |
| TMF1 | 4 | 0.13119 | 0.26961 | 0.998561 | 5963 | 3 | -1.1458 | 0.90239 | 0.90558 | 1 | 19822 | 1 | -1.1458 |
| ABCA3 | 6 | 0.083203 | 0.21386 | 0.979587 | 4349 | 3 | -0.65048 | 0.90241 | 0.9465 | 1 | 19823 | 1 | -0.65048 |
| GPR161 | 5 | 0.090052 | 0.21067 | 0.979587 | 4595 | 3 | -1.7403 | 0.90242 | 0.92478 | 1 | 19824 | 1 | -1.7403 |
| NonTargetingControlG | 1 | 0.097571 | 0.097539 | 0.945253 | 4873 | 1 | -4.8461 | 0.90243 | 0.90279 | 1 | 19825 | 0 | -4.8461 |
| hsa-mir-3170 | 4 | 0.16189 | 0.30143 | 0.999994 | 6787 | 2 | -1.0923 | 0.90246 | 0.90564 | 1 | 19826 | 1 | -1.0923 |
| C1orf116 | 5 | 0.057556 | 0.15802 | 0.967126 | 3318 | 2 | -0.14138 | 0.90247 | 0.9248 | 1 | 19827 | 1 | -0.14138 |
| RGS22 | 5 | 0.039179 | 0.11384 | 0.951052 | 2474 | 4 | -1.5009 | 0.9025 | 0.92481 | 1 | 19828 | 1 | -1.5009 |
| COL7A1 | 5 | 0.32018 | 0.49746 | 0.999994 | 10514 | 2 | -0.072768 | 0.9026 | 0.92485 | 1 | 19829 | 1 | -0.072768 |
| SLC9C2 | 5 | 0.03236 | 0.096514 | 0.945031 | 2124 | 4 | -1.8439 | 0.90264 | 0.92487 | 1 | 19830 | 1 | -1.8439 |
| hsa-mir-548at | 2 | 0.097253 | 0.16581 | 0.967126 | 4869 | 2 | -1.3878 | 0.90275 | 0.90266 | 1 | 19831 | 0 | -1.3878 |
| ENKUR | 6 | 0.30954 | 0.51782 | 0.999994 | 10317 | 3 | -0.53882 | 0.90281 | 0.9466 | 1 | 19832 | 1 | -0.53882 |
| IL17D | 5 | 0.051349 | 0.14366 | 0.961453 | 3041 | 3 | -2.4935 | 0.90285 | 0.92498 | 1 | 19833 | 1 | -2.4935 |
| EMID1 | 4 | 0.22611 | 0.36748 | 0.999994 | 8428 | 2 | -1.0805 | 0.90288 | 0.90601 | 1 | 19834 | 1 | -1.0805 |
| C9orf173 | 2 | 0.096918 | 0.1653 | 0.967126 | 4854 | 2 | -1.6208 | 0.90308 | 0.90301 | 1 | 19835 | 0 | -1.6208 |
| hsa-mir-3146 | 3 | 0.55297 | 0.58238 | 0.999994 | 14258 | 1 | -0.44123 | 0.90313 | 0.90315 | 1 | 19836 | 0 | -0.44123 |
| LOC100287399 | 6 | 0.34085 | 0.55362 | 0.999994 | 10958 | 3 | -0.63059 | 0.90314 | 0.94668 | 1 | 19837 | 1 | -0.63059 |
| IL17RC | 4 | 0.082054 | 0.1825 | 0.968526 | 4299 | 2 | -1.7572 | 0.90321 | 0.90625 | 1 | 19838 | 1 | -1.7572 |
| MUC2 | 5 | 0.89354 | 0.89368 | 0.999994 | 19096 | 0 | -0.36845 | 0.90322 | 0.92512 | 1 | 19839 | 1 | -0.36845 |
| ICAM3 | 5 | 0.09347 | 0.21613 | 0.980699 | 4738 | 2 | -0.42586 | 0.90332 | 0.92517 | 1 | 19840 | 1 | -0.42586 |
| hsa-mir-5590 | 2 | 0.096654 | 0.1649 | 0.967126 | 4843 | 2 | -1.0651 | 0.90335 | 0.90328 | 1 | 19841 | 0 | -1.0651 |
| CEBPZ | 3 | 0.24537 | 0.36924 | 0.999994 | 8909 | 2 | -0.89549 | 0.90335 | 0.90338 | 1 | 19842 | 0 | -0.89549 |
| PUF60 | 2 | 0.066988 | 0.11747 | 0.951052 | 3720 | 2 | -4.9435 | 0.90336 | 0.90328 | 1 | 19843 | 0 | -4.9435 |
| LIN7A | 5 | 0.083675 | 0.20051 | 0.977164 | 4364 | 1 | -0.20713 | 0.90339 | 0.9252 | 1 | 19844 | 1 | -0.20713 |
| NonTargetingControlG | 2 | 0.096507 | 0.16466 | 0.967126 | 4838 | 2 | -1.1134 | 0.90349 | 0.90343 | 1 | 19845 | 0 | -1.1134 |
| NonTargetingControlG | 2 | 0.096443 | 0.16457 | 0.967126 | 4834 | 2 | -0.99071 | 0.90356 | 0.90349 | 1 | 19846 | 0 | -0.99071 |
| ALDH1L1 | 6 | 0.025997 | 0.089351 | 0.94395 | 1806 | 4 | -2.4221 | 0.90357 | 0.94679 | 1 | 19847 | 1 | -2.4221 |
| NonTargetingControlG | 2 | 0.096422 | 0.16454 | 0.967126 | 4832 | 2 | -1.2221 | 0.90358 | 0.90352 | 1 | 19848 | 0 | -1.2221 |
| TM7SF3 | 6 | 0.2116 | 0.40202 | 0.999994 | 8059 | 3 | -0.64197 | 0.90394 | 0.94688 | 1 | 19849 | 1 | -0.64197 |
| NonTargetingControlG | 2 | 0.09601 | 0.16389 | 0.967126 | 4817 | 2 | -1.6249 | 0.90399 | 0.90392 | 1 | 19850 | 0 | -1.6249 |
| TRIM42 | 6 | 0.013523 | 0.050864 | 0.909101 | 1102 | 5 | -1.0786 | 0.90399 | 0.94689 | 1 | 19851 | 1 | -1.0786 |
| PLEKHG4B | 5 | 0.22402 | 0.40887 | 0.999994 | 8380 | 3 | -0.69749 | 0.90419 | 0.92557 | 1 | 19852 | 1 | -0.69749 |
| hsa-mir-6728 | 4 | 0.38371 | 0.52894 | 0.999994 | 11699 | 2 | -0.63459 | 0.90422 | 0.90706 | 1 | 19853 | 1 | -0.63459 |
| MAGEB6 | 6 | 0.27032 | 0.47211 | 0.999994 | 9484 | 3 | -0.68116 | 0.90427 | 0.94695 | 1 | 19854 | 1 | -0.68116 |
| ASIC4 | 6 | 0.030576 | 0.10013 | 0.946394 | 2034 | 3 | -1.5085 | 0.90428 | 0.94696 | 1 | 19855 | 1 | -1.5085 |
| hsa-mir-4458 | 4 | 0.10505 | 0.22415 | 0.983354 | 5135 | 3 | -1.8277 | 0.90439 | 0.90719 | 1 | 19856 | 1 | -1.8277 |
| FAM76B | 4 | 0.015569 | 0.05083 | 0.909101 | 1221 | 3 | -3.5004 | 0.90455 | 0.90733 | 1 | 19857 | 1 | -3.5004 |
| NRIP2 | 4 | 0.0079005 | 0.027069 | 0.848412 | 723 | 3 | -3.6655 | 0.90463 | 0.90739 | 1 | 19858 | 1 | -3.6655 |
| PLXDC2 | 5 | 0.15368 | 0.30828 | 0.999994 | 6554 | 3 | -1.1706 | 0.90468 | 0.92578 | 1 | 19859 | 1 | -1.1706 |
| C9orf3 | 4 | 0.77007 | 0.77584 | 0.999994 | 17082 | 1 | -0.10416 | 0.90481 | 0.90753 | 1 | 19860 | 1 | -0.10416 |
| hsa-mir-499a | 4 | 0.10954 | 0.23206 | 0.984502 | 5290 | 3 | -0.99043 | 0.90484 | 0.90755 | 1 | 19861 | 1 | -0.99043 |
| NPTX1 | 4 | 0.09982 | 0.21479 | 0.979587 | 4971 | 3 | -1.6376 | 0.90501 | 0.90768 | 1 | 19862 | 1 | -1.6376 |
| SP100 | 6 | 0.0011406 | 0.0057602 | 0.623123 | 182 | 5 | -2.5598 | 0.9052 | 0.94718 | 1 | 19863 | 1 | -2.5598 |

| | | | | | | | | | | | | | |
|---|---|---|---|---|---|---|---|---|---|---|---|---|---|
| MBLAC2 | 5 | 0.14576 | 0.29642 | 0.999994 | 6344 | 3 | -1.1193 | 0.90545 | 0.92614 | 1 | 19864 | 1 | -1.1193 |
| IFI27L1 | 5 | 0.67863 | 0.75697 | 0.999994 | 16070 | 1 | -0.25631 | 0.90555 | 0.92619 | 1 | 19865 | 1 | -0.25631 |
| NRN1L | 6 | 0.02018 | 0.072553 | 0.93914 | 1469 | 4 | -2.6092 | 0.90572 | 0.94731 | 1 | 19866 | 1 | -2.6092 |
| EPPIN | 6 | 0.022563 | 0.080179 | 0.93914 | 1622 | 5 | -1.8185 | 0.90573 | 0.94732 | 1 | 19867 | 1 | -1.8185 |
| MUSTN1 | 3 | 0.13796 | 0.23292 | 0.984502 | 6132 | 2 | -1.6801 | 0.90575 | 0.90581 | 1 | 19868 | 0 | -1.6801 |
| hsa-mir-598 | 3 | 0.16218 | 0.26431 | 0.996734 | 6798 | 2 | -1.4475 | 0.906 | 0.90607 | 1 | 19869 | 0 | -1.4475 |
| LOXHD1 | 6 | 0.037748 | 0.11682 | 0.951052 | 2417 | 4 | -1.6532 | 0.90609 | 0.94741 | 1 | 19870 | 1 | -1.6532 |
| KRTAP22-2 | 6 | 0.019831 | 0.071445 | 0.93914 | 1449 | 4 | -1.8303 | 0.90616 | 0.94742 | 1 | 19871 | 1 | -1.8303 |
| NonTargetingControlG | 1 | 0.093739 | 0.09369 | 0.945031 | 4743 | 1 | -6.4254 | 0.90626 | 0.9066 | 1 | 19872 | 0 | -6.4254 |
| C10orf90 | 5 | 0.22539 | 0.41083 | 0.999994 | 8409 | 3 | -0.70493 | 0.90637 | 0.92656 | 1 | 19873 | 1 | -0.70493 |
| FAM209B | 5 | 0.047151 | 0.13351 | 0.95318 | 2858 | 4 | -2.0878 | 0.90644 | 0.92659 | 1 | 19874 | 1 | -2.0878 |
| CECR1 | 6 | 0.17237 | 0.35365 | 0.999994 | 7046 | 3 | -0.88298 | 0.90654 | 0.94752 | 1 | 19875 | 1 | -0.88298 |
| WDR12 | 4 | 0.243 | 0.38472 | 0.999994 | 8863 | 2 | -0.62609 | 0.90657 | 0.90895 | 1 | 19876 | 1 | -0.62609 |
| hsa-mir-6752 | 2 | 0.093318 | 0.15969 | 0.967126 | 4732 | 2 | -3.9741 | 0.90668 | 0.90668 | 1 | 19877 | 0 | -3.9741 |
| FCER2 | 5 | 0.000047026 | 0.00021466 | 0.177301 | 25 | 4 | -2.3015 | 0.90673 | 0.92672 | 1 | 19878 | 1 | -2.3015 |
| SLC22A31 | 5 | 0.50518 | 0.65246 | 0.999994 | 13514 | 2 | -0.52758 | 0.90687 | 0.92678 | 1 | 19879 | 1 | -0.52758 |
| LAMA1 | 6 | 0.025111 | 0.087259 | 0.94395 | 1759 | 4 | -2.1144 | 0.90689 | 0.9476 | 1 | 19880 | 1 | -2.1144 |
| hsa-mir-7154 | 4 | 0.14171 | 0.28066 | 0.999632 | 6234 | 2 | -1.1001 | 0.90693 | 0.90926 | 1 | 19881 | 1 | -1.1001 |
| TIMELESS | 5 | 0.04155 | 0.11978 | 0.951052 | 2593 | 4 | -1.1051 | 0.90719 | 0.92692 | 1 | 19882 | 1 | -1.1051 |
| SNRPD2 | 3 | 0.30455 | 0.43095 | 0.999994 | 10217 | 1 | -0.25623 | 0.90728 | 0.90731 | 1 | 19883 | 0 | -0.25623 |
| SDR42E1 | 5 | 0.92131 | 0.92149 | 0.999994 | 19662 | 0 | 0.51989 | 0.90731 | 0.92698 | 1 | 19884 | 1 | 0.51989 |
| CLEC1A | 6 | 0.22401 | 0.41694 | 0.999994 | 8379 | 2 | -0.19795 | 0.90736 | 0.94774 | 1 | 19885 | 1 | -0.19795 |
| NonTargetingControlG | 2 | 0.092571 | 0.15854 | 0.967126 | 4707 | 2 | -2.1121 | 0.90743 | 0.90741 | 1 | 19886 | 0 | -2.1121 |
| CDK5R2 | 5 | 0.034533 | 0.10209 | 0.946414 | 2243 | 4 | -1.6772 | 0.9075 | 0.92707 | 1 | 19887 | 1 | -1.6772 |
| HIST1H1A | 6 | 0.044722 | 0.13248 | 0.95318 | 2748 | 4 | -1.6086 | 0.90751 | 0.94777 | 1 | 19888 | 1 | -1.6086 |
| CFI | 4 | 0.016527 | 0.053208 | 0.916684 | 1277 | 3 | -4.3465 | 0.90751 | 0.90974 | 1 | 19889 | 1 | -4.3465 |
| RSPH1 | 6 | 0.20629 | 0.39554 | 0.999994 | 7940 | 3 | -0.53827 | 0.90784 | 0.94786 | 1 | 19890 | 1 | -0.53827 |
| TTC18 | 5 | 0.10027 | 0.22684 | 0.983354 | 4981 | 3 | -1.5277 | 0.90785 | 0.92724 | 1 | 19891 | 1 | -1.5277 |
| FAM72A | 6 | 0.061034 | 0.16798 | 0.967126 | 3470 | 4 | -1.2865 | 0.90802 | 0.9479 | 1 | 19892 | 1 | -1.2865 |
| ATP9A | 6 | 0.066194 | 0.17879 | 0.968171 | 3694 | 4 | -1.7096 | 0.90818 | 0.94794 | 1 | 19893 | 1 | -1.7096 |
| NR0B2 | 5 | 0.31849 | 0.49611 | 0.999994 | 10486 | 2 | -0.53054 | 0.90838 | 0.92747 | 1 | 19894 | 1 | -0.53054 |
| TTC17 | 5 | 0.49053 | 0.63967 | 0.999994 | 13285 | 2 | 0.038751 | 0.90853 | 0.92754 | 1 | 19895 | 1 | 0.038751 |
| SLC5A7 | 4 | 0.20877 | 0.34972 | 0.999994 | 7996 | 2 | -1.0146 | 0.90854 | 0.91053 | 1 | 19896 | 1 | -1.0146 |
| hsa-mir-4277 | 4 | 0.74953 | 0.75972 | 0.999994 | 16854 | 1 | 0.41495 | 0.90862 | 0.91059 | 1 | 19897 | 1 | 0.41495 |
| hsa-mir-621 | 3 | 0.29036 | 0.42343 | 0.999994 | 9915 | 1 | 0.024526 | 0.90882 | 0.90882 | 1 | 19898 | 0 | 0.024526 |
| NonTargetingControlG | 2 | 0.091085 | 0.15623 | 0.967126 | 4640 | 2 | -1.1346 | 0.90892 | 0.9089 | 1 | 19899 | 0 | -1.1346 |
| UBP1 | 5 | 0.034279 | 0.10143 | 0.946414 | 2228 | 4 | -1.0204 | 0.90911 | 0.92781 | 1 | 19900 | 1 | -1.0204 |
| RAB44 | 4 | 0.077749 | 0.17456 | 0.968171 | 4137 | 3 | -2.1964 | 0.90916 | 0.91104 | 1 | 19901 | 1 | -2.1964 |
| INHA | 6 | 0.20112 | 0.38922 | 0.999994 | 7793 | 3 | -0.85667 | 0.90921 | 0.94822 | 1 | 19902 | 1 | -0.85667 |
| NonTargetingControlG | 2 | 0.090715 | 0.15564 | 0.967126 | 4626 | 2 | -1.004 | 0.90928 | 0.9093 | 1 | 19903 | 0 | -1.004 |
| CTNNBL1 | 6 | 0.0092607 | 0.036421 | 0.880243 | 813 | 3 | -3.7401 | 0.90931 | 0.94824 | 1 | 19904 | 1 | -3.7401 |
| TNNC1 | 5 | 0.0013 | 0.0057826 | 0.623123 | 193 | 4 | -5.5417 | 0.90932 | 0.92791 | 1 | 19905 | 1 | -5.5417 |
| FBP1 | 6 | 0.52642 | 0.68701 | 0.999994 | 13835 | 2 | 0.096996 | 0.9096 | 0.94831 | 1 | 19906 | 1 | 0.096996 |
| TMEM164 | 6 | 0.13323 | 0.30444 | 0.999994 | 6003 | 3 | -1.0149 | 0.90972 | 0.94834 | 1 | 19907 | 1 | -1.0149 |
| PRSS55 | 4 | 0.27544 | 0.4179 | 0.999994 | 9604 | 1 | -0.26507 | 0.90981 | 0.91158 | 1 | 19908 | 1 | -0.26507 |
| BCL7B | 6 | 0.090627 | 0.22874 | 0.983354 | 4622 | 2 | -0.024535 | 0.90985 | 0.94838 | 1 | 19909 | 1 | -0.024535 |
| hsa-mir-5701-1 | 4 | 0.5895 | 0.65969 | 0.999994 | 14814 | 1 | -0.54598 | 0.90989 | 0.91164 | 1 | 19910 | 1 | -0.54598 |
| GGT6 | 5 | 0.008959 | 0.033054 | 0.867348 | 789 | 4 | -2.5534 | 0.91002 | 0.92824 | 1 | 19911 | 1 | -2.5534 |
| hsa-mir-4534 | 3 | 0.1598 | 0.2612 | 0.994943 | 6726 | 2 | -1.5796 | 0.91005 | 0.91009 | 1 | 19912 | 0 | -1.5796 |
| hsa-mir-206 | 4 | 0.44271 | 0.58956 | 0.999994 | 12620 | 2 | -0.15797 | 0.91021 | 0.91192 | 1 | 19913 | 1 | -0.15797 |
| RQCD1 | 6 | 0.00028394 | 0.0015177 | 0.388244 | 76 | 4 | -5.181 | 0.91021 | 0.94847 | 1 | 19914 | 1 | -5.181 |
| CHRNA1 | 5 | 0.5542 | 0.69549 | 0.999994 | 14272 | 2 | 0.17187 | 0.91025 | 0.92834 | 1 | 19915 | 1 | 0.17187 |

| | | | | | | | | | | | | | |
|---|---|---|---|---|---|---|---|---|---|---|---|---|---|
| hsa-mir-4757 | 4 | 0.25881 | 0.40092 | 0.999994 | 9211 | 2 | -0.81263 | 0.91029 | 0.91199 | 1 | 19916 | 1 | -0.81263 |
| NonTargetingControlG | 2 | 0.024606 | 0.045426 | 0.891621 | 1737 | 2 | -5.5806 | 0.9106 | 0.91064 | 1 | 19917 | 0 | -5.5806 |
| hsa-mir-3197 | 2 | 0.08929 | 0.15338 | 0.967126 | 4568 | 2 | -1.2671 | 0.91071 | 0.91075 | 1 | 19918 | 0 | -1.2671 |
| NonTargetingControlG | 2 | 0.089233 | 0.15329 | 0.967126 | 4562 | 2 | -1.3797 | 0.91077 | 0.9108 | 1 | 19919 | 0 | -1.3797 |
| BDH1 | 5 | 0.11362 | 0.24759 | 0.990682 | 5425 | 3 | -1.4011 | 0.91084 | 0.92863 | 1 | 19920 | 1 | -1.4011 |
| hsa-mir-514a-2 | 1 | 0.089086 | 0.089047 | 0.94395 | 4556 | 1 | -3.8265 | 0.91091 | 0.9112 | 1 | 19921 | 0 | -3.8265 |
| FBXO36 | 5 | 0.062231 | 0.16577 | 0.967126 | 3520 | 3 | -1.838 | 0.91105 | 0.92874 | 1 | 19922 | 1 | -1.838 |
| NonTargetingControlG | 2 | 0.088935 | 0.15282 | 0.967126 | 4551 | 2 | -1.3043 | 0.91106 | 0.91112 | 1 | 19923 | 0 | -1.3043 |
| TCL1B | 6 | 0.073663 | 0.19437 | 0.975797 | 3979 | 2 | -0.46666 | 0.91108 | 0.9487 | 1 | 19924 | 1 | -0.46666 |
| NonTargetingControlG | 2 | 0.033479 | 0.060986 | 0.932932 | 2190 | 2 | -4.4892 | 0.91124 | 0.9113 | 1 | 19925 | 0 | -4.4892 |
| DNPEP | 5 | 0.16749 | 0.32842 | 0.999994 | 6923 | 3 | -0.96973 | 0.91125 | 0.92883 | 1 | 19926 | 1 | -0.96973 |
| APITD1 | 6 | 0.081554 | 0.21049 | 0.979587 | 4284 | 4 | -1.055 | 0.91134 | 0.94876 | 1 | 19927 | 1 | -1.055 |
| hsa-mir-1278 | 3 | 0.6041 | 0.62032 | 0.999994 | 14997 | 1 | 0.066147 | 0.91134 | 0.91136 | 1 | 19928 | 0 | 0.066147 |
| HSPA12A | 5 | 0.087567 | 0.20669 | 0.978744 | 4496 | 3 | -1.886 | 0.91166 | 0.92902 | 1 | 19929 | 1 | -1.886 |
| RAC1 | 6 | 0.0066176 | 0.027072 | 0.848412 | 635 | 5 | -3.129 | 0.91168 | 0.94886 | 1 | 19930 | 1 | -3.129 |
| ACTR5 | 6 | 0.55403 | 0.70654 | 0.999994 | 14269 | 2 | -0.019764 | 0.91169 | 0.94886 | 1 | 19931 | 1 | -0.019764 |
| NonTargetingControlG | 2 | 0.088208 | 0.15165 | 0.967126 | 4521 | 2 | -1.2249 | 0.91179 | 0.91185 | 1 | 19932 | 0 | -1.2249 |
| hsa-mir-924 | 4 | 0.028884 | 0.079253 | 0.93914 | 1950 | 3 | -4.3137 | 0.91201 | 0.91348 | 1 | 19933 | 1 | -4.3137 |
| NCOA4 | 5 | 0.091778 | 0.21342 | 0.979587 | 4670 | 2 | -0.49945 | 0.91203 | 0.9292 | 1 | 19934 | 1 | -0.49945 |
| DNAH7 | 6 | 0.12181 | 0.28918 | 0.999632 | 5695 | 2 | -0.21487 | 0.91205 | 0.94897 | 1 | 19935 | 1 | -0.21487 |
| BET1 | 5 | 0.16198 | 0.32044 | 0.999994 | 6796 | 2 | -0.56214 | 0.91207 | 0.92921 | 1 | 19936 | 1 | -0.56214 |
| OR1A1 | 6 | 0.34184 | 0.55476 | 0.999994 | 10978 | 2 | -0.088994 | 0.91207 | 0.94897 | 1 | 19937 | 1 | -0.088994 |
| UBA2 | 3 | 0.036226 | 0.089275 | 0.94395 | 2329 | 2 | -0.94169 | 0.91214 | 0.91215 | 1 | 19938 | 0 | -0.94169 |
| KLK10 | 5 | 0.18853 | 0.35882 | 0.999994 | 7437 | 2 | -0.18507 | 0.91226 | 0.92932 | 1 | 19939 | 1 | -0.18507 |
| EFNA4 | 4 | 0.046862 | 0.1153 | 0.951052 | 2845 | 2 | -0.52676 | 0.91229 | 0.91369 | 1 | 19940 | 1 | -0.52676 |
| NonTargetingControlG | 2 | 0.08768 | 0.15083 | 0.966359 | 4502 | 2 | -2.4356 | 0.91232 | 0.91237 | 1 | 19941 | 0 | -2.4356 |
| DCAF17 | 4 | 0.099559 | 0.21432 | 0.979587 | 4959 | 3 | -1.3527 | 0.91235 | 0.91375 | 1 | 19942 | 1 | -1.3527 |
| CDK13 | 5 | 0.085027 | 0.2026 | 0.977973 | 4408 | 2 | -0.49407 | 0.91252 | 0.92942 | 1 | 19943 | 1 | -0.49407 |
| OR6C4 | 4 | 0.26866 | 0.41087 | 0.999994 | 9436 | 2 | -0.58889 | 0.91283 | 0.91417 | 1 | 19944 | 1 | -0.58889 |
| ACAD9 | 4 | 0.23674 | 0.37837 | 0.999994 | 8688 | 2 | -0.88729 | 0.91287 | 0.9142 | 1 | 19945 | 1 | -0.88729 |
| MTM1 | 5 | 0.29677 | 0.47844 | 0.999994 | 10060 | 2 | -0.30516 | 0.91303 | 0.9297 | 1 | 19946 | 1 | -0.30516 |
| GIT2 | 4 | 0.068539 | 0.15708 | 0.967126 | 3776 | 2 | -0.95148 | 0.91313 | 0.91441 | 1 | 19947 | 1 | -0.95148 |
| MKL1 | 6 | 0.0089364 | 0.03529 | 0.86993 | 787 | 5 | -3.8917 | 0.91315 | 0.94927 | 1 | 19948 | 1 | -3.8917 |
| CEP41 | 6 | 0.28945 | 0.49444 | 0.999994 | 9900 | 3 | -0.51374 | 0.91316 | 0.94927 | 1 | 19949 | 1 | -0.51374 |
| PRPS1L1 | 4 | 0.076754 | 0.1727 | 0.96793 | 4101 | 3 | -1.2448 | 0.91324 | 0.9145 | 1 | 19950 | 1 | -1.2448 |
| BMP8A | 5 | 0.18264 | 0.35041 | 0.999994 | 7280 | 3 | -0.95071 | 0.91325 | 0.92981 | 1 | 19951 | 1 | -0.95071 |
| hsa-mir-4771-2 | 1 | 0.086662 | 0.086627 | 0.94395 | 4471 | 1 | -4.9332 | 0.91334 | 0.91368 | 1 | 19952 | 0 | -4.9332 |
| PPM1B | 5 | 0.62147 | 0.73409 | 0.999994 | 15252 | 1 | -0.28844 | 0.91355 | 0.92997 | 1 | 19953 | 1 | -0.28844 |
| SLC27A3 | 6 | 0.013996 | 0.052454 | 0.914561 | 1128 | 4 | -3.2294 | 0.91357 | 0.94938 | 1 | 19954 | 1 | -3.2294 |
| ELK4 | 6 | 0.22291 | 0.41567 | 0.999994 | 8354 | 2 | 0.087396 | 0.91361 | 0.9494 | 1 | 19955 | 1 | 0.087396 |
| CDK12 | 5 | 0.060749 | 0.16331 | 0.967126 | 3455 | 3 | -3.5179 | 0.91377 | 0.93009 | 1 | 19956 | 1 | -3.5179 |
| KCNJ13 | 6 | 0.023122 | 0.081966 | 0.93914 | 1659 | 5 | -1.6473 | 0.91379 | 0.94944 | 1 | 19957 | 1 | -1.6473 |
| NPB | 5 | 0.056418 | 0.15578 | 0.967126 | 3266 | 4 | -2.0546 | 0.91382 | 0.93012 | 1 | 19958 | 1 | -2.0546 |
| C14orf39 | 6 | 0.10864 | 0.2641 | 0.996734 | 5262 | 4 | -0.78119 | 0.91398 | 0.9495 | 1 | 19959 | 1 | -0.78119 |
| RNF24 | 5 | 0.0065591 | 0.02612 | 0.848412 | 629 | 4 | -2.1138 | 0.91409 | 0.93025 | 1 | 19960 | 1 | -2.1138 |
| ZSCAN5A | 4 | 0.10921 | 0.23145 | 0.984304 | 5275 | 3 | -1.148 | 0.91412 | 0.91522 | 1 | 19961 | 1 | -1.148 |
| NonTargetingControlG | 2 | 0.085808 | 0.14779 | 0.965554 | 4438 | 2 | -1.2748 | 0.91419 | 0.91425 | 1 | 19962 | 0 | -1.2748 |
| PLP1 | 5 | 0.043196 | 0.12381 | 0.951052 | 2684 | 4 | -1.1076 | 0.91424 | 0.93032 | 1 | 19963 | 1 | -1.1076 |
| NOC3L | 4 | 0.1262 | 0.26103 | 0.994943 | 5827 | 2 | -1.5889 | 0.91425 | 0.91532 | 1 | 19964 | 1 | -1.5889 |
| PITPNB | 5 | 0.12703 | 0.2681 | 0.998561 | 5850 | 3 | -1.2699 | 0.91438 | 0.93038 | 1 | 19965 | 1 | -1.2699 |
| hsa-mir-4489 | 2 | 0.085433 | 0.14721 | 0.965554 | 4426 | 2 | -1.5409 | 0.91457 | 0.91461 | 1 | 19966 | 0 | -1.5409 |
| MLH1 | 5 | 0.51059 | 0.65714 | 0.999994 | 13583 | 2 | -0.16879 | 0.91501 | 0.93069 | 1 | 19967 | 1 | -0.16879 |

| | | | | | | | | | | | | | |
|---|---|---|---|---|---|---|---|---|---|---|---|---|---|
| NonTargetingControlG | 1 | 0.084871 | 0.084825 | 0.94395 | 4405 | 1 | -6.8084 | 0.91513 | 0.9155 | 1 | 19968 | 0 | -6.8084 |
| TRIQK | 4 | 0.31171 | 0.45502 | 0.999994 | 10353 | 2 | -0.81644 | 0.91524 | 0.91616 | 1 | 19969 | 1 | -0.81644 |
| CYS1 | 4 | 0.37302 | 0.51797 | 0.999994 | 11558 | 2 | -0.33333 | 0.91527 | 0.91618 | 1 | 19970 | 1 | -0.33333 |
| DUS1L | 4 | 0.084402 | 0.18681 | 0.969409 | 4390 | 3 | -1.2092 | 0.91527 | 0.91618 | 1 | 19971 | 1 | -1.2092 |
| FAM210B | 5 | 0.012906 | 0.044324 | 0.891621 | 1056 | 3 | -2.0604 | 0.9156 | 0.93101 | 1 | 19972 | 1 | -2.0604 |
| XPC | 5 | 0.22167 | 0.40557 | 0.999994 | 8324 | 2 | -0.28688 | 0.9157 | 0.93107 | 1 | 19973 | 1 | -0.28688 |
| DDX58 | 4 | 0.78304 | 0.78643 | 0.999994 | 17252 | 1 | 0.035475 | 0.91587 | 0.91671 | 1 | 19974 | 1 | 0.035475 |
| FLJ22184 | 4 | 0.14716 | 0.28624 | 0.999632 | 6379 | 2 | -0.20308 | 0.91603 | 0.91685 | 1 | 19975 | 1 | -0.20308 |
| SORCS2 | 4 | 0.0031829 | 0.011376 | 0.720297 | 373 | 3 | -4.5881 | 0.91648 | 0.91722 | 1 | 19976 | 1 | -4.5881 |
| NUP133 | 5 | 0.021288 | 0.067569 | 0.93914 | 1534 | 3 | -1.1779 | 0.91662 | 0.93152 | 1 | 19977 | 1 | -1.1779 |
| hsa-mir-4648 | 4 | 0.028134 | 0.077698 | 0.93914 | 1912 | 3 | -2.2044 | 0.91673 | 0.91745 | 1 | 19978 | 1 | -2.2044 |
| ERLIN2 | 6 | 0.068704 | 0.18406 | 0.968526 | 3780 | 4 | -1.142 | 0.91693 | 0.9503 | 1 | 19979 | 1 | -1.142 |
| HIST1H1C | 4 | 0.1002 | 0.21546 | 0.980236 | 4978 | 3 | -1.7073 | 0.917 | 0.91769 | 1 | 19980 | 1 | -1.7073 |
| RELN | 5 | 0.12769 | 0.26909 | 0.998561 | 5865 | 3 | -1.3001 | 0.917 | 0.93171 | 1 | 19981 | 1 | -1.3001 |
| KDELR3 | 5 | 0.22435 | 0.40934 | 0.999994 | 8387 | 3 | -0.59596 | 0.91703 | 0.93173 | 1 | 19982 | 1 | -0.59596 |
| SPRY1 | 4 | 0.33385 | 0.47762 | 0.999994 | 10808 | 1 | -0.10576 | 0.91714 | 0.91782 | 1 | 19983 | 1 | -0.10576 |
| TTC21B | 5 | 0.21527 | 0.3967 | 0.999994 | 8149 | 2 | -0.58449 | 0.91719 | 0.93181 | 1 | 19984 | 1 | -0.58449 |
| YES1 | 6 | 0.17258 | 0.35393 | 0.999994 | 7050 | 3 | -0.92563 | 0.91748 | 0.95045 | 1 | 19985 | 1 | -0.92563 |
| hsa-mir-3682 | 4 | 0.1121 | 0.2366 | 0.985449 | 5373 | 3 | -1.8707 | 0.91754 | 0.91819 | 1 | 19986 | 1 | -1.8707 |
| MTMR14 | 4 | 0.044051 | 0.10985 | 0.951052 | 2712 | 3 | -1.9432 | 0.91762 | 0.91827 | 1 | 19987 | 1 | -1.9432 |
| BCL6 | 2 | 0.055093 | 0.097761 | 0.945253 | 3203 | 2 | -5.2291 | 0.91766 | 0.91774 | 1 | 19988 | 0 | -5.2291 |
| GTF3C1 | 3 | 0.12916 | 0.22147 | 0.983093 | 5918 | 2 | -1.7408 | 0.91766 | 0.91769 | 1 | 19989 | 0 | -1.7408 |
| LCE3E | 3 | 0.10813 | 0.19378 | 0.975797 | 5242 | 1 | -1.2059 | 0.91766 | 0.91769 | 1 | 19990 | 0 | -1.2059 |
| CHRNB4 | 5 | 0.12396 | 0.26339 | 0.996734 | 5767 | 3 | -1.0436 | 0.91777 | 0.93211 | 1 | 19991 | 1 | -1.0436 |
| XAB2 | 4 | 0.19667 | 0.3373 | 0.999994 | 7668 | 2 | -0.93094 | 0.91785 | 0.91845 | 1 | 19992 | 1 | -0.93094 |
| APAF1 | 5 | 0.023581 | 0.073731 | 0.93914 | 1691 | 3 | -0.92134 | 0.91787 | 0.93216 | 1 | 19993 | 1 | -0.92134 |
| NonTargetingControlG | 2 | 0.082088 | 0.14191 | 0.959146 | 4301 | 2 | -1.5376 | 0.91791 | 0.91799 | 1 | 19994 | 0 | -1.5376 |
| ECE1 | 6 | 0.60113 | 0.74078 | 0.999994 | 14962 | 1 | 0.065105 | 0.91793 | 0.9506 | 1 | 19995 | 1 | 0.065105 |
| LIFR | 4 | 0.1868 | 0.32709 | 0.999994 | 7382 | 2 | -0.87448 | 0.91797 | 0.91855 | 1 | 19996 | 1 | -0.87448 |
| RBM41 | 4 | 0.10702 | 0.2276 | 0.983354 | 5212 | 2 | -1.8866 | 0.91818 | 0.91874 | 1 | 19997 | 1 | -1.8866 |
| hsa-mir-6796 | 4 | 0.67422 | 0.70798 | 0.999994 | 15991 | 1 | 0.18982 | 0.9183 | 0.91884 | 1 | 19998 | 1 | 0.18982 |
| NonTargetingControlG | 2 | 0.08164 | 0.14119 | 0.959146 | 4287 | 2 | -2.6038 | 0.91836 | 0.91843 | 1 | 19999 | 0 | -2.6038 |
| SHC4 | 5 | 0.10197 | 0.22949 | 0.983354 | 5038 | 2 | -0.37635 | 0.91839 | 0.93243 | 1 | 20000 | 1 | -0.37635 |
| PTPLB | 5 | 0.010755 | 0.038228 | 0.888237 | 916 | 3 | -4.0637 | 0.91848 | 0.93247 | 1 | 20001 | 1 | -4.0637 |
| hsa-mir-3155a | 4 | 0.88269 | 0.88232 | 0.999994 | 18885 | 0 | 0.10836 | 0.91849 | 0.919 | 1 | 20002 | 1 | 0.10836 |
| CALCOCO1 | 5 | 0.1871 | 0.35682 | 0.999994 | 7393 | 3 | -0.87997 | 0.91861 | 0.93255 | 1 | 20003 | 1 | -0.87997 |
| ADARB2 | 5 | 0.030063 | 0.090641 | 0.94395 | 2011 | 4 | -1.6592 | 0.91862 | 0.93256 | 1 | 20004 | 1 | -1.6592 |
| hsa-mir-646 | 3 | 0.13542 | 0.22966 | 0.983354 | 6063 | 2 | -1.6393 | 0.91868 | 0.91871 | 1 | 20005 | 0 | -1.6393 |
| HLF | 5 | 0.01384 | 0.046938 | 0.896157 | 1119 | 4 | -2.4711 | 0.91879 | 0.93265 | 1 | 20006 | 1 | -2.4711 |
| IQCD | 4 | 0.20276 | 0.3435 | 0.999994 | 7852 | 2 | -0.98156 | 0.91882 | 0.91933 | 1 | 20007 | 1 | -0.98156 |
| PLK5 | 5 | 0.27764 | 0.46295 | 0.999994 | 9648 | 2 | -0.21263 | 0.91885 | 0.93268 | 1 | 20008 | 1 | -0.21263 |
| ACSF2 | 5 | 0.047343 | 0.134 | 0.95318 | 2872 | 4 | -1.5174 | 0.91917 | 0.93286 | 1 | 20009 | 1 | -1.5174 |
| ORC1 | 3 | 0.22898 | 0.3488 | 0.999994 | 8504 | 2 | -1.0432 | 0.9192 | 0.91925 | 1 | 20010 | 0 | -1.0432 |
| SPECC1 | 6 | 0.053768 | 0.15225 | 0.967126 | 3153 | 4 | -1.466 | 0.91925 | 0.95099 | 1 | 20011 | 1 | -1.466 |
| MPDZ | 5 | 0.15508 | 0.31033 | 0.999994 | 6596 | 3 | -1.0383 | 0.91932 | 0.93294 | 1 | 20012 | 1 | -1.0383 |
| PDE6H | 5 | 0.39724 | 0.56126 | 0.999994 | 11909 | 2 | 0.077131 | 0.91934 | 0.93295 | 1 | 20013 | 1 | 0.077131 |
| hsa-mir-4769 | 1 | 0.080506 | 0.080542 | 0.93914 | 4250 | 1 | -6.9974 | 0.91949 | 0.9199 | 1 | 20014 | 0 | -6.9974 |
| C16orf87 | 5 | 0.12824 | 0.26991 | 0.998561 | 5886 | 3 | -1.3607 | 0.91959 | 0.93309 | 1 | 20015 | 1 | -1.3607 |
| hsa-mir-1302-3 | 3 | 0.057348 | 0.12583 | 0.951052 | 3308 | 1 | -0.31165 | 0.91964 | 0.91966 | 1 | 20016 | 0 | -0.31165 |
| OGFR | 5 | 0.15043 | 0.30343 | 0.999994 | 6466 | 3 | -1.1455 | 0.91971 | 0.93314 | 1 | 20017 | 1 | -1.1455 |
| WDR17 | 4 | 0.034981 | 0.091684 | 0.94395 | 2267 | 3 | -3.9563 | 0.91985 | 0.92023 | 1 | 20018 | 1 | -3.9563 |
| SOBP | 5 | 0.0125 | 0.043207 | 0.891621 | 1030 | 4 | -4.1447 | 0.91999 | 0.9333 | 1 | 20019 | 1 | -4.1447 |

| | | | | | | | | | | | | | |
|---|---|---|---|---|---|---|---|---|---|---|---|---|---|
| CDK5RAP1 | 5 | 0.098715 | 0.22441 | 0.983354 | 4922 | 3 | -1.6529 | 0.92017 | 0.93338 | 1 | 20020 | 1 | -1.6529 |
| EPPIN-WFDC6 | 2 | 0.079829 | 0.13823 | 0.957317 | 4228 | 2 | -1.3119 | 0.92017 | 0.92026 | 1 | 20021 | 0 | -1.3119 |
| LOC100129636 | 6 | 0.02129 | 0.076122 | 0.93914 | 1535 | 5 | -2.0757 | 0.92019 | 0.95127 | 1 | 20022 | 1 | -2.0757 |
| NonTargetingControlG | 2 | 0.066742 | 0.11707 | 0.951052 | 3715 | 2 | -5.0908 | 0.92023 | 0.92032 | 1 | 20023 | 0 | -5.0908 |
| hsa-mir-885 | 3 | 0.27599 | 0.40708 | 0.999994 | 9616 | 2 | -0.67678 | 0.9203 | 0.92035 | 1 | 20024 | 0 | -0.67678 |
| GALNT2 | 5 | 0.69902 | 0.76604 | 0.999994 | 16295 | 1 | 0.20794 | 0.92047 | 0.93356 | 1 | 20025 | 1 | 0.20794 |
| DGCR2 | 6 | 0.29714 | 0.50339 | 0.999994 | 10070 | 3 | -0.98755 | 0.92047 | 0.95135 | 1 | 20026 | 1 | -0.98755 |
| TTI1 | 5 | 0.03241 | 0.09665 | 0.945031 | 2130 | 3 | -2.96 | 0.92049 | 0.93358 | 1 | 20027 | 1 | -2.96 |
| IL21 | 4 | 0.25202 | 0.39398 | 0.999994 | 9057 | 2 | -0.95303 | 0.92054 | 0.92084 | 1 | 20028 | 1 | -0.95303 |
| NonTargetingControlG | 2 | 0.079441 | 0.13763 | 0.957317 | 4204 | 2 | -1.1937 | 0.92056 | 0.92064 | 1 | 20029 | 0 | -1.1937 |
| AKR7A2 | 4 | 0.14065 | 0.27961 | 0.999632 | 6206 | 2 | -0.84877 | 0.92067 | 0.92096 | 1 | 20030 | 1 | -0.84877 |
| NXPH2 | 5 | 0.19011 | 0.3611 | 0.999994 | 7488 | 3 | -0.76152 | 0.92071 | 0.93369 | 1 | 20031 | 1 | -0.76152 |
| LRRC40 | 6 | 0.18722 | 0.37206 | 0.999994 | 7399 | 3 | -0.7895 | 0.9209 | 0.95149 | 1 | 20032 | 1 | -0.7895 |
| NonTargetingControlG | 2 | 0.079037 | 0.137 | 0.957161 | 4190 | 2 | -1.6726 | 0.92096 | 0.92104 | 1 | 20033 | 0 | -1.6726 |
| PIGP | 4 | 0.094959 | 0.20603 | 0.978744 | 4785 | 3 | -1.4847 | 0.921 | 0.92124 | 1 | 20034 | 1 | -1.4847 |
| NonTargetingControlG | 1 | 0.078931 | 0.078952 | 0.93914 | 4186 | 1 | -5.5348 | 0.92107 | 0.92141 | 1 | 20035 | 0 | -5.5348 |
| SLC35C1 | 5 | 0.21924 | 0.40221 | 0.999994 | 8269 | 3 | -0.76321 | 0.92125 | 0.93397 | 1 | 20036 | 1 | -0.76321 |
| CCDC121 | 6 | 0.0015399 | 0.007702 | 0.672529 | 226 | 5 | -5.8071 | 0.92134 | 0.95161 | 1 | 20037 | 1 | -5.8071 |
| SEC14L4 | 6 | 0.016541 | 0.060774 | 0.932932 | 1279 | 4 | -2.9207 | 0.92135 | 0.95161 | 1 | 20038 | 1 | -2.9207 |
| C9orf163 | 5 | 0.086178 | 0.20445 | 0.978186 | 4458 | 2 | -1.2059 | 0.92147 | 0.93409 | 1 | 20039 | 1 | -1.2059 |
| ZNF282 | 6 | 0.21579 | 0.4071 | 0.999994 | 8166 | 3 | -0.40509 | 0.92157 | 0.95168 | 1 | 20040 | 1 | -0.40509 |
| PDGFB | 6 | 0.35259 | 0.56692 | 0.999994 | 11207 | 2 | -0.35305 | 0.9216 | 0.95168 | 1 | 20041 | 1 | -0.35305 |
| CYP7B1 | 5 | 0.025638 | 0.079187 | 0.93914 | 1782 | 3 | -1.6097 | 0.92163 | 0.93416 | 1 | 20042 | 1 | -1.6097 |
| TAB3 | 3 | 0.18674 | 0.2959 | 0.999994 | 7381 | 2 | -1.5651 | 0.92183 | 0.92188 | 1 | 20043 | 0 | -1.5651 |
| NonTargetingControlG | 2 | 0.07808 | 0.13546 | 0.95468 | 4152 | 2 | -1.2884 | 0.92192 | 0.92197 | 1 | 20044 | 0 | -1.2884 |
| PPAN | 6 | 0.015164 | 0.056252 | 0.9235 | 1198 | 5 | -1.1168 | 0.92197 | 0.95179 | 1 | 20045 | 1 | -1.1168 |
| FXYD2 | 6 | 0.11392 | 0.27419 | 0.998561 | 5445 | 4 | -1.0127 | 0.92203 | 0.95181 | 1 | 20046 | 1 | -1.0127 |
| CCL14 | 6 | 0.19283 | 0.37906 | 0.999994 | 7569 | 3 | -0.72517 | 0.9221 | 0.95184 | 1 | 20047 | 1 | -0.72517 |
| PAX2 | 5 | 0.33683 | 0.51113 | 0.999994 | 10869 | 2 | -0.12583 | 0.92214 | 0.93444 | 1 | 20048 | 1 | -0.12583 |
| CHD5 | 4 | 0.78781 | 0.7905 | 0.999994 | 17317 | 1 | -0.062936 | 0.92215 | 0.92232 | 1 | 20049 | 1 | -0.062936 |
| TOX | 5 | 0.20395 | 0.38084 | 0.999994 | 7878 | 3 | -0.792 | 0.92217 | 0.93446 | 1 | 20050 | 1 | -0.792 |
| OPN1MW | 6 | 0.058811 | 0.16321 | 0.967126 | 3374 | 4 | -1.3924 | 0.92218 | 0.95186 | 1 | 20051 | 1 | -1.3924 |
| GZMA | 6 | 0.045601 | 0.13438 | 0.953565 | 2793 | 4 | -1.9101 | 0.9223 | 0.95191 | 1 | 20052 | 1 | -1.9101 |
| SLC5A11 | 5 | 0.22645 | 0.41231 | 0.999994 | 8436 | 3 | -0.67489 | 0.92241 | 0.93459 | 1 | 20053 | 1 | -0.67489 |
| NonTargetingControlG | 2 | 0.077575 | 0.13467 | 0.953952 | 4134 | 2 | -1.4882 | 0.92242 | 0.9225 | 1 | 20054 | 0 | -1.4882 |
| PPP2R5C | 6 | 0.0052632 | 0.022126 | 0.819701 | 539 | 5 | -3.7865 | 0.92245 | 0.95196 | 1 | 20055 | 1 | -3.7865 |
| SPIN4 | 5 | 0.042043 | 0.121 | 0.951052 | 2615 | 4 | -0.99395 | 0.92249 | 0.93464 | 1 | 20056 | 1 | -0.99395 |
| NonTargetingControlG | 2 | 0.077502 | 0.13454 | 0.953565 | 4130 | 2 | -2.2354 | 0.9225 | 0.92257 | 1 | 20057 | 0 | -2.2354 |
| NonTargetingControlG | 2 | 0.077439 | 0.13444 | 0.953565 | 4126 | 2 | -2.8567 | 0.92256 | 0.92263 | 1 | 20058 | 0 | -2.8567 |
| hsa-mir-320b-2 | 4 | 0.058953 | 0.13877 | 0.957514 | 3379 | 3 | -1.5648 | 0.92292 | 0.92307 | 1 | 20059 | 1 | -1.5648 |
| PPFIBP1 | 4 | 0.19882 | 0.33946 | 0.999994 | 7730 | 2 | -1.2341 | 0.923 | 0.92315 | 1 | 20060 | 1 | -1.2341 |
| ZBTB20 | 4 | 0.0017149 | 0.0062325 | 0.636218 | 240 | 3 | -1.9129 | 0.92301 | 0.92316 | 1 | 20061 | 1 | -1.9129 |
| KAT6B | 5 | 0.10538 | 0.23478 | 0.984971 | 5148 | 3 | -1.1744 | 0.92302 | 0.93493 | 1 | 20062 | 1 | -1.1744 |
| TUT1 | 3 | 0.15769 | 0.25844 | 0.994943 | 6665 | 1 | -0.5657 | 0.92307 | 0.92312 | 1 | 20063 | 0 | -0.5657 |
| hsa-mir-195 | 4 | 0.11568 | 0.24275 | 0.988088 | 5509 | 3 | -0.98292 | 0.9232 | 0.92332 | 1 | 20064 | 1 | -0.98292 |
| hsa-mir-922 | 4 | 0.21335 | 0.35445 | 0.999994 | 8097 | 2 | -0.90004 | 0.9232 | 0.92333 | 1 | 20065 | 1 | -0.90004 |
| ESD | 5 | 0.060192 | 0.16243 | 0.967126 | 3429 | 3 | -1.7721 | 0.9232 | 0.93503 | 1 | 20066 | 1 | -1.7721 |
| SUN5 | 5 | 0.092149 | 0.21404 | 0.979587 | 4685 | 3 | -1.166 | 0.92328 | 0.93507 | 1 | 20067 | 1 | -1.166 |
| PARVA | 3 | 0.19663 | 0.30839 | 0.999994 | 7665 | 1 | -0.14175 | 0.9234 | 0.92343 | 1 | 20068 | 0 | -0.14175 |
| hsa-mir-576 | 4 | 0.47643 | 0.60592 | 0.999994 | 13088 | 1 | -0.081478 | 0.9234 | 0.92351 | 1 | 20069 | 1 | -0.081478 |
| ANXA8 | 6 | 0.26802 | 0.46936 | 0.999994 | 9421 | 3 | -0.32402 | 0.92341 | 0.95224 | 1 | 20070 | 1 | -0.32402 |
| STRN3 | 5 | 0.13697 | 0.28318 | 0.999632 | 6107 | 2 | -0.30652 | 0.92342 | 0.93514 | 1 | 20071 | 1 | -0.30652 |

| | | | | | | | | | | | | | |
|---|---|---|---|---|---|---|---|---|---|---|---|---|---|
| CDRT15L2 | 5 | 0.55422 | 0.6955 | 0.999994 | 14273 | 2 | 0.28447 | 0.92359 | 0.93524 | 1 | 20072 | 1 | 0.28447 |
| DYNAP | 4 | 0.092099 | 0.20086 | 0.977164 | 4684 | 2 | -1.5351 | 0.92362 | 0.92372 | 1 | 20073 | 1 | -1.5351 |
| LRRC48 | 6 | 0.10786 | 0.26258 | 0.996166 | 5234 | 3 | -0.99497 | 0.92363 | 0.95231 | 1 | 20074 | 1 | -0.99497 |
| HAVCR1 | 6 | 0.35494 | 0.56959 | 0.999994 | 11251 | 3 | -0.31392 | 0.92363 | 0.95231 | 1 | 20075 | 1 | -0.31392 |
| GFM2 | 5 | 0.89787 | 0.89799 | 0.999994 | 19176 | 0 | -0.061157 | 0.92364 | 0.93527 | 1 | 20076 | 1 | -0.061157 |
| TREML1 | 5 | 0.042776 | 0.12276 | 0.951052 | 2666 | 4 | -1.4848 | 0.92369 | 0.9353 | 1 | 20077 | 1 | -1.4848 |
| STAMBP | 6 | 0.22423 | 0.41721 | 0.999994 | 8385 | 3 | -0.80444 | 0.92375 | 0.95234 | 1 | 20078 | 1 | -0.80444 |
| NonTargetingControlG | 1 | 0.076159 | 0.076159 | 0.93914 | 4081 | 1 | -5.744 | 0.92384 | 0.92418 | 1 | 20079 | 0 | -5.744 |
| PPP1R13B | 4 | 0.28913 | 0.43191 | 0.999994 | 9895 | 2 | -0.21488 | 0.9239 | 0.92397 | 1 | 20080 | 1 | -0.21488 |
| SH2D6 | 4 | 0.0068807 | 0.023798 | 0.832699 | 651 | 3 | -3.59 | 0.92391 | 0.92397 | 1 | 20081 | 1 | -3.59 |
| hsa-mir-6888 | 4 | 0.27803 | 0.42056 | 0.999994 | 9655 | 1 | -0.44914 | 0.92404 | 0.92409 | 1 | 20082 | 1 | -0.44914 |
| KCNK2 | 5 | 0.034847 | 0.10288 | 0.946414 | 2260 | 4 | -1.0129 | 0.92422 | 0.9356 | 1 | 20083 | 1 | -1.0129 |
| CD160 | 4 | 0.099575 | 0.21435 | 0.979587 | 4960 | 2 | -1.1757 | 0.92423 | 0.92426 | 1 | 20084 | 1 | -1.1757 |
| OR2W1 | 3 | 0.045952 | 0.11006 | 0.951052 | 2806 | 2 | -3.0583 | 0.9243 | 0.92434 | 1 | 20085 | 0 | -3.0583 |
| CACNB2 | 4 | 0.10666 | 0.22696 | 0.983354 | 5191 | 3 | -1.3556 | 0.9245 | 0.9245 | 1 | 20086 | 1 | -1.3556 |
| NonTargetingControlG | 2 | 0.075458 | 0.13125 | 0.95318 | 4053 | 2 | -3.2348 | 0.92454 | 0.92466 | 1 | 20087 | 0 | -3.2348 |
| PAX1 | 3 | 0.16074 | 0.26241 | 0.995875 | 6759 | 2 | -1.4486 | 0.92455 | 0.9246 | 1 | 20088 | 0 | -1.4486 |
| LAPTM5 | 5 | 0.45277 | 0.60756 | 0.999994 | 12745 | 1 | -0.17051 | 0.92465 | 0.93582 | 1 | 20089 | 1 | -0.17051 |
| GPBP1L1 | 5 | 0.024 | 0.074825 | 0.93914 | 1709 | 3 | -3.3175 | 0.92467 | 0.93584 | 1 | 20090 | 1 | -3.3175 |
| DNAH1 | 6 | 0.0038391 | 0.016896 | 0.785648 | 430 | 5 | -1.7714 | 0.92475 | 0.95263 | 1 | 20091 | 1 | -1.7714 |
| PRAMEF11 | 5 | 0.0049186 | 0.020496 | 0.81027 | 514 | 4 | -4.0183 | 0.92479 | 0.9359 | 1 | 20092 | 1 | -4.0183 |
| OR4A5 | 6 | 0.20175 | 0.39 | 0.999994 | 7817 | 3 | -0.47993 | 0.92483 | 0.95265 | 1 | 20093 | 1 | -0.47993 |
| PRKDC | 4 | 0.10055 | 0.21608 | 0.980699 | 4993 | 2 | -1.1369 | 0.92489 | 0.92487 | 1 | 20094 | 1 | -1.1369 |
| INCENP | 3 | 0.21945 | 0.33699 | 0.999994 | 8275 | 2 | -1.1138 | 0.92492 | 0.92496 | 1 | 20095 | 0 | -1.1138 |
| SLX4 | 6 | 0.060403 | 0.1666 | 0.967126 | 3439 | 3 | -1.1638 | 0.92494 | 0.95268 | 1 | 20096 | 1 | -1.1638 |
| NXPH1 | 6 | 0.1612 | 0.33977 | 0.999994 | 6771 | 3 | -0.89451 | 0.92501 | 0.9527 | 1 | 20097 | 1 | -0.89451 |
| hsa-mir-4478 | 3 | 0.52143 | 0.56049 | 0.999994 | 13765 | 1 | -0.20183 | 0.92522 | 0.92527 | 1 | 20098 | 0 | -0.20183 |
| NonTargetingControlG | 1 | 0.074735 | 0.074739 | 0.93914 | 4025 | 1 | -7.2311 | 0.92526 | 0.9256 | 1 | 20099 | 0 | -7.2311 |
| VAMP2 | 5 | 0.0054163 | 0.022446 | 0.819701 | 552 | 4 | -2.2601 | 0.92532 | 0.9362 | 1 | 20100 | 1 | -2.2601 |
| RBM39 | 4 | 0.0095324 | 0.032217 | 0.865899 | 839 | 2 | -3.9889 | 0.92539 | 0.92534 | 1 | 20101 | 1 | -3.9889 |
| NonTargetingControlG | 1 | 0.074603 | 0.074611 | 0.93914 | 4019 | 1 | -5.687 | 0.9254 | 0.92572 | 1 | 20102 | 0 | -5.687 |
| IQCF6 | 4 | 0.039829 | 0.10146 | 0.946414 | 2506 | 3 | -2.5356 | 0.92544 | 0.9254 | 1 | 20103 | 1 | -2.5356 |
| LAMTOR1 | 5 | 0.5224 | 0.66739 | 0.999994 | 13785 | 2 | -0.45375 | 0.92549 | 0.93629 | 1 | 20104 | 1 | -0.45375 |
| CST4 | 4 | 0.11612 | 0.2435 | 0.988088 | 5518 | 3 | -1.182 | 0.92558 | 0.92553 | 1 | 20105 | 1 | -1.182 |
| NonTargetingControlG | 1 | 0.074264 | 0.074274 | 0.93914 | 4002 | 1 | -5.6761 | 0.92574 | 0.92602 | 1 | 20106 | 0 | -5.6761 |
| CCNI2 | 5 | 0.11565 | 0.25074 | 0.992497 | 5502 | 3 | -0.89422 | 0.92577 | 0.93643 | 1 | 20107 | 1 | -0.89422 |
| hsa-mir-4665 | 3 | 0.15897 | 0.26014 | 0.994943 | 6700 | 2 | -1.5454 | 0.92578 | 0.92581 | 1 | 20108 | 0 | -1.5454 |
| EP400 | 4 | 0.0092267 | 0.031284 | 0.862223 | 809 | 3 | -1.8143 | 0.92593 | 0.92588 | 1 | 20109 | 1 | -1.8143 |
| HLA-DQB2 | 6 | 0.29292 | 0.49845 | 0.999994 | 9974 | 3 | -0.60005 | 0.92599 | 0.95299 | 1 | 20110 | 1 | -0.60005 |
| RAE1 | 5 | 0.040592 | 0.11736 | 0.951052 | 2541 | 4 | -2.2897 | 0.92608 | 0.93662 | 1 | 20111 | 1 | -2.2897 |
| EXD2 | 6 | 0.29153 | 0.49682 | 0.999994 | 9938 | 1 | 0.30723 | 0.92612 | 0.95304 | 1 | 20112 | 1 | 0.30723 |
| LOC100288332 | 5 | 0.21959 | 0.4027 | 0.999994 | 8280 | 3 | -0.69468 | 0.92617 | 0.93666 | 1 | 20113 | 1 | -0.69468 |
| IMMP1L | 5 | 0.059477 | 0.1612 | 0.967126 | 3400 | 3 | -1.7255 | 0.9262 | 0.93668 | 1 | 20114 | 1 | -1.7255 |
| GPX8 | 5 | 0.080512 | 0.19542 | 0.975797 | 4251 | 2 | -0.036391 | 0.92629 | 0.93673 | 1 | 20115 | 1 | -0.036391 |
| IQCJ-SCHIP1 | 2 | 0.073653 | 0.12831 | 0.951777 | 3977 | 2 | -2.3517 | 0.92635 | 0.92648 | 1 | 20116 | 0 | -2.3517 |
| TMC6 | 6 | 0.033043 | 0.10597 | 0.950444 | 2160 | 4 | -1.8209 | 0.92636 | 0.95312 | 1 | 20117 | 1 | -1.8209 |
| IQCF2 | 5 | 0.063207 | 0.16736 | 0.967126 | 3568 | 2 | -0.10824 | 0.92647 | 0.93683 | 1 | 20118 | 1 | -0.10824 |
| PIP5K1C | 3 | 0.2693 | 0.39882 | 0.999994 | 9451 | 2 | -0.74022 | 0.92664 | 0.9267 | 1 | 20119 | 0 | -0.74022 |
| GNA11 | 5 | 0.56986 | 0.7094 | 0.999994 | 14535 | 2 | -0.43394 | 0.92671 | 0.93697 | 1 | 20120 | 1 | -0.43394 |
| hsa-mir-194-1 | 4 | 0.70398 | 0.72707 | 0.999994 | 16352 | 1 | 0.33453 | 0.92672 | 0.9266 | 1 | 20121 | 1 | 0.33453 |
| CD79B | 3 | 0.039173 | 0.09597 | 0.945031 | 2473 | 2 | -4.0553 | 0.92681 | 0.92687 | 1 | 20122 | 0 | -4.0553 |
| DAGLA | 5 | 0.029658 | 0.089574 | 0.94395 | 1991 | 3 | -2.8558 | 0.92684 | 0.93704 | 1 | 20123 | 1 | -2.8558 |

| | | | | | | | | | | | | | |
|---|---|---|---|---|---|---|---|---|---|---|---|---|---|
| C15orf38-AP3S2 | 4 | 0.3659 | 0.51057 | 0.999994 | 11441 | 2 | -0.42157 | 0.92687 | 0.92674 | 1 | 20124 | 1 | -0.42157 |
| NonTargetingControlG | 2 | 0.073129 | 0.12742 | 0.951052 | 3963 | 2 | -1.7913 | 0.92687 | 0.92701 | 1 | 20125 | 0 | -1.7913 |
| HBB | 6 | 0.11516 | 0.27658 | 0.999379 | 5482 | 3 | -0.94737 | 0.92691 | 0.95329 | 1 | 20126 | 1 | -0.94737 |
| TWF1 | 5 | 0.022656 | 0.071263 | 0.93914 | 1629 | 1 | -0.081548 | 0.92704 | 0.93715 | 1 | 20127 | 1 | -0.081548 |
| CHIA | 4 | 0.24819 | 0.39006 | 0.999994 | 8974 | 1 | -0.12218 | 0.92704 | 0.9269 | 1 | 20128 | 1 | -0.12218 |
| NUPL1 | 3 | 0.037802 | 0.092839 | 0.943968 | 2418 | 2 | -4.0888 | 0.9271 | 0.92715 | 1 | 20129 | 0 | -4.0888 |
| hsa-mir-5703 | 4 | 0.10779 | 0.22894 | 0.983354 | 5232 | 3 | -2.0413 | 0.92727 | 0.92711 | 1 | 20130 | 1 | -2.0413 |
| SERPINI2 | 5 | 0.22105 | 0.4047 | 0.999994 | 8309 | 3 | -0.7118 | 0.9273 | 0.9373 | 1 | 20131 | 1 | -0.7118 |
| hsa-mir-589 | 3 | 0.27739 | 0.40881 | 0.999994 | 9643 | 2 | -0.63353 | 0.92733 | 0.92739 | 1 | 20132 | 0 | -0.63353 |
| BEND2 | 5 | 0.024494 | 0.076127 | 0.93914 | 1730 | 4 | -2.0776 | 0.92745 | 0.93739 | 1 | 20133 | 1 | -2.0776 |
| SYCP1 | 4 | 0.46982 | 0.60303 | 0.999994 | 13002 | 1 | 0.14557 | 0.92747 | 0.9273 | 1 | 20134 | 1 | 0.14557 |
| LONRF1 | 5 | 0.084859 | 0.20233 | 0.977973 | 4403 | 3 | -1.0667 | 0.92752 | 0.93743 | 1 | 20135 | 1 | -1.0667 |
| RNF148 | 4 | 0.030771 | 0.083031 | 0.94336 | 2050 | 2 | -0.51962 | 0.92752 | 0.92735 | 1 | 20136 | 1 | -0.51962 |
| FNBP4 | 4 | 0.026602 | 0.074543 | 0.93914 | 1833 | 3 | -2.2494 | 0.92755 | 0.92737 | 1 | 20137 | 1 | -2.2494 |
| LDB1 | 6 | 0.12096 | 0.28754 | 0.999632 | 5661 | 4 | -1.032 | 0.92757 | 0.95352 | 1 | 20138 | 1 | -1.032 |
| hsa-mir-6814 | 4 | 0.074572 | 0.16853 | 0.967126 | 4018 | 3 | -2.2214 | 0.92761 | 0.92744 | 1 | 20139 | 1 | -2.2214 |
| NonTargetingControlG | 1 | 0.072364 | 0.072381 | 0.93914 | 3936 | 1 | -7.3259 | 0.92764 | 0.92786 | 1 | 20140 | 0 | -7.3259 |
| hsa-mir-548ak | 1 | 0.072364 | 0.072381 | 0.93914 | 3935 | 1 | -7.3259 | 0.92764 | 0.92786 | 1 | 20141 | 0 | -7.3259 |
| PAQR9 | 5 | 0.10173 | 0.22914 | 0.983354 | 5027 | 3 | -0.91493 | 0.92771 | 0.93753 | 1 | 20142 | 1 | -0.91493 |
| RIMBP3B | 6 | 0.060578 | 0.16697 | 0.967126 | 3448 | 4 | -1.5913 | 0.92774 | 0.95358 | 1 | 20143 | 1 | -1.5913 |
| hsa-mir-4513 | 4 | 0.80642 | 0.80702 | 0.999994 | 17576 | 1 | 0.21774 | 0.92779 | 0.92761 | 1 | 20144 | 1 | 0.21774 |
| RFTN2 | 6 | 0.23836 | 0.43403 | 0.999994 | 8734 | 2 | -0.32259 | 0.9278 | 0.9536 | 1 | 20145 | 1 | -0.32259 |
| NonTargetingControlG | 2 | 0.072174 | 0.12588 | 0.951052 | 3924 | 2 | -1.5876 | 0.92783 | 0.92796 | 1 | 20146 | 0 | -1.5876 |
| VASH2 | 6 | 0.14571 | 0.32027 | 0.999994 | 6342 | 2 | -0.14338 | 0.92791 | 0.95363 | 1 | 20147 | 1 | -0.14338 |
| NonTargetingControlG | 2 | 0.071976 | 0.12557 | 0.951052 | 3917 | 2 | -1.8548 | 0.92802 | 0.92816 | 1 | 20148 | 0 | -1.8548 |
| CCDC42 | 5 | 0.45781 | 0.61181 | 0.999994 | 12818 | 2 | 0.064452 | 0.92817 | 0.93781 | 1 | 20149 | 1 | 0.064452 |
| MON1A | 6 | 0.0092361 | 0.036336 | 0.879988 | 810 | 5 | -1.5274 | 0.92825 | 0.95374 | 1 | 20150 | 1 | -1.5274 |
| NonTargetingControlG | 2 | 0.071739 | 0.12522 | 0.951052 | 3908 | 2 | -2.2926 | 0.92826 | 0.92841 | 1 | 20151 | 0 | -2.2926 |
| PPP4R2 | 3 | 0.047303 | 0.11194 | 0.951052 | 2868 | 2 | -5.0735 | 0.92836 | 0.92845 | 1 | 20152 | 0 | -5.0735 |
| PRPF39 | 4 | 0.1189 | 0.24832 | 0.990682 | 5599 | 2 | -1.4216 | 0.92843 | 0.92824 | 1 | 20153 | 1 | -1.4216 |
| TRMT10B | 5 | 0.33647 | 0.51084 | 0.999994 | 10857 | 2 | -0.57281 | 0.92844 | 0.93798 | 1 | 20154 | 1 | -0.57281 |
| HOXA2 | 5 | 0.017373 | 0.056895 | 0.925104 | 1329 | 4 | -1.8352 | 0.92881 | 0.93819 | 1 | 20155 | 1 | -1.8352 |
| ARL2 | 4 | 0.060697 | 0.1421 | 0.959216 | 3451 | 3 | -2.0379 | 0.92886 | 0.92866 | 1 | 20156 | 1 | -2.0379 |
| SULF2 | 5 | 0.029459 | 0.089064 | 0.94395 | 1982 | 4 | -1.1499 | 0.92903 | 0.93833 | 1 | 20157 | 1 | -1.1499 |
| FREM2 | 3 | 0.24409 | 0.36763 | 0.999994 | 8884 | 2 | -0.9031 | 0.92913 | 0.92923 | 1 | 20158 | 0 | -0.9031 |
| FASLG | 5 | 0.14926 | 0.30164 | 0.999994 | 6436 | 2 | 0.071193 | 0.92915 | 0.93839 | 1 | 20159 | 1 | 0.071193 |
| KRTAP27-1 | 6 | 0.021255 | 0.076013 | 0.93914 | 1532 | 5 | -1.5777 | 0.92916 | 0.95404 | 1 | 20160 | 1 | -1.5777 |
| hsa-mir-6770-1 | 4 | 0.082438 | 0.18323 | 0.968526 | 4315 | 3 | -1.1159 | 0.9292 | 0.92898 | 1 | 20161 | 1 | -1.1159 |
| hsa-mir-4286 | 4 | 0.023659 | 0.068391 | 0.93914 | 1693 | 3 | -2.8377 | 0.92926 | 0.92904 | 1 | 20162 | 1 | -2.8377 |
| NAA50 | 4 | 0.27911 | 0.42165 | 0.999994 | 9679 | 2 | -0.56287 | 0.92927 | 0.92905 | 1 | 20163 | 1 | -0.56287 |
| B3GNT3 | 4 | 0.056293 | 0.13373 | 0.95318 | 3257 | 3 | -2.2206 | 0.9294 | 0.92918 | 1 | 20164 | 1 | -2.2206 |
| NUP153 | 4 | 0.15484 | 0.29419 | 0.999994 | 6591 | 1 | 0.36629 | 0.92943 | 0.92921 | 1 | 20165 | 1 | 0.36629 |
| hsa-mir-1183 | 4 | 0.31292 | 0.45625 | 0.999994 | 10375 | 2 | -0.6429 | 0.92946 | 0.92925 | 1 | 20166 | 1 | -0.6429 |
| GTF3C3 | 6 | 0.00026727 | 0.0014368 | 0.384267 | 73 | 5 | -4.3781 | 0.92947 | 0.95414 | 1 | 20167 | 1 | -4.3781 |
| HIST1H3B | 6 | 0.12143 | 0.28844 | 0.999632 | 5680 | 4 | -0.72523 | 0.92952 | 0.95416 | 1 | 20168 | 1 | -0.72523 |
| PABPC1 | 5 | 0.47442 | 0.62587 | 0.999994 | 13054 | 2 | -0.13343 | 0.92953 | 0.9386 | 1 | 20169 | 1 | -0.13343 |
| ADM | 6 | 0.027208 | 0.09224 | 0.943968 | 1865 | 2 | -0.1355 | 0.92963 | 0.9542 | 1 | 20170 | 1 | -0.1355 |
| GJB2 | 4 | 0.19436 | 0.33486 | 0.999994 | 7617 | 1 | -0.32679 | 0.92979 | 0.92957 | 1 | 20171 | 1 | -0.32679 |
| GRK5 | 6 | 0.021173 | 0.075742 | 0.93914 | 1525 | 5 | -1.2457 | 0.92985 | 0.95427 | 1 | 20172 | 1 | -1.2457 |
| PGAP1 | 6 | 0.038602 | 0.11875 | 0.951052 | 2444 | 4 | -1.7483 | 0.92986 | 0.95428 | 1 | 20173 | 1 | -1.7483 |
| CPED1 | 4 | 0.3801 | 0.52524 | 0.999994 | 11649 | 2 | -0.67871 | 0.92993 | 0.9297 | 1 | 20174 | 1 | -0.67871 |
| ANKIB1 | 6 | 0.072592 | 0.19215 | 0.974607 | 3944 | 3 | -1.323 | 0.92994 | 0.9543 | 1 | 20175 | 1 | -1.323 |

| | | | | | | | | | | | | | |
|---|---|---|---|---|---|---|---|---|---|---|---|---|---|
| RCE1 | 4 | 0.25389 | 0.39592 | 0.999994 | 9101 | 2 | -0.67347 | 0.92997 | 0.92975 | 1 | 20176 | 1 | -0.67347 |
| ANGPT4 | 5 | 0.15947 | 0.3168 | 0.999994 | 6718 | 2 | -0.21306 | 0.93004 | 0.93891 | 1 | 20177 | 1 | -0.21306 |
| PRSS33 | 5 | 0.082003 | 0.19784 | 0.977164 | 4298 | 3 | -0.64425 | 0.93011 | 0.93896 | 1 | 20178 | 1 | -0.64425 |
| FGF18 | 5 | 0.51368 | 0.65981 | 0.999994 | 13637 | 1 | 0.22505 | 0.93024 | 0.93903 | 1 | 20179 | 1 | 0.22505 |
| PCDHB4 | 6 | 0.095908 | 0.2393 | 0.987253 | 4809 | 4 | -1.0269 | 0.93037 | 0.95444 | 1 | 20180 | 1 | -1.0269 |
| ADC | 6 | 0.098383 | 0.24416 | 0.988309 | 4909 | 3 | -0.752 | 0.9304 | 0.95445 | 1 | 20181 | 1 | -0.752 |
| H2BFWT | 6 | 0.088086 | 0.2237 | 0.983354 | 4518 | 4 | -1.2752 | 0.93053 | 0.9545 | 1 | 20182 | 1 | -1.2752 |
| VAT1 | 5 | 0.051393 | 0.14377 | 0.961477 | 3042 | 4 | -0.75631 | 0.93068 | 0.93929 | 1 | 20183 | 1 | -0.75631 |
| HLA-DRB5 | 5 | 0.57325 | 0.71241 | 0.999994 | 14595 | 2 | -0.32667 | 0.93102 | 0.93949 | 1 | 20184 | 1 | -0.32667 |
| LDLRAD2 | 6 | 0.0070536 | 0.028615 | 0.858757 | 663 | 4 | -2.9699 | 0.93104 | 0.95468 | 1 | 20185 | 1 | -2.9699 |
| RPF1 | 5 | 0.20927 | 0.38836 | 0.999994 | 8013 | 2 | -0.011901 | 0.93105 | 0.93951 | 1 | 20186 | 1 | -0.011901 |
| ACTB | 3 | 0.15146 | 0.25038 | 0.992497 | 6494 | 2 | -1.3504 | 0.93112 | 0.93118 | 1 | 20187 | 0 | -1.3504 |
| NonTargetingControlG | 1 | 0.068734 | 0.068764 | 0.93914 | 3782 | 1 | -7.4616 | 0.93127 | 0.93143 | 1 | 20188 | 0 | -7.4616 |
| CYP24A1 | 6 | 0.0029703 | 0.013624 | 0.75005 | 352 | 5 | -4.1839 | 0.9313 | 0.95476 | 1 | 20189 | 1 | -4.1839 |
| SETDB2 | 5 | 0.29153 | 0.47419 | 0.999994 | 9939 | 2 | 0.14564 | 0.93147 | 0.93975 | 1 | 20190 | 1 | 0.14564 |
| ZIC3 | 6 | 0.21224 | 0.40281 | 0.999994 | 8067 | 3 | -0.48863 | 0.9315 | 0.95483 | 1 | 20191 | 1 | -0.48863 |
| TNRC18 | 3 | 0.25647 | 0.38299 | 0.999994 | 9161 | 2 | -0.78128 | 0.93151 | 0.93156 | 1 | 20192 | 0 | -0.78128 |
| LALBA | 6 | 0.72091 | 0.81861 | 0.999994 | 16505 | 1 | 0.0072831 | 0.93154 | 0.95485 | 1 | 20193 | 1 | 0.0072831 |
| LYSMD4 | 6 | 0.042222 | 0.1269 | 0.951052 | 2627 | 4 | -1.7131 | 0.93155 | 0.95485 | 1 | 20194 | 1 | -1.7131 |
| PCDHGB3 | 2 | 0.06837 | 0.11974 | 0.951052 | 3761 | 2 | -3.6202 | 0.93163 | 0.93175 | 1 | 20195 | 0 | -3.6202 |
| DENND2C | 6 | 0.086077 | 0.21965 | 0.980699 | 4450 | 4 | -1.55 | 0.9317 | 0.9549 | 1 | 20196 | 1 | -1.55 |
| C12orf54 | 4 | 0.3834 | 0.52863 | 0.999994 | 11696 | 2 | -0.1956 | 0.93175 | 0.9315 | 1 | 20197 | 0 | -0.1956 |
| OGFOD3 | 5 | 0.0086729 | 0.032249 | 0.865899 | 772 | 4 | -3.2633 | 0.93178 | 0.93993 | 1 | 20198 | 1 | -3.2633 |
| VSTM4 | 5 | 0.029093 | 0.088151 | 0.94395 | 1961 | 4 | -2.2257 | 0.93181 | 0.93995 | 1 | 20199 | 1 | -2.2257 |
| hsa-mir-7641-1 | 1 | 0.068149 | 0.068175 | 0.93914 | 3755 | 1 | -7.4817 | 0.93185 | 0.93202 | 1 | 20200 | 0 | -7.4817 |
| NonTargetingControlG | 2 | 0.068001 | 0.11912 | 0.951052 | 3750 | 2 | -4.7199 | 0.932 | 0.93214 | 1 | 20201 | 0 | -4.7199 |
| DEFB1 | 5 | 0.020723 | 0.06602 | 0.93914 | 1491 | 3 | -2.9776 | 0.93209 | 0.94011 | 1 | 20202 | 1 | -2.9776 |
| LIN7C | 4 | 0.48903 | 0.61146 | 0.999994 | 13267 | 1 | -0.061789 | 0.93219 | 0.93193 | 1 | 20203 | 0 | -0.061789 |
| C22orf43 | 4 | 0.106 | 0.22583 | 0.983354 | 5175 | 2 | -1.1009 | 0.9322 | 0.93194 | 1 | 20204 | 0 | -1.1009 |
| KRTAP1-4 | 6 | 0.19772 | 0.38512 | 0.999994 | 7700 | 2 | -0.12743 | 0.93223 | 0.95507 | 1 | 20205 | 1 | -0.12743 |
| RBMY1A1 | 6 | 0.62753 | 0.76049 | 0.999994 | 15344 | 2 | 0.047662 | 0.93224 | 0.95507 | 1 | 20206 | 1 | 0.047662 |
| SZT2 | 4 | 0.032465 | 0.08653 | 0.94395 | 2133 | 2 | -0.35418 | 0.93228 | 0.93202 | 1 | 20207 | 0 | -0.35418 |
| GEMIN6 | 3 | 0.024713 | 0.06283 | 0.937389 | 1743 | 2 | -5.3627 | 0.93228 | 0.93233 | 1 | 20208 | 0 | -5.3627 |
| CASP3 | 5 | 0.1279 | 0.26939 | 0.998561 | 5868 | 3 | -1.1803 | 0.93233 | 0.94025 | 1 | 20209 | 1 | -1.1803 |
| FOXN2 | 6 | 0.091676 | 0.2309 | 0.983933 | 4664 | 4 | -0.95154 | 0.93234 | 0.9551 | 1 | 20210 | 1 | -0.95154 |
| SIRPA | 5 | 0.026342 | 0.080994 | 0.93914 | 1820 | 4 | -2.5986 | 0.93257 | 0.9404 | 1 | 20211 | 1 | -2.5986 |
| hsa-mir-140 | 4 | 0.83881 | 0.83826 | 0.999994 | 18068 | 0 | 0.046386 | 0.93261 | 0.93235 | 1 | 20212 | 0 | 0.046386 |
| CWF19L2 | 4 | 0.077183 | 0.1735 | 0.96793 | 4114 | 3 | -1.5971 | 0.93266 | 0.93241 | 1 | 20213 | 0 | -1.5971 |
| MYBL1 | 4 | 0.14035 | 0.2793 | 0.999632 | 6195 | 1 | 0.22936 | 0.93269 | 0.93244 | 1 | 20214 | 0 | 0.22936 |
| MTA1 | 5 | 0.012459 | 0.0431 | 0.891621 | 1026 | 4 | -2.3331 | 0.93273 | 0.9405 | 1 | 20215 | 1 | -2.3331 |
| CBWD2 | 1 | 0.067155 | 0.06721 | 0.93914 | 3724 | 1 | -5.7094 | 0.93285 | 0.93303 | 1 | 20216 | 0 | -5.7094 |
| FOXD4L3 | 3 | 0.15917 | 0.2604 | 0.994943 | 6709 | 2 | -1.5352 | 0.93287 | 0.93295 | 1 | 20217 | 0 | -1.5352 |
| NPFFR1 | 5 | 0.39093 | 0.55591 | 0.999994 | 11814 | 2 | -0.45567 | 0.93308 | 0.9407 | 1 | 20218 | 1 | -0.45567 |
| BARHL1 | 4 | 0.034226 | 0.090173 | 0.94395 | 2226 | 3 | -4.3535 | 0.93309 | 0.93284 | 1 | 20219 | 0 | -4.3535 |
| hsa-mir-4781 | 3 | 0.28277 | 0.41537 | 0.999994 | 9754 | 2 | -0.63017 | 0.93309 | 0.93318 | 1 | 20220 | 0 | -0.63017 |
| LCE1F | 4 | 0.56087 | 0.64511 | 0.999994 | 14385 | 1 | -0.048404 | 0.93326 | 0.93302 | 1 | 20221 | 0 | -0.048404 |
| RNF112 | 6 | 0.33646 | 0.54862 | 0.999994 | 10856 | 3 | -0.50292 | 0.93328 | 0.9554 | 1 | 20222 | 1 | -0.50292 |
| NonTargetingControlG | 2 | 0.066497 | 0.11665 | 0.951052 | 3703 | 2 | -4.1942 | 0.9335 | 0.93359 | 1 | 20223 | 0 | -4.1942 |
| ALKBH4 | 5 | 0.77191 | 0.80279 | 0.999994 | 17110 | 1 | -0.13482 | 0.93351 | 0.94094 | 1 | 20224 | 1 | -0.13482 |
| NonTargetingControlG | 2 | 0.066407 | 0.11648 | 0.951052 | 3701 | 2 | -4.6083 | 0.93359 | 0.93368 | 1 | 20225 | 0 | -4.6083 |
| LCE1C | 4 | 0.33754 | 0.48143 | 0.999994 | 10888 | 2 | -0.34574 | 0.93365 | 0.93342 | 1 | 20226 | 0 | -0.34574 |
| hsa-mir-758 | 4 | 0.64561 | 0.69075 | 0.999994 | 15594 | 1 | 0.28807 | 0.93373 | 0.9335 | 1 | 20227 | 0 | 0.28807 |

| | | | | | | | | | | | | | |
|---|---|---|---|---|---|---|---|---|---|---|---|---|---|
| PCED1A | 6 | 0.073268 | 0.19355 | 0.975797 | 3969 | 4 | -1.5201 | 0.93388 | 0.95559 | 1 | 20228 | 1 | -1.5201 |
| hsa-mir-6819 | 2 | 0.066111 | 0.11602 | 0.951052 | 3690 | 2 | -3.2049 | 0.93389 | 0.93397 | 1 | 20229 | 0 | -3.2049 |
| TRMT44 | 4 | 0.057531 | 0.13604 | 0.95468 | 3315 | 3 | -1.7304 | 0.93403 | 0.93382 | 1 | 20230 | 0 | -1.7304 |
| TCEA1 | 4 | 0.048543 | 0.11858 | 0.951052 | 2910 | 3 | -1.9508 | 0.93404 | 0.93383 | 1 | 20231 | 0 | -1.9508 |
| SCN10A | 5 | 0.022489 | 0.070817 | 0.93914 | 1618 | 4 | -2.4663 | 0.93408 | 0.94129 | 1 | 20232 | 1 | -2.4663 |
| MYL12B | 4 | 0.13092 | 0.26916 | 0.998561 | 5957 | 2 | -0.26304 | 0.93413 | 0.93392 | 1 | 20233 | 0 | -0.26304 |
| MICB | 5 | 0.2157 | 0.39726 | 0.999994 | 8159 | 3 | -0.70722 | 0.93419 | 0.94135 | 1 | 20234 | 1 | -0.70722 |
| AK9 | 5 | 0.47706 | 0.6281 | 0.999994 | 13100 | 2 | -0.25967 | 0.93433 | 0.94144 | 1 | 20235 | 1 | -0.25967 |
| LIMS1 | 6 | 0.059213 | 0.16406 | 0.967126 | 3388 | 3 | -1.0637 | 0.9344 | 0.95578 | 1 | 20236 | 1 | -1.0637 |
| ZAR1 | 4 | 0.30749 | 0.45065 | 0.999994 | 10277 | 2 | -0.9009 | 0.93441 | 0.93422 | 1 | 20237 | 0 | -0.9009 |
| DNAJC17 | 3 | 0.017212 | 0.044742 | 0.891621 | 1313 | 2 | -7.1108 | 0.93465 | 0.93476 | 1 | 20238 | 0 | -7.1108 |
| DUSP22 | 6 | 0.006209 | 0.025578 | 0.845628 | 606 | 5 | -3.5933 | 0.93467 | 0.95587 | 1 | 20239 | 1 | -3.5933 |
| ZFYVE27 | 6 | 0.048192 | 0.14012 | 0.958716 | 2896 | 4 | -1.9339 | 0.93469 | 0.95588 | 1 | 20240 | 1 | -1.9339 |
| TNFAIP8L2-SCNM1 | 1 | 0.065288 | 0.065327 | 0.93914 | 3665 | 1 | -5.5222 | 0.93471 | 0.93492 | 1 | 20241 | 0 | -5.5222 |
| GPC3 | 6 | 0.011413 | 0.043807 | 0.891621 | 950 | 5 | -1.9661 | 0.93484 | 0.95593 | 1 | 20242 | 1 | -1.9661 |
| SLC30A4 | 6 | 0.54316 | 0.69878 | 0.999994 | 14114 | 2 | -0.042308 | 0.9349 | 0.95594 | 1 | 20243 | 1 | -0.042308 |
| hsa-mir-4771-1 | 4 | 0.41371 | 0.55975 | 0.999994 | 12164 | 2 | -0.19138 | 0.9349 | 0.93472 | 1 | 20244 | 0 | -0.19138 |
| NLRP14 | 6 | 0.011153 | 0.042918 | 0.891621 | 936 | 5 | -1.8298 | 0.9352 | 0.95604 | 1 | 20245 | 1 | -1.8298 |
| BBOX1 | 5 | 0.05406 | 0.15019 | 0.966359 | 3160 | 4 | -1.5464 | 0.93527 | 0.94203 | 1 | 20246 | 1 | -1.5464 |
| NonTargetingControlG | 2 | 0.060209 | 0.10626 | 0.950812 | 3431 | 2 | -4.4142 | 0.93543 | 0.93554 | 1 | 20247 | 0 | -4.4142 |
| NONO | 6 | 0.033474 | 0.10695 | 0.950995 | 2189 | 4 | -1.7606 | 0.9355 | 0.95614 | 1 | 20248 | 1 | -1.7606 |
| hsa-mir-4710 | 2 | 0.064478 | 0.11335 | 0.951052 | 3628 | 2 | -1.9446 | 0.93552 | 0.93564 | 1 | 20249 | 0 | -1.9446 |
| IRS1 | 6 | 0.042522 | 0.12758 | 0.951052 | 2647 | 4 | -1.8054 | 0.93554 | 0.95616 | 1 | 20250 | 1 | -1.8054 |
| TPST1 | 4 | 0.25359 | 0.3956 | 0.999994 | 9094 | 2 | -0.48895 | 0.93559 | 0.93539 | 1 | 20251 | 0 | -0.48895 |
| NonTargetingControlG | 2 | 0.064296 | 0.11305 | 0.951052 | 3620 | 2 | -4.2149 | 0.9357 | 0.93582 | 1 | 20252 | 0 | -4.2149 |
| LGALS3BP | 2 | 0.06427 | 0.11301 | 0.951052 | 3619 | 2 | -4.977 | 0.93573 | 0.93585 | 1 | 20253 | 0 | -4.977 |
| GLIPR1 | 5 | 0.01922 | 0.06197 | 0.937033 | 1419 | 4 | -1.4603 | 0.93586 | 0.94241 | 1 | 20254 | 1 | -1.4603 |
| RENBP | 6 | 0.17694 | 0.35926 | 0.999994 | 7155 | 3 | -0.2899 | 0.9359 | 0.95628 | 1 | 20255 | 1 | -0.2899 |
| DNAAF1 | 5 | 0.38659 | 0.55229 | 0.999994 | 11744 | 2 | -0.41352 | 0.93607 | 0.94254 | 1 | 20256 | 1 | -0.41352 |
| MAPK13 | 5 | 0.29624 | 0.47801 | 0.999994 | 10041 | 1 | -0.11194 | 0.9361 | 0.94256 | 1 | 20257 | 1 | -0.11194 |
| DPP10 | 6 | 0.27171 | 0.47372 | 0.999994 | 9514 | 3 | -0.51206 | 0.93615 | 0.95637 | 1 | 20258 | 1 | -0.51206 |
| OGG1 | 4 | 0.05099 | 0.12343 | 0.951052 | 3019 | 3 | -2.1892 | 0.93616 | 0.93596 | 1 | 20259 | 0 | -2.1892 |
| RALGAPB | 5 | 0.21292 | 0.39347 | 0.999994 | 8091 | 3 | -0.68202 | 0.93617 | 0.94261 | 1 | 20260 | 1 | -0.68202 |
| SH3PXD2B | 4 | 0.12149 | 0.25289 | 0.994501 | 5686 | 2 | -0.39026 | 0.93623 | 0.93603 | 1 | 20261 | 0 | -0.39026 |
| C8orf42 | 4 | 0.62306 | 0.6781 | 0.999994 | 15274 | 1 | 0.098762 | 0.93631 | 0.93613 | 1 | 20262 | 0 | 0.098762 |
| F11R | 5 | 0.16357 | 0.32273 | 0.999994 | 6830 | 3 | -1.0602 | 0.93641 | 0.94275 | 1 | 20263 | 1 | -1.0602 |
| PRR20A | 6 | 0.34625 | 0.55979 | 0.999994 | 11080 | 3 | -0.32757 | 0.93642 | 0.95645 | 1 | 20264 | 1 | -0.32757 |
| MRPL53 | 5 | 0.0088459 | 0.032725 | 0.867348 | 782 | 3 | -5.9335 | 0.93657 | 0.94285 | 1 | 20265 | 1 | -5.9335 |
| hsa-mir-3661 | 3 | 0.049875 | 0.11552 | 0.951052 | 2969 | 2 | -3.5379 | 0.93676 | 0.93688 | 1 | 20266 | 0 | -3.5379 |
| NSDHL | 4 | 0.085367 | 0.18858 | 0.969543 | 4425 | 2 | -1.4701 | 0.93678 | 0.93663 | 1 | 20267 | 0 | -1.4701 |
| KHDRBS3 | 5 | 0.17329 | 0.33694 | 0.999994 | 7070 | 3 | -0.97021 | 0.93684 | 0.943 | 1 | 20268 | 1 | -0.97021 |
| AKNAD1 | 6 | 0.05485 | 0.1546 | 0.967126 | 3193 | 4 | -1.7208 | 0.93692 | 0.95662 | 1 | 20269 | 1 | -1.7208 |
| DICER1 | 5 | 0.006745 | 0.026631 | 0.848412 | 643 | 3 | -1.7264 | 0.93698 | 0.94307 | 1 | 20270 | 1 | -1.7264 |
| hsa-mir-6134 | 4 | 0.18763 | 0.32794 | 0.999994 | 7412 | 2 | -1.3697 | 0.93698 | 0.93683 | 1 | 20271 | 0 | -1.3697 |
| SEL1L | 4 | 0.13087 | 0.26906 | 0.998561 | 5954 | 3 | -1.1292 | 0.93724 | 0.93709 | 1 | 20272 | 0 | -1.1292 |
| ACVR2B | 4 | 0.0091144 | 0.03091 | 0.862223 | 805 | 3 | -2.7242 | 0.93725 | 0.93711 | 1 | 20273 | 0 | -2.7242 |
| hsa-mir-548h-3 | 3 | 0.23811 | 0.36019 | 0.999994 | 8729 | 2 | -0.8845 | 0.93727 | 0.93741 | 1 | 20274 | 0 | -0.8845 |
| ASB15 | 6 | 0.22907 | 0.42295 | 0.999994 | 8507 | 3 | -0.72092 | 0.93737 | 0.95677 | 1 | 20275 | 1 | -0.72092 |
| KPNA6 | 4 | 0.1109 | 0.23446 | 0.984971 | 5328 | 2 | -1.3865 | 0.93743 | 0.93729 | 1 | 20276 | 0 | -1.3865 |
| GP2 | 5 | 0.20039 | 0.37578 | 0.999994 | 7770 | 3 | -0.81892 | 0.93749 | 0.94339 | 1 | 20277 | 1 | -0.81892 |
| hsa-mir-510 | 4 | 0.23506 | 0.3767 | 0.999994 | 8646 | 2 | -1.0076 | 0.93756 | 0.93742 | 1 | 20278 | 0 | -1.0076 |
| hsa-mir-625 | 3 | 0.44679 | 0.51226 | 0.999994 | 12672 | 1 | -0.26566 | 0.9376 | 0.93773 | 1 | 20279 | 0 | -0.26566 |

| | | | | | | | | | | | | | |
|---|---|---|---|---|---|---|---|---|---|---|---|---|---|
| MS4A13 | 6 | 0.012672 | 0.047999 | 0.900643 | 1043 | 5 | -1.7627 | 0.93761 | 0.95685 | 1 | 20280 | 1 | -1.7627 |
| NAALADL2 | 4 | 0.27244 | 0.41484 | 0.999994 | 9525 | 2 | -0.92848 | 0.93764 | 0.9375 | 1 | 20281 | 0 | -0.92848 |
| NonTargetingControlG | 2 | 0.062322 | 0.10977 | 0.951052 | 3526 | 2 | -5.0242 | 0.93768 | 0.93782 | 1 | 20282 | 0 | -5.0242 |
| GPR173 | 2 | 0.062252 | 0.10966 | 0.951052 | 3522 | 2 | -1.7216 | 0.93775 | 0.93789 | 1 | 20283 | 0 | -1.7216 |
| TMEM19 | 6 | 0.53183 | 0.69085 | 0.999994 | 13931 | 2 | -0.2889 | 0.93778 | 0.95691 | 1 | 20284 | 1 | -0.2889 |
| SSBP2 | 5 | 0.092419 | 0.21446 | 0.979587 | 4698 | 3 | -1.697 | 0.93778 | 0.94356 | 1 | 20285 | 1 | -1.697 |
| NPAS2 | 3 | 0.023768 | 0.060608 | 0.932932 | 1697 | 2 | -4.6458 | 0.93783 | 0.93797 | 1 | 20286 | 0 | -4.6458 |
| OR6Y1 | 4 | 0.085849 | 0.18944 | 0.970473 | 4440 | 2 | -1.8286 | 0.93791 | 0.93776 | 1 | 20287 | 0 | -1.8286 |
| USH2A | 5 | 0.79989 | 0.81944 | 0.999994 | 17477 | 1 | -0.39643 | 0.93794 | 0.94365 | 1 | 20288 | 1 | -0.39643 |
| RUFY2 | 6 | 0.19175 | 0.3777 | 0.999994 | 7535 | 3 | -0.98268 | 0.93796 | 0.95696 | 1 | 20289 | 1 | -0.98268 |
| UBE2Z | 5 | 0.082728 | 0.19902 | 0.977164 | 4327 | 3 | -1.8261 | 0.93802 | 0.9437 | 1 | 20290 | 1 | -1.8261 |
| SHC1 | 5 | 0.16186 | 0.32028 | 0.999994 | 6786 | 3 | -1.0701 | 0.93806 | 0.94373 | 1 | 20291 | 1 | -1.0701 |
| ACSBG2 | 4 | 0.0091314 | 0.030965 | 0.862223 | 806 | 3 | -4.9014 | 0.93808 | 0.93795 | 1 | 20292 | 0 | -4.9014 |
| HIST3H2BB | 6 | 0.0030861 | 0.014062 | 0.75278 | 365 | 5 | -2.9183 | 0.93825 | 0.95705 | 1 | 20293 | 1 | -2.9183 |
| SLC4A11 | 5 | 0.18516 | 0.35404 | 0.999994 | 7345 | 3 | -0.85411 | 0.93826 | 0.94387 | 1 | 20294 | 1 | -0.85411 |
| NonTargetingControlG | 2 | 0.06173 | 0.1088 | 0.951052 | 3501 | 2 | -1.6311 | 0.93827 | 0.93841 | 1 | 20295 | 0 | -1.6311 |
| CDC20 | 2 | 0.061599 | 0.10858 | 0.951052 | 3496 | 2 | -2.7719 | 0.9384 | 0.93853 | 1 | 20296 | 0 | -2.7719 |
| TRMT2A | 4 | 0.078351 | 0.17568 | 0.968171 | 4166 | 3 | -1.3801 | 0.93873 | 0.9386 | 1 | 20297 | 0 | -1.3801 |
| hsa-mir-4258 | 4 | 0.51074 | 0.62112 | 0.999994 | 13588 | 1 | 0.42974 | 0.93876 | 0.93864 | 1 | 20298 | 0 | 0.42974 |
| DDTL | 4 | 0.26427 | 0.40643 | 0.999994 | 9338 | 2 | -0.82776 | 0.93879 | 0.93867 | 1 | 20299 | 0 | -0.82776 |
| PLEKHM3 | 5 | 0.12712 | 0.26823 | 0.998561 | 5851 | 3 | -1.2949 | 0.93888 | 0.94425 | 1 | 20300 | 1 | -1.2949 |
| NBPF4 | 5 | 0.0096499 | 0.035067 | 0.867348 | 850 | 4 | -1.9237 | 0.9389 | 0.94428 | 1 | 20301 | 1 | -1.9237 |
| C8orf44 | 4 | 0.060781 | 0.14227 | 0.959553 | 3459 | 3 | -2.5805 | 0.93895 | 0.93885 | 1 | 20302 | 0 | -2.5805 |
| MGAT2 | 5 | 0.17588 | 0.34067 | 0.999994 | 7131 | 3 | -1.0188 | 0.93898 | 0.94433 | 1 | 20303 | 1 | -1.0188 |
| NonTargetingControlG | 2 | 0.060881 | 0.10736 | 0.950995 | 3463 | 2 | -1.7136 | 0.93912 | 0.93926 | 1 | 20304 | 0 | -1.7136 |
| B3GALNT2 | 5 | 0.4652 | 0.61804 | 0.999994 | 12935 | 1 | 0.13785 | 0.93984 | 0.94486 | 1 | 20305 | 1 | 0.13785 |
| PLA2G10 | 5 | 0.11738 | 0.25335 | 0.994501 | 5554 | 2 | -0.070163 | 0.93985 | 0.94487 | 1 | 20306 | 1 | -0.070163 |
| hsa-mir-105-2 | 2 | 0.060041 | 0.10597 | 0.950444 | 3425 | 2 | -2.224 | 0.93996 | 0.94008 | 1 | 20307 | 0 | -2.224 |
| ZNF273 | 5 | 0.0093205 | 0.034113 | 0.867348 | 816 | 4 | -2.2199 | 0.93996 | 0.94493 | 1 | 20308 | 1 | -2.2199 |
| INPP5F | 6 | 0.068326 | 0.18328 | 0.968526 | 3758 | 4 | -1.6724 | 0.94001 | 0.95771 | 1 | 20309 | 1 | -1.6724 |
| PRSS38 | 6 | 0.090665 | 0.22883 | 0.983354 | 4624 | 4 | -0.97075 | 0.94002 | 0.95771 | 1 | 20310 | 1 | -0.97075 |
| OR51I1 | 5 | 0.05805 | 0.15885 | 0.967126 | 3341 | 3 | -0.88169 | 0.94003 | 0.94498 | 1 | 20311 | 1 | -0.88169 |
| CRISP3 | 5 | 0.10253 | 0.23035 | 0.983933 | 5055 | 3 | -1.4479 | 0.94015 | 0.94506 | 1 | 20312 | 1 | -1.4479 |
| ZNF142 | 5 | 0.14213 | 0.29105 | 0.999632 | 6242 | 3 | -1.1307 | 0.94023 | 0.94511 | 1 | 20313 | 1 | -1.1307 |
| CUL9 | 4 | 0.10409 | 0.22239 | 0.983354 | 5099 | 2 | -1.7046 | 0.94024 | 0.94014 | 1 | 20314 | 0 | -1.7046 |
| RGMA | 4 | 0.34533 | 0.48954 | 0.999994 | 11057 | 2 | -0.80306 | 0.94034 | 0.94023 | 1 | 20315 | 0 | -0.80306 |
| OLFML2B | 5 | 0.44034 | 0.59709 | 0.999994 | 12582 | 1 | 0.42418 | 0.94042 | 0.94524 | 1 | 20316 | 1 | 0.42418 |
| IL36G | 6 | 0.003721 | 0.016435 | 0.772095 | 423 | 5 | -2.0297 | 0.94043 | 0.95785 | 1 | 20317 | 1 | -2.0297 |
| PARD6B | 6 | 0.029528 | 0.097676 | 0.945253 | 1985 | 4 | -0.84603 | 0.94043 | 0.95785 | 1 | 20318 | 1 | -0.84603 |
| hsa-mir-4326 | 2 | 0.059284 | 0.10472 | 0.948754 | 3393 | 2 | -2.7764 | 0.94072 | 0.94082 | 1 | 20319 | 0 | -2.7764 |
| PDE6B | 6 | 0.022064 | 0.078576 | 0.93914 | 1589 | 5 | -1.155 | 0.94072 | 0.95798 | 1 | 20320 | 1 | -1.155 |
| FAM198A | 4 | 0.34711 | 0.49131 | 0.999994 | 11095 | 2 | -0.7913 | 0.94073 | 0.94062 | 1 | 20321 | 0 | -0.7913 |
| TNFRSF10A | 6 | 0.81357 | 0.85387 | 0.999994 | 17684 | 1 | 0.20375 | 0.94086 | 0.95803 | 1 | 20322 | 1 | 0.20375 |
| ZDHHC4 | 5 | 0.4706 | 0.62266 | 0.999994 | 13012 | 2 | -0.52731 | 0.941 | 0.94564 | 1 | 20323 | 1 | -0.52731 |
| PAPLN | 5 | 0.037441 | 0.10944 | 0.951052 | 2394 | 4 | -1.9055 | 0.94106 | 0.94569 | 1 | 20324 | 1 | -1.9055 |
| PLCG2 | 4 | 0.66468 | 0.70211 | 0.999994 | 15852 | 1 | -0.2092 | 0.9411 | 0.941 | 1 | 20325 | 0 | -0.2092 |
| LAMC2 | 4 | 0.011118 | 0.037241 | 0.881473 | 934 | 2 | -4.4627 | 0.9411 | 0.941 | 1 | 20326 | 0 | -4.4627 |
| LARS | 4 | 0.05567 | 0.13251 | 0.95318 | 3230 | 2 | -3.2594 | 0.9411 | 0.941 | 1 | 20327 | 0 | -3.2594 |
| hsa-mir-6866 | 3 | 0.23131 | 0.35172 | 0.999994 | 8551 | 2 | -0.84457 | 0.94124 | 0.94135 | 1 | 20328 | 0 | -0.84457 |
| NYNRIN | 6 | 0.056374 | 0.1579 | 0.967126 | 3263 | 4 | -1.3393 | 0.94138 | 0.95823 | 1 | 20329 | 1 | -1.3393 |
| COL23A1 | 4 | 0.73197 | 0.74661 | 0.999994 | 16627 | 1 | 0.18137 | 0.94146 | 0.94134 | 1 | 20330 | 0 | 0.18137 |
| TSGA10 | 3 | 0.57563 | 0.59878 | 0.999994 | 14625 | 1 | -0.59553 | 0.94151 | 0.94161 | 1 | 20331 | 0 | -0.59553 |

| | | | | | | | | | | | | | |
|---|---|---|---|---|---|---|---|---|---|---|---|---|---|
| ANKRD17 | 4 | 0.34313 | 0.48723 | 0.999994 | 11009 | 2 | -0.50622 | 0.94153 | 0.94141 | 1 | 20332 | 0 | -0.50622 |
| RAET1G | 6 | 0.20546 | 0.39451 | 0.999994 | 7907 | 2 | -1.0109 | 0.94157 | 0.95829 | 1 | 20333 | 1 | -1.0109 |
| PAPOLG | 5 | 0.018595 | 0.060279 | 0.932932 | 1392 | 3 | -3.4478 | 0.9417 | 0.9461 | 1 | 20334 | 1 | -3.4478 |
| GSAP | 5 | 0.14768 | 0.29928 | 0.999994 | 6396 | 3 | -1.0554 | 0.94171 | 0.94611 | 1 | 20335 | 1 | -1.0554 |
| UQCRC2 | 4 | 0.22529 | 0.36664 | 0.999994 | 8407 | 2 | -0.65799 | 0.94178 | 0.94166 | 1 | 20336 | 0 | -0.65799 |
| PRMT1 | 6 | 0.0035674 | 0.015844 | 0.766918 | 412 | 5 | -3.438 | 0.94187 | 0.95841 | 1 | 20337 | 1 | -3.438 |
| PRAMEF20 | 4 | 0.26253 | 0.40465 | 0.999994 | 9298 | 2 | -0.98289 | 0.94206 | 0.94193 | 1 | 20338 | 0 | -0.98289 |
| hsa-mir-429 | 4 | 0.062747 | 0.14602 | 0.964852 | 3546 | 3 | -1.4441 | 0.94207 | 0.94194 | 1 | 20339 | 0 | -1.4441 |
| LANCL2 | 6 | 0.0048834 | 0.020716 | 0.81027 | 511 | 5 | -1.6165 | 0.94221 | 0.95855 | 1 | 20340 | 1 | -1.6165 |
| ACVRL1 | 5 | 0.039709 | 0.11518 | 0.951052 | 2500 | 4 | -1.4929 | 0.94222 | 0.94646 | 1 | 20341 | 1 | -1.4929 |
| TMEM176B | 6 | 0.0022193 | 0.010788 | 0.707936 | 294 | 5 | -5.0327 | 0.94223 | 0.95855 | 1 | 20342 | 1 | -5.0327 |
| LYPLAL1 | 6 | 0.043769 | 0.13039 | 0.95318 | 2701 | 4 | -1.7525 | 0.94245 | 0.95863 | 1 | 20343 | 1 | -1.7525 |
| OR5H2 | 5 | 0.070593 | 0.17935 | 0.968171 | 3859 | 3 | -1.8312 | 0.94255 | 0.94667 | 1 | 20344 | 1 | -1.8312 |
| CNRIP1 | 5 | 0.26087 | 0.44945 | 0.999994 | 9266 | 2 | -0.29428 | 0.94261 | 0.94671 | 1 | 20345 | 1 | -0.29428 |
| CCKAR | 4 | 0.24664 | 0.38845 | 0.999994 | 8937 | 2 | -0.67238 | 0.94271 | 0.94257 | 1 | 20346 | 0 | -0.67238 |
| MORN4 | 5 | 0.038064 | 0.11102 | 0.951052 | 2423 | 3 | -1.4814 | 0.94277 | 0.94683 | 1 | 20347 | 1 | -1.4814 |
| ANKRD61 | 5 | 0.14152 | 0.29012 | 0.999632 | 6232 | 3 | -0.69549 | 0.94282 | 0.94685 | 1 | 20348 | 1 | -0.69549 |
| TJP3 | 6 | 0.017465 | 0.063753 | 0.937389 | 1338 | 5 | -1.5257 | 0.94291 | 0.95882 | 1 | 20349 | 1 | -1.5257 |
| INTS10 | 5 | 0.12109 | 0.25902 | 0.994943 | 5666 | 3 | -1.3905 | 0.94308 | 0.94704 | 1 | 20350 | 1 | -1.3905 |
| YKT6 | 4 | 0.0074653 | 0.025688 | 0.845768 | 690 | 3 | -1.5364 | 0.94316 | 0.94301 | 1 | 20351 | 0 | -1.5364 |
| TLDC2 | 6 | 0.20334 | 0.39192 | 0.999994 | 7862 | 3 | -0.5465 | 0.94325 | 0.95895 | 1 | 20352 | 1 | -0.5465 |
| SLCO1A2 | 6 | 0.032652 | 0.105 | 0.95013 | 2142 | 4 | -1.7128 | 0.94326 | 0.95895 | 1 | 20353 | 1 | -1.7128 |
| FRG1 | 5 | 0.18035 | 0.34713 | 0.999994 | 7227 | 3 | -0.9502 | 0.94329 | 0.94719 | 1 | 20354 | 1 | -0.9502 |
| NPSR1 | 5 | 0.018749 | 0.060692 | 0.932932 | 1400 | 4 | -2.0283 | 0.94333 | 0.94721 | 1 | 20355 | 1 | -2.0283 |
| OR8J1 | 4 | 0.3558 | 0.50017 | 0.999994 | 11264 | 2 | -0.67977 | 0.94337 | 0.94321 | 1 | 20356 | 0 | -0.67977 |
| RNF34 | 6 | 0.23882 | 0.43458 | 0.999994 | 8742 | 3 | -0.58245 | 0.94339 | 0.959 | 1 | 20357 | 1 | -0.58245 |
| XKR4 | 6 | 0.14922 | 0.32463 | 0.999994 | 6434 | 3 | -1.05 | 0.94364 | 0.95911 | 1 | 20358 | 1 | -1.05 |
| KIF20B | 4 | 0.60364 | 0.66731 | 0.999994 | 14991 | 1 | -0.10304 | 0.9438 | 0.94366 | 1 | 20359 | 0 | -0.10304 |
| NonTargetingControlG | 1 | 0.056076 | 0.05606 | 0.9235 | 3251 | 1 | -7.9542 | 0.94392 | 0.94411 | 1 | 20360 | 0 | -7.9542 |
| LOH12CR1 | 5 | 0.051177 | 0.14323 | 0.960247 | 3032 | 4 | -0.90055 | 0.94409 | 0.94773 | 1 | 20361 | 1 | -0.90055 |
| IPO8 | 5 | 0.024866 | 0.077111 | 0.93914 | 1749 | 4 | -1.8282 | 0.94411 | 0.94775 | 1 | 20362 | 1 | -1.8282 |
| MB | 6 | 0.11358 | 0.27358 | 0.998561 | 5424 | 3 | -1.0342 | 0.94415 | 0.9593 | 1 | 20363 | 1 | -1.0342 |
| PSMF1 | 6 | 0.063479 | 0.17312 | 0.96793 | 3576 | 3 | -0.26038 | 0.94423 | 0.95933 | 1 | 20364 | 1 | -0.26038 |
| hsa-mir-3179-3 | 2 | 0.055714 | 0.098787 | 0.945253 | 3232 | 2 | -3.5365 | 0.94429 | 0.9443 | 1 | 20365 | 0 | -3.5365 |
| ZFYVE16 | 5 | 0.039922 | 0.11569 | 0.951052 | 2513 | 4 | -1.227 | 0.94433 | 0.9479 | 1 | 20366 | 1 | -1.227 |
| GREB1L | 6 | 0.037062 | 0.11528 | 0.951052 | 2379 | 4 | -0.69843 | 0.94442 | 0.95941 | 1 | 20367 | 1 | -0.69843 |
| OR2A5 | 5 | 0.61875 | 0.7331 | 0.999994 | 15199 | 1 | -0.18692 | 0.94443 | 0.94797 | 1 | 20368 | 1 | -0.18692 |
| ZNF883 | 5 | 0.2205 | 0.40395 | 0.999994 | 8300 | 3 | -0.72668 | 0.94448 | 0.948 | 1 | 20369 | 1 | -0.72668 |
| CEP192 | 3 | 0.15384 | 0.25346 | 0.994501 | 6562 | 2 | -0.92925 | 0.94448 | 0.94462 | 1 | 20370 | 0 | -0.92925 |
| PDC | 6 | 0.079573 | 0.20644 | 0.978744 | 4212 | 3 | -1.2709 | 0.94451 | 0.95944 | 1 | 20371 | 1 | -1.2709 |
| BAIAP2 | 5 | 0.15368 | 0.30828 | 0.999994 | 6555 | 3 | -1.0379 | 0.94457 | 0.94807 | 1 | 20372 | 1 | -1.0379 |
| CAD | 6 | 0.0395 | 0.12077 | 0.951052 | 2490 | 4 | -1.9217 | 0.9446 | 0.95948 | 1 | 20373 | 1 | -1.9217 |
| RGL1 | 6 | 0.0017978 | 0.0089621 | 0.680529 | 251 | 5 | -5.8191 | 0.94461 | 0.95948 | 1 | 20374 | 1 | -5.8191 |
| NonTargetingControlG | 2 | 0.055354 | 0.098203 | 0.945253 | 3213 | 2 | -3.273 | 0.94465 | 0.94467 | 1 | 20375 | 0 | -3.273 |
| REV3L | 6 | 0.09257 | 0.23264 | 0.984502 | 4706 | 4 | -1.081 | 0.94466 | 0.9595 | 1 | 20376 | 1 | -1.081 |
| FOS | 6 | 0.024449 | 0.085695 | 0.94395 | 1729 | 2 | -0.38954 | 0.94489 | 0.95959 | 1 | 20377 | 1 | -0.38954 |
| USP17L8 | 5 | 0.17009 | 0.33229 | 0.999994 | 6986 | 3 | -1.0272 | 0.94498 | 0.94834 | 1 | 20378 | 1 | -1.0272 |
| STAT4 | 5 | 0.77785 | 0.8062 | 0.999994 | 17182 | 1 | -0.083376 | 0.94501 | 0.94835 | 1 | 20379 | 1 | -0.083376 |
| hsa-mir-2392 | 3 | 0.64254 | 0.65089 | 0.999994 | 15555 | 1 | -0.56927 | 0.94505 | 0.94518 | 1 | 20380 | 0 | -0.56927 |
| PAQR8 | 5 | 0.12163 | 0.25984 | 0.994943 | 5690 | 3 | -0.89327 | 0.9451 | 0.94842 | 1 | 20381 | 1 | -0.89327 |
| SSH2 | 5 | 0.12233 | 0.26089 | 0.994943 | 5714 | 3 | -0.57252 | 0.94522 | 0.94852 | 1 | 20382 | 1 | -0.57252 |
| SLITRK5 | 4 | 0.083246 | 0.18472 | 0.968526 | 4350 | 3 | -2.1812 | 0.94528 | 0.94514 | 1 | 20383 | 0 | -2.1812 |

| | | | | | | | | | | | | | |
|---|---|---|---|---|---|---|---|---|---|---|---|---|---|
| MC4R | 5 | 0.23556 | 0.42488 | 0.999994 | 8661 | 2 | -0.60603 | 0.9453 | 0.94857 | 1 | 20384 | 1 | -0.60603 |
| CXXC11 | 6 | 0.0094021 | 0.036926 | 0.881473 | 819 | 5 | -2.0598 | 0.94541 | 0.95981 | 1 | 20385 | 1 | -2.0598 |
| hsa-mir-4321 | 3 | 0.05727 | 0.12572 | 0.951052 | 3303 | 2 | -3.0146 | 0.94543 | 0.94558 | 1 | 20386 | 0 | -3.0146 |
| ITGAV | 5 | 0.20424 | 0.38126 | 0.999994 | 7881 | 3 | -0.77061 | 0.94543 | 0.94867 | 1 | 20387 | 1 | -0.77061 |
| hsa-mir-1244-2 | 2 | 0.054497 | 0.096745 | 0.945031 | 3176 | 2 | -2.6993 | 0.9455 | 0.94551 | 1 | 20388 | 0 | -2.6993 |
| GPX7 | 5 | 0.053208 | 0.14811 | 0.965554 | 3124 | 4 | -1.1211 | 0.94554 | 0.94875 | 1 | 20389 | 1 | -1.1211 |
| DEPDC4 | 6 | 0.16559 | 0.34519 | 0.999994 | 6875 | 3 | -0.97922 | 0.94566 | 0.9599 | 1 | 20390 | 1 | -0.97922 |
| SMO | 5 | 0.5196 | 0.66496 | 0.999994 | 13740 | 2 | -0.29739 | 0.94582 | 0.94897 | 1 | 20391 | 1 | -0.29739 |
| KLHL17 | 6 | 0.17306 | 0.3545 | 0.999994 | 7066 | 3 | -0.57747 | 0.94584 | 0.95997 | 1 | 20392 | 1 | -0.57747 |
| C7orf57 | 5 | 0.0080514 | 0.030439 | 0.862223 | 736 | 4 | -2.5953 | 0.94585 | 0.949 | 1 | 20393 | 1 | -2.5953 |
| SYK | 4 | 0.046839 | 0.11525 | 0.951052 | 2844 | 2 | -3.0567 | 0.94586 | 0.94575 | 1 | 20394 | 0 | -3.0567 |
| SNX15 | 6 | 0.052247 | 0.149 | 0.965566 | 3080 | 4 | -1.5071 | 0.94588 | 0.95998 | 1 | 20395 | 1 | -1.5071 |
| SEMA4C | 3 | 0.27953 | 0.41143 | 0.999994 | 9694 | 2 | -0.66952 | 0.94596 | 0.94612 | 1 | 20396 | 0 | -0.66952 |
| hsa-mir-23b | 4 | 0.10745 | 0.22834 | 0.983354 | 5224 | 3 | -0.85399 | 0.94599 | 0.94589 | 1 | 20397 | 0 | -0.85399 |
| FAM162B | 5 | 0.0132 | 0.04515 | 0.891621 | 1079 | 3 | -1.7234 | 0.946 | 0.94909 | 1 | 20398 | 1 | -1.7234 |
| SAMD11 | 5 | 0.42407 | 0.58349 | 0.999994 | 12310 | 1 | -0.16006 | 0.94613 | 0.94919 | 1 | 20399 | 1 | -0.16006 |
| NCOA6 | 4 | 0.62709 | 0.68026 | 0.999994 | 15339 | 1 | 0.083459 | 0.94616 | 0.94608 | 1 | 20400 | 0 | 0.083459 |
| CDH5 | 4 | 0.73565 | 0.74926 | 0.999994 | 16670 | 1 | -0.018088 | 0.94622 | 0.94612 | 1 | 20401 | 0 | -0.018088 |
| CD300LF | 4 | 0.38233 | 0.52755 | 0.999994 | 11683 | 2 | -0.64856 | 0.94629 | 0.9462 | 1 | 20402 | 0 | -0.64856 |
| hsa-mir-548w | 2 | 0.05369 | 0.09535 | 0.945031 | 3150 | 2 | -4.866 | 0.94631 | 0.94632 | 1 | 20403 | 0 | -4.866 |
| SPATA7 | 5 | 0.021185 | 0.067288 | 0.93914 | 1526 | 3 | -3.2899 | 0.94632 | 0.94932 | 1 | 20404 | 1 | -3.2899 |
| hsa-mir-4502 | 4 | 0.1385 | 0.27738 | 0.999379 | 6152 | 2 | -1.4433 | 0.94632 | 0.94624 | 1 | 20405 | 0 | -1.4433 |
| NonTargetingControlG | 2 | 0.053673 | 0.09532 | 0.945031 | 3149 | 2 | -3.7524 | 0.94633 | 0.94634 | 1 | 20406 | 0 | -3.7524 |
| FZD8 | 5 | 0.044008 | 0.12584 | 0.951052 | 2711 | 4 | -1.0116 | 0.94634 | 0.94934 | 1 | 20407 | 1 | -1.0116 |
| NonTargetingControlG | 1 | 0.053625 | 0.053657 | 0.919981 | 3148 | 1 | -8.0498 | 0.94638 | 0.94656 | 1 | 20408 | 0 | -8.0498 |
| AGAP6 | 2 | 0.053548 | 0.095111 | 0.945031 | 3142 | 2 | -4.2549 | 0.94645 | 0.94646 | 1 | 20409 | 0 | -4.2549 |
| LGR5 | 6 | 0.27328 | 0.47558 | 0.999994 | 9547 | 3 | -0.66371 | 0.94647 | 0.96021 | 1 | 20410 | 1 | -0.66371 |
| NDUFA6 | 5 | 0.46023 | 0.61384 | 0.999994 | 12847 | 2 | -0.42193 | 0.94654 | 0.94947 | 1 | 20411 | 1 | -0.42193 |
| FABP4 | 6 | 0.26762 | 0.46887 | 0.999994 | 9412 | 3 | -0.49217 | 0.94666 | 0.96029 | 1 | 20412 | 1 | -0.49217 |
| NonTargetingControlG | 2 | 0.053315 | 0.094721 | 0.945031 | 3128 | 2 | -1.5968 | 0.94669 | 0.9467 | 1 | 20413 | 0 | -1.5968 |
| C4BPB | 4 | 0.22196 | 0.36325 | 0.999994 | 8329 | 2 | -1.1061 | 0.9467 | 0.94665 | 1 | 20414 | 0 | -1.1061 |
| CLIC6 | 6 | 0.05712 | 0.15955 | 0.967126 | 3298 | 4 | -1.6709 | 0.94679 | 0.96036 | 1 | 20415 | 1 | -1.6709 |
| SLC25A53 | 6 | 0.071512 | 0.18993 | 0.970473 | 3895 | 3 | -0.38378 | 0.94682 | 0.96037 | 1 | 20416 | 1 | -0.38378 |
| KRT2 | 4 | 0.029328 | 0.080165 | 0.93914 | 1974 | 2 | -3.5384 | 0.94686 | 0.94681 | 1 | 20417 | 0 | -3.5384 |
| TMEM110-MUSTN1 | 4 | 0.016842 | 0.05389 | 0.921374 | 1295 | 3 | -3.8067 | 0.94697 | 0.94693 | 1 | 20418 | 0 | -3.8067 |
| STATH | 4 | 0.07794 | 0.17491 | 0.968171 | 4148 | 3 | -1.6916 | 0.94703 | 0.94699 | 1 | 20419 | 0 | -1.6916 |
| hsa-mir-6821 | 3 | 0.28394 | 0.41679 | 0.999994 | 9781 | 2 | -0.64467 | 0.94704 | 0.94722 | 1 | 20420 | 0 | -0.64467 |
| SPCS1 | 5 | 0.030053 | 0.090617 | 0.94395 | 2010 | 4 | -2.495 | 0.94706 | 0.94986 | 1 | 20421 | 1 | -2.495 |
| EDNRB | 5 | 0.040598 | 0.11738 | 0.951052 | 2543 | 1 | -0.13048 | 0.94706 | 0.94986 | 1 | 20422 | 1 | -0.13048 |
| OR13J1 | 5 | 0.03767 | 0.11003 | 0.951052 | 2413 | 3 | -2.5727 | 0.94707 | 0.94987 | 1 | 20423 | 1 | -2.5727 |
| IMP3 | 2 | 0.044118 | 0.079329 | 0.93914 | 2716 | 2 | -5.5866 | 0.94718 | 0.9472 | 1 | 20424 | 0 | -5.5866 |
| IQSEC1 | 5 | 0.14556 | 0.29612 | 0.999994 | 6338 | 3 | -0.70727 | 0.94719 | 0.94995 | 1 | 20425 | 1 | -0.70727 |
| RPL4 | 3 | 0.15954 | 0.26086 | 0.994943 | 6721 | 2 | -1.1251 | 0.94721 | 0.9474 | 1 | 20426 | 0 | -1.1251 |
| NonTargetingControlG | 2 | 0.052781 | 0.093818 | 0.945031 | 3104 | 2 | -2.3414 | 0.94722 | 0.94724 | 1 | 20427 | 0 | -2.3414 |
| hsa-mir-5583-1 | 4 | 0.25448 | 0.39652 | 0.999994 | 9112 | 2 | -0.44602 | 0.94724 | 0.94722 | 1 | 20428 | 0 | -0.44602 |
| ZNF572 | 5 | 0.26905 | 0.45608 | 0.999994 | 9445 | 2 | -0.44548 | 0.94728 | 0.95001 | 1 | 20429 | 1 | -0.44548 |
| POU6F2 | 4 | 0.018586 | 0.057646 | 0.929232 | 1391 | 3 | -2.648 | 0.9473 | 0.94728 | 1 | 20430 | 0 | -2.648 |
| C1orf204 | 6 | 0.19908 | 0.38675 | 0.999994 | 7739 | 2 | 0.39849 | 0.94735 | 0.9606 | 1 | 20431 | 1 | 0.39849 |
| ZNF676 | 3 | 0.24409 | 0.36763 | 0.999994 | 8885 | 2 | -1.6165 | 0.9475 | 0.9477 | 1 | 20432 | 0 | -1.6165 |
| hsa-mir-7854 | 4 | 0.082626 | 0.18357 | 0.968526 | 4323 | 3 | -1.9456 | 0.9476 | 0.94755 | 1 | 20433 | 0 | -1.9456 |
| RNF19B | 6 | 0.011606 | 0.044431 | 0.891621 | 966 | 5 | -2.571 | 0.94762 | 0.96072 | 1 | 20434 | 1 | -2.571 |
| EIF4E2 | 6 | 0.042569 | 0.12768 | 0.951464 | 2652 | 3 | -2.0623 | 0.94763 | 0.96072 | 1 | 20435 | 1 | -2.0623 |

| | | | | | | | | | | | | |
|---|---|---|---|---|---|---|---|---|---|---|---|---|
| KLK7 | 5 | 0.53048 | 0.67448 | 0.999994 | 13904 | 2 | -0.37978 | 0.94763 | 0.95027 | 1 | 20436 | 1 | -0.37978 |
| NonTargetingControlG | 2 | 0.052335 | 0.093067 | 0.943968 | 3085 | 2 | -1.95 | 0.94766 | 0.94768 | 1 | 20437 | 0 | -1.95 |
| PHF5A | 5 | 0.17013 | 0.33234 | 0.999994 | 6988 | 2 | -0.60015 | 0.9477 | 0.95032 | 1 | 20438 | 1 | -0.60015 |
| NCLN | 4 | 0.055455 | 0.13208 | 0.95318 | 3219 | 3 | -2.2487 | 0.94782 | 0.94777 | 1 | 20439 | 0 | -2.2487 |
| TYW5 | 5 | 0.045106 | 0.12857 | 0.951777 | 2762 | 4 | -0.75043 | 0.94782 | 0.95041 | 1 | 20440 | 1 | -0.75043 |
| SMIM8 | 5 | 0.3226 | 0.4994 | 0.999994 | 10567 | 1 | -0.39115 | 0.94789 | 0.95046 | 1 | 20441 | 1 | -0.39115 |
| hsa-mir-1468 | 4 | 0.030624 | 0.082736 | 0.942973 | 2037 | 2 | -0.97057 | 0.94793 | 0.94788 | 1 | 20442 | 0 | -0.97057 |
| FIGN | 6 | 0.06214 | 0.1703 | 0.967126 | 3516 | 4 | -1.2633 | 0.94795 | 0.96084 | 1 | 20443 | 1 | -1.2633 |
| NDUFA3 | 6 | 0.063399 | 0.17296 | 0.96793 | 3572 | 3 | -0.52388 | 0.94796 | 0.96085 | 1 | 20444 | 1 | -0.52388 |
| TOMM22 | 5 | 0.38508 | 0.551 | 0.999994 | 11720 | 2 | -0.31303 | 0.94797 | 0.95052 | 1 | 20445 | 1 | -0.31303 |
| ZNF287 | 4 | 0.065546 | 0.15137 | 0.967126 | 3675 | 3 | -1.4237 | 0.948 | 0.94795 | 1 | 20446 | 0 | -1.4237 |
| CLCC1 | 4 | 0.72722 | 0.74315 | 0.999994 | 16570 | 1 | -0.855 | 0.9481 | 0.94805 | 1 | 20447 | 0 | -0.855 |
| hsa-mir-5685 | 4 | 0.41003 | 0.55598 | 0.999994 | 12109 | 2 | -0.33941 | 0.94816 | 0.9481 | 1 | 20448 | 0 | -0.33941 |
| CEPT1 | 6 | 0.086434 | 0.22036 | 0.980699 | 4466 | 3 | -0.77959 | 0.94819 | 0.96095 | 1 | 20449 | 1 | -0.77959 |
| WBSCR16 | 5 | 0.009528 | 0.034715 | 0.867348 | 837 | 4 | -3.9158 | 0.94822 | 0.9507 | 1 | 20450 | 1 | -3.9158 |
| PAICS | 3 | 0.048332 | 0.1134 | 0.951052 | 2900 | 2 | -0.66134 | 0.94823 | 0.94839 | 1 | 20451 | 0 | -0.66134 |
| CPXM1 | 5 | 0.0067882 | 0.026759 | 0.848412 | 645 | 4 | -3.9772 | 0.94825 | 0.95073 | 1 | 20452 | 1 | -3.9772 |
| ALG3 | 5 | 0.043772 | 0.12525 | 0.951052 | 2702 | 3 | -2.0643 | 0.94825 | 0.95073 | 1 | 20453 | 1 | -2.0643 |
| hsa-mir-4263 | 4 | 0.025473 | 0.072212 | 0.93914 | 1776 | 3 | -2.3999 | 0.94861 | 0.94855 | 1 | 20454 | 0 | -2.3999 |
| SEMA5A | 5 | 0.13354 | 0.27807 | 0.999536 | 6010 | 3 | -1.1668 | 0.94864 | 0.95103 | 1 | 20455 | 1 | -1.1668 |
| GAB1 | 6 | 0.0069662 | 0.028289 | 0.858439 | 658 | 4 | -1.5805 | 0.94865 | 0.96113 | 1 | 20456 | 1 | -1.5805 |
| C2orf80 | 5 | 0.037314 | 0.10911 | 0.951052 | 2389 | 3 | -2.8206 | 0.94869 | 0.95106 | 1 | 20457 | 1 | -2.8206 |
| CPSF7 | 5 | 0.0090544 | 0.033313 | 0.867348 | 798 | 4 | -2.5143 | 0.9487 | 0.95106 | 1 | 20458 | 1 | -2.5143 |
| FAM98A | 5 | 0.014383 | 0.048445 | 0.90205 | 1158 | 4 | -1.6149 | 0.94873 | 0.95109 | 1 | 20459 | 1 | -1.6149 |
| CNOT3 | 5 | 0.18118 | 0.34835 | 0.999994 | 7253 | 3 | -0.61191 | 0.94876 | 0.95112 | 1 | 20460 | 1 | -0.61191 |
| hsa-mir-6838 | 4 | 0.82746 | 0.82703 | 0.999994 | 17889 | 0 | -0.20252 | 0.94882 | 0.94875 | 1 | 20461 | 0 | -0.20252 |
| USH1G | 4 | 0.19378 | 0.33426 | 0.999994 | 7592 | 2 | -1.0927 | 0.94882 | 0.94875 | 1 | 20462 | 0 | -1.0927 |
| NonTargetingControlG | 2 | 0.051156 | 0.091114 | 0.94395 | 3030 | 2 | -2.6277 | 0.94884 | 0.94886 | 1 | 20463 | 0 | -2.6277 |
| CD19 | 4 | 0.03416 | 0.090035 | 0.94395 | 2223 | 3 | -3.1025 | 0.94885 | 0.94878 | 1 | 20464 | 0 | -3.1025 |
| NonTargetingControlG | 1 | 0.051145 | 0.051206 | 0.909241 | 3028 | 1 | -8.1527 | 0.94885 | 0.94907 | 1 | 20465 | 0 | -8.1527 |
| hsa-mir-571 | 4 | 0.64893 | 0.69265 | 0.999994 | 15641 | 1 | -0.059301 | 0.94892 | 0.94885 | 1 | 20466 | 0 | -0.059301 |
| SLC38A3 | 5 | 0.0078885 | 0.029936 | 0.862223 | 722 | 3 | -4.1789 | 0.94897 | 0.95127 | 1 | 20467 | 1 | -4.1789 |
| hsa-mir-3685 | 3 | 0.021803 | 0.055956 | 0.9235 | 1565 | 2 | -0.8112 | 0.94902 | 0.94919 | 1 | 20468 | 0 | -0.8112 |
| NonTargetingControlG | 2 | 0.050516 | 0.090032 | 0.94395 | 2996 | 2 | -1.8752 | 0.94948 | 0.94952 | 1 | 20469 | 0 | -1.8752 |
| hsa-mir-4322 | 3 | 0.28774 | 0.42137 | 0.999994 | 9870 | 2 | -0.62289 | 0.94949 | 0.94967 | 1 | 20470 | 0 | -0.62289 |
| CKB | 4 | 0.084771 | 0.18748 | 0.969409 | 4398 | 3 | -1.2706 | 0.9495 | 0.94942 | 1 | 20471 | 0 | -1.2706 |
| ZMAT1 | 5 | 0.56974 | 0.70929 | 0.999994 | 14534 | 2 | -0.42325 | 0.9495 | 0.95166 | 1 | 20472 | 1 | -0.42325 |
| AGAP9 | 2 | 0.050442 | 0.089907 | 0.94395 | 2992 | 2 | -5.4777 | 0.94956 | 0.9496 | 1 | 20473 | 0 | -5.4777 |
| ZNF19 | 6 | 0.014312 | 0.05349 | 0.919407 | 1151 | 4 | -3.2075 | 0.94963 | 0.96154 | 1 | 20474 | 1 | -3.2075 |
| NonTargetingControlG | 2 | 0.050319 | 0.089684 | 0.94395 | 2989 | 2 | -4.0828 | 0.94968 | 0.94973 | 1 | 20475 | 0 | -4.0828 |
| SMR3B | 4 | 0.37003 | 0.5149 | 0.999994 | 11510 | 2 | -0.44357 | 0.94976 | 0.94967 | 1 | 20476 | 0 | -0.44357 |
| KIF11 | 5 | 0.0013939 | 0.0061914 | 0.636218 | 210 | 4 | -8.0674 | 0.94988 | 0.95194 | 1 | 20477 | 1 | -8.0674 |
| CCDC173 | 5 | 0.041435 | 0.1195 | 0.951052 | 2585 | 3 | -2.4023 | 0.94988 | 0.95194 | 1 | 20478 | 1 | -2.4023 |
| NonTargetingControlG | 2 | 0.050015 | 0.089184 | 0.94395 | 2981 | 2 | -1.9624 | 0.94999 | 0.95004 | 1 | 20479 | 0 | -1.9624 |
| STAT5B | 6 | 0.11211 | 0.27075 | 0.998561 | 5374 | 3 | -1.1675 | 0.95005 | 0.96172 | 1 | 20480 | 1 | -1.1675 |
| FAHD1 | 5 | 0.44448 | 0.60057 | 0.999994 | 12640 | 2 | -0.42499 | 0.95024 | 0.95221 | 1 | 20481 | 1 | -0.42499 |
| GTSE1 | 5 | 0.14267 | 0.29186 | 0.999632 | 6261 | 2 | -0.51689 | 0.95026 | 0.95222 | 1 | 20482 | 1 | -0.51689 |
| TAS2R4 | 5 | 0.81321 | 0.82806 | 0.999994 | 17678 | 1 | 0.2242 | 0.9504 | 0.95233 | 1 | 20483 | 1 | 0.2242 |
| ICMT | 6 | 0.046932 | 0.13735 | 0.957317 | 2848 | 4 | -2.2223 | 0.95041 | 0.96187 | 1 | 20484 | 1 | -2.2223 |
| WEE2 | 5 | 0.0024462 | 0.010636 | 0.706752 | 312 | 2 | -0.086021 | 0.95043 | 0.95235 | 1 | 20485 | 1 | -0.086021 |
| hsa-mir-5190 | 3 | 0.25739 | 0.38411 | 0.999994 | 9175 | 2 | -0.69987 | 0.95055 | 0.95075 | 1 | 20486 | 0 | -0.69987 |
| LSP1 | 5 | 0.054716 | 0.15178 | 0.967126 | 3182 | 3 | -1.8535 | 0.95082 | 0.95266 | 1 | 20487 | 1 | -1.8535 |

| | | | | | | | | | | | | | |
|---|---|---|---|---|---|---|---|---|---|---|---|---|---|
| ETNK2 | 4 | 0.023681 | 0.068442 | 0.93914 | 1694 | 3 | -3.36 | 0.95092 | 0.95083 | 1 | 20488 | 0 | -3.36 |
| NUDCD1 | 5 | 0.032996 | 0.098227 | 0.945253 | 2157 | 2 | 0.35554 | 0.95095 | 0.95277 | 1 | 20489 | 1 | 0.35554 |
| FAM78B | 4 | 0.022437 | 0.065756 | 0.93914 | 1612 | 3 | -2.0806 | 0.95098 | 0.95087 | 1 | 20490 | 0 | -2.0806 |
| OR2B6 | 6 | 0.011588 | 0.044367 | 0.891621 | 963 | 3 | -2.3752 | 0.95106 | 0.96217 | 1 | 20491 | 1 | -2.3752 |
| RTP2 | 5 | 0.18048 | 0.34731 | 0.999994 | 7229 | 3 | -0.98944 | 0.9511 | 0.95288 | 1 | 20492 | 1 | -0.98944 |
| CAMP | 5 | 0.16159 | 0.31989 | 0.999994 | 6779 | 2 | -0.010468 | 0.95119 | 0.95296 | 1 | 20493 | 1 | -0.010468 |
| CADPS | 6 | 0.02188 | 0.078006 | 0.93914 | 1574 | 4 | -2.0373 | 0.95121 | 0.96223 | 1 | 20494 | 1 | -2.0373 |
| LOC401052 | 5 | 0.022516 | 0.070894 | 0.93914 | 1620 | 3 | -2.6624 | 0.95123 | 0.95299 | 1 | 20495 | 1 | -2.6624 |
| LOC100132146 | 5 | 0.030256 | 0.091146 | 0.94395 | 2023 | 4 | -1.674 | 0.95123 | 0.95299 | 1 | 20496 | 1 | -1.674 |
| hsa-mir-887 | 4 | 0.1364 | 0.2752 | 0.999239 | 6094 | 2 | -1.2563 | 0.95126 | 0.95116 | 1 | 20497 | 0 | -1.2563 |
| ASAP2 | 4 | 0.31194 | 0.45525 | 0.999994 | 10358 | 2 | -0.67322 | 0.95129 | 0.9512 | 1 | 20498 | 0 | -0.67322 |
| hsa-mir-4714 | 4 | 0.049337 | 0.12016 | 0.951052 | 2945 | 3 | -1.6596 | 0.95131 | 0.95121 | 1 | 20499 | 0 | -1.6596 |
| CYFIP2 | 5 | 0.031604 | 0.094596 | 0.945031 | 2084 | 4 | -2.8472 | 0.95145 | 0.95316 | 1 | 20500 | 1 | -2.8472 |
| SYCE3 | 5 | 0.02567 | 0.079266 | 0.93914 | 1784 | 4 | -1.5604 | 0.95154 | 0.95323 | 1 | 20501 | 1 | -1.5604 |
| NonTargetingControlG | 2 | 0.048454 | 0.086558 | 0.94395 | 2906 | 2 | -2.3933 | 0.95155 | 0.95155 | 1 | 20502 | 0 | -2.3933 |
| CXorf40B | 4 | 0.14897 | 0.28811 | 0.999632 | 6426 | 2 | -1.3545 | 0.95181 | 0.95169 | 1 | 20503 | 0 | -1.3545 |
| UCK1 | 5 | 0.0091111 | 0.033472 | 0.867348 | 802 | 3 | -2.6619 | 0.95187 | 0.95348 | 1 | 20504 | 1 | -2.6619 |
| KLHL30 | 5 | 0.02708 | 0.082946 | 0.94336 | 1861 | 3 | -2.682 | 0.95189 | 0.9535 | 1 | 20505 | 1 | -2.682 |
| ZFHX4 | 5 | 0.083982 | 0.20094 | 0.977164 | 4375 | 3 | -1.0612 | 0.95211 | 0.95366 | 1 | 20506 | 1 | -1.0612 |
| POLR2A | 4 | 0.019789 | 0.060178 | 0.932932 | 1446 | 3 | -2.7259 | 0.95217 | 0.95204 | 1 | 20507 | 0 | -2.7259 |
| hsa-mir-6871 | 3 | 0.16826 | 0.27215 | 0.998561 | 6945 | 2 | -1.3401 | 0.95223 | 0.95245 | 1 | 20508 | 0 | -1.3401 |
| FBXO21 | 6 | 0.005339 | 0.022392 | 0.819701 | 544 | 3 | -1.0161 | 0.95227 | 0.96269 | 1 | 20509 | 1 | -1.0161 |
| hsa-mir-1286 | 4 | 0.41422 | 0.56027 | 0.999994 | 12168 | 2 | -0.64673 | 0.95235 | 0.95223 | 1 | 20510 | 0 | -0.64673 |
| PBLD | 6 | 0.0073917 | 0.029793 | 0.862223 | 685 | 4 | -3.6859 | 0.95241 | 0.96276 | 1 | 20511 | 1 | -3.6859 |
| DNASE1L3 | 6 | 0.014216 | 0.053175 | 0.916684 | 1144 | 4 | -2.3036 | 0.95243 | 0.96278 | 1 | 20512 | 1 | -2.3036 |
| ATG7 | 5 | 0.037026 | 0.10841 | 0.951052 | 2376 | 4 | -1.1085 | 0.95249 | 0.95397 | 1 | 20513 | 1 | -1.1085 |
| FAM72B | 4 | 0.36055 | 0.50511 | 0.999994 | 11342 | 2 | -0.60387 | 0.95257 | 0.95244 | 1 | 20514 | 0 | -0.60387 |
| VPS33B | 6 | 0.031287 | 0.10184 | 0.946414 | 2073 | 4 | -1.5569 | 0.95258 | 0.96286 | 1 | 20515 | 1 | -1.5569 |
| NIF3L1 | 6 | 0.29519 | 0.5011 | 0.999994 | 10019 | 3 | -0.28674 | 0.95264 | 0.96288 | 1 | 20516 | 1 | -0.28674 |
| NonTargetingControlG | 2 | 0.047229 | 0.084528 | 0.94395 | 2866 | 2 | -2.2714 | 0.95277 | 0.95278 | 1 | 20517 | 0 | -2.2714 |
| SYS1 | 5 | 0.013038 | 0.044691 | 0.891621 | 1065 | 3 | -3.9753 | 0.95282 | 0.95424 | 1 | 20518 | 1 | -3.9753 |
| C1orf68 | 3 | 0.2765 | 0.40771 | 0.999994 | 9631 | 2 | -0.69771 | 0.95285 | 0.95308 | 1 | 20519 | 0 | -0.69771 |
| ZMYND10 | 4 | 0.1155 | 0.24242 | 0.988088 | 5497 | 3 | -0.74757 | 0.95288 | 0.95273 | 1 | 20520 | 0 | -0.74757 |
| FAM220A | 5 | 0.20606 | 0.38382 | 0.999994 | 7930 | 3 | -0.76922 | 0.95292 | 0.95432 | 1 | 20521 | 1 | -0.76922 |
| CNDP2 | 4 | 0.86725 | 0.86682 | 0.999994 | 18579 | 0 | 0.0072502 | 0.95296 | 0.95281 | 1 | 20522 | 0 | 0.0072502 |
| DAXX | 6 | 0.00076802 | 0.003941 | 0.560963 | 137 | 4 | -6.1842 | 0.95304 | 0.96306 | 1 | 20523 | 1 | -6.1842 |
| LOC440243 | 4 | 0.387 | 0.53228 | 0.999994 | 11753 | 2 | -1.2196 | 0.95308 | 0.95294 | 1 | 20524 | 0 | -1.2196 |
| NonTargetingControlG | 2 | 0.046886 | 0.083957 | 0.94395 | 2847 | 2 | -2.162 | 0.95311 | 0.9531 | 1 | 20525 | 0 | -2.162 |
| ITIH4 | 6 | 0.0044241 | 0.019027 | 0.804634 | 473 | 5 | -2.7584 | 0.9532 | 0.96313 | 1 | 20526 | 1 | -2.7584 |
| RBMS2 | 4 | 0.11863 | 0.24785 | 0.990682 | 5592 | 2 | -1.3642 | 0.95322 | 0.95308 | 1 | 20527 | 0 | -1.3642 |
| TFEC | 5 | 0.0014492 | 0.0064287 | 0.636218 | 217 | 3 | -2.3658 | 0.95331 | 0.95463 | 1 | 20528 | 1 | -2.3658 |
| SCN4B | 4 | 0.83428 | 0.83382 | 0.999994 | 17992 | 0 | -0.43057 | 0.95332 | 0.95318 | 1 | 20529 | 0 | -0.43057 |
| CYYR1 | 5 | 0.14265 | 0.29183 | 0.999632 | 6260 | 3 | -1.1702 | 0.95334 | 0.95466 | 1 | 20530 | 1 | -1.1702 |
| LTA4H | 6 | 0.23795 | 0.43354 | 0.999994 | 8728 | 3 | -0.56616 | 0.95338 | 0.96322 | 1 | 20531 | 1 | -0.56616 |
| TP53INP2 | 5 | 0.054993 | 0.15246 | 0.967126 | 3198 | 3 | -1.04 | 0.9534 | 0.9547 | 1 | 20532 | 1 | -1.04 |
| SERTAD4 | 5 | 0.67558 | 0.75568 | 0.999994 | 16013 | 1 | 0.084253 | 0.9534 | 0.9547 | 1 | 20533 | 1 | 0.084253 |
| PRPH | 6 | 0.074512 | 0.19615 | 0.975797 | 4015 | 4 | -1.2952 | 0.95344 | 0.96324 | 1 | 20534 | 1 | -1.2952 |
| DEFB121 | 3 | 0.15483 | 0.2547 | 0.994835 | 6590 | 2 | -1.1957 | 0.95344 | 0.95369 | 1 | 20535 | 0 | -1.1957 |
| VGF | 4 | 0.085964 | 0.18964 | 0.970473 | 4445 | 3 | -1.4483 | 0.95347 | 0.95334 | 1 | 20536 | 0 | -1.4483 |
| GREM2 | 4 | 0.36661 | 0.51132 | 0.999994 | 11451 | 2 | -0.72673 | 0.95349 | 0.95336 | 1 | 20537 | 0 | -0.72673 |
| CCL1 | 5 | 0.14605 | 0.29686 | 0.999994 | 6352 | 3 | -1.0858 | 0.95352 | 0.9548 | 1 | 20538 | 1 | -1.0858 |
| AGBL5 | 6 | 0.19398 | 0.3805 | 0.999994 | 7602 | 3 | -0.61238 | 0.95353 | 0.96329 | 1 | 20539 | 1 | -0.61238 |

| | | | | | | | | | | | | | |
|---|---|---|---|---|---|---|---|---|---|---|---|---|---|
| MAGEA1 | 6 | 0.10978 | 0.26626 | 0.998254 | 5296 | 4 | -0.8906 | 0.95357 | 0.96331 | 1 | 20540 | 1 | -0.8906 |
| CSGALNACT1 | 6 | 0.19287 | 0.37911 | 0.999994 | 7572 | 3 | -0.88753 | 0.95358 | 0.96331 | 1 | 20541 | 1 | -0.88753 |
| LRIF1 | 5 | 0.014431 | 0.048577 | 0.902886 | 1160 | 4 | -1.5576 | 0.95367 | 0.95491 | 1 | 20542 | 1 | -1.5576 |
| MAPRE3 | 6 | 0.083538 | 0.21456 | 0.979587 | 4355 | 4 | -0.98783 | 0.95369 | 0.96336 | 1 | 20543 | 1 | -0.98783 |
| CARNS1 | 5 | 0.15046 | 0.30347 | 0.999994 | 6470 | 3 | -0.79533 | 0.9538 | 0.95504 | 1 | 20544 | 1 | -0.79533 |
| hsa-mir-4687 | 4 | 0.17282 | 0.31274 | 0.999994 | 7058 | 2 | -1.1315 | 0.95383 | 0.9537 | 1 | 20545 | 0 | -1.1315 |
| HNRNPLL | 6 | 0.0063794 | 0.02619 | 0.848412 | 619 | 5 | -1.8065 | 0.95402 | 0.96353 | 1 | 20546 | 1 | -1.8065 |
| NonTargetingControlG | 2 | 0.045947 | 0.082372 | 0.940526 | 2805 | 2 | -3.0032 | 0.95405 | 0.9541 | 1 | 20547 | 0 | -3.0032 |
| KHK | 5 | 0.13522 | 0.28057 | 0.999632 | 6058 | 3 | -1.1675 | 0.95414 | 0.9553 | 1 | 20548 | 1 | -1.1675 |
| MRPL45 | 4 | 0.02 | 0.06062 | 0.932932 | 1460 | 3 | -4.2286 | 0.95414 | 0.954 | 1 | 20549 | 0 | -4.2286 |
| LY6G6F | 6 | 0.23681 | 0.43219 | 0.999994 | 8690 | 3 | -0.81626 | 0.9542 | 0.96361 | 1 | 20550 | 1 | -0.81626 |
| UCHL5 | 5 | 0.070087 | 0.17852 | 0.968171 | 3830 | 3 | -0.81635 | 0.95423 | 0.95537 | 1 | 20551 | 1 | -0.81635 |
| FEZF1 | 5 | 0.029841 | 0.090058 | 0.94395 | 2000 | 2 | -0.28284 | 0.95444 | 0.95555 | 1 | 20552 | 1 | -0.28284 |
| hsa-mir-548al | 2 | 0.045528 | 0.081647 | 0.93914 | 2792 | 2 | -3.5985 | 0.95447 | 0.95453 | 1 | 20553 | 0 | -3.5985 |
| hsa-mir-135a-1 | 4 | 0.033869 | 0.089401 | 0.94395 | 2210 | 3 | -2.7934 | 0.9545 | 0.95436 | 1 | 20554 | 0 | -2.7934 |
| DMAP1 | 6 | 0.0064662 | 0.02653 | 0.848412 | 623 | 5 | -1.7419 | 0.9545 | 0.96377 | 1 | 20555 | 1 | -1.7419 |
| HSPA6 | 6 | 0.58756 | 0.73079 | 0.999994 | 14783 | 2 | -0.23286 | 0.95456 | 0.96379 | 1 | 20556 | 1 | -0.23286 |
| MMP12 | 5 | 0.15876 | 0.31572 | 0.999994 | 6690 | 3 | -0.97785 | 0.9546 | 0.95568 | 1 | 20557 | 1 | -0.97785 |
| ZMYM5 | 5 | 0.036006 | 0.10579 | 0.950444 | 2313 | 4 | -1.2183 | 0.95463 | 0.9557 | 1 | 20558 | 1 | -1.2183 |
| MAMLD1 | 6 | 0.080305 | 0.20791 | 0.979108 | 4239 | 4 | -1.2083 | 0.95471 | 0.96386 | 1 | 20559 | 1 | -1.2083 |
| RAB3IP | 6 | 0.27417 | 0.47662 | 0.999994 | 9568 | 3 | -0.4184 | 0.95479 | 0.96391 | 1 | 20560 | 1 | -0.4184 |
| FGFBP2 | 4 | 0.016356 | 0.052841 | 0.916123 | 1264 | 2 | -4.1475 | 0.95486 | 0.95471 | 1 | 20561 | 0 | -4.1475 |
| CHRNG | 5 | 0.028794 | 0.087372 | 0.94395 | 1942 | 4 | -3.145 | 0.95503 | 0.95603 | 1 | 20562 | 1 | -3.145 |
| C18orf8 | 6 | 0.083711 | 0.21489 | 0.979587 | 4365 | 4 | -1.3639 | 0.95504 | 0.96403 | 1 | 20563 | 1 | -1.3639 |
| SLC25A27 | 5 | 0.2576 | 0.44678 | 0.999994 | 9183 | 2 | -0.44934 | 0.95512 | 0.9561 | 1 | 20564 | 1 | -0.44934 |
| ECT2 | 4 | 0.12584 | 0.26045 | 0.994943 | 5815 | 3 | -0.77714 | 0.95529 | 0.95518 | 1 | 20565 | 0 | -0.77714 |
| FLYWCH1 | 4 | 0.10937 | 0.23177 | 0.984479 | 5284 | 3 | -0.92161 | 0.95541 | 0.95531 | 1 | 20566 | 0 | -0.92161 |
| NonTargetingControlG | 2 | 0.044555 | 0.080086 | 0.93914 | 2738 | 2 | -2.8391 | 0.95544 | 0.95553 | 1 | 20567 | 0 | -2.8391 |
| NonTargetingControlG | 2 | 0.044432 | 0.079874 | 0.93914 | 2730 | 2 | -2.7035 | 0.95557 | 0.95566 | 1 | 20568 | 0 | -2.7035 |
| hsa-mir-548i-1 | 3 | 0.044369 | 0.10757 | 0.951052 | 2728 | 3 | -1.8206 | 0.95563 | 0.95584 | 1 | 20569 | 0 | -1.8206 |
| TMOD1 | 5 | 0.034022 | 0.10078 | 0.946414 | 2218 | 4 | -1.1331 | 0.95564 | 0.95654 | 1 | 20570 | 1 | -1.1331 |
| DENND6A | 5 | 0.0015357 | 0.0068018 | 0.643671 | 225 | 2 | -0.046358 | 0.95574 | 0.95663 | 1 | 20571 | 1 | -0.046358 |
| OTUB1 | 5 | 0.49179 | 0.64081 | 0.999994 | 13298 | 2 | 0.13165 | 0.9558 | 0.95668 | 1 | 20572 | 1 | 0.13165 |
| TAF4 | 4 | 0.036712 | 0.095196 | 0.945031 | 2357 | 3 | -1.8043 | 0.95581 | 0.95571 | 1 | 20573 | 0 | -1.8043 |
| PLEK | 5 | 0.091443 | 0.21289 | 0.979587 | 4653 | 3 | -1.6946 | 0.95596 | 0.95681 | 1 | 20574 | 1 | -1.6946 |
| GCNT4 | 6 | 0.010377 | 0.040306 | 0.891621 | 892 | 5 | -2.4466 | 0.95596 | 0.96447 | 1 | 20575 | 1 | -2.4466 |
| INPPL1 | 6 | 0.38336 | 0.59168 | 0.999994 | 11695 | 2 | 0.020969 | 0.956 | 0.96448 | 1 | 20576 | 1 | 0.020969 |
| SLC25A40 | 6 | 0.38841 | 0.59493 | 0.999994 | 11774 | 2 | -0.14884 | 0.95606 | 0.96452 | 1 | 20577 | 1 | -0.14884 |
| NonTargetingControlG | 2 | 0.043901 | 0.078952 | 0.93914 | 2708 | 2 | -2.2173 | 0.9561 | 0.95621 | 1 | 20578 | 0 | -2.2173 |
| USP17L1P | 5 | 0.02642 | 0.081206 | 0.93914 | 1825 | 4 | -1.9063 | 0.95622 | 0.95704 | 1 | 20579 | 1 | -1.9063 |
| FBXL13 | 5 | 0.53358 | 0.67722 | 0.999994 | 13960 | 1 | -0.10397 | 0.95627 | 0.95708 | 1 | 20580 | 1 | -0.10397 |
| ADPRH | 3 | 0.043713 | 0.10613 | 0.950452 | 2699 | 3 | -1.7368 | 0.95629 | 0.95649 | 1 | 20581 | 0 | -1.7368 |
| RNF216 | 5 | 0.16596 | 0.32619 | 0.999994 | 6883 | 2 | -0.41375 | 0.95638 | 0.95717 | 1 | 20582 | 1 | -0.41375 |
| NonTargetingControlG | 2 | 0.043538 | 0.07832 | 0.93914 | 2696 | 2 | -2.0542 | 0.95646 | 0.95655 | 1 | 20583 | 0 | -2.0542 |
| SPEG | 5 | 0.077637 | 0.19078 | 0.972248 | 4135 | 3 | -1.0678 | 0.95653 | 0.95729 | 1 | 20584 | 1 | -1.0678 |
| NonTargetingControlG | 1 | 0.043466 | 0.043438 | 0.891621 | 2693 | 1 | -8.4829 | 0.95653 | 0.95668 | 1 | 20585 | 0 | -8.4829 |
| TMEM68 | 6 | 0.034843 | 0.11016 | 0.951052 | 2258 | 4 | -1.9862 | 0.95655 | 0.96474 | 1 | 20586 | 1 | -1.9862 |
| FBLN2 | 5 | 0.21596 | 0.39762 | 0.999994 | 8168 | 3 | -0.71529 | 0.95657 | 0.95732 | 1 | 20587 | 1 | -0.71529 |
| TUBA4A | 6 | 0.00023492 | 0.0012498 | 0.377819 | 66 | 5 | -4.0829 | 0.95658 | 0.96475 | 1 | 20588 | 1 | -4.0829 |
| CFC1 | 5 | 0.052 | 0.14523 | 0.963012 | 3066 | 4 | -0.75216 | 0.95674 | 0.95746 | 1 | 20589 | 1 | -0.75216 |
| PHF7 | 5 | 0.0078255 | 0.029749 | 0.862223 | 717 | 4 | -2.5816 | 0.95676 | 0.95748 | 1 | 20590 | 1 | -2.5816 |
| KCTD8 | 4 | 0.027378 | 0.076151 | 0.93914 | 1875 | 2 | -0.93058 | 0.95684 | 0.95676 | 1 | 20591 | 0 | -0.93058 |

| | | | | | | | | | | | | | |
|---|---|---|---|---|---|---|---|---|---|---|---|---|---|
| FAM168B | 5 | 0.011078 | 0.039183 | 0.891621 | 931 | 4 | -2.3987 | 0.95686 | 0.95756 | 1 | 20592 | 1 | -2.3987 |
| MT1X | 5 | 0.18737 | 0.3572 | 0.999994 | 7403 | 3 | -0.77504 | 0.95687 | 0.95757 | 1 | 20593 | 1 | -0.77504 |
| XPNPEP2 | 6 | 0.68331 | 0.80355 | 0.999994 | 16134 | 1 | 0.12785 | 0.95701 | 0.96498 | 1 | 20594 | 1 | 0.12785 |
| WDR45 | 4 | 0.37727 | 0.52236 | 0.999994 | 11609 | 2 | -0.76791 | 0.9571 | 0.95703 | 1 | 20595 | 0 | -0.76791 |
| ISY1-RAB43 | 1 | 0.042843 | 0.042804 | 0.891621 | 2668 | 1 | -8.5066 | 0.95716 | 0.95732 | 1 | 20596 | 0 | -8.5066 |
| CDK4 | 5 | 0.54168 | 0.68439 | 0.999994 | 14082 | 2 | -0.20655 | 0.95717 | 0.95784 | 1 | 20597 | 1 | -0.20655 |
| hsa-mir-4268 | 4 | 0.43838 | 0.58508 | 0.999994 | 12544 | 2 | -0.61845 | 0.95723 | 0.95717 | 1 | 20598 | 0 | -0.61845 |
| C8orf44-SGK3 | 5 | 0.7029 | 0.76781 | 0.999994 | 16336 | 1 | -0.029157 | 0.95728 | 0.95793 | 1 | 20599 | 1 | -0.029157 |
| hsa-mir-4267 | 3 | 0.042678 | 0.10377 | 0.947284 | 2659 | 3 | -1.0011 | 0.95732 | 0.95753 | 1 | 20600 | 0 | -1.0011 |
| KDM1A | 6 | 0.018664 | 0.067684 | 0.93914 | 1399 | 5 | -1.6572 | 0.95733 | 0.96514 | 1 | 20601 | 1 | -1.6572 |
| EPHA10 | 4 | 0.29157 | 0.4344 | 0.999994 | 9940 | 2 | -0.6139 | 0.95735 | 0.9573 | 1 | 20602 | 0 | -0.6139 |
| CALHM2 | 2 | 0.042615 | 0.076774 | 0.93914 | 2654 | 2 | -5.4368 | 0.95739 | 0.95747 | 1 | 20603 | 0 | -5.4368 |
| SKP2 | 6 | 0.077858 | 0.20302 | 0.97798 | 4145 | 4 | -1.1562 | 0.95742 | 0.96519 | 1 | 20604 | 1 | -1.1562 |
| TRIM61 | 5 | 0.196 | 0.3695 | 0.999994 | 7655 | 3 | -0.80291 | 0.95742 | 0.95805 | 1 | 20605 | 1 | -0.80291 |
| PCSK6 | 5 | 0.41548 | 0.57634 | 0.999994 | 12184 | 2 | -0.48184 | 0.95747 | 0.9581 | 1 | 20606 | 1 | -0.48184 |
| hsa-mir-711 | 4 | 0.38867 | 0.53401 | 0.999994 | 11776 | 2 | -0.65933 | 0.95749 | 0.95745 | 1 | 20607 | 0 | -0.65933 |
| PRM2 | 3 | 0.042509 | 0.10339 | 0.946414 | 2643 | 3 | -2.5778 | 0.95749 | 0.95768 | 1 | 20608 | 0 | -2.5778 |
| PTMA | 5 | 0.00094566 | 0.0042382 | 0.566526 | 157 | 4 | -4.0333 | 0.95752 | 0.95814 | 1 | 20609 | 1 | -4.0333 |
| MYL6 | 5 | 0.016479 | 0.054405 | 0.922677 | 1275 | 4 | -1.7454 | 0.95755 | 0.95816 | 1 | 20610 | 1 | -1.7454 |
| NonTargetingControlG | 1 | 0.04241 | 0.042366 | 0.891621 | 2636 | 1 | -8.5338 | 0.95759 | 0.95777 | 1 | 20611 | 0 | -8.5338 |
| G6PC2 | 5 | 0.032874 | 0.097886 | 0.945253 | 2152 | 4 | -1.3733 | 0.95762 | 0.95823 | 1 | 20612 | 1 | -1.3733 |
| LOC100130705 | 6 | 0.072537 | 0.19205 | 0.974607 | 3942 | 3 | -0.66333 | 0.95763 | 0.96529 | 1 | 20613 | 1 | -0.66333 |
| B3GALT1 | 6 | 0.0057846 | 0.024005 | 0.834563 | 577 | 5 | -1.8463 | 0.95766 | 0.96531 | 1 | 20614 | 1 | -1.8463 |
| MT2A | 4 | 0.01159 | 0.038696 | 0.890863 | 964 | 3 | -3.1078 | 0.95769 | 0.95766 | 1 | 20615 | 0 | -3.1078 |
| CEP350 | 5 | 0.000014872 | 0.000060352 | 0.087459 | 16 | 4 | -5.1399 | 0.95769 | 0.95829 | 1 | 20616 | 1 | -5.1399 |
| TIMM9 | 5 | 0.022077 | 0.069704 | 0.93914 | 1590 | 2 | -0.52507 | 0.95774 | 0.95832 | 1 | 20617 | 1 | -0.52507 |
| ESPL1 | 4 | 0.045959 | 0.11356 | 0.951052 | 2807 | 3 | -3.0213 | 0.95805 | 0.95801 | 1 | 20618 | 0 | -3.0213 |
| NUP214 | 5 | 0.098198 | 0.22359 | 0.983354 | 4901 | 2 | -0.47752 | 0.95818 | 0.95872 | 1 | 20619 | 1 | -0.47752 |
| HORMAD2 | 5 | 0.076338 | 0.18869 | 0.969543 | 4084 | 2 | -0.531 | 0.95821 | 0.95874 | 1 | 20620 | 1 | -0.531 |
| CT47A1 | 5 | 0.24264 | 0.43452 | 0.999994 | 8852 | 3 | -0.51777 | 0.95823 | 0.95876 | 1 | 20621 | 1 | -0.51777 |
| ZNF469 | 5 | 0.021123 | 0.067122 | 0.93914 | 1519 | 3 | -0.70608 | 0.95827 | 0.95879 | 1 | 20622 | 1 | -0.70608 |
| EXOC3L2 | 6 | 0.21848 | 0.41029 | 0.999994 | 8239 | 3 | -0.85114 | 0.95827 | 0.96562 | 1 | 20623 | 1 | -0.85114 |
| CCDC169-SOHLH2 | 4 | 0.12609 | 0.26085 | 0.994943 | 5823 | 2 | -0.55144 | 0.95835 | 0.95829 | 1 | 20624 | 0 | -0.55144 |
| CTAGE5 | 4 | 0.092618 | 0.20181 | 0.977164 | 4708 | 2 | -0.72262 | 0.95836 | 0.9583 | 1 | 20625 | 0 | -0.72262 |
| NGEF | 5 | 0.012127 | 0.042168 | 0.891621 | 1001 | 2 | -0.072332 | 0.9584 | 0.95891 | 1 | 20626 | 1 | -0.072332 |
| ELF4 | 6 | 0.091333 | 0.2302 | 0.983933 | 4649 | 4 | -1.306 | 0.95845 | 0.96571 | 1 | 20627 | 1 | -1.306 |
| FBXO22 | 5 | 0.033464 | 0.099392 | 0.945253 | 2188 | 4 | -2.222 | 0.95846 | 0.95897 | 1 | 20628 | 1 | -2.222 |
| MFSD2A | 6 | 0.46304 | 0.64378 | 0.999994 | 12906 | 2 | -0.1689 | 0.9585 | 0.96574 | 1 | 20629 | 1 | -0.1689 |
| hsa-mir-3690-1 | 4 | 0.12827 | 0.26462 | 0.996734 | 5887 | 2 | -0.79674 | 0.95852 | 0.95847 | 1 | 20630 | 0 | -0.79674 |
| CSB-PGBD3 | 4 | 0.64915 | 0.69279 | 0.999994 | 15644 | 1 | -0.27365 | 0.95853 | 0.95848 | 1 | 20631 | 0 | -0.27365 |
| FOXA1 | 5 | 0.023722 | 0.074082 | 0.93914 | 1695 | 4 | -2.1484 | 0.95853 | 0.95902 | 1 | 20632 | 1 | -2.1484 |
| FAM153A | 5 | 0.19555 | 0.36886 | 0.999994 | 7652 | 3 | -0.75077 | 0.95855 | 0.95904 | 1 | 20633 | 1 | -0.75077 |
| NonTargetingControlG | 2 | 0.041451 | 0.07477 | 0.93914 | 2587 | 2 | -2.7318 | 0.95855 | 0.95862 | 1 | 20634 | 0 | -2.7318 |
| CLIC1 | 5 | 0.15702 | 0.31318 | 0.999994 | 6650 | 3 | -1.117 | 0.95859 | 0.95908 | 1 | 20635 | 1 | -1.117 |
| KCNK15 | 6 | 0.034431 | 0.10917 | 0.951052 | 2235 | 4 | -1.6125 | 0.95859 | 0.96579 | 1 | 20636 | 1 | -1.6125 |
| DDX4 | 6 | 0.24472 | 0.44163 | 0.999994 | 8895 | 3 | -0.76264 | 0.95864 | 0.96581 | 1 | 20637 | 1 | -0.76264 |
| hsa-mir-6837 | 4 | 0.12279 | 0.25513 | 0.994943 | 5733 | 3 | -2.2877 | 0.95864 | 0.95859 | 1 | 20638 | 0 | -2.2877 |
| OTOA | 4 | 0.064044 | 0.14849 | 0.965554 | 3612 | 3 | -2.729 | 0.95867 | 0.95862 | 1 | 20639 | 0 | -2.729 |
| COPS4 | 6 | 0.065659 | 0.17768 | 0.968171 | 3679 | 4 | -1.7246 | 0.95867 | 0.96583 | 1 | 20640 | 1 | -1.7246 |
| USP38 | 4 | 0.1268 | 0.26205 | 0.995544 | 5840 | 2 | -1.3767 | 0.95867 | 0.95862 | 1 | 20641 | 0 | -1.3767 |
| MROH1 | 4 | 0.1289 | 0.2657 | 0.997535 | 5913 | 3 | -1.3158 | 0.95869 | 0.95865 | 1 | 20642 | 0 | -1.3158 |
| PNRC2 | 4 | 0.079734 | 0.17824 | 0.968171 | 4221 | 1 | -0.3432 | 0.95871 | 0.95867 | 1 | 20643 | 0 | -0.3432 |

| | | | | | | | | | | | | | |
|---|---|---|---|---|---|---|---|---|---|---|---|---|---|
| BARHL2 | 4 | 0.026547 | 0.074425 | 0.93914 | 1831 | 3 | -3.1959 | 0.95878 | 0.95873 | 1 | 20644 | 0 | -3.1959 |
| MORN1 | 6 | 0.087176 | 0.22182 | 0.983139 | 4485 | 4 | -1.0059 | 0.95879 | 0.96588 | 1 | 20645 | 1 | -1.0059 |
| ARPC5 | 5 | 0.85684 | 0.85979 | 0.999994 | 18386 | 1 | 0.45908 | 0.95883 | 0.9593 | 1 | 20646 | 1 | 0.45908 |
| NonTargetingControlG | 2 | 0.041129 | 0.074235 | 0.93914 | 2567 | 2 | -2.3586 | 0.95887 | 0.95894 | 1 | 20647 | 0 | -2.3586 |
| NonTargetingControlG | 2 | 0.041121 | 0.074223 | 0.93914 | 2566 | 2 | -5.4285 | 0.95888 | 0.95895 | 1 | 20648 | 0 | -5.4285 |
| hsa-mir-6834 | 4 | 0.83855 | 0.83801 | 0.999994 | 18062 | 0 | -0.40534 | 0.95904 | 0.95896 | 1 | 20649 | 0 | -0.40534 |
| hsa-mir-4676 | 3 | 0.040837 | 0.099718 | 0.946394 | 2551 | 3 | -1.7133 | 0.95916 | 0.95933 | 1 | 20650 | 0 | -1.7133 |
| NonTargetingControlG | 2 | 0.040805 | 0.073681 | 0.93914 | 2550 | 2 | -2.1403 | 0.9592 | 0.95926 | 1 | 20651 | 0 | -2.1403 |
| CD55 | 5 | 0.81729 | 0.83079 | 0.999994 | 17732 | 1 | -0.28818 | 0.9592 | 0.95961 | 1 | 20652 | 1 | -0.28818 |
| GNG7 | 5 | 0.048083 | 0.1358 | 0.95468 | 2891 | 4 | -2.294 | 0.95925 | 0.95966 | 1 | 20653 | 1 | -2.294 |
| GLOD4 | 6 | 0.038372 | 0.11824 | 0.951052 | 2438 | 4 | -2.2103 | 0.95932 | 0.96614 | 1 | 20654 | 1 | -2.2103 |
| FOXD1 | 5 | 0.33843 | 0.51245 | 0.999994 | 10906 | 2 | 0.55729 | 0.95938 | 0.95978 | 1 | 20655 | 1 | 0.55729 |
| hsa-mir-431 | 4 | 0.08304 | 0.18435 | 0.968526 | 4340 | 3 | -1.4935 | 0.95944 | 0.95936 | 1 | 20656 | 0 | -1.4935 |
| KLK15 | 6 | 0.022486 | 0.079931 | 0.93914 | 1616 | 4 | -1.9964 | 0.95946 | 0.96621 | 1 | 20657 | 1 | -1.9964 |
| hsa-mir-1292 | 3 | 0.040509 | 0.098992 | 0.945253 | 2536 | 3 | -1.8554 | 0.95949 | 0.95966 | 1 | 20658 | 0 | -1.8554 |
| TFAP2C | 5 | 0.32628 | 0.50246 | 0.999994 | 10650 | 2 | -0.49554 | 0.95951 | 0.9599 | 1 | 20659 | 1 | -0.49554 |
| GLRX2 | 5 | 0.33421 | 0.50898 | 0.999994 | 10818 | 2 | 0.33113 | 0.95952 | 0.95991 | 1 | 20660 | 1 | 0.33113 |
| CD2AP | 6 | 0.10833 | 0.2635 | 0.996734 | 5249 | 4 | -0.83692 | 0.95953 | 0.96624 | 1 | 20661 | 1 | -0.83692 |
| INTS9 | 3 | 0.0053912 | 0.014828 | 0.755578 | 550 | 3 | -8.4997 | 0.95954 | 0.95971 | 1 | 20662 | 0 | -8.4997 |
| DMRTC1B | 4 | 0.064605 | 0.14958 | 0.966317 | 3635 | 3 | -1.6009 | 0.9596 | 0.95952 | 1 | 20663 | 0 | -1.6009 |
| HRNR | 5 | 0.37747 | 0.54463 | 0.999994 | 11617 | 2 | -0.29427 | 0.9597 | 0.96007 | 1 | 20664 | 1 | -0.29427 |
| hsa-mir-4642 | 4 | 0.34606 | 0.49028 | 0.999994 | 11067 | 2 | -0.75679 | 0.95973 | 0.95965 | 1 | 20665 | 0 | -0.75679 |
| RNF166 | 5 | 0.87059 | 0.87159 | 0.999994 | 18647 | 1 | -0.28565 | 0.95981 | 0.96016 | 1 | 20666 | 1 | -0.28565 |
| NonTargetingControlG | 1 | 0.040151 | 0.040074 | 0.891621 | 2527 | 1 | -8.6191 | 0.95985 | 0.96001 | 1 | 20667 | 0 | -8.6191 |
| SCAP | 3 | 0.040113 | 0.098113 | 0.945253 | 2524 | 3 | -2.6386 | 0.95989 | 0.96006 | 1 | 20668 | 0 | -2.6386 |
| SLC35F1 | 6 | 0.15732 | 0.33485 | 0.999994 | 6656 | 3 | -0.54756 | 0.95994 | 0.96646 | 1 | 20669 | 1 | -0.54756 |
| PRKCE | 5 | 0.17892 | 0.34506 | 0.999994 | 7195 | 3 | -0.98018 | 0.95997 | 0.96031 | 1 | 20670 | 1 | -0.98018 |
| LOC649330 | 1 | 0.039991 | 0.039911 | 0.891621 | 2514 | 1 | -8.6254 | 0.96001 | 0.96016 | 1 | 20671 | 0 | -8.6254 |
| POLDIP2 | 4 | 0.061769 | 0.14417 | 0.961983 | 3502 | 3 | -2.435 | 0.96007 | 0.95997 | 1 | 20672 | 0 | -2.435 |
| ANXA11 | 6 | 0.10293 | 0.25303 | 0.994501 | 5065 | 4 | -0.87087 | 0.96008 | 0.96653 | 1 | 20673 | 1 | -0.87087 |
| PRSS53 | 5 | 0.71088 | 0.77148 | 0.999994 | 16407 | 1 | -0.24864 | 0.96014 | 0.96046 | 1 | 20674 | 1 | -0.24864 |
| EIF1B | 6 | 0.046324 | 0.136 | 0.95468 | 2822 | 4 | -1.6369 | 0.96025 | 0.96663 | 1 | 20675 | 1 | -1.6369 |
| GATSL2 | 4 | 0.1246 | 0.25829 | 0.994943 | 5785 | 3 | -1.109 | 0.96034 | 0.96024 | 1 | 20676 | 0 | -1.109 |
| SLC35F5 | 5 | 0.12448 | 0.26419 | 0.996734 | 5781 | 3 | -1.2292 | 0.96038 | 0.96067 | 1 | 20677 | 1 | -1.2292 |
| PEA15 | 6 | 0.0043298 | 0.018693 | 0.801598 | 467 | 5 | -2.7756 | 0.96039 | 0.96669 | 1 | 20678 | 1 | -2.7756 |
| DLGAP1 | 6 | 0.014798 | 0.055062 | 0.922677 | 1176 | 5 | -1.3194 | 0.96039 | 0.96669 | 1 | 20679 | 1 | -1.3194 |
| CRYBA4 | 4 | 0.021993 | 0.06482 | 0.93914 | 1583 | 3 | -4.4931 | 0.96041 | 0.96031 | 1 | 20680 | 0 | -4.4931 |
| PDSS1 | 4 | 0.077708 | 0.17447 | 0.968171 | 4136 | 3 | -1.513 | 0.96054 | 0.96044 | 1 | 20681 | 0 | -1.513 |
| GALNT13 | 6 | 0.16645 | 0.34629 | 0.999994 | 6897 | 3 | -0.81675 | 0.96058 | 0.96679 | 1 | 20682 | 1 | -0.81675 |
| PILRA | 5 | 0.034975 | 0.1032 | 0.946414 | 2266 | 4 | -1.3955 | 0.96059 | 0.96086 | 1 | 20683 | 1 | -1.3955 |
| CDC42BPA | 5 | 0.19202 | 0.3638 | 0.999994 | 7547 | 3 | -0.88212 | 0.96073 | 0.961 | 1 | 20684 | 1 | -0.88212 |
| NonTargetingControlG | 2 | 0.01831 | 0.034101 | 0.867348 | 1376 | 2 | -6.3353 | 0.96079 | 0.96088 | 1 | 20685 | 0 | -6.3353 |
| NPPC | 4 | 0.017199 | 0.054679 | 0.922677 | 1311 | 2 | -4.1395 | 0.96082 | 0.96073 | 1 | 20686 | 0 | -4.1395 |
| OR8B12 | 5 | 0.71498 | 0.77347 | 0.999994 | 16448 | 1 | -0.14052 | 0.96085 | 0.96112 | 1 | 20687 | 1 | -0.14052 |
| ZEB1 | 5 | 0.18102 | 0.34811 | 0.999994 | 7247 | 3 | -0.79516 | 0.96088 | 0.96115 | 1 | 20688 | 1 | -0.79516 |
| SIRPG | 5 | 0.068515 | 0.17602 | 0.968171 | 3772 | 3 | -1.9295 | 0.96092 | 0.96118 | 1 | 20689 | 1 | -1.9295 |
| GTF3C5 | 4 | 0.082071 | 0.18254 | 0.968526 | 4300 | 2 | -0.75904 | 0.96097 | 0.96087 | 1 | 20690 | 0 | -0.75904 |
| MIR205HG | 6 | 0.11829 | 0.28248 | 0.999632 | 5582 | 4 | -1.1393 | 0.96107 | 0.96705 | 1 | 20691 | 1 | -1.1393 |
| IMPAD1 | 4 | 0.24263 | 0.38434 | 0.999994 | 8851 | 2 | -0.92541 | 0.96114 | 0.96103 | 1 | 20692 | 0 | -0.92541 |
| MORC3 | 6 | 0.00020407 | 0.0010921 | 0.352868 | 57 | 5 | -7.2248 | 0.96115 | 0.96709 | 1 | 20693 | 1 | -7.2248 |
| DFNB31 | 6 | 0.015875 | 0.058635 | 0.931872 | 1237 | 3 | -0.63846 | 0.96136 | 0.9672 | 1 | 20694 | 1 | -0.63846 |
| REM1 | 6 | 0.0218 | 0.077753 | 0.93914 | 1564 | 5 | -1.5736 | 0.96148 | 0.96727 | 1 | 20695 | 1 | -1.5736 |

| | | | | | | | | | | | | | |
|---|---|---|---|---|---|---|---|---|---|---|---|---|---|
| AMELY | 4 | 0.26192 | 0.40402 | 0.999994 | 9284 | 1 | -0.011746 | 0.9615 | 0.9614 | 1 | 20696 | 0 | -0.011746 |
| SPDYE5 | 3 | 0.029442 | 0.073883 | 0.93914 | 1981 | 3 | -6.0519 | 0.96157 | 0.96174 | 1 | 20697 | 0 | -6.0519 |
| hsa-mir-6754 | 3 | 0.038424 | 0.094241 | 0.945031 | 2439 | 3 | -3.0622 | 0.96158 | 0.96174 | 1 | 20698 | 0 | -3.0622 |
| hsa-mir-5195 | 4 | 0.094378 | 0.20498 | 0.978186 | 4763 | 3 | -1.5981 | 0.96165 | 0.96154 | 1 | 20699 | 0 | -1.5981 |
| NonTargetingControlG | 2 | 0.038289 | 0.069301 | 0.93914 | 2430 | 2 | -2.6222 | 0.96171 | 0.9618 | 1 | 20700 | 0 | -2.6222 |
| PYDC1 | 5 | 0.37234 | 0.54035 | 0.999994 | 11545 | 2 | -0.55203 | 0.9618 | 0.96203 | 1 | 20701 | 1 | -0.55203 |
| SYT6 | 5 | 0.27795 | 0.46319 | 0.999994 | 9651 | 2 | 0.25382 | 0.96182 | 0.96205 | 1 | 20702 | 1 | 0.25382 |
| SNX2 | 6 | 0.07635 | 0.19996 | 0.977164 | 4086 | 4 | -1.4155 | 0.96192 | 0.96751 | 1 | 20703 | 1 | -1.4155 |
| OR52N2 | 5 | 0.35844 | 0.52877 | 0.999994 | 11309 | 2 | -0.1598 | 0.96193 | 0.96215 | 1 | 20704 | 1 | -0.1598 |
| MPLKIP | 4 | 0.0013061 | 0.0047859 | 0.588856 | 194 | 3 | -5.1305 | 0.96211 | 0.962 | 1 | 20705 | 0 | -5.1305 |
| FAM57B | 5 | 0.0042747 | 0.017922 | 0.795727 | 462 | 4 | -3.6786 | 0.96216 | 0.96237 | 1 | 20706 | 1 | -3.6786 |
| DNAH5 | 6 | 0.00025378 | 0.0013486 | 0.384267 | 70 | 5 | -4.2868 | 0.96216 | 0.96765 | 1 | 20707 | 1 | -4.2868 |
| BCLAF1 | 6 | 0.00099123 | 0.005036 | 0.589484 | 166 | 4 | -5.7933 | 0.96217 | 0.96765 | 1 | 20708 | 1 | -5.7933 |
| SCYL2 | 5 | 0.23213 | 0.42018 | 0.999994 | 8572 | 3 | -0.66181 | 0.96226 | 0.96246 | 1 | 20709 | 1 | -0.66181 |
| PLK1S1 | 4 | 0.15044 | 0.28964 | 0.999632 | 6468 | 2 | -1.375 | 0.96226 | 0.96215 | 1 | 20710 | 0 | -1.375 |
| N6AMT2 | 6 | 0.0024583 | 0.011694 | 0.727553 | 314 | 3 | -0.64035 | 0.9623 | 0.96773 | 1 | 20711 | 1 | -0.64035 |
| APOBEC3G | 5 | 0.04353 | 0.12464 | 0.951052 | 2695 | 4 | -2.1203 | 0.96246 | 0.96265 | 1 | 20712 | 1 | -2.1203 |
| SLC9A8 | 5 | 0.065594 | 0.17121 | 0.967541 | 3677 | 3 | -1.7869 | 0.96253 | 0.96272 | 1 | 20713 | 1 | -1.7869 |
| NGFR | 3 | 0.037415 | 0.091974 | 0.943968 | 2393 | 3 | -1.0322 | 0.96259 | 0.96272 | 1 | 20714 | 0 | -1.0322 |
| hsa-mir-3613 | 4 | 0.093069 | 0.20266 | 0.977973 | 4721 | 3 | -1.3086 | 0.96259 | 0.9625 | 1 | 20715 | 0 | -1.3086 |
| RNF40 | 4 | 0.017529 | 0.05539 | 0.922677 | 1341 | 3 | -2.8487 | 0.96263 | 0.96254 | 1 | 20716 | 0 | -2.8487 |
| WIF1 | 6 | 0.027816 | 0.093656 | 0.945031 | 1896 | 4 | -1.9896 | 0.96272 | 0.96793 | 1 | 20717 | 1 | -1.9896 |
| TP73 | 5 | 0.6321 | 0.73811 | 0.999994 | 15401 | 1 | 0.38426 | 0.96282 | 0.96301 | 1 | 20718 | 1 | 0.38426 |
| EXOC7 | 5 | 0.040574 | 0.11732 | 0.951052 | 2540 | 4 | -1.2775 | 0.96287 | 0.96306 | 1 | 20719 | 1 | -1.2775 |
| EVC | 6 | 0.23447 | 0.42942 | 0.999994 | 8627 | 3 | -0.79789 | 0.96288 | 0.96802 | 1 | 20720 | 1 | -0.79789 |
| SCARB2 | 6 | 0.35753 | 0.57252 | 0.999994 | 11296 | 2 | -0.05953 | 0.9629 | 0.96803 | 1 | 20721 | 1 | -0.05953 |
| FAM47E-STBD1 | 2 | 0.037081 | 0.06723 | 0.93914 | 2380 | 2 | -2.6114 | 0.96292 | 0.96299 | 1 | 20722 | 0 | -2.6114 |
| CIRBP | 4 | 0.027138 | 0.075641 | 0.93914 | 1864 | 2 | -1.33 | 0.96294 | 0.96283 | 1 | 20723 | 0 | -1.33 |
| hsa-mir-3680-1 | 3 | 0.037049 | 0.091113 | 0.94395 | 2378 | 3 | -1.0424 | 0.96295 | 0.96309 | 1 | 20724 | 0 | -1.0424 |
| MS4A2 | 6 | 0.21053 | 0.40074 | 0.999994 | 8033 | 3 | -0.51179 | 0.96301 | 0.9681 | 1 | 20725 | 1 | -0.51179 |
| HMGCS1 | 4 | 0.10115 | 0.21719 | 0.980699 | 5010 | 2 | -1.7913 | 0.96307 | 0.96296 | 1 | 20726 | 0 | -1.7913 |
| hsa-mir-1178 | 4 | 0.42277 | 0.56903 | 0.999994 | 12293 | 2 | -0.44317 | 0.96311 | 0.96301 | 1 | 20727 | 0 | -0.44317 |
| LRRC47 | 5 | 0.22539 | 0.41083 | 0.999994 | 8410 | 1 | 0.17113 | 0.96312 | 0.9633 | 1 | 20728 | 1 | 0.17113 |
| SPANXA1 | 6 | 0.18124 | 0.36471 | 0.999994 | 7259 | 3 | -0.92107 | 0.96318 | 0.96819 | 1 | 20729 | 1 | -0.92107 |
| NonTargetingControlG | 2 | 0.036784 | 0.066689 | 0.93914 | 2362 | 2 | -5.8025 | 0.96322 | 0.96329 | 1 | 20730 | 0 | -5.8025 |
| HIST2H4A | 3 | 0.036717 | 0.09036 | 0.94395 | 2359 | 3 | -1.3062 | 0.96328 | 0.96343 | 1 | 20731 | 0 | -1.3062 |
| hsa-mir-219b | 4 | 0.39529 | 0.54088 | 0.999994 | 11875 | 2 | -0.60376 | 0.9633 | 0.96319 | 1 | 20732 | 0 | -0.60376 |
| TRIB2 | 5 | 0.061682 | 0.16486 | 0.967126 | 3500 | 3 | -1.9725 | 0.96333 | 0.96349 | 1 | 20733 | 1 | -1.9725 |
| hsa-mir-4682 | 4 | 0.26632 | 0.40852 | 0.999994 | 9375 | 1 | -0.54017 | 0.96337 | 0.96327 | 1 | 20734 | 0 | -0.54017 |
| hsa-mir-4518 | 4 | 0.12429 | 0.25775 | 0.994943 | 5775 | 3 | -0.75376 | 0.96341 | 0.9633 | 1 | 20735 | 0 | -0.75376 |
| RHOT1 | 4 | 0.068755 | 0.15753 | 0.967126 | 3785 | 3 | -3.2346 | 0.96358 | 0.96347 | 1 | 20736 | 0 | -3.2346 |
| SENP2 | 6 | 0.020546 | 0.073718 | 0.93914 | 1486 | 5 | -1.6753 | 0.96377 | 0.96851 | 1 | 20737 | 1 | -1.6753 |
| DNAJC10 | 5 | 0.19961 | 0.37468 | 0.999994 | 7752 | 3 | -0.81919 | 0.96382 | 0.96397 | 1 | 20738 | 1 | -0.81919 |
| hsa-mir-3689e | 2 | 0.036139 | 0.065552 | 0.93914 | 2327 | 2 | -4.0099 | 0.96386 | 0.96394 | 1 | 20739 | 0 | -4.0099 |
| hsa-mir-548aq | 2 | 0.036136 | 0.065546 | 0.93914 | 2326 | 2 | -5.0113 | 0.96386 | 0.96394 | 1 | 20740 | 0 | -5.0113 |
| hsa-mir-574 | 4 | 0.031048 | 0.083623 | 0.94395 | 2063 | 2 | -0.90827 | 0.9639 | 0.96378 | 1 | 20741 | 0 | -0.90827 |
| OR4D5 | 5 | 0.0035025 | 0.014845 | 0.755578 | 406 | 4 | -2.8574 | 0.9639 | 0.96406 | 1 | 20742 | 1 | -2.8574 |
| SPATA31D3 | 6 | 0.033576 | 0.10718 | 0.950995 | 2194 | 3 | -0.82197 | 0.96391 | 0.9686 | 1 | 20743 | 1 | -0.82197 |
| hsa-mir-6836 | 3 | 0.036066 | 0.088918 | 0.94395 | 2323 | 3 | -1.3837 | 0.96393 | 0.96405 | 1 | 20744 | 0 | -1.3837 |
| LAD1 | 6 | 0.020182 | 0.072562 | 0.93914 | 1471 | 4 | -2.9091 | 0.9641 | 0.96871 | 1 | 20745 | 1 | -2.9091 |
| CYP4F3 | 6 | 0.11298 | 0.27242 | 0.998561 | 5402 | 3 | -0.86098 | 0.9641 | 0.96871 | 1 | 20746 | 1 | -0.86098 |
| OXNAD1 | 4 | 0.3751 | 0.52011 | 0.999994 | 11581 | 2 | -1.0934 | 0.96418 | 0.96408 | 1 | 20747 | 0 | -1.0934 |

| | | | | | | | | | | | | | |
|---|---|---|---|---|---|---|---|---|---|---|---|---|---|
| RNF215 | 6 | 0.048439 | 0.14067 | 0.959146 | 2905 | 4 | -1.7425 | 0.96435 | 0.96884 | 1 | 20748 | 1 | -1.7425 |
| SLC26A6 | 6 | 0.0038637 | 0.016977 | 0.785648 | 433 | 5 | -3.2122 | 0.96448 | 0.96893 | 1 | 20749 | 1 | -3.2122 |
| FLJ27352 | 4 | 0.035488 | 0.092711 | 0.943968 | 2293 | 2 | -2.5505 | 0.96448 | 0.96437 | 1 | 20750 | 0 | -2.5505 |
| RBMX2 | 3 | 0.035481 | 0.087611 | 0.94395 | 2292 | 3 | -2.495 | 0.96452 | 0.96463 | 1 | 20751 | 0 | -2.495 |
| ATP8A2 | 5 | 0.36423 | 0.53358 | 0.999994 | 11406 | 1 | -0.22907 | 0.96455 | 0.96469 | 1 | 20752 | 0 | -0.22907 |
| ATP2B1 | 5 | 0.0072616 | 0.028103 | 0.855594 | 675 | 4 | -2.3012 | 0.96461 | 0.96476 | 1 | 20753 | 0 | -2.3012 |
| SMR3A | 5 | 0.22248 | 0.40674 | 0.999994 | 8341 | 2 | 0.34278 | 0.96461 | 0.96477 | 1 | 20754 | 0 | 0.34278 |
| FAM46B | 6 | 0.0042193 | 0.018277 | 0.795727 | 459 | 4 | -3.6406 | 0.9647 | 0.96905 | 1 | 20755 | 1 | -3.6406 |
| RER1 | 5 | 0.053338 | 0.14842 | 0.965554 | 3130 | 4 | -1.645 | 0.96473 | 0.96489 | 1 | 20756 | 0 | -1.645 |
| PRR18 | 3 | 0.035155 | 0.086871 | 0.94395 | 2277 | 3 | -1.2478 | 0.96484 | 0.96495 | 1 | 20757 | 0 | -1.2478 |
| hsa-mir-4443 | 3 | 0.035131 | 0.086817 | 0.94395 | 2275 | 3 | -1.0836 | 0.96487 | 0.96497 | 1 | 20758 | 0 | -1.0836 |
| NRTN | 5 | 0.019804 | 0.063566 | 0.937389 | 1447 | 4 | -1.3616 | 0.96492 | 0.96509 | 1 | 20759 | 0 | -1.3616 |
| KIF13A | 6 | 0.060259 | 0.16629 | 0.967126 | 3432 | 4 | -1.321 | 0.96495 | 0.96919 | 1 | 20760 | 1 | -1.321 |
| PPIP5K2 | 5 | 0.62874 | 0.73682 | 0.999994 | 15358 | 1 | -0.12699 | 0.96496 | 0.96512 | 1 | 20761 | 0 | -0.12699 |
| PDK4 | 5 | 0.027458 | 0.083901 | 0.94395 | 1876 | 3 | -2.8532 | 0.96499 | 0.96515 | 1 | 20762 | 0 | -2.8532 |
| hsa-mir-196b | 4 | 0.034801 | 0.091313 | 0.94395 | 2257 | 3 | -2.5158 | 0.965 | 0.96489 | 1 | 20763 | 0 | -2.5158 |
| BRD7 | 6 | 0.030245 | 0.099353 | 0.945253 | 2022 | 1 | 0.27088 | 0.96502 | 0.96923 | 1 | 20764 | 1 | 0.27088 |
| GGH | 5 | 0.57725 | 0.7159 | 0.999994 | 14647 | 2 | -0.10271 | 0.96511 | 0.96526 | 1 | 20765 | 0 | -0.10271 |
| USP42 | 6 | 0.10993 | 0.26657 | 0.998525 | 5302 | 4 | -0.81823 | 0.96513 | 0.96929 | 1 | 20766 | 1 | -0.81823 |
| VPS39 | 5 | 0.044628 | 0.12739 | 0.951052 | 2744 | 4 | -2.5815 | 0.96516 | 0.9653 | 1 | 20767 | 0 | -2.5815 |
| USP17L11 | 6 | 0.11422 | 0.27479 | 0.998561 | 5454 | 4 | -0.60622 | 0.9652 | 0.96933 | 1 | 20768 | 1 | -0.60622 |
| CAP2 | 6 | 0.87532 | 0.88665 | 0.999994 | 18742 | 1 | -0.10596 | 0.96524 | 0.96935 | 1 | 20769 | 1 | -0.10596 |
| PRMT10 | 5 | 0.015637 | 0.052019 | 0.911383 | 1226 | 4 | -1.712 | 0.96524 | 0.96539 | 1 | 20770 | 0 | -1.712 |
| OTUD5 | 5 | 0.27799 | 0.46323 | 0.999994 | 9654 | 2 | -0.57493 | 0.96525 | 0.9654 | 1 | 20771 | 0 | -0.57493 |
| C20orf85 | 5 | 0.37747 | 0.54463 | 0.999994 | 11618 | 2 | -0.31733 | 0.96531 | 0.96546 | 1 | 20772 | 0 | -0.31733 |
| FAM13C | 6 | 0.02385 | 0.084246 | 0.94395 | 1703 | 5 | -1.2103 | 0.96552 | 0.96954 | 1 | 20773 | 1 | -1.2103 |
| FGFR1OP | 6 | 0.0079456 | 0.031779 | 0.865899 | 727 | 3 | -0.76856 | 0.96553 | 0.96954 | 1 | 20774 | 1 | -0.76856 |
| DPY19L1 | 5 | 0.14939 | 0.30187 | 0.999994 | 6440 | 2 | 0.19363 | 0.96559 | 0.96574 | 1 | 20775 | 0 | 0.19363 |
| CHRM1 | 4 | 0.15804 | 0.29748 | 0.999994 | 6671 | 2 | -1.2847 | 0.96562 | 0.9655 | 1 | 20776 | 0 | -1.2847 |
| NECAP2 | 6 | 0.41929 | 0.61491 | 0.999994 | 12239 | 2 | -0.13003 | 0.96562 | 0.9696 | 1 | 20777 | 1 | -0.13003 |
| hsa-mir-3609 | 4 | 0.044942 | 0.11161 | 0.951052 | 2754 | 3 | -2.2455 | 0.96564 | 0.96552 | 1 | 20778 | 0 | -2.2455 |
| hsa-mir-7113 | 2 | 0.034135 | 0.062112 | 0.937033 | 2221 | 2 | -5.3906 | 0.96586 | 0.96588 | 1 | 20779 | 0 | -5.3906 |
| PCNT | 6 | 0.23601 | 0.43126 | 0.999994 | 8671 | 3 | -0.87492 | 0.96588 | 0.96976 | 1 | 20780 | 1 | -0.87492 |
| FABP2 | 6 | 0.010056 | 0.039199 | 0.891621 | 872 | 4 | -0.79731 | 0.96594 | 0.96979 | 1 | 20781 | 1 | -0.79731 |
| B4GALNT4 | 6 | 0.20906 | 0.39896 | 0.999994 | 8002 | 3 | -0.64806 | 0.96594 | 0.96979 | 1 | 20782 | 1 | -0.64806 |
| NonTargetingControlG | 2 | 0.034017 | 0.061891 | 0.937033 | 2217 | 2 | -3.1096 | 0.96598 | 0.966 | 1 | 20783 | 0 | -3.1096 |
| OR51B2 | 5 | 0.064105 | 0.16881 | 0.967126 | 3618 | 3 | -2.4124 | 0.96598 | 0.96613 | 1 | 20784 | 0 | -2.4124 |
| LRP4 | 5 | 0.51406 | 0.66014 | 0.999994 | 13642 | 2 | -0.12984 | 0.966 | 0.96615 | 1 | 20785 | 0 | -0.12984 |
| TMEM240 | 6 | 0.055173 | 0.1553 | 0.967126 | 3207 | 3 | -0.6822 | 0.96604 | 0.96986 | 1 | 20786 | 1 | -0.6822 |
| ADCY10 | 6 | 0.038078 | 0.11757 | 0.951052 | 2424 | 4 | -2.408 | 0.96607 | 0.96987 | 1 | 20787 | 1 | -2.408 |
| CBX7 | 5 | 0.12393 | 0.26334 | 0.996734 | 5764 | 3 | -1.3872 | 0.9661 | 0.96626 | 1 | 20788 | 0 | -1.3872 |
| NonTargetingControlG | 2 | 0.033876 | 0.061648 | 0.937033 | 2211 | 2 | -5.3175 | 0.96612 | 0.96613 | 1 | 20789 | 0 | -5.3175 |
| ZNF474 | 6 | 0.12123 | 0.28806 | 0.999632 | 5672 | 4 | -0.86151 | 0.96618 | 0.96993 | 1 | 20790 | 1 | -0.86151 |
| SAA1 | 5 | 0.53289 | 0.6766 | 0.999994 | 13951 | 2 | -0.37681 | 0.96622 | 0.96638 | 1 | 20791 | 0 | -0.37681 |
| HIF1AN | 5 | 0.064347 | 0.16923 | 0.967126 | 3625 | 3 | -0.60955 | 0.96629 | 0.96645 | 1 | 20792 | 0 | -0.60955 |
| CLPTM1L | 5 | 0.21909 | 0.40201 | 0.999994 | 8264 | 3 | -0.74983 | 0.96638 | 0.96655 | 1 | 20793 | 0 | -0.74983 |
| ZC3H12B | 4 | 0.27084 | 0.41314 | 0.999994 | 9496 | 1 | -0.23064 | 0.96642 | 0.9663 | 1 | 20794 | 0 | -0.23064 |
| SLC29A4 | 5 | 0.022991 | 0.072184 | 0.93914 | 1649 | 4 | -1.1486 | 0.96651 | 0.96671 | 1 | 20795 | 0 | -1.1486 |
| DDX41 | 5 | 0.09579 | 0.21977 | 0.980699 | 4806 | 3 | -1.7238 | 0.96653 | 0.96673 | 1 | 20796 | 0 | -1.7238 |
| MFSD12 | 5 | 0.0015783 | 0.0069723 | 0.645102 | 232 | 4 | -2.4616 | 0.96655 | 0.96674 | 1 | 20797 | 0 | -2.4616 |
| RBM14-RBM4 | 2 | 0.033397 | 0.06085 | 0.932932 | 2181 | 2 | -3.2395 | 0.9666 | 0.96661 | 1 | 20798 | 0 | -3.2395 |
| GAS8 | 5 | 0.19384 | 0.36641 | 0.999994 | 7594 | 2 | 0.21982 | 0.96669 | 0.96688 | 1 | 20799 | 0 | 0.21982 |

| | | | | | | | | | | | | |
|---|---|---|---|---|---|---|---|---|---|---|---|---|
| NELFCD | 4 | 1.2737E-06 | 3.4291E-06 | 0.008251 | 10 | 3 | -11.058 | 0.96672 | 0.96661 | 1 | 20800 | 0 | -11.058 |
| SLC12A4 | 5 | 0.53536 | 0.67876 | 0.999994 | 13986 | 2 | -0.27662 | 0.96675 | 0.96693 | 1 | 20801 | 0 | -0.27662 |
| SLC16A14 | 5 | 0.104 | 0.23262 | 0.984502 | 5095 | 3 | -1.3831 | 0.96679 | 0.96697 | 1 | 20802 | 0 | -1.3831 |
| TIMMDC1 | 5 | 0.56128 | 0.70176 | 0.999994 | 14393 | 1 | -0.5321 | 0.96682 | 0.967 | 1 | 20803 | 0 | -0.5321 |
| DDC | 6 | 0.17961 | 0.36267 | 0.999994 | 7214 | 3 | -0.51939 | 0.96688 | 0.97033 | 1 | 20804 | 1 | -0.51939 |
| IFIT2 | 6 | 0.13476 | 0.30636 | 0.999994 | 6044 | 2 | 0.060435 | 0.96691 | 0.97034 | 1 | 20805 | 1 | 0.060435 |
| KPNA3 | 5 | 0.04989 | 0.14007 | 0.958716 | 2970 | 4 | -0.99804 | 0.96696 | 0.96713 | 1 | 20806 | 0 | -0.99804 |
| CHD4 | 6 | 0.0089902 | 0.035487 | 0.870441 | 793 | 4 | -3.7859 | 0.96696 | 0.97037 | 1 | 20807 | 1 | -3.7859 |
| EIF4E3 | 5 | 0.050144 | 0.14072 | 0.959146 | 2984 | 4 | -0.93072 | 0.96702 | 0.9672 | 1 | 20808 | 0 | -0.93072 |
| BTAF1 | 6 | 0.001346 | 0.0067341 | 0.643671 | 202 | 5 | -3.3252 | 0.96706 | 0.97044 | 1 | 20809 | 1 | -3.3252 |
| FLRT1 | 6 | 0.097671 | 0.24276 | 0.988088 | 4876 | 3 | -1.1715 | 0.96708 | 0.97045 | 1 | 20810 | 1 | -1.1715 |
| DIEXF | 5 | 0.002608 | 0.011277 | 0.716138 | 325 | 4 | -7.3738 | 0.96714 | 0.96733 | 1 | 20811 | 0 | -7.3738 |
| IVNS1ABP | 5 | 0.075381 | 0.18714 | 0.969409 | 4050 | 3 | -2.9135 | 0.96715 | 0.96734 | 1 | 20812 | 0 | -2.9135 |
| hsa-mir-4435-1 | 3 | 0.032751 | 0.081436 | 0.93914 | 2146 | 3 | -1.5963 | 0.96725 | 0.96733 | 1 | 20813 | 0 | -1.5963 |
| ADCY1 | 6 | 0.045147 | 0.13339 | 0.95318 | 2767 | 4 | -0.96244 | 0.96725 | 0.97055 | 1 | 20814 | 1 | -0.96244 |
| C5orf47 | 5 | 0.34002 | 0.51372 | 0.999994 | 10934 | 2 | -0.49326 | 0.96726 | 0.96744 | 1 | 20815 | 0 | -0.49326 |
| CYP4F22 | 5 | 0.8718 | 0.87268 | 0.999994 | 18674 | 1 | -0.041636 | 0.9673 | 0.96748 | 1 | 20816 | 0 | -0.041636 |
| MBOAT1 | 5 | 0.22364 | 0.40836 | 0.999994 | 8371 | 3 | -0.73736 | 0.96739 | 0.96755 | 1 | 20817 | 0 | -0.73736 |
| BTNL8 | 5 | 0.00091894 | 0.0041097 | 0.560963 | 153 | 4 | -3.5694 | 0.96739 | 0.96756 | 1 | 20818 | 0 | -3.5694 |
| LRP12 | 6 | 0.028368 | 0.094993 | 0.945031 | 1918 | 4 | -1.8682 | 0.9674 | 0.97064 | 1 | 20819 | 1 | -1.8682 |
| SPRY3_X | 5 | 0.26962 | 0.45653 | 0.999994 | 9466 | 1 | -0.2048 | 0.96743 | 0.96759 | 1 | 20820 | 0 | -0.2048 |
| DMRT3 | 5 | 0.00043723 | 0.0020284 | 0.439257 | 102 | 4 | -8.1133 | 0.96747 | 0.96763 | 1 | 20821 | 0 | -8.1133 |
| SULT1A2 | 6 | 0.17207 | 0.3533 | 0.999994 | 7039 | 3 | -0.57137 | 0.96751 | 0.9707 | 1 | 20822 | 1 | -0.57137 |
| CDC40 | 6 | 0.035142 | 0.11086 | 0.951052 | 2276 | 3 | -0.99875 | 0.96754 | 0.97072 | 1 | 20823 | 1 | -0.99875 |
| HBA1 | 6 | 0.034562 | 0.10947 | 0.951052 | 2244 | 4 | -1.5868 | 0.96754 | 0.97072 | 1 | 20824 | 1 | -1.5868 |
| SLC26A1 | 4 | 0.07606 | 0.17138 | 0.967658 | 4073 | 3 | -1.3543 | 0.96762 | 0.96751 | 1 | 20825 | 0 | -1.3543 |
| KRTAP5-8 | 5 | 0.069009 | 0.17681 | 0.968171 | 3799 | 3 | -2.0016 | 0.96766 | 0.96782 | 1 | 20826 | 0 | -2.0016 |
| PLA2G4E | 5 | 0.85472 | 0.85806 | 0.999994 | 18346 | 1 | 0.092178 | 0.96772 | 0.96787 | 1 | 20827 | 0 | 0.092178 |
| TRNAU1AP | 4 | 0.035405 | 0.092558 | 0.943968 | 2287 | 3 | -1.8384 | 0.96783 | 0.96772 | 1 | 20828 | 0 | -1.8384 |
| FIBP | 3 | 0.016442 | 0.042875 | 0.891621 | 1273 | 3 | -2.1672 | 0.9679 | 0.96797 | 1 | 20829 | 0 | -2.1672 |
| MGST3 | 5 | 0.0090391 | 0.033272 | 0.867348 | 796 | 4 | -2.3096 | 0.9679 | 0.96805 | 1 | 20830 | 0 | -2.3096 |
| PFN3 | 6 | 0.022288 | 0.079313 | 0.93914 | 1598 | 3 | -2.2488 | 0.96792 | 0.97096 | 1 | 20831 | 1 | -2.2488 |
| ANKRD35 | 5 | 0.02928 | 0.088619 | 0.94395 | 1972 | 2 | -0.40862 | 0.96806 | 0.9682 | 1 | 20832 | 0 | -0.40862 |
| NISCH | 5 | 0.025703 | 0.079346 | 0.93914 | 1785 | 3 | -2.2962 | 0.9681 | 0.96823 | 1 | 20833 | 0 | -2.2962 |
| C9orf129 | 5 | 0.4969 | 0.64518 | 0.999994 | 13374 | 1 | 0.27017 | 0.96812 | 0.96826 | 1 | 20834 | 0 | 0.27017 |
| NAA16 | 6 | 0.047452 | 0.13848 | 0.957336 | 2878 | 4 | -1.5176 | 0.9682 | 0.97114 | 1 | 20835 | 1 | -1.5176 |
| SRCRB4D | 6 | 0.11718 | 0.28041 | 0.999632 | 5543 | 4 | -0.7983 | 0.96828 | 0.97119 | 1 | 20836 | 1 | -0.7983 |
| CAPZA2 | 6 | 0.070225 | 0.18724 | 0.969409 | 3842 | 4 | -1.2035 | 0.9683 | 0.9712 | 1 | 20837 | 1 | -1.2035 |
| CUL2 | 6 | 0.22605 | 0.41935 | 0.999994 | 8426 | 3 | -0.79274 | 0.96833 | 0.97122 | 1 | 20838 | 1 | -0.79274 |
| BOLA3 | 5 | 0.15607 | 0.31179 | 0.999994 | 6630 | 3 | -1.1691 | 0.96835 | 0.96849 | 1 | 20839 | 0 | -1.1691 |
| DNAJB6 | 5 | 0.49384 | 0.64259 | 0.999994 | 13323 | 2 | -0.2243 | 0.96839 | 0.96853 | 1 | 20840 | 0 | -0.2243 |
| C4orf45 | 4 | 0.29242 | 0.43526 | 0.999994 | 9959 | 2 | -0.79894 | 0.96847 | 0.96838 | 1 | 20841 | 0 | -0.79894 |
| OR2A25 | 5 | 0.013074 | 0.044794 | 0.891621 | 1069 | 4 | -1.9509 | 0.96869 | 0.96881 | 1 | 20842 | 0 | -1.9509 |
| GOLT1B | 6 | 0.022976 | 0.081511 | 0.93914 | 1648 | 4 | -1.7661 | 0.96871 | 0.97146 | 1 | 20843 | 1 | -1.7661 |
| NonTargetingControlG | 2 | 0.031268 | 0.057153 | 0.92665 | 2072 | 2 | -2.2662 | 0.96873 | 0.96875 | 1 | 20844 | 0 | -2.2662 |
| GRK4 | 5 | 0.039468 | 0.11457 | 0.951052 | 2488 | 3 | -2.3744 | 0.96875 | 0.96888 | 1 | 20845 | 0 | -2.3744 |
| ACTG1 | 5 | 0.021914 | 0.069259 | 0.93914 | 1576 | 3 | -2.5122 | 0.96888 | 0.96899 | 1 | 20846 | 0 | -2.5122 |
| SLC35B3 | 5 | 0.071087 | 0.18014 | 0.968171 | 3878 | 3 | -1.7134 | 0.96892 | 0.96903 | 1 | 20847 | 0 | -1.7134 |
| STON1-GTF2A1L | 4 | 0.28439 | 0.42703 | 0.999994 | 9795 | 2 | -0.8288 | 0.96901 | 0.96889 | 1 | 20848 | 0 | -0.8288 |
| MAMDC2 | 5 | 0.1771 | 0.34244 | 0.999994 | 7160 | 3 | -0.98515 | 0.96903 | 0.96912 | 1 | 20849 | 0 | -0.98515 |
| DAG1 | 5 | 0.30411 | 0.48436 | 0.999994 | 10211 | 2 | -0.32262 | 0.96903 | 0.96913 | 1 | 20850 | 0 | -0.32262 |
| CES2 | 6 | 0.020335 | 0.073055 | 0.93914 | 1479 | 4 | -2.3009 | 0.96904 | 0.97166 | 1 | 20851 | 1 | -2.3009 |

| | | | | | | | | | | | | | |
|---|---|---|---|---|---|---|---|---|---|---|---|---|---|
| USP40 | 6 | 0.1193 | 0.28441 | 0.999632 | 5608 | 3 | -0.94786 | 0.96907 | 0.97168 | 1 | 20852 | 1 | -0.94786 |
| hsa-mir-3929 | 1 | 0.030916 | 0.030813 | 0.862223 | 2056 | 1 | -9.011 | 0.96908 | 0.96918 | 1 | 20853 | 0 | -9.011 |
| TLR8 | 4 | 0.10592 | 0.22571 | 0.983354 | 5173 | 3 | -1.8929 | 0.96911 | 0.96899 | 1 | 20854 | 0 | -1.8929 |
| NonTargetingControlG | 2 | 0.030876 | 0.05648 | 0.9235 | 2055 | 2 | -2.3115 | 0.96912 | 0.96915 | 1 | 20855 | 0 | -2.3115 |
| RTL1 | 5 | 0.03187 | 0.095291 | 0.945031 | 2098 | 4 | -1.3376 | 0.96917 | 0.96926 | 1 | 20856 | 0 | -1.3376 |
| FAM86B2 | 1 | 0.030812 | 0.030706 | 0.862223 | 2052 | 1 | -9.0172 | 0.96919 | 0.96929 | 1 | 20857 | 0 | -9.0172 |
| APOBEC2 | 6 | 0.33782 | 0.55019 | 0.999994 | 10892 | 3 | -0.30403 | 0.96923 | 0.97178 | 1 | 20858 | 1 | -0.30403 |
| AMIGO3 | 6 | 0.10673 | 0.26041 | 0.994943 | 5200 | 3 | -0.44416 | 0.96925 | 0.9718 | 1 | 20859 | 1 | -0.44416 |
| TRAM2 | 5 | 0.27276 | 0.45902 | 0.999994 | 9532 | 1 | -0.039362 | 0.96932 | 0.96942 | 1 | 20860 | 0 | -0.039362 |
| POC1A | 6 | 0.03121 | 0.10167 | 0.946414 | 2069 | 4 | -1.2766 | 0.96938 | 0.97187 | 1 | 20861 | 1 | -1.2766 |
| TP53I11 | 5 | 0.0034405 | 0.0146 | 0.755578 | 399 | 4 | -2.4913 | 0.96944 | 0.96952 | 1 | 20862 | 0 | -2.4913 |
| HIST1H3F | 5 | 0.0038474 | 0.016211 | 0.772095 | 431 | 4 | -7.1974 | 0.96955 | 0.96962 | 1 | 20863 | 0 | -7.1974 |
| HIST1H2BE | 4 | 0.020174 | 0.061004 | 0.932932 | 1467 | 3 | -3.3562 | 0.96957 | 0.96945 | 1 | 20864 | 0 | -3.3562 |
| IPMK | 5 | 0.2392 | 0.42984 | 0.999994 | 8752 | 3 | -0.64762 | 0.96957 | 0.96964 | 1 | 20865 | 0 | -0.64762 |
| IQCF3 | 6 | 0.018076 | 0.06575 | 0.93914 | 1364 | 5 | -1.5195 | 0.96958 | 0.972 | 1 | 20866 | 1 | -1.5195 |
| SNAPC4 | 3 | 0.0094483 | 0.025337 | 0.84412 | 827 | 3 | -7.9078 | 0.9696 | 0.9697 | 1 | 20867 | 0 | -7.9078 |
| ZNF232 | 4 | 0.056737 | 0.13457 | 0.953565 | 3282 | 3 | -1.9516 | 0.9696 | 0.96949 | 1 | 20868 | 0 | -1.9516 |
| DAAM1 | 6 | 0.028953 | 0.096342 | 0.945031 | 1951 | 4 | -2.2623 | 0.96962 | 0.97202 | 1 | 20869 | 1 | -2.2623 |
| MBD3 | 5 | 0.10854 | 0.23973 | 0.987253 | 5258 | 3 | -1.2723 | 0.96965 | 0.96971 | 1 | 20870 | 0 | -1.2723 |
| SLC6A9 | 5 | 0.42987 | 0.58831 | 0.999994 | 12412 | 2 | -0.52705 | 0.96967 | 0.96973 | 1 | 20871 | 0 | -0.52705 |
| hsa-mir-188 | 4 | 0.077852 | 0.17474 | 0.968171 | 4144 | 2 | -2.0448 | 0.96974 | 0.96964 | 1 | 20872 | 0 | -2.0448 |
| THOC3 | 4 | 0.05659 | 0.13428 | 0.953565 | 3277 | 3 | -3.552 | 0.96975 | 0.96965 | 1 | 20873 | 0 | -3.552 |
| KCNS1 | 4 | 0.040531 | 0.10285 | 0.946414 | 2537 | 3 | -2.4438 | 0.96976 | 0.96965 | 1 | 20874 | 0 | -2.4438 |
| NonTargetingControlG | 2 | 0.03023 | 0.055337 | 0.922677 | 2021 | 2 | -3.2595 | 0.96977 | 0.96979 | 1 | 20875 | 0 | -3.2595 |
| NR4A3 | 6 | 0.33273 | 0.54439 | 0.999994 | 10786 | 3 | -0.43634 | 0.96982 | 0.97216 | 1 | 20876 | 1 | -0.43634 |
| EPC2 | 5 | 0.0033768 | 0.014342 | 0.753239 | 392 | 3 | -0.79739 | 0.96982 | 0.96988 | 1 | 20877 | 0 | -0.79739 |
| CDK14 | 5 | 0.14904 | 0.3013 | 0.999994 | 6430 | 3 | -0.96543 | 0.96991 | 0.96997 | 1 | 20878 | 0 | -0.96543 |
| NonTargetingControlG | 1 | 0.030082 | 0.030004 | 0.862223 | 2012 | 1 | -7.0964 | 0.96992 | 0.97002 | 1 | 20879 | 0 | -7.0964 |
| OR2B11 | 4 | 0.083923 | 0.18595 | 0.969409 | 4374 | 3 | -1.9143 | 0.96993 | 0.96981 | 1 | 20880 | 0 | -1.9143 |
| SLC13A5 | 5 | 0.027599 | 0.084255 | 0.94395 | 1882 | 3 | -0.85074 | 0.96994 | 0.97 | 1 | 20881 | 0 | -0.85074 |
| ETV5 | 5 | 0.08989 | 0.21043 | 0.979587 | 4589 | 3 | -1.612 | 0.96995 | 0.97 | 1 | 20882 | 0 | -1.612 |
| IFNAR2 | 6 | 0.036011 | 0.11288 | 0.951052 | 2315 | 3 | -0.35188 | 0.97001 | 0.97227 | 1 | 20883 | 1 | -0.35188 |
| HMX2 | 6 | 0.018652 | 0.067648 | 0.93914 | 1396 | 5 | -1.4578 | 0.97007 | 0.97232 | 1 | 20884 | 1 | -1.4578 |
| HTRA1 | 5 | 0.035757 | 0.10516 | 0.95013 | 2304 | 4 | -1.9064 | 0.9701 | 0.97015 | 1 | 20885 | 0 | -1.9064 |
| STRAP | 5 | 0.68174 | 0.75831 | 0.999994 | 16113 | 1 | -0.51691 | 0.97016 | 0.97021 | 1 | 20886 | 0 | -0.51691 |
| NonTargetingControlG | 2 | 0.029789 | 0.054599 | 0.922677 | 1999 | 2 | -4.4596 | 0.97021 | 0.97022 | 1 | 20887 | 0 | -4.4596 |
| BOP1 | 4 | 0.0047063 | 0.016647 | 0.77694 | 498 | 3 | -6.5105 | 0.97038 | 0.97027 | 1 | 20888 | 0 | -6.5105 |
| SHISA4 | 5 | 0.064332 | 0.1692 | 0.967126 | 3623 | 3 | -1.8643 | 0.9704 | 0.97046 | 1 | 20889 | 0 | -1.8643 |
| TRPM1 | 6 | 0.31045 | 0.51887 | 0.999994 | 10334 | 3 | -0.61484 | 0.97042 | 0.97254 | 1 | 20890 | 1 | -0.61484 |
| SMC2 | 4 | 0.012357 | 0.04105 | 0.891621 | 1018 | 2 | -4.2956 | 0.97058 | 0.97046 | 1 | 20891 | 0 | -4.2956 |
| CRIP1 | 6 | 0.0077091 | 0.030934 | 0.862223 | 708 | 5 | -1.8806 | 0.97062 | 0.97267 | 1 | 20892 | 1 | -1.8806 |
| MMP21 | 5 | 0.029777 | 0.089893 | 0.94395 | 1998 | 4 | -2.3781 | 0.97062 | 0.97068 | 1 | 20893 | 0 | -2.3781 |
| ZNF496 | 6 | 0.024809 | 0.086556 | 0.94395 | 1747 | 4 | -2.1586 | 0.97076 | 0.97278 | 1 | 20894 | 1 | -2.1586 |
| hsa-mir-92a-1 | 4 | 0.022223 | 0.06533 | 0.93914 | 1595 | 3 | -4.2397 | 0.97082 | 0.97071 | 1 | 20895 | 0 | -4.2397 |
| ITGA4 | 6 | 0.048474 | 0.14075 | 0.959146 | 2907 | 4 | -1.387 | 0.97086 | 0.97284 | 1 | 20896 | 1 | -1.387 |
| NPM1 | 3 | 0.029065 | 0.073019 | 0.93914 | 1959 | 3 | -4.2231 | 0.97094 | 0.97099 | 1 | 20897 | 0 | -4.2231 |
| PREPL | 5 | 0.47791 | 0.62886 | 0.999994 | 13112 | 2 | 0.073946 | 0.97098 | 0.97103 | 1 | 20898 | 0 | 0.073946 |
| UTP6 | 5 | 0.070399 | 0.17904 | 0.968171 | 3850 | 2 | 0.25349 | 0.97116 | 0.97121 | 1 | 20899 | 0 | 0.25349 |
| hsa-mir-18b | 4 | 0.0094523 | 0.031976 | 0.865899 | 828 | 3 | -3.1912 | 0.97118 | 0.97106 | 1 | 20900 | 0 | -3.1912 |
| CDH13 | 6 | 0.046867 | 0.13721 | 0.957161 | 2846 | 3 | -0.58807 | 0.97121 | 0.97308 | 1 | 20901 | 1 | -0.58807 |
| MON1B | 6 | 0.28595 | 0.4904 | 0.999994 | 9832 | 2 | 0.26236 | 0.97123 | 0.97309 | 1 | 20902 | 1 | 0.26236 |
| TM9SF3 | 5 | 0.012578 | 0.043411 | 0.891621 | 1034 | 4 | -2.0169 | 0.97125 | 0.9713 | 1 | 20903 | 0 | -2.0169 |

| | | | | | | | | | | | | | |
|---|---|---|---|---|---|---|---|---|---|---|---|---|---|
| DVL1 | 5 | 0.083808 | 0.2007 | 0.977164 | 4369 | 3 | -1.6588 | 0.9713 | 0.97136 | 1 | 20904 | 0 | -1.6588 |
| NonTargetingControlG | 2 | 0.028641 | 0.052566 | 0.914561 | 1934 | 2 | -5.1222 | 0.97136 | 0.97133 | 1 | 20905 | 0 | -5.1222 |
| PPP1R17 | 6 | 0.037968 | 0.11731 | 0.951052 | 2422 | 4 | -1.6924 | 0.97148 | 0.97326 | 1 | 20906 | 1 | -1.6924 |
| SNTB2 | 5 | 0.12173 | 0.25999 | 0.994943 | 5692 | 2 | -0.40244 | 0.97151 | 0.97156 | 1 | 20907 | 0 | -0.40244 |
| hsa-mir-548h-5 | 2 | 0.028466 | 0.052251 | 0.913966 | 1923 | 2 | -4.7988 | 0.97153 | 0.9715 | 1 | 20908 | 0 | -4.7988 |
| COBL | 6 | 0.015505 | 0.057369 | 0.927806 | 1216 | 5 | -1.1527 | 0.97156 | 0.97331 | 1 | 20909 | 1 | -1.1527 |
| PLXNB1 | 6 | 0.0031356 | 0.014234 | 0.753188 | 370 | 4 | -1.5031 | 0.97164 | 0.97336 | 1 | 20910 | 1 | -1.5031 |
| PRR14 | 6 | 0.022824 | 0.081029 | 0.93914 | 1637 | 5 | -1.7595 | 0.97166 | 0.97337 | 1 | 20911 | 1 | -1.7595 |
| SLC26A3 | 5 | 0.042762 | 0.12273 | 0.951052 | 2665 | 4 | -1.0896 | 0.97167 | 0.97172 | 1 | 20912 | 0 | -1.0896 |
| ARFGAP2 | 6 | 0.016864 | 0.061844 | 0.937033 | 1296 | 4 | -2.7884 | 0.97167 | 0.97338 | 1 | 20913 | 1 | -2.7884 |
| PTN | 4 | 0.3232 | 0.4667 | 0.999994 | 10579 | 2 | -0.71946 | 0.97172 | 0.97162 | 1 | 20914 | 0 | -0.71946 |
| SMCHD1 | 6 | 0.12148 | 0.28854 | 0.999632 | 5684 | 4 | -0.73726 | 0.97173 | 0.97342 | 1 | 20915 | 1 | -0.73726 |
| ALOXE3 | 6 | 0.11619 | 0.27858 | 0.999632 | 5519 | 3 | -1.0555 | 0.97174 | 0.97343 | 1 | 20916 | 1 | -1.0555 |
| NonTargetingControlG | 2 | 0.012735 | 0.024047 | 0.834563 | 1048 | 2 | -6.6643 | 0.97177 | 0.97173 | 1 | 20917 | 0 | -6.6643 |
| OR8K1 | 6 | 0.30993 | 0.51828 | 0.999994 | 10324 | 3 | -0.48772 | 0.97181 | 0.97348 | 1 | 20918 | 1 | -0.48772 |
| SLC6A16 | 5 | 0.052684 | 0.14685 | 0.964852 | 3099 | 4 | -0.9626 | 0.97191 | 0.97195 | 1 | 20919 | 0 | -0.9626 |
| SPANXN3 | 6 | 0.071959 | 0.19086 | 0.972248 | 3916 | 3 | -0.93947 | 0.97192 | 0.97354 | 1 | 20920 | 1 | -0.93947 |
| TMC2 | 6 | 0.089187 | 0.22588 | 0.983354 | 4561 | 4 | -2.0733 | 0.97193 | 0.97355 | 1 | 20921 | 1 | -2.0733 |
| SYT10 | 5 | 0.081104 | 0.19639 | 0.976541 | 4272 | 3 | -1.6809 | 0.97204 | 0.97208 | 1 | 20922 | 0 | -1.6809 |
| CES5A | 5 | 0.12189 | 0.26021 | 0.994943 | 5699 | 3 | -1.4637 | 0.9721 | 0.97214 | 1 | 20923 | 0 | -1.4637 |
| MBD4 | 6 | 0.013162 | 0.049658 | 0.907155 | 1074 | 3 | -3.518 | 0.97222 | 0.97374 | 1 | 20924 | 1 | -3.518 |
| FGFR4 | 5 | 0.036028 | 0.10585 | 0.950444 | 2318 | 3 | -0.72119 | 0.97228 | 0.97235 | 1 | 20925 | 0 | -0.72119 |
| TRAIP | 4 | 0.092064 | 0.20079 | 0.977164 | 4683 | 3 | -1.5653 | 0.97242 | 0.9723 | 1 | 20926 | 0 | -1.5653 |
| ADAMTSL5 | 6 | 0.015937 | 0.058839 | 0.931872 | 1241 | 4 | -2.863 | 0.97259 | 0.974 | 1 | 20927 | 1 | -2.863 |
| PLEKHS1 | 4 | 0.093821 | 0.20399 | 0.978186 | 4745 | 3 | -1.7092 | 0.97259 | 0.97247 | 1 | 20928 | 0 | -1.7092 |
| PPP1R12A | 6 | 0.002176 | 0.010623 | 0.706752 | 286 | 4 | -5.2173 | 0.97261 | 0.97402 | 1 | 20929 | 1 | -5.2173 |
| hsa-mir-8057 | 2 | 0.027342 | 0.050244 | 0.909101 | 1874 | 2 | -4.9674 | 0.97266 | 0.97265 | 1 | 20930 | 0 | -4.9674 |
| VWA3B | 6 | 0.086963 | 0.22141 | 0.983093 | 4477 | 4 | -1.0038 | 0.97269 | 0.97407 | 1 | 20931 | 1 | -1.0038 |
| SLC25A33 | 6 | 0.22321 | 0.416 | 0.999994 | 8362 | 3 | -0.85087 | 0.9727 | 0.97407 | 1 | 20932 | 1 | -0.85087 |
| POLE | 2 | 0.027301 | 0.050172 | 0.909101 | 1872 | 2 | -2.9135 | 0.9727 | 0.97269 | 1 | 20933 | 0 | -2.9135 |
| NonTargetingControlG | 2 | 0.02727 | 0.050119 | 0.909101 | 1870 | 2 | -5.574 | 0.97273 | 0.97272 | 1 | 20934 | 0 | -5.574 |
| ITSN1 | 6 | 0.029708 | 0.098086 | 0.945253 | 1993 | 3 | -2.0246 | 0.9729 | 0.97421 | 1 | 20935 | 1 | -2.0246 |
| WFDC1 | 5 | 0.14622 | 0.29709 | 0.999994 | 6360 | 3 | -1.062 | 0.9729 | 0.97295 | 1 | 20936 | 0 | -1.062 |
| GPR158 | 5 | 0.12062 | 0.2583 | 0.994943 | 5648 | 3 | -1.1308 | 0.97292 | 0.97297 | 1 | 20937 | 0 | -1.1308 |
| KIF2B | 5 | 0.39567 | 0.55995 | 0.999994 | 11886 | 2 | -0.31337 | 0.97297 | 0.97302 | 1 | 20938 | 0 | -0.31337 |
| GATA5 | 4 | 0.13978 | 0.27868 | 0.999632 | 6179 | 2 | -2.0938 | 0.97298 | 0.97286 | 1 | 20939 | 0 | -2.0938 |
| NDUFA13 | 4 | 0.099796 | 0.21475 | 0.979587 | 4970 | 3 | -1.5569 | 0.97298 | 0.97286 | 1 | 20940 | 0 | -1.5569 |
| TEN1 | 4 | 0.013421 | 0.044288 | 0.891621 | 1095 | 3 | -3.9887 | 0.97298 | 0.97286 | 1 | 20941 | 0 | -3.9887 |
| FGFR1OP2 | 6 | 0.52489 | 0.68593 | 0.999994 | 13814 | 2 | 0.1944 | 0.97306 | 0.97432 | 1 | 20942 | 1 | 0.1944 |
| GIGYF1 | 5 | 0.018374 | 0.059663 | 0.932932 | 1382 | 4 | -1.5993 | 0.97307 | 0.97313 | 1 | 20943 | 0 | -1.5993 |
| LIMS3 | 6 | 0.043775 | 0.1304 | 0.95318 | 2703 | 4 | -1.4806 | 0.97321 | 0.97442 | 1 | 20944 | 1 | -1.4806 |
| hsa-mir-1260b | 4 | 0.16148 | 0.301 | 0.999994 | 6774 | 2 | -1.3498 | 0.97326 | 0.97315 | 1 | 20945 | 0 | -1.3498 |
| SPIN2A | 6 | 0.10991 | 0.26652 | 0.998525 | 5300 | 4 | -1.0586 | 0.9734 | 0.97457 | 1 | 20946 | 1 | -1.0586 |
| FUNDC1 | 6 | 0.078157 | 0.20362 | 0.978186 | 4155 | 2 | -0.051528 | 0.97352 | 0.97465 | 1 | 20947 | 1 | -0.051528 |
| C9orf172 | 5 | 0.030927 | 0.092876 | 0.943968 | 2058 | 4 | -2.2372 | 0.97356 | 0.9736 | 1 | 20948 | 0 | -2.2372 |
| EGFL6 | 6 | 0.079038 | 0.20539 | 0.978342 | 4191 | 4 | -1.0802 | 0.97367 | 0.97476 | 1 | 20949 | 1 | -1.0802 |
| RUNX1 | 4 | 0.32527 | 0.46887 | 0.999994 | 10628 | 2 | -0.86053 | 0.97384 | 0.97371 | 1 | 20950 | 0 | -0.86053 |
| SUPT5H | 3 | 0.026147 | 0.066202 | 0.93914 | 1811 | 3 | -4.7622 | 0.97385 | 0.97393 | 1 | 20951 | 0 | -4.7622 |
| TAF10 | 4 | 0.029432 | 0.080363 | 0.93914 | 1979 | 3 | -3.6824 | 0.97388 | 0.97376 | 1 | 20952 | 0 | -3.6824 |
| TADA1 | 6 | 0.098229 | 0.24387 | 0.988214 | 4904 | 4 | -1.181 | 0.97392 | 0.97494 | 1 | 20953 | 1 | -1.181 |
| hsa-mir-654 | 4 | 0.11329 | 0.23867 | 0.987253 | 5417 | 3 | -1.6677 | 0.97395 | 0.97381 | 1 | 20954 | 0 | -1.6677 |
| AGO1 | 4 | 0.014734 | 0.048303 | 0.90085 | 1174 | 3 | -4.5865 | 0.97403 | 0.97389 | 1 | 20955 | 0 | -4.5865 |

| | | | | | | | | | | | | | |
|---|---|---|---|---|---|---|---|---|---|---|---|---|---|
| hsa-mir-9-1 | 3 | 0.025911 | 0.065645 | 0.93914 | 1799 | 3 | -1.1483 | 0.97409 | 0.97416 | 1 | 20956 | 0 | -1.1483 |
| FZD9 | 5 | 0.0052336 | 0.021726 | 0.819701 | 535 | 4 | -6.8417 | 0.97411 | 0.97415 | 1 | 20957 | 0 | -6.8417 |
| hsa-mir-644a | 3 | 0.025886 | 0.065592 | 0.93914 | 1797 | 3 | -1.9183 | 0.97411 | 0.97418 | 1 | 20958 | 0 | -1.9183 |
| PLEKHJ1 | 5 | 0.030421 | 0.09155 | 0.94395 | 2028 | 3 | -2.6383 | 0.97414 | 0.97418 | 1 | 20959 | 0 | -2.6383 |
| MTFR1L | 6 | 0.0098217 | 0.03841 | 0.889591 | 862 | 4 | -2.8122 | 0.97417 | 0.97512 | 1 | 20960 | 1 | -2.8122 |
| WDR31 | 6 | 0.20909 | 0.39899 | 0.999994 | 8005 | 1 | -0.33812 | 0.9743 | 0.97522 | 1 | 20961 | 1 | -0.33812 |
| HAAO | 4 | 0.10522 | 0.22446 | 0.983354 | 5140 | 3 | -1.4452 | 0.97432 | 0.97418 | 1 | 20962 | 0 | -1.4452 |
| SPANXN1 | 6 | 0.4408 | 0.62896 | 0.999994 | 12589 | 2 | -0.32933 | 0.97433 | 0.97525 | 1 | 20963 | 1 | -0.32933 |
| GORASP2 | 5 | 0.1684 | 0.32978 | 0.999994 | 6946 | 3 | -1.0571 | 0.97435 | 0.97439 | 1 | 20964 | 0 | -1.0571 |
| HAPLN2 | 6 | 0.058272 | 0.16206 | 0.967126 | 3351 | 3 | -1.5036 | 0.97437 | 0.97527 | 1 | 20965 | 1 | -1.5036 |
| ANKRD24 | 4 | 0.0055638 | 0.019506 | 0.808847 | 561 | 3 | -4.8473 | 0.97446 | 0.97434 | 1 | 20966 | 0 | -4.8473 |
| FAM71F1 | 6 | 0.0011792 | 0.0059422 | 0.623123 | 185 | 5 | -5.9255 | 0.97448 | 0.97536 | 1 | 20967 | 1 | -5.9255 |
| IL7 | 6 | 0.11008 | 0.26684 | 0.998555 | 5308 | 4 | -0.87102 | 0.97454 | 0.9754 | 1 | 20968 | 1 | -0.87102 |
| GRTP1 | 5 | 0.018506 | 0.060026 | 0.932932 | 1386 | 4 | -1.3712 | 0.97456 | 0.97462 | 1 | 20969 | 0 | -1.3712 |
| C2orf61 | 4 | 0.15834 | 0.2978 | 0.999994 | 6681 | 2 | -0.72725 | 0.97457 | 0.97446 | 1 | 20970 | 0 | -0.72725 |
| C10orf95 | 6 | 0.030563 | 0.1001 | 0.946394 | 2033 | 4 | -1.7483 | 0.97458 | 0.97543 | 1 | 20971 | 1 | -1.7483 |
| NonTargetingControlG | 2 | 0.025336 | 0.046696 | 0.896157 | 1768 | 2 | -5.7112 | 0.97466 | 0.97466 | 1 | 20972 | 0 | -5.7112 |
| APOBEC1 | 5 | 0.21382 | 0.39472 | 0.999994 | 8108 | 2 | -0.44007 | 0.97472 | 0.97478 | 1 | 20973 | 0 | -0.44007 |
| NUPL2 | 6 | 0.32232 | 0.53257 | 0.999994 | 10563 | 3 | -0.49585 | 0.97482 | 0.97562 | 1 | 20974 | 1 | -0.49585 |
| ITPK1 | 6 | 0.17396 | 0.35559 | 0.999994 | 7083 | 3 | -0.68264 | 0.97488 | 0.97567 | 1 | 20975 | 1 | -0.68264 |
| SLC27A1 | 5 | 0.090871 | 0.21199 | 0.979587 | 4633 | 3 | -1.844 | 0.97505 | 0.97512 | 1 | 20976 | 0 | -1.844 |
| CCL26 | 6 | 0.037274 | 0.11577 | 0.951052 | 2387 | 4 | -1.8552 | 0.97506 | 0.97582 | 1 | 20977 | 1 | -1.8552 |
| BEND7 | 6 | 0.077981 | 0.20327 | 0.97798 | 4150 | 4 | -1.3227 | 0.97506 | 0.97582 | 1 | 20978 | 1 | -1.3227 |
| FBXO28 | 5 | 0.056711 | 0.1565 | 0.967126 | 3281 | 4 | -0.81606 | 0.97507 | 0.97514 | 1 | 20979 | 0 | -0.81606 |
| ISM2 | 5 | 0.21803 | 0.40054 | 0.999994 | 8229 | 3 | -0.74383 | 0.97512 | 0.97519 | 1 | 20980 | 0 | -0.74383 |
| ARHGAP30 | 5 | 0.82649 | 0.83699 | 0.999994 | 17875 | 1 | 0.057042 | 0.97512 | 0.97519 | 1 | 20981 | 0 | 0.057042 |
| NonTargetingControlG | 2 | 0.024797 | 0.045738 | 0.892434 | 1745 | 2 | -2.949 | 0.9752 | 0.9752 | 1 | 20982 | 0 | -2.949 |
| CYB5R1 | 5 | 0.29539 | 0.47731 | 0.999994 | 10023 | 2 | -0.53187 | 0.9752 | 0.97527 | 1 | 20983 | 0 | -0.53187 |
| ZSWIM2 | 6 | 0.11397 | 0.2743 | 0.998561 | 5446 | 4 | -1.1782 | 0.97528 | 0.97599 | 1 | 20984 | 1 | -1.1782 |
| CACNA2D2 | 6 | 0.038284 | 0.11805 | 0.951052 | 2429 | 4 | -1.6347 | 0.97535 | 0.97605 | 1 | 20985 | 1 | -1.6347 |
| CD244 | 6 | 0.019227 | 0.069491 | 0.93914 | 1420 | 5 | -1.1004 | 0.97551 | 0.97617 | 1 | 20986 | 1 | -1.1004 |
| CAV2 | 5 | 0.023901 | 0.074566 | 0.93914 | 1705 | 4 | -3.0721 | 0.9756 | 0.97566 | 1 | 20987 | 0 | -3.0721 |
| OXGR1 | 4 | 0.80933 | 0.80967 | 0.999994 | 17621 | 1 | -0.41111 | 0.9757 | 0.97557 | 1 | 20988 | 0 | -0.41111 |
| GNAL | 5 | 0.062662 | 0.16645 | 0.967126 | 3542 | 3 | -2.202 | 0.97576 | 0.97582 | 1 | 20989 | 0 | -2.202 |
| TFF1 | 6 | 0.0019535 | 0.0097471 | 0.680529 | 268 | 4 | -4.5591 | 0.97577 | 0.97638 | 1 | 20990 | 1 | -4.5591 |
| SRSF7 | 3 | 0.011292 | 0.030067 | 0.862223 | 943 | 3 | -7.6877 | 0.97583 | 0.97592 | 1 | 20991 | 0 | -7.6877 |
| TMCC2 | 6 | 0.064048 | 0.17433 | 0.968171 | 3613 | 3 | -0.21949 | 0.97583 | 0.97642 | 1 | 20992 | 1 | -0.21949 |
| MMP13 | 6 | 0.24441 | 0.44125 | 0.999994 | 8889 | 3 | -0.47632 | 0.97584 | 0.97644 | 1 | 20993 | 1 | -0.47632 |
| ATF3 | 5 | 0.10781 | 0.23859 | 0.987253 | 5233 | 3 | -1.3864 | 0.97591 | 0.97598 | 1 | 20994 | 0 | -1.3864 |
| TMEM50B | 5 | 0.1082 | 0.2392 | 0.987253 | 5244 | 3 | -1.5293 | 0.97595 | 0.97601 | 1 | 20995 | 0 | -1.5293 |
| CCZ1 | 5 | 0.047194 | 0.13363 | 0.95318 | 2863 | 4 | -0.98002 | 0.97595 | 0.97602 | 1 | 20996 | 0 | -0.98002 |
| TBX20 | 6 | 0.079671 | 0.20663 | 0.978744 | 4218 | 3 | -0.87269 | 0.97601 | 0.97658 | 1 | 20997 | 1 | -0.87269 |
| MOV10 | 6 | 0.048001 | 0.1397 | 0.958716 | 2890 | 2 | -0.52497 | 0.97605 | 0.97661 | 1 | 20998 | 1 | -0.52497 |
| hsa-mir-513c | 4 | 0.14718 | 0.28626 | 0.999632 | 6381 | 2 | -1.4437 | 0.97605 | 0.97594 | 1 | 20999 | 0 | -1.4437 |
| CLDN15 | 6 | 0.014169 | 0.05302 | 0.916684 | 1139 | 3 | -2.4618 | 0.97608 | 0.97663 | 1 | 21000 | 1 | -2.4618 |
| MT1G | 6 | 0.07655 | 0.20037 | 0.977164 | 4094 | 4 | -1.5455 | 0.97609 | 0.97664 | 1 | 21001 | 1 | -1.5455 |
| BMPER | 6 | 0.27537 | 0.47802 | 0.999994 | 9598 | 3 | -0.53169 | 0.9761 | 0.97664 | 1 | 21002 | 1 | -0.53169 |
| HMP19 | 6 | 0.034476 | 0.10927 | 0.951052 | 2239 | 4 | -2.3754 | 0.97628 | 0.97678 | 1 | 21003 | 1 | -2.3754 |
| hsa-mir-7150 | 4 | 0.66967 | 0.70517 | 0.999994 | 15925 | 1 | -0.46027 | 0.97635 | 0.97623 | 1 | 21004 | 0 | -0.46027 |
| hsa-mir-1199 | 2 | 0.023502 | 0.043459 | 0.891621 | 1686 | 2 | -3.1335 | 0.9765 | 0.97648 | 1 | 21005 | 0 | -3.1335 |
| BHLHA9 | 6 | 0.032803 | 0.10536 | 0.95013 | 2149 | 4 | -2.495 | 0.97654 | 0.97699 | 1 | 21006 | 1 | -2.495 |
| OAZ1 | 5 | 0.0038487 | 0.016218 | 0.772095 | 432 | 4 | -4.9239 | 0.97664 | 0.9767 | 1 | 21007 | 0 | -4.9239 |

| | | | | | | | | | | | | | |
|---|---|---|---|---|---|---|---|---|---|---|---|---|---|
| SPATA31C2 | 5 | 0.21947 | 0.40255 | 0.999994 | 8277 | 1 | 0.18077 | 0.97664 | 0.9767 | 1 | 21008 | 0 | 0.18077 |
| hsa-mir-326 | 3 | 0.023361 | 0.059625 | 0.932932 | 1672 | 3 | -1.453 | 0.97664 | 0.97674 | 1 | 21009 | 0 | -1.453 |
| THSD7B | 6 | 0.27513 | 0.47774 | 0.999994 | 9595 | 3 | -0.62665 | 0.97669 | 0.97712 | 1 | 21010 | 1 | -0.62665 |
| hsa-mir-6823 | 3 | 0.023267 | 0.059416 | 0.932932 | 1668 | 3 | -1.9769 | 0.97673 | 0.97684 | 1 | 21011 | 0 | -1.9769 |
| NFU1 | 6 | 0.18771 | 0.37269 | 0.999994 | 7413 | 2 | 0.031695 | 0.97674 | 0.97716 | 1 | 21012 | 1 | 0.031695 |
| TSPAN13 | 6 | 0.23217 | 0.42669 | 0.999994 | 8573 | 3 | -0.56702 | 0.97683 | 0.97724 | 1 | 21013 | 1 | -0.56702 |
| HUS1B | 6 | 0.2992 | 0.50576 | 0.999994 | 10117 | 3 | -0.59054 | 0.97683 | 0.97724 | 1 | 21014 | 1 | -0.59054 |
| PCDHB12 | 6 | 0.085251 | 0.218 | 0.980699 | 4420 | 3 | -1.2474 | 0.9769 | 0.97729 | 1 | 21015 | 1 | -1.2474 |
| UBE3C | 6 | 0.00054514 | 0.0028546 | 0.519469 | 114 | 5 | -4.0257 | 0.97695 | 0.97732 | 1 | 21016 | 1 | -4.0257 |
| IL1R2 | 5 | 0.2529 | 0.44308 | 0.999994 | 9082 | 2 | -0.53182 | 0.97695 | 0.97703 | 1 | 21017 | 0 | -0.53182 |
| hsa-mir-4678 | 4 | 0.072719 | 0.16507 | 0.967126 | 3949 | 3 | -1.3329 | 0.97695 | 0.97689 | 1 | 21018 | 0 | -1.3329 |
| RBBP4 | 6 | 0.0064841 | 0.026594 | 0.848412 | 625 | 4 | -3.0082 | 0.977 | 0.97737 | 1 | 21019 | 1 | -3.0082 |
| FAM8A1 | 6 | 0.24271 | 0.43923 | 0.999994 | 8856 | 3 | -0.60583 | 0.97703 | 0.97739 | 1 | 21020 | 1 | -0.60583 |
| SNX17 | 5 | 0.23794 | 0.42812 | 0.999994 | 8726 | 3 | -0.63164 | 0.97705 | 0.97713 | 1 | 21021 | 0 | -0.63164 |
| LRFN5 | 5 | 0.28854 | 0.47173 | 0.999994 | 9883 | 2 | -0.24955 | 0.97709 | 0.97717 | 1 | 21022 | 0 | -0.24955 |
| KNG1 | 5 | 0.4758 | 0.62699 | 0.999994 | 13074 | 2 | -0.56361 | 0.97713 | 0.97721 | 1 | 21023 | 0 | -0.56361 |
| IFNGR2 | 6 | 0.017549 | 0.064016 | 0.93914 | 1343 | 4 | -0.90391 | 0.97722 | 0.97755 | 1 | 21024 | 1 | -0.90391 |
| INO80B | 5 | 0.013654 | 0.046401 | 0.893965 | 1109 | 4 | -3.9698 | 0.97723 | 0.9773 | 1 | 21025 | 0 | -3.9698 |
| SPNS3 | 4 | 0.06817 | 0.15638 | 0.967126 | 3756 | 3 | -1.4404 | 0.97737 | 0.97732 | 1 | 21026 | 0 | -1.4404 |
| hsa-let-7g | 3 | 0.022629 | 0.057878 | 0.929685 | 1625 | 3 | -2.5572 | 0.97737 | 0.97745 | 1 | 21027 | 0 | -2.5572 |
| PDE12 | 4 | 0.050518 | 0.1225 | 0.951052 | 2997 | 3 | -1.7274 | 0.97741 | 0.97736 | 1 | 21028 | 0 | -1.7274 |
| STPG1 | 6 | 0.26913 | 0.47071 | 0.999994 | 9446 | 3 | -0.62092 | 0.97743 | 0.97774 | 1 | 21029 | 1 | -0.62092 |
| NonTargetingControlG | 2 | 0.022558 | 0.041799 | 0.891621 | 1621 | 2 | -2.9077 | 0.97744 | 0.97742 | 1 | 21030 | 0 | -2.9077 |
| DAGLB | 6 | 0.015987 | 0.058992 | 0.931872 | 1245 | 3 | -0.88466 | 0.9775 | 0.97779 | 1 | 21031 | 1 | -0.88466 |
| GNAI2 | 6 | 0.00026758 | 0.0014373 | 0.384267 | 74 | 5 | -2.9164 | 0.97753 | 0.97782 | 1 | 21032 | 1 | -2.9164 |
| YAE1D1 | 4 | 0.11803 | 0.2468 | 0.98999 | 5573 | 3 | -1.2922 | 0.97754 | 0.9775 | 1 | 21033 | 0 | -1.2922 |
| NonTargetingControlG | 1 | 0.022426 | 0.022385 | 0.819701 | 1610 | 1 | -7.4727 | 0.97757 | 0.9776 | 1 | 21034 | 0 | -7.4727 |
| SRSF3 | 2 | 0.022411 | 0.041528 | 0.891621 | 1609 | 2 | -4.7875 | 0.97759 | 0.97757 | 1 | 21035 | 0 | -4.7875 |
| DDN | 6 | 0.12892 | 0.29891 | 0.999994 | 5914 | 3 | -1.0078 | 0.97759 | 0.97787 | 1 | 21036 | 1 | -1.0078 |
| CFHR4 | 6 | 0.23436 | 0.42928 | 0.999994 | 8626 | 2 | -0.21051 | 0.97778 | 0.97803 | 1 | 21037 | 1 | -0.21051 |
| SNX10 | 6 | 0.035367 | 0.1114 | 0.951052 | 2285 | 4 | -1.7141 | 0.97788 | 0.97811 | 1 | 21038 | 1 | -1.7141 |
| CASP14 | 5 | 0.32773 | 0.50367 | 0.999994 | 10678 | 1 | -0.062267 | 0.9779 | 0.97797 | 1 | 21039 | 0 | -0.062267 |
| CREBL2 | 6 | 0.59046 | 0.73294 | 0.999994 | 14830 | 2 | 0.28655 | 0.97792 | 0.97815 | 1 | 21040 | 1 | 0.28655 |
| SIRT4 | 6 | 0.0077005 | 0.030911 | 0.862223 | 707 | 5 | -2.4399 | 0.9781 | 0.97831 | 1 | 21041 | 1 | -2.4399 |
| FKTN | 6 | 0.13643 | 0.30854 | 0.999994 | 6095 | 2 | -0.51211 | 0.97818 | 0.97838 | 1 | 21042 | 1 | -0.51211 |
| SLC2A11 | 5 | 0.022864 | 0.071809 | 0.93914 | 1641 | 4 | -1.2306 | 0.9782 | 0.97828 | 1 | 21043 | 0 | -1.2306 |
| TMEM42 | 5 | 0.69071 | 0.76231 | 0.999994 | 16216 | 1 | -0.14171 | 0.97834 | 0.97842 | 1 | 21044 | 0 | -0.14171 |
| CTIF | 6 | 0.051675 | 0.14775 | 0.965554 | 3053 | 4 | -1.735 | 0.97838 | 0.97856 | 1 | 21045 | 1 | -1.735 |
| NDUFA1 | 5 | 0.5463 | 0.68845 | 0.999994 | 14155 | 2 | -0.12578 | 0.97841 | 0.97848 | 1 | 21046 | 0 | -0.12578 |
| CALM2 | 5 | 0.10342 | 0.23173 | 0.984479 | 5081 | 3 | -1.0232 | 0.97852 | 0.9786 | 1 | 21047 | 0 | -1.0232 |
| NonTargetingControlG | 2 | 0.019861 | 0.036927 | 0.881473 | 1451 | 2 | -6.4917 | 0.97853 | 0.97851 | 1 | 21048 | 0 | -6.4917 |
| BMP2 | 6 | 0.23748 | 0.43298 | 0.999994 | 8715 | 3 | -0.73803 | 0.97858 | 0.97871 | 1 | 21049 | 1 | -0.73803 |
| NonTargetingControlG | 2 | 0.021384 | 0.03966 | 0.891621 | 1540 | 2 | -3.5027 | 0.97862 | 0.9786 | 1 | 21050 | 0 | -3.5027 |
| KISS1 | 5 | 0.030667 | 0.092211 | 0.943968 | 2039 | 4 | -2.0439 | 0.97863 | 0.9787 | 1 | 21051 | 0 | -2.0439 |
| SNAP29 | 6 | 0.10683 | 0.26062 | 0.994943 | 5205 | 4 | -0.85405 | 0.97866 | 0.97878 | 1 | 21052 | 1 | -0.85405 |
| hsa-mir-181d | 4 | 0.023394 | 0.067821 | 0.93914 | 1676 | 3 | -2.7005 | 0.97869 | 0.97867 | 1 | 21053 | 0 | -2.7005 |
| ADTRP | 6 | 0.003927 | 0.017217 | 0.785648 | 437 | 2 | -0.2066 | 0.97871 | 0.97882 | 1 | 21054 | 1 | -0.2066 |
| IARS2 | 3 | 0.021153 | 0.054403 | 0.922677 | 1523 | 3 | -5.0735 | 0.97885 | 0.97893 | 1 | 21055 | 0 | -5.0735 |
| hsa-mir-933 | 4 | 0.047709 | 0.11697 | 0.951052 | 2882 | 3 | -2.6656 | 0.97885 | 0.97882 | 1 | 21056 | 0 | -2.6656 |
| HAS3 | 5 | 0.09796 | 0.22324 | 0.983354 | 4892 | 3 | -1.573 | 0.97886 | 0.97894 | 1 | 21057 | 0 | -1.573 |
| ATP5E | 1 | 0.021139 | 0.02109 | 0.812607 | 1522 | 1 | -9.5251 | 0.97886 | 0.97888 | 1 | 21058 | 0 | -9.5251 |
| PAPOLA | 4 | 0.020778 | 0.062248 | 0.937033 | 1495 | 3 | -2.8842 | 0.97888 | 0.97885 | 1 | 21059 | 0 | -2.8842 |

| | | | | | | | | | | | | | |
|---|---|---|---|---|---|---|---|---|---|---|---|---|---|
| CHFR | 4 | 0.051341 | 0.12413 | 0.951052 | 3040 | 3 | -1.4474 | 0.97893 | 0.97891 | 1 | 21060 | 0 | -1.4474 |
| AP1B1 | 6 | 0.086205 | 0.2199 | 0.980699 | 4460 | 4 | -1.2987 | 0.97895 | 0.97903 | 1 | 21061 | 1 | -1.2987 |
| PSMD7 | 2 | 0.020969 | 0.038942 | 0.891621 | 1513 | 2 | -3.6217 | 0.97903 | 0.97901 | 1 | 21062 | 0 | -3.6217 |
| TRIM67 | 6 | 0.052009 | 0.14851 | 0.965554 | 3068 | 4 | -1.7361 | 0.97904 | 0.9791 | 1 | 21063 | 1 | -1.7361 |
| APOA5 | 6 | 0.29385 | 0.49953 | 0.999994 | 9993 | 3 | -0.51892 | 0.97914 | 0.97919 | 1 | 21064 | 1 | -0.51892 |
| PARP4 | 6 | 0.096453 | 0.24038 | 0.987253 | 4835 | 4 | -0.91895 | 0.97915 | 0.9792 | 1 | 21065 | 1 | -0.91895 |
| ZFX | 5 | 0.21892 | 0.40176 | 0.999994 | 8258 | 2 | -0.18025 | 0.97919 | 0.97929 | 1 | 21066 | 0 | -0.18025 |
| GABRR2 | 6 | 0.11271 | 0.27192 | 0.998561 | 5393 | 3 | -0.61124 | 0.9792 | 0.97924 | 1 | 21067 | 1 | -0.61124 |
| NUPR1L | 5 | 0.12329 | 0.26234 | 0.995875 | 5747 | 2 | -0.097947 | 0.97922 | 0.97932 | 1 | 21068 | 0 | -0.097947 |
| EZH2 | 5 | 0.0066716 | 0.026433 | 0.848412 | 640 | 4 | -2.9823 | 0.97923 | 0.97933 | 1 | 21069 | 0 | -2.9823 |
| TAF12 | 5 | 0.036074 | 0.10597 | 0.950444 | 2324 | 4 | -1.0036 | 0.97926 | 0.97935 | 1 | 21070 | 0 | -1.0036 |
| ZNF354A | 6 | 0.052031 | 0.14856 | 0.965554 | 3070 | 4 | -1.5336 | 0.97926 | 0.9793 | 1 | 21071 | 1 | -1.5336 |
| UCN | 6 | 0.06115 | 0.16823 | 0.967126 | 3475 | 4 | -1.3278 | 0.9795 | 0.97953 | 1 | 21072 | 1 | -1.3278 |
| ATRN | 6 | 0.088121 | 0.22377 | 0.983354 | 4520 | 3 | -1.3232 | 0.97957 | 0.97959 | 1 | 21073 | 1 | -1.3232 |
| GPR123 | 6 | 0.016369 | 0.060213 | 0.932932 | 1267 | 5 | -1.52 | 0.97958 | 0.9796 | 1 | 21074 | 1 | -1.52 |
| ACSL4 | 5 | 0.42065 | 0.58061 | 0.999994 | 12259 | 2 | -0.30797 | 0.97964 | 0.97975 | 1 | 21075 | 0 | -0.30797 |
| MIDN | 4 | 0.0022042 | 0.0079873 | 0.680529 | 290 | 3 | -6.8157 | 0.97971 | 0.97969 | 1 | 21076 | 0 | -6.8157 |
| TRIM25 | 5 | 0.014226 | 0.048012 | 0.900643 | 1145 | 3 | -0.77654 | 0.97973 | 0.97983 | 1 | 21077 | 0 | -0.77654 |
| hsa-mir-892c | 4 | 0.10489 | 0.22387 | 0.983354 | 5128 | 3 | -1.8784 | 0.97973 | 0.97973 | 1 | 21078 | 0 | -1.8784 |
| ABCA13 | 5 | 0.35451 | 0.52557 | 0.999994 | 11239 | 2 | -0.20889 | 0.97975 | 0.97986 | 1 | 21079 | 0 | -0.20889 |
| GABRG3 | 6 | 0.91703 | 0.91768 | 0.999994 | 19562 | 1 | 0.0016039 | 0.97979 | 0.97979 | 1 | 21080 | 1 | 0.0016039 |
| TRIM40 | 5 | 0.4093 | 0.57131 | 0.999994 | 12098 | 2 | -0.34502 | 0.97981 | 0.97991 | 1 | 21081 | 0 | -0.34502 |
| GLIPR2 | 4 | 0.03921 | 0.10019 | 0.946394 | 2477 | 3 | -3.7007 | 0.97982 | 0.97982 | 1 | 21082 | 0 | -3.7007 |
| GNL1 | 5 | 0.0336 | 0.099736 | 0.946394 | 2195 | 4 | -1.0372 | 0.97983 | 0.97994 | 1 | 21083 | 0 | -1.0372 |
| BTF3 | 5 | 0.012803 | 0.044034 | 0.891621 | 1052 | 3 | -1.2059 | 0.97987 | 0.97998 | 1 | 21084 | 0 | -1.2059 |
| ITPRIPL1 | 5 | 0.0038026 | 0.016027 | 0.771254 | 429 | 4 | -3.4242 | 0.9799 | 0.98001 | 1 | 21085 | 0 | -3.4242 |
| C1orf94 | 4 | 0.033823 | 0.089313 | 0.94395 | 2206 | 3 | -1.9223 | 0.98004 | 0.98003 | 1 | 21086 | 0 | -1.9223 |
| TNFAIP6 | 6 | 0.11143 | 0.26945 | 0.998561 | 5347 | 4 | -0.85156 | 0.98006 | 0.98003 | 1 | 21087 | 1 | -0.85156 |
| PHOSPHO2 | 5 | 0.051094 | 0.14303 | 0.95983 | 3022 | 4 | -0.72571 | 0.98009 | 0.98019 | 1 | 21088 | 0 | -0.72571 |
| KLRB1 | 5 | 0.47621 | 0.62735 | 0.999994 | 13085 | 2 | -0.45308 | 0.98021 | 0.98031 | 1 | 21089 | 0 | -0.45308 |
| PDP2 | 5 | 0.20778 | 0.38625 | 0.999994 | 7968 | 3 | -0.66437 | 0.98027 | 0.98037 | 1 | 21090 | 0 | -0.66437 |
| HEYL | 5 | 0.048149 | 0.13598 | 0.95468 | 2894 | 4 | -1.5975 | 0.98028 | 0.98038 | 1 | 21091 | 0 | -1.5975 |
| SI | 3 | 0.019673 | 0.050745 | 0.909101 | 1436 | 3 | -1.7107 | 0.98033 | 0.9804 | 1 | 21092 | 0 | -1.7107 |
| CD80 | 5 | 0.15911 | 0.31624 | 0.999994 | 6705 | 3 | -1.1189 | 0.98035 | 0.98044 | 1 | 21093 | 0 | -1.1189 |
| TNFSF14 | 4 | 0.40662 | 0.55248 | 0.999994 | 12054 | 2 | -0.67509 | 0.9804 | 0.98038 | 1 | 21094 | 0 | -0.67509 |
| NLRP1 | 4 | 0.064904 | 0.15017 | 0.966359 | 3648 | 3 | -0.99634 | 0.9805 | 0.98047 | 1 | 21095 | 0 | -0.99634 |
| UGT2A1 | 5 | 0.19704 | 0.37103 | 0.999994 | 7674 | 3 | -0.84288 | 0.98052 | 0.98062 | 1 | 21096 | 0 | -0.84288 |
| UNK | 6 | 0.29896 | 0.50549 | 0.999994 | 10112 | 2 | -0.405 | 0.98054 | 0.98049 | 1 | 21097 | 1 | -0.405 |
| CCDC138 | 6 | 0.019587 | 0.070656 | 0.93914 | 1433 | 5 | -1.4063 | 0.98058 | 0.98054 | 1 | 21098 | 1 | -1.4063 |
| ORAI3 | 5 | 0.056156 | 0.15519 | 0.967126 | 3254 | 3 | -2.0957 | 0.98079 | 0.98089 | 1 | 21099 | 0 | -2.0957 |
| ZNF510 | 5 | 0.41595 | 0.57675 | 0.999994 | 12192 | 2 | -0.35643 | 0.98083 | 0.98093 | 1 | 21100 | 0 | -0.35643 |
| CHAC1 | 5 | 0.041387 | 0.11937 | 0.951052 | 2583 | 4 | -1.0022 | 0.98084 | 0.98094 | 1 | 21101 | 0 | -1.0022 |
| PKP1 | 6 | 0.0066953 | 0.027343 | 0.853176 | 642 | 4 | -3.0008 | 0.98089 | 0.98083 | 1 | 21102 | 1 | -3.0008 |
| LIMS2 | 5 | 0.025807 | 0.079605 | 0.93914 | 1791 | 4 | -2.8812 | 0.9809 | 0.98101 | 1 | 21103 | 0 | -2.8812 |
| AGXT2 | 6 | 0.010769 | 0.041621 | 0.891621 | 917 | 4 | -2.9961 | 0.98091 | 0.98085 | 1 | 21104 | 1 | -2.9961 |
| EMC8 | 6 | 0.50247 | 0.67044 | 0.999994 | 13469 | 1 | 0.083987 | 0.98092 | 0.98085 | 1 | 21105 | 1 | 0.083987 |
| NOP2 | 6 | 0.016976 | 0.062188 | 0.937033 | 1301 | 4 | -2.74 | 0.98095 | 0.98089 | 1 | 21106 | 1 | -2.74 |
| CHD1 | 6 | 0.0091015 | 0.035881 | 0.875006 | 801 | 2 | -0.094074 | 0.98097 | 0.9809 | 1 | 21107 | 1 | -0.094074 |
| DYNLL1 | 2 | 0.0022898 | 0.0044402 | 0.56977 | 304 | 2 | -7.6137 | 0.98109 | 0.9811 | 1 | 21108 | 0 | -7.6137 |
| RBAK | 6 | 0.14987 | 0.32545 | 0.999994 | 6456 | 3 | -0.94898 | 0.9811 | 0.98102 | 1 | 21109 | 1 | -0.94898 |
| NPM3 | 5 | 0.17391 | 0.33781 | 0.999994 | 7082 | 3 | -0.90842 | 0.98112 | 0.98123 | 1 | 21110 | 0 | -0.90842 |
| AIP | 6 | 0.12623 | 0.29553 | 0.999994 | 5831 | 2 | -0.25934 | 0.98127 | 0.98118 | 1 | 21111 | 1 | -0.25934 |

| | | | | | | | | | | | | |
|---|---|---|---|---|---|---|---|---|---|---|---|---|
| URB2 | 6 | 0.60507 | 0.74374 | 0.999994 | 15011 | 2 | 0.071125 | 0.98133 | 0.98124 | 1 | 21112 | 1 | 0.071125 |
| PIGF | 6 | 0.11204 | 0.27061 | 0.998561 | 5365 | 3 | -1.1594 | 0.98134 | 0.98125 | 1 | 21113 | 1 | -1.1594 |
| hsa-mir-1204 | 3 | 0.013187 | 0.034797 | 0.867348 | 1076 | 3 | -7.4922 | 0.98134 | 0.98143 | 1 | 21114 | 0 | -7.4922 |
| AGPAT5 | 6 | 0.65969 | 0.78509 | 0.999994 | 15773 | 2 | -0.28659 | 0.98136 | 0.98127 | 1 | 21115 | 1 | -0.28659 |
| OR8G2 | 6 | 0.019763 | 0.071232 | 0.93914 | 1444 | 4 | -2.7223 | 0.9814 | 0.98132 | 1 | 21116 | 1 | -2.7223 |
| KAT2A | 5 | 0.0046505 | 0.019417 | 0.808616 | 492 | 4 | -3.2775 | 0.98144 | 0.98157 | 1 | 21117 | 0 | -3.2775 |
| RBAK-LOC389458 | 2 | 0.018556 | 0.034551 | 0.867348 | 1389 | 2 | -5.0853 | 0.98144 | 0.98147 | 1 | 21118 | 0 | -5.0853 |
| ATXN1 | 4 | 0.12131 | 0.25259 | 0.994481 | 5675 | 3 | -0.90448 | 0.98145 | 0.98142 | 1 | 21119 | 0 | -0.90448 |
| TULP4 | 5 | 0.057929 | 0.15865 | 0.967126 | 3334 | 3 | -2.1038 | 0.98149 | 0.9816 | 1 | 21120 | 0 | -2.1038 |
| QRICH1 | 5 | 0.06984 | 0.17813 | 0.968171 | 3824 | 2 | -0.4673 | 0.98155 | 0.98166 | 1 | 21121 | 0 | -0.4673 |
| C10orf54 | 6 | 0.56299 | 0.71295 | 0.999994 | 14430 | 2 | -0.1955 | 0.98155 | 0.98147 | 1 | 21122 | 1 | -0.1955 |
| LYG2 | 6 | 0.20909 | 0.39899 | 0.999994 | 8006 | 3 | -0.77823 | 0.98158 | 0.9815 | 1 | 21123 | 1 | -0.77823 |
| CHRAC1 | 4 | 0.096515 | 0.20884 | 0.979587 | 4840 | 3 | -0.96656 | 0.98167 | 0.98165 | 1 | 21124 | 0 | -0.96656 |
| KPNA2 | 6 | 0.1045 | 0.25612 | 0.994943 | 5113 | 4 | -1.1041 | 0.98167 | 0.98159 | 1 | 21125 | 1 | -1.1041 |
| FBXL7 | 6 | 0.11453 | 0.27536 | 0.999239 | 5463 | 4 | -1.1627 | 0.98168 | 0.9816 | 1 | 21126 | 1 | -1.1627 |
| TSPAN31 | 5 | 0.18311 | 0.35111 | 0.999994 | 7296 | 3 | -0.67784 | 0.98169 | 0.98181 | 1 | 21127 | 0 | -0.67784 |
| ANO7 | 5 | 0.026183 | 0.080595 | 0.93914 | 1813 | 3 | -2.7216 | 0.98176 | 0.98187 | 1 | 21128 | 0 | -2.7216 |
| TOP2B | 6 | 0.30517 | 0.51275 | 0.999994 | 10235 | 3 | -0.55216 | 0.98183 | 0.98175 | 1 | 21129 | 0 | -0.55216 |
| ADSL | 6 | 0.020175 | 0.072537 | 0.93914 | 1468 | 4 | -2.6915 | 0.98191 | 0.98184 | 1 | 21130 | 0 | -2.6915 |
| BCL11A | 5 | 0.063079 | 0.16715 | 0.967126 | 3558 | 3 | -2.2474 | 0.98196 | 0.98207 | 1 | 21131 | 0 | -2.2474 |
| PPP2R3C | 6 | 0.019152 | 0.069257 | 0.93914 | 1417 | 3 | -1.9599 | 0.98199 | 0.98192 | 1 | 21132 | 0 | -1.9599 |
| CTDSPL2 | 6 | 0.003047 | 0.013912 | 0.75278 | 361 | 5 | -2.3394 | 0.98208 | 0.98201 | 1 | 21133 | 0 | -2.3394 |
| CGN | 5 | 0.055444 | 0.15352 | 0.967126 | 3218 | 4 | -1.3063 | 0.98209 | 0.98221 | 1 | 21134 | 0 | -1.3063 |
| ZNF77 | 6 | 0.070651 | 0.18816 | 0.969543 | 3861 | 4 | -1.7342 | 0.98211 | 0.98203 | 1 | 21135 | 0 | -1.7342 |
| TBX3 | 5 | 0.035713 | 0.10505 | 0.95013 | 2301 | 4 | -1.8399 | 0.98215 | 0.98226 | 1 | 21136 | 0 | -1.8399 |
| HSFX1 | 5 | 0.061053 | 0.16384 | 0.967126 | 3471 | 3 | -1.8767 | 0.98217 | 0.98228 | 1 | 21137 | 0 | -1.8767 |
| KLRC4-KLRK1 | 2 | 0.017816 | 0.033232 | 0.867348 | 1353 | 2 | -6.1749 | 0.98218 | 0.98217 | 1 | 21138 | 0 | -6.1749 |
| E2F4 | 4 | 0.00037726 | 0.0014272 | 0.384267 | 89 | 3 | -7.3618 | 0.9822 | 0.98216 | 1 | 21139 | 0 | -7.3618 |
| hsa-mir-548ad | 4 | 0.12679 | 0.26202 | 0.995544 | 5839 | 3 | -1.4407 | 0.98225 | 0.98222 | 1 | 21140 | 0 | -1.4407 |
| TFCP2L1 | 6 | 0.085688 | 0.21889 | 0.980699 | 4434 | 4 | -0.97679 | 0.98226 | 0.98219 | 1 | 21141 | 0 | -0.97679 |
| DNAJC28 | 6 | 0.021194 | 0.075809 | 0.93914 | 1527 | 5 | -1.6722 | 0.9823 | 0.98223 | 1 | 21142 | 0 | -1.6722 |
| CDC42BPB | 5 | 0.15981 | 0.3173 | 0.999994 | 6728 | 3 | -1.1777 | 0.98236 | 0.98246 | 1 | 21143 | 0 | -1.1777 |
| GOT2 | 5 | 0.02997 | 0.090399 | 0.94395 | 2007 | 3 | -1.1181 | 0.98243 | 0.98251 | 1 | 21144 | 0 | -1.1181 |
| NCSTN | 4 | 0.030845 | 0.083189 | 0.94383 | 2054 | 2 | -1.1509 | 0.98251 | 0.98247 | 1 | 21145 | 0 | -1.1509 |
| PTBP1 | 5 | 0.021974 | 0.069421 | 0.93914 | 1581 | 4 | -2.0415 | 0.98259 | 0.98266 | 1 | 21146 | 0 | -2.0415 |
| RNFT1 | 5 | 0.51332 | 0.65955 | 0.999994 | 13626 | 2 | -0.22498 | 0.9826 | 0.98266 | 1 | 21147 | 0 | -0.22498 |
| RXFP3 | 6 | 0.046344 | 0.13605 | 0.95468 | 2825 | 4 | -1.2723 | 0.9827 | 0.98264 | 1 | 21148 | 0 | -1.2723 |
| hsa-mir-4717 | 4 | 0.04045 | 0.10268 | 0.946414 | 2533 | 3 | -1.6641 | 0.98271 | 0.98267 | 1 | 21149 | 0 | -1.6641 |
| ANGPTL4 | 6 | 0.036514 | 0.11405 | 0.951052 | 2346 | 3 | -1.8872 | 0.98289 | 0.98284 | 1 | 21150 | 0 | -1.8872 |
| DSG1 | 5 | 0.50182 | 0.64951 | 0.999994 | 13455 | 2 | -0.48792 | 0.9829 | 0.98297 | 1 | 21151 | 0 | -0.48792 |
| UBFD1 | 4 | 0.11497 | 0.24152 | 0.987253 | 5476 | 3 | -0.7874 | 0.98291 | 0.98288 | 1 | 21152 | 0 | -0.7874 |
| CEP78 | 6 | 0.24706 | 0.44443 | 0.999994 | 8945 | 3 | -0.72579 | 0.98294 | 0.9829 | 1 | 21153 | 0 | -0.72579 |
| RLBP1 | 5 | 0.0032343 | 0.013773 | 0.751291 | 378 | 3 | -7.3721 | 0.98302 | 0.98308 | 1 | 21154 | 0 | -7.3721 |
| TP53I3 | 6 | 0.14443 | 0.31866 | 0.999994 | 6313 | 3 | -1.0024 | 0.98302 | 0.98297 | 1 | 21155 | 0 | -1.0024 |
| HOMER1 | 6 | 0.0078176 | 0.031335 | 0.862223 | 716 | 5 | -2.227 | 0.98304 | 0.98299 | 1 | 21156 | 0 | -2.227 |
| PAX8 | 5 | 0.0032946 | 0.01401 | 0.75278 | 384 | 3 | -7.3492 | 0.9831 | 0.98317 | 1 | 21157 | 0 | -7.3492 |
| TBR1 | 5 | 0.027692 | 0.084489 | 0.94395 | 1888 | 4 | -2.1116 | 0.98311 | 0.98318 | 1 | 21158 | 0 | -2.1116 |
| PIK3R4 | 6 | 0.21809 | 0.40983 | 0.999994 | 8230 | 3 | -0.48277 | 0.98318 | 0.98312 | 1 | 21159 | 0 | -0.48277 |
| TTC7A | 6 | 0.14265 | 0.31643 | 0.999994 | 6259 | 3 | -0.4697 | 0.98324 | 0.98319 | 1 | 21160 | 0 | -0.4697 |
| RPS12 | 2 | 0.016689 | 0.031174 | 0.862223 | 1289 | 2 | -6.0794 | 0.98331 | 0.98328 | 1 | 21161 | 0 | -6.0794 |
| OR10A7 | 5 | 0.0054702 | 0.022649 | 0.820184 | 557 | 4 | -2.7279 | 0.98336 | 0.98343 | 1 | 21162 | 0 | -2.7279 |
| OR5A1 | 6 | 0.083849 | 0.21519 | 0.979587 | 4372 | 4 | -1.543 | 0.98338 | 0.98333 | 1 | 21163 | 0 | -1.543 |

| | | | | | | | | | | | | |
|---|---|---|---|---|---|---|---|---|---|---|---|---|
| RAB19 | 5 | 0.12129 | 0.25934 | 0.994943 | 5674 | 3 | -0.61191 | 0.9834 | 0.98347 | 1 | 21164 | 0 | -0.61191 |
| GPATCH2 | 6 | 0.11225 | 0.271 | 0.998561 | 5377 | 4 | -0.83408 | 0.9834 | 0.98335 | 1 | 21165 | 0 | -0.83408 |
| POTEJ | 2 | 0.016596 | 0.03103 | 0.862223 | 1282 | 2 | -4.183 | 0.9834 | 0.98338 | 1 | 21166 | 0 | -4.183 |
| ZNF433 | 5 | 0.035095 | 0.10348 | 0.946414 | 2273 | 4 | -2.7 | 0.98349 | 0.98356 | 1 | 21167 | 0 | -2.7 |
| ICAM2 | 6 | 0.10402 | 0.25516 | 0.994943 | 5097 | 4 | -1.0196 | 0.98349 | 0.98343 | 1 | 21168 | 0 | -1.0196 |
| APEX1 | 6 | 0.026877 | 0.091459 | 0.94395 | 1849 | 4 | -3.1381 | 0.9835 | 0.98345 | 1 | 21169 | 0 | -3.1381 |
| DNAJA3 | 4 | 0.010087 | 0.033996 | 0.867348 | 875 | 3 | -3.917 | 0.98353 | 0.9835 | 1 | 21170 | 0 | -3.917 |
| ASF1A | 5 | 0.071522 | 0.18085 | 0.968171 | 3898 | 2 | -0.40879 | 0.98357 | 0.98365 | 1 | 21171 | 0 | -0.40879 |
| hsa-mir-4468 | 4 | 0.11384 | 0.23959 | 0.987253 | 5442 | 3 | -0.77108 | 0.98363 | 0.98359 | 1 | 21172 | 0 | -0.77108 |
| PMPCA | 1 | 0.016363 | 0.016292 | 0.772095 | 1266 | 1 | -9.8202 | 0.98364 | 0.98366 | 1 | 21173 | 0 | -9.8202 |
| TMED4 | 6 | 0.13761 | 0.31002 | 0.999994 | 6124 | 3 | -0.86347 | 0.98366 | 0.98361 | 1 | 21174 | 0 | -0.86347 |
| hsa-mir-4793 | 4 | 0.027667 | 0.076759 | 0.93914 | 1886 | 3 | -2.1809 | 0.98372 | 0.98367 | 1 | 21175 | 0 | -2.1809 |
| RAB2A | 6 | 0.1034 | 0.25397 | 0.994632 | 5078 | 4 | -0.81043 | 0.98372 | 0.98367 | 1 | 21176 | 0 | -0.81043 |
| DHFRL1 | 5 | 0.039505 | 0.11466 | 0.951052 | 2491 | 4 | -1.0165 | 0.98378 | 0.98384 | 1 | 21177 | 0 | -1.0165 |
| MAP9 | 6 | 0.081073 | 0.20948 | 0.979587 | 4271 | 3 | -1.3267 | 0.98378 | 0.98373 | 1 | 21178 | 0 | -1.3267 |
| STX6 | 5 | 0.014681 | 0.04929 | 0.906088 | 1171 | 4 | -2.4048 | 0.98383 | 0.9839 | 1 | 21179 | 0 | -2.4048 |
| SYT13 | 6 | 0.040073 | 0.12207 | 0.951052 | 2519 | 4 | -1.4379 | 0.98388 | 0.98383 | 1 | 21180 | 0 | -1.4379 |
| MID1IP1 | 5 | 0.039611 | 0.11493 | 0.951052 | 2495 | 4 | -1.1874 | 0.98389 | 0.98397 | 1 | 21181 | 0 | -1.1874 |
| hsa-mir-5692c-2 | 4 | 0.023397 | 0.067829 | 0.93914 | 1677 | 3 | -3.1091 | 0.98389 | 0.98386 | 1 | 21182 | 0 | -3.1091 |
| GRPR | 6 | 0.026988 | 0.091727 | 0.94395 | 1856 | 4 | -1.8706 | 0.98391 | 0.98386 | 1 | 21183 | 0 | -1.8706 |
| HIST1H2BA | 5 | 0.01166 | 0.040815 | 0.891621 | 968 | 3 | -2.4996 | 0.98392 | 0.98399 | 1 | 21184 | 0 | -2.4996 |
| TBCA | 3 | 0.016036 | 0.041853 | 0.891621 | 1247 | 3 | -6.263 | 0.98396 | 0.98409 | 1 | 21185 | 0 | -6.263 |
| ANKRD52 | 6 | 0.013017 | 0.049175 | 0.905507 | 1062 | 5 | -1.5824 | 0.98403 | 0.98397 | 1 | 21186 | 0 | -1.5824 |
| ZNF32 | 5 | 0.12435 | 0.26398 | 0.996734 | 5776 | 3 | -1.4077 | 0.98403 | 0.98411 | 1 | 21187 | 0 | -1.4077 |
| GNG2 | 4 | 0.059646 | 0.14011 | 0.958716 | 3407 | 3 | -1.9526 | 0.98405 | 0.98402 | 1 | 21188 | 0 | -1.9526 |
| DOPEY2 | 6 | 0.33192 | 0.54347 | 0.999994 | 10765 | 3 | -0.62033 | 0.98405 | 0.984 | 1 | 21189 | 0 | -0.62033 |
| MLN | 6 | 0.014329 | 0.053546 | 0.919407 | 1154 | 5 | -1.4882 | 0.98409 | 0.98403 | 1 | 21190 | 0 | -1.4882 |
| ROBO4 | 5 | 0.012471 | 0.043133 | 0.891621 | 1028 | 3 | -3.7856 | 0.98418 | 0.98426 | 1 | 21191 | 0 | -3.7856 |
| NEUROG2 | 6 | 0.28741 | 0.49208 | 0.999994 | 9862 | 3 | -0.57115 | 0.98423 | 0.98416 | 1 | 21192 | 0 | -0.57115 |
| GLTPD1 | 6 | 0.0036703 | 0.016235 | 0.772095 | 417 | 5 | -1.7447 | 0.98429 | 0.98423 | 1 | 21193 | 0 | -1.7447 |
| STAM | 6 | 0.30672 | 0.51456 | 0.999994 | 10268 | 3 | -0.62033 | 0.98429 | 0.98423 | 1 | 21194 | 0 | -0.62033 |
| CCDC84 | 3 | 0.012464 | 0.032978 | 0.867348 | 1027 | 3 | -6.5116 | 0.9843 | 0.98443 | 1 | 21195 | 0 | -6.5116 |
| TACO1 | 6 | 0.042413 | 0.12732 | 0.951052 | 2637 | 4 | -2.037 | 0.98432 | 0.98427 | 1 | 21196 | 0 | -2.037 |
| C1orf85 | 6 | 0.022248 | 0.079173 | 0.93914 | 1596 | 5 | -2.0691 | 0.98436 | 0.98429 | 1 | 21197 | 0 | -2.0691 |
| RACGAP1 | 4 | 0.015614 | 0.050965 | 0.909101 | 1224 | 4 | -2.6251 | 0.98439 | 0.98435 | 1 | 21198 | 0 | -2.6251 |
| FYTTD1 | 5 | 0.17957 | 0.34601 | 0.999994 | 7211 | 3 | -0.62802 | 0.98447 | 0.98455 | 1 | 21199 | 0 | -0.62802 |
| HIST1H2AC | 5 | 0.14954 | 0.30209 | 0.999994 | 6446 | 3 | -1.2112 | 0.9845 | 0.98457 | 1 | 21200 | 0 | -1.2112 |
| NonTargetingControlG | 2 | 0.015477 | 0.029023 | 0.859137 | 1213 | 2 | -6.3437 | 0.98452 | 0.9845 | 1 | 21201 | 0 | -6.3437 |
| hsa-mir-4304 | 2 | 0.015472 | 0.029018 | 0.859137 | 1212 | 2 | -4.1031 | 0.98453 | 0.98451 | 1 | 21202 | 0 | -4.1031 |
| LOC100289561 | 4 | 0.01543 | 0.050406 | 0.909101 | 1211 | 4 | -1.3425 | 0.98457 | 0.98453 | 1 | 21203 | 0 | -1.3425 |
| LOC100130357 | 5 | 0.046938 | 0.13301 | 0.95318 | 2849 | 3 | -2.0384 | 0.98467 | 0.98474 | 1 | 21204 | 0 | -2.0384 |
| HRAS | 6 | 0.10497 | 0.25701 | 0.994943 | 5133 | 4 | -1.0894 | 0.98467 | 0.98461 | 1 | 21205 | 0 | -1.0894 |
| ETNK1 | 6 | 0.12072 | 0.28709 | 0.999632 | 5653 | 4 | -0.79358 | 0.9847 | 0.98463 | 1 | 21206 | 0 | -0.79358 |
| PLEKHN1 | 6 | 0.15309 | 0.32955 | 0.999994 | 6537 | 3 | -0.7703 | 0.98478 | 0.98473 | 1 | 21207 | 0 | -0.7703 |
| ITIH3 | 5 | 0.000033435 | 0.00014608 | 0.157291 | 21 | 4 | -9.7911 | 0.98483 | 0.9849 | 1 | 21208 | 0 | -9.7911 |
| NT5DC2 | 5 | 0.0030255 | 0.012937 | 0.733879 | 359 | 4 | -2.8903 | 0.98484 | 0.9849 | 1 | 21209 | 0 | -2.8903 |
| GTF2IRD2 | 6 | 0.013881 | 0.052061 | 0.911383 | 1121 | 5 | -2.2025 | 0.98491 | 0.98486 | 1 | 21210 | 0 | -2.2025 |
| hsa-mir-5090 | 4 | 0.015079 | 0.049376 | 0.906138 | 1192 | 4 | -1.1742 | 0.98492 | 0.98488 | 1 | 21211 | 0 | -1.1742 |
| ARHGDIB | 6 | 0.0095357 | 0.037404 | 0.881473 | 841 | 5 | -1.3985 | 0.98501 | 0.98497 | 1 | 21212 | 0 | -1.3985 |
| NonTargetingControlG | 2 | 0.01499 | 0.028131 | 0.855594 | 1188 | 2 | -3.7754 | 0.98501 | 0.98497 | 1 | 21213 | 0 | -3.7754 |
| hsa-mir-450b | 4 | 0.014956 | 0.049006 | 0.904562 | 1185 | 4 | -2.0184 | 0.98504 | 0.98501 | 1 | 21214 | 0 | -2.0184 |
| XPO7 | 6 | 0.0037624 | 0.016584 | 0.775667 | 426 | 5 | -1.9698 | 0.98508 | 0.98504 | 1 | 21215 | 0 | -1.9698 |

| | | | | | | | | | | | | | |
|---|---|---|---|---|---|---|---|---|---|---|---|---|---|
| IWS1 | 6 | 0.051504 | 0.14737 | 0.965554 | 3047 | 4 | -1.7727 | 0.98508 | 0.98505 | 1 | 21216 | 0 | -1.7727 |
| GPX4 | 2 | 0.01487 | 0.027909 | 0.855011 | 1182 | 2 | -6.1556 | 0.98513 | 0.98509 | 1 | 21217 | 0 | -6.1556 |
| hsa-mir-3652 | 2 | 0.01485 | 0.027867 | 0.855011 | 1181 | 2 | -5.6768 | 0.98515 | 0.98512 | 1 | 21218 | 0 | -5.6768 |
| GTDC1 | 6 | 0.03415 | 0.10851 | 0.951052 | 2222 | 4 | -1.8937 | 0.98515 | 0.98511 | 1 | 21219 | 0 | -1.8937 |
| COL6A6 | 5 | 0.14385 | 0.29361 | 0.999994 | 6294 | 3 | -1.3025 | 0.98517 | 0.98523 | 1 | 21220 | 0 | -1.3025 |
| CAND2 | 4 | 0.014804 | 0.048525 | 0.902764 | 1178 | 4 | -1.5665 | 0.9852 | 0.98517 | 1 | 21221 | 0 | -1.5665 |
| C10orf111 | 4 | 0.014649 | 0.048037 | 0.900643 | 1170 | 4 | -1.4379 | 0.98535 | 0.98532 | 1 | 21222 | 0 | -1.4379 |
| PPME1 | 6 | 0.022952 | 0.081433 | 0.93914 | 1646 | 5 | -1.556 | 0.98548 | 0.98544 | 1 | 21223 | 0 | -1.556 |
| ZCCHC12 | 6 | 0.15286 | 0.32926 | 0.999994 | 6530 | 3 | -0.7298 | 0.98549 | 0.98546 | 1 | 21224 | 0 | -0.7298 |
| IRF2BP1 | 6 | 0.035126 | 0.11083 | 0.951052 | 2274 | 3 | -1.5921 | 0.98556 | 0.98553 | 1 | 21225 | 0 | -1.5921 |
| THPO | 5 | 0.23323 | 0.42167 | 0.999994 | 8600 | 3 | -0.58026 | 0.98561 | 0.98565 | 1 | 21226 | 0 | -0.58026 |
| NonTargetingControlG | 1 | 0.014336 | 0.014246 | 0.753188 | 1155 | 1 | -9.9641 | 0.98566 | 0.98564 | 1 | 21227 | 0 | -9.9641 |
| hsa-mir-548av | 4 | 0.014311 | 0.046961 | 0.896157 | 1150 | 4 | -0.86293 | 0.98569 | 0.98566 | 1 | 21228 | 0 | -0.86293 |
| NRM | 6 | 0.39227 | 0.5974 | 0.999994 | 11838 | 2 | -0.45579 | 0.9857 | 0.98567 | 1 | 21229 | 0 | -0.45579 |
| LCA5L | 5 | 0.0092519 | 0.033895 | 0.867348 | 811 | 4 | -1.9033 | 0.98578 | 0.9858 | 1 | 21230 | 0 | -1.9033 |
| PPA2 | 4 | 0.014176 | 0.046545 | 0.895138 | 1141 | 4 | -1.4006 | 0.98582 | 0.9858 | 1 | 21231 | 0 | -1.4006 |
| PPP1R3D | 6 | 0.07849 | 0.20427 | 0.978186 | 4171 | 4 | -1.4548 | 0.98585 | 0.98581 | 1 | 21232 | 0 | -1.4548 |
| TFAP4 | 6 | 3.8674E-06 | 0.00001806 | 0.030084 | 13 | 5 | -9.2412 | 0.98588 | 0.98584 | 1 | 21233 | 0 | -9.2412 |
| CDK5 | 5 | 0.0013077 | 0.0058201 | 0.623123 | 195 | 4 | -3.7994 | 0.98589 | 0.98591 | 1 | 21234 | 0 | -3.7994 |
| CLUH | 5 | 0.023142 | 0.072586 | 0.93914 | 1660 | 4 | -1.8513 | 0.9859 | 0.98591 | 1 | 21235 | 0 | -1.8513 |
| OR6C70 | 5 | 0.19751 | 0.37167 | 0.999994 | 7688 | 3 | -0.86667 | 0.98595 | 0.98597 | 1 | 21236 | 0 | -0.86667 |
| PQLC3 | 6 | 0.12085 | 0.28731 | 0.999632 | 5655 | 3 | -1.1672 | 0.98596 | 0.98592 | 1 | 21237 | 0 | -1.1672 |
| NRBP1 | 6 | 0.21248 | 0.40311 | 0.999994 | 8073 | 3 | -0.7731 | 0.98597 | 0.98593 | 1 | 21238 | 0 | -0.7731 |
| HTR2C | 5 | 0.0020605 | 0.0090156 | 0.680529 | 279 | 4 | -4.6608 | 0.98597 | 0.98599 | 1 | 21239 | 0 | -4.6608 |
| WBSCR27 | 5 | 0.53109 | 0.67504 | 0.999994 | 13913 | 2 | -0.58737 | 0.98599 | 0.98601 | 1 | 21240 | 0 | -0.58737 |
| STAB1 | 6 | 0.00030462 | 0.0016197 | 0.398571 | 81 | 5 | -4.6653 | 0.98604 | 0.986 | 1 | 21241 | 0 | -4.6653 |
| DCTN3 | 4 | 0.013937 | 0.045839 | 0.892434 | 1123 | 4 | -2.0049 | 0.98606 | 0.98604 | 1 | 21242 | 0 | -2.0049 |
| DEFB126 | 5 | 0.074877 | 0.18631 | 0.969409 | 4030 | 3 | -1.7332 | 0.98613 | 0.98614 | 1 | 21243 | 0 | -1.7332 |
| ERGIC1 | 6 | 0.022466 | 0.07987 | 0.93914 | 1615 | 5 | -1.8235 | 0.98617 | 0.98613 | 1 | 21244 | 0 | -1.8235 |
| LAMTOR2 | 5 | 0.26962 | 0.45653 | 0.999994 | 9465 | 2 | -0.51956 | 0.98618 | 0.98619 | 1 | 21245 | 0 | -0.51956 |
| LTC4S | 5 | 0.045807 | 0.13029 | 0.95318 | 2797 | 4 | -1.8885 | 0.9862 | 0.98621 | 1 | 21246 | 0 | -1.8885 |
| MERTK | 5 | 0.0023677 | 0.010326 | 0.69657 | 311 | 4 | -2.9884 | 0.98623 | 0.98624 | 1 | 21247 | 0 | -2.9884 |
| HMHA1 | 5 | 0.0034827 | 0.014775 | 0.755578 | 404 | 4 | -3.0995 | 0.98635 | 0.98637 | 1 | 21248 | 0 | -3.0995 |
| IL13 | 6 | 0.42284 | 0.61727 | 0.999994 | 12294 | 2 | -0.11491 | 0.98639 | 0.98635 | 1 | 21249 | 0 | -0.11491 |
| CCDC17 | 5 | 0.04162 | 0.11995 | 0.951052 | 2596 | 4 | -1.2204 | 0.9864 | 0.98642 | 1 | 21250 | 0 | -1.2204 |
| EVX2 | 4 | 0.013543 | 0.044638 | 0.891621 | 1104 | 4 | -1.309 | 0.98646 | 0.98641 | 1 | 21251 | 0 | -1.309 |
| DNAJC9 | 5 | 0.014634 | 0.049146 | 0.905507 | 1168 | 4 | -2.7475 | 0.98647 | 0.98648 | 1 | 21252 | 0 | -2.7475 |
| C5orf38 | 5 | 0.29198 | 0.47454 | 0.999994 | 9947 | 2 | -0.51942 | 0.9865 | 0.98651 | 1 | 21253 | 0 | -0.51942 |
| CBWD1 | 5 | 0.018663 | 0.060468 | 0.932932 | 1398 | 4 | -1.4201 | 0.98652 | 0.98652 | 1 | 21254 | 0 | -1.4201 |
| NonTargetingControlG | 2 | 0.013448 | 0.025306 | 0.84412 | 1097 | 2 | -6.9466 | 0.98655 | 0.98651 | 1 | 21255 | 0 | -6.9466 |
| ACTR8 | 4 | 0.01339 | 0.044193 | 0.891621 | 1092 | 4 | -2.3665 | 0.98661 | 0.98655 | 1 | 21256 | 0 | -2.3665 |
| RPL15 | 2 | 0.013313 | 0.025058 | 0.844082 | 1083 | 2 | -6.5433 | 0.98669 | 0.98665 | 1 | 21257 | 0 | -6.5433 |
| COL6A1 | 4 | 0.013141 | 0.043429 | 0.891621 | 1073 | 4 | -1.5806 | 0.98686 | 0.98681 | 1 | 21258 | 0 | -1.5806 |
| TWF2 | 6 | 0.00019749 | 0.0010601 | 0.352868 | 55 | 5 | -4.7776 | 0.98686 | 0.9868 | 1 | 21259 | 0 | -4.7776 |
| CD2 | 6 | 0.18212 | 0.36581 | 0.999994 | 7274 | 2 | -0.5349 | 0.98693 | 0.98687 | 1 | 21260 | 0 | -0.5349 |
| ZSCAN21 | 6 | 0.0065557 | 0.026856 | 0.848412 | 628 | 5 | -1.821 | 0.98707 | 0.98702 | 1 | 21261 | 0 | -1.821 |
| KCNA5 | 5 | 0.016964 | 0.05576 | 0.9235 | 1300 | 4 | -3.4884 | 0.98716 | 0.98713 | 1 | 21262 | 0 | -3.4884 |
| MPV17L2 | 4 | 0.012826 | 0.042473 | 0.891621 | 1054 | 4 | -3.7942 | 0.98717 | 0.98713 | 1 | 21263 | 0 | -3.7942 |
| hsa-mir-4692 | 4 | 0.012822 | 0.042462 | 0.891621 | 1053 | 4 | -1.8791 | 0.98718 | 0.98713 | 1 | 21264 | 0 | -1.8791 |
| YWHAE | 5 | 0.42486 | 0.58412 | 0.999994 | 12334 | 2 | -0.51582 | 0.9872 | 0.98717 | 1 | 21265 | 0 | -0.51582 |
| HDAC5 | 6 | 0.017851 | 0.065022 | 0.93914 | 1354 | 5 | -1.3377 | 0.98727 | 0.98723 | 1 | 21266 | 0 | -1.3377 |
| GJA8 | 6 | 0.0078166 | 0.031334 | 0.862223 | 715 | 4 | -3.0343 | 0.9873 | 0.98726 | 1 | 21267 | 0 | -3.0343 |

| | | | | | | | | | | | | | |
|---|---|---|---|---|---|---|---|---|---|---|---|---|---|
| hsa-mir-3622a | 4 | 0.012673 | 0.04199 | 0.891621 | 1044 | 4 | -1.4262 | 0.98733 | 0.9873 | 1 | 21268 | 0 | -1.4262 |
| NOD1 | 6 | 0.094481 | 0.23647 | 0.985449 | 4767 | 4 | -0.79985 | 0.98736 | 0.98731 | 1 | 21269 | 0 | -0.79985 |
| COG7 | 5 | 0.025589 | 0.079061 | 0.93914 | 1780 | 4 | -1.7041 | 0.98737 | 0.98734 | 1 | 21270 | 0 | -1.7041 |
| C3orf30 | 6 | 0.003516 | 0.015641 | 0.766139 | 408 | 5 | -2.4083 | 0.98748 | 0.98743 | 1 | 21271 | 0 | -2.4083 |
| FAM13B | 4 | 0.012184 | 0.04052 | 0.891621 | 1006 | 4 | -3.0408 | 0.98753 | 0.9875 | 1 | 21272 | 0 | -3.0408 |
| CHODL | 5 | 0.014817 | 0.049683 | 0.907155 | 1180 | 4 | -1.56 | 0.98756 | 0.98754 | 1 | 21273 | 0 | -1.56 |
| YBX1 | 4 | 0.00005972 | 0.00021512 | 0.177301 | 31 | 4 | -9.4407 | 0.98756 | 0.98753 | 1 | 21274 | 0 | -9.4407 |
| KLC2 | 6 | 0.0028246 | 0.013083 | 0.733879 | 342 | 5 | -4.7133 | 0.98757 | 0.98753 | 1 | 21275 | 0 | -4.7133 |
| TMEM155 | 6 | 0.0035498 | 0.015779 | 0.766139 | 411 | 3 | -4.0487 | 0.98757 | 0.98753 | 1 | 21276 | 0 | -4.0487 |
| TMEM63A | 6 | 0.012523 | 0.047492 | 0.899064 | 1031 | 4 | -2.3464 | 0.98763 | 0.9876 | 1 | 21277 | 0 | -2.3464 |
| GIPC3 | 6 | 0.011605 | 0.044424 | 0.891621 | 965 | 5 | -2.1054 | 0.98766 | 0.98763 | 1 | 21278 | 0 | -2.1054 |
| SSPO | 4 | 0.012308 | 0.040902 | 0.891621 | 1013 | 4 | -2.2793 | 0.98769 | 0.98766 | 1 | 21279 | 0 | -2.2793 |
| KLHL15 | 6 | 0.011345 | 0.043571 | 0.891621 | 946 | 4 | -2.8863 | 0.98776 | 0.98773 | 1 | 21280 | 0 | -2.8863 |
| PLCB3 | 6 | 0.017414 | 0.063581 | 0.937389 | 1334 | 4 | -2.2625 | 0.98777 | 0.98774 | 1 | 21281 | 0 | -2.2625 |
| DOCK2 | 6 | 0.00019471 | 0.0010436 | 0.352868 | 54 | 5 | -5.8423 | 0.98782 | 0.98778 | 1 | 21282 | 0 | -5.8423 |
| MICA | 5 | 0.051251 | 0.14342 | 0.960973 | 3035 | 4 | -1.7162 | 0.98793 | 0.98792 | 1 | 21283 | 0 | -1.7162 |
| NOTCH1 | 6 | 0.24772 | 0.44521 | 0.999994 | 8964 | 3 | -0.49141 | 0.98801 | 0.98797 | 1 | 21284 | 0 | -0.49141 |
| LOC554223 | 6 | 0.20231 | 0.3907 | 0.999994 | 7839 | 3 | -0.66648 | 0.98802 | 0.98798 | 1 | 21285 | 0 | -0.66648 |
| LOC284385 | 4 | 0.011944 | 0.039781 | 0.891621 | 987 | 4 | -2.468 | 0.98806 | 0.98803 | 1 | 21286 | 0 | -2.468 |
| NonTargetingControlG | 2 | 0.01179 | 0.022297 | 0.819701 | 979 | 2 | -4.905 | 0.98821 | 0.98811 | 1 | 21287 | 0 | -4.905 |
| NonTargetingControlG | 2 | 0.01178 | 0.02228 | 0.819701 | 977 | 2 | -4.4334 | 0.98822 | 0.98812 | 1 | 21288 | 0 | -4.4334 |
| TAS1R1 | 6 | 0.020232 | 0.072718 | 0.93914 | 1474 | 5 | -0.94088 | 0.98826 | 0.98823 | 1 | 21289 | 0 | -0.94088 |
| SLC39A6 | 4 | 0.011709 | 0.039056 | 0.891621 | 973 | 4 | -1.8039 | 0.98829 | 0.98826 | 1 | 21290 | 0 | -1.8039 |
| hsa-mir-6740 | 3 | 0.011681 | 0.031028 | 0.862223 | 970 | 3 | -2.9713 | 0.98832 | 0.98848 | 1 | 21291 | 0 | -2.9713 |
| DGCR8 | 4 | 0.011563 | 0.038616 | 0.890863 | 960 | 4 | -3.1294 | 0.98844 | 0.98841 | 1 | 21292 | 0 | -3.1294 |
| CRYGN | 4 | 0.0020653 | 0.0074954 | 0.665213 | 280 | 4 | -7.1136 | 0.98846 | 0.98844 | 1 | 21293 | 0 | -7.1136 |
| CCDC28B | 6 | 0.26176 | 0.46198 | 0.999994 | 9281 | 3 | -0.71049 | 0.98847 | 0.98843 | 1 | 21294 | 0 | -0.71049 |
| RNF20 | 6 | 0.020741 | 0.074355 | 0.93914 | 1493 | 5 | -2.5608 | 0.98851 | 0.98847 | 1 | 21295 | 0 | -2.5608 |
| ATP13A3 | 5 | 0.020265 | 0.064785 | 0.93914 | 1475 | 4 | -2.3193 | 0.98851 | 0.98853 | 1 | 21296 | 0 | -2.3193 |
| CDKN2D | 3 | 0.011433 | 0.030411 | 0.862223 | 952 | 3 | -2.8318 | 0.98857 | 0.98872 | 1 | 21297 | 0 | -2.8318 |
| CLDN20 | 5 | 0.085069 | 0.20266 | 0.977973 | 4411 | 3 | -1.0425 | 0.98881 | 0.98885 | 1 | 21298 | 0 | -1.0425 |
| NKX3-1 | 5 | 0.28993 | 0.47286 | 0.999994 | 9909 | 2 | -0.38439 | 0.98882 | 0.98886 | 1 | 21299 | 0 | -0.38439 |
| CYTIP | 6 | 0.026274 | 0.089998 | 0.94395 | 1817 | 3 | -2.1344 | 0.98884 | 0.9888 | 1 | 21300 | 0 | -2.1344 |
| STX12 | 6 | 0.020158 | 0.072485 | 0.93914 | 1465 | 4 | -2.281 | 0.98885 | 0.98881 | 1 | 21301 | 0 | -2.281 |
| ARPC1A | 6 | 0.020872 | 0.074786 | 0.93914 | 1502 | 5 | -0.78591 | 0.98887 | 0.98882 | 1 | 21302 | 0 | -0.78591 |
| NonTargetingControlG | 1 | 0.011064 | 0.010999 | 0.71022 | 929 | 1 | -10.225 | 0.98894 | 0.98898 | 1 | 21303 | 0 | -10.225 |
| hsa-mir-4804 | 4 | 0.011057 | 0.037037 | 0.881473 | 927 | 4 | -0.91414 | 0.98894 | 0.98892 | 1 | 21304 | 0 | -0.91414 |
| DLL1 | 4 | 0.011023 | 0.036928 | 0.881473 | 925 | 4 | -1.6084 | 0.98898 | 0.98896 | 1 | 21305 | 0 | -1.6084 |
| KCNMB4 | 6 | 0.020371 | 0.073173 | 0.93914 | 1481 | 4 | -1.9858 | 0.98898 | 0.98894 | 1 | 21306 | 0 | -1.9858 |
| GTF2A2 | 6 | 0.004454 | 0.01915 | 0.805137 | 477 | 5 | -4.1766 | 0.98898 | 0.98894 | 1 | 21307 | 0 | -4.1766 |
| CNBP | 6 | 0.035832 | 0.11248 | 0.951052 | 2307 | 3 | -1.0549 | 0.98898 | 0.98894 | 1 | 21308 | 0 | -1.0549 |
| DBR1 | 5 | 0.00030795 | 0.0014551 | 0.384267 | 82 | 4 | -6.5713 | 0.98904 | 0.98909 | 1 | 21309 | 0 | -6.5713 |
| PTGR1 | 5 | 0.032766 | 0.097587 | 0.945253 | 2148 | 4 | -1.4517 | 0.98904 | 0.98909 | 1 | 21310 | 0 | -1.4517 |
| S100PBP | 3 | 0.010917 | 0.029171 | 0.859467 | 922 | 3 | -2.6943 | 0.98908 | 0.98922 | 1 | 21311 | 0 | -2.6943 |
| PWP1 | 5 | 0.18844 | 0.35871 | 0.999994 | 7433 | 3 | -0.87232 | 0.9892 | 0.98923 | 1 | 21312 | 0 | -0.87232 |
| STIP1 | 5 | 0.052565 | 0.14657 | 0.964852 | 3095 | 4 | -1.1377 | 0.98921 | 0.98923 | 1 | 21313 | 0 | -1.1377 |
| NSMCE2 | 6 | 0.0098126 | 0.038384 | 0.889591 | 860 | 3 | -3.6629 | 0.9893 | 0.98926 | 1 | 21314 | 0 | -3.6629 |
| GRIK4 | 5 | 0.19502 | 0.36811 | 0.999994 | 7633 | 3 | -0.89648 | 0.98932 | 0.98936 | 1 | 21315 | 0 | -0.89648 |
| NonTargetingControlG | 2 | 0.010649 | 0.020159 | 0.808847 | 908 | 2 | -7.3625 | 0.98935 | 0.98927 | 1 | 21316 | 0 | -7.3625 |
| NKTR | 5 | 0.010441 | 0.037324 | 0.881473 | 899 | 3 | -3.1133 | 0.98941 | 0.98945 | 1 | 21317 | 0 | -3.1133 |
| SNTG1 | 5 | 0.034754 | 0.10263 | 0.946414 | 2254 | 4 | -2.6399 | 0.98943 | 0.98947 | 1 | 21318 | 0 | -2.6399 |
| AKAP11 | 6 | 0.014589 | 0.054377 | 0.922677 | 1167 | 5 | -3.0969 | 0.9895 | 0.98947 | 1 | 21319 | 0 | -3.0969 |

| | | | | | | | | | | | | | |
|---|---|---|---|---|---|---|---|---|---|---|---|---|---|
| TAF5L | 6 | 0.01375 | 0.05162 | 0.910734 | 1115 | 4 | -2.2694 | 0.98962 | 0.98958 | 1 | 21320 | 0 | -2.2694 |
| SMIM21 | 6 | 0.32587 | 0.53653 | 0.999994 | 10639 | 3 | -0.50888 | 0.98965 | 0.98961 | 1 | 21321 | 0 | -0.50888 |
| LLPH | 5 | 0.0030511 | 0.013045 | 0.733879 | 362 | 4 | -4.1784 | 0.98966 | 0.9897 | 1 | 21322 | 0 | -4.1784 |
| MS4A7 | 5 | 0.012185 | 0.042327 | 0.891621 | 1007 | 4 | -2.386 | 0.98969 | 0.98974 | 1 | 21323 | 0 | -2.386 |
| PTPRB | 6 | 0.10571 | 0.25844 | 0.994943 | 5164 | 4 | -0.83074 | 0.98969 | 0.98964 | 1 | 21324 | 0 | -0.83074 |
| ZNF638 | 6 | 0.00066147 | 0.0034248 | 0.541338 | 127 | 5 | -4.1686 | 0.9897 | 0.98965 | 1 | 21325 | 0 | -4.1686 |
| VRK1 | 6 | 0.061137 | 0.1682 | 0.967126 | 3473 | 4 | -1.5559 | 0.98974 | 0.98969 | 1 | 21326 | 0 | -1.5559 |
| hsa-mir-6511a-4 | 4 | 0.01022 | 0.034409 | 0.867348 | 884 | 4 | -2.7403 | 0.98978 | 0.98972 | 1 | 21327 | 0 | -2.7403 |
| ZNF560 | 4 | 0.010202 | 0.034358 | 0.867348 | 883 | 4 | -4.6736 | 0.9898 | 0.98974 | 1 | 21328 | 0 | -4.6736 |
| TMEM115 | 4 | 0.010185 | 0.034304 | 0.867348 | 882 | 4 | -1.7014 | 0.98981 | 0.98976 | 1 | 21329 | 0 | -1.7014 |
| SCAF8 | 6 | 0.00043599 | 0.0022941 | 0.45324 | 101 | 5 | -6.9362 | 0.98989 | 0.98984 | 1 | 21330 | 0 | -6.9362 |
| SP4 | 6 | 0.022308 | 0.079373 | 0.93914 | 1602 | 5 | -1.3109 | 0.98993 | 0.98988 | 1 | 21331 | 0 | -1.3109 |
| IRF6 | 6 | 0.31565 | 0.52491 | 0.999994 | 10434 | 3 | -0.61503 | 0.98999 | 0.98995 | 1 | 21332 | 0 | -0.61503 |
| THBS4 | 5 | 0.14039 | 0.28842 | 0.999632 | 6196 | 3 | -1.1563 | 0.99 | 0.99003 | 1 | 21333 | 0 | -1.1563 |
| TCF20 | 6 | 0.000015321 | 0.000092129 | 0.114686 | 17 | 5 | -7.1135 | 0.99003 | 0.99 | 1 | 21334 | 0 | -7.1135 |
| CDIP1 | 5 | 0.044191 | 0.12628 | 0.951052 | 2718 | 4 | -1.8901 | 0.99008 | 0.99011 | 1 | 21335 | 0 | -1.8901 |
| NIPA2 | 6 | 0.059982 | 0.16571 | 0.967126 | 3420 | 4 | -1.2183 | 0.99009 | 0.99006 | 1 | 21336 | 0 | -1.2183 |
| TP63 | 6 | 0.32603 | 0.53672 | 0.999994 | 10643 | 3 | -0.4749 | 0.99012 | 0.99009 | 1 | 21337 | 0 | -0.4749 |
| MMP28 | 6 | 0.017467 | 0.06376 | 0.937389 | 1340 | 5 | -1.1633 | 0.99015 | 0.99012 | 1 | 21338 | 0 | -1.1633 |
| SMAD5 | 6 | 0.056412 | 0.158 | 0.967126 | 3265 | 4 | -1.8823 | 0.99019 | 0.99015 | 1 | 21339 | 0 | -1.8823 |
| UQCC2 | 5 | 0.0055604 | 0.022999 | 0.827176 | 560 | 4 | -2.5146 | 0.99021 | 0.99025 | 1 | 21340 | 0 | -2.5146 |
| RHOF | 5 | 0.014086 | 0.047618 | 0.899802 | 1134 | 4 | -3.0427 | 0.99024 | 0.99027 | 1 | 21341 | 0 | -3.0427 |
| SEPT5 | 6 | 0.18849 | 0.37363 | 0.999994 | 7435 | 3 | -0.92486 | 0.99029 | 0.99025 | 1 | 21342 | 0 | -0.92486 |
| TASP1 | 6 | 0.071471 | 0.18985 | 0.970473 | 3890 | 4 | -1.4999 | 0.99038 | 0.99035 | 1 | 21343 | 0 | -1.4999 |
| DTX3L | 6 | 0.014889 | 0.055365 | 0.922677 | 1183 | 5 | -0.856 | 0.99042 | 0.99039 | 1 | 21344 | 0 | -0.856 |
| EEPD1 | 5 | 0.030731 | 0.092369 | 0.943968 | 2045 | 4 | -1.4514 | 0.99049 | 0.99052 | 1 | 21345 | 0 | -1.4514 |
| PSPC1 | 5 | 0.032395 | 0.096612 | 0.945031 | 2129 | 4 | -2.3814 | 0.99051 | 0.99054 | 1 | 21346 | 0 | -2.3814 |
| hsa-mir-3134 | 3 | 0.0094895 | 0.025447 | 0.845628 | 833 | 3 | -1.9371 | 0.99051 | 0.99062 | 1 | 21347 | 0 | -1.9371 |
| hsa-mir-1299 | 2 | 0.0094779 | 0.017989 | 0.795727 | 830 | 2 | -7.4574 | 0.99052 | 0.9904 | 1 | 21348 | 0 | -7.4574 |
| ADAMTS1 | 6 | 0.25129 | 0.44948 | 0.999994 | 9044 | 3 | -0.77316 | 0.99053 | 0.99049 | 1 | 21349 | 0 | -0.77316 |
| CR2 | 5 | 0.010865 | 0.03853 | 0.89046 | 919 | 4 | -2.9194 | 0.99059 | 0.99062 | 1 | 21350 | 0 | -2.9194 |
| GMNC | 6 | 0.022381 | 0.079608 | 0.93914 | 1608 | 4 | -3.557 | 0.99061 | 0.99056 | 1 | 21351 | 0 | -3.557 |
| PTGFR | 6 | 0.036555 | 0.11414 | 0.951052 | 2348 | 4 | -1.7421 | 0.99069 | 0.99065 | 1 | 21352 | 0 | -1.7421 |
| OR6C75 | 6 | 0.017214 | 0.062951 | 0.937389 | 1314 | 3 | -0.7097 | 0.99072 | 0.99068 | 1 | 21353 | 0 | -0.7097 |
| SLC7A14 | 5 | 0.028064 | 0.085476 | 0.94395 | 1905 | 4 | -1.9861 | 0.99072 | 0.99077 | 1 | 21354 | 0 | -1.9861 |
| RBMX | 4 | 0.0077653 | 0.026653 | 0.848412 | 712 | 4 | -4.9086 | 0.99073 | 0.99068 | 1 | 21355 | 0 | -4.9086 |
| TM4SF20 | 5 | 0.057734 | 0.15832 | 0.967126 | 3325 | 3 | -2.0621 | 0.99078 | 0.99083 | 1 | 21356 | 0 | -2.0621 |
| LOC100505679 | 6 | 0.24905 | 0.4468 | 0.999994 | 8994 | 3 | -0.76607 | 0.99086 | 0.99081 | 1 | 21357 | 0 | -0.76607 |
| PTS | 6 | 0.018041 | 0.065627 | 0.93914 | 1362 | 5 | -2.2393 | 0.99087 | 0.99083 | 1 | 21358 | 0 | -2.2393 |
| URB1 | 5 | 0.045206 | 0.12882 | 0.951777 | 2774 | 4 | -1.6411 | 0.99091 | 0.99097 | 1 | 21359 | 0 | -1.6411 |
| TNS3 | 5 | 0.056024 | 0.15487 | 0.967126 | 3248 | 4 | -0.73808 | 0.99094 | 0.991 | 1 | 21360 | 0 | -0.73808 |
| NEMF | 5 | 0.041721 | 0.1202 | 0.951052 | 2602 | 4 | -0.99972 | 0.991 | 0.99106 | 1 | 21361 | 0 | -0.99972 |
| LPXN | 5 | 0.15234 | 0.30625 | 0.999994 | 6516 | 3 | -1.0618 | 0.99101 | 0.99106 | 1 | 21362 | 0 | -1.0618 |
| TOMM7 | 6 | 0.18962 | 0.37501 | 0.999994 | 7473 | 3 | -0.66539 | 0.99101 | 0.99098 | 1 | 21363 | 0 | -0.66539 |
| WDR67 | 5 | 0.024339 | 0.075716 | 0.93914 | 1725 | 3 | -3.1719 | 0.99121 | 0.99127 | 1 | 21364 | 0 | -3.1719 |
| TMEM247 | 6 | 0.017954 | 0.065353 | 0.93914 | 1359 | 5 | -2.0343 | 0.99123 | 0.99119 | 1 | 21365 | 0 | -2.0343 |
| UBE2D4 | 6 | 0.0097265 | 0.038076 | 0.887545 | 855 | 3 | -1.1647 | 0.99125 | 0.99122 | 1 | 21366 | 0 | -1.1647 |
| MPV17 | 6 | 0.013815 | 0.051836 | 0.911128 | 1118 | 5 | -2.3601 | 0.99126 | 0.99122 | 1 | 21367 | 0 | -2.3601 |
| SUCLG1 | 6 | 0.017144 | 0.062729 | 0.937389 | 1309 | 4 | -2.3834 | 0.99133 | 0.99129 | 1 | 21368 | 0 | -2.3834 |
| TUBA1B | 3 | 0.0086234 | 0.023208 | 0.83009 | 769 | 3 | -6.7744 | 0.99138 | 0.99146 | 1 | 21369 | 0 | -6.7744 |
| GDPD4 | 6 | 0.51858 | 0.68155 | 0.999994 | 13718 | 2 | -0.39319 | 0.99142 | 0.99138 | 1 | 21370 | 0 | -0.39319 |
| FAM208A | 6 | 0.000023333 | 0.00013831 | 0.157291 | 19 | 5 | -5.8849 | 0.99144 | 0.99139 | 1 | 21371 | 0 | -5.8849 |

| | | | | | | | | | | | | | |
|---|---|---|---|---|---|---|---|---|---|---|---|---|---|
| LYST | 6 | 0.0028621 | 0.013217 | 0.733879 | 346 | 5 | -5.3699 | 0.99146 | 0.99141 | 1 | 21372 | 0 | -5.3699 |
| IGFBP5 | 6 | 0.10201 | 0.25125 | 0.993254 | 5040 | 4 | -1.1241 | 0.9915 | 0.99145 | 1 | 21373 | 0 | -1.1241 |
| ABCB9 | 5 | 0.10432 | 0.23313 | 0.984913 | 5108 | 3 | -0.68841 | 0.99151 | 0.99156 | 1 | 21374 | 0 | -0.68841 |
| PI4KB | 6 | 0.012678 | 0.048017 | 0.900643 | 1045 | 4 | -2.5157 | 0.99152 | 0.99148 | 1 | 21375 | 0 | -2.5157 |
| HSCB | 4 | 0.0084762 | 0.0289 | 0.859137 | 758 | 4 | -1.6196 | 0.99152 | 0.99148 | 1 | 21376 | 0 | -1.6196 |
| NonTargetingControlG | 1 | 0.0084008 | 0.0083755 | 0.680529 | 755 | 1 | -10.472 | 0.9916 | 0.99157 | 1 | 21377 | 0 | -10.472 |
| EHBP1 | 6 | 0.035472 | 0.11165 | 0.951052 | 2291 | 4 | -0.90522 | 0.99162 | 0.99157 | 1 | 21378 | 0 | -0.90522 |
| PTPN11 | 5 | 0.0029438 | 0.012623 | 0.733879 | 350 | 4 | -3.4649 | 0.99162 | 0.99166 | 1 | 21379 | 0 | -3.4649 |
| PPDPF | 4 | 0.0083422 | 0.028476 | 0.858757 | 753 | 4 | -1.9075 | 0.99166 | 0.99161 | 1 | 21380 | 0 | -1.9075 |
| ARMC5 | 5 | 0.022449 | 0.070694 | 0.93914 | 1614 | 4 | -1.4064 | 0.99167 | 0.99171 | 1 | 21381 | 0 | -1.4064 |
| FAM53A | 5 | 0.040734 | 0.11769 | 0.951052 | 2547 | 4 | -1.3132 | 0.99174 | 0.99178 | 1 | 21382 | 0 | -1.3132 |
| NEFH | 6 | 0.077452 | 0.20219 | 0.977973 | 4127 | 3 | -0.85841 | 0.99175 | 0.99172 | 1 | 21383 | 0 | -0.85841 |
| EDC4 | 5 | 0.039758 | 0.1153 | 0.951052 | 2502 | 4 | -1.0739 | 0.99176 | 0.99179 | 1 | 21384 | 0 | -1.0739 |
| LRP2 | 6 | 0.2861 | 0.49057 | 0.999994 | 9834 | 3 | -0.73719 | 0.99177 | 0.99173 | 1 | 21385 | 0 | -0.73719 |
| SKIV2L2 | 4 | 0.0082226 | 0.028088 | 0.855594 | 746 | 4 | -1.7622 | 0.99178 | 0.99174 | 1 | 21386 | 0 | -1.7622 |
| ETV7 | 4 | 0.008182 | 0.02796 | 0.855011 | 741 | 4 | -1.1773 | 0.99182 | 0.9918 | 1 | 21387 | 0 | -1.1773 |
| DEFB108B | 4 | 0.007451 | 0.02564 | 0.845768 | 687 | 4 | -5.3564 | 0.9919 | 0.99187 | 1 | 21388 | 0 | -5.3564 |
| SERPINB7 | 4 | 0.0080982 | 0.027692 | 0.855011 | 738 | 4 | -1.9442 | 0.9919 | 0.99187 | 1 | 21389 | 0 | -1.9442 |
| CENPM | 4 | 0.0052784 | 0.018575 | 0.801598 | 540 | 4 | -5.1086 | 0.99192 | 0.99189 | 1 | 21390 | 0 | -5.1086 |
| SLC12A7 | 4 | 0.0080342 | 0.027478 | 0.854935 | 732 | 4 | -2.0873 | 0.99197 | 0.99193 | 1 | 21391 | 0 | -2.0873 |
| PRF1 | 6 | 0.0052495 | 0.022069 | 0.819701 | 537 | 5 | -2.6762 | 0.99197 | 0.99192 | 1 | 21392 | 0 | -2.6762 |
| CASP6 | 6 | 0.33418 | 0.54604 | 0.999994 | 10817 | 3 | -0.39761 | 0.99198 | 0.99193 | 1 | 21393 | 0 | -0.39761 |
| SELL | 5 | 0.11309 | 0.24676 | 0.98999 | 5408 | 3 | -1.2982 | 0.992 | 0.99202 | 1 | 21394 | 0 | -1.2982 |
| ANKRD20A3 | 2 | 0.0079623 | 0.015151 | 0.757255 | 729 | 2 | -7.3546 | 0.99204 | 0.99196 | 1 | 21395 | 0 | -7.3546 |
| DGUOK | 5 | 0.093672 | 0.21644 | 0.980699 | 4742 | 3 | -1.1636 | 0.99209 | 0.99212 | 1 | 21396 | 0 | -1.1636 |
| SP5 | 5 | 0.0060664 | 0.024615 | 0.843409 | 598 | 4 | -3.2712 | 0.9921 | 0.99212 | 1 | 21397 | 0 | -3.2712 |
| EXOC8 | 5 | 0.0053854 | 0.02233 | 0.819701 | 548 | 4 | -2.5769 | 0.9921 | 0.99212 | 1 | 21398 | 0 | -2.5769 |
| FAM53C | 5 | 0.034913 | 0.10305 | 0.946414 | 2264 | 4 | -1.4326 | 0.99211 | 0.99213 | 1 | 21399 | 0 | -1.4326 |
| NonTargetingControlG | 2 | 0.0078525 | 0.014939 | 0.755841 | 718 | 2 | -5.7373 | 0.99215 | 0.99207 | 1 | 21400 | 0 | -5.7373 |
| NonTargetingControlG | 2 | 0.0077924 | 0.014845 | 0.755578 | 713 | 2 | -7.8827 | 0.99221 | 0.99212 | 1 | 21401 | 0 | -7.8827 |
| ARL6IP6 | 5 | 0.0029791 | 0.012756 | 0.733879 | 354 | 4 | -3.3835 | 0.99226 | 0.99227 | 1 | 21402 | 0 | -3.3835 |
| BEND5 | 6 | 0.11308 | 0.27262 | 0.998561 | 5407 | 4 | -1.0783 | 0.99232 | 0.99228 | 1 | 21403 | 0 | -1.0783 |
| LOC100652824 | 5 | 0.042901 | 0.12308 | 0.951052 | 2674 | 4 | -2.4693 | 0.99233 | 0.99235 | 1 | 21404 | 0 | -2.4693 |
| ATG3 | 5 | 0.00014195 | 0.00065222 | 0.309052 | 46 | 4 | -9.3327 | 0.99236 | 0.99238 | 1 | 21405 | 0 | -9.3327 |
| MAPK4 | 5 | 0.14189 | 0.29069 | 0.999632 | 6236 | 3 | -1.287 | 0.99241 | 0.99243 | 1 | 21406 | 0 | -1.287 |
| ARHGAP40 | 6 | 0.040893 | 0.12393 | 0.951052 | 2556 | 4 | -1.3407 | 0.99241 | 0.99237 | 1 | 21407 | 0 | -1.3407 |
| RCN1 | 6 | 0.034431 | 0.10917 | 0.951052 | 2236 | 4 | -2.1226 | 0.99248 | 0.99244 | 1 | 21408 | 0 | -2.1226 |
| CDK6 | 6 | 0.0017829 | 0.0088889 | 0.680529 | 248 | 4 | -4.2607 | 0.9925 | 0.99245 | 1 | 21409 | 0 | -4.2607 |
| GRXCR2 | 5 | 0.045865 | 0.13045 | 0.95318 | 2799 | 4 | -1.1274 | 0.9925 | 0.99252 | 1 | 21410 | 0 | -1.1274 |
| PPM1A | 4 | 0.0049999 | 0.017629 | 0.795727 | 520 | 4 | -5.2423 | 0.9925 | 0.99248 | 1 | 21411 | 0 | -5.2423 |
| LSM12 | 5 | 0.003549 | 0.015026 | 0.757255 | 410 | 4 | -4.3372 | 0.99251 | 0.99253 | 1 | 21412 | 0 | -4.3372 |
| ANKRD36 | 6 | 0.020358 | 0.073128 | 0.93914 | 1480 | 2 | -0.40646 | 0.9926 | 0.99256 | 1 | 21413 | 0 | -0.40646 |
| hsa-mir-4457 | 3 | 0.0073544 | 0.019963 | 0.808847 | 682 | 3 | -2.2474 | 0.99265 | 0.99274 | 1 | 21414 | 0 | -2.2474 |
| hsa-mir-17 | 4 | 0.0073243 | 0.025216 | 0.84412 | 681 | 4 | -2.0072 | 0.99268 | 0.99264 | 1 | 21415 | 0 | -2.0072 |
| GPR62 | 6 | 0.023031 | 0.081683 | 0.93914 | 1652 | 5 | -1.265 | 0.99271 | 0.99267 | 1 | 21416 | 0 | -1.265 |
| WDR37 | 4 | 0.0072687 | 0.02503 | 0.844082 | 677 | 4 | -4.2445 | 0.99273 | 0.9927 | 1 | 21417 | 0 | -4.2445 |
| POTEI | 3 | 0.0072638 | 0.019739 | 0.808847 | 676 | 3 | -2.5212 | 0.99274 | 0.99283 | 1 | 21418 | 0 | -2.5212 |
| hsa-mir-3116-1 | 4 | 0.0072172 | 0.02487 | 0.844082 | 671 | 4 | -1.4303 | 0.99278 | 0.99275 | 1 | 21419 | 0 | -1.4303 |
| C2orf15 | 6 | 0.27462 | 0.47715 | 0.999994 | 9586 | 3 | -0.60649 | 0.9928 | 0.99277 | 1 | 21420 | 0 | -0.60649 |
| hsa-mir-4683 | 3 | 0.0071593 | 0.019465 | 0.808847 | 668 | 3 | -2.8956 | 0.99284 | 0.99294 | 1 | 21421 | 0 | -2.8956 |
| NDRG3 | 6 | 0.0028127 | 0.01304 | 0.733879 | 341 | 5 | -3.4263 | 0.99294 | 0.99293 | 1 | 21422 | 0 | -3.4263 |
| ZNF664 | 6 | 0.026877 | 0.091459 | 0.94395 | 1850 | 4 | -2.2975 | 0.99302 | 0.99298 | 1 | 21423 | 0 | -2.2975 |

| | | | | | | | | | | | | | |
|---|---|---|---|---|---|---|---|---|---|---|---|---|---|
| TUBB1 | 6 | 0.026728 | 0.091086 | 0.94395 | 1843 | 3 | -1.3928 | 0.99304 | 0.99299 | 1 | 21424 | 0 | -1.3928 |
| GNPTAB | 6 | 0.0018326 | 0.0091294 | 0.680529 | 255 | 5 | -4.4151 | 0.99305 | 0.99301 | 1 | 21425 | 0 | -4.4151 |
| TRRAP | 3 | 0.0069502 | 0.018921 | 0.803792 | 657 | 3 | -3.2234 | 0.99305 | 0.99317 | 1 | 21426 | 0 | -3.2234 |
| SVIL | 6 | 0.084597 | 0.2167 | 0.980699 | 4391 | 4 | -1.5595 | 0.99316 | 0.99312 | 1 | 21427 | 0 | -1.5595 |
| ZNF212 | 5 | 0.045894 | 0.13051 | 0.95318 | 2802 | 4 | -1.1435 | 0.99318 | 0.99317 | 1 | 21428 | 0 | -1.1435 |
| OSBPL10 | 6 | 0.009993 | 0.038982 | 0.891621 | 870 | 4 | -4.1194 | 0.99326 | 0.99321 | 1 | 21429 | 0 | -4.1194 |
| hsa-mir-128-2 | 4 | 0.006688 | 0.023184 | 0.83009 | 641 | 4 | -5.5293 | 0.99331 | 0.99328 | 1 | 21430 | 0 | -5.5293 |
| LRRC24 | 2 | 0.0066354 | 0.012686 | 0.733879 | 636 | 2 | -6.7551 | 0.99336 | 0.99328 | 1 | 21431 | 0 | -6.7551 |
| ATP6V1E2 | 5 | 0.025103 | 0.077732 | 0.93914 | 1758 | 4 | -1.4937 | 0.9934 | 0.99338 | 1 | 21432 | 0 | -1.4937 |
| GLYCTK | 6 | 0.020606 | 0.073922 | 0.93914 | 1488 | 5 | -1.8866 | 0.9934 | 0.99334 | 1 | 21433 | 0 | -1.8866 |
| NUP37 | 6 | 0.0019122 | 0.0095345 | 0.680529 | 262 | 5 | -3.1291 | 0.99345 | 0.99339 | 1 | 21434 | 0 | -3.1291 |
| XRRA1 | 5 | 0.0026655 | 0.011498 | 0.723794 | 328 | 4 | -3.5369 | 0.99347 | 0.99345 | 1 | 21435 | 0 | -3.5369 |
| DUSP14 | 4 | 0.0065107 | 0.022598 | 0.819701 | 627 | 4 | -1.5386 | 0.99349 | 0.99345 | 1 | 21436 | 0 | -1.5386 |
| CBX5 | 6 | 0.0094467 | 0.037074 | 0.881473 | 826 | 4 | -1.4721 | 0.99351 | 0.99345 | 1 | 21437 | 0 | -1.4721 |
| EPS8L2 | 6 | 0.049005 | 0.14191 | 0.959146 | 2930 | 4 | -1.5337 | 0.99351 | 0.99345 | 1 | 21438 | 0 | -1.5337 |
| C7orf60 | 5 | 0.042212 | 0.1214 | 0.951052 | 2626 | 4 | -1.7245 | 0.99361 | 0.99358 | 1 | 21439 | 0 | -1.7245 |
| PARN | 5 | 0.006566 | 0.026143 | 0.848412 | 630 | 4 | -4.6387 | 0.99363 | 0.99361 | 1 | 21440 | 0 | -4.6387 |
| TOX3 | 6 | 0.07454 | 0.1962 | 0.975833 | 4017 | 4 | -1.1088 | 0.99364 | 0.99359 | 1 | 21441 | 0 | -1.1088 |
| C2CD5 | 5 | 0.00078829 | 0.0035752 | 0.553006 | 140 | 4 | -8.4717 | 0.99365 | 0.99363 | 1 | 21442 | 0 | -8.4717 |
| FBXW11 | 5 | 0.053745 | 0.14942 | 0.966183 | 3152 | 4 | -0.76946 | 0.99369 | 0.99368 | 1 | 21443 | 0 | -0.76946 |
| PPP1R2 | 6 | 0.055497 | 0.15601 | 0.967126 | 3220 | 4 | -1.4209 | 0.99373 | 0.99368 | 1 | 21444 | 0 | -1.4209 |
| MCM4 | 4 | 0.0062561 | 0.021771 | 0.819701 | 610 | 4 | -2.8846 | 0.99374 | 0.99371 | 1 | 21445 | 0 | -2.8846 |
| VIMP | 6 | 0.0010471 | 0.0053186 | 0.60466 | 175 | 5 | -5.8486 | 0.99374 | 0.99369 | 1 | 21446 | 0 | -5.8486 |
| NonTargetingControlG | 2 | 0.0062547 | 0.011966 | 0.729405 | 609 | 2 | -8.0386 | 0.99375 | 0.99365 | 1 | 21447 | 0 | -8.0386 |
| STX5 | 6 | 0.090412 | 0.22831 | 0.983354 | 4610 | 3 | -0.5737 | 0.99376 | 0.99371 | 1 | 21448 | 0 | -0.5737 |
| NonTargetingControlG | 2 | 0.0062182 | 0.011895 | 0.729405 | 607 | 2 | -8.5316 | 0.99378 | 0.9937 | 1 | 21449 | 0 | -8.5316 |
| KDM4A | 6 | 0.0078673 | 0.031505 | 0.864134 | 719 | 5 | -1.619 | 0.99379 | 0.99374 | 1 | 21450 | 0 | -1.619 |
| GAL3ST4 | 4 | 0.0061448 | 0.021403 | 0.81457 | 602 | 4 | -4.5633 | 0.99386 | 0.99383 | 1 | 21451 | 0 | -4.5633 |
| SLCO5A1 | 6 | 0.015552 | 0.057549 | 0.929232 | 1218 | 5 | -2.0093 | 0.99389 | 0.99384 | 1 | 21452 | 0 | -2.0093 |
| AQP1 | 6 | 0.004165 | 0.018087 | 0.795727 | 455 | 5 | -4.8019 | 0.99391 | 0.99386 | 1 | 21453 | 0 | -4.8019 |
| NPPA | 5 | 0.0087325 | 0.032414 | 0.865899 | 774 | 4 | -2.8211 | 0.99394 | 0.99392 | 1 | 21454 | 0 | -2.8211 |
| SIGMAR1 | 5 | 0.0032343 | 0.013773 | 0.751291 | 377 | 4 | -7.3721 | 0.99398 | 0.99396 | 1 | 21455 | 0 | -7.3721 |
| MFSD9 | 3 | 0.0060093 | 0.016452 | 0.772095 | 593 | 3 | -2.782 | 0.99399 | 0.99409 | 1 | 21456 | 0 | -2.782 |
| XPO5 | 5 | 0.00016161 | 0.00076561 | 0.325083 | 49 | 4 | -8.0951 | 0.994 | 0.99399 | 1 | 21457 | 0 | -8.0951 |
| MESP1 | 4 | 0.0059587 | 0.020792 | 0.81027 | 590 | 4 | -2.1692 | 0.99404 | 0.99402 | 1 | 21458 | 0 | -2.1692 |
| F8 | 4 | 0.0059256 | 0.020682 | 0.81027 | 586 | 4 | -2.9982 | 0.99407 | 0.99406 | 1 | 21459 | 0 | -2.9982 |
| TREM2 | 6 | 0.034222 | 0.10866 | 0.951052 | 2225 | 4 | -2.0033 | 0.99408 | 0.99405 | 1 | 21460 | 0 | -2.0033 |
| PPIAL4A | 6 | 0.016777 | 0.06156 | 0.937033 | 1293 | 5 | -2.8007 | 0.99409 | 0.99405 | 1 | 21461 | 0 | -2.8007 |
| hsa-mir-5692b | 4 | 0.0059063 | 0.020626 | 0.81027 | 585 | 4 | -1.7346 | 0.99409 | 0.99408 | 1 | 21462 | 0 | -1.7346 |
| E2F3 | 4 | 0.005883 | 0.020549 | 0.81027 | 582 | 4 | -5.3726 | 0.99412 | 0.9941 | 1 | 21463 | 0 | -5.3726 |
| WDR78 | 5 | 0.028426 | 0.086431 | 0.94395 | 1921 | 4 | -1.2773 | 0.99417 | 0.99415 | 1 | 21464 | 0 | -1.2773 |
| TMEM110 | 6 | 0.00036304 | 0.0019294 | 0.439257 | 87 | 5 | -3.9474 | 0.99422 | 0.99419 | 1 | 21465 | 0 | -3.9474 |
| ZP2 | 6 | 0.0040614 | 0.017689 | 0.795727 | 446 | 5 | -4.3121 | 0.99422 | 0.9942 | 1 | 21466 | 0 | -4.3121 |
| hsa-mir-550a-1 | 4 | 0.0057213 | 0.020027 | 0.808847 | 573 | 4 | -2.3532 | 0.99428 | 0.99426 | 1 | 21467 | 0 | -2.3532 |
| SHQ1 | 6 | 0.015683 | 0.058001 | 0.929685 | 1228 | 4 | -2.375 | 0.99429 | 0.99427 | 1 | 21468 | 0 | -2.375 |
| PTGES | 3 | 0.0056952 | 0.015617 | 0.766139 | 570 | 3 | -2.315 | 0.9943 | 0.99439 | 1 | 21469 | 0 | -2.315 |
| ABHD14B | 6 | 0.0075566 | 0.030407 | 0.862223 | 695 | 5 | -2.9116 | 0.99437 | 0.99435 | 1 | 21470 | 0 | -2.9116 |
| SAT1 | 5 | 0.0055688 | 0.023033 | 0.827176 | 562 | 5 | -1.1134 | 0.99443 | 0.99441 | 1 | 21471 | 0 | -1.1134 |
| RUSC1 | 4 | 0.005533 | 0.019407 | 0.808616 | 559 | 4 | -1.6977 | 0.99447 | 0.99444 | 1 | 21472 | 0 | -1.6977 |
| SEMA3A | 6 | 0.30285 | 0.51005 | 0.999994 | 10191 | 3 | -0.54719 | 0.99448 | 0.99445 | 1 | 21473 | 0 | -0.54719 |
| CCDC67 | 6 | 0.23859 | 0.4343 | 0.999994 | 8736 | 3 | -0.55661 | 0.99448 | 0.99445 | 1 | 21474 | 0 | -0.55661 |
| TADA3 | 6 | 0.027535 | 0.093024 | 0.943968 | 1878 | 4 | -1.619 | 0.99456 | 0.99454 | 1 | 21475 | 0 | -1.619 |

| | | | | | | | | | | | | | |
|---|---|---|---|---|---|---|---|---|---|---|---|---|---|
| PTK2 | 5 | 0.0027673 | 0.011909 | 0.729405 | 335 | 5 | -5.2221 | 0.99464 | 0.99462 | 1 | 21476 | 0 | -5.2221 |
| HEMK1 | 6 | 0.14247 | 0.3162 | 0.999994 | 6251 | 3 | -1.079 | 0.99476 | 0.99473 | 1 | 21477 | 0 | -1.079 |
| NFYB | 6 | 0.0060721 | 0.025071 | 0.844082 | 599 | 5 | -4.5634 | 0.99482 | 0.99479 | 1 | 21478 | 0 | -4.5634 |
| MEA1 | 6 | 0.034526 | 0.10939 | 0.951052 | 2242 | 3 | -1.2336 | 0.99487 | 0.99485 | 1 | 21479 | 0 | -1.2336 |
| UBASH3A | 6 | 0.067868 | 0.18229 | 0.968526 | 3746 | 4 | -1.3753 | 0.99495 | 0.99493 | 1 | 21480 | 0 | -1.3753 |
| PRRT4 | 6 | 0.011563 | 0.044274 | 0.891621 | 959 | 5 | -1.8111 | 0.99502 | 0.99501 | 1 | 21481 | 0 | -1.8111 |
| GTF2A1 | 6 | 0.0084395 | 0.033468 | 0.867348 | 757 | 5 | -2.3761 | 0.99507 | 0.99505 | 1 | 21482 | 0 | -2.3761 |
| C7orf66 | 4 | 0.0048916 | 0.017274 | 0.785648 | 512 | 4 | -1.5997 | 0.99511 | 0.9951 | 1 | 21483 | 0 | -1.5997 |
| BAZ1B | 6 | 0.0013996 | 0.0070006 | 0.645102 | 211 | 5 | -3.9961 | 0.99512 | 0.9951 | 1 | 21484 | 0 | -3.9961 |
| OR2G6 | 5 | 0.0047375 | 0.019771 | 0.808847 | 501 | 5 | -1.2299 | 0.99526 | 0.99524 | 1 | 21485 | 0 | -1.2299 |
| hsa-mir-8064 | 4 | 0.0046523 | 0.016444 | 0.772095 | 493 | 4 | -3.4536 | 0.99535 | 0.99535 | 1 | 21486 | 0 | -3.4536 |
| STIM1 | 5 | 0.0046476 | 0.019406 | 0.808616 | 491 | 5 | -2.1956 | 0.99535 | 0.99533 | 1 | 21487 | 0 | -2.1956 |
| CNOT10 | 3 | 0.0046475 | 0.012826 | 0.733879 | 490 | 3 | -3.2157 | 0.99535 | 0.99546 | 1 | 21488 | 0 | -3.2157 |
| EPGN | 6 | 0.0080344 | 0.032079 | 0.865899 | 733 | 5 | -2.1943 | 0.99535 | 0.99535 | 1 | 21489 | 0 | -2.1943 |
| PSMB5 | 6 | 0.00086929 | 0.0044526 | 0.56977 | 147 | 5 | -4.8107 | 0.99536 | 0.99535 | 1 | 21490 | 0 | -4.8107 |
| TREH | 4 | 0.0023403 | 0.0084541 | 0.680529 | 308 | 4 | -6.6211 | 0.99539 | 0.99539 | 1 | 21491 | 0 | -6.6211 |
| TUBGCP2 | 6 | 0.0022928 | 0.011066 | 0.71022 | 305 | 4 | -3.8379 | 0.9954 | 0.99539 | 1 | 21492 | 0 | -3.8379 |
| POLD2 | 2 | 0.0045886 | 0.008797 | 0.680529 | 485 | 2 | -7.0265 | 0.99541 | 0.99533 | 1 | 21493 | 0 | -7.0265 |
| NDUFB10 | 3 | 0.0045769 | 0.012653 | 0.733879 | 484 | 3 | -3.3626 | 0.99542 | 0.99553 | 1 | 21494 | 0 | -3.3626 |
| NME9 | 6 | 0.022711 | 0.080657 | 0.93914 | 1632 | 5 | -1.6026 | 0.99543 | 0.99542 | 1 | 21495 | 0 | -1.6026 |
| hsa-mir-7109 | 3 | 0.0045249 | 0.0125 | 0.733879 | 483 | 3 | -2.9707 | 0.99548 | 0.99558 | 1 | 21496 | 0 | -2.9707 |
| C1GALT1 | 5 | 0.0044982 | 0.01882 | 0.803792 | 482 | 5 | -1.0809 | 0.9955 | 0.99547 | 1 | 21497 | 0 | -1.0809 |
| TRPS1 | 6 | 0.009966 | 0.03889 | 0.891621 | 867 | 5 | -1.8502 | 0.99551 | 0.9955 | 1 | 21498 | 0 | -1.8502 |
| PDE7A | 5 | 0.0044677 | 0.018693 | 0.801598 | 480 | 5 | -1.543 | 0.99553 | 0.9955 | 1 | 21499 | 0 | -1.543 |
| CCM2 | 4 | 0.0044351 | 0.015695 | 0.766139 | 475 | 4 | -3.7105 | 0.99556 | 0.99557 | 1 | 21500 | 0 | -3.7105 |
| CCDC7 | 5 | 0.0043667 | 0.018299 | 0.795727 | 470 | 5 | -1.3944 | 0.99563 | 0.9956 | 1 | 21501 | 0 | -1.3944 |
| ASPRV1 | 6 | 0.00058245 | 0.0030508 | 0.537109 | 117 | 5 | -7.4666 | 0.99569 | 0.99568 | 1 | 21502 | 0 | -7.4666 |
| C2orf83 | 6 | 0.021418 | 0.076559 | 0.93914 | 1542 | 5 | -0.86208 | 0.99573 | 0.99571 | 1 | 21503 | 0 | -0.86208 |
| SLC39A9 | 6 | 0.00041743 | 0.0022049 | 0.45324 | 98 | 5 | -6.4643 | 0.99584 | 0.99584 | 1 | 21504 | 0 | -6.4643 |
| URGCP-MRPS24 | 3 | 0.00018299 | 0.00053746 | 0.285361 | 53 | 3 | -10.523 | 0.99584 | 0.99592 | 1 | 21505 | 0 | -10.523 |
| NDUFS4 | 4 | 0.0041229 | 0.014625 | 0.755578 | 451 | 4 | -2.7078 | 0.99588 | 0.99589 | 1 | 21506 | 0 | -2.7078 |
| GOLGA4 | 4 | 0.004057 | 0.014401 | 0.753239 | 445 | 4 | -1.6914 | 0.99594 | 0.99595 | 1 | 21507 | 0 | -1.6914 |
| CT45A4 | 2 | 0.0039983 | 0.0076842 | 0.672529 | 442 | 2 | -8.2455 | 0.996 | 0.99595 | 1 | 21508 | 0 | -8.2455 |
| GRHPR | 5 | 0.0023542 | 0.010272 | 0.695096 | 310 | 5 | -2.4563 | 0.99612 | 0.99609 | 1 | 21509 | 0 | -2.4563 |
| SFRP4 | 6 | 0.0025458 | 0.012026 | 0.729498 | 320 | 5 | -2.4895 | 0.99615 | 0.99614 | 1 | 21510 | 0 | -2.4895 |
| KLHL41 | 6 | 0.0041833 | 0.018146 | 0.795727 | 456 | 5 | -2.393 | 0.99617 | 0.99616 | 1 | 21511 | 0 | -2.393 |
| FMOD | 6 | 0.083074 | 0.2136 | 0.979587 | 4341 | 3 | -1.4885 | 0.99625 | 0.99624 | 1 | 21512 | 0 | -1.4885 |
| SMS | 6 | 0.0066045 | 0.027026 | 0.848412 | 634 | 5 | -4.9337 | 0.99627 | 0.99626 | 1 | 21513 | 0 | -4.9337 |
| CARS2 | 6 | 0.021477 | 0.07674 | 0.93914 | 1546 | 5 | -0.89235 | 0.99628 | 0.99626 | 1 | 21514 | 0 | -0.89235 |
| hsa-mir-3180-4 | 3 | 0.0037042 | 0.010241 | 0.695096 | 422 | 3 | -3.6086 | 0.9963 | 0.99636 | 1 | 21515 | 0 | -3.6086 |
| AASS | 4 | 0.0037039 | 0.013212 | 0.733879 | 421 | 4 | -2.5423 | 0.9963 | 0.9963 | 1 | 21516 | 0 | -2.5423 |
| ZNF20 | 6 | 0.00012837 | 0.00069931 | 0.309052 | 43 | 5 | -6.1921 | 0.99633 | 0.99632 | 1 | 21517 | 0 | -6.1921 |
| CNOT11 | 4 | 0.0036484 | 0.013024 | 0.733879 | 414 | 4 | -2.9428 | 0.99635 | 0.99636 | 1 | 21518 | 0 | -2.9428 |
| NFE2L3 | 6 | 0.011917 | 0.045432 | 0.891621 | 986 | 5 | -1.3895 | 0.99635 | 0.99634 | 1 | 21519 | 0 | -1.3895 |
| FEV | 6 | 0.005255 | 0.022097 | 0.819701 | 538 | 5 | -1.8279 | 0.9964 | 0.99638 | 1 | 21520 | 0 | -1.8279 |
| hsa-mir-6872 | 4 | 0.0034583 | 0.012359 | 0.733879 | 401 | 4 | -3.7481 | 0.99654 | 0.99653 | 1 | 21521 | 0 | -3.7481 |
| DARS2 | 5 | 0.0034227 | 0.014534 | 0.754766 | 398 | 5 | -3.1453 | 0.99658 | 0.99654 | 1 | 21522 | 0 | -3.1453 |
| C11orf35 | 6 | 0.021396 | 0.076484 | 0.93914 | 1541 | 5 | -1.6024 | 0.99658 | 0.99655 | 1 | 21523 | 0 | -1.6024 |
| HUNK | 5 | 0.0033942 | 0.014421 | 0.753239 | 394 | 5 | -2.1605 | 0.99661 | 0.99658 | 1 | 21524 | 0 | -2.1605 |
| SOX12 | 6 | 0.0078823 | 0.031564 | 0.864134 | 721 | 5 | -4.1492 | 0.99662 | 0.9966 | 1 | 21525 | 0 | -4.1492 |
| CCDC101 | 5 | 0.0033141 | 0.014089 | 0.75278 | 387 | 5 | -3.7127 | 0.99669 | 0.99665 | 1 | 21526 | 0 | -3.7127 |
| SLC35F2 | 5 | 0.0033029 | 0.014041 | 0.75278 | 386 | 5 | -1.9765 | 0.9967 | 0.99667 | 1 | 21527 | 0 | -1.9765 |

| | | | | | | | | | | | | | |
|---|---|---|---|---|---|---|---|---|---|---|---|---|---|
| PRIM2 | 3 | 0.0032725 | 0.0090595 | 0.680529 | 381 | 3 | -6.5105 | 0.99673 | 0.99678 | 1 | 21528 | 0 | -6.5105 |
| NCKAP1 | 5 | 0.0032621 | 0.013895 | 0.75278 | 379 | 5 | -2.3895 | 0.99674 | 0.99672 | 1 | 21529 | 0 | -2.3895 |
| IRAK1 | 4 | 0.003205 | 0.011455 | 0.72322 | 375 | 4 | -4.7846 | 0.9968 | 0.99679 | 1 | 21530 | 0 | -4.7846 |
| GATAD2A | 5 | 0.0031196 | 0.013304 | 0.736813 | 369 | 5 | -3.1154 | 0.99688 | 0.99686 | 1 | 21531 | 0 | -3.1154 |
| PCK1 | 4 | 0.0030991 | 0.011085 | 0.71022 | 368 | 4 | -3.0454 | 0.9969 | 0.99689 | 1 | 21532 | 0 | -3.0454 |
| MARCH3 | 5 | 0.0030881 | 0.013182 | 0.733879 | 366 | 5 | -1.6244 | 0.99691 | 0.99689 | 1 | 21533 | 0 | -1.6244 |
| USP48 | 6 | 0.023398 | 0.082848 | 0.943262 | 1678 | 5 | -1.1472 | 0.99691 | 0.99691 | 1 | 21534 | 0 | -1.1472 |
| RNF139 | 4 | 0.0030813 | 0.01102 | 0.71022 | 364 | 4 | -4.2949 | 0.99692 | 0.99691 | 1 | 21535 | 0 | -4.2949 |
| ARTN | 4 | 0.0029594 | 0.0106 | 0.706752 | 351 | 4 | -4.0879 | 0.99704 | 0.99703 | 1 | 21536 | 0 | -4.0879 |
| VWDE | 6 | 0.02078 | 0.074489 | 0.93914 | 1496 | 5 | -1.6499 | 0.99714 | 0.99713 | 1 | 21537 | 0 | -1.6499 |
| SLC30A1 | 5 | 0.00077755 | 0.0035281 | 0.553006 | 138 | 5 | -6.0959 | 0.99716 | 0.99715 | 1 | 21538 | 0 | -6.0959 |
| hsa-mir-381 | 4 | 0.0027762 | 0.0099588 | 0.680529 | 338 | 4 | -2.8521 | 0.99722 | 0.99722 | 1 | 21539 | 0 | -2.8521 |
| PM20D1 | 6 | 0.022436 | 0.079778 | 0.93914 | 1611 | 4 | -0.82046 | 0.99725 | 0.99723 | 1 | 21540 | 0 | -0.82046 |
| TPR | 3 | 0.0026867 | 0.0074643 | 0.665213 | 330 | 3 | -7.3888 | 0.99731 | 0.99732 | 1 | 21541 | 0 | -7.3888 |
| MYCT1 | 6 | 0.076062 | 0.19937 | 0.977164 | 4074 | 4 | -1.1264 | 0.99735 | 0.99733 | 1 | 21542 | 0 | -1.1264 |
| PKD1L1 | 6 | 0.046307 | 0.13597 | 0.95468 | 2821 | 4 | -1.5716 | 0.99737 | 0.99735 | 1 | 21543 | 0 | -1.5716 |
| KRTAP19-7 | 4 | 0.0025991 | 0.009342 | 0.680529 | 323 | 4 | -2.3781 | 0.9974 | 0.9974 | 1 | 21544 | 0 | -2.3781 |
| FAM134B | 6 | 0.010137 | 0.039488 | 0.891621 | 880 | 5 | -1.4922 | 0.99745 | 0.99743 | 1 | 21545 | 0 | -1.4922 |
| JMJD6 | 6 | 0.00497 | 0.02104 | 0.812607 | 515 | 5 | -2.2989 | 0.99748 | 0.99748 | 1 | 21546 | 0 | -2.2989 |
| C10orf76 | 6 | 0.015219 | 0.056435 | 0.9235 | 1204 | 4 | -2.7185 | 0.99754 | 0.99754 | 1 | 21547 | 0 | -2.7185 |
| XKRX | 6 | 0.0074632 | 0.030042 | 0.862223 | 689 | 5 | -1.9438 | 0.99755 | 0.99754 | 1 | 21548 | 0 | -1.9438 |
| CALCRL | 6 | 0.0095097 | 0.037315 | 0.881473 | 834 | 5 | -2.238 | 0.99755 | 0.99755 | 1 | 21549 | 0 | -2.238 |
| DCPS | 6 | 0.0049796 | 0.021076 | 0.812607 | 516 | 5 | -2.1916 | 0.99756 | 0.99756 | 1 | 21550 | 0 | -2.1916 |
| LDLR | 6 | 0.0044647 | 0.019185 | 0.805137 | 479 | 5 | -3.3074 | 0.99765 | 0.99765 | 1 | 21551 | 0 | -3.3074 |
| POM121L12 | 5 | 0.00228 | 0.0099556 | 0.680529 | 302 | 5 | -2.4911 | 0.99772 | 0.99771 | 1 | 21552 | 0 | -2.4911 |
| HNMT | 6 | 0.010672 | 0.0413 | 0.891621 | 909 | 5 | -2.2304 | 0.99774 | 0.99772 | 1 | 21553 | 0 | -2.2304 |
| PDE5A | 4 | 0.0022477 | 0.0081391 | 0.680529 | 297 | 4 | -2.4992 | 0.99775 | 0.99777 | 1 | 21554 | 0 | -2.4992 |
| SP140L | 5 | 0.0022052 | 0.0096351 | 0.680529 | 292 | 5 | -2.2397 | 0.99779 | 0.9978 | 1 | 21555 | 0 | -2.2397 |
| NVL | 5 | 0.002182 | 0.0095391 | 0.680529 | 289 | 5 | -5.964 | 0.99782 | 0.99782 | 1 | 21556 | 0 | -5.964 |
| C1orf234 | 5 | 0.0021795 | 0.0095277 | 0.680529 | 287 | 5 | -1.88 | 0.99782 | 0.99783 | 1 | 21557 | 0 | -1.88 |
| MFF | 6 | 0.021985 | 0.078326 | 0.93914 | 1582 | 5 | -1.4866 | 0.99783 | 0.99782 | 1 | 21558 | 0 | -1.4866 |
| PDK1 | 6 | 0.027061 | 0.091906 | 0.943968 | 1860 | 4 | -1.6325 | 0.99786 | 0.99784 | 1 | 21559 | 0 | -1.6325 |
| ORC6 | 5 | 0.0021359 | 0.0093425 | 0.680529 | 283 | 5 | -6.0232 | 0.99786 | 0.99788 | 1 | 21560 | 0 | -6.0232 |
| HEPN1 | 6 | 0.014341 | 0.053581 | 0.919407 | 1157 | 5 | -3.5626 | 0.99793 | 0.9979 | 1 | 21561 | 0 | -3.5626 |
| SPDYE2L | 2 | 0.0020371 | 0.0039469 | 0.560963 | 278 | 2 | -8.6597 | 0.99796 | 0.99792 | 1 | 21562 | 0 | -8.6597 |
| EIF3D | 3 | 0.0020088 | 0.0056372 | 0.623123 | 276 | 3 | -5.7203 | 0.99799 | 0.99801 | 1 | 21563 | 0 | -5.7203 |
| NELFA | 5 | 0.001976 | 0.0086594 | 0.680529 | 272 | 5 | -5.5887 | 0.99802 | 0.99804 | 1 | 21564 | 0 | -5.5887 |
| LOC100132247 | 5 | 0.0019616 | 0.0085995 | 0.680529 | 271 | 5 | -1.6587 | 0.99804 | 0.99805 | 1 | 21565 | 0 | -1.6587 |
| ARL16 | 5 | 0.0019459 | 0.0085351 | 0.680529 | 265 | 5 | -1.7621 | 0.99805 | 0.99807 | 1 | 21566 | 0 | -1.7621 |
| BCL2L11 | 6 | 0.0019293 | 0.0096155 | 0.680529 | 263 | 6 | -1.8208 | 0.99807 | 0.99805 | 1 | 21567 | 0 | -1.8208 |
| hsa-mir-8072 | 4 | 0.00021674 | 0.00082504 | 0.343583 | 60 | 4 | -8.748 | 0.9981 | 0.99812 | 1 | 21568 | 0 | -8.748 |
| SLC10A1 | 6 | 0.0018912 | 0.009433 | 0.680529 | 260 | 6 | -3.5499 | 0.99811 | 0.99808 | 1 | 21569 | 0 | -3.5499 |
| LRRC72 | 6 | 0.0018782 | 0.0093686 | 0.680529 | 258 | 6 | -2.6545 | 0.99812 | 0.9981 | 1 | 21570 | 0 | -2.6545 |
| SMC6 | 6 | 0.0017954 | 0.0089479 | 0.680529 | 250 | 6 | -5.397 | 0.9982 | 0.99818 | 1 | 21571 | 0 | -5.397 |
| ENPP3 | 6 | 0.0017538 | 0.0087445 | 0.680529 | 244 | 6 | -2.0607 | 0.99825 | 0.99822 | 1 | 21572 | 0 | -2.0607 |
| ELL3 | 6 | 0.0017266 | 0.008605 | 0.680529 | 241 | 6 | -1.7146 | 0.99827 | 0.99825 | 1 | 21573 | 0 | -1.7146 |
| SLC6A8 | 3 | 0.0015913 | 0.0044915 | 0.56977 | 234 | 3 | -4.6381 | 0.99841 | 0.99842 | 1 | 21574 | 0 | -4.6381 |
| KLHDC8B | 4 | 0.0015911 | 0.0058142 | 0.623123 | 233 | 4 | -3.2207 | 0.99841 | 0.99842 | 1 | 21575 | 0 | -3.2207 |
| ARID4A | 4 | 0.0015597 | 0.0057049 | 0.623123 | 228 | 4 | -2.508 | 0.99844 | 0.99845 | 1 | 21576 | 0 | -2.508 |
| GBAS | 5 | 0.0015467 | 0.0068525 | 0.643671 | 227 | 5 | -2.4454 | 0.99845 | 0.99848 | 1 | 21577 | 0 | -2.4454 |
| SLC37A1 | 5 | 0.0015101 | 0.006692 | 0.643671 | 224 | 5 | -3.6153 | 0.99849 | 0.99852 | 1 | 21578 | 0 | -3.6153 |
| PLAA | 6 | 0.0014843 | 0.0074163 | 0.665213 | 222 | 6 | -1.1099 | 0.99852 | 0.9985 | 1 | 21579 | 0 | -1.1099 |

| | | | | | | | | | | | | | |
|---|---|---|---|---|---|---|---|---|---|---|---|---|---|
| C17orf97 | 5 | 0.0014762 | 0.006543 | 0.643671 | 220 | 5 | -2.0453 | 0.99852 | 0.99854 | 1 | 21580 | 0 | -2.0453 |
| LGMN | 6 | 0.0014633 | 0.0073061 | 0.661979 | 219 | 6 | -1.9477 | 0.99854 | 0.99852 | 1 | 21581 | 0 | -1.9477 |
| PSMA5 | 3 | 0.0014327 | 0.0040301 | 0.560963 | 213 | 3 | -6.2382 | 0.99857 | 0.99858 | 1 | 21582 | 0 | -6.2382 |
| PPM1M | 6 | 0.0013834 | 0.0069229 | 0.643671 | 209 | 6 | -2.8988 | 0.99862 | 0.9986 | 1 | 21583 | 0 | -2.8988 |
| ALDH1B1 | 6 | 0.0013652 | 0.0068315 | 0.643671 | 206 | 6 | -2.5459 | 0.99863 | 0.99862 | 1 | 21584 | 0 | -2.5459 |
| hsa-mir-3193 | 3 | 0.0013634 | 0.0038509 | 0.560963 | 205 | 3 | -5.9844 | 0.99864 | 0.99865 | 1 | 21585 | 0 | -5.9844 |
| SOX9 | 5 | 0.0013348 | 0.0059349 | 0.623123 | 198 | 5 | -2.4859 | 0.99867 | 0.99868 | 1 | 21586 | 0 | -2.4859 |
| GMNN | 5 | 0.0013296 | 0.0059134 | 0.623123 | 197 | 5 | -3.0718 | 0.99867 | 0.99869 | 1 | 21587 | 0 | -3.0718 |
| hsa-mir-4706 | 3 | 0.001329 | 0.0037595 | 0.560963 | 196 | 3 | -7.5736 | 0.99867 | 0.99868 | 1 | 21588 | 0 | -7.5736 |
| TBRG4 | 5 | 0.00029638 | 0.0013966 | 0.384267 | 79 | 5 | -8.392 | 0.99876 | 0.99878 | 1 | 21589 | 0 | -8.392 |
| SENP3 | 6 | 0.00010867 | 0.00060055 | 0.302318 | 37 | 6 | -3.7177 | 0.99883 | 0.99882 | 1 | 21590 | 0 | -3.7177 |
| CCT4 | 4 | 0.0011437 | 0.0041979 | 0.564633 | 183 | 4 | -5.6925 | 0.99886 | 0.99885 | 1 | 21591 | 0 | -5.6925 |
| M1AP | 4 | 0.0011184 | 0.0041115 | 0.560963 | 181 | 4 | -3.8114 | 0.99888 | 0.99887 | 1 | 21592 | 0 | -3.8114 |
| RANBP3 | 5 | 0.0011026 | 0.0049199 | 0.589484 | 179 | 5 | -3.8238 | 0.9989 | 0.99891 | 1 | 21593 | 0 | -3.8238 |
| LHX9 | 6 | 0.00014047 | 0.00076515 | 0.325083 | 45 | 6 | -5.3743 | 0.99895 | 0.99894 | 1 | 21594 | 0 | -5.3743 |
| METTL6 | 6 | 0.0010115 | 0.0051357 | 0.594721 | 171 | 6 | -2.1797 | 0.99899 | 0.99897 | 1 | 21595 | 0 | -2.1797 |
| TLR9 | 5 | 0.00099891 | 0.0044562 | 0.56977 | 170 | 5 | -4.7065 | 0.999 | 0.99902 | 1 | 21596 | 0 | -4.7065 |
| TMEM260 | 6 | 0.00097874 | 0.0049752 | 0.589484 | 160 | 6 | -3.0233 | 0.99902 | 0.999 | 1 | 21597 | 0 | -3.0233 |
| USP5 | 5 | 0.00097514 | 0.0043666 | 0.56977 | 159 | 5 | -6.0424 | 0.99902 | 0.99904 | 1 | 21598 | 0 | -6.0424 |
| COASY | 5 | 0.0009567 | 0.0042871 | 0.56955 | 158 | 5 | -2.2157 | 0.99904 | 0.99906 | 1 | 21599 | 0 | -2.2157 |
| DEPDC5 | 5 | 0.00092066 | 0.0041188 | 0.560963 | 154 | 5 | -5.9481 | 0.99908 | 0.9991 | 1 | 21600 | 0 | -5.9481 |
| TMIGD2 | 6 | 0.00091483 | 0.0046826 | 0.586133 | 152 | 6 | -3.8129 | 0.99909 | 0.99906 | 1 | 21601 | 0 | -3.8129 |
| URGCP | 5 | 0.00089118 | 0.0039958 | 0.560963 | 151 | 5 | -2.6305 | 0.99911 | 0.99913 | 1 | 21602 | 0 | -2.6305 |
| SMARCAL1 | 6 | 0.00087041 | 0.0044581 | 0.56977 | 148 | 6 | -4.111 | 0.99913 | 0.99911 | 1 | 21603 | 0 | -4.111 |
| RAB3IL1 | 4 | 0.00084211 | 0.0031093 | 0.537554 | 146 | 4 | -5.7026 | 0.99916 | 0.99915 | 1 | 21604 | 0 | -5.7026 |
| C4orf21 | 5 | 0.00082287 | 0.0037224 | 0.560963 | 145 | 5 | -2.2492 | 0.99918 | 0.9992 | 1 | 21605 | 0 | -2.2492 |
| C19orf43 | 6 | 0.00022367 | 0.0011899 | 0.368105 | 63 | 6 | -4.8288 | 0.99921 | 0.99919 | 1 | 21606 | 0 | -4.8288 |
| ACY1 | 4 | 0.00048214 | 0.0018122 | 0.435802 | 106 | 4 | -6.8927 | 0.99921 | 0.99921 | 1 | 21607 | 0 | -6.8927 |
| SNRNP25 | 5 | 0.00071196 | 0.0032542 | 0.541338 | 132 | 5 | -6.7135 | 0.99924 | 0.99924 | 1 | 21608 | 0 | -6.7135 |
| DNAJB13 | 6 | 0.00075314 | 0.0038728 | 0.560963 | 134 | 6 | -1.2326 | 0.99925 | 0.99923 | 1 | 21609 | 0 | -1.2326 |
| RTN4R | 6 | 0.00017956 | 0.00097684 | 0.352558 | 52 | 6 | -5.7228 | 0.99927 | 0.99924 | 1 | 21610 | 0 | -5.7228 |
| TMEM39B | 5 | 0.00069276 | 0.0031774 | 0.537554 | 130 | 5 | -6.7395 | 0.99931 | 0.99931 | 1 | 21611 | 0 | -6.7395 |
| NFS1 | 4 | 0.00067658 | 0.0025213 | 0.483177 | 128 | 4 | -7.6914 | 0.99932 | 0.99933 | 1 | 21612 | 0 | -7.6914 |
| PPIH | 6 | 0.00065989 | 0.0034165 | 0.541338 | 126 | 6 | -1.9908 | 0.99934 | 0.99931 | 1 | 21613 | 0 | -1.9908 |
| NRXN1 | 6 | 0.00065447 | 0.0033882 | 0.541338 | 124 | 6 | -2.1302 | 0.99935 | 0.99932 | 1 | 21614 | 0 | -2.1302 |
| EPDR1 | 6 | 0.00064028 | 0.0033265 | 0.541338 | 122 | 6 | -2.3858 | 0.99936 | 0.99933 | 1 | 21615 | 0 | -2.3858 |
| SLC25A20 | 5 | 0.00061685 | 0.0028203 | 0.517578 | 121 | 5 | -4.0532 | 0.99938 | 0.99938 | 1 | 21616 | 0 | -4.0532 |
| TMEM150C | 5 | 0.00060608 | 0.0027737 | 0.517578 | 119 | 5 | -2.5324 | 0.99939 | 0.99939 | 1 | 21617 | 0 | -2.5324 |
| hsa-mir-320a | 3 | 0.00050913 | 0.0014784 | 0.385721 | 113 | 3 | -8.9627 | 0.99949 | 0.99949 | 1 | 21618 | 0 | -8.9627 |
| ATP6V0D1 | 2 | 0.00050397 | 0.00095215 | 0.352558 | 111 | 2 | -9.7657 | 0.9995 | 0.99948 | 1 | 21619 | 0 | -9.7657 |
| WDHD1 | 6 | 0.0004602 | 0.0024139 | 0.466717 | 105 | 6 | -4.4883 | 0.99954 | 0.99952 | 1 | 21620 | 0 | -4.4883 |
| NonTargetingControlG | 1 | 0.00044786 | 0.0004515 | 0.27159 | 104 | 1 | -12.624 | 0.99955 | 0.99958 | 1 | 21621 | 0 | -12.624 |
| MANEAL | 6 | 0.00041405 | 0.002188 | 0.45324 | 97 | 6 | -2.2003 | 0.99959 | 0.99957 | 1 | 21622 | 0 | -2.2003 |
| GPC4 | 4 | 0.00041004 | 0.0015438 | 0.388244 | 96 | 4 | -6.8041 | 0.99959 | 0.9996 | 1 | 21623 | 0 | -6.8041 |
| SUMO2 | 6 | 0.00040428 | 0.0021345 | 0.448765 | 95 | 6 | -2.9633 | 0.9996 | 0.99958 | 1 | 21624 | 0 | -2.9633 |
| SHH | 6 | 0.00039125 | 0.0020728 | 0.44006 | 94 | 6 | -4.7478 | 0.99961 | 0.99959 | 1 | 21625 | 0 | -4.7478 |
| hsa-mir-3655 | 4 | 0.0003824 | 0.0014437 | 0.384267 | 92 | 4 | -5.7007 | 0.99962 | 0.99963 | 1 | 21626 | 0 | -5.7007 |
| SMIM10 | 6 | 0.00038073 | 0.0020216 | 0.439257 | 91 | 6 | -3.0903 | 0.99962 | 0.9996 | 1 | 21627 | 0 | -3.0903 |
| SHB | 6 | 0.00034605 | 0.0018401 | 0.435802 | 83 | 6 | -3.3874 | 0.99965 | 0.99963 | 1 | 21628 | 0 | -3.3874 |
| SENP1 | 6 | 0.000059043 | 0.00031662 | 0.207771 | 30 | 6 | -4.6602 | 0.99967 | 0.99965 | 1 | 21629 | 0 | -4.6602 |
| HNRNPF | 6 | 0.00029195 | 0.0015598 | 0.388244 | 78 | 6 | -3.2549 | 0.99971 | 0.99969 | 1 | 21630 | 0 | -3.2549 |
| TAF3 | 5 | 0.00022789 | 0.0010797 | 0.352868 | 64 | 5 | -9.1332 | 0.99972 | 0.99972 | 1 | 21631 | 0 | -9.1332 |

| | | | | | | | | | | | | | |
|---|---|---|---|---|---|---|---|---|---|---|---|---|---|
| DHX36 | 4 | 0.00025621 | 0.00096998 | 0.352558 | 71 | 4 | -6.7887 | 0.99974 | 0.99975 | 1 | 21632 | 0 | -6.7887 |
| hsa-mir-943 | 4 | 0.0002291 | 0.00087671 | 0.352558 | 65 | 4 | -5.2147 | 0.99977 | 0.99977 | 1 | 21633 | 0 | -5.2147 |
| CCDC115 | 5 | 0.00020589 | 0.00097227 | 0.352558 | 58 | 5 | -3.2518 | 0.99979 | 0.9998 | 1 | 21634 | 0 | -3.2518 |
| ZNF395 | 6 | 0.00016537 | 0.00090094 | 0.352558 | 51 | 6 | -2.1948 | 0.99983 | 0.99982 | 1 | 21635 | 0 | -2.1948 |
| CDKN2AIP | 5 | 0.00014926 | 0.00069565 | 0.309052 | 48 | 5 | -4.0187 | 0.99985 | 0.99986 | 1 | 21636 | 0 | -4.0187 |
| CEP97 | 5 | 0.00014377 | 0.00066456 | 0.309052 | 47 | 5 | -5.3699 | 0.99986 | 0.99987 | 1 | 21637 | 0 | -5.3699 |
| SCAPER | 4 | 0.0001272 | 0.00049585 | 0.275324 | 41 | 4 | -5.615 | 0.99987 | 0.99988 | 1 | 21638 | 0 | -5.615 |
| LOXL3 | 5 | 0.00012148 | 0.00055346 | 0.285361 | 39 | 5 | -2.8118 | 0.99988 | 0.99989 | 1 | 21639 | 0 | -2.8118 |
| COPS8 | 5 | 0.00011968 | 0.0005466 | 0.285361 | 38 | 5 | -7.6318 | 0.99988 | 0.99989 | 1 | 21640 | 0 | -7.6318 |
| SRSF6 | 2 | 0.000072568 | 0.00015294 | 0.157291 | 35 | 2 | -10.674 | 0.99993 | 0.99992 | 1 | 21641 | 0 | -10.674 |
| PPP2R1A | 5 | 0.000048203 | 0.00022106 | 0.177301 | 26 | 5 | -8.5245 | 0.99993 | 0.99994 | 1 | 21642 | 0 | -8.5245 |
| MORC2 | 6 | 1.1126E-09 | 2.2861E-07 | 0.001238 | 2 | 6 | -10.035 | 0.99994 | 0.99993 | 1 | 21643 | 0 | -10.035 |
| NEK4 | 6 | 0.000062802 | 0.00033445 | 0.213017 | 32 | 6 | -4.8495 | 0.99994 | 0.99993 | 1 | 21644 | 0 | -4.8495 |
| U2AF1 | 5 | 0.000051696 | 0.00023798 | 0.18112 | 28 | 5 | -6.9974 | 0.99995 | 0.99996 | 1 | 21645 | 0 | -6.9974 |
| SP3 | 6 | 0.000050345 | 0.00027456 | 0.198185 | 27 | 6 | -4.5931 | 0.99995 | 0.99995 | 1 | 21646 | 0 | -4.5931 |
| CNOT1 | 6 | 0.000034351 | 0.00019775 | 0.177301 | 22 | 6 | -5.1293 | 0.99997 | 0.99996 | 1 | 21647 | 0 | -5.1293 |
| NELFE | 6 | 0.000031996 | 0.00018494 | 0.174128 | 20 | 6 | -6.3119 | 0.99997 | 0.99996 | 1 | 21648 | 0 | -6.3119 |
| TRIM24 | 6 | 7.9992E-07 | 3.4291E-06 | 0.008251 | 8 | 6 | -7.262 | 0.99998 | 0.99998 | 1 | 21649 | 0 | -7.262 |
| ACTR1A | 6 | 0.000010038 | 0.000060581 | 0.087459 | 14 | 6 | -7.3599 | 0.99999 | 0.99999 | 1 | 21650 | 0 | -7.3599 |
| DHX29 | 5 | 2.1351E-07 | 6.8582E-07 | 0.00297 | 6 | 5 | -8.1382 | 1 | 1 | 1 | 21651 | 0 | -8.1382 |
| WDR82 | 6 | 1.6323E-09 | 2.2861E-07 | 0.001238 | 3 | 6 | -11.387 | 1 | 1 | 1 | 21652 | 0 | -11.387 |
| TRIM33 | 6 | 2.7824E-08 | 2.2861E-07 | 0.001238 | 4 | 6 | -10.338 | 1 | 1 | 1 | 21653 | 0 | -10.338 |
| ATF7IP | 6 | 1.6724E-07 | 0.000001143 | 0.004125 | 5 | 6 | -6.855 | 1 | 1 | 1 | 21654 | 0 | -6.855 |
| MPHOSPH8 | 6 | 1.0961E-15 | 2.2861E-07 | 0.001238 | 1 | 6 | -12.339 | 1 | 1 | 1 | 21655 | 0 | -12.339 |
